# Oleic acid Induces Tissue Resident FoxP3 Regulatory T cell Lineage Stability and Suppressive Functions

**DOI:** 10.1101/2020.04.14.041525

**Authors:** Saige L. Pompura, Allon Wagner, Alexandra Kitz, Jacob LaPerche, Nir Yosef, Margarita Dominguez-Villar, David A. Hafler

## Abstract

FoxP3 positive regulatory T cells (T_regs_) rely on fatty acid β-oxidation (FAO)-driven OXPHOS for differentiation and function. Recent data have demonstrated a role for T_regs_ in the maintenance of tissue homeostasis with tissue-resident T_regs_ possessing tissue-specific transcriptomes. However, specific signals that establish these tissue-resident T_regs_ programs are largely unknown. As T_regs_ metabolically rely on FAO, and considering the lipid-rich environments of tissues, we hypothesized that environmental lipids drive T_reg_ homeostasis. Using human adipose tissue as a model for tissue residency, we identify oleic acid as the most prevalent free fatty acid in human adipose tissue. Mechanistically, oleic acid amplifies T_reg_ FAO-driven OXPHOS metabolism, creating a positive feedback mechanism that induces the expression of Foxp3 and enhances phosphorylation of STAT5, which acts to stabilize the T_reg_ lineage and increase suppressive function. Comparing the transcriptomic program induced by oleic acid to that of the pro-inflammatory arachidonic acid, we find that T_regs_ sorted from peripheral blood and adipose of healthy donors transcriptomically resemble the oleic acid *in vitro* treated T_regs_, whereas T_regs_ obtained from the adipose tissue of relapsing-remitting MS patients more closely resemble an arachidonic acid profile. Finally, we find that oleic acid concentrations are reduced in the fat tissue of MS patients, and exposure of dysfunctional MS T_regs_ to oleic acid restores defects in their suppressive function. These data demonstrate the importance of fatty acids in regulating tissue inflammatory signals.

## Introduction

Metabolic signatures of T cells are intricately linked to their differentiation and activation status. Metabolic pathways facilitate cellular functions, therefore linking metabolic remodeling to the development, activation, differentiation, and survival of T cells. Upon activation, quiescent, naïve T cells become rapidly dividing effector T (T_eff_) cells and switch their metabolic program from oxidative phosphorylation (OXPHOS) to aerobic glycolysis, known as the Warburg effect, in order to meet the increase in demand for cellular energy and biomass (*1, 2*). However, despite having similar developmental origins, regulatory T cells (T_regs_) rely predominantly on a fatty acid β-oxidation (FAO)-driven OXPHOS metabolic program to maintain their suppressive phenotype, which is further promoted by the expression of FoxP3 (*3–5*). Forced expression of FoxP3 in T cells suppresses glycolysis-related genes while inducing lipid and oxidative metabolic-related genes that are required for maximum suppression (*4*). In addition, cytokines that promote T_reg_ differentiation, such as TGF-β (*6*), activate AMPK (*7*) and promote FAO to skew naïve T cells into a T_reg_ phenotype (*3, 8*). Furthermore, T_reg_ differentiation and suppression are reduced by inhibiting FAO (*3*), highlighting the importance of FAO-driven OXPHOS in the initiation and maintenance of the T_reg_ phenotype.

The suppressive function of T_reg_ cells is critical for controlling immune responses and preventing autoimmunity. In this regard, we have identified functional T_reg_ defects in patients with autoimmune disease (*9*). However, a second critical function for T_regs_ is the regulation of tissue homeostasis that can secondarily impact organismal biology. Studies have shown that in mice and humans, T_regs_ infiltrate tissues not only during inflammatory conditions or injury, but also during homeostasis (*10–12*), and reside within tumors (*13–16*). Resident T_regs_ adapt to perform tissue-specific functions in order to regulate inflammation and perform homeostatic functions, such as wound repair and maintain metabolic indices (*10, 12, 17–20*). Specifically, in the visceral adipose tissue (VAT) which comprises fat depots that surrounds internal organs and is a site of molecular crosstalk between the immune and metabolic systems, VAT-resident T_regs_ possess a unique epigenetic and transcriptional profile. This allows T_reg_ survival in lipotoxic environments and enables the utilization of FAO as metabolic fuel (*10, 21*). Importantly, VAT-T_regs_ acquire expression of PPARγ, a master regulator of adipocytes, which is critical for VAT-T_reg_ function and has been shown to interact with FoxP3 (*22, 23*). Despite these observations, the signals that act to balance canonical T_reg_ function and T_reg_ adaptations of tissue-specific signals remain unknown.

Dissecting the signals that act to either promote or inhibit T_reg_ adaptation in tissues are essential to our understanding of tissue-T_reg_ biology. An example are signals that prevent loss of FoxP3 or promote generation of Th-like T_regs_ (*9, 24–28*). While both local antigens and cytokine milieu play a role in tissue adaptation (*29*), we hypothesized that environmental cues, such as lipids, may be critical considering their role in shaping the T_reg_ metabolic-functional axis. Here we identify oleic acid as the most prevalent free fatty acid in human adipose tissue and dissect its involvement in the maintenance of T_reg_ function. Oleic acid amplifies T_reg_ FAO-driven OXPHOS metabolism, creating a positive feedback mechanism that induces the expression of Foxp3 and enhances phosphorylation of STAT5, which acts to stabilize the T_reg_ lineage and increase suppressive function (*30–38*). We compare the transcriptomic program induced by oleic acid to that of the pro-inflammatory arachidonic acid (*39, 40*), and derive a computational transcriptome signature to quantify the similarity of the T_reg_ RNA profile to either state. Interestingly, we find that T_regs_ sorted from peripheral blood and adipose of healthy donors transcriptomically resemble the oleic acid *in vitro* treated T_regs_, whereas T_regs_ obtained from the adipose tissue of relapsing-remitting MS patients more closely resemble the arachidonic acid treated profile. A similar trend is observed in the comparison of treated and untreated MS groups. Finally, we find that oleic acid concentrations are reduced in the fat tissue of MS patients, and that exposure of dysfunctional MS T_regs_ to oleic acid partially restores their suppressive function, highlighting the importance of fatty acids in regulating tissue inflammatory signals.

## Results

### Oleic acid upregulates fatty acid β-oxidation in T_regs_

In order to determine which free fatty acids T_regs_ may be responding to in human adipose tissue, we performed mass spectrometry on the supernatant from healthy human adipose tissue and paired serum samples. In agreement with previous reports (*41–43*), we found that oleic acid is the most abundant free fatty acid in adipose tissue (Figure 1A). Oleic acid is a monounsaturated, omega-9 long chain fatty acid (LCFA) that is found in most animal and vegetable sources. In animal tissues, oleic acid is one of the most abundant free fatty acids regardless of tissue or species (*41–44*), implying that tissue-resident T_regs_ are sensing oleic acid, and that oleic acid may be an important signal for tissue-resident T_regs_. LCFAs can alter cellular proliferation and viability in a dose dependent manner (*3, 45–51*). However, at a concentration of 10μM of oleic acid, we observe increases in *FOXP3* expression (Supplemental Figure 1A-B), but do not find any differences in T_reg_ viability (Supplemental Figure 1C) or proliferation (Supplemental Figure 1D) in the presence of either oleic acid or arachidonic acid compared to vehicle. We thus compared the effects of oleic acid with those of arachidonic acid, since arachidonic acid is known to induce a pro-inflammatory phenotype in T cells (*39, 40*). Furthermore, LCFA treatment does not affect *FoxP3* demethylation (Supplemental Figure 1E), suggesting that the stability of FoxP3 expression is unchanged under these conditions.

**Figure 1.**
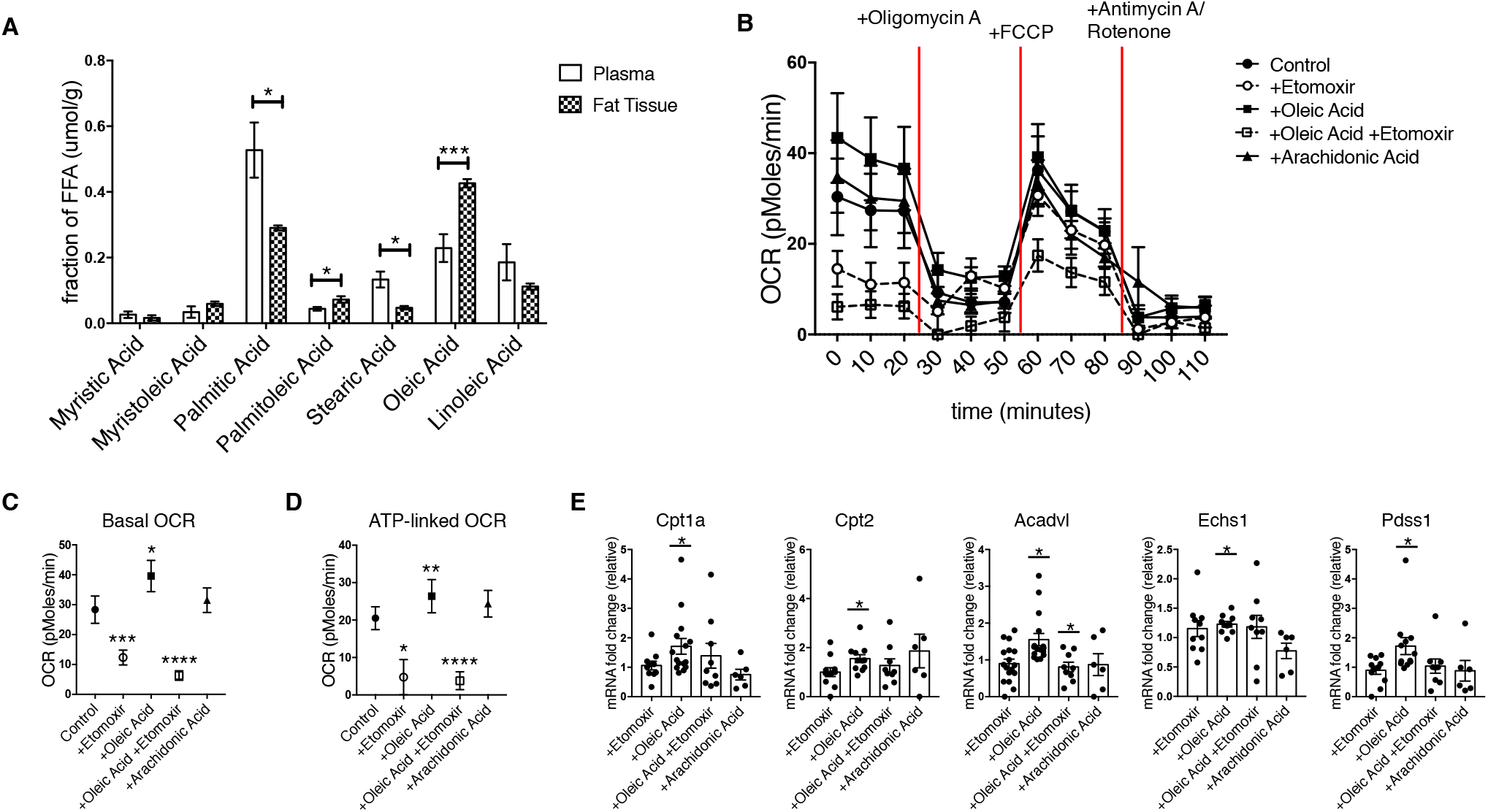
Oleic acid increases fatty acid β-oxidation in T_regs_. (**A**) Mass spectrometry analysis of long chain fatty acids in supernatant from human adipose samples compared to blood plasma (n=6) Paired t-test; \**P* < 0.05; \*\**P* < 0.01; \*\*\**P* < 0.001. (**B**) Oxygen Consumption Rate of T_regs_ stimulated in the presence or absence of Etomoxir (50μM), oleic acid (10μM), oleic acid and Etomoxir, or arachidonic acid (10μM) for 72hrs. n=12. Summary of (**C**) Basal oxygen consumption rate and (**D**) ATP-linked oxygen consumption rate of T_regs_ stimulated with same conditions as in (**B**). n=12. Paired t-test; \**P* < 0.05; \*\**P* < 0.01; \*\*\**P* < 0.001; **** P < 0.0001. (**E**) Fold change of mRNA expression in T_regs_ stimulated with conditions described in (**C**) for 72hrs. Paired t-test; \**P* < 0.05.

To determine whether FAO drives T_reg_ differentiation and function (*3–5*), we examined the metabolic phenotype of CD25^+^;CD127^lo/-^ T_regs_ in the presence of LCFAs. We first measured whether oleic acid is used as a metabolic substrate by T_regs_. We measured oxygen consumption rate (OCR) by Seahorse analysis as a measurement of OXPHOS, and found that oleic acid specifically increases OCR in T_regs_ (Figure 1B). T_regs_ cultured with oleic acid showed elevated basal respiration (Figure 1C) and ATP-linked oxygen consumption (Figure 1D), effects that were blocked by the inhibition of Cpt1a via Etomoxir, suggesting that the effects of oleic acid are mediated by its entry into FAO. Importantly, we also observed increases in mitochondrial mass (Supplemental Figure 2A) but not reactive oxygen species in the presence of oleic acid (Supplemental Figure 2B). Thus, the observed increases in OCR could be attributed to increases in total numbers of mitochondria rather than individual mitochondria increasing their respiratory rates. Basal respiration was found to be greater than or equal to maximal respiration rates (Figure 1B-C and Supplemental Figure 3A) in most donors, which is consistent with previous reports in human and murine T_regs_ (*48, 52–54*). We also observed a decrease in the overall spare respiratory capacity in the presence of oleic acid (Supplemental Figure 3B). However, the addition of Etomoxir to stimulated T_regs_ increased spare respiratory capacity (Supplemental Figure 3B), indicating that oleic acid-driven FAO might act to deplete this capacity in T_regs_.

We compared oleic acid and arachidonic acid and observed that while both increased basal respiration (Supplemental Figure 3C-D), only arachidonic acid increased ATP-linked respiration in CD127^+^;CD25^lo/-^ T_eff_ cells (Supplemental Figure 3E), both of which are negated by the addition of Etomoxir. Unlike T_regs_, the maximum oxygen consumption rate and spare respiratory capacity of T_eff_ cells was increased with oleic acid, also in a Cpt1a-dependent manner (Supplemental Figure 3F-G), while we observed no differences in mitochondrial mass (Supplemental Figure 2C) or reactive oxygen species (Supplemental Figure 2D). These data demonstrate that while both T_regs_ and T_eff_ are targeting lipids into FAO, they exhibit different capacities to respond to an excess of lipid uptake, and FAO inhibition by Etomoxir. These differences might reflect unique metabolic requirements of effector versus regulatory functions, or the inherent ability of T_reg_ versus T_eff_ cells to metabolically adapt to niche tissue environments.

We then measured the expression of mitochondrial metabolic and integrity genes in T_regs_ by quantitative PCR analysis and confirmed *CPT1A, CPT2, ACADVL, ECHS1, and PDSS1* are all upregulated in the presence of oleic acid (Figure 1E). In contrast, expression of *ACADVL and PDSS1* are dependent on FAO, as their expression was downregulated by the addition of Etomoxir (Figure 1E). Conversely, no significant effects were observed in T_eff_ cells (Supplemental Figure 4). These data demonstrate that free fatty acids in the tissues may have differential effects on T_regs_ as compared to T_eff_ cells due to T_reg_ reliance on FAO-driven OXPHOS, which is supported by the uptake of extracellular lipids (*3, 55*). Of note, expression of *PPARG* and *ACACB*, which are measurements of global lipid metabolism and storage and fatty acid synthesis, respectively, trended upward in T_regs_ (Supplemental Figure 5) but downward in T_eff_ cells (Supplemental Figure 4). These data provide further evidence that the two cell types are utilizing oleic acid differently. Taken together, our data show that oleic acid drives a unique gene expression profile in T_regs_, characterized by the upregulation of FAO-driven OXPHOS.

Previous reports have found human T_regs_ to be engaged in both FAO and glycolysis to support their expansion (*52, 56*). To examine whether LCFAs are specifically upregulating FAO and not simply increasing activation status of cells, we measured glycolysis activity via extracellular acidification rate by Seahorse analysis. We did not find increases in T_reg_ (Supplemental Figure 6A-B) or T_eff_ (Supplemental Figure 6C-D) cells cultured with LCFAs suggesting that exposure to LCFAs is not directly affecting cellular activation or expansion.

### Oleic acid drives T_reg_ suppression by upregulating fatty acid β-oxidation-dependent T_reg_ genes

In order to determine whether oleic acid influences T_reg_ function, we performed a suppression assay with T_regs_ pre-incubated with oleic acid, arachidonic acid, or IL-12 as a negative control (*9*). We found that oleic acid specifically increased T_reg_ suppression (Figure 2A-B), which is dependent on oleic acid-driven FAO, as pre-incubation with oleic acid and Etomoxir inhibited the increase in T_reg_ suppression as compared to the addition of oleic acid alone (Figure 2C-D).

**Figure 2.**
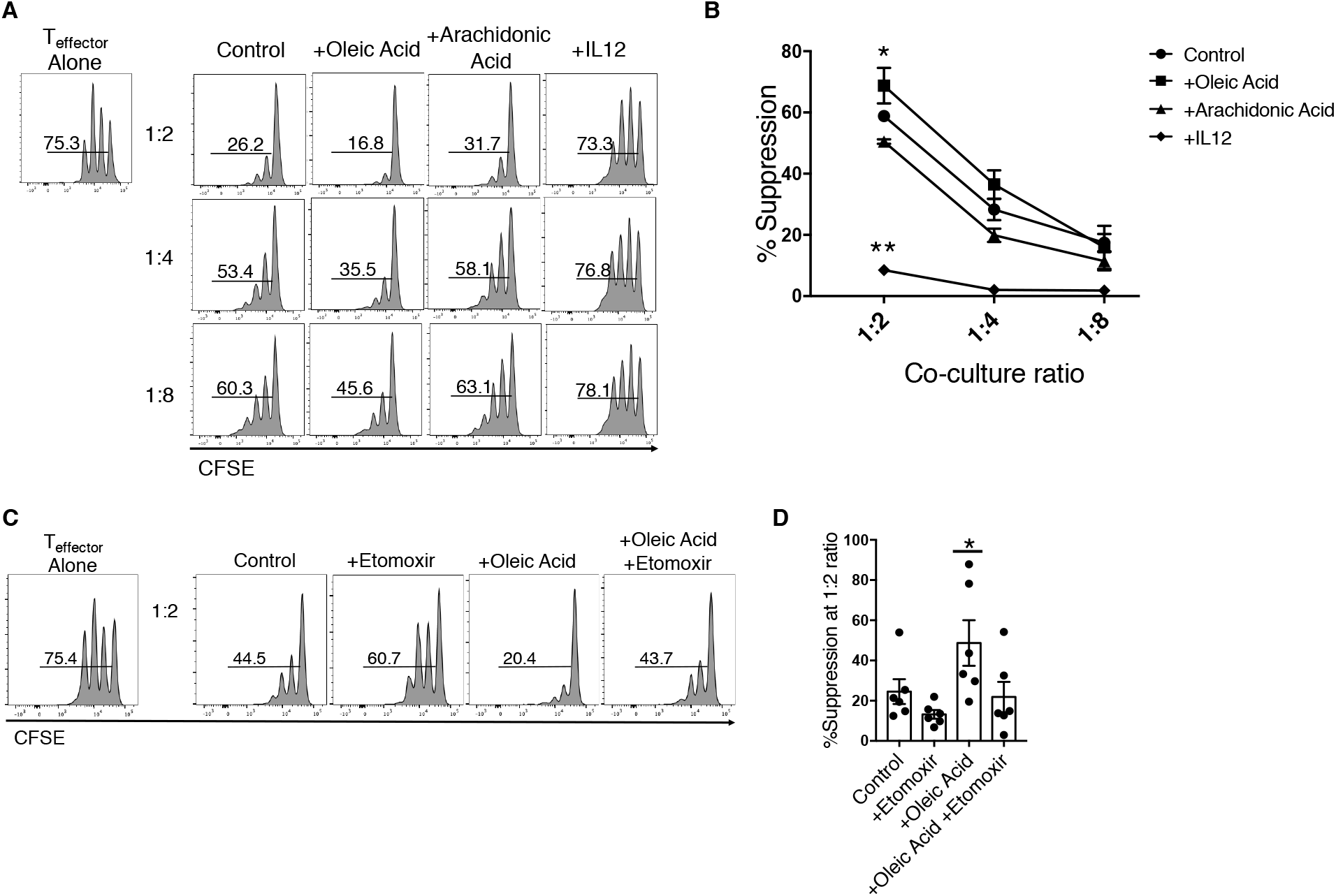
Oleic acid increases suppressive capacity of T_regs_. (**A**) Proliferation of T_eff_ cells, as measured by CFSE, that were cultured with sorted T_regs_ pre-incubated with vehicle (first column), 10μM oleic acid (second column), 10μM arachidonic acid (third column), or 25ng/mL IL-12 (fourth column) for 72hrs. Proliferation was measured after 4 days. n=10. Histograms are one representative experiment. (**B**) Summary of suppression assay described in (**A**), of 5 independent experiments. Paired t-test; \**P* < 0.05; \*\**P* < 0.01. (**C**) Proliferation T_eff_ cells, as measured by CFSE, that were cultured with T_regs_ pre-incubated with vehicle (first column), 50μM Etomoxir (second column), 10μM oleic acid (third column), Etomoxir and oleic acid (fourth column) for 72hrs. Proliferation was measured after 4 days. n=8. Histograms are one representative experiment. (**D**) Summary data of T_eff_ proliferation co-cultured with T_regs_ at 1:2 ratio. Paired t-test; \**P* < 0.05.

T_reg_ differentiation requires an intrinsic metabolic switch to FAO, and expression of FoxP3 promotes lipid and OXPHOS related gene expression (*3–5*). In order to understand how oleic acid-driven FAO enhances T_reg_ suppression, we measured protein expression of FoxP3 and phosphorylation status of STAT5 (p-STAT5), which promotes T_reg_ lineage stability via FoxP3 demethylation (*36–38*). We found that expression of *FoxP3*, and specifically *Foxp3* exon 2 (*57–59*) (Figure 3A and Supplemental Figure 7A), as well as p-STAT5, were increased in the presence of oleic acid (Figure 3B and Supplemental Figure 7B-C). These increases are dependent on oleic acid-driven FAO as the addition of Etomoxir reversed oleic acid-specific effects. Again, oleic acid did not increase expression of FoxP3 (Supplemental Figure 8A-B) or increase p-STAT5 (Supplemental Figure 8C-D) in T_eff_ cells. Furthermore, lentiviral knockdown of CPT1A (Figure 3C) and ACADVL (Figure 3D) both reduced the expression of FoxP3, further demonstrating that oleic acid-driven FAO is driving tissue resident T_reg_ function.

**Figure 3.**
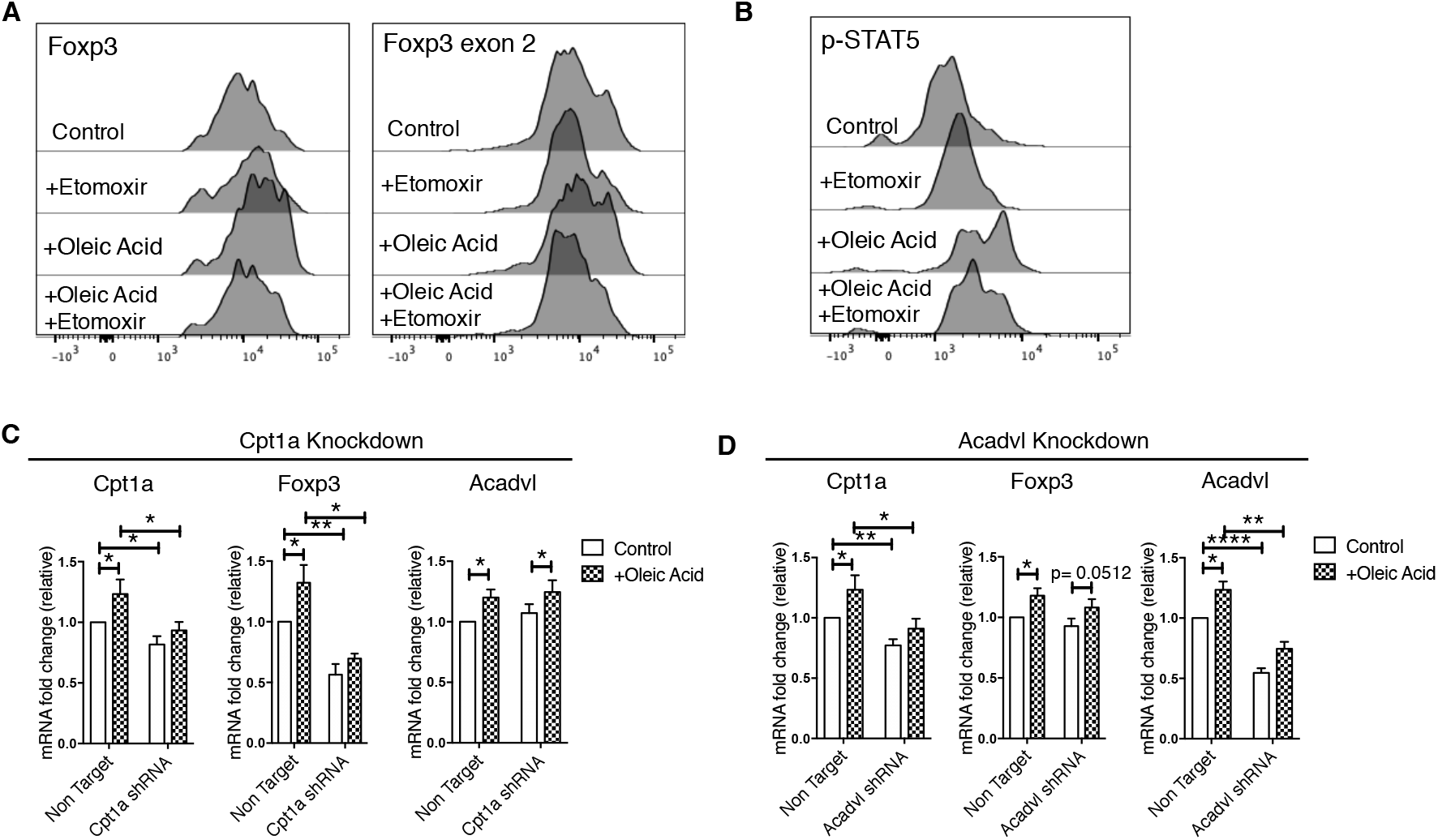
Oleic acid-driven fatty acid β-oxidation drives FoxP3 expression in T_regs_. (**A**) One representative experiment of FoxP3 protein expression in T_regs_ treated with indicated conditions after 72hrs. n=12. (**B**) One representative experiment of phosphorylation of STAT5 in T_regs_ after 6hrs of stimulation in the presence of indicated conditions. n=10. (**C**) mRNA expression of indicated genes in T_regs_ cultured in the presence of absence of 10μM oleic acid after lentiviral knockdown of either *Cptla* (top) or *Acadvl* (bottom). n=8. Paired t-test; \**P* < 0.05; \*\**P* < 0.01; \*\*\**P* < 0.001.

The enhanced p-STAT5 driven by oleic acid stimulation led us to investigate the expression of CD25 on T_regs_. We found at day three, oleic acid increased the total expression of CD25 on T_regs_, which could explain the increases observed in p-STAT5 (Supplemental Figure 9A-B), as IL-2 drives p-STAT5 (*36, 38*). Further, inhibition of different components of the electron transport chain inhibited the oleic acid-driven increase, and overall expression of CD25 (Supplemental Figure 9A-B) providing evidence that oleic acid-driven mitochondrial respiration drives the CD25-STAT5 axis and provides an additional layer of stabilization to FoxP3 and the T_reg_ lineage (*36, 38, 60*). In total, oleic acid-driven increases in FAO and mitochondrial metabolism drives T_reg_ suppressive functions through stabilization of the T_reg_ lineage by increasing p-STAT5 and upregulating FoxP3 expression.

### Global transcriptomic effects of oleic acid vs. arachidonic acid

Given the observed changes in mitochondrial respiration and suppressive function in T_regs_ specifically induced by oleic acid, we studied the effects of oleic acid on the T_reg_ transcriptional state. To do so, we isolated T_regs_ from peripheral blood of nine healthy donors, cultured them *in vitro* in the presence or absence of oleic or arachidonic acid, and performed bulk RNA sequencing (Supplemental Figure 10 and Supplementary Table 1). We observed considerable donor-specific effects that obstructed attempts to infer differentially expressed genes between the control and treatment groups through inclusion of the donor as a nuisance covariate of a generalized linear model (data not shown). We therefore took advantage of the experimental paired design by subtracting the high-dimensional gene expression vector of the vehicle state from either treatment, producing two vectors for each patient that capture the differential effect of oleic or arachidonic acid treatment (Methods). The first principal component (PC1) computed over this set of vectors captures the differential effect of the two fatty acids (Figure 4A; PC1 accounts for 24.7% of the variance). While the dynamic range as well as the magnitude of the PC1 effect are donor-specific, the overall trend captured by PC1 is robust for the entire donor group (Figure 4B). Reassuringly, the gene loadings on PC1 were also consistent with a gene-wise linear model designed to directly compare the effects of oleic vs. arachidonic acid (Supplemental Figure 11, Methods, Supplementary Table 2). Overall, this analysis indicates that despite the considerable donor-specificity in T_reg_ response to fatty acids, aggregation of the signal across multiple genes can successfully discern transcriptomic effect that is consistent across donors and is a primary source of variability in data.

**Figure 4.**
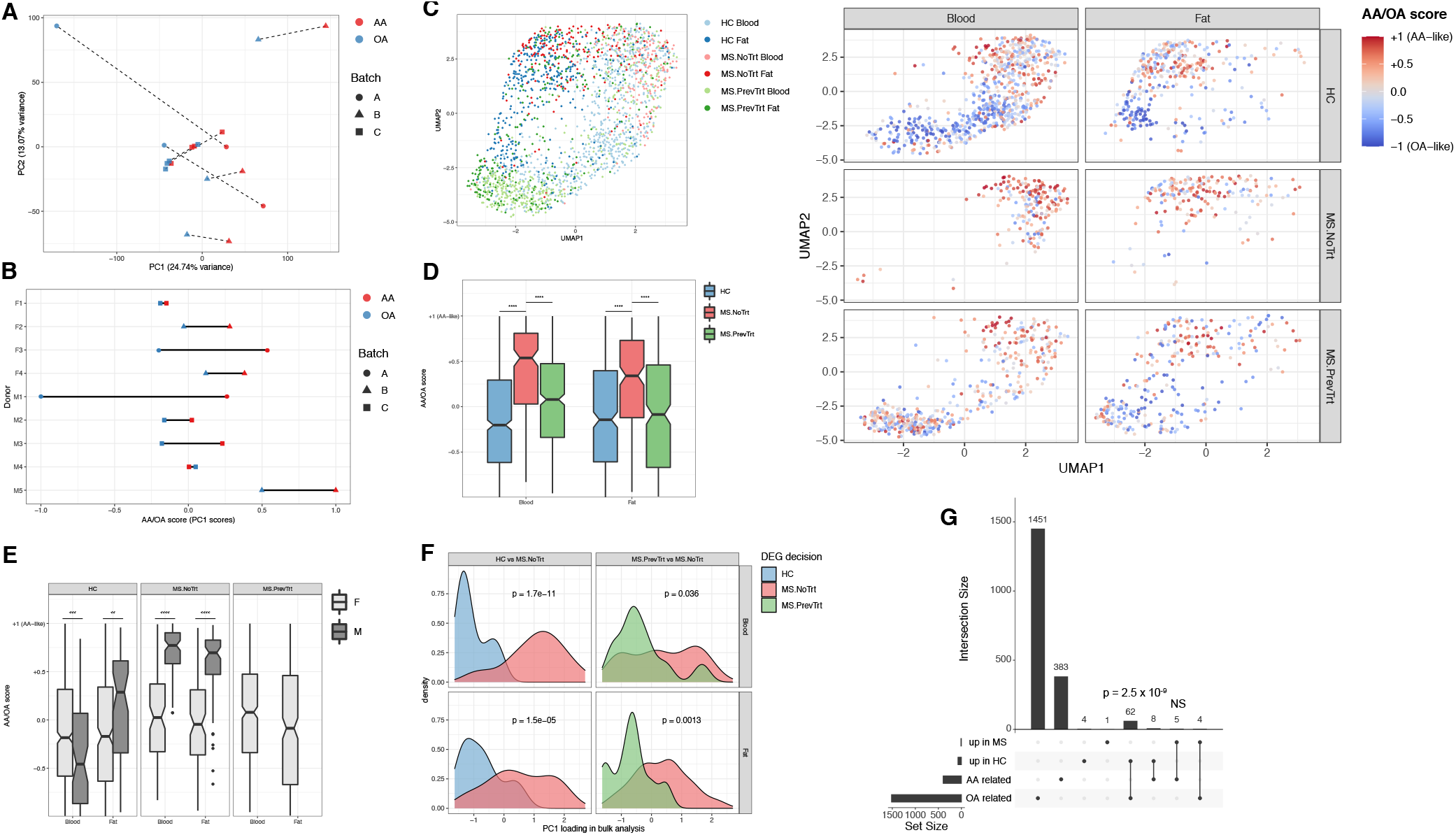
Oleic acid transcriptomic signature characterizes healthy but not MS T_regs_. (**A-B**) T_regs_ from peripheral blood of healthy donors were stimulated in the presence or absence of 10μM of oleic or arachidonic acid. Every donor is represented by two dots, corresponding to the difference between the oleic acid (OA) and control or arachidonic acid (AA) and control. (**A**) Principal component analysis, the two dots associated with each patient are connected by a dashed line. Donors were partitioned into 3 separate dates for tissue collection, denoted as batches (batch A took place during two consecutive days). (**B**) PC1 is scaled to [-1,1], every line represents one donor (donor’s sex indicated on the left). (**C**) Singlecell transcriptomes of T_regs_ were sorted from peripheral blood or adipose of healthy donors or MS patients and projected to a bi-dimensional space with UMAP (left). The same data is shown on the right, stratified by tissue and donor group. Colors indicate a computational signature of similarity to oleic or arachidonic acid stimulated blood T_regs_ (Main text and Methods; HC = healthy controls, MS.NoTrt and MS.PrevTrt = untreated and previously treated patients with MS as defined in the main text). (**D-E**) The computational signatures of single cells are aggregated and stratified by tissue, treatment group, and donor sex. Significance is indicated for a two-sided Welch t-test. (**F**) Left: densities of genes upregulated in healthy donors vs. untreated MS patients with respect to the loadings of PC1 shown in panel A. Right: similarly for a comparison of previously treated and untreated MS patients. Significance is indicated for a two-sided Welch t-test. (**G**) An upset plot (*127*) showing the overlap of differentially expressed genes upregulated in healthy or MS (aggregating previously treated and untreated patients) states by the in vivo single-cell RNA-Seq, and genes belonging to the oleic or arachidonic acid modules in the in vitro bulk RNA. P-value is for a hypergeometric enrichment test (NS = non-significant).

The loadings of PC1 rank genes with respect to their importance in capturing treatment effects. We defined the 250 genes with the lowest loading as the module of oleic acid related genes, and similarly, the 250 with the highest loadings as the module of arachidonic acid related genes. Notably, the oleic acid module was enriched for genes that related to long chain FAO, the enforcement of mitochondrial integrity, and promotion of T_reg_ generation. For example, *ACBD7, ACOT11, and ACSL6* have been shown to bind and convert long chain fatty acids to acyl-CoA for degradation by FAO (*61–64*). *LPIN3* has a general role in regulating fatty acid metabolism (*65*) and *LETM1* is thought to regulate mitochondrial dynamics and maintain normal mitochondrial morphology (*66*). Other genes upregulated in the oleic acid module included *SYAP1* and zinc finger protein, *ZFP30*, that have roles in adipogenesis (*67, 68*). We also observed upregulation of C5AR2, which has been shown to promote induced T_regs_ in human and murine models (*69*).

On the other hand, in the arachidonic acid module we observed an enrichment of genes related to both glycolysis and mitochondrial respiration, as well as genes that promote pro-inflammatory T_eff_ subsets. For instance, *PGK1* and *PGM1* catalyze the breakdown of glucose (*70–72*) and *SLC2A7* is a known glucose transporter (*73*). *ACAA2* catalyzes steps in FAO (*74*) and there were multiple genes upregulated (*NDUFA4, NDUFAF4, NDUFB11*, and *NDUFS5*) that correspond to accessory subunits of mitochondrial complex I (*75, 76*). Importantly, SIRT4, which regulates FAO via inhibition of PPARA transcription (*77*) and FABP5, which is a known intracellular lipid chaperone (*78, 79*) were upregulated as well. Furthermore, enrichment of CD69 and costimulatory factor CD9 would suggest arachidonic acid treated T_regs_ are more activated (*80–82*). Of note, we also detected genes that promote pro-inflammatory T cells. For example, CKS2, LDHA, and LDHB, promote Th17 differentiation (*83, 84*), and Th1 T_eff_ cells (*85*), respectively. LDHA is induced by T cell activation to support aerobic glycolysis, however, it also promotes IFNγ production via enhanced histone acetylation through acetyl-CoA (*85*). In contrast to genes enriched in the oleic acid module, arachidonic acid-related genes imply more activated T_regs_ that are potentially adopting a more pro-inflammatory phenotype.

### Adipose-derived T_regs_ from healthy subjects, not MS patients, are more similar to oleic acid treated T_reg_ profiles

To study the relevance of fatty acid milieu *in vivo*, we wanted to examine the transcriptional profile of T_regs_ isolated from the adipose tissue of healthy donors and patients diagnosed with an inflammatory autoimmune disease associated with altered T_reg_ function. Our lab has previously shown that T_regs_ isolated from patients with MS express IFNγ^+^ and are less suppressive *in vitro* (*9*). However, Th1-like T_regs_ have also been described in models of chronic infection (*86–88*) and patients with type one diabetes (T1D) (*26*). Furthermore, childhood obesity has been linked as a risk factor for the development of MS (*89–92*). Therefore, using MS as our model, we used single cell RNA-sequencing to profile 1,334 T_regs_ from peripheral blood and 805 T_regs_ from adipose tissue of eight healthy donors and eight MS patients, two of whom are untreated (> ~1.5yrs at the time of the procedure), and the rest had previously received disease modifying treatments (DMTs) (> 6 months prior to the procedure, but were currently off treatment) (Supplemental Figure 10, Supplemental Figure 12, and Supplementary Table 3). We assigned a quantitative score to each cell based on the PC1 axis we inferred from the *in vitro* data above (Figure 4C-D, Methods), that represents the similarity between the single cell’s transcriptome and the bulk dataset from either oleic or arachidonic acid treated blood T_regs_. The score ranges between −1 and +1, which represents more similarity to oleic acid or arachidonic acid, respectively.

T_regs_ collected from the adipose tissue of healthy donors had a significantly lower quantitative score than T_regs_ collected from the adipose of previously treated MS patients, which themselves had a significantly lower score than T_regs_ isolated from untreated MS patients (Figure 4D; two-sided Welch t-test). This suggests that oleic acid might be more prevalent in the lipid milieu encountered by T_regs_ in healthy individuals compared to subjects with MS. Moreover, immunotherapy may partially restore the transcriptomic T_reg_ state that is characteristic of a healthy donor, consistent with previous observations regarding the link between transcriptome restoration in response to treatment (*93*). However, these observations alone do not necessarily indicate a causative role for fatty acids in development of autoimmunity since the MS patients in our sample are on average older and have higher BMIs than the healthy donors. While the distribution of scores reflecting the similarity to oleic- or arachidonic-acid-treated T_regs_ was sex-specific (Figure 4E), the generalizability of these particular data is unclear due to the small number of donors in the current study.

### Significant overlap of oleic acid responsive genes and genes suppressed in MS

We further examined the relevance of the *in vitro-derived* oleic acid similarity score to human autoimmune disease by computing the set of differentially expressed genes between T_regs_ isolated from the blood and adipose tissues of healthy donors and T_regs_ isolated from the same tissues of untreated MS patients based on the single cell RNA-Seq (Supplementary Table 4, Methods). Indeed, we observed the genes that are upregulated in the healthy donors had significantly lower PC1 loadings with respect to the *in vitro* PC1 derived above (Figure 4F, left). A similar result was obtained comparing the previously treated and untreated MS patients (Figure 4F, right). Further, when comparing the healthy donors to the patients with MS (both previously treated and untreated), there existed a significant overlap between genes upregulated in the healthy state and genes belonging to the oleic acid gene module defined above based on the *in vitro* PC1 (Figure 4G; 1.65 fold, hypergeometric p = 2.5 x 10^−9^). In contrast, there was no significant overlap between genes upregulated in the MS state and the arachidonic acid module (hypergeometric p = 0.4). These data corroborate our findings that, in comparison to patients with MS, the transcriptional signature observed in T_regs_ isolated from the blood and adipose tissue of healthy donors is significantly more similar to blood T_regs_ stimulated with oleic acid than to those stimulated with arachidonic acid. This again suggests that T_regs_ in the blood and adipose of healthy donors might be exposed to a lipid milieu that contains greater amounts of oleic acid than in patients with MS. It also provides evidence that the exposure of peripherally-derived T_regs_ to oleic acid can, to a certain degree, recapitulate clinically-relevant aspects of the adipose-derived phenotype. Thus, oleic acid may counteract inflammatory signals in the tissues by reinforcing the canonical T_reg_ program and suppressive function.

### Oleic acid driven-fatty acid β-oxidation partially restores defective T_reg_ function

In order to address the physiological importance of oleic acid driving T_reg_ stability and function in the tissues and to address the hypothesis that lipids, as a metabolic cue, are able to buffer inflammatory signals, we cultured T_regs_ isolated from untreated (> ~1.5yrs at the time of the blood draw) MS patients with oleic acid. We found that oleic acid was able to partially restore T_reg_ suppression in comparison to T_regs_ from healthy, aged-matched controls (Figure 5A-B). Furthermore, oleic acid decreased the percentage of IFNγ^+^ while increasing the percentage of IL-10^+^ T_regs_ from MS subjects (Figure 5C-D). We also found that exposure to oleic acid upregulated *FOXP3* and *CPT1A* transcripts in MS T_regs_ (Figure 5E), suggesting oleic acid-driven FAO is negating T_reg_ dysfunction.

**Figure 5.**
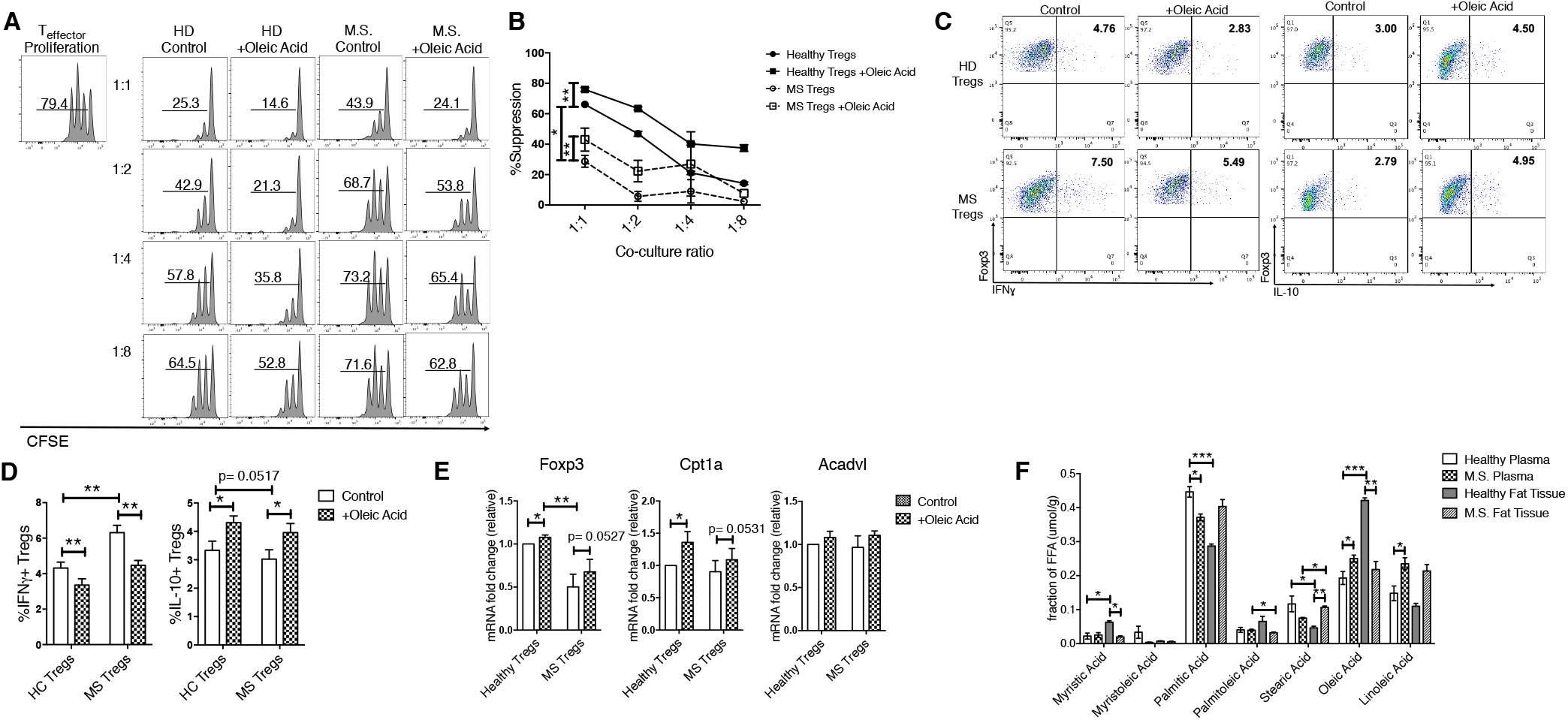
Oleic acid partially restores suppressive defects in MS T_regs_. (**A**) Proliferation of T_eff_ cells, as measured by CFSE, cultured T_regs_ sorted from frozen PBMC’s from either healthy or MS subjects. T_regs_ were pre-incubated with vehicle or 10μM of oleic acid for 72hrs and proliferation was measured after 4 days in co-culture. n=8. Histograms are one representative experiment. (**B**) Summary of 4 independent experiments, n=8. Paired t-test; \**P* < 0.05; \*\**P* < 0.01. (**C**) One representative dot plot of healthy or MS T_regs_ sorted from frozen PBMC’s, and stimulated in the presence or absence of 10μM oleic acid for 72hrs. Intracellular staining of IFNγ (left) or IL-10 (right) was measured after a 4hr stimulation of PMA and ionomycin in the presence of Golgistop. (**D**) Summary of IFNγ^+^ (left) and IL-10^+^ (right) healthy and MS T_regs_ after treatment with 10μM oleic acid for 3 days. n=8. Paired t-test; \**P* < 0.05; \*\**P* < 0.01. (**E**) mRNA expression of indicated genes measured in healthy or MS T_regs_ sorted from frozen PBMC’s, and stimulated in the presence or absence of 10μM oleic acid for 72hrs. n=10. Paired t-test; \**P* < 0.05; \*\**P* < 0.01. (**F**) Mass spectrometry analysis of long chain fatty acids in supernatant from human adipose samples compared to blood plasma in healthy or MS patients (n=6) Paired t-test; \**P* < 0.05; \*\**P* < 0.01; \*\*\**P* < 0.001.

To determine whether FAO-driven OXPHOS can influence T_reg_ IFNγ and IL-10 protein expression, we measured the percentage of IFNγ^+^ and IL-10^+^ T_regs_ isolated from healthy donors in the presence of electron transport chain inhibitors. We found that in the presence of Etomoxir, the percentage of IFNγ^+^ T_regs_ increase, suggesting FAO normally acts to inhibit IFNγ expression (Supplemental Figure 13A-B). Conversely, the percentage of IL-10^+^ T_regs_ decreases when Cpt1a and Complex I are inhibited by Etomoxir and Rotenone, respectively, indicating that FAO and the electron transport chain promote IL-10 expression (Supplemental Figure 13C-D). Together, these data provide evidence that the regulation of pro-versus anti-inflammatory cytokine expression is regulated in a FAO-dependent manner in T_regs_. This adds to previous reports showing specific electron transport chain complexes differentially regulate proliferation and cytokine expression in Th1 T_eff_ cells and provides evidence these two processes can be uncoupled metabolically (*94*).

To directly address whether T_regs_ resident in healthy adipose tissue are exposed to different concentrations of oleic acid compared to adipose-resident T_regs_ in MS patients, we measured the concentrations of LCFAs in the plasma and adipose supernatant of MS donors. We found that the fraction of oleic acid present in the adipose supernatant of MS patients was strikingly reduced compared to healthy donors, whereas in the plasma, oleic acid was increased in MS patients (Figure 5F and Supplemental Figure 14). Overall, we observed inverse trends in the fraction of oleic acid present in healthy versus MS tissue compartments, that is, in healthy donors more oleic acid is detected in adipose supernatant relative to plasma, but in MS subjects there are no significant differences. This could suggest either the existence of a basal state of inflammation or an inherent dysregulation of LCFA uptake and storage in MS patients. Further, we observed an overall change in LCFA composition between healthy and MS subjects (Figure 5F), consistent with an intrinsic defect in the regulation of LCFAs in the autoimmune, and in this case, the MS disease state.

## Discussion

Since the discovery of T_regs_ by Sakaguchi and co-workers in mice (*95*) and their identification in humans by our lab and others (*96–99*), this T cell lineage driven by the FoxP3 transcription factor has been thought of in the context of autoimmunity. That is, genetic deletion of FoxP3 results in spontaneous autoimmune disease in mice and humans (*100*). Additionally, defects in T_reg_ function have been observed in several common human autoimmune diseases (*9, 26, 101, 102*). However, more recent data has unveiled a new role for T_regs_ in the regulation of tissue homeostasis at multiple tissue sites, including skeletal muscle, intestines, skin, lung, adipose, and other organs (*103*).

Tissue-resident T_reg_ populations possess unique epigenetic and transcriptional profiles that allows them to fine-tune their tissue-specific functions (*10, 12, 104–106*). For example, T_regs_ in muscle expand upon injury, and play a role in the maintenance of homeostasis and tissue regeneration (*12, 107, 108*). In skin, T_regs_ colonize the skin barrier during neonatal development, and are necessary for preventing skin lesions, hypersensitivity and atopic dermatitis (*109–112*). In adipose tissue, VAT-resident T_regs_ are highly enriched within the VAT CD4^+^ T cell compartment (*10*) and possess a distinct repertoire of antigen-specific TCRs that exhibit clonal expansion in lean B6 mice (*113*), suggesting response to local antigens. Differentially expressed genes of VAT-T_regs_ encode for transcription factors, chemokines/chemokine receptors, cytokines/cytokine receptors, and genes related to lipid metabolism that allow survival in lipotoxic environments and enable the utilization of fatty acid oxidation as metabolic fuel (*21*). However, modes of T_reg_ metabolic adaptation to different tissue environments, and the signals that act to balance adaptation and the canonical T_reg_ program are unknown.

Here we show that oleic acid is the most prevalent LCFA in human adipose tissue and serves as a critical environmental signal that stabilizes FoxP3 and drives T_reg_ suppression by enhancing the FAO-OXPHOS metabolic program. Moreover, oleic acid-specific enhancement of FAO can partially restore T_reg_ suppression in patients with MS. LCFAs exert specific transcriptional effects in T_regs_, and the oleic acid-derived signature more closely resembles the expression profile of blood- and adipose-derived T_regs_ from healthy donors compared to donors diagnosed with MS. Interestingly, the transcriptomic analysis suggests that the balance of oleic and arachidonic acids in the extracellular environment modulates the T_reg_ phenotype. The high degree of variability with respect to both the range and magnitude of the LCFA effect that is donor-specific may be attributed to the highly variable nature of human donors, such as dietary or life-style patterns that might affect lipid uptake or metabolic adaptations of T_regs_.

We provide new evidence regarding the influence of environmental factors on tissue-resident immune cell function and how LCFA composition fluctuates between healthy and autoimmune disease states. First, we found that the gene expression profile of T_regs_ isolated from the adipose tissue of healthy donors is different from that of MS patients. Moreover, when we compare these signatures to data derived from peripheral blood T_regs_ exposed to oleic acid, we find that the oleic acid signature is more reflective of healthy adipose-resident T_regs_ than MS adipose-resident T_regs_. In contrast the pro-inflammatory arachidonic acid more closely resembles the T_regs_ isolated from MS adipose tissue. This is further supported by the significant overlap observed between genes upregulated in the healthy state and genes belonging to oleic acid treatment. These data demonstrate that exposure of peripherally-derived T_regs_ to oleic acid can partially recapitulate the transcriptional profile of adipose-resident T_regs_, and perhaps identify a new signal necessary for maintenance of the canonical T_reg_ program in tissue-resident T_regs_, especially during inflammation. In comparison, we did not observe the same trends with arachidonic acid, supporting our *in vitro* data that LCFAs might be metabolized differently, resulting in different functional effects. Arachidonic acid is known to be metabolized into pro-inflammatory lipid mediators, and further experiments are needed to understand how this lipid functions in T_regs_.

We hypothesized that tissue specific environmental signals allow both adaptation and acquisition of unique, tissue-specific functions, and stabilize and promote canonical T_reg_ functions. This is of importance as without this balance, T_regs_ can acquire Th-effector properties and lose their suppressive functions as seen in environments of chronic inflammation (*9, 26, 86–88*). Considering the lipid-rich environment of most tissues, we posit that environmental lipids, specifically oleic acid, are an important signal in striking a functional balance in tissue-resident T_regs_. As mentioned above, we demonstrate that healthy subjects and patients with MS have distinct lipid composition profiles in both the plasma and fat supernatant tissue compartments, suggesting that there could be an inherent defect in the ability to regulate LCFAs in MS adipose, or that there is a basal state of inflammation that contributes to the contrasting lipid profiles. However, the MS donors had a higher average BMI than healthy donors, so undoubtedly, dietary and other lifestyle choices could explain these differences as well.

We find that oleic acid is the most prevalent LCFA in healthy fat supernatant, but is reduced in patients with MS. Oleic acid engages a FAO-driven OXPHOS metabolic program in T_regs_ that reinforces canonical regulators of the T_reg_ lineage and T_reg_ suppressive function. The reduction of oleic acid might provide one mechanism by which T_regs_ in MS patients are more susceptible to dysfunction in environments of chronic inflammation, as the exposure to oleic acid partially restores suppressive function. Taken together, our data serve as a model for how environmental lipids acts to support T_reg_ function and phenotype during homeostatic and inflammatory conditions within tissues.

These investigations show the importance of FAO and OXPHOS in promoting T_reg_ survival and functions (*3*). However, recent reports revealed different metabolic requirements of T_regs_ during development versus those needed in established T_regs_ for proper function. In human T_regs_, OXPHOS and glycolytic engagement upon activation have been reported (*52, 56*), and it is known that glycolysis must be engaged to prevent enolase-1 suppression of FoxP3 (*114*). However, T_regs_ have lower measured ECAR relative to other Th-effector subsets (*115*), and CD45RO^+^ T_regs_ have greater mitochondrial mass relative to CD45RO^+^ T_eff_ cells (*56*). It can be considered that glycolytic restriction or FAO engagement favors T_reg_ development, then once the T_reg_ program is established glycolysis is re-engaged in order to preserve the phenotype. In this regard, enolase-1 has been shown to suppress FoxP3, and acetylation, a by-product of FAO and OXPHOS, enhances FoxP3 stability (*114, 116*). However, FAO-driven OXPHOS is still, nevertheless, the major metabolic program as we have provided evidence that it reinforces T_reg_ stability in existing T_reg_ populations.

Our data show that oleic acid drives FAO-driven OXPHOS in T_regs_, acting as a stabilizing factor of T_reg_ lineage, as it drives the expression of FoxP3, CD25, and p-STAT5, all of which act to reinforce the T_reg_ lineage by inducing demethylation of the *FoxP3* TSDR region. CD25 expression is also critical for T_regs_ development, allowing differentiation of thymic T_regs_, with recruitment of demethylation enzymes to CNS1 and CNS2 regions, creating a positive feedback loop that ensures FoxP3 expression and stability, especially under inflammatory conditions (*36, 38, 117–119*). CD25 also increases sensitivity to IL-2, which acts to increase T_reg_ lineage stability via STAT5 occupancy at the FoxP3 enhancer region and is a critical mechanism by which a stable T_reg_ phenotype is established, especially in the periphery (*36, 38*). Importantly, the increase of CD25 expression might also serve as a suppressive mechanism as loss of environmental IL-2 deprives T_eff_ cells of a crucial growth factor while simultaneously driving the T_reg_ lineage (*15, 120–122*).

In summary, we define a new mechanism by which environmental lipids drive cellular-specific metabolic programs that establish a positive feedback loop designed to enhance the stability and function of T_regs_ via the CD25-STAT5-FoxP3 axis. These signals act to balance inflammatory and tissue-specific cues so that the canonical T_reg_ phenotype and function can be maintained as these cells acquire tissue-specific plasticity. Moreover, oleic acid partially restores defects in suppressive function of T_regs_ isolated from patients with MS, which further suggests the importance of fatty acid species in counteracting inflammatory signals in the tissue. Investigating cross-talk between T_regs_ and tissue-resident cell populations, and the mechanisms by which T_regs_ metabolic programs orchestrate unique tissue-resident phenotypes and functions will impact our understanding of tissue resident T_regs_ development and perhaps treatment of autoimmune disorders associated with T_reg_ dysfunction.

## Materials & Methods

### Study Design

The objective of this study was to interrogate the role of environmental lipids in shaping the tissue-resident regulatory T cell phenotype. To do this, we used a combination of ex vivo computational analyses and in vitro experimental assays with human regulatory T cells isolated from peripheral blood and adipose tissue. We designed and performed the experiments mainly in the fields of cellular immunology and computational biology. The number of replicates for each experiment is indicated in the figure legends.

### Study subjects

Peripheral blood mononuclear cells (PBMCs) for suppression assays were cryopreserved from 8 Relapsing-Remitting MS (MS) patients (average age, 39 years; minimum 32 years; maximum 55 years) and 8 healthy individuals (average age, 29 years; minimum 21 years; maximum 35 years). The patients were diagnosed with either Clinically Isolated Syndrome (CIS) or MS by 2010 MacDonald Criteria and were not treated with any immunomodulatory therapy at the time of the blood draw (Supplementary Table 5). For bulk RNA sequencing, PBMCs were drawn for healthy individuals (average age, 25.4 years; minimum 21 years; maximum 33 years) (Supplementary Table 1). Adipose tissue biopsy healthy subjects were of a median age of 40.8 ± 12 years and BMI of 24.75 ± 3.16 Adipose tissue biopsies from MS subjects were a median age of 49.1 ± 9 years and a BMI of 29.5 ± 5.42, and off modulatory therapy for at least six months (Supplementary Table 3). These same samples were used to quantify long chain fatty acid concentrations by mass spectrometry. All experiments conformed to the principles set out in the WMA Declaration of Helsinki and the Department of Health and Human Services Belmont Report. Healthy individual PBMCs for experiments not matched with MS subjects PBMCs were freshly isolated.

### Human T Cell Isolation and Culture

Peripheral blood mononuclear cells (PBMCs) were isolated from donors by Ficoll-Paque PLUS (GE Healthcare) or Lymphoprep (Stemcell) gradient centrifugation. Total CD4^+^ T cells were isolated by negative magnetic selection using a CD4 T cell isolation kit (Stemcell) and CD4^+^CD25^hi^CD127^lo-neg^ T_reg_ cells were sorted on a FACS Aria (BD Biosciences). T_reg_ cells were cultured in RPMI 1640 medium supplemented with 5% Human serum, 2 nM L-glutamine, 5 mM HEPES, and 100 U/ml penicillin, 100 μg/ml streptomycin, 0.5 mM sodium pyruvate, 0.05 mM nonessential amino acids, and 5% human AB serum (Gemini Bio-Products). 96-well round bottom plates (Corning) were pre-coated with anti-human CD3 (UCHT1) (1 μg/ml) and used for T_reg_ in vitro culture with soluble anti-human CD28 (28.2) (1 μg/ml) (BD Bioscience) and human IL-2 (50 U/ml). Human IL-2 was obtained through the AIDS Research and Reference Reagent Program, Division of AIDS, National Institute of Allergy and Infectious Diseases (NIAID), National Institutes of Health (NIH). T_H_1-T_reg_ cells were induced with human recombinant IL-12 (20 ng/ml) (R&D). 10μM of free fatty acids solubilized by DMSO and conjugated in 250μM BSA-lipid free RPMI (Gibco 27016021): Oleic Acid (Sigma, O1008), Arachidonic Acid (Sigma, A3611). Metabolic inhibitors used were Etomoxir 50μM (Sigma, E1905), 2-Deoxy-D-glucose (2DG) 250μM (Sigma, D6134), 5-Aminoimidazole-4-carboxamide 1-β-D-ribofuranoside, Acadesine, N^1^-(β-D-Ribofuranosyl)-5-aminoimidazole-4-carboxamide (AICAR) 250μM (Sigma, A9978), Oligomycin A 1 μM (Sigma, 75351), 5-(Tetradecyloxy)-2-furoic acid (TOFA) 5μg/mL (Sigma, T6575), Rotenone 1 μM (Sigma, R8875), Antimycin A 1 μM (Sigma, A8674), and Dimethyl Malonate 10mM (Sigma, 136441)

### Suppression Assay

CD4^+^CD25^+^ T_reg_ cells were sorted from peripheral blood on a FACS Aria (BD Biosciences) and stimulated with anti-CD3 and anti-CD28 in the presence or absence of IL-12, Oleic Acid, Arachidonic Acid, or Etomoxir for 3 days, washed, and co-cultured with 10^4^ CFSE-labeled responder CD4^+^CD25^dim/low^CD127^+^ T cells at different T_reg_:Tresp ratios. The stimulus used was T_reg_ Inspector Beads (Miltenyi) at a 1:2 cell:bead ratio. At day 4, co-cultures were stained for viability with LIVE/DEAD^™^ Far Red Fixable Viability dye from (ThermoFischer) and fixed using FoxP3 staining buffer (eBioscience), and proliferation of viable responder T cells was analyzed on a Fortessa flow cytometer (BD Biosciences).

### Quantitative PCR

Total RNA was extracted using RNeasy Micro Kit (QIAGEN). RNA was treated with DNase and reverse transcribed using TaqMan Reverse Transcription Reagents (Applied Biosystems). cDNAs were amplified with Taqman probes (Taqman Gene Expression Arrays) and TaqMan Fast Advanced Master Mix on a StepOne Real-Time PCR System (Applied Biosystems) according to the manufacturer’s instructions. mRNA expression was measured relative to *B2M* expression. Values are represented as the difference in *C* _t_ values normalized to β2-microglobulin for each sample as per the following formula: Relative RNA expression = (2^−d*C*^_t_) × 1,000.

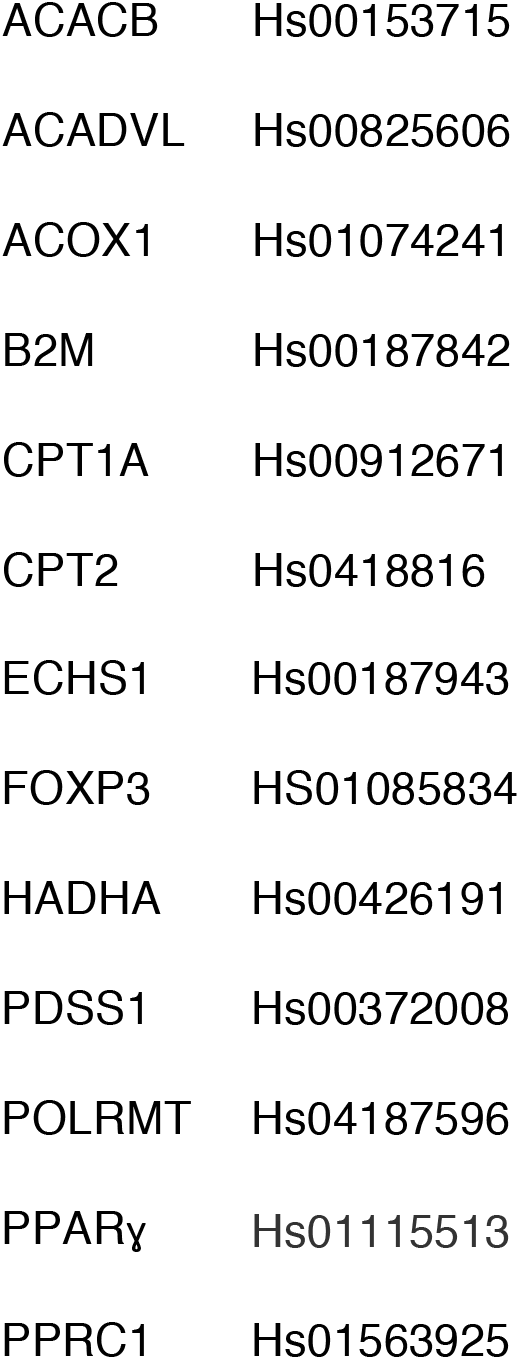

|  |  |
| --- | --- |
| ECHS1 | Hs00187943 |
| FOXP3 | HS01085834 |
| HADHA | Hs00426191 |
| PDSS1 | Hs00372008 |
| POLRMT | Hs04187596 |
| PPARγ | Hs01115513 |
| PPRC1 | Hs01563925 |

### Flow Cytometry Analysis

Cells were prepared from PBMCs and stained with fixable viability dye for 20 min at room temperature, followed by staining with surface antibodies for 30 min also at room temperature. For intracellular staining, cells were fixed and permeabilized with the Foxp3 Fix/Perm buffer set (eBioscience) for 20 min at room temperature, followed by staining with intracellular antibodies. For cytokine staining, cells were stimulated with phorbol-12-myristate-13-acetate (PMA) (50 ng/ml) and ionomycin (250 ng/ml) in the presence of GolgiStop (BD Bioscience) for 4 h at 37°C. Antibodies and reagents used for flow cytometric analysis are listed as follows: anti-CD25 PE (M-A251) BD Biosciences, anti-CD127 APC (HIL-7R-M21) BD Biosciences, anti-Foxp3 (PCH101) from eBioscience, anti-Foxp3 Exon 2 (AD2) Biolegend, anti-IFNγ (4S.B3) Biolegend, anti-IL-10 (JES3-9D7) Biolegend, anti-Ctla4 (L3D10) Biolegend, anti-Pd1 (NAT105) Biolegend, anti-Tigit (A15153G) Biolegend, anti-CD73 (HIL7R-M21) BD Bioscience, and LIVE/DEAD^™^ Far Red Fixable Viability dye from ThermoFischer. Stained samples were analyzed with a BD Fortessa flow cytometer (BD Bioscience). Data were analyzed with FlowJo software (Treestar).

### Phosphorylation Staining

Freshly isolated T_reg_ and T_eff_ cells were stimulated for indicated time points with 50 nM phorbol-12-myristate-13-acetate (PMA) (MilliporeSigma) and 250 nM ionomycin (MilliporeSigma) in specified conditions. After indicated time, cells were fixed with BD Cytofix buffer (BD Biosciences, 554655), followed by a 10-minute incubation at 37°C. Fixed cells were permeabilized with ice-cold BD PhosphoFlow Perm buffer III (BD Biosciences, 558050) and stained with the monoclonal antibodies αSTAT5 Tyr694 (eBioscience 11-9010-42). After 45-minute incubation, cells were washed and acquired with a BD Fortessa flow cytometer.

### Metabolic Assays

For ROS and mitochondrial mass, freshly isolated T_regs_ and T_eff_ cells were stimulated for 3 days as described above; in specified conditions. Cells were then stained with LIVE/DEAD^™^ Far Red Fixable Viability dye from ThermoFischer at 1:10,000x for 20 minutes at room temperature. Cells were then incubated with MitoTracker Green FM and MitoTracker^®^ Orange CM-H_2_TMRos at 250nM (Life Technologies) for 20 minutes at 37 °C for 20 minutes and then analyzed by flow cytometry. For the Seahorse assay, 400,000 freshly isolated T_regs_ or T_eff_ cells were stimulated for 3 days as described above; in specified conditions, and OCR and ECAR were measured on a Seahorse XF96 analyzer (Agilent) in the presence of the mitochondrial inhibitor oligomycin (1.5μM), mitochondrial uncoupler FCCP (1μM), and respiratory chain inhibitor antimycin A/rotenone (0.5μM).

### Proliferation Assay

Cells were sorted on a FACS Aria (BD Biosciences), spun down, and re-suspended at 10^6^ cells/ml in room temperature PBS, 0.1%BSA. CFSE was added to the cells at 1 μ L/mL, inverted to mix, and immediately incubated at 37°C for 5 min. Cells were then incubated in ice-cold RPMI complete media for 10 minutes. After, cells were spun, counted and re-suspended to desired numbers.

### FoxP3 Demethylation

The methylation status of the *FoxP3* gene was determined from DNA purified from frozen aliquots of sorted CD25^hi^;CD127^lo/neg^ T_regs_ cultured in the presence of anti-human CD3, soluble anti-human CD28, and IL-2 in the presence or absence of Oleic Acid and Arachidonic Acid as described above. Samples were sent to Epiontis (Berlin, Germany) for bisulfite modification and quantification of TSDR methylation by epigenetic human FoxP3 qPCR Assay.

### Lentiviral Transduction for shRNA Gene Silencing

Lentiviral plasmids encoding shRNAs were obtained from Sigma-Aldrich and all-in-one vectors carrying *CTNNB1* sgRNA/Cas9 with GFP reporter were obtained from Applied Biological Materials. Each plasmid was transformed into One Shot^®^ Stbl3^™^ chemically competent cells (Invitrogen) and purified by ZymoPURE plasmid Maxiprep kit (Zymo research). Lentiviral pseudoparticles were obtained after plasmid transfection of 293FT cells using Lipofectamine 2000 (Invitrogen). The lentivirus-containing media was harvested 48 or 72 h after transfection and concentrated 40 – 50 times using Lenti-X concentrator (Takara Clontech). Sorted T_reg_ cells were stimulated with plate-bound anti-CD3 (1 μg/ml) and soluble anti-CD28 (1 μg/ml) for 24 h and transduced with lentiviral particles by spinfection (1000 x *g* for 90 min at 32°C) in the presence of Polybrene (5 μg/ml) on the plates coated with Retronectin (50 μg/ml) (Takara/Clontech) and anti-CD3 (1–2 μg/ml). Human T_reg_ cells were directly transduced with lentiviral particles by spinfection. 24hrs after transduction, 10uM of oleic acid was added to culture. Five days after transduction, cells were sorted on the basis of expression of GFP and gene expression was measured using qPCR methods described above.

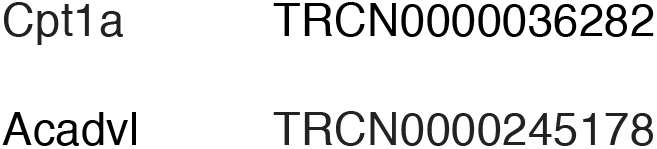

### Mass Spectrometry of Free Fatty Acids

Fatty Acid Concentrations: NEFA concentration in plasma and WAT (chloroform:methanol extraction 2:1) were determined using the WAKO HR Series NEFA-HR2 in vitro enzymatic colorimetric assay (FUJIFILM Wako Diagnostics, USA).

Fatty Acid Profiles: Plasma and WAT were extracted into chloroform:methanol (2:1) and NEFA were isolated with the use of a weak anion exchange column (Kinesis TELOS aminopropyl-NH2, 200mg, 3ml) for determination of fatty acid profile in the samples. (1) Neutral lipids were eluted with chloroform-2-propanol (2:1), and the NEFA fraction was then eluted with 2% formic acid in diethyl ether, dried under N2 gas, and derivatized to the fatty acid methyl ester with boron trifluoride methanol 14% (Sigma). Fatty acid methyl esters (C14 to C18) were determined by GC-MS analysis (CI mode), selective ion monitoring of masses 241 to 299 with an HP-1 column (25m, 0.2mm ID, 0.33μm film), with temperature gradient from 100 to 220 °C.

### Statistical analysis of non-transcriptomic assays

Significance was determined by a paired two-tailed Students’ *t* test. All statistical tests were performed with GraphPad Prism V8 software. Data are shown as means ± SEM. \**P* < 0.05; \*\**P* < 0.01; \*\*\**P* < 0.001; \*\*\*\**P* < 0.0001. Values of *P* <0.05 or less were considered significant.

### RNA-seq Library Preparation and Data Analysis

#### Preparation of cells for RNA-seq

For T_reg_ populations cultured with 10μM oleic acid, 10μM arachidonic acid, CD4^+^CD25^hi^CD127^lo/neg^ T_reg_ cells from healthy donors were sorted and cultured with specified conditions for three days as described above. Cells were harvested and RNA was isolated using RNeasy Micro Kit (QIAGEN), and immediately processed for cDNA preparation. Samples were collected from nine healthy subjects for identification free fatty acid-induced signatures.

#### cDNA and Library Preparation and Sequencing

Illumina TruSeq Stranded mRNA kit was used for library prep, and sequenced with a 2 × 100 bp paired-end protocol on the HiSeq 2000 Sequencing System (Illumina).

#### Bulk RNA-seq Data Analysis

Libraries were pseuoaligned with Kallisto v0.46.0 (*123*)to the Ensembl v96 human transcripome. Transcript TPMs were aggregated and summed for gene-level analysis, and then log2-transformed; non-protein-coding transcripts were excluded. This resulted in 3 gene expression vectors per donor, of T_regs_ stimulated in the presence of oleic acid, arachidonic acid, or none (Control). To mitigate donor-specific effects, we transformed them into two vectors by subtracting the control vector from either treatment (note that subtraction was done in log space, and it therefore represents the ratio between the original signals). Coordinates of these vectors were z-scaled for the purpose of PCA computation. An alternative analysis was performed by subtracting a donor’s arachidonic acid vector from his/her oleic acid vector. The limma R package (*124*) was used to fit a linear regression model, with the vector as the dependent variable, an intercept, and two sum-to-zero nuisance covariates to regress out the division of the donors into the 3 collection batches, followed by an empirical Bayes moderation of the gene-wise sample variances with mean-variance trend (the limma-trend method, as described in (*125*)). A moderated t-test was used to determine whether the intercept coefficient (which is supposed to capture the differential effect on that gene, comparing between oleic acid and arachidonic acid) was significantly different than 0.

#### Single-cell RNA-seq data analysis

Single-cell libraries were pseudoaligned as described above for bulk libraries and processed with the sctransform workflow of Seurat v3 (*126*). We identified non-T_regs_ cell clusters in the data, excluded them, and repeated the processing to eliminate their effect on the subsequent workflow. Differentially expressed genes were called with a Wilcoxon rank sum test as implemented in Seurat’s FindAllMarkers function with default parameters.

We defined a transcriptomic signature based on the in vitro bulk RNA data as follows. As described in the main text, we defined the top 250 genes with lowest (most negative) loadings with respect to PC1 of the bulk data the oleic acid related module, and the top 250 genes with highest loadings as the arachidonic acid module. The score of a single-cell is the dot product of its normalized gene expression vector with a vector that contains −1 for the oleic acid module genes, +1 for arachidonic acid module genes, and 0 otherwise. The resulting scores were rank-transformed and then scaled to a [−1,+1] range, with −1 representing the most oleic-acid-like and +1 the most arachidonic-acid-like cell in the data. Statistical significance is encoded by asterisks same as described for the non-transcriptomic assays.

## Acknowledgments

We would like to thank Gary Cline and the Mouse Metabolic Phenotyping Center at Yale University, School of Medicine for mass spectrometry analysis of fat supernatant and serum samples, and the Yale Center for Genome Analysis for cDNA library preparation and sequencing of samples. This work was supported by grants to D.A.H. from the National Institutes of Health (U19 AI089992, R25 NS079193, P01 AI073748, U24 AI11867, R01 AI22220, UM 1HG009390, P01 AI039671, P50 CA121974, R01 CA227473), the National Multiple Sclerosis Society (NMSS) (CA 1061-A-18, RG-1802-30153), the Nancy Taylor Foundation for Chronic Diseases, and Erase MS. This investigation was supported by NIH Training Grant from T32AI07019, Yale Interdisciplinary Immunobiology Training Program.

## Author Contributions

S.L.P performed in vitro experiments with the help of M.D.V.; A.W. and N.Y. analyzed the RNA-sequencing data; J.L. and A.K. designed protocols, collected, and sequenced human adipose tissue samples; S.L.P. performed data analysis and wrote the manuscript with A.W. under the supervision of N.Y., M.D.V and D.A.H.; and M.D.V. and D.A.H. supervised the overall study.

