## Supplementary Figures and Tables for "Oleic acid Induces Tissue Resident FoxP3 Regulatory T cell Lineage Stability and Suppressive Functions"

**Supplementary Figure 1. Oleic acid does not affect $T_{reg}$ viability or proliferation.**

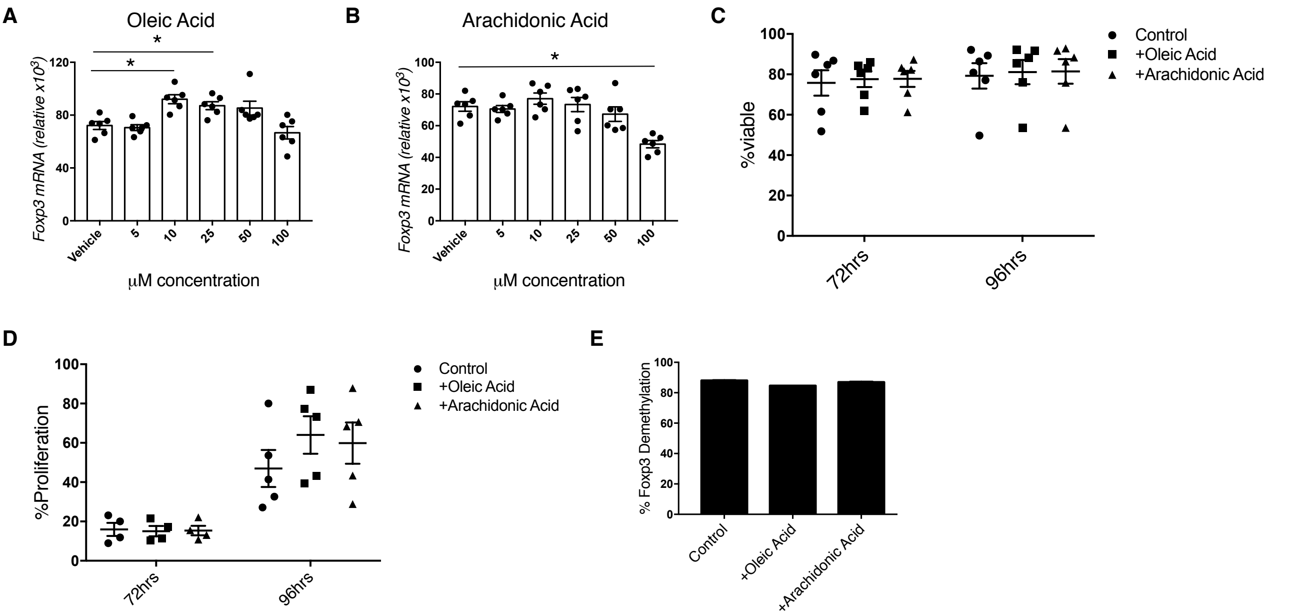

**Supplementary Figure 1. Oleic acid does not affect $T_{reg}$ viability or proliferation.** (**A**) $T_{regs}$ stimulated in the presence of increasing concentrations of Oleic Acid. Paired t-test; $*P < 0.05$. (**B**) $T_{regs}$ stimulated in the presence of increasing concentrations of arachidonic acid. Paired t-test; $*P < 0.05$. $T_{eff}$ and $T_{regs}$ stimulated in the presence or absence of 10μM Oleic or Arachidonic Acid. (**C**) Viability and (**D**) proliferation was measured after 72 and 96hrs. (**E**) $T_{regs}$ stimulated in the presence or absence of 10μM oleic or arachidonic acid and analyzed for bisulfite modification and quantification of TSDR methylation by epigenetic human FoxP3 qPCR Assay (Epiontis). n=6.

**Supplementary Figure 2. Oleic acid increases mitochondrial mass in $T_{regs}$.**

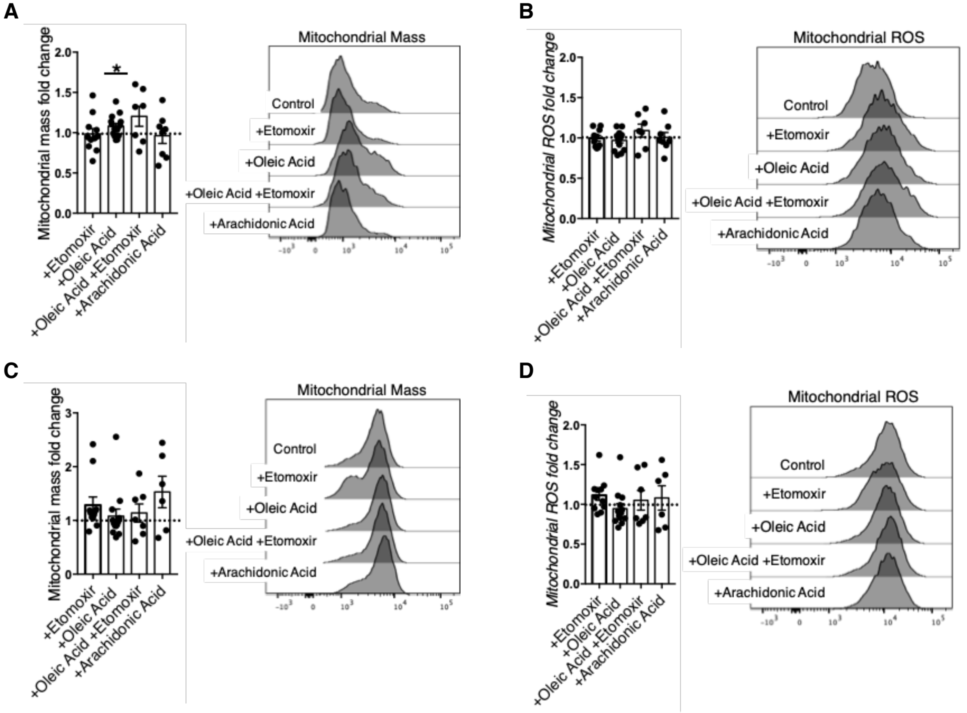

**Supplementary Figure 2. Oleic acid increases mitochondrial mass in $T_{regs}$.** (**A**) Mitochondrial mass and (**B**) reactive oxygen species measured in $T_{regs}$ stimulated in the presence or absence of Etomoxir (50μM), oleic acid (10μM), oleic acid and Etomoxir, or arachidonic acid (10μM) for 72hrs. Left is summary data, and right is one representative histogram. Paired t-test; $^*P < 0.05$. (**C**) Mitochondrial mass and (**D**) reactive oxygen species measured in $T_{eff}$ stimulated in the presence or absence of Etomoxir (50μM), oleic acid (10μM), oleic acid and Etomoxir, or arachidonic acid (10μM) for 72hrs. Left is summary data, and right is one representative histogram.

**Supplementary Figure 3. Oleic acid does not affect mitochondrial respiration in $T_{eff}$ cells.**

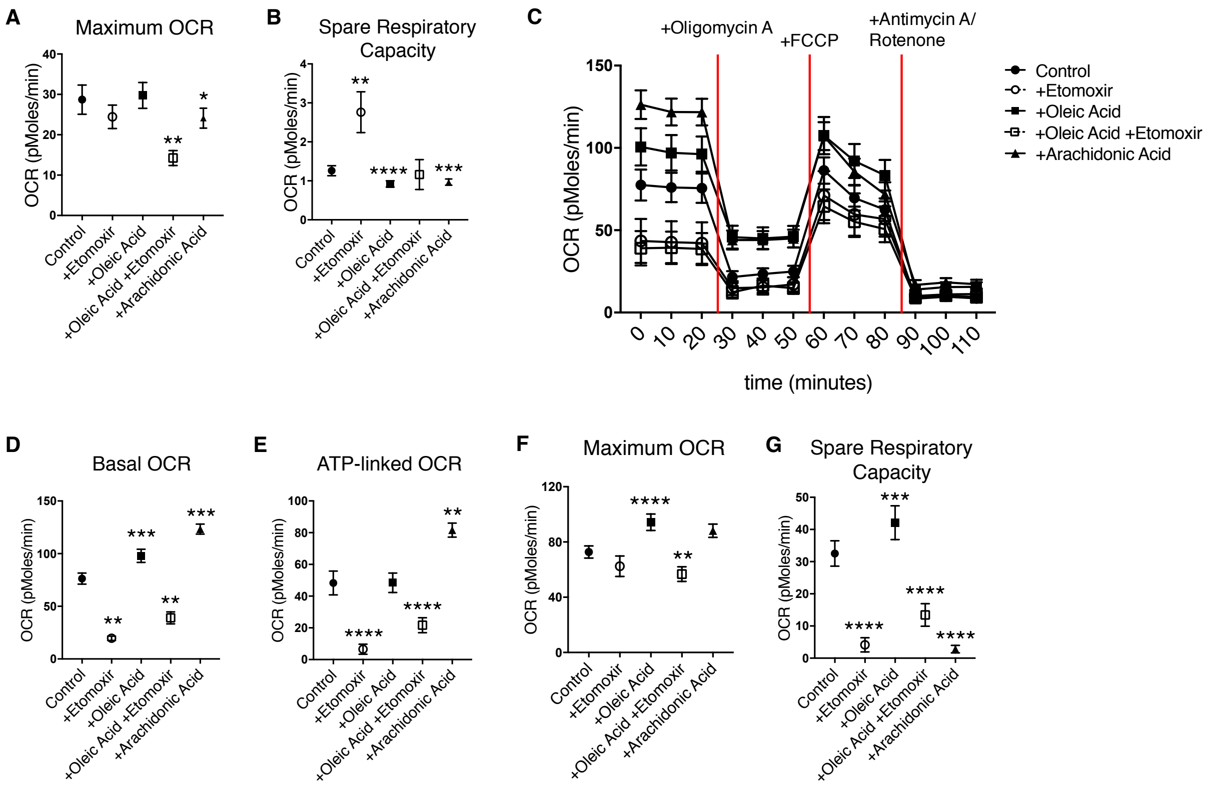

**Supplementary Figure 3. Oleic acid does not affect mitochondrial respiration in $T_{eff}$ cells.** (**A**) Maximum oxygen consumption rate and (**B**) spare respiratory capacity of $T_{regs}$ stimulated in the presence or absence of Etomoxir (50μM), oleic acid (10μM), oleic acid and Etomoxir, or arachidonic acid (10μM) for 72hrs. Graphs are a summary of n=12. Paired t-test; *$P < 0.05$; **$P < 0.01$; ***$P < 0.001$; **** $P < 0.0001$. (**C**) Oxygen consumption rate of sorted $T_{eff}$ stimulated in the presence or absence of Etomoxir (50μM), oleic acid (10μM), oleic acid and Etomoxir, or arachidonic acid (10μM) for 72hrs. n=12. Summary data of (**D**) Basal oxygen consumption, (**E**) ATP-linked oxygen consumption, (**F**) Maximum oxygen consumption, and (**G**) spare respiratory capacity of $T_{eff}$ stimulated with the same conditions as (**C**). n=12. Paired t-test; *$P < 0.05$; **$P < 0.01$; ***$P < 0.001$; **** $P < 0.0001$.

**Supplementary Figure 4. Oleic acid-induced expression of fatty acid oxidation-related genes in $T_{eff}$ cells.**

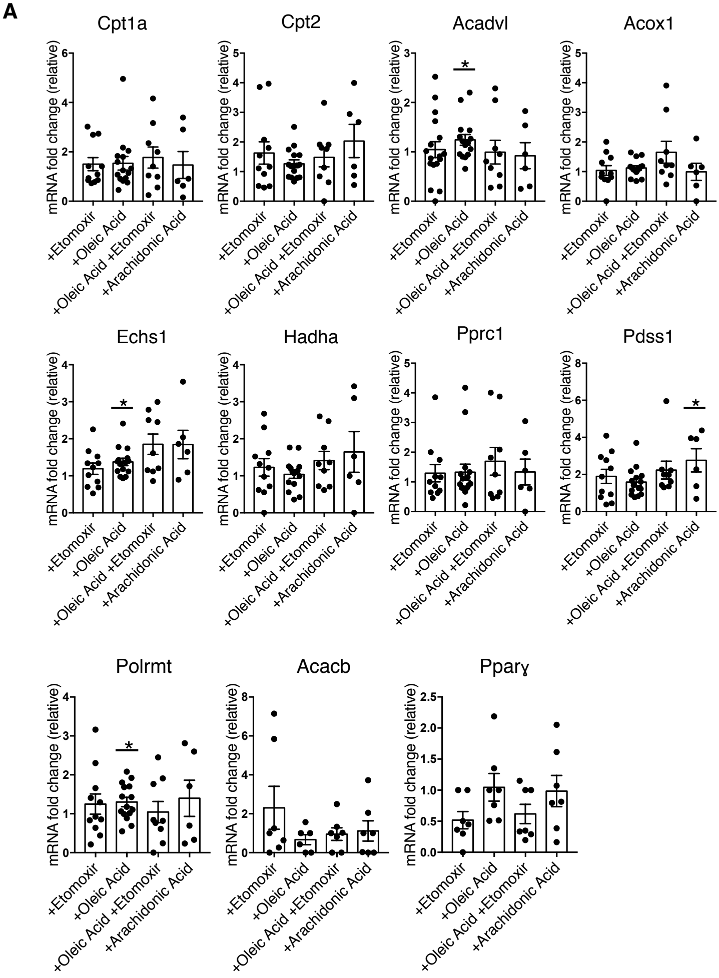

**Supplementary Figure 4. Oleic acid-induced expression of fatty acid oxidation-related genes in $T_{eff}$ cells.** (**A**) Fold change of mRNA expression of indicated genes in $T_{effs}$ stimulated in the presence or absence of Etomoxir (50μM), oleic acid (10μM), oleic acid and Etomoxir, or arachidonic acid (10μM) for 72hrs. Paired t-test; $^{*}P < 0.05$.

**Supplementary Figure 5. Oleic acid-induced expression of fatty acid oxidation-related genes in $T_{reg}$ cells.**

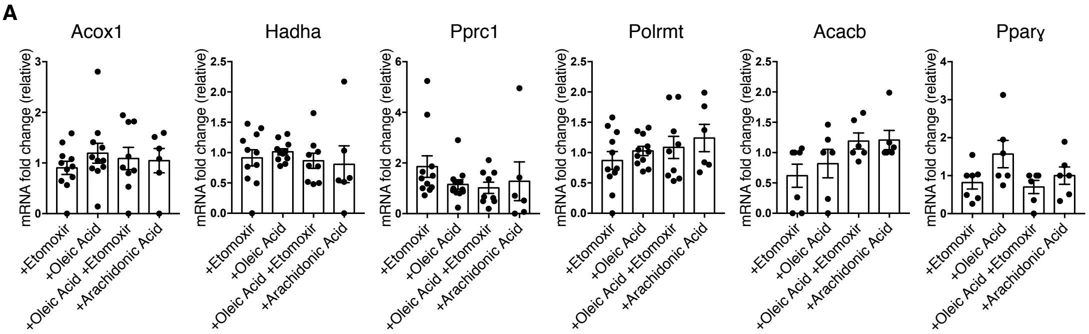

**Supplementary Figure 5. Oleic acid-induced expression of fatty acid oxidation-related genes in $T_{reg}$ cells.** (**A**) Fold change of mRNA expression of indicated genes in $T_{regs}$ stimulated in the presence or absence of Etomoxir (50μM), oleic acid (10μM), oleic acid and Etomoxir, or arachidonic acid (10μM) for 72hrs. Paired t-test; $*P < 0.05$.

**Supplementary Figure 6. Oleic acid does not drive ECAR in $T_{regs}$ or $T_{eff}$ cells.**

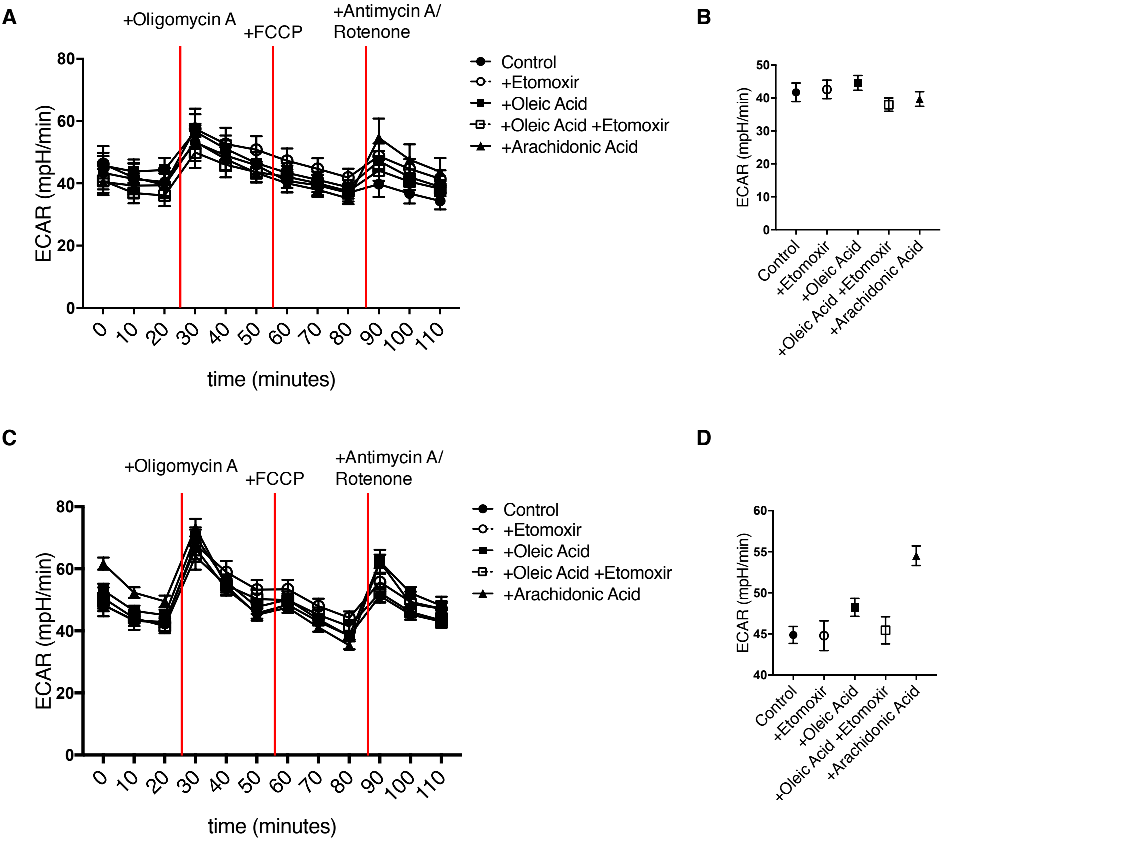

**Supplementary Figure 6. Oleic acid does not drive ECAR in $T_{regs}$ or $T_{eff}$ cells.** (**A**) Extracellular acidification rate of $T_{regs}$ stimulated in the presence or absence of Etomoxir (50μM), oleic acid (10μM), oleic acid and Etomoxir, or arachidonic acid (10μM) for 72hrs. n=12. Summary data of (**B**) Summary data of basal extracellular acidification rate in $T_{regs}$ stimulated with the same conditions as (**A**). (**C**) Extracellular acidification rate of $T_{eff}$ stimulated in the presence or absence of Etomoxir (50μM), oleic acid (10μM), oleic acid and Etomoxir, or arachidonic acid (10μM) for 72hrs. n=12. (**D**) Summary data of basal extracellular acidification rate in $T_{eff}$ stimulated with the same conditions as (**C**). Paired t-test; $*P < 0.05$; $**P < 0.01$; $***P < 0.001$.

**Supplementary Figure 7. Oleic acid-induced expression of Foxp3 and STAT5 phosphorylation in $T_{reg}$ cells.**

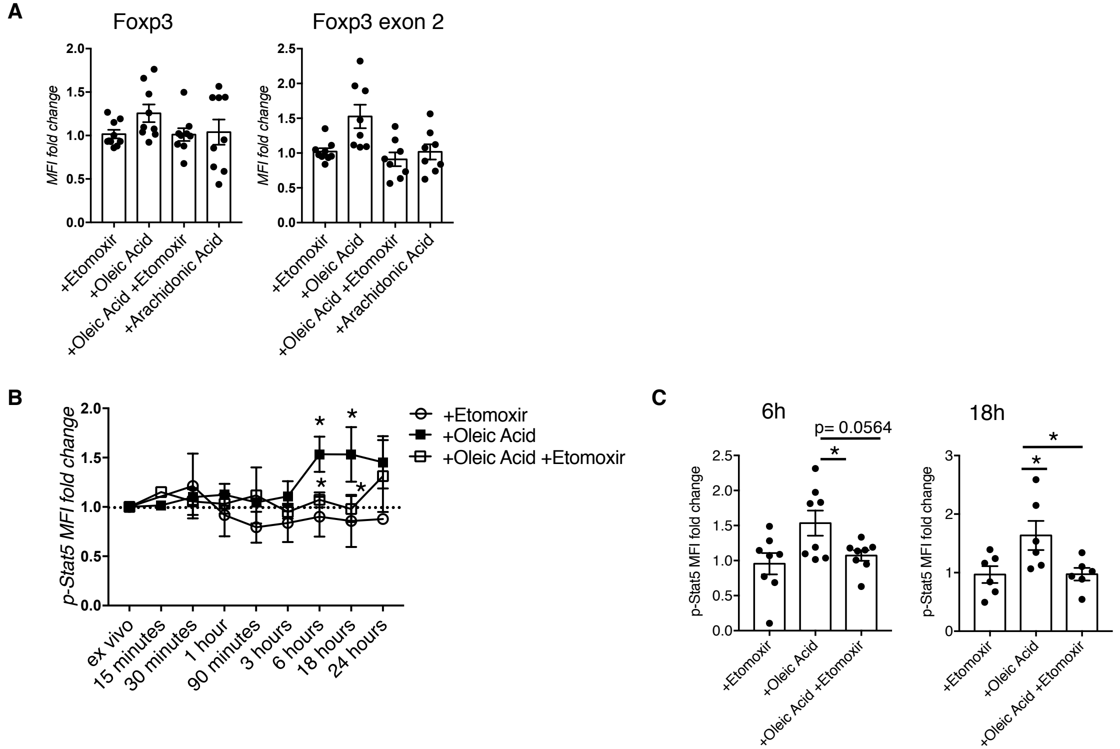

**Supplementary Figure 7. Oleic acid-induced expression of Foxp3 and STAT5 phosphorylation in $T_{reg}$ cells.** (**A**) Summary data for expression of FoxP3 or FoxP3 exon 2 mean fluorescence intensity in $T_{reg}$ cells stimulated in the presence or absence of Etomoxir (50μM), oleic acid (10μM), oleic acid and Etomoxir, or arachidonic acid (10μM) for 72hrs. Paired t-test; **P* < 0.05; ***P* < 0.01. (**B**) Kinetics of STAT5 phosphorylation in $T_{regs}$ stimulated with conditions described in (**A**). n=8 Paired t-test; **P* < 0.05. (**C**) One representative experiment of STAT5 phosphorylation in $T_{regs}$ after 6hrs (left) and 18hrs (right) of stimulation in the presence of conditions described in (**A**) Paired t-test; **P* < 0.05.

**Supplementary Figure 8. Expression of Foxp3 and phosphorylation of STAT5 are not upregulated by Oleic Acid in $T_{eff.}$**

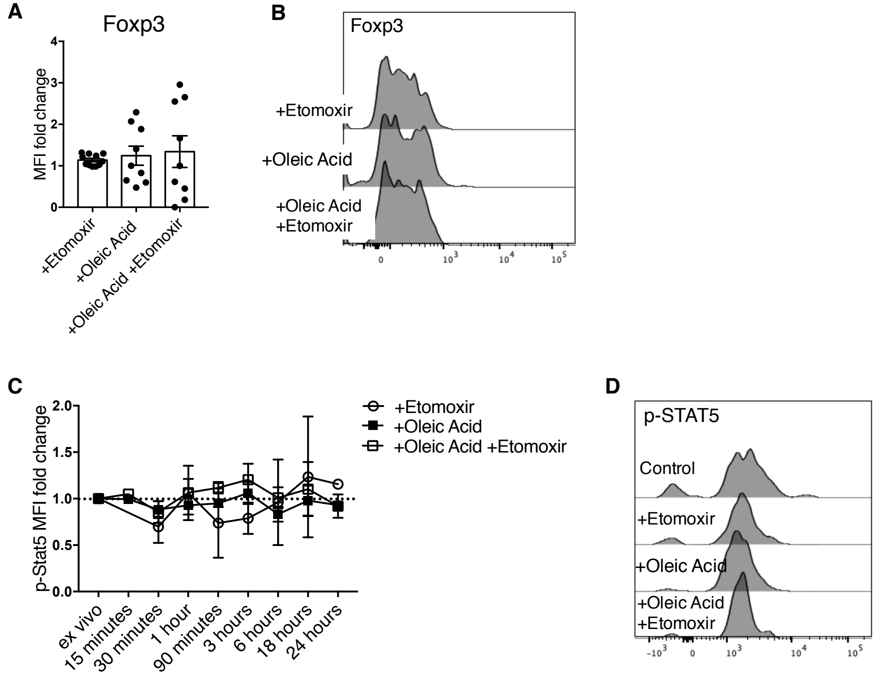

**Supplementary Figure 8. Expression of Foxp3 and phosphorylation of STAT5 are not upregulated by Oleic Acid in $T_{eff.}$** (**A**) Summary data for expression of Foxp3 mean fluorescence intensity in $T_{eff}$ cells stimulated in the presence or absence of Etomoxir (50$\mu$M), oleic acid (10$\mu$M), oleic acid and Etomoxir, or arachidonic acid (10$\mu$M) for 72hrs. (**B**) One representative experiment of protein expression in $T_{eff}$ treated with conditions specified in (**A**), after 72hrs. (**C**) Kinetics of STAT5 phosphorylation in $T_{eff}$ stimulated with conditions described in (**A)**. n=8 (**D**) One representative experiment of STAT5 phosphorylation in $T_{eff}$ after 6hrs of stimulation in the presence of conditions described in (**A**).

**Supplementary Figure 9. CD25 expression is dependent on mitochondrial respiration.**

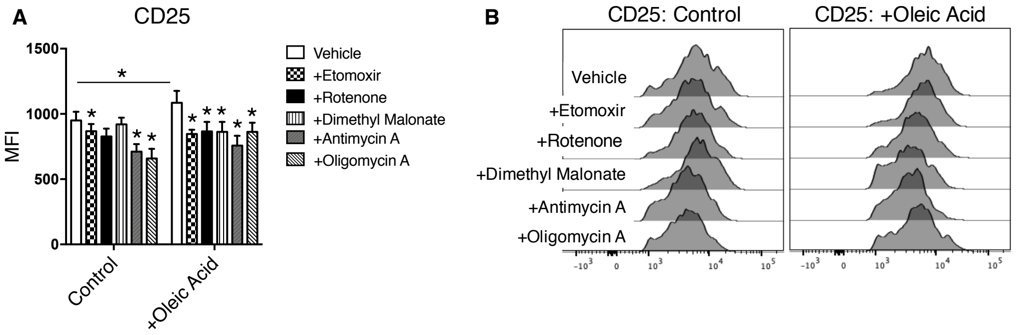

**Supplementary Figure 9. CD25 expression is dependent on mitochondrial respiration.** (**A**) Mean fluorescence intensity of CD25 $T_{regs}$ stimulated in the presence or absence of 50μM Etomoxir, 1μM Rotenone, 10mM Dimethyl Malonate, 1μM Antimycin A, or 1μM Oligomycin A for 72hrs. On the left are summary plots from 4 independent experiments (n=8) Paired t-test; *$P < 0.05$. (**B**) One representative histogram plot from summary data in (**A**).

### Supplementary Figure 10. Donor Information

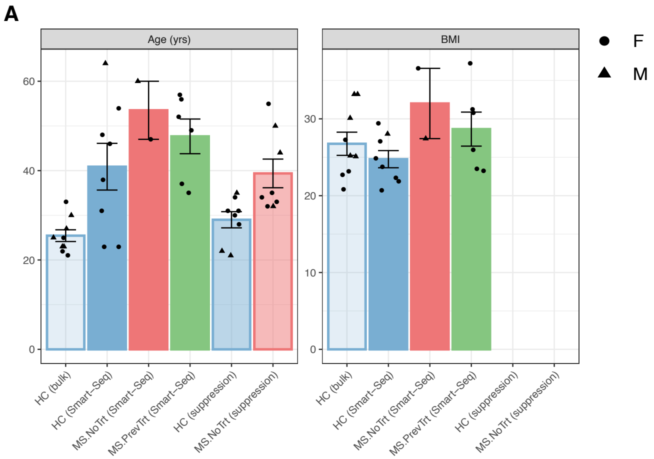

**Supplementary Figure 10. Donor Information.** (**A**) Mean and SEM error bars for age and BMI of the donors whose samples were used in the bulk RNA-Seq, single-cell Smart-Seq, and suppression assays. Self-reported height and weight (hence, BMI) were recorded only for the RNA assays. HC = healthy controls, MS.NoTrt and MS.PrevTrt = untreated and previously treated patients with MS as defined in the main text.

**Supplementary Figure 11. PC1 correlates with the intercept of a linear model that infers oleic versus arachidonic acid effects**

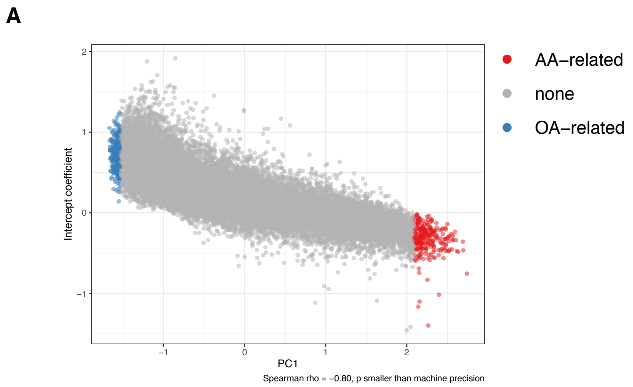

**Supplementary Figure 11. PC1 correlates with intercept of linear models to infer oleic versus arachidonic specific effects.** (**A**) Correlation of PC1 loadings and intercept of a direct model comparison (Methods) per gene, colored dots are the bottom and top 250 genes by PC1 loading defined as the oleic- and arachidonic acid related genes.

**Supplementary Figure 12. Number of cells sequenced per donor.**

A

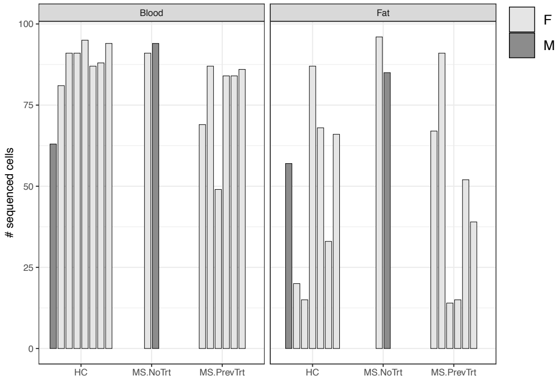

**Supplementary Figure 12. Number of cells sequenced per donor.** (**A**) Number of cells that were sequenced and satisfied QC criteria per donor. Bars represent donors, shown in the same order in both panels.

**Supplementary Figure 13. Mitochondrial respiration influences expression of cytokines in $T_{regs}$.**

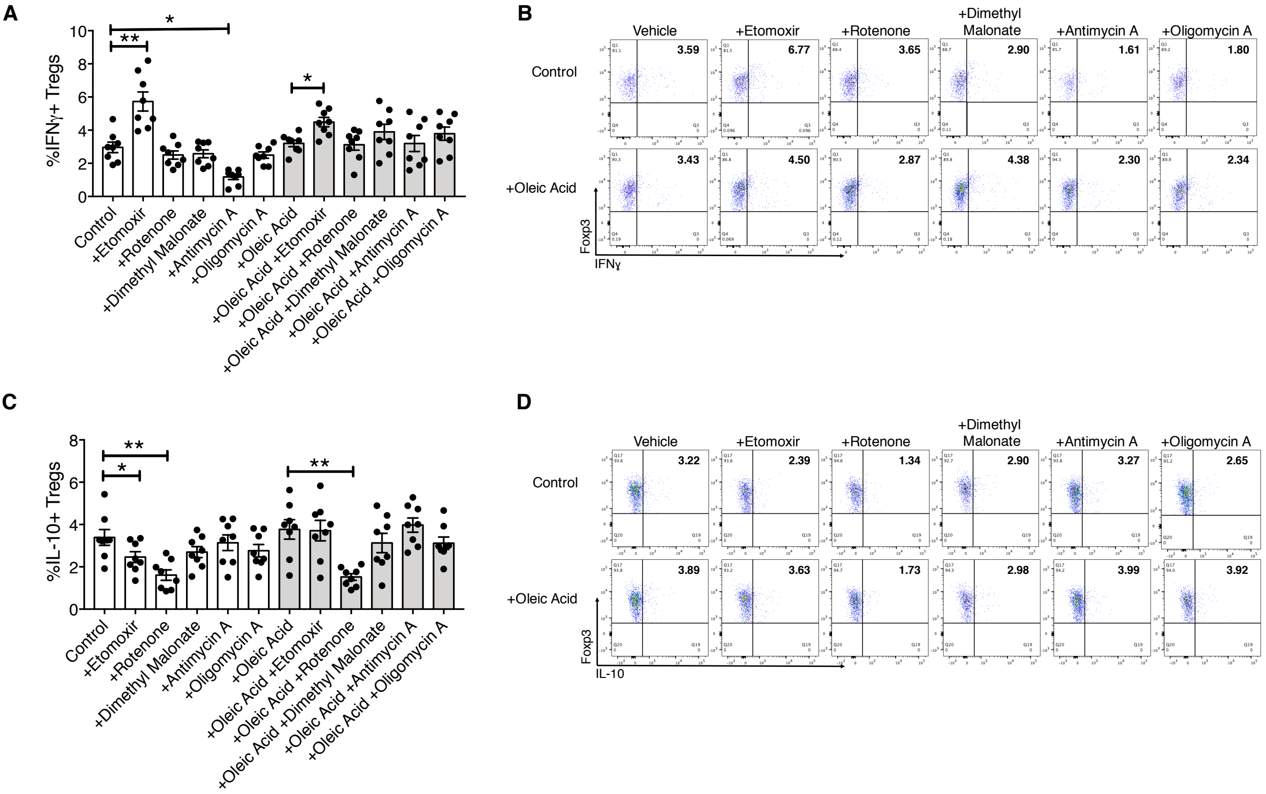

**Supplementary Figure 13. Mitochondrial respiration influences expression of cytokines in $T_{regs}$.** (**A**) Percentage of IFNγ positive in $T_{regs}$ stimulated in the presence or absence of 50μM Etomoxir, 1μM Rotenone, 10mM Dimethyl Malonate, 1μM Antimycin A, or 1μM Oligomycin A for 72hrs. Prior to fixation and antibody staining, $T_{regs}$ were stimulated for 4 hours with PMA and ionomycin in the presence of Golgistop (n=8). Paired t-test; $^{*}P < 0.05$. (**B**) One representative dot plot of summary data in (**A**). (**C**) Percentage of IL-10 positive in $T_{regs}$ stimulated in the presence or absence of 50μM Etomoxir, 1μM Rotenone, 10mM Dimethyl Malonate, 1μM Antimycin A, or 1μM Oligomycin A for 72hrs. Prior to fixation and antibody staining, $T_{regs}$ were stimulated for 4 hours with PMA and ionomycin in the presence of Golgistop (n=8). Paired t-test; $^{*}P < 0.05$. (**D**) One representative dot plot of summary data in (**C**).

**Supplementary Figure 14. PCA plot for the concentrations of LCFAs in the plasma and adipose supernatant of healthy donors and MS donors.**

A

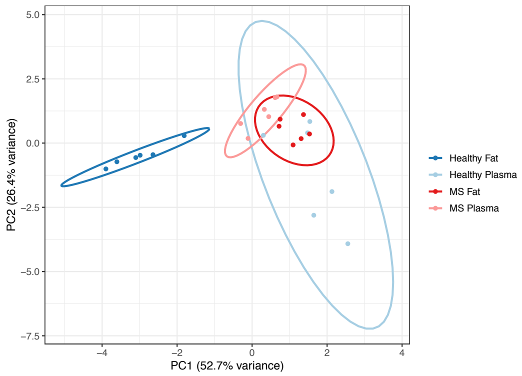

**Supplementary Figure 14. PCA plot for the concentrations of LCFAs in the plasma and adipose supernatant of healthy donors and MS donors.** Dots represent donors, with a 95% confidence ellipses for group means based on a bivariate t-distribution.

**Supplementary Table 1. Donor demographics for bulk RNA sequencing of $T_{regs}$ treated with oleic or arachidonic acid**

| Age | Height (cm) | Weight (kg) | BMI | Gender | Date Collected | Date Processed | Date Sequenced |
|---|---|---|---|---|---|---|---|
| 21 | 170.1 | 65.8 | 22.7 | F | 6-18-18 | 6-24-18 | 6-28-18 |
| 30 | 193.0 | 93.9 | 25.2 | M | 6-19-18 | 6-24-18 | 6-28-18 |
| 23 | 175.3 | 102.1 | 33.2 | M | 8-6-18 | 8-13-18 | 8-20-18 |
| 33 | 162.6 | 61.2 | 23.2 | F | 8-6-18 | 8-13-18 | 8-20-18 |
| 22 | 154.9 | 49.9 | 20.8 | F | 8-6-18 | 8-13-18 | 8-20-18 |
| 25 | 160.0 | 69.9 | 27.3 | F | 1-7-19 | 1-7-19 | 1-25-19 |
| 25 | 182.9 | 83.9 | 25.1 | M | 1-7-19 | 1-7-19 | 1-25-19 |
| 27 | 177.8 | 95.3 | 30.1 | M | 1-7-19 | 1-7-19 | 1-25-19 |
| 23 | 175.3 | 102.1 | 33.2 | M | 1-7-19 | 1-7-19 | 1-25-19 |

**Supplementary Table 1. Donor demographics for bulk RNA sequencing of $T_{regs}$ treated with oleic or arachidonic acid.** Age, height, weight, body mass index (BMI), gender, and processing dates for peripheral blood $T_{regs}$ from healthy subjects used for bulk sequencing after stimulated in the presence of oleic or arachidonic acid.

**Supplementary Table 2. Transcriptomic signature of oleic- vs. arachidonic stimuli**

| | limma-based analysis | | | | | | PC1-based analysis | |
|---|---|---|---|---|---|---|---|---|
| symbol | logFC | AveExpr | t | P.Value | adj.P.Va | B | PC1 | PC1 gene module |
| GDA | 1.382 | 1.0695 | 10.653 | 1.19E-06 | 0.003 | 5.91 | -1.4859 | none |
| EPHB4 | 1.211 | 0.9936 | 9.7736 | 2.56E-06 | 0.003 | 5.23 | -1.4756 | none |
| IGDCC3 | 1.402 | 1.1349 | 9.4921 | 3.32E-06 | 0.003 | 4.99 | -1.2976 | none |
| A4GALT | 1.423 | 1.2497 | 9.3523 | 3.78E-06 | 0.003 | 4.87 | -1.2301 | none |
| FAM117A | 1.168 | 0.9322 | 9.2526 | 4.15E-06 | 0.003 | 4.79 | -1.0209 | none |
| TUBB4A | 1.412 | 0.9808 | 9.2047 | 4.34E-06 | 0.003 | 4.75 | -1.2581 | none |
| GINS3 | 1.154 | 0.9533 | 9.0167 | 5.20E-06 | 0.003 | 4.58 | -1.1929 | none |
| ZSCAN2 | 1.131 | 0.8166 | 8.9607 | 5.49E-06 | 0.003 | 4.53 | -1.2315 | none |
| GPR173 | 1.162 | 0.9122 | 8.9532 | 5.53E-06 | 0.003 | 4.52 | -1.1335 | none |
| CSF2RA | 1.18 | 0.9632 | 8.95 | 5.55E-06 | 0.003 | 4.52 | -1.28 | none |
| BTNL9 | 1.196 | 0.947 | 8.9473 | 5.56E-06 | 0.003 | 4.52 | -1.3063 | none |
| RIMS2 | 1.155 | 0.9478 | 8.9143 | 5.74E-06 | 0.003 | 4.49 | -1.2264 | none |
| SLC19A3 | 1.389 | 1.1266 | 8.8942 | 5.86E-06 | 0.003 | 4.47 | -1.2381 | none |
| ARSE | 1.353 | 1.095 | 8.8918 | 5.87E-06 | 0.003 | 4.47 | -0.9005 | none |
| DMWD | 1.128 | 0.9113 | 8.8768 | 5.96E-06 | 0.003 | 4.45 | -1.3384 | none |
| PCGF6 | 1.054 | 0.7806 | 8.8508 | 6.11E-06 | 0.003 | 4.43 | -1.2517 | none |
| COPS7B | 1.17 | 0.9709 | 8.7965 | 6.45E-06 | 0.003 | 4.38 | -1.3272 | none |
| WNK4 | 1.042 | 0.8473 | 8.7817 | 6.54E-06 | 0.003 | 4.37 | -1.5055 | none |
| ZNF726 | 1.564 | 1.2707 | 8.7793 | 6.56E-06 | 0.003 | 4.37 | -0.951 | none |
| SMPD3 | 1.119 | 0.9216 | 8.7616 | 6.67E-06 | 0.003 | 4.35 | -1.3286 | none |
| MAPK3 | 1.004 | 0.7429 | 8.7358 | 6.85E-06 | 0.003 | 4.33 | -0.7326 | none |
| REPS1 | 1.241 | 1.0258 | 8.693 | 7.14E-06 | 0.003 | 4.29 | -1.1675 | none |
| ZNF90 | 1.185 | 0.9639 | 8.69 | 7.16E-06 | 0.003 | 4.28 | -1.3105 | none |
| RAB5B | 1.054 | 0.8049 | 8.6853 | 7.20E-06 | 0.003 | 4.28 | -1.5151 | none |
| ZNF169 | 1.045 | 0.8424 | 8.6768 | 7.26E-06 | 0.003 | 4.27 | -1.5196 | none |
| PANO1 | 1.172 | 0.8659 | 8.6512 | 7.45E-06 | 0.003 | 4.25 | -1.147 | none |
| MYH7B | 1.048 | 0.887 | 8.5909 | 7.91E-06 | 0.003 | 4.19 | -1.365 | none |
| SEMA6B | 1.26 | 1.0597 | 8.5749 | 8.04E-06 | 0.003 | 4.18 | -1.1662 | none |
| TSEN2 | 1.222 | 0.9777 | 8.5738 | 8.05E-06 | 0.003 | 4.17 | -1.2215 | none |
| GAB3 | 1.13 | 1.0347 | 8.5084 | 8.60E-06 | 0.003 | 4.11 | -1.0922 | none |
| PADI4 | 1.058 | 0.7957 | 8.5051 | 8.63E-06 | 0.003 | 4.11 | -1.4457 | none |
| AP3B2 | 1.147 | 0.8956 | 8.4999 | 8.67E-06 | 0.003 | 4.1 | -1.4958 | none |
| ARNT2 | 1.1 | 0.8447 | 8.4908 | 8.75E-06 | 0.003 | 4.1 | -1.3079 | none |
| PHLDB1 | 1.13 | 0.9314 | 8.4644 | 8.99E-06 | 0.003 | 4.07 | -1.4808 | none |
| CYP3A43 | 0.992 | 0.7851 | 8.4553 | 9.07E-06 | 0.003 | 4.06 | -1.0946 | none |
| ABCA8 | 0.954 | 0.7071 | 8.4469 | 9.15E-06 | 0.003 | 4.05 | -1.1935 | none |
| COG7 | 1.509 | 1.2902 | 8.4046 | 9.55E-06 | 0.003 | 4.01 | -1.1816 | none |
| SIRT5 | 1.056 | 0.8016 | 8.4025 | 9.58E-06 | 0.003 | 4.01 | -1.2781 | none |

| | | | | | | | | |
|---|---|---|---|---|---|---|---|---|
| CNTNAP3E | 1.002 | 0.7953 | 8.4007 | 9.59E-06 | 0.003 | 4.01 | -1.4494 | none |
| ZNF674 | 1.301 | 1.0325 | 8.3881 | 9.72E-06 | 0.003 | 4 | -1.5118 | none |
| AS3MT | 1.049 | 0.9075 | 8.3599 | 1.00E-05 | 0.003 | 3.97 | -1.5306 | none |
| CHMP1B | 0.961 | 0.7105 | 8.3586 | 1.00E-05 | 0.003 | 3.97 | -1.1124 | none |
| PLA2G4C | 1.247 | 1.0659 | 8.3272 | 1.03E-05 | 0.003 | 3.94 | -1.4285 | none |
| SLC1A2 | 0.928 | 0.7001 | 8.3123 | 1.05E-05 | 0.003 | 3.92 | -1.443 | none |
| FAM221B | 1.026 | 0.8067 | 8.2892 | 1.08E-05 | 0.003 | 3.9 | -0.9206 | none |
| PIWIL2 | 0.907 | 0.632 | 8.2324 | 1.14E-05 | 0.003 | 3.85 | -0.836 | none |
| ZNF527 | 1.12 | 0.9059 | 8.2099 | 1.17E-05 | 0.003 | 3.82 | -0.9552 | none |
| ITGA10 | 0.972 | 0.7904 | 8.2076 | 1.17E-05 | 0.003 | 3.82 | -1.291 | none |
| FAM178B | 1.087 | 0.8647 | 8.204 | 1.18E-05 | 0.003 | 3.82 | -1.1762 | none |
| SUDS3 | 1.042 | 0.8515 | 8.1891 | 1.19E-05 | 0.003 | 3.8 | -1.1361 | none |
| DLST | 1.16 | 0.9146 | 8.1624 | 1.23E-05 | 0.003 | 3.78 | -1.383 | none |
| RASGEF1B | 1.35 | 1.1745 | 8.1391 | 1.26E-05 | 0.003 | 3.75 | -1.1252 | none |
| LRRC49 | 1.33 | 1.0191 | 8.1335 | 1.27E-05 | 0.003 | 3.75 | -1.2779 | none |
| LRRC8E | 0.973 | 0.7705 | 8.081 | 1.34E-05 | 0.003 | 3.7 | -1.5287 | none |
| HECW1 | 1.118 | 0.8601 | 8.0758 | 1.34E-05 | 0.003 | 3.69 | -1.4642 | none |
| TCF23 | 0.957 | 0.7457 | 8.0728 | 1.35E-05 | 0.003 | 3.69 | -1.6057 | OA-related |
| TLE2 | 1.229 | 0.9549 | 8.0618 | 1.36E-05 | 0.003 | 3.68 | -1.3341 | none |
| PCNX2 | 1.029 | 0.8807 | 8.0431 | 1.39E-05 | 0.003 | 3.66 | -1.2594 | none |
| TMEM145 | 1.192 | 1.0017 | 8.031 | 1.41E-05 | 0.003 | 3.65 | -1.2669 | none |
| WDR59 | 1.227 | 1.0095 | 8.0214 | 1.42E-05 | 0.003 | 3.64 | -1.2873 | none |
| KLC4 | 1.281 | 1.0821 | 8.0203 | 1.43E-05 | 0.003 | 3.64 | -1.2248 | none |
| FN3K | 1.125 | 0.8946 | 8.0078 | 1.44E-05 | 0.003 | 3.62 | -1.1952 | none |
| ZNF684 | 0.959 | 0.7163 | 7.9913 | 1.47E-05 | 0.003 | 3.61 | -1.2584 | none |
| TBC1D30 | 1.114 | 0.8754 | 7.9754 | 1.50E-05 | 0.003 | 3.59 | -1.3227 | none |
| EVI5 | 1.055 | 0.9232 | 7.964 | 1.51E-05 | 0.003 | 3.58 | -1.0819 | none |
| TVP23C | 1.098 | 0.8788 | 7.963 | 1.52E-05 | 0.003 | 3.58 | -1.2603 | none |
| VASH2 | 1.309 | 1.1433 | 7.932 | 1.57E-05 | 0.003 | 3.55 | -1.1479 | none |
| ZNF556 | 0.916 | 0.6976 | 7.9251 | 1.58E-05 | 0.003 | 3.54 | -1.5733 | OA-related |
| FRMPD1 | 1.065 | 0.8317 | 7.9243 | 1.58E-05 | 0.003 | 3.54 | -1.2668 | none |
| IL31RA | 0.97 | 0.7734 | 7.9225 | 1.58E-05 | 0.003 | 3.54 | -1.3996 | none |
| MOB3B | 1.055 | 0.8222 | 7.9143 | 1.60E-05 | 0.003 | 3.53 | -1.531 | none |
| FMO3 | 1.149 | 0.9374 | 7.9143 | 1.60E-05 | 0.003 | 3.53 | -1.4726 | none |
| ATL2 | 1.112 | 0.8472 | 7.8892 | 1.64E-05 | 0.003 | 3.5 | -1.4119 | none |
| MROH8 | 1.074 | 0.7536 | 7.8833 | 1.65E-05 | 0.003 | 3.5 | -1.1909 | none |
| SLC9A6 | 0.96 | 0.7719 | 7.8806 | 1.65E-05 | 0.003 | 3.49 | -1.0024 | none |
| INMT | 0.987 | 0.7624 | 7.8802 | 1.66E-05 | 0.003 | 3.49 | -1.5664 | OA-related |
| RASSF6 | 1.05 | 0.8464 | 7.8726 | 1.67E-05 | 0.003 | 3.49 | -1.295 | none |
| TLCD2 | 0.916 | 0.7042 | 7.859 | 1.69E-05 | 0.003 | 3.47 | -1.6334 | OA-related |

| | | | | | | | | |
|---|---|---|---|---|---|---|---|---|
| ATP8B1 | 1.384 | 1.2569 | 7.8565 | 1.70E-05 | 0.003 | 3.47 | -0.697 | none |
| ACE | 1.083 | 0.8805 | 7.8246 | 1.76E-05 | 0.003 | 3.44 | -1.5817 | OA-related |
| PSD3 | 1.423 | 1.1777 | 7.8233 | 1.76E-05 | 0.003 | 3.44 | -0.8647 | none |
| NMT2 | 1.016 | 0.7854 | 7.8188 | 1.77E-05 | 0.003 | 3.43 | -1.5005 | none |
| CCDC40 | 1.121 | 0.9208 | 7.8107 | 1.78E-05 | 0.003 | 3.42 | -1.3009 | none |
| FAM129C | 0.971 | 0.7882 | 7.8063 | 1.79E-05 | 0.003 | 3.42 | -1.5963 | OA-related |
| EPB41L4A | 0.918 | 0.75 | 7.8053 | 1.79E-05 | 0.003 | 3.42 | -0.7739 | none |
| CD22 | 1.117 | 0.858 | 7.8031 | 1.80E-05 | 0.003 | 3.41 | -1.3248 | none |
| ETNPPL | 0.902 | 0.7254 | 7.7693 | 1.87E-05 | 0.003 | 3.38 | -0.8541 | none |
| SLC4A5 | 0.974 | 0.717 | 7.7589 | 1.89E-05 | 0.003 | 3.37 | -1.5056 | none |
| CALCRL | 1.023 | 0.8321 | 7.7534 | 1.90E-05 | 0.003 | 3.36 | -1.3451 | none |
| CERCAM | 1.004 | 0.8309 | 7.7481 | 1.91E-05 | 0.003 | 3.36 | -1.3596 | none |
| SFI1 | 1.036 | 0.8341 | 7.7384 | 1.93E-05 | 0.003 | 3.35 | -1.3638 | none |
| RASL10B | 1.171 | 0.9501 | 7.7374 | 1.93E-05 | 0.003 | 3.35 | -1.107 | none |
| PRKAA1 | 1.046 | 0.813 | 7.7314 | 1.94E-05 | 0.003 | 3.34 | -1.1533 | none |
| DEFB133 | 0.816 | 0.5534 | 7.7175 | 1.97E-05 | 0.003 | 3.33 | -0.8891 | none |
| MS4A7 | 1.172 | 0.8629 | 7.7173 | 1.97E-05 | 0.003 | 3.33 | -1.3517 | none |
| MILR1 | 1.302 | 1.0556 | 7.7074 | 2.00E-05 | 0.003 | 3.32 | -1.4239 | none |
| LHFPL5 | 0.959 | 0.8284 | 7.6979 | 2.02E-05 | 0.003 | 3.31 | -1.4104 | none |
| CDH3 | 1.227 | 0.9193 | 7.6738 | 2.07E-05 | 0.003 | 3.28 | -1.0937 | none |
| CLEC4C | 0.839 | 0.6441 | 7.6684 | 2.08E-05 | 0.003 | 3.27 | -1.5918 | OA-related |
| SCIN | 1.12 | 0.8632 | 7.6682 | 2.08E-05 | 0.003 | 3.27 | -1.4616 | none |
| OSBPL6 | 1.016 | 0.8079 | 7.6641 | 2.09E-05 | 0.003 | 3.27 | -1.212 | none |
| MYT1L | 1.097 | 0.8918 | 7.6533 | 2.12E-05 | 0.003 | 3.26 | -1.3765 | none |
| NR5A2 | 1.025 | 0.8082 | 7.6398 | 2.15E-05 | 0.003 | 3.24 | -1.5343 | OA-related |
| TP63 | 1.08 | 0.8787 | 7.6352 | 2.16E-05 | 0.003 | 3.24 | -1.064 | none |
| WDR73 | 1.272 | 1.0453 | 7.6315 | 2.17E-05 | 0.003 | 3.24 | -1.3367 | none |
| NPAS3 | 1.876 | 1.6086 | 7.6276 | 2.18E-05 | 0.003 | 3.23 | -1.2034 | none |
| DYNC2LI1 | 1.007 | 0.7447 | 7.626 | 2.18E-05 | 0.003 | 3.23 | -0.9972 | none |
| TMEM98 | 1.411 | 1.0366 | 7.6002 | 2.25E-05 | 0.003 | 3.2 | -0.9995 | none |
| NSUN3 | 1.054 | 0.7757 | 7.5823 | 2.29E-05 | 0.003 | 3.18 | -1.0756 | none |
| TMEM213 | 0.953 | 0.7285 | 7.5752 | 2.31E-05 | 0.003 | 3.18 | -1.2309 | none |
| PHYHD1 | 1.498 | 1.2676 | 7.571 | 2.32E-05 | 0.003 | 3.17 | -1.165 | none |
| ECD | 0.991 | 0.7752 | 7.5694 | 2.32E-05 | 0.003 | 3.17 | -1.5102 | none |
| MORN4 | 0.938 | 0.7263 | 7.5652 | 2.33E-05 | 0.003 | 3.17 | -1.3962 | none |
| ZNF567 | 1.026 | 0.8783 | 7.5652 | 2.33E-05 | 0.003 | 3.17 | -1.1231 | none |
| IFT80 | 0.941 | 0.7553 | 7.561 | 2.34E-05 | 0.003 | 3.16 | -1.4736 | none |
| SNUPN | 0.944 | 0.7297 | 7.5518 | 2.37E-05 | 0.003 | 3.15 | -1.4555 | none |
| IDO1 | 0.829 | 0.5731 | 7.5456 | 2.39E-05 | 0.003 | 3.15 | -1.5159 | none |
| AACS | 0.929 | 0.7392 | 7.5311 | 2.42E-05 | 0.003 | 3.13 | -1.5224 | none |

| | | | | | | | | |
|---|---|---|---|---|---|---|---|---|
| RNF212 | 1.549 | 1.2432 | 7.5263 | 2.44E-05 | 0.003 | 3.12 | -0.8615 | none |
| SLC4A1 | 0.915 | 0.7007 | 7.5225 | 2.45E-05 | 0.003 | 3.12 | -1.3974 | none |
| CLN3 | 0.958 | 0.7077 | 7.5175 | 2.46E-05 | 0.003 | 3.12 | -1.3704 | none |
| ALDH7A1 | 1.051 | 0.8493 | 7.504 | 2.50E-05 | 0.003 | 3.1 | -1.6688 | OA-related |
| PIP5KL1 | 0.98 | 0.8037 | 7.5019 | 2.50E-05 | 0.003 | 3.1 | -0.4353 | none |
| UGT2B4 | 1.01 | 0.7262 | 7.4935 | 2.53E-05 | 0.003 | 3.09 | -1.1892 | none |
| GDPD1 | 1.105 | 0.9405 | 7.468 | 2.60E-05 | 0.003 | 3.06 | -1.3071 | none |
| CXXC5 | 1.408 | 1.1984 | 7.4607 | 2.62E-05 | 0.003 | 3.05 | -1.3075 | none |
| SHROOM4 | 1.005 | 0.7822 | 7.4529 | 2.64E-05 | 0.003 | 3.05 | -1.3959 | none |
| DNAAF1 | 0.964 | 0.7711 | 7.4442 | 2.67E-05 | 0.003 | 3.04 | -0.635 | none |
| ZNF554 | 0.912 | 0.7491 | 7.44 | 2.68E-05 | 0.003 | 3.03 | -1.5096 | none |
| PLCE1 | 1.358 | 1.1043 | 7.4311 | 2.71E-05 | 0.003 | 3.02 | -1.3022 | none |
| CNN1 | 1.166 | 0.9314 | 7.431 | 2.71E-05 | 0.003 | 3.02 | -1.1719 | none |
| LCTL | 0.79 | 0.6023 | 7.4274 | 2.72E-05 | 0.003 | 3.02 | -1.2767 | none |
| CCDC81 | 0.931 | 0.7442 | 7.4155 | 2.76E-05 | 0.003 | 3.01 | -1.4821 | none |
| ZNF675 | 1.085 | 0.9315 | 7.411 | 2.77E-05 | 0.003 | 3 | -1.4849 | none |
| FAM57B | 0.909 | 0.7394 | 7.4008 | 2.80E-05 | 0.003 | 2.99 | -1.4349 | none |
| MYCBPAP | 1.339 | 1.1393 | 7.3992 | 2.81E-05 | 0.003 | 2.99 | -0.9533 | none |
| TRAF3IP2 | 0.881 | 0.6924 | 7.3943 | 2.82E-05 | 0.003 | 2.98 | -1.4657 | none |
| VSTM4 | 0.867 | 0.6884 | 7.3901 | 2.84E-05 | 0.003 | 2.98 | -1.4111 | none |
| MTHFR | 1.072 | 0.937 | 7.388 | 2.84E-05 | 0.003 | 2.98 | -1.387 | none |
| ZNF429 | 1.237 | 1.0245 | 7.3854 | 2.85E-05 | 0.003 | 2.97 | -1.1082 | none |
| VSNL1 | 1.008 | 0.8115 | 7.3775 | 2.88E-05 | 0.003 | 2.96 | -0.776 | none |
| APLP1 | 0.985 | 0.7697 | 7.3662 | 2.92E-05 | 0.003 | 2.95 | -1.4646 | none |
| HCAR1 | 0.803 | 0.611 | 7.3573 | 2.95E-05 | 0.003 | 2.94 | -1.3618 | none |
| EPN2 | 1.048 | 0.8695 | 7.3561 | 2.95E-05 | 0.003 | 2.94 | -1.3613 | none |
| CNTNAP3 | 1.194 | 0.9504 | 7.3496 | 2.97E-05 | 0.003 | 2.93 | -1.1895 | none |
| DDB2 | 0.945 | 0.7518 | 7.3401 | 3.00E-05 | 0.003 | 2.92 | -1.5084 | none |
| FXR2 | 1.087 | 0.9349 | 7.331 | 3.03E-05 | 0.003 | 2.91 | -1.3227 | none |
| ZNF587 | 1.105 | 0.9212 | 7.3187 | 3.08E-05 | 0.003 | 2.9 | -1.1024 | none |
| EXTL3 | 1.417 | 1.1896 | 7.3067 | 3.12E-05 | 0.003 | 2.89 | -1.0903 | none |
| CD151 | 0.879 | 0.6809 | 7.3058 | 3.12E-05 | 0.003 | 2.89 | -1.3486 | none |
| ZFHX3 | 1.031 | 0.8228 | 7.3025 | 3.13E-05 | 0.003 | 2.88 | -1.1732 | none |
| LTBR | 0.971 | 0.7706 | 7.2946 | 3.16E-05 | 0.003 | 2.87 | -1.4571 | none |
| ZNF385A | 1.527 | 1.2563 | 7.2926 | 3.17E-05 | 0.003 | 2.87 | -1.2842 | none |
| PYGL | 1.351 | 1.0972 | 7.2915 | 3.17E-05 | 0.003 | 2.87 | -1.4154 | none |
| ATG16L2 | 0.971 | 0.7771 | 7.2902 | 3.18E-05 | 0.003 | 2.87 | -1.4546 | none |
| ADGRV1 | 1.102 | 0.8497 | 7.2825 | 3.21E-05 | 0.003 | 2.86 | -1.2398 | none |
| COMP | 1.111 | 0.8898 | 7.2821 | 3.21E-05 | 0.003 | 2.86 | -1.2737 | none |
| FHL2 | 1.054 | 0.7884 | 7.2745 | 3.24E-05 | 0.003 | 2.85 | -1.0978 | none |

| | | | | | | | | |
|---|---|---|---|---|---|---|---|---|
| CFAP74 | 0.952 | 0.8113 | 7.264 | 3.27E-05 | 0.003 | 2.84 | -1.3963 | none |
| LRFN3 | 1.301 | 1.0267 | 7.2627 | 3.28E-05 | 0.003 | 2.84 | -1.2557 | none |
| RAB21 | 1.044 | 0.8277 | 7.2539 | 3.31E-05 | 0.003 | 2.83 | -1.3376 | none |
| NRP2 | 0.983 | 0.849 | 7.2535 | 3.31E-05 | 0.003 | 2.83 | -1.2237 | none |
| GULP1 | 0.968 | 0.7609 | 7.2457 | 3.34E-05 | 0.003 | 2.82 | -1.4278 | none |
| DACT3 | 0.882 | 0.7066 | 7.2409 | 3.36E-05 | 0.003 | 2.81 | -1.478 | none |
| USP6NL | 0.763 | 0.5673 | 7.2404 | 3.36E-05 | 0.003 | 2.81 | -1.0094 | none |
| ZBTB8A | 0.913 | 0.6941 | 7.2377 | 3.37E-05 | 0.003 | 2.81 | -1.6501 | OA-related |
| PDILT | 1.016 | 0.816 | 7.2206 | 3.44E-05 | 0.003 | 2.79 | -0.7203 | none |
| OR10G3 | 0.935 | 0.7231 | 7.2165 | 3.46E-05 | 0.003 | 2.79 | -1.479 | none |
| AKT3 | 1.055 | 0.8574 | 7.2092 | 3.49E-05 | 0.003 | 2.78 | -1.4145 | none |
| CREG2 | 1.042 | 0.7646 | 7.2088 | 3.49E-05 | 0.003 | 2.78 | -1.4861 | none |
| SHROOM3 | 0.992 | 0.771 | 7.2072 | 3.49E-05 | 0.003 | 2.78 | -1.0102 | none |
| CALCOCO2 | 0.989 | 0.7818 | 7.2037 | 3.51E-05 | 0.003 | 2.77 | -1.4122 | none |
| FRS3 | 1.18 | 0.9984 | 7.2003 | 3.52E-05 | 0.003 | 2.77 | -1.0342 | none |
| TMEM130 | 0.911 | 0.68 | 7.1993 | 3.53E-05 | 0.003 | 2.77 | -1.2679 | none |
| SLC25A42 | 0.878 | 0.6607 | 7.1919 | 3.56E-05 | 0.003 | 2.76 | -1.2846 | none |
| EARS2 | 1.051 | 0.8326 | 7.1833 | 3.59E-05 | 0.003 | 2.75 | -1.2395 | none |
| EFHB | 0.806 | 0.5707 | 7.1763 | 3.62E-05 | 0.003 | 2.74 | -0.0765 | none |
| ITGB8 | 0.808 | 0.6553 | 7.1714 | 3.64E-05 | 0.003 | 2.74 | -1.4321 | none |
| BAG1 | 1.06 | 0.9097 | 7.1665 | 3.66E-05 | 0.003 | 2.73 | -1.5433 | OA-related |
| ZNF724 | 1.421 | 1.2046 | 7.1591 | 3.69E-05 | 0.003 | 2.72 | -1.4151 | none |
| ALS2CR12 | 1.421 | 1.0772 | 7.1565 | 3.70E-05 | 0.003 | 2.72 | -1.1278 | none |
| PLA2G16 | 1.026 | 0.8101 | 7.1547 | 3.71E-05 | 0.003 | 2.72 | -1.3307 | none |
| SHOX | 0.876 | 0.6712 | 7.1489 | 3.74E-05 | 0.003 | 2.71 | -1.4807 | none |
| TMEM186 | 0.992 | 0.7224 | 7.1388 | 3.78E-05 | 0.003 | 2.7 | -1.2425 | none |
| DRAXIN | 0.769 | 0.5781 | 7.1214 | 3.86E-05 | 0.003 | 2.68 | -1.5575 | OA-related |
| DIRAS2 | 1.003 | 0.787 | 7.1186 | 3.87E-05 | 0.003 | 2.68 | -1.3281 | none |
| SPOP | 0.814 | 0.6108 | 7.1118 | 3.90E-05 | 0.003 | 2.67 | -1.3626 | none |
| ZNF157 | 0.819 | 0.6402 | 7.1113 | 3.90E-05 | 0.003 | 2.67 | -1.607 | OA-related |
| KBTBD2 | 0.856 | 0.6471 | 7.1081 | 3.92E-05 | 0.003 | 2.67 | -1.1636 | none |
| PBX1 | 1.261 | 1.0668 | 7.1067 | 3.92E-05 | 0.003 | 2.67 | -1.2947 | none |
| ECT2 | 1.266 | 1.0953 | 7.0957 | 3.97E-05 | 0.003 | 2.65 | -1.3234 | none |
| RAB42 | 0.727 | 0.565 | 7.0935 | 3.98E-05 | 0.003 | 2.65 | -1.223 | none |
| PHKA2 | 1.371 | 1.1702 | 7.0894 | 4.00E-05 | 0.003 | 2.65 | -1.3881 | none |
| ELAVL2 | 0.721 | 0.5502 | 7.0877 | 4.01E-05 | 0.003 | 2.64 | -1.5121 | none |
| PEX11A | 1.015 | 0.7979 | 7.0873 | 4.01E-05 | 0.003 | 2.64 | -1.3601 | none |
| PRDM16 | 1.318 | 1.0519 | 7.0797 | 4.05E-05 | 0.003 | 2.64 | -1.1733 | none |
| SCIMP | 1.002 | 0.7221 | 7.0751 | 4.07E-05 | 0.003 | 2.63 | -1.2383 | none |
| DUOX1 | 1.216 | 0.9838 | 7.0736 | 4.08E-05 | 0.003 | 2.63 | -1.5537 | OA-related |

| | | | | | | | | |
|---|---|---|---|---|---|---|---|---|
| FUT6 | 0.803 | 0.6211 | 7.0643 | 4.12E-05 | 0.003 | 2.62 | -1.3794 | none |
| IVD | 1.086 | 0.9338 | 7.0623 | 4.13E-05 | 0.003 | 2.62 | -1.1744 | none |
| PXDN | 1.207 | 0.9948 | 7.0615 | 4.14E-05 | 0.003 | 2.61 | -1.049 | none |
| PIDD1 | 0.83 | 0.6603 | 7.059 | 4.15E-05 | 0.003 | 2.61 | -1.476 | none |
| ZNF208 | 0.946 | 0.747 | 7.0575 | 4.15E-05 | 0.003 | 2.61 | -0.8984 | none |
| TMEM52B | 0.968 | 0.7764 | 7.0566 | 4.16E-05 | 0.003 | 2.61 | -1.4171 | none |
| FAM76A | 0.937 | 0.813 | 7.0395 | 4.24E-05 | 0.003 | 2.59 | -1.2741 | none |
| ZNF677 | 0.866 | 0.6746 | 7.0349 | 4.26E-05 | 0.003 | 2.58 | -1.1011 | none |
| TRIM72 | 0.807 | 0.6212 | 7.0323 | 4.28E-05 | 0.003 | 2.58 | -1.5898 | OA-related |
| MIPOL1 | 0.774 | 0.6008 | 7.0315 | 4.28E-05 | 0.003 | 2.58 | -1.5008 | none |
| RHBDF2 | 1.037 | 0.8429 | 7.0281 | 4.30E-05 | 0.003 | 2.58 | -1.4289 | none |
| ZNF596 | 0.902 | 0.7203 | 7.0187 | 4.35E-05 | 0.003 | 2.57 | -1.3104 | none |
| SIRPB2 | 0.965 | 0.7255 | 7.018 | 4.35E-05 | 0.003 | 2.57 | -1.4614 | none |
| PRICKLE2 | 0.831 | 0.632 | 7.0143 | 4.37E-05 | 0.003 | 2.56 | -1.5363 | OA-related |
| CDHR4 | 0.872 | 0.6407 | 7.012 | 4.38E-05 | 0.003 | 2.56 | -1.3867 | none |
| STRA6 | 0.996 | 0.7744 | 7.0062 | 4.41E-05 | 0.003 | 2.55 | -1.3557 | none |
| HPN | 0.947 | 0.785 | 7.003 | 4.43E-05 | 0.003 | 2.55 | -0.4095 | none |
| RIPPLY3 | 0.845 | 0.6406 | 6.9861 | 4.52E-05 | 0.003 | 2.53 | -1.1668 | none |
| ENAM | 1.02 | 0.7186 | 6.9856 | 4.52E-05 | 0.003 | 2.53 | -1.4022 | none |
| CLNK | 0.864 | 0.6501 | 6.9821 | 4.54E-05 | 0.003 | 2.52 | -1.478 | none |
| SMARCAL1 | 1.32 | 1.0612 | 6.9816 | 4.54E-05 | 0.003 | 2.52 | -1.4487 | none |
| SLC37A3 | 0.913 | 0.6688 | 6.9799 | 4.55E-05 | 0.003 | 2.52 | -1.2808 | none |
| SLC11A1 | 1.093 | 0.8709 | 6.9778 | 4.56E-05 | 0.003 | 2.52 | -1.3026 | none |
| SPACA6 | 1.109 | 0.9237 | 6.9728 | 4.59E-05 | 0.003 | 2.51 | -1.4119 | none |
| ZNF347 | 1.082 | 0.8692 | 6.9725 | 4.59E-05 | 0.003 | 2.51 | -1.3599 | none |
| C21ORF62 | 0.685 | 0.4782 | 6.9717 | 4.59E-05 | 0.003 | 2.51 | -1.4146 | none |
| OR7D4 | 0.725 | 0.5357 | 6.9689 | 4.61E-05 | 0.003 | 2.51 | -1.5919 | OA-related |
| RHAG | 1.358 | 1.1047 | 6.9677 | 4.61E-05 | 0.003 | 2.51 | -1.1003 | none |
| PRSS27 | 0.961 | 0.7294 | 6.9671 | 4.62E-05 | 0.003 | 2.51 | -1.5108 | none |
| MED11 | 0.879 | 0.6914 | 6.9588 | 4.66E-05 | 0.003 | 2.5 | -1.583 | OA-related |
| SLC36A2 | 0.791 | 0.5912 | 6.9548 | 4.68E-05 | 0.003 | 2.49 | -1.2975 | none |
| ADGRG1 | 1.028 | 0.8522 | 6.9447 | 4.74E-05 | 0.003 | 2.48 | -0.9769 | none |
| DNAJC14 | 1.063 | 0.8711 | 6.9444 | 4.74E-05 | 0.003 | 2.48 | -1.4088 | none |
| SEPSECS | 0.988 | 0.7993 | 6.9444 | 4.74E-05 | 0.003 | 2.48 | -1.3854 | none |
| FBLIM1 | 0.813 | 0.629 | 6.9441 | 4.74E-05 | 0.003 | 2.48 | -1.4158 | none |
| GIPR | 0.926 | 0.7064 | 6.9424 | 4.75E-05 | 0.003 | 2.48 | -1.3904 | none |
| MLXIPL | 1.078 | 0.8241 | 6.9422 | 4.75E-05 | 0.003 | 2.48 | -1.2293 | none |
| OR7D2 | 0.856 | 0.6234 | 6.94 | 4.77E-05 | 0.003 | 2.48 | -1.6061 | OA-related |
| BCAM | 1.068 | 0.8571 | 6.9232 | 4.86E-05 | 0.003 | 2.46 | -1.2888 | none |
| HECW2 | 0.973 | 0.7091 | 6.9198 | 4.88E-05 | 0.003 | 2.45 | -1.0035 | none |

| | | | | | | | | |
|---|---|---|---|---|---|---|---|---|
| TINF2 | 0.91 | 0.6945 | 6.9191 | 4.89E-05 | 0.003 | 2.45 | -1.2268 | none |
| ITSN1 | 1.223 | 1.0567 | 6.9174 | 4.90E-05 | 0.003 | 2.45 | -1.5201 | none |
| TERF2 | 1.027 | 0.8628 | 6.9138 | 4.92E-05 | 0.003 | 2.45 | -1.6123 | OA-related |
| SMIM14 | 1.079 | 0.8443 | 6.9103 | 4.94E-05 | 0.003 | 2.44 | -1.2157 | none |
| INTS8 | 1.088 | 0.8319 | 6.9035 | 4.98E-05 | 0.003 | 2.43 | -1.3425 | none |
| ZNF707 | 1.28 | 0.921 | 6.9026 | 4.98E-05 | 0.003 | 2.43 | -1.33 | none |
| ASAP3 | 0.84 | 0.6449 | 6.8897 | 5.06E-05 | 0.003 | 2.42 | -1.2558 | none |
| ACACB | 1.025 | 0.8515 | 6.8878 | 5.07E-05 | 0.003 | 2.42 | -1.4322 | none |
| CDH13 | 1.11 | 0.9045 | 6.8853 | 5.09E-05 | 0.003 | 2.41 | -1.102 | none |
| INSL6 | 1.061 | 0.8639 | 6.8847 | 5.09E-05 | 0.003 | 2.41 | -1.4657 | none |
| ZNF70 | 0.802 | 0.6239 | 6.8807 | 5.11E-05 | 0.003 | 2.41 | -1.489 | none |
| ZNF843 | 0.685 | 0.5055 | 6.8772 | 5.13E-05 | 0.003 | 2.4 | -1.5126 | none |
| CLDN4 | 1.065 | 0.8489 | 6.876 | 5.14E-05 | 0.003 | 2.4 | -1.1286 | none |
| CYSLTR1 | 0.827 | 0.6313 | 6.8752 | 5.15E-05 | 0.003 | 2.4 | -1.3816 | none |
| WDR93 | 0.962 | 0.7513 | 6.8697 | 5.18E-05 | 0.003 | 2.4 | -1.5775 | OA-related |
| ZBTB8B | 0.764 | 0.5642 | 6.8631 | 5.22E-05 | 0.003 | 2.39 | -1.582 | OA-related |
| CNR2 | 0.794 | 0.5848 | 6.86 | 5.24E-05 | 0.003 | 2.38 | -1.5638 | OA-related |
| ABCC9 | 0.768 | 0.6235 | 6.8491 | 5.31E-05 | 0.003 | 2.37 | -1.4869 | none |
| ZNF605 | 1.096 | 0.916 | 6.84 | 5.37E-05 | 0.003 | 2.36 | -1.2209 | none |
| MAGI2 | 1.096 | 0.8767 | 6.8395 | 5.37E-05 | 0.003 | 2.36 | -1.431 | none |
| HIST3H2BI | 0.723 | 0.52 | 6.8394 | 5.37E-05 | 0.003 | 2.36 | -1.5189 | none |
| C11ORF80 | 0.889 | 0.6296 | 6.8329 | 5.41E-05 | 0.003 | 2.35 | -1.4317 | none |
| ZNF212 | 1.223 | 1.0128 | 6.8308 | 5.43E-05 | 0.003 | 2.35 | -1.1066 | none |
| FAM122C | 1.062 | 0.7895 | 6.8291 | 5.44E-05 | 0.003 | 2.35 | -1.2669 | none |
| IAPP | 0.771 | 0.5637 | 6.8255 | 5.46E-05 | 0.003 | 2.34 | -1.2568 | none |
| TPD52 | 0.923 | 0.7668 | 6.8232 | 5.48E-05 | 0.003 | 2.34 | -1.2307 | none |
| ZNF320 | 1.267 | 1.0082 | 6.8229 | 5.48E-05 | 0.003 | 2.34 | -1.2944 | none |
| FPR2 | 1.348 | 1.0638 | 6.8189 | 5.50E-05 | 0.003 | 2.34 | -1.3117 | none |
| POMT2 | 0.983 | 0.7863 | 6.8143 | 5.53E-05 | 0.003 | 2.33 | -1.2079 | none |
| EYS | 0.727 | 0.5468 | 6.8113 | 5.55E-05 | 0.003 | 2.33 | -1.3626 | none |
| BHMT2 | 0.803 | 0.6197 | 6.8074 | 5.58E-05 | 0.003 | 2.32 | -1.5222 | none |
| MS4A10 | 0.967 | 0.715 | 6.8021 | 5.61E-05 | 0.003 | 2.32 | -1.4934 | none |
| LIAS | 0.745 | 0.6003 | 6.801 | 5.62E-05 | 0.003 | 2.32 | -1.4264 | none |
| CATIP | 0.993 | 0.8054 | 6.7959 | 5.66E-05 | 0.003 | 2.31 | -1.6346 | OA-related |
| C12ORF76 | 0.993 | 0.7468 | 6.7949 | 5.66E-05 | 0.003 | 2.31 | -1.3753 | none |
| PDE4C | 0.847 | 0.6727 | 6.7944 | 5.67E-05 | 0.003 | 2.31 | -1.5043 | none |
| WDR31 | 0.877 | 0.6955 | 6.7934 | 5.67E-05 | 0.003 | 2.31 | -1.5638 | OA-related |
| TAPT1 | 0.983 | 0.7414 | 6.7903 | 5.69E-05 | 0.003 | 2.3 | -1.2938 | none |
| KLHDC1 | 0.824 | 0.6804 | 6.7818 | 5.75E-05 | 0.004 | 2.29 | -0.4664 | none |
| KLK10 | 0.895 | 0.705 | 6.771 | 5.83E-05 | 0.004 | 2.28 | -1.5028 | none |

| | | | | | | | | |
|---|---|---|---|---|---|---|---|---|
| LTBP4 | 0.676 | 0.4863 | 6.7709 | 5.83E-05 | 0.004 | 2.28 | -1.2657 | none |
| SIGLEC10 | 0.844 | 0.6801 | 6.7691 | 5.84E-05 | 0.004 | 2.28 | -1.3887 | none |
| ZNF331 | 1.179 | 0.9618 | 6.7545 | 5.94E-05 | 0.004 | 2.26 | -1.5768 | OA-related |
| PLA2G15 | 0.833 | 0.6622 | 6.7516 | 5.97E-05 | 0.004 | 2.26 | -1.4727 | none |
| FAM160B2 | 0.959 | 0.7857 | 6.7512 | 5.97E-05 | 0.004 | 2.26 | -1.4453 | none |
| RSPH10B2 | 0.827 | 0.6649 | 6.7468 | 6.00E-05 | 0.004 | 2.25 | -1.0495 | none |
| GTF2E2 | 0.742 | 0.5359 | 6.7312 | 6.11E-05 | 0.004 | 2.23 | -1.4313 | none |
| ABCB11 | 1.238 | 1.0125 | 6.7304 | 6.12E-05 | 0.004 | 2.23 | -1.1552 | none |
| FAXDC2 | 1.122 | 0.9082 | 6.7302 | 6.12E-05 | 0.004 | 2.23 | -1.0366 | none |
| AFMID | 0.848 | 0.6711 | 6.7278 | 6.14E-05 | 0.004 | 2.23 | -1.454 | none |
| TNFRSF10I | 1.02 | 0.8448 | 6.7242 | 6.16E-05 | 0.004 | 2.23 | -1.4262 | none |
| AKR1D1 | 0.764 | 0.546 | 6.7127 | 6.25E-05 | 0.004 | 2.21 | -1.2249 | none |
| CFAP54 | 0.691 | 0.5175 | 6.7125 | 6.25E-05 | 0.004 | 2.21 | -1.4341 | none |
| ZNF546 | 1.207 | 0.9615 | 6.7107 | 6.27E-05 | 0.004 | 2.21 | -1.3894 | none |
| GALC | 1.136 | 0.8385 | 6.7099 | 6.27E-05 | 0.004 | 2.21 | -1.0924 | none |
| SCRG1 | 0.747 | 0.5667 | 6.7026 | 6.33E-05 | 0.004 | 2.2 | -1.4794 | none |
| KREMEN1 | 0.644 | 0.4362 | 6.7003 | 6.35E-05 | 0.004 | 2.2 | -1.0774 | none |
| PHF7 | 1.061 | 0.8291 | 6.6909 | 6.42E-05 | 0.004 | 2.19 | -1.4375 | none |
| ATCAY | 0.93 | 0.7219 | 6.6903 | 6.42E-05 | 0.004 | 2.19 | -1.3192 | none |
| HIF1A | 1.055 | 0.8703 | 6.6888 | 6.43E-05 | 0.004 | 2.18 | -1.4768 | none |
| MAGT1 | 0.712 | 0.5343 | 6.6882 | 6.44E-05 | 0.004 | 2.18 | -1.5347 | OA-related |
| CACNA1A | 0.795 | 0.5946 | 6.6868 | 6.45E-05 | 0.004 | 2.18 | -1.4698 | none |
| ITGB3BP | 1.192 | 0.8384 | 6.6862 | 6.45E-05 | 0.004 | 2.18 | -1.0735 | none |
| SAMD12 | 0.755 | 0.6198 | 6.6782 | 6.52E-05 | 0.004 | 2.17 | -1.4754 | none |
| DENND2A | 0.795 | 0.6042 | 6.6771 | 6.52E-05 | 0.004 | 2.17 | -1.4196 | none |
| PTK2 | 0.997 | 0.8105 | 6.6767 | 6.53E-05 | 0.004 | 2.17 | -1.2597 | none |
| GSK3B | 1.177 | 1.0025 | 6.6723 | 6.56E-05 | 0.004 | 2.16 | -1.4445 | none |
| SYTL4 | 0.705 | 0.5206 | 6.6717 | 6.57E-05 | 0.004 | 2.16 | -1.6209 | OA-related |
| NSD1 | 1.085 | 0.9124 | 6.6703 | 6.58E-05 | 0.004 | 2.16 | -1.3504 | none |
| SMIM22 | 0.921 | 0.7883 | 6.6699 | 6.58E-05 | 0.004 | 2.16 | -1.3285 | none |
| PNLIPRP1 | 1.057 | 0.8432 | 6.6685 | 6.59E-05 | 0.004 | 2.16 | -1.0293 | none |
| CFHR1 | 0.835 | 0.6517 | 6.6594 | 6.67E-05 | 0.004 | 2.15 | -0.8412 | none |
| THAP9 | 1.169 | 0.9355 | 6.6557 | 6.70E-05 | 0.004 | 2.15 | -1.2737 | none |
| SSTR2 | 0.753 | 0.6033 | 6.6528 | 6.72E-05 | 0.004 | 2.14 | -1.5614 | OA-related |
| SETDB2 | 1.237 | 1.0408 | 6.6513 | 6.73E-05 | 0.004 | 2.14 | -1.4691 | none |
| P2RX3 | 0.792 | 0.5653 | 6.6484 | 6.76E-05 | 0.004 | 2.14 | -1.0819 | none |
| RSPH10B | 0.788 | 0.6299 | 6.6466 | 6.77E-05 | 0.004 | 2.13 | -1.0034 | none |
| PKNOX1 | 0.875 | 0.7425 | 6.6434 | 6.80E-05 | 0.004 | 2.13 | -1.5141 | none |
| PTCD3 | 0.905 | 0.6993 | 6.6369 | 6.85E-05 | 0.004 | 2.12 | -1.4476 | none |
| SAMD4B | 1.211 | 1.0625 | 6.6337 | 6.88E-05 | 0.004 | 2.12 | -1.2326 | none |

| | | | | | | | | |
|---|---|---|---|---|---|---|---|---|
| SH3GL1 | 1.277 | 1.0816 | 6.6321 | 6.89E-05 | 0.004 | 2.12 | -1.4637 | none |
| SYT5 | 0.662 | 0.4867 | 6.6315 | 6.90E-05 | 0.004 | 2.12 | -1.425 | none |
| PLIN5 | 0.828 | 0.6078 | 6.6295 | 6.91E-05 | 0.004 | 2.11 | -1.4117 | none |
| CNTN2 | 0.742 | 0.5526 | 6.6291 | 6.92E-05 | 0.004 | 2.11 | -1.4796 | none |
| TARSL2 | 1.177 | 1.0411 | 6.6276 | 6.93E-05 | 0.004 | 2.11 | -1.4831 | none |
| ERMAP | 0.824 | 0.6229 | 6.6255 | 6.95E-05 | 0.004 | 2.11 | -1.4355 | none |
| PGBD4 | 0.767 | 0.5929 | 6.6242 | 6.96E-05 | 0.004 | 2.11 | -1.5983 | OA-related |
| ZNF506 | 1.004 | 0.8154 | 6.6191 | 7.00E-05 | 0.004 | 2.1 | -1.1632 | none |
| STEAP2 | 0.713 | 0.5518 | 6.6126 | 7.06E-05 | 0.004 | 2.09 | -1.5926 | OA-related |
| SLC31A1 | 0.936 | 0.7012 | 6.6095 | 7.08E-05 | 0.004 | 2.09 | -1.3698 | none |
| C5AR1 | 0.892 | 0.6821 | 6.608 | 7.10E-05 | 0.004 | 2.09 | -1.2894 | none |
| MGST1 | 1.272 | 1.0655 | 6.6039 | 7.13E-05 | 0.004 | 2.08 | -1.0187 | none |
| C1ORF109 | 0.784 | 0.6236 | 6.6019 | 7.15E-05 | 0.004 | 2.08 | -1.5782 | OA-related |
| OAF | 0.937 | 0.6958 | 6.6009 | 7.16E-05 | 0.004 | 2.08 | -1.3039 | none |
| ST18 | 1.149 | 0.8635 | 6.5985 | 7.18E-05 | 0.004 | 2.08 | -0.9963 | none |
| SENP5 | 1.079 | 0.906 | 6.5985 | 7.18E-05 | 0.004 | 2.08 | -1.4391 | none |
| LPO | 0.798 | 0.6079 | 6.5966 | 7.20E-05 | 0.004 | 2.08 | -1.3979 | none |
| KCNK6 | 1.181 | 1.0195 | 6.5957 | 7.20E-05 | 0.004 | 2.07 | -1.3135 | none |
| NUGGC | 0.934 | 0.7431 | 6.59 | 7.25E-05 | 0.004 | 2.07 | -1.557 | OA-related |
| CENPBD1 | 0.771 | 0.6337 | 6.5874 | 7.28E-05 | 0.004 | 2.06 | -1.4958 | none |
| ZNF606 | 1.147 | 0.93 | 6.5869 | 7.28E-05 | 0.004 | 2.06 | -1.0741 | none |
| PPEF2 | 1.453 | 1.3053 | 6.5795 | 7.35E-05 | 0.004 | 2.05 | -0.3786 | none |
| FAM9C | 0.716 | 0.5107 | 6.5699 | 7.43E-05 | 0.004 | 2.04 | -1.4068 | none |
| WDR90 | 1.124 | 0.9084 | 6.5699 | 7.43E-05 | 0.004 | 2.04 | -1.2777 | none |
| BRDT | 0.755 | 0.5321 | 6.5688 | 7.44E-05 | 0.004 | 2.04 | -0.6098 | none |
| CRYZL1 | 0.943 | 0.7005 | 6.5685 | 7.45E-05 | 0.004 | 2.04 | -1.2849 | none |
| ADAMTSL3 | 0.877 | 0.7177 | 6.5605 | 7.52E-05 | 0.004 | 2.03 | -1.3888 | none |
| FBXO36 | 1.14 | 0.905 | 6.5589 | 7.53E-05 | 0.004 | 2.03 | -1.6142 | OA-related |
| MARVELD: | 0.757 | 0.6065 | 6.5486 | 7.63E-05 | 0.004 | 2.02 | -1.5016 | none |
| TMC8 | 0.965 | 0.8603 | 6.5484 | 7.63E-05 | 0.004 | 2.02 | -1.5625 | OA-related |
| CC2D2A | 1.066 | 0.8099 | 6.5482 | 7.63E-05 | 0.004 | 2.02 | -1.469 | none |
| CPT1C | 0.933 | 0.6997 | 6.5464 | 7.65E-05 | 0.004 | 2.01 | -1.2726 | none |
| C1ORF216 | 0.874 | 0.6695 | 6.5378 | 7.73E-05 | 0.004 | 2 | -1.3831 | none |
| 9-Mar | 0.939 | 0.7518 | 6.536 | 7.75E-05 | 0.004 | 2 | -0.8534 | none |
| CASC10 | 0.671 | 0.4852 | 6.5308 | 7.80E-05 | 0.004 | 2 | -1.4862 | none |
| ZNF561 | 0.892 | 0.6936 | 6.5285 | 7.82E-05 | 0.004 | 1.99 | -0.9881 | none |
| CEP89 | 0.986 | 0.8805 | 6.5284 | 7.82E-05 | 0.004 | 1.99 | -1.3662 | none |
| PLEKHG1 | 0.956 | 0.7695 | 6.5284 | 7.82E-05 | 0.004 | 1.99 | -1.5689 | OA-related |
| MS4A4E | 0.818 | 0.6783 | 6.5279 | 7.83E-05 | 0.004 | 1.99 | -0.677 | none |
| KRT8 | 1.012 | 0.7828 | 6.5254 | 7.85E-05 | 0.004 | 1.99 | -1.3195 | none |

| | | | | | | | | |
|---|---|---|---|---|---|---|---|---|
| RPUSD4 | 0.904 | 0.6625 | 6.522 | 7.88E-05 | 0.004 | 1.99 | -1.286 | none |
| TMEM210 | 0.767 | 0.5902 | 6.521 | 7.89E-05 | 0.004 | 1.98 | -1.1865 | none |
| NMNAT1 | 1.181 | 1.0154 | 6.5127 | 7.98E-05 | 0.004 | 1.97 | -1.4172 | none |
| FBXO44 | 1.028 | 0.8194 | 6.5047 | 8.05E-05 | 0.004 | 1.96 | -1.376 | none |
| ARIH2OS | 1.037 | 0.8199 | 6.5003 | 8.10E-05 | 0.004 | 1.96 | -1.469 | none |
| FBXO24 | 0.925 | 0.7229 | 6.4911 | 8.19E-05 | 0.004 | 1.95 | -1.0598 | none |
| TBXA2R | 0.797 | 0.5126 | 6.4803 | 8.30E-05 | 0.004 | 1.94 | -1.1017 | none |
| TRIM52 | 1.012 | 0.8249 | 6.4801 | 8.30E-05 | 0.004 | 1.93 | -1.0888 | none |
| LINC0067[illegible] | 0.774 | 0.5957 | 6.4783 | 8.32E-05 | 0.004 | 1.93 | -1.6608 | OA-related |
| ERC1 | 1.235 | 1.1033 | 6.4726 | 8.38E-05 | 0.004 | 1.93 | -1.358 | none |
| TSPAN10 | 1.061 | 0.8536 | 6.47 | 8.41E-05 | 0.004 | 1.92 | -1.3735 | none |
| PDLIM5 | 1.145 | 0.8675 | 6.4684 | 8.42E-05 | 0.004 | 1.92 | -1.2604 | none |
| OR10H5 | 0.793 | 0.5939 | 6.466 | 8.45E-05 | 0.004 | 1.92 | -1.5327 | OA-related |
| CHST6 | 0.669 | 0.5097 | 6.4627 | 8.48E-05 | 0.004 | 1.91 | -1.5298 | none |
| MKNK1 | 0.79 | 0.6183 | 6.4611 | 8.50E-05 | 0.004 | 1.91 | -1.461 | none |
| UTP4 | 0.693 | 0.5097 | 6.4605 | 8.51E-05 | 0.004 | 1.91 | -1.3778 | none |
| NIM1K | 0.842 | 0.6082 | 6.4563 | 8.55E-05 | 0.004 | 1.91 | -0.8644 | none |
| SDHAF4 | 0.862 | 0.6103 | 6.4545 | 8.57E-05 | 0.004 | 1.9 | -1.3055 | none |
| WNT10A | 0.995 | 0.7095 | 6.4529 | 8.59E-05 | 0.004 | 1.9 | -0.9893 | none |
| HLCS | 0.841 | 0.6692 | 6.4497 | 8.62E-05 | 0.004 | 1.9 | -1.4753 | none |
| MYOM1 | 0.687 | 0.4989 | 6.4493 | 8.62E-05 | 0.004 | 1.9 | -1.2341 | none |
| SPG7 | 0.912 | 0.7633 | 6.4489 | 8.63E-05 | 0.004 | 1.9 | -1.4984 | none |
| HPS5 | 0.963 | 0.7125 | 6.4479 | 8.64E-05 | 0.004 | 1.9 | -1.3857 | none |
| SH2D3A | 0.756 | 0.529 | 6.4358 | 8.77E-05 | 0.004 | 1.88 | -1.4658 | none |
| RHBDL2 | 0.754 | 0.5807 | 6.4348 | 8.78E-05 | 0.004 | 1.88 | -1.3506 | none |
| MYZAP | 0.806 | 0.6185 | 6.43 | 8.83E-05 | 0.004 | 1.87 | -0.0924 | none |
| TRPV1 | 0.81 | 0.6688 | 6.4286 | 8.85E-05 | 0.004 | 1.87 | -1.4086 | none |
| POMGNT1 | 1.08 | 0.8751 | 6.4275 | 8.86E-05 | 0.004 | 1.87 | -1.0738 | none |
| ARID3A | 0.945 | 0.8178 | 6.4271 | 8.86E-05 | 0.004 | 1.87 | -1.4006 | none |
| FARP1 | 0.989 | 0.7873 | 6.4268 | 8.87E-05 | 0.004 | 1.87 | -1.2305 | none |
| AQP6 | 0.877 | 0.6741 | 6.4256 | 8.88E-05 | 0.004 | 1.87 | -1.3718 | none |
| THAP6 | 0.933 | 0.7472 | 6.424 | 8.90E-05 | 0.004 | 1.87 | -1.4833 | none |
| TATDN3 | 0.832 | 0.6004 | 6.423 | 8.91E-05 | 0.004 | 1.87 | -1.2657 | none |
| IZUMO2 | 1.035 | 0.7955 | 6.416 | 8.99E-05 | 0.004 | 1.86 | -1.309 | none |
| OR51M1 | 0.942 | 0.7044 | 6.4153 | 9.00E-05 | 0.004 | 1.86 | -1.4953 | none |
| LIPT1 | 0.906 | 0.6806 | 6.4067 | 9.09E-05 | 0.004 | 1.85 | -1.168 | none |
| MAGEA10 | 0.892 | 0.6766 | 6.4067 | 9.09E-05 | 0.004 | 1.85 | -1.5667 | OA-related |
| HOOK2 | 1.206 | 0.9443 | 6.4062 | 9.10E-05 | 0.004 | 1.85 | -1.1205 | none |
| ZNF793 | 0.942 | 0.7722 | 6.4049 | 9.11E-05 | 0.004 | 1.84 | -1.535 | OA-related |
| DAG1 | 1.277 | 1.059 | 6.3995 | 9.17E-05 | 0.004 | 1.84 | -1.277 | none |

| | | | | | | | | |
|---|---|---|---|---|---|---|---|---|
| RPAP1 | 1.12 | 0.9998 | 6.3985 | 9.19E-05 | 0.004 | 1.84 | -1.2352 | none |
| SPIRE2 | 0.759 | 0.5678 | 6.3982 | 9.19E-05 | 0.004 | 1.84 | -1.534 | OA-related |
| NDRG2 | 1.029 | 0.7865 | 6.3966 | 9.21E-05 | 0.004 | 1.83 | -0.8324 | none |
| PNMA2 | 0.794 | 0.5964 | 6.3959 | 9.22E-05 | 0.004 | 1.83 | -1.5285 | none |
| PKIB | 1.164 | 0.9241 | 6.3937 | 9.24E-05 | 0.004 | 1.83 | -1.3311 | none |
| RRP12 | 0.993 | 0.8426 | 6.3913 | 9.27E-05 | 0.004 | 1.83 | -1.5296 | none |
| ZBTB11 | 1.124 | 0.9975 | 6.39 | 9.28E-05 | 0.004 | 1.83 | -1.5038 | none |
| FGD2 | 0.87 | 0.6297 | 6.3868 | 9.32E-05 | 0.004 | 1.82 | -1.2332 | none |
| HINFP | 0.897 | 0.6778 | 6.3844 | 9.35E-05 | 0.004 | 1.82 | -1.4563 | none |
| CCNB1IP1 | 0.791 | 0.613 | 6.3836 | 9.36E-05 | 0.004 | 1.82 | -1.4535 | none |
| MMP11 | 0.847 | 0.6753 | 6.3832 | 9.36E-05 | 0.004 | 1.82 | -1.0251 | none |
| VGLL4 | 0.862 | 0.6934 | 6.3775 | 9.43E-05 | 0.004 | 1.81 | -1.3386 | none |
| ZNF383 | 0.822 | 0.5997 | 6.3744 | 9.47E-05 | 0.004 | 1.81 | -1.2663 | none |
| EML1 | 0.833 | 0.6512 | 6.3739 | 9.47E-05 | 0.004 | 1.81 | -1.1925 | none |
| ZNF532 | 1.108 | 0.8917 | 6.3721 | 9.49E-05 | 0.004 | 1.8 | -1.3274 | none |
| RNF185 | 0.74 | 0.5426 | 6.367 | 9.55E-05 | 0.004 | 1.8 | -1.3295 | none |
| RAB29 | 0.694 | 0.5181 | 6.3609 | 9.63E-05 | 0.004 | 1.79 | -1.4324 | none |
| OTUD4 | 1.214 | 1.004 | 6.3607 | 9.63E-05 | 0.004 | 1.79 | -1.5174 | none |
| C1ORF167 | 0.889 | 0.6578 | 6.3551 | 9.70E-05 | 0.004 | 1.78 | -1.4904 | none |
| ABHD11 | 0.756 | 0.6191 | 6.3547 | 9.70E-05 | 0.004 | 1.78 | -1.35 | none |
| AGXT | 0.617 | 0.4518 | 6.35 | 9.76E-05 | 0.004 | 1.78 | -1.2105 | none |
| SYNE2 | 1.034 | 0.937 | 6.3497 | 9.76E-05 | 0.004 | 1.78 | -1.0472 | none |
| KANK2 | 1.524 | 1.2351 | 6.3491 | 9.77E-05 | 0.004 | 1.78 | -1.1046 | none |
| 2-Sep | 0.605 | 0.434 | 6.3471 | 9.79E-05 | 0.004 | 1.77 | -0.7535 | none |
| PIH1D3 | 0.579 | 0.377 | 6.3433 | 9.84E-05 | 0.004 | 1.77 | -0.7294 | none |
| TMEM266 | 0.897 | 0.7024 | 6.3395 | 9.89E-05 | 0.004 | 1.76 | -1.4405 | none |
| KCNH6 | 0.703 | 0.5292 | 6.3352 | 9.94E-05 | 0.004 | 1.76 | -1.5239 | none |
| COLEC11 | 1.036 | 0.8625 | 6.3335 | 9.96E-05 | 0.004 | 1.76 | -1.3847 | none |
| CTRC | 0.673 | 0.5033 | 6.3324 | 9.98E-05 | 0.004 | 1.75 | -1.6169 | OA-related |
| GYPA | 0.976 | 0.7895 | 6.3307 | 1.00E-04 | 0.004 | 1.75 | -1.0496 | none |
| SWSAP1 | 0.83 | 0.6527 | 6.3298 | 1.00E-04 | 0.004 | 1.75 | -1.1127 | none |
| C1ORF210 | 0.738 | 0.5547 | 6.3292 | 1.00E-04 | 0.004 | 1.75 | -1.4903 | none |
| FER1L5 | 0.72 | 0.5767 | 6.3289 | 1.00E-04 | 0.004 | 1.75 | -1.5037 | none |
| OR4D9 | 0.813 | 0.616 | 6.3289 | 1.00E-04 | 0.004 | 1.75 | -1.4348 | none |
| DUSP3 | 0.878 | 0.6915 | 6.3232 | 1.01E-04 | 0.004 | 1.74 | -1.2918 | none |
| SLC15A1 | 0.886 | 0.676 | 6.323 | 1.01E-04 | 0.004 | 1.74 | -1.4587 | none |
| HMGA2 | 0.929 | 0.723 | 6.3227 | 1.01E-04 | 0.004 | 1.74 | -1.1553 | none |
| C10ORF82 | 0.922 | 0.7164 | 6.3193 | 1.01E-04 | 0.004 | 1.74 | -1.3662 | none |
| DST | 1.27 | 1.075 | 6.3166 | 1.02E-04 | 0.004 | 1.74 | -0.8763 | none |
| XYLT2 | 1.27 | 1.1169 | 6.3164 | 1.02E-04 | 0.004 | 1.73 | -1.3351 | none |

| | | | | | | | | |
|---|---|---|---|---|---|---|---|---|
| ANO10 | 0.961 | 0.7157 | 6.3121 | 1.02E-04 | 0.004 | 1.73 | -1.3411 | none |
| TNFSF15 | 0.636 | 0.4726 | 6.3106 | 1.03E-04 | 0.004 | 1.73 | -1.4941 | none |
| MFSD4B | 0.673 | 0.5358 | 6.3096 | 1.03E-04 | 0.004 | 1.73 | -1.5319 | OA-related |
| ARIH2 | 0.722 | 0.5329 | 6.3069 | 1.03E-04 | 0.004 | 1.72 | -1.2724 | none |
| LAT2 | 0.772 | 0.6049 | 6.301 | 1.04E-04 | 0.004 | 1.72 | -1.6596 | OA-related |
| LRRC36 | 1.085 | 0.8402 | 6.2995 | 1.04E-04 | 0.004 | 1.71 | -0.738 | none |
| NEK5 | 0.678 | 0.5274 | 6.2964 | 1.04E-04 | 0.004 | 1.71 | -1.4636 | none |
| ASTN2 | 0.718 | 0.5595 | 6.2947 | 1.05E-04 | 0.004 | 1.71 | -1.3724 | none |
| WDR62 | 1.133 | 0.9779 | 6.2861 | 1.06E-04 | 0.004 | 1.7 | -1.4181 | none |
| KCNA7 | 0.594 | 0.4403 | 6.2856 | 1.06E-04 | 0.004 | 1.7 | -1.3132 | none |
| ZNF16 | 0.844 | 0.6814 | 6.2839 | 1.06E-04 | 0.004 | 1.69 | -1.5874 | OA-related |
| PRKAG2 | 0.666 | 0.5054 | 6.2812 | 1.06E-04 | 0.004 | 1.69 | -1.012 | none |
| EXD1 | 0.649 | 0.4505 | 6.2797 | 1.07E-04 | 0.004 | 1.69 | -1.3354 | none |
| GREB1 | 0.791 | 0.6443 | 6.2791 | 1.07E-04 | 0.004 | 1.69 | -0.8371 | none |
| ZNF615 | 1.538 | 1.2896 | 6.2775 | 1.07E-04 | 0.004 | 1.69 | -1.2649 | none |
| TXNRD2 | 0.708 | 0.5959 | 6.2739 | 1.07E-04 | 0.004 | 1.68 | -1.2623 | none |
| PDE11A | 0.617 | 0.4608 | 6.2731 | 1.07E-04 | 0.004 | 1.68 | -1.4313 | none |
| DAGLB | 0.854 | 0.6799 | 6.2697 | 1.08E-04 | 0.004 | 1.68 | -1.1634 | none |
| SIGLEC8 | 0.658 | 0.5102 | 6.2656 | 1.09E-04 | 0.004 | 1.67 | -1.487 | none |
| ZNF566 | 0.823 | 0.5957 | 6.2637 | 1.09E-04 | 0.004 | 1.67 | -1.2652 | none |
| SLC47A1 | 0.849 | 0.6523 | 6.2632 | 1.09E-04 | 0.004 | 1.67 | -1.196 | none |
| FUT1 | 0.768 | 0.6384 | 6.2612 | 1.09E-04 | 0.004 | 1.67 | -1.0512 | none |
| CDH1 | 0.803 | 0.5749 | 6.2577 | 1.10E-04 | 0.004 | 1.66 | -1.4178 | none |
| ACBD7 | 0.973 | 0.7639 | 6.2536 | 1.10E-04 | 0.004 | 1.66 | -1.5428 | OA-related |
| CCDC171 | 1.031 | 0.9048 | 6.2528 | 1.10E-04 | 0.004 | 1.66 | -1.2913 | none |
| PDHX | 0.869 | 0.6639 | 6.2438 | 1.12E-04 | 0.004 | 1.64 | -1.3401 | none |
| PYGM | 0.892 | 0.7366 | 6.2437 | 1.12E-04 | 0.004 | 1.64 | -1.3693 | none |
| TYRO3 | 0.948 | 0.7631 | 6.2431 | 1.12E-04 | 0.004 | 1.64 | -1.4496 | none |
| PTPRK | 1.104 | 0.8244 | 6.2414 | 1.12E-04 | 0.004 | 1.64 | -1.4638 | none |
| WDR41 | 0.784 | 0.553 | 6.2405 | 1.12E-04 | 0.004 | 1.64 | -0.8761 | none |
| PLEKHH2 | 1.147 | 0.92 | 6.2397 | 1.12E-04 | 0.004 | 1.64 | -1.3739 | none |
| DNAJA3 | 0.712 | 0.567 | 6.2357 | 1.13E-04 | 0.004 | 1.63 | -1.4699 | none |
| SLC1A6 | 0.722 | 0.5746 | 6.2355 | 1.13E-04 | 0.004 | 1.63 | -1.5309 | none |
| TCF4 | 1.035 | 0.7666 | 6.2335 | 1.13E-04 | 0.004 | 1.63 | -0.8872 | none |
| CRLF3 | 0.907 | 0.7052 | 6.2331 | 1.13E-04 | 0.004 | 1.63 | -1.2502 | none |
| TNFRSF13( | 0.78 | 0.5886 | 6.2327 | 1.13E-04 | 0.004 | 1.63 | -1.6437 | OA-related |
| OR7G2 | 0.665 | 0.4916 | 6.2312 | 1.13E-04 | 0.004 | 1.63 | -1.5681 | OA-related |
| DZANK1 | 0.832 | 0.6912 | 6.2278 | 1.14E-04 | 0.004 | 1.62 | -1.4511 | none |
| OPHN1 | 0.784 | 0.6464 | 6.2262 | 1.14E-04 | 0.004 | 1.62 | -1.5077 | none |
| ABCB9 | 0.869 | 0.7111 | 6.2239 | 1.14E-04 | 0.004 | 1.62 | -1.5091 | none |

| | | | | | | | | |
|---|---|---|---|---|---|---|---|---|
| ZNF141 | 0.931 | 0.7211 | 6.2231 | 1.15E-04 | 0.004 | 1.62 | -1.2787 | none |
| TAF8 | 0.882 | 0.7548 | 6.2218 | 1.15E-04 | 0.004 | 1.62 | -1.3802 | none |
| ARHGAP8 | 1.012 | 0.8218 | 6.2185 | 1.15E-04 | 0.004 | 1.61 | -1.4548 | none |
| DAND5 | 0.864 | 0.6461 | 6.2184 | 1.15E-04 | 0.004 | 1.61 | -1.3853 | none |
| GBA2 | 1.077 | 0.9097 | 6.2166 | 1.15E-04 | 0.004 | 1.61 | -1.1126 | none |
| ABCC10 | 0.959 | 0.7999 | 6.216 | 1.16E-04 | 0.004 | 1.61 | -1.468 | none |
| AIPL1 | 0.632 | 0.4733 | 6.2116 | 1.16E-04 | 0.004 | 1.6 | -1.0886 | none |
| GCM1 | 0.728 | 0.5143 | 6.2097 | 1.16E-04 | 0.004 | 1.6 | -1.3713 | none |
| SLC26A8 | 0.711 | 0.5385 | 6.2018 | 1.18E-04 | 0.004 | 1.59 | -1.368 | none |
| OR1I1 | 0.717 | 0.5347 | 6.2016 | 1.18E-04 | 0.004 | 1.59 | -1.59 | OA-related |
| CSMD3 | 0.668 | 0.5237 | 6.2003 | 1.18E-04 | 0.004 | 1.59 | -0.5 | none |
| ANKRD40 | 1.004 | 0.7633 | 6.2 | 1.18E-04 | 0.004 | 1.59 | -1.3782 | none |
| TRIQK | 1.497 | 1.2202 | 6.198 | 1.18E-04 | 0.004 | 1.59 | -0.9765 | none |
| ZNF844 | 0.761 | 0.6048 | 6.1965 | 1.18E-04 | 0.004 | 1.59 | -1.5328 | OA-related |
| CYP4F8 | 0.82 | 0.6336 | 6.196 | 1.19E-04 | 0.004 | 1.58 | -1.6056 | OA-related |
| OR52I1 | 0.652 | 0.5013 | 6.194 | 1.19E-04 | 0.004 | 1.58 | -1.0983 | none |
| CREB3L1 | 0.71 | 0.5474 | 6.1905 | 1.19E-04 | 0.004 | 1.58 | -0.8403 | none |
| NPIPB15 | 1.494 | 1.2004 | 6.1888 | 1.20E-04 | 0.004 | 1.58 | -0.8776 | none |
| CEP112 | 0.962 | 0.7478 | 6.1864 | 1.20E-04 | 0.004 | 1.57 | -1.2829 | none |
| MYO15A | 0.724 | 0.5329 | 6.183 | 1.20E-04 | 0.004 | 1.57 | -1.4536 | none |
| C5AR2 | 0.749 | 0.5603 | 6.183 | 1.20E-04 | 0.004 | 1.57 | -1.6276 | OA-related |
| SCN3B | 0.604 | 0.4334 | 6.1824 | 1.21E-04 | 0.004 | 1.57 | -1.2256 | none |
| ZNF385C | 1.029 | 0.7602 | 6.1818 | 1.21E-04 | 0.004 | 1.57 | -1.5452 | OA-related |
| HLA-DOA | 0.9 | 0.6684 | 6.1813 | 1.21E-04 | 0.004 | 1.57 | -0.878 | none |
| SCLY | 0.771 | 0.6178 | 6.1797 | 1.21E-04 | 0.004 | 1.56 | -1.3263 | none |
| PLEKHA5 | 0.961 | 0.7525 | 6.1765 | 1.21E-04 | 0.004 | 1.56 | -1.3439 | none |
| GPALPP1 | 1.027 | 0.8557 | 6.1757 | 1.22E-04 | 0.004 | 1.56 | -1.5666 | OA-related |
| MOB1B | 0.851 | 0.6805 | 6.1709 | 1.22E-04 | 0.004 | 1.55 | -1.3779 | none |
| FUT8 | 1.075 | 0.926 | 6.1706 | 1.22E-04 | 0.004 | 1.55 | -1.5162 | none |
| TSGA13 | 0.677 | 0.4967 | 6.1693 | 1.23E-04 | 0.004 | 1.55 | -1.4205 | none |
| CSNK1D | 1.013 | 0.8597 | 6.1688 | 1.23E-04 | 0.004 | 1.55 | -1.6311 | OA-related |
| POU2AF1 | 0.933 | 0.8316 | 6.1682 | 1.23E-04 | 0.004 | 1.55 | -1.3231 | none |
| KLB | 0.636 | 0.486 | 6.167 | 1.23E-04 | 0.004 | 1.55 | -1.5673 | OA-related |
| ZNF492 | 0.701 | 0.5353 | 6.1668 | 1.23E-04 | 0.004 | 1.55 | -1.4691 | none |
| KLC3 | 0.993 | 0.8188 | 6.1557 | 1.25E-04 | 0.004 | 1.53 | -0.9953 | none |
| TRAPPC2 | 0.704 | 0.5072 | 6.1552 | 1.25E-04 | 0.004 | 1.53 | -1.5022 | none |
| ELAVL1 | 0.866 | 0.7087 | 6.1537 | 1.25E-04 | 0.004 | 1.53 | -1.6281 | OA-related |
| TINCR | 1.13 | 0.8937 | 6.1534 | 1.25E-04 | 0.004 | 1.53 | -1.4333 | none |
| C19ORF25 | 0.79 | 0.5734 | 6.1526 | 1.25E-04 | 0.004 | 1.53 | -1.4097 | none |
| FCHO2 | 0.697 | 0.5464 | 6.1513 | 1.25E-04 | 0.004 | 1.53 | -1.3019 | none |

| | | | | | | | | |
|---|---|---|---|---|---|---|---|---|
| USHBP1 | 0.69 | 0.53 | 6.1487 | 1.26E-04 | 0.004 | 1.53 | -1.055 | none |
| TFPI | 0.778 | 0.5887 | 6.1426 | 1.27E-04 | 0.004 | 1.52 | -1.4371 | none |
| COBLL1 | 0.794 | 0.6152 | 6.1381 | 1.28E-04 | 0.004 | 1.51 | -1.5457 | OA-related |
| HDAC6 | 1.158 | 1.0105 | 6.1381 | 1.28E-04 | 0.004 | 1.51 | -1.3555 | none |
| FRRS1L | 0.749 | 0.5726 | 6.1364 | 1.28E-04 | 0.004 | 1.51 | -1.5537 | OA-related |
| CCDC24 | 0.815 | 0.6251 | 6.1358 | 1.28E-04 | 0.004 | 1.51 | -1.5047 | none |
| ZNF430 | 1.227 | 1.0889 | 6.1322 | 1.29E-04 | 0.004 | 1.5 | -1.4485 | none |
| TMEM185 | 0.863 | 0.6392 | 6.1316 | 1.29E-04 | 0.004 | 1.5 | -1.3138 | none |
| TERB1 | 1.557 | 1.254 | 6.1315 | 1.29E-04 | 0.004 | 1.5 | -0.8746 | none |
| C12ORF50 | 0.75 | 0.5744 | 6.1294 | 1.29E-04 | 0.004 | 1.5 | -1.4048 | none |
| CHRNB1 | 0.76 | 0.5441 | 6.1267 | 1.29E-04 | 0.004 | 1.5 | -1.2808 | none |
| DYDC1 | 1.582 | 1.2867 | 6.1236 | 1.30E-04 | 0.004 | 1.49 | -1.2707 | none |
| ETFDH | 0.617 | 0.4912 | 6.123 | 1.30E-04 | 0.004 | 1.49 | -1.4488 | none |
| PGS1 | 0.745 | 0.5899 | 6.1191 | 1.31E-04 | 0.004 | 1.49 | -1.4837 | none |
| CYP1A2 | 0.772 | 0.6126 | 6.1162 | 1.31E-04 | 0.004 | 1.48 | -1.6119 | OA-related |
| SPACA9 | 0.682 | 0.4884 | 6.1151 | 1.31E-04 | 0.004 | 1.48 | -1.2342 | none |
| SLC52A1 | 0.846 | 0.6352 | 6.1146 | 1.31E-04 | 0.004 | 1.48 | -1.473 | none |
| C9ORF85 | 0.829 | 0.6256 | 6.1141 | 1.32E-04 | 0.004 | 1.48 | -1.5001 | none |
| ZIM3 | 0.66 | 0.4896 | 6.1134 | 1.32E-04 | 0.004 | 1.48 | -1.5733 | OA-related |
| NRROS | 0.658 | 0.5723 | 6.1129 | 1.32E-04 | 0.004 | 1.48 | -0.8137 | none |
| MPRIP | 1.233 | 1.0951 | 6.1114 | 1.32E-04 | 0.004 | 1.48 | -1.3025 | none |
| GAL3ST1 | 0.738 | 0.564 | 6.1098 | 1.32E-04 | 0.004 | 1.48 | -1.2527 | none |
| CCDC36 | 0.819 | 0.6643 | 6.109 | 1.32E-04 | 0.004 | 1.48 | -0.955 | none |
| ARID1B | 1.58 | 1.4346 | 6.1045 | 1.33E-04 | 0.004 | 1.47 | -0.9199 | none |
| UEVLD | 0.773 | 0.5471 | 6.1022 | 1.34E-04 | 0.004 | 1.47 | -1.3998 | none |
| TAS2R4 | 0.63 | 0.5173 | 6.101 | 1.34E-04 | 0.004 | 1.47 | -1.4989 | none |
| CEACAM1! | 1.142 | 0.957 | 6.1005 | 1.34E-04 | 0.004 | 1.46 | -0.8188 | none |
| SFMBT1 | 0.983 | 0.7956 | 6.0944 | 1.35E-04 | 0.004 | 1.46 | -1.2259 | none |
| BBIP1 | 0.703 | 0.4292 | 6.0943 | 1.35E-04 | 0.004 | 1.46 | -0.2578 | none |
| LAMC2 | 0.635 | 0.4772 | 6.094 | 1.35E-04 | 0.004 | 1.46 | -1.2085 | none |
| RASEF | 0.673 | 0.5398 | 6.0933 | 1.35E-04 | 0.004 | 1.46 | -1.5487 | OA-related |
| CNKSR3 | 0.621 | 0.4518 | 6.0923 | 1.35E-04 | 0.004 | 1.45 | -1.4711 | none |
| ZNF846 | 1.058 | 0.8883 | 6.0915 | 1.35E-04 | 0.004 | 1.45 | -0.7219 | none |
| CDH18 | 0.943 | 0.7456 | 6.0913 | 1.35E-04 | 0.004 | 1.45 | -1.1485 | none |
| SLC50A1 | 0.697 | 0.4742 | 6.0864 | 1.36E-04 | 0.004 | 1.45 | -1.3709 | none |
| CLEC3A | 0.874 | 0.6665 | 6.0861 | 1.36E-04 | 0.004 | 1.45 | -1.4558 | none |
| ADAMTS4 | 0.721 | 0.5815 | 6.0854 | 1.37E-04 | 0.004 | 1.45 | -1.3359 | none |
| NXN | 0.786 | 0.5919 | 6.0832 | 1.37E-04 | 0.004 | 1.44 | -0.8436 | none |
| COG1 | 1.028 | 0.8868 | 6.0826 | 1.37E-04 | 0.004 | 1.44 | -1.2424 | none |
| EFCAB2 | 0.973 | 0.7132 | 6.0823 | 1.37E-04 | 0.004 | 1.44 | -1.4014 | none |

| | | | | | | | | |
|---|---|---|---|---|---|---|---|---|
| MARK3 | 1.22 | 1.0489 | 6.0804 | 1.37E-04 | 0.004 | 1.44 | -1.528 | none |
| COL8A1 | 0.766 | 0.6083 | 6.0781 | 1.38E-04 | 0.004 | 1.44 | -0.7549 | none |
| GSDMB | 0.765 | 0.5914 | 6.077 | 1.38E-04 | 0.004 | 1.43 | -1.2225 | none |
| EFCAB11 | 1.171 | 0.8735 | 6.074 | 1.39E-04 | 0.004 | 1.43 | -1.1881 | none |
| MEFV | 0.784 | 0.5631 | 6.0633 | 1.40E-04 | 0.004 | 1.42 | -1.4252 | none |
| ZNF586 | 0.78 | 0.5908 | 6.0604 | 1.41E-04 | 0.004 | 1.41 | -1.3969 | none |
| GAB1 | 1.03 | 0.8156 | 6.0544 | 1.42E-04 | 0.004 | 1.41 | -1.5167 | none |
| IFT122 | 1.128 | 0.9399 | 6.0485 | 1.43E-04 | 0.004 | 1.4 | -1.416 | none |
| OR7A17 | 0.768 | 0.5701 | 6.0484 | 1.43E-04 | 0.004 | 1.4 | -1.521 | none |
| ZNF114 | 0.745 | 0.5448 | 6.048 | 1.43E-04 | 0.004 | 1.4 | -1.4348 | none |
| S1PR3 | 0.635 | 0.4607 | 6.0472 | 1.43E-04 | 0.004 | 1.4 | -1.5542 | OA-related |
| ABCG8 | 0.707 | 0.5627 | 6.0428 | 1.44E-04 | 0.004 | 1.39 | -1.5053 | none |
| KCNC1 | 1.164 | 0.9981 | 6.0383 | 1.45E-04 | 0.004 | 1.39 | -1.1027 | none |
| STAP2 | 0.897 | 0.704 | 6.0376 | 1.45E-04 | 0.004 | 1.38 | -1.3874 | none |
| FGFR1 | 0.798 | 0.6107 | 6.0367 | 1.45E-04 | 0.004 | 1.38 | -1.3878 | none |
| LGMN | 0.88 | 0.7044 | 6.0359 | 1.46E-04 | 0.004 | 1.38 | -1.4653 | none |
| KIAA1191 | 0.797 | 0.603 | 6.0344 | 1.46E-04 | 0.004 | 1.38 | -0.9041 | none |
| RPRD1A | 0.765 | 0.5444 | 6.0292 | 1.47E-04 | 0.004 | 1.37 | -0.4985 | none |
| SLC24A4 | 0.639 | 0.4818 | 6.0272 | 1.47E-04 | 0.004 | 1.37 | -1.621 | OA-related |
| RASGRP3 | 1.008 | 0.8334 | 6.0271 | 1.47E-04 | 0.004 | 1.37 | -1.4525 | none |
| C6ORF141 | 1.213 | 1.0085 | 6.0233 | 1.48E-04 | 0.004 | 1.37 | -1.0781 | none |
| UTP14C | 0.77 | 0.5131 | 6.0223 | 1.48E-04 | 0.004 | 1.37 | 0.0645 | none |
| SGCE | 0.85 | 0.6604 | 6.0207 | 1.48E-04 | 0.004 | 1.36 | -1.567 | OA-related |
| ASB9 | 0.981 | 0.7875 | 6.0171 | 1.49E-04 | 0.004 | 1.36 | 0.22548 | none |
| TADA2A | 1.267 | 1.0475 | 6.0169 | 1.49E-04 | 0.004 | 1.36 | -1.3285 | none |
| GP2 | 0.63 | 0.491 | 6.0141 | 1.50E-04 | 0.004 | 1.35 | -0.1776 | none |
| ZNF3 | 1.067 | 0.8993 | 6.0136 | 1.50E-04 | 0.004 | 1.35 | -1.5245 | none |
| FBXO43 | 0.774 | 0.5837 | 6.0132 | 1.50E-04 | 0.004 | 1.35 | -1.422 | none |
| HVCN1 | 0.79 | 0.552 | 6.0129 | 1.50E-04 | 0.004 | 1.35 | -1.0141 | none |
| METTL2B | 0.831 | 0.6624 | 6.011 | 1.50E-04 | 0.004 | 1.35 | -1.3083 | none |
| RUVBL1 | 0.802 | 0.6446 | 6.0093 | 1.51E-04 | 0.004 | 1.35 | -1.4965 | none |
| KCTD21 | 1.047 | 0.814 | 6.0035 | 1.52E-04 | 0.004 | 1.34 | -1.3817 | none |
| STK38L | 0.752 | 0.5691 | 6.0031 | 1.52E-04 | 0.004 | 1.34 | -1.2979 | none |
| ZNF713 | 0.849 | 0.705 | 5.9999 | 1.52E-04 | 0.004 | 1.34 | -1.5386 | OA-related |
| SUOX | 0.727 | 0.5603 | 5.9994 | 1.53E-04 | 0.004 | 1.34 | -1.5022 | none |
| PTPN2 | 0.728 | 0.5507 | 5.9973 | 1.53E-04 | 0.004 | 1.33 | -1.2591 | none |
| PTGR2 | 0.706 | 0.5868 | 5.996 | 1.53E-04 | 0.004 | 1.33 | -1.3507 | none |
| LAMB4 | 0.828 | 0.6101 | 5.9912 | 1.54E-04 | 0.004 | 1.33 | -1.2028 | none |
| GHR | 0.95 | 0.7458 | 5.9907 | 1.54E-04 | 0.004 | 1.32 | -1.4194 | none |
| COG4 | 0.737 | 0.582 | 5.9901 | 1.54E-04 | 0.004 | 1.32 | -1.3415 | none |

| | | | | | | | | |
|---|---|---|---|---|---|---|---|---|
| TTLL9 | 0.579 | 0.4421 | 5.9899 | 1.54E-04 | 0.004 | 1.32 | -1.4198 | none |
| C8ORF44 | 0.982 | 0.6776 | 5.9897 | 1.54E-04 | 0.004 | 1.32 | -1.4439 | none |
| NFS1 | 0.799 | 0.6485 | 5.9879 | 1.55E-04 | 0.004 | 1.32 | -1.461 | none |
| MOGAT3 | 0.707 | 0.5305 | 5.9835 | 1.56E-04 | 0.004 | 1.32 | -1.5928 | OA-related |
| SLC11A2 | 0.648 | 0.4509 | 5.9817 | 1.56E-04 | 0.004 | 1.31 | -1.5199 | none |
| GNPTG | 0.814 | 0.7143 | 5.9808 | 1.56E-04 | 0.004 | 1.31 | -1.0513 | none |
| CLEC4M | 0.847 | 0.6681 | 5.9793 | 1.57E-04 | 0.004 | 1.31 | -1.1588 | none |
| FBXL20 | 0.617 | 0.4432 | 5.9789 | 1.57E-04 | 0.004 | 1.31 | -1.3216 | none |
| TANGO6 | 1.018 | 0.8406 | 5.9787 | 1.57E-04 | 0.004 | 1.31 | -1.0497 | none |
| SIRT2 | 0.798 | 0.6022 | 5.9743 | 1.58E-04 | 0.004 | 1.3 | -1.2058 | none |
| KDM1B | 0.977 | 0.8271 | 5.9723 | 1.58E-04 | 0.004 | 1.3 | -1.0823 | none |
| TRIM65 | 0.978 | 0.8144 | 5.9722 | 1.58E-04 | 0.004 | 1.3 | -1.2938 | none |
| NUPL2 | 0.771 | 0.561 | 5.9684 | 1.59E-04 | 0.004 | 1.3 | -1.3478 | none |
| PDZD2 | 0.849 | 0.6667 | 5.9682 | 1.59E-04 | 0.004 | 1.3 | -0.6122 | none |
| GJC1 | 0.744 | 0.5283 | 5.9675 | 1.59E-04 | 0.004 | 1.29 | -1.5043 | none |
| MSS51 | 0.769 | 0.5913 | 5.9673 | 1.59E-04 | 0.004 | 1.29 | -1.5043 | none |
| TRPM4 | 0.706 | 0.5441 | 5.9672 | 1.59E-04 | 0.004 | 1.29 | -1.4277 | none |
| MUTYH | 0.894 | 0.7157 | 5.9654 | 1.59E-04 | 0.004 | 1.29 | -1.5911 | OA-related |
| MME | 0.789 | 0.5918 | 5.9635 | 1.60E-04 | 0.004 | 1.29 | -1.3289 | none |
| PAWR | 0.824 | 0.707 | 5.9622 | 1.60E-04 | 0.004 | 1.29 | -1.2639 | none |
| BOLL | 0.612 | 0.4556 | 5.9619 | 1.60E-04 | 0.004 | 1.29 | -1.0938 | none |
| AAMDC | 0.925 | 0.6401 | 5.9619 | 1.60E-04 | 0.004 | 1.29 | -1.2324 | none |
| SULT2A1 | 0.777 | 0.5759 | 5.9618 | 1.60E-04 | 0.004 | 1.29 | -1.5421 | OA-related |
| SF3B3 | 0.614 | 0.5066 | 5.9602 | 1.61E-04 | 0.004 | 1.29 | -1.0856 | none |
| ATF7IP2 | 0.601 | 0.4378 | 5.9593 | 1.61E-04 | 0.004 | 1.28 | -0.8742 | none |
| C15ORF41 | 0.8 | 0.5663 | 5.956 | 1.61E-04 | 0.004 | 1.28 | -0.9202 | none |
| IRGQ | 0.889 | 0.7295 | 5.9551 | 1.62E-04 | 0.004 | 1.28 | -1.5003 | none |
| PPIL6 | 0.858 | 0.6919 | 5.9542 | 1.62E-04 | 0.004 | 1.28 | -1.1666 | none |
| CHRNA6 | 1.536 | 1.1863 | 5.952 | 1.62E-04 | 0.004 | 1.27 | -1.3544 | none |
| OSTM1 | 0.766 | 0.5505 | 5.9507 | 1.63E-04 | 0.004 | 1.27 | -0.8885 | none |
| ARHGDIG | 1.2 | 0.9882 | 5.9501 | 1.63E-04 | 0.004 | 1.27 | -1.1903 | none |
| FCN1 | 0.567 | 0.4488 | 5.9473 | 1.63E-04 | 0.004 | 1.27 | -1.4598 | none |
| ZNF564 | 0.861 | 0.621 | 5.9428 | 1.64E-04 | 0.004 | 1.26 | -1.03 | none |
| NAP1L6 | 0.858 | 0.6793 | 5.9402 | 1.65E-04 | 0.004 | 1.26 | -0.9036 | none |
| TM7SF3 | 0.914 | 0.6384 | 5.9382 | 1.65E-04 | 0.004 | 1.26 | -1.1723 | none |
| LYPD5 | 0.744 | 0.547 | 5.9371 | 1.65E-04 | 0.004 | 1.26 | -1.5748 | OA-related |
| OR2L2 | 0.646 | 0.4945 | 5.9358 | 1.66E-04 | 0.004 | 1.25 | -1.5336 | OA-related |
| SLC35E3 | 0.884 | 0.6945 | 5.9354 | 1.66E-04 | 0.004 | 1.25 | -0.6225 | none |
| CNTN1 | 0.846 | 0.6517 | 5.9334 | 1.66E-04 | 0.004 | 1.25 | -1.0128 | none |
| ZNF397 | 0.836 | 0.6419 | 5.9333 | 1.66E-04 | 0.004 | 1.25 | -1.39 | none |

| | | | | | | | | |
|---|---|---|---|---|---|---|---|---|
| FAM228B | 0.942 | 0.8086 | 5.9331 | 1.66E-04 | 0.004 | 1.25 | -1.3903 | none |
| NOVA2 | 0.991 | 0.7445 | 5.9322 | 1.66E-04 | 0.004 | 1.25 | -1.2108 | none |
| MYH11 | 0.803 | 0.6364 | 5.932 | 1.67E-04 | 0.004 | 1.25 | -1.3422 | none |
| AC007325 | 0.61 | 0.4481 | 5.9319 | 1.67E-04 | 0.004 | 1.25 | -1.1803 | none |
| CYP3A5 | 0.691 | 0.5656 | 5.9313 | 1.67E-04 | 0.004 | 1.25 | -1.5291 | none |
| SPATA21 | 0.651 | 0.4985 | 5.9304 | 1.67E-04 | 0.004 | 1.25 | -1.3576 | none |
| GSG1 | 0.706 | 0.5655 | 5.9288 | 1.67E-04 | 0.004 | 1.25 | -1.5378 | OA-related |
| ZNF839 | 0.747 | 0.5989 | 5.9266 | 1.68E-04 | 0.004 | 1.24 | -0.9592 | none |
| ZNF805 | 0.685 | 0.4752 | 5.9189 | 1.69E-04 | 0.004 | 1.23 | -1.3362 | none |
| CNPY4 | 0.921 | 0.7424 | 5.9184 | 1.70E-04 | 0.004 | 1.23 | -1.5359 | OA-related |
| ELF5 | 0.817 | 0.6504 | 5.9172 | 1.70E-04 | 0.004 | 1.23 | -1.5509 | OA-related |
| DEPDC5 | 0.934 | 0.7688 | 5.9168 | 1.70E-04 | 0.004 | 1.23 | -1.2565 | none |
| TLR3 | 0.532 | 0.3794 | 5.9142 | 1.70E-04 | 0.004 | 1.23 | -1.2206 | none |
| FRK | 0.658 | 0.5039 | 5.9134 | 1.71E-04 | 0.004 | 1.23 | -1.4513 | none |
| SMURF2 | 0.9 | 0.7818 | 5.9129 | 1.71E-04 | 0.004 | 1.22 | -1.5338 | OA-related |
| CASK | 0.971 | 0.8018 | 5.9127 | 1.71E-04 | 0.004 | 1.22 | -1.0156 | none |
| TRPM8 | 0.677 | 0.4699 | 5.9122 | 1.71E-04 | 0.004 | 1.22 | -1.3541 | none |
| PDE12 | 0.568 | 0.4426 | 5.91 | 1.71E-04 | 0.004 | 1.22 | -1.0111 | none |
| ZNF396 | 0.81 | 0.5726 | 5.9088 | 1.72E-04 | 0.004 | 1.22 | -0.9842 | none |
| BACE1 | 1.037 | 0.937 | 5.9076 | 1.72E-04 | 0.004 | 1.22 | -1.4648 | none |
| SEC14L6 | 0.757 | 0.5708 | 5.9074 | 1.72E-04 | 0.004 | 1.22 | -1.6092 | OA-related |
| SLC23A1 | 1.013 | 0.8972 | 5.9049 | 1.73E-04 | 0.004 | 1.21 | -0.4876 | none |
| ZNF12 | 1.036 | 0.8459 | 5.9041 | 1.73E-04 | 0.004 | 1.21 | -1.5375 | OA-related |
| RILPL1 | 0.801 | 0.6704 | 5.9036 | 1.73E-04 | 0.004 | 1.21 | -1.6272 | OA-related |
| NSRP1 | 1.289 | 1.1558 | 5.9027 | 1.73E-04 | 0.004 | 1.21 | -1.4265 | none |
| TNFAIP8L3 | 0.703 | 0.5228 | 5.9025 | 1.73E-04 | 0.004 | 1.21 | -1.3811 | none |
| SYT12 | 0.555 | 0.4295 | 5.901 | 1.73E-04 | 0.004 | 1.21 | -1.4201 | none |
| DNAH3 | 0.843 | 0.6764 | 5.9007 | 1.73E-04 | 0.004 | 1.21 | -1.3489 | none |
| ANKRD16 | 0.593 | 0.436 | 5.9004 | 1.74E-04 | 0.004 | 1.21 | -1.4894 | none |
| SFXN5 | 0.858 | 0.5843 | 5.8998 | 1.74E-04 | 0.004 | 1.21 | -1.1413 | none |
| CPM | 0.891 | 0.7217 | 5.897 | 1.74E-04 | 0.004 | 1.2 | -1.0769 | none |
| ABCA6 | 0.729 | 0.5621 | 5.8952 | 1.75E-04 | 0.004 | 1.2 | -1.221 | none |
| ACOT11 | 0.748 | 0.6034 | 5.895 | 1.75E-04 | 0.004 | 1.2 | -1.5563 | OA-related |
| ZNF283 | 0.966 | 0.75 | 5.8906 | 1.76E-04 | 0.004 | 1.2 | -1.4471 | none |
| ATP6V0A1 | 0.665 | 0.4877 | 5.8875 | 1.77E-04 | 0.004 | 1.19 | -1.4741 | none |
| CSNK2A1 | 0.576 | 0.4351 | 5.8818 | 1.78E-04 | 0.004 | 1.18 | -1.0028 | none |
| STK25 | 0.802 | 0.659 | 5.8799 | 1.78E-04 | 0.004 | 1.18 | -1.2932 | none |
| GALNT9 | 0.708 | 0.572 | 5.8759 | 1.79E-04 | 0.004 | 1.18 | -0.8726 | none |
| SCPEP1 | 0.643 | 0.4309 | 5.8742 | 1.80E-04 | 0.004 | 1.17 | -1.3134 | none |
| ANKLE2 | 1.049 | 0.9472 | 5.8718 | 1.80E-04 | 0.004 | 1.17 | -1.4423 | none |

| | | | | | | | | |
|---|---|---|---|---|---|---|---|---|
| SPINK1 | 1.037 | 0.7518 | 5.8683 | 1.81E-04 | 0.005 | 1.17 | -1.3864 | none |
| KIF6 | 0.895 | 0.7225 | 5.8656 | 1.82E-04 | 0.005 | 1.16 | -1.4666 | none |
| CHN2 | 1.629 | 1.3654 | 5.8637 | 1.82E-04 | 0.005 | 1.16 | -1.1337 | none |
| BRAF | 0.74 | 0.5959 | 5.8625 | 1.82E-04 | 0.005 | 1.16 | -1.5515 | OA-related |
| SNTG1 | 0.913 | 0.7402 | 5.8597 | 1.83E-04 | 0.005 | 1.16 | -1.2972 | none |
| NPC1 | 0.577 | 0.4767 | 5.8586 | 1.83E-04 | 0.005 | 1.15 | -1.3919 | none |
| ARHGAP2: | 0.848 | 0.6619 | 5.8582 | 1.83E-04 | 0.005 | 1.15 | -0.7805 | none |
| C9ORF66 | 0.713 | 0.5531 | 5.8544 | 1.84E-04 | 0.005 | 1.15 | -1.4295 | none |
| NEK10 | 0.778 | 0.6125 | 5.8542 | 1.84E-04 | 0.005 | 1.15 | -1.3314 | none |
| FRMPD3 | 0.719 | 0.5454 | 5.8521 | 1.85E-04 | 0.005 | 1.15 | -1.4397 | none |
| HSD17B13 | 0.73 | 0.5704 | 5.8438 | 1.87E-04 | 0.005 | 1.13 | -1.5002 | none |
| PCDHB1 | 0.691 | 0.5147 | 5.843 | 1.87E-04 | 0.005 | 1.13 | -1.5646 | OA-related |
| PLBD1 | 0.871 | 0.6886 | 5.8416 | 1.88E-04 | 0.005 | 1.13 | -1.1155 | none |
| GALK2 | 0.726 | 0.5012 | 5.8406 | 1.88E-04 | 0.005 | 1.13 | -1.4737 | none |
| STX8 | 0.881 | 0.6618 | 5.8404 | 1.88E-04 | 0.005 | 1.13 | -1.0563 | none |
| GFI1B | 0.662 | 0.5074 | 5.8387 | 1.88E-04 | 0.005 | 1.13 | -1.2996 | none |
| L3MBTL1 | 0.776 | 0.5703 | 5.8383 | 0.00019 | 0.005 | 1.13 | -1.5154 | none |
| PRKG1 | 0.893 | 0.6717 | 5.8376 | 1.88E-04 | 0.005 | 1.13 | -1.0669 | none |
| DAPK2 | 0.782 | 0.6174 | 5.8376 | 1.88E-04 | 0.005 | 1.13 | -1.3803 | none |
| GPR82 | 0.985 | 0.7497 | 5.8343 | 1.89E-04 | 0.005 | 1.12 | -0.9649 | none |
| ZNF487 | 0.845 | 0.6308 | 5.83 | 1.90E-04 | 0.005 | 1.12 | -0.7032 | none |
| GTF2IRD2 | 0.927 | 0.7365 | 5.8283 | 1.91E-04 | 0.005 | 1.11 | -1.3855 | none |
| INHBE | 0.535 | 0.3938 | 5.8225 | 1.92E-04 | 0.005 | 1.11 | -1.5082 | none |
| FUT2 | 0.702 | 0.5512 | 5.8209 | 1.93E-04 | 0.005 | 1.1 | -1.5575 | OA-related |
| OR56A4 | 0.729 | 0.5372 | 5.8192 | 1.93E-04 | 0.005 | 1.1 | -1.1095 | none |
| CERS5 | 0.849 | 0.6106 | 5.816 | 1.94E-04 | 0.005 | 1.1 | -1.1234 | none |
| GATAD1 | 0.81 | 0.703 | 5.8156 | 1.94E-04 | 0.005 | 1.1 | -1.5065 | none |
| KANSL3 | 1.075 | 0.8973 | 5.815 | 1.94E-04 | 0.005 | 1.1 | -1.5278 | none |
| TPM1 | 0.788 | 0.6075 | 5.813 | 1.95E-04 | 0.005 | 1.09 | -1.5828 | OA-related |
| PUS10 | 0.911 | 0.725 | 5.8127 | 1.95E-04 | 0.005 | 1.09 | -1.3377 | none |
| PFKM | 0.968 | 0.8037 | 5.8081 | 1.96E-04 | 0.005 | 1.09 | -1.4857 | none |
| SPP2 | 0.861 | 0.6577 | 5.807 | 1.96E-04 | 0.005 | 1.09 | -0.8926 | none |
| PPIAL4G | 0.594 | 0.4624 | 5.8067 | 1.96E-04 | 0.005 | 1.09 | -1.4716 | none |
| P3H4 | 0.814 | 0.6203 | 5.8061 | 1.97E-04 | 0.005 | 1.09 | -1.3322 | none |
| FCER1G | 0.861 | 0.6181 | 5.8058 | 1.97E-04 | 0.005 | 1.08 | -0.9957 | none |
| TMEM67 | 1.016 | 0.7554 | 5.8031 | 1.97E-04 | 0.005 | 1.08 | -1.1904 | none |
| GFM1 | 0.832 | 0.6698 | 5.8025 | 1.97E-04 | 0.005 | 1.08 | -0.955 | none |
| PCDHGA6 | 0.586 | 0.4254 | 5.8023 | 1.97E-04 | 0.005 | 1.08 | -1.5227 | none |
| DCN | 1.023 | 0.8452 | 5.8017 | 1.98E-04 | 0.005 | 1.08 | -1.1938 | none |
| SOX6 | 0.939 | 0.7977 | 5.8016 | 1.98E-04 | 0.005 | 1.08 | -0.5472 | none |

| | | | | | | | | |
|---|---|---|---|---|---|---|---|---|
| ZSWIM1 | 0.844 | 0.7458 | 5.8011 | 1.98E-04 | 0.005 | 1.08 | -1.5657 | OA-related |
| C20ORF14 | 0.798 | 0.5979 | 5.7993 | 1.98E-04 | 0.005 | 1.08 | -1.5683 | OA-related |
| SNIP1 | 0.651 | 0.5232 | 5.7984 | 1.99E-04 | 0.005 | 1.08 | -1.5273 | none |
| S100PBP | 0.742 | 0.4983 | 5.797 | 1.99E-04 | 0.005 | 1.07 | -1.3943 | none |
| ABCG1 | 1.159 | 0.8757 | 5.7966 | 1.99E-04 | 0.005 | 1.07 | -1.0044 | none |
| CELF5 | 0.75 | 0.5854 | 5.7965 | 1.99E-04 | 0.005 | 1.07 | -1.2937 | none |
| SOX13 | 0.598 | 0.4263 | 5.7958 | 1.99E-04 | 0.005 | 1.07 | -1.4881 | none |
| NRG4 | 1.408 | 1.2299 | 5.7951 | 1.99E-04 | 0.005 | 1.07 | -1.089 | none |
| RNF220 | 0.836 | 0.6996 | 5.7944 | 2.00E-04 | 0.005 | 1.07 | -1.2126 | none |
| CSMD1 | 0.894 | 0.8411 | 5.7923 | 2.00E-04 | 0.005 | 1.07 | -0.4422 | none |
| DAAM2 | 0.824 | 0.6049 | 5.7908 | 2.01E-04 | 0.005 | 1.07 | -1.1831 | none |
| ZNF665 | 1.023 | 0.7701 | 5.7907 | 2.01E-04 | 0.005 | 1.07 | -1.2771 | none |
| ELN | 1.094 | 0.8225 | 5.7886 | 2.01E-04 | 0.005 | 1.06 | -0.5786 | none |
| TMEM212 | 0.703 | 0.5375 | 5.7879 | 2.01E-04 | 0.005 | 1.06 | -1.34 | none |
| DNAJC5G | 0.778 | 0.6017 | 5.7843 | 2.02E-04 | 0.005 | 1.06 | -0.3891 | none |
| OR1A1 | 0.632 | 0.4984 | 5.784 | 2.02E-04 | 0.005 | 1.06 | -1.6414 | OA-related |
| CEBPZOS | 0.694 | 0.5141 | 5.7823 | 2.03E-04 | 0.005 | 1.05 | -1.3594 | none |
| PNPO | 0.552 | 0.3929 | 5.7788 | 2.04E-04 | 0.005 | 1.05 | -1.1689 | none |
| ERO1A | 0.774 | 0.5286 | 5.7776 | 2.04E-04 | 0.005 | 1.05 | -1.5479 | OA-related |
| BLOC1S6 | 0.595 | 0.4454 | 5.7772 | 2.04E-04 | 0.005 | 1.05 | -1.2355 | none |
| CBFA2T2 | 0.892 | 0.7361 | 5.777 | 2.04E-04 | 0.005 | 1.05 | -1.5974 | OA-related |
| FITM2 | 0.662 | 0.5231 | 5.7751 | 2.05E-04 | 0.005 | 1.04 | -1.5401 | OA-related |
| UQCRHL | 0.62 | 0.4484 | 5.775 | 2.05E-04 | 0.005 | 1.04 | -1.2546 | none |
| PPP2R2D | 0.705 | 0.5621 | 5.7749 | 2.05E-04 | 0.005 | 1.04 | -0.8953 | none |
| PIGN | 0.577 | 0.3477 | 5.7745 | 2.05E-04 | 0.005 | 1.04 | -1.1367 | none |
| PROCR | 1.015 | 0.7693 | 5.7739 | 2.05E-04 | 0.005 | 1.04 | -1.2134 | none |
| TIAM2 | 1.428 | 1.1828 | 5.7711 | 2.06E-04 | 0.005 | 1.04 | -1.3935 | none |
| OR51B5 | 0.639 | 0.4522 | 5.7699 | 2.06E-04 | 0.005 | 1.04 | -0.9761 | none |
| FAM71F2 | 0.954 | 0.632 | 5.7696 | 2.06E-04 | 0.005 | 1.04 | -1.2023 | none |
| ZNF544 | 0.855 | 0.6638 | 5.769 | 2.06E-04 | 0.005 | 1.04 | -1.4164 | none |
| A2ML1 | 0.652 | 0.5428 | 5.7658 | 2.07E-04 | 0.005 | 1.03 | -1.2933 | none |
| CLCN6 | 0.826 | 0.6976 | 5.7636 | 2.08E-04 | 0.005 | 1.03 | -1.2529 | none |
| OPRM1 | 0.576 | 0.4445 | 5.7604 | 2.09E-04 | 0.005 | 1.03 | -1.6403 | OA-related |
| CDC20B | 0.677 | 0.5057 | 5.7586 | 2.09E-04 | 0.005 | 1.02 | -1.3443 | none |
| FAM189B | 0.612 | 0.4618 | 5.7582 | 2.09E-04 | 0.005 | 1.02 | -1.5899 | OA-related |
| TMEM40 | 0.793 | 0.5974 | 5.7563 | 2.10E-04 | 0.005 | 1.02 | -1.2453 | none |
| SRSF12 | 0.933 | 0.6406 | 5.7559 | 2.10E-04 | 0.005 | 1.02 | -1.4303 | none |
| GTF2I | 1.002 | 0.8758 | 5.7545 | 2.10E-04 | 0.005 | 1.02 | -1.5524 | OA-related |
| TNFRSF1A | 0.625 | 0.4765 | 5.7534 | 2.11E-04 | 0.005 | 1.02 | -1.1816 | none |
| C5 | 1.046 | 0.8396 | 5.7519 | 2.11E-04 | 0.005 | 1.01 | -1.3821 | none |

| | | | | | | | | |
|---|---|---|---|---|---|---|---|---|
| OR2B11 | 0.521 | 0.3902 | 5.7477 | 2.12E-04 | 0.005 | 1.01 | -1.6537 | OA-related |
| ZFP42 | 0.652 | 0.4917 | 5.7477 | 2.12E-04 | 0.005 | 1.01 | -1.555 | OA-related |
| TRAF6 | 0.837 | 0.6396 | 5.7468 | 2.13E-04 | 0.005 | 1.01 | -0.8714 | none |
| KCNN1 | 0.745 | 0.5601 | 5.7462 | 2.13E-04 | 0.005 | 1.01 | -1.4383 | none |
| CA5A | 0.768 | 0.5646 | 5.7449 | 2.13E-04 | 0.005 | 1 | -1.5911 | OA-related |
| ABCC2 | 0.891 | 0.6845 | 5.7445 | 2.13E-04 | 0.005 | 1 | -0.8125 | none |
| BHMT | 0.836 | 0.7136 | 5.7434 | 2.14E-04 | 0.005 | 1 | -0.6839 | none |
| USP54 | 1.184 | 1.0341 | 5.7415 | 2.14E-04 | 0.005 | 1 | -1.2409 | none |
| IL10 | 0.966 | 0.8171 | 5.7367 | 2.15E-04 | 0.005 | 0.99 | -1.0211 | none |
| PQLC1 | 0.879 | 0.6702 | 5.7353 | 2.16E-04 | 0.005 | 0.99 | -1.071 | none |
| RBM23 | 0.713 | 0.5621 | 5.7349 | 2.16E-04 | 0.005 | 0.99 | -1.1371 | none |
| GIGYF2 | 1.15 | 0.9682 | 5.7331 | 2.17E-04 | 0.005 | 0.99 | -1.2678 | none |
| SHC1 | 0.719 | 0.5986 | 5.7326 | 2.17E-04 | 0.005 | 0.99 | -1.2061 | none |
| ZMAT3 | 0.597 | 0.4122 | 5.7315 | 2.17E-04 | 0.005 | 0.99 | -1.3396 | none |
| MAP3K10 | 0.925 | 0.794 | 5.7314 | 2.17E-04 | 0.005 | 0.99 | -1.4845 | none |
| SPATA31A | 0.951 | 0.8268 | 5.7304 | 2.17E-04 | 0.005 | 0.99 | -1.4923 | none |
| CAPN10 | 0.567 | 0.4907 | 5.7273 | 2.18E-04 | 0.005 | 0.98 | -0.7109 | none |
| HSD11B1L | 1.297 | 0.9646 | 5.7243 | 2.19E-04 | 0.005 | 0.98 | -1.1802 | none |
| HMGCS1 | 0.621 | 0.4819 | 5.7236 | 2.19E-04 | 0.005 | 0.98 | -0.9093 | none |
| ZNF785 | 0.789 | 0.6092 | 5.7218 | 2.20E-04 | 0.005 | 0.97 | -1.2867 | none |
| NUP133 | 0.604 | 0.4789 | 5.7212 | 2.20E-04 | 0.005 | 0.97 | -1.0085 | none |
| MTFMT | 0.562 | 0.3793 | 5.7207 | 2.20E-04 | 0.005 | 0.97 | -1.2717 | none |
| RDH16 | 0.534 | 0.387 | 5.72 | 2.20E-04 | 0.005 | 0.97 | -1.6027 | OA-related |
| AKAP9 | 1.016 | 0.8647 | 5.7188 | 2.21E-04 | 0.005 | 0.97 | -1.5057 | none |
| OXR1 | 1.061 | 0.9066 | 5.7176 | 2.21E-04 | 0.005 | 0.97 | -0.8545 | none |
| ST5 | 1.364 | 1.1947 | 5.7162 | 2.21E-04 | 0.005 | 0.97 | -1.2426 | none |
| SERPINF1 | 0.744 | 0.6448 | 5.7155 | 2.22E-04 | 0.005 | 0.97 | -1.1708 | none |
| LRRTM4 | 0.944 | 0.7951 | 5.7154 | 2.22E-04 | 0.005 | 0.97 | -1.0879 | none |
| HOGA1 | 0.753 | 0.5709 | 5.7133 | 2.22E-04 | 0.005 | 0.96 | -1.0532 | none |
| ANKRD62 | 0.494 | 0.3589 | 5.7129 | 2.22E-04 | 0.005 | 0.96 | -1.4833 | none |
| CRK | 0.716 | 0.5781 | 5.7127 | 2.22E-04 | 0.005 | 0.96 | -1.4394 | none |
| MATN2 | 0.765 | 0.5971 | 5.7099 | 2.23E-04 | 0.005 | 0.96 | -1.4048 | none |
| FHAD1 | 0.773 | 0.6372 | 5.7079 | 2.24E-04 | 0.005 | 0.96 | -0.8665 | none |
| KCP | 0.619 | 0.4547 | 5.7066 | 2.24E-04 | 0.005 | 0.95 | -1.3138 | none |
| PAPLN | 0.575 | 0.4504 | 5.7054 | 2.25E-04 | 0.005 | 0.95 | -1.5209 | none |
| RBBP5 | 0.725 | 0.5372 | 5.7032 | 2.25E-04 | 0.005 | 0.95 | -1.6175 | OA-related |
| HERC2 | 0.956 | 0.7779 | 5.7001 | 2.26E-04 | 0.005 | 0.95 | -1.2527 | none |
| UGGT2 | 0.801 | 0.5714 | 5.6978 | 2.27E-04 | 0.005 | 0.94 | -1.2155 | none |
| ADGRD2 | 0.808 | 0.6302 | 5.6931 | 2.28E-04 | 0.005 | 0.94 | -1.3532 | none |
| NVL | 0.858 | 0.7227 | 5.6919 | 2.29E-04 | 0.005 | 0.93 | -1.2996 | none |

| | | | | | | | | |
|---|---|---|---|---|---|---|---|---|
| CLDN16 | 0.68 | 0.4879 | 5.6914 | 2.29E-04 | 0.005 | 0.93 | -1.3275 | none |
| ELMOD3 | 0.607 | 0.462 | 5.6913 | 2.29E-04 | 0.005 | 0.93 | -1.4792 | none |
| FAAP24 | 0.797 | 0.6701 | 5.6901 | 2.29E-04 | 0.005 | 0.93 | -1.3874 | none |
| LETM2 | 0.728 | 0.5806 | 5.6892 | 2.30E-04 | 0.005 | 0.93 | -1.0278 | none |
| RPS6KL1 | 0.867 | 0.7033 | 5.6862 | 2.31E-04 | 0.005 | 0.93 | -0.8781 | none |
| TAF1B | 0.841 | 0.6505 | 5.6824 | 2.32E-04 | 0.005 | 0.92 | -1.1189 | none |
| OR14J1 | 0.525 | 0.4026 | 5.6809 | 2.32E-04 | 0.005 | 0.92 | -1.3686 | none |
| ST3GAL3 | 0.817 | 0.6361 | 5.6798 | 2.33E-04 | 0.005 | 0.92 | -1.6637 | OA-related |
| SLC25A21 | 0.717 | 0.5032 | 5.6777 | 2.33E-04 | 0.005 | 0.92 | -1.2287 | none |
| AC087651 | 0.644 | 0.4815 | 5.6771 | 2.33E-04 | 0.005 | 0.91 | -1.5646 | OA-related |
| OR5AS1 | 0.648 | 0.4982 | 5.6766 | 2.33E-04 | 0.005 | 0.91 | -1.3985 | none |
| CTF1 | 0.655 | 0.4878 | 5.6761 | 2.34E-04 | 0.005 | 0.91 | -1.039 | none |
| PTGES3L | 0.987 | 0.7413 | 5.6727 | 2.35E-04 | 0.005 | 0.91 | -1.2764 | none |
| EFCAB12 | 0.577 | 0.4482 | 5.6719 | 2.35E-04 | 0.005 | 0.91 | -1.4974 | none |
| DCBLD2 | 1.301 | 1.0184 | 5.6716 | 2.35E-04 | 0.005 | 0.91 | -1.1769 | none |
| TMED4 | 0.711 | 0.5067 | 5.6692 | 2.36E-04 | 0.005 | 0.9 | -1.5265 | none |
| TTLL3 | 0.988 | 0.8195 | 5.6692 | 2.36E-04 | 0.005 | 0.9 | -1.2686 | none |
| RICTOR | 0.989 | 0.8279 | 5.6676 | 2.36E-04 | 0.005 | 0.9 | -1.5858 | OA-related |
| LPIN1 | 0.923 | 0.7613 | 5.664 | 2.37E-04 | 0.005 | 0.9 | -1.3412 | none |
| GSE1 | 0.965 | 0.7996 | 5.6626 | 2.38E-04 | 0.005 | 0.9 | -1.2914 | none |
| TTC23 | 0.874 | 0.6941 | 5.6622 | 2.38E-04 | 0.005 | 0.89 | -1.3977 | none |
| EIF2D | 0.603 | 0.4447 | 5.6621 | 2.38E-04 | 0.005 | 0.89 | -1.2188 | none |
| POMZP3 | 0.732 | 0.5222 | 5.6617 | 2.38E-04 | 0.005 | 0.89 | -1.3539 | none |
| C3ORF70 | 0.63 | 0.4694 | 5.6609 | 2.38E-04 | 0.005 | 0.89 | -1.4527 | none |
| NAV1 | 0.648 | 0.4952 | 5.6601 | 2.39E-04 | 0.005 | 0.89 | -1.5413 | OA-related |
| PLEKHD1 | 0.629 | 0.493 | 5.6598 | 2.39E-04 | 0.005 | 0.89 | -1.5619 | OA-related |
| SEC23A | 0.923 | 0.7438 | 5.6587 | 2.39E-04 | 0.005 | 0.89 | -1.4592 | none |
| ETFBKMT | 1.226 | 0.9219 | 5.6582 | 2.39E-04 | 0.005 | 0.89 | -1.1257 | none |
| CRTAM | 0.658 | 0.5004 | 5.6579 | 2.39E-04 | 0.005 | 0.89 | -0.7682 | none |
| VPS11 | 0.756 | 0.6425 | 5.6574 | 2.40E-04 | 0.005 | 0.89 | -1.2985 | none |
| KCNMB1 | 0.633 | 0.4934 | 5.6555 | 2.40E-04 | 0.005 | 0.89 | -1.5347 | OA-related |
| ULK2 | 0.896 | 0.6783 | 5.6532 | 2.41E-04 | 0.005 | 0.88 | -0.9906 | none |
| RENBP | 1.162 | 0.9419 | 5.653 | 2.41E-04 | 0.005 | 0.88 | -1.4403 | none |
| CDH15 | 0.693 | 0.5088 | 5.6526 | 2.41E-04 | 0.005 | 0.88 | -1.0392 | none |
| SPDYE5 | 1.131 | 0.9839 | 5.6525 | 2.41E-04 | 0.005 | 0.88 | -1.4073 | none |
| ATP6V0A2 | 0.765 | 0.5491 | 5.6523 | 2.41E-04 | 0.005 | 0.88 | -1.3874 | none |
| PHACTR1 | 0.938 | 0.7526 | 5.6513 | 2.42E-04 | 0.005 | 0.88 | -1.0711 | none |
| AICDA | 0.64 | 0.4618 | 5.6512 | 2.42E-04 | 0.005 | 0.88 | -1.4777 | none |
| CDH12 | 0.814 | 0.63 | 5.6503 | 2.42E-04 | 0.005 | 0.88 | -1.4189 | none |
| SPECC1 | 1.244 | 1.0625 | 5.6503 | 2.42E-04 | 0.005 | 0.88 | -1.5442 | OA-related |

| | | | | | | | | |
|---|---|---|---|---|---|---|---|---|
| CHST12 | 0.674 | 0.4995 | 5.6494 | 2.42E-04 | 0.005 | 0.88 | -1.126 | none |
| MYL10 | 0.922 | 0.7429 | 5.649 | 2.42E-04 | 0.005 | 0.88 | -1.5264 | none |
| STIL | 1.078 | 0.8449 | 5.6478 | 2.43E-04 | 0.005 | 0.88 | -0.854 | none |
| DYNLT3 | 0.743 | 0.5088 | 5.6459 | 2.43E-04 | 0.005 | 0.87 | -0.9524 | none |
| TRIM16L | 0.761 | 0.5007 | 5.6444 | 2.44E-04 | 0.005 | 0.87 | -1.2619 | none |
| MAGEB10 | 0.711 | 0.5557 | 5.6435 | 2.44E-04 | 0.005 | 0.87 | -1.257 | none |
| APLP2 | 1.046 | 0.9039 | 5.6417 | 2.45E-04 | 0.005 | 0.87 | -1.3515 | none |
| BEND2 | 0.631 | 0.479 | 5.6411 | 2.45E-04 | 0.005 | 0.87 | -1.5853 | OA-related |
| KCTD19 | 1.62 | 1.3571 | 5.6401 | 2.45E-04 | 0.005 | 0.87 | -1.1594 | none |
| ACSS3 | 0.672 | 0.4691 | 5.6384 | 2.46E-04 | 0.005 | 0.86 | -1.2453 | none |
| ULBP3 | 0.86 | 0.7176 | 5.638 | 2.46E-04 | 0.005 | 0.86 | -1.1618 | none |
| SPAG7 | 0.682 | 0.5613 | 5.6349 | 2.47E-04 | 0.005 | 0.86 | -1.1521 | none |
| MLLT10 | 1.059 | 0.8399 | 5.629 | 2.49E-04 | 0.005 | 0.85 | -1.2613 | none |
| CCDC191 | 0.889 | 0.6646 | 5.6278 | 2.49E-04 | 0.005 | 0.85 | -0.6325 | none |
| ZNF8 | 0.686 | 0.55 | 5.6245 | 2.50E-04 | 0.005 | 0.84 | -1.6358 | OA-related |
| RNF217 | 0.946 | 0.8761 | 5.6224 | 2.51E-04 | 0.005 | 0.84 | -0.9062 | none |
| AXL | 0.675 | 0.4869 | 5.6215 | 2.51E-04 | 0.005 | 0.84 | -1.2165 | none |
| UPB1 | 0.635 | 0.4897 | 5.6194 | 2.52E-04 | 0.005 | 0.84 | -1.397 | none |
| LEMD3 | 1.109 | 0.9238 | 5.6188 | 2.52E-04 | 0.005 | 0.84 | -1.6003 | OA-related |
| NR6A1 | 0.581 | 0.4496 | 5.6186 | 2.52E-04 | 0.005 | 0.84 | -1.5099 | none |
| SLC48A1 | 0.625 | 0.4354 | 5.6099 | 2.55E-04 | 0.005 | 0.82 | -1.391 | none |
| GABRA1 | 0.721 | 0.5248 | 5.609 | 2.56E-04 | 0.005 | 0.82 | -1.5343 | OA-related |
| CAPS | 0.644 | 0.5022 | 5.6087 | 2.56E-04 | 0.005 | 0.82 | -1.4981 | none |
| UGT1A1 | 0.545 | 0.4159 | 5.6084 | 2.56E-04 | 0.005 | 0.82 | -1.1683 | none |
| TMC1 | 0.77 | 0.5848 | 5.6045 | 2.57E-04 | 0.005 | 0.82 | -0.9861 | none |
| UBE2O | 1.435 | 1.3061 | 5.6038 | 2.58E-04 | 0.005 | 0.82 | -1.276 | none |
| THRA | 0.693 | 0.6215 | 5.6025 | 2.58E-04 | 0.005 | 0.81 | -1.356 | none |
| HTR3B | 0.656 | 0.4892 | 5.6024 | 2.58E-04 | 0.005 | 0.81 | -1.5587 | OA-related |
| GRK4 | 0.84 | 0.6476 | 5.6023 | 2.58E-04 | 0.005 | 0.81 | -1.4317 | none |
| TMEM143 | 0.583 | 0.4723 | 5.6021 | 2.58E-04 | 0.005 | 0.81 | -1.1579 | none |
| MFSD13A | 0.966 | 0.7604 | 5.5978 | 2.60E-04 | 0.005 | 0.81 | -1.3384 | none |
| FAM186A | 0.611 | 0.4876 | 5.5954 | 2.60E-04 | 0.005 | 0.81 | -1.4253 | none |
| CYP2W1 | 0.676 | 0.4605 | 5.595 | 2.61E-04 | 0.005 | 0.8 | -1.4287 | none |
| PPP1R13L | 1.019 | 0.8586 | 5.5943 | 2.61E-04 | 0.005 | 0.8 | -1.3172 | none |
| DNM2 | 1.257 | 1.0872 | 5.591 | 2.62E-04 | 0.005 | 0.8 | -1.4356 | none |
| UBE2S | 1.06 | 0.8189 | 5.5896 | 2.63E-04 | 0.005 | 0.8 | -1.0548 | none |
| BST1 | 0.874 | 0.6803 | 5.5895 | 2.63E-04 | 0.005 | 0.8 | -1.5453 | OA-related |
| TRNAU1AI | 0.894 | 0.645 | 5.5888 | 2.63E-04 | 0.005 | 0.8 | -1.0141 | none |
| TMEM119 | 0.887 | 0.6226 | 5.5883 | 2.63E-04 | 0.005 | 0.8 | -1.1882 | none |
| HTR1D | 0.662 | 0.5263 | 5.5881 | 2.63E-04 | 0.005 | 0.8 | -1.4943 | none |

| | | | | | | | | |
|---|---|---|---|---|---|---|---|---|
| OR10A3 | 0.529 | 0.3834 | 5.5878 | 2.63E-04 | 0.005 | 0.8 | -1.535 | OA-related |
| SENP7 | 1.101 | 0.9222 | 5.587 | 2.63E-04 | 0.005 | 0.79 | -1.3346 | none |
| CYTH4 | 0.784 | 0.6336 | 5.5796 | 2.66E-04 | 0.005 | 0.78 | -1.028 | none |
| GOSR1 | 0.628 | 0.4758 | 5.5755 | 2.68E-04 | 0.005 | 0.78 | -1.2209 | none |
| SDCCAG8 | 0.801 | 0.5438 | 5.5742 | 2.68E-04 | 0.005 | 0.78 | -0.9257 | none |
| ZNF343 | 0.828 | 0.7328 | 5.5724 | 2.69E-04 | 0.005 | 0.77 | -1.1783 | none |
| MCU | 0.633 | 0.4123 | 5.5718 | 2.69E-04 | 0.005 | 0.77 | -0.3193 | none |
| DPPA4 | 0.733 | 0.511 | 5.5707 | 2.69E-04 | 0.005 | 0.77 | -1.5185 | none |
| ZNF529 | 0.984 | 0.728 | 5.5669 | 2.71E-04 | 0.005 | 0.77 | -1.388 | none |
| ZNF670-ZN | 0.599 | 0.4869 | 5.566 | 2.71E-04 | 0.005 | 0.77 | -1.318 | none |
| ZNF799 | 1.069 | 0.8248 | 5.5622 | 2.72E-04 | 0.005 | 0.76 | -1.3358 | none |
| ICA1L | 0.974 | 0.8507 | 5.5611 | 2.73E-04 | 0.005 | 0.76 | -1.0447 | none |
| ZNF776 | 1.059 | 0.8495 | 5.5599 | 2.73E-04 | 0.005 | 0.76 | -1.285 | none |
| MFSD14C | 0.941 | 0.8223 | 5.5595 | 2.73E-04 | 0.005 | 0.76 | -1.5033 | none |
| GGA1 | 0.795 | 0.6709 | 5.5586 | 2.74E-04 | 0.005 | 0.76 | -1.5521 | OA-related |
| GTPBP2 | 1.128 | 0.8998 | 5.557 | 2.74E-04 | 0.005 | 0.75 | -1.1032 | none |
| C8ORF86 | 0.798 | 0.621 | 5.5568 | 2.74E-04 | 0.005 | 0.75 | -1.2726 | none |
| NEK9 | 0.738 | 0.5951 | 5.5556 | 2.75E-04 | 0.005 | 0.75 | -1.2935 | none |
| RUBCN | 1.31 | 1.1494 | 5.5551 | 2.75E-04 | 0.005 | 0.75 | -1.1965 | none |
| OR7C1 | 0.58 | 0.4459 | 5.5541 | 2.76E-04 | 0.005 | 0.75 | -1.4625 | none |
| ZPR1 | 0.67 | 0.5368 | 5.5528 | 2.76E-04 | 0.005 | 0.75 | -1.3823 | none |
| MAEL | 0.661 | 0.4677 | 5.5524 | 2.76E-04 | 0.005 | 0.75 | -1.4485 | none |
| IQCF3 | 0.729 | 0.6414 | 5.5524 | 2.76E-04 | 0.005 | 0.75 | 0.46327 | none |
| WFDC1 | 0.664 | 0.5226 | 5.5521 | 2.76E-04 | 0.005 | 0.75 | -1.4764 | none |
| CATSPERG | 0.923 | 0.7031 | 5.5509 | 2.77E-04 | 0.005 | 0.75 | -1.1993 | none |
| FKBP6 | 0.793 | 0.6018 | 5.5494 | 2.77E-04 | 0.005 | 0.74 | -1.3782 | none |
| PLA2G6 | 1.178 | 0.9817 | 5.5462 | 2.78E-04 | 0.005 | 0.74 | -1.0128 | none |
| C6ORF132 | 0.991 | 0.7571 | 5.5443 | 2.79E-04 | 0.005 | 0.74 | -1.3625 | none |
| SIGLEC6 | 0.564 | 0.4477 | 5.5435 | 2.80E-04 | 0.005 | 0.74 | -1.4175 | none |
| TRIP4 | 0.99 | 0.7663 | 5.5434 | 2.80E-04 | 0.005 | 0.74 | -1.0387 | none |
| NT5DC3 | 0.738 | 0.6268 | 5.5421 | 2.80E-04 | 0.005 | 0.73 | -1.5534 | OA-related |
| FAM153C | 0.776 | 0.559 | 5.5418 | 2.80E-04 | 0.005 | 0.73 | -1.1305 | none |
| KCNE4 | 0.937 | 0.7236 | 5.5396 | 2.81E-04 | 0.005 | 0.73 | -1.2698 | none |
| VMAC | 0.799 | 0.6187 | 5.5368 | 2.82E-04 | 0.005 | 0.73 | -1.3246 | none |
| TACC2 | 0.672 | 0.4686 | 5.5367 | 2.82E-04 | 0.005 | 0.73 | -1.2972 | none |
| ARIH1 | 1.04 | 0.787 | 5.5367 | 2.82E-04 | 0.005 | 0.73 | -1.2281 | none |
| MIER2 | 1.06 | 0.9221 | 5.5361 | 2.82E-04 | 0.005 | 0.73 | -1.2998 | none |
| KCTD6 | 0.66 | 0.429 | 5.5361 | 2.82E-04 | 0.005 | 0.73 | -1.0366 | none |
| RNF4 | 0.49 | 0.3606 | 5.5357 | 2.82E-04 | 0.005 | 0.72 | -1.2391 | none |
| ZNF563 | 0.483 | 0.3512 | 5.5353 | 2.83E-04 | 0.005 | 0.72 | -0.9393 | none |

| | | | | | | | | |
|---|---|---|---|---|---|---|---|---|
| TMEM156 | 0.775 | 0.5711 | 5.5347 | 2.83E-04 | 0.005 | 0.72 | -1.257 | none |
| OPRD1 | 0.576 | 0.4116 | 5.5306 | 2.84E-04 | 0.005 | 0.72 | -1.5261 | none |
| ELF3 | 1.261 | 1.05 | 5.5267 | 2.86E-04 | 0.005 | 0.71 | -1.381 | none |
| ZNF490 | 0.872 | 0.7122 | 5.5253 | 2.87E-04 | 0.005 | 0.71 | -1.3032 | none |
| PPFIA1 | 1.432 | 1.3241 | 5.5237 | 2.87E-04 | 0.005 | 0.71 | -1.2214 | none |
| SLC51A | 1.058 | 0.8914 | 5.5208 | 2.88E-04 | 0.005 | 0.7 | -0.915 | none |
| PPP1R27 | 0.638 | 0.4264 | 5.5205 | 2.88E-04 | 0.005 | 0.7 | -0.9255 | none |
| CCDC158 | 0.739 | 0.5169 | 5.5137 | 2.91E-04 | 0.005 | 0.7 | -1.5629 | OA-related |
| MED17 | 0.566 | 0.4118 | 5.5132 | 2.91E-04 | 0.005 | 0.69 | -1.4757 | none |
| TESMIN | 1.138 | 0.8157 | 5.5113 | 2.92E-04 | 0.005 | 0.69 | -1.461 | none |
| KDM2A | 1.153 | 1.03 | 5.5111 | 2.92E-04 | 0.005 | 0.69 | -1.4369 | none |
| FAM13A | 0.626 | 0.5427 | 5.5103 | 2.92E-04 | 0.005 | 0.69 | -0.9158 | none |
| PKD2L2 | 1.212 | 1.0036 | 5.5102 | 2.92E-04 | 0.005 | 0.69 | -1.2224 | none |
| ALDH6A1 | 0.692 | 0.5297 | 5.5081 | 2.93E-04 | 0.005 | 0.69 | -1.5778 | OA-related |
| EPS15L1 | 0.931 | 0.8106 | 5.5064 | 2.94E-04 | 0.005 | 0.69 | -1.2239 | none |
| ZFAND3 | 1.001 | 0.8436 | 5.5064 | 2.94E-04 | 0.005 | 0.69 | -1.1269 | none |
| APOBEC3A | 0.829 | 0.6518 | 5.5063 | 2.94E-04 | 0.005 | 0.69 | -1.4302 | none |
| C8ORF34 | 0.777 | 0.5878 | 5.5048 | 2.95E-04 | 0.005 | 0.68 | -1.4136 | none |
| NUP160 | 0.631 | 0.4671 | 5.504 | 2.95E-04 | 0.005 | 0.68 | -1.4964 | none |
| PAEP | 0.937 | 0.7102 | 5.5022 | 2.96E-04 | 0.005 | 0.68 | -1.1798 | none |
| ZSCAN5A | 0.92 | 0.7283 | 5.4995 | 2.97E-04 | 0.005 | 0.68 | -1.5819 | OA-related |
| ARNT | 1.054 | 0.8933 | 5.4962 | 2.98E-04 | 0.005 | 0.67 | -1.1071 | none |
| NME6 | 0.531 | 0.358 | 5.4904 | 3.00E-04 | 0.005 | 0.66 | -1.2983 | none |
| LRRC37A3 | 1.175 | 1.0013 | 5.4883 | 3.01E-04 | 0.005 | 0.66 | -1.043 | none |
| DOCK7 | 0.871 | 0.6817 | 5.4876 | 3.02E-04 | 0.005 | 0.66 | -1.1471 | none |
| EMP2 | 0.639 | 0.4917 | 5.4875 | 3.02E-04 | 0.005 | 0.66 | -1.5015 | none |
| TANGO2 | 1.062 | 0.8478 | 5.4866 | 3.02E-04 | 0.005 | 0.66 | -1.4343 | none |
| PROSER3 | 0.887 | 0.7839 | 5.486 | 3.02E-04 | 0.005 | 0.66 | -1.4001 | none |
| IZUMO1 | 0.574 | 0.4144 | 5.4848 | 3.03E-04 | 0.005 | 0.66 | -1.4265 | none |
| MYO3B | 0.556 | 0.3859 | 5.4835 | 3.03E-04 | 0.005 | 0.65 | -1.3893 | none |
| IGF2BP1 | 1.016 | 0.8548 | 5.4819 | 3.04E-04 | 0.005 | 0.65 | -1.0436 | none |
| RBSN | 0.916 | 0.7232 | 5.4795 | 3.05E-04 | 0.005 | 0.65 | -1.5407 | OA-related |
| P4HTM | 0.761 | 0.5883 | 5.4791 | 3.05E-04 | 0.005 | 0.65 | -1.2491 | none |
| SGK3 | 0.766 | 0.5745 | 5.4789 | 3.05E-04 | 0.005 | 0.65 | -1.2292 | none |
| ZNF483 | 0.724 | 0.496 | 5.4774 | 3.06E-04 | 0.005 | 0.65 | -1.3699 | none |
| AKR1B10 | 0.73 | 0.4994 | 5.4757 | 3.07E-04 | 0.005 | 0.64 | -1.0954 | none |
| CHD2 | 0.933 | 0.7123 | 5.4752 | 3.07E-04 | 0.005 | 0.64 | -1.2912 | none |
| ZNF81 | 1.092 | 0.9348 | 5.4728 | 3.08E-04 | 0.005 | 0.64 | -1.2911 | none |
| MBD5 | 0.592 | 0.4251 | 5.4696 | 3.09E-04 | 0.005 | 0.64 | -1.1357 | none |
| CDH8 | 0.64 | 0.4816 | 5.4694 | 3.09E-04 | 0.005 | 0.63 | -1.4641 | none |

| | | | | | | | | |
|---|---|---|---|---|---|---|---|---|
| GNAL | 0.722 | 0.5923 | 5.4693 | 3.09E-04 | 0.005 | 0.63 | -1.5096 | none |
| ZDHHC17 | 0.631 | 0.5169 | 5.4669 | 3.10E-04 | 0.005 | 0.63 | -0.7815 | none |
| PCBD2 | 0.798 | 0.6153 | 5.4668 | 3.10E-04 | 0.005 | 0.63 | -1.4217 | none |
| MAPK1IP1 | 0.625 | 0.4307 | 5.4657 | 3.11E-04 | 0.005 | 0.63 | -1.2488 | none |
| PGA3 | 0.726 | 0.5808 | 5.4642 | 3.11E-04 | 0.005 | 0.63 | -1.1381 | none |
| CCDC69 | 0.766 | 0.6034 | 5.4612 | 3.13E-04 | 0.005 | 0.62 | -1.299 | none |
| ZNF611 | 1.168 | 0.9709 | 5.461 | 3.13E-04 | 0.005 | 0.62 | -1.2405 | none |
| MED13 | 1.11 | 0.9019 | 5.4529 | 3.16E-04 | 0.006 | 0.61 | -1.417 | none |
| TTPAL | 0.611 | 0.4995 | 5.4525 | 3.16E-04 | 0.006 | 0.61 | -1.4902 | none |
| ABCD4 | 0.558 | 0.4373 | 5.4514 | 3.17E-04 | 0.006 | 0.61 | -1.5226 | none |
| CNKSR1 | 0.921 | 0.6543 | 5.4492 | 3.18E-04 | 0.006 | 0.61 | -0.9522 | none |
| TMSB15B | 0.629 | 0.4668 | 5.4489 | 3.18E-04 | 0.006 | 0.61 | -0.8504 | none |
| BRWD1 | 0.791 | 0.5935 | 5.4446 | 3.20E-04 | 0.006 | 0.6 | -1.3759 | none |
| PLPP7 | 0.575 | 0.4159 | 5.4413 | 3.21E-04 | 0.006 | 0.6 | -1.2215 | none |
| TCFL5 | 0.963 | 0.7963 | 5.4404 | 3.22E-04 | 0.006 | 0.6 | -1.6076 | OA-related |
| ANKS4B | 0.492 | 0.3487 | 5.4401 | 3.22E-04 | 0.006 | 0.59 | -1.5568 | OA-related |
| EML2 | 0.713 | 0.6647 | 5.4385 | 3.23E-04 | 0.006 | 0.59 | -0.7926 | none |
| BCL2L14 | 0.638 | 0.4905 | 5.4375 | 3.23E-04 | 0.006 | 0.59 | -1.2575 | none |
| DNAH14 | 1.097 | 0.9846 | 5.4359 | 3.24E-04 | 0.006 | 0.59 | -0.348 | none |
| SCN1B | 0.722 | 0.5501 | 5.4341 | 3.25E-04 | 0.006 | 0.59 | -1.3605 | none |
| GFRAL | 0.69 | 0.4966 | 5.433 | 3.25E-04 | 0.006 | 0.59 | -0.9633 | none |
| ING5 | 0.906 | 0.7836 | 5.4302 | 3.26E-04 | 0.006 | 0.58 | -1.4546 | none |
| SLC2A13 | 0.684 | 0.4869 | 5.4299 | 3.26E-04 | 0.006 | 0.58 | -0.838 | none |
| ZNF891 | 0.71 | 0.5654 | 5.4288 | 3.27E-04 | 0.006 | 0.58 | -1.517 | none |
| METTL6 | 0.583 | 0.3747 | 5.4286 | 3.27E-04 | 0.006 | 0.58 | -1.33 | none |
| AL589743. | 1.151 | 0.85 | 5.4271 | 3.28E-04 | 0.006 | 0.58 | -1.1162 | none |
| LIPC | 0.712 | 0.5384 | 5.4254 | 3.28E-04 | 0.006 | 0.57 | -1.0703 | none |
| SLC13A5 | 0.753 | 0.5859 | 5.425 | 3.29E-04 | 0.006 | 0.57 | -1.4901 | none |
| SNX11 | 0.821 | 0.5665 | 5.424 | 3.29E-04 | 0.006 | 0.57 | -1.3565 | none |
| MACC1 | 0.889 | 0.6319 | 5.423 | 3.30E-04 | 0.006 | 0.57 | -1.09 | none |
| BMP1 | 0.967 | 0.8205 | 5.423 | 3.30E-04 | 0.006 | 0.57 | -1.4622 | none |
| IDH3A | 0.472 | 0.2626 | 5.4226 | 3.30E-04 | 0.006 | 0.57 | -0.7793 | none |
| FAT1 | 1.017 | 0.845 | 5.4215 | 3.30E-04 | 0.006 | 0.57 | -0.54 | none |
| RUFY3 | 0.903 | 0.7732 | 5.42 | 3.31E-04 | 0.006 | 0.57 | -1.1305 | none |
| NANOG | 0.61 | 0.4744 | 5.417 | 3.32E-04 | 0.006 | 0.56 | -1.2151 | none |
| SAG | 0.692 | 0.5003 | 5.4168 | 3.32E-04 | 0.006 | 0.56 | -1.5169 | none |
| NPLOC4 | 0.642 | 0.4767 | 5.4165 | 3.33E-04 | 0.006 | 0.56 | -1.4287 | none |
| BEST1 | 0.695 | 0.5109 | 5.4128 | 3.34E-04 | 0.006 | 0.56 | -1.2944 | none |
| ZNF714 | 0.896 | 0.7375 | 5.4121 | 3.35E-04 | 0.006 | 0.56 | -1.6662 | OA-related |
| C2CD3 | 1.041 | 0.9415 | 5.4103 | 3.35E-04 | 0.006 | 0.55 | -1.2527 | none |

| | | | | | | | | |
|---|---|---|---|---|---|---|---|---|
| HNF1A | 0.482 | 0.3806 | 5.409 | 3.36E-04 | 0.006 | 0.55 | -0.7203 | none |
| LRRC57 | 0.77 | 0.6263 | 5.4084 | 3.36E-04 | 0.006 | 0.55 | -1.5308 | none |
| IL18 | 0.726 | 0.6189 | 5.4083 | 3.36E-04 | 0.006 | 0.55 | -0.4811 | none |
| OR3A3 | 0.529 | 0.3967 | 5.408 | 3.36E-04 | 0.006 | 0.55 | -1.529 | none |
| NUPR1 | 0.814 | 0.6281 | 5.4058 | 3.37E-04 | 0.006 | 0.55 | -1.5191 | none |
| SSH2 | 0.68 | 0.5298 | 5.4046 | 3.38E-04 | 0.006 | 0.55 | -1.5324 | OA-related |
| GPLD1 | 0.929 | 0.7693 | 5.4038 | 3.38E-04 | 0.006 | 0.55 | -1.1501 | none |
| C2ORF88 | 0.625 | 0.4635 | 5.4038 | 3.38E-04 | 0.006 | 0.54 | -0.6578 | none |
| PRKCZ | 0.915 | 0.7828 | 5.4034 | 3.39E-04 | 0.006 | 0.54 | -1.0177 | none |
| ZWILCH | 0.764 | 0.5196 | 5.4018 | 3.39E-04 | 0.006 | 0.54 | -1.033 | none |
| CCDC27 | 0.673 | 0.5109 | 5.3992 | 3.41E-04 | 0.006 | 0.54 | -1.427 | none |
| SRGAP3 | 0.751 | 0.601 | 5.3956 | 3.42E-04 | 0.006 | 0.53 | -1.4239 | none |
| NDST1 | 0.917 | 0.7954 | 5.394 | 3.43E-04 | 0.006 | 0.53 | -1.2988 | none |
| CSTL1 | 1.278 | 1.0664 | 5.3932 | 3.43E-04 | 0.006 | 0.53 | -0.6395 | none |
| MNT | 1.025 | 0.9168 | 5.3921 | 3.44E-04 | 0.006 | 0.53 | -0.974 | none |
| DNAJC22 | 0.674 | 0.4921 | 5.3898 | 3.45E-04 | 0.006 | 0.53 | -1.3143 | none |
| HEATR6 | 1.081 | 0.8793 | 5.3884 | 3.46E-04 | 0.006 | 0.52 | -1.5534 | OA-related |
| CDKL5 | 0.744 | 0.5796 | 5.3876 | 3.46E-04 | 0.006 | 0.52 | -1.4973 | none |
| PIF1 | 0.837 | 0.7071 | 5.3876 | 3.46E-04 | 0.006 | 0.52 | -1.2197 | none |
| KLHDC10 | 1.359 | 1.1382 | 5.387 | 3.46E-04 | 0.006 | 0.52 | -0.6061 | none |
| CHFR | 0.831 | 0.6801 | 5.3854 | 3.47E-04 | 0.006 | 0.52 | -1.484 | none |
| SLC35E1 | 0.912 | 0.7864 | 5.3843 | 3.48E-04 | 0.006 | 0.52 | -1.1719 | none |
| MANSC1 | 0.537 | 0.3851 | 5.3825 | 3.49E-04 | 0.006 | 0.52 | -1.5696 | OA-related |
| MXRA7 | 0.666 | 0.41 | 5.3824 | 3.49E-04 | 0.006 | 0.52 | -1.221 | none |
| VWA2 | 0.648 | 0.486 | 5.3823 | 3.49E-04 | 0.006 | 0.52 | -1.5221 | none |
| RBFOX3 | 0.705 | 0.5686 | 5.3821 | 3.49E-04 | 0.006 | 0.52 | -1.4335 | none |
| FEZ1 | 0.964 | 0.7893 | 5.3817 | 3.49E-04 | 0.006 | 0.51 | -1.3266 | none |
| ARGFX | 0.703 | 0.5362 | 5.3797 | 3.50E-04 | 0.006 | 0.51 | -1.6092 | OA-related |
| COL1A1 | 1.002 | 0.7216 | 5.3797 | 3.50E-04 | 0.006 | 0.51 | -1.1237 | none |
| P2RX5-TA) | 0.679 | 0.5293 | 5.3786 | 3.50E-04 | 0.006 | 0.51 | -1.5793 | OA-related |
| DHTKD1 | 0.832 | 0.7065 | 5.3766 | 3.51E-04 | 0.006 | 0.51 | -0.8344 | none |
| IDO2 | 0.635 | 0.4546 | 5.3756 | 3.52E-04 | 0.006 | 0.51 | -1.5105 | none |
| IRF9 | 0.596 | 0.4391 | 5.3714 | 3.54E-04 | 0.006 | 0.5 | -1.1007 | none |
| CCPG1 | 1.136 | 0.957 | 5.3713 | 3.54E-04 | 0.006 | 0.5 | -1.161 | none |
| ALPP | 0.508 | 0.3985 | 5.3706 | 3.54E-04 | 0.006 | 0.5 | -0.5727 | none |
| MIGA1 | 0.647 | 0.4848 | 5.3674 | 3.56E-04 | 0.006 | 0.49 | -1.4479 | none |
| PAQR7 | 0.608 | 0.4569 | 5.3652 | 3.57E-04 | 0.006 | 0.49 | -1.4952 | none |
| PLXDC1 | 0.863 | 0.6622 | 5.3643 | 3.57E-04 | 0.006 | 0.49 | -0.8094 | none |
| PLPP2 | 0.718 | 0.5279 | 5.3637 | 3.58E-04 | 0.006 | 0.49 | -1.4533 | none |
| EOGT | 0.673 | 0.5472 | 5.3636 | 3.58E-04 | 0.006 | 0.49 | -0.5667 | none |

| | | | | | | | | |
|---|---|---|---|---|---|---|---|---|
| PRKAB1 | 0.578 | 0.46 | 5.3619 | 3.59E-04 | 0.006 | 0.49 | -1.3913 | none |
| 14-Sep | 1.019 | 0.7642 | 5.361 | 3.59E-04 | 0.006 | 0.49 | -1.1425 | none |
| RHBG | 0.842 | 0.6657 | 5.3604 | 3.59E-04 | 0.006 | 0.49 | -1.3882 | none |
| PCDH11Y | 0.47 | 0.3438 | 5.3603 | 3.59E-04 | 0.006 | 0.49 | -1.3884 | none |
| TRIM58 | 0.586 | 0.4201 | 5.3596 | 3.60E-04 | 0.006 | 0.48 | -1.5862 | OA-related |
| SLC7A7 | 1.222 | 0.9944 | 5.3593 | 3.60E-04 | 0.006 | 0.48 | -1.1648 | none |
| HKR1 | 1.004 | 0.8503 | 5.358 | 3.61E-04 | 0.006 | 0.48 | -1.535 | OA-related |
| CABYR | 1.024 | 0.8202 | 5.3536 | 3.63E-04 | 0.006 | 0.48 | -1.0482 | none |
| ATP9B | 0.802 | 0.7224 | 5.3536 | 3.63E-04 | 0.006 | 0.48 | -1.1766 | none |
| TBX6 | 0.715 | 0.5276 | 5.352 | 3.64E-04 | 0.006 | 0.47 | -1.4699 | none |
| KIRREL3 | 0.752 | 0.5515 | 5.3517 | 3.64E-04 | 0.006 | 0.47 | -1.2694 | none |
| ERBB2 | 1.305 | 1.005 | 5.3512 | 3.64E-04 | 0.006 | 0.47 | -1.3338 | none |
| TRPV3 | 0.624 | 0.5198 | 5.3507 | 3.64E-04 | 0.006 | 0.47 | -1.4994 | none |
| CD34 | 0.553 | 0.4056 | 5.3505 | 3.64E-04 | 0.006 | 0.47 | -1.4653 | none |
| ANGPT2 | 0.725 | 0.5578 | 5.3497 | 3.65E-04 | 0.006 | 0.47 | -1.4994 | none |
| TCTN1 | 0.638 | 0.4819 | 5.348 | 3.66E-04 | 0.006 | 0.47 | -1.0925 | none |
| CPT1B | 0.779 | 0.75 | 5.3447 | 3.67E-04 | 0.006 | 0.46 | -0.1266 | none |
| IFT20 | 0.682 | 0.4587 | 5.3399 | 3.70E-04 | 0.006 | 0.46 | -1.0988 | none |
| ATP8B3 | 0.663 | 0.5043 | 5.3398 | 3.70E-04 | 0.006 | 0.46 | -0.8703 | none |
| SLC4A8 | 0.936 | 0.6679 | 5.3393 | 3.70E-04 | 0.006 | 0.46 | -1.2722 | none |
| EMCN | 0.82 | 0.6895 | 5.3392 | 3.70E-04 | 0.006 | 0.46 | -0.9119 | none |
| RELN | 0.965 | 0.6761 | 5.332 | 3.74E-04 | 0.006 | 0.45 | -1.2532 | none |
| SIPA1L3 | 1.317 | 1.1995 | 5.3307 | 3.74E-04 | 0.006 | 0.44 | -1.4183 | none |
| NRBP1 | 0.695 | 0.4904 | 5.3288 | 3.75E-04 | 0.006 | 0.44 | -1.4094 | none |
| GGT5 | 0.615 | 0.4777 | 5.3276 | 3.76E-04 | 0.006 | 0.44 | -0.9369 | none |
| SLC1A4 | 0.731 | 0.5499 | 5.3245 | 3.78E-04 | 0.006 | 0.44 | -1.3279 | none |
| HACD2 | 0.796 | 0.568 | 5.3239 | 3.78E-04 | 0.006 | 0.43 | -0.9403 | none |
| DDX25 | 0.736 | 0.5394 | 5.3231 | 3.78E-04 | 0.006 | 0.43 | -1.2619 | none |
| TYW5 | 0.483 | 0.3511 | 5.3212 | 3.79E-04 | 0.006 | 0.43 | -1.4314 | none |
| ACSL6 | 0.728 | 0.571 | 5.3208 | 3.80E-04 | 0.006 | 0.43 | -1.5685 | OA-related |
| ETS1 | 0.868 | 0.7174 | 5.3199 | 3.80E-04 | 0.006 | 0.43 | -1.46 | none |
| GOLM1 | 1.162 | 1.0401 | 5.3195 | 3.80E-04 | 0.006 | 0.43 | -0.992 | none |
| PSPH | 0.828 | 0.5947 | 5.318 | 3.81E-04 | 0.006 | 0.43 | -1.28 | none |
| DDHD1 | 1.128 | 0.9453 | 5.315 | 3.83E-04 | 0.006 | 0.42 | -1.5848 | OA-related |
| PDLIM3 | 0.756 | 0.6112 | 5.3149 | 3.83E-04 | 0.006 | 0.42 | -1.5037 | none |
| PSPN | 0.67 | 0.5569 | 5.3141 | 3.83E-04 | 0.006 | 0.42 | -1.6061 | OA-related |
| LGALS4 | 0.535 | 0.3531 | 5.3104 | 3.85E-04 | 0.006 | 0.42 | -1.3168 | none |
| ALDOB | 0.653 | 0.514 | 5.3098 | 3.85E-04 | 0.006 | 0.42 | -1.0999 | none |
| ARMC9 | 0.8 | 0.6161 | 5.3094 | 3.86E-04 | 0.006 | 0.41 | -1.0924 | none |
| SGIP1 | 0.623 | 0.4253 | 5.3092 | 3.86E-04 | 0.006 | 0.41 | 0.44284 | none |

| | | | | | | | | |
|---|---|---|---|---|---|---|---|---|
| TRMT12 | 0.765 | 0.5542 | 5.308 | 3.86E-04 | 0.006 | 0.41 | -0.2928 | none |
| ZNF382 | 0.959 | 0.8212 | 5.308 | 3.86E-04 | 0.006 | 0.41 | -1.2958 | none |
| ZNF667 | 0.648 | 0.5239 | 5.3073 | 3.87E-04 | 0.006 | 0.41 | -1.4806 | none |
| RBMS2 | 1.017 | 0.8099 | 5.3065 | 3.87E-04 | 0.006 | 0.41 | -1.0379 | none |
| ENTPD1 | 0.765 | 0.5799 | 5.3034 | 3.89E-04 | 0.006 | 0.41 | -1.1545 | none |
| AK1 | 0.648 | 0.5923 | 5.3022 | 3.90E-04 | 0.006 | 0.4 | -0.3204 | none |
| NDEL1 | 0.639 | 0.5017 | 5.3016 | 3.90E-04 | 0.006 | 0.4 | -1.0586 | none |
| TTC19 | 0.832 | 0.6731 | 5.2991 | 3.91E-04 | 0.006 | 0.4 | -1.1982 | none |
| RHOBTB3 | 0.574 | 0.4246 | 5.2958 | 3.93E-04 | 0.006 | 0.4 | -0.3489 | none |
| MTBP | 0.772 | 0.593 | 5.2933 | 3.94E-04 | 0.006 | 0.39 | -1.414 | none |
| CCDC88A | 0.93 | 0.7614 | 5.2889 | 3.97E-04 | 0.006 | 0.39 | -1.2016 | none |
| IFT74 | 1.106 | 0.9501 | 5.2848 | 3.99E-04 | 0.006 | 0.38 | -1.481 | none |
| SLC25A30 | 0.687 | 0.4439 | 5.2831 | 4.00E-04 | 0.006 | 0.38 | -1.0697 | none |
| CHEK2 | 0.635 | 0.4003 | 5.2826 | 4.00E-04 | 0.006 | 0.38 | -1.0337 | none |
| SLC38A6 | 0.889 | 0.6562 | 5.2819 | 4.01E-04 | 0.006 | 0.38 | -0.7817 | none |
| QPRT | 0.713 | 0.5482 | 5.2807 | 4.01E-04 | 0.006 | 0.37 | -1.6385 | OA-related |
| MOAP1 | 0.524 | 0.3884 | 5.2747 | 4.05E-04 | 0.006 | 0.37 | -0.8285 | none |
| LRTOMT | 1.015 | 0.8266 | 5.2701 | 4.07E-04 | 0.006 | 0.36 | -0.7502 | none |
| TMEM191 | 0.75 | 0.5195 | 5.2701 | 4.07E-04 | 0.006 | 0.36 | -1.3387 | none |
| CHI3L1 | 0.511 | 0.407 | 5.2698 | 4.08E-04 | 0.006 | 0.36 | -1.3726 | none |
| TRMT2B | 0.538 | 0.4141 | 5.2689 | 4.08E-04 | 0.006 | 0.36 | -1.2727 | none |
| PKD1L1 | 0.676 | 0.5025 | 5.2657 | 4.10E-04 | 0.006 | 0.35 | -1.6526 | OA-related |
| CHMP4C | 0.851 | 0.6804 | 5.2656 | 4.10E-04 | 0.006 | 0.35 | -1.3923 | none |
| XRRA1 | 1.057 | 0.9049 | 5.2654 | 4.10E-04 | 0.006 | 0.35 | -1.2552 | none |
| RNF222 | 0.609 | 0.4459 | 5.2645 | 4.11E-04 | 0.006 | 0.35 | -1.4003 | none |
| ARL9 | 0.63 | 0.4636 | 5.2633 | 4.11E-04 | 0.006 | 0.35 | -1.5268 | none |
| TLR5 | 1.049 | 0.808 | 5.2607 | 4.13E-04 | 0.006 | 0.35 | -0.963 | none |
| CYP4B1 | 1.074 | 0.8649 | 5.2567 | 4.15E-04 | 0.006 | 0.34 | -1.3867 | none |
| VSIG10 | 0.945 | 0.6805 | 5.2557 | 4.16E-04 | 0.006 | 0.34 | -1.3157 | none |
| VTI1A | 0.771 | 0.5334 | 5.2553 | 4.16E-04 | 0.006 | 0.34 | -1.333 | none |
| RBMS1 | 0.818 | 0.6759 | 5.2531 | 4.17E-04 | 0.006 | 0.34 | -1.2031 | none |
| C19ORF54 | 0.618 | 0.5574 | 5.2467 | 4.21E-04 | 0.006 | 0.33 | -1.3309 | none |
| NPHP1 | 0.746 | 0.607 | 5.2446 | 4.22E-04 | 0.006 | 0.32 | -1.2939 | none |
| LPCAT3 | 0.72 | 0.5372 | 5.2445 | 4.22E-04 | 0.006 | 0.32 | -0.898 | none |
| ARHGEF1( | 0.657 | 0.5039 | 5.2442 | 4.23E-04 | 0.006 | 0.32 | -1.4499 | none |
| CETP | 0.625 | 0.5283 | 5.2433 | 4.23E-04 | 0.006 | 0.32 | -1.2769 | none |
| OR2A25 | 0.833 | 0.5935 | 5.2431 | 4.23E-04 | 0.006 | 0.32 | -1.1399 | none |
| FADS1 | 0.789 | 0.6961 | 5.2428 | 4.23E-04 | 0.006 | 0.32 | -0.6932 | none |
| BLMH | 0.823 | 0.6413 | 5.2421 | 4.24E-04 | 0.006 | 0.32 | -1.3717 | none |
| CACNB2 | 0.682 | 0.5101 | 5.2397 | 4.25E-04 | 0.006 | 0.32 | -1.1453 | none |

| | | | | | | | | |
|---|---|---|---|---|---|---|---|---|
| EXOSC6 | 0.779 | 0.597 | 5.2396 | 4.25E-04 | 0.006 | 0.32 | -1.6636 | OA-related |
| ANXA13 | 0.623 | 0.4431 | 5.2355 | 4.28E-04 | 0.006 | 0.31 | -1.5062 | none |
| SPIN3 | 0.653 | 0.5219 | 5.2348 | 4.28E-04 | 0.006 | 0.31 | -1.4735 | none |
| MEIOC | 0.814 | 0.6729 | 5.2326 | 0.00043 | 0.006 | 0.31 | -1.4758 | none |
| LDLR | 1.061 | 1.0148 | 5.2319 | 4.30E-04 | 0.006 | 0.31 | -1.2113 | none |
| ARFGAP2 | 0.623 | 0.4711 | 5.2294 | 4.31E-04 | 0.006 | 0.3 | -1.2369 | none |
| DNASE1 | 0.941 | 0.8348 | 5.2291 | 4.32E-04 | 0.006 | 0.3 | -1.2879 | none |
| TRAPPC8 | 1.184 | 1.0341 | 5.2291 | 4.32E-04 | 0.006 | 0.3 | -1.3002 | none |
| LRRC34 | 0.689 | 0.5252 | 5.229 | 4.32E-04 | 0.006 | 0.3 | -1.1954 | none |
| PRELP | 0.517 | 0.4056 | 5.229 | 4.32E-04 | 0.006 | 0.3 | -1.6521 | OA-related |
| AGBL4 | 0.506 | 0.3776 | 5.2264 | 4.33E-04 | 0.006 | 0.3 | -1.3412 | none |
| CRX | 0.708 | 0.5156 | 5.225 | 4.34E-04 | 0.006 | 0.3 | -1.5057 | none |
| BEST2 | 1.112 | 0.8668 | 5.2214 | 4.36E-04 | 0.007 | 0.29 | -1.4229 | none |
| PTPN3 | 0.892 | 0.6933 | 5.2207 | 4.37E-04 | 0.007 | 0.29 | -1.4831 | none |
| SYTL2 | 0.908 | 0.6771 | 5.2204 | 4.37E-04 | 0.007 | 0.29 | -1.1067 | none |
| ATXN3 | 0.845 | 0.6632 | 5.2201 | 4.37E-04 | 0.007 | 0.29 | -1.4956 | none |
| ISPD | 0.814 | 0.6529 | 5.2197 | 4.37E-04 | 0.007 | 0.29 | -1.2943 | none |
| BCL11A | 0.483 | 0.334 | 5.2195 | 4.37E-04 | 0.007 | 0.29 | -1.577 | OA-related |
| ARSK | 0.863 | 0.6649 | 5.2168 | 4.39E-04 | 0.007 | 0.29 | -1.2771 | none |
| STPG1 | 0.754 | 0.6159 | 5.2166 | 4.39E-04 | 0.007 | 0.29 | -1.3377 | none |
| PODN | 0.738 | 0.5527 | 5.2153 | 4.40E-04 | 0.007 | 0.28 | -1.0514 | none |
| BOD1L1 | 1.062 | 0.9784 | 5.2092 | 4.44E-04 | 0.007 | 0.27 | -1.3298 | none |
| TLL2 | 0.789 | 0.6053 | 5.2059 | 4.46E-04 | 0.007 | 0.27 | -0.9487 | none |
| ZNF154 | 0.657 | 0.4939 | 5.205 | 4.46E-04 | 0.007 | 0.27 | -1.4966 | none |
| SLC2A5 | 0.546 | 0.4241 | 5.2048 | 4.47E-04 | 0.007 | 0.27 | -1.2132 | none |
| C4ORF19 | 1.023 | 0.8276 | 5.2044 | 4.47E-04 | 0.007 | 0.27 | -1.2565 | none |
| SEC14L3 | 0.442 | 0.3238 | 5.2024 | 4.48E-04 | 0.007 | 0.27 | -1.3412 | none |
| SZT2 | 0.736 | 0.6329 | 5.2011 | 4.49E-04 | 0.007 | 0.26 | -1.4688 | none |
| GVQW2 | 1.916 | 1.5815 | 5.2001 | 4.50E-04 | 0.007 | 0.26 | -0.8564 | none |
| PBX3 | 0.62 | 0.4461 | 5.1991 | 4.50E-04 | 0.007 | 0.26 | -0.6384 | none |
| ASB3 | 0.499 | 0.3187 | 5.199 | 4.50E-04 | 0.007 | 0.26 | -1.3399 | none |
| CCDC157 | 0.583 | 0.4037 | 5.1987 | 4.50E-04 | 0.007 | 0.26 | -1.3567 | none |
| ATP1B2 | 0.51 | 0.443 | 5.1986 | 4.50E-04 | 0.007 | 0.26 | -0.038 | none |
| PUS3 | 0.67 | 0.5257 | 5.1981 | 4.51E-04 | 0.007 | 0.26 | -0.9749 | none |
| CCDC57 | 0.909 | 0.6515 | 5.1977 | 4.51E-04 | 0.007 | 0.26 | -0.9726 | none |
| PDE6A | 0.633 | 0.4682 | 5.1968 | 4.52E-04 | 0.007 | 0.26 | -1.4172 | none |
| NAV2 | 1.339 | 1.0086 | 5.1958 | 4.52E-04 | 0.007 | 0.26 | -1.0616 | none |
| ARHGEF12 | 0.771 | 0.5639 | 5.1952 | 4.53E-04 | 0.007 | 0.25 | -1.0309 | none |
| ZNF431 | 1.112 | 0.9077 | 5.1945 | 4.53E-04 | 0.007 | 0.25 | -1.3122 | none |
| LRCH3 | 1.022 | 0.8971 | 5.1943 | 4.53E-04 | 0.007 | 0.25 | -1.406 | none |

| | | | | | | | | |
|---|---|---|---|---|---|---|---|---|
| PUS1 | 0.675 | 0.5775 | 5.1915 | 4.55E-04 | 0.007 | 0.25 | -1.5373 | OA-related |
| SVOP | 0.588 | 0.4452 | 5.1907 | 4.55E-04 | 0.007 | 0.25 | -1.1886 | none |
| NAA16 | 0.922 | 0.8142 | 5.1878 | 4.57E-04 | 0.007 | 0.24 | -0.9027 | none |
| ARHGAP28 | 0.693 | 0.5092 | 5.1869 | 4.58E-04 | 0.007 | 0.24 | -1.404 | none |
| SLC38A3 | 1.259 | 0.9267 | 5.1868 | 4.58E-04 | 0.007 | 0.24 | -0.0073 | none |
| C2ORF74 | 0.628 | 0.5026 | 5.1859 | 4.59E-04 | 0.007 | 0.24 | -1.2029 | none |
| NCKAP5L | 0.866 | 0.7162 | 5.1844 | 4.60E-04 | 0.007 | 0.24 | -1.3634 | none |
| ENO3 | 0.964 | 0.8884 | 5.1832 | 4.60E-04 | 0.007 | 0.24 | -0.9771 | none |
| SLC6A1 | 0.825 | 0.6041 | 5.1807 | 4.62E-04 | 0.007 | 0.23 | -1.3183 | none |
| HSD17B2 | 0.449 | 0.3109 | 5.18 | 4.62E-04 | 0.007 | 0.23 | -0.8006 | none |
| TMEM120 | 0.834 | 0.5851 | 5.1791 | 4.63E-04 | 0.007 | 0.23 | -1.1676 | none |
| CARS2 | 0.698 | 0.5803 | 5.1788 | 4.63E-04 | 0.007 | 0.23 | -1.3749 | none |
| STARD10 | 0.874 | 0.6512 | 5.178 | 4.64E-04 | 0.007 | 0.23 | -1.2549 | none |
| HOPX | 0.731 | 0.5162 | 5.1772 | 4.64E-04 | 0.007 | 0.23 | -1.2298 | none |
| RPS6KA3 | 0.788 | 0.6392 | 5.1736 | 4.67E-04 | 0.007 | 0.22 | -1.5114 | none |
| TMUB2 | 0.66 | 0.4559 | 5.1724 | 4.67E-04 | 0.007 | 0.22 | -1.2427 | none |
| OR2M4 | 0.531 | 0.3799 | 5.1723 | 4.68E-04 | 0.007 | 0.22 | -1.3593 | none |
| PIWIL3 | 0.642 | 0.459 | 5.1714 | 4.68E-04 | 0.007 | 0.22 | -1.2994 | none |
| ARSG | 0.819 | 0.6235 | 5.1684 | 4.70E-04 | 0.007 | 0.22 | -1.0295 | none |
| MORN3 | 0.738 | 0.5261 | 5.1658 | 4.72E-04 | 0.007 | 0.21 | -1.5389 | OA-related |
| CD200 | 0.695 | 0.5125 | 5.1638 | 4.73E-04 | 0.007 | 0.21 | -1.4153 | none |
| CABP4 | 0.979 | 0.7524 | 5.1605 | 4.75E-04 | 0.007 | 0.21 | -1.4452 | none |
| AKT2 | 0.646 | 0.5513 | 5.1588 | 4.77E-04 | 0.007 | 0.2 | -0.9463 | none |
| VWCE | 0.818 | 0.6594 | 5.1584 | 4.77E-04 | 0.007 | 0.2 | -1.6472 | OA-related |
| OLFML2A | 0.507 | 0.359 | 5.1582 | 4.77E-04 | 0.007 | 0.2 | -1.4756 | none |
| NUMA1 | 1.116 | 1.0442 | 5.158 | 4.77E-04 | 0.007 | 0.2 | -0.9338 | none |
| STEAP4 | 0.833 | 0.5963 | 5.1554 | 4.79E-04 | 0.007 | 0.2 | -1.3944 | none |
| KYNU | 0.471 | 0.3501 | 5.1533 | 4.80E-04 | 0.007 | 0.2 | -1.5247 | none |
| GRIK2 | 0.691 | 0.518 | 5.152 | 4.81E-04 | 0.007 | 0.19 | -1.3121 | none |
| PPP2CB | 0.962 | 0.778 | 5.1511 | 4.82E-04 | 0.007 | 0.19 | -1.0657 | none |
| PCDHA9 | 0.531 | 0.3894 | 5.1509 | 4.82E-04 | 0.007 | 0.19 | -1.6026 | OA-related |
| FCHO1 | 0.874 | 0.77 | 5.1508 | 4.82E-04 | 0.007 | 0.19 | -1.1352 | none |
| HNMT | 0.994 | 0.746 | 5.1499 | 4.83E-04 | 0.007 | 0.19 | -1.0997 | none |
| C1ORF158 | 0.518 | 0.3672 | 5.1477 | 4.84E-04 | 0.007 | 0.19 | -1.0977 | none |
| GLS | 0.595 | 0.4919 | 5.147 | 4.85E-04 | 0.007 | 0.19 | -1.1171 | none |
| NMNAT3 | 0.692 | 0.5124 | 5.1444 | 4.86E-04 | 0.007 | 0.18 | -1.2065 | none |
| DBT | 0.732 | 0.633 | 5.1443 | 4.86E-04 | 0.007 | 0.18 | -1.4493 | none |
| SLCO2B1 | 0.733 | 0.6294 | 5.1415 | 4.88E-04 | 0.007 | 0.18 | -0.8908 | none |
| DUSP18 | 0.66 | 0.5815 | 5.1356 | 4.92E-04 | 0.007 | 0.17 | -1.1913 | none |
| TMEM45B | 0.579 | 0.3989 | 5.1351 | 4.93E-04 | 0.007 | 0.17 | -1.2074 | none |

| | | | | | | | | |
|---|---|---|---|---|---|---|---|---|
| PACRG | 1.221 | 0.9947 | 5.1329 | 4.94E-04 | 0.007 | 0.17 | -0.8942 | none |
| DAPL1 | 0.698 | 0.5393 | 5.1328 | 4.94E-04 | 0.007 | 0.17 | 0.0721 | none |
| SCNN1A | 0.805 | 0.619 | 5.1327 | 4.94E-04 | 0.007 | 0.17 | -1.0906 | none |
| GPT2 | 0.733 | 0.5721 | 5.132 | 4.95E-04 | 0.007 | 0.17 | -1.4535 | none |
| ZNF136 | 0.965 | 0.7404 | 5.1299 | 4.96E-04 | 0.007 | 0.16 | -1.2038 | none |
| KMT2D | 1.306 | 1.2134 | 5.1298 | 4.96E-04 | 0.007 | 0.16 | -1.3647 | none |
| ZNF333 | 0.708 | 0.5759 | 5.1239 | 5.01E-04 | 0.007 | 0.15 | -1.5126 | none |
| DDX51 | 0.868 | 0.7886 | 5.1216 | 5.02E-04 | 0.007 | 0.15 | -1.567 | OA-related |
| TCTN2 | 0.588 | 0.4134 | 5.1184 | 5.05E-04 | 0.007 | 0.15 | -1.5491 | OA-related |
| PLCXD1 | 0.811 | 0.6681 | 5.1174 | 5.05E-04 | 0.007 | 0.15 | -1.2765 | none |
| OR8A1 | 0.508 | 0.3779 | 5.1171 | 5.06E-04 | 0.007 | 0.14 | -1.3687 | none |
| BMP8A | 0.59 | 0.4569 | 5.1168 | 5.06E-04 | 0.007 | 0.14 | -1.4075 | none |
| DPY19L3 | 0.898 | 0.7618 | 5.1166 | 5.06E-04 | 0.007 | 0.14 | -1.4292 | none |
| ZNF557 | 0.677 | 0.544 | 5.1161 | 5.06E-04 | 0.007 | 0.14 | -1.6608 | OA-related |
| LRRN4CL | 0.646 | 0.4848 | 5.1139 | 5.08E-04 | 0.007 | 0.14 | -1.4537 | none |
| DGKH | 0.9 | 0.7986 | 5.1136 | 5.08E-04 | 0.007 | 0.14 | -1.6416 | OA-related |
| TRIM73 | 0.824 | 0.6893 | 5.1135 | 5.08E-04 | 0.007 | 0.14 | -1.2763 | none |
| ABHD12B | 1.027 | 0.8885 | 5.113 | 5.08E-04 | 0.007 | 0.14 | -0.8366 | none |
| BCL2L11 | 0.643 | 0.474 | 5.1128 | 5.09E-04 | 0.007 | 0.14 | -0.7219 | none |
| DSG2 | 0.537 | 0.386 | 5.1111 | 5.10E-04 | 0.007 | 0.14 | -1.5621 | OA-related |
| NALCN | 0.932 | 0.6882 | 5.11 | 5.11E-04 | 0.007 | 0.13 | -1.3345 | none |
| MAP4K1 | 0.632 | 0.5229 | 5.1094 | 5.11E-04 | 0.007 | 0.13 | -1.4515 | none |
| COPB2 | 0.582 | 0.4331 | 5.1092 | 5.11E-04 | 0.007 | 0.13 | -0.9857 | none |
| C3 | 1.122 | 0.8748 | 5.1079 | 5.12E-04 | 0.007 | 0.13 | -1.2749 | none |
| EFCAB7 | 0.76 | 0.5659 | 5.1065 | 5.13E-04 | 0.007 | 0.13 | -0.6634 | none |
| ST3GAL2 | 0.843 | 0.7182 | 5.1064 | 5.13E-04 | 0.007 | 0.13 | -1.3989 | none |
| AC019206 | 1.011 | 0.7499 | 5.1045 | 5.15E-04 | 0.007 | 0.13 | -0.6609 | none |
| TBC1D12 | 0.631 | 0.476 | 5.1012 | 5.17E-04 | 0.007 | 0.12 | -0.879 | none |
| CBX7 | 0.539 | 0.4434 | 5.0984 | 5.19E-04 | 0.007 | 0.12 | -1.4028 | none |
| MALL | 0.6 | 0.4562 | 5.0977 | 5.20E-04 | 0.007 | 0.12 | -1.0828 | none |
| EPS8 | 0.95 | 0.8165 | 5.0974 | 5.20E-04 | 0.007 | 0.12 | -1.1539 | none |
| APCDD1 | 0.851 | 0.608 | 5.0942 | 5.22E-04 | 0.007 | 0.11 | -1.4981 | none |
| FGFR4 | 0.717 | 0.577 | 5.0935 | 5.23E-04 | 0.007 | 0.11 | -1.147 | none |
| ABCB5 | 0.588 | 0.4247 | 5.0928 | 5.23E-04 | 0.007 | 0.11 | -1.3552 | none |
| OCLN | 0.466 | 0.3695 | 5.0915 | 5.24E-04 | 0.007 | 0.11 | -1.2839 | none |
| TPD52L1 | 0.638 | 0.4201 | 5.0908 | 5.25E-04 | 0.007 | 0.11 | -1.4204 | none |
| CYP2C19 | 0.558 | 0.4146 | 5.0884 | 5.27E-04 | 0.007 | 0.1 | -1.2954 | none |
| C2ORF50 | 0.479 | 0.3727 | 5.0864 | 5.28E-04 | 0.007 | 0.1 | -1.3408 | none |
| BTBD19 | 0.829 | 0.6709 | 5.0826 | 5.31E-04 | 0.007 | 0.1 | -1.5352 | OA-related |
| SLC29A4 | 0.681 | 0.5048 | 5.0821 | 5.31E-04 | 0.007 | 0.09 | -1.396 | none |

| | | | | | | | | |
|---|---|---|---|---|---|---|---|---|
| PQLC2 | 0.624 | 0.5092 | 5.0795 | 5.33E-04 | 0.007 | 0.09 | -0.2378 | none |
| ROBO1 | 0.647 | 0.4933 | 5.079 | 5.34E-04 | 0.007 | 0.09 | -1.2213 | none |
| RTBDN | 0.605 | 0.4767 | 5.079 | 5.34E-04 | 0.007 | 0.09 | -0.033 | none |
| TNRC6A | 1.034 | 0.995 | 5.0782 | 5.34E-04 | 0.007 | 0.09 | -1.2199 | none |
| ADGRE2 | 0.623 | 0.4946 | 5.0761 | 5.36E-04 | 0.007 | 0.09 | -1.1862 | none |
| SLC25A33 | 0.621 | 0.4529 | 5.073 | 5.38E-04 | 0.007 | 0.08 | -1.1774 | none |
| GGCX | 0.986 | 0.7003 | 5.0697 | 5.41E-04 | 0.007 | 0.08 | -1.3514 | none |
| MAB21L3 | 0.511 | 0.428 | 5.0683 | 5.42E-04 | 0.007 | 0.08 | -1.2016 | none |
| PCYOX1 | 0.71 | 0.5814 | 5.0681 | 5.42E-04 | 0.007 | 0.07 | -1.3086 | none |
| SYCP3 | 0.684 | 0.4929 | 5.0651 | 5.44E-04 | 0.007 | 0.07 | -1.4869 | none |
| NOCT | 0.945 | 0.8316 | 5.0641 | 5.45E-04 | 0.007 | 0.07 | -1.5446 | OA-related |
| ZNF814 | 0.798 | 0.6165 | 5.0613 | 5.47E-04 | 0.007 | 0.07 | -1.1017 | none |
| ORC4 | 0.62 | 0.4185 | 5.0575 | 5.50E-04 | 0.007 | 0.06 | -1.2391 | none |
| GDPGP1 | 0.998 | 0.7704 | 5.0568 | 5.51E-04 | 0.007 | 0.06 | -1.5492 | OA-related |
| KCNE3 | 0.603 | 0.4808 | 5.0548 | 5.52E-04 | 0.007 | 0.06 | -1.2704 | none |
| MORC4 | 0.885 | 0.8046 | 5.0516 | 5.55E-04 | 0.007 | 0.05 | -1.3962 | none |
| ANKRD54 | 0.613 | 0.5074 | 5.0514 | 5.55E-04 | 0.007 | 0.05 | -1.2905 | none |
| PIGR | 0.847 | 0.6893 | 5.051 | 5.55E-04 | 0.007 | 0.05 | -0.9774 | none |
| RBFOX1 | 0.696 | 0.5424 | 5.0484 | 5.58E-04 | 0.007 | 0.05 | -1.4512 | none |
| DHRSX | 0.681 | 0.4413 | 5.0471 | 5.59E-04 | 0.008 | 0.04 | -0.8953 | none |
| INTU | 0.521 | 0.4149 | 5.0465 | 5.59E-04 | 0.008 | 0.04 | -1.3896 | none |
| HAP1 | 0.586 | 0.4196 | 5.0442 | 5.61E-04 | 0.008 | 0.04 | -1.6227 | OA-related |
| WIPI1 | 0.714 | 0.5173 | 5.0438 | 5.61E-04 | 0.008 | 0.04 | -1.3671 | none |
| HERC4 | 0.713 | 0.5467 | 5.0431 | 5.62E-04 | 0.008 | 0.04 | -1.342 | none |
| FER | 0.684 | 0.5946 | 5.0418 | 5.63E-04 | 0.008 | 0.04 | -1.204 | none |
| EXPH5 | 0.547 | 0.385 | 5.0406 | 5.64E-04 | 0.008 | 0.04 | -1.4937 | none |
| PLLP | 0.673 | 0.5447 | 5.0402 | 5.64E-04 | 0.008 | 0.04 | -1.3821 | none |
| COX18 | 0.776 | 0.6292 | 5.0388 | 5.65E-04 | 0.008 | 0.03 | -1.0049 | none |
| TNNT1 | 0.957 | 0.7653 | 5.0384 | 5.66E-04 | 0.008 | 0.03 | -0.97 | none |
| ZNF83 | 0.979 | 0.8118 | 5.0373 | 5.66E-04 | 0.008 | 0.03 | -1.4116 | none |
| ENPP1 | 0.639 | 0.4569 | 5.0369 | 5.67E-04 | 0.008 | 0.03 | -1.3211 | none |
| NOS1 | 0.611 | 0.4846 | 5.0362 | 5.67E-04 | 0.008 | 0.03 | -1.3287 | none |
| ZNF660 | 0.438 | 0.3299 | 5.035 | 5.68E-04 | 0.008 | 0.03 | -1.4298 | none |
| CDC42BPG | 0.643 | 0.5109 | 5.0338 | 5.69E-04 | 0.008 | 0.03 | -1.0507 | none |
| ACOT2 | 0.564 | 0.3345 | 5.0312 | 5.71E-04 | 0.008 | 0.02 | -1.1194 | none |
| IFT88 | 0.773 | 0.6308 | 5.0296 | 5.73E-04 | 0.008 | 0.02 | -0.9284 | none |
| OR10K1 | 0.454 | 0.3426 | 5.0272 | 5.75E-04 | 0.008 | 0.02 | -0.8692 | none |
| C1RL | 0.952 | 0.8286 | 5.0265 | 5.75E-04 | 0.008 | 0.02 | -1.4152 | none |
| IGSF6 | 0.84 | 0.7137 | 5.0254 | 5.76E-04 | 0.008 | 0.01 | -1.3824 | none |
| SPTB | 0.682 | 0.5661 | 5.0252 | 5.76E-04 | 0.008 | 0.01 | -1.1968 | none |

| | | | | | | | | |
|---|---|---|---|---|---|---|---|---|
| ZNF789 | 0.749 | 0.5703 | 5.0247 | 5.77E-04 | 0.008 | 0.01 | -1.3147 | none |
| YBX3 | 1.006 | 0.8909 | 5.0241 | 5.77E-04 | 0.008 | 0.01 | -1.3964 | none |
| CCBE1 | 0.569 | 0.381 | 5.0206 | 5.80E-04 | 0.008 | 0.01 | -1.0427 | none |
| SBSPON | 0.571 | 0.443 | 5.0197 | 5.81E-04 | 0.008 | 0.01 | -1.2461 | none |
| DCAF17 | 0.903 | 0.6685 | 5.0188 | 5.82E-04 | 0.008 | 0 | -1.137 | none |
| CFAP221 | 0.599 | 0.4931 | 5.0179 | 5.82E-04 | 0.008 | 0 | -0.9849 | none |
| SLC3A1 | 0.83 | 0.6176 | 5.0178 | 5.82E-04 | 0.008 | 0 | -1.4872 | none |
| FILIP1L | 0.95 | 0.8309 | 5.0176 | 5.83E-04 | 0.008 | 0 | -0.9919 | none |
| TRIM5 | 0.87 | 0.6567 | 5.0175 | 5.83E-04 | 0.008 | 0 | -0.8844 | none |
| NEIL1 | 0.771 | 0.6056 | 5.0174 | 5.83E-04 | 0.008 | 0 | -1.5045 | none |
| KCNK3 | 0.542 | 0.4093 | 5.0167 | 5.83E-04 | 0.008 | 0 | -1.449 | none |
| COL1A2 | 0.526 | 0.3804 | 5.0158 | 5.84E-04 | 0.008 | ##### | -0.9916 | none |
| SLC2A14 | 0.82 | 0.6325 | 5.0153 | 5.85E-04 | 0.008 | ##### | -1.2063 | none |
| GLB1L | 0.838 | 0.6407 | 5.0145 | 5.85E-04 | 0.008 | -0 | -0.7651 | none |
| NPFFR1 | 0.527 | 0.3873 | 5.0136 | 5.86E-04 | 0.008 | -0 | -1.5775 | OA-related |
| GCNT3 | 0.583 | 0.4224 | 5.0125 | 5.87E-04 | 0.008 | -0 | -1.3362 | none |
| GABPB2 | 0.724 | 0.5932 | 5.0119 | 5.87E-04 | 0.008 | -0.01 | -1.4883 | none |
| PIGBOS1 | 0.743 | 0.5573 | 5.0113 | 5.88E-04 | 0.008 | -0.01 | -1.0587 | none |
| P3H1 | 0.827 | 0.674 | 5.0109 | 5.88E-04 | 0.008 | -0.01 | -1.1939 | none |
| FAM180A | 0.762 | 0.5459 | 5.0107 | 5.88E-04 | 0.008 | -0.01 | -1.2635 | none |
| WDR26 | 0.754 | 0.6275 | 5.0075 | 5.91E-04 | 0.008 | -0.01 | -1.4547 | none |
| SATL1 | 0.558 | 0.4266 | 5.0073 | 5.91E-04 | 0.008 | -0.01 | -1.1955 | none |
| KIAA1328 | 0.692 | 0.5537 | 5.0071 | 5.92E-04 | 0.008 | -0.01 | -1.2425 | none |
| SNX29 | 1.167 | 0.9433 | 5.0068 | 5.92E-04 | 0.008 | -0.01 | -1.356 | none |
| CADPS | 0.557 | 0.4132 | 5.0067 | 5.92E-04 | 0.008 | -0.01 | -1.3387 | none |
| POU2F3 | 0.552 | 0.4382 | 5.0041 | 5.94E-04 | 0.008 | -0.02 | -1.4092 | none |
| ZNF595 | 0.862 | 0.6895 | 5.0036 | 5.95E-04 | 0.008 | -0.02 | -1.3051 | none |
| THEM4 | 0.657 | 0.5044 | 5.0026 | 5.95E-04 | 0.008 | -0.02 | -0.9282 | none |
| APOBEC3F | 0.531 | 0.4305 | 5.0002 | 5.97E-04 | 0.008 | -0.02 | -1.5061 | none |
| CEP76 | 0.659 | 0.4837 | 4.9979 | 5.99E-04 | 0.008 | -0.03 | -1.0492 | none |
| TMEM135 | 1.054 | 0.8241 | 4.9966 | 6.00E-04 | 0.008 | -0.03 | -0.9149 | none |
| DMD | 0.871 | 0.7154 | 4.9941 | 6.03E-04 | 0.008 | -0.03 | -1.3151 | none |
| FRMD4B | 1.174 | 1.0303 | 4.9918 | 6.05E-04 | 0.008 | -0.03 | -1.4167 | none |
| AHDC1 | 1.021 | 0.9495 | 4.9873 | 6.09E-04 | 0.008 | -0.04 | -1.3551 | none |
| TIE1 | 0.631 | 0.4799 | 4.9872 | 6.09E-04 | 0.008 | -0.04 | -1.1708 | none |
| ABCC3 | 0.968 | 0.7302 | 4.9857 | 6.10E-04 | 0.008 | -0.04 | -1.5267 | none |
| C1S | 0.6 | 0.4889 | 4.9843 | 6.11E-04 | 0.008 | -0.05 | -1.4408 | none |
| ANKS3 | 0.741 | 0.6111 | 4.9839 | 6.12E-04 | 0.008 | -0.05 | -1.4941 | none |
| FAM9B | 0.498 | 0.3404 | 4.9835 | 6.12E-04 | 0.008 | -0.05 | -1.3659 | none |
| ZNF641 | 0.634 | 0.5196 | 4.9823 | 6.13E-04 | 0.008 | -0.05 | -0.5973 | none |

| | | | | | | | | |
|---|---|---|---|---|---|---|---|---|
| ANKLE1 | 0.536 | 0.434 | 4.9789 | 6.16E-04 | 0.008 | -0.05 | -0.7624 | none |
| AFAP1L1 | 0.552 | 0.4091 | 4.978 | 6.17E-04 | 0.008 | -0.05 | -1.5576 | OA-related |
| RHD | 0.766 | 0.6192 | 4.9776 | 6.17E-04 | 0.008 | -0.05 | -1.5337 | OA-related |
| LILRA2 | 0.68 | 0.5161 | 4.9776 | 6.17E-04 | 0.008 | -0.05 | -1.5988 | OA-related |
| SIAE | 0.575 | 0.4744 | 4.9769 | 6.18E-04 | 0.008 | -0.06 | -0.9552 | none |
| SMIM5 | 0.469 | 0.3582 | 4.9759 | 6.19E-04 | 0.008 | -0.06 | -1.3243 | none |
| ZNF662 | 0.561 | 0.3814 | 4.9759 | 6.19E-04 | 0.008 | -0.06 | -1.4049 | none |
| VRK3 | 0.508 | 0.3672 | 4.9757 | 6.19E-04 | 0.008 | -0.06 | -1.3962 | none |
| LIMK1 | 0.888 | 0.736 | 4.9737 | 6.21E-04 | 0.008 | -0.06 | -1.4345 | none |
| PATJ | 0.929 | 0.7895 | 4.9716 | 6.23E-04 | 0.008 | -0.06 | -1.4928 | none |
| MTPAP | 0.596 | 0.4975 | 4.9715 | 6.23E-04 | 0.008 | -0.06 | -1.2004 | none |
| SLC35F5 | 0.745 | 0.5029 | 4.9715 | 6.23E-04 | 0.008 | -0.06 | -1.2127 | none |
| STARD9 | 0.786 | 0.6162 | 4.9713 | 6.23E-04 | 0.008 | -0.06 | -1.2763 | none |
| KAT8 | 0.767 | 0.6173 | 4.9703 | 6.24E-04 | 0.008 | -0.07 | -1.4386 | none |
| SLC25A23 | 1.224 | 0.9857 | 4.9689 | 6.25E-04 | 0.008 | -0.07 | -1.0253 | none |
| C15ORF40 | 0.831 | 0.6184 | 4.9667 | 6.27E-04 | 0.008 | -0.07 | -1.1492 | none |
| ZNF7 | 0.784 | 0.6764 | 4.9667 | 6.27E-04 | 0.008 | -0.07 | -1.2297 | none |
| VPS50 | 0.776 | 0.6428 | 4.9657 | 6.28E-04 | 0.008 | -0.07 | -1.0612 | none |
| SLC30A8 | 1.15 | 0.953 | 4.9652 | 6.28E-04 | 0.008 | -0.07 | -0.5096 | none |
| CEP19 | 0.7 | 0.5393 | 4.9626 | 6.31E-04 | 0.008 | -0.08 | -1.3126 | none |
| AKAP10 | 0.794 | 0.6538 | 4.9614 | 6.32E-04 | 0.008 | -0.08 | -1.4537 | none |
| CFLAR | 0.52 | 0.3436 | 4.9612 | 6.32E-04 | 0.008 | -0.08 | -1.2252 | none |
| PPIP5K2 | 0.877 | 0.7064 | 4.9599 | 6.33E-04 | 0.008 | -0.08 | -1.0995 | none |
| MTFR1 | 0.998 | 0.8387 | 4.9591 | 6.34E-04 | 0.008 | -0.08 | -0.7788 | none |
| PRR23C | 0.546 | 0.3923 | 4.9581 | 6.35E-04 | 0.008 | -0.08 | -1.3362 | none |
| HTRA4 | 0.58 | 0.4105 | 4.9536 | 6.39E-04 | 0.008 | -0.09 | -1.0777 | none |
| SLC25A45 | 0.897 | 0.7218 | 4.9528 | 6.40E-04 | 0.008 | -0.09 | -1.4091 | none |
| ZC3HAV1L | 0.47 | 0.3604 | 4.9517 | 6.41E-04 | 0.008 | -0.09 | -0.7814 | none |
| PCDHB11 | 0.675 | 0.4994 | 4.9506 | 6.42E-04 | 0.008 | -0.09 | -1.4688 | none |
| NT5C3B | 0.59 | 0.4369 | 4.9496 | 6.43E-04 | 0.008 | -0.1 | -0.9833 | none |
| DSEL | 0.429 | 0.3163 | 4.9492 | 6.43E-04 | 0.008 | -0.1 | -1.5623 | OA-related |
| BARX2 | 0.543 | 0.395 | 4.9474 | 6.45E-04 | 0.008 | -0.1 | -1.4938 | none |
| ZSCAN18 | 1.173 | 0.9925 | 4.9463 | 6.46E-04 | 0.008 | -0.1 | -0.6317 | none |
| CLCC1 | 0.572 | 0.4808 | 4.9459 | 6.46E-04 | 0.008 | -0.1 | -1.4477 | none |
| RALGPS2 | 0.659 | 0.4552 | 4.9453 | 6.47E-04 | 0.008 | -0.1 | -1.1826 | none |
| TEPP | 0.916 | 0.6653 | 4.9447 | 6.47E-04 | 0.008 | -0.1 | -1.3826 | none |
| UBXN2A | 0.763 | 0.6417 | 4.9436 | 6.48E-04 | 0.008 | -0.1 | -1.5344 | OA-related |
| DIP2A | 0.687 | 0.6044 | 4.943 | 6.49E-04 | 0.008 | -0.1 | -1.3801 | none |
| EREG | 0.738 | 0.5929 | 4.9424 | 6.49E-04 | 0.008 | -0.11 | -1.3824 | none |
| KLHL24 | 0.645 | 0.4572 | 4.9402 | 6.51E-04 | 0.008 | -0.11 | -1.0931 | none |

| | | | | | | | | |
|---|---|---|---|---|---|---|---|---|
| CARF | 0.608 | 0.4882 | 4.9401 | 6.51E-04 | 0.008 | -0.11 | -1.4878 | none |
| SEC23B | 0.514 | 0.3527 | 4.9385 | 6.53E-04 | 0.008 | -0.11 | -0.6292 | none |
| CCDC122 | 0.513 | 0.3674 | 4.9378 | 6.54E-04 | 0.008 | -0.11 | -1.4921 | none |
| CYP27C1 | 0.49 | 0.3496 | 4.9362 | 6.55E-04 | 0.008 | -0.11 | -1.4319 | none |
| CENPV | 0.735 | 0.5226 | 4.9356 | 6.56E-04 | 0.008 | -0.12 | -1.3434 | none |
| ASB18 | 0.724 | 0.5777 | 4.9323 | 6.59E-04 | 0.008 | -0.12 | -1.3016 | none |
| ZNF559 | 0.791 | 0.5997 | 4.9316 | 6.60E-04 | 0.008 | -0.12 | -1.1503 | none |
| SETDB1 | 0.772 | 0.6349 | 4.9295 | 6.62E-04 | 0.008 | -0.12 | -1.5883 | OA-related |
| TFR2 | 0.693 | 0.5416 | 4.9283 | 6.63E-04 | 0.008 | -0.13 | -0.9515 | none |
| SAMD5 | 0.479 | 0.3734 | 4.928 | 6.63E-04 | 0.008 | -0.13 | -1.43 | none |
| SLC22A25 | 0.477 | 0.3564 | 4.9278 | 6.63E-04 | 0.008 | -0.13 | -1.4663 | none |
| PTGIS | 0.657 | 0.5223 | 4.925 | 6.66E-04 | 0.008 | -0.13 | -1.5523 | OA-related |
| CROCC | 0.846 | 0.8223 | 4.9244 | 6.66E-04 | 0.008 | -0.13 | -0.3654 | none |
| STK33 | 0.813 | 0.7014 | 4.9232 | 6.68E-04 | 0.008 | -0.13 | -1.1551 | none |
| VWA7 | 0.497 | 0.3612 | 4.9215 | 6.69E-04 | 0.008 | -0.14 | -1.4601 | none |
| COL9A1 | 0.684 | 0.4918 | 4.9209 | 6.70E-04 | 0.008 | -0.14 | -0.9888 | none |
| CLVS1 | 0.65 | 0.4555 | 4.9208 | 6.70E-04 | 0.008 | -0.14 | -1.0611 | none |
| AKAP5 | 0.709 | 0.5863 | 4.9194 | 6.71E-04 | 0.008 | -0.14 | -1.5368 | OA-related |
| MEF2A | 1.149 | 1.0034 | 4.9191 | 6.72E-04 | 0.008 | -0.14 | -1.0392 | none |
| TBCE | 0.644 | 0.4736 | 4.9189 | 6.72E-04 | 0.008 | -0.14 | -0.9708 | none |
| SERPINB6 | 0.596 | 0.4278 | 4.9189 | 6.72E-04 | 0.008 | -0.14 | -1.3607 | none |
| P2RX4 | 0.788 | 0.5924 | 4.9184 | 6.72E-04 | 0.008 | -0.14 | -1.3794 | none |
| H6PD | 0.717 | 0.6626 | 4.9159 | 6.75E-04 | 0.008 | -0.14 | -1.4761 | none |
| GAREM1 | 1.007 | 0.7839 | 4.9158 | 6.75E-04 | 0.008 | -0.14 | -1.0115 | none |
| YTHDF3 | 0.862 | 0.6922 | 4.9153 | 6.75E-04 | 0.008 | -0.14 | -1.1527 | none |
| OR2AT4 | 0.412 | 0.3147 | 4.9152 | 6.75E-04 | 0.008 | -0.14 | -1.2082 | none |
| MRPL48 | 0.592 | 0.4092 | 4.914 | 6.76E-04 | 0.008 | -0.15 | -1.3777 | none |
| L2HGDH | 0.515 | 0.4056 | 4.9132 | 6.77E-04 | 0.008 | -0.15 | -0.9943 | none |
| KCNG4 | 0.496 | 0.3881 | 4.9125 | 6.78E-04 | 0.008 | -0.15 | -1.6206 | OA-related |
| SSBP2 | 0.609 | 0.4679 | 4.9124 | 6.78E-04 | 0.008 | -0.15 | -1.142 | none |
| ZNF135 | 0.644 | 0.4858 | 4.9116 | 6.79E-04 | 0.008 | -0.15 | -1.5008 | none |
| WHAMM | 0.872 | 0.7406 | 4.9113 | 6.79E-04 | 0.008 | -0.15 | -1.6278 | OA-related |
| CDAN1 | 0.767 | 0.662 | 4.9108 | 6.80E-04 | 0.008 | -0.15 | -1.2591 | none |
| SS18 | 0.71 | 0.5302 | 4.9107 | 6.80E-04 | 0.008 | -0.15 | -1.2243 | none |
| ZFP14 | 0.575 | 0.4583 | 4.9101 | 6.80E-04 | 0.008 | -0.15 | -1.5732 | OA-related |
| AGMAT | 0.582 | 0.4517 | 4.9096 | 6.81E-04 | 0.008 | -0.15 | -0.935 | none |
| SPICE1 | 0.836 | 0.78 | 4.908 | 6.82E-04 | 0.008 | -0.16 | -0.5515 | none |
| OR10G4 | 0.538 | 0.3955 | 4.9077 | 6.83E-04 | 0.008 | -0.16 | -0.8245 | none |
| DNASE1L3 | 0.619 | 0.4779 | 4.9072 | 6.83E-04 | 0.008 | -0.16 | -1.3504 | none |
| CLK4 | 0.549 | 0.4245 | 4.907 | 6.83E-04 | 0.008 | -0.16 | -1.299 | none |

| | | | | | | | | |
|---|---|---|---|---|---|---|---|---|
| FGD5 | 0.714 | 0.5097 | 4.904 | 6.86E-04 | 0.008 | -0.16 | -1.525 | none |
| THAP5 | 0.862 | 0.6952 | 4.9017 | 6.89E-04 | 0.008 | -0.16 | -1.3721 | none |
| PALM | 0.598 | 0.4189 | 4.9001 | 6.90E-04 | 0.008 | -0.17 | -1.4669 | none |
| FAM177A1 | 0.76 | 0.5483 | 4.8991 | 6.91E-04 | 0.008 | -0.17 | -1.3414 | none |
| SDE2 | 0.527 | 0.3527 | 4.8971 | 6.93E-04 | 0.008 | -0.17 | -1.321 | none |
| PLEKHB2 | 0.925 | 0.675 | 4.8962 | 6.94E-04 | 0.008 | -0.17 | -1.2512 | none |
| PTGR1 | 0.545 | 0.4328 | 4.8956 | 6.95E-04 | 0.008 | -0.17 | -1.3512 | none |
| CDC73 | 0.564 | 0.4101 | 4.8943 | 6.96E-04 | 0.008 | -0.18 | -1.1867 | none |
| NAIF1 | 0.643 | 0.4983 | 4.8927 | 6.98E-04 | 0.008 | -0.18 | -1.4102 | none |
| PM20D2 | 0.595 | 0.453 | 4.8919 | 6.99E-04 | 0.008 | -0.18 | -0.8174 | none |
| TNS4 | 0.72 | 0.573 | 4.8916 | 6.99E-04 | 0.008 | -0.18 | -1.4631 | none |
| SPATA6L | 0.752 | 0.5531 | 4.8908 | 7.00E-04 | 0.008 | -0.18 | -1.2044 | none |
| ADAMTS1 | 0.493 | 0.3518 | 4.89 | 7.01E-04 | 0.008 | -0.18 | -1.5894 | OA-related |
| APOE | 1.159 | 0.8947 | 4.8889 | 7.02E-04 | 0.008 | -0.18 | -1.344 | none |
| CYBRD1 | 0.97 | 0.714 | 4.8886 | 7.02E-04 | 0.008 | -0.18 | -0.8604 | none |
| SPC24 | 0.619 | 0.4689 | 4.8873 | 7.03E-04 | 0.008 | -0.19 | -1.5259 | none |
| GLMP | 1.252 | 0.9145 | 4.8863 | 7.04E-04 | 0.008 | -0.19 | -1.1556 | none |
| MSANTD1 | 0.656 | 0.4939 | 4.8835 | 7.07E-04 | 0.008 | -0.19 | -1.0389 | none |
| ZNF354B | 0.673 | 0.5217 | 4.8822 | 7.09E-04 | 0.008 | -0.19 | -0.5263 | none |
| PLPP5 | 0.729 | 0.5193 | 4.8815 | 7.09E-04 | 0.008 | -0.19 | -0.7186 | none |
| RSRP1 | 0.648 | 0.514 | 4.8795 | 7.11E-04 | 0.008 | -0.2 | -1.1619 | none |
| CCDC9 | 1.031 | 0.8635 | 4.8787 | 7.12E-04 | 0.008 | -0.2 | -1.4591 | none |
| SYTL1 | 0.584 | 0.4314 | 4.8769 | 7.14E-04 | 0.008 | -0.2 | -1.4733 | none |
| SDSL | 0.777 | 0.5818 | 4.8767 | 7.14E-04 | 0.008 | -0.2 | -1.5489 | OA-related |
| SPRED3 | 0.603 | 0.5031 | 4.8766 | 7.14E-04 | 0.008 | -0.2 | -1.4041 | none |
| PRRG4 | 0.692 | 0.5434 | 4.8757 | 7.15E-04 | 0.008 | -0.2 | -1.6108 | OA-related |
| TNNT2 | 0.823 | 0.6743 | 4.8741 | 7.17E-04 | 0.009 | -0.2 | -1.3636 | none |
| FCRL2 | 0.921 | 0.7079 | 4.873 | 7.18E-04 | 0.009 | -0.21 | -1.5131 | none |
| TAF1L | 0.416 | 0.2978 | 4.8729 | 7.18E-04 | 0.009 | -0.21 | -1.4587 | none |
| PTPRS | 0.684 | 0.5094 | 4.8725 | 7.19E-04 | 0.009 | -0.21 | -1.1077 | none |
| DLG4 | 0.786 | 0.6353 | 4.8703 | 7.21E-04 | 0.009 | -0.21 | -1.3951 | none |
| SYNJ2 | 0.771 | 0.6097 | 4.8699 | 7.21E-04 | 0.009 | -0.21 | -1.4004 | none |
| ASH1L | 0.937 | 0.8203 | 4.869 | 7.22E-04 | 0.009 | -0.21 | -1.0695 | none |
| ASGR1 | 1.049 | 0.8077 | 4.868 | 7.23E-04 | 0.009 | -0.21 | -1.1304 | none |
| SLC30A4 | 0.591 | 0.4777 | 4.8659 | 7.26E-04 | 0.009 | -0.22 | -1.3835 | none |
| CRLF2 | 0.621 | 0.4223 | 4.8622 | 7.29E-04 | 0.009 | -0.22 | -1.4724 | none |
| GABRG2 | 0.694 | 0.5483 | 4.8621 | 0.00073 | 0.009 | -0.22 | -1.4194 | none |
| ZNF716 | 0.634 | 0.4575 | 4.8597 | 7.32E-04 | 0.009 | -0.23 | -1.5761 | OA-related |
| ZKSCAN3 | 0.629 | 0.4794 | 4.8588 | 7.33E-04 | 0.009 | -0.23 | -1.4254 | none |
| TUSC3 | 0.73 | 0.5227 | 4.8584 | 7.33E-04 | 0.009 | -0.23 | -0.897 | none |

| | | | | | | | | |
|---|---|---|---|---|---|---|---|---|
| ACAD8 | 0.471 | 0.3887 | 4.8579 | 7.34E-04 | 0.009 | -0.23 | -1.4429 | none |
| ANKRD18/ | 1.047 | 0.8105 | 4.8568 | 7.35E-04 | 0.009 | -0.23 | -0.4682 | none |
| LARP6 | 0.525 | 0.4197 | 4.8567 | 7.35E-04 | 0.009 | -0.23 | -1.3321 | none |
| SCD5 | 0.628 | 0.4486 | 4.8554 | 7.37E-04 | 0.009 | -0.23 | -1.4493 | none |
| NELFA | 0.807 | 0.6951 | 4.8553 | 7.37E-04 | 0.009 | -0.23 | -1.5404 | OA-related |
| PPP1R37 | 0.839 | 0.7003 | 4.8552 | 7.37E-04 | 0.009 | -0.23 | -1.4087 | none |
| WTIP | 0.466 | 0.3473 | 4.8538 | 7.38E-04 | 0.009 | -0.23 | -1.4378 | none |
| MYH3 | 1.186 | 0.9656 | 4.8531 | 7.39E-04 | 0.009 | -0.24 | -1.0452 | none |
| CA6 | 0.803 | 0.5922 | 4.8522 | 7.40E-04 | 0.009 | -0.24 | -1.2631 | none |
| PCDHGA5 | 0.431 | 0.3306 | 4.8499 | 7.43E-04 | 0.009 | -0.24 | -1.4156 | none |
| ZNF215 | 0.929 | 0.719 | 4.8495 | 7.43E-04 | 0.009 | -0.24 | -1.2323 | none |
| GAN | 0.628 | 0.5442 | 4.8484 | 7.44E-04 | 0.009 | -0.24 | -1.1909 | none |
| ZKSCAN7 | 0.655 | 0.4654 | 4.8472 | 7.46E-04 | 0.009 | -0.24 | -1.4178 | none |
| PHF11 | 0.614 | 0.4539 | 4.8455 | 7.47E-04 | 0.009 | -0.25 | -1.3351 | none |
| STKLD1 | 0.752 | 0.5673 | 4.8446 | 7.48E-04 | 0.009 | -0.25 | -1.1999 | none |
| DENND2C | 0.735 | 0.5675 | 4.8436 | 7.50E-04 | 0.009 | -0.25 | -1.3174 | none |
| ITGA3 | 0.765 | 0.6482 | 4.8429 | 7.50E-04 | 0.009 | -0.25 | -1.3045 | none |
| CIART | 0.746 | 0.5544 | 4.8428 | 7.50E-04 | 0.009 | -0.25 | -1.3565 | none |
| RSPH3 | 0.628 | 0.499 | 4.8416 | 7.52E-04 | 0.009 | -0.25 | -1.128 | none |
| FAM153A | 1.2 | 0.8865 | 4.8411 | 7.52E-04 | 0.009 | -0.25 | -0.8169 | none |
| CHST5 | 0.604 | 0.4499 | 4.8408 | 7.53E-04 | 0.009 | -0.25 | -1.5572 | OA-related |
| DOCK5 | 0.902 | 0.7414 | 4.8406 | 7.53E-04 | 0.009 | -0.25 | -1.1801 | none |
| SLC16A4 | 0.624 | 0.4885 | 4.8401 | 7.53E-04 | 0.009 | -0.25 | -1.4332 | none |
| PLA2G2D | 0.712 | 0.5031 | 4.84 | 7.53E-04 | 0.009 | -0.25 | -1.5005 | none |
| AJUBA | 0.718 | 0.4909 | 4.8372 | 7.57E-04 | 0.009 | -0.26 | -1.3345 | none |
| MX1 | 0.969 | 0.7818 | 4.8337 | 7.60E-04 | 0.009 | -0.26 | -1.4891 | none |
| IREB2 | 0.714 | 0.5723 | 4.8306 | 7.64E-04 | 0.009 | -0.27 | -0.7266 | none |
| B3GALT1 | 0.733 | 0.5472 | 4.8284 | 7.66E-04 | 0.009 | -0.27 | -1.5793 | OA-related |
| LGR6 | 0.58 | 0.4078 | 4.8278 | 7.67E-04 | 0.009 | -0.27 | -1.3833 | none |
| CCDC120 | 0.506 | 0.4035 | 4.8266 | 7.68E-04 | 0.009 | -0.27 | -1.3879 | none |
| RAD9A | 0.842 | 0.6332 | 4.8242 | 7.71E-04 | 0.009 | -0.28 | -1.5817 | OA-related |
| ZNF225 | 0.863 | 0.7504 | 4.8239 | 7.71E-04 | 0.009 | -0.28 | -1.3403 | none |
| MACF1 | 1.209 | 1.1145 | 4.8233 | 7.72E-04 | 0.009 | -0.28 | -1.0679 | none |
| NEURL1 | 0.833 | 0.629 | 4.8229 | 7.72E-04 | 0.009 | -0.28 | -0.7871 | none |
| MSI2 | 0.614 | 0.4457 | 4.8228 | 7.73E-04 | 0.009 | -0.28 | -1.438 | none |
| DNAH17 | 0.768 | 0.765 | 4.8216 | 7.74E-04 | 0.009 | -0.28 | -1.0153 | none |
| HAUS3 | 0.608 | 0.5203 | 4.82 | 7.76E-04 | 0.009 | -0.28 | -0.707 | none |
| ADIPOQ | 0.606 | 0.4143 | 4.8163 | 7.80E-04 | 0.009 | -0.29 | -1.4078 | none |
| ATP1A3 | 0.752 | 0.5347 | 4.8151 | 7.81E-04 | 0.009 | -0.29 | -0.8054 | none |
| HINT3 | 0.469 | 0.3447 | 4.8136 | 7.83E-04 | 0.009 | -0.29 | -0.5419 | none |

| | | | | | | | | |
|---|---|---|---|---|---|---|---|---|
| GLOD4 | 0.596 | 0.385 | 4.8132 | 7.84E-04 | 0.009 | -0.29 | -1.2894 | none |
| BTBD1 | 0.64 | 0.5045 | 4.8128 | 7.84E-04 | 0.009 | -0.29 | -0.9918 | none |
| LEP | 0.456 | 0.3354 | 4.8123 | 7.85E-04 | 0.009 | -0.29 | -1.1234 | none |
| EDC3 | 0.395 | 0.2554 | 4.811 | 7.86E-04 | 0.009 | -0.3 | -0.3759 | none |
| CCDC125 | 0.671 | 0.591 | 4.8099 | 7.87E-04 | 0.009 | -0.3 | -0.9056 | none |
| DCST1 | 0.632 | 0.4385 | 4.8071 | 7.91E-04 | 0.009 | -0.3 | -0.7565 | none |
| VGLL3 | 0.808 | 0.593 | 4.8057 | 7.92E-04 | 0.009 | -0.3 | -1.3886 | none |
| ZNF69 | 1.126 | 0.8784 | 4.8041 | 7.94E-04 | 0.009 | -0.31 | -1.3493 | none |
| TARS2 | 0.749 | 0.6419 | 4.803 | 7.95E-04 | 0.009 | -0.31 | -1.4101 | none |
| SULT1B1 | 0.637 | 0.4682 | 4.8026 | 7.96E-04 | 0.009 | -0.31 | -1.256 | none |
| ISG20L2 | 0.542 | 0.4453 | 4.8019 | 7.97E-04 | 0.009 | -0.31 | -1.4072 | none |
| BPIFB1 | 0.671 | 0.5152 | 4.8014 | 7.97E-04 | 0.009 | -0.31 | -1.2795 | none |
| LPIN3 | 0.792 | 0.6423 | 4.8013 | 7.97E-04 | 0.009 | -0.31 | -1.5548 | OA-related |
| ADAM29 | 0.415 | 0.2587 | 4.7996 | 7.99E-04 | 0.009 | -0.31 | -1.2986 | none |
| GTPBP3 | 0.546 | 0.4713 | 4.7994 | 8.00E-04 | 0.009 | -0.31 | -1.0173 | none |
| SLC9A4 | 0.506 | 0.3492 | 4.7985 | 8.01E-04 | 0.009 | -0.32 | -1.6144 | OA-related |
| TNFRSF11I | 0.55 | 0.3971 | 4.7985 | 8.01E-04 | 0.009 | -0.32 | -0.5132 | none |
| CHRNA7 | 0.67 | 0.4723 | 4.798 | 8.01E-04 | 0.009 | -0.32 | -1.2524 | none |
| LMOD3 | 0.521 | 0.3738 | 4.7966 | 8.03E-04 | 0.009 | -0.32 | -1.478 | none |
| ZNF883 | 0.895 | 0.7125 | 4.7934 | 8.07E-04 | 0.009 | -0.32 | -1.2672 | none |
| ZNF274 | 0.826 | 0.7155 | 4.7917 | 8.09E-04 | 0.009 | -0.33 | -0.5911 | none |
| TRIM16 | 0.707 | 0.5623 | 4.791 | 8.09E-04 | 0.009 | -0.33 | -1.3483 | none |
| GTF2IRD2I | 0.784 | 0.5857 | 4.7896 | 8.11E-04 | 0.009 | -0.33 | -0.6871 | none |
| USP42 | 1.068 | 0.9472 | 4.7877 | 8.13E-04 | 0.009 | -0.33 | -1.4775 | none |
| PODNL1 | 1.103 | 0.8698 | 4.7854 | 8.16E-04 | 0.009 | -0.33 | -0.8196 | none |
| SMYD4 | 1.106 | 0.9447 | 4.7847 | 8.17E-04 | 0.009 | -0.34 | -1.4837 | none |
| DHODH | 0.873 | 0.7652 | 4.7845 | 8.17E-04 | 0.009 | -0.34 | -0.8486 | none |
| C12ORF49 | 0.542 | 0.3323 | 4.783 | 8.19E-04 | 0.009 | -0.34 | -1.4283 | none |
| C3ORF67 | 0.671 | 0.4702 | 4.7817 | 8.21E-04 | 0.009 | -0.34 | -1.0206 | none |
| ZNF346 | 0.511 | 0.4467 | 4.7815 | 8.21E-04 | 0.009 | -0.34 | -0.4237 | none |
| LIN52 | 0.712 | 0.5114 | 4.7779 | 8.25E-04 | 0.009 | -0.35 | -1.009 | none |
| SH2D6 | 1.023 | 0.9541 | 4.7777 | 8.25E-04 | 0.009 | -0.35 | -0.7434 | none |
| EFTUD2 | 0.588 | 0.4524 | 4.7772 | 8.26E-04 | 0.009 | -0.35 | -1.0474 | none |
| ECHDC1 | 0.511 | 0.3608 | 4.777 | 8.26E-04 | 0.009 | -0.35 | -1.2079 | none |
| C3ORF62 | 0.881 | 0.79 | 4.7764 | 8.27E-04 | 0.009 | -0.35 | -1.4568 | none |
| SPDYE2 | 0.618 | 0.559 | 4.774 | 8.30E-04 | 0.009 | -0.35 | -0.4814 | none |
| SMOC1 | 0.986 | 0.7774 | 4.773 | 8.31E-04 | 0.009 | -0.35 | -1.3287 | none |
| SYAP1 | 0.655 | 0.5475 | 4.7714 | 8.33E-04 | 0.009 | -0.35 | -1.615 | OA-related |
| C1ORF50 | 0.779 | 0.6654 | 4.7698 | 8.35E-04 | 0.009 | -0.36 | -1.0708 | none |
| APC | 0.938 | 0.8297 | 4.7695 | 8.35E-04 | 0.009 | -0.36 | -1.3725 | none |

| | | | | | | | | |
|---|---|---|---|---|---|---|---|---|
| PBLD | 0.566 | 0.4137 | 4.7693 | 8.36E-04 | 0.009 | -0.36 | -0.8115 | none |
| NMUR1 | 0.487 | 0.342 | 4.7685 | 8.37E-04 | 0.009 | -0.36 | -1.4401 | none |
| MRPL10 | 0.618 | 0.3853 | 4.7667 | 8.39E-04 | 0.009 | -0.36 | -1.0283 | none |
| FKRP | 0.863 | 0.7657 | 4.7663 | 8.39E-04 | 0.009 | -0.36 | -1.4118 | none |
| CCDC77 | 1.065 | 0.9027 | 4.7648 | 8.41E-04 | 0.009 | -0.36 | -1.2445 | none |
| DBF4B | 0.887 | 0.7578 | 4.7646 | 8.41E-04 | 0.009 | -0.36 | -1.6391 | OA-related |
| TRIM59 | 0.62 | 0.4623 | 4.7644 | 8.42E-04 | 0.009 | -0.37 | -1.3328 | none |
| FAM221A | 0.952 | 0.7689 | 4.7633 | 8.43E-04 | 0.009 | -0.37 | -1.356 | none |
| PPIAL4E | 0.476 | 0.3246 | 4.7632 | 8.43E-04 | 0.009 | -0.37 | 0.31 | none |
| PLEKHG4B | 0.657 | 0.4667 | 4.7625 | 8.44E-04 | 0.009 | -0.37 | -1.1808 | none |
| ATP7A | 0.79 | 0.6604 | 4.7608 | 8.46E-04 | 0.009 | -0.37 | -1.1388 | none |
| PPARA | 0.633 | 0.4849 | 4.7597 | 8.48E-04 | 0.009 | -0.37 | -1.1852 | none |
| TOR1AIP2 | 0.785 | 0.6223 | 4.7597 | 8.48E-04 | 0.009 | -0.37 | -1.6307 | OA-related |
| KCNJ16 | 0.611 | 0.4554 | 4.7578 | 8.50E-04 | 0.009 | -0.37 | -1.4489 | none |
| CAMTA2 | 0.528 | 0.4393 | 4.7577 | 8.50E-04 | 0.009 | -0.38 | -1.4701 | none |
| TRNT1 | 0.401 | 0.3129 | 4.7577 | 8.50E-04 | 0.009 | -0.38 | -0.8724 | none |
| TAC4 | 1.315 | 1.0912 | 4.7558 | 8.52E-04 | 0.009 | -0.38 | -1.4146 | none |
| DIS3L | 1.096 | 0.9547 | 4.7548 | 8.54E-04 | 0.009 | -0.38 | -1.2589 | none |
| NSG1 | 1.228 | 1.087 | 4.7523 | 8.57E-04 | 0.009 | -0.38 | -1.1751 | none |
| SIDT1 | 0.96 | 0.78 | 4.7522 | 8.57E-04 | 0.009 | -0.38 | -1.5724 | OA-related |
| CANX | 0.641 | 0.5057 | 4.752 | 8.57E-04 | 0.009 | -0.38 | 0.00615 | none |
| LRRD1 | 0.558 | 0.4035 | 4.7483 | 8.62E-04 | 0.009 | -0.39 | -1.4548 | none |
| MRM3 | 0.43 | 0.3468 | 4.748 | 8.62E-04 | 0.009 | -0.39 | -1.5365 | OA-related |
| MYLK3 | 0.577 | 0.4373 | 4.745 | 8.66E-04 | 0.009 | -0.39 | -1.4619 | none |
| SLC26A4 | 0.641 | 0.5007 | 4.7441 | 8.67E-04 | 0.009 | -0.4 | -0.7819 | none |
| UIMC1 | 1.001 | 0.8505 | 4.7427 | 8.69E-04 | 0.01 | -0.4 | -1.2342 | none |
| BRIP1 | 0.896 | 0.7068 | 4.7418 | 8.70E-04 | 0.01 | -0.4 | -0.8218 | none |
| GOLGA8R | 0.517 | 0.4254 | 4.7409 | 8.71E-04 | 0.01 | -0.4 | -0.9933 | none |
| NPIPB5 | 1.183 | 1.0836 | 4.7395 | 8.73E-04 | 0.01 | -0.4 | -0.9619 | none |
| GPSM2 | 0.977 | 0.8195 | 4.7393 | 8.73E-04 | 0.01 | -0.4 | -1.264 | none |
| NCOA7 | 1.249 | 1.0932 | 4.7376 | 8.76E-04 | 0.01 | -0.4 | -1.3679 | none |
| LOXL2 | 1.072 | 0.813 | 4.7372 | 8.76E-04 | 0.01 | -0.41 | -1.1673 | none |
| CPSF4 | 0.644 | 0.5457 | 4.7359 | 8.78E-04 | 0.01 | -0.41 | -0.996 | none |
| SHISA9 | 0.615 | 0.4371 | 4.7346 | 8.80E-04 | 0.01 | -0.41 | -1.4199 | none |
| C11ORF49 | 0.454 | 0.3538 | 4.7331 | 8.81E-04 | 0.01 | -0.41 | -1.4349 | none |
| PEX16 | 0.737 | 0.5663 | 4.7329 | 8.82E-04 | 0.01 | -0.41 | -1.2827 | none |
| ZFAND4 | 0.528 | 0.405 | 4.7329 | 8.82E-04 | 0.01 | -0.41 | -1.098 | none |
| FTO | 0.724 | 0.5972 | 4.7328 | 8.82E-04 | 0.01 | -0.41 | -1.4608 | none |
| NUP107 | 0.498 | 0.3826 | 4.7321 | 8.83E-04 | 0.01 | -0.41 | -1.1013 | none |
| OR2AG2 | 0.478 | 0.3422 | 4.7306 | 8.85E-04 | 0.01 | -0.42 | -1.5814 | OA-related |

| | | | | | | | | |
|---|---|---|---|---|---|---|---|---|
| BCO2 | 0.769 | 0.5861 | 4.7297 | 8.86E-04 | 0.01 | -0.42 | -1.4276 | none |
| TRIM22 | 0.576 | 0.4177 | 4.7294 | 8.86E-04 | 0.01 | -0.42 | -0.7422 | none |
| PPIAL4A | -0.399 | -0.2657 | -4.727 | 8.90E-04 | 0.01 | -0.42 | 1.25335 | none |
| FAM160B1 | 0.65 | 0.4657 | 4.7264 | 8.90E-04 | 0.01 | -0.42 | 0.03582 | none |
| SERINC5 | 0.692 | 0.5179 | 4.7262 | 8.91E-04 | 0.01 | -0.42 | -1.4902 | none |
| PRDM15 | 0.911 | 0.711 | 4.7246 | 8.93E-04 | 0.01 | -0.42 | -1.4437 | none |
| SAFB2 | 1.047 | 0.9574 | 4.7242 | 8.93E-04 | 0.01 | -0.42 | -1.3379 | none |
| GRAMD1A | 0.756 | 0.6741 | 4.7234 | 8.94E-04 | 0.01 | -0.43 | -1.4384 | none |
| CEP57L1 | 0.909 | 0.8129 | 4.7231 | 8.94E-04 | 0.01 | -0.43 | -1.0245 | none |
| S1PR2 | 0.549 | 0.3932 | 4.7223 | 8.96E-04 | 0.01 | -0.43 | -1.6253 | OA-related |
| MAP2K7 | 0.847 | 0.6902 | 4.7221 | 8.96E-04 | 0.01 | -0.43 | -0.9624 | none |
| FBXW10 | 1.024 | 0.7767 | 4.7216 | 8.97E-04 | 0.01 | -0.43 | -1.2735 | none |
| PARD6B | 0.635 | 0.4433 | 4.7201 | 8.99E-04 | 0.01 | -0.43 | -1.5846 | OA-related |
| FKBP9 | 0.604 | 0.5163 | 4.7165 | 9.03E-04 | 0.01 | -0.44 | -0.1004 | none |
| PROM2 | 0.475 | 0.3384 | 4.7098 | 9.12E-04 | 0.01 | -0.45 | -0.9403 | none |
| ACKR2 | 0.662 | 0.5239 | 4.7076 | 9.15E-04 | 0.01 | -0.45 | -1.0218 | none |
| ASPH | 0.874 | 0.7682 | 4.707 | 9.16E-04 | 0.01 | -0.45 | -1.2662 | none |
| FOXK1 | 0.702 | 0.6348 | 4.7066 | 9.17E-04 | 0.01 | -0.45 | -1.3472 | none |
| POU6F2 | 0.629 | 0.524 | 4.7064 | 9.17E-04 | 0.01 | -0.45 | -0.9948 | none |
| MOK | 0.884 | 0.6383 | 4.7047 | 9.19E-04 | 0.01 | -0.45 | -1.2985 | none |
| ZNF687 | 0.722 | 0.6138 | 4.7043 | 9.20E-04 | 0.01 | -0.45 | -1.4654 | none |
| PCSK6 | 0.603 | 0.4388 | 4.7042 | 9.20E-04 | 0.01 | -0.45 | -1.3522 | none |
| IYD | 0.646 | 0.5102 | 4.7036 | 9.21E-04 | 0.01 | -0.46 | -1.4636 | none |
| AP4M1 | 0.74 | 0.5935 | 4.702 | 9.23E-04 | 0.01 | -0.46 | -0.7372 | none |
| ZFP30 | 1.096 | 0.9619 | 4.7011 | 9.24E-04 | 0.01 | -0.46 | -1.5618 | OA-related |
| TIMM23B | 0.692 | 0.5929 | 4.6992 | 9.27E-04 | 0.01 | -0.46 | -0.6604 | none |
| VAV3 | 0.73 | 0.5509 | 4.6992 | 9.27E-04 | 0.01 | -0.46 | -0.7596 | none |
| OSBPL1A | 1.207 | 0.9894 | 4.6934 | 9.35E-04 | 0.01 | -0.47 | -1.1408 | none |
| GRM6 | 0.624 | 0.4747 | 4.6934 | 9.35E-04 | 0.01 | -0.47 | -1.4438 | none |
| RNF207 | 0.544 | 0.4851 | 4.692 | 9.37E-04 | 0.01 | -0.47 | -0.6841 | none |
| LRRFIP2 | 0.843 | 0.6432 | 4.6917 | 9.37E-04 | 0.01 | -0.47 | -0.7808 | none |
| ZNF780B | 0.586 | 0.4566 | 4.6917 | 9.37E-04 | 0.01 | -0.47 | -1.1495 | none |
| KIAA0586 | 1.177 | 1.0665 | 4.6913 | 9.38E-04 | 0.01 | -0.47 | -1.2711 | none |
| SHE | 0.529 | 0.3772 | 4.6912 | 9.38E-04 | 0.01 | -0.47 | -1.349 | none |
| GPR83 | 0.414 | 0.3143 | 4.6892 | 9.41E-04 | 0.01 | -0.48 | -1.4775 | none |
| RWDD2A | 0.455 | 0.301 | 4.6892 | 9.41E-04 | 0.01 | -0.48 | -0.7293 | none |
| GIMAP2 | 0.652 | 0.3798 | 4.6879 | 9.42E-04 | 0.01 | -0.48 | -1.1435 | none |
| TUBGCP4 | 0.52 | 0.4305 | 4.6861 | 9.45E-04 | 0.01 | -0.48 | -0.3032 | none |
| ZNF747 | 0.599 | 0.4521 | 4.6849 | 9.47E-04 | 0.01 | -0.48 | -1.3145 | none |
| SEMA4F | 0.53 | 0.3669 | 4.6847 | 9.47E-04 | 0.01 | -0.48 | -0.026 | none |

| | | | | | | | | |
|---|---|---|---|---|---|---|---|---|
| ABCA13 | 0.503 | 0.3609 | 4.6823 | 9.50E-04 | 0.01 | -0.49 | -1.492 | none |
| STOM | 0.457 | 0.3277 | 4.6818 | 9.51E-04 | 0.01 | -0.49 | -1.2439 | none |
| SH3BP5 | 0.791 | 0.6786 | 4.6811 | 9.52E-04 | 0.01 | -0.49 | -1.4325 | none |
| MCF2L | 0.777 | 0.6727 | 4.6807 | 9.52E-04 | 0.01 | -0.49 | -0.7174 | none |
| TFDP2 | 0.678 | 0.5295 | 4.6805 | 9.53E-04 | 0.01 | -0.49 | -0.9407 | none |
| ZNF345 | 1.033 | 0.7851 | 4.6797 | 9.54E-04 | 0.01 | -0.49 | -1.2165 | none |
| R3HDM4 | 0.611 | 0.4342 | 4.6793 | 9.54E-04 | 0.01 | -0.49 | -0.1632 | none |
| MGP | 0.82 | 0.5953 | 4.6785 | 9.56E-04 | 0.01 | -0.49 | -1.465 | none |
| POLQ | 1.201 | 1.0832 | 4.676 | 9.59E-04 | 0.01 | -0.5 | -1.3949 | none |
| RABL2B | 0.567 | 0.4367 | 4.6747 | 9.61E-04 | 0.01 | -0.5 | -1.075 | none |
| METTL7A | 0.692 | 0.5241 | 4.6735 | 9.63E-04 | 0.01 | -0.5 | -1.4966 | none |
| RFC2 | 0.53 | 0.4085 | 4.6728 | 9.64E-04 | 0.01 | -0.5 | -1.4081 | none |
| CEP135 | 0.882 | 0.7812 | 4.6694 | 9.69E-04 | 0.01 | -0.51 | -1.3494 | none |
| AARSD1 | 0.483 | 0.3573 | 4.6665 | 9.73E-04 | 0.01 | -0.51 | -1.2661 | none |
| CNOT10 | 0.907 | 0.7408 | 4.6645 | 9.76E-04 | 0.01 | -0.51 | -1.1622 | none |
| USP33 | 0.874 | 0.7167 | 4.6644 | 9.76E-04 | 0.01 | -0.51 | -1.2789 | none |
| GBA | 0.45 | 0.3143 | 4.6641 | 9.76E-04 | 0.01 | -0.51 | -1.0925 | none |
| SLC13A3 | 0.445 | 0.2938 | 4.664 | 9.76E-04 | 0.01 | -0.51 | -1.468 | none |
| SLC5A5 | 0.457 | 0.3244 | 4.6639 | 9.77E-04 | 0.01 | -0.51 | -1.3215 | none |
| SCAMP4 | 0.558 | 0.4317 | 4.6636 | 9.77E-04 | 0.01 | -0.51 | -1.3361 | none |
| LMO3 | 0.429 | 0.3131 | 4.6635 | 9.77E-04 | 0.01 | -0.51 | -0.5183 | none |
| MYO19 | 0.734 | 0.6394 | 4.6632 | 9.78E-04 | 0.01 | -0.51 | -1.4049 | none |
| KLHL5 | 0.665 | 0.5017 | 4.6626 | 9.78E-04 | 0.01 | -0.52 | -1.4817 | none |
| RBL2 | 0.596 | 0.461 | 4.6622 | 9.79E-04 | 0.01 | -0.52 | -0.6855 | none |
| ARL16 | 0.564 | 0.4163 | 4.6609 | 9.81E-04 | 0.01 | -0.52 | -1.3412 | none |
| TIGAR | 0.643 | 0.4787 | 4.6595 | 9.83E-04 | 0.01 | -0.52 | -0.9759 | none |
| AK9 | 0.654 | 0.5022 | 4.6582 | 9.85E-04 | 0.01 | -0.52 | -1.3427 | none |
| CR1 | 0.391 | 0.2545 | 4.6577 | 9.86E-04 | 0.01 | -0.52 | -1.5053 | none |
| DFFB | 0.598 | 0.3987 | 4.6568 | 9.87E-04 | 0.01 | -0.52 | -1.1593 | none |
| B3GALNT1 | 0.651 | 0.4568 | 4.6529 | 9.93E-04 | 0.01 | -0.53 | -1.175 | none |
| NOX4 | 0.382 | 0.2784 | 4.6519 | 9.94E-04 | 0.01 | -0.53 | -1.4127 | none |
| CTRB1 | 0.481 | 0.3347 | 4.6492 | 9.98E-04 | 0.01 | -0.54 | -1.3367 | none |
| CRTC2 | 0.703 | 0.632 | 4.6492 | 9.98E-04 | 0.01 | -0.54 | -1.3875 | none |
| APPBP2 | 0.539 | 0.3787 | 4.6471 | 0.001 | 0.01 | -0.54 | -0.1086 | none |
| TBL2 | 0.566 | 0.4499 | 4.646 | 0.001 | 0.01 | -0.54 | -0.8453 | none |
| CYP51A1 | 0.494 | 0.3506 | 4.645 | 0.001 | 0.01 | -0.54 | -1.2514 | none |
| ZNF619 | 0.543 | 0.4122 | 4.6408 | 0.00101 | 0.01 | -0.55 | -1.4552 | none |
| SLC25A51 | 0.458 | 0.3366 | 4.6396 | 0.00101 | 0.01 | -0.55 | -1.5052 | none |
| MITD1 | 0.648 | 0.4614 | 4.6395 | 0.00101 | 0.01 | -0.55 | -1.2299 | none |
| SLC28A2 | 0.708 | 0.5112 | 4.6394 | 0.00101 | 0.01 | -0.55 | -1.4273 | none |

| | | | | | | | | |
|---|---|---|---|---|---|---|---|---|
| PCDHA4 | 0.527 | 0.4043 | 4.6383 | 0.00101 | 0.01 | -0.55 | -1.5637 | OA-related |
| GOLGB1 | 0.831 | 0.721 | 4.6362 | 0.00102 | 0.01 | -0.56 | -1.0283 | none |
| ORAI2 | 0.557 | 0.4515 | 4.6352 | 0.00102 | 0.01 | -0.56 | -1.2452 | none |
| MAP3K15 | 0.481 | 0.3631 | 4.6349 | 0.00102 | 0.01 | -0.56 | -1.4423 | none |
| KLHL26 | 0.758 | 0.5293 | 4.6349 | 0.00102 | 0.01 | -0.56 | -1.557 | OA-related |
| PAICS | 0.49 | 0.3769 | 4.6349 | 0.00102 | 0.01 | -0.56 | -0.4215 | none |
| ACTR3C | 0.666 | 0.5356 | 4.6313 | 0.00103 | 0.011 | -0.56 | 0.06861 | none |
| KLLN | 0.564 | 0.3968 | 4.6312 | 0.00103 | 0.011 | -0.56 | -1.444 | none |
| CCDC151 | 0.439 | 0.3158 | 4.6294 | 0.00103 | 0.011 | -0.57 | -1.5212 | none |
| ZNF525 | 0.899 | 0.7225 | 4.6293 | 0.00103 | 0.011 | -0.57 | -1.1933 | none |
| DCAF16 | 0.742 | 0.6151 | 4.6263 | 0.00103 | 0.011 | -0.57 | -1.3364 | none |
| DTWD2 | 0.61 | 0.4675 | 4.6262 | 0.00103 | 0.011 | -0.57 | -1.4888 | none |
| FBXL22 | 0.678 | 0.5127 | 4.6262 | 0.00103 | 0.011 | -0.57 | -1.1172 | none |
| SPN | 0.685 | 0.552 | 4.6262 | 0.00103 | 0.011 | -0.57 | -1.2991 | none |
| D2HGDH | 0.659 | 0.5301 | 4.6246 | 0.00104 | 0.011 | -0.57 | -1.0574 | none |
| PCMTD2 | 0.491 | 0.3782 | 4.6237 | 0.00104 | 0.011 | -0.57 | -1.317 | none |
| HES2 | 0.636 | 0.5084 | 4.6234 | 0.00104 | 0.011 | -0.57 | -0.9493 | none |
| CLIP2 | 0.872 | 0.6568 | 4.6227 | 0.00104 | 0.011 | -0.58 | -1.3106 | none |
| MOCOS | 0.554 | 0.3945 | 4.6221 | 0.00104 | 0.011 | -0.58 | -1.2349 | none |
| ARPIN | 0.513 | 0.3979 | 4.622 | 0.00104 | 0.011 | -0.58 | -1.5239 | none |
| LETM1 | 0.881 | 0.8096 | 4.622 | 0.00104 | 0.011 | -0.58 | -1.5623 | OA-related |
| DISP2 | 0.655 | 0.4691 | 4.6214 | 0.00104 | 0.011 | -0.58 | -1.4897 | none |
| TRIM13 | 0.587 | 0.3867 | 4.6208 | 0.00104 | 0.011 | -0.58 | -1.3691 | none |
| BAZ2B | 0.771 | 0.7007 | 4.6208 | 0.00104 | 0.011 | -0.58 | -1.1373 | none |
| TSPAN6 | 0.598 | 0.4326 | 4.6192 | 0.00104 | 0.011 | -0.58 | -1.4503 | none |
| VPS53 | 0.722 | 0.6485 | 4.6188 | 0.00104 | 0.011 | -0.58 | -1.506 | none |
| ECHDC2 | 0.52 | 0.3751 | 4.6185 | 0.00104 | 0.011 | -0.58 | -1.2469 | none |
| BRI3 | 0.859 | 0.5934 | 4.6176 | 0.00105 | 0.011 | -0.58 | -1.1648 | none |
| STARD6 | 0.841 | 0.6372 | 4.617 | 0.00105 | 0.011 | -0.58 | -1.2391 | none |
| PARD6G | 0.512 | 0.3887 | 4.6165 | 0.00105 | 0.011 | -0.58 | -1.1452 | none |
| WDPCP | 0.675 | 0.5364 | 4.6153 | 0.00105 | 0.011 | -0.59 | -1.5464 | OA-related |
| HSPA4L | 0.74 | 0.6032 | 4.6141 | 0.00105 | 0.011 | -0.59 | -1.3855 | none |
| C2ORF15 | 0.564 | 0.3587 | 4.6119 | 0.00106 | 0.011 | -0.59 | -0.7195 | none |
| KIAA1958 | 0.642 | 0.5714 | 4.6117 | 0.00106 | 0.011 | -0.59 | -1.4172 | none |
| ZSCAN22 | 0.588 | 0.4725 | 4.6089 | 0.00106 | 0.011 | -0.6 | -1.4824 | none |
| ZNF233 | 0.704 | 0.5881 | 4.6078 | 0.00106 | 0.011 | -0.6 | -0.8246 | none |
| CDON | 0.56 | 0.4147 | 4.6073 | 0.00106 | 0.011 | -0.6 | -0.5225 | none |
| PDZD7 | 0.662 | 0.5359 | 4.6073 | 0.00106 | 0.011 | -0.6 | -1.1838 | none |
| KCNJ11 | 0.472 | 0.3567 | 4.6061 | 0.00106 | 0.011 | -0.6 | -1.2565 | none |
| RPS6KB1 | 0.985 | 0.8529 | 4.6047 | 0.00107 | 0.011 | -0.6 | -1.2785 | none |

| | | | | | | | | |
|---|---|---|---|---|---|---|---|---|
| CXORF58 | 0.41 | 0.2822 | 4.6044 | 0.00107 | 0.011 | -0.6 | -1.4831 | none |
| ELOVL7 | 0.536 | 0.3847 | 4.6044 | 0.00107 | 0.011 | -0.6 | -1.2616 | none |
| TRPM6 | 0.707 | 0.5716 | 4.6042 | 0.00107 | 0.011 | -0.6 | -1.3156 | none |
| ANGPT1 | 0.656 | 0.486 | 4.6037 | 0.00107 | 0.011 | -0.6 | -0.9993 | none |
| CASP10 | 1.154 | 0.9272 | 4.6018 | 0.00107 | 0.011 | -0.61 | -1.4589 | none |
| CA12 | 0.469 | 0.358 | 4.6017 | 0.00107 | 0.011 | -0.61 | -1.1425 | none |
| APBB3 | 0.489 | 0.4385 | 4.6007 | 0.00107 | 0.011 | -0.61 | 0.14157 | none |
| ZNF180 | 0.597 | 0.4813 | 4.6001 | 0.00107 | 0.011 | -0.61 | -1.2205 | none |
| PPM1F | 0.788 | 0.6165 | 4.599 | 0.00108 | 0.011 | -0.61 | -1.2377 | none |
| NIPAL3 | 0.502 | 0.3847 | 4.598 | 0.00108 | 0.011 | -0.61 | -1.5869 | OA-related |
| C2ORF83 | 0.667 | 0.5023 | 4.5971 | 0.00108 | 0.011 | -0.61 | -1.5864 | OA-related |
| CAMK2A | 0.445 | 0.3081 | 4.597 | 0.00108 | 0.011 | -0.61 | -1.2105 | none |
| ITPRIPL2 | 0.497 | 0.4197 | 4.5948 | 0.00108 | 0.011 | -0.62 | -1.4285 | none |
| ZNF85 | 0.628 | 0.43 | 4.5934 | 0.00108 | 0.011 | -0.62 | -1.0555 | none |
| STAT5B | 0.743 | 0.63 | 4.593 | 0.00109 | 0.011 | -0.62 | -1.3752 | none |
| LRRC27 | 0.619 | 0.5174 | 4.5929 | 0.00109 | 0.011 | -0.62 | -1.2328 | none |
| FAM49B | 0.486 | 0.3038 | 4.5881 | 0.00109 | 0.011 | -0.63 | -1.2311 | none |
| MFSD8 | 0.907 | 0.6317 | 4.5877 | 0.00109 | 0.011 | -0.63 | -1.17 | none |
| SEC14L4 | 0.413 | 0.3164 | 4.5861 | 0.0011 | 0.011 | -0.63 | -1.4338 | none |
| HSBP1L1 | 0.565 | 0.3929 | 4.5852 | 0.0011 | 0.011 | -0.63 | -0.079 | none |
| NICN1 | 0.572 | 0.3754 | 4.5837 | 0.0011 | 0.011 | -0.63 | -1.3546 | none |
| CLCN3 | 0.739 | 0.6037 | 4.5825 | 0.0011 | 0.011 | -0.64 | -0.2296 | none |
| HP | 0.61 | 0.3932 | 4.5822 | 0.0011 | 0.011 | -0.64 | -1.3554 | none |
| NUDT19 | 0.622 | 0.4457 | 4.5811 | 0.0011 | 0.011 | -0.64 | -1.5548 | OA-related |
| FAM83F | 0.451 | 0.3402 | 4.5809 | 0.00111 | 0.011 | -0.64 | -1.3596 | none |
| PNPLA3 | 0.619 | 0.4642 | 4.5806 | 0.00111 | 0.011 | -0.64 | -1.3663 | none |
| MLIP | 0.564 | 0.388 | 4.5797 | 0.00111 | 0.011 | -0.64 | -1.4626 | none |
| ALDH3B1 | 0.935 | 0.7334 | 4.5795 | 0.00111 | 0.011 | -0.64 | -1.2149 | none |
| SLC38A7 | 0.741 | 0.6423 | 4.5782 | 0.00111 | 0.011 | -0.64 | -1.416 | none |
| SEMA3C | 0.885 | 0.6485 | 4.5782 | 0.00111 | 0.011 | -0.64 | -1.0533 | none |
| PPP1R16A | 0.771 | 0.6674 | 4.5775 | 0.00111 | 0.011 | -0.64 | -1.4282 | none |
| AP000769 | 0.893 | 0.64 | 4.5757 | 0.00111 | 0.011 | -0.65 | -1.2226 | none |
| STARD8 | 0.523 | 0.4316 | 4.575 | 0.00112 | 0.011 | -0.65 | -0.9528 | none |
| TNKS | 1.192 | 1.0258 | 4.5721 | 0.00112 | 0.011 | -0.65 | -1.4505 | none |
| SEMA4C | 0.747 | 0.6499 | 4.572 | 0.00112 | 0.011 | -0.65 | -1.3868 | none |
| SPDYE3 | 0.998 | 0.8521 | 4.5718 | 0.00112 | 0.011 | -0.65 | -1.5008 | none |
| PRR5 | 0.59 | 0.4314 | 4.5714 | 0.00112 | 0.011 | -0.65 | -1.1133 | none |
| UVSSA | 0.716 | 0.5979 | 4.5695 | 0.00112 | 0.011 | -0.66 | -0.9583 | none |
| CARD18 | 0.566 | 0.4223 | 4.5694 | 0.00112 | 0.011 | -0.66 | -1.1458 | none |
| LGI1 | 0.766 | 0.5859 | 4.5694 | 0.00112 | 0.011 | -0.66 | -1.399 | none |

| | | | | | | | | |
|---|---|---|---|---|---|---|---|---|
| CAPRIN2 | 1.001 | 0.852 | 4.5686 | 0.00113 | 0.011 | -0.66 | -1.4227 | none |
| CDH6 | 0.396 | 0.3012 | 4.5666 | 0.00113 | 0.011 | -0.66 | -0.9051 | none |
| ANAPC5 | 0.434 | 0.3026 | 4.5662 | 0.00113 | 0.011 | -0.66 | -1.2418 | none |
| MSH5 | 0.865 | 0.6607 | 4.5655 | 0.00113 | 0.011 | -0.66 | -0.9125 | none |
| PTAFR | 0.492 | 0.395 | 4.5643 | 0.00113 | 0.011 | -0.66 | -1.0611 | none |
| CARD14 | 0.696 | 0.5226 | 4.5641 | 0.00113 | 0.011 | -0.66 | -1.2265 | none |
| PLA2G7 | 0.468 | 0.3861 | 4.564 | 0.00113 | 0.011 | -0.66 | -1.1769 | none |
| ARHGAP2( | 0.942 | 0.7922 | 4.5637 | 0.00113 | 0.011 | -0.66 | -1.2624 | none |
| CASP8 | 0.796 | 0.6386 | 4.5626 | 0.00114 | 0.011 | -0.67 | -1.4813 | none |
| LIN28A | 0.726 | 0.5654 | 4.5613 | 0.00114 | 0.011 | -0.67 | -1.5846 | OA-related |
| APOPT1 | 0.699 | 0.545 | 4.5593 | 0.00114 | 0.011 | -0.67 | -1.2077 | none |
| GAS2L3 | 0.588 | 0.4376 | 4.5589 | 0.00114 | 0.011 | -0.67 | -1.3319 | none |
| SPATS2 | 0.889 | 0.7545 | 4.5584 | 0.00114 | 0.011 | -0.67 | -1.479 | none |
| TJP2 | 1.032 | 0.9268 | 4.5565 | 0.00115 | 0.011 | -0.67 | -0.7844 | none |
| POLK | 0.905 | 0.7724 | 4.5563 | 0.00115 | 0.011 | -0.67 | -1.4017 | none |
| AK7 | 1.45 | 1.2051 | 4.5563 | 0.00115 | 0.011 | -0.67 | -1.0721 | none |
| TBC1D8B | 0.65 | 0.5314 | 4.5558 | 0.00115 | 0.011 | -0.68 | -1.0091 | none |
| KIAA2012 | 0.422 | 0.3256 | 4.5535 | 0.00115 | 0.011 | -0.68 | -1.6479 | OA-related |
| CLDN11 | 0.477 | 0.4384 | 4.5527 | 0.00115 | 0.011 | -0.68 | -0.9328 | none |
| ATM | 0.99 | 0.8195 | 4.5516 | 0.00115 | 0.011 | -0.68 | -1.5685 | OA-related |
| C1ORF174 | 0.489 | 0.3433 | 4.5506 | 0.00116 | 0.011 | -0.68 | -1.5683 | OA-related |
| DTD2 | 0.637 | 0.442 | 4.5504 | 0.00116 | 0.011 | -0.68 | -1.1276 | none |
| ZBTB21 | 0.721 | 0.5499 | 4.5503 | 0.00116 | 0.011 | -0.68 | -1.0916 | none |
| ZNF415 | 1.019 | 0.8116 | 4.5502 | 0.00116 | 0.011 | -0.68 | -1.2006 | none |
| ARFGEF2 | 1.15 | 1.0395 | 4.5502 | 0.00116 | 0.011 | -0.68 | -1.3455 | none |
| UCN | 0.574 | 0.4907 | 4.5474 | 0.00116 | 0.011 | -0.69 | -1.0106 | none |
| NAA60 | 0.68 | 0.5666 | 4.5474 | 0.00116 | 0.011 | -0.69 | -1.3182 | none |
| TBC1D24 | 0.665 | 0.5701 | 4.5472 | 0.00116 | 0.011 | -0.69 | -1.2881 | none |
| IQCC | 0.723 | 0.6396 | 4.5471 | 0.00116 | 0.011 | -0.69 | -1.5717 | OA-related |
| FCAR | 0.415 | 0.304 | 4.5471 | 0.00116 | 0.011 | -0.69 | -1.5988 | OA-related |
| FAM120A( | 0.581 | 0.4265 | 4.546 | 0.00116 | 0.011 | -0.69 | -1.2308 | none |
| IQCG | 0.913 | 0.801 | 4.545 | 0.00117 | 0.011 | -0.69 | -1.1775 | none |
| GRTP1 | 0.618 | 0.4196 | 4.5443 | 0.00117 | 0.011 | -0.69 | -1.2902 | none |
| TXK | 0.652 | 0.4767 | 4.5443 | 0.00117 | 0.011 | -0.69 | -1.386 | none |
| GGT6 | 0.554 | 0.4431 | 4.5424 | 0.00117 | 0.011 | -0.7 | -1.2283 | none |
| USP29 | 1.235 | 0.9606 | 4.5409 | 0.00117 | 0.011 | -0.7 | -1.3359 | none |
| SIGLEC11 | 0.669 | 0.4855 | 4.5406 | 0.00117 | 0.011 | -0.7 | -1.5107 | none |
| SLC43A2 | 0.65 | 0.4848 | 4.5396 | 0.00118 | 0.011 | -0.7 | -1.064 | none |
| TOP3A | 0.895 | 0.7801 | 4.5386 | 0.00118 | 0.011 | -0.7 | -1.3046 | none |
| OR2A5 | 0.823 | 0.6442 | 4.5379 | 0.00118 | 0.011 | -0.7 | -1.3829 | none |

| | | | | | | | | |
|---|---|---|---|---|---|---|---|---|
| KIF19 | 0.729 | 0.5278 | 4.5374 | 0.00118 | 0.011 | -0.7 | -1.4279 | none |
| ZNF318 | 1.012 | 0.8854 | 4.5364 | 0.00118 | 0.011 | -0.7 | -1.3729 | none |
| AP1S3 | 0.842 | 0.7036 | 4.5352 | 0.00118 | 0.011 | -0.71 | -1.4699 | none |
| TRABD2A | 0.488 | 0.3624 | 4.5345 | 0.00118 | 0.011 | -0.71 | -1.183 | none |
| OR2G6 | 0.411 | 0.3017 | 4.5321 | 0.00119 | 0.011 | -0.71 | -1.4277 | none |
| KCTD1 | 0.957 | 0.851 | 4.5315 | 0.00119 | 0.011 | -0.71 | -1.4286 | none |
| RNF115 | 0.547 | 0.3724 | 4.5315 | 0.00119 | 0.011 | -0.71 | -1.2342 | none |
| HSD17B4 | 0.425 | 0.33 | 4.531 | 0.00119 | 0.011 | -0.71 | 1.00776 | none |
| POLB | 0.644 | 0.4266 | 4.5293 | 0.00119 | 0.011 | -0.72 | -1.1306 | none |
| CAGE1 | 0.683 | 0.4955 | 4.5291 | 0.00119 | 0.011 | -0.72 | -1.4361 | none |
| METTL15 | 0.472 | 0.3191 | 4.5282 | 0.0012 | 0.011 | -0.72 | -0.1507 | none |
| KLRK1 | 0.556 | 0.3904 | 4.5271 | 0.0012 | 0.011 | -0.72 | -1.2558 | none |
| APAF1 | 0.731 | 0.6271 | 4.5249 | 0.0012 | 0.011 | -0.72 | -0.1692 | none |
| ZNF701 | 0.847 | 0.6952 | 4.5248 | 0.0012 | 0.011 | -0.72 | -1.3163 | none |
| RAB22A | 0.467 | 0.3438 | 4.5246 | 0.0012 | 0.011 | -0.72 | -1.2674 | none |
| HTN1 | 0.593 | 0.4586 | 4.5243 | 0.0012 | 0.011 | -0.72 | -0.8758 | none |
| MOCS3 | 0.558 | 0.361 | 4.5232 | 0.00121 | 0.011 | -0.72 | -1.0417 | none |
| ZNF442 | 0.534 | 0.4282 | 4.523 | 0.00121 | 0.011 | -0.73 | -1.5837 | OA-related |
| FBN1 | 0.514 | 0.3919 | 4.5211 | 0.00121 | 0.011 | -0.73 | -1.4495 | none |
| KRIT1 | 0.944 | 0.8584 | 4.5205 | 0.00121 | 0.012 | -0.73 | -1.2463 | none |
| NLRP12 | 0.658 | 0.4904 | 4.5182 | 0.00121 | 0.012 | -0.73 | -1.3707 | none |
| MCOLN3 | 0.399 | 0.3041 | 4.5176 | 0.00122 | 0.012 | -0.73 | -0.8179 | none |
| DPH1 | 0.43 | 0.3092 | 4.517 | 0.00122 | 0.012 | -0.73 | -0.9153 | none |
| RPP14 | 0.644 | 0.5418 | 4.517 | 0.00122 | 0.012 | -0.73 | -1.2295 | none |
| OR52A1 | 0.479 | 0.3514 | 4.5152 | 0.00122 | 0.012 | -0.74 | -1.5621 | OA-related |
| WNT9B | 0.491 | 0.3926 | 4.5148 | 0.00122 | 0.012 | -0.74 | -1.2173 | none |
| TNNI1 | 0.447 | 0.317 | 4.5126 | 0.00122 | 0.012 | -0.74 | -1.5567 | OA-related |
| TMEM192 | 0.504 | 0.3476 | 4.512 | 0.00123 | 0.012 | -0.74 | -0.8999 | none |
| CASP2 | 0.529 | 0.4107 | 4.5097 | 0.00123 | 0.012 | -0.75 | -1.584 | OA-related |
| UBA3 | 0.604 | 0.3862 | 4.5088 | 0.00123 | 0.012 | -0.75 | -1.1416 | none |
| ABCA1 | 0.877 | 0.9274 | 4.507 | 0.00124 | 0.012 | -0.75 | 0.28462 | none |
| HERC1 | 0.97 | 0.8848 | 4.5069 | 0.00124 | 0.012 | -0.75 | -1.4059 | none |
| DPYSL5 | 0.468 | 0.3046 | 4.5067 | 0.00124 | 0.012 | -0.75 | -1.5128 | none |
| RHBDD1 | 0.724 | 0.4971 | 4.5035 | 0.00124 | 0.012 | -0.75 | -1.0001 | none |
| CDK17 | 0.5 | 0.4206 | 4.5031 | 0.00124 | 0.012 | -0.76 | -0.3743 | none |
| HIST4H4 | 1.016 | 0.7377 | 4.5027 | 0.00124 | 0.012 | -0.76 | -1.1734 | none |
| SLC22A13 | 0.537 | 0.4024 | 4.5023 | 0.00124 | 0.012 | -0.76 | 0.25982 | none |
| FOXL2NB | 0.376 | 0.281 | 4.5012 | 0.00125 | 0.012 | -0.76 | -1.18 | none |
| SYNPO2 | 0.446 | 0.3551 | 4.5011 | 0.00125 | 0.012 | -0.76 | -1.4091 | none |
| PDXK | 0.505 | 0.4207 | 4.5004 | 0.00125 | 0.012 | -0.76 | -0.0808 | none |

| | | | | | | | | |
|---|---|---|---|---|---|---|---|---|
| MLH1 | 0.605 | 0.5029 | 4.499 | 0.00125 | 0.012 | -0.76 | -1.1937 | none |
| ERN1 | 0.63 | 0.5153 | 4.498 | 0.00125 | 0.012 | -0.76 | -0.7234 | none |
| XRN1 | 1.145 | 1.0101 | 4.4969 | 0.00125 | 0.012 | -0.76 | -1.4666 | none |
| ZNF20 | 1.106 | 0.9501 | 4.4968 | 0.00125 | 0.012 | -0.76 | -0.7659 | none |
| SPAST | 0.576 | 0.4107 | 4.496 | 0.00126 | 0.012 | -0.77 | -1.2718 | none |
| TIMM44 | 0.607 | 0.557 | 4.4948 | 0.00126 | 0.012 | -0.77 | -1.3556 | none |
| CASQ1 | 1.242 | 0.9393 | 4.4921 | 0.00126 | 0.012 | -0.77 | -0.7175 | none |
| AP1G1 | 0.513 | 0.4095 | 4.4917 | 0.00126 | 0.012 | -0.77 | -1.5508 | OA-related |
| C21ORF58 | 0.613 | 0.5412 | 4.4896 | 0.00127 | 0.012 | -0.78 | -1.3513 | none |
| TRPM3 | 0.529 | 0.3822 | 4.4872 | 0.00127 | 0.012 | -0.78 | -1.4517 | none |
| GSDMC | 0.495 | 0.3548 | 4.487 | 0.00127 | 0.012 | -0.78 | -1.3349 | none |
| ZNF791 | 0.735 | 0.6153 | 4.4853 | 0.00128 | 0.012 | -0.78 | -1.6231 | OA-related |
| PRR19 | 0.688 | 0.5436 | 4.4825 | 0.00128 | 0.012 | -0.79 | -1.4158 | none |
| DCAF8 | 0.648 | 0.6088 | 4.4805 | 0.00129 | 0.012 | -0.79 | -1.3154 | none |
| ART4 | 0.486 | 0.3535 | 4.4802 | 0.00129 | 0.012 | -0.79 | -1.4704 | none |
| C19ORF44 | 0.903 | 0.734 | 4.4795 | 0.00129 | 0.012 | -0.79 | -1.183 | none |
| DOT1L | 1.008 | 0.8996 | 4.4783 | 0.00129 | 0.012 | -0.79 | -1.4045 | none |
| HSCB | 0.918 | 0.67 | 4.4765 | 0.00129 | 0.012 | -0.8 | -0.9174 | none |
| C3ORF33 | 0.666 | 0.5532 | 4.4762 | 0.00129 | 0.012 | -0.8 | -1.2125 | none |
| ARRDC5 | 0.538 | 0.3743 | 4.4759 | 0.00129 | 0.012 | -0.8 | -1.4116 | none |
| RAB33B | 0.702 | 0.5575 | 4.4739 | 0.0013 | 0.012 | -0.8 | -0.9391 | none |
| HMX3 | 0.472 | 0.3679 | 4.4738 | 0.0013 | 0.012 | -0.8 | -1.0892 | none |
| VAV1 | 0.937 | 0.8186 | 4.4697 | 0.00131 | 0.012 | -0.81 | -1.4058 | none |
| NRG3 | 0.484 | 0.391 | 4.4689 | 0.00131 | 0.012 | -0.81 | -0.2876 | none |
| ANKRD13/ | 0.799 | 0.6636 | 4.4673 | 0.00131 | 0.012 | -0.81 | -1.3673 | none |
| SRL | 0.54 | 0.3823 | 4.4661 | 0.00131 | 0.012 | -0.81 | -1.3606 | none |
| LIX1L | 0.478 | 0.2773 | 4.465 | 0.00132 | 0.012 | -0.81 | -1.2886 | none |
| ALDH8A1 | 0.913 | 0.6676 | 4.4634 | 0.00132 | 0.012 | -0.82 | -1.0881 | none |
| LCN15 | 0.507 | 0.3598 | 4.4633 | 0.00132 | 0.012 | -0.82 | -1.1728 | none |
| FOXI3 | 0.457 | 0.3113 | 4.4633 | 0.00132 | 0.012 | -0.82 | -1.4869 | none |
| GNA11 | 1.361 | 1.1437 | 4.4632 | 0.00132 | 0.012 | -0.82 | -0.7652 | none |
| PPP5D1 | 0.608 | 0.4537 | 4.4616 | 0.00132 | 0.012 | -0.82 | -1.0157 | none |
| ISLR2 | 0.559 | 0.4049 | 4.4594 | 0.00133 | 0.012 | -0.82 | -1.0564 | none |
| SYT15 | 0.923 | 0.7192 | 4.4589 | 0.00133 | 0.012 | -0.82 | -1.2581 | none |
| UBXN7 | 1.021 | 0.8421 | 4.4549 | 0.00134 | 0.012 | -0.83 | -1.3857 | none |
| SPINK9 | 0.465 | 0.3904 | 4.4532 | 0.00134 | 0.012 | -0.83 | -0.7572 | none |
| GDF5 | 0.481 | 0.3694 | 4.4499 | 0.00135 | 0.012 | -0.84 | -1.2322 | none |
| NLRP2 | 0.88 | 0.8675 | 4.4489 | 0.00135 | 0.012 | -0.84 | -1.0209 | none |
| FGD4 | 0.608 | 0.4733 | 4.4462 | 0.00135 | 0.012 | -0.84 | -1.2688 | none |
| SV2C | 0.429 | 0.3099 | 4.4459 | 0.00135 | 0.012 | -0.84 | -1.3938 | none |

| | | | | | | | | |
|---|---|---|---|---|---|---|---|---|
| CAMKK2 | 0.537 | 0.5181 | 4.4454 | 0.00136 | 0.012 | -0.84 | -0.2215 | none |
| ZNF394 | 0.508 | 0.3707 | 4.4427 | 0.00136 | 0.012 | -0.85 | -1.5624 | OA-related |
| APOBEC3D | 0.744 | 0.6253 | 4.4426 | 0.00136 | 0.012 | -0.85 | -1.5233 | none |
| ZNF227 | 1.063 | 0.851 | 4.4425 | 0.00136 | 0.012 | -0.85 | -1.166 | none |
| RABL2A | 0.641 | 0.5952 | 4.4424 | 0.00136 | 0.012 | -0.85 | -0.6489 | none |
| ZNF774 | 0.754 | 0.5806 | 4.4421 | 0.00136 | 0.012 | -0.85 | -1.0139 | none |
| ORC6 | 0.609 | 0.5042 | 4.442 | 0.00136 | 0.012 | -0.85 | -0.9919 | none |
| SPAG9 | 1.39 | 1.2283 | 4.4415 | 0.00136 | 0.012 | -0.85 | -1.5177 | none |
| MIXL1 | 0.473 | 0.3669 | 4.4413 | 0.00136 | 0.012 | -0.85 | -1.3563 | none |
| CEACAM5 | 0.593 | 0.4305 | 4.4409 | 0.00136 | 0.012 | -0.85 | -0.6663 | none |
| VSIG1 | 0.656 | 0.5264 | 4.4386 | 0.00137 | 0.013 | -0.85 | -1.3155 | none |
| WSB2 | 0.687 | 0.4606 | 4.4379 | 0.00137 | 0.013 | -0.85 | -1.365 | none |
| HAMP | 0.74 | 0.536 | 4.4371 | 0.00137 | 0.013 | -0.86 | -1.4054 | none |
| CACNB4 | 0.629 | 0.4767 | 4.437 | 0.00137 | 0.013 | -0.86 | -1.6216 | OA-related |
| WDR76 | 0.58 | 0.4344 | 4.4359 | 0.00138 | 0.013 | -0.86 | -1.1236 | none |
| UNC80 | 0.812 | 0.6287 | 4.4356 | 0.00138 | 0.013 | -0.86 | -0.988 | none |
| SHF | 0.655 | 0.4919 | 4.4335 | 0.00138 | 0.013 | -0.86 | -1.1208 | none |
| B3GALNT2 | 0.46 | 0.4155 | 4.433 | 0.00138 | 0.013 | -0.86 | -0.9373 | none |
| ZNF616 | 0.915 | 0.7469 | 4.4325 | 0.00138 | 0.013 | -0.86 | -1.0722 | none |
| CHAMP1 | 0.687 | 0.5759 | 4.4324 | 0.00138 | 0.013 | -0.86 | -1.1669 | none |
| CHGB | 0.846 | 0.6372 | 4.432 | 0.00138 | 0.013 | -0.86 | -1.363 | none |
| C12ORF4 | 0.807 | 0.7217 | 4.4311 | 0.00139 | 0.013 | -0.86 | -0.8795 | none |
| PDPK1 | 0.801 | 0.6398 | 4.4306 | 0.00139 | 0.013 | -0.87 | -1.1896 | none |
| TMEM209 | 1.039 | 0.8214 | 4.4294 | 0.00139 | 0.013 | -0.87 | -1.3636 | none |
| CDH7 | 0.399 | 0.3132 | 4.4288 | 0.00139 | 0.013 | -0.87 | -1.2928 | none |
| TMOD2 | 0.383 | 0.2619 | 4.4287 | 0.00139 | 0.013 | -0.87 | -0.5912 | none |
| TMEM41A | 0.505 | 0.3489 | 4.4283 | 0.00139 | 0.013 | -0.87 | -1.0717 | none |
| C6ORF203 | 0.634 | 0.4748 | 4.4281 | 0.00139 | 0.013 | -0.87 | -1.3736 | none |
| SMG1 | 0.904 | 0.8295 | 4.4276 | 0.00139 | 0.013 | -0.87 | -1.4496 | none |
| ZNF230 | 0.775 | 0.5428 | 4.4275 | 0.00139 | 0.013 | -0.87 | -1.2608 | none |
| FBXL18 | 0.772 | 0.6252 | 4.4272 | 0.00139 | 0.013 | -0.87 | -1.5464 | OA-related |
| NIPA1 | 0.549 | 0.4211 | 4.4266 | 0.00139 | 0.013 | -0.87 | -1.3273 | none |
| CDH10 | 0.648 | 0.4934 | 4.4264 | 0.0014 | 0.013 | -0.87 | -1.3245 | none |
| SLC12A4 | 0.692 | 0.5566 | 4.4257 | 0.0014 | 0.013 | -0.87 | -1.0614 | none |
| XIAP | 0.741 | 0.6425 | 4.4256 | 0.0014 | 0.013 | -0.87 | -1.038 | none |
| GART | 0.403 | 0.3103 | 4.4256 | 0.0014 | 0.013 | -0.87 | -0.7357 | none |
| EYA1 | 0.76 | 0.5829 | 4.4256 | 0.0014 | 0.013 | -0.87 | -1.4106 | none |
| FAM193B | 0.851 | 0.7782 | 4.4249 | 0.0014 | 0.013 | -0.87 | -1.3667 | none |
| MLXIP | 0.892 | 0.7889 | 4.4248 | 0.0014 | 0.013 | -0.87 | -1.5042 | none |
| GLYR1 | 0.821 | 0.7239 | 4.4246 | 0.0014 | 0.013 | -0.87 | -0.8321 | none |

| | | | | | | | | |
|---|---|---|---|---|---|---|---|---|
| ZNF578 | 0.715 | 0.5769 | 4.4244 | 0.0014 | 0.013 | -0.87 | -1.6009 | OA-related |
| G6PC | 0.505 | 0.3959 | 4.4241 | 0.0014 | 0.013 | -0.87 | -1.3642 | none |
| PHKB | 0.869 | 0.7196 | 4.4237 | 0.0014 | 0.013 | -0.88 | -1.1139 | none |
| CAMK1D | 0.779 | 0.6136 | 4.4235 | 0.0014 | 0.013 | -0.88 | -1.4009 | none |
| CNPY1 | 0.496 | 0.393 | 4.423 | 0.0014 | 0.013 | -0.88 | -0.6934 | none |
| DOCK6 | 0.704 | 0.5647 | 4.4225 | 0.0014 | 0.013 | -0.88 | 0.19843 | none |
| TOR3A | 0.574 | 0.4156 | 4.422 | 0.0014 | 0.013 | -0.88 | -1.179 | none |
| KIAA0100 | 0.803 | 0.7071 | 4.4218 | 0.00141 | 0.013 | -0.88 | -1.2817 | none |
| PEAK1 | 0.992 | 0.8393 | 4.4215 | 0.00141 | 0.013 | -0.88 | -1.1349 | none |
| ADCY7 | 0.793 | 0.6797 | 4.4171 | 0.00142 | 0.013 | -0.89 | -1.1946 | none |
| ZNF766 | 1.178 | 1.0164 | 4.4164 | 0.00142 | 0.013 | -0.89 | -1.2114 | none |
| NKIRAS2 | 0.545 | 0.398 | 4.4134 | 0.00142 | 0.013 | -0.89 | -1.2057 | none |
| FUT11 | 0.695 | 0.6299 | 4.4134 | 0.00142 | 0.013 | -0.89 | -1.1543 | none |
| SLIT3 | 0.447 | 0.3122 | 4.4131 | 0.00142 | 0.013 | -0.89 | -1.524 | none |
| SPNS1 | 0.493 | 0.3416 | 4.4129 | 0.00142 | 0.013 | -0.89 | -0.8645 | none |
| RCC1L | 0.455 | 0.368 | 4.4126 | 0.00142 | 0.013 | -0.89 | -1.0101 | none |
| RTN2 | 0.811 | 0.6173 | 4.4123 | 0.00143 | 0.013 | -0.89 | -0.3808 | none |
| VENTX | 0.422 | 0.3264 | 4.4118 | 0.00143 | 0.013 | -0.89 | -1.2875 | none |
| CACNA1D | 0.669 | 0.5597 | 4.4106 | 0.00143 | 0.013 | -0.9 | -1.081 | none |
| HRG | 0.939 | 0.7175 | 4.4091 | 0.00143 | 0.013 | -0.9 | -1.3232 | none |
| KAT6B | 0.785 | 0.688 | 4.4081 | 0.00143 | 0.013 | -0.9 | -1.3898 | none |
| LRRC8B | 0.483 | 0.4004 | 4.4072 | 0.00144 | 0.013 | -0.9 | -0.4317 | none |
| NEK4 | 1.091 | 0.9609 | 4.4064 | 0.00144 | 0.013 | -0.9 | -1.0592 | none |
| TCTA | 0.59 | 0.429 | 4.4061 | 0.00144 | 0.013 | -0.9 | -1.2492 | none |
| NANOS1 | 0.492 | 0.3694 | 4.405 | 0.00144 | 0.013 | -0.9 | -1.4512 | none |
| FAM53C | 0.574 | 0.4141 | 4.4049 | 0.00144 | 0.013 | -0.9 | -1.241 | none |
| CRISPLD2 | 0.498 | 0.3617 | 4.4047 | 0.00144 | 0.013 | -0.9 | -1.3748 | none |
| IFT22 | 0.563 | 0.3725 | 4.4047 | 0.00144 | 0.013 | -0.9 | -1.4408 | none |
| HMG20B | 0.6 | 0.4427 | 4.4045 | 0.00144 | 0.013 | -0.9 | -1.1446 | none |
| SERPINA1( | 0.475 | 0.3757 | 4.4039 | 0.00144 | 0.013 | -0.91 | -1.3135 | none |
| AK3 | 0.432 | 0.3245 | 4.4033 | 0.00145 | 0.013 | -0.91 | 0.56581 | none |
| IGFL2 | 0.461 | 0.3806 | 4.4026 | 0.00145 | 0.013 | -0.91 | -0.6477 | none |
| PTGES | 0.745 | 0.5744 | 4.4004 | 0.00145 | 0.013 | -0.91 | -1.3032 | none |
| ZNF749 | 0.687 | 0.6602 | 4.3999 | 0.00145 | 0.013 | -0.91 | -0.1857 | none |
| SRGAP2 | 1.2 | 1.0905 | 4.3992 | 0.00145 | 0.013 | -0.91 | -1.1301 | none |
| TRIP11 | 0.74 | 0.7286 | 4.3985 | 0.00146 | 0.013 | -0.91 | -1.0284 | none |
| GBP6 | 0.585 | 0.4029 | 4.3978 | 0.00146 | 0.013 | -0.92 | -1.0324 | none |
| STAM2 | 0.84 | 0.6785 | 4.3969 | 0.00146 | 0.013 | -0.92 | -1.3204 | none |
| SECISBP2L | 1.123 | 0.9523 | 4.3968 | 0.00146 | 0.013 | -0.92 | -1.284 | none |
| TSC2 | 0.752 | 0.7128 | 4.3955 | 0.00146 | 0.013 | -0.92 | -1.0114 | none |

| | | | | | | | | |
|---|---|---|---|---|---|---|---|---|
| C14ORF17 | 1.367 | 1.1237 | 4.395 | 0.00146 | 0.013 | -0.92 | -1.0428 | none |
| VKORC1L1 | 0.533 | 0.3576 | 4.3935 | 0.00147 | 0.013 | -0.92 | -1.0594 | none |
| ADAMTS2 | 0.344 | 0.263 | 4.3919 | 0.00147 | 0.013 | -0.92 | -0.2096 | none |
| SCUBE3 | 0.478 | 0.3549 | 4.3918 | 0.00147 | 0.013 | -0.92 | -1.3277 | none |
| CELA2A | 0.859 | 0.6602 | 4.3915 | 0.00147 | 0.013 | -0.92 | -1.215 | none |
| PIGB | 0.812 | 0.5708 | 4.3914 | 0.00147 | 0.013 | -0.92 | -1.1006 | none |
| SLC25A16 | 0.695 | 0.4616 | 4.3906 | 0.00147 | 0.013 | -0.93 | -1.1362 | none |
| HDAC4 | 0.915 | 0.8195 | 4.389 | 0.00148 | 0.013 | -0.93 | -0.9895 | none |
| ZNF253 | 0.729 | 0.6044 | 4.3884 | 0.00148 | 0.013 | -0.93 | -0.6026 | none |
| PLK3 | 0.678 | 0.5776 | 4.3864 | 0.00148 | 0.013 | -0.93 | -1.2394 | none |
| SLC6A14 | 0.36 | 0.2585 | 4.3855 | 0.00148 | 0.013 | -0.93 | -1.1807 | none |
| REV3L | 1.122 | 0.9937 | 4.3854 | 0.00149 | 0.013 | -0.93 | -1.3124 | none |
| REEP2 | 0.563 | 0.4981 | 4.3829 | 0.00149 | 0.013 | -0.94 | -1.2506 | none |
| SOGA1 | 0.424 | 0.3559 | 4.3807 | 0.0015 | 0.013 | -0.94 | -1.3963 | none |
| NLGN4X | 0.731 | 0.5203 | 4.3806 | 0.0015 | 0.013 | -0.94 | -1.3099 | none |
| NUP43 | 0.896 | 0.7642 | 4.38 | 0.0015 | 0.013 | -0.94 | -1.1921 | none |
| NLN | 0.645 | 0.5517 | 4.3798 | 0.0015 | 0.013 | -0.94 | -1.0986 | none |
| MGRN1 | 0.557 | 0.4653 | 4.3796 | 0.0015 | 0.013 | -0.94 | -1.078 | none |
| POU5F2 | 0.466 | 0.344 | 4.3796 | 0.0015 | 0.013 | -0.94 | -1.5476 | OA-related |
| ATXN7L1 | 0.526 | 0.4098 | 4.379 | 0.0015 | 0.013 | -0.94 | -0.8804 | none |
| SUN1 | 0.483 | 0.4004 | 4.3789 | 0.0015 | 0.013 | -0.94 | -0.6371 | none |
| ZNF765 | 0.859 | 0.7404 | 4.3786 | 0.0015 | 0.013 | -0.94 | -1.548 | OA-related |
| ZNF669 | 0.861 | 0.6922 | 4.3779 | 0.0015 | 0.013 | -0.95 | -1.5703 | OA-related |
| RFX5 | 0.597 | 0.4966 | 4.3763 | 0.00151 | 0.013 | -0.95 | -1.208 | none |
| OSBPL9 | 0.516 | 0.3797 | 4.3759 | 0.00151 | 0.013 | -0.95 | -0.5673 | none |
| GALNT16 | 0.497 | 0.354 | 4.3743 | 0.00151 | 0.013 | -0.95 | -1.5567 | OA-related |
| C1ORF61 | 0.677 | 0.4025 | 4.3732 | 0.00151 | 0.013 | -0.95 | -1.3859 | none |
| SLC2A4 | 0.5 | 0.3657 | 4.3732 | 0.00151 | 0.013 | -0.95 | -1.1792 | none |
| SDS | 0.507 | 0.4633 | 4.3723 | 0.00152 | 0.013 | -0.95 | -1.0638 | none |
| TYK2 | 0.785 | 0.682 | 4.3705 | 0.00152 | 0.013 | -0.96 | -1.1798 | none |
| FAM83B | 0.478 | 0.3483 | 4.3697 | 0.00152 | 0.013 | -0.96 | -1.4169 | none |
| SLC25A40 | 0.684 | 0.4959 | 4.3694 | 0.00152 | 0.013 | -0.96 | -0.3576 | none |
| C1QTNF1 | 0.612 | 0.5087 | 4.3674 | 0.00153 | 0.013 | -0.96 | -1.3012 | none |
| P2RX6 | 0.672 | 0.5063 | 4.3659 | 0.00153 | 0.013 | -0.96 | -1.4347 | none |
| DOC2A | 1.027 | 0.7851 | 4.3658 | 0.00153 | 0.013 | -0.96 | -1.2344 | none |
| LIPH | 0.544 | 0.4297 | 4.3649 | 0.00153 | 0.013 | -0.97 | -1.5789 | OA-related |
| EBF2 | 0.652 | 0.5102 | 4.3647 | 0.00153 | 0.013 | -0.97 | -1.284 | none |
| ABHD1 | 0.677 | 0.5003 | 4.3643 | 0.00153 | 0.013 | -0.97 | -1.05 | none |
| MECP2 | 1.1 | 0.9874 | 4.3643 | 0.00153 | 0.013 | -0.97 | -1.2536 | none |
| TBX4 | 0.614 | 0.4571 | 4.3635 | 0.00154 | 0.013 | -0.97 | -1.2251 | none |

| | | | | | | | | |
|---|---|---|---|---|---|---|---|---|
| ASPA | 0.394 | 0.2882 | 4.3625 | 0.00154 | 0.013 | -0.97 | -1.4867 | none |
| OTOA | 0.425 | 0.2945 | 4.3622 | 0.00154 | 0.013 | -0.97 | -1.2157 | none |
| ZNF131 | 0.753 | 0.6003 | 4.362 | 0.00154 | 0.013 | -0.97 | -1.4593 | none |
| TPH1 | 0.412 | 0.3056 | 4.3616 | 0.00154 | 0.013 | -0.97 | -1.417 | none |
| CFL1 | -0.467 | -0.3924 | -4.361 | 0.00154 | 0.013 | -0.97 | 2.69753 | AA-related |
| RAB40B | 0.685 | 0.4313 | 4.3611 | 0.00154 | 0.013 | -0.97 | -1.0657 | none |
| NUMB | 0.647 | 0.5465 | 4.3607 | 0.00154 | 0.013 | -0.97 | -0.4546 | none |
| SIN3B | 0.844 | 0.7572 | 4.3603 | 0.00154 | 0.013 | -0.97 | -1.2493 | none |
| TIMM17A | 0.56 | 0.3663 | 4.3591 | 0.00155 | 0.013 | -0.97 | -1.3573 | none |
| GUCA1B | 0.457 | 0.3444 | 4.3582 | 0.00155 | 0.013 | -0.98 | -1.5571 | OA-related |
| PRELID3A | 0.813 | 0.7075 | 4.3574 | 0.00155 | 0.013 | -0.98 | -1.136 | none |
| TMEM220 | 0.786 | 0.5866 | 4.3568 | 0.00155 | 0.013 | -0.98 | -1.0827 | none |
| SLC4A7 | 0.461 | 0.3791 | 4.3561 | 0.00155 | 0.013 | -0.98 | -0.7544 | none |
| RND1 | 0.638 | 0.4833 | 4.3557 | 0.00155 | 0.013 | -0.98 | -0.8653 | none |
| TMEM17 | 0.657 | 0.4983 | 4.3555 | 0.00155 | 0.013 | -0.98 | -1.3914 | none |
| UNKL | 0.61 | 0.5377 | 4.3542 | 0.00156 | 0.013 | -0.98 | -1.1414 | none |
| ARHGEF15 | 0.719 | 0.5384 | 4.354 | 0.00156 | 0.013 | -0.98 | -1.2067 | none |
| CCL5 | 0.675 | 0.5712 | 4.3534 | 0.00156 | 0.013 | -0.98 | -1.1018 | none |
| SYCP1 | 1.276 | 0.8801 | 4.3529 | 0.00156 | 0.013 | -0.98 | -0.0128 | none |
| COQ5 | 0.557 | 0.3923 | 4.3528 | 0.00156 | 0.013 | -0.98 | -1.3634 | none |
| NPIPA5 | 0.627 | 0.426 | 4.3527 | 0.00156 | 0.013 | -0.98 | -0.8546 | none |
| CSNK1E | 0.695 | 0.6034 | 4.3515 | 0.00156 | 0.013 | -0.99 | -1.5227 | none |
| KLHL8 | 0.497 | 0.3526 | 4.3505 | 0.00157 | 0.013 | -0.99 | -1.1994 | none |
| NEDD1 | 0.651 | 0.4607 | 4.35 | 0.00157 | 0.013 | -0.99 | -0.6581 | none |
| ETV5 | 0.857 | 0.7437 | 4.3488 | 0.00157 | 0.013 | -0.99 | -0.7355 | none |
| GMPR | 0.7 | 0.4467 | 4.3483 | 0.00157 | 0.013 | -0.99 | -1.3819 | none |
| CGN | 0.573 | 0.531 | 4.3475 | 0.00157 | 0.013 | -0.99 | -0.4312 | none |
| COX6B2 | 0.664 | 0.5032 | 4.3469 | 0.00158 | 0.013 | -0.99 | -1.3129 | none |
| TMPRSS12 | 0.489 | 0.3692 | 4.3466 | 0.00158 | 0.013 | -0.99 | -1.1527 | none |
| SCAF8 | 1.256 | 1.1208 | 4.3458 | 0.00158 | 0.013 | -0.99 | -0.5128 | none |
| KLF17 | 0.424 | 0.2973 | 4.3458 | 0.00158 | 0.013 | -0.99 | -1.0861 | none |
| BANP | 0.553 | 0.5251 | 4.3457 | 0.00158 | 0.013 | -0.99 | -1.1701 | none |
| TUBGCP6 | 0.701 | 0.6319 | 4.3453 | 0.00158 | 0.013 | -1 | -1.2037 | none |
| KBTBD6 | 0.578 | 0.4248 | 4.3445 | 0.00158 | 0.013 | -1 | -0.8246 | none |
| ASZ1 | 0.435 | 0.311 | 4.3443 | 0.00158 | 0.013 | -1 | -1.0057 | none |
| BCORL1 | 0.756 | 0.6085 | 4.344 | 0.00158 | 0.013 | -1 | -0.8891 | none |
| ACSF2 | 0.642 | 0.5315 | 4.3426 | 0.00159 | 0.013 | -1 | -1.502 | none |
| MTRF1 | 0.85 | 0.695 | 4.3425 | 0.00159 | 0.013 | -1 | -1.0353 | none |
| TNS3 | 0.797 | 0.8018 | 4.3411 | 0.00159 | 0.013 | -1 | -0.4242 | none |
| MIER1 | 0.627 | 0.4443 | 4.3406 | 0.00159 | 0.013 | -1 | -1.3812 | none |

| | | | | | | | | |
|---|---|---|---|---|---|---|---|---|
| NUP98 | 0.792 | 0.7064 | 4.3403 | 0.00159 | 0.013 | -1 | -1.3198 | none |
| USP22 | 0.632 | 0.5559 | 4.3391 | 0.00159 | 0.013 | -1.01 | -1.2961 | none |
| GPHN | 0.841 | 0.7425 | 4.339 | 0.00159 | 0.013 | -1.01 | -1.3502 | none |
| RP1 | 0.673 | 0.5069 | 4.339 | 0.00159 | 0.013 | -1.01 | -1.3397 | none |
| ADRB2 | 0.363 | 0.2671 | 4.3389 | 0.00159 | 0.013 | -1.01 | -1.403 | none |
| OR2T33 | 0.381 | 0.2803 | 4.3386 | 0.0016 | 0.013 | -1.01 | -1.1031 | none |
| MIA3 | 1.03 | 0.9091 | 4.3372 | 0.0016 | 0.013 | -1.01 | -1.1919 | none |
| ARL11 | 0.459 | 0.295 | 4.3369 | 0.0016 | 0.013 | -1.01 | -1.3223 | none |
| PSG5 | 1.068 | 0.8022 | 4.3338 | 0.00161 | 0.014 | -1.01 | -1.2664 | none |
| FFAR2 | 0.87 | 0.6343 | 4.3322 | 0.00161 | 0.014 | -1.02 | -0.9232 | none |
| ZNF200 | 0.462 | 0.3411 | 4.3302 | 0.00162 | 0.014 | -1.02 | -1.4969 | none |
| JKAMP | 0.651 | 0.4073 | 4.3293 | 0.00162 | 0.014 | -1.02 | -0.9702 | none |
| GRIFIN | -0.37 | -0.2384 | -4.329 | 0.00162 | 0.014 | -1.02 | 1.93287 | none |
| CASC1 | 0.484 | 0.2688 | 4.3279 | 0.00162 | 0.014 | -1.02 | -1.0545 | none |
| TTC38 | 0.782 | 0.6018 | 4.3275 | 0.00162 | 0.014 | -1.02 | -1.1706 | none |
| BLOC1S3 | 0.723 | 0.5377 | 4.3269 | 0.00162 | 0.014 | -1.02 | -1.5821 | OA-related |
| MFAP3 | 0.725 | 0.5178 | 4.3261 | 0.00163 | 0.014 | -1.02 | -1.437 | none |
| MPP4 | 0.517 | 0.3654 | 4.3254 | 0.00163 | 0.014 | -1.03 | -1.0575 | none |
| SORT1 | 0.692 | 0.5329 | 4.3251 | 0.00163 | 0.014 | -1.03 | -1.3786 | none |
| TNFRSF10/ | 0.589 | 0.4913 | 4.3247 | 0.00163 | 0.014 | -1.03 | -1.3398 | none |
| OPA3 | 0.685 | 0.6055 | 4.3246 | 0.00163 | 0.014 | -1.03 | -1.2805 | none |
| STXBP3 | 0.608 | 0.4576 | 4.3234 | 0.00163 | 0.014 | -1.03 | -1.4744 | none |
| MARCKSL1 | -0.485 | -0.3921 | -4.323 | 0.00163 | 0.014 | -1.03 | 2.08002 | none |
| NDUFAF6 | 0.506 | 0.3548 | 4.3228 | 0.00163 | 0.014 | -1.03 | -0.7243 | none |
| DSG3 | 0.364 | 0.2617 | 4.3227 | 0.00163 | 0.014 | -1.03 | -1.3104 | none |
| DOK4 | 0.392 | 0.3344 | 4.3226 | 0.00163 | 0.014 | -1.03 | -0.9271 | none |
| USF3 | 0.854 | 0.7395 | 4.32 | 0.00164 | 0.014 | -1.03 | -1.4686 | none |
| KCTD5 | -0.566 | -0.3923 | -4.32 | 0.00164 | 0.014 | -1.03 | 1.70968 | none |
| CAD | 0.874 | 0.7661 | 4.319 | 0.00164 | 0.014 | -1.04 | -1.039 | none |
| ST8SIA6 | 0.689 | 0.487 | 4.319 | 0.00164 | 0.014 | -1.04 | -0.9825 | none |
| TIAF1 | 0.409 | 0.3792 | 4.3183 | 0.00165 | 0.014 | -1.04 | -1.2185 | none |
| HPS4 | 0.625 | 0.4727 | 4.3176 | 0.00165 | 0.014 | -1.04 | -0.9872 | none |
| C16ORF89 | 0.612 | 0.4358 | 4.3174 | 0.00165 | 0.014 | -1.04 | -1.401 | none |
| CXORF38 | 0.648 | 0.5697 | 4.3174 | 0.00165 | 0.014 | -1.04 | -1.2466 | none |
| SPON2 | 0.664 | 0.5035 | 4.3163 | 0.00165 | 0.014 | -1.04 | -0.7007 | none |
| SEMA4B | 0.813 | 0.6601 | 4.3161 | 0.00165 | 0.014 | -1.04 | -1.2047 | none |
| C16ORF72 | 0.493 | 0.3858 | 4.3156 | 0.00165 | 0.014 | -1.04 | -1.6181 | OA-related |
| ZYG11A | 0.69 | 0.5194 | 4.3154 | 0.00165 | 0.014 | -1.04 | -1.4285 | none |
| OR2AG1 | 0.335 | 0.2467 | 4.3153 | 0.00165 | 0.014 | -1.04 | -1.3745 | none |
| IKBKB | 0.843 | 0.7568 | 4.3141 | 0.00166 | 0.014 | -1.04 | -1.4488 | none |

| | | | | | | | | |
|---|---|---|---|---|---|---|---|---|
| RFFL | 0.528 | 0.425 | 4.3139 | 0.00166 | 0.014 | -1.04 | -0.5877 | none |
| SLC6A6 | 0.633 | 0.5714 | 4.3129 | 0.00166 | 0.014 | -1.05 | -0.2178 | none |
| PTK6 | 0.384 | 0.255 | 4.3127 | 0.00166 | 0.014 | -1.05 | -1.3498 | none |
| ASB8 | 0.357 | 0.2446 | 4.3122 | 0.00166 | 0.014 | -1.05 | -1.0017 | none |
| DMBX1 | 0.542 | 0.3829 | 4.3109 | 0.00166 | 0.014 | -1.05 | -1.521 | none |
| TBC1D17 | 0.673 | 0.5659 | 4.3101 | 0.00167 | 0.014 | -1.05 | -1.5045 | none |
| DNAJC24 | 0.77 | 0.5949 | 4.309 | 0.00167 | 0.014 | -1.05 | -1.0679 | none |
| BORCS8 | 0.687 | 0.516 | 4.3086 | 0.00167 | 0.014 | -1.05 | -0.9626 | none |
| VPS37B | 0.579 | 0.4391 | 4.3085 | 0.00167 | 0.014 | -1.05 | -1.3991 | none |
| PKP2 | 0.63 | 0.4899 | 4.3083 | 0.00167 | 0.014 | -1.05 | -1.1975 | none |
| CTSB | 0.554 | 0.43 | 4.3083 | 0.00167 | 0.014 | -1.05 | -1.0669 | none |
| ADSSL1 | 0.575 | 0.485 | 4.3069 | 0.00167 | 0.014 | -1.05 | -0.5559 | none |
| TUBD1 | 0.724 | 0.5976 | 4.3036 | 0.00168 | 0.014 | -1.06 | -1.3499 | none |
| CENPO | 0.516 | 0.4152 | 4.3033 | 0.00168 | 0.014 | -1.06 | 0.10423 | none |
| VPS33A | 0.52 | 0.3853 | 4.3008 | 0.00169 | 0.014 | -1.06 | -1.3144 | none |
| LYRM7 | 0.795 | 0.6345 | 4.3005 | 0.00169 | 0.014 | -1.06 | -0.905 | none |
| LSM11 | 0.442 | 0.3418 | 4.3001 | 0.00169 | 0.014 | -1.06 | -1.6353 | OA-related |
| NATD1 | 0.543 | 0.443 | 4.2996 | 0.00169 | 0.014 | -1.07 | -1.5137 | none |
| ESYT3 | 0.533 | 0.3946 | 4.2994 | 0.00169 | 0.014 | -1.07 | -1.3472 | none |
| AFAP1 | 0.922 | 0.7898 | 4.299 | 0.0017 | 0.014 | -1.07 | -1.0893 | none |
| FOXN3 | 0.994 | 0.8335 | 4.297 | 0.0017 | 0.014 | -1.07 | -1.3109 | none |
| TBC1D20 | 0.472 | 0.3394 | 4.297 | 0.0017 | 0.014 | -1.07 | -1.1479 | none |
| ZNF552 | 0.718 | 0.6359 | 4.2969 | 0.0017 | 0.014 | -1.07 | -1.0286 | none |
| FANCA | 0.483 | 0.4426 | 4.2965 | 0.0017 | 0.014 | -1.07 | -1.2808 | none |
| CEP126 | 0.546 | 0.423 | 4.2955 | 0.0017 | 0.014 | -1.07 | -1.4301 | none |
| ZSCAN10 | 0.949 | 0.7783 | 4.2952 | 0.00171 | 0.014 | -1.07 | -0.9448 | none |
| TDRD3 | 0.811 | 0.6654 | 4.2946 | 0.00171 | 0.014 | -1.07 | -1.6056 | OA-related |
| ZNF273 | 0.678 | 0.5347 | 4.2946 | 0.00171 | 0.014 | -1.07 | -1.4213 | none |
| TNFSF13 | 0.57 | 0.3801 | 4.2928 | 0.00171 | 0.014 | -1.08 | -0.3231 | none |
| DDHD2 | 0.542 | 0.4035 | 4.2927 | 0.00171 | 0.014 | -1.08 | -0.3117 | none |
| OR4F17 | 0.595 | 0.4419 | 4.2916 | 0.00171 | 0.014 | -1.08 | -1.0919 | none |
| SGPL1 | 0.54 | 0.4294 | 4.2916 | 0.00171 | 0.014 | -1.08 | -1.4327 | none |
| KPNA5 | 0.526 | 0.4402 | 4.2909 | 0.00172 | 0.014 | -1.08 | -0.8968 | none |
| BSND | 0.365 | 0.2581 | 4.2898 | 0.00172 | 0.014 | -1.08 | -1.4748 | none |
| WDR92 | 0.468 | 0.3642 | 4.2896 | 0.00172 | 0.014 | -1.08 | -1.27 | none |
| GRK7 | 0.495 | 0.3775 | 4.2893 | 0.00172 | 0.014 | -1.08 | -0.8889 | none |
| KANSL1L | 0.772 | 0.5446 | 4.2887 | 0.00172 | 0.014 | -1.08 | -1.4967 | none |
| ANKRD30I | 0.532 | 0.4056 | 4.2868 | 0.00173 | 0.014 | -1.09 | -1.1599 | none |
| AP4E1 | 0.662 | 0.5808 | 4.2867 | 0.00173 | 0.014 | -1.09 | -0.929 | none |
| KLK7 | 0.417 | 0.3413 | 4.286 | 0.00173 | 0.014 | -1.09 | -0.3784 | none |

| | | | | | | | | |
|---|---|---|---|---|---|---|---|---|
| FGF5 | 0.417 | 0.283 | 4.2856 | 0.00173 | 0.014 | -1.09 | -1.4474 | none |
| HAUS5 | 0.492 | 0.433 | 4.2846 | 0.00173 | 0.014 | -1.09 | -1.363 | none |
| ITFG2 | 0.529 | 0.4382 | 4.2843 | 0.00173 | 0.014 | -1.09 | -0.1522 | none |
| KCNN2 | 0.688 | 0.57 | 4.2839 | 0.00173 | 0.014 | -1.09 | 0.0111 | none |
| ZKSCAN1 | 0.973 | 0.8734 | 4.2837 | 0.00174 | 0.014 | -1.09 | -1.263 | none |
| OR2V2 | 0.435 | 0.3025 | 4.2832 | 0.00174 | 0.014 | -1.09 | -1.243 | none |
| OR6K3 | 0.577 | 0.4 | 4.2821 | 0.00174 | 0.014 | -1.09 | -1.565 | OA-related |
| MBTPS1 | 0.645 | 0.5628 | 4.2819 | 0.00174 | 0.014 | -1.09 | -1.3101 | none |
| POU5F1 | 0.633 | 0.4837 | 4.2818 | 0.00174 | 0.014 | -1.09 | -1.3141 | none |
| SOX7 | 0.406 | 0.3009 | 4.2806 | 0.00174 | 0.014 | -1.1 | -1.4824 | none |
| ZCCHC4 | 0.51 | 0.3386 | 4.2797 | 0.00175 | 0.014 | -1.1 | -1.0924 | none |
| ZGRF1 | 1.106 | 0.9572 | 4.2772 | 0.00175 | 0.014 | -1.1 | -1.37 | none |
| DNAJC18 | 0.543 | 0.3389 | 4.2764 | 0.00176 | 0.014 | -1.1 | -1.0921 | none |
| WDR55 | 0.494 | 0.3946 | 4.2747 | 0.00176 | 0.014 | -1.1 | -1.3991 | none |
| NCCRP1 | 0.537 | 0.4127 | 4.2737 | 0.00176 | 0.014 | -1.11 | -1.3112 | none |
| FAAH | 0.754 | 0.6653 | 4.2721 | 0.00177 | 0.014 | -1.11 | -0.0774 | none |
| PRKCA | 1.012 | 0.9145 | 4.2701 | 0.00177 | 0.014 | -1.11 | -1.3523 | none |
| KLHDC4 | 0.688 | 0.6742 | 4.2697 | 0.00177 | 0.014 | -1.11 | -1.1048 | none |
| BPNT1 | 0.604 | 0.5115 | 4.2694 | 0.00177 | 0.014 | -1.11 | -0.7169 | none |
| IL17RE | 0.617 | 0.5068 | 4.269 | 0.00178 | 0.014 | -1.11 | -0.729 | none |
| NOL11 | 0.594 | 0.4364 | 4.2685 | 0.00178 | 0.014 | -1.11 | -1.3891 | none |
| BCAS1 | 0.505 | 0.3449 | 4.2685 | 0.00178 | 0.014 | -1.11 | -0.9021 | none |
| INPP5B | 1.006 | 0.9207 | 4.2685 | 0.00178 | 0.014 | -1.11 | -1.3373 | none |
| ZNF391 | 0.48 | 0.3949 | 4.2662 | 0.00178 | 0.014 | -1.12 | -0.2355 | none |
| SYMPK | 1.179 | 1.0371 | 4.2644 | 0.00179 | 0.014 | -1.12 | -1.3012 | none |
| MFAP5 | 0.515 | 0.4193 | 4.2638 | 0.00179 | 0.014 | -1.12 | -1.302 | none |
| CHST1 | 0.445 | 0.3027 | 4.2637 | 0.00179 | 0.014 | -1.12 | -1.316 | none |
| NRG1 | 0.695 | 0.496 | 4.2617 | 0.0018 | 0.014 | -1.12 | -1.3434 | none |
| LILRB5 | 0.416 | 0.2886 | 4.2616 | 0.0018 | 0.014 | -1.12 | -1.3979 | none |
| ATXN2 | 1.081 | 0.9904 | 4.26 | 0.0018 | 0.014 | -1.13 | -1.1058 | none |
| SPRYD4 | 0.503 | 0.3377 | 4.2597 | 0.0018 | 0.014 | -1.13 | -1.3497 | none |
| SNX27 | 0.806 | 0.664 | 4.2595 | 0.0018 | 0.014 | -1.13 | -1.5949 | OA-related |
| PCDHGA8 | 0.453 | 0.3325 | 4.2589 | 0.0018 | 0.014 | -1.13 | -1.2652 | none |
| MCOLN1 | 0.579 | 0.4501 | 4.2584 | 0.0018 | 0.014 | -1.13 | -1.2382 | none |
| EZH1 | 0.76 | 0.6856 | 4.2553 | 0.00181 | 0.015 | -1.13 | -1.3177 | none |
| ZMYM6 | 0.782 | 0.6469 | 4.2549 | 0.00181 | 0.015 | -1.13 | -1.0579 | none |
| QPCTL | 0.769 | 0.6111 | 4.2544 | 0.00182 | 0.015 | -1.14 | -1.2253 | none |
| CCDC150 | 0.815 | 0.6687 | 4.2543 | 0.00182 | 0.015 | -1.14 | -1.4656 | none |
| NBPF12 | 0.681 | 0.6043 | 4.2535 | 0.00182 | 0.015 | -1.14 | -1.2196 | none |
| RFX2 | 0.417 | 0.359 | 4.2522 | 0.00182 | 0.015 | -1.14 | -0.5434 | none |

| | | | | | | | | |
|---|---|---|---|---|---|---|---|---|
| MS4A2 | 0.513 | 0.4062 | 4.2515 | 0.00182 | 0.015 | -1.14 | -1.1069 | none |
| STMND1 | 0.879 | 0.7106 | 4.2512 | 0.00182 | 0.015 | -1.14 | -1.1552 | none |
| MAPK10 | 0.664 | 0.5337 | 4.2507 | 0.00183 | 0.015 | -1.14 | -1.3391 | none |
| NOL9 | 0.684 | 0.616 | 4.2481 | 0.00183 | 0.015 | -1.15 | -1.4277 | none |
| RNF44 | 0.469 | 0.3899 | 4.2476 | 0.00183 | 0.015 | -1.15 | -1.3304 | none |
| NSUN6 | 0.75 | 0.6088 | 4.2465 | 0.00184 | 0.015 | -1.15 | -1.204 | none |
| NELL1 | 0.585 | 0.4848 | 4.2461 | 0.00184 | 0.015 | -1.15 | -1.3707 | none |
| CTCFL | 0.491 | 0.3636 | 4.2456 | 0.00184 | 0.015 | -1.15 | -1.3975 | none |
| ANKRD10 | 0.646 | 0.6279 | 4.2454 | 0.00184 | 0.015 | -1.15 | -1.149 | none |
| PHLPP2 | 0.535 | 0.4438 | 4.2452 | 0.00184 | 0.015 | -1.15 | -1.4796 | none |
| ZC3H12B | 0.589 | 0.4325 | 4.2446 | 0.00184 | 0.015 | -1.15 | -1.5917 | OA-related |
| TNFRSF17 | 0.864 | 0.7284 | 4.2443 | 0.00184 | 0.015 | -1.15 | -0.5561 | none |
| LRRIQ1 | 0.622 | 0.4408 | 4.2443 | 0.00184 | 0.015 | -1.15 | -1.463 | none |
| RPAP3 | 0.945 | 0.7571 | 4.2414 | 0.00185 | 0.015 | -1.16 | -0.9308 | none |
| DEPDC4 | 0.524 | 0.3629 | 4.2409 | 0.00185 | 0.015 | -1.16 | -1.3773 | none |
| MYOZ3 | 0.613 | 0.4522 | 4.2394 | 0.00186 | 0.015 | -1.16 | -1.3616 | none |
| LDHB | -0.39 | -0.3259 | -4.239 | 0.00186 | 0.015 | -1.16 | 2.31012 | AA-related |
| IMPA2 | 0.683 | 0.5097 | 4.2386 | 0.00186 | 0.015 | -1.16 | -1.3501 | none |
| PARP15 | 0.573 | 0.4366 | 4.2368 | 0.00187 | 0.015 | -1.16 | -1.5241 | none |
| MPPE1 | 0.677 | 0.4903 | 4.2364 | 0.00187 | 0.015 | -1.16 | -1.1145 | none |
| ADAT1 | 0.679 | 0.5511 | 4.2342 | 0.00187 | 0.015 | -1.17 | -1.0088 | none |
| WWOX | 0.493 | 0.4531 | 4.234 | 0.00187 | 0.015 | -1.17 | -0.1039 | none |
| TRIM6 | 0.874 | 0.6481 | 4.2336 | 0.00187 | 0.015 | -1.17 | -1.0983 | none |
| SLC25A34 | 0.623 | 0.4673 | 4.2331 | 0.00188 | 0.015 | -1.17 | -1.5056 | none |
| OLAH | 0.534 | 0.3296 | 4.2322 | 0.00188 | 0.015 | -1.17 | -1.2205 | none |
| ZNF627 | 0.916 | 0.7085 | 4.2304 | 0.00188 | 0.015 | -1.17 | -1.5286 | none |
| PIGA | 0.631 | 0.3701 | 4.2296 | 0.00189 | 0.015 | -1.17 | -0.7923 | none |
| OR10J1 | 0.574 | 0.4211 | 4.2285 | 0.00189 | 0.015 | -1.18 | -1.626 | OA-related |
| PON2 | 0.598 | 0.3962 | 4.2282 | 0.00189 | 0.015 | -1.18 | -0.9313 | none |
| WDR97 | 0.383 | 0.2915 | 4.2273 | 0.00189 | 0.015 | -1.18 | -1.2774 | none |
| NYAP2 | 0.419 | 0.3053 | 4.2264 | 0.0019 | 0.015 | -1.18 | -1.5313 | none |
| MICALL1 | 0.681 | 0.6393 | 4.2252 | 0.0019 | 0.015 | -1.18 | -1.1781 | none |
| OR10A2 | 0.468 | 0.349 | 4.2247 | 0.0019 | 0.015 | -1.18 | -1.4019 | none |
| PFAS | 0.813 | 0.7372 | 4.2236 | 0.0019 | 0.015 | -1.18 | -1.3485 | none |
| TMEM116 | 0.665 | 0.4122 | 4.2229 | 0.00191 | 0.015 | -1.18 | -0.821 | none |
| CCDC113 | 0.721 | 0.5859 | 4.2222 | 0.00191 | 0.015 | -1.19 | -1.1005 | none |
| CD209 | 0.452 | 0.3084 | 4.2215 | 0.00191 | 0.015 | -1.19 | -1.117 | none |
| TRIM7 | 0.436 | 0.3059 | 4.2199 | 0.00192 | 0.015 | -1.19 | -1.191 | none |
| NAT10 | 0.691 | 0.6126 | 4.2184 | 0.00192 | 0.015 | -1.19 | -1.2801 | none |
| RTTN | 0.459 | 0.3626 | 4.2138 | 0.00193 | 0.015 | -1.2 | -1.2823 | none |

| | | | | | | | | |
|---|---|---|---|---|---|---|---|---|
| CLN6 | 0.462 | 0.353 | 4.2121 | 0.00194 | 0.015 | -1.2 | -0.6074 | none |
| ARSB | 0.429 | 0.377 | 4.2119 | 0.00194 | 0.015 | -1.2 | -1.5503 | OA-related |
| TMCC3 | 0.916 | 0.7032 | 4.2114 | 0.00194 | 0.015 | -1.2 | -0.9608 | none |
| ZC3H6 | 0.545 | 0.4381 | 4.211 | 0.00194 | 0.015 | -1.2 | -1.2762 | none |
| GSS | 0.506 | 0.3829 | 4.2103 | 0.00194 | 0.015 | -1.2 | -1.244 | none |
| ACADL | 0.581 | 0.4592 | 4.21 | 0.00194 | 0.015 | -1.2 | -1.5143 | none |
| AKIRIN1 | 0.41 | 0.313 | 4.2098 | 0.00195 | 0.015 | -1.2 | 0.71305 | none |
| SLC30A5 | 0.722 | 0.4768 | 4.2089 | 0.00195 | 0.015 | -1.21 | -1.1201 | none |
| NBPF15 | 0.722 | 0.6728 | 4.206 | 0.00196 | 0.015 | -1.21 | -0.9146 | none |
| SIN3A | 1.006 | 0.7992 | 4.2047 | 0.00196 | 0.015 | -1.21 | -0.7985 | none |
| TFRC | 0.649 | 0.4953 | 4.2013 | 0.00197 | 0.015 | -1.22 | -1.4186 | none |
| PLEKHM3 | 0.514 | 0.4463 | 4.2005 | 0.00197 | 0.015 | -1.22 | -1.4007 | none |
| DGKD | 0.647 | 0.5482 | 4.2003 | 0.00197 | 0.015 | -1.22 | -1.3468 | none |
| HMG20A | 0.732 | 0.5124 | 4.1957 | 0.00199 | 0.015 | -1.23 | -1.2012 | none |
| TRAF5 | 0.454 | 0.391 | 4.1953 | 0.00199 | 0.015 | -1.23 | -0.4914 | none |
| BCL2L15 | 0.59 | 0.3735 | 4.1952 | 0.00199 | 0.015 | -1.23 | -1.2494 | none |
| NFAM1 | 0.496 | 0.3664 | 4.1946 | 0.00199 | 0.015 | -1.23 | -1.2083 | none |
| SHPK | 0.389 | 0.3174 | 4.1923 | 0.002 | 0.016 | -1.23 | -0.5754 | none |
| NUSAP1 | 0.78 | 0.6738 | 4.1916 | 0.002 | 0.016 | -1.23 | -1.3969 | none |
| SYP | 0.485 | 0.3293 | 4.1916 | 0.002 | 0.016 | -1.23 | -1.3765 | none |
| CAMK1 | 0.577 | 0.4233 | 4.1908 | 0.002 | 0.016 | -1.23 | -1.2319 | none |
| EXD2 | 0.642 | 0.4464 | 4.1905 | 0.002 | 0.016 | -1.23 | -1.1149 | none |
| ELMSAN1 | 0.856 | 0.772 | 4.1904 | 0.002 | 0.016 | -1.23 | -1.4041 | none |
| FAM110D | 0.582 | 0.4375 | 4.1902 | 0.00201 | 0.016 | -1.24 | -1.3338 | none |
| COL15A1 | 0.719 | 0.5597 | 4.1897 | 0.00201 | 0.016 | -1.24 | -0.9051 | none |
| MTX3 | 0.565 | 0.448 | 4.189 | 0.00201 | 0.016 | -1.24 | -1.6195 | OA-related |
| MCM9 | 0.616 | 0.5753 | 4.1881 | 0.00201 | 0.016 | -1.24 | -0.4845 | none |
| TRMT10B | 0.639 | 0.4762 | 4.1866 | 0.00202 | 0.016 | -1.24 | -1.3213 | none |
| TDGF1 | 0.673 | 0.4701 | 4.1852 | 0.00202 | 0.016 | -1.24 | -1.3216 | none |
| HELZ2 | 0.894 | 0.764 | 4.1847 | 0.00202 | 0.016 | -1.24 | -1.1149 | none |
| PHGDH | 0.688 | 0.5548 | 4.1845 | 0.00202 | 0.016 | -1.24 | -1.3175 | none |
| EBPL | 0.878 | 0.5975 | 4.1839 | 0.00203 | 0.016 | -1.24 | -0.9983 | none |
| AUTS2 | 0.642 | 0.5048 | 4.1838 | 0.00203 | 0.016 | -1.24 | -1.4427 | none |
| GSTO2 | 0.658 | 0.5208 | 4.1835 | 0.00203 | 0.016 | -1.25 | -1.4851 | none |
| SIDT2 | 0.687 | 0.5376 | 4.1826 | 0.00203 | 0.016 | -1.25 | -1.4182 | none |
| MIOX | 0.5 | 0.3855 | 4.1825 | 0.00203 | 0.016 | -1.25 | -1.0021 | none |
| DCTN5 | 0.497 | 0.3194 | 4.1813 | 0.00203 | 0.016 | -1.25 | -1.2691 | none |
| PHACTR4 | 1.019 | 0.9454 | 4.1802 | 0.00204 | 0.016 | -1.25 | -1.4054 | none |
| FRMD4A | 0.827 | 0.7166 | 4.1798 | 0.00204 | 0.016 | -1.25 | -1.2884 | none |
| SLC29A2 | 0.354 | 0.3087 | 4.1797 | 0.00204 | 0.016 | -1.25 | -0.1362 | none |

| | | | | | | | | |
|---|---|---|---|---|---|---|---|---|
| ARL10 | 0.571 | 0.5075 | 4.1795 | 0.00204 | 0.016 | -1.25 | -1.4995 | none |
| SLC7A6 | 0.569 | 0.4193 | 4.1795 | 0.00204 | 0.016 | -1.25 | -1.2794 | none |
| LAMB1 | 0.652 | 0.593 | 4.1785 | 0.00204 | 0.016 | -1.25 | -0.7913 | none |
| CFHR2 | 0.586 | 0.4406 | 4.1782 | 0.00204 | 0.016 | -1.25 | -1.3486 | none |
| KIAA0895 | 0.569 | 0.3524 | 4.1775 | 0.00205 | 0.016 | -1.25 | -0.8264 | none |
| ZNF26 | 0.586 | 0.5046 | 4.1769 | 0.00205 | 0.016 | -1.26 | -1.0285 | none |
| SGK2 | 0.75 | 0.5852 | 4.1753 | 0.00205 | 0.016 | -1.26 | -0.8559 | none |
| MICAL2 | 0.671 | 0.5886 | 4.1738 | 0.00206 | 0.016 | -1.26 | -1.2877 | none |
| ARRB1 | 0.629 | 0.4588 | 4.1735 | 0.00206 | 0.016 | -1.26 | -1.4421 | none |
| LIN9 | -0.473 | -0.3949 | -4.173 | 0.00206 | 0.016 | -1.26 | 2.61752 | AA-related |
| GID4 | 0.648 | 0.4681 | 4.1713 | 0.00207 | 0.016 | -1.26 | -1.1722 | none |
| RPRD1B | 0.792 | 0.5884 | 4.1695 | 0.00207 | 0.016 | -1.27 | -0.9473 | none |
| DYRK3 | 0.423 | 0.3369 | 4.1679 | 0.00208 | 0.016 | -1.27 | -1.17 | none |
| DRC3 | 1.178 | 1.0512 | 4.1674 | 0.00208 | 0.016 | -1.27 | -0.9742 | none |
| SGPP2 | 0.419 | 0.3612 | 4.1666 | 0.00208 | 0.016 | -1.27 | -1.1778 | none |
| ZKSCAN4 | 0.386 | 0.2796 | 4.165 | 0.00209 | 0.016 | -1.27 | -0.8611 | none |
| FEM1A | 0.734 | 0.6384 | 4.1648 | 0.00209 | 0.016 | -1.27 | -1.2268 | none |
| VWA5B1 | 0.59 | 0.4346 | 4.1643 | 0.00209 | 0.016 | -1.28 | -1.2627 | none |
| FAM161B | 0.548 | 0.4258 | 4.1639 | 0.00209 | 0.016 | -1.28 | -1.483 | none |
| C17ORF99 | 0.743 | 0.5385 | 4.1636 | 0.00209 | 0.016 | -1.28 | -0.7287 | none |
| TTLL7 | 0.615 | 0.3607 | 4.1621 | 0.0021 | 0.016 | -1.28 | -1.0054 | none |
| PLD6 | 0.63 | 0.5032 | 4.1608 | 0.0021 | 0.016 | -1.28 | -1.1268 | none |
| TTC22 | 0.519 | 0.439 | 4.1599 | 0.0021 | 0.016 | -1.28 | -0.5319 | none |
| ZNF808 | 0.924 | 0.7999 | 4.1598 | 0.0021 | 0.016 | -1.28 | -1.4802 | none |
| TMEM108 | 0.433 | 0.3132 | 4.1583 | 0.00211 | 0.016 | -1.28 | -1.4601 | none |
| PPP1R3G | 0.369 | 0.2972 | 4.1572 | 0.00211 | 0.016 | -1.29 | -1.1548 | none |
| ITIH1 | 1.069 | 0.8489 | 4.1569 | 0.00211 | 0.016 | -1.29 | -1.0152 | none |
| ZNF117 | 0.498 | 0.3819 | 4.1568 | 0.00211 | 0.016 | -1.29 | -1.3834 | none |
| C1ORF122 | -0.692 | -0.5093 | -4.154 | 0.00212 | 0.016 | -1.29 | 2.14833 | AA-related |
| OTUD6A | 0.694 | 0.5254 | 4.1521 | 0.00213 | 0.016 | -1.29 | -1.2861 | none |
| SLC47A2 | 0.527 | 0.3726 | 4.1517 | 0.00213 | 0.016 | -1.3 | -1.2256 | none |
| HDAC9 | 0.855 | 0.7055 | 4.1496 | 0.00214 | 0.016 | -1.3 | -0.836 | none |
| COLEC12 | 0.762 | 0.5673 | 4.1494 | 0.00214 | 0.016 | -1.3 | -1.2549 | none |
| STRN3 | 0.922 | 0.7727 | 4.1486 | 0.00214 | 0.016 | -1.3 | -1.415 | none |
| NPHP3 | 0.832 | 0.7917 | 4.1473 | 0.00214 | 0.016 | -1.3 | -1.0518 | none |
| ALG14 | 0.74 | 0.57 | 4.1466 | 0.00215 | 0.016 | -1.3 | -0.5459 | none |
| SLCO1A2 | 0.671 | 0.4637 | 4.1462 | 0.00215 | 0.016 | -1.3 | -1.1275 | none |
| ZNF445 | 0.63 | 0.537 | 4.1447 | 0.00215 | 0.016 | -1.31 | -1.4713 | none |
| ACAD10 | 0.855 | 0.7153 | 4.1444 | 0.00215 | 0.016 | -1.31 | -1.1283 | none |
| TMEM169 | 0.314 | 0.137 | 4.1435 | 0.00216 | 0.016 | -1.31 | 0.04085 | none |

| | | | | | | | | |
|---|---|---|---|---|---|---|---|---|
| RAB3B | 0.386 | 0.2725 | 4.1396 | 0.00217 | 0.016 | -1.31 | -1.5992 | OA-related |
| FLYWCH1 | 0.724 | 0.5856 | 4.1385 | 0.00217 | 0.016 | -1.32 | -1.5733 | OA-related |
| LDB3 | 0.582 | 0.4508 | 4.1382 | 0.00217 | 0.016 | -1.32 | -1.1742 | none |
| PLEKHH3 | 0.489 | 0.3088 | 4.1365 | 0.00218 | 0.016 | -1.32 | -1.1871 | none |
| BTN3A1 | 0.631 | 0.5418 | 4.1362 | 0.00218 | 0.016 | -1.32 | -0.9962 | none |
| SLC25A41 | 0.558 | 0.423 | 4.1356 | 0.00218 | 0.016 | -1.32 | -1.11 | none |
| HYLS1 | 0.492 | 0.3584 | 4.1338 | 0.00219 | 0.016 | -1.32 | -0.7651 | none |
| SYNPO2L | 0.527 | 0.4027 | 4.1321 | 0.0022 | 0.017 | -1.33 | -1.2704 | none |
| KLK11 | 0.613 | 0.4654 | 4.1309 | 0.0022 | 0.017 | -1.33 | -0.5542 | none |
| ALDH2 | 0.587 | 0.4136 | 4.1308 | 0.0022 | 0.017 | -1.33 | -1.5232 | none |
| GNG4 | 0.564 | 0.4482 | 4.1305 | 0.0022 | 0.017 | -1.33 | -1.2373 | none |
| LZTFL1 | 0.658 | 0.4639 | 4.1298 | 0.0022 | 0.017 | -1.33 | -0.8686 | none |
| ZNF493 | 0.715 | 0.594 | 4.1294 | 0.0022 | 0.017 | -1.33 | -1.195 | none |
| LARS2 | 0.842 | 0.7539 | 4.1294 | 0.0022 | 0.017 | -1.33 | -0.8157 | none |
| LRRC37B | 0.793 | 0.6987 | 4.1282 | 0.00221 | 0.017 | -1.33 | -1.2086 | none |
| SENP1 | 0.553 | 0.4428 | 4.1267 | 0.00221 | 0.017 | -1.33 | -1.0336 | none |
| DEGS1 | 0.414 | 0.264 | 4.1259 | 0.00222 | 0.017 | -1.34 | -0.2188 | none |
| ASB16 | 0.473 | 0.4158 | 4.1258 | 0.00222 | 0.017 | -1.34 | -0.6893 | none |
| ANKRD45 | 0.74 | 0.5562 | 4.1256 | 0.00222 | 0.017 | -1.34 | -1.5698 | OA-related |
| ZNF584 | 0.632 | 0.4888 | 4.1254 | 0.00222 | 0.017 | -1.34 | -1.3237 | none |
| MKRN1 | 0.712 | 0.5214 | 4.1254 | 0.00222 | 0.017 | -1.34 | -1.3612 | none |
| CYB5RL | 0.646 | 0.5164 | 4.1253 | 0.00222 | 0.017 | -1.34 | -1.3666 | none |
| RNFT2 | 0.484 | 0.3595 | 4.1245 | 0.00222 | 0.017 | -1.34 | -1.3443 | none |
| SMUG1 | 0.771 | 0.5165 | 4.1235 | 0.00222 | 0.017 | -1.34 | -1.2316 | none |
| RAB3D | 0.425 | 0.2794 | 4.1232 | 0.00223 | 0.017 | -1.34 | -1.4484 | none |
| BICDL2 | 0.403 | 0.3071 | 4.1226 | 0.00223 | 0.017 | -1.34 | -0.9282 | none |
| TMEM170 | 0.369 | 0.2439 | 4.1224 | 0.00223 | 0.017 | -1.34 | -0.8024 | none |
| TNS2 | 0.78 | 0.6288 | 4.1214 | 0.00223 | 0.017 | -1.34 | -1.0613 | none |
| SIM2 | 0.679 | 0.4822 | 4.1204 | 0.00224 | 0.017 | -1.34 | -1.1264 | none |
| NRIP3 | 0.59 | 0.4804 | 4.1202 | 0.00224 | 0.017 | -1.34 | -1.1111 | none |
| PLD1 | 0.844 | 0.733 | 4.119 | 0.00224 | 0.017 | -1.35 | -0.9328 | none |
| SENP8 | 0.492 | 0.3277 | 4.117 | 0.00225 | 0.017 | -1.35 | -0.954 | none |
| BICDL1 | 0.917 | 0.8473 | 4.1168 | 0.00225 | 0.017 | -1.35 | -1.3983 | none |
| ZNF550 | 0.637 | 0.6137 | 4.1145 | 0.00226 | 0.017 | -1.35 | -0.6518 | none |
| ZNF589 | 0.82 | 0.7683 | 4.1119 | 0.00227 | 0.017 | -1.36 | -0.8963 | none |
| DNAJC5B | 0.572 | 0.4308 | 4.1118 | 0.00227 | 0.017 | -1.36 | -1.2319 | none |
| OR6C75 | 0.64 | 0.4538 | 4.1115 | 0.00227 | 0.017 | -1.36 | -1.4302 | none |
| GTF2H3 | 0.509 | 0.3276 | 4.1086 | 0.00228 | 0.017 | -1.36 | -0.6309 | none |
| ACADVL | 0.447 | 0.4829 | 4.1079 | 0.00228 | 0.017 | -1.36 | 0.95719 | none |
| SIPA1 | 0.827 | 0.7739 | 4.1072 | 0.00228 | 0.017 | -1.36 | -1.1294 | none |

| | | | | | | | | |
|---|---|---|---|---|---|---|---|---|
| DNAL1 | 0.578 | 0.4434 | 4.1049 | 0.00229 | 0.017 | -1.37 | -1.2422 | none |
| SHCBP1L | 0.737 | 0.5757 | 4.1034 | 0.0023 | 0.017 | -1.37 | -0.7725 | none |
| UBA7 | 0.433 | 0.3796 | 4.1025 | 0.0023 | 0.017 | -1.37 | -0.4962 | none |
| RPS6KA5 | 0.618 | 0.4856 | 4.1023 | 0.0023 | 0.017 | -1.37 | -1.4375 | none |
| LYRM4 | 0.565 | 0.3193 | 4.1004 | 0.00231 | 0.017 | -1.38 | -1.0806 | none |
| GCC1 | 0.97 | 0.8347 | 4.0993 | 0.00231 | 0.017 | -1.38 | -1.1532 | none |
| GINS4 | 0.874 | 0.7122 | 4.0986 | 0.00231 | 0.017 | -1.38 | -1.2552 | none |
| POLR2J2 | 0.623 | 0.487 | 4.0983 | 0.00231 | 0.017 | -1.38 | -1.1566 | none |
| ATG2A | 0.702 | 0.6532 | 4.0972 | 0.00232 | 0.017 | -1.38 | -0.8216 | none |
| C6ORF163 | 0.514 | 0.3942 | 4.0969 | 0.00232 | 0.017 | -1.38 | -0.6594 | none |
| DHCR24 | 0.466 | 0.4481 | 4.0952 | 0.00233 | 0.017 | -1.38 | 0.10868 | none |
| ATXN1 | 0.489 | 0.3389 | 4.0948 | 0.00233 | 0.017 | -1.38 | -1.4672 | none |
| MROH7 | 0.859 | 0.6431 | 4.0948 | 0.00233 | 0.017 | -1.38 | -1.1523 | none |
| TMEM181 | 0.488 | 0.3904 | 4.0942 | 0.00233 | 0.017 | -1.38 | -1.1653 | none |
| KCNB1 | 0.665 | 0.5447 | 4.0941 | 0.00233 | 0.017 | -1.39 | -1.304 | none |
| BMPER | 0.687 | 0.4852 | 4.0932 | 0.00233 | 0.017 | -1.39 | -0.9048 | none |
| PRAP1 | 0.645 | 0.4516 | 4.0925 | 0.00234 | 0.017 | -1.39 | -1.376 | none |
| PTGIR | 0.516 | 0.4368 | 4.0924 | 0.00234 | 0.017 | -1.39 | -1.1696 | none |
| PCYT1B | 0.387 | 0.2481 | 4.0923 | 0.00234 | 0.017 | -1.39 | -1.3435 | none |
| NPIPB4 | 1.175 | 1.0689 | 4.0921 | 0.00234 | 0.017 | -1.39 | -1.1832 | none |
| ATP6V0D2 | 0.871 | 0.7337 | 4.0915 | 0.00234 | 0.017 | -1.39 | -0.9713 | none |
| RUNDC1 | 0.685 | 0.571 | 4.0914 | 0.00234 | 0.017 | -1.39 | -1.507 | none |
| SPECC1L-A | 0.528 | 0.4261 | 4.0913 | 0.00234 | 0.017 | -1.39 | -0.5122 | none |
| MCF2L2 | 0.58 | 0.471 | 4.0905 | 0.00234 | 0.017 | -1.39 | -0.9033 | none |
| GPAT3 | -0.501 | -0.3865 | -4.09 | 0.00234 | 0.017 | -1.39 | 1.92547 | none |
| ARL5C | 0.96 | 0.7566 | 4.09 | 0.00234 | 0.017 | -1.39 | -0.9406 | none |
| KCNH1 | 0.606 | 0.4046 | 4.09 | 0.00234 | 0.017 | -1.39 | -1.378 | none |
| TMEM182 | 0.657 | 0.5147 | 4.089 | 0.00235 | 0.017 | -1.39 | -0.7098 | none |
| BTBD7 | 0.706 | 0.5671 | 4.088 | 0.00235 | 0.017 | -1.39 | -1.2718 | none |
| FAM98C | 0.976 | 0.7833 | 4.0871 | 0.00236 | 0.017 | -1.4 | -0.8992 | none |
| FKBP14 | 0.628 | 0.5834 | 4.0864 | 0.00236 | 0.017 | -1.4 | -1.1544 | none |
| DNAH9 | 0.626 | 0.4389 | 4.0859 | 0.00236 | 0.017 | -1.4 | -0.6148 | none |
| PNPLA1 | 0.47 | 0.3596 | 4.0855 | 0.00236 | 0.017 | -1.4 | -1.1332 | none |
| IRGM | 0.474 | 0.3884 | 4.0851 | 0.00236 | 0.017 | -1.4 | -1.1869 | none |
| CCDC88C | 0.528 | 0.4497 | 4.0848 | 0.00236 | 0.017 | -1.4 | -1.2932 | none |
| CFTR | 1.111 | 0.8367 | 4.0836 | 0.00237 | 0.017 | -1.4 | -0.6858 | none |
| SEC31A | 0.647 | 0.5282 | 4.0835 | 0.00237 | 0.017 | -1.4 | -1.472 | none |
| OR4P4 | 0.478 | 0.3661 | 4.0817 | 0.00238 | 0.017 | -1.4 | -1.0228 | none |
| NAPB | 0.469 | 0.3142 | 4.081 | 0.00238 | 0.017 | -1.41 | 0.13869 | none |
| ITGAD | 0.736 | 0.5122 | 4.0798 | 0.00238 | 0.017 | -1.41 | -1.1332 | none |

| | | | | | | | | |
|---|---|---|---|---|---|---|---|---|
| HSPBAP1 | 0.462 | 0.3808 | 4.0798 | 0.00238 | 0.017 | -1.41 | -1.0149 | none |
| HSD17B1 | 0.495 | 0.4478 | 4.0781 | 0.00239 | 0.017 | -1.41 | -0.2404 | none |
| ATF7 | 0.637 | 0.4781 | 4.0781 | 0.00239 | 0.017 | -1.41 | -1.4165 | none |
| XPO5 | 0.531 | 0.4646 | 4.0778 | 0.00239 | 0.017 | -1.41 | -1.2212 | none |
| C22ORF39 | 0.364 | 0.295 | 4.0765 | 0.00239 | 0.017 | -1.41 | -0.149 | none |
| PAX8 | 0.667 | 0.5535 | 4.0747 | 0.0024 | 0.017 | -1.42 | -0.4725 | none |
| ZMYM1 | 0.768 | 0.6445 | 4.0741 | 0.0024 | 0.017 | -1.42 | -0.9251 | none |
| SLC14A2 | 0.32 | 0.2352 | 4.0718 | 0.00241 | 0.018 | -1.42 | -1.4999 | none |
| NCMAP | 0.504 | 0.3513 | 4.0709 | 0.00242 | 0.018 | -1.42 | -1.45 | none |
| CHRAC1 | 0.416 | 0.2844 | 4.0694 | 0.00242 | 0.018 | -1.42 | -0.7989 | none |
| ACADSB | 0.548 | 0.4126 | 4.0682 | 0.00243 | 0.018 | -1.43 | -0.4381 | none |
| MTMR12 | 0.401 | 0.3202 | 4.0673 | 0.00243 | 0.018 | -1.43 | -0.5898 | none |
| RASGRP4 | 0.711 | 0.6114 | 4.066 | 0.00243 | 0.018 | -1.43 | -1.2334 | none |
| FAM161A | 0.582 | 0.5186 | 4.0648 | 0.00244 | 0.018 | -1.43 | -1.1875 | none |
| L3MBTL2 | 0.615 | 0.5194 | 4.0646 | 0.00244 | 0.018 | -1.43 | -1.1699 | none |
| HIVEP2 | 1.174 | 1.1145 | 4.0644 | 0.00244 | 0.018 | -1.43 | -1.2957 | none |
| IQSEC2 | 0.66 | 0.525 | 4.063 | 0.00245 | 0.018 | -1.43 | -0.8781 | none |
| FSIP2 | 0.372 | 0.2977 | 4.0618 | 0.00245 | 0.018 | -1.44 | -0.975 | none |
| DECR2 | 0.637 | 0.4332 | 4.059 | 0.00246 | 0.018 | -1.44 | -0.9697 | none |
| RASL10A | -0.467 | -0.328 | -4.056 | 0.00247 | 0.018 | -1.44 | 0.87494 | none |
| FOXJ2 | 0.663 | 0.571 | 4.0555 | 0.00247 | 0.018 | -1.45 | -1.481 | none |
| PITPNM2 | 0.628 | 0.5423 | 4.0551 | 0.00248 | 0.018 | -1.45 | -1.2024 | none |
| CNDP1 | 0.957 | 0.8203 | 4.0551 | 0.00248 | 0.018 | -1.45 | -0.97 | none |
| CCDC163 | 0.622 | 0.4186 | 4.0546 | 0.00248 | 0.018 | -1.45 | -1.108 | none |
| NRIP2 | 0.636 | 0.4676 | 4.054 | 0.00248 | 0.018 | -1.45 | -1.194 | none |
| SLC35E4 | 0.353 | 0.2759 | 4.0538 | 0.00248 | 0.018 | -1.45 | -0.5296 | none |
| COL28A1 | 0.66 | 0.4937 | 4.0519 | 0.00249 | 0.018 | -1.45 | -0.2794 | none |
| FAM129A | 1.172 | 1.054 | 4.0517 | 0.00249 | 0.018 | -1.45 | -1.355 | none |
| C9ORF3 | 0.436 | 0.328 | 4.0504 | 0.00249 | 0.018 | -1.45 | -1.0935 | none |
| MELK | 0.522 | 0.3596 | 4.0501 | 0.0025 | 0.018 | -1.45 | -1.2785 | none |
| ZNRF1 | 0.87 | 0.6524 | 4.0501 | 0.0025 | 0.018 | -1.45 | -1.3278 | none |
| RNF103 | 0.415 | 0.2845 | 4.0493 | 0.0025 | 0.018 | -1.46 | -0.7878 | none |
| GTF2IRD1 | 0.62 | 0.4639 | 4.0479 | 0.0025 | 0.018 | -1.46 | -1.3235 | none |
| SLC16A12 | 0.723 | 0.5196 | 4.0476 | 0.00251 | 0.018 | -1.46 | -1.4727 | none |
| ATPAF2 | 0.483 | 0.3777 | 4.0472 | 0.00251 | 0.018 | -1.46 | -1.1756 | none |
| HBP1 | 0.508 | 0.3089 | 4.046 | 0.00251 | 0.018 | -1.46 | -0.7362 | none |
| L1TD1 | 0.372 | 0.2796 | 4.0451 | 0.00252 | 0.018 | -1.46 | -1.5689 | OA-related |
| CASC3 | 0.709 | 0.6454 | 4.0449 | 0.00252 | 0.018 | -1.46 | -1.2838 | none |
| MTHFD1 | 0.76 | 0.6584 | 4.0422 | 0.00253 | 0.018 | -1.47 | -0.9694 | none |
| NUB1 | 0.671 | 0.5083 | 4.0419 | 0.00253 | 0.018 | -1.47 | -1.3721 | none |

| | | | | | | | | |
|---|---|---|---|---|---|---|---|---|
| LIX1 | 0.356 | 0.2595 | 4.0416 | 0.00253 | 0.018 | -1.47 | -0.8244 | none |
| RPH3A | 0.656 | 0.5838 | 4.041 | 0.00253 | 0.018 | -1.47 | -1.052 | none |
| KLHL2 | 0.678 | 0.4895 | 4.0397 | 0.00254 | 0.018 | -1.47 | -1.2642 | none |
| DACH2 | 0.448 | 0.3436 | 4.0376 | 0.00255 | 0.018 | -1.47 | -1.5283 | none |
| CFAP69 | 0.543 | 0.4345 | 4.0374 | 0.00255 | 0.018 | -1.47 | -1.2465 | none |
| SRGAP1 | 0.428 | 0.2854 | 4.0366 | 0.00255 | 0.018 | -1.48 | -1.2678 | none |
| FRMD8 | 0.422 | 0.3124 | 4.0364 | 0.00255 | 0.018 | -1.48 | 0.03394 | none |
| CPT1A | 0.829 | 0.8463 | 4.0364 | 0.00255 | 0.018 | -1.48 | -1.0394 | none |
| SLC9A7 | 0.458 | 0.3628 | 4.0356 | 0.00255 | 0.018 | -1.48 | -1.0238 | none |
| RBP7 | 0.675 | 0.5146 | 4.0336 | 0.00256 | 0.018 | -1.48 | -1.2369 | none |
| PLPP3 | 0.461 | 0.333 | 4.0328 | 0.00256 | 0.018 | -1.48 | -0.9746 | none |
| KCNRG | 0.53 | 0.4773 | 4.0325 | 0.00257 | 0.018 | -1.48 | -0.8662 | none |
| ARNTL2 | 0.542 | 0.4706 | 4.0324 | 0.00257 | 0.018 | -1.48 | -1.2249 | none |
| CEP170 | 1.067 | 1.025 | 4.0324 | 0.00257 | 0.018 | -1.48 | -0.8846 | none |
| XKR6 | 0.579 | 0.4369 | 4.0322 | 0.00257 | 0.018 | -1.48 | -0.5378 | none |
| GAPVD1 | 0.975 | 0.8505 | 4.0317 | 0.00257 | 0.018 | -1.48 | -1.0299 | none |
| LRRC69 | 0.961 | 0.7149 | 4.0299 | 0.00258 | 0.018 | -1.49 | -1.2092 | none |
| SLC25A37 | 0.444 | 0.335 | 4.0289 | 0.00258 | 0.018 | -1.49 | -1.3044 | none |
| XAF1 | 0.529 | 0.4881 | 4.0287 | 0.00258 | 0.018 | -1.49 | -0.6568 | none |
| TBCK | 0.414 | 0.304 | 4.0285 | 0.00258 | 0.018 | -1.49 | -0.9563 | none |
| FERMT2 | 0.521 | 0.4233 | 4.0284 | 0.00258 | 0.018 | -1.49 | -1.3814 | none |
| PRR16 | 0.414 | 0.2803 | 4.0277 | 0.00259 | 0.018 | -1.49 | -1.2497 | none |
| SULT2B1 | 0.536 | 0.4263 | 4.0256 | 0.00259 | 0.018 | -1.49 | -1.1178 | none |
| PRDM7 | 0.674 | 0.5367 | 4.0256 | 0.00259 | 0.018 | -1.49 | -1.06 | none |
| CATSPERD | 0.47 | 0.3822 | 4.0249 | 0.0026 | 0.018 | -1.49 | -0.7721 | none |
| DDX19A | 0.906 | 0.8619 | 4.0241 | 0.0026 | 0.018 | -1.5 | -1.0405 | none |
| CLIP1 | 1.125 | 1.0947 | 4.0235 | 0.0026 | 0.018 | -1.5 | -1.1731 | none |
| PRND | 0.539 | 0.3947 | 4.022 | 0.00261 | 0.018 | -1.5 | -1.3276 | none |
| LMNTD1 | 0.831 | 0.6257 | 4.0219 | 0.00261 | 0.018 | -1.5 | -1.0455 | none |
| PNPT1 | 0.583 | 0.4847 | 4.0217 | 0.00261 | 0.018 | -1.5 | -0.9214 | none |
| TNFAIP8L1 | 0.61 | 0.5434 | 4.0214 | 0.00261 | 0.018 | -1.5 | -1.3843 | none |
| PLEKHM1 | 0.702 | 0.5482 | 4.0199 | 0.00262 | 0.019 | -1.5 | -0.9638 | none |
| XAGE5 | 0.562 | 0.4465 | 4.0194 | 0.00262 | 0.019 | -1.5 | -1.328 | none |
| INTS4 | 0.524 | 0.3995 | 4.0188 | 0.00262 | 0.019 | -1.5 | -1.2267 | none |
| UNC79 | 0.389 | 0.2709 | 4.0181 | 0.00262 | 0.019 | -1.5 | -1.3364 | none |
| MIA2 | 0.801 | 0.6571 | 4.0181 | 0.00262 | 0.019 | -1.5 | -1.3109 | none |
| NIPSNAP1 | 0.473 | 0.37 | 4.0169 | 0.00263 | 0.019 | -1.51 | -1.2455 | none |
| SLCO2A1 | 0.342 | 0.2848 | 4.0168 | 0.00263 | 0.019 | -1.51 | -1.4692 | none |
| FGF11 | 0.574 | 0.4331 | 4.0162 | 0.00263 | 0.019 | -1.51 | -1.4022 | none |
| NUP88 | 0.509 | 0.324 | 4.0152 | 0.00264 | 0.019 | -1.51 | -1.322 | none |

| | | | | | | | | |
|---|---|---|---|---|---|---|---|---|
| RAF1 | 0.457 | 0.3858 | 4.0143 | 0.00264 | 0.019 | -1.51 | -0.6929 | none |
| POLG2 | 0.616 | 0.4887 | 4.0134 | 0.00264 | 0.019 | -1.51 | -1.1974 | none |
| MRI1 | 0.634 | 0.535 | 4.013 | 0.00265 | 0.019 | -1.51 | -0.5573 | none |
| SCN9A | 0.392 | 0.2715 | 4.0122 | 0.00265 | 0.019 | -1.51 | -1.3823 | none |
| FKTN | 0.638 | 0.44 | 4.0048 | 0.00268 | 0.019 | -1.53 | -0.2744 | none |
| ODF2L | 0.423 | 0.2573 | 4.0037 | 0.00269 | 0.019 | -1.53 | -1.2459 | none |
| PDP2 | 0.762 | 0.6344 | 4.0005 | 0.0027 | 0.019 | -1.53 | -0.4195 | none |
| SNRNP35 | 0.652 | 0.5113 | 3.9991 | 0.0027 | 0.019 | -1.53 | -1.3883 | none |
| SULT1C4 | 0.493 | 0.3941 | 3.9991 | 0.0027 | 0.019 | -1.53 | -1.4606 | none |
| ANXA4 | 0.453 | 0.3327 | 3.9976 | 0.00271 | 0.019 | -1.54 | -0.0791 | none |
| RASSF9 | 0.525 | 0.3673 | 3.9973 | 0.00271 | 0.019 | -1.54 | -1.3915 | none |
| ADGRF1 | 0.558 | 0.4302 | 3.9967 | 0.00272 | 0.019 | -1.54 | -1.47 | none |
| ARPC3 | -0.45 | -0.4154 | -3.996 | 0.00272 | 0.019 | -1.54 | 2.12672 | AA-related |
| AZIN2 | 0.534 | 0.4015 | 3.9954 | 0.00272 | 0.019 | -1.54 | -0.89 | none |
| MOV10L1 | 0.584 | 0.4423 | 3.9945 | 0.00272 | 0.019 | -1.54 | -1.5099 | none |
| DCC | 1.004 | 0.8201 | 3.9938 | 0.00273 | 0.019 | -1.54 | -1.0399 | none |
| FAM216A | 0.423 | 0.3144 | 3.992 | 0.00274 | 0.019 | -1.55 | -1.0998 | none |
| GIPC1 | 0.696 | 0.5165 | 3.9918 | 0.00274 | 0.019 | -1.55 | -0.8392 | none |
| NLRC5 | 0.756 | 0.6649 | 3.9915 | 0.00274 | 0.019 | -1.55 | -1.4257 | none |
| CCL15 | 0.777 | 0.6507 | 3.9911 | 0.00274 | 0.019 | -1.55 | -0.5215 | none |
| POGZ | 0.836 | 0.7178 | 3.99 | 0.00274 | 0.019 | -1.55 | -1.1856 | none |
| SCFD1 | 0.494 | 0.3175 | 3.9898 | 0.00274 | 0.019 | -1.55 | -1.2351 | none |
| RUFY4 | 0.608 | 0.4883 | 3.9884 | 0.00275 | 0.019 | -1.55 | -0.3688 | none |
| RBM48 | 0.421 | 0.3565 | 3.9864 | 0.00276 | 0.019 | -1.55 | -1.1686 | none |
| ARID1A | 1.108 | 1.0597 | 3.9861 | 0.00276 | 0.019 | -1.56 | -1.185 | none |
| DLK2 | 0.57 | 0.4131 | 3.986 | 0.00276 | 0.019 | -1.56 | -1.1985 | none |
| FBXL12 | 0.555 | 0.4338 | 3.9858 | 0.00276 | 0.019 | -1.56 | -1.2874 | none |
| FNTA | 0.567 | 0.3846 | 3.9844 | 0.00277 | 0.019 | -1.56 | -1.139 | none |
| ZNF568 | 0.659 | 0.4825 | 3.9837 | 0.00277 | 0.019 | -1.56 | -1.6447 | OA-related |
| SRRD | 0.728 | 0.5494 | 3.9835 | 0.00277 | 0.019 | -1.56 | -1.244 | none |
| NUBPL | 0.636 | 0.4644 | 3.9832 | 0.00277 | 0.019 | -1.56 | -1.1128 | none |
| CREG1 | 0.46 | 0.3857 | 3.9831 | 0.00277 | 0.019 | -1.56 | 0.89919 | none |
| MAML1 | 0.804 | 0.7669 | 3.982 | 0.00278 | 0.019 | -1.56 | -1.2727 | none |
| AEBP2 | 1.137 | 1.0185 | 3.9811 | 0.00278 | 0.019 | -1.56 | -1.4146 | none |
| ENDOV | 0.668 | 0.478 | 3.9806 | 0.00278 | 0.019 | -1.56 | -0.9402 | none |
| DUSP19 | 0.471 | 0.3179 | 3.9792 | 0.00279 | 0.019 | -1.57 | -0.9387 | none |
| KBTBD12 | 0.519 | 0.382 | 3.979 | 0.00279 | 0.019 | -1.57 | -1.3046 | none |
| KCNJ14 | 0.401 | 0.3299 | 3.9786 | 0.00279 | 0.019 | -1.57 | -1.4494 | none |
| ANKMY1 | 0.654 | 0.6037 | 3.9762 | 0.0028 | 0.019 | -1.57 | -0.3273 | none |
| MAST1 | 0.688 | 0.5335 | 3.9741 | 0.00281 | 0.02 | -1.57 | -1.4047 | none |

| | | | | | | | | |
|---|---|---|---|---|---|---|---|---|
| MRO | 0.444 | 0.3439 | 3.9732 | 0.00282 | 0.02 | -1.58 | -0.852 | none |
| WDR83 | 0.645 | 0.5663 | 3.9729 | 0.00282 | 0.02 | -1.58 | -1.5492 | OA-related |
| KIAA0319I | 0.747 | 0.6727 | 3.9716 | 0.00282 | 0.02 | -1.58 | -1.3111 | none |
| ZNF528 | 0.989 | 0.8461 | 3.9707 | 0.00283 | 0.02 | -1.58 | -1.0365 | none |
| GDF5OS | 0.584 | 0.4559 | 3.9703 | 0.00283 | 0.02 | -1.58 | -0.9089 | none |
| CACNB3 | 0.51 | 0.4054 | 3.9702 | 0.00283 | 0.02 | -1.58 | -1.0168 | none |
| WDR19 | 0.45 | 0.3794 | 3.9693 | 0.00284 | 0.02 | -1.58 | -1.0107 | none |
| ZNF19 | 0.57 | 0.4091 | 3.9691 | 0.00284 | 0.02 | -1.58 | -1.2913 | none |
| RORB | 0.416 | 0.3098 | 3.969 | 0.00284 | 0.02 | -1.58 | -1.4028 | none |
| SLIT1 | 0.947 | 0.6662 | 3.9682 | 0.00284 | 0.02 | -1.58 | -0.9946 | none |
| SNCAIP | 1.144 | 0.9677 | 3.9677 | 0.00284 | 0.02 | -1.58 | -0.7448 | none |
| GAPDH | -0.483 | -0.4037 | -3.967 | 0.00284 | 0.02 | -1.58 | 2.03704 | none |
| CCDC33 | -0.444 | -0.438 | -3.967 | 0.00284 | 0.02 | -1.58 | 1.06481 | none |
| USP39 | 0.356 | 0.2525 | 3.965 | 0.00285 | 0.02 | -1.59 | -1.0528 | none |
| PIAS4 | 0.612 | 0.5728 | 3.965 | 0.00285 | 0.02 | -1.59 | -0.8623 | none |
| METTL21A | 0.517 | 0.3551 | 3.9637 | 0.00286 | 0.02 | -1.59 | -0.6104 | none |
| PABPC4 | 0.65 | 0.5575 | 3.9621 | 0.00287 | 0.02 | -1.59 | -0.8744 | none |
| RBM12B | 0.475 | 0.4572 | 3.962 | 0.00287 | 0.02 | -1.59 | -0.0548 | none |
| ATAD2B | 0.739 | 0.6004 | 3.9612 | 0.00287 | 0.02 | -1.59 | -1.2167 | none |
| HMBOX1 | 0.851 | 0.717 | 3.9609 | 0.00287 | 0.02 | -1.59 | -0.9457 | none |
| ZER1 | 0.521 | 0.4481 | 3.9599 | 0.00288 | 0.02 | -1.6 | -0.0086 | none |
| ALG1L | 0.928 | 0.7503 | 3.9597 | 0.00288 | 0.02 | -1.6 | -1.2288 | none |
| TPSB2 | 0.587 | 0.4186 | 3.9591 | 0.00288 | 0.02 | -1.6 | -0.5367 | none |
| SUMO4 | 0.471 | 0.3144 | 3.9591 | 0.00288 | 0.02 | -1.6 | -0.6061 | none |
| OR8D4 | 0.421 | 0.3248 | 3.9578 | 0.00289 | 0.02 | -1.6 | -1.0899 | none |
| SLC12A9 | 0.739 | 0.5987 | 3.9577 | 0.00289 | 0.02 | -1.6 | -0.9775 | none |
| GK5 | 0.61 | 0.3857 | 3.9577 | 0.00289 | 0.02 | -1.6 | -0.8548 | none |
| GON4L | 0.692 | 0.5938 | 3.9573 | 0.00289 | 0.02 | -1.6 | -1.4548 | none |
| TMOD4 | 0.909 | 0.7281 | 3.9567 | 0.00289 | 0.02 | -1.6 | -0.9547 | none |
| PPP2R3A | 0.797 | 0.6298 | 3.9551 | 0.0029 | 0.02 | -1.6 | -1.1822 | none |
| FBXL13 | 0.567 | 0.3901 | 3.955 | 0.0029 | 0.02 | -1.6 | -1.0897 | none |
| TSACC | 0.504 | 0.3793 | 3.9535 | 0.00291 | 0.02 | -1.61 | -0.0703 | none |
| PIAS1 | 0.416 | 0.3138 | 3.9529 | 0.00291 | 0.02 | -1.61 | -0.2787 | none |
| GPRC5A | 0.484 | 0.3415 | 3.9527 | 0.00291 | 0.02 | -1.61 | -1.169 | none |
| SLC22A9 | 0.591 | 0.456 | 3.9518 | 0.00292 | 0.02 | -1.61 | -1.3691 | none |
| OPN3 | 0.614 | 0.4218 | 3.9503 | 0.00292 | 0.02 | -1.61 | -1.0541 | none |
| PINLYP | 0.557 | 0.4359 | 3.9501 | 0.00292 | 0.02 | -1.61 | -0.8096 | none |
| TKFC | 0.5 | 0.4153 | 3.9498 | 0.00292 | 0.02 | -1.61 | -0.9636 | none |
| RIOK3 | 0.647 | 0.5152 | 3.9493 | 0.00293 | 0.02 | -1.61 | -0.9531 | none |
| ENG | 0.791 | 0.6646 | 3.9485 | 0.00293 | 0.02 | -1.61 | -1.1497 | none |

| | | | | | | | | |
|---|---|---|---|---|---|---|---|---|
| TOX4 | 0.491 | 0.372 | 3.948 | 0.00293 | 0.02 | -1.62 | -0.7086 | none |
| ARID2 | 1.035 | 0.9144 | 3.9479 | 0.00293 | 0.02 | -1.62 | -1.2562 | none |
| STMN1 | -0.559 | -0.5034 | -3.946 | 0.00294 | 0.02 | -1.62 | 2.26644 | AA-related |
| CCZ1B | 0.861 | 0.7125 | 3.945 | 0.00295 | 0.02 | -1.62 | -1.5591 | OA-related |
| FOXM1 | 0.844 | 0.7284 | 3.9448 | 0.00295 | 0.02 | -1.62 | -1.3816 | none |
| PCCA | 0.604 | 0.44 | 3.9447 | 0.00295 | 0.02 | -1.62 | -1.3612 | none |
| ANAPC1 | 0.46 | 0.3666 | 3.9441 | 0.00295 | 0.02 | -1.62 | -0.5486 | none |
| ZNF440 | 0.885 | 0.7431 | 3.9441 | 0.00295 | 0.02 | -1.62 | -1.3047 | none |
| PTPN22 | 0.534 | 0.4005 | 3.9439 | 0.00295 | 0.02 | -1.62 | -0.9587 | none |
| MTMR3 | 0.585 | 0.5553 | 3.9437 | 0.00295 | 0.02 | -1.62 | -0.8131 | none |
| PCDH11X | 0.418 | 0.2925 | 3.9434 | 0.00295 | 0.02 | -1.62 | -1.4152 | none |
| ANKRD46 | 0.733 | 0.4698 | 3.9431 | 0.00296 | 0.02 | -1.62 | -0.8279 | none |
| ERCC4 | 0.727 | 0.56 | 3.9431 | 0.00296 | 0.02 | -1.62 | -1.0824 | none |
| ELP4 | 0.513 | 0.3246 | 3.9421 | 0.00296 | 0.02 | -1.62 | -0.71 | none |
| PDIA5 | 0.818 | 0.6774 | 3.9415 | 0.00296 | 0.02 | -1.63 | -1.1018 | none |
| PRPF3 | 1.122 | 0.9984 | 3.9406 | 0.00297 | 0.02 | -1.63 | -1.1552 | none |
| HYAL1 | 0.916 | 0.7629 | 3.939 | 0.00297 | 0.02 | -1.63 | -0.0572 | none |
| METTL16 | 0.919 | 0.8284 | 3.9384 | 0.00298 | 0.02 | -1.63 | -1.2524 | none |
| CDC23 | 0.379 | 0.245 | 3.9377 | 0.00298 | 0.02 | -1.63 | -0.8731 | none |
| CNOT1 | 0.816 | 0.7255 | 3.9375 | 0.00298 | 0.02 | -1.63 | -1.1896 | none |
| ATG10 | 0.713 | 0.5289 | 3.9369 | 0.00298 | 0.02 | -1.63 | -1.416 | none |
| CAMK2G | 0.466 | 0.3874 | 3.9365 | 0.00299 | 0.02 | -1.63 | -0.8164 | none |
| OR52N1 | 0.353 | 0.2143 | 3.9354 | 0.00299 | 0.02 | -1.64 | -0.8397 | none |
| NAV3 | 0.772 | 0.6252 | 3.9335 | 0.003 | 0.02 | -1.64 | -1.1343 | none |
| PCSK7 | 0.898 | 0.8115 | 3.9332 | 0.003 | 0.02 | -1.64 | -1.2459 | none |
| KANK4 | 0.437 | 0.2988 | 3.9317 | 0.00301 | 0.02 | -1.64 | -1.4523 | none |
| NUDT5 | -0.379 | -0.3241 | -3.931 | 0.00301 | 0.02 | -1.64 | 2.13276 | AA-related |
| IL20RB | 0.609 | 0.4254 | 3.9313 | 0.00301 | 0.02 | -1.64 | -0.9346 | none |
| GLRA1 | 0.638 | 0.4844 | 3.93 | 0.00302 | 0.02 | -1.64 | -1.2942 | none |
| KLF3 | 0.934 | 0.737 | 3.9296 | 0.00302 | 0.02 | -1.64 | -1.451 | none |
| CHAF1B | 0.454 | 0.3409 | 3.9289 | 0.00302 | 0.02 | -1.65 | -1.2828 | none |
| CSTF3 | 0.639 | 0.5188 | 3.9285 | 0.00302 | 0.02 | -1.65 | -1.3531 | none |
| WEE1 | 0.506 | 0.4056 | 3.9282 | 0.00303 | 0.02 | -1.65 | -1.188 | none |
| BORCS7 | 0.479 | 0.3093 | 3.9272 | 0.00303 | 0.02 | -1.65 | -0.8885 | none |
| C11ORF94 | -0.437 | -0.3417 | -3.925 | 0.00304 | 0.02 | -1.65 | 0.60014 | none |
| FAM124A | 0.705 | 0.6389 | 3.9246 | 0.00304 | 0.02 | -1.65 | -1.3404 | none |
| IL4R | 0.749 | 0.6987 | 3.9245 | 0.00304 | 0.02 | -1.65 | -1.3003 | none |
| CLN5 | 0.417 | 0.2614 | 3.9244 | 0.00304 | 0.02 | -1.65 | -1.2599 | none |
| GNAI1 | 0.508 | 0.3643 | 3.9226 | 0.00305 | 0.02 | -1.66 | -1.397 | none |
| UBE3C | 0.447 | 0.314 | 3.9221 | 0.00306 | 0.021 | -1.66 | -1.0217 | none |

| | | | | | | | | |
|---|---|---|---|---|---|---|---|---|
| ARID3B | 0.571 | 0.5456 | 3.9217 | 0.00306 | 0.021 | -1.66 | -0.118 | none |
| STAT6 | 0.684 | 0.611 | 3.9212 | 0.00306 | 0.021 | -1.66 | -1.1115 | none |
| ACSM5 | 0.395 | 0.3489 | 3.9207 | 0.00306 | 0.021 | -1.66 | 0.65824 | none |
| RAPGEF6 | 0.819 | 0.6595 | 3.9204 | 0.00306 | 0.021 | -1.66 | -0.6251 | none |
| RHBDD3 | 0.433 | 0.3703 | 3.9195 | 0.00307 | 0.021 | -1.66 | -1.1553 | none |
| TMEM236 | 0.474 | 0.3507 | 3.9169 | 0.00308 | 0.021 | -1.66 | -1.507 | none |
| KIAA1841 | 0.764 | 0.6587 | 3.9166 | 0.00308 | 0.021 | -1.67 | -1.209 | none |
| PARP2 | 0.548 | 0.4604 | 3.9165 | 0.00308 | 0.021 | -1.67 | -0.7101 | none |
| HECTD1 | 1.006 | 0.9138 | 3.9156 | 0.00309 | 0.021 | -1.67 | -0.8854 | none |
| SLC26A11 | 0.617 | 0.4573 | 3.9153 | 0.00309 | 0.021 | -1.67 | -1.2844 | none |
| EPCAM | -0.481 | -0.4063 | -3.915 | 0.00309 | 0.021 | -1.67 | 2.03496 | none |
| RAD23B | 0.432 | 0.3512 | 3.9151 | 0.00309 | 0.021 | -1.67 | 0.73717 | none |
| HMGCR | 1.707 | 1.5536 | 3.9135 | 0.0031 | 0.021 | -1.67 | -1.0801 | none |
| FFAR4 | 0.529 | 0.4072 | 3.9125 | 0.0031 | 0.021 | -1.67 | -1.1667 | none |
| FBXW12 | 0.632 | 0.4498 | 3.9124 | 0.0031 | 0.021 | -1.67 | -1.0048 | none |
| SMAD1 | 0.301 | 0.1696 | 3.9123 | 0.0031 | 0.021 | -1.67 | -0.557 | none |
| EFEMP2 | -0.657 | -0.529 | -3.912 | 0.00311 | 0.021 | -1.67 | -0.0737 | none |
| SEC24D | 0.555 | 0.4796 | 3.9115 | 0.00311 | 0.021 | -1.67 | -0.2216 | none |
| TREX2 | 0.308 | 0.2388 | 3.9112 | 0.00311 | 0.021 | -1.67 | -1.0932 | none |
| PKNOX2 | -0.315 | -0.238 | -3.911 | 0.00311 | 0.021 | -1.67 | 1.91317 | none |
| ABAT | 0.501 | 0.4736 | 3.9099 | 0.00312 | 0.021 | -1.68 | -0.3235 | none |
| SLC12A3 | 0.505 | 0.3634 | 3.9099 | 0.00312 | 0.021 | -1.68 | -1.4082 | none |
| ZNF543 | 0.583 | 0.4749 | 3.9096 | 0.00312 | 0.021 | -1.68 | -1.5099 | none |
| QRSL1 | 0.498 | 0.3776 | 3.9091 | 0.00312 | 0.021 | -1.68 | -0.725 | none |
| ZNF284 | 0.595 | 0.473 | 3.9087 | 0.00312 | 0.021 | -1.68 | -1.4086 | none |
| JAK3 | 0.632 | 0.565 | 3.9087 | 0.00312 | 0.021 | -1.68 | -1.5225 | none |
| BRI3BP | 0.662 | 0.5688 | 3.9084 | 0.00312 | 0.021 | -1.68 | -1.4921 | none |
| CACNG8 | 0.372 | 0.2604 | 3.9082 | 0.00312 | 0.021 | -1.68 | -1.2623 | none |
| DPY19L4 | 0.407 | 0.2736 | 3.908 | 0.00312 | 0.021 | -1.68 | -0.4979 | none |
| RNF223 | 0.296 | 0.1982 | 3.9078 | 0.00313 | 0.021 | -1.68 | -0.7639 | none |
| ZNF23 | 0.487 | 0.4452 | 3.9078 | 0.00313 | 0.021 | -1.68 | -0.8629 | none |
| OR2AP1 | 0.723 | 0.5411 | 3.9067 | 0.00313 | 0.021 | -1.68 | -1.41 | none |
| CYB561 | 0.783 | 0.5513 | 3.9039 | 0.00315 | 0.021 | -1.69 | -1.0962 | none |
| CXCL16 | 0.588 | 0.4227 | 3.9037 | 0.00315 | 0.021 | -1.69 | -1.3044 | none |
| EIF4A1 | -0.377 | -0.3086 | -3.903 | 0.00315 | 0.021 | -1.69 | 2.25717 | AA-related |
| ZNF639 | 0.545 | 0.3975 | 3.9024 | 0.00315 | 0.021 | -1.69 | -1.4107 | none |
| GSDMA | 0.49 | 0.4139 | 3.9022 | 0.00315 | 0.021 | -1.69 | -1.1577 | none |
| ZMPSTE24 | 0.7 | 0.4411 | 3.9018 | 0.00316 | 0.021 | -1.69 | -0.9027 | none |
| EYA3 | 0.623 | 0.5022 | 3.9016 | 0.00316 | 0.021 | -1.69 | -1.069 | none |
| CNGA1 | 0.743 | 0.5822 | 3.8994 | 0.00317 | 0.021 | -1.69 | -0.4538 | none |

| | | | | | | | | |
|---|---|---|---|---|---|---|---|---|
| TMEM150 | 0.775 | 0.5599 | 3.8993 | 0.00317 | 0.021 | -1.69 | -1.1543 | none |
| BVES | 0.379 | 0.2734 | 3.8992 | 0.00317 | 0.021 | -1.69 | -0.9407 | none |
| ZHX3 | 0.87 | 0.8083 | 3.8982 | 0.00317 | 0.021 | -1.69 | -0.6009 | none |
| PIK3R1 | 0.89 | 0.7452 | 3.8977 | 0.00318 | 0.021 | -1.7 | -0.8084 | none |
| TSPAN5 | 0.428 | 0.3321 | 3.8975 | 0.00318 | 0.021 | -1.7 | -0.232 | none |
| TRPC6 | 0.733 | 0.4963 | 3.8972 | 0.00318 | 0.021 | -1.7 | -0.4213 | none |
| KCNJ15 | 0.532 | 0.406 | 3.8955 | 0.00319 | 0.021 | -1.7 | -0.9563 | none |
| CD163 | 0.602 | 0.4494 | 3.8951 | 0.00319 | 0.021 | -1.7 | -0.9271 | none |
| OR4K13 | 0.409 | 0.2896 | 3.8949 | 0.00319 | 0.021 | -1.7 | -1.5017 | none |
| DCAF10 | 0.533 | 0.3985 | 3.8948 | 0.00319 | 0.021 | -1.7 | -0.5675 | none |
| ZFP2 | 0.567 | 0.4324 | 3.8947 | 0.00319 | 0.021 | -1.7 | -1.4522 | none |
| CUX1 | 0.876 | 0.8359 | 3.8937 | 0.0032 | 0.021 | -1.7 | -0.5925 | none |
| C12ORF65 | 0.631 | 0.5428 | 3.8936 | 0.0032 | 0.021 | -1.7 | -1.1807 | none |
| PIK3C2B | 0.768 | 0.7376 | 3.8922 | 0.0032 | 0.021 | -1.7 | -0.3665 | none |
| NADK2 | 0.561 | 0.4121 | 3.8909 | 0.00321 | 0.021 | -1.71 | -1.1262 | none |
| DNAJC12 | 0.732 | 0.5537 | 3.8901 | 0.00321 | 0.021 | -1.71 | -1.3543 | none |
| CSF3R | 1.092 | 0.8276 | 3.8899 | 0.00322 | 0.021 | -1.71 | -1.1454 | none |
| ZNF763 | 0.393 | 0.2762 | 3.8898 | 0.00322 | 0.021 | -1.71 | -1.1175 | none |
| ZNF441 | 0.618 | 0.4873 | 3.8895 | 0.00322 | 0.021 | -1.71 | -0.3417 | none |
| PDE8A | 0.981 | 0.798 | 3.8891 | 0.00322 | 0.021 | -1.71 | -1.3441 | none |
| ABCC1 | 0.861 | 0.7484 | 3.8891 | 0.00322 | 0.021 | -1.71 | -1.5078 | none |
| ATG13 | 0.614 | 0.5122 | 3.8871 | 0.00323 | 0.021 | -1.71 | -1.4696 | none |
| FRRS1 | 0.684 | 0.5223 | 3.8862 | 0.00323 | 0.021 | -1.71 | -1.2056 | none |
| ENO1 | -0.372 | -0.29 | -3.886 | 0.00324 | 0.021 | -1.71 | 2.06627 | none |
| MED18 | 0.755 | 0.6518 | 3.8859 | 0.00324 | 0.021 | -1.71 | -1.0222 | none |
| PRR27 | 0.357 | 0.2618 | 3.8853 | 0.00324 | 0.021 | -1.72 | -0.8028 | none |
| ITGAL | 0.768 | 0.6558 | 3.8853 | 0.00324 | 0.021 | -1.72 | -1.4163 | none |
| PAFAH1B2 | 0.562 | 0.3498 | 3.8826 | 0.00325 | 0.021 | -1.72 | -1.4745 | none |
| NDUFV3 | 0.568 | 0.4869 | 3.8822 | 0.00326 | 0.021 | -1.72 | -1.2725 | none |
| PRKACA | 0.713 | 0.6844 | 3.8821 | 0.00326 | 0.021 | -1.72 | -0.6948 | none |
| CMBL | 0.476 | 0.3465 | 3.8815 | 0.00326 | 0.021 | -1.72 | -1.331 | none |
| ARMC8 | 0.452 | 0.345 | 3.8814 | 0.00326 | 0.021 | -1.72 | -1.0855 | none |
| APOOL | 0.871 | 0.738 | 3.8804 | 0.00327 | 0.021 | -1.72 | -1.4428 | none |
| NASP | 0.761 | 0.6563 | 3.8799 | 0.00327 | 0.021 | -1.72 | -1.2676 | none |
| KNG1 | 0.368 | 0.2628 | 3.8793 | 0.00327 | 0.021 | -1.72 | -1.2013 | none |
| NOX5 | 0.421 | 0.2941 | 3.8779 | 0.00328 | 0.021 | -1.73 | -1.2882 | none |
| USE1 | 0.533 | 0.3676 | 3.8779 | 0.00328 | 0.021 | -1.73 | -1.2873 | none |
| ACSS2 | 0.377 | 0.247 | 3.8779 | 0.00328 | 0.021 | -1.73 | -0.3973 | none |
| OR1L8 | 0.569 | 0.402 | 3.8779 | 0.00328 | 0.021 | -1.73 | -0.939 | none |
| TRAF3 | 0.666 | 0.6014 | 3.8777 | 0.00328 | 0.021 | -1.73 | -1.3332 | none |

| | | | | | | | | |
|---|---|---|---|---|---|---|---|---|
| MICAL3 | 1.028 | 0.9576 | 3.8773 | 0.00328 | 0.021 | -1.73 | -0.9399 | none |
| ITGA1 | 0.647 | 0.5116 | 3.8765 | 0.00329 | 0.021 | -1.73 | -0.9049 | none |
| ACBD5 | 0.609 | 0.4879 | 3.8753 | 0.00329 | 0.021 | -1.73 | -0.5505 | none |
| ATP1A4 | 0.616 | 0.4566 | 3.8746 | 0.0033 | 0.021 | -1.73 | -1.0012 | none |
| GPR78 | 0.448 | 0.3547 | 3.8733 | 0.0033 | 0.021 | -1.73 | -0.5611 | none |
| PCDHB16 | 0.33 | 0.2495 | 3.8725 | 0.00331 | 0.021 | -1.74 | -1.4817 | none |
| CNEP1R1 | 0.53 | 0.3642 | 3.8716 | 0.00331 | 0.021 | -1.74 | -0.4445 | none |
| LGALS8 | 0.614 | 0.4328 | 3.8716 | 0.00331 | 0.021 | -1.74 | -1.1226 | none |
| ABHD16A | 1.295 | 0.9305 | 3.8714 | 0.00331 | 0.021 | -1.74 | -0.6719 | none |
| PRPF19 | 0.436 | 0.4086 | 3.8685 | 0.00333 | 0.022 | -1.74 | -0.6547 | none |
| ZNF829 | 0.739 | 0.6141 | 3.8674 | 0.00333 | 0.022 | -1.74 | -1.607 | OA-related |
| TNRC18 | 1.484 | 1.3627 | 3.8668 | 0.00334 | 0.022 | -1.74 | -1.1014 | none |
| THOC5 | 0.646 | 0.5376 | 3.8667 | 0.00334 | 0.022 | -1.74 | -0.8249 | none |
| ENTPD4 | 0.681 | 0.6033 | 3.8665 | 0.00334 | 0.022 | -1.75 | -1.0569 | none |
| CYP8B1 | 0.388 | 0.283 | 3.8664 | 0.00334 | 0.022 | -1.75 | -1.2191 | none |
| IRAK1 | 0.554 | 0.5686 | 3.8644 | 0.00335 | 0.022 | -1.75 | -0.6881 | none |
| MAPK6 | 0.758 | 0.5762 | 3.8635 | 0.00335 | 0.022 | -1.75 | -1.1606 | none |
| USP9Y | 0.435 | 0.3116 | 3.8633 | 0.00336 | 0.022 | -1.75 | -1.2497 | none |
| TUFT1 | 0.782 | 0.5914 | 3.863 | 0.00336 | 0.022 | -1.75 | -1.4169 | none |
| SMU1 | 0.368 | 0.2478 | 3.8626 | 0.00336 | 0.022 | -1.75 | -1.3668 | none |
| SLC15A4 | 0.593 | 0.4414 | 3.8618 | 0.00336 | 0.022 | -1.75 | -1.0427 | none |
| C12ORF60 | 0.618 | 0.5025 | 3.8618 | 0.00336 | 0.022 | -1.75 | -1.1918 | none |
| ZNF99 | 0.416 | 0.299 | 3.8612 | 0.00337 | 0.022 | -1.75 | -1.1164 | none |
| PLXNB1 | -0.477 | -0.3151 | -3.861 | 0.00337 | 0.022 | -1.75 | 1.98502 | none |
| ART3 | 0.605 | 0.4416 | 3.8589 | 0.00338 | 0.022 | -1.76 | -0.9936 | none |
| NISCH | 0.843 | 0.786 | 3.858 | 0.00338 | 0.022 | -1.76 | -1.338 | none |
| GATAD2A | 1.497 | 1.4088 | 3.8577 | 0.00339 | 0.022 | -1.76 | -1.0995 | none |
| DTX3L | 0.61 | 0.5073 | 3.8563 | 0.00339 | 0.022 | -1.76 | -1.3184 | none |
| SPATS2L | 1.094 | 0.9487 | 3.8563 | 0.00339 | 0.022 | -1.76 | -1.3115 | none |
| NFASC | 0.541 | 0.4175 | 3.8559 | 0.0034 | 0.022 | -1.76 | -1.4192 | none |
| OR2T12 | 0.485 | 0.3822 | 3.8551 | 0.0034 | 0.022 | -1.76 | -1.4027 | none |
| CASR | 0.3 | 0.2323 | 3.8548 | 0.0034 | 0.022 | -1.76 | -1.417 | none |
| PTK7 | 0.831 | 0.64 | 3.8527 | 0.00341 | 0.022 | -1.77 | -1.469 | none |
| RGS10 | -0.359 | -0.2874 | -3.851 | 0.00342 | 0.022 | -1.77 | 1.99007 | none |
| SCAI | 0.412 | 0.3265 | 3.8512 | 0.00342 | 0.022 | -1.77 | 0.28622 | none |
| SETD7 | 0.581 | 0.4692 | 3.8512 | 0.00342 | 0.022 | -1.77 | -1.4683 | none |
| LEAP2 | 0.475 | 0.4339 | 3.851 | 0.00342 | 0.022 | -1.77 | -1.2852 | none |
| ZNF783 | 0.553 | 0.473 | 3.8505 | 0.00342 | 0.022 | -1.77 | -0.5881 | none |
| NSUN2 | 0.874 | 0.803 | 3.8499 | 0.00343 | 0.022 | -1.77 | -0.8442 | none |
| DCBLD1 | 0.654 | 0.5151 | 3.8496 | 0.00343 | 0.022 | -1.77 | -1.2932 | none |

| | | | | | | | | |
|---|---|---|---|---|---|---|---|---|
| KIAA1324 | 0.648 | 0.5552 | 3.8495 | 0.00343 | 0.022 | -1.77 | -0.9372 | none |
| MRPS31 | 0.6 | 0.4461 | 3.8495 | 0.00343 | 0.022 | -1.77 | -1.4573 | none |
| RAD9B | 0.662 | 0.5403 | 3.8489 | 0.00343 | 0.022 | -1.77 | -0.794 | none |
| GPR161 | 1.5 | 1.2081 | 3.8484 | 0.00344 | 0.022 | -1.77 | -1.2161 | none |
| TMEM63A | 0.552 | 0.4907 | 3.8476 | 0.00344 | 0.022 | -1.78 | -0.82 | none |
| ATG14 | 0.49 | 0.4076 | 3.8476 | 0.00344 | 0.022 | -1.78 | -1.3723 | none |
| PPIP5K1 | 0.627 | 0.5213 | 3.8471 | 0.00344 | 0.022 | -1.78 | -1.2265 | none |
| PLCD3 | 0.68 | 0.4978 | 3.8469 | 0.00344 | 0.022 | -1.78 | -0.7463 | none |
| ZBED5 | 0.499 | 0.396 | 3.8464 | 0.00345 | 0.022 | -1.78 | -0.5371 | none |
| GPATCH1 | 0.825 | 0.6963 | 3.8447 | 0.00346 | 0.022 | -1.78 | -1.1654 | none |
| CLPB | 0.584 | 0.5036 | 3.8447 | 0.00346 | 0.022 | -1.78 | -0.708 | none |
| LRRC19 | 0.339 | 0.2453 | 3.8445 | 0.00346 | 0.022 | -1.78 | -1.1702 | none |
| SMAGP | 0.683 | 0.499 | 3.8436 | 0.00346 | 0.022 | -1.78 | -1.0131 | none |
| C2CD4A | 0.42 | 0.2994 | 3.8433 | 0.00346 | 0.022 | -1.78 | -1.3281 | none |
| KLF8 | 0.347 | 0.2633 | 3.8433 | 0.00346 | 0.022 | -1.78 | -1.2102 | none |
| ARPC4-TTI | -0.673 | -0.5477 | -3.843 | 0.00347 | 0.022 | -1.78 | 2.07115 | none |
| SCARF1 | 0.579 | 0.5127 | 3.8413 | 0.00348 | 0.022 | -1.79 | -1.2277 | none |
| L3MBTL4 | 0.474 | 0.3751 | 3.8412 | 0.00348 | 0.022 | -1.79 | -1.4156 | none |
| ZNF486 | 0.614 | 0.4666 | 3.8406 | 0.00348 | 0.022 | -1.79 | -0.932 | none |
| TCAIM | 0.487 | 0.3513 | 3.8399 | 0.00348 | 0.022 | -1.79 | -0.1688 | none |
| MAP1S | 0.588 | 0.4809 | 3.8398 | 0.00348 | 0.022 | -1.79 | -1.4152 | none |
| ITGA2B | 0.551 | 0.4029 | 3.8392 | 0.00349 | 0.022 | -1.79 | -1.1453 | none |
| PPARG | 0.571 | 0.427 | 3.8391 | 0.00349 | 0.022 | -1.79 | -1.0365 | none |
| NDUFA7 | 0.652 | 0.4002 | 3.8389 | 0.00349 | 0.022 | -1.79 | -1.2014 | none |
| ZNF570 | 0.598 | 0.487 | 3.8387 | 0.00349 | 0.022 | -1.79 | -1.4209 | none |
| TBCEL | 0.82 | 0.6457 | 3.8386 | 0.00349 | 0.022 | -1.79 | -0.6322 | none |
| PCNX1 | 0.924 | 0.6947 | 3.8384 | 0.00349 | 0.022 | -1.79 | -0.67 | none |
| DSTYK | 0.603 | 0.4989 | 3.8376 | 0.0035 | 0.022 | -1.79 | -1.3216 | none |
| AARD | 0.707 | 0.6017 | 3.8375 | 0.0035 | 0.022 | -1.79 | -0.9748 | none |
| SLC9B1 | 0.646 | 0.4167 | 3.8365 | 0.0035 | 0.022 | -1.79 | -1.136 | none |
| CERKL | 0.84 | 0.6803 | 3.8361 | 0.0035 | 0.022 | -1.79 | -1.086 | none |
| DLG5 | 0.446 | 0.3722 | 3.8348 | 0.00351 | 0.022 | -1.8 | -1.1587 | none |
| HFE | 0.513 | 0.3876 | 3.8338 | 0.00352 | 0.022 | -1.8 | -1.2738 | none |
| KCNJ6 | 0.323 | 0.2321 | 3.8334 | 0.00352 | 0.022 | -1.8 | -1.3811 | none |
| XPO1 | 0.69 | 0.5578 | 3.8329 | 0.00352 | 0.022 | -1.8 | -0.8547 | none |
| RNASE10 | 0.386 | 0.3147 | 3.8322 | 0.00353 | 0.022 | -1.8 | -0.8263 | none |
| COG5 | 0.549 | 0.4079 | 3.8322 | 0.00353 | 0.022 | -1.8 | -0.735 | none |
| RUVBL2 | 0.368 | 0.3227 | 3.8319 | 0.00353 | 0.022 | -1.8 | -1.2035 | none |
| POLR3A | 0.82 | 0.7383 | 3.8311 | 0.00353 | 0.022 | -1.8 | -1.1272 | none |
| LCAT | 0.796 | 0.7083 | 3.8308 | 0.00353 | 0.022 | -1.8 | -1.1741 | none |

| | | | | | | | | |
|---|---|---|---|---|---|---|---|---|
| LIPG | 0.475 | 0.3601 | 3.8308 | 0.00353 | 0.022 | -1.8 | -1.452 | none |
| GLT8D1 | 0.531 | 0.3445 | 3.8304 | 0.00354 | 0.022 | -1.8 | -1.3091 | none |
| YTHDC2 | 0.85 | 0.7454 | 3.8301 | 0.00354 | 0.022 | -1.8 | -0.4864 | none |
| GNLY | 0.61 | 0.4504 | 3.8301 | 0.00354 | 0.022 | -1.8 | -0.5079 | none |
| SH2D1B | 0.717 | 0.4661 | 3.8298 | 0.00354 | 0.022 | -1.8 | -0.0922 | none |
| CYP2F1 | 0.95 | 0.7922 | 3.8297 | 0.00354 | 0.022 | -1.8 | -0.9183 | none |
| TBC1D32 | 0.585 | 0.4496 | 3.8295 | 0.00354 | 0.022 | -1.8 | -0.9817 | none |
| CD36 | 0.64 | 0.4707 | 3.8282 | 0.00355 | 0.022 | -1.81 | -1.3808 | none |
| ABCA5 | 0.553 | 0.4186 | 3.8277 | 0.00355 | 0.022 | -1.81 | -1.0004 | none |
| ATP10B | 0.612 | 0.4742 | 3.8261 | 0.00356 | 0.022 | -1.81 | -0.5829 | none |
| TNK1 | 0.372 | 0.3763 | 3.8248 | 0.00357 | 0.022 | -1.81 | 0.587 | none |
| KIF1BP | 0.476 | 0.3649 | 3.8236 | 0.00358 | 0.022 | -1.81 | -1.2554 | none |
| ZC3HC1 | 0.511 | 0.3272 | 3.8234 | 0.00358 | 0.022 | -1.81 | -0.7726 | none |
| WDR20 | 0.409 | 0.2967 | 3.8233 | 0.00358 | 0.022 | -1.81 | -0.9988 | none |
| F3 | 0.509 | 0.3687 | 3.8226 | 0.00358 | 0.022 | -1.82 | -1.3455 | none |
| ME2 | 0.4 | 0.3111 | 3.8225 | 0.00358 | 0.022 | -1.82 | 0.35518 | none |
| SRPK3 | 0.515 | 0.3511 | 3.8219 | 0.00358 | 0.022 | -1.82 | -1.1548 | none |
| ERAP1 | 0.599 | 0.4922 | 3.8213 | 0.00359 | 0.022 | -1.82 | -0.8332 | none |
| RSAD2 | 0.488 | 0.3663 | 3.8209 | 0.00359 | 0.022 | -1.82 | -1.0569 | none |
| LRRC25 | 0.524 | 0.3859 | 3.8202 | 0.00359 | 0.023 | -1.82 | -1.4765 | none |
| FBXO38 | 0.581 | 0.4675 | 3.8175 | 0.00361 | 0.023 | -1.82 | -1.4925 | none |
| CCDC159 | 0.631 | 0.4613 | 3.8175 | 0.00361 | 0.023 | -1.82 | -1.3779 | none |
| MCPH1 | 0.83 | 0.7199 | 3.8174 | 0.00361 | 0.023 | -1.82 | -0.988 | none |
| KIAA0408 | 0.508 | 0.4293 | 3.8163 | 0.00362 | 0.023 | -1.83 | -1.5034 | none |
| LBX2 | 0.656 | 0.4096 | 3.8155 | 0.00362 | 0.023 | -1.83 | -1.4223 | none |
| POLR2A | 1.029 | 0.9754 | 3.8154 | 0.00362 | 0.023 | -1.83 | -1.2249 | none |
| SUSD6 | 0.367 | 0.2997 | 3.8143 | 0.00363 | 0.023 | -1.83 | -0.2365 | none |
| FSD2 | 0.427 | 0.3045 | 3.8143 | 0.00363 | 0.023 | -1.83 | -1.0481 | none |
| DPF2 | 0.358 | 0.2437 | 3.8142 | 0.00363 | 0.023 | -1.83 | -0.9811 | none |
| EHMT2 | 0.493 | 0.4194 | 3.8138 | 0.00363 | 0.023 | -1.83 | -0.8412 | none |
| PAPOLG | 0.867 | 0.7042 | 3.8137 | 0.00363 | 0.023 | -1.83 | -0.8079 | none |
| HIF3A | 0.603 | 0.4378 | 3.8137 | 0.00363 | 0.023 | -1.83 | -1.4546 | none |
| REV1 | 0.685 | 0.6005 | 3.8129 | 0.00364 | 0.023 | -1.83 | -1.1519 | none |
| TAF2 | 0.692 | 0.5951 | 3.8119 | 0.00364 | 0.023 | -1.83 | -0.8466 | none |
| SLC25A15 | 0.489 | 0.4019 | 3.8116 | 0.00364 | 0.023 | -1.83 | -1.1077 | none |
| C9ORF78 | 0.583 | 0.489 | 3.8114 | 0.00365 | 0.023 | -1.83 | -1.3085 | none |
| GLTP | 0.573 | 0.4696 | 3.8113 | 0.00365 | 0.023 | -1.83 | -1.5724 | OA-related |
| MAVS | 0.612 | 0.5826 | 3.8112 | 0.00365 | 0.023 | -1.83 | -0.9922 | none |
| FBXW8 | 0.955 | 0.7722 | 3.8105 | 0.00365 | 0.023 | -1.83 | -1.5249 | none |
| TPMT | 0.449 | 0.2715 | 3.8103 | 0.00365 | 0.023 | -1.83 | -1.0969 | none |

| | | | | | | | | |
|---|---|---|---|---|---|---|---|---|
| NFAT5 | 0.842 | 0.7504 | 3.8093 | 0.00366 | 0.023 | -1.84 | -1.2281 | none |
| EFCAB3 | 0.707 | 0.6088 | 3.8084 | 0.00366 | 0.023 | -1.84 | -1.3556 | none |
| OR2J2 | 0.487 | 0.3366 | 3.808 | 0.00367 | 0.023 | -1.84 | -1.4331 | none |
| ZNF480 | 0.942 | 0.7957 | 3.8078 | 0.00367 | 0.023 | -1.84 | -1.3758 | none |
| EHBP1 | 0.673 | 0.5544 | 3.807 | 0.00367 | 0.023 | -1.84 | -1.3517 | none |
| ROBO3 | -0.445 | -0.2359 | -3.807 | 0.00367 | 0.023 | -1.84 | 1.26489 | none |
| CERS3 | 0.78 | 0.5929 | 3.8066 | 0.00367 | 0.023 | -1.84 | -1.4005 | none |
| DDX47 | 0.42 | 0.2802 | 3.805 | 0.00368 | 0.023 | -1.84 | -1.4464 | none |
| DNAJC16 | 0.843 | 0.6433 | 3.8029 | 0.0037 | 0.023 | -1.85 | -1.417 | none |
| CTSW | 0.566 | 0.3972 | 3.8027 | 0.0037 | 0.023 | -1.85 | -0.9758 | none |
| EIF4E | 0.44 | 0.231 | 3.8023 | 0.0037 | 0.023 | -1.85 | -0.7709 | none |
| ZNF93 | 0.511 | 0.3487 | 3.8014 | 0.0037 | 0.023 | -1.85 | -1.1882 | none |
| NBPF11 | 0.453 | 0.3715 | 3.8011 | 0.00371 | 0.023 | -1.85 | 0.13532 | none |
| DUSP28 | 0.558 | 0.3953 | 3.8004 | 0.00371 | 0.023 | -1.85 | -1.2103 | none |
| C8ORF76 | 0.348 | 0.1651 | 3.8003 | 0.00371 | 0.023 | -1.85 | -0.3822 | none |
| UBE3A | 0.519 | 0.4592 | 3.8001 | 0.00371 | 0.023 | -1.85 | -0.5569 | none |
| INCA1 | 0.557 | 0.4651 | 3.7997 | 0.00371 | 0.023 | -1.85 | -0.7765 | none |
| TM4SF19 | 0.582 | 0.4296 | 3.7989 | 0.00372 | 0.023 | -1.85 | -1.4757 | none |
| WDFY3 | 0.602 | 0.4813 | 3.7983 | 0.00372 | 0.023 | -1.85 | -1.4488 | none |
| SBF2 | 0.68 | 0.5329 | 3.7974 | 0.00373 | 0.023 | -1.86 | -1.1354 | none |
| R3HDM2 | 1.007 | 0.952 | 3.7962 | 0.00374 | 0.023 | -1.86 | -1.3471 | none |
| PTPN14 | 0.491 | 0.3937 | 3.7923 | 0.00376 | 0.023 | -1.86 | -1.273 | none |
| UBIAD1 | 0.492 | 0.3662 | 3.7919 | 0.00376 | 0.023 | -1.86 | -0.9126 | none |
| RAP1GAP2 | 0.451 | 0.3387 | 3.7916 | 0.00376 | 0.023 | -1.86 | -1.0097 | none |
| EIF4E1B | 0.466 | 0.3377 | 3.7913 | 0.00376 | 0.023 | -1.87 | -1.4669 | none |
| AFF3 | 0.565 | 0.4365 | 3.7913 | 0.00376 | 0.023 | -1.87 | -0.9883 | none |
| ZHX1 | 1.068 | 0.9753 | 3.7911 | 0.00377 | 0.023 | -1.87 | -0.9341 | none |
| OGFRL1 | 0.546 | 0.418 | 3.7889 | 0.00378 | 0.023 | -1.87 | -0.946 | none |
| LURAP1 | 0.523 | 0.3824 | 3.7886 | 0.00378 | 0.023 | -1.87 | -1.0813 | none |
| HRH4 | 0.479 | 0.3201 | 3.7872 | 0.00379 | 0.023 | -1.87 | -1.4853 | none |
| MLPH | 0.866 | 0.6729 | 3.7855 | 0.0038 | 0.023 | -1.87 | -0.7137 | none |
| UBN2 | 0.754 | 0.6405 | 3.7845 | 0.00381 | 0.023 | -1.88 | -1.5191 | none |
| CCDC170 | 0.299 | 0.1913 | 3.7843 | 0.00381 | 0.023 | -1.88 | -1.302 | none |
| CA5B | 0.547 | 0.3863 | 3.7836 | 0.00381 | 0.023 | -1.88 | -1.1419 | none |
| CLASP2 | 0.719 | 0.652 | 3.783 | 0.00382 | 0.023 | -1.88 | -0.9263 | none |
| BTBD2 | 0.474 | 0.4281 | 3.7826 | 0.00382 | 0.023 | -1.88 | -0.9974 | none |
| KCNJ5 | 0.444 | 0.3332 | 3.7824 | 0.00382 | 0.023 | -1.88 | -1.5067 | none |
| CD3G | 0.458 | 0.293 | 3.7824 | 0.00382 | 0.023 | -1.88 | -1.3416 | none |
| COL4A1 | 0.681 | 0.5517 | 3.7823 | 0.00382 | 0.023 | -1.88 | -1.2467 | none |
| SFXN3 | 0.428 | 0.3339 | 3.7822 | 0.00382 | 0.023 | -1.88 | -1.3139 | none |

| | | | | | | | | |
|---|---|---|---|---|---|---|---|---|
| IL12RB1 | 0.415 | 0.2638 | 3.7817 | 0.00382 | 0.023 | -1.88 | -1.3989 | none |
| GLDC | 0.718 | 0.5343 | 3.7808 | 0.00383 | 0.023 | -1.88 | -0.9681 | none |
| GTPBP1 | 0.63 | 0.587 | 3.7795 | 0.00384 | 0.023 | -1.88 | -1.4506 | none |
| B3GNT6 | 0.463 | 0.3592 | 3.7795 | 0.00384 | 0.023 | -1.88 | -1.1851 | none |
| MYO5A | 1.442 | 1.3417 | 3.7788 | 0.00384 | 0.023 | -1.89 | -1.2455 | none |
| SLC13A4 | 0.338 | 0.283 | 3.7779 | 0.00385 | 0.024 | -1.89 | -1.168 | none |
| DVL2 | 0.436 | 0.3816 | 3.7753 | 0.00386 | 0.024 | -1.89 | -1.0216 | none |
| MLYCD | 0.356 | 0.2447 | 3.7744 | 0.00387 | 0.024 | -1.89 | -0.709 | none |
| SNF8 | 0.553 | 0.4075 | 3.7739 | 0.00387 | 0.024 | -1.89 | -1.229 | none |
| PLA2G2C | 0.478 | 0.3857 | 3.7739 | 0.00387 | 0.024 | -1.89 | -1.3605 | none |
| CDK13 | 1.004 | 0.948 | 3.7737 | 0.00387 | 0.024 | -1.89 | -1.1732 | none |
| HIP1R | 0.538 | 0.5098 | 3.773 | 0.00388 | 0.024 | -1.89 | -0.4677 | none |
| TMC2 | 0.475 | 0.3844 | 3.773 | 0.00388 | 0.024 | -1.89 | -1.2717 | none |
| ALDH5A1 | 0.461 | 0.355 | 3.7719 | 0.00388 | 0.024 | -1.9 | -0.8158 | none |
| RAB11A | -0.436 | -0.3706 | -3.771 | 0.00389 | 0.024 | -1.9 | 2.18129 | AA-related |
| ZNF234 | 0.577 | 0.4734 | 3.771 | 0.00389 | 0.024 | -1.9 | -1.4303 | none |
| TBXAS1 | 0.605 | 0.4611 | 3.7698 | 0.0039 | 0.024 | -1.9 | 0.34288 | none |
| ZNF835 | 0.607 | 0.4817 | 3.7687 | 0.0039 | 0.024 | -1.9 | -0.9976 | none |
| PTCHD3 | 0.692 | 0.5208 | 3.7687 | 0.0039 | 0.024 | -1.9 | -1.2028 | none |
| CDC42EP1 | 0.588 | 0.4434 | 3.7674 | 0.00391 | 0.024 | -1.9 | -1.4367 | none |
| EXOC2 | 0.449 | 0.4003 | 3.7673 | 0.00391 | 0.024 | -1.9 | -0.4709 | none |
| MAPKAPK | 0.564 | 0.4696 | 3.7669 | 0.00392 | 0.024 | -1.9 | -0.1344 | none |
| ZNF738 | 0.614 | 0.4312 | 3.7666 | 0.00392 | 0.024 | -1.9 | -0.787 | none |
| FABP2 | 0.285 | 0.1991 | 3.7657 | 0.00392 | 0.024 | -1.91 | 0.64058 | none |
| ZNF579 | 0.465 | 0.2809 | 3.7654 | 0.00392 | 0.024 | -1.91 | 0.54512 | none |
| MRPS27 | 0.499 | 0.3951 | 3.7653 | 0.00393 | 0.024 | -1.91 | -0.8544 | none |
| AKAP6 | 0.587 | 0.3949 | 3.7646 | 0.00393 | 0.024 | -1.91 | -1.2087 | none |
| MCCC2 | 0.314 | 0.2199 | 3.7645 | 0.00393 | 0.024 | -1.91 | -0.5717 | none |
| ZNF329 | 0.843 | 0.7271 | 3.7645 | 0.00393 | 0.024 | -1.91 | -1.2165 | none |
| ZNF414 | 0.629 | 0.5015 | 3.7644 | 0.00393 | 0.024 | -1.91 | -1.265 | none |
| HELB | 0.682 | 0.5942 | 3.7636 | 0.00394 | 0.024 | -1.91 | -1.295 | none |
| SUCLA2 | 0.401 | 0.3215 | 3.7636 | 0.00394 | 0.024 | -1.91 | -1.4191 | none |
| ABHD15 | 0.351 | 0.2827 | 3.7629 | 0.00394 | 0.024 | -1.91 | -1.0873 | none |
| XYLB | 0.394 | 0.2918 | 3.7627 | 0.00394 | 0.024 | -1.91 | -1.0022 | none |
| ZNF689 | 0.367 | 0.3286 | 3.7626 | 0.00394 | 0.024 | -1.91 | -0.5426 | none |
| ESR1 | 0.622 | 0.4666 | 3.7622 | 0.00394 | 0.024 | -1.91 | -0.812 | none |
| FBXO42 | 0.408 | 0.3717 | 3.7613 | 0.00395 | 0.024 | -1.91 | -0.9457 | none |
| ADGRG2 | 0.486 | 0.3538 | 3.7613 | 0.00395 | 0.024 | -1.91 | -1.4822 | none |
| ZBTB20 | 0.817 | 0.6064 | 3.7613 | 0.00395 | 0.024 | -1.91 | -0.6678 | none |
| UBFD1 | 0.479 | 0.3831 | 3.7612 | 0.00395 | 0.024 | -1.91 | -0.9738 | none |

| | | | | | | | | |
|---|---|---|---|---|---|---|---|---|
| IARS | 0.634 | 0.517 | 3.7611 | 0.00395 | 0.024 | -1.91 | -1.1038 | none |
| NPAP1 | 0.27 | 0.1747 | 3.7609 | 0.00395 | 0.024 | -1.91 | -1.0501 | none |
| ADH1B | 0.381 | 0.2776 | 3.7602 | 0.00396 | 0.024 | -1.92 | -1.4169 | none |
| CYP2B6 | 0.6 | 0.4056 | 3.759 | 0.00397 | 0.024 | -1.92 | -1.2076 | none |
| CA14 | 0.567 | 0.4616 | 3.7587 | 0.00397 | 0.024 | -1.92 | -1.3442 | none |
| ASB11 | 0.498 | 0.4061 | 3.7585 | 0.00397 | 0.024 | -1.92 | -0.989 | none |
| TBC1D23 | 0.327 | 0.2572 | 3.7583 | 0.00397 | 0.024 | -1.92 | -0.8258 | none |
| OLR1 | 0.686 | 0.5059 | 3.7567 | 0.00398 | 0.024 | -1.92 | -1.1954 | none |
| PAFAH2 | 0.482 | 0.3543 | 3.756 | 0.00398 | 0.024 | -1.92 | -1.307 | none |
| PSEN1 | 0.484 | 0.3304 | 3.7547 | 0.00399 | 0.024 | -1.92 | -0.8557 | none |
| ICAM4 | 0.584 | 0.4219 | 3.754 | 0.004 | 0.024 | -1.93 | -1.4492 | none |
| EIF2S3 | 0.525 | 0.3915 | 3.7533 | 0.004 | 0.024 | -1.93 | -1.4768 | none |
| NUP155 | 0.683 | 0.5678 | 3.7532 | 0.004 | 0.024 | -1.93 | -0.9827 | none |
| SYDE2 | 0.566 | 0.444 | 3.7527 | 0.00401 | 0.024 | -1.93 | -1.3937 | none |
| EFR3B | 0.55 | 0.4356 | 3.7517 | 0.00401 | 0.024 | -1.93 | -1.6175 | OA-related |
| OTUD7B | 0.693 | 0.6377 | 3.7517 | 0.00401 | 0.024 | -1.93 | -1.427 | none |
| CAMKMT | 0.48 | 0.324 | 3.7511 | 0.00402 | 0.024 | -1.93 | -1.0074 | none |
| CNTNAP1 | 0.754 | 0.4963 | 3.7502 | 0.00402 | 0.024 | -1.93 | -0.9822 | none |
| ZCWPW2 | 0.913 | 0.6649 | 3.7501 | 0.00402 | 0.024 | -1.93 | -1.4726 | none |
| SIPA1L1 | 1.254 | 1.1838 | 3.7496 | 0.00403 | 0.024 | -1.93 | -1.2356 | none |
| LRRC2 | 0.484 | 0.3376 | 3.7493 | 0.00403 | 0.024 | -1.93 | -1.2067 | none |
| HYKK | 0.608 | 0.4207 | 3.7492 | 0.00403 | 0.024 | -1.93 | -1.5733 | OA-related |
| RRP36 | 0.678 | 0.5674 | 3.7489 | 0.00403 | 0.024 | -1.93 | -1.337 | none |
| KLK2 | 0.866 | 0.6349 | 3.7479 | 0.00404 | 0.024 | -1.93 | -1.3917 | none |
| CSAD | 0.827 | 0.6501 | 3.7479 | 0.00404 | 0.024 | -1.94 | -1.5464 | OA-related |
| ADD3 | 0.634 | 0.5245 | 3.7476 | 0.00404 | 0.024 | -1.94 | -0.7711 | none |
| NPIPB11 | 0.57 | 0.4222 | 3.7475 | 0.00404 | 0.024 | -1.94 | -1.5433 | OA-related |
| STX16 | 0.58 | 0.5057 | 3.7474 | 0.00404 | 0.024 | -1.94 | -0.3354 | none |
| ZNF276 | 0.706 | 0.7054 | 3.7467 | 0.00404 | 0.024 | -1.94 | -0.8294 | none |
| CHEK1 | 0.476 | 0.3047 | 3.7462 | 0.00405 | 0.024 | -1.94 | -1.2373 | none |
| ZNF470 | 0.659 | 0.5452 | 3.7451 | 0.00406 | 0.024 | -1.94 | -1.6196 | OA-related |
| MID1IP1 | 0.49 | 0.3585 | 3.745 | 0.00406 | 0.024 | -1.94 | -1.0939 | none |
| ZNF426 | 0.802 | 0.6799 | 3.7448 | 0.00406 | 0.024 | -1.94 | -1.4111 | none |
| TTLL10 | 0.969 | 0.7401 | 3.7445 | 0.00406 | 0.024 | -1.94 | -1.0457 | none |
| TMEM44 | 0.849 | 0.6817 | 3.7442 | 0.00406 | 0.024 | -1.94 | -1.3428 | none |
| PDK3 | 0.42 | 0.2473 | 3.7427 | 0.00407 | 0.024 | -1.94 | -1.2247 | none |
| GPX8 | 0.533 | 0.4018 | 3.7418 | 0.00408 | 0.024 | -1.94 | -1.4097 | none |
| ADTRP | 0.407 | 0.2762 | 3.7409 | 0.00408 | 0.024 | -1.95 | -0.2492 | none |
| ZNF850 | 0.74 | 0.6005 | 3.7407 | 0.00408 | 0.024 | -1.95 | -0.808 | none |
| TMEM68 | 0.661 | 0.4836 | 3.7407 | 0.00408 | 0.024 | -1.95 | -0.7753 | none |

| | | | | | | | | |
|---|---|---|---|---|---|---|---|---|
| ENO2 | 0.397 | 0.3128 | 3.7401 | 0.00409 | 0.024 | -1.95 | 0.35518 | none |
| MICU1 | 0.455 | 0.3452 | 3.7395 | 0.00409 | 0.024 | -1.95 | -1.375 | none |
| CECR2 | 0.449 | 0.325 | 3.7387 | 0.0041 | 0.024 | -1.95 | -0.8703 | none |
| LACTB | 0.548 | 0.4231 | 3.7384 | 0.0041 | 0.024 | -1.95 | -0.0759 | none |
| OR8H1 | 0.431 | 0.2919 | 3.7379 | 0.0041 | 0.024 | -1.95 | -1.3048 | none |
| ANK3 | 0.703 | 0.605 | 3.7371 | 0.00411 | 0.024 | -1.95 | -1.3135 | none |
| KANSL1 | 0.78 | 0.668 | 3.7365 | 0.00411 | 0.024 | -1.95 | -1.17 | none |
| PRMT8 | 0.491 | 0.3382 | 3.7361 | 0.00411 | 0.024 | -1.95 | -1.031 | none |
| CD226 | -0.436 | -0.3467 | -3.736 | 0.00412 | 0.024 | -1.95 | 1.16927 | none |
| TIMP1 | -0.367 | -0.2852 | -3.735 | 0.00412 | 0.024 | -1.96 | 1.46016 | none |
| NT5E | -0.558 | -0.4374 | -3.732 | 0.00414 | 0.025 | -1.96 | 2.01981 | none |
| ZBTB8OS | 0.519 | 0.3269 | 3.7312 | 0.00415 | 0.025 | -1.96 | -1.1935 | none |
| CFAP206 | 0.448 | 0.3056 | 3.731 | 0.00415 | 0.025 | -1.96 | -1.3736 | none |
| NCOR1 | 1.198 | 1.1522 | 3.7306 | 0.00415 | 0.025 | -1.96 | -1.1664 | none |
| HMGN2 | -0.282 | -0.2034 | -3.731 | 0.00415 | 0.025 | -1.96 | 0.91686 | none |
| C22ORF46 | 0.74 | 0.694 | 3.7305 | 0.00415 | 0.025 | -1.96 | -1.1264 | none |
| CDCA8 | -0.3 | -0.2109 | -3.73 | 0.00415 | 0.025 | -1.96 | 1.77358 | none |
| ESYT2 | 0.45 | 0.3592 | 3.7298 | 0.00416 | 0.025 | -1.96 | -1.444 | none |
| CCDC47 | 0.436 | 0.3623 | 3.7296 | 0.00416 | 0.025 | -1.96 | -0.9586 | none |
| FAM131A | 0.54 | 0.3281 | 3.7288 | 0.00416 | 0.025 | -1.97 | -1.1549 | none |
| TMPRSS11 | 0.393 | 0.2929 | 3.7274 | 0.00417 | 0.025 | -1.97 | -1.0331 | none |
| BBS2 | 0.899 | 0.6856 | 3.7273 | 0.00417 | 0.025 | -1.97 | -0.8276 | none |
| FRMD1 | 0.463 | 0.3351 | 3.7257 | 0.00418 | 0.025 | -1.97 | -1.1698 | none |
| ZNF100 | 0.876 | 0.7128 | 3.7248 | 0.00419 | 0.025 | -1.97 | -1.0759 | none |
| MSL1 | 0.986 | 0.8104 | 3.7243 | 0.00419 | 0.025 | -1.97 | -1.2445 | none |
| ZNF121 | 0.812 | 0.7297 | 3.7235 | 0.0042 | 0.025 | -1.97 | -1.4251 | none |
| PARN | 0.569 | 0.4732 | 3.7228 | 0.0042 | 0.025 | -1.98 | -0.6043 | none |
| TTC16 | 0.437 | 0.3228 | 3.7215 | 0.00421 | 0.025 | -1.98 | -1.2568 | none |
| MYH14 | 0.306 | 0.2221 | 3.7211 | 0.00422 | 0.025 | -1.98 | -1.3183 | none |
| STX1B | 0.373 | 0.2929 | 3.7207 | 0.00422 | 0.025 | -1.98 | -1.2943 | none |
| NTRK3 | 0.684 | 0.5486 | 3.7203 | 0.00422 | 0.025 | -1.98 | -1.0564 | none |
| CAV2 | 0.683 | 0.4743 | 3.72 | 0.00422 | 0.025 | -1.98 | -1.1621 | none |
| CAMSAP1 | 0.957 | 0.9728 | 3.7198 | 0.00422 | 0.025 | -1.98 | -0.9974 | none |
| ITIH5 | 0.81 | 0.547 | 3.7197 | 0.00422 | 0.025 | -1.98 | -1.1475 | none |
| NKAP | 0.914 | 0.8083 | 3.7192 | 0.00423 | 0.025 | -1.98 | -1.1174 | none |
| LETMD1 | 0.525 | 0.3663 | 3.7188 | 0.00423 | 0.025 | -1.98 | -0.9929 | none |
| KDM5C | 0.955 | 0.8932 | 3.7188 | 0.00423 | 0.025 | -1.98 | -1.0998 | none |
| HIST1H2B( | 0.65 | 0.485 | 3.7181 | 0.00424 | 0.025 | -1.98 | -1.0876 | none |
| RNF10 | 0.484 | 0.3872 | 3.7179 | 0.00424 | 0.025 | -1.98 | -0.416 | none |
| ZRSR2 | 0.55 | 0.4381 | 3.7172 | 0.00424 | 0.025 | -1.98 | -1.4133 | none |

| | | | | | | | | |
|---|---|---|---|---|---|---|---|---|
| IGFBPL1 | 0.463 | 0.3555 | 3.7163 | 0.00425 | 0.025 | -1.99 | -1.4375 | none |
| XDH | 0.435 | 0.3215 | 3.7153 | 0.00425 | 0.025 | -1.99 | -0.9262 | none |
| SLC6A4 | 0.307 | 0.213 | 3.7149 | 0.00426 | 0.025 | -1.99 | -1.3019 | none |
| TCEANC2 | 0.697 | 0.4836 | 3.7136 | 0.00427 | 0.025 | -1.99 | -1.2472 | none |
| ZSCAN25 | 0.495 | 0.4577 | 3.7123 | 0.00428 | 0.025 | -1.99 | -0.3983 | none |
| ASPSCR1 | 0.431 | 0.3029 | 3.7119 | 0.00428 | 0.025 | -1.99 | -1.4688 | none |
| RPL3L | 0.536 | 0.3931 | 3.7108 | 0.00429 | 0.025 | -1.99 | -0.9795 | none |
| MICALL2 | 0.421 | 0.4107 | 3.7101 | 0.00429 | 0.025 | -2 | 0.0463 | none |
| 8-Mar | 0.739 | 0.5622 | 3.71 | 0.00429 | 0.025 | -2 | -0.9702 | none |
| TCF25 | 0.515 | 0.4398 | 3.7098 | 0.00429 | 0.025 | -2 | -1.4983 | none |
| CEACAM1 | 0.583 | 0.4861 | 3.7084 | 0.0043 | 0.025 | -2 | -0.8035 | none |
| DENND5B | 0.478 | 0.3787 | 3.708 | 0.00431 | 0.025 | -2 | -0.8559 | none |
| C4BPA | 1.181 | 0.9786 | 3.7078 | 0.00431 | 0.025 | -2 | -1.2916 | none |
| MTRNR2L! | 0.717 | 0.5636 | 3.7076 | 0.00431 | 0.025 | -2 | -1.0332 | none |
| ANXA3 | 0.574 | 0.4622 | 3.7051 | 0.00433 | 0.025 | -2 | -1.0221 | none |
| SPG21 | 0.544 | 0.3386 | 3.7043 | 0.00433 | 0.025 | -2.01 | -0.901 | none |
| CREBRF | 0.681 | 0.5857 | 3.7037 | 0.00434 | 0.025 | -2.01 | -0.7202 | none |
| ZNF101 | 0.772 | 0.6492 | 3.703 | 0.00434 | 0.025 | -2.01 | -1.491 | none |
| ERVW-1 | 0.402 | 0.3875 | 3.7025 | 0.00434 | 0.025 | -2.01 | -0.4765 | none |
| RARRES1 | 0.9 | 0.5687 | 3.7018 | 0.00435 | 0.025 | -2.01 | -1.0993 | none |
| PNKD | -0.404 | -0.3279 | -3.701 | 0.00435 | 0.025 | -2.01 | 1.31448 | none |
| ZNF45 | 0.953 | 0.8179 | 3.7001 | 0.00436 | 0.025 | -2.01 | -1.3857 | none |
| NPIPB6 | 0.514 | 0.4489 | 3.6996 | 0.00436 | 0.025 | -2.01 | -0.6897 | none |
| CDH11 | 0.946 | 0.7308 | 3.6992 | 0.00437 | 0.025 | -2.01 | -0.919 | none |
| C4BPB | 0.557 | 0.4299 | 3.6982 | 0.00437 | 0.025 | -2.01 | -1.0154 | none |
| CDH17 | 0.81 | 0.628 | 3.698 | 0.00438 | 0.025 | -2.02 | -1.3758 | none |
| SENP6 | 1.101 | 1.022 | 3.6978 | 0.00438 | 0.025 | -2.02 | -1.3022 | none |
| PTPN9 | 0.636 | 0.5142 | 3.6977 | 0.00438 | 0.025 | -2.02 | -1.2941 | none |
| TTC7B | 1.188 | 1.0796 | 3.6976 | 0.00438 | 0.025 | -2.02 | -0.9931 | none |
| WDR24 | 0.386 | 0.3426 | 3.6974 | 0.00438 | 0.025 | -2.02 | -0.6367 | none |
| ITGB6 | 0.589 | 0.4432 | 3.6957 | 0.00439 | 0.026 | -2.02 | -0.9574 | none |
| TRIM41 | 0.602 | 0.5401 | 3.6947 | 0.0044 | 0.026 | -2.02 | -1.3581 | none |
| PGAM5 | 0.51 | 0.4176 | 3.6942 | 0.0044 | 0.026 | -2.02 | -1.3869 | none |
| GABRD | 0.515 | 0.3727 | 3.6937 | 0.00441 | 0.026 | -2.02 | -1.1749 | none |
| CCNL2 | 0.6 | 0.5107 | 3.6934 | 0.00441 | 0.026 | -2.02 | -1.3014 | none |
| ALDH1B1 | 0.546 | 0.4798 | 3.6926 | 0.00441 | 0.026 | -2.02 | -0.9782 | none |
| ZNF468 | 0.723 | 0.6001 | 3.6912 | 0.00442 | 0.026 | -2.03 | -1.2424 | none |
| WIPF2 | 0.674 | 0.5199 | 3.6893 | 0.00444 | 0.026 | -2.03 | -0.986 | none |
| INSC | 0.435 | 0.324 | 3.6892 | 0.00444 | 0.026 | -2.03 | -1.2378 | none |
| CD276 | 0.861 | 0.7235 | 3.6883 | 0.00444 | 0.026 | -2.03 | -1.213 | none |

| | | | | | | | | |
|---|---|---|---|---|---|---|---|---|
| TRIM68 | 0.49 | 0.3777 | 3.6871 | 0.00445 | 0.026 | -2.03 | -1.2793 | none |
| NPHS1 | 0.318 | 0.214 | 3.6857 | 0.00446 | 0.026 | -2.04 | -1.2549 | none |
| USP40 | 0.675 | 0.557 | 3.6843 | 0.00447 | 0.026 | -2.04 | -0.8854 | none |
| PRELID2 | 0.558 | 0.4336 | 3.6838 | 0.00448 | 0.026 | -2.04 | -1.1497 | none |
| FAM149A | 1.248 | 1.0185 | 3.6827 | 0.00449 | 0.026 | -2.04 | -1.1427 | none |
| TMEM45A | 0.615 | 0.4635 | 3.6826 | 0.00449 | 0.026 | -2.04 | -1.0406 | none |
| AC124312 | 0.522 | 0.3757 | 3.6818 | 0.00449 | 0.026 | -2.04 | -1.02 | none |
| IPO7 | 0.73 | 0.5647 | 3.6806 | 0.0045 | 0.026 | -2.04 | -0.9345 | none |
| HIST1H2AI | 1.012 | 0.7305 | 3.6803 | 0.0045 | 0.026 | -2.04 | -0.0735 | none |
| GCLC | 0.576 | 0.4798 | 3.6769 | 0.00453 | 0.026 | -2.05 | -1.3982 | none |
| MYO10 | 0.508 | 0.4122 | 3.6765 | 0.00453 | 0.026 | -2.05 | -0.7153 | none |
| EHMT1 | 1.024 | 1.0305 | 3.6765 | 0.00453 | 0.026 | -2.05 | -0.9083 | none |
| SOCS7 | 0.464 | 0.4012 | 3.6756 | 0.00454 | 0.026 | -2.05 | -1.3354 | none |
| BTF3 | -0.354 | -0.3308 | -3.675 | 0.00454 | 0.026 | -2.05 | 1.64388 | none |
| LAMC3 | 0.513 | 0.3827 | 3.6754 | 0.00454 | 0.026 | -2.05 | -1.2112 | none |
| F2RL3 | 0.55 | 0.4176 | 3.6753 | 0.00454 | 0.026 | -2.05 | -1.5865 | OA-related |
| ZNF816 | 0.828 | 0.6691 | 3.6749 | 0.00454 | 0.026 | -2.05 | -1.5021 | none |
| SUPV3L1 | 0.456 | 0.4044 | 3.6749 | 0.00454 | 0.026 | -2.05 | -0.0534 | none |
| USP4 | 0.741 | 0.6735 | 3.6748 | 0.00454 | 0.026 | -2.05 | -1.3434 | none |
| BROX | 0.597 | 0.4663 | 3.674 | 0.00455 | 0.026 | -2.05 | -1.5589 | OA-related |
| SLC35F1 | 0.389 | 0.2726 | 3.6715 | 0.00457 | 0.026 | -2.06 | -0.5766 | none |
| RHOB | -0.379 | -0.291 | -3.67 | 0.00458 | 0.026 | -2.06 | 1.32121 | none |
| SGF29 | 0.33 | 0.2945 | 3.6697 | 0.00458 | 0.026 | -2.06 | -0.3866 | none |
| ZSWIM7 | 0.525 | 0.3018 | 3.6697 | 0.00458 | 0.026 | -2.06 | -0.9981 | none |
| VPS13B | 0.566 | 0.4965 | 3.6692 | 0.00458 | 0.026 | -2.06 | -1.3 | none |
| MAPK9 | 0.638 | 0.4195 | 3.6689 | 0.00459 | 0.026 | -2.06 | -1.1283 | none |
| AMN1 | 0.662 | 0.4359 | 3.6688 | 0.00459 | 0.026 | -2.06 | -0.921 | none |
| ARHGEF38 | 0.701 | 0.662 | 3.6679 | 0.00459 | 0.026 | -2.06 | -0.6259 | none |
| MAS1 | 0.296 | 0.214 | 3.6674 | 0.0046 | 0.026 | -2.06 | -1.3676 | none |
| WDR82 | 0.313 | 0.2166 | 3.6665 | 0.0046 | 0.026 | -2.07 | -0.78 | none |
| ZNF609 | 0.86 | 0.8546 | 3.6662 | 0.00461 | 0.026 | -2.07 | -0.9235 | none |
| UBE4B | 0.687 | 0.6011 | 3.664 | 0.00462 | 0.026 | -2.07 | -1.453 | none |
| INO80C | 0.439 | 0.3294 | 3.6638 | 0.00462 | 0.026 | -2.07 | -0.7474 | none |
| IL13RA1 | 0.381 | 0.2883 | 3.6637 | 0.00462 | 0.026 | -2.07 | -0.9834 | none |
| KMT5B | 0.642 | 0.554 | 3.6622 | 0.00464 | 0.027 | -2.07 | -1.2189 | none |
| SLC52A3 | 0.294 | 0.2182 | 3.6618 | 0.00464 | 0.027 | -2.07 | -1.3559 | none |
| ATL1 | 0.966 | 0.6871 | 3.661 | 0.00465 | 0.027 | -2.08 | -1.1109 | none |
| PDXDC1 | 0.653 | 0.6449 | 3.6591 | 0.00466 | 0.027 | -2.08 | -0.8834 | none |
| TMPRSS7 | 0.33 | 0.245 | 3.6581 | 0.00467 | 0.027 | -2.08 | -1.0278 | none |
| TRIM36 | 0.866 | 0.6721 | 3.6578 | 0.00467 | 0.027 | -2.08 | -0.8772 | none |

| | | | | | | | | |
|---|---|---|---|---|---|---|---|---|
| UQCC1 | 0.535 | 0.4261 | 3.6577 | 0.00467 | 0.027 | -2.08 | -1.0305 | none |
| MSH6 | 0.58 | 0.5132 | 3.6569 | 0.00468 | 0.027 | -2.08 | -0.4111 | none |
| MLLT6 | 0.803 | 0.786 | 3.6565 | 0.00468 | 0.027 | -2.08 | -0.9615 | none |
| SLC35E2B | 0.608 | 0.5327 | 3.6564 | 0.00468 | 0.027 | -2.08 | -0.3572 | none |
| RPAP2 | 0.663 | 0.5315 | 3.6559 | 0.00468 | 0.027 | -2.08 | -1.2222 | none |
| NDRG3 | 0.309 | 0.2287 | 3.6558 | 0.00468 | 0.027 | -2.08 | 0.15691 | none |
| SETD6 | 0.499 | 0.4615 | 3.6544 | 0.00469 | 0.027 | -2.09 | -1.1446 | none |
| LRRC1 | 0.458 | 0.3002 | 3.6543 | 0.0047 | 0.027 | -2.09 | -1.3788 | none |
| MYADM | 0.605 | 0.4885 | 3.6525 | 0.00471 | 0.027 | -2.09 | -0.5739 | none |
| HCK | 0.356 | 0.2454 | 3.6524 | 0.00471 | 0.027 | -2.09 | -1.278 | none |
| MAP3K12 | 0.729 | 0.5797 | 3.6523 | 0.00471 | 0.027 | -2.09 | -1.3581 | none |
| KIF25 | 0.614 | 0.526 | 3.6513 | 0.00472 | 0.027 | -2.09 | -0.9872 | none |
| IL17RC | 0.769 | 0.6438 | 3.6502 | 0.00473 | 0.027 | -2.09 | -0.8838 | none |
| TTLL2 | 0.352 | 0.2394 | 3.6491 | 0.00474 | 0.027 | -2.09 | -0.5833 | none |
| NLRP10 | 0.36 | 0.2472 | 3.6468 | 0.00475 | 0.027 | -2.1 | -1.2647 | none |
| GXYLT2 | 0.358 | 0.2619 | 3.6467 | 0.00475 | 0.027 | -2.1 | -1.348 | none |
| SMOX | 0.751 | 0.7494 | 3.6464 | 0.00476 | 0.027 | -2.1 | -1.0098 | none |
| BCS1L | 0.372 | 0.2845 | 3.6462 | 0.00476 | 0.027 | -2.1 | -0.6763 | none |
| RP1L1 | 0.543 | 0.4589 | 3.6458 | 0.00476 | 0.027 | -2.1 | -0.5839 | none |
| GAA | 0.4 | 0.3595 | 3.6422 | 0.00479 | 0.027 | -2.11 | 0.64758 | none |
| TRIM56 | 0.798 | 0.7626 | 3.642 | 0.00479 | 0.027 | -2.11 | -1.1385 | none |
| ABCC4 | 0.955 | 0.7366 | 3.6413 | 0.0048 | 0.027 | -2.11 | -1.2918 | none |
| UBOX5 | 0.444 | 0.3507 | 3.64 | 0.00481 | 0.027 | -2.11 | -0.8105 | none |
| C15ORF53 | 1.161 | 0.9568 | 3.6387 | 0.00482 | 0.027 | -2.11 | -0.8029 | none |
| SPDYE1 | 0.979 | 0.8388 | 3.6383 | 0.00482 | 0.027 | -2.11 | -1.415 | none |
| CTH | 0.648 | 0.4971 | 3.6382 | 0.00482 | 0.027 | -2.11 | -0.7886 | none |
| NOL6 | 0.725 | 0.6685 | 3.6377 | 0.00482 | 0.027 | -2.11 | -1.1163 | none |
| SLC26A1 | 0.7 | 0.5758 | 3.6371 | 0.00483 | 0.027 | -2.11 | 0.03944 | none |
| TWSG1 | 0.42 | 0.2924 | 3.6369 | 0.00483 | 0.027 | -2.11 | -1.2485 | none |
| RINL | 0.8 | 0.7154 | 3.6365 | 0.00483 | 0.027 | -2.11 | -1.0083 | none |
| TMEM65 | 0.369 | 0.2249 | 3.6364 | 0.00483 | 0.027 | -2.11 | -0.7268 | none |
| ANKIB1 | 0.886 | 0.7349 | 3.6356 | 0.00484 | 0.027 | -2.12 | -1.3609 | none |
| SNAPC3 | 0.493 | 0.4004 | 3.6349 | 0.00485 | 0.027 | -2.12 | -0.4791 | none |
| CTNNA1 | 0.973 | 0.8608 | 3.6342 | 0.00485 | 0.027 | -2.12 | -1.29 | none |
| DDX11 | 0.599 | 0.6116 | 3.634 | 0.00485 | 0.027 | -2.12 | -0.6086 | none |
| CAPN12 | 0.99 | 0.7761 | 3.6334 | 0.00486 | 0.027 | -2.12 | -0.8561 | none |
| MC1R | 0.42 | 0.3085 | 3.6322 | 0.00487 | 0.027 | -2.12 | -0.8509 | none |
| WFDC8 | 0.754 | 0.5107 | 3.6321 | 0.00487 | 0.027 | -2.12 | -1.3851 | none |
| TFIP11 | 0.716 | 0.654 | 3.6314 | 0.00487 | 0.027 | -2.12 | -0.4731 | none |
| IL7R | 0.971 | 0.7866 | 3.6313 | 0.00487 | 0.027 | -2.12 | -1.2537 | none |

| | | | | | | | | |
|---|---|---|---|---|---|---|---|---|
| IFI30 | -0.433 | -0.3991 | -3.631 | 0.00487 | 0.027 | -2.12 | 2.09932 | AA-related |
| PLK5 | 0.441 | 0.3197 | 3.6307 | 0.00488 | 0.027 | -2.12 | -1.4091 | none |
| ASCC2 | 0.722 | 0.6907 | 3.6301 | 0.00488 | 0.027 | -2.13 | -0.9653 | none |
| TPGS1 | 0.331 | 0.1806 | 3.6297 | 0.00489 | 0.028 | -2.13 | -1.0148 | none |
| SAA2 | 0.726 | 0.632 | 3.6291 | 0.00489 | 0.028 | -2.13 | -1.3588 | none |
| GOLGA3 | 0.745 | 0.6439 | 3.6285 | 0.0049 | 0.028 | -2.13 | -1.2502 | none |
| CLMN | 0.394 | 0.3089 | 3.6284 | 0.0049 | 0.028 | -2.13 | -1.5498 | OA-related |
| MAPK12 | 0.61 | 0.4884 | 3.6282 | 0.0049 | 0.028 | -2.13 | -1.3523 | none |
| SRCAP | 1.042 | 1.0071 | 3.6279 | 0.0049 | 0.028 | -2.13 | -0.8069 | none |
| UBAP1L | 0.443 | 0.3826 | 3.6277 | 0.0049 | 0.028 | -2.13 | -0.8573 | none |
| CCDC60 | 0.896 | 0.7124 | 3.6275 | 0.0049 | 0.028 | -2.13 | -1.3024 | none |
| STK3 | 0.926 | 0.7703 | 3.6264 | 0.00491 | 0.028 | -2.13 | -0.7745 | none |
| ERMN | 0.636 | 0.4962 | 3.6261 | 0.00492 | 0.028 | -2.13 | -1.2265 | none |
| DDB1 | 0.402 | 0.2916 | 3.6255 | 0.00492 | 0.028 | -2.13 | -0.7493 | none |
| TMEM106 | 0.614 | 0.4946 | 3.6249 | 0.00493 | 0.028 | -2.13 | -0.8386 | none |
| ANKDD1A | 0.501 | 0.4106 | 3.6242 | 0.00493 | 0.028 | -2.13 | -1.4884 | none |
| CCR4 | 0.757 | 0.6309 | 3.623 | 0.00494 | 0.028 | -2.14 | -1.3594 | none |
| RNF43 | 0.354 | 0.2947 | 3.6218 | 0.00495 | 0.028 | -2.14 | -1.2933 | none |
| CD82 | 0.403 | 0.35 | 3.6217 | 0.00495 | 0.028 | -2.14 | -0.9043 | none |
| ACP2 | 0.399 | 0.309 | 3.6212 | 0.00495 | 0.028 | -2.14 | -1.0764 | none |
| SNAPC1 | 0.396 | 0.3005 | 3.6205 | 0.00496 | 0.028 | -2.14 | -1.1225 | none |
| RGS9 | 0.453 | 0.3031 | 3.6205 | 0.00496 | 0.028 | -2.14 | -1.3238 | none |
| HOOK3 | 0.97 | 0.8092 | 3.6204 | 0.00496 | 0.028 | -2.14 | -1.4118 | none |
| MISP | 0.874 | 0.6564 | 3.6198 | 0.00497 | 0.028 | -2.14 | -1.0187 | none |
| ZFPM2 | 0.652 | 0.5196 | 3.6195 | 0.00497 | 0.028 | -2.14 | -1.2076 | none |
| TMEM150 | 0.627 | 0.496 | 3.618 | 0.00498 | 0.028 | -2.14 | -1.1711 | none |
| CHRNB4 | 0.423 | 0.2916 | 3.6172 | 0.00499 | 0.028 | -2.15 | -0.719 | none |
| GNB4 | 0.62 | 0.4131 | 3.6159 | 0.005 | 0.028 | -2.15 | -0.3066 | none |
| FAM227A | 0.479 | 0.342 | 3.6156 | 0.005 | 0.028 | -2.15 | -1.5247 | none |
| TBC1D22A | 0.594 | 0.4855 | 3.6155 | 0.005 | 0.028 | -2.15 | -1.268 | none |
| LDLRAD3 | 0.598 | 0.4736 | 3.6138 | 0.00501 | 0.028 | -2.15 | -1.1686 | none |
| SCARB1 | 0.468 | 0.4787 | 3.6134 | 0.00502 | 0.028 | -2.15 | 1.14553 | none |
| SMTNL1 | 0.509 | 0.3932 | 3.6133 | 0.00502 | 0.028 | -2.15 | -1.4037 | none |
| ZCCHC8 | 0.655 | 0.5835 | 3.6128 | 0.00502 | 0.028 | -2.15 | -0.908 | none |
| 1-Dec | 0.545 | 0.4307 | 3.6126 | 0.00502 | 0.028 | -2.15 | -1.2176 | none |
| SERPINB5 | 0.483 | 0.3912 | 3.6126 | 0.00502 | 0.028 | -2.15 | -0.4282 | none |
| MRVI1 | 0.832 | 0.617 | 3.6114 | 0.00503 | 0.028 | -2.16 | -1.1227 | none |
| FUT4 | 0.422 | 0.2932 | 3.6102 | 0.00504 | 0.028 | -2.16 | -1.5379 | OA-related |
| PROS1 | 0.586 | 0.3839 | 3.6102 | 0.00504 | 0.028 | -2.16 | -1.1533 | none |
| ZNF696 | 0.58 | 0.4725 | 3.6096 | 0.00505 | 0.028 | -2.16 | -0.5549 | none |

| | | | | | | | | |
|---|---|---|---|---|---|---|---|---|
| MARVELD: | 0.402 | 0.31 | 3.6089 | 0.00506 | 0.028 | -2.16 | -1.3729 | none |
| ZBTB3 | 0.957 | 0.7016 | 3.6088 | 0.00506 | 0.028 | -2.16 | -1.1457 | none |
| ZNF98 | 0.746 | 0.5316 | 3.6086 | 0.00506 | 0.028 | -2.16 | -1.3123 | none |
| INPP5K | 0.335 | 0.2296 | 3.6083 | 0.00506 | 0.028 | -2.16 | -0.0148 | none |
| NEK2 | 0.357 | 0.2202 | 3.6077 | 0.00507 | 0.028 | -2.16 | -1.0918 | none |
| DNAJC28 | 0.326 | 0.2298 | 3.6074 | 0.00507 | 0.028 | -2.16 | -1.4315 | none |
| KPNA1 | 0.29 | 0.2105 | 3.6072 | 0.00507 | 0.028 | -2.16 | 0.04489 | none |
| MIS18A | -0.446 | -0.3627 | -3.607 | 0.00507 | 0.028 | -2.16 | 2.00124 | none |
| SUMO3 | 0.519 | 0.3401 | 3.606 | 0.00508 | 0.028 | -2.16 | -0.8536 | none |
| AP5S1 | 0.405 | 0.3109 | 3.6042 | 0.00509 | 0.028 | -2.17 | -1.1475 | none |
| PARP6 | 0.532 | 0.5529 | 3.6035 | 0.0051 | 0.028 | -2.17 | -0.2923 | none |
| HAPLN4 | 0.449 | 0.3381 | 3.5994 | 0.00513 | 0.028 | -2.17 | -1.0912 | none |
| FANCD2 | 0.622 | 0.6002 | 3.5993 | 0.00513 | 0.028 | -2.18 | -1.0899 | none |
| NPL | 0.743 | 0.5432 | 3.5992 | 0.00514 | 0.028 | -2.18 | -0.8229 | none |
| WSB1 | 0.353 | 0.228 | 3.5984 | 0.00514 | 0.028 | -2.18 | -0.4434 | none |
| EPHB2 | 0.373 | 0.2962 | 3.5981 | 0.00514 | 0.028 | -2.18 | -0.7792 | none |
| TRIML2 | 1.054 | 0.8823 | 3.5975 | 0.00515 | 0.028 | -2.18 | -1.2792 | none |
| RGMA | 0.335 | 0.2427 | 3.5975 | 0.00515 | 0.028 | -2.18 | -1.4109 | none |
| CHP1 | 0.532 | 0.336 | 3.5973 | 0.00515 | 0.028 | -2.18 | -1.4714 | none |
| ANKRD42 | 0.429 | 0.3283 | 3.5968 | 0.00516 | 0.028 | -2.18 | -1.1521 | none |
| ZNF254 | 0.707 | 0.5125 | 3.5967 | 0.00516 | 0.028 | -2.18 | -0.9818 | none |
| RRAS2 | 0.419 | 0.3239 | 3.5962 | 0.00516 | 0.028 | -2.18 | -0.9086 | none |
| FOCAD | 0.889 | 0.5936 | 3.5959 | 0.00516 | 0.028 | -2.18 | -1.134 | none |
| MIEF2 | 0.378 | 0.2128 | 3.5958 | 0.00516 | 0.028 | -2.18 | -0.4884 | none |
| NLRP8 | 0.377 | 0.2709 | 3.5948 | 0.00517 | 0.029 | -2.18 | -1.2857 | none |
| TAOK1 | 1.026 | 0.9884 | 3.5947 | 0.00517 | 0.029 | -2.18 | -0.9637 | none |
| CLASRP | 0.671 | 0.7174 | 3.5945 | 0.00517 | 0.029 | -2.18 | 0.1255 | none |
| POU5F1B | 0.436 | 0.3124 | 3.594 | 0.00518 | 0.029 | -2.18 | -1.3333 | none |
| NONO | 0.59 | 0.4894 | 3.5932 | 0.00519 | 0.029 | -2.18 | -1.1671 | none |
| SLCO3A1 | 0.522 | 0.3904 | 3.5931 | 0.00519 | 0.029 | -2.19 | -0.891 | none |
| STAG2 | 0.877 | 0.7498 | 3.5926 | 0.00519 | 0.029 | -2.19 | -1.3994 | none |
| RCSD1 | 0.574 | 0.4756 | 3.5923 | 0.00519 | 0.029 | -2.19 | -0.4906 | none |
| TIGD1 | 0.371 | 0.3469 | 3.5917 | 0.0052 | 0.029 | -2.19 | -0.8626 | none |
| CORO2A | 0.589 | 0.4934 | 3.591 | 0.0052 | 0.029 | -2.19 | -1.5508 | OA-related |
| ZNF496 | 0.522 | 0.4842 | 3.5905 | 0.00521 | 0.029 | -2.19 | -0.2697 | none |
| VPS18 | 0.393 | 0.3504 | 3.5899 | 0.00521 | 0.029 | -2.19 | -1.3039 | none |
| TAB2 | 0.976 | 0.9018 | 3.5897 | 0.00522 | 0.029 | -2.19 | -1.3195 | none |
| MYH10 | 0.782 | 0.7512 | 3.589 | 0.00522 | 0.029 | -2.19 | -0.3737 | none |
| DCTN6 | 0.517 | 0.2948 | 3.588 | 0.00523 | 0.029 | -2.19 | -0.9004 | none |
| SH3TC2 | 0.367 | 0.2643 | 3.588 | 0.00523 | 0.029 | -2.19 | -0.5346 | none |

| | | | | | | | | |
|---|---|---|---|---|---|---|---|---|
| TCEANC | 0.817 | 0.6276 | 3.5872 | 0.00524 | 0.029 | -2.19 | -1.1398 | none |
| TMOD3 | 0.581 | 0.4656 | 3.5864 | 0.00524 | 0.029 | -2.2 | -0.7101 | none |
| MDM4 | 0.497 | 0.4065 | 3.5862 | 0.00525 | 0.029 | -2.2 | -1.0362 | none |
| IFFO1 | 0.472 | 0.4136 | 3.5861 | 0.00525 | 0.029 | -2.2 | -0.6117 | none |
| DMKN | 0.439 | 0.3701 | 3.5859 | 0.00525 | 0.029 | -2.2 | -0.8445 | none |
| CCDC84 | 0.569 | 0.4148 | 3.5858 | 0.00525 | 0.029 | -2.2 | -1.316 | none |
| TAF6 | 0.606 | 0.5164 | 3.5857 | 0.00525 | 0.029 | -2.2 | -0.8409 | none |
| MAN2C1 | 0.531 | 0.4927 | 3.5834 | 0.00527 | 0.029 | -2.2 | -0.9196 | none |
| SNX22 | 0.522 | 0.3941 | 3.5833 | 0.00527 | 0.029 | -2.2 | -1.4914 | none |
| SYNJ2BP | 0.484 | 0.3045 | 3.5811 | 0.00529 | 0.029 | -2.2 | -1.1099 | none |
| ZNF292 | 0.74 | 0.7199 | 3.5806 | 0.00529 | 0.029 | -2.21 | -1.1611 | none |
| DGKA | 0.589 | 0.5239 | 3.58 | 0.0053 | 0.029 | -2.21 | -0.8047 | none |
| PLOD3 | 0.492 | 0.378 | 3.5795 | 0.0053 | 0.029 | -2.21 | -1.4053 | none |
| PALM2 | 0.31 | 0.2573 | 3.5791 | 0.00531 | 0.029 | -2.21 | -1.1216 | none |
| HIVEP1 | 0.817 | 0.7382 | 3.5788 | 0.00531 | 0.029 | -2.21 | -1.2387 | none |
| STAT2 | 0.64 | 0.5299 | 3.5784 | 0.00531 | 0.029 | -2.21 | -1.3867 | none |
| TMIGD2 | 0.601 | 0.4432 | 3.5775 | 0.00532 | 0.029 | -2.21 | -0.6337 | none |
| KIAA0513 | 0.587 | 0.5307 | 3.577 | 0.00532 | 0.029 | -2.21 | -1.4917 | none |
| RPGRIP1L | 0.658 | 0.5037 | 3.5763 | 0.00533 | 0.029 | -2.21 | -1.1213 | none |
| MAK16 | 0.668 | 0.5764 | 3.5749 | 0.00534 | 0.029 | -2.21 | -0.877 | none |
| TM2D1 | 0.559 | 0.3587 | 3.5727 | 0.00536 | 0.029 | -2.22 | -1.0944 | none |
| IL17D | 0.502 | 0.3894 | 3.5719 | 0.00537 | 0.029 | -2.22 | -0.9023 | none |
| TONSL | 0.555 | 0.5048 | 3.5699 | 0.00539 | 0.029 | -2.22 | -0.7023 | none |
| IGFL4 | 0.5 | 0.3804 | 3.5677 | 0.00541 | 0.029 | -2.23 | -0.0876 | none |
| LIMS1 | 0.306 | 0.2094 | 3.5676 | 0.00541 | 0.029 | -2.23 | -0.3344 | none |
| SOWAHC | 0.419 | 0.3697 | 3.5675 | 0.00541 | 0.029 | -2.23 | -1.3714 | none |
| MYT1 | 0.336 | 0.2679 | 3.5657 | 0.00542 | 0.03 | -2.23 | -0.8945 | none |
| BTRC | 0.514 | 0.3405 | 3.5654 | 0.00543 | 0.03 | -2.23 | -0.1245 | none |
| RFTN2 | 0.48 | 0.3762 | 3.5635 | 0.00544 | 0.03 | -2.23 | -1.4745 | none |
| MRS2 | 0.428 | 0.285 | 3.5624 | 0.00545 | 0.03 | -2.23 | -0.9265 | none |
| GINS1 | 0.38 | 0.2647 | 3.562 | 0.00546 | 0.03 | -2.24 | -1.1732 | none |
| POLR2B | 0.516 | 0.4053 | 3.5619 | 0.00546 | 0.03 | -2.24 | -1.3908 | none |
| FOXK2 | 0.871 | 0.8176 | 3.5612 | 0.00546 | 0.03 | -2.24 | -0.9148 | none |
| USP10 | 0.535 | 0.448 | 3.5603 | 0.00547 | 0.03 | -2.24 | -0.3137 | none |
| ORC2 | 0.457 | 0.4054 | 3.5602 | 0.00547 | 0.03 | -2.24 | -0.7055 | none |
| SAMD3 | 0.68 | 0.5827 | 3.5598 | 0.00548 | 0.03 | -2.24 | -0.3742 | none |
| C7 | 0.437 | 0.3026 | 3.5592 | 0.00548 | 0.03 | -2.24 | -1.2276 | none |
| MEGF11 | 0.36 | 0.2524 | 3.5587 | 0.00549 | 0.03 | -2.24 | -1.0858 | none |
| ZFYVE16 | 0.67 | 0.59 | 3.5572 | 0.0055 | 0.03 | -2.24 | -0.0805 | none |
| USP53 | 0.924 | 0.808 | 3.5571 | 0.0055 | 0.03 | -2.24 | -0.8777 | none |

| | | | | | | | | |
|---|---|---|---|---|---|---|---|---|
| ERN2 | 0.687 | 0.4458 | 3.5564 | 0.00551 | 0.03 | -2.24 | -0.8291 | none |
| U2AF1L5 | -0.317 | -0.2306 | -3.556 | 0.00551 | 0.03 | -2.25 | 1.41339 | none |
| ZDHHC21 | 0.611 | 0.4299 | 3.5554 | 0.00552 | 0.03 | -2.25 | -0.726 | none |
| ZNF471 | 0.345 | 0.2706 | 3.5552 | 0.00552 | 0.03 | -2.25 | -0.9764 | none |
| TGM2 | 0.474 | 0.3737 | 3.5549 | 0.00552 | 0.03 | -2.25 | -1.0788 | none |
| WDR45 | 0.48 | 0.3247 | 3.5548 | 0.00552 | 0.03 | -2.25 | -1.0909 | none |
| SRGAP2C | 0.577 | 0.5093 | 3.5547 | 0.00552 | 0.03 | -2.25 | -1.1071 | none |
| REPS2 | 0.313 | 0.2195 | 3.5541 | 0.00553 | 0.03 | -2.25 | -1.2974 | none |
| TP53 | 0.448 | 0.331 | 3.5529 | 0.00554 | 0.03 | -2.25 | -1.511 | none |
| FBXL2 | 0.491 | 0.3276 | 3.5529 | 0.00554 | 0.03 | -2.25 | -1.1746 | none |
| IDE | 0.615 | 0.5537 | 3.5527 | 0.00554 | 0.03 | -2.25 | -1.0618 | none |
| CCDC90B | 0.395 | 0.2401 | 3.5523 | 0.00554 | 0.03 | -2.25 | -1.005 | none |
| KNTC1 | 0.737 | 0.6413 | 3.5518 | 0.00555 | 0.03 | -2.25 | -1.4509 | none |
| ACVRL1 | 0.507 | 0.4735 | 3.5511 | 0.00555 | 0.03 | -2.25 | 0.26937 | none |
| MTG2 | 0.559 | 0.4773 | 3.5508 | 0.00556 | 0.03 | -2.25 | -1.1721 | none |
| NF2 | 0.737 | 0.6583 | 3.5505 | 0.00556 | 0.03 | -2.25 | -1.502 | none |
| RNF168 | 0.7 | 0.6452 | 3.5502 | 0.00556 | 0.03 | -2.25 | -1.346 | none |
| TRIM66 | 0.856 | 0.8199 | 3.5491 | 0.00557 | 0.03 | -2.26 | -0.548 | none |
| HNRNPA1 | -0.276 | -0.2371 | -3.547 | 0.00559 | 0.03 | -2.26 | 1.88162 | none |
| CLDN15 | 0.63 | 0.543 | 3.5473 | 0.00559 | 0.03 | -2.26 | -0.8258 | none |
| DBN1 | 0.46 | 0.3497 | 3.5472 | 0.00559 | 0.03 | -2.26 | -0.2761 | none |
| SLAMF6 | 0.405 | 0.2917 | 3.5465 | 0.0056 | 0.03 | -2.26 | -1.1245 | none |
| TPCN1 | 0.663 | 0.5906 | 3.5448 | 0.00561 | 0.03 | -2.26 | -0.8611 | none |
| TMEM185 | 0.42 | 0.3068 | 3.5437 | 0.00562 | 0.03 | -2.27 | -1.0575 | none |
| ACER1 | 0.72 | 0.538 | 3.5435 | 0.00562 | 0.03 | -2.27 | -0.6429 | none |
| SLC30A2 | 0.548 | 0.3954 | 3.5429 | 0.00563 | 0.03 | -2.27 | -1.5848 | OA-related |
| MED25 | 0.438 | 0.438 | 3.5429 | 0.00563 | 0.03 | -2.27 | -0.657 | none |
| KLHL36 | 0.462 | 0.3865 | 3.5426 | 0.00563 | 0.03 | -2.27 | -0.8331 | none |
| TMPRSS3 | 0.465 | 0.36 | 3.5424 | 0.00563 | 0.03 | -2.27 | -0.9555 | none |
| ISY1 | 0.309 | 0.2245 | 3.5416 | 0.00564 | 0.03 | -2.27 | -1.2267 | none |
| IRAK1BP1 | 0.482 | 0.3657 | 3.5407 | 0.00565 | 0.03 | -2.27 | -0.7201 | none |
| FLVCR1 | 0.431 | 0.2854 | 3.5392 | 0.00566 | 0.03 | -2.27 | -0.5945 | none |
| MEN1 | 0.66 | 0.5905 | 3.5352 | 0.0057 | 0.031 | -2.28 | -0.8163 | none |
| CGNL1 | 0.424 | 0.3811 | 3.5348 | 0.0057 | 0.031 | -2.28 | -0.6756 | none |
| PTCD1 | 0.664 | 0.6649 | 3.5344 | 0.00571 | 0.031 | -2.28 | -1.0349 | none |
| CHD1L | 0.385 | 0.3239 | 3.534 | 0.00571 | 0.031 | -2.28 | 0.08662 | none |
| SLC39A14 | 0.512 | 0.4201 | 3.5338 | 0.00571 | 0.031 | -2.28 | -0.3338 | none |
| RUFY1 | 0.805 | 0.7143 | 3.5338 | 0.00571 | 0.031 | -2.28 | -1.2475 | none |
| LMBR1L | 0.681 | 0.4931 | 3.5333 | 0.00572 | 0.031 | -2.28 | -0.9959 | none |
| SCHIP1 | 0.888 | 0.6535 | 3.5324 | 0.00573 | 0.031 | -2.28 | -1.1432 | none |

| | | | | | | | | |
|---|---|---|---|---|---|---|---|---|
| URB1 | 0.719 | 0.6971 | 3.5324 | 0.00573 | 0.031 | -2.28 | -0.9657 | none |
| FLCN | 0.638 | 0.5063 | 3.5308 | 0.00574 | 0.031 | -2.29 | -1.3016 | none |
| CYTIP | -0.411 | -0.4015 | -3.53 | 0.00575 | 0.031 | -2.29 | 1.92361 | none |
| LZIC | 0.423 | 0.2656 | 3.5302 | 0.00575 | 0.031 | -2.29 | -0.8034 | none |
| NCF1 | 0.835 | 0.7192 | 3.5301 | 0.00575 | 0.031 | -2.29 | -0.6793 | none |
| DISC1 | 0.713 | 0.5509 | 3.5294 | 0.00575 | 0.031 | -2.29 | -1.5979 | OA-related |
| KSR1 | 0.767 | 0.7174 | 3.5289 | 0.00576 | 0.031 | -2.29 | -1.1021 | none |
| WDR5B | 0.463 | 0.3373 | 3.5279 | 0.00577 | 0.031 | -2.29 | -0.5688 | none |
| MAU2 | 0.633 | 0.6068 | 3.5276 | 0.00577 | 0.031 | -2.29 | -0.175 | none |
| ZRANB3 | 0.536 | 0.3734 | 3.527 | 0.00578 | 0.031 | -2.29 | -0.8285 | none |
| VIL1 | 0.403 | 0.3502 | 3.5269 | 0.00578 | 0.031 | -2.29 | -1.029 | none |
| PRSS53 | 0.344 | 0.2776 | 3.526 | 0.00579 | 0.031 | -2.29 | -0.3056 | none |
| TULP4 | 0.918 | 0.8095 | 3.5255 | 0.00579 | 0.031 | -2.29 | -0.9794 | none |
| CATSPERB | 0.716 | 0.5427 | 3.5238 | 0.00581 | 0.031 | -2.3 | -0.3848 | none |
| DHRS7B | 0.378 | 0.2531 | 3.5238 | 0.00581 | 0.031 | -2.3 | -1.0787 | none |
| SELENBP1 | 0.426 | 0.261 | 3.523 | 0.00581 | 0.031 | -2.3 | -0.5677 | none |
| RHPN2 | 0.533 | 0.3896 | 3.5208 | 0.00584 | 0.031 | -2.3 | -0.512 | none |
| REL | 0.86 | 0.8273 | 3.5203 | 0.00584 | 0.031 | -2.3 | -1.22 | none |
| DENND4B | 0.666 | 0.6414 | 3.52 | 0.00584 | 0.031 | -2.3 | -1.1263 | none |
| HMCN2 | 0.408 | 0.2795 | 3.519 | 0.00585 | 0.031 | -2.31 | -0.9716 | none |
| CENPL | 0.345 | 0.1976 | 3.518 | 0.00586 | 0.031 | -2.31 | -1.3387 | none |
| DCUN1D2 | 0.81 | 0.6887 | 3.518 | 0.00586 | 0.031 | -2.31 | -0.8001 | none |
| PLS1 | 0.611 | 0.4854 | 3.5177 | 0.00587 | 0.031 | -2.31 | -0.3514 | none |
| RGPD5 | 0.803 | 0.6596 | 3.5176 | 0.00587 | 0.031 | -2.31 | -0.9261 | none |
| C17ORF51 | 0.57 | 0.4409 | 3.5176 | 0.00587 | 0.031 | -2.31 | -1.2538 | none |
| LRG1 | 0.697 | 0.5701 | 3.5168 | 0.00587 | 0.031 | -2.31 | -1.4588 | none |
| GATM | 0.619 | 0.4374 | 3.5168 | 0.00587 | 0.031 | -2.31 | -0.7639 | none |
| ITPR1 | 0.603 | 0.552 | 3.5159 | 0.00588 | 0.031 | -2.31 | -0.9983 | none |
| FOXP1 | 0.772 | 0.696 | 3.5155 | 0.00589 | 0.031 | -2.31 | -1.4151 | none |
| TPI1 | -0.357 | -0.2653 | -3.514 | 0.0059 | 0.031 | -2.31 | 1.84854 | none |
| MAMLD1 | 0.826 | 0.6369 | 3.511 | 0.00593 | 0.031 | -2.32 | -1.0373 | none |
| CLIC1 | -0.379 | -0.319 | -3.511 | 0.00594 | 0.031 | -2.32 | 2.21923 | AA-related |
| TM4SF18 | 0.721 | 0.5079 | 3.5093 | 0.00595 | 0.031 | -2.32 | -1.2747 | none |
| CBS | 0.502 | 0.3355 | 3.5089 | 0.00595 | 0.031 | -2.32 | -0.5283 | none |
| EEF1A1 | -0.428 | -0.4046 | -3.509 | 0.00595 | 0.031 | -2.32 | 1.40704 | none |
| DAP3 | 0.248 | 0.128 | 3.5077 | 0.00596 | 0.032 | -2.32 | -0.766 | none |
| C16ORF58 | 0.393 | 0.3603 | 3.5075 | 0.00596 | 0.032 | -2.32 | -0.7693 | none |
| DPF1 | 0.415 | 0.2858 | 3.5074 | 0.00597 | 0.032 | -2.32 | -1.3754 | none |
| PPARD | 0.561 | 0.5121 | 3.5069 | 0.00597 | 0.032 | -2.33 | -1.305 | none |
| FAM234B | 0.421 | 0.3234 | 3.5066 | 0.00597 | 0.032 | -2.33 | -1.3143 | none |

| | | | | | | | | |
|---|---|---|---|---|---|---|---|---|
| MACROD2 | 0.558 | 0.4475 | 3.5064 | 0.00598 | 0.032 | -2.33 | -0.9775 | none |
| CPA4 | 0.543 | 0.4335 | 3.5064 | 0.00598 | 0.032 | -2.33 | -1.1546 | none |
| MPP5 | 0.687 | 0.5755 | 3.5063 | 0.00598 | 0.032 | -2.33 | -0.6931 | none |
| STX2 | 0.706 | 0.5608 | 3.5056 | 0.00598 | 0.032 | -2.33 | -0.7401 | none |
| OR5A1 | 0.274 | 0.1973 | 3.5055 | 0.00598 | 0.032 | -2.33 | -1.5245 | none |
| TFEB | 0.517 | 0.4485 | 3.5032 | 0.00601 | 0.032 | -2.33 | 0.11422 | none |
| CCDC7 | 0.553 | 0.4224 | 3.503 | 0.00601 | 0.032 | -2.33 | 0.2125 | none |
| GAS7 | 0.372 | 0.2666 | 3.5027 | 0.00601 | 0.032 | -2.33 | -0.5357 | none |
| BTBD18 | 0.275 | 0.2347 | 3.5023 | 0.00602 | 0.032 | -2.33 | -0.7617 | none |
| SKP1 | -0.375 | -0.367 | -3.501 | 0.00603 | 0.032 | -2.33 | 1.95626 | none |
| MCM4 | 0.837 | 0.7627 | 3.5012 | 0.00603 | 0.032 | -2.33 | -0.6333 | none |
| HERC6 | 0.554 | 0.4793 | 3.4982 | 0.00606 | 0.032 | -2.34 | -0.2437 | none |
| TYR | 0.456 | 0.2933 | 3.4979 | 0.00606 | 0.032 | -2.34 | -0.9817 | none |
| GRB2 | 0.35 | 0.275 | 3.4977 | 0.00606 | 0.032 | -2.34 | -1.269 | none |
| SPIB | 0.598 | 0.4028 | 3.4973 | 0.00606 | 0.032 | -2.34 | -1.3857 | none |
| ZNF124 | 0.488 | 0.3642 | 3.4968 | 0.00607 | 0.032 | -2.34 | -0.8937 | none |
| PFDN6 | -0.303 | -0.2302 | -3.497 | 0.00607 | 0.032 | -2.34 | 2.14829 | AA-related |
| ERI1 | 0.389 | 0.2938 | 3.4961 | 0.00608 | 0.032 | -2.34 | -0.6509 | none |
| ST7L | 0.89 | 0.7186 | 3.4959 | 0.00608 | 0.032 | -2.34 | -0.6344 | none |
| ZNF540 | 0.478 | 0.3517 | 3.4938 | 0.0061 | 0.032 | -2.35 | -0.7397 | none |
| FASTKD1 | 0.401 | 0.276 | 3.4922 | 0.00612 | 0.032 | -2.35 | 0.27409 | none |
| SCN1A | 0.427 | 0.3032 | 3.4903 | 0.00614 | 0.032 | -2.35 | -1.2217 | none |
| 6-Mar | 0.436 | 0.2987 | 3.4891 | 0.00615 | 0.032 | -2.35 | -1.3491 | none |
| GSPT1 | 0.526 | 0.4456 | 3.4884 | 0.00615 | 0.032 | -2.36 | -1.3095 | none |
| KCTD18 | 0.488 | 0.3462 | 3.4879 | 0.00616 | 0.032 | -2.36 | -1.1702 | none |
| GGA2 | 0.567 | 0.5012 | 3.4878 | 0.00616 | 0.032 | -2.36 | -0.6189 | none |
| PPP1R15B | 0.37 | 0.3069 | 3.4878 | 0.00616 | 0.032 | -2.36 | -1.0777 | none |
| PACSIN2 | 0.931 | 0.8511 | 3.4877 | 0.00616 | 0.032 | -2.36 | -1.1829 | none |
| SLC30A7 | 0.592 | 0.4108 | 3.4876 | 0.00616 | 0.032 | -2.36 | -1.309 | none |
| GFPT1 | 0.363 | 0.2269 | 3.4861 | 0.00618 | 0.032 | -2.36 | -1.2041 | none |
| METTL22 | 0.35 | 0.2928 | 3.4855 | 0.00618 | 0.032 | -2.36 | -0.6772 | none |
| ASAH1 | 0.373 | 0.199 | 3.4849 | 0.00619 | 0.032 | -2.36 | -0.2135 | none |
| PDE4DIP | 0.747 | 0.6969 | 3.4847 | 0.00619 | 0.032 | -2.36 | -1.1041 | none |
| RUFY2 | 0.721 | 0.6184 | 3.484 | 0.0062 | 0.032 | -2.36 | -0.8822 | none |
| MAP3K13 | 0.275 | 0.1384 | 3.4835 | 0.0062 | 0.032 | -2.36 | -0.4304 | none |
| KLRD1 | 0.576 | 0.4458 | 3.4825 | 0.00621 | 0.032 | -2.37 | -0.8622 | none |
| PRDX1 | -0.483 | -0.4588 | -3.483 | 0.00621 | 0.032 | -2.37 | 2.33016 | AA-related |
| ARRDC3 | 0.508 | 0.3183 | 3.4816 | 0.00622 | 0.032 | -2.37 | -1.0004 | none |
| ESPL1 | 0.643 | 0.6392 | 3.4814 | 0.00623 | 0.032 | -2.37 | -1.0376 | none |
| TROAP | 0.364 | 0.299 | 3.4808 | 0.00623 | 0.033 | -2.37 | -1.3587 | none |

| | | | | | | | | |
|---|---|---|---|---|---|---|---|---|
| TTN | 0.466 | 0.3086 | 3.4801 | 0.00624 | 0.033 | -2.37 | 0.0301 | none |
| LRRFIP1 | 1.099 | 1.0377 | 3.4798 | 0.00624 | 0.033 | -2.37 | -1.1016 | none |
| ACP7 | 0.454 | 0.3042 | 3.4795 | 0.00624 | 0.033 | -2.37 | -1.1359 | none |
| SLC23A3 | 0.51 | 0.4518 | 3.4791 | 0.00625 | 0.033 | -2.37 | -0.353 | none |
| SMARCB1 | 0.839 | 0.6759 | 3.4784 | 0.00626 | 0.033 | -2.37 | -1.181 | none |
| ZNF718 | 0.412 | 0.2836 | 3.4776 | 0.00626 | 0.033 | -2.37 | -1.0849 | none |
| UGT8 | 0.42 | 0.3012 | 3.4764 | 0.00628 | 0.033 | -2.38 | -1.3255 | none |
| TNFSF14 | 0.815 | 0.7418 | 3.4741 | 0.0063 | 0.033 | -2.38 | -1.0588 | none |
| RCOR3 | 0.512 | 0.4323 | 3.4728 | 0.00631 | 0.033 | -2.38 | -1.2832 | none |
| ZBED1 | 0.67 | 0.5752 | 3.4712 | 0.00633 | 0.033 | -2.38 | -0.967 | none |
| SEC31B | 0.471 | 0.4234 | 3.4712 | 0.00633 | 0.033 | -2.38 | -1.2067 | none |
| STRN4 | 0.691 | 0.6318 | 3.4707 | 0.00634 | 0.033 | -2.38 | -1.5136 | none |
| ARHGAP2 | 0.552 | 0.5331 | 3.4701 | 0.00634 | 0.033 | -2.39 | -0.2214 | none |
| PPIL3 | 0.52 | 0.3083 | 3.4698 | 0.00635 | 0.033 | -2.39 | -0.8169 | none |
| FAF1 | 0.499 | 0.4113 | 3.4684 | 0.00636 | 0.033 | -2.39 | -1.1044 | none |
| CLUAP1 | 0.757 | 0.6514 | 3.4682 | 0.00636 | 0.033 | -2.39 | -1.0328 | none |
| CNOT2 | 0.601 | 0.4853 | 3.468 | 0.00636 | 0.033 | -2.39 | -1.2736 | none |
| CAPN2 | 0.794 | 0.741 | 3.4679 | 0.00636 | 0.033 | -2.39 | -0.1017 | none |
| CD302 | 0.378 | 0.2764 | 3.4674 | 0.00637 | 0.033 | -2.39 | -0.7664 | none |
| COG3 | 0.644 | 0.5667 | 3.467 | 0.00637 | 0.033 | -2.39 | -0.4124 | none |
| ACBD4 | 0.615 | 0.4619 | 3.4666 | 0.00638 | 0.033 | -2.39 | -0.3079 | none |
| FUZ | 0.537 | 0.4051 | 3.466 | 0.00638 | 0.033 | -2.39 | -1.1707 | none |
| BTBD8 | 0.642 | 0.5364 | 3.4658 | 0.00639 | 0.033 | -2.39 | -0.8535 | none |
| POLR2J3 | 0.647 | 0.4891 | 3.465 | 0.00639 | 0.033 | -2.39 | -1.2501 | none |
| TMEM138 | 0.383 | 0.2616 | 3.4647 | 0.0064 | 0.033 | -2.39 | -1.3745 | none |
| MFN1 | 0.347 | 0.2393 | 3.4638 | 0.00641 | 0.033 | -2.4 | -0.6125 | none |
| MIB2 | 0.403 | 0.3151 | 3.4631 | 0.00641 | 0.033 | -2.4 | 0.28767 | none |
| DCTN4 | 0.579 | 0.4368 | 3.4629 | 0.00642 | 0.033 | -2.4 | -0.9506 | none |
| ZNF57 | 0.456 | 0.3622 | 3.4628 | 0.00642 | 0.033 | -2.4 | -0.3052 | none |
| CAPN7 | 0.734 | 0.6356 | 3.4623 | 0.00642 | 0.033 | -2.4 | -0.9417 | none |
| ZNF610 | 0.551 | 0.4761 | 3.4604 | 0.00644 | 0.033 | -2.4 | -0.7823 | none |
| AKAP7 | 0.684 | 0.5336 | 3.4596 | 0.00645 | 0.033 | -2.4 | -0.8553 | none |
| RIMBP3C | 0.323 | 0.2608 | 3.4591 | 0.00646 | 0.033 | -2.4 | -1.3523 | none |
| SLX4 | 0.67 | 0.647 | 3.4572 | 0.00648 | 0.033 | -2.41 | -1.0079 | none |
| RPL7L1 | 0.765 | 0.6963 | 3.4561 | 0.00649 | 0.034 | -2.41 | -1.0145 | none |
| CCNJL | 0.75 | 0.5727 | 3.4555 | 0.0065 | 0.034 | -2.41 | -1.2457 | none |
| VAMP4 | -0.505 | -0.4835 | -3.455 | 0.0065 | 0.034 | -2.41 | 1.41059 | none |
| PAG1 | 0.792 | 0.6346 | 3.455 | 0.0065 | 0.034 | -2.41 | -0.922 | none |
| PECAM1 | 0.539 | 0.4278 | 3.4537 | 0.00651 | 0.034 | -2.41 | -0.4763 | none |
| IRS2 | 0.346 | 0.2471 | 3.4537 | 0.00652 | 0.034 | -2.41 | -1.3533 | none |

| | | | | | | | | |
|---|---|---|---|---|---|---|---|---|
| CYB5D1 | 0.544 | 0.4772 | 3.4534 | 0.00652 | 0.034 | -2.41 | -1.3255 | none |
| CHMP7 | 0.337 | 0.2834 | 3.4532 | 0.00652 | 0.034 | -2.41 | -1.0661 | none |
| FGF13 | 0.467 | 0.3454 | 3.453 | 0.00652 | 0.034 | -2.41 | -1.3277 | none |
| NLGN4Y | 0.408 | 0.2863 | 3.4529 | 0.00652 | 0.034 | -2.41 | 0.20554 | none |
| GMEB1 | 0.571 | 0.4522 | 3.4523 | 0.00653 | 0.034 | -2.41 | -1.1719 | none |
| TNFRSF13I | 0.391 | 0.325 | 3.4511 | 0.00654 | 0.034 | -2.42 | -0.5374 | none |
| PMS2 | 0.629 | 0.5806 | 3.4511 | 0.00654 | 0.034 | -2.42 | -0.905 | none |
| NFKBIB | 0.458 | 0.3957 | 3.4507 | 0.00655 | 0.034 | -2.42 | -1.401 | none |
| SERPINB9 | 0.326 | 0.2147 | 3.4498 | 0.00656 | 0.034 | -2.42 | -1.537 | OA-related |
| PHKG1 | 0.639 | 0.4765 | 3.4494 | 0.00656 | 0.034 | -2.42 | -0.0438 | none |
| ATP13A1 | 0.554 | 0.5607 | 3.4488 | 0.00657 | 0.034 | -2.42 | -1.0693 | none |
| EIF6 | -0.403 | -0.3505 | -3.447 | 0.00658 | 0.034 | -2.42 | 2.4786 | AA-related |
| CENPI | 0.577 | 0.4714 | 3.4452 | 0.00661 | 0.034 | -2.43 | -1.0363 | none |
| FOXRED2 | 0.653 | 0.6119 | 3.445 | 0.00661 | 0.034 | -2.43 | -0.8041 | none |
| OGT | 0.551 | 0.4391 | 3.4444 | 0.00661 | 0.034 | -2.43 | -0.9937 | none |
| DBI | -0.367 | -0.3553 | -3.444 | 0.00662 | 0.034 | -2.43 | 2.20389 | AA-related |
| ICMT | 0.382 | 0.3165 | 3.4431 | 0.00663 | 0.034 | -2.43 | 0.69302 | none |
| CTDSPL2 | 0.469 | 0.3182 | 3.4427 | 0.00663 | 0.034 | -2.43 | -0.6671 | none |
| AP2A2 | 0.835 | 0.7455 | 3.4412 | 0.00665 | 0.034 | -2.43 | -1.364 | none |
| PNKP | 0.382 | 0.298 | 3.441 | 0.00665 | 0.034 | -2.43 | -0.9828 | none |
| KAT7 | 0.546 | 0.46 | 3.4406 | 0.00666 | 0.034 | -2.43 | -1.3955 | none |
| CRELD1 | 0.461 | 0.3709 | 3.4402 | 0.00666 | 0.034 | -2.43 | -0.3246 | none |
| TRRAP | 1.134 | 1.0895 | 3.4395 | 0.00667 | 0.034 | -2.44 | -1.0881 | none |
| KLHL30 | 0.291 | 0.208 | 3.4376 | 0.00669 | 0.034 | -2.44 | -1.3532 | none |
| NEU3 | 0.349 | 0.3388 | 3.4376 | 0.00669 | 0.034 | -2.44 | -0.4133 | none |
| HLTF | 1.001 | 0.8931 | 3.4364 | 0.0067 | 0.034 | -2.44 | -1.2837 | none |
| VHL | 0.713 | 0.6511 | 3.4362 | 0.00671 | 0.034 | -2.44 | -0.7946 | none |
| RANBP3 | 0.815 | 0.747 | 3.436 | 0.00671 | 0.034 | -2.44 | -1.2939 | none |
| TMEM38A | 0.535 | 0.3783 | 3.4357 | 0.00671 | 0.034 | -2.44 | -0.4929 | none |
| BUB1B | 0.818 | 0.7283 | 3.4347 | 0.00672 | 0.034 | -2.44 | -0.9985 | none |
| HSH2D | 0.424 | 0.3983 | 3.4336 | 0.00673 | 0.034 | -2.44 | -1.2306 | none |
| JADE1 | 0.383 | 0.3395 | 3.4315 | 0.00676 | 0.034 | -2.45 | -0.8586 | none |
| RPP38 | 0.334 | 0.2881 | 3.4311 | 0.00676 | 0.035 | -2.45 | -0.19 | none |
| NUMBL | 0.616 | 0.5666 | 3.4283 | 0.00679 | 0.035 | -2.45 | -0.6748 | none |
| INSL3 | 0.6 | 0.469 | 3.4282 | 0.00679 | 0.035 | -2.45 | -0.5981 | none |
| BIVM | 0.547 | 0.4968 | 3.428 | 0.0068 | 0.035 | -2.45 | -0.6042 | none |
| LONRF2 | 0.331 | 0.2405 | 3.4269 | 0.00681 | 0.035 | -2.46 | -1.323 | none |
| STRBP | 0.898 | 0.8092 | 3.4265 | 0.00681 | 0.035 | -2.46 | -1.098 | none |
| OR8H2 | 0.397 | 0.2838 | 3.4264 | 0.00681 | 0.035 | -2.46 | -1.2427 | none |
| PEX11G | -0.626 | -0.5186 | -3.426 | 0.00682 | 0.035 | -2.46 | 0.17983 | none |

| | | | | | | | | |
|---|---|---|---|---|---|---|---|---|
| PCDHGA1 | 0.297 | 0.2202 | 3.4253 | 0.00683 | 0.035 | -2.46 | -1.2011 | none |
| NDUFS1 | 0.257 | 0.1721 | 3.4247 | 0.00683 | 0.035 | -2.46 | 0.33311 | none |
| NACA | -0.275 | -0.2291 | -3.424 | 0.00684 | 0.035 | -2.46 | 2.04356 | none |
| PPARGC1E | 0.523 | 0.5026 | 3.4239 | 0.00684 | 0.035 | -2.46 | -0.8926 | none |
| NCR1 | 0.327 | 0.2108 | 3.4239 | 0.00684 | 0.035 | -2.46 | -1.1223 | none |
| ATP1A2 | 0.531 | 0.426 | 3.4237 | 0.00684 | 0.035 | -2.46 | -0.9078 | none |
| RASSF8 | 0.627 | 0.506 | 3.4236 | 0.00685 | 0.035 | -2.46 | -1.2523 | none |
| TSPYL1 | 0.496 | 0.3619 | 3.4231 | 0.00685 | 0.035 | -2.46 | -1.5314 | none |
| MRPS17 | 0.593 | 0.411 | 3.4218 | 0.00687 | 0.035 | -2.46 | -1.2158 | none |
| CHRNA5 | 0.518 | 0.3279 | 3.421 | 0.00687 | 0.035 | -2.47 | -1.0343 | none |
| EEF1AKMT | -0.388 | -0.3678 | -3.421 | 0.00688 | 0.035 | -2.47 | 1.1453 | none |
| SAMD4A | 0.549 | 0.3537 | 3.4207 | 0.00688 | 0.035 | -2.47 | -1.486 | none |
| MATK | 0.38 | 0.2935 | 3.4202 | 0.00688 | 0.035 | -2.47 | -1.204 | none |
| HIST1H4E | 0.303 | 0.1853 | 3.4201 | 0.00688 | 0.035 | -2.47 | -0.4683 | none |
| MON2 | 0.67 | 0.5905 | 3.4198 | 0.00689 | 0.035 | -2.47 | -1.0284 | none |
| CCT7 | -0.337 | -0.2845 | -3.419 | 0.0069 | 0.035 | -2.47 | 2.13855 | AA-related |
| SCO1 | 0.307 | 0.262 | 3.4178 | 0.00691 | 0.035 | -2.47 | -1.003 | none |
| SGO2 | 1.014 | 0.8924 | 3.415 | 0.00694 | 0.035 | -2.48 | -1.2026 | none |
| ABHD2 | 0.456 | 0.2934 | 3.4147 | 0.00695 | 0.035 | -2.48 | -0.4922 | none |
| RGR | 0.387 | 0.2893 | 3.4145 | 0.00695 | 0.035 | -2.48 | -0.7712 | none |
| NBEAL2 | 0.821 | 0.8804 | 3.4135 | 0.00696 | 0.035 | -2.48 | -0.7531 | none |
| TNNC1 | 0.741 | 0.6476 | 3.412 | 0.00698 | 0.035 | -2.48 | -0.0313 | none |
| GPR141 | 0.565 | 0.388 | 3.412 | 0.00698 | 0.035 | -2.48 | 0.28158 | none |
| DEF8 | 0.428 | 0.3381 | 3.4117 | 0.00698 | 0.035 | -2.48 | -1.1658 | none |
| VPS39 | 0.769 | 0.722 | 3.4105 | 0.00699 | 0.035 | -2.48 | -1.1279 | none |
| HIST2H3A | 0.331 | 0.2571 | 3.4061 | 0.00704 | 0.036 | -2.49 | -1.3184 | none |
| HIST2H3C | 0.331 | 0.2571 | 3.4061 | 0.00704 | 0.036 | -2.49 | -1.3184 | none |
| CYLD | 0.391 | 0.2568 | 3.4058 | 0.00705 | 0.036 | -2.49 | -0.6147 | none |
| KLHL11 | 0.481 | 0.4027 | 3.4055 | 0.00705 | 0.036 | -2.49 | -1.1466 | none |
| PKM | -0.288 | -0.1753 | -3.404 | 0.00707 | 0.036 | -2.49 | 2.01049 | none |
| UGDH | 0.327 | 0.2164 | 3.4034 | 0.00708 | 0.036 | -2.49 | -0.9694 | none |
| SUZ12 | 0.857 | 0.7544 | 3.4031 | 0.00708 | 0.036 | -2.49 | -1.2167 | none |
| FABP5 | -0.506 | -0.4523 | -3.403 | 0.00709 | 0.036 | -2.5 | 2.23443 | AA-related |
| POMT1 | 0.321 | 0.1978 | 3.4014 | 0.0071 | 0.036 | -2.5 | -1.0279 | none |
| DGKB | 0.77 | 0.5471 | 3.401 | 0.00711 | 0.036 | -2.5 | -1.4286 | none |
| STX7 | 0.524 | 0.3491 | 3.4009 | 0.00711 | 0.036 | -2.5 | -0.8917 | none |
| DCLK1 | 0.404 | 0.3389 | 3.3986 | 0.00713 | 0.036 | -2.5 | -0.2892 | none |
| C9ORF50 | 0.448 | 0.3252 | 3.3979 | 0.00714 | 0.036 | -2.5 | -1.2214 | none |
| AP4S1 | 0.547 | 0.3764 | 3.3978 | 0.00714 | 0.036 | -2.5 | -1.0252 | none |
| CENPK | 0.577 | 0.3992 | 3.3973 | 0.00715 | 0.036 | -2.5 | -0.8257 | none |

| | | | | | | | | |
|---|---|---|---|---|---|---|---|---|
| FOXJ3 | 0.49 | 0.3617 | 3.3965 | 0.00716 | 0.036 | -2.51 | -0.5553 | none |
| DMC1 | 0.53 | 0.3238 | 3.3955 | 0.00717 | 0.036 | -2.51 | -0.8634 | none |
| PRRT3 | 0.531 | 0.5026 | 3.3944 | 0.00718 | 0.036 | -2.51 | -0.8227 | none |
| C16ORF70 | 0.4 | 0.2731 | 3.3942 | 0.00719 | 0.036 | -2.51 | -1.0898 | none |
| MTRNR2L1 | -0.252 | -0.1635 | -3.394 | 0.00719 | 0.036 | -2.51 | 1.00509 | none |
| KCNC2 | 0.792 | 0.66 | 3.3929 | 0.0072 | 0.036 | -2.51 | 0.36157 | none |
| CLDN5 | 0.385 | 0.3163 | 3.3929 | 0.0072 | 0.036 | -2.51 | -0.4368 | none |
| ZNF417 | 0.59 | 0.5419 | 3.3919 | 0.00721 | 0.036 | -2.51 | -0.3538 | none |
| RAD51 | 0.65 | 0.4925 | 3.3916 | 0.00722 | 0.036 | -2.51 | -1.3986 | none |
| CSGALNAC | 0.661 | 0.5731 | 3.391 | 0.00722 | 0.036 | -2.51 | -1.1106 | none |
| PTPRZ1 | 0.765 | 0.6152 | 3.3907 | 0.00723 | 0.036 | -2.52 | -0.8192 | none |
| AKIP1 | 0.457 | 0.2817 | 3.3905 | 0.00723 | 0.036 | -2.52 | -1.3363 | none |
| GNE | 0.757 | 0.649 | 3.3904 | 0.00723 | 0.036 | -2.52 | -0.4279 | none |
| SEMA3B | 0.306 | 0.1804 | 3.3889 | 0.00725 | 0.036 | -2.52 | -0.6754 | none |
| NUP188 | 0.744 | 0.7028 | 3.3888 | 0.00725 | 0.036 | -2.52 | -1.047 | none |
| NFE2L2 | 0.465 | 0.4169 | 3.3887 | 0.00725 | 0.036 | -2.52 | -0.4814 | none |
| SCUBE2 | 0.699 | 0.5765 | 3.3879 | 0.00726 | 0.036 | -2.52 | 0.47672 | none |
| RINT1 | 0.767 | 0.5439 | 3.3875 | 0.00726 | 0.036 | -2.52 | -0.6133 | none |
| ALDH1L1 | 0.627 | 0.4856 | 3.3867 | 0.00727 | 0.036 | -2.52 | -0.9355 | none |
| DYNLRB1 | -0.391 | -0.3936 | -3.387 | 0.00728 | 0.036 | -2.52 | 2.12252 | AA-related |
| SULT1A4 | 0.623 | 0.4495 | 3.3863 | 0.00728 | 0.036 | -2.52 | -0.65 | none |
| BRCA1 | 1.066 | 1.0199 | 3.3859 | 0.00728 | 0.036 | -2.52 | -1.1712 | none |
| TEX9 | 0.639 | 0.4933 | 3.3843 | 0.0073 | 0.036 | -2.53 | -1.4599 | none |
| RNF135 | 0.372 | 0.2769 | 3.3839 | 0.00731 | 0.037 | -2.53 | -0.0818 | none |
| AC010287 | 1.046 | 0.8163 | 3.3833 | 0.00732 | 0.037 | -2.53 | -1.0849 | none |
| SEC61A2 | 0.443 | 0.301 | 3.3833 | 0.00732 | 0.037 | -2.53 | -0.1712 | none |
| RMDN1 | 0.455 | 0.2972 | 3.3832 | 0.00732 | 0.037 | -2.53 | -0.8447 | none |
| MUC1 | 0.546 | 0.6151 | 3.3819 | 0.00733 | 0.037 | -2.53 | 0.82845 | none |
| ZYG11B | 0.464 | 0.3422 | 3.3808 | 0.00735 | 0.037 | -2.53 | -0.9675 | none |
| VEGFA | 0.376 | 0.4307 | 3.3808 | 0.00735 | 0.037 | -2.53 | 0.60034 | none |
| SLC2A11 | 0.439 | 0.2203 | 3.3802 | 0.00735 | 0.037 | -2.53 | -1.0495 | none |
| EPM2AIP1 | 0.525 | 0.3615 | 3.3797 | 0.00736 | 0.037 | -2.53 | -1.444 | none |
| NR1I3 | 0.697 | 0.5828 | 3.379 | 0.00737 | 0.037 | -2.53 | -0.8323 | none |
| ENTPD3 | 1.019 | 0.702 | 3.3786 | 0.00737 | 0.037 | -2.53 | -0.3099 | none |
| PLCB1 | 0.789 | 0.5787 | 3.3784 | 0.00738 | 0.037 | -2.54 | -0.5538 | none |
| TMEM241 | 0.378 | 0.2315 | 3.3782 | 0.00738 | 0.037 | -2.54 | -0.5259 | none |
| PLBD2 | 0.642 | 0.5809 | 3.3781 | 0.00738 | 0.037 | -2.54 | -0.7482 | none |
| FBXO48 | 0.38 | 0.2816 | 3.378 | 0.00738 | 0.037 | -2.54 | -1.4031 | none |
| PIGG | 0.541 | 0.4274 | 3.3775 | 0.00739 | 0.037 | -2.54 | -1.1468 | none |
| CRKL | 0.441 | 0.3638 | 3.3768 | 0.00739 | 0.037 | -2.54 | 0.15089 | none |

| | | | | | | | | |
|---|---|---|---|---|---|---|---|---|
| ELL | 0.47 | 0.425 | 3.3764 | 0.0074 | 0.037 | -2.54 | 0.24802 | none |
| RNF149 | 0.364 | 0.2169 | 3.3763 | 0.0074 | 0.037 | -2.54 | -0.6258 | none |
| AMER3 | 0.291 | 0.2195 | 3.3744 | 0.00742 | 0.037 | -2.54 | -1.1084 | none |
| NPIPB3 | 1.006 | 1.0493 | 3.3744 | 0.00742 | 0.037 | -2.54 | -0.7325 | none |
| GLCCI1 | 0.491 | 0.3715 | 3.3743 | 0.00742 | 0.037 | -2.54 | -1.0114 | none |
| METTL2A | 0.388 | 0.266 | 3.3741 | 0.00743 | 0.037 | -2.54 | -0.7499 | none |
| OR4F15 | 0.492 | 0.3167 | 3.3738 | 0.00743 | 0.037 | -2.54 | -1.2413 | none |
| DDX3X | 0.573 | 0.4881 | 3.3736 | 0.00743 | 0.037 | -2.54 | -1.1008 | none |
| C10ORF95 | 0.418 | 0.302 | 3.3735 | 0.00743 | 0.037 | -2.54 | -1.431 | none |
| DNAJA4 | 0.484 | 0.4636 | 3.3728 | 0.00744 | 0.037 | -2.54 | -1.1793 | none |
| SV2A | 0.477 | 0.3025 | 3.3726 | 0.00745 | 0.037 | -2.54 | -1.4179 | none |
| IBTK | 0.398 | 0.2959 | 3.3726 | 0.00745 | 0.037 | -2.54 | -1.3614 | none |
| PRPH2 | 0.335 | 0.2125 | 3.3723 | 0.00745 | 0.037 | -2.55 | -0.6873 | none |
| GGT1 | -0.341 | -0.2647 | -3.372 | 0.00745 | 0.037 | -2.55 | 2.56722 | AA-related |
| STK10 | 0.87 | 0.8492 | 3.3717 | 0.00746 | 0.037 | -2.55 | -0.9953 | none |
| C12ORF66 | 0.404 | 0.2627 | 3.3715 | 0.00746 | 0.037 | -2.55 | -0.51 | none |
| RASA4B | 0.509 | 0.5035 | 3.3712 | 0.00746 | 0.037 | -2.55 | -0.6273 | none |
| ZNF354A | 0.646 | 0.599 | 3.3702 | 0.00748 | 0.037 | -2.55 | -0.4401 | none |
| ZNF266 | 0.502 | 0.4408 | 3.37 | 0.00748 | 0.037 | -2.55 | -0.9908 | none |
| GRPEL1 | -0.283 | -0.2407 | -3.37 | 0.00748 | 0.037 | -2.55 | 2.56947 | AA-related |
| TTF1 | 0.954 | 0.8815 | 3.3699 | 0.00748 | 0.037 | -2.55 | -1.1403 | none |
| GUF1 | 0.454 | 0.3582 | 3.3698 | 0.00748 | 0.037 | -2.55 | -0.7018 | none |
| LRAT | 0.318 | 0.2272 | 3.3692 | 0.00749 | 0.037 | -2.55 | -1.139 | none |
| NBPF19 | 0.672 | 0.7281 | 3.3689 | 0.00749 | 0.037 | -2.55 | -0.7196 | none |
| UBE2G2 | 0.41 | 0.3224 | 3.3683 | 0.0075 | 0.037 | -2.55 | -0.5038 | none |
| PER2 | 0.749 | 0.6649 | 3.3679 | 0.0075 | 0.037 | -2.55 | -1.1944 | none |
| MPLKIP | 0.457 | 0.2859 | 3.3663 | 0.00752 | 0.037 | -2.56 | -1.0443 | none |
| CHDH | 0.329 | 0.208 | 3.3646 | 0.00754 | 0.037 | -2.56 | -1.3284 | none |
| CPS1 | 0.557 | 0.417 | 3.3636 | 0.00756 | 0.037 | -2.56 | -1.0124 | none |
| XPO4 | 0.418 | 0.2884 | 3.3636 | 0.00756 | 0.037 | -2.56 | 0.27736 | none |
| DUSP2 | -0.638 | -0.4856 | -3.363 | 0.00756 | 0.037 | -2.56 | 1.23014 | none |
| BSDC1 | 0.418 | 0.3875 | 3.363 | 0.00756 | 0.037 | -2.56 | -1.0358 | none |
| NOSTRIN | 0.67 | 0.4591 | 3.3627 | 0.00757 | 0.037 | -2.56 | -0.2448 | none |
| TESC | 0.413 | 0.2959 | 3.3626 | 0.00757 | 0.037 | -2.56 | -0.6673 | none |
| C9ORF57 | 0.291 | 0.2037 | 3.3621 | 0.00758 | 0.037 | -2.56 | -1.3289 | none |
| KYAT1 | 0.903 | 0.7406 | 3.362 | 0.00758 | 0.037 | -2.56 | -0.223 | none |
| CLASP1 | 0.976 | 0.9474 | 3.3612 | 0.00759 | 0.037 | -2.56 | -0.7627 | none |
| EHD2 | 0.627 | 0.4011 | 3.3611 | 0.00759 | 0.037 | -2.56 | -1.1108 | none |
| ZNF256 | 0.35 | 0.2661 | 3.3606 | 0.00759 | 0.037 | -2.56 | 0.71933 | none |
| ZNF813 | 0.597 | 0.5139 | 3.3596 | 0.00761 | 0.037 | -2.57 | -1.4386 | none |

| | | | | | | | | |
|---|---|---|---|---|---|---|---|---|
| EPPIN | 0.318 | 0.2178 | 3.3589 | 0.00762 | 0.037 | -2.57 | -0.6803 | none |
| PEX5L | 0.331 | 0.2783 | 3.3578 | 0.00763 | 0.037 | -2.57 | -1.2571 | none |
| PREPL | 0.856 | 0.6766 | 3.3573 | 0.00764 | 0.038 | -2.57 | -1.1365 | none |
| P2RX7 | 0.68 | 0.63 | 3.3567 | 0.00764 | 0.038 | -2.57 | -1.2422 | none |
| SLC9A1 | 0.645 | 0.6178 | 3.3565 | 0.00765 | 0.038 | -2.57 | -1.1104 | none |
| MYO1F | 0.801 | 0.7135 | 3.3564 | 0.00765 | 0.038 | -2.57 | -1.42 | none |
| ESPN | -0.33 | -0.2186 | -3.356 | 0.00765 | 0.038 | -2.57 | 2.56567 | AA-related |
| PLIN2 | 0.381 | 0.3279 | 3.3549 | 0.00767 | 0.038 | -2.57 | 0.45113 | none |
| YWHAZ | -0.278 | -0.2665 | -3.355 | 0.00767 | 0.038 | -2.57 | 1.5326 | none |
| ZNF860 | 0.282 | 0.1988 | 3.3548 | 0.00767 | 0.038 | -2.57 | -1.4644 | none |
| WDTC1 | 0.397 | 0.36 | 3.3542 | 0.00768 | 0.038 | -2.58 | -0.8334 | none |
| EID1 | -0.352 | -0.3219 | -3.353 | 0.00769 | 0.038 | -2.58 | 2.30876 | AA-related |
| SNRPD1 | -0.473 | -0.4344 | -3.353 | 0.0077 | 0.038 | -2.58 | 2.51144 | AA-related |
| GJB4 | 0.279 | 0.202 | 3.3517 | 0.00771 | 0.038 | -2.58 | -1.5827 | OA-related |
| SPACA7 | 0.364 | 0.2836 | 3.3516 | 0.00771 | 0.038 | -2.58 | -0.8178 | none |
| PDK1 | 0.409 | 0.3442 | 3.3513 | 0.00771 | 0.038 | -2.58 | 0.15638 | none |
| EMC4 | 0.439 | 0.2735 | 3.351 | 0.00772 | 0.038 | -2.58 | -1.2936 | none |
| MCFD2 | 0.556 | 0.4393 | 3.3503 | 0.00772 | 0.038 | -2.58 | -1.4264 | none |
| ATF6B | 0.782 | 0.5861 | 3.35 | 0.00773 | 0.038 | -2.58 | -1.1114 | none |
| COPS2 | 0.58 | 0.4537 | 3.35 | 0.00773 | 0.038 | -2.58 | -0.7168 | none |
| EXOSC2 | 0.384 | 0.2943 | 3.3495 | 0.00774 | 0.038 | -2.58 | -1.1749 | none |
| ADAM32 | 0.635 | 0.5278 | 3.3495 | 0.00774 | 0.038 | -2.58 | -0.2171 | none |
| SP140L | 0.869 | 0.7877 | 3.3477 | 0.00776 | 0.038 | -2.59 | -1.1274 | none |
| P3H2 | 0.873 | 0.7253 | 3.347 | 0.00777 | 0.038 | -2.59 | -1.3577 | none |
| NAP1L1 | -0.269 | -0.2363 | -3.347 | 0.00777 | 0.038 | -2.59 | 1.85865 | none |
| ST8SIA4 | 0.462 | 0.3368 | 3.3466 | 0.00777 | 0.038 | -2.59 | -0.2619 | none |
| IFI35 | 0.413 | 0.2447 | 3.3465 | 0.00777 | 0.038 | -2.59 | -1.1304 | none |
| EGFL7 | -0.354 | -0.315 | -3.346 | 0.00777 | 0.038 | -2.59 | 1.73783 | none |
| XRCC2 | 0.508 | 0.471 | 3.3463 | 0.00778 | 0.038 | -2.59 | -1.3273 | none |
| KCNAB3 | 0.594 | 0.5227 | 3.3444 | 0.0078 | 0.038 | -2.59 | -0.9586 | none |
| ANKRD9 | 0.539 | 0.4264 | 3.3439 | 0.00781 | 0.038 | -2.59 | -1.1352 | none |
| SLC2A8 | 0.578 | 0.4403 | 3.3436 | 0.00781 | 0.038 | -2.59 | -0.2691 | none |
| ZDBF2 | 0.456 | 0.4298 | 3.3433 | 0.00782 | 0.038 | -2.59 | -0.9008 | none |
| EPS8L2 | -0.709 | -0.4705 | -3.343 | 0.00782 | 0.038 | -2.59 | 1.8698 | none |
| BAHCC1 | -0.28 | -0.1958 | -3.342 | 0.00783 | 0.038 | -2.59 | 1.26297 | none |
| TMSB4X | -0.442 | -0.4214 | -3.342 | 0.00783 | 0.038 | -2.6 | 2.04448 | none |
| DRAM1 | 0.535 | 0.3373 | 3.3417 | 0.00784 | 0.038 | -2.6 | -0.9848 | none |
| SAMHD1 | 0.656 | 0.5206 | 3.341 | 0.00785 | 0.038 | -2.6 | -1.2153 | none |
| ERGIC1 | 0.293 | 0.2144 | 3.3389 | 0.00787 | 0.038 | -2.6 | -1.1261 | none |
| GRM7 | 0.514 | 0.4089 | 3.3383 | 0.00788 | 0.038 | -2.6 | -1.2804 | none |

| | | | | | | | | |
|---|---|---|---|---|---|---|---|---|
| AVIL | 0.599 | 0.4743 | 3.3378 | 0.00789 | 0.038 | -2.6 | -0.7591 | none |
| IFNAR1 | 0.632 | 0.5233 | 3.3373 | 0.00789 | 0.038 | -2.6 | -1.0185 | none |
| CPT2 | 0.53 | 0.4371 | 3.3373 | 0.00789 | 0.038 | -2.6 | -0.457 | none |
| HLA-G | 0.746 | 0.4802 | 3.3362 | 0.00791 | 0.038 | -2.6 | -0.3641 | none |
| ZC3H10 | 0.491 | 0.3989 | 3.3359 | 0.00791 | 0.038 | -2.6 | -0.843 | none |
| ADRA1A | 0.613 | 0.3853 | 3.3348 | 0.00793 | 0.038 | -2.61 | -0.9272 | none |
| TMEM231 | 0.456 | 0.2621 | 3.334 | 0.00794 | 0.038 | -2.61 | -0.9272 | none |
| ZNF736 | 0.483 | 0.2999 | 3.3334 | 0.00794 | 0.039 | -2.61 | -0.7708 | none |
| GALR1 | 0.267 | 0.192 | 3.3331 | 0.00795 | 0.039 | -2.61 | -0.8349 | none |
| BDKRB1 | 0.594 | 0.44 | 3.3324 | 0.00796 | 0.039 | -2.61 | -0.9104 | none |
| C3ORF49 | 1.258 | 0.9439 | 3.3322 | 0.00796 | 0.039 | -2.61 | -1.4252 | none |
| QKI | 0.66 | 0.5491 | 3.3322 | 0.00796 | 0.039 | -2.61 | -1.473 | none |
| SLC14A1 | 0.581 | 0.4999 | 3.3306 | 0.00798 | 0.039 | -2.61 | -1.2378 | none |
| GPATCH8 | 0.979 | 0.9762 | 3.3304 | 0.00798 | 0.039 | -2.61 | -0.8007 | none |
| CLEC7A | 0.338 | 0.1838 | 3.3303 | 0.00798 | 0.039 | -2.61 | -0.9946 | none |
| GNAI3 | 0.278 | 0.1561 | 3.3293 | 0.008 | 0.039 | -2.62 | -0.8184 | none |
| C3ORF22 | 0.274 | 0.274 | 3.3288 | 0.008 | 0.039 | -2.62 | -0.2214 | none |
| TRPM7 | 0.934 | 0.8215 | 3.3287 | 0.00801 | 0.039 | -2.62 | -1.3666 | none |
| RRNAD1 | 0.663 | 0.5131 | 3.3282 | 0.00801 | 0.039 | -2.62 | -1.0399 | none |
| FBXO45 | 0.377 | 0.2592 | 3.3281 | 0.00801 | 0.039 | -2.62 | -1.1896 | none |
| ESR2 | 0.421 | 0.3122 | 3.3273 | 0.00802 | 0.039 | -2.62 | -1.1269 | none |
| AAK1 | 0.617 | 0.5613 | 3.3273 | 0.00803 | 0.039 | -2.62 | -1.0067 | none |
| MBNL3 | 0.498 | 0.3872 | 3.327 | 0.00803 | 0.039 | -2.62 | -1.2094 | none |
| CD27 | -0.383 | -0.333 | -3.327 | 0.00803 | 0.039 | -2.62 | 1.75282 | none |
| DPP9 | 0.645 | 0.6084 | 3.3266 | 0.00803 | 0.039 | -2.62 | -0.9692 | none |
| NR2C1 | 0.571 | 0.4854 | 3.3255 | 0.00805 | 0.039 | -2.62 | -0.9745 | none |
| TUBGCP5 | 0.671 | 0.5674 | 3.3251 | 0.00805 | 0.039 | -2.62 | -0.2 | none |
| GLYATL2 | 0.307 | 0.2377 | 3.3248 | 0.00806 | 0.039 | -2.62 | -0.5341 | none |
| ANKRD36( | 0.588 | 0.5488 | 3.3237 | 0.00807 | 0.039 | -2.62 | -0.4197 | none |
| RNASEL | 0.432 | 0.3755 | 3.3235 | 0.00808 | 0.039 | -2.63 | -1.1063 | none |
| CCDC8 | 0.267 | 0.2115 | 3.3234 | 0.00808 | 0.039 | -2.63 | -1.2499 | none |
| PAFAH1B1 | 0.376 | 0.2602 | 3.3233 | 0.00808 | 0.039 | -2.63 | -1.5066 | none |
| FPR1 | 0.437 | 0.3356 | 3.3232 | 0.00808 | 0.039 | -2.63 | -0.2606 | none |
| G6PC3 | 0.548 | 0.4433 | 3.3231 | 0.00808 | 0.039 | -2.63 | -1.0956 | none |
| ZNF614 | 0.74 | 0.6013 | 3.3222 | 0.00809 | 0.039 | -2.63 | -1.4515 | none |
| EDA | 0.551 | 0.4423 | 3.322 | 0.0081 | 0.039 | -2.63 | -0.2186 | none |
| WBP1L | 0.46 | 0.3535 | 3.3218 | 0.0081 | 0.039 | -2.63 | -1.1911 | none |
| C14ORF93 | 0.579 | 0.5022 | 3.3209 | 0.00811 | 0.039 | -2.63 | -0.8133 | none |
| DUSP4 | -0.345 | -0.2352 | -3.32 | 0.00812 | 0.039 | -2.63 | 0.972 | none |
| PGK1 | -0.451 | -0.4075 | -3.32 | 0.00812 | 0.039 | -2.63 | 2.42384 | AA-related |

| | | | | | | | | |
|---|---|---|---|---|---|---|---|---|
| SPDYE2B | 0.631 | 0.5525 | 3.3197 | 0.00813 | 0.039 | -2.63 | -1.2749 | none |
| LRRC74B | 0.479 | 0.3678 | 3.3195 | 0.00813 | 0.039 | -2.63 | -1.4475 | none |
| IPO5 | 0.493 | 0.4264 | 3.3195 | 0.00813 | 0.039 | -2.63 | -0.8906 | none |
| ARAF | 0.459 | 0.337 | 3.3187 | 0.00814 | 0.039 | -2.63 | -1.1035 | none |
| NETO2 | 0.483 | 0.3505 | 3.3184 | 0.00814 | 0.039 | -2.63 | -0.7356 | none |
| ZNF75A | 0.627 | 0.5573 | 3.3182 | 0.00815 | 0.039 | -2.63 | -1.425 | none |
| ZNF574 | 0.411 | 0.3683 | 3.3177 | 0.00815 | 0.039 | -2.63 | -0.8001 | none |
| RBM43 | 0.5 | 0.3948 | 3.317 | 0.00816 | 0.039 | -2.64 | -1.5855 | OA-related |
| PRPSAP2 | 0.387 | 0.2289 | 3.3167 | 0.00817 | 0.039 | -2.64 | -0.9047 | none |
| AIM2 | 0.548 | 0.4317 | 3.3166 | 0.00817 | 0.039 | -2.64 | -0.9908 | none |
| RTCB | -0.302 | -0.2615 | -3.316 | 0.00817 | 0.039 | -2.64 | 2.51638 | AA-related |
| L3MBTL3 | 0.683 | 0.5503 | 3.3161 | 0.00818 | 0.039 | -2.64 | 0.06919 | none |
| MTHFD1L | 0.465 | 0.4004 | 3.3155 | 0.00818 | 0.039 | -2.64 | -1.0785 | none |
| PIPOX | 0.76 | 0.5391 | 3.3138 | 0.00821 | 0.039 | -2.64 | -0.6519 | none |
| TLR7 | 0.385 | 0.2796 | 3.3132 | 0.00821 | 0.039 | -2.64 | -1.4464 | none |
| DHX40 | 0.326 | 0.1561 | 3.3129 | 0.00822 | 0.039 | -2.64 | -0.6078 | none |
| HHLA2 | 0.765 | 0.6074 | 3.3129 | 0.00822 | 0.039 | -2.64 | -0.6034 | none |
| TMEM41B | 0.488 | 0.2984 | 3.3125 | 0.00822 | 0.039 | -2.64 | -1.0976 | none |
| RND2 | 0.543 | 0.4572 | 3.3122 | 0.00823 | 0.039 | -2.64 | -0.5789 | none |
| FANCI | 0.603 | 0.5488 | 3.3121 | 0.00823 | 0.039 | -2.64 | -0.7135 | none |
| ARFIP1 | 0.666 | 0.493 | 3.3115 | 0.00824 | 0.039 | -2.65 | -0.4533 | none |
| ZNF594 | 0.566 | 0.4844 | 3.311 | 0.00824 | 0.039 | -2.65 | -1.3089 | none |
| C7ORF43 | -0.38 | -0.2352 | -3.311 | 0.00825 | 0.039 | -2.65 | 1.29894 | none |
| TBL1X | 0.782 | 0.7111 | 3.3095 | 0.00827 | 0.039 | -2.65 | -1.0961 | none |
| FAM151B | 0.657 | 0.5452 | 3.3086 | 0.00828 | 0.039 | -2.65 | -0.9907 | none |
| SOD2 | 0.335 | 0.2039 | 3.3061 | 0.00831 | 0.04 | -2.65 | -1.2877 | none |
| PRKCD | 1.005 | 0.8463 | 3.3058 | 0.00832 | 0.04 | -2.65 | -1.1473 | none |
| CDC27 | 0.516 | 0.4863 | 3.3057 | 0.00832 | 0.04 | -2.65 | -0.7062 | none |
| CCNL1 | 0.535 | 0.5646 | 3.3055 | 0.00832 | 0.04 | -2.65 | -0.3093 | none |
| TBC1D8 | 0.793 | 0.7513 | 3.3051 | 0.00832 | 0.04 | -2.66 | -1.286 | none |
| OR3A2 | 0.496 | 0.4241 | 3.3048 | 0.00833 | 0.04 | -2.66 | -0.8731 | none |
| ELMOD1 | 0.593 | 0.4918 | 3.3045 | 0.00833 | 0.04 | -2.66 | -1.0518 | none |
| SMIM17 | 0.582 | 0.3911 | 3.3044 | 0.00833 | 0.04 | -2.66 | -0.7846 | none |
| SCLT1 | 0.322 | 0.2256 | 3.3029 | 0.00836 | 0.04 | -2.66 | -1.0959 | none |
| ZNF37A | 0.674 | 0.6218 | 3.3024 | 0.00836 | 0.04 | -2.66 | -1.3332 | none |
| COQ6 | 0.415 | 0.2748 | 3.3019 | 0.00837 | 0.04 | -2.66 | -1.0561 | none |
| GALNT10 | 0.59 | 0.5273 | 3.3009 | 0.00838 | 0.04 | -2.66 | -0.8304 | none |
| ABCA9 | 0.714 | 0.574 | 3.3006 | 0.00839 | 0.04 | -2.66 | -1.1457 | none |
| KANSL2 | 0.286 | 0.2206 | 3.3004 | 0.00839 | 0.04 | -2.66 | 0.86694 | none |
| AARS2 | 0.547 | 0.5512 | 3.3001 | 0.00839 | 0.04 | -2.66 | -0.5193 | none |

| | | | | | | | | |
|---|---|---|---|---|---|---|---|---|
| NEDD4 | 0.722 | 0.6655 | 3.2992 | 0.00841 | 0.04 | -2.67 | -1.1943 | none |
| ZFP82 | 0.368 | 0.2658 | 3.2984 | 0.00842 | 0.04 | -2.67 | -1.4142 | none |
| R3HDM1 | 1.124 | 1.0546 | 3.2976 | 0.00843 | 0.04 | -2.67 | -0.9896 | none |
| ZNF621 | 0.708 | 0.6123 | 3.2973 | 0.00843 | 0.04 | -2.67 | -1.0089 | none |
| TOR4A | 0.303 | 0.2264 | 3.2968 | 0.00844 | 0.04 | -2.67 | -1.1027 | none |
| NOTO | 0.312 | 0.2321 | 3.2956 | 0.00846 | 0.04 | -2.67 | -1.4305 | none |
| HPS3 | 0.571 | 0.4347 | 3.2954 | 0.00846 | 0.04 | -2.67 | -0.6954 | none |
| ZNF821 | 0.65 | 0.4605 | 3.2951 | 0.00846 | 0.04 | -2.67 | -0.4678 | none |
| GNB5 | 0.477 | 0.2948 | 3.2949 | 0.00847 | 0.04 | -2.67 | -1.2448 | none |
| ELL2 | 0.637 | 0.673 | 3.2947 | 0.00847 | 0.04 | -2.67 | 0.03122 | none |
| BOC | 0.265 | 0.2292 | 3.2946 | 0.00847 | 0.04 | -2.67 | 0.2988 | none |
| SAMD15 | 0.426 | 0.3119 | 3.2941 | 0.00848 | 0.04 | -2.67 | -0.9814 | none |
| ANKRD11 | 0.729 | 0.7659 | 3.294 | 0.00848 | 0.04 | -2.67 | -0.9835 | none |
| ZNF624 | 1.074 | 0.8794 | 3.2937 | 0.00848 | 0.04 | -2.67 | -0.5882 | none |
| XKR9 | 0.471 | 0.2949 | 3.2936 | 0.00849 | 0.04 | -2.67 | -1.2048 | none |
| CLK2 | 0.302 | 0.2889 | 3.2931 | 0.00849 | 0.04 | -2.68 | -0.2084 | none |
| ZFP90 | 0.823 | 0.7747 | 3.2927 | 0.0085 | 0.04 | -2.68 | -0.9506 | none |
| NEB | 0.531 | 0.4124 | 3.292 | 0.00851 | 0.04 | -2.68 | -1.4978 | none |
| ZSCAN32 | 0.543 | 0.4495 | 3.292 | 0.00851 | 0.04 | -2.68 | -1.2592 | none |
| TOP3B | 0.781 | 0.7094 | 3.2919 | 0.00851 | 0.04 | -2.68 | -1.3145 | none |
| SSBP3 | 0.772 | 0.7017 | 3.2917 | 0.00851 | 0.04 | -2.68 | -0.5398 | none |
| WDR17 | 0.69 | 0.5888 | 3.2914 | 0.00852 | 0.04 | -2.68 | -0.8972 | none |
| HNRNPUL1 | 1.059 | 1.0118 | 3.2914 | 0.00852 | 0.04 | -2.68 | -1.0991 | none |
| CALB1 | 0.477 | 0.4235 | 3.2895 | 0.00854 | 0.04 | -2.68 | 0.22532 | none |
| SNRPB2 | -0.371 | -0.3705 | -3.288 | 0.00856 | 0.04 | -2.68 | 1.77823 | none |
| RD3 | 0.358 | 0.268 | 3.2876 | 0.00857 | 0.04 | -2.68 | -1.543 | OA-related |
| SPPL3 | 0.435 | 0.3091 | 3.2874 | 0.00857 | 0.04 | -2.68 | -0.9313 | none |
| FBXL8 | 0.603 | 0.4439 | 3.2874 | 0.00857 | 0.04 | -2.68 | -1.2624 | none |
| NNT | 0.439 | 0.3499 | 3.2873 | 0.00857 | 0.04 | -2.68 | -0.1655 | none |
| MEGF9 | 0.475 | 0.3872 | 3.2868 | 0.00858 | 0.04 | -2.69 | -1.2021 | none |
| NKTR | 0.652 | 0.6003 | 3.2864 | 0.00859 | 0.04 | -2.69 | -1.0903 | none |
| KCND3 | 0.26 | 0.1886 | 3.2859 | 0.00859 | 0.04 | -2.69 | -1.4833 | none |
| ANTXR2 | 0.437 | 0.3201 | 3.2842 | 0.00862 | 0.041 | -2.69 | -0.2297 | none |
| SLC30A6 | 0.558 | 0.3181 | 3.284 | 0.00862 | 0.041 | -2.69 | -0.9484 | none |
| PLXDC2 | 0.296 | 0.202 | 3.2831 | 0.00863 | 0.041 | -2.69 | -1.5528 | OA-related |
| LRP1B | 0.351 | 0.2411 | 3.2827 | 0.00864 | 0.041 | -2.69 | -0.1858 | none |
| PLEKHH1 | 0.529 | 0.4023 | 3.2825 | 0.00864 | 0.041 | -2.69 | -1.2801 | none |
| ZBED9 | 0.362 | 0.2703 | 3.2818 | 0.00865 | 0.041 | -2.69 | -1.415 | none |
| PEX26 | 0.541 | 0.4562 | 3.2813 | 0.00866 | 0.041 | -2.69 | 0.02294 | none |
| CAPN8 | 0.871 | 0.7264 | 3.2804 | 0.00867 | 0.041 | -2.7 | -0.2069 | none |

| | | | | | | | | |
|---|---|---|---|---|---|---|---|---|
| APBB2 | 0.869 | 0.7388 | 3.2801 | 0.00868 | 0.041 | -2.7 | -0.9882 | none |
| TMCO6 | 0.776 | 0.6895 | 3.2789 | 0.0087 | 0.041 | -2.7 | -0.2185 | none |
| EPG5 | 0.585 | 0.4801 | 3.2788 | 0.0087 | 0.041 | -2.7 | -0.9344 | none |
| SEMA3D | 0.598 | 0.481 | 3.2787 | 0.0087 | 0.041 | -2.7 | -0.9339 | none |
| PBRM1 | 0.859 | 0.7734 | 3.2786 | 0.0087 | 0.041 | -2.7 | -1.2145 | none |
| CFAP73 | 0.556 | 0.443 | 3.2768 | 0.00873 | 0.041 | -2.7 | -0.7372 | none |
| PRKX | 0.441 | 0.3216 | 3.2759 | 0.00874 | 0.041 | -2.7 | -1.0144 | none |
| ITK | 0.532 | 0.4335 | 3.2746 | 0.00876 | 0.041 | -2.71 | -1.0796 | none |
| TRPC1 | 0.636 | 0.5978 | 3.2736 | 0.00877 | 0.041 | -2.71 | -0.0128 | none |
| PTCD2 | 0.271 | 0.1795 | 3.2735 | 0.00877 | 0.041 | -2.71 | -0.377 | none |
| MYLK4 | 0.285 | 0.1837 | 3.2729 | 0.00878 | 0.041 | -2.71 | -1.1195 | none |
| PPHLN1 | 0.371 | 0.2531 | 3.2727 | 0.00879 | 0.041 | -2.71 | -0.492 | none |
| FBXO27 | 0.551 | 0.4037 | 3.2723 | 0.00879 | 0.041 | -2.71 | -1.1436 | none |
| ARRB2 | 0.561 | 0.4604 | 3.2704 | 0.00882 | 0.041 | -2.71 | -0.9519 | none |
| TGFBR3L | -0.433 | -0.3306 | -3.27 | 0.00883 | 0.041 | -2.71 | 0.65388 | none |
| XPOT | 0.41 | 0.3787 | 3.268 | 0.00885 | 0.041 | -2.72 | 0.09274 | none |
| PTTG1 | -0.46 | -0.4152 | -3.267 | 0.00887 | 0.041 | -2.72 | 2.3342 | AA-related |
| OR4E2 | 0.322 | 0.2223 | 3.2654 | 0.00889 | 0.042 | -2.72 | -1.2193 | none |
| THOC3 | 0.438 | 0.2467 | 3.2653 | 0.00889 | 0.042 | -2.72 | -0.9808 | none |
| HSF1 | 0.468 | 0.4107 | 3.2649 | 0.0089 | 0.042 | -2.72 | -0.0533 | none |
| KLHL22 | 0.396 | 0.3349 | 3.2646 | 0.0089 | 0.042 | -2.72 | -0.7863 | none |
| ZNF519 | 0.68 | 0.5569 | 3.2644 | 0.00891 | 0.042 | -2.72 | -0.9705 | none |
| ARHGEF39 | 0.323 | 0.2195 | 3.2637 | 0.00892 | 0.042 | -2.72 | -0.4865 | none |
| NUCB2 | 0.743 | 0.6125 | 3.2635 | 0.00892 | 0.042 | -2.72 | -1.3583 | none |
| SLC16A11 | -0.274 | -0.2115 | -3.263 | 0.00893 | 0.042 | -2.72 | 1.15084 | none |
| APOD | 0.676 | 0.539 | 3.2626 | 0.00893 | 0.042 | -2.73 | -0.9463 | none |
| FAM222B | 0.698 | 0.67 | 3.2611 | 0.00896 | 0.042 | -2.73 | -0.2792 | none |
| CRTC3 | 0.558 | 0.5483 | 3.261 | 0.00896 | 0.042 | -2.73 | -1.2506 | none |
| AKNAD1 | 0.38 | 0.3114 | 3.2607 | 0.00896 | 0.042 | -2.73 | -0.7262 | none |
| CCNB2 | -0.447 | -0.4245 | -3.261 | 0.00896 | 0.042 | -2.73 | 2.4439 | AA-related |
| PFKFB2 | 0.428 | 0.2582 | 3.2599 | 0.00897 | 0.042 | -2.73 | -0.2682 | none |
| SPRYD7 | 0.541 | 0.4199 | 3.2584 | 0.009 | 0.042 | -2.73 | -0.5172 | none |
| PTPN12 | 1.002 | 0.8763 | 3.2583 | 0.009 | 0.042 | -2.73 | -1.3143 | none |
| ATP6AP1L | 0.544 | 0.4195 | 3.2571 | 0.00902 | 0.042 | -2.73 | -1.3126 | none |
| CTCF | 0.863 | 0.8006 | 3.2555 | 0.00904 | 0.042 | -2.74 | -1.1788 | none |
| GPR157 | 0.786 | 0.6765 | 3.2553 | 0.00904 | 0.042 | -2.74 | -0.8063 | none |
| CBL | 0.402 | 0.3563 | 3.2548 | 0.00905 | 0.042 | -2.74 | 0.08777 | none |
| SLC35C1 | 0.33 | 0.211 | 3.2545 | 0.00905 | 0.042 | -2.74 | -0.1888 | none |
| APBA2 | -0.492 | -0.3156 | -3.253 | 0.00907 | 0.042 | -2.74 | 0.97934 | none |
| CLPX | 0.572 | 0.4951 | 3.2532 | 0.00907 | 0.042 | -2.74 | -1.0744 | none |

| | | | | | | | | |
|---|---|---|---|---|---|---|---|---|
| TARDBP | 0.299 | 0.1891 | 3.2529 | 0.00908 | 0.042 | -2.74 | -0.4236 | none |
| ST6GAL2 | 0.402 | 0.3496 | 3.2529 | 0.00908 | 0.042 | -2.74 | -1.3423 | none |
| SFXN1 | 0.245 | 0.1511 | 3.2526 | 0.00908 | 0.042 | -2.74 | -0.4817 | none |
| ZDHHC18 | 0.402 | 0.3119 | 3.2522 | 0.00909 | 0.042 | -2.74 | -1.0563 | none |
| FOXP4 | 0.785 | 0.7225 | 3.2521 | 0.00909 | 0.042 | -2.74 | -1.1434 | none |
| FMO5 | 0.533 | 0.3751 | 3.2514 | 0.0091 | 0.042 | -2.74 | -1.4268 | none |
| BAIAP2L1 | 0.635 | 0.5684 | 3.2514 | 0.0091 | 0.042 | -2.74 | -1.1898 | none |
| FUT10 | -0.341 | -0.289 | -3.251 | 0.00911 | 0.042 | -2.74 | 1.31901 | none |
| EIF1AD | 0.523 | 0.3651 | 3.2493 | 0.00913 | 0.042 | -2.75 | -1.4138 | none |
| USP37 | 0.717 | 0.6503 | 3.2476 | 0.00916 | 0.042 | -2.75 | -0.4271 | none |
| PACS1 | 0.701 | 0.642 | 3.2467 | 0.00917 | 0.042 | -2.75 | -1.1274 | none |
| KBTBD8 | 0.448 | 0.3101 | 3.2447 | 0.0092 | 0.043 | -2.75 | 0.07764 | none |
| FBRSL1 | 0.707 | 0.7618 | 3.2439 | 0.00921 | 0.043 | -2.76 | -0.1581 | none |
| TAF1 | 0.95 | 0.8246 | 3.2439 | 0.00922 | 0.043 | -2.76 | -1.0831 | none |
| GDF11 | 0.457 | 0.4036 | 3.2415 | 0.00925 | 0.043 | -2.76 | -0.8715 | none |
| DENND4C | 0.657 | 0.5806 | 3.2403 | 0.00927 | 0.043 | -2.76 | -1.19 | none |
| INPP5F | -0.307 | -0.1959 | -3.24 | 0.00927 | 0.043 | -2.76 | 1.65733 | none |
| TCF7L1 | 0.843 | 0.6036 | 3.2401 | 0.00927 | 0.043 | -2.76 | -0.5434 | none |
| CAMK2D | 0.265 | 0.199 | 3.2401 | 0.00927 | 0.043 | -2.76 | 0.33899 | none |
| PWWP2A | 0.753 | 0.6442 | 3.2391 | 0.00929 | 0.043 | -2.76 | -1.2197 | none |
| GCLM | 0.371 | 0.1734 | 3.2389 | 0.00929 | 0.043 | -2.76 | -1.0753 | none |
| SLFN12L | 0.576 | 0.4781 | 3.2383 | 0.0093 | 0.043 | -2.77 | -1.2368 | none |
| PPM1K | 0.41 | 0.3134 | 3.2376 | 0.00931 | 0.043 | -2.77 | -1.3095 | none |
| FAM98B | 0.624 | 0.5279 | 3.2375 | 0.00931 | 0.043 | -2.77 | -1.0653 | none |
| FAM47E-S | 0.242 | 0.1358 | 3.2375 | 0.00931 | 0.043 | -2.77 | -0.135 | none |
| PDZK1 | 0.515 | 0.4303 | 3.237 | 0.00932 | 0.043 | -2.77 | -1.0736 | none |
| DZIP1 | 0.294 | 0.2108 | 3.2367 | 0.00933 | 0.043 | -2.77 | -1.6189 | OA-related |
| LDHAL6A | 0.544 | 0.3922 | 3.2363 | 0.00933 | 0.043 | -2.77 | -1.2952 | none |
| C2CD2L | 0.593 | 0.5477 | 3.2349 | 0.00935 | 0.043 | -2.77 | -0.651 | none |
| LRP11 | 0.312 | 0.1722 | 3.2347 | 0.00936 | 0.043 | -2.77 | -1.2198 | none |
| GADD45A | 0.388 | 0.3628 | 3.2344 | 0.00936 | 0.043 | -2.77 | 0.08492 | none |
| BCL7A | 0.467 | 0.402 | 3.2343 | 0.00936 | 0.043 | -2.77 | -1.1374 | none |
| NBPF3 | 0.563 | 0.5105 | 3.2339 | 0.00937 | 0.043 | -2.77 | -1.1278 | none |
| AGO4 | 0.547 | 0.5487 | 3.2338 | 0.00937 | 0.043 | -2.77 | -0.5211 | none |
| AMPD3 | 0.626 | 0.5102 | 3.2337 | 0.00937 | 0.043 | -2.77 | -0.393 | none |
| GNG7 | 0.302 | 0.2469 | 3.2335 | 0.00938 | 0.043 | -2.77 | -1.4302 | none |
| NFIC | 0.803 | 0.776 | 3.2323 | 0.0094 | 0.043 | -2.78 | -0.4443 | none |
| PTPN23 | 0.703 | 0.7163 | 3.2322 | 0.0094 | 0.043 | -2.78 | -0.4517 | none |
| SLC37A2 | 0.425 | 0.3968 | 3.2318 | 0.0094 | 0.043 | -2.78 | -0.0856 | none |
| DDX6 | 0.716 | 0.5778 | 3.2311 | 0.00941 | 0.043 | -2.78 | -1.1133 | none |

| | | | | | | | | |
|---|---|---|---|---|---|---|---|---|
| FAM78A | 0.535 | 0.5191 | 3.2306 | 0.00942 | 0.043 | -2.78 | -0.9651 | none |
| FAM91A1 | 0.425 | 0.3757 | 3.2293 | 0.00944 | 0.043 | -2.78 | -0.6592 | none |
| AMIGO2 | -0.332 | -0.3293 | -3.228 | 0.00946 | 0.043 | -2.78 | 0.76938 | none |
| CCDC65 | 0.512 | 0.4888 | 3.2265 | 0.00949 | 0.044 | -2.78 | -0.261 | none |
| TTBK2 | 0.638 | 0.4906 | 3.226 | 0.00949 | 0.044 | -2.79 | -1.0171 | none |
| STK17B | 0.391 | 0.2323 | 3.2247 | 0.00951 | 0.044 | -2.79 | -1.2267 | none |
| EXD3 | 0.554 | 0.418 | 3.2246 | 0.00952 | 0.044 | -2.79 | -1.381 | none |
| PGM1 | -0.273 | -0.1946 | -3.224 | 0.00952 | 0.044 | -2.79 | 2.26992 | AA-related |
| YPEL2 | 0.458 | 0.3065 | 3.2236 | 0.00953 | 0.044 | -2.79 | -0.3544 | none |
| SLC35F6 | 0.36 | 0.2683 | 3.2235 | 0.00953 | 0.044 | -2.79 | -1.4259 | none |
| ZNF160 | 0.723 | 0.6143 | 3.2228 | 0.00954 | 0.044 | -2.79 | -0.7276 | none |
| WDR7 | 0.496 | 0.4636 | 3.2226 | 0.00955 | 0.044 | -2.79 | -0.0602 | none |
| FKBP10 | 0.865 | 0.7134 | 3.2212 | 0.00957 | 0.044 | -2.79 | -0.7235 | none |
| SYNRG | 0.895 | 0.804 | 3.2206 | 0.00958 | 0.044 | -2.79 | -1.3413 | none |
| AKTIP | 0.402 | 0.2605 | 3.2202 | 0.00959 | 0.044 | -2.8 | -0.8262 | none |
| SNRK | 0.467 | 0.397 | 3.2201 | 0.00959 | 0.044 | -2.8 | -1.3927 | none |
| AMMECR1 | 0.465 | 0.3445 | 3.2199 | 0.00959 | 0.044 | -2.8 | 0.37426 | none |
| KATNBL1 | 0.39 | 0.2282 | 3.2196 | 0.0096 | 0.044 | -2.8 | -1.0895 | none |
| OR51E2 | 0.443 | 0.3326 | 3.2195 | 0.0096 | 0.044 | -2.8 | -1.1161 | none |
| TEFM | 0.575 | 0.4143 | 3.2186 | 0.00961 | 0.044 | -2.8 | -0.9724 | none |
| FAM192A | 0.484 | 0.4316 | 3.2186 | 0.00961 | 0.044 | -2.8 | -1.2453 | none |
| CHRNA3 | 0.523 | 0.3984 | 3.2183 | 0.00962 | 0.044 | -2.8 | -0.735 | none |
| SMARCA4 | 0.774 | 0.7957 | 3.2183 | 0.00962 | 0.044 | -2.8 | -0.9646 | none |
| OR1M1 | 0.522 | 0.3557 | 3.2183 | 0.00962 | 0.044 | -2.8 | -1.4493 | none |
| PLA2G10 | 0.417 | 0.2736 | 3.2168 | 0.00964 | 0.044 | -2.8 | -0.1952 | none |
| SDK2 | 0.831 | 0.6123 | 3.2166 | 0.00964 | 0.044 | -2.8 | -1.222 | none |
| TRIM46 | 0.503 | 0.3666 | 3.2166 | 0.00964 | 0.044 | -2.8 | -1.26 | none |
| TLR10 | 0.472 | 0.328 | 3.216 | 0.00965 | 0.044 | -2.8 | -1.384 | none |
| RALBP1 | 0.868 | 0.8071 | 3.2148 | 0.00967 | 0.044 | -2.8 | -0.869 | none |
| C16ORF54 | 0.434 | 0.2725 | 3.2146 | 0.00968 | 0.044 | -2.8 | -1.3889 | none |
| TMEM189 | 0.347 | 0.2753 | 3.2146 | 0.00968 | 0.044 | -2.8 | -0.0536 | none |
| CRTAP | 0.446 | 0.2701 | 3.2145 | 0.00968 | 0.044 | -2.8 | -0.4507 | none |
| DNAJC30 | 0.304 | 0.2051 | 3.2141 | 0.00968 | 0.044 | -2.81 | -0.8875 | none |
| ESD | -0.469 | -0.4419 | -3.214 | 0.00969 | 0.044 | -2.81 | 2.47881 | AA-related |
| FES | -0.327 | -0.0607 | -3.214 | 0.00969 | 0.044 | -2.81 | 0.06587 | none |
| AFG3L2 | 0.484 | 0.5019 | 3.2133 | 0.0097 | 0.044 | -2.81 | -0.0253 | none |
| EIF3G | 0.308 | 0.2397 | 3.2133 | 0.0097 | 0.044 | -2.81 | -1.0928 | none |
| CCNT2 | 0.79 | 0.6816 | 3.212 | 0.00972 | 0.044 | -2.81 | -0.9232 | none |
| DPYSL2 | 0.35 | 0.2785 | 3.2116 | 0.00972 | 0.044 | -2.81 | -0.9132 | none |
| ZNF197 | 0.424 | 0.3528 | 3.2109 | 0.00974 | 0.044 | -2.81 | -1.301 | none |

| | | | | | | | | |
|---|---|---|---|---|---|---|---|---|
| ZNF708 | 0.661 | 0.5112 | 3.2099 | 0.00975 | 0.044 | -2.81 | -0.9979 | none |
| HIST1H4I | 0.286 | 0.1144 | 3.2096 | 0.00976 | 0.044 | -2.81 | -0.4726 | none |
| GPNMB | 0.69 | 0.5977 | 3.2093 | 0.00976 | 0.044 | -2.81 | -0.7457 | none |
| PYCR2 | 0.405 | 0.3329 | 3.209 | 0.00977 | 0.044 | -2.81 | -0.9265 | none |
| KMT2A | 1.028 | 0.9847 | 3.209 | 0.00977 | 0.044 | -2.81 | -1.2794 | none |
| HECA | 0.41 | 0.3251 | 3.2089 | 0.00977 | 0.044 | -2.81 | -1.1189 | none |
| DENND1A | 0.428 | 0.4291 | 3.2083 | 0.00978 | 0.044 | -2.81 | -0.9925 | none |
| LRRC37A | 0.418 | 0.3006 | 3.2082 | 0.00978 | 0.044 | -2.81 | -0.7163 | none |
| RELL1 | 0.508 | 0.3826 | 3.2079 | 0.00978 | 0.044 | -2.82 | -1.4624 | none |
| RPL7A | -0.432 | -0.4152 | -3.207 | 0.00979 | 0.044 | -2.82 | 1.916 | none |
| KDELC2 | 0.606 | 0.4584 | 3.207 | 0.0098 | 0.044 | -2.82 | -0.9615 | none |
| CYP17A1 | 0.464 | 0.3635 | 3.2052 | 0.00983 | 0.045 | -2.82 | -0.8242 | none |
| DMPK | 0.582 | 0.477 | 3.2052 | 0.00983 | 0.045 | -2.82 | -0.6986 | none |
| RUNX2 | 0.819 | 0.7082 | 3.2044 | 0.00984 | 0.045 | -2.82 | -1.0526 | none |
| ZNF845 | 0.583 | 0.4991 | 3.2042 | 0.00984 | 0.045 | -2.82 | -1.2039 | none |
| CORIN | 0.369 | 0.2652 | 3.2032 | 0.00986 | 0.045 | -2.82 | -1.4845 | none |
| STAT3 | 0.698 | 0.6441 | 3.203 | 0.00986 | 0.045 | -2.82 | -0.7522 | none |
| DENR | 0.353 | 0.2692 | 3.2017 | 0.00989 | 0.045 | -2.83 | -1.1811 | none |
| ARPC2 | -0.273 | -0.2263 | -3.202 | 0.00989 | 0.045 | -2.83 | 1.8266 | none |
| IL1R1 | 0.312 | 0.223 | 3.2007 | 0.0099 | 0.045 | -2.83 | -1.433 | none |
| RILPL2 | 0.348 | 0.2761 | 3.2003 | 0.00991 | 0.045 | -2.83 | -1.5059 | none |
| ITPKB | 0.605 | 0.5418 | 3.1995 | 0.00992 | 0.045 | -2.83 | -1.3825 | none |
| ZHX1-C8O | -0.707 | -0.5736 | -3.199 | 0.00993 | 0.045 | -2.83 | 1.52517 | none |
| PTPRE | 0.615 | 0.5732 | 3.1985 | 0.00994 | 0.045 | -2.83 | 0.24328 | none |
| RABGAP1L | 0.307 | 0.1753 | 3.1982 | 0.00994 | 0.045 | -2.83 | -1.1808 | none |
| CEP83 | 0.735 | 0.6328 | 3.1982 | 0.00994 | 0.045 | -2.83 | -1.1674 | none |
| FBXO15 | 0.485 | 0.3576 | 3.1976 | 0.00995 | 0.045 | -2.83 | -0.7999 | none |
| ADCY2 | 0.416 | 0.2881 | 3.1968 | 0.00997 | 0.045 | -2.83 | -0.8199 | none |
| ASF1B | -0.387 | -0.2994 | -3.196 | 0.00998 | 0.045 | -2.84 | 2.48276 | AA-related |
| ZNF786 | 0.536 | 0.4639 | 3.1942 | 0.01001 | 0.045 | -2.84 | -0.9893 | none |
| SEC14L1 | 0.271 | 0.2188 | 3.1937 | 0.01002 | 0.045 | -2.84 | -0.8627 | none |
| HERPUD2 | 0.486 | 0.3443 | 3.1934 | 0.01002 | 0.045 | -2.84 | -1.1834 | none |
| GSTCD | 0.706 | 0.5861 | 3.1932 | 0.01003 | 0.045 | -2.84 | -1.298 | none |
| HP1BP3 | 0.638 | 0.6099 | 3.1932 | 0.01003 | 0.045 | -2.84 | -1 | none |
| MAPK8IP2 | 0.35 | 0.2539 | 3.1925 | 0.01004 | 0.045 | -2.84 | -1.1809 | none |
| TMEM239 | 0.268 | 0.1895 | 3.1907 | 0.01007 | 0.045 | -2.84 | -1.3594 | none |
| HID1 | 0.53 | 0.3667 | 3.1901 | 0.01008 | 0.045 | -2.84 | -0.3705 | none |
| SHROOM1 | 0.556 | 0.3541 | 3.189 | 0.0101 | 0.045 | -2.85 | -1.165 | none |
| MPHOSPH | 1.038 | 1.0207 | 3.1883 | 0.01011 | 0.045 | -2.85 | -0.9559 | none |
| RPL19 | -0.322 | -0.2793 | -3.188 | 0.01011 | 0.045 | -2.85 | 2.09855 | AA-related |

| | | | | | | | | |
|---|---|---|---|---|---|---|---|---|
| KCNJ3 | 0.414 | 0.2824 | 3.1877 | 0.01012 | 0.045 | -2.85 | -1.2311 | none |
| MUC5B | -0.208 | -0.1303 | -3.187 | 0.01012 | 0.045 | -2.85 | 0.84415 | none |
| LRRK1 | 0.372 | 0.3414 | 3.1865 | 0.01014 | 0.046 | -2.85 | -0.983 | none |
| CDKN2C | -0.31 | -0.2235 | -3.186 | 0.01014 | 0.046 | -2.85 | 0.16602 | none |
| TSHR | 0.608 | 0.4322 | 3.1855 | 0.01016 | 0.046 | -2.85 | -1.2487 | none |
| GABRP | 0.848 | 0.8241 | 3.185 | 0.01017 | 0.046 | -2.85 | -0.4959 | none |
| KIF27 | 0.665 | 0.475 | 3.1844 | 0.01017 | 0.046 | -2.85 | -1.1391 | none |
| ZNF461 | 0.785 | 0.5186 | 3.1835 | 0.01019 | 0.046 | -2.86 | -0.7922 | none |
| ST8SIA5 | 0.349 | 0.2793 | 3.1831 | 0.0102 | 0.046 | -2.86 | -1.2477 | none |
| CEACAM2: | 0.407 | 0.3045 | 3.1828 | 0.0102 | 0.046 | -2.86 | -0.3422 | none |
| FAM126B | 0.641 | 0.6153 | 3.1819 | 0.01022 | 0.046 | -2.86 | -1.0572 | none |
| TRIM69 | 0.527 | 0.4617 | 3.181 | 0.01023 | 0.046 | -2.86 | -1.3114 | none |
| PPIA | -0.42 | -0.4009 | -3.178 | 0.01028 | 0.046 | -2.86 | 1.59169 | none |
| CASC4 | 0.549 | 0.4043 | 3.1783 | 0.01028 | 0.046 | -2.86 | -0.835 | none |
| PROCA1 | 0.684 | 0.5985 | 3.1765 | 0.01031 | 0.046 | -2.87 | -1.1564 | none |
| ODC1 | -0.376 | -0.3076 | -3.175 | 0.01033 | 0.046 | -2.87 | 1.07365 | none |
| ROGDI | -0.603 | -0.4758 | -3.175 | 0.01033 | 0.046 | -2.87 | 1.02323 | none |
| ABLIM3 | 0.512 | 0.5629 | 3.1735 | 0.01036 | 0.046 | -2.87 | -0.5368 | none |
| TFDP1 | -0.315 | -0.2197 | -3.173 | 0.01037 | 0.046 | -2.87 | 1.25615 | none |
| FFAR1 | 0.33 | 0.2488 | 3.1732 | 0.01037 | 0.046 | -2.87 | -1.1462 | none |
| NPIPA1 | 0.696 | 0.5975 | 3.1725 | 0.01038 | 0.046 | -2.87 | -1.1373 | none |
| SH2B1 | 1.294 | 1.1175 | 3.1723 | 0.01038 | 0.046 | -2.87 | -0.47 | none |
| DYM | 0.511 | 0.3756 | 3.1707 | 0.01041 | 0.047 | -2.88 | 0.04157 | none |
| PLA2G5 | 0.391 | 0.3297 | 3.1696 | 0.01043 | 0.047 | -2.88 | -0.204 | none |
| GOLPH3L | 0.345 | 0.1514 | 3.1685 | 0.01045 | 0.047 | -2.88 | 0.541 | none |
| LDHA | -0.527 | -0.476 | -3.168 | 0.01046 | 0.047 | -2.88 | 2.09457 | AA-related |
| LIF | 0.416 | 0.4757 | 3.1678 | 0.01046 | 0.047 | -2.88 | 0.00805 | none |
| NMD3 | 0.275 | 0.2027 | 3.1677 | 0.01046 | 0.047 | -2.88 | 0.21095 | none |
| NFATC2IP | 0.459 | 0.4218 | 3.1676 | 0.01046 | 0.047 | -2.88 | -0.7032 | none |
| CWF19L1 | 0.659 | 0.5717 | 3.167 | 0.01047 | 0.047 | -2.88 | -0.9869 | none |
| ZC3H13 | 0.489 | 0.5433 | 3.1661 | 0.01049 | 0.047 | -2.88 | 0.10737 | none |
| ATRNL1 | 0.869 | 0.5955 | 3.166 | 0.01049 | 0.047 | -2.88 | -1.2253 | none |
| USP35 | 0.42 | 0.3962 | 3.1658 | 0.0105 | 0.047 | -2.88 | -0.1827 | none |
| PCDH9 | 0.27 | 0.2009 | 3.1652 | 0.01051 | 0.047 | -2.89 | -1.5135 | none |
| SERAC1 | 0.266 | 0.147 | 3.165 | 0.01051 | 0.047 | -2.89 | -0.4126 | none |
| FBXO6 | 0.404 | 0.2262 | 3.1649 | 0.01051 | 0.047 | -2.89 | -0.859 | none |
| PYGO1 | 0.264 | 0.1856 | 3.1647 | 0.01052 | 0.047 | -2.89 | -0.5699 | none |
| MAP6D1 | 0.344 | 0.3966 | 3.1646 | 0.01052 | 0.047 | -2.89 | 0.46288 | none |
| STAC3 | 0.673 | 0.5638 | 3.164 | 0.01053 | 0.047 | -2.89 | -0.4202 | none |
| ANKRD20/ | 0.56 | 0.4138 | 3.1637 | 0.01053 | 0.047 | -2.89 | -1.1276 | none |

| | | | | | | | | |
|---|---|---|---|---|---|---|---|---|
| ACSL4 | 0.398 | 0.3145 | 3.1636 | 0.01053 | 0.047 | -2.89 | -0.877 | none |
| AP1G2 | 0.53 | 0.5274 | 3.1631 | 0.01054 | 0.047 | -2.89 | -0.6803 | none |
| STAC2 | 0.452 | 0.3464 | 3.1629 | 0.01055 | 0.047 | -2.89 | -1.2468 | none |
| RAB11FIP3 | 0.547 | 0.5232 | 3.1625 | 0.01055 | 0.047 | -2.89 | -0.1119 | none |
| ALKBH4 | 0.381 | 0.2861 | 3.162 | 0.01056 | 0.047 | -2.89 | -1.317 | none |
| RNF144A | 0.567 | 0.3781 | 3.1606 | 0.01059 | 0.047 | -2.89 | -0.0138 | none |
| PON3 | 0.677 | 0.464 | 3.1599 | 0.0106 | 0.047 | -2.89 | -0.1377 | none |
| RPAIN | 0.414 | 0.2379 | 3.159 | 0.01061 | 0.047 | -2.9 | -1.0925 | none |
| KDM2B | 0.531 | 0.5141 | 3.158 | 0.01063 | 0.047 | -2.9 | -0.6572 | none |
| CDK10 | 0.431 | 0.4336 | 3.1569 | 0.01065 | 0.047 | -2.9 | -0.6079 | none |
| KCNMB4 | 0.446 | 0.3181 | 3.155 | 0.01069 | 0.047 | -2.9 | -1.0722 | none |
| NPRL3 | 0.443 | 0.3917 | 3.1543 | 0.0107 | 0.047 | -2.9 | -1.2323 | none |
| TSPYL2 | 0.659 | 0.6715 | 3.1542 | 0.0107 | 0.047 | -2.9 | -0.8933 | none |
| ITGB1 | 0.467 | 0.3824 | 3.1531 | 0.01072 | 0.048 | -2.91 | -0.5101 | none |
| ERC2 | 0.337 | 0.2986 | 3.1529 | 0.01072 | 0.048 | -2.91 | -0.29 | none |
| CLCN2 | 0.556 | 0.3785 | 3.1523 | 0.01074 | 0.048 | -2.91 | -1.155 | none |
| POLM | 0.452 | 0.3876 | 3.1508 | 0.01076 | 0.048 | -2.91 | -0.7669 | none |
| MBIP | 0.317 | 0.1877 | 3.1507 | 0.01076 | 0.048 | -2.91 | -0.4045 | none |
| ABCC8 | 0.333 | 0.282 | 3.1505 | 0.01077 | 0.048 | -2.91 | -1.0129 | none |
| THYN1 | -0.359 | -0.3278 | -3.15 | 0.01077 | 0.048 | -2.91 | 2.69156 | AA-related |
| CXORF56 | 0.792 | 0.6616 | 3.15 | 0.01078 | 0.048 | -2.91 | -0.9614 | none |
| CRELD2 | 0.314 | 0.2636 | 3.1494 | 0.01079 | 0.048 | -2.91 | -0.2088 | none |
| GNL3L | 0.887 | 0.839 | 3.1494 | 0.01079 | 0.048 | -2.91 | -1.0518 | none |
| ECT2L | 0.398 | 0.3016 | 3.1487 | 0.0108 | 0.048 | -2.91 | -1.3042 | none |
| ABCB7 | 0.3 | 0.2017 | 3.1483 | 0.01081 | 0.048 | -2.91 | 0.7191 | none |
| BANF1 | -0.382 | -0.3672 | -3.148 | 0.01081 | 0.048 | -2.91 | 2.01201 | none |
| C9ORF64 | 0.585 | 0.3946 | 3.1479 | 0.01081 | 0.048 | -2.91 | -0.6298 | none |
| MAP3K11 | 0.719 | 0.6125 | 3.1478 | 0.01082 | 0.048 | -2.91 | -1.0719 | none |
| UBR5 | 0.958 | 0.8762 | 3.1476 | 0.01082 | 0.048 | -2.91 | -1.0638 | none |
| CD180 | 0.244 | 0.204 | 3.1472 | 0.01083 | 0.048 | -2.92 | -1.1022 | none |
| GRAP2 | -0.356 | -0.312 | -3.146 | 0.01085 | 0.048 | -2.92 | 1.88765 | none |
| PHC3 | 0.557 | 0.4428 | 3.1459 | 0.01085 | 0.048 | -2.92 | -0.9778 | none |
| PSTPIP2 | 0.431 | 0.3455 | 3.1454 | 0.01086 | 0.048 | -2.92 | -0.8101 | none |
| PUM1 | 0.534 | 0.4941 | 3.1451 | 0.01086 | 0.048 | -2.92 | -0.8678 | none |
| EI24 | 0.536 | 0.3598 | 3.1449 | 0.01087 | 0.048 | -2.92 | -1.272 | none |
| FGF22 | 0.364 | 0.291 | 3.1442 | 0.01088 | 0.048 | -2.92 | -0.9952 | none |
| SYNE4 | 0.464 | 0.3618 | 3.144 | 0.01088 | 0.048 | -2.92 | -0.6547 | none |
| SNAP29 | 0.421 | 0.3544 | 3.1439 | 0.01089 | 0.048 | -2.92 | -0.7768 | none |
| RMND5B | 0.461 | 0.3778 | 3.1439 | 0.01089 | 0.048 | -2.92 | -0.6817 | none |
| PRDX6 | -0.484 | -0.4576 | -3.142 | 0.01092 | 0.048 | -2.92 | 2.63294 | AA-related |

| | | | | | | | | |
|---|---|---|---|---|---|---|---|---|
| ENSA | 0.356 | 0.292 | 3.1418 | 0.01092 | 0.048 | -2.92 | -1.1161 | none |
| OR6C4 | 0.331 | 0.2369 | 3.1405 | 0.01095 | 0.048 | -2.93 | -1.4223 | none |
| IBA57 | 0.509 | 0.3687 | 3.1397 | 0.01096 | 0.048 | -2.93 | -0.6837 | none |
| ZNF418 | 0.711 | 0.5487 | 3.1395 | 0.01097 | 0.048 | -2.93 | -0.1246 | none |
| INTS2 | 0.493 | 0.4154 | 3.1387 | 0.01098 | 0.048 | -2.93 | -0.691 | none |
| FADS2 | 0.662 | 0.5726 | 3.1384 | 0.01099 | 0.048 | -2.93 | -0.6564 | none |
| RASAL2 | 0.635 | 0.5267 | 3.1381 | 0.01099 | 0.048 | -2.93 | -1.2768 | none |
| RAPGEF2 | 0.756 | 0.7549 | 3.1379 | 0.011 | 0.048 | -2.93 | -0.4767 | none |
| C17ORF75 | 0.28 | 0.1238 | 3.1374 | 0.011 | 0.048 | -2.93 | -0.6766 | none |
| CCNYL1 | 0.657 | 0.5009 | 3.1367 | 0.01102 | 0.048 | -2.93 | -1.3486 | none |
| TMSB10 | -0.563 | -0.5163 | -3.136 | 0.01103 | 0.048 | -2.93 | 2.45279 | AA-related |
| UNC13C | 0.305 | 0.1961 | 3.1358 | 0.01103 | 0.048 | -2.93 | -1.3768 | none |
| RABGAP1 | 0.761 | 0.7359 | 3.1337 | 0.01107 | 0.049 | -2.94 | -0.8846 | none |
| JOSD1 | 0.328 | 0.2703 | 3.1335 | 0.01108 | 0.049 | -2.94 | -1.3743 | none |
| SAE1 | -0.28 | -0.1827 | -3.133 | 0.01109 | 0.049 | -2.94 | 1.18092 | none |
| ARHGEF26 | 0.289 | 0.218 | 3.1324 | 0.0111 | 0.049 | -2.94 | -1.053 | none |
| SRGAP2B | 0.397 | 0.4008 | 3.1323 | 0.0111 | 0.049 | -2.94 | -0.5468 | none |
| CHPT1 | 0.571 | 0.4114 | 3.1322 | 0.0111 | 0.049 | -2.94 | -0.5633 | none |
| C12ORF43 | 0.684 | 0.6345 | 3.1322 | 0.0111 | 0.049 | -2.94 | -1.0298 | none |
| HOXA13 | -0.225 | -0.161 | -3.132 | 0.0111 | 0.049 | -2.94 | 1.07738 | none |
| CPAMD8 | 0.572 | 0.399 | 3.1317 | 0.01111 | 0.049 | -2.94 | -0.7977 | none |
| MTRNR2L3 | 0.332 | 0.2612 | 3.1315 | 0.01111 | 0.049 | -2.94 | -0.3255 | none |
| AOC1 | 0.319 | 0.2289 | 3.1305 | 0.01113 | 0.049 | -2.94 | -1.1566 | none |
| FLG2 | 0.218 | 0.1489 | 3.1304 | 0.01113 | 0.049 | -2.94 | -1.218 | none |
| ULK4 | 0.521 | 0.4374 | 3.13 | 0.01114 | 0.049 | -2.94 | -1.2819 | none |
| TSFM | -0.306 | -0.2679 | -3.129 | 0.01115 | 0.049 | -2.94 | 2.37716 | AA-related |
| SLC25A32 | 0.322 | 0.1952 | 3.1291 | 0.01116 | 0.049 | -2.95 | -0.8315 | none |
| VPS9D1 | 0.477 | 0.3871 | 3.129 | 0.01116 | 0.049 | -2.95 | -1.1315 | none |
| CA7 | -0.277 | -0.2047 | -3.128 | 0.01118 | 0.049 | -2.95 | 0.24436 | none |
| TXNDC12 | -0.5 | -0.4731 | -3.128 | 0.01119 | 0.049 | -2.95 | 2.18243 | AA-related |
| RAB2B | 0.31 | 0.188 | 3.1275 | 0.01119 | 0.049 | -2.95 | -0.8137 | none |
| PTCHD1 | 0.27 | 0.1835 | 3.1271 | 0.0112 | 0.049 | -2.95 | -1.47 | none |
| ZNF526 | 0.511 | 0.4519 | 3.1268 | 0.0112 | 0.049 | -2.95 | -1.0298 | none |
| PKD1 | 0.545 | 0.5184 | 3.1266 | 0.01121 | 0.049 | -2.95 | -0.8514 | none |
| CSPP1 | 0.674 | 0.6452 | 3.1265 | 0.01121 | 0.049 | -2.95 | -0.233 | none |
| ATP8B2 | 0.696 | 0.6334 | 3.1231 | 0.01127 | 0.049 | -2.96 | -0.6554 | none |
| DBR1 | 0.425 | 0.239 | 3.123 | 0.01127 | 0.049 | -2.96 | -1.2438 | none |
| CCL28 | 0.484 | 0.3294 | 3.1224 | 0.01128 | 0.049 | -2.96 | -1.2191 | none |
| ZNF425 | 0.394 | 0.3186 | 3.1213 | 0.01131 | 0.049 | -2.96 | -0.4159 | none |
| FAM72A | 0.401 | 0.3601 | 3.1212 | 0.01131 | 0.049 | -2.96 | -0.3914 | none |

| | | | | | | | | |
|---|---|---|---|---|---|---|---|---|
| DCAF4L1 | 0.276 | 0.2006 | 3.1211 | 0.01131 | 0.049 | -2.96 | -1.0774 | none |
| ZNF681 | 0.55 | 0.4058 | 3.1201 | 0.01133 | 0.049 | -2.96 | -0.9679 | none |
| PPP1R14D | 0.549 | 0.439 | 3.12 | 0.01133 | 0.049 | -2.96 | -1.325 | none |
| GTF3C2 | 0.451 | 0.385 | 3.1196 | 0.01134 | 0.049 | -2.96 | -1.3554 | none |
| WNT2B | 0.23 | 0.1799 | 3.1196 | 0.01134 | 0.049 | -2.96 | -1.3908 | none |
| RAP1B | 0.304 | 0.136 | 3.1184 | 0.01136 | 0.049 | -2.96 | -0.5502 | none |
| HNF4A | 0.228 | 0.1837 | 3.1182 | 0.01136 | 0.049 | -2.96 | -0.688 | none |
| TBK1 | 0.319 | 0.2548 | 3.1179 | 0.01137 | 0.049 | -2.96 | -0.0676 | none |
| EMC2 | -0.48 | -0.4576 | -3.118 | 0.01137 | 0.049 | -2.96 | 0.75655 | none |
| EMX2 | 0.229 | 0.18 | 3.1167 | 0.01139 | 0.05 | -2.97 | -0.8248 | none |
| OR6C6 | 0.447 | 0.2684 | 3.1145 | 0.01143 | 0.05 | -2.97 | -1.1855 | none |
| MED22 | 0.426 | 0.4087 | 3.114 | 0.01144 | 0.05 | -2.97 | -0.1538 | none |
| RPS6KC1 | 0.423 | 0.3711 | 3.1139 | 0.01145 | 0.05 | -2.97 | -0.4671 | none |
| GREM1 | 0.262 | 0.1871 | 3.1137 | 0.01145 | 0.05 | -2.97 | -1.5308 | none |
| GPANK1 | 0.36 | 0.2406 | 3.1137 | 0.01145 | 0.05 | -2.97 | -0.7702 | none |
| MOB3A | 0.415 | 0.3644 | 3.1125 | 0.01147 | 0.05 | -2.97 | -0.9581 | none |
| MED13L | 0.681 | 0.6261 | 3.1125 | 0.01147 | 0.05 | -2.97 | -1.0804 | none |
| FOXD1 | 0.462 | 0.4538 | 3.1122 | 0.01148 | 0.05 | -2.97 | 0.94744 | none |
| CYP7B1 | 0.538 | 0.4 | 3.1117 | 0.01149 | 0.05 | -2.97 | -1.1463 | none |
| CDIP1 | 0.367 | 0.3086 | 3.1117 | 0.01149 | 0.05 | -2.97 | -0.6163 | none |
| SUMF2 | 0.447 | 0.3156 | 3.1117 | 0.01149 | 0.05 | -2.97 | -1.4787 | none |
| PNRC2 | 0.261 | 0.1442 | 3.1115 | 0.01149 | 0.05 | -2.97 | -0.5612 | none |
| LY75 | 0.4 | 0.3306 | 3.1115 | 0.01149 | 0.05 | -2.97 | -0.5164 | none |
| FNDC3A | 0.717 | 0.6241 | 3.1104 | 0.01151 | 0.05 | -2.98 | -1.2952 | none |
| LRP2BP | 0.375 | 0.2573 | 3.1103 | 0.01152 | 0.05 | -2.98 | -1.2764 | none |
| H2AFZ | -0.505 | -0.454 | -3.11 | 0.01152 | 0.05 | -2.98 | 1.82981 | none |
| TYW3 | 0.421 | 0.3012 | 3.1095 | 0.01153 | 0.05 | -2.98 | -0.1853 | none |
| ZNF517 | 0.625 | 0.479 | 3.1085 | 0.01155 | 0.05 | -2.98 | -1.2823 | none |
| CALD1 | 1.058 | 0.8863 | 3.1075 | 0.01157 | 0.05 | -2.98 | -0.3026 | none |
| STRADA | 0.574 | 0.4448 | 3.107 | 0.01158 | 0.05 | -2.98 | -0.6524 | none |
| AASDHPP1 | -0.469 | -0.3943 | -3.107 | 0.01158 | 0.05 | -2.98 | 2.23238 | AA-related |
| RGP1 | 0.231 | 0.1597 | 3.1061 | 0.0116 | 0.05 | -2.98 | -0.0416 | none |
| MAP4K5 | 0.53 | 0.4777 | 3.1061 | 0.0116 | 0.05 | -2.98 | -0.4228 | none |
| ZNF33B | 0.478 | 0.3362 | 3.1057 | 0.0116 | 0.05 | -2.98 | -0.7993 | none |
| EPS15 | 0.758 | 0.6387 | 3.1057 | 0.0116 | 0.05 | -2.98 | -0.7555 | none |
| MAP2K2 | 0.493 | 0.4582 | 3.1043 | 0.01163 | 0.05 | -2.99 | -1.3537 | none |
| ZEB2 | 0.386 | 0.3299 | 3.1041 | 0.01163 | 0.05 | -2.99 | -0.9338 | none |
| AP3M1 | 0.285 | 0.1538 | 3.1033 | 0.01165 | 0.05 | -2.99 | 0.55284 | none |
| NCKAP1L | 0.418 | 0.352 | 3.1029 | 0.01166 | 0.05 | -2.99 | 0.06597 | none |
| SPTLC1 | 0.458 | 0.2733 | 3.1018 | 0.01168 | 0.05 | -2.99 | -0.9719 | none |

| | | | | | | | | |
|---|---|---|---|---|---|---|---|---|
| CMC1 | -0.489 | -0.4591 | -3.101 | 0.0117 | 0.05 | -2.99 | 1.22942 | none |
| DYNC1I1 | 0.487 | 0.3869 | 3.1005 | 0.01171 | 0.05 | -2.99 | -1.1026 | none |
| AP4B1 | 0.277 | 0.1812 | 3.1005 | 0.01171 | 0.05 | -2.99 | -0.9314 | none |
| NCBP3 | 0.421 | 0.361 | 3.1004 | 0.01171 | 0.05 | -2.99 | -0.3751 | none |
| CREB3L2 | 0.444 | 0.3708 | 3.0999 | 0.01172 | 0.05 | -2.99 | -1.324 | none |
| NHLRC3 | 0.445 | 0.2853 | 3.0998 | 0.01172 | 0.05 | -2.99 | -0.6741 | none |
| TSEN54 | 0.317 | 0.2316 | 3.0997 | 0.01172 | 0.05 | -2.99 | -0.3892 | none |
| ESRP1 | 0.89 | 0.6595 | 3.0997 | 0.01172 | 0.05 | -2.99 | -0.9654 | none |
| BIK | -0.535 | -0.4755 | -3.1 | 0.01172 | 0.05 | -2.99 | 0.83138 | none |
| LSM12 | 0.322 | 0.2471 | 3.0991 | 0.01173 | 0.05 | -2.99 | -1.3993 | none |
| SULF1 | 1.03 | 0.8787 | 3.0991 | 0.01173 | 0.05 | -2.99 | -0.1905 | none |
| CD40 | 0.284 | 0.1188 | 3.0984 | 0.01175 | 0.051 | -3 | -0.9743 | none |
| JMJD7-PLA | 0.506 | 0.4308 | 3.0981 | 0.01175 | 0.051 | -3 | -1.2637 | none |
| HCN3 | 0.316 | 0.2207 | 3.0955 | 0.0118 | 0.051 | -3 | -0.8049 | none |
| ZNF79 | 0.493 | 0.4183 | 3.0955 | 0.0118 | 0.051 | -3 | -0.1365 | none |
| CUTC | -0.329 | -0.3368 | -3.095 | 0.01181 | 0.051 | -3 | 2.28682 | AA-related |
| IRF3 | 0.423 | 0.3148 | 3.0945 | 0.01182 | 0.051 | -3 | -0.7155 | none |
| ZNF280D | -0.758 | -0.6868 | -3.094 | 0.01184 | 0.051 | -3 | 1.6027 | none |
| GRIP1 | 0.484 | 0.3967 | 3.0934 | 0.01185 | 0.051 | -3 | -0.3186 | none |
| LRRC29 | 0.683 | 0.5235 | 3.0928 | 0.01186 | 0.051 | -3.01 | -0.682 | none |
| HECTD4 | 0.829 | 0.769 | 3.0927 | 0.01186 | 0.051 | -3.01 | -0.9824 | none |
| DPY19L1 | -0.462 | -0.4519 | -3.092 | 0.01187 | 0.051 | -3.01 | 0.14977 | none |
| ZNF558 | 0.626 | 0.4321 | 3.0917 | 0.01188 | 0.051 | -3.01 | -1.0395 | none |
| EPAS1 | 0.715 | 0.6791 | 3.0914 | 0.01189 | 0.051 | -3.01 | -0.8894 | none |
| RNF2 | 0.315 | 0.2117 | 3.0913 | 0.01189 | 0.051 | -3.01 | 0.24047 | none |
| 10-Sep | 0.799 | 0.6828 | 3.0912 | 0.01189 | 0.051 | -3.01 | -1.3236 | none |
| TUBE1 | 0.399 | 0.2652 | 3.0911 | 0.01189 | 0.051 | -3.01 | 0.12129 | none |
| MEIS3 | 0.489 | 0.365 | 3.091 | 0.01189 | 0.051 | -3.01 | -1.3113 | none |
| NWD1 | 0.273 | 0.1832 | 3.0906 | 0.0119 | 0.051 | -3.01 | -1.3122 | none |
| HEATR1 | 0.761 | 0.6421 | 3.0905 | 0.0119 | 0.051 | -3.01 | -0.875 | none |
| LRRN2 | 0.372 | 0.271 | 3.0903 | 0.01191 | 0.051 | -3.01 | -1.4314 | none |
| USP26 | 0.338 | 0.2597 | 3.0898 | 0.01192 | 0.051 | -3.01 | -0.2025 | none |
| SP3 | 0.497 | 0.4271 | 3.0896 | 0.01192 | 0.051 | -3.01 | -0.0708 | none |
| PARP16 | 0.456 | 0.3555 | 3.0894 | 0.01192 | 0.051 | -3.01 | -0.6998 | none |
| IGLL5 | -0.528 | -0.4902 | -3.089 | 0.01192 | 0.051 | -3.01 | 1.48694 | none |
| GSTP1 | -0.517 | -0.4932 | -3.089 | 0.01194 | 0.051 | -3.01 | 2.17157 | AA-related |
| DMGDH | 0.751 | 0.5945 | 3.0884 | 0.01194 | 0.051 | -3.01 | -0.8126 | none |
| MDM2 | 0.626 | 0.5286 | 3.0884 | 0.01195 | 0.051 | -3.01 | -1.2369 | none |
| RRM2B | 0.501 | 0.3686 | 3.0881 | 0.01195 | 0.051 | -3.01 | -1.1901 | none |
| ABI3 | -0.366 | -0.2888 | -3.087 | 0.01197 | 0.051 | -3.01 | 2.44846 | AA-related |

| | | | | | | | | |
|---|---|---|---|---|---|---|---|---|
| EID2B | 1.081 | 0.84 | 3.0868 | 0.01198 | 0.051 | -3.02 | -0.5271 | none |
| CCR6 | 0.457 | 0.27 | 3.0863 | 0.01199 | 0.051 | -3.02 | -0.4033 | none |
| GALR2 | 0.414 | 0.2794 | 3.0861 | 0.01199 | 0.051 | -3.02 | -0.7104 | none |
| CD300LD | 0.436 | 0.3358 | 3.0861 | 0.01199 | 0.051 | -3.02 | -1.1414 | none |
| CREB3L4 | 0.652 | 0.4923 | 3.0857 | 0.012 | 0.051 | -3.02 | -0.9488 | none |
| MLLT11 | 0.258 | 0.1794 | 3.0856 | 0.012 | 0.051 | -3.02 | -0.8733 | none |
| ABCF3 | 0.391 | 0.3937 | 3.0855 | 0.012 | 0.051 | -3.02 | -0.6943 | none |
| BAG5 | 0.433 | 0.3487 | 3.0846 | 0.01202 | 0.051 | -3.02 | -0.5642 | none |
| PHF6 | 0.426 | 0.3425 | 3.0841 | 0.01203 | 0.051 | -3.02 | -0.9539 | none |
| GPR137C | 0.542 | 0.4145 | 3.084 | 0.01203 | 0.051 | -3.02 | -0.5194 | none |
| NABP2 | 0.307 | 0.1999 | 3.084 | 0.01203 | 0.051 | -3.02 | -1.1391 | none |
| DENND6A | 0.58 | 0.4468 | 3.0824 | 0.01207 | 0.051 | -3.02 | -0.3991 | none |
| FAM169B | 0.679 | 0.4931 | 3.0822 | 0.01207 | 0.051 | -3.02 | -1.3628 | none |
| CFI | 0.527 | 0.3648 | 3.0813 | 0.01209 | 0.051 | -3.02 | -1.402 | none |
| TAGLN2 | -0.327 | -0.302 | -3.08 | 0.01211 | 0.052 | -3.03 | 2.11727 | AA-related |
| CDKAL1 | 0.449 | 0.347 | 3.0796 | 0.01212 | 0.052 | -3.03 | -1.5713 | OA-related |
| CHN1 | 0.521 | 0.3801 | 3.0792 | 0.01213 | 0.052 | -3.03 | -0.8068 | none |
| ZNF778 | 0.588 | 0.6075 | 3.079 | 0.01214 | 0.052 | -3.03 | 0.99909 | none |
| ZP3 | 0.503 | 0.3375 | 3.078 | 0.01215 | 0.052 | -3.03 | -1.2212 | none |
| RBM8A | -0.26 | -0.1909 | -3.078 | 0.01216 | 0.052 | -3.03 | 1.25778 | none |
| RAB11FIP1 | 0.612 | 0.5469 | 3.0773 | 0.01217 | 0.052 | -3.03 | -1.2586 | none |
| UBE2L6 | -0.352 | -0.3454 | -3.077 | 0.01218 | 0.052 | -3.03 | 2.03812 | none |
| CDHR3 | 0.478 | 0.3358 | 3.0764 | 0.01219 | 0.052 | -3.03 | -0.92 | none |
| RFESD | -0.635 | -0.5728 | -3.075 | 0.01222 | 0.052 | -3.04 | 1.61095 | none |
| EPHA4 | 0.617 | 0.5353 | 3.0741 | 0.01223 | 0.052 | -3.04 | -0.926 | none |
| MARCO | 0.321 | 0.3215 | 3.0739 | 0.01224 | 0.052 | -3.04 | 0.53088 | none |
| ATP13A4 | 0.485 | 0.3443 | 3.0735 | 0.01225 | 0.052 | -3.04 | -1.3237 | none |
| KIF18B | 0.665 | 0.6417 | 3.0732 | 0.01225 | 0.052 | -3.04 | -1.0754 | none |
| CA13 | 0.508 | 0.3832 | 3.073 | 0.01226 | 0.052 | -3.04 | -1.2671 | none |
| DHX58 | 0.614 | 0.5767 | 3.0721 | 0.01228 | 0.052 | -3.04 | -0.5977 | none |
| LONP1 | 0.675 | 0.6812 | 3.072 | 0.01228 | 0.052 | -3.04 | -1.1183 | none |
| PNP | -0.36 | -0.3612 | -3.072 | 0.01228 | 0.052 | -3.04 | 1.63485 | none |
| SLC16A8 | 0.273 | 0.2429 | 3.0719 | 0.01228 | 0.052 | -3.04 | -0.3861 | none |
| COASY | 0.432 | 0.3148 | 3.0712 | 0.01229 | 0.052 | -3.04 | -0.8566 | none |
| TGM5 | 0.281 | 0.2375 | 3.0709 | 0.0123 | 0.052 | -3.04 | -0.349 | none |
| ZNF43 | 0.776 | 0.6502 | 3.0707 | 0.01231 | 0.052 | -3.04 | -1.3886 | none |
| IPO8 | 0.741 | 0.6479 | 3.0706 | 0.01231 | 0.052 | -3.04 | -1.059 | none |
| ZNF600 | 0.784 | 0.5904 | 3.0702 | 0.01231 | 0.052 | -3.04 | -0.0771 | none |
| PECR | 0.461 | 0.308 | 3.0678 | 0.01236 | 0.052 | -3.05 | -0.3113 | none |
| PPIB | -0.393 | -0.3508 | -3.068 | 0.01237 | 0.052 | -3.05 | 2.09331 | AA-related |

| | | | | | | | | |
|---|---|---|---|---|---|---|---|---|
| PICK1 | 0.588 | 0.501 | 3.0672 | 0.01238 | 0.052 | -3.05 | -1.0241 | none |
| FAM167B | 0.566 | 0.3984 | 3.0667 | 0.01239 | 0.052 | -3.05 | -1.1194 | none |
| TCF7 | 0.557 | 0.4548 | 3.0662 | 0.0124 | 0.052 | -3.05 | -1.3123 | none |
| USP24 | 0.84 | 0.7772 | 3.0661 | 0.0124 | 0.052 | -3.05 | -1.0387 | none |
| MTMR10 | 0.46 | 0.4003 | 3.0655 | 0.01241 | 0.052 | -3.05 | -0.2699 | none |
| INPP1 | 0.248 | 0.1999 | 3.0653 | 0.01242 | 0.052 | -3.05 | -0.3338 | none |
| WDR83OS | -0.477 | -0.4458 | -3.065 | 0.01242 | 0.052 | -3.05 | 2.51314 | AA-related |
| SCN11A | 0.415 | 0.2905 | 3.0651 | 0.01242 | 0.052 | -3.05 | -1.2328 | none |
| SLC27A5 | 0.469 | 0.3264 | 3.065 | 0.01242 | 0.052 | -3.05 | -0.4883 | none |
| AC018638 | 1.058 | 0.9464 | 3.0639 | 0.01244 | 0.052 | -3.05 | -1.1004 | none |
| RPS7 | -0.557 | -0.5346 | -3.063 | 0.01246 | 0.053 | -3.05 | 2.34343 | AA-related |
| COL27A1 | 0.511 | 0.4984 | 3.0626 | 0.01247 | 0.053 | -3.06 | -0.7441 | none |
| DDX19B | 0.344 | 0.2603 | 3.0624 | 0.01248 | 0.053 | -3.06 | -0.3644 | none |
| RAMP1 | -0.382 | -0.3228 | -3.061 | 0.01251 | 0.053 | -3.06 | 0.46864 | none |
| FBXW4 | 0.559 | 0.5071 | 3.0605 | 0.01252 | 0.053 | -3.06 | -1.261 | none |
| WFDC11 | 0.45 | 0.3309 | 3.0602 | 0.01252 | 0.053 | -3.06 | -0.5413 | none |
| PCBP1 | -0.246 | -0.222 | -3.06 | 0.01253 | 0.053 | -3.06 | 1.74092 | none |
| ZNF587B | 0.72 | 0.6818 | 3.0597 | 0.01253 | 0.053 | -3.06 | -1.1345 | none |
| G3BP2 | 0.359 | 0.2275 | 3.0593 | 0.01254 | 0.053 | -3.06 | -0.9792 | none |
| BMPR1A | 0.452 | 0.3247 | 3.058 | 0.01257 | 0.053 | -3.06 | -1.2247 | none |
| FAM84B | 0.397 | 0.2611 | 3.0562 | 0.01261 | 0.053 | -3.07 | -0.6589 | none |
| SGSM2 | 0.529 | 0.59 | 3.0538 | 0.01266 | 0.053 | -3.07 | 0.56417 | none |
| ADARB2 | 0.352 | 0.2813 | 3.0533 | 0.01267 | 0.053 | -3.07 | -1.371 | none |
| ZMYND19 | 0.387 | 0.4596 | 3.0522 | 0.01269 | 0.053 | -3.07 | -0.3084 | none |
| HYOU1 | 0.593 | 0.5762 | 3.0517 | 0.0127 | 0.053 | -3.07 | -1.033 | none |
| TRAM2 | 0.398 | 0.3677 | 3.0515 | 0.01271 | 0.053 | -3.07 | -0.8997 | none |
| TSPAN19 | 0.6 | 0.398 | 3.0515 | 0.01271 | 0.053 | -3.07 | 0.04258 | none |
| TEDDM1 | 0.331 | 0.2155 | 3.0507 | 0.01272 | 0.053 | -3.07 | -0.5929 | none |
| CHST11 | 0.655 | 0.5337 | 3.0488 | 0.01277 | 0.054 | -3.08 | -0.2181 | none |
| FCER2 | 0.477 | 0.4979 | 3.0487 | 0.01277 | 0.054 | -3.08 | -0.564 | none |
| CYP46A1 | 0.475 | 0.3032 | 3.0481 | 0.01278 | 0.054 | -3.08 | -0.7197 | none |
| IRAK4 | 0.483 | 0.4176 | 3.048 | 0.01278 | 0.054 | -3.08 | -1.0837 | none |
| RAET1E | 0.275 | 0.2102 | 3.0478 | 0.01279 | 0.054 | -3.08 | -1.4014 | none |
| RHOF | 0.279 | 0.2244 | 3.046 | 0.01283 | 0.054 | -3.08 | -0.7437 | none |
| UBR4 | 0.755 | 0.7855 | 3.0454 | 0.01284 | 0.054 | -3.08 | -0.6765 | none |
| SNRPC | -0.323 | -0.2975 | -3.044 | 0.01287 | 0.054 | -3.09 | 2.47327 | AA-related |
| PTMA | -0.311 | -0.2162 | -3.044 | 0.01287 | 0.054 | -3.09 | 1.36398 | none |
| PIP5K1A | 0.368 | 0.2904 | 3.0435 | 0.01288 | 0.054 | -3.09 | -1.3601 | none |
| ATAD2 | 1.01 | 0.9456 | 3.0424 | 0.0129 | 0.054 | -3.09 | -1.3503 | none |
| ZSCAN29 | 0.55 | 0.5505 | 3.0417 | 0.01292 | 0.054 | -3.09 | -1.046 | none |

| | | | | | | | | |
|---|---|---|---|---|---|---|---|---|
| ATP2A2 | 0.589 | 0.5073 | 3.0413 | 0.01293 | 0.054 | -3.09 | -1.4545 | none |
| RUNX1 | 0.664 | 0.5997 | 3.0411 | 0.01293 | 0.054 | -3.09 | -0.5723 | none |
| TMX3 | 0.566 | 0.3579 | 3.0405 | 0.01294 | 0.054 | -3.09 | -1.1936 | none |
| PIK3R2 | 0.613 | 0.5445 | 3.0399 | 0.01296 | 0.054 | -3.09 | -1.236 | none |
| OR11G2 | 0.248 | 0.1718 | 3.0396 | 0.01296 | 0.054 | -3.09 | -1.3676 | none |
| ACOX1 | 0.397 | 0.2508 | 3.0384 | 0.01299 | 0.054 | -3.1 | -1.0056 | none |
| MTHFD2L | 0.397 | 0.2795 | 3.0379 | 0.013 | 0.054 | -3.1 | -0.7002 | none |
| CARMIL1 | 0.487 | 0.4516 | 3.0376 | 0.01301 | 0.054 | -3.1 | -1.2664 | none |
| RAB38 | -0.755 | -0.6438 | -3.037 | 0.01301 | 0.054 | -3.1 | 2.74101 | AA-related |
| ABRA | 0.31 | 0.2121 | 3.037 | 0.01302 | 0.054 | -3.1 | -1.4253 | none |
| EBI3 | 0.441 | 0.3596 | 3.0365 | 0.01303 | 0.054 | -3.1 | -0.0883 | none |
| CD33 | 0.446 | 0.2963 | 3.0362 | 0.01304 | 0.054 | -3.1 | -0.7872 | none |
| DNM1 | 0.434 | 0.3317 | 3.0352 | 0.01306 | 0.054 | -3.1 | -0.6513 | none |
| ABCC6 | 0.547 | 0.43 | 3.0351 | 0.01306 | 0.054 | -3.1 | -1.0192 | none |
| NAMPT | 0.307 | 0.2296 | 3.035 | 0.01306 | 0.054 | -3.1 | -0.7774 | none |
| GLIPR1L2 | 0.42 | 0.311 | 3.0347 | 0.01307 | 0.054 | -3.1 | -1.5455 | OA-related |
| RARS2 | 0.299 | 0.1834 | 3.0347 | 0.01307 | 0.054 | -3.1 | -0.6595 | none |
| COX16 | -0.538 | -0.5167 | -3.034 | 0.01308 | 0.054 | -3.1 | 1.12559 | none |
| TCP11L2 | 0.459 | 0.355 | 3.0339 | 0.01309 | 0.055 | -3.1 | -0.2462 | none |
| PLEKHA2 | 0.631 | 0.4882 | 3.0339 | 0.01309 | 0.055 | -3.1 | -1.2834 | none |
| VCL | 0.415 | 0.4183 | 3.0334 | 0.0131 | 0.055 | -3.1 | -0.4303 | none |
| COX6C | -0.461 | -0.4522 | -3.033 | 0.01311 | 0.055 | -3.1 | 2.38783 | AA-related |
| RAN | -0.455 | -0.4066 | -3.033 | 0.01311 | 0.055 | -3.1 | 2.09659 | AA-related |
| ABRACL | -0.479 | -0.4806 | -3.032 | 0.01312 | 0.055 | -3.11 | 1.58103 | none |
| BORCS5 | -0.35 | -0.3024 | -3.032 | 0.01313 | 0.055 | -3.11 | 0.60287 | none |
| TIGD7 | 0.821 | 0.6162 | 3.031 | 0.01315 | 0.055 | -3.11 | -1.2628 | none |
| TRIOBP | 0.395 | 0.4275 | 3.0301 | 0.01317 | 0.055 | -3.11 | -0.8478 | none |
| SPSB1 | 0.583 | 0.5107 | 3.0298 | 0.01318 | 0.055 | -3.11 | 0.34035 | none |
| DNAJB4 | 0.419 | 0.3465 | 3.0297 | 0.01318 | 0.055 | -3.11 | -0.8072 | none |
| LYRM2 | 0.735 | 0.6067 | 3.0278 | 0.01322 | 0.055 | -3.11 | -0.947 | none |
| DDIT4 | -0.384 | -0.2638 | -3.027 | 0.01323 | 0.055 | -3.11 | 2.01917 | none |
| CISD1 | -0.329 | -0.3223 | -3.027 | 0.01324 | 0.055 | -3.11 | 1.84934 | none |
| CCDC30 | 0.524 | 0.3768 | 3.0267 | 0.01325 | 0.055 | -3.11 | -1.2626 | none |
| NDUFA8 | -0.359 | -0.3322 | -3.026 | 0.01325 | 0.055 | -3.12 | 1.75517 | none |
| XKR4 | 0.281 | 0.2006 | 3.026 | 0.01326 | 0.055 | -3.12 | -1.3598 | none |
| LGSN | 0.381 | 0.2726 | 3.0249 | 0.01329 | 0.055 | -3.12 | -1.1804 | none |
| FGL2 | 0.428 | 0.3056 | 3.0243 | 0.0133 | 0.055 | -3.12 | -1.1385 | none |
| MAN2A2 | 0.712 | 0.6725 | 3.0236 | 0.01332 | 0.055 | -3.12 | -0.4301 | none |
| KDM5B | 0.713 | 0.6462 | 3.0232 | 0.01333 | 0.055 | -3.12 | -1.1953 | none |
| SLC12A2 | 0.571 | 0.4761 | 3.0226 | 0.01334 | 0.055 | -3.12 | -1.5365 | OA-related |

| | | | | | | | | |
|---|---|---|---|---|---|---|---|---|
| SCN7A | 0.566 | 0.4102 | 3.0224 | 0.01334 | 0.055 | -3.12 | -1.1405 | none |
| YWHAB | -0.336 | -0.293 | -3.022 | 0.01336 | 0.055 | -3.12 | 1.20521 | none |
| SH3TC1 | 0.359 | 0.3264 | 3.0216 | 0.01336 | 0.055 | -3.12 | -0.5207 | none |
| MX2 | 0.575 | 0.4886 | 3.0212 | 0.01337 | 0.055 | -3.12 | -0.9857 | none |
| YEATS2 | 0.898 | 0.9106 | 3.0198 | 0.0134 | 0.055 | -3.13 | -1.0223 | none |
| B3GNT8 | 0.396 | 0.2754 | 3.0198 | 0.0134 | 0.055 | -3.13 | -0.9387 | none |
| GLB1L3 | 0.402 | 0.2155 | 3.0194 | 0.01341 | 0.055 | -3.13 | -1.1978 | none |
| SH2D4A | 0.752 | 0.7278 | 3.0187 | 0.01343 | 0.056 | -3.13 | -0.5877 | none |
| ADM2 | 0.363 | 0.2695 | 3.018 | 0.01344 | 0.056 | -3.13 | -1.095 | none |
| TMEM232 | 0.564 | 0.4307 | 3.0179 | 0.01344 | 0.056 | -3.13 | -0.7666 | none |
| ZC3HAV1 | 0.364 | 0.3252 | 3.0177 | 0.01345 | 0.056 | -3.13 | -1.3499 | none |
| CHRM5 | 0.253 | 0.2031 | 3.0171 | 0.01346 | 0.056 | -3.13 | -0.6985 | none |
| FMC1 | 0.713 | 0.4556 | 3.0168 | 0.01347 | 0.056 | -3.13 | -1.0579 | none |
| NDNF | 0.473 | 0.276 | 3.016 | 0.01349 | 0.056 | -3.13 | -1.0883 | none |
| PRCC | 0.408 | 0.3923 | 3.0158 | 0.01349 | 0.056 | -3.13 | -1.1217 | none |
| PAQR3 | 0.768 | 0.5269 | 3.0158 | 0.01349 | 0.056 | -3.13 | -0.7708 | none |
| ACSS1 | 0.371 | 0.2771 | 3.0153 | 0.0135 | 0.056 | -3.13 | -1.058 | none |
| ELK4 | 0.384 | 0.302 | 3.0147 | 0.01352 | 0.056 | -3.13 | -0.9628 | none |
| HAUS8 | 0.399 | 0.3719 | 3.0146 | 0.01352 | 0.056 | -3.13 | -0.8015 | none |
| PRICKLE1 | 0.741 | 0.4845 | 3.0128 | 0.01356 | 0.056 | -3.14 | -1.1092 | none |
| FAM47E | 0.562 | 0.3517 | 3.0117 | 0.01359 | 0.056 | -3.14 | -1.0211 | none |
| MYPN | 0.502 | 0.3753 | 3.0115 | 0.01359 | 0.056 | -3.14 | -1.0525 | none |
| MDM1 | 0.413 | 0.3534 | 3.0113 | 0.01359 | 0.056 | -3.14 | 0.61123 | none |
| NAIP | 0.646 | 0.4988 | 3.0113 | 0.01359 | 0.056 | -3.14 | -0.8896 | none |
| ARHGAP1 | 0.574 | 0.5125 | 3.0104 | 0.01361 | 0.056 | -3.14 | -1.234 | none |
| RNF152 | 0.293 | 0.1918 | 3.0093 | 0.01364 | 0.056 | -3.14 | -1.4594 | none |
| TMEM268 | 0.406 | 0.3264 | 3.0086 | 0.01366 | 0.056 | -3.14 | -1.1478 | none |
| TMEM94 | 0.825 | 0.764 | 3.0084 | 0.01366 | 0.056 | -3.14 | -1.1037 | none |
| AGAP6 | 0.625 | 0.6437 | 3.0081 | 0.01367 | 0.056 | -3.15 | -0.2083 | none |
| RPGRIP1 | 0.573 | 0.5717 | 3.0071 | 0.01369 | 0.056 | -3.15 | -0.8894 | none |
| TEPSIN | 0.594 | 0.5186 | 3.0069 | 0.01369 | 0.056 | -3.15 | -0.8907 | none |
| SF3B6 | -0.473 | -0.4673 | -3.007 | 0.0137 | 0.056 | -3.15 | 1.77353 | none |
| RGPD6 | 0.858 | 0.7472 | 3.0061 | 0.01371 | 0.056 | -3.15 | -1.1306 | none |
| CCM2 | 0.384 | 0.3167 | 3.0061 | 0.01371 | 0.056 | -3.15 | -1.0369 | none |
| LMLN | 0.457 | 0.3837 | 3.0056 | 0.01373 | 0.056 | -3.15 | 0.03601 | none |
| FXYD5 | -0.36 | -0.3004 | -3.006 | 0.01373 | 0.056 | -3.15 | 1.76134 | none |
| RC3H1 | 0.803 | 0.7695 | 3.0053 | 0.01373 | 0.056 | -3.15 | -1.2726 | none |
| C1ORF56 | 0.459 | 0.3434 | 3.0053 | 0.01373 | 0.056 | -3.15 | -0.8057 | none |
| HAUS1 | 0.411 | 0.2553 | 3.0035 | 0.01377 | 0.057 | -3.15 | -1.1454 | none |
| PHTF2 | 0.404 | 0.3346 | 3.0034 | 0.01378 | 0.057 | -3.15 | -0.6915 | none |

| | | | | | | | | |
|---|---|---|---|---|---|---|---|---|
| CCDC130 | 0.317 | 0.2997 | 3.0031 | 0.01378 | 0.057 | -3.15 | -0.8336 | none |
| DYRK1A | 0.66 | 0.5659 | 3.0029 | 0.01379 | 0.057 | -3.15 | -1.2497 | none |
| CFDP1 | 0.546 | 0.4807 | 3.0025 | 0.0138 | 0.057 | -3.15 | -0.9241 | none |
| DSC3 | 0.243 | 0.1532 | 3.0024 | 0.0138 | 0.057 | -3.15 | -1.3241 | none |
| AUNIP | 0.39 | 0.3299 | 3.0021 | 0.01381 | 0.057 | -3.16 | -0.7689 | none |
| NRDE2 | 0.602 | 0.5978 | 3.0019 | 0.01381 | 0.057 | -3.16 | -0.6595 | none |
| IL6R | 0.358 | 0.313 | 3.0009 | 0.01383 | 0.057 | -3.16 | -0.1097 | none |
| LCMT2 | 0.315 | 0.2061 | 3.0004 | 0.01384 | 0.057 | -3.16 | 0.07775 | none |
| MCUR1 | 0.341 | 0.2433 | 3.0002 | 0.01385 | 0.057 | -3.16 | -0.1666 | none |
| BNIPL | 0.638 | 0.4307 | 2.9997 | 0.01386 | 0.057 | -3.16 | -0.9577 | none |
| CNNM2 | 0.295 | 0.2308 | 2.9992 | 0.01387 | 0.057 | -3.16 | -1.2616 | none |
| NDST3 | 0.248 | 0.1704 | 2.997 | 0.01392 | 0.057 | -3.16 | -1.5224 | none |
| AC074141 | 0.392 | 0.2728 | 2.9968 | 0.01393 | 0.057 | -3.16 | -1.3191 | none |
| HARBI1 | 0.328 | 0.2129 | 2.9964 | 0.01394 | 0.057 | -3.16 | -1.0848 | none |
| DTNA | 0.754 | 0.7195 | 2.9955 | 0.01396 | 0.057 | -3.17 | -0.6075 | none |
| MCM3AP | 0.72 | 0.7005 | 2.994 | 0.01399 | 0.057 | -3.17 | -1.4326 | none |
| CCDC14 | 0.503 | 0.4556 | 2.9937 | 0.014 | 0.057 | -3.17 | -0.6037 | none |
| PDE7B | 0.347 | 0.2697 | 2.9936 | 0.014 | 0.057 | -3.17 | -0.7331 | none |
| SKIDA1 | 0.51 | 0.4218 | 2.9933 | 0.01401 | 0.057 | -3.17 | -0.9558 | none |
| RSF1 | 0.499 | 0.4658 | 2.9931 | 0.01402 | 0.057 | -3.17 | -0.272 | none |
| CIITA | 0.588 | 0.4778 | 2.9931 | 0.01402 | 0.057 | -3.17 | -0.7657 | none |
| C11ORF1 | 0.375 | 0.1479 | 2.9917 | 0.01405 | 0.057 | -3.17 | 0.34635 | none |
| TRIM61 | 0.277 | 0.2103 | 2.9916 | 0.01405 | 0.057 | -3.17 | 0.81573 | none |
| MRLN | 0.688 | 0.4719 | 2.9912 | 0.01406 | 0.057 | -3.17 | -0.4 | none |
| ESRRG | 0.411 | 0.3248 | 2.9911 | 0.01406 | 0.057 | -3.17 | -0.0701 | none |
| SMARCD3 | -0.284 | -0.1089 | -2.991 | 0.01406 | 0.057 | -3.17 | 0.2455 | none |
| MAGI3 | 0.285 | 0.2182 | 2.9904 | 0.01408 | 0.057 | -3.17 | 0.4947 | none |
| ATRIP | 0.509 | 0.5183 | 2.9902 | 0.01409 | 0.057 | -3.17 | -0.4139 | none |
| SRSF7 | -0.484 | -0.4514 | -2.99 | 0.0141 | 0.057 | -3.18 | 2.06063 | none |
| NAA35 | 0.277 | 0.2316 | 2.9896 | 0.0141 | 0.057 | -3.18 | -0.1862 | none |
| FUK | 0.394 | 0.2986 | 2.9894 | 0.0141 | 0.057 | -3.18 | -1.2785 | none |
| EML6 | 0.522 | 0.3601 | 2.9891 | 0.01411 | 0.057 | -3.18 | 0.04583 | none |
| CD248 | -0.216 | -0.1602 | -2.989 | 0.01412 | 0.057 | -3.18 | 0.92573 | none |
| FTH1 | -0.456 | -0.4178 | -2.988 | 0.01413 | 0.058 | -3.18 | 1.91723 | none |
| GPR85 | 0.679 | 0.4703 | 2.9878 | 0.01414 | 0.058 | -3.18 | -1.1827 | none |
| GPR22 | 0.281 | 0.2122 | 2.9875 | 0.01415 | 0.058 | -3.18 | -0.9687 | none |
| GCC2 | 0.694 | 0.6402 | 2.9875 | 0.01415 | 0.058 | -3.18 | -1.0947 | none |
| FAM3A | 0.435 | 0.3369 | 2.9871 | 0.01416 | 0.058 | -3.18 | -1.4522 | none |
| HSF2BP | -0.242 | -0.1791 | -2.986 | 0.01418 | 0.058 | -3.18 | 0.69251 | none |
| SRCIN1 | 0.322 | 0.2371 | 2.9859 | 0.01419 | 0.058 | -3.18 | -1.3485 | none |

| | | | | | | | | |
|---|---|---|---|---|---|---|---|---|
| RSPH9 | -0.23 | -0.1236 | -2.986 | 0.01419 | 0.058 | -3.18 | 1.53088 | none |
| NR2C2 | 0.698 | 0.6261 | 2.9857 | 0.01419 | 0.058 | -3.18 | -0.9795 | none |
| OFD1 | 0.678 | 0.6075 | 2.9852 | 0.0142 | 0.058 | -3.18 | -0.2041 | none |
| LPAR4 | 0.686 | 0.4944 | 2.9849 | 0.01421 | 0.058 | -3.18 | -0.5962 | none |
| H3F3A | -0.575 | -0.5915 | -2.985 | 0.01421 | 0.058 | -3.18 | 1.93271 | none |
| COG8 | 0.354 | 0.2886 | 2.9848 | 0.01421 | 0.058 | -3.18 | -1.2952 | none |
| FYCO1 | 0.632 | 0.6268 | 2.9842 | 0.01423 | 0.058 | -3.18 | -1.0375 | none |
| ZNF473 | 0.667 | 0.5483 | 2.9837 | 0.01424 | 0.058 | -3.19 | -0.7405 | none |
| CNDP2 | 0.408 | 0.3723 | 2.9834 | 0.01425 | 0.058 | -3.19 | -0.4335 | none |
| KMT5C | 0.683 | 0.5846 | 2.9831 | 0.01425 | 0.058 | -3.19 | -1.3978 | none |
| SMAD2 | 0.356 | 0.2277 | 2.9825 | 0.01427 | 0.058 | -3.19 | -0.3231 | none |
| FAM149B1 | 0.603 | 0.4416 | 2.9824 | 0.01427 | 0.058 | -3.19 | -0.911 | none |
| ERAP2 | 0.549 | 0.4953 | 2.9822 | 0.01428 | 0.058 | -3.19 | -1.1659 | none |
| BZW2 | -0.276 | -0.2094 | -2.982 | 0.01428 | 0.058 | -3.19 | 2.05323 | none |
| TBC1D9B | 0.585 | 0.549 | 2.982 | 0.01428 | 0.058 | -3.19 | -0.8909 | none |
| CPQ | 0.507 | 0.4331 | 2.981 | 0.0143 | 0.058 | -3.19 | -0.4825 | none |
| RGS17 | 0.332 | 0.2221 | 2.9788 | 0.01436 | 0.058 | -3.19 | -1.3744 | none |
| CLYBL | 0.405 | 0.3392 | 2.9786 | 0.01436 | 0.058 | -3.19 | -0.4771 | none |
| CLIC2 | 0.577 | 0.4283 | 2.9763 | 0.01442 | 0.058 | -3.2 | -0.6229 | none |
| LSM4 | -0.31 | -0.235 | -2.976 | 0.01442 | 0.058 | -3.2 | 1.29918 | none |
| C19ORF47 | 0.487 | 0.3321 | 2.9761 | 0.01442 | 0.058 | -3.2 | -0.613 | none |
| PMAIP1 | -0.343 | -0.3418 | -2.975 | 0.01444 | 0.058 | -3.2 | 2.11649 | AA-related |
| RMND1 | 0.431 | 0.327 | 2.9753 | 0.01444 | 0.058 | -3.2 | -0.8781 | none |
| WWP2 | 0.634 | 0.5647 | 2.9749 | 0.01445 | 0.058 | -3.2 | -1.2103 | none |
| SF3A2 | 0.671 | 0.6828 | 2.9728 | 0.0145 | 0.059 | -3.2 | -1.1177 | none |
| BCAR1 | 0.289 | 0.2945 | 2.9725 | 0.01451 | 0.059 | -3.2 | -0.9496 | none |
| EIF2A | 0.396 | 0.3161 | 2.9724 | 0.01451 | 0.059 | -3.2 | -0.902 | none |
| FAM118A | 0.496 | 0.443 | 2.9719 | 0.01452 | 0.059 | -3.2 | -1.0737 | none |
| VILL | 0.464 | 0.4356 | 2.9717 | 0.01453 | 0.059 | -3.21 | -0.2409 | none |
| EML3 | 0.36 | 0.3469 | 2.9712 | 0.01454 | 0.059 | -3.21 | -1.0725 | none |
| ANKS1B | 0.509 | 0.3188 | 2.9683 | 0.01461 | 0.059 | -3.21 | -1.1855 | none |
| MREG | 0.652 | 0.5485 | 2.9679 | 0.01462 | 0.059 | -3.21 | -1.0028 | none |
| RAB33A | -0.431 | -0.3861 | -2.968 | 0.01462 | 0.059 | -3.21 | 2.04025 | none |
| HSPA12A | 0.509 | 0.3979 | 2.9677 | 0.01463 | 0.059 | -3.21 | -1.3671 | none |
| UVRAG | 0.53 | 0.4099 | 2.9677 | 0.01463 | 0.059 | -3.21 | -1.2156 | none |
| IGFLR1 | 0.528 | 0.3217 | 2.9671 | 0.01464 | 0.059 | -3.21 | -0.6348 | none |
| BLOC1S1 | -0.406 | -0.3999 | -2.967 | 0.01465 | 0.059 | -3.21 | 1.79124 | none |
| CHD5 | 0.374 | 0.3066 | 2.9655 | 0.01468 | 0.059 | -3.22 | -0.6477 | none |
| HSBP1 | -0.499 | -0.4631 | -2.965 | 0.01468 | 0.059 | -3.22 | 1.57163 | none |
| ZNF549 | 0.485 | 0.3962 | 2.9643 | 0.01471 | 0.059 | -3.22 | -1.0467 | none |

| | | | | | | | | |
|---|---|---|---|---|---|---|---|---|
| OR5AU1 | 0.226 | 0.1934 | 2.9636 | 0.01473 | 0.059 | -3.22 | -0.6376 | none |
| 1-Mar | 0.351 | 0.2638 | 2.9629 | 0.01475 | 0.059 | -3.22 | -1.1747 | none |
| REM2 | -0.263 | -0.179 | -2.963 | 0.01475 | 0.059 | -3.22 | 2.44456 | AA-related |
| SIGLEC15 | 0.519 | 0.3757 | 2.9624 | 0.01476 | 0.059 | -3.22 | -0.7698 | none |
| MOB1A | 0.299 | 0.1609 | 2.9624 | 0.01476 | 0.059 | -3.22 | -1.0669 | none |
| OXCT2 | -0.263 | -0.2069 | -2.962 | 0.01476 | 0.059 | -3.22 | 2.08827 | none |
| PIK3C3 | 0.489 | 0.3391 | 2.9622 | 0.01476 | 0.059 | -3.22 | -0.7382 | none |
| FAM156A | 0.423 | 0.3997 | 2.9622 | 0.01476 | 0.059 | -3.22 | 0.45655 | none |
| C16ORF95 | 0.514 | 0.3728 | 2.9618 | 0.01477 | 0.059 | -3.22 | -0.642 | none |
| C11ORF98 | -0.262 | -0.1975 | -2.961 | 0.01478 | 0.059 | -3.22 | 1.09775 | none |
| PHLDA1 | 0.398 | 0.393 | 2.9615 | 0.01478 | 0.059 | -3.22 | -0.0663 | none |
| FXYD7 | -0.361 | -0.3112 | -2.961 | 0.01478 | 0.059 | -3.22 | 1.5391 | none |
| TMEM8B | 0.36 | 0.2508 | 2.9612 | 0.01479 | 0.059 | -3.22 | -0.7476 | none |
| TAT | 0.387 | 0.2742 | 2.961 | 0.01479 | 0.059 | -3.22 | -1.3951 | none |
| CX3CL1 | 0.232 | 0.146 | 2.961 | 0.01479 | 0.059 | -3.22 | -1.0872 | none |
| MCTS1 | -0.525 | -0.5091 | -2.96 | 0.01483 | 0.059 | -3.23 | 2.39289 | AA-related |
| PLOD1 | 0.53 | 0.4673 | 2.9593 | 0.01483 | 0.059 | -3.23 | -0.5321 | none |
| MTMR4 | 0.502 | 0.4893 | 2.9592 | 0.01484 | 0.059 | -3.23 | -1.0965 | none |
| GLCE | 0.354 | 0.2159 | 2.9592 | 0.01484 | 0.059 | -3.23 | -1.0351 | none |
| TRAF3IP1 | 0.619 | 0.5454 | 2.9587 | 0.01485 | 0.059 | -3.23 | -0.7709 | none |
| NAGA | -0.261 | -0.1723 | -2.959 | 0.01485 | 0.059 | -3.23 | 1.88626 | none |
| MSMP | 0.348 | 0.1953 | 2.9587 | 0.01485 | 0.059 | -3.23 | -0.4588 | none |
| LIPE | 0.664 | 0.583 | 2.9587 | 0.01485 | 0.059 | -3.23 | -1.2363 | none |
| ALDH4A1 | -0.286 | -0.1897 | -2.959 | 0.01485 | 0.059 | -3.23 | 2.15939 | AA-related |
| BCAS3 | 0.458 | 0.3023 | 2.9579 | 0.01487 | 0.059 | -3.23 | -0.8586 | none |
| ZNF264 | 0.372 | 0.2378 | 2.9577 | 0.01488 | 0.06 | -3.23 | -1.1176 | none |
| C16ORF45 | 0.4 | 0.3312 | 2.9567 | 0.0149 | 0.06 | -3.23 | -0.9199 | none |
| TBX19 | 0.424 | 0.3134 | 2.9565 | 0.01491 | 0.06 | -3.23 | -0.6312 | none |
| WSCD2 | 0.307 | 0.3346 | 2.9565 | 0.01491 | 0.06 | -3.23 | 1.23423 | none |
| NLRC3 | 0.634 | 0.6197 | 2.956 | 0.01492 | 0.06 | -3.23 | -0.5554 | none |
| EPHX2 | 0.36 | 0.3069 | 2.956 | 0.01492 | 0.06 | -3.23 | -0.4062 | none |
| MOCS2 | -0.397 | -0.3717 | -2.956 | 0.01492 | 0.06 | -3.23 | 1.90832 | none |
| RSBN1L | 0.613 | 0.5362 | 2.9555 | 0.01493 | 0.06 | -3.23 | -0.9371 | none |
| CD300LG | 0.614 | 0.4202 | 2.9551 | 0.01494 | 0.06 | -3.23 | -0.6954 | none |
| OR10K2 | 0.335 | 0.2613 | 2.9545 | 0.01496 | 0.06 | -3.23 | -1.2607 | none |
| ELP3 | 0.336 | 0.2071 | 2.9541 | 0.01497 | 0.06 | -3.23 | -0.7552 | none |
| TXNDC11 | 0.308 | 0.2679 | 2.9536 | 0.01498 | 0.06 | -3.24 | -0.8811 | none |
| HIST1H2BI | 0.436 | 0.2786 | 2.9529 | 0.015 | 0.06 | -3.24 | -1.2819 | none |
| CHRNG | 0.45 | 0.3553 | 2.9524 | 0.01501 | 0.06 | -3.24 | -1.1253 | none |
| LANCL3 | 0.315 | 0.2269 | 2.9519 | 0.01502 | 0.06 | -3.24 | -1.2157 | none |

| | | | | | | | | |
|---|---|---|---|---|---|---|---|---|
| BMP8B | 0.239 | 0.202 | 2.9518 | 0.01502 | 0.06 | -3.24 | -0.9968 | none |
| RAD54L2 | 0.694 | 0.6727 | 2.9516 | 0.01503 | 0.06 | -3.24 | -1.2235 | none |
| ITGA2 | 0.373 | 0.3031 | 2.9516 | 0.01503 | 0.06 | -3.24 | -1.0927 | none |
| CTHRC1 | -0.352 | -0.2436 | -2.951 | 0.01505 | 0.06 | -3.24 | 1.45693 | none |
| STRIP2 | 0.42 | 0.4365 | 2.9504 | 0.01506 | 0.06 | -3.24 | -0.6286 | none |
| PSIP1 | 0.555 | 0.5309 | 2.9499 | 0.01507 | 0.06 | -3.24 | -1.0309 | none |
| NHSL1 | 0.596 | 0.4655 | 2.9495 | 0.01508 | 0.06 | -3.24 | -0.5105 | none |
| DDX23 | 0.792 | 0.741 | 2.9494 | 0.01509 | 0.06 | -3.24 | -0.9486 | none |
| ZC3H7B | 0.789 | 0.7981 | 2.9488 | 0.0151 | 0.06 | -3.24 | -0.9694 | none |
| TMEM30B | 0.258 | 0.2026 | 2.9487 | 0.0151 | 0.06 | -3.24 | -0.118 | none |
| TRMT5 | -0.401 | -0.3155 | -2.948 | 0.01511 | 0.06 | -3.24 | 0.65492 | none |
| TTL | 0.531 | 0.4791 | 2.948 | 0.01512 | 0.06 | -3.24 | -0.7652 | none |
| EPHX4 | 0.391 | 0.2167 | 2.9476 | 0.01513 | 0.06 | -3.25 | -0.1079 | none |
| IDNK | 0.253 | 0.1001 | 2.9474 | 0.01514 | 0.06 | -3.25 | -0.8516 | none |
| SLC8B1 | 0.229 | 0.1601 | 2.9466 | 0.01516 | 0.06 | -3.25 | 0.08983 | none |
| ZC3H14 | 0.719 | 0.6237 | 2.9461 | 0.01517 | 0.06 | -3.25 | -0.625 | none |
| CDC42SE1 | -0.215 | -0.1869 | -2.945 | 0.01519 | 0.06 | -3.25 | 2.30195 | AA-related |
| TPRN | -0.418 | -0.2946 | -2.945 | 0.01519 | 0.06 | -3.25 | 0.43839 | none |
| UBE2C | -0.348 | -0.325 | -2.945 | 0.01519 | 0.06 | -3.25 | 2.15082 | AA-related |
| BBS1 | 0.784 | 0.6728 | 2.9452 | 0.01519 | 0.06 | -3.25 | -1.1527 | none |
| SAMD7 | 0.44 | 0.294 | 2.9451 | 0.01519 | 0.06 | -3.25 | -0.7748 | none |
| DFFA | 0.352 | 0.2317 | 2.9446 | 0.01521 | 0.06 | -3.25 | -1.2131 | none |
| CNTROB | 0.627 | 0.6239 | 2.9444 | 0.01521 | 0.06 | -3.25 | -0.646 | none |
| ANAPC7 | 0.344 | 0.2771 | 2.9434 | 0.01524 | 0.06 | -3.25 | -0.7656 | none |
| CIPC | 0.381 | 0.3051 | 2.9422 | 0.01527 | 0.06 | -3.25 | -1.0592 | none |
| PDIK1L | 0.368 | 0.2723 | 2.9422 | 0.01527 | 0.06 | -3.25 | -1.0646 | none |
| FBXO46 | 0.523 | 0.442 | 2.9422 | 0.01527 | 0.06 | -3.25 | -0.901 | none |
| CCNJ | 0.355 | 0.2753 | 2.9417 | 0.01528 | 0.061 | -3.25 | -0.51 | none |
| ERCC8 | 0.451 | 0.2751 | 2.9411 | 0.0153 | 0.061 | -3.26 | -0.454 | none |
| NDUFA3 | 0.63 | 0.4078 | 2.9409 | 0.0153 | 0.061 | -3.26 | -0.7625 | none |
| AVEN | 0.44 | 0.3755 | 2.9406 | 0.01531 | 0.061 | -3.26 | -0.8074 | none |
| IPO13 | 0.425 | 0.4921 | 2.9398 | 0.01533 | 0.061 | -3.26 | -0.1169 | none |
| HPRT1 | -0.496 | -0.4603 | -2.94 | 0.01533 | 0.061 | -3.26 | 1.89178 | none |
| TMEM245 | 0.38 | 0.2982 | 2.9388 | 0.01535 | 0.061 | -3.26 | -0.2255 | none |
| HS3ST3B1 | 0.455 | 0.394 | 2.9385 | 0.01536 | 0.061 | -3.26 | -1.4242 | none |
| PSME2 | -0.37 | -0.3145 | -2.938 | 0.01537 | 0.061 | -3.26 | 1.2408 | none |
| HELZ | 0.536 | 0.4889 | 2.9379 | 0.01538 | 0.061 | -3.26 | -1.0266 | none |
| VPS36 | 0.403 | 0.3133 | 2.937 | 0.0154 | 0.061 | -3.26 | -0.6242 | none |
| MYOM3 | 0.318 | 0.2398 | 2.9368 | 0.01541 | 0.061 | -3.26 | -1.2568 | none |
| AGO3 | 0.297 | 0.2712 | 2.9363 | 0.01542 | 0.061 | -3.26 | 0.43637 | none |

| | | | | | | | | |
|---|---|---|---|---|---|---|---|---|
| B3GNT5 | 0.509 | 0.4529 | 2.9359 | 0.01543 | 0.061 | -3.26 | -0.5007 | none |
| SUPT20H | 0.353 | 0.3205 | 2.9356 | 0.01544 | 0.061 | -3.26 | 0.17806 | none |
| HSF2 | -0.566 | -0.4598 | -2.935 | 0.01544 | 0.061 | -3.27 | 2.13756 | AA-related |
| NUP50 | 0.7 | 0.6149 | 2.9351 | 0.01545 | 0.061 | -3.27 | -1.1079 | none |
| PEX1 | 0.405 | 0.3924 | 2.933 | 0.01551 | 0.061 | -3.27 | -0.0708 | none |
| NEGR1 | 0.324 | 0.2627 | 2.9328 | 0.01551 | 0.061 | -3.27 | -1.3043 | none |
| SNTA1 | -0.224 | -0.1888 | -2.933 | 0.01551 | 0.061 | -3.27 | 1.1977 | none |
| MACROD1 | -0.643 | -0.3686 | -2.933 | 0.01552 | 0.061 | -3.27 | 0.89643 | none |
| PLEKHA7 | 0.682 | 0.6629 | 2.9326 | 0.01552 | 0.061 | -3.27 | -0.087 | none |
| FAM133B | 0.566 | 0.4507 | 2.9318 | 0.01554 | 0.061 | -3.27 | -1.1863 | none |
| HTN3 | 0.453 | 0.328 | 2.9312 | 0.01555 | 0.061 | -3.27 | -0.8826 | none |
| NECAB3 | 0.477 | 0.3383 | 2.9312 | 0.01555 | 0.061 | -3.27 | -1.3507 | none |
| EMC1 | 0.428 | 0.4132 | 2.9308 | 0.01556 | 0.061 | -3.27 | -0.8419 | none |
| SIMC1 | 0.594 | 0.5662 | 2.9302 | 0.01558 | 0.061 | -3.27 | -0.4109 | none |
| MSR1 | 0.617 | 0.488 | 2.929 | 0.01561 | 0.061 | -3.28 | -0.3571 | none |
| EPB41 | 0.905 | 0.8471 | 2.928 | 0.01564 | 0.062 | -3.28 | -0.951 | none |
| MZT2A | 0.404 | 0.3146 | 2.9273 | 0.01566 | 0.062 | -3.28 | -1.1476 | none |
| HDGF | -0.324 | -0.1991 | -2.927 | 0.01566 | 0.062 | -3.28 | 1.07444 | none |
| SLC40A1 | 0.302 | 0.2406 | 2.9268 | 0.01567 | 0.062 | -3.28 | -0.6484 | none |
| SUMO2 | -0.478 | -0.487 | -2.927 | 0.01567 | 0.062 | -3.28 | 1.66083 | none |
| HILPDA | 0.461 | 0.3603 | 2.9268 | 0.01567 | 0.062 | -3.28 | -0.1381 | none |
| TTC39C | 0.417 | 0.2259 | 2.9264 | 0.01568 | 0.062 | -3.28 | -1.0937 | none |
| BTBD9 | 0.407 | 0.4558 | 2.9256 | 0.0157 | 0.062 | -3.28 | -0.5318 | none |
| TBCCD1 | 0.549 | 0.3799 | 2.9255 | 0.0157 | 0.062 | -3.28 | -1.3625 | none |
| PNCK | 0.431 | 0.407 | 2.9251 | 0.01571 | 0.062 | -3.28 | -0.3364 | none |
| RPS10-NU | 0.511 | 0.3701 | 2.9247 | 0.01572 | 0.062 | -3.28 | -1.1272 | none |
| NBPF10 | 0.366 | 0.3971 | 2.9246 | 0.01573 | 0.062 | -3.28 | 0.09595 | none |
| HGSNAT | 0.582 | 0.3359 | 2.9245 | 0.01573 | 0.062 | -3.28 | -1.0778 | none |
| R3HCC1L | 0.778 | 0.6794 | 2.9245 | 0.01573 | 0.062 | -3.28 | -1.3828 | none |
| CARD8 | 0.397 | 0.2844 | 2.9242 | 0.01574 | 0.062 | -3.28 | -1.041 | none |
| TUBA1B | -0.596 | -0.5735 | -2.924 | 0.01575 | 0.062 | -3.28 | 2.10906 | AA-related |
| FAM135A | 0.604 | 0.4045 | 2.9231 | 0.01577 | 0.062 | -3.29 | -0.7757 | none |
| CXCL12 | -0.268 | -0.1528 | -2.923 | 0.01577 | 0.062 | -3.29 | 1.20617 | none |
| ZPBP2 | 0.416 | 0.269 | 2.9229 | 0.01577 | 0.062 | -3.29 | -1.1828 | none |
| ESCO1 | 0.596 | 0.5418 | 2.9216 | 0.01581 | 0.062 | -3.29 | -0.6619 | none |
| CCP110 | 0.82 | 0.859 | 2.9212 | 0.01582 | 0.062 | -3.29 | -0.4451 | none |
| GALNT7 | 0.355 | 0.3374 | 2.9212 | 0.01582 | 0.062 | -3.29 | -0.4576 | none |
| USP49 | 0.577 | 0.4884 | 2.9211 | 0.01582 | 0.062 | -3.29 | -1.4895 | none |
| ATP6V1B2 | 0.264 | 0.1527 | 2.921 | 0.01582 | 0.062 | -3.29 | -0.8301 | none |
| DNAJB14 | 0.427 | 0.3494 | 2.9196 | 0.01586 | 0.062 | -3.29 | -0.7685 | none |

| | | | | | | | | |
|---|---|---|---|---|---|---|---|---|
| TUBA1C | -0.513 | -0.5095 | -2.918 | 0.0159 | 0.062 | -3.29 | 1.55699 | none |
| ARL2 | 0.315 | 0.2516 | 2.9178 | 0.01591 | 0.062 | -3.29 | -1.1686 | none |
| ELF2 | 0.434 | 0.4263 | 2.9174 | 0.01592 | 0.062 | -3.29 | -0.6441 | none |
| TMEM175 | 0.37 | 0.3015 | 2.9171 | 0.01593 | 0.062 | -3.3 | -0.7579 | none |
| NAPSA | -0.422 | -0.3213 | -2.917 | 0.01593 | 0.062 | -3.3 | 2.33528 | AA-related |
| HOMER3 | 0.323 | 0.196 | 2.9165 | 0.01594 | 0.062 | -3.3 | -0.2968 | none |
| BCAT1 | 0.38 | 0.2451 | 2.9161 | 0.01595 | 0.062 | -3.3 | -0.9603 | none |
| ACVR1 | 0.36 | 0.2694 | 2.9161 | 0.01595 | 0.062 | -3.3 | 0.70557 | none |
| ARSD | 0.664 | 0.5028 | 2.9149 | 0.01599 | 0.062 | -3.3 | -1.1521 | none |
| MMRN2 | 0.433 | 0.3298 | 2.9148 | 0.01599 | 0.062 | -3.3 | -1.1046 | none |
| PLP2 | -0.353 | -0.3372 | -2.915 | 0.01599 | 0.062 | -3.3 | 2.51941 | AA-related |
| FDFT1 | -0.289 | -0.2342 | -2.914 | 0.01601 | 0.062 | -3.3 | 2.26655 | AA-related |
| AOAH | 0.758 | 0.6702 | 2.914 | 0.01601 | 0.062 | -3.3 | -0.6871 | none |
| DDX17 | 0.258 | 0.2393 | 2.9136 | 0.01602 | 0.062 | -3.3 | -0.2346 | none |
| CHST2 | -0.308 | -0.2383 | -2.913 | 0.01603 | 0.062 | -3.3 | 1.89055 | none |
| GCK | 0.252 | 0.2371 | 2.9133 | 0.01603 | 0.062 | -3.3 | 0.15561 | none |
| FRMPD2 | 0.378 | 0.2663 | 2.9128 | 0.01604 | 0.063 | -3.3 | -1.2008 | none |
| ABCE1 | 0.241 | 0.1978 | 2.9117 | 0.01607 | 0.063 | -3.3 | -0.1365 | none |
| FNIP1 | 0.448 | 0.3841 | 2.9102 | 0.01611 | 0.063 | -3.31 | -1.1785 | none |
| VPS8 | 0.435 | 0.3694 | 2.9091 | 0.01614 | 0.063 | -3.31 | -0.7976 | none |
| WFS1 | 0.347 | 0.2727 | 2.909 | 0.01615 | 0.063 | -3.31 | 0.21558 | none |
| CHD1 | 0.981 | 0.9293 | 2.9086 | 0.01616 | 0.063 | -3.31 | -0.9469 | none |
| HSD11B1 | -0.305 | -0.3396 | -2.908 | 0.01618 | 0.063 | -3.31 | 1.91892 | none |
| PPP1R3B | 0.475 | 0.3953 | 2.9078 | 0.01618 | 0.063 | -3.31 | -0.6492 | none |
| TNFAIP8L2 | -0.394 | -0.2662 | -2.907 | 0.01621 | 0.063 | -3.31 | 2.32394 | AA-related |
| GAS6 | 0.45 | 0.3058 | 2.9061 | 0.01622 | 0.063 | -3.31 | 0.40404 | none |
| LCORL | 0.425 | 0.3474 | 2.9054 | 0.01624 | 0.063 | -3.31 | -0.3929 | none |
| PKHD1L1 | 0.35 | 0.2736 | 2.9048 | 0.01626 | 0.063 | -3.32 | -0.8122 | none |
| PPP2R5C | 0.231 | 0.1231 | 2.9048 | 0.01626 | 0.063 | -3.32 | -1.2045 | none |
| C1QTNF2 | 0.414 | 0.2784 | 2.9047 | 0.01626 | 0.063 | -3.32 | -1.1731 | none |
| SYNE3 | 0.582 | 0.5914 | 2.9038 | 0.01629 | 0.063 | -3.32 | -0.8066 | none |
| CCDC167 | -0.403 | -0.3976 | -2.904 | 0.01629 | 0.063 | -3.32 | 1.16472 | none |
| ZFP64 | 0.556 | 0.5229 | 2.9031 | 0.01631 | 0.063 | -3.32 | -1.1618 | none |
| CCDC62 | 0.383 | 0.2685 | 2.903 | 0.01631 | 0.063 | -3.32 | -0.8163 | none |
| MCMBP | 0.253 | 0.1641 | 2.9028 | 0.01632 | 0.063 | -3.32 | -0.4254 | none |
| MSL2 | 0.302 | 0.2175 | 2.9017 | 0.01634 | 0.063 | -3.32 | -0.5256 | none |
| OVCA2 | -0.83 | -0.6737 | -2.902 | 0.01634 | 0.063 | -3.32 | 2.25464 | AA-related |
| CHIT1 | -0.474 | -0.3505 | -2.901 | 0.01636 | 0.063 | -3.32 | 2.41806 | AA-related |
| CELF1 | 0.345 | 0.3513 | 2.8989 | 0.01642 | 0.064 | -3.33 | 0.04376 | none |
| ZSCAN16 | -0.552 | -0.3859 | -2.899 | 0.01643 | 0.064 | -3.33 | 2.08578 | none |

| | | | | | | | | |
|---|---|---|---|---|---|---|---|---|
| SLC12A1 | 0.301 | 0.2281 | 2.8986 | 0.01643 | 0.064 | -3.33 | -1.2394 | none |
| WDR34 | -0.319 | -0.2718 | -2.899 | 0.01643 | 0.064 | -3.33 | 2.27137 | AA-related |
| ADK | -0.304 | -0.2513 | -2.899 | 0.01643 | 0.064 | -3.33 | 2.35004 | AA-related |
| SASH1 | 0.27 | 0.2091 | 2.8979 | 0.01645 | 0.064 | -3.33 | 0.20351 | none |
| AGK | 0.352 | 0.3101 | 2.8978 | 0.01645 | 0.064 | -3.33 | -0.0418 | none |
| NUDT9 | 0.546 | 0.4357 | 2.8975 | 0.01646 | 0.064 | -3.33 | -1.3628 | none |
| PRDX2 | -0.351 | -0.2994 | -2.897 | 0.01647 | 0.064 | -3.33 | 0.6446 | none |
| PHF3 | 0.955 | 0.9343 | 2.8968 | 0.01648 | 0.064 | -3.33 | -0.8511 | none |
| CARM1 | 0.531 | 0.5418 | 2.896 | 0.0165 | 0.064 | -3.33 | 0.20215 | none |
| RBBP9 | 0.556 | 0.4181 | 2.8957 | 0.01651 | 0.064 | -3.33 | 0.29515 | none |
| FAM83G | 0.397 | 0.3919 | 2.8948 | 0.01654 | 0.064 | -3.33 | -0.5975 | none |
| OXCT1 | 0.339 | 0.2592 | 2.8946 | 0.01654 | 0.064 | -3.33 | -0.0373 | none |
| BRD8 | 0.795 | 0.7722 | 2.8933 | 0.01658 | 0.064 | -3.33 | -1.1804 | none |
| IFIT3 | 0.307 | 0.1961 | 2.8932 | 0.01658 | 0.064 | -3.33 | -0.7935 | none |
| SMC6 | 0.857 | 0.8264 | 2.8928 | 0.01659 | 0.064 | -3.34 | -0.1009 | none |
| ARID4A | 0.647 | 0.5239 | 2.8923 | 0.01661 | 0.064 | -3.34 | -1.1596 | none |
| CD160 | 0.552 | 0.4765 | 2.892 | 0.01661 | 0.064 | -3.34 | -0.5469 | none |
| CINP | 0.418 | 0.278 | 2.8917 | 0.01662 | 0.064 | -3.34 | -0.208 | none |
| ZNF419 | 0.37 | 0.2685 | 2.8916 | 0.01663 | 0.064 | -3.34 | -1.0855 | none |
| BRICD5 | 0.689 | 0.6538 | 2.8911 | 0.01664 | 0.064 | -3.34 | -1.2731 | none |
| CCDC39 | 0.372 | 0.2641 | 2.8896 | 0.01668 | 0.064 | -3.34 | -0.7502 | none |
| VPS4B | 0.346 | 0.2224 | 2.8888 | 0.0167 | 0.064 | -3.34 | -1.0759 | none |
| HSPA1A | -0.362 | -0.2748 | -2.888 | 0.01672 | 0.064 | -3.34 | 0.50508 | none |
| MCOLN2 | 0.511 | 0.3292 | 2.8873 | 0.01675 | 0.065 | -3.34 | -1.0199 | none |
| POP5 | 0.333 | 0.213 | 2.8871 | 0.01675 | 0.065 | -3.34 | -1.2188 | none |
| UAP1L1 | 0.367 | 0.3039 | 2.8869 | 0.01676 | 0.065 | -3.35 | -0.9107 | none |
| POLR3GL | -0.254 | -0.2284 | -2.887 | 0.01676 | 0.065 | -3.35 | 1.22837 | none |
| PSMA2 | -0.44 | -0.3891 | -2.886 | 0.01679 | 0.065 | -3.35 | 1.07435 | none |
| GTDC1 | 0.386 | 0.305 | 2.8855 | 0.0168 | 0.065 | -3.35 | -0.6433 | none |
| APOF | 0.315 | 0.2289 | 2.8853 | 0.0168 | 0.065 | -3.35 | -1.1847 | none |
| TDRD9 | 0.492 | 0.3779 | 2.885 | 0.01681 | 0.065 | -3.35 | -0.9332 | none |
| PHLPP1 | 0.764 | 0.6765 | 2.8844 | 0.01683 | 0.065 | -3.35 | -1.0676 | none |
| UBQLN2 | 0.395 | 0.3355 | 2.8833 | 0.01686 | 0.065 | -3.35 | 0.10741 | none |
| SLCO4A1 | 0.283 | 0.2889 | 2.8831 | 0.01687 | 0.065 | -3.35 | 0.26743 | none |
| TAF1A | 0.398 | 0.2306 | 2.8831 | 0.01687 | 0.065 | -3.35 | -0.6838 | none |
| GALNT4 | 0.347 | 0.2324 | 2.883 | 0.01687 | 0.065 | -3.35 | -0.8258 | none |
| MTERF2 | 0.41 | 0.3428 | 2.8827 | 0.01688 | 0.065 | -3.35 | 0.87578 | none |
| KLHL6 | 0.54 | 0.4918 | 2.8825 | 0.01688 | 0.065 | -3.35 | -1.3664 | none |
| TEX2 | 0.526 | 0.5267 | 2.8821 | 0.01689 | 0.065 | -3.35 | -1.1521 | none |
| STK31 | 0.442 | 0.3661 | 2.8813 | 0.01692 | 0.065 | -3.35 | -1.0115 | none |

| | | | | | | | | |
|---|---|---|---|---|---|---|---|---|
| BEAN1 | 0.466 | 0.3765 | 2.881 | 0.01693 | 0.065 | -3.35 | -1.2383 | none |
| SLC22A4 | 0.258 | 0.1376 | 2.8807 | 0.01693 | 0.065 | -3.36 | -0.4467 | none |
| WDR61 | -0.446 | -0.4146 | -2.881 | 0.01694 | 0.065 | -3.36 | 2.18562 | AA-related |
| CACTIN | 0.594 | 0.5924 | 2.8794 | 0.01697 | 0.065 | -3.36 | -1.1592 | none |
| ECHS1 | -0.299 | -0.2303 | -2.879 | 0.01697 | 0.065 | -3.36 | 2.05634 | none |
| MKRN3 | 0.503 | 0.3691 | 2.8794 | 0.01697 | 0.065 | -3.36 | -1.0622 | none |
| SPCS3 | -0.475 | -0.4234 | -2.879 | 0.01698 | 0.065 | -3.36 | 2.1637 | AA-related |
| CYFIP2 | 0.609 | 0.5753 | 2.8789 | 0.01698 | 0.065 | -3.36 | -0.8359 | none |
| MTA3 | 0.398 | 0.2632 | 2.8786 | 0.01699 | 0.065 | -3.36 | -0.8606 | none |
| CRCP | 0.729 | 0.6759 | 2.8786 | 0.01699 | 0.065 | -3.36 | -0.6566 | none |
| PCNX4 | 0.359 | 0.2775 | 2.8783 | 0.017 | 0.065 | -3.36 | -0.981 | none |
| NPPA | -0.324 | -0.257 | -2.878 | 0.01702 | 0.065 | -3.36 | 1.30793 | none |
| HAVCR1 | 0.602 | 0.4327 | 2.8768 | 0.01705 | 0.065 | -3.36 | -1.0928 | none |
| OMD | 0.386 | 0.3011 | 2.8765 | 0.01706 | 0.065 | -3.36 | -1.0132 | none |
| PIK3CB | 0.649 | 0.592 | 2.8761 | 0.01707 | 0.065 | -3.36 | -1.2738 | none |
| STAM | 0.336 | 0.1928 | 2.876 | 0.01707 | 0.065 | -3.36 | -1.2005 | none |
| SARDH | 0.317 | 0.227 | 2.8752 | 0.01709 | 0.065 | -3.36 | -0.9272 | none |
| IKZF3 | 0.798 | 0.8238 | 2.8746 | 0.01711 | 0.065 | -3.37 | -0.8267 | none |
| LUC7L | 0.357 | 0.3409 | 2.8737 | 0.01714 | 0.066 | -3.37 | -1.3586 | none |
| FANCL | -0.267 | -0.21 | -2.873 | 0.01715 | 0.066 | -3.37 | 2.31114 | AA-related |
| UPF3A | 0.526 | 0.4749 | 2.8732 | 0.01715 | 0.066 | -3.37 | -0.7865 | none |
| KCNMB3 | 0.21 | 0.1536 | 2.8722 | 0.01718 | 0.066 | -3.37 | -1.3407 | none |
| MORC3 | 0.568 | 0.5255 | 2.8715 | 0.0172 | 0.066 | -3.37 | -1.0108 | none |
| IL17RA | 0.48 | 0.447 | 2.8713 | 0.0172 | 0.066 | -3.37 | 0.04421 | none |
| TMEM159 | 0.441 | 0.2525 | 2.8711 | 0.01721 | 0.066 | -3.37 | -1.1149 | none |
| APOC1 | 0.855 | 0.6653 | 2.871 | 0.01721 | 0.066 | -3.37 | -0.439 | none |
| PSMG2 | -0.526 | -0.5069 | -2.871 | 0.01721 | 0.066 | -3.37 | 2.14741 | AA-related |
| TBC1D15 | 0.339 | 0.2311 | 2.8708 | 0.01722 | 0.066 | -3.37 | -0.5073 | none |
| BRCA2 | 0.665 | 0.6967 | 2.8707 | 0.01722 | 0.066 | -3.37 | -0.9011 | none |
| COLGALT1 | 0.425 | 0.39 | 2.8701 | 0.01724 | 0.066 | -3.37 | -1.3612 | none |
| RNF145 | 0.239 | 0.2333 | 2.8699 | 0.01724 | 0.066 | -3.37 | 0.4475 | none |
| TLL1 | 0.232 | 0.2326 | 2.8693 | 0.01726 | 0.066 | -3.37 | 0.89998 | none |
| PAQR8 | 0.357 | 0.2472 | 2.8689 | 0.01727 | 0.066 | -3.37 | -0.5977 | none |
| ZNF623 | 0.477 | 0.4219 | 2.8687 | 0.01728 | 0.066 | -3.38 | -1.116 | none |
| KDM4B | 0.676 | 0.6738 | 2.8683 | 0.01729 | 0.066 | -3.38 | -0.8004 | none |
| ZBED2 | -0.298 | -0.1929 | -2.868 | 0.01729 | 0.066 | -3.38 | 2.09228 | AA-related |
| TSHZ2 | 0.57 | 0.5472 | 2.8679 | 0.0173 | 0.066 | -3.38 | -1.2357 | none |
| CBX5 | 0.347 | 0.2653 | 2.8675 | 0.01732 | 0.066 | -3.38 | -1.2667 | none |
| ARHGEF4 | 0.558 | 0.4363 | 2.8672 | 0.01732 | 0.066 | -3.38 | -1.0415 | none |
| CNOT6L | 0.251 | 0.181 | 2.8671 | 0.01733 | 0.066 | -3.38 | -0.6 | none |

| | | | | | | | | |
|---|---|---|---|---|---|---|---|---|
| PARP8 | 0.501 | 0.3883 | 2.8669 | 0.01733 | 0.066 | -3.38 | -0.8148 | none |
| ZNF18 | 0.676 | 0.5017 | 2.8664 | 0.01735 | 0.066 | -3.38 | -1.489 | none |
| SRGN | -0.609 | -0.5885 | -2.866 | 0.01735 | 0.066 | -3.38 | 1.62087 | none |
| RFWD3 | 0.539 | 0.5008 | 2.8657 | 0.01737 | 0.066 | -3.38 | -1.1183 | none |
| SGCB | 0.48 | 0.3423 | 2.8656 | 0.01737 | 0.066 | -3.38 | -0.0876 | none |
| TGIF2 | 0.364 | 0.2849 | 2.8656 | 0.01737 | 0.066 | -3.38 | -0.7588 | none |
| SUMF1 | 0.374 | 0.2479 | 2.8648 | 0.0174 | 0.066 | -3.38 | -0.9489 | none |
| EXOC3L2 | 0.431 | 0.3209 | 2.8644 | 0.01741 | 0.066 | -3.38 | -1.028 | none |
| PSPC1 | 0.431 | 0.4437 | 2.8636 | 0.01743 | 0.066 | -3.38 | -0.8192 | none |
| CYP27B1 | 0.806 | 0.579 | 2.863 | 0.01745 | 0.066 | -3.38 | -1.0215 | none |
| FRMD5 | 0.466 | 0.4545 | 2.8627 | 0.01746 | 0.066 | -3.39 | -0.5034 | none |
| CACNA1F | 0.657 | 0.5142 | 2.8622 | 0.01747 | 0.066 | -3.39 | -1.101 | none |
| FAM210B | 0.373 | 0.2614 | 2.8619 | 0.01748 | 0.066 | -3.39 | -0.0492 | none |
| ATP1B4 | 0.413 | 0.3029 | 2.8618 | 0.01748 | 0.066 | -3.39 | -1.4093 | none |
| THUMPD3 | 0.409 | 0.3574 | 2.8613 | 0.0175 | 0.066 | -3.39 | -0.5417 | none |
| CUL9 | 0.395 | 0.3727 | 2.8591 | 0.01756 | 0.067 | -3.39 | -1.0552 | none |
| EEF1A2 | 0.511 | 0.3453 | 2.8577 | 0.01761 | 0.067 | -3.39 | -1.1573 | none |
| DHRS1 | 0.384 | 0.2505 | 2.8566 | 0.01763 | 0.067 | -3.4 | -0.6006 | none |
| ZC3H8 | 0.261 | 0.196 | 2.8564 | 0.01764 | 0.067 | -3.4 | -0.7114 | none |
| NXNL2 | 0.383 | 0.289 | 2.856 | 0.01765 | 0.067 | -3.4 | -1.2892 | none |
| FKBP11 | 0.315 | 0.1827 | 2.856 | 0.01765 | 0.067 | -3.4 | -0.6295 | none |
| GLRA3 | 0.236 | 0.1684 | 2.8549 | 0.01769 | 0.067 | -3.4 | -1.3492 | none |
| TXN | -0.556 | -0.5566 | -2.855 | 0.0177 | 0.067 | -3.4 | 2.16813 | AA-related |
| PRAF2 | -0.657 | -0.523 | -2.854 | 0.01773 | 0.067 | -3.4 | 1.98804 | none |
| ZNF773 | 0.513 | 0.3761 | 2.8532 | 0.01774 | 0.067 | -3.4 | -1.1523 | none |
| KIAA1257 | 0.582 | 0.4773 | 2.8519 | 0.01778 | 0.067 | -3.4 | -1.0706 | none |
| L1CAM | -0.327 | -0.2507 | -2.852 | 0.01778 | 0.067 | -3.4 | 0.29134 | none |
| GPR155 | 0.482 | 0.3299 | 2.8511 | 0.0178 | 0.067 | -3.4 | -0.6948 | none |
| C1QTNF6 | 0.401 | 0.2985 | 2.8509 | 0.01781 | 0.067 | -3.4 | -0.9056 | none |
| TMEM91 | 0.377 | 0.2067 | 2.8509 | 0.01781 | 0.067 | -3.4 | -0.8661 | none |
| POLD4 | 0.361 | 0.2631 | 2.8507 | 0.01781 | 0.067 | -3.4 | -1.0587 | none |
| OR4D1 | 0.315 | 0.2297 | 2.8498 | 0.01784 | 0.067 | -3.41 | -1.285 | none |
| PDCD7 | 0.424 | 0.346 | 2.8497 | 0.01784 | 0.067 | -3.41 | -1.5027 | none |
| MTIF2 | 0.555 | 0.4661 | 2.849 | 0.01786 | 0.067 | -3.41 | -0.9053 | none |
| CLEC16A | 0.509 | 0.5337 | 2.849 | 0.01786 | 0.067 | -3.41 | -0.3827 | none |
| SULT1C2 | 0.383 | 0.2596 | 2.8487 | 0.01787 | 0.068 | -3.41 | -1.4049 | none |
| RPS24 | -0.378 | -0.3808 | -2.849 | 0.01788 | 0.068 | -3.41 | 2.12533 | AA-related |
| NHP2 | -0.327 | -0.2984 | -2.848 | 0.01789 | 0.068 | -3.41 | 1.72965 | none |
| OTUD1 | -0.278 | -0.193 | -2.848 | 0.01789 | 0.068 | -3.41 | 1.93371 | none |
| SHOC2 | 0.351 | 0.1706 | 2.8478 | 0.0179 | 0.068 | -3.41 | -0.3366 | none |

| | | | | | | | | |
|---|---|---|---|---|---|---|---|---|
| SH2D1A | -0.592 | -0.5488 | -2.848 | 0.0179 | 0.068 | -3.41 | 0.72843 | none |
| ICOSLG | 0.448 | 0.402 | 2.8471 | 0.01792 | 0.068 | -3.41 | -1.4249 | none |
| OR8J3 | 0.264 | 0.1816 | 2.8463 | 0.01795 | 0.068 | -3.41 | -1.4047 | none |
| TMEM80 | 0.35 | 0.2885 | 2.8451 | 0.01798 | 0.068 | -3.41 | -0.3805 | none |
| KPNA4 | 0.489 | 0.4395 | 2.8447 | 0.01799 | 0.068 | -3.41 | -0.5604 | none |
| E2F2 | 0.353 | 0.3062 | 2.8441 | 0.01801 | 0.068 | -3.42 | -1.4854 | none |
| MUSTN1 | -0.554 | -0.4057 | -2.844 | 0.01802 | 0.068 | -3.42 | 1.55122 | none |
| GRIA1 | -0.533 | -0.4175 | -2.844 | 0.01803 | 0.068 | -3.42 | -0.0875 | none |
| ZNF236 | 0.95 | 0.8503 | 2.8434 | 0.01803 | 0.068 | -3.42 | -0.9721 | none |
| PLEK2 | -0.663 | -0.4936 | -2.843 | 0.01803 | 0.068 | -3.42 | 1.98708 | none |
| ZNF524 | -0.296 | -0.3327 | -2.843 | 0.01804 | 0.068 | -3.42 | 1.35051 | none |
| FRA10AC1 | 0.566 | 0.5763 | 2.843 | 0.01805 | 0.068 | -3.42 | -0.6098 | none |
| C11ORF52 | 0.464 | 0.317 | 2.8428 | 0.01805 | 0.068 | -3.42 | -1.0902 | none |
| LILRA6 | 0.216 | 0.1589 | 2.8421 | 0.01807 | 0.068 | -3.42 | -1.4884 | none |
| DHX30 | 0.412 | 0.3956 | 2.8414 | 0.01809 | 0.068 | -3.42 | -0.26 | none |
| RPSA | -0.53 | -0.5123 | -2.841 | 0.01809 | 0.068 | -3.42 | 1.75425 | none |
| HACE1 | 0.411 | 0.2688 | 2.841 | 0.01811 | 0.068 | -3.42 | -0.9975 | none |
| ZNF823 | 0.436 | 0.4305 | 2.8404 | 0.01812 | 0.068 | -3.42 | 0.30872 | none |
| PSG4 | -0.21 | -0.1346 | -2.84 | 0.01813 | 0.068 | -3.42 | -0.2092 | none |
| TRIB3 | -0.275 | -0.2025 | -2.839 | 0.01816 | 0.068 | -3.42 | 1.82888 | none |
| COA5 | 0.302 | 0.1617 | 2.8385 | 0.01818 | 0.068 | -3.43 | 0.33515 | none |
| NRF1 | 0.308 | 0.2634 | 2.8384 | 0.01818 | 0.068 | -3.43 | -0.2494 | none |
| CNST | 0.537 | 0.5087 | 2.8373 | 0.01822 | 0.068 | -3.43 | 0.15633 | none |
| CDCA7 | -0.339 | -0.2962 | -2.837 | 0.01823 | 0.068 | -3.43 | 2.6076 | AA-related |
| ARL17B | 0.764 | 0.4825 | 2.837 | 0.01823 | 0.068 | -3.43 | -1.3714 | none |
| LEPR | 0.257 | 0.1475 | 2.8369 | 0.01823 | 0.068 | -3.43 | 0.58279 | none |
| POLE | 0.745 | 0.7668 | 2.8367 | 0.01824 | 0.068 | -3.43 | -1.0846 | none |
| HCFC2 | 0.345 | 0.2817 | 2.8367 | 0.01824 | 0.068 | -3.43 | -1.2753 | none |
| SNURF | 0.887 | 0.5252 | 2.8365 | 0.01825 | 0.068 | -3.43 | -0.709 | none |
| STK11 | 0.496 | 0.4876 | 2.8351 | 0.01829 | 0.069 | -3.43 | -0.8687 | none |
| IRAK2 | 0.35 | 0.3029 | 2.8347 | 0.0183 | 0.069 | -3.43 | -0.7403 | none |
| LLGL1 | 0.319 | 0.2952 | 2.834 | 0.01832 | 0.069 | -3.43 | -1.1355 | none |
| MADD | 0.636 | 0.6319 | 2.8334 | 0.01834 | 0.069 | -3.43 | -1.2416 | none |
| ZNF432 | 0.488 | 0.4215 | 2.8333 | 0.01834 | 0.069 | -3.43 | -0.8921 | none |
| FAM120C | 0.249 | 0.1926 | 2.8332 | 0.01835 | 0.069 | -3.43 | 0.20117 | none |
| SATB2 | 0.521 | 0.3942 | 2.8324 | 0.01837 | 0.069 | -3.44 | -0.7462 | none |
| CCDC74A | 0.338 | 0.3035 | 2.8319 | 0.01839 | 0.069 | -3.44 | -0.9523 | none |
| POM121C | 0.673 | 0.7277 | 2.8316 | 0.0184 | 0.069 | -3.44 | -0.9241 | none |
| NDUFS5 | -0.39 | -0.3993 | -2.83 | 0.01844 | 0.069 | -3.44 | 2.19897 | AA-related |
| PJA2 | 0.391 | 0.2877 | 2.83 | 0.01845 | 0.069 | -3.44 | -1.3139 | none |

| | | | | | | | | |
|---|---|---|---|---|---|---|---|---|
| TMEM132 | 0.279 | 0.1918 | 2.8292 | 0.01847 | 0.069 | -3.44 | -1.3801 | none |
| SRP14 | -0.237 | -0.2215 | -2.829 | 0.01849 | 0.069 | -3.44 | 1.57968 | none |
| PDF | 0.594 | 0.5457 | 2.8284 | 0.0185 | 0.069 | -3.44 | 0.38007 | none |
| PRKRA | 0.242 | 0.1353 | 2.8283 | 0.0185 | 0.069 | -3.44 | -0.3396 | none |
| ZZEF1 | 0.644 | 0.6513 | 2.8283 | 0.0185 | 0.069 | -3.44 | -0.6845 | none |
| CREB3L3 | 0.418 | 0.4413 | 2.8283 | 0.0185 | 0.069 | -3.44 | -0.3496 | none |
| FAM13B | 0.644 | 0.5627 | 2.8283 | 0.0185 | 0.069 | -3.44 | -0.907 | none |
| PGD | -0.214 | -0.158 | -2.826 | 0.01856 | 0.069 | -3.44 | 2.2267 | AA-related |
| SIK2 | 0.594 | 0.5445 | 2.8262 | 0.01857 | 0.069 | -3.45 | -0.9134 | none |
| NBR1 | 0.562 | 0.501 | 2.8255 | 0.01859 | 0.069 | -3.45 | -1.0854 | none |
| SFTPB | 0.472 | 0.3486 | 2.825 | 0.0186 | 0.069 | -3.45 | -1.1291 | none |
| QTRT2 | 0.386 | 0.291 | 2.8248 | 0.01861 | 0.069 | -3.45 | -0.2667 | none |
| GPR137 | 0.488 | 0.3743 | 2.824 | 0.01864 | 0.069 | -3.45 | -0.0162 | none |
| HOXB7 | 0.961 | 0.8688 | 2.8236 | 0.01865 | 0.07 | -3.45 | -0.8988 | none |
| ZNF257 | 0.792 | 0.5603 | 2.8232 | 0.01866 | 0.07 | -3.45 | -0.8442 | none |
| HLA-E | 0.295 | 0.1568 | 2.8228 | 0.01867 | 0.07 | -3.45 | 0.00791 | none |
| STAU2 | 0.48 | 0.3527 | 2.8217 | 0.01871 | 0.07 | -3.45 | -0.0722 | none |
| EXT2 | 0.265 | 0.1681 | 2.8216 | 0.01871 | 0.07 | -3.45 | 0.40009 | none |
| TCAF2 | 0.373 | 0.3342 | 2.8215 | 0.01871 | 0.07 | -3.45 | -0.8082 | none |
| C10ORF11 | 0.563 | 0.4287 | 2.8213 | 0.01872 | 0.07 | -3.45 | -0.9255 | none |
| SUB1 | -0.385 | -0.3712 | -2.821 | 0.01872 | 0.07 | -3.45 | 1.83375 | none |
| SIRPG | -0.232 | -0.2086 | -2.821 | 0.01873 | 0.07 | -3.45 | 1.62802 | none |
| MBTD1 | 0.474 | 0.4022 | 2.8207 | 0.01874 | 0.07 | -3.45 | -0.8286 | none |
| AVPR1A | 0.24 | 0.1578 | 2.8201 | 0.01876 | 0.07 | -3.46 | -0.2033 | none |
| TBCA | -0.352 | -0.3557 | -2.819 | 0.0188 | 0.07 | -3.46 | 1.85352 | none |
| DDX10 | 0.663 | 0.6421 | 2.8187 | 0.0188 | 0.07 | -3.46 | -0.298 | none |
| GOLGA2 | 0.614 | 0.5483 | 2.8186 | 0.0188 | 0.07 | -3.46 | -0.7017 | none |
| NDUFA4 | -0.444 | -0.4504 | -2.818 | 0.01881 | 0.07 | -3.46 | 2.24306 | AA-related |
| ZNF106 | 0.91 | 0.895 | 2.8183 | 0.01881 | 0.07 | -3.46 | -0.8424 | none |
| SNX15 | 0.556 | 0.4162 | 2.8181 | 0.01882 | 0.07 | -3.46 | -0.5333 | none |
| NXPE3 | 0.421 | 0.3033 | 2.8179 | 0.01883 | 0.07 | -3.46 | -0.3408 | none |
| QRICH1 | 0.692 | 0.7022 | 2.8174 | 0.01884 | 0.07 | -3.46 | -0.6979 | none |
| POLE3 | -0.276 | -0.256 | -2.817 | 0.01885 | 0.07 | -3.46 | 2.29567 | AA-related |
| RRM2 | -0.355 | -0.3107 | -2.817 | 0.01886 | 0.07 | -3.46 | 2.10634 | AA-related |
| ZNF678 | 0.386 | 0.2876 | 2.8164 | 0.01887 | 0.07 | -3.46 | -0.4971 | none |
| RDH13 | 0.637 | 0.584 | 2.8164 | 0.01888 | 0.07 | -3.46 | -0.2773 | none |
| FAM184A | 0.668 | 0.5937 | 2.8148 | 0.01892 | 0.07 | -3.46 | -0.091 | none |
| FAM172A | 0.47 | 0.3431 | 2.8146 | 0.01893 | 0.07 | -3.46 | 0.58631 | none |
| CCDC146 | 0.411 | 0.2452 | 2.8142 | 0.01894 | 0.07 | -3.47 | -0.6023 | none |
| GPX3 | 0.809 | 0.7954 | 2.8142 | 0.01895 | 0.07 | -3.47 | -1.0203 | none |

| | | | | | | | | |
|---|---|---|---|---|---|---|---|---|
| MPDU1 | 0.412 | 0.2149 | 2.8139 | 0.01896 | 0.07 | -3.47 | -0.8093 | none |
| SHMT2 | -0.242 | -0.1787 | -2.814 | 0.01896 | 0.07 | -3.47 | 0.55244 | none |
| TSNARE1 | 0.518 | 0.4334 | 2.813 | 0.01898 | 0.07 | -3.47 | -1.0008 | none |
| HIST1H1D | -0.381 | -0.3525 | -2.813 | 0.01899 | 0.07 | -3.47 | 0.93128 | none |
| MGAT4B | 0.309 | 0.3032 | 2.8121 | 0.01901 | 0.07 | -3.47 | -0.2704 | none |
| DCDC2 | 0.375 | 0.2707 | 2.8119 | 0.01902 | 0.07 | -3.47 | -1.3808 | none |
| U2AF1L4 | 0.375 | 0.2299 | 2.8114 | 0.01903 | 0.07 | -3.47 | -1.1741 | none |
| DNAJC25 | 0.32 | 0.2602 | 2.8114 | 0.01903 | 0.07 | -3.47 | -0.6939 | none |
| SCARB2 | 0.449 | 0.3171 | 2.8104 | 0.01907 | 0.071 | -3.47 | -1.1373 | none |
| SULF2 | 0.495 | 0.512 | 2.8096 | 0.01909 | 0.071 | -3.47 | -0.1314 | none |
| FAM227B | 0.568 | 0.4187 | 2.8091 | 0.01911 | 0.071 | -3.47 | 0.09682 | none |
| PRPF39 | 0.507 | 0.4324 | 2.809 | 0.01911 | 0.071 | -3.47 | -0.1398 | none |
| C4ORF36 | 0.406 | 0.3694 | 2.8084 | 0.01913 | 0.071 | -3.47 | -0.5665 | none |
| ACSM3 | 0.545 | 0.4891 | 2.8083 | 0.01913 | 0.071 | -3.47 | -0.6307 | none |
| RPL22L1 | -0.587 | -0.5732 | -2.807 | 0.01916 | 0.071 | -3.48 | 2.28251 | AA-related |
| WDR72 | 0.822 | 0.6861 | 2.8073 | 0.01917 | 0.071 | -3.48 | -0.6163 | none |
| RPL22 | -0.408 | -0.4355 | -2.807 | 0.01918 | 0.071 | -3.48 | 1.97159 | none |
| CYB5D2 | 0.469 | 0.3184 | 2.8064 | 0.01919 | 0.071 | -3.48 | -1.0377 | none |
| CLDN12 | 0.542 | 0.4253 | 2.8063 | 0.0192 | 0.071 | -3.48 | -1.3191 | none |
| PCCB | 0.226 | 0.1535 | 2.8063 | 0.0192 | 0.071 | -3.48 | -0.5891 | none |
| ADAT2 | 0.365 | 0.2757 | 2.8057 | 0.01922 | 0.071 | -3.48 | -1.0211 | none |
| LMO7 | 0.422 | 0.3686 | 2.8056 | 0.01922 | 0.071 | -3.48 | -0.1592 | none |
| TP53AIP1 | 0.492 | 0.3446 | 2.8043 | 0.01926 | 0.071 | -3.48 | -1.2701 | none |
| B4GALNT2 | 0.231 | 0.2031 | 2.8043 | 0.01926 | 0.071 | -3.48 | -0.9176 | none |
| NINJ2 | -0.381 | -0.4006 | -2.804 | 0.01928 | 0.071 | -3.48 | 1.32467 | none |
| CCL22 | 0.662 | 0.5264 | 2.8037 | 0.01928 | 0.071 | -3.48 | -1.2979 | none |
| MRPL30 | 0.339 | 0.2235 | 2.8035 | 0.01929 | 0.071 | -3.48 | -1.0887 | none |
| INTS6L | 0.426 | 0.3546 | 2.8029 | 0.01931 | 0.071 | -3.48 | 0.29687 | none |
| KMT2E | 0.653 | 0.6167 | 2.8028 | 0.01931 | 0.071 | -3.48 | -0.809 | none |
| SYT13 | 0.273 | 0.181 | 2.8026 | 0.01932 | 0.071 | -3.48 | -1.3056 | none |
| RAB15 | 0.455 | 0.3998 | 2.8023 | 0.01933 | 0.071 | -3.48 | -1.2535 | none |
| RAI1 | 0.386 | 0.4055 | 2.802 | 0.01934 | 0.071 | -3.49 | -0.5013 | none |
| PRLHR | 0.278 | 0.2002 | 2.8018 | 0.01935 | 0.071 | -3.49 | -1.1788 | none |
| GNS | 0.328 | 0.1969 | 2.8011 | 0.01937 | 0.071 | -3.49 | -0.9698 | none |
| VEZT | 0.549 | 0.4715 | 2.8006 | 0.01938 | 0.071 | -3.49 | -0.8834 | none |
| CHCHD2 | -0.456 | -0.422 | -2.8 | 0.0194 | 0.071 | -3.49 | 2.12148 | AA-related |
| FAM168A | 0.766 | 0.7129 | 2.8 | 0.0194 | 0.071 | -3.49 | -1.1366 | none |
| DDX42 | 0.916 | 0.8983 | 2.7999 | 0.01941 | 0.071 | -3.49 | -1.0387 | none |
| MED10 | -0.208 | -0.1727 | -2.799 | 0.01944 | 0.071 | -3.49 | 1.74367 | none |
| AMT | 0.338 | 0.3154 | 2.798 | 0.01947 | 0.072 | -3.49 | -0.6922 | none |

| | | | | | | | | |
|---|---|---|---|---|---|---|---|---|
| XPO6 | 0.416 | 0.3953 | 2.7978 | 0.01948 | 0.072 | -3.49 | -1.0328 | none |
| 12-Sep | 0.747 | 0.5805 | 2.7977 | 0.01948 | 0.072 | -3.49 | -0.8982 | none |
| C14ORF28 | 0.387 | 0.2035 | 2.7971 | 0.0195 | 0.072 | -3.49 | -0.588 | none |
| ZNF385D | 0.314 | 0.2502 | 2.7969 | 0.01951 | 0.072 | -3.49 | -0.8675 | none |
| DNASE2 | 0.338 | 0.2868 | 2.7961 | 0.01953 | 0.072 | -3.49 | -0.1565 | none |
| LGALS7B | -0.194 | -0.1292 | -2.794 | 0.01961 | 0.072 | -3.5 | -0.2163 | none |
| USP7 | 0.884 | 0.8185 | 2.7936 | 0.01962 | 0.072 | -3.5 | -0.7851 | none |
| PYHIN1 | 0.573 | 0.4311 | 2.7934 | 0.01962 | 0.072 | -3.5 | -0.9861 | none |
| PRDM5 | 0.898 | 0.7208 | 2.7929 | 0.01964 | 0.072 | -3.5 | -0.9138 | none |
| STON1 | 0.24 | 0.2019 | 2.7924 | 0.01966 | 0.072 | -3.5 | -1.2368 | none |
| ADM | -0.56 | -0.3998 | -2.792 | 0.01968 | 0.072 | -3.5 | 1.76375 | none |
| TDRKH | 0.58 | 0.4371 | 2.7914 | 0.01969 | 0.072 | -3.5 | -0.8738 | none |
| NQO1 | -0.377 | -0.4287 | -2.791 | 0.01969 | 0.072 | -3.5 | 1.98918 | none |
| IL34 | 0.474 | 0.3374 | 2.7911 | 0.0197 | 0.072 | -3.5 | -0.2395 | none |
| EIF2AK4 | 0.602 | 0.573 | 2.7911 | 0.0197 | 0.072 | -3.5 | -1.279 | none |
| C2ORF80 | 0.618 | 0.407 | 2.7909 | 0.0197 | 0.072 | -3.5 | -1.3268 | none |
| FAHD1 | 0.327 | 0.2565 | 2.7904 | 0.01972 | 0.072 | -3.5 | -1.0741 | none |
| PPP1CA | -0.339 | -0.2978 | -2.79 | 0.01975 | 0.072 | -3.51 | 1.34758 | none |
| FAM153B | 0.827 | 0.6591 | 2.7879 | 0.0198 | 0.072 | -3.51 | -0.9507 | none |
| GJB3 | -0.283 | -0.1767 | -2.787 | 0.01983 | 0.073 | -3.51 | 1.13124 | none |
| HELLS | 0.501 | 0.5012 | 2.7871 | 0.01983 | 0.073 | -3.51 | -1.2712 | none |
| MAST3 | 0.532 | 0.4821 | 2.7871 | 0.01983 | 0.073 | -3.51 | -1.1303 | none |
| RPGR | 0.397 | 0.2877 | 2.787 | 0.01983 | 0.073 | -3.51 | -1.094 | none |
| MAP3K19 | -0.31 | -0.219 | -2.787 | 0.01984 | 0.073 | -3.51 | 0.17736 | none |
| ZSCAN12 | 0.572 | 0.4808 | 2.7867 | 0.01985 | 0.073 | -3.51 | -1.2388 | none |
| TRIM45 | 0.317 | 0.2349 | 2.786 | 0.01987 | 0.073 | -3.51 | -1.1369 | none |
| EPHA10 | 0.506 | 0.3208 | 2.7857 | 0.01988 | 0.073 | -3.51 | -1.2265 | none |
| NDUFA5 | 0.313 | 0.1519 | 2.7856 | 0.01988 | 0.073 | -3.51 | -0.9497 | none |
| OSMR | 0.24 | 0.1348 | 2.784 | 0.01994 | 0.073 | -3.51 | -0.7811 | none |
| BMPR2 | 0.547 | 0.5544 | 2.7837 | 0.01994 | 0.073 | -3.52 | -0.0402 | none |
| UFSP2 | -0.335 | -0.3092 | -2.783 | 0.01996 | 0.073 | -3.52 | 1.93346 | none |
| MAST4 | 0.556 | 0.5843 | 2.7827 | 0.01998 | 0.073 | -3.52 | -0.6045 | none |
| CREM | -0.363 | -0.3605 | -2.781 | 0.02004 | 0.073 | -3.52 | 1.17424 | none |
| EPC1 | 0.252 | 0.2351 | 2.7807 | 0.02005 | 0.073 | -3.52 | 0.28717 | none |
| GPR89A | 0.326 | 0.2105 | 2.7807 | 0.02005 | 0.073 | -3.52 | -0.3955 | none |
| IL18RAP | -0.285 | -0.2572 | -2.78 | 0.02007 | 0.073 | -3.52 | 1.28849 | none |
| DNA2 | 0.585 | 0.4954 | 2.7796 | 0.02009 | 0.073 | -3.52 | -1.3027 | none |
| UTS2B | 0.347 | 0.2645 | 2.7795 | 0.02009 | 0.073 | -3.52 | -0.6898 | none |
| RANBP9 | 0.617 | 0.5464 | 2.779 | 0.02011 | 0.073 | -3.52 | 0.19421 | none |
| GLMN | 0.356 | 0.2626 | 2.7781 | 0.02014 | 0.073 | -3.52 | -0.2496 | none |

| | | | | | | | | |
|---|---|---|---|---|---|---|---|---|
| ARHGDIB | -0.258 | -0.2114 | -2.776 | 0.02022 | 0.074 | -3.53 | 1.9976 | none |
| NDUFAF4 | -0.454 | -0.4045 | -2.774 | 0.02026 | 0.074 | -3.53 | 2.55481 | AA-related |
| FAM199X | 0.252 | 0.1457 | 2.7735 | 0.02029 | 0.074 | -3.53 | -0.428 | none |
| NKIRAS1 | 0.419 | 0.276 | 2.773 | 0.02031 | 0.074 | -3.53 | -0.7817 | none |
| HSPA8 | -0.306 | -0.2623 | -2.773 | 0.02031 | 0.074 | -3.53 | 2.18412 | AA-related |
| APOL2 | 0.654 | 0.6107 | 2.7725 | 0.02033 | 0.074 | -3.53 | -1.3865 | none |
| ID3 | -0.437 | -0.3741 | -2.772 | 0.02035 | 0.074 | -3.53 | 2.22865 | AA-related |
| MAP7D1 | 0.68 | 0.6734 | 2.7716 | 0.02036 | 0.074 | -3.54 | -0.9755 | none |
| UBAP2L | 0.62 | 0.6324 | 2.7716 | 0.02036 | 0.074 | -3.54 | -0.7058 | none |
| ACSL3 | 0.408 | 0.3519 | 2.7715 | 0.02036 | 0.074 | -3.54 | -0.2707 | none |
| ZNF334 | 0.274 | 0.2062 | 2.7707 | 0.02039 | 0.074 | -3.54 | -1.247 | none |
| LSM5 | -0.544 | -0.5531 | -2.771 | 0.02039 | 0.074 | -3.54 | 2.13291 | AA-related |
| PDE1B | 0.416 | 0.3558 | 2.7707 | 0.02039 | 0.074 | -3.54 | -1.118 | none |
| GCNT4 | 0.252 | 0.1124 | 2.7706 | 0.02039 | 0.074 | -3.54 | -0.6236 | none |
| UBA6 | 0.56 | 0.475 | 2.7695 | 0.02043 | 0.074 | -3.54 | -1.2776 | none |
| CABP1 | 0.563 | 0.4318 | 2.7684 | 0.02047 | 0.074 | -3.54 | -0.9576 | none |
| KPNA2 | -0.3 | -0.2308 | -2.768 | 0.02049 | 0.074 | -3.54 | 1.37468 | none |
| TTC39B | 0.326 | 0.1834 | 2.7679 | 0.02049 | 0.074 | -3.54 | -1.0255 | none |
| C8ORF37 | 0.401 | 0.3445 | 2.7672 | 0.02051 | 0.074 | -3.54 | -0.3346 | none |
| SLC39A13 | 0.382 | 0.2869 | 2.7669 | 0.02052 | 0.074 | -3.54 | -0.3105 | none |
| MFF | -0.355 | -0.3103 | -2.766 | 0.02054 | 0.074 | -3.54 | 2.32114 | AA-related |
| ZNF880 | 0.431 | 0.3318 | 2.7659 | 0.02055 | 0.075 | -3.54 | -0.5645 | none |
| NDC80 | 0.644 | 0.5853 | 2.7657 | 0.02056 | 0.075 | -3.55 | -1.2772 | none |
| PIH1D2 | 0.838 | 0.6432 | 2.7652 | 0.02058 | 0.075 | -3.55 | -1.3055 | none |
| MRPS6 | -0.456 | -0.465 | -2.764 | 0.02061 | 0.075 | -3.55 | 1.6634 | none |
| CSNK1G3 | 0.41 | 0.4173 | 2.7635 | 0.02064 | 0.075 | -3.55 | 0.7655 | none |
| SEMA5B | 0.26 | 0.1726 | 2.7635 | 0.02064 | 0.075 | -3.55 | -1.206 | none |
| CD300A | 0.38 | 0.2813 | 2.7632 | 0.02065 | 0.075 | -3.55 | -1.0338 | none |
| DHX37 | 0.707 | 0.6758 | 2.7625 | 0.02067 | 0.075 | -3.55 | -1.1518 | none |
| OR6J1 | 0.292 | 0.2213 | 2.7619 | 0.02069 | 0.075 | -3.55 | -0.7799 | none |
| NUP37 | -0.351 | -0.3577 | -2.762 | 0.0207 | 0.075 | -3.55 | 1.88946 | none |
| SH3YL1 | 0.353 | 0.1713 | 2.7618 | 0.0207 | 0.075 | -3.55 | -0.8545 | none |
| PAM | 0.342 | 0.3451 | 2.7612 | 0.02072 | 0.075 | -3.55 | 0.22862 | none |
| C2ORF91 | 0.331 | 0.2905 | 2.7611 | 0.02072 | 0.075 | -3.55 | -0.3689 | none |
| SENP2 | 0.299 | 0.152 | 2.761 | 0.02073 | 0.075 | -3.55 | -1.2188 | none |
| CLSTN2 | 0.214 | 0.1458 | 2.7608 | 0.02073 | 0.075 | -3.55 | -1.3951 | none |
| KDM4C | 0.635 | 0.6131 | 2.7602 | 0.02075 | 0.075 | -3.55 | -0.3997 | none |
| LRRC37A2 | 0.523 | 0.506 | 2.7598 | 0.02077 | 0.075 | -3.55 | -0.7299 | none |
| MRPL51 | -0.411 | -0.3909 | -2.76 | 0.02078 | 0.075 | -3.56 | 1.39708 | none |
| NPM3 | -0.517 | -0.4724 | -2.759 | 0.02078 | 0.075 | -3.56 | 2.29971 | AA-related |

| | | | | | | | | |
|---|---|---|---|---|---|---|---|---|
| CCT4 | -0.29 | -0.2527 | -2.759 | 0.02079 | 0.075 | -3.56 | 1.94383 | none |
| ACOXL | 0.441 | 0.3551 | 2.7582 | 0.02082 | 0.075 | -3.56 | -1.1041 | none |
| PALB2 | 0.659 | 0.6406 | 2.7577 | 0.02084 | 0.075 | -3.56 | -0.1244 | none |
| STIM1 | 0.308 | 0.2643 | 2.7573 | 0.02086 | 0.075 | -3.56 | 0.01025 | none |
| AL358113. | 0.677 | 0.4481 | 2.757 | 0.02086 | 0.075 | -3.56 | -0.4883 | none |
| TGM1 | 0.342 | 0.3238 | 2.7569 | 0.02087 | 0.075 | -3.56 | -1.0211 | none |
| FHIT | 0.546 | 0.3148 | 2.7568 | 0.02087 | 0.075 | -3.56 | -0.5764 | none |
| GPR12 | 0.291 | 0.225 | 2.7559 | 0.0209 | 0.075 | -3.56 | -0.5165 | none |
| C7ORF61 | 0.326 | 0.3571 | 2.7553 | 0.02092 | 0.075 | -3.56 | -0.1613 | none |
| ERO1B | -0.337 | -0.3146 | -2.755 | 0.02094 | 0.075 | -3.56 | 1.61697 | none |
| INPP5E | 0.286 | 0.2663 | 2.7547 | 0.02095 | 0.075 | -3.56 | -0.9184 | none |
| CTTNBP2N | 0.556 | 0.5271 | 2.7546 | 0.02095 | 0.075 | -3.56 | -1.0395 | none |
| MYL12B | -0.291 | -0.318 | -2.754 | 0.02096 | 0.075 | -3.56 | 1.85049 | none |
| PRKAG1 | 0.348 | 0.1927 | 2.7539 | 0.02098 | 0.076 | -3.56 | -0.7718 | none |
| ZNF260 | 0.525 | 0.4827 | 2.7537 | 0.02098 | 0.076 | -3.56 | -0.6987 | none |
| LRIG2 | 0.611 | 0.5685 | 2.7534 | 0.02099 | 0.076 | -3.57 | -1.4381 | none |
| PPP4C | 0.303 | 0.2152 | 2.7531 | 0.021 | 0.076 | -3.57 | -0.126 | none |
| RPS6KA6 | 0.316 | 0.232 | 2.7528 | 0.02102 | 0.076 | -3.57 | -1.4293 | none |
| TMEM233 | 0.36 | 0.2942 | 2.7528 | 0.02102 | 0.076 | -3.57 | 0.0668 | none |
| NUTM2D | 0.328 | 0.2943 | 2.7527 | 0.02102 | 0.076 | -3.57 | -1.4345 | none |
| DCLRE1C | 0.783 | 0.7582 | 2.7527 | 0.02102 | 0.076 | -3.57 | -1.1215 | none |
| DHRS12 | -0.294 | -0.2413 | -2.752 | 0.02103 | 0.076 | -3.57 | 0.50268 | none |
| ATG2B | 0.705 | 0.6344 | 2.7522 | 0.02104 | 0.076 | -3.57 | -0.6321 | none |
| PRICKLE3 | 0.306 | 0.2499 | 2.752 | 0.02104 | 0.076 | -3.57 | 0.07352 | none |
| KDM6A | 0.47 | 0.4316 | 2.7515 | 0.02106 | 0.076 | -3.57 | 0.14633 | none |
| FHDC1 | 0.257 | 0.1991 | 2.7509 | 0.02108 | 0.076 | -3.57 | -1.5574 | OA-related |
| DERL2 | -0.422 | -0.3824 | -2.75 | 0.02112 | 0.076 | -3.57 | 1.23252 | none |
| ZNF551 | 0.339 | 0.3076 | 2.7491 | 0.02114 | 0.076 | -3.57 | -0.861 | none |
| IFRD1 | 0.397 | 0.3714 | 2.7476 | 0.0212 | 0.076 | -3.57 | -0.4323 | none |
| CNNM3 | 0.373 | 0.3096 | 2.7476 | 0.0212 | 0.076 | -3.57 | -0.4822 | none |
| SHANK1 | 0.302 | 0.2585 | 2.7476 | 0.0212 | 0.076 | -3.57 | 0.11376 | none |
| RPL14 | -0.36 | -0.3459 | -2.747 | 0.02124 | 0.076 | -3.58 | 1.82513 | none |
| CDKN2AIP | 0.357 | 0.2038 | 2.7463 | 0.02125 | 0.076 | -3.58 | -0.8294 | none |
| ADAL | 0.439 | 0.3855 | 2.746 | 0.02126 | 0.076 | -3.58 | -1.2984 | none |
| ACSM2A | 0.384 | 0.3454 | 2.746 | 0.02126 | 0.076 | -3.58 | -0.5765 | none |
| CD79B | -0.237 | -0.1921 | -2.745 | 0.02128 | 0.076 | -3.58 | 1.41173 | none |
| GPC2 | 0.487 | 0.4164 | 2.7454 | 0.02128 | 0.076 | -3.58 | -1.2233 | none |
| SAXO2 | 0.818 | 0.6456 | 2.7445 | 0.02131 | 0.076 | -3.58 | -0.8881 | none |
| HBS1L | 0.412 | 0.3656 | 2.7445 | 0.02131 | 0.076 | -3.58 | -0.9257 | none |
| MYL6 | -0.371 | -0.3247 | -2.744 | 0.02134 | 0.076 | -3.58 | 0.50183 | none |

| | | | | | | | | |
|---|---|---|---|---|---|---|---|---|
| MOBP | 0.41 | 0.304 | 2.7435 | 0.02135 | 0.076 | -3.58 | -1.4079 | none |
| PRDM2 | 0.551 | 0.4845 | 2.7433 | 0.02135 | 0.076 | -3.58 | -1.2404 | none |
| CCAR1 | 0.91 | 0.8819 | 2.7409 | 0.02144 | 0.077 | -3.59 | -0.5222 | none |
| NUP210L | 0.195 | 0.1517 | 2.7404 | 0.02146 | 0.077 | -3.59 | -1.1486 | none |
| VN1R1 | 0.229 | 0.1315 | 2.74 | 0.02148 | 0.077 | -3.59 | -1.254 | none |
| SMAD9 | 0.297 | 0.235 | 2.7399 | 0.02148 | 0.077 | -3.59 | -0.9721 | none |
| FLT1 | 0.734 | 0.6924 | 2.7381 | 0.02154 | 0.077 | -3.59 | -0.2724 | none |
| SULT1A3 | 0.495 | 0.4698 | 2.7376 | 0.02156 | 0.077 | -3.59 | -0.2496 | none |
| RBM6 | 0.745 | 0.7001 | 2.7373 | 0.02157 | 0.077 | -3.59 | -1.3205 | none |
| MUC17 | 0.568 | 0.3425 | 2.737 | 0.02158 | 0.077 | -3.59 | -0.0097 | none |
| TOMM6 | -0.404 | -0.3932 | -2.737 | 0.02158 | 0.077 | -3.59 | 2.00937 | none |
| GDF15 | 0.668 | 0.4983 | 2.737 | 0.02158 | 0.077 | -3.59 | -1.4328 | none |
| PIK3CD | 0.568 | 0.5756 | 2.7363 | 0.02161 | 0.077 | -3.59 | -0.2413 | none |
| PIGO | 0.416 | 0.3233 | 2.7362 | 0.02161 | 0.077 | -3.59 | -0.6842 | none |
| VAC14 | 0.549 | 0.5483 | 2.7361 | 0.02161 | 0.077 | -3.59 | -0.622 | none |
| SNX21 | 0.626 | 0.4686 | 2.7353 | 0.02164 | 0.077 | -3.59 | -1.3373 | none |
| MFGE8 | 0.377 | 0.2675 | 2.7351 | 0.02165 | 0.077 | -3.6 | -1.0303 | none |
| ALDOA | -0.318 | -0.2442 | -2.735 | 0.02165 | 0.077 | -3.6 | 1.40942 | none |
| SLITRK5 | 0.248 | 0.1731 | 2.7349 | 0.02166 | 0.077 | -3.6 | -1.412 | none |
| ZNF211 | 0.355 | 0.3107 | 2.734 | 0.02169 | 0.077 | -3.6 | -0.1138 | none |
| NEIL3 | 0.237 | 0.1382 | 2.7339 | 0.0217 | 0.077 | -3.6 | 0.22623 | none |
| LARGE1 | 0.36 | 0.3332 | 2.7337 | 0.0217 | 0.077 | -3.6 | -0.8839 | none |
| MRPL21 | -0.409 | -0.3981 | -2.734 | 0.02171 | 0.077 | -3.6 | 1.9044 | none |
| ENAH | 0.35 | 0.2673 | 2.7321 | 0.02176 | 0.078 | -3.6 | -1.1786 | none |
| TBX3 | -0.205 | -0.1477 | -2.732 | 0.02178 | 0.078 | -3.6 | 1.19534 | none |
| ATPAF1 | 0.381 | 0.3315 | 2.7311 | 0.0218 | 0.078 | -3.6 | 0.78617 | none |
| HPD | 0.261 | 0.2615 | 2.7307 | 0.02181 | 0.078 | -3.6 | 0.96786 | none |
| UBR3 | 0.583 | 0.5781 | 2.7306 | 0.02182 | 0.078 | -3.6 | -0.0875 | none |
| SPG11 | 0.575 | 0.4753 | 2.73 | 0.02184 | 0.078 | -3.6 | -0.9396 | none |
| FDXACB1 | 0.26 | 0.1058 | 2.7298 | 0.02185 | 0.078 | -3.6 | -0.8165 | none |
| JAML | -0.259 | -0.244 | -2.729 | 0.02186 | 0.078 | -3.6 | 0.52625 | none |
| POLR1A | 0.735 | 0.8074 | 2.7287 | 0.02189 | 0.078 | -3.61 | -0.2639 | none |
| RAB12 | 0.725 | 0.5761 | 2.7286 | 0.02189 | 0.078 | -3.61 | -1.3973 | none |
| XPNPEP3 | 0.4 | 0.3112 | 2.7282 | 0.02191 | 0.078 | -3.61 | -1.4104 | none |
| CEP68 | 0.478 | 0.3903 | 2.7276 | 0.02193 | 0.078 | -3.61 | -0.6111 | none |
| KLC2 | 0.535 | 0.5208 | 2.7273 | 0.02194 | 0.078 | -3.61 | 0.2489 | none |
| ZNF548 | 0.55 | 0.3942 | 2.7273 | 0.02194 | 0.078 | -3.61 | -0.3164 | none |
| SOS1 | 0.763 | 0.6975 | 2.7268 | 0.02196 | 0.078 | -3.61 | -0.4675 | none |
| SLC7A11 | 0.282 | 0.2112 | 2.7265 | 0.02197 | 0.078 | -3.61 | -0.845 | none |
| EFCAB5 | 0.89 | 0.6615 | 2.7252 | 0.02202 | 0.078 | -3.61 | -0.6518 | none |

| | | | | | | | | |
|---|---|---|---|---|---|---|---|---|
| ERH | -0.396 | -0.3902 | -2.725 | 0.02203 | 0.078 | -3.61 | 1.63414 | none |
| DGAT1 | -0.491 | -0.3606 | -2.724 | 0.02206 | 0.078 | -3.61 | 1.14148 | none |
| AKR1B15 | 0.283 | 0.2695 | 2.7238 | 0.02207 | 0.078 | -3.61 | -0.6633 | none |
| SLC44A5 | 0.383 | 0.3122 | 2.7234 | 0.02208 | 0.078 | -3.61 | -1.2235 | none |
| PFDN1 | -0.214 | -0.2295 | -2.723 | 0.02208 | 0.078 | -3.61 | 1.72373 | none |
| PLA2G4E | 0.2 | 0.1528 | 2.7231 | 0.0221 | 0.078 | -3.62 | -0.9906 | none |
| ZWINT | -0.208 | -0.159 | -2.723 | 0.02211 | 0.078 | -3.62 | 1.69709 | none |
| TMC7 | 0.576 | 0.4124 | 2.7216 | 0.02215 | 0.079 | -3.62 | -1.2738 | none |
| RAC2 | -0.29 | -0.2401 | -2.721 | 0.02217 | 0.079 | -3.62 | 2.09859 | AA-related |
| LRPPRC | 0.658 | 0.6148 | 2.7203 | 0.0222 | 0.079 | -3.62 | -0.4419 | none |
| LILRA5 | 0.372 | 0.265 | 2.7201 | 0.02221 | 0.079 | -3.62 | -0.8625 | none |
| PKN1 | 0.435 | 0.37 | 2.72 | 0.02221 | 0.079 | -3.62 | -1.0364 | none |
| TBX1 | 0.526 | 0.418 | 2.7195 | 0.02223 | 0.079 | -3.62 | -0.0757 | none |
| F2RL2 | 0.455 | 0.3153 | 2.7183 | 0.02228 | 0.079 | -3.62 | -0.8601 | none |
| LCLAT1 | 0.332 | 0.2919 | 2.7178 | 0.02229 | 0.079 | -3.62 | -0.8108 | none |
| SLF2 | 0.693 | 0.6516 | 2.7175 | 0.02231 | 0.079 | -3.62 | -0.745 | none |
| BIRC6 | 0.922 | 0.8892 | 2.7165 | 0.02234 | 0.079 | -3.63 | -0.9853 | none |
| ABCA7 | 0.388 | 0.3991 | 2.7165 | 0.02235 | 0.079 | -3.63 | -0.1362 | none |
| TMEM249 | 0.207 | 0.1559 | 2.7164 | 0.02235 | 0.079 | -3.63 | -1.0549 | none |
| NME9 | 0.369 | 0.3218 | 2.7159 | 0.02237 | 0.079 | -3.63 | -0.7001 | none |
| RHPN1 | 0.347 | 0.3047 | 2.7154 | 0.02238 | 0.079 | -3.63 | -0.3431 | none |
| BST2 | -0.396 | -0.365 | -2.715 | 0.02239 | 0.079 | -3.63 | 2.17598 | AA-related |
| IGSF1 | 0.298 | 0.2048 | 2.7145 | 0.02242 | 0.079 | -3.63 | -0.6851 | none |
| KRT15 | 0.37 | 0.2345 | 2.7144 | 0.02242 | 0.079 | -3.63 | -0.7825 | none |
| SEC24C | 0.258 | 0.2649 | 2.7126 | 0.02249 | 0.079 | -3.63 | 0.01807 | none |
| RAE1 | -0.214 | -0.199 | -2.712 | 0.0225 | 0.079 | -3.63 | 2.26098 | AA-related |
| FTL | -0.428 | -0.4018 | -2.712 | 0.0225 | 0.079 | -3.63 | 2.14515 | AA-related |
| NAA25 | 0.4 | 0.3178 | 2.7115 | 0.02253 | 0.08 | -3.63 | -0.8311 | none |
| PAN3 | 0.467 | 0.3932 | 2.7115 | 0.02253 | 0.08 | -3.63 | -1.2385 | none |
| ZSCAN20 | 0.519 | 0.4232 | 2.7114 | 0.02254 | 0.08 | -3.63 | -1.3717 | none |
| CLCN5 | 0.325 | 0.1886 | 2.7109 | 0.02256 | 0.08 | -3.64 | -1.3174 | none |
| TTLL6 | 0.448 | 0.378 | 2.7103 | 0.02258 | 0.08 | -3.64 | -0.8885 | none |
| RERG | 0.572 | 0.5139 | 2.7101 | 0.02259 | 0.08 | -3.64 | -0.7309 | none |
| SEMA6A | 0.643 | 0.4847 | 2.7098 | 0.0226 | 0.08 | -3.64 | -0.957 | none |
| TLK2 | 0.571 | 0.4703 | 2.7095 | 0.02261 | 0.08 | -3.64 | -0.8266 | none |
| BAP1 | 0.555 | 0.5746 | 2.7091 | 0.02262 | 0.08 | -3.64 | -0.8897 | none |
| CHD7 | 0.486 | 0.4598 | 2.7089 | 0.02263 | 0.08 | -3.64 | 0.66789 | none |
| ONECUT3 | 0.298 | 0.2048 | 2.7087 | 0.02264 | 0.08 | -3.64 | -1.2614 | none |
| DHX34 | 0.344 | 0.3583 | 2.7068 | 0.02271 | 0.08 | -3.64 | -0.1912 | none |
| CTC1 | 0.471 | 0.4873 | 2.7067 | 0.02272 | 0.08 | -3.64 | -0.8082 | none |

| | | | | | | | | |
|---|---|---|---|---|---|---|---|---|
| OAS3 | 0.588 | 0.5577 | 2.7067 | 0.02272 | 0.08 | -3.64 | -1.1325 | none |
| GZMB | -0.396 | -0.1616 | -2.707 | 0.02272 | 0.08 | -3.64 | 1.58142 | none |
| RPE | 0.45 | 0.2543 | 2.7063 | 0.02273 | 0.08 | -3.64 | -0.9428 | none |
| SLC39A5 | 0.484 | 0.3167 | 2.7058 | 0.02275 | 0.08 | -3.64 | -1.0564 | none |
| ZNF341 | 0.431 | 0.3559 | 2.7057 | 0.02276 | 0.08 | -3.64 | -1.0447 | none |
| WNT7B | 0.375 | 0.2502 | 2.7056 | 0.02276 | 0.08 | -3.64 | -0.4776 | none |
| SNRNP25 | -0.273 | -0.2464 | -2.705 | 0.02278 | 0.08 | -3.64 | 1.84231 | none |
| COQ10A | 0.325 | 0.2837 | 2.7044 | 0.02281 | 0.08 | -3.65 | -0.557 | none |
| MR1 | 0.217 | 0.1095 | 2.7039 | 0.02282 | 0.08 | -3.65 | -0.4083 | none |
| VPS54 | 0.456 | 0.4087 | 2.7035 | 0.02284 | 0.08 | -3.65 | -0.9037 | none |
| DECR1 | -0.282 | -0.3183 | -2.703 | 0.02284 | 0.08 | -3.65 | 1.66751 | none |
| PGM2L1 | 0.29 | 0.2655 | 2.7032 | 0.02285 | 0.08 | -3.65 | -1.1789 | none |
| TMEM50B | 0.487 | 0.3045 | 2.703 | 0.02286 | 0.08 | -3.65 | -0.3044 | none |
| RABEP2 | 0.426 | 0.4047 | 2.7028 | 0.02287 | 0.08 | -3.65 | -1.174 | none |
| LMNA | 0.519 | 0.4774 | 2.7028 | 0.02287 | 0.08 | -3.65 | -1.0998 | none |
| ZNF446 | -0.337 | -0.2483 | -2.703 | 0.02287 | 0.08 | -3.65 | 1.91656 | none |
| INTS3 | 0.829 | 0.801 | 2.7026 | 0.02287 | 0.08 | -3.65 | -0.6095 | none |
| DIAPH3 | 0.687 | 0.6087 | 2.7019 | 0.0229 | 0.08 | -3.65 | -0.9112 | none |
| DNAJC25-( | -0.41 | -0.3366 | -2.701 | 0.02295 | 0.08 | -3.65 | 2.03187 | none |
| SLC33A1 | 0.342 | 0.1517 | 2.7007 | 0.02295 | 0.08 | -3.65 | -0.2823 | none |
| AHR | 0.693 | 0.6743 | 2.7007 | 0.02295 | 0.08 | -3.65 | -0.1453 | none |
| RCAN1 | 0.352 | 0.2705 | 2.6999 | 0.02298 | 0.081 | -3.65 | -0.7092 | none |
| SEZ6L2 | 0.27 | 0.1787 | 2.6998 | 0.02298 | 0.081 | -3.65 | 0.00791 | none |
| MMAB | 0.386 | 0.317 | 2.6996 | 0.02299 | 0.081 | -3.65 | -1.2237 | none |
| CNGB1 | 0.449 | 0.3497 | 2.6994 | 0.023 | 0.081 | -3.65 | -0.4339 | none |
| ZNF625 | 0.414 | 0.3289 | 2.6989 | 0.02302 | 0.081 | -3.65 | -1.3265 | none |
| SFN | 0.315 | 0.2594 | 2.6985 | 0.02303 | 0.081 | -3.66 | -0.7737 | none |
| CYP4V2 | 0.625 | 0.4329 | 2.6984 | 0.02304 | 0.081 | -3.66 | -0.9729 | none |
| MBD3 | 0.416 | 0.414 | 2.6982 | 0.02305 | 0.081 | -3.66 | -0.9792 | none |
| TMEM154 | 0.589 | 0.4514 | 2.6976 | 0.02307 | 0.081 | -3.66 | -0.2705 | none |
| ABTB2 | 0.344 | 0.4191 | 2.6974 | 0.02308 | 0.081 | -3.66 | 0.30327 | none |
| FAM3B | 0.315 | 0.2268 | 2.6974 | 0.02308 | 0.081 | -3.66 | -0.7184 | none |
| GLG1 | 0.584 | 0.5882 | 2.6973 | 0.02308 | 0.081 | -3.66 | 0.2075 | none |
| LARP4B | 0.719 | 0.7405 | 2.6967 | 0.0231 | 0.081 | -3.66 | -1.1743 | none |
| WDFY2 | 0.615 | 0.5331 | 2.6964 | 0.02311 | 0.081 | -3.66 | -0.972 | none |
| HSPBP1 | -0.246 | -0.1799 | -2.696 | 0.02312 | 0.081 | -3.66 | 1.94518 | none |
| TMC6 | 0.351 | 0.2523 | 2.6957 | 0.02314 | 0.081 | -3.66 | -1.2402 | none |
| DHDH | 0.504 | 0.3246 | 2.6953 | 0.02316 | 0.081 | -3.66 | 0.23175 | none |
| C10ORF53 | 0.314 | 0.25 | 2.6949 | 0.02318 | 0.081 | -3.66 | -0.567 | none |
| ZFP41 | 0.606 | 0.5635 | 2.6945 | 0.02319 | 0.081 | -3.66 | -0.923 | none |

| | | | | | | | | |
|---|---|---|---|---|---|---|---|---|
| KRTAP9-6 | 0.278 | 0.186 | 2.6939 | 0.02321 | 0.081 | -3.66 | -0.7245 | none |
| ERFE | 0.401 | 0.324 | 2.6939 | 0.02322 | 0.081 | -3.66 | -0.5975 | none |
| ZNF132 | 0.354 | 0.3479 | 2.6924 | 0.02327 | 0.081 | -3.67 | -0.4981 | none |
| HINT1 | -0.489 | -0.4991 | -2.692 | 0.02327 | 0.081 | -3.67 | 1.75881 | none |
| ADAP2 | 0.452 | 0.2946 | 2.692 | 0.02329 | 0.081 | -3.67 | -1.0958 | none |
| MED8 | -0.249 | -0.2333 | -2.691 | 0.02331 | 0.081 | -3.67 | 1.20927 | none |
| TBC1D4 | 0.365 | 0.3117 | 2.6912 | 0.02332 | 0.081 | -3.67 | 0.15151 | none |
| COMMD1( | 0.352 | 0.1945 | 2.691 | 0.02333 | 0.081 | -3.67 | -0.1192 | none |
| ALG11 | 0.397 | 0.3344 | 2.6904 | 0.02335 | 0.081 | -3.67 | -1.061 | none |
| SVIP | -0.53 | -0.4604 | -2.69 | 0.02335 | 0.081 | -3.67 | 0.82021 | none |
| ISCA1 | 0.328 | 0.1874 | 2.6904 | 0.02335 | 0.081 | -3.67 | -0.4636 | none |
| INTS1 | 0.615 | 0.6773 | 2.6898 | 0.02337 | 0.081 | -3.67 | -0.9613 | none |
| OAZ1 | -0.359 | -0.2918 | -2.689 | 0.02339 | 0.081 | -3.67 | 1.76376 | none |
| CDK9 | 0.508 | 0.4403 | 2.6894 | 0.02339 | 0.081 | -3.67 | -0.2796 | none |
| CEP350 | 0.752 | 0.824 | 2.6892 | 0.0234 | 0.081 | -3.67 | -0.6724 | none |
| CCDC142 | 0.368 | 0.2557 | 2.6886 | 0.02342 | 0.082 | -3.67 | -1.2888 | none |
| ANKUB1 | 0.345 | 0.2648 | 2.688 | 0.02345 | 0.082 | -3.67 | -1.1275 | none |
| OR51D1 | 0.302 | 0.212 | 2.688 | 0.02345 | 0.082 | -3.67 | -0.7724 | none |
| CYB561D1 | 0.339 | 0.2743 | 2.6866 | 0.0235 | 0.082 | -3.67 | -0.567 | none |
| PSMB7 | -0.288 | -0.2685 | -2.686 | 0.02351 | 0.082 | -3.68 | 2.00672 | none |
| PCDHA10 | 0.19 | 0.1515 | 2.6862 | 0.02352 | 0.082 | -3.68 | -1.0626 | none |
| LRRC58 | 0.301 | 0.2313 | 2.6859 | 0.02353 | 0.082 | -3.68 | -0.0869 | none |
| SLC27A1 | 0.442 | 0.3466 | 2.6857 | 0.02354 | 0.082 | -3.68 | -1.3033 | none |
| ACIN1 | 0.875 | 0.929 | 2.6856 | 0.02354 | 0.082 | -3.68 | -0.9956 | none |
| OR6Y1 | 0.254 | 0.1835 | 2.685 | 0.02357 | 0.082 | -3.68 | -0.8117 | none |
| ICAM1 | 0.694 | 0.7191 | 2.6849 | 0.02357 | 0.082 | -3.68 | -0.78 | none |
| CNOT4 | 0.637 | 0.5597 | 2.6849 | 0.02357 | 0.082 | -3.68 | -1.3225 | none |
| NPIPB8 | 0.425 | 0.3202 | 2.6847 | 0.02358 | 0.082 | -3.68 | -1.3155 | none |
| SPEF2 | 0.416 | 0.3023 | 2.6841 | 0.0236 | 0.082 | -3.68 | -0.6182 | none |
| WWP1 | 0.423 | 0.425 | 2.684 | 0.02361 | 0.082 | -3.68 | -0.8233 | none |
| ZNF407 | 0.518 | 0.4942 | 2.684 | 0.02361 | 0.082 | -3.68 | -1.1541 | none |
| GOLGA8B | 0.526 | 0.537 | 2.684 | 0.02361 | 0.082 | -3.68 | -0.7019 | none |
| MED24 | 0.403 | 0.408 | 2.6837 | 0.02362 | 0.082 | -3.68 | -1.0188 | none |
| MRPL16 | -0.211 | -0.158 | -2.683 | 0.02363 | 0.082 | -3.68 | 0.55318 | none |
| COX20 | -0.383 | -0.3992 | -2.683 | 0.02364 | 0.082 | -3.68 | 1.03035 | none |
| KLHL9 | 0.265 | 0.1703 | 2.6828 | 0.02365 | 0.082 | -3.68 | -0.609 | none |
| PCDHGB6 | 0.302 | 0.224 | 2.6828 | 0.02365 | 0.082 | -3.68 | -1.4702 | none |
| ATAD5 | 0.607 | 0.6384 | 2.6824 | 0.02367 | 0.082 | -3.68 | -0.8426 | none |
| SPATA17 | 0.684 | 0.7027 | 2.6822 | 0.02368 | 0.082 | -3.68 | -1.0428 | none |
| PSMD13 | -0.3 | -0.2786 | -2.682 | 0.02369 | 0.082 | -3.68 | 2.12508 | AA-related |

| | | | | | | | | |
|---|---|---|---|---|---|---|---|---|
| IPP | 0.402 | 0.3862 | 2.6813 | 0.02371 | 0.082 | -3.68 | -0.1792 | none |
| SHKBP1 | 0.313 | 0.2353 | 2.6812 | 0.02372 | 0.082 | -3.68 | -0.106 | none |
| SEC24B | 0.472 | 0.4417 | 2.6809 | 0.02373 | 0.082 | -3.68 | 0.09066 | none |
| DNAJB11 | -0.231 | -0.182 | -2.679 | 0.02379 | 0.082 | -3.69 | 2.29798 | AA-related |
| DGKE | 0.288 | 0.2431 | 2.6794 | 0.02379 | 0.082 | -3.69 | -0.0973 | none |
| ANKRD13[ | 0.317 | 0.2325 | 2.6785 | 0.02383 | 0.082 | -3.69 | -1.2472 | none |
| USP19 | 0.561 | 0.5755 | 2.6784 | 0.02383 | 0.082 | -3.69 | -0.4391 | none |
| SHISA2 | 0.211 | 0.1791 | 2.6779 | 0.02385 | 0.082 | -3.69 | 0.26686 | none |
| PLEKHG2 | 0.764 | 0.6828 | 2.6777 | 0.02386 | 0.082 | -3.69 | -0.6903 | none |
| SNTN | 0.254 | 0.1889 | 2.6768 | 0.02389 | 0.083 | -3.69 | -0.6998 | none |
| RECQL5 | 0.577 | 0.5765 | 2.6764 | 0.02391 | 0.083 | -3.69 | -0.8051 | none |
| SEC61G | -0.595 | -0.594 | -2.676 | 0.02391 | 0.083 | -3.69 | 2.04981 | none |
| SLC8A1 | -0.298 | -0.3009 | -2.676 | 0.02392 | 0.083 | -3.69 | 0.65398 | none |
| PFN1 | -0.318 | -0.2354 | -2.675 | 0.02395 | 0.083 | -3.69 | 1.47672 | none |
| WDR48 | 0.421 | 0.3441 | 2.6749 | 0.02397 | 0.083 | -3.69 | -0.1983 | none |
| IP6K2 | 0.302 | 0.2702 | 2.6746 | 0.02399 | 0.083 | -3.69 | -0.3995 | none |
| MFSD6 | 0.608 | 0.5334 | 2.6744 | 0.02399 | 0.083 | -3.69 | -0.7463 | none |
| PSMA7 | -0.319 | -0.2692 | -2.673 | 0.02403 | 0.083 | -3.7 | 1.58573 | none |
| UBXN11 | 0.407 | 0.3681 | 2.6728 | 0.02406 | 0.083 | -3.7 | -0.4001 | none |
| DYNLL1 | -0.431 | -0.4478 | -2.673 | 0.02407 | 0.083 | -3.7 | 1.3613 | none |
| KCTD20 | 0.379 | 0.3094 | 2.6724 | 0.02407 | 0.083 | -3.7 | -0.9287 | none |
| HMGB1 | -0.204 | -0.1478 | -2.672 | 0.02408 | 0.083 | -3.7 | 1.92993 | none |
| ACCS | 0.46 | 0.2578 | 2.6719 | 0.02409 | 0.083 | -3.7 | -0.1548 | none |
| VPS45 | 0.454 | 0.3688 | 2.6715 | 0.02411 | 0.083 | -3.7 | -1.3203 | none |
| GP1BB | 0.364 | 0.2857 | 2.6711 | 0.02412 | 0.083 | -3.7 | -0.5452 | none |
| MROH1 | 0.445 | 0.3546 | 2.6706 | 0.02415 | 0.083 | -3.7 | -0.9577 | none |
| NEK1 | 0.476 | 0.4532 | 2.6705 | 0.02415 | 0.083 | -3.7 | -0.391 | none |
| SHQ1 | 0.313 | 0.221 | 2.6702 | 0.02416 | 0.083 | -3.7 | -1.4295 | none |
| RBM20 | 0.489 | 0.3299 | 2.6697 | 0.02418 | 0.083 | -3.7 | -0.887 | none |
| TMED8 | 0.302 | 0.2698 | 2.6697 | 0.02418 | 0.083 | -3.7 | -1.3718 | none |
| RAB14 | -0.227 | -0.1642 | -2.669 | 0.02421 | 0.083 | -3.7 | 1.17695 | none |
| ZNF652 | 0.357 | 0.2874 | 2.6688 | 0.02422 | 0.083 | -3.7 | -1.3768 | none |
| RP2 | 0.321 | 0.1208 | 2.6686 | 0.02423 | 0.083 | -3.7 | -0.7402 | none |
| NAAA | 0.394 | 0.263 | 2.6683 | 0.02424 | 0.083 | -3.7 | -0.6312 | none |
| TMC5 | 0.26 | 0.2164 | 2.6682 | 0.02425 | 0.083 | -3.71 | -1.0834 | none |
| PXN | 0.342 | 0.2957 | 2.6677 | 0.02427 | 0.083 | -3.71 | -0.2924 | none |
| PPP1R21 | 0.453 | 0.4162 | 2.6673 | 0.02428 | 0.083 | -3.71 | 0.45762 | none |
| CFAP161 | 0.429 | 0.3607 | 2.6673 | 0.02428 | 0.083 | -3.71 | 0.81303 | none |
| ZNF317 | 0.52 | 0.5048 | 2.6672 | 0.02428 | 0.083 | -3.71 | -0.5388 | none |
| PTPN4 | 0.629 | 0.529 | 2.6664 | 0.02432 | 0.084 | -3.71 | -0.8271 | none |

| | | | | | | | | |
|---|---|---|---|---|---|---|---|---|
| PIGT | 0.258 | 0.1825 | 2.6657 | 0.02435 | 0.084 | -3.71 | -0.7258 | none |
| ARFGAP3 | 0.36 | 0.2556 | 2.6656 | 0.02435 | 0.084 | -3.71 | -0.4728 | none |
| SYT14 | 0.247 | 0.1622 | 2.6642 | 0.02441 | 0.084 | -3.71 | -1.1386 | none |
| TES | 0.335 | 0.2395 | 2.6636 | 0.02444 | 0.084 | -3.71 | -1.4012 | none |
| POLR2G | -0.404 | -0.4015 | -2.663 | 0.02444 | 0.084 | -3.71 | 1.92958 | none |
| PITPNA | -0.427 | -0.2695 | -2.663 | 0.02445 | 0.084 | -3.71 | 0.25511 | none |
| SNRPF | -0.438 | -0.4174 | -2.663 | 0.02446 | 0.084 | -3.71 | 1.59105 | none |
| RBM14-RE | 0.435 | 0.5175 | 2.6621 | 0.02449 | 0.084 | -3.72 | -0.5692 | none |
| RARS | 0.266 | 0.1814 | 2.6621 | 0.0245 | 0.084 | -3.72 | -0.636 | none |
| RETSAT | 0.239 | 0.1826 | 2.6614 | 0.02452 | 0.084 | -3.72 | -0.5627 | none |
| PSMB2 | -0.285 | -0.252 | -2.661 | 0.02455 | 0.084 | -3.72 | 2.52701 | AA-related |
| PSMA5 | -0.459 | -0.4569 | -2.661 | 0.02456 | 0.084 | -3.72 | 1.97969 | none |
| MT-CO3 | -0.376 | -0.3512 | -2.661 | 0.02456 | 0.084 | -3.72 | 1.17693 | none |
| OR7A5 | 0.184 | 0.1437 | 2.6605 | 0.02456 | 0.084 | -3.72 | -0.9019 | none |
| TLCD1 | 0.728 | 0.5135 | 2.6603 | 0.02457 | 0.084 | -3.72 | -0.9137 | none |
| GOLGA8A | 0.506 | 0.5373 | 2.6594 | 0.02461 | 0.084 | -3.72 | -0.7318 | none |
| IL21R | 0.248 | 0.2421 | 2.6589 | 0.02463 | 0.084 | -3.72 | 0.66404 | none |
| ARPC4 | -0.445 | -0.4039 | -2.659 | 0.02463 | 0.084 | -3.72 | 0.98394 | none |
| BAZ1A | 0.864 | 0.8656 | 2.6587 | 0.02464 | 0.084 | -3.72 | -0.9068 | none |
| RBM39 | 0.337 | 0.3157 | 2.6586 | 0.02464 | 0.084 | -3.72 | -1.3108 | none |
| GABARAP | -0.479 | -0.4945 | -2.658 | 0.02467 | 0.084 | -3.72 | 1.8725 | none |
| SCNN1G | 0.192 | 0.1382 | 2.6579 | 0.02467 | 0.084 | -3.72 | 0.10611 | none |
| HIST1H2A( | 0.442 | 0.3729 | 2.6574 | 0.02469 | 0.084 | -3.72 | -0.4018 | none |
| FRYL | 0.752 | 0.7334 | 2.6573 | 0.02469 | 0.084 | -3.72 | -0.8286 | none |
| GSKIP | -0.39 | -0.4382 | -2.657 | 0.02473 | 0.085 | -3.72 | 1.20558 | none |
| CAST | 1.182 | 1.182 | 2.6562 | 0.02474 | 0.085 | -3.72 | -0.6923 | none |
| TMEM167 | 0.253 | 0.1373 | 2.6555 | 0.02477 | 0.085 | -3.73 | -0.8686 | none |
| SAAL1 | -0.304 | -0.2602 | -2.655 | 0.02478 | 0.085 | -3.73 | 1.93546 | none |
| GORAB | 0.514 | 0.4467 | 2.6531 | 0.02487 | 0.085 | -3.73 | -0.9718 | none |
| PGGHG | 0.322 | 0.2914 | 2.6524 | 0.0249 | 0.085 | -3.73 | 0.02265 | none |
| POMP | -0.422 | -0.4165 | -2.652 | 0.0249 | 0.085 | -3.73 | 1.27617 | none |
| OGDH | 0.427 | 0.4041 | 2.6523 | 0.02491 | 0.085 | -3.73 | -0.676 | none |
| ATXN7 | 0.528 | 0.533 | 2.6518 | 0.02493 | 0.085 | -3.73 | -1.1435 | none |
| APP | 0.485 | 0.4408 | 2.6517 | 0.02493 | 0.085 | -3.73 | -0.8697 | none |
| VDAC3 | -0.348 | -0.3562 | -2.651 | 0.02495 | 0.085 | -3.73 | 1.82504 | none |
| NRIP1 | 0.992 | 0.8905 | 2.6511 | 0.02496 | 0.085 | -3.73 | -0.9954 | none |
| GALNT5 | 0.213 | 0.1452 | 2.651 | 0.02496 | 0.085 | -3.73 | -1.3461 | none |
| DDX56 | -0.318 | -0.2169 | -2.65 | 0.025 | 0.085 | -3.73 | 2.2244 | AA-related |
| SART3 | 0.67 | 0.6898 | 2.6492 | 0.02504 | 0.085 | -3.74 | -0.3913 | none |
| 8-Sep | 0.313 | 0.2877 | 2.6481 | 0.02508 | 0.085 | -3.74 | -0.5467 | none |

| | | | | | | | | |
|---|---|---|---|---|---|---|---|---|
| SAP18 | -0.322 | -0.3184 | -2.648 | 0.02509 | 0.085 | -3.74 | 1.4247 | none |
| AURKA | 0.255 | 0.1696 | 2.6478 | 0.0251 | 0.085 | -3.74 | -0.2062 | none |
| SUSD5 | 0.23 | 0.1653 | 2.6476 | 0.0251 | 0.086 | -3.74 | -0.9378 | none |
| KIF1B | 0.654 | 0.6172 | 2.6475 | 0.02511 | 0.086 | -3.74 | -0.9648 | none |
| TYW1 | 0.449 | 0.4225 | 2.6467 | 0.02514 | 0.086 | -3.74 | -0.8826 | none |
| CASS4 | 0.27 | 0.1977 | 2.6466 | 0.02515 | 0.086 | -3.74 | -1.3788 | none |
| POC1A | -0.38 | -0.3314 | -2.646 | 0.02518 | 0.086 | -3.74 | 1.70372 | none |
| CTBP2 | 0.497 | 0.4844 | 2.6457 | 0.02518 | 0.086 | -3.74 | -0.7224 | none |
| BDP1 | 0.573 | 0.5167 | 2.6454 | 0.0252 | 0.086 | -3.74 | -0.8914 | none |
| NADK | 0.348 | 0.3061 | 2.6448 | 0.02522 | 0.086 | -3.74 | -0.1075 | none |
| ZCCHC18 | 0.659 | 0.5212 | 2.6446 | 0.02523 | 0.086 | -3.74 | -0.6181 | none |
| EPS8L1 | -0.532 | -0.4669 | -2.644 | 0.02525 | 0.086 | -3.74 | 1.07139 | none |
| CRY2 | 0.369 | 0.3194 | 2.6431 | 0.0253 | 0.086 | -3.75 | -1.1535 | none |
| TAS1R3 | 0.345 | 0.2376 | 2.6428 | 0.02531 | 0.086 | -3.75 | -0.7149 | none |
| GPBP1L1 | 0.436 | 0.3397 | 2.6424 | 0.02533 | 0.086 | -3.75 | -1.1853 | none |
| ADIRF | 0.478 | 0.4315 | 2.6416 | 0.02536 | 0.086 | -3.75 | 0.9918 | none |
| ETF1 | -0.305 | -0.2355 | -2.642 | 0.02536 | 0.086 | -3.75 | 2.5045 | AA-related |
| ZNF541 | -0.277 | -0.2325 | -2.641 | 0.02537 | 0.086 | -3.75 | 1.01319 | none |
| ACAD11 | 0.35 | 0.2806 | 2.641 | 0.02538 | 0.086 | -3.75 | -0.1403 | none |
| GTPBP8 | 0.23 | 0.1269 | 2.641 | 0.02539 | 0.086 | -3.75 | -0.4221 | none |
| AZU1 | 0.327 | 0.2106 | 2.6405 | 0.02541 | 0.086 | -3.75 | -1.2621 | none |
| KDM5A | 0.725 | 0.7182 | 2.6396 | 0.02545 | 0.086 | -3.75 | -0.5021 | none |
| PLGLB1 | 0.718 | 0.596 | 2.6392 | 0.02546 | 0.086 | -3.75 | -1.2699 | none |
| SLC12A8 | 0.449 | 0.3148 | 2.6392 | 0.02546 | 0.086 | -3.75 | -1.1126 | none |
| ITPK1 | 0.353 | 0.307 | 2.6391 | 0.02547 | 0.086 | -3.75 | -0.4452 | none |
| HS2ST1 | -0.367 | -0.2562 | -2.638 | 0.0255 | 0.086 | -3.75 | 0.48569 | none |
| CMTR1 | 0.327 | 0.276 | 2.6383 | 0.0255 | 0.086 | -3.75 | -1.21 | none |
| SPOCD1 | 0.454 | 0.3326 | 2.6382 | 0.02551 | 0.086 | -3.75 | -1.2345 | none |
| TNIP3 | 0.462 | 0.375 | 2.638 | 0.02551 | 0.086 | -3.75 | -1.349 | none |
| CSF2 | -0.385 | -0.1884 | -2.637 | 0.02554 | 0.087 | -3.76 | 0.80249 | none |
| HIP1 | 0.658 | 0.667 | 2.6367 | 0.02557 | 0.087 | -3.76 | -1.1298 | none |
| LPAR2 | 0.212 | 0.1394 | 2.6362 | 0.02559 | 0.087 | -3.76 | -0.8906 | none |
| SPATA33 | 0.463 | 0.4229 | 2.6358 | 0.02561 | 0.087 | -3.76 | -0.4293 | none |
| ZNF66 | 0.578 | 0.3999 | 2.6353 | 0.02563 | 0.087 | -3.76 | 0.00362 | none |
| CACYBP | -0.199 | -0.143 | -2.634 | 0.02567 | 0.087 | -3.76 | 1.02115 | none |
| APH1B | 0.67 | 0.4307 | 2.6342 | 0.02568 | 0.087 | -3.76 | -0.7238 | none |
| METRNL | 0.425 | 0.3833 | 2.6341 | 0.02569 | 0.087 | -3.76 | 1.13357 | none |
| SYNGAP1 | 1.167 | 1.231 | 2.6338 | 0.0257 | 0.087 | -3.76 | 0.44628 | none |
| CHCHD6 | 0.362 | 0.2921 | 2.6336 | 0.0257 | 0.087 | -3.76 | -0.1257 | none |
| TGOLN2 | 0.529 | 0.5451 | 2.6335 | 0.02571 | 0.087 | -3.76 | -0.3682 | none |

| | | | | | | | | |
|---|---|---|---|---|---|---|---|---|
| TEX13D | 0.39 | 0.2458 | 2.6335 | 0.02571 | 0.087 | -3.76 | -0.8896 | none |
| TRAPPC3L | 0.252 | 0.1998 | 2.6327 | 0.02574 | 0.087 | -3.76 | -1.1033 | none |
| GSTO1 | -0.549 | -0.5543 | -2.632 | 0.02576 | 0.087 | -3.76 | 1.91211 | none |
| C1D | -0.498 | -0.5441 | -2.632 | 0.02576 | 0.087 | -3.76 | 1.82326 | none |
| DISP1 | -0.297 | -0.1714 | -2.631 | 0.02581 | 0.087 | -3.77 | 0.96348 | none |
| FAM81A | -0.378 | -0.2476 | -2.631 | 0.02583 | 0.087 | -3.77 | 0.32057 | none |
| COMT | -0.286 | -0.2247 | -2.631 | 0.02583 | 0.087 | -3.77 | 2.02636 | none |
| ARVCF | 0.455 | 0.3385 | 2.6306 | 0.02584 | 0.087 | -3.77 | 0.07411 | none |
| TVP23A | 0.489 | 0.3356 | 2.6305 | 0.02584 | 0.087 | -3.77 | -1.2985 | none |
| EXOC1 | 0.394 | 0.3194 | 2.6296 | 0.02588 | 0.087 | -3.77 | -0.1551 | none |
| TAF4B | 0.503 | 0.5282 | 2.629 | 0.02591 | 0.087 | -3.77 | 0.88134 | none |
| GFAP | 0.394 | 0.2385 | 2.6289 | 0.02591 | 0.087 | -3.77 | -0.3845 | none |
| EIF2S2 | -0.27 | -0.1814 | -2.628 | 0.02594 | 0.087 | -3.77 | 2.23469 | AA-related |
| SMNDC1 | -0.349 | -0.3577 | -2.628 | 0.02595 | 0.087 | -3.77 | 0.53062 | none |
| LEO1 | 0.841 | 0.8036 | 2.627 | 0.02599 | 0.088 | -3.77 | -1.0383 | none |
| CCDC144A | 0.381 | 0.2665 | 2.6264 | 0.02602 | 0.088 | -3.77 | -1.3876 | none |
| PSMD14 | -0.295 | -0.2865 | -2.626 | 0.02603 | 0.088 | -3.77 | 2.02982 | none |
| WDR27 | 0.44 | 0.3467 | 2.6252 | 0.02607 | 0.088 | -3.78 | 0.15485 | none |
| SPCS1 | -0.432 | -0.4439 | -2.624 | 0.02612 | 0.088 | -3.78 | 2.16556 | AA-related |
| DDX55 | 0.44 | 0.4429 | 2.6239 | 0.02613 | 0.088 | -3.78 | -1.1155 | none |
| NHLRC1 | 0.239 | 0.173 | 2.6232 | 0.02616 | 0.088 | -3.78 | -0.7475 | none |
| RPL24 | -0.476 | -0.4744 | -2.623 | 0.02616 | 0.088 | -3.78 | 1.42408 | none |
| OGFOD1 | 0.341 | 0.297 | 2.6232 | 0.02616 | 0.088 | -3.78 | -0.8498 | none |
| STT3B | 0.389 | 0.2456 | 2.6231 | 0.02617 | 0.088 | -3.78 | -1.1198 | none |
| CC2D1B | 0.559 | 0.5923 | 2.623 | 0.02617 | 0.088 | -3.78 | -0.2659 | none |
| MED4 | 0.308 | 0.2377 | 2.6229 | 0.02617 | 0.088 | -3.78 | -1.0212 | none |
| DLG1 | 0.662 | 0.5659 | 2.6228 | 0.02618 | 0.088 | -3.78 | -0.861 | none |
| HUNK | 0.327 | 0.2686 | 2.6225 | 0.02619 | 0.088 | -3.78 | -0.3874 | none |
| TOMM22 | -0.41 | -0.4167 | -2.623 | 0.02619 | 0.088 | -3.78 | 2.32968 | AA-related |
| DUT | -0.285 | -0.2608 | -2.621 | 0.02626 | 0.088 | -3.78 | 2.00488 | none |
| COX7B2 | 0.388 | 0.3418 | 2.6203 | 0.02629 | 0.088 | -3.78 | 1.3191 | none |
| MYH15 | 0.255 | 0.18 | 2.6198 | 0.02631 | 0.088 | -3.78 | -0.9682 | none |
| LMAN1 | -0.324 | -0.2386 | -2.62 | 0.02632 | 0.088 | -3.78 | 2.19224 | AA-related |
| MGA | 0.809 | 0.7921 | 2.6196 | 0.02632 | 0.088 | -3.78 | -1.0147 | none |
| GMFG | -0.361 | -0.3693 | -2.62 | 0.02632 | 0.088 | -3.78 | 1.35066 | none |
| RBM5 | 0.456 | 0.4922 | 2.6194 | 0.02633 | 0.088 | -3.79 | -0.7473 | none |
| ADCK1 | 0.462 | 0.3516 | 2.6191 | 0.02634 | 0.088 | -3.79 | -0.3279 | none |
| DOCK8 | 0.767 | 0.7231 | 2.6191 | 0.02635 | 0.088 | -3.79 | -0.8905 | none |
| PRKD3 | 0.534 | 0.3864 | 2.6185 | 0.02637 | 0.088 | -3.79 | -1.2191 | none |
| TMEM255 | -0.202 | -0.1148 | -2.618 | 0.0264 | 0.089 | -3.79 | -0.0608 | none |

| | | | | | | | | |
|---|---|---|---|---|---|---|---|---|
| KCNQ2 | -0.909 | -0.6884 | -2.617 | 0.02642 | 0.089 | -3.79 | 0.98172 | none |
| FEZ2 | 0.38 | 0.2568 | 2.6172 | 0.02643 | 0.089 | -3.79 | -1.1483 | none |
| FNTB | 0.279 | 0.2555 | 2.6168 | 0.02645 | 0.089 | -3.79 | -1.3214 | none |
| C11ORF54 | 0.373 | 0.1812 | 2.6163 | 0.02647 | 0.089 | -3.79 | -1.0881 | none |
| OCRL | 0.254 | 0.2111 | 2.6163 | 0.02647 | 0.089 | -3.79 | -0.5563 | none |
| CSMD2 | 0.256 | 0.1556 | 2.6153 | 0.02651 | 0.089 | -3.79 | -1.3352 | none |
| RFC1 | 0.844 | 0.8897 | 2.6153 | 0.02651 | 0.089 | -3.79 | -0.4589 | none |
| CSRNP1 | -0.385 | -0.264 | -2.615 | 0.02653 | 0.089 | -3.79 | 1.80596 | none |
| DLG3 | 0.508 | 0.4206 | 2.6148 | 0.02653 | 0.089 | -3.79 | -0.9517 | none |
| MMADHC | -0.43 | -0.444 | -2.615 | 0.02655 | 0.089 | -3.79 | 1.55083 | none |
| PLA2R1 | 0.22 | 0.1685 | 2.6141 | 0.02657 | 0.089 | -3.79 | -1.459 | none |
| IL11 | 0.323 | 0.2379 | 2.6136 | 0.02659 | 0.089 | -3.79 | -1.0666 | none |
| VNN2 | 0.261 | 0.0975 | 2.6134 | 0.0266 | 0.089 | -3.79 | 0.62504 | none |
| PHOSPHO: | 0.463 | 0.3059 | 2.6127 | 0.02663 | 0.089 | -3.8 | -0.6993 | none |
| TUBGCP2 | 0.417 | 0.3861 | 2.6124 | 0.02664 | 0.089 | -3.8 | -1.1907 | none |
| PRAM1 | 0.533 | 0.3536 | 2.6113 | 0.02669 | 0.089 | -3.8 | -1.4356 | none |
| LYPLAL1 | -0.501 | -0.4694 | -2.611 | 0.0267 | 0.089 | -3.8 | 1.72 | none |
| SNRPB | -0.37 | -0.3052 | -2.611 | 0.02672 | 0.089 | -3.8 | 1.34227 | none |
| CD52 | -0.526 | -0.5451 | -2.611 | 0.02673 | 0.089 | -3.8 | 1.84605 | none |
| CHST9 | 0.188 | 0.1374 | 2.6099 | 0.02675 | 0.089 | -3.8 | -1.4566 | none |
| COL6A3 | 0.567 | 0.6308 | 2.6099 | 0.02675 | 0.089 | -3.8 | -0.4917 | none |
| CYP2J2 | -0.273 | -0.2537 | -2.609 | 0.02679 | 0.089 | -3.8 | 0.07828 | none |
| MBOAT7 | 0.24 | 0.2443 | 2.609 | 0.0268 | 0.089 | -3.8 | -0.1019 | none |
| XBP1 | 0.281 | 0.2019 | 2.6084 | 0.02682 | 0.09 | -3.8 | -0.3261 | none |
| ARPC5 | -0.314 | -0.3146 | -2.608 | 0.02683 | 0.09 | -3.8 | 1.51831 | none |
| MSMO1 | 0.36 | 0.2028 | 2.6083 | 0.02683 | 0.09 | -3.8 | -0.4917 | none |
| ABCB4 | -0.485 | -0.3952 | -2.608 | 0.02683 | 0.09 | -3.8 | 1.40564 | none |
| NAA15 | 0.814 | 0.7694 | 2.6078 | 0.02685 | 0.09 | -3.8 | -0.6925 | none |
| ADI1 | -0.288 | -0.2551 | -2.607 | 0.02688 | 0.09 | -3.81 | 1.73275 | none |
| NME2 | -0.34 | -0.3133 | -2.607 | 0.0269 | 0.09 | -3.81 | 1.94616 | none |
| PRLR | -0.359 | -0.2841 | -2.606 | 0.02691 | 0.09 | -3.81 | 0.4949 | none |
| OR8B8 | 0.252 | 0.1732 | 2.6058 | 0.02694 | 0.09 | -3.81 | -1.4119 | none |
| N4BP2L1 | 0.498 | 0.356 | 2.6057 | 0.02694 | 0.09 | -3.81 | -0.6104 | none |
| HCFC1R1 | -0.425 | -0.4189 | -2.606 | 0.02695 | 0.09 | -3.81 | 0.9919 | none |
| EIF5A | -0.319 | -0.2356 | -2.604 | 0.02702 | 0.09 | -3.81 | 1.51271 | none |
| NPW | 0.473 | 0.4166 | 2.6035 | 0.02705 | 0.09 | -3.81 | -0.9846 | none |
| GJD4 | 0.262 | 0.2203 | 2.6025 | 0.02709 | 0.09 | -3.81 | -1.1586 | none |
| MON1B | 0.403 | 0.3386 | 2.6021 | 0.02711 | 0.09 | -3.81 | -0.3942 | none |
| SLFN13 | 0.507 | 0.4336 | 2.6016 | 0.02713 | 0.09 | -3.81 | 0.07817 | none |
| RAPGEF3 | 0.265 | 0.1518 | 2.6014 | 0.02715 | 0.09 | -3.81 | -0.7387 | none |

| | | | | | | | | |
|---|---|---|---|---|---|---|---|---|
| SLCO5A1 | 0.237 | 0.2325 | 2.6013 | 0.02715 | 0.09 | -3.81 | -0.78 | none |
| ARFGEF3 | 0.288 | 0.244 | 2.6012 | 0.02715 | 0.09 | -3.81 | -1.3756 | none |
| GPATCH2L | 0.252 | 0.2437 | 2.6008 | 0.02717 | 0.09 | -3.82 | -0.8565 | none |
| NOP10 | -0.406 | -0.3801 | -2.599 | 0.02724 | 0.091 | -3.82 | 1.56471 | none |
| FAM78B | 0.268 | 0.2534 | 2.5991 | 0.02725 | 0.091 | -3.82 | -0.4797 | none |
| SORL1 | 0.602 | 0.5333 | 2.5989 | 0.02726 | 0.091 | -3.82 | -0.6972 | none |
| PASK | 0.312 | 0.2313 | 2.5988 | 0.02726 | 0.091 | -3.82 | -0.6976 | none |
| MTSS1 | 0.56 | 0.3971 | 2.5982 | 0.02729 | 0.091 | -3.82 | -1.0938 | none |
| ENTPD5 | 0.272 | 0.1833 | 2.5981 | 0.02729 | 0.091 | -3.82 | 1.2347 | none |
| PPP1R14B | -0.299 | -0.2307 | -2.597 | 0.02733 | 0.091 | -3.82 | 0.01686 | none |
| MAP1LC3C | 0.839 | 0.5451 | 2.5973 | 0.02733 | 0.091 | -3.82 | -0.6506 | none |
| ZNF646 | 0.392 | 0.4479 | 2.5972 | 0.02734 | 0.091 | -3.82 | -0.3116 | none |
| ATG9A | 0.334 | 0.3416 | 2.5965 | 0.02737 | 0.091 | -3.82 | 0.66317 | none |
| TANC2 | 0.327 | 0.3042 | 2.5964 | 0.02737 | 0.091 | -3.82 | -0.3862 | none |
| BTN2A2 | -0.35 | -0.2271 | -2.596 | 0.02739 | 0.091 | -3.82 | 1.96084 | none |
| TMEM253 | -0.237 | -0.2167 | -2.596 | 0.0274 | 0.091 | -3.82 | 0.69777 | none |
| HSF4 | 0.581 | 0.5372 | 2.5957 | 0.02741 | 0.091 | -3.82 | -0.3359 | none |
| RBMS3 | 0.328 | 0.2465 | 2.5956 | 0.02741 | 0.091 | -3.82 | -0.8368 | none |
| UGGT1 | 0.771 | 0.7509 | 2.5952 | 0.02743 | 0.091 | -3.82 | -0.5835 | none |
| TK1 | -0.457 | -0.4011 | -2.595 | 0.02745 | 0.091 | -3.83 | 2.52957 | AA-related |
| ZNHIT6 | 0.522 | 0.4815 | 2.5946 | 0.02746 | 0.091 | -3.83 | -0.745 | none |
| VAMP8 | -0.353 | -0.3626 | -2.594 | 0.02747 | 0.091 | -3.83 | 1.78297 | none |
| TMEM79 | 0.329 | 0.3158 | 2.5943 | 0.02747 | 0.091 | -3.83 | -0.6599 | none |
| CEP63 | 0.629 | 0.5976 | 2.5939 | 0.02749 | 0.091 | -3.83 | -0.8628 | none |
| ADH1C | 0.24 | 0.1684 | 2.5935 | 0.02751 | 0.091 | -3.83 | -0.6233 | none |
| LIMD1 | 0.651 | 0.6036 | 2.593 | 0.02753 | 0.091 | -3.83 | -0.737 | none |
| FANCE | 0.239 | 0.2506 | 2.5925 | 0.02756 | 0.091 | -3.83 | -0.5485 | none |
| MAEA | 0.275 | 0.2391 | 2.5924 | 0.02756 | 0.091 | -3.83 | -0.8045 | none |
| FAM117B | 0.442 | 0.3687 | 2.592 | 0.02758 | 0.091 | -3.83 | -1.2873 | none |
| SMCHD1 | 0.798 | 0.769 | 2.5918 | 0.02759 | 0.091 | -3.83 | -0.4184 | none |
| TMEM123 | 0.303 | 0.2309 | 2.5916 | 0.0276 | 0.091 | -3.83 | 0.8304 | none |
| SMARCC2 | 0.747 | 0.729 | 2.5906 | 0.02765 | 0.091 | -3.83 | -0.8651 | none |
| EDEM3 | 0.55 | 0.4666 | 2.5906 | 0.02765 | 0.091 | -3.83 | -1.0404 | none |
| CLMP | 0.286 | 0.1939 | 2.5905 | 0.02765 | 0.091 | -3.83 | -1.0681 | none |
| CNTLN | 0.42 | 0.3033 | 2.5904 | 0.02765 | 0.091 | -3.83 | -1.1751 | none |
| CCDC61 | 0.307 | 0.2525 | 2.59 | 0.02767 | 0.091 | -3.83 | 0.82348 | none |
| ADGRA3 | 0.479 | 0.5759 | 2.5899 | 0.02768 | 0.091 | -3.83 | 0.45167 | none |
| SDK1 | -0.409 | -0.3593 | -2.59 | 0.02769 | 0.091 | -3.83 | 0.09335 | none |
| RAD54L | 0.221 | 0.2027 | 2.5883 | 0.02775 | 0.092 | -3.84 | -1.153 | none |
| FANCC | 0.431 | 0.3602 | 2.5879 | 0.02777 | 0.092 | -3.84 | -0.9596 | none |

| | | | | | | | | |
|---|---|---|---|---|---|---|---|---|
| DENND3 | 0.392 | 0.3846 | 2.5879 | 0.02777 | 0.092 | -3.84 | -1.0351 | none |
| ATP13A2 | 0.479 | 0.439 | 2.5876 | 0.02778 | 0.092 | -3.84 | -1.1615 | none |
| HHLA3 | 0.276 | 0.1925 | 2.5862 | 0.02785 | 0.092 | -3.84 | -0.2498 | none |
| N4BP2L2 | 0.382 | 0.3022 | 2.5861 | 0.02785 | 0.092 | -3.84 | -0.9324 | none |
| NKX3-1 | -0.255 | -0.2083 | -2.586 | 0.02786 | 0.092 | -3.84 | 1.26928 | none |
| XKR5 | 0.275 | 0.1856 | 2.5861 | 0.02786 | 0.092 | -3.84 | -1.4168 | none |
| LAIR1 | 0.567 | 0.4238 | 2.5859 | 0.02787 | 0.092 | -3.84 | -0.8893 | none |
| RPA2 | -0.212 | -0.1981 | -2.584 | 0.02795 | 0.092 | -3.84 | 0.64551 | none |
| ATF7IP | 0.75 | 0.7473 | 2.584 | 0.02795 | 0.092 | -3.84 | -0.5803 | none |
| RPS23 | -0.518 | -0.5445 | -2.584 | 0.02796 | 0.092 | -3.84 | 2.03509 | none |
| DARS | -0.229 | -0.2237 | -2.583 | 0.02802 | 0.092 | -3.85 | 1.79486 | none |
| STX16-NPI | 0.398 | 0.3219 | 2.5822 | 0.02804 | 0.092 | -3.85 | -0.7118 | none |
| UQCRC2 | 0.326 | 0.1439 | 2.5821 | 0.02804 | 0.092 | -3.85 | 0.42606 | none |
| MGST2 | -0.471 | -0.4922 | -2.581 | 0.0281 | 0.093 | -3.85 | 1.51531 | none |
| TUBG1 | 0.281 | 0.227 | 2.5807 | 0.02811 | 0.093 | -3.85 | -0.8609 | none |
| MYO1E | 0.736 | 0.7519 | 2.5804 | 0.02812 | 0.093 | -3.85 | -0.4623 | none |
| WDR36 | 0.376 | 0.2608 | 2.5799 | 0.02815 | 0.093 | -3.85 | -0.6975 | none |
| NUDT2 | -0.263 | -0.2468 | -2.579 | 0.02819 | 0.093 | -3.85 | 1.33753 | none |
| COL4A3BP | 0.506 | 0.4578 | 2.5782 | 0.02823 | 0.093 | -3.85 | -0.9199 | none |
| TALDO1 | -0.322 | -0.2991 | -2.578 | 0.02824 | 0.093 | -3.85 | 1.85362 | none |
| ZSCAN5B | 0.462 | 0.3674 | 2.5775 | 0.02826 | 0.093 | -3.85 | -1.2193 | none |
| FSD1 | 0.406 | 0.3147 | 2.5764 | 0.02831 | 0.093 | -3.86 | -0.7302 | none |
| ATP11A | 0.739 | 0.7908 | 2.5746 | 0.0284 | 0.093 | -3.86 | -0.7709 | none |
| ATP6V1E1 | 0.297 | 0.2058 | 2.5746 | 0.0284 | 0.093 | -3.86 | -1.3266 | none |
| RNASEH2A | 0.362 | 0.2276 | 2.5744 | 0.02841 | 0.093 | -3.86 | -0.8866 | none |
| MAN1C1 | 0.476 | 0.2976 | 2.5742 | 0.02842 | 0.093 | -3.86 | -0.8992 | none |
| RRAGA | -0.374 | -0.3265 | -2.574 | 0.02842 | 0.093 | -3.86 | 2.42083 | AA-related |
| BOLA3 | -0.373 | -0.3586 | -2.574 | 0.02843 | 0.093 | -3.86 | 2.18453 | AA-related |
| SLC7A9 | 0.423 | 0.3433 | 2.5738 | 0.02844 | 0.093 | -3.86 | -1.33 | none |
| HEATR4 | 0.349 | 0.2883 | 2.5735 | 0.02845 | 0.093 | -3.86 | 0.6305 | none |
| BCL2A1 | -0.443 | -0.3999 | -2.574 | 0.02845 | 0.093 | -3.86 | 1.69986 | none |
| POLR3C | 0.319 | 0.2957 | 2.5734 | 0.02846 | 0.093 | -3.86 | -0.4211 | none |
| CUL7 | 0.344 | 0.3611 | 2.5733 | 0.02846 | 0.093 | -3.86 | 0.24596 | none |
| ASPG | -0.582 | -0.4866 | -2.573 | 0.02847 | 0.093 | -3.86 | 0.8402 | none |
| ARSJ | 0.43 | 0.3693 | 2.573 | 0.02848 | 0.093 | -3.86 | 0.11312 | none |
| RASSF4 | 0.651 | 0.5486 | 2.573 | 0.02848 | 0.093 | -3.86 | 0.26532 | none |
| VARS2 | 0.553 | 0.4397 | 2.572 | 0.02853 | 0.094 | -3.86 | -0.645 | none |
| PREX2 | 0.35 | 0.2506 | 2.5714 | 0.02856 | 0.094 | -3.86 | -1.2156 | none |
| ADAD2 | 0.331 | 0.2232 | 2.5708 | 0.02858 | 0.094 | -3.86 | -0.3357 | none |
| PFKFB3 | 0.502 | 0.4614 | 2.5708 | 0.02859 | 0.094 | -3.86 | -0.6704 | none |

| | | | | | | | | |
|---|---|---|---|---|---|---|---|---|
| MRPL47 | -0.34 | -0.3164 | -2.571 | 0.02859 | 0.094 | -3.86 | 1.71732 | none |
| IKZF1 | 0.436 | 0.4338 | 2.5705 | 0.0286 | 0.094 | -3.86 | -0.5908 | none |
| KIAA1109 | 0.69 | 0.6832 | 2.5705 | 0.0286 | 0.094 | -3.86 | -0.6911 | none |
| LOX | 0.419 | 0.2816 | 2.5702 | 0.02861 | 0.094 | -3.87 | -1.4124 | none |
| SOD1 | -0.385 | -0.3988 | -2.57 | 0.02864 | 0.094 | -3.87 | 1.91346 | none |
| CCDC66 | 0.504 | 0.3844 | 2.5696 | 0.02864 | 0.094 | -3.87 | -0.7255 | none |
| BASP1 | 0.461 | 0.4813 | 2.5689 | 0.02868 | 0.094 | -3.87 | -1.0417 | none |
| VCPKMT | 0.494 | 0.3743 | 2.5688 | 0.02868 | 0.094 | -3.87 | -1.1097 | none |
| KAT6A | 0.773 | 0.7331 | 2.5685 | 0.0287 | 0.094 | -3.87 | -0.9785 | none |
| APC2 | 0.351 | 0.2805 | 2.5685 | 0.0287 | 0.094 | -3.87 | -1.5102 | none |
| TNRC6B | 0.819 | 0.8356 | 2.5685 | 0.0287 | 0.094 | -3.87 | -1.057 | none |
| CRBN | 0.317 | 0.1636 | 2.5676 | 0.02874 | 0.094 | -3.87 | -0.5772 | none |
| C3AR1 | -0.319 | -0.3373 | -2.567 | 0.02876 | 0.094 | -3.87 | 1.52741 | none |
| USP14 | 0.221 | 0.171 | 2.5672 | 0.02876 | 0.094 | -3.87 | 0.77523 | none |
| RFX8 | -0.398 | -0.3706 | -2.566 | 0.0288 | 0.094 | -3.87 | 1.18852 | none |
| UTP23 | -0.297 | -0.1926 | -2.566 | 0.02884 | 0.094 | -3.87 | 0.78682 | none |
| DNAAF5 | 0.432 | 0.4239 | 2.5654 | 0.02884 | 0.094 | -3.87 | 0.21846 | none |
| GBP3 | 0.683 | 0.6242 | 2.5647 | 0.02888 | 0.094 | -3.87 | -0.7805 | none |
| ZNF175 | 0.487 | 0.4164 | 2.5644 | 0.0289 | 0.094 | -3.87 | -0.2585 | none |
| PGAM1 | -0.286 | -0.2167 | -2.564 | 0.0289 | 0.094 | -3.87 | 1.41125 | none |
| ALDH1L2 | -0.269 | -0.2771 | -2.564 | 0.02892 | 0.094 | -3.88 | 0.53609 | none |
| TRIM11 | 0.253 | 0.1641 | 2.5633 | 0.02895 | 0.094 | -3.88 | -0.0608 | none |
| PLEKHG3 | 0.424 | 0.3967 | 2.5628 | 0.02897 | 0.095 | -3.88 | 0.01292 | none |
| TXNDC9 | -0.327 | -0.3546 | -2.561 | 0.02904 | 0.095 | -3.88 | 2.03911 | none |
| CLDN18 | 0.352 | 0.2451 | 2.5603 | 0.0291 | 0.095 | -3.88 | -1.1509 | none |
| RAB27A | 0.339 | 0.2311 | 2.5601 | 0.0291 | 0.095 | -3.88 | -0.9039 | none |
| ZNF620 | 0.526 | 0.3786 | 2.5598 | 0.02912 | 0.095 | -3.88 | -1.0712 | none |
| FBF1 | 0.362 | 0.4099 | 2.5594 | 0.02914 | 0.095 | -3.88 | -0.6119 | none |
| HSPA2 | 0.482 | 0.4251 | 2.5589 | 0.02917 | 0.095 | -3.88 | -0.7566 | none |
| DPP4 | 0.262 | 0.173 | 2.5588 | 0.02917 | 0.095 | -3.88 | 0.54842 | none |
| MAP2K4 | 0.413 | 0.3647 | 2.5576 | 0.02923 | 0.095 | -3.89 | -0.2098 | none |
| DARS2 | 0.248 | 0.1912 | 2.5572 | 0.02925 | 0.095 | -3.89 | -0.2241 | none |
| SNX1 | 0.353 | 0.3766 | 2.5569 | 0.02926 | 0.095 | -3.89 | -1.1121 | none |
| ROCK1 | 0.906 | 0.9122 | 2.5558 | 0.02932 | 0.095 | -3.89 | -0.7971 | none |
| POMK | 0.365 | 0.2132 | 2.5557 | 0.02932 | 0.095 | -3.89 | -0.9983 | none |
| GJA10 | 0.562 | 0.4335 | 2.5556 | 0.02933 | 0.095 | -3.89 | -1.0285 | none |
| MOSPD2 | 0.522 | 0.4234 | 2.5555 | 0.02933 | 0.095 | -3.89 | -0.7619 | none |
| TMEM19 | 0.307 | 0.1285 | 2.5555 | 0.02934 | 0.095 | -3.89 | -0.6748 | none |
| POLH | 0.464 | 0.3851 | 2.5544 | 0.02939 | 0.096 | -3.89 | -0.5582 | none |
| ZNF585B | 1.043 | 0.821 | 2.5538 | 0.02942 | 0.096 | -3.89 | 0.28983 | none |

| | | | | | | | | |
|---|---|---|---|---|---|---|---|---|
| DSTN | -0.316 | -0.3526 | -2.554 | 0.02943 | 0.096 | -3.89 | 1.78207 | none |
| PTCHD4 | 0.23 | 0.158 | 2.5532 | 0.02945 | 0.096 | -3.89 | -0.2415 | none |
| C12ORF71 | -0.173 | -0.1304 | -2.553 | 0.02948 | 0.096 | -3.89 | 0.91931 | none |
| ELP2 | 0.375 | 0.2549 | 2.5518 | 0.02952 | 0.096 | -3.9 | -0.6293 | none |
| PCSK2 | 0.234 | 0.1903 | 2.5514 | 0.02954 | 0.096 | -3.9 | 0.80593 | none |
| TAF9 | -0.263 | -0.2898 | -2.551 | 0.02954 | 0.096 | -3.9 | 1.16867 | none |
| PTGES3 | -0.327 | -0.3299 | -2.551 | 0.02954 | 0.096 | -3.9 | 1.72286 | none |
| NBEAL1 | 0.359 | 0.2889 | 2.5512 | 0.02955 | 0.096 | -3.9 | -0.5583 | none |
| TIRAP | 0.383 | 0.3136 | 2.5511 | 0.02955 | 0.096 | -3.9 | -0.3955 | none |
| FBXW11 | 0.335 | 0.2353 | 2.5506 | 0.02958 | 0.096 | -3.9 | 0.07508 | none |
| UTP20 | 0.77 | 0.7718 | 2.5503 | 0.02959 | 0.096 | -3.9 | -0.4994 | none |
| ZDHHC24 | 0.341 | 0.2057 | 2.55 | 0.02961 | 0.096 | -3.9 | -0.9816 | none |
| EIF3H | -0.259 | -0.2322 | -2.55 | 0.02962 | 0.096 | -3.9 | 2.15744 | AA-related |
| ANKRD13( | 0.375 | 0.3377 | 2.5497 | 0.02962 | 0.096 | -3.9 | -1.0911 | none |
| ASMT | 0.407 | 0.3454 | 2.5497 | 0.02962 | 0.096 | -3.9 | -0.5472 | none |
| PSMB8 | -0.495 | -0.4832 | -2.55 | 0.02963 | 0.096 | -3.9 | 1.38747 | none |
| EVA1B | -0.337 | -0.2933 | -2.549 | 0.02964 | 0.096 | -3.9 | 1.58418 | none |
| EP300 | 0.807 | 0.824 | 2.5487 | 0.02967 | 0.096 | -3.9 | -0.7677 | none |
| SNCG | 0.372 | 0.3003 | 2.5487 | 0.02967 | 0.096 | -3.9 | -0.4396 | none |
| TSPOAP1 | -0.54 | -0.4332 | -2.549 | 0.02968 | 0.096 | -3.9 | 1.03788 | none |
| EVL | 0.454 | 0.4372 | 2.5469 | 0.02976 | 0.096 | -3.9 | -0.6539 | none |
| GTF3C3 | 0.303 | 0.1867 | 2.5468 | 0.02977 | 0.096 | -3.9 | -0.0279 | none |
| NDUFA1 | -0.432 | -0.4686 | -2.546 | 0.0298 | 0.096 | -3.9 | 1.3283 | none |
| PFN4 | 0.341 | 0.3213 | 2.5461 | 0.0298 | 0.096 | -3.9 | -0.6041 | none |
| CARMIL2 | 0.451 | 0.5282 | 2.546 | 0.02981 | 0.096 | -3.9 | -0.8129 | none |
| ZNF74 | 0.382 | 0.3359 | 2.5453 | 0.02984 | 0.097 | -3.91 | 0.36908 | none |
| IL17RD | 0.396 | 0.2454 | 2.5453 | 0.02984 | 0.097 | -3.91 | -0.8854 | none |
| C12ORF10 | -0.313 | -0.2677 | -2.545 | 0.02986 | 0.097 | -3.91 | 1.76935 | none |
| CD2BP2 | -0.389 | -0.3013 | -2.545 | 0.02987 | 0.097 | -3.91 | 2.43993 | AA-related |
| ZDHHC13 | 0.314 | 0.1398 | 2.5433 | 0.02994 | 0.097 | -3.91 | -0.628 | none |
| IRF2BP1 | -0.245 | -0.1415 | -2.543 | 0.02996 | 0.097 | -3.91 | 1.95389 | none |
| SUGP2 | 0.537 | 0.5365 | 2.5428 | 0.02997 | 0.097 | -3.91 | -0.5901 | none |
| EXOC5 | 0.364 | 0.3099 | 2.5424 | 0.02999 | 0.097 | -3.91 | 0.46717 | none |
| RHEBL1 | -0.392 | -0.3239 | -2.542 | 0.03 | 0.097 | -3.91 | 0.83187 | none |
| IPO11 | 0.449 | 0.3157 | 2.542 | 0.03001 | 0.097 | -3.91 | -0.3072 | none |
| SRPX2 | 0.242 | 0.154 | 2.5413 | 0.03005 | 0.097 | -3.91 | 0.21387 | none |
| PRR3 | 0.763 | 0.5485 | 2.5409 | 0.03007 | 0.097 | -3.91 | -1.0769 | none |
| PAQR5 | 0.314 | 0.2441 | 2.5403 | 0.0301 | 0.097 | -3.91 | -0.9355 | none |
| FPGT | -0.299 | -0.1868 | -2.54 | 0.0301 | 0.097 | -3.91 | 0.41372 | none |
| VWA3B | 0.427 | 0.3639 | 2.5399 | 0.03012 | 0.097 | -3.91 | 0.00455 | none |

| | | | | | | | | |
|---|---|---|---|---|---|---|---|---|
| CIR1 | 0.818 | 0.8095 | 2.5386 | 0.03018 | 0.097 | -3.92 | -1.0195 | none |
| COX10 | 0.232 | 0.1803 | 2.5382 | 0.0302 | 0.097 | -3.92 | -0.8958 | none |
| CREB1 | 0.425 | 0.3382 | 2.5382 | 0.0302 | 0.097 | -3.92 | 0.15401 | none |
| URGCP-MI | -0.44 | -0.4849 | -2.538 | 0.03021 | 0.097 | -3.92 | 1.37922 | none |
| CEP72 | 0.343 | 0.3694 | 2.5378 | 0.03023 | 0.097 | -3.92 | -0.7216 | none |
| CHADL | -0.27 | -0.1981 | -2.537 | 0.03024 | 0.097 | -3.92 | 0.05132 | none |
| NCOA1 | 0.809 | 0.739 | 2.5352 | 0.03036 | 0.098 | -3.92 | -0.4944 | none |
| CTSC | -0.319 | -0.3112 | -2.535 | 0.03037 | 0.098 | -3.92 | 2.37256 | AA-related |
| FAM219A | 0.509 | 0.4844 | 2.5346 | 0.03039 | 0.098 | -3.92 | -0.2664 | none |
| B4GALT4 | 0.454 | 0.3077 | 2.5346 | 0.03039 | 0.098 | -3.92 | 0.38038 | none |
| LGALS1 | -0.58 | -0.547 | -2.535 | 0.03039 | 0.098 | -3.92 | 2.05338 | none |
| ANKHD1 | 0.483 | 0.4635 | 2.5344 | 0.0304 | 0.098 | -3.92 | -0.6644 | none |
| AP5Z1 | 0.378 | 0.2998 | 2.5343 | 0.0304 | 0.098 | -3.92 | -1.3195 | none |
| WAPL | 0.734 | 0.6979 | 2.5339 | 0.03043 | 0.098 | -3.92 | -0.4738 | none |
| ANGEL2 | 0.258 | 0.1668 | 2.5337 | 0.03043 | 0.098 | -3.92 | -0.763 | none |
| SMIM11B | 0.388 | 0.2262 | 2.5336 | 0.03044 | 0.098 | -3.92 | -0.5168 | none |
| TMEM132 | 0.431 | 0.3581 | 2.5331 | 0.03046 | 0.098 | -3.93 | -1.2923 | none |
| ZNF224 | 0.418 | 0.3505 | 2.5327 | 0.03048 | 0.098 | -3.93 | -0.4269 | none |
| PHYH | -0.3 | -0.2703 | -2.533 | 0.03049 | 0.098 | -3.93 | 1.03309 | none |
| KLKB1 | 0.211 | 0.1289 | 2.5324 | 0.0305 | 0.098 | -3.93 | -0.3316 | none |
| FCRL3 | 0.579 | 0.4602 | 2.5311 | 0.03057 | 0.098 | -3.93 | -1.1481 | none |
| PAX6 | 0.336 | 0.2811 | 2.5304 | 0.03061 | 0.098 | -3.93 | -0.1762 | none |
| SHISA7 | 0.211 | 0.1599 | 2.5304 | 0.03061 | 0.098 | -3.93 | -0.894 | none |
| SLC5A10 | 0.307 | 0.2381 | 2.5298 | 0.03064 | 0.098 | -3.93 | -1.0764 | none |
| CD53 | -0.347 | -0.3942 | -2.53 | 0.03064 | 0.098 | -3.93 | 1.92477 | none |
| SIAH2 | -0.234 | -0.2051 | -2.529 | 0.03067 | 0.098 | -3.93 | 0.19383 | none |
| ASH2L | 0.328 | 0.2505 | 2.5289 | 0.03068 | 0.098 | -3.93 | -0.0646 | none |
| NEMF | 0.612 | 0.5609 | 2.5277 | 0.03075 | 0.099 | -3.93 | -1.0898 | none |
| KIF1C | 0.711 | 0.6954 | 2.5276 | 0.03075 | 0.099 | -3.93 | -0.8976 | none |
| FOXO3 | 0.457 | 0.4481 | 2.5276 | 0.03075 | 0.099 | -3.93 | -0.8089 | none |
| GEMIN2 | 0.374 | 0.2016 | 2.5275 | 0.03076 | 0.099 | -3.93 | -1.0757 | none |
| PRDM14 | 0.334 | 0.2334 | 2.5271 | 0.03078 | 0.099 | -3.94 | -1.4868 | none |
| AC091180 | 0.903 | 0.7048 | 2.5265 | 0.03081 | 0.099 | -3.94 | -1.3961 | none |
| PCDHB9 | 0.227 | 0.1717 | 2.5262 | 0.03082 | 0.099 | -3.94 | -0.8845 | none |
| KDM3B | 0.738 | 0.7224 | 2.526 | 0.03083 | 0.099 | -3.94 | -0.663 | none |
| PRUNE2 | 0.299 | 0.2591 | 2.5258 | 0.03084 | 0.099 | -3.94 | 0.10199 | none |
| C12ORF40 | 0.275 | 0.2149 | 2.5239 | 0.03095 | 0.099 | -3.94 | -1.0417 | none |
| PKP4 | 0.685 | 0.7453 | 2.5236 | 0.03096 | 0.099 | -3.94 | -0.7735 | none |
| MYO6 | 0.227 | 0.1665 | 2.5235 | 0.03096 | 0.099 | -3.94 | 0.05325 | none |
| PLP1 | 0.223 | 0.2292 | 2.5231 | 0.03098 | 0.099 | -3.94 | -0.576 | none |

| | | | | | | | | |
|---|---|---|---|---|---|---|---|---|
| RIDA | -0.465 | -0.3886 | -2.523 | 0.03099 | 0.099 | -3.94 | 1.15701 | none |
| NGDN | -0.3 | -0.2777 | -2.523 | 0.03099 | 0.099 | -3.94 | 1.15571 | none |
| NPIPA2 | 0.504 | 0.3594 | 2.5226 | 0.03101 | 0.099 | -3.94 | -0.4995 | none |
| MTMR2 | 0.231 | 0.1729 | 2.5223 | 0.03103 | 0.099 | -3.94 | 0.15249 | none |
| RNF157 | 0.391 | 0.2577 | 2.5222 | 0.03103 | 0.099 | -3.94 | -1.1836 | none |
| PDZD11 | -0.393 | -0.4188 | -2.522 | 0.03105 | 0.099 | -3.94 | 0.94119 | none |
| NDUFB4 | -0.339 | -0.3305 | -2.522 | 0.03106 | 0.099 | -3.94 | 2.00568 | none |
| TSPAN3 | -0.328 | -0.2616 | -2.522 | 0.03107 | 0.099 | -3.94 | 1.82352 | none |
| LDB1 | 0.601 | 0.5357 | 2.5215 | 0.03107 | 0.099 | -3.94 | -1.1908 | none |
| CDC26 | -0.378 | -0.3918 | -2.521 | 0.03107 | 0.099 | -3.94 | 1.35368 | none |
| BTLA | -0.596 | -0.5324 | -2.521 | 0.0311 | 0.099 | -3.95 | 0.71874 | none |
| PSME1 | -0.256 | -0.2688 | -2.521 | 0.03111 | 0.099 | -3.95 | 2.31974 | AA-related |
| AP2B1 | 0.557 | 0.4669 | 2.5206 | 0.03112 | 0.099 | -3.95 | -0.6432 | none |
| KLRB1 | -0.484 | -0.5152 | -2.519 | 0.03118 | 0.1 | -3.95 | 1.52199 | none |
| GLDN | 0.317 | 0.2585 | 2.5194 | 0.03118 | 0.1 | -3.95 | -0.9673 | none |
| FUT9 | 0.198 | 0.1442 | 2.519 | 0.0312 | 0.1 | -3.95 | -1.442 | none |
| PAK6 | 1.04 | 0.7937 | 2.5188 | 0.03121 | 0.1 | -3.95 | 0.24391 | none |
| FMNL3 | 0.468 | 0.5322 | 2.5187 | 0.03122 | 0.1 | -3.95 | -0.961 | none |
| TANC1 | 0.194 | 0.1746 | 2.5181 | 0.03125 | 0.1 | -3.95 | -1.217 | none |
| PHF1 | 0.69 | 0.5395 | 2.5174 | 0.03129 | 0.1 | -3.95 | -0.6797 | none |
| NPEPPS | 0.327 | 0.288 | 2.5173 | 0.03129 | 0.1 | -3.95 | -0.0608 | none |
| GTF2H2 | 0.481 | 0.3167 | 2.517 | 0.0313 | 0.1 | -3.95 | -0.7837 | none |
| PRSS23 | 0.491 | 0.391 | 2.5169 | 0.03131 | 0.1 | -3.95 | -0.1313 | none |
| RPA1 | 0.256 | 0.1921 | 2.5166 | 0.03133 | 0.1 | -3.95 | 0.07399 | none |
| ITGA6 | 0.547 | 0.5243 | 2.5165 | 0.03133 | 0.1 | -3.95 | 0.04528 | none |
| COL10A1 | -0.248 | -0.1697 | -2.516 | 0.03134 | 0.1 | -3.95 | 0.4618 | none |
| NOC2L | 0.472 | 0.469 | 2.5163 | 0.03134 | 0.1 | -3.95 | -0.6849 | none |
| UHRF1BP1 | 0.491 | 0.5099 | 2.5163 | 0.03134 | 0.1 | -3.95 | -0.5757 | none |
| CHCHD1 | -0.344 | -0.3344 | -2.516 | 0.03135 | 0.1 | -3.95 | 0.79828 | none |
| RWDD2B | 0.388 | 0.3211 | 2.5162 | 0.03135 | 0.1 | -3.95 | -0.6472 | none |
| B3GALT5 | 0.265 | 0.2074 | 2.5161 | 0.03136 | 0.1 | -3.95 | -1.0931 | none |
| BCAR3 | 0.548 | 0.4001 | 2.5159 | 0.03136 | 0.1 | -3.95 | -1.3289 | none |
| COX4I1 | -0.416 | -0.396 | -2.516 | 0.03138 | 0.1 | -3.95 | 1.82711 | none |
| MBNL1 | 0.473 | 0.4179 | 2.5153 | 0.0314 | 0.1 | -3.95 | -0.3282 | none |
| DNAJC7 | 0.588 | 0.5387 | 2.5152 | 0.0314 | 0.1 | -3.95 | -0.8439 | none |
| WRAP53 | -0.255 | -0.1728 | -2.515 | 0.03142 | 0.1 | -3.96 | 1.27829 | none |
| ACSL1 | 0.346 | 0.3332 | 2.5146 | 0.03143 | 0.1 | -3.96 | -0.4801 | none |
| FKBP3 | 0.242 | 0.1618 | 2.5145 | 0.03144 | 0.1 | -3.96 | -0.553 | none |
| RREB1 | 0.531 | 0.5788 | 2.5142 | 0.03145 | 0.1 | -3.96 | -0.6012 | none |
| MED6 | 0.201 | 0.0697 | 2.5141 | 0.03146 | 0.1 | -3.96 | -0.4568 | none |

| | | | | | | | | |
|---|---|---|---|---|---|---|---|---|
| BCAP31 | -0.319 | -0.301 | -2.514 | 0.03146 | 0.1 | -3.96 | 1.9761 | none |
| ADD2 | 0.275 | 0.2034 | 2.5129 | 0.03153 | 0.1 | -3.96 | -0.3138 | none |
| PDE3B | 0.472 | 0.3782 | 2.5128 | 0.03153 | 0.1 | -3.96 | 0.08766 | none |
| ENKUR | 0.216 | 0.2195 | 2.5123 | 0.03156 | 0.1 | -3.96 | 0.48374 | none |
| SGSH | 0.437 | 0.4395 | 2.5123 | 0.03156 | 0.1 | -3.96 | -0.5804 | none |
| PBDC1 | -0.229 | -0.1781 | -2.512 | 0.03159 | 0.1 | -3.96 | 0.99727 | none |
| MYDGF | -0.241 | -0.2088 | -2.511 | 0.03162 | 0.1 | -3.96 | 2.33201 | AA-related |
| ODF2 | 0.759 | 0.7363 | 2.5099 | 0.03169 | 0.101 | -3.96 | -0.6314 | none |
| HARS2 | 0.196 | 0.1454 | 2.5097 | 0.0317 | 0.101 | -3.96 | -0.7157 | none |
| MAN2B2 | 0.644 | 0.5788 | 2.5096 | 0.0317 | 0.101 | -3.96 | -0.884 | none |
| SRPRB | -0.354 | -0.3346 | -2.509 | 0.03172 | 0.101 | -3.96 | 2.40855 | AA-related |
| AIFM2 | 0.294 | 0.2686 | 2.5086 | 0.03175 | 0.101 | -3.97 | -1.3284 | none |
| PILRB | 0.384 | 0.3878 | 2.5085 | 0.03176 | 0.101 | -3.97 | -0.4626 | none |
| TMEM25 | 0.369 | 0.2668 | 2.5072 | 0.03183 | 0.101 | -3.97 | -0.3625 | none |
| ANO7 | 0.424 | 0.3498 | 2.5069 | 0.03184 | 0.101 | -3.97 | -0.843 | none |
| COL17A1 | 0.28 | 0.2472 | 2.5068 | 0.03185 | 0.101 | -3.97 | -0.6185 | none |
| CLCN7 | 0.447 | 0.3997 | 2.5067 | 0.03186 | 0.101 | -3.97 | -0.4739 | none |
| ACPP | 0.386 | 0.3638 | 2.5065 | 0.03187 | 0.101 | -3.97 | -0.9566 | none |
| PCNA | -0.357 | -0.3411 | -2.506 | 0.03187 | 0.101 | -3.97 | 1.73821 | none |
| RPL5 | -0.218 | -0.2244 | -2.506 | 0.03188 | 0.101 | -3.97 | 1.42014 | none |
| USP9X | 0.645 | 0.6512 | 2.5052 | 0.03194 | 0.101 | -3.97 | -0.4441 | none |
| MYCT1 | 0.343 | 0.2221 | 2.5047 | 0.03196 | 0.101 | -3.97 | 0.27373 | none |
| FAM193A | 0.584 | 0.6366 | 2.5041 | 0.032 | 0.101 | -3.97 | -0.3649 | none |
| NUDT3 | 0.396 | 0.3579 | 2.5032 | 0.03204 | 0.101 | -3.97 | -0.8046 | none |
| PTPRF | 0.603 | 0.4892 | 2.5031 | 0.03205 | 0.101 | -3.97 | -1.1893 | none |
| PCNT | 0.554 | 0.6727 | 2.5031 | 0.03205 | 0.101 | -3.97 | -0.8062 | none |
| DCUN1D5 | -0.328 | -0.2886 | -2.503 | 0.03206 | 0.101 | -3.97 | 1.6774 | none |
| CTSA | 0.346 | 0.3272 | 2.5028 | 0.03207 | 0.101 | -3.98 | -0.5418 | none |
| ZNF771 | 0.313 | 0.1916 | 2.5027 | 0.03207 | 0.101 | -3.98 | -0.998 | none |
| PRKCH | 0.355 | 0.3334 | 2.5026 | 0.03208 | 0.101 | -3.98 | 0.22653 | none |
| RAB26 | 0.331 | 0.2287 | 2.5024 | 0.03209 | 0.101 | -3.98 | -1.1851 | none |
| ATIC | -0.197 | -0.1629 | -2.502 | 0.03213 | 0.101 | -3.98 | 2.28433 | AA-related |
| BARD1 | 0.223 | 0.1175 | 2.5014 | 0.03214 | 0.101 | -3.98 | 0.08563 | none |
| MEIG1 | -0.323 | -0.3122 | -2.501 | 0.03214 | 0.101 | -3.98 | 1.7452 | none |
| BRK1 | -0.478 | -0.5302 | -2.501 | 0.03218 | 0.102 | -3.98 | 1.82862 | none |
| SMAD5 | 0.411 | 0.2502 | 2.5002 | 0.03221 | 0.102 | -3.98 | -0.6763 | none |
| ZNF451 | 0.531 | 0.3837 | 2.5001 | 0.03221 | 0.102 | -3.98 | -1.05 | none |
| TNIK | 0.353 | 0.2949 | 2.4996 | 0.03224 | 0.102 | -3.98 | -0.9575 | none |
| MNS1 | 0.546 | 0.4271 | 2.4996 | 0.03224 | 0.102 | -3.98 | -0.9688 | none |
| NLRP1 | 0.432 | 0.4443 | 2.4995 | 0.03224 | 0.102 | -3.98 | -0.8405 | none |

| | | | | | | | | |
|---|---|---|---|---|---|---|---|---|
| PCDHB13 | 0.227 | 0.1923 | 2.4988 | 0.03228 | 0.102 | -3.98 | -1.1629 | none |
| NME1 | -0.308 | -0.2809 | -2.498 | 0.03232 | 0.102 | -3.98 | 1.59091 | none |
| DNAH5 | 0.711 | 0.5695 | 2.498 | 0.03232 | 0.102 | -3.98 | -0.7344 | none |
| LY86 | -0.289 | -0.2701 | -2.498 | 0.03233 | 0.102 | -3.98 | 1.57342 | none |
| PSMB1 | -0.377 | -0.3621 | -2.498 | 0.03233 | 0.102 | -3.98 | 1.13806 | none |
| SMURF1 | 0.365 | 0.3737 | 2.4975 | 0.03235 | 0.102 | -3.98 | -0.3891 | none |
| MSTO1 | 0.348 | 0.4141 | 2.4966 | 0.0324 | 0.102 | -3.99 | 0.15693 | none |
| PTPA | 0.398 | 0.2978 | 2.4965 | 0.03241 | 0.102 | -3.99 | -1.1342 | none |
| PARP12 | 0.28 | 0.221 | 2.4965 | 0.03241 | 0.102 | -3.99 | -0.6155 | none |
| PDE10A | 0.446 | 0.3699 | 2.4965 | 0.03241 | 0.102 | -3.99 | 0.44509 | none |
| ERI3 | -0.398 | -0.3421 | -2.496 | 0.03244 | 0.102 | -3.99 | 1.01087 | none |
| UBXN6 | -0.264 | -0.1925 | -2.495 | 0.03247 | 0.102 | -3.99 | 1.17203 | none |
| QTRT1 | 0.33 | 0.2732 | 2.4953 | 0.03247 | 0.102 | -3.99 | -0.5251 | none |
| NAP1L2 | -0.232 | -0.1787 | -2.495 | 0.0325 | 0.102 | -3.99 | 1.61277 | none |
| LPXN | -0.324 | -0.3454 | -2.495 | 0.03251 | 0.102 | -3.99 | 2.00174 | none |
| ACOX2 | -0.229 | -0.2287 | -2.494 | 0.03252 | 0.102 | -3.99 | 1.38067 | none |
| PPIL1 | -0.37 | -0.3579 | -2.494 | 0.03253 | 0.102 | -3.99 | 2.16067 | AA-related |
| SMG7 | 0.639 | 0.6216 | 2.494 | 0.03255 | 0.102 | -3.99 | -0.909 | none |
| NDOR1 | 0.319 | 0.299 | 2.4939 | 0.03255 | 0.102 | -3.99 | -0.4028 | none |
| ZNF676 | 0.488 | 0.3 | 2.4939 | 0.03255 | 0.102 | -3.99 | -1.0457 | none |
| LRFN1 | 0.257 | 0.2335 | 2.4939 | 0.03255 | 0.102 | -3.99 | -1.0738 | none |
| CST7 | -0.267 | -0.2465 | -2.494 | 0.03256 | 0.102 | -3.99 | 0.47687 | none |
| TMEM243 | 0.372 | 0.2342 | 2.4937 | 0.03256 | 0.102 | -3.99 | -0.2659 | none |
| LCOR | 0.57 | 0.4994 | 2.4935 | 0.03257 | 0.102 | -3.99 | -0.8108 | none |
| ZNF630 | 0.367 | 0.3074 | 2.4935 | 0.03257 | 0.102 | -3.99 | -0.7089 | none |
| TTC28 | 0.532 | 0.4133 | 2.4935 | 0.03258 | 0.102 | -3.99 | -0.8169 | none |
| MED1 | 0.654 | 0.6667 | 2.4934 | 0.03258 | 0.102 | -3.99 | -0.4398 | none |
| ERCC1 | 0.344 | 0.2311 | 2.493 | 0.0326 | 0.102 | -3.99 | -1.2972 | none |
| TRIM14 | 0.558 | 0.5433 | 2.4919 | 0.03266 | 0.102 | -3.99 | 0.45807 | none |
| TMEM184 | 0.288 | 0.1875 | 2.4919 | 0.03266 | 0.102 | -3.99 | 0.83694 | none |
| ZNF691 | 0.287 | 0.203 | 2.4917 | 0.03267 | 0.102 | -3.99 | -0.4259 | none |
| FAM210A | 0.243 | 0.172 | 2.4916 | 0.03268 | 0.102 | -3.99 | -0.9435 | none |
| NRSN2 | -0.336 | -0.2341 | -2.491 | 0.03273 | 0.103 | -3.99 | 1.61892 | none |
| PSMA4 | -0.253 | -0.2361 | -2.49 | 0.03279 | 0.103 | -4 | 2.04472 | none |
| TCF12 | 0.555 | 0.5722 | 2.4891 | 0.03282 | 0.103 | -4 | -0.2823 | none |
| ANGPT4 | 0.253 | 0.1869 | 2.4889 | 0.03283 | 0.103 | -4 | -1.3277 | none |
| MAP2K3 | -0.17 | -0.0763 | -2.489 | 0.03284 | 0.103 | -4 | 1.3559 | none |
| NXF1 | 0.453 | 0.3908 | 2.4886 | 0.03284 | 0.103 | -4 | -0.9804 | none |
| RNF31 | 0.458 | 0.4021 | 2.4885 | 0.03285 | 0.103 | -4 | -1.181 | none |
| AP2A1 | 0.448 | 0.4269 | 2.4883 | 0.03286 | 0.103 | -4 | -0.886 | none |

| | | | | | | | | |
|---|---|---|---|---|---|---|---|---|
| CKS2 | -0.535 | -0.5382 | -2.488 | 0.03287 | 0.103 | -4 | 2.1611 | AA-related |
| SCFD2 | 0.248 | 0.1949 | 2.4878 | 0.03289 | 0.103 | -4 | -1.0215 | none |
| POLL | 0.381 | 0.3313 | 2.4876 | 0.0329 | 0.103 | -4 | -0.7781 | none |
| NUP214 | 0.613 | 0.6512 | 2.4875 | 0.03291 | 0.103 | -4 | -0.8178 | none |
| PCYT1A | 0.229 | 0.154 | 2.487 | 0.03293 | 0.103 | -4 | -0.1856 | none |
| KCNK2 | 0.344 | 0.2968 | 2.4863 | 0.03297 | 0.103 | -4 | -0.2439 | none |
| EDRF1 | 0.517 | 0.5202 | 2.4862 | 0.03298 | 0.103 | -4 | -1.1184 | none |
| KCNG1 | 0.514 | 0.4225 | 2.486 | 0.03299 | 0.103 | -4 | -0.442 | none |
| SLC17A7 | 0.312 | 0.2424 | 2.4857 | 0.03301 | 0.103 | -4 | -1.0039 | none |
| N4BP2 | 0.652 | 0.6415 | 2.4855 | 0.03302 | 0.103 | -4 | -0.5654 | none |
| ACOT4 | -0.412 | -0.345 | -2.485 | 0.03303 | 0.103 | -4 | 1.63801 | none |
| PHF13 | -0.291 | -0.2075 | -2.485 | 0.03306 | 0.103 | -4 | 2.17665 | AA-related |
| MAATS1 | 0.422 | 0.2974 | 2.4845 | 0.03307 | 0.103 | -4 | -0.4928 | none |
| ATF2 | 0.202 | 0.1096 | 2.4844 | 0.03308 | 0.103 | -4 | -0.8348 | none |
| TM4SF20 | 0.208 | 0.1234 | 2.4838 | 0.03311 | 0.103 | -4.01 | -0.9428 | none |
| SGO1 | 0.442 | 0.3549 | 2.4835 | 0.03313 | 0.103 | -4.01 | -1.1255 | none |
| RELA | 0.345 | 0.3203 | 2.483 | 0.03316 | 0.103 | -4.01 | -1.0645 | none |
| TSC1 | 0.491 | 0.4439 | 2.4828 | 0.03317 | 0.103 | -4.01 | -1.3052 | none |
| USB1 | 0.277 | 0.2352 | 2.4818 | 0.03322 | 0.104 | -4.01 | 0.37479 | none |
| PSMB5 | -0.344 | -0.3248 | -2.482 | 0.03323 | 0.104 | -4.01 | 1.94743 | none |
| HSFY1 | 0.481 | 0.3498 | 2.4809 | 0.03327 | 0.104 | -4.01 | -0.8321 | none |
| AGBL1 | 0.194 | 0.1441 | 2.4803 | 0.03331 | 0.104 | -4.01 | -1.3755 | none |
| FOSB | 0.269 | 0.2077 | 2.4803 | 0.03331 | 0.104 | -4.01 | -0.2894 | none |
| TTC25 | 0.546 | 0.4357 | 2.48 | 0.03333 | 0.104 | -4.01 | -0.2875 | none |
| SNTB2 | 0.42 | 0.3658 | 2.4798 | 0.03334 | 0.104 | -4.01 | -1.0633 | none |
| PPP4R3B | 0.434 | 0.4134 | 2.4796 | 0.03335 | 0.104 | -4.01 | -0.2384 | none |
| ASB6 | 0.319 | 0.2734 | 2.4795 | 0.03335 | 0.104 | -4.01 | -0.6927 | none |
| PLIN3 | 0.324 | 0.2546 | 2.4791 | 0.03337 | 0.104 | -4.01 | -0.9322 | none |
| CDKL1 | 0.52 | 0.4083 | 2.4791 | 0.03338 | 0.104 | -4.01 | -0.9785 | none |
| NCKIPSD | 0.477 | 0.4615 | 2.4789 | 0.03339 | 0.104 | -4.01 | 0.04342 | none |
| RECK | 0.364 | 0.2863 | 2.4788 | 0.03339 | 0.104 | -4.01 | -1.169 | none |
| DCAF11 | 0.285 | 0.2241 | 2.4778 | 0.03345 | 0.104 | -4.02 | -0.2859 | none |
| UBR1 | 0.488 | 0.4222 | 2.4774 | 0.03347 | 0.104 | -4.02 | -1.1804 | none |
| ERP44 | -0.268 | -0.2735 | -2.477 | 0.03347 | 0.104 | -4.02 | 2.07252 | none |
| YARS | 0.357 | 0.3381 | 2.4772 | 0.03348 | 0.104 | -4.02 | -0.2848 | none |
| SDHB | -0.274 | -0.2566 | -2.477 | 0.03352 | 0.104 | -4.02 | 1.97486 | none |
| PCGF3 | 0.209 | 0.144 | 2.4759 | 0.03356 | 0.104 | -4.02 | -0.6109 | none |
| HLA-DQB2 | 0.262 | 0.1916 | 2.4749 | 0.03362 | 0.104 | -4.02 | -0.2372 | none |
| OSBP2 | 0.784 | 0.5418 | 2.4747 | 0.03363 | 0.105 | -4.02 | -1.0683 | none |
| PLS3 | 0.299 | 0.2515 | 2.4742 | 0.03365 | 0.105 | -4.02 | -0.3266 | none |

| | | | | | | | | |
|---|---|---|---|---|---|---|---|---|
| ST3GAL5 | 0.248 | 0.1668 | 2.4735 | 0.03369 | 0.105 | -4.02 | -1.0023 | none |
| SPSB2 | -0.21 | -0.173 | -2.473 | 0.0337 | 0.105 | -4.02 | 0.69852 | none |
| B3GAT3 | 0.399 | 0.3319 | 2.4734 | 0.0337 | 0.105 | -4.02 | -1.059 | none |
| RHOT2 | 0.246 | 0.2392 | 2.4733 | 0.0337 | 0.105 | -4.02 | 0.08796 | none |
| MRPS35 | -0.2 | -0.1814 | -2.473 | 0.03372 | 0.105 | -4.02 | 2.12974 | AA-related |
| FAN1 | 0.897 | 0.776 | 2.4727 | 0.03374 | 0.105 | -4.02 | -0.1456 | none |
| CD3EAP | 0.734 | 0.7858 | 2.4725 | 0.03375 | 0.105 | -4.02 | -1.0898 | none |
| OSGIN1 | 0.315 | 0.2777 | 2.47 | 0.03389 | 0.105 | -4.03 | -0.0855 | none |
| YY1AP1 | 0.528 | 0.5052 | 2.4696 | 0.03392 | 0.105 | -4.03 | -0.4684 | none |
| CACNB1 | 0.312 | 0.3547 | 2.4694 | 0.03393 | 0.105 | -4.03 | 0.30229 | none |
| RALA | -0.29 | -0.2445 | -2.469 | 0.03393 | 0.105 | -4.03 | 2.35644 | AA-related |
| CES3 | 0.272 | 0.2197 | 2.4693 | 0.03393 | 0.105 | -4.03 | -0.4562 | none |
| SERPINA5 | 0.483 | 0.3642 | 2.4691 | 0.03394 | 0.105 | -4.03 | -0.7586 | none |
| AADAT | 0.245 | 0.1937 | 2.469 | 0.03395 | 0.105 | -4.03 | -0.9956 | none |
| TSPAN17 | -0.274 | -0.2458 | -2.469 | 0.03397 | 0.105 | -4.03 | 2.36746 | AA-related |
| SMIM7 | 0.347 | 0.1831 | 2.4685 | 0.03398 | 0.105 | -4.03 | -0.4747 | none |
| DMTF1 | 0.43 | 0.3356 | 2.4683 | 0.03399 | 0.105 | -4.03 | -0.5433 | none |
| PHTF1 | 0.517 | 0.3195 | 2.4682 | 0.034 | 0.105 | -4.03 | -0.8647 | none |
| SCAMP5 | 0.518 | 0.3159 | 2.468 | 0.03401 | 0.105 | -4.03 | -0.7399 | none |
| NDUFB6 | -0.461 | -0.4699 | -2.467 | 0.03405 | 0.105 | -4.03 | 1.24363 | none |
| GHRL | 0.751 | 0.7184 | 2.4671 | 0.03406 | 0.105 | -4.03 | 0.2648 | none |
| CHD4 | 0.954 | 0.9637 | 2.4663 | 0.0341 | 0.106 | -4.03 | -0.9233 | none |
| SEC61B | -0.445 | -0.4512 | -2.466 | 0.03411 | 0.106 | -4.03 | 1.83606 | none |
| ASAH2B | 0.342 | 0.2522 | 2.4661 | 0.03412 | 0.106 | -4.03 | -0.1556 | none |
| DVL3 | 0.365 | 0.3274 | 2.4657 | 0.03414 | 0.106 | -4.04 | -0.5257 | none |
| UBE2D4 | 0.235 | 0.1168 | 2.4657 | 0.03414 | 0.106 | -4.04 | -0.9031 | none |
| RPUSD3 | 0.26 | 0.1783 | 2.4655 | 0.03415 | 0.106 | -4.04 | -0.6635 | none |
| TPP2 | 0.669 | 0.6106 | 2.4653 | 0.03416 | 0.106 | -4.04 | -0.1956 | none |
| RHOA | -0.401 | -0.4203 | -2.465 | 0.0342 | 0.106 | -4.04 | 1.92017 | none |
| ZFYVE26 | 0.552 | 0.4408 | 2.4646 | 0.0342 | 0.106 | -4.04 | -1.3865 | none |
| DDIAS | 0.56 | 0.4733 | 2.4642 | 0.03423 | 0.106 | -4.04 | -0.8212 | none |
| IFT57 | -0.334 | -0.2904 | -2.463 | 0.03428 | 0.106 | -4.04 | 1.59686 | none |
| LYZ | 0.643 | 0.4578 | 2.4627 | 0.03431 | 0.106 | -4.04 | -1.0078 | none |
| GBP4 | 0.271 | 0.1842 | 2.4625 | 0.03432 | 0.106 | -4.04 | -1.0162 | none |
| MTRNR2L4 | 0.195 | 0.1384 | 2.4624 | 0.03433 | 0.106 | -4.04 | 0.19669 | none |
| UQCRH | -0.416 | -0.4449 | -2.462 | 0.03434 | 0.106 | -4.04 | 1.5486 | none |
| RPL15 | -0.393 | -0.4014 | -2.462 | 0.03437 | 0.106 | -4.04 | 2.05844 | none |
| COX6B1 | -0.376 | -0.3601 | -2.461 | 0.03439 | 0.106 | -4.04 | 1.66291 | none |
| TPP1 | 0.371 | 0.3389 | 2.4608 | 0.03443 | 0.106 | -4.04 | -0.6642 | none |
| PSMA1 | -0.33 | -0.3382 | -2.46 | 0.03445 | 0.106 | -4.04 | 1.92081 | none |

| | | | | | | | | |
|---|---|---|---|---|---|---|---|---|
| RUNDC3A | 0.395 | 0.2479 | 2.4592 | 0.03452 | 0.106 | -4.05 | -1.047 | none |
| UQCR10 | -0.311 | -0.3135 | -2.459 | 0.03453 | 0.106 | -4.05 | 0.63934 | none |
| PRKAR2A | 0.57 | 0.555 | 2.4587 | 0.03455 | 0.107 | -4.05 | -0.5003 | none |
| TUBB4B | -0.262 | -0.2501 | -2.457 | 0.03466 | 0.107 | -4.05 | 1.57577 | none |
| NCOA3 | 0.664 | 0.6576 | 2.4565 | 0.03467 | 0.107 | -4.05 | -0.5883 | none |
| SIRT4 | -0.274 | -0.2595 | -2.457 | 0.03467 | 0.107 | -4.05 | 2.19194 | AA-related |
| URB2 | 0.265 | 0.236 | 2.4563 | 0.03468 | 0.107 | -4.05 | 0.28093 | none |
| CNKSR2 | 0.496 | 0.4408 | 2.4559 | 0.03471 | 0.107 | -4.05 | -0.3753 | none |
| ANKRD7 | 0.348 | 0.2316 | 2.4557 | 0.03472 | 0.107 | -4.05 | -0.3809 | none |
| IL32 | -0.281 | -0.2653 | -2.456 | 0.03473 | 0.107 | -4.05 | 1.92073 | none |
| SPAG4 | 0.468 | 0.3923 | 2.4555 | 0.03473 | 0.107 | -4.05 | -0.7503 | none |
| NUP210 | 0.588 | 0.6463 | 2.4554 | 0.03474 | 0.107 | -4.05 | -0.5061 | none |
| PUS7 | 0.395 | 0.3974 | 2.4548 | 0.03478 | 0.107 | -4.05 | -0.0363 | none |
| CSTB | -0.328 | -0.3265 | -2.454 | 0.03483 | 0.107 | -4.05 | 1.9723 | none |
| CLEC17A | 0.3 | 0.2409 | 2.4536 | 0.03485 | 0.107 | -4.05 | -0.7773 | none |
| KIFC2 | 0.468 | 0.4297 | 2.4532 | 0.03487 | 0.107 | -4.06 | -0.0203 | none |
| DCX | 0.194 | 0.134 | 2.4523 | 0.03492 | 0.107 | -4.06 | -1.453 | none |
| RB1 | 0.359 | 0.3191 | 2.452 | 0.03494 | 0.107 | -4.06 | 0.02281 | none |
| S100A11 | -0.256 | -0.2656 | -2.452 | 0.03495 | 0.107 | -4.06 | 0.90683 | none |
| TTBK1 | 0.208 | 0.2021 | 2.4509 | 0.035 | 0.108 | -4.06 | -1.2253 | none |
| LGALS7 | 0.303 | 0.25 | 2.4508 | 0.03501 | 0.108 | -4.06 | -0.327 | none |
| HSPB1 | -0.322 | -0.2627 | -2.451 | 0.03501 | 0.108 | -4.06 | 1.58535 | none |
| DHX33 | 0.375 | 0.4299 | 2.4501 | 0.03505 | 0.108 | -4.06 | -0.4606 | none |
| CHMP5 | -0.432 | -0.4456 | -2.449 | 0.03512 | 0.108 | -4.06 | 1.51269 | none |
| ZNF148 | 0.433 | 0.4212 | 2.4488 | 0.03513 | 0.108 | -4.06 | -0.3727 | none |
| UMOD | -0.236 | -0.1772 | -2.449 | 0.03513 | 0.108 | -4.06 | 0.04135 | none |
| ZNF195 | 0.361 | 0.3042 | 2.4483 | 0.03516 | 0.108 | -4.06 | -1.1774 | none |
| LY6E | -0.444 | -0.4053 | -2.448 | 0.03517 | 0.108 | -4.06 | 1.96734 | none |
| AIF1L | -0.397 | -0.3674 | -2.447 | 0.03524 | 0.108 | -4.07 | 2.07553 | none |
| BTNL8 | 0.272 | 0.1951 | 2.4463 | 0.03527 | 0.108 | -4.07 | -0.0795 | none |
| GXYLT1 | 0.47 | 0.3955 | 2.4463 | 0.03528 | 0.108 | -4.07 | -0.4467 | none |
| CSF2RB | 0.255 | 0.2537 | 2.446 | 0.03529 | 0.108 | -4.07 | -0.7943 | none |
| MED15 | 0.615 | 0.6542 | 2.4449 | 0.03536 | 0.108 | -4.07 | -0.4986 | none |
| MYLK | 0.332 | 0.2933 | 2.4444 | 0.03539 | 0.108 | -4.07 | -0.8891 | none |
| P4HA3 | 0.469 | 0.2521 | 2.4439 | 0.03542 | 0.109 | -4.07 | -0.9819 | none |
| CLSTN3 | 0.389 | 0.4664 | 2.4438 | 0.03542 | 0.109 | -4.07 | 0.42872 | none |
| ACOT13 | -0.287 | -0.2544 | -2.444 | 0.03543 | 0.109 | -4.07 | 1.90335 | none |
| EVA1C | -0.458 | -0.3373 | -2.444 | 0.03543 | 0.109 | -4.07 | 2.03719 | none |
| MLF1 | 0.335 | 0.2284 | 2.4436 | 0.03544 | 0.109 | -4.07 | -0.8875 | none |
| GTF2E1 | 0.225 | 0.0597 | 2.4432 | 0.03546 | 0.109 | -4.07 | -0.6677 | none |

| | | | | | | | | |
|---|---|---|---|---|---|---|---|---|
| SEL1L3 | 0.295 | 0.2655 | 2.443 | 0.03547 | 0.109 | -4.07 | -0.4794 | none |
| PRPF4 | 0.212 | 0.1847 | 2.4429 | 0.03548 | 0.109 | -4.07 | -0.934 | none |
| F10 | 0.205 | 0.1963 | 2.4424 | 0.03551 | 0.109 | -4.07 | 0.43733 | none |
| EXOSC10 | 0.674 | 0.6603 | 2.4423 | 0.03551 | 0.109 | -4.07 | -0.8904 | none |
| NEXN | 0.351 | 0.2378 | 2.4421 | 0.03553 | 0.109 | -4.07 | -0.6934 | none |
| SRFBP1 | 0.496 | 0.4759 | 2.4419 | 0.03554 | 0.109 | -4.07 | -0.4899 | none |
| TECPR1 | 0.493 | 0.4682 | 2.4418 | 0.03555 | 0.109 | -4.07 | -0.7073 | none |
| DCDC1 | 0.268 | 0.2019 | 2.4413 | 0.03557 | 0.109 | -4.07 | -1.3381 | none |
| ACTA2 | 0.367 | 0.264 | 2.4413 | 0.03557 | 0.109 | -4.07 | -1.0878 | none |
| GOSR2 | 0.297 | 0.2063 | 2.4412 | 0.03558 | 0.109 | -4.07 | -0.0543 | none |
| KIDINS220 | 0.614 | 0.6268 | 2.4402 | 0.03564 | 0.109 | -4.08 | -0.7498 | none |
| TMEM265 | 0.317 | 0.2821 | 2.44 | 0.03565 | 0.109 | -4.08 | -0.2633 | none |
| UBAP2 | 0.525 | 0.4769 | 2.4398 | 0.03566 | 0.109 | -4.08 | -0.5727 | none |
| SLC18A1 | 0.242 | 0.2149 | 2.4397 | 0.03567 | 0.109 | -4.08 | 0.22271 | none |
| ANKFY1 | 0.493 | 0.4773 | 2.4391 | 0.03571 | 0.109 | -4.08 | -0.8079 | none |
| SBNO2 | 0.446 | 0.4396 | 2.4385 | 0.03574 | 0.109 | -4.08 | -0.4173 | none |
| SLC1A5 | -0.25 | -0.2095 | -2.438 | 0.03576 | 0.109 | -4.08 | 1.89479 | none |
| SIRT3 | 0.396 | 0.3091 | 2.438 | 0.03577 | 0.109 | -4.08 | -1.0267 | none |
| COA6 | -0.425 | -0.425 | -2.437 | 0.03583 | 0.109 | -4.08 | 1.73256 | none |
| B3GNTL1 | 0.42 | 0.3399 | 2.437 | 0.03583 | 0.109 | -4.08 | -0.1428 | none |
| RCN3 | 0.32 | 0.2536 | 2.4369 | 0.03584 | 0.109 | -4.08 | -1.1386 | none |
| HNF4G | 0.871 | 0.636 | 2.4367 | 0.03585 | 0.109 | -4.08 | -0.9476 | none |
| KANK3 | 0.325 | 0.2543 | 2.4366 | 0.03586 | 0.109 | -4.08 | -0.4487 | none |
| LASP1 | 0.324 | 0.2672 | 2.4364 | 0.03587 | 0.109 | -4.08 | -0.0653 | none |
| SLC44A2 | 0.377 | 0.2526 | 2.4363 | 0.03588 | 0.109 | -4.08 | -0.7221 | none |
| COQ10B | 0.249 | 0.0913 | 2.436 | 0.0359 | 0.109 | -4.08 | -0.21 | none |
| PHYKPL | 0.247 | 0.1851 | 2.4334 | 0.03605 | 0.11 | -4.09 | 0.036 | none |
| UQCRFS1 | -0.415 | -0.3923 | -2.433 | 0.03606 | 0.11 | -4.09 | 2.03752 | none |
| CCT8 | -0.248 | -0.1826 | -2.433 | 0.03607 | 0.11 | -4.09 | 2.18123 | AA-related |
| LRRC47 | 0.294 | 0.2545 | 2.4331 | 0.03607 | 0.11 | -4.09 | -0.401 | none |
| DPP6 | 0.204 | 0.1537 | 2.433 | 0.03608 | 0.11 | -4.09 | -1.0571 | none |
| GLI4 | 0.384 | 0.2548 | 2.4328 | 0.03609 | 0.11 | -4.09 | 0.16239 | none |
| ABHD18 | 0.3 | 0.1512 | 2.4323 | 0.03612 | 0.11 | -4.09 | -1.1272 | none |
| UBTD1 | -0.395 | -0.3132 | -2.432 | 0.03614 | 0.11 | -4.09 | 0.77582 | none |
| HAUS2 | 0.299 | 0.1761 | 2.4312 | 0.03619 | 0.11 | -4.09 | -1.082 | none |
| CHCHD4 | -0.271 | -0.2619 | -2.431 | 0.03622 | 0.11 | -4.09 | 2.26083 | AA-related |
| RAP2B | 0.216 | 0.0884 | 2.4307 | 0.03622 | 0.11 | -4.09 | 0.17105 | none |
| USP28 | 0.419 | 0.3474 | 2.4306 | 0.03623 | 0.11 | -4.09 | 0.40531 | none |
| INSR | 0.354 | 0.2277 | 2.4302 | 0.03624 | 0.11 | -4.09 | -0.8719 | none |
| NHSL2 | 0.192 | 0.157 | 2.4302 | 0.03625 | 0.11 | -4.09 | -0.8924 | none |

| | | | | | | | | |
|---|---|---|---|---|---|---|---|---|
| NDUFAB1 | -0.4 | -0.3961 | -2.43 | 0.03628 | 0.11 | -4.09 | 1.83869 | none |
| MCCC1 | 0.268 | 0.2129 | 2.4297 | 0.03628 | 0.11 | -4.09 | -0.5339 | none |
| ZNF384 | 0.594 | 0.572 | 2.4293 | 0.0363 | 0.11 | -4.09 | -0.857 | none |
| YIPF3 | 0.369 | 0.2205 | 2.4288 | 0.03633 | 0.11 | -4.09 | -0.6338 | none |
| FNBP4 | 0.621 | 0.5865 | 2.4287 | 0.03634 | 0.11 | -4.09 | -0.203 | none |
| SLC41A1 | 0.237 | 0.2592 | 2.4284 | 0.03636 | 0.11 | -4.1 | -0.4907 | none |
| STOML2 | -0.324 | -0.2954 | -2.428 | 0.03641 | 0.111 | -4.1 | 2.48889 | AA-related |
| RAB19 | 0.746 | 0.5805 | 2.4275 | 0.03641 | 0.111 | -4.1 | -1.2175 | none |
| RBAK | 0.621 | 0.5508 | 2.4275 | 0.03641 | 0.111 | -4.1 | -1.0836 | none |
| CHRNA2 | 0.34 | 0.2936 | 2.4271 | 0.03644 | 0.111 | -4.1 | 0.06722 | none |
| PIGL | 0.368 | 0.2867 | 2.4268 | 0.03645 | 0.111 | -4.1 | -1.3573 | none |
| BORCS7-A | -0.181 | -0.1813 | -2.427 | 0.03647 | 0.111 | -4.1 | 0.6785 | none |
| OVGP1 | 0.362 | 0.3129 | 2.4262 | 0.03649 | 0.111 | -4.1 | -0.5822 | none |
| FMNL2 | 0.458 | 0.3844 | 2.426 | 0.03651 | 0.111 | -4.1 | -0.0067 | none |
| NFX1 | 0.602 | 0.5886 | 2.426 | 0.03651 | 0.111 | -4.1 | -1.0595 | none |
| IQGAP2 | 0.882 | 0.7995 | 2.4259 | 0.03651 | 0.111 | -4.1 | -0.9845 | none |
| RCC1 | 0.268 | 0.2432 | 2.4257 | 0.03652 | 0.111 | -4.1 | 0.3354 | none |
| PIK3C2A | 0.693 | 0.7099 | 2.4253 | 0.03655 | 0.111 | -4.1 | -0.0157 | none |
| ARL3 | -0.277 | -0.2612 | -2.425 | 0.03655 | 0.111 | -4.1 | 1.36561 | none |
| LSM1 | -0.341 | -0.337 | -2.425 | 0.03656 | 0.111 | -4.1 | 1.02289 | none |
| CHMP4A | 0.362 | 0.3271 | 2.4251 | 0.03656 | 0.111 | -4.1 | -1.3328 | none |
| ZNF134 | 0.388 | 0.3021 | 2.4248 | 0.03658 | 0.111 | -4.1 | -0.4968 | none |
| MTRNR2L( | -0.392 | -0.3427 | -2.425 | 0.0366 | 0.111 | -4.1 | 0.89187 | none |
| TACC1 | 0.514 | 0.4994 | 2.4244 | 0.0366 | 0.111 | -4.1 | -0.0539 | none |
| TRIP10 | 0.284 | 0.3122 | 2.4236 | 0.03665 | 0.111 | -4.1 | -0.4097 | none |
| SFXN2 | 0.274 | 0.1899 | 2.4236 | 0.03666 | 0.111 | -4.1 | -0.0343 | none |
| PRKCE | 0.541 | 0.5004 | 2.4232 | 0.03668 | 0.111 | -4.1 | -0.9168 | none |
| PIAS2 | 0.362 | 0.2648 | 2.4229 | 0.0367 | 0.111 | -4.1 | 0.32182 | none |
| TFAP4 | 0.412 | 0.4158 | 2.4228 | 0.0367 | 0.111 | -4.1 | -0.2396 | none |
| LAMP1 | 0.337 | 0.2687 | 2.4224 | 0.03673 | 0.111 | -4.11 | -1.1695 | none |
| WDR45B | 0.218 | 0.1061 | 2.4221 | 0.03675 | 0.111 | -4.11 | -0.2759 | none |
| RBX1 | -0.392 | -0.3903 | -2.421 | 0.03678 | 0.111 | -4.11 | 1.58599 | none |
| TPH2 | 0.399 | 0.3141 | 2.4214 | 0.03679 | 0.111 | -4.11 | -0.7546 | none |
| MOSPD3 | 0.284 | 0.1774 | 2.4211 | 0.03681 | 0.111 | -4.11 | -0.4102 | none |
| WDR89 | 0.207 | 0.1095 | 2.421 | 0.03682 | 0.111 | -4.11 | -0.5359 | none |
| SCYL1 | 0.406 | 0.4045 | 2.42 | 0.03688 | 0.111 | -4.11 | -0.1248 | none |
| AKT1S1 | 0.522 | 0.403 | 2.4199 | 0.03689 | 0.111 | -4.11 | -1.1841 | none |
| CD81 | -0.378 | -0.2882 | -2.42 | 0.03691 | 0.111 | -4.11 | 0.98785 | none |
| CENPN | -0.205 | -0.1683 | -2.419 | 0.03691 | 0.111 | -4.11 | 1.12526 | none |
| FMNL1 | 0.599 | 0.6307 | 2.4194 | 0.03692 | 0.111 | -4.11 | -0.9847 | none |

| | | | | | | | | |
|---|---|---|---|---|---|---|---|---|
| IRF5 | 0.465 | 0.3133 | 2.4189 | 0.03694 | 0.111 | -4.11 | -0.9581 | none |
| SRSF6 | 0.579 | 0.5588 | 2.4184 | 0.03698 | 0.112 | -4.11 | -1.104 | none |
| CTNNB1 | 0.283 | 0.2868 | 2.4183 | 0.03699 | 0.112 | -4.11 | -0.735 | none |
| NF1 | 0.594 | 0.5815 | 2.4181 | 0.03699 | 0.112 | -4.11 | -0.781 | none |
| JAZF1 | 0.389 | 0.2885 | 2.4179 | 0.03701 | 0.112 | -4.11 | -0.5991 | none |
| CD83 | -0.235 | -0.1726 | -2.418 | 0.03702 | 0.112 | -4.11 | 2.59559 | AA-related |
| NCOA6 | 0.599 | 0.6069 | 2.4173 | 0.03704 | 0.112 | -4.11 | -0.9774 | none |
| SEC11C | -0.348 | -0.3773 | -2.417 | 0.03706 | 0.112 | -4.11 | 1.34324 | none |
| CBX2 | 0.289 | 0.3139 | 2.4159 | 0.03713 | 0.112 | -4.12 | -0.2713 | none |
| SCYL3 | 0.297 | 0.2299 | 2.415 | 0.03719 | 0.112 | -4.12 | -1.0665 | none |
| SLC26A2 | 0.397 | 0.3229 | 2.4149 | 0.03719 | 0.112 | -4.12 | -1.2217 | none |
| ZC3H12D | 0.619 | 0.6689 | 2.4145 | 0.03722 | 0.112 | -4.12 | -0.4444 | none |
| RPS27A | -0.443 | -0.4651 | -2.414 | 0.03724 | 0.112 | -4.12 | 1.84806 | none |
| OLFM2 | 0.33 | 0.2361 | 2.4142 | 0.03724 | 0.112 | -4.12 | -0.9588 | none |
| ELAC1 | 0.257 | 0.1875 | 2.4136 | 0.03728 | 0.112 | -4.12 | -0.2384 | none |
| CLIP3 | 0.518 | 0.5157 | 2.4133 | 0.03729 | 0.112 | -4.12 | 0.05481 | none |
| SUPT16H | 0.742 | 0.7506 | 2.4129 | 0.03732 | 0.112 | -4.12 | -0.8666 | none |
| AKAP13 | 0.633 | 0.6147 | 2.4127 | 0.03734 | 0.112 | -4.12 | -0.8261 | none |
| ZSCAN21 | 0.328 | 0.2135 | 2.4119 | 0.03739 | 0.112 | -4.12 | -0.562 | none |
| CLSPN | 0.713 | 0.7257 | 2.4107 | 0.03746 | 0.113 | -4.12 | -1.014 | none |
| SFXN4 | -0.272 | -0.2757 | -2.411 | 0.03746 | 0.113 | -4.12 | 1.83069 | none |
| PTPN20 | 0.32 | 0.239 | 2.4091 | 0.03756 | 0.113 | -4.13 | -0.7508 | none |
| DHX35 | 0.226 | 0.2107 | 2.4089 | 0.03757 | 0.113 | -4.13 | 0.14996 | none |
| C2ORF69 | 0.279 | 0.2159 | 2.4087 | 0.03759 | 0.113 | -4.13 | 1.06411 | none |
| GSR | 0.458 | 0.428 | 2.4082 | 0.03762 | 0.113 | -4.13 | -0.721 | none |
| SERHL2 | 0.422 | 0.2586 | 2.4082 | 0.03762 | 0.113 | -4.13 | -0.9664 | none |
| CTIF | 0.35 | 0.288 | 2.4081 | 0.03762 | 0.113 | -4.13 | -0.521 | none |
| PRKCSH | 0.534 | 0.5501 | 2.4075 | 0.03766 | 0.113 | -4.13 | -0.6029 | none |
| KIZ | 0.407 | 0.3653 | 2.4075 | 0.03766 | 0.113 | -4.13 | -0.6064 | none |
| ANKRD29 | 0.213 | 0.1649 | 2.4075 | 0.03766 | 0.113 | -4.13 | -0.3005 | none |
| MRPS2 | -0.254 | -0.19 | -2.407 | 0.03769 | 0.113 | -4.13 | 1.921 | none |
| SIX4 | 0.217 | 0.182 | 2.4065 | 0.03772 | 0.113 | -4.13 | -1.0041 | none |
| CELA3B | 0.529 | 0.2694 | 2.4063 | 0.03774 | 0.113 | -4.13 | -1.1584 | none |
| PHF19 | -0.266 | -0.2141 | -2.406 | 0.03775 | 0.113 | -4.13 | 2.25199 | AA-related |
| ZC3H18 | 0.762 | 0.824 | 2.4061 | 0.03775 | 0.113 | -4.13 | -0.4284 | none |
| PIGW | 0.343 | 0.2095 | 2.4054 | 0.03779 | 0.113 | -4.13 | 0.20203 | none |
| RPA4 | -0.19 | -0.1284 | -2.405 | 0.03783 | 0.113 | -4.13 | -0.1138 | none |
| C1ORF100 | 0.354 | 0.2799 | 2.4049 | 0.03783 | 0.113 | -4.13 | -0.8191 | none |
| DZIP3 | 0.453 | 0.3687 | 2.4038 | 0.0379 | 0.114 | -4.13 | -0.7345 | none |
| DDX60L | 0.549 | 0.4723 | 2.4037 | 0.03791 | 0.114 | -4.14 | -1.4173 | none |

| | | | | | | | | |
|---|---|---|---|---|---|---|---|---|
| WBP2 | 0.229 | 0.1843 | 2.4032 | 0.03793 | 0.114 | -4.14 | -0.0411 | none |
| BCKDHB | 0.279 | 0.1481 | 2.4026 | 0.03798 | 0.114 | -4.14 | -0.2678 | none |
| IMMP2L | 0.517 | 0.32 | 2.4018 | 0.03803 | 0.114 | -4.14 | -0.9317 | none |
| SCRN2 | 0.338 | 0.2882 | 2.4017 | 0.03803 | 0.114 | -4.14 | -0.6964 | none |
| TRO | -0.28 | -0.213 | -2.401 | 0.03805 | 0.114 | -4.14 | 1.52361 | none |
| CDRT4 | 0.363 | 0.2549 | 2.4014 | 0.03805 | 0.114 | -4.14 | -0.5709 | none |
| MAPKAP1 | 0.377 | 0.3549 | 2.4009 | 0.03808 | 0.114 | -4.14 | -0.8737 | none |
| KCNE1 | 0.276 | 0.1563 | 2.3997 | 0.03816 | 0.114 | -4.14 | -1.0838 | none |
| EEF1G | -0.271 | -0.2476 | -2.4 | 0.03817 | 0.114 | -4.14 | 0.70757 | none |
| NOTCH3 | 0.561 | 0.4397 | 2.3995 | 0.03818 | 0.114 | -4.14 | 0.11729 | none |
| TBC1D25 | 0.378 | 0.2608 | 2.3988 | 0.03822 | 0.114 | -4.14 | -0.8049 | none |
| DDX39A | -0.212 | -0.1294 | -2.399 | 0.03822 | 0.114 | -4.14 | 1.66807 | none |
| GOLGA4 | 0.39 | 0.4239 | 2.3987 | 0.03822 | 0.114 | -4.14 | -0.0786 | none |
| ZNF622 | -0.195 | -0.1128 | -2.399 | 0.03824 | 0.114 | -4.14 | 1.7575 | none |
| TAX1BP3 | -0.308 | -0.2896 | -2.398 | 0.03825 | 0.114 | -4.14 | 1.761 | none |
| EVC | 0.292 | 0.2445 | 2.3974 | 0.03831 | 0.114 | -4.15 | -0.8746 | none |
| PRRC2A | 0.833 | 0.7263 | 2.3972 | 0.03832 | 0.115 | -4.15 | -1.1943 | none |
| ITPRIPL1 | 0.251 | 0.1809 | 2.3969 | 0.03834 | 0.115 | -4.15 | -0.5797 | none |
| TMBIM6 | -0.418 | -0.4185 | -2.397 | 0.03834 | 0.115 | -4.15 | 2.29402 | AA-related |
| THOC1 | 0.477 | 0.4332 | 2.3963 | 0.03838 | 0.115 | -4.15 | 0.08044 | none |
| JUN | 0.363 | 0.4 | 2.396 | 0.0384 | 0.115 | -4.15 | 0.66921 | none |
| CDK1 | -0.394 | -0.3524 | -2.396 | 0.03842 | 0.115 | -4.15 | 1.96737 | none |
| CCDC117 | 0.403 | 0.2734 | 2.3955 | 0.03843 | 0.115 | -4.15 | 0.28487 | none |
| PRG4 | 0.497 | 0.4729 | 2.3953 | 0.03845 | 0.115 | -4.15 | -0.3601 | none |
| FBXW2 | 0.21 | 0.1456 | 2.395 | 0.03846 | 0.115 | -4.15 | -0.9588 | none |
| THNSL2 | 0.377 | 0.2733 | 2.3947 | 0.03848 | 0.115 | -4.15 | -0.9139 | none |
| PIKFYVE | 0.561 | 0.6033 | 2.3947 | 0.03848 | 0.115 | -4.15 | -0.2075 | none |
| ANGPTL4 | 0.528 | 0.4259 | 2.3941 | 0.03852 | 0.115 | -4.15 | -1.0445 | none |
| FDX2 | 0.41 | 0.3283 | 2.3929 | 0.0386 | 0.115 | -4.15 | -0.5473 | none |
| SUN3 | 0.635 | 0.4703 | 2.3922 | 0.03865 | 0.115 | -4.15 | -1.3165 | none |
| IGF2R | 0.693 | 0.7863 | 2.3921 | 0.03866 | 0.115 | -4.15 | -0.3976 | none |
| CCZ1 | -0.235 | -0.219 | -2.392 | 0.03866 | 0.115 | -4.15 | 1.30118 | none |
| MTCP1 | -0.813 | -0.7288 | -2.392 | 0.03867 | 0.115 | -4.15 | 1.54589 | none |
| MSANTD3 | 0.286 | 0.2773 | 2.3918 | 0.03867 | 0.115 | -4.15 | -0.9563 | none |
| SRSF3 | -0.328 | -0.316 | -2.392 | 0.03868 | 0.115 | -4.15 | 1.61559 | none |
| TNPO1 | 0.24 | 0.1717 | 2.3916 | 0.03868 | 0.115 | -4.15 | -0.254 | none |
| POLR1B | 0.298 | 0.2659 | 2.3915 | 0.03869 | 0.115 | -4.15 | 0.15815 | none |
| CDC25A | 0.234 | 0.2736 | 2.3914 | 0.0387 | 0.115 | -4.16 | -0.5821 | none |
| PBOV1 | 0.208 | 0.1491 | 2.3903 | 0.03877 | 0.115 | -4.16 | -0.4887 | none |
| RPL17 | -0.353 | -0.3622 | -2.39 | 0.03877 | 0.115 | -4.16 | 2.04426 | none |

| | | | | | | | | |
|---|---|---|---|---|---|---|---|---|
| TNKS1BP1 | 0.436 | 0.442 | 2.3896 | 0.03882 | 0.115 | -4.16 | -0.117 | none |
| MAPK8 | 0.419 | 0.2577 | 2.3893 | 0.03884 | 0.115 | -4.16 | -0.5102 | none |
| WRN | 0.504 | 0.4688 | 2.3892 | 0.03884 | 0.115 | -4.16 | -0.6101 | none |
| EFHC1 | 0.392 | 0.2849 | 2.3892 | 0.03884 | 0.115 | -4.16 | -1.3679 | none |
| SDCBP | -0.369 | -0.3731 | -2.389 | 0.03885 | 0.115 | -4.16 | 1.77346 | none |
| ATAD3C | 0.624 | 0.5218 | 2.389 | 0.03886 | 0.115 | -4.16 | -0.9056 | none |
| NAE1 | 0.197 | 0.095 | 2.3889 | 0.03886 | 0.115 | -4.16 | -0.1888 | none |
| AVL9 | 0.27 | 0.207 | 2.3889 | 0.03886 | 0.115 | -4.16 | -1.2464 | none |
| NEDD4L | 0.488 | 0.3999 | 2.3888 | 0.03887 | 0.115 | -4.16 | -0.892 | none |
| BHLHB9 | 0.265 | 0.171 | 2.3886 | 0.03888 | 0.116 | -4.16 | -0.4904 | none |
| SP4 | 0.423 | 0.3515 | 2.3883 | 0.0389 | 0.116 | -4.16 | -0.9461 | none |
| SLC2A10 | 0.362 | 0.2566 | 2.3871 | 0.03898 | 0.116 | -4.16 | -0.8705 | none |
| SOCS3 | 0.193 | 0.181 | 2.3868 | 0.039 | 0.116 | -4.16 | 0.66836 | none |
| PRC1 | 0.413 | 0.3879 | 2.3866 | 0.03901 | 0.116 | -4.16 | -0.7189 | none |
| AMZ2 | 0.193 | 0.1453 | 2.3865 | 0.03902 | 0.116 | -4.16 | 0.68627 | none |
| TMA16 | 0.255 | 0.1319 | 2.3865 | 0.03902 | 0.116 | -4.16 | -0.9205 | none |
| SRA1 | -0.333 | -0.2938 | -2.386 | 0.03904 | 0.116 | -4.16 | 1.43095 | none |
| ST3GAL6 | 0.444 | 0.3231 | 2.3859 | 0.03906 | 0.116 | -4.16 | -0.9747 | none |
| THOC7 | -0.285 | -0.3081 | -2.386 | 0.03908 | 0.116 | -4.16 | 1.75662 | none |
| HIST2H2BI | 0.327 | 0.3811 | 2.3845 | 0.03915 | 0.116 | -4.17 | 0.27605 | none |
| MED20 | 0.29 | 0.1727 | 2.3843 | 0.03916 | 0.116 | -4.17 | -1.0222 | none |
| PPARGC1A | 0.433 | 0.3354 | 2.3838 | 0.0392 | 0.116 | -4.17 | 1.049 | none |
| SLFNL1 | 0.275 | 0.2823 | 2.3828 | 0.03926 | 0.116 | -4.17 | -0.4336 | none |
| KCTD3 | 0.578 | 0.4068 | 2.3827 | 0.03927 | 0.116 | -4.17 | -1.0076 | none |
| SYNE1 | 0.553 | 0.5155 | 2.3827 | 0.03927 | 0.116 | -4.17 | -1.0616 | none |
| ITGAE | -0.348 | -0.345 | -2.382 | 0.03929 | 0.116 | -4.17 | 1.34284 | none |
| CNN2 | 0.283 | 0.1816 | 2.3822 | 0.0393 | 0.116 | -4.17 | -0.2593 | none |
| CR2 | -0.312 | -0.2906 | -2.382 | 0.03931 | 0.116 | -4.17 | 0.39531 | none |
| DUS4L | 0.269 | 0.2238 | 2.3814 | 0.03936 | 0.117 | -4.17 | 1.42903 | none |
| NRG2 | -0.195 | -0.1501 | -2.381 | 0.03937 | 0.117 | -4.17 | 0.20422 | none |
| LAMTOR2 | -0.189 | -0.1355 | -2.381 | 0.03938 | 0.117 | -4.17 | 0.95839 | none |
| MXD1 | 0.293 | 0.2385 | 2.3804 | 0.03942 | 0.117 | -4.17 | 0.59638 | none |
| HPS6 | -0.223 | -0.1474 | -2.38 | 0.03944 | 0.117 | -4.17 | 1.38152 | none |
| OTUD5 | 0.487 | 0.4137 | 2.3801 | 0.03944 | 0.117 | -4.17 | -1.0275 | none |
| ACVR2B | 0.209 | 0.1954 | 2.38 | 0.03945 | 0.117 | -4.17 | -0.7015 | none |
| FAM20B | 0.45 | 0.3789 | 2.379 | 0.03952 | 0.117 | -4.17 | -1.2954 | none |
| ADAM12 | 0.238 | 0.194 | 2.3787 | 0.03954 | 0.117 | -4.18 | -0.1435 | none |
| SLC39A3 | 0.311 | 0.1745 | 2.3785 | 0.03955 | 0.117 | -4.18 | -0.6363 | none |
| LYN | 0.311 | 0.2233 | 2.3784 | 0.03955 | 0.117 | -4.18 | -0.4705 | none |
| GDI1 | 0.313 | 0.2792 | 2.3778 | 0.0396 | 0.117 | -4.18 | -0.302 | none |

| | | | | | | | | |
|---|---|---|---|---|---|---|---|---|
| NCOR2 | 0.572 | 0.6329 | 2.3777 | 0.0396 | 0.117 | -4.18 | -0.7997 | none |
| ZNRF3 | 0.291 | 0.2382 | 2.3771 | 0.03964 | 0.117 | -4.18 | -1.3896 | none |
| NELL2 | 0.672 | 0.449 | 2.3769 | 0.03965 | 0.117 | -4.18 | -0.7809 | none |
| FCGBP | 0.415 | 0.3432 | 2.3768 | 0.03966 | 0.117 | -4.18 | 0.70356 | none |
| COMMD3 | -0.326 | -0.353 | -2.377 | 0.03966 | 0.117 | -4.18 | 1.47217 | none |
| PSMG4 | 0.441 | 0.2508 | 2.3759 | 0.03972 | 0.117 | -4.18 | -0.7077 | none |
| SPTAN1 | 0.752 | 0.8613 | 2.3759 | 0.03972 | 0.117 | -4.18 | -0.5289 | none |
| TSNAX | 0.249 | 0.1057 | 2.3746 | 0.03981 | 0.117 | -4.18 | -0.5348 | none |
| SH3BP1 | 0.45 | 0.4356 | 2.3745 | 0.03981 | 0.117 | -4.18 | -0.4036 | none |
| GON7 | 1.084 | 0.9419 | 2.3743 | 0.03983 | 0.118 | -4.18 | 0.55958 | none |
| SUPT4H1 | -0.408 | -0.443 | -2.374 | 0.03986 | 0.118 | -4.18 | 1.65275 | none |
| CNTNAP2 | 0.618 | 0.5297 | 2.3735 | 0.03988 | 0.118 | -4.18 | -0.5533 | none |
| H1F0 | 0.301 | 0.2802 | 2.373 | 0.03991 | 0.118 | -4.18 | 0.3095 | none |
| NRDC | 0.384 | 0.3502 | 2.3718 | 0.04 | 0.118 | -4.19 | -0.1072 | none |
| C12ORF45 | 0.34 | 0.1883 | 2.3715 | 0.04002 | 0.118 | -4.19 | -0.9753 | none |
| PEX12 | 0.22 | 0.1544 | 2.3701 | 0.04012 | 0.118 | -4.19 | -1.0796 | none |
| TMEM229 | 0.251 | 0.2036 | 2.3697 | 0.04014 | 0.118 | -4.19 | -0.4632 | none |
| FILIP1 | 0.186 | 0.1386 | 2.3695 | 0.04015 | 0.118 | -4.19 | -1.2926 | none |
| GALNT15 | 0.534 | 0.3809 | 2.3694 | 0.04016 | 0.118 | -4.19 | -1.105 | none |
| ATP1B1 | 0.272 | 0.2583 | 2.3681 | 0.04025 | 0.119 | -4.19 | -0.4202 | none |
| HEMGN | 0.286 | 0.231 | 2.3681 | 0.04025 | 0.119 | -4.19 | -1.4258 | none |
| IL18BP | 0.451 | 0.3872 | 2.3672 | 0.04031 | 0.119 | -4.19 | -0.6472 | none |
| HBQ1 | 0.355 | 0.4039 | 2.367 | 0.04033 | 0.119 | -4.19 | 1.48792 | none |
| SSBP1 | -0.311 | -0.3084 | -2.367 | 0.04033 | 0.119 | -4.19 | 1.65143 | none |
| RNF113A | -0.222 | -0.2242 | -2.367 | 0.04035 | 0.119 | -4.19 | 2.35228 | AA-related |
| TPT1 | -0.452 | -0.4656 | -2.367 | 0.04035 | 0.119 | -4.19 | 1.54772 | none |
| SSR2 | -0.376 | -0.3993 | -2.366 | 0.04038 | 0.119 | -4.2 | 2.2508 | AA-related |
| ASNA1 | -0.318 | -0.264 | -2.366 | 0.04038 | 0.119 | -4.2 | 1.15927 | none |
| INO80D | 0.28 | 0.2453 | 2.3661 | 0.04038 | 0.119 | -4.2 | -0.9885 | none |
| ST8SIA1 | 0.37 | 0.2396 | 2.3661 | 0.04038 | 0.119 | -4.2 | -0.5565 | none |
| PLEKHA1 | 0.364 | 0.2404 | 2.3651 | 0.04045 | 0.119 | -4.2 | -0.9451 | none |
| PPWD1 | 0.34 | 0.2696 | 2.3649 | 0.04046 | 0.119 | -4.2 | -0.8714 | none |
| ZNF91 | 0.673 | 0.5421 | 2.3643 | 0.04051 | 0.119 | -4.2 | -0.8529 | none |
| ZNF322 | 0.427 | 0.4406 | 2.3643 | 0.04051 | 0.119 | -4.2 | 0.27924 | none |
| TAF5L | 0.295 | 0.2565 | 2.3634 | 0.04057 | 0.119 | -4.2 | -1.0431 | none |
| WFDC2 | -0.273 | -0.2265 | -2.363 | 0.0406 | 0.119 | -4.2 | 1.09138 | none |
| MMACHC | 0.402 | 0.3277 | 2.3627 | 0.04062 | 0.119 | -4.2 | -1.079 | none |
| KRBA2 | 0.363 | 0.2248 | 2.3623 | 0.04064 | 0.119 | -4.2 | -1.2538 | none |
| LNX2 | 0.233 | 0.1476 | 2.3622 | 0.04065 | 0.119 | -4.2 | -1.2993 | none |
| ARHGAP1: | 0.315 | 0.4152 | 2.3618 | 0.04068 | 0.119 | -4.2 | 1.06275 | none |

| | | | | | | | | |
|---|---|---|---|---|---|---|---|---|
| IGF1R | 0.485 | 0.5221 | 2.3617 | 0.04069 | 0.119 | -4.2 | -0.282 | none |
| ECE1 | 0.737 | 0.7523 | 2.3612 | 0.04072 | 0.119 | -4.2 | -0.2593 | none |
| NANOGNE | 0.591 | 0.4622 | 2.3611 | 0.04072 | 0.119 | -4.2 | -0.3512 | none |
| ZCCHC2 | 0.48 | 0.4641 | 2.3611 | 0.04072 | 0.119 | -4.2 | -0.3243 | none |
| B4GALT2 | 0.186 | 0.1919 | 2.3607 | 0.04076 | 0.12 | -4.2 | -0.1663 | none |
| SLC24A2 | 0.201 | 0.144 | 2.36 | 0.0408 | 0.12 | -4.21 | -1.3723 | none |
| ZFP28 | 0.456 | 0.383 | 2.3598 | 0.04081 | 0.12 | -4.21 | 0.28081 | none |
| NR1H3 | 0.353 | 0.2053 | 2.3596 | 0.04083 | 0.12 | -4.21 | -1.0458 | none |
| PRRG2 | -0.258 | -0.2113 | -2.359 | 0.04089 | 0.12 | -4.21 | 1.40989 | none |
| CEACAM4 | 0.286 | 0.1844 | 2.3584 | 0.04091 | 0.12 | -4.21 | -1.1258 | none |
| IGSF9B | -0.289 | -0.1448 | -2.358 | 0.04091 | 0.12 | -4.21 | 0.83338 | none |
| MMS22L | 0.31 | 0.1969 | 2.3583 | 0.04092 | 0.12 | -4.21 | -1.2584 | none |
| UGT2B7 | 0.408 | 0.2877 | 2.3582 | 0.04092 | 0.12 | -4.21 | -0.6027 | none |
| HOMEZ | 0.325 | 0.2686 | 2.3581 | 0.04093 | 0.12 | -4.21 | 0.53439 | none |
| KLHDC8B | -0.333 | -0.2603 | -2.358 | 0.04094 | 0.12 | -4.21 | 0.9794 | none |
| DIABLO | 0.286 | 0.1428 | 2.3579 | 0.04094 | 0.12 | -4.21 | -0.9426 | none |
| CTSS | 0.237 | 0.1049 | 2.3571 | 0.041 | 0.12 | -4.21 | -0.3628 | none |
| ZNF644 | 0.47 | 0.496 | 2.3562 | 0.04106 | 0.12 | -4.21 | -0.2709 | none |
| UQCRQ | -0.357 | -0.3485 | -2.356 | 0.04108 | 0.12 | -4.21 | 1.48604 | none |
| PMM2 | 0.354 | 0.3346 | 2.3559 | 0.04108 | 0.12 | -4.21 | -0.3134 | none |
| SPATA5L1 | 0.273 | 0.2551 | 2.3548 | 0.04116 | 0.12 | -4.21 | 0.18222 | none |
| SND1 | 0.386 | 0.3594 | 2.3546 | 0.04117 | 0.12 | -4.21 | 0.02285 | none |
| EPC2 | 0.474 | 0.4255 | 2.3544 | 0.04118 | 0.12 | -4.21 | -0.3929 | none |
| ATP8A1 | 0.339 | 0.3453 | 2.3542 | 0.0412 | 0.12 | -4.21 | 0.32446 | none |
| MGAT4A | 0.326 | 0.1708 | 2.354 | 0.04121 | 0.12 | -4.21 | -0.1739 | none |
| TRAPPC1 | -0.186 | -0.1573 | -2.354 | 0.04122 | 0.12 | -4.22 | 2.29541 | AA-related |
| USP48 | 0.351 | 0.3453 | 2.3536 | 0.04125 | 0.12 | -4.22 | 0.29191 | none |
| MANF | -0.291 | -0.2282 | -2.353 | 0.0413 | 0.121 | -4.22 | 1.29602 | none |
| PYGB | 0.267 | 0.2785 | 2.3527 | 0.0413 | 0.121 | -4.22 | 0.13907 | none |
| RPL12 | -0.385 | -0.4073 | -2.353 | 0.04131 | 0.121 | -4.22 | 2.11179 | AA-related |
| IKBKG | 0.357 | 0.3284 | 2.3521 | 0.04135 | 0.121 | -4.22 | -0.1305 | none |
| MAP3K14 | 0.487 | 0.4532 | 2.352 | 0.04136 | 0.121 | -4.22 | -0.4804 | none |
| DGCR8 | 0.441 | 0.491 | 2.3519 | 0.04136 | 0.121 | -4.22 | -0.8876 | none |
| TWISTNB | -0.408 | -0.3291 | -2.352 | 0.04137 | 0.121 | -4.22 | 1.72728 | none |
| CD99 | -0.303 | -0.3082 | -2.352 | 0.04138 | 0.121 | -4.22 | 1.85659 | none |
| TMA7 | -0.31 | -0.2967 | -2.351 | 0.04145 | 0.121 | -4.22 | 0.95459 | none |
| PROSER1 | 0.31 | 0.3112 | 2.35 | 0.04149 | 0.121 | -4.22 | -0.7278 | none |
| NOD1 | 0.317 | 0.2581 | 2.35 | 0.04149 | 0.121 | -4.22 | 0.35846 | none |
| ZNF177 | -0.267 | -0.2358 | -2.35 | 0.04151 | 0.121 | -4.22 | 1.57403 | none |
| TAS2R5 | 0.293 | 0.2437 | 2.3494 | 0.04153 | 0.121 | -4.22 | -0.6659 | none |

| | | | | | | | | |
|---|---|---|---|---|---|---|---|---|
| METAP2 | 0.372 | 0.3194 | 2.3491 | 0.04156 | 0.121 | -4.22 | -0.314 | none |
| MIP | -0.296 | -0.2389 | -2.349 | 0.04156 | 0.121 | -4.22 | 0.59734 | none |
| YLPM1 | 0.695 | 0.7153 | 2.349 | 0.04156 | 0.121 | -4.22 | -0.662 | none |
| COX8A | -0.467 | -0.4496 | -2.349 | 0.04157 | 0.121 | -4.22 | 1.94743 | none |
| TNNT3 | 0.385 | 0.2938 | 2.3486 | 0.04159 | 0.121 | -4.22 | -0.7914 | none |
| SAP30 | -0.4 | -0.3398 | -2.348 | 0.04163 | 0.121 | -4.22 | 1.6637 | none |
| LEPROT | 0.465 | 0.2862 | 2.3478 | 0.04164 | 0.121 | -4.22 | -0.1857 | none |
| GTSF1 | -0.26 | -0.2839 | -2.348 | 0.04166 | 0.121 | -4.23 | 1.37479 | none |
| ZNF562 | 0.429 | 0.3806 | 2.3469 | 0.04171 | 0.121 | -4.23 | -1.3874 | none |
| CALML6 | -0.154 | -0.1015 | -2.347 | 0.04173 | 0.121 | -4.23 | 1.21453 | none |
| SVBP | 0.26 | 0.1197 | 2.3464 | 0.04175 | 0.121 | -4.23 | 0.88493 | none |
| MTCH2 | -0.371 | -0.4034 | -2.346 | 0.04176 | 0.121 | -4.23 | 1.68397 | none |
| FHOD1 | 0.431 | 0.4407 | 2.346 | 0.04177 | 0.121 | -4.23 | -0.5982 | none |
| STYX | 0.36 | 0.1804 | 2.346 | 0.04177 | 0.121 | -4.23 | -0.7527 | none |
| RPL35A | -0.406 | -0.4403 | -2.346 | 0.04179 | 0.121 | -4.23 | 2.07965 | none |
| ABL2 | 0.639 | 0.6149 | 2.3458 | 0.04179 | 0.121 | -4.23 | -1.3006 | none |
| RDM1 | -0.448 | -0.4056 | -2.346 | 0.04179 | 0.121 | -4.23 | 1.1848 | none |
| MRPS10 | 0.209 | 0.0833 | 2.3454 | 0.04182 | 0.122 | -4.23 | -0.381 | none |
| KCTD2 | 0.392 | 0.4145 | 2.3451 | 0.04183 | 0.122 | -4.23 | -0.7899 | none |
| SFSWAP | 0.587 | 0.5899 | 2.345 | 0.04184 | 0.122 | -4.23 | -0.9806 | none |
| LCA5L | 0.483 | 0.3872 | 2.3444 | 0.04188 | 0.122 | -4.23 | -1.2809 | none |
| SNX20 | 0.433 | 0.4347 | 2.3439 | 0.04192 | 0.122 | -4.23 | 0.48926 | none |
| TLK1 | 0.637 | 0.599 | 2.3435 | 0.04195 | 0.122 | -4.23 | -0.891 | none |
| BCAS4 | 0.387 | 0.2543 | 2.3433 | 0.04197 | 0.122 | -4.23 | -0.0182 | none |
| STEAP1B | 0.26 | 0.2167 | 2.3429 | 0.04199 | 0.122 | -4.23 | -1.209 | none |
| PRR11 | 0.901 | 0.9327 | 2.3426 | 0.04201 | 0.122 | -4.23 | -0.8555 | none |
| EIF5AL1 | -0.327 | -0.2268 | -2.342 | 0.04205 | 0.122 | -4.23 | 0.39023 | none |
| CARD6 | 0.208 | 0.1582 | 2.3417 | 0.04208 | 0.122 | -4.23 | -1.4711 | none |
| KBTBD3 | -0.547 | -0.4914 | -2.342 | 0.04209 | 0.122 | -4.23 | 1.24224 | none |
| C1ORF53 | 0.48 | 0.3653 | 2.3412 | 0.04211 | 0.122 | -4.24 | 0.87904 | none |
| ARL8A | 0.414 | 0.3454 | 2.341 | 0.04212 | 0.122 | -4.24 | 0.6251 | none |
| CEP120 | 0.432 | 0.4119 | 2.3405 | 0.04216 | 0.122 | -4.24 | -0.2564 | none |
| CSNK1G1 | 0.557 | 0.4674 | 2.3401 | 0.04219 | 0.122 | -4.24 | -1.3401 | none |
| CRYBG3 | 0.388 | 0.4575 | 2.34 | 0.0422 | 0.122 | -4.24 | -0.4716 | none |
| C17ORF50 | 0.301 | 0.2393 | 2.3397 | 0.04222 | 0.122 | -4.24 | -0.2323 | none |
| FCF1 | 0.435 | 0.3335 | 2.3395 | 0.04224 | 0.122 | -4.24 | -0.846 | none |
| PHF20L1 | 0.439 | 0.456 | 2.3391 | 0.04226 | 0.122 | -4.24 | -0.5808 | none |
| C2ORF66 | 0.193 | 0.1339 | 2.3386 | 0.0423 | 0.122 | -4.24 | -0.6341 | none |
| DHX15 | 0.193 | 0.1173 | 2.3382 | 0.04233 | 0.123 | -4.24 | 0.14207 | none |
| SRD5A1 | 0.262 | 0.1532 | 2.3378 | 0.04235 | 0.123 | -4.24 | -0.0286 | none |

| | | | | | | | | |
|---|---|---|---|---|---|---|---|---|
| KLHL20 | -0.313 | -0.2489 | -2.338 | 0.04238 | 0.123 | -4.24 | 1.47217 | none |
| KLF7 | 0.471 | 0.4239 | 2.3375 | 0.04238 | 0.123 | -4.24 | -0.4336 | none |
| PHB2 | -0.282 | -0.2355 | -2.337 | 0.04244 | 0.123 | -4.24 | 2.29572 | AA-related |
| CD69 | -0.42 | -0.4215 | -2.336 | 0.04248 | 0.123 | -4.24 | 2.16656 | AA-related |
| ERV3-1 | 0.542 | 0.5276 | 2.3357 | 0.0425 | 0.123 | -4.24 | 0.03132 | none |
| AGAP5 | 0.331 | 0.2972 | 2.3356 | 0.04251 | 0.123 | -4.24 | -0.2263 | none |
| PM20D1 | 0.501 | 0.4297 | 2.3349 | 0.04256 | 0.123 | -4.25 | -1.175 | none |
| PRPF8 | 0.84 | 0.8565 | 2.3347 | 0.04258 | 0.123 | -4.25 | -0.5407 | none |
| MRPL15 | -0.26 | -0.2647 | -2.334 | 0.04259 | 0.123 | -4.25 | 1.91069 | none |
| USP12 | -0.337 | -0.3045 | -2.334 | 0.04263 | 0.123 | -4.25 | 1.93513 | none |
| RAD18 | 0.379 | 0.352 | 2.3337 | 0.04265 | 0.123 | -4.25 | -0.0319 | none |
| SWAP70 | 0.461 | 0.4037 | 2.3336 | 0.04266 | 0.123 | -4.25 | -0.8178 | none |
| GJA9 | 0.193 | 0.1544 | 2.3331 | 0.04269 | 0.123 | -4.25 | -1.1858 | none |
| LITAF | -0.241 | -0.2681 | -2.333 | 0.04272 | 0.123 | -4.25 | 1.76514 | none |
| DNM1L | 0.396 | 0.3506 | 2.332 | 0.04277 | 0.123 | -4.25 | -0.0777 | none |
| MRPS23 | -0.309 | -0.3293 | -2.332 | 0.04279 | 0.124 | -4.25 | 0.2299 | none |
| PSD4 | 0.522 | 0.5617 | 2.3314 | 0.04281 | 0.124 | -4.25 | -0.6228 | none |
| CLOCK | 0.356 | 0.3713 | 2.331 | 0.04284 | 0.124 | -4.25 | -0.5917 | none |
| UBE2T | -0.356 | -0.3618 | -2.331 | 0.04286 | 0.124 | -4.25 | 1.51408 | none |
| ZNF2 | 0.205 | 0.1077 | 2.3306 | 0.04287 | 0.124 | -4.25 | -0.7748 | none |
| DENND2D | -0.206 | -0.1991 | -2.33 | 0.04288 | 0.124 | -4.25 | 2.15668 | AA-related |
| ADAP1 | -0.493 | -0.3771 | -2.33 | 0.0429 | 0.124 | -4.25 | 1.95833 | none |
| OCIAD2 | -0.275 | -0.3078 | -2.33 | 0.04293 | 0.124 | -4.25 | -0.1169 | none |
| PLEKHB1 | 0.307 | 0.2833 | 2.3298 | 0.04293 | 0.124 | -4.25 | 0.06298 | none |
| EPB41L2 | 0.753 | 0.8101 | 2.3298 | 0.04293 | 0.124 | -4.25 | -0.6191 | none |
| POTEC | 0.253 | 0.1792 | 2.3297 | 0.04294 | 0.124 | -4.25 | -1.234 | none |
| SCMH1 | 0.482 | 0.488 | 2.3293 | 0.04296 | 0.124 | -4.25 | -0.2847 | none |
| CETN3 | -0.554 | -0.4944 | -2.329 | 0.04299 | 0.124 | -4.26 | 2.32789 | AA-related |
| TNIP2 | 0.243 | 0.2804 | 2.3287 | 0.04301 | 0.124 | -4.26 | 0.017 | none |
| TSR3 | -0.267 | -0.2203 | -2.328 | 0.04303 | 0.124 | -4.26 | 0.96883 | none |
| UNC5CL | 0.233 | 0.2287 | 2.3283 | 0.04303 | 0.124 | -4.26 | -0.5938 | none |
| PRRC2B | 0.693 | 0.8022 | 2.3283 | 0.04304 | 0.124 | -4.26 | -0.6377 | none |
| ZNF44 | 0.34 | 0.2656 | 2.3282 | 0.04304 | 0.124 | -4.26 | -0.9563 | none |
| MYO1D | 0.464 | 0.3983 | 2.3272 | 0.04312 | 0.124 | -4.26 | -0.338 | none |
| KRT86 | -0.237 | -0.2316 | -2.326 | 0.04319 | 0.124 | -4.26 | 1.32035 | none |
| SPATA25 | -0.283 | -0.166 | -2.326 | 0.0432 | 0.124 | -4.26 | 2.23454 | AA-related |
| NPC1L1 | 0.19 | 0.1312 | 2.3257 | 0.04323 | 0.124 | -4.26 | -1.2932 | none |
| IGF2BP3 | 0.335 | 0.247 | 2.3254 | 0.04325 | 0.124 | -4.26 | -0.7125 | none |
| SPTY2D1 | 0.508 | 0.5174 | 2.3253 | 0.04325 | 0.124 | -4.26 | -1.1063 | none |
| PMVK | -0.298 | -0.2747 | -2.324 | 0.04333 | 0.125 | -4.26 | 0.92783 | none |

| | | | | | | | | |
|---|---|---|---|---|---|---|---|---|
| STAB1 | -0.491 | -0.3635 | -2.324 | 0.04335 | 0.125 | -4.26 | 0.26248 | none |
| PCDHA5 | 0.154 | 0.1097 | 2.3239 | 0.04336 | 0.125 | -4.26 | -1.1537 | none |
| LAMTOR1 | -0.323 | -0.3062 | -2.324 | 0.04336 | 0.125 | -4.26 | 1.68373 | none |
| MFSD11 | 0.346 | 0.1614 | 2.3237 | 0.04337 | 0.125 | -4.26 | 0.90491 | none |
| CCDC50 | 0.45 | 0.357 | 2.3237 | 0.04337 | 0.125 | -4.26 | 0.16949 | none |
| NGRN | 0.229 | 0.1475 | 2.3236 | 0.04337 | 0.125 | -4.26 | -0.9715 | none |
| C15ORF48 | -0.552 | -0.5719 | -2.323 | 0.04346 | 0.125 | -4.27 | 1.02141 | none |
| RPL29 | -0.283 | -0.2788 | -2.322 | 0.04348 | 0.125 | -4.27 | 1.98119 | none |
| LCN10 | -0.461 | -0.452 | -2.322 | 0.04353 | 0.125 | -4.27 | 1.28444 | none |
| RORC | 0.347 | 0.3244 | 2.3209 | 0.04358 | 0.125 | -4.27 | -1.2534 | none |
| GTF3C4 | 0.313 | 0.2634 | 2.3208 | 0.04359 | 0.125 | -4.27 | -1.2465 | none |
| ZBTB16 | 0.285 | 0.1829 | 2.3204 | 0.04361 | 0.125 | -4.27 | -1.2819 | none |
| CHAF1A | 0.718 | 0.8172 | 2.3201 | 0.04364 | 0.125 | -4.27 | -0.7039 | none |
| ITGAX | 0.534 | 0.4436 | 2.3195 | 0.04368 | 0.125 | -4.27 | -1.1215 | none |
| PCSK5 | 0.558 | 0.5229 | 2.319 | 0.04371 | 0.125 | -4.27 | -0.0315 | none |
| TMEM168 | 0.481 | 0.3507 | 2.3179 | 0.04379 | 0.126 | -4.27 | -0.9259 | none |
| RPS17 | -0.442 | -0.4609 | -2.318 | 0.04382 | 0.126 | -4.27 | 2.18977 | AA-related |
| PPM1L | 0.393 | 0.3011 | 2.3171 | 0.04386 | 0.126 | -4.27 | 0.077 | none |
| FAM19A5 | -0.277 | -0.2176 | -2.317 | 0.04387 | 0.126 | -4.27 | 1.02254 | none |
| SLFN11 | 0.531 | 0.4937 | 2.3168 | 0.04387 | 0.126 | -4.27 | -0.3159 | none |
| UBL5 | -0.357 | -0.3689 | -2.317 | 0.04389 | 0.126 | -4.27 | 1.794 | none |
| PYROXD1 | 0.404 | 0.2775 | 2.3166 | 0.04389 | 0.126 | -4.27 | -0.4137 | none |
| RRAGC | 0.237 | 0.1347 | 2.3147 | 0.04403 | 0.126 | -4.28 | -0.1869 | none |
| CD86 | 0.276 | 0.2024 | 2.3146 | 0.04404 | 0.126 | -4.28 | 0.14936 | none |
| CXXC1 | 0.402 | 0.3323 | 2.3144 | 0.04405 | 0.126 | -4.28 | -0.4959 | none |
| DDI2 | 0.654 | 0.6585 | 2.3142 | 0.04407 | 0.126 | -4.28 | -0.3958 | none |
| RPL23A | -0.359 | -0.3808 | -2.314 | 0.04409 | 0.126 | -4.28 | 2.01977 | none |
| SLC17A5 | 0.29 | 0.1309 | 2.3134 | 0.04413 | 0.126 | -4.28 | -0.8327 | none |
| OSBPL2 | 0.361 | 0.31 | 2.312 | 0.04423 | 0.127 | -4.28 | -0.741 | none |
| FREM2 | 0.217 | 0.1581 | 2.3106 | 0.04433 | 0.127 | -4.28 | -0.847 | none |
| MT-ND4L | 0.572 | 0.5939 | 2.3105 | 0.04434 | 0.127 | -4.28 | -0.4981 | none |
| B2M | -0.368 | -0.398 | -2.31 | 0.04441 | 0.127 | -4.29 | 1.66944 | none |
| STX12 | -0.359 | -0.3113 | -2.31 | 0.04442 | 0.127 | -4.29 | 1.283 | none |
| CD48 | -0.525 | -0.5547 | -2.309 | 0.04443 | 0.127 | -4.29 | 1.78837 | none |
| NRN1 | -0.278 | -0.2647 | -2.309 | 0.04444 | 0.127 | -4.29 | 1.92254 | none |
| AMBRA1 | 0.619 | 0.5639 | 2.3089 | 0.04446 | 0.127 | -4.29 | -1.2374 | none |
| TTC36 | -0.278 | -0.2313 | -2.309 | 0.04448 | 0.127 | -4.29 | 1.73331 | none |
| FSTL1 | 0.217 | 0.2306 | 2.3084 | 0.0445 | 0.127 | -4.29 | 0.90055 | none |
| DLGAP2 | 0.28 | 0.2877 | 2.308 | 0.04453 | 0.127 | -4.29 | 0.94274 | none |
| TMEM217 | 0.336 | 0.331 | 2.3076 | 0.04456 | 0.127 | -4.29 | 0.38995 | none |

| | | | | | | | | |
|---|---|---|---|---|---|---|---|---|
| NPFFR2 | 0.159 | 0.1002 | 2.3074 | 0.04458 | 0.127 | -4.29 | -1.0935 | none |
| NDUFB11 | -0.44 | -0.4258 | -2.307 | 0.04458 | 0.127 | -4.29 | 2.46859 | AA-related |
| EFNB1 | 0.294 | 0.2669 | 2.3072 | 0.04459 | 0.127 | -4.29 | 0.02049 | none |
| AC240274 | 0.364 | 0.2985 | 2.3071 | 0.0446 | 0.127 | -4.29 | -0.7017 | none |
| ACTR3B | 0.267 | 0.1873 | 2.3071 | 0.0446 | 0.127 | -4.29 | -0.6971 | none |
| FAM83E | 0.606 | 0.5796 | 2.3065 | 0.04464 | 0.127 | -4.29 | 1.01796 | none |
| LSM14A | 0.583 | 0.4743 | 2.3063 | 0.04466 | 0.127 | -4.29 | -0.2382 | none |
| PELP1 | 0.404 | 0.4156 | 2.3063 | 0.04466 | 0.127 | -4.29 | -0.6822 | none |
| NKPD1 | 0.233 | 0.1682 | 2.3058 | 0.0447 | 0.127 | -4.29 | -0.9215 | none |
| PNPLA6 | 0.45 | 0.4253 | 2.3058 | 0.0447 | 0.127 | -4.29 | -0.1424 | none |
| UPK1A | 0.459 | 0.3293 | 2.3057 | 0.0447 | 0.127 | -4.29 | -1.2767 | none |
| TSPAN31 | 0.431 | 0.1575 | 2.3051 | 0.04475 | 0.128 | -4.29 | -0.6214 | none |
| TTC7A | 0.318 | 0.3507 | 2.304 | 0.04483 | 0.128 | -4.29 | -0.8087 | none |
| RPL11 | -0.367 | -0.3925 | -2.304 | 0.04485 | 0.128 | -4.3 | 2.14877 | AA-related |
| SLC12A7 | 0.527 | 0.5443 | 2.3026 | 0.04493 | 0.128 | -4.3 | 0.05634 | none |
| ZNF404 | 0.398 | 0.3195 | 2.3024 | 0.04495 | 0.128 | -4.3 | -0.8315 | none |
| GABPB1 | 0.234 | 0.1218 | 2.3022 | 0.04497 | 0.128 | -4.3 | -0.7525 | none |
| EDF1 | -0.227 | -0.1598 | -2.302 | 0.04501 | 0.128 | -4.3 | 1.87658 | none |
| ERCC6 | 0.53 | 0.5283 | 2.301 | 0.04506 | 0.128 | -4.3 | 0.10929 | none |
| MAGOH | -0.241 | -0.2552 | -2.301 | 0.04508 | 0.128 | -4.3 | 1.27157 | none |
| RCOR1 | 0.741 | 0.6752 | 2.2998 | 0.04515 | 0.128 | -4.3 | -0.7236 | none |
| CDC123 | -0.292 | -0.2607 | -2.299 | 0.04517 | 0.129 | -4.3 | 1.06113 | none |
| IER3 | 0.979 | 0.7517 | 2.2991 | 0.0452 | 0.129 | -4.3 | -1.1868 | none |
| TMIE | 0.291 | 0.203 | 2.2988 | 0.04523 | 0.129 | -4.3 | -0.4598 | none |
| TTC21B | 0.409 | 0.3673 | 2.2987 | 0.04523 | 0.129 | -4.3 | -0.7742 | none |
| CCDC180 | 0.297 | 0.1967 | 2.2985 | 0.04525 | 0.129 | -4.3 | -1.1552 | none |
| LMAN1L | 0.391 | 0.2895 | 2.2983 | 0.04526 | 0.129 | -4.3 | 1.01191 | none |
| DCAF15 | 0.374 | 0.3988 | 2.298 | 0.04529 | 0.129 | -4.3 | 0.25564 | none |
| INTS6 | 0.576 | 0.5177 | 2.2978 | 0.0453 | 0.129 | -4.3 | -0.3122 | none |
| RHNO1 | 0.283 | 0.1582 | 2.2972 | 0.04535 | 0.129 | -4.31 | -1.0183 | none |
| TMEM136 | 0.305 | 0.1882 | 2.2969 | 0.04537 | 0.129 | -4.31 | -0.0792 | none |
| CAMK4 | 0.285 | 0.1892 | 2.2967 | 0.04538 | 0.129 | -4.31 | -0.0805 | none |
| ZNF226 | 0.512 | 0.3527 | 2.2962 | 0.04542 | 0.129 | -4.31 | -0.9128 | none |
| UQCRB | -0.318 | -0.3411 | -2.295 | 0.04548 | 0.129 | -4.31 | 1.67846 | none |
| NARS | 0.265 | 0.2396 | 2.2951 | 0.04551 | 0.129 | -4.31 | 0.38499 | none |
| BCR | 0.578 | 0.5983 | 2.295 | 0.04551 | 0.129 | -4.31 | -0.4059 | none |
| TJAP1 | 0.364 | 0.3456 | 2.295 | 0.04551 | 0.129 | -4.31 | -1.0106 | none |
| MLF2 | -0.262 | -0.1964 | -2.295 | 0.04552 | 0.129 | -4.31 | 1.41537 | none |
| SCTR | 0.311 | 0.2834 | 2.2948 | 0.04553 | 0.129 | -4.31 | -0.0461 | none |
| SKA3 | 0.311 | 0.2182 | 2.2945 | 0.04555 | 0.129 | -4.31 | -0.3534 | none |

| | | | | | | | | |
|---|---|---|---|---|---|---|---|---|
| ZGLP1 | -0.262 | -0.2052 | -2.294 | 0.04556 | 0.129 | -4.31 | 0.34635 | none |
| CLEC19A | 0.31 | 0.2528 | 2.2943 | 0.04557 | 0.129 | -4.31 | -1.003 | none |
| FBXO10 | 0.42 | 0.4788 | 2.2937 | 0.04561 | 0.129 | -4.31 | 0.40548 | none |
| GNPTAB | 0.616 | 0.6434 | 2.2932 | 0.04565 | 0.129 | -4.31 | 0.21222 | none |
| SAP30L | -0.561 | -0.3515 | -2.293 | 0.04569 | 0.129 | -4.31 | 1.34641 | none |
| ALOX5AP | -0.575 | -0.6532 | -2.293 | 0.0457 | 0.129 | -4.31 | 1.44181 | none |
| NSUN4 | 0.226 | 0.1391 | 2.2922 | 0.04573 | 0.13 | -4.31 | -0.9685 | none |
| TULP1 | 0.285 | 0.2678 | 2.291 | 0.04582 | 0.13 | -4.32 | -0.4496 | none |
| SLC4A1AP | 0.54 | 0.5405 | 2.2908 | 0.04584 | 0.13 | -4.32 | -0.5039 | none |
| GTF2H1 | 0.545 | 0.5311 | 2.2905 | 0.04586 | 0.13 | -4.32 | -0.572 | none |
| LY6G5C | 0.496 | 0.1655 | 2.2901 | 0.04589 | 0.13 | -4.32 | -0.3458 | none |
| SRRM2 | 0.576 | 0.6879 | 2.2901 | 0.04589 | 0.13 | -4.32 | -0.4187 | none |
| RARRES3 | -0.411 | -0.5046 | -2.29 | 0.04591 | 0.13 | -4.32 | 1.35517 | none |
| RASGRF1 | 0.28 | 0.1652 | 2.2898 | 0.04591 | 0.13 | -4.32 | -1.1894 | none |
| VPS33B | 0.379 | 0.248 | 2.2888 | 0.04599 | 0.13 | -4.32 | -1.207 | none |
| PAXBP1 | 0.603 | 0.5533 | 2.2881 | 0.04605 | 0.13 | -4.32 | -0.9101 | none |
| RPL27 | -0.434 | -0.444 | -2.288 | 0.04605 | 0.13 | -4.32 | 1.43587 | none |
| ZBTB33 | -0.243 | -0.1649 | -2.288 | 0.04608 | 0.13 | -4.32 | 2.06762 | none |
| GLRX | -0.316 | -0.3811 | -2.288 | 0.04609 | 0.13 | -4.32 | 1.23497 | none |
| KAT2B | 0.502 | 0.5097 | 2.2875 | 0.04609 | 0.13 | -4.32 | -0.83 | none |
| PLD3 | 0.313 | 0.3072 | 2.2864 | 0.04618 | 0.131 | -4.32 | -1.0069 | none |
| TAF1D | 0.545 | 0.4435 | 2.2862 | 0.04619 | 0.131 | -4.32 | -1.0973 | none |
| FAM135B | 0.545 | 0.3474 | 2.2861 | 0.0462 | 0.131 | -4.32 | -1.0905 | none |
| ATP6V0E1 | -0.366 | -0.4037 | -2.286 | 0.04621 | 0.131 | -4.32 | 1.74027 | none |
| SLC38A1 | 0.199 | 0.1523 | 2.2854 | 0.04626 | 0.131 | -4.32 | -0.3845 | none |
| RC3H2 | 0.369 | 0.382 | 2.2851 | 0.04628 | 0.131 | -4.32 | -0.0116 | none |
| EIF4ENIF1 | 0.482 | 0.5014 | 2.2849 | 0.04629 | 0.131 | -4.33 | -0.4929 | none |
| PPRC1 | 0.511 | 0.5791 | 2.2849 | 0.04629 | 0.131 | -4.33 | -0.1978 | none |
| ZNF286A | 0.363 | 0.3458 | 2.2848 | 0.0463 | 0.131 | -4.33 | 0.04248 | none |
| CIZ1 | 0.66 | 0.7972 | 2.2843 | 0.04634 | 0.131 | -4.33 | -0.2512 | none |
| PAMR1 | 0.16 | 0.0961 | 2.2842 | 0.04635 | 0.131 | -4.33 | -1.2401 | none |
| ROCK2 | 0.58 | 0.6035 | 2.2839 | 0.04637 | 0.131 | -4.33 | -0.8693 | none |
| LATS1 | 0.457 | 0.4615 | 2.2828 | 0.04646 | 0.131 | -4.33 | -0.94 | none |
| VDAC2 | -0.258 | -0.2449 | -2.283 | 0.04646 | 0.131 | -4.33 | 1.82438 | none |
| PPA1 | -0.269 | -0.2938 | -2.283 | 0.04646 | 0.131 | -4.33 | 1.58217 | none |
| NUTF2 | -0.337 | -0.3447 | -2.282 | 0.04648 | 0.131 | -4.33 | 1.78654 | none |
| CPSF2 | 0.385 | 0.34 | 2.2823 | 0.04649 | 0.131 | -4.33 | -1.4069 | none |
| KCTD17 | -0.532 | -0.3528 | -2.282 | 0.04652 | 0.131 | -4.33 | 1.01692 | none |
| OR4K2 | 0.226 | 0.1654 | 2.281 | 0.0466 | 0.131 | -4.33 | -0.9166 | none |
| OAZ3 | 0.353 | 0.2469 | 2.2805 | 0.04664 | 0.131 | -4.33 | -0.1078 | none |

| | | | | | | | | |
|---|---|---|---|---|---|---|---|---|
| C8ORF58 | -0.324 | -0.2423 | -2.28 | 0.04667 | 0.131 | -4.33 | 2.18293 | AA-related |
| JRK | 0.311 | 0.3472 | 2.2801 | 0.04667 | 0.131 | -4.33 | -0.1412 | none |
| SLC25A13 | 0.212 | 0.1301 | 2.2798 | 0.04669 | 0.131 | -4.33 | -0.2125 | none |
| PARVB | 0.27 | 0.232 | 2.2795 | 0.04671 | 0.132 | -4.33 | -0.5943 | none |
| TEX14 | 0.18 | 0.1701 | 2.2794 | 0.04672 | 0.132 | -4.33 | 0.74825 | none |
| LAX1 | 0.604 | 0.6055 | 2.2792 | 0.04674 | 0.132 | -4.33 | -0.9277 | none |
| RIPK2 | 0.242 | 0.2222 | 2.2791 | 0.04675 | 0.132 | -4.33 | -0.1937 | none |
| HRH1 | 0.287 | 0.2529 | 2.2786 | 0.04678 | 0.132 | -4.34 | -0.8663 | none |
| CCDC127 | 0.533 | 0.5036 | 2.2785 | 0.0468 | 0.132 | -4.34 | 0.01795 | none |
| CLEC2D | 0.338 | 0.2661 | 2.2784 | 0.0468 | 0.132 | -4.34 | -1.1251 | none |
| PPID | 0.214 | 0.1523 | 2.2783 | 0.04681 | 0.132 | -4.34 | -1.1089 | none |
| ZNF33A | 0.503 | 0.4619 | 2.2782 | 0.04682 | 0.132 | -4.34 | -1.0096 | none |
| UBE2V2 | 0.223 | 0.1697 | 2.2781 | 0.04683 | 0.132 | -4.34 | -1.1424 | none |
| CEP95 | 0.521 | 0.4637 | 2.278 | 0.04683 | 0.132 | -4.34 | -1.2236 | none |
| PSMD10 | -0.323 | -0.298 | -2.278 | 0.04683 | 0.132 | -4.34 | 1.58402 | none |
| GCNT2 | 0.706 | 0.5528 | 2.278 | 0.04683 | 0.132 | -4.34 | -1.2436 | none |
| ZNF287 | 0.38 | 0.3863 | 2.2779 | 0.04684 | 0.132 | -4.34 | 0.21342 | none |
| ATG12 | 0.269 | 0.1326 | 2.2777 | 0.04686 | 0.132 | -4.34 | -0.9952 | none |
| WDR35 | 0.291 | 0.2722 | 2.2777 | 0.04686 | 0.132 | -4.34 | -1.161 | none |
| AKR1C1 | 0.466 | 0.244 | 2.2775 | 0.04687 | 0.132 | -4.34 | -0.806 | none |
| TSGA10IP | 0.313 | 0.285 | 2.277 | 0.04691 | 0.132 | -4.34 | -0.8502 | none |
| RIF1 | 0.756 | 0.7955 | 2.2769 | 0.04691 | 0.132 | -4.34 | -0.1135 | none |
| ARHGEF1 | 0.397 | 0.3958 | 2.2766 | 0.04694 | 0.132 | -4.34 | -0.2016 | none |
| PAXIP1 | 0.434 | 0.4245 | 2.2761 | 0.04698 | 0.132 | -4.34 | -0.8554 | none |
| CIT | 0.413 | 0.4394 | 2.276 | 0.04699 | 0.132 | -4.34 | -0.6897 | none |
| FLT3LG | 0.363 | 0.2331 | 2.2756 | 0.04702 | 0.132 | -4.34 | -1.01 | none |
| CCND2 | 0.272 | 0.2558 | 2.2743 | 0.04713 | 0.132 | -4.34 | -0.1692 | none |
| CYP3A7 | -0.273 | -0.2353 | -2.274 | 0.04715 | 0.132 | -4.34 | 1.93614 | none |
| ARHGAP2: | 0.682 | 0.6866 | 2.2734 | 0.0472 | 0.132 | -4.34 | -0.9505 | none |
| SETD5 | 0.56 | 0.6327 | 2.2733 | 0.04721 | 0.132 | -4.34 | -0.0862 | none |
| ARHGEF2 | 0.421 | 0.4755 | 2.2732 | 0.04721 | 0.132 | -4.34 | -0.3122 | none |
| PRKG2 | 0.314 | 0.2347 | 2.2732 | 0.04721 | 0.132 | -4.34 | -0.2091 | none |
| TAPBP | 0.488 | 0.5102 | 2.2731 | 0.04722 | 0.132 | -4.34 | 1.35147 | none |
| GCFC2 | 0.219 | 0.1883 | 2.2724 | 0.04727 | 0.132 | -4.34 | -0.2028 | none |
| SNRNP20( | 0.543 | 0.5684 | 2.2724 | 0.04728 | 0.132 | -4.35 | -0.805 | none |
| SYBU | 0.2 | 0.1557 | 2.2722 | 0.04729 | 0.132 | -4.35 | -1.3847 | none |
| MAPK11 | -0.225 | -0.1053 | -2.272 | 0.04731 | 0.132 | -4.35 | 1.05848 | none |
| PARP9 | 0.373 | 0.3335 | 2.2715 | 0.04735 | 0.133 | -4.35 | -0.4977 | none |
| JRKL | 0.223 | 0.19 | 2.2712 | 0.04737 | 0.133 | -4.35 | 0.50541 | none |
| SNRNP48 | -0.433 | -0.3226 | -2.271 | 0.04739 | 0.133 | -4.35 | 2.05542 | none |

| | | | | | | | | |
|---|---|---|---|---|---|---|---|---|
| C2ORF81 | 0.32 | 0.2076 | 2.2708 | 0.0474 | 0.133 | -4.35 | -0.1974 | none |
| ACAD9 | 0.259 | 0.2351 | 2.2706 | 0.04742 | 0.133 | -4.35 | 0.13857 | none |
| DNAJC13 | 0.723 | 0.7164 | 2.2704 | 0.04743 | 0.133 | -4.35 | -0.871 | none |
| RALGPS1 | 0.374 | 0.2498 | 2.2704 | 0.04744 | 0.133 | -4.35 | -0.0256 | none |
| IL33 | -0.175 | -0.0372 | -2.27 | 0.04745 | 0.133 | -4.35 | 0.86454 | none |
| GCSAM | -0.179 | -0.1643 | -2.27 | 0.04745 | 0.133 | -4.35 | 0.79419 | none |
| CDK11A | 0.497 | 0.5038 | 2.2702 | 0.04745 | 0.133 | -4.35 | -0.6063 | none |
| KAT5 | 0.374 | 0.3589 | 2.27 | 0.04746 | 0.133 | -4.35 | -0.6932 | none |
| GNPAT | -0.191 | -0.1676 | -2.269 | 0.04751 | 0.133 | -4.35 | 1.74804 | none |
| EFCAB10 | 0.316 | 0.2142 | 2.2692 | 0.04753 | 0.133 | -4.35 | -0.3218 | none |
| RNF6 | 0.554 | 0.512 | 2.2691 | 0.04754 | 0.133 | -4.35 | -0.4431 | none |
| PLCG1 | 0.555 | 0.5968 | 2.2684 | 0.0476 | 0.133 | -4.35 | -0.3748 | none |
| SREBF2 | 0.47 | 0.5472 | 2.2682 | 0.04761 | 0.133 | -4.35 | 0.36208 | none |
| MVB12A | 0.356 | 0.2402 | 2.2677 | 0.04765 | 0.133 | -4.35 | -0.5977 | none |
| IRX5 | 0.238 | 0.2102 | 2.2674 | 0.04768 | 0.133 | -4.35 | 0.42933 | none |
| KCNT1 | 0.426 | 0.2735 | 2.2673 | 0.04768 | 0.133 | -4.35 | 0.05188 | none |
| LINGO4 | 0.165 | 0.1376 | 2.2667 | 0.04773 | 0.133 | -4.35 | -0.5289 | none |
| DNHD1 | 0.368 | 0.3902 | 2.2666 | 0.04774 | 0.133 | -4.35 | -0.4672 | none |
| RASA4 | 0.495 | 0.4632 | 2.2666 | 0.04774 | 0.133 | -4.35 | -1.0763 | none |
| SERPINH1 | 0.395 | 0.3744 | 2.2666 | 0.04774 | 0.133 | -4.35 | -0.7077 | none |
| RPS11 | -0.406 | -0.4204 | -2.266 | 0.04776 | 0.133 | -4.35 | 1.54988 | none |
| FAM98A | 0.357 | 0.332 | 2.2662 | 0.04777 | 0.133 | -4.35 | -0.5402 | none |
| TMEM161 | 0.501 | 0.3967 | 2.2658 | 0.0478 | 0.133 | -4.36 | -0.8238 | none |
| CCDC28A | -0.246 | -0.2739 | -2.265 | 0.04785 | 0.133 | -4.36 | 0.95363 | none |
| HADH | -0.221 | -0.2176 | -2.265 | 0.04789 | 0.133 | -4.36 | 1.62977 | none |
| SH3BGRL3 | -0.288 | -0.2728 | -2.264 | 0.04792 | 0.134 | -4.36 | 1.61905 | none |
| SLC16A10 | 0.464 | 0.2901 | 2.2642 | 0.04793 | 0.134 | -4.36 | -0.9982 | none |
| BTBD11 | 0.281 | 0.2217 | 2.2637 | 0.04797 | 0.134 | -4.36 | -1.0129 | none |
| ANKZF1 | 0.38 | 0.3512 | 2.2627 | 0.04805 | 0.134 | -4.36 | -0.9459 | none |
| IRAK3 | 0.274 | 0.2061 | 2.2624 | 0.04807 | 0.134 | -4.36 | -0.6312 | none |
| LCN12 | 0.329 | 0.1693 | 2.2624 | 0.04808 | 0.134 | -4.36 | 1.21385 | none |
| ZNF395 | 0.32 | 0.2951 | 2.2621 | 0.0481 | 0.134 | -4.36 | -0.045 | none |
| NBPF14 | 0.432 | 0.4727 | 2.2618 | 0.04812 | 0.134 | -4.36 | -0.3496 | none |
| ATOX1 | -0.275 | -0.283 | -2.261 | 0.04817 | 0.134 | -4.36 | 1.97404 | none |
| AC129492 | 0.391 | 0.2907 | 2.2611 | 0.04818 | 0.134 | -4.36 | -0.825 | none |
| GZMH | -0.311 | -0.1905 | -2.261 | 0.04821 | 0.134 | -4.36 | 1.81884 | none |
| CD68 | 0.334 | 0.2568 | 2.2604 | 0.04824 | 0.134 | -4.36 | -0.9552 | none |
| BACH2 | 0.356 | 0.3988 | 2.2603 | 0.04824 | 0.134 | -4.36 | 0.06656 | none |
| TAF3 | 0.595 | 0.666 | 2.2598 | 0.04829 | 0.134 | -4.37 | -1.0229 | none |
| C1GALT1C | -0.244 | -0.2834 | -2.259 | 0.04831 | 0.134 | -4.37 | 0.88864 | none |

| | | | | | | | | |
|---|---|---|---|---|---|---|---|---|
| FAM155A | 0.167 | 0.1211 | 2.2587 | 0.04838 | 0.134 | -4.37 | -1.3455 | none |
| FAM32A | 0.261 | 0.2069 | 2.2586 | 0.04838 | 0.134 | -4.37 | -1.3329 | none |
| IQCB1 | 0.271 | 0.1777 | 2.2585 | 0.04839 | 0.134 | -4.37 | -0.5081 | none |
| FBXL14 | -0.276 | -0.1656 | -2.258 | 0.04844 | 0.135 | -4.37 | 1.8118 | none |
| ITGB7 | -0.301 | -0.2704 | -2.258 | 0.04845 | 0.135 | -4.37 | 1.22012 | none |
| WDR33 | 0.351 | 0.2757 | 2.2577 | 0.04846 | 0.135 | -4.37 | -0.249 | none |
| DAD1 | -0.394 | -0.4063 | -2.258 | 0.04847 | 0.135 | -4.37 | 1.73605 | none |
| BCL10 | 0.256 | 0.0927 | 2.2568 | 0.04852 | 0.135 | -4.37 | -1.2258 | none |
| HACD1 | 0.318 | 0.2157 | 2.2568 | 0.04853 | 0.135 | -4.37 | 0.5597 | none |
| DNMT1 | 0.789 | 0.8312 | 2.2565 | 0.04855 | 0.135 | -4.37 | -0.7101 | none |
| GATAD2B | 0.752 | 0.7205 | 2.2562 | 0.04858 | 0.135 | -4.37 | -0.199 | none |
| CHMP3 | -0.259 | -0.2802 | -2.256 | 0.04859 | 0.135 | -4.37 | 0.59797 | none |
| ANKK1 | -0.193 | -0.1856 | -2.256 | 0.04863 | 0.135 | -4.37 | 0.06743 | none |
| DIDO1 | 0.32 | 0.3597 | 2.254 | 0.04875 | 0.135 | -4.37 | -0.0576 | none |
| SPRY4 | -0.259 | -0.2054 | -2.254 | 0.04877 | 0.135 | -4.37 | 0.86393 | none |
| FRAT1 | -0.223 | -0.1759 | -2.254 | 0.04877 | 0.135 | -4.37 | 0.83633 | none |
| GSTA4 | -0.531 | -0.4749 | -2.254 | 0.04879 | 0.135 | -4.37 | 2.02489 | none |
| TAOK3 | 0.716 | 0.6989 | 2.2533 | 0.04881 | 0.135 | -4.38 | -0.9193 | none |
| FAM209A | 0.31 | 0.2839 | 2.2527 | 0.04886 | 0.135 | -4.38 | 0.02313 | none |
| ASPHD2 | -0.194 | -0.1156 | -2.252 | 0.04891 | 0.136 | -4.38 | 0.99364 | none |
| SLC22A31 | 0.291 | 0.2555 | 2.2511 | 0.04899 | 0.136 | -4.38 | 0.84284 | none |
| INO80 | 0.546 | 0.6142 | 2.2509 | 0.04901 | 0.136 | -4.38 | -0.4259 | none |
| RPL38 | -0.437 | -0.456 | -2.251 | 0.04901 | 0.136 | -4.38 | 1.48837 | none |
| CDK12 | 0.82 | 0.888 | 2.2502 | 0.04907 | 0.136 | -4.38 | -0.866 | none |
| CTU2 | -0.257 | -0.2 | -2.25 | 0.04907 | 0.136 | -4.38 | 0.45889 | none |
| STAG1 | 0.561 | 0.465 | 2.249 | 0.04917 | 0.136 | -4.38 | -0.8895 | none |
| ZFYVE21 | 0.285 | 0.1774 | 2.2487 | 0.04919 | 0.136 | -4.38 | -0.9098 | none |
| ETAA1 | 0.416 | 0.4114 | 2.2487 | 0.04919 | 0.136 | -4.38 | -0.7946 | none |
| C11ORF21 | 0.482 | 0.4126 | 2.2485 | 0.04921 | 0.136 | -4.38 | 0.03483 | none |
| GMCL1 | 0.197 | 0.1448 | 2.2485 | 0.04921 | 0.136 | -4.38 | -0.9168 | none |
| SCARF2 | 0.235 | 0.2073 | 2.2485 | 0.04921 | 0.136 | -4.38 | -1.0789 | none |
| DLGAP1 | 0.738 | 0.5794 | 2.2481 | 0.04924 | 0.136 | -4.38 | -0.7413 | none |
| HMGB2 | -0.2 | -0.1311 | -2.247 | 0.0493 | 0.136 | -4.38 | 0.80545 | none |
| OR2M3 | 0.169 | 0.1353 | 2.2468 | 0.04934 | 0.136 | -4.39 | -1.0122 | none |
| SPATA5 | 0.363 | 0.3269 | 2.2468 | 0.04934 | 0.136 | -4.39 | -0.8848 | none |
| BPTF | 0.745 | 0.7741 | 2.2465 | 0.04937 | 0.136 | -4.39 | -0.9219 | none |
| UBE2A | 0.276 | 0.1019 | 2.2463 | 0.04939 | 0.136 | -4.39 | -0.5362 | none |
| ARAP1 | 0.459 | 0.485 | 2.2456 | 0.04945 | 0.137 | -4.39 | -0.673 | none |
| CGB5 | 0.35 | 0.3007 | 2.2456 | 0.04945 | 0.137 | -4.39 | 0.22393 | none |
| IGLL1 | -0.234 | -0.1578 | -2.245 | 0.04948 | 0.137 | -4.39 | 1.28261 | none |

| | | | | | | | | |
|---|---|---|---|---|---|---|---|---|
| RPL37 | -0.426 | -0.4313 | -2.245 | 0.04948 | 0.137 | -4.39 | 1.0285 | none |
| ARHGAP4! | 0.441 | 0.4716 | 2.245 | 0.04949 | 0.137 | -4.39 | 0.05421 | none |
| LACTB2 | -0.348 | -0.3252 | -2.244 | 0.04956 | 0.137 | -4.39 | 1.77859 | none |
| HCN1 | 0.178 | 0.1295 | 2.2439 | 0.04959 | 0.137 | -4.39 | 0.11626 | none |
| ZNF547 | 0.589 | 0.4574 | 2.2438 | 0.04959 | 0.137 | -4.39 | -0.84 | none |
| SGTA | 0.291 | 0.2625 | 2.2429 | 0.04967 | 0.137 | -4.39 | -1.1161 | none |
| MGAT3 | 0.286 | 0.1548 | 2.2427 | 0.04969 | 0.137 | -4.39 | -0.4054 | none |
| MGAT4C | 0.177 | 0.1346 | 2.2423 | 0.04972 | 0.137 | -4.39 | -0.7897 | none |
| SLC25A28 | 0.423 | 0.3953 | 2.2423 | 0.04972 | 0.137 | -4.39 | -0.147 | none |
| RTEL1 | 0.517 | 0.5174 | 2.2419 | 0.04975 | 0.137 | -4.39 | -0.768 | none |
| UBE2L3 | -0.157 | -0.1411 | -2.242 | 0.04976 | 0.137 | -4.39 | 0.50333 | none |
| SRRM1 | 0.791 | 0.826 | 2.2413 | 0.0498 | 0.137 | -4.39 | -0.7741 | none |
| SNU13 | -0.348 | -0.3362 | -2.241 | 0.04985 | 0.137 | -4.4 | 2.05858 | none |
| THAP12 | 0.213 | 0.1637 | 2.2407 | 0.04986 | 0.137 | -4.4 | 0.75159 | none |
| RBM4 | 0.342 | 0.1854 | 2.2404 | 0.04988 | 0.137 | -4.4 | -0.7694 | none |
| PPP1R11 | -0.542 | -0.5153 | -2.24 | 0.04988 | 0.137 | -4.4 | 1.48913 | none |
| HSPB6 | -0.253 | -0.2383 | -2.24 | 0.04992 | 0.137 | -4.4 | 2.05245 | none |
| SCN2B | 0.251 | 0.1928 | 2.2399 | 0.04992 | 0.137 | -4.4 | -1.1571 | none |
| BBC3 | -0.25 | -0.1486 | -2.24 | 0.04992 | 0.137 | -4.4 | 1.97594 | none |
| GARNL3 | 0.368 | 0.311 | 2.2397 | 0.04994 | 0.137 | -4.4 | 0.31431 | none |
| LSM2 | -0.558 | -0.5689 | -2.239 | 0.04999 | 0.138 | -4.4 | 2.03818 | none |
| GIT2 | 0.387 | 0.3668 | 2.2387 | 0.05002 | 0.138 | -4.4 | 0.12884 | none |
| SLC44A4 | 0.203 | 0.1377 | 2.2385 | 0.05004 | 0.138 | -4.4 | -0.9486 | none |
| ARHGEF37 | 0.669 | 0.5119 | 2.2384 | 0.05005 | 0.138 | -4.4 | -0.2718 | none |
| RB1CC1 | 0.78 | 0.739 | 2.2376 | 0.05012 | 0.138 | -4.4 | -0.4107 | none |
| TOM1L1 | 0.801 | 0.6877 | 2.2376 | 0.05012 | 0.138 | -4.4 | -1.0057 | none |
| MYO9B | 0.652 | 0.7497 | 2.2375 | 0.05012 | 0.138 | -4.4 | -0.7184 | none |
| PCMT1 | -0.28 | -0.2752 | -2.237 | 0.05016 | 0.138 | -4.4 | 1.49921 | none |
| 2-Mar | 0.32 | 0.1716 | 2.237 | 0.05016 | 0.138 | -4.4 | -0.6192 | none |
| NUAK2 | 0.399 | 0.3206 | 2.2369 | 0.05018 | 0.138 | -4.4 | -0.4956 | none |
| NIPAL2 | 0.513 | 0.4709 | 2.2359 | 0.05025 | 0.138 | -4.4 | 0.22424 | none |
| LARS | 0.476 | 0.389 | 2.2359 | 0.05026 | 0.138 | -4.4 | -1.1247 | none |
| CGRRF1 | 0.371 | 0.2171 | 2.2354 | 0.0503 | 0.138 | -4.4 | -0.017 | none |
| RBM44 | 0.159 | 0.1063 | 2.2354 | 0.0503 | 0.138 | -4.4 | -0.4434 | none |
| ZNF879 | 0.367 | 0.3233 | 2.2349 | 0.05034 | 0.138 | -4.4 | 0.06989 | none |
| WDR66 | 0.534 | 0.2973 | 2.2348 | 0.05035 | 0.138 | -4.4 | -1.0526 | none |
| JAKMIP2 | 0.396 | 0.3497 | 2.234 | 0.05042 | 0.138 | -4.41 | -1.0111 | none |
| EIF1 | -0.294 | -0.2832 | -2.233 | 0.05049 | 0.139 | -4.41 | 2.20311 | AA-related |
| EPHB1 | -0.269 | -0.1958 | -2.233 | 0.05051 | 0.139 | -4.41 | 0.53566 | none |
| CD63 | -0.485 | -0.4778 | -2.233 | 0.05053 | 0.139 | -4.41 | 1.95637 | none |

| | | | | | | | | |
|---|---|---|---|---|---|---|---|---|
| FYTTD1 | 0.259 | 0.1654 | 2.2319 | 0.05059 | 0.139 | -4.41 | -0.6457 | none |
| RADIL | 0.268 | 0.211 | 2.2311 | 0.05066 | 0.139 | -4.41 | -1.0509 | none |
| TIAM1 | 0.373 | 0.3608 | 2.2306 | 0.05071 | 0.139 | -4.41 | -0.1734 | none |
| CCL3L1 | 0.33 | 0.2535 | 2.2305 | 0.05071 | 0.139 | -4.41 | -0.483 | none |
| MRPL12 | -0.233 | -0.1805 | -2.231 | 0.05071 | 0.139 | -4.41 | 1.80947 | none |
| APOA2 | 0.335 | 0.2141 | 2.2303 | 0.05073 | 0.139 | -4.41 | -1.171 | none |
| ZMAT1 | 0.264 | 0.179 | 2.2301 | 0.05075 | 0.139 | -4.41 | -1.1123 | none |
| CHID1 | 0.198 | 0.1513 | 2.2291 | 0.05083 | 0.139 | -4.41 | 0.00898 | none |
| CBX1 | 0.425 | 0.4055 | 2.2284 | 0.0509 | 0.139 | -4.41 | -0.2704 | none |
| GDI2 | -0.197 | -0.2227 | -2.227 | 0.051 | 0.14 | -4.42 | 1.4341 | none |
| POLG | 0.472 | 0.5864 | 2.2271 | 0.05101 | 0.14 | -4.42 | 0.85772 | none |
| FMN2 | 0.21 | 0.1507 | 2.2269 | 0.05102 | 0.14 | -4.42 | -1.3561 | none |
| EIF2S1 | -0.379 | -0.2919 | -2.227 | 0.05104 | 0.14 | -4.42 | 1.72691 | none |
| ZNF827 | 0.609 | 0.5773 | 2.2259 | 0.05111 | 0.14 | -4.42 | -0.8422 | none |
| ETV1 | 0.325 | 0.2756 | 2.2251 | 0.05118 | 0.14 | -4.42 | 0.49054 | none |
| HOMER2 | 0.568 | 0.558 | 2.225 | 0.05118 | 0.14 | -4.42 | 0.04145 | none |
| FBXO9 | 0.365 | 0.2861 | 2.2245 | 0.05122 | 0.14 | -4.42 | -0.8914 | none |
| SMAD4 | 0.223 | 0.1758 | 2.2243 | 0.05124 | 0.14 | -4.42 | 0.88021 | none |
| CD70 | 0.36 | 0.2058 | 2.2243 | 0.05124 | 0.14 | -4.42 | -0.4521 | none |
| PRPF38B | 0.544 | 0.6022 | 2.2241 | 0.05126 | 0.14 | -4.42 | -0.4242 | none |
| MANEA | 0.164 | 0.0606 | 2.224 | 0.05127 | 0.14 | -4.42 | 1.10803 | none |
| GRHL1 | 0.334 | 0.2647 | 2.2238 | 0.05129 | 0.14 | -4.42 | -0.1768 | none |
| CC2D1A | 0.566 | 0.5999 | 2.2237 | 0.0513 | 0.14 | -4.42 | -1.065 | none |
| CDC20 | -0.26 | -0.2001 | -2.223 | 0.05135 | 0.14 | -4.42 | 1.85394 | none |
| ZFP37 | 0.273 | 0.1889 | 2.2231 | 0.05135 | 0.14 | -4.42 | -0.6505 | none |
| CYP24A1 | 0.424 | 0.4091 | 2.2229 | 0.05136 | 0.14 | -4.42 | 1.4577 | none |
| C11ORF88 | 0.287 | 0.1633 | 2.2229 | 0.05137 | 0.14 | -4.42 | -0.5738 | none |
| ARMC6 | 0.254 | 0.1761 | 2.2226 | 0.05139 | 0.14 | -4.42 | 0.05932 | none |
| HSPE1 | -0.303 | -0.2945 | -2.222 | 0.05142 | 0.14 | -4.42 | 2.01868 | none |
| RMND5A | 0.334 | 0.2719 | 2.2206 | 0.05156 | 0.141 | -4.43 | -0.8254 | none |
| APOL3 | 0.385 | 0.3578 | 2.2205 | 0.05157 | 0.141 | -4.43 | -0.2726 | none |
| ARHGAP17 | 0.652 | 0.6682 | 2.2199 | 0.05162 | 0.141 | -4.43 | 0.03365 | none |
| PSMB3 | -0.365 | -0.3462 | -2.22 | 0.05164 | 0.141 | -4.43 | 1.75987 | none |
| FAM169A | 0.442 | 0.3619 | 2.2195 | 0.05166 | 0.141 | -4.43 | 0.22005 | none |
| ZSCAN4 | 0.262 | 0.1914 | 2.2192 | 0.05168 | 0.141 | -4.43 | -1.4252 | none |
| DUS3L | 0.224 | 0.1812 | 2.2189 | 0.05171 | 0.141 | -4.43 | 0.10018 | none |
| RNASEK | -0.422 | -0.4236 | -2.219 | 0.05172 | 0.141 | -4.43 | 2.07841 | none |
| NEK11 | -0.213 | -0.1797 | -2.217 | 0.05186 | 0.141 | -4.43 | 1.0309 | none |
| SLC5A1 | 0.178 | 0.1383 | 2.2171 | 0.05186 | 0.141 | -4.43 | -1.0596 | none |
| FASN | 0.435 | 0.5677 | 2.217 | 0.05187 | 0.141 | -4.43 | -0.0873 | none |

| | | | | | | | | |
|---|---|---|---|---|---|---|---|---|
| SERPIND1 | 0.166 | 0.1125 | 2.2166 | 0.05191 | 0.142 | -4.43 | -0.5639 | none |
| NRK | 0.161 | 0.1057 | 2.2161 | 0.05195 | 0.142 | -4.43 | -1.3177 | none |
| SLC35F2 | 0.293 | 0.2027 | 2.216 | 0.05196 | 0.142 | -4.43 | -1.2177 | none |
| CAND1 | 0.295 | 0.2815 | 2.2156 | 0.052 | 0.142 | -4.43 | -0.358 | none |
| APPL1 | 0.222 | 0.1023 | 2.2151 | 0.05204 | 0.142 | -4.44 | 0.49672 | none |
| TMEM139 | 0.294 | 0.2818 | 2.2149 | 0.05206 | 0.142 | -4.44 | -1.2825 | none |
| ACAP2 | 0.352 | 0.3437 | 2.2146 | 0.05208 | 0.142 | -4.44 | 0.2294 | none |
| ATP13A5 | 0.446 | 0.3201 | 2.2146 | 0.05208 | 0.142 | -4.44 | -0.8482 | none |
| CDK5RAP3 | 0.233 | 0.2297 | 2.2143 | 0.05211 | 0.142 | -4.44 | -1.0816 | none |
| BCKDHA | 0.258 | 0.227 | 2.2143 | 0.05211 | 0.142 | -4.44 | 0.1799 | none |
| DNAJB1 | 0.261 | 0.1851 | 2.2133 | 0.05219 | 0.142 | -4.44 | -1.2616 | none |
| TMEM59 | -0.343 | -0.4002 | -2.213 | 0.0522 | 0.142 | -4.44 | 0.88809 | none |
| ABCA3 | 0.356 | 0.272 | 2.213 | 0.05222 | 0.142 | -4.44 | -1.0246 | none |
| DOCK4 | 0.333 | 0.2369 | 2.213 | 0.05222 | 0.142 | -4.44 | -0.6817 | none |
| ARHGEF18 | 0.661 | 0.6241 | 2.2127 | 0.05225 | 0.142 | -4.44 | -0.9187 | none |
| SIGLECL1 | -0.163 | -0.1392 | -2.212 | 0.05227 | 0.142 | -4.44 | 0.6297 | none |
| AGER | 0.199 | 0.1886 | 2.2124 | 0.05228 | 0.142 | -4.44 | 0.71271 | none |
| HNF1B | 0.261 | 0.1789 | 2.2122 | 0.05229 | 0.142 | -4.44 | -1.0813 | none |
| DYNC1H1 | 0.772 | 0.8132 | 2.212 | 0.05231 | 0.142 | -4.44 | -0.5656 | none |
| TTC14 | 0.434 | 0.4358 | 2.2106 | 0.05243 | 0.142 | -4.44 | 0.33201 | none |
| CABIN1 | 0.566 | 0.649 | 2.2105 | 0.05244 | 0.142 | -4.44 | -0.7997 | none |
| SYK | 0.234 | 0.1727 | 2.2104 | 0.05245 | 0.142 | -4.44 | -1.1411 | none |
| EHD4 | 0.408 | 0.4065 | 2.2103 | 0.05246 | 0.142 | -4.44 | -0.6461 | none |
| NPAT | 0.559 | 0.5458 | 2.2102 | 0.05247 | 0.142 | -4.44 | -0.9155 | none |
| ADAM15 | 0.2 | 0.1728 | 2.2094 | 0.05254 | 0.143 | -4.44 | 0.51709 | none |
| ZSWIM8 | 0.46 | 0.4878 | 2.2092 | 0.05256 | 0.143 | -4.44 | -1.0706 | none |
| TPRG1L | 0.305 | 0.1808 | 2.2091 | 0.05257 | 0.143 | -4.45 | -0.9866 | none |
| TRUB2 | 0.262 | 0.2196 | 2.209 | 0.05257 | 0.143 | -4.45 | 0.53988 | none |
| GOPC | 0.213 | 0.2136 | 2.2081 | 0.05265 | 0.143 | -4.45 | 0.67205 | none |
| EIF3M | -0.277 | -0.2756 | -2.207 | 0.05275 | 0.143 | -4.45 | 0.95509 | none |
| TMEM56 | 0.189 | 0.1391 | 2.2069 | 0.05276 | 0.143 | -4.45 | -1.2472 | none |
| MT1G | -0.466 | -0.2217 | -2.207 | 0.05279 | 0.143 | -4.45 | 0.73975 | none |
| SLC22A1 | 0.337 | 0.3301 | 2.2063 | 0.05281 | 0.143 | -4.45 | 0.12821 | none |
| MT2A | -0.339 | -0.3389 | -2.206 | 0.05284 | 0.143 | -4.45 | 2.12749 | AA-related |
| TP73 | 0.443 | 0.3706 | 2.2057 | 0.05286 | 0.143 | -4.45 | -1.201 | none |
| SLC7A13 | 0.288 | 0.1832 | 2.2057 | 0.05286 | 0.143 | -4.45 | -1.3652 | none |
| SLC35A3 | 0.33 | 0.2099 | 2.2057 | 0.05286 | 0.143 | -4.45 | -0.3827 | none |
| FAM163A | 0.215 | 0.1837 | 2.2056 | 0.05287 | 0.143 | -4.45 | -0.6675 | none |
| PINK1 | 0.298 | 0.272 | 2.2056 | 0.05287 | 0.143 | -4.45 | -0.4004 | none |
| CCDC28B | 0.428 | 0.3342 | 2.2049 | 0.05293 | 0.143 | -4.45 | 0.11074 | none |

| | | | | | | | | |
|---|---|---|---|---|---|---|---|---|
| PCOLCE2 | 0.413 | 0.3645 | 2.2046 | 0.05296 | 0.143 | -4.45 | 0.42784 | none |
| SYT17 | 0.26 | 0.2046 | 2.2046 | 0.05296 | 0.143 | -4.45 | -1.1525 | none |
| CFAP157 | 0.311 | 0.2769 | 2.2041 | 0.05301 | 0.144 | -4.45 | 1.28997 | none |
| FADS3 | 0.381 | 0.3462 | 2.2029 | 0.05311 | 0.144 | -4.45 | 0.30657 | none |
| PSEN2 | 0.208 | 0.2025 | 2.2028 | 0.05312 | 0.144 | -4.45 | -0.0096 | none |
| ZNF316 | 0.597 | 0.5684 | 2.2027 | 0.05313 | 0.144 | -4.46 | -1.185 | none |
| ABHD14A- | 0.509 | 0.4331 | 2.2025 | 0.05315 | 0.144 | -4.46 | -0.2559 | none |
| WDR88 | 0.333 | 0.2353 | 2.2015 | 0.05323 | 0.144 | -4.46 | -0.9126 | none |
| BRD7 | 0.635 | 0.6249 | 2.2014 | 0.05324 | 0.144 | -4.46 | -0.7473 | none |
| SFPQ | 0.407 | 0.4118 | 2.2014 | 0.05325 | 0.144 | -4.46 | -0.4573 | none |
| SMCR8 | 0.381 | 0.3817 | 2.2011 | 0.05328 | 0.144 | -4.46 | -0.7217 | none |
| GABRG3 | 0.242 | 0.144 | 2.1991 | 0.05345 | 0.145 | -4.46 | -1.3496 | none |
| S100A10 | -0.297 | -0.3147 | -2.199 | 0.05346 | 0.145 | -4.46 | 1.75423 | none |
| SLC35B4 | 0.243 | 0.1723 | 2.1986 | 0.0535 | 0.145 | -4.46 | -0.354 | none |
| IL9 | -0.359 | -0.2143 | -2.198 | 0.05351 | 0.145 | -4.46 | 1.43522 | none |
| BTAF1 | 0.422 | 0.4629 | 2.1984 | 0.05351 | 0.145 | -4.46 | -0.1286 | none |
| PARP14 | 0.691 | 0.6829 | 2.1984 | 0.05351 | 0.145 | -4.46 | -0.7621 | none |
| LYRM9 | 0.701 | 0.5435 | 2.1981 | 0.05354 | 0.145 | -4.46 | -0.9275 | none |
| URI1 | 0.479 | 0.3792 | 2.198 | 0.05355 | 0.145 | -4.46 | -0.5814 | none |
| TRMT10C | -0.213 | -0.1576 | -2.198 | 0.05358 | 0.145 | -4.46 | 2.17653 | AA-related |
| NIPSNAP3. | -0.266 | -0.3208 | -2.198 | 0.05359 | 0.145 | -4.46 | 2.103 | AA-related |
| XPA | 0.377 | 0.3056 | 2.1975 | 0.05359 | 0.145 | -4.46 | 0.37656 | none |
| RBM26 | 0.642 | 0.6761 | 2.1968 | 0.05366 | 0.145 | -4.46 | -0.6657 | none |
| PMPCA | 0.216 | 0.2155 | 2.1967 | 0.05366 | 0.145 | -4.46 | -0.8413 | none |
| RPL17-C18 | -0.244 | -0.2759 | -2.197 | 0.05367 | 0.145 | -4.46 | 0.21127 | none |
| RPS6 | -0.219 | -0.2015 | -2.195 | 0.05379 | 0.145 | -4.47 | 2.01699 | none |
| MRPS36 | -0.281 | -0.3558 | -2.195 | 0.05379 | 0.145 | -4.47 | 0.3627 | none |
| F2R | 0.312 | 0.2006 | 2.1952 | 0.0538 | 0.145 | -4.47 | 0.44046 | none |
| OPRK1 | 0.216 | 0.137 | 2.1951 | 0.05381 | 0.145 | -4.47 | -0.827 | none |
| VWA5A | -0.18 | -0.1661 | -2.195 | 0.05382 | 0.145 | -4.47 | 1.10506 | none |
| OR4K17 | 0.2 | 0.1314 | 2.1947 | 0.05384 | 0.145 | -4.47 | -1.4084 | none |
| NOS3 | 0.318 | 0.3287 | 2.1943 | 0.05388 | 0.145 | -4.47 | 0.99925 | none |
| KIR3DX1 | -0.228 | -0.165 | -2.194 | 0.05395 | 0.145 | -4.47 | 1.26451 | none |
| C14ORF11 | -0.278 | -0.2783 | -2.194 | 0.05395 | 0.145 | -4.47 | 1.44167 | none |
| ENY2 | -0.255 | -0.2461 | -2.193 | 0.054 | 0.145 | -4.47 | 1.17486 | none |
| HIGD2A | -0.265 | -0.2981 | -2.193 | 0.054 | 0.145 | -4.47 | 1.08232 | none |
| TTC13 | 0.559 | 0.5129 | 2.1926 | 0.05404 | 0.145 | -4.47 | -0.6739 | none |
| NDUFB9 | -0.429 | -0.4346 | -2.192 | 0.05405 | 0.145 | -4.47 | 1.73273 | none |
| MTFR1L | 0.198 | 0.111 | 2.1925 | 0.05405 | 0.145 | -4.47 | 0.44034 | none |
| APOL1 | 0.587 | 0.5898 | 2.1919 | 0.0541 | 0.146 | -4.47 | -0.8454 | none |

| | | | | | | | | |
|---|---|---|---|---|---|---|---|---|
| LAP3 | -0.211 | -0.1759 | -2.192 | 0.0541 | 0.146 | -4.47 | 1.76098 | none |
| BRF1 | 0.333 | 0.358 | 2.1918 | 0.05411 | 0.146 | -4.47 | -0.7668 | none |
| CCDC102B | 0.328 | 0.1874 | 2.1916 | 0.05412 | 0.146 | -4.47 | -0.8101 | none |
| ASTE1 | 0.261 | 0.1991 | 2.1915 | 0.05413 | 0.146 | -4.47 | -0.3903 | none |
| ADHFE1 | 0.608 | 0.4782 | 2.1915 | 0.05413 | 0.146 | -4.47 | -0.9519 | none |
| PRKD2 | 0.356 | 0.3865 | 2.1912 | 0.05416 | 0.146 | -4.47 | -0.6559 | none |
| OST4 | -0.383 | -0.4139 | -2.191 | 0.05421 | 0.146 | -4.47 | 1.76196 | none |
| IFT172 | 0.342 | 0.2712 | 2.1899 | 0.05428 | 0.146 | -4.48 | -0.347 | none |
| USP38 | 0.203 | 0.1432 | 2.189 | 0.05436 | 0.146 | -4.48 | 1.73283 | none |
| COPA | 0.31 | 0.2902 | 2.189 | 0.05436 | 0.146 | -4.48 | -0.7307 | none |
| PDPR | 0.341 | 0.3303 | 2.1884 | 0.05442 | 0.146 | -4.48 | -0.4892 | none |
| MRAS | 0.385 | 0.1851 | 2.1875 | 0.0545 | 0.146 | -4.48 | -1.0893 | none |
| TMEM131 | 0.627 | 0.7251 | 2.1874 | 0.05451 | 0.146 | -4.48 | 0.40533 | none |
| NCALD | 0.431 | 0.2719 | 2.1871 | 0.05453 | 0.146 | -4.48 | -1.1788 | none |
| UTY | 0.349 | 0.2863 | 2.187 | 0.05454 | 0.146 | -4.48 | -0.6147 | none |
| IDS | 0.229 | 0.0906 | 2.187 | 0.05455 | 0.146 | -4.48 | 0.15062 | none |
| MRPL54 | -0.296 | -0.2611 | -2.187 | 0.05456 | 0.146 | -4.48 | 2.18505 | AA-related |
| KCNH5 | 0.22 | 0.1742 | 2.1867 | 0.05457 | 0.146 | -4.48 | -0.1677 | none |
| TF | 0.421 | 0.2867 | 2.1866 | 0.05458 | 0.146 | -4.48 | -0.6385 | none |
| CHKA | 0.344 | 0.2744 | 2.1864 | 0.0546 | 0.146 | -4.48 | -0.1355 | none |
| ELAC2 | 0.385 | 0.4096 | 2.1861 | 0.05463 | 0.147 | -4.48 | -0.8056 | none |
| CD3D | -0.401 | -0.4448 | -2.186 | 0.05463 | 0.147 | -4.48 | 1.62654 | none |
| DRC7 | 0.204 | 0.1739 | 2.1848 | 0.05474 | 0.147 | -4.48 | -0.9566 | none |
| GTPBP6 | 0.167 | 0.1408 | 2.1846 | 0.05476 | 0.147 | -4.48 | -0.4868 | none |
| STK16 | -0.375 | -0.351 | -2.184 | 0.05483 | 0.147 | -4.48 | 1.77877 | none |
| GGA3 | 0.433 | 0.459 | 2.1833 | 0.05488 | 0.147 | -4.49 | -0.5652 | none |
| CPED1 | 0.263 | 0.2023 | 2.1833 | 0.05488 | 0.147 | -4.49 | -0.0428 | none |
| ARID4B | 0.572 | 0.632 | 2.1831 | 0.0549 | 0.147 | -4.49 | -0.2165 | none |
| UBA52 | -0.336 | -0.3459 | -2.183 | 0.05494 | 0.147 | -4.49 | 1.83709 | none |
| ELK3 | 0.496 | 0.469 | 2.1824 | 0.05496 | 0.147 | -4.49 | -0.9205 | none |
| MIER3 | -0.287 | -0.2812 | -2.182 | 0.055 | 0.147 | -4.49 | 0.76087 | none |
| C11ORF65 | 0.372 | 0.4219 | 2.1814 | 0.05505 | 0.147 | -4.49 | 0.80989 | none |
| ATL3 | 0.223 | 0.0243 | 2.181 | 0.05509 | 0.148 | -4.49 | 0.20958 | none |
| WDR91 | 0.458 | 0.3955 | 2.1807 | 0.05512 | 0.148 | -4.49 | -0.9413 | none |
| RPL13A | -0.333 | -0.3208 | -2.18 | 0.05516 | 0.148 | -4.49 | 1.65322 | none |
| PTGER4 | 0.248 | 0.2395 | 2.1795 | 0.05523 | 0.148 | -4.49 | 1.35589 | none |
| ENOSF1 | 0.363 | 0.3281 | 2.179 | 0.05527 | 0.148 | -4.49 | -0.0942 | none |
| CES4A | 0.319 | 0.2412 | 2.1787 | 0.0553 | 0.148 | -4.49 | -0.4434 | none |
| GABRA6 | 0.411 | 0.3153 | 2.1785 | 0.05532 | 0.148 | -4.49 | 0.11635 | none |
| SEC11A | -0.349 | -0.3998 | -2.178 | 0.05534 | 0.148 | -4.49 | 1.77045 | none |

| | | | | | | | | |
|---|---|---|---|---|---|---|---|---|
| NFATC1 | 0.344 | 0.3086 | 2.178 | 0.05537 | 0.148 | -4.49 | 0.14872 | none |
| TAZ | 0.22 | 0.1601 | 2.1779 | 0.05538 | 0.148 | -4.49 | 0.75776 | none |
| GCSH | 0.231 | 0.1253 | 2.1778 | 0.05539 | 0.148 | -4.49 | 0.09191 | none |
| CNTD1 | 0.341 | 0.2622 | 2.1778 | 0.05539 | 0.148 | -4.49 | -0.4316 | none |
| CCDC137 | 0.453 | 0.5297 | 2.1776 | 0.05541 | 0.148 | -4.49 | -0.3652 | none |
| RARA | 0.378 | 0.3517 | 2.1755 | 0.0556 | 0.149 | -4.5 | -0.6238 | none |
| SNX19 | 0.462 | 0.4599 | 2.1741 | 0.05573 | 0.149 | -4.5 | -0.2324 | none |
| RPUSD1 | 0.219 | 0.1966 | 2.174 | 0.05574 | 0.149 | -4.5 | 0.85241 | none |
| ALPK1 | 0.578 | 0.3913 | 2.1737 | 0.05577 | 0.149 | -4.5 | -0.9071 | none |
| MYB | 0.481 | 0.5094 | 2.1736 | 0.05577 | 0.149 | -4.5 | -0.6831 | none |
| CENPE | 0.282 | 0.1901 | 2.1735 | 0.05578 | 0.149 | -4.5 | 0.02051 | none |
| FUBP3 | 0.42 | 0.3753 | 2.1732 | 0.05582 | 0.149 | -4.5 | -0.9422 | none |
| IQGAP1 | 0.765 | 0.7932 | 2.173 | 0.05583 | 0.149 | -4.5 | -0.9274 | none |
| PAX1 | 0.378 | 0.2967 | 2.1729 | 0.05585 | 0.149 | -4.5 | 0.24585 | none |
| PTPN11 | 0.571 | 0.6096 | 2.1715 | 0.05598 | 0.149 | -4.5 | -0.4973 | none |
| GPR26 | 0.153 | 0.1064 | 2.1714 | 0.05599 | 0.149 | -4.5 | -1.2235 | none |
| AGA | -0.541 | -0.4909 | -2.171 | 0.05601 | 0.149 | -4.5 | 1.74018 | none |
| RPS10 | -0.43 | -0.4368 | -2.171 | 0.05602 | 0.149 | -4.5 | 1.48129 | none |
| PLSCR1 | -0.743 | -0.6423 | -2.171 | 0.05603 | 0.149 | -4.51 | 2.15406 | AA-related |
| LRRC43 | 0.42 | 0.3738 | 2.1707 | 0.05605 | 0.149 | -4.51 | -0.6386 | none |
| IMPDH2 | -0.222 | -0.1699 | -2.17 | 0.05611 | 0.15 | -4.51 | 2.21283 | AA-related |
| SPATA3 | 0.262 | 0.2862 | 2.1695 | 0.05616 | 0.15 | -4.51 | 0.01167 | none |
| CFL2 | 0.326 | 0.1781 | 2.1694 | 0.05617 | 0.15 | -4.51 | -0.1246 | none |
| SULT6B1 | -0.491 | -0.3835 | -2.169 | 0.0562 | 0.15 | -4.51 | 2.07141 | none |
| MPPED1 | 0.26 | 0.1955 | 2.1682 | 0.05629 | 0.15 | -4.51 | -0.334 | none |
| IMP3 | -0.327 | -0.3211 | -2.168 | 0.05631 | 0.15 | -4.51 | 2.47358 | AA-related |
| ZNF133 | 0.335 | 0.2941 | 2.1676 | 0.05634 | 0.15 | -4.51 | -1.0429 | none |
| RPL37A | -0.443 | -0.4608 | -2.168 | 0.05635 | 0.15 | -4.51 | 1.44832 | none |
| ERVK3-1 | -0.178 | -0.1495 | -2.167 | 0.05635 | 0.15 | -4.51 | 0.7819 | none |
| APBA1 | 0.238 | 0.1955 | 2.1674 | 0.05636 | 0.15 | -4.51 | -1.295 | none |
| DCUN1D4 | 0.424 | 0.2931 | 2.1673 | 0.05637 | 0.15 | -4.51 | -0.0605 | none |
| SEC22B | 0.201 | 0.0835 | 2.1671 | 0.05639 | 0.15 | -4.51 | -0.9293 | none |
| EGR3 | 0.252 | 0.2823 | 2.1669 | 0.05641 | 0.15 | -4.51 | -0.5817 | none |
| RACK1 | -0.36 | -0.3708 | -2.167 | 0.05642 | 0.15 | -4.51 | 1.93437 | none |
| HGS | 0.468 | 0.5322 | 2.1662 | 0.05647 | 0.15 | -4.51 | -0.3573 | none |
| HIST1H4G | 0.467 | 0.467 | 2.1661 | 0.05648 | 0.15 | -4.51 | 1.10751 | none |
| DCAF4 | 0.364 | 0.2995 | 2.1657 | 0.05652 | 0.15 | -4.51 | -0.8578 | none |
| UGT2A1 | 0.323 | 0.2539 | 2.165 | 0.05659 | 0.15 | -4.51 | -1.0008 | none |
| DPF3 | 0.278 | 0.3119 | 2.1646 | 0.05662 | 0.151 | -4.51 | -0.1838 | none |
| CREBBP | 0.542 | 0.6435 | 2.1644 | 0.05664 | 0.151 | -4.52 | -0.398 | none |

| | | | | | | | | |
|---|---|---|---|---|---|---|---|---|
| SCNM1 | 0.242 | 0.1466 | 2.1643 | 0.05665 | 0.151 | -4.52 | -0.8243 | none |
| GP1BA | 0.262 | 0.1725 | 2.1641 | 0.05667 | 0.151 | -4.52 | -0.8313 | none |
| KCNK12 | -0.161 | -0.1335 | -2.164 | 0.05671 | 0.151 | -4.52 | 1.84419 | none |
| SNX30 | 0.334 | 0.3007 | 2.1633 | 0.05674 | 0.151 | -4.52 | -0.8904 | none |
| TRAF4 | -0.362 | -0.19 | -2.163 | 0.05676 | 0.151 | -4.52 | 1.57006 | none |
| RPS6KA2 | 0.225 | 0.2359 | 2.1622 | 0.05685 | 0.151 | -4.52 | 0.77052 | none |
| KRT7 | -0.544 | -0.3943 | -2.162 | 0.05687 | 0.151 | -4.52 | ######## | none |
| INPP5D | 0.487 | 0.4919 | 2.1619 | 0.05688 | 0.151 | -4.52 | -0.5628 | none |
| ZNF583 | 0.515 | 0.3415 | 2.1617 | 0.05689 | 0.151 | -4.52 | -0.836 | none |
| SPX | 0.171 | 0.1585 | 2.1613 | 0.05693 | 0.151 | -4.52 | 0.89985 | none |
| CYTH1 | 0.328 | 0.3447 | 2.1611 | 0.05695 | 0.151 | -4.52 | -0.3347 | none |
| CDC45 | -0.252 | -0.2375 | -2.16 | 0.05707 | 0.151 | -4.52 | 2.26225 | AA-related |
| SOX11 | 0.169 | 0.1216 | 2.1598 | 0.05707 | 0.151 | -4.52 | -1.106 | none |
| C4ORF33 | 0.456 | 0.2627 | 2.1598 | 0.05708 | 0.151 | -4.52 | -0.446 | none |
| EEF1B2 | -0.361 | -0.39 | -2.16 | 0.05709 | 0.151 | -4.52 | 1.48566 | none |
| C6ORF136 | -0.945 | -0.8351 | -2.159 | 0.05715 | 0.152 | -4.52 | 1.0346 | none |
| RPTOR | 0.608 | 0.6475 | 2.1589 | 0.05717 | 0.152 | -4.52 | -1.1596 | none |
| ZMYM5 | 0.393 | 0.2626 | 2.1574 | 0.05731 | 0.152 | -4.53 | -0.5027 | none |
| FBL | -0.228 | -0.198 | -2.157 | 0.05738 | 0.152 | -4.53 | 2.22328 | AA-related |
| ELMO2 | 0.38 | 0.3588 | 2.1565 | 0.0574 | 0.152 | -4.53 | -0.7548 | none |
| RPL27A | -0.464 | -0.5001 | -2.156 | 0.05741 | 0.152 | -4.53 | 1.29722 | none |
| GPATCH2 | 0.289 | 0.2245 | 2.1563 | 0.05741 | 0.152 | -4.53 | -0.3082 | none |
| FAM3D | 0.196 | 0.1956 | 2.1559 | 0.05745 | 0.152 | -4.53 | -0.5356 | none |
| LPAR3 | 0.236 | 0.1194 | 2.1556 | 0.05748 | 0.152 | -4.53 | -0.6698 | none |
| ATP2B3 | -0.193 | -0.147 | -2.155 | 0.05751 | 0.152 | -4.53 | 0.55353 | none |
| NRSN1 | 0.318 | 0.1993 | 2.155 | 0.05753 | 0.152 | -4.53 | -0.8653 | none |
| ROMO1 | -0.444 | -0.4258 | -2.155 | 0.05756 | 0.152 | -4.53 | 1.80376 | none |
| SKAP2 | 0.352 | 0.3669 | 2.1545 | 0.05758 | 0.152 | -4.53 | -0.035 | none |
| NR1H2 | 0.236 | 0.176 | 2.1542 | 0.05761 | 0.152 | -4.53 | -1.4014 | none |
| GSTM1 | 0.229 | 0.1346 | 2.1541 | 0.05763 | 0.152 | -4.53 | -0.2825 | none |
| FCHSD1 | 0.421 | 0.4408 | 2.1541 | 0.05763 | 0.152 | -4.53 | -0.7679 | none |
| TEX35 | 0.44 | 0.3884 | 2.1531 | 0.05772 | 0.153 | -4.53 | -0.8091 | none |
| ANKRD20/ | 0.188 | 0.1571 | 2.1529 | 0.05774 | 0.153 | -4.53 | -0.8586 | none |
| KCTD13 | 0.284 | 0.2599 | 2.1528 | 0.05774 | 0.153 | -4.53 | -0.7384 | none |
| RNF166 | 0.251 | 0.2234 | 2.1526 | 0.05776 | 0.153 | -4.53 | 1.07208 | none |
| PEF1 | -0.265 | -0.2665 | -2.152 | 0.05779 | 0.153 | -4.53 | 2.07731 | none |
| RNF125 | 0.507 | 0.4378 | 2.1523 | 0.0578 | 0.153 | -4.53 | -1.0735 | none |
| SNRPD2 | -0.226 | -0.198 | -2.152 | 0.05783 | 0.153 | -4.53 | 0.82923 | none |
| SNED1 | 0.456 | 0.3631 | 2.1518 | 0.05784 | 0.153 | -4.53 | -0.0948 | none |
| FOLR3 | 0.357 | 0.2832 | 2.1517 | 0.05785 | 0.153 | -4.53 | -0.5748 | none |

| | | | | | | | | |
|---|---|---|---|---|---|---|---|---|
| LTK | -0.198 | -0.1539 | -2.152 | 0.05785 | 0.153 | -4.53 | 0.185 | none |
| TIAL1 | 0.236 | 0.1104 | 2.1512 | 0.0579 | 0.153 | -4.54 | -0.8694 | none |
| SLC7A14 | 0.642 | 0.4423 | 2.151 | 0.05792 | 0.153 | -4.54 | -0.85 | none |
| PML | 0.399 | 0.4015 | 2.1507 | 0.05795 | 0.153 | -4.54 | -0.2156 | none |
| DENND4A | 0.647 | 0.5884 | 2.1506 | 0.05796 | 0.153 | -4.54 | -0.6308 | none |
| C17ORF53 | 0.315 | 0.3337 | 2.1505 | 0.05797 | 0.153 | -4.54 | 0.52435 | none |
| HCST | -0.306 | -0.3214 | -2.15 | 0.05797 | 0.153 | -4.54 | 1.71992 | none |
| TM9SF3 | 0.339 | 0.2463 | 2.1504 | 0.05797 | 0.153 | -4.54 | 1.0456 | none |
| LNPEP | 0.276 | 0.1984 | 2.1497 | 0.05805 | 0.153 | -4.54 | 0.65061 | none |
| SFT2D2 | 0.336 | 0.1976 | 2.1495 | 0.05807 | 0.153 | -4.54 | -0.6356 | none |
| VPS29 | -0.288 | -0.3213 | -2.149 | 0.05809 | 0.153 | -4.54 | 0.73014 | none |
| BLOC1S4 | -0.342 | -0.2251 | -2.149 | 0.05813 | 0.153 | -4.54 | 0.91389 | none |
| CBR1 | -0.244 | -0.2247 | -2.149 | 0.05814 | 0.153 | -4.54 | 2.16765 | AA-related |
| DISP3 | 0.235 | 0.2005 | 2.1484 | 0.05817 | 0.153 | -4.54 | -1.1804 | none |
| SRI | -0.326 | -0.3369 | -2.148 | 0.05818 | 0.153 | -4.54 | 1.46925 | none |
| COMMD6 | -0.452 | -0.4547 | -2.148 | 0.0582 | 0.153 | -4.54 | 2.13066 | AA-related |
| ZFYVE28 | 0.311 | 0.2333 | 2.148 | 0.05821 | 0.153 | -4.54 | -0.2824 | none |
| NCF2 | 0.521 | 0.4411 | 2.1478 | 0.05823 | 0.153 | -4.54 | -0.8528 | none |
| SLC27A2 | -0.273 | -0.2291 | -2.148 | 0.05823 | 0.153 | -4.54 | 0.81369 | none |
| KIAA2026 | -0.331 | -0.1869 | -2.147 | 0.05828 | 0.153 | -4.54 | 1.37131 | none |
| ADAMTS6 | 0.196 | 0.1743 | 2.1473 | 0.05828 | 0.153 | -4.54 | -0.4077 | none |
| BFSP1 | 0.195 | 0.1793 | 2.1472 | 0.05829 | 0.153 | -4.54 | -0.4682 | none |
| PTPN18 | 0.545 | 0.5542 | 2.1467 | 0.05834 | 0.154 | -4.54 | -0.4995 | none |
| PPP2R3C | 0.211 | 0.1419 | 2.1466 | 0.05835 | 0.154 | -4.54 | 0.49237 | none |
| HYAL4 | 0.504 | 0.3323 | 2.1465 | 0.05836 | 0.154 | -4.54 | -0.6496 | none |
| ZBTB37 | 0.286 | 0.2248 | 2.1462 | 0.05839 | 0.154 | -4.54 | -0.0605 | none |
| POLR2D | 0.184 | 0.135 | 2.1451 | 0.0585 | 0.154 | -4.55 | -1.3576 | none |
| CALCB | -0.249 | -0.2674 | -2.145 | 0.05852 | 0.154 | -4.55 | 0.07782 | none |
| ALMS1 | 0.443 | 0.4904 | 2.1436 | 0.05864 | 0.154 | -4.55 | -0.2328 | none |
| MORN2 | -0.513 | -0.5876 | -2.144 | 0.05864 | 0.154 | -4.55 | 0.9337 | none |
| SPIN4 | 0.213 | 0.1517 | 2.1435 | 0.05865 | 0.154 | -4.55 | -0.6256 | none |
| TMEM38B | -0.321 | -0.3341 | -2.143 | 0.05867 | 0.154 | -4.55 | 0.68823 | none |
| MAP4 | 0.692 | 0.7765 | 2.1432 | 0.05868 | 0.154 | -4.55 | -0.4265 | none |
| ZNF841 | 0.547 | 0.4367 | 2.143 | 0.0587 | 0.154 | -4.55 | 0.10635 | none |
| RPS6KA4 | 0.233 | 0.2346 | 2.1428 | 0.05871 | 0.154 | -4.55 | -0.9364 | none |
| ARL5A | 0.311 | 0.194 | 2.1427 | 0.05873 | 0.154 | -4.55 | 0.17974 | none |
| LMX1B | 0.246 | 0.1612 | 2.1424 | 0.05875 | 0.154 | -4.55 | -1.2058 | none |
| S100A5 | -0.385 | -0.2699 | -2.142 | 0.05878 | 0.154 | -4.55 | 0.74118 | none |
| OCEL1 | 0.351 | 0.1537 | 2.1419 | 0.0588 | 0.154 | -4.55 | -0.6597 | none |
| LDAH | 0.325 | 0.2651 | 2.1417 | 0.05883 | 0.154 | -4.55 | -0.8479 | none |

| | | | | | | | | |
|---|---|---|---|---|---|---|---|---|
| UBA1 | 0.287 | 0.2807 | 2.1416 | 0.05884 | 0.154 | -4.55 | -0.8034 | none |
| PCSK4 | -0.295 | -0.2525 | -2.141 | 0.05893 | 0.155 | -4.55 | 0.77695 | none |
| RAB11B | 0.295 | 0.2119 | 2.1405 | 0.05895 | 0.155 | -4.55 | -0.2469 | none |
| ADD1 | 0.366 | 0.3675 | 2.1401 | 0.05898 | 0.155 | -4.55 | -0.9735 | none |
| ARGLU1 | 0.306 | 0.2779 | 2.1398 | 0.05901 | 0.155 | -4.55 | 0.35732 | none |
| MUC13 | 0.263 | 0.2698 | 2.1391 | 0.05909 | 0.155 | -4.55 | -0.6999 | none |
| SAT1 | -0.193 | -0.1995 | -2.139 | 0.05909 | 0.155 | -4.55 | 0.11322 | none |
| HUS1B | -0.165 | -0.1136 | -2.139 | 0.0591 | 0.155 | -4.55 | 1.2031 | none |
| LHFPL4 | 0.335 | 0.332 | 2.1385 | 0.05915 | 0.155 | -4.56 | 0.56722 | none |
| PAK1 | 0.365 | 0.3655 | 2.1381 | 0.05918 | 0.155 | -4.56 | -0.0982 | none |
| ZNF444 | 0.443 | 0.3542 | 2.1375 | 0.05924 | 0.155 | -4.56 | -0.902 | none |
| PDGFA | 0.416 | 0.4263 | 2.1367 | 0.05932 | 0.155 | -4.56 | -0.1762 | none |
| GNB1 | 0.264 | 0.2365 | 2.1366 | 0.05933 | 0.155 | -4.56 | -0.3322 | none |
| CRLS1 | 0.3 | 0.2349 | 2.1364 | 0.05935 | 0.155 | -4.56 | -0.1194 | none |
| ALS2CL | 0.291 | 0.2547 | 2.1364 | 0.05935 | 0.155 | -4.56 | 0.08944 | none |
| PDPN | 0.244 | 0.1419 | 2.1362 | 0.05936 | 0.155 | -4.56 | -0.4731 | none |
| ZNF165 | 0.422 | 0.3928 | 2.1358 | 0.05941 | 0.156 | -4.56 | -0.6932 | none |
| LPGAT1 | 0.213 | 0.1499 | 2.1354 | 0.05945 | 0.156 | -4.56 | 0.27467 | none |
| ZKSCAN2 | 0.324 | 0.2902 | 2.1352 | 0.05947 | 0.156 | -4.56 | -1.0345 | none |
| GNA12 | -0.198 | -0.0273 | -2.135 | 0.05951 | 0.156 | -4.56 | 0.79085 | none |
| SHBG | -0.645 | -0.5008 | -2.134 | 0.05955 | 0.156 | -4.56 | 0.71545 | none |
| RDH10 | 0.279 | 0.1334 | 2.1343 | 0.05956 | 0.156 | -4.56 | -1.0721 | none |
| UTP11 | 0.196 | 0.1699 | 2.1339 | 0.0596 | 0.156 | -4.56 | -0.7117 | none |
| HIKESHI | -0.502 | -0.5219 | -2.134 | 0.05964 | 0.156 | -4.56 | 1.59739 | none |
| GOLIM4 | 0.334 | 0.3989 | 2.1334 | 0.05965 | 0.156 | -4.56 | -0.0568 | none |
| ST3GAL1 | 0.306 | 0.3107 | 2.1333 | 0.05966 | 0.156 | -4.56 | 0.53799 | none |
| ACAP1 | 0.24 | 0.1741 | 2.1327 | 0.05972 | 0.156 | -4.56 | -0.6402 | none |
| FAAP100 | 0.283 | 0.2777 | 2.1324 | 0.05974 | 0.156 | -4.57 | 0.21087 | none |
| GYPC | -0.317 | -0.2879 | -2.132 | 0.05978 | 0.156 | -4.57 | 1.03672 | none |
| TSSK3 | 0.294 | 0.239 | 2.1317 | 0.05982 | 0.156 | -4.57 | -0.4438 | none |
| CLINT1 | 0.384 | 0.3876 | 2.1315 | 0.05984 | 0.156 | -4.57 | -0.2494 | none |
| RAB8B | -0.293 | -0.3167 | -2.131 | 0.0599 | 0.156 | -4.57 | 0.6769 | none |
| APIP | -0.368 | -0.3743 | -2.131 | 0.05991 | 0.156 | -4.57 | 1.22795 | none |
| BAZ2A | 0.575 | 0.5885 | 2.1307 | 0.05991 | 0.156 | -4.57 | -0.4376 | none |
| MYO9A | 0.662 | 0.5958 | 2.1305 | 0.05993 | 0.156 | -4.57 | 0.03445 | none |
| SLC46A1 | 0.67 | 0.5445 | 2.1299 | 0.05999 | 0.157 | -4.57 | -1.1189 | none |
| CEACAM8 | 0.332 | 0.218 | 2.1299 | 0.06 | 0.157 | -4.57 | -0.9278 | none |
| ADAR | 0.729 | 0.7471 | 2.1297 | 0.06001 | 0.157 | -4.57 | -0.7898 | none |
| IMPG2 | 0.232 | 0.1726 | 2.1294 | 0.06004 | 0.157 | -4.57 | -1.1861 | none |
| GPR160 | 0.268 | 0.1696 | 2.1289 | 0.06009 | 0.157 | -4.57 | -0.5951 | none |

| | | | | | | | | |
|---|---|---|---|---|---|---|---|---|
| TRAP1 | 0.225 | 0.2008 | 2.1288 | 0.06011 | 0.157 | -4.57 | -0.2798 | none |
| CKLF | -0.377 | -0.3952 | -2.129 | 0.06012 | 0.157 | -4.57 | 0.95216 | none |
| NMRK2 | 0.191 | 0.191 | 2.1284 | 0.06014 | 0.157 | -4.57 | 1.38882 | none |
| PPM1B | 0.3 | 0.2195 | 2.1284 | 0.06015 | 0.157 | -4.57 | -0.1212 | none |
| GFM2 | 0.36 | 0.2536 | 2.1282 | 0.06016 | 0.157 | -4.57 | 0.56695 | none |
| MORN5 | 0.355 | 0.3947 | 2.1279 | 0.06019 | 0.157 | -4.57 | 0.12302 | none |
| ERBIN | 0.6 | 0.6531 | 2.1278 | 0.06021 | 0.157 | -4.57 | 0.1944 | none |
| MBD1 | 0.349 | 0.3778 | 2.1277 | 0.06022 | 0.157 | -4.57 | 0.25639 | none |
| TMEM129 | 0.211 | 0.236 | 2.1276 | 0.06023 | 0.157 | -4.57 | -1.0288 | none |
| HLA-DOB | 0.217 | 0.202 | 2.1275 | 0.06023 | 0.157 | -4.57 | 0.3002 | none |
| GOT1 | -0.197 | -0.1864 | -2.127 | 0.06025 | 0.157 | -4.57 | 1.28349 | none |
| MDH1 | -0.212 | -0.2067 | -2.127 | 0.06029 | 0.157 | -4.57 | 0.45269 | none |
| SLC9A9 | -0.385 | -0.5018 | -2.126 | 0.06036 | 0.157 | -4.57 | 0.6018 | none |
| TGFBR3 | 0.338 | 0.293 | 2.1262 | 0.06036 | 0.157 | -4.57 | 1.22098 | none |
| QSOX2 | 0.396 | 0.3643 | 2.1261 | 0.06037 | 0.157 | -4.57 | -1.1345 | none |
| NCSTN | 0.22 | 0.1444 | 2.126 | 0.06038 | 0.157 | -4.57 | -0.7779 | none |
| SEL1L | 0.419 | 0.3743 | 2.1258 | 0.0604 | 0.157 | -4.58 | -0.4991 | none |
| CPNE8 | 0.404 | 0.3131 | 2.1257 | 0.06041 | 0.157 | -4.58 | -1.168 | none |
| RPL7 | -0.24 | -0.2363 | -2.126 | 0.06043 | 0.157 | -4.58 | 1.77237 | none |
| PXMP4 | 0.36 | 0.2004 | 2.1255 | 0.06044 | 0.157 | -4.58 | -0.2192 | none |
| MSLN | -0.222 | -0.1381 | -2.125 | 0.06049 | 0.157 | -4.58 | 0.70739 | none |
| LRRC42 | -0.189 | -0.0902 | -2.125 | 0.06052 | 0.157 | -4.58 | 2.47487 | AA-related |
| F11R | 0.381 | 0.2379 | 2.1244 | 0.06055 | 0.157 | -4.58 | -0.8956 | none |
| NDUFC2 | -0.42 | -0.402 | -2.124 | 0.06055 | 0.157 | -4.58 | 1.72285 | none |
| SLC13A2 | -0.189 | -0.132 | -2.124 | 0.06057 | 0.157 | -4.58 | 0.7205 | none |
| SCML4 | 0.302 | 0.2435 | 2.1242 | 0.06057 | 0.157 | -4.58 | 0.04249 | none |
| BRMS1L | -0.247 | -0.2527 | -2.124 | 0.06061 | 0.157 | -4.58 | 0.7189 | none |
| ZNF286B | 0.503 | 0.491 | 2.1233 | 0.06066 | 0.158 | -4.58 | -0.8805 | none |
| TMEM87A | 0.201 | 0.0706 | 2.123 | 0.06069 | 0.158 | -4.58 | 0.30893 | none |
| OPTC | 0.182 | 0.1824 | 2.1227 | 0.06072 | 0.158 | -4.58 | 0.87864 | none |
| TRIM33 | 0.524 | 0.5041 | 2.1224 | 0.06075 | 0.158 | -4.58 | -0.2667 | none |
| RPL10A | -0.299 | -0.2918 | -2.122 | 0.06077 | 0.158 | -4.58 | 2.04521 | none |
| FBXO5 | 0.198 | 0.1753 | 2.122 | 0.06079 | 0.158 | -4.58 | -0.1322 | none |
| PANK1 | 0.412 | 0.2961 | 2.1217 | 0.06082 | 0.158 | -4.58 | 0.0406 | none |
| PFDN4 | -0.246 | -0.3225 | -2.122 | 0.06082 | 0.158 | -4.58 | 0.92347 | none |
| SMARCD2 | -0.215 | -0.105 | -2.121 | 0.06093 | 0.158 | -4.58 | 1.52237 | none |
| MFSD2B | 0.211 | 0.1664 | 2.1204 | 0.06095 | 0.158 | -4.58 | -1.0926 | none |
| ARHGEF40 | 0.221 | 0.1822 | 2.1203 | 0.06096 | 0.158 | -4.58 | -1.1152 | none |
| UCK2 | -0.241 | -0.1048 | -2.12 | 0.06098 | 0.158 | -4.58 | 0.82849 | none |
| E2F6 | 0.26 | 0.1831 | 2.1199 | 0.061 | 0.158 | -4.58 | 0.9591 | none |

| | | | | | | | | |
|---|---|---|---|---|---|---|---|---|
| MYBL1 | 0.266 | 0.2275 | 2.1199 | 0.061 | 0.158 | -4.58 | 0.34426 | none |
| ZNF213 | 0.318 | 0.323 | 2.1195 | 0.06105 | 0.158 | -4.59 | 0.82633 | none |
| NHLRC2 | 0.263 | 0.1862 | 2.1186 | 0.06113 | 0.158 | -4.59 | -0.6539 | none |
| SMIM2 | 0.358 | 0.2897 | 2.1186 | 0.06114 | 0.158 | -4.59 | 0.02598 | none |
| AFF1 | 0.556 | 0.5699 | 2.1182 | 0.06117 | 0.158 | -4.59 | -0.5864 | none |
| RPS8 | -0.281 | -0.2832 | -2.118 | 0.06118 | 0.158 | -4.59 | 1.74349 | none |
| CPEB1 | 0.273 | 0.2346 | 2.1181 | 0.06118 | 0.158 | -4.59 | -0.8127 | none |
| SERPINI2 | 0.34 | 0.2471 | 2.1181 | 0.06119 | 0.158 | -4.59 | -0.5524 | none |
| EBP | -0.285 | -0.281 | -2.118 | 0.06123 | 0.159 | -4.59 | 1.03087 | none |
| PMEPA1 | 0.312 | 0.2776 | 2.1175 | 0.06124 | 0.159 | -4.59 | 0.60112 | none |
| CMKLR1 | 0.314 | 0.2772 | 2.1174 | 0.06126 | 0.159 | -4.59 | 0.94506 | none |
| DAW1 | 0.308 | 0.1924 | 2.1174 | 0.06126 | 0.159 | -4.59 | -0.0219 | none |
| GMEB2 | 0.271 | 0.2654 | 2.1173 | 0.06126 | 0.159 | -4.59 | -0.5073 | none |
| GFOD2 | 0.318 | 0.2132 | 2.1159 | 0.06141 | 0.159 | -4.59 | -1.168 | none |
| NRN1L | 0.238 | 0.2492 | 2.1156 | 0.06144 | 0.159 | -4.59 | -0.0366 | none |
| FLOT1 | -0.478 | -0.2959 | -2.115 | 0.06155 | 0.159 | -4.59 | 1.93451 | none |
| DSC2 | 0.268 | 0.1919 | 2.1143 | 0.06158 | 0.159 | -4.59 | -1.0206 | none |
| KIF15 | 0.439 | 0.5193 | 2.1138 | 0.06162 | 0.159 | -4.59 | -0.1485 | none |
| CALN1 | 0.214 | 0.1968 | 2.1138 | 0.06162 | 0.159 | -4.59 | 0.4143 | none |
| RBM41 | 0.471 | 0.4064 | 2.1136 | 0.06165 | 0.159 | -4.59 | -0.4086 | none |
| TMEM127 | 0.311 | 0.2236 | 2.1132 | 0.06168 | 0.159 | -4.59 | -0.4001 | none |
| CKB | -0.56 | -0.5385 | -2.113 | 0.06169 | 0.159 | -4.6 | 0.15246 | none |
| SEH1L | 0.295 | 0.1556 | 2.113 | 0.0617 | 0.159 | -4.6 | -0.3188 | none |
| EIF2AK1 | 0.37 | 0.3246 | 2.1118 | 0.06183 | 0.16 | -4.6 | 0.22443 | none |
| RFX1 | 0.628 | 0.6047 | 2.1114 | 0.06187 | 0.16 | -4.6 | -1.0916 | none |
| ERCC5 | 0.62 | 0.6322 | 2.1114 | 0.06187 | 0.16 | -4.6 | -0.6544 | none |
| HYDIN | 0.565 | 0.4576 | 2.1107 | 0.06194 | 0.16 | -4.6 | -1.0902 | none |
| ARL17A | 0.308 | 0.3099 | 2.1105 | 0.06196 | 0.16 | -4.6 | -0.7095 | none |
| SVEP1 | 0.162 | 0.1196 | 2.1099 | 0.06202 | 0.16 | -4.6 | -1.3851 | none |
| DHFR2 | 0.376 | 0.2846 | 2.1096 | 0.06205 | 0.16 | -4.6 | -1.1603 | none |
| MSX1 | 0.163 | 0.1827 | 2.1094 | 0.06207 | 0.16 | -4.6 | 0.75046 | none |
| SH3BP2 | 0.286 | 0.3429 | 2.1088 | 0.06214 | 0.16 | -4.6 | -0.0501 | none |
| HAGH | 0.357 | 0.2348 | 2.1083 | 0.06219 | 0.16 | -4.6 | -0.7654 | none |
| PCM1 | 0.799 | 0.8494 | 2.1082 | 0.0622 | 0.16 | -4.6 | -0.5681 | none |
| PURB | 0.372 | 0.3411 | 2.1078 | 0.06224 | 0.16 | -4.6 | -0.9425 | none |
| EEA1 | 0.438 | 0.4896 | 2.1077 | 0.06225 | 0.161 | -4.6 | -0.7671 | none |
| NEDD8 | -0.232 | -0.2262 | -2.107 | 0.06229 | 0.161 | -4.6 | 0.94845 | none |
| NPIPA8 | 0.455 | 0.3903 | 2.1071 | 0.06232 | 0.161 | -4.6 | -0.9807 | none |
| XCL2 | -0.338 | -0.2626 | -2.107 | 0.06234 | 0.161 | -4.6 | 0.49524 | none |
| TMEM163 | -0.149 | -0.121 | -2.107 | 0.06236 | 0.161 | -4.6 | 0.05229 | none |

| | | | | | | | | |
|---|---|---|---|---|---|---|---|---|
| SORCS2 | -0.338 | -0.2488 | -2.107 | 0.06237 | 0.161 | -4.61 | 1.24618 | none |
| OMA1 | 0.227 | 0.0675 | 2.1066 | 0.06237 | 0.161 | -4.61 | 0.22648 | none |
| RPL26 | -0.313 | -0.3479 | -2.107 | 0.06237 | 0.161 | -4.61 | 1.44013 | none |
| LRCOL1 | 0.316 | 0.2347 | 2.1065 | 0.06238 | 0.161 | -4.61 | 0.35996 | none |
| CLTA | -0.26 | -0.2386 | -2.106 | 0.0624 | 0.161 | -4.61 | 2.10395 | AA-related |
| GLO1 | -0.239 | -0.2746 | -2.106 | 0.0624 | 0.161 | -4.61 | 1.2974 | none |
| CREBZF | 0.216 | 0.2269 | 2.1062 | 0.06241 | 0.161 | -4.61 | 0.03819 | none |
| IQGAP3 | 0.224 | 0.213 | 2.1059 | 0.06244 | 0.161 | -4.61 | -1.0067 | none |
| ATP6V1A | 0.183 | 0.0943 | 2.1058 | 0.06244 | 0.161 | -4.61 | -0.5644 | none |
| TUBB3 | 0.321 | 0.2359 | 2.1057 | 0.06245 | 0.161 | -4.61 | -0.7659 | none |
| ERLIN1 | -0.222 | -0.1578 | -2.106 | 0.06247 | 0.161 | -4.61 | 2.34133 | AA-related |
| S100A6 | -0.422 | -0.4318 | -2.105 | 0.06249 | 0.161 | -4.61 | 1.80985 | none |
| SYCE2 | 0.493 | 0.2785 | 2.1054 | 0.06249 | 0.161 | -4.61 | 0.0988 | none |
| RPL39 | -0.543 | -0.6106 | -2.104 | 0.06266 | 0.161 | -4.61 | 1.97363 | none |
| DNAJB13 | 0.457 | 0.3581 | 2.1035 | 0.06269 | 0.161 | -4.61 | -1.3004 | none |
| DPYD | 0.25 | 0.1943 | 2.1028 | 0.06276 | 0.161 | -4.61 | 0.19732 | none |
| IGSF22 | 0.218 | 0.1379 | 2.1027 | 0.06277 | 0.161 | -4.61 | 0.3916 | none |
| ARL6IP1 | -0.318 | -0.3579 | -2.102 | 0.06283 | 0.161 | -4.61 | 1.9152 | none |
| AGRN | 0.351 | 0.2839 | 2.102 | 0.06284 | 0.161 | -4.61 | -0.8785 | none |
| GPR3 | 0.275 | 0.2027 | 2.1017 | 0.06287 | 0.162 | -4.61 | -0.3484 | none |
| THOC2 | 0.621 | 0.6926 | 2.1017 | 0.06288 | 0.162 | -4.61 | -0.2841 | none |
| ZNF174 | 0.241 | 0.1859 | 2.1014 | 0.06291 | 0.162 | -4.61 | -1.0946 | none |
| PAAF1 | 0.271 | 0.1171 | 2.101 | 0.06294 | 0.162 | -4.61 | -0.6523 | none |
| DAZAP1 | 0.299 | 0.3162 | 2.101 | 0.06295 | 0.162 | -4.61 | -0.9127 | none |
| NOTCH1 | 0.484 | 0.5438 | 2.1007 | 0.06298 | 0.162 | -4.61 | -0.7581 | none |
| TUB | 0.213 | 0.2049 | 2.1005 | 0.063 | 0.162 | -4.61 | -0.1497 | none |
| CD80 | 0.383 | 0.217 | 2.0996 | 0.0631 | 0.162 | -4.62 | 0.23039 | none |
| CTNND2 | -0.256 | -0.2053 | -2.099 | 0.06311 | 0.162 | -4.62 | 2.22677 | AA-related |
| TMBIM4 | -0.327 | -0.3772 | -2.099 | 0.06314 | 0.162 | -4.62 | 1.12894 | none |
| MAL2 | -0.254 | -0.2632 | -2.099 | 0.06315 | 0.162 | -4.62 | 1.59001 | none |
| GATC | 0.352 | 0.3104 | 2.099 | 0.06315 | 0.162 | -4.62 | -1.2109 | none |
| CAPN3 | 0.362 | 0.3144 | 2.099 | 0.06316 | 0.162 | -4.62 | 0.50215 | none |
| IRF2BP2 | 0.334 | 0.2701 | 2.0984 | 0.06322 | 0.162 | -4.62 | -1.0636 | none |
| ATP2C1 | 0.242 | 0.1645 | 2.0983 | 0.06324 | 0.162 | -4.62 | 0.56782 | none |
| AL354822. | 0.24 | 0.2111 | 2.0981 | 0.06326 | 0.162 | -4.62 | -1.4135 | none |
| NLE1 | 0.233 | 0.2407 | 2.097 | 0.06337 | 0.162 | -4.62 | 0.68911 | none |
| GEN1 | 0.485 | 0.5526 | 2.0964 | 0.06343 | 0.163 | -4.62 | 0.64337 | none |
| MBLAC2 | 0.234 | 0.2026 | 2.0964 | 0.06343 | 0.163 | -4.62 | 0.36678 | none |
| C1ORF54 | 0.474 | 0.3855 | 2.0961 | 0.06346 | 0.163 | -4.62 | -0.558 | none |
| CCDC114 | 0.312 | 0.2817 | 2.0959 | 0.06348 | 0.163 | -4.62 | -0.7244 | none |

| | | | | | | | | |
|---|---|---|---|---|---|---|---|---|
| FBXL5 | -0.297 | -0.3067 | -2.096 | 0.06348 | 0.163 | -4.62 | 1.33421 | none |
| S100A4 | -0.642 | -0.7168 | -2.096 | 0.0635 | 0.163 | -4.62 | 1.6017 | none |
| SLC41A2 | 0.259 | 0.1335 | 2.0953 | 0.06355 | 0.163 | -4.62 | -0.4411 | none |
| SHCBP1 | 0.254 | 0.1962 | 2.0949 | 0.06359 | 0.163 | -4.62 | -0.4068 | none |
| NDUFV2 | -0.287 | -0.2926 | -2.095 | 0.0636 | 0.163 | -4.62 | 0.47728 | none |
| P4HA2 | 0.632 | 0.5536 | 2.0944 | 0.06364 | 0.163 | -4.62 | -0.9712 | none |
| CYP1B1 | -0.307 | -0.2418 | -2.094 | 0.06367 | 0.163 | -4.62 | 0.13729 | none |
| CDC16 | 0.26 | 0.2095 | 2.0939 | 0.0637 | 0.163 | -4.62 | 0.15703 | none |
| KCNA3 | 0.271 | 0.2293 | 2.0935 | 0.06374 | 0.163 | -4.63 | 0.00855 | none |
| HNRNPC | -0.176 | -0.1434 | -2.093 | 0.06374 | 0.163 | -4.63 | 1.17381 | none |
| RPLP0 | -0.333 | -0.3277 | -2.093 | 0.06375 | 0.163 | -4.63 | 1.07481 | none |
| RHOJ | 0.225 | 0.1604 | 2.0929 | 0.0638 | 0.163 | -4.63 | -0.9907 | none |
| RDH12 | 0.33 | 0.2542 | 2.0919 | 0.06391 | 0.163 | -4.63 | -1.1136 | none |
| MED23 | 0.397 | 0.3181 | 2.0917 | 0.06393 | 0.163 | -4.63 | -0.8213 | none |
| KMT2B | 0.449 | 0.5554 | 2.0911 | 0.064 | 0.164 | -4.63 | 0.06542 | none |
| DNMT3A | 0.306 | 0.3194 | 2.0911 | 0.064 | 0.164 | -4.63 | -0.0301 | none |
| BCAN | 0.522 | 0.4571 | 2.0909 | 0.06401 | 0.164 | -4.63 | 0.71275 | none |
| GPR4 | 0.235 | 0.1534 | 2.0904 | 0.06406 | 0.164 | -4.63 | -1.4157 | none |
| PRRG1 | 0.236 | 0.1824 | 2.0904 | 0.06407 | 0.164 | -4.63 | -0.5597 | none |
| MCM8 | 0.438 | 0.4406 | 2.0901 | 0.0641 | 0.164 | -4.63 | -1.0777 | none |
| EXOC3L1 | -0.172 | -0.1683 | -2.09 | 0.0641 | 0.164 | -4.63 | -0.2289 | none |
| OSBPL3 | 0.357 | 0.3278 | 2.0894 | 0.06417 | 0.164 | -4.63 | -0.8218 | none |
| RBPJL | -0.189 | -0.1245 | -2.089 | 0.06418 | 0.164 | -4.63 | 0.25699 | none |
| SEMA4G | 0.239 | 0.2506 | 2.0892 | 0.06419 | 0.164 | -4.63 | 1.29656 | none |
| AGMO | 0.497 | 0.3185 | 2.089 | 0.06422 | 0.164 | -4.63 | -1.1602 | none |
| COX5A | -0.296 | -0.2925 | -2.088 | 0.06428 | 0.164 | -4.63 | 1.77543 | none |
| DONSON | 0.393 | 0.3159 | 2.0879 | 0.06433 | 0.164 | -4.63 | 0.34288 | none |
| KRTCAP2 | -0.472 | -0.4931 | -2.088 | 0.06436 | 0.164 | -4.63 | 1.5658 | none |
| ASIC4 | 0.151 | 0.1207 | 2.0876 | 0.06436 | 0.164 | -4.63 | -1.1571 | none |
| MRPL33 | -0.32 | -0.3306 | -2.088 | 0.06437 | 0.164 | -4.63 | 1.21277 | none |
| ZNF491 | 0.28 | 0.1888 | 2.0875 | 0.06438 | 0.164 | -4.63 | 0.07178 | none |
| TMIGD3 | -0.258 | -0.2526 | -2.087 | 0.06442 | 0.164 | -4.64 | 0.54856 | none |
| FHOD3 | 0.284 | 0.2096 | 2.0865 | 0.06448 | 0.164 | -4.64 | -1.014 | none |
| NDN | -0.225 | -0.1769 | -2.086 | 0.06449 | 0.164 | -4.64 | 1.48309 | none |
| HEATR3 | 0.455 | 0.4594 | 2.0864 | 0.0645 | 0.164 | -4.64 | 0.12676 | none |
| MAK | 0.254 | 0.2416 | 2.0858 | 0.06456 | 0.164 | -4.64 | -0.6478 | none |
| FKBP1A | -0.292 | -0.2949 | -2.086 | 0.06458 | 0.164 | -4.64 | 1.94065 | none |
| PDS5A | 0.455 | 0.4471 | 2.0851 | 0.06463 | 0.165 | -4.64 | -0.8863 | none |
| DHH | 0.241 | 0.1679 | 2.0849 | 0.06465 | 0.165 | -4.64 | -1.2552 | none |
| ADAMTSL: | 0.268 | 0.189 | 2.0849 | 0.06465 | 0.165 | -4.64 | -1.4607 | none |

| | | | | | | | | |
|---|---|---|---|---|---|---|---|---|
| CAB39L | 0.294 | 0.1647 | 2.0849 | 0.06465 | 0.165 | -4.64 | -0.1794 | none |
| C1ORF194 | 0.218 | 0.1793 | 2.0847 | 0.06467 | 0.165 | -4.64 | 0.82531 | none |
| CEP152 | 0.685 | 0.5625 | 2.0846 | 0.06469 | 0.165 | -4.64 | -0.9776 | none |
| INCENP | 0.498 | 0.6415 | 2.0844 | 0.06471 | 0.165 | -4.64 | -0.1942 | none |
| PKN2 | 0.447 | 0.4223 | 2.0843 | 0.06472 | 0.165 | -4.64 | -0.8098 | none |
| FOXO1 | 0.396 | 0.3379 | 2.0841 | 0.06474 | 0.165 | -4.64 | -0.3727 | none |
| RPS20 | -0.456 | -0.4816 | -2.084 | 0.06479 | 0.165 | -4.64 | 1.06939 | none |
| MSRB3 | 0.581 | 0.3108 | 2.0833 | 0.06483 | 0.165 | -4.64 | -1.0118 | none |
| CHMP2B | -0.289 | -0.251 | -2.083 | 0.06489 | 0.165 | -4.64 | 1.37117 | none |
| SRSF10 | 0.173 | 0.0871 | 2.0825 | 0.06491 | 0.165 | -4.64 | -0.0836 | none |
| CDHR2 | 0.254 | 0.2214 | 2.0823 | 0.06493 | 0.165 | -4.64 | 0.3994 | none |
| RAP1A | -0.283 | -0.3049 | -2.082 | 0.06493 | 0.165 | -4.64 | 1.38843 | none |
| GALNT11 | 0.217 | 0.1713 | 2.0822 | 0.06494 | 0.165 | -4.64 | 0.6371 | none |
| ZIK1 | 0.212 | 0.1334 | 2.0816 | 0.06501 | 0.165 | -4.64 | 0.54842 | none |
| PPP1R36 | 0.261 | 0.1327 | 2.0815 | 0.06502 | 0.165 | -4.64 | -0.5348 | none |
| PSMA3 | -0.289 | -0.3288 | -2.081 | 0.06512 | 0.165 | -4.65 | 1.03407 | none |
| TMEM206 | -0.457 | -0.3571 | -2.08 | 0.06513 | 0.165 | -4.65 | 2.07118 | none |
| TCHP | 0.543 | 0.5523 | 2.0804 | 0.06514 | 0.165 | -4.65 | -0.8535 | none |
| NLRP11 | 0.276 | 0.2064 | 2.0804 | 0.06514 | 0.165 | -4.65 | -1.2603 | none |
| CREB5 | 0.493 | 0.3602 | 2.0799 | 0.06519 | 0.165 | -4.65 | -1.1756 | none |
| ZBTB38 | 0.631 | 0.6312 | 2.0798 | 0.06521 | 0.165 | -4.65 | 0.11609 | none |
| ING4 | -0.356 | -0.3375 | -2.08 | 0.06522 | 0.165 | -4.65 | 0.18386 | none |
| METTL3 | 0.229 | 0.1675 | 2.0794 | 0.06524 | 0.165 | -4.65 | -0.6322 | none |
| TMEM50A | -0.337 | -0.3808 | -2.079 | 0.06524 | 0.165 | -4.65 | 1.06339 | none |
| NDUFA12 | -0.251 | -0.2602 | -2.079 | 0.06526 | 0.165 | -4.65 | 0.15961 | none |
| GCH1 | -0.313 | -0.3127 | -2.079 | 0.06526 | 0.165 | -4.65 | 1.7481 | none |
| TCEA1 | -0.191 | -0.1797 | -2.079 | 0.06528 | 0.166 | -4.65 | 1.7279 | none |
| ERVMER34 | 0.344 | 0.2537 | 2.0789 | 0.0653 | 0.166 | -4.65 | 0.71737 | none |
| CLC | -0.29 | -0.3936 | -2.079 | 0.06531 | 0.166 | -4.65 | 0.17406 | none |
| HECTD3 | 0.311 | 0.3145 | 2.0785 | 0.06534 | 0.166 | -4.65 | 0.31412 | none |
| POLR2J | -0.3 | -0.2676 | -2.079 | 0.06534 | 0.166 | -4.65 | 2.20959 | AA-related |
| IFIH1 | 0.558 | 0.5877 | 2.0783 | 0.06536 | 0.166 | -4.65 | -0.0593 | none |
| POLR1E | 0.298 | 0.3157 | 2.0769 | 0.06552 | 0.166 | -4.65 | -0.3654 | none |
| ZNF92 | 0.397 | 0.3501 | 2.0765 | 0.06556 | 0.166 | -4.65 | -0.0747 | none |
| SAR1B | 0.269 | 0.0802 | 2.0763 | 0.06558 | 0.166 | -4.65 | -0.6236 | none |
| TRAF2 | 0.254 | 0.2407 | 2.0763 | 0.06558 | 0.166 | -4.65 | -0.9203 | none |
| INPP4A | 0.383 | 0.3495 | 2.0761 | 0.0656 | 0.166 | -4.65 | -0.423 | none |
| CXORF21 | 0.182 | 0.1125 | 2.0755 | 0.06567 | 0.166 | -4.65 | 0.14826 | none |
| ERAS | 0.161 | 0.1661 | 2.0754 | 0.06568 | 0.166 | -4.65 | 0.05986 | none |
| RNF111 | 0.667 | 0.5871 | 2.0749 | 0.06573 | 0.166 | -4.65 | -0.6494 | none |

| | | | | | | | | |
|---|---|---|---|---|---|---|---|---|
| SNTG2 | 0.487 | 0.3627 | 2.0747 | 0.06576 | 0.166 | -4.65 | -1.186 | none |
| PELI1 | 0.3 | 0.174 | 2.0746 | 0.06577 | 0.166 | -4.65 | -0.1336 | none |
| UBE2Q1 | -0.309 | -0.1658 | -2.074 | 0.06588 | 0.167 | -4.66 | 1.53623 | none |
| F7 | 0.305 | 0.3024 | 2.0735 | 0.06589 | 0.167 | -4.66 | -0.2873 | none |
| MAST2 | 0.397 | 0.4272 | 2.073 | 0.06594 | 0.167 | -4.66 | -0.2375 | none |
| KLF4 | 0.316 | 0.2227 | 2.0729 | 0.06596 | 0.167 | -4.66 | -0.8173 | none |
| CDC40 | 0.703 | 0.5402 | 2.0719 | 0.06606 | 0.167 | -4.66 | -0.8025 | none |
| PATE2 | 0.351 | 0.2451 | 2.0719 | 0.06607 | 0.167 | -4.66 | -1.3521 | none |
| FAM102B | 0.222 | 0.2298 | 2.0717 | 0.06609 | 0.167 | -4.66 | 0.21674 | none |
| GDNF | 0.181 | 0.1292 | 2.0713 | 0.06613 | 0.167 | -4.66 | -1.0683 | none |
| SSR4 | -0.449 | -0.4552 | -2.071 | 0.06613 | 0.167 | -4.66 | 1.4453 | none |
| STAU1 | 0.408 | 0.4292 | 2.0712 | 0.06614 | 0.167 | -4.66 | -0.9117 | none |
| SSX5 | -0.459 | -0.3761 | -2.071 | 0.06616 | 0.167 | -4.66 | 1.67207 | none |
| CCL27 | 0.225 | 0.221 | 2.0707 | 0.06619 | 0.167 | -4.66 | 0.10719 | none |
| SCAF11 | 0.563 | 0.6212 | 2.0706 | 0.06621 | 0.167 | -4.66 | -0.5828 | none |
| PCF11 | 0.416 | 0.419 | 2.0701 | 0.06626 | 0.167 | -4.66 | -0.3504 | none |
| KLHL21 | 0.238 | 0.2237 | 2.0697 | 0.06631 | 0.167 | -4.66 | -0.3737 | none |
| XIRP1 | 0.567 | 0.7489 | 2.0694 | 0.06634 | 0.167 | -4.66 | -0.4769 | none |
| GDPD2 | 0.219 | 0.1729 | 2.0688 | 0.0664 | 0.168 | -4.66 | 0.26649 | none |
| DOHH | 0.267 | 0.2676 | 2.068 | 0.06649 | 0.168 | -4.66 | 0.20489 | none |
| IL3RA | -0.208 | -0.2335 | -2.068 | 0.06652 | 0.168 | -4.67 | 1.27494 | none |
| PCGF1 | 0.264 | 0.1878 | 2.0677 | 0.06653 | 0.168 | -4.67 | -0.5447 | none |
| MTR | 0.638 | 0.6441 | 2.0676 | 0.06653 | 0.168 | -4.67 | -0.4813 | none |
| SPEN | 0.594 | 0.7684 | 2.0676 | 0.06654 | 0.168 | -4.67 | -0.4606 | none |
| ERVV-2 | 0.249 | 0.1888 | 2.067 | 0.0666 | 0.168 | -4.67 | -0.8816 | none |
| FAM89B | -0.451 | -0.3482 | -2.067 | 0.06661 | 0.168 | -4.67 | 1.99656 | none |
| CYSLTR2 | -0.334 | -0.2595 | -2.067 | 0.06664 | 0.168 | -4.67 | 0.23564 | none |
| TMEM11 | -0.207 | -0.1813 | -2.067 | 0.06665 | 0.168 | -4.67 | 1.54342 | none |
| HLA-DPA1 | 0.26 | 0.1452 | 2.0665 | 0.06666 | 0.168 | -4.67 | -0.6282 | none |
| RPL35 | -0.312 | -0.2942 | -2.066 | 0.06672 | 0.168 | -4.67 | 0.75998 | none |
| BIRC3 | 0.365 | 0.3689 | 2.0658 | 0.06674 | 0.168 | -4.67 | -0.9341 | none |
| RNF216 | 0.429 | 0.4759 | 2.0655 | 0.06677 | 0.168 | -4.67 | -0.7071 | none |
| OGFOD2 | 0.291 | 0.2507 | 2.0652 | 0.0668 | 0.168 | -4.67 | -1.0305 | none |
| PTRH1 | 0.322 | 0.1959 | 2.0651 | 0.06681 | 0.168 | -4.67 | -0.9683 | none |
| MAGEA4 | -0.184 | -0.1632 | -2.065 | 0.06682 | 0.168 | -4.67 | 1.69427 | none |
| RPL28 | -0.414 | -0.4412 | -2.065 | 0.06686 | 0.168 | -4.67 | 2.08357 | none |
| SP110 | 0.68 | 0.707 | 2.0641 | 0.06693 | 0.168 | -4.67 | -0.274 | none |
| IK | 0.626 | 0.6112 | 2.064 | 0.06694 | 0.168 | -4.67 | -0.5784 | none |
| MYO18A | 0.506 | 0.557 | 2.0636 | 0.06698 | 0.168 | -4.67 | -0.3198 | none |
| N6AMT1 | -0.286 | -0.3303 | -2.063 | 0.06699 | 0.169 | -4.67 | 1.54196 | none |

| | | | | | | | | |
|---|---|---|---|---|---|---|---|---|
| IST1 | 0.183 | 0.1028 | 2.0634 | 0.067 | 0.169 | -4.67 | -0.7255 | none |
| TUBB2A | 0.314 | 0.2013 | 2.063 | 0.06705 | 0.169 | -4.67 | -0.0762 | none |
| AK4 | -0.309 | -0.2215 | -2.063 | 0.06708 | 0.169 | -4.67 | 1.70516 | none |
| SVIL | 0.741 | 0.7229 | 2.0626 | 0.06709 | 0.169 | -4.67 | 0.12413 | none |
| TMEM258 | -0.41 | -0.4069 | -2.062 | 0.06714 | 0.169 | -4.67 | 1.41791 | none |
| PODXL | 0.313 | 0.2751 | 2.0619 | 0.06717 | 0.169 | -4.67 | 1.3517 | none |
| PEBP1 | -0.189 | -0.1612 | -2.061 | 0.06722 | 0.169 | -4.67 | 2.03455 | none |
| WNT10B | -0.209 | -0.1896 | -2.06 | 0.06733 | 0.169 | -4.68 | 1.82699 | none |
| UNC13D | 0.33 | 0.3355 | 2.0601 | 0.06737 | 0.169 | -4.68 | -0.15 | none |
| FAM126A | -0.213 | -0.2369 | -2.06 | 0.06741 | 0.169 | -4.68 | 1.3547 | none |
| CRYBB3 | 0.26 | 0.2712 | 2.0591 | 0.06748 | 0.169 | -4.68 | 0.65427 | none |
| TXNDC17 | -0.354 | -0.3411 | -2.059 | 0.0675 | 0.169 | -4.68 | 1.56954 | none |
| TAF4 | -0.548 | -0.3963 | -2.058 | 0.06756 | 0.17 | -4.68 | 0.78324 | none |
| RBMX | 0.216 | 0.179 | 2.0579 | 0.06762 | 0.17 | -4.68 | 0.83223 | none |
| HKDC1 | 0.184 | 0.1887 | 2.0571 | 0.0677 | 0.17 | -4.68 | -0.4808 | none |
| IFT140 | 0.527 | 0.4543 | 2.0571 | 0.0677 | 0.17 | -4.68 | -0.3164 | none |
| SERF2 | -0.287 | -0.2628 | -2.056 | 0.06777 | 0.17 | -4.68 | 0.77846 | none |
| RBP1 | 0.475 | 0.3505 | 2.0563 | 0.06779 | 0.17 | -4.68 | -0.1908 | none |
| DUOX2 | -0.224 | -0.1662 | -2.055 | 0.06791 | 0.17 | -4.68 | 0.50322 | none |
| ADAT3 | 0.203 | 0.1159 | 2.0552 | 0.06792 | 0.17 | -4.68 | 0.28232 | none |
| ANKRD33I | 0.275 | 0.3429 | 2.0552 | 0.06792 | 0.17 | -4.68 | 0.05776 | none |
| NCR3 | -1.015 | -0.8317 | -2.055 | 0.06794 | 0.17 | -4.68 | 2.39725 | AA-related |
| FH | -0.151 | -0.1433 | -2.055 | 0.06796 | 0.17 | -4.68 | 2.12034 | AA-related |
| MDN1 | 0.54 | 0.5918 | 2.0538 | 0.06807 | 0.171 | -4.69 | 0.01914 | none |
| KLHL35 | 0.313 | 0.1558 | 2.0538 | 0.06807 | 0.171 | -4.69 | -0.7702 | none |
| ASB14 | -0.135 | -0.0572 | -2.054 | 0.06809 | 0.171 | -4.69 | 1.29654 | none |
| HSD17B10 | -0.232 | -0.2219 | -2.054 | 0.06809 | 0.171 | -4.69 | 2.03025 | none |
| UBASH3A | 0.234 | 0.1955 | 2.0533 | 0.06813 | 0.171 | -4.69 | -0.176 | none |
| SLC6A11 | 0.204 | 0.1649 | 2.0532 | 0.06814 | 0.171 | -4.69 | 1.14947 | none |
| TUBG2 | 0.19 | 0.2362 | 2.0531 | 0.06816 | 0.171 | -4.69 | 0.50005 | none |
| AC023491 | 0.247 | 0.1585 | 2.0528 | 0.06819 | 0.171 | -4.69 | -0.3892 | none |
| SAP130 | 0.572 | 0.5895 | 2.0523 | 0.06825 | 0.171 | -4.69 | -0.2619 | none |
| LZTS1 | 0.193 | 0.1707 | 2.0516 | 0.06833 | 0.171 | -4.69 | -0.5991 | none |
| TMTC3 | 0.27 | 0.2679 | 2.051 | 0.06839 | 0.171 | -4.69 | -0.747 | none |
| TNFRSF10 | 0.211 | 0.1707 | 2.051 | 0.0684 | 0.171 | -4.69 | 1.26964 | none |
| RAG1 | -0.276 | -0.2323 | -2.051 | 0.06845 | 0.171 | -4.69 | -0.3741 | none |
| HEPH | -0.15 | -0.1406 | -2.05 | 0.0685 | 0.171 | -4.69 | 1.52926 | none |
| ATP6V0E2 | -0.157 | -0.1724 | -2.05 | 0.06853 | 0.171 | -4.69 | 1.88746 | none |
| ACSM2B | 0.503 | 0.3953 | 2.0497 | 0.06854 | 0.171 | -4.69 | 0.36021 | none |
| FAM86C1 | 0.253 | 0.2358 | 2.0494 | 0.06858 | 0.171 | -4.69 | -0.0111 | none |

| | | | | | | | | |
|---|---|---|---|---|---|---|---|---|
| ZNF32 | -0.208 | -0.2072 | -2.049 | 0.06858 | 0.171 | -4.69 | 2.42846 | AA-related |
| PGBD5 | 0.195 | 0.1168 | 2.0492 | 0.0686 | 0.171 | -4.69 | -0.6537 | none |
| CKS1B | -0.263 | -0.2738 | -2.049 | 0.06864 | 0.172 | -4.69 | 1.79771 | none |
| DCTD | -0.21 | -0.1919 | -2.048 | 0.06868 | 0.172 | -4.69 | 2.40391 | AA-related |
| UXT | -0.401 | -0.4243 | -2.048 | 0.06869 | 0.172 | -4.69 | 1.84449 | none |
| OARD1 | -0.213 | -0.222 | -2.048 | 0.06876 | 0.172 | -4.7 | 0.95157 | none |
| GIN1 | 0.415 | 0.3013 | 2.0473 | 0.06881 | 0.172 | -4.7 | 0.12187 | none |
| PARP11 | 0.205 | 0.0891 | 2.0472 | 0.06882 | 0.172 | -4.7 | -0.4333 | none |
| DGCR2 | 0.325 | 0.3114 | 2.0472 | 0.06882 | 0.172 | -4.7 | 0.73923 | none |
| RAVER1 | 0.381 | 0.3668 | 2.0469 | 0.06886 | 0.172 | -4.7 | 0.21596 | none |
| ATF6 | 0.296 | 0.2563 | 2.0459 | 0.06897 | 0.172 | -4.7 | -0.509 | none |
| KIF23 | 0.559 | 0.5333 | 2.0452 | 0.06906 | 0.172 | -4.7 | -0.5487 | none |
| ZEB1 | 0.639 | 0.6831 | 2.0451 | 0.06906 | 0.172 | -4.7 | -0.6024 | none |
| ICA1 | 0.284 | 0.2006 | 2.0448 | 0.06909 | 0.172 | -4.7 | -0.2375 | none |
| EFNA4 | -0.226 | -0.2638 | -2.045 | 0.06911 | 0.172 | -4.7 | 1.05635 | none |
| TFF2 | 0.391 | 0.3994 | 2.0446 | 0.06912 | 0.172 | -4.7 | -0.0312 | none |
| TAP1 | -0.298 | -0.2678 | -2.045 | 0.06912 | 0.172 | -4.7 | 1.77166 | none |
| COQ8A | 0.25 | 0.2389 | 2.0443 | 0.06916 | 0.172 | -4.7 | 0.21582 | none |
| RP9 | -0.27 | -0.1055 | -2.044 | 0.06924 | 0.173 | -4.7 | 1.01511 | none |
| RPS28 | -0.49 | -0.522 | -2.043 | 0.06925 | 0.173 | -4.7 | 1.86863 | none |
| CUL4A | 0.21 | 0.2213 | 2.0434 | 0.06925 | 0.173 | -4.7 | 0.68849 | none |
| GIMAP5 | -0.193 | -0.2178 | -2.043 | 0.06926 | 0.173 | -4.7 | 0.44916 | none |
| MEI1 | 0.504 | 0.2941 | 2.0422 | 0.06939 | 0.173 | -4.7 | -0.6933 | none |
| GOLGA6L6 | 0.364 | 0.2514 | 2.0421 | 0.0694 | 0.173 | -4.7 | -0.6303 | none |
| KRTAP4-7 | 0.19 | 0.1515 | 2.0419 | 0.06943 | 0.173 | -4.7 | -0.6791 | none |
| MMP15 | 0.249 | 0.2534 | 2.0419 | 0.06944 | 0.173 | -4.7 | 1.09527 | none |
| RNF130 | 0.195 | 0.1507 | 2.0415 | 0.06947 | 0.173 | -4.71 | -0.0554 | none |
| RPL23 | -0.378 | -0.4057 | -2.041 | 0.06951 | 0.173 | -4.71 | 1.48084 | none |
| CENPH | 0.315 | 0.1974 | 2.0412 | 0.06951 | 0.173 | -4.71 | -0.4321 | none |
| SUN2 | 0.312 | 0.3505 | 2.041 | 0.06954 | 0.173 | -4.71 | -0.6406 | none |
| CAPN15 | 0.364 | 0.4649 | 2.0395 | 0.0697 | 0.173 | -4.71 | -0.2323 | none |
| KPTN | 0.464 | 0.4283 | 2.0392 | 0.06974 | 0.174 | -4.71 | -0.1193 | none |
| PPIH | -0.302 | -0.3125 | -2.039 | 0.06976 | 0.174 | -4.71 | 1.54974 | none |
| PPM1N | 0.517 | 0.3081 | 2.0377 | 0.06991 | 0.174 | -4.71 | -0.8557 | none |
| ZNF304 | 0.158 | 0.1171 | 2.0369 | 0.07001 | 0.174 | -4.71 | 0.53582 | none |
| RAB20 | -0.375 | -0.3453 | -2.037 | 0.07005 | 0.174 | -4.71 | 1.55407 | none |
| GINS2 | -0.202 | -0.1707 | -2.036 | 0.07007 | 0.174 | -4.71 | 1.47064 | none |
| GUCD1 | 0.285 | 0.2946 | 2.0359 | 0.07012 | 0.174 | -4.71 | -0.5572 | none |
| CCDC175 | 0.471 | 0.3851 | 2.0359 | 0.07012 | 0.174 | -4.71 | 0.08509 | none |
| RPL32 | -0.351 | -0.3743 | -2.036 | 0.07014 | 0.174 | -4.71 | 2.26519 | AA-related |

| | | | | | | | | |
|---|---|---|---|---|---|---|---|---|
| NUDT15 | 0.285 | 0.1407 | 2.0357 | 0.07015 | 0.174 | -4.71 | -0.1337 | none |
| GOLGA6L4 | 0.455 | 0.3877 | 2.0355 | 0.07017 | 0.174 | -4.71 | -0.5784 | none |
| ANKRD27 | 0.585 | 0.6067 | 2.035 | 0.07022 | 0.174 | -4.72 | -0.1492 | none |
| MIEF1 | 0.209 | 0.2046 | 2.0348 | 0.07025 | 0.175 | -4.72 | 0.14336 | none |
| VRK2 | 0.364 | 0.359 | 2.0347 | 0.07026 | 0.175 | -4.72 | -0.3375 | none |
| RSRC1 | 0.444 | 0.3848 | 2.0343 | 0.07031 | 0.175 | -4.72 | -0.6485 | none |
| RALB | 0.24 | 0.1128 | 2.0337 | 0.07037 | 0.175 | -4.72 | -0.7248 | none |
| ANKRD18B | 0.432 | 0.2956 | 2.0335 | 0.07041 | 0.175 | -4.72 | -0.7636 | none |
| ZIC4 | 0.223 | 0.2226 | 2.0335 | 0.07041 | 0.175 | -4.72 | 0.08485 | none |
| ZKSCAN5 | 0.447 | 0.4484 | 2.0334 | 0.07041 | 0.175 | -4.72 | -0.9196 | none |
| RPS13 | -0.408 | -0.4482 | -2.033 | 0.07043 | 0.175 | -4.72 | 1.36386 | none |
| GPX2 | 0.385 | 0.2346 | 2.0326 | 0.07051 | 0.175 | -4.72 | -1.0476 | none |
| SCGB3A1 | -0.518 | -0.4601 | -2.033 | 0.07052 | 0.175 | -4.72 | 2.27702 | AA-related |
| VAMP2 | 0.22 | 0.158 | 2.0325 | 0.07052 | 0.175 | -4.72 | -0.3086 | none |
| NBPF1 | 0.339 | 0.3175 | 2.032 | 0.07057 | 0.175 | -4.72 | -0.4532 | none |
| HSD17B7 | -0.264 | -0.2406 | -2.032 | 0.07059 | 0.175 | -4.72 | 0.74455 | none |
| MVP | 0.352 | 0.3497 | 2.0316 | 0.07062 | 0.175 | -4.72 | -0.9865 | none |
| CSNK2A2 | 0.448 | 0.4477 | 2.0315 | 0.07063 | 0.175 | -4.72 | -0.9599 | none |
| BCL7B | 0.313 | 0.2049 | 2.0307 | 0.07073 | 0.175 | -4.72 | -0.6581 | none |
| TBC1D19 | 0.261 | 0.0949 | 2.0306 | 0.07074 | 0.175 | -4.72 | -0.2224 | none |
| GPR61 | -0.239 | -0.1483 | -2.031 | 0.07075 | 0.175 | -4.72 | 2.2177 | AA-related |
| TUBA1A | -0.278 | -0.3729 | -2.031 | 0.07075 | 0.175 | -4.72 | 1.07845 | none |
| SLC2A6 | 0.253 | 0.2065 | 2.03 | 0.07081 | 0.175 | -4.72 | 0.64197 | none |
| GPR182 | 0.274 | 0.2507 | 2.0299 | 0.07083 | 0.175 | -4.72 | 0.04892 | none |
| SRRM5 | 0.165 | 0.1482 | 2.0295 | 0.07087 | 0.175 | -4.72 | -1.3125 | none |
| REEP5 | -0.278 | -0.3024 | -2.03 | 0.07087 | 0.175 | -4.72 | 1.39639 | none |
| PDCD6IP | 0.36 | 0.324 | 2.0295 | 0.07087 | 0.175 | -4.72 | 0.32275 | none |
| SPATA9 | -0.167 | -0.1591 | -2.029 | 0.07093 | 0.176 | -4.72 | -0.2759 | none |
| NPIPB7 | 0.214 | 0.1779 | 2.0289 | 0.07094 | 0.176 | -4.72 | 0.89348 | none |
| COX19 | 0.296 | 0.2361 | 2.0286 | 0.07097 | 0.176 | -4.73 | -0.2992 | none |
| UBASH3B | 0.264 | 0.278 | 2.0283 | 0.07101 | 0.176 | -4.73 | 0.79109 | none |
| NFATC3 | 0.305 | 0.2922 | 2.0281 | 0.07103 | 0.176 | -4.73 | -0.3278 | none |
| OR52K1 | 0.18 | 0.1424 | 2.0279 | 0.07105 | 0.176 | -4.73 | -1.0896 | none |
| HOOK1 | 0.53 | 0.4449 | 2.0279 | 0.07106 | 0.176 | -4.73 | -0.3153 | none |
| BTN2A1 | 0.263 | 0.2344 | 2.0277 | 0.07108 | 0.176 | -4.73 | -0.1264 | none |
| RPS4Y2 | 0.251 | 0.1947 | 2.0276 | 0.07109 | 0.176 | -4.73 | -1.297 | none |
| DLGAP4 | 0.569 | 0.6064 | 2.0261 | 0.07127 | 0.176 | -4.73 | -0.1571 | none |
| RPS12 | -0.495 | -0.549 | -2.026 | 0.07128 | 0.176 | -4.73 | 1.89743 | none |
| PDCD5 | -0.157 | -0.1354 | -2.026 | 0.07128 | 0.176 | -4.73 | 1.81052 | none |
| SLC39A9 | -0.249 | -0.2287 | -2.026 | 0.07132 | 0.176 | -4.73 | 1.99065 | none |

| | | | | | | | | |
|---|---|---|---|---|---|---|---|---|
| CLIC4 | -0.302 | -0.231 | -2.025 | 0.07139 | 0.176 | -4.73 | 0.916 | none |
| STRIP1 | 0.229 | 0.2547 | 2.0251 | 0.07139 | 0.176 | -4.73 | -0.6214 | none |
| RNF121 | 0.286 | 0.2023 | 2.0249 | 0.07141 | 0.176 | -4.73 | -0.392 | none |
| INPP5J | -0.16 | -0.1064 | -2.024 | 0.07146 | 0.176 | -4.73 | 0.22784 | none |
| LAPTM5 | -0.198 | -0.1948 | -2.024 | 0.07152 | 0.177 | -4.73 | 1.70339 | none |
| SH3GLB1 | -0.344 | -0.2602 | -2.023 | 0.07164 | 0.177 | -4.73 | 2.32764 | AA-related |
| IFITM3 | 0.249 | 0.146 | 2.0229 | 0.07164 | 0.177 | -4.73 | -0.7449 | none |
| TEN1 | 0.416 | 0.3321 | 2.0228 | 0.07165 | 0.177 | -4.73 | -0.7714 | none |
| ROR2 | -0.275 | -0.2721 | -2.023 | 0.07166 | 0.177 | -4.73 | 0.09323 | none |
| CALM2 | -0.233 | -0.25 | -2.023 | 0.07168 | 0.177 | -4.73 | 1.4569 | none |
| CCER2 | -0.176 | -0.1134 | -2.023 | 0.07169 | 0.177 | -4.73 | 0.637 | none |
| SRP68 | 0.359 | 0.3739 | 2.0225 | 0.07169 | 0.177 | -4.73 | -0.0616 | none |
| PARP10 | 0.195 | 0.1584 | 2.0225 | 0.0717 | 0.177 | -4.73 | 0.9961 | none |
| GAR1 | 0.223 | 0.178 | 2.0223 | 0.07172 | 0.177 | -4.73 | 0.31187 | none |
| KIF3A | 0.402 | 0.3342 | 2.0221 | 0.07174 | 0.177 | -4.73 | -0.3289 | none |
| SLC25A43 | 0.273 | 0.2181 | 2.0219 | 0.07176 | 0.177 | -4.74 | 0.036 | none |
| DENND1B | 0.16 | 0.1395 | 2.0209 | 0.07188 | 0.177 | -4.74 | 0.22619 | none |
| SH2B3 | 0.402 | 0.4196 | 2.0208 | 0.0719 | 0.177 | -4.74 | -0.0292 | none |
| C19ORF66 | 0.205 | 0.1502 | 2.0202 | 0.07196 | 0.177 | -4.74 | 0.41747 | none |
| RAB6B | 0.177 | 0.1583 | 2.0198 | 0.07201 | 0.177 | -4.74 | -0.61 | none |
| APOL6 | 0.591 | 0.5866 | 2.0195 | 0.07205 | 0.177 | -4.74 | -0.77 | none |
| TMCO3 | 0.358 | 0.2211 | 2.0193 | 0.07207 | 0.178 | -4.74 | -0.4933 | none |
| RPS4X | -0.332 | -0.3423 | -2.019 | 0.07209 | 0.178 | -4.74 | 2.12062 | AA-related |
| TXNIP | -0.306 | -0.3229 | -2.019 | 0.07214 | 0.178 | -4.74 | 0.73858 | none |
| GOLT1B | -0.462 | -0.4705 | -2.018 | 0.07217 | 0.178 | -4.74 | 1.16655 | none |
| MVD | 0.208 | 0.2038 | 2.0181 | 0.07221 | 0.178 | -4.74 | -0.4401 | none |
| PRAC1 | -0.301 | -0.2406 | -2.018 | 0.07224 | 0.178 | -4.74 | 2.01689 | none |
| DHDDS | 0.181 | 0.173 | 2.0178 | 0.07225 | 0.178 | -4.74 | -0.8886 | none |
| NFKBIA | -0.267 | -0.1503 | -2.017 | 0.0723 | 0.178 | -4.74 | 1.40545 | none |
| UNC119 | -0.182 | -0.1656 | -2.017 | 0.07231 | 0.178 | -4.74 | 1.28102 | none |
| NDUFA11 | -0.41 | -0.4118 | -2.017 | 0.07233 | 0.178 | -4.74 | 1.96207 | none |
| PNPLA8 | 0.449 | 0.4005 | 2.0171 | 0.07234 | 0.178 | -4.74 | -0.4817 | none |
| USF2 | 0.35 | 0.3367 | 2.0169 | 0.07236 | 0.178 | -4.74 | -0.4848 | none |
| SEBOX | 0.269 | 0.2764 | 2.0169 | 0.07236 | 0.178 | -4.74 | 0.99877 | none |
| ORM1 | 0.298 | 0.2314 | 2.0167 | 0.07238 | 0.178 | -4.74 | 0.37999 | none |
| CYB5R3 | 0.273 | 0.2739 | 2.0166 | 0.07239 | 0.178 | -4.74 | -0.3797 | none |
| C19ORF53 | -0.193 | -0.154 | -2.017 | 0.0724 | 0.178 | -4.74 | 0.46643 | none |
| FKBPL | -0.153 | -0.1034 | -2.017 | 0.0724 | 0.178 | -4.74 | 0.55885 | none |
| TNK2 | 0.368 | 0.4618 | 2.0163 | 0.07243 | 0.178 | -4.74 | -0.2719 | none |
| NT5C2 | 0.305 | 0.2262 | 2.0163 | 0.07243 | 0.178 | -4.74 | -0.9698 | none |

| | | | | | | | | |
|---|---|---|---|---|---|---|---|---|
| GPAT4 | 0.239 | 0.1908 | 2.0159 | 0.07248 | 0.178 | -4.74 | 0.19476 | none |
| DHPS | 0.173 | 0.105 | 2.0159 | 0.07248 | 0.178 | -4.74 | 0.18568 | none |
| EXOC7 | 0.404 | 0.4339 | 2.0151 | 0.07258 | 0.178 | -4.75 | -0.1836 | none |
| DMXL2 | 0.355 | 0.315 | 2.015 | 0.07259 | 0.178 | -4.75 | -0.7133 | none |
| PRMT7 | 0.214 | 0.2215 | 2.0149 | 0.0726 | 0.178 | -4.75 | -0.7926 | none |
| ELP6 | 0.267 | 0.17 | 2.0149 | 0.0726 | 0.178 | -4.75 | -0.7409 | none |
| ZNF862 | 0.279 | 0.2882 | 2.0147 | 0.07262 | 0.178 | -4.75 | 0.79504 | none |
| MGLL | 0.316 | 0.2759 | 2.0147 | 0.07262 | 0.178 | -4.75 | -0.4707 | none |
| AHI1 | 0.525 | 0.541 | 2.0139 | 0.07272 | 0.178 | -4.75 | -0.8905 | none |
| PHLDB3 | 0.415 | 0.3718 | 2.0136 | 0.07275 | 0.178 | -4.75 | -0.7266 | none |
| RAD23A | 0.231 | 0.2174 | 2.0136 | 0.07275 | 0.178 | -4.75 | -1.254 | none |
| E2F5 | 0.209 | 0.0712 | 2.0135 | 0.07276 | 0.178 | -4.75 | -0.3999 | none |
| TBC1D22B | 0.172 | 0.1675 | 2.0131 | 0.07282 | 0.179 | -4.75 | -0.3924 | none |
| C10ORF67 | 0.263 | 0.1975 | 2.013 | 0.07282 | 0.179 | -4.75 | -1.2628 | none |
| PMS1 | 0.239 | 0.2159 | 2.013 | 0.07283 | 0.179 | -4.75 | -0.5261 | none |
| RFXAP | 0.25 | 0.1377 | 2.0129 | 0.07283 | 0.179 | -4.75 | -0.4528 | none |
| GEMIN5 | 0.462 | 0.5312 | 2.0119 | 0.07296 | 0.179 | -4.75 | 0.30493 | none |
| MTTP | -0.208 | -0.17 | -2.012 | 0.07296 | 0.179 | -4.75 | 1.16327 | none |
| ZNF439 | 0.354 | 0.2615 | 2.0115 | 0.073 | 0.179 | -4.75 | -1.1967 | none |
| NR3C2 | -0.333 | -0.1787 | -2.011 | 0.07301 | 0.179 | -4.75 | 1.21182 | none |
| RABL6 | 0.45 | 0.4943 | 2.0114 | 0.07301 | 0.179 | -4.75 | -0.443 | none |
| PHACTR2 | 0.725 | 0.7693 | 2.0112 | 0.07304 | 0.179 | -4.75 | -0.5649 | none |
| RAB32 | -0.223 | -0.2238 | -2.01 | 0.07317 | 0.179 | -4.75 | 0.71154 | none |
| CLDND2 | -0.335 | -0.2728 | -2.01 | 0.07323 | 0.179 | -4.75 | 1.14982 | none |
| CEP85 | 0.306 | 0.3127 | 2.0089 | 0.07332 | 0.179 | -4.76 | -0.5502 | none |
| FAU | -0.396 | -0.4222 | -2.008 | 0.0734 | 0.18 | -4.76 | 2.26514 | AA-related |
| HAND2 | 0.256 | 0.1943 | 2.0078 | 0.07345 | 0.18 | -4.76 | -0.6799 | none |
| SNRPE | -0.288 | -0.3401 | -2.008 | 0.07349 | 0.18 | -4.76 | 1.18545 | none |
| MT1B | -0.282 | -0.282 | -2.007 | 0.07349 | 0.18 | -4.76 | -0.1514 | none |
| SPIN2A | 0.169 | 0.1058 | 2.0065 | 0.07361 | 0.18 | -4.76 | -0.0376 | none |
| IL2RG | -0.248 | -0.2158 | -2.006 | 0.07364 | 0.18 | -4.76 | 1.49555 | none |
| URM1 | -0.317 | -0.2199 | -2.006 | 0.07365 | 0.18 | -4.76 | 2.41789 | AA-related |
| DAO | 0.239 | 0.2392 | 2.0055 | 0.07373 | 0.18 | -4.76 | -0.3603 | none |
| SMC5 | 0.705 | 0.7091 | 2.0055 | 0.07373 | 0.18 | -4.76 | -0.2613 | none |
| FCGR1B | 0.159 | 0.1671 | 2.0054 | 0.07374 | 0.18 | -4.76 | 0.60808 | none |
| GNG5 | -0.279 | -0.2934 | -2.005 | 0.07375 | 0.18 | -4.76 | 1.46912 | none |
| CFAP77 | 0.271 | 0.1937 | 2.0053 | 0.07376 | 0.18 | -4.76 | -0.7837 | none |
| GGH | -0.297 | -0.3324 | -2.005 | 0.07376 | 0.18 | -4.76 | 1.47758 | none |
| EXOC4 | 0.32 | 0.3725 | 2.0049 | 0.07381 | 0.18 | -4.76 | 0.3196 | none |
| C1QTNF3 | 0.41 | 0.3105 | 2.0045 | 0.07385 | 0.18 | -4.76 | 0.48363 | none |

| | | | | | | | | |
|---|---|---|---|---|---|---|---|---|
| ZNF668 | 0.228 | 0.2448 | 2.0044 | 0.07387 | 0.18 | -4.76 | -0.4922 | none |
| BPHL | 0.268 | 0.193 | 2.0041 | 0.0739 | 0.18 | -4.76 | -1.2461 | none |
| RPLP2 | -0.435 | -0.4532 | -2.004 | 0.07392 | 0.18 | -4.76 | 1.76891 | none |
| ERICH5 | -0.233 | -0.1551 | -2.004 | 0.07395 | 0.181 | -4.76 | 0.94479 | none |
| LINGO1 | -0.221 | -0.1583 | -2.004 | 0.07395 | 0.181 | -4.76 | 0.83893 | none |
| SBNO1 | 0.482 | 0.5016 | 2.0035 | 0.07398 | 0.181 | -4.76 | -1.0095 | none |
| NUP205 | 0.419 | 0.3275 | 2.0034 | 0.07399 | 0.181 | -4.76 | -1.1803 | none |
| HBA2 | -0.264 | -0.1529 | -2.003 | 0.074 | 0.181 | -4.76 | 1.24251 | none |
| WDR78 | 0.355 | 0.2256 | 2.0033 | 0.074 | 0.181 | -4.76 | -0.9644 | none |
| NBPF20 | 0.383 | 0.4053 | 2.003 | 0.07403 | 0.181 | -4.76 | -0.7407 | none |
| MLKL | 0.515 | 0.4694 | 2.0029 | 0.07405 | 0.181 | -4.76 | -0.5785 | none |
| C12ORF29 | -0.191 | -0.1833 | -2.003 | 0.07408 | 0.181 | -4.76 | 1.45314 | none |
| SIAH1 | 0.292 | 0.255 | 2.0027 | 0.07408 | 0.181 | -4.76 | -0.3805 | none |
| MRPL11 | -0.304 | -0.3105 | -2.002 | 0.07413 | 0.181 | -4.77 | 1.3649 | none |
| PRKD1 | 0.215 | 0.1869 | 2.0021 | 0.07415 | 0.181 | -4.77 | -0.9261 | none |
| AP2S1 | -0.299 | -0.2828 | -2.002 | 0.07416 | 0.181 | -4.77 | 1.38425 | none |
| TYROBP | -0.648 | -0.7062 | -2.002 | 0.07417 | 0.181 | -4.77 | 1.94398 | none |
| SRRT | 0.456 | 0.5112 | 2.0017 | 0.07419 | 0.181 | -4.77 | -0.0449 | none |
| LRSAM1 | 0.345 | 0.343 | 2.0015 | 0.07422 | 0.181 | -4.77 | -0.7769 | none |
| SCRN3 | 0.241 | 0.2171 | 2.0015 | 0.07422 | 0.181 | -4.77 | 0.34267 | none |
| IL6ST | 0.486 | 0.3995 | 2.0015 | 0.07423 | 0.181 | -4.77 | -1.3208 | none |
| TSTD3 | 0.242 | 0.1315 | 2.0014 | 0.07423 | 0.181 | -4.77 | -1.0729 | none |
| F8 | -0.179 | -0.1585 | -2.001 | 0.07428 | 0.181 | -4.77 | 0.50879 | none |
| PAPOLA | 0.202 | 0.1914 | 2.0009 | 0.07429 | 0.181 | -4.77 | 0.40007 | none |
| DDX60 | 0.242 | 0.1183 | 2.0006 | 0.07433 | 0.181 | -4.77 | -0.7577 | none |
| KXD1 | 0.242 | 0.1947 | 2.0006 | 0.07433 | 0.181 | -4.77 | -1.0091 | none |
| MRGPRX3 | 0.141 | 0.1015 | 2.0006 | 0.07433 | 0.181 | -4.77 | -1.0309 | none |
| COL19A1 | 0.334 | 0.3385 | 2.0003 | 0.07437 | 0.181 | -4.77 | 1.01062 | none |
| TBC1D3B | 0.619 | 0.5956 | 1.9999 | 0.07441 | 0.181 | -4.77 | -0.5888 | none |
| KRTAP19-4 | 0.283 | 0.2834 | 1.9999 | 0.07441 | 0.181 | -4.77 | 0.87693 | none |
| GNG3 | 0.349 | 0.2957 | 1.9995 | 0.07446 | 0.181 | -4.77 | -0.4963 | none |
| MSH3 | 0.504 | 0.4799 | 1.9994 | 0.07448 | 0.181 | -4.77 | -0.0124 | none |
| NAXD | 0.205 | 0.1715 | 1.9992 | 0.0745 | 0.181 | -4.77 | -0.7338 | none |
| LGI4 | -0.225 | -0.1757 | -1.999 | 0.07451 | 0.181 | -4.77 | 0.32059 | none |
| HIST1H2AI | -0.739 | -0.6492 | -1.999 | 0.07453 | 0.181 | -4.77 | 1.15254 | none |
| KNSTRN | 0.233 | 0.2112 | 1.9989 | 0.07453 | 0.181 | -4.77 | -1.1735 | none |
| LSM6 | -0.315 | -0.3502 | -1.999 | 0.07456 | 0.181 | -4.77 | 1.45931 | none |
| GPX4 | -0.26 | -0.2232 | -1.999 | 0.07458 | 0.181 | -4.77 | 2.54987 | AA-related |
| LAMTOR3 | -0.406 | -0.4517 | -1.998 | 0.0746 | 0.181 | -4.77 | 1.75589 | none |
| PTCH2 | 0.181 | 0.1425 | 1.998 | 0.07465 | 0.181 | -4.77 | 0.43836 | none |

| | | | | | | | | |
|---|---|---|---|---|---|---|---|---|
| MTRNR2L1 | -0.195 | -0.1704 | -1.998 | 0.07469 | 0.181 | -4.77 | 1.47144 | none |
| MRPL13 | -0.294 | -0.288 | -1.998 | 0.07471 | 0.181 | -4.77 | 1.14459 | none |
| ZCCHC10 | 0.353 | 0.2454 | 1.9975 | 0.07471 | 0.181 | -4.77 | -0.9333 | none |
| DHRS11 | 0.243 | 0.2319 | 1.9974 | 0.07472 | 0.181 | -4.77 | -0.2382 | none |
| PRDM9 | -0.229 | -0.256 | -1.997 | 0.07475 | 0.181 | -4.77 | 1.67459 | none |
| PSMB4 | -0.34 | -0.3256 | -1.997 | 0.07481 | 0.182 | -4.77 | 0.30128 | none |
| TCP1 | -0.195 | -0.173 | -1.996 | 0.07484 | 0.182 | -4.77 | 1.97392 | none |
| GMPR2 | 0.329 | 0.2395 | 1.9962 | 0.07487 | 0.182 | -4.77 | 0.72704 | none |
| SLC7A1 | 0.277 | 0.3472 | 1.9962 | 0.07488 | 0.182 | -4.77 | 0.07222 | none |
| CHMP2A | -0.175 | -0.1684 | -1.996 | 0.07496 | 0.182 | -4.78 | 1.21313 | none |
| LIN7B | 0.383 | 0.266 | 1.9951 | 0.075 | 0.182 | -4.78 | -0.6445 | none |
| PPFIA4 | 0.613 | 0.5592 | 1.9949 | 0.07504 | 0.182 | -4.78 | -1.0031 | none |
| FGF2 | 0.157 | 0.1088 | 1.9948 | 0.07504 | 0.182 | -4.78 | -1.3369 | none |
| ING1 | 0.418 | 0.2989 | 1.9945 | 0.07509 | 0.182 | -4.78 | -0.7274 | none |
| TCAP | 0.339 | 0.2204 | 1.994 | 0.07514 | 0.182 | -4.78 | -0.2222 | none |
| ENOX2 | 0.579 | 0.4881 | 1.994 | 0.07514 | 0.182 | -4.78 | -0.38 | none |
| PSME4 | 0.472 | 0.4133 | 1.9937 | 0.07519 | 0.182 | -4.78 | -1.0667 | none |
| RER1 | -0.259 | -0.2355 | -1.993 | 0.07522 | 0.182 | -4.78 | 1.07863 | none |
| LRR1 | -0.301 | -0.3493 | -1.993 | 0.07522 | 0.182 | -4.78 | 0.92338 | none |
| ARID5B | 0.518 | 0.5509 | 1.9931 | 0.07525 | 0.182 | -4.78 | -0.9443 | none |
| RASL11B | 0.233 | 0.2061 | 1.993 | 0.07527 | 0.182 | -4.78 | -0.8808 | none |
| WNT4 | 0.31 | 0.2808 | 1.9923 | 0.07535 | 0.182 | -4.78 | -0.9542 | none |
| IGFBP4 | -0.209 | -0.1742 | -1.992 | 0.07538 | 0.182 | -4.78 | 1.10288 | none |
| MORC2 | 0.412 | 0.468 | 1.9918 | 0.07542 | 0.183 | -4.78 | -0.4083 | none |
| CHML | 0.462 | 0.4673 | 1.9914 | 0.07547 | 0.183 | -4.78 | -0.958 | none |
| TTC26 | 0.305 | 0.1809 | 1.9911 | 0.0755 | 0.183 | -4.78 | 0.18587 | none |
| MRPL35 | -0.282 | -0.3002 | -1.99 | 0.07559 | 0.183 | -4.78 | 1.05457 | none |
| TRIM37 | 0.452 | 0.4083 | 1.9899 | 0.07565 | 0.183 | -4.78 | 0.63988 | none |
| ADGRA2 | 0.199 | 0.1728 | 1.9894 | 0.07572 | 0.183 | -4.78 | -1.0013 | none |
| GADL1 | 0.276 | 0.2164 | 1.989 | 0.07577 | 0.183 | -4.79 | -1.1488 | none |
| YIPF6 | 0.283 | 0.1116 | 1.9887 | 0.07581 | 0.183 | -4.79 | -0.9347 | none |
| MRPL18 | -0.305 | -0.3024 | -1.988 | 0.07584 | 0.183 | -4.79 | 1.19736 | none |
| RGS1 | -0.318 | -0.34 | -1.988 | 0.07587 | 0.183 | -4.79 | 0.81108 | none |
| TNP1 | 0.246 | 0.251 | 1.9871 | 0.076 | 0.184 | -4.79 | 0.82459 | none |
| CYSRT1 | 0.311 | 0.1899 | 1.9863 | 0.07611 | 0.184 | -4.79 | 0.09401 | none |
| PIGK | -0.324 | -0.3726 | -1.986 | 0.07611 | 0.184 | -4.79 | 1.70261 | none |
| NUP153 | 0.413 | 0.4491 | 1.9857 | 0.07618 | 0.184 | -4.79 | 0.56862 | none |
| ACSF3 | 0.263 | 0.227 | 1.9855 | 0.0762 | 0.184 | -4.79 | 0.13216 | none |
| APEX1 | -0.184 | -0.1533 | -1.985 | 0.07626 | 0.184 | -4.79 | 1.74106 | none |
| RPS5 | -0.421 | -0.4587 | -1.984 | 0.07638 | 0.184 | -4.79 | 1.9676 | none |

| | | | | | | | | |
|---|---|---|---|---|---|---|---|---|
| TRMT2A | 0.237 | 0.2362 | 1.9836 | 0.07644 | 0.184 | -4.79 | 0.0333 | none |
| HBB | -0.328 | -0.2801 | -1.984 | 0.07645 | 0.184 | -4.79 | 0.71207 | none |
| CD24 | 0.162 | 0.1415 | 1.9835 | 0.07645 | 0.184 | -4.79 | -0.9932 | none |
| ABHD17B | 0.261 | 0.1603 | 1.9835 | 0.07645 | 0.184 | -4.79 | 0.29545 | none |
| PIBF1 | 0.438 | 0.5183 | 1.9834 | 0.07647 | 0.185 | -4.79 | 0.50593 | none |
| UQCC3 | 0.306 | 0.2044 | 1.9833 | 0.07647 | 0.185 | -4.79 | -0.6849 | none |
| SULT1A1 | 0.237 | 0.2254 | 1.9825 | 0.07659 | 0.185 | -4.8 | -0.1232 | none |
| MNAT1 | -0.216 | -0.2276 | -1.982 | 0.07662 | 0.185 | -4.8 | 1.98596 | none |
| CPD | 0.461 | 0.3917 | 1.9821 | 0.07663 | 0.185 | -4.8 | -0.6391 | none |
| CCT5 | -0.169 | -0.1288 | -1.982 | 0.07664 | 0.185 | -4.8 | 0.64314 | none |
| ATG9B | 0.259 | 0.1965 | 1.9819 | 0.07666 | 0.185 | -4.8 | 0.31172 | none |
| ACTR3 | -0.16 | -0.1599 | -1.982 | 0.0767 | 0.185 | -4.8 | 1.63773 | none |
| OR5A2 | 0.138 | 0.1153 | 1.9815 | 0.07671 | 0.185 | -4.8 | -0.8185 | none |
| RUSC2 | -0.191 | -0.0754 | -1.981 | 0.07673 | 0.185 | -4.8 | 1.93818 | none |
| ZNF740 | 0.322 | 0.3371 | 1.9813 | 0.07674 | 0.185 | -4.8 | 0.49944 | none |
| NEFH | 0.438 | 0.5248 | 1.981 | 0.07677 | 0.185 | -4.8 | 0.47838 | none |
| OTOG | 0.343 | 0.2617 | 1.9809 | 0.07678 | 0.185 | -4.8 | -0.1763 | none |
| ITPR3 | 0.411 | 0.4905 | 1.9806 | 0.07683 | 0.185 | -4.8 | -0.4315 | none |
| SNX16 | -0.281 | -0.2273 | -1.981 | 0.07683 | 0.185 | -4.8 | 1.26224 | none |
| AC096949 | 0.468 | 0.3121 | 1.9802 | 0.07688 | 0.185 | -4.8 | 0.10073 | none |
| LDLRAD4 | 0.198 | 0.2028 | 1.9798 | 0.07693 | 0.185 | -4.8 | -0.2717 | none |
| MORF4L1 | 0.213 | 0.1319 | 1.9795 | 0.07696 | 0.185 | -4.8 | -1.1375 | none |
| ADAMTS1: | 0.198 | 0.1519 | 1.979 | 0.07703 | 0.185 | -4.8 | -0.3501 | none |
| GGNBP2 | 0.379 | 0.3936 | 1.9785 | 0.07708 | 0.185 | -4.8 | -0.0664 | none |
| PRDX3 | -0.31 | -0.305 | -1.978 | 0.07718 | 0.186 | -4.8 | 1.01874 | none |
| GRIPAP1 | 0.613 | 0.6041 | 1.9763 | 0.07737 | 0.186 | -4.8 | -0.981 | none |
| CST6 | -0.309 | -0.2894 | -1.976 | 0.07737 | 0.186 | -4.8 | 1.13039 | none |
| RPS2 | -0.356 | -0.3009 | -1.976 | 0.07739 | 0.186 | -4.8 | 0.91914 | none |
| RNF165 | 0.265 | 0.1905 | 1.9761 | 0.0774 | 0.186 | -4.8 | -0.6061 | none |
| EIF4G3 | 0.606 | 0.6618 | 1.9758 | 0.07743 | 0.186 | -4.81 | -0.6443 | none |
| TBC1D10B | 0.228 | 0.2278 | 1.9758 | 0.07743 | 0.186 | -4.81 | -0.3954 | none |
| GPX1 | -0.246 | -0.2182 | -1.975 | 0.07748 | 0.186 | -4.81 | 1.59328 | none |
| IQCE | 0.304 | 0.3313 | 1.9754 | 0.07748 | 0.186 | -4.81 | 0.49459 | none |
| API5 | 0.3 | 0.2755 | 1.9754 | 0.07749 | 0.186 | -4.81 | -0.0641 | none |
| ILF2 | -0.169 | -0.1426 | -1.975 | 0.07752 | 0.186 | -4.81 | 1.25289 | none |
| LSS | 0.241 | 0.251 | 1.9748 | 0.07755 | 0.186 | -4.81 | 0.44265 | none |
| KLF13 | 0.29 | 0.2603 | 1.9746 | 0.07758 | 0.186 | -4.81 | 0.39165 | none |
| RCBTB1 | 0.211 | 0.1184 | 1.9741 | 0.07764 | 0.186 | -4.81 | 0.1233 | none |
| CAPZA2 | 0.203 | 0.0393 | 1.974 | 0.07766 | 0.186 | -4.81 | -0.7364 | none |
| TMCO1 | -0.283 | -0.3236 | -1.974 | 0.07766 | 0.186 | -4.81 | 1.76691 | none |

| | | | | | | | | |
|---|---|---|---|---|---|---|---|---|
| LSM3 | -0.259 | -0.2606 | -1.974 | 0.07768 | 0.186 | -4.81 | 0.64555 | none |
| ARFGEF1 | 0.489 | 0.5143 | 1.9731 | 0.07777 | 0.187 | -4.81 | -0.5066 | none |
| SLC25A3 | -0.267 | -0.2733 | -1.973 | 0.07783 | 0.187 | -4.81 | 1.77375 | none |
| AL513122. | -0.125 | -0.0813 | -1.973 | 0.07784 | 0.187 | -4.81 | 0.19337 | none |
| PPP3CC | 0.275 | 0.1644 | 1.9724 | 0.07786 | 0.187 | -4.81 | 0.2351 | none |
| RTCA | 0.172 | 0.0395 | 1.9724 | 0.07787 | 0.187 | -4.81 | -0.0385 | none |
| SZRD1 | 0.169 | 0.1819 | 1.9722 | 0.07789 | 0.187 | -4.81 | -0.3846 | none |
| ODF3B | 0.274 | 0.1296 | 1.9721 | 0.07791 | 0.187 | -4.81 | 0.58951 | none |
| DGUOK | -0.251 | -0.2753 | -1.972 | 0.07791 | 0.187 | -4.81 | 1.99656 | none |
| GPR6 | 0.189 | 0.0926 | 1.9717 | 0.07795 | 0.187 | -4.81 | -0.708 | none |
| RAB17 | 0.231 | 0.2268 | 1.9716 | 0.07797 | 0.187 | -4.81 | 0.26526 | none |
| SEZ6 | 0.395 | 0.3397 | 1.9716 | 0.07797 | 0.187 | -4.81 | -0.8152 | none |
| APH1A | -0.342 | -0.3602 | -1.971 | 0.07799 | 0.187 | -4.81 | 1.95879 | none |
| PRRG3 | 0.265 | 0.2206 | 1.9714 | 0.07799 | 0.187 | -4.81 | -0.722 | none |
| MANBA | 0.198 | 0.1257 | 1.9704 | 0.07812 | 0.187 | -4.81 | 0.52519 | none |
| ZNF577 | 0.214 | 0.1269 | 1.9702 | 0.07814 | 0.187 | -4.81 | -0.5004 | none |
| BEND7 | 0.462 | 0.3487 | 1.9695 | 0.07823 | 0.187 | -4.81 | -0.0625 | none |
| TRMT112 | -0.307 | -0.3065 | -1.969 | 0.07828 | 0.187 | -4.82 | 1.64625 | none |
| PHGR1 | -0.278 | -0.2188 | -1.969 | 0.07828 | 0.187 | -4.82 | 0.5042 | none |
| PTBP2 | 0.287 | 0.2154 | 1.9682 | 0.0784 | 0.188 | -4.82 | 0.19605 | none |
| POLR1D | -0.318 | -0.3295 | -1.968 | 0.07846 | 0.188 | -4.82 | 1.91451 | none |
| RAB1B | -0.254 | -0.1899 | -1.966 | 0.07867 | 0.188 | -4.82 | 1.42115 | none |
| GNL3 | 0.363 | 0.367 | 1.9659 | 0.07869 | 0.188 | -4.82 | 0.1748 | none |
| HIST1H3B | -0.514 | -0.491 | -1.966 | 0.07872 | 0.188 | -4.82 | 0.8431 | none |
| RIMS1 | 0.205 | 0.1433 | 1.9657 | 0.07873 | 0.188 | -4.82 | -0.8782 | none |
| MYEOV | 0.486 | 0.332 | 1.9656 | 0.07874 | 0.188 | -4.82 | -0.496 | none |
| AEBP1 | 0.389 | 0.4456 | 1.9656 | 0.07874 | 0.188 | -4.82 | 0.26253 | none |
| ZNF530 | 0.401 | 0.3345 | 1.9651 | 0.0788 | 0.188 | -4.82 | 0.19717 | none |
| TADA2B | 0.326 | 0.2714 | 1.9644 | 0.0789 | 0.189 | -4.82 | -0.9972 | none |
| TMEM184 | 0.199 | 0.1307 | 1.9637 | 0.07898 | 0.189 | -4.82 | -0.7013 | none |
| SDC3 | 0.195 | 0.1287 | 1.9633 | 0.07904 | 0.189 | -4.82 | -1.2285 | none |
| SARS2 | 0.175 | 0.1289 | 1.9623 | 0.07917 | 0.189 | -4.83 | 0.90264 | none |
| LMTK2 | 0.479 | 0.5208 | 1.9623 | 0.07917 | 0.189 | -4.83 | -0.6294 | none |
| ZNF285 | 0.269 | 0.2169 | 1.9608 | 0.07936 | 0.19 | -4.83 | -0.1695 | none |
| OBSL1 | 0.208 | 0.1387 | 1.96 | 0.07947 | 0.19 | -4.83 | -0.9645 | none |
| ZNF613 | 0.672 | 0.5582 | 1.9599 | 0.07948 | 0.19 | -4.83 | -1.1936 | none |
| GBGT1 | 0.378 | 0.2161 | 1.9599 | 0.07949 | 0.19 | -4.83 | -1.1196 | none |
| SSX2 | 0.169 | 0.1624 | 1.9596 | 0.07951 | 0.19 | -4.83 | -0.6965 | none |
| SEMA7A | 0.386 | 0.3348 | 1.958 | 0.07973 | 0.19 | -4.83 | -0.2271 | none |
| LIMK2 | 0.243 | 0.2884 | 1.9578 | 0.07975 | 0.19 | -4.83 | 0.08188 | none |

| | | | | | | | | |
|---|---|---|---|---|---|---|---|---|
| SCAF1 | 0.662 | 0.7183 | 1.9578 | 0.07976 | 0.19 | -4.83 | -0.6739 | none |
| GRM5 | 0.218 | 0.1569 | 1.9572 | 0.07983 | 0.191 | -4.83 | 0.0141 | none |
| H3F3B | -0.246 | -0.1932 | -1.957 | 0.07986 | 0.191 | -4.83 | 0.22374 | none |
| RPS29 | -0.553 | -0.6013 | -1.957 | 0.07988 | 0.191 | -4.83 | 1.19948 | none |
| ITIH2 | -0.256 | -0.2337 | -1.957 | 0.07989 | 0.191 | -4.83 | 0.01395 | none |
| FLYWCH2 | 0.218 | 0.2001 | 1.9567 | 0.0799 | 0.191 | -4.83 | 0.16526 | none |
| DLX3 | 0.208 | 0.1595 | 1.9567 | 0.0799 | 0.191 | -4.83 | 1.54237 | none |
| GOLGA8Q | 0.23 | 0.1747 | 1.9564 | 0.07993 | 0.191 | -4.83 | 0.41493 | none |
| IQCJ-SCHIF | 0.378 | 0.4043 | 1.9562 | 0.07996 | 0.191 | -4.83 | -0.2815 | none |
| ZNF599 | 0.281 | 0.1403 | 1.9559 | 0.08 | 0.191 | -4.84 | -0.7717 | none |
| TBL1XR1 | 0.255 | 0.1569 | 1.9553 | 0.08008 | 0.191 | -4.84 | -0.945 | none |
| SORCS1 | -0.231 | -0.1805 | -1.955 | 0.08009 | 0.191 | -4.84 | -0.1214 | none |
| SETBP1 | 0.237 | 0.2321 | 1.9548 | 0.08015 | 0.191 | -4.84 | -0.061 | none |
| PRR29 | 0.22 | 0.2286 | 1.9544 | 0.0802 | 0.191 | -4.84 | -1.0851 | none |
| PRCD | 0.241 | 0.2751 | 1.9542 | 0.08023 | 0.191 | -4.84 | 0.71319 | none |
| C1ORF141 | 0.434 | 0.3413 | 1.9542 | 0.08023 | 0.191 | -4.84 | 0.09852 | none |
| ZG16 | 0.219 | 0.1749 | 1.9537 | 0.0803 | 0.191 | -4.84 | -0.7023 | none |
| MEA1 | -0.21 | -0.2033 | -1.954 | 0.0803 | 0.191 | -4.84 | 1.58213 | none |
| C3ORF52 | 0.18 | 0.1262 | 1.9535 | 0.08032 | 0.191 | -4.84 | 0.56696 | none |
| RPS21 | -0.423 | -0.4554 | -1.953 | 0.08034 | 0.191 | -4.84 | 1.72949 | none |
| AGFG2 | 0.288 | 0.2372 | 1.9533 | 0.08035 | 0.191 | -4.84 | -0.2591 | none |
| FLT4 | -0.248 | -0.1234 | -1.953 | 0.08036 | 0.191 | -4.84 | 1.17546 | none |
| IGF1 | 0.326 | 0.2872 | 1.9522 | 0.08049 | 0.192 | -4.84 | -0.4551 | none |
| ACTC1 | -0.266 | -0.1757 | -1.952 | 0.08049 | 0.192 | -4.84 | 1.54978 | none |
| SFTPC | -0.283 | -0.1715 | -1.952 | 0.0805 | 0.192 | -4.84 | 0.05698 | none |
| DYNAP | -0.155 | -0.0946 | -1.952 | 0.08052 | 0.192 | -4.84 | -0.2389 | none |
| PVR | 0.368 | 0.3916 | 1.9519 | 0.08053 | 0.192 | -4.84 | -0.0593 | none |
| CYP2R1 | -0.507 | -0.3856 | -1.952 | 0.08058 | 0.192 | -4.84 | 2.03199 | none |
| SFT2D1 | -0.337 | -0.3884 | -1.951 | 0.08065 | 0.192 | -4.84 | 1.12033 | none |
| FABP6 | -0.351 | -0.3037 | -1.951 | 0.08066 | 0.192 | -4.84 | 1.34779 | none |
| IFNL2 | -0.168 | -0.1529 | -1.951 | 0.08069 | 0.192 | -4.84 | 1.84221 | none |
| TMEM170 | 0.145 | 0.0993 | 1.9502 | 0.08075 | 0.192 | -4.84 | -0.8839 | none |
| TSR2 | 0.214 | 0.2023 | 1.9501 | 0.08077 | 0.192 | -4.84 | -1.2386 | none |
| DYNLT1 | -0.306 | -0.3713 | -1.95 | 0.08079 | 0.192 | -4.84 | 1.45182 | none |
| ZMIZ1 | 0.505 | 0.5885 | 1.9499 | 0.0808 | 0.192 | -4.84 | -0.6361 | none |
| RPS16 | -0.402 | -0.4249 | -1.95 | 0.0808 | 0.192 | -4.84 | 1.4165 | none |
| HIGD1B | 0.505 | 0.3593 | 1.9496 | 0.08083 | 0.192 | -4.84 | -0.9985 | none |
| DCAKD | 0.204 | 0.1415 | 1.9495 | 0.08085 | 0.192 | -4.84 | -0.5408 | none |
| CDNF | 0.226 | 0.1517 | 1.9493 | 0.08088 | 0.192 | -4.85 | -0.0733 | none |
| NUDT4 | 0.208 | 0.1457 | 1.9491 | 0.0809 | 0.192 | -4.85 | -0.3839 | none |

| | | | | | | | | |
|---|---|---|---|---|---|---|---|---|
| MIF | -0.352 | -0.317 | -1.949 | 0.08094 | 0.192 | -4.85 | 1.463 | none |
| ARCN1 | 0.231 | 0.1725 | 1.9485 | 0.08098 | 0.192 | -4.85 | 0.07625 | none |
| GPN3 | 0.251 | 0.1194 | 1.9477 | 0.08108 | 0.192 | -4.85 | -0.2997 | none |
| KLHDC3 | -0.319 | -0.3386 | -1.947 | 0.08114 | 0.193 | -4.85 | 1.30857 | none |
| PPP1R12A | 0.731 | 0.773 | 1.9471 | 0.08117 | 0.193 | -4.85 | -0.2361 | none |
| CD300LF | 0.196 | 0.1462 | 1.947 | 0.08118 | 0.193 | -4.85 | -0.3268 | none |
| SFT2D3 | 0.388 | 0.2588 | 1.9461 | 0.08129 | 0.193 | -4.85 | -0.9201 | none |
| JPH4 | 0.21 | 0.1502 | 1.9459 | 0.08133 | 0.193 | -4.85 | -0.0113 | none |
| ZMYM2 | 0.466 | 0.459 | 1.9456 | 0.08137 | 0.193 | -4.85 | -0.0408 | none |
| 11-Sep | -0.245 | -0.1015 | -1.946 | 0.08137 | 0.193 | -4.85 | 1.24096 | none |
| M6PR | -0.229 | -0.2357 | -1.945 | 0.08139 | 0.193 | -4.85 | 1.68184 | none |
| DNTTIP1 | 0.141 | 0.1426 | 1.9447 | 0.08149 | 0.193 | -4.85 | 0.66929 | none |
| UBE2R2 | -0.155 | -0.1278 | -1.944 | 0.08156 | 0.193 | -4.85 | 1.07594 | none |
| HDAC5 | 0.356 | 0.2726 | 1.9437 | 0.08162 | 0.193 | -4.85 | -0.9923 | none |
| RIN1 | 0.529 | 0.4861 | 1.9435 | 0.08165 | 0.193 | -4.85 | -0.6532 | none |
| PSMD5 | 0.151 | 0.1384 | 1.9434 | 0.08166 | 0.193 | -4.85 | -0.054 | none |
| KRT14 | 0.19 | 0.1996 | 1.9434 | 0.08167 | 0.193 | -4.85 | 1.15997 | none |
| KIF4A | 0.66 | 0.6968 | 1.9429 | 0.08172 | 0.193 | -4.85 | -0.2691 | none |
| TEAD2 | -0.174 | -0.1588 | -1.943 | 0.08174 | 0.193 | -4.86 | 0.30584 | none |
| GLIPR1 | -0.406 | -0.4837 | -1.943 | 0.08175 | 0.193 | -4.86 | 1.82812 | none |
| CSRP2 | 0.266 | 0.1952 | 1.9427 | 0.08175 | 0.193 | -4.86 | -1.1578 | none |
| CABP5 | 0.24 | 0.24 | 1.9426 | 0.08176 | 0.193 | -4.86 | 0.56439 | none |
| CCNT1 | 0.414 | 0.3428 | 1.9426 | 0.08177 | 0.193 | -4.86 | -0.1958 | none |
| FAM162A | -0.264 | -0.2763 | -1.942 | 0.08179 | 0.193 | -4.86 | 1.39837 | none |
| PTGDR2 | 0.141 | 0.0919 | 1.9424 | 0.08179 | 0.193 | -4.86 | -1.1921 | none |
| PLEKHG4 | 0.215 | 0.2421 | 1.9418 | 0.08188 | 0.194 | -4.86 | 0.76827 | none |
| HR | -0.25 | -0.147 | -1.942 | 0.08191 | 0.194 | -4.86 | -0.2544 | none |
| KIF11 | 0.608 | 0.628 | 1.9413 | 0.08194 | 0.194 | -4.86 | -0.5215 | none |
| KIAA1211l | 0.262 | 0.2747 | 1.9402 | 0.08209 | 0.194 | -4.86 | 0.6937 | none |
| LAIR2 | 0.711 | 0.4966 | 1.94 | 0.08211 | 0.194 | -4.86 | 0.07514 | none |
| ZC3H7A | 0.348 | 0.3368 | 1.9397 | 0.08216 | 0.194 | -4.86 | -0.8127 | none |
| HELQ | -0.332 | -0.2639 | -1.939 | 0.08223 | 0.194 | -4.86 | 0.19945 | none |
| IRF8 | 0.268 | 0.2339 | 1.9389 | 0.08226 | 0.194 | -4.86 | -0.5398 | none |
| C11ORF68 | 0.307 | 0.1392 | 1.9387 | 0.0823 | 0.194 | -4.86 | -1.1727 | none |
| NOXA1 | -0.145 | -0.1095 | -1.938 | 0.08233 | 0.194 | -4.86 | 0.31713 | none |
| SLC12A6 | 0.476 | 0.473 | 1.9381 | 0.08238 | 0.195 | -4.86 | -0.9473 | none |
| NAT1 | 0.266 | 0.0486 | 1.9379 | 0.0824 | 0.195 | -4.86 | 0.01905 | none |
| NT5DC1 | 0.275 | 0.1294 | 1.9379 | 0.08241 | 0.195 | -4.86 | 0.30921 | none |
| PDS5B | 0.582 | 0.5925 | 1.9377 | 0.08243 | 0.195 | -4.86 | -0.207 | none |
| ZNF182 | 0.298 | 0.3186 | 1.9376 | 0.08244 | 0.195 | -4.86 | -0.8197 | none |

| | | | | | | | | |
|---|---|---|---|---|---|---|---|---|
| SCGB1D2 | 0.266 | 0.2657 | 1.9375 | 0.08245 | 0.195 | -4.86 | 1.00034 | none |
| SKA1 | 0.49 | 0.4303 | 1.9375 | 0.08246 | 0.195 | -4.86 | -0.1928 | none |
| NECAP1 | 0.285 | 0.1575 | 1.9375 | 0.08246 | 0.195 | -4.86 | -0.2713 | none |
| NDUFC1 | -0.329 | -0.34 | -1.937 | 0.08247 | 0.195 | -4.86 | 1.52596 | none |
| INO80B-W | -0.25 | -0.3339 | -1.937 | 0.08249 | 0.195 | -4.86 | 0.75644 | none |
| BTN3A2 | 0.391 | 0.3432 | 1.9371 | 0.08251 | 0.195 | -4.86 | -0.6206 | none |
| MTG1 | 0.193 | 0.157 | 1.9371 | 0.08251 | 0.195 | -4.86 | -0.3059 | none |
| MCC | 0.322 | 0.2817 | 1.937 | 0.08252 | 0.195 | -4.86 | -0.6483 | none |
| ZNF831 | 0.173 | 0.1788 | 1.9369 | 0.08254 | 0.195 | -4.86 | -0.3137 | none |
| OR7C2 | 0.17 | 0.145 | 1.9368 | 0.08255 | 0.195 | -4.86 | -0.3257 | none |
| SLC22A11 | 0.336 | 0.2442 | 1.9366 | 0.08258 | 0.195 | -4.86 | -0.1166 | none |
| GALNT1 | 0.194 | 0.1702 | 1.9364 | 0.0826 | 0.195 | -4.86 | 0.09305 | none |
| RHOT1 | 0.382 | 0.3133 | 1.9362 | 0.08263 | 0.195 | -4.86 | -0.5811 | none |
| ASNSD1 | -0.196 | -0.1967 | -1.936 | 0.08264 | 0.195 | -4.86 | 1.52749 | none |
| EVI5L | 0.278 | 0.2283 | 1.9355 | 0.08273 | 0.195 | -4.87 | -0.9597 | none |
| ITPR2 | 0.575 | 0.6501 | 1.9352 | 0.08277 | 0.195 | -4.87 | -0.2632 | none |
| GABBR1 | -0.206 | -0.1389 | -1.935 | 0.08279 | 0.195 | -4.87 | 0.24789 | none |
| SPATA45 | -0.247 | -0.2996 | -1.935 | 0.08282 | 0.195 | -4.87 | -0.4168 | none |
| ABCA2 | 0.432 | 0.4242 | 1.9346 | 0.08284 | 0.195 | -4.87 | -0.4243 | none |
| OOEP | 0.316 | 0.1984 | 1.9343 | 0.08289 | 0.195 | -4.87 | -0.1275 | none |
| FKBP5 | 0.326 | 0.3259 | 1.9341 | 0.08291 | 0.195 | -4.87 | 0.18893 | none |
| ZBP1 | 0.316 | 0.2168 | 1.9335 | 0.08299 | 0.195 | -4.87 | -0.5049 | none |
| PLEKHF1 | 0.182 | 0.1504 | 1.9333 | 0.08303 | 0.195 | -4.87 | 0.56425 | none |
| NOP2 | 0.418 | 0.4853 | 1.9332 | 0.08304 | 0.195 | -4.87 | 0.12913 | none |
| YWHAE | -0.189 | -0.1542 | -1.933 | 0.08304 | 0.195 | -4.87 | 0.26249 | none |
| RPA3 | -0.455 | -0.5149 | -1.932 | 0.08314 | 0.196 | -4.87 | 1.32091 | none |
| AGO2 | 0.302 | 0.3543 | 1.9321 | 0.08319 | 0.196 | -4.87 | 0.27768 | none |
| EPM2A | 0.154 | 0.1127 | 1.932 | 0.0832 | 0.196 | -4.87 | -0.9014 | none |
| GALNT6 | 0.493 | 0.5167 | 1.9314 | 0.08328 | 0.196 | -4.87 | -0.6576 | none |
| ANO9 | 0.235 | 0.2681 | 1.9306 | 0.08339 | 0.196 | -4.87 | 0.22865 | none |
| C2ORF68 | 0.318 | 0.2107 | 1.9304 | 0.08342 | 0.196 | -4.87 | -0.9574 | none |
| NAPG | -0.194 | -0.1822 | -1.93 | 0.08343 | 0.196 | -4.87 | 1.09504 | none |
| SNAPIN | -0.214 | -0.2381 | -1.93 | 0.08349 | 0.196 | -4.87 | 2.31416 | AA-related |
| ZNF500 | 0.323 | 0.3216 | 1.9298 | 0.0835 | 0.196 | -4.87 | -0.4919 | none |
| RIBC1 | -0.335 | -0.1945 | -1.93 | 0.08351 | 0.196 | -4.87 | 1.19374 | none |
| STRC | 0.231 | 0.2161 | 1.9292 | 0.08359 | 0.196 | -4.88 | -0.3373 | none |
| CEP128 | 0.357 | 0.3359 | 1.9289 | 0.08363 | 0.196 | -4.88 | -0.275 | none |
| SLITRK2 | 0.152 | 0.1029 | 1.9287 | 0.08365 | 0.196 | -4.88 | -1.2355 | none |
| CLK1 | 0.209 | 0.2015 | 1.9287 | 0.08365 | 0.196 | -4.88 | 0.39109 | none |
| WDR47 | 0.178 | 0.1929 | 1.9285 | 0.08369 | 0.196 | -4.88 | -0.4545 | none |

| | | | | | | | | |
|---|---|---|---|---|---|---|---|---|
| ASTL | -0.201 | -0.1572 | -1.928 | 0.08372 | 0.197 | -4.88 | 1.9907 | none |
| RNASEH2C | -0.406 | -0.3676 | -1.928 | 0.08377 | 0.197 | -4.88 | 2.04649 | none |
| SUPT5H | 0.449 | 0.4706 | 1.9266 | 0.08394 | 0.197 | -4.88 | -0.5162 | none |
| FCRL1 | 0.322 | 0.2483 | 1.9264 | 0.08396 | 0.197 | -4.88 | -0.2313 | none |
| THEM5 | -0.483 | -0.2635 | -1.926 | 0.08397 | 0.197 | -4.88 | 1.59352 | none |
| LRP2 | 0.198 | 0.1924 | 1.9263 | 0.08399 | 0.197 | -4.88 | 0.48378 | none |
| PCP2 | -0.304 | -0.2324 | -1.926 | 0.08401 | 0.197 | -4.88 | 1.07521 | none |
| CSGALNAC | 0.199 | 0.1739 | 1.926 | 0.08402 | 0.197 | -4.88 | -0.3792 | none |
| SLC16A2 | 0.156 | 0.1154 | 1.926 | 0.08402 | 0.197 | -4.88 | -1.353 | none |
| RRP1 | 0.286 | 0.3306 | 1.9256 | 0.08408 | 0.197 | -4.88 | 0.12343 | none |
| ITM2A | -0.293 | -0.3757 | -1.926 | 0.08408 | 0.197 | -4.88 | 1.25182 | none |
| PTPRB | 0.326 | 0.2299 | 1.9255 | 0.08409 | 0.197 | -4.88 | -1.3892 | none |
| SNRPN | -0.232 | -0.2628 | -1.925 | 0.08414 | 0.197 | -4.88 | 2.23552 | AA-related |
| PPP4R2 | 0.312 | 0.2812 | 1.925 | 0.08416 | 0.197 | -4.88 | -1.1492 | none |
| TAF13 | 0.373 | 0.4118 | 1.9247 | 0.08421 | 0.197 | -4.88 | -0.5157 | none |
| IKBIP | 0.329 | 0.2826 | 1.9246 | 0.08421 | 0.197 | -4.88 | -0.7103 | none |
| RPL10 | -0.335 | -0.3565 | -1.925 | 0.08422 | 0.197 | -4.88 | 2.08177 | none |
| CENPVL1 | 0.136 | 0.1055 | 1.9245 | 0.08424 | 0.197 | -4.88 | -0.3228 | none |
| CENPVL2 | 0.136 | 0.1055 | 1.9245 | 0.08424 | 0.197 | -4.88 | -0.3228 | none |
| TDP2 | -0.29 | -0.2829 | -1.924 | 0.08425 | 0.197 | -4.88 | 1.70706 | none |
| RGS6 | -0.271 | -0.2176 | -1.924 | 0.08428 | 0.197 | -4.88 | 1.8267 | none |
| GGCT | -0.134 | -0.1005 | -1.924 | 0.0843 | 0.197 | -4.88 | 0.76315 | none |
| CLCNKA | 0.189 | 0.189 | 1.924 | 0.0843 | 0.197 | -4.88 | 0.55525 | none |
| SMPD1 | -0.262 | -0.2445 | -1.924 | 0.08432 | 0.197 | -4.88 | -0.0737 | none |
| TGFBR1 | 0.174 | 0.0935 | 1.9232 | 0.08441 | 0.197 | -4.88 | 1.00557 | none |
| CAPRIN1 | 0.354 | 0.358 | 1.9227 | 0.08448 | 0.198 | -4.89 | 0.64313 | none |
| SMG6 | 0.29 | 0.2725 | 1.9225 | 0.0845 | 0.198 | -4.89 | 0.03867 | none |
| SNRNP40 | -0.164 | -0.148 | -1.922 | 0.08451 | 0.198 | -4.89 | 0.05821 | none |
| CEP290 | -0.317 | -0.2711 | -1.922 | 0.08452 | 0.198 | -4.89 | 1.70678 | none |
| WIPF1 | 0.328 | 0.3036 | 1.9223 | 0.08453 | 0.198 | -4.89 | -0.6303 | none |
| RAB30 | 0.402 | 0.2513 | 1.9218 | 0.0846 | 0.198 | -4.89 | -0.6053 | none |
| KCTD7 | 0.28 | 0.2283 | 1.9216 | 0.08463 | 0.198 | -4.89 | 0.61865 | none |
| CALCOCO1 | 0.476 | 0.4306 | 1.9214 | 0.08465 | 0.198 | -4.89 | -1.3942 | none |
| CD38 | -0.264 | -0.288 | -1.921 | 0.08467 | 0.198 | -4.89 | 2.00681 | none |
| NELFE | -0.155 | -0.123 | -1.921 | 0.08467 | 0.198 | -4.89 | 1.94655 | none |
| JAG2 | 0.339 | 0.3093 | 1.9202 | 0.08482 | 0.198 | -4.89 | 0.77115 | none |
| ATXN7L3B | 0.177 | 0.124 | 1.92 | 0.08485 | 0.198 | -4.89 | 0.49501 | none |
| PRTFDC1 | -0.247 | -0.2124 | -1.92 | 0.08487 | 0.198 | -4.89 | 0.07047 | none |
| EIF1AX | -0.153 | -0.1193 | -1.919 | 0.08494 | 0.198 | -4.89 | 1.13298 | none |
| PRTG | -0.225 | -0.175 | -1.919 | 0.08494 | 0.198 | -4.89 | 0.60489 | none |

| | | | | | | | | |
|---|---|---|---|---|---|---|---|---|
| SULT1E1 | 0.191 | 0.1809 | 1.9193 | 0.08495 | 0.198 | -4.89 | 1.12657 | none |
| ALDH3A1 | 0.405 | 0.4143 | 1.9192 | 0.08497 | 0.198 | -4.89 | -0.2545 | none |
| SPATA8 | 0.344 | 0.2291 | 1.9185 | 0.08506 | 0.198 | -4.89 | -0.9201 | none |
| SUFU | 0.274 | 0.2982 | 1.9184 | 0.08507 | 0.198 | -4.89 | -1.1325 | none |
| C5ORF24 | 0.313 | 0.3439 | 1.9184 | 0.08507 | 0.198 | -4.89 | 0.89652 | none |
| CEP55 | 0.497 | 0.4654 | 1.9184 | 0.08508 | 0.198 | -4.89 | -1.0519 | none |
| DDX59 | 0.314 | 0.2887 | 1.9183 | 0.08509 | 0.198 | -4.89 | 1.03413 | none |
| ADGRD1 | 0.207 | 0.2124 | 1.9182 | 0.0851 | 0.198 | -4.89 | -0.4331 | none |
| SLMAP | 0.559 | 0.5647 | 1.9182 | 0.0851 | 0.198 | -4.89 | 0.3609 | none |
| MAPKBP1 | 0.315 | 0.318 | 1.9177 | 0.08517 | 0.199 | -4.89 | -0.6698 | none |
| TJP3 | 0.367 | 0.2475 | 1.9176 | 0.08518 | 0.199 | -4.89 | -0.0951 | none |
| NDST4 | 0.213 | 0.1418 | 1.9173 | 0.08523 | 0.199 | -4.89 | -0.2241 | none |
| LRRC75A | -0.326 | -0.3677 | -1.917 | 0.08524 | 0.199 | -4.89 | 1.23406 | none |
| AFF4 | 0.664 | 0.6627 | 1.9171 | 0.08525 | 0.199 | -4.89 | -0.8473 | none |
| SLC35B1 | -0.321 | -0.3272 | -1.917 | 0.08526 | 0.199 | -4.89 | 1.51691 | none |
| FBXO2 | 0.394 | 0.3555 | 1.9168 | 0.0853 | 0.199 | -4.89 | -0.3605 | none |
| RPS15A | -0.428 | -0.4993 | -1.917 | 0.08534 | 0.199 | -4.89 | 2.01025 | none |
| SMIM11A | 0.287 | 0.1237 | 1.9163 | 0.08537 | 0.199 | -4.89 | 0.55511 | none |
| RNF24 | 0.263 | 0.2081 | 1.9159 | 0.08542 | 0.199 | -4.9 | -0.4162 | none |
| MYO7B | -0.231 | -0.1937 | -1.916 | 0.08543 | 0.199 | -4.9 | 2.337 | AA-related |
| UBR2 | 0.366 | 0.3458 | 1.9159 | 0.08543 | 0.199 | -4.9 | -0.7066 | none |
| GLA | -0.29 | -0.3305 | -1.916 | 0.08547 | 0.199 | -4.9 | 1.31708 | none |
| ACVR1C | 0.139 | 0.1261 | 1.9154 | 0.0855 | 0.199 | -4.9 | -0.2929 | none |
| PIM3 | -0.25 | -0.1857 | -1.915 | 0.08553 | 0.199 | -4.9 | 0.55224 | none |
| SPIN2B | 0.398 | 0.2801 | 1.9148 | 0.08558 | 0.199 | -4.9 | -0.6053 | none |
| NUP62 | 0.44 | 0.4917 | 1.9146 | 0.0856 | 0.199 | -4.9 | -0.6558 | none |
| TDRD1 | 0.205 | 0.1883 | 1.9145 | 0.08563 | 0.199 | -4.9 | -0.7146 | none |
| HIC2 | 0.198 | 0.2137 | 1.9143 | 0.08565 | 0.199 | -4.9 | -0.2961 | none |
| MOSPD1 | 0.306 | 0.1299 | 1.9142 | 0.08566 | 0.199 | -4.9 | 0.31218 | none |
| VWA8 | 0.494 | 0.4192 | 1.9139 | 0.0857 | 0.199 | -4.9 | 1.18655 | none |
| SYCE1L | -0.351 | -0.2475 | -1.914 | 0.08572 | 0.199 | -4.9 | 0.76568 | none |
| ARHGAP3: | 0.481 | 0.4774 | 1.9124 | 0.08591 | 0.2 | -4.9 | -0.8334 | none |
| DCAF6 | 0.338 | 0.2804 | 1.9118 | 0.086 | 0.2 | -4.9 | -0.0254 | none |
| STX3 | 0.37 | 0.2899 | 1.9117 | 0.08601 | 0.2 | -4.9 | -0.7094 | none |
| ZBTB40 | 0.409 | 0.4365 | 1.9117 | 0.08601 | 0.2 | -4.9 | -0.1767 | none |
| KIF21B | 0.49 | 0.5937 | 1.9117 | 0.08602 | 0.2 | -4.9 | -0.3611 | none |
| PXYLP1 | 0.406 | 0.2101 | 1.9115 | 0.08605 | 0.2 | -4.9 | -0.5884 | none |
| HSPB9 | 0.203 | 0.1744 | 1.9109 | 0.08613 | 0.2 | -4.9 | -1.0318 | none |
| PCDHB2 | 0.352 | 0.2597 | 1.9108 | 0.08615 | 0.2 | -4.9 | -1.2798 | none |
| RAB40C | 0.263 | 0.2498 | 1.9107 | 0.08616 | 0.2 | -4.9 | 0.27851 | none |

| | | | | | | | | |
|---|---|---|---|---|---|---|---|---|
| KIAA1549 | 0.178 | 0.1574 | 1.9106 | 0.08617 | 0.2 | -4.9 | -1.0247 | none |
| ZNF10 | 0.313 | 0.2455 | 1.9106 | 0.08617 | 0.2 | -4.9 | 0.72671 | none |
| ZFY | 0.266 | 0.2279 | 1.9106 | 0.08617 | 0.2 | -4.9 | -1.2669 | none |
| KIAA1210 | 0.169 | 0.12 | 1.9103 | 0.08621 | 0.2 | -4.9 | -1.3647 | none |
| TEP1 | 0.465 | 0.5591 | 1.9103 | 0.08621 | 0.2 | -4.9 | -0.1493 | none |
| SSX1 | -0.19 | -0.1816 | -1.91 | 0.08623 | 0.2 | -4.9 | 0.89986 | none |
| TMTC2 | 0.177 | 0.1428 | 1.9102 | 0.08623 | 0.2 | -4.9 | -0.3813 | none |
| APOH | 0.761 | 0.4526 | 1.91 | 0.08626 | 0.2 | -4.9 | -0.9593 | none |
| ACTR1A | -0.168 | -0.1132 | -1.91 | 0.08626 | 0.2 | -4.9 | 1.01629 | none |
| SORCS3 | -0.224 | -0.2243 | -1.91 | 0.08626 | 0.2 | -4.9 | 1.51352 | none |
| IDH1 | -0.216 | -0.2658 | -1.91 | 0.08627 | 0.2 | -4.9 | 0.50195 | none |
| PWP2 | 0.494 | 0.4583 | 1.9081 | 0.08651 | 0.2 | -4.91 | -0.0598 | none |
| LONP2 | 0.302 | 0.3485 | 1.9081 | 0.08653 | 0.2 | -4.91 | -0.4481 | none |
| AHCY | -0.206 | -0.1783 | -1.908 | 0.08653 | 0.2 | -4.91 | 2.48093 | AA-related |
| MRPL58 | -0.201 | -0.2 | -1.908 | 0.08655 | 0.2 | -4.91 | 1.99611 | none |
| CNPPD1 | 0.168 | 0.1553 | 1.9079 | 0.08655 | 0.2 | -4.91 | 0.22532 | none |
| RPUSD2 | 0.223 | 0.2151 | 1.9079 | 0.08655 | 0.2 | -4.91 | -0.6534 | none |
| CACNA2D3 | 0.384 | 0.3092 | 1.9078 | 0.08656 | 0.2 | -4.91 | -1.1828 | none |
| SERPINB7 | -0.23 | -0.2299 | -1.908 | 0.08658 | 0.201 | -4.91 | 1.65197 | none |
| MRPL41 | -0.331 | -0.2908 | -1.908 | 0.0866 | 0.201 | -4.91 | 1.31077 | none |
| ANK1 | 0.251 | 0.3385 | 1.9074 | 0.08662 | 0.201 | -4.91 | -0.0553 | none |
| RTN4RL2 | 0.265 | 0.1943 | 1.9073 | 0.08664 | 0.201 | -4.91 | 0.61214 | none |
| DHX57 | 0.368 | 0.3782 | 1.9064 | 0.08676 | 0.201 | -4.91 | -1.0223 | none |
| MISP3 | -0.155 | -0.1722 | -1.906 | 0.08686 | 0.201 | -4.91 | 1.78611 | none |
| INTS9 | 0.3 | 0.2555 | 1.9054 | 0.0869 | 0.201 | -4.91 | -0.4045 | none |
| NMRAL1 | 0.242 | 0.1403 | 1.9049 | 0.08698 | 0.201 | -4.91 | -0.1076 | none |
| DCTPP1 | -0.307 | -0.2793 | -1.905 | 0.08699 | 0.201 | -4.91 | 1.19517 | none |
| RPS3A | -0.256 | -0.2606 | -1.905 | 0.08701 | 0.201 | -4.91 | 0.53306 | none |
| TMEM87B | -0.33 | -0.281 | -1.904 | 0.08712 | 0.201 | -4.91 | 1.64213 | none |
| SLC35G2 | 0.193 | 0.0673 | 1.9038 | 0.08713 | 0.201 | -4.91 | -0.0103 | none |
| ZNF777 | 0.347 | 0.4123 | 1.9035 | 0.08717 | 0.202 | -4.91 | -0.3332 | none |
| CTTNBP2 | 0.362 | 0.2362 | 1.9035 | 0.08718 | 0.202 | -4.91 | -1.3693 | none |
| AKR1B1 | 0.192 | 0.084 | 1.9028 | 0.08728 | 0.202 | -4.91 | -0.0963 | none |
| BANF2 | 0.219 | 0.1572 | 1.9024 | 0.08733 | 0.202 | -4.92 | 0.80366 | none |
| EMC7 | -0.337 | -0.3414 | -1.902 | 0.08735 | 0.202 | -4.92 | 1.35126 | none |
| CDC42 | -0.313 | -0.3818 | -1.902 | 0.08737 | 0.202 | -4.92 | 0.56091 | none |
| GPKOW | 0.252 | 0.3007 | 1.9021 | 0.08737 | 0.202 | -4.92 | -0.6205 | none |
| TRIP12 | 0.66 | 0.725 | 1.9017 | 0.08743 | 0.202 | -4.92 | -0.449 | none |
| CD6 | 0.236 | 0.2031 | 1.9016 | 0.08745 | 0.202 | -4.92 | -0.3119 | none |
| LUC7L2 | 0.344 | 0.4183 | 1.9016 | 0.08745 | 0.202 | -4.92 | 0.27759 | none |

| | | | | | | | | |
|---|---|---|---|---|---|---|---|---|
| ZNF770 | -0.263 | -0.2514 | -1.902 | 0.08745 | 0.202 | -4.92 | 1.49254 | none |
| SLC23A2 | 0.234 | 0.2221 | 1.9015 | 0.08746 | 0.202 | -4.92 | 0.72713 | none |
| FOPNL | -0.272 | -0.2436 | -1.901 | 0.08749 | 0.202 | -4.92 | 0.89087 | none |
| ZNF76 | 0.299 | 0.3248 | 1.9011 | 0.08752 | 0.202 | -4.92 | -0.3282 | none |
| CDC42EP3 | 0.345 | 0.2845 | 1.9008 | 0.08756 | 0.202 | -4.92 | -1.0623 | none |
| PAIP2B | 0.146 | 0.094 | 1.9008 | 0.08756 | 0.202 | -4.92 | -1.2358 | none |
| ZC3H3 | 0.283 | 0.2693 | 1.9004 | 0.08761 | 0.202 | -4.92 | -0.3629 | none |
| ITFG1 | 0.194 | 0.1594 | 1.9001 | 0.08766 | 0.202 | -4.92 | -0.4608 | none |
| LYST | 0.269 | 0.1409 | 1.9 | 0.08768 | 0.202 | -4.92 | -0.5447 | none |
| CGA | 0.385 | 0.2552 | 1.8998 | 0.0877 | 0.202 | -4.92 | -0.107 | none |
| FBRS | 0.403 | 0.4658 | 1.8997 | 0.08771 | 0.202 | -4.92 | -0.0142 | none |
| SPSB4 | 0.278 | 0.2137 | 1.8997 | 0.08772 | 0.202 | -4.92 | 0.41917 | none |
| PARG | 0.504 | 0.5148 | 1.8997 | 0.08772 | 0.202 | -4.92 | -0.28 | none |
| PDE4D | 0.234 | 0.1925 | 1.8995 | 0.08775 | 0.202 | -4.92 | -0.4037 | none |
| ACO2 | 0.322 | 0.3415 | 1.8994 | 0.08776 | 0.202 | -4.92 | -0.2397 | none |
| NUP85 | 0.201 | 0.1539 | 1.8991 | 0.0878 | 0.202 | -4.92 | -0.7868 | none |
| ZNF790 | 0.309 | 0.2503 | 1.899 | 0.08782 | 0.202 | -4.92 | -0.6164 | none |
| CRTC1 | 0.329 | 0.4029 | 1.8989 | 0.08783 | 0.202 | -4.92 | -0.4756 | none |
| TMEM52 | 0.309 | 0.1704 | 1.8989 | 0.08784 | 0.202 | -4.92 | -0.9151 | none |
| COMMD4 | 0.284 | 0.1646 | 1.8988 | 0.08784 | 0.202 | -4.92 | -0.8199 | none |
| RNF19A | 0.477 | 0.4147 | 1.8987 | 0.08786 | 0.202 | -4.92 | -0.8392 | none |
| LIN54 | 0.492 | 0.444 | 1.8985 | 0.08789 | 0.202 | -4.92 | -0.6975 | none |
| GAGE12B | -1.164 | -0.8074 | -1.898 | 0.08793 | 0.202 | -4.92 | 2.14239 | AA-related |
| UBE2I | -0.217 | -0.2033 | -1.898 | 0.08795 | 0.202 | -4.92 | 1.66439 | none |
| PLD2 | 0.462 | 0.3492 | 1.8974 | 0.08804 | 0.203 | -4.92 | -1.0551 | none |
| TBC1D2 | 0.264 | 0.2474 | 1.8972 | 0.08808 | 0.203 | -4.92 | 0.51838 | none |
| RAB9A | -0.334 | -0.2708 | -1.897 | 0.08808 | 0.203 | -4.92 | 1.90755 | none |
| REST | 0.699 | 0.657 | 1.8969 | 0.08812 | 0.203 | -4.92 | -0.6427 | none |
| TMEM33 | 0.332 | 0.1617 | 1.8968 | 0.08813 | 0.203 | -4.92 | -0.7812 | none |
| SACS | 0.422 | 0.4403 | 1.8966 | 0.08817 | 0.203 | -4.92 | 0.59195 | none |
| PCBP3 | 0.389 | 0.3816 | 1.8965 | 0.08818 | 0.203 | -4.92 | 0.2658 | none |
| GID8 | -0.181 | -0.1011 | -1.896 | 0.08819 | 0.203 | -4.92 | 2.18408 | AA-related |
| CNN3 | 0.307 | 0.2393 | 1.896 | 0.08825 | 0.203 | -4.92 | -0.0462 | none |
| SMN2 | 0.697 | 0.4613 | 1.8958 | 0.08828 | 0.203 | -4.93 | -0.5636 | none |
| PADI2 | 0.168 | 0.1217 | 1.8956 | 0.08831 | 0.203 | -4.93 | -1.0781 | none |
| SLC22A17 | 0.425 | 0.3773 | 1.8955 | 0.08832 | 0.203 | -4.93 | -0.1129 | none |
| RBM19 | 0.68 | 0.8063 | 1.8955 | 0.08832 | 0.203 | -4.93 | -0.4893 | none |
| GCKR | 0.312 | 0.3227 | 1.8955 | 0.08833 | 0.203 | -4.93 | -0.3923 | none |
| RGS12 | 0.339 | 0.2725 | 1.8951 | 0.08837 | 0.203 | -4.93 | -0.9957 | none |
| HIST1H3I | -1.116 | -0.9245 | -1.895 | 0.08843 | 0.203 | -4.93 | 0.86908 | none |

| | | | | | | | | |
|---|---|---|---|---|---|---|---|---|
| GPR37L1 | 0.221 | 0.1501 | 1.8944 | 0.08848 | 0.203 | -4.93 | -1.3644 | none |
| HMGN1 | -0.155 | -0.1223 | -1.894 | 0.08851 | 0.203 | -4.93 | 1.89373 | none |
| NAA40 | 0.238 | 0.1825 | 1.8937 | 0.08859 | 0.203 | -4.93 | -0.7221 | none |
| WAC | 0.436 | 0.4337 | 1.8936 | 0.08859 | 0.203 | -4.93 | -0.4073 | none |
| MIS18BP1 | 0.604 | 0.6779 | 1.8933 | 0.08863 | 0.203 | -4.93 | -0.5498 | none |
| TBCB | 0.234 | 0.1351 | 1.8933 | 0.08864 | 0.203 | -4.93 | -0.5216 | none |
| ZMYM3 | 0.283 | 0.3268 | 1.8932 | 0.08865 | 0.203 | -4.93 | 0.40247 | none |
| UACA | 0.292 | 0.2071 | 1.8931 | 0.08866 | 0.203 | -4.93 | 0.59992 | none |
| IQCF2 | 0.355 | 0.2341 | 1.8931 | 0.08867 | 0.203 | -4.93 | -0.9413 | none |
| OXNAD1 | -0.25 | -0.197 | -1.893 | 0.0887 | 0.203 | -4.93 | 2.05005 | none |
| LNPK | -0.233 | -0.295 | -1.892 | 0.08877 | 0.204 | -4.93 | 1.034 | none |
| RNF167 | -0.178 | -0.1806 | -1.892 | 0.08882 | 0.204 | -4.93 | 0.92426 | none |
| CDK19 | 0.377 | 0.3276 | 1.892 | 0.08883 | 0.204 | -4.93 | -1.125 | none |
| WDR6 | 0.179 | 0.1937 | 1.8919 | 0.08884 | 0.204 | -4.93 | 0.64966 | none |
| STAG3 | 0.253 | 0.188 | 1.8916 | 0.08888 | 0.204 | -4.93 | 0.98613 | none |
| EXOC3 | 0.329 | 0.3333 | 1.8916 | 0.08889 | 0.204 | -4.93 | -0.0601 | none |
| GPATCH11 | 0.315 | 0.2831 | 1.8913 | 0.08892 | 0.204 | -4.93 | -0.7418 | none |
| DSN1 | 0.185 | 0.0709 | 1.8911 | 0.08895 | 0.204 | -4.93 | -0.164 | none |
| ZBTB25 | 0.217 | 0.1799 | 1.8908 | 0.089 | 0.204 | -4.93 | -0.3101 | none |
| KCNJ13 | 0.47 | 0.3936 | 1.8898 | 0.08915 | 0.204 | -4.93 | -0.9125 | none |
| KLHL33 | 0.155 | 0.1162 | 1.8897 | 0.08916 | 0.204 | -4.93 | 0.1446 | none |
| RAB23 | -0.243 | -0.2029 | -1.889 | 0.08927 | 0.204 | -4.94 | 1.20071 | none |
| MYO1C | 0.369 | 0.406 | 1.8888 | 0.08929 | 0.204 | -4.94 | -0.6152 | none |
| KTN1 | 0.648 | 0.7172 | 1.8884 | 0.08935 | 0.205 | -4.94 | 0.21473 | none |
| MRM1 | 0.274 | 0.2899 | 1.8883 | 0.08936 | 0.205 | -4.94 | 0.73808 | none |
| PKN3 | 0.191 | 0.2244 | 1.8872 | 0.08952 | 0.205 | -4.94 | 0.73462 | none |
| CHI3L2 | -0.385 | -0.3704 | -1.887 | 0.08955 | 0.205 | -4.94 | 0.09508 | none |
| FAR1 | 0.418 | 0.3148 | 1.887 | 0.08956 | 0.205 | -4.94 | -0.3529 | none |
| PLXNC1 | 0.586 | 0.5248 | 1.8869 | 0.08957 | 0.205 | -4.94 | -0.7277 | none |
| CCDC152 | 0.274 | 0.2215 | 1.8865 | 0.08963 | 0.205 | -4.94 | -0.9423 | none |
| PPDPF | 0.34 | 0.2418 | 1.8862 | 0.08967 | 0.205 | -4.94 | -0.333 | none |
| NDUFS4 | -0.309 | -0.3898 | -1.886 | 0.08969 | 0.205 | -4.94 | 0.46399 | none |
| CHD8 | 0.532 | 0.5933 | 1.8856 | 0.08976 | 0.205 | -4.94 | -0.6501 | none |
| DOLPP1 | 0.173 | 0.1597 | 1.8851 | 0.08983 | 0.205 | -4.94 | -0.4031 | none |
| SETD2 | 0.558 | 0.6504 | 1.8849 | 0.08986 | 0.205 | -4.94 | -0.0247 | none |
| TRPC3 | 0.227 | 0.1474 | 1.8848 | 0.08987 | 0.205 | -4.94 | -1.2522 | none |
| CD55 | 0.18 | 0.0811 | 1.8847 | 0.08989 | 0.205 | -4.94 | -0.7788 | none |
| ELK1 | -0.247 | -0.2152 | -1.885 | 0.08989 | 0.205 | -4.94 | 2.31183 | AA-related |
| P2RY14 | -0.247 | -0.267 | -1.884 | 0.08996 | 0.205 | -4.94 | 0.32043 | none |
| SETD1B | 0.422 | 0.4764 | 1.884 | 0.08999 | 0.206 | -4.94 | -0.4365 | none |

| | | | | | | | | |
|---|---|---|---|---|---|---|---|---|
| SERPINB1 | 0.15 | 0.0485 | 1.8835 | 0.09007 | 0.206 | -4.94 | 0.28588 | none |
| CERS1 | 0.217 | 0.1643 | 1.8832 | 0.0901 | 0.206 | -4.94 | -0.387 | none |
| KIAA0825 | 0.202 | 0.1593 | 1.8827 | 0.09018 | 0.206 | -4.94 | -0.6417 | none |
| SEC63 | 0.427 | 0.4944 | 1.8825 | 0.09022 | 0.206 | -4.94 | -0.5929 | none |
| IL22 | -0.357 | -0.2621 | -1.882 | 0.09024 | 0.206 | -4.95 | 1.18411 | none |
| RNASEK-C | -0.371 | -0.3735 | -1.882 | 0.09026 | 0.206 | -4.95 | -0.5086 | none |
| GIMAP6 | -0.18 | -0.2029 | -1.881 | 0.09036 | 0.206 | -4.95 | 1.8041 | none |
| RGS11 | 0.213 | 0.1166 | 1.8815 | 0.09036 | 0.206 | -4.95 | 0.4077 | none |
| HERPUD1 | 0.149 | 0.0877 | 1.8814 | 0.09038 | 0.206 | -4.95 | -0.9397 | none |
| ETNK1 | 0.294 | 0.2038 | 1.8813 | 0.09038 | 0.206 | -4.95 | -0.6047 | none |
| PPP1R13B | 0.487 | 0.4279 | 1.881 | 0.09044 | 0.206 | -4.95 | -0.9299 | none |
| EDEM1 | 0.212 | 0.2116 | 1.8808 | 0.09047 | 0.206 | -4.95 | -0.7451 | none |
| HHIP | 0.328 | 0.2471 | 1.8802 | 0.09055 | 0.206 | -4.95 | -0.019 | none |
| FGD3 | 0.444 | 0.4369 | 1.88 | 0.09058 | 0.207 | -4.95 | -0.4562 | none |
| UBE2F-SCI | 0.207 | 0.2525 | 1.8799 | 0.09059 | 0.207 | -4.95 | -0.4125 | none |
| ZNF555 | 0.365 | 0.3321 | 1.8795 | 0.09066 | 0.207 | -4.95 | -0.3944 | none |
| TTPA | 0.14 | 0.1184 | 1.8794 | 0.09067 | 0.207 | -4.95 | -0.6178 | none |
| GTF3C1 | 0.437 | 0.5024 | 1.8791 | 0.09071 | 0.207 | -4.95 | -0.4674 | none |
| IFI6 | -0.363 | -0.4298 | -1.879 | 0.09076 | 0.207 | -4.95 | 0.99915 | none |
| C1QBP | -0.298 | -0.2668 | -1.879 | 0.09079 | 0.207 | -4.95 | 1.42553 | none |
| PDZD8 | 0.483 | 0.4739 | 1.8782 | 0.09085 | 0.207 | -4.95 | -0.9939 | none |
| C14ORF13 | 0.238 | 0.1997 | 1.8774 | 0.09097 | 0.207 | -4.95 | -0.2781 | none |
| PAQR4 | -0.164 | -0.0427 | -1.877 | 0.09098 | 0.207 | -4.95 | 1.59318 | none |
| SLC25A4 | -0.243 | -0.2334 | -1.877 | 0.09103 | 0.207 | -4.95 | 2.22516 | AA-related |
| CASP6 | 0.249 | 0.1429 | 1.8767 | 0.09106 | 0.207 | -4.95 | -0.5697 | none |
| NFKB2 | 0.37 | 0.4017 | 1.8765 | 0.0911 | 0.207 | -4.95 | -0.9917 | none |
| SLC7A5 | -0.176 | -0.0522 | -1.876 | 0.09112 | 0.207 | -4.95 | 0.93593 | none |
| LY6K | 0.281 | 0.2235 | 1.8763 | 0.09112 | 0.207 | -4.95 | -1.2576 | none |
| PRODH | 0.333 | 0.2642 | 1.8762 | 0.09114 | 0.207 | -4.95 | -1.0997 | none |
| NUBP1 | -0.142 | -0.0867 | -1.875 | 0.09128 | 0.208 | -4.96 | 0.27799 | none |
| TM4SF5 | -0.153 | -0.085 | -1.875 | 0.09136 | 0.208 | -4.96 | 0.67925 | none |
| CRYBB1 | -0.293 | -0.2029 | -1.875 | 0.09137 | 0.208 | -4.96 | 0.59095 | none |
| ANXA5 | -0.187 | -0.2247 | -1.875 | 0.09137 | 0.208 | -4.96 | 1.66709 | none |
| PSAT1 | -0.191 | -0.2297 | -1.875 | 0.09139 | 0.208 | -4.96 | 1.55352 | none |
| SETX | 0.589 | 0.6591 | 1.8745 | 0.09139 | 0.208 | -4.96 | -0.2117 | none |
| AGR3 | 0.217 | 0.2171 | 1.8743 | 0.09142 | 0.208 | -4.96 | 0.94619 | none |
| RPL6 | -0.192 | -0.1124 | -1.874 | 0.09143 | 0.208 | -4.96 | 2.14451 | AA-related |
| CALM1 | -0.262 | -0.1899 | -1.873 | 0.09157 | 0.208 | -4.96 | 0.99037 | none |
| TIGD5 | 0.154 | 0.1212 | 1.8733 | 0.09157 | 0.208 | -4.96 | -1.1976 | none |
| RECQL4 | 0.406 | 0.4647 | 1.873 | 0.09162 | 0.208 | -4.96 | -0.0798 | none |

| | | | | | | | | |
|---|---|---|---|---|---|---|---|---|
| SEMA3A | -0.398 | -0.2692 | -1.873 | 0.09164 | 0.208 | -4.96 | 1.10395 | none |
| DMXL1 | 0.406 | 0.4135 | 1.8728 | 0.09165 | 0.208 | -4.96 | -0.131 | none |
| PGF | 0.183 | 0.1182 | 1.8726 | 0.09168 | 0.208 | -4.96 | -0.6483 | none |
| RGL2 | 0.216 | 0.1875 | 1.8724 | 0.0917 | 0.208 | -4.96 | 0.46647 | none |
| CENPC | 0.502 | 0.4632 | 1.8722 | 0.09174 | 0.208 | -4.96 | -0.2112 | none |
| BCL2L2-PA | -0.343 | -0.3541 | -1.872 | 0.09177 | 0.208 | -4.96 | 0.97302 | none |
| WTAP | 0.156 | 0.0926 | 1.8713 | 0.09188 | 0.209 | -4.96 | -0.0297 | none |
| MYCBP2 | 0.543 | 0.5903 | 1.8712 | 0.09188 | 0.209 | -4.96 | -0.4069 | none |
| ASPN | -0.125 | -0.0855 | -1.871 | 0.09191 | 0.209 | -4.96 | 0.99173 | none |
| EPO | 0.18 | 0.1346 | 1.8708 | 0.09195 | 0.209 | -4.96 | -0.3859 | none |
| ZNF350 | 0.293 | 0.1879 | 1.8707 | 0.09196 | 0.209 | -4.96 | -0.8339 | none |
| YARS2 | 0.164 | 0.102 | 1.8706 | 0.09198 | 0.209 | -4.96 | 0.83596 | none |
| MAFF | 0.291 | 0.2914 | 1.8699 | 0.09208 | 0.209 | -4.96 | -0.0822 | none |
| KLF5 | 0.298 | 0.3906 | 1.8697 | 0.09212 | 0.209 | -4.96 | -0.3332 | none |
| TBC1D2B | 0.362 | 0.2837 | 1.8696 | 0.09213 | 0.209 | -4.96 | 0.19404 | none |
| DEAF1 | 0.334 | 0.3313 | 1.8694 | 0.09216 | 0.209 | -4.96 | -0.2803 | none |
| WBP2NL | 0.417 | 0.2371 | 1.8691 | 0.09221 | 0.209 | -4.96 | -1.1171 | none |
| NFRKB | 0.442 | 0.5382 | 1.8689 | 0.09223 | 0.209 | -4.96 | -0.2294 | none |
| FAM72B | 0.386 | 0.3825 | 1.8681 | 0.09234 | 0.209 | -4.97 | -0.9937 | none |
| STK4 | 0.449 | 0.4244 | 1.868 | 0.09237 | 0.209 | -4.97 | -0.2784 | none |
| SLX4IP | 0.274 | 0.2309 | 1.8678 | 0.0924 | 0.209 | -4.97 | -1.1245 | none |
| SECISBP2 | 0.455 | 0.4775 | 1.8676 | 0.09242 | 0.209 | -4.97 | -0.5193 | none |
| COA3 | -0.244 | -0.2689 | -1.868 | 0.09243 | 0.209 | -4.97 | 0.41757 | none |
| COL4A2 | 0.284 | 0.2699 | 1.8676 | 0.09243 | 0.209 | -4.97 | -0.5279 | none |
| FARP2 | 0.444 | 0.4754 | 1.867 | 0.09251 | 0.209 | -4.97 | -1.0789 | none |
| VWC2 | 0.145 | 0.1138 | 1.8667 | 0.09257 | 0.21 | -4.97 | -1.2093 | none |
| CTLA4 | -0.186 | -0.1468 | -1.866 | 0.0926 | 0.21 | -4.97 | 0.80358 | none |
| LAG3 | 0.327 | 0.3974 | 1.8664 | 0.09261 | 0.21 | -4.97 | -0.1787 | none |
| NXT1 | -0.307 | -0.2626 | -1.865 | 0.09275 | 0.21 | -4.97 | 1.49183 | none |
| RNF25 | 0.35 | 0.3268 | 1.8654 | 0.09276 | 0.21 | -4.97 | -1.2061 | none |
| NR0B1 | -0.279 | -0.2322 | -1.865 | 0.09277 | 0.21 | -4.97 | 0.88317 | none |
| TAS2R38 | 0.213 | 0.1506 | 1.8651 | 0.0928 | 0.21 | -4.97 | -0.7915 | none |
| RASSF2 | 0.346 | 0.3089 | 1.8649 | 0.09284 | 0.21 | -4.97 | -1.0529 | none |
| PCDH15 | 0.161 | 0.1064 | 1.8648 | 0.09285 | 0.21 | -4.97 | -0.1864 | none |
| GRAMD4 | 0.371 | 0.4347 | 1.8646 | 0.09288 | 0.21 | -4.97 | -0.4909 | none |
| BABAM1 | 0.208 | 0.1 | 1.8643 | 0.09292 | 0.21 | -4.97 | -0.3614 | none |
| MT-ND4 | 0.485 | 0.5272 | 1.8642 | 0.09294 | 0.21 | -4.97 | -0.2379 | none |
| NOC3L | 0.443 | 0.4588 | 1.8642 | 0.09295 | 0.21 | -4.97 | -0.0657 | none |
| ZNF800 | 0.553 | 0.4942 | 1.8637 | 0.09301 | 0.21 | -4.97 | 0.40577 | none |
| PI4KA | 0.457 | 0.5223 | 1.8637 | 0.09302 | 0.21 | -4.97 | 0.01984 | none |

| | | | | | | | | |
|---|---|---|---|---|---|---|---|---|
| RNF175 | 0.33 | 0.1756 | 1.8637 | 0.09302 | 0.21 | -4.97 | -0.1758 | none |
| AC005042 | 0.263 | 0.1644 | 1.8634 | 0.09307 | 0.21 | -4.97 | -0.1272 | none |
| GPR21 | 0.19 | 0.1175 | 1.8632 | 0.09309 | 0.21 | -4.97 | -0.6288 | none |
| ZNF682 | 0.326 | 0.1476 | 1.8631 | 0.0931 | 0.21 | -4.97 | 0.38149 | none |
| TCOF1 | 0.6 | 0.7349 | 1.8631 | 0.0931 | 0.21 | -4.97 | -0.3121 | none |
| AC008074 | -0.369 | -0.3159 | -1.863 | 0.09312 | 0.21 | -4.97 | 0.09796 | none |
| GDPD5 | 0.392 | 0.4326 | 1.8626 | 0.09319 | 0.21 | -4.97 | -0.4023 | none |
| YIPF7 | -0.249 | -0.169 | -1.862 | 0.09322 | 0.21 | -4.97 | 0.51509 | none |
| EFHD1 | 0.721 | 0.6555 | 1.8622 | 0.09324 | 0.21 | -4.97 | -0.9059 | none |
| TAF7L | 0.152 | 0.1007 | 1.862 | 0.09327 | 0.21 | -4.98 | 0.50266 | none |
| B9D1 | -0.227 | -0.1707 | -1.862 | 0.0933 | 0.211 | -4.98 | 2.18656 | AA-related |
| TMPO | 0.384 | 0.4272 | 1.8615 | 0.09335 | 0.211 | -4.98 | 0.52334 | none |
| NEURL3 | 0.201 | 0.2138 | 1.8614 | 0.09336 | 0.211 | -4.98 | -0.4565 | none |
| TRIAP1 | -0.249 | -0.2794 | -1.861 | 0.09342 | 0.211 | -4.98 | 1.89655 | none |
| NUDT10 | -0.189 | -0.1426 | -1.861 | 0.09342 | 0.211 | -4.98 | 1.73877 | none |
| GTF2A2 | -0.283 | -0.3203 | -1.861 | 0.09347 | 0.211 | -4.98 | 1.13344 | none |
| VMP1 | 0.227 | 0.1279 | 1.8607 | 0.09347 | 0.211 | -4.98 | -0.7781 | none |
| CUTA | -0.358 | -0.3884 | -1.861 | 0.09348 | 0.211 | -4.98 | 2.1523 | AA-related |
| KRTAP12-3 | -0.264 | -0.1763 | -1.86 | 0.09356 | 0.211 | -4.98 | 0.47004 | none |
| BGN | 0.466 | 0.3174 | 1.8601 | 0.09357 | 0.211 | -4.98 | -0.8092 | none |
| ZNF575 | 0.292 | 0.2651 | 1.86 | 0.09357 | 0.211 | -4.98 | -0.2809 | none |
| EXOG | 0.265 | 0.1376 | 1.8599 | 0.09359 | 0.211 | -4.98 | 0.32134 | none |
| FAH | -0.188 | -0.1856 | -1.86 | 0.09362 | 0.211 | -4.98 | 0.55835 | none |
| ACTN1 | 0.443 | 0.4693 | 1.8597 | 0.09362 | 0.211 | -4.98 | 0.28629 | none |
| RPL18 | -0.321 | -0.3409 | -1.859 | 0.09372 | 0.211 | -4.98 | 0.90502 | none |
| TTYH1 | 0.301 | 0.1987 | 1.859 | 0.09373 | 0.211 | -4.98 | 0.7979 | none |
| RBP5 | 0.375 | 0.3538 | 1.8586 | 0.09379 | 0.211 | -4.98 | -0.76 | none |
| TDP1 | 0.322 | 0.2536 | 1.8586 | 0.09379 | 0.211 | -4.98 | -1.0148 | none |
| DPH7 | 0.27 | 0.2081 | 1.8583 | 0.09383 | 0.211 | -4.98 | 0.10867 | none |
| KNL1 | 0.643 | 0.726 | 1.8582 | 0.09384 | 0.211 | -4.98 | 0.42253 | none |
| NUDT17 | -0.304 | -0.2862 | -1.858 | 0.09389 | 0.211 | -4.98 | 2.20161 | AA-related |
| NPC2 | -0.394 | -0.4006 | -1.858 | 0.09393 | 0.211 | -4.98 | 0.56984 | none |
| FAM120B | 0.261 | 0.2717 | 1.8576 | 0.09395 | 0.211 | -4.98 | -0.416 | none |
| ASCC1 | -0.219 | -0.2739 | -1.857 | 0.09403 | 0.212 | -4.98 | -0.1022 | none |
| ZP1 | -0.195 | -0.1337 | -1.857 | 0.09405 | 0.212 | -4.98 | -0.5076 | none |
| CRYAB | 0.265 | 0.1986 | 1.8561 | 0.09417 | 0.212 | -4.98 | -0.1293 | none |
| RPL8 | -0.283 | -0.2692 | -1.856 | 0.09418 | 0.212 | -4.98 | 1.95631 | none |
| CIB1 | -0.27 | -0.2548 | -1.856 | 0.09419 | 0.212 | -4.98 | 1.16014 | none |
| NPIPB9 | -0.201 | -0.1517 | -1.856 | 0.09421 | 0.212 | -4.98 | 1.04469 | none |
| KYAT3 | -0.198 | -0.1912 | -1.856 | 0.09424 | 0.212 | -4.98 | 2.28409 | AA-related |

| | | | | | | | | |
|---|---|---|---|---|---|---|---|---|
| HDLBP | 0.664 | 0.7592 | 1.8556 | 0.09424 | 0.212 | -4.98 | -0.7252 | none |
| SELP | 0.207 | 0.145 | 1.8555 | 0.09426 | 0.212 | -4.98 | 0.28182 | none |
| SDC4 | -0.228 | -0.1933 | -1.855 | 0.09431 | 0.212 | -4.99 | -0.0375 | none |
| STX5 | 0.235 | 0.1253 | 1.8549 | 0.09436 | 0.212 | -4.99 | -0.5863 | none |
| CEP164 | 0.369 | 0.3849 | 1.8544 | 0.09443 | 0.212 | -4.99 | 0.22433 | none |
| QRICH2 | 0.246 | 0.2032 | 1.8541 | 0.09448 | 0.212 | -4.99 | 0.45378 | none |
| TMPRSS11 | -0.351 | -0.2389 | -1.854 | 0.09448 | 0.212 | -4.99 | 0.90439 | none |
| GAK | 0.416 | 0.4588 | 1.8528 | 0.09468 | 0.213 | -4.99 | -0.1541 | none |
| ANAPC13 | -0.325 | -0.3379 | -1.853 | 0.09471 | 0.213 | -4.99 | 1.96416 | none |
| MPZL3 | 0.245 | 0.1095 | 1.8525 | 0.09472 | 0.213 | -4.99 | -1.1102 | none |
| CT47A12 | -0.173 | -0.1631 | -1.853 | 0.09472 | 0.213 | -4.99 | -0.2612 | none |
| HS3ST2 | 0.273 | 0.1934 | 1.8523 | 0.09475 | 0.213 | -4.99 | -1.2006 | none |
| CARD9 | 0.284 | 0.2738 | 1.8523 | 0.09475 | 0.213 | -4.99 | -0.355 | none |
| RPL31 | -0.355 | -0.4119 | -1.852 | 0.09479 | 0.213 | -4.99 | 1.64318 | none |
| RPS27L | -0.337 | -0.4159 | -1.852 | 0.09487 | 0.213 | -4.99 | 1.05374 | none |
| TERF1 | 0.327 | 0.2504 | 1.8511 | 0.09493 | 0.213 | -4.99 | -0.0504 | none |
| ACYP2 | 0.373 | 0.2184 | 1.8511 | 0.09495 | 0.213 | -4.99 | -0.5309 | none |
| RPS25 | -0.328 | -0.3682 | -1.851 | 0.09496 | 0.213 | -4.99 | 1.14876 | none |
| ZSCAN9 | 0.155 | 0.1056 | 1.8507 | 0.09499 | 0.213 | -4.99 | 0.53358 | none |
| GPD1L | 0.245 | 0.2029 | 1.8502 | 0.09507 | 0.213 | -4.99 | 0.07011 | none |
| FGF6 | -0.233 | -0.1576 | -1.85 | 0.09509 | 0.213 | -4.99 | 1.71112 | none |
| NARF | 0.292 | 0.1769 | 1.8501 | 0.0951 | 0.213 | -4.99 | -0.4726 | none |
| PARVA | 0.413 | 0.3379 | 1.8499 | 0.09512 | 0.213 | -4.99 | -0.0661 | none |
| OR51L1 | 0.168 | 0.1183 | 1.8498 | 0.09513 | 0.213 | -4.99 | -1.0685 | none |
| MAP3K9 | 0.256 | 0.26 | 1.8495 | 0.09519 | 0.213 | -4.99 | -1.0561 | none |
| ATP2B1 | 0.413 | 0.4665 | 1.8493 | 0.09522 | 0.213 | -4.99 | -0.7381 | none |
| HS6ST1 | 0.435 | 0.5093 | 1.8491 | 0.09525 | 0.213 | -4.99 | -0.4774 | none |
| AKAP1 | 0.43 | 0.4347 | 1.8485 | 0.09534 | 0.213 | -5 | -0.6304 | none |
| DNAJC3 | 0.399 | 0.3774 | 1.8484 | 0.09535 | 0.213 | -5 | -1.2396 | none |
| ST6GALNA | 0.485 | 0.4543 | 1.848 | 0.09541 | 0.214 | -5 | -0.0136 | none |
| ANKRD44 | 0.372 | 0.3063 | 1.8479 | 0.09544 | 0.214 | -5 | -1.0257 | none |
| S100Z | -0.24 | -0.2226 | -1.848 | 0.09545 | 0.214 | -5 | 1.58536 | none |
| COPS9 | -0.355 | -0.3486 | -1.847 | 0.09562 | 0.214 | -5 | 1.00982 | none |
| SGSM1 | 0.345 | 0.2394 | 1.8466 | 0.09563 | 0.214 | -5 | -0.0199 | none |
| MEGF8 | 0.279 | 0.2686 | 1.8463 | 0.09568 | 0.214 | -5 | -1.116 | none |
| MAPK8IP3 | 0.371 | 0.4541 | 1.846 | 0.09572 | 0.214 | -5 | -0.0688 | none |
| LRRC3 | 0.183 | 0.121 | 1.8459 | 0.09574 | 0.214 | -5 | -1.0676 | none |
| SPI1 | -0.154 | -0.1291 | -1.846 | 0.09575 | 0.214 | -5 | 0.7773 | none |
| SF3B1 | 0.374 | 0.3356 | 1.8457 | 0.09577 | 0.214 | -5 | -0.619 | none |
| HIVEP3 | 0.436 | 0.5942 | 1.8457 | 0.09578 | 0.214 | -5 | -0.5768 | none |

| | | | | | | | | |
|---|---|---|---|---|---|---|---|---|
| PTGES2 | 0.159 | 0.1721 | 1.8454 | 0.09582 | 0.214 | -5 | 0.21886 | none |
| LAMB3 | 0.2 | 0.1718 | 1.8454 | 0.09582 | 0.214 | -5 | -0.0902 | none |
| MEX3C | 0.153 | 0.1185 | 1.8445 | 0.09595 | 0.214 | -5 | -0.1375 | none |
| RPLP1 | -0.34 | -0.3598 | -1.844 | 0.09597 | 0.214 | -5 | 1.86529 | none |
| TMTC1 | 0.142 | 0.1203 | 1.8444 | 0.09598 | 0.214 | -5 | -0.6325 | none |
| FZD4 | 0.134 | 0.0933 | 1.8441 | 0.09602 | 0.215 | -5 | -1.2499 | none |
| JAK1 | 0.512 | 0.5875 | 1.8434 | 0.09613 | 0.215 | -5 | 0.48527 | none |
| RPL36 | -0.29 | -0.2882 | -1.843 | 0.09618 | 0.215 | -5 | 1.76492 | none |
| RPL30 | -0.341 | -0.4087 | -1.843 | 0.09618 | 0.215 | -5 | 1.55266 | none |
| SF3B5 | -0.3 | -0.3179 | -1.843 | 0.09623 | 0.215 | -5 | 1.2139 | none |
| SLC6A7 | 0.156 | 0.1562 | 1.8426 | 0.09625 | 0.215 | -5 | 1.01859 | none |
| SEC13 | -0.144 | -0.1162 | -1.842 | 0.09631 | 0.215 | -5 | 2.22001 | AA-related |
| ACAT2 | -0.262 | -0.2505 | -1.842 | 0.09632 | 0.215 | -5 | 0.76925 | none |
| BHLHE40 | 0.343 | 0.4044 | 1.8412 | 0.09647 | 0.215 | -5.01 | -0.6291 | none |
| CHODL | 0.169 | 0.1539 | 1.8406 | 0.09657 | 0.215 | -5.01 | -1.1172 | none |
| POLR2F | -0.319 | -0.3689 | -1.84 | 0.09661 | 0.216 | -5.01 | 0.92638 | none |
| MASP2 | 0.36 | 0.2827 | 1.84 | 0.09666 | 0.216 | -5.01 | -0.8 | none |
| NUDCD3 | 0.199 | 0.1606 | 1.8396 | 0.09673 | 0.216 | -5.01 | 0.11346 | none |
| AIG1 | 0.497 | 0.3014 | 1.8395 | 0.09674 | 0.216 | -5.01 | -0.9998 | none |
| C6 | -0.188 | -0.1465 | -1.839 | 0.09676 | 0.216 | -5.01 | -0.1108 | none |
| CPLX4 | 0.156 | 0.1177 | 1.8394 | 0.09676 | 0.216 | -5.01 | -1.1243 | none |
| G2E3 | 0.221 | 0.1247 | 1.839 | 0.09682 | 0.216 | -5.01 | -0.8211 | none |
| PPP6R1 | 0.382 | 0.406 | 1.8387 | 0.09687 | 0.216 | -5.01 | -0.494 | none |
| NGLY1 | 0.139 | 0.069 | 1.8383 | 0.09693 | 0.216 | -5.01 | 0.49421 | none |
| AMY2A | -0.167 | -0.1403 | -1.838 | 0.09695 | 0.216 | -5.01 | 1.35324 | none |
| IDH3G | -0.16 | -0.1172 | -1.838 | 0.09698 | 0.216 | -5.01 | 1.07896 | none |
| NFE2L3 | 0.353 | 0.3893 | 1.838 | 0.09698 | 0.216 | -5.01 | -0.6172 | none |
| RPS3 | -0.372 | -0.3962 | -1.838 | 0.09703 | 0.216 | -5.01 | 0.9619 | none |
| PRPF40B | 0.342 | 0.3817 | 1.8373 | 0.09709 | 0.216 | -5.01 | -1.0909 | none |
| TMEM211 | -0.162 | -0.1479 | -1.837 | 0.09715 | 0.216 | -5.01 | -0.0217 | none |
| ADAM10 | 0.406 | 0.3973 | 1.8364 | 0.09723 | 0.217 | -5.01 | -0.1378 | none |
| TXNRD1 | 0.147 | 0.0826 | 1.8362 | 0.09725 | 0.217 | -5.01 | 0.23374 | none |
| MAGOHB | -0.371 | -0.3728 | -1.836 | 0.09729 | 0.217 | -5.01 | 1.18547 | none |
| RNF112 | 0.212 | 0.1478 | 1.8358 | 0.09732 | 0.217 | -5.01 | 0.1841 | none |
| DIS3L2 | 0.471 | 0.4297 | 1.8354 | 0.09739 | 0.217 | -5.01 | -0.7858 | none |
| PIGU | -0.432 | -0.3873 | -1.835 | 0.09751 | 0.217 | -5.02 | 2.22944 | AA-related |
| MAN2B1 | 0.318 | 0.2788 | 1.8346 | 0.09752 | 0.217 | -5.02 | -0.1922 | none |
| FRY | -0.412 | -0.3047 | -1.834 | 0.09767 | 0.217 | -5.02 | 0.63239 | none |
| RPP40 | -0.359 | -0.3569 | -1.833 | 0.0977 | 0.217 | -5.02 | 1.9077 | none |
| INPP5A | 0.304 | 0.2996 | 1.8331 | 0.09775 | 0.217 | -5.02 | 0.54467 | none |

| | | | | | | | | |
|---|---|---|---|---|---|---|---|---|
| TYW1B | 0.334 | 0.2429 | 1.833 | 0.09776 | 0.217 | -5.02 | -0.5077 | none |
| CACNA2D2 | 0.169 | 0.1408 | 1.8323 | 0.09788 | 0.218 | -5.02 | 0.17358 | none |
| VAMP1 | 0.229 | 0.2013 | 1.8321 | 0.09791 | 0.218 | -5.02 | 0.0906 | none |
| PAK4 | 0.428 | 0.4129 | 1.8314 | 0.09802 | 0.218 | -5.02 | -0.3892 | none |
| RIC8B | 0.364 | 0.3353 | 1.8314 | 0.09803 | 0.218 | -5.02 | -0.2104 | none |
| PGR | -0.179 | -0.1225 | -1.83 | 0.09822 | 0.218 | -5.02 | 0.82582 | none |
| ECI2 | -0.217 | -0.2269 | -1.83 | 0.09822 | 0.218 | -5.02 | 0.91644 | none |
| TERB2 | 0.632 | 0.5536 | 1.8297 | 0.09829 | 0.218 | -5.02 | 1.17226 | none |
| TAL2 | 0.185 | 0.1475 | 1.8296 | 0.0983 | 0.218 | -5.02 | 0.69671 | none |
| CENPW | -0.338 | -0.3477 | -1.829 | 0.09834 | 0.218 | -5.02 | 2.0921 | AA-related |
| ARMCX6 | -0.188 | -0.1571 | -1.829 | 0.09834 | 0.218 | -5.02 | 1.55829 | none |
| ELOVL5 | -0.266 | -0.293 | -1.829 | 0.09839 | 0.218 | -5.02 | 1.90952 | none |
| PLEC | 0.423 | 0.5051 | 1.829 | 0.0984 | 0.218 | -5.02 | -0.0132 | none |
| PHF21A | 0.361 | 0.454 | 1.8288 | 0.09844 | 0.219 | -5.02 | 0.42875 | none |
| CACNA1E | 0.255 | 0.1715 | 1.8286 | 0.09846 | 0.219 | -5.02 | -0.1459 | none |
| CCDC85C | 0.229 | 0.1404 | 1.8283 | 0.09851 | 0.219 | -5.02 | -0.3051 | none |
| TP53I3 | -0.22 | -0.1929 | -1.828 | 0.09851 | 0.219 | -5.02 | 1.9929 | none |
| SCGB1D4 | 0.2 | 0.1998 | 1.8277 | 0.0986 | 0.219 | -5.03 | 1.07091 | none |
| PLAUR | 0.26 | 0.1903 | 1.8272 | 0.09869 | 0.219 | -5.03 | -0.8034 | none |
| OR52N4 | 0.277 | 0.1596 | 1.827 | 0.09873 | 0.219 | -5.03 | -0.7787 | none |
| TBC1D14 | 0.301 | 0.3433 | 1.8262 | 0.09885 | 0.219 | -5.03 | -0.227 | none |
| DCTN1 | 0.485 | 0.554 | 1.826 | 0.09889 | 0.219 | -5.03 | 0.11525 | none |
| MRPS28 | 0.287 | 0.1104 | 1.8255 | 0.09896 | 0.219 | -5.03 | -0.9965 | none |
| YAP1 | 0.202 | 0.204 | 1.8251 | 0.09903 | 0.22 | -5.03 | 0.18148 | none |
| HNRNPH1 | 0.343 | 0.3983 | 1.825 | 0.09905 | 0.22 | -5.03 | -0.7496 | none |
| ZNF692 | 0.296 | 0.3523 | 1.8246 | 0.09911 | 0.22 | -5.03 | -0.1856 | none |
| SMYD5 | 0.21 | 0.1784 | 1.824 | 0.0992 | 0.22 | -5.03 | -0.0769 | none |
| PCDHGA9 | 0.142 | 0.1458 | 1.8234 | 0.0993 | 0.22 | -5.03 | 0.75431 | none |
| PRRC1 | 0.336 | 0.2283 | 1.8233 | 0.09931 | 0.22 | -5.03 | -0.6564 | none |
| AMOTL2 | -0.208 | -0.1654 | -1.823 | 0.09933 | 0.22 | -5.03 | -0.157 | none |
| CLDN9 | 0.148 | 0.0958 | 1.8225 | 0.09945 | 0.22 | -5.03 | 0.18068 | none |
| TNFRSF25 | 0.296 | 0.2035 | 1.8223 | 0.09947 | 0.22 | -5.03 | -0.7989 | none |
| SNAI3 | 0.194 | 0.1007 | 1.8223 | 0.09948 | 0.22 | -5.03 | -0.1904 | none |
| LMAN2L | -0.338 | -0.3347 | -1.822 | 0.09949 | 0.22 | -5.03 | 1.77547 | none |
| RWDD1 | -0.175 | -0.1647 | -1.822 | 0.09949 | 0.22 | -5.03 | 1.50151 | none |
| ITGA4 | 0.291 | 0.2958 | 1.8221 | 0.0995 | 0.22 | -5.03 | 0.93796 | none |
| FBXO40 | 0.207 | 0.153 | 1.8219 | 0.09954 | 0.22 | -5.03 | -0.2505 | none |
| ERBB3 | 0.54 | 0.3551 | 1.8216 | 0.09959 | 0.22 | -5.03 | -0.6028 | none |
| CDK5 | -0.265 | -0.2711 | -1.821 | 0.09964 | 0.22 | -5.03 | 0.86676 | none |
| CLPTM1 | 0.2 | 0.2082 | 1.8212 | 0.09965 | 0.22 | -5.03 | 0.78396 | none |

| | | | | | | | | |
|---|---|---|---|---|---|---|---|---|
| RNF41 | 0.278 | 0.1532 | 1.8211 | 0.09966 | 0.22 | -5.04 | -0.3036 | none |
| ODF3 | 0.28 | 0.1946 | 1.8208 | 0.09971 | 0.221 | -5.04 | -0.4359 | none |
| ZBTB24 | 0.331 | 0.3468 | 1.8208 | 0.09971 | 0.221 | -5.04 | -0.9913 | none |
| KIAA1755 | 0.151 | 0.1139 | 1.8206 | 0.09974 | 0.221 | -5.04 | -0.8873 | none |
| GRHL2 | -0.168 | -0.1265 | -1.821 | 0.09976 | 0.221 | -5.04 | -0.4503 | none |
| PLEKHA4 | 0.323 | 0.2229 | 1.8199 | 0.09986 | 0.221 | -5.04 | -0.4871 | none |
| TRAPPC12 | -0.365 | -0.134 | -1.82 | 0.09987 | 0.221 | -5.04 | 1.14554 | none |
| DNAJC27 | 0.318 | 0.3597 | 1.8194 | 0.09994 | 0.221 | -5.04 | 1.38717 | none |
| KIAA0391 | -0.202 | -0.2418 | -1.819 | 0.09995 | 0.221 | -5.04 | 0.99795 | none |
| POLN | 0.425 | 0.271 | 1.8191 | 0.09999 | 0.221 | -5.04 | -0.935 | none |
| CHD9 | 0.289 | 0.3141 | 1.8185 | 0.10009 | 0.221 | -5.04 | 0.13648 | none |
| FAM20C | -0.132 | -0.1245 | -1.818 | 0.10015 | 0.221 | -5.04 | -0.1307 | none |
| TBC1D7 | 0.281 | 0.1744 | 1.818 | 0.10016 | 0.221 | -5.04 | -0.3232 | none |
| RGL4 | 0.423 | 0.2226 | 1.8177 | 0.10022 | 0.221 | -5.04 | 0.10895 | none |
| TIMM13 | -0.236 | -0.2126 | -1.817 | 0.10025 | 0.221 | -5.04 | 1.22443 | none |
| GNPNAT1 | 0.213 | 0.0825 | 1.8173 | 0.10028 | 0.221 | -5.04 | 0.64515 | none |
| APOBR | 0.346 | 0.3536 | 1.817 | 0.10034 | 0.221 | -5.04 | -0.5621 | none |
| VWA1 | 0.271 | 0.2935 | 1.8169 | 0.10035 | 0.221 | -5.04 | -0.0576 | none |
| ATP10D | 0.491 | 0.3829 | 1.8167 | 0.10037 | 0.222 | -5.04 | -0.5423 | none |
| SHMT1 | 0.189 | 0.1224 | 1.8163 | 0.10045 | 0.222 | -5.04 | -0.3966 | none |
| RANBP1 | -0.153 | -0.0432 | -1.816 | 0.10045 | 0.222 | -5.04 | 0.30885 | none |
| BEX1 | -0.183 | -0.1503 | -1.816 | 0.10057 | 0.222 | -5.04 | 0.5215 | none |
| GKAP1 | 0.646 | 0.4339 | 1.8154 | 0.10059 | 0.222 | -5.04 | -0.6227 | none |
| MAGI1 | 0.169 | 0.1699 | 1.8153 | 0.10061 | 0.222 | -5.04 | -0.2342 | none |
| AGAP4 | 0.724 | 0.6722 | 1.8153 | 0.10061 | 0.222 | -5.04 | -0.5423 | none |
| NNAT | -0.181 | -0.1337 | -1.815 | 0.10066 | 0.222 | -5.04 | 0.73771 | none |
| NGEF | 0.202 | 0.1429 | 1.8144 | 0.10076 | 0.222 | -5.04 | 0.14946 | none |
| CCDC189 | -0.153 | -0.1345 | -1.814 | 0.10078 | 0.222 | -5.05 | 0.32545 | none |
| UBE2Z | 0.361 | 0.2968 | 1.8142 | 0.10079 | 0.222 | -5.05 | 0.31777 | none |
| APOBEC2 | 0.354 | 0.2075 | 1.8141 | 0.1008 | 0.222 | -5.05 | -0.9435 | none |
| ENPP2 | 0.164 | 0.1459 | 1.8141 | 0.10081 | 0.222 | -5.05 | 0.72161 | none |
| PRDM11 | 0.177 | 0.1461 | 1.8139 | 0.10084 | 0.222 | -5.05 | -0.4399 | none |
| AMHR2 | 0.24 | 0.2413 | 1.8134 | 0.10092 | 0.222 | -5.05 | -0.868 | none |
| CYP21A2 | 0.235 | 0.1823 | 1.8134 | 0.10092 | 0.222 | -5.05 | -0.3118 | none |
| UBXN8 | 0.203 | 0.0616 | 1.8133 | 0.10093 | 0.222 | -5.05 | -0.5247 | none |
| NDUFA13 | -0.293 | -0.2771 | -1.813 | 0.10099 | 0.222 | -5.05 | 0.708 | none |
| LIN28B | 0.338 | 0.2199 | 1.8128 | 0.10102 | 0.222 | -5.05 | -0.9111 | none |
| ZNF337 | 0.301 | 0.2883 | 1.8127 | 0.10104 | 0.222 | -5.05 | -1.0176 | none |
| BTBD3 | -0.204 | -0.219 | -1.813 | 0.10104 | 0.222 | -5.05 | 0.35159 | none |
| NRL | 0.158 | 0.1265 | 1.8126 | 0.10105 | 0.222 | -5.05 | -0.6253 | none |

| | | | | | | | | |
|---|---|---|---|---|---|---|---|---|
| LSR | 0.337 | 0.1965 | 1.8121 | 0.10113 | 0.223 | -5.05 | -0.6348 | none |
| ARL6IP4 | -0.165 | -0.102 | -1.812 | 0.10113 | 0.223 | -5.05 | 1.24011 | none |
| REG4 | -0.204 | -0.2188 | -1.812 | 0.10117 | 0.223 | -5.05 | 1.33829 | none |
| GMNN | -0.156 | -0.1713 | -1.812 | 0.10121 | 0.223 | -5.05 | 1.34051 | none |
| CIB2 | 0.555 | 0.2759 | 1.8106 | 0.10137 | 0.223 | -5.05 | -0.4832 | none |
| SPPL2B | 0.231 | 0.2608 | 1.8105 | 0.10139 | 0.223 | -5.05 | 1.17878 | none |
| NUDCD2 | -0.273 | -0.2955 | -1.81 | 0.10143 | 0.223 | -5.05 | 1.29534 | none |
| JMJD8 | 0.244 | 0.2603 | 1.8098 | 0.1015 | 0.223 | -5.05 | -0.2722 | none |
| CXORF40A | -0.227 | -0.2111 | -1.81 | 0.10154 | 0.223 | -5.05 | 1.31166 | none |
| SMS | -0.129 | -0.1285 | -1.809 | 0.10158 | 0.223 | -5.05 | 1.6192 | none |
| JMY | 0.264 | 0.2805 | 1.8093 | 0.10159 | 0.223 | -5.05 | -0.545 | none |
| STK40 | 0.229 | 0.2546 | 1.809 | 0.10164 | 0.223 | -5.05 | 0.09168 | none |
| THPO | -0.494 | -0.3729 | -1.809 | 0.10167 | 0.223 | -5.05 | 0.41223 | none |
| TMEM208 | -0.348 | -0.3621 | -1.809 | 0.10167 | 0.223 | -5.05 | 1.62837 | none |
| MAMDC4 | 0.224 | 0.2367 | 1.8088 | 0.10168 | 0.223 | -5.05 | -0.7916 | none |
| ARPC1A | -0.169 | -0.1746 | -1.809 | 0.10169 | 0.223 | -5.05 | 1.22473 | none |
| ARHGAP5 | 0.425 | 0.3983 | 1.8086 | 0.1017 | 0.223 | -5.05 | 0.34016 | none |
| SH3GLB2 | 0.225 | 0.2537 | 1.808 | 0.10181 | 0.224 | -5.05 | 0.40458 | none |
| LBHD1 | 0.259 | 0.1948 | 1.808 | 0.10181 | 0.224 | -5.05 | -0.6412 | none |
| ZFYVE1 | 0.2 | 0.2148 | 1.8078 | 0.10183 | 0.224 | -5.05 | 0.67749 | none |
| PRSS21 | 0.377 | 0.3002 | 1.8076 | 0.10187 | 0.224 | -5.05 | -0.6423 | none |
| TGIF1 | 0.218 | 0.1685 | 1.8074 | 0.1019 | 0.224 | -5.05 | -0.4278 | none |
| TBCC | 0.206 | 0.1655 | 1.8068 | 0.102 | 0.224 | -5.06 | -0.9794 | none |
| AC021106 | 0.301 | 0.3111 | 1.8066 | 0.10204 | 0.224 | -5.06 | -0.038 | none |
| PLCB2 | 0.452 | 0.3959 | 1.8066 | 0.10204 | 0.224 | -5.06 | -0.3983 | none |
| OSBPL11 | 0.329 | 0.3842 | 1.8061 | 0.10212 | 0.224 | -5.06 | 0.50071 | none |
| SDR39U1 | 0.163 | 0.1114 | 1.8059 | 0.10215 | 0.224 | -5.06 | 0.74519 | none |
| SSPO | 0.369 | 0.3599 | 1.8055 | 0.10221 | 0.224 | -5.06 | -0.5364 | none |
| TMEM201 | 0.182 | 0.1589 | 1.8055 | 0.10222 | 0.224 | -5.06 | 0.47462 | none |
| XCL1 | -0.199 | -0.2073 | -1.805 | 0.10225 | 0.224 | -5.06 | 0.23006 | none |
| FNDC5 | 0.274 | 0.198 | 1.8052 | 0.10226 | 0.224 | -5.06 | 0.41627 | none |
| C9ORF153 | 0.182 | 0.168 | 1.805 | 0.10229 | 0.224 | -5.06 | -0.6039 | none |
| FAM71F1 | 0.212 | 0.2116 | 1.8046 | 0.10236 | 0.224 | -5.06 | -0.2744 | none |
| PDGFC | 0.208 | 0.1648 | 1.8045 | 0.10238 | 0.224 | -5.06 | -0.7504 | none |
| AMER2 | 0.119 | 0.0932 | 1.8043 | 0.10242 | 0.224 | -5.06 | -0.1509 | none |
| ZSCAN30 | 0.304 | 0.3715 | 1.8036 | 0.10253 | 0.225 | -5.06 | -0.5131 | none |
| HFM1 | -0.235 | -0.16 | -1.804 | 0.10253 | 0.225 | -5.06 | -0.129 | none |
| PRDX4 | -0.336 | -0.3478 | -1.804 | 0.10254 | 0.225 | -5.06 | 1.02433 | none |
| SOX12 | 0.162 | 0.1615 | 1.8032 | 0.1026 | 0.225 | -5.06 | -1.1058 | none |
| ALDH9A1 | -0.187 | -0.2073 | -1.803 | 0.10263 | 0.225 | -5.06 | 1.08192 | none |

| | | | | | | | | |
|---|---|---|---|---|---|---|---|---|
| HOXB2 | 0.187 | 0.1459 | 1.803 | 0.10263 | 0.225 | -5.06 | 0.64454 | none |
| FKBP15 | 0.325 | 0.3746 | 1.8029 | 0.10265 | 0.225 | -5.06 | 0.22543 | none |
| PRCP | 0.308 | 0.1467 | 1.8028 | 0.10267 | 0.225 | -5.06 | -0.8327 | none |
| CLEC4E | 0.151 | 0.1049 | 1.8026 | 0.10269 | 0.225 | -5.06 | 0.58634 | none |
| PEX13 | 0.218 | 0.0856 | 1.8023 | 0.10274 | 0.225 | -5.06 | -0.2754 | none |
| OLA1 | -0.158 | -0.1931 | -1.802 | 0.10276 | 0.225 | -5.06 | 1.84557 | none |
| ZDHHC14 | 0.303 | 0.1882 | 1.802 | 0.10279 | 0.225 | -5.06 | 0.1746 | none |
| SLC30A1 | 0.187 | 0.1396 | 1.8018 | 0.10283 | 0.225 | -5.06 | 0.10571 | none |
| GAL3ST4 | 0.225 | 0.1323 | 1.8017 | 0.10284 | 0.225 | -5.06 | -0.3645 | none |
| PWP1 | -0.151 | -0.1467 | -1.802 | 0.10287 | 0.225 | -5.06 | 1.07526 | none |
| RFT1 | -0.252 | -0.2721 | -1.801 | 0.103 | 0.225 | -5.06 | 0.73777 | none |
| HIST1H4A | -0.333 | -0.2942 | -1.801 | 0.10301 | 0.225 | -5.06 | 0.77133 | none |
| MATR3 | 0.362 | 0.3337 | 1.8 | 0.10312 | 0.225 | -5.07 | 0.095 | none |
| RFC4 | -0.157 | -0.1337 | -1.8 | 0.10321 | 0.226 | -5.07 | 1.28758 | none |
| CIAO1 | -0.166 | -0.1011 | -1.799 | 0.10336 | 0.226 | -5.07 | 0.89947 | none |
| ATXN10 | -0.137 | -0.126 | -1.798 | 0.10347 | 0.226 | -5.07 | 1.38022 | none |
| PCED1A | 0.358 | 0.2212 | 1.7972 | 0.10359 | 0.226 | -5.07 | 0.88946 | none |
| PLCXD2 | 0.187 | 0.1668 | 1.797 | 0.10363 | 0.226 | -5.07 | -1.1824 | none |
| C16ORF74 | 0.225 | 0.2032 | 1.7966 | 0.1037 | 0.226 | -5.07 | -0.4072 | none |
| EPHA1 | 0.274 | 0.1509 | 1.7963 | 0.10374 | 0.227 | -5.07 | -1.2271 | none |
| ISYNA1 | -0.197 | -0.1688 | -1.796 | 0.10378 | 0.227 | -5.07 | 1.70452 | none |
| TIMELESS | 0.306 | 0.3556 | 1.796 | 0.10379 | 0.227 | -5.07 | -0.0365 | none |
| MOXD1 | 0.194 | 0.1725 | 1.796 | 0.10379 | 0.227 | -5.07 | 0.44894 | none |
| WDR44 | 0.285 | 0.3408 | 1.796 | 0.1038 | 0.227 | -5.07 | 0.53308 | none |
| CCDC106 | -0.26 | -0.1524 | -1.796 | 0.10382 | 0.227 | -5.07 | 1.6528 | none |
| LZTR1 | 0.243 | 0.2675 | 1.7958 | 0.10383 | 0.227 | -5.07 | 0.04756 | none |
| TRDMT1 | -0.283 | -0.3133 | -1.796 | 0.10385 | 0.227 | -5.07 | -0.0286 | none |
| TUT1 | 0.199 | 0.1959 | 1.7955 | 0.10387 | 0.227 | -5.07 | 0.19857 | none |
| MBTPS2 | 0.265 | 0.1547 | 1.7955 | 0.10388 | 0.227 | -5.07 | -0.839 | none |
| NRXN1 | 0.294 | 0.2551 | 1.7954 | 0.1039 | 0.227 | -5.07 | -0.1581 | none |
| SH3BP4 | 0.262 | 0.2718 | 1.7952 | 0.10392 | 0.227 | -5.07 | -0.0851 | none |
| CLP1 | -0.256 | -0.2329 | -1.795 | 0.10393 | 0.227 | -5.07 | 2.04744 | none |
| TEX11 | 0.216 | 0.1571 | 1.7948 | 0.10399 | 0.227 | -5.07 | 0.1012 | none |
| TXNL1 | -0.189 | -0.2203 | -1.793 | 0.10424 | 0.227 | -5.08 | 1.59193 | none |
| TBC1D3L | 0.342 | 0.3553 | 1.7933 | 0.10424 | 0.227 | -5.08 | -0.3168 | none |
| AQR | 0.337 | 0.3054 | 1.793 | 0.10429 | 0.227 | -5.08 | -0.8192 | none |
| COX7B | -0.272 | -0.3329 | -1.793 | 0.1043 | 0.227 | -5.08 | 1.48988 | none |
| CHD6 | 0.395 | 0.3644 | 1.793 | 0.1043 | 0.227 | -5.08 | -0.6742 | none |
| CD84 | 0.397 | 0.3281 | 1.7929 | 0.10431 | 0.227 | -5.08 | -0.9783 | none |
| PIEZO1 | 0.513 | 0.6148 | 1.7929 | 0.10432 | 0.227 | -5.08 | -0.3754 | none |

| | | | | | | | | |
|---|---|---|---|---|---|---|---|---|
| RNF19B | 0.176 | 0.1652 | 1.7927 | 0.10435 | 0.227 | -5.08 | 0.11962 | none |
| PTH1R | 0.151 | 0.1401 | 1.7927 | 0.10435 | 0.227 | -5.08 | 0.40905 | none |
| PEX7 | 0.366 | 0.1611 | 1.7921 | 0.10444 | 0.227 | -5.08 | -0.6027 | none |
| UNC45A | 0.277 | 0.2916 | 1.7921 | 0.10444 | 0.227 | -5.08 | -0.4553 | none |
| TTI1 | 0.18 | 0.2445 | 1.792 | 0.10446 | 0.227 | -5.08 | -0.2314 | none |
| STOML1 | 0.221 | 0.2027 | 1.7916 | 0.10453 | 0.228 | -5.08 | 0.08465 | none |
| TWF2 | -0.188 | -0.1564 | -1.791 | 0.10459 | 0.228 | -5.08 | 1.86596 | none |
| SYT11 | 0.282 | 0.2734 | 1.7908 | 0.10466 | 0.228 | -5.08 | -0.6322 | none |
| FGF7 | 0.176 | 0.1274 | 1.7906 | 0.1047 | 0.228 | -5.08 | -0.746 | none |
| MKS1 | -0.202 | -0.1665 | -1.791 | 0.10471 | 0.228 | -5.08 | 0.96835 | none |
| MSL3 | 0.206 | 0.1268 | 1.7897 | 0.10486 | 0.228 | -5.08 | -1.063 | none |
| ANGEL1 | 0.196 | 0.1165 | 1.7896 | 0.10487 | 0.228 | -5.08 | -0.5519 | none |
| KCTD12 | 0.155 | 0.171 | 1.7889 | 0.10499 | 0.228 | -5.08 | -0.0424 | none |
| F11 | -0.16 | -0.1312 | -1.789 | 0.10499 | 0.228 | -5.08 | 0.50993 | none |
| SBF1 | 0.335 | 0.3914 | 1.7889 | 0.10499 | 0.228 | -5.08 | -0.1664 | none |
| NSMCE3 | 0.183 | 0.1269 | 1.7887 | 0.10502 | 0.228 | -5.08 | -1.2423 | none |
| HEG1 | 0.265 | 0.3305 | 1.7885 | 0.10505 | 0.228 | -5.08 | 0.91916 | none |
| HJURP | 0.321 | 0.3306 | 1.7883 | 0.10509 | 0.228 | -5.08 | 0.3918 | none |
| PTGDS | 0.358 | 0.2757 | 1.7881 | 0.10512 | 0.228 | -5.08 | 0.08212 | none |
| PNOC | 0.356 | 0.2912 | 1.7872 | 0.10527 | 0.229 | -5.08 | 0.71987 | none |
| KIF3C | -0.295 | -0.1563 | -1.787 | 0.10539 | 0.229 | -5.09 | 0.71336 | none |
| BLZF1 | 0.391 | 0.313 | 1.7863 | 0.10543 | 0.229 | -5.09 | -0.0889 | none |
| ZDHHC1 | 0.148 | 0.2051 | 1.7862 | 0.10545 | 0.229 | -5.09 | 0.86579 | none |
| SETD4 | 0.388 | 0.241 | 1.7861 | 0.10546 | 0.229 | -5.09 | -0.274 | none |
| CCDC88B | 0.567 | 0.6289 | 1.7859 | 0.1055 | 0.229 | -5.09 | -0.4599 | none |
| MTRNR2L1 | -0.19 | -0.1851 | -1.786 | 0.10551 | 0.229 | -5.09 | 1.16715 | none |
| PDIA6 | -0.186 | -0.1698 | -1.786 | 0.10556 | 0.229 | -5.09 | 1.45473 | none |
| PRR12 | 0.351 | 0.4017 | 1.7852 | 0.10561 | 0.229 | -5.09 | -0.1077 | none |
| ANKRD53 | -0.263 | -0.3146 | -1.785 | 0.10561 | 0.229 | -5.09 | 0.6135 | none |
| ZNF420 | 0.337 | 0.1345 | 1.785 | 0.10565 | 0.229 | -5.09 | -0.1308 | none |
| KCTD9 | 0.184 | 0.1095 | 1.7849 | 0.10566 | 0.229 | -5.09 | -1.0406 | none |
| DNMT3L | -0.15 | -0.0997 | -1.785 | 0.10569 | 0.229 | -5.09 | -0.1475 | none |
| RPL36A | -0.298 | -0.3305 | -1.784 | 0.10585 | 0.23 | -5.09 | 1.22179 | none |
| FGFR2 | 0.374 | 0.2366 | 1.7832 | 0.10595 | 0.23 | -5.09 | -0.6963 | none |
| PPP3CB | -0.394 | -0.3805 | -1.783 | 0.10601 | 0.23 | -5.09 | 1.68152 | none |
| MRPL43 | -0.271 | -0.2606 | -1.783 | 0.10604 | 0.23 | -5.09 | 1.4427 | none |
| RNF114 | 0.325 | 0.2095 | 1.782 | 0.10616 | 0.23 | -5.09 | -0.7085 | none |
| REN | 0.132 | 0.1319 | 1.782 | 0.10616 | 0.23 | -5.09 | 0.75649 | none |
| ISCU | -0.222 | -0.2085 | -1.781 | 0.10626 | 0.23 | -5.09 | 1.82684 | none |
| KISS1R | -0.193 | -0.1316 | -1.781 | 0.10633 | 0.23 | -5.09 | 0.71241 | none |

| | | | | | | | | |
|---|---|---|---|---|---|---|---|---|
| LDB2 | 0.184 | 0.1861 | 1.7809 | 0.10635 | 0.23 | -5.09 | 0.97429 | none |
| OR2J3 | 0.141 | 0.0963 | 1.7808 | 0.10637 | 0.23 | -5.09 | -1.5539 | OA-related |
| RPS15 | -0.328 | -0.3307 | -1.781 | 0.10637 | 0.23 | -5.09 | 2.2341 | AA-related |
| OSR2 | -0.224 | -0.1965 | -1.78 | 0.10644 | 0.231 | -5.09 | 2.00256 | none |
| RASGRP2 | 0.374 | 0.2659 | 1.7795 | 0.10659 | 0.231 | -5.1 | 0.89894 | none |
| GAL | 0.308 | 0.3316 | 1.7795 | 0.1066 | 0.231 | -5.1 | 1.67067 | none |
| KMT2C | 0.392 | 0.5318 | 1.7793 | 0.10662 | 0.231 | -5.1 | -0.5526 | none |
| ZCCHC17 | -0.187 | -0.2188 | -1.779 | 0.10663 | 0.231 | -5.1 | 0.78159 | none |
| RET | 0.15 | 0.1977 | 1.779 | 0.10668 | 0.231 | -5.1 | 0.65381 | none |
| SURF4 | -0.236 | -0.231 | -1.779 | 0.1067 | 0.231 | -5.1 | 2.09788 | AA-related |
| COPS6 | 0.208 | 0.136 | 1.7787 | 0.10673 | 0.231 | -5.1 | -0.8837 | none |
| SLC6A13 | -0.336 | -0.3341 | -1.778 | 0.10683 | 0.231 | -5.1 | 1.15831 | none |
| PANX2 | 0.253 | 0.2441 | 1.7773 | 0.10696 | 0.231 | -5.1 | -1.269 | none |
| ZMAT2 | -0.195 | -0.1643 | -1.777 | 0.10698 | 0.231 | -5.1 | 2.08511 | none |
| THOP1 | 0.26 | 0.308 | 1.7771 | 0.107 | 0.231 | -5.1 | -0.2452 | none |
| SLTM | 0.496 | 0.6022 | 1.7769 | 0.10704 | 0.231 | -5.1 | 0.05052 | none |
| ACTG2 | 0.246 | 0.0763 | 1.7765 | 0.10711 | 0.232 | -5.1 | 0.0424 | none |
| DGAT2L6 | -0.16 | -0.1553 | -1.776 | 0.10712 | 0.232 | -5.1 | 1.34828 | none |
| RTF1 | 0.495 | 0.484 | 1.7764 | 0.10713 | 0.232 | -5.1 | -0.6813 | none |
| TYMP | 0.278 | 0.226 | 1.7762 | 0.10716 | 0.232 | -5.1 | -0.1043 | none |
| NSMCE1 | 0.236 | 0.0768 | 1.7759 | 0.10722 | 0.232 | -5.1 | -0.2351 | none |
| SCN4B | 0.174 | 0.129 | 1.7754 | 0.10729 | 0.232 | -5.1 | 0.06303 | none |
| RTP3 | -0.179 | -0.1793 | -1.775 | 0.10734 | 0.232 | -5.1 | 0.8507 | none |
| UHRF1 | 0.506 | 0.5775 | 1.7749 | 0.10738 | 0.232 | -5.1 | 0.29336 | none |
| IGBP1 | -0.222 | -0.2079 | -1.775 | 0.1074 | 0.232 | -5.1 | 1.14314 | none |
| FAM200B | 0.375 | 0.1538 | 1.7748 | 0.1074 | 0.232 | -5.1 | 0.22895 | none |
| GGTLC2 | 0.258 | 0.265 | 1.7746 | 0.10743 | 0.232 | -5.1 | 1.58762 | none |
| CDK2 | -0.197 | -0.2004 | -1.774 | 0.10751 | 0.232 | -5.1 | 2.09478 | AA-related |
| RPS19 | -0.256 | -0.228 | -1.774 | 0.10754 | 0.232 | -5.1 | 0.98425 | none |
| CEP250 | 0.378 | 0.4211 | 1.7739 | 0.10755 | 0.232 | -5.1 | 0.18069 | none |
| SLPI | -0.213 | -0.1422 | -1.774 | 0.10756 | 0.232 | -5.1 | 1.24897 | none |
| RHBDF1 | -0.226 | -0.1569 | -1.774 | 0.10761 | 0.232 | -5.1 | 0.17075 | none |
| XKR8 | 0.266 | 0.1241 | 1.773 | 0.10771 | 0.232 | -5.1 | -0.7451 | none |
| PSD | 0.211 | 0.1377 | 1.773 | 0.10772 | 0.232 | -5.1 | 1.36275 | none |
| OTX1 | -0.161 | -0.1412 | -1.773 | 0.10774 | 0.232 | -5.1 | -0.1018 | none |
| TGFBRAP1 | 0.319 | 0.3704 | 1.7727 | 0.10776 | 0.232 | -5.11 | -0.2057 | none |
| HRAS | 0.155 | 0.1501 | 1.7726 | 0.10779 | 0.232 | -5.11 | -0.2508 | none |
| NACC1 | -0.332 | -0.2081 | -1.772 | 0.10781 | 0.232 | -5.11 | 1.39217 | none |
| TIMM10 | -0.309 | -0.3294 | -1.772 | 0.10797 | 0.233 | -5.11 | 1.01645 | none |
| IL10RA | 0.365 | 0.3311 | 1.7715 | 0.10797 | 0.233 | -5.11 | -0.1904 | none |

| | | | | | | | | |
|---|---|---|---|---|---|---|---|---|
| TEX264 | -0.288 | -0.2701 | -1.771 | 0.10798 | 0.233 | -5.11 | 1.7018 | none |
| MRPS33 | 0.25 | 0.0842 | 1.7714 | 0.10798 | 0.233 | -5.11 | -0.8738 | none |
| UROD | -0.229 | -0.1933 | -1.771 | 0.108 | 0.233 | -5.11 | 2.21448 | AA-related |
| MB | 0.549 | 0.4659 | 1.7711 | 0.10805 | 0.233 | -5.11 | -0.292 | none |
| PRKCB | 0.34 | 0.3885 | 1.771 | 0.10805 | 0.233 | -5.11 | 0.72747 | none |
| PLCD1 | 0.338 | 0.2606 | 1.7709 | 0.10807 | 0.233 | -5.11 | -0.7285 | none |
| RNASE7 | -0.242 | -0.2037 | -1.771 | 0.10814 | 0.233 | -5.11 | 1.25473 | none |
| MGAT1 | -0.209 | -0.2122 | -1.77 | 0.10823 | 0.233 | -5.11 | 1.97408 | none |
| PIK3R5 | 0.415 | 0.4128 | 1.7697 | 0.10828 | 0.233 | -5.11 | 0.19409 | none |
| BAG3 | 0.351 | 0.411 | 1.7694 | 0.10833 | 0.233 | -5.11 | -0.2634 | none |
| PIP5K1B | 0.173 | 0.1383 | 1.7692 | 0.10836 | 0.233 | -5.11 | -0.3791 | none |
| LY9 | 0.305 | 0.133 | 1.7692 | 0.10838 | 0.233 | -5.11 | -0.7295 | none |
| DRAM2 | -0.335 | -0.3532 | -1.768 | 0.1085 | 0.234 | -5.11 | 1.21324 | none |
| FLNB | 0.576 | 0.5772 | 1.7679 | 0.1086 | 0.234 | -5.11 | -0.3986 | none |
| HIBCH | 0.168 | 0.0987 | 1.7678 | 0.10862 | 0.234 | -5.11 | 0.53997 | none |
| PARPBP | 0.296 | 0.1905 | 1.7677 | 0.10864 | 0.234 | -5.11 | -0.49 | none |
| RPL41 | -0.341 | -0.3877 | -1.767 | 0.10872 | 0.234 | -5.11 | 2.1759 | AA-related |
| CTBP1 | 0.22 | 0.2786 | 1.7672 | 0.10873 | 0.234 | -5.11 | -0.195 | none |
| ERP27 | 0.328 | 0.2715 | 1.7671 | 0.10874 | 0.234 | -5.11 | -0.8554 | none |
| YPEL5 | -0.242 | -0.2677 | -1.767 | 0.10875 | 0.234 | -5.11 | 1.77329 | none |
| TNFRSF12/ | 0.386 | 0.3273 | 1.7666 | 0.10882 | 0.234 | -5.11 | -0.1531 | none |
| TMEM86B | 0.339 | 0.3377 | 1.7665 | 0.10883 | 0.234 | -5.11 | -0.9608 | none |
| HCFC1 | 0.532 | 0.6722 | 1.7662 | 0.1089 | 0.234 | -5.11 | -0.0993 | none |
| KCTD15 | -0.243 | -0.1581 | -1.766 | 0.10891 | 0.234 | -5.11 | 1.19645 | none |
| SPON1 | -0.181 | -0.1439 | -1.766 | 0.10893 | 0.234 | -5.11 | 0.17318 | none |
| CMPK1 | -0.186 | -0.2143 | -1.766 | 0.10896 | 0.234 | -5.12 | 1.99098 | none |
| SNRPG | -0.292 | -0.3149 | -1.766 | 0.10901 | 0.234 | -5.12 | 0.94955 | none |
| IL23R | 0.322 | 0.2673 | 1.7652 | 0.10907 | 0.234 | -5.12 | -0.6349 | none |
| PTPRO | -0.175 | -0.1556 | -1.765 | 0.10909 | 0.234 | -5.12 | 1.92345 | none |
| PSTK | 0.311 | 0.2629 | 1.7651 | 0.10909 | 0.234 | -5.12 | -0.8415 | none |
| CHCHD7 | -0.261 | -0.2958 | -1.765 | 0.10912 | 0.234 | -5.12 | 1.08775 | none |
| CCL14 | -0.183 | -0.1292 | -1.764 | 0.10926 | 0.235 | -5.12 | 0.72952 | none |
| PGAP3 | 0.221 | 0.186 | 1.7634 | 0.10938 | 0.235 | -5.12 | 1.01615 | none |
| FAM229A | -0.266 | -0.2148 | -1.763 | 0.10938 | 0.235 | -5.12 | 0.70377 | none |
| VRTN | -0.17 | -0.1291 | -1.763 | 0.10952 | 0.235 | -5.12 | 0.2379 | none |
| NAT9 | 0.258 | 0.2287 | 1.7623 | 0.10959 | 0.235 | -5.12 | -0.272 | none |
| PBK | -0.332 | -0.311 | -1.762 | 0.10961 | 0.235 | -5.12 | 2.06613 | none |
| STX11 | 0.134 | 0.1344 | 1.7621 | 0.10962 | 0.235 | -5.12 | 0.40145 | none |
| BCL9L | 0.546 | 0.6091 | 1.762 | 0.10963 | 0.235 | -5.12 | -0.5194 | none |
| KIF14 | 0.475 | 0.5296 | 1.7613 | 0.10976 | 0.235 | -5.12 | -0.1056 | none |

| | | | | | | | | |
|---|---|---|---|---|---|---|---|---|
| FBXO11 | 0.465 | 0.4529 | 1.7607 | 0.10986 | 0.236 | -5.12 | -0.7017 | none |
| ECSCR | -0.248 | -0.2495 | -1.761 | 0.10989 | 0.236 | -5.12 | -0.561 | none |
| IVNS1ABP | 0.126 | 0.1182 | 1.7603 | 0.10993 | 0.236 | -5.12 | 0.30383 | none |
| KRBA1 | 0.264 | 0.3149 | 1.7597 | 0.11003 | 0.236 | -5.12 | 0.33906 | none |
| CFH | 0.287 | 0.1817 | 1.7595 | 0.11007 | 0.236 | -5.12 | -0.832 | none |
| TRIM62 | 0.39 | 0.3019 | 1.7593 | 0.11011 | 0.236 | -5.12 | -0.4767 | none |
| MAP7 | 0.356 | 0.457 | 1.7592 | 0.11013 | 0.236 | -5.12 | -0.2268 | none |
| AASS | 0.29 | 0.2571 | 1.7585 | 0.11024 | 0.236 | -5.13 | -1.0893 | none |
| AGTPBP1 | 0.269 | 0.2552 | 1.7585 | 0.11025 | 0.236 | -5.13 | 0.63401 | none |
| PRDM4 | 0.305 | 0.3673 | 1.7579 | 0.11035 | 0.236 | -5.13 | 0.59274 | none |
| SLC28A3 | -0.253 | -0.1552 | -1.758 | 0.11037 | 0.236 | -5.13 | 0.5414 | none |
| ZNF280B | 0.207 | 0.173 | 1.7575 | 0.11043 | 0.237 | -5.13 | 0.15294 | none |
| 5-Mar | -0.571 | -0.4693 | -1.757 | 0.11045 | 0.237 | -5.13 | 2.20606 | AA-related |
| JMJD1C | 0.487 | 0.5668 | 1.7573 | 0.11046 | 0.237 | -5.13 | -0.2005 | none |
| TOMM7 | -0.343 | -0.3805 | -1.757 | 0.1105 | 0.237 | -5.13 | 0.73533 | none |
| SPTBN4 | 0.201 | 0.1806 | 1.7569 | 0.11052 | 0.237 | -5.13 | -0.6729 | none |
| SLC6A12 | 0.272 | 0.221 | 1.7567 | 0.11057 | 0.237 | -5.13 | 0.05647 | none |
| MUL1 | -0.184 | -0.1674 | -1.757 | 0.11057 | 0.237 | -5.13 | 1.11437 | none |
| PPP1R35 | -0.436 | -0.4337 | -1.756 | 0.11065 | 0.237 | -5.13 | 0.98412 | none |
| IL13 | -0.291 | -0.2104 | -1.756 | 0.11065 | 0.237 | -5.13 | 0.85546 | none |
| COPB1 | 0.187 | 0.0785 | 1.756 | 0.11069 | 0.237 | -5.13 | 0.9123 | none |
| GINM1 | 0.204 | 0.0959 | 1.7559 | 0.11071 | 0.237 | -5.13 | -0.6817 | none |
| DNAH2 | 0.243 | 0.1432 | 1.7555 | 0.11077 | 0.237 | -5.13 | -1.0823 | none |
| BPGM | -0.182 | -0.2047 | -1.755 | 0.11079 | 0.237 | -5.13 | 0.32611 | none |
| ANO8 | -0.148 | -0.0938 | -1.755 | 0.11081 | 0.237 | -5.13 | -0.1456 | none |
| SLC22A8 | 0.164 | 0.1644 | 1.7551 | 0.11086 | 0.237 | -5.13 | 1.02679 | none |
| ADGRE5 | 0.342 | 0.2721 | 1.7549 | 0.11088 | 0.237 | -5.13 | -0.6744 | none |
| ARG2 | -0.35 | -0.2743 | -1.755 | 0.11093 | 0.237 | -5.13 | 2.1524 | AA-related |
| STYXL1 | -0.163 | -0.2269 | -1.755 | 0.11095 | 0.237 | -5.13 | 0.18904 | none |
| THUMPD1 | 0.315 | 0.3089 | 1.7544 | 0.11099 | 0.237 | -5.13 | -0.416 | none |
| MTFP1 | -0.231 | -0.1838 | -1.754 | 0.11099 | 0.237 | -5.13 | 1.64871 | none |
| RPL26L1 | 0.218 | 0.0669 | 1.7542 | 0.11102 | 0.237 | -5.13 | -0.2342 | none |
| MRPL37 | -0.245 | -0.2083 | -1.754 | 0.11106 | 0.237 | -5.13 | 0.90883 | none |
| SPEG | 0.157 | 0.0634 | 1.7539 | 0.11107 | 0.237 | -5.13 | -0.4519 | none |
| SHD | 0.296 | 0.231 | 1.7537 | 0.1111 | 0.237 | -5.13 | 0.40111 | none |
| B4GALT5 | 0.166 | 0.1466 | 1.7522 | 0.11137 | 0.238 | -5.13 | -0.9673 | none |
| ITGAV | 0.301 | 0.385 | 1.7522 | 0.11138 | 0.238 | -5.13 | -0.5518 | none |
| RRN3 | 0.173 | 0.1452 | 1.7521 | 0.11139 | 0.238 | -5.13 | -0.2947 | none |
| STX17 | 0.237 | 0.1915 | 1.7514 | 0.11152 | 0.238 | -5.14 | 1.01099 | none |
| WDHD1 | 0.389 | 0.4047 | 1.7512 | 0.11155 | 0.238 | -5.14 | -0.0838 | none |

| | | | | | | | | |
|---|---|---|---|---|---|---|---|---|
| YES1 | 0.34 | 0.2982 | 1.7511 | 0.11156 | 0.238 | -5.14 | 0.18293 | none |
| ZFC3H1 | 0.439 | 0.4797 | 1.751 | 0.11159 | 0.238 | -5.14 | -0.5919 | none |
| HMX1 | -0.164 | -0.1261 | -1.751 | 0.11166 | 0.238 | -5.14 | 1.20213 | none |
| KATNAL1 | 0.142 | 0.0713 | 1.7501 | 0.11174 | 0.238 | -5.14 | -0.2733 | none |
| GSTK1 | -0.225 | -0.2346 | -1.75 | 0.11175 | 0.238 | -5.14 | 1.06192 | none |
| ZDHHC22 | -0.165 | -0.1508 | -1.75 | 0.11179 | 0.238 | -5.14 | 1.36672 | none |
| INTS5 | -0.301 | -0.3051 | -1.75 | 0.11179 | 0.238 | -5.14 | 2.1559 | AA-related |
| CYP4F22 | -0.124 | -0.0905 | -1.75 | 0.11183 | 0.238 | -5.14 | 1.65279 | none |
| CCDC6 | 0.372 | 0.3759 | 1.7496 | 0.11184 | 0.238 | -5.14 | -0.0792 | none |
| MED12 | 0.346 | 0.3781 | 1.7495 | 0.11186 | 0.238 | -5.14 | -0.5182 | none |
| PITPNM1 | 0.364 | 0.4214 | 1.7492 | 0.1119 | 0.238 | -5.14 | -0.729 | none |
| PRDM10 | 0.274 | 0.2934 | 1.749 | 0.11194 | 0.239 | -5.14 | 0.15396 | none |
| RELL2 | 0.199 | 0.2157 | 1.7488 | 0.11197 | 0.239 | -5.14 | 0.30135 | none |
| MT1X | -0.377 | -0.3739 | -1.749 | 0.11203 | 0.239 | -5.14 | 2.28873 | AA-related |
| PPP1R9A | 0.14 | 0.1155 | 1.7484 | 0.11205 | 0.239 | -5.14 | -1.4172 | none |
| GPA33 | 0.29 | 0.1499 | 1.7483 | 0.11206 | 0.239 | -5.14 | 1.04376 | none |
| BTK | -0.315 | -0.1952 | -1.748 | 0.11206 | 0.239 | -5.14 | 0.77171 | none |
| DPY30 | -0.254 | -0.2949 | -1.748 | 0.11212 | 0.239 | -5.14 | 1.31371 | none |
| DDA1 | -0.221 | -0.1732 | -1.748 | 0.11213 | 0.239 | -5.14 | 0.65841 | none |
| TCERG1 | 0.541 | 0.6372 | 1.7479 | 0.11214 | 0.239 | -5.14 | 0.14584 | none |
| ASXL2 | 0.539 | 0.5781 | 1.7479 | 0.11214 | 0.239 | -5.14 | -0.4851 | none |
| SLC25A26 | 0.228 | 0.1026 | 1.7478 | 0.11217 | 0.239 | -5.14 | -0.2307 | none |
| BLVRA | -0.264 | -0.2241 | -1.747 | 0.11222 | 0.239 | -5.14 | 1.96939 | none |
| DDR1 | 0.376 | 0.3256 | 1.7474 | 0.11223 | 0.239 | -5.14 | -0.3436 | none |
| POLR2L | -0.234 | -0.2269 | -1.747 | 0.11228 | 0.239 | -5.14 | 0.68096 | none |
| ATXN2L | 0.406 | 0.4462 | 1.747 | 0.11231 | 0.239 | -5.14 | -0.4838 | none |
| BCDIN3D | 0.24 | 0.1519 | 1.7466 | 0.11238 | 0.239 | -5.14 | -0.098 | none |
| DCST2 | -0.159 | -0.1039 | -1.746 | 0.11241 | 0.239 | -5.14 | 1.97408 | none |
| ANKRD1 | -0.119 | -0.0834 | -1.746 | 0.11242 | 0.239 | -5.14 | 0.81131 | none |
| FAM72D | -0.372 | -0.2358 | -1.746 | 0.11246 | 0.239 | -5.14 | 0.53665 | none |
| KMT5A | 0.352 | 0.4289 | 1.7461 | 0.11246 | 0.239 | -5.14 | -0.3667 | none |
| SNAPC5 | 0.205 | 0.1166 | 1.746 | 0.11248 | 0.239 | -5.14 | -0.294 | none |
| POLR3K | 0.21 | 0.1127 | 1.7456 | 0.11255 | 0.239 | -5.14 | -0.9709 | none |
| CDCA5 | -0.163 | -0.1214 | -1.746 | 0.11257 | 0.239 | -5.14 | 2.08448 | none |
| UTP6 | 0.137 | 0.0795 | 1.7454 | 0.11258 | 0.239 | -5.14 | 0.60106 | none |
| MRPL28 | -0.199 | -0.1637 | -1.745 | 0.11265 | 0.239 | -5.14 | 1.34256 | none |
| CLEC18C | -0.214 | -0.1323 | -1.744 | 0.11279 | 0.24 | -5.15 | -0.1931 | none |
| PACRGL | 0.511 | 0.3789 | 1.7443 | 0.1128 | 0.24 | -5.15 | 0.15433 | none |
| IL3 | 0.321 | 0.4617 | 1.7441 | 0.11283 | 0.24 | -5.15 | 0.63795 | none |
| ANKRD28 | 0.447 | 0.4811 | 1.744 | 0.11284 | 0.24 | -5.15 | 0.40238 | none |

| | | | | | | | | |
|---|---|---|---|---|---|---|---|---|
| PRKACB | -0.225 | -0.248 | -1.744 | 0.11286 | 0.24 | -5.15 | 1.28377 | none |
| RELB | 0.207 | 0.1914 | 1.7435 | 0.11293 | 0.24 | -5.15 | 0.04734 | none |
| ETV6 | 0.352 | 0.3654 | 1.743 | 0.11303 | 0.24 | -5.15 | -1.0489 | none |
| COL20A1 | 0.187 | 0.1637 | 1.7427 | 0.11308 | 0.24 | -5.15 | 0.68625 | none |
| RAX | -0.183 | -0.1686 | -1.743 | 0.11311 | 0.24 | -5.15 | 1.31308 | none |
| VPS13C | 0.361 | 0.3108 | 1.7423 | 0.11315 | 0.24 | -5.15 | -0.7409 | none |
| SF3B4 | 0.168 | 0.1871 | 1.7419 | 0.11322 | 0.24 | -5.15 | 0.33597 | none |
| NIT2 | -0.222 | -0.218 | -1.742 | 0.11323 | 0.24 | -5.15 | 1.2059 | none |
| SPATA13 | 0.339 | 0.3538 | 1.7416 | 0.11327 | 0.24 | -5.15 | -0.645 | none |
| STIP1 | 0.507 | 0.5694 | 1.7415 | 0.11329 | 0.24 | -5.15 | -0.4568 | none |
| ASRGL1 | 0.33 | 0.2809 | 1.7415 | 0.1133 | 0.24 | -5.15 | -0.1542 | none |
| NDUFS2 | -0.152 | -0.1488 | -1.741 | 0.11343 | 0.24 | -5.15 | 1.24691 | none |
| TMEM18 | -0.234 | -0.2313 | -1.74 | 0.11362 | 0.241 | -5.15 | 1.88612 | none |
| ZNF700 | 0.3 | 0.269 | 1.7395 | 0.11366 | 0.241 | -5.15 | -0.0884 | none |
| ZFP62 | 0.322 | 0.3084 | 1.7393 | 0.11369 | 0.241 | -5.15 | 0.93931 | none |
| COLQ | 0.213 | 0.1787 | 1.7393 | 0.1137 | 0.241 | -5.15 | -0.1192 | none |
| AC233755 | -0.516 | -0.4828 | -1.739 | 0.11373 | 0.241 | -5.15 | 0.83202 | none |
| KSR2 | 0.127 | 0.1107 | 1.7386 | 0.11383 | 0.241 | -5.15 | -1.2832 | none |
| SPATC1 | 0.169 | 0.1298 | 1.7379 | 0.11394 | 0.241 | -5.15 | -0.1969 | none |
| FAM186B | -0.137 | -0.0736 | -1.738 | 0.11398 | 0.241 | -5.16 | 0.87241 | none |
| CBLN3 | -0.184 | -0.0662 | -1.738 | 0.114 | 0.241 | -5.16 | 0.27652 | none |
| SOS2 | 0.252 | 0.2215 | 1.7373 | 0.11406 | 0.242 | -5.16 | -0.4368 | none |
| PYCR1 | 0.163 | 0.1385 | 1.7369 | 0.11414 | 0.242 | -5.16 | -0.0901 | none |
| GPR35 | 0.251 | 0.2531 | 1.7356 | 0.11437 | 0.242 | -5.16 | -0.2066 | none |
| GCHFR | -0.304 | -0.2818 | -1.735 | 0.11441 | 0.242 | -5.16 | 0.7918 | none |
| SS18L1 | 0.25 | 0.3187 | 1.7354 | 0.11441 | 0.242 | -5.16 | -0.4137 | none |
| LTBP3 | 0.366 | 0.296 | 1.7352 | 0.11444 | 0.242 | -5.16 | -0.8274 | none |
| KIF5C | 0.229 | 0.2752 | 1.735 | 0.11448 | 0.242 | -5.16 | -0.0953 | none |
| NKAPL | 0.149 | 0.1207 | 1.735 | 0.11448 | 0.242 | -5.16 | -0.6686 | none |
| TRIM38 | 0.286 | 0.2682 | 1.7348 | 0.11452 | 0.242 | -5.16 | -0.9075 | none |
| OVOL3 | -0.201 | -0.1472 | -1.734 | 0.11464 | 0.242 | -5.16 | 1.67977 | none |
| UBE2N | -0.269 | -0.2883 | -1.734 | 0.11467 | 0.242 | -5.16 | 1.05874 | none |
| BMP4 | 0.205 | 0.1358 | 1.734 | 0.11467 | 0.242 | -5.16 | -0.2226 | none |
| PRX | 0.225 | 0.1998 | 1.7337 | 0.11473 | 0.243 | -5.16 | -0.1444 | none |
| DDO | -0.13 | -0.0877 | -1.734 | 0.11474 | 0.243 | -5.16 | 0.45659 | none |
| COMMD7 | -0.22 | -0.1986 | -1.733 | 0.11486 | 0.243 | -5.16 | 1.24719 | none |
| NCBP2-AS | 0.208 | 0.1602 | 1.7328 | 0.11488 | 0.243 | -5.16 | -0.475 | none |
| MYO1H | 0.234 | 0.2884 | 1.7325 | 0.11494 | 0.243 | -5.16 | 0.64996 | none |
| TMEM71 | 0.347 | 0.2039 | 1.7324 | 0.11496 | 0.243 | -5.16 | -0.832 | none |
| NPAS2 | 0.3 | 0.2291 | 1.7324 | 0.11496 | 0.243 | -5.16 | -0.9638 | none |

| | | | | | | | | |
|---|---|---|---|---|---|---|---|---|
| NPIPA7 | 0.806 | 0.6313 | 1.7323 | 0.11497 | 0.243 | -5.16 | -0.1604 | none |
| ARHGAP1( | 0.243 | 0.2894 | 1.732 | 0.11503 | 0.243 | -5.16 | 0.29802 | none |
| IFT81 | 0.422 | 0.3101 | 1.7318 | 0.11507 | 0.243 | -5.16 | -0.7412 | none |
| RPS18 | -0.335 | -0.3786 | -1.732 | 0.11507 | 0.243 | -5.16 | 1.92288 | none |
| COX15 | 0.177 | 0.1125 | 1.7317 | 0.11509 | 0.243 | -5.16 | -0.7192 | none |
| INS | -0.196 | -0.1304 | -1.732 | 0.1151 | 0.243 | -5.16 | -0.0086 | none |
| C17ORF49 | -0.267 | -0.2568 | -1.732 | 0.11511 | 0.243 | -5.16 | 1.70626 | none |
| VPS13A | 0.588 | 0.5863 | 1.7307 | 0.11527 | 0.243 | -5.17 | -0.8485 | none |
| PDE6B | 0.317 | 0.2006 | 1.7306 | 0.11528 | 0.243 | -5.17 | 0.60921 | none |
| MTMR9 | 0.149 | 0.1242 | 1.7303 | 0.11534 | 0.243 | -5.17 | 0.12676 | none |
| HAPLN1 | 0.439 | 0.2825 | 1.7302 | 0.11537 | 0.243 | -5.17 | -0.8921 | none |
| CDK6 | 0.163 | 0.1849 | 1.73 | 0.11541 | 0.243 | -5.17 | 0.79354 | none |
| SLC2A7 | -0.127 | -0.1092 | -1.73 | 0.11545 | 0.244 | -5.17 | 2.3646 | AA-related |
| HNRNPA0 | -0.15 | -0.1422 | -1.729 | 0.11549 | 0.244 | -5.17 | 1.27637 | none |
| RBL1 | 0.311 | 0.3279 | 1.7294 | 0.11552 | 0.244 | -5.17 | -0.0802 | none |
| CRB3 | -0.275 | -0.1633 | -1.729 | 0.11553 | 0.244 | -5.17 | 1.75797 | none |
| OPTN | 0.427 | 0.4459 | 1.7292 | 0.11555 | 0.244 | -5.17 | -0.3807 | none |
| HK2 | 0.383 | 0.4763 | 1.7283 | 0.11571 | 0.244 | -5.17 | 0.3627 | none |
| PLOD2 | 0.161 | 0.1495 | 1.7281 | 0.11575 | 0.244 | -5.17 | -0.1163 | none |
| TEN1-CDK | 0.287 | 0.2518 | 1.7279 | 0.11579 | 0.244 | -5.17 | -0.6774 | none |
| CNIH4 | -0.349 | -0.4019 | -1.728 | 0.11581 | 0.244 | -5.17 | 0.95088 | none |
| UBALD2 | -0.414 | -0.3557 | -1.728 | 0.11583 | 0.244 | -5.17 | 1.47827 | none |
| ANAPC15 | 0.132 | 0.1069 | 1.7273 | 0.1159 | 0.244 | -5.17 | -0.1272 | none |
| MAGIX | 0.339 | 0.2155 | 1.7272 | 0.11592 | 0.244 | -5.17 | -0.4802 | none |
| CCHCR1 | 0.228 | 0.1823 | 1.7271 | 0.11593 | 0.244 | -5.17 | -0.4545 | none |
| SPAG5 | 0.564 | 0.6495 | 1.7267 | 0.11601 | 0.244 | -5.17 | -0.4638 | none |
| ABO | 0.255 | 0.2195 | 1.7265 | 0.11605 | 0.244 | -5.17 | -0.7861 | none |
| USP8 | 0.617 | 0.6032 | 1.7264 | 0.11606 | 0.244 | -5.17 | -0.3935 | none |
| ANKRD36 | 0.298 | 0.1441 | 1.7261 | 0.11612 | 0.244 | -5.17 | -0.5754 | none |
| POLRMT | -0.305 | -0.1352 | -1.726 | 0.11613 | 0.244 | -5.17 | 1.09007 | none |
| OTP | 0.135 | 0.1363 | 1.7252 | 0.11629 | 0.245 | -5.17 | 0.96251 | none |
| GOLGA6B | 0.358 | 0.2276 | 1.7251 | 0.1163 | 0.245 | -5.17 | -0.3459 | none |
| PCK2 | 0.148 | 0.1028 | 1.7243 | 0.11646 | 0.245 | -5.17 | -0.3886 | none |
| PSG1 | 0.184 | 0.2087 | 1.724 | 0.11651 | 0.245 | -5.17 | -0.4702 | none |
| CTPS1 | 0.124 | 0.1104 | 1.724 | 0.11652 | 0.245 | -5.17 | 0.04143 | none |
| PPA2 | -0.183 | -0.2228 | -1.723 | 0.11661 | 0.245 | -5.18 | 1.1201 | none |
| TMEM43 | 0.198 | 0.1998 | 1.7234 | 0.11662 | 0.245 | -5.18 | 0.14946 | none |
| IP6K1 | -0.224 | -0.1896 | -1.723 | 0.11665 | 0.245 | -5.18 | 1.58736 | none |
| HIST2H2A( | -0.593 | -0.6505 | -1.723 | 0.11678 | 0.246 | -5.18 | 0.65201 | none |
| PDCD10 | -0.22 | -0.2339 | -1.722 | 0.11685 | 0.246 | -5.18 | 1.1103 | none |

| | | | | | | | | |
|---|---|---|---|---|---|---|---|---|
| IL1RN | -0.308 | -0.2444 | -1.722 | 0.11692 | 0.246 | -5.18 | 1.36114 | none |
| PEX3 | -0.224 | -0.1892 | -1.722 | 0.11693 | 0.246 | -5.18 | 1.10138 | none |
| SNAP23 | 0.246 | 0.0907 | 1.7217 | 0.11694 | 0.246 | -5.18 | 0.24469 | none |
| ALOX15B | 0.242 | 0.1593 | 1.7217 | 0.11694 | 0.246 | -5.18 | -1.0055 | none |
| LMNB1 | 0.418 | 0.5126 | 1.7217 | 0.11694 | 0.246 | -5.18 | 0.6444 | none |
| USP46 | 0.37 | 0.1964 | 1.7214 | 0.11699 | 0.246 | -5.18 | -0.9955 | none |
| POGLUT1 | 0.176 | 0.169 | 1.7213 | 0.11702 | 0.246 | -5.18 | -0.7126 | none |
| PXDC1 | -0.169 | -0.1669 | -1.721 | 0.11709 | 0.246 | -5.18 | 0.02749 | none |
| CTDSPL | 0.605 | 0.4101 | 1.7204 | 0.11718 | 0.246 | -5.18 | -0.2419 | none |
| MRPL27 | -0.208 | -0.1959 | -1.72 | 0.11721 | 0.246 | -5.18 | 0.83386 | none |
| ANKRD61 | 0.128 | 0.1443 | 1.7199 | 0.11728 | 0.246 | -5.18 | -0.1234 | none |
| CRIP2 | -0.295 | -0.1946 | -1.72 | 0.11732 | 0.246 | -5.18 | 1.8286 | none |
| CCNH | -0.215 | -0.216 | -1.72 | 0.11732 | 0.246 | -5.18 | 2.08239 | none |
| ARHGDIA | 0.189 | 0.2081 | 1.7195 | 0.11735 | 0.246 | -5.18 | 0.99908 | none |
| GPRC5B | -0.176 | -0.1468 | -1.72 | 0.11735 | 0.246 | -5.18 | 1.58652 | none |
| PRKAA2 | 0.122 | 0.0824 | 1.7192 | 0.1174 | 0.246 | -5.18 | -1.2447 | none |
| SSX7 | 0.199 | 0.1158 | 1.7174 | 0.11774 | 0.247 | -5.18 | -1.0749 | none |
| SYN3 | 0.408 | 0.34 | 1.7168 | 0.11787 | 0.247 | -5.19 | -0.5525 | none |
| CBY3 | 0.145 | 0.1452 | 1.7166 | 0.11789 | 0.247 | -5.19 | -0.3557 | none |
| MDK | -0.158 | -0.102 | -1.717 | 0.1179 | 0.247 | -5.19 | -0.1812 | none |
| PSMD11 | -0.159 | -0.0865 | -1.716 | 0.11796 | 0.247 | -5.19 | 0.36268 | none |
| KIF17 | 0.325 | 0.2418 | 1.7161 | 0.11799 | 0.247 | -5.19 | 0.21639 | none |
| C18ORF54 | 0.233 | 0.2531 | 1.716 | 0.11801 | 0.247 | -5.19 | 0.47363 | none |
| DCTN2 | 0.211 | 0.1628 | 1.7156 | 0.11809 | 0.248 | -5.19 | 0.61924 | none |
| GHRHR | 0.244 | 0.2376 | 1.7155 | 0.11811 | 0.248 | -5.19 | -0.5038 | none |
| MYPOP | 0.149 | 0.1764 | 1.7153 | 0.11813 | 0.248 | -5.19 | 0.13752 | none |
| HSD3B7 | 0.235 | 0.1572 | 1.7151 | 0.11818 | 0.248 | -5.19 | -0.2512 | none |
| LTBP2 | -0.171 | -0.1538 | -1.715 | 0.11821 | 0.248 | -5.19 | 1.0703 | none |
| MIA | 0.464 | 0.382 | 1.7147 | 0.11826 | 0.248 | -5.19 | -1.0047 | none |
| PSG6 | -0.168 | -0.0989 | -1.714 | 0.1183 | 0.248 | -5.19 | 2.19802 | AA-related |
| NECAP2 | -0.189 | -0.2034 | -1.714 | 0.1184 | 0.248 | -5.19 | 1.14324 | none |
| SLC4A10 | -0.183 | -0.0962 | -1.714 | 0.11846 | 0.248 | -5.19 | 2.36738 | AA-related |
| ZNF335 | 0.369 | 0.4417 | 1.7129 | 0.1186 | 0.248 | -5.19 | -0.6581 | none |
| JUNB | -0.108 | -0.0499 | -1.713 | 0.1186 | 0.248 | -5.19 | 1.57848 | none |
| ARHGAP1: | 0.238 | 0.2009 | 1.7128 | 0.11862 | 0.248 | -5.19 | 1.20964 | none |
| PRDM1 | 0.381 | 0.4341 | 1.7127 | 0.11863 | 0.248 | -5.19 | -0.7132 | none |
| TFB2M | -0.331 | -0.3053 | -1.712 | 0.11867 | 0.248 | -5.19 | 1.79267 | none |
| NFYC | 0.243 | 0.1908 | 1.7124 | 0.11868 | 0.248 | -5.19 | -0.7654 | none |
| PHF14 | 0.476 | 0.4749 | 1.7119 | 0.11878 | 0.248 | -5.19 | 0.08977 | none |
| SERTAD1 | -0.221 | -0.1924 | -1.712 | 0.11881 | 0.249 | -5.19 | 1.23729 | none |

| | | | | | | | | |
|---|---|---|---|---|---|---|---|---|
| NCAPD2 | 0.444 | 0.4901 | 1.7117 | 0.11882 | 0.249 | -5.19 | -0.3695 | none |
| P2RY10 | -0.252 | -0.2905 | -1.712 | 0.11885 | 0.249 | -5.19 | 2.11196 | AA-related |
| UBL7 | -0.203 | -0.1922 | -1.711 | 0.1189 | 0.249 | -5.19 | 1.8725 | none |
| FBXO22 | 0.299 | 0.089 | 1.7112 | 0.11892 | 0.249 | -5.19 | -0.5795 | none |
| ADPRM | 0.262 | 0.0799 | 1.7108 | 0.11899 | 0.249 | -5.19 | 0.68775 | none |
| DDX27 | 0.492 | 0.6103 | 1.7107 | 0.11901 | 0.249 | -5.19 | -0.4555 | none |
| CLHC1 | 0.323 | 0.2509 | 1.7103 | 0.11909 | 0.249 | -5.19 | -0.6383 | none |
| TINAGL1 | 0.246 | 0.1969 | 1.71 | 0.11915 | 0.249 | -5.19 | -0.9571 | none |
| SATB1 | 0.32 | 0.3474 | 1.7098 | 0.11918 | 0.249 | -5.19 | -0.0431 | none |
| RPS26 | -0.339 | -0.3601 | -1.709 | 0.11927 | 0.249 | -5.2 | 1.99767 | none |
| HPGD | 0.175 | 0.0276 | 1.7092 | 0.1193 | 0.249 | -5.2 | 0.84064 | none |
| DAZAP2 | -0.213 | -0.2541 | -1.709 | 0.11943 | 0.249 | -5.2 | 0.60293 | none |
| ADAMTSL4 | 0.212 | 0.1757 | 1.7084 | 0.11944 | 0.249 | -5.2 | -0.8315 | none |
| FOXR1 | -0.209 | -0.1759 | -1.708 | 0.11945 | 0.249 | -5.2 | 0.89101 | none |
| CXCL8 | -0.559 | -0.4937 | -1.708 | 0.11953 | 0.25 | -5.2 | 0.99565 | none |
| FIBCD1 | 0.23 | 0.1869 | 1.7079 | 0.11954 | 0.25 | -5.2 | -0.1852 | none |
| CSK | -0.198 | -0.1205 | -1.708 | 0.11962 | 0.25 | -5.2 | 1.67592 | none |
| PRIM2 | 0.368 | 0.2028 | 1.7069 | 0.11973 | 0.25 | -5.2 | -0.2909 | none |
| HNRNPUL1 | 0.652 | 0.7313 | 1.706 | 0.1199 | 0.25 | -5.2 | 0.03093 | none |
| ATF4 | -0.175 | -0.0758 | -1.706 | 0.11995 | 0.25 | -5.2 | 1.52463 | none |
| LCA5 | 0.156 | 0.1039 | 1.7055 | 0.12 | 0.25 | -5.2 | -0.9755 | none |
| MRPL49 | 0.253 | 0.1558 | 1.7045 | 0.1202 | 0.251 | -5.2 | -1.1534 | none |
| ZFP91 | 0.231 | 0.2437 | 1.7043 | 0.12024 | 0.251 | -5.2 | 0.22324 | none |
| SNX5 | 0.165 | 0.1481 | 1.7042 | 0.12025 | 0.251 | -5.2 | -0.9376 | none |
| COQ2 | 0.278 | 0.206 | 1.7042 | 0.12026 | 0.251 | -5.2 | 0.73775 | none |
| USP11 | 0.254 | 0.2055 | 1.7041 | 0.12026 | 0.251 | -5.2 | 0.04538 | none |
| ARHGAP11 | 0.486 | 0.5791 | 1.7039 | 0.12031 | 0.251 | -5.2 | 0.81292 | none |
| DIS3 | 0.359 | 0.4135 | 1.7034 | 0.1204 | 0.251 | -5.2 | -0.0877 | none |
| MLEC | 0.358 | 0.3534 | 1.7033 | 0.12042 | 0.251 | -5.2 | 0.1328 | none |
| C8ORF74 | 0.32 | 0.2408 | 1.7028 | 0.12051 | 0.251 | -5.2 | -0.7238 | none |
| KIAA1671 | 0.276 | 0.256 | 1.7027 | 0.12054 | 0.251 | -5.2 | -0.384 | none |
| GPR63 | 0.151 | 0.0969 | 1.7027 | 0.12054 | 0.251 | -5.2 | 0.24285 | none |
| DUSP6 | -0.185 | -0.0599 | -1.702 | 0.12063 | 0.251 | -5.21 | 1.5779 | none |
| CNIH2 | 0.369 | 0.3048 | 1.7021 | 0.12065 | 0.251 | -5.21 | -0.4575 | none |
| NUDT16L1 | -0.224 | -0.2029 | -1.701 | 0.12083 | 0.252 | -5.21 | 2.0547 | none |
| RNF122 | -0.142 | -0.1268 | -1.701 | 0.12084 | 0.252 | -5.21 | 1.22138 | none |
| LRRC56 | 0.26 | 0.1423 | 1.7011 | 0.12084 | 0.252 | -5.21 | -1.0847 | none |
| SMC1A | 0.653 | 0.7491 | 1.7005 | 0.12095 | 0.252 | -5.21 | 0.21109 | none |
| THBS3 | 0.278 | 0.2226 | 1.6999 | 0.12108 | 0.252 | -5.21 | -0.531 | none |
| HIST1H4H | 0.178 | -0.0244 | 1.6996 | 0.12113 | 0.252 | -5.21 | -0.5885 | none |

| | | | | | | | | |
|---|---|---|---|---|---|---|---|---|
| KIF12 | 0.187 | 0.152 | 1.6996 | 0.12114 | 0.252 | -5.21 | 0.72857 | none |
| CHURC1 | -0.259 | -0.3237 | -1.699 | 0.12118 | 0.252 | -5.21 | 0.71614 | none |
| PABPC1L | 0.25 | 0.2615 | 1.6992 | 0.12122 | 0.252 | -5.21 | -0.6154 | none |
| SEPHS1 | 0.255 | 0.2189 | 1.6987 | 0.12131 | 0.252 | -5.21 | -0.6453 | none |
| EPX | 0.144 | 0.1009 | 1.6985 | 0.12134 | 0.252 | -5.21 | -1.323 | none |
| GM2A | 0.142 | 0.0644 | 1.6984 | 0.12137 | 0.252 | -5.21 | -0.7036 | none |
| ZNF460 | 0.225 | 0.1954 | 1.698 | 0.12144 | 0.253 | -5.21 | -0.7466 | none |
| MRPS34 | -0.169 | -0.1103 | -1.698 | 0.12147 | 0.253 | -5.21 | 1.91033 | none |
| NOC4L | -0.155 | -0.1243 | -1.698 | 0.12152 | 0.253 | -5.21 | 1.92078 | none |
| MRPL19 | -0.19 | -0.1662 | -1.697 | 0.1217 | 0.253 | -5.21 | 2.1971 | AA-related |
| GCG | 0.191 | 0.1275 | 1.6964 | 0.12175 | 0.253 | -5.21 | -0.6396 | none |
| UFSP1 | 0.25 | 0.1293 | 1.6956 | 0.1219 | 0.253 | -5.21 | 0.09701 | none |
| STAB2 | -0.375 | -0.2263 | -1.695 | 0.12194 | 0.253 | -5.22 | 1.12791 | none |
| HADHB | -0.145 | -0.1793 | -1.695 | 0.12194 | 0.253 | -5.22 | 1.27333 | none |
| ARHGEF25 | 0.248 | 0.2539 | 1.695 | 0.12203 | 0.253 | -5.22 | 1.33643 | none |
| HIST1H4J | -0.526 | -0.4882 | -1.695 | 0.12204 | 0.253 | -5.22 | 0.22845 | none |
| AKR1A1 | 0.238 | 0.1155 | 1.6949 | 0.12205 | 0.253 | -5.22 | -0.285 | none |
| DOCK9 | 0.504 | 0.6702 | 1.6948 | 0.12206 | 0.253 | -5.22 | ######## | none |
| APELA | 0.591 | 0.4685 | 1.6948 | 0.12206 | 0.253 | -5.22 | -1.0839 | none |
| DEF6 | 0.329 | 0.3712 | 1.6948 | 0.12206 | 0.253 | -5.22 | 0.64408 | none |
| SOCS4 | -0.364 | -0.2748 | -1.694 | 0.12223 | 0.254 | -5.22 | 1.94928 | none |
| MORN1 | 0.361 | 0.3101 | 1.6937 | 0.12228 | 0.254 | -5.22 | -0.585 | none |
| GTPBP10 | 0.247 | 0.1295 | 1.6933 | 0.12235 | 0.254 | -5.22 | 0.48601 | none |
| SNX33 | 0.194 | 0.2072 | 1.6933 | 0.12236 | 0.254 | -5.22 | -0.3875 | none |
| NIF3L1 | -0.214 | -0.2479 | -1.693 | 0.12238 | 0.254 | -5.22 | 1.60239 | none |
| ZNF410 | 0.191 | 0.0712 | 1.6925 | 0.1225 | 0.254 | -5.22 | -0.2722 | none |
| PPP2R3B | 0.229 | 0.2027 | 1.6923 | 0.12254 | 0.254 | -5.22 | -0.8472 | none |
| ZNF454 | 0.203 | 0.1204 | 1.6913 | 0.12274 | 0.255 | -5.22 | -0.3864 | none |
| PELI2 | 0.21 | 0.162 | 1.6913 | 0.12274 | 0.255 | -5.22 | -0.2963 | none |
| SLC38A10 | 0.156 | 0.1547 | 1.6913 | 0.12275 | 0.255 | -5.22 | -0.2405 | none |
| NFKBIZ | 0.46 | 0.5545 | 1.691 | 0.1228 | 0.255 | -5.22 | 0.21086 | none |
| ARID5A | 0.244 | 0.2766 | 1.691 | 0.12281 | 0.255 | -5.22 | 0.13374 | none |
| TM4SF4 | -0.161 | -0.112 | -1.691 | 0.12288 | 0.255 | -5.22 | -0.6168 | none |
| C1ORF162 | 0.276 | 0.1546 | 1.69 | 0.123 | 0.255 | -5.22 | -0.3319 | none |
| ETV4 | 0.417 | 0.3466 | 1.6899 | 0.12301 | 0.255 | -5.22 | -0.4697 | none |
| MTHFD2 | -0.146 | -0.1226 | -1.69 | 0.12308 | 0.255 | -5.22 | 1.56648 | none |
| SMIM19 | -0.231 | -0.3156 | -1.689 | 0.12316 | 0.255 | -5.22 | 0.50026 | none |
| CENPM | -0.281 | -0.263 | -1.689 | 0.12321 | 0.255 | -5.22 | 1.21233 | none |
| CNPY2 | 0.241 | 0.1108 | 1.6887 | 0.12325 | 0.255 | -5.22 | -0.7539 | none |
| PPP6R3 | 0.396 | 0.4365 | 1.6886 | 0.12328 | 0.255 | -5.22 | -0.883 | none |

| | | | | | | | | |
|---|---|---|---|---|---|---|---|---|
| CNIH1 | -0.349 | -0.4113 | -1.688 | 0.12333 | 0.255 | -5.23 | 1.65801 | none |
| RANBP10 | 0.277 | 0.3852 | 1.6882 | 0.12334 | 0.255 | -5.23 | 0.28177 | none |
| ZMIZ2 | 0.446 | 0.4655 | 1.6882 | 0.12335 | 0.255 | -5.23 | -0.3607 | none |
| ERCC6L | 0.433 | 0.4689 | 1.688 | 0.12338 | 0.255 | -5.23 | -0.6395 | none |
| NMT1 | 0.246 | 0.2297 | 1.6876 | 0.12346 | 0.256 | -5.23 | 0.77751 | none |
| MVK | 0.141 | 0.13 | 1.6869 | 0.1236 | 0.256 | -5.23 | -0.6198 | none |
| TMEM184 | 0.511 | 0.51 | 1.6861 | 0.12376 | 0.256 | -5.23 | 0.16875 | none |
| GEMIN4 | 0.188 | 0.1635 | 1.685 | 0.12399 | 0.257 | -5.23 | 0.61281 | none |
| ATP1A1 | 0.329 | 0.3343 | 1.6848 | 0.12402 | 0.257 | -5.23 | -0.201 | none |
| ZC2HC1A | -0.191 | -0.2255 | -1.685 | 0.12403 | 0.257 | -5.23 | 0.06559 | none |
| GGN | 0.356 | 0.2596 | 1.6846 | 0.12406 | 0.257 | -5.23 | -0.1185 | none |
| HBM | -0.179 | -0.1194 | -1.684 | 0.12418 | 0.257 | -5.23 | 1.24897 | none |
| USP32 | 0.42 | 0.4816 | 1.6839 | 0.12419 | 0.257 | -5.23 | -0.023 | none |
| LEMD2 | 0.288 | 0.2529 | 1.6836 | 0.12426 | 0.257 | -5.23 | -0.3458 | none |
| PHAX | 0.411 | 0.4356 | 1.6835 | 0.12428 | 0.257 | -5.23 | -0.8225 | none |
| ZNF653 | 0.34 | 0.2988 | 1.6832 | 0.12432 | 0.257 | -5.23 | -0.6612 | none |
| C6ORF58 | 0.124 | 0.1569 | 1.6827 | 0.12444 | 0.257 | -5.23 | -0.8067 | none |
| NSMAF | 0.245 | 0.2372 | 1.6824 | 0.12449 | 0.257 | -5.23 | -0.7712 | none |
| ELOVL6 | 0.523 | 0.5016 | 1.6823 | 0.1245 | 0.257 | -5.23 | -0.5305 | none |
| ZBTB49 | 0.357 | 0.3168 | 1.6822 | 0.12452 | 0.257 | -5.23 | 0.53239 | none |
| NTMT1 | -0.244 | -0.2042 | -1.682 | 0.12459 | 0.257 | -5.23 | 1.13798 | none |
| RGS20 | -0.225 | -0.1407 | -1.682 | 0.12461 | 0.257 | -5.23 | -0.3422 | none |
| USP31 | 0.15 | 0.1471 | 1.6818 | 0.12462 | 0.257 | -5.23 | -0.0423 | none |
| JMJD4 | 0.174 | 0.1313 | 1.6815 | 0.12467 | 0.257 | -5.23 | -0.4266 | none |
| NAT16 | 0.213 | 0.141 | 1.6815 | 0.12467 | 0.257 | -5.23 | 0.04378 | none |
| GABRR1 | 0.223 | 0.205 | 1.6811 | 0.12474 | 0.258 | -5.24 | -0.4809 | none |
| UBE2E3 | -0.145 | -0.1288 | -1.68 | 0.12499 | 0.258 | -5.24 | 1.19065 | none |
| ENKD1 | 0.395 | 0.3854 | 1.6798 | 0.125 | 0.258 | -5.24 | -0.5918 | none |
| COX7C | -0.267 | -0.3044 | -1.68 | 0.125 | 0.258 | -5.24 | 1.49214 | none |
| METRN | -0.244 | -0.1896 | -1.68 | 0.12503 | 0.258 | -5.24 | 1.45784 | none |
| TRIM2 | 0.354 | 0.3343 | 1.6797 | 0.12503 | 0.258 | -5.24 | 0.88544 | none |
| ESYT1 | 0.147 | 0.1322 | 1.6795 | 0.12506 | 0.258 | -5.24 | 1.02114 | none |
| MICU2 | 0.247 | 0.1921 | 1.6794 | 0.12508 | 0.258 | -5.24 | 0.85881 | none |
| PRAMEF1 | 0.203 | 0.1629 | 1.6794 | 0.12508 | 0.258 | -5.24 | 0.30873 | none |
| COL24A1 | 0.224 | 0.2268 | 1.6792 | 0.12512 | 0.258 | -5.24 | 1.36184 | none |
| VMA21 | -0.3 | -0.3063 | -1.679 | 0.12514 | 0.258 | -5.24 | 1.51273 | none |
| CXORF51A | -0.199 | -0.1683 | -1.679 | 0.12514 | 0.258 | -5.24 | 1.76991 | none |
| HLA-DMB | 0.154 | 0.1374 | 1.6791 | 0.12514 | 0.258 | -5.24 | 0.45786 | none |
| AC141272 | -0.177 | -0.1183 | -1.679 | 0.12516 | 0.258 | -5.24 | 1.24897 | none |
| RETNLB | 0.171 | 0.1211 | 1.679 | 0.12517 | 0.258 | -5.24 | 0.18447 | none |

| | | | | | | | | |
|---|---|---|---|---|---|---|---|---|
| SLC34A1 | 0.114 | 0.1333 | 1.6789 | 0.12518 | 0.258 | -5.24 | -0.7854 | none |
| BMF | 0.368 | 0.3517 | 1.6789 | 0.12518 | 0.258 | -5.24 | 0.40435 | none |
| METTL8 | 0.254 | 0.1762 | 1.6789 | 0.12519 | 0.258 | -5.24 | 0.344 | none |
| CDC34 | 0.325 | 0.2259 | 1.6788 | 0.12521 | 0.258 | -5.24 | -0.6257 | none |
| PATL1 | 0.341 | 0.3989 | 1.6787 | 0.12522 | 0.258 | -5.24 | -0.1651 | none |
| OSTF1 | -0.117 | -0.1441 | -1.679 | 0.12525 | 0.258 | -5.24 | 1.31305 | none |
| STAR | 0.223 | 0.115 | 1.6783 | 0.1253 | 0.258 | -5.24 | 0.42243 | none |
| OPN5 | 0.207 | 0.1472 | 1.6783 | 0.12531 | 0.258 | -5.24 | -1.2028 | none |
| PSMD7 | -0.115 | -0.075 | -1.678 | 0.12535 | 0.258 | -5.24 | -0.1349 | none |
| CHD3 | 0.688 | 0.7847 | 1.6779 | 0.12538 | 0.258 | -5.24 | -0.5293 | none |
| TBX21 | 0.225 | 0.286 | 1.6779 | 0.12538 | 0.258 | -5.24 | -0.5626 | none |
| HNRNPF | -0.206 | -0.2068 | -1.678 | 0.1254 | 0.258 | -5.24 | 1.82385 | none |
| C18ORF21 | -0.347 | -0.2749 | -1.678 | 0.12545 | 0.258 | -5.24 | 1.78205 | none |
| DES | 0.251 | 0.2549 | 1.6775 | 0.12547 | 0.258 | -5.24 | -0.32 | none |
| RNPEP | 0.127 | 0.1219 | 1.6768 | 0.12561 | 0.258 | -5.24 | 0.10144 | none |
| HMGN3 | -0.147 | -0.1795 | -1.676 | 0.12568 | 0.259 | -5.24 | 0.55 | none |
| ERGIC2 | -0.235 | -0.1997 | -1.676 | 0.12578 | 0.259 | -5.24 | 1.6728 | none |
| SMTN | 0.377 | 0.3519 | 1.6758 | 0.12581 | 0.259 | -5.24 | -0.1732 | none |
| BET1L | 0.197 | 0.2398 | 1.6757 | 0.12582 | 0.259 | -5.24 | 1.1053 | none |
| CORO1A | -0.164 | -0.1306 | -1.676 | 0.12584 | 0.259 | -5.24 | 1.43635 | none |
| NAA11 | 0.204 | 0.1356 | 1.6754 | 0.12588 | 0.259 | -5.24 | -1.155 | none |
| NT5C1A | -0.168 | -0.1273 | -1.675 | 0.1259 | 0.259 | -5.24 | 2.04812 | none |
| PTGER2 | 0.176 | 0.0638 | 1.6751 | 0.12594 | 0.259 | -5.24 | 0.67806 | none |
| WFDC3 | 0.591 | 0.3985 | 1.6737 | 0.12622 | 0.259 | -5.25 | -0.7499 | none |
| HS3ST3A1 | 0.298 | 0.3046 | 1.6736 | 0.12624 | 0.259 | -5.25 | -0.679 | none |
| ST6GALNA | 0.167 | 0.1256 | 1.6732 | 0.12632 | 0.26 | -5.25 | 0.18521 | none |
| SUPT6H | 0.567 | 0.7103 | 1.6726 | 0.12645 | 0.26 | -5.25 | -0.5052 | none |
| RPS19BP1 | -0.137 | -0.0595 | -1.672 | 0.12647 | 0.26 | -5.25 | 1.37993 | none |
| PDCD2L | -0.367 | -0.3795 | -1.672 | 0.12651 | 0.26 | -5.25 | 1.11627 | none |
| KITLG | 0.128 | 0.1621 | 1.6718 | 0.12661 | 0.26 | -5.25 | -0.5724 | none |
| HIC1 | 0.343 | 0.2812 | 1.6717 | 0.12662 | 0.26 | -5.25 | -0.8682 | none |
| SOX15 | 0.183 | 0.2149 | 1.6716 | 0.12665 | 0.26 | -5.25 | 0.44248 | none |
| FUS | 0.223 | 0.2434 | 1.6711 | 0.12675 | 0.26 | -5.25 | -0.0239 | none |
| SLC24A1 | 0.616 | 0.5913 | 1.671 | 0.12677 | 0.26 | -5.25 | -0.7592 | none |
| BCL2L13 | 0.172 | 0.0869 | 1.6706 | 0.12684 | 0.26 | -5.25 | -1.2194 | none |
| B3GALT2 | 0.165 | 0.1177 | 1.6706 | 0.12684 | 0.26 | -5.25 | -0.8527 | none |
| TCEAL8 | -0.305 | -0.2798 | -1.67 | 0.12692 | 0.26 | -5.25 | 1.77877 | none |
| RPL4 | -0.162 | -0.0966 | -1.67 | 0.12697 | 0.26 | -5.25 | 1.69024 | none |
| ZGPAT | 0.19 | 0.2037 | 1.67 | 0.12697 | 0.26 | -5.25 | -0.7016 | none |
| IL1RL2 | -0.215 | -0.1791 | -1.669 | 0.12709 | 0.261 | -5.25 | -0.0587 | none |

| | | | | | | | | |
|---|---|---|---|---|---|---|---|---|
| STK26 | 0.149 | 0.1076 | 1.6693 | 0.1271 | 0.261 | -5.25 | 1.05724 | none |
| IGFBP2 | 0.261 | 0.1999 | 1.6691 | 0.12714 | 0.261 | -5.25 | 0.02834 | none |
| TRIM3 | 0.234 | 0.1929 | 1.669 | 0.12717 | 0.261 | -5.25 | 0.18623 | none |
| TTC6 | 0.464 | 0.3956 | 1.6688 | 0.1272 | 0.261 | -5.25 | -0.8227 | none |
| KRT77 | 0.247 | 0.159 | 1.6686 | 0.12725 | 0.261 | -5.25 | -1.295 | none |
| SLC6A18 | -0.129 | -0.1009 | -1.668 | 0.12728 | 0.261 | -5.25 | 0.08072 | none |
| NEDD9 | 0.371 | 0.4111 | 1.668 | 0.12737 | 0.261 | -5.25 | -0.2194 | none |
| CYSTM1 | 0.343 | 0.2206 | 1.6676 | 0.12744 | 0.261 | -5.25 | -0.3017 | none |
| BOLA2B | -0.339 | -0.3543 | -1.668 | 0.12746 | 0.261 | -5.25 | 1.24999 | none |
| BLM | 0.415 | 0.5004 | 1.666 | 0.12777 | 0.262 | -5.26 | 0.05369 | none |
| BOLA2 | -0.322 | -0.3028 | -1.666 | 0.12778 | 0.262 | -5.26 | 0.89827 | none |
| MON1A | 0.261 | 0.2084 | 1.6658 | 0.12782 | 0.262 | -5.26 | -0.7138 | none |
| ZFYVE9 | 0.156 | 0.1485 | 1.6655 | 0.12787 | 0.262 | -5.26 | -0.5222 | none |
| RAB13 | 0.325 | 0.2268 | 1.6651 | 0.12795 | 0.262 | -5.26 | -1.1291 | none |
| ADAM9 | 0.262 | 0.2653 | 1.665 | 0.12797 | 0.262 | -5.26 | 0.35627 | none |
| STK32C | 0.244 | 0.2331 | 1.6649 | 0.128 | 0.262 | -5.26 | 0.34071 | none |
| SSH1 | 0.512 | 0.525 | 1.6642 | 0.12813 | 0.262 | -5.26 | -0.3009 | none |
| ARF1 | -0.163 | -0.1305 | -1.664 | 0.12821 | 0.262 | -5.26 | 1.53534 | none |
| TXLNB | 0.207 | 0.1754 | 1.6638 | 0.12822 | 0.262 | -5.26 | -0.1474 | none |
| TRAPPC4 | 0.233 | 0.0925 | 1.6638 | 0.12822 | 0.262 | -5.26 | -0.847 | none |
| PPAT | 0.213 | 0.211 | 1.6635 | 0.12829 | 0.262 | -5.26 | -0.1084 | none |
| BCLAF1 | 0.658 | 0.7598 | 1.6634 | 0.12829 | 0.262 | -5.26 | -0.1862 | none |
| ZFR | 0.381 | 0.4232 | 1.6632 | 0.12833 | 0.262 | -5.26 | -0.2474 | none |
| FAT3 | 0.215 | 0.171 | 1.663 | 0.12839 | 0.263 | -5.26 | -0.2741 | none |
| C5ORF63 | 0.223 | 0.1104 | 1.6628 | 0.12841 | 0.263 | -5.26 | 0.05278 | none |
| LAMA5 | 0.288 | 0.2368 | 1.6626 | 0.12846 | 0.263 | -5.26 | -0.3521 | none |
| ZBTB4 | 0.214 | 0.2265 | 1.6626 | 0.12847 | 0.263 | -5.26 | -0.425 | none |
| BRD4 | 0.556 | 0.6729 | 1.6625 | 0.12849 | 0.263 | -5.26 | 0.29201 | none |
| C7ORF26 | 0.432 | 0.3564 | 1.6623 | 0.12851 | 0.263 | -5.26 | -1.0884 | none |
| NFYA | 0.3 | 0.3092 | 1.6622 | 0.12854 | 0.263 | -5.26 | -0.4721 | none |
| LRBA | 0.277 | 0.2523 | 1.662 | 0.12859 | 0.263 | -5.26 | 0.20541 | none |
| NRP1 | 0.415 | 0.4624 | 1.6616 | 0.12866 | 0.263 | -5.26 | -0.3752 | none |
| NDUFB8 | -0.222 | -0.2078 | -1.661 | 0.12876 | 0.263 | -5.26 | -0.058 | none |
| DBNDD1 | 0.239 | 0.2074 | 1.6609 | 0.12881 | 0.263 | -5.26 | 0.21661 | none |
| TMEM39B | 0.255 | 0.1828 | 1.6607 | 0.12885 | 0.263 | -5.26 | -0.1744 | none |
| TM9SF4 | 0.18 | 0.2065 | 1.6601 | 0.12898 | 0.263 | -5.26 | 0.07511 | none |
| MBD6 | 0.355 | 0.4236 | 1.6599 | 0.12901 | 0.263 | -5.26 | 0.90756 | none |
| CD1E | -0.205 | -0.1747 | -1.66 | 0.12902 | 0.263 | -5.26 | 0.93543 | none |
| IL4 | -0.448 | -0.4373 | -1.66 | 0.12903 | 0.263 | -5.26 | 1.30889 | none |
| GLIS3 | 0.422 | 0.3476 | 1.6597 | 0.12905 | 0.263 | -5.27 | -1.0438 | none |

| | | | | | | | | |
|---|---|---|---|---|---|---|---|---|
| CD9 | -0.386 | -0.3256 | -1.66 | 0.12909 | 0.263 | -5.27 | 2.17402 | AA-related |
| ABHD3 | 0.193 | 0.07 | 1.6593 | 0.12913 | 0.264 | -5.27 | 0.57044 | none |
| RETN | -0.172 | -0.1144 | -1.659 | 0.12914 | 0.264 | -5.27 | 0.65738 | none |
| RASA3 | 0.278 | 0.2796 | 1.6591 | 0.12916 | 0.264 | -5.27 | 0.42069 | none |
| TMEM8A | 0.148 | 0.1458 | 1.6591 | 0.12917 | 0.264 | -5.27 | -0.4087 | none |
| OTC | -0.172 | -0.1145 | -1.659 | 0.12921 | 0.264 | -5.27 | -0.1926 | none |
| SEMA4D | 0.445 | 0.5067 | 1.6589 | 0.12921 | 0.264 | -5.27 | -0.8613 | none |
| IDH2 | -0.256 | -0.223 | -1.659 | 0.12927 | 0.264 | -5.27 | 2.08407 | none |
| PC | 0.274 | 0.3313 | 1.6586 | 0.12928 | 0.264 | -5.27 | 0.324 | none |
| MYL12A | -0.187 | -0.2541 | -1.659 | 0.12929 | 0.264 | -5.27 | 0.71871 | none |
| UBLCP1 | -0.193 | -0.1641 | -1.658 | 0.1293 | 0.264 | -5.27 | 2.15085 | AA-related |
| NBL1 | -0.243 | -0.207 | -1.658 | 0.12934 | 0.264 | -5.27 | -0.0537 | none |
| CHMP6 | -0.152 | -0.0775 | -1.658 | 0.12937 | 0.264 | -5.27 | 1.70688 | none |
| SPRN | 0.203 | 0.1282 | 1.6576 | 0.12949 | 0.264 | -5.27 | -1.2542 | none |
| THAP4 | 0.206 | 0.2494 | 1.6572 | 0.12957 | 0.264 | -5.27 | -0.3115 | none |
| CEP295 | 0.412 | 0.4069 | 1.657 | 0.1296 | 0.264 | -5.27 | -0.1356 | none |
| BAG2 | 0.114 | 0.0551 | 1.6569 | 0.12963 | 0.264 | -5.27 | 0.53549 | none |
| USP34 | 0.41 | 0.4196 | 1.6567 | 0.12967 | 0.264 | -5.27 | -0.5719 | none |
| FN3KRP | 0.203 | 0.1541 | 1.6566 | 0.12969 | 0.264 | -5.27 | -0.5582 | none |
| KRT5 | -0.177 | -0.1567 | -1.655 | 0.12991 | 0.265 | -5.27 | 0.70152 | none |
| RALGDS | 0.343 | 0.419 | 1.6549 | 0.13004 | 0.265 | -5.27 | -0.0506 | none |
| EIF4EBP3 | -0.62 | -0.524 | -1.655 | 0.1301 | 0.265 | -5.27 | 0.48719 | none |
| S1PR5 | 0.303 | 0.2149 | 1.6539 | 0.13025 | 0.265 | -5.27 | 0.07663 | none |
| CNP | 0.26 | 0.2386 | 1.6534 | 0.13034 | 0.265 | -5.27 | -0.7404 | none |
| CDK4 | -0.211 | -0.218 | -1.652 | 0.13054 | 0.266 | -5.28 | 1.3517 | none |
| UPF1 | 0.492 | 0.61 | 1.652 | 0.13064 | 0.266 | -5.28 | -0.1872 | none |
| HUWE1 | 0.493 | 0.6418 | 1.6517 | 0.1307 | 0.266 | -5.28 | -0.6889 | none |
| PSORS1C1 | -0.147 | -0.1006 | -1.651 | 0.13077 | 0.266 | -5.28 | 0.9826 | none |
| NHLH1 | 0.129 | 0.1143 | 1.6509 | 0.13085 | 0.266 | -5.28 | -0.7888 | none |
| TSKU | 0.158 | 0.0988 | 1.6507 | 0.13091 | 0.266 | -5.28 | 0.28329 | none |
| MAP7D3 | 0.471 | 0.5007 | 1.6502 | 0.131 | 0.266 | -5.28 | -0.7725 | none |
| TOPORS | 0.399 | 0.4563 | 1.6501 | 0.13103 | 0.266 | -5.28 | 0.07854 | none |
| NDUFB3 | -0.217 | -0.2579 | -1.65 | 0.13112 | 0.267 | -5.28 | 0.89413 | none |
| PRELID1 | -0.173 | -0.1346 | -1.649 | 0.13115 | 0.267 | -5.28 | 1.59035 | none |
| SLFN5 | 0.418 | 0.4495 | 1.6493 | 0.13118 | 0.267 | -5.28 | 0.38667 | none |
| TRAPPC3 | -0.208 | -0.2202 | -1.649 | 0.13133 | 0.267 | -5.28 | 1.36396 | none |
| EP400 | 0.442 | 0.5607 | 1.6483 | 0.13139 | 0.267 | -5.28 | -0.378 | none |
| ARL4A | -0.235 | -0.1931 | -1.648 | 0.13151 | 0.267 | -5.28 | 1.68763 | none |
| SLC39A8 | 0.212 | 0.1214 | 1.6474 | 0.13159 | 0.267 | -5.28 | 1.41367 | none |
| PSMC5 | -0.137 | -0.094 | -1.647 | 0.13167 | 0.267 | -5.28 | 1.3159 | none |

| | | | | | | | | |
|---|---|---|---|---|---|---|---|---|
| CXCR5 | 0.342 | 0.4212 | 1.6469 | 0.13168 | 0.267 | -5.28 | -0.5403 | none |
| HABP4 | 0.271 | 0.249 | 1.6466 | 0.13176 | 0.268 | -5.28 | -0.9719 | none |
| TBKBP1 | 0.246 | 0.2607 | 1.6462 | 0.13183 | 0.268 | -5.28 | -0.5564 | none |
| HAX1 | -0.303 | -0.3585 | -1.646 | 0.13187 | 0.268 | -5.28 | 1.74946 | none |
| NDUFAF7 | -0.171 | -0.2007 | -1.646 | 0.13194 | 0.268 | -5.28 | 1.12183 | none |
| NPEPL1 | 0.17 | 0.134 | 1.6456 | 0.13196 | 0.268 | -5.28 | 0.1314 | none |
| DMRT1 | 0.21 | 0.1237 | 1.6455 | 0.13199 | 0.268 | -5.28 | -1.2437 | none |
| RFX4 | 0.136 | 0.0893 | 1.6453 | 0.13203 | 0.268 | -5.29 | -0.4127 | none |
| SYPL2 | 0.14 | 0.1297 | 1.6453 | 0.13203 | 0.268 | -5.29 | 0.2591 | none |
| MRPL20 | -0.215 | -0.2434 | -1.645 | 0.13206 | 0.268 | -5.29 | 0.49796 | none |
| ZNF488 | -0.112 | -0.0674 | -1.645 | 0.13209 | 0.268 | -5.29 | 0.24113 | none |
| SSTR3 | 0.205 | 0.227 | 1.6449 | 0.1321 | 0.268 | -5.29 | 1.02329 | none |
| F9 | -0.137 | -0.0997 | -1.645 | 0.13212 | 0.268 | -5.29 | 1.84901 | none |
| ALAD | 0.211 | 0.2266 | 1.6448 | 0.13213 | 0.268 | -5.29 | -0.2133 | none |
| TSPAN2 | -0.154 | -0.1509 | -1.645 | 0.13217 | 0.268 | -5.29 | 0.71973 | none |
| NPM2 | 0.375 | 0.2428 | 1.6443 | 0.13222 | 0.268 | -5.29 | -1.2131 | none |
| DNAJC21 | 0.508 | 0.5558 | 1.6442 | 0.13225 | 0.268 | -5.29 | -0.3866 | none |
| SCD | 0.242 | 0.3586 | 1.644 | 0.1323 | 0.268 | -5.29 | 0.02055 | none |
| GADD45G | 0.371 | 0.3868 | 1.644 | 0.1323 | 0.268 | -5.29 | -0.8704 | none |
| DTHD1 | -0.159 | -0.1144 | -1.644 | 0.13231 | 0.268 | -5.29 | 0.28288 | none |
| CERS4 | 0.216 | 0.1439 | 1.6435 | 0.1324 | 0.268 | -5.29 | 0.53471 | none |
| HIPK3 | 0.167 | 0.1473 | 1.6434 | 0.13242 | 0.268 | -5.29 | 0.41982 | none |
| NAA30 | 0.212 | 0.1054 | 1.6432 | 0.13245 | 0.268 | -5.29 | 0.02687 | none |
| CPSF4L | 0.202 | 0.1816 | 1.6432 | 0.13246 | 0.268 | -5.29 | 0.69015 | none |
| ZDHHC20 | 0.293 | 0.2699 | 1.6427 | 0.13257 | 0.268 | -5.29 | 0.28333 | none |
| TMCO2 | 0.213 | 0.1093 | 1.6423 | 0.13264 | 0.269 | -5.29 | -0.5344 | none |
| GHITM | -0.183 | -0.2003 | -1.642 | 0.13272 | 0.269 | -5.29 | 0.64216 | none |
| AGPAT3 | 0.373 | 0.3461 | 1.6416 | 0.13279 | 0.269 | -5.29 | -1.0347 | none |
| ACOT1 | 0.333 | 0.2814 | 1.6416 | 0.13279 | 0.269 | -5.29 | -0.0252 | none |
| EFL1 | 0.448 | 0.4366 | 1.6414 | 0.13283 | 0.269 | -5.29 | -0.1867 | none |
| KLK5 | -0.145 | -0.1309 | -1.641 | 0.13293 | 0.269 | -5.29 | 1.33192 | none |
| SSX2B | 0.208 | 0.1823 | 1.6408 | 0.13297 | 0.269 | -5.29 | 0.89495 | none |
| WARS | 0.202 | 0.1892 | 1.6403 | 0.13306 | 0.269 | -5.29 | 0.09051 | none |
| CT45A7 | -0.166 | -0.1108 | -1.64 | 0.13311 | 0.269 | -5.29 | 0.23066 | none |
| ZNF654 | 0.245 | 0.2577 | 1.6399 | 0.13314 | 0.269 | -5.29 | 0.16739 | none |
| PCP4L1 | 0.131 | 0.1261 | 1.6398 | 0.13318 | 0.269 | -5.29 | -0.0239 | none |
| ZNF571 | 0.271 | 0.2466 | 1.6397 | 0.13319 | 0.269 | -5.29 | -0.0606 | none |
| GANC | 0.26 | 0.2433 | 1.6396 | 0.13322 | 0.269 | -5.29 | -0.7825 | none |
| NACC2 | 0.191 | 0.206 | 1.6396 | 0.13322 | 0.269 | -5.29 | -0.6585 | none |
| SYT6 | -0.16 | -0.148 | -1.639 | 0.13333 | 0.27 | -5.29 | 1.04285 | none |

| | | | | | | | | |
|---|---|---|---|---|---|---|---|---|
| CNOT3 | 0.434 | 0.4978 | 1.639 | 0.13334 | 0.27 | -5.29 | 0.0222 | none |
| IQCF1 | 0.31 | 0.2252 | 1.639 | 0.13335 | 0.27 | -5.29 | -1.0159 | none |
| TTLL11 | -0.207 | -0.1396 | -1.639 | 0.13339 | 0.27 | -5.29 | -0.4446 | none |
| KIAA1614 | 0.429 | 0.3975 | 1.638 | 0.13355 | 0.27 | -5.3 | 0.1093 | none |
| TMEM256 | -0.317 | -0.383 | -1.638 | 0.1336 | 0.27 | -5.3 | 1.87655 | none |
| EIF3D | -0.16 | -0.098 | -1.637 | 0.13371 | 0.27 | -5.3 | 1.96197 | none |
| RBBP7 | -0.124 | -0.138 | -1.637 | 0.13374 | 0.27 | -5.3 | 1.20004 | none |
| AP3B1 | 0.435 | 0.4411 | 1.6371 | 0.13374 | 0.27 | -5.3 | -0.5653 | none |
| ESAM | 0.223 | 0.1284 | 1.637 | 0.13376 | 0.27 | -5.3 | -0.5414 | none |
| CHPF2 | 0.178 | 0.1244 | 1.6367 | 0.13383 | 0.27 | -5.3 | 0.6502 | none |
| RIN3 | 0.366 | 0.4192 | 1.6366 | 0.13385 | 0.27 | -5.3 | -0.3288 | none |
| PHF12 | 0.494 | 0.5409 | 1.6362 | 0.13392 | 0.27 | -5.3 | -0.7945 | none |
| ETFB | -0.165 | -0.1252 | -1.636 | 0.13398 | 0.27 | -5.3 | 0.37807 | none |
| POU2F2 | 0.432 | 0.549 | 1.6357 | 0.13404 | 0.271 | -5.3 | 0.54629 | none |
| ZBTB39 | 0.167 | 0.1958 | 1.6355 | 0.13408 | 0.271 | -5.3 | -0.4175 | none |
| PLXNA4 | 0.166 | 0.1578 | 1.6352 | 0.13414 | 0.271 | -5.3 | 0.51548 | none |
| SPANXN5 | 0.113 | 0.1128 | 1.6346 | 0.13427 | 0.271 | -5.3 | -0.0749 | none |
| QDPR | -0.212 | -0.1888 | -1.634 | 0.13447 | 0.271 | -5.3 | 2.16259 | AA-related |
| CYB561A3 | 0.278 | 0.0916 | 1.6328 | 0.13465 | 0.272 | -5.3 | -0.1754 | none |
| OPA1 | 0.47 | 0.4969 | 1.6322 | 0.13479 | 0.272 | -5.3 | 0.15643 | none |
| FOSL2 | 0.335 | 0.3367 | 1.6321 | 0.13481 | 0.272 | -5.3 | -0.8341 | none |
| PIP4K2C | 0.129 | 0.084 | 1.6317 | 0.13488 | 0.272 | -5.3 | -0.1493 | none |
| SLC36A3 | 0.161 | 0.0915 | 1.6316 | 0.1349 | 0.272 | -5.3 | -0.9085 | none |
| HORMAD2 | 0.171 | 0.1651 | 1.6316 | 0.13491 | 0.272 | -5.3 | -0.1003 | none |
| USP16 | 0.561 | 0.5407 | 1.6315 | 0.13492 | 0.272 | -5.3 | 0.4405 | none |
| JAKMIP1 | 0.253 | 0.2627 | 1.6314 | 0.13494 | 0.272 | -5.3 | 0.79704 | none |
| TTLL1 | 0.186 | 0.1222 | 1.6312 | 0.135 | 0.272 | -5.3 | 0.65544 | none |
| SDHC | -0.275 | -0.3161 | -1.631 | 0.13503 | 0.272 | -5.3 | 0.97954 | none |
| BEX4 | -0.14 | -0.1459 | -1.631 | 0.13504 | 0.272 | -5.3 | 2.0962 | AA-related |
| HMGXB3 | 0.277 | 0.3633 | 1.6309 | 0.13506 | 0.272 | -5.3 | 0.35387 | none |
| CBX4 | 0.337 | 0.3301 | 1.6308 | 0.13507 | 0.272 | -5.31 | 0.23708 | none |
| CREB3 | -0.137 | -0.1044 | -1.631 | 0.13507 | 0.272 | -5.31 | 1.68206 | none |
| ZNF780A | 0.259 | 0.2548 | 1.6306 | 0.13513 | 0.272 | -5.31 | 0.7339 | none |
| HLA-DMA | 0.435 | 0.3132 | 1.6304 | 0.13516 | 0.272 | -5.31 | 0.10301 | none |
| SMDT1 | -0.26 | -0.2608 | -1.63 | 0.13519 | 0.272 | -5.31 | 1.64246 | none |
| LANCL2 | -0.223 | -0.1622 | -1.63 | 0.13522 | 0.272 | -5.31 | -0.34 | none |
| NDUFA4L2 | -0.296 | -0.1499 | -1.63 | 0.13526 | 0.272 | -5.31 | 1.06953 | none |
| CLEC4A | -0.394 | -0.287 | -1.63 | 0.13527 | 0.272 | -5.31 | 1.34232 | none |
| AHSA1 | 0.144 | 0.1127 | 1.6293 | 0.13539 | 0.272 | -5.31 | -0.7087 | none |
| LIG4 | 0.251 | 0.1868 | 1.629 | 0.13546 | 0.273 | -5.31 | 0.18379 | none |

| | | | | | | | | |
|---|---|---|---|---|---|---|---|---|
| UQCR11 | -0.255 | -0.2674 | -1.628 | 0.13558 | 0.273 | -5.31 | 1.65658 | none |
| FANCG | 0.169 | 0.1552 | 1.6282 | 0.13563 | 0.273 | -5.31 | 0.12795 | none |
| EIF3K | -0.27 | -0.314 | -1.628 | 0.13563 | 0.273 | -5.31 | 1.37061 | none |
| TWIST1 | 0.217 | 0.2105 | 1.6279 | 0.13569 | 0.273 | -5.31 | -0.8942 | none |
| HAVCR2 | 0.188 | 0.0729 | 1.6278 | 0.13572 | 0.273 | -5.31 | -0.6712 | none |
| ATP6V0B | -0.347 | -0.3591 | -1.628 | 0.13573 | 0.273 | -5.31 | 1.44922 | none |
| HOXC9 | 0.15 | 0.1519 | 1.6267 | 0.13594 | 0.273 | -5.31 | 0.66468 | none |
| LARP1 | 0.523 | 0.6134 | 1.6266 | 0.13597 | 0.273 | -5.31 | -0.3104 | none |
| SNW1 | 0.382 | 0.4182 | 1.6265 | 0.136 | 0.273 | -5.31 | -0.3863 | none |
| ITSN2 | 0.454 | 0.5084 | 1.6263 | 0.13603 | 0.273 | -5.31 | -0.539 | none |
| ARL4D | 0.235 | 0.1661 | 1.6262 | 0.13606 | 0.273 | -5.31 | -0.3572 | none |
| C12ORF57 | -0.287 | -0.3371 | -1.626 | 0.13608 | 0.273 | -5.31 | 0.83769 | none |
| RNASE4 | -0.279 | -0.2579 | -1.626 | 0.13617 | 0.274 | -5.31 | 1.87438 | none |
| TAB3 | 0.273 | 0.2959 | 1.6253 | 0.13626 | 0.274 | -5.31 | -0.2896 | none |
| SLA2 | 0.153 | 0.1187 | 1.6244 | 0.13644 | 0.274 | -5.31 | -0.6271 | none |
| C1QTNF9E | -0.256 | -0.2677 | -1.623 | 0.13669 | 0.274 | -5.32 | 1.01171 | none |
| MCMDC2 | 0.141 | 0.0577 | 1.6233 | 0.13669 | 0.274 | -5.32 | 0.03875 | none |
| RASA2 | 0.369 | 0.2879 | 1.6232 | 0.13671 | 0.274 | -5.32 | 0.23688 | none |
| NCOA2 | -0.15 | -0.0879 | -1.622 | 0.13694 | 0.275 | -5.32 | 0.14789 | none |
| ERAL1 | -0.141 | -0.0895 | -1.622 | 0.13699 | 0.275 | -5.32 | 2.50097 | AA-related |
| ARSF | 0.113 | 0.0826 | 1.6218 | 0.137 | 0.275 | -5.32 | 0 | none |
| SRSF1 | -0.163 | -0.1544 | -1.622 | 0.13703 | 0.275 | -5.32 | 1.72986 | none |
| USP44 | 0.163 | 0.1483 | 1.6215 | 0.13707 | 0.275 | -5.32 | 0.00486 | none |
| SP1 | 0.365 | 0.3272 | 1.6214 | 0.13709 | 0.275 | -5.32 | 0.66329 | none |
| SLC45A3 | -0.125 | -0.1055 | -1.621 | 0.13713 | 0.275 | -5.32 | 0.51562 | none |
| FZR1 | 0.356 | 0.3902 | 1.6211 | 0.13716 | 0.275 | -5.32 | -0.2391 | none |
| ACTL10 | -0.149 | -0.1522 | -1.621 | 0.13724 | 0.275 | -5.32 | -0.1459 | none |
| PRR14L | 0.428 | 0.4754 | 1.6206 | 0.13726 | 0.275 | -5.32 | 0.07914 | none |
| PKD2L1 | 0.15 | 0.1071 | 1.6205 | 0.13729 | 0.275 | -5.32 | -0.7868 | none |
| PDCD11 | 0.366 | 0.4593 | 1.6198 | 0.13743 | 0.276 | -5.32 | 0.15002 | none |
| NUCB1 | 0.262 | 0.3007 | 1.6196 | 0.13747 | 0.276 | -5.32 | -0.9136 | none |
| ZNF326 | 0.28 | 0.2888 | 1.6196 | 0.13748 | 0.276 | -5.32 | -0.3195 | none |
| CLK3 | 0.22 | 0.2623 | 1.6193 | 0.13754 | 0.276 | -5.32 | -0.1598 | none |
| MUC12 | 0.321 | 0.3451 | 1.6193 | 0.13754 | 0.276 | -5.32 | 0.36148 | none |
| ZCCHC7 | 0.256 | 0.17 | 1.6191 | 0.13759 | 0.276 | -5.32 | -0.1512 | none |
| TMED1 | 0.136 | 0.119 | 1.6189 | 0.13763 | 0.276 | -5.32 | -0.1259 | none |
| TRIM63 | 0.211 | 0.1942 | 1.6184 | 0.13773 | 0.276 | -5.32 | 1.15769 | none |
| ANGPTL2 | -0.18 | -0.1151 | -1.618 | 0.13774 | 0.276 | -5.32 | -0.5916 | none |
| MCL1 | 0.16 | 0.1277 | 1.6183 | 0.13775 | 0.276 | -5.32 | 0.24624 | none |
| MMP19 | 0.218 | 0.2226 | 1.6181 | 0.1378 | 0.276 | -5.32 | 1.10671 | none |

| | | | | | | | | |
|---|---|---|---|---|---|---|---|---|
| CALCA | -0.181 | -0.1898 | -1.618 | 0.1378 | 0.276 | -5.32 | 0.08974 | none |
| CCDC187 | 0.126 | 0.1024 | 1.6179 | 0.13785 | 0.276 | -5.32 | -1.2454 | none |
| MYO18B | 0.206 | 0.1817 | 1.6179 | 0.13785 | 0.276 | -5.32 | -0.9864 | none |
| ATG101 | -0.244 | -0.2342 | -1.618 | 0.13786 | 0.276 | -5.32 | 1.21412 | none |
| NPAS1 | -0.227 | -0.2079 | -1.618 | 0.13793 | 0.276 | -5.32 | 0.0819 | none |
| SHISA8 | -0.193 | -0.1066 | -1.617 | 0.13798 | 0.276 | -5.32 | 0.58692 | none |
| DDX43 | 0.296 | 0.1509 | 1.6172 | 0.13799 | 0.276 | -5.32 | -0.1723 | none |
| COPS5 | -0.135 | -0.1719 | -1.617 | 0.13801 | 0.276 | -5.32 | 1.68977 | none |
| MPDZ | -0.12 | -0.0853 | -1.617 | 0.1381 | 0.276 | -5.32 | 0.80752 | none |
| ASIP | 0.155 | 0.17 | 1.6167 | 0.13811 | 0.276 | -5.32 | -0.5444 | none |
| CNNM4 | -0.181 | -0.0743 | -1.616 | 0.13818 | 0.276 | -5.32 | 1.8767 | none |
| NBAS | 0.415 | 0.3951 | 1.6161 | 0.13824 | 0.276 | -5.33 | -0.3572 | none |
| AC007326 | 0.168 | 0.2214 | 1.6159 | 0.13828 | 0.276 | -5.33 | 1.1581 | none |
| DNAH7 | 0.291 | 0.2901 | 1.6158 | 0.13831 | 0.277 | -5.33 | -0.1987 | none |
| TMED3 | -0.222 | -0.184 | -1.616 | 0.13834 | 0.277 | -5.33 | 1.64956 | none |
| RAVER2 | 0.129 | 0.0916 | 1.6154 | 0.13839 | 0.277 | -5.33 | -0.6347 | none |
| SLC20A1 | 0.154 | 0.1112 | 1.615 | 0.13847 | 0.277 | -5.33 | 0.65781 | none |
| SNX31 | 0.673 | 0.4812 | 1.6149 | 0.1385 | 0.277 | -5.33 | -1.0186 | none |
| SLC35F4 | 0.218 | 0.2392 | 1.6137 | 0.13875 | 0.277 | -5.33 | 0.04258 | none |
| FAM216B | 0.123 | 0.0932 | 1.6136 | 0.13878 | 0.277 | -5.33 | -1.2576 | none |
| FXYD6-FXY | -0.159 | -0.1061 | -1.613 | 0.13886 | 0.277 | -5.33 | 0.86195 | none |
| MPP2 | 0.125 | 0.0933 | 1.6121 | 0.13911 | 0.278 | -5.33 | -0.007 | none |
| VMO1 | -0.135 | -0.1376 | -1.611 | 0.13931 | 0.278 | -5.33 | -0.1619 | none |
| DDRGK1 | 0.202 | 0.217 | 1.6108 | 0.13938 | 0.278 | -5.33 | 0.35925 | none |
| OXA1L | 0.15 | 0.0821 | 1.6107 | 0.13941 | 0.278 | -5.33 | 0.1467 | none |
| KRCC1 | -0.283 | -0.2658 | -1.611 | 0.13943 | 0.278 | -5.33 | 2.18614 | AA-related |
| COQ3 | 0.247 | 0.134 | 1.6102 | 0.13951 | 0.278 | -5.33 | -0.6183 | none |
| OR13C5 | -0.136 | -0.0999 | -1.61 | 0.13952 | 0.278 | -5.33 | 0.87773 | none |
| RAPSN | 0.246 | 0.1628 | 1.6094 | 0.13969 | 0.279 | -5.33 | -0.7329 | none |
| TMEM121 | 0.174 | 0.1704 | 1.6087 | 0.13986 | 0.279 | -5.34 | -0.0484 | none |
| TXLNA | 0.367 | 0.4495 | 1.6083 | 0.13994 | 0.279 | -5.34 | -0.5461 | none |
| NME4 | 0.18 | 0.0711 | 1.6082 | 0.13997 | 0.279 | -5.34 | 0.40889 | none |
| UBAP1 | 0.125 | 0.0929 | 1.6081 | 0.13998 | 0.279 | -5.34 | -1.0656 | none |
| DOCK11 | 0.548 | 0.5055 | 1.6081 | 0.13999 | 0.279 | -5.34 | -0.2294 | none |
| TLN2 | 0.523 | 0.5858 | 1.6077 | 0.14007 | 0.279 | -5.34 | 0.04354 | none |
| TMEM125 | -0.159 | -0.1335 | -1.607 | 0.14017 | 0.28 | -5.34 | -0.1476 | none |
| SLC41A3 | 0.231 | 0.1248 | 1.6071 | 0.1402 | 0.28 | -5.34 | -0.7185 | none |
| RHEB | -0.185 | -0.2013 | -1.607 | 0.14021 | 0.28 | -5.34 | 0.75727 | none |
| ABT1 | -0.168 | -0.144 | -1.607 | 0.14023 | 0.28 | -5.34 | 0.9625 | none |
| MC2R | 0.109 | 0.0804 | 1.6069 | 0.14025 | 0.28 | -5.34 | 0.14857 | none |

| | | | | | | | | |
|---|---|---|---|---|---|---|---|---|
| MTA1 | 0.432 | 0.5349 | 1.6063 | 0.14037 | 0.28 | -5.34 | -0.4944 | none |
| SMARCA5 | 0.513 | 0.5967 | 1.6062 | 0.1404 | 0.28 | -5.34 | -0.2089 | none |
| UBQLN1 | 0.261 | 0.2798 | 1.6048 | 0.14071 | 0.28 | -5.34 | 0.24789 | none |
| MBP | 0.175 | 0.1114 | 1.6047 | 0.14074 | 0.28 | -5.34 | -0.0284 | none |
| LGALS14 | -0.174 | -0.2068 | -1.605 | 0.14074 | 0.28 | -5.34 | 1.25013 | none |
| ZSCAN5C | 0.191 | 0.1292 | 1.6044 | 0.14081 | 0.28 | -5.34 | -0.9346 | none |
| SPHK2 | 0.144 | 0.1085 | 1.6038 | 0.14093 | 0.281 | -5.34 | -0.8446 | none |
| SLC25A5 | -0.221 | -0.213 | -1.604 | 0.14099 | 0.281 | -5.34 | 0.25591 | none |
| NOM1 | 0.431 | 0.4767 | 1.6033 | 0.14103 | 0.281 | -5.34 | -0.4756 | none |
| PI4KB | 0.169 | 0.1702 | 1.6031 | 0.14109 | 0.281 | -5.34 | 0.57065 | none |
| HIST3H3 | 0.3 | 0.237 | 1.6027 | 0.14118 | 0.281 | -5.34 | -1.2638 | none |
| KCNN4 | 0.163 | 0.2246 | 1.6027 | 0.14118 | 0.281 | -5.34 | 0.2878 | none |
| ZNF720 | -0.38 | -0.3292 | -1.603 | 0.14119 | 0.281 | -5.34 | 0.16935 | none |
| KAZN | 0.132 | 0.0784 | 1.6026 | 0.1412 | 0.281 | -5.34 | -0.5414 | none |
| GABRE | -0.117 | -0.0797 | -1.602 | 0.14127 | 0.281 | -5.34 | -0.3036 | none |
| SLC30A10 | 0.124 | 0.0917 | 1.6018 | 0.14138 | 0.281 | -5.34 | -0.7064 | none |
| VBP1 | -0.217 | -0.2387 | -1.601 | 0.14156 | 0.282 | -5.35 | 1.51693 | none |
| BZW1 | -0.18 | -0.1827 | -1.601 | 0.14164 | 0.282 | -5.35 | 1.25506 | none |
| AP001888 | 0.151 | 0.1512 | 1.6004 | 0.14169 | 0.282 | -5.35 | -0.4344 | none |
| NELFCD | 0.13 | 0.1011 | 1.6001 | 0.14174 | 0.282 | -5.35 | 0.34425 | none |
| MED27 | 0.238 | 0.1003 | 1.6 | 0.14178 | 0.282 | -5.35 | -0.3034 | none |
| APEX2 | 0.145 | 0.1577 | 1.6 | 0.14178 | 0.282 | -5.35 | -0.8226 | none |
| SRBD1 | 0.213 | 0.1765 | 1.599 | 0.142 | 0.282 | -5.35 | 0.15591 | none |
| NDUFS8 | 0.253 | 0.1481 | 1.5983 | 0.14216 | 0.283 | -5.35 | -0.8587 | none |
| COQ9 | 0.147 | 0.1535 | 1.5981 | 0.14219 | 0.283 | -5.35 | 0.16853 | none |
| FYN | -0.372 | -0.2154 | -1.598 | 0.14226 | 0.283 | -5.35 | 1.61899 | none |
| RRBP1 | 0.432 | 0.5506 | 1.5974 | 0.14234 | 0.283 | -5.35 | -0.1784 | none |
| TMEM106 | -0.186 | -0.1918 | -1.596 | 0.14256 | 0.283 | -5.35 | 1.96214 | none |
| TNFRSF4 | -0.198 | -0.1626 | -1.596 | 0.14259 | 0.283 | -5.35 | 1.3063 | none |
| DPPA2 | 0.167 | 0.1208 | 1.5962 | 0.14262 | 0.283 | -5.35 | -0.8497 | none |
| GPM6B | 0.21 | 0.1456 | 1.596 | 0.14267 | 0.283 | -5.35 | -0.9967 | none |
| ZNF398 | 0.304 | 0.332 | 1.5957 | 0.14274 | 0.283 | -5.35 | -0.232 | none |
| PSENEN | -0.199 | -0.2815 | -1.596 | 0.14275 | 0.283 | -5.35 | 1.66201 | none |
| TMEM198 | 0.265 | 0.1983 | 1.5953 | 0.14281 | 0.284 | -5.35 | -0.4046 | none |
| ZFX | 0.337 | 0.3195 | 1.5947 | 0.14295 | 0.284 | -5.35 | 0.26833 | none |
| NUDT14 | 0.291 | 0.1291 | 1.5947 | 0.14295 | 0.284 | -5.35 | -0.28 | none |
| PARD6A | -0.276 | -0.2603 | -1.595 | 0.14299 | 0.284 | -5.35 | 1.75543 | none |
| R3HCC1 | -0.15 | -0.115 | -1.594 | 0.143 | 0.284 | -5.35 | 0.88453 | none |
| MRPL34 | -0.249 | -0.2137 | -1.594 | 0.14301 | 0.284 | -5.35 | 1.8276 | none |
| LHPP | 0.203 | 0.2109 | 1.5939 | 0.14314 | 0.284 | -5.36 | -0.3795 | none |

| | | | | | | | | |
|---|---|---|---|---|---|---|---|---|
| PIH1D1 | -0.184 | -0.1936 | -1.593 | 0.14326 | 0.284 | -5.36 | 0.37071 | none |
| ZHX2 | 0.323 | 0.3743 | 1.5932 | 0.14329 | 0.284 | -5.36 | -0.2336 | none |
| LARP4 | 0.285 | 0.2924 | 1.593 | 0.14332 | 0.284 | -5.36 | -0.7864 | none |
| MFSD4A | 0.224 | 0.1161 | 1.5929 | 0.14336 | 0.284 | -5.36 | -0.6379 | none |
| MT-ND2 | 0.309 | 0.2781 | 1.5925 | 0.14344 | 0.284 | -5.36 | 0.01004 | none |
| HIPK1 | 0.226 | 0.2318 | 1.5923 | 0.14348 | 0.284 | -5.36 | 0.23928 | none |
| KLHL7 | 0.263 | 0.1698 | 1.5922 | 0.1435 | 0.284 | -5.36 | -0.4102 | none |
| BLOC1S5 | 0.252 | 0.1432 | 1.592 | 0.14356 | 0.285 | -5.36 | 0.53705 | none |
| C10ORF90 | -0.154 | -0.1399 | -1.592 | 0.14358 | 0.285 | -5.36 | 0.49803 | none |
| RIIAD1 | 0.283 | 0.2928 | 1.5916 | 0.14364 | 0.285 | -5.36 | 0.02034 | none |
| TUBA4B | 0.147 | 0.0241 | 1.5911 | 0.14375 | 0.285 | -5.36 | -0.8766 | none |
| C4ORF3 | -0.228 | -0.2705 | -1.591 | 0.14376 | 0.285 | -5.36 | 1.43519 | none |
| EMX1 | 0.199 | 0.1893 | 1.5908 | 0.14383 | 0.285 | -5.36 | 0.7682 | none |
| ATP9A | 0.186 | 0.2023 | 1.5908 | 0.14383 | 0.285 | -5.36 | 0.18918 | none |
| ZBTB17 | 0.275 | 0.3539 | 1.5907 | 0.14384 | 0.285 | -5.36 | -0.3155 | none |
| TMEM97 | 0.159 | 0.0943 | 1.5903 | 0.14394 | 0.285 | -5.36 | -0.1969 | none |
| CHAC1 | -0.23 | -0.1982 | -1.59 | 0.14408 | 0.285 | -5.36 | 1.25261 | none |
| DBP | 0.358 | 0.1957 | 1.5896 | 0.14409 | 0.285 | -5.36 | -0.905 | none |
| FAM200A | 0.162 | 0.1026 | 1.5893 | 0.14416 | 0.285 | -5.36 | -0.5338 | none |
| ZNF787 | -0.279 | -0.1806 | -1.588 | 0.14435 | 0.286 | -5.36 | 1.48057 | none |
| LIN7C | 0.255 | 0.1494 | 1.5884 | 0.14436 | 0.286 | -5.36 | -0.5532 | none |
| EIF4G2 | 0.333 | 0.3774 | 1.5882 | 0.14442 | 0.286 | -5.36 | -0.1573 | none |
| AKAP8 | 0.309 | 0.3982 | 1.5878 | 0.1445 | 0.286 | -5.36 | -0.0322 | none |
| SSPN | 0.222 | 0.1917 | 1.5872 | 0.14464 | 0.286 | -5.36 | -1.1733 | none |
| SHISA4 | 0.127 | 0.1205 | 1.5869 | 0.1447 | 0.286 | -5.36 | 0.46856 | none |
| RAX2 | 0.165 | 0.1118 | 1.5869 | 0.14471 | 0.286 | -5.37 | -1.0876 | none |
| MAML2 | 0.301 | 0.3498 | 1.5867 | 0.14475 | 0.286 | -5.37 | -0.1396 | none |
| SMAD3 | -0.277 | -0.2024 | -1.587 | 0.14478 | 0.286 | -5.37 | 0.59551 | none |
| TTLL5 | 0.366 | 0.4188 | 1.5863 | 0.14483 | 0.286 | -5.37 | -0.0122 | none |
| KIF16B | 0.309 | 0.3703 | 1.5862 | 0.14487 | 0.286 | -5.37 | -0.0972 | none |
| OLFM1 | 0.15 | 0.1122 | 1.586 | 0.14492 | 0.286 | -5.37 | 1.27165 | none |
| CLDN7 | 0.346 | 0.2172 | 1.586 | 0.14492 | 0.286 | -5.37 | -0.5601 | none |
| SART1 | 0.511 | 0.5562 | 1.5853 | 0.14506 | 0.287 | -5.37 | -0.8253 | none |
| NEK6 | -0.177 | -0.1079 | -1.585 | 0.1451 | 0.287 | -5.37 | 1.54266 | none |
| GTF3C6 | -0.145 | -0.1335 | -1.585 | 0.14513 | 0.287 | -5.37 | 0.18504 | none |
| NAT8 | -0.152 | -0.1014 | -1.585 | 0.14523 | 0.287 | -5.37 | 0.86195 | none |
| ANP32E | 0.282 | 0.2696 | 1.5846 | 0.14524 | 0.287 | -5.37 | -0.9754 | none |
| DIP2B | 0.31 | 0.2818 | 1.5844 | 0.14528 | 0.287 | -5.37 | -1.0537 | none |
| METTL25 | 0.29 | 0.2203 | 1.5842 | 0.14532 | 0.287 | -5.37 | 0.39005 | none |
| ZNF146 | 0.479 | 0.5423 | 1.5835 | 0.14547 | 0.287 | -5.37 | 0.18638 | none |

| | | | | | | | | |
|---|---|---|---|---|---|---|---|---|
| AC010655 | -0.152 | -0.1012 | -1.583 | 0.14551 | 0.287 | -5.37 | 1.24897 | none |
| SPATA18 | -0.153 | -0.159 | -1.583 | 0.14551 | 0.287 | -5.37 | 1.98912 | none |
| FAM76B | 0.273 | 0.2125 | 1.5833 | 0.14552 | 0.287 | -5.37 | -0.2879 | none |
| SYT8 | -0.196 | -0.1637 | -1.583 | 0.14552 | 0.287 | -5.37 | 1.00409 | none |
| DERL3 | -0.157 | -0.0677 | -1.583 | 0.14553 | 0.287 | -5.37 | 1.20935 | none |
| GALNT8 | -0.106 | -0.0659 | -1.583 | 0.14557 | 0.287 | -5.37 | 1.43205 | none |
| LAMA1 | 0.13 | 0.0786 | 1.583 | 0.14558 | 0.287 | -5.37 | -1.2269 | none |
| ANXA2 | -0.135 | -0.1293 | -1.583 | 0.14559 | 0.287 | -5.37 | 1.26573 | none |
| RIPPLY2 | -0.13 | -0.0629 | -1.583 | 0.14561 | 0.287 | -5.37 | 0.99292 | none |
| EIF2AK2 | 0.337 | 0.3145 | 1.5826 | 0.14568 | 0.287 | -5.37 | -0.2056 | none |
| MS4A5 | 0.4 | 0.2808 | 1.5826 | 0.14569 | 0.287 | -5.37 | -0.4236 | none |
| CCNF | 0.275 | 0.3224 | 1.5825 | 0.1457 | 0.287 | -5.37 | -0.1808 | none |
| NPTN | 0.174 | 0.1473 | 1.582 | 0.14583 | 0.288 | -5.37 | -0.5878 | none |
| TNKS2 | 0.178 | 0.1687 | 1.5818 | 0.14588 | 0.288 | -5.37 | -0.1639 | none |
| TSPAN18 | 0.152 | 0.1497 | 1.5817 | 0.14589 | 0.288 | -5.37 | -0.1511 | none |
| AHCYL2 | -0.144 | -0.1074 | -1.582 | 0.1459 | 0.288 | -5.37 | 1.22514 | none |
| NKAIN4 | -0.16 | -0.1341 | -1.581 | 0.14595 | 0.288 | -5.37 | -0.1935 | none |
| TMCC2 | 0.222 | 0.224 | 1.5812 | 0.146 | 0.288 | -5.37 | -0.5208 | none |
| SGTB | 0.229 | 0.2478 | 1.5809 | 0.14606 | 0.288 | -5.37 | 0.4387 | none |
| PDE5A | 0.238 | 0.152 | 1.5808 | 0.14609 | 0.288 | -5.37 | 0.01674 | none |
| CPLX2 | -0.27 | -0.1489 | -1.58 | 0.14623 | 0.288 | -5.37 | 0.26856 | none |
| CRYL1 | -0.209 | -0.203 | -1.58 | 0.14625 | 0.288 | -5.37 | 0.71818 | none |
| SLC5A12 | -0.11 | -0.069 | -1.58 | 0.14625 | 0.288 | -5.37 | 2.29375 | AA-related |
| STEAP3 | 0.166 | 0.1471 | 1.5801 | 0.14625 | 0.288 | -5.37 | -0.4597 | none |
| KLF16 | -0.171 | -0.1131 | -1.58 | 0.14632 | 0.288 | -5.37 | 1.83943 | none |
| RCL1 | 0.168 | 0.1038 | 1.5797 | 0.14633 | 0.288 | -5.37 | 0.56993 | none |
| BATF | -0.138 | -0.062 | -1.58 | 0.14635 | 0.288 | -5.37 | 0.51719 | none |
| PLEKHF2 | 0.147 | 0.0549 | 1.5794 | 0.14642 | 0.288 | -5.38 | 0.13324 | none |
| GRB7 | 0.195 | 0.1178 | 1.5793 | 0.14642 | 0.288 | -5.38 | 0.03098 | none |
| SORBS2 | 0.248 | 0.1618 | 1.5792 | 0.14647 | 0.288 | -5.38 | -0.7844 | none |
| USP5 | 0.257 | 0.3032 | 1.5789 | 0.14653 | 0.288 | -5.38 | -0.4304 | none |
| DCAF5 | 0.268 | 0.3279 | 1.5787 | 0.14658 | 0.288 | -5.38 | -0.6179 | none |
| CWC25 | 0.306 | 0.3072 | 1.5785 | 0.14661 | 0.288 | -5.38 | -0.2399 | none |
| QSER1 | 0.403 | 0.4993 | 1.5783 | 0.14666 | 0.288 | -5.38 | 0.23719 | none |
| SLC30A3 | 0.227 | 0.1543 | 1.5779 | 0.14675 | 0.289 | -5.38 | -0.3447 | none |
| MMD | 0.305 | 0.216 | 1.5778 | 0.14678 | 0.289 | -5.38 | 0.77584 | none |
| NIFK | -0.143 | -0.1644 | -1.578 | 0.14679 | 0.289 | -5.38 | 0.76593 | none |
| VWF | 0.183 | 0.1542 | 1.5773 | 0.14689 | 0.289 | -5.38 | -0.2083 | none |
| BRD1 | 0.345 | 0.4371 | 1.5773 | 0.1469 | 0.289 | -5.38 | 0.14689 | none |
| LRRC3C | 0.246 | 0.1503 | 1.5771 | 0.14694 | 0.289 | -5.38 | -1.1999 | none |

| | | | | | | | | |
|---|---|---|---|---|---|---|---|---|
| GTF2A1 | -0.179 | -0.1996 | -1.577 | 0.14697 | 0.289 | -5.38 | 1.31625 | none |
| ST8SIA3 | -0.151 | -0.1257 | -1.577 | 0.147 | 0.289 | -5.38 | 0.2123 | none |
| ANGPTL6 | 0.225 | 0.2082 | 1.5768 | 0.14702 | 0.289 | -5.38 | -0.1974 | none |
| METTL11B | -0.15 | -0.1002 | -1.576 | 0.14712 | 0.289 | -5.38 | 0.63037 | none |
| PLA1A | -0.217 | -0.0791 | -1.576 | 0.14721 | 0.289 | -5.38 | -0.1225 | none |
| ATP6V1H | -0.201 | -0.1841 | -1.576 | 0.14724 | 0.289 | -5.38 | 1.66374 | none |
| NCOA5 | -0.144 | -0.0938 | -1.576 | 0.14725 | 0.289 | -5.38 | -0.011 | none |
| BRAP | 0.35 | 0.4165 | 1.5755 | 0.1473 | 0.289 | -5.38 | -0.598 | none |
| GRK2 | 0.258 | 0.2934 | 1.5747 | 0.14748 | 0.29 | -5.38 | -0.1911 | none |
| STEAP1 | -0.182 | -0.2059 | -1.574 | 0.14754 | 0.29 | -5.38 | 0.6842 | none |
| L3HYPDH | 0.19 | 0.1677 | 1.5741 | 0.14764 | 0.29 | -5.38 | 0.34144 | none |
| TM6SF1 | 0.158 | 0.0648 | 1.5735 | 0.14778 | 0.29 | -5.38 | 0.57502 | none |
| SLC2A4RG | 0.235 | 0.1763 | 1.571 | 0.14834 | 0.291 | -5.39 | 0.46743 | none |
| KRTAP2-4 | 0.118 | 0.0996 | 1.571 | 0.14835 | 0.291 | -5.39 | -0.062 | none |
| NLK | -0.224 | -0.2437 | -1.571 | 0.14838 | 0.291 | -5.39 | 0.40893 | none |
| FAM19A1 | 0.18 | 0.1088 | 1.5706 | 0.14843 | 0.291 | -5.39 | -1.1259 | none |
| CYP20A1 | 0.216 | 0.01 | 1.5704 | 0.14849 | 0.291 | -5.39 | -0.5046 | none |
| NKD1 | 0.181 | 0.1105 | 1.5701 | 0.14854 | 0.291 | -5.39 | -0.8569 | none |
| MPV17L | 0.21 | 0.0854 | 1.5699 | 0.14859 | 0.291 | -5.39 | 0.20845 | none |
| USP25 | 0.372 | 0.3666 | 1.5698 | 0.14861 | 0.291 | -5.39 | 0.22775 | none |
| SEC16A | 0.4 | 0.5182 | 1.5698 | 0.14862 | 0.291 | -5.39 | -0.3492 | none |
| PLA2G4D | -0.189 | -0.11 | -1.57 | 0.14864 | 0.291 | -5.39 | 1.77994 | none |
| ENHO | -0.158 | -0.1312 | -1.569 | 0.14872 | 0.291 | -5.39 | 0.26761 | none |
| POLD1 | 0.209 | 0.2538 | 1.5694 | 0.14872 | 0.291 | -5.39 | 0.35459 | none |
| CCDC34 | 0.376 | 0.4267 | 1.5693 | 0.14873 | 0.291 | -5.39 | 0.17074 | none |
| SLC35A4 | 0.144 | 0.1529 | 1.5692 | 0.14876 | 0.291 | -5.39 | 0.64136 | none |
| ARHGAP29 | 0.21 | 0.2004 | 1.5691 | 0.14879 | 0.292 | -5.39 | -0.5879 | none |
| PCMTD1 | 0.188 | 0.0746 | 1.5687 | 0.14887 | 0.292 | -5.39 | 0.13388 | none |
| ANKRD26 | 0.35 | 0.3194 | 1.5686 | 0.14891 | 0.292 | -5.39 | 0.15086 | none |
| DBNDD2 | -0.241 | -0.3297 | -1.569 | 0.14891 | 0.292 | -5.39 | 0.30885 | none |
| CCDC124 | 0.246 | 0.2896 | 1.5684 | 0.14895 | 0.292 | -5.39 | -0.6974 | none |
| FANCD2OS | 0.85 | 0.6029 | 1.5673 | 0.1492 | 0.292 | -5.39 | -0.8921 | none |
| ERP29 | 0.183 | 0.1184 | 1.5669 | 0.1493 | 0.292 | -5.39 | -0.9134 | none |
| SP2 | 0.561 | 0.5539 | 1.5668 | 0.14932 | 0.292 | -5.39 | -0.658 | none |
| ZNF250 | 0.246 | 0.2224 | 1.5668 | 0.14933 | 0.292 | -5.39 | 0.30118 | none |
| S100A16 | 0.137 | 0.1389 | 1.5667 | 0.14934 | 0.292 | -5.39 | 0.60516 | none |
| RAB5C | 0.193 | 0.0932 | 1.5667 | 0.14934 | 0.292 | -5.39 | -0.0333 | none |
| MAP3K5 | 0.388 | 0.4468 | 1.5667 | 0.14934 | 0.292 | -5.39 | 0.11564 | none |
| SPDYE16 | -1.397 | -1.0166 | -1.567 | 0.14938 | 0.292 | -5.39 | 2.2645 | AA-related |
| ARSA | 0.22 | 0.0854 | 1.5661 | 0.14948 | 0.292 | -5.39 | -0.0239 | none |

| | | | | | | | | |
|---|---|---|---|---|---|---|---|---|
| KEAP1 | 0.271 | 0.2476 | 1.566 | 0.1495 | 0.292 | -5.39 | -0.2277 | none |
| AKR1E2 | 0.544 | 0.5015 | 1.5657 | 0.14957 | 0.293 | -5.39 | 0.83035 | none |
| CNTN5 | 0.146 | 0.0995 | 1.5657 | 0.14958 | 0.293 | -5.39 | -1.3245 | none |
| GABARAPI | -0.217 | -0.2041 | -1.566 | 0.1496 | 0.293 | -5.39 | -0.2262 | none |
| SPACA4 | -0.106 | -0.0917 | -1.565 | 0.14967 | 0.293 | -5.39 | -0.5051 | none |
| C19ORF71 | -0.204 | -0.0856 | -1.565 | 0.14969 | 0.293 | -5.39 | 0.00963 | none |
| GFOD1 | 0.217 | 0.2178 | 1.5652 | 0.14969 | 0.293 | -5.39 | -0.3837 | none |
| KPNB1 | 0.526 | 0.652 | 1.5633 | 0.15013 | 0.293 | -5.4 | -0.582 | none |
| GBF1 | 0.412 | 0.5475 | 1.5631 | 0.15018 | 0.293 | -5.4 | -0.3233 | none |
| TTK | 0.35 | 0.3163 | 1.5629 | 0.15023 | 0.294 | -5.4 | -0.309 | none |
| RABGEF1 | 0.186 | 0.1643 | 1.5628 | 0.15025 | 0.294 | -5.4 | -0.5003 | none |
| ADAM22 | 0.234 | 0.1775 | 1.5627 | 0.15027 | 0.294 | -5.4 | 0.04994 | none |
| GSTT2B | 0.148 | 0.1179 | 1.5626 | 0.1503 | 0.294 | -5.4 | 0.05897 | none |
| LPAR6 | 0.213 | 0.1423 | 1.5622 | 0.15039 | 0.294 | -5.4 | -0.3728 | none |
| SIT1 | -0.217 | -0.2152 | -1.562 | 0.1504 | 0.294 | -5.4 | 2.10714 | AA-related |
| PJA1 | -0.174 | -0.102 | -1.562 | 0.15052 | 0.294 | -5.4 | 1.38251 | none |
| PITX1 | -0.162 | -0.1513 | -1.561 | 0.15057 | 0.294 | -5.4 | -0.1636 | none |
| PPP1R3F | -0.351 | -0.298 | -1.561 | 0.15059 | 0.294 | -5.4 | 0.80279 | none |
| PLEKHS1 | 0.232 | 0.1336 | 1.5611 | 0.15065 | 0.294 | -5.4 | -1.1914 | none |
| CEP57 | 0.301 | 0.2847 | 1.5609 | 0.15069 | 0.294 | -5.4 | 0.42851 | none |
| TNFRSF10I | 0.159 | 0.221 | 1.5603 | 0.15084 | 0.294 | -5.4 | 0.68927 | none |
| PRKDC | 0.489 | 0.5712 | 1.5601 | 0.15089 | 0.294 | -5.4 | -0.0681 | none |
| CORO7 | 0.294 | 0.2729 | 1.5591 | 0.15113 | 0.295 | -5.4 | -0.4557 | none |
| CCDC183 | -0.31 | -0.2039 | -1.559 | 0.15113 | 0.295 | -5.4 | 1.10098 | none |
| BCOR | 0.348 | 0.5057 | 1.559 | 0.15115 | 0.295 | -5.4 | -0.6071 | none |
| RPL18A | -0.273 | -0.2798 | -1.559 | 0.15123 | 0.295 | -5.4 | 0.76148 | none |
| KCNK1 | -0.2 | -0.21 | -1.558 | 0.15128 | 0.295 | -5.4 | 1.94844 | none |
| AKAP14 | 0.487 | 0.2853 | 1.5583 | 0.15131 | 0.295 | -5.4 | -0.5179 | none |
| KCNMB2 | 0.15 | 0.1692 | 1.5581 | 0.15136 | 0.295 | -5.4 | 0.2893 | none |
| NTAN1 | 0.243 | 0.062 | 1.5577 | 0.15144 | 0.295 | -5.4 | -0.7509 | none |
| PSMF1 | -0.123 | -0.115 | -1.557 | 0.15162 | 0.296 | -5.41 | 1.32348 | none |
| DNMBP | 0.272 | 0.2672 | 1.5564 | 0.15176 | 0.296 | -5.41 | -0.8568 | none |
| PLK1 | -0.156 | -0.086 | -1.556 | 0.15177 | 0.296 | -5.41 | 0.7551 | none |
| MRPL24 | -0.17 | -0.1391 | -1.556 | 0.1518 | 0.296 | -5.41 | 0.86255 | none |
| FLRT2 | 0.248 | 0.1517 | 1.5559 | 0.15186 | 0.296 | -5.41 | -0.9284 | none |
| SLFN12 | -0.172 | -0.1288 | -1.556 | 0.15194 | 0.296 | -5.41 | 1.68333 | none |
| MTMR7 | 0.209 | 0.1218 | 1.5556 | 0.15194 | 0.296 | -5.41 | -0.8387 | none |
| GNG8 | -0.58 | -0.3462 | -1.555 | 0.15207 | 0.296 | -5.41 | 2.09782 | AA-related |
| KIAA1324I | 0.459 | 0.3875 | 1.5548 | 0.15213 | 0.296 | -5.41 | -0.6985 | none |
| CIC | 0.387 | 0.4714 | 1.5544 | 0.15222 | 0.296 | -5.41 | -0.2187 | none |

| | | | | | | | | |
|---|---|---|---|---|---|---|---|---|
| ALAS2 | -0.139 | -0.112 | -1.554 | 0.1523 | 0.297 | -5.41 | -0.1996 | none |
| RIPK3 | 0.203 | 0.1862 | 1.5539 | 0.15235 | 0.297 | -5.41 | 0.57027 | none |
| SVOPL | 0.152 | 0.156 | 1.5538 | 0.15237 | 0.297 | -5.41 | 0.66656 | none |
| PI4K2A | -0.119 | -0.0441 | -1.554 | 0.15239 | 0.297 | -5.41 | 0.32248 | none |
| DDX58 | 0.203 | 0.1536 | 1.5535 | 0.15245 | 0.297 | -5.41 | 0.96759 | none |
| NECTIN3 | -0.26 | -0.2605 | -1.553 | 0.15249 | 0.297 | -5.41 | 1.94519 | none |
| PGBD1 | -0.168 | -0.096 | -1.552 | 0.15268 | 0.297 | -5.41 | 0.93417 | none |
| EMC10 | 0.219 | 0.0735 | 1.5524 | 0.15271 | 0.297 | -5.41 | -0.2103 | none |
| NHLRC4 | -0.172 | -0.1251 | -1.552 | 0.15284 | 0.297 | -5.41 | 1.38071 | none |
| RCCD1 | 0.192 | 0.072 | 1.5515 | 0.15292 | 0.297 | -5.41 | 0.62863 | none |
| UBA5 | 0.157 | 0.1563 | 1.5508 | 0.15308 | 0.298 | -5.41 | 0.62305 | none |
| LAMTOR5 | -0.216 | -0.242 | -1.551 | 0.1531 | 0.298 | -5.41 | 1.0979 | none |
| NOG | 0.28 | 0.1764 | 1.5507 | 0.1531 | 0.298 | -5.41 | -0.6919 | none |
| WEE2 | 0.167 | 0.1923 | 1.5506 | 0.15313 | 0.298 | -5.41 | 0.77004 | none |
| PLEKHA8 | 0.23 | 0.2131 | 1.5504 | 0.15318 | 0.298 | -5.41 | -1.2176 | none |
| TMEM82 | -0.132 | -0.1321 | -1.55 | 0.15325 | 0.298 | -5.41 | 1.17573 | none |
| FARS2 | 0.144 | 0.0814 | 1.5499 | 0.15329 | 0.298 | -5.41 | 0.9347 | none |
| UQCC2 | -0.265 | -0.2321 | -1.55 | 0.15334 | 0.298 | -5.42 | 1.36637 | none |
| GRINA | 0.188 | 0.1943 | 1.5496 | 0.15335 | 0.298 | -5.42 | -0.3589 | none |
| H2BFM | 0.333 | 0.2508 | 1.5494 | 0.15341 | 0.298 | -5.42 | -0.7757 | none |
| LOXHD1 | 0.256 | 0.1717 | 1.5481 | 0.15373 | 0.299 | -5.42 | -0.1874 | none |
| UCP3 | 0.189 | 0.1533 | 1.548 | 0.15375 | 0.299 | -5.42 | -0.174 | none |
| PRKAR1B | 0.147 | 0.145 | 1.5479 | 0.15376 | 0.299 | -5.42 | -0.6293 | none |
| ATF1 | 0.231 | 0.1417 | 1.5478 | 0.1538 | 0.299 | -5.42 | 1.48494 | none |
| NTRK1 | 0.231 | 0.2245 | 1.5477 | 0.15382 | 0.299 | -5.42 | 0.66292 | none |
| SRP72 | 0.366 | 0.3948 | 1.5471 | 0.15396 | 0.299 | -5.42 | 0.27428 | none |
| ZNF672 | 0.16 | 0.1441 | 1.5471 | 0.15397 | 0.299 | -5.42 | 0.72566 | none |
| NOMO3 | 0.209 | 0.193 | 1.5464 | 0.15414 | 0.299 | -5.42 | -0.7286 | none |
| DAPP1 | 0.207 | 0.1821 | 1.5454 | 0.15437 | 0.3 | -5.42 | -0.8999 | none |
| ANAPC2 | 0.243 | 0.281 | 1.5454 | 0.15438 | 0.3 | -5.42 | -0.5049 | none |
| KCNQ5 | -0.146 | -0.0387 | -1.545 | 0.15451 | 0.3 | -5.42 | 0.68108 | none |
| KNDC1 | 0.222 | 0.2304 | 1.5447 | 0.15453 | 0.3 | -5.42 | 1.08084 | none |
| TOMM20L | 0.431 | 0.3168 | 1.5442 | 0.15465 | 0.3 | -5.42 | -0.7156 | none |
| ATF5 | -0.312 | -0.127 | -1.544 | 0.15465 | 0.3 | -5.42 | 1.7454 | none |
| PIK3IP1 | 0.169 | 0.1287 | 1.5437 | 0.15479 | 0.3 | -5.42 | 0.16926 | none |
| HIST1H3G | -0.175 | -0.1612 | -1.543 | 0.15492 | 0.3 | -5.42 | -0.1712 | none |
| STRN | 0.256 | 0.301 | 1.5427 | 0.15501 | 0.301 | -5.42 | 0.82444 | none |
| S1PR4 | -0.192 | -0.2089 | -1.543 | 0.15503 | 0.301 | -5.42 | 0.65629 | none |
| DNAH11 | 0.45 | 0.5441 | 1.5423 | 0.15512 | 0.301 | -5.43 | -0.4364 | none |
| GNPDA1 | -0.187 | -0.1878 | -1.542 | 0.15527 | 0.301 | -5.43 | 1.54118 | none |

| | | | | | | | | |
|---|---|---|---|---|---|---|---|---|
| SCGB1A1 | -0.372 | -0.2949 | -1.542 | 0.1553 | 0.301 | -5.43 | 1.99986 | none |
| SKIL | 0.317 | 0.3386 | 1.5412 | 0.15537 | 0.301 | -5.43 | -0.6793 | none |
| PPAN | 0.201 | 0.2916 | 1.5408 | 0.15548 | 0.301 | -5.43 | -0.2186 | none |
| RNF40 | 0.52 | 0.5806 | 1.5406 | 0.15553 | 0.301 | -5.43 | 0.06265 | none |
| HIGD1C | -0.228 | -0.2627 | -1.541 | 0.15554 | 0.301 | -5.43 | 0.05513 | none |
| HMGCL | 0.179 | 0.1093 | 1.5403 | 0.15561 | 0.301 | -5.43 | -0.8207 | none |
| AKAP2 | 0.35 | 0.3945 | 1.5402 | 0.15561 | 0.301 | -5.43 | 0.4692 | none |
| NDST2 | 0.221 | 0.2019 | 1.5402 | 0.15561 | 0.301 | -5.43 | -0.4762 | none |
| C5ORF38 | 0.195 | 0.1304 | 1.5402 | 0.15561 | 0.301 | -5.43 | -0.3495 | none |
| GNRH1 | 0.237 | 0.232 | 1.5396 | 0.15575 | 0.302 | -5.43 | 0.55469 | none |
| GCDH | 0.147 | 0.136 | 1.5394 | 0.1558 | 0.302 | -5.43 | 0.18651 | none |
| HSPE1-MC | -0.395 | -0.4384 | -1.539 | 0.15583 | 0.302 | -5.43 | 1.76372 | none |
| COX7A2L | -0.275 | -0.3056 | -1.539 | 0.15585 | 0.302 | -5.43 | 1.24584 | none |
| SH2D3C | 0.297 | 0.3539 | 1.5392 | 0.15587 | 0.302 | -5.43 | -0.7169 | none |
| MMP25 | 0.246 | 0.1928 | 1.5391 | 0.15589 | 0.302 | -5.43 | -0.4044 | none |
| ADPGK | 0.167 | 0.1023 | 1.5388 | 0.15597 | 0.302 | -5.43 | 0.79095 | none |
| HS1BP3 | 0.2 | 0.1576 | 1.5386 | 0.15599 | 0.302 | -5.43 | 1.09155 | none |
| MYO1G | 0.291 | 0.3018 | 1.5381 | 0.15611 | 0.302 | -5.43 | -0.1672 | none |
| SQLE | 0.179 | 0.1443 | 1.5375 | 0.15628 | 0.302 | -5.43 | 0.52537 | none |
| LEKR1 | -0.278 | -0.3409 | -1.537 | 0.15637 | 0.302 | -5.43 | 1.52853 | none |
| ANAPC11 | -0.28 | -0.2638 | -1.537 | 0.15639 | 0.302 | -5.43 | 1.41718 | none |
| KRT72 | 0.223 | 0.1741 | 1.5368 | 0.15643 | 0.302 | -5.43 | -1.0803 | none |
| FN1 | 0.327 | 0.217 | 1.5367 | 0.15646 | 0.302 | -5.43 | -0.828 | none |
| RNF215 | 0.168 | 0.2123 | 1.5359 | 0.15665 | 0.303 | -5.43 | -0.3016 | none |
| ACAA2 | -0.196 | -0.1509 | -1.536 | 0.15667 | 0.303 | -5.43 | 2.44305 | AA-related |
| PGRMC2 | 0.267 | 0.1812 | 1.5357 | 0.1567 | 0.303 | -5.43 | -0.4541 | none |
| SOCS1 | -0.224 | -0.1958 | -1.535 | 0.15684 | 0.303 | -5.43 | 0.96286 | none |
| ZDHHC5 | 0.214 | 0.271 | 1.5348 | 0.15693 | 0.303 | -5.43 | 0.02557 | none |
| CEP85L | 0.196 | 0.2079 | 1.5347 | 0.15696 | 0.303 | -5.44 | -0.425 | none |
| C2ORF49 | -0.199 | -0.1752 | -1.534 | 0.15705 | 0.303 | -5.44 | 0.47001 | none |
| ALG10B | 0.378 | 0.2495 | 1.5342 | 0.15707 | 0.303 | -5.44 | 0.03545 | none |
| AAGAB | -0.245 | -0.2402 | -1.534 | 0.15708 | 0.303 | -5.44 | 1.9938 | none |
| FRG2C | -0.131 | -0.1051 | -1.534 | 0.1571 | 0.303 | -5.44 | 0.59826 | none |
| TOMM20 | -0.249 | -0.2964 | -1.534 | 0.15713 | 0.303 | -5.44 | 1.52292 | none |
| TMEM203 | 0.31 | 0.2487 | 1.5336 | 0.15721 | 0.304 | -5.44 | 0.13822 | none |
| FEM1B | 0.314 | 0.1505 | 1.5335 | 0.15725 | 0.304 | -5.44 | -0.268 | none |
| LRP4 | 0.252 | 0.265 | 1.5333 | 0.15728 | 0.304 | -5.44 | 0.5888 | none |
| AREG | -0.146 | -0.1051 | -1.533 | 0.15733 | 0.304 | -5.44 | 1.9519 | none |
| NTN1 | -0.147 | -0.1386 | -1.533 | 0.15746 | 0.304 | -5.44 | 1.24347 | none |
| AGRP | 0.283 | 0.2077 | 1.5324 | 0.1575 | 0.304 | -5.44 | 0.13814 | none |

| | | | | | | | | |
|---|---|---|---|---|---|---|---|---|
| TBC1D31 | 0.298 | 0.1985 | 1.5324 | 0.15751 | 0.304 | -5.44 | -0.6646 | none |
| MATN4 | 0.14 | 0.1353 | 1.5321 | 0.15758 | 0.304 | -5.44 | 0.77575 | none |
| SMARCA2 | 0.369 | 0.4723 | 1.532 | 0.1576 | 0.304 | -5.44 | 0.32155 | none |
| HIST1H1B | -0.19 | -0.0768 | -1.532 | 0.15763 | 0.304 | -5.44 | 1.97538 | none |
| POT1 | -0.223 | -0.214 | -1.532 | 0.15766 | 0.304 | -5.44 | -0.0271 | none |
| GPRIN3 | 0.419 | 0.4535 | 1.5317 | 0.15768 | 0.304 | -5.44 | -0.1849 | none |
| CEP192 | 0.314 | 0.3864 | 1.5316 | 0.15771 | 0.304 | -5.44 | -0.3834 | none |
| S100A7A | 0.17 | 0.1805 | 1.5314 | 0.15775 | 0.304 | -5.44 | -0.7173 | none |
| EXOC8 | 0.245 | 0.259 | 1.5314 | 0.15775 | 0.304 | -5.44 | -0.6746 | none |
| RPN2 | -0.15 | -0.1042 | -1.531 | 0.15775 | 0.304 | -5.44 | 1.59377 | none |
| NAGS | -0.146 | -0.1344 | -1.531 | 0.15787 | 0.304 | -5.44 | 1.39832 | none |
| NAGLU | 0.225 | 0.148 | 1.5305 | 0.15798 | 0.304 | -5.44 | -0.3939 | none |
| STAMBPL1 | 0.174 | 0.2099 | 1.53 | 0.15808 | 0.305 | -5.44 | 0.01703 | none |
| ITGB1BP1 | -0.171 | -0.1844 | -1.53 | 0.1581 | 0.305 | -5.44 | 1.25782 | none |
| MMP12 | -0.135 | -0.1375 | -1.53 | 0.15821 | 0.305 | -5.44 | 0.67564 | none |
| VPS37A | 0.177 | 0.1688 | 1.5289 | 0.15837 | 0.305 | -5.44 | 0.77533 | none |
| KLC1 | 0.352 | 0.4337 | 1.5284 | 0.15848 | 0.305 | -5.44 | -0.26 | none |
| ECEL1 | -0.104 | -0.0843 | -1.528 | 0.15848 | 0.305 | -5.44 | 0.78088 | none |
| FPGT-TNN | 0.275 | 0.2617 | 1.5283 | 0.15851 | 0.305 | -5.44 | -0.0507 | none |
| PPCDC | 0.201 | 0.223 | 1.5272 | 0.15878 | 0.306 | -5.45 | 1.03767 | none |
| TMEM219 | -0.303 | -0.3401 | -1.527 | 0.15879 | 0.306 | -5.45 | 1.76115 | none |
| UBL3 | 0.137 | 0.0883 | 1.5266 | 0.15894 | 0.306 | -5.45 | 0.27538 | none |
| JARID2 | 0.368 | 0.4736 | 1.5263 | 0.15899 | 0.306 | -5.45 | 0.07078 | none |
| DSCC1 | -0.166 | -0.1437 | -1.526 | 0.15915 | 0.306 | -5.45 | 1.72749 | none |
| CDYL | 0.309 | 0.3642 | 1.5255 | 0.15919 | 0.306 | -5.45 | -1.0492 | none |
| AURKAIP1 | -0.211 | -0.1736 | -1.525 | 0.15925 | 0.306 | -5.45 | 1.00398 | none |
| DNAJC17 | -0.211 | -0.1493 | -1.525 | 0.15926 | 0.306 | -5.45 | -0.0147 | none |
| PIGQ | 0.2 | 0.1799 | 1.5252 | 0.15926 | 0.306 | -5.45 | -0.4449 | none |
| CHST14 | -0.168 | -0.1359 | -1.525 | 0.15934 | 0.306 | -5.45 | 1.37781 | none |
| OTUD3 | 0.292 | 0.3159 | 1.5249 | 0.15935 | 0.306 | -5.45 | -0.5563 | none |
| RSPH1 | 0.203 | 0.1292 | 1.5247 | 0.15941 | 0.306 | -5.45 | 0.38271 | none |
| FKBP1B | -0.154 | -0.1141 | -1.524 | 0.15967 | 0.307 | -5.45 | 0.99094 | none |
| ARHGAP4 | 0.21 | 0.2366 | 1.5235 | 0.1597 | 0.307 | -5.45 | 0.1923 | none |
| CTNS | 0.134 | 0.0861 | 1.5232 | 0.15976 | 0.307 | -5.45 | 0.38451 | none |
| DNAJB5 | 0.21 | 0.2018 | 1.5232 | 0.15976 | 0.307 | -5.45 | 0.61171 | none |
| FOLR2 | -0.174 | -0.0873 | -1.523 | 0.15978 | 0.307 | -5.45 | 0.31145 | none |
| TNNC2 | 0.287 | 0.1827 | 1.5228 | 0.15986 | 0.307 | -5.45 | -0.5504 | none |
| GRIN3A | -0.155 | -0.1274 | -1.522 | 0.16 | 0.307 | -5.45 | 1.89958 | none |
| PPP2R5A | 0.147 | 0.1093 | 1.5221 | 0.16005 | 0.307 | -5.45 | -0.4668 | none |
| PACSIN3 | -0.264 | -0.2313 | -1.522 | 0.16012 | 0.308 | -5.45 | 0.88471 | none |

| | | | | | | | | |
|---|---|---|---|---|---|---|---|---|
| PIK3R3 | 0.2 | 0.0916 | 1.5211 | 0.16028 | 0.308 | -5.45 | 0.42701 | none |
| CBSL | 0.241 | 0.2462 | 1.521 | 0.16031 | 0.308 | -5.45 | 1.06255 | none |
| AGBL5 | 0.231 | 0.228 | 1.5207 | 0.16039 | 0.308 | -5.45 | -0.6014 | none |
| RRP15 | -0.165 | -0.1257 | -1.52 | 0.1605 | 0.308 | -5.45 | 0.91727 | none |
| PDE7A | 0.181 | 0.1718 | 1.5202 | 0.16051 | 0.308 | -5.45 | -0.6403 | none |
| LMO2 | -0.162 | -0.1047 | -1.52 | 0.16057 | 0.308 | -5.45 | 0.19882 | none |
| TMEM155 | 0.182 | 0.1738 | 1.5197 | 0.16063 | 0.308 | -5.46 | 0.00161 | none |
| KIAA0355 | 0.415 | 0.3912 | 1.5196 | 0.16067 | 0.308 | -5.46 | -0.7932 | none |
| YWHAQ | -0.199 | -0.1737 | -1.52 | 0.16067 | 0.308 | -5.46 | 1.03412 | none |
| MLANA | 0.384 | 0.3187 | 1.5195 | 0.16069 | 0.308 | -5.46 | -0.9035 | none |
| ABHD16B | 0.212 | 0.1941 | 1.5193 | 0.16073 | 0.308 | -5.46 | 0.5708 | none |
| MAGED1 | 0.241 | 0.2457 | 1.5192 | 0.16076 | 0.308 | -5.46 | -0.2843 | none |
| CYFIP1 | 0.215 | 0.2752 | 1.5182 | 0.161 | 0.309 | -5.46 | -0.3029 | none |
| HIST1H2BI | -0.354 | -0.3517 | -1.518 | 0.1611 | 0.309 | -5.46 | 0.83973 | none |
| TM2D3 | -0.176 | -0.1935 | -1.518 | 0.16116 | 0.309 | -5.46 | 1.48543 | none |
| VASP | 0.168 | 0.1084 | 1.5175 | 0.16119 | 0.309 | -5.46 | -0.9316 | none |
| PINX1 | -0.252 | -0.0497 | -1.517 | 0.16128 | 0.309 | -5.46 | 1.6541 | none |
| TTC17 | 0.341 | 0.3869 | 1.5169 | 0.16134 | 0.309 | -5.46 | -0.352 | none |
| TOPBP1 | 0.542 | 0.6068 | 1.5165 | 0.16143 | 0.309 | -5.46 | 0.09827 | none |
| CCNB1 | -0.149 | -0.1368 | -1.516 | 0.16146 | 0.309 | -5.46 | 0.28197 | none |
| KLK15 | 0.212 | 0.1586 | 1.5163 | 0.16148 | 0.309 | -5.46 | 0.74326 | none |
| CAPN11 | 0.116 | 0.1245 | 1.5161 | 0.16152 | 0.309 | -5.46 | -0.6293 | none |
| KRT85 | 0.167 | 0.1079 | 1.5161 | 0.16152 | 0.309 | -5.46 | -0.8096 | none |
| NOP14 | 0.563 | 0.7127 | 1.5159 | 0.16157 | 0.309 | -5.46 | 0.14573 | none |
| OGDHL | -0.244 | -0.2426 | -1.516 | 0.16166 | 0.31 | -5.46 | 0.30767 | none |
| RPS6KB2 | -0.178 | -0.1546 | -1.515 | 0.16176 | 0.31 | -5.46 | 1.76447 | none |
| NDUFB7 | -0.247 | -0.2528 | -1.515 | 0.16176 | 0.31 | -5.46 | 1.49253 | none |
| CELSR3 | 0.273 | 0.319 | 1.5145 | 0.16192 | 0.31 | -5.46 | -0.5688 | none |
| VTA1 | -0.133 | -0.1547 | -1.514 | 0.16198 | 0.31 | -5.46 | 0.74204 | none |
| CFAP100 | 0.402 | 0.3464 | 1.5142 | 0.162 | 0.31 | -5.46 | 0.29247 | none |
| SLC16A6 | 0.228 | 0.0975 | 1.514 | 0.16205 | 0.31 | -5.46 | -0.4597 | none |
| ST7 | 0.169 | 0.1108 | 1.5139 | 0.16208 | 0.31 | -5.46 | -0.7294 | none |
| TENM4 | 0.15 | 0.1368 | 1.5138 | 0.16209 | 0.31 | -5.46 | 0.9154 | none |
| DIAPH1 | 0.333 | 0.3792 | 1.5138 | 0.1621 | 0.31 | -5.46 | -0.7298 | none |
| MRPS15 | -0.18 | -0.1854 | -1.514 | 0.16212 | 0.31 | -5.46 | 0.81487 | none |
| TIPIN | 0.168 | 0.0736 | 1.5136 | 0.16215 | 0.31 | -5.46 | 0.3681 | none |
| PPIL4 | 0.256 | 0.2604 | 1.5134 | 0.16219 | 0.31 | -5.46 | 0.1496 | none |
| SAFB | 0.452 | 0.5565 | 1.5131 | 0.16227 | 0.31 | -5.46 | -0.2519 | none |
| GTF2H2C | 0.452 | 0.2333 | 1.5131 | 0.16227 | 0.31 | -5.46 | -0.6895 | none |
| SUCLG1 | -0.157 | -0.2056 | -1.513 | 0.16227 | 0.31 | -5.46 | 0.32778 | none |

| | | | | | | | | |
|---|---|---|---|---|---|---|---|---|
| POLE2 | 0.279 | 0.103 | 1.5127 | 0.16238 | 0.31 | -5.46 | -0.2574 | none |
| PDGFRB | -0.174 | -0.149 | -1.513 | 0.16241 | 0.31 | -5.46 | -0.2555 | none |
| DCP1A | 0.223 | 0.244 | 1.5123 | 0.16249 | 0.311 | -5.46 | 0.27539 | none |
| PABPC1L2. | 0.114 | 0.0908 | 1.5121 | 0.16254 | 0.311 | -5.47 | 0.88165 | none |
| MRPS18A | -0.141 | -0.1511 | -1.512 | 0.16257 | 0.311 | -5.47 | 1.5181 | none |
| KLHL3 | -0.289 | -0.2181 | -1.511 | 0.1627 | 0.311 | -5.47 | 0.0197 | none |
| MAPRE1 | -0.151 | -0.1641 | -1.511 | 0.16276 | 0.311 | -5.47 | 0.88942 | none |
| FNDC8 | 0.165 | 0.1408 | 1.5107 | 0.16289 | 0.311 | -5.47 | 0.56387 | none |
| TNF | 0.721 | 0.6417 | 1.5103 | 0.16297 | 0.311 | -5.47 | 0.79868 | none |
| KMO | 0.289 | 0.2443 | 1.5096 | 0.16315 | 0.312 | -5.47 | 0.00365 | none |
| FUT7 | 0.208 | 0.0798 | 1.5094 | 0.16321 | 0.312 | -5.47 | -0.6571 | none |
| PARP4 | 0.339 | 0.3667 | 1.5093 | 0.16324 | 0.312 | -5.47 | -0.314 | none |
| RPL13 | -0.242 | -0.2414 | -1.509 | 0.16329 | 0.312 | -5.47 | 1.60768 | none |
| GPR1 | 0.158 | 0.1144 | 1.5088 | 0.16335 | 0.312 | -5.47 | 0.1628 | none |
| MROH7-TT | -0.361 | -0.2935 | -1.508 | 0.16345 | 0.312 | -5.47 | 1.50043 | none |
| MALSU1 | -0.261 | -0.2457 | -1.508 | 0.16349 | 0.312 | -5.47 | 1.00488 | none |
| NEK7 | 0.218 | 0.1516 | 1.5081 | 0.16353 | 0.312 | -5.47 | 0.6487 | none |
| BTD | -0.246 | -0.3098 | -1.508 | 0.1636 | 0.312 | -5.47 | 1.81904 | none |
| IFNG | -0.282 | -0.1905 | -1.507 | 0.1637 | 0.312 | -5.47 | 1.85424 | none |
| FBXO7 | -0.202 | -0.2154 | -1.507 | 0.16373 | 0.312 | -5.47 | 2.22898 | AA-related |
| SPATC1L | 0.134 | 0.0937 | 1.5072 | 0.16377 | 0.312 | -5.47 | 0.39013 | none |
| NSUN7 | 0.219 | 0.2115 | 1.5071 | 0.16379 | 0.312 | -5.47 | -0.4016 | none |
| PKD2 | 0.313 | 0.277 | 1.5067 | 0.16389 | 0.313 | -5.47 | -0.926 | none |
| DNAJC2 | 0.366 | 0.4777 | 1.5065 | 0.16394 | 0.313 | -5.47 | -0.0749 | none |
| AIMP2 | -0.194 | -0.2243 | -1.506 | 0.16394 | 0.313 | -5.47 | 1.39834 | none |
| DTD1 | -0.136 | -0.1549 | -1.506 | 0.16395 | 0.313 | -5.47 | 0.91891 | none |
| ARF6 | 0.181 | 0.1204 | 1.5064 | 0.16396 | 0.313 | -5.47 | -0.9667 | none |
| GRIN2C | -0.168 | -0.1332 | -1.506 | 0.16397 | 0.313 | -5.47 | 1.11577 | none |
| GPR75-ASI | -0.295 | -0.1622 | -1.506 | 0.16402 | 0.313 | -5.47 | 0.36498 | none |
| NOLC1 | 0.595 | 0.7087 | 1.5056 | 0.16415 | 0.313 | -5.47 | -0.1608 | none |
| CCDC141 | 0.244 | 0.2444 | 1.5056 | 0.16416 | 0.313 | -5.47 | -0.3381 | none |
| SERBP1 | -0.15 | -0.0571 | -1.506 | 0.16418 | 0.313 | -5.47 | 1.6944 | none |
| TXN2 | -0.148 | -0.1411 | -1.506 | 0.16418 | 0.313 | -5.47 | 1.18619 | none |
| ZDHHC7 | 0.407 | 0.5002 | 1.5053 | 0.16423 | 0.313 | -5.47 | 0.62806 | none |
| TCAF1 | 0.347 | 0.4039 | 1.5053 | 0.16425 | 0.313 | -5.47 | -0.0499 | none |
| DPM2 | -0.123 | -0.0796 | -1.505 | 0.16429 | 0.313 | -5.47 | 0.47134 | none |
| MRPS30 | 0.117 | 0.076 | 1.5051 | 0.16429 | 0.313 | -5.47 | -0.8658 | none |
| GPM6A | 0.207 | 0.1378 | 1.5047 | 0.16439 | 0.313 | -5.47 | -0.1292 | none |
| CALB2 | 0.13 | 0.1296 | 1.5045 | 0.16443 | 0.313 | -5.47 | 0.80662 | none |
| SFR1 | 0.208 | 0.1102 | 1.5042 | 0.16452 | 0.313 | -5.48 | -0.6456 | none |

| | | | | | | | | |
|---|---|---|---|---|---|---|---|---|
| TMEM237 | -0.167 | -0.1858 | -1.504 | 0.16459 | 0.313 | -5.48 | 0.76346 | none |
| COX6A1 | -0.185 | -0.2054 | -1.504 | 0.16463 | 0.313 | -5.48 | 1.85654 | none |
| GLB1L2 | 0.258 | 0.2546 | 1.5037 | 0.16463 | 0.313 | -5.48 | 0.57756 | none |
| AGPS | 0.227 | 0.1871 | 1.5037 | 0.16464 | 0.313 | -5.48 | -0.5611 | none |
| TPM4 | 0.257 | 0.2296 | 1.5036 | 0.16467 | 0.313 | -5.48 | -0.3479 | none |
| GATB | -0.137 | -0.0645 | -1.503 | 0.16474 | 0.313 | -5.48 | 1.10957 | none |
| CADM1 | 0.42 | 0.3451 | 1.5028 | 0.16487 | 0.314 | -5.48 | -1.1046 | none |
| MXI1 | 0.257 | 0.2329 | 1.5028 | 0.16487 | 0.314 | -5.48 | 1.01788 | none |
| LSG1 | 0.359 | 0.3895 | 1.5026 | 0.16493 | 0.314 | -5.48 | -0.5254 | none |
| HIST1H2BI | 0.194 | 0.015 | 1.5024 | 0.16497 | 0.314 | -5.48 | -0.5787 | none |
| RASSF3 | 0.18 | 0.1126 | 1.502 | 0.16507 | 0.314 | -5.48 | 1.09267 | none |
| MED29 | 0.23 | 0.217 | 1.5017 | 0.16515 | 0.314 | -5.48 | -0.8599 | none |
| EIF1B | -0.215 | -0.2848 | -1.501 | 0.16524 | 0.314 | -5.48 | 1.70033 | none |
| FGFBP3 | 0.127 | 0.1007 | 1.501 | 0.16534 | 0.314 | -5.48 | 1.21869 | none |
| SLC10A7 | 0.158 | 0.0928 | 1.5009 | 0.16535 | 0.314 | -5.48 | -0.3499 | none |
| HIST1H3F | -0.434 | -0.4803 | -1.501 | 0.16537 | 0.314 | -5.48 | 1.72867 | none |
| PHKG2 | 0.167 | 0.1552 | 1.5008 | 0.16538 | 0.314 | -5.48 | -0.2936 | none |
| TRMU | 0.142 | 0.163 | 1.5003 | 0.16551 | 0.314 | -5.48 | -0.17 | none |
| SERPING1 | 0.137 | 0.1458 | 1.5 | 0.16557 | 0.314 | -5.48 | 1.46374 | none |
| OCM | -0.18 | -0.1691 | -1.5 | 0.16558 | 0.314 | -5.48 | 1.6827 | none |
| MYOZ1 | 0.142 | 0.0927 | 1.5 | 0.16559 | 0.314 | -5.48 | -1.264 | none |
| SLC30A9 | -0.115 | -0.1106 | -1.5 | 0.1656 | 0.314 | -5.48 | 1.92307 | none |
| PROC | -0.187 | -0.1314 | -1.5 | 0.16561 | 0.314 | -5.48 | 0.78772 | none |
| UBQLNL | 0.122 | 0.0892 | 1.4999 | 0.16562 | 0.314 | -5.48 | -0.5937 | none |
| APPL2 | 0.299 | 0.3664 | 1.4997 | 0.16566 | 0.314 | -5.48 | 0.2402 | none |
| CYP4F2 | -0.178 | -0.1392 | -1.499 | 0.16584 | 0.315 | -5.48 | -0.1178 | none |
| SHH | -0.162 | -0.1677 | -1.499 | 0.1659 | 0.315 | -5.48 | 1.20019 | none |
| TMEM35A | 0.125 | 0.1245 | 1.4982 | 0.16604 | 0.315 | -5.48 | 0.31786 | none |
| TADA1 | -0.174 | -0.1937 | -1.498 | 0.16616 | 0.315 | -5.48 | 0.59329 | none |
| NOL8 | 0.429 | 0.498 | 1.4975 | 0.16622 | 0.315 | -5.48 | -0.5386 | none |
| SLC2A3 | 0.17 | 0.0859 | 1.4975 | 0.16623 | 0.315 | -5.48 | 0.23888 | none |
| CDKN3 | -0.234 | -0.2809 | -1.497 | 0.1663 | 0.315 | -5.48 | 0.45602 | none |
| RTKN | 0.204 | 0.1707 | 1.4968 | 0.16639 | 0.316 | -5.49 | -0.6066 | none |
| PRH1 | 0.352 | 0.245 | 1.4966 | 0.16646 | 0.316 | -5.49 | 0.12211 | none |
| PLK2 | 0.22 | 0.3072 | 1.4964 | 0.1665 | 0.316 | -5.49 | 1.37889 | none |
| OR6C76 | 0.127 | 0.1273 | 1.4959 | 0.16663 | 0.316 | -5.49 | -0.0353 | none |
| POC1B-GA | -0.2 | -0.191 | -1.496 | 0.16672 | 0.316 | -5.49 | 1.09506 | none |
| HRK | 0.176 | 0.0884 | 1.4952 | 0.16682 | 0.316 | -5.49 | 0.04965 | none |
| NUDT12 | 0.166 | 0.1208 | 1.495 | 0.16685 | 0.316 | -5.49 | 0.26709 | none |
| SLC37A1 | -0.258 | -0.1548 | -1.495 | 0.16687 | 0.316 | -5.49 | 1.07739 | none |

| | | | | | | | | |
|---|---|---|---|---|---|---|---|---|
| ULBP1 | 0.104 | 0.0723 | 1.4949 | 0.16689 | 0.316 | -5.49 | -0.5767 | none |
| AHCYL1 | 0.21 | 0.1815 | 1.4947 | 0.16694 | 0.316 | -5.49 | -0.4109 | none |
| CYP26B1 | 0.13 | 0.1305 | 1.4945 | 0.16698 | 0.316 | -5.49 | -0.5049 | none |
| ERMP1 | -0.133 | -0.1034 | -1.495 | 0.16699 | 0.316 | -5.49 | 1.52215 | none |
| KRI1 | 0.423 | 0.5234 | 1.4942 | 0.16707 | 0.316 | -5.49 | -0.2659 | none |
| SFMBT2 | 0.272 | 0.2782 | 1.4931 | 0.16734 | 0.317 | -5.49 | 0.0069 | none |
| BEST3 | -0.309 | -0.2211 | -1.492 | 0.16759 | 0.317 | -5.49 | 1.11096 | none |
| PPP5C | 0.209 | 0.2415 | 1.4917 | 0.1677 | 0.317 | -5.49 | 0.18845 | none |
| ZNF184 | 0.23 | 0.2008 | 1.4915 | 0.16777 | 0.318 | -5.49 | -0.2845 | none |
| DHRS4L2 | -0.15 | -0.1806 | -1.491 | 0.16785 | 0.318 | -5.49 | 0.75731 | none |
| PARP3 | 0.283 | 0.2834 | 1.4907 | 0.16796 | 0.318 | -5.49 | -0.0805 | none |
| CCR7 | 0.171 | 0.1682 | 1.4906 | 0.168 | 0.318 | -5.49 | -0.046 | none |
| MT1F | -0.373 | -0.4121 | -1.49 | 0.16802 | 0.318 | -5.49 | 1.0846 | none |
| TLR2 | 0.175 | 0.1543 | 1.4903 | 0.16807 | 0.318 | -5.49 | 0.2516 | none |
| ATP6V1G2 | 0.162 | 0.1293 | 1.4897 | 0.16823 | 0.318 | -5.49 | -0.5088 | none |
| ANAPC16 | 0.182 | 0.0303 | 1.4895 | 0.16829 | 0.318 | -5.49 | -0.0925 | none |
| NDE1 | 0.449 | 0.4658 | 1.489 | 0.16841 | 0.318 | -5.5 | 0.0346 | none |
| MTO1 | 0.221 | 0.1578 | 1.489 | 0.16842 | 0.318 | -5.5 | -0.9232 | none |
| C5ORF34 | -0.174 | -0.1522 | -1.487 | 0.16881 | 0.319 | -5.5 | 1.66666 | none |
| ATRX | 0.445 | 0.5526 | 1.4872 | 0.16888 | 0.319 | -5.5 | -0.1329 | none |
| MSH2 | 0.257 | 0.2542 | 1.487 | 0.16892 | 0.319 | -5.5 | 1.19553 | none |
| HIST1H2A. | 0.508 | 0.3782 | 1.4869 | 0.16894 | 0.319 | -5.5 | 0.5601 | none |
| ILDR2 | 0.207 | 0.2043 | 1.4869 | 0.16895 | 0.319 | -5.5 | 0.77043 | none |
| OR1Q1 | 0.119 | 0.0968 | 1.4867 | 0.16901 | 0.319 | -5.5 | 0.59825 | none |
| PSRC1 | -0.222 | -0.1314 | -1.487 | 0.16905 | 0.319 | -5.5 | 0.64618 | none |
| FHL1 | 0.198 | 0.1254 | 1.4857 | 0.16926 | 0.32 | -5.5 | 0.97195 | none |
| SPRED1 | 0.232 | 0.1809 | 1.4851 | 0.16941 | 0.32 | -5.5 | -0.3481 | none |
| SPIN1 | 0.166 | 0.0502 | 1.485 | 0.16944 | 0.32 | -5.5 | 0.06565 | none |
| TRMT1L | 0.234 | 0.2631 | 1.4847 | 0.16952 | 0.32 | -5.5 | -0.0658 | none |
| KLF9 | 0.168 | 0.1388 | 1.4846 | 0.16955 | 0.32 | -5.5 | -0.4528 | none |
| HRASLS | -0.17 | -0.1958 | -1.484 | 0.16981 | 0.321 | -5.5 | 1.68861 | none |
| CNIH3 | 0.179 | 0.1566 | 1.4831 | 0.16993 | 0.321 | -5.5 | -0.4124 | none |
| MRPL45 | -0.518 | -0.3775 | -1.483 | 0.17007 | 0.321 | -5.5 | 0.33533 | none |
| CLEC3B | -0.123 | -0.0901 | -1.482 | 0.17013 | 0.321 | -5.5 | 1.03225 | none |
| BICD1 | 0.241 | 0.2665 | 1.4822 | 0.17018 | 0.321 | -5.5 | 0.34532 | none |
| PPP2R2B | 0.221 | 0.168 | 1.4822 | 0.17018 | 0.321 | -5.5 | -0.6243 | none |
| TBC1D10A | -0.156 | -0.1456 | -1.482 | 0.1702 | 0.321 | -5.5 | 2.07367 | none |
| KLHL12 | -0.12 | -0.1205 | -1.482 | 0.17027 | 0.321 | -5.5 | 1.86251 | none |
| FUCA1 | -0.159 | -0.14 | -1.482 | 0.17029 | 0.321 | -5.5 | 0.47888 | none |
| MT-CO1 | 0.142 | 0.1079 | 1.4817 | 0.17031 | 0.321 | -5.5 | 0.65161 | none |

| | | | | | | | | |
|---|---|---|---|---|---|---|---|---|
| DNAI2 | -0.096 | -0.0682 | -1.482 | 0.17035 | 0.321 | -5.51 | 1.88228 | none |
| APRT | -0.227 | -0.2249 | -1.481 | 0.17039 | 0.321 | -5.51 | 2.00154 | none |
| SPANXN3 | 0.135 | 0.0901 | 1.4814 | 0.17039 | 0.321 | -5.51 | 0.20876 | none |
| PWWP2B | 0.164 | 0.1346 | 1.4812 | 0.17042 | 0.321 | -5.51 | -1.1408 | none |
| MPHOSPH | 0.447 | 0.4965 | 1.481 | 0.17049 | 0.321 | -5.51 | -0.6169 | none |
| UNC93B1 | 0.25 | 0.1751 | 1.4798 | 0.17079 | 0.322 | -5.51 | 0.10283 | none |
| CCNA2 | -0.157 | -0.1644 | -1.48 | 0.17081 | 0.322 | -5.51 | 1.6836 | none |
| BRD3 | 0.287 | 0.2701 | 1.4797 | 0.17082 | 0.322 | -5.51 | -0.7407 | none |
| STON2 | 0.263 | 0.2385 | 1.4796 | 0.17085 | 0.322 | -5.51 | -0.9818 | none |
| HSPB7 | -0.155 | -0.1173 | -1.48 | 0.17086 | 0.322 | -5.51 | 1.09674 | none |
| BAK1 | -0.159 | -0.1775 | -1.479 | 0.17089 | 0.322 | -5.51 | 1.8048 | none |
| GLIPR2 | -0.162 | -0.2097 | -1.479 | 0.17099 | 0.322 | -5.51 | 0.25158 | none |
| KLK6 | -0.117 | -0.0868 | -1.479 | 0.17102 | 0.322 | -5.51 | 0.49785 | none |
| PDAP1 | 0.27 | 0.3285 | 1.4784 | 0.17118 | 0.322 | -5.51 | 0.09268 | none |
| NOSIP | -0.167 | -0.1511 | -1.477 | 0.17141 | 0.323 | -5.51 | 1.54993 | none |
| POTEG | 0.177 | 0.1456 | 1.477 | 0.17155 | 0.323 | -5.51 | -0.9753 | none |
| LRP1 | 0.235 | 0.1743 | 1.4766 | 0.17165 | 0.323 | -5.51 | -0.766 | none |
| VPS25 | -0.199 | -0.219 | -1.476 | 0.17167 | 0.323 | -5.51 | 1.32198 | none |
| ZNF80 | 0.182 | 0.1309 | 1.476 | 0.17178 | 0.323 | -5.51 | 0.94554 | none |
| GPR143 | -0.118 | -0.0928 | -1.476 | 0.17189 | 0.323 | -5.51 | -0.0779 | none |
| MAP2K1 | -0.19 | -0.1419 | -1.476 | 0.17191 | 0.323 | -5.51 | 1.28497 | none |
| PIGX | 0.289 | 0.164 | 1.4756 | 0.17191 | 0.323 | -5.51 | -0.7389 | none |
| WBP4 | -0.224 | -0.1139 | -1.475 | 0.172 | 0.324 | -5.51 | 1.90607 | none |
| GNB1L | -0.207 | -0.2187 | -1.475 | 0.17218 | 0.324 | -5.51 | 1.33022 | none |
| TMEM126 | -0.25 | -0.3221 | -1.475 | 0.17218 | 0.324 | -5.51 | 0.66063 | none |
| GPATCH4 | 0.409 | 0.5637 | 1.4744 | 0.17221 | 0.324 | -5.51 | -0.6047 | none |
| GADD45G | -0.183 | -0.1683 | -1.474 | 0.17224 | 0.324 | -5.51 | 0.93179 | none |
| PGM2 | -0.14 | -0.1053 | -1.473 | 0.17259 | 0.325 | -5.52 | 1.91224 | none |
| POM121 | 0.328 | 0.4009 | 1.4729 | 0.17262 | 0.325 | -5.52 | -0.5199 | none |
| FAM131B | -0.134 | -0.1145 | -1.473 | 0.17263 | 0.325 | -5.52 | 0.24012 | none |
| KCNK7 | 0.22 | 0.2378 | 1.4724 | 0.17274 | 0.325 | -5.52 | 0.94567 | none |
| ZNF30 | 0.264 | 0.2533 | 1.472 | 0.17284 | 0.325 | -5.52 | 0.89373 | none |
| NSF | 0.148 | 0.1405 | 1.4719 | 0.17289 | 0.325 | -5.52 | -0.5803 | none |
| SLC18B1 | 0.262 | 0.1476 | 1.4718 | 0.1729 | 0.325 | -5.52 | 0.81543 | none |
| CHAC2 | -0.367 | -0.3465 | -1.472 | 0.1729 | 0.325 | -5.52 | 0.98174 | none |
| YTHDF2 | 0.177 | 0.1859 | 1.4716 | 0.17295 | 0.325 | -5.52 | 0.74591 | none |
| ZFP1 | 0.132 | 0.1682 | 1.4716 | 0.17295 | 0.325 | -5.52 | 1.4641 | none |
| VPS37C | -0.249 | -0.1434 | -1.471 | 0.17316 | 0.325 | -5.52 | 1.53036 | none |
| PITPNB | 0.2 | 0.1663 | 1.4708 | 0.17317 | 0.325 | -5.52 | 0.33325 | none |
| SKAP1 | -0.124 | -0.1021 | -1.471 | 0.17317 | 0.325 | -5.52 | 1.01384 | none |

| | | | | | | | | |
|---|---|---|---|---|---|---|---|---|
| EIF1AY | 0.241 | 0.1385 | 1.4702 | 0.17333 | 0.325 | -5.52 | -0.2173 | none |
| KARS | -0.125 | -0.1054 | -1.47 | 0.17334 | 0.325 | -5.52 | 1.17782 | none |
| SNX7 | 0.358 | 0.2166 | 1.47 | 0.17338 | 0.326 | -5.52 | -0.8132 | none |
| EXOC6B | 0.349 | 0.3191 | 1.4699 | 0.17341 | 0.326 | -5.52 | -0.9402 | none |
| HLA-A | 0.138 | 0.0159 | 1.4698 | 0.17343 | 0.326 | -5.52 | 0.06733 | none |
| EAF1 | 0.156 | 0.0949 | 1.4696 | 0.17349 | 0.326 | -5.52 | -1.0284 | none |
| WDR11 | 0.234 | 0.1963 | 1.4696 | 0.17349 | 0.326 | -5.52 | -0.8047 | none |
| NKAIN1 | -0.246 | -0.2187 | -1.469 | 0.17356 | 0.326 | -5.52 | 2.00384 | none |
| IL1RL1 | -0.26 | -0.1774 | -1.469 | 0.17358 | 0.326 | -5.52 | 2.11452 | AA-related |
| CD164 | -0.168 | -0.1574 | -1.469 | 0.17363 | 0.326 | -5.52 | 1.34529 | none |
| PNISR | 0.297 | 0.3037 | 1.4689 | 0.17367 | 0.326 | -5.52 | 0.57652 | none |
| NXPH1 | -0.134 | -0.1345 | -1.469 | 0.1737 | 0.326 | -5.52 | 1.93047 | none |
| LCN8 | 0.177 | 0.1392 | 1.4688 | 0.17371 | 0.326 | -5.52 | 0.25834 | none |
| CHPF | 0.125 | 0.117 | 1.4683 | 0.17383 | 0.326 | -5.52 | 0.63789 | none |
| GET4 | -0.154 | -0.1006 | -1.468 | 0.17384 | 0.326 | -5.52 | 1.44592 | none |
| EPRS | 0.513 | 0.5787 | 1.4681 | 0.17388 | 0.326 | -5.52 | -0.1688 | none |
| TIMP2 | 0.214 | 0.1935 | 1.4678 | 0.17398 | 0.326 | -5.52 | -0.5035 | none |
| RABGGTA | -0.173 | -0.1199 | -1.468 | 0.17399 | 0.326 | -5.52 | 0.9111 | none |
| FSTL4 | 0.358 | 0.2687 | 1.4672 | 0.17412 | 0.326 | -5.52 | 0.00207 | none |
| ALS2 | 0.228 | 0.2068 | 1.4669 | 0.1742 | 0.326 | -5.52 | 0.45775 | none |
| RNF8 | 0.247 | 0.263 | 1.4669 | 0.17422 | 0.326 | -5.52 | 0.10744 | none |
| PHC2 | -0.292 | -0.2276 | -1.467 | 0.17428 | 0.326 | -5.52 | 0.39158 | none |
| ANKS1A | 0.203 | 0.2796 | 1.4666 | 0.17429 | 0.326 | -5.52 | -0.2085 | none |
| MPZL1 | 0.148 | 0.0853 | 1.4665 | 0.17432 | 0.326 | -5.52 | 0.33272 | none |
| NTF3 | -0.131 | -0.1313 | -1.466 | 0.17436 | 0.327 | -5.52 | 1.92427 | none |
| INA | 0.141 | 0.1434 | 1.466 | 0.17444 | 0.327 | -5.53 | 0.33708 | none |
| FDXR | 0.221 | 0.1807 | 1.4659 | 0.17446 | 0.327 | -5.53 | -0.9591 | none |
| IPO4 | 0.191 | 0.1866 | 1.4653 | 0.17464 | 0.327 | -5.53 | -0.5916 | none |
| CAPN1 | 0.168 | 0.118 | 1.4652 | 0.17464 | 0.327 | -5.53 | 0.00281 | none |
| SLC17A3 | -0.153 | -0.0972 | -1.465 | 0.17465 | 0.327 | -5.53 | 0.04485 | none |
| RNF180 | 0.231 | 0.2028 | 1.4652 | 0.17467 | 0.327 | -5.53 | -0.1493 | none |
| PSG11 | 0.224 | 0.1961 | 1.4651 | 0.17469 | 0.327 | -5.53 | -0.6603 | none |
| NECTIN4 | 0.123 | 0.0883 | 1.4649 | 0.17473 | 0.327 | -5.53 | -1.0449 | none |
| MUC2 | -0.159 | -0.1058 | -1.464 | 0.17485 | 0.327 | -5.53 | 0.96489 | none |
| SLC4A9 | 0.13 | 0.1269 | 1.4643 | 0.17489 | 0.327 | -5.53 | -0.3392 | none |
| GAPT | 0.19 | 0.1704 | 1.464 | 0.17498 | 0.327 | -5.53 | -0.2087 | none |
| VSTM1 | -0.128 | -0.0853 | -1.464 | 0.175 | 0.327 | -5.53 | 0.35444 | none |
| ZDHHC11E | -0.2 | -0.225 | -1.464 | 0.17504 | 0.327 | -5.53 | 1.36071 | none |
| AC007325 | -0.23 | -0.2 | -1.463 | 0.17521 | 0.328 | -5.53 | 0.65628 | none |
| IL1R2 | -0.163 | -0.1431 | -1.463 | 0.17526 | 0.328 | -5.53 | 2.00444 | none |

| | | | | | | | | |
|---|---|---|---|---|---|---|---|---|
| LMBRD2 | -0.213 | -0.2073 | -1.463 | 0.17533 | 0.328 | -5.53 | 0.00281 | none |
| OR7G1 | -0.114 | -0.0646 | -1.462 | 0.17549 | 0.328 | -5.53 | 1.0039 | none |
| UBE3D | 0.259 | 0.1042 | 1.4621 | 0.1755 | 0.328 | -5.53 | -0.7217 | none |
| ELF1 | 0.171 | 0.1563 | 1.4618 | 0.17555 | 0.328 | -5.53 | -0.2233 | none |
| ZMYND8 | 0.397 | 0.4418 | 1.4612 | 0.17572 | 0.328 | -5.53 | -0.8633 | none |
| RAB11FIP4 | 0.432 | 0.3359 | 1.4609 | 0.1758 | 0.328 | -5.53 | -0.5212 | none |
| HIST1H4F | 0.546 | 0.552 | 1.4607 | 0.17587 | 0.329 | -5.53 | -0.3194 | none |
| OR2A1 | 0.117 | 0.0873 | 1.4605 | 0.17592 | 0.329 | -5.53 | -0.0287 | none |
| BIN2 | 0.45 | 0.4726 | 1.4605 | 0.17593 | 0.329 | -5.53 | -0.3343 | none |
| TRAFD1 | 0.195 | 0.0963 | 1.4601 | 0.17603 | 0.329 | -5.53 | -0.8077 | none |
| LDOC1 | -0.313 | -0.2411 | -1.46 | 0.17615 | 0.329 | -5.53 | 1.84563 | none |
| RBM34 | 0.292 | 0.305 | 1.4593 | 0.17623 | 0.329 | -5.53 | -0.2823 | none |
| LRRC14 | 0.242 | 0.1837 | 1.4593 | 0.17624 | 0.329 | -5.53 | -0.2672 | none |
| PPFIBP1 | 0.349 | 0.3298 | 1.4592 | 0.17627 | 0.329 | -5.53 | -0.0747 | none |
| ZBED3 | 0.121 | 0.1123 | 1.459 | 0.17632 | 0.329 | -5.53 | -0.3827 | none |
| PAOX | -0.285 | -0.2139 | -1.459 | 0.17639 | 0.329 | -5.53 | 1.04402 | none |
| NIPBL | 0.386 | 0.52 | 1.4586 | 0.17643 | 0.329 | -5.53 | -0.1671 | none |
| CDK2AP1 | 0.193 | 0.113 | 1.4586 | 0.17643 | 0.329 | -5.53 | -0.1849 | none |
| PRPF6 | 0.367 | 0.445 | 1.4585 | 0.17644 | 0.329 | -5.53 | -0.2306 | none |
| MMP9 | -0.131 | -0.0933 | -1.458 | 0.17646 | 0.329 | -5.53 | 0.38347 | none |
| HPCAL4 | 0.155 | 0.1325 | 1.4571 | 0.17682 | 0.33 | -5.54 | -0.1174 | none |
| AQP7 | 0.558 | 0.4532 | 1.4569 | 0.17689 | 0.33 | -5.54 | -0.8697 | none |
| PARS2 | -0.126 | -0.0567 | -1.457 | 0.17694 | 0.33 | -5.54 | 1.54435 | none |
| DTNB | -0.18 | -0.0429 | -1.457 | 0.17697 | 0.33 | -5.54 | 1.40236 | none |
| RLN3 | -0.121 | -0.0866 | -1.457 | 0.17697 | 0.33 | -5.54 | 1.41439 | none |
| SLC6A8 | 0.486 | 0.2867 | 1.4562 | 0.17707 | 0.33 | -5.54 | -0.4919 | none |
| CDK11B | 0.386 | 0.4611 | 1.4559 | 0.17716 | 0.33 | -5.54 | -0.465 | none |
| ETHE1 | -0.239 | -0.2657 | -1.456 | 0.17718 | 0.33 | -5.54 | 1.32708 | none |
| ZNF705B | 0.121 | 0.1211 | 1.4557 | 0.17721 | 0.33 | -5.54 | 1.0648 | none |
| FAM217B | 0.144 | 0.1352 | 1.4548 | 0.17746 | 0.331 | -5.54 | 0.52224 | none |
| EXT1 | 0.236 | 0.2189 | 1.4546 | 0.1775 | 0.331 | -5.54 | -0.1226 | none |
| EMC6 | -0.239 | -0.2237 | -1.455 | 0.17751 | 0.331 | -5.54 | 2.00071 | none |
| RNASET2 | 0.16 | 0.0623 | 1.4536 | 0.17779 | 0.331 | -5.54 | 0.35075 | none |
| TRMT11 | -0.154 | -0.1473 | -1.454 | 0.17779 | 0.331 | -5.54 | 1.43823 | none |
| ETV3 | 0.148 | 0.1376 | 1.4534 | 0.17783 | 0.331 | -5.54 | -0.7862 | none |
| ADAM28 | 0.119 | 0.115 | 1.4529 | 0.17795 | 0.331 | -5.54 | 1.4224 | none |
| GPRIN1 | 0.211 | 0.2628 | 1.4526 | 0.17805 | 0.332 | -5.54 | 0.13615 | none |
| DAAM1 | 0.359 | 0.3285 | 1.4521 | 0.17817 | 0.332 | -5.54 | -0.5498 | none |
| MRPL14 | -0.241 | -0.2507 | -1.452 | 0.1783 | 0.332 | -5.54 | 0.80972 | none |
| IMMT | 0.403 | 0.4459 | 1.4511 | 0.17845 | 0.332 | -5.54 | 0.2414 | none |

| | | | | | | | | |
|---|---|---|---|---|---|---|---|---|
| EIF4G1 | 0.476 | 0.6511 | 1.4511 | 0.17846 | 0.332 | -5.54 | -0.3864 | none |
| RAB28 | -0.135 | -0.187 | -1.451 | 0.17853 | 0.332 | -5.54 | -0.1388 | none |
| CRAMP1 | 0.301 | 0.3396 | 1.4506 | 0.17859 | 0.332 | -5.55 | -0.2439 | none |
| UROS | 0.197 | 0.1347 | 1.4502 | 0.1787 | 0.332 | -5.55 | -0.0946 | none |
| ZFPL1 | -0.299 | -0.1652 | -1.449 | 0.1789 | 0.333 | -5.55 | 1.56761 | none |
| PRR7 | -0.158 | -0.1413 | -1.449 | 0.17891 | 0.333 | -5.55 | 1.7178 | none |
| TAP2 | 0.346 | 0.3096 | 1.4494 | 0.17891 | 0.333 | -5.55 | 0.31043 | none |
| KATNA1 | -0.177 | -0.1419 | -1.449 | 0.17904 | 0.333 | -5.55 | 1.36189 | none |
| ARL14EP | -0.191 | -0.1552 | -1.449 | 0.17904 | 0.333 | -5.55 | 2.18701 | AA-related |
| TRAPPC2B | 0.215 | 0.0327 | 1.4487 | 0.1791 | 0.333 | -5.55 | -0.8145 | none |
| SH3BP5L | -0.239 | -0.0705 | -1.449 | 0.1791 | 0.333 | -5.55 | 1.81721 | none |
| FAM207A | 0.139 | 0.1546 | 1.4483 | 0.1792 | 0.333 | -5.55 | 0.6316 | none |
| CNBD2 | 0.312 | 0.1442 | 1.4483 | 0.17923 | 0.333 | -5.55 | -0.813 | none |
| C9ORF40 | -0.274 | -0.2082 | -1.448 | 0.17925 | 0.333 | -5.55 | 2.02704 | none |
| EPN1 | 0.191 | 0.1761 | 1.4482 | 0.17926 | 0.333 | -5.55 | 0.26159 | none |
| GRB14 | -0.12 | -0.1204 | -1.447 | 0.17955 | 0.334 | -5.55 | 1.36588 | none |
| CABP2 | -0.114 | -0.0886 | -1.447 | 0.1796 | 0.334 | -5.55 | 0.83869 | none |
| NRCAM | 0.183 | 0.2165 | 1.4468 | 0.17964 | 0.334 | -5.55 | 0.01377 | none |
| RCAN3 | 0.177 | 0.084 | 1.4465 | 0.1797 | 0.334 | -5.55 | -0.3316 | none |
| TFB1M | 0.162 | 0.0662 | 1.446 | 0.17983 | 0.334 | -5.55 | -0.5295 | none |
| DDX54 | 0.386 | 0.4544 | 1.446 | 0.17984 | 0.334 | -5.55 | -0.5504 | none |
| SAMD13 | -0.141 | -0.1036 | -1.446 | 0.17984 | 0.334 | -5.55 | -0.3773 | none |
| NPR1 | 0.123 | 0.1281 | 1.446 | 0.17985 | 0.334 | -5.55 | -0.7533 | none |
| ZSWIM6 | 0.219 | 0.2482 | 1.4459 | 0.17988 | 0.334 | -5.55 | 0.12849 | none |
| FOXD4 | 0.13 | 0.0969 | 1.4458 | 0.17989 | 0.334 | -5.55 | -1.1597 | none |
| RERE | 0.502 | 0.5715 | 1.4458 | 0.17991 | 0.334 | -5.55 | -0.612 | none |
| ARMCX3 | 0.141 | 0.0757 | 1.4457 | 0.17993 | 0.334 | -5.55 | 0.72218 | none |
| OR8G1 | 0.122 | 0.093 | 1.4456 | 0.17995 | 0.334 | -5.55 | -1.259 | none |
| AGAP2 | 0.424 | 0.5133 | 1.4449 | 0.18014 | 0.334 | -5.55 | 0.02899 | none |
| AHRR | 0.309 | 0.2556 | 1.4448 | 0.18018 | 0.334 | -5.55 | -0.5327 | none |
| TSHZ1 | 0.277 | 0.306 | 1.4448 | 0.18018 | 0.334 | -5.55 | -0.3271 | none |
| SCNN1D | -0.153 | -0.1493 | -1.445 | 0.18025 | 0.334 | -5.55 | 0.28512 | none |
| HK1 | 0.232 | 0.2792 | 1.4443 | 0.18032 | 0.334 | -5.55 | 0.54166 | none |
| SAYSD1 | 0.171 | 0.066 | 1.4436 | 0.18051 | 0.335 | -5.55 | 0.05822 | none |
| NEURL4 | 0.345 | 0.3963 | 1.4435 | 0.18052 | 0.335 | -5.55 | -0.5803 | none |
| CLPSL1 | -0.132 | -0.1119 | -1.444 | 0.18052 | 0.335 | -5.55 | -0.5689 | none |
| TP53TG3D | 0.162 | 0.1064 | 1.4434 | 0.18056 | 0.335 | -5.55 | 0.34025 | none |
| TMX2-CTN | -0.104 | -0.1385 | -1.443 | 0.18058 | 0.335 | -5.55 | 0.57316 | none |
| ZFHX2 | 0.272 | 0.2754 | 1.4433 | 0.18059 | 0.335 | -5.55 | -0.4398 | none |
| UMODL1 | -0.128 | -0.1231 | -1.443 | 0.18067 | 0.335 | -5.55 | 0.13477 | none |

| | | | | | | | | |
|---|---|---|---|---|---|---|---|---|
| KLF12 | 0.244 | 0.295 | 1.4429 | 0.18068 | 0.335 | -5.55 | 0.41099 | none |
| SAMM50 | -0.157 | -0.1944 | -1.443 | 0.18073 | 0.335 | -5.56 | 0.6537 | none |
| PPP2CA | -0.176 | -0.1371 | -1.443 | 0.18079 | 0.335 | -5.56 | 1.90332 | none |
| ESRRB | 0.169 | 0.1288 | 1.4425 | 0.1808 | 0.335 | -5.56 | -1.0807 | none |
| NEO1 | 0.688 | 0.4944 | 1.4423 | 0.18085 | 0.335 | -5.56 | 0.16371 | none |
| NDUFB1 | -0.261 | -0.3203 | -1.442 | 0.18092 | 0.335 | -5.56 | 0.78451 | none |
| LCN6 | -0.124 | -0.0827 | -1.441 | 0.18112 | 0.335 | -5.56 | 0.86893 | none |
| SLC36A1 | 0.294 | 0.3491 | 1.4412 | 0.18115 | 0.335 | -5.56 | 1.07086 | none |
| GAS2L1 | 0.134 | 0.1434 | 1.4409 | 0.18125 | 0.336 | -5.56 | 0.34856 | none |
| GEMIN6 | 0.235 | 0.0634 | 1.4405 | 0.18135 | 0.336 | -5.56 | -0.7592 | none |
| ANKHD1-E | 0.382 | 0.3981 | 1.4404 | 0.18138 | 0.336 | -5.56 | -0.4426 | none |
| ADAMTS1( | 0.202 | 0.1396 | 1.4404 | 0.18139 | 0.336 | -5.56 | -0.6428 | none |
| SMG5 | 0.293 | 0.4144 | 1.4403 | 0.18142 | 0.336 | -5.56 | 0.86172 | none |
| TMEM235 | 0.159 | 0.1561 | 1.44 | 0.1815 | 0.336 | -5.56 | 0.5237 | none |
| LUZP2 | 0.123 | 0.0707 | 1.4398 | 0.18156 | 0.336 | -5.56 | -0.5713 | none |
| KIF9 | 0.315 | 0.1985 | 1.4392 | 0.18172 | 0.336 | -5.56 | 0.88866 | none |
| SERPINI1 | 0.376 | 0.2315 | 1.4388 | 0.18184 | 0.336 | -5.56 | -0.6614 | none |
| 3-Sep | -0.138 | -0.0764 | -1.439 | 0.18186 | 0.336 | -5.56 | 0.93965 | none |
| ITGB3 | 0.141 | 0.0992 | 1.4383 | 0.18197 | 0.337 | -5.56 | -1.2899 | none |
| LCN9 | 0.137 | 0.1374 | 1.4382 | 0.18198 | 0.337 | -5.56 | 0.7069 | none |
| HHIPL1 | 0.141 | 0.1222 | 1.4382 | 0.18199 | 0.337 | -5.56 | -0.6795 | none |
| GJB7 | 0.321 | 0.2714 | 1.438 | 0.18205 | 0.337 | -5.56 | -0.0951 | none |
| KAT2A | 0.265 | 0.2815 | 1.4377 | 0.18213 | 0.337 | -5.56 | -0.3437 | none |
| BUB1 | 0.285 | 0.3198 | 1.4375 | 0.18217 | 0.337 | -5.56 | 0.54107 | none |
| ICAM3 | -0.167 | -0.1918 | -1.437 | 0.18245 | 0.337 | -5.56 | 1.99788 | none |
| YTHDC1 | 0.38 | 0.4903 | 1.4365 | 0.18245 | 0.337 | -5.56 | 0.59176 | none |
| SNRPD3 | -0.167 | -0.1501 | -1.436 | 0.18257 | 0.337 | -5.56 | 0.17173 | none |
| ENPP6 | 0.134 | 0.092 | 1.4355 | 0.18273 | 0.338 | -5.56 | -0.7377 | none |
| MYH9 | 0.493 | 0.6438 | 1.4354 | 0.18276 | 0.338 | -5.56 | -0.2555 | none |
| NET1 | 0.164 | 0.1638 | 1.4351 | 0.18284 | 0.338 | -5.56 | -0.3626 | none |
| SNTB1 | 0.231 | 0.2634 | 1.4351 | 0.18284 | 0.338 | -5.56 | -0.0668 | none |
| ZNF565 | 0.124 | 0.1192 | 1.4342 | 0.1831 | 0.338 | -5.57 | -0.123 | none |
| ZNF14 | 0.228 | 0.2403 | 1.4342 | 0.18311 | 0.338 | -5.57 | -0.2351 | none |
| SPTLC2 | 0.203 | 0.151 | 1.4337 | 0.18323 | 0.338 | -5.57 | 0.98819 | none |
| KIF13B | 0.402 | 0.4249 | 1.4335 | 0.1833 | 0.338 | -5.57 | 0.46236 | none |
| C4ORF45 | 0.154 | 0.1346 | 1.4332 | 0.18339 | 0.339 | -5.57 | 0.67903 | none |
| SAMD10 | 0.207 | 0.1432 | 1.4329 | 0.18345 | 0.339 | -5.57 | 0.38319 | none |
| DALRD3 | 0.206 | 0.1031 | 1.4329 | 0.18347 | 0.339 | -5.57 | 0.87298 | none |
| BRSK2 | 0.273 | 0.2299 | 1.4321 | 0.18368 | 0.339 | -5.57 | -0.5711 | none |
| UBE4A | 0.231 | 0.2407 | 1.4316 | 0.18382 | 0.339 | -5.57 | -0.0425 | none |

| | | | | | | | | |
|---|---|---|---|---|---|---|---|---|
| SUV39H2 | 0.223 | 0.0931 | 1.4306 | 0.18409 | 0.34 | -5.57 | -0.2291 | none |
| NDFIP1 | -0.193 | -0.2401 | -1.43 | 0.18426 | 0.34 | -5.57 | 0.81322 | none |
| DSC1 | 0.104 | 0.0832 | 1.4298 | 0.18432 | 0.34 | -5.57 | -0.3511 | none |
| EIF2AK3 | 0.212 | 0.2287 | 1.4298 | 0.18433 | 0.34 | -5.57 | -0.5566 | none |
| KRT6A | -0.244 | -0.2245 | -1.43 | 0.18438 | 0.34 | -5.57 | 0.22449 | none |
| NDUFA9 | -0.219 | -0.235 | -1.429 | 0.18446 | 0.34 | -5.57 | 0.04156 | none |
| CTNNA2 | 0.153 | 0.1537 | 1.429 | 0.18456 | 0.34 | -5.57 | -0.5508 | none |
| TMEM221 | 0.303 | 0.2444 | 1.429 | 0.18456 | 0.34 | -5.57 | -0.8086 | none |
| ASPM | 0.394 | 0.4959 | 1.4289 | 0.18459 | 0.34 | -5.57 | 0.23943 | none |
| TEX38 | -0.138 | -0.0912 | -1.428 | 0.18471 | 0.34 | -5.57 | 0.31088 | none |
| PTP4A2 | 0.119 | 0.0176 | 1.4283 | 0.18473 | 0.34 | -5.57 | 0.6178 | none |
| STK35 | 0.184 | 0.237 | 1.4283 | 0.18474 | 0.34 | -5.57 | -0.0082 | none |
| CACNG5 | 0.13 | 0.1297 | 1.4282 | 0.18478 | 0.34 | -5.57 | 1.17005 | none |
| FBXO34 | 0.309 | 0.3165 | 1.4277 | 0.1849 | 0.341 | -5.57 | -0.1991 | none |
| FCHSD2 | 0.283 | 0.2948 | 1.4274 | 0.185 | 0.341 | -5.57 | 0.96267 | none |
| NNMT | -0.129 | -0.0996 | -1.427 | 0.18513 | 0.341 | -5.58 | 0.27456 | none |
| PIP4K2A | 0.275 | 0.3007 | 1.4269 | 0.18513 | 0.341 | -5.58 | -0.9943 | none |
| HSD17B14 | 0.262 | 0.2055 | 1.4266 | 0.18521 | 0.341 | -5.58 | -0.3507 | none |
| NHS | 0.276 | 0.3633 | 1.4264 | 0.18529 | 0.341 | -5.58 | 0.99864 | none |
| DET1 | 0.256 | 0.1514 | 1.425 | 0.18568 | 0.342 | -5.58 | -0.2971 | none |
| TRMT1 | 0.123 | 0.1491 | 1.4248 | 0.18572 | 0.342 | -5.58 | 0.49513 | none |
| PCP4 | -0.125 | -0.1247 | -1.425 | 0.1858 | 0.342 | -5.58 | 1.04028 | none |
| SLC9A8 | 0.185 | 0.1988 | 1.4242 | 0.1859 | 0.342 | -5.58 | -0.9473 | none |
| TPO | 0.1 | 0.0763 | 1.4239 | 0.18597 | 0.342 | -5.58 | -0.4985 | none |
| ACAN | 0.122 | 0.0965 | 1.4239 | 0.18599 | 0.342 | -5.58 | -0.0843 | none |
| RGS19 | -0.169 | -0.1971 | -1.424 | 0.18603 | 0.342 | -5.58 | 1.06986 | none |
| MT-ND6 | 0.452 | 0.4907 | 1.4237 | 0.18603 | 0.342 | -5.58 | -0.2514 | none |
| NMU | 0.35 | 0.1898 | 1.4236 | 0.18607 | 0.342 | -5.58 | 0.4515 | none |
| HSF5 | -0.155 | -0.1539 | -1.423 | 0.18609 | 0.342 | -5.58 | 1.19339 | none |
| ASCL3 | 0.125 | 0.0831 | 1.4235 | 0.18611 | 0.342 | -5.58 | 1.11751 | none |
| WNT5A | 0.276 | 0.2761 | 1.4234 | 0.18611 | 0.342 | -5.58 | 1.11927 | none |
| CEBPB | -0.115 | -0.0688 | -1.423 | 0.18611 | 0.342 | -5.58 | 0.75461 | none |
| VDR | 0.33 | 0.4479 | 1.4234 | 0.18612 | 0.342 | -5.58 | 0.38143 | none |
| ATP23 | 0.233 | 0.1794 | 1.4231 | 0.18619 | 0.342 | -5.58 | -0.0703 | none |
| CTNNAL1 | 0.162 | 0.1574 | 1.423 | 0.18623 | 0.342 | -5.58 | 0.99033 | none |
| PGAP1 | 0.166 | 0.1058 | 1.4228 | 0.18629 | 0.342 | -5.58 | -0.1929 | none |
| CPSF1 | 0.246 | 0.3142 | 1.4227 | 0.18631 | 0.342 | -5.58 | 0.15665 | none |
| PREP | 0.174 | 0.1718 | 1.4227 | 0.18631 | 0.342 | -5.58 | 0.75697 | none |
| PARVG | -0.127 | -0.0584 | -1.423 | 0.18631 | 0.342 | -5.58 | 1.56015 | none |
| KCNA5 | 0.135 | 0.0867 | 1.4226 | 0.18634 | 0.342 | -5.58 | -1.0658 | none |

| | | | | | | | | |
|---|---|---|---|---|---|---|---|---|
| RGS22 | 0.166 | 0.1166 | 1.4225 | 0.18636 | 0.342 | -5.58 | 0.43535 | none |
| TFEC | -0.168 | -0.0904 | -1.423 | 0.18636 | 0.342 | -5.58 | 0.87259 | none |
| ARHGAP3! | 0.37 | 0.519 | 1.4225 | 0.18637 | 0.342 | -5.58 | -0.1056 | none |
| PTPN7 | 0.214 | 0.2818 | 1.4225 | 0.18639 | 0.342 | -5.58 | -0.0287 | none |
| ANKRD12 | 0.201 | 0.2274 | 1.4224 | 0.1864 | 0.342 | -5.58 | -0.0836 | none |
| RANBP6 | 0.211 | 0.1597 | 1.4223 | 0.18644 | 0.342 | -5.58 | 0.32611 | none |
| IMPAD1 | 0.305 | 0.1823 | 1.4222 | 0.18646 | 0.342 | -5.58 | -0.7993 | none |
| CCNE1 | 0.214 | 0.1913 | 1.4221 | 0.18649 | 0.342 | -5.58 | 0.07754 | none |
| KIAA1586 | 0.327 | 0.2741 | 1.422 | 0.18652 | 0.342 | -5.58 | -0.1325 | none |
| F12 | 0.407 | 0.3098 | 1.4216 | 0.18662 | 0.342 | -5.58 | -0.0543 | none |
| SH3KBP1 | 0.42 | 0.5022 | 1.421 | 0.1868 | 0.343 | -5.58 | 0.57178 | none |
| NME7 | 0.213 | 0.0552 | 1.421 | 0.18681 | 0.343 | -5.58 | -0.432 | none |
| NDUFA10 | -0.103 | -0.0537 | -1.421 | 0.18687 | 0.343 | -5.58 | 1.68996 | none |
| AREL1 | 0.208 | 0.1611 | 1.4206 | 0.1869 | 0.343 | -5.58 | -1.2668 | none |
| BAHD1 | 0.311 | 0.341 | 1.4206 | 0.18691 | 0.343 | -5.58 | 0.42194 | none |
| MT-ATP8 | 0.382 | 0.3601 | 1.4205 | 0.18693 | 0.343 | -5.58 | 0.35852 | none |
| AARS | 0.25 | 0.2927 | 1.4204 | 0.18697 | 0.343 | -5.58 | 0.3369 | none |
| NAGPA | 0.153 | 0.1155 | 1.4202 | 0.18702 | 0.343 | -5.58 | -0.0766 | none |
| ZNF781 | 0.504 | 0.5076 | 1.4202 | 0.18702 | 0.343 | -5.58 | -0.4497 | none |
| DCAF12 | 0.147 | 0.0593 | 1.4202 | 0.18703 | 0.343 | -5.58 | 1.02301 | none |
| SOGA3 | 0.158 | 0.1932 | 1.4201 | 0.18704 | 0.343 | -5.58 | -0.3399 | none |
| SMCO4 | -0.243 | -0.2006 | -1.42 | 0.18707 | 0.343 | -5.58 | 1.67563 | none |
| TRIM49D2 | -0.126 | -0.1347 | -1.42 | 0.18718 | 0.343 | -5.58 | 0.81413 | none |
| RPL39L | -0.238 | -0.2992 | -1.42 | 0.1872 | 0.343 | -5.58 | 1.48546 | none |
| ZNF626 | 0.205 | 0.0791 | 1.4195 | 0.18722 | 0.343 | -5.58 | 0.19679 | none |
| TRA2B | -0.157 | -0.1808 | -1.419 | 0.18731 | 0.343 | -5.59 | 2.11396 | AA-related |
| IGFL1 | -0.113 | -0.113 | -1.419 | 0.18732 | 0.343 | -5.59 | -0.2788 | none |
| LUZP1 | 0.416 | 0.4876 | 1.4191 | 0.18735 | 0.343 | -5.59 | -0.2283 | none |
| SLC25A27 | 0.334 | 0.2569 | 1.4191 | 0.18735 | 0.343 | -5.59 | -0.4242 | none |
| MYL6B | 0.174 | 0.1064 | 1.419 | 0.18737 | 0.343 | -5.59 | 0.44476 | none |
| CEMP1 | 0.397 | 0.2825 | 1.4186 | 0.18747 | 0.343 | -5.59 | 0.2999 | none |
| GOT1L1 | 0.16 | 0.1746 | 1.4186 | 0.18748 | 0.343 | -5.59 | -0.0275 | none |
| ERRFI1 | -0.157 | -0.1447 | -1.419 | 0.18749 | 0.343 | -5.59 | 1.4724 | none |
| BRIX1 | 0.174 | 0.1698 | 1.4183 | 0.18757 | 0.343 | -5.59 | -0.7288 | none |
| IL5RA | 0.152 | 0.1368 | 1.4182 | 0.18759 | 0.343 | -5.59 | -1.1164 | none |
| TREH | 0.231 | 0.2308 | 1.4182 | 0.1876 | 0.343 | -5.59 | 0.43847 | none |
| VRK1 | -0.146 | -0.1268 | -1.418 | 0.18761 | 0.343 | -5.59 | 1.40319 | none |
| HDX | -0.221 | -0.1821 | -1.418 | 0.18761 | 0.343 | -5.59 | 1.32715 | none |
| EVA1A | 0.143 | 0.1514 | 1.4178 | 0.18769 | 0.343 | -5.59 | 0.5925 | none |
| ZNF792 | 0.125 | 0.1449 | 1.4175 | 0.18779 | 0.343 | -5.59 | 1.47722 | none |

| | | | | | | | | |
|---|---|---|---|---|---|---|---|---|
| CDKN1C | -0.153 | -0.1802 | -1.418 | 0.18779 | 0.343 | -5.59 | 1.44231 | none |
| HDAC2 | -0.182 | -0.0991 | -1.417 | 0.1879 | 0.344 | -5.59 | 2.00813 | none |
| KIF24 | 0.246 | 0.2706 | 1.4171 | 0.18791 | 0.344 | -5.59 | -0.4051 | none |
| EEF2KMT | 0.25 | 0.2102 | 1.4171 | 0.18792 | 0.344 | -5.59 | 0.16104 | none |
| ZBTB41 | 0.217 | 0.2227 | 1.417 | 0.18792 | 0.344 | -5.59 | 0.55922 | none |
| ATP1B3 | -0.127 | -0.1372 | -1.416 | 0.18814 | 0.344 | -5.59 | 1.05854 | none |
| BEST4 | 0.148 | 0.1248 | 1.4161 | 0.18818 | 0.344 | -5.59 | 0.72592 | none |
| COL6A2 | 0.42 | 0.3584 | 1.416 | 0.1882 | 0.344 | -5.59 | -0.6089 | none |
| PCED1B | 0.213 | 0.1731 | 1.4158 | 0.18828 | 0.344 | -5.59 | -0.458 | none |
| OR5H14 | 0.105 | 0.0782 | 1.4157 | 0.18831 | 0.344 | -5.59 | -0.4496 | none |
| BICD2 | 0.354 | 0.4706 | 1.4156 | 0.18834 | 0.344 | -5.59 | 0.07054 | none |
| MYOF | 0.435 | 0.5631 | 1.4154 | 0.18838 | 0.344 | -5.59 | 0.07046 | none |
| PLA2G2A | -0.179 | -0.12 | -1.415 | 0.18847 | 0.344 | -5.59 | 1.0021 | none |
| SYCP2L | -0.288 | -0.2266 | -1.415 | 0.18848 | 0.344 | -5.59 | 1.52929 | none |
| CEND1 | 0.359 | 0.3908 | 1.415 | 0.1885 | 0.344 | -5.59 | 1.06506 | none |
| ACAP3 | 0.323 | 0.2907 | 1.4141 | 0.18877 | 0.345 | -5.59 | -0.1777 | none |
| PGLYRP3 | -0.116 | -0.0829 | -1.414 | 0.18887 | 0.345 | -5.59 | 0.94332 | none |
| KCNH2 | 0.124 | 0.1063 | 1.4133 | 0.189 | 0.345 | -5.59 | -0.1468 | none |
| KCNJ2 | -0.099 | -0.0796 | -1.413 | 0.18902 | 0.345 | -5.59 | 2.30605 | AA-related |
| REG1B | 0.169 | 0.1691 | 1.4131 | 0.18904 | 0.345 | -5.59 | -0.5233 | none |
| TIGIT | -0.175 | -0.238 | -1.413 | 0.18908 | 0.345 | -5.59 | 1.49783 | none |
| STK32A | 0.116 | 0.0976 | 1.413 | 0.18908 | 0.345 | -5.59 | -0.9077 | none |
| AFF2 | 0.144 | 0.1138 | 1.4127 | 0.18914 | 0.345 | -5.59 | -1.1666 | none |
| RNF170 | 0.221 | 0.0701 | 1.4125 | 0.18923 | 0.345 | -5.59 | 0.07265 | none |
| PON1 | 0.29 | 0.1818 | 1.4122 | 0.1893 | 0.345 | -5.59 | -0.6391 | none |
| KRTAP13-3 | 0.116 | 0.116 | 1.4121 | 0.18933 | 0.345 | -5.59 | 0.86214 | none |
| FXYD4 | -0.125 | -0.0795 | -1.412 | 0.1894 | 0.345 | -5.59 | 1.2992 | none |
| GPN2 | 0.179 | 0.0935 | 1.4116 | 0.18947 | 0.346 | -5.59 | 0.73087 | none |
| HLF | 0.194 | 0.1047 | 1.4111 | 0.18961 | 0.346 | -5.6 | 0.44903 | none |
| AK8 | -0.364 | -0.328 | -1.411 | 0.18961 | 0.346 | -5.6 | 0.8354 | none |
| GRK3 | 0.277 | 0.2781 | 1.4111 | 0.18961 | 0.346 | -5.6 | -0.4221 | none |
| MAPKAPK | -0.124 | -0.0171 | -1.411 | 0.18971 | 0.346 | -5.6 | 1.72055 | none |
| HNRNPU | 0.496 | 0.6282 | 1.4105 | 0.18978 | 0.346 | -5.6 | -0.1074 | none |
| C1ORF21 | -0.223 | -0.2203 | -1.41 | 0.18979 | 0.346 | -5.6 | 1.7252 | none |
| AUH | 0.109 | 0.1192 | 1.4099 | 0.18996 | 0.346 | -5.6 | 1.7425 | none |
| OR10V1 | 0.112 | 0.0862 | 1.4098 | 0.19 | 0.346 | -5.6 | 1.2838 | none |
| CEP104 | 0.389 | 0.3631 | 1.4097 | 0.19002 | 0.346 | -5.6 | -0.6758 | none |
| LYSMD3 | 0.172 | 0.048 | 1.4094 | 0.1901 | 0.346 | -5.6 | 1.26126 | none |
| GPRC5C | -0.174 | -0.119 | -1.409 | 0.19012 | 0.346 | -5.6 | 0.31139 | none |
| CRISPLD1 | -0.123 | -0.1273 | -1.409 | 0.19013 | 0.346 | -5.6 | 2.06316 | none |

| | | | | | | | | |
|---|---|---|---|---|---|---|---|---|
| OC90 | 0.124 | 0.082 | 1.409 | 0.19023 | 0.346 | -5.6 | -0.5465 | none |
| NEMP2 | 0.15 | 0.0917 | 1.4089 | 0.19024 | 0.346 | -5.6 | 0.68727 | none |
| HTR3A | -0.107 | -0.0972 | -1.409 | 0.19033 | 0.346 | -5.6 | 2.13398 | AA-related |
| THUMPD2 | 0.155 | 0.153 | 1.4086 | 0.19033 | 0.346 | -5.6 | 1.00681 | none |
| MCM7 | -0.119 | -0.0673 | -1.409 | 0.19033 | 0.346 | -5.6 | 1.4246 | none |
| CDRT15 | 0.343 | 0.3586 | 1.4086 | 0.19034 | 0.346 | -5.6 | -0.6679 | none |
| ADAM17 | 0.308 | 0.238 | 1.4085 | 0.19035 | 0.346 | -5.6 | -0.4085 | none |
| PAK2 | 0.156 | 0.1315 | 1.4085 | 0.19036 | 0.346 | -5.6 | -0.2391 | none |
| TNRC6C | 0.231 | 0.2244 | 1.4084 | 0.19039 | 0.346 | -5.6 | 0.22388 | none |
| VPS4A | 0.187 | 0.2296 | 1.4083 | 0.1904 | 0.346 | -5.6 | 1.40818 | none |
| SPIDR | 0.172 | 0.118 | 1.4082 | 0.19045 | 0.346 | -5.6 | -0.4478 | none |
| KRTAP10-1 | -0.119 | -0.0791 | -1.408 | 0.19054 | 0.347 | -5.6 | 1.67587 | none |
| EFCAB6 | 0.304 | 0.2373 | 1.4078 | 0.19058 | 0.347 | -5.6 | 0.10699 | none |
| RAB4B | 0.185 | 0.0682 | 1.4074 | 0.19068 | 0.347 | -5.6 | -0.4761 | none |
| CES2 | 0.212 | 0.2863 | 1.4065 | 0.19093 | 0.347 | -5.6 | -0.1496 | none |
| GP6 | 0.171 | 0.1064 | 1.4065 | 0.19093 | 0.347 | -5.6 | -1.0196 | none |
| RPS14 | -0.291 | -0.3262 | -1.406 | 0.19108 | 0.347 | -5.6 | 0.86495 | none |
| RBKS | -0.256 | -0.2577 | -1.406 | 0.19119 | 0.348 | -5.6 | 1.26617 | none |
| CSE1L | 0.136 | 0.1286 | 1.4055 | 0.19122 | 0.348 | -5.6 | 0.47939 | none |
| S100B | 0.145 | 0.0692 | 1.4051 | 0.19134 | 0.348 | -5.6 | -0.3195 | none |
| IL4I1 | -0.196 | -0.0301 | -1.405 | 0.19139 | 0.348 | -5.6 | 1.0505 | none |
| EEFSEC | 0.209 | 0.2715 | 1.4049 | 0.19141 | 0.348 | -5.6 | 1.15261 | none |
| ARHGEF28 | 0.416 | 0.4087 | 1.4045 | 0.19152 | 0.348 | -5.6 | 0.52441 | none |
| BCL3 | 0.178 | 0.1724 | 1.404 | 0.19166 | 0.348 | -5.6 | -0.3083 | none |
| ERCC6L2 | 0.386 | 0.4871 | 1.4038 | 0.19173 | 0.348 | -5.6 | 0.0467 | none |
| CATSPER1 | -0.133 | -0.1035 | -1.404 | 0.19176 | 0.348 | -5.6 | 1.11599 | none |
| SIRPB1 | -0.201 | -0.1755 | -1.404 | 0.19177 | 0.348 | -5.6 | 0.00469 | none |
| NTF4 | 0.141 | 0.1413 | 1.4035 | 0.19179 | 0.348 | -5.6 | 0.24563 | none |
| PLEKHA6 | -0.141 | -0.1389 | -1.403 | 0.19187 | 0.348 | -5.61 | 2.03013 | none |
| TMEM134 | 0.234 | 0.1436 | 1.4033 | 0.19187 | 0.348 | -5.61 | -0.0538 | none |
| FFAR3 | -0.095 | -0.0733 | -1.403 | 0.19194 | 0.348 | -5.61 | 0.89399 | none |
| IER5L | -0.112 | -0.1084 | -1.403 | 0.19195 | 0.348 | -5.61 | 0.68661 | none |
| KPNA6 | 0.141 | 0.0982 | 1.403 | 0.19196 | 0.348 | -5.61 | -0.1812 | none |
| UPK3B | -0.139 | -0.0836 | -1.403 | 0.19207 | 0.349 | -5.61 | 1.22049 | none |
| WNT8B | -0.096 | -0.0735 | -1.402 | 0.19214 | 0.349 | -5.61 | 1.2281 | none |
| ZBTB44 | 0.185 | 0.1768 | 1.4019 | 0.19226 | 0.349 | -5.61 | -0.5892 | none |
| SERF1A | 0.378 | 0.3578 | 1.4014 | 0.19242 | 0.349 | -5.61 | 0.93592 | none |
| HBG1 | 0.223 | 0.1677 | 1.4011 | 0.19249 | 0.349 | -5.61 | -0.7374 | none |
| TSSK6 | -0.117 | -0.031 | -1.401 | 0.19255 | 0.349 | -5.61 | 0.40205 | none |
| CD28 | -0.117 | -0.1151 | -1.401 | 0.19264 | 0.349 | -5.61 | 1.20557 | none |

| | | | | | | | | |
|---|---|---|---|---|---|---|---|---|
| ZBTB6 | -0.123 | -0.1366 | -1.4 | 0.19273 | 0.349 | -5.61 | 0.64104 | none |
| GIGYF1 | 0.259 | 0.2139 | 1.3999 | 0.19284 | 0.35 | -5.61 | -0.5905 | none |
| USP47 | 0.353 | 0.4502 | 1.3999 | 0.19284 | 0.35 | -5.61 | 0.66471 | none |
| PPP1R15A | 0.281 | 0.324 | 1.3997 | 0.19289 | 0.35 | -5.61 | -0.8759 | none |
| CELA3A | 0.113 | 0.1133 | 1.3996 | 0.19293 | 0.35 | -5.61 | 1.67968 | none |
| NME5 | 0.135 | 0.1084 | 1.3989 | 0.19312 | 0.35 | -5.61 | 0.40814 | none |
| PIN4 | 0.222 | 0.1061 | 1.3986 | 0.19322 | 0.35 | -5.61 | -0.7445 | none |
| RAD52 | 0.28 | 0.3121 | 1.3986 | 0.19323 | 0.35 | -5.61 | -0.3825 | none |
| FNBP1 | 0.42 | 0.5034 | 1.3981 | 0.19336 | 0.35 | -5.61 | -0.5215 | none |
| PTAR1 | -0.185 | -0.086 | -1.398 | 0.19349 | 0.351 | -5.61 | 1.93693 | none |
| WIPF3 | 0.178 | 0.1083 | 1.3973 | 0.1936 | 0.351 | -5.61 | -1.3523 | none |
| GSN | 0.373 | 0.2661 | 1.3972 | 0.19364 | 0.351 | -5.61 | -0.3995 | none |
| ATP11B | 0.239 | 0.2704 | 1.3968 | 0.19376 | 0.351 | -5.61 | -0.8924 | none |
| ANKDD1B | 0.132 | 0.1322 | 1.3967 | 0.19378 | 0.351 | -5.61 | 0.76294 | none |
| CD1B | -0.093 | -0.0939 | -1.397 | 0.19379 | 0.351 | -5.61 | 1.01106 | none |
| REXO4 | 0.3 | 0.3594 | 1.3957 | 0.19407 | 0.351 | -5.61 | -0.1109 | none |
| TCIRG1 | 0.125 | 0.1317 | 1.3956 | 0.19411 | 0.351 | -5.61 | 1.38573 | none |
| H2AFX | -0.135 | -0.1053 | -1.396 | 0.19413 | 0.351 | -5.61 | 1.05761 | none |
| MSN | 0.265 | 0.3032 | 1.3955 | 0.19414 | 0.351 | -5.61 | 0.08916 | none |
| HLA-DRA | -0.198 | -0.1418 | -1.395 | 0.19421 | 0.351 | -5.62 | -0.1084 | none |
| CAP2 | 0.136 | 0.1005 | 1.395 | 0.19428 | 0.352 | -5.62 | 0.42222 | none |
| TMEM165 | 0.183 | 0.1184 | 1.395 | 0.19429 | 0.352 | -5.62 | 0.98551 | none |
| CDC42BPB | 0.155 | 0.2407 | 1.3947 | 0.19436 | 0.352 | -5.62 | 1.3493 | none |
| PUDP | -0.224 | -0.2898 | -1.394 | 0.19461 | 0.352 | -5.62 | 1.75625 | none |
| UBE2Q2 | -0.223 | -0.2011 | -1.394 | 0.19461 | 0.352 | -5.62 | 0.99529 | none |
| GCN1 | 0.368 | 0.4612 | 1.3937 | 0.19465 | 0.352 | -5.62 | 0.29102 | none |
| TMCO4 | 0.212 | 0.1425 | 1.3936 | 0.19469 | 0.352 | -5.62 | -0.5121 | none |
| NRBP2 | 0.119 | 0.1118 | 1.3935 | 0.19472 | 0.352 | -5.62 | -0.5289 | none |
| PROK2 | -0.215 | -0.2446 | -1.393 | 0.19479 | 0.352 | -5.62 | 0.56697 | none |
| PET117 | -0.373 | -0.2246 | -1.393 | 0.19484 | 0.352 | -5.62 | 1.88904 | none |
| OR6M1 | -0.141 | -0.0963 | -1.393 | 0.19488 | 0.352 | -5.62 | 1.29587 | none |
| ABHD6 | 0.374 | 0.3078 | 1.3928 | 0.19491 | 0.352 | -5.62 | -0.4538 | none |
| GDPD4 | -0.172 | -0.0363 | -1.393 | 0.19493 | 0.352 | -5.62 | -0.045 | none |
| PPP6C | -0.184 | -0.2118 | -1.392 | 0.19515 | 0.353 | -5.62 | 1.8931 | none |
| WDR3 | 0.104 | 0.1015 | 1.3915 | 0.19529 | 0.353 | -5.62 | -0.032 | none |
| FOXP3 | 0.333 | 0.4273 | 1.3914 | 0.19534 | 0.353 | -5.62 | -0.651 | none |
| TPR | 0.248 | 0.1987 | 1.3912 | 0.19539 | 0.353 | -5.62 | -0.5276 | none |
| MTCL1 | -0.292 | -0.2822 | -1.391 | 0.19541 | 0.353 | -5.62 | -0.211 | none |
| EMG1 | -0.179 | -0.2075 | -1.391 | 0.1955 | 0.353 | -5.62 | 2.19655 | AA-related |
| HINT2 | -0.194 | -0.1846 | -1.391 | 0.19551 | 0.353 | -5.62 | 1.49791 | none |

| | | | | | | | | |
|---|---|---|---|---|---|---|---|---|
| APOC4 | 0.423 | 0.3698 | 1.3905 | 0.1956 | 0.353 | -5.62 | -0.7055 | none |
| SMYD2 | 0.121 | 0.0637 | 1.3903 | 0.19566 | 0.353 | -5.62 | 0.12868 | none |
| GPAT2 | 0.187 | 0.1911 | 1.3902 | 0.19567 | 0.353 | -5.62 | -0.415 | none |
| RSC1A1 | 0.351 | 0.4464 | 1.3902 | 0.19568 | 0.353 | -5.62 | 1.34575 | none |
| ABCG2 | 0.244 | 0.1518 | 1.3901 | 0.19571 | 0.353 | -5.62 | -0.394 | none |
| PPP4R3A | 0.284 | 0.3658 | 1.39 | 0.19575 | 0.353 | -5.62 | 0.38542 | none |
| DRP2 | 0.116 | 0.1267 | 1.3899 | 0.19578 | 0.353 | -5.62 | 0.497 | none |
| NLRP7 | 0.124 | 0.0994 | 1.3895 | 0.1959 | 0.353 | -5.62 | -1.062 | none |
| MOV10 | 0.258 | 0.238 | 1.3889 | 0.19606 | 0.354 | -5.62 | -0.3921 | none |
| RSBN1 | 0.256 | 0.189 | 1.3889 | 0.19607 | 0.354 | -5.62 | -0.3636 | none |
| NIPAL1 | 0.202 | 0.1694 | 1.3884 | 0.1962 | 0.354 | -5.62 | -0.161 | none |
| LRRN3 | -0.329 | -0.2347 | -1.388 | 0.19621 | 0.354 | -5.62 | 1.41089 | none |
| ERICH2 | 0.184 | 0.2016 | 1.3883 | 0.19624 | 0.354 | -5.62 | 1.50707 | none |
| RELT | 0.156 | 0.1766 | 1.3882 | 0.19627 | 0.354 | -5.62 | 0.11865 | none |
| LYRM1 | 0.198 | 0.1233 | 1.388 | 0.19633 | 0.354 | -5.62 | -0.1158 | none |
| GIPC2 | -0.133 | -0.1632 | -1.388 | 0.19636 | 0.354 | -5.62 | 1.15935 | none |
| MAD2L1BI | -0.174 | -0.1759 | -1.388 | 0.19645 | 0.354 | -5.62 | 2.28616 | AA-related |
| STYK1 | 0.315 | 0.2799 | 1.3871 | 0.1966 | 0.354 | -5.63 | 0.99621 | none |
| EIF3A | 0.477 | 0.6077 | 1.3866 | 0.19673 | 0.355 | -5.63 | 0.05341 | none |
| ZBTB7C | -0.137 | -0.145 | -1.386 | 0.19683 | 0.355 | -5.63 | 0.83513 | none |
| VCPIP1 | 0.248 | 0.2436 | 1.3862 | 0.19685 | 0.355 | -5.63 | -0.0411 | none |
| FAM47C | 0.102 | 0.0775 | 1.3862 | 0.19687 | 0.355 | -5.63 | -1.1738 | none |
| SNX3 | -0.172 | -0.2156 | -1.386 | 0.19689 | 0.355 | -5.63 | 0.84667 | none |
| PFDN2 | -0.17 | -0.1059 | -1.386 | 0.19689 | 0.355 | -5.63 | 0.6002 | none |
| SLC17A2 | 0.112 | 0.0782 | 1.386 | 0.19692 | 0.355 | -5.63 | -0.7929 | none |
| MTRF1L | 0.246 | 0.2777 | 1.3858 | 0.19698 | 0.355 | -5.63 | 1.15185 | none |
| C19ORF57 | 0.183 | 0.2421 | 1.3857 | 0.197 | 0.355 | -5.63 | 0.32113 | none |
| MAN1B1 | 0.254 | 0.2108 | 1.3855 | 0.19708 | 0.355 | -5.63 | -0.608 | none |
| WDFY4 | -0.144 | -0.1102 | -1.385 | 0.19716 | 0.355 | -5.63 | 0.59437 | none |
| IL1B | -0.134 | -0.0307 | -1.385 | 0.19717 | 0.355 | -5.63 | 0.33249 | none |
| RAD50 | 0.398 | 0.4981 | 1.3851 | 0.19718 | 0.355 | -5.63 | -0.1719 | none |
| SIPA1L2 | 0.321 | 0.2142 | 1.3847 | 0.1973 | 0.355 | -5.63 | 0.49537 | none |
| VPS13D | 0.314 | 0.3828 | 1.3846 | 0.19734 | 0.355 | -5.63 | 0.31674 | none |
| C1R | 0.245 | 0.2137 | 1.3845 | 0.19736 | 0.355 | -5.63 | 0.55649 | none |
| GNL1 | -0.152 | -0.1293 | -1.384 | 0.19749 | 0.355 | -5.63 | 0.16412 | none |
| ABHD10 | -0.166 | -0.1773 | -1.384 | 0.1975 | 0.355 | -5.63 | 1.41553 | none |
| TOR1A | 0.139 | 0.0618 | 1.3838 | 0.19756 | 0.355 | -5.63 | -0.1485 | none |
| CSRNP2 | 0.123 | 0.1456 | 1.3836 | 0.19764 | 0.355 | -5.63 | 0.71164 | none |
| P3H3 | 0.423 | 0.3083 | 1.3834 | 0.19769 | 0.355 | -5.63 | -1.0084 | none |
| IDH3B | -0.119 | -0.1001 | -1.383 | 0.19772 | 0.355 | -5.63 | -0.3095 | none |

| | | | | | | | | |
|---|---|---|---|---|---|---|---|---|
| MYL3 | 0.133 | 0.1326 | 1.3833 | 0.19772 | 0.355 | -5.63 | 1.64295 | none |
| CHURC1-F | -1.09 | -0.931 | -1.383 | 0.19772 | 0.355 | -5.63 | 1.63039 | none |
| CUL1 | 0.334 | 0.3983 | 1.3833 | 0.19773 | 0.355 | -5.63 | 0.28977 | none |
| DHX38 | 0.377 | 0.4639 | 1.3832 | 0.19774 | 0.355 | -5.63 | -0.3167 | none |
| VANGL1 | 0.177 | 0.108 | 1.383 | 0.19781 | 0.355 | -5.63 | 1.50149 | none |
| SRRM3 | 0.198 | 0.1977 | 1.3829 | 0.19784 | 0.356 | -5.63 | -0.4836 | none |
| ZNF853 | 0.098 | 0.1095 | 1.3825 | 0.19794 | 0.356 | -5.63 | -0.0191 | none |
| FAM53B | 0.191 | 0.2586 | 1.3824 | 0.19798 | 0.356 | -5.63 | 0.2214 | none |
| ZNF680 | 0.301 | 0.2629 | 1.3822 | 0.19806 | 0.356 | -5.63 | 1.03778 | none |
| COPZ1 | -0.169 | -0.2434 | -1.382 | 0.19814 | 0.356 | -5.63 | 1.81404 | none |
| TRMT44 | 0.256 | 0.3461 | 1.3818 | 0.19816 | 0.356 | -5.63 | 0.0623 | none |
| PATL2 | 0.18 | 0.1095 | 1.3815 | 0.19824 | 0.356 | -5.63 | 0.11926 | none |
| ULK1 | 0.173 | 0.1218 | 1.3815 | 0.19825 | 0.356 | -5.63 | -0.8272 | none |
| TMEM230 | 0.197 | 0.0235 | 1.3812 | 0.19834 | 0.356 | -5.63 | -0.2021 | none |
| C15ORF62 | 0.148 | 0.1852 | 1.3812 | 0.19834 | 0.356 | -5.63 | 0.75952 | none |
| THADA | 0.135 | 0.1826 | 1.3812 | 0.19835 | 0.356 | -5.63 | 0.55974 | none |
| ACE2 | -0.115 | -0.0764 | -1.381 | 0.19839 | 0.356 | -5.63 | 1.06169 | none |
| ZNF569 | 0.308 | 0.2845 | 1.3807 | 0.19848 | 0.356 | -5.63 | 0.69584 | none |
| PTPN1 | 0.215 | 0.272 | 1.3802 | 0.19865 | 0.356 | -5.63 | -0.0654 | none |
| AP1S2 | -0.123 | -0.1332 | -1.38 | 0.19869 | 0.356 | -5.63 | 1.67776 | none |
| REC114 | 0.12 | 0.0865 | 1.38 | 0.19869 | 0.356 | -5.63 | 0.26663 | none |
| BBOF1 | -0.277 | -0.2071 | -1.38 | 0.19878 | 0.357 | -5.63 | 0.29956 | none |
| MAGED4B | -0.307 | -0.3192 | -1.379 | 0.19887 | 0.357 | -5.63 | 1.99708 | none |
| DSCAML1 | -0.102 | -0.0879 | -1.379 | 0.19899 | 0.357 | -5.64 | 0.98639 | none |
| TUBA3E | 0.139 | 0.0976 | 1.3787 | 0.19907 | 0.357 | -5.64 | 1.06436 | none |
| SEMA3F | 0.227 | 0.1158 | 1.3785 | 0.19916 | 0.357 | -5.64 | 0.24954 | none |
| OAS1 | 0.164 | 0.0905 | 1.378 | 0.1993 | 0.357 | -5.64 | -0.604 | none |
| SMARCE1 | 0.303 | 0.334 | 1.3778 | 0.19936 | 0.357 | -5.64 | -0.2077 | none |
| EXOSC8 | -0.148 | -0.1309 | -1.378 | 0.19941 | 0.357 | -5.64 | 0.75509 | none |
| SPTBN1 | 0.462 | 0.6228 | 1.3775 | 0.19943 | 0.357 | -5.64 | -0.1095 | none |
| LAPTM4A | -0.19 | -0.26 | -1.377 | 0.19948 | 0.358 | -5.64 | 1.37884 | none |
| DHX8 | 0.496 | 0.5155 | 1.3765 | 0.19974 | 0.358 | -5.64 | 0.82463 | none |
| MTX2 | -0.189 | -0.2141 | -1.376 | 0.19987 | 0.358 | -5.64 | 0.95053 | none |
| LRCH4 | -0.156 | -0.0555 | -1.376 | 0.1999 | 0.358 | -5.64 | 1.23536 | none |
| PTEN | 0.345 | 0.3578 | 1.3759 | 0.19992 | 0.358 | -5.64 | -0.4116 | none |
| RNASE6 | 0.13 | 0.1502 | 1.3759 | 0.19993 | 0.358 | -5.64 | 0.88503 | none |
| UGCG | 0.172 | 0.1415 | 1.3758 | 0.19995 | 0.358 | -5.64 | 0.65081 | none |
| KNOP1 | 0.353 | 0.3764 | 1.3757 | 0.19999 | 0.358 | -5.64 | -0.7083 | none |
| MPZL2 | -0.169 | -0.1969 | -1.376 | 0.20002 | 0.358 | -5.64 | 1.03547 | none |
| OTUB2 | -0.177 | -0.1681 | -1.376 | 0.20003 | 0.358 | -5.64 | 1.55217 | none |

| | | | | | | | | |
|---|---|---|---|---|---|---|---|---|
| TMOD1 | -0.118 | -0.0741 | -1.375 | 0.20006 | 0.358 | -5.64 | 1.04203 | none |
| RHCE | 0.127 | 0.1306 | 1.3752 | 0.20014 | 0.358 | -5.64 | 0.22265 | none |
| ATP2B4 | 0.383 | 0.3757 | 1.3752 | 0.20015 | 0.358 | -5.64 | -0.0277 | none |
| ADAM11 | -0.18 | -0.1636 | -1.375 | 0.20018 | 0.358 | -5.64 | 2.03456 | none |
| HDAC11 | 0.262 | 0.187 | 1.375 | 0.2002 | 0.358 | -5.64 | 0.55837 | none |
| GPRASP1 | 0.21 | 0.1732 | 1.3749 | 0.20023 | 0.358 | -5.64 | 0.18292 | none |
| RNF214 | -0.198 | -0.0476 | -1.375 | 0.20026 | 0.358 | -5.64 | 1.64226 | none |
| NUFIP2 | 0.286 | 0.3349 | 1.3746 | 0.20031 | 0.358 | -5.64 | -0.7949 | none |
| SMARCC1 | 0.406 | 0.4905 | 1.3746 | 0.20032 | 0.358 | -5.64 | -0.2173 | none |
| KLRG2 | 0.211 | 0.1637 | 1.3742 | 0.20045 | 0.359 | -5.64 | -0.6087 | none |
| ATOH8 | 0.154 | 0.1454 | 1.374 | 0.20049 | 0.359 | -5.64 | -0.5928 | none |
| METTL4 | -0.332 | -0.399 | -1.373 | 0.20067 | 0.359 | -5.64 | 0.85215 | none |
| PTCH1 | 0.357 | 0.3255 | 1.3733 | 0.2007 | 0.359 | -5.64 | -0.2994 | none |
| NPAS4 | 0.18 | 0.2057 | 1.3731 | 0.20076 | 0.359 | -5.64 | -0.3527 | none |
| DYSF | 0.147 | 0.1916 | 1.3727 | 0.20088 | 0.359 | -5.64 | -0.2852 | none |
| ADNP | 0.343 | 0.3871 | 1.3726 | 0.2009 | 0.359 | -5.64 | -0.446 | none |
| ST6GAL1 | 0.113 | -0.0061 | 1.3725 | 0.20095 | 0.359 | -5.64 | -0.6921 | none |
| TBC1D3C | 0.253 | 0.262 | 1.3724 | 0.20099 | 0.359 | -5.64 | 1.21861 | none |
| SMCO2 | 0.126 | 0.1274 | 1.3724 | 0.20099 | 0.359 | -5.64 | -0.8396 | none |
| CCDC25 | -0.121 | -0.0963 | -1.372 | 0.20103 | 0.359 | -5.64 | 1.81606 | none |
| MAGEH1 | -0.197 | -0.1964 | -1.372 | 0.20104 | 0.359 | -5.64 | 1.77697 | none |
| GLB1 | 0.152 | 0.0371 | 1.3719 | 0.20114 | 0.359 | -5.64 | 0.18363 | none |
| BMP2K | 0.227 | 0.1877 | 1.3714 | 0.20128 | 0.359 | -5.64 | 0.65266 | none |
| CCDC22 | 0.189 | 0.1666 | 1.3714 | 0.20128 | 0.359 | -5.64 | 0.62332 | none |
| CETN2 | -0.256 | -0.2572 | -1.371 | 0.20129 | 0.359 | -5.64 | 1.8451 | none |
| TESK2 | -0.153 | -0.1022 | -1.371 | 0.20131 | 0.359 | -5.65 | 0.30328 | none |
| DNTTIP2 | 0.495 | 0.6215 | 1.3712 | 0.20134 | 0.359 | -5.65 | 0.04931 | none |
| TUFM | -0.127 | -0.0804 | -1.371 | 0.20144 | 0.36 | -5.65 | 1.57 | none |
| PRPSAP1 | 0.174 | 0.1379 | 1.3707 | 0.20148 | 0.36 | -5.65 | -0.1272 | none |
| LMF2 | -0.14 | -0.133 | -1.371 | 0.20153 | 0.36 | -5.65 | 1.08542 | none |
| FAM173A | 0.293 | 0.2266 | 1.3706 | 0.20153 | 0.36 | -5.65 | 1.02733 | none |
| CNNM1 | 0.121 | 0.1303 | 1.3703 | 0.20159 | 0.36 | -5.65 | -0.7421 | none |
| PEA15 | 0.242 | 0.3254 | 1.3703 | 0.2016 | 0.36 | -5.65 | 0.02833 | none |
| CEP97 | 0.339 | 0.3086 | 1.3701 | 0.20168 | 0.36 | -5.65 | -0.3764 | none |
| RBM14 | 0.121 | 0.0953 | 1.37 | 0.2017 | 0.36 | -5.65 | 0.16686 | none |
| GANAB | 0.186 | 0.1715 | 1.3697 | 0.2018 | 0.36 | -5.65 | -0.3986 | none |
| RBM24 | 0.125 | 0.1515 | 1.3695 | 0.20185 | 0.36 | -5.65 | 0.8345 | none |
| TRIM24 | 0.399 | 0.4866 | 1.3691 | 0.20197 | 0.36 | -5.65 | 0.33521 | none |
| MEF2C | -0.398 | -0.3587 | -1.369 | 0.20198 | 0.36 | -5.65 | 1.27254 | none |
| LPCAT4 | 0.242 | 0.2239 | 1.3691 | 0.20198 | 0.36 | -5.65 | -0.557 | none |

| | | | | | | | | |
|---|---|---|---|---|---|---|---|---|
| DENND6B | 0.235 | 0.1929 | 1.369 | 0.20199 | 0.36 | -5.65 | -0.0793 | none |
| PLCG2 | 0.306 | 0.3213 | 1.3685 | 0.20214 | 0.36 | -5.65 | -0.019 | none |
| DEXI | 0.206 | 0.2128 | 1.368 | 0.20232 | 0.361 | -5.65 | 1.74513 | none |
| CHRDL2 | 0.09 | 0.0892 | 1.3678 | 0.20238 | 0.361 | -5.65 | -0.8356 | none |
| CAPSL | 0.135 | 0.1351 | 1.3677 | 0.20239 | 0.361 | -5.65 | 1.27608 | none |
| NAPEPLD | 0.163 | 0.1931 | 1.3675 | 0.20244 | 0.361 | -5.65 | -0.4272 | none |
| MASTL | 0.208 | 0.2101 | 1.3669 | 0.20263 | 0.361 | -5.65 | 0.25101 | none |
| SCGB1C1 | 0.138 | 0.1384 | 1.3666 | 0.20272 | 0.361 | -5.65 | 1.04011 | none |
| LRP10 | 0.101 | 0.0614 | 1.3666 | 0.20274 | 0.361 | -5.65 | -0.3684 | none |
| TMEM74B | 0.315 | 0.215 | 1.3665 | 0.20275 | 0.361 | -5.65 | -0.901 | none |
| BARX1 | -0.149 | -0.1123 | -1.366 | 0.20278 | 0.361 | -5.65 | 0.82041 | none |
| ABHD8 | -0.184 | -0.1119 | -1.366 | 0.20287 | 0.361 | -5.65 | 1.09133 | none |
| ZNF251 | 0.182 | 0.1531 | 1.3661 | 0.20289 | 0.361 | -5.65 | -0.2535 | none |
| OR2M7 | -0.112 | -0.0749 | -1.366 | 0.20299 | 0.361 | -5.65 | 0.86195 | none |
| OR7G3 | -0.112 | -0.0749 | -1.366 | 0.20299 | 0.361 | -5.65 | 1.39014 | none |
| BSCL2 | 0.258 | 0.1164 | 1.3655 | 0.20305 | 0.361 | -5.65 | -0.4077 | none |
| C19ORF48 | 0.124 | 0.1295 | 1.3653 | 0.20313 | 0.362 | -5.65 | -1.0653 | none |
| CEP41 | 0.128 | 0.0908 | 1.3651 | 0.20318 | 0.362 | -5.65 | -1.1168 | none |
| ARMT1 | 0.139 | 0.0685 | 1.3648 | 0.20327 | 0.362 | -5.65 | 0.6948 | none |
| CARS | -0.246 | -0.138 | -1.364 | 0.20339 | 0.362 | -5.65 | 1.74175 | none |
| OFCC1 | 0.25 | 0.1968 | 1.3642 | 0.20346 | 0.362 | -5.65 | -0.9503 | none |
| PPP6R2 | 0.281 | 0.3345 | 1.3642 | 0.20346 | 0.362 | -5.65 | 0.58835 | none |
| TMEM173 | 0.136 | 0.0764 | 1.3641 | 0.20349 | 0.362 | -5.65 | -1.0757 | none |
| CYP4F12 | 0.208 | 0.143 | 1.3636 | 0.20365 | 0.362 | -5.65 | -0.9757 | none |
| TCF20 | 0.177 | 0.2293 | 1.3635 | 0.20367 | 0.362 | -5.65 | -0.1272 | none |
| JTB | -0.229 | -0.2279 | -1.363 | 0.20374 | 0.362 | -5.65 | 1.12433 | none |
| AXIN1 | 0.323 | 0.3381 | 1.3629 | 0.20384 | 0.362 | -5.66 | -1.0148 | none |
| MRPS18B | 0.15 | 0.0581 | 1.3627 | 0.2039 | 0.362 | -5.66 | -0.7927 | none |
| GPD2 | -0.153 | -0.0904 | -1.362 | 0.204 | 0.363 | -5.66 | 0.78374 | none |
| MEIKIN | -0.25 | -0.2382 | -1.362 | 0.2041 | 0.363 | -5.66 | 0.9378 | none |
| TIMM9 | -0.218 | -0.2516 | -1.362 | 0.2041 | 0.363 | -5.66 | 0.90454 | none |
| PSMD1 | 0.3 | 0.3425 | 1.362 | 0.20413 | 0.363 | -5.66 | 0.25871 | none |
| SLC39A6 | 0.204 | 0.2017 | 1.3614 | 0.2043 | 0.363 | -5.66 | 0.30899 | none |
| FOXO4 | 0.286 | 0.2657 | 1.3613 | 0.20433 | 0.363 | -5.66 | -0.8507 | none |
| TIMM50 | 0.171 | 0.099 | 1.3612 | 0.20438 | 0.363 | -5.66 | -0.7582 | none |
| IRF4 | 0.278 | 0.3228 | 1.3608 | 0.20449 | 0.363 | -5.66 | -0.7843 | none |
| PEX11B | -0.177 | -0.2186 | -1.361 | 0.2045 | 0.363 | -5.66 | 1.34529 | none |
| SREBF1 | 0.166 | 0.2648 | 1.3608 | 0.2045 | 0.363 | -5.66 | -0.6112 | none |
| PNPLA4 | -0.306 | -0.2878 | -1.361 | 0.20452 | 0.363 | -5.66 | 0.8446 | none |
| TREM1 | -0.131 | -0.1102 | -1.361 | 0.20457 | 0.363 | -5.66 | 1.45407 | none |

| | | | | | | | | |
|---|---|---|---|---|---|---|---|---|
| BSPRY | 0.157 | 0.1304 | 1.3602 | 0.20467 | 0.363 | -5.66 | 0.66977 | none |
| DKKL1 | 0.131 | 0.1307 | 1.3599 | 0.20476 | 0.363 | -5.66 | -0.6614 | none |
| ANP32B | -0.137 | -0.0516 | -1.36 | 0.20477 | 0.363 | -5.66 | 1.65203 | none |
| P4HB | -0.141 | -0.0844 | -1.359 | 0.20495 | 0.364 | -5.66 | 1.68334 | none |
| NKX6-2 | -0.146 | -0.0864 | -1.359 | 0.20502 | 0.364 | -5.66 | 2.26338 | AA-related |
| LIPT2 | -0.195 | -0.1336 | -1.359 | 0.20505 | 0.364 | -5.66 | 1.54292 | none |
| NCOA4 | 0.114 | 0.0621 | 1.3585 | 0.2052 | 0.364 | -5.66 | 0.67982 | none |
| NAF1 | 0.232 | 0.1851 | 1.3585 | 0.20521 | 0.364 | -5.66 | 0.21824 | none |
| GYG1 | 0.155 | 0.0288 | 1.3583 | 0.20524 | 0.364 | -5.66 | 0.32873 | none |
| ST20-MTH | 0.509 | 0.4032 | 1.358 | 0.20535 | 0.364 | -5.66 | -0.6676 | none |
| SMC4 | 0.52 | 0.6967 | 1.3578 | 0.20541 | 0.364 | -5.66 | -0.1209 | none |
| EGLN1 | 0.164 | 0.1058 | 1.3578 | 0.20542 | 0.364 | -5.66 | -0.4781 | none |
| TRA2A | 0.136 | 0.0793 | 1.3577 | 0.20545 | 0.364 | -5.66 | -0.385 | none |
| PPM1J | -0.115 | -0.0549 | -1.357 | 0.20566 | 0.365 | -5.66 | 0.97409 | none |
| MEF2D | 0.207 | 0.1958 | 1.3569 | 0.20568 | 0.365 | -5.66 | -0.336 | none |
| CRACR2A | 0.221 | 0.3332 | 1.3568 | 0.20571 | 0.365 | -5.66 | 0.41926 | none |
| DPYSL4 | 0.16 | 0.1805 | 1.3568 | 0.20572 | 0.365 | -5.66 | 0.2219 | none |
| PHOSPHO: | -0.166 | -0.1451 | -1.356 | 0.20583 | 0.365 | -5.66 | 0.85053 | none |
| ZNF41 | 0.275 | 0.3661 | 1.3562 | 0.20589 | 0.365 | -5.66 | 0.2997 | none |
| FOXA1 | 0.13 | 0.176 | 1.356 | 0.20597 | 0.365 | -5.66 | 0.12838 | none |
| CCDC134 | 0.15 | 0.1297 | 1.3553 | 0.20617 | 0.365 | -5.66 | 0.45159 | none |
| MPP1 | 0.133 | 0.1335 | 1.3552 | 0.2062 | 0.365 | -5.66 | 0.34223 | none |
| HEBP1 | 0.16 | 0.1453 | 1.3551 | 0.20625 | 0.365 | -5.67 | 1.12054 | none |
| POLR3F | -0.262 | -0.2362 | -1.355 | 0.20634 | 0.365 | -5.67 | 1.01058 | none |
| SPOPL | -0.227 | -0.1919 | -1.355 | 0.20639 | 0.365 | -5.67 | 0.2005 | none |
| SPAG1 | 0.344 | 0.2672 | 1.3544 | 0.20647 | 0.366 | -5.67 | -0.8675 | none |
| GRIA4 | -0.184 | -0.146 | -1.354 | 0.20647 | 0.366 | -5.67 | 1.37981 | none |
| REPIN1 | 0.172 | 0.2039 | 1.3542 | 0.20652 | 0.366 | -5.67 | ######## | none |
| MDH2 | -0.117 | -0.0963 | -1.354 | 0.20654 | 0.366 | -5.67 | 1.79169 | none |
| GPR89B | -0.31 | -0.2629 | -1.354 | 0.20659 | 0.366 | -5.67 | 2.09272 | AA-related |
| GCA | 0.186 | 0.0239 | 1.3539 | 0.20662 | 0.366 | -5.67 | 0.35601 | none |
| TEX30 | -0.242 | -0.2748 | -1.354 | 0.20669 | 0.366 | -5.67 | 0.84732 | none |
| C1QA | -0.111 | -0.0737 | -1.354 | 0.20672 | 0.366 | -5.67 | -0.029 | none |
| RNFT1 | -0.292 | -0.3377 | -1.353 | 0.20675 | 0.366 | -5.67 | 1.40775 | none |
| IFI27L2 | -0.262 | -0.2506 | -1.353 | 0.20678 | 0.366 | -5.67 | 1.59715 | none |
| SLC25A38 | -0.152 | -0.127 | -1.353 | 0.20683 | 0.366 | -5.67 | 1.08497 | none |
| CENPF | 0.202 | 0.3272 | 1.3531 | 0.20685 | 0.366 | -5.67 | 0.1286 | none |
| KCNK16 | -0.108 | -0.0877 | -1.353 | 0.20688 | 0.366 | -5.67 | 0.7405 | none |
| ELMO3 | 0.172 | 0.2064 | 1.3525 | 0.20704 | 0.366 | -5.67 | 0.04998 | none |
| PAQR6 | 0.142 | 0.1134 | 1.352 | 0.20718 | 0.366 | -5.67 | -0.2669 | none |

| | | | | | | | | |
|---|---|---|---|---|---|---|---|---|
| REEP3 | 0.192 | 0.0624 | 1.3519 | 0.20724 | 0.366 | -5.67 | -0.9083 | none |
| CDK18 | 0.439 | 0.391 | 1.3512 | 0.20743 | 0.367 | -5.67 | -0.1997 | none |
| CEP131 | 0.281 | 0.285 | 1.3511 | 0.20747 | 0.367 | -5.67 | -0.7835 | none |
| KIF5B | 0.468 | 0.5793 | 1.3506 | 0.20761 | 0.367 | -5.67 | 0.19187 | none |
| TCTE3 | 0.093 | 0.0651 | 1.3504 | 0.20768 | 0.367 | -5.67 | -0.4132 | none |
| TOMM5 | -0.187 | -0.208 | -1.35 | 0.20769 | 0.367 | -5.67 | 0.40623 | none |
| ETNK2 | 0.276 | 0.272 | 1.3504 | 0.2077 | 0.367 | -5.67 | -0.5767 | none |
| CBFB | 0.191 | 0.1271 | 1.3503 | 0.20771 | 0.367 | -5.67 | -0.2851 | none |
| NLRC4 | -0.095 | -0.069 | -1.35 | 0.20779 | 0.367 | -5.67 | 1.663 | none |
| ADCY10 | 0.141 | 0.0974 | 1.3496 | 0.20795 | 0.367 | -5.67 | -0.7117 | none |
| ZNF202 | 0.227 | 0.2489 | 1.3494 | 0.20799 | 0.367 | -5.67 | 0.74046 | none |
| IQCA1 | 0.12 | 0.1154 | 1.3493 | 0.20801 | 0.367 | -5.67 | 0.28803 | none |
| ATP10A | 0.408 | 0.3789 | 1.349 | 0.20812 | 0.367 | -5.67 | -0.4104 | none |
| PRMT1 | 0.148 | 0.0883 | 1.3489 | 0.20814 | 0.367 | -5.67 | -0.6261 | none |
| C1ORF131 | 0.187 | 0.195 | 1.3489 | 0.20814 | 0.367 | -5.67 | -0.6899 | none |
| CBLN2 | -0.127 | -0.1268 | -1.349 | 0.20819 | 0.367 | -5.67 | 0.14088 | none |
| PLVAP | 0.173 | 0.1594 | 1.3487 | 0.2082 | 0.367 | -5.67 | -0.8096 | none |
| MTERF3 | 0.176 | 0.1061 | 1.3484 | 0.20832 | 0.368 | -5.67 | -0.4812 | none |
| MYO5C | 0.18 | 0.0884 | 1.3476 | 0.20856 | 0.368 | -5.67 | 0.80926 | none |
| PDRG1 | -0.115 | -0.0565 | -1.347 | 0.20865 | 0.368 | -5.67 | 1.43956 | none |
| GALNS | 0.165 | 0.114 | 1.3473 | 0.20866 | 0.368 | -5.67 | -0.4658 | none |
| LDLRAP1 | 0.133 | 0.0781 | 1.346 | 0.20906 | 0.369 | -5.68 | -0.5125 | none |
| ZDHHC19 | -0.114 | -0.085 | -1.346 | 0.20906 | 0.369 | -5.68 | 0.84676 | none |
| CNRIP1 | 0.104 | 0.0968 | 1.3457 | 0.20916 | 0.369 | -5.68 | -0.2304 | none |
| RFX6 | -0.105 | -0.0725 | -1.345 | 0.20944 | 0.369 | -5.68 | 0.8909 | none |
| TMEM242 | 0.1 | 0.095 | 1.3443 | 0.20958 | 0.37 | -5.68 | -0.0972 | none |
| POF1B | -0.092 | -0.0732 | -1.344 | 0.20962 | 0.37 | -5.68 | 0.72809 | none |
| KRTAP4-3 | -0.109 | -0.0728 | -1.344 | 0.20968 | 0.37 | -5.68 | 0.86195 | none |
| MFHAS1 | 0.192 | 0.2128 | 1.3436 | 0.20981 | 0.37 | -5.68 | 0.80585 | none |
| TTYH2 | 0.269 | 0.1665 | 1.3435 | 0.20983 | 0.37 | -5.68 | -0.1108 | none |
| CYP2C9 | 0.196 | 0.1363 | 1.3432 | 0.20992 | 0.37 | -5.68 | -0.5916 | none |
| NLRX1 | 0.331 | 0.3037 | 1.3432 | 0.20993 | 0.37 | -5.68 | -0.4861 | none |
| XRCC3 | 0.167 | 0.1657 | 1.3431 | 0.20997 | 0.37 | -5.68 | -0.3724 | none |
| PATE1 | 0.148 | 0.1587 | 1.3429 | 0.21001 | 0.37 | -5.68 | 0.23544 | none |
| DIO2 | 0.19 | 0.238 | 1.3428 | 0.21004 | 0.37 | -5.68 | 0.90001 | none |
| FANK1 | 0.217 | 0.0826 | 1.3426 | 0.21012 | 0.37 | -5.68 | -1.0008 | none |
| PDZD4 | 0.126 | 0.136 | 1.3418 | 0.21035 | 0.37 | -5.68 | 0.38524 | none |
| TECR | -0.143 | -0.1813 | -1.342 | 0.21035 | 0.37 | -5.68 | 1.75867 | none |
| CAMP | 0.107 | 0.1067 | 1.3418 | 0.21037 | 0.37 | -5.68 | -0.7439 | none |
| NAA10 | -0.124 | -0.0714 | -1.342 | 0.21039 | 0.37 | -5.68 | 1.90188 | none |

| | | | | | | | | |
|---|---|---|---|---|---|---|---|---|
| NXNL1 | 0.262 | 0.2007 | 1.3417 | 0.2104 | 0.37 | -5.68 | 0.30997 | none |
| ZNF281 | 0.336 | 0.3375 | 1.3413 | 0.21051 | 0.371 | -5.68 | 0.5609 | none |
| MSC | 0.237 | 0.2603 | 1.3413 | 0.21052 | 0.371 | -5.68 | 1.5101 | none |
| ZNF443 | -0.195 | -0.1323 | -1.341 | 0.21063 | 0.371 | -5.68 | 1.82823 | none |
| CBWD3 | -0.208 | -0.2538 | -1.341 | 0.21064 | 0.371 | -5.68 | -0.1705 | none |
| EIF3L | -0.105 | -0.1135 | -1.341 | 0.21066 | 0.371 | -5.68 | 1.69793 | none |
| XCR1 | 0.73 | 0.5515 | 1.3405 | 0.21078 | 0.371 | -5.68 | 0.24042 | none |
| U2SURP | 0.336 | 0.4381 | 1.34 | 0.21093 | 0.371 | -5.68 | -0.3047 | none |
| ICK | 0.258 | 0.2865 | 1.3397 | 0.21103 | 0.371 | -5.68 | 0.31446 | none |
| C6ORF62 | 0.181 | 0.171 | 1.3394 | 0.2111 | 0.371 | -5.68 | -0.8722 | none |
| DLEC1 | 0.373 | 0.293 | 1.3393 | 0.21114 | 0.371 | -5.68 | -0.713 | none |
| SECTM1 | -0.36 | -0.1848 | -1.339 | 0.21114 | 0.371 | -5.68 | 0.88415 | none |
| GRAMD1B | 0.291 | 0.3909 | 1.3392 | 0.21116 | 0.371 | -5.68 | -0.3248 | none |
| BCKDK | -0.151 | -0.1068 | -1.339 | 0.21122 | 0.371 | -5.68 | 1.81757 | none |
| FIGN | -0.105 | -0.0851 | -1.339 | 0.21122 | 0.371 | -5.68 | 2.13142 | AA-related |
| SMIM24 | -0.25 | -0.2279 | -1.339 | 0.21122 | 0.371 | -5.68 | 1.63391 | none |
| IFITM2 | 0.123 | 0.0576 | 1.3385 | 0.2114 | 0.372 | -5.69 | 0.1 | none |
| USP2 | 0.21 | 0.2276 | 1.338 | 0.21155 | 0.372 | -5.69 | 0.02054 | none |
| ST20 | 0.276 | 0.1528 | 1.338 | 0.21155 | 0.372 | -5.69 | 0.17519 | none |
| RRP1B | 0.337 | 0.4661 | 1.338 | 0.21155 | 0.372 | -5.69 | 0.35753 | none |
| USP45 | 0.254 | 0.1359 | 1.3372 | 0.2118 | 0.372 | -5.69 | -0.2525 | none |
| PHB | -0.138 | -0.1133 | -1.337 | 0.21189 | 0.372 | -5.69 | 0.6655 | none |
| CSHL1 | 0.112 | 0.1116 | 1.3367 | 0.21196 | 0.372 | -5.69 | 0.20808 | none |
| MYBBP1A | 0.314 | 0.42 | 1.3367 | 0.21197 | 0.372 | -5.69 | -0.1223 | none |
| PSMA6 | -0.166 | -0.2145 | -1.337 | 0.21199 | 0.372 | -5.69 | 0.89078 | none |
| MAGEF1 | -0.15 | -0.0566 | -1.337 | 0.21199 | 0.372 | -5.69 | 1.31783 | none |
| COLCA2 | -0.1 | -0.082 | -1.337 | 0.21202 | 0.372 | -5.69 | 1.59588 | none |
| RRP7A | -0.146 | -0.0449 | -1.336 | 0.2121 | 0.372 | -5.69 | 0.04505 | none |
| ANKRD52 | 0.439 | 0.5082 | 1.3357 | 0.21227 | 0.373 | -5.69 | -0.1201 | none |
| VWDE | -0.119 | -0.0795 | -1.336 | 0.21227 | 0.373 | -5.69 | 1.90725 | none |
| PLA2G4B | 0.229 | 0.199 | 1.3352 | 0.21244 | 0.373 | -5.69 | -0.6131 | none |
| ADGRE1 | -0.213 | -0.2305 | -1.335 | 0.21246 | 0.373 | -5.69 | 0.7833 | none |
| FBXO25 | -0.156 | -0.185 | -1.335 | 0.21251 | 0.373 | -5.69 | 1.40839 | none |
| ZNF280C | 0.262 | 0.2346 | 1.335 | 0.21251 | 0.373 | -5.69 | -0.525 | none |
| PLEKHG5 | 0.241 | 0.1711 | 1.3349 | 0.21254 | 0.373 | -5.69 | -1.0223 | none |
| CAAP1 | 0.201 | 0.1697 | 1.3348 | 0.21257 | 0.373 | -5.69 | 0.44424 | none |
| ACTN2 | -0.182 | -0.2294 | -1.335 | 0.21264 | 0.373 | -5.69 | 0.21121 | none |
| TLN1 | 0.481 | 0.6297 | 1.3343 | 0.2127 | 0.373 | -5.69 | -0.5042 | none |
| CAPZB | -0.135 | -0.0679 | -1.334 | 0.21279 | 0.373 | -5.69 | 1.83818 | none |
| CCAR2 | 0.207 | 0.2393 | 1.3341 | 0.2128 | 0.373 | -5.69 | 0.14629 | none |

| | | | | | | | | |
|---|---|---|---|---|---|---|---|---|
| NR4A3 | 0.223 | 0.2658 | 1.333 | 0.21314 | 0.374 | -5.69 | -0.4799 | none |
| ZNF71 | 0.126 | 0.1039 | 1.3329 | 0.21317 | 0.374 | -5.69 | 0.30209 | none |
| E2F4 | 0.119 | 0.1482 | 1.3326 | 0.21324 | 0.374 | -5.69 | 0.33776 | none |
| TACO1 | 0.124 | 0.0657 | 1.3325 | 0.21327 | 0.374 | -5.69 | 1.00818 | none |
| CYP3A4 | 0.156 | 0.1185 | 1.3324 | 0.2133 | 0.374 | -5.69 | 0.51041 | none |
| FAM118B | -0.344 | -0.3037 | -1.332 | 0.21334 | 0.374 | -5.69 | 2.13181 | AA-related |
| SSR3 | -0.241 | -0.3105 | -1.332 | 0.21341 | 0.374 | -5.69 | 1.14385 | none |
| GOLGA8N | 0.122 | 0.1348 | 1.3321 | 0.21343 | 0.374 | -5.69 | -0.0099 | none |
| AC011043 | 0.11 | 0.0831 | 1.3318 | 0.21351 | 0.374 | -5.69 | -0.0687 | none |
| AC011841 | 0.11 | 0.0831 | 1.3318 | 0.21351 | 0.374 | -5.69 | -0.0687 | none |
| KIAA0930 | 0.324 | 0.3218 | 1.3318 | 0.21352 | 0.374 | -5.69 | 0.51889 | none |
| SIK3 | 0.299 | 0.2475 | 1.3318 | 0.21352 | 0.374 | -5.69 | -0.7342 | none |
| RBM3 | -0.193 | -0.2652 | -1.331 | 0.21364 | 0.374 | -5.69 | 0.81708 | none |
| PCDHA11 | 0.112 | 0.1185 | 1.3313 | 0.21366 | 0.374 | -5.69 | -0.2036 | none |
| MOG | 0.37 | 0.2893 | 1.3313 | 0.21368 | 0.374 | -5.69 | 1.00811 | none |
| FGF3 | -0.096 | -0.0729 | -1.331 | 0.21373 | 0.374 | -5.69 | 0.0339 | none |
| TEX29 | 0.242 | 0.1626 | 1.3308 | 0.21384 | 0.374 | -5.69 | -0.566 | none |
| CDC5L | 0.353 | 0.3805 | 1.3306 | 0.2139 | 0.374 | -5.69 | -0.1871 | none |
| C22ORF31 | 0.147 | 0.1038 | 1.3304 | 0.21396 | 0.374 | -5.7 | -0.1693 | none |
| ZNF362 | 0.321 | 0.3432 | 1.33 | 0.21407 | 0.375 | -5.7 | 0.6951 | none |
| CHAT | -0.095 | -0.091 | -1.33 | 0.21417 | 0.375 | -5.7 | 1.80806 | none |
| ZNF784 | 0.1 | 0.1131 | 1.3297 | 0.21419 | 0.375 | -5.7 | 0.90507 | none |
| OCIAD1 | 0.146 | 0.066 | 1.3295 | 0.21423 | 0.375 | -5.7 | -0.3728 | none |
| COX5B | -0.215 | -0.2465 | -1.329 | 0.21438 | 0.375 | -5.7 | 0.90216 | none |
| TAF1C | 0.165 | 0.1715 | 1.3288 | 0.21445 | 0.375 | -5.7 | 0.09505 | none |
| ACY1 | 0.19 | 0.1204 | 1.3285 | 0.21456 | 0.375 | -5.7 | 0.25487 | none |
| TAX1BP1 | 0.496 | 0.5792 | 1.3285 | 0.21457 | 0.375 | -5.7 | -0.0301 | none |
| CA9 | 0.209 | 0.1266 | 1.3283 | 0.21462 | 0.375 | -5.7 | 0.91896 | none |
| EPHX1 | -0.224 | -0.1986 | -1.328 | 0.21479 | 0.375 | -5.7 | 0.18548 | none |
| ZNF709 | 0.171 | 0.1725 | 1.3277 | 0.21481 | 0.375 | -5.7 | 1.49993 | none |
| PRR4 | -0.31 | -0.232 | -1.327 | 0.21494 | 0.376 | -5.7 | 1.5613 | none |
| STARD4 | -0.199 | -0.2402 | -1.327 | 0.21498 | 0.376 | -5.7 | -0.1193 | none |
| FNDC3B | 0.269 | 0.2238 | 1.3271 | 0.215 | 0.376 | -5.7 | -0.3269 | none |
| PLPPR2 | 0.122 | -0.0194 | 1.327 | 0.21504 | 0.376 | -5.7 | -0.639 | none |
| ITM2B | -0.196 | -0.2788 | -1.327 | 0.21506 | 0.376 | -5.7 | 1.09639 | none |
| ACTL6A | -0.124 | -0.1576 | -1.327 | 0.21514 | 0.376 | -5.7 | 0.7015 | none |
| CYP2U1 | 0.126 | 0.0724 | 1.3261 | 0.21531 | 0.376 | -5.7 | -0.1171 | none |
| ZSCAN31 | -0.24 | -0.2381 | -1.326 | 0.21534 | 0.376 | -5.7 | 0.35409 | none |
| NXF2B | -0.101 | -0.0801 | -1.326 | 0.21534 | 0.376 | -5.7 | 1.3115 | none |
| LENEP | -0.12 | -0.0692 | -1.326 | 0.21547 | 0.376 | -5.7 | 1.07509 | none |

| | | | | | | | | |
|---|---|---|---|---|---|---|---|---|
| ODF3L2 | 0.173 | 0.1938 | 1.3255 | 0.2155 | 0.376 | -5.7 | 0.43568 | none |
| MROH6 | 0.116 | 0.1493 | 1.3252 | 0.21561 | 0.376 | -5.7 | 1.02596 | none |
| ARHGEF5 | 0.323 | 0.2235 | 1.3251 | 0.21563 | 0.376 | -5.7 | -0.8953 | none |
| SDC2 | 0.163 | 0.1538 | 1.3248 | 0.21573 | 0.377 | -5.7 | 0.50579 | none |
| POLR2I | -0.205 | -0.2154 | -1.325 | 0.21581 | 0.377 | -5.7 | 0.37588 | none |
| ACYP1 | 0.193 | 0.139 | 1.3241 | 0.21594 | 0.377 | -5.7 | -0.2174 | none |
| PYY | 0.119 | 0.1219 | 1.3241 | 0.21596 | 0.377 | -5.7 | 0.35764 | none |
| GJA1 | 0.094 | 0.0836 | 1.3241 | 0.21597 | 0.377 | -5.7 | -0.005 | none |
| PORCN | 0.289 | 0.1854 | 1.3239 | 0.21602 | 0.377 | -5.7 | 0.20879 | none |
| RASGEF1C | -0.246 | -0.1718 | -1.323 | 0.2163 | 0.377 | -5.7 | 1.92758 | none |
| PHKA1 | 0.173 | 0.2437 | 1.3229 | 0.21636 | 0.377 | -5.7 | 0.63155 | none |
| SNX32 | -0.123 | -0.0202 | -1.323 | 0.2164 | 0.377 | -5.7 | 1.02741 | none |
| PFDN5 | -0.177 | -0.2312 | -1.322 | 0.21657 | 0.378 | -5.71 | 0.97732 | none |
| CABP7 | -0.109 | -0.0728 | -1.322 | 0.21658 | 0.378 | -5.71 | 1.56517 | none |
| CPNE9 | 0.172 | 0.1799 | 1.3216 | 0.21675 | 0.378 | -5.71 | 0.365 | none |
| MFNG | -0.143 | -0.1101 | -1.322 | 0.21679 | 0.378 | -5.71 | 1.47435 | none |
| DLX4 | -0.129 | -0.1269 | -1.321 | 0.2169 | 0.378 | -5.71 | 1.46844 | none |
| BTG2 | -0.143 | -0.0873 | -1.321 | 0.21708 | 0.378 | -5.71 | 1.33754 | none |
| MRPS14 | -0.166 | -0.2162 | -1.32 | 0.21713 | 0.378 | -5.71 | 1.34971 | none |
| CFAP20 | -0.121 | -0.1484 | -1.32 | 0.21715 | 0.378 | -5.71 | 1.27241 | none |
| UPF3B | 0.33 | 0.4047 | 1.3201 | 0.21725 | 0.379 | -5.71 | -0.1283 | none |
| PLK4 | 0.37 | 0.3286 | 1.32 | 0.21728 | 0.379 | -5.71 | -0.8891 | none |
| PSMB6 | -0.21 | -0.2281 | -1.32 | 0.21735 | 0.379 | -5.71 | 0.17873 | none |
| GYG2 | -0.099 | -0.0697 | -1.32 | 0.21738 | 0.379 | -5.71 | 0.55601 | none |
| SYNGR4 | 0.184 | 0.2247 | 1.319 | 0.2176 | 0.379 | -5.71 | 0.40947 | none |
| TSEN34 | 0.267 | 0.1246 | 1.3188 | 0.21765 | 0.379 | -5.71 | 0.33111 | none |
| TPSG1 | -0.121 | -0.1205 | -1.319 | 0.21769 | 0.379 | -5.71 | 0.71194 | none |
| RASGRP1 | 0.279 | 0.3359 | 1.3185 | 0.21774 | 0.379 | -5.71 | 0.05802 | none |
| LRP5L | 0.146 | 0.1532 | 1.3183 | 0.21782 | 0.379 | -5.71 | -0.5823 | none |
| SLC52A2 | -0.181 | -0.1656 | -1.318 | 0.21787 | 0.379 | -5.71 | 0.96825 | none |
| HM13 | 0.118 | 0.0878 | 1.3178 | 0.21799 | 0.379 | -5.71 | -1.1285 | none |
| PRKAG3 | -0.141 | -0.1235 | -1.318 | 0.21806 | 0.379 | -5.71 | -0.251 | none |
| FLII | 0.339 | 0.3832 | 1.3172 | 0.21818 | 0.38 | -5.71 | 0.24005 | none |
| AHNAK | 0.321 | 0.4298 | 1.3171 | 0.21822 | 0.38 | -5.71 | -0.3669 | none |
| MS4A6A | -0.232 | -0.1866 | -1.317 | 0.21832 | 0.38 | -5.71 | 1.76788 | none |
| A3GALT2 | -0.112 | -0.1122 | -1.316 | 0.21843 | 0.38 | -5.71 | 1.38436 | none |
| MARS | 0.108 | 0.0832 | 1.3164 | 0.21843 | 0.38 | -5.71 | 0.40403 | none |
| RNF186 | 0.106 | 0.1063 | 1.3164 | 0.21844 | 0.38 | -5.71 | -0.143 | none |
| CD14 | 0.163 | 0.1128 | 1.3164 | 0.21844 | 0.38 | -5.71 | 1.36652 | none |
| GJB5 | 0.093 | 0.0763 | 1.3162 | 0.21851 | 0.38 | -5.71 | -1.0813 | none |

| | | | | | | | | |
|---|---|---|---|---|---|---|---|---|
| GZMA | -0.261 | -0.2816 | -1.316 | 0.21855 | 0.38 | -5.71 | 1.8467 | none |
| ZNF699 | 0.157 | 0.1473 | 1.3158 | 0.21864 | 0.38 | -5.71 | -0.547 | none |
| ELOVL1 | -0.149 | -0.1774 | -1.316 | 0.21868 | 0.38 | -5.71 | 1.09265 | none |
| ADCK5 | 0.185 | 0.1262 | 1.3156 | 0.2187 | 0.38 | -5.71 | -0.9075 | none |
| ATP6V1F | -0.179 | -0.2263 | -1.315 | 0.2188 | 0.38 | -5.71 | 1.68228 | none |
| PTPMT1 | -0.153 | -0.1303 | -1.315 | 0.21883 | 0.38 | -5.71 | 2.25217 | AA-related |
| EVPL | 0.31 | 0.3266 | 1.315 | 0.21887 | 0.38 | -5.71 | 0.90527 | none |
| CUL5 | 0.235 | 0.2462 | 1.3149 | 0.21891 | 0.38 | -5.71 | 1.01116 | none |
| RIOK2 | 0.141 | 0.0719 | 1.3148 | 0.21893 | 0.38 | -5.71 | -0.2548 | none |
| MS4A4A | 0.203 | 0.1398 | 1.3148 | 0.21895 | 0.38 | -5.71 | -1.0358 | none |
| SLC19A1 | 0.13 | 0.0963 | 1.3148 | 0.21895 | 0.38 | -5.71 | 1.38314 | none |
| PLXNB3 | -0.095 | -0.0929 | -1.314 | 0.21922 | 0.381 | -5.71 | 0.08636 | none |
| MAT2B | -0.141 | -0.1906 | -1.314 | 0.21937 | 0.381 | -5.72 | 0.82015 | none |
| ZNF585A | 0.201 | 0.0743 | 1.3134 | 0.2194 | 0.381 | -5.72 | -0.5857 | none |
| KCTD8 | 0.129 | 0.0774 | 1.3134 | 0.21941 | 0.381 | -5.72 | -1.1473 | none |
| ANKRD55 | 0.229 | 0.1201 | 1.3131 | 0.21949 | 0.381 | -5.72 | -0.5999 | none |
| CCDC93 | 0.298 | 0.3392 | 1.3129 | 0.21955 | 0.381 | -5.72 | -0.422 | none |
| TMEM106 | -0.144 | -0.1556 | -1.313 | 0.21963 | 0.381 | -5.72 | 1.31645 | none |
| ACR | 0.121 | 0.121 | 1.3125 | 0.21971 | 0.381 | -5.72 | 1.2947 | none |
| ZNF17 | 0.253 | 0.1814 | 1.3122 | 0.21981 | 0.381 | -5.72 | -0.7042 | none |
| DHX29 | 0.345 | 0.3839 | 1.312 | 0.21987 | 0.381 | -5.72 | -0.1883 | none |
| C15ORF61 | 0.25 | 0.1765 | 1.3119 | 0.21988 | 0.381 | -5.72 | -0.2919 | none |
| RPN1 | -0.129 | -0.134 | -1.312 | 0.21992 | 0.381 | -5.72 | 1.54426 | none |
| CHMP4B | 0.214 | 0.2886 | 1.3117 | 0.21994 | 0.381 | -5.72 | -0.7384 | none |
| WWC3 | 0.239 | 0.2598 | 1.3117 | 0.21996 | 0.381 | -5.72 | -0.0763 | none |
| ZNF268 | 0.288 | 0.2413 | 1.311 | 0.22017 | 0.382 | -5.72 | -0.6697 | none |
| TUBB6 | 0.229 | 0.1755 | 1.3103 | 0.22039 | 0.382 | -5.72 | -0.4046 | none |
| ARL5B | 0.1 | 0.0216 | 1.3103 | 0.2204 | 0.382 | -5.72 | 0.76314 | none |
| PKP3 | -0.258 | -0.1927 | -1.31 | 0.22047 | 0.382 | -5.72 | -0.3148 | none |
| ACRBP | 0.428 | 0.2963 | 1.31 | 0.22051 | 0.382 | -5.72 | -0.2065 | none |
| CASP4 | -0.144 | -0.1372 | -1.31 | 0.22057 | 0.382 | -5.72 | 0.03796 | none |
| KDM5D | 0.262 | 0.2455 | 1.3096 | 0.22063 | 0.382 | -5.72 | 0.02957 | none |
| WRB | -0.194 | -0.2317 | -1.309 | 0.2207 | 0.382 | -5.72 | 0.75971 | none |
| EBF1 | 0.151 | 0.1147 | 1.3086 | 0.22096 | 0.383 | -5.72 | -0.8127 | none |
| ZBED6CL | 0.14 | 0.2376 | 1.3085 | 0.22099 | 0.383 | -5.72 | 0.41737 | none |
| MGAT5B | 0.102 | 0.0603 | 1.3083 | 0.22107 | 0.383 | -5.72 | -0.8939 | none |
| TRADD | 0.125 | 0.0912 | 1.3081 | 0.22114 | 0.383 | -5.72 | 0.81239 | none |
| MEOX1 | 0.133 | 0.0348 | 1.3074 | 0.22136 | 0.383 | -5.72 | 0.26653 | none |
| QPCT | -0.3 | -0.332 | -1.307 | 0.2216 | 0.384 | -5.72 | -0.1681 | none |
| MORF4L2 | 0.145 | 0.0547 | 1.3065 | 0.22165 | 0.384 | -5.72 | 0.10599 | none |

| | | | | | | | | |
|---|---|---|---|---|---|---|---|---|
| RAB11FIP2 | 0.278 | 0.2244 | 1.3063 | 0.22171 | 0.384 | -5.72 | 0.24801 | none |
| ZBBX | 0.122 | 0.0755 | 1.3062 | 0.22175 | 0.384 | -5.72 | 0.29973 | none |
| CD2 | -0.124 | -0.0661 | -1.306 | 0.22177 | 0.384 | -5.72 | 0.96041 | none |
| TTC12 | 0.289 | 0.2677 | 1.3061 | 0.22177 | 0.384 | -5.72 | -0.9349 | none |
| POTEJ | 0.107 | 0.071 | 1.3057 | 0.22189 | 0.384 | -5.72 | 0.07437 | none |
| ZXDA | -0.098 | -0.0678 | -1.306 | 0.2219 | 0.384 | -5.72 | 0.79104 | none |
| TSPYL4 | -0.16 | -0.1119 | -1.306 | 0.22194 | 0.384 | -5.73 | 1.80174 | none |
| COX11 | 0.203 | 0.0938 | 1.3052 | 0.22208 | 0.384 | -5.73 | -0.368 | none |
| GTF2F2 | -0.131 | -0.0715 | -1.305 | 0.22209 | 0.384 | -5.73 | 1.10561 | none |
| ILF3 | 0.387 | 0.4912 | 1.3048 | 0.22219 | 0.384 | -5.73 | 0.21972 | none |
| PPP1R12C | -0.291 | -0.1318 | -1.305 | 0.22221 | 0.384 | -5.73 | 0.5323 | none |
| LSM10 | -0.183 | -0.2009 | -1.305 | 0.22226 | 0.384 | -5.73 | 1.77018 | none |
| RLIM | 0.245 | 0.2312 | 1.3046 | 0.22227 | 0.384 | -5.73 | -0.3417 | none |
| PPM1G | 0.185 | 0.2023 | 1.304 | 0.22245 | 0.385 | -5.73 | 0.45209 | none |
| TMEM81 | -0.261 | -0.1965 | -1.304 | 0.2225 | 0.385 | -5.73 | 0.3529 | none |
| ITPKC | 0.204 | 0.2756 | 1.3038 | 0.22254 | 0.385 | -5.73 | 0.18588 | none |
| AKNA | 0.433 | 0.5603 | 1.3036 | 0.22258 | 0.385 | -5.73 | -0.1033 | none |
| METTL1 | 0.105 | 0.1078 | 1.3036 | 0.22261 | 0.385 | -5.73 | 0.22455 | none |
| PANK4 | 0.253 | 0.2668 | 1.3031 | 0.22277 | 0.385 | -5.73 | -0.4564 | none |
| CLDN19 | 0.123 | 0.0917 | 1.3027 | 0.22287 | 0.385 | -5.73 | -1.014 | none |
| IL15RA | 0.145 | 0.1407 | 1.3025 | 0.22297 | 0.385 | -5.73 | 0.51156 | none |
| PSMD2 | 0.155 | 0.2068 | 1.3023 | 0.22302 | 0.385 | -5.73 | 0.55475 | none |
| DNAJC15 | -0.254 | -0.2758 | -1.302 | 0.22316 | 0.386 | -5.73 | 1.95209 | none |
| FAM71E2 | -0.09 | -0.0703 | -1.301 | 0.22329 | 0.386 | -5.73 | 0 | none |
| FUT3 | 0.194 | 0.2698 | 1.3014 | 0.22333 | 0.386 | -5.73 | 0.56081 | none |
| CPXM2 | -0.175 | -0.1634 | -1.301 | 0.22334 | 0.386 | -5.73 | 0.60301 | none |
| NID2 | 0.38 | 0.1985 | 1.3013 | 0.22335 | 0.386 | -5.73 | -0.0039 | none |
| RBM47 | 0.247 | 0.1622 | 1.301 | 0.22345 | 0.386 | -5.73 | -0.0776 | none |
| MYO1B | 0.295 | 0.3372 | 1.3008 | 0.2235 | 0.386 | -5.73 | -0.6477 | none |
| JAM2 | 0.283 | 0.1901 | 1.3006 | 0.22358 | 0.386 | -5.73 | -0.8861 | none |
| ACTRT2 | 0.092 | 0.0758 | 1.3006 | 0.22359 | 0.386 | -5.73 | -0.5539 | none |
| ATP6AP2 | 0.133 | 0.029 | 1.3003 | 0.22367 | 0.386 | -5.73 | -0.4269 | none |
| CPB2 | 0.143 | 0.1038 | 1.3002 | 0.22372 | 0.386 | -5.73 | -0.4564 | none |
| DNAJB7 | 0.118 | 0.0961 | 1.2999 | 0.22382 | 0.386 | -5.73 | -0.4001 | none |
| OR10S1 | -0.105 | -0.0878 | -1.3 | 0.22388 | 0.386 | -5.73 | 1.8745 | none |
| PRR14 | 0.217 | 0.2408 | 1.2991 | 0.22406 | 0.387 | -5.73 | 0.85663 | none |
| CNTD2 | -0.109 | -0.1037 | -1.299 | 0.22413 | 0.387 | -5.73 | 1.24883 | none |
| PHOX2A | 0.113 | 0.1127 | 1.2984 | 0.22431 | 0.387 | -5.73 | -0.7616 | none |
| RTN3 | -0.132 | -0.173 | -1.298 | 0.22437 | 0.387 | -5.73 | 1.76699 | none |
| LRRC20 | 0.145 | 0.1872 | 1.2979 | 0.22446 | 0.387 | -5.73 | 0.72578 | none |

| | | | | | | | | |
|---|---|---|---|---|---|---|---|---|
| PHF2 | 0.296 | 0.3895 | 1.2978 | 0.2245 | 0.387 | -5.73 | -0.3375 | none |
| OR2A12 | 0.156 | 0.0959 | 1.2975 | 0.2246 | 0.387 | -5.73 | -0.6044 | none |
| MRPL32 | -0.155 | -0.1678 | -1.297 | 0.22478 | 0.388 | -5.74 | 0.40101 | none |
| SRSF5 | 0.112 | 0.0997 | 1.2967 | 0.22488 | 0.388 | -5.74 | 0.61478 | none |
| DLX2 | 0.1 | 0.0754 | 1.2966 | 0.22489 | 0.388 | -5.74 | -0.1272 | none |
| PDE1A | 0.106 | 0.1044 | 1.2964 | 0.22495 | 0.388 | -5.74 | -0.5637 | none |
| HUS1 | 0.16 | 0.0718 | 1.2964 | 0.22497 | 0.388 | -5.74 | 0.54409 | none |
| LLGL2 | 0.227 | 0.2007 | 1.2962 | 0.22502 | 0.388 | -5.74 | -0.3666 | none |
| NIPSNAP3 | -0.137 | -0.1249 | -1.296 | 0.22504 | 0.388 | -5.74 | 1.1013 | none |
| TARBP1 | 0.232 | 0.2302 | 1.296 | 0.22509 | 0.388 | -5.74 | -1.1503 | none |
| IL16 | -0.166 | -0.0801 | -1.296 | 0.22509 | 0.388 | -5.74 | 0.69524 | none |
| ZRANB1 | 0.164 | 0.1481 | 1.2957 | 0.2252 | 0.388 | -5.74 | -0.4098 | none |
| CCDC74B | 0.297 | 0.2378 | 1.2956 | 0.22523 | 0.388 | -5.74 | -0.517 | none |
| UNG | -0.132 | -0.141 | -1.295 | 0.22538 | 0.388 | -5.74 | 2.18956 | AA-related |
| H2AFB2 | -0.188 | -0.1769 | -1.295 | 0.22539 | 0.388 | -5.74 | 0.03723 | none |
| PABPC3 | 0.094 | 0.0546 | 1.2948 | 0.22548 | 0.388 | -5.74 | -1.009 | none |
| INTS10 | 0.11 | 0.1116 | 1.2941 | 0.22572 | 0.389 | -5.74 | -0.0779 | none |
| DNAJC19 | -0.216 | -0.2757 | -1.294 | 0.2258 | 0.389 | -5.74 | 1.31783 | none |
| SOX5 | -0.129 | -0.1245 | -1.294 | 0.22582 | 0.389 | -5.74 | -0.6509 | none |
| WNK3 | 0.108 | 0.0505 | 1.2935 | 0.22592 | 0.389 | -5.74 | -1.0884 | none |
| CLEC2L | 0.122 | 0.0921 | 1.2932 | 0.22602 | 0.389 | -5.74 | -0.2994 | none |
| TLE4 | 0.261 | 0.2919 | 1.2923 | 0.22632 | 0.389 | -5.74 | -0.2966 | none |
| PSMD6 | -0.096 | -0.0969 | -1.292 | 0.22633 | 0.389 | -5.74 | 0.71507 | none |
| BOP1 | 0.129 | 0.1534 | 1.2923 | 0.22634 | 0.389 | -5.74 | ######## | none |
| POLD3 | 0.311 | 0.4445 | 1.2915 | 0.2266 | 0.39 | -5.74 | 0.05243 | none |
| SIVA1 | -0.271 | -0.2909 | -1.291 | 0.22672 | 0.39 | -5.74 | 1.55354 | none |
| MYBPC3 | 0.088 | 0.0826 | 1.2911 | 0.22673 | 0.39 | -5.74 | -0.9267 | none |
| ATP11C | 0.222 | 0.2465 | 1.2907 | 0.22685 | 0.39 | -5.74 | 0.17106 | none |
| HBG2 | 0.193 | 0.2398 | 1.2906 | 0.22688 | 0.39 | -5.74 | 0.79074 | none |
| ADPRHL1 | 0.188 | 0.1403 | 1.2906 | 0.2269 | 0.39 | -5.74 | -0.1499 | none |
| MOGS | 0.216 | 0.1496 | 1.2903 | 0.227 | 0.39 | -5.74 | -0.7848 | none |
| COL13A1 | -0.088 | -0.0641 | -1.29 | 0.22703 | 0.39 | -5.74 | -0.6211 | none |
| ABCB8 | 0.148 | 0.1522 | 1.2898 | 0.22714 | 0.39 | -5.74 | 1.06189 | none |
| MDC1 | 0.369 | 0.5214 | 1.2897 | 0.22718 | 0.39 | -5.74 | 0.04603 | none |
| NDUFS6 | -0.197 | -0.1695 | -1.289 | 0.22726 | 0.391 | -5.74 | 0.16516 | none |
| UBTF | 0.394 | 0.5281 | 1.2894 | 0.22729 | 0.391 | -5.74 | 0.11595 | none |
| OCLM | 0.251 | 0.2298 | 1.2893 | 0.22733 | 0.391 | -5.74 | 0.16612 | none |
| H3F3C | 0.211 | 0.165 | 1.2893 | 0.22733 | 0.391 | -5.74 | -0.5702 | none |
| TRPS1 | 0.264 | 0.3806 | 1.2892 | 0.22735 | 0.391 | -5.74 | 0.02625 | none |
| FBP1 | -0.291 | -0.2451 | -1.289 | 0.22738 | 0.391 | -5.74 | 1.68197 | none |

| | | | | | | | | |
|---|---|---|---|---|---|---|---|---|
| DPH3 | -0.228 | -0.2563 | -1.289 | 0.22738 | 0.391 | -5.74 | 1.29421 | none |
| WDR86 | 0.102 | 0.1277 | 1.2889 | 0.22745 | 0.391 | -5.74 | 0.27224 | none |
| ITIH6 | -0.102 | -0.0683 | -1.289 | 0.2275 | 0.391 | -5.75 | 1.63349 | none |
| PDK2 | 0.155 | 0.156 | 1.2888 | 0.2275 | 0.391 | -5.75 | 0.3974 | none |
| LY6G5B | 0.66 | 0.6103 | 1.2886 | 0.22756 | 0.391 | -5.75 | 0.08317 | none |
| CCNO | -0.102 | -0.0994 | -1.288 | 0.2277 | 0.391 | -5.75 | 0.99885 | none |
| FBXL17 | 0.17 | 0.1021 | 1.2879 | 0.2278 | 0.391 | -5.75 | 0.96938 | none |
| PCDHGB5 | -0.087 | -0.0682 | -1.288 | 0.22784 | 0.391 | -5.75 | 1.11272 | none |
| LRRC28 | 0.142 | -0.0147 | 1.2876 | 0.22788 | 0.391 | -5.75 | -0.1352 | none |
| CMIP | 0.255 | 0.3624 | 1.2869 | 0.22813 | 0.391 | -5.75 | 0.02418 | none |
| CASKIN1 | 0.133 | 0.1476 | 1.2864 | 0.22829 | 0.392 | -5.75 | 1.56865 | none |
| DOK1 | -0.128 | -0.0406 | -1.286 | 0.22836 | 0.392 | -5.75 | 1.57079 | none |
| FAM45A | 0.175 | 0.0587 | 1.2862 | 0.22836 | 0.392 | -5.75 | -0.4824 | none |
| CD99L2 | 0.112 | 0.0834 | 1.2861 | 0.2284 | 0.392 | -5.75 | 0.56987 | none |
| AKR1C4 | 0.105 | 0.0733 | 1.2857 | 0.22853 | 0.392 | -5.75 | 0.74411 | none |
| PPFIA3 | -0.101 | -0.0541 | -1.286 | 0.22855 | 0.392 | -5.75 | 1.53961 | none |
| C16ORF91 | -0.19 | -0.1251 | -1.286 | 0.22857 | 0.392 | -5.75 | 0.97524 | none |
| TMEM37 | -0.152 | -0.1175 | -1.285 | 0.2286 | 0.392 | -5.75 | 0.92103 | none |
| ELP5 | -0.21 | -0.266 | -1.285 | 0.22867 | 0.392 | -5.75 | 1.01938 | none |
| ADRM1 | -0.129 | -0.0653 | -1.285 | 0.2287 | 0.392 | -5.75 | 0.66817 | none |
| DPYSL3 | -0.184 | -0.1428 | -1.285 | 0.22875 | 0.392 | -5.75 | -0.2685 | none |
| PRELID3B | -0.183 | -0.2058 | -1.285 | 0.22878 | 0.392 | -5.75 | 1.50013 | none |
| EMC8 | 0.185 | 0.0889 | 1.2846 | 0.2289 | 0.392 | -5.75 | 0.78858 | none |
| MED12L | 0.244 | 0.2584 | 1.2845 | 0.22894 | 0.392 | -5.75 | -0.0094 | none |
| RNASEH2E | -0.117 | -0.1571 | -1.284 | 0.22902 | 0.392 | -5.75 | 1.07883 | none |
| S1PR1 | -0.132 | -0.1579 | -1.284 | 0.22908 | 0.392 | -5.75 | 0.03711 | none |
| IARS2 | 0.141 | 0.1801 | 1.284 | 0.2291 | 0.392 | -5.75 | 0.9071 | none |
| TFE3 | 0.219 | 0.2595 | 1.2836 | 0.22922 | 0.393 | -5.75 | 0.12411 | none |
| KIAA0753 | 0.339 | 0.3662 | 1.2836 | 0.22922 | 0.393 | -5.75 | 0.40733 | none |
| SH3BGRL | -0.144 | -0.1724 | -1.284 | 0.22926 | 0.393 | -5.75 | 1.54205 | none |
| CDKN2D | -0.159 | -0.0935 | -1.284 | 0.22926 | 0.393 | -5.75 | 1.32276 | none |
| BNIP3 | -0.139 | -0.1147 | -1.283 | 0.22946 | 0.393 | -5.75 | 1.31986 | none |
| SMAP2 | -0.09 | -0.0507 | -1.283 | 0.22956 | 0.393 | -5.75 | 0.74369 | none |
| SLC25A25 | 0.13 | 0.1556 | 1.2826 | 0.22958 | 0.393 | -5.75 | 1.30882 | none |
| RPF1 | 0.105 | 0.0502 | 1.2824 | 0.22965 | 0.393 | -5.75 | -0.2036 | none |
| DQX1 | -0.226 | -0.2093 | -1.282 | 0.22966 | 0.393 | -5.75 | 1.64358 | none |
| COQ8B | 0.147 | 0.1239 | 1.2823 | 0.22966 | 0.393 | -5.75 | 1.0327 | none |
| USF1 | 0.398 | 0.4582 | 1.2818 | 0.22984 | 0.393 | -5.75 | -0.4607 | none |
| SOX30 | -0.104 | -0.0638 | -1.282 | 0.22991 | 0.393 | -5.75 | -0.1653 | none |
| GCNT1 | 0.2 | 0.1734 | 1.2815 | 0.22993 | 0.393 | -5.75 | 0.11264 | none |

| | | | | | | | | |
|---|---|---|---|---|---|---|---|---|
| OXTR | -0.147 | -0.1437 | -1.281 | 0.22998 | 0.393 | -5.75 | 0.74759 | none |
| ZDHHC23 | 0.12 | 0.1349 | 1.2813 | 0.23001 | 0.393 | -5.75 | -1.0003 | none |
| PEX2 | -0.23 | -0.2509 | -1.281 | 0.23011 | 0.394 | -5.75 | 1.4776 | none |
| SRSF11 | 0.22 | 0.2457 | 1.2809 | 0.23015 | 0.394 | -5.75 | 0.15121 | none |
| CD93 | 0.106 | 0.116 | 1.2808 | 0.23017 | 0.394 | -5.75 | -0.6613 | none |
| RPS9 | -0.281 | -0.2337 | -1.28 | 0.23058 | 0.394 | -5.76 | 0.26967 | none |
| MTHFS | -0.144 | -0.1642 | -1.279 | 0.23061 | 0.394 | -5.76 | 1.22717 | none |
| POMGNT2 | -0.111 | -0.0416 | -1.279 | 0.23075 | 0.394 | -5.76 | 1.16635 | none |
| KANK1 | 0.258 | 0.1971 | 1.2789 | 0.23081 | 0.394 | -5.76 | -0.1526 | none |
| NGFR | 0.109 | 0.0912 | 1.2786 | 0.23091 | 0.395 | -5.76 | 0.67844 | none |
| NCR3LG1 | 0.099 | 0.0961 | 1.2783 | 0.23101 | 0.395 | -5.76 | -0.0438 | none |
| TBC1D3D | -0.167 | -0.0537 | -1.278 | 0.23109 | 0.395 | -5.76 | 0.85261 | none |
| HEY1 | -0.086 | -0.0471 | -1.278 | 0.23113 | 0.395 | -5.76 | 1.42796 | none |
| HSPD1 | -0.132 | -0.0528 | -1.278 | 0.23122 | 0.395 | -5.76 | 1.75892 | none |
| UBAC1 | 0.122 | 0.1296 | 1.2776 | 0.23125 | 0.395 | -5.76 | 0.45314 | none |
| APLF | -0.113 | 0.0067 | -1.277 | 0.23132 | 0.395 | -5.76 | 1.56616 | none |
| DNM3 | -0.177 | -0.1298 | -1.277 | 0.23138 | 0.395 | -5.76 | 1.65123 | none |
| ZFP69 | 0.16 | 0.116 | 1.2772 | 0.23139 | 0.395 | -5.76 | -0.8495 | none |
| METTL26 | -0.166 | -0.1537 | -1.277 | 0.23162 | 0.395 | -5.76 | 1.43277 | none |
| ALKBH8 | 0.171 | 0.1554 | 1.2764 | 0.23166 | 0.395 | -5.76 | 1.02612 | none |
| TSPAN13 | -0.173 | -0.1668 | -1.276 | 0.23186 | 0.396 | -5.76 | 2.02059 | none |
| IFT27 | -0.196 | -0.2284 | -1.276 | 0.23196 | 0.396 | -5.76 | 0.63974 | none |
| MAD2L1 | -0.202 | -0.2304 | -1.275 | 0.23202 | 0.396 | -5.76 | 0.82457 | none |
| SQSTM1 | -0.125 | -0.1291 | -1.275 | 0.23218 | 0.396 | -5.76 | 1.5937 | none |
| TOM1L2 | 0.17 | 0.2328 | 1.2749 | 0.23218 | 0.396 | -5.76 | -0.045 | none |
| YME1L1 | -0.207 | -0.1388 | -1.275 | 0.23228 | 0.396 | -5.76 | 0.55919 | none |
| KRTAP3-1 | -0.134 | -0.1279 | -1.274 | 0.23232 | 0.396 | -5.76 | 1.11089 | none |
| NCKAP1 | 0.225 | 0.1595 | 1.2744 | 0.23233 | 0.396 | -5.76 | -0.5519 | none |
| PRRC2C | 0.428 | 0.591 | 1.2743 | 0.23236 | 0.396 | -5.76 | 0.03982 | none |
| H2AFB3 | -0.1 | -0.0666 | -1.274 | 0.2325 | 0.397 | -5.76 | 1.24898 | none |
| MAF1 | 0.102 | 0.0952 | 1.2736 | 0.23263 | 0.397 | -5.76 | 0.97225 | none |
| C9ORF24 | 0.18 | 0.162 | 1.2733 | 0.23271 | 0.397 | -5.76 | 0.7926 | none |
| TC2N | 0.169 | 0.0527 | 1.2733 | 0.23273 | 0.397 | -5.76 | 1.22308 | none |
| RPL21 | -0.148 | -0.1595 | -1.273 | 0.23277 | 0.397 | -5.76 | 0.83259 | none |
| BAZ1B | 0.474 | 0.6436 | 1.2728 | 0.23287 | 0.397 | -5.76 | -0.0797 | none |
| MRAP | -0.175 | -0.1048 | -1.273 | 0.23289 | 0.397 | -5.76 | 0.91002 | none |
| ZNF277 | 0.167 | 0.0587 | 1.2728 | 0.23289 | 0.397 | -5.76 | 0.39506 | none |
| TRIM25 | 0.186 | 0.2479 | 1.2725 | 0.23297 | 0.397 | -5.76 | -0.2725 | none |
| MTHFSD | 0.191 | 0.166 | 1.2723 | 0.23307 | 0.397 | -5.76 | -0.1637 | none |
| VSIG2 | 0.148 | 0.0659 | 1.2722 | 0.23308 | 0.397 | -5.76 | -0.4505 | none |

| | | | | | | | | |
|---|---|---|---|---|---|---|---|---|
| KRT74 | 0.111 | 0.1148 | 1.2718 | 0.23322 | 0.397 | -5.77 | -0.3854 | none |
| LRRC41 | 0.158 | 0.12 | 1.2712 | 0.23342 | 0.398 | -5.77 | -0.6777 | none |
| TP53BP1 | 0.574 | 0.6738 | 1.2707 | 0.23361 | 0.398 | -5.77 | -0.1815 | none |
| MTCH1 | -0.17 | -0.1221 | -1.271 | 0.23366 | 0.398 | -5.77 | 1.18887 | none |
| SIGLEC12 | 0.161 | 0.2108 | 1.2704 | 0.23368 | 0.398 | -5.77 | -0.4921 | none |
| RAB3A | 0.324 | 0.1838 | 1.27 | 0.23385 | 0.398 | -5.77 | -0.28 | none |
| SEC23IP | 0.236 | 0.2113 | 1.2698 | 0.2339 | 0.398 | -5.77 | 0.65899 | none |
| MRTO4 | 0.198 | 0.2146 | 1.2698 | 0.2339 | 0.398 | -5.77 | -0.0492 | none |
| GGT7 | -0.231 | -0.173 | -1.27 | 0.23393 | 0.398 | -5.77 | 1.38746 | none |
| C19ORF81 | 0.185 | 0.0709 | 1.2693 | 0.23409 | 0.398 | -5.77 | -0.5077 | none |
| INO80E | -0.19 | -0.1195 | -1.269 | 0.23411 | 0.398 | -5.77 | 1.86998 | none |
| GGT2 | -0.102 | -0.0653 | -1.269 | 0.2342 | 0.399 | -5.77 | 1.57462 | none |
| CCT8L2 | -0.082 | -0.0533 | -1.269 | 0.23428 | 0.399 | -5.77 | 1.01512 | none |
| FCN3 | 0.137 | 0.1372 | 1.2684 | 0.23439 | 0.399 | -5.77 | 0.8562 | none |
| ACHE | -0.144 | -0.0819 | -1.268 | 0.23456 | 0.399 | -5.77 | 1.29094 | none |
| VAX1 | 0.083 | 0.0868 | 1.2678 | 0.2346 | 0.399 | -5.77 | -0.5032 | none |
| CD96 | 0.186 | 0.026 | 1.2676 | 0.23467 | 0.399 | -5.77 | -0.6883 | none |
| LRP5 | -0.395 | -0.1284 | -1.267 | 0.23481 | 0.399 | -5.77 | 0.62209 | none |
| PIK3AP1 | -0.212 | -0.1255 | -1.267 | 0.23481 | 0.399 | -5.77 | 1.12504 | none |
| OSGEP | 0.147 | 0.0287 | 1.2668 | 0.23493 | 0.399 | -5.77 | -0.6634 | none |
| HAPLN2 | 0.1 | 0.1005 | 1.2667 | 0.23497 | 0.399 | -5.77 | 1.23506 | none |
| RGS13 | 0.087 | 0.0692 | 1.2666 | 0.23498 | 0.399 | -5.77 | -0.2094 | none |
| NCBP1 | 0.179 | 0.1236 | 1.2666 | 0.235 | 0.399 | -5.77 | -0.436 | none |
| SCAMP1 | 0.159 | 0.0333 | 1.2665 | 0.23502 | 0.399 | -5.77 | -0.232 | none |
| ZNF302 | 0.134 | 0.0286 | 1.2661 | 0.23516 | 0.4 | -5.77 | 0.95662 | none |
| ANO6 | 0.242 | 0.2136 | 1.2661 | 0.23516 | 0.4 | -5.77 | -0.9002 | none |
| SNX25 | 0.235 | 0.2744 | 1.2658 | 0.23527 | 0.4 | -5.77 | 0.02697 | none |
| IRF7 | -0.537 | -0.3862 | -1.266 | 0.23533 | 0.4 | -5.77 | 2.01348 | none |
| MCHR2 | -0.091 | -0.069 | -1.266 | 0.23535 | 0.4 | -5.77 | 1.31599 | none |
| SMOC2 | 0.134 | 0.1225 | 1.2654 | 0.23542 | 0.4 | -5.77 | 0.47927 | none |
| KLHL42 | 0.17 | 0.1998 | 1.2652 | 0.23548 | 0.4 | -5.77 | 0.57049 | none |
| DNAJB12 | 0.201 | 0.2577 | 1.265 | 0.23554 | 0.4 | -5.77 | -0.2872 | none |
| TMEM214 | -0.169 | 0.021 | -1.265 | 0.23554 | 0.4 | -5.77 | 0.43598 | none |
| PTPRM | 0.251 | 0.2425 | 1.265 | 0.23556 | 0.4 | -5.77 | 0.48634 | none |
| TMEM115 | 0.131 | 0.1016 | 1.2649 | 0.23558 | 0.4 | -5.77 | 1.09847 | none |
| WBP11 | 0.237 | 0.2546 | 1.2647 | 0.23566 | 0.4 | -5.77 | -0.3907 | none |
| LTA | -0.236 | -0.2283 | -1.265 | 0.2357 | 0.4 | -5.77 | 1.46341 | none |
| SYN2 | -0.137 | -0.1102 | -1.264 | 0.2358 | 0.4 | -5.77 | 1.61144 | none |
| SCO2 | -0.245 | -0.2284 | -1.264 | 0.23585 | 0.4 | -5.77 | 1.72266 | none |
| TRIB1 | 0.133 | 0.1346 | 1.2641 | 0.23586 | 0.4 | -5.77 | 0.43914 | none |

| | | | | | | | | |
|---|---|---|---|---|---|---|---|---|
| CLDN25 | 0.097 | 0.0966 | 1.264 | 0.23589 | 0.4 | -5.77 | 1.06121 | none |
| INAFM1 | 0.166 | 0.1649 | 1.2639 | 0.23593 | 0.4 | -5.77 | 1.34849 | none |
| CCDC12 | 0.124 | 0.1096 | 1.2633 | 0.23614 | 0.401 | -5.78 | -0.51 | none |
| MRPL39 | -0.142 | -0.2038 | -1.263 | 0.23625 | 0.401 | -5.78 | 1.79476 | none |
| CRAT | 0.333 | 0.277 | 1.2627 | 0.23633 | 0.401 | -5.78 | 0.09166 | none |
| DESI2 | -0.239 | -0.2345 | -1.262 | 0.23643 | 0.401 | -5.78 | 1.21677 | none |
| GABARAPI | -0.206 | -0.2818 | -1.262 | 0.23655 | 0.401 | -5.78 | 1.0019 | none |
| COPG1 | 0.274 | 0.3513 | 1.2619 | 0.23662 | 0.401 | -5.78 | -0.0195 | none |
| PDIA2 | 0.098 | 0.0982 | 1.2613 | 0.23682 | 0.402 | -5.78 | 0.6961 | none |
| RIT1 | -0.161 | -0.2015 | -1.261 | 0.23687 | 0.402 | -5.78 | 1.84404 | none |
| IFNK | 0.105 | 0.1055 | 1.2611 | 0.23688 | 0.402 | -5.78 | 0.5892 | none |
| ANO2 | -0.159 | -0.1585 | -1.261 | 0.2369 | 0.402 | -5.78 | 1.78783 | none |
| MARK2 | 0.514 | 0.5903 | 1.2608 | 0.23699 | 0.402 | -5.78 | 0.49004 | none |
| CMA1 | 0.141 | 0.1609 | 1.2608 | 0.237 | 0.402 | -5.78 | 1.86153 | none |
| ZDHHC8 | 0.173 | 0.2289 | 1.2607 | 0.23703 | 0.402 | -5.78 | -0.4859 | none |
| PITPNM3 | -0.092 | -0.0581 | -1.261 | 0.23709 | 0.402 | -5.78 | 1.65401 | none |
| DDC | -0.153 | -0.1551 | -1.26 | 0.23712 | 0.402 | -5.78 | 0.81648 | none |
| ERICH1 | 0.21 | 0.2709 | 1.2603 | 0.23715 | 0.402 | -5.78 | -0.5339 | none |
| CORO1B | -0.183 | -0.1598 | -1.259 | 0.23768 | 0.403 | -5.78 | 1.08185 | none |
| EID2 | 0.133 | 0.0695 | 1.2587 | 0.23772 | 0.403 | -5.78 | -0.7414 | none |
| NFIA | -0.183 | -0.1679 | -1.259 | 0.23773 | 0.403 | -5.78 | 1.16719 | none |
| DENND5A | 0.171 | 0.1846 | 1.2584 | 0.23782 | 0.403 | -5.78 | 0.58197 | none |
| RRAGB | 0.284 | 0.1979 | 1.2582 | 0.23789 | 0.403 | -5.78 | -0.1557 | none |
| HES3 | -0.095 | -0.0653 | -1.258 | 0.23803 | 0.403 | -5.78 | 2.16608 | AA-related |
| GLI1 | -0.208 | -0.1715 | -1.258 | 0.23804 | 0.403 | -5.78 | 1.65401 | none |
| ACLY | 0.245 | 0.3142 | 1.2577 | 0.23806 | 0.403 | -5.78 | 0.17235 | none |
| GNG10 | -0.219 | -0.2757 | -1.258 | 0.23808 | 0.403 | -5.78 | 0.58754 | none |
| SSTR4 | -0.094 | -0.0939 | -1.258 | 0.23812 | 0.403 | -5.78 | 0.79024 | none |
| GPR15 | -0.19 | -0.2735 | -1.258 | 0.23813 | 0.403 | -5.78 | 0.25162 | none |
| PER3 | -0.189 | -0.1086 | -1.257 | 0.23814 | 0.403 | -5.78 | 0.71464 | none |
| IFNW1 | 0.161 | 0.1022 | 1.2574 | 0.23818 | 0.403 | -5.78 | -1.0198 | none |
| ACOT7 | -0.138 | -0.0818 | -1.257 | 0.23825 | 0.403 | -5.78 | 1.91109 | none |
| ZNF219 | 0.33 | 0.2742 | 1.2569 | 0.23833 | 0.403 | -5.78 | 0.63673 | none |
| TCTE1 | 0.109 | 0.09 | 1.2566 | 0.23843 | 0.403 | -5.78 | -0.2133 | none |
| DUSP26 | 0.122 | 0.1234 | 1.2564 | 0.23851 | 0.403 | -5.78 | -0.3012 | none |
| S100A3 | 0.137 | 0.1315 | 1.2563 | 0.23856 | 0.403 | -5.78 | 0.83308 | none |
| DPCD | 0.193 | 0.157 | 1.2561 | 0.23863 | 0.403 | -5.78 | -0.8262 | none |
| PRDX5 | -0.178 | -0.1914 | -1.256 | 0.23864 | 0.403 | -5.78 | -0.1253 | none |
| CHMP1A | -0.14 | -0.1201 | -1.256 | 0.23872 | 0.404 | -5.78 | 1.59614 | none |
| METAP1 | -0.207 | -0.1999 | -1.255 | 0.23883 | 0.404 | -5.78 | 1.87847 | none |

| | | | | | | | | |
|---|---|---|---|---|---|---|---|---|
| C11ORF91 | -0.205 | -0.2155 | -1.255 | 0.23885 | 0.404 | -5.78 | 0.70019 | none |
| ANGPTL8 | -0.108 | -0.1186 | -1.255 | 0.23886 | 0.404 | -5.78 | 1.72802 | none |
| SLC1A7 | 0.091 | 0.0927 | 1.2554 | 0.23887 | 0.404 | -5.78 | 0.40323 | none |
| ZBTB1 | 0.169 | 0.162 | 1.2554 | 0.23888 | 0.404 | -5.78 | 0.65562 | none |
| SLC25A6 | -0.155 | -0.1652 | -1.255 | 0.23889 | 0.404 | -5.78 | 1.58917 | none |
| TRAPPC2L | -0.18 | -0.2307 | -1.255 | 0.23893 | 0.404 | -5.78 | 0.54233 | none |
| UNC5A | -0.086 | -0.0572 | -1.255 | 0.23895 | 0.404 | -5.78 | 0.5047 | none |
| PROM1 | -0.118 | -0.0935 | -1.255 | 0.23902 | 0.404 | -5.78 | 1.31906 | none |
| TRIM29 | 0.168 | 0.1495 | 1.2548 | 0.23908 | 0.404 | -5.79 | -0.8094 | none |
| COX7A1 | -0.184 | -0.2019 | -1.254 | 0.23929 | 0.404 | -5.79 | 0.81395 | none |
| GIMAP1 | 0.21 | 0.1412 | 1.2539 | 0.23937 | 0.404 | -5.79 | -0.1481 | none |
| HIBADH | -0.146 | -0.1104 | -1.254 | 0.23946 | 0.404 | -5.79 | 1.66934 | none |
| PNMT | 0.098 | 0.106 | 1.2536 | 0.23948 | 0.404 | -5.79 | -0.0135 | none |
| TMEM205 | -0.229 | -0.2688 | -1.253 | 0.23956 | 0.404 | -5.79 | 0.74208 | none |
| RHBDD2 | -0.152 | -0.154 | -1.253 | 0.23962 | 0.404 | -5.79 | 1.352 | none |
| GFRA2 | 0.114 | 0.1233 | 1.2526 | 0.23983 | 0.405 | -5.79 | -0.8507 | none |
| CHST3 | 0.172 | 0.2692 | 1.2526 | 0.23984 | 0.405 | -5.79 | 0.25872 | none |
| PRSS22 | 0.142 | 0.1557 | 1.2525 | 0.23987 | 0.405 | -5.79 | 1.33988 | none |
| UMPS | -0.126 | -0.1193 | -1.252 | 0.23993 | 0.405 | -5.79 | 0.40411 | none |
| UBE2Q2L | -0.318 | -0.2774 | -1.252 | 0.23998 | 0.405 | -5.79 | 0.00709 | none |
| AFDN | 0.342 | 0.4371 | 1.2521 | 0.24001 | 0.405 | -5.79 | -0.1523 | none |
| GPR132 | 0.323 | 0.4249 | 1.2514 | 0.24025 | 0.405 | -5.79 | 0.23099 | none |
| TELO2 | 0.144 | 0.2074 | 1.2512 | 0.24033 | 0.405 | -5.79 | 0.60842 | none |
| FOXA3 | -0.138 | -0.1449 | -1.251 | 0.24039 | 0.405 | -5.79 | 0.07943 | none |
| ZNFX1 | 0.366 | 0.4376 | 1.2494 | 0.24097 | 0.406 | -5.79 | -0.0132 | none |
| SNCA | -0.181 | -0.1114 | -1.249 | 0.24097 | 0.406 | -5.79 | 0.53066 | none |
| PCDH7 | 0.16 | 0.1559 | 1.2492 | 0.24103 | 0.406 | -5.79 | -0.61 | none |
| AGAP1 | 0.205 | 0.2207 | 1.2491 | 0.24107 | 0.406 | -5.79 | 0.49139 | none |
| MRPL40 | 0.136 | 0.0524 | 1.2487 | 0.24122 | 0.407 | -5.79 | -0.7913 | none |
| WRNIP1 | -0.143 | -0.0589 | -1.249 | 0.24124 | 0.407 | -5.79 | 1.26394 | none |
| BEX5 | -1.1 | -1.0204 | -1.248 | 0.2413 | 0.407 | -5.79 | 2.15724 | AA-related |
| ZIC1 | 0.152 | 0.1694 | 1.2483 | 0.24135 | 0.407 | -5.79 | 0.33514 | none |
| JAM3 | -0.105 | -0.0182 | -1.248 | 0.24136 | 0.407 | -5.79 | 1.18421 | none |
| MLX | 0.147 | 0.0492 | 1.2483 | 0.24136 | 0.407 | -5.79 | -0.5419 | none |
| NDUFA6 | -0.262 | -0.254 | -1.248 | 0.24143 | 0.407 | -5.79 | 0.52389 | none |
| DSP | -0.094 | -0.0724 | -1.248 | 0.24146 | 0.407 | -5.79 | 0.04228 | none |
| POFUT1 | 0.108 | 0.0807 | 1.2479 | 0.24148 | 0.407 | -5.79 | -0.4893 | none |
| NLRP6 | 0.117 | 0.1128 | 1.2479 | 0.24148 | 0.407 | -5.79 | -0.66 | none |
| CYHR1 | 0.106 | 0.1361 | 1.2477 | 0.24154 | 0.407 | -5.79 | 0.39083 | none |
| ADH1A | 0.151 | 0.151 | 1.2474 | 0.24166 | 0.407 | -5.79 | 1.40909 | none |

| | | | | | | | | |
|---|---|---|---|---|---|---|---|---|
| TBP | 0.132 | 0.0755 | 1.2468 | 0.24187 | 0.407 | -5.79 | -0.4535 | none |
| AMMECR1 | 0.103 | 0.1173 | 1.2467 | 0.2419 | 0.407 | -5.79 | 0.36028 | none |
| EIF4H | -0.099 | -0.0709 | -1.247 | 0.24193 | 0.407 | -5.79 | 1.05207 | none |
| SLC6A16 | 0.525 | 0.4533 | 1.2463 | 0.24203 | 0.407 | -5.79 | 0.67315 | none |
| OGG1 | 0.126 | 0.0863 | 1.2463 | 0.24205 | 0.407 | -5.79 | 1.0092 | none |
| TTLL4 | 0.228 | 0.2299 | 1.2463 | 0.24205 | 0.407 | -5.79 | -0.7289 | none |
| RAB36 | 0.115 | 0.1201 | 1.2459 | 0.24219 | 0.407 | -5.8 | -0.1364 | none |
| COPS8 | -0.135 | -0.1569 | -1.246 | 0.24221 | 0.407 | -5.8 | 1.78698 | none |
| SLIRP | -0.212 | -0.2889 | -1.245 | 0.24243 | 0.408 | -5.8 | 0.82437 | none |
| ATMIN | 0.148 | 0.1966 | 1.2451 | 0.24248 | 0.408 | -5.8 | -0.3391 | none |
| ARMS2 | 0.111 | 0.1107 | 1.245 | 0.24251 | 0.408 | -5.8 | 1.82213 | none |
| STARD3NL | 0.185 | 0.0607 | 1.2447 | 0.24261 | 0.408 | -5.8 | 0.16512 | none |
| ATP6V1B1 | 0.198 | 0.1984 | 1.2446 | 0.24264 | 0.408 | -5.8 | 1.02907 | none |
| FOXD4L4 | -0.078 | -0.0508 | -1.244 | 0.24283 | 0.408 | -5.8 | -0.0151 | none |
| CARNMT1 | 0.215 | 0.1973 | 1.2439 | 0.24289 | 0.408 | -5.8 | 0.44356 | none |
| CCDC97 | 0.161 | 0.1485 | 1.2431 | 0.24316 | 0.409 | -5.8 | -0.9961 | none |
| MOGAT2 | 0.121 | 0.0812 | 1.2426 | 0.24335 | 0.409 | -5.8 | -1.3106 | none |
| LIG1 | 0.306 | 0.4272 | 1.2426 | 0.24336 | 0.409 | -5.8 | 0.32818 | none |
| PHLDB2 | 0.174 | 0.087 | 1.2426 | 0.24336 | 0.409 | -5.8 | 0.13694 | none |
| ZBTB26 | 0.253 | 0.1824 | 1.2425 | 0.24337 | 0.409 | -5.8 | 0.00596 | none |
| STRAP | -0.141 | -0.1785 | -1.242 | 0.24344 | 0.409 | -5.8 | 0.43689 | none |
| PPIF | -0.123 | -0.0815 | -1.242 | 0.24345 | 0.409 | -5.8 | 1.28995 | none |
| B3GNT3 | 0.108 | 0.097 | 1.2423 | 0.24347 | 0.409 | -5.8 | -0.1679 | none |
| CHRNA10 | 0.14 | 0.1291 | 1.2421 | 0.24351 | 0.409 | -5.8 | -0.1142 | none |
| FNDC9 | 0.23 | 0.2299 | 1.242 | 0.24356 | 0.409 | -5.8 | -0.8402 | none |
| RSRC2 | 0.401 | 0.4463 | 1.2418 | 0.24364 | 0.409 | -5.8 | -0.2015 | none |
| SLC39A1 | -0.129 | -0.0806 | -1.242 | 0.24369 | 0.409 | -5.8 | 1.52075 | none |
| ZP2 | 0.143 | 0.0839 | 1.2415 | 0.24373 | 0.409 | -5.8 | -0.6305 | none |
| HNRNPR | 0.153 | 0.1629 | 1.2412 | 0.24384 | 0.409 | -5.8 | 0.32767 | none |
| CCDC86 | 0.275 | 0.3538 | 1.241 | 0.24391 | 0.409 | -5.8 | 0.24979 | none |
| FGD6 | 0.281 | 0.1929 | 1.241 | 0.24392 | 0.409 | -5.8 | -0.0108 | none |
| TYSND1 | 0.116 | 0.1008 | 1.2408 | 0.24399 | 0.409 | -5.8 | 0.94138 | none |
| CEP170B | 0.225 | 0.2867 | 1.2407 | 0.24401 | 0.409 | -5.8 | -0.9598 | none |
| EVI2A | -0.156 | -0.2317 | -1.241 | 0.24405 | 0.409 | -5.8 | 0.95806 | none |
| CASP14 | 0.331 | 0.2342 | 1.2404 | 0.24411 | 0.41 | -5.8 | -0.8144 | none |
| DLGAP3 | -0.092 | -0.0431 | -1.24 | 0.2443 | 0.41 | -5.8 | -0.3886 | none |
| PSMC6 | 0.101 | 0.0021 | 1.2398 | 0.24434 | 0.41 | -5.8 | 0.70086 | none |
| ACRV1 | -0.193 | -0.1115 | -1.24 | 0.24439 | 0.41 | -5.8 | 0.48149 | none |
| CBR3 | -0.169 | -0.1868 | -1.239 | 0.24451 | 0.41 | -5.8 | 1.24585 | none |
| SNX13 | 0.226 | 0.2411 | 1.2392 | 0.24455 | 0.41 | -5.8 | -0.0921 | none |

| | | | | | | | | |
|---|---|---|---|---|---|---|---|---|
| RHOBTB2 | 0.168 | 0.2521 | 1.2391 | 0.24458 | 0.41 | -5.8 | 0.45464 | none |
| MRPL55 | 0.181 | 0.1353 | 1.2391 | 0.24459 | 0.41 | -5.8 | -0.0123 | none |
| ATR | 0.271 | 0.2787 | 1.2389 | 0.24466 | 0.41 | -5.8 | 0.61315 | none |
| CYP3A7-C\ | 0.098 | 0.0858 | 1.2379 | 0.245 | 0.411 | -5.8 | -0.6407 | none |
| ZNF775 | 0.132 | 0.1225 | 1.2377 | 0.24507 | 0.411 | -5.8 | -0.5181 | none |
| BAD | -0.227 | -0.2241 | -1.237 | 0.24529 | 0.411 | -5.81 | 1.64628 | none |
| EWSR1 | 0.226 | 0.3007 | 1.2369 | 0.24536 | 0.411 | -5.81 | -0.4506 | none |
| USP36 | 0.267 | 0.365 | 1.2362 | 0.2456 | 0.411 | -5.81 | -0.3877 | none |
| MAFG | 0.215 | 0.1792 | 1.2362 | 0.24562 | 0.411 | -5.81 | -0.548 | none |
| TBRG1 | 0.145 | 0.13 | 1.2359 | 0.24572 | 0.412 | -5.81 | 0.0318 | none |
| RPS27 | -0.26 | -0.3529 | -1.236 | 0.24574 | 0.412 | -5.81 | 1.2086 | none |
| UTP14A | 0.437 | 0.5372 | 1.2355 | 0.24586 | 0.412 | -5.81 | 0.18796 | none |
| SCAPER | 0.318 | 0.4063 | 1.2351 | 0.24599 | 0.412 | -5.81 | 0.12166 | none |
| TAC3 | 0.119 | 0.1368 | 1.2349 | 0.24606 | 0.412 | -5.81 | 0.91188 | none |
| ZBTB10 | 0.126 | 0.1208 | 1.2347 | 0.24616 | 0.412 | -5.81 | -0.9829 | none |
| OVCH1 | -0.132 | -0.1319 | -1.234 | 0.24631 | 0.412 | -5.81 | 1.37371 | none |
| CWC22 | 0.375 | 0.4414 | 1.234 | 0.24639 | 0.412 | -5.81 | 0.31714 | none |
| DMRT3 | 0.123 | 0.1296 | 1.2332 | 0.24668 | 0.413 | -5.81 | 1.42669 | none |
| ACAT1 | 0.141 | 0.1148 | 1.233 | 0.24675 | 0.413 | -5.81 | -0.1341 | none |
| TBC1D3 | 0.21 | 0.2226 | 1.2329 | 0.24677 | 0.413 | -5.81 | 0.09037 | none |
| ANKH | 0.152 | 0.1546 | 1.2329 | 0.24679 | 0.413 | -5.81 | 0.73768 | none |
| RBPMS | -0.133 | -0.0923 | -1.233 | 0.24689 | 0.413 | -5.81 | -0.6725 | none |
| TMEM132 | 0.089 | 0.0892 | 1.2325 | 0.24694 | 0.413 | -5.81 | -0.9319 | none |
| LGALS9C | 0.155 | 0.1653 | 1.232 | 0.24711 | 0.413 | -5.81 | 0.60229 | none |
| MLLT3 | 0.323 | 0.3117 | 1.2312 | 0.24738 | 0.414 | -5.81 | 0.21284 | none |
| SLAMF7 | 0.154 | 0.1422 | 1.231 | 0.24746 | 0.414 | -5.81 | 0.84628 | none |
| CD3E | -0.095 | -0.0689 | -1.23 | 0.24765 | 0.414 | -5.81 | 0.61668 | none |
| CCT3 | -0.111 | -0.0557 | -1.23 | 0.24767 | 0.414 | -5.81 | 1.47565 | none |
| HPCA | -0.238 | -0.1788 | -1.23 | 0.24773 | 0.414 | -5.81 | 0.75152 | none |
| EML4 | 0.287 | 0.3361 | 1.2298 | 0.2479 | 0.414 | -5.81 | 0.97623 | none |
| RILP | -0.219 | -0.1239 | -1.23 | 0.24794 | 0.414 | -5.81 | 1.76248 | none |
| GTF2B | -0.18 | -0.2354 | -1.229 | 0.24802 | 0.415 | -5.81 | 1.69086 | none |
| 7-Mar | 0.124 | 0.0527 | 1.229 | 0.24819 | 0.415 | -5.81 | -0.0067 | none |
| RSPO2 | -0.103 | -0.0644 | -1.229 | 0.24828 | 0.415 | -5.82 | 0.4238 | none |
| SERTAD3 | -0.106 | -0.1635 | -1.229 | 0.24828 | 0.415 | -5.82 | 0.53549 | none |
| KNCN | 0.112 | 0.1159 | 1.2287 | 0.2483 | 0.415 | -5.82 | 0.57257 | none |
| KRT4 | 0.193 | 0.1935 | 1.2285 | 0.24835 | 0.415 | -5.82 | 1.21354 | none |
| HENMT1 | 0.121 | 0.0135 | 1.2285 | 0.24837 | 0.415 | -5.82 | 0.60693 | none |
| POTEH | 0.189 | 0.1049 | 1.2284 | 0.2484 | 0.415 | -5.82 | -0.6518 | none |
| XRCC4 | -0.257 | -0.2467 | -1.228 | 0.24844 | 0.415 | -5.82 | 0.72226 | none |

| | | | | | | | | |
|---|---|---|---|---|---|---|---|---|
| ATG3 | -0.099 | -0.0947 | -1.228 | 0.24845 | 0.415 | -5.82 | 1.89567 | none |
| TGFA | -0.145 | -0.1264 | -1.228 | 0.24851 | 0.415 | -5.82 | 0.1144 | none |
| SYPL1 | -0.186 | -0.2629 | -1.228 | 0.24856 | 0.415 | -5.82 | 1.43767 | none |
| SH3PXD2A | 0.218 | 0.2671 | 1.2278 | 0.24861 | 0.415 | -5.82 | -0.122 | none |
| BAG6 | 0.332 | 0.4036 | 1.2274 | 0.24875 | 0.415 | -5.82 | 0.10107 | none |
| C5ORF51 | 0.16 | 0.0123 | 1.2272 | 0.24881 | 0.415 | -5.82 | -0.1715 | none |
| PEX10 | 0.178 | 0.0789 | 1.2269 | 0.24895 | 0.415 | -5.82 | -0.1748 | none |
| SLC25A18 | 0.103 | 0.1056 | 1.2269 | 0.24895 | 0.415 | -5.82 | -0.6156 | none |
| DHRS2 | -0.162 | -0.0502 | -1.227 | 0.24901 | 0.415 | -5.82 | 0.78893 | none |
| CALR | -0.145 | -0.087 | -1.226 | 0.24909 | 0.416 | -5.82 | 1.08997 | none |
| TESPA1 | 0.29 | 0.3266 | 1.2264 | 0.24913 | 0.416 | -5.82 | -0.2375 | none |
| STX4 | 0.201 | 0.2226 | 1.2263 | 0.24914 | 0.416 | -5.82 | -1.1619 | none |
| RPRD2 | 0.333 | 0.3914 | 1.2262 | 0.24918 | 0.416 | -5.82 | 0.53668 | none |
| TMEM190 | -0.125 | -0.1437 | -1.226 | 0.24919 | 0.416 | -5.82 | -0.0744 | none |
| C11ORF86 | -0.118 | -0.0745 | -1.226 | 0.24922 | 0.416 | -5.82 | 0.41824 | none |
| ANKFN1 | -0.178 | -0.0703 | -1.226 | 0.24924 | 0.416 | -5.82 | 0.60525 | none |
| TOP1MT | 0.188 | 0.2856 | 1.2258 | 0.24934 | 0.416 | -5.82 | -0.389 | none |
| ADCY3 | -0.165 | 0.0283 | -1.226 | 0.24935 | 0.416 | -5.82 | 0.27263 | none |
| C18ORF25 | 0.135 | 0.0617 | 1.2256 | 0.2494 | 0.416 | -5.82 | 0.3041 | none |
| DRAP1 | 0.211 | 0.1898 | 1.2255 | 0.24945 | 0.416 | -5.82 | 0.49734 | none |
| BTG3 | -0.107 | -0.1102 | -1.225 | 0.24946 | 0.416 | -5.82 | 0.99812 | none |
| B4GALNT1 | 0.101 | 0.031 | 1.2252 | 0.24956 | 0.416 | -5.82 | 0.96956 | none |
| TMC4 | 0.095 | 0.0946 | 1.225 | 0.24962 | 0.416 | -5.82 | -0.5691 | none |
| NEPRO | -0.11 | -0.0938 | -1.224 | 0.24983 | 0.416 | -5.82 | 1.08583 | none |
| DNAH10 | 0.101 | 0.0958 | 1.2241 | 0.24994 | 0.416 | -5.82 | 0.08783 | none |
| PGPEP1 | 0.133 | ######## | 1.224 | 0.24997 | 0.416 | -5.82 | -0.4894 | none |
| KIF3B | 0.33 | 0.4113 | 1.2238 | 0.25005 | 0.416 | -5.82 | -0.4557 | none |
| BLVRB | -0.188 | -0.18 | -1.224 | 0.25008 | 0.416 | -5.82 | 1.48334 | none |
| ZNRF2 | 0.197 | 0.1537 | 1.2237 | 0.25009 | 0.416 | -5.82 | 0.93778 | none |
| S100A9 | -0.133 | -0.1102 | -1.224 | 0.25013 | 0.417 | -5.82 | -0.3165 | none |
| DCAF1 | 0.156 | 0.1796 | 1.2233 | 0.25023 | 0.417 | -5.82 | -0.3322 | none |
| ZFP36L2 | 0.14 | 0.1304 | 1.2231 | 0.25031 | 0.417 | -5.82 | -0.3473 | none |
| UBE2D3 | -0.164 | -0.2034 | -1.223 | 0.25034 | 0.417 | -5.82 | 1.36651 | none |
| PANK2 | 0.208 | 0.1611 | 1.2228 | 0.2504 | 0.417 | -5.82 | 1.13316 | none |
| RPL10L | 0.257 | 0.1312 | 1.2227 | 0.25044 | 0.417 | -5.82 | -0.6996 | none |
| POC1B | 0.131 | 0.103 | 1.2224 | 0.25055 | 0.417 | -5.82 | 1.368 | none |
| NPR2 | 0.328 | 0.2764 | 1.2222 | 0.25061 | 0.417 | -5.82 | -0.459 | none |
| GRAP | 0.227 | 0.1905 | 1.2221 | 0.25065 | 0.417 | -5.82 | 0.54412 | none |
| MSANTD4 | 0.201 | 0.1369 | 1.2221 | 0.25066 | 0.417 | -5.82 | -0.0259 | none |
| MMP7 | 0.151 | 0.0862 | 1.2221 | 0.25066 | 0.417 | -5.82 | 0.65484 | none |

| | | | | | | | | |
|---|---|---|---|---|---|---|---|---|
| HIST1H3H | 0.211 | 0.1008 | 1.2212 | 0.25098 | 0.417 | -5.82 | -0.5777 | none |
| MPHOSPH | 0.425 | 0.482 | 1.2212 | 0.25099 | 0.417 | -5.82 | 0.15908 | none |
| PLEKHG6 | 0.237 | 0.1953 | 1.2212 | 0.25099 | 0.417 | -5.82 | -0.3167 | none |
| DEFB103A | 0.095 | 0.0788 | 1.2209 | 0.25108 | 0.417 | -5.82 | 0.85479 | none |
| DEFB103B | 0.095 | 0.0788 | 1.2209 | 0.25108 | 0.417 | -5.82 | 0.85479 | none |
| RAB3GAP1 | 0.212 | 0.1895 | 1.2209 | 0.2511 | 0.417 | -5.82 | -0.2032 | none |
| PRDM8 | 0.126 | 0.1615 | 1.2207 | 0.25115 | 0.418 | -5.82 | 1.24004 | none |
| SAP30BP | 0.148 | 0.1621 | 1.2206 | 0.25122 | 0.418 | -5.82 | -0.0018 | none |
| PIM2 | -0.138 | -0.1458 | -1.22 | 0.25127 | 0.418 | -5.82 | 1.98585 | none |
| POLR3E | 0.199 | 0.2153 | 1.2203 | 0.25132 | 0.418 | -5.82 | -0.3308 | none |
| GTF2F1 | 0.412 | 0.5169 | 1.2202 | 0.25136 | 0.418 | -5.82 | -0.0938 | none |
| TRPM5 | 0.096 | 0.0764 | 1.2201 | 0.25138 | 0.418 | -5.82 | -0.9812 | none |
| DCTN3 | -0.174 | -0.2142 | -1.22 | 0.25143 | 0.418 | -5.83 | 0.14409 | none |
| RPIA | -0.117 | -0.0844 | -1.22 | 0.25149 | 0.418 | -5.83 | 1.75981 | none |
| TMEM63C | 0.159 | 0.1818 | 1.2198 | 0.2515 | 0.418 | -5.83 | 0.60832 | none |
| DOCK2 | 0.351 | 0.5073 | 1.2193 | 0.25167 | 0.418 | -5.83 | -0.1783 | none |
| POLA2 | 0.152 | 0.1508 | 1.2192 | 0.25173 | 0.418 | -5.83 | -0.4461 | none |
| MTFR2 | -0.192 | -0.2421 | -1.219 | 0.25181 | 0.418 | -5.83 | 0.98942 | none |
| DNAJC9 | -0.161 | -0.1394 | -1.219 | 0.25181 | 0.418 | -5.83 | 1.57424 | none |
| MKI67 | 0.423 | 0.614 | 1.2187 | 0.25191 | 0.418 | -5.83 | -0.0576 | none |
| GAPDHS | 0.244 | 0.286 | 1.2185 | 0.25196 | 0.418 | -5.83 | 1.68195 | none |
| TTC32 | 0.258 | 0.2109 | 1.2185 | 0.25199 | 0.418 | -5.83 | 0.24701 | none |
| COMMD8 | -0.138 | -0.1957 | -1.218 | 0.25199 | 0.418 | -5.83 | 1.18219 | none |
| DHRS4 | 0.154 | 0.1061 | 1.2184 | 0.252 | 0.418 | -5.83 | 0.58021 | none |
| UCHL3 | -0.15 | -0.1897 | -1.218 | 0.25202 | 0.418 | -5.83 | 0.64508 | none |
| RNF14 | 0.154 | 0.0485 | 1.2183 | 0.25205 | 0.418 | -5.83 | -0.4799 | none |
| UNK | 0.173 | 0.1221 | 1.2181 | 0.2521 | 0.418 | -5.83 | -0.5781 | none |
| CPO | -0.091 | -0.0667 | -1.218 | 0.25212 | 0.418 | -5.83 | 2.08514 | none |
| EMB | 0.203 | 0.1319 | 1.2181 | 0.25213 | 0.418 | -5.83 | 1.32578 | none |
| H1FX | 0.085 | 0.0928 | 1.218 | 0.25216 | 0.418 | -5.83 | 0.13876 | none |
| ZNF25 | 0.279 | 0.274 | 1.2171 | 0.25248 | 0.419 | -5.83 | -0.1785 | none |
| CAV1 | -0.1 | -0.0821 | -1.217 | 0.25249 | 0.419 | -5.83 | 0.96891 | none |
| SLC5A6 | 0.171 | 0.1152 | 1.2169 | 0.25255 | 0.419 | -5.83 | 0.99168 | none |
| CIRBP | 0.157 | 0.1314 | 1.2162 | 0.25279 | 0.419 | -5.83 | 0.95224 | none |
| C8ORF59 | -0.196 | -0.2911 | -1.216 | 0.25293 | 0.419 | -5.83 | 0.6828 | none |
| COQ4 | 0.171 | 0.095 | 1.2158 | 0.25295 | 0.419 | -5.83 | -0.334 | none |
| RAB5A | 0.119 | 0.0863 | 1.2157 | 0.25299 | 0.419 | -5.83 | 0.58244 | none |
| GLT8D2 | 0.243 | 0.1788 | 1.2156 | 0.25301 | 0.419 | -5.83 | 1.05848 | none |
| FGF12 | -0.27 | -0.1988 | -1.215 | 0.25338 | 0.42 | -5.83 | 1.62093 | none |
| PCDH1 | -0.12 | -0.0874 | -1.214 | 0.25347 | 0.42 | -5.83 | 0.86643 | none |

| | | | | | | | | |
|---|---|---|---|---|---|---|---|---|
| CCDC121 | -0.125 | -0.0871 | -1.214 | 0.2535 | 0.42 | -5.83 | 1.33964 | none |
| GIMAP1-G | -0.352 | -0.2209 | -1.214 | 0.25358 | 0.42 | -5.83 | 1.52512 | none |
| AGPAT1 | 0.217 | 0.0246 | 1.2139 | 0.25364 | 0.42 | -5.83 | -0.061 | none |
| PRSS3 | 0.146 | 0.1352 | 1.2138 | 0.25368 | 0.42 | -5.83 | -0.5756 | none |
| PIGY | 0.379 | 0.5059 | 1.2137 | 0.25373 | 0.42 | -5.83 | -0.1427 | none |
| KCNQ4 | -0.081 | -0.039 | -1.213 | 0.25386 | 0.42 | -5.83 | 0.06021 | none |
| VNN3 | -0.118 | -0.087 | -1.213 | 0.25391 | 0.42 | -5.83 | 0.1691 | none |
| OPRL1 | 0.094 | 0.0624 | 1.213 | 0.25395 | 0.42 | -5.83 | 0.21225 | none |
| C4A | 0.312 | 0.2228 | 1.2129 | 0.25401 | 0.421 | -5.83 | 0.92463 | none |
| EIF3B | 0.214 | 0.2897 | 1.2125 | 0.25416 | 0.421 | -5.83 | -0.3697 | none |
| ZNF217 | 0.389 | 0.4046 | 1.2123 | 0.25424 | 0.421 | -5.83 | -0.2522 | none |
| VSTM2B | 0.079 | 0.08 | 1.2122 | 0.25425 | 0.421 | -5.83 | 0 | none |
| CYP27A1 | -0.104 | -0.0815 | -1.212 | 0.25427 | 0.421 | -5.83 | 1.57104 | none |
| CPVL | 0.262 | 0.2144 | 1.2121 | 0.2543 | 0.421 | -5.83 | -0.7828 | none |
| RHOQ | 0.182 | 0.2186 | 1.212 | 0.25434 | 0.421 | -5.83 | -0.3592 | none |
| MROH2A | 0.139 | 0.1399 | 1.2115 | 0.25453 | 0.421 | -5.83 | 1.63779 | none |
| GAREM2 | 0.115 | 0.1076 | 1.2115 | 0.25453 | 0.421 | -5.83 | 0.0677 | none |
| RAD51B | 0.218 | 0.1547 | 1.2115 | 0.25453 | 0.421 | -5.83 | -0.7059 | none |
| WNK1 | 0.455 | 0.5913 | 1.2114 | 0.25456 | 0.421 | -5.83 | -0.3333 | none |
| TNS1 | 0.196 | 0.2032 | 1.2112 | 0.25464 | 0.421 | -5.84 | 0.57872 | none |
| ENPEP | -0.131 | -0.1397 | -1.211 | 0.25469 | 0.421 | -5.84 | 0.46044 | none |
| ARPC1B | -0.136 | -0.0832 | -1.211 | 0.25477 | 0.421 | -5.84 | 0.97807 | none |
| INPPL1 | 0.24 | 0.3497 | 1.2106 | 0.25484 | 0.421 | -5.84 | 0.52861 | none |
| PROX1 | 0.255 | 0.2596 | 1.2106 | 0.25486 | 0.421 | -5.84 | -0.7883 | none |
| IQCH | 0.144 | 0.0985 | 1.2105 | 0.25488 | 0.421 | -5.84 | -0.2284 | none |
| CKAP5 | 0.49 | 0.6483 | 1.2105 | 0.2549 | 0.421 | -5.84 | -0.195 | none |
| WDR60 | -0.192 | -0.1076 | -1.21 | 0.25492 | 0.421 | -5.84 | 1.25927 | none |
| TOLLIP | 0.152 | 0.1294 | 1.2101 | 0.25504 | 0.421 | -5.84 | 0.15678 | none |
| DIMT1 | 0.152 | 0.0258 | 1.21 | 0.25505 | 0.421 | -5.84 | 0.74561 | none |
| ASB13 | -0.158 | -0.1196 | -1.21 | 0.25509 | 0.421 | -5.84 | 1.79431 | none |
| TMEM167 | -0.2 | -0.275 | -1.21 | 0.2551 | 0.421 | -5.84 | 0.71763 | none |
| CRNKL1 | 0.173 | 0.0887 | 1.2096 | 0.25521 | 0.422 | -5.84 | -0.3487 | none |
| TMEM150 | 0.2 | 0.1085 | 1.2096 | 0.25523 | 0.422 | -5.84 | -0.0436 | none |
| MXRA8 | 0.22 | 0.1899 | 1.2096 | 0.25523 | 0.422 | -5.84 | 0.19935 | none |
| MT3 | -0.218 | -0.2197 | -1.21 | 0.25525 | 0.422 | -5.84 | 1.13212 | none |
| LAMB2 | 0.224 | 0.3107 | 1.2094 | 0.25528 | 0.422 | -5.84 | 0.12605 | none |
| IL17RB | 0.105 | 0.1178 | 1.209 | 0.25544 | 0.422 | -5.84 | 1.1035 | none |
| SF1 | 0.23 | 0.2729 | 1.2089 | 0.25548 | 0.422 | -5.84 | 0.11336 | none |
| WDR4 | 0.11 | 0.0815 | 1.2086 | 0.25557 | 0.422 | -5.84 | 0.19293 | none |
| PSMC2 | -0.102 | -0.1129 | -1.208 | 0.25564 | 0.422 | -5.84 | 1.97531 | none |

| | | | | | | | | |
|---|---|---|---|---|---|---|---|---|
| C17ORF78 | 0.107 | 0.0937 | 1.2084 | 0.25566 | 0.422 | -5.84 | -0.1957 | none |
| NMNAT2 | -0.149 | -0.1327 | -1.208 | 0.25585 | 0.422 | -5.84 | 2.11599 | AA-related |
| SLC29A3 | 0.192 | 0.1405 | 1.2078 | 0.25587 | 0.422 | -5.84 | -0.8579 | none |
| MAP4K4 | 0.362 | 0.4309 | 1.2075 | 0.25599 | 0.422 | -5.84 | -0.5104 | none |
| C1QTNF4 | -0.085 | -0.0761 | -1.207 | 0.25601 | 0.422 | -5.84 | 2.31129 | AA-related |
| IKBKE | 0.138 | 0.116 | 1.2074 | 0.25601 | 0.422 | -5.84 | 0.52291 | none |
| FAM177B | -0.092 | -0.0606 | -1.207 | 0.2561 | 0.422 | -5.84 | 1.37395 | none |
| LARP1B | 0.132 | 0.1247 | 1.2072 | 0.2561 | 0.422 | -5.84 | -0.465 | none |
| ULK3 | 0.132 | 0.156 | 1.2071 | 0.25614 | 0.422 | -5.84 | 0.66262 | none |
| SPATS1 | -0.135 | -0.1268 | -1.207 | 0.25619 | 0.422 | -5.84 | 1.32606 | none |
| PANX3 | 0.117 | 0.0778 | 1.2068 | 0.25623 | 0.422 | -5.84 | -1.2762 | none |
| FTCDNL1 | -0.275 | -0.2256 | -1.206 | 0.25642 | 0.423 | -5.84 | 1.92309 | none |
| KLK9 | -0.092 | -0.0612 | -1.206 | 0.25643 | 0.423 | -5.84 | 0.86195 | none |
| FOXH1 | -0.111 | -0.0657 | -1.206 | 0.25643 | 0.423 | -5.84 | 1.02853 | none |
| MED26 | 0.213 | 0.2922 | 1.2062 | 0.25647 | 0.423 | -5.84 | -0.2306 | none |
| PSMA8 | -0.191 | -0.1708 | -1.206 | 0.25652 | 0.423 | -5.84 | 1.4096 | none |
| FAM217A | 0.244 | 0.3351 | 1.2052 | 0.25682 | 0.423 | -5.84 | 0.29132 | none |
| TMCC1 | 0.272 | 0.3038 | 1.2052 | 0.25684 | 0.423 | -5.84 | -0.0888 | none |
| RHOG | -0.146 | -0.1359 | -1.205 | 0.25692 | 0.423 | -5.84 | 1.80811 | none |
| TMEM179 | -0.247 | -0.2842 | -1.205 | 0.25696 | 0.423 | -5.84 | 1.84028 | none |
| RTN4IP1 | 0.122 | -0.0121 | 1.2047 | 0.25701 | 0.423 | -5.84 | -0.2637 | none |
| BEND4 | -0.128 | -0.1358 | -1.205 | 0.25701 | 0.423 | -5.84 | 1.48484 | none |
| FAM136A | 0.09 | 0.0452 | 1.2045 | 0.25709 | 0.423 | -5.84 | -0.4923 | none |
| ZNF35 | -0.149 | -0.0753 | -1.204 | 0.25711 | 0.423 | -5.84 | 1.9149 | none |
| PRR22 | 0.181 | 0.166 | 1.2041 | 0.25724 | 0.423 | -5.84 | 0.42332 | none |
| CHRNE | 0.176 | 0.1556 | 1.2041 | 0.25725 | 0.423 | -5.84 | 0.0824 | none |
| YWHAH | -0.081 | -0.0394 | -1.204 | 0.25726 | 0.423 | -5.84 | 1.11962 | none |
| HIST1H2BI | 0.357 | 0.1008 | 1.204 | 0.25726 | 0.423 | -5.84 | -0.3727 | none |
| PTRH2 | -0.116 | -0.1238 | -1.204 | 0.25727 | 0.423 | -5.84 | 0.38349 | none |
| C11ORF45 | -0.171 | -0.1493 | -1.204 | 0.25742 | 0.424 | -5.84 | 1.63328 | none |
| SPAG17 | 0.187 | 0.1956 | 1.2036 | 0.25742 | 0.424 | -5.84 | 1.1623 | none |
| IL9R | -0.236 | -0.2696 | -1.204 | 0.25746 | 0.424 | -5.84 | 1.01086 | none |
| NCR2 | -0.12 | -0.1283 | -1.203 | 0.25751 | 0.424 | -5.84 | 0.17126 | none |
| UPK1B | 0.222 | 0.2031 | 1.2033 | 0.25753 | 0.424 | -5.84 | 0.01473 | none |
| RNF181 | -0.2 | -0.2222 | -1.203 | 0.25758 | 0.424 | -5.84 | 0.63138 | none |
| RSL1D1 | 0.504 | 0.6478 | 1.203 | 0.25763 | 0.424 | -5.84 | -0.077 | none |
| C2CD4C | -0.077 | -0.051 | -1.203 | 0.25773 | 0.424 | -5.84 | 1.60727 | none |
| DCSTAMP | 0.112 | 0.1342 | 1.2028 | 0.25774 | 0.424 | -5.84 | -0.793 | none |
| ADAMTS1' | -0.128 | -0.1607 | -1.203 | 0.25777 | 0.424 | -5.84 | 1.06185 | none |
| COCH | 0.101 | 0.0228 | 1.2016 | 0.25817 | 0.424 | -5.85 | 0.67004 | none |

| | | | | | | | | |
|---|---|---|---|---|---|---|---|---|
| OCSTAMP | -0.094 | -0.0658 | -1.202 | 0.25818 | 0.424 | -5.85 | 1.24697 | none |
| THBS4 | 0.303 | 0.1983 | 1.2014 | 0.25824 | 0.424 | -5.85 | -1.0264 | none |
| KIF20A | -0.27 | -0.0621 | -1.201 | 0.25834 | 0.425 | -5.85 | 1.66792 | none |
| LYPD6B | -0.179 | -0.1395 | -1.201 | 0.25839 | 0.425 | -5.85 | 0.04119 | none |
| GNAT1 | -0.135 | -0.1257 | -1.201 | 0.25842 | 0.425 | -5.85 | 1.77766 | none |
| SPNS3 | -0.173 | -0.1756 | -1.2 | 0.25859 | 0.425 | -5.85 | 1.51279 | none |
| PAPPA | -0.148 | -0.1334 | -1.2 | 0.25865 | 0.425 | -5.85 | 1.60951 | none |
| TRIM35 | 0.184 | 0.2434 | 1.1985 | 0.25929 | 0.426 | -5.85 | 1.19462 | none |
| BRCC3 | 0.347 | 0.4486 | 1.1978 | 0.25959 | 0.426 | -5.85 | -0.4266 | none |
| GUSB | 0.136 | 0.0629 | 1.1973 | 0.25975 | 0.427 | -5.85 | 0.32691 | none |
| VAMP7 | -0.148 | -0.2204 | -1.197 | 0.25984 | 0.427 | -5.85 | 0.91334 | none |
| RNF182 | 0.101 | 0.0923 | 1.1969 | 0.25992 | 0.427 | -5.85 | -0.741 | none |
| CKAP2L | 0.329 | 0.4017 | 1.1968 | 0.25995 | 0.427 | -5.85 | -0.2026 | none |
| RNF103-CI | -0.244 | -0.3204 | -1.197 | 0.25998 | 0.427 | -5.85 | 0.51447 | none |
| LRRC10 | 0.106 | 0.0875 | 1.1966 | 0.26001 | 0.427 | -5.85 | 0.75148 | none |
| IRF2BPL | 0.242 | 0.2832 | 1.1959 | 0.26027 | 0.427 | -5.85 | -0.3386 | none |
| PPP1R3D | -0.121 | -0.0901 | -1.196 | 0.26029 | 0.427 | -5.85 | 1.47094 | none |
| MRPL50 | -0.201 | -0.2248 | -1.196 | 0.2603 | 0.427 | -5.85 | 0.41034 | none |
| RABL3 | -0.128 | -0.1376 | -1.196 | 0.26034 | 0.427 | -5.85 | 0.34268 | none |
| OAZ2 | -0.143 | -0.2083 | -1.195 | 0.26054 | 0.428 | -5.85 | -0.0227 | none |
| ELMO1 | 0.18 | 0.1798 | 1.1951 | 0.26059 | 0.428 | -5.85 | -0.0721 | none |
| WFDC12 | 0.215 | 0.1287 | 1.1945 | 0.2608 | 0.428 | -5.85 | -0.9845 | none |
| KLHL17 | 0.321 | 0.3453 | 1.1943 | 0.26089 | 0.428 | -5.85 | -0.8495 | none |
| RACGAP1 | 0.105 | 0.0655 | 1.1942 | 0.26091 | 0.428 | -5.85 | 0.38627 | none |
| P2RY6 | 0.139 | 0.1448 | 1.1941 | 0.26093 | 0.428 | -5.85 | 0.19538 | none |
| BID | 0.16 | 0.067 | 1.1938 | 0.26107 | 0.428 | -5.85 | -0.2692 | none |
| SAMD8 | 0.184 | 0.0976 | 1.1935 | 0.26119 | 0.428 | -5.86 | 0.01411 | none |
| TGS1 | 0.277 | 0.3379 | 1.1933 | 0.26125 | 0.428 | -5.86 | 0.52371 | none |
| LRRC8A | 0.237 | 0.3321 | 1.1933 | 0.26125 | 0.428 | -5.86 | -0.0529 | none |
| ACTR8 | 0.158 | 0.1367 | 1.1931 | 0.26133 | 0.428 | -5.86 | 0.59993 | none |
| RGN | -0.101 | -0.0681 | -1.193 | 0.26136 | 0.428 | -5.86 | 1.62648 | none |
| NPY1R | 0.094 | 0.0947 | 1.1929 | 0.26139 | 0.428 | -5.86 | 0.71364 | none |
| NMI | 0.201 | 0.0921 | 1.1922 | 0.26165 | 0.429 | -5.86 | -0.6768 | none |
| NDUFB2 | -0.202 | -0.2269 | -1.192 | 0.26172 | 0.429 | -5.86 | 0.61833 | none |
| UFM1 | -0.16 | -0.1386 | -1.192 | 0.26178 | 0.429 | -5.86 | 1.60037 | none |
| HIST1H2AI | 0.355 | 0.2747 | 1.1918 | 0.26181 | 0.429 | -5.86 | 0.80256 | none |
| NABP1 | 0.184 | 0.1322 | 1.1915 | 0.26191 | 0.429 | -5.86 | -0.5722 | none |
| MFSD9 | 0.158 | 0.0693 | 1.1915 | 0.26193 | 0.429 | -5.86 | 0.23295 | none |
| FAM124B | 0.148 | 0.0967 | 1.1915 | 0.26193 | 0.429 | -5.86 | 0.34537 | none |
| GMIP | 0.21 | 0.3039 | 1.1909 | 0.26216 | 0.429 | -5.86 | 0.80588 | none |

| | | | | | | | | |
|---|---|---|---|---|---|---|---|---|
| RIPK1 | 0.269 | 0.2921 | 1.1908 | 0.26217 | 0.429 | -5.86 | 0.10812 | none |
| GSTZ1 | -0.15 | -0.1269 | -1.191 | 0.2622 | 0.429 | -5.86 | 0.91169 | none |
| SNAI2 | -0.102 | -0.0974 | -1.191 | 0.26222 | 0.429 | -5.86 | 1.26543 | none |
| SNAP91 | -0.12 | -0.112 | -1.19 | 0.26234 | 0.429 | -5.86 | -0.1319 | none |
| PHF5A | -0.151 | -0.1949 | -1.19 | 0.26236 | 0.429 | -5.86 | -0.2325 | none |
| 2-Mar | -0.097 | -0.0998 | -1.19 | 0.26237 | 0.429 | -5.86 | -0.601 | none |
| DPEP2 | 0.164 | 0.093 | 1.1901 | 0.26243 | 0.429 | -5.86 | -0.144 | none |
| CCER1 | 0.089 | 0.0623 | 1.1901 | 0.26244 | 0.429 | -5.86 | -0.9612 | none |
| UBE2W | 0.211 | 0.0036 | 1.1899 | 0.26252 | 0.43 | -5.86 | -0.3059 | none |
| FKBP7 | 0.2 | 0.1078 | 1.1892 | 0.26279 | 0.43 | -5.86 | 1.48625 | none |
| ILDR1 | 0.113 | 0.1056 | 1.1891 | 0.26282 | 0.43 | -5.86 | 0.98807 | none |
| AC016734 | 0.242 | 0.2423 | 1.1891 | 0.26283 | 0.43 | -5.86 | -0.2407 | none |
| IRF2 | -0.097 | -0.0842 | -1.188 | 0.2631 | 0.43 | -5.86 | 2.12091 | AA-related |
| FAM71A | -0.081 | -0.0653 | -1.188 | 0.26329 | 0.431 | -5.86 | 1.44676 | none |
| GOLGA6L9 | 0.148 | 0.1208 | 1.1875 | 0.26344 | 0.431 | -5.86 | -0.3837 | none |
| RNF113B | 0.087 | 0.0686 | 1.1874 | 0.26345 | 0.431 | -5.86 | 0 | none |
| IRS1 | -0.137 | -0.0741 | -1.187 | 0.26346 | 0.431 | -5.86 | 0.19106 | none |
| TVP23B | -0.147 | -0.2383 | -1.187 | 0.2635 | 0.431 | -5.86 | 0.41961 | none |
| WRAP73 | -0.115 | -0.0812 | -1.187 | 0.2635 | 0.431 | -5.86 | 1.27868 | none |
| C2ORF42 | -0.214 | -0.1255 | -1.187 | 0.26375 | 0.431 | -5.86 | 0.75064 | none |
| TMEM51 | 0.106 | 0.1015 | 1.1862 | 0.26391 | 0.431 | -5.86 | 1.54144 | none |
| LYPD8 | -0.095 | -0.0947 | -1.186 | 0.26399 | 0.431 | -5.86 | 0.68574 | none |
| CHERP | 0.281 | 0.3845 | 1.1853 | 0.26426 | 0.432 | -5.86 | -0.0684 | none |
| DRD4 | -0.213 | -0.1748 | -1.185 | 0.2644 | 0.432 | -5.86 | 0.42575 | none |
| HAGHL | 0.291 | 0.1985 | 1.1848 | 0.26444 | 0.432 | -5.86 | -0.3621 | none |
| PRDM6 | -0.084 | -0.0567 | -1.185 | 0.26447 | 0.432 | -5.86 | 1.04178 | none |
| CRADD | -0.229 | -0.2737 | -1.185 | 0.26447 | 0.432 | -5.86 | 0.9084 | none |
| KCNMA1 | 0.162 | 0.1875 | 1.1846 | 0.26453 | 0.432 | -5.87 | 0.22206 | none |
| EPHB6 | 0.286 | 0.2358 | 1.1843 | 0.26461 | 0.432 | -5.87 | 0.51033 | none |
| FUBP1 | 0.249 | 0.2908 | 1.1842 | 0.26468 | 0.432 | -5.87 | -0.1565 | none |
| HTR4 | 0.093 | 0.1046 | 1.1841 | 0.26469 | 0.432 | -5.87 | -0.8317 | none |
| UBE2J2 | 0.147 | 0.1094 | 1.1839 | 0.26476 | 0.432 | -5.87 | -0.7987 | none |
| AP5B1 | 0.106 | 0.1086 | 1.1837 | 0.26484 | 0.432 | -5.87 | -0.2546 | none |
| IRX2 | 0.084 | 0.0886 | 1.1835 | 0.26494 | 0.432 | -5.87 | -0.4808 | none |
| SPATA2L | 0.096 | 0.0301 | 1.1833 | 0.26499 | 0.433 | -5.87 | 0.39875 | none |
| NRXN3 | 0.152 | 0.1164 | 1.1833 | 0.26502 | 0.433 | -5.87 | -0.4206 | none |
| ANK2 | 0.229 | 0.2077 | 1.1829 | 0.26517 | 0.433 | -5.87 | 0.21154 | none |
| RAB39A | 0.161 | 0.1036 | 1.1825 | 0.2653 | 0.433 | -5.87 | -0.5929 | none |
| NAP1L4 | 0.153 | 0.1512 | 1.1825 | 0.26532 | 0.433 | -5.87 | 0.3661 | none |
| THRAP3 | 0.401 | 0.5279 | 1.1822 | 0.26541 | 0.433 | -5.87 | -0.1122 | none |

| | | | | | | | | |
|---|---|---|---|---|---|---|---|---|
| GOLGA7B | -0.087 | -0.0638 | -1.182 | 0.26546 | 0.433 | -5.87 | 1.46065 | none |
| MPC2 | -0.195 | -0.2142 | -1.181 | 0.26571 | 0.433 | -5.87 | 0.38336 | none |
| DDX3Y | 0.105 | 0.1281 | 1.1814 | 0.26573 | 0.433 | -5.87 | 1.17436 | none |
| SPDL1 | 0.345 | 0.4078 | 1.1811 | 0.26585 | 0.434 | -5.87 | 0.78169 | none |
| GUK1 | -0.175 | -0.1754 | -1.181 | 0.26587 | 0.434 | -5.87 | 1.51532 | none |
| PLCL1 | 0.176 | 0.1866 | 1.1809 | 0.2659 | 0.434 | -5.87 | -0.1288 | none |
| HTATIP2 | -0.142 | -0.2059 | -1.181 | 0.26596 | 0.434 | -5.87 | 1.21671 | none |
| RAC1 | -0.225 | -0.208 | -1.181 | 0.26603 | 0.434 | -5.87 | 0.9442 | none |
| CCNY | -0.14 | -0.2059 | -1.181 | 0.26605 | 0.434 | -5.87 | 0.48163 | none |
| SCGB2B2 | 0.172 | 0.1442 | 1.1795 | 0.26645 | 0.434 | -5.87 | -0.0963 | none |
| POLE4 | -0.149 | -0.1803 | -1.179 | 0.26648 | 0.434 | -5.87 | 2.08849 | none |
| MAP3K6 | -0.295 | -0.2802 | -1.179 | 0.26652 | 0.434 | -5.87 | 0.50824 | none |
| SESN3 | 0.181 | 0.091 | 1.1793 | 0.26654 | 0.434 | -5.87 | 0.8737 | none |
| BEX2 | -0.28 | -0.3981 | -1.179 | 0.26655 | 0.434 | -5.87 | 0.61848 | none |
| MAGEA9B | -0.086 | -0.0596 | -1.179 | 0.26656 | 0.434 | -5.87 | -0.5771 | none |
| GPRASP2 | 0.127 | 0.1048 | 1.1788 | 0.26671 | 0.434 | -5.87 | -0.4525 | none |
| ARL4C | -0.177 | -0.1596 | -1.179 | 0.26678 | 0.435 | -5.87 | 1.71145 | none |
| IL6 | 0.133 | 0.1249 | 1.1779 | 0.26704 | 0.435 | -5.87 | -0.8494 | none |
| FURIN | 0.228 | 0.3143 | 1.1778 | 0.26708 | 0.435 | -5.87 | -0.2226 | none |
| DUPD1 | 0.09 | 0.068 | 1.1778 | 0.26709 | 0.435 | -5.87 | 0.10136 | none |
| WIF1 | 0.13 | 0.1387 | 1.1775 | 0.26721 | 0.435 | -5.87 | 0.53554 | none |
| GDAP1L1 | -0.12 | -0.0972 | -1.177 | 0.26725 | 0.435 | -5.87 | 0.20605 | none |
| CYP1A1 | -0.15 | -0.1104 | -1.177 | 0.26735 | 0.435 | -5.87 | 0.4653 | none |
| SLC25A39 | -0.168 | -0.1651 | -1.177 | 0.26737 | 0.435 | -5.87 | 1.55347 | none |
| GTF2H5 | -0.219 | -0.2548 | -1.177 | 0.26743 | 0.435 | -5.87 | 0.97159 | none |
| ADH7 | 0.181 | 0.194 | 1.1762 | 0.26772 | 0.436 | -5.87 | 0.89896 | none |
| PNMA3 | 0.164 | 0.106 | 1.1757 | 0.26791 | 0.436 | -5.88 | -0.4134 | none |
| ASCC3 | 0.302 | 0.3732 | 1.1752 | 0.26806 | 0.436 | -5.88 | 0.27604 | none |
| CRIPT | -0.21 | -0.3164 | -1.175 | 0.26811 | 0.436 | -5.88 | 1.31614 | none |
| WWTR1 | 0.184 | 0.1265 | 1.1749 | 0.26819 | 0.436 | -5.88 | -0.2689 | none |
| PLAT | 0.207 | 0.2259 | 1.1748 | 0.26822 | 0.436 | -5.88 | -0.5785 | none |
| EID3 | 0.202 | 0.1643 | 1.1748 | 0.26824 | 0.436 | -5.88 | 0.43995 | none |
| GRIP2 | -0.078 | -0.0598 | -1.173 | 0.26896 | 0.437 | -5.88 | -0.155 | none |
| SURF6 | 0.29 | 0.3592 | 1.1719 | 0.26933 | 0.438 | -5.88 | -0.293 | none |
| SIGLEC14 | 0.093 | 0.058 | 1.1718 | 0.26936 | 0.438 | -5.88 | 0.78112 | none |
| SP5 | -0.096 | -0.0564 | -1.172 | 0.2694 | 0.438 | -5.88 | 1.11127 | none |
| SUMO1 | -0.115 | -0.1667 | -1.172 | 0.26941 | 0.438 | -5.88 | 0.45164 | none |
| NADSYN1 | 0.107 | 0.1035 | 1.1717 | 0.26943 | 0.438 | -5.88 | -0.8031 | none |
| CCDC186 | 0.314 | 0.3152 | 1.1716 | 0.26945 | 0.438 | -5.88 | -0.1418 | none |
| GLRX3 | 0.102 | 0.0065 | 1.1711 | 0.26964 | 0.438 | -5.88 | -0.3929 | none |

| | | | | | | | | |
|---|---|---|---|---|---|---|---|---|
| NAA20 | -0.14 | -0.1868 | -1.171 | 0.26978 | 0.438 | -5.88 | 0.89231 | none |
| TCF24 | -0.134 | -0.1064 | -1.17 | 0.26993 | 0.439 | -5.88 | 0.85401 | none |
| OXER1 | 0.117 | 0.0839 | 1.1702 | 0.26998 | 0.439 | -5.88 | 0.34923 | none |
| AIFM1 | 0.122 | 0.08 | 1.1702 | 0.26998 | 0.439 | -5.88 | -0.1821 | none |
| TAMM41 | 0.094 | -0.0033 | 1.1701 | 0.27003 | 0.439 | -5.88 | -0.6643 | none |
| NME8 | -0.131 | -0.1395 | -1.169 | 0.27032 | 0.439 | -5.88 | -0.2082 | none |
| TBATA | -0.08 | -0.0738 | -1.169 | 0.27037 | 0.439 | -5.88 | 1.67922 | none |
| ROBO4 | -0.078 | -0.046 | -1.169 | 0.27039 | 0.439 | -5.88 | 0.64386 | none |
| KCNN3 | 0.092 | 0.0518 | 1.169 | 0.27046 | 0.439 | -5.88 | 0.46316 | none |
| MAFK | 0.158 | 0.2235 | 1.1687 | 0.27057 | 0.439 | -5.88 | -0.4312 | none |
| TTI2 | -0.133 | -0.1472 | -1.168 | 0.27085 | 0.44 | -5.88 | 0.27743 | none |
| KLHL23 | 0.131 | 0.1336 | 1.1679 | 0.27088 | 0.44 | -5.88 | -0.2161 | none |
| EPB41L5 | -0.218 | -0.2363 | -1.168 | 0.27102 | 0.44 | -5.88 | 0.4057 | none |
| UQCRC1 | -0.126 | -0.1105 | -1.167 | 0.27118 | 0.44 | -5.88 | 0.94804 | none |
| COL9A3 | 0.329 | 0.3116 | 1.167 | 0.27122 | 0.44 | -5.88 | -0.0426 | none |
| G0S2 | 0.163 | 0.1442 | 1.1669 | 0.27124 | 0.44 | -5.88 | -0.4439 | none |
| FAR2 | -0.244 | -0.3057 | -1.166 | 0.27145 | 0.44 | -5.89 | 0.26737 | none |
| SERP1 | -0.176 | -0.2012 | -1.166 | 0.27149 | 0.44 | -5.89 | 0.92044 | none |
| CAMKK1 | -0.099 | -0.025 | -1.166 | 0.27179 | 0.441 | -5.89 | 1.39527 | none |
| VASH1 | 0.214 | 0.1872 | 1.1654 | 0.27185 | 0.441 | -5.89 | 0.26635 | none |
| TEAD4 | 0.119 | 0.1748 | 1.1652 | 0.27192 | 0.441 | -5.89 | 1.89686 | none |
| KRT19 | 0.092 | 0.0884 | 1.1651 | 0.27194 | 0.441 | -5.89 | -0.0282 | none |
| ORMDL3 | 0.202 | 0.2027 | 1.165 | 0.272 | 0.441 | -5.89 | -0.5312 | none |
| MARK4 | 0.264 | 0.3503 | 1.1648 | 0.27208 | 0.441 | -5.89 | -0.3357 | none |
| OR2A4 | -0.093 | -0.0692 | -1.165 | 0.27208 | 0.441 | -5.89 | 0.31285 | none |
| TFAM | -0.088 | -0.0974 | -1.165 | 0.2721 | 0.441 | -5.89 | 1.89986 | none |
| SYN1 | -0.101 | -0.0786 | -1.164 | 0.27224 | 0.441 | -5.89 | 1.29738 | none |
| NFATC2 | 0.259 | 0.3269 | 1.1642 | 0.2723 | 0.441 | -5.89 | -0.2426 | none |
| ZCCHC24 | 0.095 | 0.1092 | 1.164 | 0.27236 | 0.441 | -5.89 | 1.00548 | none |
| AMPD2 | 0.109 | 0.1717 | 1.1634 | 0.2726 | 0.442 | -5.89 | 1.08642 | none |
| HMHB1 | 0.2 | 0.2078 | 1.1629 | 0.27281 | 0.442 | -5.89 | 1.23597 | none |
| MAP4K2 | 0.117 | 0.2011 | 1.1627 | 0.27288 | 0.442 | -5.89 | 0.42989 | none |
| DPM1 | -0.154 | -0.2388 | -1.162 | 0.27307 | 0.442 | -5.89 | 0.7847 | none |
| MICU3 | -0.173 | -0.0768 | -1.162 | 0.27312 | 0.442 | -5.89 | 1.6303 | none |
| CTSE | 0.136 | 0.1373 | 1.1617 | 0.27327 | 0.443 | -5.89 | 0.48971 | none |
| OSR1 | 0.129 | 0.092 | 1.1616 | 0.27333 | 0.443 | -5.89 | -0.9842 | none |
| KLHDC8A | 0.096 | 0.0965 | 1.1614 | 0.2734 | 0.443 | -5.89 | -0.5242 | none |
| SLC9C2 | -0.206 | -0.2058 | -1.161 | 0.27341 | 0.443 | -5.89 | 0.51327 | none |
| SLC31A2 | -0.217 | -0.3077 | -1.161 | 0.27344 | 0.443 | -5.89 | 0.08206 | none |
| NAGK | 0.148 | 0.0789 | 1.1606 | 0.27371 | 0.443 | -5.89 | -0.3155 | none |

| | | | | | | | | |
|---|---|---|---|---|---|---|---|---|
| P2RX5 | 0.181 | 0.2248 | 1.1603 | 0.2738 | 0.443 | -5.89 | 0.28198 | none |
| GJA4 | -0.094 | -0.0819 | -1.16 | 0.27386 | 0.443 | -5.89 | 1.57402 | none |
| OASL | -0.164 | -0.118 | -1.16 | 0.27391 | 0.443 | -5.89 | 1.29139 | none |
| TRUB1 | -0.249 | -0.1648 | -1.16 | 0.27399 | 0.443 | -5.89 | 1.57184 | none |
| MT-ND5 | 0.37 | 0.4642 | 1.1596 | 0.27407 | 0.443 | -5.89 | -0.2419 | none |
| KRTAP4-9 | 0.168 | 0.1619 | 1.1593 | 0.2742 | 0.444 | -5.89 | 1.29901 | none |
| CCDC144N | -0.168 | -0.0761 | -1.159 | 0.27421 | 0.444 | -5.89 | 0.48543 | none |
| TP53BP2 | 0.322 | 0.4066 | 1.1587 | 0.27442 | 0.444 | -5.89 | -0.3276 | none |
| MYO15B | 0.215 | 0.169 | 1.1582 | 0.27462 | 0.444 | -5.89 | 0.28663 | none |
| PLXNA3 | 0.23 | 0.2608 | 1.1581 | 0.27468 | 0.444 | -5.89 | -0.4568 | none |
| CYP4A22 | 0.089 | 0.0809 | 1.158 | 0.27469 | 0.444 | -5.89 | -1.0473 | none |
| OSBPL10 | 0.29 | 0.2521 | 1.1575 | 0.27489 | 0.445 | -5.9 | -0.0741 | none |
| DGCR6 | 0.376 | 0.3036 | 1.1573 | 0.27496 | 0.445 | -5.9 | -0.4297 | none |
| CHST10 | 0.202 | 0.2765 | 1.157 | 0.27508 | 0.445 | -5.9 | 0.18629 | none |
| SRPX | 0.085 | 0.0603 | 1.157 | 0.27511 | 0.445 | -5.9 | 0.78011 | none |
| CNTF | 0.095 | 0.1234 | 1.1569 | 0.27513 | 0.445 | -5.9 | -0.4192 | none |
| DIAPH2 | 0.191 | 0.2383 | 1.1568 | 0.27516 | 0.445 | -5.9 | -0.0416 | none |
| PTMS | 0.217 | 0.1263 | 1.1564 | 0.27533 | 0.445 | -5.9 | -0.8719 | none |
| CLEC18A | 0.081 | 0.0843 | 1.1561 | 0.27543 | 0.445 | -5.9 | -0.3653 | none |
| TRIM34 | 0.21 | 0.0273 | 1.1561 | 0.27546 | 0.445 | -5.9 | -0.7121 | none |
| PAGE2B | -0.146 | -0.1531 | -1.156 | 0.27551 | 0.445 | -5.9 | 0.93899 | none |
| FCAMR | -0.091 | -0.0825 | -1.156 | 0.27552 | 0.445 | -5.9 | 0.24837 | none |
| TNNI2 | -0.1 | -0.1133 | -1.156 | 0.27555 | 0.445 | -5.9 | -0.1499 | none |
| FZD6 | 0.164 | 0.0764 | 1.1557 | 0.2756 | 0.445 | -5.9 | -0.4789 | none |
| TNFAIP3 | 0.26 | 0.3699 | 1.1554 | 0.27572 | 0.445 | -5.9 | 0.34152 | none |
| GALNT18 | 0.136 | 0.1752 | 1.1553 | 0.27577 | 0.445 | -5.9 | 1.43611 | none |
| CHRNB3 | 0.13 | 0.0786 | 1.155 | 0.27588 | 0.445 | -5.9 | 0.98785 | none |
| TEX13C | 0.085 | 0.0598 | 1.1544 | 0.27611 | 0.446 | -5.9 | -1.247 | none |
| CTRL | 0.276 | 0.2924 | 1.1543 | 0.27614 | 0.446 | -5.9 | -0.6269 | none |
| UBE2B | -0.183 | -0.2595 | -1.154 | 0.27627 | 0.446 | -5.9 | 1.20877 | none |
| CUZD1 | 0.112 | 0.0892 | 1.1538 | 0.27633 | 0.446 | -5.9 | -0.5526 | none |
| GIT1 | 0.297 | 0.4232 | 1.1533 | 0.27656 | 0.446 | -5.9 | -0.4462 | none |
| MBOAT1 | -0.11 | -0.1232 | -1.153 | 0.27657 | 0.446 | -5.9 | 1.07664 | none |
| GAL3ST3 | 0.118 | 0.1059 | 1.153 | 0.27665 | 0.446 | -5.9 | -0.8618 | none |
| TIMM21 | 0.144 | 0.0814 | 1.1526 | 0.27683 | 0.447 | -5.9 | -0.3339 | none |
| KDELR2 | -0.178 | -0.2293 | -1.152 | 0.27687 | 0.447 | -5.9 | 1.57122 | none |
| IGSF23 | -0.123 | -0.1319 | -1.152 | 0.27703 | 0.447 | -5.9 | 1.09881 | none |
| HS3ST5 | 0.165 | 0.1647 | 1.1518 | 0.27713 | 0.447 | -5.9 | 0.48003 | none |
| CUEDC2 | 0.1 | 0.1074 | 1.1516 | 0.27719 | 0.447 | -5.9 | -0.9704 | none |
| ALDH1A2 | 0.193 | 0.1711 | 1.1516 | 0.27721 | 0.447 | -5.9 | -0.9229 | none |

| | | | | | | | | |
|---|---|---|---|---|---|---|---|---|
| PXT1 | 0.114 | 0.0687 | 1.1516 | 0.27722 | 0.447 | -5.9 | 0.07266 | none |
| FBXO8 | -0.235 | -0.2937 | -1.151 | 0.27728 | 0.447 | -5.9 | 1.16667 | none |
| FMR1 | 0.278 | 0.3328 | 1.1511 | 0.27738 | 0.447 | -5.9 | 0.39611 | none |
| TRHDE | -0.123 | -0.103 | -1.151 | 0.27741 | 0.447 | -5.9 | 2.0499 | none |
| TMTC4 | 0.21 | 0.2338 | 1.151 | 0.27744 | 0.447 | -5.9 | 1.11599 | none |
| CMC2 | -0.153 | -0.1973 | -1.151 | 0.27747 | 0.447 | -5.9 | 1.36067 | none |
| GOT2 | 0.108 | 0.0705 | 1.1506 | 0.27759 | 0.447 | -5.9 | 1.12822 | none |
| UBE2K | 0.092 | 0.0305 | 1.1505 | 0.27765 | 0.447 | -5.9 | -0.4293 | none |
| TNP2 | 0.099 | 0.1089 | 1.1501 | 0.27778 | 0.448 | -5.9 | 1.11911 | none |
| BTBD6 | -0.154 | -0.1289 | -1.149 | 0.27809 | 0.448 | -5.9 | 0.45349 | none |
| ZNF267 | 0.208 | 0.2085 | 1.149 | 0.27822 | 0.448 | -5.9 | 0.57948 | none |
| PBX2 | 0.137 | 0.1051 | 1.149 | 0.27824 | 0.448 | -5.9 | -0.3968 | none |
| RPL34 | -0.211 | -0.2745 | -1.149 | 0.27834 | 0.448 | -5.9 | 0.75859 | none |
| THAP1 | 0.196 | 0.1977 | 1.1484 | 0.27846 | 0.448 | -5.91 | 0.54365 | none |
| CSNK1A1 | 0.11 | 0.094 | 1.1483 | 0.27851 | 0.449 | -5.91 | 0.61367 | none |
| CXCL1 | -0.116 | -0.1295 | -1.148 | 0.27855 | 0.449 | -5.91 | 1.42844 | none |
| CDK20 | -0.201 | -0.1277 | -1.148 | 0.27856 | 0.449 | -5.91 | 1.76809 | none |
| KHDC1 | 0.254 | 0.0651 | 1.148 | 0.27862 | 0.449 | -5.91 | -0.2249 | none |
| CCL25 | 0.104 | 0.1706 | 1.1478 | 0.27868 | 0.449 | -5.91 | 0.9886 | none |
| NAXE | -0.188 | -0.2069 | -1.148 | 0.27869 | 0.449 | -5.91 | 0.50202 | none |
| MUC6 | 0.093 | 0.0832 | 1.1476 | 0.27877 | 0.449 | -5.91 | -0.3951 | none |
| ENTPD7 | 0.176 | 0.1716 | 1.1475 | 0.2788 | 0.449 | -5.91 | -0.209 | none |
| PRPF4B | 0.443 | 0.5452 | 1.1475 | 0.27881 | 0.449 | -5.91 | -0.1559 | none |
| RHBDL1 | 0.109 | 0.0935 | 1.1471 | 0.27896 | 0.449 | -5.91 | 1.66228 | none |
| AQP10 | 0.123 | 0.1302 | 1.1468 | 0.27908 | 0.449 | -5.91 | -0.273 | none |
| EFNA5 | -0.105 | -0.0814 | -1.147 | 0.27915 | 0.449 | -5.91 | 0.45808 | none |
| SPAG11B | 0.119 | 0.1188 | 1.1465 | 0.27919 | 0.449 | -5.91 | 0.77714 | none |
| GEM | -0.286 | -0.1676 | -1.146 | 0.27922 | 0.449 | -5.91 | 1.27786 | none |
| ANAPC4 | 0.113 | 0.0603 | 1.1464 | 0.27923 | 0.449 | -5.91 | -0.5272 | none |
| NR1D1 | 0.16 | 0.2082 | 1.1453 | 0.27967 | 0.45 | -5.91 | -0.2609 | none |
| BDH1 | 0.232 | 0.2026 | 1.1448 | 0.27988 | 0.45 | -5.91 | 0.21983 | none |
| GHSR | 0.151 | 0.0741 | 1.1445 | 0.27998 | 0.45 | -5.91 | -0.5485 | none |
| NPBWR2 | 0.084 | 0.0591 | 1.1445 | 0.27999 | 0.45 | -5.91 | 0 | none |
| DHX9 | 0.198 | 0.2726 | 1.1442 | 0.28012 | 0.45 | -5.91 | 0.33181 | none |
| CCDC38 | 0.174 | 0.0562 | 1.1441 | 0.28015 | 0.45 | -5.91 | -1.1181 | none |
| ACTL6B | -0.241 | -0.2407 | -1.144 | 0.28025 | 0.45 | -5.91 | 1.31926 | none |
| MMGT1 | -0.189 | -0.2297 | -1.144 | 0.28029 | 0.45 | -5.91 | 1.47106 | none |
| SGK1 | 0.142 | 0.0665 | 1.1435 | 0.28041 | 0.451 | -5.91 | 0.29934 | none |
| REEP6 | 0.249 | 0.2092 | 1.1432 | 0.28051 | 0.451 | -5.91 | -0.2921 | none |
| MKLN1 | 0.133 | 0.1611 | 1.143 | 0.2806 | 0.451 | -5.91 | 0.74161 | none |

| | | | | | | | | |
|---|---|---|---|---|---|---|---|---|
| CRIP3 | -0.211 | -0.1448 | -1.143 | 0.28069 | 0.451 | -5.91 | 0.39343 | none |
| PERP | -0.149 | -0.1912 | -1.143 | 0.28078 | 0.451 | -5.91 | 1.57301 | none |
| HGD | -0.217 | -0.1909 | -1.142 | 0.28081 | 0.451 | -5.91 | 1.63813 | none |
| AC011043 | 0.162 | 0.0782 | 1.1424 | 0.28084 | 0.451 | -5.91 | -0.0989 | none |
| ZBTB7B | 0.167 | 0.2417 | 1.1423 | 0.28088 | 0.451 | -5.91 | 0.74801 | none |
| ZNHIT3 | 0.135 | -0.0441 | 1.1417 | 0.2811 | 0.451 | -5.91 | -0.0694 | none |
| HAUS4 | -0.144 | -0.1371 | -1.141 | 0.28124 | 0.452 | -5.91 | 1.96136 | none |
| SRSF2 | -0.156 | -0.1702 | -1.141 | 0.28147 | 0.452 | -5.91 | 1.86099 | none |
| PPP3CA | 0.177 | 0.1428 | 1.1407 | 0.28151 | 0.452 | -5.91 | -0.3725 | none |
| IFNL3 | 0.093 | 0.1251 | 1.1405 | 0.28157 | 0.452 | -5.91 | -0.1543 | none |
| MYOCD | -0.082 | -0.0571 | -1.14 | 0.2816 | 0.452 | -5.91 | -0.3269 | none |
| CDK5RAP1 | 0.095 | 0.0258 | 1.1403 | 0.28165 | 0.452 | -5.91 | -0.5649 | none |
| SLC25A46 | 0.151 | 0.0707 | 1.1403 | 0.28166 | 0.452 | -5.91 | 0.5433 | none |
| TAS2R50 | -0.077 | -0.0699 | -1.14 | 0.28172 | 0.452 | -5.91 | 0 | none |
| PRSS37 | 0.086 | 0.0842 | 1.1399 | 0.2818 | 0.452 | -5.91 | 1.40684 | none |
| SLC1A1 | 0.176 | 0.2017 | 1.1399 | 0.28183 | 0.452 | -5.91 | 1.17776 | none |
| SPINT1 | 0.168 | 0.2022 | 1.1398 | 0.28187 | 0.452 | -5.91 | 0.5544 | none |
| CKAP4 | 0.209 | 0.2354 | 1.1398 | 0.28187 | 0.452 | -5.91 | -0.5873 | none |
| CCND3 | -0.118 | -0.1249 | -1.14 | 0.28187 | 0.452 | -5.91 | 2.06309 | none |
| HTRA2 | -0.127 | -0.1073 | -1.139 | 0.28201 | 0.452 | -5.91 | 1.83902 | none |
| ABCD2 | -0.105 | -0.0757 | -1.139 | 0.28206 | 0.452 | -5.91 | 1.45999 | none |
| EZR | 0.244 | 0.3124 | 1.1389 | 0.28221 | 0.452 | -5.92 | 0.04057 | none |
| USP15 | 0.235 | 0.2458 | 1.1385 | 0.28239 | 0.453 | -5.92 | -0.5681 | none |
| SHTN1 | 0.231 | 0.1936 | 1.1384 | 0.28243 | 0.453 | -5.92 | 0.2456 | none |
| CCDC15 | 0.179 | 0.1907 | 1.1382 | 0.28248 | 0.453 | -5.92 | -0.7771 | none |
| GTPBP4 | 0.186 | 0.206 | 1.138 | 0.28256 | 0.453 | -5.92 | 0.23128 | none |
| GPER1 | -0.155 | -0.0914 | -1.138 | 0.28258 | 0.453 | -5.92 | 1.79988 | none |
| TMED10 | -0.143 | -0.1767 | -1.138 | 0.28266 | 0.453 | -5.92 | -0.2093 | none |
| HIST1H2AI | -0.248 | -0.2237 | -1.138 | 0.28276 | 0.453 | -5.92 | 1.55807 | none |
| PHF21B | 0.182 | 0.1465 | 1.1372 | 0.28287 | 0.453 | -5.92 | -0.4275 | none |
| ZMYM4 | 0.372 | 0.4937 | 1.137 | 0.28297 | 0.453 | -5.92 | 0.95223 | none |
| CAMKV | 0.096 | 0.0688 | 1.1369 | 0.283 | 0.453 | -5.92 | -0.8744 | none |
| TSPAN14 | 0.125 | 0.0617 | 1.1365 | 0.28316 | 0.454 | -5.92 | 0.12001 | none |
| SKIV2L | -0.267 | -0.264 | -1.136 | 0.28329 | 0.454 | -5.92 | 1.28873 | none |
| KLRF1 | -0.137 | -0.1156 | -1.136 | 0.2833 | 0.454 | -5.92 | 0.61342 | none |
| TSSK2 | -0.122 | -0.1202 | -1.136 | 0.28333 | 0.454 | -5.92 | 0.24163 | none |
| ATG4B | 0.132 | 0.1265 | 1.1359 | 0.28341 | 0.454 | -5.92 | 0.76648 | none |
| PCNX3 | 0.241 | 0.348 | 1.1357 | 0.2835 | 0.454 | -5.92 | -0.4365 | none |
| TMEM117 | 0.238 | 0.1779 | 1.1355 | 0.28355 | 0.454 | -5.92 | -0.4363 | none |
| A1BG | 0.26 | 0.1102 | 1.1355 | 0.28357 | 0.454 | -5.92 | -0.7324 | none |

| | | | | | | | | |
|---|---|---|---|---|---|---|---|---|
| CDH23 | 0.182 | 0.1396 | 1.1354 | 0.2836 | 0.454 | -5.92 | -0.1388 | none |
| R3HDML | 0.121 | 0.0959 | 1.1348 | 0.28386 | 0.454 | -5.92 | -0.928 | none |
| REEP1 | 0.104 | 0.1208 | 1.1344 | 0.28401 | 0.454 | -5.92 | 0.41097 | none |
| MOB2 | 0.246 | 0.2434 | 1.1339 | 0.28419 | 0.455 | -5.92 | 0.73956 | none |
| SNX8 | 0.138 | 0.1054 | 1.1339 | 0.2842 | 0.455 | -5.92 | 0.80815 | none |
| KRTAP21-3 | 0.231 | 0.2315 | 1.1338 | 0.28426 | 0.455 | -5.92 | 0.66005 | none |
| DKK3 | 0.09 | 0.0881 | 1.1337 | 0.28427 | 0.455 | -5.92 | 1.52019 | none |
| C6ORF47 | 0.225 | 0.1608 | 1.1337 | 0.28428 | 0.455 | -5.92 | -0.6994 | none |
| ZCWPW1 | 0.113 | 0.1017 | 1.1337 | 0.2843 | 0.455 | -5.92 | 0.35312 | none |
| CEP78 | 0.177 | 0.2298 | 1.1335 | 0.28435 | 0.455 | -5.92 | 0.33673 | none |
| FOXB1 | 0.142 | 0.1518 | 1.1335 | 0.28436 | 0.455 | -5.92 | -0.146 | none |
| PPP1R14A | -0.145 | -0.1582 | -1.133 | 0.28437 | 0.455 | -5.92 | 0.45078 | none |
| SLC16A13 | -0.115 | -0.0577 | -1.133 | 0.28454 | 0.455 | -5.92 | 2.04542 | none |
| PIGH | 0.138 | 0.0588 | 1.1324 | 0.28479 | 0.455 | -5.92 | -0.0611 | none |
| ZMYND10 | -0.19 | -0.1328 | -1.132 | 0.28483 | 0.455 | -5.92 | 1.026 | none |
| C19ORF38 | -0.164 | -0.1776 | -1.132 | 0.28485 | 0.455 | -5.92 | 0.74151 | none |
| SNX14 | 0.183 | 0.0965 | 1.1322 | 0.28487 | 0.455 | -5.92 | 1.29336 | none |
| ZNF143 | 0.205 | 0.2124 | 1.1319 | 0.28499 | 0.455 | -5.92 | -0.4507 | none |
| NT5C3A | 0.163 | 0.0208 | 1.1317 | 0.28508 | 0.455 | -5.92 | 0.29786 | none |
| OR4C3 | -0.081 | -0.081 | -1.132 | 0.28508 | 0.455 | -5.92 | 1.63945 | none |
| FGFRL1 | 0.185 | 0.201 | 1.1317 | 0.28508 | 0.455 | -5.92 | 0.19477 | none |
| MT1E | -0.241 | -0.2438 | -1.131 | 0.28532 | 0.456 | -5.92 | 1.65293 | none |
| ABR | 0.5 | 0.3389 | 1.1306 | 0.28553 | 0.456 | -5.92 | -0.9425 | none |
| LOXL1 | 0.25 | 0.2564 | 1.1304 | 0.28562 | 0.456 | -5.92 | 0.97923 | none |
| STARD13 | -0.09 | -0.0698 | -1.13 | 0.28563 | 0.456 | -5.92 | -0.4192 | none |
| SLC39A7 | -0.268 | -0.1438 | -1.13 | 0.28571 | 0.456 | -5.92 | 1.01817 | none |
| P2RY11 | 0.248 | 0.2869 | 1.1301 | 0.28572 | 0.456 | -5.92 | -0.6146 | none |
| CPEB3 | 0.093 | 0.085 | 1.13 | 0.28575 | 0.456 | -5.92 | -0.6614 | none |
| DNAJC1 | 0.333 | 0.3895 | 1.1299 | 0.28582 | 0.456 | -5.93 | 0.70979 | none |
| FOXN2 | -0.204 | -0.1431 | -1.13 | 0.28585 | 0.456 | -5.93 | 2.2156 | AA-related |
| ARPP19 | 0.17 | 0.0781 | 1.1297 | 0.28591 | 0.456 | -5.93 | -0.9672 | none |
| LCE1E | -0.187 | -0.1877 | -1.129 | 0.28602 | 0.456 | -5.93 | 0.88699 | none |
| RNF208 | 0.194 | 0.171 | 1.1293 | 0.28606 | 0.456 | -5.93 | -0.4871 | none |
| MDP1 | -0.194 | -0.2145 | -1.129 | 0.28609 | 0.456 | -5.93 | 0.64959 | none |
| VIM | -0.159 | -0.1423 | -1.129 | 0.28621 | 0.457 | -5.93 | 1.05541 | none |
| FAM25C | -0.208 | -0.2083 | -1.129 | 0.28629 | 0.457 | -5.93 | 1.9054 | none |
| MTMR14 | 0.118 | 0.0499 | 1.1286 | 0.28632 | 0.457 | -5.93 | -0.0935 | none |
| SEL1L2 | -0.155 | -0.1768 | -1.128 | 0.2864 | 0.457 | -5.93 | 0.69722 | none |
| RPP25 | 0.1 | 0.1292 | 1.1282 | 0.2865 | 0.457 | -5.93 | 1.13186 | none |
| SDR9C7 | 0.082 | 0.0824 | 1.128 | 0.28656 | 0.457 | -5.93 | -0.7589 | none |

| | | | | | | | | |
|---|---|---|---|---|---|---|---|---|
| MEAF6 | -0.133 | -0.1621 | -1.128 | 0.2866 | 0.457 | -5.93 | 2.00981 | none |
| TMED9 | -0.132 | -0.0959 | -1.128 | 0.28673 | 0.457 | -5.93 | 1.01031 | none |
| RTKN2 | 0.124 | 0.007 | 1.1276 | 0.28674 | 0.457 | -5.93 | 0.10412 | none |
| CHGA | 0.126 | 0.15 | 1.1275 | 0.28679 | 0.457 | -5.93 | 0.83816 | none |
| MYO7A | -0.338 | -0.2935 | -1.127 | 0.28687 | 0.457 | -5.93 | -0.4529 | none |
| ABI3BP | 0.304 | 0.2629 | 1.1272 | 0.28691 | 0.457 | -5.93 | -0.1567 | none |
| MPP3 | -0.132 | -0.1137 | -1.127 | 0.28696 | 0.457 | -5.93 | 1.57831 | none |
| RALYL | -0.194 | -0.1947 | -1.127 | 0.287 | 0.457 | -5.93 | -0.1806 | none |
| KCNAB2 | 0.192 | 0.2748 | 1.1269 | 0.28703 | 0.457 | -5.93 | 0.74744 | none |
| DOK5 | 0.17 | 0.045 | 1.1268 | 0.28705 | 0.457 | -5.93 | 0.27963 | none |
| PPP1R7 | 0.195 | 0.2025 | 1.1266 | 0.28715 | 0.457 | -5.93 | -0.8181 | none |
| HLA-DQB1 | 0.134 | 0.0628 | 1.1266 | 0.28715 | 0.457 | -5.93 | 0.32075 | none |
| FAXC | 0.125 | 0.079 | 1.1265 | 0.28719 | 0.457 | -5.93 | -0.8507 | none |
| AKAP8L | 0.237 | 0.36 | 1.1262 | 0.28728 | 0.457 | -5.93 | -0.0384 | none |
| LAS1L | 0.151 | 0.2112 | 1.1261 | 0.28732 | 0.457 | -5.93 | 0.09312 | none |
| PRSS45 | -0.091 | -0.0803 | -1.126 | 0.28753 | 0.458 | -5.93 | 0.34552 | none |
| LTN1 | 0.183 | 0.1771 | 1.1255 | 0.28757 | 0.458 | -5.93 | 1.20516 | none |
| CASP8AP2 | 0.301 | 0.3386 | 1.1252 | 0.2877 | 0.458 | -5.93 | 0.06758 | none |
| TMEM110 | 0.146 | 0.0722 | 1.1251 | 0.28774 | 0.458 | -5.93 | 1.05873 | none |
| XXYLT1 | 0.181 | 0.2173 | 1.1248 | 0.28784 | 0.458 | -5.93 | 0.89108 | none |
| MAGED4 | -0.152 | -0.0393 | -1.125 | 0.28797 | 0.458 | -5.93 | 1.25099 | none |
| MRPS7 | -0.115 | -0.1087 | -1.124 | 0.28804 | 0.458 | -5.93 | 0.10657 | none |
| ORC1 | 0.204 | 0.2411 | 1.1243 | 0.28808 | 0.458 | -5.93 | 0.0836 | none |
| LONRF1 | 0.157 | 0.1208 | 1.1234 | 0.28844 | 0.459 | -5.93 | -0.7947 | none |
| PHIP | 0.275 | 0.3199 | 1.1226 | 0.28876 | 0.459 | -5.93 | 0.31777 | none |
| ZBED6 | 0.208 | 0.2731 | 1.1223 | 0.28886 | 0.459 | -5.93 | -0.2543 | none |
| AC008622 | 0.21 | 0.2104 | 1.1222 | 0.28892 | 0.46 | -5.93 | 1.19688 | none |
| GNRH2 | 0.108 | 0.1264 | 1.1219 | 0.28903 | 0.46 | -5.93 | 0.09752 | none |
| LGR4 | 0.093 | 0.0545 | 1.1218 | 0.28906 | 0.46 | -5.93 | 1.18659 | none |
| IRF1 | 0.14 | 0.1067 | 1.1214 | 0.28923 | 0.46 | -5.93 | -0.093 | none |
| USP18 | -0.131 | -0.1686 | -1.121 | 0.2893 | 0.46 | -5.93 | 1.04487 | none |
| FUNDC1 | 0.121 | 0.0277 | 1.1209 | 0.28944 | 0.46 | -5.93 | 0.88443 | none |
| SGMS1 | 0.169 | 0.1103 | 1.1205 | 0.28961 | 0.46 | -5.94 | -0.2557 | none |
| FLNA | 0.361 | 0.5643 | 1.1204 | 0.28966 | 0.46 | -5.94 | -0.0349 | none |
| ERMARD | 0.283 | 0.2794 | 1.1202 | 0.28972 | 0.46 | -5.94 | -0.2325 | none |
| PRKCI | 0.157 | 0.1236 | 1.1202 | 0.28974 | 0.46 | -5.94 | 1.24566 | none |
| RLN2 | -0.296 | -0.2856 | -1.12 | 0.28978 | 0.46 | -5.94 | 1.63853 | none |
| IL2 | -0.212 | -0.2432 | -1.12 | 0.28984 | 0.46 | -5.94 | 0.88602 | none |
| OSER1 | 0.076 | -0.0032 | 1.1198 | 0.28986 | 0.46 | -5.94 | -0.2013 | none |
| CCDC96 | 0.111 | 0.1123 | 1.1197 | 0.28991 | 0.46 | -5.94 | 0.20499 | none |

| | | | | | | | | |
|---|---|---|---|---|---|---|---|---|
| CIDEC | 0.101 | 0.0971 | 1.1197 | 0.28994 | 0.46 | -5.94 | 0.60614 | none |
| SAC3D1 | -0.166 | -0.1672 | -1.12 | 0.28995 | 0.46 | -5.94 | -0.2121 | none |
| TMEM54 | 0.104 | 0.18 | 1.1196 | 0.28996 | 0.46 | -5.94 | 1.11078 | none |
| CAPS2 | -0.163 | -0.171 | -1.12 | 0.28996 | 0.46 | -5.94 | 0.88294 | none |
| IFITM1 | -0.185 | -0.283 | -1.12 | 0.28999 | 0.46 | -5.94 | 0.94325 | none |
| OVOL1 | -0.08 | -0.0516 | -1.119 | 0.29002 | 0.46 | -5.94 | -0.768 | none |
| CTR9 | 0.29 | 0.3419 | 1.1188 | 0.2903 | 0.461 | -5.94 | 0.18128 | none |
| SIM1 | 0.085 | 0.0725 | 1.1187 | 0.29031 | 0.461 | -5.94 | -0.2329 | none |
| DDX5 | 0.153 | 0.1921 | 1.1185 | 0.29039 | 0.461 | -5.94 | 1.32188 | none |
| ARHGEF19 | 0.131 | 0.1353 | 1.1185 | 0.29041 | 0.461 | -5.94 | 0.28923 | none |
| ASMTL | 0.138 | 0.1288 | 1.1171 | 0.29097 | 0.462 | -5.94 | 0.05406 | none |
| RBM11 | 0.137 | 0.0801 | 1.117 | 0.29101 | 0.462 | -5.94 | 0.19912 | none |
| KCNS3 | -0.133 | -0.1373 | -1.117 | 0.29101 | 0.462 | -5.94 | 0.50269 | none |
| CNBP | -0.093 | -0.0859 | -1.116 | 0.29123 | 0.462 | -5.94 | 1.07005 | none |
| FAM25G | -0.19 | -0.1897 | -1.116 | 0.29128 | 0.462 | -5.94 | 0.38565 | none |
| CDC42BPA | 0.133 | 0.1411 | 1.1162 | 0.29136 | 0.462 | -5.94 | 0.55659 | none |
| SLC8A2 | 0.215 | 0.228 | 1.1159 | 0.29145 | 0.462 | -5.94 | 0.0079 | none |
| MMP28 | 0.088 | 0.085 | 1.1159 | 0.29148 | 0.462 | -5.94 | 0.05739 | none |
| OCM2 | 0.2 | 0.2002 | 1.1154 | 0.29166 | 0.462 | -5.94 | -0.7522 | none |
| HPF1 | -0.118 | -0.1608 | -1.115 | 0.29169 | 0.462 | -5.94 | -0.0523 | none |
| DEFB128 | -0.188 | -0.1881 | -1.115 | 0.29177 | 0.463 | -5.94 | 1.03077 | none |
| CHM | 0.197 | 0.1876 | 1.115 | 0.29182 | 0.463 | -5.94 | 0.40297 | none |
| ANO4 | -0.122 | -0.0654 | -1.115 | 0.29185 | 0.463 | -5.94 | 1.36794 | none |
| EDDM3B | 0.081 | 0.0814 | 1.1146 | 0.29199 | 0.463 | -5.94 | -0.7502 | none |
| MDH1B | -0.113 | -0.0672 | -1.115 | 0.29201 | 0.463 | -5.94 | 0.16016 | none |
| TRAIP | 0.266 | 0.2681 | 1.1144 | 0.29207 | 0.463 | -5.94 | 0.18733 | none |
| CYCS | -0.158 | -0.1788 | -1.114 | 0.29207 | 0.463 | -5.94 | 1.81729 | none |
| TMEM26 | -0.102 | -0.1019 | -1.114 | 0.29209 | 0.463 | -5.94 | 1.94783 | none |
| HIST1H2B( | -0.189 | -0.2523 | -1.114 | 0.29212 | 0.463 | -5.94 | 0.55986 | none |
| HIGD1A | -0.152 | -0.2179 | -1.114 | 0.2922 | 0.463 | -5.94 | 0.79229 | none |
| ADNP2 | 0.089 | 0.0808 | 1.1141 | 0.29221 | 0.463 | -5.94 | 0.67942 | none |
| ZCCHC3 | 0.133 | 0.064 | 1.1141 | 0.29222 | 0.463 | -5.94 | 0.00151 | none |
| SCARA5 | 0.079 | 0.0786 | 1.1139 | 0.29228 | 0.463 | -5.94 | 0.80665 | none |
| ESCO2 | 0.267 | 0.2697 | 1.1138 | 0.29233 | 0.463 | -5.94 | 0.58941 | none |
| HOXB3 | 0.284 | 0.3094 | 1.1136 | 0.29239 | 0.463 | -5.94 | 0.44552 | none |
| RIMKLA | 0.163 | 0.0778 | 1.1136 | 0.2924 | 0.463 | -5.94 | 0.61047 | none |
| SRPK2 | -0.156 | -0.0856 | -1.113 | 0.29254 | 0.463 | -5.94 | 0.12557 | none |
| MMAA | -0.334 | -0.2912 | -1.113 | 0.2927 | 0.463 | -5.94 | 1.27436 | none |
| PPP1R10 | 0.556 | 0.5382 | 1.1128 | 0.29272 | 0.463 | -5.94 | 0.46168 | none |
| PALLD | -0.25 | -0.073 | -1.113 | 0.2928 | 0.463 | -5.94 | 0.12811 | none |

| | | | | | | | | |
|---|---|---|---|---|---|---|---|---|
| ZNF48 | 0.168 | 0.2185 | 1.1125 | 0.29283 | 0.463 | -5.94 | 0.44481 | none |
| C2ORF72 | -0.111 | -0.1125 | -1.112 | 0.2931 | 0.464 | -5.94 | 1.58819 | none |
| BMS1 | 0.327 | 0.4751 | 1.1118 | 0.29315 | 0.464 | -5.94 | -0.0188 | none |
| LYL1 | 0.14 | 0.1249 | 1.1114 | 0.29331 | 0.464 | -5.94 | -0.1492 | none |
| NSUN5 | -0.121 | -0.09 | -1.111 | 0.29335 | 0.464 | -5.95 | 1.32223 | none |
| NOA1 | -0.09 | -0.0478 | -1.111 | 0.29342 | 0.464 | -5.95 | 0.82414 | none |
| FAM19A3 | 0.08 | 0.0875 | 1.111 | 0.29349 | 0.464 | -5.95 | -0.355 | none |
| LEMD1 | 0.104 | 0.0997 | 1.1108 | 0.29357 | 0.464 | -5.95 | -0.0738 | none |
| KIF21A | 0.3 | 0.4631 | 1.1106 | 0.29364 | 0.464 | -5.95 | 0.08296 | none |
| PRKAR2B | 0.082 | 0.0589 | 1.1103 | 0.29376 | 0.464 | -5.95 | -0.4987 | none |
| DNAJA1 | -0.123 | -0.0818 | -1.11 | 0.29395 | 0.465 | -5.95 | 1.42125 | none |
| DYNC1I2 | -0.136 | -0.1023 | -1.11 | 0.29405 | 0.465 | -5.95 | 1.35964 | none |
| OR8D1 | -0.086 | -0.0477 | -1.11 | 0.29408 | 0.465 | -5.95 | 0.22522 | none |
| TAC1 | 0.096 | 0.1282 | 1.1095 | 0.29409 | 0.465 | -5.95 | 0.11709 | none |
| EEF1D | -0.136 | -0.0928 | -1.109 | 0.29411 | 0.465 | -5.95 | 0.95845 | none |
| ZNF232 | 0.149 | 0.1496 | 1.1094 | 0.29412 | 0.465 | -5.95 | -0.143 | none |
| LTC4S | -0.113 | -0.1196 | -1.109 | 0.29424 | 0.465 | -5.95 | 0.23576 | none |
| DCP1B | 0.165 | 0.1544 | 1.1089 | 0.29432 | 0.465 | -5.95 | 0.89724 | none |
| IGSF10 | 0.114 | 0.1144 | 1.1089 | 0.29434 | 0.465 | -5.95 | 1.42493 | none |
| MRPL36 | -0.137 | -0.1381 | -1.109 | 0.29434 | 0.465 | -5.95 | 1.00354 | none |
| SPINK2 | 0.232 | 0.161 | 1.1087 | 0.29439 | 0.465 | -5.95 | 0.70457 | none |
| PDHA1 | 0.094 | 0.1254 | 1.1082 | 0.29461 | 0.465 | -5.95 | 1.0324 | none |
| LBR | 0.138 | 0.0567 | 1.1082 | 0.29463 | 0.465 | -5.95 | 0.94694 | none |
| PIP4K2B | 0.179 | 0.1832 | 1.1077 | 0.29482 | 0.466 | -5.95 | -0.201 | none |
| FAM83H | 0.221 | 0.1442 | 1.1075 | 0.29491 | 0.466 | -5.95 | -0.8581 | none |
| FAS | 0.112 | 0.0567 | 1.1073 | 0.29498 | 0.466 | -5.95 | 0.37022 | none |
| MKKS | -0.152 | -0.1329 | -1.107 | 0.2952 | 0.466 | -5.95 | 1.18528 | none |
| ALOX5 | -0.084 | -0.0947 | -1.106 | 0.29535 | 0.466 | -5.95 | -0.0359 | none |
| ZNF436 | 0.085 | 0.0961 | 1.1059 | 0.29554 | 0.466 | -5.95 | -0.3281 | none |
| SLC2A9 | 0.179 | 0.2726 | 1.1059 | 0.29555 | 0.466 | -5.95 | 0.82954 | none |
| RAET1L | 0.102 | 0.1016 | 1.1059 | 0.29558 | 0.466 | -5.95 | -0.2435 | none |
| C7ORF31 | -0.154 | -0.1006 | -1.106 | 0.29559 | 0.466 | -5.95 | 1.29966 | none |
| EFNA1 | 0.157 | 0.1473 | 1.1051 | 0.29588 | 0.467 | -5.95 | 0.03763 | none |
| SPANXN2 | -0.1 | -0.0927 | -1.105 | 0.29605 | 0.467 | -5.95 | 1.00702 | none |
| TMEM42 | 0.181 | 0.1256 | 1.1047 | 0.29607 | 0.467 | -5.95 | 1.14656 | none |
| RWDD4 | -0.133 | -0.0916 | -1.105 | 0.29608 | 0.467 | -5.95 | 0.27365 | none |
| PEG10 | -0.095 | -0.0648 | -1.104 | 0.29618 | 0.467 | -5.95 | 1.50828 | none |
| B4GAT1 | -0.161 | -0.1117 | -1.104 | 0.29638 | 0.467 | -5.95 | 1.72701 | none |
| KTI12 | 0.116 | 0.0772 | 1.1039 | 0.29638 | 0.467 | -5.95 | -0.4158 | none |
| SLC38A9 | -0.244 | -0.2348 | -1.104 | 0.29639 | 0.467 | -5.95 | 1.04754 | none |

| | | | | | | | | |
|---|---|---|---|---|---|---|---|---|
| PRR15L | 0.081 | 0.0831 | 1.1039 | 0.29639 | 0.467 | -5.95 | -0.3156 | none |
| AIFM3 | -0.09 | -0.0252 | -1.104 | 0.29642 | 0.467 | -5.95 | 0.58156 | none |
| KLHDC9 | -0.208 | -0.2018 | -1.104 | 0.29644 | 0.467 | -5.95 | 0.76012 | none |
| GALM | 0.151 | 0.1384 | 1.1036 | 0.29652 | 0.467 | -5.95 | -0.7363 | none |
| SREK1IP1 | 0.2 | 0.1252 | 1.1035 | 0.29654 | 0.467 | -5.95 | 0.27296 | none |
| CNOT7 | -0.146 | -0.1613 | -1.104 | 0.29655 | 0.467 | -5.95 | 1.98457 | none |
| GNAS | 0.3 | 0.4004 | 1.1034 | 0.29659 | 0.467 | -5.95 | -0.6888 | none |
| APOBEC4 | 0.107 | 0.1072 | 1.1029 | 0.29678 | 0.468 | -5.95 | 0.68084 | none |
| FOXL2 | 0.082 | 0.0557 | 1.1026 | 0.29691 | 0.468 | -5.95 | -1.215 | none |
| AC110602 | -0.198 | -0.1134 | -1.102 | 0.29697 | 0.468 | -5.95 | 1.44705 | none |
| GPCPD1 | 0.27 | 0.2536 | 1.1025 | 0.29698 | 0.468 | -5.95 | -0.6468 | none |
| PROB1 | 0.173 | 0.2227 | 1.1024 | 0.29702 | 0.468 | -5.95 | -0.653 | none |
| TBC1D3I | -0.193 | -0.1878 | -1.102 | 0.29702 | 0.468 | -5.95 | 0.51185 | none |
| THAP2 | 0.174 | 0.1306 | 1.1024 | 0.29702 | 0.468 | -5.95 | 1.27073 | none |
| CDC25B | 0.136 | 0.0982 | 1.1023 | 0.29704 | 0.468 | -5.95 | -0.6496 | none |
| GSG1L | 0.09 | 0.043 | 1.102 | 0.29716 | 0.468 | -5.95 | -0.6895 | none |
| MRPL57 | 0.152 | 0.0759 | 1.1017 | 0.29731 | 0.468 | -5.96 | -0.5118 | none |
| USP50 | -0.085 | -0.051 | -1.102 | 0.29731 | 0.468 | -5.96 | -0.4621 | none |
| NCDN | 0.131 | 0.2117 | 1.1017 | 0.29731 | 0.468 | -5.96 | 0.79097 | none |
| SYF2 | 0.135 | 0.1054 | 1.1014 | 0.2974 | 0.468 | -5.96 | -0.8216 | none |
| ZNF484 | 0.156 | 0.1652 | 1.1011 | 0.29756 | 0.468 | -5.96 | -0.7463 | none |
| MMD2 | 0.137 | 0.1367 | 1.101 | 0.29759 | 0.468 | -5.96 | 0.91134 | none |
| KIAA1217 | 0.114 | 0.1205 | 1.101 | 0.29761 | 0.468 | -5.96 | 1.26087 | none |
| ROM1 | 0.46 | 0.3253 | 1.1007 | 0.2977 | 0.468 | -5.96 | -0.7027 | none |
| TNR | 0.079 | 0.0503 | 1.1003 | 0.29787 | 0.469 | -5.96 | 0 | none |
| OR8S1 | -0.079 | -0.0775 | -1.1 | 0.29791 | 0.469 | -5.96 | -0.1107 | none |
| MSANTD2 | -0.108 | -0.1195 | -1.1 | 0.29794 | 0.469 | -5.96 | 1.59791 | none |
| UBE2M | -0.158 | -0.1215 | -1.1 | 0.29813 | 0.469 | -5.96 | 1.0626 | none |
| ZNF705A | -0.128 | -0.1399 | -1.1 | 0.29819 | 0.469 | -5.96 | 1.48163 | none |
| TIMM10B | 0.09 | 0.0341 | 1.0995 | 0.29823 | 0.469 | -5.96 | -0.0553 | none |
| NOL12 | 0.127 | 0.1597 | 1.0992 | 0.29835 | 0.469 | -5.96 | -0.0688 | none |
| CAMSAP2 | 0.187 | 0.248 | 1.099 | 0.29842 | 0.469 | -5.96 | -0.4662 | none |
| DCD | 0.207 | 0.1059 | 1.0989 | 0.29845 | 0.469 | -5.96 | -0.6041 | none |
| PHLDA3 | 0.182 | 0.1623 | 1.0988 | 0.29851 | 0.469 | -5.96 | -0.552 | none |
| AL356585. | -0.08 | -0.0757 | -1.099 | 0.2986 | 0.469 | -5.96 | 1.61434 | none |
| HSP90AB1 | -0.14 | -0.0342 | -1.099 | 0.29861 | 0.469 | -5.96 | 1.78357 | none |
| ACTL8 | -0.083 | -0.0962 | -1.098 | 0.29869 | 0.469 | -5.96 | 1.42652 | none |
| FZD7 | 0.073 | 0.0387 | 1.0982 | 0.29874 | 0.469 | -5.96 | -0.8559 | none |
| LIPF | -0.169 | -0.169 | -1.098 | 0.29875 | 0.469 | -5.96 | 0.82838 | none |
| DUSP15 | -0.114 | -0.0948 | -1.098 | 0.29875 | 0.469 | -5.96 | 0.87388 | none |

| | | | | | | | | |
|---|---|---|---|---|---|---|---|---|
| NECAB2 | -0.157 | -0.133 | -1.098 | 0.29879 | 0.469 | -5.96 | 0.47239 | none |
| FAM214B | 0.106 | 0.0966 | 1.0978 | 0.29893 | 0.469 | -5.96 | -0.0648 | none |
| CKMT1B | 0.112 | 0.111 | 1.0978 | 0.29893 | 0.469 | -5.96 | -0.0328 | none |
| C1QTNF3-. | 0.094 | 0.0636 | 1.0977 | 0.29894 | 0.469 | -5.96 | 0.18261 | none |
| RBM33 | 0.171 | 0.2535 | 1.0977 | 0.29895 | 0.469 | -5.96 | 0.77978 | none |
| TECPR2 | 0.195 | 0.2431 | 1.0976 | 0.29898 | 0.469 | -5.96 | 0.6444 | none |
| CCDC149 | 0.16 | 0.0955 | 1.0976 | 0.29898 | 0.469 | -5.96 | -0.8416 | none |
| P2RY8 | 0.211 | 0.2342 | 1.0975 | 0.29904 | 0.469 | -5.96 | 0.18081 | none |
| BBS5 | 0.232 | 0.2739 | 1.0974 | 0.29909 | 0.469 | -5.96 | -0.0262 | none |
| HLA-DRB5 | -0.104 | -0.0894 | -1.097 | 0.29913 | 0.469 | -5.96 | 1.07978 | none |
| FAM168B | 0.102 | 0.117 | 1.0971 | 0.2992 | 0.469 | -5.96 | 0.52782 | none |
| C5ORF67 | 0.131 | 0.1354 | 1.0971 | 0.29921 | 0.469 | -5.96 | -0.0563 | none |
| COL21A1 | -0.08 | -0.0533 | -1.097 | 0.29929 | 0.469 | -5.96 | -0.2506 | none |
| IDI2 | -0.084 | -0.0446 | -1.097 | 0.2993 | 0.469 | -5.96 | 2.09167 | AA-related |
| DGKG | -0.238 | -0.194 | -1.097 | 0.29931 | 0.469 | -5.96 | 1.44392 | none |
| ARFGAP1 | 0.134 | 0.1715 | 1.0967 | 0.29937 | 0.47 | -5.96 | 0.9951 | none |
| PDGFRA | 0.317 | 0.3476 | 1.0967 | 0.29938 | 0.47 | -5.96 | -0.0261 | none |
| WDR77 | -0.149 | -0.1275 | -1.097 | 0.29945 | 0.47 | -5.96 | 1.85127 | none |
| BPIFB4 | -0.101 | -0.1013 | -1.096 | 0.29946 | 0.47 | -5.96 | 1.76804 | none |
| SH2D5 | -0.104 | -0.0961 | -1.096 | 0.2996 | 0.47 | -5.96 | 0.33608 | none |
| CNFN | -0.085 | -0.0629 | -1.096 | 0.29965 | 0.47 | -5.96 | 0.53257 | none |
| SPINT4 | 0.177 | 0.1765 | 1.096 | 0.29967 | 0.47 | -5.96 | 1.19688 | none |
| IMPACT | 0.118 | 0.0688 | 1.0959 | 0.29971 | 0.47 | -5.96 | -0.2846 | none |
| DDX52 | 0.215 | 0.1882 | 1.0957 | 0.29979 | 0.47 | -5.96 | 0.25178 | none |
| SENP3-EIF | 0.16 | 0.2053 | 1.0954 | 0.29991 | 0.47 | -5.96 | 0.18584 | none |
| EPS8L3 | -0.114 | -0.1082 | -1.095 | 0.29995 | 0.47 | -5.96 | -0.3991 | none |
| NLRP3 | 0.13 | 0.0989 | 1.0953 | 0.29996 | 0.47 | -5.96 | 0.81164 | none |
| DHX36 | 0.243 | 0.2538 | 1.0951 | 0.30003 | 0.47 | -5.96 | -0.074 | none |
| EGR2 | 0.128 | 0.2002 | 1.095 | 0.3001 | 0.47 | -5.96 | 0.38029 | none |
| SEMA3G | 0.139 | 0.1329 | 1.0947 | 0.30021 | 0.47 | -5.96 | -1.177 | none |
| PTPRC | 0.289 | 0.359 | 1.0944 | 0.30033 | 0.47 | -5.96 | 0.31892 | none |
| RAB41 | -0.149 | -0.1082 | -1.094 | 0.30038 | 0.47 | -5.96 | 1.38746 | none |
| HEXB | -0.154 | -0.198 | -1.094 | 0.30046 | 0.471 | -5.96 | 2.0378 | none |
| ZBED4 | 0.159 | 0.2258 | 1.0938 | 0.30057 | 0.471 | -5.96 | -0.5732 | none |
| SCML2 | 0.235 | 0.1787 | 1.0936 | 0.30066 | 0.471 | -5.96 | -0.9295 | none |
| ANXA10 | 0.099 | 0.1081 | 1.0929 | 0.30094 | 0.471 | -5.96 | 0.84925 | none |
| RAD51D | 0.157 | 0.1363 | 1.0928 | 0.301 | 0.471 | -5.96 | -0.6458 | none |
| CAPNS1 | 0.157 | 0.1119 | 1.0926 | 0.30108 | 0.471 | -5.96 | -0.8047 | none |
| FGFBP2 | 0.11 | 0.1119 | 1.0926 | 0.3011 | 0.471 | -5.96 | 1.14804 | none |
| SCAND1 | -0.17 | -0.2195 | -1.092 | 0.30118 | 0.471 | -5.97 | 1.56751 | none |

| | | | | | | | | |
|---|---|---|---|---|---|---|---|---|
| WDCP | -0.173 | -0.1286 | -1.092 | 0.30132 | 0.471 | -5.97 | 0.31273 | none |
| TRMT10A | -0.209 | -0.205 | -1.092 | 0.30153 | 0.472 | -5.97 | 0.17476 | none |
| LACC1 | 0.136 | 0.1581 | 1.0915 | 0.30153 | 0.472 | -5.97 | 0.00761 | none |
| MMP13 | 0.09 | 0.0657 | 1.0913 | 0.30161 | 0.472 | -5.97 | -0.6001 | none |
| HSDL1 | 0.133 | 0.0409 | 1.091 | 0.30177 | 0.472 | -5.97 | 0.5869 | none |
| IL18R1 | 0.202 | 0.2424 | 1.0909 | 0.30178 | 0.472 | -5.97 | 0.18526 | none |
| FNDC1 | -0.079 | -0.0839 | -1.091 | 0.30184 | 0.472 | -5.97 | -0.0473 | none |
| ANKRD20/ | 0.164 | 0.1278 | 1.0908 | 0.30185 | 0.472 | -5.97 | 0.64284 | none |
| MFN2 | 0.135 | 0.2032 | 1.0907 | 0.30187 | 0.472 | -5.97 | 0.61451 | none |
| CD274 | -0.169 | -0.1726 | -1.091 | 0.30189 | 0.472 | -5.97 | 0.56538 | none |
| CPB1 | 0.783 | 0.7166 | 1.0902 | 0.30209 | 0.472 | -5.97 | -0.2537 | none |
| VKORC1 | -0.179 | -0.2094 | -1.09 | 0.30219 | 0.472 | -5.97 | 0.88039 | none |
| PMCH | -0.138 | -0.1998 | -1.09 | 0.30235 | 0.473 | -5.97 | 0.71898 | none |
| EBAG9 | -0.102 | -0.123 | -1.09 | 0.30235 | 0.473 | -5.97 | 1.71014 | none |
| NXT2 | -0.171 | -0.2216 | -1.089 | 0.30259 | 0.473 | -5.97 | 1.20103 | none |
| COPZ2 | -0.204 | -0.1959 | -1.089 | 0.30264 | 0.473 | -5.97 | 0.17845 | none |
| TASP1 | -0.111 | -0.0986 | -1.089 | 0.30266 | 0.473 | -5.97 | -0.3773 | none |
| ACY3 | -0.095 | -0.085 | -1.089 | 0.30276 | 0.473 | -5.97 | 0.57399 | none |
| USP13 | 0.196 | 0.2213 | 1.0879 | 0.30304 | 0.473 | -5.97 | 0.6019 | none |
| OLFML3 | 0.087 | 0.076 | 1.0875 | 0.30322 | 0.474 | -5.97 | -0.9506 | none |
| ENPP4 | 0.168 | 0.104 | 1.0871 | 0.30339 | 0.474 | -5.97 | -0.1898 | none |
| CLN8 | 0.17 | 0.1265 | 1.087 | 0.30344 | 0.474 | -5.97 | 0.99749 | none |
| ART5 | -0.089 | -0.08 | -1.087 | 0.30356 | 0.474 | -5.97 | -0.5832 | none |
| TBL3 | 0.122 | 0.1728 | 1.0867 | 0.30357 | 0.474 | -5.97 | 0.38247 | none |
| DEFB115 | 0.167 | 0.1673 | 1.0865 | 0.30364 | 0.474 | -5.97 | 0.44852 | none |
| MYNN | 0.127 | 0.0854 | 1.0864 | 0.30368 | 0.474 | -5.97 | -0.3699 | none |
| ZNF559-ZN | 0.243 | 0.2188 | 1.0862 | 0.30378 | 0.474 | -5.97 | 1.55737 | none |
| KHDRBS1 | 0.144 | 0.187 | 1.0858 | 0.30393 | 0.474 | -5.97 | 0.28446 | none |
| PQLC2L | -0.078 | -0.0782 | -1.086 | 0.30402 | 0.474 | -5.97 | -0.3985 | none |
| HNRNPLL | -0.1 | -0.0766 | -1.085 | 0.3041 | 0.475 | -5.97 | 1.17516 | none |
| KLHL18 | 0.14 | 0.1682 | 1.0853 | 0.30415 | 0.475 | -5.97 | 0.53656 | none |
| CDC14A | 0.167 | 0.1696 | 1.0852 | 0.30417 | 0.475 | -5.97 | 0.66136 | none |
| HDAC7 | 0.181 | 0.2674 | 1.0852 | 0.30418 | 0.475 | -5.97 | -0.3565 | none |
| LIPA | 0.1 | -0.0035 | 1.0848 | 0.30436 | 0.475 | -5.97 | -0.4962 | none |
| SAP25 | 0.272 | 0.2493 | 1.0846 | 0.30445 | 0.475 | -5.97 | -0.4762 | none |
| PNLIP | 0.085 | 0.0848 | 1.0845 | 0.30447 | 0.475 | -5.97 | 1.28988 | none |
| VPREB3 | -0.097 | -0.086 | -1.084 | 0.30454 | 0.475 | -5.97 | 0.67267 | none |
| GPBAR1 | 0.086 | 0.0766 | 1.0842 | 0.30461 | 0.475 | -5.97 | -0.5706 | none |
| CCSER1 | 0.175 | 0.1784 | 1.0839 | 0.30474 | 0.475 | -5.97 | -0.4746 | none |
| PODXL2 | 0.117 | 0.1265 | 1.0835 | 0.30489 | 0.475 | -5.97 | 0.91244 | none |

| | | | | | | | | |
|---|---|---|---|---|---|---|---|---|
| OR5M3 | -0.077 | -0.0528 | -1.083 | 0.30513 | 0.476 | -5.97 | 1.75388 | none |
| 5-Sep | -0.251 | -0.0778 | -1.083 | 0.30518 | 0.476 | -5.97 | 0.00795 | none |
| ZNF28 | 0.194 | 0.1003 | 1.0826 | 0.3053 | 0.476 | -5.98 | 0.88794 | none |
| MZT1 | -0.206 | -0.2588 | -1.083 | 0.3053 | 0.476 | -5.98 | 1.25297 | none |
| C20ORF27 | -0.146 | -0.1391 | -1.082 | 0.30534 | 0.476 | -5.98 | 1.80044 | none |
| C8B | 0.164 | 0.1636 | 1.0823 | 0.3054 | 0.476 | -5.98 | 1.02049 | none |
| TCF19 | -0.347 | -0.2095 | -1.082 | 0.30545 | 0.476 | -5.98 | 1.63335 | none |
| AP3D1 | 0.347 | 0.5407 | 1.082 | 0.30552 | 0.476 | -5.98 | -0.2789 | none |
| MFAP3L | 0.154 | 0.1689 | 1.0806 | 0.30613 | 0.477 | -5.98 | 0.37969 | none |
| GALNT12 | -0.149 | -0.1658 | -1.08 | 0.30632 | 0.477 | -5.98 | 0.64095 | none |
| AGR2 | 0.13 | 0.078 | 1.08 | 0.3064 | 0.477 | -5.98 | 0.42263 | none |
| ASPDH | -0.136 | -0.0901 | -1.08 | 0.30642 | 0.477 | -5.98 | 0.24702 | none |
| TMEM56-I | -0.088 | -0.0097 | -1.08 | 0.30647 | 0.477 | -5.98 | -0.0445 | none |
| MEMO1 | -0.122 | -0.1506 | -1.08 | 0.30648 | 0.477 | -5.98 | 1.46397 | none |
| KRTDAP | -0.153 | -0.153 | -1.08 | 0.30656 | 0.477 | -5.98 | 0.21342 | none |
| PCDHGC4 | -0.098 | -0.1018 | -1.079 | 0.3067 | 0.477 | -5.98 | 1.73467 | none |
| C1ORF43 | -0.146 | -0.2149 | -1.079 | 0.30674 | 0.477 | -5.98 | 1.38779 | none |
| HMBS | -0.155 | -0.1403 | -1.079 | 0.30686 | 0.478 | -5.98 | 1.64107 | none |
| ACP1 | -0.123 | -0.1439 | -1.079 | 0.30698 | 0.478 | -5.98 | 0.09469 | none |
| PLAC8L1 | 0.166 | 0.1009 | 1.0785 | 0.30702 | 0.478 | -5.98 | 0.56752 | none |
| ISL1 | 0.077 | 0.0787 | 1.0783 | 0.3071 | 0.478 | -5.98 | -0.8379 | none |
| CEBPZ | 0.274 | 0.3322 | 1.0782 | 0.30716 | 0.478 | -5.98 | 0.17166 | none |
| C1ORF146 | -0.189 | -0.1692 | -1.078 | 0.30717 | 0.478 | -5.98 | 0.64183 | none |
| TRAM1 | -0.154 | -0.2124 | -1.078 | 0.30732 | 0.478 | -5.98 | 1.48442 | none |
| AKAP17A | 0.297 | 0.4461 | 1.0776 | 0.3074 | 0.478 | -5.98 | 0.51255 | none |
| CC2D2B | 0.301 | 0.2655 | 1.0774 | 0.30748 | 0.478 | -5.98 | 0.60171 | none |
| EIF4B | 0.189 | 0.235 | 1.0773 | 0.30754 | 0.478 | -5.98 | -0.0362 | none |
| ONECUT2 | 0.11 | 0.1058 | 1.0772 | 0.30755 | 0.478 | -5.98 | 0.58568 | none |
| MIB1 | 0.134 | 0.1355 | 1.0771 | 0.30759 | 0.478 | -5.98 | 0.0097 | none |
| ARL13B | 0.247 | 0.2425 | 1.077 | 0.30763 | 0.478 | -5.98 | 0.51965 | none |
| FAM220A | 0.367 | 0.2554 | 1.0769 | 0.30772 | 0.478 | -5.98 | 0.37774 | none |
| TBPL2 | 0.199 | 0.15 | 1.0768 | 0.30775 | 0.478 | -5.98 | -0.6727 | none |
| SALL1 | -0.112 | -0.1066 | -1.077 | 0.30777 | 0.478 | -5.98 | 1.03887 | none |
| DNAJB9 | -0.174 | -0.2073 | -1.077 | 0.30782 | 0.478 | -5.98 | 1.08886 | none |
| COX17 | -0.18 | -0.2009 | -1.076 | 0.30796 | 0.479 | -5.98 | 0.35642 | none |
| 1-Mar | 0.189 | 0.1125 | 1.0759 | 0.3081 | 0.479 | -5.98 | -0.6713 | none |
| AHCTF1 | 0.494 | 0.5556 | 1.0758 | 0.30818 | 0.479 | -5.98 | -0.5756 | none |
| ZNF688 | 0.162 | 0.1118 | 1.0756 | 0.30824 | 0.479 | -5.98 | -0.0829 | none |
| CMAS | -0.099 | -0.0594 | -1.076 | 0.30825 | 0.479 | -5.98 | 1.22498 | none |
| COL23A1 | -0.081 | -0.0542 | -1.075 | 0.30831 | 0.479 | -5.98 | 1.99598 | none |

| | | | | | | | | |
|---|---|---|---|---|---|---|---|---|
| AP1B1 | 0.232 | 0.3296 | 1.0754 | 0.30834 | 0.479 | -5.98 | 0.56094 | none |
| TSPY2 | 0.158 | 0.1578 | 1.0752 | 0.3084 | 0.479 | -5.98 | -0.2399 | none |
| FAM71E1 | -0.125 | -0.1166 | -1.075 | 0.30851 | 0.479 | -5.98 | 1.70359 | none |
| LILRB2 | 0.119 | 0.1231 | 1.0747 | 0.30864 | 0.479 | -5.98 | 0.59894 | none |
| CARHSP1 | 0.184 | 0.083 | 1.074 | 0.30892 | 0.48 | -5.98 | -0.482 | none |
| GPR33 | 0.081 | 0.0806 | 1.074 | 0.30894 | 0.48 | -5.98 | 1.07626 | none |
| ZNF513 | 0.185 | 0.1654 | 1.0739 | 0.30897 | 0.48 | -5.98 | -0.2325 | none |
| USP17L4 | 0.079 | 0.0527 | 1.0736 | 0.30909 | 0.48 | -5.98 | 0.10877 | none |
| CLSTN1 | 0.294 | 0.4187 | 1.0735 | 0.30914 | 0.48 | -5.98 | -0.3251 | none |
| XRCC6 | -0.116 | -0.0423 | -1.073 | 0.30924 | 0.48 | -5.98 | 1.53386 | none |
| GTF3C5 | 0.132 | 0.1464 | 1.0732 | 0.30926 | 0.48 | -5.98 | 1.18921 | none |
| KDM4A | 0.261 | 0.3308 | 1.0732 | 0.30928 | 0.48 | -5.99 | -0.3384 | none |
| ITIH3 | -0.207 | -0.1754 | -1.073 | 0.30931 | 0.48 | -5.99 | 0.62808 | none |
| RFPL3 | -0.103 | -0.0876 | -1.073 | 0.30932 | 0.48 | -5.99 | 0.4433 | none |
| BCAP29 | 0.104 | 0.0389 | 1.0727 | 0.30949 | 0.48 | -5.99 | 0.16221 | none |
| YPEL4 | 0.084 | 0.0862 | 1.0727 | 0.3095 | 0.48 | -5.99 | 0.2166 | none |
| IFRD2 | -0.119 | -0.0743 | -1.073 | 0.3095 | 0.48 | -5.99 | 2.02546 | none |
| AC136616 | 0.156 | 0.1556 | 1.0725 | 0.30958 | 0.48 | -5.99 | 1.4938 | none |
| CD244 | -0.116 | -0.1345 | -1.072 | 0.30973 | 0.48 | -5.99 | 1.11964 | none |
| CCRL2 | -0.14 | -0.1488 | -1.072 | 0.30978 | 0.48 | -5.99 | 0.22462 | none |
| KIF18A | 0.305 | 0.3623 | 1.0719 | 0.30981 | 0.48 | -5.99 | -0.1735 | none |
| CCR1 | -0.104 | -0.1457 | -1.072 | 0.30989 | 0.48 | -5.99 | 0.14924 | none |
| FMO4 | -0.211 | -0.1297 | -1.072 | 0.3099 | 0.48 | -5.99 | 0.82041 | none |
| ZNF721 | 0.214 | 0.1876 | 1.0717 | 0.30991 | 0.48 | -5.99 | 0.09288 | none |
| ENO4 | 0.077 | 0.0551 | 1.0714 | 0.31005 | 0.48 | -5.99 | 0.71267 | none |
| DEFB106A | 0.155 | 0.1548 | 1.0714 | 0.31005 | 0.48 | -5.99 | 0.44852 | none |
| SPANXB1 | 0.155 | 0.1547 | 1.0712 | 0.31012 | 0.481 | -5.99 | 0.66005 | none |
| DCLK3 | 0.137 | 0.1365 | 1.0706 | 0.31038 | 0.481 | -5.99 | 1.22583 | none |
| ZNF416 | -0.114 | -0.1039 | -1.07 | 0.31064 | 0.481 | -5.99 | 2.12563 | AA-related |
| NBPF6 | 0.116 | 0.0743 | 1.0697 | 0.31078 | 0.481 | -5.99 | -0.0169 | none |
| EDARADD | 0.216 | 0.21 | 1.0694 | 0.31089 | 0.482 | -5.99 | -0.5762 | none |
| CYP2E1 | 0.14 | 0.0944 | 1.0694 | 0.31091 | 0.482 | -5.99 | -0.4791 | none |
| GPRC5D | 0.299 | 0.2035 | 1.0693 | 0.31094 | 0.482 | -5.99 | -0.9537 | none |
| PTF1A | 0.153 | 0.1532 | 1.0692 | 0.31096 | 0.482 | -5.99 | 1.19688 | none |
| SF3B2 | 0.233 | 0.3695 | 1.0691 | 0.31104 | 0.482 | -5.99 | -0.1476 | none |
| FAM160A2 | 0.136 | 0.1629 | 1.069 | 0.31107 | 0.482 | -5.99 | -0.1004 | none |
| GSX2 | -0.145 | -0.1454 | -1.069 | 0.31109 | 0.482 | -5.99 | 0.57507 | none |
| GTF2A1L | 0.153 | 0.1528 | 1.0687 | 0.31118 | 0.482 | -5.99 | 0.87018 | none |
| CKAP2 | 0.341 | 0.4197 | 1.0683 | 0.31138 | 0.482 | -5.99 | 0.20086 | none |
| AKR7A2 | -0.149 | -0.1613 | -1.068 | 0.31147 | 0.482 | -5.99 | 1.43261 | none |

| | | | | | | | | |
|---|---|---|---|---|---|---|---|---|
| MFSD3 | 0.116 | 0.089 | 1.0679 | 0.31152 | 0.482 | -5.99 | 0.99589 | none |
| PPL | -0.212 | -0.1373 | -1.068 | 0.31156 | 0.482 | -5.99 | 0.98531 | none |
| RRM1 | 0.163 | 0.1668 | 1.0676 | 0.31165 | 0.482 | -5.99 | 0.18335 | none |
| CYP2D6 | 0.126 | 0.0966 | 1.0673 | 0.31178 | 0.482 | -5.99 | 0.15169 | none |
| TPST2 | -0.095 | -0.1148 | -1.067 | 0.31195 | 0.483 | -5.99 | 1.95643 | none |
| LRRC17 | 0.078 | 0.0896 | 1.0669 | 0.31196 | 0.483 | -5.99 | 1.91741 | none |
| RBCK1 | 0.094 | 0.0749 | 1.0666 | 0.31208 | 0.483 | -5.99 | -0.6169 | none |
| CCDC103 | 0.203 | 0.149 | 1.0666 | 0.31209 | 0.483 | -5.99 | -0.3341 | none |
| SMG9 | 0.175 | 0.3031 | 1.0665 | 0.31215 | 0.483 | -5.99 | -0.394 | none |
| SLC16A5 | 0.405 | 0.3396 | 1.0664 | 0.31217 | 0.483 | -5.99 | -0.74 | none |
| HIST1H2B/ | -0.143 | -0.1434 | -1.066 | 0.31242 | 0.483 | -5.99 | 1.9054 | none |
| FAM111A | 0.246 | 0.2629 | 1.0658 | 0.31244 | 0.483 | -5.99 | 0.33192 | none |
| ANKMY2 | -0.172 | -0.0696 | -1.066 | 0.31246 | 0.483 | -5.99 | 1.72633 | none |
| BGLAP | -0.267 | -0.1541 | -1.065 | 0.31263 | 0.483 | -5.99 | 0.98708 | none |
| CCDC63 | -0.123 | -0.1112 | -1.064 | 0.31305 | 0.484 | -5.99 | -0.2931 | none |
| PLGRKT | -0.16 | -0.2145 | -1.064 | 0.31312 | 0.484 | -5.99 | 0.92525 | none |
| GOLGA8M | -0.142 | -0.1418 | -1.063 | 0.31348 | 0.484 | -6 | 1.9054 | none |
| CCDC78 | 0.117 | 0.1277 | 1.0633 | 0.31353 | 0.484 | -6 | 0.49487 | none |
| TIPARP | -0.163 | -0.1728 | -1.063 | 0.31359 | 0.484 | -6 | 2.01804 | none |
| SYCE3 | -0.17 | -0.1039 | -1.063 | 0.31372 | 0.485 | -6 | 0.56238 | none |
| TNNI3K | 0.139 | 0.1403 | 1.0625 | 0.31388 | 0.485 | -6 | -0.7364 | none |
| PSG7 | -0.166 | -0.0849 | -1.062 | 0.31391 | 0.485 | -6 | 0.1833 | none |
| LENG8 | -0.353 | -0.0805 | -1.062 | 0.31397 | 0.485 | -6 | 1.22896 | none |
| FAIM | 0.14 | 0.0543 | 1.0622 | 0.31401 | 0.485 | -6 | 0.77318 | none |
| SCML1 | -0.19 | -0.1734 | -1.062 | 0.31413 | 0.485 | -6 | 0.89019 | none |
| TTC39A | 0.213 | 0.1198 | 1.0618 | 0.31417 | 0.485 | -6 | -0.1308 | none |
| PVRIG | -0.111 | -0.1238 | -1.062 | 0.31424 | 0.485 | -6 | 0.7644 | none |
| MYH6 | -0.072 | -0.0671 | -1.062 | 0.31427 | 0.485 | -6 | 1.22307 | none |
| CDC14B | 0.235 | 0.1407 | 1.0612 | 0.31442 | 0.485 | -6 | -0.3818 | none |
| TGFB2 | 0.13 | 0.1335 | 1.0608 | 0.31458 | 0.486 | -6 | 0.1498 | none |
| SIRPD | -0.088 | -0.0533 | -1.061 | 0.31465 | 0.486 | -6 | -0.2604 | none |
| ULBP2 | 0.172 | 0.1378 | 1.0606 | 0.31466 | 0.486 | -6 | 0.1682 | none |
| TSC22D4 | -0.09 | 0.0089 | -1.06 | 0.31476 | 0.486 | -6 | 1.10669 | none |
| GPC3 | -0.132 | -0.166 | -1.06 | 0.31481 | 0.486 | -6 | 0.79609 | none |
| WARS2 | -0.147 | -0.1066 | -1.06 | 0.31488 | 0.486 | -6 | 1.58239 | none |
| AQP4 | 0.147 | 0.1468 | 1.06 | 0.31492 | 0.486 | -6 | -0.3407 | none |
| TRIB2 | 0.145 | 0.0311 | 1.0599 | 0.31497 | 0.486 | -6 | -0.9286 | none |
| GPX6 | 0.11 | 0.1205 | 1.0599 | 0.315 | 0.486 | -6 | 0.327 | none |
| FLNC | -0.148 | -0.1437 | -1.059 | 0.31533 | 0.486 | -6 | 0.91949 | none |
| NDUFB5 | -0.171 | -0.2807 | -1.059 | 0.31538 | 0.486 | -6 | 1.32062 | none |

| | | | | | | | | |
|---|---|---|---|---|---|---|---|---|
| TPST1 | 0.211 | 0.1414 | 1.0589 | 0.31543 | 0.486 | -6 | 1.77561 | none |
| AMOTL1 | -0.09 | -0.0572 | -1.059 | 0.31547 | 0.486 | -6 | -0.5832 | none |
| PHF8 | 0.19 | 0.315 | 1.0586 | 0.31557 | 0.486 | -6 | 0.63146 | none |
| SPTA1 | -0.079 | -0.0721 | -1.058 | 0.31578 | 0.487 | -6 | 0.52786 | none |
| SPATA7 | 0.284 | 0.2468 | 1.058 | 0.31582 | 0.487 | -6 | -0.7479 | none |
| WDR74 | 0.078 | 0.0983 | 1.0578 | 0.3159 | 0.487 | -6 | 1.40667 | none |
| TPCN2 | 0.103 | 0.0655 | 1.057 | 0.31624 | 0.487 | -6 | -0.4678 | none |
| CITED4 | -0.096 | -0.0907 | -1.057 | 0.31624 | 0.487 | -6 | 1.0209 | none |
| DYRK1B | 0.133 | 0.1603 | 1.057 | 0.31625 | 0.487 | -6 | -0.074 | none |
| WDR53 | 0.158 | 0.1206 | 1.0569 | 0.31627 | 0.487 | -6 | 0.30669 | none |
| CXCL3 | -0.1 | -0.0791 | -1.057 | 0.31634 | 0.487 | -6 | 2.08918 | none |
| AIRE | 0.137 | 0.107 | 1.0566 | 0.31642 | 0.487 | -6 | -0.4882 | none |
| SEZ6L | -0.099 | -0.0977 | -1.057 | 0.31645 | 0.487 | -6 | 1.51812 | none |
| SSC4D | 0.13 | 0.0961 | 1.0565 | 0.31647 | 0.487 | -6 | 1.06049 | none |
| ACOX3 | 0.085 | 0.1173 | 1.0563 | 0.31656 | 0.487 | -6 | 0.44887 | none |
| SLF1 | 0.168 | 0.1372 | 1.0559 | 0.31672 | 0.488 | -6 | -0.6 | none |
| PRLH | 0.144 | 0.144 | 1.0558 | 0.31675 | 0.488 | -6 | 1.19688 | none |
| DEPDC7 | 0.104 | 0.0585 | 1.0555 | 0.31688 | 0.488 | -6 | -0.0535 | none |
| PAK3 | 0.158 | 0.0834 | 1.0552 | 0.31702 | 0.488 | -6 | -0.4309 | none |
| C10ORF55 | 0.085 | 0.0882 | 1.055 | 0.31711 | 0.488 | -6 | 0.13565 | none |
| LRMP | 0.21 | 0.1218 | 1.055 | 0.31712 | 0.488 | -6 | 0.07225 | none |
| ABL1 | 0.239 | 0.4283 | 1.0547 | 0.31722 | 0.488 | -6 | 0.52698 | none |
| LKAAEAR1 | 0.075 | 0.0756 | 1.0547 | 0.31724 | 0.488 | -6 | 0.67078 | none |
| BHLHE41 | 0.115 | 0.121 | 1.0547 | 0.31725 | 0.488 | -6 | -0.7042 | none |
| KRTAP19-8 | -0.136 | -0.1361 | -1.054 | 0.31758 | 0.489 | -6 | 1.9054 | none |
| PRICKLE4 | 0.235 | 0.1895 | 1.0537 | 0.31768 | 0.489 | -6.01 | -1.1891 | none |
| MAP1LC3E | -0.192 | -0.1715 | -1.054 | 0.31773 | 0.489 | -6.01 | 0.47518 | none |
| NUCKS1 | 0.221 | 0.2528 | 1.0536 | 0.31773 | 0.489 | -6.01 | -0.4605 | none |
| LHFPL2 | -0.171 | -0.0687 | -1.053 | 0.31779 | 0.489 | -6.01 | 1.17288 | none |
| E2F8 | 0.217 | 0.2691 | 1.0534 | 0.31782 | 0.489 | -6.01 | -0.1194 | none |
| HIST1H4C | -0.328 | -0.2681 | -1.053 | 0.31789 | 0.489 | -6.01 | 1.13426 | none |
| GTSE1 | 0.157 | 0.1852 | 1.053 | 0.31797 | 0.489 | -6.01 | -0.6196 | none |
| LRIT1 | 0.084 | 0.0843 | 1.0529 | 0.31803 | 0.489 | -6.01 | 0.75515 | none |
| SPRR1A | 0.142 | 0.142 | 1.0526 | 0.31814 | 0.489 | -6.01 | 1.19688 | none |
| C12ORF56 | -0.109 | -0.0841 | -1.053 | 0.3182 | 0.489 | -6.01 | 1.4843 | none |
| HNRNPAB | -0.116 | -0.024 | -1.052 | 0.31825 | 0.489 | -6.01 | 0.68576 | none |
| RBM12 | 0.113 | 0.1273 | 1.0523 | 0.31828 | 0.489 | -6.01 | -0.2172 | none |
| NFXL1 | 0.198 | 0.2061 | 1.0522 | 0.31833 | 0.489 | -6.01 | 0.10146 | none |
| SYDE1 | -0.163 | -0.0818 | -1.052 | 0.31833 | 0.489 | -6.01 | 0.20251 | none |
| BAG4 | 0.212 | 0.1837 | 1.052 | 0.31841 | 0.489 | -6.01 | -0.4847 | none |

| | | | | | | | | |
|---|---|---|---|---|---|---|---|---|
| CFAP61 | -0.136 | -0.0649 | -1.052 | 0.31861 | 0.489 | -6.01 | -0.0342 | none |
| LCE3D | 0.141 | 0.1413 | 1.0515 | 0.31862 | 0.489 | -6.01 | 1.19688 | none |
| MLST8 | 0.13 | 0.0963 | 1.0511 | 0.3188 | 0.49 | -6.01 | -0.0313 | none |
| WDR75 | 0.136 | 0.1102 | 1.0511 | 0.31882 | 0.49 | -6.01 | 1.28576 | none |
| CPTP | 0.152 | 0.1426 | 1.0509 | 0.3189 | 0.49 | -6.01 | 0.14772 | none |
| RABIF | -0.137 | -0.1756 | -1.051 | 0.31891 | 0.49 | -6.01 | 0.8865 | none |
| RAD17 | 0.304 | 0.1689 | 1.0506 | 0.31901 | 0.49 | -6.01 | -0.9524 | none |
| IFIT5 | 0.103 | 0.0514 | 1.0506 | 0.31904 | 0.49 | -6.01 | 1.08178 | none |
| APOA5 | -0.082 | -0.0586 | -1.05 | 0.31908 | 0.49 | -6.01 | 0.66201 | none |
| GPIHBP1 | 0.069 | 0.0689 | 1.0503 | 0.31915 | 0.49 | -6.01 | 0.65061 | none |
| MRPL4 | 0.11 | 0.0819 | 1.05 | 0.3193 | 0.49 | -6.01 | -0.3319 | none |
| METTL18 | -0.12 | -0.164 | -1.05 | 0.31932 | 0.49 | -6.01 | 0.99819 | none |
| MSLNL | -0.072 | -0.034 | -1.05 | 0.31936 | 0.49 | -6.01 | 1.95957 | none |
| DEFB107A | -0.092 | -0.0519 | -1.05 | 0.31943 | 0.49 | -6.01 | 0.17954 | none |
| DEFB107B | -0.092 | -0.0519 | -1.05 | 0.31943 | 0.49 | -6.01 | 0.17954 | none |
| NXF3 | 0.078 | 0.1083 | 1.0497 | 0.31943 | 0.49 | -6.01 | -0.58 | none |
| GRHL3 | -0.115 | -0.1258 | -1.049 | 0.31956 | 0.49 | -6.01 | 0.63837 | none |
| KRTAP22-1 | 0.14 | 0.14 | 1.0493 | 0.31958 | 0.49 | -6.01 | -0.511 | none |
| GNB2 | -0.231 | -0.1954 | -1.049 | 0.31979 | 0.49 | -6.01 | 0.94654 | none |
| HSD17B12 | 0.162 | 0.0506 | 1.0488 | 0.31981 | 0.49 | -6.01 | -0.3017 | none |
| DEFB123 | 0.14 | 0.1397 | 1.0488 | 0.31983 | 0.49 | -6.01 | 0.44852 | none |
| FABP12 | -0.075 | -0.0502 | -1.049 | 0.31992 | 0.49 | -6.01 | 1.24897 | none |
| HEPACAM | -0.08 | -0.0708 | -1.049 | 0.31994 | 0.49 | -6.01 | 0.49575 | none |
| ZNF24 | -0.147 | -0.0967 | -1.048 | 0.31995 | 0.49 | -6.01 | 1.85106 | none |
| RLF | 0.265 | 0.3404 | 1.0485 | 0.31997 | 0.49 | -6.01 | 0.08349 | none |
| SLC6A15 | 0.098 | 0.1018 | 1.0482 | 0.32006 | 0.491 | -6.01 | 0.63154 | none |
| UCHL1 | 0.213 | 0.2672 | 1.0482 | 0.32006 | 0.491 | -6.01 | 0.46696 | none |
| BATF2 | 0.083 | 0.0722 | 1.0482 | 0.32008 | 0.491 | -6.01 | 0.16655 | none |
| C1GALT1 | 0.127 | 0.0839 | 1.0481 | 0.32011 | 0.491 | -6.01 | 0.63999 | none |
| SPRR3 | -0.133 | -0.1328 | -1.048 | 0.32019 | 0.491 | -6.01 | 1.55345 | none |
| PHOX2B | -0.174 | -0.1736 | -1.048 | 0.32019 | 0.491 | -6.01 | 1.57654 | none |
| NUDT6 | 0.121 | 0.0666 | 1.0475 | 0.32037 | 0.491 | -6.01 | -0.186 | none |
| BCL11B | 0.194 | 0.1774 | 1.0473 | 0.32046 | 0.491 | -6.01 | 0.08143 | none |
| RFX7 | 0.28 | 0.3695 | 1.0472 | 0.32052 | 0.491 | -6.01 | -0.2526 | none |
| CFAP65 | -0.098 | -0.0547 | -1.046 | 0.32087 | 0.491 | -6.01 | 1.06404 | none |
| CRIM1 | 0.236 | 0.3139 | 1.0463 | 0.32091 | 0.491 | -6.01 | 0.67161 | none |
| CRYM | -0.078 | -0.0474 | -1.046 | 0.32093 | 0.491 | -6.01 | 1.34597 | none |
| STK39 | -0.162 | -0.0575 | -1.046 | 0.32116 | 0.492 | -6.01 | 0.12207 | none |
| DNAJC6 | -0.124 | -0.0135 | -1.046 | 0.32119 | 0.492 | -6.01 | 1.68465 | none |
| C2CD5 | 0.241 | 0.2151 | 1.0457 | 0.32119 | 0.492 | -6.01 | -0.3825 | none |

| | | | | | | | | |
|---|---|---|---|---|---|---|---|---|
| DYNC1LI1 | -0.134 | -0.0759 | -1.045 | 0.32127 | 0.492 | -6.01 | 1.98195 | none |
| TRHR | 0.076 | 0.0758 | 1.0453 | 0.32136 | 0.492 | -6.01 | 0.54573 | none |
| INF2 | 0.139 | 0.2045 | 1.0451 | 0.32144 | 0.492 | -6.01 | 0.2081 | none |
| ANXA1 | 0.126 | 0.062 | 1.0449 | 0.32151 | 0.492 | -6.01 | 1.528 | none |
| CACNA1I | 0.159 | 0.161 | 1.0448 | 0.32155 | 0.492 | -6.01 | -0.6978 | none |
| KIF26B | -0.084 | -0.0723 | -1.045 | 0.32158 | 0.492 | -6.01 | -0.3803 | none |
| ZNF837 | 0.101 | 0.118 | 1.0447 | 0.3216 | 0.492 | -6.01 | 0.69791 | none |
| TAF10 | -0.124 | -0.0826 | -1.045 | 0.32164 | 0.492 | -6.01 | 1.7523 | none |
| MTPN | -0.144 | -0.1946 | -1.045 | 0.32169 | 0.492 | -6.01 | 1.23771 | none |
| PAFAH1B3 | -0.136 | -0.1646 | -1.044 | 0.32176 | 0.492 | -6.01 | 0.09084 | none |
| HNRNPCL4 | 0.081 | 0.0701 | 1.0441 | 0.32188 | 0.492 | -6.01 | 0.09135 | none |
| TMEM240 | 0.088 | 0.1004 | 1.044 | 0.32194 | 0.492 | -6.01 | 0.85572 | none |
| JAK2 | 0.209 | 0.3249 | 1.0438 | 0.32202 | 0.492 | -6.02 | 0.83136 | none |
| CDK8 | 0.12 | 0.0883 | 1.0436 | 0.32211 | 0.492 | -6.02 | 1.05444 | none |
| PLIN1 | 0.137 | 0.1526 | 1.0434 | 0.32217 | 0.493 | -6.02 | 0.86691 | none |
| ADAMTS7 | -0.123 | -0.0673 | -1.043 | 0.32223 | 0.493 | -6.02 | -0.2733 | none |
| FOXD2 | 0.077 | 0.0938 | 1.0431 | 0.32233 | 0.493 | -6.02 | -0.3531 | none |
| CSTF1 | 0.093 | -0.0246 | 1.0429 | 0.3224 | 0.493 | -6.02 | -0.2962 | none |
| BBS4 | 0.294 | 0.2735 | 1.0429 | 0.32242 | 0.493 | -6.02 | -0.5029 | none |
| ZNF140 | 0.128 | 0.1078 | 1.0426 | 0.32252 | 0.493 | -6.02 | 0.262 | none |
| HEBP2 | 0.127 | 0.084 | 1.0425 | 0.32258 | 0.493 | -6.02 | 1.52875 | none |
| TET2 | 0.378 | 0.4822 | 1.0423 | 0.32265 | 0.493 | -6.02 | 0.55382 | none |
| MROH9 | -0.071 | -0.0556 | -1.042 | 0.32265 | 0.493 | -6.02 | 0 | none |
| PPP1R17 | 0.087 | 0.0873 | 1.0423 | 0.32266 | 0.493 | -6.02 | -0.7793 | none |
| 3-Mar | 0.19 | 0.1202 | 1.0421 | 0.32277 | 0.493 | -6.02 | -0.4345 | none |
| SPIC | -0.075 | -0.0497 | -1.042 | 0.32281 | 0.493 | -6.02 | 0.24793 | none |
| IER5 | -0.12 | -0.06 | -1.041 | 0.32307 | 0.493 | -6.02 | 0.1799 | none |
| KRTAP22-2 | 0.135 | 0.1352 | 1.041 | 0.32324 | 0.494 | -6.02 | 1.19688 | none |
| PCDHAC2 | -0.129 | -0.1292 | -1.041 | 0.32324 | 0.494 | -6.02 | 1.05226 | none |
| MRPL9 | -0.104 | -0.1319 | -1.041 | 0.32328 | 0.494 | -6.02 | 0.6579 | none |
| KCNJ9 | -0.069 | -0.0416 | -1.041 | 0.32328 | 0.494 | -6.02 | -0.0653 | none |
| POSTN | -0.125 | -0.1257 | -1.04 | 0.32358 | 0.494 | -6.02 | 1.53529 | none |
| AVPI1 | -0.085 | 0.0049 | -1.04 | 0.32368 | 0.494 | -6.02 | 0.83059 | none |
| GSAP | 0.137 | 0.1241 | 1.0399 | 0.32373 | 0.494 | -6.02 | 1.26372 | none |
| HHIPL2 | 0.135 | 0.1171 | 1.0398 | 0.32377 | 0.494 | -6.02 | 0.72611 | none |
| FANCB | 0.114 | 0.1197 | 1.0396 | 0.32388 | 0.494 | -6.02 | 0.91092 | none |
| TPM3 | -0.094 | -0.0447 | -1.039 | 0.32391 | 0.494 | -6.02 | 0.26797 | none |
| DNAJB6 | 0.102 | 0.0801 | 1.0392 | 0.32402 | 0.494 | -6.02 | 0.1424 | none |
| ADIG | -0.128 | -0.1282 | -1.039 | 0.32413 | 0.494 | -6.02 | 0.88916 | none |
| MRPL53 | -0.149 | -0.1655 | -1.039 | 0.32421 | 0.495 | -6.02 | 1.55349 | none |

| | | | | | | | | |
|---|---|---|---|---|---|---|---|---|
| MUC3A | 0.098 | 0.0768 | 1.0387 | 0.32427 | 0.495 | -6.02 | 1.09553 | none |
| ZNF502 | -0.081 | -0.06 | -1.038 | 0.3244 | 0.495 | -6.02 | 0.34409 | none |
| CTSH | -0.2 | -0.1757 | -1.038 | 0.32445 | 0.495 | -6.02 | 1.32482 | none |
| TNFSF11 | 0.17 | 0.0961 | 1.038 | 0.32455 | 0.495 | -6.02 | 0.17586 | none |
| MBD3L2 | 0.134 | 0.1336 | 1.038 | 0.32456 | 0.495 | -6.02 | -0.7798 | none |
| FAM13C | 0.085 | 0.0854 | 1.0376 | 0.32474 | 0.495 | -6.02 | -0.4811 | none |
| COL5A3 | 0.09 | 0.1294 | 1.0375 | 0.3248 | 0.495 | -6.02 | 0.26707 | none |
| ARHGAP4; | 0.081 | 0.0586 | 1.0374 | 0.32483 | 0.495 | -6.02 | -1.247 | none |
| MED16 | 0.164 | 0.1621 | 1.0373 | 0.32487 | 0.495 | -6.02 | 0.38135 | none |
| TNFRSF18 | -0.11 | -0.0985 | -1.037 | 0.32496 | 0.495 | -6.02 | 1.12138 | none |
| KDF1 | 0.107 | 0.0817 | 1.0367 | 0.32513 | 0.495 | -6.02 | -1.1139 | none |
| SLC25A52 | 0.111 | 0.1411 | 1.0367 | 0.32514 | 0.495 | -6.02 | 1.23816 | none |
| HIPK2 | 0.265 | 0.3921 | 1.0366 | 0.32517 | 0.495 | -6.02 | -0.3283 | none |
| RNF38 | 0.128 | 0.0851 | 1.0362 | 0.32538 | 0.496 | -6.02 | -0.3896 | none |
| CPSF6 | 0.118 | 0.1403 | 1.0361 | 0.32539 | 0.496 | -6.02 | -0.0537 | none |
| MFSD10 | -0.121 | -0.1344 | -1.036 | 0.32539 | 0.496 | -6.02 | 1.6341 | none |
| CCR8 | 0.289 | 0.3772 | 1.0361 | 0.32543 | 0.496 | -6.02 | -0.0813 | none |
| PPFIBP2 | 0.327 | 0.4001 | 1.036 | 0.32544 | 0.496 | -6.02 | -0.2093 | none |
| PCBP2 | -0.084 | -0.1065 | -1.036 | 0.32556 | 0.496 | -6.02 | 1.70848 | none |
| NYNRIN | -0.081 | -0.0852 | -1.036 | 0.32567 | 0.496 | -6.02 | -0.7074 | none |
| TRPM1 | -0.094 | -0.0825 | -1.035 | 0.32595 | 0.496 | -6.02 | -0.4408 | none |
| ZNF142 | 0.278 | 0.437 | 1.0348 | 0.326 | 0.496 | -6.02 | 0.24207 | none |
| TMEM88 | -0.153 | -0.1198 | -1.035 | 0.326 | 0.496 | -6.02 | 0.8529 | none |
| TNIP1 | 0.252 | 0.3847 | 1.0346 | 0.32607 | 0.496 | -6.02 | 0.44634 | none |
| TOP1 | 0.417 | 0.5821 | 1.0345 | 0.3261 | 0.496 | -6.02 | -0.0929 | none |
| ABCA4 | 0.146 | 0.1554 | 1.0344 | 0.32614 | 0.496 | -6.02 | 0.58594 | none |
| NSFL1C | 0.094 | 0.1031 | 1.0344 | 0.32617 | 0.496 | -6.02 | -0.7243 | none |
| MYF6 | 0.073 | 0.0552 | 1.0343 | 0.3262 | 0.496 | -6.02 | 0.38883 | none |
| OR5K3 | 0.075 | 0.0501 | 1.0341 | 0.32631 | 0.496 | -6.02 | 0.20876 | none |
| AP1S1 | -0.145 | -0.1689 | -1.034 | 0.32646 | 0.497 | -6.03 | 0.39214 | none |
| ZNF510 | 0.269 | 0.3743 | 1.0337 | 0.32648 | 0.497 | -6.03 | 1.21274 | none |
| HLA-DRB1 | -0.125 | -0.1338 | -1.033 | 0.32658 | 0.497 | -6.03 | 0.89155 | none |
| ACTR5 | -0.158 | -0.0772 | -1.033 | 0.32672 | 0.497 | -6.03 | 1.85054 | none |
| RNASEH1 | 0.124 | 0.1021 | 1.0331 | 0.32675 | 0.497 | -6.03 | -0.3566 | none |
| FXYD1 | -0.125 | -0.1662 | -1.033 | 0.3269 | 0.497 | -6.03 | 0.3438 | none |
| THY1 | 0.111 | 0.1812 | 1.0326 | 0.32695 | 0.497 | -6.03 | 0.80369 | none |
| IGDCC4 | 0.071 | 0.0805 | 1.0326 | 0.32696 | 0.497 | -6.03 | -0.4963 | none |
| GPR146 | 0.284 | 0.2051 | 1.0323 | 0.32708 | 0.497 | -6.03 | 0.53184 | none |
| TSR1 | 0.147 | 0.1746 | 1.0323 | 0.32711 | 0.497 | -6.03 | 0.4892 | none |
| DCPS | -0.134 | -0.1095 | -1.032 | 0.32713 | 0.497 | -6.03 | 1.65028 | none |

| | | | | | | | | |
|---|---|---|---|---|---|---|---|---|
| RSL24D1 | -0.132 | -0.1652 | -1.032 | 0.32713 | 0.497 | -6.03 | 0.35563 | none |
| ZMYND11 | 0.197 | 0.2336 | 1.0318 | 0.3273 | 0.497 | -6.03 | 0.45336 | none |
| SRP54 | -0.106 | -0.0838 | -1.031 | 0.32745 | 0.498 | -6.03 | 1.34856 | none |
| DOK7 | 0.098 | 0.0727 | 1.031 | 0.32766 | 0.498 | -6.03 | -0.6189 | none |
| YPEL1 | 0.156 | 0.0853 | 1.031 | 0.32769 | 0.498 | -6.03 | 0.24165 | none |
| NUS1 | 0.115 | 0.0737 | 1.0309 | 0.32773 | 0.498 | -6.03 | 0.33245 | none |
| TMEM61 | -0.086 | -0.0768 | -1.031 | 0.32774 | 0.498 | -6.03 | 2.03347 | none |
| FBXL6 | -0.128 | -0.0464 | -1.031 | 0.32776 | 0.498 | -6.03 | 0.84675 | none |
| GNG12 | 0.1 | 0.1036 | 1.0305 | 0.32788 | 0.498 | -6.03 | 1.2488 | none |
| NRTN | 0.105 | 0.0628 | 1.0305 | 0.32788 | 0.498 | -6.03 | -0.7194 | none |
| RING1 | -0.097 | -0.0378 | -1.03 | 0.3279 | 0.498 | -6.03 | 1.30127 | none |
| RIMS4 | 0.082 | 0.0832 | 1.0303 | 0.328 | 0.498 | -6.03 | 0.74543 | none |
| RAB24 | 0.112 | 0.0947 | 1.0301 | 0.3281 | 0.498 | -6.03 | -0.0665 | none |
| NME1-NM | 0.113 | 0.0359 | 1.03 | 0.32812 | 0.498 | -6.03 | -0.2809 | none |
| CBLN1 | 0.098 | 0.0729 | 1.0299 | 0.32815 | 0.498 | -6.03 | 1.46465 | none |
| LYG2 | 0.23 | 0.1656 | 1.0296 | 0.3283 | 0.498 | -6.03 | -0.7097 | none |
| PLAC1 | -0.245 | -0.2436 | -1.029 | 0.32848 | 0.498 | -6.03 | 0.48123 | none |
| EVC2 | 0.079 | 0.0972 | 1.0291 | 0.32851 | 0.498 | -6.03 | 0.62064 | none |
| KALRN | 0.202 | 0.1526 | 1.0291 | 0.32854 | 0.498 | -6.03 | -0.9369 | none |
| PNPLA7 | 0.085 | 0.0728 | 1.0289 | 0.32859 | 0.499 | -6.03 | -0.1125 | none |
| GABRA4 | 0.109 | 0.109 | 1.0287 | 0.32872 | 0.499 | -6.03 | 0.4387 | none |
| HEXIM1 | 0.202 | 0.2698 | 1.0283 | 0.32889 | 0.499 | -6.03 | 0.57111 | none |
| ZNF761 | 0.248 | 0.2961 | 1.0283 | 0.3289 | 0.499 | -6.03 | 0.62771 | none |
| KIF20B | 0.33 | 0.4025 | 1.028 | 0.32899 | 0.499 | -6.03 | -0.3423 | none |
| FDPS | -0.11 | -0.1158 | -1.028 | 0.329 | 0.499 | -6.03 | 0.59289 | none |
| MAF | 0.164 | 0.2085 | 1.028 | 0.32903 | 0.499 | -6.03 | -0.2359 | none |
| MMP21 | 0.096 | 0.1204 | 1.0272 | 0.32936 | 0.499 | -6.03 | 0.58728 | none |
| GALNT3 | 0.144 | 0.1412 | 1.0266 | 0.32963 | 0.5 | -6.03 | -0.1168 | none |
| AKR7A3 | 0.083 | 0.0969 | 1.0266 | 0.32966 | 0.5 | -6.03 | 1.13938 | none |
| UBB | -0.123 | -0.1417 | -1.026 | 0.32978 | 0.5 | -6.03 | 0.84143 | none |
| TMF1 | 0.229 | 0.2638 | 1.0254 | 0.33018 | 0.5 | -6.03 | -0.2869 | none |
| CD47 | -0.129 | -0.1737 | -1.025 | 0.33023 | 0.5 | -6.03 | 1.37377 | none |
| LYZL1 | 0.127 | 0.127 | 1.025 | 0.33034 | 0.501 | -6.03 | 0.7508 | none |
| ENPP3 | -0.1 | -0.0643 | -1.025 | 0.33051 | 0.501 | -6.03 | 0.57792 | none |
| MTMR1 | 0.165 | 0.1944 | 1.0247 | 0.33051 | 0.501 | -6.03 | -0.5619 | none |
| TSC22D1 | 0.115 | 0.1382 | 1.0247 | 0.33051 | 0.501 | -6.03 | -0.1671 | none |
| MESP1 | -0.206 | -0.1777 | -1.024 | 0.33064 | 0.501 | -6.03 | 0.90511 | none |
| CASP7 | 0.103 | 0.117 | 1.0244 | 0.33065 | 0.501 | -6.03 | 0.78685 | none |
| EPHA2 | -0.133 | -0.0418 | -1.024 | 0.33081 | 0.501 | -6.04 | 0.58 | none |
| ZBED8 | -0.096 | -0.091 | -1.024 | 0.33086 | 0.501 | -6.04 | 0.20017 | none |

| | | | | | | | | |
|---|---|---|---|---|---|---|---|---|
| TMEM53 | 0.161 | 0.156 | 1.0235 | 0.33102 | 0.501 | -6.04 | 0.73519 | none |
| IL5 | -0.196 | -0.0504 | -1.024 | 0.33103 | 0.501 | -6.04 | 0.64862 | none |
| TBRG4 | 0.12 | 0.1075 | 1.0234 | 0.33107 | 0.501 | -6.04 | 0.2658 | none |
| ACTR1B | 0.12 | 0.1216 | 1.0225 | 0.33148 | 0.502 | -6.04 | 1.32834 | none |
| PPP1R2 | -0.11 | -0.1394 | -1.022 | 0.33156 | 0.502 | -6.04 | 1.49132 | none |
| PA2G4 | -0.128 | -0.031 | -1.022 | 0.33169 | 0.502 | -6.04 | 1.59566 | none |
| AATK | -0.111 | -0.1007 | -1.022 | 0.33169 | 0.502 | -6.04 | 1.49902 | none |
| CCDC153 | 0.102 | 0.1242 | 1.022 | 0.33171 | 0.502 | -6.04 | 0.91733 | none |
| UBE2F | 0.121 | 0.0329 | 1.0217 | 0.33185 | 0.502 | -6.04 | -0.4992 | none |
| BNIP1 | -0.097 | -0.1146 | -1.021 | 0.33193 | 0.502 | -6.04 | 0.9937 | none |
| SIRT7 | 0.189 | 0.1699 | 1.0214 | 0.33195 | 0.502 | -6.04 | -0.6933 | none |
| SRPK1 | 0.107 | 0.0884 | 1.0214 | 0.33196 | 0.502 | -6.04 | -0.3412 | none |
| AMFR | 0.239 | 0.1675 | 1.0214 | 0.33199 | 0.502 | -6.04 | 0.86259 | none |
| SHC3 | -0.104 | -0.1017 | -1.021 | 0.33205 | 0.502 | -6.04 | 0.83391 | none |
| EIF2B2 | -0.075 | -0.071 | -1.021 | 0.33213 | 0.502 | -6.04 | 1.33052 | none |
| CT62 | -0.071 | -0.0502 | -1.021 | 0.33226 | 0.502 | -6.04 | 2.06384 | none |
| LYPD4 | 0.082 | 0.0573 | 1.0208 | 0.33227 | 0.502 | -6.04 | -1.1505 | none |
| AC241377 | 0.125 | 0.125 | 1.0206 | 0.33232 | 0.502 | -6.04 | -0.5847 | none |
| SLC22A23 | 0.096 | 0.0935 | 1.0204 | 0.3324 | 0.503 | -6.04 | 1.24376 | none |
| MCM10 | 0.301 | 0.4231 | 1.0204 | 0.33243 | 0.503 | -6.04 | 0.6824 | none |
| ASAH2 | 0.189 | -0.0014 | 1.0202 | 0.33251 | 0.503 | -6.04 | -0.5143 | none |
| RPL3 | -0.101 | -0.0371 | -1.02 | 0.33255 | 0.503 | -6.04 | 0.60208 | none |
| GDF3 | -0.082 | -0.0596 | -1.02 | 0.3326 | 0.503 | -6.04 | 1.19215 | none |
| NREP | 0.217 | 0.2426 | 1.02 | 0.33261 | 0.503 | -6.04 | -0.1497 | none |
| HIST1H3E | 0.237 | 0.129 | 1.0198 | 0.3327 | 0.503 | -6.04 | -0.8045 | none |
| TMEM60 | -0.178 | -0.2624 | -1.02 | 0.33279 | 0.503 | -6.04 | 1.65819 | none |
| MEPCE | -0.147 | -0.014 | -1.019 | 0.33287 | 0.503 | -6.04 | 0.23544 | none |
| AMER1 | 0.099 | 0.1346 | 1.0194 | 0.33289 | 0.503 | -6.04 | 0.55482 | none |
| PYROXD2 | 0.326 | 0.2232 | 1.0194 | 0.3329 | 0.503 | -6.04 | -0.4427 | none |
| AAR2 | -0.078 | -0.0555 | -1.019 | 0.333 | 0.503 | -6.04 | 1.24309 | none |
| ARHGAP2: | 0.177 | 0.2029 | 1.0191 | 0.33303 | 0.503 | -6.04 | 0.89552 | none |
| LY75-CD3( | 0.162 | 0.2066 | 1.019 | 0.33304 | 0.503 | -6.04 | 0.29233 | none |
| TEX43 | 0.124 | 0.1242 | 1.019 | 0.33305 | 0.503 | -6.04 | 1.66721 | none |
| MFAP2 | 0.116 | 0.1177 | 1.019 | 0.33307 | 0.503 | -6.04 | 1.21943 | none |
| DERL1 | -0.151 | -0.2307 | -1.019 | 0.33307 | 0.503 | -6.04 | 0.00733 | none |
| C2ORF70 | -0.103 | -0.1012 | -1.019 | 0.33322 | 0.503 | -6.04 | -0.4792 | none |
| LRTM1 | -0.086 | -0.0845 | -1.019 | 0.33323 | 0.503 | -6.04 | 1.32879 | none |
| SERP2 | -0.124 | -0.1054 | -1.019 | 0.33327 | 0.503 | -6.04 | 0.36887 | none |
| VPS41 | 0.132 | 0.062 | 1.018 | 0.33351 | 0.503 | -6.04 | 0.63943 | none |
| TRIM21 | -0.102 | -0.0866 | -1.018 | 0.33358 | 0.503 | -6.04 | 1.32425 | none |

| | | | | | | | | |
|---|---|---|---|---|---|---|---|---|
| GDAP2 | 0.13 | 0.0261 | 1.0175 | 0.33371 | 0.504 | -6.04 | 0.84997 | none |
| TFG | 0.078 | 0.0668 | 1.0175 | 0.33375 | 0.504 | -6.04 | 0.73868 | none |
| COPS4 | -0.107 | -0.1202 | -1.017 | 0.33377 | 0.504 | -6.04 | 0.70865 | none |
| MYOT | 0.245 | 0.0888 | 1.0173 | 0.33381 | 0.504 | -6.04 | -0.9719 | none |
| ATP6V0C | 0.12 | 0.0291 | 1.0172 | 0.33387 | 0.504 | -6.04 | -0.2716 | none |
| NFU1 | -0.208 | -0.2879 | -1.017 | 0.33397 | 0.504 | -6.04 | 0.64922 | none |
| IDI1 | -0.125 | -0.1187 | -1.017 | 0.33398 | 0.504 | -6.04 | 2.14599 | AA-related |
| CHORDC1 | 0.093 | 0.0438 | 1.0165 | 0.3342 | 0.504 | -6.04 | -0.949 | none |
| BHLHA15 | 0.113 | 0.0755 | 1.0164 | 0.33423 | 0.504 | -6.04 | 1.49017 | none |
| NANOS3 | 0.188 | 0.1058 | 1.0163 | 0.33426 | 0.504 | -6.04 | 1.50487 | none |
| SPRY1 | 0.141 | 0.1518 | 1.0162 | 0.33433 | 0.504 | -6.04 | 0.84474 | none |
| ERCC2 | 0.174 | 0.1649 | 1.0161 | 0.33439 | 0.504 | -6.04 | -0.1661 | none |
| WDR25 | 0.18 | 0.079 | 1.016 | 0.33439 | 0.504 | -6.04 | -0.8226 | none |
| GNPDA2 | 0.111 | -0.0145 | 1.016 | 0.3344 | 0.504 | -6.04 | 1.45839 | none |
| TMEM14B | -0.181 | -0.2801 | -1.016 | 0.33443 | 0.504 | -6.04 | 0.79536 | none |
| PPIL2 | 0.202 | 0.2903 | 1.0159 | 0.33446 | 0.504 | -6.04 | 0.83598 | none |
| FECH | 0.089 | 0.0724 | 1.0156 | 0.33461 | 0.504 | -6.04 | 1.55203 | none |
| CD44 | 0.079 | 0.0162 | 1.0156 | 0.33461 | 0.504 | -6.04 | 0.03875 | none |
| TAGLN | 0.196 | 0.2315 | 1.0154 | 0.33469 | 0.504 | -6.04 | 1.01772 | none |
| ZBTB2 | 0.185 | 0.2488 | 1.0154 | 0.33469 | 0.504 | -6.04 | 0.33641 | none |
| CTSV | 0.106 | 0.0065 | 1.0148 | 0.33497 | 0.505 | -6.04 | -0.5079 | none |
| ADCY4 | -0.092 | -0.0628 | -1.015 | 0.33501 | 0.505 | -6.04 | 0.95898 | none |
| SELPLG | 0.231 | 0.24 | 1.0147 | 0.33501 | 0.505 | -6.04 | 0.12154 | none |
| ATP2A3 | 0.166 | 0.2312 | 1.0143 | 0.33518 | 0.505 | -6.04 | -0.0074 | none |
| ZNF518B | 0.236 | 0.2011 | 1.0141 | 0.33527 | 0.505 | -6.04 | 0.07849 | none |
| HSPA1B | 0.301 | 0.2598 | 1.0141 | 0.33529 | 0.505 | -6.04 | 0.56569 | none |
| MT-ATP6 | 0.22 | 0.2462 | 1.014 | 0.33532 | 0.505 | -6.04 | 0.4636 | none |
| TXNDC2 | -0.071 | -0.0498 | -1.014 | 0.33532 | 0.505 | -6.04 | 1.06194 | none |
| RYR1 | -0.205 | -0.0316 | -1.014 | 0.33534 | 0.505 | -6.05 | 0.32883 | none |
| MDGA1 | 0.238 | 0.1943 | 1.0139 | 0.33536 | 0.505 | -6.05 | -0.6505 | none |
| C1ORF127 | -0.15 | -0.138 | -1.014 | 0.33553 | 0.505 | -6.05 | 1.204 | none |
| CALY | -0.077 | -0.0784 | -1.013 | 0.33556 | 0.505 | -6.05 | -0.7692 | none |
| ABCF1 | 0.455 | 0.5335 | 1.0134 | 0.33558 | 0.505 | -6.05 | -0.9732 | none |
| S100A12 | -0.117 | -0.1168 | -1.013 | 0.33559 | 0.505 | -6.05 | 0.88916 | none |
| CD4 | 0.202 | 0.2632 | 1.0134 | 0.3356 | 0.505 | -6.05 | -0.343 | none |
| DOK3 | -0.119 | 0.0614 | -1.013 | 0.33571 | 0.505 | -6.05 | 0.91709 | none |
| CDC42EP2 | 0.152 | 0.1677 | 1.0129 | 0.33583 | 0.505 | -6.05 | 0.40971 | none |
| TTC3 | 0.312 | 0.3823 | 1.0124 | 0.33604 | 0.505 | -6.05 | 0.52317 | none |
| PRPS1 | -0.132 | -0.1403 | -1.012 | 0.3362 | 0.506 | -6.05 | 1.72646 | none |
| LY6G6E | 0.127 | 0.0837 | 1.012 | 0.33623 | 0.506 | -6.05 | 0.80262 | none |

| | | | | | | | | |
|---|---|---|---|---|---|---|---|---|
| STAT4 | 0.203 | 0.1883 | 1.0116 | 0.33642 | 0.506 | -6.05 | 0.11462 | none |
| KEL | -0.088 | -0.0642 | -1.011 | 0.33653 | 0.506 | -6.05 | 0.82374 | none |
| ZSCAN23 | -0.109 | -0.0438 | -1.011 | 0.33661 | 0.506 | -6.05 | 1.49365 | none |
| MAL | -0.191 | -0.2827 | -1.011 | 0.3368 | 0.506 | -6.05 | 1.079 | none |
| TRIM49D1 | 0.079 | 0.0817 | 1.0105 | 0.33693 | 0.506 | -6.05 | -0.6216 | none |
| CMTM5 | 0.081 | 0.0815 | 1.0104 | 0.33698 | 0.506 | -6.05 | 1.43614 | none |
| SKOR2 | -0.104 | -0.1094 | -1.01 | 0.337 | 0.506 | -6.05 | -0.3543 | none |
| ACTR2 | 0.094 | 0.0428 | 1.0101 | 0.33711 | 0.507 | -6.05 | 0.25406 | none |
| SLAMF1 | 0.17 | 0.2018 | 1.0098 | 0.33722 | 0.507 | -6.05 | -0.3595 | none |
| GAGE2A | -0.173 | -0.1674 | -1.01 | 0.33723 | 0.507 | -6.05 | 0.52517 | none |
| MSRA | 0.228 | 0.234 | 1.0097 | 0.33729 | 0.507 | -6.05 | 0.79717 | none |
| NKX6-3 | 0.07 | 0.0735 | 1.0096 | 0.33732 | 0.507 | -6.05 | -0.4771 | none |
| MT1M | -0.115 | -0.1151 | -1.009 | 0.33746 | 0.507 | -6.05 | 1.9054 | none |
| ZNF155 | 0.279 | 0.2681 | 1.0092 | 0.33749 | 0.507 | -6.05 | -0.1324 | none |
| ZNF746 | 0.174 | 0.273 | 1.0092 | 0.33751 | 0.507 | -6.05 | 1.02559 | none |
| SYNGR3 | 0.292 | 0.2647 | 1.0091 | 0.33755 | 0.507 | -6.05 | -0.0984 | none |
| TBX5 | 0.071 | 0.0715 | 1.0089 | 0.33765 | 0.507 | -6.05 | 0.76095 | none |
| NCEH1 | 0.103 | 0.0107 | 1.0089 | 0.33765 | 0.507 | -6.05 | -0.7349 | none |
| GALNT14 | 0.159 | 0.1158 | 1.0088 | 0.33767 | 0.507 | -6.05 | -0.9004 | none |
| SLC22A3 | 0.079 | 0.0531 | 1.0084 | 0.33785 | 0.507 | -6.05 | 0.7242 | none |
| FAHD2A | 0.1 | 0.0443 | 1.0079 | 0.33812 | 0.507 | -6.05 | -0.4977 | none |
| DNAJC10 | -0.132 | -0.033 | -1.008 | 0.33815 | 0.507 | -6.05 | 0.93809 | none |
| CA1 | 0.249 | 0.169 | 1.0075 | 0.33826 | 0.507 | -6.05 | -0.6213 | none |
| UGT2B10 | -0.066 | -0.0456 | -1.007 | 0.33828 | 0.507 | -6.05 | 0 | none |
| JADE2 | 0.25 | 0.33 | 1.0075 | 0.33829 | 0.507 | -6.05 | 0.32031 | none |
| MDFIC | 0.139 | 0.0632 | 1.0075 | 0.33829 | 0.507 | -6.05 | -0.5075 | none |
| DLGAP5 | 0.384 | 0.4825 | 1.0073 | 0.33838 | 0.507 | -6.05 | -0.112 | none |
| C5ORF46 | 0.119 | 0.1191 | 1.0073 | 0.33838 | 0.507 | -6.05 | -0.4058 | none |
| TRANK1 | -0.152 | -0.0331 | -1.007 | 0.33842 | 0.507 | -6.05 | 0.90637 | none |
| TOMM70 | 0.151 | 0.194 | 1.0071 | 0.33845 | 0.507 | -6.05 | 0.95621 | none |
| RUSC1 | -0.102 | -0.0801 | -1.007 | 0.33848 | 0.507 | -6.05 | 1.97173 | none |
| FAM50B | -0.093 | -0.0672 | -1.007 | 0.3385 | 0.507 | -6.05 | 1.441 | none |
| PDE8B | 0.122 | 0.1332 | 1.007 | 0.3385 | 0.507 | -6.05 | -0.3063 | none |
| LPCAT1 | 0.123 | 0.1738 | 1.0068 | 0.33858 | 0.507 | -6.05 | 0.11841 | none |
| STON1-GT | 0.127 | 0.1268 | 1.0067 | 0.33862 | 0.507 | -6.05 | 1.24571 | none |
| CDC6 | 0.155 | 0.1858 | 1.0067 | 0.33865 | 0.507 | -6.05 | 0.50984 | none |
| FA2H | -0.244 | -0.1082 | -1.007 | 0.33866 | 0.507 | -6.05 | 1.73456 | none |
| GOLGA7 | -0.14 | -0.1947 | -1.007 | 0.33867 | 0.507 | -6.05 | 1.30333 | none |
| HEYL | -0.078 | -0.0467 | -1.007 | 0.33867 | 0.507 | -6.05 | 0.90179 | none |
| TUSC1 | -0.073 | -0.0661 | -1.006 | 0.3391 | 0.508 | -6.05 | 1.8667 | none |

| | | | | | | | | |
|---|---|---|---|---|---|---|---|---|
| ZNF514 | 0.155 | 0.1228 | 1.0056 | 0.33914 | 0.508 | -6.05 | -0.2524 | none |
| RFPL2 | 0.084 | 0.0807 | 1.0055 | 0.33919 | 0.508 | -6.05 | -0.6256 | none |
| TTC31 | 0.11 | 0.1236 | 1.0054 | 0.33922 | 0.508 | -6.05 | 1.22381 | none |
| TTC33 | -0.177 | -0.2093 | -1.005 | 0.33923 | 0.508 | -6.05 | 1.27824 | none |
| TTC8 | 0.192 | 0.1962 | 1.0053 | 0.33927 | 0.508 | -6.05 | -0.3051 | none |
| NDUFC2-K | -1.413 | -1.1234 | -1.005 | 0.33932 | 0.508 | -6.05 | 2.04631 | none |
| TBX10 | 0.07 | 0.0713 | 1.0052 | 0.33934 | 0.508 | -6.05 | 0 | none |
| FKBP1C | 0.091 | 0.1004 | 1.005 | 0.33944 | 0.508 | -6.05 | -0.4638 | none |
| LRFN4 | 0.125 | 0.0696 | 1.0049 | 0.33948 | 0.508 | -6.05 | 0.70321 | none |
| TCEAL9 | -0.168 | -0.087 | -1.005 | 0.33961 | 0.508 | -6.05 | 1.98692 | none |
| DYNC1LI2 | 0.116 | 0.1763 | 1.0045 | 0.33964 | 0.508 | -6.05 | 1.02691 | none |
| TMEM263 | 0.108 | -0.0029 | 1.0041 | 0.33985 | 0.509 | -6.05 | -0.2332 | none |
| GRIK1 | 0.086 | 0.0555 | 1.004 | 0.33986 | 0.509 | -6.05 | 0.08617 | none |
| CRYBA2 | 0.094 | 0.0664 | 1.0037 | 0.34 | 0.509 | -6.06 | 1.6563 | none |
| CERS2 | -0.119 | -0.1516 | -1.004 | 0.34002 | 0.509 | -6.06 | 0.91743 | none |
| TAS2R40 | -0.074 | -0.0634 | -1.004 | 0.34003 | 0.509 | -6.06 | 0 | none |
| GLI2 | 0.073 | 0.0807 | 1.0034 | 0.34015 | 0.509 | -6.06 | 1.31073 | none |
| ISL2 | -0.091 | -0.0865 | -1.003 | 0.34034 | 0.509 | -6.06 | 1.67951 | none |
| ASPRV1 | -0.094 | -0.0678 | -1.003 | 0.34034 | 0.509 | -6.06 | 0.46299 | none |
| CWF19L2 | 0.27 | 0.3217 | 1.0028 | 0.34041 | 0.509 | -6.06 | -0.4056 | none |
| HNRNPA1l | 0.131 | 0.1953 | 1.0028 | 0.34044 | 0.509 | -6.06 | 0.54655 | none |
| HSPA13 | 0.12 | 0.0057 | 1.0026 | 0.34054 | 0.509 | -6.06 | 0.08529 | none |
| RBM25 | 0.307 | 0.4423 | 1.0025 | 0.34056 | 0.509 | -6.06 | 0.19668 | none |
| FZD3 | 0.226 | 0.1094 | 1.0025 | 0.34057 | 0.509 | -6.06 | 0.3198 | none |
| TJP1 | 0.135 | 0.1501 | 1.0025 | 0.34058 | 0.509 | -6.06 | -0.6082 | none |
| ARF4 | -0.172 | -0.262 | -1.002 | 0.34067 | 0.509 | -6.06 | 0.76075 | none |
| ACTN4 | 0.344 | 0.4662 | 1.0022 | 0.34072 | 0.509 | -6.06 | 0.24171 | none |
| IGSF9 | -0.082 | -0.0651 | -1.002 | 0.34079 | 0.509 | -6.06 | 0.99267 | none |
| KIF1A | 0.195 | 0.1573 | 1.0019 | 0.34086 | 0.509 | -6.06 | -0.668 | none |
| POLR2K | -0.174 | -0.2525 | -1.002 | 0.3409 | 0.509 | -6.06 | 0.21078 | none |
| METTL5 | -0.112 | -0.1771 | -1.002 | 0.34093 | 0.509 | -6.06 | 0.63566 | none |
| DGKZ | 0.167 | 0.1718 | 1.0017 | 0.34094 | 0.509 | -6.06 | -0.2041 | none |
| SLC35A5 | 0.182 | 0.0324 | 1.0015 | 0.34102 | 0.509 | -6.06 | 0.30551 | none |
| FOXI1 | 0.067 | 0.0672 | 1.0013 | 0.3411 | 0.509 | -6.06 | -0.0401 | none |
| RNF20 | 0.319 | 0.4287 | 1.0013 | 0.3411 | 0.509 | -6.06 | 0.03719 | none |
| LRRCC1 | 0.228 | 0.2998 | 1.0013 | 0.34113 | 0.509 | -6.06 | 0.58388 | none |
| PYCARD | -0.207 | -0.286 | -1.001 | 0.34116 | 0.509 | -6.06 | 0.34611 | none |
| PPM1D | 0.108 | 0.0676 | 1.0005 | 0.34148 | 0.51 | -6.06 | 0.67785 | none |
| GLUD1 | 0.119 | 0.1222 | 1.0003 | 0.34159 | 0.51 | -6.06 | 0.08558 | none |
| E2F7 | -0.151 | -0.0091 | -1 | 0.34161 | 0.51 | -6.06 | 1.56495 | none |

| | | | | | | | | |
|---|---|---|---|---|---|---|---|---|
| AP3S1 | -0.119 | -0.1726 | -1 | 0.34172 | 0.51 | -6.06 | 0.68519 | none |
| RASGRF2 | -0.098 | -0.0555 | -1 | 0.34188 | 0.51 | -6.06 | 1.15187 | none |
| TICRR | 0.23 | 0.337 | 0.9995 | 0.34193 | 0.51 | -6.06 | -0.5478 | none |
| TTC27 | 0.1 | 0.0275 | 0.9995 | 0.34194 | 0.51 | -6.06 | -0.2834 | none |
| TSPY8 | -0.111 | -0.1114 | -0.999 | 0.34198 | 0.51 | -6.06 | 0.12942 | none |
| LMBRD1 | 0.144 | -0.0178 | 0.9993 | 0.34202 | 0.51 | -6.06 | 0.19731 | none |
| SLC7A8 | 0.183 | 0.1848 | 0.9993 | 0.34203 | 0.51 | -6.06 | 0.85799 | none |
| NENF | -0.187 | -0.1806 | -0.999 | 0.34213 | 0.51 | -6.06 | -0.0302 | none |
| AMPD1 | 0.072 | 0.0733 | 0.999 | 0.34219 | 0.51 | -6.06 | 1.16625 | none |
| C19ORF67 | -0.109 | -0.0529 | -0.998 | 0.34255 | 0.511 | -6.06 | -0.3199 | none |
| SPINK8 | 0.115 | 0.1153 | 0.9979 | 0.3427 | 0.511 | -6.06 | 0.51333 | none |
| KCNK4 | -0.12 | -0.0803 | -0.998 | 0.34281 | 0.511 | -6.06 | 0.83625 | none |
| KRTAP12-2 | 0.115 | 0.1151 | 0.9973 | 0.34297 | 0.511 | -6.06 | -0.1375 | none |
| SPRED2 | 0.135 | 0.1428 | 0.9972 | 0.34298 | 0.511 | -6.06 | 0.06742 | none |
| GTSF1L | 0.135 | 0.1174 | 0.9972 | 0.34298 | 0.511 | -6.06 | 0.17453 | none |
| VEGFB | 0.132 | 0.1243 | 0.9969 | 0.34316 | 0.512 | -6.06 | 0.25496 | none |
| CYC1 | -0.161 | -0.1613 | -0.996 | 0.34339 | 0.512 | -6.06 | 0.64848 | none |
| SH3RF2 | 0.118 | 0.0792 | 0.9963 | 0.34342 | 0.512 | -6.06 | -0.0521 | none |
| WNT1 | 0.073 | 0.0494 | 0.9962 | 0.34344 | 0.512 | -6.06 | 0.00644 | none |
| TXLNG | 0.242 | 0.3272 | 0.9962 | 0.34346 | 0.512 | -6.06 | 0.43209 | none |
| SPOUT1 | 0.125 | 0.172 | 0.9961 | 0.34353 | 0.512 | -6.06 | -0.568 | none |
| SMPDL3B | 0.177 | 0.1437 | 0.9959 | 0.34359 | 0.512 | -6.06 | -0.756 | none |
| U2AF1 | 0.21 | 0.1821 | 0.9958 | 0.34365 | 0.512 | -6.06 | 0.24589 | none |
| PSORS1C2 | -0.07 | -0.0469 | -0.996 | 0.34371 | 0.512 | -6.06 | 0 | none |
| ARHGEF10 | -0.101 | -0.1013 | -0.996 | 0.34378 | 0.512 | -6.06 | -0.1743 | none |
| ADGRL1 | -0.174 | 0.0202 | -0.995 | 0.34379 | 0.512 | -6.06 | -0.0392 | none |
| DUSP5 | -0.116 | -0.1086 | -0.995 | 0.344 | 0.512 | -6.06 | -0.3764 | none |
| FSCN3 | -0.125 | -0.0523 | -0.995 | 0.34401 | 0.512 | -6.06 | 0.14825 | none |
| PTP4A1 | -0.132 | -0.1545 | -0.994 | 0.34436 | 0.513 | -6.06 | 1.18125 | none |
| UBXN4 | 0.171 | 0.2054 | 0.9936 | 0.34469 | 0.513 | -6.07 | 0.5248 | none |
| TSHZ3 | -0.095 | 0.0122 | -0.993 | 0.34484 | 0.513 | -6.07 | -0.3155 | none |
| HNRNPDL | -0.101 | -0.0616 | -0.993 | 0.34488 | 0.513 | -6.07 | 1.8843 | none |
| NUTM2E | -0.069 | -0.0507 | -0.993 | 0.34513 | 0.514 | -6.07 | 0.58425 | none |
| PLEKHN1 | -0.232 | -0.1197 | -0.992 | 0.34522 | 0.514 | -6.07 | 1.97458 | none |
| COL18A1 | 0.211 | 0.2689 | 0.9922 | 0.34531 | 0.514 | -6.07 | 0.26205 | none |
| CCDC188 | -0.069 | -0.0467 | -0.992 | 0.34534 | 0.514 | -6.07 | 0 | none |
| AIF1 | 0.35 | 0.2787 | 0.9921 | 0.34536 | 0.514 | -6.07 | -0.1773 | none |
| ACTRT3 | -0.12 | -0.0864 | -0.992 | 0.34544 | 0.514 | -6.07 | 1.05807 | none |
| CDH22 | -0.069 | -0.0567 | -0.992 | 0.34551 | 0.514 | -6.07 | -0.4876 | none |
| PAX5 | -0.253 | -0.1815 | -0.992 | 0.34557 | 0.514 | -6.07 | 0.24573 | none |

| | | | | | | | | |
|---|---|---|---|---|---|---|---|---|
| SCGB1D1 | 0.113 | 0.1129 | 0.9916 | 0.34561 | 0.514 | -6.07 | -0.5846 | none |
| BBS12 | 0.116 | 0.0856 | 0.9915 | 0.34565 | 0.514 | -6.07 | 0.30907 | none |
| TRIP13 | 0.079 | 0.0618 | 0.991 | 0.34587 | 0.514 | -6.07 | 0.36485 | none |
| VWA5B2 | -0.132 | -0.1229 | -0.991 | 0.34602 | 0.515 | -6.07 | 0.72391 | none |
| ZSWIM4 | 0.146 | 0.2252 | 0.9906 | 0.34607 | 0.515 | -6.07 | 0.17181 | none |
| DNAAF3 | -0.123 | -0.0855 | -0.991 | 0.34608 | 0.515 | -6.07 | 0.26112 | none |
| OS9 | 0.197 | 0.2703 | 0.9901 | 0.34631 | 0.515 | -6.07 | -0.1961 | none |
| CCDC70 | 0.136 | 0.122 | 0.99 | 0.34632 | 0.515 | -6.07 | 0.52546 | none |
| KCNV2 | -0.077 | -0.0392 | -0.99 | 0.34635 | 0.515 | -6.07 | 1.36394 | none |
| S100G | 0.112 | 0.1122 | 0.9898 | 0.34643 | 0.515 | -6.07 | 0.38115 | none |
| ACOT6 | -0.113 | -0.0557 | -0.989 | 0.34661 | 0.515 | -6.07 | 0.92929 | none |
| DSE | 0.144 | 0.1341 | 0.9891 | 0.34675 | 0.515 | -6.07 | -0.931 | none |
| ARHGAP24 | 0.08 | 0.0809 | 0.9891 | 0.34676 | 0.515 | -6.07 | 0.32509 | none |
| PPFIA2 | 0.094 | 0.0942 | 0.989 | 0.34679 | 0.515 | -6.07 | 1.66596 | none |
| BCO1 | 0.077 | 0.0781 | 0.989 | 0.34679 | 0.515 | -6.07 | -0.8465 | none |
| LCN2 | -0.108 | -0.1077 | -0.989 | 0.34682 | 0.515 | -6.07 | 0.93889 | none |
| C2 | 0.147 | 0.0683 | 0.9886 | 0.34696 | 0.515 | -6.07 | -0.8743 | none |
| TPTE2 | 0.09 | 0.0787 | 0.9885 | 0.34705 | 0.516 | -6.07 | -1.0165 | none |
| CBX8 | 0.193 | 0.2271 | 0.9882 | 0.34717 | 0.516 | -6.07 | -0.6615 | none |
| SYS1 | -0.205 | -0.2116 | -0.988 | 0.34725 | 0.516 | -6.07 | 1.02799 | none |
| TAF15 | 0.318 | 0.4576 | 0.9879 | 0.3473 | 0.516 | -6.07 | -0.1179 | none |
| ECM2 | 0.14 | 0.1007 | 0.9876 | 0.34743 | 0.516 | -6.07 | -0.1255 | none |
| CHL1 | 0.108 | 0.1121 | 0.9876 | 0.34745 | 0.516 | -6.07 | 1.15842 | none |
| KRT78 | -0.124 | -0.1141 | -0.987 | 0.34754 | 0.516 | -6.07 | 1.63323 | none |
| MYOC | -0.074 | -0.0677 | -0.987 | 0.34761 | 0.516 | -6.07 | 1.12074 | none |
| MAMSTR | -0.131 | -0.0995 | -0.987 | 0.34764 | 0.516 | -6.07 | 1.93831 | none |
| PIGS | 0.114 | 0.0285 | 0.9869 | 0.34778 | 0.516 | -6.07 | -0.1952 | none |
| CHST8 | 0.068 | 0.0683 | 0.9867 | 0.34786 | 0.516 | -6.07 | 0.5601 | none |
| STXBP1 | -0.164 | -0.0482 | -0.987 | 0.34787 | 0.516 | -6.07 | 1.38006 | none |
| C9 | 0.084 | 0.0841 | 0.9865 | 0.34795 | 0.516 | -6.07 | 1.06079 | none |
| RBM27 | 0.234 | 0.3059 | 0.9864 | 0.34802 | 0.516 | -6.07 | 0.24328 | none |
| NUTM2F | 0.077 | 0.0542 | 0.9861 | 0.34816 | 0.517 | -6.07 | 0.35145 | none |
| DEFB110 | 0.111 | 0.1109 | 0.986 | 0.34819 | 0.517 | -6.07 | 1.19688 | none |
| SH3PXD2B | 0.067 | 0.0714 | 0.9859 | 0.34823 | 0.517 | -6.07 | -0.8862 | none |
| CUL3 | 0.25 | 0.3086 | 0.9856 | 0.34839 | 0.517 | -6.07 | 0.12287 | none |
| IGFBP5 | -0.085 | -0.0889 | -0.985 | 0.34852 | 0.517 | -6.07 | 1.97304 | none |
| RASSF5 | 0.157 | 0.2194 | 0.9852 | 0.34855 | 0.517 | -6.07 | 0.90268 | none |
| SP140 | 0.15 | 0.2038 | 0.9852 | 0.34858 | 0.517 | -6.07 | -0.1612 | none |
| DRG1 | -0.11 | -0.1414 | -0.985 | 0.34874 | 0.517 | -6.07 | 0.9055 | none |
| SLC37A4 | 0.16 | 0.0704 | 0.9848 | 0.34875 | 0.517 | -6.07 | 0.30667 | none |

| | | | | | | | | |
|---|---|---|---|---|---|---|---|---|
| COL5A1 | 0.114 | 0.1401 | 0.9844 | 0.34894 | 0.517 | -6.07 | 0.0189 | none |
| ARHGEF6 | 0.201 | 0.2259 | 0.9843 | 0.34899 | 0.517 | -6.07 | 0.1517 | none |
| FCRLA | 0.076 | 0.0848 | 0.9839 | 0.34919 | 0.518 | -6.07 | 0.36921 | none |
| LRRTM3 | -0.117 | -0.1174 | -0.984 | 0.34927 | 0.518 | -6.07 | 0.43974 | none |
| ALDH18A1 | -0.169 | -0.0738 | -0.984 | 0.34929 | 0.518 | -6.07 | 1.58638 | none |
| PIK3C2G | 0.126 | 0.1321 | 0.9835 | 0.34934 | 0.518 | -6.07 | 0.98959 | none |
| CCT6A | -0.099 | -0.0664 | -0.983 | 0.34936 | 0.518 | -6.07 | 1.88172 | none |
| RAB8A | 0.179 | 0.25 | 0.9833 | 0.34948 | 0.518 | -6.08 | -0.0176 | none |
| ARL6IP5 | 0.11 | 0.0392 | 0.9829 | 0.34963 | 0.518 | -6.08 | -0.268 | none |
| BORCS6 | -0.078 | -0.0857 | -0.983 | 0.34965 | 0.518 | -6.08 | 1.0152 | none |
| GREB1L | 0.123 | 0.135 | 0.9828 | 0.3497 | 0.518 | -6.08 | -0.7429 | none |
| PATE4 | 0.072 | 0.0551 | 0.9827 | 0.34975 | 0.518 | -6.08 | 0.55179 | none |
| LRRC4C | 0.209 | 0.1121 | 0.9826 | 0.3498 | 0.518 | -6.08 | -0.2293 | none |
| ZRANB2 | 0.1 | 0.1024 | 0.9824 | 0.34987 | 0.518 | -6.08 | 1.17112 | none |
| TRERF1 | 0.129 | 0.1866 | 0.9824 | 0.3499 | 0.518 | -6.08 | -0.5199 | none |
| LYG1 | 0.114 | 0.0554 | 0.9823 | 0.34992 | 0.518 | -6.08 | 0.83335 | none |
| YRDC | -0.156 | -0.1503 | -0.982 | 0.34997 | 0.518 | -6.08 | 0.15771 | none |
| SOCS5 | 0.094 | 0.0564 | 0.982 | 0.35006 | 0.518 | -6.08 | -1.1653 | none |
| AP5M1 | 0.188 | 0.0678 | 0.982 | 0.35007 | 0.518 | -6.08 | 0.54057 | none |
| MT1A | -0.119 | -0.1202 | -0.982 | 0.3503 | 0.518 | -6.08 | 0.08613 | none |
| RAB25 | 0.276 | 0.1804 | 0.9814 | 0.35035 | 0.518 | -6.08 | -0.5593 | none |
| PGC | -0.253 | -0.1717 | -0.981 | 0.35042 | 0.519 | -6.08 | 0.45535 | none |
| ACSBG1 | 0.323 | 0.4276 | 0.9811 | 0.35049 | 0.519 | -6.08 | 0.41208 | none |
| ILK | -0.091 | -0.0872 | -0.981 | 0.35058 | 0.519 | -6.08 | 0.64563 | none |
| KRT18 | 0.17 | 0.1448 | 0.9805 | 0.35077 | 0.519 | -6.08 | -0.7479 | none |
| FEM1C | 0.093 | 0.0447 | 0.9803 | 0.35086 | 0.519 | -6.08 | 0.83461 | none |
| MINK1 | 0.258 | 0.359 | 0.9803 | 0.35088 | 0.519 | -6.08 | -0.1117 | none |
| C12ORF42 | 0.346 | 0.2355 | 0.9802 | 0.3509 | 0.519 | -6.08 | 0.29562 | none |
| OR14I1 | 0.069 | 0.0689 | 0.9802 | 0.3509 | 0.519 | -6.08 | 0.60476 | none |
| NRGN | 0.108 | 0.1388 | 0.9801 | 0.35095 | 0.519 | -6.08 | 0.91475 | none |
| ZNF507 | 0.209 | 0.2888 | 0.98 | 0.35099 | 0.519 | -6.08 | -0.1064 | none |
| PPIC | 0.071 | 0.0516 | 0.98 | 0.35102 | 0.519 | -6.08 | 0.07846 | none |
| ZNF683 | 0.311 | 0.1771 | 0.9799 | 0.35103 | 0.519 | -6.08 | -0.1749 | none |
| CRABP1 | -0.091 | -0.0971 | -0.98 | 0.35108 | 0.519 | -6.08 | 1.99808 | none |
| GFRA3 | 0.068 | 0.0713 | 0.9798 | 0.35109 | 0.519 | -6.08 | -0.291 | none |
| FTMT | 0.069 | 0.0685 | 0.9797 | 0.35113 | 0.519 | -6.08 | -0.8746 | none |
| FAM120A | 0.215 | 0.292 | 0.9797 | 0.35116 | 0.519 | -6.08 | -0.1738 | none |
| SLC38A2 | -0.098 | -0.1632 | -0.98 | 0.35116 | 0.519 | -6.08 | 0.6388 | none |
| PNN | 0.274 | 0.4287 | 0.9796 | 0.35119 | 0.519 | -6.08 | 0.3455 | none |
| PRR13 | -0.098 | -0.1315 | -0.979 | 0.35126 | 0.519 | -6.08 | 0.70301 | none |

| | | | | | | | | |
|---|---|---|---|---|---|---|---|---|
| GPHB5 | 0.109 | 0.1085 | 0.9794 | 0.3513 | 0.519 | -6.08 | 1.19688 | none |
| UBE2J1 | -0.078 | -0.0664 | -0.979 | 0.35142 | 0.519 | -6.08 | 0.99225 | none |
| DLX1 | 0.086 | 0.0846 | 0.9789 | 0.35154 | 0.519 | -6.08 | -0.5588 | none |
| XAGE1A | 0.108 | 0.1082 | 0.9785 | 0.35172 | 0.519 | -6.08 | -0.7569 | none |
| CDHR5 | -0.072 | -0.0451 | -0.978 | 0.35183 | 0.52 | -6.08 | 0.94955 | none |
| POP4 | -0.109 | -0.0882 | -0.978 | 0.35195 | 0.52 | -6.08 | 1.63879 | none |
| KCNIP4 | -0.088 | -0.0729 | -0.978 | 0.35197 | 0.52 | -6.08 | 1.16316 | none |
| PCBP4 | 0.217 | 0.2561 | 0.9775 | 0.35218 | 0.52 | -6.08 | 0.33577 | none |
| MIEN1 | -0.149 | -0.1877 | -0.977 | 0.35228 | 0.52 | -6.08 | 1.22792 | none |
| FUCA2 | -0.11 | -0.1421 | -0.977 | 0.35231 | 0.52 | -6.08 | 0.15935 | none |
| GLS2 | -0.153 | -0.0817 | -0.977 | 0.3526 | 0.52 | -6.08 | 1.12991 | none |
| C11ORF40 | -0.083 | -0.0892 | -0.976 | 0.35287 | 0.521 | -6.08 | 1.79234 | none |
| HIST1H2AI | 0.175 | 0.082 | 0.9757 | 0.35302 | 0.521 | -6.08 | 1.1358 | none |
| EIF5B | 0.359 | 0.5285 | 0.9756 | 0.35305 | 0.521 | -6.08 | -0.0221 | none |
| PKIA | 0.13 | 0.1066 | 0.9756 | 0.35305 | 0.521 | -6.08 | 1.11434 | none |
| HAAO | 0.123 | 0.1143 | 0.9756 | 0.35307 | 0.521 | -6.08 | -0.2546 | none |
| SEMA6C | 0.103 | 0.1329 | 0.9751 | 0.35332 | 0.521 | -6.08 | -0.3831 | none |
| KCTD14 | -0.082 | -0.1003 | -0.975 | 0.35351 | 0.521 | -6.08 | 0.04134 | none |
| POR | 0.222 | 0.1938 | 0.9745 | 0.35358 | 0.521 | -6.08 | -0.357 | none |
| NOMO2 | -0.16 | -0.0495 | -0.974 | 0.35375 | 0.522 | -6.08 | 0.46203 | none |
| LIMA1 | 0.295 | 0.3512 | 0.9741 | 0.35379 | 0.522 | -6.08 | 0.29265 | none |
| OSTN | 0.107 | 0.1067 | 0.9739 | 0.35386 | 0.522 | -6.08 | -0.714 | none |
| AGT | -0.094 | -0.1073 | -0.974 | 0.35404 | 0.522 | -6.08 | 1.11903 | none |
| RIT2 | 0.071 | 0.0914 | 0.9732 | 0.35423 | 0.522 | -6.08 | 0.79603 | none |
| SERINC1 | 0.12 | -0.0076 | 0.9731 | 0.35427 | 0.522 | -6.08 | -0.1371 | none |
| THTPA | 0.12 | 0.0536 | 0.973 | 0.35429 | 0.522 | -6.08 | 0.73997 | none |
| NT5DC2 | 0.143 | 0.1834 | 0.9727 | 0.35443 | 0.522 | -6.09 | -0.2123 | none |
| LILRA4 | 0.106 | 0.1062 | 0.9726 | 0.35447 | 0.522 | -6.09 | -0.3748 | none |
| CCSER2 | 0.231 | 0.3188 | 0.9726 | 0.35448 | 0.522 | -6.09 | 0.86103 | none |
| OR4B1 | 0.106 | 0.1062 | 0.9725 | 0.35452 | 0.522 | -6.09 | 0.861 | none |
| RBMX2 | 0.116 | 0.1466 | 0.9721 | 0.35472 | 0.523 | -6.09 | 1.01719 | none |
| NPSR1 | -0.071 | -0.0633 | -0.972 | 0.35477 | 0.523 | -6.09 | -0.6262 | none |
| ATP6V1D | 0.129 | 0.0486 | 0.972 | 0.35477 | 0.523 | -6.09 | -0.4436 | none |
| SBDS | 0.08 | 0.0925 | 0.9717 | 0.35493 | 0.523 | -6.09 | 0.86417 | none |
| ALG3 | -0.116 | -0.1067 | -0.972 | 0.35493 | 0.523 | -6.09 | 0.77049 | none |
| BLNK | -0.071 | -0.0689 | -0.972 | 0.35494 | 0.523 | -6.09 | 1.90337 | none |
| IFITM5 | -0.119 | -0.1167 | -0.971 | 0.3553 | 0.523 | -6.09 | 1.82983 | none |
| CXORF67 | -0.071 | -0.0411 | -0.971 | 0.35531 | 0.523 | -6.09 | 1.65861 | none |
| GPR62 | -0.067 | -0.0593 | -0.971 | 0.35532 | 0.523 | -6.09 | 0 | none |
| MTRNR2L8 | -0.084 | -0.1068 | -0.971 | 0.35537 | 0.523 | -6.09 | 0.9992 | none |

| | | | | | | | | |
|---|---|---|---|---|---|---|---|---|
| COX7A2 | -0.143 | -0.2211 | -0.971 | 0.35545 | 0.523 | -6.09 | 0.24967 | none |
| TCP11 | 0.111 | 0.1231 | 0.9705 | 0.3555 | 0.523 | -6.09 | 0.76994 | none |
| TNFSF12 | 0.276 | 0.0944 | 0.9703 | 0.35559 | 0.523 | -6.09 | -0.4952 | none |
| ZNF222 | -0.13 | -0.0789 | -0.97 | 0.35565 | 0.523 | -6.09 | 1.91062 | none |
| AMDHD2 | 0.185 | 0.0924 | 0.9698 | 0.3558 | 0.524 | -6.09 | -0.1844 | none |
| COL22A1 | 0.105 | 0.1053 | 0.9698 | 0.3558 | 0.524 | -6.09 | -0.8743 | none |
| GIMAP4 | -0.123 | -0.1413 | -0.97 | 0.35581 | 0.524 | -6.09 | 0.57354 | none |
| FERMT1 | 0.112 | 0.1233 | 0.9687 | 0.35635 | 0.524 | -6.09 | -0.8655 | none |
| OXGR1 | -0.071 | -0.0709 | -0.968 | 0.35645 | 0.524 | -6.09 | 1.38879 | none |
| ZFAT | 0.177 | 0.2318 | 0.968 | 0.35667 | 0.525 | -6.09 | -0.2391 | none |
| AC145212 | 0.135 | 0.1565 | 0.9678 | 0.35678 | 0.525 | -6.09 | -0.1623 | none |
| TTC24 | 0.192 | 0.1716 | 0.9674 | 0.35695 | 0.525 | -6.09 | -0.2149 | none |
| MCAT | 0.088 | 0.1214 | 0.9674 | 0.35696 | 0.525 | -6.09 | 1.32267 | none |
| CEP295NL | -0.144 | -0.0904 | -0.967 | 0.357 | 0.525 | -6.09 | 1.59751 | none |
| PDLIM2 | 0.127 | 0.0805 | 0.967 | 0.35714 | 0.525 | -6.09 | -0.0113 | none |
| FBXO21 | 0.157 | 0.1321 | 0.9667 | 0.3573 | 0.525 | -6.09 | 0.39725 | none |
| SMIM13 | 0.14 | 0.0664 | 0.9665 | 0.35737 | 0.525 | -6.09 | -0.0589 | none |
| JMJD7 | -0.318 | -0.2655 | -0.966 | 0.35749 | 0.525 | -6.09 | 0.9203 | none |
| HPS1 | 0.117 | 0.0753 | 0.9659 | 0.35766 | 0.526 | -6.09 | 0.90776 | none |
| COQ7 | 0.138 | 0.0776 | 0.9658 | 0.35771 | 0.526 | -6.09 | -0.2757 | none |
| FAM122A | 0.092 | 0.0255 | 0.9656 | 0.35782 | 0.526 | -6.09 | 0.40998 | none |
| HYAL3 | -0.188 | -0.1282 | -0.965 | 0.35808 | 0.526 | -6.09 | 1.60398 | none |
| NAB2 | 0.1 | 0.1231 | 0.9649 | 0.35814 | 0.526 | -6.09 | 1.05044 | none |
| HMOX2 | 0.119 | 0.1496 | 0.9647 | 0.35822 | 0.526 | -6.09 | -0.1149 | none |
| SLC7A6OS | 0.216 | 0.1957 | 0.9641 | 0.35852 | 0.527 | -6.09 | 0.58487 | none |
| NBEA | 0.073 | 0.052 | 0.9641 | 0.35853 | 0.527 | -6.09 | -1.0415 | none |
| YIPF2 | 0.165 | 0.0953 | 0.964 | 0.35855 | 0.527 | -6.09 | -0.7065 | none |
| NFE2L1 | 0.3 | 0.4024 | 0.964 | 0.35857 | 0.527 | -6.09 | 0.71314 | none |
| BOLA1 | 0.157 | 0.0783 | 0.9635 | 0.35883 | 0.527 | -6.09 | 0.23558 | none |
| MRPS24 | -0.156 | -0.1797 | -0.963 | 0.35884 | 0.527 | -6.09 | 0.76265 | none |
| FIG4 | -0.14 | -0.1577 | -0.963 | 0.35895 | 0.527 | -6.09 | 0.59777 | none |
| TNPO2 | 0.159 | 0.2769 | 0.9631 | 0.359 | 0.527 | -6.09 | -0.4243 | none |
| NPTXR | 0.09 | 0.0802 | 0.9628 | 0.35912 | 0.527 | -6.09 | -0.5491 | none |
| SLAMF8 | -0.146 | -0.1769 | -0.963 | 0.35918 | 0.527 | -6.09 | 1.49394 | none |
| RNF13 | -0.209 | -0.2339 | -0.963 | 0.35921 | 0.527 | -6.09 | 1.32105 | none |
| RBBP6 | -0.156 | -0.1025 | -0.962 | 0.35946 | 0.528 | -6.1 | 1.00855 | none |
| POLR2C | 0.149 | 0.0283 | 0.9621 | 0.35946 | 0.528 | -6.1 | 0.2095 | none |
| NUDT11 | -0.134 | -0.1493 | -0.962 | 0.35953 | 0.528 | -6.1 | 1.42214 | none |
| ADGRF3 | 0.231 | 0.2393 | 0.9619 | 0.35957 | 0.528 | -6.1 | 0.10194 | none |
| SLK | 0.242 | 0.4454 | 0.9619 | 0.35958 | 0.528 | -6.1 | 0.39528 | none |

| | | | | | | | | |
|---|---|---|---|---|---|---|---|---|
| RBFA | -0.089 | -0.069 | -0.962 | 0.35968 | 0.528 | -6.1 | 1.76154 | none |
| C9ORF116 | -0.231 | -0.162 | -0.962 | 0.35968 | 0.528 | -6.1 | 0.6458 | none |
| NOXRED1 | -0.1 | -0.0835 | -0.961 | 0.35979 | 0.528 | -6.1 | 0.01936 | none |
| EME1 | -0.239 | -0.0364 | -0.961 | 0.35979 | 0.528 | -6.1 | 1.4751 | none |
| CXORF51B | 0.103 | 0.1026 | 0.9613 | 0.35988 | 0.528 | -6.1 | 0.66005 | none |
| LRRC63 | 0.076 | 0.08 | 0.9611 | 0.35994 | 0.528 | -6.1 | 1.3432 | none |
| FBXO33 | 0.082 | 0.032 | 0.9611 | 0.35996 | 0.528 | -6.1 | 0.8386 | none |
| OR5B3 | -0.066 | -0.047 | -0.961 | 0.36 | 0.528 | -6.1 | 0 | none |
| TMEM14C | -0.144 | -0.2121 | -0.961 | 0.36002 | 0.528 | -6.1 | 0.63522 | none |
| SELL | -0.107 | -0.1164 | -0.961 | 0.36003 | 0.528 | -6.1 | 1.69334 | none |
| WFDC10A | 0.102 | 0.1025 | 0.9608 | 0.36008 | 0.528 | -6.1 | 0.44852 | none |
| ASB4 | 0.074 | 0.0841 | 0.9604 | 0.36026 | 0.528 | -6.1 | -0.4354 | none |
| CITED1 | 0.268 | 0.088 | 0.96 | 0.36046 | 0.528 | -6.1 | -0.2395 | none |
| RNF151 | 0.195 | 0.1486 | 0.9599 | 0.36055 | 0.528 | -6.1 | -1.0507 | none |
| GAST | -0.099 | -0.0986 | -0.96 | 0.36063 | 0.528 | -6.1 | -0.2316 | none |
| RNF212B | -0.171 | -0.0984 | -0.96 | 0.36069 | 0.528 | -6.1 | 1.17209 | none |
| TAS2R19 | -0.08 | -0.0452 | -0.959 | 0.36099 | 0.529 | -6.1 | 0.68906 | none |
| ERLEC1 | 0.147 | 0.1261 | 0.9587 | 0.36109 | 0.529 | -6.1 | -0.6641 | none |
| GAS8 | 0.262 | 0.2641 | 0.9587 | 0.36109 | 0.529 | -6.1 | 0.16708 | none |
| FANCM | 0.211 | 0.2309 | 0.9585 | 0.3612 | 0.529 | -6.1 | -0.5681 | none |
| RMI1 | -0.094 | -0.1327 | -0.958 | 0.36126 | 0.529 | -6.1 | 0.96889 | none |
| TAGLN3 | -0.105 | -0.1137 | -0.958 | 0.36127 | 0.529 | -6.1 | -0.6876 | none |
| MFRP | -0.123 | -0.0793 | -0.958 | 0.36134 | 0.529 | -6.1 | 0.00804 | none |
| RAD51AP1 | -0.13 | -0.1114 | -0.958 | 0.3614 | 0.529 | -6.1 | 0.24235 | none |
| ICE1 | 0.256 | 0.3607 | 0.958 | 0.36142 | 0.529 | -6.1 | 0.24872 | none |
| ANKRD6 | -0.157 | -0.0772 | -0.958 | 0.3616 | 0.529 | -6.1 | 0.81832 | none |
| ITGB1BP2 | -0.161 | -0.0426 | -0.958 | 0.36163 | 0.529 | -6.1 | 1.38224 | none |
| MNX1 | 0.136 | 0.2089 | 0.9576 | 0.36164 | 0.529 | -6.1 | 0.74043 | none |
| HMGCS2 | 0.135 | 0.1175 | 0.9576 | 0.36165 | 0.529 | -6.1 | 0.49376 | none |
| IFT43 | -0.174 | -0.1073 | -0.957 | 0.36176 | 0.529 | -6.1 | 0.9846 | none |
| ZNF330 | -0.116 | -0.1457 | -0.957 | 0.36178 | 0.529 | -6.1 | 1.29873 | none |
| SCYL2 | 0.122 | 0.0969 | 0.9572 | 0.36183 | 0.529 | -6.1 | 0.70617 | none |
| CUL4B | 0.146 | 0.1611 | 0.9572 | 0.36183 | 0.529 | -6.1 | 0.55831 | none |
| AGO1 | 0.153 | 0.2046 | 0.9569 | 0.36197 | 0.529 | -6.1 | -0.7447 | none |
| SPRR4 | 0.101 | 0.1012 | 0.9567 | 0.36206 | 0.53 | -6.1 | -0.4 | none |
| PITX2 | 0.073 | 0.0669 | 0.9563 | 0.36224 | 0.53 | -6.1 | -0.7671 | none |
| COL12A1 | -0.08 | -0.0418 | -0.956 | 0.36234 | 0.53 | -6.1 | 0.3882 | none |
| ANKRD24 | -0.076 | -0.0286 | -0.956 | 0.3624 | 0.53 | -6.1 | 1.4464 | none |
| CNTRL | 0.17 | 0.297 | 0.9559 | 0.36242 | 0.53 | -6.1 | 0.12119 | none |
| PLPP4 | 0.121 | 0.1181 | 0.9559 | 0.36244 | 0.53 | -6.1 | 1.18623 | none |

| | | | | | | | | |
|---|---|---|---|---|---|---|---|---|
| PLCD4 | 0.264 | 0.3388 | 0.9558 | 0.3625 | 0.53 | -6.1 | 0.06877 | none |
| AKT1 | 0.126 | 0.1372 | 0.9558 | 0.36251 | 0.53 | -6.1 | -0.0941 | none |
| ACP5 | -0.179 | -0.1792 | -0.956 | 0.36255 | 0.53 | -6.1 | 1.64971 | none |
| TPPP2 | -0.097 | -0.0974 | -0.955 | 0.36274 | 0.53 | -6.1 | 0.79493 | none |
| SYCP2 | 0.139 | 0.0715 | 0.9552 | 0.36279 | 0.53 | -6.1 | -0.5016 | none |
| NOXO1 | 0.182 | 0.1685 | 0.955 | 0.36288 | 0.53 | -6.1 | 0.95464 | none |
| TEKT3 | 0.231 | 0.1922 | 0.9546 | 0.36305 | 0.53 | -6.1 | 0.72399 | none |
| CMC4 | 0.218 | 0.1479 | 0.9545 | 0.36314 | 0.531 | -6.1 | 0.61117 | none |
| KLF11 | -0.066 | -0.0154 | -0.954 | 0.36317 | 0.531 | -6.1 | 0.58261 | none |
| TPM2 | 0.194 | 0.1314 | 0.9543 | 0.36322 | 0.531 | -6.1 | -0.8337 | none |
| AQP2 | 0.079 | 0.083 | 0.9543 | 0.36323 | 0.531 | -6.1 | 0.36474 | none |
| TFAP2A | 0.096 | 0.113 | 0.9541 | 0.36332 | 0.531 | -6.1 | 0.92157 | none |
| ATG4A | -0.134 | -0.112 | -0.954 | 0.36335 | 0.531 | -6.1 | 1.97407 | none |
| TMEM102 | -0.087 | -0.0577 | -0.954 | 0.36335 | 0.531 | -6.1 | 1.01121 | none |
| TMPRSS5 | 0.136 | 0.156 | 0.9533 | 0.3637 | 0.531 | -6.1 | 1.55781 | none |
| HSPH1 | 0.343 | 0.5105 | 0.9532 | 0.36373 | 0.531 | -6.1 | 0.61306 | none |
| ZNF865 | 0.13 | 0.2099 | 0.953 | 0.36383 | 0.531 | -6.1 | -0.2756 | none |
| SPINK7 | -0.066 | -0.0443 | -0.953 | 0.36394 | 0.531 | -6.1 | -0.6074 | none |
| RAB18 | -0.144 | -0.1919 | -0.953 | 0.36398 | 0.531 | -6.1 | 1.53765 | none |
| OR13J1 | -0.065 | -0.0509 | -0.953 | 0.36404 | 0.531 | -6.1 | 0.12838 | none |
| CUX2 | 0.085 | 0.0964 | 0.9524 | 0.36412 | 0.531 | -6.1 | -0.2278 | none |
| SPARCL1 | 0.069 | 0.0863 | 0.9524 | 0.36412 | 0.531 | -6.1 | -0.3358 | none |
| C4ORF47 | -0.132 | -0.1542 | -0.952 | 0.36417 | 0.531 | -6.1 | 1.20884 | none |
| GRIK3 | 0.105 | 0.1084 | 0.9522 | 0.36422 | 0.531 | -6.1 | -0.4514 | none |
| FTSJ3 | 0.248 | 0.3852 | 0.9521 | 0.36427 | 0.531 | -6.1 | 0.23759 | none |
| TNFSF13B | -0.174 | -0.1116 | -0.952 | 0.36434 | 0.531 | -6.1 | 1.25253 | none |
| CPNE2 | 0.11 | 0.0554 | 0.9518 | 0.36441 | 0.532 | -6.1 | 1.393 | none |
| MSH5-SAF | 0.075 | 0.0853 | 0.9515 | 0.36458 | 0.532 | -6.11 | 0.78767 | none |
| LRRK2 | 0.135 | 0.1346 | 0.951 | 0.36481 | 0.532 | -6.11 | 1.21341 | none |
| RTN4RL1 | 0.082 | 0.0674 | 0.9506 | 0.36497 | 0.532 | -6.11 | -0.5447 | none |
| SYTL3 | -0.157 | -0.0546 | -0.951 | 0.365 | 0.532 | -6.11 | 1.35389 | none |
| RANGRF | -0.175 | -0.2005 | -0.951 | 0.36501 | 0.532 | -6.11 | 0.96847 | none |
| SCAF4 | 0.279 | 0.4163 | 0.9505 | 0.36505 | 0.532 | -6.11 | 0.79817 | none |
| GPR107 | 0.081 | 0.1206 | 0.9496 | 0.36547 | 0.533 | -6.11 | 0.77381 | none |
| PPP2R2A | 0.103 | 0.1026 | 0.9494 | 0.36558 | 0.533 | -6.11 | 0.00194 | none |
| EIF5 | 0.144 | 0.2126 | 0.9493 | 0.36561 | 0.533 | -6.11 | -0.1389 | none |
| VPS16 | 0.107 | 0.1678 | 0.9491 | 0.36573 | 0.533 | -6.11 | 0.84428 | none |
| EXO5 | -0.099 | -0.1685 | -0.949 | 0.3658 | 0.533 | -6.11 | -0.6852 | none |
| GPR17 | 0.116 | 0.116 | 0.9485 | 0.36599 | 0.533 | -6.11 | 1.01342 | none |
| MAGEE1 | 0.112 | 0.1189 | 0.9483 | 0.36612 | 0.533 | -6.11 | -0.1542 | none |

| | | | | | | | | |
|---|---|---|---|---|---|---|---|---|
| MT-ND3 | -0.148 | -0.1861 | -0.948 | 0.36623 | 0.534 | -6.11 | 0.69041 | none |
| TSPO | -0.136 | -0.1451 | -0.947 | 0.36667 | 0.534 | -6.11 | 0.88127 | none |
| STPG3 | -0.123 | -0.1035 | -0.947 | 0.36668 | 0.534 | -6.11 | 1.64791 | none |
| MTDH | 0.241 | 0.3264 | 0.9466 | 0.36693 | 0.534 | -6.11 | 0.53604 | none |
| CYP39A1 | 0.082 | 0.0812 | 0.9466 | 0.36694 | 0.534 | -6.11 | 0.74491 | none |
| MRPL3 | -0.093 | -0.1336 | -0.946 | 0.36708 | 0.535 | -6.11 | 1.12175 | none |
| RABGGTB | -0.123 | -0.1908 | -0.946 | 0.36709 | 0.535 | -6.11 | 0.23467 | none |
| CPOX | -0.173 | -0.1339 | -0.946 | 0.36715 | 0.535 | -6.11 | 1.02009 | none |
| HOXB6 | 0.079 | 0.0571 | 0.946 | 0.36721 | 0.535 | -6.11 | 0.35788 | none |
| SIGMAR1 | 0.096 | 0.0768 | 0.9459 | 0.36725 | 0.535 | -6.11 | 0.61041 | none |
| NCAPD3 | 0.192 | 0.2811 | 0.9459 | 0.36726 | 0.535 | -6.11 | 0.95982 | none |
| DCT | -0.06 | -0.0343 | -0.946 | 0.36735 | 0.535 | -6.11 | 0 | none |
| PAGR1 | -0.073 | -0.0882 | -0.946 | 0.36745 | 0.535 | -6.11 | 1.47899 | none |
| KIAA1147 | 0.126 | 0.1641 | 0.9452 | 0.36759 | 0.535 | -6.11 | 0.24487 | none |
| STS | 0.093 | 0.1166 | 0.9451 | 0.36766 | 0.535 | -6.11 | 0.19953 | none |
| ARPP21 | -0.182 | -0.0756 | -0.945 | 0.36786 | 0.535 | -6.11 | 0.41796 | none |
| GALK1 | -0.168 | -0.189 | -0.944 | 0.36798 | 0.535 | -6.11 | 0.80055 | none |
| KIAA0556 | 0.209 | 0.2109 | 0.9443 | 0.36804 | 0.535 | -6.11 | -0.8035 | none |
| GAMT | -0.115 | -0.1407 | -0.944 | 0.36823 | 0.536 | -6.11 | 0.71002 | none |
| PSG2 | 0.308 | 0.261 | 0.9439 | 0.36824 | 0.536 | -6.11 | 0.62269 | none |
| PLCL2 | 0.147 | 0.2018 | 0.9438 | 0.36829 | 0.536 | -6.11 | 0.22687 | none |
| FLOT2 | -0.12 | -0.1321 | -0.944 | 0.36835 | 0.536 | -6.11 | 1.68054 | none |
| TACC3 | 0.227 | 0.3297 | 0.9434 | 0.36847 | 0.536 | -6.11 | -0.0164 | none |
| CCL3 | -0.526 | -0.1277 | -0.943 | 0.36859 | 0.536 | -6.11 | 0.65427 | none |
| FBXO28 | 0.118 | 0.1116 | 0.9432 | 0.36859 | 0.536 | -6.11 | -0.118 | none |
| EEF2K | -0.249 | -0.1392 | -0.943 | 0.36869 | 0.536 | -6.11 | -0.6134 | none |
| CRYBB2 | 0.092 | 0.093 | 0.9428 | 0.36879 | 0.536 | -6.11 | 0.46526 | none |
| KCNE2 | 0.075 | 0.0825 | 0.9427 | 0.36882 | 0.536 | -6.11 | 0.88671 | none |
| MSMB | -0.094 | -0.0938 | -0.942 | 0.36912 | 0.536 | -6.11 | 1.23273 | none |
| PRAMEF2( | -0.094 | -0.0938 | -0.942 | 0.36914 | 0.536 | -6.11 | 0.85875 | none |
| MEIS2 | -0.102 | -0.0788 | -0.942 | 0.3693 | 0.537 | -6.11 | 0.90863 | none |
| APCDD1L | -0.081 | -0.0535 | -0.942 | 0.36936 | 0.537 | -6.11 | 1.08564 | none |
| POLI | 0.14 | 0.1175 | 0.9413 | 0.36949 | 0.537 | -6.11 | 0.17005 | none |
| KAZALD1 | -0.072 | -0.0884 | -0.941 | 0.36962 | 0.537 | -6.12 | 1.80404 | none |
| ZDHHC12 | -0.12 | -0.0863 | -0.941 | 0.36971 | 0.537 | -6.12 | 2.24396 | AA-related |
| PDSS2 | 0.131 | 0.0565 | 0.9406 | 0.36982 | 0.537 | -6.12 | -0.4698 | none |
| CCSAP | 0.139 | 0.1493 | 0.9405 | 0.3699 | 0.537 | -6.12 | 0.31584 | none |
| TRPC5 | 0.065 | 0.0384 | 0.9402 | 0.37004 | 0.537 | -6.12 | -0.7667 | none |
| CCT6B | 0.108 | 0.0615 | 0.9401 | 0.37009 | 0.537 | -6.12 | -0.6194 | none |
| EPPIN-WFI | -0.064 | -0.0465 | -0.94 | 0.37009 | 0.537 | -6.12 | 1.0871 | none |

| | | | | | | | | |
|---|---|---|---|---|---|---|---|---|
| SNAPC4 | 0.228 | 0.3066 | 0.9399 | 0.37018 | 0.537 | -6.12 | 0.25407 | none |
| ZNF764 | 0.089 | 0.0389 | 0.9398 | 0.37023 | 0.537 | -6.12 | 0.35391 | none |
| ORMDL2 | -0.221 | -0.3399 | -0.94 | 0.37026 | 0.537 | -6.12 | 0.9472 | none |
| NUDT1 | -0.119 | -0.1847 | -0.94 | 0.37027 | 0.537 | -6.12 | 0.16236 | none |
| CLEC12A | 0.083 | 0.0248 | 0.9396 | 0.37035 | 0.538 | -6.12 | -0.4366 | none |
| RPS6KA1 | -0.088 | -0.025 | -0.939 | 0.37041 | 0.538 | -6.12 | 1.40482 | none |
| AC022210 | -0.173 | -0.0973 | -0.939 | 0.37054 | 0.538 | -6.12 | 0.92351 | none |
| WWC2 | 0.105 | 0.0912 | 0.9392 | 0.37055 | 0.538 | -6.12 | 0.06 | none |
| OR4N5 | -0.065 | -0.0435 | -0.939 | 0.37058 | 0.538 | -6.12 | 0 | none |
| MYEF2 | 0.209 | 0.1114 | 0.9391 | 0.3706 | 0.538 | -6.12 | -0.7558 | none |
| BOK | 0.084 | 0.0757 | 0.9388 | 0.37074 | 0.538 | -6.12 | 0.37749 | none |
| SEC14L5 | -0.062 | -0.0344 | -0.939 | 0.37077 | 0.538 | -6.12 | 0.96085 | none |
| GABBR2 | 0.11 | 0.0877 | 0.9384 | 0.37093 | 0.538 | -6.12 | -0.0776 | none |
| SLC1A3 | -0.103 | -0.0804 | -0.938 | 0.37096 | 0.538 | -6.12 | 1.02978 | none |
| CTXN1 | -0.072 | -0.0956 | -0.938 | 0.37108 | 0.538 | -6.12 | 0.93162 | none |
| PLD5 | 0.067 | 0.0426 | 0.9379 | 0.37115 | 0.538 | -6.12 | 1.6923 | none |
| FAM71D | -0.186 | 0.0157 | -0.937 | 0.37138 | 0.538 | -6.12 | 0.01789 | none |
| MYD88 | -0.094 | -0.1023 | -0.937 | 0.37143 | 0.538 | -6.12 | 0.8347 | none |
| AC006445 | 0.439 | 0.2772 | 0.9373 | 0.37145 | 0.538 | -6.12 | 0.47873 | none |
| CENPJ | 0.22 | 0.3381 | 0.937 | 0.37159 | 0.539 | -6.12 | 0.55494 | none |
| CRYGC | -0.092 | -0.0925 | -0.937 | 0.37171 | 0.539 | -6.12 | 1.9054 | none |
| DNAL4 | -0.167 | -0.1632 | -0.937 | 0.37179 | 0.539 | -6.12 | 0.81817 | none |
| MRPS11 | -0.077 | -0.0711 | -0.937 | 0.3718 | 0.539 | -6.12 | -0.2083 | none |
| PCDHGC5 | 0.067 | 0.0668 | 0.9365 | 0.37183 | 0.539 | -6.12 | -0.3646 | none |
| CHRNB2 | 0.16 | 0.1537 | 0.9364 | 0.37189 | 0.539 | -6.12 | -0.2365 | none |
| APMAP | -0.104 | -0.1375 | -0.936 | 0.37204 | 0.539 | -6.12 | 1.29681 | none |
| ENDOD1 | -0.084 | -0.0344 | -0.936 | 0.37213 | 0.539 | -6.12 | 1.6292 | none |
| SARS | -0.067 | -0.0483 | -0.936 | 0.37213 | 0.539 | -6.12 | 0.29505 | none |
| CLRN1 | 0.081 | 0.0424 | 0.9357 | 0.37226 | 0.539 | -6.12 | -0.6057 | none |
| CDIPT | -0.113 | -0.137 | -0.935 | 0.37234 | 0.539 | -6.12 | 0.52984 | none |
| ANKAR | -0.164 | -0.0969 | -0.935 | 0.37235 | 0.539 | -6.12 | 0.20801 | none |
| CABLES2 | 0.165 | 0.2449 | 0.9354 | 0.37237 | 0.539 | -6.12 | 0.7047 | none |
| TNFRSF1B | 0.134 | 0.166 | 0.9352 | 0.37249 | 0.539 | -6.12 | 0.32632 | none |
| HTATSF1 | 0.223 | 0.3261 | 0.935 | 0.37259 | 0.539 | -6.12 | 0.68512 | none |
| GNA15 | -0.099 | -0.0995 | -0.935 | 0.37265 | 0.539 | -6.12 | 1.05058 | none |
| FLI1 | 0.16 | 0.169 | 0.9349 | 0.37265 | 0.539 | -6.12 | -0.8754 | none |
| LILRB3 | 0.13 | 0.0601 | 0.9347 | 0.37275 | 0.54 | -6.12 | 1.28775 | none |
| MAP3K7 | 0.129 | 0.1651 | 0.9346 | 0.37278 | 0.54 | -6.12 | 0.64501 | none |
| CNTFR | -0.069 | -0.0426 | -0.934 | 0.37289 | 0.54 | -6.12 | 0.75286 | none |
| NPIPA3 | 0.127 | 0.1281 | 0.9344 | 0.37289 | 0.54 | -6.12 | 1.47931 | none |

| | | | | | | | | |
|---|---|---|---|---|---|---|---|---|
| FAM24A | -0.092 | -0.0918 | -0.934 | 0.3731 | 0.54 | -6.12 | -0.5011 | none |
| HSPB8 | -0.079 | -0.0394 | -0.933 | 0.37335 | 0.54 | -6.12 | -0.0323 | none |
| CXORF49 | 0.102 | 0.1144 | 0.9333 | 0.3734 | 0.54 | -6.12 | 1.11704 | none |
| CXORF49B | 0.102 | 0.1144 | 0.9333 | 0.3734 | 0.54 | -6.12 | 1.11704 | none |
| MED7 | 0.12 | 0.0653 | 0.9331 | 0.37352 | 0.54 | -6.12 | -0.3731 | none |
| MAP10 | -0.097 | -0.0359 | -0.933 | 0.37356 | 0.54 | -6.12 | 0.77295 | none |
| POFUT2 | -0.18 | -0.1287 | -0.933 | 0.37363 | 0.54 | -6.12 | 0.17041 | none |
| RAB3C | 0.089 | 0.0873 | 0.9328 | 0.37366 | 0.54 | -6.12 | 1.75078 | none |
| CAPN14 | -0.066 | -0.0461 | -0.933 | 0.37376 | 0.54 | -6.12 | 0 | none |
| TBC1D10C | -0.102 | -0.1062 | -0.933 | 0.37379 | 0.54 | -6.12 | 0.41133 | none |
| GABPA | 0.142 | 0.1558 | 0.9325 | 0.3738 | 0.54 | -6.12 | -0.2592 | none |
| SLC45A2 | -0.103 | -0.1029 | -0.932 | 0.37388 | 0.54 | -6.12 | 1.17337 | none |
| CTSZ | -0.087 | -0.0319 | -0.932 | 0.37391 | 0.54 | -6.12 | 1.58549 | none |
| STPG2 | -0.064 | -0.0412 | -0.932 | 0.37397 | 0.54 | -6.12 | 1.64801 | none |
| CAT | 0.083 | -0.0179 | 0.9321 | 0.374 | 0.54 | -6.12 | -0.6442 | none |
| OR1D2 | 0.096 | 0.0921 | 0.9321 | 0.37402 | 0.54 | -6.12 | 0.24304 | none |
| SLBP | 0.113 | 0.1361 | 0.9319 | 0.37408 | 0.541 | -6.12 | 1.38233 | none |
| TMEM238 | -0.115 | -0.1382 | -0.932 | 0.37418 | 0.541 | -6.12 | 0.51224 | none |
| FBXO30 | 0.092 | 0.0464 | 0.9312 | 0.37443 | 0.541 | -6.12 | 0.75165 | none |
| ITM2C | -0.085 | -0.1357 | -0.931 | 0.37446 | 0.541 | -6.12 | 0.74437 | none |
| P2RX2 | 0.093 | 0.0611 | 0.9311 | 0.37451 | 0.541 | -6.12 | -0.1898 | none |
| BBOX1 | 0.094 | 0.094 | 0.9309 | 0.37461 | 0.541 | -6.12 | -0.8888 | none |
| RBM15 | 0.093 | 0.1173 | 0.9308 | 0.37466 | 0.541 | -6.12 | 0.37795 | none |
| ALG9 | 0.133 | 0.0022 | 0.9305 | 0.37479 | 0.541 | -6.12 | 0.53439 | none |
| JPH1 | 0.088 | 0.1177 | 0.9304 | 0.37483 | 0.541 | -6.12 | -0.0316 | none |
| PRR23A | 0.094 | 0.0939 | 0.9304 | 0.37485 | 0.541 | -6.12 | 0.99131 | none |
| SNAPC2 | -0.099 | -0.0797 | -0.93 | 0.3749 | 0.541 | -6.13 | 0.39253 | none |
| C22ORF42 | 0.084 | 0.0912 | 0.9303 | 0.37491 | 0.541 | -6.13 | 0.59266 | none |
| ZNF598 | 0.18 | 0.3178 | 0.9296 | 0.37523 | 0.542 | -6.13 | 0.18608 | none |
| MIGA2 | 0.129 | 0.1063 | 0.9295 | 0.3753 | 0.542 | -6.13 | 0.44386 | none |
| PDLIM4 | 0.152 | 0.1676 | 0.9293 | 0.37538 | 0.542 | -6.13 | 0.88628 | none |
| STMN3 | 0.114 | 0.0741 | 0.9292 | 0.37542 | 0.542 | -6.13 | -0.3813 | none |
| OSBPL5 | -0.089 | -0.0483 | -0.929 | 0.37569 | 0.542 | -6.13 | 0.70259 | none |
| CXORF66 | 0.066 | 0.0663 | 0.9282 | 0.37592 | 0.542 | -6.13 | -0.129 | none |
| CBWD5 | 0.09 | 0.011 | 0.928 | 0.37603 | 0.542 | -6.13 | 0.1967 | none |
| RNGTT | 0.143 | 0.1031 | 0.9279 | 0.37607 | 0.543 | -6.13 | -0.8002 | none |
| CFAP52 | 0.3 | 0.2763 | 0.9278 | 0.37615 | 0.543 | -6.13 | 0.4904 | none |
| GRIN1 | 0.115 | 0.1171 | 0.9275 | 0.37628 | 0.543 | -6.13 | 0.35447 | none |
| ENTPD6 | 0.171 | 0.1205 | 0.9275 | 0.37628 | 0.543 | -6.13 | 0.33997 | none |
| NOL10 | 0.22 | 0.2662 | 0.9273 | 0.37639 | 0.543 | -6.13 | -0.0143 | none |

| | | | | | | | | |
|---|---|---|---|---|---|---|---|---|
| HIST1H3C | 0.483 | 0.2801 | 0.9272 | 0.37642 | 0.543 | -6.13 | -0.1693 | none |
| NOP56 | 0.16 | 0.2861 | 0.9271 | 0.37645 | 0.543 | -6.13 | 0.51476 | none |
| MYCL | 0.079 | 0.0926 | 0.9271 | 0.37646 | 0.543 | -6.13 | 0.52499 | none |
| COX4I2 | 0.093 | 0.093 | 0.9271 | 0.37647 | 0.543 | -6.13 | 1.19688 | none |
| ISX | 0.068 | 0.0423 | 0.927 | 0.3765 | 0.543 | -6.13 | -1.317 | none |
| MYLPF | 0.333 | 0.2159 | 0.9269 | 0.37656 | 0.543 | -6.13 | -0.182 | none |
| UPK2 | 0.078 | 0.0974 | 0.9258 | 0.3771 | 0.543 | -6.13 | 1.5828 | none |
| OR1J2 | -0.064 | -0.0427 | -0.926 | 0.37719 | 0.544 | -6.13 | 0.49761 | none |
| PMEL | 0.163 | 0.2582 | 0.9256 | 0.37722 | 0.544 | -6.13 | 1.22934 | none |
| GRAMD1C | 0.143 | 0.1598 | 0.9255 | 0.37724 | 0.544 | -6.13 | 0.02599 | none |
| MTUS1 | 0.358 | 0.4025 | 0.9252 | 0.37742 | 0.544 | -6.13 | 0.79564 | none |
| NOS2 | 0.068 | 0.0524 | 0.9249 | 0.37755 | 0.544 | -6.13 | -0.5364 | none |
| CYBA | -0.188 | -0.2404 | -0.925 | 0.37769 | 0.544 | -6.13 | 0.28056 | none |
| FBLN7 | -0.112 | -0.0903 | -0.925 | 0.3777 | 0.544 | -6.13 | 1.42536 | none |
| LINS1 | 0.096 | 0.0955 | 0.9246 | 0.3777 | 0.544 | -6.13 | 0.32117 | none |
| GRIA2 | -0.088 | -0.0884 | -0.924 | 0.37779 | 0.544 | -6.13 | 1.58675 | none |
| ARHGEF33 | -0.192 | -0.1457 | -0.924 | 0.37805 | 0.544 | -6.13 | 0.35461 | none |
| MMP3 | 0.09 | 0.0902 | 0.9239 | 0.37807 | 0.544 | -6.13 | 0.04014 | none |
| ITGB2 | -0.099 | -0.0403 | -0.924 | 0.3781 | 0.544 | -6.13 | 0.97685 | none |
| TMEM255 | 0.072 | 0.0744 | 0.9236 | 0.37819 | 0.544 | -6.13 | 0.21466 | none |
| CLIC3 | -0.202 | -0.1523 | -0.923 | 0.37845 | 0.545 | -6.13 | 1.64428 | none |
| PPP1R1B | 0.093 | 0.0817 | 0.923 | 0.37851 | 0.545 | -6.13 | -0.1276 | none |
| AOC3 | 0.172 | 0.1941 | 0.9225 | 0.37874 | 0.545 | -6.13 | -0.3458 | none |
| FAM204A | 0.079 | 0.0389 | 0.9224 | 0.37877 | 0.545 | -6.13 | 0.17839 | none |
| C20ORF14 | 0.092 | 0.0918 | 0.9223 | 0.37887 | 0.545 | -6.13 | 0.54171 | none |
| SERPINA6 | 0.068 | 0.066 | 0.9215 | 0.37923 | 0.546 | -6.13 | 1.52692 | none |
| SLC25A12 | 0.111 | 0.0587 | 0.9211 | 0.37942 | 0.546 | -6.13 | 0.1067 | none |
| AC015688 | -0.089 | -0.0887 | -0.921 | 0.37944 | 0.546 | -6.13 | 0.92737 | none |
| TSPAN4 | 0.174 | 0.2355 | 0.9211 | 0.37945 | 0.546 | -6.13 | -0.1908 | none |
| MRFAP1 | -0.096 | -0.1061 | -0.921 | 0.37947 | 0.546 | -6.13 | 0.64748 | none |
| SRR | 0.098 | -0.0107 | 0.9204 | 0.37977 | 0.546 | -6.13 | -0.3196 | none |
| ATXN7L2 | 0.129 | 0.2307 | 0.9202 | 0.37986 | 0.546 | -6.13 | -0.4545 | none |
| ACTR6 | 0.181 | 0.0822 | 0.9202 | 0.37987 | 0.546 | -6.13 | 0.60302 | none |
| SLC39A2 | 0.154 | 0.0787 | 0.9195 | 0.38024 | 0.547 | -6.13 | -0.4938 | none |
| GSPT2 | 0.122 | 0.1643 | 0.919 | 0.3805 | 0.547 | -6.14 | -0.4318 | none |
| POTEF | 0.116 | 0.0911 | 0.9188 | 0.3806 | 0.547 | -6.14 | -0.2479 | none |
| KHK | 0.086 | 0.0703 | 0.9186 | 0.38068 | 0.547 | -6.14 | 1.14526 | none |
| RFX3 | 0.205 | 0.2726 | 0.9181 | 0.38093 | 0.548 | -6.14 | -0.5658 | none |
| MATN3 | 0.101 | 0.1005 | 0.9179 | 0.38104 | 0.548 | -6.14 | -0.7204 | none |
| PKP1 | 0.072 | 0.0762 | 0.9179 | 0.38105 | 0.548 | -6.14 | -0.6636 | none |

| | | | | | | | | |
|---|---|---|---|---|---|---|---|---|
| ENDOG | -0.162 | -0.148 | -0.918 | 0.38107 | 0.548 | -6.14 | 0.95275 | none |
| TOB2 | 0.11 | 0.1648 | 0.9177 | 0.38111 | 0.548 | -6.14 | 0.71713 | none |
| ACADS | 0.111 | 0.0929 | 0.9172 | 0.38139 | 0.548 | -6.14 | 0.66368 | none |
| CHKB | 0.108 | 0.1024 | 0.917 | 0.38146 | 0.548 | -6.14 | 0.6287 | none |
| GPR108 | -0.098 | -0.0613 | -0.917 | 0.3817 | 0.548 | -6.14 | 0.29607 | none |
| UBE3B | 0.099 | 0.0456 | 0.9164 | 0.38179 | 0.548 | -6.14 | 0.09373 | none |
| STRA8 | -0.063 | -0.0429 | -0.916 | 0.38184 | 0.548 | -6.14 | 1.88681 | none |
| TSC22D3 | 0.256 | 0.1455 | 0.9159 | 0.38202 | 0.549 | -6.14 | 0.25675 | none |
| COL2A1 | -0.086 | -0.0991 | -0.916 | 0.38214 | 0.549 | -6.14 | 0.23047 | none |
| ARF5 | -0.138 | -0.1629 | -0.915 | 0.38223 | 0.549 | -6.14 | 1.53276 | none |
| ZFHX4 | -0.182 | -0.1806 | -0.915 | 0.38238 | 0.549 | -6.14 | 1.40875 | none |
| TOP2A | 0.323 | 0.476 | 0.9151 | 0.38241 | 0.549 | -6.14 | 0.28544 | none |
| SCRN1 | -0.116 | -0.0528 | -0.915 | 0.38258 | 0.549 | -6.14 | 0.27899 | none |
| ZC3H12A | 0.186 | 0.2877 | 0.9148 | 0.38259 | 0.549 | -6.14 | 0.35558 | none |
| TSKS | 0.179 | 0.1323 | 0.9141 | 0.3829 | 0.55 | -6.14 | 0.36152 | none |
| OR52W1 | 0.078 | 0.0817 | 0.9139 | 0.38302 | 0.55 | -6.14 | 0.93667 | none |
| PTPRH | 0.091 | 0.0648 | 0.9134 | 0.38327 | 0.55 | -6.14 | 0.22636 | none |
| ADAMTSL! | 0.179 | 0.1609 | 0.9129 | 0.38352 | 0.55 | -6.14 | -0.121 | none |
| PAQR9 | 0.073 | 0.0785 | 0.9129 | 0.38352 | 0.55 | -6.14 | -0.76 | none |
| PAX2 | 0.086 | 0.0915 | 0.9126 | 0.38366 | 0.55 | -6.14 | 1.0248 | none |
| BPI | -0.071 | -0.0652 | -0.913 | 0.38368 | 0.55 | -6.14 | 1.10458 | none |
| MOB3C | 0.098 | 0.1519 | 0.9125 | 0.38374 | 0.55 | -6.14 | -0.3759 | none |
| RBM17 | 0.127 | 0.2074 | 0.9124 | 0.38377 | 0.55 | -6.14 | -0.0631 | none |
| HAUS6 | 0.21 | 0.2456 | 0.9119 | 0.38405 | 0.551 | -6.14 | 0.72788 | none |
| RGS8 | -0.113 | -0.1149 | -0.912 | 0.38412 | 0.551 | -6.14 | 0.04322 | none |
| CLCN4 | -0.064 | -0.0413 | -0.912 | 0.38415 | 0.551 | -6.14 | 1.43582 | none |
| TTC9C | -0.121 | -0.1633 | -0.912 | 0.38421 | 0.551 | -6.14 | 1.84487 | none |
| CLEC5A | -0.063 | -0.0419 | -0.911 | 0.38427 | 0.551 | -6.14 | -0.7456 | none |
| HIST1H2AI | -0.173 | -0.2509 | -0.911 | 0.38447 | 0.551 | -6.14 | 0.32053 | none |
| CCDC107 | -0.103 | -0.1219 | -0.911 | 0.38465 | 0.551 | -6.14 | 1.78212 | none |
| GIP | 0.089 | 0.089 | 0.9106 | 0.38467 | 0.551 | -6.14 | 0.83946 | none |
| MRGPRE | 0.064 | 0.0471 | 0.9101 | 0.38492 | 0.552 | -6.14 | -1.2832 | none |
| OTUD7A | 0.155 | 0.1653 | 0.9101 | 0.38495 | 0.552 | -6.14 | 0.9063 | none |
| TMEM30A | 0.118 | 0.0682 | 0.91 | 0.38496 | 0.552 | -6.14 | 1.18465 | none |
| OLFML1 | 0.061 | 0.0426 | 0.9099 | 0.38502 | 0.552 | -6.14 | 0.47524 | none |
| F2RL1 | 0.101 | 0.022 | 0.9098 | 0.38507 | 0.552 | -6.14 | 0.22653 | none |
| PLB1 | -0.119 | -0.165 | -0.909 | 0.38527 | 0.552 | -6.14 | 1.67372 | none |
| CYP11A1 | 0.153 | 0.1177 | 0.909 | 0.38547 | 0.552 | -6.14 | -0.3246 | none |
| NKX2-6 | 0.088 | 0.0885 | 0.9086 | 0.3857 | 0.552 | -6.14 | -0.71 | none |
| PPP1R8 | 0.159 | 0.0871 | 0.9084 | 0.38576 | 0.553 | -6.15 | -0.2836 | none |

| | | | | | | | | |
|---|---|---|---|---|---|---|---|---|
| FSIP1 | 0.076 | 0.0288 | 0.9082 | 0.38587 | 0.553 | -6.15 | 0.02324 | none |
| RPL9 | -0.165 | -0.2445 | -0.908 | 0.3859 | 0.553 | -6.15 | 0.82223 | none |
| DRICH1 | 0.081 | 0.0451 | 0.908 | 0.38598 | 0.553 | -6.15 | 0.98676 | none |
| NFKB1 | 0.176 | 0.2769 | 0.9078 | 0.38608 | 0.553 | -6.15 | -0.0887 | none |
| LRRC45 | 0.135 | 0.1805 | 0.9077 | 0.38614 | 0.553 | -6.15 | -0.0385 | none |
| FAM86B2 | 0.263 | 0.1904 | 0.9074 | 0.38629 | 0.553 | -6.15 | 0.74523 | none |
| GCGR | -0.077 | -0.056 | -0.907 | 0.38629 | 0.553 | -6.15 | 1.34322 | none |
| CMTR2 | 0.139 | 0.0791 | 0.9073 | 0.38635 | 0.553 | -6.15 | 0.72713 | none |
| NYX | -0.081 | -0.0556 | -0.907 | 0.38637 | 0.553 | -6.15 | 0.88075 | none |
| CSRNP3 | -0.108 | -0.1076 | -0.906 | 0.38682 | 0.554 | -6.15 | -0.2684 | none |
| PUM3 | 0.218 | 0.3324 | 0.9062 | 0.38691 | 0.554 | -6.15 | 0.49596 | none |
| ARL2-SNX1 | 0.075 | 0.0803 | 0.9055 | 0.38725 | 0.554 | -6.15 | 0.46135 | none |
| CCL26 | -0.085 | -0.0851 | -0.905 | 0.3874 | 0.554 | -6.15 | -0.2281 | none |
| TMEM35B | -0.122 | -0.234 | -0.905 | 0.38751 | 0.554 | -6.15 | 1.21771 | none |
| HHLA1 | 0.161 | 0.1824 | 0.9048 | 0.38758 | 0.554 | -6.15 | 0.11558 | none |
| LTB | 0.143 | 0.1342 | 0.9046 | 0.3877 | 0.555 | -6.15 | 0.25319 | none |
| QSOX1 | -0.092 | -0.0331 | -0.904 | 0.3878 | 0.555 | -6.15 | 1.36941 | none |
| EPHX3 | 0.064 | 0.0721 | 0.9042 | 0.38789 | 0.555 | -6.15 | 0.58211 | none |
| NIP7 | 0.097 | -0.0031 | 0.9038 | 0.38809 | 0.555 | -6.15 | -0.9283 | none |
| IL23A | -0.08 | -0.0303 | -0.903 | 0.38829 | 0.555 | -6.15 | 0.62411 | none |
| ZNF706 | -0.126 | -0.1736 | -0.903 | 0.38837 | 0.555 | -6.15 | 0.30656 | none |
| ANXA9 | 0.102 | 0.1483 | 0.9032 | 0.38841 | 0.555 | -6.15 | -0.2037 | none |
| UGT1A7 | 0.075 | 0.0809 | 0.903 | 0.38852 | 0.555 | -6.15 | 0.84138 | none |
| IFI16 | 0.313 | 0.4087 | 0.9028 | 0.38859 | 0.555 | -6.15 | 0.48107 | none |
| PPP2R1B | 0.239 | 0.1137 | 0.9027 | 0.38863 | 0.555 | -6.15 | -0.8067 | none |
| SORBS3 | 0.114 | 0.0803 | 0.9026 | 0.38869 | 0.555 | -6.15 | 0.56721 | none |
| HTRA1 | 0.062 | 0.0645 | 0.9024 | 0.38882 | 0.556 | -6.15 | 1.2718 | none |
| GOLGA6A | 0.062 | 0.0644 | 0.9014 | 0.3893 | 0.556 | -6.15 | -0.1228 | none |
| TMX4 | 0.168 | 0.0687 | 0.9011 | 0.38946 | 0.556 | -6.15 | 1.12528 | none |
| PDCD6 | -0.109 | -0.1183 | -0.901 | 0.38967 | 0.557 | -6.15 | 0.55982 | none |
| CYB561D2 | -0.129 | -0.1236 | -0.9 | 0.38992 | 0.557 | -6.15 | 0.59615 | none |
| ZFAND6 | -0.137 | -0.2027 | -0.9 | 0.38995 | 0.557 | -6.15 | 0.84663 | none |
| VCP | -0.098 | -0.0529 | -0.9 | 0.38997 | 0.557 | -6.15 | 1.47558 | none |
| REG1A | 0.086 | 0.0865 | 0.8998 | 0.39013 | 0.557 | -6.15 | 0.28046 | none |
| MT-ND1 | 0.164 | 0.1871 | 0.8996 | 0.39021 | 0.557 | -6.15 | 0.18992 | none |
| ORAI3 | 0.089 | 0.0741 | 0.8991 | 0.39049 | 0.558 | -6.15 | -0.2725 | none |
| IFITM10 | 0.086 | 0.1184 | 0.8989 | 0.39055 | 0.558 | -6.15 | 0.92068 | none |
| NAALAD2 | 0.065 | 0.0748 | 0.8984 | 0.39081 | 0.558 | -6.15 | 0.75975 | none |
| B4GALNT4 | -0.116 | -0.0986 | -0.898 | 0.39103 | 0.558 | -6.15 | 1.87243 | none |
| RBP2 | 0.096 | 0.094 | 0.8979 | 0.39107 | 0.558 | -6.15 | 0.44088 | none |

| | | | | | | | | |
|---|---|---|---|---|---|---|---|---|
| RFPL1 | -0.069 | -0.0048 | -0.898 | 0.3911 | 0.558 | -6.15 | 0.13082 | none |
| PLAC8 | -0.154 | -0.2306 | -0.898 | 0.39124 | 0.558 | -6.15 | 0.51917 | none |
| FRG1 | -0.096 | -0.0852 | -0.898 | 0.39127 | 0.558 | -6.15 | 0.71692 | none |
| ADAMTSL1 | -0.071 | -0.0726 | -0.897 | 0.3914 | 0.558 | -6.16 | 0.71239 | none |
| IFI27L1 | -0.204 | -0.2233 | -0.897 | 0.39142 | 0.558 | -6.16 | 0.12348 | none |
| IRX1 | -0.063 | -0.0547 | -0.897 | 0.39143 | 0.558 | -6.16 | 0 | none |
| DUSP13 | -0.091 | -0.0978 | -0.897 | 0.39148 | 0.558 | -6.16 | 0.60746 | none |
| SLC15A2 | 0.121 | 0.0759 | 0.897 | 0.39154 | 0.559 | -6.16 | 0.47448 | none |
| CPNE1 | 0.08 | 0.0501 | 0.8968 | 0.39163 | 0.559 | -6.16 | -0.8242 | none |
| TMX2 | -0.117 | -0.1477 | -0.897 | 0.39164 | 0.559 | -6.16 | 1.79856 | none |
| CARD17 | 0.178 | -0.0592 | 0.8966 | 0.39172 | 0.559 | -6.16 | 0.57062 | none |
| C20ORF96 | -0.161 | -0.0892 | -0.896 | 0.39182 | 0.559 | -6.16 | 1.85057 | none |
| ARHGEF9 | 0.219 | 0.2795 | 0.8964 | 0.39182 | 0.559 | -6.16 | -0.4517 | none |
| ASB1 | 0.181 | 0.1294 | 0.8963 | 0.3919 | 0.559 | -6.16 | 0.28787 | none |
| COL16A1 | 0.121 | 0.1585 | 0.8959 | 0.3921 | 0.559 | -6.16 | -0.582 | none |
| GBP1 | 0.235 | 0.2744 | 0.8954 | 0.39234 | 0.559 | -6.16 | 0.36649 | none |
| TRNP1 | 0.075 | 0.0719 | 0.8953 | 0.39241 | 0.559 | -6.16 | -0.5868 | none |
| LHX6 | 0.266 | 0.2132 | 0.8951 | 0.39249 | 0.559 | -6.16 | -0.8792 | none |
| ARL6IP6 | -0.182 | -0.2235 | -0.895 | 0.39262 | 0.56 | -6.16 | 1.11681 | none |
| CFHR3 | 0.08 | 0.0692 | 0.8946 | 0.39275 | 0.56 | -6.16 | 0.88077 | none |
| ZNF618 | 0.068 | 0.0731 | 0.8944 | 0.39286 | 0.56 | -6.16 | 0.46139 | none |
| CA11 | 0.099 | 0.1472 | 0.8941 | 0.39299 | 0.56 | -6.16 | 0.52765 | none |
| PDC | -0.063 | -0.0545 | -0.894 | 0.39299 | 0.56 | -6.16 | 0 | none |
| AURKC | 0.126 | 0.1322 | 0.8938 | 0.39316 | 0.56 | -6.16 | 0.26524 | none |
| BBX | 0.226 | 0.3093 | 0.8937 | 0.3932 | 0.56 | -6.16 | 0.18089 | none |
| SARNP | -0.074 | -0.0577 | -0.894 | 0.39323 | 0.56 | -6.16 | -0.0493 | none |
| DDIT3 | 0.188 | 0.149 | 0.8937 | 0.39324 | 0.56 | -6.16 | 0.09244 | none |
| ECHDC3 | 0.155 | 0.115 | 0.8936 | 0.39329 | 0.56 | -6.16 | 1.2043 | none |
| SCGN | 0.088 | 0.0876 | 0.8935 | 0.39331 | 0.56 | -6.16 | 1.07554 | none |
| FEV | -0.106 | -0.0831 | -0.894 | 0.39331 | 0.56 | -6.16 | -0.134 | none |
| CXADR | 0.157 | 0.1679 | 0.8935 | 0.39332 | 0.56 | -6.16 | -0.0063 | none |
| MYCBP | -0.179 | -0.1369 | -0.893 | 0.39337 | 0.56 | -6.16 | 1.43901 | none |
| PRAMEF14 | 0.088 | 0.0867 | 0.8934 | 0.39339 | 0.56 | -6.16 | 0.00397 | none |
| IKZF4 | 0.14 | 0.1266 | 0.8931 | 0.39354 | 0.56 | -6.16 | -0.5744 | none |
| PTRHD1 | -0.16 | -0.2037 | -0.893 | 0.39368 | 0.56 | -6.16 | 0.46235 | none |
| CDK5R1 | -0.081 | -0.0815 | -0.892 | 0.39387 | 0.561 | -6.16 | -0.1611 | none |
| PTK2B | 0.154 | 0.2138 | 0.892 | 0.39409 | 0.561 | -6.16 | -0.2574 | none |
| FAM114A2 | -0.114 | -0.0879 | -0.892 | 0.39418 | 0.561 | -6.16 | 0.42727 | none |
| EFR3A | 0.125 | 0.0619 | 0.8917 | 0.39421 | 0.561 | -6.16 | 0.46834 | none |
| GPC1 | -0.162 | -0.0734 | -0.892 | 0.39429 | 0.561 | -6.16 | 0.29118 | none |

| | | | | | | | | |
|---|---|---|---|---|---|---|---|---|
| UPRT | -0.131 | -0.1252 | -0.891 | 0.39439 | 0.561 | -6.16 | 1.32833 | none |
| CDC7 | 0.159 | 0.03 | 0.8913 | 0.39445 | 0.561 | -6.16 | -0.7909 | none |
| STX19 | 0.063 | 0.0437 | 0.8912 | 0.39449 | 0.561 | -6.16 | 1.02951 | none |
| PYM1 | -0.074 | -0.0107 | -0.891 | 0.39454 | 0.561 | -6.16 | 0.231 | none |
| CXCL6 | -0.082 | -0.0821 | -0.891 | 0.39454 | 0.561 | -6.16 | 1.9054 | none |
| SMPD4 | 0.115 | 0.2007 | 0.8911 | 0.39455 | 0.561 | -6.16 | 0.88259 | none |
| TRPV2 | 0.121 | 0.0853 | 0.8909 | 0.39465 | 0.561 | -6.16 | -0.6036 | none |
| ACKR3 | -0.164 | -0.1159 | -0.891 | 0.39473 | 0.561 | -6.16 | 1.51076 | none |
| TNFSF9 | -0.133 | -0.1561 | -0.891 | 0.3948 | 0.561 | -6.16 | 0.87372 | none |
| SMARCAD | 0.169 | 0.2103 | 0.8904 | 0.39489 | 0.561 | -6.16 | 0.11419 | none |
| RASL11A | -0.207 | -0.2861 | -0.89 | 0.39495 | 0.561 | -6.16 | -0.1838 | none |
| NEBL | 0.2 | 0.3134 | 0.8902 | 0.395 | 0.561 | -6.16 | -0.0285 | none |
| HAUS7 | 0.14 | 0.0948 | 0.8902 | 0.39501 | 0.561 | -6.16 | -0.9935 | none |
| ZFR2 | 0.16 | 0.0697 | 0.8902 | 0.39502 | 0.561 | -6.16 | 0.13799 | none |
| AC118758 | -0.314 | -0.1932 | -0.89 | 0.39505 | 0.561 | -6.16 | 1.83154 | none |
| EDN3 | 0.06 | 0.0599 | 0.8899 | 0.39515 | 0.561 | -6.16 | 1.4015 | none |
| FNBP1L | 0.117 | 0.1688 | 0.8897 | 0.39525 | 0.561 | -6.16 | 0.64734 | none |
| CSF1R | 0.204 | 0.215 | 0.8896 | 0.3953 | 0.562 | -6.16 | -0.6402 | none |
| NTHL1 | -0.088 | -0.0758 | -0.889 | 0.39542 | 0.562 | -6.16 | 0.54549 | none |
| CCR2 | -0.116 | -0.1022 | -0.889 | 0.39565 | 0.562 | -6.16 | 0.58695 | none |
| PHF24 | 0.083 | 0.0887 | 0.8885 | 0.39587 | 0.562 | -6.16 | 1.40049 | none |
| LRIG3 | 0.084 | 0.0885 | 0.8883 | 0.39596 | 0.562 | -6.16 | -0.6186 | none |
| CHAD | -0.068 | -0.0373 | -0.888 | 0.396 | 0.562 | -6.16 | 0.85085 | none |
| RAI2 | -0.105 | -0.0598 | -0.888 | 0.39601 | 0.562 | -6.16 | 0.79909 | none |
| MZB1 | 0.115 | 0.0528 | 0.8881 | 0.39607 | 0.562 | -6.16 | 0.03253 | none |
| LINGO3 | -0.093 | -0.0398 | -0.888 | 0.39629 | 0.563 | -6.16 | 1.71951 | none |
| PRMT6 | 0.09 | 0.0353 | 0.8875 | 0.39638 | 0.563 | -6.16 | -0.1005 | none |
| PPTC7 | 0.076 | 0.0132 | 0.8873 | 0.39649 | 0.563 | -6.16 | 0.0855 | none |
| UGP2 | -0.064 | -0.0791 | -0.887 | 0.39655 | 0.563 | -6.16 | 1.35243 | none |
| TMEM86A | -0.127 | -0.1009 | -0.887 | 0.39656 | 0.563 | -6.16 | -0.3577 | none |
| LRIT2 | 0.061 | 0.061 | 0.8871 | 0.39657 | 0.563 | -6.16 | 0.99008 | none |
| SNPH | 0.11 | 0.1201 | 0.8867 | 0.3968 | 0.563 | -6.16 | 0.10348 | none |
| POLA1 | 0.26 | 0.3504 | 0.8867 | 0.3968 | 0.563 | -6.16 | 0.46997 | none |
| CPNE6 | -0.081 | -0.0604 | -0.887 | 0.39687 | 0.563 | -6.16 | 2.2982 | AA-related |
| CHRND | -0.058 | -0.0342 | -0.886 | 0.39703 | 0.563 | -6.16 | 0 | none |
| PPIE | 0.078 | 0.0408 | 0.8862 | 0.39705 | 0.563 | -6.16 | -0.5122 | none |
| ZNF592 | 0.228 | 0.3563 | 0.886 | 0.39713 | 0.563 | -6.17 | -0.3539 | none |
| PTPRCAP | -0.132 | -0.1706 | -0.886 | 0.39721 | 0.563 | -6.17 | 1.41237 | none |
| CCR9 | -0.073 | -0.0768 | -0.886 | 0.39723 | 0.563 | -6.17 | 1.2634 | none |
| LYPD2 | -0.061 | -0.0404 | -0.886 | 0.39735 | 0.563 | -6.17 | 0 | none |

| | | | | | | | | |
|---|---|---|---|---|---|---|---|---|
| NACA2 | 0.168 | 0.1375 | 0.8854 | 0.39744 | 0.563 | -6.17 | 1.25226 | none |
| HAL | -0.134 | -0.0411 | -0.885 | 0.39745 | 0.563 | -6.17 | 0.46421 | none |
| TGDS | -0.105 | -0.1469 | -0.885 | 0.39753 | 0.564 | -6.17 | 1.18174 | none |
| TMEM196 | 0.081 | 0.083 | 0.8852 | 0.39758 | 0.564 | -6.17 | 1.29681 | none |
| MAT2A | -0.094 | -0.0696 | -0.885 | 0.39762 | 0.564 | -6.17 | 0.71372 | none |
| JAGN1 | -0.139 | -0.1803 | -0.885 | 0.39791 | 0.564 | -6.17 | 1.67914 | none |
| HORMAD1 | -0.129 | -0.0983 | -0.884 | 0.398 | 0.564 | -6.17 | 0.61052 | none |
| SPANXC | -0.167 | -0.1562 | -0.884 | 0.39801 | 0.564 | -6.17 | 0.35013 | none |
| MTUS2 | -0.108 | -0.0837 | -0.884 | 0.39812 | 0.564 | -6.17 | 1.6632 | none |
| PPP2R5B | 0.129 | 0.1606 | 0.8841 | 0.39813 | 0.564 | -6.17 | -0.0481 | none |
| STARD5 | -0.221 | -0.1074 | -0.884 | 0.39813 | 0.564 | -6.17 | 1.40177 | none |
| TULP3 | 0.161 | 0.147 | 0.8839 | 0.39824 | 0.564 | -6.17 | 0.39195 | none |
| TYRP1 | -0.081 | -0.0806 | -0.883 | 0.39847 | 0.564 | -6.17 | 1.14241 | none |
| FAM83A | -0.21 | -0.2304 | -0.883 | 0.39879 | 0.565 | -6.17 | 0.52485 | none |
| ADIPOR2 | 0.132 | 0.1268 | 0.8827 | 0.39884 | 0.565 | -6.17 | -0.4067 | none |
| ID1 | -0.118 | -0.1022 | -0.883 | 0.39893 | 0.565 | -6.17 | 1.43431 | none |
| HOXC10 | 0.083 | 0.0827 | 0.8825 | 0.39895 | 0.565 | -6.17 | 0.63623 | none |
| ABHD14A | 0.135 | -0.0031 | 0.8825 | 0.39896 | 0.565 | -6.17 | 0.46321 | none |
| CCL13 | -0.08 | -0.0804 | -0.882 | 0.39897 | 0.565 | -6.17 | 1.9054 | none |
| ATAD3B | 0.12 | 0.1819 | 0.8824 | 0.39898 | 0.565 | -6.17 | 0.20447 | none |
| ODF4 | 0.06 | 0.0604 | 0.8823 | 0.39903 | 0.565 | -6.17 | 1.54859 | none |
| LAT | -0.105 | -0.0991 | -0.882 | 0.39905 | 0.565 | -6.17 | 0.44139 | none |
| BCL2 | -0.107 | -0.0655 | -0.882 | 0.39909 | 0.565 | -6.17 | 1.00333 | none |
| SMARCD1 | -0.167 | -0.1568 | -0.882 | 0.39913 | 0.565 | -6.17 | 0.56381 | none |
| ASIC1 | 0.151 | 0.1742 | 0.882 | 0.39921 | 0.565 | -6.17 | 1.0464 | none |
| PAF1 | 0.16 | 0.2407 | 0.882 | 0.39921 | 0.565 | -6.17 | 0.61562 | none |
| TRIM54 | -0.071 | -0.0359 | -0.882 | 0.39923 | 0.565 | -6.17 | 0.11852 | none |
| IL20RA | 0.083 | 0.0825 | 0.8816 | 0.39942 | 0.565 | -6.17 | -0.8671 | none |
| CCDC116 | 0.106 | 0.0987 | 0.8815 | 0.39947 | 0.565 | -6.17 | -0.6504 | none |
| VAPA | -0.109 | -0.1305 | -0.881 | 0.39961 | 0.565 | -6.17 | 1.37446 | none |
| MEX3D | 0.096 | 0.1402 | 0.8812 | 0.39961 | 0.565 | -6.17 | 0.54302 | none |
| ADGRE3 | -0.112 | -0.1124 | -0.881 | 0.39971 | 0.565 | -6.17 | 2.15035 | AA-related |
| METTL17 | 0.095 | 0.0686 | 0.8808 | 0.39984 | 0.565 | -6.17 | 0.64788 | none |
| TRIM17 | -0.059 | -0.0271 | -0.881 | 0.39984 | 0.565 | -6.17 | 1.07357 | none |
| AKAP3 | -0.116 | -0.0323 | -0.881 | 0.39995 | 0.565 | -6.17 | 0.39782 | none |
| HTR1A | -0.08 | -0.0797 | -0.879 | 0.40064 | 0.566 | -6.17 | 2.05429 | none |
| KRT1 | 0.09 | 0.0978 | 0.8787 | 0.40089 | 0.567 | -6.17 | 1.64245 | none |
| IFT46 | 0.101 | 0.0271 | 0.8786 | 0.40096 | 0.567 | -6.17 | 1.23215 | none |
| IGSF11 | -0.16 | -0.1311 | -0.878 | 0.40103 | 0.567 | -6.17 | -0.4135 | none |
| DKK2 | 0.061 | 0.0552 | 0.8783 | 0.40109 | 0.567 | -6.17 | 0 | none |

| | | | | | | | | |
|---|---|---|---|---|---|---|---|---|
| GJB6 | 0.152 | 0.1048 | 0.8781 | 0.40122 | 0.567 | -6.17 | 0.87636 | none |
| SYNDIG1L | 0.063 | 0.0746 | 0.8781 | 0.40124 | 0.567 | -6.17 | -0.7199 | none |
| BMX | 0.082 | 0.0817 | 0.8779 | 0.40131 | 0.567 | -6.17 | 1.19688 | none |
| POGK | 0.125 | 0.1655 | 0.8778 | 0.40138 | 0.567 | -6.17 | 1.02123 | none |
| SPDYE6 | 0.239 | 0.2685 | 0.8774 | 0.40159 | 0.567 | -6.17 | 0.60262 | none |
| AC107081 | 0.088 | 0.1175 | 0.8773 | 0.40162 | 0.567 | -6.17 | 0.40726 | none |
| ANKRD13I | 0.229 | 0.2751 | 0.8773 | 0.40163 | 0.567 | -6.17 | -0.0013 | none |
| GORASP1 | 0.083 | 0.1677 | 0.8772 | 0.40169 | 0.567 | -6.17 | 0.72794 | none |
| C19ORF12 | 0.065 | 0.0249 | 0.8771 | 0.40171 | 0.567 | -6.17 | 0.04813 | none |
| UBN1 | 0.205 | 0.335 | 0.8771 | 0.40173 | 0.567 | -6.17 | 0.24407 | none |
| GK | 0.079 | 0.0064 | 0.8769 | 0.40181 | 0.567 | -6.17 | -0.0473 | none |
| TRIT1 | -0.128 | -0.0923 | -0.877 | 0.40188 | 0.567 | -6.17 | 0.32361 | none |
| ZNF534 | 0.117 | 0.0683 | 0.8766 | 0.40198 | 0.567 | -6.17 | -0.3451 | none |
| CCDC102A | -0.122 | -0.0247 | -0.877 | 0.40201 | 0.567 | -6.17 | 0.88333 | none |
| CXCL17 | 0.108 | 0.0777 | 0.8765 | 0.40206 | 0.567 | -6.17 | 0.03167 | none |
| NEUROD2 | 0.148 | 0.0794 | 0.8764 | 0.40209 | 0.567 | -6.17 | -0.0697 | none |
| HCRT | 0.134 | 0.0938 | 0.8763 | 0.40215 | 0.567 | -6.17 | 0.50621 | none |
| OR11L1 | -0.079 | -0.0791 | -0.876 | 0.4023 | 0.568 | -6.17 | 0.64435 | none |
| EMC9 | 0.166 | 0.0948 | 0.8758 | 0.4024 | 0.568 | -6.17 | -0.8112 | none |
| OR56A3 | -0.067 | -0.0556 | -0.876 | 0.40248 | 0.568 | -6.17 | 0.81278 | none |
| GOLGA6L1 | 0.082 | 0.0549 | 0.8755 | 0.40254 | 0.568 | -6.17 | 0.28888 | none |
| KIN | 0.094 | 0.0571 | 0.8753 | 0.40267 | 0.568 | -6.17 | 0.6775 | none |
| SPOCK1 | 0.103 | 0.1684 | 0.8749 | 0.40285 | 0.568 | -6.17 | -0.2214 | none |
| CCNA1 | -0.073 | -0.0667 | -0.874 | 0.40346 | 0.569 | -6.18 | 0.88832 | none |
| CCL11 | -0.079 | -0.0787 | -0.874 | 0.40353 | 0.569 | -6.18 | 2.13744 | AA-related |
| TMPRSS2 | -0.155 | -0.1266 | -0.874 | 0.40354 | 0.569 | -6.18 | 2.08895 | none |
| ARAP3 | -0.089 | -0.0534 | -0.874 | 0.40357 | 0.569 | -6.18 | -0.1791 | none |
| TMEM147 | -0.157 | -0.1698 | -0.873 | 0.40368 | 0.569 | -6.18 | 1.43293 | none |
| PLAA | -0.104 | -0.045 | -0.873 | 0.40377 | 0.569 | -6.18 | 1.98196 | none |
| TRPC5OS | -0.069 | -0.0658 | -0.873 | 0.4039 | 0.569 | -6.18 | 1.9792 | none |
| DCXR | -0.096 | -0.112 | -0.873 | 0.40403 | 0.569 | -6.18 | 1.53633 | none |
| PGA4 | 0.076 | 0.1187 | 0.8727 | 0.40403 | 0.569 | -6.18 | -0.3144 | none |
| TSC22D2 | 0.183 | 0.225 | 0.8726 | 0.40406 | 0.569 | -6.18 | 0.47002 | none |
| SSBP4 | -0.145 | -0.1169 | -0.872 | 0.40412 | 0.569 | -6.18 | 0.63529 | none |
| KRT10 | -0.093 | -0.0778 | -0.872 | 0.40415 | 0.569 | -6.18 | 1.22987 | none |
| OXSR1 | 0.157 | 0.2241 | 0.8722 | 0.40426 | 0.57 | -6.18 | -0.0343 | none |
| ANTXR1 | 0.107 | 0.1122 | 0.872 | 0.40439 | 0.57 | -6.18 | 0.63746 | none |
| MBLAC1 | 0.12 | 0.0047 | 0.8715 | 0.40466 | 0.57 | -6.18 | 0.00662 | none |
| SLC5A2 | 0.136 | 0.1009 | 0.8714 | 0.4047 | 0.57 | -6.18 | 1.32434 | none |
| AL513523. | -0.059 | -0.0311 | -0.871 | 0.40475 | 0.57 | -6.18 | 1.61352 | none |

| | | | | | | | | |
|---|---|---|---|---|---|---|---|---|
| GAGE10 | 0.08 | 0.0804 | 0.8712 | 0.40479 | 0.57 | -6.18 | 1.52574 | none |
| PRL | -0.08 | -0.0516 | -0.871 | 0.40485 | 0.57 | -6.18 | -0.4674 | none |
| SERINC4 | -0.102 | -0.0745 | -0.871 | 0.40487 | 0.57 | -6.18 | 1.1658 | none |
| CT45A6 | 0.08 | 0.0803 | 0.8708 | 0.40499 | 0.57 | -6.18 | 1.00727 | none |
| KRT27 | 0.08 | 0.0803 | 0.8708 | 0.40499 | 0.57 | -6.18 | 1.19688 | none |
| OR3A1 | 0.11 | 0.0952 | 0.8707 | 0.40504 | 0.57 | -6.18 | -0.7339 | none |
| MPC1 | -0.174 | -0.2508 | -0.87 | 0.40525 | 0.57 | -6.18 | 0.39964 | none |
| ISOC1 | -0.121 | -0.124 | -0.87 | 0.40528 | 0.57 | -6.18 | 0.60313 | none |
| CYP19A1 | -0.199 | -0.227 | -0.87 | 0.40531 | 0.57 | -6.18 | -0.447 | none |
| PSME3 | -0.125 | -0.1066 | -0.87 | 0.4054 | 0.57 | -6.18 | 1.16847 | none |
| EVX1 | 0.091 | 0.1037 | 0.869 | 0.40594 | 0.571 | -6.18 | -0.8091 | none |
| ALG13 | -0.149 | -0.2459 | -0.868 | 0.40622 | 0.572 | -6.18 | 0.33939 | none |
| GLYATL1 | 0.168 | 0.0761 | 0.8683 | 0.40628 | 0.572 | -6.18 | -0.2654 | none |
| FMR1NB | -0.1 | -0.0898 | -0.868 | 0.40653 | 0.572 | -6.18 | 1.29506 | none |
| USP17L7 | 0.059 | 0.0418 | 0.8678 | 0.40656 | 0.572 | -6.18 | 0 | none |
| SETD1A | 0.223 | 0.384 | 0.8677 | 0.40658 | 0.572 | -6.18 | 0.07955 | none |
| C1QTNF7 | 0.065 | 0.0684 | 0.8674 | 0.40677 | 0.572 | -6.18 | 1.10778 | none |
| TPPP3 | -0.104 | -0.0355 | -0.867 | 0.40686 | 0.572 | -6.18 | 0.58715 | none |
| XPNPEP2 | -0.067 | -0.0438 | -0.867 | 0.40691 | 0.572 | -6.18 | 0.61382 | none |
| PCBD1 | -0.154 | -0.1475 | -0.867 | 0.40694 | 0.572 | -6.18 | 1.37026 | none |
| FDCSP | -0.077 | -0.0773 | -0.867 | 0.40722 | 0.573 | -6.18 | 1.9054 | none |
| OAT | 0.085 | -0.034 | 0.8664 | 0.40729 | 0.573 | -6.18 | 0.23808 | none |
| VAV2 | 0.138 | 0.2173 | 0.8663 | 0.40731 | 0.573 | -6.18 | -0.2444 | none |
| APOC3 | 0.079 | 0.0802 | 0.8662 | 0.4074 | 0.573 | -6.18 | 0.8025 | none |
| POLR3G | -0.145 | -0.1053 | -0.866 | 0.40756 | 0.573 | -6.18 | 1.06982 | none |
| ARMC1 | -0.133 | -0.1681 | -0.866 | 0.40757 | 0.573 | -6.18 | 0.52924 | none |
| SDAD1 | 0.264 | 0.3041 | 0.8653 | 0.40787 | 0.573 | -6.18 | -0.2936 | none |
| CKMT2 | 0.245 | 0.1941 | 0.8651 | 0.40797 | 0.573 | -6.18 | -0.8787 | none |
| PIANP | -0.065 | -0.0392 | -0.865 | 0.40801 | 0.573 | -6.18 | 1.37377 | none |
| IGSF3 | 0.124 | 0.1135 | 0.8648 | 0.40813 | 0.573 | -6.18 | -0.0899 | none |
| PRPF40A | 0.238 | 0.3508 | 0.8647 | 0.40818 | 0.573 | -6.18 | 0.47809 | none |
| LGALS9 | -0.178 | -0.2069 | -0.865 | 0.40826 | 0.573 | -6.18 | 0.61491 | none |
| HIST1H3J | -0.3 | -0.3381 | -0.864 | 0.40832 | 0.573 | -6.18 | 1.5622 | none |
| TG | -0.123 | -0.0504 | -0.864 | 0.40835 | 0.573 | -6.18 | 0.27824 | none |
| NUDT18 | -0.118 | -0.1113 | -0.864 | 0.4084 | 0.574 | -6.18 | 0.4842 | none |
| PPT1 | -0.13 | -0.1493 | -0.864 | 0.40853 | 0.574 | -6.18 | 1.34765 | none |
| GSTM2 | 0.126 | 0.0473 | 0.8637 | 0.40867 | 0.574 | -6.18 | -0.6188 | none |
| LEFTY1 | -0.147 | -0.0822 | -0.864 | 0.40868 | 0.574 | -6.18 | 0.59395 | none |
| GAB4 | -0.062 | -0.0615 | -0.864 | 0.40872 | 0.574 | -6.18 | 0.57238 | none |
| GUCA2B | 0.079 | 0.0789 | 0.8636 | 0.40873 | 0.574 | -6.18 | 0.44852 | none |

| | | | | | | | | |
|---|---|---|---|---|---|---|---|---|
| CADM4 | -0.084 | -0.083 | -0.864 | 0.40879 | 0.574 | -6.18 | 0.95784 | none |
| KIAA0040 | 0.147 | 0.1931 | 0.8635 | 0.40881 | 0.574 | -6.18 | 0.03906 | none |
| GYS1 | 0.133 | 0.2297 | 0.8635 | 0.40882 | 0.574 | -6.18 | 0.92698 | none |
| CPA6 | 0.062 | 0.062 | 0.8633 | 0.40888 | 0.574 | -6.18 | 0.76829 | none |
| HOXA11 | 0.062 | 0.0622 | 0.8633 | 0.40892 | 0.574 | -6.18 | 0 | none |
| HIST1H2AI | -0.195 | -0.1816 | -0.863 | 0.40893 | 0.574 | -6.18 | 1.42628 | none |
| S100P | 0.169 | 0.0317 | 0.8632 | 0.40898 | 0.574 | -6.19 | 1.28066 | none |
| INTS7 | 0.109 | 0.0755 | 0.8631 | 0.40899 | 0.574 | -6.19 | -0.2639 | none |
| HMOX1 | 0.082 | 0.0871 | 0.8631 | 0.40902 | 0.574 | -6.19 | 0.55482 | none |
| IL21 | -0.158 | -0.167 | -0.863 | 0.40918 | 0.574 | -6.19 | 1.4096 | none |
| A1CF | 0.078 | 0.0499 | 0.8627 | 0.40921 | 0.574 | -6.19 | 0.12959 | none |
| TST | -0.267 | -0.2897 | -0.863 | 0.40926 | 0.574 | -6.19 | 1.94894 | none |
| NSMCE2 | 0.111 | 0.0496 | 0.8626 | 0.40928 | 0.574 | -6.19 | -0.428 | none |
| SLX1B | 0.115 | 0.0024 | 0.8624 | 0.40938 | 0.574 | -6.19 | -0.8176 | none |
| RRS1 | 0.129 | 0.174 | 0.862 | 0.4096 | 0.574 | -6.19 | 0.52336 | none |
| IQCF5 | 0.079 | 0.0785 | 0.8618 | 0.40967 | 0.574 | -6.19 | -0.9162 | none |
| BCL6 | 0.169 | 0.2535 | 0.8618 | 0.40969 | 0.574 | -6.19 | -0.3477 | none |
| CBX6 | 0.17 | 0.3205 | 0.8615 | 0.40985 | 0.574 | -6.19 | -0.4196 | none |
| CDRT1 | 0.113 | 0.1244 | 0.8611 | 0.41008 | 0.575 | -6.19 | -0.1255 | none |
| LGALS13 | 0.078 | 0.0784 | 0.861 | 0.41009 | 0.575 | -6.19 | 0.66005 | none |
| SPIRE1 | -0.12 | -0.018 | -0.861 | 0.41015 | 0.575 | -6.19 | -0.0632 | none |
| FIGNL1 | -0.126 | -0.102 | -0.861 | 0.4103 | 0.575 | -6.19 | -0.0854 | none |
| SLC25A19 | -0.085 | -0.0516 | -0.86 | 0.41039 | 0.575 | -6.19 | 0.69729 | none |
| NOP58 | 0.185 | 0.2487 | 0.8603 | 0.41049 | 0.575 | -6.19 | 0.53492 | none |
| ALDOC | -0.111 | -0.0792 | -0.86 | 0.41062 | 0.575 | -6.19 | 0.90752 | none |
| USO1 | 0.167 | 0.1956 | 0.8599 | 0.41066 | 0.575 | -6.19 | 0.7035 | none |
| SUSD3 | 0.095 | 0.0419 | 0.8598 | 0.41074 | 0.575 | -6.19 | 0.08007 | none |
| NFIL3 | -0.085 | -0.0142 | -0.86 | 0.41076 | 0.575 | -6.19 | 1.69043 | none |
| YIF1A | -0.147 | -0.1908 | -0.86 | 0.41077 | 0.575 | -6.19 | 0.62849 | none |
| EMSY | 0.138 | 0.2427 | 0.8595 | 0.41087 | 0.575 | -6.19 | 0.55793 | none |
| KDM3A | 0.213 | 0.2821 | 0.8591 | 0.41113 | 0.576 | -6.19 | -0.7291 | none |
| GAS2L2 | -0.059 | -0.0488 | -0.859 | 0.41122 | 0.576 | -6.19 | 1.29326 | none |
| FGR | 0.106 | 0.1276 | 0.8587 | 0.41129 | 0.576 | -6.19 | 0.21867 | none |
| FBXW7 | 0.243 | 0.3539 | 0.8584 | 0.41148 | 0.576 | -6.19 | 0.84091 | none |
| TMEM70 | -0.125 | -0.144 | -0.858 | 0.41152 | 0.576 | -6.19 | 1.35697 | none |
| NT5C | 0.09 | 0.0571 | 0.8582 | 0.41156 | 0.576 | -6.19 | 1.41802 | none |
| TMPRSS4 | 0.172 | 0.0819 | 0.8578 | 0.41179 | 0.576 | -6.19 | -0.5486 | none |
| PDLIM7 | 0.096 | 0.1394 | 0.8577 | 0.41186 | 0.576 | -6.19 | 0.14681 | none |
| MALT1 | 0.184 | 0.1054 | 0.8576 | 0.41189 | 0.576 | -6.19 | 0.01033 | none |
| SLC12A5 | -0.283 | -0.2133 | -0.858 | 0.41191 | 0.576 | -6.19 | 0.85946 | none |

| | | | | | | | | |
|---|---|---|---|---|---|---|---|---|
| PLA2G1B | -0.111 | -0.104 | -0.857 | 0.41195 | 0.576 | -6.19 | 0.62456 | none |
| ADAD1 | -0.101 | -0.1014 | -0.857 | 0.41201 | 0.576 | -6.19 | 0.63836 | none |
| HOXA7 | 0.091 | 0.0946 | 0.8568 | 0.41229 | 0.577 | -6.19 | -0.3275 | none |
| PKDCC | 0.171 | 0.1707 | 0.8568 | 0.41232 | 0.577 | -6.19 | 0.57062 | none |
| HIST2H2A/ | -0.539 | -0.4339 | -0.857 | 0.41233 | 0.577 | -6.19 | 1.24856 | none |
| EXOSC9 | -0.126 | -0.0859 | -0.857 | 0.4124 | 0.577 | -6.19 | 0.70195 | none |
| GRK5 | 0.173 | 0.2413 | 0.8564 | 0.41254 | 0.577 | -6.19 | -0.3017 | none |
| OR52K2 | 0.058 | 0.0582 | 0.8563 | 0.41257 | 0.577 | -6.19 | 0.59355 | none |
| SPAG6 | 0.081 | 0.0786 | 0.856 | 0.41273 | 0.577 | -6.19 | 0.44486 | none |
| FABP1 | -0.113 | -0.1237 | -0.856 | 0.41274 | 0.577 | -6.19 | -0.0121 | none |
| FRMD6 | -0.138 | -0.132 | -0.856 | 0.41291 | 0.577 | -6.19 | 1.63002 | none |
| NPHP3-AC | 0.07 | 0.0928 | 0.8556 | 0.41293 | 0.577 | -6.19 | 0.32129 | none |
| CLDN3 | -0.071 | -0.0546 | -0.856 | 0.41296 | 0.577 | -6.19 | 0.33276 | none |
| TRAT1 | -0.127 | -0.2378 | -0.855 | 0.41305 | 0.577 | -6.19 | 0.7302 | none |
| LIMS2 | 0.109 | 0.0935 | 0.8554 | 0.41305 | 0.577 | -6.19 | -0.5724 | none |
| ADARB1 | 0.273 | 0.24 | 0.8552 | 0.41317 | 0.577 | -6.19 | -0.1539 | none |
| GBX1 | 0.071 | 0.0801 | 0.855 | 0.41327 | 0.577 | -6.19 | 0.24643 | none |
| SULT4A1 | 0.106 | 0.114 | 0.8549 | 0.41329 | 0.577 | -6.19 | -0.624 | none |
| SP100 | 0.15 | 0.1812 | 0.8549 | 0.41331 | 0.577 | -6.19 | 0.29251 | none |
| TPTE | -0.07 | -0.0696 | -0.855 | 0.41347 | 0.578 | -6.19 | 1.1485 | none |
| MAX | -0.098 | -0.1467 | -0.855 | 0.4135 | 0.578 | -6.19 | -0.0712 | none |
| STXBP4 | 0.144 | 0.0477 | 0.8544 | 0.41359 | 0.578 | -6.19 | 0.17394 | none |
| RBBP8 | 0.295 | 0.4172 | 0.8542 | 0.41368 | 0.578 | -6.19 | 0.42608 | none |
| HSPB11 | -0.124 | -0.2253 | -0.854 | 0.41391 | 0.578 | -6.19 | 0.36776 | none |
| LCP2 | 0.168 | 0.1923 | 0.8537 | 0.41396 | 0.578 | -6.19 | -0.8116 | none |
| DLL3 | -0.143 | -0.1036 | -0.854 | 0.414 | 0.578 | -6.19 | 1.79883 | none |
| SRSF4 | 0.17 | 0.2516 | 0.8531 | 0.41426 | 0.578 | -6.19 | 1.00272 | none |
| ING3 | 0.105 | 0.0899 | 0.853 | 0.4143 | 0.578 | -6.19 | 0.4272 | none |
| LSM7 | 0.14 | 0.078 | 0.852 | 0.41482 | 0.579 | -6.19 | -0.5989 | none |
| KIF5A | 0.071 | 0.0654 | 0.852 | 0.41483 | 0.579 | -6.19 | 0.52389 | none |
| LAMA4 | -0.127 | -0.0873 | -0.852 | 0.41496 | 0.579 | -6.19 | 0.94734 | none |
| TMPRSS11 | -0.062 | -0.0643 | -0.852 | 0.41511 | 0.579 | -6.2 | 0 | none |
| MRPL52 | -0.127 | -0.1546 | -0.851 | 0.41522 | 0.579 | -6.2 | 1.20722 | none |
| BCL2L12 | -0.132 | -0.1382 | -0.851 | 0.41537 | 0.58 | -6.2 | 1.08948 | none |
| MAOB | -0.128 | -0.0883 | -0.851 | 0.41548 | 0.58 | -6.2 | 0.21675 | none |
| PRPF18 | -0.109 | -0.1369 | -0.851 | 0.41562 | 0.58 | -6.2 | 0.45293 | none |
| SDHA | 0.09 | 0.105 | 0.8503 | 0.41573 | 0.58 | -6.2 | 0.87324 | none |
| MXD4 | -0.083 | -0.0736 | -0.85 | 0.4159 | 0.58 | -6.2 | 0.72365 | none |
| FOLH1 | 0.058 | 0.039 | 0.85 | 0.41592 | 0.58 | -6.2 | -0.3176 | none |
| GGTLC1 | 0.062 | 0.0337 | 0.8496 | 0.41611 | 0.58 | -6.2 | 0.20297 | none |

| | | | | | | | | |
|---|---|---|---|---|---|---|---|---|
| KRTAP19-2 | -0.074 | -0.0743 | -0.849 | 0.41617 | 0.58 | -6.2 | 2.06233 | none |
| HTT | 0.244 | 0.2985 | 0.8492 | 0.41633 | 0.58 | -6.2 | 0.93788 | none |
| TUBGCP3 | 0.096 | 0.0985 | 0.8492 | 0.41633 | 0.58 | -6.2 | 1.69208 | none |
| TMEM109 | -0.102 | -0.0772 | -0.849 | 0.41633 | 0.58 | -6.2 | 1.09979 | none |
| MEIOB | 0.064 | 0.0638 | 0.849 | 0.41645 | 0.58 | -6.2 | 1.23395 | none |
| IQCA1L | 0.086 | 0.0995 | 0.8486 | 0.41663 | 0.581 | -6.2 | -0.0882 | none |
| C17ORF58 | -0.172 | -0.0798 | -0.849 | 0.41663 | 0.581 | -6.2 | 1.12499 | none |
| GALNT2 | 0.133 | 0.168 | 0.8486 | 0.41664 | 0.581 | -6.2 | 0.35039 | none |
| ANKRD36B | 0.138 | 0.0348 | 0.8485 | 0.41671 | 0.581 | -6.2 | 0.11074 | none |
| BRWD3 | 0.176 | 0.2666 | 0.8481 | 0.41693 | 0.581 | -6.2 | 0.77058 | none |
| LDHC | 0.151 | 0.0298 | 0.8474 | 0.4173 | 0.581 | -6.2 | -0.7064 | none |
| ERICH4 | 0.076 | 0.0757 | 0.8468 | 0.41761 | 0.582 | -6.2 | -0.4532 | none |
| SLC28A1 | -0.09 | -0.0981 | -0.847 | 0.41767 | 0.582 | -6.2 | -0.2481 | none |
| DRG2 | 0.072 | 0.0801 | 0.8466 | 0.41769 | 0.582 | -6.2 | 1.22965 | none |
| GH2 | -0.074 | -0.0738 | -0.847 | 0.41771 | 0.582 | -6.2 | 0.54647 | none |
| ARMCX4 | -0.126 | -0.1275 | -0.846 | 0.41782 | 0.582 | -6.2 | 0.61106 | none |
| PSKH2 | 0.076 | 0.0756 | 0.8463 | 0.41784 | 0.582 | -6.2 | 0.20449 | none |
| ZNF282 | -0.136 | 0.0561 | -0.846 | 0.41807 | 0.582 | -6.2 | 0.87759 | none |
| TFPT | -0.142 | -0.1015 | -0.846 | 0.41811 | 0.582 | -6.2 | 0.89575 | none |
| LEF1 | 0.108 | 0.0853 | 0.8455 | 0.41828 | 0.582 | -6.2 | 0.828 | none |
| CWH43 | 0.11 | 0.1178 | 0.8454 | 0.41835 | 0.582 | -6.2 | 0.44006 | none |
| DLL1 | 0.083 | 0.1135 | 0.8452 | 0.41844 | 0.582 | -6.2 | -0.2292 | none |
| KLHL15 | 0.074 | 0.0043 | 0.8451 | 0.41852 | 0.583 | -6.2 | -0.1547 | none |
| CACNG7 | 0.092 | 0.0342 | 0.845 | 0.41857 | 0.583 | -6.2 | 0.1585 | none |
| SPATA20 | 0.104 | 0.1306 | 0.8448 | 0.41869 | 0.583 | -6.2 | 1.14419 | none |
| DACT1 | 0.061 | 0.0364 | 0.8447 | 0.41874 | 0.583 | -6.2 | 1.22627 | none |
| DYDC2 | 0.11 | 0.0661 | 0.8446 | 0.41877 | 0.583 | -6.2 | -0.7302 | none |
| MYC | 0.113 | 0.1986 | 0.8445 | 0.41881 | 0.583 | -6.2 | 1.02606 | none |
| ALG10 | -0.141 | -0.1585 | -0.844 | 0.41898 | 0.583 | -6.2 | 1.66887 | none |
| TNFSF12-T | -0.094 | -0.1153 | -0.844 | 0.4191 | 0.583 | -6.2 | 1.12064 | none |
| THBS1 | 0.095 | 0.1041 | 0.8439 | 0.41912 | 0.583 | -6.2 | 0.65511 | none |
| SLC9B2 | 0.084 | 0.0461 | 0.8439 | 0.41913 | 0.583 | -6.2 | -0.2132 | none |
| CXCL13 | -0.153 | -0.1642 | -0.844 | 0.41931 | 0.583 | -6.2 | 0.50744 | none |
| TAS2R14 | -0.061 | -0.0507 | -0.843 | 0.41936 | 0.583 | -6.2 | 1.65485 | none |
| TUBAL3 | -0.072 | -0.0598 | -0.843 | 0.41937 | 0.583 | -6.2 | 0.10949 | none |
| ARHGAP30 | 0.191 | 0.2771 | 0.8435 | 0.41937 | 0.583 | -6.2 | 0.44409 | none |
| SSU72 | -0.082 | -0.1147 | -0.843 | 0.41938 | 0.583 | -6.2 | 0.18389 | none |
| NSA2 | 0.125 | 0.0744 | 0.8435 | 0.41938 | 0.583 | -6.2 | -0.6973 | none |
| MS4A6E | 0.075 | 0.0751 | 0.8435 | 0.41938 | 0.583 | -6.2 | 0.66005 | none |
| NDUFAF2 | 0.203 | 0.1287 | 0.8434 | 0.41943 | 0.583 | -6.2 | -0.0458 | none |

| | | | | | | | | |
|---|---|---|---|---|---|---|---|---|
| KRT16 | 0.1 | 0.1108 | 0.8433 | 0.41945 | 0.583 | -6.2 | 0.16877 | none |
| KCNC4 | 0.251 | 0.1805 | 0.8433 | 0.41947 | 0.583 | -6.2 | 0.41535 | none |
| CHRM1 | 0.071 | 0.0792 | 0.8431 | 0.41959 | 0.583 | -6.2 | 0.56499 | none |
| IL15 | -0.075 | -0.0077 | -0.843 | 0.41965 | 0.583 | -6.2 | 1.59178 | none |
| OR9I1 | 0.059 | 0.0594 | 0.8428 | 0.41971 | 0.583 | -6.2 | -0.2365 | none |
| ACD | 0.094 | 0.1124 | 0.8426 | 0.41983 | 0.583 | -6.2 | -0.0765 | none |
| FSTL5 | 0.089 | 0.0829 | 0.8423 | 0.41999 | 0.583 | -6.2 | 1.15012 | none |
| F5 | 0.323 | 0.5375 | 0.842 | 0.42013 | 0.583 | -6.2 | 0.31248 | none |
| PKLR | 0.058 | 0.0582 | 0.842 | 0.42014 | 0.583 | -6.2 | 0 | none |
| WDR81 | 0.135 | 0.2339 | 0.842 | 0.42018 | 0.583 | -6.2 | 0.39984 | none |
| PID1 | 0.208 | 0.1519 | 0.8419 | 0.4202 | 0.583 | -6.2 | 1.46357 | none |
| LRRC9 | 0.07 | 0.0721 | 0.8415 | 0.42043 | 0.584 | -6.2 | -0.6112 | none |
| CRYBA1 | 0.083 | 0.0822 | 0.8415 | 0.42044 | 0.584 | -6.2 | 1.28545 | none |
| SLC25A11 | -0.106 | -0.0845 | -0.841 | 0.42045 | 0.584 | -6.2 | 1.37057 | none |
| EMP3 | -0.116 | -0.1568 | -0.841 | 0.42063 | 0.584 | -6.2 | 1.4972 | none |
| INSRR | 0.062 | 0.0542 | 0.8409 | 0.42075 | 0.584 | -6.2 | -1.0215 | none |
| WDR63 | 0.075 | 0.0748 | 0.8408 | 0.42078 | 0.584 | -6.2 | -0.8743 | none |
| TMEM151 | 0.061 | 0.0608 | 0.8408 | 0.4208 | 0.584 | -6.2 | 0.15445 | none |
| C10ORF12 | 0.075 | 0.0746 | 0.8406 | 0.42092 | 0.584 | -6.2 | -0.4083 | none |
| ZNF207 | -0.082 | -0.0927 | -0.84 | 0.421 | 0.584 | -6.2 | 1.98633 | none |
| STXBP2 | 0.115 | 0.1382 | 0.8403 | 0.42105 | 0.584 | -6.2 | -0.2775 | none |
| ILVBL | -0.126 | -0.1371 | -0.84 | 0.42131 | 0.584 | -6.21 | 0.85497 | none |
| SFRP2 | -0.058 | -0.0552 | -0.84 | 0.42132 | 0.584 | -6.21 | 0.05211 | none |
| SLC26A10 | -0.116 | -0.1164 | -0.84 | 0.42136 | 0.584 | -6.21 | 0.41959 | none |
| SYT1 | -0.078 | -0.0779 | -0.84 | 0.42138 | 0.584 | -6.21 | 0.53951 | none |
| JMJD6 | -0.101 | 0.0017 | -0.84 | 0.42143 | 0.584 | -6.21 | 1.65039 | none |
| ADGRF2 | 0.071 | 0.0508 | 0.8395 | 0.42151 | 0.584 | -6.21 | -1.1142 | none |
| TPD52L2 | -0.059 | -0.0151 | -0.839 | 0.42152 | 0.584 | -6.21 | 0.81304 | none |
| DPEP3 | 0.068 | 0.0817 | 0.8393 | 0.42157 | 0.584 | -6.21 | 0.56527 | none |
| CD40LG | -0.109 | -0.1053 | -0.839 | 0.42158 | 0.584 | -6.21 | 0.17888 | none |
| CXCL10 | 0.086 | 0.0981 | 0.8393 | 0.42159 | 0.584 | -6.21 | 0.92767 | none |
| TMEM100 | 0.074 | 0.0743 | 0.8391 | 0.42169 | 0.585 | -6.21 | -0.5512 | none |
| LHX5 | -0.073 | -0.0732 | -0.839 | 0.42171 | 0.585 | -6.21 | 0.94872 | none |
| AKR1C2 | 0.142 | 0.0912 | 0.839 | 0.42176 | 0.585 | -6.21 | -0.11 | none |
| GLRX2 | -0.142 | -0.183 | -0.839 | 0.42178 | 0.585 | -6.21 | 0.54592 | none |
| PAGE2 | 0.074 | 0.0743 | 0.8389 | 0.4218 | 0.585 | -6.21 | 1.19688 | none |
| FIBIN | 0.057 | 0.057 | 0.8387 | 0.4219 | 0.585 | -6.21 | 0 | none |
| PRSS57 | 0.069 | 0.0402 | 0.8386 | 0.42197 | 0.585 | -6.21 | -0.5976 | none |
| OR2T10 | -0.057 | -0.0379 | -0.839 | 0.422 | 0.585 | -6.21 | 0.311 | none |
| ACTR10 | 0.142 | 0.022 | 0.8383 | 0.42214 | 0.585 | -6.21 | -0.7482 | none |

| | | | | | | | | |
|---|---|---|---|---|---|---|---|---|
| KRT13 | -0.07 | -0.0704 | -0.838 | 0.42216 | 0.585 | -6.21 | 1.54273 | none |
| FAM72C | 0.227 | 0.237 | 0.8381 | 0.42222 | 0.585 | -6.21 | 1.12447 | none |
| PLAU | 0.07 | 0.1398 | 0.8379 | 0.42236 | 0.585 | -6.21 | 0.06148 | none |
| C11ORF53 | -0.059 | -0.0504 | -0.838 | 0.42237 | 0.585 | -6.21 | 0 | none |
| HIST1H2AI | -0.119 | -0.123 | -0.838 | 0.4224 | 0.585 | -6.21 | 1.26822 | none |
| LYSMD2 | 0.165 | 0.1473 | 0.8377 | 0.42244 | 0.585 | -6.21 | 0.60821 | none |
| CLPS | -0.291 | -0.2615 | -0.838 | 0.42253 | 0.585 | -6.21 | 2.16009 | AA-related |
| KIF7 | 0.125 | 0.1404 | 0.8375 | 0.42256 | 0.585 | -6.21 | 1.07954 | none |
| SUSD4 | 0.147 | 0.0767 | 0.8374 | 0.42259 | 0.585 | -6.21 | 0.20577 | none |
| RESP18 | -0.057 | -0.0378 | -0.837 | 0.42259 | 0.585 | -6.21 | 0 | none |
| FBXL16 | 0.089 | 0.1183 | 0.8374 | 0.42262 | 0.585 | -6.21 | 0.11337 | none |
| DMRTC2 | 0.125 | 0.1248 | 0.8372 | 0.42272 | 0.585 | -6.21 | -0.0128 | none |
| TNFAIP6 | -0.126 | -0.0369 | -0.837 | 0.4228 | 0.585 | -6.21 | 0.74295 | none |
| MED21 | -0.144 | -0.2475 | -0.837 | 0.42293 | 0.585 | -6.21 | -0.275 | none |
| BCL2L2 | 0.079 | 0.0815 | 0.8364 | 0.42314 | 0.585 | -6.21 | -0.2218 | none |
| FKBP8 | -0.107 | -0.0249 | -0.836 | 0.42328 | 0.586 | -6.21 | 0.1474 | none |
| ADAMTS1! | -0.089 | -0.0373 | -0.836 | 0.42352 | 0.586 | -6.21 | 0.63198 | none |
| TGFBI | 0.076 | 0.0737 | 0.8354 | 0.42366 | 0.586 | -6.21 | -0.3877 | none |
| DRD2 | -0.129 | -0.1242 | -0.835 | 0.42373 | 0.586 | -6.21 | -0.6467 | none |
| TAF5 | 0.097 | 0.1399 | 0.8352 | 0.42378 | 0.586 | -6.21 | 0.17213 | none |
| WDR43 | 0.179 | 0.2598 | 0.8349 | 0.42396 | 0.586 | -6.21 | 0.4082 | none |
| ACSM6 | 0.108 | 0.0797 | 0.8348 | 0.42402 | 0.586 | -6.21 | 0.06393 | none |
| TBC1D13 | 0.093 | 0.1133 | 0.8346 | 0.42411 | 0.586 | -6.21 | 0.72896 | none |
| TTC4 | -0.07 | -0.0578 | -0.835 | 0.42412 | 0.586 | -6.21 | 0.3885 | none |
| KRTAP5-1: | -0.164 | -0.047 | -0.834 | 0.42448 | 0.587 | -6.21 | 0.64607 | none |
| SUGT1 | -0.063 | -0.0729 | -0.834 | 0.42452 | 0.587 | -6.21 | 0.75911 | none |
| CMTM6 | -0.109 | -0.1907 | -0.834 | 0.42459 | 0.587 | -6.21 | 1.34791 | none |
| KRTAP8-1 | 0.073 | 0.0734 | 0.8337 | 0.42461 | 0.587 | -6.21 | 1.19688 | none |
| HMGB4 | -0.072 | -0.0715 | -0.833 | 0.42481 | 0.587 | -6.21 | 0.86537 | none |
| OR1S2 | -0.072 | -0.0715 | -0.833 | 0.42485 | 0.587 | -6.21 | -0.3212 | none |
| ZNF365 | 0.108 | 0.1 | 0.8331 | 0.42493 | 0.587 | -6.21 | 1.67828 | none |
| TNNI3 | -0.068 | -0.0675 | -0.833 | 0.42501 | 0.587 | -6.21 | 1.15491 | none |
| FBXW9 | -0.083 | -0.042 | -0.833 | 0.42504 | 0.587 | -6.21 | 1.54048 | none |
| GPR162 | -0.056 | -0.0384 | -0.833 | 0.42509 | 0.587 | -6.21 | 1.49424 | none |
| KLF15 | -0.081 | -0.0851 | -0.833 | 0.42511 | 0.587 | -6.21 | -0.3472 | none |
| DAPK1 | -0.171 | -0.0946 | -0.833 | 0.42516 | 0.587 | -6.21 | 0.63592 | none |
| CCL2 | -0.07 | -0.0932 | -0.832 | 0.42526 | 0.587 | -6.21 | 0.88525 | none |
| FGA | -0.092 | -0.1005 | -0.832 | 0.42564 | 0.588 | -6.21 | 0.71948 | none |
| OR5B21 | -0.114 | -0.08 | -0.831 | 0.42591 | 0.588 | -6.21 | -0.5298 | none |
| MPC1L | -0.065 | -0.0683 | -0.831 | 0.42592 | 0.588 | -6.21 | 1.5055 | none |

| | | | | | | | | |
|---|---|---|---|---|---|---|---|---|
| PARP1 | 0.225 | 0.3826 | 0.8309 | 0.42608 | 0.588 | -6.21 | 0.32191 | none |
| CPEB4 | 0.192 | 0.1784 | 0.8309 | 0.42611 | 0.588 | -6.21 | 0.41587 | none |
| XPNPEP1 | 0.088 | 0.1245 | 0.8307 | 0.4262 | 0.588 | -6.21 | 0.42783 | none |
| ASGR2 | -0.1 | -0.0563 | -0.831 | 0.4262 | 0.588 | -6.21 | 1.45651 | none |
| ZNF576 | 0.118 | 0.0314 | 0.8304 | 0.42635 | 0.588 | -6.21 | -0.0103 | none |
| CRTAC1 | 0.101 | 0.1009 | 0.8301 | 0.42651 | 0.589 | -6.21 | 0.13446 | none |
| SWI5 | -0.113 | -0.1083 | -0.83 | 0.42664 | 0.589 | -6.21 | 1.10291 | none |
| XAGE1B | -0.071 | -0.071 | -0.83 | 0.42664 | 0.589 | -6.21 | -0.1025 | none |
| JPH3 | -0.071 | -0.076 | -0.83 | 0.42671 | 0.589 | -6.21 | -0.194 | none |
| HDDC2 | 0.072 | 0.0426 | 0.8295 | 0.42686 | 0.589 | -6.21 | -0.8884 | none |
| PSG9 | -0.056 | -0.0558 | -0.829 | 0.42688 | 0.589 | -6.21 | -0.271 | none |
| MTM1 | -0.165 | -0.1516 | -0.829 | 0.42698 | 0.589 | -6.21 | 0.85614 | none |
| U2AF2 | -0.112 | -0.0256 | -0.829 | 0.42704 | 0.589 | -6.21 | 0.09909 | none |
| FANCF | 0.064 | 0.0065 | 0.8291 | 0.4271 | 0.589 | -6.21 | 0.12751 | none |
| FASTKD3 | -0.133 | -0.2023 | -0.829 | 0.42721 | 0.589 | -6.21 | 0.80695 | none |
| AQP3 | -0.127 | -0.1809 | -0.829 | 0.42727 | 0.589 | -6.21 | 1.75153 | none |
| MS4A8 | 0.083 | 0.0827 | 0.8285 | 0.42739 | 0.589 | -6.21 | -0.0605 | none |
| CPN2 | 0.061 | 0.0618 | 0.8285 | 0.42739 | 0.589 | -6.21 | 1.39219 | none |
| IGFBP1 | -0.127 | -0.1545 | -0.828 | 0.42739 | 0.589 | -6.21 | 0.37427 | none |
| HMGN4 | 0.099 | -0.016 | 0.8283 | 0.42752 | 0.589 | -6.21 | 0.8431 | none |
| CACNA1B | -0.094 | -0.0946 | -0.828 | 0.42762 | 0.589 | -6.21 | 0.40959 | none |
| PNLDC1 | -0.081 | -0.0386 | -0.828 | 0.42767 | 0.589 | -6.22 | 0.80142 | none |
| COPE | -0.09 | -0.0511 | -0.828 | 0.42782 | 0.59 | -6.22 | 0.33683 | none |
| ZNF658 | -0.216 | -0.0843 | -0.828 | 0.42787 | 0.59 | -6.22 | 0.50457 | none |
| SSR1 | 0.084 | -0.0194 | 0.8273 | 0.42802 | 0.59 | -6.22 | -0.4987 | none |
| BMI1 | 0.139 | 0.0891 | 0.8269 | 0.42826 | 0.59 | -6.22 | -0.5928 | none |
| RHOXF1 | -0.069 | -0.0704 | -0.827 | 0.42831 | 0.59 | -6.22 | 0.88887 | none |
| TK2 | -0.075 | -0.067 | -0.827 | 0.42832 | 0.59 | -6.22 | 0.89034 | none |
| PSAP | -0.086 | -0.1044 | -0.827 | 0.4284 | 0.59 | -6.22 | 1.21503 | none |
| BEX3 | -0.156 | -0.1704 | -0.827 | 0.42841 | 0.59 | -6.22 | 0.40882 | none |
| TMEM14A | -0.153 | -0.2578 | -0.827 | 0.42845 | 0.59 | -6.22 | 0.89367 | none |
| TEX26 | -0.07 | -0.0703 | -0.826 | 0.42866 | 0.59 | -6.22 | 0.88916 | none |
| FAM89A | -0.139 | -0.1306 | -0.826 | 0.42869 | 0.59 | -6.22 | -0.1694 | none |
| CARNS1 | 0.14 | 0.1183 | 0.826 | 0.42873 | 0.59 | -6.22 | 0.00443 | none |
| SEC16B | -0.055 | -0.0309 | -0.826 | 0.42878 | 0.59 | -6.22 | -0.2364 | none |
| CDCA4 | -0.125 | -0.0645 | -0.826 | 0.42886 | 0.59 | -6.22 | 1.96944 | none |
| KHSRP | 0.176 | 0.2597 | 0.8253 | 0.42914 | 0.591 | -6.22 | 0.37302 | none |
| SERPINF2 | 0.074 | 0.0781 | 0.8248 | 0.4294 | 0.591 | -6.22 | 1.29741 | none |
| SLC18A3 | -0.057 | -0.0389 | -0.825 | 0.42948 | 0.591 | -6.22 | 0.03385 | none |
| FICD | 0.131 | 0.1816 | 0.8246 | 0.42952 | 0.591 | -6.22 | 0.46424 | none |

| | | | | | | | | |
|---|---|---|---|---|---|---|---|---|
| NUDT21 | 0.115 | -0.0202 | 0.8243 | 0.42968 | 0.591 | -6.22 | 0.51329 | none |
| MCF2 | -0.062 | -0.0422 | -0.824 | 0.42973 | 0.591 | -6.22 | -0.442 | none |
| PLXNA1 | -0.166 | ######## | -0.824 | 0.42997 | 0.592 | -6.22 | 0.88062 | none |
| DIRAS3 | -0.071 | -0.0263 | -0.824 | 0.43 | 0.592 | -6.22 | 1.59485 | none |
| TIMM17B | 0.133 | 0.0256 | 0.8237 | 0.43 | 0.592 | -6.22 | -0.3681 | none |
| RAD54B | -0.144 | -0.0888 | -0.824 | 0.43006 | 0.592 | -6.22 | 0.3674 | none |
| TREML4 | 0.078 | 0.0757 | 0.8229 | 0.43042 | 0.592 | -6.22 | 1.62775 | none |
| USP17L20 | 0.08 | 0.084 | 0.8225 | 0.43065 | 0.592 | -6.22 | 1.41237 | none |
| DLAT | -0.14 | -0.0631 | -0.822 | 0.43091 | 0.593 | -6.22 | 2.06144 | none |
| PPIAL4C | -0.105 | -0.1296 | -0.822 | 0.43102 | 0.593 | -6.22 | 0.84053 | none |
| TMED5 | 0.122 | 0.0476 | 0.8215 | 0.43119 | 0.593 | -6.22 | 1.09899 | none |
| STK38 | 0.11 | 0.1259 | 0.8212 | 0.43131 | 0.593 | -6.22 | 1.22498 | none |
| YIPF1 | -0.112 | -0.1297 | -0.821 | 0.43132 | 0.593 | -6.22 | 0.29825 | none |
| NKX2-1 | 0.07 | 0.0666 | 0.821 | 0.43144 | 0.593 | -6.22 | 0.5764 | none |
| STAC | 0.113 | 0.1169 | 0.821 | 0.43148 | 0.593 | -6.22 | 0.37861 | none |
| ACVR2A | 0.216 | 0.1665 | 0.8209 | 0.43153 | 0.593 | -6.22 | -0.765 | none |
| SLX1A | 0.149 | 0.0729 | 0.8207 | 0.43161 | 0.593 | -6.22 | 0.33332 | none |
| TP53TG3C | -0.059 | -0.0349 | -0.821 | 0.43169 | 0.593 | -6.22 | 1.27997 | none |
| CKLF-CMT | 0.24 | 0.2298 | 0.8202 | 0.43189 | 0.593 | -6.22 | 0.36639 | none |
| RGS9BP | 0.056 | 0.0443 | 0.8201 | 0.43191 | 0.593 | -6.22 | -1.1162 | none |
| AGAP3 | 0.116 | 0.2738 | 0.8199 | 0.43203 | 0.594 | -6.22 | 0.59389 | none |
| DRD5 | 0.057 | 0.039 | 0.8196 | 0.43219 | 0.594 | -6.22 | 1.5649 | none |
| ASIC2 | -0.096 | -0.0742 | -0.819 | 0.4323 | 0.594 | -6.22 | 0.07474 | none |
| ZDHHC16 | -0.114 | -0.1383 | -0.819 | 0.4325 | 0.594 | -6.22 | 1.53243 | none |
| WASF2 | 0.118 | 0.1778 | 0.819 | 0.43256 | 0.594 | -6.22 | -0.1071 | none |
| CIB4 | 0.071 | 0.0709 | 0.8189 | 0.43261 | 0.594 | -6.22 | -0.6397 | none |
| TIGD2 | 0.094 | 0.0444 | 0.8185 | 0.43278 | 0.594 | -6.22 | 1.17142 | none |
| RNF138 | -0.25 | -0.2441 | -0.818 | 0.43282 | 0.594 | -6.22 | 1.8461 | none |
| AL513523. | 0.062 | 0.0632 | 0.8183 | 0.4329 | 0.594 | -6.22 | -0.0092 | none |
| MALRD1 | -0.07 | -0.0682 | -0.818 | 0.43297 | 0.594 | -6.22 | 2.10187 | AA-related |
| DCLRE1B | 0.086 | 0.0616 | 0.8176 | 0.43327 | 0.595 | -6.22 | 1.39292 | none |
| CELA1 | 0.069 | 0.0707 | 0.8176 | 0.43328 | 0.595 | -6.22 | -0.6538 | none |
| SNAP25 | -0.07 | -0.0698 | -0.817 | 0.43342 | 0.595 | -6.22 | 2.14624 | AA-related |
| HNRNPUL: | 0.182 | 0.258 | 0.8174 | 0.43342 | 0.595 | -6.22 | 0.56954 | none |
| LRP6 | -0.115 | 0.0109 | -0.817 | 0.43352 | 0.595 | -6.22 | -0.0296 | none |
| CXORF65 | 0.134 | 0.2413 | 0.8171 | 0.43357 | 0.595 | -6.22 | 0.78536 | none |
| TFCP2 | 0.125 | 0.1554 | 0.8168 | 0.43373 | 0.595 | -6.22 | -0.0153 | none |
| LRRC75B | 0.092 | 0.1451 | 0.8166 | 0.43383 | 0.595 | -6.22 | 1.20164 | none |
| AUP1 | -0.106 | -0.0796 | -0.817 | 0.43389 | 0.595 | -6.22 | 0.43033 | none |
| MYL7 | 0.092 | 0.086 | 0.8157 | 0.43431 | 0.596 | -6.23 | 0.32408 | none |

| | | | | | | | | |
|---|---|---|---|---|---|---|---|---|
| PIGC | -0.107 | -0.1691 | -0.815 | 0.43448 | 0.596 | -6.23 | 1.20797 | none |
| ANXA8L1 | 0.06 | 0.0669 | 0.8153 | 0.43453 | 0.596 | -6.23 | -0.0602 | none |
| LENG9 | 0.102 | 0.0796 | 0.8153 | 0.43457 | 0.596 | -6.23 | 0.30813 | none |
| LGALS12 | 0.064 | 0.0388 | 0.8147 | 0.43486 | 0.596 | -6.23 | -0.9235 | none |
| FMO1 | 0.069 | 0.075 | 0.8144 | 0.43506 | 0.597 | -6.23 | -0.6417 | none |
| OR10J5 | -0.068 | -0.0684 | -0.814 | 0.43519 | 0.597 | -6.23 | 0.47745 | none |
| TMC3 | 0.067 | 0.0515 | 0.8137 | 0.4354 | 0.597 | -6.23 | -0.6032 | none |
| CREBL2 | -0.079 | -0.1134 | -0.813 | 0.43559 | 0.597 | -6.23 | 1.27258 | none |
| SSH3 | 0.111 | 0.1488 | 0.8128 | 0.43592 | 0.598 | -6.23 | 0.50861 | none |
| STIM2 | 0.164 | 0.192 | 0.8125 | 0.43609 | 0.598 | -6.23 | 0.79499 | none |
| ZC3H11A | 0.206 | 0.2963 | 0.8124 | 0.43613 | 0.598 | -6.23 | -0.1239 | none |
| MET | 0.149 | 0.1275 | 0.8123 | 0.43621 | 0.598 | -6.23 | -0.2525 | none |
| ITGBL1 | 0.08 | 0.0838 | 0.8119 | 0.43639 | 0.598 | -6.23 | 1.30831 | none |
| MAOA | 0.105 | 0.1579 | 0.8118 | 0.43645 | 0.598 | -6.23 | 0.44433 | none |
| BHLHE23 | -0.055 | -0.0364 | -0.811 | 0.43709 | 0.599 | -6.23 | 0 | none |
| FST | 0.08 | 0.083 | 0.8104 | 0.43722 | 0.599 | -6.23 | -0.5016 | none |
| GDF9 | 0.14 | 0.0838 | 0.8103 | 0.43728 | 0.599 | -6.23 | -0.3939 | none |
| EHD3 | 0.088 | 0.0882 | 0.8102 | 0.43733 | 0.599 | -6.23 | 0.41456 | none |
| RIC1 | 0.12 | 0.1552 | 0.8099 | 0.43751 | 0.599 | -6.23 | -0.086 | none |
| KCNIP3 | -0.097 | -0.0789 | -0.81 | 0.4377 | 0.599 | -6.23 | 1.57578 | none |
| TSLP | 0.055 | 0.0563 | 0.8092 | 0.43789 | 0.6 | -6.23 | -0.4061 | none |
| CCDC160 | -0.068 | -0.0567 | -0.809 | 0.43792 | 0.6 | -6.23 | 0.2096 | none |
| PARL | -0.096 | -0.0948 | -0.809 | 0.43793 | 0.6 | -6.23 | 0.49258 | none |
| UBE2H | 0.188 | 0.1194 | 0.8089 | 0.43808 | 0.6 | -6.23 | -0.1209 | none |
| ZW10 | -0.08 | -0.0432 | -0.808 | 0.43829 | 0.6 | -6.23 | 1.53464 | none |
| IMPDH1 | 0.077 | 0.1209 | 0.8078 | 0.43868 | 0.601 | -6.23 | 0.58356 | none |
| ALCAM | 0.198 | 0.2856 | 0.8076 | 0.43879 | 0.601 | -6.23 | -0.3397 | none |
| TOB1 | 0.076 | 0.019 | 0.8075 | 0.43884 | 0.601 | -6.23 | -1.0309 | none |
| SPTLC3 | 0.145 | 0.1742 | 0.8073 | 0.43894 | 0.601 | -6.23 | 0.25676 | none |
| ZNF704 | 0.101 | 0.1004 | 0.8071 | 0.43906 | 0.601 | -6.23 | -0.6482 | none |
| PLN | -0.054 | -0.0406 | -0.807 | 0.43931 | 0.601 | -6.23 | 1.77964 | none |
| CD109 | 0.128 | 0.0645 | 0.8066 | 0.43932 | 0.601 | -6.23 | -0.4654 | none |
| LRRC39 | -0.06 | -0.0538 | -0.807 | 0.43932 | 0.601 | -6.23 | 0.31497 | none |
| POP7 | -0.101 | -0.0943 | -0.806 | 0.43942 | 0.601 | -6.23 | 0.64345 | none |
| ITPA | -0.091 | -0.0953 | -0.806 | 0.43945 | 0.601 | -6.23 | 0.70392 | none |
| CYGB | 0.084 | 0.1034 | 0.8063 | 0.43948 | 0.601 | -6.23 | 1.39319 | none |
| HBE1 | 0.069 | 0.0688 | 0.8058 | 0.43975 | 0.601 | -6.23 | -0.3054 | none |
| VPS35 | 0.077 | 0.0283 | 0.8057 | 0.4398 | 0.601 | -6.23 | 0.67499 | none |
| ZNHIT1 | -0.123 | -0.1638 | -0.806 | 0.43988 | 0.602 | -6.23 | 1.27754 | none |
| RUNX1T1 | -0.067 | -0.065 | -0.805 | 0.44011 | 0.602 | -6.23 | 1.7084 | none |

| | | | | | | | | |
|---|---|---|---|---|---|---|---|---|
| SF3A1 | 0.2 | 0.3537 | 0.8051 | 0.44014 | 0.602 | -6.23 | 0.86163 | none |
| MINPP1 | -0.095 | -0.0821 | -0.805 | 0.4402 | 0.602 | -6.23 | 2.1967 | AA-related |
| NRM | 0.091 | 0.0585 | 0.8049 | 0.44023 | 0.602 | -6.23 | 0.39288 | none |
| ZNF324B | 0.104 | 0.1939 | 0.8048 | 0.44031 | 0.602 | -6.23 | 1.63124 | none |
| C1QTNF5 | 0.069 | 0.0686 | 0.8047 | 0.44036 | 0.602 | -6.23 | 0.24127 | none |
| PDCD4 | 0.118 | 0.0925 | 0.8044 | 0.44052 | 0.602 | -6.23 | 0.62486 | none |
| GLI3 | -0.118 | -0.1174 | -0.804 | 0.44054 | 0.602 | -6.23 | -0.4616 | none |
| LRRC23 | 0.234 | 0.1277 | 0.8042 | 0.44062 | 0.602 | -6.23 | -0.3394 | none |
| MYRFL | 0.059 | 0.06 | 0.8039 | 0.44082 | 0.602 | -6.23 | -0.7584 | none |
| TLR1 | -0.129 | -0.0887 | -0.804 | 0.44086 | 0.602 | -6.23 | 0.43771 | none |
| RHOBTB1 | -0.072 | -0.0911 | -0.804 | 0.44088 | 0.602 | -6.24 | 0.7922 | none |
| CLEC4D | 0.069 | 0.0685 | 0.8037 | 0.44089 | 0.602 | -6.24 | 1.19405 | none |
| SMYD3 | 0.128 | 0.1293 | 0.8036 | 0.44096 | 0.602 | -6.24 | 0.88771 | none |
| PAH | 0.098 | 0.151 | 0.8035 | 0.44103 | 0.602 | -6.24 | -0.2084 | none |
| BTG1 | 0.1 | 0.0573 | 0.803 | 0.44127 | 0.603 | -6.24 | -0.7139 | none |
| WAS | 0.121 | 0.0974 | 0.8029 | 0.44138 | 0.603 | -6.24 | -0.2052 | none |
| ZYX | -0.084 | -0.0567 | -0.803 | 0.44144 | 0.603 | -6.24 | 0.36924 | none |
| MLLT1 | 0.202 | 0.3428 | 0.8026 | 0.44149 | 0.603 | -6.24 | -0.0674 | none |
| CAMLG | -0.102 | -0.1557 | -0.802 | 0.44174 | 0.603 | -6.24 | 1.10051 | none |
| KRTAP10-1 | -0.054 | -0.0359 | -0.802 | 0.44182 | 0.603 | -6.24 | 0 | none |
| SLC27A6 | 0.061 | 0.0664 | 0.8019 | 0.44188 | 0.603 | -6.24 | 0 | none |
| MANBAL | 0.081 | 0.0434 | 0.8019 | 0.4419 | 0.603 | -6.24 | 0.04138 | none |
| ANKRD2 | -0.066 | -0.0149 | -0.802 | 0.4419 | 0.603 | -6.24 | 0.12555 | none |
| RAPGEF1 | 0.2 | 0.3652 | 0.8019 | 0.44192 | 0.603 | -6.24 | 0.0042 | none |
| ARHGAP3 | 0.106 | 0.0669 | 0.8018 | 0.44195 | 0.603 | -6.24 | 0.35199 | none |
| ASB15 | -0.068 | -0.0671 | -0.802 | 0.44202 | 0.603 | -6.24 | 1.83911 | none |
| CRYGB | -0.067 | -0.0665 | -0.802 | 0.44205 | 0.603 | -6.24 | 1.9054 | none |
| TTC37 | 0.272 | 0.3316 | 0.8016 | 0.44207 | 0.603 | -6.24 | 0.28181 | none |
| SERGEF | -0.109 | -0.14 | -0.802 | 0.44212 | 0.603 | -6.24 | 1.09497 | none |
| FAIM2 | 0.191 | 0.1787 | 0.8013 | 0.44224 | 0.603 | -6.24 | 0.43205 | none |
| COMMD3- | -0.1 | -0.0995 | -0.801 | 0.44232 | 0.603 | -6.24 | 0.9439 | none |
| PLPP1 | 0.117 | 0.0618 | 0.8011 | 0.44235 | 0.603 | -6.24 | 0.47981 | none |
| NFIX | 0.074 | 0.1018 | 0.8008 | 0.4425 | 0.603 | -6.24 | -0.2694 | none |
| GLRX5 | -0.135 | -0.1455 | -0.801 | 0.44253 | 0.603 | -6.24 | 0.79736 | none |
| PPY | 0.144 | 0.0712 | 0.8007 | 0.44256 | 0.603 | -6.24 | -0.1477 | none |
| GOLGA8J | 0.098 | 0.061 | 0.8006 | 0.44262 | 0.603 | -6.24 | -0.6537 | none |
| APOBEC3H | -0.157 | -0.1336 | -0.8 | 0.44296 | 0.604 | -6.24 | 0.82868 | none |
| SLC35G5 | -0.06 | -0.0134 | -0.799 | 0.44334 | 0.604 | -6.24 | 1.64961 | none |
| HIST3H2A | -0.12 | -0.0897 | -0.799 | 0.44337 | 0.604 | -6.24 | 1.40935 | none |
| CANT1 | 0.167 | 0.1494 | 0.7991 | 0.44344 | 0.604 | -6.24 | -0.6028 | none |

| | | | | | | | | |
|---|---|---|---|---|---|---|---|---|
| KRTAP12-1 | 0.068 | 0.0677 | 0.7991 | 0.44345 | 0.604 | -6.24 | 1.19688 | none |
| ADGB | 0.086 | 0.067 | 0.7991 | 0.44346 | 0.604 | -6.24 | 0.06546 | none |
| AMTN | -0.066 | -0.0662 | -0.799 | 0.44351 | 0.604 | -6.24 | 0.88916 | none |
| IFNA13 | -0.066 | -0.0661 | -0.799 | 0.44362 | 0.604 | -6.24 | 1.9054 | none |
| TSSK1B | 0.06 | 0.0477 | 0.7987 | 0.44365 | 0.604 | -6.24 | -0.3303 | none |
| NR5A1 | -0.057 | -0.0291 | -0.799 | 0.4437 | 0.604 | -6.24 | -0.2196 | none |
| KLHL34 | 0.08 | 0.041 | 0.7985 | 0.44377 | 0.604 | -6.24 | 0.77154 | none |
| MYL5 | 0.122 | -0.0099 | 0.7983 | 0.4439 | 0.605 | -6.24 | -0.0684 | none |
| MRPS9 | -0.122 | -0.0357 | -0.798 | 0.44403 | 0.605 | -6.24 | 1.77435 | none |
| OR2T27 | 0.054 | 0.0362 | 0.798 | 0.44404 | 0.605 | -6.24 | 0 | none |
| UBC | 0.183 | 0.2199 | 0.798 | 0.44407 | 0.605 | -6.24 | -0.07 | none |
| COA7 | 0.118 | 0.1496 | 0.7977 | 0.4442 | 0.605 | -6.24 | -0.3249 | none |
| EFHD2 | -0.077 | -0.0168 | -0.797 | 0.44434 | 0.605 | -6.24 | 0.58182 | none |
| MCEMP1 | -0.066 | -0.0659 | -0.797 | 0.44441 | 0.605 | -6.24 | 0.1193 | none |
| GLRB | -0.068 | -0.067 | -0.797 | 0.44448 | 0.605 | -6.24 | 0.88257 | none |
| SPPL2A | -0.119 | -0.1701 | -0.797 | 0.4445 | 0.605 | -6.24 | 1.55117 | none |
| KCNQ1 | 0.088 | 0.0544 | 0.7972 | 0.4445 | 0.605 | -6.24 | -0.8441 | none |
| SLC35F3 | -0.084 | -0.1205 | -0.797 | 0.44454 | 0.605 | -6.24 | 1.41126 | none |
| EXOSC7 | -0.1 | -0.1507 | -0.797 | 0.44454 | 0.605 | -6.24 | 1.37365 | none |
| ATP4A | -0.079 | -0.0241 | -0.797 | 0.44455 | 0.605 | -6.24 | -0.0531 | none |
| PDE6D | 0.096 | 0.0409 | 0.7971 | 0.44456 | 0.605 | -6.24 | -0.6741 | none |
| KRTAP1-4 | -0.066 | -0.0659 | -0.797 | 0.44461 | 0.605 | -6.24 | -0.2536 | none |
| VDAC1 | -0.065 | -0.0955 | -0.797 | 0.44462 | 0.605 | -6.24 | 1.48922 | none |
| SFTPA1 | -0.189 | -0.1809 | -0.797 | 0.44463 | 0.605 | -6.24 | 0.84644 | none |
| HCRTR1 | -0.063 | -0.0339 | -0.797 | 0.44479 | 0.605 | -6.24 | 0.18079 | none |
| TDRP | -0.119 | -0.1075 | -0.796 | 0.44506 | 0.605 | -6.24 | 1.69978 | none |
| IGFBP6 | -0.16 | -0.0628 | -0.796 | 0.44529 | 0.606 | -6.24 | 1.92487 | none |
| NFKBIL1 | 0.23 | 0.1921 | 0.7956 | 0.44539 | 0.606 | -6.24 | 0.33715 | none |
| OR13F1 | -0.066 | -0.0657 | -0.796 | 0.44539 | 0.606 | -6.24 | 0.47965 | none |
| PTS | -0.137 | -0.2002 | -0.795 | 0.44569 | 0.606 | -6.24 | 0.70802 | none |
| SLC7A4 | -0.055 | -0.0467 | -0.795 | 0.4458 | 0.606 | -6.24 | -0.5825 | none |
| TIMM8B | -0.127 | -0.1477 | -0.795 | 0.44587 | 0.606 | -6.24 | -0.0363 | none |
| PLET1 | 0.055 | 0.0353 | 0.7947 | 0.44588 | 0.606 | -6.24 | 0 | none |
| MED14 | 0.211 | 0.2972 | 0.7946 | 0.44594 | 0.606 | -6.24 | -0.1502 | none |
| DDX41 | 0.167 | 0.2461 | 0.7946 | 0.44595 | 0.606 | -6.24 | 0.38735 | none |
| OR10G7 | 0.067 | 0.067 | 0.7945 | 0.44597 | 0.606 | -6.24 | 1.01178 | none |
| CCDC91 | 0.179 | 0.23 | 0.7945 | 0.44601 | 0.606 | -6.24 | -0.4673 | none |
| EPB41L3 | -0.148 | -0.1176 | -0.794 | 0.44602 | 0.606 | -6.24 | -0.3824 | none |
| SLC22A7 | 0.066 | 0.0782 | 0.7943 | 0.4461 | 0.606 | -6.24 | 1.68334 | none |
| PTGER1 | -0.058 | -0.0497 | -0.794 | 0.44613 | 0.606 | -6.24 | 0.2674 | none |

| | | | | | | | | |
|---|---|---|---|---|---|---|---|---|
| LRRC71 | 0.065 | 0.0662 | 0.7942 | 0.44614 | 0.606 | -6.24 | 0.55263 | none |
| M1AP | 0.097 | 0.0051 | 0.7942 | 0.44617 | 0.606 | -6.24 | -0.4912 | none |
| RECQL | 0.136 | 0.1511 | 0.794 | 0.44625 | 0.606 | -6.24 | 0.80958 | none |
| LYZL4 | -0.065 | -0.0654 | -0.794 | 0.44626 | 0.606 | -6.24 | -0.2672 | none |
| ZNF597 | 0.064 | 0.0436 | 0.794 | 0.44627 | 0.606 | -6.24 | -0.5664 | none |
| ELAVL3 | -0.075 | -0.0379 | -0.794 | 0.44639 | 0.606 | -6.24 | -0.3877 | none |
| OR6P1 | 0.067 | 0.0669 | 0.7937 | 0.44646 | 0.606 | -6.24 | 1.44028 | none |
| MCTP2 | -0.127 | -0.1237 | -0.793 | 0.44656 | 0.606 | -6.24 | 0.01867 | none |
| KCNE5 | 0.054 | 0.0726 | 0.7929 | 0.44687 | 0.607 | -6.24 | 0.20361 | none |
| GRIA3 | -0.061 | -0.0344 | -0.793 | 0.44689 | 0.607 | -6.24 | 1.31688 | none |
| NBPF26 | -0.163 | 0.0301 | -0.792 | 0.44712 | 0.607 | -6.24 | 0.93903 | none |
| PLPPR5 | 0.078 | 0.0783 | 0.7923 | 0.44719 | 0.607 | -6.24 | 1.24893 | none |
| SARM1 | 0.113 | 0.1652 | 0.7922 | 0.44729 | 0.607 | -6.24 | 0.90502 | none |
| DEPDC1B | 0.092 | 0.0306 | 0.792 | 0.44738 | 0.607 | -6.24 | 0.25499 | none |
| GRIN2B | -0.174 | -0.0399 | -0.792 | 0.44755 | 0.607 | -6.24 | 1.5264 | none |
| EFCAB14 | 0.148 | 0.2221 | 0.7916 | 0.44759 | 0.607 | -6.24 | 0.58589 | none |
| C4ORF51 | 0.166 | 0.1671 | 0.7914 | 0.44769 | 0.607 | -6.24 | 1.06812 | none |
| REC8 | 0.08 | 0.0728 | 0.7914 | 0.44773 | 0.607 | -6.25 | -0.4077 | none |
| NDUFA2 | -0.133 | -0.1869 | -0.791 | 0.44793 | 0.608 | -6.25 | 1.36181 | none |
| MLC1 | -0.109 | -0.068 | -0.791 | 0.44805 | 0.608 | -6.25 | 0.57463 | none |
| YBX2 | 0.066 | 0.0801 | 0.7906 | 0.44816 | 0.608 | -6.25 | 0.91664 | none |
| KLHL13 | 0.078 | 0.0845 | 0.7902 | 0.44838 | 0.608 | -6.25 | 1.1652 | none |
| NCAN | 0.074 | 0.0779 | 0.7902 | 0.44841 | 0.608 | -6.25 | 1 | none |
| MPP6 | 0.148 | 0.1821 | 0.7901 | 0.44843 | 0.608 | -6.25 | 1.10494 | none |
| CCDC169 | 0.086 | 0.1113 | 0.79 | 0.44851 | 0.608 | -6.25 | -0.2181 | none |
| RNLS | 0.197 | 0.1136 | 0.7898 | 0.44863 | 0.608 | -6.25 | -0.001 | none |
| RASAL1 | -0.095 | -0.0468 | -0.79 | 0.44864 | 0.608 | -6.25 | 1.63283 | none |
| ZNF479 | 0.056 | 0.0557 | 0.7897 | 0.44868 | 0.608 | -6.25 | 0.98551 | none |
| EXOSC5 | -0.085 | -0.0858 | -0.789 | 0.44888 | 0.608 | -6.25 | 0.86873 | none |
| RIMBP3 | -0.066 | -0.0342 | -0.789 | 0.44889 | 0.608 | -6.25 | 1.00751 | none |
| HSPA5 | 0.103 | 0.1717 | 0.7893 | 0.44891 | 0.608 | -6.25 | 0.54534 | none |
| SAPCD1 | -0.105 | -0.1027 | -0.789 | 0.44898 | 0.608 | -6.25 | 0.57659 | none |
| PGA5 | -0.065 | -0.0647 | -0.789 | 0.44901 | 0.608 | -6.25 | 0.21041 | none |
| SERPINB12 | 0.053 | 0.0531 | 0.7891 | 0.44902 | 0.608 | -6.25 | -0.5163 | none |
| SARAF | -0.122 | -0.2167 | -0.789 | 0.44906 | 0.608 | -6.25 | 0.928 | none |
| PDLIM1 | 0.098 | 0.1064 | 0.7888 | 0.44917 | 0.608 | -6.25 | -0.1172 | none |
| EIF2B1 | 0.068 | 0.0652 | 0.7888 | 0.44917 | 0.608 | -6.25 | 0.06499 | none |
| POTEE | -0.069 | -0.0406 | -0.789 | 0.44933 | 0.609 | -6.25 | 1.85224 | none |
| MGAM | 0.08 | 0.0803 | 0.7884 | 0.44936 | 0.609 | -6.25 | -0.6101 | none |
| RDH11 | 0.107 | 0.1212 | 0.7884 | 0.44938 | 0.609 | -6.25 | 0.65031 | none |

| | | | | | | | | |
|---|---|---|---|---|---|---|---|---|
| CELF2 | 0.11 | 0.116 | 0.7879 | 0.44965 | 0.609 | -6.25 | 0.65486 | none |
| PIGP | -0.135 | -0.2214 | -0.788 | 0.44976 | 0.609 | -6.25 | 0.87041 | none |
| PTPRN | -0.138 | -0.0627 | -0.788 | 0.44983 | 0.609 | -6.25 | 0.91296 | none |
| MRPL44 | 0.07 | -0.0527 | 0.7874 | 0.44993 | 0.609 | -6.25 | 0.52355 | none |
| SIGLEC9 | 0.094 | 0.0835 | 0.7873 | 0.44998 | 0.609 | -6.25 | 1.11401 | none |
| PDP1 | 0.187 | 0.1507 | 0.7873 | 0.45 | 0.609 | -6.25 | 0.05142 | none |
| ARAP2 | 0.2 | 0.2132 | 0.7872 | 0.45003 | 0.609 | -6.25 | 0.16708 | none |
| DKK1 | 0.085 | 0.0905 | 0.7872 | 0.45004 | 0.609 | -6.25 | 1.1712 | none |
| CAMK2N1 | 0.087 | 0.0752 | 0.787 | 0.45016 | 0.609 | -6.25 | 0.71638 | none |
| MICAL1 | 0.186 | 0.285 | 0.7868 | 0.45026 | 0.609 | -6.25 | 0.6877 | none |
| REXO2 | 0.068 | 0.0286 | 0.7862 | 0.45062 | 0.61 | -6.25 | 1.43489 | none |
| CNGA4 | 0.054 | 0.0556 | 0.786 | 0.4507 | 0.61 | -6.25 | -0.7226 | none |
| PRAMEF19 | 0.068 | 0.0683 | 0.7858 | 0.45083 | 0.61 | -6.25 | 1.29653 | none |
| GUCA1C | -0.086 | -0.0856 | -0.786 | 0.45085 | 0.61 | -6.25 | -0.0019 | none |
| TAPBPL | -0.069 | -0.0285 | -0.786 | 0.45092 | 0.61 | -6.25 | 0.60941 | none |
| CDK16 | 0.154 | 0.3059 | 0.7855 | 0.45102 | 0.61 | -6.25 | 0.95208 | none |
| GTF3A | -0.067 | -0.0517 | -0.785 | 0.45109 | 0.61 | -6.25 | 1.60759 | none |
| MEP1A | -0.052 | -0.0293 | -0.785 | 0.45112 | 0.61 | -6.25 | 0 | none |
| NAT2 | -0.064 | -0.0641 | -0.785 | 0.45137 | 0.61 | -6.25 | 1.9054 | none |
| HTR1E | -0.052 | -0.0519 | -0.785 | 0.45148 | 0.61 | -6.25 | 0 | none |
| SMIM12 | -0.114 | -0.1749 | -0.784 | 0.45157 | 0.611 | -6.25 | 0.71855 | none |
| RBAK-RBA | 0.077 | 0.0376 | 0.7842 | 0.45173 | 0.611 | -6.25 | 0.67452 | none |
| EAF2 | 0.109 | 0.0161 | 0.7842 | 0.45173 | 0.611 | -6.25 | -0.7194 | none |
| HHAT | -0.065 | -0.0428 | -0.784 | 0.45179 | 0.611 | -6.25 | 1.36904 | none |
| ORAI1 | 0.118 | 0.0438 | 0.784 | 0.45184 | 0.611 | -6.25 | -0.0863 | none |
| CCDC42 | 0.076 | 0.0414 | 0.7839 | 0.45191 | 0.611 | -6.25 | 1.03004 | none |
| QRFP | 0.054 | 0.0673 | 0.7839 | 0.45191 | 0.611 | -6.25 | -0.9455 | none |
| ATG4D | 0.14 | 0.1341 | 0.7837 | 0.452 | 0.611 | -6.25 | -0.9201 | none |
| PTPRR | 0.061 | 0.0634 | 0.7837 | 0.45203 | 0.611 | -6.25 | 0.31829 | none |
| TH | -0.103 | -0.0645 | -0.783 | 0.45216 | 0.611 | -6.25 | 1.78675 | none |
| CCDC92 | 0.242 | 0.2511 | 0.7834 | 0.45216 | 0.611 | -6.25 | 0.17146 | none |
| HES7 | -0.061 | -0.0768 | -0.783 | 0.45221 | 0.611 | -6.25 | 1.6718 | none |
| VIPAS39 | 0.062 | 0.059 | 0.7833 | 0.45222 | 0.611 | -6.25 | -0.6255 | none |
| HIST1H1A | -0.13 | -0.1618 | -0.783 | 0.45223 | 0.611 | -6.25 | 0.51258 | none |
| RMDN3 | -0.063 | 0.0184 | -0.783 | 0.45231 | 0.611 | -6.25 | 1.3124 | none |
| AIMP1 | -0.113 | -0.1328 | -0.783 | 0.4524 | 0.611 | -6.25 | 1.20453 | none |
| SULT1A2 | 0.074 | 0.0887 | 0.783 | 0.45243 | 0.611 | -6.25 | -0.9603 | none |
| ZNF497 | 0.077 | 0.0396 | 0.7829 | 0.45246 | 0.611 | -6.25 | -0.7128 | none |
| KRT222 | -0.063 | -0.0305 | -0.783 | 0.45259 | 0.611 | -6.25 | 1.00369 | none |
| TRAF3IP3 | 0.149 | 0.1845 | 0.7826 | 0.45265 | 0.611 | -6.25 | -0.1942 | none |

| | | | | | | | | |
|---|---|---|---|---|---|---|---|---|
| GABRB3 | 0.177 | 0.0543 | 0.7825 | 0.45266 | 0.611 | -6.25 | -0.8358 | none |
| BRD2 | 0.188 | 0.3635 | 0.7819 | 0.453 | 0.611 | -6.25 | 0.06534 | none |
| HCCS | -0.106 | -0.1333 | -0.782 | 0.45305 | 0.611 | -6.25 | 1.69139 | none |
| SLC6A17 | -0.079 | -0.061 | -0.782 | 0.45321 | 0.612 | -6.25 | 1.24072 | none |
| C20ORF19 | 0.1 | 0.1302 | 0.7814 | 0.45332 | 0.612 | -6.25 | -0.0231 | none |
| EGFR | -0.076 | -0.0704 | -0.781 | 0.4534 | 0.612 | -6.25 | 1.25002 | none |
| PACS2 | 0.147 | 0.2217 | 0.7811 | 0.45349 | 0.612 | -6.25 | -0.0837 | none |
| GPR87 | -0.097 | -0.1122 | -0.781 | 0.45352 | 0.612 | -6.25 | 1.1952 | none |
| BRPF1 | 0.224 | 0.3603 | 0.7807 | 0.45371 | 0.612 | -6.25 | -0.2553 | none |
| C5ORF52 | 0.088 | 0.1017 | 0.7804 | 0.45387 | 0.612 | -6.25 | -0.0304 | none |
| BEND6 | 0.056 | 0.0543 | 0.7804 | 0.45388 | 0.612 | -6.25 | -0.8686 | none |
| ASB5 | -0.063 | -0.0634 | -0.78 | 0.45398 | 0.612 | -6.25 | 1.96868 | none |
| CSRP1 | -0.08 | -0.0668 | -0.78 | 0.45412 | 0.612 | -6.25 | 1.56543 | none |
| C17ORF10 | 0.064 | 0.0566 | 0.7799 | 0.45417 | 0.612 | -6.25 | 1.5569 | none |
| BRPF3 | 0.169 | 0.2478 | 0.7798 | 0.45419 | 0.612 | -6.25 | -0.452 | none |
| CCDC140 | 0.065 | 0.0648 | 0.7798 | 0.45419 | 0.612 | -6.25 | -0.9329 | none |
| ZSCAN26 | 0.177 | 0.193 | 0.7798 | 0.45421 | 0.612 | -6.25 | 0.25366 | none |
| TLX3 | -0.061 | -0.0517 | -0.779 | 0.45458 | 0.613 | -6.25 | 0.52268 | none |
| KLHL28 | 0.117 | 0.0291 | 0.779 | 0.45464 | 0.613 | -6.25 | 0.61147 | none |
| OR52B2 | 0.062 | 0.0721 | 0.7789 | 0.45471 | 0.613 | -6.25 | 0.86922 | none |
| USPL1 | 0.143 | 0.1774 | 0.7787 | 0.45482 | 0.613 | -6.26 | 0.39802 | none |
| RNF5 | -0.126 | -0.2026 | -0.779 | 0.45484 | 0.613 | -6.26 | 0.98498 | none |
| CXCR6 | -0.099 | -0.1579 | -0.778 | 0.45498 | 0.613 | -6.26 | 1.08963 | none |
| CDH9 | 0.121 | 0.1255 | 0.7783 | 0.45502 | 0.613 | -6.26 | 0.97741 | none |
| NOVA1 | -0.069 | -0.0603 | -0.778 | 0.45504 | 0.613 | -6.26 | 0.82445 | none |
| CASKIN2 | -0.164 | -0.1036 | -0.778 | 0.45506 | 0.613 | -6.26 | 1.41356 | none |
| KLHL32 | -0.069 | -0.0457 | -0.778 | 0.45517 | 0.613 | -6.26 | 1.80086 | none |
| DEFB108B | 0.065 | 0.0645 | 0.7779 | 0.45525 | 0.613 | -6.26 | 1.19688 | none |
| ABCA10 | -0.118 | -0.0711 | -0.778 | 0.45528 | 0.613 | -6.26 | 1.29987 | none |
| DOK2 | -0.084 | -0.0514 | -0.778 | 0.45532 | 0.613 | -6.26 | 0.60787 | none |
| ZSWIM3 | -0.101 | -0.1085 | -0.778 | 0.45533 | 0.613 | -6.26 | 0.55491 | none |
| MATN1 | 0.116 | 0.0841 | 0.7777 | 0.45537 | 0.613 | -6.26 | -0.1621 | none |
| CACUL1 | 0.107 | 0.1434 | 0.7775 | 0.45549 | 0.613 | -6.26 | -0.5758 | none |
| HES4 | -0.06 | -0.0802 | -0.777 | 0.45558 | 0.613 | -6.26 | 0.79957 | none |
| HAT1 | 0.075 | 0.001 | 0.7773 | 0.45562 | 0.613 | -6.26 | -0.0713 | none |
| GRIN2A | -0.095 | -0.0934 | -0.777 | 0.45566 | 0.613 | -6.26 | -0.234 | none |
| AC213203 | -0.209 | -0.2095 | -0.777 | 0.45569 | 0.613 | -6.26 | 1.08356 | none |
| GRHPR | 0.113 | 0.0688 | 0.7771 | 0.45569 | 0.613 | -6.26 | -0.2606 | none |
| H2AFV | -0.075 | -0.112 | -0.777 | 0.45578 | 0.613 | -6.26 | 0.22152 | none |
| DNLZ | -0.109 | -0.1037 | -0.777 | 0.45583 | 0.613 | -6.26 | 0.54465 | none |

| | | | | | | | | |
|---|---|---|---|---|---|---|---|---|
| PLXNA2 | 0.062 | 0.0825 | 0.7766 | 0.456 | 0.614 | -6.26 | -0.4259 | none |
| CADM3 | -0.062 | -0.0563 | -0.777 | 0.45601 | 0.614 | -6.26 | 1.45123 | none |
| SYNGR1 | 0.094 | 0.082 | 0.776 | 0.45631 | 0.614 | -6.26 | -0.1039 | none |
| GCNT7 | -0.13 | -0.0896 | -0.776 | 0.4564 | 0.614 | -6.26 | 1.33487 | none |
| CLDN14 | 0.066 | 0.0638 | 0.7758 | 0.45643 | 0.614 | -6.26 | 0.62521 | none |
| UTRN | 0.158 | 0.2565 | 0.7756 | 0.45655 | 0.614 | -6.26 | -0.3069 | none |
| FABP7 | 0.05 | 0.0213 | 0.7754 | 0.45667 | 0.614 | -6.26 | 0.11438 | none |
| NDUFB10 | -0.101 | -0.1045 | -0.775 | 0.45681 | 0.614 | -6.26 | 1.12511 | none |
| RNASE8 | 0.064 | 0.0641 | 0.7751 | 0.45684 | 0.614 | -6.26 | 0.1843 | none |
| TTF2 | 0.241 | 0.3762 | 0.7751 | 0.45685 | 0.614 | -6.26 | 0.2071 | none |
| ANTXRL | 0.297 | 0.1566 | 0.7751 | 0.45685 | 0.614 | -6.26 | 0.33061 | none |
| DPH2 | 0.077 | 0.0662 | 0.7747 | 0.4571 | 0.614 | -6.26 | 0.61697 | none |
| C16ORF46 | 0.076 | 0.0763 | 0.7745 | 0.45717 | 0.615 | -6.26 | 1.15822 | none |
| CLECL1 | -0.176 | -0.1744 | -0.774 | 0.45722 | 0.615 | -6.26 | 1.21298 | none |
| NPR3 | -0.06 | -0.0118 | -0.774 | 0.45726 | 0.615 | -6.26 | 1.26874 | none |
| SERPINC1 | 0.058 | 0.0676 | 0.7742 | 0.45733 | 0.615 | -6.26 | 1.35416 | none |
| DPPA5 | -0.063 | -0.0626 | -0.774 | 0.45736 | 0.615 | -6.26 | 1.9054 | none |
| VPS52 | 0.212 | 0.1119 | 0.7741 | 0.4574 | 0.615 | -6.26 | 0.54487 | none |
| HRH2 | 0.131 | 0.2341 | 0.7739 | 0.45752 | 0.615 | -6.26 | 0.62772 | none |
| NT5C1B | 0.123 | 0.1508 | 0.7739 | 0.45753 | 0.615 | -6.26 | 0.27373 | none |
| PRAMEF2! | 0.053 | 0.0359 | 0.7738 | 0.45757 | 0.615 | -6.26 | 0 | none |
| PIGZ | -0.06 | -0.0326 | -0.774 | 0.45758 | 0.615 | -6.26 | 0.85343 | none |
| FBXL15 | 0.117 | 0.0802 | 0.7737 | 0.45764 | 0.615 | -6.26 | 0.58292 | none |
| MYO16 | -0.067 | -0.0834 | -0.774 | 0.45769 | 0.615 | -6.26 | 1.09756 | none |
| DNPEP | 0.107 | 0.0715 | 0.7735 | 0.45774 | 0.615 | -6.26 | -0.564 | none |
| OTX2 | 0.061 | 0.0672 | 0.7735 | 0.45774 | 0.615 | -6.26 | 0.91633 | none |
| RIC3 | 0.226 | 0.1902 | 0.7735 | 0.45775 | 0.615 | -6.26 | 1.07586 | none |
| MDFI | 0.071 | 0.0939 | 0.7725 | 0.45831 | 0.615 | -6.26 | 1.3695 | none |
| LACRT | 0.064 | 0.0637 | 0.7723 | 0.4584 | 0.615 | -6.26 | 1.19688 | none |
| RPF2 | 0.095 | 0.0603 | 0.7723 | 0.4584 | 0.615 | -6.26 | -0.137 | none |
| MEGF10 | -0.086 | -0.0867 | -0.772 | 0.45872 | 0.616 | -6.26 | 0.63464 | none |
| GNL2 | 0.233 | 0.3422 | 0.7715 | 0.45887 | 0.616 | -6.26 | 0.31328 | none |
| DTX3 | 0.158 | 0.1719 | 0.7715 | 0.45888 | 0.616 | -6.26 | 0.2242 | none |
| ARHGAP4 | 0.083 | 0.0324 | 0.7711 | 0.4591 | 0.616 | -6.26 | -0.8042 | none |
| BMP7 | 0.127 | 0.1774 | 0.7708 | 0.45929 | 0.616 | -6.26 | 0.58749 | none |
| TP53INP2 | 0.081 | 0.1197 | 0.7704 | 0.45949 | 0.617 | -6.26 | 0.14209 | none |
| SERPINB4 | 0.063 | 0.0634 | 0.7704 | 0.45952 | 0.617 | -6.26 | 0.36521 | none |
| HIST1H1E | -0.151 | 0.0209 | -0.77 | 0.45956 | 0.617 | -6.26 | 2.0461 | none |
| HIST1H2BI | 0.148 | 0.0337 | 0.7702 | 0.45961 | 0.617 | -6.26 | -0.5664 | none |
| SPTBN5 | 0.054 | 0.054 | 0.7699 | 0.4598 | 0.617 | -6.26 | -0.0482 | none |

| | | | | | | | | |
|---|---|---|---|---|---|---|---|---|
| VOPP1 | 0.088 | 0.0237 | 0.7696 | 0.45994 | 0.617 | -6.26 | 1.13094 | none |
| ONECUT1 | -0.068 | -0.0593 | -0.77 | 0.46 | 0.617 | -6.26 | -0.1836 | none |
| PTPRQ | 0.066 | 0.0669 | 0.7695 | 0.46 | 0.617 | -6.26 | -0.2605 | none |
| C1ORF74 | -0.073 | -0.0766 | -0.769 | 0.46016 | 0.617 | -6.26 | 1.18791 | none |
| GPR19 | -0.122 | -0.1928 | -0.769 | 0.46024 | 0.617 | -6.26 | 1.03158 | none |
| RBM22 | 0.075 | 0.065 | 0.7688 | 0.4604 | 0.617 | -6.26 | 0.70639 | none |
| PAX4 | 0.073 | 0.091 | 0.7685 | 0.46056 | 0.617 | -6.26 | 0.08219 | none |
| PCDHA13 | -0.054 | -0.0391 | -0.768 | 0.46067 | 0.618 | -6.26 | 0 | none |
| THAP8 | -0.182 | -0.1376 | -0.768 | 0.46074 | 0.618 | -6.26 | 0.06673 | none |
| FIS1 | -0.136 | -0.1871 | -0.768 | 0.46078 | 0.618 | -6.26 | 1.06212 | none |
| EIF4EBP1 | -0.09 | -0.0395 | -0.768 | 0.46102 | 0.618 | -6.26 | 0.32532 | none |
| PLA2G2F | 0.054 | 0.0498 | 0.7677 | 0.46104 | 0.618 | -6.26 | 0 | none |
| CTAGE4 | -0.054 | -0.0667 | -0.767 | 0.46122 | 0.618 | -6.26 | 1.16849 | none |
| FXYD2 | 0.274 | 0.1531 | 0.767 | 0.4614 | 0.618 | -6.26 | -0.3674 | none |
| H2AFJ | -0.112 | -0.1723 | -0.767 | 0.46158 | 0.618 | -6.26 | 0.4989 | none |
| HACL1 | -0.077 | -0.1332 | -0.767 | 0.46171 | 0.619 | -6.26 | 0.83652 | none |
| GJD3 | -0.058 | -0.0635 | -0.766 | 0.46173 | 0.619 | -6.26 | 1.50067 | none |
| STAT1 | 0.112 | 0.1351 | 0.7664 | 0.46178 | 0.619 | -6.26 | 0.33524 | none |
| SLC27A3 | 0.129 | 0.181 | 0.7661 | 0.46196 | 0.619 | -6.27 | 0.01677 | none |
| PSMB10 | -0.123 | -0.1568 | -0.766 | 0.46202 | 0.619 | -6.27 | 1.46803 | none |
| ENDOU | -0.108 | -0.074 | -0.765 | 0.46232 | 0.619 | -6.27 | 1.12799 | none |
| SFTA3 | -0.091 | -0.0591 | -0.765 | 0.46232 | 0.619 | -6.27 | 0.67467 | none |
| BAIAP2L2 | -0.058 | -0.0441 | -0.765 | 0.46235 | 0.619 | -6.27 | 1.61804 | none |
| LIME1 | -0.119 | -0.2753 | -0.765 | 0.46246 | 0.619 | -6.27 | 0.32842 | none |
| COTL1 | -0.114 | -0.1393 | -0.765 | 0.46247 | 0.619 | -6.27 | 0.24338 | none |
| PCYOX1L | 0.093 | 0.0889 | 0.7651 | 0.46251 | 0.619 | -6.27 | 0.95635 | none |
| ARHGAP19 | -0.167 | -0.0507 | -0.765 | 0.46254 | 0.619 | -6.27 | 1.27732 | none |
| SLC4A2 | -0.097 | -0.0456 | -0.765 | 0.46276 | 0.619 | -6.27 | 1.02244 | none |
| FBXO17 | 0.067 | 0.0394 | 0.7645 | 0.46282 | 0.619 | -6.27 | -0.4265 | none |
| LRP12 | 0.099 | 0.1206 | 0.7645 | 0.46285 | 0.619 | -6.27 | 0.21218 | none |
| B9D2 | 0.17 | 0.0864 | 0.7645 | 0.46287 | 0.619 | -6.27 | -0.6012 | none |
| TMEM187 | 0.156 | 0.0625 | 0.7644 | 0.46288 | 0.619 | -6.27 | -0.1663 | none |
| GALE | 0.065 | 0.0358 | 0.7642 | 0.46301 | 0.619 | -6.27 | 0.5802 | none |
| NOL4 | 0.063 | 0.0626 | 0.7641 | 0.46305 | 0.619 | -6.27 | -0.6951 | none |
| CEACAM3 | 0.085 | 0.1002 | 0.7638 | 0.46326 | 0.62 | -6.27 | -0.6509 | none |
| PMF1-BGL | -0.124 | -0.1779 | -0.764 | 0.46329 | 0.62 | -6.27 | 0.36722 | none |
| LDHD | 0.114 | 0.0848 | 0.7635 | 0.46342 | 0.62 | -6.27 | -0.3273 | none |
| CSH2 | 0.064 | 0.0632 | 0.7634 | 0.46345 | 0.62 | -6.27 | 0.92852 | none |
| TTC34 | 0.049 | 0.0281 | 0.7627 | 0.46385 | 0.62 | -6.27 | 0 | none |
| C6ORF120 | -0.067 | -0.0508 | -0.763 | 0.46395 | 0.62 | -6.27 | 2.15265 | AA-related |

| | | | | | | | | |
|---|---|---|---|---|---|---|---|---|
| SV2B | 0.091 | 0.0352 | 0.7624 | 0.46402 | 0.62 | -6.27 | -0.799 | none |
| GPR55 | 0.142 | 0.0851 | 0.7621 | 0.4642 | 0.621 | -6.27 | 0.38007 | none |
| SDHAF2 | 0.114 | 0.0395 | 0.7616 | 0.46451 | 0.621 | -6.27 | 0.11805 | none |
| SYNGR2 | -0.125 | -0.1514 | -0.761 | 0.46461 | 0.621 | -6.27 | 0.62172 | none |
| SAMD1 | 0.091 | 0.1603 | 0.761 | 0.46483 | 0.621 | -6.27 | 0.94222 | none |
| SEMA6D | 0.053 | 0.0521 | 0.7609 | 0.4649 | 0.621 | -6.27 | -0.4481 | none |
| HIST1H2BJ | 0.142 | 0.077 | 0.7609 | 0.46491 | 0.621 | -6.27 | 0.22092 | none |
| COMMD5 | -0.1 | -0.0647 | -0.76 | 0.46516 | 0.622 | -6.27 | 1.28255 | none |
| KRTCAP3 | -0.098 | 0.0328 | -0.76 | 0.46518 | 0.622 | -6.27 | 1.44919 | none |
| EDDM3A | 0.062 | 0.062 | 0.7604 | 0.46519 | 0.622 | -6.27 | 1.19688 | none |
| MAD2L2 | -0.075 | -0.0733 | -0.76 | 0.46543 | 0.622 | -6.27 | 0.95532 | none |
| AASDH | -0.151 | -0.1336 | -0.76 | 0.46546 | 0.622 | -6.27 | 1.28499 | none |
| ELL3 | 0.109 | 0.0517 | 0.7593 | 0.4658 | 0.622 | -6.27 | -0.9494 | none |
| PARD3 | 0.112 | 0.1461 | 0.7592 | 0.46589 | 0.622 | -6.27 | -0.2576 | none |
| MFSD6L | -0.048 | -0.0277 | -0.759 | 0.46592 | 0.622 | -6.27 | 1.30958 | none |
| CA4 | -0.057 | -0.0087 | -0.759 | 0.46592 | 0.622 | -6.27 | 2.09051 | none |
| NCS1 | -0.064 | -0.0323 | -0.759 | 0.46601 | 0.622 | -6.27 | 0.1194 | none |
| DLX5 | 0.051 | 0.0509 | 0.7588 | 0.46609 | 0.622 | -6.27 | 0 | none |
| OR4M2 | -0.074 | -0.0312 | -0.758 | 0.46635 | 0.623 | -6.27 | 1.48825 | none |
| PLEK | 0.132 | 0.0745 | 0.7582 | 0.46645 | 0.623 | -6.27 | -0.681 | none |
| SPARC | 0.096 | 0.0645 | 0.758 | 0.46653 | 0.623 | -6.27 | 1.42914 | none |
| EBLN2 | 0.078 | 0.0533 | 0.7577 | 0.46675 | 0.623 | -6.27 | 0.56527 | none |
| SH3GL2 | -0.052 | -0.0419 | -0.757 | 0.46688 | 0.623 | -6.27 | 1.12675 | none |
| ZNF878 | -0.062 | -0.0053 | -0.757 | 0.46694 | 0.623 | -6.27 | 0.84291 | none |
| UTP3 | 0.155 | 0.2538 | 0.7573 | 0.46694 | 0.623 | -6.27 | 0.01549 | none |
| CEL | 0.057 | 0.0651 | 0.7573 | 0.46695 | 0.623 | -6.27 | -0.5147 | none |
| ABHD12 | 0.119 | 0.1138 | 0.7572 | 0.467 | 0.623 | -6.27 | -0.2367 | none |
| GEMIN8 | -0.058 | -0.0619 | -0.757 | 0.46701 | 0.623 | -6.27 | 0.78034 | none |
| PPIG | 0.198 | 0.3056 | 0.7569 | 0.46717 | 0.623 | -6.27 | 0.36561 | none |
| TGM6 | -0.087 | -0.0967 | -0.756 | 0.46748 | 0.624 | -6.27 | 1.34552 | none |
| TMEM72 | 0.066 | 0.0862 | 0.7561 | 0.46763 | 0.624 | -6.27 | 1.40251 | none |
| TMEM126 | -0.093 | -0.1345 | -0.756 | 0.4677 | 0.624 | -6.27 | 0.37966 | none |
| SYT7 | -0.081 | -0.042 | -0.756 | 0.46774 | 0.624 | -6.27 | -0.0546 | none |
| BICC1 | -0.066 | -0.0687 | -0.756 | 0.46781 | 0.624 | -6.27 | 1.81406 | none |
| MGMT | -0.101 | -0.0941 | -0.756 | 0.46785 | 0.624 | -6.27 | 1.6285 | none |
| OR4C13 | -0.06 | -0.06 | -0.755 | 0.46831 | 0.624 | -6.27 | -0.2807 | none |
| DOCK10 | 0.224 | 0.3265 | 0.7549 | 0.46833 | 0.624 | -6.27 | -0.3037 | none |
| MEDAG | -0.078 | -0.0779 | -0.755 | 0.46837 | 0.624 | -6.27 | 1.94317 | none |
| PTPRA | 0.196 | 0.2714 | 0.7546 | 0.46852 | 0.625 | -6.27 | 0.39051 | none |
| OR4D11 | 0.061 | 0.0612 | 0.7545 | 0.46857 | 0.625 | -6.27 | 0.84279 | none |

| | | | | | | | | |
|---|---|---|---|---|---|---|---|---|
| FSCN2 | 0.065 | 0.0797 | 0.7543 | 0.46866 | 0.625 | -6.27 | 0.37922 | none |
| NAT14 | 0.175 | 0.1411 | 0.7543 | 0.46869 | 0.625 | -6.27 | 0.37437 | none |
| HSPB3 | -0.06 | -0.0599 | -0.754 | 0.4687 | 0.625 | -6.27 | 1.9054 | none |
| DNAJC4 | 0.102 | 0.0648 | 0.7542 | 0.46872 | 0.625 | -6.27 | 0.56901 | none |
| ITPRIP | 0.13 | 0.2157 | 0.7541 | 0.46876 | 0.625 | -6.27 | 0.6142 | none |
| TMSB15A | -0.112 | -0.1114 | -0.754 | 0.46882 | 0.625 | -6.27 | 0.23288 | none |
| TPSAB1 | -0.069 | -0.0321 | -0.754 | 0.46884 | 0.625 | -6.27 | 1.12912 | none |
| PSMG3 | -0.113 | -0.1241 | -0.754 | 0.469 | 0.625 | -6.27 | -0.3287 | none |
| FAM222A | -0.084 | -0.0307 | -0.753 | 0.46919 | 0.625 | -6.27 | 0.31072 | none |
| COX8C | -0.061 | -0.0607 | -0.753 | 0.46925 | 0.625 | -6.27 | 0.5687 | none |
| CORO7-PA | -0.081 | -0.0515 | -0.753 | 0.46933 | 0.625 | -6.28 | 0.58415 | none |
| RTN4 | 0.069 | 0.0452 | 0.7531 | 0.46934 | 0.625 | -6.28 | -0.1379 | none |
| TOX2 | 0.139 | 0.2181 | 0.753 | 0.46943 | 0.625 | -6.28 | 0.036 | none |
| CEBPG | 0.06 | 0.0809 | 0.753 | 0.46944 | 0.625 | -6.28 | 0.56824 | none |
| TUBA8 | 0.166 | 0.0968 | 0.7529 | 0.46947 | 0.625 | -6.28 | -0.1974 | none |
| CFAP43 | -0.056 | -0.0277 | -0.753 | 0.46947 | 0.625 | -6.28 | -0.3015 | none |
| SLC22A14 | 0.077 | 0.0812 | 0.7529 | 0.4695 | 0.625 | -6.28 | 0.02849 | none |
| MAGEA2 | -0.06 | -0.0597 | -0.753 | 0.46966 | 0.625 | -6.28 | 1.75405 | none |
| OR2M5 | 0.061 | 0.0609 | 0.7525 | 0.46972 | 0.625 | -6.28 | 1.19688 | none |
| PLCH2 | 0.175 | 0.1222 | 0.7524 | 0.46976 | 0.625 | -6.28 | -0.2078 | none |
| SMAD6 | 0.055 | 0.0564 | 0.7517 | 0.47016 | 0.626 | -6.28 | -0.348 | none |
| DYNC2H1 | -0.051 | -0.0262 | -0.752 | 0.47024 | 0.626 | -6.28 | 0.58259 | none |
| CAPN13 | 0.082 | 0.0822 | 0.7509 | 0.47063 | 0.626 | -6.28 | 1.58251 | none |
| TRIM67 | 0.053 | 0.0409 | 0.7508 | 0.47067 | 0.626 | -6.28 | 0.04936 | none |
| ARFRP1 | 0.065 | 0.0748 | 0.7507 | 0.47072 | 0.626 | -6.28 | 0.80529 | none |
| PQLC3 | 0.101 | -0.073 | 0.7504 | 0.47088 | 0.626 | -6.28 | 0.4934 | none |
| EPHB3 | 0.088 | 0.0991 | 0.7504 | 0.4709 | 0.626 | -6.28 | 0.46418 | none |
| ZNF354C | 0.087 | 0.1368 | 0.7501 | 0.4711 | 0.627 | -6.28 | 0.73509 | none |
| PITRM1 | 0.17 | 0.2455 | 0.7499 | 0.47117 | 0.627 | -6.28 | 0.97518 | none |
| MTSS1L | -0.188 | -0.1922 | -0.75 | 0.47123 | 0.627 | -6.28 | 1.28226 | none |
| FCN2 | 0.061 | 0.0606 | 0.7498 | 0.47124 | 0.627 | -6.28 | 0.98208 | none |
| CDC37 | 0.145 | 0.2296 | 0.7495 | 0.47143 | 0.627 | -6.28 | 0.17455 | none |
| NAA50 | 0.102 | 0.0783 | 0.7494 | 0.47146 | 0.627 | -6.28 | 0.86787 | none |
| CHIA | -0.06 | -0.0602 | -0.749 | 0.47164 | 0.627 | -6.28 | 1.98404 | none |
| SLC38A8 | 0.179 | 0.0888 | 0.749 | 0.4717 | 0.627 | -6.28 | -0.2752 | none |
| BANK1 | 0.268 | 0.2068 | 0.749 | 0.47171 | 0.627 | -6.28 | 0.77135 | none |
| C10ORF62 | -0.065 | -0.0498 | -0.749 | 0.47184 | 0.627 | -6.28 | -0.4463 | none |
| KRT17 | 0.123 | 0.0394 | 0.7488 | 0.47185 | 0.627 | -6.28 | 0.08768 | none |
| GNA13 | 0.071 | 0.0246 | 0.7486 | 0.47196 | 0.627 | -6.28 | 0.77123 | none |
| GCSAML | 0.092 | 0.0933 | 0.7483 | 0.47209 | 0.627 | -6.28 | -0.332 | none |

| | | | | | | | | |
|---|---|---|---|---|---|---|---|---|
| TUBA3D | -0.107 | -0.1044 | -0.748 | 0.47221 | 0.627 | -6.28 | 0.75953 | none |
| TSPY3 | 0.055 | 0.0599 | 0.748 | 0.47227 | 0.627 | -6.28 | 1.13029 | none |
| FBXO39 | -0.076 | -0.0555 | -0.748 | 0.47231 | 0.627 | -6.28 | 1.9354 | none |
| HMGB3 | -0.105 | -0.1346 | -0.748 | 0.47232 | 0.627 | -6.28 | 0.18816 | none |
| RIMBP2 | 0.071 | 0.0837 | 0.7474 | 0.47262 | 0.628 | -6.28 | -0.385 | none |
| SLC4A3 | -0.068 | -0.0425 | -0.747 | 0.47268 | 0.628 | -6.28 | 0.37443 | none |
| DHX32 | 0.105 | 0.1807 | 0.7469 | 0.47293 | 0.628 | -6.28 | 1.23549 | none |
| BTN3A3 | 0.082 | -0.0081 | 0.7466 | 0.47309 | 0.628 | -6.28 | 0.64354 | none |
| NCAPG | 0.131 | 0.1635 | 0.7466 | 0.47311 | 0.628 | -6.28 | 0.59747 | none |
| PTHLH | -0.11 | -0.0735 | -0.746 | 0.47322 | 0.628 | -6.28 | 0.6519 | none |
| SLU7 | 0.233 | 0.3531 | 0.7464 | 0.47322 | 0.628 | -6.28 | 0.48322 | none |
| LDLRAD1 | 0.053 | 0.054 | 0.7464 | 0.47322 | 0.628 | -6.28 | -0.0731 | none |
| IGLON5 | 0.052 | 0.0532 | 0.7462 | 0.47332 | 0.628 | -6.28 | -0.5149 | none |
| YPEL3 | -0.115 | -0.1311 | -0.746 | 0.47337 | 0.628 | -6.28 | -0.4618 | none |
| KATNAL2 | 0.16 | 0.1995 | 0.7459 | 0.47351 | 0.628 | -6.28 | 1.15425 | none |
| DDT | -0.12 | -0.1613 | -0.745 | 0.47383 | 0.629 | -6.28 | 1.46118 | none |
| RASSF7 | -0.076 | -0.0834 | -0.745 | 0.47397 | 0.629 | -6.28 | 2.08997 | none |
| RBM10 | 0.18 | 0.304 | 0.745 | 0.47402 | 0.629 | -6.28 | 0.37807 | none |
| TYMS | -0.093 | -0.0391 | -0.744 | 0.47446 | 0.629 | -6.28 | 1.59861 | none |
| FERD3L | 0.06 | 0.0598 | 0.7441 | 0.47454 | 0.63 | -6.28 | 1.19688 | none |
| LSAMP | -0.157 | -0.1162 | -0.744 | 0.47457 | 0.63 | -6.28 | 0.63393 | none |
| CST1 | 0.065 | 0.0654 | 0.7439 | 0.47466 | 0.63 | -6.28 | -0.0788 | none |
| SLC35A2 | -0.145 | -0.1524 | -0.744 | 0.47468 | 0.63 | -6.28 | 1.2532 | none |
| FKBP2 | -0.069 | -0.0091 | -0.744 | 0.47483 | 0.63 | -6.28 | 0.97608 | none |
| CX3CR1 | -0.073 | -0.0481 | -0.744 | 0.47487 | 0.63 | -6.28 | 0.00898 | none |
| DXO | -0.214 | -0.1215 | -0.743 | 0.47502 | 0.63 | -6.28 | 0.69809 | none |
| GRASP | -0.166 | -0.1563 | -0.743 | 0.47503 | 0.63 | -6.28 | 0.06022 | none |
| RAB3GAP2 | 0.167 | 0.2052 | 0.7432 | 0.47508 | 0.63 | -6.28 | 0.21529 | none |
| ABCA12 | 0.07 | 0.0785 | 0.7431 | 0.4751 | 0.63 | -6.28 | 1.19418 | none |
| IL1RAP | 0.111 | 0.1591 | 0.7431 | 0.47513 | 0.63 | -6.28 | -0.153 | none |
| ACTL7A | -0.059 | -0.0363 | -0.743 | 0.47527 | 0.63 | -6.28 | 0.04755 | none |
| CLEC4G | -0.054 | -0.0297 | -0.743 | 0.4753 | 0.63 | -6.28 | 1.04562 | none |
| SSRP1 | 0.161 | 0.2588 | 0.7428 | 0.4753 | 0.63 | -6.28 | 0.36494 | none |
| PLA2G12B | 0.083 | 0.0321 | 0.7426 | 0.47542 | 0.63 | -6.28 | 0.883 | none |
| SOD3 | -0.058 | -0.0584 | -0.743 | 0.47543 | 0.63 | -6.28 | 2.02693 | none |
| FBXO32 | 0.126 | 0.1643 | 0.7425 | 0.47546 | 0.63 | -6.28 | 0.88932 | none |
| DNALI1 | -0.055 | -0.0474 | -0.742 | 0.47557 | 0.63 | -6.28 | 0.41893 | none |
| RAB43 | 0.113 | 0.1099 | 0.7421 | 0.47569 | 0.63 | -6.28 | -0.1867 | none |
| ZBTB48 | 0.16 | 0.1665 | 0.742 | 0.47574 | 0.63 | -6.28 | -0.3758 | none |
| TOP2B | 0.273 | 0.3763 | 0.7417 | 0.47592 | 0.63 | -6.28 | 0.13894 | none |

| | | | | | | | | |
|---|---|---|---|---|---|---|---|---|
| ATG16L1 | 0.084 | 0.1174 | 0.7416 | 0.47601 | 0.63 | -6.28 | 0.79843 | none |
| PPM1H | 0.053 | 0.0621 | 0.7415 | 0.47606 | 0.63 | -6.28 | 0.09451 | none |
| DTWD1 | 0.088 | 0.0552 | 0.7411 | 0.47629 | 0.631 | -6.28 | 0.637 | none |
| VEGFC | 0.055 | 0.0583 | 0.741 | 0.47635 | 0.631 | -6.28 | 1.453 | none |
| TCEA2 | -0.077 | -0.0244 | -0.741 | 0.47635 | 0.631 | -6.28 | 0.49628 | none |
| RNPEPL1 | 0.131 | 0.1516 | 0.7405 | 0.47665 | 0.631 | -6.28 | 0.34465 | none |
| IL11RA | 0.12 | 0.1295 | 0.7404 | 0.47666 | 0.631 | -6.28 | 0.00851 | none |
| CXCR4 | -0.099 | -0.2083 | -0.74 | 0.47679 | 0.631 | -6.28 | 1.20528 | none |
| TRAF7 | -0.072 | 0.0073 | -0.74 | 0.47697 | 0.631 | -6.29 | 0.72025 | none |
| STX6 | 0.106 | 0.1862 | 0.7396 | 0.47714 | 0.631 | -6.29 | 0.26267 | none |
| NOP16 | -0.079 | -0.0834 | -0.739 | 0.47732 | 0.632 | -6.29 | 0.08472 | none |
| SLC35B3 | -0.142 | -0.2207 | -0.739 | 0.47737 | 0.632 | -6.29 | 1.03131 | none |
| YKT6 | -0.083 | -0.0215 | -0.739 | 0.4774 | 0.632 | -6.29 | 0.95087 | none |
| DTX4 | 0.134 | 0.1243 | 0.739 | 0.47746 | 0.632 | -6.29 | -0.5547 | none |
| TMEM164 | -0.137 | -0.035 | -0.739 | 0.47754 | 0.632 | -6.29 | -0.0301 | none |
| KRTAP9-8 | 0.059 | 0.0591 | 0.7388 | 0.47763 | 0.632 | -6.29 | -0.5246 | none |
| PTP4A3 | -0.101 | -0.1018 | -0.739 | 0.47763 | 0.632 | -6.29 | 1.20197 | none |
| PGLYRP1 | -0.079 | -0.0531 | -0.739 | 0.47764 | 0.632 | -6.29 | -0.1957 | none |
| PPP1R12B | 0.167 | 0.1718 | 0.7386 | 0.4777 | 0.632 | -6.29 | -0.5858 | none |
| PGM5 | 0.105 | 0.1329 | 0.7385 | 0.47777 | 0.632 | -6.29 | -0.3023 | none |
| FAM107A | 0.049 | 0.0546 | 0.7382 | 0.47793 | 0.632 | -6.29 | -0.6434 | none |
| FBLN5 | 0.08 | 0.0188 | 0.7382 | 0.47793 | 0.632 | -6.29 | -0.1967 | none |
| MTMR6 | 0.109 | 0.0641 | 0.7379 | 0.4781 | 0.632 | -6.29 | 1.40532 | none |
| IL7 | -0.12 | -0.2215 | -0.737 | 0.47844 | 0.632 | -6.29 | -0.1736 | none |
| LGI3 | 0.05 | 0.053 | 0.7369 | 0.47868 | 0.633 | -6.29 | -0.8506 | none |
| MYBPC2 | 0.116 | 0.1205 | 0.7369 | 0.47873 | 0.633 | -6.29 | 0.9051 | none |
| RAB2A | 0.062 | 0.0384 | 0.7367 | 0.47882 | 0.633 | -6.29 | 0.59749 | none |
| CGB7 | -0.049 | -0.031 | -0.737 | 0.47885 | 0.633 | -6.29 | 0 | none |
| SMPDL3A | -0.088 | -0.0454 | -0.737 | 0.47889 | 0.633 | -6.29 | 0.57936 | none |
| GLUD2 | -0.049 | -0.0482 | -0.736 | 0.47899 | 0.633 | -6.29 | 0.70545 | none |
| C6ORF226 | -0.167 | -0.1271 | -0.736 | 0.47899 | 0.633 | -6.29 | 1.34741 | none |
| NAA38 | -0.129 | -0.1733 | -0.736 | 0.47899 | 0.633 | -6.29 | 0.53054 | none |
| MRPS25 | -0.083 | -0.0714 | -0.736 | 0.47901 | 0.633 | -6.29 | 0.0067 | none |
| RIPK4 | -0.05 | -0.033 | -0.736 | 0.4791 | 0.633 | -6.29 | 0.21716 | none |
| MGAT5 | 0.149 | 0.19 | 0.7361 | 0.47916 | 0.633 | -6.29 | 0.66354 | none |
| ASAP1 | -0.19 | 0.011 | -0.736 | 0.4793 | 0.633 | -6.29 | -0.0165 | none |
| LNX1 | 0.143 | 0.1128 | 0.7357 | 0.4794 | 0.633 | -6.29 | 0.98842 | none |
| CLEC9A | 0.059 | 0.0587 | 0.7357 | 0.47942 | 0.633 | -6.29 | 1.2124 | none |
| LST1 | -0.151 | -0.2936 | -0.735 | 0.47955 | 0.633 | -6.29 | 1.15711 | none |
| FAM24B | 0.124 | 0.0909 | 0.7354 | 0.47956 | 0.633 | -6.29 | 0.0546 | none |

| | | | | | | | | |
|---|---|---|---|---|---|---|---|---|
| MST1R | -0.087 | -0.0132 | -0.735 | 0.47969 | 0.633 | -6.29 | 0.08133 | none |
| GPR52 | -0.049 | -0.0306 | -0.735 | 0.47981 | 0.633 | -6.29 | 2.25038 | AA-related |
| KCNC3 | 0.058 | 0.094 | 0.735 | 0.47981 | 0.633 | -6.29 | 0.54314 | none |
| FLVCR2 | 0.112 | 0.0962 | 0.7349 | 0.47989 | 0.633 | -6.29 | 0.97361 | none |
| RBM42 | -0.064 | -0.0269 | -0.735 | 0.48005 | 0.634 | -6.29 | 0.52099 | none |
| OR52D1 | -0.057 | -0.0574 | -0.735 | 0.48005 | 0.634 | -6.29 | 1.9054 | none |
| ELF4 | 0.093 | 0.1193 | 0.7346 | 0.48007 | 0.634 | -6.29 | 0.58143 | none |
| FGF23 | 0.095 | 0.0607 | 0.7345 | 0.48012 | 0.634 | -6.29 | -0.1772 | none |
| FBLN2 | -0.123 | -0.0334 | -0.734 | 0.48024 | 0.634 | -6.29 | -0.3839 | none |
| PRPF31 | 0.096 | 0.1633 | 0.7342 | 0.48027 | 0.634 | -6.29 | 0.54348 | none |
| SSC5D | 0.062 | 0.0898 | 0.734 | 0.48038 | 0.634 | -6.29 | 1.59834 | none |
| XPC | 0.134 | 0.2637 | 0.7336 | 0.48063 | 0.634 | -6.29 | 0.24447 | none |
| MCTP1 | -0.092 | -0.0494 | -0.733 | 0.48077 | 0.634 | -6.29 | -0.2425 | none |
| EFCC1 | -0.049 | -0.0242 | -0.733 | 0.48084 | 0.634 | -6.29 | 0 | none |
| ZNF239 | -0.079 | -0.0745 | -0.733 | 0.48091 | 0.634 | -6.29 | 0.21068 | none |
| NQO2 | 0.087 | -0.0014 | 0.733 | 0.48098 | 0.634 | -6.29 | -0.1438 | none |
| ADAM8 | 0.087 | 0.1764 | 0.7329 | 0.48102 | 0.634 | -6.29 | 1.31558 | none |
| PRF1 | 0.072 | 0.0811 | 0.7328 | 0.4811 | 0.634 | -6.29 | -0.2251 | none |
| TRPM2 | 0.085 | 0.0939 | 0.7325 | 0.48128 | 0.635 | -6.29 | 0.15241 | none |
| ADCY1 | -0.083 | -0.0555 | -0.732 | 0.48144 | 0.635 | -6.29 | 0.2631 | none |
| KBTBD11 | 0.107 | 0.1427 | 0.732 | 0.48155 | 0.635 | -6.29 | 0.30114 | none |
| RAB34 | -0.129 | -0.1754 | -0.732 | 0.48173 | 0.635 | -6.29 | 0.54646 | none |
| TEKT2 | 0.085 | 0.0414 | 0.7317 | 0.48175 | 0.635 | -6.29 | -0.9254 | none |
| USP1 | 0.162 | 0.2339 | 0.7315 | 0.48184 | 0.635 | -6.29 | 0.70035 | none |
| SPEF1 | 0.054 | 0.0619 | 0.7315 | 0.48184 | 0.635 | -6.29 | -0.0862 | none |
| ARHGAP25 | 0.177 | 0.2168 | 0.7311 | 0.48208 | 0.635 | -6.29 | -0.1437 | none |
| SLC9A5 | 0.099 | 0.114 | 0.731 | 0.48215 | 0.635 | -6.29 | 0.59526 | none |
| CDCA7L | -0.097 | -0.0333 | -0.731 | 0.48216 | 0.635 | -6.29 | 2.09793 | AA-related |
| UTP18 | -0.064 | -0.0343 | -0.731 | 0.48235 | 0.635 | -6.29 | 1.99077 | none |
| FAM228A | 0.076 | 0.0784 | 0.7305 | 0.48241 | 0.635 | -6.29 | 0.08996 | none |
| WNT8A | -0.049 | -0.0494 | -0.731 | 0.48242 | 0.635 | -6.29 | -0.2181 | none |
| CTSL | -0.124 | -0.0858 | -0.73 | 0.48257 | 0.636 | -6.29 | 1.27315 | none |
| PSMD12 | 0.083 | 0.1113 | 0.7302 | 0.4826 | 0.636 | -6.29 | 0.21092 | none |
| GADD45B | -0.19 | 0.0073 | -0.73 | 0.48281 | 0.636 | -6.29 | 1.89098 | none |
| TIMD4 | 0.121 | 0.0235 | 0.7298 | 0.48286 | 0.636 | -6.29 | -0.0253 | none |
| FBXL4 | 0.077 | -0.0167 | 0.7295 | 0.48302 | 0.636 | -6.29 | 0.44963 | none |
| IL27 | -0.062 | -0.0171 | -0.729 | 0.48318 | 0.636 | -6.29 | 0.33711 | none |
| TSPAN32 | 0.226 | 0.1214 | 0.729 | 0.4833 | 0.636 | -6.29 | 0.08057 | none |
| BORA | -0.131 | -0.1159 | -0.729 | 0.48333 | 0.636 | -6.29 | 1.1274 | none |
| MAP3K8 | 0.086 | 0.0345 | 0.729 | 0.48333 | 0.636 | -6.29 | -0.6545 | none |

| | | | | | | | | |
|---|---|---|---|---|---|---|---|---|
| DAXX | -0.245 | 0.0319 | -0.728 | 0.4837 | 0.637 | -6.29 | 0.4282 | none |
| ALKBH5 | 0.071 | 0.1004 | 0.728 | 0.48388 | 0.637 | -6.29 | 0.65908 | none |
| C5ORF15 | -0.117 | -0.1659 | -0.728 | 0.48395 | 0.637 | -6.29 | 0.5445 | none |
| CLCNKB | -0.08 | -0.0799 | -0.728 | 0.48403 | 0.637 | -6.29 | -0.2456 | none |
| CHCHD5 | 0.143 | 0.0883 | 0.7277 | 0.48409 | 0.637 | -6.29 | -0.4345 | none |
| MRFAP1L1 | -0.059 | -0.1142 | -0.727 | 0.48429 | 0.637 | -6.29 | 0.4767 | none |
| EPOR | -0.095 | -0.1074 | -0.727 | 0.48448 | 0.637 | -6.29 | 0.43055 | none |
| MMP2 | 0.062 | 0.0708 | 0.727 | 0.48448 | 0.637 | -6.29 | -0.5022 | none |
| TANK | -0.134 | -0.0677 | -0.727 | 0.4847 | 0.638 | -6.3 | 1.54945 | none |
| PPP1CC | 0.077 | -0.0192 | 0.7258 | 0.4852 | 0.638 | -6.3 | -0.0201 | none |
| RXRA | 0.085 | 0.1373 | 0.7255 | 0.48533 | 0.638 | -6.3 | 0.30825 | none |
| SH3RF1 | 0.147 | 0.1812 | 0.7254 | 0.48543 | 0.638 | -6.3 | -0.5279 | none |
| IPO9 | 0.154 | 0.1893 | 0.7253 | 0.48547 | 0.638 | -6.3 | 0.58715 | none |
| CBARP | -0.055 | -0.0345 | -0.725 | 0.48588 | 0.639 | -6.3 | 1.42756 | none |
| GNA14 | 0.062 | 0.0569 | 0.7246 | 0.48589 | 0.639 | -6.3 | 1.08718 | none |
| ANKRD63 | 0.055 | 0.0567 | 0.7244 | 0.48602 | 0.639 | -6.3 | 0.62656 | none |
| DLD | -0.073 | -0.09 | -0.724 | 0.48618 | 0.639 | -6.3 | 0.77286 | none |
| TRPT1 | -0.111 | -0.1731 | -0.724 | 0.48619 | 0.639 | -6.3 | 1.00933 | none |
| NR3C1 | 0.101 | 0.115 | 0.7238 | 0.48635 | 0.639 | -6.3 | -0.1846 | none |
| SDR42E1 | -0.072 | -0.0778 | -0.724 | 0.4864 | 0.639 | -6.3 | -0.41 | none |
| TPRG1 | 0.202 | 0.036 | 0.7237 | 0.48641 | 0.639 | -6.3 | -0.7991 | none |
| SMC3 | 0.268 | 0.4291 | 0.7237 | 0.48641 | 0.639 | -6.3 | 0.41847 | none |
| CDR2L | -0.077 | 0.0114 | -0.723 | 0.48654 | 0.639 | -6.3 | -0.0192 | none |
| FIZ1 | 0.135 | 0.1429 | 0.7232 | 0.48673 | 0.639 | -6.3 | 0.37516 | none |
| SHISA5 | 0.072 | 0.0119 | 0.7231 | 0.48676 | 0.639 | -6.3 | -0.4449 | none |
| LCN1 | 0.057 | 0.0571 | 0.7229 | 0.48688 | 0.64 | -6.3 | 0.44852 | none |
| FGG | -0.071 | -0.0854 | -0.723 | 0.48689 | 0.64 | -6.3 | 0.56939 | none |
| LMCD1 | -0.081 | -0.0642 | -0.722 | 0.48727 | 0.64 | -6.3 | -0.2905 | none |
| LRRN4 | 0.069 | 0.0786 | 0.7221 | 0.48734 | 0.64 | -6.3 | -0.9418 | none |
| NKG7 | 0.182 | 0.0774 | 0.7221 | 0.48735 | 0.64 | -6.3 | 0.48655 | none |
| SDC1 | -0.052 | -0.052 | -0.722 | 0.48759 | 0.64 | -6.3 | 0.08681 | none |
| MICB | -0.127 | -0.0458 | -0.722 | 0.48765 | 0.64 | -6.3 | 0.16539 | none |
| RANBP17 | -0.118 | -0.1342 | -0.721 | 0.48783 | 0.64 | -6.3 | 0.63478 | none |
| CDH24 | -0.054 | -0.0478 | -0.721 | 0.48805 | 0.641 | -6.3 | 0.32273 | none |
| GSC | -0.053 | -0.0551 | -0.721 | 0.48816 | 0.641 | -6.3 | 0 | none |
| FMN1 | 0.062 | 0.026 | 0.7207 | 0.48817 | 0.641 | -6.3 | -0.8786 | none |
| SNX6 | 0.066 | 0.0043 | 0.7207 | 0.48817 | 0.641 | -6.3 | 0.76761 | none |
| IFT52 | 0.087 | -0.0532 | 0.7202 | 0.48849 | 0.641 | -6.3 | 0.17396 | none |
| FBXL19 | 0.188 | 0.2944 | 0.7201 | 0.48851 | 0.641 | -6.3 | 0.49924 | none |
| HMGXB4 | 0.12 | 0.1697 | 0.7201 | 0.48853 | 0.641 | -6.3 | 0.77531 | none |

| | | | | | | | | |
|---|---|---|---|---|---|---|---|---|
| UPF2 | 0.203 | 0.2741 | 0.7197 | 0.48874 | 0.641 | -6.3 | -0.2147 | none |
| ADIPOR1 | -0.094 | -0.1265 | -0.72 | 0.4888 | 0.641 | -6.3 | 0.23871 | none |
| PPP1R26 | -0.113 | -0.0327 | -0.719 | 0.48907 | 0.642 | -6.3 | 0.34642 | none |
| RGS7BP | 0.057 | 0.0566 | 0.7191 | 0.48911 | 0.642 | -6.3 | 1.16258 | none |
| HIST1H1T | -0.055 | -0.0555 | -0.719 | 0.4893 | 0.642 | -6.3 | -0.3521 | none |
| INSL5 | -0.055 | -0.0555 | -0.719 | 0.48932 | 0.642 | -6.3 | 1.9054 | none |
| FAHD2B | -0.119 | -0.0421 | -0.719 | 0.48932 | 0.642 | -6.3 | 1.49957 | none |
| GPR148 | 0.062 | 0.0438 | 0.7187 | 0.48933 | 0.642 | -6.3 | 0.67502 | none |
| GBP2 | 0.195 | 0.28 | 0.7187 | 0.48937 | 0.642 | -6.3 | 0.851 | none |
| GBP5 | 0.16 | 0.184 | 0.7184 | 0.48955 | 0.642 | -6.3 | 0.7275 | none |
| BCL7C | 0.084 | 0.0458 | 0.7183 | 0.48958 | 0.642 | -6.3 | -0.6513 | none |
| MAP1LC3A | 0.108 | -0.0064 | 0.718 | 0.48977 | 0.642 | -6.3 | -0.1745 | none |
| SRMS | 0.055 | 0.0388 | 0.7179 | 0.4898 | 0.642 | -6.3 | 0 | none |
| PHLDA2 | 0.106 | 0.152 | 0.7179 | 0.48982 | 0.642 | -6.3 | 1.44226 | none |
| OSTC | -0.116 | -0.2358 | -0.718 | 0.48982 | 0.642 | -6.3 | 0.4089 | none |
| SMIM15 | -0.096 | -0.1083 | -0.718 | 0.48986 | 0.642 | -6.3 | 1.83371 | none |
| UXS1 | 0.078 | 0.0915 | 0.7166 | 0.49058 | 0.643 | -6.3 | -0.6552 | none |
| SMIM18 | -0.367 | -0.288 | -0.716 | 0.49073 | 0.643 | -6.3 | 1.70328 | none |
| OTUD6B | 0.158 | 0.1308 | 0.7161 | 0.4909 | 0.643 | -6.3 | -0.0808 | none |
| IL12A | 0.157 | 0.1758 | 0.716 | 0.49097 | 0.643 | -6.3 | 0.68817 | none |
| SLC22A18 | -0.149 | -0.3053 | -0.716 | 0.491 | 0.643 | -6.3 | 0.36134 | none |
| CCDC115 | 0.107 | 0.0049 | 0.7159 | 0.49103 | 0.643 | -6.3 | 0.08179 | none |
| ANKRD31 | 0.05 | 0.0402 | 0.7158 | 0.49108 | 0.643 | -6.3 | -0.7578 | none |
| TLE1 | -0.095 | -0.0877 | -0.716 | 0.49121 | 0.643 | -6.3 | -0.2579 | none |
| ALDH3B2 | -0.096 | -0.0508 | -0.715 | 0.49135 | 0.643 | -6.3 | 0.47464 | none |
| C6ORF201 | -0.208 | -0.1228 | -0.715 | 0.49149 | 0.644 | -6.3 | 1.61679 | none |
| ADPRHL2 | -0.077 | -0.037 | -0.715 | 0.49155 | 0.644 | -6.3 | 0.82948 | none |
| DNER | 0.068 | 0.0488 | 0.7149 | 0.49161 | 0.644 | -6.3 | 0.86371 | none |
| CTSF | -0.08 | -0.0846 | -0.715 | 0.49164 | 0.644 | -6.3 | 1.76422 | none |
| ING2 | 0.11 | 0.1094 | 0.7148 | 0.49166 | 0.644 | -6.3 | 1.38101 | none |
| PGGT1B | -0.165 | -0.1424 | -0.715 | 0.4918 | 0.644 | -6.3 | 0.55976 | none |
| HIST1H3A | 0.216 | 0.2178 | 0.7144 | 0.49188 | 0.644 | -6.3 | 0.3258 | none |
| PIGM | 0.067 | 0.0164 | 0.7139 | 0.49219 | 0.644 | -6.3 | -0.7157 | none |
| CD300C | 0.082 | 0.1416 | 0.7139 | 0.49221 | 0.644 | -6.3 | 0.51352 | none |
| GRM1 | 0.053 | 0.022 | 0.7132 | 0.49259 | 0.645 | -6.31 | -0.614 | none |
| SNX24 | -0.12 | -0.0979 | -0.713 | 0.49265 | 0.645 | -6.31 | 0.07572 | none |
| OR4Q3 | 0.1 | 0.0819 | 0.7131 | 0.49267 | 0.645 | -6.31 | -0.2337 | none |
| TMEM171 | 0.049 | 0.0647 | 0.7131 | 0.49267 | 0.645 | -6.31 | 0.07128 | none |
| MAZ | -0.048 | -0.0145 | -0.713 | 0.49268 | 0.645 | -6.31 | 1.58169 | none |
| XPO7 | 0.15 | 0.2382 | 0.7128 | 0.49284 | 0.645 | -6.31 | 0.4544 | none |

| | | | | | | | | |
|---|---|---|---|---|---|---|---|---|
| AIDA | 0.058 | 0.0252 | 0.7127 | 0.49287 | 0.645 | -6.31 | 0.59135 | none |
| OIT3 | -0.059 | -0.0591 | -0.713 | 0.49288 | 0.645 | -6.31 | 0.74937 | none |
| DNAH1 | 0.096 | 0.1974 | 0.7125 | 0.493 | 0.645 | -6.31 | 1.54026 | none |
| HARS | 0.08 | 0.0792 | 0.712 | 0.49329 | 0.645 | -6.31 | 0.07937 | none |
| PDCL3 | -0.064 | -0.0887 | -0.712 | 0.49346 | 0.645 | -6.31 | -0.5849 | none |
| ARRDC4 | 0.051 | 0.0475 | 0.7117 | 0.49348 | 0.645 | -6.31 | 1.18718 | none |
| WDYHV1 | -0.094 | -0.1665 | -0.712 | 0.49348 | 0.645 | -6.31 | 0.80654 | none |
| FAM174A | -0.123 | -0.117 | -0.711 | 0.49376 | 0.645 | -6.31 | 1.61181 | none |
| GJC3 | -0.05 | -0.065 | -0.711 | 0.49384 | 0.646 | -6.31 | 1.65341 | none |
| FCRL4 | 0.06 | 0.0764 | 0.7111 | 0.49385 | 0.646 | -6.31 | -0.1498 | none |
| ITGA9 | 0.08 | 0.0989 | 0.7109 | 0.49399 | 0.646 | -6.31 | 0.42551 | none |
| SDF2L1 | -0.088 | -0.0976 | -0.711 | 0.49412 | 0.646 | -6.31 | 0.50497 | none |
| PLEKHO2 | 0.068 | 0.0382 | 0.7104 | 0.49426 | 0.646 | -6.31 | 0.36955 | none |
| BATF3 | -0.122 | -0.1414 | -0.71 | 0.49445 | 0.646 | -6.31 | 1.1065 | none |
| RDH14 | -0.091 | -0.0644 | -0.71 | 0.49464 | 0.646 | -6.31 | 1.92763 | none |
| PPP1R3C | 0.065 | 0.0152 | 0.7097 | 0.49467 | 0.646 | -6.31 | -0.8385 | none |
| UAP1 | 0.102 | 0.0231 | 0.7094 | 0.49485 | 0.646 | -6.31 | 0.45737 | none |
| OR4A16 | 0.055 | 0.0554 | 0.7094 | 0.49487 | 0.646 | -6.31 | 0.66005 | none |
| RPL36A-HI | -0.157 | -0.1395 | -0.709 | 0.49488 | 0.646 | -6.31 | 1.41152 | none |
| GPR65 | 0.071 | -0.0575 | 0.709 | 0.49507 | 0.647 | -6.31 | -0.032 | none |
| BRAT1 | 0.05 | 0.0649 | 0.7087 | 0.49524 | 0.647 | -6.31 | 0.62442 | none |
| MYBPH | 0.046 | 0.0511 | 0.7085 | 0.49537 | 0.647 | -6.31 | 0 | none |
| CD74 | 0.131 | 0.1558 | 0.7083 | 0.49548 | 0.647 | -6.31 | 0.34875 | none |
| PPP1CB | 0.093 | 0.0108 | 0.7082 | 0.49555 | 0.647 | -6.31 | -0.681 | none |
| OR52B6 | 0.055 | 0.0553 | 0.7082 | 0.49556 | 0.647 | -6.31 | -0.5265 | none |
| C20ORF17 | 0.099 | 0.0988 | 0.7076 | 0.49591 | 0.647 | -6.31 | 0.67856 | none |
| ZNF485 | 0.114 | 0.1303 | 0.7075 | 0.49599 | 0.647 | -6.31 | 0.01106 | none |
| SREK1 | 0.178 | 0.2913 | 0.7074 | 0.49601 | 0.647 | -6.31 | -0.2886 | none |
| SMR3B | 0.055 | 0.0552 | 0.7074 | 0.49604 | 0.647 | -6.31 | 0.08915 | none |
| VHLL | 0.055 | 0.0552 | 0.7074 | 0.49604 | 0.647 | -6.31 | -0.7211 | none |
| SS18L2 | -0.102 | -0.1974 | -0.707 | 0.49611 | 0.647 | -6.31 | 0.1322 | none |
| SLC3A2 | -0.066 | -0.006 | -0.707 | 0.49638 | 0.648 | -6.31 | 0.84603 | none |
| DKK4 | 0.05 | 0.0547 | 0.7067 | 0.49647 | 0.648 | -6.31 | 1.58387 | none |
| CBFA2T3 | 0.078 | 0.1232 | 0.7066 | 0.49649 | 0.648 | -6.31 | -0.7356 | none |
| MPZ | -0.08 | -0.0203 | -0.707 | 0.49653 | 0.648 | -6.31 | 1.57719 | none |
| SRSF9 | -0.09 | -0.0611 | -0.707 | 0.49654 | 0.648 | -6.31 | 0.3103 | none |
| C11ORF71 | 0.135 | 0.0598 | 0.706 | 0.49686 | 0.648 | -6.31 | 0.15593 | none |
| APOBEC3E | -0.137 | -0.0677 | -0.706 | 0.497 | 0.648 | -6.31 | 1.37454 | none |
| OR2Z1 | -0.056 | -0.0171 | -0.706 | 0.49701 | 0.648 | -6.31 | 0.10458 | none |
| TXNDC15 | 0.087 | -0.0384 | 0.7058 | 0.49701 | 0.648 | -6.31 | -0.7707 | none |

| | | | | | | | | |
|---|---|---|---|---|---|---|---|---|
| TFPI2 | 0.052 | 0.0546 | 0.7058 | 0.49702 | 0.648 | -6.31 | -0.8452 | none |
| TVP23C-CI | 0.213 | 0.0833 | 0.7056 | 0.49709 | 0.648 | -6.31 | 0.18226 | none |
| KRBOX4 | 0.132 | 0.1178 | 0.7056 | 0.49709 | 0.648 | -6.31 | 0.06928 | none |
| SLC38A4 | -0.053 | -0.0536 | -0.706 | 0.49709 | 0.648 | -6.31 | 0.5382 | none |
| C19ORF24 | -0.144 | -0.0917 | -0.706 | 0.49714 | 0.648 | -6.31 | 1.29642 | none |
| TNFRSF8 | 0.093 | 0.1689 | 0.7053 | 0.49727 | 0.648 | -6.31 | 0.08581 | none |
| ITLN2 | -0.054 | -0.0539 | -0.705 | 0.49728 | 0.648 | -6.31 | -0.7212 | none |
| LIG3 | 0.097 | 0.2006 | 0.7051 | 0.49742 | 0.648 | -6.31 | 1.3002 | none |
| NUP54 | 0.096 | 0.0257 | 0.7049 | 0.49753 | 0.648 | -6.31 | -0.6411 | none |
| DEFA6 | 0.055 | 0.0548 | 0.7042 | 0.49792 | 0.649 | -6.31 | 0.66005 | none |
| KDR | -0.074 | -0.0511 | -0.704 | 0.49808 | 0.649 | -6.31 | 0.0539 | none |
| NPRL2 | -0.065 | -0.0987 | -0.704 | 0.49832 | 0.649 | -6.31 | 1.23873 | none |
| SRD5A3 | 0.098 | 0.0022 | 0.7035 | 0.49834 | 0.649 | -6.31 | 1.01971 | none |
| SLC35G6 | 0.052 | 0.0374 | 0.7035 | 0.49838 | 0.649 | -6.31 | 0.97443 | none |
| COX14 | -0.115 | -0.1732 | -0.703 | 0.4984 | 0.649 | -6.31 | 0.89668 | none |
| UBR7 | -0.069 | -0.0678 | -0.703 | 0.4985 | 0.649 | -6.31 | 1.69837 | none |
| OBSCN | 0.106 | 0.1426 | 0.7033 | 0.4985 | 0.649 | -6.31 | -0.6322 | none |
| DHX16 | 0.136 | 0.2582 | 0.7027 | 0.49886 | 0.65 | -6.31 | 0.79826 | none |
| ATP12A | 0.047 | 0.0473 | 0.7025 | 0.49895 | 0.65 | -6.31 | 0.46778 | none |
| CBY1 | 0.104 | 0.069 | 0.7025 | 0.49898 | 0.65 | -6.31 | 0.9977 | none |
| DCP2 | 0.179 | 0.1014 | 0.7024 | 0.49904 | 0.65 | -6.31 | 0.49758 | none |
| SENP3 | -0.084 | 0.0102 | -0.702 | 0.49904 | 0.65 | -6.31 | 0.61259 | none |
| SPAG8 | 0.089 | 0.0865 | 0.7021 | 0.4992 | 0.65 | -6.31 | 0.97526 | none |
| MCCD1 | 0.055 | 0.0545 | 0.7019 | 0.4993 | 0.65 | -6.31 | 0.66005 | none |
| GPS1 | 0.095 | 0.0818 | 0.7019 | 0.49933 | 0.65 | -6.31 | -0.4148 | none |
| KRT2 | 0.062 | 0.069 | 0.7016 | 0.49947 | 0.65 | -6.31 | 0.48712 | none |
| SEC61A1 | 0.073 | 0.0521 | 0.7016 | 0.49949 | 0.65 | -6.31 | 1.08782 | none |
| SLC43A3 | -0.084 | 0.0074 | -0.701 | 0.49958 | 0.65 | -6.31 | 1.0456 | none |
| ANGPTL7 | -0.055 | -0.057 | -0.701 | 0.49992 | 0.65 | -6.31 | 1.14986 | none |
| ATXN7L3 | -0.074 | 0.0238 | -0.701 | 0.49997 | 0.65 | -6.31 | 1.42989 | none |
| LECT2 | -0.107 | -0.0803 | -0.701 | 0.49999 | 0.65 | -6.31 | 0.00909 | none |
| BSPH1 | 0.056 | 0.056 | 0.7007 | 0.50003 | 0.65 | -6.31 | -0.1117 | none |
| PRR26 | 0.051 | 0.021 | 0.7006 | 0.5001 | 0.65 | -6.31 | -0.1462 | none |
| GBE1 | -0.147 | -0.1113 | -0.7 | 0.50017 | 0.65 | -6.31 | 2.14384 | AA-related |
| ADORA2B | -0.073 | -0.071 | -0.7 | 0.50018 | 0.65 | -6.31 | 0.12115 | none |
| CNOT8 | -0.062 | -0.1149 | -0.7 | 0.50021 | 0.65 | -6.31 | 0.20466 | none |
| TOMM34 | 0.095 | 0.1565 | 0.7004 | 0.50023 | 0.65 | -6.31 | -0.1258 | none |
| TIFA | 0.09 | -0.0211 | 0.7 | 0.50046 | 0.651 | -6.31 | 1.39403 | none |
| AGBL2 | 0.115 | 0.0854 | 0.6998 | 0.50053 | 0.651 | -6.31 | -0.0649 | none |
| NKRF | 0.127 | 0.2081 | 0.6997 | 0.50061 | 0.651 | -6.31 | 1.15101 | none |

| | | | | | | | | |
|---|---|---|---|---|---|---|---|---|
| SPATA32 | 0.164 | 0.1356 | 0.6997 | 0.50061 | 0.651 | -6.31 | -0.048 | none |
| KLF2 | -0.189 | -0.275 | -0.7 | 0.50069 | 0.651 | -6.31 | 0.06724 | none |
| ZNF695 | 0.125 | 0.117 | 0.6995 | 0.50075 | 0.651 | -6.32 | 1.32841 | none |
| UBQLN4 | 0.112 | 0.1666 | 0.6994 | 0.50079 | 0.651 | -6.32 | 0.04161 | none |
| MBNL2 | 0.099 | 0.1332 | 0.6992 | 0.5009 | 0.651 | -6.32 | 0.43693 | none |
| ADAMTS1! | 0.054 | 0.0542 | 0.6992 | 0.50092 | 0.651 | -6.32 | 1.19688 | none |
| RCC2 | 0.079 | 0.1416 | 0.6991 | 0.50099 | 0.651 | -6.32 | -0.379 | none |
| C6ORF223 | 0.052 | 0.0075 | 0.6984 | 0.50143 | 0.651 | -6.32 | -0.507 | none |
| SMAD7 | 0.107 | 0.1626 | 0.6983 | 0.50149 | 0.651 | -6.32 | 0.47313 | none |
| ABI1 | -0.076 | -0.1488 | -0.698 | 0.5015 | 0.651 | -6.32 | 1.15941 | none |
| CYB5A | -0.095 | -0.1874 | -0.698 | 0.50151 | 0.651 | -6.32 | 0.48546 | none |
| PIK3R6 | -0.06 | -0.0273 | -0.698 | 0.50172 | 0.652 | -6.32 | 1.47315 | none |
| TATDN1 | 0.102 | -0.0284 | 0.6978 | 0.50176 | 0.652 | -6.32 | 0.46979 | none |
| MEGF6 | -0.106 | -0.0419 | -0.698 | 0.50177 | 0.652 | -6.32 | 0.75355 | none |
| C19ORF73 | -0.104 | ######## | -0.698 | 0.50179 | 0.652 | -6.32 | 0.42971 | none |
| ADGRB1 | 0.086 | 0.0949 | 0.6976 | 0.50185 | 0.652 | -6.32 | 0.90835 | none |
| E4F1 | -0.12 | 0.0467 | -0.698 | 0.50187 | 0.652 | -6.32 | 1.46018 | none |
| NR2C2AP | -0.086 | -0.0732 | -0.698 | 0.5019 | 0.652 | -6.32 | 0.62666 | none |
| USP30 | 0.132 | 0.08 | 0.6975 | 0.50191 | 0.652 | -6.32 | 0.96463 | none |
| OR56A5 | -0.053 | -0.053 | -0.697 | 0.5021 | 0.652 | -6.32 | 1.85525 | none |
| INO80B | -0.148 | -0.0184 | -0.697 | 0.50233 | 0.652 | -6.32 | 1.29619 | none |
| CLDN10 | 0.049 | 0.0482 | 0.6967 | 0.5024 | 0.652 | -6.32 | 1.49129 | none |
| DDAH2 | -0.147 | -0.1559 | -0.697 | 0.5024 | 0.652 | -6.32 | 0.7959 | none |
| MAD1L1 | 0.086 | 0.0455 | 0.6966 | 0.5025 | 0.652 | -6.32 | -0.6499 | none |
| CAPG | 0.086 | 0.0757 | 0.6957 | 0.50301 | 0.653 | -6.32 | 1.09877 | none |
| KCNE1B | -0.079 | -0.0533 | -0.696 | 0.5031 | 0.653 | -6.32 | 1.28781 | none |
| NUDCD1 | -0.101 | -0.1163 | -0.695 | 0.50316 | 0.653 | -6.32 | 1.24507 | none |
| BRF2 | -0.112 | -0.0353 | -0.695 | 0.50327 | 0.653 | -6.32 | 0.91954 | none |
| CCDC136 | 0.289 | 0.2177 | 0.6952 | 0.5033 | 0.653 | -6.32 | -0.8542 | none |
| OSBP | 0.139 | 0.1925 | 0.6952 | 0.50333 | 0.653 | -6.32 | -0.0755 | none |
| FOLR1 | -0.077 | -0.0601 | -0.695 | 0.50336 | 0.653 | -6.32 | 1.05875 | none |
| INVS | 0.114 | 0.1714 | 0.6948 | 0.50356 | 0.653 | -6.32 | 0.91129 | none |
| GNB3 | 0.171 | 0.1006 | 0.6948 | 0.50356 | 0.653 | -6.32 | -0.1138 | none |
| STOX2 | 0.052 | 0.0559 | 0.6947 | 0.50359 | 0.653 | -6.32 | 0.70046 | none |
| DIXDC1 | 0.191 | 0.3294 | 0.6947 | 0.50359 | 0.653 | -6.32 | -0.4064 | none |
| COL4A6 | 0.052 | 0.051 | 0.6946 | 0.50365 | 0.653 | -6.32 | -0.7515 | none |
| C10ORF14 | 0.054 | 0.0536 | 0.6943 | 0.50387 | 0.653 | -6.32 | -0.665 | none |
| LIPM | 0.052 | 0.0532 | 0.694 | 0.50401 | 0.653 | -6.32 | 0.53627 | none |
| EDC4 | 0.132 | 0.2509 | 0.6937 | 0.50419 | 0.654 | -6.32 | 0.6643 | none |
| CDHR1 | -0.09 | -0.0597 | -0.694 | 0.50426 | 0.654 | -6.32 | -0.0878 | none |

| | | | | | | | | |
|---|---|---|---|---|---|---|---|---|
| GRK6 | 0.081 | 0.1251 | 0.6935 | 0.50432 | 0.654 | -6.32 | -0.314 | none |
| GJA5 | -0.053 | -0.0535 | -0.693 | 0.50435 | 0.654 | -6.32 | 0.83551 | none |
| FAM162B | 0.054 | 0.0535 | 0.6935 | 0.50436 | 0.654 | -6.32 | -0.5288 | none |
| EGLN3 | 0.125 | 0.0252 | 0.6933 | 0.50445 | 0.654 | -6.32 | 0.17957 | none |
| ACOT9 | 0.081 | -0.0174 | 0.6932 | 0.50453 | 0.654 | -6.32 | -0.1539 | none |
| PERM1 | -0.083 | -0.0133 | -0.693 | 0.5046 | 0.654 | -6.32 | 1.16565 | none |
| AL049872. | -0.233 | -0.2243 | -0.693 | 0.50476 | 0.654 | -6.32 | 1.10672 | none |
| CORO6 | -0.121 | -0.0632 | -0.693 | 0.50487 | 0.654 | -6.32 | 1.28857 | none |
| LAMC1 | 0.079 | 0.0762 | 0.6926 | 0.50488 | 0.654 | -6.32 | -0.527 | none |
| CPSF7 | 0.078 | 0.1206 | 0.6915 | 0.50553 | 0.655 | -6.32 | -0.1449 | none |
| SGSM3 | 0.065 | 0.0563 | 0.6914 | 0.50559 | 0.655 | -6.32 | 0.63858 | none |
| FASTKD2 | 0.077 | 0.0311 | 0.6907 | 0.506 | 0.655 | -6.32 | 1.17327 | none |
| POLD2 | -0.078 | -0.0344 | -0.69 | 0.50629 | 0.656 | -6.32 | 0.8678 | none |
| TBCD | 0.133 | 0.1607 | 0.69 | 0.50641 | 0.656 | -6.32 | 1.13916 | none |
| SLC35G4 | 0.054 | 0.0537 | 0.6896 | 0.50671 | 0.656 | -6.32 | -0.8278 | none |
| CST9L | 0.053 | 0.053 | 0.6894 | 0.50678 | 0.656 | -6.32 | 1.19688 | none |
| SMIM20 | -0.121 | -0.1231 | -0.689 | 0.50683 | 0.656 | -6.32 | 1.50234 | none |
| TPK1 | 0.094 | 0.0361 | 0.6893 | 0.50687 | 0.656 | -6.32 | 1.35785 | none |
| CEMIP | 0.083 | 0.1062 | 0.6892 | 0.50695 | 0.656 | -6.32 | -0.8169 | none |
| C2CD4B | -0.058 | -0.0471 | -0.689 | 0.50704 | 0.656 | -6.32 | 0.22443 | none |
| KLK8 | -0.121 | -0.1205 | -0.689 | 0.5071 | 0.656 | -6.32 | 0.28585 | none |
| AGPAT2 | 0.099 | 0.054 | 0.6889 | 0.50712 | 0.656 | -6.32 | -0.053 | none |
| RAP2A | 0.053 | 0.0694 | 0.6886 | 0.5073 | 0.656 | -6.32 | 0.87589 | none |
| PVALB | 0.05 | 0.0436 | 0.6877 | 0.5078 | 0.657 | -6.32 | 0.61885 | none |
| MS4A3 | 0.048 | 0.0613 | 0.6876 | 0.50791 | 0.657 | -6.32 | 1.93555 | none |
| IRF6 | -0.08 | -0.0574 | -0.687 | 0.50804 | 0.657 | -6.32 | -0.1306 | none |
| CALHM1 | -0.049 | -0.0368 | -0.687 | 0.50816 | 0.657 | -6.32 | -0.1361 | none |
| RBM38 | -0.088 | -0.0479 | -0.687 | 0.50852 | 0.658 | -6.32 | -0.1485 | none |
| CCL20 | 0.153 | 0.0248 | 0.6864 | 0.50863 | 0.658 | -6.32 | 0.76134 | none |
| TRIM32 | 0.069 | 0.0179 | 0.6863 | 0.50867 | 0.658 | -6.32 | 0.0975 | none |
| NRBF2 | -0.087 | -0.049 | -0.686 | 0.50872 | 0.658 | -6.32 | 0.56392 | none |
| GOLGA8F | 0.045 | 0.055 | 0.6859 | 0.50889 | 0.658 | -6.32 | -0.076 | none |
| DLG2 | -0.098 | -0.0667 | -0.686 | 0.50894 | 0.658 | -6.32 | 1.34994 | none |
| CYP4F11 | -0.051 | -0.0314 | -0.686 | 0.50905 | 0.658 | -6.32 | 0.67906 | none |
| SASS6 | 0.127 | 0.1844 | 0.6856 | 0.50907 | 0.658 | -6.32 | 0.42254 | none |
| KIFAP3 | -0.075 | -0.099 | -0.686 | 0.50911 | 0.658 | -6.32 | 1.54262 | none |
| PTBP1 | 0.082 | 0.1662 | 0.6856 | 0.50911 | 0.658 | -6.32 | 1.46179 | none |
| NXF5 | 0.053 | 0.065 | 0.6854 | 0.50919 | 0.658 | -6.33 | 0.66409 | none |
| BDH2 | 0.135 | 0.019 | 0.6849 | 0.5095 | 0.658 | -6.33 | 0.41451 | none |
| CHST15 | 0.073 | 0.1006 | 0.6849 | 0.5095 | 0.658 | -6.33 | 1.36879 | none |

| | | | | | | | | |
|---|---|---|---|---|---|---|---|---|
| SUV39H1 | -0.062 | 0.0103 | -0.685 | 0.50961 | 0.658 | -6.33 | 1.25136 | none |
| DRD3 | -0.048 | -0.0524 | -0.685 | 0.50965 | 0.658 | -6.33 | 1.48116 | none |
| IFNGR2 | -0.108 | -0.1261 | -0.684 | 0.50979 | 0.659 | -6.33 | 0.47372 | none |
| CSDE1 | 0.091 | 0.0954 | 0.6844 | 0.5098 | 0.659 | -6.33 | 1.08289 | none |
| CASZ1 | 0.075 | 0.0477 | 0.684 | 0.51006 | 0.659 | -6.33 | 0.76241 | none |
| SNX17 | -0.086 | -0.1024 | -0.684 | 0.5101 | 0.659 | -6.33 | 0.49164 | none |
| WDR87 | 0.062 | 0.0625 | 0.6839 | 0.51013 | 0.659 | -6.33 | -0.7553 | none |
| TUBB | -0.066 | -0.039 | -0.684 | 0.51019 | 0.659 | -6.33 | 1.39929 | none |
| AGTR1 | -0.051 | -0.0515 | -0.684 | 0.51021 | 0.659 | -6.33 | 0 | none |
| CACNA1G | 0.124 | 0.1027 | 0.6829 | 0.51073 | 0.659 | -6.33 | 0.48119 | none |
| EXO1 | 0.136 | 0.1816 | 0.6827 | 0.51083 | 0.659 | -6.33 | 0.06219 | none |
| ITPKA | -0.073 | -0.0261 | -0.683 | 0.51086 | 0.659 | -6.33 | 1.31552 | none |
| ABTB1 | 0.068 | 0.0204 | 0.6827 | 0.51086 | 0.659 | -6.33 | -0.4539 | none |
| A2M | -0.21 | -0.2045 | -0.683 | 0.51095 | 0.659 | -6.33 | 0.42102 | none |
| THBS2 | 0.108 | 0.0926 | 0.6821 | 0.51121 | 0.66 | -6.33 | -0.1409 | none |
| CASD1 | -0.161 | -0.1643 | -0.682 | 0.51142 | 0.66 | -6.33 | 1.27678 | none |
| CDO1 | 0.053 | 0.0529 | 0.6816 | 0.51148 | 0.66 | -6.33 | 0.02895 | none |
| IGFBP3 | 0.14 | 0.1174 | 0.6815 | 0.51159 | 0.66 | -6.33 | -0.1291 | none |
| COBL | 0.064 | 0.0599 | 0.6812 | 0.51175 | 0.66 | -6.33 | 0.5923 | none |
| OR51F2 | 0.052 | 0.0521 | 0.6812 | 0.51178 | 0.66 | -6.33 | 0.66005 | none |
| SMIM3 | -0.092 | -0.1536 | -0.681 | 0.51199 | 0.66 | -6.33 | 0.13819 | none |
| FAAP20 | 0.107 | 0.0768 | 0.6804 | 0.51223 | 0.661 | -6.33 | -0.7704 | none |
| NEK3 | -0.105 | -0.1012 | -0.68 | 0.51227 | 0.661 | -6.33 | 0.14409 | none |
| CPSF3 | -0.061 | -0.065 | -0.68 | 0.51243 | 0.661 | -6.33 | 0.20185 | none |
| FAM219B | 0.089 | 0.0966 | 0.6801 | 0.51244 | 0.661 | -6.33 | 0.23664 | none |
| ALG12 | -0.133 | -0.079 | -0.68 | 0.51244 | 0.661 | -6.33 | 0.50005 | none |
| ANKRD23 | -0.083 | -0.0543 | -0.68 | 0.51245 | 0.661 | -6.33 | 0.10641 | none |
| SIX5 | -0.063 | -0.0012 | -0.679 | 0.51292 | 0.661 | -6.33 | 0.96595 | none |
| ZNF638 | 0.113 | 0.183 | 0.679 | 0.51305 | 0.661 | -6.33 | 0.86022 | none |
| TRIM77 | 0.052 | 0.0518 | 0.6786 | 0.51335 | 0.662 | -6.33 | 0.89752 | none |
| KRT73 | 0.049 | 0.0571 | 0.678 | 0.5137 | 0.662 | -6.33 | 0.26433 | none |
| KRTAP1-3 | -0.051 | -0.0507 | -0.677 | 0.51415 | 0.663 | -6.33 | 1.9054 | none |
| AJAP1 | 0.049 | 0.0596 | 0.6764 | 0.51468 | 0.663 | -6.33 | -0.3595 | none |
| STX18 | 0.148 | 0.1633 | 0.6763 | 0.51474 | 0.663 | -6.33 | 1.14371 | none |
| TGFB1I1 | 0.126 | 0.1097 | 0.676 | 0.5149 | 0.663 | -6.33 | 0.02796 | none |
| SLC34A2 | 0.181 | 0.169 | 0.676 | 0.5149 | 0.663 | -6.33 | 0.46888 | none |
| HLX | -0.051 | 0.003 | -0.676 | 0.51498 | 0.664 | -6.33 | 1.46011 | none |
| DPP3 | 0.073 | 0.0752 | 0.6757 | 0.51506 | 0.664 | -6.33 | 0.37497 | none |
| MMP8 | 0.071 | 0.0642 | 0.6757 | 0.51511 | 0.664 | -6.33 | 0.05616 | none |
| PPP4R1 | 0.101 | 0.0783 | 0.6753 | 0.51535 | 0.664 | -6.33 | -0.8322 | none |

| | | | | | | | | |
|---|---|---|---|---|---|---|---|---|
| SURF2 | 0.093 | 0.1121 | 0.6753 | 0.51536 | 0.664 | -6.33 | 0.30584 | none |
| IGFBP7 | 0.086 | 0.053 | 0.6751 | 0.51543 | 0.664 | -6.33 | -0.7284 | none |
| MZF1 | 0.082 | 0.0865 | 0.6747 | 0.51569 | 0.664 | -6.33 | 0.95641 | none |
| MRPS12 | -0.092 | -0.0968 | -0.674 | 0.51582 | 0.664 | -6.33 | 0.79505 | none |
| OSBPL8 | 0.155 | 0.26 | 0.6744 | 0.51589 | 0.664 | -6.33 | 0.35902 | none |
| SAMD9 | 0.192 | 0.3011 | 0.6736 | 0.51634 | 0.665 | -6.33 | 0.58483 | none |
| STK11IP | 0.096 | 0.1494 | 0.6734 | 0.51647 | 0.665 | -6.33 | 0.71915 | none |
| IQSEC1 | 0.179 | 0.2823 | 0.6734 | 0.51647 | 0.665 | -6.33 | -0.1041 | none |
| EBNA1BP2 | -0.061 | -0.0192 | -0.673 | 0.51666 | 0.665 | -6.33 | 2.90E-04 | none |
| TKTL1 | -0.161 | -0.1371 | -0.673 | 0.5167 | 0.665 | -6.33 | 1.72842 | none |
| DPEP1 | 0.095 | 0.1573 | 0.6729 | 0.51678 | 0.665 | -6.33 | 0.13819 | none |
| ALOX12 | -0.062 | 0.0677 | -0.673 | 0.51679 | 0.665 | -6.33 | 0.23409 | none |
| C8ORF88 | -0.115 | -0.077 | -0.673 | 0.5169 | 0.665 | -6.33 | 1.86027 | none |
| PAGE4 | 0.06 | 0.056 | 0.6726 | 0.51699 | 0.665 | -6.33 | 1.6921 | none |
| SLC22A16 | -0.054 | -0.0759 | -0.672 | 0.51743 | 0.666 | -6.33 | -0.3688 | none |
| ARHGEF11 | 0.111 | 0.113 | 0.6717 | 0.51753 | 0.666 | -6.33 | 0.22716 | none |
| C6ORF48 | 0.076 | 0.0713 | 0.6716 | 0.51761 | 0.666 | -6.33 | 0.10419 | none |
| BTNL2 | 0.066 | 0.0664 | 0.6715 | 0.51762 | 0.666 | -6.33 | 1.13082 | none |
| HIST1H2BI | 0.11 | 0.0127 | 0.6714 | 0.51771 | 0.666 | -6.33 | 0.38804 | none |
| SESTD1 | 0.202 | 0.2849 | 0.6713 | 0.51774 | 0.666 | -6.33 | 0.07479 | none |
| PPP1R42 | 0.175 | 0.1162 | 0.6713 | 0.51777 | 0.666 | -6.33 | 1.18614 | none |
| TREML2 | 0.101 | 0.1559 | 0.6713 | 0.51777 | 0.666 | -6.33 | -0.2247 | none |
| HNRNPA3 | 0.097 | 0.1008 | 0.6712 | 0.51782 | 0.666 | -6.34 | 0.65435 | none |
| METTL23 | -0.09 | -0.1556 | -0.671 | 0.518 | 0.666 | -6.34 | 0.79794 | none |
| RHBDL3 | 0.064 | 0.068 | 0.6708 | 0.51808 | 0.666 | -6.34 | 1.17066 | none |
| HIST1H2B( | -0.165 | -0.1612 | -0.671 | 0.51815 | 0.666 | -6.34 | 1.07102 | none |
| ORM2 | -0.062 | -0.0685 | -0.671 | 0.5182 | 0.666 | -6.34 | 0.64577 | none |
| SLC10A1 | -0.055 | -0.0366 | -0.67 | 0.51851 | 0.667 | -6.34 | 1.53032 | none |
| RNF17 | 0.056 | 0.0442 | 0.6691 | 0.51909 | 0.667 | -6.34 | -0.5722 | none |
| ALOX12B | 0.106 | 0.0966 | 0.6688 | 0.51931 | 0.667 | -6.34 | -0.6714 | none |
| PNMA1 | 0.072 | 0.056 | 0.6687 | 0.51933 | 0.667 | -6.34 | 0.45013 | none |
| PDYN | 0.181 | 0.1814 | 0.6683 | 0.51959 | 0.668 | -6.34 | 0.86642 | none |
| OR6C68 | -0.05 | -0.0498 | -0.668 | 0.5197 | 0.668 | -6.34 | -0.2081 | none |
| PANX1 | 0.078 | 0.0939 | 0.6679 | 0.51984 | 0.668 | -6.34 | 0.50638 | none |
| XG | -0.05 | -0.0497 | -0.668 | 0.51986 | 0.668 | -6.34 | 1.9054 | none |
| SCAMP3 | -0.072 | -0.0659 | -0.667 | 0.52012 | 0.668 | -6.34 | -0.0329 | none |
| MUCL1 | 0.047 | 0.0567 | 0.6674 | 0.52013 | 0.668 | -6.34 | 0.55246 | none |
| H2BFS | -0.088 | -0.1357 | -0.667 | 0.52015 | 0.668 | -6.34 | 1.0612 | none |
| MKRN2OS | 0.094 | 0.1222 | 0.6673 | 0.52022 | 0.668 | -6.34 | 3.63E-04 | none |
| GPAM | 0.12 | 0.1236 | 0.6666 | 0.52062 | 0.669 | -6.34 | 0.75113 | none |

| | | | | | | | | |
|---|---|---|---|---|---|---|---|---|
| RDH8 | 0.179 | 0.193 | 0.6665 | 0.52069 | 0.669 | -6.34 | 0.44928 | none |
| TAF6L | -0.114 | -0.0212 | -0.666 | 0.52073 | 0.669 | -6.34 | -0.1453 | none |
| CXORF40B | -0.074 | -0.1052 | -0.666 | 0.52074 | 0.669 | -6.34 | 1.19158 | none |
| BEGAIN | 0.104 | 0.2011 | 0.6659 | 0.52105 | 0.669 | -6.34 | 1.04173 | none |
| FSD1L | -0.081 | -0.0212 | -0.666 | 0.52107 | 0.669 | -6.34 | -0.0956 | none |
| TCEA3 | 0.108 | 0.0747 | 0.6657 | 0.52117 | 0.669 | -6.34 | -0.1183 | none |
| RFC5 | -0.078 | -0.106 | -0.665 | 0.52135 | 0.669 | -6.34 | 1.67155 | none |
| LEPROTL1 | 0.11 | -0.0711 | 0.6653 | 0.52142 | 0.669 | -6.34 | -0.2325 | none |
| ISM2 | -0.05 | -0.0488 | -0.665 | 0.52146 | 0.669 | -6.34 | -0.4142 | none |
| ALG5 | -0.091 | -0.1803 | -0.665 | 0.5216 | 0.669 | -6.34 | 0.22795 | none |
| IWS1 | 0.214 | 0.3951 | 0.6648 | 0.52173 | 0.669 | -6.34 | 0.23733 | none |
| WASF3 | 0.051 | 0.054 | 0.6647 | 0.5218 | 0.67 | -6.34 | 0.87432 | none |
| ME3 | 0.143 | 0.1344 | 0.6645 | 0.52193 | 0.67 | -6.34 | 0.43042 | none |
| EIF3C | 0.202 | 0.3651 | 0.6644 | 0.52201 | 0.67 | -6.34 | 0.73601 | none |
| FEZF2 | -0.048 | -0.0499 | -0.664 | 0.52201 | 0.67 | -6.34 | 0 | none |
| OR4A15 | -0.049 | -0.0493 | -0.664 | 0.52211 | 0.67 | -6.34 | 0.88916 | none |
| OAS2 | 0.126 | 0.1679 | 0.664 | 0.52225 | 0.67 | -6.34 | 0.14395 | none |
| KCNJ8 | -0.053 | -0.0504 | -0.664 | 0.52239 | 0.67 | -6.34 | 1.97179 | none |
| P2RY12 | -0.047 | -0.0256 | -0.663 | 0.5226 | 0.67 | -6.34 | 0 | none |
| HIST2H2AI | 0.088 | 0.0891 | 0.6634 | 0.5226 | 0.67 | -6.34 | -0.4419 | none |
| HDHD3 | 0.08 | 0.0446 | 0.6632 | 0.52271 | 0.67 | -6.34 | 1.17218 | none |
| CKM | 0.048 | 0.053 | 0.6625 | 0.52315 | 0.671 | -6.34 | -0.9827 | none |
| MRPL46 | 0.072 | 0.015 | 0.6622 | 0.52331 | 0.671 | -6.34 | -0.155 | none |
| ME1 | -0.048 | -0.0423 | -0.662 | 0.52332 | 0.671 | -6.34 | 1.99806 | none |
| CLTB | -0.051 | -0.0131 | -0.662 | 0.52333 | 0.671 | -6.34 | -0.2438 | none |
| ZNF772 | 0.089 | -0.0282 | 0.6622 | 0.52334 | 0.671 | -6.34 | -0.9777 | none |
| CST11 | -0.065 | -0.0869 | -0.662 | 0.52335 | 0.671 | -6.34 | -0.4838 | none |
| ZADH2 | 0.117 | 0.0785 | 0.6621 | 0.52337 | 0.671 | -6.34 | 0.80753 | none |
| TMEM92 | 0.045 | 0.0502 | 0.6621 | 0.52338 | 0.671 | -6.34 | -0.4713 | none |
| LPIN2 | 0.126 | 0.1264 | 0.6621 | 0.52341 | 0.671 | -6.34 | -0.3623 | none |
| S100A1 | -0.111 | -0.0956 | -0.662 | 0.52362 | 0.671 | -6.34 | 0.68479 | none |
| CTAGE15 | 0.053 | 0.0433 | 0.6612 | 0.52396 | 0.671 | -6.34 | 0.84605 | none |
| FAM171A2 | -0.048 | -0.0429 | -0.661 | 0.524 | 0.671 | -6.34 | 1.25377 | none |
| IGFN1 | 0.049 | 0.0533 | 0.661 | 0.52408 | 0.671 | -6.34 | 1.33707 | none |
| APOBEC3C | 0.111 | 0.1428 | 0.6607 | 0.52427 | 0.671 | -6.34 | 0.12614 | none |
| IMPA1 | -0.093 | -0.1654 | -0.661 | 0.52432 | 0.671 | -6.34 | 1.03003 | none |
| B3GNT4 | 0.143 | 0.1508 | 0.6606 | 0.52432 | 0.671 | -6.34 | 0.08709 | none |
| HOMER1 | 0.183 | 0.2877 | 0.6604 | 0.52446 | 0.672 | -6.34 | 0.64332 | none |
| GSTM5 | 0.081 | 0.0727 | 0.6601 | 0.5246 | 0.672 | -6.34 | -0.6114 | none |
| CACNA1H | -0.048 | -0.0686 | -0.66 | 0.52466 | 0.672 | -6.34 | 1.64765 | none |

| | | | | | | | | |
|---|---|---|---|---|---|---|---|---|
| CACNG2 | 0.051 | 0.0489 | 0.6594 | 0.52503 | 0.672 | -6.34 | -0.6729 | none |
| EQTN | 0.05 | 0.0496 | 0.6587 | 0.52548 | 0.673 | -6.34 | 1.66579 | none |
| EGFL8 | 0.085 | 0.0928 | 0.6585 | 0.5256 | 0.673 | -6.34 | 1.17298 | none |
| LY6G6C | 0.052 | 0.0503 | 0.6584 | 0.52569 | 0.673 | -6.34 | 1.32715 | none |
| GSTA5 | 0.05 | 0.0495 | 0.6583 | 0.52574 | 0.673 | -6.34 | 0.66005 | none |
| ZBTB43 | 0.129 | 0.0898 | 0.6581 | 0.52583 | 0.673 | -6.34 | -0.5486 | none |
| PIP5K1C | -0.138 | 0.0252 | -0.658 | 0.52583 | 0.673 | -6.34 | 0.97375 | none |
| CSF3 | -0.054 | -0.0818 | -0.658 | 0.52587 | 0.673 | -6.34 | -0.4937 | none |
| STUM | -0.078 | -0.1105 | -0.658 | 0.5259 | 0.673 | -6.34 | 0.35201 | none |
| OR6C3 | 0.049 | 0.0495 | 0.658 | 0.5259 | 0.673 | -6.34 | 1.19688 | none |
| FAM92B | 0.073 | 0.0391 | 0.6579 | 0.52595 | 0.673 | -6.34 | -0.739 | none |
| ALOX15 | 0.092 | 0.0458 | 0.6579 | 0.52597 | 0.673 | -6.34 | -0.3607 | none |
| AHSG | -0.095 | -0.1199 | -0.658 | 0.52617 | 0.673 | -6.34 | 0.04013 | none |
| GPD1 | -0.09 | -0.0499 | -0.657 | 0.5263 | 0.673 | -6.34 | 1.47593 | none |
| TMPRSS9 | 0.061 | 0.0655 | 0.6572 | 0.52638 | 0.673 | -6.34 | 1.61723 | none |
| KRTAP4-8 | 0.049 | 0.0494 | 0.6572 | 0.52639 | 0.673 | -6.34 | -0.8261 | none |
| HNRNPCL3 | 0.045 | 0.0451 | 0.6572 | 0.5264 | 0.673 | -6.34 | 0 | none |
| ALLC | 0.051 | 0.0508 | 0.657 | 0.52653 | 0.673 | -6.34 | 0.37625 | none |
| GSTA2 | -0.047 | -0.0485 | -0.657 | 0.52662 | 0.673 | -6.34 | 1.00139 | none |
| PRKAR1A | 0.087 | 0.05 | 0.6566 | 0.5268 | 0.673 | -6.35 | 1.12652 | none |
| HSPA4 | 0.148 | 0.2372 | 0.6564 | 0.52687 | 0.673 | -6.35 | 0.87415 | none |
| CDKN1B | -0.139 | -0.15 | -0.656 | 0.5269 | 0.673 | -6.35 | 1.58444 | none |
| ZNF503 | -0.048 | -0.0276 | -0.656 | 0.52692 | 0.673 | -6.35 | 1.95204 | none |
| NAP1L5 | 0.125 | 0.1411 | 0.6563 | 0.52696 | 0.673 | -6.35 | 1.49633 | none |
| PARD3B | -0.112 | -0.1203 | -0.656 | 0.52706 | 0.674 | -6.35 | 0.30363 | none |
| TDRD6 | -0.051 | -0.0483 | -0.656 | 0.52725 | 0.674 | -6.35 | 0 | none |
| TXNL4B | 0.099 | 0.0081 | 0.6555 | 0.52746 | 0.674 | -6.35 | -0.5193 | none |
| ZNF408 | 0.078 | 0.0982 | 0.6554 | 0.52754 | 0.674 | -6.35 | 0.12723 | none |
| SMARCA1 | -0.058 | -0.0627 | -0.655 | 0.52762 | 0.674 | -6.35 | 1.71717 | none |
| ABHD14B | 0.083 | 0.0898 | 0.6552 | 0.52762 | 0.674 | -6.35 | 1.38017 | none |
| LGALS3BP | 0.067 | 0.0425 | 0.6551 | 0.5277 | 0.674 | -6.35 | 0.65375 | none |
| APEH | 0.094 | 0.0512 | 0.6549 | 0.52782 | 0.674 | -6.35 | 0.02655 | none |
| SH3D21 | 0.133 | 0.1808 | 0.6547 | 0.52794 | 0.674 | -6.35 | -0.0672 | none |
| S100A7 | -0.056 | -0.0516 | -0.654 | 0.52808 | 0.674 | -6.35 | 2.03412 | none |
| DUOXA1 | -0.1 | -0.0675 | -0.654 | 0.52828 | 0.675 | -6.35 | 1.69342 | none |
| TRAPPC11 | -0.069 | -0.0017 | -0.654 | 0.52831 | 0.675 | -6.35 | 1.2574 | none |
| UCN2 | -0.048 | -0.0223 | -0.654 | 0.52841 | 0.675 | -6.35 | 1.10503 | none |
| CYTH3 | 0.159 | 0.0746 | 0.6538 | 0.52852 | 0.675 | -6.35 | -0.5953 | none |
| ERICH6 | -0.046 | -0.048 | -0.653 | 0.52875 | 0.675 | -6.35 | 1.39928 | none |
| FCGR2A | -0.076 | -0.0773 | -0.653 | 0.52878 | 0.675 | -6.35 | 0.68268 | none |

| | | | | | | | | |
|---|---|---|---|---|---|---|---|---|
| DDX21 | 0.175 | 0.2904 | 0.653 | 0.52898 | 0.675 | -6.35 | 0.25631 | none |
| STK36 | 0.089 | 0.2146 | 0.653 | 0.52899 | 0.675 | -6.35 | 0.42774 | none |
| EIF3F | -0.069 | -0.072 | -0.653 | 0.52915 | 0.675 | -6.35 | -0.1902 | none |
| WDR12 | 0.078 | 0.0522 | 0.6527 | 0.52918 | 0.675 | -6.35 | 0.29297 | none |
| AC136612 | -0.048 | -0.0239 | -0.653 | 0.5293 | 0.675 | -6.35 | 0.89723 | none |
| ISOC2 | -0.081 | -0.0759 | -0.652 | 0.52944 | 0.676 | -6.35 | 1.55494 | none |
| DUS1L | 0.114 | 0.1475 | 0.6522 | 0.52951 | 0.676 | -6.35 | 0.01709 | none |
| C10ORF88 | -0.088 | -0.172 | -0.652 | 0.5296 | 0.676 | -6.35 | 0.7983 | none |
| ZNF275 | -0.062 | -0.0594 | -0.652 | 0.5299 | 0.676 | -6.35 | 1.53165 | none |
| TSTD2 | 0.122 | 0.1022 | 0.6515 | 0.52993 | 0.676 | -6.35 | -0.3428 | none |
| H2AFY | -0.059 | 0.0044 | -0.651 | 0.52993 | 0.676 | -6.35 | 0.99464 | none |
| IFNGR1 | 0.092 | 0.0152 | 0.6514 | 0.53 | 0.676 | -6.35 | 0.2976 | none |
| EXOSC4 | -0.086 | -0.0989 | -0.651 | 0.53006 | 0.676 | -6.35 | 0.1476 | none |
| TMEM244 | -0.069 | -0.1206 | -0.651 | 0.5301 | 0.676 | -6.35 | 0.29349 | none |
| PCOLCE | 0.093 | 0.009 | 0.651 | 0.53021 | 0.676 | -6.35 | -0.7549 | none |
| NEIL2 | 0.059 | 0.0201 | 0.651 | 0.53025 | 0.676 | -6.35 | -0.3021 | none |
| FAF2 | -0.11 | -0.0778 | -0.651 | 0.5303 | 0.676 | -6.35 | 1.27355 | none |
| CLLU1OS | -0.052 | -0.0492 | -0.651 | 0.53036 | 0.676 | -6.35 | -0.1742 | none |
| FAM166B | 0.087 | 0.0464 | 0.6504 | 0.53062 | 0.676 | -6.35 | 0.23975 | none |
| ALG6 | 0.071 | -0.029 | 0.6501 | 0.5308 | 0.677 | -6.35 | 0.85034 | none |
| SMIM4 | 0.201 | 0.0145 | 0.65 | 0.53083 | 0.677 | -6.35 | -0.2159 | none |
| PABPC1 | 0.137 | 0.2053 | 0.6499 | 0.53092 | 0.677 | -6.35 | 0.55131 | none |
| C3ORF84 | 0.07 | 0.1024 | 0.6495 | 0.53116 | 0.677 | -6.35 | 0.91166 | none |
| ALK | -0.074 | -0.0277 | -0.649 | 0.53116 | 0.677 | -6.35 | -0.2885 | none |
| CATSPER2 | -0.065 | -0.0281 | -0.649 | 0.53119 | 0.677 | -6.35 | 0.6138 | none |
| NCL | 0.238 | 0.4354 | 0.6487 | 0.53163 | 0.677 | -6.35 | 0.28302 | none |
| TMEM9B | 0.072 | -0.0346 | 0.6486 | 0.53172 | 0.677 | -6.35 | -0.0528 | none |
| B3GLCT | 0.063 | 0.0439 | 0.6484 | 0.53181 | 0.677 | -6.35 | 0.00751 | none |
| ZDHHC4 | -0.088 | -0.1476 | -0.648 | 0.53183 | 0.677 | -6.35 | 1.29382 | none |
| RMDN2 | 0.121 | 0.1756 | 0.6481 | 0.53203 | 0.678 | -6.35 | 0.75313 | none |
| ACTB | -0.076 | -0.0394 | -0.648 | 0.53207 | 0.678 | -6.35 | 1.43499 | none |
| MYOM2 | -0.111 | -0.0737 | -0.648 | 0.53231 | 0.678 | -6.35 | 1.52325 | none |
| SERTAD4 | -0.049 | -0.0488 | -0.647 | 0.5325 | 0.678 | -6.35 | 1.19237 | none |
| ANKEF1 | 0.072 | 0.0588 | 0.6459 | 0.5334 | 0.679 | -6.35 | -0.262 | none |
| C19ORF33 | 0.106 | 0.1123 | 0.6459 | 0.53341 | 0.679 | -6.35 | 1.61806 | none |
| ADH5 | -0.075 | -0.1447 | -0.646 | 0.53345 | 0.679 | -6.35 | -0.4052 | none |
| PDZD3 | -0.045 | -0.0251 | -0.646 | 0.53355 | 0.679 | -6.35 | 0.35106 | none |
| CSTF2 | 0.059 | 0.0862 | 0.6455 | 0.53364 | 0.679 | -6.35 | 0.05695 | none |
| LYPLA2 | -0.08 | -0.0704 | -0.645 | 0.5337 | 0.679 | -6.35 | 0.692 | none |
| TTC29 | -0.047 | -0.0474 | -0.645 | 0.53375 | 0.679 | -6.35 | -0.6726 | none |

| | | | | | | | | |
|---|---|---|---|---|---|---|---|---|
| USP17L17 | 0.048 | 0.0481 | 0.645 | 0.53393 | 0.679 | -6.35 | -0.6841 | none |
| OR2T8 | 0.047 | 0.0481 | 0.645 | 0.53397 | 0.679 | -6.35 | 0 | none |
| MAP3K1 | 0.136 | 0.1736 | 0.6449 | 0.53399 | 0.679 | -6.35 | 0.67607 | none |
| SLC9A3 | 0.062 | 0.043 | 0.6445 | 0.53423 | 0.68 | -6.35 | 1.79449 | none |
| LRRC4B | -0.062 | -0.0371 | -0.644 | 0.53427 | 0.68 | -6.35 | 1.49857 | none |
| CLNS1A | -0.065 | -0.1275 | -0.644 | 0.53431 | 0.68 | -6.35 | 1.25136 | none |
| SALL4 | 0.085 | 0.0108 | 0.6443 | 0.53442 | 0.68 | -6.35 | 0.64706 | none |
| MED30 | -0.084 | -0.1408 | -0.644 | 0.53446 | 0.68 | -6.35 | 0.4983 | none |
| GJB2 | -0.108 | -0.1389 | -0.644 | 0.53448 | 0.68 | -6.35 | 0.41389 | none |
| MZT2B | -0.114 | -0.1052 | -0.644 | 0.5347 | 0.68 | -6.35 | 0.03078 | none |
| HPDL | -0.086 | -0.019 | -0.644 | 0.53473 | 0.68 | -6.35 | 2.25695 | AA-related |
| PDSS1 | -0.058 | 0.0044 | -0.644 | 0.5348 | 0.68 | -6.35 | 1.42631 | none |
| TUBA4A | -0.067 | -0.1484 | -0.643 | 0.53496 | 0.68 | -6.35 | 0.99355 | none |
| DDN | 0.052 | 0.0837 | 0.6433 | 0.535 | 0.68 | -6.35 | 0.39219 | none |
| SPACA5 | -0.093 | -0.09 | -0.643 | 0.53501 | 0.68 | -6.35 | 0.08374 | none |
| ARL13A | -0.11 | -0.0798 | -0.643 | 0.53503 | 0.68 | -6.35 | -0.1143 | none |
| NUFIP1 | 0.084 | 0.1573 | 0.6429 | 0.53526 | 0.68 | -6.35 | 0.26209 | none |
| TOX | 0.064 | 0.0717 | 0.6428 | 0.53531 | 0.68 | -6.35 | -0.3856 | none |
| HBZ | 0.065 | 0.0579 | 0.6424 | 0.5356 | 0.681 | -6.35 | 0.74388 | none |
| C2ORF16 | -0.047 | -0.0619 | -0.642 | 0.53575 | 0.681 | -6.35 | 1.03291 | none |
| HGFAC | -0.069 | -0.0296 | -0.642 | 0.53595 | 0.681 | -6.36 | 1.03296 | none |
| AC105052 | 0.07 | 0.0384 | 0.6416 | 0.53607 | 0.681 | -6.36 | 0.03741 | none |
| CYP2A7 | -0.047 | -0.047 | -0.642 | 0.53611 | 0.681 | -6.36 | 0.7865 | none |
| GFI1 | 0.104 | 0.1733 | 0.6415 | 0.53611 | 0.681 | -6.36 | -0.6366 | none |
| CDX2 | -0.044 | -0.0336 | -0.641 | 0.5362 | 0.681 | -6.36 | 0.53669 | none |
| WDSUB1 | 0.072 | -0.0094 | 0.6413 | 0.53626 | 0.681 | -6.36 | 1.00559 | none |
| NFATC4 | -0.041 | -0.0183 | -0.641 | 0.53634 | 0.681 | -6.36 | 0.22191 | none |
| PRSS16 | 0.134 | 0.0459 | 0.641 | 0.53645 | 0.681 | -6.36 | 0.10156 | none |
| LIFR | 0.181 | 0.1486 | 0.641 | 0.53646 | 0.681 | -6.36 | -0.127 | none |
| COA1 | 0.103 | -0.0308 | 0.6409 | 0.5365 | 0.681 | -6.36 | 0.43828 | none |
| AC004556 | -0.071 | -0.1361 | -0.641 | 0.53662 | 0.681 | -6.36 | 0.77453 | none |
| CWC15 | 0.067 | 0.0528 | 0.6404 | 0.53682 | 0.682 | -6.36 | 0.42536 | none |
| PDCL | 0.084 | 0.0595 | 0.6403 | 0.53686 | 0.682 | -6.36 | 0.3539 | none |
| NCK1 | -0.073 | -0.1419 | -0.64 | 0.53701 | 0.682 | -6.36 | 1.24065 | none |
| SLC35D1 | -0.055 | -0.0819 | -0.64 | 0.53707 | 0.682 | -6.36 | 1.15062 | none |
| CDC42EP4 | 0.082 | 0.1187 | 0.64 | 0.53708 | 0.682 | -6.36 | 1.56669 | none |
| OXLD1 | 0.083 | 0.0479 | 0.64 | 0.53708 | 0.682 | -6.36 | -0.0146 | none |
| SIRT1 | -0.121 | -0.0998 | -0.64 | 0.53732 | 0.682 | -6.36 | 1.13575 | none |
| CUL2 | 0.163 | 0.2176 | 0.6393 | 0.53749 | 0.682 | -6.36 | 0.41828 | none |
| SERPINE3 | 0.126 | 0.1396 | 0.6393 | 0.53749 | 0.682 | -6.36 | 1.43981 | none |

| | | | | | | | | |
|---|---|---|---|---|---|---|---|---|
| PAIP2 | -0.07 | -0.1588 | -0.639 | 0.53753 | 0.682 | -6.36 | -0.2391 | none |
| PHF23 | -0.091 | -0.043 | -0.639 | 0.53768 | 0.682 | -6.36 | 0.94686 | none |
| AFTPH | 0.075 | 0.1109 | 0.6389 | 0.53774 | 0.682 | -6.36 | 1.45093 | none |
| MFAP4 | 0.05 | 0.0524 | 0.6388 | 0.5378 | 0.682 | -6.36 | 1.20882 | none |
| PNO1 | 0.087 | 0.0646 | 0.6387 | 0.53788 | 0.682 | -6.36 | 0.15573 | none |
| NCAPH | 0.12 | 0.2063 | 0.6387 | 0.53789 | 0.682 | -6.36 | 0.75843 | none |
| KCMF1 | 0.073 | -0.0474 | 0.6384 | 0.53803 | 0.682 | -6.36 | 0.31155 | none |
| TMEM179 | 0.055 | 0.066 | 0.6382 | 0.53817 | 0.682 | -6.36 | 0.5441 | none |
| FGB | -0.053 | -0.0673 | -0.638 | 0.53818 | 0.682 | -6.36 | 1.29809 | none |
| NDUFV1 | 0.077 | 0.0635 | 0.6382 | 0.53819 | 0.682 | -6.36 | 0.7338 | none |
| PCDH12 | -0.057 | -0.0451 | -0.638 | 0.53823 | 0.682 | -6.36 | -0.6937 | none |
| AFAP1L2 | -0.098 | -0.0489 | -0.638 | 0.53824 | 0.682 | -6.36 | 0.0858 | none |
| CDH2 | 0.057 | 0.0548 | 0.6379 | 0.53839 | 0.682 | -6.36 | -0.7311 | none |
| TMEM140 | 0.073 | 0.0531 | 0.6377 | 0.53847 | 0.682 | -6.36 | -0.0044 | none |
| ALG1L2 | -0.176 | -0.0986 | -0.637 | 0.53891 | 0.683 | -6.36 | 0.1229 | none |
| SIGIRR | -0.137 | -0.2227 | -0.637 | 0.53896 | 0.683 | -6.36 | 0.44521 | none |
| LZTS2 | 0.05 | 0.0281 | 0.6369 | 0.53897 | 0.683 | -6.36 | 0.98908 | none |
| ASIC3 | 0.119 | 0.0621 | 0.6369 | 0.53901 | 0.683 | -6.36 | 1.57364 | none |
| ZNF107 | 0.144 | 0.2703 | 0.6367 | 0.53915 | 0.683 | -6.36 | 0.615 | none |
| USP17L2 | 0.044 | 0.0494 | 0.6364 | 0.53928 | 0.683 | -6.36 | 1.70709 | none |
| SLC5A3 | -0.06 | -0.0807 | -0.636 | 0.53933 | 0.683 | -6.36 | 1.01039 | none |
| INHBC | -0.044 | -0.0288 | -0.636 | 0.53935 | 0.683 | -6.36 | 0 | none |
| METAP1D | 0.139 | 0.0988 | 0.6363 | 0.53935 | 0.683 | -6.36 | 0.07416 | none |
| LTA4H | -0.054 | -0.0814 | -0.636 | 0.53958 | 0.683 | -6.36 | 1.60755 | none |
| GHDC | 0.102 | 0.1336 | 0.6356 | 0.53979 | 0.684 | -6.36 | 1.66322 | none |
| TMEM204 | -0.047 | -0.0649 | -0.635 | 0.54004 | 0.684 | -6.36 | 0.39474 | none |
| SMO | 0.07 | 0.065 | 0.6351 | 0.54014 | 0.684 | -6.36 | -0.4911 | none |
| PTX4 | 0.07 | 0.0521 | 0.6351 | 0.54014 | 0.684 | -6.36 | 0.941 | none |
| PUM2 | 0.117 | 0.1007 | 0.6348 | 0.54033 | 0.684 | -6.36 | -0.5692 | none |
| CATSPER3 | -0.078 | -0.0501 | -0.634 | 0.5408 | 0.685 | -6.36 | -0.0028 | none |
| GALNTL5 | -0.078 | -0.0236 | -0.634 | 0.54084 | 0.685 | -6.36 | 1.67783 | none |
| TMEM69 | -0.09 | -0.1715 | -0.634 | 0.54087 | 0.685 | -6.36 | 1.74339 | none |
| CACNA2D4 | -0.088 | -0.1262 | -0.634 | 0.54089 | 0.685 | -6.36 | 0.09197 | none |
| CTSG | -0.046 | -0.0462 | -0.634 | 0.541 | 0.685 | -6.36 | -0.0349 | none |
| GGACT | 0.148 | 0.1497 | 0.6336 | 0.54104 | 0.685 | -6.36 | 0.58126 | none |
| OTULIN | 0.091 | ######## | 0.6336 | 0.54107 | 0.685 | -6.36 | -0.5154 | none |
| CYP4F3 | 0.18 | 0.0255 | 0.6335 | 0.54114 | 0.685 | -6.36 | -1.1849 | none |
| FBXO16 | -0.23 | -0.1152 | -0.633 | 0.54124 | 0.685 | -6.36 | 1.52288 | none |
| RBFOX2 | 0.078 | 0.1375 | 0.6331 | 0.54135 | 0.685 | -6.36 | 0.04169 | none |
| RASL12 | 0.054 | 0.0747 | 0.633 | 0.54142 | 0.685 | -6.36 | 1.47012 | none |

| | | | | | | | | |
|---|---|---|---|---|---|---|---|---|
| ZNF511 | -0.104 | -0.1293 | -0.633 | 0.54152 | 0.685 | -6.36 | 1.33047 | none |
| ALB | -0.169 | -0.2256 | -0.633 | 0.54154 | 0.685 | -6.36 | 0.39065 | none |
| LYSMD4 | 0.118 | 0.179 | 0.6328 | 0.54158 | 0.685 | -6.36 | 0.31466 | none |
| ERCC3 | 0.092 | 0.1856 | 0.6327 | 0.54161 | 0.685 | -6.36 | 0.69883 | none |
| NFIB | 0.109 | 0.1291 | 0.6325 | 0.54177 | 0.685 | -6.36 | -0.518 | none |
| FAM102A | 0.076 | 0.0531 | 0.6323 | 0.54189 | 0.685 | -6.36 | -0.5217 | none |
| CHTOP | 0.049 | 0.0734 | 0.6323 | 0.54189 | 0.685 | -6.36 | -0.5323 | none |
| HSD17B6 | -0.217 | -0.2007 | -0.632 | 0.5421 | 0.685 | -6.36 | 0.45147 | none |
| ADCYAP1F | 0.051 | 0.0506 | 0.6319 | 0.54215 | 0.685 | -6.36 | 0.26365 | none |
| OR6B3 | -0.046 | -0.046 | -0.632 | 0.54216 | 0.685 | -6.36 | -0.4862 | none |
| N4BP3 | 0.05 | 0.0611 | 0.6318 | 0.54218 | 0.685 | -6.36 | 0.17853 | none |
| KAT14 | 0.058 | 0.0359 | 0.6318 | 0.5422 | 0.685 | -6.36 | -0.6097 | none |
| CDYL2 | 0.074 | 0.0842 | 0.6318 | 0.5422 | 0.685 | -6.36 | 1.29884 | none |
| HTR3C | 0.049 | 0.049 | 0.6318 | 0.54222 | 0.685 | -6.36 | 0.20764 | none |
| CASP3 | -0.077 | -0.1208 | -0.632 | 0.54222 | 0.685 | -6.36 | 1.43466 | none |
| H1FOO | 0.047 | 0.0467 | 0.6317 | 0.54227 | 0.685 | -6.36 | -0.5168 | none |
| PPM1M | -0.061 | -0.0228 | -0.632 | 0.54227 | 0.685 | -6.36 | 1.34922 | none |
| CLCF1 | -0.078 | -0.0311 | -0.632 | 0.54231 | 0.685 | -6.36 | 1.13958 | none |
| TRIM64C | 0.047 | 0.0467 | 0.6314 | 0.54242 | 0.685 | -6.36 | 0 | none |
| FXN | -0.073 | -0.0752 | -0.631 | 0.5427 | 0.685 | -6.36 | 0.62712 | none |
| SDCBP2 | 0.08 | 0.1485 | 0.631 | 0.54272 | 0.685 | -6.36 | 0.67644 | none |
| JAKMIP3 | 0.044 | 0.0368 | 0.6307 | 0.54286 | 0.686 | -6.36 | -0.7618 | none |
| MVB12B | 0.123 | 0.0046 | 0.6304 | 0.54305 | 0.686 | -6.36 | -0.8828 | none |
| SLC17A4 | -0.043 | -0.0257 | -0.63 | 0.54306 | 0.686 | -6.36 | 0 | none |
| FP325317. | 0.047 | 0.0466 | 0.6301 | 0.54324 | 0.686 | -6.36 | 0.16654 | none |
| VPS37D | 0.048 | 0.0549 | 0.6295 | 0.54362 | 0.686 | -6.36 | 0.02191 | none |
| RXFP1 | 0.095 | 0.1964 | 0.6291 | 0.54385 | 0.687 | -6.36 | 1.02363 | none |
| CDKL2 | 0.044 | 0.0614 | 0.6291 | 0.54386 | 0.687 | -6.36 | 1.31296 | none |
| WDR64 | 0.066 | 0.0649 | 0.629 | 0.54391 | 0.687 | -6.36 | -0.8214 | none |
| TSPY10 | 0.046 | 0.0464 | 0.6286 | 0.54422 | 0.687 | -6.36 | 1.39467 | none |
| PKIG | 0.058 | 0.0485 | 0.6285 | 0.54424 | 0.687 | -6.36 | 0.48469 | none |
| USP17L22 | 0.147 | 0.1427 | 0.6285 | 0.54426 | 0.687 | -6.36 | 1.45888 | none |
| TRIM26 | 0.094 | 0.1584 | 0.6281 | 0.54454 | 0.687 | -6.36 | 1.23047 | none |
| ABHD4 | 0.105 | 0.0375 | 0.6279 | 0.54464 | 0.687 | -6.36 | -1.1452 | none |
| PAPSS1 | -0.055 | -0.093 | -0.627 | 0.54504 | 0.688 | -6.36 | 1.07152 | none |
| GLYAT | -0.057 | -0.0957 | -0.627 | 0.54516 | 0.688 | -6.36 | -0.0577 | none |
| LRRC38 | -0.042 | -0.025 | -0.627 | 0.54516 | 0.688 | -6.36 | 0 | none |
| SLC10A3 | 0.079 | 0.0684 | 0.6269 | 0.54526 | 0.688 | -6.36 | 0.37208 | none |
| MAPK14 | 0.069 | 0.0493 | 0.6266 | 0.54543 | 0.688 | -6.37 | 0.76575 | none |
| SNX2 | -0.07 | -0.0429 | -0.627 | 0.5455 | 0.688 | -6.37 | 1.66933 | none |

| | | | | | | | | |
|---|---|---|---|---|---|---|---|---|
| CCDC73 | -0.043 | -0.0351 | -0.626 | 0.54574 | 0.688 | -6.37 | 0.53341 | none |
| 6-Sep | 0.136 | 0.1979 | 0.6258 | 0.54593 | 0.688 | -6.37 | 0.54814 | none |
| BMT2 | 0.138 | 0.013 | 0.6258 | 0.54598 | 0.688 | -6.37 | -0.3993 | none |
| THSD7A | 0.07 | 0.0225 | 0.6255 | 0.54615 | 0.689 | -6.37 | 0.97214 | none |
| SAPCD2 | 0.091 | 0.168 | 0.6252 | 0.54634 | 0.689 | -6.37 | -0.1265 | none |
| MMP23B | -0.075 | -0.002 | -0.625 | 0.54644 | 0.689 | -6.37 | 1.68567 | none |
| IER3IP1 | 0.112 | -0.0421 | 0.625 | 0.54644 | 0.689 | -6.37 | 0.08571 | none |
| GPHA2 | 0.064 | 0.0914 | 0.625 | 0.54646 | 0.689 | -6.37 | 1.63851 | none |
| MARCKS | 0.045 | 0.0454 | 0.6249 | 0.54653 | 0.689 | -6.37 | -0.8245 | none |
| MED28 | -0.11 | -0.1836 | -0.625 | 0.54666 | 0.689 | -6.37 | 0.40964 | none |
| BUB3 | -0.056 | -0.0599 | -0.624 | 0.54678 | 0.689 | -6.37 | -0.3245 | none |
| RAB37 | -0.123 | -0.175 | -0.624 | 0.5469 | 0.689 | -6.37 | 1.00927 | none |
| ASB7 | 0.067 | 0.0592 | 0.6243 | 0.54693 | 0.689 | -6.37 | 0.55624 | none |
| ETFA | -0.059 | -0.1111 | -0.624 | 0.54697 | 0.689 | -6.37 | -0.4527 | none |
| CARTPT | 0.046 | 0.0459 | 0.6241 | 0.54703 | 0.689 | -6.37 | -0.4725 | none |
| GPR137B | -0.141 | -0.1912 | -0.624 | 0.54726 | 0.689 | -6.37 | 0.39137 | none |
| SLCO1C1 | 0.109 | 0.1297 | 0.6231 | 0.54763 | 0.69 | -6.37 | 1.36334 | none |
| SHPRH | 0.135 | 0.1734 | 0.623 | 0.54773 | 0.69 | -6.37 | 0.46318 | none |
| HERC5 | 0.121 | 0.1473 | 0.6229 | 0.54778 | 0.69 | -6.37 | 1.28389 | none |
| CPA2 | -0.046 | -0.013 | -0.623 | 0.54786 | 0.69 | -6.37 | 0.9513 | none |
| SLC25A20 | 0.075 | 0.0663 | 0.6226 | 0.54797 | 0.69 | -6.37 | 0.01096 | none |
| GPSM3 | -0.121 | -0.142 | -0.622 | 0.54815 | 0.69 | -6.37 | 0.62323 | none |
| ROPN1B | 0.053 | 0.0632 | 0.6221 | 0.54826 | 0.69 | -6.37 | -0.8684 | none |
| NUF2 | 0.118 | 0.1439 | 0.6221 | 0.54829 | 0.69 | -6.37 | 0.78169 | none |
| ZNF433 | 0.13 | 0.1569 | 0.6221 | 0.54829 | 0.69 | -6.37 | -0.0324 | none |
| ALKBH6 | 0.142 | 0.1645 | 0.622 | 0.54837 | 0.69 | -6.37 | 1.22017 | none |
| LANCL1 | 0.113 | ######## | 0.6219 | 0.54839 | 0.69 | -6.37 | -0.2334 | none |
| MGARP | -0.053 | -0.0329 | -0.622 | 0.54847 | 0.69 | -6.37 | -0.3672 | none |
| RUNX3 | 0.104 | 0.1901 | 0.6216 | 0.54858 | 0.69 | -6.37 | 0.00353 | none |
| PRPS2 | -0.085 | -0.0879 | -0.622 | 0.54859 | 0.69 | -6.37 | 1.60404 | none |
| CDH26 | 0.119 | 0.1587 | 0.6213 | 0.54877 | 0.691 | -6.37 | 0.37986 | none |
| DTL | 0.064 | 0.0864 | 0.6209 | 0.54905 | 0.691 | -6.37 | 1.16982 | none |
| ALKBH3 | -0.1 | -0.1435 | -0.621 | 0.54908 | 0.691 | -6.37 | -0.2801 | none |
| C22ORF23 | -0.085 | -0.0339 | -0.621 | 0.54912 | 0.691 | -6.37 | 1.06407 | none |
| DMTN | 0.096 | 0.0991 | 0.6203 | 0.54943 | 0.691 | -6.37 | 0.31726 | none |
| S100A13 | -0.176 | -0.1246 | -0.62 | 0.54954 | 0.691 | -6.37 | 1.85347 | none |
| E2F1 | -0.069 | -0.0173 | -0.62 | 0.54961 | 0.691 | -6.37 | 1.48286 | none |
| FAM129B | -0.159 | -0.0672 | -0.62 | 0.54965 | 0.691 | -6.37 | -0.1584 | none |
| FAM110C | 0.092 | 0.0928 | 0.6197 | 0.54977 | 0.691 | -6.37 | 1.82538 | none |
| HOXA1 | -0.053 | -0.012 | -0.619 | 0.55008 | 0.692 | -6.37 | 1.57918 | none |

| | | | | | | | | |
|---|---|---|---|---|---|---|---|---|
| ZG16B | -0.098 | -0.0868 | -0.619 | 0.5505 | 0.692 | -6.37 | 1.3793 | none |
| TARS | -0.086 | -0.0126 | -0.618 | 0.5506 | 0.692 | -6.37 | 1.72907 | none |
| NACAD | -0.043 | -0.0352 | -0.618 | 0.55065 | 0.692 | -6.37 | 0 | none |
| BBS9 | -0.102 | -0.0874 | -0.618 | 0.55068 | 0.692 | -6.37 | 0.47234 | none |
| OSGIN2 | -0.102 | -0.1405 | -0.618 | 0.55074 | 0.692 | -6.37 | 0.7849 | none |
| NKX6-1 | 0.045 | 0.0263 | 0.618 | 0.55087 | 0.692 | -6.37 | 1.54885 | none |
| VTI1B | 0.061 | 0.0454 | 0.618 | 0.55088 | 0.692 | -6.37 | -0.7703 | none |
| THRB | 0.054 | 0.0542 | 0.6179 | 0.55091 | 0.692 | -6.37 | 0.80584 | none |
| SLC25A29 | -0.124 | -0.1954 | -0.618 | 0.55116 | 0.693 | -6.37 | -0.6686 | none |
| PNMA6A | -0.054 | -0.0231 | -0.617 | 0.55124 | 0.693 | -6.37 | 1.2961 | none |
| SLC44A3 | 0.064 | 0.0576 | 0.6174 | 0.55127 | 0.693 | -6.37 | 1.13399 | none |
| OR5M11 | -0.045 | -0.0446 | -0.617 | 0.5513 | 0.693 | -6.37 | 1.50474 | none |
| PAPSS2 | -0.057 | -0.0056 | -0.617 | 0.55131 | 0.693 | -6.37 | 1.08437 | none |
| TRIM60 | 0.116 | 0.066 | 0.6172 | 0.55138 | 0.693 | -6.37 | -0.761 | none |
| EPSTI1 | 0.097 | 0.106 | 0.6171 | 0.55142 | 0.693 | -6.37 | 0.34181 | none |
| MRPS26 | -0.052 | -0.0408 | -0.617 | 0.55143 | 0.693 | -6.37 | 1.03455 | none |
| OR2AE1 | 0.047 | 0.0455 | 0.617 | 0.55149 | 0.693 | -6.37 | -0.1632 | none |
| IL17C | -0.077 | -0.0497 | -0.617 | 0.55156 | 0.693 | -6.37 | 1.15185 | none |
| WLS | -0.108 | -0.1342 | -0.617 | 0.55162 | 0.693 | -6.37 | 0.36459 | none |
| C5ORF22 | 0.092 | 0.0169 | 0.6168 | 0.55166 | 0.693 | -6.37 | 0.20875 | none |
| LGALS9B | -0.096 | -0.036 | -0.617 | 0.55179 | 0.693 | -6.37 | 1.18472 | none |
| SMCP | 0.045 | 0.0451 | 0.6163 | 0.55194 | 0.693 | -6.37 | -0.846 | none |
| GOLGA8G | 0.042 | 0.0478 | 0.6162 | 0.55202 | 0.693 | -6.37 | -0.3005 | none |
| APOA1 | -0.073 | -0.0656 | -0.616 | 0.55206 | 0.693 | -6.37 | 0.40063 | none |
| PSG8 | -0.073 | -0.073 | -0.616 | 0.5523 | 0.693 | -6.37 | 2.16217 | AA-related |
| OR6N2 | 0.042 | 0.0423 | 0.6155 | 0.55243 | 0.693 | -6.37 | 1.52134 | none |
| CBLB | 0.098 | 0.1262 | 0.6155 | 0.55248 | 0.693 | -6.37 | 0.57386 | none |
| SSB | 0.186 | 0.3121 | 0.6152 | 0.55262 | 0.693 | -6.37 | -0.2298 | none |
| CRYBA4 | -0.068 | -0.0439 | -0.615 | 0.5527 | 0.694 | -6.37 | 1.35065 | none |
| GRM8 | -0.051 | -0.0511 | -0.615 | 0.55299 | 0.694 | -6.37 | 1.06361 | none |
| SLC16A1 | 0.063 | 0.0468 | 0.6145 | 0.55306 | 0.694 | -6.37 | 0.02435 | none |
| PSD2 | -0.042 | -0.0403 | -0.614 | 0.55311 | 0.694 | -6.37 | 0 | none |
| C22ORF15 | 0.06 | 0.0558 | 0.6144 | 0.55316 | 0.694 | -6.37 | 0.44846 | none |
| NLGN2 | -0.054 | -0.0068 | -0.614 | 0.55348 | 0.694 | -6.37 | 0.64123 | none |
| IGF2 | -0.096 | -0.1086 | -0.614 | 0.55349 | 0.694 | -6.37 | -0.3064 | none |
| DLK1 | -0.07 | -0.0718 | -0.614 | 0.55367 | 0.694 | -6.37 | 1.28421 | none |
| OR5T2 | -0.042 | -0.0161 | -0.614 | 0.55372 | 0.694 | -6.37 | 0.74998 | none |
| AC136352 | 0.045 | 0.0448 | 0.6135 | 0.55373 | 0.694 | -6.37 | 0 | none |
| MANEAL | 0.062 | 0.0675 | 0.6135 | 0.55373 | 0.694 | -6.37 | 0.66639 | none |
| DOLK | 0.084 | 0.0136 | 0.6131 | 0.55398 | 0.695 | -6.37 | -0.4038 | none |

| | | | | | | | | |
|---|---|---|---|---|---|---|---|---|
| TESK1 | 0.077 | 0.1304 | 0.6131 | 0.55399 | 0.695 | -6.37 | -0.5488 | none |
| SRM | 0.077 | 0.0616 | 0.6124 | 0.55441 | 0.695 | -6.37 | -0.7025 | none |
| TERT | -0.142 | -0.094 | -0.612 | 0.55462 | 0.695 | -6.37 | -0.107 | none |
| AADACL4 | 0.045 | 0.0447 | 0.6119 | 0.55471 | 0.695 | -6.37 | 0 | none |
| HYAL2 | 0.111 | 0.122 | 0.6115 | 0.55498 | 0.696 | -6.37 | 0.05704 | none |
| TET3 | 0.107 | 0.2013 | 0.6115 | 0.555 | 0.696 | -6.37 | 1.32618 | none |
| IKZF5 | 0.155 | 0.1762 | 0.6114 | 0.55506 | 0.696 | -6.37 | 0.6185 | none |
| SIL1 | -0.062 | 0.0044 | -0.611 | 0.55528 | 0.696 | -6.38 | 0.64842 | none |
| CTBS | -0.075 | -0.1014 | -0.611 | 0.55538 | 0.696 | -6.38 | 0.82354 | none |
| MFSD1 | 0.071 | -0.0014 | 0.6108 | 0.55543 | 0.696 | -6.38 | 0.60715 | none |
| BCL6B | 0.067 | 0.0983 | 0.6107 | 0.55549 | 0.696 | -6.38 | 1.39544 | none |
| FAM111B | 0.248 | 0.3706 | 0.6105 | 0.55561 | 0.696 | -6.38 | -0.0785 | none |
| SMIM6 | -0.047 | -0.0376 | -0.61 | 0.55614 | 0.697 | -6.38 | 0.71349 | none |
| ACMSD | 0.107 | 0.074 | 0.6095 | 0.55624 | 0.697 | -6.38 | -0.6649 | none |
| IFNE | 0.044 | 0.0444 | 0.6094 | 0.55631 | 0.697 | -6.38 | 0 | none |
| B4GALNT3 | 0.075 | 0.1697 | 0.6092 | 0.55646 | 0.697 | -6.38 | -0.0092 | none |
| ARMCX5 | 0.11 | 0.0626 | 0.6092 | 0.55647 | 0.697 | -6.38 | -0.2119 | none |
| KCNH3 | -0.095 | -0.0158 | -0.609 | 0.5567 | 0.697 | -6.38 | 1.1753 | none |
| CDH16 | -0.069 | -0.061 | -0.609 | 0.55676 | 0.697 | -6.38 | 1.9259 | none |
| TAOK2 | 0.14 | 0.2685 | 0.6082 | 0.55706 | 0.697 | -6.38 | 0.25363 | none |
| C1ORF159 | 0.066 | 0.1009 | 0.6082 | 0.55708 | 0.697 | -6.38 | 0.89495 | none |
| HRASLS5 | -0.047 | -0.0591 | -0.608 | 0.55712 | 0.697 | -6.38 | 1.53994 | none |
| ZNF582 | 0.102 | 0.2125 | 0.6081 | 0.55716 | 0.697 | -6.38 | 1.56403 | none |
| MGAT4D | 0.078 | 0.0782 | 0.6078 | 0.55737 | 0.697 | -6.38 | 1.05585 | none |
| SAR1A | 0.081 | 0.003 | 0.6078 | 0.55737 | 0.697 | -6.38 | -0.2911 | none |
| HSD3B2 | 0.044 | 0.0443 | 0.6077 | 0.55739 | 0.697 | -6.38 | 1.68353 | none |
| TRMT13 | 0.101 | 0.0572 | 0.6077 | 0.55742 | 0.697 | -6.38 | 1.03839 | none |
| SLC27A4 | -0.053 | -0.0116 | -0.608 | 0.55743 | 0.697 | -6.38 | 1.42098 | none |
| CGREF1 | 0.092 | 0.2165 | 0.6069 | 0.55794 | 0.698 | -6.38 | 0.43025 | none |
| PRSS55 | 0.05 | 0.036 | 0.6064 | 0.55825 | 0.698 | -6.38 | 0.40683 | none |
| IFNA8 | -0.043 | -0.0435 | -0.606 | 0.55834 | 0.698 | -6.38 | 0 | none |
| INTS12 | -0.076 | -0.0042 | -0.606 | 0.55843 | 0.698 | -6.38 | 0.74468 | none |
| BCL9 | 0.188 | 0.3614 | 0.6059 | 0.55856 | 0.699 | -6.38 | 0.21411 | none |
| CIDEA | 0.047 | 0.0465 | 0.6055 | 0.5588 | 0.699 | -6.38 | 0.78245 | none |
| ZNHIT2 | -0.069 | -0.0509 | -0.605 | 0.55903 | 0.699 | -6.38 | 0.11418 | none |
| ANP32A | -0.088 | -0.0085 | -0.605 | 0.55921 | 0.699 | -6.38 | 0.93063 | none |
| FTCD | -0.07 | -0.0695 | -0.605 | 0.55923 | 0.699 | -6.38 | 1.92315 | none |
| SHB | 0.044 | 0.058 | 0.6048 | 0.55928 | 0.699 | -6.38 | -0.1555 | none |
| METTL14 | 0.141 | 0.1353 | 0.6042 | 0.55962 | 0.7 | -6.38 | -0.0199 | none |
| DHRS13 | -0.057 | -0.0405 | -0.604 | 0.55968 | 0.7 | -6.38 | 1.20255 | none |

| | | | | | | | | |
|---|---|---|---|---|---|---|---|---|
| IL13RA2 | -0.043 | -0.0433 | -0.604 | 0.55969 | 0.7 | -6.38 | 1.51533 | none |
| TRIM15 | -0.043 | -0.0341 | -0.604 | 0.55976 | 0.7 | -6.38 | 0.35918 | none |
| IL2RB | 0.095 | 0.1388 | 0.604 | 0.55977 | 0.7 | -6.38 | 0.70827 | none |
| MXD3 | 0.072 | 0.0385 | 0.6037 | 0.55993 | 0.7 | -6.38 | 0.6562 | none |
| S100A8 | -0.053 | -0.0706 | -0.604 | 0.55996 | 0.7 | -6.38 | 0.64998 | none |
| CRYAA | 0.044 | 0.0439 | 0.6035 | 0.56006 | 0.7 | -6.38 | 0 | none |
| GOLGA6D | -0.043 | -0.0432 | -0.603 | 0.56027 | 0.7 | -6.38 | 1.22662 | none |
| CA2 | -0.086 | 0.0283 | -0.603 | 0.56067 | 0.7 | -6.38 | 1.58245 | none |
| ARID3C | 0.042 | 0.0313 | 0.6025 | 0.56075 | 0.7 | -6.38 | 0 | none |
| TIGD6 | -0.074 | -0.0309 | -0.602 | 0.56081 | 0.701 | -6.38 | 0.46811 | none |
| HAS1 | -0.051 | -0.0523 | -0.602 | 0.56088 | 0.701 | -6.38 | -0.0212 | none |
| GLUL | -0.111 | -0.1273 | -0.602 | 0.56099 | 0.701 | -6.38 | 1.46624 | none |
| CHCHD3 | -0.077 | -0.0308 | -0.602 | 0.56103 | 0.701 | -6.38 | 1.61549 | none |
| ACOT8 | -0.074 | -0.1176 | -0.601 | 0.56137 | 0.701 | -6.38 | 0.71967 | none |
| DUSP14 | 0.258 | 0.2485 | 0.6013 | 0.56151 | 0.701 | -6.38 | 0.10592 | none |
| GAB2 | 0.143 | 0.1617 | 0.6012 | 0.56155 | 0.701 | -6.38 | 0.35689 | none |
| CD8B | -0.074 | -0.0127 | -0.601 | 0.56167 | 0.701 | -6.38 | 0.29113 | none |
| FGF4 | -0.042 | -0.0422 | -0.601 | 0.56171 | 0.701 | -6.38 | 1.1314 | none |
| TEX101 | 0.08 | 0.0619 | 0.6009 | 0.56176 | 0.701 | -6.38 | 0.85941 | none |
| B3GAT2 | 0.041 | 0.0142 | 0.6008 | 0.56179 | 0.701 | -6.38 | 0.54834 | none |
| DAB2 | -0.118 | -0.1048 | -0.601 | 0.5618 | 0.701 | -6.38 | 0.35806 | none |
| LSM14B | 0.08 | 0.1295 | 0.6006 | 0.56195 | 0.701 | -6.38 | 0.63443 | none |
| ZNF717 | 0.073 | 0.0071 | 0.6002 | 0.56219 | 0.702 | -6.38 | 0.3865 | none |
| ZNF84 | -0.076 | -0.075 | -0.6 | 0.56223 | 0.702 | -6.38 | 0.61269 | none |
| ATF3 | -0.128 | -0.06 | -0.6 | 0.5623 | 0.702 | -6.38 | 1.6849 | none |
| OR1N1 | -0.042 | -0.0431 | -0.6 | 0.56242 | 0.702 | -6.38 | 0 | none |
| THNSL1 | -0.088 | -0.0456 | -0.6 | 0.56256 | 0.702 | -6.38 | 1.79659 | none |
| ASS1 | -0.131 | -0.1577 | -0.6 | 0.56261 | 0.702 | -6.38 | 0.21123 | none |
| CALML4 | 0.072 | 0.1208 | 0.599 | 0.56297 | 0.702 | -6.38 | 0.57716 | none |
| ZNF248 | -0.138 | -0.1165 | -0.599 | 0.56304 | 0.702 | -6.38 | 1.17592 | none |
| MCUB | 0.106 | 0.0013 | 0.5984 | 0.56334 | 0.703 | -6.38 | 0.41507 | none |
| CST3 | -0.074 | -0.0391 | -0.598 | 0.56381 | 0.703 | -6.38 | 0.89437 | none |
| SERPINA3 | -0.093 | -0.0914 | -0.597 | 0.56399 | 0.703 | -6.38 | 1.07129 | none |
| UST | 0.166 | 0.155 | 0.5974 | 0.56401 | 0.703 | -6.38 | 0.93241 | none |
| FOS | 0.112 | 0.0919 | 0.597 | 0.56423 | 0.703 | -6.38 | 0.54091 | none |
| FOXD4L5 | -0.044 | -0.0353 | -0.597 | 0.56433 | 0.704 | -6.38 | 0.39054 | none |
| AGL | 0.068 | 0.0872 | 0.5968 | 0.56436 | 0.704 | -6.38 | 0.86883 | none |
| PENK | 0.062 | 0.085 | 0.5966 | 0.5645 | 0.704 | -6.38 | -0.31 | none |
| TAAR2 | -0.043 | -0.0426 | -0.597 | 0.56451 | 0.704 | -6.38 | 0 | none |
| SMIM10 | 0.093 | 0.0936 | 0.5963 | 0.56468 | 0.704 | -6.38 | -0.4119 | none |

| | | | | | | | | |
|---|---|---|---|---|---|---|---|---|
| GRWD1 | 0.063 | 0.0529 | 0.5962 | 0.56474 | 0.704 | -6.38 | -0.0456 | none |
| PRSS54 | -0.049 | -0.0148 | -0.596 | 0.56487 | 0.704 | -6.38 | 1.06067 | none |
| OR4K14 | 0.043 | 0.0431 | 0.5959 | 0.56493 | 0.704 | -6.38 | 0 | none |
| MUS81 | 0.05 | 0.066 | 0.5959 | 0.56495 | 0.704 | -6.38 | -0.408 | none |
| HOXD12 | 0.045 | 0.0455 | 0.5959 | 0.56495 | 0.704 | -6.38 | -0.1265 | none |
| TMEM202 | 0.043 | 0.0431 | 0.5958 | 0.56499 | 0.704 | -6.38 | 0 | none |
| CD177 | 0.057 | 0.0421 | 0.5957 | 0.56508 | 0.704 | -6.38 | -0.9413 | none |
| MTRNR2L1 | 0.061 | 0.0587 | 0.5956 | 0.56511 | 0.704 | -6.38 | 0.87654 | none |
| CAV3 | -0.108 | -0.0794 | -0.596 | 0.56516 | 0.704 | -6.38 | 0.4035 | none |
| OGN | -0.054 | -0.0514 | -0.596 | 0.56517 | 0.704 | -6.38 | -0.1422 | none |
| NOD2 | -0.05 | -0.0224 | -0.595 | 0.56527 | 0.704 | -6.38 | 0.13236 | none |
| ENOPH1 | -0.059 | -0.0881 | -0.595 | 0.56535 | 0.704 | -6.38 | 1.28654 | none |
| EVI2B | 0.103 | 0.0878 | 0.5951 | 0.56549 | 0.704 | -6.39 | 1.04788 | none |
| ADORA3 | 0.041 | 0.0371 | 0.595 | 0.5655 | 0.704 | -6.39 | 0.68944 | none |
| AOC2 | 0.058 | 0.1143 | 0.5947 | 0.56569 | 0.704 | -6.39 | -0.3066 | none |
| LONRF3 | -0.061 | 0.0225 | -0.595 | 0.56569 | 0.704 | -6.39 | 0.63524 | none |
| LCE5A | -0.042 | -0.0424 | -0.594 | 0.56591 | 0.704 | -6.39 | 1.04303 | none |
| TREM2 | -0.042 | -0.0424 | -0.594 | 0.56591 | 0.704 | -6.39 | -0.5763 | none |
| GPN1 | -0.053 | -0.097 | -0.594 | 0.56597 | 0.704 | -6.39 | 0.35258 | none |
| PCGF5 | 0.051 | 0.0222 | 0.5942 | 0.56601 | 0.704 | -6.39 | 1.90414 | none |
| WWC1 | 0.118 | 0.1648 | 0.5942 | 0.56604 | 0.704 | -6.39 | 0.84581 | none |
| ANXA8 | 0.055 | 0.0633 | 0.5941 | 0.56611 | 0.704 | -6.39 | 0.02547 | none |
| LMO4 | -0.086 | -0.0919 | -0.594 | 0.56611 | 0.704 | -6.39 | 1.25816 | none |
| SLC25A36 | 0.064 | 0.0147 | 0.594 | 0.56617 | 0.704 | -6.39 | 1.30542 | none |
| NKX2-8 | -0.048 | -0.0251 | -0.594 | 0.56636 | 0.704 | -6.39 | 1.52557 | none |
| RFTN1 | 0.076 | 0.1424 | 0.5937 | 0.56639 | 0.704 | -6.39 | 1.07439 | none |
| ADCY9 | 0.056 | 0.1182 | 0.5937 | 0.56639 | 0.704 | -6.39 | 0.78397 | none |
| PRRX2 | -0.062 | -0.0251 | -0.594 | 0.56644 | 0.704 | -6.39 | 1.11632 | none |
| ABCD3 | 0.06 | 0.08 | 0.5936 | 0.56645 | 0.704 | -6.39 | 1.91909 | none |
| PTPRJ | 0.183 | 0.2587 | 0.5932 | 0.56668 | 0.705 | -6.39 | -0.1075 | none |
| RNF169 | 0.139 | 0.2695 | 0.5932 | 0.56668 | 0.705 | -6.39 | 0.59505 | none |
| CTNNA3 | -0.054 | -0.0496 | -0.593 | 0.56677 | 0.705 | -6.39 | 1.71784 | none |
| EPB41L1 | -0.126 | -0.1561 | -0.593 | 0.56695 | 0.705 | -6.39 | 0.63463 | none |
| MSRB1 | -0.087 | -0.101 | -0.593 | 0.56705 | 0.705 | -6.39 | 0.8088 | none |
| KRTAP13-4 | 0.043 | 0.0428 | 0.5924 | 0.56719 | 0.705 | -6.39 | -0.832 | none |
| POC5 | -0.1 | -0.1019 | -0.592 | 0.56724 | 0.705 | -6.39 | 0.46696 | none |
| RBMXL3 | 0.046 | 0.0229 | 0.592 | 0.56744 | 0.705 | -6.39 | -0.8884 | none |
| OR6C2 | 0.043 | 0.0427 | 0.5919 | 0.56749 | 0.705 | -6.39 | 0.39601 | none |
| USP17L23 | -0.125 | -0.0498 | -0.592 | 0.56766 | 0.705 | -6.39 | -0.3349 | none |
| MFSD14A | -0.092 | -0.1197 | -0.592 | 0.56766 | 0.705 | -6.39 | 1.42712 | none |

| | | | | | | | | |
|---|---|---|---|---|---|---|---|---|
| TARBP2 | -0.073 | -0.0926 | -0.592 | 0.56768 | 0.705 | -6.39 | 0.64828 | none |
| IZUMO1R | 0.043 | 0.0427 | 0.5916 | 0.56772 | 0.705 | -6.39 | 0 | none |
| SPECC1L | 0.195 | 0.3107 | 0.5915 | 0.56777 | 0.705 | -6.39 | 0.5103 | none |
| NLGN3 | 0.073 | 0.0775 | 0.5915 | 0.56778 | 0.705 | -6.39 | -0.0414 | none |
| ARC | -0.059 | -0.0389 | -0.591 | 0.5678 | 0.705 | -6.39 | 1.2913 | none |
| NR2F2 | 0.074 | 0.0811 | 0.5915 | 0.56781 | 0.705 | -6.39 | 1.01743 | none |
| GAL3ST2 | -0.042 | -0.0421 | -0.591 | 0.56791 | 0.705 | -6.39 | 1.30781 | none |
| OR10J3 | 0.043 | 0.0427 | 0.5913 | 0.56792 | 0.705 | -6.39 | 0 | none |
| MYBL2 | 0.107 | 0.24 | 0.5912 | 0.568 | 0.705 | -6.39 | 0.04325 | none |
| SLIT2 | 0.089 | 0.0887 | 0.5911 | 0.56803 | 0.705 | -6.39 | -0.6813 | none |
| ENPP5 | -0.071 | -0.0653 | -0.591 | 0.56803 | 0.705 | -6.39 | 0.16218 | none |
| DUSP16 | 0.118 | 0.1585 | 0.5907 | 0.56829 | 0.706 | -6.39 | -0.3645 | none |
| TECRL | 0.113 | 0.1113 | 0.5906 | 0.56836 | 0.706 | -6.39 | 1.65904 | none |
| ABHD5 | 0.056 | -0.033 | 0.5906 | 0.56836 | 0.706 | -6.39 | -0.3024 | none |
| RCOR2 | 0.047 | 0.0474 | 0.5903 | 0.56856 | 0.706 | -6.39 | 1.59297 | none |
| DAOA | 0.1 | 0.05 | 0.5903 | 0.56857 | 0.706 | -6.39 | -0.4762 | none |
| RIC8A | -0.07 | -0.0465 | -0.59 | 0.56868 | 0.706 | -6.39 | 1.36103 | none |
| UNCX | -0.04 | -0.0192 | -0.59 | 0.56884 | 0.706 | -6.39 | 0 | none |
| ACKR4 | 0.042 | 0.0582 | 0.5898 | 0.56889 | 0.706 | -6.39 | 0.99439 | none |
| CDK15 | -0.061 | -0.0668 | -0.59 | 0.56894 | 0.706 | -6.39 | 0.87926 | none |
| PRSS56 | 0.042 | 0.0459 | 0.5896 | 0.569 | 0.706 | -6.39 | 0 | none |
| CCS | -0.063 | -0.0488 | -0.59 | 0.56905 | 0.706 | -6.39 | 0.09497 | none |
| DTX2 | 0.062 | 0.0349 | 0.5894 | 0.56914 | 0.706 | -6.39 | -0.3719 | none |
| MUC20 | -0.057 | -0.0172 | -0.589 | 0.56918 | 0.706 | -6.39 | 0.90755 | none |
| KRTAP5-6 | 0.042 | 0.0425 | 0.5893 | 0.5692 | 0.706 | -6.39 | -0.5385 | none |
| EXOC3L4 | -0.112 | -0.0795 | -0.589 | 0.56925 | 0.706 | -6.39 | -0.0389 | none |
| BPIFB2 | -0.042 | -0.0512 | -0.589 | 0.5693 | 0.706 | -6.39 | 1.50573 | none |
| PTPDC1 | -0.052 | 0.0194 | -0.589 | 0.5697 | 0.706 | -6.39 | 1.25946 | none |
| CARD10 | -0.072 | -0.0589 | -0.588 | 0.56971 | 0.706 | -6.39 | 0.61124 | none |
| CXCR2 | 0.089 | 0.165 | 0.5884 | 0.56977 | 0.706 | -6.39 | 0.47826 | none |
| SGCA | -0.063 | -0.0531 | -0.588 | 0.56983 | 0.707 | -6.39 | 1.62983 | none |
| NECTIN2 | -0.067 | -0.0448 | -0.588 | 0.57 | 0.707 | -6.39 | 0.15914 | none |
| OR5F1 | 0.042 | 0.0424 | 0.588 | 0.57001 | 0.707 | -6.39 | -0.599 | none |
| RCVRN | 0.069 | 0.0661 | 0.5879 | 0.57007 | 0.707 | -6.39 | -0.4249 | none |
| ALDH16A1 | 0.051 | 0.0169 | 0.5879 | 0.5701 | 0.707 | -6.39 | 0.46467 | none |
| ARHGEF3 | 0.103 | 0.0982 | 0.5879 | 0.57011 | 0.707 | -6.39 | 0.43422 | none |
| SLC35B2 | -0.087 | -0.1227 | -0.588 | 0.57018 | 0.707 | -6.39 | 1.27842 | none |
| MAGEB17 | -0.042 | -0.0417 | -0.588 | 0.57018 | 0.707 | -6.39 | 1.52319 | none |
| ENTPD8 | 0.046 | 0.0494 | 0.5877 | 0.57022 | 0.707 | -6.39 | 0.70123 | none |
| CT47A7 | 0.07 | 0.0704 | 0.5876 | 0.57031 | 0.707 | -6.39 | 0.74555 | none |

| | | | | | | | | |
|---|---|---|---|---|---|---|---|---|
| SFRP4 | -0.052 | -0.0296 | -0.588 | 0.57034 | 0.707 | -6.39 | 0.87122 | none |
| SLC25A10 | -0.053 | -0.0149 | -0.587 | 0.57039 | 0.707 | -6.39 | 1.37752 | none |
| LAMTOR4 | -0.075 | -0.0811 | -0.587 | 0.57044 | 0.707 | -6.39 | 0.70824 | none |
| ADAMTS9 | -0.122 | -0.0986 | -0.587 | 0.57046 | 0.707 | -6.39 | 0.17821 | none |
| HIST1H2BI | 0.066 | -0.0477 | 0.5871 | 0.5706 | 0.707 | -6.39 | 0.0832 | none |
| TSPAN1 | -0.092 | -0.0703 | -0.587 | 0.57085 | 0.707 | -6.39 | -0.1632 | none |
| ENTPD2 | -0.075 | -0.0365 | -0.586 | 0.57128 | 0.707 | -6.39 | 0.66473 | none |
| RYBP | -0.071 | -0.046 | -0.586 | 0.57136 | 0.707 | -6.39 | 1.0421 | none |
| NETO1 | -0.14 | -0.1064 | -0.586 | 0.57136 | 0.707 | -6.39 | 1.63974 | none |
| EPB42 | 0.099 | 0.1013 | 0.5858 | 0.57144 | 0.708 | -6.39 | -0.2802 | none |
| SOCS2 | 0.078 | 0.0204 | 0.5856 | 0.57156 | 0.708 | -6.39 | 0.51279 | none |
| TMEM262 | 0.05 | 0.0882 | 0.5856 | 0.57159 | 0.708 | -6.39 | 0.13256 | none |
| KCTD11 | 0.062 | 0.0782 | 0.5855 | 0.57164 | 0.708 | -6.39 | -0.3184 | none |
| PRSS2 | -0.105 | -0.127 | -0.585 | 0.57169 | 0.708 | -6.39 | 0.77704 | none |
| ETV2 | 0.058 | 0.0346 | 0.5853 | 0.57177 | 0.708 | -6.39 | -0.0248 | none |
| NYAP1 | -0.054 | -0.0585 | -0.585 | 0.57178 | 0.708 | -6.39 | 0.3978 | none |
| BORCS8-M | 0.119 | 0.1449 | 0.5852 | 0.57184 | 0.708 | -6.39 | 0.90592 | none |
| CCR5 | 0.05 | -0.0268 | 0.5847 | 0.57215 | 0.708 | -6.39 | -0.7586 | none |
| GDF10 | -0.046 | -0.0294 | -0.585 | 0.57215 | 0.708 | -6.39 | 0.84794 | none |
| MAN1A2 | 0.076 | 0.024 | 0.5846 | 0.57221 | 0.708 | -6.39 | 0.06313 | none |
| NINL | 0.134 | 0.1837 | 0.5846 | 0.57221 | 0.708 | -6.39 | -0.3174 | none |
| WDR37 | 0.072 | 0.1656 | 0.5845 | 0.57229 | 0.708 | -6.39 | -0.4198 | none |
| SLC25A14 | 0.075 | 0.0355 | 0.5844 | 0.57233 | 0.708 | -6.39 | 0.06229 | none |
| MRPS5 | 0.044 | 0.0581 | 0.5843 | 0.57244 | 0.708 | -6.39 | 1.04769 | none |
| FOXS1 | -0.043 | -0.0286 | -0.584 | 0.57244 | 0.708 | -6.39 | 0.81398 | none |
| MBD3L1 | -0.094 | -0.0922 | -0.584 | 0.57246 | 0.708 | -6.39 | 0.25337 | none |
| TMEM223 | 0.095 | 0.054 | 0.5841 | 0.57256 | 0.708 | -6.39 | 0.21738 | none |
| WBP1 | 0.147 | 0.1258 | 0.584 | 0.57262 | 0.708 | -6.39 | 0.36559 | none |
| CABLES1 | -0.118 | -0.2264 | -0.584 | 0.57266 | 0.708 | -6.39 | 0.15989 | none |
| VIPR1 | -0.077 | 0.0081 | -0.584 | 0.57266 | 0.708 | -6.39 | 0.24199 | none |
| FAP | 0.058 | 0.0624 | 0.5839 | 0.5727 | 0.708 | -6.39 | -0.7832 | none |
| ZFAND5 | 0.076 | 0.124 | 0.5837 | 0.57278 | 0.708 | -6.39 | 1.25581 | none |
| THRSP | -0.046 | -0.0292 | -0.584 | 0.57287 | 0.708 | -6.39 | 0.34236 | none |
| MAPK15 | -0.048 | 0.0266 | -0.583 | 0.57318 | 0.708 | -6.39 | 0.14309 | none |
| CHRFAM7. | -0.053 | -0.0145 | -0.582 | 0.57367 | 0.709 | -6.39 | 0.10182 | none |
| HMCES | -0.044 | -0.0603 | -0.582 | 0.5737 | 0.709 | -6.39 | -0.208 | none |
| YAF2 | 0.058 | -0.0217 | 0.5823 | 0.57372 | 0.709 | -6.39 | 0.71366 | none |
| CCDC190 | -0.041 | -0.0412 | -0.582 | 0.57372 | 0.709 | -6.39 | 0.11503 | none |
| GRM4 | 0.081 | 0.0234 | 0.5822 | 0.57375 | 0.709 | -6.39 | -0.2281 | none |
| FUT5 | 0.041 | 0.0419 | 0.5822 | 0.57376 | 0.709 | -6.39 | 0 | none |

| | | | | | | | | |
|---|---|---|---|---|---|---|---|---|
| CCNB3 | -0.079 | -0.0827 | -0.582 | 0.5739 | 0.709 | -6.39 | -0.2466 | none |
| ENC1 | -0.061 | -0.0582 | -0.582 | 0.57391 | 0.709 | -6.39 | 1.58472 | none |
| FTSJ1 | -0.058 | -0.0798 | -0.582 | 0.5742 | 0.709 | -6.39 | 0.87777 | none |
| MAPRE3 | 0.045 | 0.05 | 0.5814 | 0.57431 | 0.709 | -6.39 | 0.72538 | none |
| TMED2 | -0.108 | -0.1983 | -0.581 | 0.57441 | 0.709 | -6.39 | 1.18468 | none |
| GRM2 | 0.051 | 0.0832 | 0.581 | 0.57454 | 0.709 | -6.39 | 1.44406 | none |
| LRRC8D | -0.065 | -0.011 | -0.581 | 0.57483 | 0.71 | -6.39 | 0.6117 | none |
| NBPF4 | 0.04 | 0.042 | 0.5802 | 0.57504 | 0.71 | -6.39 | 0 | none |
| RARRES2 | -0.087 | -0.0396 | -0.58 | 0.57505 | 0.71 | -6.39 | 1.1821 | none |
| LRRC59 | 0.066 | 0.1228 | 0.5798 | 0.57536 | 0.71 | -6.39 | 0.54514 | none |
| SDF2 | 0.07 | -0.0139 | 0.5795 | 0.57553 | 0.71 | -6.39 | -0.3829 | none |
| KBTBD13 | 0.041 | 0.0502 | 0.5793 | 0.57568 | 0.711 | -6.39 | 1.80324 | none |
| RXRG | -0.043 | -0.041 | -0.579 | 0.57582 | 0.711 | -6.39 | -0.6333 | none |
| RAD51C | 0.08 | -0.0731 | 0.5788 | 0.57598 | 0.711 | -6.4 | -0.216 | none |
| EXOC6 | 0.111 | 0.1305 | 0.5788 | 0.57598 | 0.711 | -6.4 | 0.45012 | none |
| CD101 | -0.122 | -0.0937 | -0.579 | 0.57612 | 0.711 | -6.4 | 0.1761 | none |
| AKAP12 | 0.061 | 0.1032 | 0.5785 | 0.5762 | 0.711 | -6.4 | 0.41452 | none |
| CDX1 | -0.041 | -0.0409 | -0.578 | 0.5762 | 0.711 | -6.4 | 0 | none |
| GSTA3 | -0.041 | -0.0409 | -0.578 | 0.57635 | 0.711 | -6.4 | 1.00425 | none |
| TRIM4 | -0.079 | -0.067 | -0.578 | 0.57637 | 0.711 | -6.4 | 1.21224 | none |
| ZNF300 | 0.043 | 0.0353 | 0.5777 | 0.5767 | 0.711 | -6.4 | 0.49812 | none |
| SLC39A10 | 0.15 | 0.2101 | 0.5774 | 0.57689 | 0.712 | -6.4 | 1.3306 | none |
| RNF148 | 0.041 | 0.0413 | 0.5769 | 0.57722 | 0.712 | -6.4 | 0 | none |
| DNAJA2 | 0.045 | 0.02 | 0.5767 | 0.57737 | 0.712 | -6.4 | 1.33447 | none |
| RAPGEF4 | 0.08 | 0.0672 | 0.5765 | 0.57748 | 0.712 | -6.4 | 0.10476 | none |
| TMEM63B | -0.093 | 0.0133 | -0.576 | 0.57787 | 0.713 | -6.4 | 1.22609 | none |
| MND1 | -0.068 | -0.0619 | -0.576 | 0.57803 | 0.713 | -6.4 | 1.00346 | none |
| SMIM1 | -0.09 | -0.0411 | -0.575 | 0.57812 | 0.713 | -6.4 | 1.46679 | none |
| C6ORF89 | -0.093 | -0.0759 | -0.575 | 0.57817 | 0.713 | -6.4 | 0.99553 | none |
| HOXD10 | -0.051 | -0.0509 | -0.575 | 0.57828 | 0.713 | -6.4 | -0.5339 | none |
| IQCD | -0.043 | -0.0164 | -0.575 | 0.57852 | 0.713 | -6.4 | 0.4781 | none |
| RHOD | -0.121 | -0.0876 | -0.575 | 0.57872 | 0.713 | -6.4 | 1.93708 | none |
| ASB2 | -0.08 | -0.0147 | -0.574 | 0.57879 | 0.713 | -6.4 | 0.61275 | none |
| C8ORF33 | 0.086 | 0.0591 | 0.5743 | 0.57892 | 0.713 | -6.4 | 0.77267 | none |
| DNAAF2 | 0.063 | 0.0609 | 0.5741 | 0.579 | 0.713 | -6.4 | 0.93509 | none |
| ZNRD1 | -0.212 | -0.3112 | -0.574 | 0.57927 | 0.714 | -6.4 | 1.28622 | none |
| DDX18 | 0.094 | 0.1198 | 0.5731 | 0.57966 | 0.714 | -6.4 | 0.78098 | none |
| COG6 | 0.125 | 0.1685 | 0.5726 | 0.57999 | 0.715 | -6.4 | 1.06036 | none |
| TSPAN9 | 0.069 | 0.0552 | 0.5725 | 0.58006 | 0.715 | -6.4 | -0.7719 | none |
| FGF21 | 0.058 | 0.0577 | 0.5722 | 0.58029 | 0.715 | -6.4 | 0.75344 | none |

| | | | | | | | | |
|---|---|---|---|---|---|---|---|---|
| AHSP | 0.049 | 0.0651 | 0.5721 | 0.58035 | 0.715 | -6.4 | -0.7714 | none |
| ARSI | 0.046 | 0.0455 | 0.5715 | 0.58075 | 0.715 | -6.4 | 0.20959 | none |
| FAM8A1 | 0.05 | 0.0094 | 0.5713 | 0.58084 | 0.715 | -6.4 | 0.38823 | none |
| PCDHB5 | -0.046 | -0.0473 | -0.571 | 0.5809 | 0.715 | -6.4 | -0.0304 | none |
| CD8A | 0.065 | -0.0013 | 0.5709 | 0.5811 | 0.716 | -6.4 | 0.3318 | none |
| C12ORF74 | 0.04 | 0.0281 | 0.5707 | 0.58124 | 0.716 | -6.4 | -0.1801 | none |
| RMI2 | -0.133 | -0.0858 | -0.571 | 0.58126 | 0.716 | -6.4 | 1.0273 | none |
| BFSP2 | -0.096 | 0.0133 | -0.571 | 0.58127 | 0.716 | -6.4 | 0.45036 | none |
| TMEM248 | 0.058 | 0.0752 | 0.5704 | 0.58141 | 0.716 | -6.4 | 1.56796 | none |
| AP1M2 | 0.229 | 0.1864 | 0.5701 | 0.58163 | 0.716 | -6.4 | -0.5049 | none |
| CAND2 | 0.064 | 0.1367 | 0.5701 | 0.58164 | 0.716 | -6.4 | 0.51799 | none |
| ANKRD30E | 0.095 | -0.0093 | 0.5699 | 0.58178 | 0.716 | -6.4 | -0.0395 | none |
| C10ORF10 | 0.04 | 0.0338 | 0.5698 | 0.58181 | 0.716 | -6.4 | -0.781 | none |
| C5ORF58 | 0.102 | 0.1268 | 0.5695 | 0.58199 | 0.716 | -6.4 | 1.0945 | none |
| BECN2 | 0.038 | 0.0416 | 0.5694 | 0.58206 | 0.716 | -6.4 | 0.77936 | none |
| SCCPDH | 0.072 | -0.0145 | 0.5694 | 0.5821 | 0.716 | -6.4 | 1.17477 | none |
| RALGAPB | 0.092 | 0.1504 | 0.5692 | 0.58225 | 0.716 | -6.4 | 0.19725 | none |
| CEP70 | 0.144 | 0.0968 | 0.569 | 0.58234 | 0.716 | -6.4 | -0.1503 | none |
| DUXA | -0.042 | -0.0265 | -0.569 | 0.58249 | 0.717 | -6.4 | 0.70421 | none |
| RSPRY1 | 0.069 | 0.068 | 0.5685 | 0.58268 | 0.717 | -6.4 | 0.55457 | none |
| TREX1 | -0.064 | -0.0454 | -0.568 | 0.58272 | 0.717 | -6.4 | 1.96592 | none |
| PSG3 | 0.04 | 0.0405 | 0.5682 | 0.58288 | 0.717 | -6.4 | -0.9207 | none |
| MT-CYB | 0.091 | 0.122 | 0.5679 | 0.58309 | 0.717 | -6.4 | 0.11246 | none |
| PPP1R9B | 0.145 | 0.2485 | 0.5678 | 0.58312 | 0.717 | -6.4 | 0.04711 | none |
| CSNK1G2 | 0.106 | 0.1979 | 0.5677 | 0.58317 | 0.717 | -6.4 | -0.0283 | none |
| BECN1 | 0.055 | 0.0312 | 0.5677 | 0.5832 | 0.717 | -6.4 | -0.9435 | none |
| GPR174 | -0.078 | -0.1958 | -0.568 | 0.58325 | 0.717 | -6.4 | -0.0278 | none |
| FZD5 | 0.043 | 0.0363 | 0.5676 | 0.58326 | 0.717 | -6.4 | 0.19178 | none |
| PHF20 | 0.078 | 0.0275 | 0.5675 | 0.58336 | 0.717 | -6.4 | 0.04846 | none |
| ZNF679 | 0.04 | 0.0404 | 0.5674 | 0.58337 | 0.717 | -6.4 | 0 | none |
| SNRNP27 | 0.065 | 0.0147 | 0.5671 | 0.58357 | 0.717 | -6.4 | -0.7916 | none |
| ZXDC | 0.11 | 0.2033 | 0.5667 | 0.58384 | 0.718 | -6.4 | 0.32468 | none |
| PI16 | 0.167 | -0.0342 | 0.5666 | 0.58389 | 0.718 | -6.4 | -0.2486 | none |
| SYVN1 | 0.066 | 0.1515 | 0.5665 | 0.58396 | 0.718 | -6.4 | 0.52892 | none |
| UHRF1BP1 | 0.13 | 0.2102 | 0.5664 | 0.58407 | 0.718 | -6.4 | 1.31296 | none |
| PEX6 | -0.053 | -0.0271 | -0.566 | 0.58411 | 0.718 | -6.4 | 1.12337 | none |
| IL12B | -0.041 | -0.0188 | -0.566 | 0.58415 | 0.718 | -6.4 | -0.3633 | none |
| TCP11L1 | 0.093 | 0.1219 | 0.5659 | 0.58437 | 0.718 | -6.4 | -0.0998 | none |
| CACNG1 | -0.045 | -0.0304 | -0.566 | 0.58438 | 0.718 | -6.4 | 1.93674 | none |
| TUBB8 | -0.061 | -0.0423 | -0.566 | 0.58444 | 0.718 | -6.4 | 1.20678 | none |

| | | | | | | | | |
|---|---|---|---|---|---|---|---|---|
| WDR54 | 0.116 | -0.002 | 0.5651 | 0.58487 | 0.718 | -6.4 | 0.14372 | none |
| PDCL2 | -0.04 | -0.0397 | -0.565 | 0.58492 | 0.718 | -6.4 | 0 | none |
| CLU | -0.121 | -0.1776 | -0.565 | 0.58505 | 0.718 | -6.4 | -0.1253 | none |
| BAIAP3 | 0.064 | 0.1179 | 0.5645 | 0.58526 | 0.719 | -6.4 | 1.10706 | none |
| CCR10 | -0.092 | -0.2294 | -0.565 | 0.58527 | 0.719 | -6.4 | 0.7708 | none |
| TMEM104 | 0.105 | 0.0844 | 0.5645 | 0.58528 | 0.719 | -6.4 | 0.34048 | none |
| ZNF735 | 0.042 | 0.0475 | 0.5643 | 0.5854 | 0.719 | -6.4 | 0.70873 | none |
| MYBPHL | 0.042 | 0.0531 | 0.5643 | 0.5854 | 0.719 | -6.4 | 0.03576 | none |
| LXN | -0.083 | -0.0904 | -0.564 | 0.58546 | 0.719 | -6.4 | 0.11097 | none |
| CLTCL1 | 0.085 | 0.1017 | 0.5637 | 0.58583 | 0.719 | -6.4 | 0.40271 | none |
| ACBD6 | -0.045 | 0.0094 | -0.564 | 0.58592 | 0.719 | -6.4 | 2.09406 | AA-related |
| NTPCR | -0.078 | -0.2102 | -0.563 | 0.586 | 0.719 | -6.4 | 1.28793 | none |
| XRCC5 | -0.06 | -0.0149 | -0.563 | 0.58601 | 0.719 | -6.4 | 0.17508 | none |
| MS4A13 | 0.083 | 0.083 | 0.5633 | 0.58608 | 0.719 | -6.4 | -0.4698 | none |
| HEATR5B | 0.203 | 0.2313 | 0.5631 | 0.58619 | 0.719 | -6.4 | 1.05239 | none |
| CTDSP1 | 0.05 | 0.0991 | 0.563 | 0.58625 | 0.719 | -6.4 | 1.01128 | none |
| METTL9 | -0.089 | -0.152 | -0.563 | 0.58636 | 0.719 | -6.4 | 0.70221 | none |
| BPY2 | 0.04 | 0.04 | 0.5626 | 0.58655 | 0.719 | -6.4 | -0.841 | none |
| BPY2B | 0.04 | 0.04 | 0.5626 | 0.58655 | 0.719 | -6.4 | -0.841 | none |
| BPY2C | 0.04 | 0.04 | 0.5626 | 0.58655 | 0.719 | -6.4 | -0.841 | none |
| DZIP1L | 0.08 | 0.1196 | 0.5623 | 0.58675 | 0.72 | -6.4 | 0.19136 | none |
| TNFRSF21 | 0.048 | 0.0658 | 0.562 | 0.58689 | 0.72 | -6.41 | -0.1495 | none |
| GALNTL6 | 0.039 | 0.0419 | 0.5618 | 0.58708 | 0.72 | -6.41 | 0 | none |
| ACER2 | 0.052 | 0.0017 | 0.5615 | 0.58723 | 0.72 | -6.41 | -0.2997 | none |
| TOE1 | -0.052 | 0.0074 | -0.562 | 0.58724 | 0.72 | -6.41 | 1.98229 | none |
| ATN1 | 0.138 | 0.1949 | 0.5614 | 0.58729 | 0.72 | -6.41 | -0.8366 | none |
| PDHB | -0.056 | -0.088 | -0.561 | 0.58745 | 0.72 | -6.41 | 0.69836 | none |
| ADO | 0.042 | 0.0246 | 0.5609 | 0.58763 | 0.72 | -6.41 | -0.13 | none |
| RASSF1 | 0.051 | 0.0825 | 0.5609 | 0.58767 | 0.72 | -6.41 | 1.21146 | none |
| HNRNPCL2 | -0.047 | -0.0473 | -0.561 | 0.58774 | 0.72 | -6.41 | 0.85531 | none |
| GZMM | -0.109 | -0.1623 | -0.561 | 0.58782 | 0.72 | -6.41 | 1.33527 | none |
| PLXNB2 | 0.06 | 0.0383 | 0.5606 | 0.58786 | 0.72 | -6.41 | -0.3914 | none |
| FSBP | 0.087 | 0.0581 | 0.5605 | 0.58789 | 0.72 | -6.41 | 0.07482 | none |
| RGS5 | 0.082 | -0.0623 | 0.5604 | 0.58795 | 0.72 | -6.41 | 1.47317 | none |
| ACSBG2 | 0.11 | 0.1011 | 0.5604 | 0.58798 | 0.72 | -6.41 | 1.29656 | none |
| TRMT61B | 0.08 | -0.0111 | 0.5604 | 0.58799 | 0.72 | -6.41 | 0.25511 | none |
| TEX28 | 0.039 | 0.0401 | 0.5603 | 0.58801 | 0.72 | -6.41 | 0.37129 | none |
| TPPP | 0.039 | 0.0672 | 0.5602 | 0.5881 | 0.72 | -6.41 | 0.67563 | none |
| SCGB3A2 | 0.111 | 0.2957 | 0.56 | 0.58821 | 0.72 | -6.41 | 1.12433 | none |
| PRB4 | 0.04 | 0.0397 | 0.56 | 0.58826 | 0.72 | -6.41 | 0 | none |

| | | | | | | | | |
|---|---|---|---|---|---|---|---|---|
| INS-IGF2 | 0.04 | 0.04 | 0.5599 | 0.58831 | 0.72 | -6.41 | 1.18529 | none |
| SERF1B | 0.103 | 0.0235 | 0.5599 | 0.58831 | 0.72 | -6.41 | -0.4596 | none |
| FASTK | -0.089 | -0.1299 | -0.56 | 0.5884 | 0.72 | -6.41 | 1.54898 | none |
| SGCD | 0.046 | 0.0454 | 0.5597 | 0.58841 | 0.72 | -6.41 | 0.80964 | none |
| SNRPA1 | -0.041 | -0.0384 | -0.56 | 0.58846 | 0.72 | -6.41 | 0.57242 | none |
| RHCG | 0.06 | 0.055 | 0.5596 | 0.58851 | 0.72 | -6.41 | 0.91729 | none |
| DNAJC11 | 0.115 | 0.2715 | 0.5596 | 0.58851 | 0.72 | -6.41 | -0.0126 | none |
| C11ORF96 | -0.046 | -0.0318 | -0.559 | 0.58879 | 0.721 | -6.41 | 0.31117 | none |
| GNAI2 | -0.104 | -0.0244 | -0.559 | 0.58886 | 0.721 | -6.41 | 0.54452 | none |
| EED | 0.066 | 0.0599 | 0.5587 | 0.5891 | 0.721 | -6.41 | -0.2181 | none |
| TNFRSF19 | 0.075 | 0.0749 | 0.5587 | 0.58911 | 0.721 | -6.41 | 0.39695 | none |
| IGSF21 | -0.043 | -0.0175 | -0.559 | 0.58912 | 0.721 | -6.41 | 1.80627 | none |
| HEMK1 | 0.095 | 0.0221 | 0.5585 | 0.58924 | 0.721 | -6.41 | 0.99853 | none |
| MSI1 | -0.045 | -0.0289 | -0.558 | 0.58925 | 0.721 | -6.41 | 0.41234 | none |
| GOLGA1 | -0.105 | -0.0022 | -0.558 | 0.58961 | 0.721 | -6.41 | 1.53715 | none |
| NAB1 | 0.112 | 0.1779 | 0.5579 | 0.58963 | 0.721 | -6.41 | 0.2009 | none |
| OBP2A | -0.074 | -0.0509 | -0.558 | 0.58963 | 0.721 | -6.41 | 1.56204 | none |
| TRIM39-RI | 0.04 | 0.0395 | 0.5578 | 0.58968 | 0.721 | -6.41 | 0 | none |
| CLDN34 | 0.039 | 0.0395 | 0.5571 | 0.59012 | 0.722 | -6.41 | 0.39591 | none |
| TMEM260 | 0.107 | 0.0671 | 0.557 | 0.5902 | 0.722 | -6.41 | 0.47618 | none |
| KHDC1L | -0.037 | -0.0159 | -0.557 | 0.59048 | 0.722 | -6.41 | -0.5982 | none |
| STT3A | 0.066 | -0.0226 | 0.5565 | 0.5905 | 0.722 | -6.41 | 0.73315 | none |
| ADSL | -0.076 | -0.1631 | -0.556 | 0.59072 | 0.722 | -6.41 | 1.38717 | none |
| PANK3 | 0.076 | 0.1265 | 0.5561 | 0.59081 | 0.722 | -6.41 | -0.0486 | none |
| IL26 | -0.047 | -0.029 | -0.556 | 0.59105 | 0.723 | -6.41 | 1.58142 | none |
| SET | 0.071 | 0.1063 | 0.5556 | 0.59113 | 0.723 | -6.41 | -0.1888 | none |
| HLA-F | 0.061 | -0.0661 | 0.5555 | 0.59117 | 0.723 | -6.41 | -0.3434 | none |
| FXYD6 | -0.119 | -0.0642 | -0.555 | 0.59125 | 0.723 | -6.41 | 0.92883 | none |
| SRPRA | 0.09 | 0.1795 | 0.5552 | 0.59138 | 0.723 | -6.41 | 0.69794 | none |
| MFSD14B | 0.089 | -0.0083 | 0.5552 | 0.59139 | 0.723 | -6.41 | -0.4007 | none |
| ADRB3 | -0.058 | -0.0481 | -0.555 | 0.59154 | 0.723 | -6.41 | 0.85725 | none |
| PPP1R18 | -0.044 | -0.0728 | -0.555 | 0.59166 | 0.723 | -6.41 | 1.35995 | none |
| EPYC | 0.04 | 0.0391 | 0.5548 | 0.59166 | 0.723 | -6.41 | -0.7614 | none |
| TMEM267 | 0.104 | -0.0034 | 0.5547 | 0.59169 | 0.723 | -6.41 | 0.78135 | none |
| SULT1C3 | -0.039 | -0.0387 | -0.555 | 0.59173 | 0.723 | -6.41 | 0 | none |
| ATP6V1C1 | -0.069 | -0.1099 | -0.554 | 0.59186 | 0.723 | -6.41 | 0.83296 | none |
| KCNIP1 | -0.038 | -0.0377 | -0.554 | 0.59191 | 0.723 | -6.41 | -0.693 | none |
| GLIPR1L1 | 0.045 | 0.0259 | 0.5544 | 0.59192 | 0.723 | -6.41 | 0.66407 | none |
| ACTRT1 | 0.039 | 0.0392 | 0.5544 | 0.59193 | 0.723 | -6.41 | 0 | none |
| DCAF7 | -0.041 | -0.0642 | -0.554 | 0.59195 | 0.723 | -6.41 | 1.20695 | none |

| | | | | | | | | |
|---|---|---|---|---|---|---|---|---|
| CLTC | -0.075 | -0.0236 | -0.554 | 0.59212 | 0.723 | -6.41 | 0.02232 | none |
| CCDC174 | 0.132 | 0.2115 | 0.5536 | 0.59247 | 0.723 | -6.41 | 0.62556 | none |
| BNC2 | 0.048 | 0.069 | 0.5532 | 0.59267 | 0.724 | -6.41 | -0.5683 | none |
| ANPEP | -0.12 | -0.0868 | -0.553 | 0.59302 | 0.724 | -6.41 | 0.31924 | none |
| ABCD1 | 0.087 | 0.1201 | 0.5527 | 0.59304 | 0.724 | -6.41 | 1.394 | none |
| RFC3 | -0.072 | -0.11 | -0.552 | 0.59322 | 0.724 | -6.41 | 1.58732 | none |
| AMIGO3 | -0.053 | -0.0627 | -0.552 | 0.59323 | 0.724 | -6.41 | 0.48259 | none |
| HLA-C | 0.101 | -0.0113 | 0.5522 | 0.59334 | 0.724 | -6.41 | 0.24808 | none |
| RNH1 | 0.048 | 0.062 | 0.5521 | 0.5934 | 0.724 | -6.41 | 0.73462 | none |
| VAMP3 | -0.055 | -0.0959 | -0.552 | 0.59347 | 0.724 | -6.41 | 0.42722 | none |
| CGB1 | 0.04 | 0.0389 | 0.552 | 0.59349 | 0.724 | -6.41 | 0.54348 | none |
| AXIN2 | 0.068 | 0.0719 | 0.5519 | 0.59357 | 0.724 | -6.41 | -0.4139 | none |
| LAGE3 | 0.088 | 0.0852 | 0.5518 | 0.59363 | 0.724 | -6.41 | 0.83909 | none |
| NAALADL1 | 0.07 | 0.0468 | 0.5518 | 0.59364 | 0.724 | -6.41 | 0.21773 | none |
| PGAM4 | -0.137 | -0.053 | -0.552 | 0.59368 | 0.724 | -6.41 | -0.0808 | none |
| PDIA4 | 0.112 | 0.2263 | 0.5511 | 0.59406 | 0.724 | -6.41 | 0.94296 | none |
| GRAPL | 0.084 | 0.1696 | 0.551 | 0.59413 | 0.724 | -6.41 | 0.99826 | none |
| OR2T35 | 0.04 | 0.0505 | 0.551 | 0.59415 | 0.724 | -6.41 | 0.65405 | none |
| GPC5 | 0.076 | 0.0417 | 0.5509 | 0.5942 | 0.724 | -6.41 | 0.11779 | none |
| CLDND1 | -0.095 | -0.1645 | -0.551 | 0.59422 | 0.724 | -6.41 | 0.4431 | none |
| FAM122B | 0.084 | -0.0222 | 0.5509 | 0.59424 | 0.724 | -6.41 | -0.7874 | none |
| VASN | -0.041 | -0.0388 | -0.551 | 0.59424 | 0.724 | -6.41 | 0.70724 | none |
| C1ORF185 | 0.039 | 0.0389 | 0.5509 | 0.59425 | 0.724 | -6.41 | 0.18059 | none |
| LHFPL3 | 0.037 | 0.0337 | 0.5507 | 0.59432 | 0.724 | -6.41 | -0.7712 | none |
| VAT1L | 0.052 | 0.055 | 0.5507 | 0.59434 | 0.724 | -6.41 | 1.00246 | none |
| PPCS | 0.078 | ######## | 0.5505 | 0.5945 | 0.725 | -6.41 | -0.4073 | none |
| PAX3 | 0.059 | 0.0549 | 0.5499 | 0.59488 | 0.725 | -6.41 | -0.7616 | none |
| ELOVL4 | -0.068 | -0.0804 | -0.55 | 0.59504 | 0.725 | -6.41 | 1.06922 | none |
| PLCB3 | 0.104 | 0.2032 | 0.5496 | 0.59506 | 0.725 | -6.41 | -0.2322 | none |
| ZNF625-ZN | 0.077 | 0.0509 | 0.5492 | 0.59536 | 0.725 | -6.41 | 0.61378 | none |
| PIFO | -0.049 | -0.0497 | -0.549 | 0.59549 | 0.726 | -6.41 | 0.04747 | none |
| BAMBI | -0.04 | -0.0439 | -0.549 | 0.5955 | 0.726 | -6.41 | 0.38472 | none |
| PNRC1 | 0.075 | -0.0269 | 0.5488 | 0.5956 | 0.726 | -6.41 | 0.13872 | none |
| BHLHA9 | 0.039 | 0.0387 | 0.5488 | 0.59563 | 0.726 | -6.41 | 0.20016 | none |
| EGLN2 | -0.042 | 0.0014 | -0.549 | 0.59567 | 0.726 | -6.41 | 1.10974 | none |
| CSDC2 | -0.08 | -0.0884 | -0.548 | 0.59592 | 0.726 | -6.41 | -0.3203 | none |
| LRIT3 | 0.039 | 0.0261 | 0.5482 | 0.596 | 0.726 | -6.41 | 0 | none |
| CFAP46 | -0.097 | -0.0873 | -0.548 | 0.59609 | 0.726 | -6.41 | 1.95025 | none |
| SPRY2 | 0.041 | 0.0299 | 0.5481 | 0.5961 | 0.726 | -6.41 | -0.4393 | none |
| AGAP9 | -0.061 | -0.0171 | -0.548 | 0.59619 | 0.726 | -6.41 | 0.88322 | none |

| | | | | | | | | |
|---|---|---|---|---|---|---|---|---|
| RBM28 | 0.129 | 0.2551 | 0.5477 | 0.59631 | 0.726 | -6.41 | 1.16326 | none |
| ECH1 | 0.053 | 0.0697 | 0.5477 | 0.59635 | 0.726 | -6.41 | -0.1184 | none |
| ATP6V1G3 | 0.05 | 0.0072 | 0.5473 | 0.59658 | 0.726 | -6.41 | 1.44766 | none |
| PDZRN3 | -0.1 | -0.0999 | -0.547 | 0.59661 | 0.726 | -6.41 | 0.05158 | none |
| XKR3 | 0.038 | 0.0386 | 0.5471 | 0.59672 | 0.726 | -6.41 | 0 | none |
| PSCA | 0.046 | 0.0554 | 0.547 | 0.5968 | 0.726 | -6.41 | -0.321 | none |
| DEFA4 | -0.038 | -0.038 | -0.547 | 0.59699 | 0.727 | -6.41 | -0.0522 | none |
| SEMA5A | 0.062 | 0.0208 | 0.5466 | 0.59706 | 0.727 | -6.41 | -0.702 | none |
| BCAS2 | -0.044 | -0.0505 | -0.547 | 0.59708 | 0.727 | -6.41 | 0.3495 | none |
| TMEM222 | -0.054 | -0.0352 | -0.546 | 0.59718 | 0.727 | -6.41 | 1.4368 | none |
| CST9 | -0.038 | -0.0262 | -0.546 | 0.59722 | 0.727 | -6.41 | 0 | none |
| LIMD2 | 0.066 | 0.0732 | 0.5462 | 0.5973 | 0.727 | -6.41 | 0.19892 | none |
| AGPAT4 | 0.071 | 0.0755 | 0.5462 | 0.59735 | 0.727 | -6.41 | 1.17786 | none |
| LSM8 | -0.125 | -0.1731 | -0.546 | 0.59751 | 0.727 | -6.41 | 1.02358 | none |
| ZNF768 | 0.07 | 0.1292 | 0.5458 | 0.59757 | 0.727 | -6.41 | 0.86419 | none |
| PRAMEF8 | 0.038 | 0.0386 | 0.5457 | 0.59764 | 0.727 | -6.41 | 0 | none |
| ANLN | 0.156 | 0.3295 | 0.5455 | 0.59781 | 0.727 | -6.41 | 0.73903 | none |
| LGALS3 | -0.092 | -0.18 | -0.545 | 0.59818 | 0.727 | -6.42 | 1.02702 | none |
| KIAA2013 | -0.082 | -0.0624 | -0.545 | 0.59821 | 0.727 | -6.42 | -0.2427 | none |
| ZNF235 | 0.11 | 0.0679 | 0.5445 | 0.59844 | 0.728 | -6.42 | -0.4391 | none |
| RLBP1 | -0.075 | -0.0571 | -0.544 | 0.59849 | 0.728 | -6.42 | -0.0267 | none |
| FAM110A | 0.072 | 0.0296 | 0.5444 | 0.59852 | 0.728 | -6.42 | 0.48327 | none |
| PLPPR3 | 0.105 | 0.1002 | 0.544 | 0.59879 | 0.728 | -6.42 | 0.36784 | none |
| TMEM254 | -0.069 | -0.1727 | -0.544 | 0.59883 | 0.728 | -6.42 | 0.61781 | none |
| TPGS2 | -0.063 | -0.0526 | -0.544 | 0.59904 | 0.728 | -6.42 | 0.66671 | none |
| P2RY1 | 0.042 | 0.0564 | 0.5433 | 0.59921 | 0.728 | -6.42 | 0.39563 | none |
| UHMK1 | 0.079 | -0.0264 | 0.5433 | 0.59922 | 0.728 | -6.42 | 1.07828 | none |
| ATP6V0A4 | 0.152 | 0.0542 | 0.5433 | 0.59922 | 0.728 | -6.42 | -0.2773 | none |
| PREB | 0.059 | 0.0215 | 0.5433 | 0.59925 | 0.728 | -6.42 | 1.15515 | none |
| PSMD4 | 0.051 | 0.0455 | 0.5432 | 0.59931 | 0.728 | -6.42 | -0.8701 | none |
| TAGAP | 0.07 | 0.0254 | 0.5431 | 0.59936 | 0.728 | -6.42 | 0.39681 | none |
| SESN1 | 0.075 | 0.0423 | 0.543 | 0.59944 | 0.728 | -6.42 | 0.78972 | none |
| GH1 | 0.04 | 0.038 | 0.5428 | 0.59961 | 0.728 | -6.42 | -0.1976 | none |
| LIMCH1 | 0.054 | 0.0519 | 0.5425 | 0.59977 | 0.728 | -6.42 | -0.1492 | none |
| TRPV4 | 0.052 | 0.0644 | 0.5425 | 0.5998 | 0.728 | -6.42 | -0.8484 | none |
| SPZ1 | 0.038 | 0.0381 | 0.5423 | 0.59993 | 0.729 | -6.42 | -0.9277 | none |
| SON | 0.14 | 0.2536 | 0.542 | 0.60013 | 0.729 | -6.42 | -0.1323 | none |
| NLRP5 | 0.09 | 0.0513 | 0.5417 | 0.60033 | 0.729 | -6.42 | 0.60867 | none |
| POLDIP3 | 0.048 | 0.0532 | 0.5416 | 0.60038 | 0.729 | -6.42 | 1.24716 | none |
| ACACA | 0.087 | 0.1188 | 0.5416 | 0.6004 | 0.729 | -6.42 | -0.2958 | none |

| | | | | | | | | |
|---|---|---|---|---|---|---|---|---|
| CD5L | 0.041 | 0.0521 | 0.5413 | 0.60055 | 0.729 | -6.42 | 1.84921 | none |
| ENOX1 | -0.048 | -0.0425 | -0.541 | 0.6006 | 0.729 | -6.42 | 0.53781 | none |
| RNF141 | -0.066 | -0.1089 | -0.541 | 0.60102 | 0.73 | -6.42 | 0.65662 | none |
| PSMG1 | -0.051 | -0.1135 | -0.54 | 0.60122 | 0.73 | -6.42 | 0.55057 | none |
| YJEFN3 | 0.068 | 0.1141 | 0.5402 | 0.60132 | 0.73 | -6.42 | -0.3477 | none |
| C1ORF115 | 0.039 | 0.0438 | 0.5401 | 0.60139 | 0.73 | -6.42 | 0.55665 | none |
| XYLT1 | 0.057 | 0.1114 | 0.5399 | 0.60149 | 0.73 | -6.42 | 0.70994 | none |
| FAM53A | -0.036 | -0.0231 | -0.54 | 0.60164 | 0.73 | -6.42 | 1.14919 | none |
| LATS2 | 0.051 | 0.1006 | 0.5393 | 0.60187 | 0.73 | -6.42 | -0.8133 | none |
| ZNF728 | -0.036 | -0.0157 | -0.539 | 0.60195 | 0.73 | -6.42 | 0.84376 | none |
| OVOL2 | -0.049 | -0.0208 | -0.539 | 0.60224 | 0.731 | -6.42 | -0.4903 | none |
| NOL7 | -0.053 | 0.0044 | -0.539 | 0.60234 | 0.731 | -6.42 | 1.18311 | none |
| OR5J2 | -0.037 | -0.0373 | -0.539 | 0.60235 | 0.731 | -6.42 | -0.2977 | none |
| OR6C70 | -0.037 | -0.0373 | -0.539 | 0.60235 | 0.731 | -6.42 | -0.5817 | none |
| CCL21 | 0.038 | 0.0378 | 0.5385 | 0.60246 | 0.731 | -6.42 | -0.8467 | none |
| AC233755 | 0.045 | 0.0594 | 0.5382 | 0.60265 | 0.731 | -6.42 | 1.19517 | none |
| PRRX1 | -0.062 | -0.0535 | -0.538 | 0.60266 | 0.731 | -6.42 | 0.87027 | none |
| IGHMBP2 | 0.112 | 0.2149 | 0.5378 | 0.60291 | 0.731 | -6.42 | 1.03565 | none |
| CCDC18 | 0.135 | 0.213 | 0.5375 | 0.60308 | 0.731 | -6.42 | 0.76354 | none |
| TEX13A | 0.038 | 0.0377 | 0.5372 | 0.60327 | 0.731 | -6.42 | 0 | none |
| KLHDC2 | 0.059 | -0.0227 | 0.537 | 0.60345 | 0.731 | -6.42 | 0.9949 | none |
| ACSM1 | -0.037 | -0.0396 | -0.537 | 0.60346 | 0.731 | -6.42 | -0.0859 | none |
| APOB | 0.054 | 0.0172 | 0.5369 | 0.60346 | 0.731 | -6.42 | -0.2766 | none |
| BRD9 | 0.065 | 0.1298 | 0.5359 | 0.60412 | 0.732 | -6.42 | 0.4818 | none |
| URGCP | 0.116 | 0.2224 | 0.5359 | 0.60414 | 0.732 | -6.42 | 0.63763 | none |
| ZNF385B | -0.058 | -0.0239 | -0.536 | 0.60416 | 0.732 | -6.42 | 0.65805 | none |
| METTL24 | -0.048 | -0.0305 | -0.536 | 0.60433 | 0.732 | -6.42 | -0.559 | none |
| CCDC138 | 0.065 | 0.086 | 0.5356 | 0.60436 | 0.732 | -6.42 | 0.38688 | none |
| ZDHHC11 | 0.112 | 0.1557 | 0.5354 | 0.6045 | 0.732 | -6.42 | 1.00466 | none |
| RPH3AL | 0.129 | 0.0385 | 0.5353 | 0.60458 | 0.732 | -6.42 | -0.4423 | none |
| H1FNT | 0.064 | 0.0794 | 0.535 | 0.60475 | 0.733 | -6.42 | 0.97052 | none |
| RTN4R | 0.094 | 0.0339 | 0.5348 | 0.60491 | 0.733 | -6.42 | 0.07208 | none |
| BIRC2 | 0.063 | 0.0219 | 0.5347 | 0.60493 | 0.733 | -6.42 | 0.58682 | none |
| ZNF830 | 0.082 | 0.0991 | 0.5347 | 0.60493 | 0.733 | -6.42 | 0.9556 | none |
| DEPTOR | -0.043 | -0.0305 | -0.535 | 0.60504 | 0.733 | -6.42 | 0.72538 | none |
| NCBP2L | 0.037 | 0.0377 | 0.5344 | 0.60518 | 0.733 | -6.42 | 0.72753 | none |
| GRIK5 | -0.039 | -0.0168 | -0.534 | 0.60525 | 0.733 | -6.42 | 1.60311 | none |
| ARL2BP | 0.087 | 0.0191 | 0.5338 | 0.60552 | 0.733 | -6.42 | 0.22917 | none |
| TEAD1 | -0.04 | -0.0349 | -0.533 | 0.60588 | 0.733 | -6.42 | -0.3773 | none |
| FGFBP1 | 0.037 | 0.0373 | 0.5333 | 0.60589 | 0.733 | -6.42 | 0 | none |

| | | | | | | | | |
|---|---|---|---|---|---|---|---|---|
| FMOD | -0.048 | -0.0656 | -0.533 | 0.60592 | 0.733 | -6.42 | 0.04663 | none |
| MED9 | 0.046 | 0.0284 | 0.5331 | 0.606 | 0.734 | -6.42 | -0.2655 | none |
| OR2B6 | 0.037 | 0.0384 | 0.5331 | 0.60604 | 0.734 | -6.42 | 0 | none |
| GEMIN7 | 0.051 | -0.0013 | 0.5329 | 0.60617 | 0.734 | -6.42 | -0.8237 | none |
| GPR101 | 0.038 | 0.04 | 0.5327 | 0.60626 | 0.734 | -6.42 | -0.2757 | none |
| RIMKLB | -0.133 | -0.1335 | -0.533 | 0.6063 | 0.734 | -6.42 | 1.03314 | none |
| TADA3 | 0.082 | 0.1044 | 0.5325 | 0.60642 | 0.734 | -6.42 | -0.0551 | none |
| SPOCK2 | -0.081 | -0.0233 | -0.532 | 0.60705 | 0.734 | -6.42 | 0.93197 | none |
| UGT1A6 | -0.055 | -0.0576 | -0.532 | 0.60706 | 0.734 | -6.42 | 1.20406 | none |
| BTBD17 | 0.037 | 0.0372 | 0.5313 | 0.60719 | 0.735 | -6.42 | -0.3669 | none |
| C7ORF25 | 0.124 | 0.0562 | 0.5313 | 0.60725 | 0.735 | -6.42 | -0.5162 | none |
| SORBS1 | 0.092 | 0.0824 | 0.5309 | 0.60748 | 0.735 | -6.42 | 0.04286 | none |
| SUCLG2 | -0.051 | -0.0429 | -0.531 | 0.60754 | 0.735 | -6.42 | 0.93453 | none |
| MOS | 0.037 | 0.0371 | 0.5308 | 0.60757 | 0.735 | -6.42 | -0.5995 | none |
| LMAN2 | -0.066 | -0.0623 | -0.53 | 0.60778 | 0.735 | -6.42 | -0.2463 | none |
| GP5 | 0.039 | 0.0408 | 0.5299 | 0.60814 | 0.735 | -6.42 | 0.97646 | none |
| SLC25A35 | -0.17 | -0.1086 | -0.529 | 0.60852 | 0.736 | -6.42 | 0.94345 | none |
| ZFP92 | 0.053 | 0.0938 | 0.5292 | 0.60864 | 0.736 | -6.42 | -0.156 | none |
| YBEY | -0.071 | -0.1511 | -0.529 | 0.60865 | 0.736 | -6.42 | 0.35093 | none |
| CYBB | -0.037 | -0.0463 | -0.529 | 0.60884 | 0.736 | -6.42 | 0 | none |
| OR2G2 | -0.036 | -0.0365 | -0.529 | 0.60887 | 0.736 | -6.42 | 0 | none |
| SEMA3E | -0.212 | -0.1506 | -0.529 | 0.60899 | 0.736 | -6.42 | 1.58348 | none |
| SOWAHD | 0.04 | 0.0403 | 0.5284 | 0.60913 | 0.736 | -6.42 | -0.2449 | none |
| KRTAP10-8 | 0.043 | 0.015 | 0.5284 | 0.60914 | 0.736 | -6.42 | 1.06669 | none |
| RAB1A | -0.069 | -0.1286 | -0.528 | 0.60924 | 0.736 | -6.42 | 0.91938 | none |
| OR51H1 | -0.036 | -0.0364 | -0.528 | 0.60931 | 0.736 | -6.42 | 0 | none |
| TCEAL6 | 0.037 | 0.0369 | 0.528 | 0.60944 | 0.736 | -6.42 | -0.1349 | none |
| BFAR | 0.097 | 0.0176 | 0.5279 | 0.6095 | 0.736 | -6.42 | 0.36482 | none |
| GMPPA | -0.062 | -0.0711 | -0.528 | 0.60957 | 0.736 | -6.42 | 0.08398 | none |
| TTYH3 | -0.07 | 0.0188 | -0.528 | 0.60969 | 0.737 | -6.42 | 0.97457 | none |
| KRTAP4-5 | 0.037 | 0.0368 | 0.5273 | 0.60987 | 0.737 | -6.42 | 1.61682 | none |
| OGFR | 0.095 | 0.2171 | 0.5273 | 0.60987 | 0.737 | -6.42 | 0.20554 | none |
| GMPPB | 0.067 | 0.049 | 0.5273 | 0.6099 | 0.737 | -6.42 | 1.03064 | none |
| C17ORF47 | -0.037 | -0.022 | -0.527 | 0.61015 | 0.737 | -6.43 | 0 | none |
| EPHA8 | -0.037 | -0.019 | -0.527 | 0.6102 | 0.737 | -6.43 | 1.31985 | none |
| CATSPER4 | 0.034 | 0.04 | 0.5268 | 0.61023 | 0.737 | -6.43 | -0.2432 | none |
| ATP6V0D1 | -0.065 | -0.1319 | -0.527 | 0.61026 | 0.737 | -6.43 | 1.15616 | none |
| GATA3 | 0.073 | 0.0919 | 0.5261 | 0.61066 | 0.737 | -6.43 | 0.25681 | none |
| DNASE1L2 | -0.038 | -0.0033 | -0.526 | 0.61071 | 0.737 | -6.43 | 0.20117 | none |
| KLK4 | 0.056 | 0.0797 | 0.5259 | 0.61086 | 0.737 | -6.43 | 0.99901 | none |

| | | | | | | | | |
|---|---|---|---|---|---|---|---|---|
| CCDC173 | 0.042 | 0.0608 | 0.5257 | 0.61097 | 0.737 | -6.43 | 0.75282 | none |
| CHKB-CPT: | 0.117 | 0.0789 | 0.5256 | 0.61105 | 0.738 | -6.43 | -0.0372 | none |
| TMEM234 | -0.096 | -0.0074 | -0.525 | 0.61123 | 0.738 | -6.43 | 0.97226 | none |
| CGB3 | 0.075 | 0.0816 | 0.5252 | 0.6113 | 0.738 | -6.43 | -0.5962 | none |
| RNF187 | 0.045 | 0.0465 | 0.5251 | 0.61136 | 0.738 | -6.43 | 0.51516 | none |
| ARSH | 0.037 | 0.0366 | 0.5249 | 0.61147 | 0.738 | -6.43 | 0 | none |
| SFRP1 | 0.038 | 0.0487 | 0.5246 | 0.61171 | 0.738 | -6.43 | 0 | none |
| PIK3R4 | -0.082 | 0.0452 | -0.525 | 0.61175 | 0.738 | -6.43 | 1.83515 | none |
| MT-CO2 | 0.053 | 0.0647 | 0.5245 | 0.61175 | 0.738 | -6.43 | 0.47333 | none |
| MARVELD: | -0.072 | -0.0296 | -0.524 | 0.61221 | 0.739 | -6.43 | -0.2336 | none |
| COL6A6 | 0.038 | 0.0385 | 0.5234 | 0.61248 | 0.739 | -6.43 | 0.91899 | none |
| ZSWIM5 | 0.069 | 0.0818 | 0.5233 | 0.61259 | 0.739 | -6.43 | 1.26255 | none |
| CCDC126 | -0.092 | -0.1297 | -0.523 | 0.6126 | 0.739 | -6.43 | -0.3819 | none |
| CD200R1 | 0.045 | 0.004 | 0.5231 | 0.61271 | 0.739 | -6.43 | 0.90692 | none |
| HSD3B1 | 0.091 | 0.1182 | 0.523 | 0.6128 | 0.739 | -6.43 | 1.48764 | none |
| TCF7L2 | -0.175 | -0.1372 | -0.523 | 0.61283 | 0.739 | -6.43 | 0.46187 | none |
| RGSL1 | 0.095 | 0.1239 | 0.5227 | 0.61296 | 0.739 | -6.43 | 0.99497 | none |
| PTGER3 | -0.056 | -0.0845 | -0.523 | 0.613 | 0.739 | -6.43 | 0.47129 | none |
| ZNF469 | 0.035 | 0.0576 | 0.5225 | 0.61309 | 0.739 | -6.43 | -0.0051 | none |
| KRT82 | 0.035 | 0.0411 | 0.5221 | 0.61339 | 0.739 | -6.43 | 0 | none |
| AC104389 | 0.036 | 0.0363 | 0.5217 | 0.61362 | 0.74 | -6.43 | 0 | none |
| BIN3 | -0.071 | -0.0172 | -0.522 | 0.61374 | 0.74 | -6.43 | 1.40544 | none |
| SRRM4 | 0.138 | 0.1169 | 0.5211 | 0.61402 | 0.74 | -6.43 | -0.0022 | none |
| HLA-B | -0.059 | -0.1202 | -0.521 | 0.61408 | 0.74 | -6.43 | 0.49025 | none |
| OR51T1 | -0.036 | -0.0358 | -0.52 | 0.61452 | 0.74 | -6.43 | 0 | none |
| GP9 | 0.036 | 0.0362 | 0.5204 | 0.61453 | 0.74 | -6.43 | -0.6649 | none |
| LEXM | -0.039 | -0.0444 | -0.52 | 0.61461 | 0.74 | -6.43 | 1.97131 | none |
| EOMES | 0.042 | 0.0487 | 0.5202 | 0.61462 | 0.74 | -6.43 | 0.83261 | none |
| OR52E8 | 0.036 | 0.0362 | 0.52 | 0.61479 | 0.741 | -6.43 | 0 | none |
| CCDC17 | -0.09 | 0.0153 | -0.52 | 0.61485 | 0.741 | -6.43 | 1.70873 | none |
| UTS2 | -0.113 | -0.0989 | -0.52 | 0.61485 | 0.741 | -6.43 | 0.89973 | none |
| ZFP91-CN1 | 0.091 | 0.1963 | 0.5197 | 0.61501 | 0.741 | -6.43 | 0.0707 | none |
| CST8 | 0.036 | 0.0361 | 0.5196 | 0.61502 | 0.741 | -6.43 | 0 | none |
| CLEC18B | 0.121 | 0.1092 | 0.5195 | 0.6151 | 0.741 | -6.43 | 0.99591 | none |
| WDFY1 | 0.051 | 0.1215 | 0.5193 | 0.61525 | 0.741 | -6.43 | 0.60936 | none |
| FOXD4L3 | -0.034 | -0.0116 | -0.519 | 0.61528 | 0.741 | -6.43 | 0 | none |
| KLHL29 | -0.035 | -0.0226 | -0.519 | 0.61541 | 0.741 | -6.43 | 0 | none |
| ITGA8 | -0.072 | -0.0648 | -0.519 | 0.61549 | 0.741 | -6.43 | 1.33749 | none |
| SIGLEC7 | 0.083 | 0.033 | 0.5189 | 0.6155 | 0.741 | -6.43 | 0.89368 | none |
| NEU1 | -0.102 | -0.1368 | -0.519 | 0.61551 | 0.741 | -6.43 | 0.19437 | none |

| | | | | | | | | |
|---|---|---|---|---|---|---|---|---|
| GAS2 | -0.072 | -0.0474 | -0.519 | 0.61557 | 0.741 | -6.43 | -0.1346 | none |
| UCKL1 | 0.053 | 0.0473 | 0.5188 | 0.61559 | 0.741 | -6.43 | 0.355 | none |
| IL12RB2 | 0.11 | 0.1863 | 0.5188 | 0.6156 | 0.741 | -6.43 | -0.0532 | none |
| C15ORF65 | -0.051 | 0.0116 | -0.518 | 0.61613 | 0.741 | -6.43 | 0.60808 | none |
| CT45A2 | 0.036 | 0.036 | 0.5175 | 0.61648 | 0.742 | -6.43 | 1.26503 | none |
| TRIM44 | -0.09 | 0.0249 | -0.517 | 0.61648 | 0.742 | -6.43 | 1.00697 | none |
| PISD | 0.044 | 0.0922 | 0.5174 | 0.6165 | 0.742 | -6.43 | -0.0582 | none |
| AP2M1 | -0.054 | 0.0107 | -0.517 | 0.61654 | 0.742 | -6.43 | 1.43168 | none |
| IMMP1L | 0.065 | -0.0779 | 0.5173 | 0.6166 | 0.742 | -6.43 | -0.2789 | none |
| TGFB3 | 0.096 | 0.1011 | 0.5173 | 0.61661 | 0.742 | -6.43 | 0.81479 | none |
| LMNB2 | 0.096 | 0.2022 | 0.5172 | 0.61668 | 0.742 | -6.43 | 0.32229 | none |
| VIP | 0.041 | 0.0564 | 0.5169 | 0.61687 | 0.742 | -6.43 | -0.0049 | none |
| RIPPLY1 | -0.04 | -0.0146 | -0.517 | 0.61702 | 0.742 | -6.43 | -0.3799 | none |
| ATP6V1G1 | -0.058 | -0.1414 | -0.517 | 0.61706 | 0.742 | -6.43 | -0.3164 | none |
| GATA4 | 0.053 | 0.0534 | 0.5164 | 0.61721 | 0.742 | -6.43 | -0.1723 | none |
| ARNTL | 0.056 | 0.0704 | 0.5161 | 0.61739 | 0.742 | -6.43 | 0.94594 | none |
| PRTN3 | 0.038 | 0.0486 | 0.516 | 0.61747 | 0.742 | -6.43 | 1.65452 | none |
| LSMEM1 | 0.084 | 0.0651 | 0.5159 | 0.61752 | 0.742 | -6.43 | 0.56936 | none |
| DEGS2 | 0.058 | 0.0522 | 0.5159 | 0.61752 | 0.742 | -6.43 | 1.74492 | none |
| RAB11FIP5 | -0.047 | -0.0074 | -0.516 | 0.61757 | 0.742 | -6.43 | 0.98603 | none |
| TRAK1 | 0.109 | 0.2531 | 0.5157 | 0.61764 | 0.742 | -6.43 | -0.2138 | none |
| CLUH | 0.129 | 0.2905 | 0.5156 | 0.61772 | 0.742 | -6.43 | -0.2249 | none |
| TRAPPC13 | -0.082 | -0.0534 | -0.516 | 0.61779 | 0.742 | -6.43 | 1.26043 | none |
| AANAT | 0.083 | 0.0342 | 0.5148 | 0.61829 | 0.743 | -6.43 | -0.1393 | none |
| WNT11 | 0.103 | 0.121 | 0.5147 | 0.61831 | 0.743 | -6.43 | 0.63143 | none |
| LRIG1 | 0.105 | 0.2752 | 0.5147 | 0.61833 | 0.743 | -6.43 | 0.11528 | none |
| ANP32D | 0.036 | 0.0357 | 0.5147 | 0.61834 | 0.743 | -6.43 | 0.46363 | none |
| SLC10A4 | 0.043 | 0.0582 | 0.5147 | 0.61836 | 0.743 | -6.43 | -0.601 | none |
| MPV17 | 0.064 | -0.02 | 0.5145 | 0.61849 | 0.743 | -6.43 | 0.02488 | none |
| IL19 | -0.039 | -0.0503 | -0.514 | 0.61855 | 0.743 | -6.43 | 1.2551 | none |
| IFNA4 | -0.035 | -0.0352 | -0.514 | 0.61874 | 0.743 | -6.43 | 0 | none |
| FETUB | 0.036 | 0.0356 | 0.5134 | 0.61924 | 0.744 | -6.43 | -0.7372 | none |
| RS1 | 0.05 | 0.0502 | 0.5133 | 0.61925 | 0.744 | -6.43 | -0.0707 | none |
| OR13C4 | 0.036 | 0.0356 | 0.5133 | 0.61931 | 0.744 | -6.43 | 0 | none |
| ARL8B | -0.052 | -0.0733 | -0.513 | 0.6195 | 0.744 | -6.43 | 0.68595 | none |
| DACT2 | 0.035 | 0.0178 | 0.5129 | 0.61956 | 0.744 | -6.43 | 0.52905 | none |
| TIMMDC1 | -0.048 | -0.1064 | -0.513 | 0.61973 | 0.744 | -6.43 | 0.21846 | none |
| ATP2B2 | 0.035 | 0.0363 | 0.5124 | 0.61988 | 0.744 | -6.43 | -0.0794 | none |
| ATAD3A | 0.074 | 0.1726 | 0.5123 | 0.61993 | 0.744 | -6.43 | 0.01303 | none |
| SLA | 0.09 | 0.1864 | 0.5123 | 0.61995 | 0.744 | -6.43 | -0.4948 | none |

| | | | | | | | | |
|---|---|---|---|---|---|---|---|---|
| CCDC181 | 0.038 | 0.0465 | 0.5123 | 0.61996 | 0.744 | -6.43 | 0.32171 | none |
| MADCAM1 | 0.04 | 0.0589 | 0.5121 | 0.62006 | 0.744 | -6.43 | 0.73885 | none |
| MSANTD3 | 0.046 | 0.0541 | 0.5117 | 0.62038 | 0.744 | -6.43 | 1.25279 | none |
| AMPH | -0.122 | -0.1271 | -0.512 | 0.62039 | 0.744 | -6.43 | 0.75205 | none |
| GRSF1 | -0.049 | -0.051 | -0.512 | 0.62045 | 0.744 | -6.43 | 1.16584 | none |
| ARMCX1 | -0.063 | -0.0388 | -0.511 | 0.62052 | 0.745 | -6.43 | 1.3525 | none |
| SAT2 | -0.076 | -0.112 | -0.511 | 0.62067 | 0.745 | -6.43 | 0.61906 | none |
| RERGL | -0.039 | -0.023 | -0.511 | 0.62086 | 0.745 | -6.43 | 1.59989 | none |
| ZNF518A | -0.118 | 0.0048 | -0.511 | 0.62103 | 0.745 | -6.43 | 0.78889 | none |
| GALNT13 | 0.059 | 0.0447 | 0.5107 | 0.62103 | 0.745 | -6.43 | -0.5707 | none |
| ZBTB7A | 0.113 | 0.269 | 0.5104 | 0.62121 | 0.745 | -6.43 | 0.67358 | none |
| LRRC26 | 0.041 | 0.0273 | 0.51 | 0.62154 | 0.745 | -6.43 | 0.52775 | none |
| CP | -0.076 | -0.1098 | -0.51 | 0.62162 | 0.745 | -6.43 | 0.24027 | none |
| UNC119B | 0.069 | 0.0677 | 0.5097 | 0.62174 | 0.746 | -6.43 | 0.12751 | none |
| OR13H1 | 0.035 | 0.0353 | 0.5096 | 0.62179 | 0.746 | -6.43 | 0 | none |
| OR1K1 | 0.04 | 0.0144 | 0.5096 | 0.6218 | 0.746 | -6.43 | 1.08183 | none |
| FASTKD5 | -0.035 | -0.0396 | -0.509 | 0.62206 | 0.746 | -6.43 | 1.08037 | none |
| CACNA1C | -0.048 | -0.0355 | -0.509 | 0.62216 | 0.746 | -6.43 | 1.84687 | none |
| SLC39A4 | 0.05 | 0.0446 | 0.509 | 0.62216 | 0.746 | -6.43 | -0.2898 | none |
| DDX39B | -0.069 | -0.0706 | -0.509 | 0.62223 | 0.746 | -6.43 | 1.82217 | none |
| IL36RN | 0.076 | 0.0884 | 0.5088 | 0.62229 | 0.746 | -6.43 | 0.55974 | none |
| PPP1R1C | 0.099 | 0.1178 | 0.5086 | 0.62243 | 0.746 | -6.44 | 0.21546 | none |
| FHL3 | 0.056 | 0.1297 | 0.5085 | 0.62249 | 0.746 | -6.44 | 0.6623 | none |
| CHST7 | 0.062 | 0.0753 | 0.5084 | 0.62259 | 0.746 | -6.44 | 0.50134 | none |
| FAM155B | 0.037 | 0.0609 | 0.5084 | 0.62262 | 0.746 | -6.44 | 0 | none |
| DVL1 | -0.099 | -0.0167 | -0.508 | 0.6227 | 0.746 | -6.44 | 0.8132 | none |
| HCN2 | -0.034 | -0.0207 | -0.508 | 0.62293 | 0.746 | -6.44 | 0 | none |
| DYRK4 | -0.068 | -0.1408 | -0.508 | 0.62302 | 0.746 | -6.44 | 0.75495 | none |
| CD79A | -0.058 | -0.0495 | -0.508 | 0.62304 | 0.746 | -6.44 | 0.59002 | none |
| MMP26 | -0.035 | -0.0347 | -0.507 | 0.6232 | 0.746 | -6.44 | 0 | none |
| OPN1SW | 0.05 | 0.0534 | 0.5075 | 0.62321 | 0.746 | -6.44 | -0.7332 | none |
| CDH19 | 0.047 | 0.0376 | 0.5067 | 0.62377 | 0.747 | -6.44 | 0.22565 | none |
| DDX49 | 0.072 | 0.1473 | 0.5064 | 0.62397 | 0.747 | -6.44 | 0.38115 | none |
| CBWD1 | 0.058 | ######## | 0.5063 | 0.62401 | 0.747 | -6.44 | -0.5501 | none |
| TNFSF10 | -0.073 | -0.0767 | -0.506 | 0.62404 | 0.747 | -6.44 | -0.3645 | none |
| ATP6V1E2 | -0.11 | -0.0456 | -0.506 | 0.62405 | 0.747 | -6.44 | 0.87381 | none |
| ZMYND15 | 0.044 | 0.035 | 0.5062 | 0.62406 | 0.747 | -6.44 | 1.36899 | none |
| ZC4H2 | 0.087 | 0.0162 | 0.5061 | 0.62414 | 0.747 | -6.44 | 0.17012 | none |
| HIST2H3D | -0.12 | -0.1344 | -0.506 | 0.62424 | 0.747 | -6.44 | 1.71303 | none |
| IGF2BP2 | 0.077 | 0.0739 | 0.5057 | 0.62439 | 0.747 | -6.44 | 1.71896 | none |

| | | | | | | | | |
|---|---|---|---|---|---|---|---|---|
| UCN3 | -0.035 | -0.0345 | -0.506 | 0.62441 | 0.747 | -6.44 | 0 | none |
| MYO3A | 0.043 | 0.0447 | 0.5056 | 0.62445 | 0.747 | -6.44 | 0.44406 | none |
| C1QC | 0.035 | 0.0349 | 0.5055 | 0.62453 | 0.747 | -6.44 | -0.6576 | none |
| TTR | 0.103 | 0.0452 | 0.5052 | 0.62478 | 0.748 | -6.44 | -0.5152 | none |
| CCDC43 | -0.058 | -0.0395 | -0.505 | 0.62509 | 0.748 | -6.44 | 0.71001 | none |
| PXDNL | -0.037 | -0.0189 | -0.505 | 0.62512 | 0.748 | -6.44 | 0.62377 | none |
| KCNF1 | 0.038 | 0.0462 | 0.5046 | 0.62513 | 0.748 | -6.44 | -0.1667 | none |
| SC5D | 0.093 | 0.0225 | 0.5045 | 0.62522 | 0.748 | -6.44 | 0.81625 | none |
| CRHBP | -0.102 | -0.0758 | -0.504 | 0.62526 | 0.748 | -6.44 | 0.1097 | none |
| KY | 0.057 | 0.0629 | 0.5043 | 0.62538 | 0.748 | -6.44 | 0.87267 | none |
| ATG5 | -0.047 | -0.069 | -0.504 | 0.62549 | 0.748 | -6.44 | 1.61267 | none |
| PSMB11 | -0.034 | -0.016 | -0.504 | 0.62559 | 0.748 | -6.44 | 1.82105 | none |
| COL8A2 | -0.035 | -0.0268 | -0.504 | 0.62562 | 0.748 | -6.44 | 0 | none |
| SBSN | 0.08 | 0.0796 | 0.5037 | 0.62579 | 0.748 | -6.44 | 0.43345 | none |
| MRPS21 | -0.063 | -0.1139 | -0.504 | 0.62587 | 0.748 | -6.44 | 0.37886 | none |
| PIGV | -0.073 | -0.1212 | -0.503 | 0.62592 | 0.748 | -6.44 | 1.73766 | none |
| FRS2 | 0.06 | 0.0548 | 0.5035 | 0.62592 | 0.748 | -6.44 | 1.51107 | none |
| POU3F1 | -0.034 | -0.0148 | -0.503 | 0.62593 | 0.748 | -6.44 | 0.54825 | none |
| RBM18 | -0.101 | -0.1276 | -0.503 | 0.62617 | 0.748 | -6.44 | 1.5969 | none |
| C17ORF10 | 0.079 | 0.0147 | 0.503 | 0.62627 | 0.749 | -6.44 | -0.3112 | none |
| ZC2HC1B | -0.037 | -0.0077 | -0.503 | 0.62644 | 0.749 | -6.44 | -0.2141 | none |
| GSC2 | 0.035 | 0.0274 | 0.5027 | 0.62647 | 0.749 | -6.44 | 0.19512 | none |
| DUSP1 | -0.061 | -0.0669 | -0.503 | 0.62647 | 0.749 | -6.44 | 0.24976 | none |
| OR2A42 | -0.035 | -0.0187 | -0.502 | 0.62706 | 0.749 | -6.44 | 0.98324 | none |
| EPHA3 | 0.036 | 0.0298 | 0.5017 | 0.62713 | 0.749 | -6.44 | 0.37349 | none |
| CEBPD | -0.037 | -0.0446 | -0.501 | 0.62739 | 0.75 | -6.44 | 1.12509 | none |
| FAM107B | 0.047 | ######## | 0.5012 | 0.62745 | 0.75 | -6.44 | 0.037 | none |
| MIDN | 0.088 | 0.1425 | 0.5011 | 0.6275 | 0.75 | -6.44 | -0.0202 | none |
| OSCP1 | 0.127 | 0.0306 | 0.5006 | 0.62784 | 0.75 | -6.44 | 0.86031 | none |
| EIF4A3 | -0.045 | -0.0194 | -0.501 | 0.62785 | 0.75 | -6.44 | 1.71553 | none |
| TMPRSS6 | 0.085 | 0.1037 | 0.5004 | 0.62797 | 0.75 | -6.44 | 0.42319 | none |
| ST13 | -0.047 | -0.0154 | -0.5 | 0.628 | 0.75 | -6.44 | 1.02332 | none |
| RNASE1 | -0.064 | -0.064 | -0.5 | 0.62826 | 0.75 | -6.44 | 0.70485 | none |
| 4-Sep | 0.097 | 0.0484 | 0.4999 | 0.62833 | 0.75 | -6.44 | -0.5616 | none |
| EPDR1 | -0.057 | -0.0548 | -0.5 | 0.62833 | 0.75 | -6.44 | 0.01656 | none |
| OR6Q1 | -0.034 | -0.0341 | -0.5 | 0.62844 | 0.75 | -6.44 | 0 | none |
| PCID2 | -0.049 | -0.0993 | -0.5 | 0.62851 | 0.75 | -6.44 | 0.29484 | none |
| IAH1 | -0.067 | -0.1395 | -0.5 | 0.62859 | 0.75 | -6.44 | 0.6149 | none |
| MCRS1 | 0.061 | 0.0392 | 0.4992 | 0.6288 | 0.751 | -6.44 | 0.54249 | none |
| EIF3I | -0.037 | -0.0833 | -0.499 | 0.62896 | 0.751 | -6.44 | 0.46827 | none |

| | | | | | | | | |
|---|---|---|---|---|---|---|---|---|
| THSD4 | 0.058 | 0.0173 | 0.4984 | 0.62937 | 0.751 | -6.44 | -0.5852 | none |
| FDX1 | -0.089 | -0.1181 | -0.498 | 0.62943 | 0.751 | -6.44 | 0.99139 | none |
| ICAM5 | 0.043 | 0.0625 | 0.4982 | 0.62953 | 0.751 | -6.44 | 0.60176 | none |
| ELAVL4 | -0.04 | -0.0404 | -0.498 | 0.62953 | 0.751 | -6.44 | 0.34731 | none |
| MRM2 | 0.05 | 0.1086 | 0.498 | 0.62964 | 0.751 | -6.44 | 0.9228 | none |
| ZNF77 | 0.069 | 0.0589 | 0.4977 | 0.62983 | 0.751 | -6.44 | -0.3642 | none |
| CMPK2 | -0.054 | -0.0735 | -0.498 | 0.6299 | 0.751 | -6.44 | 0.24293 | none |
| ST6GALNA | 0.062 | -0.0438 | 0.4974 | 0.63002 | 0.752 | -6.44 | 0.93741 | none |
| NR4A1 | 0.056 | 0.1142 | 0.497 | 0.63029 | 0.752 | -6.44 | 0.17298 | none |
| ADCY5 | -0.095 | -0.0584 | -0.497 | 0.63043 | 0.752 | -6.44 | 1.17116 | none |
| MTMR11 | 0.066 | 0.0193 | 0.4966 | 0.63059 | 0.752 | -6.44 | 1.19734 | none |
| SRP9 | -0.078 | -0.1431 | -0.497 | 0.63062 | 0.752 | -6.44 | 0.18321 | none |
| SNN | 0.042 | 0.0241 | 0.4964 | 0.63069 | 0.752 | -6.44 | 0.06947 | none |
| ICE2 | 0.088 | 0.0974 | 0.4964 | 0.63069 | 0.752 | -6.44 | 0.65192 | none |
| OR8K5 | 0.034 | 0.0342 | 0.4963 | 0.63076 | 0.752 | -6.44 | 0 | none |
| SLC25A47 | -0.032 | -0.0141 | -0.496 | 0.63081 | 0.752 | -6.44 | -0.2453 | none |
| MRPL23 | -0.088 | -0.0639 | -0.496 | 0.63082 | 0.752 | -6.44 | 0.07118 | none |
| MYL9 | 0.047 | 0.0775 | 0.4962 | 0.63083 | 0.752 | -6.44 | 1.02633 | none |
| LHX9 | -0.079 | -0.0902 | -0.496 | 0.631 | 0.752 | -6.44 | 0.53933 | none |
| THEG | -0.039 | -0.0096 | -0.496 | 0.63106 | 0.752 | -6.44 | -0.7151 | none |
| IFNA5 | 0.034 | 0.0341 | 0.4956 | 0.63125 | 0.752 | -6.44 | 0 | none |
| KCNIP2 | 0.082 | 0.1305 | 0.4954 | 0.63138 | 0.752 | -6.44 | 0.45998 | none |
| C8G | 0.133 | 0.1435 | 0.4953 | 0.63148 | 0.752 | -6.44 | -0.3016 | none |
| LARP7 | 0.124 | 0.2699 | 0.4951 | 0.63164 | 0.753 | -6.44 | 0.46819 | none |
| TNFAIP2 | 0.118 | 0.0716 | 0.4947 | 0.6319 | 0.753 | -6.44 | -0.4371 | none |
| PGM3 | 0.11 | 0.1433 | 0.4944 | 0.6321 | 0.753 | -6.44 | 1.47188 | none |
| AMOT | -0.062 | -0.0561 | -0.494 | 0.63211 | 0.753 | -6.44 | 0.91556 | none |
| C3ORF14 | -0.11 | -0.221 | -0.494 | 0.63229 | 0.753 | -6.44 | 0.19035 | none |
| TBC1D26 | 0.05 | 0.0721 | 0.4939 | 0.63239 | 0.753 | -6.44 | -0.4524 | none |
| VSTM2L | -0.036 | -0.0486 | -0.494 | 0.63243 | 0.753 | -6.44 | 0.25943 | none |
| RAB3IL1 | -0.079 | -0.0251 | -0.494 | 0.63258 | 0.753 | -6.44 | 0.0091 | none |
| C12ORF54 | 0.068 | 0.0538 | 0.4936 | 0.63264 | 0.753 | -6.44 | 1.28249 | none |
| IFI27 | -0.141 | -0.3015 | -0.493 | 0.63273 | 0.753 | -6.44 | 0.09332 | none |
| LRTM2 | -0.042 | -0.0298 | -0.493 | 0.63288 | 0.754 | -6.44 | 1.57682 | none |
| IL27RA | 0.078 | 0.159 | 0.4932 | 0.63293 | 0.754 | -6.44 | 1.13807 | none |
| C17ORF67 | -0.066 | -0.0653 | -0.493 | 0.63317 | 0.754 | -6.44 | 0.27086 | none |
| NIT1 | -0.075 | -0.0856 | -0.493 | 0.63323 | 0.754 | -6.44 | 1.43439 | none |
| OR5AC2 | 0.034 | 0.0338 | 0.4923 | 0.63349 | 0.754 | -6.44 | 0 | none |
| OR7A10 | 0.034 | 0.0338 | 0.4923 | 0.63349 | 0.754 | -6.44 | 0 | none |
| CD300LB | 0.035 | 0.035 | 0.4922 | 0.63354 | 0.754 | -6.44 | 0 | none |

| | | | | | | | | |
|---|---|---|---|---|---|---|---|---|
| CTAG1A | 0.034 | 0.0338 | 0.4922 | 0.63358 | 0.754 | -6.44 | 0 | none |
| UBL4A | -0.051 | 0.0106 | -0.492 | 0.63372 | 0.754 | -6.44 | 1.51405 | none |
| LHX3 | -0.037 | -0.0225 | -0.492 | 0.63386 | 0.754 | -6.44 | 0.67306 | none |
| CYP2A6 | 0.034 | 0.0338 | 0.4917 | 0.63392 | 0.754 | -6.44 | -0.7974 | none |
| URAD | 0.034 | 0.0338 | 0.4917 | 0.63392 | 0.754 | -6.44 | 0 | none |
| DERA | 0.075 | -0.0614 | 0.4916 | 0.634 | 0.754 | -6.44 | 0.50368 | none |
| RRP9 | -0.052 | 0.0143 | -0.492 | 0.63403 | 0.754 | -6.44 | 1.32206 | none |
| WIPI2 | -0.065 | -0.0473 | -0.491 | 0.63417 | 0.754 | -6.44 | 1.23713 | none |
| DAB1 | -0.049 | -0.0339 | -0.491 | 0.63424 | 0.754 | -6.44 | 0.74991 | none |
| ISCA2 | -0.069 | -0.1013 | -0.491 | 0.63428 | 0.754 | -6.44 | 0.51931 | none |
| HIST1H2BI | 0.119 | 0.144 | 0.4908 | 0.6345 | 0.755 | -6.44 | 0.87311 | none |
| ZNF629 | 0.036 | 0.0771 | 0.4904 | 0.63477 | 0.755 | -6.44 | 0.90355 | none |
| CCL19 | -0.068 | -0.0675 | -0.49 | 0.63493 | 0.755 | -6.44 | 0.76786 | none |
| FAM20A | 0.046 | 0.06 | 0.4896 | 0.63537 | 0.755 | -6.45 | -0.6685 | none |
| HDAC1 | 0.039 | 0.0319 | 0.4895 | 0.63545 | 0.755 | -6.45 | 0.36332 | none |
| HRCT1 | 0.034 | 0.0336 | 0.4892 | 0.63562 | 0.756 | -6.45 | -0.7237 | none |
| AQP5 | 0.036 | 0.0363 | 0.4886 | 0.636 | 0.756 | -6.45 | 0.62862 | none |
| TULP2 | -0.098 | -0.1345 | -0.489 | 0.63605 | 0.756 | -6.45 | 1.36529 | none |
| OR52J3 | 0.034 | 0.0335 | 0.4886 | 0.63606 | 0.756 | -6.45 | 0 | none |
| HEXA | -0.063 | -0.0922 | -0.488 | 0.63611 | 0.756 | -6.45 | 0.06814 | none |
| MBOAT2 | 0.067 | ######## | 0.4885 | 0.63613 | 0.756 | -6.45 | 1.03385 | none |
| TP53I13 | 0.041 | 0.0185 | 0.4883 | 0.6362 | 0.756 | -6.45 | 1.33789 | none |
| CTNNBIP1 | -0.143 | -0.2029 | -0.488 | 0.63622 | 0.756 | -6.45 | -0.4949 | none |
| AMN | -0.041 | -0.0262 | -0.488 | 0.63634 | 0.756 | -6.45 | 0.42002 | none |
| BNIP2 | 0.033 | -0.0201 | 0.4881 | 0.63636 | 0.756 | -6.45 | 0.98994 | none |
| NUDT16 | 0.037 | 0.1226 | 0.4881 | 0.63637 | 0.756 | -6.45 | 0.76992 | none |
| FBXL3 | -0.082 | -0.1264 | -0.488 | 0.63643 | 0.756 | -6.45 | 1.27075 | none |
| SLC5A11 | 0.08 | 0.0639 | 0.488 | 0.63644 | 0.756 | -6.45 | 0.68684 | none |
| NAPA | 0.066 | 0.0545 | 0.488 | 0.63646 | 0.756 | -6.45 | 0.16604 | none |
| NTSR1 | 0.054 | 0.0923 | 0.4879 | 0.63653 | 0.756 | -6.45 | 0.7771 | none |
| CD320 | -0.059 | -0.0496 | -0.488 | 0.63668 | 0.756 | -6.45 | -0.1478 | none |
| SOX10 | 0.098 | 0.1388 | 0.4876 | 0.63674 | 0.756 | -6.45 | 0.38759 | none |
| DEDD2 | -0.044 | -0.0579 | -0.488 | 0.63677 | 0.756 | -6.45 | 0.39956 | none |
| TMEM144 | 0.051 | 0.0101 | 0.4875 | 0.6368 | 0.756 | -6.45 | 0.45971 | none |
| OTOF | -0.055 | -0.0309 | -0.487 | 0.63682 | 0.756 | -6.45 | 0.13039 | none |
| KLRF2 | 0.057 | 0.0137 | 0.4874 | 0.63686 | 0.756 | -6.45 | 0.6407 | none |
| IFNA16 | 0.033 | 0.0334 | 0.4866 | 0.6374 | 0.757 | -6.45 | 0 | none |
| OR8H3 | 0.033 | 0.0334 | 0.4866 | 0.6374 | 0.757 | -6.45 | 0 | none |
| GPR39 | 0.04 | 0.0397 | 0.4865 | 0.63744 | 0.757 | -6.45 | 0 | none |
| ZIM2 | 0.033 | 0.0333 | 0.4865 | 0.63744 | 0.757 | -6.45 | -0.8173 | none |

| | | | | | | | | |
|---|---|---|---|---|---|---|---|---|
| GJB1 | 0.088 | 0.1276 | 0.4863 | 0.6376 | 0.757 | -6.45 | 1.26461 | none |
| THEMIS2 | -0.052 | 0.0039 | -0.486 | 0.63763 | 0.757 | -6.45 | 1.52384 | none |
| RAPGEF5 | -0.061 | 0.0188 | -0.486 | 0.63783 | 0.757 | -6.45 | -0.0611 | none |
| C1ORF226 | 0.037 | 0.0413 | 0.4858 | 0.63797 | 0.757 | -6.45 | 1.2349 | none |
| NXPH4 | 0.047 | 0.07 | 0.4856 | 0.63811 | 0.757 | -6.45 | 0.24266 | none |
| SEC22C | 0.099 | -0.0353 | 0.4853 | 0.63826 | 0.757 | -6.45 | -0.3897 | none |
| FAM86B1 | 0.088 | 0.0898 | 0.4852 | 0.63833 | 0.757 | -6.45 | 0.97119 | none |
| CHSY1 | 0.117 | 0.154 | 0.4852 | 0.63835 | 0.757 | -6.45 | 0.67972 | none |
| PRAMEF18 | -0.034 | -0.0209 | -0.485 | 0.63849 | 0.757 | -6.45 | 0 | none |
| SRP19 | -0.067 | -0.1585 | -0.484 | 0.63898 | 0.758 | -6.45 | 0.4111 | none |
| MIF4GD | -0.042 | -0.0845 | -0.484 | 0.63923 | 0.758 | -6.45 | 1.26255 | none |
| ESPNL | -0.072 | -0.0418 | -0.484 | 0.63939 | 0.758 | -6.45 | 1.1252 | none |
| SLAMF9 | 0.055 | 0.0552 | 0.4833 | 0.63966 | 0.758 | -6.45 | 1.21269 | none |
| DCDC2B | -0.037 | 0.0123 | -0.483 | 0.63982 | 0.759 | -6.45 | 2.27065 | AA-related |
| AP1AR | 0.051 | 0.0431 | 0.483 | 0.63984 | 0.759 | -6.45 | 0.47482 | none |
| DCK | -0.063 | -0.1476 | -0.483 | 0.63986 | 0.759 | -6.45 | 0.83635 | none |
| PITX3 | -0.035 | 0.0041 | -0.483 | 0.63987 | 0.759 | -6.45 | 0.22496 | none |
| IGSF5 | -0.049 | -0.066 | -0.483 | 0.64005 | 0.759 | -6.45 | 1.02673 | none |
| PROK1 | 0.034 | 0.0327 | 0.4827 | 0.64006 | 0.759 | -6.45 | 0 | none |
| SLC38A11 | -0.033 | -0.0146 | -0.483 | 0.64011 | 0.759 | -6.45 | 0.7074 | none |
| DLC1 | 0.033 | 0.0429 | 0.4826 | 0.64013 | 0.759 | -6.45 | -0.1027 | none |
| DROSHA | 0.118 | 0.1878 | 0.4819 | 0.64061 | 0.759 | -6.45 | -0.2146 | none |
| GOLGA8K | 0.054 | 0.0875 | 0.4816 | 0.64081 | 0.759 | -6.45 | 1.16957 | none |
| RALY | -0.053 | ######## | -0.482 | 0.64086 | 0.759 | -6.45 | 1.75401 | none |
| HNRNPD | 0.082 | 0.126 | 0.4813 | 0.64101 | 0.759 | -6.45 | -0.2198 | none |
| MYO5B | 0.046 | 0.0966 | 0.4811 | 0.64117 | 0.76 | -6.45 | 0.35722 | none |
| MCEE | 0.081 | -0.0411 | 0.481 | 0.64124 | 0.76 | -6.45 | 0.45522 | none |
| MED19 | 0.064 | 0.0494 | 0.4807 | 0.64142 | 0.76 | -6.45 | -0.2241 | none |
| ZNF812P | 0.039 | 0.029 | 0.4807 | 0.64145 | 0.76 | -6.45 | -0.4521 | none |
| NEURL1B | 0.037 | 0.0415 | 0.4805 | 0.64157 | 0.76 | -6.45 | -0.2035 | none |
| MARS2 | 0.065 | 0.0139 | 0.4805 | 0.6416 | 0.76 | -6.45 | 1.06531 | none |
| RHOC | -0.045 | -0.0951 | -0.48 | 0.64161 | 0.76 | -6.45 | 0.33261 | none |
| C21ORF91 | -0.058 | -0.1482 | -0.48 | 0.64172 | 0.76 | -6.45 | 1.70358 | none |
| OR1J1 | -0.039 | -0.0261 | -0.48 | 0.64176 | 0.76 | -6.45 | 1.30508 | none |
| MAJIN | -0.037 | -0.0473 | -0.48 | 0.6418 | 0.76 | -6.45 | 1.44693 | none |
| BX004987 | -0.049 | -0.0014 | -0.48 | 0.6419 | 0.76 | -6.45 | 0.87361 | none |
| CTSK | -0.079 | 0.0055 | -0.48 | 0.64208 | 0.76 | -6.45 | 0.33973 | none |
| CHCHD10 | -0.062 | -0.0836 | -0.48 | 0.64219 | 0.76 | -6.45 | -0.1857 | none |
| NUDT8 | 0.102 | 0.0572 | 0.4795 | 0.64226 | 0.76 | -6.45 | 0.8273 | none |
| TENM3 | 0.043 | 0.0523 | 0.4785 | 0.64298 | 0.761 | -6.45 | -0.7298 | none |

| | | | | | | | | |
|---|---|---|---|---|---|---|---|---|
| FGF14 | -0.032 | -0.0165 | -0.477 | 0.64365 | 0.762 | -6.45 | 0 | none |
| RAD1 | 0.056 | -0.0481 | 0.4773 | 0.64377 | 0.762 | -6.45 | 0.42861 | none |
| PALM3 | 0.053 | 0.0668 | 0.4772 | 0.64387 | 0.762 | -6.45 | 1.18711 | none |
| CCNI | 0.044 | -0.014 | 0.4771 | 0.64393 | 0.762 | -6.45 | 1.4007 | none |
| ITGA5 | 0.058 | 0.1222 | 0.4771 | 0.64393 | 0.762 | -6.45 | -0.2057 | none |
| OTOP2 | 0.035 | 0.0486 | 0.4768 | 0.6441 | 0.762 | -6.45 | 0.04328 | none |
| SLC6A19 | -0.035 | -0.0077 | -0.477 | 0.64424 | 0.762 | -6.45 | 0.80117 | none |
| BACH1 | -0.105 | -0.2168 | -0.476 | 0.64441 | 0.762 | -6.45 | 0.41033 | none |
| RAP1GAP | -0.064 | 0.0546 | -0.476 | 0.64441 | 0.762 | -6.45 | 0.21932 | none |
| MLH3 | 0.094 | 0.2097 | 0.4762 | 0.64454 | 0.762 | -6.45 | 0.74288 | none |
| PRKRIP1 | 0.055 | 0.1341 | 0.476 | 0.64469 | 0.762 | -6.45 | 0.3876 | none |
| FLRT1 | -0.05 | -0.0059 | -0.476 | 0.64469 | 0.762 | -6.45 | 0.97617 | none |
| IL1F10 | -0.038 | -0.0381 | -0.476 | 0.64472 | 0.762 | -6.45 | 1.32395 | none |
| NDUFS3 | -0.066 | -0.123 | -0.476 | 0.64481 | 0.762 | -6.45 | 0.82702 | none |
| STXBP5 | 0.095 | 0.1031 | 0.4757 | 0.6449 | 0.762 | -6.45 | 0.68806 | none |
| SOST | -0.037 | -0.0367 | -0.476 | 0.64495 | 0.762 | -6.45 | 1.37577 | none |
| CDC42SE2 | -0.063 | -0.188 | -0.474 | 0.64588 | 0.763 | -6.45 | 0.42385 | none |
| UFC1 | 0.049 | -0.0251 | 0.474 | 0.64607 | 0.764 | -6.45 | -0.7761 | none |
| CEPT1 | 0.051 | -0.0278 | 0.4739 | 0.6461 | 0.764 | -6.45 | 0.02407 | none |
| HOXA3 | -0.091 | -0.0295 | -0.474 | 0.6461 | 0.764 | -6.45 | 1.46401 | none |
| ELMOD2 | 0.056 | -0.0207 | 0.4732 | 0.64659 | 0.764 | -6.45 | 0.46865 | none |
| FXYD3 | 0.067 | 0.1191 | 0.4731 | 0.64665 | 0.764 | -6.45 | 1.39231 | none |
| EEF1E1 | 0.068 | -0.0305 | 0.4731 | 0.64667 | 0.764 | -6.45 | 0.00907 | none |
| BIVM-ERC( | -0.035 | -0.0469 | -0.473 | 0.64691 | 0.764 | -6.45 | 0.43137 | none |
| HMSD | -0.043 | -0.0458 | -0.473 | 0.64706 | 0.764 | -6.45 | 1.44682 | none |
| GOLGA8S | 0.035 | 0.0233 | 0.4723 | 0.64719 | 0.765 | -6.45 | -0.5119 | none |
| EPN3 | 0.09 | 0.159 | 0.4721 | 0.64734 | 0.765 | -6.45 | 0.40855 | none |
| CAPZA1 | 0.049 | 0.0226 | 0.4721 | 0.64737 | 0.765 | -6.45 | -0.741 | none |
| MPST | 0.072 | 0.0065 | 0.472 | 0.64744 | 0.765 | -6.45 | -0.5272 | none |
| GDAP1 | -0.099 | 0.0225 | -0.472 | 0.64758 | 0.765 | -6.45 | 2.09141 | none |
| DIRC1 | -0.05 | 0.0011 | -0.472 | 0.64762 | 0.765 | -6.45 | 0.48557 | none |
| CXCR1 | 0.04 | 0.0115 | 0.4715 | 0.64775 | 0.765 | -6.45 | 0.31567 | none |
| RAB9B | -0.032 | -0.0158 | -0.471 | 0.64782 | 0.765 | -6.45 | 2.1286 | AA-related |
| GALR3 | 0.054 | 0.0073 | 0.4713 | 0.64786 | 0.765 | -6.45 | -0.7543 | none |
| HIST1H4K | -0.203 | -0.1789 | -0.471 | 0.64803 | 0.765 | -6.45 | 1.03223 | none |
| PMF1 | 0.052 | -0.0076 | 0.471 | 0.64808 | 0.765 | -6.45 | -0.6457 | none |
| IMP4 | -0.052 | 0.0059 | -0.471 | 0.64823 | 0.765 | -6.45 | 0.163 | none |
| BIN1 | 0.068 | 0.1202 | 0.4707 | 0.64834 | 0.765 | -6.45 | 0.62724 | none |
| SUSD2 | 0.035 | 0.0089 | 0.4706 | 0.64841 | 0.765 | -6.45 | 0.59553 | none |
| RIMS3 | 0.039 | 0.0361 | 0.4702 | 0.64864 | 0.766 | -6.45 | 0.11144 | none |

| | | | | | | | | |
|---|---|---|---|---|---|---|---|---|
| G6PD | 0.1 | 0.1672 | 0.47 | 0.64876 | 0.766 | -6.46 | 0.5921 | none |
| ACTL9 | 0.039 | 0.047 | 0.4699 | 0.64883 | 0.766 | -6.46 | 1.84575 | none |
| AMIGO1 | 0.056 | 0.0959 | 0.4698 | 0.64891 | 0.766 | -6.46 | 1.34778 | none |
| TMEM158 | 0.038 | 0.0287 | 0.4698 | 0.64896 | 0.766 | -6.46 | 0.22101 | none |
| MRPL1 | 0.045 | -0.0426 | 0.4696 | 0.64907 | 0.766 | -6.46 | 0.42586 | none |
| IRX4 | -0.041 | -0.0305 | -0.469 | 0.64938 | 0.766 | -6.46 | 1.45344 | none |
| NDUFAF3 | -0.072 | -0.091 | -0.469 | 0.6494 | 0.766 | -6.46 | 1.60516 | none |
| SRXN1 | -0.088 | 0.0449 | -0.469 | 0.64953 | 0.766 | -6.46 | 0.81096 | none |
| SALL3 | -0.038 | -0.0282 | -0.469 | 0.64961 | 0.766 | -6.46 | -0.2811 | none |
| ATG7 | 0.049 | 0.0295 | 0.4686 | 0.64976 | 0.766 | -6.46 | 0.18183 | none |
| TNFAIP1 | 0.085 | 0.0901 | 0.4681 | 0.6501 | 0.767 | -6.46 | 0.46432 | none |
| PRR5-ARH | 0.059 | 0.0648 | 0.4676 | 0.65046 | 0.767 | -6.46 | 1.49877 | none |
| ANAPC10 | 0.118 | 0.0093 | 0.4676 | 0.65048 | 0.767 | -6.46 | 0.48022 | none |
| FAM104B | 0.11 | -0.0517 | 0.4675 | 0.65051 | 0.767 | -6.46 | -0.0033 | none |
| CDKN2B | 0.086 | 0.0132 | 0.4674 | 0.6506 | 0.767 | -6.46 | 0.3259 | none |
| WDR18 | -0.047 | -0.0316 | -0.467 | 0.65061 | 0.767 | -6.46 | 0.75559 | none |
| CILP2 | -0.061 | -0.0445 | -0.467 | 0.65067 | 0.767 | -6.46 | 0.69549 | none |
| ANXA2R | 0.086 | -0.0512 | 0.467 | 0.65083 | 0.767 | -6.46 | 0.28679 | none |
| SLC4A11 | 0.069 | 0.0808 | 0.4657 | 0.65175 | 0.768 | -6.46 | 1.00656 | none |
| WSCD1 | -0.069 | -0.0124 | -0.466 | 0.65177 | 0.768 | -6.46 | 0.64923 | none |
| C9ORF135 | -0.05 | -0.0843 | -0.465 | 0.652 | 0.768 | -6.46 | -0.56 | none |
| FAM209B | -0.048 | 0.0418 | -0.465 | 0.65218 | 0.769 | -6.46 | -0.1431 | none |
| OR56B4 | 0.032 | 0.0429 | 0.4649 | 0.65229 | 0.769 | -6.46 | 1.09028 | none |
| LRRIQ4 | 0.033 | 0.0343 | 0.4649 | 0.65231 | 0.769 | -6.46 | 1.20231 | none |
| CCT2 | -0.037 | -0.0509 | -0.465 | 0.65237 | 0.769 | -6.46 | 1.01078 | none |
| LENG1 | 0.032 | 0.0431 | 0.4646 | 0.6525 | 0.769 | -6.46 | 0.44734 | none |
| DUSP12 | -0.06 | -0.0751 | -0.465 | 0.65256 | 0.769 | -6.46 | 1.41755 | none |
| SPESP1 | -0.064 | ######## | -0.464 | 0.65262 | 0.769 | -6.46 | 0.81639 | none |
| ZFP3 | 0.034 | 0.0384 | 0.4642 | 0.65281 | 0.769 | -6.46 | 0.01746 | none |
| TMX1 | -0.069 | -0.1376 | -0.464 | 0.65288 | 0.769 | -6.46 | 0.98836 | none |
| FOSL1 | 0.097 | 0.2073 | 0.4641 | 0.65289 | 0.769 | -6.46 | 0.30969 | none |
| SOHLH2 | -0.034 | -0.0288 | -0.464 | 0.65305 | 0.769 | -6.46 | 1.05188 | none |
| HAS3 | -0.083 | 0.0137 | -0.464 | 0.65323 | 0.769 | -6.46 | 0.51941 | none |
| CT45A5 | 0.034 | 0.0132 | 0.4634 | 0.65333 | 0.769 | -6.46 | -0.2435 | none |
| ABI2 | -0.052 | -0.0649 | -0.462 | 0.65435 | 0.77 | -6.46 | 1.37548 | none |
| EMP1 | -0.08 | 0.0138 | -0.462 | 0.65442 | 0.77 | -6.46 | 0.54481 | none |
| STC1 | 0.031 | 0.0567 | 0.4618 | 0.65443 | 0.77 | -6.46 | 0.07478 | none |
| TOR2A | 0.046 | 0.0088 | 0.4618 | 0.65445 | 0.77 | -6.46 | 0.6499 | none |
| SSX2IP | -0.086 | 0.0045 | -0.462 | 0.65446 | 0.77 | -6.46 | 1.40942 | none |
| PPP1R16B | 0.097 | 0.2509 | 0.4613 | 0.6548 | 0.771 | -6.46 | -0.0304 | none |

| | | | | | | | | |
|---|---|---|---|---|---|---|---|---|
| ATP8B4 | 0.105 | 0.0068 | 0.4613 | 0.65481 | 0.771 | -6.46 | 0.44161 | none |
| LRCH1 | 0.09 | 0.1788 | 0.4612 | 0.65486 | 0.771 | -6.46 | 0.62547 | none |
| HK3 | 0.095 | 0.0068 | 0.4608 | 0.65512 | 0.771 | -6.46 | 0.36873 | none |
| PAN2 | 0.077 | 0.1058 | 0.4607 | 0.65521 | 0.771 | -6.46 | 0.71183 | none |
| UMAD1 | 0.073 | -0.0285 | 0.4605 | 0.65533 | 0.771 | -6.46 | -0.3821 | none |
| TGM4 | -0.09 | -0.084 | -0.46 | 0.65541 | 0.771 | -6.46 | 0.30141 | none |
| CCDC85B | -0.075 | -0.0573 | -0.46 | 0.65542 | 0.771 | -6.46 | 0.62872 | none |
| CEP162 | 0.06 | 0.1273 | 0.4602 | 0.65559 | 0.771 | -6.46 | 0.31227 | none |
| KBTBD7 | 0.035 | -0.0198 | 0.4601 | 0.65564 | 0.771 | -6.46 | 0.49808 | none |
| ZNF573 | 0.17 | 0.1053 | 0.46 | 0.65567 | 0.771 | -6.46 | 0.71319 | none |
| RBBP4 | 0.035 | -0.0044 | 0.46 | 0.65573 | 0.771 | -6.46 | 0.2251 | none |
| ETS2 | -0.057 | -0.0404 | -0.46 | 0.65595 | 0.771 | -6.46 | 1.5625 | none |
| CELA2B | -0.069 | -0.0366 | -0.46 | 0.65602 | 0.771 | -6.46 | 1.17566 | none |
| CCR3 | 0.084 | 0.0407 | 0.4595 | 0.65607 | 0.771 | -6.46 | -0.1981 | none |
| ZNF138 | -0.059 | -0.0611 | -0.459 | 0.65629 | 0.772 | -6.46 | 0.45196 | none |
| MITF | -0.053 | 0.0597 | -0.459 | 0.65636 | 0.772 | -6.46 | 0.55593 | none |
| TLE6 | 0.088 | 0.128 | 0.4588 | 0.6565 | 0.772 | -6.46 | 0.42228 | none |
| ARHGEF17 | 0.064 | 0.1849 | 0.4581 | 0.65703 | 0.772 | -6.46 | 0.70834 | none |
| F8A3 | -0.055 | -0.0444 | -0.458 | 0.65723 | 0.772 | -6.46 | 0.62078 | none |
| EHBP1L1 | 0.141 | 0.2182 | 0.4578 | 0.65724 | 0.772 | -6.46 | -0.2837 | none |
| SHOX2 | -0.05 | -0.0507 | -0.458 | 0.65743 | 0.773 | -6.46 | 1.61016 | none |
| 10-Mar | -0.058 | 0.0052 | -0.457 | 0.65745 | 0.773 | -6.46 | -0.0366 | none |
| SCGB2A2 | 0.049 | 0.0878 | 0.4572 | 0.65762 | 0.773 | -6.46 | 1.42595 | none |
| MBD2 | -0.118 | -0.0304 | -0.457 | 0.65776 | 0.773 | -6.46 | 0.55207 | none |
| ZNF501 | 0.039 | 0.0686 | 0.4569 | 0.65784 | 0.773 | -6.46 | -0.1606 | none |
| ZNF607 | -0.036 | 0.0291 | -0.457 | 0.6579 | 0.773 | -6.46 | 0.51288 | none |
| NEU4 | -0.036 | -0.0299 | -0.457 | 0.658 | 0.773 | -6.46 | 1.35031 | none |
| TMPRSS13 | 0.046 | 0.093 | 0.4566 | 0.65806 | 0.773 | -6.46 | 1.26825 | none |
| HEATR5A | 0.105 | -0.0593 | 0.4566 | 0.65809 | 0.773 | -6.46 | -0.4998 | none |
| SNX18 | -0.062 | -0.0131 | -0.457 | 0.65812 | 0.773 | -6.46 | 0.8941 | none |
| KIF22 | 0.063 | 0.1478 | 0.4562 | 0.6583 | 0.773 | -6.46 | 0.66578 | none |
| EPGN | 0.04 | 0.0531 | 0.4562 | 0.65832 | 0.773 | -6.46 | 0.70079 | none |
| C11ORF95 | 0.055 | 0.0984 | 0.4561 | 0.6584 | 0.773 | -6.46 | 0.78937 | none |
| POTEB | 0.061 | 0.0322 | 0.4561 | 0.65842 | 0.773 | -6.46 | -0.1013 | none |
| MPG | -0.073 | -0.134 | -0.456 | 0.65842 | 0.773 | -6.46 | 0.88452 | none |
| NUP58 | 0.069 | 0.0868 | 0.456 | 0.65844 | 0.773 | -6.46 | 1.03828 | none |
| PARK7 | -0.041 | -0.0737 | -0.456 | 0.65847 | 0.773 | -6.46 | 0.29964 | none |
| PRPH | 0.039 | 0.0614 | 0.4559 | 0.65856 | 0.773 | -6.46 | -0.2496 | none |
| DUSP9 | 0.031 | 0.0285 | 0.4556 | 0.65877 | 0.773 | -6.46 | 0.39406 | none |
| SSX4 | -0.037 | -0.0215 | -0.455 | 0.65924 | 0.774 | -6.46 | -0.1559 | none |

| | | | | | | | | |
|---|---|---|---|---|---|---|---|---|
| SSX4B | -0.037 | -0.0215 | -0.455 | 0.65924 | 0.774 | -6.46 | -0.1559 | none |
| DYNLRB2 | -0.063 | -0.0751 | -0.455 | 0.65924 | 0.774 | -6.46 | 1.66517 | none |
| CENPQ | -0.083 | -0.0621 | -0.455 | 0.65936 | 0.774 | -6.46 | 0.62367 | none |
| DUX4 | -0.155 | -0.1084 | -0.454 | 0.65952 | 0.774 | -6.46 | -0.3564 | none |
| GNAZ | -0.04 | 0.0196 | -0.454 | 0.65965 | 0.774 | -6.46 | -0.2792 | none |
| PRSS8 | 0.067 | 0.0416 | 0.4542 | 0.65972 | 0.774 | -6.46 | -0.1887 | none |
| KLHL41 | 0.076 | 0.1033 | 0.4539 | 0.65991 | 0.774 | -6.46 | 1.54461 | none |
| COA4 | -0.068 | -0.1044 | -0.454 | 0.6601 | 0.774 | -6.46 | 0.82313 | none |
| TOM1 | -0.06 | 0.035 | -0.453 | 0.66038 | 0.775 | -6.46 | 1.2463 | none |
| CCDC71L | -0.084 | -0.0528 | -0.453 | 0.66067 | 0.775 | -6.46 | 1.10855 | none |
| UBP1 | 0.067 | 0.1046 | 0.4528 | 0.66068 | 0.775 | -6.46 | 0.40613 | none |
| NTN4 | 0.074 | 0.0745 | 0.4525 | 0.66088 | 0.775 | -6.46 | -0.5397 | none |
| ZNF319 | -0.113 | -0.0775 | -0.452 | 0.66092 | 0.775 | -6.46 | 1.91799 | none |
| HEXIM2 | 0.066 | 0.1067 | 0.4523 | 0.66106 | 0.775 | -6.46 | 0.47562 | none |
| UTP15 | -0.067 | -0.0928 | -0.452 | 0.66123 | 0.775 | -6.46 | 1.21915 | none |
| SIRPA | 0.031 | 0.0386 | 0.452 | 0.66124 | 0.775 | -6.46 | 0 | none |
| DUSP22 | -0.066 | -0.0738 | -0.452 | 0.66131 | 0.775 | -6.46 | 0.84492 | none |
| UNC45B | 0.034 | 0.0257 | 0.4518 | 0.6614 | 0.775 | -6.46 | -0.9308 | none |
| SCUBE1 | 0.092 | 0.0222 | 0.4517 | 0.66146 | 0.775 | -6.46 | 0.07489 | none |
| TRAK2 | 0.09 | 0.1217 | 0.4517 | 0.66149 | 0.775 | -6.46 | 1.2887 | none |
| DDX24 | -0.046 | 0.0323 | -0.451 | 0.66167 | 0.775 | -6.46 | 0.90452 | none |
| ZNF710 | 0.161 | 0.2371 | 0.4509 | 0.66204 | 0.776 | -6.46 | -0.7447 | none |
| KLF10 | 0.038 | 0.0428 | 0.4507 | 0.66215 | 0.776 | -6.46 | 0.338 | none |
| CASP5 | -0.032 | -0.0292 | -0.451 | 0.66221 | 0.776 | -6.46 | -0.6574 | none |
| DCAF13 | -0.051 | -0.0595 | -0.451 | 0.66224 | 0.776 | -6.46 | 0.03884 | none |
| TMED6 | -0.059 | 0.019 | -0.45 | 0.66236 | 0.776 | -6.46 | -0.2099 | none |
| CYTL1 | -0.031 | -0.0307 | -0.45 | 0.66238 | 0.776 | -6.46 | 0 | none |
| ANKRD50 | 0.037 | 0.0457 | 0.4498 | 0.66278 | 0.776 | -6.46 | 0 | none |
| PIN1 | 0.055 | 0.0566 | 0.4496 | 0.66289 | 0.776 | -6.46 | 0.02432 | none |
| LRRTM2 | 0.034 | 0.0194 | 0.4495 | 0.66303 | 0.777 | -6.47 | -0.0544 | none |
| CCDC68 | 0.055 | 0.0436 | 0.4492 | 0.66318 | 0.777 | -6.47 | -0.138 | none |
| TMEM177 | -0.081 | -0.033 | -0.449 | 0.66322 | 0.777 | -6.47 | 1.8566 | none |
| SPDYE4 | -0.061 | -0.0043 | -0.449 | 0.66323 | 0.777 | -6.47 | 0.59543 | none |
| CARD16 | 0.064 | -0.0314 | 0.4491 | 0.66324 | 0.777 | -6.47 | -0.7533 | none |
| PDXP | -0.094 | -0.0921 | -0.449 | 0.66362 | 0.777 | -6.47 | -0.1562 | none |
| ABCB1 | 0.076 | 0.0719 | 0.4484 | 0.66378 | 0.777 | -6.47 | -0.6406 | none |
| IRX3 | 0.044 | 0.0611 | 0.4483 | 0.6638 | 0.777 | -6.47 | 1.50692 | none |
| FCER1A | -0.042 | -0.0638 | -0.448 | 0.66382 | 0.777 | -6.47 | 0.79459 | none |
| ANKRD33 | -0.03 | -0.0182 | -0.448 | 0.66409 | 0.777 | -6.47 | 1.68091 | none |
| JOSD2 | 0.063 | 0.0239 | 0.4478 | 0.66415 | 0.777 | -6.47 | -0.6842 | none |

| | | | | | | | | |
|---|---|---|---|---|---|---|---|---|
| GPBP1 | 0.096 | 0.1389 | 0.4476 | 0.66432 | 0.777 | -6.47 | 0.7546 | none |
| PDK4 | 0.064 | -0.0061 | 0.4474 | 0.66445 | 0.777 | -6.47 | 1.25775 | none |
| RNF139 | 0.058 | -0.0425 | 0.4473 | 0.66452 | 0.777 | -6.47 | 0.70896 | none |
| SOX4 | 0.083 | 0.056 | 0.4473 | 0.66454 | 0.777 | -6.47 | -0.5426 | none |
| XRCC1 | 0.095 | 0.2096 | 0.4472 | 0.66457 | 0.777 | -6.47 | 0.27306 | none |
| B4GALT3 | -0.067 | -0.096 | -0.447 | 0.66463 | 0.777 | -6.47 | 0.87243 | none |
| EMILIN1 | -0.034 | -0.0207 | -0.447 | 0.66463 | 0.777 | -6.47 | 1.02614 | none |
| EYA4 | 0.047 | 0.0472 | 0.4469 | 0.66477 | 0.778 | -6.47 | 0.75923 | none |
| TSPAN8 | -0.09 | -0.0032 | -0.447 | 0.66495 | 0.778 | -6.47 | 0.2374 | none |
| SIK1 | 0.116 | 0.1532 | 0.4466 | 0.66498 | 0.778 | -6.47 | -0.4537 | none |
| MTF2 | -0.047 | -0.0602 | -0.447 | 0.66507 | 0.778 | -6.47 | 0.25401 | none |
| ID2 | -0.087 | -0.14 | -0.446 | 0.6651 | 0.778 | -6.47 | 0.47915 | none |
| PRSS1 | -0.032 | -0.03 | -0.446 | 0.66513 | 0.778 | -6.47 | 1.54695 | none |
| IGIP | 0.038 | 0.022 | 0.4463 | 0.66525 | 0.778 | -6.47 | -0.3989 | none |
| SFTPA2 | -0.031 | -0.0063 | -0.446 | 0.66532 | 0.778 | -6.47 | 1.6208 | none |
| CS | 0.048 | 0.0165 | 0.446 | 0.66544 | 0.778 | -6.47 | 0.35837 | none |
| SPANXA2 | -0.106 | -0.1059 | -0.446 | 0.66549 | 0.778 | -6.47 | 1.55316 | none |
| RAB4B-EG | -0.075 | -0.0358 | -0.446 | 0.66564 | 0.778 | -6.47 | 0.82689 | none |
| NAP1L3 | 0.071 | 0.0853 | 0.4453 | 0.66595 | 0.778 | -6.47 | 0.22661 | none |
| C11ORF74 | -0.072 | -0.1677 | -0.445 | 0.66607 | 0.778 | -6.47 | 0.35944 | none |
| FBXW5 | 0.046 | 0.0703 | 0.4445 | 0.66651 | 0.779 | -6.47 | 0.58486 | none |
| POU4F1 | 0.031 | 0.0334 | 0.4444 | 0.66653 | 0.779 | -6.47 | 0.5276 | none |
| RITA1 | -0.047 | -0.0508 | -0.444 | 0.66661 | 0.779 | -6.47 | 1.77576 | none |
| TCF21 | 0.033 | 0.0422 | 0.4442 | 0.66669 | 0.779 | -6.47 | -0.582 | none |
| NOX1 | -0.07 | -0.0571 | -0.444 | 0.66716 | 0.779 | -6.47 | 0.64185 | none |
| TSSK4 | 0.06 | 0.0688 | 0.4435 | 0.6672 | 0.779 | -6.47 | -0.551 | none |
| C4ORF46 | 0.081 | -0.0137 | 0.4424 | 0.66791 | 0.78 | -6.47 | 0.97108 | none |
| SSNA1 | 0.063 | 0.0496 | 0.4424 | 0.66791 | 0.78 | -6.47 | -0.2314 | none |
| MAP3K4 | 0.119 | 0.1743 | 0.4424 | 0.66797 | 0.78 | -6.47 | 0.9682 | none |
| RGPD3 | -0.03 | -0.0428 | -0.442 | 0.66801 | 0.78 | -6.47 | 1.08499 | none |
| SAMD14 | -0.039 | -0.0296 | -0.442 | 0.66814 | 0.78 | -6.47 | 0.56241 | none |
| PROZ | -0.066 | -0.0905 | -0.442 | 0.66822 | 0.78 | -6.47 | 0.71417 | none |
| CLPTM1L | -0.077 | -0.0772 | -0.442 | 0.66831 | 0.78 | -6.47 | 1.35537 | none |
| SYNCRIP | -0.077 | 0.0316 | -0.442 | 0.66837 | 0.78 | -6.47 | 1.49926 | none |
| SHC4 | -0.093 | 0.0494 | -0.442 | 0.66841 | 0.78 | -6.47 | -0.2596 | none |
| KDM8 | 0.089 | 0.1795 | 0.4416 | 0.66849 | 0.78 | -6.47 | 1.80637 | none |
| MCIDAS | 0.038 | 0.047 | 0.4415 | 0.66855 | 0.78 | -6.47 | -0.743 | none |
| JDP2 | 0.04 | 0.0591 | 0.4413 | 0.66873 | 0.781 | -6.47 | 0.24022 | none |
| ANKRD17 | 0.126 | 0.3142 | 0.441 | 0.66893 | 0.781 | -6.47 | 0.70322 | none |
| SLC35D2 | -0.052 | -0.1529 | -0.441 | 0.66894 | 0.781 | -6.47 | -0.2327 | none |

| | | | | | | | | |
|---|---|---|---|---|---|---|---|---|
| NEFL | 0.034 | 0.0415 | 0.441 | 0.66894 | 0.781 | -6.47 | -0.6001 | none |
| CELF4 | 0.12 | 0.0949 | 0.4408 | 0.66906 | 0.781 | -6.47 | 0.96521 | none |
| AMZ1 | -0.039 | -0.0118 | -0.441 | 0.6691 | 0.781 | -6.47 | 0.5433 | none |
| MTERF1 | 0.084 | 0.1122 | 0.4406 | 0.66918 | 0.781 | -6.47 | -0.0822 | none |
| MID1 | 0.037 | 0.0154 | 0.4405 | 0.66926 | 0.781 | -6.47 | 0.00492 | none |
| INSM1 | 0.034 | 0.0339 | 0.4404 | 0.66936 | 0.781 | -6.47 | 1.26562 | none |
| BPIFA3 | 0.031 | 0.0507 | 0.44 | 0.66965 | 0.781 | -6.47 | 0.09134 | none |
| USH1G | -0.033 | -0.0163 | -0.44 | 0.66973 | 0.781 | -6.47 | 0.61422 | none |
| UGT3A1 | -0.053 | -0.0502 | -0.44 | 0.6698 | 0.781 | -6.47 | 0.60896 | none |
| PI4K2B | -0.065 | -0.058 | -0.439 | 0.67008 | 0.781 | -6.47 | 1.18604 | none |
| PGAP2 | -0.041 | -0.0657 | -0.439 | 0.67016 | 0.781 | -6.47 | 1.46604 | none |
| SCAMP2 | -0.051 | -0.0956 | -0.439 | 0.67017 | 0.781 | -6.47 | 0.52522 | none |
| IL17A | -0.111 | -0.1392 | -0.439 | 0.67018 | 0.781 | -6.47 | 0.78352 | none |
| HAPLN3 | -0.051 | -0.0781 | -0.439 | 0.6702 | 0.781 | -6.47 | 0.49396 | none |
| LMO1 | -0.055 | -0.0627 | -0.439 | 0.67038 | 0.782 | -6.47 | 0.48313 | none |
| GLTPD2 | 0.039 | 0.0213 | 0.4388 | 0.67043 | 0.782 | -6.47 | 0.52391 | none |
| SLCO1B7 | 0.032 | 0.0429 | 0.4385 | 0.67065 | 0.782 | -6.47 | -0.7347 | none |
| C12ORF73 | 0.084 | -0.0096 | 0.4385 | 0.67069 | 0.782 | -6.47 | -0.2913 | none |
| SCN4A | 0.037 | 0.0455 | 0.4384 | 0.67075 | 0.782 | -6.47 | 0.89054 | none |
| ADCY6 | 0.076 | 0.0077 | 0.4381 | 0.67096 | 0.782 | -6.47 | -0.046 | none |
| KLHL14 | -0.035 | -0.0343 | -0.438 | 0.67114 | 0.782 | -6.47 | 1.74629 | none |
| LRRC6 | -0.052 | -0.0246 | -0.438 | 0.67116 | 0.782 | -6.47 | 1.87798 | none |
| HES1 | -0.048 | -0.0468 | -0.438 | 0.67124 | 0.782 | -6.47 | 0.92111 | none |
| SDHAF3 | 0.119 | -0.0656 | 0.4375 | 0.67136 | 0.782 | -6.47 | -0.2255 | none |
| RBM15B | 0.081 | 0.1739 | 0.4373 | 0.67151 | 0.782 | -6.47 | 0.28439 | none |
| NINJ1 | -0.05 | -0.0043 | -0.437 | 0.67161 | 0.782 | -6.47 | 1.5292 | none |
| FEN1 | -0.058 | 0.0456 | -0.437 | 0.67169 | 0.783 | -6.47 | 1.24017 | none |
| ZFAND2A | 0.058 | 0.0551 | 0.4369 | 0.67178 | 0.783 | -6.47 | 0.85003 | none |
| DTX1 | -0.089 | ######## | -0.437 | 0.67193 | 0.783 | -6.47 | 0.21325 | none |
| NUTM2B | -0.054 | -0.0083 | -0.437 | 0.67203 | 0.783 | -6.47 | 0.9022 | none |
| CMTM2 | 0.048 | 0.0532 | 0.4362 | 0.67225 | 0.783 | -6.47 | 1.59291 | none |
| RDH5 | -0.175 | -0.0263 | -0.436 | 0.6723 | 0.783 | -6.47 | 0.74276 | none |
| SUSD1 | 0.091 | 0.1198 | 0.4361 | 0.67236 | 0.783 | -6.47 | -0.1491 | none |
| GNGT2 | 0.093 | -0.0256 | 0.4361 | 0.67237 | 0.783 | -6.47 | -0.7799 | none |
| TEKT1 | -0.031 | -0.0312 | -0.436 | 0.67247 | 0.783 | -6.47 | 1.0022 | none |
| CDADC1 | -0.052 | -0.0077 | -0.436 | 0.67248 | 0.783 | -6.47 | 0.94991 | none |
| LILRB4 | 0.088 | 0.0858 | 0.4355 | 0.67279 | 0.783 | -6.47 | 1.41387 | none |
| NUP93 | -0.037 | -0.01 | -0.435 | 0.67279 | 0.783 | -6.47 | 0.34791 | none |
| TKT | 0.082 | 0.1453 | 0.4354 | 0.67284 | 0.783 | -6.47 | 1.43867 | none |
| GPATCH3 | 0.037 | 0.1037 | 0.4353 | 0.67292 | 0.783 | -6.47 | 1.45837 | none |

| | | | | | | | | |
|---|---|---|---|---|---|---|---|---|
| MBD3L3 | 0.034 | 0.0512 | 0.4352 | 0.673 | 0.783 | -6.47 | 1.45851 | none |
| UCHL5 | -0.066 | -0.0398 | -0.435 | 0.67304 | 0.783 | -6.47 | 1.16449 | none |
| PILRA | -0.038 | 0.0126 | -0.435 | 0.67307 | 0.783 | -6.47 | 0.47402 | none |
| RXRB | -0.041 | -0.0084 | -0.435 | 0.6731 | 0.783 | -6.47 | 0.60746 | none |
| CDK5RAP2 | 0.11 | 0.1994 | 0.4346 | 0.67344 | 0.784 | -6.47 | 0.01384 | none |
| LCP1 | 0.086 | 0.1479 | 0.4344 | 0.67356 | 0.784 | -6.47 | 0.965 | none |
| TTC23L | -0.044 | -0.0367 | -0.434 | 0.67366 | 0.784 | -6.47 | 0.80831 | none |
| CGB2 | 0.05 | 0.0882 | 0.4341 | 0.67379 | 0.784 | -6.47 | -0.2302 | none |
| SLC25A48 | -0.035 | -0.0224 | -0.434 | 0.67379 | 0.784 | -6.47 | 0.00775 | none |
| ARHGAP4( | 0.032 | 0.0422 | 0.434 | 0.67381 | 0.784 | -6.47 | 1.58698 | none |
| OR2C1 | -0.036 | -0.0358 | -0.434 | 0.67382 | 0.784 | -6.47 | 0.89247 | none |
| HDAC3 | 0.049 | -0.0074 | 0.4336 | 0.67413 | 0.784 | -6.47 | 0.40325 | none |
| PBXIP1 | 0.079 | 0.1146 | 0.4335 | 0.67417 | 0.784 | -6.47 | -0.149 | none |
| CYB5R2 | -0.053 | -0.0795 | -0.433 | 0.67418 | 0.784 | -6.47 | 0.11044 | none |
| EMILIN2 | 0.061 | 0.0177 | 0.4334 | 0.67425 | 0.784 | -6.47 | -0.597 | none |
| NEFM | 0.047 | 0.114 | 0.433 | 0.67453 | 0.784 | -6.47 | 0.24466 | none |
| TBC1D28 | 0.072 | 0.0945 | 0.4328 | 0.67463 | 0.784 | -6.47 | -0.1449 | none |
| SLC16A9 | 0.068 | 0.1269 | 0.4328 | 0.67466 | 0.784 | -6.47 | 0.55041 | none |
| CDT1 | 0.054 | 0.0726 | 0.432 | 0.67524 | 0.785 | -6.47 | -0.0525 | none |
| STK17A | 0.043 | -0.0119 | 0.4317 | 0.67541 | 0.785 | -6.47 | 0.17651 | none |
| COL4A4 | -0.04 | -0.048 | -0.431 | 0.67558 | 0.785 | -6.47 | 0.42003 | none |
| PDE6C | -0.053 | -0.0076 | -0.431 | 0.67563 | 0.785 | -6.47 | 0.78209 | none |
| YOD1 | 0.037 | 0.0246 | 0.4314 | 0.67566 | 0.785 | -6.47 | 0.42338 | none |
| CRABP2 | -0.088 | -0.1269 | -0.431 | 0.67574 | 0.785 | -6.47 | 0.65373 | none |
| APTX | -0.044 | -0.0451 | -0.431 | 0.67582 | 0.785 | -6.47 | 0.40792 | none |
| PGLYRP2 | -0.049 | 0.0013 | -0.431 | 0.67585 | 0.785 | -6.47 | -0.1431 | none |
| COPS7A | 0.04 | 0.0129 | 0.431 | 0.6759 | 0.785 | -6.47 | 0.41367 | none |
| UBE2U | -0.03 | -0.0302 | -0.431 | 0.67599 | 0.785 | -6.47 | 1.91174 | none |
| SPATA2 | -0.048 | -0.0046 | -0.431 | 0.67608 | 0.785 | -6.47 | 0.9401 | none |
| ZFP36L1 | 0.063 | 0.1368 | 0.4307 | 0.67615 | 0.785 | -6.47 | -0.2685 | none |
| YIF1B | 0.058 | 0.0064 | 0.4305 | 0.67629 | 0.785 | -6.47 | 0.29417 | none |
| APBB1 | 0.073 | 0.0852 | 0.4303 | 0.67639 | 0.786 | -6.47 | 0.62144 | none |
| NUP35 | 0.066 | -0.0813 | 0.4302 | 0.67646 | 0.786 | -6.47 | 0.09136 | none |
| GJA8 | 0.038 | 0.0384 | 0.43 | 0.67663 | 0.786 | -6.47 | 0.90101 | none |
| EDNRA | 0.033 | 0.044 | 0.43 | 0.67666 | 0.786 | -6.47 | 1.36389 | none |
| DPAGT1 | 0.039 | -0.0035 | 0.4296 | 0.67692 | 0.786 | -6.47 | 0.73926 | none |
| C4ORF17 | -0.03 | -0.0128 | -0.429 | 0.67699 | 0.786 | -6.47 | 0 | none |
| MAGEA8 | 0.035 | 0.02 | 0.4292 | 0.67718 | 0.786 | -6.47 | -0.3871 | none |
| CPA1 | 0.036 | 0.046 | 0.429 | 0.67736 | 0.786 | -6.47 | 1.3263 | none |
| EN2 | 0.03 | 0.027 | 0.4285 | 0.67768 | 0.787 | -6.47 | 0 | none |

| | | | | | | | | |
|---|---|---|---|---|---|---|---|---|
| EGFL6 | -0.039 | -0.0108 | -0.428 | 0.67789 | 0.787 | -6.47 | 1.78453 | none |
| TFAP2D | 0.035 | 0.035 | 0.4282 | 0.67792 | 0.787 | -6.47 | 1.31366 | none |
| DGKI | 0.06 | 0.0628 | 0.428 | 0.67802 | 0.787 | -6.48 | -0.4302 | none |
| GRB10 | -0.076 | 0.0371 | -0.428 | 0.67828 | 0.787 | -6.48 | -0.0407 | none |
| PZP | -0.046 | 0.0079 | -0.427 | 0.67857 | 0.787 | -6.48 | 1.75348 | none |
| NTM | -0.06 | 0.0213 | -0.427 | 0.67872 | 0.787 | -6.48 | 0.5238 | none |
| FBXO31 | 0.125 | 0.2497 | 0.4268 | 0.6789 | 0.788 | -6.48 | -0.598 | none |
| ATAT1 | -0.053 | -0.0293 | -0.427 | 0.67906 | 0.788 | -6.48 | -0.3271 | none |
| ART1 | 0.036 | 0.0185 | 0.4263 | 0.67923 | 0.788 | -6.48 | -0.7668 | none |
| MAPRE2 | -0.078 | 0.0109 | -0.426 | 0.67925 | 0.788 | -6.48 | 1.05072 | none |
| TNFAIP8 | -0.032 | -0.0844 | -0.426 | 0.67929 | 0.788 | -6.48 | 0.7452 | none |
| SPTSSA | 0.058 | -0.0889 | 0.4261 | 0.67936 | 0.788 | -6.48 | 0.08651 | none |
| EIF3E | 0.048 | -0.0478 | 0.426 | 0.67942 | 0.788 | -6.48 | -0.3361 | none |
| ERICH6B | 0.127 | 0.2036 | 0.4254 | 0.67989 | 0.788 | -6.48 | 0.71721 | none |
| PPT2 | 0.106 | 0.0822 | 0.4252 | 0.67999 | 0.788 | -6.48 | 1.23322 | none |
| HTRA3 | -0.029 | -0.0104 | -0.425 | 0.68008 | 0.788 | -6.48 | 0.07215 | none |
| EIF4E2 | -0.062 | -0.0867 | -0.425 | 0.68008 | 0.788 | -6.48 | 0.16722 | none |
| RGL1 | 0.036 | 0.0537 | 0.4249 | 0.6802 | 0.789 | -6.48 | 1.50673 | none |
| MRPL38 | 0.05 | 0.0657 | 0.4248 | 0.68026 | 0.789 | -6.48 | 0.39828 | none |
| PTGFRN | 0.065 | 0.1253 | 0.4248 | 0.68027 | 0.789 | -6.48 | -0.6356 | none |
| SPATA4 | 0.042 | 0.0271 | 0.4248 | 0.68032 | 0.789 | -6.48 | -0.2038 | none |
| TBC1D1 | 0.058 | 0.0608 | 0.4243 | 0.68062 | 0.789 | -6.48 | -0.6722 | none |
| AQP1 | 0.044 | 0.0327 | 0.4243 | 0.68066 | 0.789 | -6.48 | 0.22162 | none |
| ZNF467 | 0.037 | 0.0319 | 0.4242 | 0.68072 | 0.789 | -6.48 | 0.4356 | none |
| CAMK2N2 | -0.03 | -0.0261 | -0.424 | 0.68111 | 0.789 | -6.48 | 0.36909 | none |
| LHX4 | -0.036 | -0.0252 | -0.423 | 0.68121 | 0.789 | -6.48 | 1.03129 | none |
| TMEM120 | -0.092 | -0.1042 | -0.423 | 0.68138 | 0.789 | -6.48 | 1.27627 | none |
| JCHAIN | -0.13 | -0.1054 | -0.423 | 0.68138 | 0.789 | -6.48 | 0.92051 | none |
| RCN2 | 0.058 | 0.0467 | 0.423 | 0.68155 | 0.79 | -6.48 | 1.33944 | none |
| HSD17B3 | -0.049 | -0.0152 | -0.423 | 0.68177 | 0.79 | -6.48 | 1.64654 | none |
| TM2D2 | -0.071 | -0.132 | -0.423 | 0.68191 | 0.79 | -6.48 | 1.03389 | none |
| SGPP1 | -0.057 | -0.0973 | -0.422 | 0.68195 | 0.79 | -6.48 | 1.06754 | none |
| SPATA24 | -0.089 | -0.0209 | -0.422 | 0.68209 | 0.79 | -6.48 | 0.92366 | none |
| AURKB | 0.04 | -0.0268 | 0.4222 | 0.68213 | 0.79 | -6.48 | 0.59081 | none |
| ESF1 | 0.117 | 0.2342 | 0.422 | 0.68224 | 0.79 | -6.48 | 1.27929 | none |
| STMN4 | -0.029 | -0.0269 | -0.422 | 0.68224 | 0.79 | -6.48 | -0.4587 | none |
| SCOC | -0.06 | -0.1591 | -0.421 | 0.68271 | 0.79 | -6.48 | 0.11771 | none |
| CRIP1 | -0.075 | -0.1333 | -0.421 | 0.68278 | 0.79 | -6.48 | 0.42272 | none |
| GATA2 | -0.066 | -0.0349 | -0.421 | 0.68289 | 0.79 | -6.48 | 0.77675 | none |
| MPPED2 | -0.043 | -0.036 | -0.421 | 0.68292 | 0.79 | -6.48 | -0.0236 | none |

| | | | | | | | | |
|---|---|---|---|---|---|---|---|---|
| OIP5 | -0.059 | -0.1016 | -0.421 | 0.68294 | 0.79 | -6.48 | 0.33961 | none |
| JUP | -0.054 | -0.0467 | -0.421 | 0.68312 | 0.791 | -6.48 | -0.7193 | none |
| ERG | 0.03 | 0.0541 | 0.4206 | 0.68326 | 0.791 | -6.48 | -0.8365 | none |
| TERF2IP | 0.035 | 0.0267 | 0.4198 | 0.68382 | 0.791 | -6.48 | 0.68608 | none |
| CISH | -0.044 | -0.0566 | -0.42 | 0.68387 | 0.791 | -6.48 | 1.73974 | none |
| KDM7A | 0.083 | 0.1101 | 0.4197 | 0.68387 | 0.791 | -6.48 | -0.1284 | none |
| PLSCR2 | 0.048 | -0.0039 | 0.4196 | 0.68394 | 0.791 | -6.48 | 0.2184 | none |
| SYCE1 | 0.045 | 0.0919 | 0.4194 | 0.68407 | 0.791 | -6.48 | 0.25377 | none |
| XRN2 | 0.051 | 0.0658 | 0.4194 | 0.68412 | 0.791 | -6.48 | 1.41227 | none |
| PFKL | 0.045 | 0.0755 | 0.4193 | 0.68418 | 0.791 | -6.48 | 1.1734 | none |
| IL24 | -0.083 | -0.1047 | -0.419 | 0.68432 | 0.792 | -6.48 | -0.2377 | none |
| OPLAH | -0.067 | -0.0784 | -0.419 | 0.68439 | 0.792 | -6.48 | 1.0576 | none |
| ENGASE | 0.073 | 0.0736 | 0.4189 | 0.68443 | 0.792 | -6.48 | -0.3679 | none |
| CSNK2B | 0.085 | -0.0143 | 0.4189 | 0.68447 | 0.792 | -6.48 | -0.0178 | none |
| LNP1 | -0.043 | -0.0187 | -0.419 | 0.68455 | 0.792 | -6.48 | 1.74229 | none |
| UBE2D2 | -0.05 | -0.112 | -0.419 | 0.68467 | 0.792 | -6.48 | 1.3361 | none |
| IP6K3 | 0.029 | 0.036 | 0.4186 | 0.68467 | 0.792 | -6.48 | -0.8455 | none |
| JADE3 | -0.055 | 0.0052 | -0.419 | 0.68473 | 0.792 | -6.48 | 0.86628 | none |
| TAS1R1 | -0.051 | -0.0122 | -0.418 | 0.68477 | 0.792 | -6.48 | 1.24935 | none |
| THEM6 | -0.042 | -0.0624 | -0.418 | 0.68508 | 0.792 | -6.48 | 1.77596 | none |
| DEUP1 | 0.03 | 0.0267 | 0.4179 | 0.68517 | 0.792 | -6.48 | -0.8842 | none |
| ARPC5L | 0.033 | 0.045 | 0.4179 | 0.68517 | 0.792 | -6.48 | 0.60716 | none |
| AWAT1 | -0.068 | -0.0169 | -0.418 | 0.68519 | 0.792 | -6.48 | 1.7235 | none |
| PPP3R1 | 0.071 | -0.0278 | 0.4178 | 0.68523 | 0.792 | -6.48 | 1.26832 | none |
| GLE1 | 0.086 | 0.1683 | 0.4178 | 0.68526 | 0.792 | -6.48 | 0.97911 | none |
| FNDC10 | 0.036 | -0.0245 | 0.4177 | 0.68533 | 0.792 | -6.48 | 0.35625 | none |
| CIB3 | -0.05 | -0.0022 | -0.417 | 0.68548 | 0.792 | -6.48 | 1.0129 | none |
| SLC22A15 | 0.037 | 0.0471 | 0.4173 | 0.68557 | 0.792 | -6.48 | 1.27404 | none |
| PDE4B | -0.072 | -0.1417 | -0.417 | 0.68567 | 0.792 | -6.48 | -0.6558 | none |
| KRTAP29-1 | 0.028 | 0.0226 | 0.4172 | 0.68567 | 0.792 | -6.48 | 0 | none |
| SESN2 | -0.049 | ######## | -0.417 | 0.68603 | 0.792 | -6.48 | 1.40997 | none |
| LIM2 | -0.03 | -0.0264 | -0.417 | 0.68612 | 0.792 | -6.48 | -0.1497 | none |
| SIX3 | 0.036 | 0.0041 | 0.4165 | 0.68613 | 0.792 | -6.48 | -0.0156 | none |
| PALM2-AK | -0.032 | -0.0149 | -0.416 | 0.68617 | 0.792 | -6.48 | 1.15957 | none |
| RDX | 0.086 | 0.1796 | 0.4162 | 0.68639 | 0.793 | -6.48 | 0.53195 | none |
| ICOS | -0.052 | -0.0445 | -0.416 | 0.68641 | 0.793 | -6.48 | 0.72846 | none |
| SFTPD | 0.039 | 0.0747 | 0.416 | 0.68652 | 0.793 | -6.48 | 0.33059 | none |
| PBX4 | -0.069 | 0.0099 | -0.416 | 0.68662 | 0.793 | -6.48 | 0.67073 | none |
| PP2D1 | 0.032 | 0.0446 | 0.4156 | 0.68676 | 0.793 | -6.48 | 0.83987 | none |
| HPGDS | -0.042 | 0.0239 | -0.415 | 0.68686 | 0.793 | -6.48 | 1.80598 | none |

| | | | | | | | | |
|---|---|---|---|---|---|---|---|---|
| LDHAL6B | -0.035 | -0.0153 | -0.415 | 0.68711 | 0.793 | -6.48 | -0.3836 | none |
| PKMYT1 | 0.04 | 0.05 | 0.415 | 0.6872 | 0.793 | -6.48 | 0.69968 | none |
| ZNF730 | 0.054 | 0.0705 | 0.4149 | 0.68725 | 0.793 | -6.48 | 1.14122 | none |
| REEP4 | -0.046 | -0.071 | -0.415 | 0.68729 | 0.793 | -6.48 | 1.51923 | none |
| PPEF1 | 0.067 | -0.0074 | 0.4149 | 0.68731 | 0.793 | -6.48 | -1.0305 | none |
| CIDEB | 0.112 | 0.1137 | 0.4147 | 0.68741 | 0.793 | -6.48 | 0.19741 | none |
| TCP10L | 0.057 | 0.0237 | 0.4145 | 0.68754 | 0.793 | -6.48 | 0.96274 | none |
| TNFSF8 | -0.076 | -0.0327 | -0.414 | 0.68762 | 0.793 | -6.48 | -0.1231 | none |
| LAMP2 | -0.063 | -0.1841 | -0.414 | 0.68764 | 0.793 | -6.48 | 0.52247 | none |
| SLAIN1 | 0.048 | -0.0396 | 0.4144 | 0.68765 | 0.793 | -6.48 | 0.78691 | none |
| B3GAT1 | 0.04 | 0.0181 | 0.4143 | 0.68772 | 0.793 | -6.48 | 0.7154 | none |
| NR2F1 | 0.028 | 0.037 | 0.414 | 0.68796 | 0.794 | -6.48 | -0.7324 | none |
| KCNJ10 | -0.029 | -0.0177 | -0.414 | 0.68824 | 0.794 | -6.48 | 0.79355 | none |
| H2AFB1 | 0.058 | 0.0385 | 0.4135 | 0.68829 | 0.794 | -6.48 | 1.29734 | none |
| LMBR1 | 0.054 | -0.052 | 0.4129 | 0.68866 | 0.794 | -6.48 | 0.35771 | none |
| MYLK2 | 0.029 | 0.0293 | 0.4127 | 0.68884 | 0.794 | -6.48 | -0.2847 | none |
| C1ORF87 | -0.048 | -0.0476 | -0.412 | 0.68906 | 0.795 | -6.48 | -0.1147 | none |
| MMS19 | -0.029 | 0.0093 | -0.412 | 0.68948 | 0.795 | -6.48 | 0.06136 | none |
| TXNL4A | -0.056 | -0.0442 | -0.411 | 0.68996 | 0.795 | -6.48 | 0.29303 | none |
| CTTN | 0.092 | 0.1747 | 0.4107 | 0.69024 | 0.796 | -6.48 | -0.5628 | none |
| DHFR | -0.063 | -0.0277 | -0.41 | 0.69058 | 0.796 | -6.48 | 0.366 | none |
| SLC16A14 | -0.05 | 0.0223 | -0.41 | 0.69063 | 0.796 | -6.48 | 1.53994 | none |
| RAPH1 | 0.058 | 0.0162 | 0.4096 | 0.69102 | 0.796 | -6.48 | 0.04364 | none |
| PXK | -0.051 | 0.0377 | -0.41 | 0.69103 | 0.796 | -6.48 | 1.23405 | none |
| PAK1IP1 | 0.055 | 0.1024 | 0.409 | 0.69143 | 0.797 | -6.48 | 0.55589 | none |
| ATRN | 0.064 | 0.154 | 0.4088 | 0.6916 | 0.797 | -6.48 | 0.36039 | none |
| ASAP2 | 0.036 | 0.05 | 0.4087 | 0.69171 | 0.797 | -6.48 | 0.8419 | none |
| NHEJ1 | -0.037 | -0.1271 | -0.409 | 0.69173 | 0.797 | -6.48 | 0.90559 | none |
| SLC22A2 | -0.041 | -0.0677 | -0.409 | 0.69178 | 0.797 | -6.48 | 0.47409 | none |
| CR1L | 0.03 | 0.0288 | 0.4085 | 0.69185 | 0.797 | -6.48 | -0.035 | none |
| HIST2H2A/ | -0.285 | -0.1525 | -0.408 | 0.69223 | 0.797 | -6.48 | 0.60113 | none |
| TCHHL1 | -0.027 | -0.0127 | -0.407 | 0.69263 | 0.798 | -6.48 | 0 | none |
| HOXA4 | 0.029 | 0.0472 | 0.4073 | 0.69265 | 0.798 | -6.48 | 0.35721 | none |
| RRAGD | -0.064 | -0.0807 | -0.407 | 0.69312 | 0.798 | -6.48 | 0.13312 | none |
| CMTM3 | -0.064 | -0.0319 | -0.406 | 0.69327 | 0.798 | -6.48 | 1.20798 | none |
| NUPR2 | -0.059 | -0.1319 | -0.406 | 0.6933 | 0.798 | -6.48 | 1.44797 | none |
| SNRNP70 | 0.078 | 0.1932 | 0.4062 | 0.69349 | 0.798 | -6.48 | 0.5864 | none |
| GNG2 | -0.048 | -0.1223 | -0.406 | 0.6935 | 0.798 | -6.48 | 1.374 | none |
| KLRG1 | -0.067 | -0.1294 | -0.406 | 0.69356 | 0.799 | -6.48 | 0.21487 | none |
| TMED7 | 0.048 | -0.0059 | 0.406 | 0.69361 | 0.799 | -6.48 | 1.53638 | none |

| | | | | | | | | |
|---|---|---|---|---|---|---|---|---|
| KCNV1 | 0.033 | 0.0439 | 0.4057 | 0.6938 | 0.799 | -6.48 | 0.27251 | none |
| TSNAX-DIS | 0.033 | 0.0371 | 0.4056 | 0.69391 | 0.799 | -6.48 | 1.34812 | none |
| TNXB | 0.049 | 0.0454 | 0.4054 | 0.69404 | 0.799 | -6.49 | 1.13566 | none |
| TRMT6 | -0.064 | 0.0063 | -0.405 | 0.69405 | 0.799 | -6.49 | 0.78209 | none |
| MRPS18C | 0.062 | -0.0464 | 0.4054 | 0.69405 | 0.799 | -6.49 | -0.4341 | none |
| TEX15 | 0.146 | 0.0961 | 0.4051 | 0.69423 | 0.799 | -6.49 | 0.84519 | none |
| GRN | -0.046 | -0.0141 | -0.405 | 0.69425 | 0.799 | -6.49 | 0.2719 | none |
| MAPK1 | 0.046 | -0.0068 | 0.405 | 0.69428 | 0.799 | -6.49 | -0.4544 | none |
| PPT2-EGFL | -0.061 | 0.0131 | -0.405 | 0.69437 | 0.799 | -6.49 | 1.89156 | none |
| TRPV6 | -0.05 | 0.0213 | -0.405 | 0.69439 | 0.799 | -6.49 | 1.39011 | none |
| ACAA1 | 0.042 | 0.0106 | 0.4047 | 0.69455 | 0.799 | -6.49 | 0.34932 | none |
| C2CD2 | 0.069 | 0.0361 | 0.4046 | 0.69458 | 0.799 | -6.49 | 0.06263 | none |
| SKP2 | 0.049 | -0.0146 | 0.4044 | 0.69472 | 0.799 | -6.49 | -0.5641 | none |
| VARS | 0.063 | 0.1112 | 0.4042 | 0.69488 | 0.799 | -6.49 | -0.0169 | none |
| CTPS2 | 0.061 | 0.1053 | 0.404 | 0.695 | 0.799 | -6.49 | 0.79612 | none |
| SLC22A18/ | -0.048 | -0.0582 | -0.404 | 0.69518 | 0.799 | -6.49 | -0.4378 | none |
| LMF1 | -0.043 | -0.0669 | -0.403 | 0.69538 | 0.8 | -6.49 | 0.09747 | none |
| CAMK1G | -0.06 | -0.0915 | -0.403 | 0.69553 | 0.8 | -6.49 | 0.36593 | none |
| NR2E3 | 0.047 | 0.0816 | 0.4026 | 0.69602 | 0.8 | -6.49 | 1.05982 | none |
| RRP8 | -0.04 | 0.0095 | -0.403 | 0.69606 | 0.8 | -6.49 | 1.26422 | none |
| GORASP2 | 0.04 | 0.0684 | 0.4021 | 0.69637 | 0.801 | -6.49 | 0.37312 | none |
| TMEM178 | -0.052 | -0.0122 | -0.402 | 0.69652 | 0.801 | -6.49 | 1.05754 | none |
| PTER | -0.04 | -0.0487 | -0.402 | 0.69663 | 0.801 | -6.49 | 0.81549 | none |
| GPR68 | -0.059 | -0.0373 | -0.402 | 0.69677 | 0.801 | -6.49 | 0.87387 | none |
| TTLL12 | 0.041 | 0.0995 | 0.4014 | 0.69688 | 0.801 | -6.49 | 0.59341 | none |
| MYRIP | 0.037 | 0.0526 | 0.4014 | 0.69689 | 0.801 | -6.49 | 0.27725 | none |
| RPL36AL | -0.048 | -0.1055 | -0.401 | 0.69697 | 0.801 | -6.49 | -0.4032 | none |
| SH3BGR | -0.055 | -0.0808 | -0.401 | 0.69707 | 0.801 | -6.49 | 1.31806 | none |
| MAGED2 | -0.046 | -0.029 | -0.401 | 0.69748 | 0.801 | -6.49 | 0.95495 | none |
| FRMD3 | 0.038 | 0.074 | 0.4005 | 0.6975 | 0.801 | -6.49 | 1.16102 | none |
| SSSCA1 | -0.052 | -0.0506 | -0.4 | 0.69796 | 0.802 | -6.49 | 0.02942 | none |
| MAGEL2 | -0.028 | -0.0095 | -0.4 | 0.69805 | 0.802 | -6.49 | 0 | none |
| AGFG1 | 0.077 | 0.045 | 0.3997 | 0.69809 | 0.802 | -6.49 | -0.2482 | none |
| WNT3 | 0.045 | 0.0689 | 0.3995 | 0.69819 | 0.802 | -6.49 | -0.3454 | none |
| PLTP | -0.05 | -0.0186 | -0.399 | 0.69831 | 0.802 | -6.49 | 0.87796 | none |
| CFD | 0.038 | 0.0084 | 0.3992 | 0.69846 | 0.802 | -6.49 | -0.4642 | none |
| ALKBH2 | 0.04 | -0.0256 | 0.3991 | 0.69848 | 0.802 | -6.49 | 0.39987 | none |
| IL2RA | 0.072 | 0.1911 | 0.399 | 0.69858 | 0.802 | -6.49 | 0.43675 | none |
| FSCN1 | 0.088 | 0.052 | 0.399 | 0.69859 | 0.802 | -6.49 | 1.04187 | none |
| C3ORF38 | -0.06 | -0.0904 | -0.399 | 0.69869 | 0.802 | -6.49 | 0.64192 | none |

| | | | | | | | | |
|---|---|---|---|---|---|---|---|---|
| TCEAL7 | 0.03 | 0.0274 | 0.3988 | 0.69872 | 0.802 | -6.49 | 0.53922 | none |
| PMP22 | -0.071 | -0.0823 | -0.399 | 0.69882 | 0.802 | -6.49 | -0.0268 | none |
| IGSF8 | 0.044 | 0.0014 | 0.3986 | 0.69889 | 0.802 | -6.49 | 0.65027 | none |
| NPM1 | -0.042 | 0.0111 | -0.398 | 0.69893 | 0.802 | -6.49 | 0.00287 | none |
| PPP1R3A | -0.032 | -0.0316 | -0.398 | 0.69911 | 0.802 | -6.49 | 0.99777 | none |
| CCM2L | 0.029 | 0.0342 | 0.3968 | 0.70018 | 0.803 | -6.49 | 0 | none |
| GIMAP7 | -0.049 | -0.1091 | -0.397 | 0.7002 | 0.803 | -6.49 | 0.51757 | none |
| GFPT2 | 0.096 | 0.1276 | 0.3967 | 0.70021 | 0.803 | -6.49 | 0.30284 | none |
| CXCL11 | 0.032 | 0.0334 | 0.3967 | 0.70023 | 0.803 | -6.49 | 1.46056 | none |
| SLC38A5 | -0.058 | -0.0702 | -0.397 | 0.70024 | 0.803 | -6.49 | 0.09993 | none |
| HYI | 0.114 | -0.0245 | 0.3965 | 0.70039 | 0.804 | -6.49 | 0.23552 | none |
| SLC35A1 | -0.034 | -0.1028 | -0.396 | 0.70056 | 0.804 | -6.49 | 1.05002 | none |
| RNF183 | 0.044 | 0.0276 | 0.3958 | 0.70085 | 0.804 | -6.49 | 1.44643 | none |
| STK32B | 0.03 | 0.0356 | 0.3957 | 0.70093 | 0.804 | -6.49 | 1.92528 | none |
| SH3RF3 | -0.028 | 0.0124 | -0.395 | 0.70114 | 0.804 | -6.49 | 0.49373 | none |
| MEP1B | 0.036 | 0.0709 | 0.3953 | 0.70119 | 0.804 | -6.49 | 0.6798 | none |
| CTU1 | 0.06 | 0.0349 | 0.3952 | 0.70129 | 0.804 | -6.49 | 0.22503 | none |
| HCLS1 | 0.048 | 0.0689 | 0.395 | 0.70144 | 0.804 | -6.49 | 0.39683 | none |
| TMEM176 | 0.03 | 0.0207 | 0.3949 | 0.70149 | 0.804 | -6.49 | 0.7208 | none |
| PREX1 | 0.064 | 0.1534 | 0.3949 | 0.70151 | 0.804 | -6.49 | 0.56206 | none |
| MCM2 | 0.054 | 0.1341 | 0.3941 | 0.70204 | 0.805 | -6.49 | 0.99155 | none |
| GNAT2 | 0.04 | 0.0434 | 0.3941 | 0.70209 | 0.805 | -6.49 | 0.87947 | none |
| PPP2R1A | 0.04 | 0.0657 | 0.394 | 0.70216 | 0.805 | -6.49 | 0.97666 | none |
| SLC51B | 0.032 | 0.0295 | 0.3938 | 0.70226 | 0.805 | -6.49 | 1.17546 | none |
| RTEL1-TNF | 0.035 | 0.08 | 0.3927 | 0.70306 | 0.806 | -6.49 | 0.78577 | none |
| FAM189A2 | 0.027 | 0.0429 | 0.3926 | 0.70311 | 0.806 | -6.49 | 0.30738 | none |
| NR1I2 | 0.04 | 0.0088 | 0.3925 | 0.70319 | 0.806 | -6.49 | 0.12553 | none |
| AC009977 | 0.042 | 0.0328 | 0.3924 | 0.70326 | 0.806 | -6.49 | -0.7269 | none |
| CPA3 | -0.044 | -0.0638 | -0.392 | 0.70369 | 0.806 | -6.49 | 0.52869 | none |
| CENPU | 0.082 | 0.1331 | 0.3917 | 0.70376 | 0.806 | -6.49 | 0.70724 | none |
| ZBTB18 | 0.035 | 0.0601 | 0.3915 | 0.70394 | 0.807 | -6.49 | 0.01715 | none |
| PYDC1 | -0.091 | -0.1174 | -0.391 | 0.70421 | 0.807 | -6.49 | 1.76417 | none |
| TRIM9 | -0.03 | -0.0357 | -0.391 | 0.70432 | 0.807 | -6.49 | 1.71273 | none |
| BBS10 | -0.031 | -0.0656 | -0.391 | 0.7046 | 0.807 | -6.49 | 0.71084 | none |
| C17ORF64 | -0.033 | -0.0217 | -0.391 | 0.70462 | 0.807 | -6.49 | -0.6339 | none |
| C17ORF97 | 0.041 | 0.0598 | 0.3901 | 0.70493 | 0.807 | -6.49 | 0.56801 | none |
| NR0B2 | 0.033 | 0.0431 | 0.39 | 0.70502 | 0.807 | -6.49 | -0.0449 | none |
| ZNF664 | 0.033 | -0.0362 | 0.39 | 0.70502 | 0.807 | -6.49 | -0.2656 | none |
| HPX | -0.123 | -0.0265 | -0.39 | 0.70511 | 0.807 | -6.49 | 0.15941 | none |
| CBR4 | 0.068 | -0.0363 | 0.3894 | 0.70547 | 0.808 | -6.49 | -0.1687 | none |

| | | | | | | | | |
|---|---|---|---|---|---|---|---|---|
| CFAP45 | 0.05 | 0.0766 | 0.3892 | 0.70556 | 0.808 | -6.49 | 0.33533 | none |
| AZI2 | 0.071 | 0.0914 | 0.3891 | 0.70564 | 0.808 | -6.49 | 1.6736 | none |
| CALU | -0.058 | -0.0458 | -0.389 | 0.70585 | 0.808 | -6.49 | 1.47369 | none |
| PRMT3 | -0.04 | -0.0523 | -0.389 | 0.70594 | 0.808 | -6.49 | 1.87514 | none |
| SLC16A7 | 0.059 | -0.0203 | 0.3885 | 0.70607 | 0.808 | -6.49 | 1.0326 | none |
| RSPH6A | 0.039 | 0.0066 | 0.3883 | 0.70625 | 0.808 | -6.49 | -1.1285 | none |
| CCDC58 | 0.046 | 0.0131 | 0.3882 | 0.70632 | 0.808 | -6.49 | 0.30574 | none |
| YIPF4 | 0.074 | -0.1003 | 0.388 | 0.70645 | 0.808 | -6.49 | 0.44606 | none |
| INSIG2 | 0.071 | -0.0061 | 0.3879 | 0.70648 | 0.808 | -6.49 | 0.29686 | none |
| USP17L11 | -0.09 | -0.0939 | -0.388 | 0.70651 | 0.808 | -6.49 | 0.77146 | none |
| TEX22 | 0.081 | 0.0586 | 0.3877 | 0.70663 | 0.808 | -6.49 | -0.6422 | none |
| RD3L | 0.031 | 0.0305 | 0.3877 | 0.70667 | 0.808 | -6.49 | -0.5388 | none |
| CCNI2 | -0.088 | -0.0205 | -0.388 | 0.70677 | 0.809 | -6.49 | 0.41891 | none |
| BAIAP2 | 0.073 | 0.0961 | 0.3871 | 0.70705 | 0.809 | -6.49 | 0.63334 | none |
| TIMM8A | 0.059 | 0.0303 | 0.3869 | 0.70719 | 0.809 | -6.49 | -0.3698 | none |
| ACER3 | 0.06 | 0.0023 | 0.3867 | 0.70733 | 0.809 | -6.49 | -0.234 | none |
| CHRNA9 | 0.035 | 0.0275 | 0.3864 | 0.70756 | 0.809 | -6.49 | 0.74028 | none |
| PSMC4 | 0.032 | 0.0123 | 0.3864 | 0.70759 | 0.809 | -6.49 | 0.80341 | none |
| MMP10 | 0.036 | 0.0249 | 0.386 | 0.7079 | 0.809 | -6.49 | 1.17321 | none |
| CRLF1 | -0.065 | 0.0243 | -0.386 | 0.70794 | 0.809 | -6.49 | -0.1087 | none |
| ALAS1 | -0.033 | 0.0099 | -0.386 | 0.70801 | 0.81 | -6.49 | 0.65225 | none |
| CLEC2B | -0.06 | -0.1985 | -0.386 | 0.7081 | 0.81 | -6.49 | 0.83977 | none |
| FGF9 | 0.054 | 0.0143 | 0.385 | 0.70857 | 0.81 | -6.49 | 0.31382 | none |
| PCGF2 | -0.026 | -0.0135 | -0.385 | 0.7086 | 0.81 | -6.49 | 1.51348 | none |
| TRIM39 | 0.055 | 0.0929 | 0.3849 | 0.70863 | 0.81 | -6.49 | 0.54615 | none |
| MAP6 | 0.026 | 0.0271 | 0.3844 | 0.70899 | 0.81 | -6.49 | -0.6441 | none |
| CLEC11A | 0.059 | 0.0314 | 0.3844 | 0.70899 | 0.81 | -6.49 | 1.21284 | none |
| VPS26A | -0.064 | -0.0762 | -0.384 | 0.70904 | 0.81 | -6.49 | 1.35033 | none |
| KIFC1 | -0.052 | 0.0312 | -0.384 | 0.70904 | 0.81 | -6.49 | 1.50888 | none |
| EZH2 | 0.063 | 0.1454 | 0.3843 | 0.70908 | 0.81 | -6.49 | 0.56726 | none |
| CTAG2 | -0.032 | -0.0385 | -0.384 | 0.70927 | 0.81 | -6.49 | 1.25984 | none |
| KIRREL2 | -0.028 | 0.0042 | -0.384 | 0.7093 | 0.81 | -6.49 | 0.18413 | none |
| BPIFA1 | -0.04 | -0.0404 | -0.384 | 0.70934 | 0.81 | -6.49 | 0.89019 | none |
| SLC2A1 | -0.045 | -0.049 | -0.384 | 0.70951 | 0.81 | -6.49 | 1.0253 | none |
| PLEKHO1 | -0.084 | -0.0324 | -0.384 | 0.70954 | 0.81 | -6.49 | 0.52638 | none |
| CHP2 | 0.03 | 0.082 | 0.3836 | 0.70956 | 0.81 | -6.49 | 1.98387 | none |
| HBEGF | -0.055 | 0.0286 | -0.384 | 0.70963 | 0.81 | -6.49 | 1.47719 | none |
| CFP | 0.044 | 0.0471 | 0.3833 | 0.70979 | 0.811 | -6.49 | 1.42735 | none |
| MMRN1 | -0.027 | -0.0263 | -0.383 | 0.70988 | 0.811 | -6.49 | 0.12253 | none |
| ALG2 | -0.06 | -0.0651 | -0.383 | 0.70988 | 0.811 | -6.49 | 1.35542 | none |

| | | | | | | | | |
|---|---|---|---|---|---|---|---|---|
| TLX1 | 0.089 | 0.0886 | 0.3831 | 0.70997 | 0.811 | -6.49 | -0.3767 | none |
| ARMC2 | -0.1 | -0.0399 | -0.383 | 0.71001 | 0.811 | -6.49 | 0.47189 | none |
| AKIRIN2 | -0.039 | -0.084 | -0.383 | 0.71002 | 0.811 | -6.49 | 1.44297 | none |
| ABCG4 | 0.034 | 0.0663 | 0.3828 | 0.71016 | 0.811 | -6.49 | 1.3165 | none |
| CARD11 | 0.126 | 0.3576 | 0.3826 | 0.71031 | 0.811 | -6.49 | 0.36343 | none |
| PACSIN1 | 0.058 | 0.0591 | 0.3825 | 0.71035 | 0.811 | -6.49 | 0.1111 | none |
| SLC10A6 | 0.038 | 0.0525 | 0.3825 | 0.71038 | 0.811 | -6.49 | -0.168 | none |
| LLPH | 0.046 | 0.0412 | 0.3824 | 0.71046 | 0.811 | -6.49 | -0.1187 | none |
| NOTCH2 | 0.083 | 0.1815 | 0.3823 | 0.7105 | 0.811 | -6.49 | -0.2092 | none |
| ARL6 | 0.062 | -0.0233 | 0.3823 | 0.71051 | 0.811 | -6.49 | -0.7312 | none |
| LAMP3 | 0.055 | -0.0059 | 0.3823 | 0.71055 | 0.811 | -6.49 | -0.0036 | none |
| AOX1 | 0.03 | 0.0445 | 0.3822 | 0.71063 | 0.811 | -6.49 | 1.45201 | none |
| HDHD2 | 0.052 | 0.0089 | 0.382 | 0.71077 | 0.811 | -6.49 | 0.98379 | none |
| HNRNPK | -0.036 | -0.0101 | -0.382 | 0.71079 | 0.811 | -6.49 | 0.10152 | none |
| TUNAR | 0.03 | 0.0438 | 0.3819 | 0.7108 | 0.811 | -6.49 | -0.2788 | none |
| CNOT6 | -0.041 | -0.0465 | -0.382 | 0.71082 | 0.811 | -6.49 | -0.2621 | none |
| CCK | 0.026 | 0.0241 | 0.3815 | 0.71108 | 0.811 | -6.5 | -0.6079 | none |
| PDZK1IP1 | -0.034 | -0.0036 | -0.381 | 0.71123 | 0.811 | -6.5 | -0.1396 | none |
| THG1L | 0.067 | 0.0118 | 0.3812 | 0.7113 | 0.811 | -6.5 | -0.7251 | none |
| BUD13 | 0.049 | 0.0898 | 0.3809 | 0.71156 | 0.811 | -6.5 | 1.13805 | none |
| CCND1 | -0.1 | -0.1158 | -0.381 | 0.7117 | 0.812 | -6.5 | 1.44553 | none |
| MKNK2 | 0.038 | 0.1096 | 0.3805 | 0.71183 | 0.812 | -6.5 | -0.116 | none |
| OTOS | 0.064 | 0.0642 | 0.38 | 0.7122 | 0.812 | -6.5 | 1.09964 | none |
| ATP2A1 | -0.06 | -0.0523 | -0.38 | 0.71224 | 0.812 | -6.5 | 0.14364 | none |
| PIWIL4 | 0.029 | 0.0374 | 0.3799 | 0.71227 | 0.812 | -6.5 | 1.28511 | none |
| AP3S2 | 0.077 | -0.0024 | 0.3798 | 0.71229 | 0.812 | -6.5 | -0.5013 | none |
| TBX20 | -0.041 | -0.0025 | -0.38 | 0.71237 | 0.812 | -6.5 | 1.04612 | none |
| OR4X1 | 0.04 | 0.0401 | 0.3793 | 0.71267 | 0.812 | -6.5 | -0.0076 | none |
| CSF1 | 0.079 | 0.2404 | 0.3792 | 0.71274 | 0.812 | -6.5 | 0.19541 | none |
| ZBTB46 | 0.041 | 0.1028 | 0.3792 | 0.71277 | 0.812 | -6.5 | 0.82132 | none |
| HEPACAM | 0.036 | 0.0419 | 0.3791 | 0.71283 | 0.812 | -6.5 | 0.8214 | none |
| RASGEF1A | -0.052 | 0.007 | -0.379 | 0.71287 | 0.812 | -6.5 | -0.0083 | none |
| EFCAB8 | -0.078 | -0.0186 | -0.379 | 0.71288 | 0.812 | -6.5 | 1.02625 | none |
| HGF | 0.027 | 0.0283 | 0.379 | 0.7129 | 0.812 | -6.5 | -0.4333 | none |
| F8A1 | -0.04 | 0.0283 | -0.378 | 0.71346 | 0.813 | -6.5 | -0.1162 | none |
| RIOK1 | 0.054 | 0.1177 | 0.378 | 0.71362 | 0.813 | -6.5 | 0.46425 | none |
| S100A14 | -0.026 | -0.0348 | -0.378 | 0.71369 | 0.813 | -6.5 | 0 | none |
| IRX6 | -0.026 | -0.0262 | -0.378 | 0.71384 | 0.813 | -6.5 | 0 | none |
| G3BP1 | 0.036 | 0.005 | 0.3774 | 0.71402 | 0.813 | -6.5 | 0.31141 | none |
| CROT | -0.039 | -0.0625 | -0.377 | 0.71409 | 0.813 | -6.5 | 1.25692 | none |

| | | | | | | | | |
|---|---|---|---|---|---|---|---|---|
| CCNG1 | 0.044 | -0.0787 | 0.3773 | 0.71412 | 0.813 | -6.5 | 1.29614 | none |
| KRT12 | -0.043 | -0.0815 | -0.377 | 0.71417 | 0.813 | -6.5 | -0.7065 | none |
| OR51B4 | 0.042 | 0.063 | 0.3767 | 0.71451 | 0.814 | -6.5 | -0.1317 | none |
| ORC3 | -0.056 | -0.0419 | -0.376 | 0.71484 | 0.814 | -6.5 | 1.16606 | none |
| TSNAXIP1 | -0.072 | 0.0304 | -0.376 | 0.71503 | 0.814 | -6.5 | 1.27549 | none |
| KIF2C | -0.048 | 0.0287 | -0.376 | 0.71526 | 0.814 | -6.5 | 0.64869 | none |
| KIF2A | 0.078 | 0.1417 | 0.3756 | 0.71531 | 0.814 | -6.5 | 0.61501 | none |
| CDKL4 | 0.047 | 0.0254 | 0.3756 | 0.71534 | 0.814 | -6.5 | 1.63253 | none |
| LTF | 0.074 | 0.0839 | 0.3748 | 0.71593 | 0.815 | -6.5 | -0.6586 | none |
| SERPINE2 | 0.034 | 0.0129 | 0.3746 | 0.71604 | 0.815 | -6.5 | 0.22001 | none |
| PEMT | 0.055 | 0.0412 | 0.3745 | 0.71613 | 0.815 | -6.5 | 1.12378 | none |
| TPRA1 | -0.042 | -0.0442 | -0.374 | 0.71614 | 0.815 | -6.5 | 0.56278 | none |
| SLC26A6 | 0.037 | 0.0676 | 0.3743 | 0.71628 | 0.815 | -6.5 | 1.94331 | none |
| ZIC2 | 0.066 | 0.0671 | 0.3741 | 0.71641 | 0.815 | -6.5 | 1.22771 | none |
| CD300E | -0.027 | -0.0177 | -0.374 | 0.71668 | 0.815 | -6.5 | 1.89954 | none |
| GPI | -0.052 | -0.0077 | -0.374 | 0.71673 | 0.815 | -6.5 | 1.26231 | none |
| ATRAID | -0.053 | -0.0975 | -0.373 | 0.71688 | 0.815 | -6.5 | 0.16927 | none |
| MAP1LC3E | -0.039 | -0.0747 | -0.373 | 0.71726 | 0.816 | -6.5 | 1.20013 | none |
| TFAP2C | 0.044 | 0.0566 | 0.3729 | 0.71731 | 0.816 | -6.5 | 0.32553 | none |
| TBC1D5 | 0.113 | 0.1775 | 0.3724 | 0.71762 | 0.816 | -6.5 | 0.32342 | none |
| CTSO | -0.084 | -0.195 | -0.372 | 0.71779 | 0.816 | -6.5 | 0.47776 | none |
| TOR1B | 0.046 | 0.0151 | 0.3719 | 0.71803 | 0.817 | -6.5 | 0.7794 | none |
| CEP44 | 0.045 | -0.0712 | 0.3717 | 0.71813 | 0.817 | -6.5 | 0.44251 | none |
| CNOT11 | -0.029 | -0.0408 | -0.371 | 0.71855 | 0.817 | -6.5 | 0.98694 | none |
| TSG101 | 0.042 | 0.024 | 0.3706 | 0.71891 | 0.817 | -6.5 | 0.29952 | none |
| HECTD2 | 0.072 | 0.068 | 0.3705 | 0.71902 | 0.817 | -6.5 | 0.92324 | none |
| TMEM64 | 0.062 | -0.0323 | 0.3702 | 0.71923 | 0.818 | -6.5 | 0.85512 | none |
| DMRTC1 | -0.03 | -0.0325 | -0.37 | 0.71925 | 0.818 | -6.5 | 1.21117 | none |
| BCCIP | -0.028 | -0.0022 | -0.37 | 0.71954 | 0.818 | -6.5 | 0.58061 | none |
| CCDC13 | 0.134 | 0.1671 | 0.3697 | 0.71962 | 0.818 | -6.5 | 0.37635 | none |
| APOC2 | 0.053 | -0.0129 | 0.3695 | 0.7197 | 0.818 | -6.5 | -0.501 | none |
| MSRB2 | 0.057 | 0.025 | 0.3694 | 0.71978 | 0.818 | -6.5 | 0.01578 | none |
| C1ORF94 | -0.026 | -0.0259 | -0.369 | 0.71982 | 0.818 | -6.5 | 1.85582 | none |
| NANP | -0.054 | -0.0426 | -0.369 | 0.71991 | 0.818 | -6.5 | 1.33189 | none |
| TRIM8 | 0.077 | 0.1527 | 0.3683 | 0.72057 | 0.819 | -6.5 | 0.18329 | none |
| TEX19 | -0.03 | -0.0254 | -0.368 | 0.72077 | 0.819 | -6.5 | 0.34774 | none |
| B3GALT6 | 0.059 | 0.0536 | 0.3677 | 0.72102 | 0.819 | -6.5 | -0.5827 | none |
| PTDSS1 | -0.052 | -0.1142 | -0.368 | 0.72103 | 0.819 | -6.5 | -0.2083 | none |
| FOXF2 | -0.025 | -0.0238 | -0.367 | 0.72134 | 0.819 | -6.5 | -0.5423 | none |
| TFAP2E | 0.032 | 0.0564 | 0.3672 | 0.72141 | 0.819 | -6.5 | 1.19666 | none |

| | | | | | | | | |
|---|---|---|---|---|---|---|---|---|
| BACE2 | 0.056 | 0.0039 | 0.3671 | 0.72145 | 0.819 | -6.5 | 0.13409 | none |
| CHRNA1 | 0.025 | 0.0309 | 0.3671 | 0.72145 | 0.819 | -6.5 | 0 | none |
| MFSD5 | 0.048 | -0.0253 | 0.3669 | 0.72159 | 0.819 | -6.5 | 0.62998 | none |
| OSM | -0.053 | -0.014 | -0.367 | 0.72188 | 0.82 | -6.5 | 0.85223 | none |
| SPINT2 | -0.067 | -0.0747 | -0.366 | 0.72191 | 0.82 | -6.5 | 1.01313 | none |
| ZBTB9 | -0.107 | -0.1014 | -0.366 | 0.72194 | 0.82 | -6.5 | 0.6226 | none |
| DNAI1 | 0.042 | 0.091 | 0.3662 | 0.72214 | 0.82 | -6.5 | 1.51118 | none |
| TPX2 | 0.131 | 0.3263 | 0.3661 | 0.72218 | 0.82 | -6.5 | 0.52038 | none |
| DPP8 | 0.069 | 0.0942 | 0.3657 | 0.72245 | 0.82 | -6.5 | -0.0669 | none |
| RGPD8 | -0.081 | 0.0767 | -0.366 | 0.72248 | 0.82 | -6.5 | 0.75863 | none |
| FAM160A1 | 0.026 | 0.031 | 0.3655 | 0.72264 | 0.82 | -6.5 | 1.81031 | none |
| DUSP23 | -0.064 | -0.0737 | -0.365 | 0.72272 | 0.82 | -6.5 | 0.67544 | none |
| RBMXL1 | 0.059 | 0.0861 | 0.3653 | 0.7228 | 0.82 | -6.5 | 1.02821 | none |
| PRSS48 | -0.039 | -0.0386 | -0.365 | 0.72288 | 0.82 | -6.5 | -0.233 | none |
| PLGLB2 | 0.056 | -0.0182 | 0.3649 | 0.72309 | 0.82 | -6.5 | 0.26226 | none |
| ST6GALNA | 0.029 | 0.0237 | 0.3648 | 0.72312 | 0.82 | -6.5 | -0.6745 | none |
| LPA | -0.029 | -0.0199 | -0.364 | 0.72339 | 0.821 | -6.5 | 0.92621 | none |
| SLC15A3 | -0.051 | -0.0527 | -0.364 | 0.72341 | 0.821 | -6.5 | 1.18289 | none |
| MEX3B | 0.027 | 0.05 | 0.3643 | 0.72348 | 0.821 | -6.5 | 1.62487 | none |
| PSMC3IP | 0.041 | 0.1272 | 0.3643 | 0.7235 | 0.821 | -6.5 | -0.4788 | none |
| PTPRN2 | -0.03 | -0.015 | -0.364 | 0.72358 | 0.821 | -6.5 | 1.30347 | none |
| CENPP | 0.037 | 0.0313 | 0.3638 | 0.72382 | 0.821 | -6.5 | 0.89907 | none |
| DESI1 | -0.047 | -0.0769 | -0.364 | 0.72385 | 0.821 | -6.5 | 1.01944 | none |
| MECOM | 0.101 | 0.1451 | 0.3638 | 0.72387 | 0.821 | -6.5 | 0.93268 | none |
| NUAK1 | -0.038 | ######## | -0.364 | 0.72393 | 0.821 | -6.5 | -0.3611 | none |
| CPE | 0.059 | 0.1016 | 0.3635 | 0.72408 | 0.821 | -6.5 | 0.07411 | none |
| EHD1 | -0.039 | -0.0299 | -0.363 | 0.72414 | 0.821 | -6.5 | 0.99643 | none |
| CEACAM2( | 0.031 | 0.0025 | 0.3626 | 0.72469 | 0.822 | -6.5 | 0.2341 | none |
| MED31 | 0.079 | -0.0424 | 0.3625 | 0.72478 | 0.822 | -6.5 | -0.304 | none |
| CBLC | 0.029 | 0.0443 | 0.3624 | 0.72486 | 0.822 | -6.5 | 0.91162 | none |
| GUCY2D | -0.025 | -0.0038 | -0.362 | 0.72492 | 0.822 | -6.5 | 0 | none |
| DMRTC1B | 0.03 | 0.0193 | 0.3621 | 0.72508 | 0.822 | -6.5 | -0.1068 | none |
| SNAP47 | 0.026 | 0.0225 | 0.3619 | 0.7252 | 0.822 | -6.5 | 0.64466 | none |
| PRRT2 | 0.123 | 0.0522 | 0.3618 | 0.7253 | 0.822 | -6.5 | 0.6186 | none |
| EDN2 | -0.044 | 0.0036 | -0.361 | 0.72561 | 0.822 | -6.5 | 0.18495 | none |
| RARG | -0.035 | -0.0044 | -0.361 | 0.72569 | 0.822 | -6.5 | 0.88498 | none |
| CISD2 | -0.055 | -0.1035 | -0.361 | 0.72569 | 0.822 | -6.5 | 1.28546 | none |
| SURF1 | -0.054 | -0.0704 | -0.361 | 0.72582 | 0.822 | -6.5 | 0.89663 | none |
| NAPRT | -0.076 | -0.0905 | -0.361 | 0.72591 | 0.822 | -6.5 | 0.23855 | none |
| CD5 | 0.09 | 0.1835 | 0.3608 | 0.72602 | 0.822 | -6.5 | 0.8014 | none |

| | | | | | | | | |
|---|---|---|---|---|---|---|---|---|
| GALT | 0.039 | 0.0127 | 0.3607 | 0.72607 | 0.822 | -6.5 | 1.20763 | none |
| ADH6 | -0.046 | -0.0653 | -0.361 | 0.72617 | 0.822 | -6.5 | 1.19163 | none |
| STARD7 | 0.041 | -0.0338 | 0.3602 | 0.72648 | 0.823 | -6.5 | 1.20143 | none |
| KCNK15 | -0.03 | -0.0322 | -0.36 | 0.72653 | 0.823 | -6.5 | 1.17693 | none |
| PSMD9 | 0.046 | -0.0176 | 0.3598 | 0.72672 | 0.823 | -6.5 | -0.7583 | none |
| ACOD1 | -0.026 | -0.028 | -0.36 | 0.72674 | 0.823 | -6.5 | 0 | none |
| IL37 | 0.025 | 0.0334 | 0.3596 | 0.72689 | 0.823 | -6.5 | 0 | none |
| STAP1 | -0.068 | -0.0502 | -0.36 | 0.72696 | 0.823 | -6.5 | 1.22086 | none |
| TAF9B | 0.037 | -0.0167 | 0.359 | 0.72735 | 0.823 | -6.5 | 0.07828 | none |
| EIF2B4 | 0.025 | 0.0587 | 0.3588 | 0.72744 | 0.823 | -6.5 | 1.01206 | none |
| FAM3C | 0.044 | 0.1094 | 0.3587 | 0.72754 | 0.823 | -6.5 | 0.85188 | none |
| CAMK2B | 0.066 | 0.0218 | 0.3582 | 0.72788 | 0.824 | -6.5 | 0.23942 | none |
| PROSER2 | -0.03 | -0.0097 | -0.358 | 0.72791 | 0.824 | -6.5 | -0.5517 | none |
| AGPAT5 | -0.07 | -0.0909 | -0.358 | 0.72796 | 0.824 | -6.5 | 0.91318 | none |
| HSFX1 | 0.038 | 0.017 | 0.3581 | 0.72801 | 0.824 | -6.5 | -0.6461 | none |
| ZNF697 | 0.025 | 0.0453 | 0.358 | 0.72806 | 0.824 | -6.5 | 0.50061 | none |
| PSKH1 | 0.052 | 0.1296 | 0.3579 | 0.72812 | 0.824 | -6.5 | 0.78316 | none |
| CNPY3 | -0.042 | -0.0162 | -0.358 | 0.72828 | 0.824 | -6.5 | 0.78379 | none |
| GPR32 | -0.032 | -0.0049 | -0.358 | 0.7283 | 0.824 | -6.5 | 0.21839 | none |
| GPS2 | 0.024 | 0.0633 | 0.3575 | 0.72844 | 0.824 | -6.5 | 0.09315 | none |
| LHX2 | -0.042 | -0.0452 | -0.357 | 0.72893 | 0.825 | -6.5 | 0.87164 | none |
| NEUROD1 | -0.024 | -0.0154 | -0.356 | 0.72914 | 0.825 | -6.5 | 0 | none |
| RABAC1 | 0.068 | -0.0243 | 0.3565 | 0.72917 | 0.825 | -6.5 | 0.48032 | none |
| CHTF18 | 0.064 | 0.162 | 0.3564 | 0.72922 | 0.825 | -6.5 | 0.98374 | none |
| LIPN | -0.025 | -0.0415 | -0.356 | 0.7295 | 0.825 | -6.51 | 1.49939 | none |
| SLC25A22 | 0.036 | 0.0313 | 0.3558 | 0.72965 | 0.825 | -6.51 | 1.40134 | none |
| SCN5A | -0.038 | 0.0071 | -0.356 | 0.72965 | 0.825 | -6.51 | 0.83989 | none |
| OPCML | -0.026 | -0.0254 | -0.356 | 0.72966 | 0.825 | -6.51 | 1.94337 | none |
| GATA1 | -0.025 | -0.0174 | -0.356 | 0.72974 | 0.825 | -6.51 | 1.09507 | none |
| STX10 | 0.056 | -0.0017 | 0.3555 | 0.72988 | 0.825 | -6.51 | 0.10256 | none |
| CENPT | 0.059 | 0.0617 | 0.3554 | 0.72994 | 0.825 | -6.51 | 0.83614 | none |
| ITLN1 | 0.026 | 0.0507 | 0.355 | 0.73022 | 0.825 | -6.51 | -0.5751 | none |
| EME2 | 0.059 | 0.0923 | 0.3549 | 0.73027 | 0.825 | -6.51 | -0.1221 | none |
| RBM4B | 0.045 | -0.0011 | 0.3545 | 0.73057 | 0.826 | -6.51 | -0.4536 | none |
| GPR183 | 0.04 | -0.0991 | 0.3544 | 0.73063 | 0.826 | -6.51 | 0.30125 | none |
| BBS7 | 0.052 | 0.008 | 0.3544 | 0.73069 | 0.826 | -6.51 | 0.89662 | none |
| HSPA9 | 0.053 | 0.1374 | 0.3542 | 0.73081 | 0.826 | -6.51 | 1.14557 | none |
| C4B | 0.106 | 0.0714 | 0.3542 | 0.73082 | 0.826 | -6.51 | 0.36241 | none |
| ADGRG6 | 0.033 | 0.0498 | 0.3541 | 0.73086 | 0.826 | -6.51 | 1.0219 | none |
| HADHA | -0.068 | 0.0318 | -0.354 | 0.73099 | 0.826 | -6.51 | 1.15739 | none |

| | | | | | | | | |
|---|---|---|---|---|---|---|---|---|
| HSD17B11 | -0.09 | -0.2184 | -0.354 | 0.73102 | 0.826 | -6.51 | 0.64268 | none |
| TM7SF2 | 0.069 | ######## | 0.3536 | 0.73123 | 0.826 | -6.51 | 1.15606 | none |
| DND1 | -0.052 | -0.0681 | -0.354 | 0.7313 | 0.826 | -6.51 | 0.48981 | none |
| WFIKKN1 | -0.024 | -0.0074 | -0.353 | 0.73137 | 0.826 | -6.51 | -0.2294 | none |
| STAMBP | -0.043 | -0.0309 | -0.353 | 0.73179 | 0.826 | -6.51 | 0.4918 | none |
| EFS | -0.027 | -0.0221 | -0.353 | 0.73183 | 0.826 | -6.51 | 1.23425 | none |
| RHOV | 0.026 | 0.0376 | 0.3526 | 0.73199 | 0.827 | -6.51 | 0.23513 | none |
| MAGEA2B | 0.038 | 0.067 | 0.3524 | 0.7321 | 0.827 | -6.51 | 1.65244 | none |
| PDE9A | -0.095 | 0.0099 | -0.352 | 0.73219 | 0.827 | -6.51 | 1.64639 | none |
| VWA3A | -0.033 | -0.0322 | -0.352 | 0.7322 | 0.827 | -6.51 | 0.22359 | none |
| MRPL17 | 0.06 | 0.0129 | 0.3522 | 0.73222 | 0.827 | -6.51 | -0.8312 | none |
| GFER | 0.054 | 0.0417 | 0.3516 | 0.73271 | 0.827 | -6.51 | 0.70076 | none |
| KIF26A | 0.025 | 0.0289 | 0.3515 | 0.73275 | 0.827 | -6.51 | 0 | none |
| IFNLR1 | 0.036 | 0.0416 | 0.3515 | 0.73278 | 0.827 | -6.51 | 1.32527 | none |
| 11-Mar | -0.044 | -0.0164 | -0.351 | 0.73283 | 0.827 | -6.51 | 2.07092 | none |
| SLC8A3 | -0.023 | -0.0192 | -0.351 | 0.7329 | 0.827 | -6.51 | 1.7151 | none |
| TCF3 | 0.066 | 0.1762 | 0.351 | 0.73309 | 0.827 | -6.51 | 0.36234 | none |
| LCK | -0.032 | -0.0423 | -0.351 | 0.73326 | 0.827 | -6.51 | -0.7312 | none |
| VGF | -0.025 | -0.0236 | -0.351 | 0.73327 | 0.827 | -6.51 | 1.64073 | none |
| PELI3 | 0.038 | 0.0883 | 0.3507 | 0.73337 | 0.827 | -6.51 | -0.1892 | none |
| MRPS16 | -0.062 | -0.145 | -0.35 | 0.73378 | 0.828 | -6.51 | 0.50907 | none |
| PEX19 | -0.036 | -0.108 | -0.35 | 0.73396 | 0.828 | -6.51 | 1.3372 | none |
| ASB12 | 0.025 | 0.0278 | 0.3496 | 0.73416 | 0.828 | -6.51 | 1.4074 | none |
| ABCB10 | 0.054 | 0.0285 | 0.3495 | 0.73421 | 0.828 | -6.51 | 0.80037 | none |
| ARV1 | 0.063 | -0.0123 | 0.349 | 0.73461 | 0.828 | -6.51 | -0.0584 | none |
| NTN5 | -0.034 | -0.0099 | -0.349 | 0.73462 | 0.828 | -6.51 | -0.1257 | none |
| MIIP | -0.047 | -0.0861 | -0.349 | 0.73474 | 0.829 | -6.51 | 1.42854 | none |
| NDUFAF5 | -0.048 | -0.0998 | -0.349 | 0.73481 | 0.829 | -6.51 | 0.28252 | none |
| VLDLR | 0.052 | 0.06 | 0.3484 | 0.73499 | 0.829 | -6.51 | 0.865 | none |
| OR1A2 | -0.026 | -0.0052 | -0.348 | 0.735 | 0.829 | -6.51 | 0.56424 | none |
| AL136097. | 0.025 | 0.0215 | 0.3483 | 0.73508 | 0.829 | -6.51 | 0.59664 | none |
| DTNBP1 | 0.038 | 0.1369 | 0.3483 | 0.73508 | 0.829 | -6.51 | 0.9312 | none |
| PTTG2 | 0.049 | 0.1152 | 0.3483 | 0.73509 | 0.829 | -6.51 | 0.49138 | none |
| TMCO5A | -0.049 | -0.0491 | -0.348 | 0.73515 | 0.829 | -6.51 | 1.06393 | none |
| NRAS | -0.06 | -0.0935 | -0.348 | 0.7352 | 0.829 | -6.51 | 0.83143 | none |
| SSUH2 | -0.079 | 0.0172 | -0.348 | 0.73521 | 0.829 | -6.51 | 1.46968 | none |
| GOLGA6L2 | 0.025 | 0.0255 | 0.3481 | 0.73522 | 0.829 | -6.51 | 1.55248 | none |
| MEPE | -0.025 | -0.0251 | -0.348 | 0.73531 | 0.829 | -6.51 | 0.6513 | none |
| TCTN3 | -0.049 | -0.0522 | -0.348 | 0.73534 | 0.829 | -6.51 | 1.44634 | none |
| NBN | 0.088 | 0.1834 | 0.3474 | 0.73577 | 0.829 | -6.51 | 0.462 | none |

| | | | | | | | | |
|---|---|---|---|---|---|---|---|---|
| MOCS1 | -0.063 | -0.067 | -0.347 | 0.73582 | 0.829 | -6.51 | 0.11399 | none |
| NT5DC4 | -0.068 | 0.0515 | -0.347 | 0.73588 | 0.829 | -6.51 | 0.55452 | none |
| BET1 | -0.048 | -0.1472 | -0.347 | 0.73592 | 0.829 | -6.51 | 0.85461 | none |
| KRT26 | 0.029 | 0.0289 | 0.347 | 0.736 | 0.829 | -6.51 | -0.1048 | none |
| WNT9A | 0.028 | 0.0521 | 0.3469 | 0.73607 | 0.829 | -6.51 | -0.0055 | none |
| IL36A | -0.04 | -0.0251 | -0.347 | 0.73627 | 0.829 | -6.51 | 1.60509 | none |
| MCM6 | -0.05 | 0.0366 | -0.347 | 0.73632 | 0.829 | -6.51 | 0.95408 | none |
| IL36G | 0.027 | 0.0352 | 0.3466 | 0.73633 | 0.829 | -6.51 | 0.27196 | none |
| MLNR | 0.026 | 0.0332 | 0.3462 | 0.73663 | 0.829 | -6.51 | -0.6913 | none |
| CTNNBL1 | 0.041 | 0.0971 | 0.3462 | 0.73665 | 0.829 | -6.51 | 0.46377 | none |
| ARL15 | -0.032 | -0.1446 | -0.346 | 0.73681 | 0.829 | -6.51 | 0.51448 | none |
| HDAC8 | 0.04 | 0.018 | 0.3459 | 0.73685 | 0.829 | -6.51 | 0.36066 | none |
| CDKL3 | -0.14 | -0.1407 | -0.346 | 0.73685 | 0.829 | -6.51 | 0.85614 | none |
| SERPINB2 | 0.03 | 0.0404 | 0.3455 | 0.73709 | 0.83 | -6.51 | 0.42446 | none |
| FAM110B | 0.066 | 0.0789 | 0.3453 | 0.73729 | 0.83 | -6.51 | -0.5462 | none |
| TIPRL | 0.046 | -0.0585 | 0.344 | 0.7382 | 0.831 | -6.51 | -0.2288 | none |
| IGFALS | 0.061 | 0.0288 | 0.344 | 0.73821 | 0.831 | -6.51 | 1.41899 | none |
| DBF4 | 0.086 | 0.1643 | 0.3439 | 0.73826 | 0.831 | -6.51 | 0.38225 | none |
| HIST1H2BI | 0.073 | 0.041 | 0.3434 | 0.73864 | 0.831 | -6.51 | -0.1677 | none |
| DENND1C | 0.079 | 0.1785 | 0.3433 | 0.73872 | 0.831 | -6.51 | 0.30214 | none |
| CALM3 | -0.028 | 0.001 | -0.343 | 0.73875 | 0.831 | -6.51 | -0.1459 | none |
| CERK | -0.031 | -0.0285 | -0.343 | 0.73877 | 0.831 | -6.51 | -0.4474 | none |
| NEURL2 | -0.04 | -0.0658 | -0.343 | 0.73882 | 0.831 | -6.51 | 1.76428 | none |
| LPP | 0.047 | 0.1267 | 0.3431 | 0.73885 | 0.831 | -6.51 | -0.1524 | none |
| HOXA9 | -0.026 | -0.022 | -0.343 | 0.73894 | 0.831 | -6.51 | 1.59515 | none |
| DPM3 | -0.076 | -0.1657 | -0.343 | 0.73896 | 0.831 | -6.51 | -0.2811 | none |
| RCBTB2 | 0.056 | -0.0453 | 0.343 | 0.73897 | 0.831 | -6.51 | -0.4444 | none |
| NUP62CL | 0.057 | 0.0818 | 0.3429 | 0.73903 | 0.831 | -6.51 | 0.5339 | none |
| CDK2AP2 | -0.039 | -0.0305 | -0.343 | 0.73909 | 0.831 | -6.51 | 1.38838 | none |
| COMMD1 | -0.054 | -0.1318 | -0.343 | 0.73921 | 0.831 | -6.51 | 0.33558 | none |
| OSGEPL1 | -0.053 | -0.1275 | -0.342 | 0.73947 | 0.831 | -6.51 | 0.30057 | none |
| RUNDC3B | 0.074 | 0.0172 | 0.3421 | 0.73958 | 0.831 | -6.51 | 0.65126 | none |
| CORT | 0.034 | 0.039 | 0.3419 | 0.73973 | 0.832 | -6.51 | -0.1174 | none |
| E2F3 | 0.051 | 0.0815 | 0.3419 | 0.73975 | 0.832 | -6.51 | 0.96065 | none |
| REXO1 | -0.076 | 0.0946 | -0.342 | 0.7398 | 0.832 | -6.51 | 0.11158 | none |
| NT5C1B-RI | 0.025 | 0.0197 | 0.3416 | 0.74 | 0.832 | -6.51 | 0.77608 | none |
| CACNA2D1 | -0.04 | -0.0604 | -0.341 | 0.74004 | 0.832 | -6.51 | 0.73895 | none |
| TAL1 | 0.032 | 0.0449 | 0.3414 | 0.74013 | 0.832 | -6.51 | 0.57633 | none |
| SCG3 | -0.025 | -0.0179 | -0.341 | 0.74019 | 0.832 | -6.51 | 1.3149 | none |
| ERLIN2 | 0.049 | -0.0484 | 0.3413 | 0.74021 | 0.832 | -6.51 | -0.6924 | none |

| | | | | | | | | |
|---|---|---|---|---|---|---|---|---|
| KDM4D | 0.03 | 0.0739 | 0.3412 | 0.74028 | 0.832 | -6.51 | 0.87949 | none |
| ISG15 | -0.062 | -0.1326 | -0.341 | 0.7406 | 0.832 | -6.51 | 0.03594 | none |
| GPR179 | 0.041 | 0.0325 | 0.3407 | 0.74064 | 0.832 | -6.51 | 0.51427 | none |
| ZNF112 | 0.023 | 0.0228 | 0.3404 | 0.74084 | 0.832 | -6.51 | 0 | none |
| ZNF729 | 0.024 | 0.024 | 0.3402 | 0.74098 | 0.832 | -6.51 | -0.4115 | none |
| MPI | 0.029 | 0.0805 | 0.3402 | 0.74102 | 0.832 | -6.51 | 0.51297 | none |
| SMIM8 | 0.049 | -0.0653 | 0.3401 | 0.74104 | 0.832 | -6.51 | -0.1997 | none |
| EGR1 | 0.066 | 0.1664 | 0.3399 | 0.7412 | 0.832 | -6.51 | -0.2132 | none |
| RTP4 | -0.044 | -0.0847 | -0.34 | 0.74123 | 0.832 | -6.51 | 1.55055 | none |
| PQBP1 | 0.025 | -0.0095 | 0.3398 | 0.74132 | 0.832 | -6.51 | 0.35226 | none |
| FIBP | -0.038 | -0.0731 | -0.339 | 0.74183 | 0.833 | -6.51 | 0.09623 | none |
| HRC | -0.071 | -0.1021 | -0.339 | 0.74196 | 0.833 | -6.51 | 1.71979 | none |
| PRSS51 | 0.026 | 0.0396 | 0.3384 | 0.74229 | 0.833 | -6.51 | -0.0453 | none |
| LAMA3 | -0.045 | -0.0288 | -0.338 | 0.74233 | 0.833 | -6.51 | 0.6074 | none |
| COPS3 | -0.043 | -0.0955 | -0.338 | 0.74262 | 0.834 | -6.51 | 1.04116 | none |
| RALGAPA1 | 0.1 | 0.1255 | 0.3375 | 0.74294 | 0.834 | -6.51 | 0.98021 | none |
| ASF1A | 0.042 | -0.0691 | 0.3373 | 0.7431 | 0.834 | -6.51 | -0.4223 | none |
| PCDHGB1 | -0.025 | -0.0104 | -0.337 | 0.74327 | 0.834 | -6.51 | 0 | none |
| RNF26 | -0.037 | -0.0387 | -0.337 | 0.74353 | 0.834 | -6.51 | 1.18903 | none |
| SOAT2 | 0.059 | 0.0591 | 0.3366 | 0.74361 | 0.834 | -6.51 | 0.42499 | none |
| CEBPA | 0.029 | 0.0724 | 0.3366 | 0.74365 | 0.834 | -6.51 | 0.72083 | none |
| SLC43A1 | -0.037 | -0.0181 | -0.336 | 0.7437 | 0.834 | -6.51 | 1.06428 | none |
| SH2D4B | 0.023 | 0.0292 | 0.3364 | 0.74375 | 0.834 | -6.51 | 0.55562 | none |
| MYBPC1 | -0.026 | -0.0257 | -0.336 | 0.74379 | 0.834 | -6.51 | -0.6861 | none |
| PNPLA2 | -0.043 | -0.0668 | -0.336 | 0.7438 | 0.834 | -6.51 | 0.66398 | none |
| CAPN5 | 0.039 | 0.0068 | 0.3363 | 0.74384 | 0.834 | -6.51 | 0.6104 | none |
| TOMM40L | -0.038 | 0.0194 | -0.336 | 0.74413 | 0.835 | -6.51 | 0.5596 | none |
| FAM181B | -0.023 | -0.0064 | -0.336 | 0.7442 | 0.835 | -6.51 | 0.01408 | none |
| PRUNE1 | 0.049 | -0.0237 | 0.3358 | 0.74422 | 0.835 | -6.51 | 0.50934 | none |
| SEC22A | 0.068 | -0.0499 | 0.3356 | 0.74435 | 0.835 | -6.51 | 0.72197 | none |
| MCM3 | 0.064 | 0.1753 | 0.3352 | 0.74462 | 0.835 | -6.51 | 0.39529 | none |
| PMM1 | 0.046 | 0.0139 | 0.3348 | 0.74496 | 0.835 | -6.51 | 0.08675 | none |
| APOBEC3G | 0.037 | -0.039 | 0.3347 | 0.74498 | 0.835 | -6.51 | 0.46805 | none |
| EXOSC1 | -0.054 | -0.1312 | -0.335 | 0.74508 | 0.835 | -6.51 | 0.72899 | none |
| PPP4R4 | 0.03 | 0.0796 | 0.3342 | 0.74538 | 0.836 | -6.51 | 0.25507 | none |
| ARHGEF16 | 0.029 | 0.0502 | 0.3342 | 0.7454 | 0.836 | -6.51 | -0.1313 | none |
| APBB1IP | 0.08 | 0.1529 | 0.3341 | 0.74544 | 0.836 | -6.51 | 0.40232 | none |
| INHA | 0.047 | 0.0477 | 0.3339 | 0.74561 | 0.836 | -6.51 | 1.69179 | none |
| ARHGAP39 | -0.033 | -0.0128 | -0.334 | 0.74564 | 0.836 | -6.51 | -0.7905 | none |
| FAM214A | 0.081 | 0.2337 | 0.3337 | 0.74573 | 0.836 | -6.51 | 0.76387 | none |

| | | | | | | | | |
|---|---|---|---|---|---|---|---|---|
| DEPDC1 | -0.054 | -0.0177 | -0.334 | 0.74577 | 0.836 | -6.51 | 1.407 | none |
| C1ORF105 | 0.076 | 0.0772 | 0.3336 | 0.7458 | 0.836 | -6.51 | 0.78806 | none |
| FAAH2 | 0.051 | 0.0246 | 0.3336 | 0.74583 | 0.836 | -6.51 | -0.0548 | none |
| TMEM189 | 0.041 | 0.0556 | 0.3334 | 0.74593 | 0.836 | -6.51 | 0.76539 | none |
| ZBTB32 | 0.033 | 0.1354 | 0.3334 | 0.74596 | 0.836 | -6.51 | 0.99595 | none |
| CNTN4 | -0.035 | -0.0588 | -0.333 | 0.74599 | 0.836 | -6.51 | -0.3238 | none |
| ARHGAP19 | 0.053 | 0.1131 | 0.3333 | 0.74602 | 0.836 | -6.51 | 0.88377 | none |
| APOL4 | 0.072 | 0.1304 | 0.3332 | 0.74608 | 0.836 | -6.51 | 0.71645 | none |
| COPG2 | 0.045 | 0.057 | 0.3332 | 0.74609 | 0.836 | -6.51 | 0.34377 | none |
| CHRNA4 | -0.152 | -0.1088 | -0.333 | 0.74634 | 0.836 | -6.51 | 1.19438 | none |
| LRRC74A | -0.049 | -0.0487 | -0.333 | 0.74639 | 0.836 | -6.51 | 0.62735 | none |
| CD72 | 0.031 | 0.0885 | 0.3326 | 0.74652 | 0.836 | -6.51 | -0.1097 | none |
| HERC3 | -0.041 | -0.0824 | -0.332 | 0.74672 | 0.836 | -6.51 | 0.08481 | none |
| ZCRB1 | 0.027 | -0.0191 | 0.3323 | 0.74674 | 0.836 | -6.51 | -0.1015 | none |
| CD247 | -0.026 | -0.0387 | -0.332 | 0.74679 | 0.836 | -6.51 | 1.14536 | none |
| TRIM28 | -0.053 | 0.0769 | -0.332 | 0.74681 | 0.836 | -6.51 | -0.1258 | none |
| ABHD13 | 0.03 | -0.0373 | 0.3322 | 0.74684 | 0.836 | -6.51 | 1.13243 | none |
| ZBTB12 | 0.026 | 0.0501 | 0.3321 | 0.7469 | 0.836 | -6.51 | -0.112 | none |
| SPINK4 | 0.038 | 0.0927 | 0.3321 | 0.74691 | 0.836 | -6.51 | -0.5131 | none |
| LARGE2 | -0.045 | -0.0432 | -0.332 | 0.74729 | 0.836 | -6.51 | 1.5111 | none |
| PLXND1 | 0.046 | 0.0562 | 0.3316 | 0.7473 | 0.836 | -6.51 | 0.18841 | none |
| HHEX | 0.028 | 0.011 | 0.3315 | 0.74734 | 0.836 | -6.51 | 0.84248 | none |
| KCNQ3 | -0.036 | -0.0472 | -0.331 | 0.74738 | 0.836 | -6.51 | -0.2263 | none |
| TCEAL3 | 0.06 | 0.0805 | 0.3312 | 0.74756 | 0.836 | -6.51 | 0.0505 | none |
| FAM174B | -0.075 | -0.1035 | -0.331 | 0.74757 | 0.836 | -6.51 | -0.2247 | none |
| RAG2 | -0.028 | -0.0184 | -0.331 | 0.7477 | 0.836 | -6.51 | 1.38788 | none |
| ARHGAP15 | 0.064 | 0.032 | 0.331 | 0.74774 | 0.836 | -6.51 | 0.32897 | none |
| ENPP7 | -0.023 | -0.0121 | -0.331 | 0.74782 | 0.837 | -6.51 | 0.50826 | none |
| CMSS1 | 0.034 | 0.0092 | 0.3307 | 0.74797 | 0.837 | -6.51 | 0.09048 | none |
| THEMIS | -0.049 | -0.0574 | -0.331 | 0.748 | 0.837 | -6.51 | 0.755 | none |
| HGH1 | -0.036 | 0.0499 | -0.33 | 0.74809 | 0.837 | -6.51 | 0.08832 | none |
| C17ORF80 | -0.085 | -0.0048 | -0.33 | 0.74829 | 0.837 | -6.51 | 1.44645 | none |
| PET100 | 0.075 | -0.0397 | 0.3301 | 0.74837 | 0.837 | -6.51 | -0.1223 | none |
| RANGAP1 | 0.052 | 0.1597 | 0.3301 | 0.74837 | 0.837 | -6.51 | 0.35669 | none |
| SCX | -0.029 | -0.002 | -0.33 | 0.74839 | 0.837 | -6.51 | 0.8632 | none |
| SUPT3H | 0.056 | 0.0287 | 0.3297 | 0.74866 | 0.837 | -6.51 | 0.07139 | none |
| NTNG1 | -0.034 | -0.0213 | -0.329 | 0.74883 | 0.837 | -6.51 | -0.1736 | none |
| PSMB9 | -0.038 | -0.0452 | -0.329 | 0.74892 | 0.837 | -6.51 | 1.35844 | none |
| FXR1 | 0.048 | 0.0693 | 0.3293 | 0.74894 | 0.837 | -6.51 | 0.91136 | none |
| TEKT5 | -0.03 | -0.0301 | -0.329 | 0.74914 | 0.837 | -6.51 | -0.2942 | none |

| | | | | | | | | |
|---|---|---|---|---|---|---|---|---|
| CGGBP1 | -0.033 | -0.0977 | -0.329 | 0.7492 | 0.837 | -6.51 | 0.04139 | none |
| TMPPE | 0.034 | -0.0062 | 0.3282 | 0.74974 | 0.838 | -6.52 | 0.41857 | none |
| RAD21 | 0.051 | 0.1086 | 0.328 | 0.74988 | 0.838 | -6.52 | 0.90759 | none |
| SPRR2D | -0.035 | -0.0349 | -0.328 | 0.75004 | 0.838 | -6.52 | 1.15163 | none |
| DNMT3B | 0.029 | 0.0541 | 0.3278 | 0.75004 | 0.838 | -6.52 | 0.16419 | none |
| CRHR2 | 0.033 | 0.0371 | 0.3278 | 0.75004 | 0.838 | -6.52 | 1.08514 | none |
| ANG | -0.065 | -0.0279 | -0.328 | 0.75014 | 0.838 | -6.52 | 1.44853 | none |
| C1GALT1C | -0.024 | -0.0285 | -0.327 | 0.75041 | 0.838 | -6.52 | 1.35295 | none |
| SLC25A24 | 0.029 | -0.0562 | 0.327 | 0.75063 | 0.839 | -6.52 | 0.26458 | none |
| GABRA5 | -0.05 | -0.0435 | -0.326 | 0.75104 | 0.839 | -6.52 | 0.02529 | none |
| 7-Sep | -0.044 | 0.0081 | -0.326 | 0.75116 | 0.839 | -6.52 | 1.29211 | none |
| KIT | -0.058 | -0.1046 | -0.326 | 0.75141 | 0.839 | -6.52 | 1.08159 | none |
| EEF2 | -0.055 | 0.0027 | -0.326 | 0.75144 | 0.839 | -6.52 | 1.41619 | none |
| CFAP99 | -0.03 | -0.0076 | -0.326 | 0.75171 | 0.839 | -6.52 | 1.2245 | none |
| TSPYL5 | 0.027 | 0.0358 | 0.3251 | 0.75203 | 0.84 | -6.52 | 1.15314 | none |
| ZBTB5 | 0.035 | 0.0506 | 0.3248 | 0.75229 | 0.84 | -6.52 | 0.9879 | none |
| AQP12A | 0.028 | 0.0536 | 0.3245 | 0.75244 | 0.84 | -6.52 | -0.0381 | none |
| TM9SF1 | -0.034 | -0.0658 | -0.325 | 0.75248 | 0.84 | -6.52 | 0.80049 | none |
| EIF3CL | 0.091 | 0.0367 | 0.3245 | 0.7525 | 0.84 | -6.52 | 0.33482 | none |
| OMG | -0.041 | -0.0292 | -0.324 | 0.75253 | 0.84 | -6.52 | 0.0455 | none |
| ADORA2A | -0.031 | 0.0183 | -0.324 | 0.75256 | 0.84 | -6.52 | 1.36964 | none |
| MCHR1 | -0.025 | -0.0269 | -0.324 | 0.75258 | 0.84 | -6.52 | 0.37772 | none |
| PTGES3L-A | 0.052 | 0.0449 | 0.3244 | 0.75258 | 0.84 | -6.52 | 1.2566 | none |
| NARS2 | 0.058 | 0.0699 | 0.3244 | 0.75258 | 0.84 | -6.52 | 0.681 | none |
| ADM5 | 0.037 | 0.096 | 0.3242 | 0.75273 | 0.84 | -6.52 | 0.55824 | none |
| JAG1 | -0.048 | -0.0137 | -0.324 | 0.75281 | 0.84 | -6.52 | 0.73329 | none |
| NIPA2 | -0.066 | -0.102 | -0.324 | 0.75286 | 0.84 | -6.52 | 1.81819 | none |
| RSU1 | 0.025 | -0.0498 | 0.3239 | 0.75292 | 0.84 | -6.52 | 0.52099 | none |
| CPEB2 | -0.032 | -0.0039 | -0.324 | 0.75306 | 0.84 | -6.52 | 1.48779 | none |
| AFP | -0.073 | -0.1679 | -0.323 | 0.75354 | 0.841 | -6.52 | 0.3842 | none |
| LPAR5 | 0.036 | 0.0563 | 0.3229 | 0.75366 | 0.841 | -6.52 | 0.27685 | none |
| COL6A1 | -0.064 | -0.0283 | -0.323 | 0.75382 | 0.841 | -6.52 | 0.84741 | none |
| MTERF4 | 0.043 | 0.0018 | 0.3227 | 0.75383 | 0.841 | -6.52 | -0.6166 | none |
| CRHR1 | -0.032 | -0.0175 | -0.323 | 0.75386 | 0.841 | -6.52 | -0.3671 | none |
| ZNF655 | -0.035 | -0.0732 | -0.323 | 0.75394 | 0.841 | -6.52 | 0.758 | none |
| RFXANK | -0.055 | -0.0692 | -0.322 | 0.75399 | 0.841 | -6.52 | 0.0484 | none |
| SPHK1 | 0.069 | 0.0656 | 0.3219 | 0.75441 | 0.841 | -6.52 | -0.5072 | none |
| SIX2 | 0.027 | 0.0257 | 0.3218 | 0.75448 | 0.841 | -6.52 | 0.72848 | none |
| EBF3 | -0.025 | ######## | -0.322 | 0.75465 | 0.841 | -6.52 | 0.40389 | none |
| CBLL1 | 0.037 | 0.052 | 0.3215 | 0.75466 | 0.841 | -6.52 | -0.637 | none |

| | | | | | | | | |
|---|---|---|---|---|---|---|---|---|
| HMGN5 | 0.091 | 0.2022 | 0.3202 | 0.75562 | 0.842 | -6.52 | 0.32596 | none |
| C3ORF18 | 0.038 | 0.0504 | 0.3199 | 0.75585 | 0.842 | -6.52 | 0.20858 | none |
| SSTR5 | -0.025 | -0.0248 | -0.32 | 0.75587 | 0.842 | -6.52 | -0.5284 | none |
| TNFRSF6B | 0.022 | 0.0036 | 0.3198 | 0.75594 | 0.842 | -6.52 | -0.3315 | none |
| PRIM1 | 0.048 | 0.0074 | 0.3197 | 0.75598 | 0.842 | -6.52 | 1.09572 | none |
| ROPN1 | 0.044 | 0.047 | 0.3188 | 0.75668 | 0.843 | -6.52 | 0.08592 | none |
| TP53I11 | -0.046 | 0.0163 | -0.318 | 0.75703 | 0.844 | -6.52 | 0.21169 | none |
| ITGB5 | -0.045 | -0.0572 | -0.318 | 0.75712 | 0.844 | -6.52 | 0.50309 | none |
| KCNK17 | 0.023 | 0.0285 | 0.3181 | 0.75716 | 0.844 | -6.52 | 0.34895 | none |
| ZP4 | -0.023 | -0.024 | -0.318 | 0.75725 | 0.844 | -6.52 | 0 | none |
| SOAT1 | 0.044 | -0.0876 | 0.3179 | 0.75732 | 0.844 | -6.52 | 0.93231 | none |
| FASLG | -0.05 | -0.0099 | -0.318 | 0.75737 | 0.844 | -6.52 | 0.74236 | none |
| SYNJ2BP-C | -1.459 | -1.3811 | -0.318 | 0.75739 | 0.844 | -6.52 | 1.99825 | none |
| AQP12B | 0.027 | 0.0353 | 0.3178 | 0.75743 | 0.844 | -6.52 | -0.3428 | none |
| TPRX1 | 0.023 | 0.0225 | 0.3176 | 0.75753 | 0.844 | -6.52 | 0.94361 | none |
| CCDC184 | -0.023 | -0.0207 | -0.318 | 0.7576 | 0.844 | -6.52 | 0 | none |
| FBLN1 | -0.055 | 0.0426 | -0.317 | 0.75773 | 0.844 | -6.52 | 0.45772 | none |
| SDHD | 0.045 | -0.0895 | 0.3173 | 0.75778 | 0.844 | -6.52 | 0.38657 | none |
| GIMAP8 | 0.044 | 0.0917 | 0.3172 | 0.75784 | 0.844 | -6.52 | 0.14996 | none |
| TMED7-TI( | -0.041 | ######## | -0.317 | 0.75786 | 0.844 | -6.52 | 1.25722 | none |
| ZKSCAN8 | -0.036 | -0.0228 | -0.317 | 0.75801 | 0.844 | -6.52 | 0.54795 | none |
| ISY1-RAB4 | 0.027 | 0.0187 | 0.3166 | 0.75832 | 0.844 | -6.52 | 0.77651 | none |
| MAP3K3 | -0.065 | 0.0312 | -0.316 | 0.75851 | 0.844 | -6.52 | 0.56759 | none |
| PTH2R | 0.1 | 0.0497 | 0.3161 | 0.75865 | 0.844 | -6.52 | -0.5654 | none |
| CRYGN | 0.022 | 0.0159 | 0.3159 | 0.75883 | 0.845 | -6.52 | 1.58362 | none |
| ZDHHC9 | -0.033 | -0.085 | -0.316 | 0.75895 | 0.845 | -6.52 | 0.91796 | none |
| MTRNR2L1 | -0.027 | -0.0649 | -0.316 | 0.75906 | 0.845 | -6.52 | 0.89478 | none |
| HIF1AN | 0.032 | -0.0013 | 0.3155 | 0.7591 | 0.845 | -6.52 | -0.6392 | none |
| PPP2R5E | 0.045 | 0.1 | 0.3145 | 0.75981 | 0.845 | -6.52 | 0.94328 | none |
| NCLN | 0.036 | 0.0756 | 0.3144 | 0.75989 | 0.845 | -6.52 | 1.4587 | none |
| DPP7 | -0.035 | -0.0336 | -0.314 | 0.75992 | 0.845 | -6.52 | 0.84152 | none |
| CASP1 | -0.024 | -0.0707 | -0.314 | 0.76004 | 0.845 | -6.52 | -0.1903 | none |
| DIRAS1 | 0.024 | 0.0134 | 0.3142 | 0.76009 | 0.845 | -6.52 | 1.5413 | none |
| DUSP7 | 0.037 | 0.074 | 0.3139 | 0.76026 | 0.846 | -6.52 | -0.1772 | none |
| BARHL1 | 0.029 | 0.0148 | 0.3137 | 0.76042 | 0.846 | -6.52 | 0.13123 | none |
| CPNE7 | -0.056 | 0.0877 | -0.314 | 0.76055 | 0.846 | -6.52 | 0.79014 | none |
| RCN1 | -0.041 | 0.0032 | -0.313 | 0.76067 | 0.846 | -6.52 | 1.44066 | none |
| COPRS | -0.069 | -0.1339 | -0.313 | 0.7608 | 0.846 | -6.52 | 0.5892 | none |
| NAALADL2 | -0.042 | -0.0532 | -0.312 | 0.76148 | 0.847 | -6.52 | 1.16088 | none |
| SPAG16 | -0.035 | -0.1411 | -0.312 | 0.76157 | 0.847 | -6.52 | 0.29378 | none |

| | | | | | | | | |
|---|---|---|---|---|---|---|---|---|
| CTSD | -0.031 | -0.0286 | -0.312 | 0.76172 | 0.847 | -6.52 | 1.60635 | none |
| AXDND1 | -0.023 | 0.0061 | -0.312 | 0.76173 | 0.847 | -6.52 | 0.26367 | none |
| DIP2C | -0.042 | ######## | -0.312 | 0.76192 | 0.847 | -6.52 | 0.96294 | none |
| VAPB | -0.034 | 0.0194 | -0.312 | 0.76193 | 0.847 | -6.52 | 1.18201 | none |
| ASPHD1 | 0.041 | 0.0539 | 0.3116 | 0.76199 | 0.847 | -6.52 | 1.4184 | none |
| HMCN1 | -0.021 | -0.0153 | -0.311 | 0.76213 | 0.847 | -6.52 | 0 | none |
| KDELR3 | -0.054 | -0.0428 | -0.311 | 0.7623 | 0.847 | -6.52 | 1.29627 | none |
| OTUB1 | -0.037 | -0.0815 | -0.311 | 0.76236 | 0.847 | -6.52 | 0.90309 | none |
| BLK | -0.053 | -0.118 | -0.311 | 0.76245 | 0.847 | -6.52 | 0.21388 | none |
| BYSL | 0.032 | 0.051 | 0.3108 | 0.76252 | 0.847 | -6.52 | 0.11234 | none |
| GCOM1 | -0.026 | 0.0099 | -0.311 | 0.76257 | 0.847 | -6.52 | 1.10823 | none |
| NPHP4 | 0.061 | 0.14 | 0.3102 | 0.76299 | 0.848 | -6.52 | 1.50878 | none |
| HSPA14 | -0.026 | -0.0705 | -0.31 | 0.76325 | 0.848 | -6.52 | 0.882 | none |
| HIST1H4L | -0.069 | -0.0928 | -0.309 | 0.76353 | 0.848 | -6.52 | 1.07801 | none |
| EIF3J | 0.053 | 0.0901 | 0.3094 | 0.76361 | 0.848 | -6.52 | 1.23043 | none |
| SPP1 | 0.065 | 0.0811 | 0.3093 | 0.76364 | 0.848 | -6.52 | 0.01623 | none |
| HSDL2 | 0.029 | -0.0451 | 0.3093 | 0.76364 | 0.848 | -6.52 | 0.31208 | none |
| TMEM141 | 0.046 | 0.0034 | 0.3091 | 0.76379 | 0.848 | -6.52 | -0.3943 | none |
| MRPL42 | -0.044 | -0.087 | -0.309 | 0.76382 | 0.848 | -6.52 | -0.1875 | none |
| CLGN | -0.026 | -0.0215 | -0.309 | 0.7639 | 0.848 | -6.52 | 1.03387 | none |
| TCP10L2 | 0.045 | 0.0887 | 0.309 | 0.76392 | 0.848 | -6.52 | -0.2044 | none |
| RGS16 | -0.026 | 0.0836 | -0.309 | 0.76395 | 0.848 | -6.52 | 0.42766 | none |
| RYR3 | -0.131 | 0.0602 | -0.309 | 0.76421 | 0.848 | -6.52 | -0.2515 | none |
| VSX2 | 0.023 | 0.0455 | 0.3082 | 0.76451 | 0.849 | -6.52 | 0.42171 | none |
| SLC25A53 | 0.064 | 0.0886 | 0.3079 | 0.76469 | 0.849 | -6.52 | 1.09431 | none |
| ALKBH1 | 0.053 | 0.0397 | 0.3078 | 0.76475 | 0.849 | -6.52 | 0.65508 | none |
| TMEM183 | -0.045 | -0.0815 | -0.308 | 0.76491 | 0.849 | -6.52 | 1.52718 | none |
| TMEM160 | -0.065 | -0.1178 | -0.307 | 0.7652 | 0.849 | -6.52 | 0.95845 | none |
| GIPC3 | -0.022 | -0.0027 | -0.307 | 0.76537 | 0.849 | -6.52 | 0.37463 | none |
| GOLGA6L1 | -0.028 | -0.0124 | -0.307 | 0.76541 | 0.849 | -6.52 | -0.224 | none |
| SPSB3 | -0.041 | -0.0421 | -0.307 | 0.76564 | 0.85 | -6.52 | 0.97475 | none |
| PRSS33 | 0.023 | 0.0231 | 0.3064 | 0.76583 | 0.85 | -6.52 | 1.49167 | none |
| DDR2 | 0.036 | 0.0358 | 0.3062 | 0.76594 | 0.85 | -6.52 | 0.51921 | none |
| CCDC110 | -0.026 | -0.018 | -0.306 | 0.76595 | 0.85 | -6.52 | 1.4035 | none |
| SEPHS2 | -0.025 | -0.0062 | -0.306 | 0.76597 | 0.85 | -6.52 | 1.29045 | none |
| PLEKHM2 | 0.043 | 0.1368 | 0.3061 | 0.76599 | 0.85 | -6.52 | 0.55834 | none |
| CACFD1 | 0.046 | -0.0046 | 0.306 | 0.76613 | 0.85 | -6.52 | 0.89024 | none |
| USP27X | 0.027 | 0.0578 | 0.3059 | 0.76616 | 0.85 | -6.52 | 0.87022 | none |
| SLC25A1 | -0.031 | 0.0181 | -0.306 | 0.76628 | 0.85 | -6.52 | 0.85451 | none |
| HIST1H1C | -0.035 | -0.0819 | -0.306 | 0.76634 | 0.85 | -6.52 | 1.29914 | none |

| | | | | | | | | |
|---|---|---|---|---|---|---|---|---|
| ADGRG5 | -0.038 | -0.066 | -0.306 | 0.76647 | 0.85 | -6.52 | 0.35485 | none |
| ANKRD22 | 0.031 | -0.0119 | 0.3054 | 0.76654 | 0.85 | -6.52 | 0.08772 | none |
| DHRS9 | 0.058 | -0.0122 | 0.3053 | 0.76664 | 0.85 | -6.52 | -0.2946 | none |
| TP53TG5 | -0.045 | -0.0033 | -0.305 | 0.76673 | 0.85 | -6.52 | 1.43444 | none |
| FGFR3 | 0.031 | 0.0756 | 0.3051 | 0.76676 | 0.85 | -6.52 | 0.76987 | none |
| RAB6A | 0.028 | 0.018 | 0.3048 | 0.76697 | 0.85 | -6.52 | 0.45822 | none |
| HOXA5 | 0.035 | 0.0305 | 0.3046 | 0.7671 | 0.85 | -6.52 | 1.53266 | none |
| PEX14 | -0.044 | -0.0296 | -0.304 | 0.76733 | 0.85 | -6.52 | -0.0391 | none |
| HIST2H4B | 0.076 | 0.0515 | 0.3043 | 0.76735 | 0.85 | -6.52 | 0.70153 | none |
| FCGR2B | -0.024 | -0.0435 | -0.304 | 0.7674 | 0.85 | -6.52 | 0.77085 | none |
| C9ORF72 | 0.047 | 0.0044 | 0.3041 | 0.76749 | 0.85 | -6.52 | 0.45013 | none |
| MAGEB3 | -0.022 | -0.0047 | -0.304 | 0.76752 | 0.85 | -6.52 | 0.86097 | none |
| CDV3 | 0.053 | 0.1002 | 0.304 | 0.76755 | 0.85 | -6.52 | -0.1094 | none |
| CFAP57 | -0.043 | 0.0102 | -0.304 | 0.76784 | 0.851 | -6.52 | 1.69394 | none |
| PDIA3 | 0.035 | 0.0994 | 0.3033 | 0.76807 | 0.851 | -6.52 | 0.8523 | none |
| PABPN1 | 0.048 | 0.1297 | 0.3032 | 0.76815 | 0.851 | -6.52 | 0.00625 | none |
| CUEDC1 | 0.082 | 0.0843 | 0.303 | 0.76829 | 0.851 | -6.52 | -0.2773 | none |
| HSP90B1 | 0.092 | 0.2386 | 0.3029 | 0.76837 | 0.851 | -6.52 | 0.67897 | none |
| PPM1A | -0.021 | -0.0057 | -0.303 | 0.7684 | 0.851 | -6.52 | 0.91089 | none |
| PRKCQ | 0.059 | 0.128 | 0.3027 | 0.76852 | 0.851 | -6.52 | 0.77544 | none |
| DUSP8 | 0.029 | 0.0402 | 0.3027 | 0.76857 | 0.851 | -6.52 | 1.45484 | none |
| LRRC32 | 0.029 | 0.068 | 0.3025 | 0.76866 | 0.851 | -6.52 | 0.083 | none |
| LZTS3 | -0.024 | -0.0206 | -0.302 | 0.76917 | 0.852 | -6.52 | 1.32728 | none |
| CHIC2 | 0.05 | -0.0267 | 0.3012 | 0.76963 | 0.852 | -6.52 | 1.66258 | none |
| CDC37L1 | 0.049 | -0.0377 | 0.3005 | 0.77015 | 0.853 | -6.52 | 0.22332 | none |
| KPNA3 | -0.04 | -0.003 | -0.3 | 0.77061 | 0.853 | -6.52 | 1.1713 | none |
| CCL23 | -0.024 | -0.0316 | -0.299 | 0.77097 | 0.853 | -6.52 | 0.96679 | none |
| ATP4B | 0.022 | 0.0367 | 0.2994 | 0.77098 | 0.853 | -6.52 | -0.0267 | none |
| TWIST2 | 0.023 | 0.023 | 0.2987 | 0.77153 | 0.854 | -6.53 | 1.8788 | none |
| PRR5L | 0.052 | 0.0871 | 0.2984 | 0.77173 | 0.854 | -6.53 | 0.88148 | none |
| PMFBP1 | 0.027 | 0.1081 | 0.2981 | 0.77198 | 0.854 | -6.53 | 0.41508 | none |
| CTAG1B | -0.023 | -0.0268 | -0.298 | 0.77222 | 0.855 | -6.53 | 0 | none |
| GNMT | -0.021 | -0.0047 | -0.298 | 0.77223 | 0.855 | -6.53 | 0.10379 | none |
| DEDD | 0.025 | -0.0026 | 0.2975 | 0.7724 | 0.855 | -6.53 | -0.8823 | none |
| MGST3 | 0.049 | -0.0651 | 0.2973 | 0.77252 | 0.855 | -6.53 | 0.73309 | none |
| RAB27B | -0.022 | -0.0285 | -0.297 | 0.77265 | 0.855 | -6.53 | 0 | none |
| ADH4 | 0.045 | -0.0161 | 0.2968 | 0.77288 | 0.855 | -6.53 | 0.04201 | none |
| CD19 | -0.033 | -0.0714 | -0.297 | 0.77292 | 0.855 | -6.53 | 1.08192 | none |
| POTEI | 0.023 | 0.0093 | 0.2967 | 0.77299 | 0.855 | -6.53 | -0.0273 | none |
| FCGRT | 0.048 | -0.113 | 0.2964 | 0.77319 | 0.855 | -6.53 | -0.2772 | none |

| | | | | | | | | |
|---|---|---|---|---|---|---|---|---|
| LRRC61 | -0.03 | ######## | -0.296 | 0.77338 | 0.855 | -6.53 | 1.65892 | none |
| CDS2 | -0.043 | -0.1462 | -0.296 | 0.77361 | 0.856 | -6.53 | 0.43072 | none |
| FADD | -0.024 | -0.0357 | -0.296 | 0.77377 | 0.856 | -6.53 | 1.14447 | none |
| ECSIT | -0.041 | -0.0574 | -0.296 | 0.77378 | 0.856 | -6.53 | 1.33981 | none |
| ICAM2 | -0.044 | -0.1186 | -0.296 | 0.77382 | 0.856 | -6.53 | -0.0276 | none |
| PTPRU | 0.025 | 0.0231 | 0.2953 | 0.77399 | 0.856 | -6.53 | 0.58178 | none |
| NCK2 | 0.031 | 0.0787 | 0.2953 | 0.774 | 0.856 | -6.53 | 1.07839 | none |
| NOP9 | -0.03 | 0.0565 | -0.295 | 0.77407 | 0.856 | -6.53 | 0.10383 | none |
| GPR152 | 0.022 | 0.0148 | 0.2952 | 0.77408 | 0.856 | -6.53 | 0.66047 | none |
| C1QTNF9 | 0.021 | 0.0241 | 0.2952 | 0.7741 | 0.856 | -6.53 | 0 | none |
| UBE2E2 | -0.045 | 0.023 | -0.295 | 0.77423 | 0.856 | -6.53 | 0.06562 | none |
| EFNA3 | -0.029 | 0.0257 | -0.295 | 0.77436 | 0.856 | -6.53 | 0.34174 | none |
| NTNG2 | 0.041 | 0.0885 | 0.2948 | 0.77438 | 0.856 | -6.53 | 0.58478 | none |
| AGBL3 | 0.049 | -0.0352 | 0.2948 | 0.77439 | 0.856 | -6.53 | -0.8944 | none |
| COMTD1 | 0.04 | 0.0137 | 0.2946 | 0.77451 | 0.856 | -6.53 | 1.51451 | none |
| LHFPL1 | 0.03 | -0.0118 | 0.2945 | 0.77463 | 0.856 | -6.53 | 1.59192 | none |
| TIGD3 | -0.03 | -0.0295 | -0.294 | 0.77468 | 0.856 | -6.53 | 0.06198 | none |
| C1QB | 0.046 | 0.0538 | 0.2944 | 0.77472 | 0.856 | -6.53 | 1.4126 | none |
| ANKRD20/ | 0.028 | 0.0258 | 0.2939 | 0.77508 | 0.856 | -6.53 | 1.76792 | none |
| CCDC155 | 0.103 | 0.1656 | 0.2937 | 0.7752 | 0.856 | -6.53 | -0.1422 | none |
| SERPINB8 | -0.054 | -0.0335 | -0.294 | 0.7753 | 0.856 | -6.53 | 0.63551 | none |
| MRGPRF | -0.041 | ######## | -0.294 | 0.7753 | 0.856 | -6.53 | 1.38705 | none |
| COL5A2 | -0.027 | 0.0114 | -0.294 | 0.77531 | 0.856 | -6.53 | 0.31063 | none |
| CLEC1A | -0.116 | -0.1554 | -0.293 | 0.77565 | 0.856 | -6.53 | -0.1267 | none |
| YDJC | -0.035 | -0.0232 | -0.293 | 0.77565 | 0.856 | -6.53 | 1.79539 | none |
| PRSS36 | -0.032 | -0.0418 | -0.293 | 0.77594 | 0.857 | -6.53 | 1.63387 | none |
| C12ORF75 | -0.048 | -0.1201 | -0.293 | 0.77595 | 0.857 | -6.53 | 1.27229 | none |
| PCDHGB2 | -0.022 | -0.023 | -0.292 | 0.77616 | 0.857 | -6.53 | 1.69853 | none |
| DMAP1 | 0.03 | 0.1393 | 0.2923 | 0.77624 | 0.857 | -6.53 | 0.58679 | none |
| SEC62 | 0.06 | 0.055 | 0.2923 | 0.77626 | 0.857 | -6.53 | 0.01659 | none |
| ZNF649 | 0.087 | 0.061 | 0.2922 | 0.77635 | 0.857 | -6.53 | 0.88164 | none |
| KREMEN2 | -0.022 | -0.0094 | -0.292 | 0.77645 | 0.857 | -6.53 | 0.43065 | none |
| CTNND1 | 0.038 | 0.0877 | 0.2919 | 0.77656 | 0.857 | -6.53 | 0.31498 | none |
| EMD | -0.05 | -0.1477 | -0.292 | 0.77675 | 0.857 | -6.53 | 1.21929 | none |
| ASL | 0.024 | 0.0503 | 0.2909 | 0.7773 | 0.858 | -6.53 | 0.73113 | none |
| CES1 | 0.054 | 0.0944 | 0.2906 | 0.77753 | 0.858 | -6.53 | 0.02019 | none |
| PRSS50 | 0.025 | 0.0104 | 0.2903 | 0.77772 | 0.858 | -6.53 | -0.2413 | none |
| CFB | -0.032 | ######## | -0.29 | 0.77785 | 0.858 | -6.53 | 0.42002 | none |
| ORMDL1 | -0.037 | -0.051 | -0.29 | 0.77786 | 0.858 | -6.53 | 1.03495 | none |
| KCTD10 | -0.032 | -0.0459 | -0.29 | 0.77787 | 0.858 | -6.53 | 0.55546 | none |

| | | | | | | | | |
|---|---|---|---|---|---|---|---|---|
| CCNE2 | 0.023 | 0.0282 | 0.2901 | 0.77787 | 0.858 | -6.53 | -0.3445 | none |
| ABLIM1 | 0.063 | 0.1038 | 0.2899 | 0.77804 | 0.858 | -6.53 | 0.60262 | none |
| VAT1 | -0.043 | -0.0737 | -0.29 | 0.77814 | 0.858 | -6.53 | 1.12593 | none |
| UBE2V1 | -0.025 | -0.0042 | -0.289 | 0.77835 | 0.858 | -6.53 | 0.88783 | none |
| CPPED1 | 0.023 | -0.0552 | 0.289 | 0.7787 | 0.859 | -6.53 | -0.5492 | none |
| ASB10 | -0.037 | -0.0317 | -0.289 | 0.77872 | 0.859 | -6.53 | 1.8263 | none |
| ST6GALNA | 0.036 | 0.0022 | 0.2888 | 0.77886 | 0.859 | -6.53 | 0.6046 | none |
| ABCG5 | 0.023 | 0.013 | 0.2887 | 0.77888 | 0.859 | -6.53 | -0.2221 | none |
| C6ORF106 | -0.035 | ######## | -0.289 | 0.77896 | 0.859 | -6.53 | 0.88914 | none |
| ALG8 | 0.041 | -0.1066 | 0.2884 | 0.77913 | 0.859 | -6.53 | -0.256 | none |
| GDF2 | 0.02 | 0.025 | 0.2884 | 0.77916 | 0.859 | -6.53 | 0 | none |
| CCL16 | -0.11 | 0.0739 | -0.288 | 0.77923 | 0.859 | -6.53 | 0.45922 | none |
| MT1H | -0.081 | -0.0969 | -0.288 | 0.77923 | 0.859 | -6.53 | 0.45498 | none |
| DPY19L2 | -0.052 | -0.0206 | -0.288 | 0.7794 | 0.859 | -6.53 | 0.06676 | none |
| AL356585. | 0.03 | 0.0653 | 0.2878 | 0.77955 | 0.859 | -6.53 | 0.48708 | none |
| OR2L5 | -0.02 | -0.0083 | -0.288 | 0.7796 | 0.859 | -6.53 | 0.05088 | none |
| MIS12 | 0.027 | -0.0742 | 0.2875 | 0.77983 | 0.859 | -6.53 | 0.3487 | none |
| FGGY | 0.05 | -0.0608 | 0.2872 | 0.78002 | 0.859 | -6.53 | -0.3931 | none |
| CBWD2 | -0.029 | -0.038 | -0.287 | 0.78005 | 0.859 | -6.53 | -0.4043 | none |
| MAPKAPK | -0.027 | -0.0272 | -0.287 | 0.78042 | 0.86 | -6.53 | 1.66354 | none |
| CPA5 | 0.078 | -0.0614 | 0.2866 | 0.78045 | 0.86 | -6.53 | 0.3674 | none |
| MAP1B | 0.027 | 0.0685 | 0.2861 | 0.78085 | 0.86 | -6.53 | -0.3592 | none |
| EXTL2 | 0.029 | -0.0053 | 0.286 | 0.78089 | 0.86 | -6.53 | 1.73565 | none |
| HSFX2 | 0.025 | 0.0042 | 0.286 | 0.78092 | 0.86 | -6.53 | -0.4876 | none |
| ZNF22 | -0.037 | -0.056 | -0.286 | 0.78095 | 0.86 | -6.53 | 0.15749 | none |
| PRM3 | 0.05 | 0.026 | 0.2859 | 0.78096 | 0.86 | -6.53 | 0.42353 | none |
| SPR | -0.027 | 0.0843 | -0.286 | 0.78098 | 0.86 | -6.53 | 1.39937 | none |
| PRR20C | 0.02 | 0.0252 | 0.2858 | 0.78106 | 0.86 | -6.53 | 0 | none |
| HPCAL1 | 0.048 | 0.0658 | 0.2858 | 0.78108 | 0.86 | -6.53 | 1.06631 | none |
| MFSD12 | 0.025 | 0.0275 | 0.2858 | 0.78109 | 0.86 | -6.53 | -0.7835 | none |
| KRAS | -0.028 | -0.0402 | -0.285 | 0.78157 | 0.86 | -6.53 | 0.96885 | none |
| USH1C | -0.044 | -0.0436 | -0.285 | 0.78184 | 0.861 | -6.53 | -0.4901 | none |
| CD7 | 0.035 | 0.0308 | 0.2843 | 0.78215 | 0.861 | -6.53 | 0.63329 | none |
| CLUL1 | 0.055 | 0.0512 | 0.2842 | 0.78227 | 0.861 | -6.53 | 1.10951 | none |
| TLDC2 | 0.029 | 0.068 | 0.2841 | 0.78231 | 0.861 | -6.53 | -0.1091 | none |
| SLC45A1 | 0.032 | -0.0054 | 0.2841 | 0.78231 | 0.861 | -6.53 | 0.70559 | none |
| USP17L15 | 0.03 | -0.0115 | 0.2841 | 0.78237 | 0.861 | -6.53 | 0.60174 | none |
| PHEX | -0.049 | -0.0542 | -0.284 | 0.7824 | 0.861 | -6.53 | 0.70016 | none |
| FJX1 | 0.031 | ######## | 0.2836 | 0.78273 | 0.861 | -6.53 | -0.523 | none |
| TRAPPC5 | -0.057 | -0.1026 | -0.283 | 0.78282 | 0.861 | -6.53 | 0.91268 | none |

| | | | | | | | | |
|---|---|---|---|---|---|---|---|---|
| PDCD1 | 0.03 | 0.1361 | 0.2832 | 0.78301 | 0.861 | -6.53 | 0.88565 | none |
| VPS28 | 0.111 | 0.0759 | 0.2832 | 0.78303 | 0.861 | -6.53 | 1.20279 | none |
| TIMM23 | -0.037 | -0.0819 | -0.283 | 0.78314 | 0.861 | -6.53 | -0.2699 | none |
| CYTH2 | -0.021 | -0.0284 | -0.283 | 0.78349 | 0.862 | -6.53 | -0.1474 | none |
| ODF1 | -0.021 | -0.0371 | -0.283 | 0.78351 | 0.862 | -6.53 | 0.54121 | none |
| EHHADH | 0.045 | 0.09 | 0.2815 | 0.78425 | 0.863 | -6.53 | 0.11335 | none |
| MTRR | 0.071 | 0.0653 | 0.2815 | 0.78426 | 0.863 | -6.53 | 0.71415 | none |
| RNF126 | -0.024 | 0.0145 | -0.281 | 0.7847 | 0.863 | -6.53 | 0.14672 | none |
| IER2 | -0.026 | 0.0322 | -0.281 | 0.78472 | 0.863 | -6.53 | 0.554 | none |
| SAXO1 | 0.028 | 0.0306 | 0.2802 | 0.7852 | 0.863 | -6.53 | 0.64061 | none |
| TMEM216 | 0.045 | -0.031 | 0.28 | 0.78537 | 0.863 | -6.53 | 0.61567 | none |
| MRGPRG | -0.023 | ######## | -0.28 | 0.7854 | 0.863 | -6.53 | 0.71676 | none |
| MIPEP | 0.035 | 0.0252 | 0.2794 | 0.78584 | 0.864 | -6.53 | 1.02119 | none |
| C11ORF24 | -0.034 | 0.0231 | -0.279 | 0.7859 | 0.864 | -6.53 | 1.38628 | none |
| LGALSL | 0.057 | 0.0577 | 0.2791 | 0.78604 | 0.864 | -6.53 | 0.48943 | none |
| ATP2C2 | 0.034 | 0.0286 | 0.279 | 0.78614 | 0.864 | -6.53 | -0.5282 | none |
| ALDH1A3 | 0.047 | 0.0468 | 0.2789 | 0.78619 | 0.864 | -6.53 | 0.58762 | none |
| PDGFB | 0.054 | 0.0279 | 0.2787 | 0.78638 | 0.864 | -6.53 | -0.1173 | none |
| KRTAP5-7 | 0.021 | 0.0232 | 0.2787 | 0.78638 | 0.864 | -6.53 | -0.473 | none |
| TSPAN12 | -0.021 | -0.0285 | -0.279 | 0.78639 | 0.864 | -6.53 | -0.2205 | none |
| WDR46 | -0.042 | 0.0117 | -0.278 | 0.78655 | 0.864 | -6.53 | 0.49745 | none |
| GTF2H4 | -0.07 | -0.1908 | -0.278 | 0.78681 | 0.864 | -6.53 | 0.76784 | none |
| ZNF34 | -0.05 | 0.0137 | -0.278 | 0.78697 | 0.865 | -6.53 | 1.32862 | none |
| SACM1L | 0.042 | -0.002 | 0.2779 | 0.78697 | 0.865 | -6.53 | 0.47179 | none |
| RRAS | -0.037 | 0.0272 | -0.278 | 0.78707 | 0.865 | -6.53 | 0.35825 | none |
| CHRM3 | -0.065 | -0.0577 | -0.277 | 0.78779 | 0.865 | -6.53 | 0.79636 | none |
| KCNK9 | 0.025 | 0.054 | 0.2766 | 0.78793 | 0.865 | -6.53 | -0.4046 | none |
| SLC6A9 | -0.043 | -0.0268 | -0.276 | 0.78809 | 0.866 | -6.53 | 0.12101 | none |
| TNC | 0.023 | 0.0118 | 0.2762 | 0.78824 | 0.866 | -6.53 | 0.63819 | none |
| VTCN1 | -0.046 | -0.0501 | -0.276 | 0.78828 | 0.866 | -6.53 | 1.15939 | none |
| TMEM101 | -0.051 | -0.0924 | -0.276 | 0.78836 | 0.866 | -6.53 | -0.2185 | none |
| STC2 | 0.044 | 0.1424 | 0.2756 | 0.78865 | 0.866 | -6.53 | 0.10148 | none |
| TAB1 | -0.05 | 0.0277 | -0.275 | 0.78876 | 0.866 | -6.53 | 1.20635 | none |
| TMEM128 | -0.051 | -0.0912 | -0.275 | 0.78889 | 0.866 | -6.53 | 0.41265 | none |
| CCNDBP1 | 0.031 | -0.0592 | 0.2752 | 0.78899 | 0.866 | -6.53 | 0.37025 | none |
| LPAR1 | -0.021 | -0.0175 | -0.275 | 0.78901 | 0.866 | -6.53 | 0 | none |
| DDX31 | 0.047 | 0.127 | 0.2749 | 0.78921 | 0.866 | -6.53 | 1.15759 | none |
| DDX50 | -0.052 | 0.017 | -0.274 | 0.78964 | 0.867 | -6.53 | 1.2618 | none |
| TM4SF1 | -0.03 | -0.0242 | -0.274 | 0.78965 | 0.867 | -6.53 | -0.4265 | none |
| DAGLA | 0.02 | 0.0142 | 0.2742 | 0.78972 | 0.867 | -6.53 | 0 | none |

| | | | | | | | | |
|---|---|---|---|---|---|---|---|---|
| SPATA19 | -0.035 | -0.0337 | -0.274 | 0.79 | 0.867 | -6.53 | -0.3287 | none |
| PUF60 | -0.051 | 0.0215 | -0.274 | 0.79014 | 0.867 | -6.53 | 0.74296 | none |
| CYP26C1 | 0.019 | 0.0314 | 0.2736 | 0.79015 | 0.867 | -6.53 | 0 | none |
| SIRT6 | 0.026 | 0.0515 | 0.2735 | 0.79022 | 0.867 | -6.53 | 0.55689 | none |
| PCDHGC3 | 0.069 | 0.1303 | 0.2735 | 0.79025 | 0.867 | -6.53 | 1.00022 | none |
| GPR42 | -0.02 | -0.0251 | -0.273 | 0.7903 | 0.867 | -6.53 | 0 | none |
| CA10 | -0.058 | -0.0917 | -0.273 | 0.79051 | 0.867 | -6.53 | 1.32825 | none |
| ZNF428 | 0.04 | 0.0565 | 0.2728 | 0.79078 | 0.867 | -6.53 | -0.0461 | none |
| BSG | -0.028 | -0.0035 | -0.272 | 0.79118 | 0.868 | -6.53 | 0.10885 | none |
| WDR13 | 0.046 | -0.0042 | 0.2721 | 0.79126 | 0.868 | -6.53 | -0.2444 | none |
| DTYMK | -0.034 | -0.0405 | -0.272 | 0.79138 | 0.868 | -6.53 | -0.1511 | none |
| PIGF | -0.064 | -0.1603 | -0.272 | 0.79153 | 0.868 | -6.53 | 1.20935 | none |
| LYPLA1 | 0.037 | -0.0833 | 0.2717 | 0.79155 | 0.868 | -6.53 | -0.0782 | none |
| C16ORF86 | -0.039 | -0.0454 | -0.272 | 0.79155 | 0.868 | -6.53 | 1.05208 | none |
| AGGF1 | 0.062 | 0.1832 | 0.2711 | 0.79202 | 0.868 | -6.53 | 1.46447 | none |
| MBOAT4 | -0.034 | -0.0322 | -0.271 | 0.79208 | 0.868 | -6.53 | 0.78176 | none |
| MRPL22 | 0.035 | -0.0054 | 0.2705 | 0.79244 | 0.869 | -6.53 | -0.5976 | none |
| TRAPPC10 | -0.04 | -0.0207 | -0.27 | 0.79304 | 0.869 | -6.53 | 0.50129 | none |
| TMEM176 | 0.03 | 0.0649 | 0.2691 | 0.79348 | 0.87 | -6.53 | 0.56927 | none |
| ADA | -0.023 | -0.0373 | -0.269 | 0.7935 | 0.87 | -6.53 | 0.16921 | none |
| CXCL14 | 0.04 | 0.0099 | 0.2688 | 0.79371 | 0.87 | -6.53 | 1.06326 | none |
| GMPS | 0.038 | 0.0533 | 0.2683 | 0.79409 | 0.87 | -6.53 | 0.94204 | none |
| FAM229B | 0.054 | -0.0616 | 0.2683 | 0.79411 | 0.87 | -6.53 | -0.2452 | none |
| HLA-DPB1 | 0.062 | -0.0135 | 0.2682 | 0.7942 | 0.87 | -6.53 | -0.0833 | none |
| SEMA4A | -0.04 | 0.0457 | -0.268 | 0.79425 | 0.87 | -6.53 | 0.19234 | none |
| PLAC9 | 0.049 | 0.1658 | 0.268 | 0.79437 | 0.87 | -6.53 | 0.97637 | none |
| SYT4 | -0.025 | 0.0157 | -0.268 | 0.7946 | 0.87 | -6.53 | -0.03 | none |
| ECI1 | -0.034 | -0.0358 | -0.268 | 0.79461 | 0.87 | -6.53 | 0.17726 | none |
| TLX2 | -0.036 | -0.0579 | -0.267 | 0.79513 | 0.871 | -6.53 | 0.26715 | none |
| MPHOSPH | -0.044 | -0.1136 | -0.267 | 0.79515 | 0.871 | -6.53 | -0.0725 | none |
| FAM167A | 0.022 | 0.0433 | 0.2661 | 0.79574 | 0.872 | -6.54 | 0.65695 | none |
| RNASE11 | -0.021 | -0.0209 | -0.266 | 0.79601 | 0.872 | -6.54 | 0.65623 | none |
| PLA2G2E | 0.02 | 0.0116 | 0.2656 | 0.79613 | 0.872 | -6.54 | 1.20093 | none |
| FMO2 | 0.018 | 0.0269 | 0.2655 | 0.79618 | 0.872 | -6.54 | 1.35013 | none |
| KIF13A | -0.026 | -0.0293 | -0.265 | 0.79636 | 0.872 | -6.54 | 0.33495 | none |
| KIAA0232 | -0.065 | 0.075 | -0.265 | 0.7964 | 0.872 | -6.54 | 0.0562 | none |
| ZNF671 | 0.024 | 0.0283 | 0.2652 | 0.79641 | 0.872 | -6.54 | 1.37969 | none |
| RAB39B | -0.035 | -0.0101 | -0.265 | 0.79642 | 0.872 | -6.54 | 1.08212 | none |
| THAP7 | 0.026 | 0.0375 | 0.265 | 0.79659 | 0.872 | -6.54 | 0.93986 | none |
| ADAM19 | -0.06 | 0.0804 | -0.265 | 0.79666 | 0.872 | -6.54 | -0.0325 | none |

| | | | | | | | | |
|---|---|---|---|---|---|---|---|---|
| HOXD8 | 0.02 | 0.0357 | 0.2645 | 0.79697 | 0.872 | -6.54 | 0.09219 | none |
| LRP8 | -0.046 | 0.109 | -0.264 | 0.79702 | 0.872 | -6.54 | 1.09731 | none |
| DIO1 | -0.038 | 0.0162 | -0.264 | 0.79725 | 0.872 | -6.54 | 1.58829 | none |
| MAP7D2 | 0.033 | 0.0132 | 0.2641 | 0.79726 | 0.872 | -6.54 | 1.47622 | none |
| PLA2G12A | -0.042 | -0.029 | -0.264 | 0.79747 | 0.873 | -6.54 | 1.51256 | none |
| CCDC80 | 0.031 | 0.017 | 0.2636 | 0.79761 | 0.873 | -6.54 | 0.02518 | none |
| TLR6 | -0.05 | -0.0752 | -0.263 | 0.79776 | 0.873 | -6.54 | -0.129 | none |
| FO538757 | 0.05 | 0.0661 | 0.2634 | 0.79778 | 0.873 | -6.54 | 0.36403 | none |
| GNGT1 | 0.023 | 0.0052 | 0.2631 | 0.79801 | 0.873 | -6.54 | 1.57739 | none |
| MPO | 0.067 | 0.1121 | 0.263 | 0.79805 | 0.873 | -6.54 | 0.15628 | none |
| UHRF2 | -0.021 | -0.0236 | -0.263 | 0.7981 | 0.873 | -6.54 | 0.30517 | none |
| CHTF8 | -0.026 | -0.0369 | -0.263 | 0.7981 | 0.873 | -6.54 | 1.65356 | none |
| ALYREF | -0.035 | ######## | -0.263 | 0.79819 | 0.873 | -6.54 | 1.29476 | none |
| SNX4 | -0.025 | -0.0629 | -0.263 | 0.79822 | 0.873 | -6.54 | -0.4303 | none |
| OGFOD3 | 0.033 | 0.0134 | 0.2626 | 0.79836 | 0.873 | -6.54 | 1.48365 | none |
| DDX46 | 0.08 | 0.2075 | 0.2624 | 0.7985 | 0.873 | -6.54 | 0.21036 | none |
| PATE3 | -0.021 | -0.0206 | -0.262 | 0.79862 | 0.873 | -6.54 | 1.81003 | none |
| AL592183. | -0.129 | -0.2012 | -0.262 | 0.79868 | 0.873 | -6.54 | 1.71067 | none |
| USP21 | 0.034 | 0.0109 | 0.2621 | 0.79877 | 0.873 | -6.54 | -0.3127 | none |
| KIAA1522 | 0.021 | 0.0338 | 0.2619 | 0.79891 | 0.873 | -6.54 | 0.86356 | none |
| CKMT1A | -0.03 | -0.0428 | -0.261 | 0.79946 | 0.874 | -6.54 | -0.8316 | none |
| RAP2C | -0.049 | -0.0875 | -0.261 | 0.79961 | 0.874 | -6.54 | 0.14359 | none |
| PPME1 | 0.018 | 0.014 | 0.2609 | 0.79962 | 0.874 | -6.54 | 1.09734 | none |
| RNASE2 | -0.028 | -0.0301 | -0.26 | 0.80008 | 0.874 | -6.54 | -0.2729 | none |
| PLEKHJ1 | 0.031 | 0.0176 | 0.2602 | 0.80016 | 0.874 | -6.54 | 0.71952 | none |
| RARB | 0.027 | 0.0353 | 0.26 | 0.80029 | 0.875 | -6.54 | 1.36669 | none |
| CD58 | -0.034 | -0.1191 | -0.26 | 0.80042 | 0.875 | -6.54 | 0.27948 | none |
| TDRD7 | -0.031 | 0.0307 | -0.26 | 0.80049 | 0.875 | -6.54 | 1.33159 | none |
| OPN4 | -0.016 | 0.0063 | -0.26 | 0.80059 | 0.875 | -6.54 | 0 | none |
| MELTF | 0.034 | 0.0028 | 0.2596 | 0.80059 | 0.875 | -6.54 | 1.36103 | none |
| SERINC2 | -0.022 | -0.0513 | -0.259 | 0.80079 | 0.875 | -6.54 | -0.1651 | none |
| NDFIP2 | 0.049 | -0.0248 | 0.259 | 0.80107 | 0.875 | -6.54 | 0.71623 | none |
| MRC1 | 0.021 | 0.061 | 0.2583 | 0.80157 | 0.875 | -6.54 | 0.66049 | none |
| TMEM251 | -0.052 | -0.0901 | -0.258 | 0.80178 | 0.876 | -6.54 | 1.04185 | none |
| HS3ST1 | 0.035 | 0.0018 | 0.258 | 0.80179 | 0.876 | -6.54 | -0.6428 | none |
| STXBP6 | -0.02 | -0.0066 | -0.258 | 0.80186 | 0.876 | -6.54 | -0.2193 | none |
| ITCH | -0.101 | 0.0708 | -0.258 | 0.80201 | 0.876 | -6.54 | 1.05186 | none |
| PUSL1 | -0.032 | -0.0701 | -0.257 | 0.80263 | 0.876 | -6.54 | 1.23767 | none |
| CMTM7 | -0.037 | -0.1019 | -0.257 | 0.80265 | 0.876 | -6.54 | 0.43987 | none |
| HOXD13 | -0.019 | -0.0181 | -0.256 | 0.80303 | 0.877 | -6.54 | 1.32223 | none |

| | | | | | | | | |
|---|---|---|---|---|---|---|---|---|
| COG2 | 0.047 | 0.0675 | 0.2562 | 0.80317 | 0.877 | -6.54 | 0.9926 | none |
| AZIN1 | 0.03 | -0.0741 | 0.2562 | 0.80318 | 0.877 | -6.54 | 1.21415 | none |
| ARMC12 | -0.019 | -0.001 | -0.256 | 0.80335 | 0.877 | -6.54 | -0.467 | none |
| CHRD | 0.031 | 0.0684 | 0.2559 | 0.80339 | 0.877 | -6.54 | 0.10295 | none |
| RWDD3 | -0.03 | -0.1447 | -0.256 | 0.80344 | 0.877 | -6.54 | -0.0078 | none |
| RAP1GDS1 | -0.029 | -0.022 | -0.255 | 0.80401 | 0.877 | -6.54 | 1.47282 | none |
| TKTL2 | 0.018 | 0.0181 | 0.2548 | 0.80423 | 0.878 | -6.54 | 0 | none |
| UPK3A | 0.025 | 0.025 | 0.2543 | 0.80461 | 0.878 | -6.54 | -0.0099 | none |
| OR52E4 | 0.018 | 0.0138 | 0.2543 | 0.80463 | 0.878 | -6.54 | 0.95298 | none |
| COMMD9 | 0.035 | -0.0325 | 0.2541 | 0.80475 | 0.878 | -6.54 | -0.0914 | none |
| TAS2R3 | 0.017 | 0.0415 | 0.2539 | 0.80486 | 0.878 | -6.54 | -0.4095 | none |
| XAB2 | -0.039 | 0.0832 | -0.254 | 0.80489 | 0.878 | -6.54 | 1.24812 | none |
| MAG | 0.04 | 0.0359 | 0.2538 | 0.80498 | 0.878 | -6.54 | 0.61175 | none |
| STK19 | 0.097 | 0.2097 | 0.2536 | 0.80509 | 0.878 | -6.54 | 0.87516 | none |
| TRABD2B | 0.042 | 0.0449 | 0.2534 | 0.80528 | 0.878 | -6.54 | 1.29537 | none |
| SNRPA | -0.05 | -0.1 | -0.253 | 0.8054 | 0.878 | -6.54 | -0.2875 | none |
| CDCA3 | 0.022 | -0.0165 | 0.2532 | 0.80545 | 0.878 | -6.54 | 0.23904 | none |
| ESRRA | -0.023 | 0.0316 | -0.253 | 0.80554 | 0.878 | -6.54 | 0.31464 | none |
| CDK7 | -0.104 | -0.2116 | -0.253 | 0.80555 | 0.878 | -6.54 | 0.35631 | none |
| HSP90AA1 | 0.085 | 0.2428 | 0.2527 | 0.80577 | 0.879 | -6.54 | 0.36081 | none |
| SP9 | -0.018 | -0.0176 | -0.253 | 0.80588 | 0.879 | -6.54 | 0 | none |
| AMY2B | 0.055 | 0.0838 | 0.2526 | 0.80589 | 0.879 | -6.54 | 0.52729 | none |
| MUC4 | 0.02 | 0.0258 | 0.2523 | 0.80608 | 0.879 | -6.54 | 1.50705 | none |
| ZNF852 | 0.023 | 0.0678 | 0.2519 | 0.80639 | 0.879 | -6.54 | 0.97607 | none |
| CD1C | -0.02 | -0.0246 | -0.252 | 0.80646 | 0.879 | -6.54 | 1.57816 | none |
| C2ORF40 | -0.043 | -0.0484 | -0.251 | 0.8068 | 0.879 | -6.54 | 0.85943 | none |
| SOHLH1 | -0.022 | -0.0122 | -0.251 | 0.80683 | 0.879 | -6.54 | 1.97996 | none |
| DGCR6L | -0.036 | -0.0669 | -0.251 | 0.80696 | 0.879 | -6.54 | 0.34345 | none |
| PSMC3 | -0.034 | 0.0529 | -0.251 | 0.80711 | 0.879 | -6.54 | -0.2313 | none |
| CTDSP2 | 0.022 | 0.0175 | 0.2509 | 0.80713 | 0.879 | -6.54 | 0.49606 | none |
| EDEM2 | -0.024 | -0.0472 | -0.251 | 0.80719 | 0.879 | -6.54 | 0.1106 | none |
| PTDSS2 | 0.06 | 0.0154 | 0.2507 | 0.8073 | 0.88 | -6.54 | -0.9334 | none |
| DRC1 | 0.042 | 0.0511 | 0.2506 | 0.8074 | 0.88 | -6.54 | 1.22983 | none |
| PCIF1 | 0.023 | 0.0733 | 0.2505 | 0.80745 | 0.88 | -6.54 | -0.1979 | none |
| HYPK | 0.045 | 0.0363 | 0.2504 | 0.80753 | 0.88 | -6.54 | 0.144 | none |
| C11ORF16 | 0.057 | -0.0419 | 0.2503 | 0.80759 | 0.88 | -6.54 | -0.1377 | none |
| TMEM99 | 0.053 | -0.1069 | 0.2499 | 0.8079 | 0.88 | -6.54 | 0.68859 | none |
| MGAT2 | -0.031 | -0.044 | -0.25 | 0.80799 | 0.88 | -6.54 | 1.07672 | none |
| ADGRB2 | -0.021 | 0.0056 | -0.249 | 0.80831 | 0.88 | -6.54 | -0.0942 | none |
| HMX2 | 0.017 | 0.0218 | 0.2492 | 0.80841 | 0.88 | -6.54 | 0 | none |

| | | | | | | | | |
|---|---|---|---|---|---|---|---|---|
| ZNF628 | 0.057 | 0.0733 | 0.2487 | 0.80884 | 0.881 | -6.54 | 0.29985 | none |
| CENPVL3 | -0.016 | -0.0103 | -0.248 | 0.80909 | 0.881 | -6.54 | -0.3239 | none |
| PTOV1 | -0.023 | 0.0561 | -0.248 | 0.80944 | 0.881 | -6.54 | 0.49319 | none |
| SNX9 | -0.054 | 0.1004 | -0.248 | 0.8095 | 0.881 | -6.54 | 0.67332 | none |
| DPPA3 | -0.018 | ######## | -0.248 | 0.80958 | 0.881 | -6.54 | 2.00441 | none |
| ZC3H15 | -0.041 | 0.0497 | -0.248 | 0.80963 | 0.881 | -6.54 | 1.43719 | none |
| MKRN2 | -0.023 | -0.0541 | -0.248 | 0.80967 | 0.881 | -6.54 | 0.24898 | none |
| FAM234A | 0.055 | 0.1313 | 0.2467 | 0.81027 | 0.882 | -6.54 | 0.70792 | none |
| PNPLA5 | 0.025 | 0.0396 | 0.2467 | 0.8103 | 0.882 | -6.54 | 0.88042 | none |
| EMID1 | 0.037 | 0.0231 | 0.2462 | 0.81067 | 0.882 | -6.54 | 0.66061 | none |
| CDKN1A | 0.027 | 0.0364 | 0.246 | 0.8108 | 0.882 | -6.54 | 0.36949 | none |
| CYP11B2 | -0.018 | -0.0185 | -0.246 | 0.81081 | 0.882 | -6.54 | 0 | none |
| RAB7B | -0.019 | -0.0181 | -0.246 | 0.81089 | 0.882 | -6.54 | -0.5811 | none |
| CPXM1 | -0.025 | 0.0425 | -0.246 | 0.81112 | 0.882 | -6.54 | 0.44792 | none |
| GSTM4 | -0.034 | -0.0262 | -0.245 | 0.81124 | 0.883 | -6.54 | 1.42489 | none |
| VPS51 | -0.019 | -0.0174 | -0.245 | 0.81127 | 0.883 | -6.54 | 1.10862 | none |
| ABHD17C | -0.034 | -0.0229 | -0.245 | 0.81144 | 0.883 | -6.54 | 0.58268 | none |
| AP1M1 | -0.032 | -0.053 | -0.245 | 0.8116 | 0.883 | -6.54 | 0.93808 | none |
| RIMBP3B | -0.017 | -0.0125 | -0.245 | 0.81173 | 0.883 | -6.54 | 1.53579 | none |
| PCDHB3 | -0.018 | -0.018 | -0.244 | 0.81196 | 0.883 | -6.54 | 0 | none |
| SLN | -0.024 | -0.0318 | -0.244 | 0.81208 | 0.883 | -6.54 | -0.4182 | none |
| TP53TG3 | -0.018 | -0.0156 | -0.244 | 0.8123 | 0.883 | -6.54 | 0.55338 | none |
| B3GNT7 | 0.022 | 0.0138 | 0.2438 | 0.81248 | 0.883 | -6.54 | -0.6161 | none |
| ZBTB14 | 0.048 | 0.1413 | 0.2437 | 0.81253 | 0.883 | -6.54 | 1.1456 | none |
| GRIN3B | -0.017 | -0.0051 | -0.243 | 0.81296 | 0.884 | -6.54 | -0.1524 | none |
| RNF34 | -0.02 | -0.0414 | -0.243 | 0.81309 | 0.884 | -6.54 | -0.3299 | none |
| CERS6 | -0.017 | 0.0156 | -0.243 | 0.81309 | 0.884 | -6.54 | 0.89264 | none |
| APBA3 | 0.025 | 0.01 | 0.243 | 0.8131 | 0.884 | -6.54 | 0.66435 | none |
| TM9SF2 | -0.031 | -0.0847 | -0.243 | 0.81315 | 0.884 | -6.54 | 1.22231 | none |
| CLPP | -0.03 | -0.0675 | -0.243 | 0.81334 | 0.884 | -6.54 | 1.16083 | none |
| RNF225 | -0.023 | -0.0234 | -0.243 | 0.81342 | 0.884 | -6.54 | -0.238 | none |
| PALMD | 0.024 | 0.0232 | 0.2425 | 0.81345 | 0.884 | -6.54 | 0.17205 | none |
| FAM9A | 0.018 | 0.0228 | 0.2425 | 0.81347 | 0.884 | -6.54 | 1.15748 | none |
| FAM43A | 0.034 | -0.0021 | 0.2423 | 0.81362 | 0.884 | -6.54 | -0.231 | none |
| FNIP2 | 0.049 | 0.1452 | 0.2423 | 0.81365 | 0.884 | -6.54 | -0.079 | none |
| FIP1L1 | 0.038 | 0.1265 | 0.2421 | 0.81378 | 0.884 | -6.54 | 1.27353 | none |
| CSRP3 | 0.024 | 0.0306 | 0.241 | 0.81462 | 0.885 | -6.54 | 0.50094 | none |
| STX1A | 0.06 | 0.2055 | 0.241 | 0.81463 | 0.885 | -6.54 | 0.92737 | none |
| EFEMP1 | 0.024 | 0.0257 | 0.2403 | 0.81513 | 0.885 | -6.54 | 1.56461 | none |
| LFNG | -0.036 | -0.043 | -0.24 | 0.81548 | 0.886 | -6.54 | -0.131 | none |

| | | | | | | | | |
|---|---|---|---|---|---|---|---|---|
| HIST1H4D | 0.042 | -0.1228 | 0.2398 | 0.81552 | 0.886 | -6.54 | 0.36302 | none |
| PATZ1 | 0.025 | 0.1053 | 0.2396 | 0.81564 | 0.886 | -6.54 | 0.59899 | none |
| NCF4 | -0.029 | -0.0776 | -0.239 | 0.81584 | 0.886 | -6.54 | 1.49284 | none |
| PLA2G4A | 0.023 | ######## | 0.2392 | 0.81595 | 0.886 | -6.54 | 0.95701 | none |
| CORO2B | -0.017 | 0.0121 | -0.239 | 0.81603 | 0.886 | -6.54 | 0.27855 | none |
| C2ORF76 | 0.065 | -0.0351 | 0.239 | 0.8161 | 0.886 | -6.54 | 0.59691 | none |
| TMEM200 | -0.018 | -0.0266 | -0.239 | 0.81617 | 0.886 | -6.54 | 1.47 | none |
| ADRA1D | -0.017 | -0.0039 | -0.239 | 0.8162 | 0.886 | -6.54 | 1.73185 | none |
| SPAG11A | 0.018 | 0.0185 | 0.2387 | 0.8163 | 0.886 | -6.54 | 0 | none |
| MST1 | -0.052 | 0.1115 | -0.239 | 0.81644 | 0.886 | -6.54 | 1.7826 | none |
| MCRIP2 | 0.03 | 0.0169 | 0.2382 | 0.81674 | 0.886 | -6.54 | 1.57912 | none |
| NIN | 0.066 | 0.1892 | 0.2378 | 0.81697 | 0.887 | -6.54 | 0.2731 | none |
| SYS1-DBNI | 0.041 | 0.059 | 0.2378 | 0.817 | 0.887 | -6.54 | 1.62907 | none |
| ADAMTS3 | 0.016 | 0.0104 | 0.2377 | 0.81705 | 0.887 | -6.54 | 0.74034 | none |
| SNX12 | 0.052 | -0.0644 | 0.2377 | 0.81706 | 0.887 | -6.54 | 0.53269 | none |
| ADCK2 | 0.036 | -0.0207 | 0.2375 | 0.81725 | 0.887 | -6.54 | 0.33438 | none |
| CLIC6 | -0.016 | -0.0021 | -0.237 | 0.8175 | 0.887 | -6.54 | 0 | none |
| HIRIP3 | 0.023 | 0.1062 | 0.237 | 0.8176 | 0.887 | -6.54 | -0.1405 | none |
| RASA1 | 0.026 | 0.0425 | 0.2368 | 0.81774 | 0.887 | -6.54 | 0.48449 | none |
| KLHL25 | 0.02 | 0.0601 | 0.2368 | 0.81779 | 0.887 | -6.54 | 1.41135 | none |
| GJC2 | -0.017 | 0.0048 | -0.236 | 0.8183 | 0.888 | -6.54 | -0.1024 | none |
| RNF11 | 0.032 | -0.0217 | 0.236 | 0.81835 | 0.888 | -6.54 | 1.75381 | none |
| BIRC5 | 0.032 | -0.0705 | 0.236 | 0.8184 | 0.888 | -6.54 | -0.1243 | none |
| THAP10 | -0.017 | -0.0167 | -0.236 | 0.8185 | 0.888 | -6.54 | 0 | none |
| SLC20A2 | -0.025 | 0.0346 | -0.236 | 0.81856 | 0.888 | -6.54 | 0.45781 | none |
| COIL | -0.03 | 0.0195 | -0.236 | 0.81856 | 0.888 | -6.54 | 0.33608 | none |
| RIN2 | -0.028 | -0.0209 | -0.236 | 0.81864 | 0.888 | -6.54 | 0.58323 | none |
| FCRL5 | 0.016 | 0.0289 | 0.2355 | 0.81874 | 0.888 | -6.54 | 0.24864 | none |
| C4ORF48 | -0.065 | -0.0402 | -0.235 | 0.81944 | 0.888 | -6.54 | 0.82792 | none |
| CXCR3 | 0.024 | 0.0175 | 0.2339 | 0.81993 | 0.889 | -6.54 | 0.2112 | none |
| KRR1 | -0.046 | 0.0498 | -0.234 | 0.82012 | 0.889 | -6.54 | 0.83273 | none |
| CCDC192 | -0.089 | -0.0476 | -0.232 | 0.82104 | 0.89 | -6.54 | 0.31255 | none |
| KRT79 | -0.026 | -0.0241 | -0.232 | 0.82108 | 0.89 | -6.54 | 1.75143 | none |
| OR10P1 | 0.03 | 0.0289 | 0.2324 | 0.82109 | 0.89 | -6.54 | 1.01309 | none |
| ATP6V1C2 | -0.035 | 0.002 | -0.232 | 0.82111 | 0.89 | -6.54 | 0.24237 | none |
| FAM57A | -0.043 | -0.0033 | -0.232 | 0.82125 | 0.89 | -6.54 | 0.94187 | none |
| ZFYVE27 | -0.028 | 0.0377 | -0.232 | 0.82134 | 0.89 | -6.54 | 0.40891 | none |
| SNX10 | -0.023 | -0.0269 | -0.232 | 0.82138 | 0.89 | -6.54 | 1.72062 | none |
| ST14 | 0.02 | 0.042 | 0.2316 | 0.82166 | 0.89 | -6.54 | 0.21232 | none |
| COMMD2 | 0.027 | -0.0577 | 0.2313 | 0.82189 | 0.89 | -6.54 | 0.22496 | none |

| | | | | | | | | |
|---|---|---|---|---|---|---|---|---|
| EVPLL | 0.024 | 0.0277 | 0.2308 | 0.82229 | 0.891 | -6.54 | 0.99277 | none |
| DOC2B | 0.021 | 0.0064 | 0.2307 | 0.82239 | 0.891 | -6.54 | 0.10109 | none |
| DGAT2 | -0.027 | -0.0117 | -0.23 | 0.82253 | 0.891 | -6.54 | 0.86417 | none |
| GARS | -0.034 | 0.029 | -0.23 | 0.82259 | 0.891 | -6.54 | 1.49613 | none |
| DDX28 | -0.024 | -0.037 | -0.23 | 0.82325 | 0.891 | -6.54 | 1.24879 | none |
| POLR3H | -0.026 | -0.0033 | -0.23 | 0.82326 | 0.891 | -6.54 | 0.62753 | none |
| GSK3A | 0.041 | 0.0553 | 0.2294 | 0.82337 | 0.891 | -6.54 | 0.91415 | none |
| ZMYND12 | -0.054 | -0.0591 | -0.229 | 0.8234 | 0.891 | -6.54 | 1.67663 | none |
| SPRTN | -0.04 | -0.0103 | -0.229 | 0.82342 | 0.891 | -6.54 | 1.39567 | none |
| SLC46A3 | -0.027 | -0.0956 | -0.229 | 0.82344 | 0.891 | -6.54 | 0.89464 | none |
| PDCD2 | -0.03 | -0.0794 | -0.229 | 0.82353 | 0.892 | -6.54 | 1.41465 | none |
| BRSK1 | -0.039 | -0.0501 | -0.229 | 0.8236 | 0.892 | -6.54 | 0.18812 | none |
| SPRYD3 | 0.031 | 0.0194 | 0.2289 | 0.82369 | 0.892 | -6.54 | 0.93114 | none |
| TIGD4 | -0.016 | -0.0019 | -0.229 | 0.82376 | 0.892 | -6.54 | 0.37369 | none |
| CDCP1 | -0.029 | -0.0328 | -0.229 | 0.82384 | 0.892 | -6.54 | 0.90344 | none |
| ZACN | -0.033 | -0.0374 | -0.229 | 0.82385 | 0.892 | -6.55 | 0.66419 | none |
| POLR2H | -0.038 | -0.1389 | -0.229 | 0.82392 | 0.892 | -6.55 | 0.42979 | none |
| TRIM23 | 0.047 | 0.0109 | 0.2286 | 0.82395 | 0.892 | -6.55 | 1.54232 | none |
| PCDHGA4 | -0.021 | -0.0216 | -0.228 | 0.82416 | 0.892 | -6.55 | 1.53809 | none |
| ZNF512 | 0.025 | 0.0457 | 0.2282 | 0.82423 | 0.892 | -6.55 | -0.4553 | none |
| ADPRH | -0.028 | -0.0526 | -0.228 | 0.82447 | 0.892 | -6.55 | 0.11832 | none |
| APLN | 0.015 | 0.0124 | 0.2277 | 0.82463 | 0.892 | -6.55 | 0.82875 | none |
| CCL4L2 | -0.029 | 0.0437 | -0.228 | 0.82463 | 0.892 | -6.55 | 0.97611 | none |
| WNT5B | -0.02 | ######## | -0.228 | 0.82475 | 0.892 | -6.55 | -0.1579 | none |
| NSDHL | -0.018 | -0.0344 | -0.227 | 0.82481 | 0.892 | -6.55 | -0.5599 | none |
| ZNF581 | -0.028 | -0.0883 | -0.227 | 0.82481 | 0.892 | -6.55 | 1.45458 | none |
| CLIP4 | 0.044 | -0.079 | 0.2274 | 0.82484 | 0.892 | -6.55 | -0.2918 | none |
| BAALC | -0.024 | ######## | -0.227 | 0.82515 | 0.892 | -6.55 | -0.0539 | none |
| NR1H4 | -0.033 | 0.0159 | -0.227 | 0.82524 | 0.892 | -6.55 | 0.89971 | none |
| ZDHHC2 | -0.037 | -0.0602 | -0.227 | 0.82543 | 0.893 | -6.55 | 0.46172 | none |
| ILKAP | 0.025 | ######## | 0.2263 | 0.82569 | 0.893 | -6.55 | 1.06349 | none |
| SEC14L2 | 0.047 | 0.0806 | 0.2262 | 0.82578 | 0.893 | -6.55 | 0.72697 | none |
| LAYN | -0.037 | -0.0306 | -0.226 | 0.82594 | 0.893 | -6.55 | 1.12291 | none |
| ADRA2B | -0.016 | -0.0047 | -0.226 | 0.82609 | 0.893 | -6.55 | 0 | none |
| ASCL2 | 0.02 | 0.0613 | 0.2257 | 0.82612 | 0.893 | -6.55 | 0.03152 | none |
| ABCC11 | -0.031 | -0.0222 | -0.226 | 0.82614 | 0.893 | -6.55 | 1.40429 | none |
| THAP11 | -0.02 | -0.0064 | -0.226 | 0.82617 | 0.893 | -6.55 | 2.12486 | AA-related |
| MAML3 | -0.026 | -0.0134 | -0.226 | 0.82624 | 0.893 | -6.55 | 0.77691 | none |
| HTR3D | 0.019 | 0.0377 | 0.2253 | 0.82647 | 0.893 | -6.55 | 0.38278 | none |
| LRRC8C | -0.032 | 0.0017 | -0.225 | 0.82649 | 0.893 | -6.55 | 1.3156 | none |

| | | | | | | | | |
|---|---|---|---|---|---|---|---|---|
| HOXB4 | -0.021 | 0.0023 | -0.225 | 0.82668 | 0.893 | -6.55 | 1.09163 | none |
| ANKRD49 | -0.034 | -0.102 | -0.225 | 0.82689 | 0.893 | -6.55 | -0.1435 | none |
| ARRDC2 | -0.025 | 0.054 | -0.224 | 0.82705 | 0.894 | -6.55 | 1.00931 | none |
| CDCA2 | 0.062 | 0.2015 | 0.2244 | 0.82713 | 0.894 | -6.55 | 0.7098 | none |
| HIST2H4A | 0.065 | 0.0468 | 0.2243 | 0.82723 | 0.894 | -6.55 | 0.18825 | none |
| IL31 | 0.029 | 0.0883 | 0.2242 | 0.82725 | 0.894 | -6.55 | 1.21639 | none |
| FPGS | 0.027 | -0.0025 | 0.2239 | 0.82749 | 0.894 | -6.55 | 0.72165 | none |
| HELT | -0.018 | -0.018 | -0.224 | 0.82751 | 0.894 | -6.55 | 1.29038 | none |
| CD46 | -0.028 | -0.0162 | -0.224 | 0.82758 | 0.894 | -6.55 | 1.68011 | none |
| RABEPK | 0.031 | -0.0484 | 0.2235 | 0.82779 | 0.894 | -6.55 | -0.0871 | none |
| C1QL1 | -0.015 | -0.0104 | -0.223 | 0.82781 | 0.894 | -6.55 | 0.85138 | none |
| ADGRG7 | 0.016 | 0.0302 | 0.2232 | 0.82803 | 0.894 | -6.55 | 0.10208 | none |
| KHNYN | 0.035 | 0.1354 | 0.2232 | 0.82806 | 0.894 | -6.55 | -0.1302 | none |
| PHF10 | 0.033 | 0.0663 | 0.223 | 0.82818 | 0.894 | -6.55 | 0.93158 | none |
| SHISA3 | 0.022 | -0.0096 | 0.223 | 0.82818 | 0.894 | -6.55 | 0.51721 | none |
| ATP7B | -0.034 | -0.0159 | -0.223 | 0.82831 | 0.894 | -6.55 | -0.2021 | none |
| ACVR1B | -0.024 | 0.071 | -0.223 | 0.82832 | 0.894 | -6.55 | 0.44044 | none |
| HCAR2 | -0.017 | -0.0117 | -0.223 | 0.82835 | 0.894 | -6.55 | 0 | none |
| POPDC2 | -0.044 | -0.0053 | -0.223 | 0.82849 | 0.894 | -6.55 | 1.00427 | none |
| IDUA | -0.056 | -0.0744 | -0.223 | 0.82852 | 0.894 | -6.55 | -0.0172 | none |
| ARMCX2 | 0.047 | 0.1283 | 0.2225 | 0.82856 | 0.894 | -6.55 | 0.71405 | none |
| PCDHGB3 | -0.015 | -0.014 | -0.222 | 0.82864 | 0.894 | -6.55 | 1.39984 | none |
| PGRMC1 | 0.025 | -0.0122 | 0.2223 | 0.8287 | 0.894 | -6.55 | 0.75014 | none |
| AMH | 0.023 | 0.0297 | 0.2222 | 0.82875 | 0.894 | -6.55 | 0.62502 | none |
| KLHL4 | -0.015 | 0.002 | -0.222 | 0.82901 | 0.894 | -6.55 | 0.34122 | none |
| LIMS3 | -0.052 | -0.1098 | -0.222 | 0.82911 | 0.894 | -6.55 | -0.5151 | none |
| ALKBH7 | 0.044 | -0.0546 | 0.2216 | 0.82926 | 0.895 | -6.55 | 0.42136 | none |
| VPS26B | -0.03 | -0.0289 | -0.221 | 0.82938 | 0.895 | -6.55 | -0.0974 | none |
| FBXO4 | -0.054 | -0.0647 | -0.221 | 0.82946 | 0.895 | -6.55 | 0.93927 | none |
| FAM180B | 0.019 | 0.0203 | 0.2212 | 0.82952 | 0.895 | -6.55 | 0.2135 | none |
| DPH6 | -0.053 | -0.2141 | -0.221 | 0.82961 | 0.895 | -6.55 | -0.1007 | none |
| LVRN | 0.028 | 0.022 | 0.2211 | 0.82962 | 0.895 | -6.55 | 0.65512 | none |
| LUZP6 | -0.061 | -0.1392 | -0.221 | 0.82989 | 0.895 | -6.55 | -0.3736 | none |
| PLAG1 | 0.029 | 0.0213 | 0.2207 | 0.82993 | 0.895 | -6.55 | 0.29897 | none |
| SOCS6 | -0.024 | -0.024 | -0.22 | 0.83018 | 0.895 | -6.55 | 0.52856 | none |
| BTF3L4 | -0.04 | -0.1456 | -0.22 | 0.83025 | 0.895 | -6.55 | 1.05518 | none |
| FOXA2 | -0.016 | -0.0161 | -0.22 | 0.83031 | 0.895 | -6.55 | 0 | none |
| ZAR1L | 0.017 | 0.0132 | 0.2198 | 0.83063 | 0.895 | -6.55 | 1.60832 | none |
| SLC5A9 | 0.016 | 0.0181 | 0.2196 | 0.83075 | 0.895 | -6.55 | -0.3929 | none |
| PRODH2 | -0.03 | 0.0381 | -0.22 | 0.83076 | 0.895 | -6.55 | 0.11013 | none |

| | | | | | | | | |
|---|---|---|---|---|---|---|---|---|
| SLC26A9 | -0.024 | -0.0234 | -0.22 | 0.83077 | 0.895 | -6.55 | -0.5473 | none |
| RBBP8NL | 0.017 | 0.0161 | 0.2195 | 0.83083 | 0.895 | -6.55 | 0.19303 | none |
| NECAB1 | -0.016 | -0.0065 | -0.219 | 0.83095 | 0.895 | -6.55 | -0.1415 | none |
| C4ORF50 | 0.014 | 0.0317 | 0.2192 | 0.83102 | 0.895 | -6.55 | 0.53253 | none |
| PRG2 | -0.024 | 0.0107 | -0.219 | 0.83108 | 0.895 | -6.55 | 0.13637 | none |
| GDE1 | -0.024 | -0.0465 | -0.219 | 0.83128 | 0.896 | -6.55 | 1.3436 | none |
| ZFPM1 | 0.048 | 0.1505 | 0.2186 | 0.8315 | 0.896 | -6.55 | 0.39836 | none |
| DAZL | -0.017 | -0.0328 | -0.219 | 0.83159 | 0.896 | -6.55 | 1.38822 | none |
| CLCA4 | -0.042 | -0.0416 | -0.218 | 0.8316 | 0.896 | -6.55 | 1.6425 | none |
| MID2 | 0.042 | -0.0045 | 0.2184 | 0.83166 | 0.896 | -6.55 | 0.58471 | none |
| MAP2K5 | -0.032 | -0.0548 | -0.218 | 0.83179 | 0.896 | -6.55 | 0.98151 | none |
| PSTPIP1 | 0.023 | 0.0563 | 0.2181 | 0.8319 | 0.896 | -6.55 | -0.1984 | none |
| RRAD | 0.038 | 0.0397 | 0.2177 | 0.83219 | 0.896 | -6.55 | 0.34531 | none |
| SRC | -0.032 | 0.0571 | -0.217 | 0.83235 | 0.896 | -6.55 | 0.80507 | none |
| PGBD2 | 0.023 | 0.0036 | 0.2171 | 0.83268 | 0.897 | -6.55 | -0.1088 | none |
| TDRD10 | 0.045 | 0.0834 | 0.2169 | 0.83282 | 0.897 | -6.55 | 0.68844 | none |
| FAM114A1 | -0.039 | -0.0142 | -0.217 | 0.83296 | 0.897 | -6.55 | -0.1643 | none |
| PITPNC1 | 0.035 | 0.0835 | 0.2166 | 0.83302 | 0.897 | -6.55 | 1.25275 | none |
| DLL4 | -0.017 | -0.0166 | -0.216 | 0.83316 | 0.897 | -6.55 | 0.32708 | none |
| DBNL | 0.019 | 0.0352 | 0.2163 | 0.83329 | 0.897 | -6.55 | 0.0707 | none |
| STARD3 | -0.033 | -0.0628 | -0.216 | 0.83357 | 0.897 | -6.55 | 0.40101 | none |
| RPRM | -0.018 | -0.0058 | -0.216 | 0.83361 | 0.897 | -6.55 | 1.08454 | none |
| LRRC4 | -0.031 | 0.0079 | -0.216 | 0.83361 | 0.897 | -6.55 | 0.42983 | none |
| HES6 | -0.069 | -0.0087 | -0.216 | 0.83364 | 0.897 | -6.55 | 0.98427 | none |
| RCAN2 | -0.024 | -0.0692 | -0.216 | 0.83368 | 0.897 | -6.55 | 0.38226 | none |
| MS4A1 | 0.036 | 0.0351 | 0.2156 | 0.83377 | 0.897 | -6.55 | 0.93998 | none |
| AAMP | -0.021 | 0.0061 | -0.215 | 0.83396 | 0.897 | -6.55 | -0.0408 | none |
| LGALS2 | -0.033 | -0.0326 | -0.215 | 0.83403 | 0.897 | -6.55 | -0.4829 | none |
| POLR1C | 0.02 | -0.0168 | 0.2152 | 0.83411 | 0.897 | -6.55 | -0.2111 | none |
| HIST2H3PS | 0.02 | 0.0375 | 0.2151 | 0.83416 | 0.897 | -6.55 | 0.7396 | none |
| KLK3 | 0.016 | 0.0157 | 0.215 | 0.83426 | 0.897 | -6.55 | 1.53512 | none |
| IL36B | 0.018 | 0.0219 | 0.2148 | 0.83438 | 0.897 | -6.55 | -0.089 | none |
| BUB1B-PA | 0.015 | ######## | 0.2146 | 0.83451 | 0.898 | -6.55 | -0.4229 | none |
| GSDMD | -0.029 | -0.0664 | -0.214 | 0.83487 | 0.898 | -6.55 | 1.34883 | none |
| ZNF181 | -0.036 | 0.0677 | -0.214 | 0.83495 | 0.898 | -6.55 | 1.43907 | none |
| NUDT22 | 0.032 | -0.0421 | 0.214 | 0.83501 | 0.898 | -6.55 | 0.40103 | none |
| TAF12 | 0.026 | -0.0063 | 0.2139 | 0.83505 | 0.898 | -6.55 | 1.13843 | none |
| PRMT5 | -0.027 | -0.0338 | -0.214 | 0.83525 | 0.898 | -6.55 | 1.09121 | none |
| GPT | -0.021 | -0.012 | -0.213 | 0.83543 | 0.898 | -6.55 | 0.70926 | none |
| RASD2 | -0.016 | -0.0042 | -0.213 | 0.8357 | 0.898 | -6.55 | 0 | none |

| | | | | | | | | |
|---|---|---|---|---|---|---|---|---|
| SLC9A3R1 | 0.022 | 0.0611 | 0.2131 | 0.83571 | 0.898 | -6.55 | -0.0235 | none |
| RGS14 | 0.025 | 0.0744 | 0.2126 | 0.83603 | 0.899 | -6.55 | 0.97308 | none |
| ARMC7 | -0.026 | 0.0216 | -0.212 | 0.83614 | 0.899 | -6.55 | 1.48761 | none |
| RGL3 | -0.053 | 0.0731 | -0.212 | 0.8362 | 0.899 | -6.55 | 1.5866 | none |
| TATDN2 | 0.034 | 0.1522 | 0.2124 | 0.8362 | 0.899 | -6.55 | 1.63721 | none |
| H2AFY2 | 0.028 | 0.0652 | 0.2123 | 0.83629 | 0.899 | -6.55 | 0.75467 | none |
| MESP2 | -0.041 | -0.0301 | -0.212 | 0.83651 | 0.899 | -6.55 | 1.95993 | none |
| FAM49A | 0.027 | 0.017 | 0.2118 | 0.83667 | 0.899 | -6.55 | 1.25174 | none |
| NME3 | 0.034 | -0.0028 | 0.2117 | 0.83673 | 0.899 | -6.55 | 1.14908 | none |
| CELF6 | -0.02 | -0.0093 | -0.211 | 0.83698 | 0.899 | -6.55 | 1.01496 | none |
| HSPG2 | 0.026 | 0.0703 | 0.2114 | 0.83701 | 0.899 | -6.55 | 1.39957 | none |
| POLR3D | 0.04 | 0.1609 | 0.2112 | 0.83713 | 0.899 | -6.55 | 0.804 | none |
| TCN2 | -0.025 | ######## | -0.211 | 0.8372 | 0.899 | -6.55 | -0.553 | none |
| AK2 | -0.014 | 0.0097 | -0.211 | 0.83738 | 0.899 | -6.55 | 0.2127 | none |
| OXSM | 0.031 | -0.032 | 0.2107 | 0.83751 | 0.9 | -6.55 | 0.36404 | none |
| PAX9 | -0.02 | 0.0384 | -0.21 | 0.83782 | 0.9 | -6.55 | 1.14585 | none |
| TRAPPC6B | 0.036 | -0.075 | 0.2103 | 0.83784 | 0.9 | -6.55 | 1.25559 | none |
| MB21D2 | -0.022 | -0.0525 | -0.21 | 0.83788 | 0.9 | -6.55 | 1.0547 | none |
| ETV7 | -0.018 | -0.0274 | -0.21 | 0.83798 | 0.9 | -6.55 | 0.24434 | none |
| GSTM3 | -0.049 | -0.1338 | -0.21 | 0.83802 | 0.9 | -6.55 | 1.16595 | none |
| CDCP2 | 0.014 | 0.0218 | 0.21 | 0.83804 | 0.9 | -6.55 | -0.2615 | none |
| CCDC3 | 0.016 | 0.0286 | 0.21 | 0.83804 | 0.9 | -6.55 | 0.49628 | none |
| RBP4 | -0.023 | -0.0417 | -0.21 | 0.83813 | 0.9 | -6.55 | 0.35894 | none |
| VSIG10L | 0.024 | 0.0865 | 0.2094 | 0.83845 | 0.9 | -6.55 | 0.45968 | none |
| SLC22A5 | -0.018 | -0.0941 | -0.209 | 0.8385 | 0.9 | -6.55 | -0.4183 | none |
| DYNLL2 | -0.022 | 0.0178 | -0.209 | 0.8385 | 0.9 | -6.55 | 0.53457 | none |
| ZDHHC15 | -0.03 | -0.0376 | -0.209 | 0.83868 | 0.9 | -6.55 | -0.3186 | none |
| SAMSN1 | -0.018 | -0.1158 | -0.209 | 0.8389 | 0.9 | -6.55 | -0.1534 | none |
| KDM1A | -0.034 | 0.1378 | -0.209 | 0.83897 | 0.9 | -6.55 | 1.34593 | none |
| RAB10 | 0.021 | -0.0094 | 0.2088 | 0.83897 | 0.9 | -6.55 | -0.2186 | none |
| ZFYVE19 | 0.036 | 0.0758 | 0.2085 | 0.83917 | 0.9 | -6.55 | 1.10195 | none |
| NELFB | -0.025 | 0.0403 | -0.208 | 0.8392 | 0.9 | -6.55 | 0.78426 | none |
| CNOT9 | -0.021 | -0.039 | -0.208 | 0.83922 | 0.9 | -6.55 | -0.5685 | none |
| EGR4 | -0.02 | 0.0244 | -0.208 | 0.83929 | 0.9 | -6.55 | 0.48579 | none |
| FZD2 | 0.015 | 0.0243 | 0.208 | 0.83958 | 0.901 | -6.55 | 1.00831 | none |
| MRPS22 | 0.025 | -0.0504 | 0.2075 | 0.83992 | 0.901 | -6.55 | 0.4121 | none |
| B4GALT7 | 0.027 | 0.0304 | 0.2071 | 0.8402 | 0.901 | -6.55 | 0.09853 | none |
| PTGDR | 0.025 | 0.0384 | 0.2071 | 0.84022 | 0.901 | -6.55 | 1.10733 | none |
| NRARP | 0.018 | 0.0271 | 0.207 | 0.84029 | 0.901 | -6.55 | 0.68276 | none |
| KCND1 | 0.025 | 0.0803 | 0.206 | 0.84105 | 0.902 | -6.55 | -0.0432 | none |

| | | | | | | | | |
|---|---|---|---|---|---|---|---|---|
| CFAP70 | 0.022 | 0.0108 | 0.206 | 0.84106 | 0.902 | -6.55 | 1.58126 | none |
| KCNK13 | 0.016 | 0.0207 | 0.2058 | 0.84126 | 0.902 | -6.55 | 0.09648 | none |
| RPP30 | -0.025 | -0.1117 | -0.206 | 0.84137 | 0.902 | -6.55 | -0.0662 | none |
| NEMP1 | 0.038 | 0.0407 | 0.2053 | 0.84162 | 0.902 | -6.55 | 1.19771 | none |
| BCL2L1 | 0.026 | 0.0386 | 0.205 | 0.84183 | 0.902 | -6.55 | 0.11688 | none |
| ARRDC1 | -0.024 | -0.0458 | -0.205 | 0.84203 | 0.903 | -6.55 | 1.5542 | none |
| IZUMO4 | 0.059 | -0.0027 | 0.2047 | 0.84205 | 0.903 | -6.55 | 0.27504 | none |
| INPP4B | 0.048 | 0.055 | 0.2045 | 0.8422 | 0.903 | -6.55 | 0.10445 | none |
| TTC5 | -0.021 | -0.0538 | -0.204 | 0.84225 | 0.903 | -6.55 | 0.21427 | none |
| ODF3L1 | 0.016 | 0.0291 | 0.2043 | 0.84235 | 0.903 | -6.55 | 0.83781 | none |
| PNMA5 | -0.017 | -0.0092 | -0.204 | 0.8424 | 0.903 | -6.55 | 0.13144 | none |
| PAIP1 | -0.025 | -0.0353 | -0.204 | 0.84245 | 0.903 | -6.55 | 1.76816 | none |
| WT1 | 0.025 | 0.0608 | 0.2041 | 0.84248 | 0.903 | -6.55 | 0.50006 | none |
| CYP4A11 | -0.029 | -0.0023 | -0.204 | 0.84253 | 0.903 | -6.55 | 1.84494 | none |
| HDDC3 | 0.027 | 0.0068 | 0.2039 | 0.84264 | 0.903 | -6.55 | 0.78846 | none |
| CCL1 | -0.054 | -0.0884 | -0.204 | 0.84266 | 0.903 | -6.55 | 1.07795 | none |
| P4HA1 | -0.016 | -0.0247 | -0.203 | 0.84317 | 0.903 | -6.55 | 1.47532 | none |
| EGFLAM | -0.027 | -0.0196 | -0.203 | 0.8433 | 0.903 | -6.55 | -0.3533 | none |
| HNRNPL | 0.027 | 0.0767 | 0.2026 | 0.84362 | 0.904 | -6.55 | -0.3154 | none |
| ANO1 | 0.035 | 0.0825 | 0.2025 | 0.84377 | 0.904 | -6.55 | 1.13295 | none |
| QRFPR | 0.016 | 0.0173 | 0.202 | 0.84411 | 0.904 | -6.55 | -0.2748 | none |
| LYNX1 | 0.029 | 0.0393 | 0.2019 | 0.84417 | 0.904 | -6.55 | -0.4568 | none |
| EFCAB13 | 0.023 | 0.0078 | 0.2018 | 0.84427 | 0.904 | -6.55 | 0.77566 | none |
| AMD1 | 0.021 | -0.0799 | 0.2017 | 0.84432 | 0.904 | -6.55 | 0.05874 | none |
| PLA2G4F | -0.041 | -0.0738 | -0.202 | 0.84439 | 0.904 | -6.55 | 0.37441 | none |
| SETMAR | 0.03 | 0.0571 | 0.2009 | 0.84496 | 0.905 | -6.55 | 0.82855 | none |
| SYT16 | 0.037 | 0.0103 | 0.2007 | 0.84511 | 0.905 | -6.55 | 0.24544 | none |
| SCAP | 0.027 | 0.0998 | 0.2003 | 0.8454 | 0.905 | -6.55 | 0.83806 | none |
| PLAGL2 | 0.02 | 0.093 | 0.2 | 0.84562 | 0.905 | -6.55 | 0.75028 | none |
| LIN37 | -0.027 | 0.0106 | -0.199 | 0.84602 | 0.905 | -6.55 | 0.08439 | none |
| RAB31 | 0.032 | -0.018 | 0.1985 | 0.84677 | 0.906 | -6.55 | 0.54753 | none |
| SH2D2A | -0.018 | 0.0722 | -0.198 | 0.84698 | 0.906 | -6.55 | 0.53993 | none |
| XKRX | 0.017 | -0.0132 | 0.1982 | 0.847 | 0.906 | -6.55 | 0.11541 | none |
| FITM1 | 0.014 | 0.0356 | 0.1982 | 0.84702 | 0.906 | -6.55 | 1.64608 | none |
| PCDHA1 | 0.014 | 0.0268 | 0.1981 | 0.84704 | 0.906 | -6.55 | 0 | none |
| DEFB129 | -0.022 | -0.0218 | -0.198 | 0.84712 | 0.906 | -6.55 | 1.19411 | none |
| SLC16A3 | -0.026 | -0.025 | -0.198 | 0.84725 | 0.906 | -6.55 | 0.70894 | none |
| PDE2A | 0.047 | 0.0742 | 0.1974 | 0.84757 | 0.907 | -6.55 | 1.12653 | none |
| GPR18 | -0.04 | -0.1494 | -0.197 | 0.84782 | 0.907 | -6.55 | 0.31267 | none |
| MAN1A1 | 0.019 | 0.0057 | 0.197 | 0.8479 | 0.907 | -6.55 | 0.88058 | none |

| | | | | | | | | |
|---|---|---|---|---|---|---|---|---|
| SERPINB13 | 0.015 | 0.0154 | 0.1968 | 0.84805 | 0.907 | -6.55 | -0.6806 | none |
| KLF1 | -0.013 | -0.014 | -0.197 | 0.84805 | 0.907 | -6.55 | 1.64649 | none |
| SYNJ1 | 0.022 | ######## | 0.1967 | 0.84814 | 0.907 | -6.55 | 0.95356 | none |
| BEND5 | 0.056 | 0.0738 | 0.1965 | 0.84826 | 0.907 | -6.55 | 1.40219 | none |
| LRP3 | -0.031 | 0.0038 | -0.197 | 0.84827 | 0.907 | -6.55 | 0.13915 | none |
| APOBEC1 | -0.018 | -0.0179 | -0.196 | 0.84861 | 0.907 | -6.55 | 0.6787 | none |
| SLC44A1 | -0.017 | -0.0703 | -0.196 | 0.84875 | 0.907 | -6.55 | -0.2123 | none |
| ANKRD65 | 0.018 | 0.045 | 0.1959 | 0.84877 | 0.907 | -6.55 | 1.77624 | none |
| FOXD4L1 | -0.014 | ######## | -0.196 | 0.8489 | 0.907 | -6.55 | 0 | none |
| MAP4K3 | 0.035 | 0.0409 | 0.1956 | 0.84895 | 0.907 | -6.55 | 0.95441 | none |
| TUBB1 | 0.014 | 0.0352 | 0.1955 | 0.84905 | 0.908 | -6.55 | 1.56501 | none |
| TMEM207 | -0.015 | -0.0151 | -0.195 | 0.84907 | 0.908 | -6.55 | -0.2598 | none |
| PFKP | 0.028 | 0.149 | 0.1954 | 0.84915 | 0.908 | -6.55 | 0.54049 | none |
| NDUFS7 | -0.034 | -0.0581 | -0.195 | 0.84946 | 0.908 | -6.55 | 1.13493 | none |
| UBE2D1 | 0.025 | -0.0207 | 0.1949 | 0.8495 | 0.908 | -6.55 | 1.09984 | none |
| SERPINA9 | 0.02 | 0.029 | 0.1943 | 0.84997 | 0.908 | -6.55 | 1.68678 | none |
| CASP9 | 0.021 | -0.0332 | 0.1943 | 0.85001 | 0.908 | -6.55 | -0.708 | none |
| CDC25C | -0.034 | -0.0388 | -0.194 | 0.85002 | 0.908 | -6.55 | 0.10872 | none |
| FCMR | -0.019 | -0.0338 | -0.194 | 0.8501 | 0.908 | -6.55 | 0.38619 | none |
| FNDC11 | 0.015 | 0.0381 | 0.194 | 0.85019 | 0.908 | -6.55 | 0.19534 | none |
| GRID2IP | -0.013 | 0.007 | -0.193 | 0.85073 | 0.909 | -6.55 | 0.88273 | none |
| BTBD10 | 0.025 | -0.0152 | 0.1932 | 0.85084 | 0.909 | -6.55 | 0.27381 | none |
| POLR3B | -0.027 | 0.0321 | -0.193 | 0.85098 | 0.909 | -6.55 | 0.51721 | none |
| CFC1 | -0.03 | -0.0159 | -0.193 | 0.85104 | 0.909 | -6.55 | 0.74998 | none |
| TRPC7 | 0.016 | 0.0159 | 0.1926 | 0.85127 | 0.909 | -6.55 | -0.6239 | none |
| RGPD1 | 0.04 | 0.0356 | 0.1925 | 0.85131 | 0.909 | -6.55 | 0.75276 | none |
| ACBD3 | 0.048 | 0.1611 | 0.1924 | 0.85142 | 0.909 | -6.55 | 0.40717 | none |
| NMB | 0.037 | 0.0128 | 0.1917 | 0.85196 | 0.91 | -6.55 | 0.33776 | none |
| GRIK4 | 0.033 | 0.0487 | 0.1916 | 0.85205 | 0.91 | -6.55 | 0.19947 | none |
| EBF4 | 0.025 | 0.0504 | 0.1915 | 0.85209 | 0.91 | -6.55 | -0.2379 | none |
| AC005779 | -0.027 | -0.0268 | -0.191 | 0.85232 | 0.91 | -6.55 | 1.59282 | none |
| ANKRD37 | -0.032 | -0.0127 | -0.191 | 0.85235 | 0.91 | -6.55 | 1.4735 | none |
| ANXA7 | -0.02 | -0.11 | -0.191 | 0.85246 | 0.91 | -6.55 | 0.16576 | none |
| RASIP1 | 0.021 | 0.033 | 0.1907 | 0.85267 | 0.91 | -6.55 | 1.24324 | none |
| CYLC2 | 0.059 | 0.1082 | 0.1906 | 0.85278 | 0.91 | -6.55 | -0.6806 | none |
| CSH1 | 0.019 | 0.0403 | 0.1905 | 0.85284 | 0.91 | -6.55 | 0.28062 | none |
| LRPAP1 | -0.028 | 0.0899 | -0.19 | 0.85306 | 0.91 | -6.55 | 0.0981 | none |
| ZBTB34 | 0.015 | 0.0201 | 0.1902 | 0.85308 | 0.91 | -6.55 | -0.1796 | none |
| TTC21A | 0.059 | 0.1052 | 0.1901 | 0.85321 | 0.91 | -6.55 | -0.2758 | none |
| ZNF516 | 0.022 | 0.0803 | 0.1899 | 0.85335 | 0.91 | -6.55 | 0.9309 | none |

| | | | | | | | | |
|---|---|---|---|---|---|---|---|---|
| LRWD1 | 0.019 | 0.0729 | 0.1898 | 0.85339 | 0.91 | -6.55 | 0.533 | none |
| ERF | -0.048 | 0.0742 | -0.19 | 0.85339 | 0.91 | -6.55 | 0.10883 | none |
| COL9A2 | 0.045 | 0.1331 | 0.1896 | 0.85358 | 0.911 | -6.55 | 1.16846 | none |
| RASD1 | 0.019 | 0.0175 | 0.1895 | 0.85361 | 0.911 | -6.55 | 1.03047 | none |
| RAB7A | 0.019 | 0.0332 | 0.1895 | 0.85366 | 0.911 | -6.55 | 0.91227 | none |
| TSEN15 | 0.029 | -0.0568 | 0.1893 | 0.8538 | 0.911 | -6.55 | 1.02567 | none |
| IPMK | 0.014 | -0.026 | 0.1891 | 0.85394 | 0.911 | -6.55 | 0.72342 | none |
| GAD2 | 0.014 | 0.0345 | 0.1891 | 0.85395 | 0.911 | -6.55 | 1.11437 | none |
| OR5C1 | -0.016 | -0.016 | -0.189 | 0.85396 | 0.911 | -6.55 | -0.6121 | none |
| HNRNPH2 | 0.019 | -0.0504 | 0.1891 | 0.85396 | 0.911 | -6.55 | 0.90545 | none |
| MAT1A | -0.016 | -0.0165 | -0.189 | 0.85427 | 0.911 | -6.55 | -0.1522 | none |
| BNIP3L | 0.016 | 0.0076 | 0.1887 | 0.85427 | 0.911 | -6.55 | -0.2898 | none |
| GABRR2 | 0.028 | 0.0278 | 0.1883 | 0.85455 | 0.911 | -6.55 | 0.41618 | none |
| B3GALT4 | 0.014 | 0.0161 | 0.1882 | 0.85463 | 0.911 | -6.55 | 0.72654 | none |
| CORO1C | -0.023 | -0.0124 | -0.188 | 0.85472 | 0.911 | -6.55 | 1.16565 | none |
| NPB | -0.015 | -0.0267 | -0.188 | 0.85491 | 0.911 | -6.55 | -0.4974 | none |
| NBPF9 | 0.033 | 0.1958 | 0.1876 | 0.85504 | 0.911 | -6.55 | 0.44533 | none |
| TSPAN11 | 0.015 | 0.0215 | 0.1875 | 0.85512 | 0.911 | -6.55 | -0.325 | none |
| MSX2 | -0.013 | -0.0052 | -0.187 | 0.85517 | 0.911 | -6.55 | -0.6895 | none |
| DGKQ | 0.026 | 0.0578 | 0.1874 | 0.85526 | 0.911 | -6.55 | -0.2384 | none |
| WNK2 | 0.032 | 0.103 | 0.1873 | 0.85533 | 0.911 | -6.55 | 1.0355 | none |
| PCDH8 | -0.013 | -0.0084 | -0.187 | 0.85535 | 0.911 | -6.55 | -0.5515 | none |
| MAP1A | 0.014 | 0.0589 | 0.1871 | 0.85548 | 0.911 | -6.55 | 0.34781 | none |
| HESX1 | -0.023 | -0.04 | -0.187 | 0.85549 | 0.911 | -6.55 | -0.4067 | none |
| NODAL | 0.014 | 0.0439 | 0.1869 | 0.85558 | 0.911 | -6.55 | 0.38287 | none |
| USP6 | -0.016 | 0.0317 | -0.187 | 0.85558 | 0.911 | -6.55 | 1.20783 | none |
| 1-Sep | 0.024 | 0.0643 | 0.1869 | 0.85564 | 0.911 | -6.55 | -0.0203 | none |
| THOC6 | -0.019 | -0.0124 | -0.187 | 0.8559 | 0.912 | -6.55 | 0.48287 | none |
| C1ORF35 | 0.023 | 0.0304 | 0.1865 | 0.85595 | 0.912 | -6.55 | -0.1125 | none |
| MTRNR2L1 | 0.02 | -0.0078 | 0.1861 | 0.85619 | 0.912 | -6.55 | 0.9718 | none |
| HYPM | -0.029 | -0.0285 | -0.186 | 0.85661 | 0.912 | -6.55 | 1.20276 | none |
| OXT | 0.023 | 0.0893 | 0.1855 | 0.85666 | 0.912 | -6.55 | 0.86115 | none |
| FAM83D | 0.017 | 0.0041 | 0.1854 | 0.85673 | 0.912 | -6.55 | 0.93034 | none |
| SYT10 | 0.012 | 0.0159 | 0.1852 | 0.85694 | 0.912 | -6.55 | 0.77493 | none |
| VTN | -0.014 | -0.026 | -0.185 | 0.85698 | 0.912 | -6.55 | -0.7647 | none |
| TMEM200 | 0.014 | 0.0161 | 0.1851 | 0.85698 | 0.912 | -6.55 | -0.6503 | none |
| CYB5R4 | -0.037 | -0.099 | -0.185 | 0.8572 | 0.912 | -6.55 | 1.06732 | none |
| FAM50A | 0.039 | 0.1775 | 0.1846 | 0.85734 | 0.912 | -6.55 | 1.25127 | none |
| TEX10 | 0.02 | -0.011 | 0.1846 | 0.85735 | 0.912 | -6.55 | 1.27186 | none |
| ZFAND1 | -0.028 | -0.14 | -0.184 | 0.85745 | 0.912 | -6.55 | 0.34923 | none |

| | | | | | | | | |
|---|---|---|---|---|---|---|---|---|
| UBE2E1 | -0.039 | -0.0778 | -0.184 | 0.85746 | 0.912 | -6.55 | 0.52914 | none |
| TMEM107 | -0.028 | -0.1285 | -0.184 | 0.85751 | 0.912 | -6.55 | 0.63227 | none |
| RCE1 | 0.024 | 0.0333 | 0.1843 | 0.85759 | 0.913 | -6.55 | 1.01189 | none |
| RANBP2 | -0.067 | 0.1971 | -0.184 | 0.85806 | 0.913 | -6.56 | 0.41386 | none |
| AEN | -0.02 | 0.0593 | -0.184 | 0.8581 | 0.913 | -6.56 | 1.5506 | none |
| GDPD3 | -0.029 | 0.0304 | -0.183 | 0.85825 | 0.913 | -6.56 | 1.11645 | none |
| NUDT13 | 0.065 | 0.0321 | 0.1834 | 0.85828 | 0.913 | -6.56 | 0.40478 | none |
| HDAC10 | 0.02 | 0.0207 | 0.1833 | 0.85837 | 0.913 | -6.56 | 1.29229 | none |
| GCAT | -0.025 | -0.036 | -0.183 | 0.85844 | 0.913 | -6.56 | 0.94289 | none |
| ADAMTS1( | 0.038 | 0.0538 | 0.1828 | 0.85876 | 0.913 | -6.56 | -0.5788 | none |
| NOB1 | -0.025 | 0.0612 | -0.182 | 0.859 | 0.913 | -6.56 | 1.53438 | none |
| GOLT1A | -0.023 | -0.0122 | -0.182 | 0.85901 | 0.913 | -6.56 | 0.31731 | none |
| B4GALT6 | 0.02 | 0.0177 | 0.1823 | 0.85909 | 0.913 | -6.56 | -0.5565 | none |
| BUD31 | 0.018 | -0.0244 | 0.1823 | 0.8591 | 0.913 | -6.56 | -0.6192 | none |
| DLEU7 | 0.015 | 0.0113 | 0.1821 | 0.85925 | 0.914 | -6.56 | 0.38296 | none |
| ZAP70 | -0.024 | 0.0121 | -0.181 | 0.86019 | 0.915 | -6.56 | 1.169 | none |
| KCNAB1 | 0.049 | ######## | 0.1806 | 0.8604 | 0.915 | -6.56 | 0.1864 | none |
| TXNDC16 | 0.042 | 0.0114 | 0.1805 | 0.86049 | 0.915 | -6.56 | 0.99161 | none |
| RAET1G | -0.014 | -0.0184 | -0.18 | 0.86097 | 0.915 | -6.56 | -0.6403 | none |
| FCGR3B | 0.018 | 0.018 | 0.1792 | 0.86151 | 0.916 | -6.56 | 0.96175 | none |
| TNPO3 | 0.028 | 0.0771 | 0.1788 | 0.86176 | 0.916 | -6.56 | 0.49318 | none |
| DEFB119 | -0.031 | -0.0306 | -0.179 | 0.86182 | 0.916 | -6.56 | 1.20526 | none |
| LRRIQ3 | -0.022 | -0.022 | -0.179 | 0.86186 | 0.916 | -6.56 | 1.48562 | none |
| C16ORF87 | -0.019 | -0.0726 | -0.178 | 0.8622 | 0.916 | -6.56 | -0.0262 | none |
| PHC1 | 0.032 | 0.0682 | 0.1781 | 0.8623 | 0.916 | -6.56 | 0.07239 | none |
| MMP24 | -0.012 | 0.0283 | -0.178 | 0.86252 | 0.916 | -6.56 | 1.23677 | none |
| UBXN2B | -0.03 | -0.0931 | -0.178 | 0.86257 | 0.916 | -6.56 | 1.08455 | none |
| PIK3CA | -0.044 | 0.1004 | -0.178 | 0.86265 | 0.916 | -6.56 | 0.86239 | none |
| ZNF263 | 0.035 | 0.1378 | 0.1777 | 0.86266 | 0.916 | -6.56 | -0.1649 | none |
| DDOST | -0.02 | -0.0782 | -0.176 | 0.86363 | 0.917 | -6.56 | 0.40023 | none |
| PELO | -0.016 | ######## | -0.176 | 0.8637 | 0.917 | -6.56 | 1.46756 | none |
| CISD3 | 0.029 | 0.028 | 0.1761 | 0.86386 | 0.918 | -6.56 | 0.10289 | none |
| POLR2M | -0.026 | -0.0534 | -0.176 | 0.86399 | 0.918 | -6.56 | 1.05678 | none |
| C1ORF189 | -0.016 | 0.0132 | -0.176 | 0.86405 | 0.918 | -6.56 | 0.0054 | none |
| TCEAL2 | 0.014 | 0.0165 | 0.1758 | 0.86407 | 0.918 | -6.56 | 0.53536 | none |
| TTC1 | 0.019 | 0.0638 | 0.1752 | 0.86452 | 0.918 | -6.56 | -0.0518 | none |
| PGPEP1L | -0.019 | 0.0164 | -0.175 | 0.86452 | 0.918 | -6.56 | -0.3845 | none |
| ELOVL3 | -0.013 | 0.0048 | -0.175 | 0.86477 | 0.918 | -6.56 | -0.1428 | none |
| PLPP6 | 0.023 | 0.0152 | 0.1748 | 0.86488 | 0.918 | -6.56 | 0.95615 | none |
| CFAP44 | 0.033 | 0.052 | 0.1746 | 0.86503 | 0.918 | -6.56 | 0.90492 | none |

| | | | | | | | | |
|---|---|---|---|---|---|---|---|---|
| GRM3 | -0.012 | 0.0029 | -0.174 | 0.86558 | 0.919 | -6.56 | -0.2622 | none |
| TMEM178 | -0.02 | -0.0025 | -0.173 | 0.86635 | 0.92 | -6.56 | -0.3159 | none |
| BLOC1S2 | 0.02 | -0.0146 | 0.1727 | 0.86648 | 0.92 | -6.56 | -0.1943 | none |
| DYRK2 | 0.028 | -0.0196 | 0.1724 | 0.86669 | 0.92 | -6.56 | 0.46542 | none |
| CYP2S1 | 0.026 | 0.0432 | 0.1721 | 0.86688 | 0.92 | -6.56 | 1.38209 | none |
| CFAP97 | -0.028 | 0.029 | -0.172 | 0.86716 | 0.92 | -6.56 | 1.71537 | none |
| MAPK7 | 0.027 | 0.1259 | 0.1718 | 0.86717 | 0.92 | -6.56 | 0.33457 | none |
| CDH5 | -0.017 | -0.0286 | -0.172 | 0.86737 | 0.92 | -6.56 | 0.66543 | none |
| MYRF | 0.023 | 0.0189 | 0.1715 | 0.8674 | 0.92 | -6.56 | -0.4668 | none |
| ZNF358 | -0.03 | ######## | -0.171 | 0.86741 | 0.92 | -6.56 | 1.0867 | none |
| CENPB | -0.022 | ######## | -0.171 | 0.86778 | 0.921 | -6.56 | -0.7112 | none |
| SIKE1 | 0.023 | -0.0898 | 0.1705 | 0.86812 | 0.921 | -6.56 | 0.04324 | none |
| KRT32 | -0.012 | -0.0067 | -0.17 | 0.86814 | 0.921 | -6.56 | 0 | none |
| C8ORF82 | 0.015 | ######## | 0.1704 | 0.86824 | 0.921 | -6.56 | -0.1344 | none |
| IPCEF1 | 0.03 | 0.0063 | 0.1702 | 0.86833 | 0.921 | -6.56 | 0.65913 | none |
| RLN1 | -0.02 | 0.0057 | -0.17 | 0.86836 | 0.921 | -6.56 | 1.22731 | none |
| ADGRG3 | 0.02 | 0.0589 | 0.1702 | 0.86838 | 0.921 | -6.56 | 0.21757 | none |
| HIST1H4B | -0.02 | -0.0397 | -0.17 | 0.86848 | 0.921 | -6.56 | 1.05785 | none |
| RNF123 | 0.034 | 0.0721 | 0.1695 | 0.86893 | 0.921 | -6.56 | 1.28094 | none |
| DUSP11 | 0.019 | -0.082 | 0.1694 | 0.86895 | 0.921 | -6.56 | 0.21781 | none |
| SPANXD | 0.019 | 0.0464 | 0.1688 | 0.86941 | 0.922 | -6.56 | 0.66625 | none |
| TTC9 | 0.021 | 0.0483 | 0.1687 | 0.86951 | 0.922 | -6.56 | 0.2255 | none |
| SWT1 | -0.027 | 0.009 | -0.168 | 0.8698 | 0.922 | -6.56 | 1.46252 | none |
| SERPINA1 | 0.039 | 0.1184 | 0.1683 | 0.86981 | 0.922 | -6.56 | 0.42341 | none |
| SERTAD2 | 0.025 | 0.0195 | 0.1682 | 0.86987 | 0.922 | -6.56 | 0.5431 | none |
| ARX | -0.056 | 0.1219 | -0.168 | 0.87002 | 0.922 | -6.56 | 0.53995 | none |
| FAM170A | -0.013 | -0.0126 | -0.168 | 0.87007 | 0.922 | -6.56 | -0.0315 | none |
| CCL17 | 0.036 | 0.1157 | 0.1679 | 0.87011 | 0.922 | -6.56 | 1.45469 | none |
| YEATS4 | 0.023 | -0.0327 | 0.1672 | 0.87063 | 0.922 | -6.56 | 0.93396 | none |
| AQP11 | -0.029 | -0.0215 | -0.167 | 0.87063 | 0.922 | -6.56 | 0.75649 | none |
| CTDNEP1 | -0.027 | 0.0807 | -0.167 | 0.87073 | 0.922 | -6.56 | 0.2514 | none |
| ANKRD34/ | -0.034 | -0.0145 | -0.167 | 0.87081 | 0.922 | -6.56 | 0.66939 | none |
| ACADM | 0.016 | 0.013 | 0.167 | 0.87084 | 0.922 | -6.56 | 0.34076 | none |
| FAM151A | 0.014 | 0.0011 | 0.167 | 0.87085 | 0.922 | -6.56 | 1.28052 | none |
| GAD1 | 0.033 | 0.0587 | 0.1668 | 0.87095 | 0.923 | -6.56 | -0.1575 | none |
| FCGR1A | 0.022 | 0.0355 | 0.1667 | 0.87101 | 0.923 | -6.56 | -0.6786 | none |
| MIOS | -0.026 | 0.0177 | -0.167 | 0.87112 | 0.923 | -6.56 | 1.53704 | none |
| CALR3 | -0.019 | 0.0087 | -0.166 | 0.87126 | 0.923 | -6.56 | 1.07585 | none |
| PLG | 0.035 | -0.0146 | 0.1664 | 0.87127 | 0.923 | -6.56 | 0.38145 | none |
| ZNF296 | -0.014 | 0.0366 | -0.166 | 0.87129 | 0.923 | -6.56 | 1.66031 | none |

| | | | | | | | | |
|---|---|---|---|---|---|---|---|---|
| SMAP1 | -0.031 | 0.0664 | -0.166 | 0.87133 | 0.923 | -6.56 | 0.96629 | none |
| UCP2 | -0.018 | -0.0365 | -0.166 | 0.87137 | 0.923 | -6.56 | 1.27315 | none |
| LUC7L3 | 0.037 | 0.1424 | 0.1657 | 0.87183 | 0.923 | -6.56 | 0.75 | none |
| NSMF | -0.037 | 0.1173 | -0.165 | 0.87197 | 0.923 | -6.56 | 0.08578 | none |
| NR4A2 | -0.024 | 0.0806 | -0.165 | 0.87198 | 0.923 | -6.56 | 0.56145 | none |
| EFCAB9 | -0.018 | -0.0184 | -0.165 | 0.87206 | 0.923 | -6.56 | 1.06067 | none |
| AMY1B | -0.016 | 0.0042 | -0.165 | 0.87258 | 0.924 | -6.56 | 1.7753 | none |
| EIF4E3 | 0.025 | -0.079 | 0.1646 | 0.87267 | 0.924 | -6.56 | 0.1093 | none |
| FCRLB | -0.015 | -0.0339 | -0.164 | 0.87316 | 0.924 | -6.56 | 1.8424 | none |
| IKZF2 | -0.018 | -0.0387 | -0.164 | 0.87325 | 0.924 | -6.56 | 1.73884 | none |
| STK24 | 0.022 | 0.0184 | 0.1636 | 0.87338 | 0.924 | -6.56 | -0.9254 | none |
| FOXE3 | 0.011 | 0.0218 | 0.1636 | 0.8734 | 0.924 | -6.56 | 0 | none |
| CITED2 | 0.038 | 0.0686 | 0.1634 | 0.87355 | 0.924 | -6.56 | 0.63732 | none |
| PAGE5 | -0.03 | -0.1134 | -0.163 | 0.87359 | 0.924 | -6.56 | 0.90061 | none |
| PCSK1N | 0.043 | 0.0946 | 0.163 | 0.8739 | 0.924 | -6.56 | -0.5698 | none |
| TCN1 | -0.018 | -0.0116 | -0.163 | 0.87392 | 0.924 | -6.56 | 1.91248 | none |
| CRYZ | -0.027 | -0.1407 | -0.163 | 0.87395 | 0.924 | -6.56 | 0.05704 | none |
| CAMTA1 | -0.03 | -0.1027 | -0.163 | 0.87397 | 0.924 | -6.56 | -0.0938 | none |
| PFKFB4 | -0.015 | 0.0638 | -0.163 | 0.87425 | 0.925 | -6.56 | 1.35049 | none |
| ROPN1L | 0.026 | -0.0437 | 0.1624 | 0.87433 | 0.925 | -6.56 | 0.96182 | none |
| EDA2R | -0.012 | 0.0062 | -0.162 | 0.87449 | 0.925 | -6.56 | 0.01022 | none |
| TTC30B | -0.011 | -0.0106 | -0.162 | 0.8746 | 0.925 | -6.56 | 0.90941 | none |
| ASXL1 | 0.023 | 0.0669 | 0.162 | 0.87467 | 0.925 | -6.56 | -0.04 | none |
| EFCAB1 | -0.014 | 0.0163 | -0.162 | 0.87467 | 0.925 | -6.56 | 0.70034 | none |
| CYB5B | -0.018 | -0.0979 | -0.162 | 0.87473 | 0.925 | -6.56 | 0.35687 | none |
| KLK12 | 0.014 | 0.0142 | 0.1614 | 0.87509 | 0.925 | -6.56 | 1.43031 | none |
| NKD2 | -0.014 | 0.0145 | -0.161 | 0.87521 | 0.925 | -6.56 | 0.61505 | none |
| SRSF8 | -0.015 | -0.0647 | -0.161 | 0.87533 | 0.925 | -6.56 | 0.30206 | none |
| A4GNT | -0.012 | -0.0115 | -0.161 | 0.87548 | 0.925 | -6.56 | 0 | none |
| PTPN5 | 0.013 | 0.0342 | 0.1608 | 0.87554 | 0.925 | -6.56 | 0.55593 | none |
| CLDN1 | -0.023 | 0.0017 | -0.161 | 0.87562 | 0.925 | -6.56 | 1.45492 | none |
| LOXL3 | 0.017 | 0.0724 | 0.1603 | 0.87593 | 0.926 | -6.56 | 0.72445 | none |
| 9-Sep | 0.031 | 0.1023 | 0.16 | 0.8762 | 0.926 | -6.56 | 0.82847 | none |
| UNC50 | 0.025 | -0.0925 | 0.1599 | 0.87626 | 0.926 | -6.56 | 0.7103 | none |
| ARHGEF35 | -0.018 | -0.0444 | -0.16 | 0.87629 | 0.926 | -6.56 | 0.64452 | none |
| ZNF536 | -0.014 | 0.0111 | -0.16 | 0.87654 | 0.926 | -6.56 | -0.3178 | none |
| TCP10 | 0.015 | 0.0313 | 0.1594 | 0.87661 | 0.926 | -6.56 | -0.5043 | none |
| MMP17 | -0.026 | -0.0343 | -0.159 | 0.87662 | 0.926 | -6.56 | 1.13134 | none |
| SASH3 | 0.027 | 0.1173 | 0.1593 | 0.87671 | 0.926 | -6.56 | 0.67291 | none |
| GPRIN2 | 0.012 | 0.0183 | 0.159 | 0.87693 | 0.926 | -6.56 | 0.28301 | none |

| | | | | | | | | |
|---|---|---|---|---|---|---|---|---|
| ASNS | 0.015 | -0.0743 | 0.1586 | 0.87728 | 0.927 | -6.56 | -0.1192 | none |
| MBD4 | -0.033 | 0.0889 | -0.158 | 0.87733 | 0.927 | -6.56 | 0.90779 | none |
| SRF | 0.022 | 0.0881 | 0.1577 | 0.87793 | 0.927 | -6.56 | 0.65851 | none |
| LRIF1 | 0.02 | 0.0442 | 0.1574 | 0.87818 | 0.927 | -6.56 | 1.56579 | none |
| COL7A1 | -0.034 | ######## | -0.157 | 0.8783 | 0.927 | -6.56 | 1.49204 | none |
| MRRF | 0.016 | -0.0308 | 0.1571 | 0.87837 | 0.927 | -6.56 | 1.55653 | none |
| TP53INP1 | 0.013 | -0.0755 | 0.1571 | 0.8784 | 0.927 | -6.56 | 0.47735 | none |
| ZNF438 | -0.02 | 0.0214 | -0.157 | 0.87846 | 0.927 | -6.56 | 1.55501 | none |
| DUSP10 | 0.019 | -0.0082 | 0.157 | 0.87847 | 0.927 | -6.56 | 0.01647 | none |
| TMLHE | -0.034 | -0.046 | -0.157 | 0.87856 | 0.927 | -6.56 | 1.2342 | none |
| NEDD8-MI | 0.029 | -0.0409 | 0.1569 | 0.87857 | 0.927 | -6.56 | 1.16694 | none |
| USP17L24 | -0.011 | 0.006 | -0.157 | 0.87858 | 0.927 | -6.56 | 0 | none |
| USP17L25 | -0.011 | 0.006 | -0.157 | 0.87858 | 0.927 | -6.56 | 0 | none |
| USP17L26 | -0.011 | 0.006 | -0.157 | 0.87858 | 0.927 | -6.56 | 0 | none |
| ANKS6 | -0.028 | -0.0226 | -0.157 | 0.87876 | 0.927 | -6.56 | 0.88133 | none |
| SLC35C2 | 0.02 | -0.0232 | 0.1566 | 0.87879 | 0.927 | -6.56 | 1.27215 | none |
| TOR1AIP1 | 0.017 | 0.021 | 0.1566 | 0.87881 | 0.927 | -6.56 | 1.04004 | none |
| TTC30A | -0.011 | -0.0061 | -0.157 | 0.87881 | 0.927 | -6.56 | -0.405 | none |
| SAV1 | -0.027 | -0.0714 | -0.156 | 0.87903 | 0.927 | -6.56 | 0.20789 | none |
| MECR | 0.017 | 0.0685 | 0.1563 | 0.87904 | 0.927 | -6.56 | -0.117 | none |
| VN1R2 | 0.011 | 0.0263 | 0.1561 | 0.87916 | 0.927 | -6.56 | 1.14937 | none |
| CTRB2 | 0.018 | 0.012 | 0.1561 | 0.87916 | 0.927 | -6.56 | 1.09092 | none |
| NIPAL4 | 0.018 | 0.0297 | 0.1555 | 0.87961 | 0.928 | -6.56 | 1.41951 | none |
| FARSA | -0.014 | 0.0272 | -0.155 | 0.87983 | 0.928 | -6.56 | 0.7266 | none |
| SETD3 | -0.016 | ######## | -0.155 | 0.88 | 0.928 | -6.56 | 1.20092 | none |
| FGL1 | -0.012 | -0.0168 | -0.155 | 0.88025 | 0.928 | -6.56 | 0.79704 | none |
| NEUROD6 | -0.011 | -0.0108 | -0.155 | 0.88037 | 0.928 | -6.56 | 0 | none |
| PTBP3 | 0.031 | -0.0259 | 0.1545 | 0.8804 | 0.928 | -6.56 | -0.1048 | none |
| C2ORF73 | -0.011 | -0.011 | -0.154 | 0.8808 | 0.929 | -6.56 | 0.50395 | none |
| AATF | 0.033 | 0.1771 | 0.1539 | 0.88086 | 0.929 | -6.56 | 0.69669 | none |
| FAM104A | 0.019 | 0.0158 | 0.1538 | 0.88096 | 0.929 | -6.56 | 0.39303 | none |
| SOBP | 0.013 | 0.0342 | 0.1535 | 0.88117 | 0.929 | -6.56 | 0.90833 | none |
| VSTM2A | -0.011 | 0.0075 | -0.153 | 0.88119 | 0.929 | -6.56 | 2.02628 | none |
| UBAC2 | 0.019 | -0.077 | 0.1535 | 0.88119 | 0.929 | -6.56 | 0.53957 | none |
| ASXL3 | 0.026 | 0.0173 | 0.1534 | 0.88124 | 0.929 | -6.56 | 0.57908 | none |
| WASF1 | -0.012 | 0.0384 | -0.153 | 0.88128 | 0.929 | -6.56 | 1.69507 | none |
| TLR9 | 0.027 | 0.0178 | 0.1533 | 0.88129 | 0.929 | -6.56 | 0.40344 | none |
| TMSB4Y | 0.012 | 0.021 | 0.1532 | 0.88141 | 0.929 | -6.56 | 1.25684 | none |
| PTPN13 | 0.012 | 0.0279 | 0.1532 | 0.88143 | 0.929 | -6.56 | 1.24575 | none |
| GOLGA6L2 | 0.023 | 0.0431 | 0.1528 | 0.88171 | 0.929 | -6.56 | -0.8125 | none |

| | | | | | | | | |
|---|---|---|---|---|---|---|---|---|
| BCAT2 | 0.014 | 0.0201 | 0.1523 | 0.88209 | 0.929 | -6.56 | 1.24791 | none |
| OR2V1 | -0.01 | 0.0112 | -0.152 | 0.88237 | 0.93 | -6.56 | 0.71908 | none |
| AMACR | 0.023 | -0.0024 | 0.1517 | 0.88254 | 0.93 | -6.56 | 0.99302 | none |
| NCAPH2 | 0.014 | 0.0337 | 0.1516 | 0.88263 | 0.93 | -6.56 | 0.02787 | none |
| RPP25L | 0.023 | -0.0364 | 0.1511 | 0.88298 | 0.93 | -6.56 | 0.60194 | none |
| TUSC2 | -0.024 | -0.0296 | -0.151 | 0.88333 | 0.93 | -6.56 | 0.55846 | none |
| PDZD9 | 0.011 | 0.0107 | 0.1505 | 0.88346 | 0.93 | -6.56 | 1.06477 | none |
| MAP3K2 | 0.032 | 0.1536 | 0.1505 | 0.88348 | 0.93 | -6.56 | 0.47163 | none |
| UBTD2 | -0.015 | -0.0104 | -0.15 | 0.88405 | 0.931 | -6.56 | 0.81601 | none |
| SLC6A5 | 0.014 | 0.0134 | 0.1496 | 0.88414 | 0.931 | -6.56 | -0.3918 | none |
| SERPINE1 | 0.016 | 0.0345 | 0.1494 | 0.88432 | 0.931 | -6.56 | 0.82088 | none |
| CCNG2 | 0.019 | -0.1149 | 0.1494 | 0.88432 | 0.931 | -6.56 | -0.0026 | none |
| GNAO1 | -0.014 | -0.028 | -0.149 | 0.8847 | 0.931 | -6.56 | -0.5972 | none |
| FGF8 | 0.019 | -0.0397 | 0.1488 | 0.88474 | 0.931 | -6.56 | -0.1914 | none |
| CMTM8 | -0.031 | -0.0738 | -0.149 | 0.88482 | 0.931 | -6.56 | 0.72465 | none |
| PSMC1 | 0.019 | 0.0745 | 0.1484 | 0.88505 | 0.932 | -6.56 | -0.5467 | none |
| SGMS2 | 0.034 | 0.0621 | 0.1484 | 0.8851 | 0.932 | -6.56 | 0.38872 | none |
| HSPA12B | 0.011 | 0.0158 | 0.1482 | 0.88523 | 0.932 | -6.56 | -0.0894 | none |
| HMMR | 0.058 | 0.2509 | 0.1482 | 0.88523 | 0.932 | -6.56 | 0.73351 | none |
| AP3M2 | 0.022 | -0.0012 | 0.1481 | 0.88533 | 0.932 | -6.56 | 1.44776 | none |
| PLEKHA3 | 0.024 | -0.0665 | 0.1478 | 0.88551 | 0.932 | -6.56 | 0.98127 | none |
| EIF4EBP2 | -0.016 | -0.096 | -0.148 | 0.88551 | 0.932 | -6.56 | 0.17989 | none |
| APOLD1 | 0.022 | 0.0444 | 0.1478 | 0.88552 | 0.932 | -6.56 | 0.32536 | none |
| KRT24 | -0.011 | -0.0052 | -0.148 | 0.88554 | 0.932 | -6.56 | 0.5088 | none |
| HNRNPA2I | -0.024 | 0.0519 | -0.147 | 0.88618 | 0.932 | -6.56 | 1.09878 | none |
| SLC7A10 | 0.012 | 0.0116 | 0.1465 | 0.88656 | 0.933 | -6.56 | 0.81301 | none |
| CD59 | 0.015 | -0.0494 | 0.1464 | 0.88661 | 0.933 | -6.56 | -0.3546 | none |
| GGPS1 | 0.017 | -0.0646 | 0.1462 | 0.88675 | 0.933 | -6.56 | -0.1371 | none |
| RXFP4 | -0.013 | 0.0112 | -0.146 | 0.88696 | 0.933 | -6.56 | -0.1907 | none |
| DMRTB1 | 0.013 | 0.0129 | 0.1459 | 0.88701 | 0.933 | -6.56 | 0.48486 | none |
| ZDHHC3 | -0.01 | -0.0087 | -0.146 | 0.88708 | 0.933 | -6.56 | 0.38149 | none |
| ANKRA2 | -0.024 | -0.0889 | -0.146 | 0.88712 | 0.933 | -6.56 | 1.57933 | none |
| CD37 | 0.016 | -0.0544 | 0.1457 | 0.88719 | 0.933 | -6.56 | 1.03445 | none |
| NDRG4 | 0.038 | 0.0623 | 0.1452 | 0.88753 | 0.933 | -6.56 | 0.58092 | none |
| RNF144B | 0.017 | 0.0457 | 0.1451 | 0.88762 | 0.933 | -6.56 | 1.34143 | none |
| VIPR2 | -0.019 | 0.0032 | -0.145 | 0.88767 | 0.933 | -6.56 | 0.83057 | none |
| CLDN20 | -0.015 | -0.0313 | -0.145 | 0.88793 | 0.933 | -6.56 | -0.5003 | none |
| YY1 | 0.029 | 0.1104 | 0.1447 | 0.88794 | 0.933 | -6.56 | 0.27906 | none |
| ANXA11 | 0.016 | 0.0388 | 0.1445 | 0.88805 | 0.933 | -6.56 | 1.03549 | none |
| C9ORF16 | -0.023 | -0.0493 | -0.144 | 0.8881 | 0.933 | -6.56 | 0.54191 | none |

| | | | | | | | | |
|---|---|---|---|---|---|---|---|---|
| MOB4 | -0.018 | -0.1102 | -0.144 | 0.88831 | 0.934 | -6.56 | 1.28042 | none |
| ATE1 | 0.025 | 0.0731 | 0.1442 | 0.88833 | 0.934 | -6.56 | 0.887 | none |
| FBN3 | -0.011 | 0.013 | -0.144 | 0.88847 | 0.934 | -6.56 | 0.77261 | none |
| ALPK2 | 0.011 | 0.0149 | 0.1439 | 0.88853 | 0.934 | -6.56 | 1.77976 | none |
| ESX1 | -0.014 | -0.0146 | -0.144 | 0.8886 | 0.934 | -6.56 | 1.31701 | none |
| NOTUM | 0.023 | 0.0028 | 0.1437 | 0.88869 | 0.934 | -6.56 | 0.56429 | none |
| SERPINB1( | -0.019 | -0.0303 | -0.143 | 0.8889 | 0.934 | -6.56 | 0.71766 | none |
| AVPR2 | -0.029 | -0.0092 | -0.143 | 0.88902 | 0.934 | -6.56 | 0.57559 | none |
| TMEM39A | -0.033 | -0.0522 | -0.143 | 0.88922 | 0.934 | -6.56 | 0.11341 | none |
| HOXB9 | -0.011 | ######## | -0.143 | 0.88952 | 0.934 | -6.56 | -0.105 | none |
| FCGR3A | -0.01 | 0.003 | -0.143 | 0.88956 | 0.934 | -6.56 | 1.06743 | none |
| HNRNPCL1 | -0.013 | -0.0077 | -0.142 | 0.88963 | 0.934 | -6.56 | 1.9268 | none |
| SCN8A | -0.019 | 0.0096 | -0.142 | 0.88972 | 0.934 | -6.56 | 1.93924 | none |
| MMEL1 | -0.054 | -0.021 | -0.142 | 0.88987 | 0.934 | -6.56 | -0.0314 | none |
| SCRIB | 0.025 | 0.1754 | 0.1421 | 0.88989 | 0.934 | -6.56 | 0.04031 | none |
| TUBB2B | -0.017 | -0.0449 | -0.142 | 0.89003 | 0.934 | -6.56 | 1.25041 | none |
| EIF4A2 | 0.016 | -0.0128 | 0.1418 | 0.89017 | 0.935 | -6.56 | 1.4336 | none |
| FAM171B | -0.012 | -0.0052 | -0.142 | 0.89018 | 0.935 | -6.56 | 0.42157 | none |
| LRRC70 | -0.019 | 0.0212 | -0.142 | 0.89023 | 0.935 | -6.56 | 0.95656 | none |
| AMDHD1 | -0.041 | 0.0727 | -0.142 | 0.89038 | 0.935 | -6.56 | 0.71786 | none |
| EIF2B3 | 0.009 | -0.0082 | 0.1412 | 0.89062 | 0.935 | -6.56 | 0.18556 | none |
| AC090958 | -0.011 | -0.0149 | -0.141 | 0.89087 | 0.935 | -6.56 | 0.25989 | none |
| MMP14 | 0.036 | 0.1118 | 0.1406 | 0.89106 | 0.935 | -6.56 | 1.03341 | none |
| DR1 | 0.014 | -0.0305 | 0.1406 | 0.8911 | 0.935 | -6.56 | 1.67286 | none |
| CAMSAP3 | -0.035 | 0.0376 | -0.14 | 0.89171 | 0.936 | -6.56 | 0.61935 | none |
| NPTX1 | -0.015 | -0.0314 | -0.139 | 0.89193 | 0.936 | -6.56 | -0.5965 | none |
| IQSEC3 | 0.011 | 0.0375 | 0.1392 | 0.89214 | 0.936 | -6.56 | -0.4981 | none |
| STUB1 | -0.018 | -0.0262 | -0.139 | 0.89232 | 0.936 | -6.56 | 1.34197 | none |
| HOXB13 | -0.016 | 0.0217 | -0.139 | 0.8924 | 0.936 | -6.56 | 0.66617 | none |
| HOXD11 | -0.01 | -0.0097 | -0.139 | 0.89241 | 0.936 | -6.56 | -0.3453 | none |
| ZNF711 | 0.013 | 0.0129 | 0.1388 | 0.89248 | 0.936 | -6.56 | 0.6135 | none |
| ACO1 | 0.012 | 0.0454 | 0.1388 | 0.89249 | 0.936 | -6.56 | 1.22094 | none |
| GPR135 | -0.011 | 0.0232 | -0.139 | 0.8925 | 0.936 | -6.56 | -0.6047 | none |
| C9ORF43 | -0.012 | 0.0222 | -0.139 | 0.89259 | 0.936 | -6.56 | 1.38932 | none |
| GPSM1 | 0.021 | 0.0942 | 0.1384 | 0.89275 | 0.936 | -6.56 | 0.50867 | none |
| OR52L1 | -0.011 | -0.011 | -0.138 | 0.89289 | 0.936 | -6.56 | 0.63284 | none |
| ARL1 | 0.016 | -0.0457 | 0.1382 | 0.89292 | 0.936 | -6.56 | 0.70659 | none |
| TCF15 | 0.01 | 0.0073 | 0.1378 | 0.89323 | 0.937 | -6.56 | 0.32807 | none |
| PCYT2 | 0.011 | 0.0168 | 0.1375 | 0.89347 | 0.937 | -6.56 | -0.5164 | none |
| ADAM21 | 0.011 | -0.0106 | 0.1374 | 0.89354 | 0.937 | -6.56 | 0.71529 | none |

| | | | | | | | | |
|---|---|---|---|---|---|---|---|---|
| FSTL3 | 0.022 | -0.0651 | 0.137 | 0.89381 | 0.937 | -6.56 | 0.1943 | none |
| SYNM | -0.01 | -0.0227 | -0.137 | 0.89384 | 0.937 | -6.56 | 0.39084 | none |
| PPP2R5D | 0.026 | 0.1334 | 0.1368 | 0.894 | 0.937 | -6.56 | 0.90782 | none |
| RCHY1 | 0.025 | -0.0631 | 0.1368 | 0.89403 | 0.937 | -6.56 | 1.25478 | none |
| SLCO6A1 | 0.019 | 0.0192 | 0.1367 | 0.89405 | 0.937 | -6.56 | 1.69849 | none |
| DNASE1L1 | -0.032 | -0.0903 | -0.136 | 0.89446 | 0.937 | -6.56 | 0.44678 | none |
| VSX1 | -0.01 | 0.0147 | -0.136 | 0.8945 | 0.937 | -6.56 | 0.53086 | none |
| GPX7 | -0.013 | -0.0407 | -0.136 | 0.89451 | 0.937 | -6.56 | 0.88195 | none |
| HNRNPM | 0.024 | 0.1218 | 0.1361 | 0.89455 | 0.937 | -6.56 | 0.3978 | none |
| HIST2H2BI | 0.011 | -0.035 | 0.1361 | 0.89458 | 0.937 | -6.56 | -0.1944 | none |
| ZBTB45 | 0.017 | 0.0259 | 0.136 | 0.89464 | 0.937 | -6.56 | 1.42043 | none |
| C7ORF50 | -0.017 | -0.056 | -0.136 | 0.89472 | 0.937 | -6.56 | 0.64592 | none |
| PGP | 0.022 | 0.0347 | 0.1358 | 0.89474 | 0.937 | -6.56 | 1.01025 | none |
| TMEM259 | -0.021 | 0.0045 | -0.136 | 0.89477 | 0.937 | -6.56 | 0.07661 | none |
| DHCR7 | 0.02 | 0.0711 | 0.1357 | 0.89483 | 0.937 | -6.56 | 0.82112 | none |
| TMBIM1 | -0.019 | -0.079 | -0.136 | 0.8949 | 0.937 | -6.56 | -0.3384 | none |
| IRS4 | 0.009 | 0.0177 | 0.135 | 0.89535 | 0.938 | -6.56 | 0 | none |
| CRY1 | 0.014 | 0.0075 | 0.1345 | 0.89577 | 0.938 | -6.56 | 0.96333 | none |
| OR4N2 | 0.02 | 0.0254 | 0.1344 | 0.89582 | 0.938 | -6.56 | 0.28641 | none |
| KIFC3 | 0.038 | 0.1747 | 0.1343 | 0.89594 | 0.938 | -6.56 | 0.59744 | none |
| LTB4R | -0.017 | 0.0109 | -0.134 | 0.89597 | 0.938 | -6.56 | 1.09999 | none |
| NDC1 | -0.025 | -0.0692 | -0.134 | 0.89601 | 0.938 | -6.56 | 0.10732 | none |
| TNFRSF14 | -0.015 | -0.0539 | -0.134 | 0.89608 | 0.938 | -6.56 | 0.98228 | none |
| MCRIP1 | -0.022 | -0.0622 | -0.134 | 0.89647 | 0.938 | -6.56 | 0.34152 | none |
| TMEM31 | -0.014 | 0.0111 | -0.133 | 0.89659 | 0.939 | -6.56 | 0.21054 | none |
| IFIT2 | -0.011 | -0.0109 | -0.133 | 0.89669 | 0.939 | -6.56 | 0.39861 | none |
| SUPT7L | -0.018 | -0.0283 | -0.133 | 0.89672 | 0.939 | -6.56 | 0.11645 | none |
| MASP1 | 0.019 | -0.0458 | 0.1332 | 0.89678 | 0.939 | -6.56 | -0.1001 | none |
| TSPAN16 | -0.01 | 0.0062 | -0.133 | 0.89709 | 0.939 | -6.56 | 0.23067 | none |
| HNRNPH3 | -0.019 | 0.0317 | -0.132 | 0.89739 | 0.939 | -6.56 | 1.28323 | none |
| MYLIP | 0.013 | 0.0176 | 0.1323 | 0.89744 | 0.939 | -6.56 | 0.09493 | none |
| TGFBR2 | -0.018 | 0.0483 | -0.132 | 0.89744 | 0.939 | -6.56 | 1.05045 | none |
| SYT3 | -0.021 | -0.0318 | -0.132 | 0.89745 | 0.939 | -6.56 | 1.45327 | none |
| YTHDF1 | -0.017 | 0.0505 | -0.132 | 0.89754 | 0.939 | -6.56 | 0.37855 | none |
| ZBTB22 | 0.018 | 0.0145 | 0.1321 | 0.89758 | 0.939 | -6.56 | 0.03646 | none |
| FCRL6 | -0.018 | -0.0321 | -0.132 | 0.89784 | 0.939 | -6.56 | 0.01552 | none |
| RPP21 | -0.044 | -0.1423 | -0.132 | 0.898 | 0.939 | -6.56 | 0.88592 | none |
| MEF2B | -0.025 | 0.0109 | -0.131 | 0.8981 | 0.939 | -6.56 | 0.68009 | none |
| C9ORF152 | -0.01 | -0.0048 | -0.131 | 0.89813 | 0.939 | -6.56 | 0.90419 | none |
| NDRG1 | -0.013 | 0.0819 | -0.131 | 0.89815 | 0.939 | -6.56 | 0.76351 | none |

| | | | | | | | | |
|---|---|---|---|---|---|---|---|---|
| FOXG1 | 0.037 | 0.0359 | 0.1314 | 0.89819 | 0.939 | -6.56 | 0.6081 | none |
| IQCK | -0.019 | -0.1202 | -0.131 | 0.89876 | 0.94 | -6.56 | -0.0292 | none |
| UNC13B | 0.023 | 0.0779 | 0.1304 | 0.8989 | 0.94 | -6.56 | 0.87817 | none |
| FABP3 | 0.017 | 0.0703 | 0.1303 | 0.89899 | 0.94 | -6.56 | 1.92109 | none |
| IFFO2 | 0.024 | 0.0774 | 0.1301 | 0.89913 | 0.94 | -6.56 | 1.24341 | none |
| HTR2B | -0.014 | -0.0546 | -0.13 | 0.89924 | 0.94 | -6.56 | 1.09061 | none |
| HSD11B2 | 0.03 | 0.019 | 0.1299 | 0.89929 | 0.94 | -6.56 | -0.0328 | none |
| FGF1 | 0.016 | 0.0061 | 0.1299 | 0.89931 | 0.94 | -6.56 | -0.5113 | none |
| SLC22A10 | -0.01 | -0.0099 | -0.13 | 0.8996 | 0.94 | -6.56 | 0 | none |
| ST3GAL4 | 0.019 | 0.0532 | 0.1294 | 0.89968 | 0.94 | -6.56 | 0.78761 | none |
| HOXA10 | -0.02 | -0.0049 | -0.129 | 0.89978 | 0.94 | -6.56 | -0.2024 | none |
| SHANK2 | 0.043 | 0.0081 | 0.1291 | 0.89994 | 0.94 | -6.56 | 0.88465 | none |
| REP15 | 0.011 | 0.0528 | 0.1291 | 0.89995 | 0.94 | -6.56 | 1.11564 | none |
| PFKFB1 | 0.014 | 0.0405 | 0.1289 | 0.9001 | 0.941 | -6.56 | 0.84689 | none |
| ARHGAP32 | -0.021 | 0.02 | -0.129 | 0.90016 | 0.941 | -6.56 | 0.6658 | none |
| GLT1D1 | -0.013 | -0.009 | -0.129 | 0.90032 | 0.941 | -6.56 | -0.3491 | none |
| FBXL7 | 0.009 | 0.0196 | 0.1284 | 0.9005 | 0.941 | -6.56 | 0 | none |
| NECTIN1 | 0.029 | 0.0751 | 0.1282 | 0.90059 | 0.941 | -6.56 | 0.70159 | none |
| MAN2A1 | 0.018 | 0.0398 | 0.1279 | 0.90086 | 0.941 | -6.56 | 0.67823 | none |
| ADGRL2 | -0.009 | 0.0113 | -0.128 | 0.90111 | 0.941 | -6.56 | -0.7104 | none |
| SCP2 | -0.018 | -0.1035 | -0.127 | 0.90117 | 0.941 | -6.56 | 0.73472 | none |
| TDG | 0.014 | -0.0247 | 0.1273 | 0.90134 | 0.941 | -6.56 | 0.58816 | none |
| C15ORF39 | 0.014 | 0.072 | 0.1272 | 0.90139 | 0.941 | -6.56 | 0.56188 | none |
| ABLIM2 | 0.018 | 0.0568 | 0.1268 | 0.90173 | 0.942 | -6.56 | -0.0625 | none |
| ARMC4 | -0.018 | 0.0423 | -0.126 | 0.90201 | 0.942 | -6.56 | 1.28512 | none |
| JSRP1 | -0.023 | -0.0574 | -0.126 | 0.90203 | 0.942 | -6.56 | 1.61748 | none |
| CARD19 | -0.02 | -0.0513 | -0.126 | 0.90243 | 0.942 | -6.56 | 0.86827 | none |
| PRKAB2 | -0.019 | 0.0642 | -0.126 | 0.90246 | 0.942 | -6.56 | 0.92252 | none |
| SIX1 | -0.021 | 0.0137 | -0.126 | 0.90253 | 0.942 | -6.56 | -0.2321 | none |
| PRAMEF17 | -0.009 | 0.0088 | -0.126 | 0.90268 | 0.942 | -6.56 | -0.4924 | none |
| FUOM | -0.022 | -0.0469 | -0.126 | 0.90269 | 0.942 | -6.56 | -0.1472 | none |
| SKI | 0.036 | 0.2211 | 0.1253 | 0.90287 | 0.942 | -6.56 | 0.53236 | none |
| EFNA2 | 0.01 | 0.0293 | 0.1253 | 0.90287 | 0.942 | -6.56 | 0.70347 | none |
| FAM163B | 0.016 | 0.0155 | 0.1251 | 0.90298 | 0.942 | -6.56 | 0.37706 | none |
| HIST1H2A( | -0.023 | -0.0708 | -0.125 | 0.90304 | 0.942 | -6.56 | 0.17261 | none |
| SNAI1 | -0.012 | ######## | -0.125 | 0.90316 | 0.942 | -6.56 | 0.45956 | none |
| ATAD1 | -0.014 | -0.1049 | -0.125 | 0.90335 | 0.943 | -6.56 | 1.23016 | none |
| DAB2IP | -0.019 | -0.0129 | -0.124 | 0.90371 | 0.943 | -6.56 | 0.61822 | none |
| SYNPR | 0.012 | 0.0414 | 0.1241 | 0.90375 | 0.943 | -6.56 | -0.5576 | none |
| C1ORF112 | -0.019 | -0.0708 | -0.124 | 0.90375 | 0.943 | -6.56 | 0.66292 | none |

| | | | | | | | | |
|---|---|---|---|---|---|---|---|---|
| POU2F1 | 0.018 | 0.0637 | 0.1236 | 0.90417 | 0.943 | -6.56 | -0.0991 | none |
| KCNK5 | -0.015 | 0.0927 | -0.123 | 0.90427 | 0.943 | -6.56 | -0.0408 | none |
| ZC2HC1C | 0.031 | 0.1007 | 0.1233 | 0.9044 | 0.943 | -6.57 | -0.3371 | none |
| DCLRE1A | -0.015 | 0.0496 | -0.123 | 0.9044 | 0.943 | -6.57 | 0.12474 | none |
| SLC36A4 | 0.022 | -0.1057 | 0.1225 | 0.90499 | 0.944 | -6.57 | 1.35146 | none |
| PAM16 | 0.036 | -0.0572 | 0.1225 | 0.90501 | 0.944 | -6.57 | 0.90952 | none |
| TMIGD1 | 0.009 | 0.0194 | 0.122 | 0.90537 | 0.944 | -6.57 | 1.6907 | none |
| SALL2 | -0.012 | 0.0344 | -0.122 | 0.90538 | 0.944 | -6.57 | 0.75759 | none |
| NUDC | -0.017 | 0.0835 | -0.122 | 0.90568 | 0.944 | -6.57 | 0.12798 | none |
| LBP | -0.009 | 0.0201 | -0.122 | 0.9057 | 0.944 | -6.57 | 1.06325 | none |
| GAP43 | -0.01 | -0.0066 | -0.122 | 0.90571 | 0.944 | -6.57 | 0.45885 | none |
| B3GNT9 | -0.011 | -0.0089 | -0.121 | 0.90593 | 0.944 | -6.57 | 0.70296 | none |
| GPR180 | -0.012 | -0.06 | -0.121 | 0.90622 | 0.945 | -6.57 | 0.42243 | none |
| ELOF1 | 0.015 | -0.0599 | 0.1209 | 0.90623 | 0.945 | -6.57 | 0.44602 | none |
| LPCAT2 | -0.02 | -0.162 | -0.121 | 0.90631 | 0.945 | -6.57 | -0.6776 | none |
| CDKN2A | 0.015 | -0.0387 | 0.1206 | 0.9065 | 0.945 | -6.57 | 0.55055 | none |
| SLC45A4 | -0.016 | 0.0405 | -0.121 | 0.90653 | 0.945 | -6.57 | 0.40541 | none |
| SH3GL3 | 0.02 | 0.0033 | 0.1205 | 0.90655 | 0.945 | -6.57 | 0.52554 | none |
| SMIM10L1 | -0.015 | 0.0582 | -0.12 | 0.9066 | 0.945 | -6.57 | 1.75159 | none |
| ACSL5 | 0.01 | -0.0431 | 0.1202 | 0.90679 | 0.945 | -6.57 | 0.77856 | none |
| RABEP1 | -0.013 | 0.115 | -0.12 | 0.9071 | 0.945 | -6.57 | 1.0376 | none |
| GAGE2E | -0.02 | -0.025 | -0.12 | 0.90717 | 0.945 | -6.57 | -0.5568 | none |
| MTURN | -0.016 | -0.0585 | -0.12 | 0.90727 | 0.945 | -6.57 | 0.74324 | none |
| DICER1 | 0.027 | 0.1607 | 0.1194 | 0.9074 | 0.945 | -6.57 | 0.57762 | none |
| AK6 | 0.013 | -0.0562 | 0.1194 | 0.90741 | 0.945 | -6.57 | 0.47857 | none |
| FAM90A1 | -0.012 | -0.0152 | -0.119 | 0.90746 | 0.945 | -6.57 | 0.95781 | none |
| CCNC | -0.016 | -0.1325 | -0.119 | 0.90758 | 0.945 | -6.57 | 0.85698 | none |
| OR6N1 | -0.016 | -0.0308 | -0.119 | 0.90758 | 0.945 | -6.57 | 0.43091 | none |
| GAS1 | 0.009 | 0.0067 | 0.1191 | 0.90764 | 0.945 | -6.57 | 0 | none |
| VPS72 | 0.014 | -0.0454 | 0.1191 | 0.90765 | 0.945 | -6.57 | 0.34445 | none |
| C17ORF98 | 0.011 | -0.0082 | 0.1189 | 0.90777 | 0.945 | -6.57 | 1.51499 | none |
| TAF7 | 0.016 | -0.0102 | 0.1188 | 0.90787 | 0.945 | -6.57 | 0.92762 | none |
| ALOXE3 | -0.009 | ######## | -0.119 | 0.90798 | 0.945 | -6.57 | 0.01161 | none |
| OR2C3 | 0.008 | 0.0139 | 0.1181 | 0.90841 | 0.946 | -6.57 | -0.6989 | none |
| AGXT2 | 0.01 | 0.0289 | 0.1181 | 0.90843 | 0.946 | -6.57 | 0.83572 | none |
| ACTG1 | -0.016 | -0.1097 | -0.118 | 0.90845 | 0.946 | -6.57 | 1.14398 | none |
| ISG20 | 0.026 | -0.0762 | 0.1176 | 0.90883 | 0.946 | -6.57 | 0.75734 | none |
| TACR1 | 0.017 | 0.0316 | 0.1173 | 0.909 | 0.946 | -6.57 | 1.06981 | none |
| HIRA | 0.023 | 0.1387 | 0.1171 | 0.90914 | 0.946 | -6.57 | -0.006 | none |
| SLC6A3 | 0.008 | 0.0087 | 0.1171 | 0.9092 | 0.946 | -6.57 | -0.0239 | none |

| | | | | | | | | |
|---|---|---|---|---|---|---|---|---|
| F2 | 0.013 | 0.0386 | 0.1169 | 0.9093 | 0.946 | -6.57 | 1.41052 | none |
| MTIF3 | -0.022 | -0.0543 | -0.117 | 0.90955 | 0.946 | -6.57 | 1.63402 | none |
| SUCO | 0.021 | 0.0553 | 0.1162 | 0.90985 | 0.947 | -6.57 | 0.83249 | none |
| YY2 | -0.01 | 0.0249 | -0.116 | 0.90989 | 0.947 | -6.57 | 0.64892 | none |
| IFI44 | -0.022 | -0.1635 | -0.116 | 0.90994 | 0.947 | -6.57 | -0.0449 | none |
| FBLL1 | -0.01 | 0.0062 | -0.116 | 0.91005 | 0.947 | -6.57 | 0.76875 | none |
| SLC6A20 | 0.018 | -0.0248 | 0.1157 | 0.91023 | 0.947 | -6.57 | -0.2931 | none |
| DNAH12 | 0.008 | 0.0024 | 0.1151 | 0.91071 | 0.947 | -6.57 | -0.3032 | none |
| ARMC5 | -0.01 | 0.0037 | -0.114 | 0.91127 | 0.948 | -6.57 | 0.12166 | none |
| PIM1 | -0.015 | -0.047 | -0.113 | 0.91198 | 0.948 | -6.57 | 1.64221 | none |
| FZD9 | 0.008 | 0.0248 | 0.1131 | 0.91227 | 0.949 | -6.57 | -0.1081 | none |
| SKOR1 | -0.008 | 0.0063 | -0.113 | 0.91232 | 0.949 | -6.57 | 0.02754 | none |
| DDIT4L | 0.017 | 0.0272 | 0.1129 | 0.91241 | 0.949 | -6.57 | 1.36465 | none |
| C1ORF198 | 0.018 | 0.1217 | 0.1127 | 0.91258 | 0.949 | -6.57 | 0.92593 | none |
| IL10RB | 0.013 | -0.1146 | 0.1127 | 0.91259 | 0.949 | -6.57 | 0.45796 | none |
| PIK3CG | -0.021 | -0.0183 | -0.113 | 0.91262 | 0.949 | -6.57 | 0.81814 | none |
| CYP26A1 | 0.008 | 0.0332 | 0.1123 | 0.91289 | 0.949 | -6.57 | -0.5039 | none |
| NCAPG2 | 0.016 | 0.0974 | 0.1123 | 0.91292 | 0.949 | -6.57 | 0.29335 | none |
| PFN2 | 0.029 | 0.0151 | 0.1122 | 0.91295 | 0.949 | -6.57 | 1.05498 | none |
| C2CD4D | 0.015 | 0.0912 | 0.1122 | 0.91296 | 0.949 | -6.57 | 0.36514 | none |
| MPP7 | 0.023 | -0.0262 | 0.112 | 0.91311 | 0.949 | -6.57 | -0.0074 | none |
| LRRN1 | -0.009 | 0.0266 | -0.112 | 0.91335 | 0.949 | -6.57 | -0.2868 | none |
| SPINK5 | -0.011 | 0.0163 | -0.111 | 0.91362 | 0.949 | -6.57 | 0.62207 | none |
| RASAL3 | 0.021 | 0.1565 | 0.1113 | 0.91364 | 0.949 | -6.57 | 1.02154 | none |
| TLE3 | 0.022 | 0.1451 | 0.111 | 0.91391 | 0.95 | -6.57 | 0.02218 | none |
| UROC1 | 0.009 | 0.0082 | 0.111 | 0.91393 | 0.95 | -6.57 | 1.25704 | none |
| C16ORF71 | -0.014 | -0.0017 | -0.111 | 0.91413 | 0.95 | -6.57 | -0.0855 | none |
| CAP1 | 0.014 | 0.0545 | 0.11 | 0.91463 | 0.95 | -6.57 | 0.82198 | none |
| SH2D7 | 0.007 | 0.0166 | 0.1099 | 0.91474 | 0.95 | -6.57 | -0.5635 | none |
| UBE2G1 | 0.015 | 0.0083 | 0.1096 | 0.91494 | 0.95 | -6.57 | 0.82248 | none |
| GLYCTK | 0.019 | 0.0569 | 0.1094 | 0.91511 | 0.95 | -6.57 | 1.24332 | none |
| IFIT1 | 0.008 | -0.0064 | 0.1094 | 0.91513 | 0.95 | -6.57 | 0.04988 | none |
| CHIC1 | 0.025 | 0.0031 | 0.1093 | 0.91522 | 0.95 | -6.57 | 1.39798 | none |
| ARFIP2 | 0.011 | 0.0119 | 0.1093 | 0.91524 | 0.95 | -6.57 | 1.62782 | none |
| DDTL | -0.023 | 0.1093 | -0.109 | 0.91533 | 0.95 | -6.57 | 0.8292 | none |
| LYSMD1 | 0.01 | 0.0563 | 0.1091 | 0.91539 | 0.95 | -6.57 | 1.22506 | none |
| OR1L4 | 0.012 | 0.0118 | 0.1087 | 0.91564 | 0.951 | -6.57 | 1.60326 | none |
| CASQ2 | -0.036 | -0.1091 | -0.109 | 0.91577 | 0.951 | -6.57 | -0.5171 | none |
| WHRN | -0.014 | 0.0908 | -0.108 | 0.91602 | 0.951 | -6.57 | 0.36337 | none |
| ARG1 | -0.012 | -0.0622 | -0.108 | 0.91653 | 0.951 | -6.57 | 1.39761 | none |

| | | | | | | | | |
|---|---|---|---|---|---|---|---|---|
| NFKBIE | -0.014 | -0.0077 | -0.107 | 0.91665 | 0.952 | -6.57 | 0.44923 | none |
| PICALM | -0.026 | ######## | -0.107 | 0.91676 | 0.952 | -6.57 | 0.88605 | none |
| PHACTR3 | 0.009 | 0.0094 | 0.1073 | 0.91679 | 0.952 | -6.57 | 1.26592 | none |
| LSP1 | 0.014 | 0.0852 | 0.107 | 0.91701 | 0.952 | -6.57 | 1.10975 | none |
| NR1D2 | -0.018 | -0.0543 | -0.107 | 0.9172 | 0.952 | -6.57 | 0.80843 | none |
| AWAT2 | 0.009 | 0.0273 | 0.1063 | 0.91754 | 0.952 | -6.57 | 0.76342 | none |
| TSPAN7 | -0.026 | -0.0339 | -0.106 | 0.91758 | 0.952 | -6.57 | 0.2852 | none |
| ZNF185 | -0.008 | 0.0229 | -0.106 | 0.91768 | 0.952 | -6.57 | 2.01189 | none |
| ARHGEF7 | 0.03 | 0.203 | 0.1061 | 0.9177 | 0.952 | -6.57 | 0.31661 | none |
| GOLPH3 | -0.013 | 0.0012 | -0.106 | 0.91798 | 0.952 | -6.57 | 1.0798 | none |
| CROCC2 | 0.011 | -0.0137 | 0.1056 | 0.91805 | 0.952 | -6.57 | 0.87021 | none |
| FAM171A1 | 0.011 | 0.0865 | 0.1056 | 0.91807 | 0.952 | -6.57 | 0.8858 | none |
| PSMD3 | 0.011 | 0.0635 | 0.1053 | 0.91833 | 0.952 | -6.57 | 1.34226 | none |
| LRRC46 | 0.011 | 0.0844 | 0.1052 | 0.91834 | 0.952 | -6.57 | 1.24564 | none |
| KRTAP5-9 | 0.01 | 0.041 | 0.1051 | 0.91844 | 0.952 | -6.57 | 0.9557 | none |
| KDSR | -0.018 | -0.0279 | -0.105 | 0.91847 | 0.952 | -6.57 | 1.04343 | none |
| RNF32 | 0.015 | 0.0383 | 0.105 | 0.91849 | 0.952 | -6.57 | 0.09668 | none |
| MPV17L2 | -0.012 | -0.0143 | -0.105 | 0.91853 | 0.952 | -6.57 | 0.26673 | none |
| PIAS3 | 0.013 | 0.0674 | 0.1049 | 0.91859 | 0.952 | -6.57 | 1.22853 | none |
| TARM1 | -0.018 | 0.0329 | -0.105 | 0.91863 | 0.952 | -6.57 | -0.1245 | none |
| TXNDC5 | -0.021 | -0.0677 | -0.105 | 0.91867 | 0.952 | -6.57 | -0.4828 | none |
| UCK1 | 0.013 | 0.069 | 0.1048 | 0.91872 | 0.952 | -6.57 | -0.1657 | none |
| ITIH4 | 0.028 | 0.0855 | 0.1047 | 0.91873 | 0.952 | -6.57 | 0.01908 | none |
| ZCCHC9 | -0.016 | -0.0644 | -0.104 | 0.91893 | 0.953 | -6.57 | 0.94458 | none |
| GPR75 | 0.014 | 0.0568 | 0.1043 | 0.91906 | 0.953 | -6.57 | 0.22877 | none |
| ZFAND2B | -0.011 | -0.031 | -0.104 | 0.91939 | 0.953 | -6.57 | -0.1098 | none |
| DPP10 | -0.014 | 0.0047 | -0.104 | 0.9196 | 0.953 | -6.57 | 1.48707 | none |
| PRADC1 | -0.016 | -0.1047 | -0.103 | 0.92007 | 0.953 | -6.57 | 0.37985 | none |
| YWHAG | 0.012 | 0.0149 | 0.1027 | 0.92033 | 0.954 | -6.57 | 1.47528 | none |
| STAT5A | 0.011 | 0.0951 | 0.1024 | 0.92051 | 0.954 | -6.57 | 1.16382 | none |
| GMFB | -0.018 | -0.1147 | -0.102 | 0.92051 | 0.954 | -6.57 | 0.65169 | none |
| RAC3 | -0.025 | ######## | -0.102 | 0.92061 | 0.954 | -6.57 | 2.12653 | AA-related |
| AGTRAP | 0.016 | -0.0478 | 0.1023 | 0.92061 | 0.954 | -6.57 | 0.54438 | none |
| TICAM1 | 0.013 | 0.0895 | 0.1021 | 0.92074 | 0.954 | -6.57 | 1.09433 | none |
| SCNN1B | 0.013 | 0.0485 | 0.1021 | 0.92078 | 0.954 | -6.57 | 0.40291 | none |
| SCG5 | -0.009 | 0.0443 | -0.102 | 0.92088 | 0.954 | -6.57 | 0.64017 | none |
| IL17B | 0.017 | 0.0679 | 0.1019 | 0.92091 | 0.954 | -6.57 | 0.02031 | none |
| CPN1 | -0.011 | -0.0388 | -0.102 | 0.92106 | 0.954 | -6.57 | 0.04383 | none |
| ABHD17A | -0.019 | -0.1121 | -0.102 | 0.9212 | 0.954 | -6.57 | 0.28089 | none |
| UBA2 | 0.009 | 0.0414 | 0.1015 | 0.92125 | 0.954 | -6.57 | -0.6354 | none |

| | | | | | | | | |
|---|---|---|---|---|---|---|---|---|
| NSL1 | -0.009 | -0.1014 | -0.101 | 0.92136 | 0.954 | -6.57 | 1.52178 | none |
| AKR1C3 | -0.018 | -0.0882 | -0.101 | 0.92159 | 0.954 | -6.57 | 0.25914 | none |
| MEX3A | 0.008 | 0.0267 | 0.1008 | 0.92176 | 0.954 | -6.57 | -0.0533 | none |
| KRT75 | -0.007 | -0.0071 | -0.101 | 0.92196 | 0.954 | -6.57 | 0 | none |
| ZCCHC14 | -0.018 | 0.1307 | -0.1 | 0.92212 | 0.955 | -6.57 | 0.71338 | none |
| DCUN1D1 | -0.007 | -0.0549 | -0.1 | 0.92224 | 0.955 | -6.57 | 0.14977 | none |
| FAM166A | -0.01 | -0.0029 | -0.1 | 0.92235 | 0.955 | -6.57 | 0.39365 | none |
| ZCCHC12 | -0.008 | 0.028 | -0.1 | 0.92244 | 0.955 | -6.57 | 0.03917 | none |
| FRAS1 | -0.007 | 0.0027 | -0.1 | 0.92269 | 0.955 | -6.57 | -0.6442 | none |
| GOLGA5 | -0.023 | 0.0892 | -0.1 | 0.92271 | 0.955 | -6.57 | 0.82353 | none |
| MAPT | -0.008 | 0.0079 | -0.1 | 0.92274 | 0.955 | -6.57 | -0.2231 | none |
| WDR49 | 0.007 | -0.0091 | 0.0989 | 0.92323 | 0.955 | -6.57 | 0.30003 | none |
| PYGO2 | -0.008 | -0.0164 | -0.099 | 0.92352 | 0.956 | -6.57 | 1.73673 | none |
| UBXN10 | -0.007 | 0.0137 | -0.099 | 0.92354 | 0.956 | -6.57 | 0.31413 | none |
| CMTM1 | 0.025 | 0.1211 | 0.0984 | 0.92367 | 0.956 | -6.57 | 0.02104 | none |
| LSMEM2 | 0.007 | 0.0169 | 0.0982 | 0.92381 | 0.956 | -6.57 | 1.17577 | none |
| VEPH1 | 0.022 | ######## | 0.0976 | 0.92424 | 0.956 | -6.57 | 1.24006 | none |
| MAMDC2 | -0.009 | -0.0114 | -0.097 | 0.92434 | 0.956 | -6.57 | -0.2612 | none |
| SMKR1 | -0.023 | -0.0719 | -0.097 | 0.92435 | 0.956 | -6.57 | 1.59354 | none |
| ZNF462 | -0.008 | -0.0126 | -0.097 | 0.92452 | 0.956 | -6.57 | 0.18109 | none |
| ITGA7 | 0.024 | 0.0659 | 0.097 | 0.92468 | 0.956 | -6.57 | 0.08437 | none |
| SH3D19 | -0.018 | -0.0503 | -0.097 | 0.9247 | 0.956 | -6.57 | 0.17167 | none |
| SLC29A1 | -0.012 | 0.0236 | -0.097 | 0.92503 | 0.957 | -6.57 | 0.4543 | none |
| RSPH14 | 0.012 | 0.0977 | 0.0964 | 0.92519 | 0.957 | -6.57 | 0.13261 | none |
| ADSS | 0.009 | -0.0063 | 0.0961 | 0.92541 | 0.957 | -6.57 | 1.76384 | none |
| PRSS12 | 0.008 | 0.0151 | 0.0961 | 0.92542 | 0.957 | -6.57 | 0.27288 | none |
| GPR25 | -0.013 | 0.0266 | -0.096 | 0.92543 | 0.957 | -6.57 | 0.13987 | none |
| TEC | -0.018 | -0.0327 | -0.096 | 0.92548 | 0.957 | -6.57 | 0.44983 | none |
| TCEAL1 | -0.015 | -0.1122 | -0.096 | 0.9256 | 0.957 | -6.57 | -0.0309 | none |
| PURA | 0.016 | 0.0331 | 0.0958 | 0.92565 | 0.957 | -6.57 | 0.51329 | none |
| ECM1 | -0.016 | -0.0737 | -0.096 | 0.92571 | 0.957 | -6.57 | -0.2028 | none |
| ABCB6 | 0.01 | -0.0573 | 0.0953 | 0.92599 | 0.957 | -6.57 | 0.68802 | none |
| MN1 | -0.007 | -0.0148 | -0.095 | 0.92628 | 0.957 | -6.57 | 1.60331 | none |
| FAM19A2 | -0.03 | -0.0797 | -0.095 | 0.92633 | 0.957 | -6.57 | 0.55411 | none |
| TRAPPC9 | 0.013 | 0.1435 | 0.0946 | 0.92657 | 0.957 | -6.57 | 0.40108 | none |
| RORA | 0.012 | 0.005 | 0.0944 | 0.92675 | 0.958 | -6.57 | 1.24983 | none |
| PTPN6 | 0.013 | 0.1122 | 0.0941 | 0.92693 | 0.958 | -6.57 | 0.4295 | none |
| MS4A15 | 0.007 | 0.0041 | 0.0936 | 0.92734 | 0.958 | -6.57 | 0.21746 | none |
| YBX1 | 0.019 | 0.1379 | 0.0929 | 0.92791 | 0.959 | -6.57 | -0.2035 | none |
| MAPK13 | -0.009 | -0.0132 | -0.093 | 0.92792 | 0.959 | -6.57 | 0.82661 | none |

| | | | | | | | | |
|---|---|---|---|---|---|---|---|---|
| KLHDC7B | 0.012 | 0.0237 | 0.0926 | 0.92808 | 0.959 | -6.57 | 1.61191 | none |
| THAP3 | -0.015 | 0.0135 | -0.092 | 0.92834 | 0.959 | -6.57 | 1.12104 | none |
| CDK3 | -0.014 | 0.0557 | -0.092 | 0.92843 | 0.959 | -6.57 | 1.19668 | none |
| TRABD | 0.009 | -0.0131 | 0.092 | 0.92854 | 0.959 | -6.57 | 1.62948 | none |
| MTA2 | 0.018 | 0.0977 | 0.0918 | 0.92876 | 0.959 | -6.57 | 1.13287 | none |
| NCAM1 | -0.018 | 0.0775 | -0.092 | 0.92896 | 0.959 | -6.57 | -0.4201 | none |
| CELSR2 | -0.008 | 0.0248 | -0.091 | 0.929 | 0.959 | -6.57 | 1.33733 | none |
| POLR2E | 0.012 | 0.045 | 0.0912 | 0.92922 | 0.959 | -6.57 | -0.4423 | none |
| SLC9A3R2 | 0.018 | 0.0059 | 0.091 | 0.92937 | 0.959 | -6.57 | 0.05395 | none |
| LRRC7 | -0.032 | 0.0508 | -0.091 | 0.9294 | 0.959 | -6.57 | 0.54869 | none |
| TBR1 | 0.006 | 0.0167 | 0.0906 | 0.92968 | 0.96 | -6.57 | 0.02522 | none |
| MEST | -0.022 | -0.1428 | -0.09 | 0.92975 | 0.96 | -6.57 | 1.39642 | none |
| TIA1 | 0.013 | -0.05 | 0.0903 | 0.92992 | 0.96 | -6.57 | -0.1783 | none |
| ATP13A3 | 0.017 | 0.0736 | 0.0901 | 0.93007 | 0.96 | -6.57 | 0.3378 | none |
| KLK1 | -0.01 | -0.0199 | -0.09 | 0.93011 | 0.96 | -6.57 | 0.28496 | none |
| CTAGE8 | -0.008 | 0.0293 | -0.09 | 0.93015 | 0.96 | -6.57 | 0.36431 | none |
| KIAA1143 | 0.013 | 0.0501 | 0.0895 | 0.93053 | 0.96 | -6.57 | -0.5786 | none |
| RAD21L1 | -0.008 | 0.0227 | -0.089 | 0.93063 | 0.96 | -6.57 | -0.3399 | none |
| TSTD1 | 0.018 | -0.0654 | 0.089 | 0.93087 | 0.96 | -6.57 | 0.3694 | none |
| WASL | 0.009 | 0.0183 | 0.0888 | 0.93107 | 0.96 | -6.57 | 0.92457 | none |
| RPS4Y1 | 0.014 | -0.1107 | 0.0887 | 0.93114 | 0.96 | -6.57 | 0.99107 | none |
| LYAR | 0.02 | 0.1615 | 0.0887 | 0.93115 | 0.96 | -6.57 | 0.54848 | none |
| DNAJC8 | 0.012 | 0.0765 | 0.0887 | 0.93117 | 0.96 | -6.57 | -0.0666 | none |
| VIT | -0.008 | 0.0171 | -0.089 | 0.93118 | 0.96 | -6.57 | -0.0843 | none |
| GPR84 | 0.009 | 0.011 | 0.0886 | 0.93119 | 0.96 | -6.57 | -0.201 | none |
| CCL4 | -0.014 | 0.0209 | -0.089 | 0.9312 | 0.96 | -6.57 | 0.50245 | none |
| MPND | -0.027 | -0.0519 | -0.088 | 0.93142 | 0.961 | -6.57 | -0.6496 | none |
| CLDN24 | 0.007 | 0.0249 | 0.0882 | 0.93154 | 0.961 | -6.57 | 0.99228 | none |
| TMEM199 | 0.011 | -0.0455 | 0.088 | 0.93169 | 0.961 | -6.57 | 1.06063 | none |
| CPNE5 | 0.007 | 0.0027 | 0.0876 | 0.93197 | 0.961 | -6.57 | 0.06927 | none |
| TRMO | 0.011 | -0.0461 | 0.0874 | 0.93215 | 0.961 | -6.57 | 1.30886 | none |
| LBH | 0.01 | -0.0258 | 0.0874 | 0.93217 | 0.961 | -6.57 | 0.54016 | none |
| CNTNAP5 | 0.014 | 0.0193 | 0.0873 | 0.93223 | 0.961 | -6.57 | 1.13402 | none |
| ABCC5 | -0.011 | 0.0566 | -0.087 | 0.93227 | 0.961 | -6.57 | -0.2047 | none |
| SLC25A44 | -0.01 | -0.0206 | -0.087 | 0.93227 | 0.961 | -6.57 | 1.18399 | none |
| GABRR3 | 0.006 | -0.001 | 0.0871 | 0.93241 | 0.961 | -6.57 | 1.36185 | none |
| TRPA1 | -0.008 | 0.0132 | -0.087 | 0.93243 | 0.961 | -6.57 | 0.29617 | none |
| CHRM4 | 0.006 | 0.0216 | 0.087 | 0.93244 | 0.961 | -6.57 | -0.0767 | none |
| DPYS | -0.013 | -0.0086 | -0.087 | 0.93255 | 0.961 | -6.57 | 0.82034 | none |
| MC5R | -0.007 | 0.0141 | -0.087 | 0.93259 | 0.961 | -6.57 | 0.36216 | none |

| | | | | | | | | |
|---|---|---|---|---|---|---|---|---|
| B3GNT2 | 0.013 | 0.0096 | 0.0865 | 0.9328 | 0.961 | -6.57 | 1.59649 | none |
| ZNF189 | -0.011 | 0.0337 | -0.087 | 0.93281 | 0.961 | -6.57 | 0.45885 | none |
| ZNF75D | 0.014 | 0.0922 | 0.0865 | 0.93282 | 0.961 | -6.57 | 1.87537 | none |
| IGFL3 | 0.009 | 0.0424 | 0.0864 | 0.93288 | 0.961 | -6.57 | 1.23377 | none |
| ARR3 | 0.009 | -0.0132 | 0.0863 | 0.93303 | 0.961 | -6.57 | 1.00829 | none |
| SLAIN2 | -0.017 | 0.0657 | -0.086 | 0.93304 | 0.961 | -6.57 | 0.80921 | none |
| XPR1 | -0.013 | ######## | -0.086 | 0.93309 | 0.961 | -6.57 | 0.53146 | none |
| MFAP1 | -0.017 | 0.078 | -0.086 | 0.9331 | 0.961 | -6.57 | 0.96409 | none |
| FAM206A | -0.012 | 0.0081 | -0.086 | 0.93328 | 0.961 | -6.57 | 0.00173 | none |
| CEACAM6 | 0.015 | 0.0061 | 0.0856 | 0.93351 | 0.961 | -6.57 | 1.45518 | none |
| CRACR2B | 0.011 | -0.0142 | 0.0856 | 0.93354 | 0.961 | -6.57 | 1.21105 | none |
| SMN1 | 0.015 | 0.101 | 0.0855 | 0.93359 | 0.961 | -6.57 | 1.82907 | none |
| PTGS2 | 0.011 | 0.0331 | 0.0855 | 0.93361 | 0.961 | -6.57 | 1.15038 | none |
| WFDC5 | 0.009 | 0.0095 | 0.0855 | 0.93364 | 0.961 | -6.57 | 0.66722 | none |
| PPP1R1A | -0.01 | -0.0074 | -0.085 | 0.93365 | 0.961 | -6.57 | 0.98675 | none |
| NFE2 | -0.009 | -0.0092 | -0.085 | 0.93374 | 0.961 | -6.57 | 0.7272 | none |
| ATOH7 | 0.009 | 0.0086 | 0.0848 | 0.93417 | 0.962 | -6.57 | -0.4546 | none |
| GABRA2 | -0.006 | 0.0037 | -0.084 | 0.93459 | 0.962 | -6.57 | -0.4209 | none |
| UNC5C | 0.007 | 0.0111 | 0.0842 | 0.93464 | 0.962 | -6.57 | 1.26835 | none |
| MTOR | 0.021 | 0.2079 | 0.084 | 0.93474 | 0.962 | -6.57 | 0.28918 | none |
| ITGAM | 0.016 | -0.0397 | 0.0837 | 0.93497 | 0.962 | -6.57 | 0.09707 | none |
| RGS3 | 0.011 | 0.0601 | 0.0837 | 0.93498 | 0.962 | -6.57 | 1.18986 | none |
| GZMK | -0.01 | -0.0211 | -0.084 | 0.93502 | 0.962 | -6.57 | 0.71141 | none |
| CFAP36 | -0.014 | -0.0346 | -0.083 | 0.93549 | 0.963 | -6.57 | 0.08153 | none |
| KBTBD4 | 0.007 | -0.0578 | 0.0829 | 0.93563 | 0.963 | -6.57 | 0.75971 | none |
| BHMG1 | 0.006 | 0.016 | 0.0828 | 0.93568 | 0.963 | -6.57 | 0.89996 | none |
| COLGALT2 | -0.006 | 0.0272 | -0.083 | 0.93576 | 0.963 | -6.57 | 0.07787 | none |
| MTAP | 0.007 | 0.0267 | 0.0824 | 0.93601 | 0.963 | -6.57 | 0.77476 | none |
| LCMT1 | -0.017 | -0.1012 | -0.082 | 0.9361 | 0.963 | -6.57 | -0.1379 | none |
| FAM71B | -0.006 | 0.0047 | -0.082 | 0.93634 | 0.963 | -6.57 | 0.73711 | none |
| TMEM62 | 0.012 | -0.0689 | 0.0817 | 0.93653 | 0.963 | -6.57 | 1.13506 | none |
| C16ORF92 | 0.02 | 0.0203 | 0.0811 | 0.93698 | 0.964 | -6.57 | -0.1693 | none |
| HLA-DQA1 | -0.014 | -0.0804 | -0.081 | 0.93704 | 0.964 | -6.57 | -0.509 | none |
| TNFSF4 | 0.007 | 0.0162 | 0.081 | 0.93712 | 0.964 | -6.57 | 0.39436 | none |
| MICA | -0.015 | -0.1098 | -0.081 | 0.93722 | 0.964 | -6.57 | 0.00146 | none |
| PIEZO2 | -0.023 | 0.0833 | -0.081 | 0.93727 | 0.964 | -6.57 | 0.23563 | none |
| C5ORF30 | -0.016 | -0.0458 | -0.081 | 0.93748 | 0.964 | -6.57 | 0.8272 | none |
| PLSCR4 | 0.007 | 0.0255 | 0.0804 | 0.93754 | 0.964 | -6.57 | -0.669 | none |
| NEU2 | 0.006 | 0.0146 | 0.0803 | 0.93763 | 0.964 | -6.57 | -0.3711 | none |
| LTV1 | -0.019 | 0.1076 | -0.08 | 0.93781 | 0.964 | -6.57 | 1.11928 | none |

| | | | | | | | | |
|---|---|---|---|---|---|---|---|---|
| MTX1 | 0.008 | -0.0137 | 0.0799 | 0.93794 | 0.964 | -6.57 | 0.21069 | none |
| PHRF1 | 0.025 | 0.2105 | 0.0799 | 0.93799 | 0.964 | -6.57 | 0.66121 | none |
| GPR34 | 0.01 | -0.0409 | 0.0797 | 0.93807 | 0.964 | -6.57 | 1.25025 | none |
| NPDC1 | 0.022 | -0.0034 | 0.0795 | 0.93828 | 0.964 | -6.57 | 0.48007 | none |
| TSTA3 | -0.009 | -0.0166 | -0.079 | 0.93865 | 0.965 | -6.57 | 1.58685 | none |
| NOMO1 | 0.021 | 0.1462 | 0.079 | 0.93868 | 0.965 | -6.57 | 0.25217 | none |
| SAMD11 | 0.006 | 0.0216 | 0.0789 | 0.93869 | 0.965 | -6.57 | -0.2607 | none |
| ALG1 | 0.009 | -0.0152 | 0.0789 | 0.93872 | 0.965 | -6.57 | -0.2914 | none |
| CCDC112 | 0.013 | -0.0048 | 0.0788 | 0.93882 | 0.965 | -6.57 | 1.43345 | none |
| COL4A3 | 0.007 | 0.0057 | 0.0784 | 0.93909 | 0.965 | -6.57 | -0.5368 | none |
| OR11H12 | -0.008 | 0.0238 | -0.078 | 0.93914 | 0.965 | -6.57 | 0.32805 | none |
| CPLX1 | 0.007 | -0.0044 | 0.0782 | 0.93927 | 0.965 | -6.57 | 1.15582 | none |
| NR2F6 | 0.016 | 0.0546 | 0.0782 | 0.93928 | 0.965 | -6.57 | 1.01099 | none |
| AC069499 | 0.022 | 0.1449 | 0.0778 | 0.93959 | 0.965 | -6.57 | 0.54886 | none |
| PSMD8 | -0.007 | -0.0457 | -0.077 | 0.94005 | 0.965 | -6.57 | -0.5311 | none |
| CD2AP | 0.02 | 0.1987 | 0.0768 | 0.94037 | 0.966 | -6.57 | 1.07685 | none |
| DEK | 0.014 | 0.1044 | 0.0767 | 0.94044 | 0.966 | -6.57 | 1.35698 | none |
| ANXA6 | -0.011 | 0.045 | -0.077 | 0.94048 | 0.966 | -6.57 | 1.07244 | none |
| OR2A7 | 0.007 | 0.065 | 0.0766 | 0.94053 | 0.966 | -6.57 | 0.94039 | none |
| PXMP2 | 0.015 | -0.0231 | 0.0765 | 0.94062 | 0.966 | -6.57 | -0.0433 | none |
| NUTM2G | -0.008 | 0.0035 | -0.076 | 0.94063 | 0.966 | -6.57 | -0.8736 | none |
| NT5M | -0.023 | -0.0679 | -0.076 | 0.94073 | 0.966 | -6.57 | -0.268 | none |
| ZNF366 | -0.009 | -0.0337 | -0.076 | 0.94086 | 0.966 | -6.57 | 0.80535 | none |
| TBC1D16 | 0.009 | 0.0497 | 0.0759 | 0.94108 | 0.966 | -6.57 | 0.56431 | none |
| ZC3H12C | -0.009 | 0.0473 | -0.076 | 0.94127 | 0.966 | -6.57 | 0.72358 | none |
| ABCF2 | 0.013 | 0.1228 | 0.0754 | 0.94145 | 0.966 | -6.57 | 1.28015 | none |
| UBXN1 | -0.008 | -0.0131 | -0.075 | 0.94159 | 0.966 | -6.57 | 0.61725 | none |
| SYCN | 0.007 | 0.028 | 0.0751 | 0.94166 | 0.966 | -6.57 | 1.46578 | none |
| TAF11 | 0.007 | -0.024 | 0.075 | 0.94174 | 0.966 | -6.57 | 0.82032 | none |
| DKC1 | 0.005 | 0.0112 | 0.0749 | 0.9418 | 0.966 | -6.57 | 0.58128 | none |
| FERMT3 | -0.008 | 0.0612 | -0.075 | 0.94196 | 0.966 | -6.57 | 0.61629 | none |
| PRPF38A | -0.005 | -0.0669 | -0.074 | 0.94218 | 0.967 | -6.57 | 0.26778 | none |
| POMC | 0.013 | -0.0222 | 0.0744 | 0.94223 | 0.967 | -6.57 | 0.52158 | none |
| ERGIC3 | -0.01 | -0.0728 | -0.074 | 0.94242 | 0.967 | -6.57 | 0.93119 | none |
| WDR38 | -0.006 | 0.029 | -0.074 | 0.94271 | 0.967 | -6.57 | -0.3059 | none |
| MYSM1 | 0.022 | 0.119 | 0.0737 | 0.94275 | 0.967 | -6.57 | 0.69073 | none |
| KDELC1 | 0.011 | 0.029 | 0.0737 | 0.94278 | 0.967 | -6.57 | 0.2611 | none |
| NUBP2 | 0.01 | -0.0217 | 0.0736 | 0.94285 | 0.967 | -6.57 | 0.4706 | none |
| TRIM27 | -0.041 | 0.0369 | -0.073 | 0.94317 | 0.967 | -6.57 | 1.24359 | none |
| CNGB3 | -0.006 | ######## | -0.073 | 0.94331 | 0.967 | -6.57 | -0.4367 | none |

| | | | | | | | | |
|---|---|---|---|---|---|---|---|---|
| ZNF521 | 0.015 | 0.023 | 0.0726 | 0.94361 | 0.968 | -6.57 | 1.33677 | none |
| GPAA1 | -0.009 | -0.0326 | -0.072 | 0.9437 | 0.968 | -6.57 | 1.64187 | none |
| UNC5B | -0.005 | 0.0136 | -0.072 | 0.94371 | 0.968 | -6.57 | 0 | none |
| CCNK | -0.019 | 0.1063 | -0.072 | 0.94393 | 0.968 | -6.57 | 0.99432 | none |
| ZNF782 | 0.01 | 0.0453 | 0.0722 | 0.94394 | 0.968 | -6.57 | -0.1483 | none |
| RFNG | -0.014 | -0.0555 | -0.072 | 0.94402 | 0.968 | -6.57 | 0.52227 | none |
| RGS2 | 0.012 | -0.0225 | 0.072 | 0.94405 | 0.968 | -6.57 | -0.0422 | none |
| GPR171 | -0.015 | -0.1011 | -0.072 | 0.94407 | 0.968 | -6.57 | -0.1383 | none |
| LYPD3 | 0.009 | -0.0172 | 0.0718 | 0.94426 | 0.968 | -6.57 | 1.58415 | none |
| SUGCT | 0.007 | ######## | 0.0716 | 0.94435 | 0.968 | -6.57 | 0.75764 | none |
| KLRC1 | 0.01 | 0.0973 | 0.0715 | 0.94447 | 0.968 | -6.57 | 1.15956 | none |
| PYURF | -0.008 | -0.0682 | -0.071 | 0.94469 | 0.968 | -6.57 | 0.28002 | none |
| RBPJ | 0.006 | -0.0358 | 0.0711 | 0.94473 | 0.968 | -6.57 | 0.03642 | none |
| CUBN | -0.008 | -0.0421 | -0.071 | 0.94477 | 0.968 | -6.57 | -0.3811 | none |
| CHUK | -0.013 | 0.0364 | -0.071 | 0.94494 | 0.968 | -6.57 | 0.85022 | none |
| HSD17B8 | -0.011 | 0.0207 | -0.071 | 0.94494 | 0.968 | -6.57 | 0.64014 | none |
| SMC2 | 0.03 | 0.2871 | 0.0704 | 0.94533 | 0.968 | -6.57 | 0.58617 | none |
| KDELR1 | -0.01 | -0.0463 | -0.07 | 0.94542 | 0.968 | -6.57 | 0.58061 | none |
| C9ORF131 | 0.019 | -0.0131 | 0.0701 | 0.94553 | 0.969 | -6.57 | -0.5496 | none |
| SPC25 | -0.03 | 0.0924 | -0.07 | 0.94574 | 0.969 | -6.57 | 0.33101 | none |
| NKX2-5 | -0.008 | -0.0397 | -0.07 | 0.9458 | 0.969 | -6.57 | 1.59014 | none |
| PITHD1 | 0.008 | -0.0253 | 0.0697 | 0.94587 | 0.969 | -6.57 | 1.45131 | none |
| IL17REL | 0.005 | 0.01 | 0.0694 | 0.9461 | 0.969 | -6.57 | -0.5592 | none |
| UPP1 | -0.014 | -0.0619 | -0.069 | 0.94616 | 0.969 | -6.57 | 0.0371 | none |
| KIAA0895l | -0.007 | -0.0201 | -0.069 | 0.94629 | 0.969 | -6.57 | -0.4977 | none |
| SYNC | 0.007 | 0.0464 | 0.0691 | 0.94631 | 0.969 | -6.57 | 0.68058 | none |
| RBM7 | -0.015 | -0.1536 | -0.069 | 0.94647 | 0.969 | -6.57 | 0.29526 | none |
| C11ORF97 | 0.007 | 0.0073 | 0.0687 | 0.94664 | 0.969 | -6.57 | 1.37194 | none |
| KRT36 | 0.005 | 0.0051 | 0.0685 | 0.94675 | 0.969 | -6.57 | 0 | none |
| CAB39 | -0.011 | 0.0609 | -0.069 | 0.94675 | 0.969 | -6.57 | 0.7411 | none |
| GRPEL2 | -0.01 | -0.0471 | -0.069 | 0.94678 | 0.969 | -6.57 | 0.86325 | none |
| KRTAP13-: | 0.007 | 0.0069 | 0.0683 | 0.94695 | 0.969 | -6.57 | -0.148 | none |
| ARHGAP18 | 0.012 | 0.0451 | 0.068 | 0.94718 | 0.969 | -6.57 | 0.0298 | none |
| ST8SIA2 | -0.005 | 0.0052 | -0.068 | 0.94723 | 0.969 | -6.57 | 0 | none |
| B4GALT1 | 0.008 | 0.0205 | 0.0678 | 0.94733 | 0.969 | -6.57 | -0.3719 | none |
| RNPS1 | -0.006 | 0.0352 | -0.068 | 0.9474 | 0.969 | -6.57 | -0.0224 | none |
| TMUB1 | 0.007 | -0.0102 | 0.0676 | 0.9475 | 0.969 | -6.57 | 0.28728 | none |
| CRYGS | -0.028 | 0.0586 | -0.067 | 0.94767 | 0.969 | -6.57 | 1.21289 | none |
| NSMCE4A | 0.007 | -0.0147 | 0.0673 | 0.94769 | 0.969 | -6.57 | 1.05768 | none |
| HPR | 0.005 | -0.0135 | 0.0672 | 0.94778 | 0.969 | -6.57 | 1.2109 | none |

| | | | | | | | | |
|---|---|---|---|---|---|---|---|---|
| SPA17 | -0.013 | -0.0259 | -0.067 | 0.9478 | 0.969 | -6.57 | -0.3041 | none |
| BRMS1 | -0.006 | ######## | -0.067 | 0.94781 | 0.969 | -6.57 | 0.53643 | none |
| DDX20 | 0.006 | 0.0624 | 0.067 | 0.94793 | 0.97 | -6.57 | 1.67102 | none |
| GMDS | -0.007 | ######## | -0.067 | 0.94809 | 0.97 | -6.57 | 1.42606 | none |
| ATXN1L | 0.009 | 0.0221 | 0.0664 | 0.94838 | 0.97 | -6.57 | 1.62094 | none |
| TIMM22 | 0.054 | -0.0467 | 0.0664 | 0.94842 | 0.97 | -6.57 | -0.4594 | none |
| GZF1 | 0.016 | 0.1122 | 0.0663 | 0.94851 | 0.97 | -6.57 | 0.77084 | none |
| C11ORF58 | 0.009 | -0.0759 | 0.0662 | 0.94858 | 0.97 | -6.57 | -0.4708 | none |
| CCDC51 | 0.009 | -0.0198 | 0.066 | 0.94874 | 0.97 | -6.57 | -0.2704 | none |
| LRRC52 | 0.005 | 0.0181 | 0.066 | 0.94876 | 0.97 | -6.57 | 1.24597 | none |
| BLOC1S5-T | -0.023 | 0.1034 | -0.066 | 0.94888 | 0.97 | -6.57 | 0.84742 | none |
| TSSC4 | -0.006 | 0.0076 | -0.066 | 0.94889 | 0.97 | -6.57 | 0.17893 | none |
| RIBC2 | -0.011 | 0.0865 | -0.066 | 0.94893 | 0.97 | -6.57 | 1.51532 | none |
| FAM185A | -0.006 | 0.0462 | -0.066 | 0.94898 | 0.97 | -6.57 | 0.90676 | none |
| TSPO2 | 0.008 | 0.0246 | 0.0656 | 0.94905 | 0.97 | -6.57 | 0.37431 | none |
| COL4A5 | 0.009 | 0.0147 | 0.0655 | 0.94913 | 0.97 | -6.57 | -0.0943 | none |
| LRRC30 | 0.006 | 0.0055 | 0.0647 | 0.94974 | 0.971 | -6.57 | -0.2947 | none |
| HMGA1 | -0.006 | 0.0161 | -0.065 | 0.94985 | 0.971 | -6.57 | 1.38879 | none |
| IL17F | 0.026 | -0.0924 | 0.0641 | 0.95017 | 0.971 | -6.57 | 0.55248 | none |
| PRIMPOL | -0.008 | -0.0521 | -0.064 | 0.95041 | 0.971 | -6.57 | -0.6641 | none |
| TPRKB | -0.008 | -0.1076 | -0.064 | 0.95055 | 0.971 | -6.57 | 0.50693 | none |
| RAPGEFL1 | 0.01 | 0.1378 | 0.0636 | 0.95062 | 0.971 | -6.57 | 0.22076 | none |
| SERINC3 | -0.009 | -0.1058 | -0.063 | 0.95067 | 0.971 | -6.57 | 0.94755 | none |
| ESRP2 | 0.012 | 0.0938 | 0.0632 | 0.95089 | 0.971 | -6.57 | 0.86638 | none |
| CXCL2 | -0.006 | 0.0204 | -0.063 | 0.951 | 0.971 | -6.57 | 0.43905 | none |
| WIZ | 0.012 | 0.1766 | 0.063 | 0.95102 | 0.971 | -6.57 | 0.71903 | none |
| DCUN1D3 | -0.009 | ######## | -0.063 | 0.95129 | 0.972 | -6.57 | 1.68064 | none |
| LMNTD2 | 0.006 | 0.0365 | 0.0626 | 0.95135 | 0.972 | -6.57 | -0.1468 | none |
| FUNDC2 | -0.007 | -0.0801 | -0.062 | 0.95165 | 0.972 | -6.57 | 0.55195 | none |
| FARSB | 0.007 | 0.0092 | 0.0622 | 0.95167 | 0.972 | -6.57 | 1.0523 | none |
| RNF146 | 0.006 | -0.0215 | 0.0621 | 0.95172 | 0.972 | -6.57 | 0.24965 | none |
| SPACA3 | -0.007 | 0.0196 | -0.062 | 0.95196 | 0.972 | -6.57 | 1.07615 | none |
| HPSE | 0.008 | -0.1052 | 0.0618 | 0.95196 | 0.972 | -6.57 | 0.41096 | none |
| CIAPIN1 | 0.006 | -0.0304 | 0.0615 | 0.95218 | 0.972 | -6.57 | 0.17196 | none |
| SHARPIN | 0.007 | 0.0065 | 0.0614 | 0.9523 | 0.972 | -6.57 | -0.024 | none |
| POLDIP2 | 0.007 | ######## | 0.0609 | 0.95266 | 0.972 | -6.57 | 0.88428 | none |
| TEX261 | -0.006 | -0.0013 | -0.061 | 0.95277 | 0.973 | -6.57 | 0.5053 | none |
| SSX3 | -0.007 | 0.0112 | -0.061 | 0.95295 | 0.973 | -6.57 | 0.56893 | none |
| TWF1 | -0.007 | -0.0167 | -0.06 | 0.95322 | 0.973 | -6.57 | 1.5032 | none |
| CA8 | 0.005 | 0.0217 | 0.0601 | 0.95334 | 0.973 | -6.57 | 1.256 | none |

| | | | | | | | | |
|---|---|---|---|---|---|---|---|---|
| DNAJC5 | -0.006 | 0.0546 | -0.06 | 0.95353 | 0.973 | -6.57 | 0.88014 | none |
| SEC24A | 0.006 | 0.0195 | 0.0597 | 0.95362 | 0.973 | -6.57 | 0.84923 | none |
| EIF5A2 | 0.011 | -0.1178 | 0.0597 | 0.95363 | 0.973 | -6.57 | 0.83784 | none |
| MRGBP | -0.006 | 0.0215 | -0.06 | 0.95364 | 0.973 | -6.57 | 0.7042 | none |
| CACNG6 | 0.005 | 0.0113 | 0.0592 | 0.95402 | 0.973 | -6.57 | 0.26554 | none |
| HDC | -0.006 | -0.0188 | -0.059 | 0.95415 | 0.973 | -6.57 | 1.6195 | none |
| PPP2R2C | -0.011 | 0.0072 | -0.059 | 0.95438 | 0.973 | -6.57 | 0.64831 | none |
| TNFRSF9 | -0.007 | 0.0473 | -0.059 | 0.95443 | 0.973 | -6.57 | 0.10587 | none |
| MTF1 | 0.013 | 0.0454 | 0.0586 | 0.95444 | 0.973 | -6.57 | 0.00574 | none |
| CWC27 | -0.015 | 0.0708 | -0.059 | 0.95449 | 0.973 | -6.57 | 1.30395 | none |
| SMPD2 | -0.007 | -0.0747 | -0.058 | 0.95457 | 0.974 | -6.57 | 0.41459 | none |
| MRPL2 | 0.007 | -0.0368 | 0.0584 | 0.95464 | 0.974 | -6.57 | -0.2125 | none |
| C1QL3 | 0.005 | 0.0383 | 0.0581 | 0.95483 | 0.974 | -6.57 | 0.7104 | none |
| MFSD2A | 0.007 | 0.0131 | 0.0578 | 0.95506 | 0.974 | -6.57 | 0.5431 | none |
| HOXD3 | 0.009 | 0.0399 | 0.0563 | 0.95624 | 0.975 | -6.57 | 0.2307 | none |
| AMY1A | -0.006 | 0.0133 | -0.056 | 0.95634 | 0.975 | -6.57 | 1.89673 | none |
| TRMT61A | 0.004 | 0.0554 | 0.0561 | 0.9564 | 0.975 | -6.57 | 0.41957 | none |
| ABCC12 | 0.005 | 0.0519 | 0.0561 | 0.95641 | 0.975 | -6.57 | -0.5518 | none |
| ACTA1 | -0.005 | ######## | -0.056 | 0.95641 | 0.975 | -6.57 | -0.3604 | none |
| ZNF593 | 0.007 | -0.0051 | 0.0554 | 0.95694 | 0.975 | -6.57 | 1.56814 | none |
| PRNP | 0.005 | -0.0406 | 0.0548 | 0.95744 | 0.976 | -6.57 | 1.24899 | none |
| INIP | 0.006 | -0.041 | 0.0547 | 0.95752 | 0.976 | -6.57 | -0.6117 | none |
| AES | 0.01 | 0.0568 | 0.0547 | 0.95753 | 0.976 | -6.57 | 1.09603 | none |
| TMEM9 | 0.009 | -0.073 | 0.0546 | 0.95757 | 0.976 | -6.57 | -0.0396 | none |
| LPL | -0.006 | 0.0122 | -0.055 | 0.95759 | 0.976 | -6.57 | 0.61384 | none |
| LTBP1 | 0.005 | 0.0328 | 0.0545 | 0.95762 | 0.976 | -6.57 | 0.22779 | none |
| WDR70 | 0.013 | 0.0587 | 0.0543 | 0.95782 | 0.976 | -6.57 | 0.62237 | none |
| TRAF1 | -0.008 | 0.1102 | -0.054 | 0.9579 | 0.976 | -6.57 | 1.36783 | none |
| AK5 | -0.018 | -0.0477 | -0.054 | 0.95794 | 0.976 | -6.57 | 0.65554 | none |
| USP20 | 0.007 | 0.1005 | 0.054 | 0.95804 | 0.976 | -6.57 | -0.003 | none |
| MERTK | 0.006 | -0.0273 | 0.0539 | 0.95808 | 0.976 | -6.57 | 1.2867 | none |
| AZGP1 | -0.007 | 0.0128 | -0.054 | 0.95816 | 0.976 | -6.57 | 0.0816 | none |
| FRAT2 | -0.008 | -0.0292 | -0.054 | 0.95819 | 0.976 | -6.57 | -0.502 | none |
| SORD | 0.006 | 0.0132 | 0.0535 | 0.95839 | 0.976 | -6.57 | 1.13224 | none |
| CEACAM1: | -0.004 | 0.0087 | -0.053 | 0.95844 | 0.976 | -6.57 | 1.05833 | none |
| TRIM74 | -0.004 | -0.0045 | -0.053 | 0.95865 | 0.976 | -6.57 | -0.8801 | none |
| ATP6AP1 | 0.006 | -0.0238 | 0.053 | 0.95879 | 0.976 | -6.57 | 0.90592 | none |
| TRAPPC6A | -0.007 | -0.065 | -0.053 | 0.95889 | 0.976 | -6.57 | 1.21548 | none |
| ZNF367 | 0.006 | 0.0427 | 0.0529 | 0.95892 | 0.976 | -6.57 | 0.18393 | none |
| RNMT | -0.01 | 0.0146 | -0.053 | 0.95893 | 0.976 | -6.57 | 0.07863 | none |

| | | | | | | | | |
|---|---|---|---|---|---|---|---|---|
| DDX1 | -0.008 | 0.0565 | -0.053 | 0.95917 | 0.976 | -6.57 | 1.31429 | none |
| PPP1R3E | -0.007 | -0.0291 | -0.052 | 0.95924 | 0.976 | -6.57 | -0.0294 | none |
| SLC35G1 | -0.01 | -0.0457 | -0.052 | 0.95957 | 0.977 | -6.57 | 0.22408 | none |
| MAPK4 | -0.004 | ######## | -0.052 | 0.95958 | 0.977 | -6.57 | -0.1593 | none |
| BEND3 | 0.008 | 0.111 | 0.0516 | 0.95988 | 0.977 | -6.57 | 0.25606 | none |
| TCTEX1D4 | 0.007 | 0.0198 | 0.0514 | 0.96004 | 0.977 | -6.57 | 0.2484 | none |
| PCNP | -0.004 | -0.0122 | -0.051 | 0.96012 | 0.977 | -6.57 | -0.8827 | none |
| APOO | -0.008 | -0.1214 | -0.051 | 0.96051 | 0.977 | -6.57 | 0.11561 | none |
| ALDH3A2 | -0.005 | -0.0485 | -0.051 | 0.96052 | 0.977 | -6.57 | -0.4549 | none |
| LY96 | -0.011 | -0.267 | -0.05 | 0.96087 | 0.978 | -6.57 | 0.01456 | none |
| MAP2K6 | 0.011 | -0.0133 | 0.0503 | 0.96089 | 0.978 | -6.57 | -0.6534 | none |
| JUND | -0.006 | 0.0161 | -0.05 | 0.96095 | 0.978 | -6.57 | -0.2291 | none |
| APOC4-AP | 0.004 | 0.0152 | 0.0501 | 0.9611 | 0.978 | -6.57 | 0 | none |
| RGPD2 | -0.011 | 0.0243 | -0.05 | 0.96115 | 0.978 | -6.57 | 0.09376 | none |
| HBD | -0.005 | 0.0105 | -0.05 | 0.96116 | 0.978 | -6.57 | 0.8233 | none |
| PI15 | -0.004 | -0.001 | -0.05 | 0.9612 | 0.978 | -6.57 | 0 | none |
| CCDC82 | -0.011 | -0.0213 | -0.05 | 0.96147 | 0.978 | -6.57 | -0.2567 | none |
| PLAGL1 | 0.007 | 0.0171 | 0.0495 | 0.96154 | 0.978 | -6.57 | -0.2035 | none |
| UBALD1 | -0.011 | 0.0251 | -0.049 | 0.96163 | 0.978 | -6.57 | 0.69634 | none |
| RAI14 | -0.011 | -0.035 | -0.049 | 0.96172 | 0.978 | -6.57 | 1.0389 | none |
| PRMT2 | -0.005 | ######## | -0.049 | 0.96173 | 0.978 | -6.57 | 0.69787 | none |
| ZFP36 | -0.006 | -0.0459 | -0.049 | 0.96178 | 0.978 | -6.57 | 1.288 | none |
| IL1A | 0.005 | 0.0619 | 0.049 | 0.96192 | 0.978 | -6.57 | -0.0109 | none |
| DNAJB2 | 0.007 | 0.0615 | 0.0488 | 0.96203 | 0.978 | -6.57 | 1.09615 | none |
| PRH2 | 0.004 | 0.0122 | 0.0488 | 0.96205 | 0.978 | -6.57 | 1.18522 | none |
| ITGB4 | 0.013 | -0.061 | 0.0487 | 0.96211 | 0.978 | -6.57 | 0.26494 | none |
| ERICH3 | -0.008 | -0.0126 | -0.049 | 0.96216 | 0.978 | -6.57 | 1.31447 | none |
| MGME1 | 0.005 | 0.0162 | 0.0486 | 0.9622 | 0.978 | -6.57 | 0.37081 | none |
| PPAN-P2R | -0.01 | 0.0851 | -0.048 | 0.9624 | 0.978 | -6.57 | 0.34624 | none |
| ACKR1 | 0.003 | 0.0125 | 0.0484 | 0.96241 | 0.978 | -6.57 | -0.4888 | none |
| TMEM191 | -0.005 | -0.0078 | -0.048 | 0.96242 | 0.978 | -6.57 | -0.2258 | none |
| CPNE3 | -0.005 | -0.0871 | -0.048 | 0.96263 | 0.978 | -6.57 | 0.16209 | none |
| FOXRED1 | -0.005 | -0.0197 | -0.048 | 0.96286 | 0.978 | -6.57 | 0.96559 | none |
| RNPC3 | -0.008 | -0.012 | -0.048 | 0.96288 | 0.978 | -6.57 | 1.31909 | none |
| SH2B2 | 0.012 | 0.15 | 0.0476 | 0.963 | 0.978 | -6.57 | 0.08109 | none |
| WNT6 | 0.005 | 0.0198 | 0.0468 | 0.96362 | 0.979 | -6.57 | -0.3005 | none |
| OR6V1 | -0.003 | 0.0154 | -0.046 | 0.96402 | 0.979 | -6.57 | -0.1122 | none |
| WDR1 | -0.006 | 0.0469 | -0.046 | 0.96404 | 0.979 | -6.57 | 1.07619 | none |
| CD207 | 0.003 | 0.01 | 0.0462 | 0.96406 | 0.979 | -6.57 | 0 | none |
| CYB5R1 | 0.005 | -0.0935 | 0.0461 | 0.96417 | 0.979 | -6.57 | -0.4599 | none |

| | | | | | | | | |
|---|---|---|---|---|---|---|---|---|
| RALGAPA2 | -0.008 | 0.0343 | -0.046 | 0.96418 | 0.979 | -6.57 | -0.1939 | none |
| LHB | 0.009 | 0.024 | 0.046 | 0.96423 | 0.979 | -6.57 | -0.7486 | none |
| FBXO3 | -0.009 | -0.0651 | -0.046 | 0.96437 | 0.979 | -6.57 | -0.4489 | none |
| TSN | 0.005 | -0.0639 | 0.0454 | 0.96471 | 0.98 | -6.57 | 0.21263 | none |
| TEAD3 | -0.004 | 0.018 | -0.045 | 0.96476 | 0.98 | -6.57 | 1.70801 | none |
| SPCS2 | -0.006 | -0.0674 | -0.045 | 0.96495 | 0.98 | -6.57 | -0.4096 | none |
| BAX | -0.005 | 0.02 | -0.045 | 0.96498 | 0.98 | -6.57 | 0.56875 | none |
| TREML1 | 0.004 | 0.0303 | 0.045 | 0.96502 | 0.98 | -6.57 | 0.39779 | none |
| TMEM161 | -0.005 | -0.0081 | -0.045 | 0.96502 | 0.98 | -6.57 | 0.67719 | none |
| RNF219 | -0.005 | -0.0216 | -0.045 | 0.96502 | 0.98 | -6.57 | 0.9533 | none |
| CELSR1 | 0.005 | 0.0529 | 0.0448 | 0.96516 | 0.98 | -6.57 | -0.0698 | none |
| SPATA22 | 0.005 | 0.0035 | 0.0448 | 0.96519 | 0.98 | -6.57 | 0.20182 | none |
| FAM47B | 0.003 | 0.0031 | 0.0448 | 0.9652 | 0.98 | -6.57 | 0 | none |
| KCNS1 | 0.004 | 0.0045 | 0.0447 | 0.96527 | 0.98 | -6.57 | 0.04782 | none |
| CXCL9 | 0.004 | 0.0254 | 0.0445 | 0.96539 | 0.98 | -6.57 | 0.03773 | none |
| FOXE1 | 0.004 | 0.0072 | 0.0445 | 0.96545 | 0.98 | -6.57 | 0.99052 | none |
| WDR5 | 0.004 | 0.0491 | 0.0444 | 0.96553 | 0.98 | -6.57 | 1.61663 | none |
| PTGS1 | -0.004 | 0.0393 | -0.044 | 0.96554 | 0.98 | -6.57 | 0.27175 | none |
| LUZP4 | 0.003 | 0.0133 | 0.0441 | 0.9657 | 0.98 | -6.57 | 0 | none |
| PUS7L | -0.006 | -0.0481 | -0.044 | 0.96598 | 0.98 | -6.57 | 0.37051 | none |
| TRIM49B | 0.005 | 0.0075 | 0.0437 | 0.966 | 0.98 | -6.57 | 1.15927 | none |
| CACHD1 | 0.003 | 0.0158 | 0.0437 | 0.96606 | 0.98 | -6.57 | 1.48108 | none |
| EEPD1 | -0.009 | -0.0089 | -0.044 | 0.96608 | 0.98 | -6.57 | 0.50496 | none |
| MAP2 | -0.003 | 0.0036 | -0.044 | 0.96608 | 0.98 | -6.57 | -0.6683 | none |
| TP53RK | -0.005 | -0.0499 | -0.043 | 0.96641 | 0.98 | -6.57 | 1.19447 | none |
| RYK | -0.011 | -0.0557 | -0.043 | 0.96657 | 0.98 | -6.57 | 1.01635 | none |
| SLC19A2 | 0.007 | 0.015 | 0.0429 | 0.96663 | 0.98 | -6.57 | 1.37488 | none |
| P2RX1 | 0.004 | 0.0166 | 0.0429 | 0.96666 | 0.98 | -6.57 | 1.3633 | none |
| VSIG4 | -0.003 | 0.0034 | -0.043 | 0.96695 | 0.98 | -6.57 | 0 | none |
| FLAD1 | 0.005 | -0.0121 | 0.0424 | 0.96706 | 0.98 | -6.57 | 0.55 | none |
| AC026348 | -0.005 | -0.0265 | -0.042 | 0.9674 | 0.981 | -6.57 | -0.0662 | none |
| KCNA2 | -0.004 | 0.0443 | -0.042 | 0.96742 | 0.981 | -6.57 | 0.58326 | none |
| HTR5A | 0.003 | 0.0076 | 0.0416 | 0.96768 | 0.981 | -6.57 | 0 | none |
| KATNB1 | -0.005 | 0.0348 | -0.041 | 0.96778 | 0.981 | -6.57 | 1.3184 | none |
| S100A2 | -0.007 | -0.0118 | -0.041 | 0.96811 | 0.981 | -6.57 | 0.70425 | none |
| DHRS3 | 0.005 | -0.0616 | 0.0409 | 0.96817 | 0.981 | -6.57 | -0.6478 | none |
| IFNAR2 | 0.005 | -0.0817 | 0.0405 | 0.96849 | 0.981 | -6.57 | 0.88523 | none |
| ST6GALNA | 0.005 | 0.0036 | 0.0404 | 0.96858 | 0.981 | -6.57 | -0.6397 | none |
| GNAQ | -0.005 | -0.0171 | -0.04 | 0.96879 | 0.982 | -6.57 | -0.2599 | none |
| ZNF205 | 0.006 | -0.018 | 0.0401 | 0.96879 | 0.982 | -6.57 | 1.06523 | none |

| | | | | | | | | |
|---|---|---|---|---|---|---|---|---|
| FKBP4 | -0.004 | 0.0375 | -0.04 | 0.96881 | 0.982 | -6.57 | 0.42223 | none |
| GALP | 0.003 | 0.0151 | 0.04 | 0.96892 | 0.982 | -6.57 | -0.314 | none |
| ZFP69B | -0.004 | 0.0232 | -0.04 | 0.96897 | 0.982 | -6.57 | 1.52245 | none |
| ZMAT5 | 0.007 | -0.0013 | 0.0398 | 0.96906 | 0.982 | -6.57 | 0.60226 | none |
| ARF3 | -0.005 | -0.0128 | -0.04 | 0.96918 | 0.982 | -6.57 | 1.6712 | none |
| MAP3K7CI | 0.006 | -0.0486 | 0.0391 | 0.96964 | 0.982 | -6.57 | -0.7152 | none |
| ARHGAP9 | 0.004 | 0.0385 | 0.039 | 0.9697 | 0.982 | -6.57 | 0.6908 | none |
| RBM45 | 0.006 | 0.0601 | 0.0388 | 0.96987 | 0.982 | -6.57 | -0.1488 | none |
| MTMR8 | 0.006 | 0.0311 | 0.0388 | 0.96987 | 0.982 | -6.57 | 0.64807 | none |
| ALX3 | 0.006 | 0.0771 | 0.0386 | 0.96997 | 0.982 | -6.57 | -0.0409 | none |
| STOX1 | 0.003 | 0.0295 | 0.0379 | 0.97055 | 0.983 | -6.57 | 1.15631 | none |
| LAPTM4B | 0.005 | -0.0734 | 0.0369 | 0.97132 | 0.983 | -6.57 | 0.71342 | none |
| KRT81 | -0.003 | -0.0075 | -0.037 | 0.97134 | 0.983 | -6.57 | 1.07175 | none |
| RNF7 | -0.006 | -0.0987 | -0.036 | 0.97164 | 0.984 | -6.57 | 0.17984 | none |
| PTPRG | 0.003 | 0.0151 | 0.0361 | 0.97195 | 0.984 | -6.57 | -0.4389 | none |
| HOXC8 | -0.003 | -0.0118 | -0.036 | 0.97197 | 0.984 | -6.57 | 0 | none |
| FBXO41 | 0.006 | 0.0702 | 0.0357 | 0.97222 | 0.984 | -6.57 | 0.82924 | none |
| ZMAT4 | 0.003 | 0.0223 | 0.0351 | 0.97272 | 0.984 | -6.57 | -0.4176 | none |
| PHPT1 | -0.005 | -0.0216 | -0.035 | 0.97286 | 0.985 | -6.57 | 0.36877 | none |
| LIMS4 | 0.003 | 0.0078 | 0.0349 | 0.97289 | 0.985 | -6.57 | 0.74567 | none |
| ZNF670 | 0.003 | -0.0545 | 0.0346 | 0.97312 | 0.985 | -6.57 | 0.56032 | none |
| IPPK | 0.003 | 0.016 | 0.0346 | 0.97312 | 0.985 | -6.57 | 1.50997 | none |
| SMG8 | 0.003 | ######## | 0.0345 | 0.97315 | 0.985 | -6.57 | 1.0133 | none |
| CCDC71 | 0.006 | 0.0906 | 0.0342 | 0.97338 | 0.985 | -6.57 | 1.3479 | none |
| TSGA10 | -0.008 | -0.0499 | -0.034 | 0.9734 | 0.985 | -6.57 | 0.52882 | none |
| ANKRD35 | -0.002 | 0.0046 | -0.034 | 0.97378 | 0.985 | -6.57 | 0.71334 | none |
| SPACA1 | 0.002 | 0.014 | 0.0336 | 0.97384 | 0.985 | -6.57 | -0.3129 | none |
| ARMC10 | -0.006 | -0.0252 | -0.033 | 0.97408 | 0.985 | -6.57 | 1.62684 | none |
| NFYB | -0.006 | -0.062 | -0.033 | 0.97418 | 0.985 | -6.57 | 0.60661 | none |
| UNC13A | 0.005 | 0.023 | 0.0332 | 0.97419 | 0.985 | -6.57 | -0.0168 | none |
| PEAR1 | 0.003 | 0.0349 | 0.0332 | 0.97419 | 0.985 | -6.57 | -0.2966 | none |
| TMEM89 | 0.007 | 0.0247 | 0.033 | 0.97437 | 0.985 | -6.57 | 0.50904 | none |
| CDKN2AIP | 0.006 | 0.103 | 0.0329 | 0.97443 | 0.985 | -6.57 | 1.8787 | none |
| DNPH1 | 0.005 | -0.0319 | 0.0326 | 0.97469 | 0.986 | -6.57 | -0.3623 | none |
| PDE1C | -0.003 | -0.01 | -0.032 | 0.97484 | 0.986 | -6.57 | 0.93525 | none |
| PCTP | 0.004 | -0.0834 | 0.0318 | 0.9753 | 0.986 | -6.57 | 1.4633 | none |
| DPH5 | -0.005 | -0.1631 | -0.032 | 0.97533 | 0.986 | -6.57 | -0.0619 | none |
| DUS2 | 0.003 | 0.0385 | 0.0316 | 0.97544 | 0.986 | -6.57 | 1.07365 | none |
| SDF4 | 0.004 | 0.07 | 0.0315 | 0.97553 | 0.986 | -6.57 | 0.55631 | none |
| RAB4A | -0.003 | -0.0187 | -0.031 | 0.97591 | 0.986 | -6.57 | 1.26466 | none |

| | | | | | | | | |
|---|---|---|---|---|---|---|---|---|
| PPOX | 0.004 | 0.0169 | 0.0308 | 0.97602 | 0.986 | -6.57 | 0.30789 | none |
| CBX3 | -0.002 | -0.0055 | -0.031 | 0.97605 | 0.986 | -6.57 | 0.23058 | none |
| KRT23 | 0.002 | 0.002 | 0.0308 | 0.97609 | 0.986 | -6.57 | 0 | none |
| ZNF836 | 0.007 | -0.0564 | 0.0307 | 0.97616 | 0.986 | -6.57 | -0.4615 | none |
| CDK14 | 0.005 | 0.0538 | 0.0306 | 0.97617 | 0.986 | -6.57 | -0.3824 | none |
| EXTL1 | 0.004 | 0.0366 | 0.0299 | 0.97677 | 0.987 | -6.57 | -0.1581 | none |
| MKX | -0.002 | ######## | -0.03 | 0.97696 | 0.987 | -6.57 | 1.21525 | none |
| RGCC | -0.007 | -0.0495 | -0.029 | 0.97738 | 0.987 | -6.57 | 1.05372 | none |
| KLF6 | -0.004 | -0.015 | -0.029 | 0.97742 | 0.987 | -6.57 | 0.73757 | none |
| KDM6B | -0.01 | 0.2515 | -0.029 | 0.97746 | 0.987 | -6.57 | 0.88742 | none |
| STRADB | 0.004 | -0.0275 | 0.029 | 0.97748 | 0.987 | -6.57 | 1.36233 | none |
| CENPA | 0.006 | -0.0226 | 0.0289 | 0.9775 | 0.987 | -6.57 | -0.0388 | none |
| HIST1H3D | 0.011 | -0.129 | 0.0289 | 0.97756 | 0.987 | -6.57 | 0.54125 | none |
| NPPC | 0.005 | 0.0027 | 0.0286 | 0.97779 | 0.987 | -6.57 | 1.23962 | none |
| USP51 | -0.004 | 0.0788 | -0.028 | 0.97792 | 0.988 | -6.57 | 1.60104 | none |
| TM6SF2 | 0.003 | 0.0103 | 0.0284 | 0.97796 | 0.988 | -6.57 | -0.1849 | none |
| CHST4 | -0.002 | 0.0091 | -0.028 | 0.97813 | 0.988 | -6.57 | -0.3286 | none |
| ANKRD39 | 0.004 | -0.0434 | 0.0275 | 0.97861 | 0.988 | -6.57 | -0.339 | none |
| SOX8 | -0.002 | -0.0042 | -0.027 | 0.97886 | 0.988 | -6.57 | 1.15341 | none |
| MEIS1 | -0.004 | 0.0081 | -0.027 | 0.9789 | 0.988 | -6.57 | -0.5973 | none |
| DBH | 0.002 | ######## | 0.027 | 0.97902 | 0.988 | -6.57 | -0.4844 | none |
| MYO1A | -0.004 | -0.0019 | -0.027 | 0.97907 | 0.988 | -6.57 | 0.54444 | none |
| TXNRD3 | -0.005 | 0.0306 | -0.027 | 0.97917 | 0.988 | -6.57 | 0.40386 | none |
| CLIC5 | 0.006 | 0.1087 | 0.0268 | 0.9792 | 0.988 | -6.57 | 0.51622 | none |
| ZNF229 | 0.003 | 0.0191 | 0.0267 | 0.97924 | 0.988 | -6.57 | 1.27085 | none |
| BOD1 | 0.004 | 0.0404 | 0.0264 | 0.97948 | 0.988 | -6.57 | 1.01438 | none |
| AIP | 0.003 | 0.0247 | 0.0264 | 0.97949 | 0.988 | -6.57 | -0.1774 | none |
| LDLRAD2 | -0.002 | -0.0034 | -0.026 | 0.9799 | 0.989 | -6.57 | -0.1938 | none |
| AC104581 | 0.003 | 0.0467 | 0.0258 | 0.97995 | 0.989 | -6.57 | 0.67516 | none |
| NANS | -0.003 | -0.0313 | -0.025 | 0.98018 | 0.989 | -6.57 | 0.83477 | none |
| NDUFAF1 | 0.003 | -0.0904 | 0.0255 | 0.9802 | 0.989 | -6.57 | 0.01378 | none |
| LRRC3B | -0.004 | -0.0157 | -0.025 | 0.98023 | 0.989 | -6.57 | -0.0305 | none |
| SKA2 | 0.004 | -0.0964 | 0.0253 | 0.98037 | 0.989 | -6.57 | -0.5623 | none |
| LRRC40 | 0.004 | ######## | 0.0252 | 0.9804 | 0.989 | -6.57 | 1.79565 | none |
| SPTBN2 | -0.005 | 0.0415 | -0.025 | 0.98059 | 0.989 | -6.57 | 0.66621 | none |
| NOL4L | 0.004 | -0.0079 | 0.0248 | 0.98071 | 0.989 | -6.57 | -0.0108 | none |
| SLC7A3 | 0.002 | 0.0031 | 0.0243 | 0.98107 | 0.989 | -6.57 | 0.50031 | none |
| EYA2 | 0.005 | 0.0162 | 0.0242 | 0.9812 | 0.989 | -6.57 | -0.2621 | none |
| CCDC59 | -0.004 | 0.046 | -0.024 | 0.98122 | 0.989 | -6.57 | -0.2668 | none |
| KHDRBS3 | -0.004 | 0.0232 | -0.024 | 0.98124 | 0.989 | -6.57 | 1.7286 | none |

| | | | | | | | | |
|---|---|---|---|---|---|---|---|---|
| PHYHIP | -0.002 | 0.0232 | -0.024 | 0.98135 | 0.989 | -6.57 | 1.15292 | none |
| APOM | 0.002 | 0.0193 | 0.0239 | 0.98145 | 0.99 | -6.57 | 0.87822 | none |
| TMEM59L | -0.003 | 0.0277 | -0.024 | 0.98165 | 0.99 | -6.57 | -0.5582 | none |
| ZNF324 | -0.003 | 0.0552 | -0.023 | 0.98196 | 0.99 | -6.57 | 0.46185 | none |
| TICAM2 | 0.002 | -0.0134 | 0.023 | 0.98208 | 0.99 | -6.57 | 0.4743 | none |
| C1ORF52 | -0.004 | -0.1309 | -0.023 | 0.98212 | 0.99 | -6.57 | 1.13734 | none |
| POPDC3 | -0.002 | 0.0065 | -0.023 | 0.98222 | 0.99 | -6.57 | 0.85469 | none |
| DHRS7 | 0.003 | -0.0574 | 0.0228 | 0.98227 | 0.99 | -6.57 | -0.1453 | none |
| PRKACG | -0.002 | ######## | -0.022 | 0.98254 | 0.99 | -6.57 | 1.53129 | none |
| PTTG1IP | -0.002 | -0.0519 | -0.022 | 0.9826 | 0.99 | -6.57 | 1.00249 | none |
| NEUROG2 | 0.001 | 0.0133 | 0.0224 | 0.98261 | 0.99 | -6.57 | -0.627 | none |
| DHRS7C | 0.002 | 0.0017 | 0.0221 | 0.9828 | 0.99 | -6.57 | 1.06571 | none |
| SLC25A17 | -0.003 | -0.1071 | -0.022 | 0.9828 | 0.99 | -6.57 | 0.53927 | none |
| SETD9 | 0.005 | -0.0964 | 0.0221 | 0.98284 | 0.99 | -6.57 | 0.87227 | none |
| ERI2 | -0.004 | 0.032 | -0.022 | 0.98305 | 0.99 | -6.57 | 1.20323 | none |
| ANHX | 0.002 | 0.0119 | 0.0216 | 0.98318 | 0.99 | -6.57 | -0.5105 | none |
| EXOSC3 | -0.002 | -0.0054 | -0.022 | 0.98325 | 0.99 | -6.57 | -0.5739 | none |
| THSD1 | 0.001 | 0.0146 | 0.0215 | 0.98332 | 0.99 | -6.57 | 1.3234 | none |
| TBPL1 | 0.003 | -0.0716 | 0.0212 | 0.98348 | 0.991 | -6.57 | -0.2419 | none |
| TEF | 0.002 | -0.0302 | 0.0212 | 0.98355 | 0.991 | -6.57 | 1.02771 | none |
| CLDN6 | 0.002 | 0.0097 | 0.021 | 0.9837 | 0.991 | -6.57 | 0.09199 | none |
| PES1 | -0.003 | 0.1464 | -0.021 | 0.98373 | 0.991 | -6.57 | 1.01639 | none |
| TSPAN33 | -0.003 | -0.0051 | -0.021 | 0.98386 | 0.991 | -6.57 | 1.21903 | none |
| TCTEX1D2 | -0.004 | -0.0611 | -0.021 | 0.98392 | 0.991 | -6.57 | 0.11626 | none |
| CCDC148 | -0.002 | -0.0151 | -0.021 | 0.98395 | 0.991 | -6.57 | 0.35411 | none |
| RSAD1 | 0.002 | 0.0312 | 0.0204 | 0.98411 | 0.991 | -6.57 | 1.36533 | none |
| NRXN2 | -0.003 | -0.0049 | -0.02 | 0.98451 | 0.991 | -6.57 | -0.3386 | none |
| CRMP1 | 0.001 | 0.0112 | 0.0198 | 0.9846 | 0.991 | -6.57 | 0.65707 | none |
| AC109486 | -0.004 | -0.0035 | -0.02 | 0.98478 | 0.991 | -6.57 | 1.60493 | none |
| KLHL10 | 0.003 | 0.0185 | 0.0195 | 0.98484 | 0.991 | -6.57 | 1.34477 | none |
| GYPE | 0.003 | -0.0547 | 0.0195 | 0.98487 | 0.991 | -6.57 | -0.2754 | none |
| CCDC154 | 0.002 | 0.0705 | 0.0193 | 0.98499 | 0.991 | -6.57 | 1.00899 | none |
| INSIG1 | 0.003 | -0.0249 | 0.0193 | 0.985 | 0.991 | -6.57 | 0.98635 | none |
| CTDP1 | -0.003 | 0.0931 | -0.019 | 0.98502 | 0.991 | -6.57 | 0.13061 | none |
| SF3A3 | 0.002 | 0.0613 | 0.0192 | 0.98511 | 0.991 | -6.57 | 0.22176 | none |
| GPR20 | -0.001 | 0.0109 | -0.019 | 0.98513 | 0.991 | -6.57 | 0 | none |
| ZNF580 | -0.002 | -0.0094 | -0.019 | 0.98528 | 0.991 | -6.57 | 1.02082 | none |
| TRPC4AP | 0.001 | 0.011 | 0.0188 | 0.98535 | 0.991 | -6.57 | 2.06835 | none |
| NCBP2 | -0.002 | -0.0025 | -0.019 | 0.98538 | 0.991 | -6.57 | 1.14479 | none |
| NUTM2A | 0.001 | 0.0298 | 0.0184 | 0.98572 | 0.992 | -6.57 | 0.02389 | none |

| | | | | | | | | |
|---|---|---|---|---|---|---|---|---|
| GPR158 | 0.001 | 0.0262 | 0.0182 | 0.98587 | 0.992 | -6.57 | 0 | none |
| LGALS16 | -0.002 | -0.0021 | -0.018 | 0.98611 | 0.992 | -6.57 | 1.87736 | none |
| QARS | -0.002 | 0.0062 | -0.018 | 0.98616 | 0.992 | -6.57 | 0.61049 | none |
| CSTF2T | -0.002 | 0.0034 | -0.018 | 0.98621 | 0.992 | -6.57 | 1.39177 | none |
| SLC39A11 | 0.002 | -0.0167 | 0.0172 | 0.98664 | 0.992 | -6.57 | 0.25001 | none |
| PEPD | 0.002 | -0.0371 | 0.0172 | 0.98664 | 0.992 | -6.57 | 1.13989 | none |
| PRMT9 | 0.001 | 0.0134 | 0.0169 | 0.98682 | 0.992 | -6.57 | 1.46307 | none |
| AAAS | 0.002 | -0.0218 | 0.0169 | 0.98686 | 0.992 | -6.57 | 1.23398 | none |
| GAGE1 | -0.001 | -0.0112 | -0.017 | 0.98692 | 0.992 | -6.57 | -0.7466 | none |
| CMTM4 | -0.003 | -0.0235 | -0.016 | 0.98725 | 0.993 | -6.57 | 0.50732 | none |
| PPP1R32 | 0.004 | 0.075 | 0.0163 | 0.98731 | 0.993 | -6.57 | -0.0491 | none |
| CCL24 | -0.002 | 0.0192 | -0.016 | 0.98733 | 0.993 | -6.57 | 0.97771 | none |
| ACP6 | 0.004 | -0.0467 | 0.0161 | 0.98749 | 0.993 | -6.57 | 0.44135 | none |
| PLRG1 | -0.002 | -0.0279 | -0.016 | 0.98766 | 0.993 | -6.57 | -0.2828 | none |
| AKAP11 | 0.003 | 0.0889 | 0.0151 | 0.98828 | 0.993 | -6.57 | 0.85468 | none |
| ORC5 | -0.002 | -0.0529 | -0.015 | 0.98843 | 0.993 | -6.57 | 0.76213 | none |
| USP43 | -0.001 | 0.034 | -0.015 | 0.98851 | 0.993 | -6.57 | 0.01797 | none |
| NMRK1 | 0.002 | -0.0426 | 0.0147 | 0.98854 | 0.993 | -6.57 | 0.23124 | none |
| LEUTX | -0.001 | -0.0011 | -0.015 | 0.9886 | 0.993 | -6.57 | 0 | none |
| SMTNL2 | -0.002 | 0.0501 | -0.015 | 0.98866 | 0.993 | -6.57 | 0.66879 | none |
| PMPCB | 0.002 | -0.0979 | 0.0145 | 0.9887 | 0.993 | -6.57 | 0.69459 | none |
| FAM189A1 | -0.004 | 0.0359 | -0.014 | 0.98875 | 0.993 | -6.57 | 1.32176 | none |
| MAP9 | 0.002 | 0.0644 | 0.0144 | 0.98878 | 0.993 | -6.57 | 0.76548 | none |
| DUOXA2 | ###### | 0.0148 | 0.0143 | 0.98886 | 0.993 | -6.57 | 0.38199 | none |
| ANKRD60 | 0.001 | ######## | 0.0142 | 0.98899 | 0.993 | -6.57 | -0.1716 | none |
| CDPF1 | -0.002 | -0.0344 | -0.014 | 0.989 | 0.993 | -6.57 | 1.05521 | none |
| RNASE9 | 0.001 | 0.001 | 0.0138 | 0.98931 | 0.994 | -6.57 | 0 | none |
| TMEM218 | -0.002 | -0.0813 | -0.014 | 0.98944 | 0.994 | -6.57 | 0.35952 | none |
| ZNF512B | -0.002 | 0.1566 | -0.013 | 0.98999 | 0.994 | -6.57 | 0.71469 | none |
| KLK14 | 0.001 | 0.0054 | 0.0129 | 0.99 | 0.994 | -6.57 | 0.84121 | none |
| CALHM2 | -0.002 | -0.0188 | -0.013 | 0.99009 | 0.994 | -6.57 | 1.52567 | none |
| HACD4 | -0.001 | -0.0816 | -0.013 | 0.99013 | 0.994 | -6.57 | 1.03369 | none |
| RGMB | -0.002 | -0.0488 | -0.013 | 0.99025 | 0.994 | -6.57 | -0.5163 | none |
| C6ORF52 | 0.002 | -0.0181 | 0.0125 | 0.99029 | 0.994 | -6.57 | 0.60372 | none |
| SDHAF1 | -0.001 | -0.0103 | -0.012 | 0.99045 | 0.994 | -6.57 | 1.12537 | none |
| FOXP2 | 0.006 | -0.0632 | 0.0122 | 0.99048 | 0.994 | -6.57 | -0.2698 | none |
| TRIO | 0.002 | 0.0847 | 0.0122 | 0.99055 | 0.994 | -6.57 | 0.34107 | none |
| EIF2B5 | 0.001 | 0.052 | 0.0115 | 0.99106 | 0.995 | -6.57 | 0.44911 | none |
| DDX4 | -0.002 | 0.0173 | -0.011 | 0.9911 | 0.995 | -6.57 | 0.23698 | none |
| PIR | 0.002 | -0.0361 | 0.0114 | 0.99112 | 0.995 | -6.57 | 0.29881 | none |

| | | | | | | | | |
|---|---|---|---|---|---|---|---|---|
| PROX2 | -0.002 | 0.0552 | -0.011 | 0.99152 | 0.995 | -6.57 | -0.3411 | none |
| PEX5 | 0.001 | -0.0014 | 0.0105 | 0.99181 | 0.995 | -6.57 | 0.48176 | none |
| SLC17A9 | 0.001 | 0.0504 | 0.01 | 0.99224 | 0.996 | -6.57 | 0.37666 | none |
| YIPF5 | ###### | -0.1108 | -0.01 | 0.99239 | 0.996 | -6.57 | -0.089 | none |
| C19ORF18 | ###### | 0.0078 | -0.01 | 0.99246 | 0.996 | -6.57 | -0.6112 | none |
| DAPK3 | 0.001 | 0.0649 | 0.0096 | 0.99255 | 0.996 | -6.57 | 0.66571 | none |
| EAPP | 0.001 | -0.0204 | 0.0094 | 0.99272 | 0.996 | -6.57 | 0.92823 | none |
| OR5B17 | ###### | ######## | -0.009 | 0.99273 | 0.996 | -6.57 | 1.86247 | none |
| VEZF1 | 0.002 | 0.1141 | 0.0092 | 0.99285 | 0.996 | -6.57 | 1.49428 | none |
| IFI44L | ###### | -0.0432 | -0.009 | 0.99294 | 0.996 | -6.57 | 0.8697 | none |
| LAD1 | -0.001 | -0.0596 | -0.009 | 0.99328 | 0.996 | -6.57 | 0.47547 | none |
| EMC3 | ###### | -0.0597 | 0.0086 | 0.99332 | 0.996 | -6.57 | 0.16298 | none |
| ITGA11 | -0.001 | -0.0433 | -0.009 | 0.99334 | 0.996 | -6.57 | -0.3356 | none |
| SPDYA | 0.002 | -0.0474 | 0.0085 | 0.9934 | 0.996 | -6.57 | 0.92049 | none |
| TNFRSF11 | -0.001 | 0.0419 | -0.008 | 0.99348 | 0.996 | -6.57 | -0.3877 | none |
| C18ORF32 | 0.001 | -0.078 | 0.0084 | 0.99348 | 0.996 | -6.57 | 0.83993 | none |
| RAB3IP | 0.001 | 0.003 | 0.0079 | 0.99387 | 0.996 | -6.57 | 0.65787 | none |
| HHATL | ###### | ######## | -0.008 | 0.99412 | 0.997 | -6.57 | 0.08831 | none |
| VAMP5 | -0.002 | -0.1049 | -0.008 | 0.99414 | 0.997 | -6.57 | 0.20287 | none |
| USP41 | -0.002 | 0.0383 | -0.007 | 0.99429 | 0.997 | -6.57 | 0.08808 | none |
| BLCAP | -0.001 | -0.0531 | -0.007 | 0.99458 | 0.997 | -6.57 | 0.51343 | none |
| RAB35 | -0.001 | 0.0035 | -0.007 | 0.99466 | 0.997 | -6.57 | 1.49945 | none |
| ELFN2 | ###### | 0.0228 | 0.0067 | 0.99481 | 0.997 | -6.57 | 0.53584 | none |
| DAP | ###### | 0.0855 | 0.0067 | 0.99483 | 0.997 | -6.57 | 1.18571 | none |
| TSPAN15 | -0.001 | -0.0167 | -0.007 | 0.99492 | 0.997 | -6.57 | 0.73594 | none |
| SAMD9L | 0.002 | 0.1398 | 0.0063 | 0.99513 | 0.997 | -6.57 | 0.90948 | none |
| RUBCNL | ###### | -0.004 | -0.006 | 0.99515 | 0.997 | -6.57 | 0.7253 | none |
| AMBP | ###### | -0.0114 | 0.0061 | 0.99529 | 0.997 | -6.57 | -0.0655 | none |
| CYP2A13 | ###### | -0.0021 | 0.006 | 0.99533 | 0.997 | -6.57 | 0.52924 | none |
| ACCSL | ###### | 0.0104 | 0.006 | 0.99533 | 0.997 | -6.57 | 0 | none |
| FGFR1OP2 | ###### | -0.0619 | 0.0059 | 0.99539 | 0.997 | -6.57 | -0.064 | none |
| DLX6 | ###### | ######## | 0.0059 | 0.99545 | 0.997 | -6.57 | 1.49553 | none |
| CPZ | ###### | ######## | 0.0055 | 0.99572 | 0.997 | -6.57 | 0.70486 | none |
| TIMP4 | ###### | -0.0083 | -0.005 | 0.99583 | 0.997 | -6.57 | -0.7972 | none |
| DNAH6 | ###### | -0.0013 | -0.005 | 0.99584 | 0.997 | -6.57 | -0.5819 | none |
| CPNE4 | ###### | 0.0063 | 0.0053 | 0.99584 | 0.997 | -6.57 | -0.3444 | none |
| CES5A | ###### | ######## | 0.0053 | 0.99591 | 0.997 | -6.57 | 1.88661 | none |
| ATG4C | ###### | -0.0506 | -0.005 | 0.99607 | 0.997 | -6.57 | 0.76732 | none |
| POP1 | 0.001 | 0.1284 | 0.0047 | 0.99636 | 0.998 | -6.57 | 1.08876 | none |
| ECE2 | ###### | -0.0015 | -0.005 | 0.99649 | 0.998 | -6.57 | 1.91482 | none |

| | | | | | | | | |
|---|---|---|---|---|---|---|---|---|
| UFL1 | ###### | 0.0777 | -0.004 | 0.9969 | 0.998 | -6.57 | 1.82433 | none |
| RHOH | ###### | -0.0903 | -0.004 | 0.99706 | 0.998 | -6.57 | 1.52253 | none |
| PDCD1LG2 | ###### | -0.0095 | 0.0033 | 0.9974 | 0.998 | -6.57 | -0.1608 | none |
| LTB4R2 | ###### | 0.0664 | -0.003 | 0.99749 | 0.998 | -6.57 | 1.02443 | none |
| HIST1H2BI | ###### | -0.0378 | -0.003 | 0.99752 | 0.998 | -6.57 | 1.31884 | none |
| FGFR1OP | ###### | -0.0251 | 0.0031 | 0.99762 | 0.998 | -6.57 | -0.3969 | none |
| NUDT7 | ###### | -0.1301 | 0.003 | 0.9977 | 0.998 | -6.57 | -0.2651 | none |
| ZDHHC6 | ###### | -0.0451 | -0.003 | 0.99795 | 0.999 | -6.57 | 1.80803 | none |
| PFN3 | ###### | 0.0217 | -0.003 | 0.99798 | 0.999 | -6.57 | 0.8171 | none |
| SPATA6 | ###### | 0.024 | 0.0025 | 0.99809 | 0.999 | -6.57 | 0.80493 | none |
| TFCP2L1 | ###### | 0.0042 | -0.002 | 0.99813 | 0.999 | -6.57 | -0.3323 | none |
| TRIP6 | ###### | 0.0477 | 0.002 | 0.99844 | 0.999 | -6.57 | 1.0276 | none |
| CFAP53 | ###### | 0.035 | -0.002 | 0.99854 | 0.999 | -6.57 | 0.05329 | none |
| TCEAL4 | ###### | 0.0784 | -0.001 | 0.99889 | 0.999 | -6.57 | 0.94028 | none |
| PLD4 | ###### | -0.0374 | -0.001 | 0.99917 | 0.999 | -6.57 | -0.2756 | none |
| RFK | ###### | -0.0851 | ####### | 0.99933 | 1 | -6.57 | 0.95739 | none |
| ALPL | ###### | ####### | ####### | 0.99943 | 1 | -6.57 | 0.531 | none |
| MCM5 | ###### | 0.0697 | ####### | 0.99957 | 1 | -6.57 | 0.52896 | none |
| EGF | ###### | -0.0099 | ####### | 0.99989 | 1 | -6.57 | 0.90435 | none |
| AC010642 | ###### | 0.0167 | ####### | 0.99996 | 1 | -6.57 | -0.7617 | none |

**Supplementary Table 2. Transcriptomic signature of oleic- vs. arachidonic acid stimuli.** Results of the two analyses described for the bulk RNA data to characterize the differential effect of the two fatty acids. PC1-based analysis determines whether genes belong to the oleic acid module, arachidonic acid module, or none (column “PC1 gene module”) based on PC1 loading described in the text. The other columns describe the results of a limma-based moderated t-test for the intercept of the described linear model (p-values adjusted with the Benjamini-Hochberg procedure).

**Supplementary Table 3. Donor demographics for single cell sequencing of blood and adipose $T_{regs}$.**

| | Age | Height (cm) | Weight (kg) | BMI | Gender | Last DMT or steroid | Date Collected | Date Sequenced |
|---|---|---|---|---|---|---|---|---|
| Healthy | 54 | 152.0 | 68.0 | 29.43 | F | N/A | 3-3-17 | 6-11-18 |
| | 48 | 157.0 | 55.0 | 22.31 | F | N/A | 7-21-17 | 2-17-18 |
| | 46 | 158.0 | 62.0 | 24.84 | F | N/A | 3-17-17 | 3-5-18 |
| | 64 | 171. | 82.0 | 28.04 | M | N/A | 10-5-16 | 4-25-18 |
| | 31 | 163.0 | 72.0 | 27.10 | F | N/A | 10-5-16 | 4-25-18 |
| | 38 | 155.0 | 57.0 | 23.73 | F | N/A | 5-19-17 | 3-8-18 |
| | 23 | 167.0 | 61.0 | 21.87 | F | N/A | 10-20-16 | 6-8-18 |
| | 23 | 163.0 | 55.0 | 20.70 | F | N/A | 10-20-16 | 6-8-18 |
| RRMS | 47 | 167.0 | 102.0 | 36.57 | F | No previous history | 3-17-17 | 6-11-18 |
| | 60 | 175.0 | 84.0 | 27.43 | M | No history since 11-2015 | 3-17-17 | 6-11-18 |
| | 57 | 161.0 | 61.0 | 23.53 | F | DMT Avonex 10-1-05, oral steroids 6-2013 | 3-3-17 | 5-15-18 |
| | 49 | 172.0 | 91.0 | 30.76 | F | DMT Tecfidera 9-1-14, oral Prednisone 1-24-17 | 3-3-17 | 5-15-18 |
| | 52 | 171.0 | 68.0 | 23.26 | F | DMT Avonex in 2000 | 3-31-17 | 3-5-18 |
| | 56 | 178.0 | 99.0 | 31.25 | F | DMT (Avonex, Copaxone, other) and IVMP in 2004 | 5-11-17 | 6-11-18 |
| | 35 | 152.0 | 60.0 | 25.97 | F | DMT Tecfidera 10-2016 | 5-19-17 | 3-8-18 |
| | 37 | 152.0 | 86.0 | 37.22 | F | DMT Tecfidera 5-2015 | 5-24-17 | 2-27-18 |

**Supplementary Table 3. Donor demographics for mass spectrometry and single cell sequencing of blood and adipose $T_{regs}$.** Age, height, weight, body mass index (BMI), gender, and disease modifying treatment (DMT) information for healthy and MS subjects. Collection and processing dates are also shown. These donors donated both blood and adipose tissue from which $T_{reg}$ cells were sorted and single cell sequenced, and blood plasma and fat supernatant were analyzed by mass spectrometry for concentrations of long chain fatty acids.

**Supplementary Table 4. Differentially expressed genes between healthy and MS donors.**

| symbol | p_val | avg_logFC | pct.1 | pct.2 | p_val_adj | comparison |
|---|---|---|---|---|---|---|
| PPDPF | 4.51E-18 | 1.39767 | 0.514 | 0.21 | 7.20E-14 | HC_vs_MS.Trt_F |
| ZFP36 | 6.30E-15 | 1.38737 | 0.673 | 0.41 | 1.01E-10 | HC_vs_MS.Trt_F |
| KPNA5 | 8.25E-15 | 0.65223 | 0.997 | 0.93 | 1.32E-10 | HC_vs_MS.Trt_F |
| CD3G | 8.43E-15 | -1.0509 | 0.931 | 0.95 | 1.35E-10 | HC_vs_MS.Trt_F |
| SAMD12 | 3.12E-14 | 0.632 | 0.994 | 0.94 | 4.98E-10 | HC_vs_MS.Trt_F |
| MORC4 | 8.59E-14 | 0.46891 | 1 | 0.99 | 1.37E-09 | HC_vs_MS.Trt_F |
| RGPD5 | 1.31E-13 | 0.84279 | 0.919 | 0.8 | 2.08E-09 | HC_vs_MS.Trt_F |
| MTRNR2L8 | 1.31E-13 | 0.43007 | 1 | 0.98 | 2.09E-09 | HC_vs_MS.Trt_F |
| C1orf56 | 5.95E-13 | -0.8723 | 0.78 | 0.84 | 9.50E-09 | HC_vs_MS.Trt_F |
| CFAP54 | 8.58E-13 | 0.66482 | 0.965 | 0.91 | 1.37E-08 | HC_vs_MS.Trt_F |
| MTRNR2L4 | 1.95E-12 | 0.46609 | 0.991 | 0.92 | 3.12E-08 | HC_vs_MS.Trt_F |
| ABLIM3 | 3.31E-12 | 2.48814 | 0.275 | 0.07 | 5.28E-08 | HC_vs_MS.Trt_F |
| MTRNR2L1 | 4.69E-12 | 0.40295 | 1 | 0.98 | 7.48E-08 | HC_vs_MS.Trt_F |
| S100A4 | 1.41E-11 | 0.68977 | 0.757 | 0.55 | 2.25E-07 | HC_vs_MS.Trt_F |
| MTRNR2L12 | 2.03E-11 | 0.35167 | 1 | 1 | 3.25E-07 | HC_vs_MS.Trt_F |
| TAC4 | 2.22E-11 | 0.89018 | 0.754 | 0.56 | 3.55E-07 | HC_vs_MS.Trt_F |
| SPATA4 | 2.62E-11 | 0.74843 | 0.72 | 0.47 | 4.18E-07 | HC_vs_MS.Trt_F |
| PBX1 | 3.70E-11 | 0.85305 | 0.934 | 0.88 | 5.91E-07 | HC_vs_MS.Trt_F |
| CYSLTR1 | 4.89E-11 | 0.57611 | 0.98 | 0.88 | 7.81E-07 | HC_vs_MS.Trt_F |
| YLPM1 | 1.21E-10 | 0.6931 | 0.749 | 0.54 | 1.94E-06 | HC_vs_MS.Trt_F |
| C4orf32 | 1.76E-10 | 0.43457 | 0.994 | 0.97 | 2.81E-06 | HC_vs_MS.Trt_F |
| HNRNPH1 | 1.98E-10 | -0.7629 | 0.991 | 0.99 | 3.16E-06 | HC_vs_MS.Trt_F |
| MYADM | 4.25E-10 | 0.96205 | 0.867 | 0.72 | 6.79E-06 | HC_vs_MS.Trt_F |
| AHNAK | 4.38E-10 | 0.7165 | 0.89 | 0.76 | 7.00E-06 | HC_vs_MS.Trt_F |
| LMNA | 4.84E-10 | 1.13869 | 0.801 | 0.65 | 7.72E-06 | HC_vs_MS.Trt_F |
| BLOC1S6 | 5.57E-10 | 0.58013 | 0.997 | 0.98 | 8.90E-06 | HC_vs_MS.Trt_F |
| C6orf62 | 6.04E-10 | -0.8343 | 0.737 | 0.84 | 9.65E-06 | HC_vs_MS.Trt_F |
| MTRNR2L10 | 8.07E-10 | 0.40013 | 0.98 | 0.89 | 1.29E-05 | HC_vs_MS.Trt_F |
| VIM | 9.05E-10 | 0.55635 | 0.893 | 0.77 | 1.45E-05 | HC_vs_MS.Trt_F |
| DDX39B | 1.14E-09 | 0.69991 | 0.783 | 0.57 | 1.83E-05 | HC_vs_MS.Trt_F |
| ZNF331 | 1.34E-09 | 1.37356 | 0.757 | 0.62 | 2.14E-05 | HC_vs_MS.Trt_F |
| RPS14 | 1.45E-09 | -0.5438 | 0.908 | 0.96 | 2.32E-05 | HC_vs_MS.Trt_F |
| MARVELD2 | 1.51E-09 | 0.75116 | 0.725 | 0.54 | 2.41E-05 | HC_vs_MS.Trt_F |
| TNFAIP3 | 1.63E-09 | 0.90209 | 0.633 | 0.42 | 2.61E-05 | HC_vs_MS.Trt_F |
| IDS | 1.65E-09 | 0.60789 | 0.879 | 0.75 | 2.64E-05 | HC_vs_MS.Trt_F |
| CLDN5 | 1.76E-09 | 0.97152 | 0.208 | 0.05 | 2.81E-05 | HC_vs_MS.Trt_F |
| LENG8 | 2.45E-09 | 0.73209 | 0.777 | 0.58 | 3.92E-05 | HC_vs_MS.Trt_F |
| MTRNR2L6 | 2.96E-09 | 0.39456 | 0.983 | 0.9 | 4.72E-05 | HC_vs_MS.Trt_F |
| TMEM45A | 3.08E-09 | 0.3743 | 1 | 0.97 | 4.91E-05 | HC_vs_MS.Trt_F |

| | | | | | | |
|---|---|---|---|---|---|---|
| TBC1D8B | 3.99E-09 | 0.54708 | 0.965 | 0.85 | 6.37E-05 | HC_vs_MS.Trt_F |
| C16orf54 | 6.16E-09 | -1.0807 | 0.737 | 0.8 | 9.84E-05 | HC_vs_MS.Trt_F |
| HACD3 | 6.32E-09 | 0.38235 | 0.821 | 0.67 | 1.01E-04 | HC_vs_MS.Trt_F |
| PCSK5 | 9.86E-09 | 1.06097 | 0.994 | 0.93 | 1.57E-04 | HC_vs_MS.Trt_F |
| EFNA5 | 1.10E-08 | 0.50042 | 0.54 | 0.32 | 1.75E-04 | HC_vs_MS.Trt_F |
| RANBP2 | 1.36E-08 | 0.67177 | 0.592 | 0.35 | 2.16E-04 | HC_vs_MS.Trt_F |
| TMSB4X | 1.37E-08 | -0.3143 | 0.974 | 1 | 2.18E-04 | HC_vs_MS.Trt_F |
| CTTN | 1.90E-08 | 0.79778 | 0.295 | 0.11 | 3.03E-04 | HC_vs_MS.Trt_F |
| KIAA1161 | 1.97E-08 | 0.50813 | 0.876 | 0.71 | 3.15E-04 | HC_vs_MS.Trt_F |
| CD69 | 1.99E-08 | 1.02523 | 0.494 | 0.28 | 3.18E-04 | HC_vs_MS.Trt_F |
| RGS1 | 2.29E-08 | 0.73001 | 0.575 | 0.36 | 3.65E-04 | HC_vs_MS.Trt_F |
| RPL3 | 2.44E-08 | -0.4 | 0.936 | 0.96 | 3.90E-04 | HC_vs_MS.Trt_F |
| DNTTIP2 | 2.89E-08 | 0.4512 | 0.96 | 0.88 | 4.61E-04 | HC_vs_MS.Trt_F |
| SULT1B1 | 2.93E-08 | 0.38038 | 0.988 | 0.9 | 4.68E-04 | HC_vs_MS.Trt_F |
| MTRNR2L11 | 3.12E-08 | 0.36627 | 0.974 | 0.89 | 4.98E-04 | HC_vs_MS.Trt_F |
| FGF13 | 3.40E-08 | 0.62617 | 0.968 | 0.92 | 5.43E-04 | HC_vs_MS.Trt_F |
| FSIP2 | 3.65E-08 | 0.53696 | 0.847 | 0.64 | 5.83E-04 | HC_vs_MS.Trt_F |
| PRRG3 | 3.81E-08 | 1.48051 | 0.737 | 0.59 | 6.08E-04 | HC_vs_MS.Trt_F |
| NR4A2 | 3.82E-08 | 0.7098 | 0.607 | 0.4 | 6.10E-04 | HC_vs_MS.Trt_F |
| CSRNP1 | 4.51E-08 | 1.53656 | 0.327 | 0.15 | 7.20E-04 | HC_vs_MS.Trt_F |
| ATXN10 | 6.12E-08 | 0.48979 | 0.587 | 0.36 | 9.77E-04 | HC_vs_MS.Trt_F |
| HLA-A | 7.95E-08 | 0.43362 | 0.899 | 0.75 | 0.001269 | HC_vs_MS.Trt_F |
| FTH1 | 7.95E-08 | 0.42519 | 0.882 | 0.74 | 0.001271 | HC_vs_MS.Trt_F |
| VWF | 8.43E-08 | 1.45704 | 0.237 | 0.08 | 0.001346 | HC_vs_MS.Trt_F |
| SAMSN1 | 8.51E-08 | 0.44882 | 0.555 | 0.33 | 0.00136 | HC_vs_MS.Trt_F |
| SPP2 | 8.83E-08 | 0.51831 | 0.809 | 0.63 | 0.00141 | HC_vs_MS.Trt_F |
| SF1 | 1.19E-07 | 0.9421 | 0.604 | 0.42 | 0.001908 | HC_vs_MS.Trt_F |
| CA4 | 1.31E-07 | 0.56693 | 0.211 | 0.07 | 0.002087 | HC_vs_MS.Trt_F |
| MT-CO2 | 1.39E-07 | -0.5155 | 0.983 | 1 | 0.00222 | HC_vs_MS.Trt_F |
| FOS | 1.43E-07 | 1.15091 | 0.451 | 0.27 | 0.002288 | HC_vs_MS.Trt_F |
| RP11-47I22. | 1.50E-07 | 0.49949 | 0.679 | 0.48 | 0.002393 | HC_vs_MS.Trt_F |
| SLTM | 1.69E-07 | 0.52324 | 0.471 | 0.27 | 0.0027 | HC_vs_MS.Trt_F |
| SP100 | 1.92E-07 | 0.42329 | 0.986 | 0.89 | 0.003071 | HC_vs_MS.Trt_F |
| SHISA9 | 1.95E-07 | 0.30713 | 0.971 | 0.91 | 0.003117 | HC_vs_MS.Trt_F |
| INTU | 2.15E-07 | 0.38785 | 0.991 | 0.98 | 0.003429 | HC_vs_MS.Trt_F |
| EPAS1 | 2.49E-07 | 0.76425 | 0.272 | 0.11 | 0.003975 | HC_vs_MS.Trt_F |
| 4-Sep | 2.50E-07 | 0.54357 | 0.529 | 0.33 | 0.004 | HC_vs_MS.Trt_F |
| MDN1 | 2.85E-07 | 0.85257 | 0.324 | 0.15 | 0.004553 | HC_vs_MS.Trt_F |
| CHMP1B | 2.89E-07 | 0.39318 | 0.855 | 0.69 | 0.004616 | HC_vs_MS.Trt_F |
| KYNU | 3.15E-07 | 0.3313 | 0.983 | 0.96 | 0.005038 | HC_vs_MS.Trt_F |

| | | | | | | |
|---|---|---|---|---|---|---|
| ITGB3BP | 3.33E-07 | 0.69368 | 0.78 | 0.67 | 0.005318 | HC_vs_MS.Trt_F |
| UGGT2 | 3.48E-07 | 0.44272 | 0.908 | 0.81 | 0.005557 | HC_vs_MS.Trt_F |
| SET | 3.48E-07 | -0.7544 | 0.829 | 0.87 | 0.005563 | HC_vs_MS.Trt_F |
| MAPK10 | 3.86E-07 | 0.82169 | 0.332 | 0.16 | 0.006172 | HC_vs_MS.Trt_F |
| NCOR2 | 3.91E-07 | 0.92636 | 0.552 | 0.36 | 0.006248 | HC_vs_MS.Trt_F |
| HSPG2 | 3.92E-07 | 1.4607 | 0.535 | 0.34 | 0.006258 | HC_vs_MS.Trt_F |
| RGPD6 | 3.94E-07 | 0.79423 | 0.685 | 0.52 | 0.006297 | HC_vs_MS.Trt_F |
| RPL34 | 4.04E-07 | -0.5355 | 0.942 | 0.95 | 0.006448 | HC_vs_MS.Trt_F |
| FMNL2 | 4.10E-07 | 0.66786 | 0.347 | 0.17 | 0.006543 | HC_vs_MS.Trt_F |
| SMCHD1 | 4.44E-07 | 0.62513 | 0.789 | 0.61 | 0.007099 | HC_vs_MS.Trt_F |
| CAPZA1 | 5.05E-07 | -0.8277 | 0.818 | 0.86 | 0.008074 | HC_vs_MS.Trt_F |
| TAGLN2 | 5.18E-07 | 0.50487 | 0.653 | 0.47 | 0.008267 | HC_vs_MS.Trt_F |
| SLAMF1 | 5.21E-07 | 0.74902 | 0.405 | 0.22 | 0.008322 | HC_vs_MS.Trt_F |
| TAP2 | 5.82E-07 | 0.31916 | 0.379 | 0.18 | 0.009302 | HC_vs_MS.Trt_F |
| MIPOL1 | 6.00E-07 | 0.35822 | 1 | 0.99 | 0.009579 | HC_vs_MS.Trt_F |
| RAPGEF5 | 8.57E-07 | 1.07041 | 0.101 | 0.01 | 0.013685 | HC_vs_MS.Trt_F |
| CRLS1 | 1.06E-06 | 0.38585 | 0.483 | 0.29 | 0.016867 | HC_vs_MS.Trt_F |
| HLA-B | 1.10E-06 | 0.43869 | 0.919 | 0.88 | 0.017624 | HC_vs_MS.Trt_F |
| SPARC | 1.30E-06 | 0.5166 | 0.249 | 0.1 | 0.02072 | HC_vs_MS.Trt_F |
| NEDD4 | 1.39E-06 | 0.47142 | 0.983 | 0.93 | 0.022188 | HC_vs_MS.Trt_F |
| LTBR | 1.40E-06 | 0.41181 | 0.951 | 0.88 | 0.022377 | HC_vs_MS.Trt_F |
| LGALS3BP | 1.53E-06 | 1.18501 | 0.711 | 0.5 | 0.02441 | HC_vs_MS.Trt_F |
| RPS3A | 1.56E-06 | -0.434 | 0.882 | 0.92 | 0.02489 | HC_vs_MS.Trt_F |
| C10orf11 | 1.66E-06 | 0.41365 | 0.986 | 0.97 | 0.026472 | HC_vs_MS.Trt_F |
| SLC2A3 | 1.90E-06 | 0.52054 | 0.908 | 0.85 | 0.030377 | HC_vs_MS.Trt_F |
| ARHGAP9 | 2.01E-06 | 0.36953 | 0.624 | 0.46 | 0.032134 | HC_vs_MS.Trt_F |
| XRCC2 | 2.14E-06 | 0.46726 | 0.858 | 0.73 | 0.034235 | HC_vs_MS.Trt_F |
| GNG11 | 2.15E-06 | 0.79775 | 0.136 | 0.03 | 0.03434 | HC_vs_MS.Trt_F |
| LPP | 2.16E-06 | 0.81429 | 0.988 | 0.96 | 0.034507 | HC_vs_MS.Trt_F |
| ADGRE2 | 2.48E-06 | 0.89021 | 0.855 | 0.72 | 0.039629 | HC_vs_MS.Trt_F |
| SLC25A44 | 2.53E-06 | -0.3452 | 0.754 | 0.59 | 0.040339 | HC_vs_MS.Trt_F |
| GPR146 | 2.57E-06 | 0.63303 | 0.237 | 0.09 | 0.041086 | HC_vs_MS.Trt_F |
| CCNH | 2.58E-06 | 0.38031 | 0.405 | 0.22 | 0.041263 | HC_vs_MS.Trt_F |
| ACIN1 | 2.67E-06 | 0.79016 | 0.633 | 0.46 | 0.042595 | HC_vs_MS.Trt_F |
| BUB1B | 2.67E-06 | -0.6924 | 0.448 | 0.27 | 0.042726 | HC_vs_MS.Trt_F |
| KIAA1462 | 2.70E-06 | 0.79406 | 0.39 | 0.22 | 0.043122 | HC_vs_MS.Trt_F |
| LDLRAD4 | 2.82E-06 | 0.78667 | 0.538 | 0.35 | 0.044973 | HC_vs_MS.Trt_F |
| USP36 | 2.84E-06 | 0.63745 | 0.535 | 0.35 | 0.045303 | HC_vs_MS.Trt_F |
| C4orf36 | 2.98E-06 | 0.39958 | 0.908 | 0.82 | 0.047554 | HC_vs_MS.Trt_F |
| EPPIN | 3.10E-06 | 0.45402 | 0.627 | 0.43 | 0.049478 | HC_vs_MS.Trt_F |

| | | | | | | |
|---|---|---|---|---|---|---|
| SRGN | 3.22E-06 | 0.50072 | 0.642 | 0.47 | 0.051396 | HC_vs_MS.Trt_F |
| ENG | 3.46E-06 | 0.72706 | 0.205 | 0.08 | 0.055328 | HC_vs_MS.Trt_F |
| RBBP6 | 3.47E-06 | 0.53152 | 0.327 | 0.15 | 0.055363 | HC_vs_MS.Trt_F |
| LRRK1 | 3.50E-06 | 0.4604 | 0.85 | 0.73 | 0.055979 | HC_vs_MS.Trt_F |
| MPZL3 | 3.87E-06 | 0.98561 | 0.743 | 0.63 | 0.06189 | HC_vs_MS.Trt_F |
| FAM213A | 4.35E-06 | 0.25107 | 0.893 | 0.77 | 0.069551 | HC_vs_MS.Trt_F |
| SLC44A5 | 4.70E-06 | -0.4655 | 0.749 | 0.82 | 0.075127 | HC_vs_MS.Trt_F |
| ANXA1 | 4.91E-06 | 0.93667 | 0.422 | 0.27 | 0.078419 | HC_vs_MS.Trt_F |
| GRB10 | 4.96E-06 | 1.25803 | 0.341 | 0.18 | 0.079196 | HC_vs_MS.Trt_F |
| PARP15 | 6.02E-06 | 0.52331 | 0.942 | 0.85 | 0.096109 | HC_vs_MS.Trt_F |
| ENTPD1 | 6.04E-06 | 0.3018 | 0.971 | 0.94 | 0.096521 | HC_vs_MS.Trt_F |
| TCF4 | 6.43E-06 | 0.40243 | 0.818 | 0.67 | 0.102785 | HC_vs_MS.Trt_F |
| SUPT5H | 6.94E-06 | 1.09069 | 0.538 | 0.36 | 0.110909 | HC_vs_MS.Trt_F |
| MECP2 | 7.20E-06 | 0.647 | 0.962 | 0.92 | 0.115046 | HC_vs_MS.Trt_F |
| PPP1CB | 7.36E-06 | -0.9109 | 0.749 | 0.85 | 0.117557 | HC_vs_MS.Trt_F |
| ARHGAP45 | 8.21E-06 | -0.6041 | 0.61 | 0.72 | 0.131183 | HC_vs_MS.Trt_F |
| KLF6 | 9.15E-06 | 0.52982 | 0.798 | 0.68 | 0.146162 | HC_vs_MS.Trt_F |
| TNFSF13B | 9.53E-06 | -0.3068 | 0.642 | 0.45 | 0.152285 | HC_vs_MS.Trt_F |
| APELA | 9.88E-06 | 0.27529 | 0.431 | 0.26 | 0.157874 | HC_vs_MS.Trt_F |
| TRERF1 | 1.01E-05 | 1.16229 | 0.251 | 0.11 | 0.16141 | HC_vs_MS.Trt_F |
| NAMPT | 1.02E-05 | 0.69761 | 0.824 | 0.72 | 0.16255 | HC_vs_MS.Trt_F |
| UBC | 1.02E-05 | 0.50061 | 0.578 | 0.38 | 0.162884 | HC_vs_MS.Trt_F |
| RABGAP1L | 1.03E-05 | 0.5176 | 0.913 | 0.84 | 0.164461 | HC_vs_MS.Trt_F |
| PER1 | 1.06E-05 | 1.13996 | 0.587 | 0.45 | 0.169791 | HC_vs_MS.Trt_F |
| NRBP1 | 1.11E-05 | 0.65105 | 0.668 | 0.5 | 0.177577 | HC_vs_MS.Trt_F |
| PLA2G7 | 1.13E-05 | 0.62962 | 0.647 | 0.48 | 0.180137 | HC_vs_MS.Trt_F |
| SIM1 | 1.16E-05 | -0.8331 | 0.312 | 0.16 | 0.184786 | HC_vs_MS.Trt_F |
| TATDN1 | 1.18E-05 | 1.28813 | 0.783 | 0.64 | 0.188682 | HC_vs_MS.Trt_F |
| TSKU | 1.27E-05 | 0.72597 | 0.587 | 0.41 | 0.202726 | HC_vs_MS.Trt_F |
| CYB5R4 | 1.28E-05 | 0.44423 | 0.922 | 0.89 | 0.203681 | HC_vs_MS.Trt_F |
| RAB12 | 1.34E-05 | 1.54462 | 0.361 | 0.23 | 0.21412 | HC_vs_MS.Trt_F |
| SFN | 1.42E-05 | 0.85476 | 0.353 | 0.21 | 0.227472 | HC_vs_MS.Trt_F |
| CRIP2 | 1.44E-05 | 0.40946 | 0.275 | 0.13 | 0.230562 | HC_vs_MS.Trt_F |
| CERCAM | 1.45E-05 | 0.34971 | 0.988 | 0.97 | 0.232175 | HC_vs_MS.Trt_F |
| REV3L | 1.47E-05 | 0.5803 | 0.74 | 0.6 | 0.234909 | HC_vs_MS.Trt_F |
| HPGD | 1.48E-05 | 0.83168 | 0.275 | 0.13 | 0.236926 | HC_vs_MS.Trt_F |
| ROBO1 | 1.60E-05 | -0.2656 | 0.512 | 0.65 | 0.255595 | HC_vs_MS.Trt_F |
| ASTN2 | 1.68E-05 | 0.38864 | 0.977 | 0.87 | 0.269136 | HC_vs_MS.Trt_F |
| NOTCH2NL | 1.69E-05 | 1.28183 | 0.292 | 0.15 | 0.270017 | HC_vs_MS.Trt_F |
| SRRM2 | 1.77E-05 | 0.4522 | 0.754 | 0.61 | 0.282329 | HC_vs_MS.Trt_F |

| | | | | | | |
|---|---|---|---|---|---|---|
| PYGL | 1.80E-05 | 0.54057 | 0.884 | 0.82 | 0.286992 | HC_vs_MS.Trt_F |
| SNRNP70 | 1.83E-05 | 1.06663 | 0.329 | 0.18 | 0.291831 | HC_vs_MS.Trt_F |
| HNRNPC | 1.90E-05 | 0.30073 | 0.809 | 0.69 | 0.303345 | HC_vs_MS.Trt_F |
| PI4KA | 1.94E-05 | 0.68604 | 0.373 | 0.22 | 0.309825 | HC_vs_MS.Trt_F |
| TNS1 | 1.95E-05 | 2.1413 | 0.39 | 0.23 | 0.311364 | HC_vs_MS.Trt_F |
| WISP2 | 1.96E-05 | -0.42 | 0.685 | 0.51 | 0.312504 | HC_vs_MS.Trt_F |
| CFAP69 | 2.00E-05 | 0.35998 | 0.873 | 0.75 | 0.319728 | HC_vs_MS.Trt_F |
| BST2 | 2.04E-05 | 0.59272 | 0.309 | 0.16 | 0.325483 | HC_vs_MS.Trt_F |
| FRYL | 2.07E-05 | 0.49264 | 0.792 | 0.67 | 0.330343 | HC_vs_MS.Trt_F |
| REL | 2.08E-05 | 0.28813 | 0.983 | 0.96 | 0.33174 | HC_vs_MS.Trt_F |
| MYCBPAP | 2.08E-05 | 0.28888 | 0.922 | 0.79 | 0.332533 | HC_vs_MS.Trt_F |
| LTBP1 | 2.17E-05 | 0.75673 | 0.107 | 0.02 | 0.34585 | HC_vs_MS.Trt_F |
| FRK | 2.24E-05 | 0.30463 | 0.997 | 0.98 | 0.357732 | HC_vs_MS.Trt_F |
| SOD2 | 2.29E-05 | 0.27071 | 1 | 1 | 0.365375 | HC_vs_MS.Trt_F |
| TF | 2.39E-05 | 0.31595 | 0.942 | 0.89 | 0.381825 | HC_vs_MS.Trt_F |
| WNK1 | 2.51E-05 | 0.66007 | 0.613 | 0.46 | 0.400697 | HC_vs_MS.Trt_F |
| VPS13C | 2.52E-05 | 0.725 | 0.838 | 0.72 | 0.401892 | HC_vs_MS.Trt_F |
| CYP4F11 | 2.63E-05 | 0.32061 | 0.46 | 0.3 | 0.420584 | HC_vs_MS.Trt_F |
| RIMS1 | 2.69E-05 | 0.52826 | 0.361 | 0.21 | 0.430432 | HC_vs_MS.Trt_F |
| MBNL1 | 2.71E-05 | 0.26728 | 0.899 | 0.75 | 0.432485 | HC_vs_MS.Trt_F |
| CATIP | 2.71E-05 | -0.3282 | 0.882 | 0.89 | 0.432969 | HC_vs_MS.Trt_F |
| ZNF670-ZNF | 2.85E-05 | 0.51463 | 0.671 | 0.51 | 0.455831 | HC_vs_MS.Trt_F |
| PTPN2 | 3.05E-05 | 0.59668 | 0.997 | 0.96 | 0.486402 | HC_vs_MS.Trt_F |
| TTLL5 | 3.10E-05 | 0.51589 | 0.749 | 0.59 | 0.495872 | HC_vs_MS.Trt_F |
| MAML2 | 3.11E-05 | 1.54641 | 0.272 | 0.14 | 0.497368 | HC_vs_MS.Trt_F |
| TSPAN3 | 3.27E-05 | 0.61905 | 0.529 | 0.35 | 0.522271 | HC_vs_MS.Trt_F |
| JAK3 | 3.51E-05 | -0.6274 | 0.731 | 0.81 | 0.560742 | HC_vs_MS.Trt_F |
| PDE4D | 3.76E-05 | 0.47707 | 0.405 | 0.25 | 0.600095 | HC_vs_MS.Trt_F |
| WTIP | 3.88E-05 | 0.35429 | 0.986 | 0.94 | 0.620344 | HC_vs_MS.Trt_F |
| BACH2 | 3.97E-05 | 1.13661 | 0.292 | 0.16 | 0.633574 | HC_vs_MS.Trt_F |
| CEP83 | 4.06E-05 | 0.83509 | 0.754 | 0.65 | 0.648773 | HC_vs_MS.Trt_F |
| IFITM3 | 4.15E-05 | 0.79529 | 0.688 | 0.53 | 0.663174 | HC_vs_MS.Trt_F |
| CABIN1 | 4.24E-05 | 0.98019 | 0.37 | 0.22 | 0.677008 | HC_vs_MS.Trt_F |
| PHTF2 | 4.25E-05 | 0.44227 | 0.552 | 0.37 | 0.679578 | HC_vs_MS.Trt_F |
| RBM12 | 4.27E-05 | 0.55281 | 0.289 | 0.15 | 0.682636 | HC_vs_MS.Trt_F |
| RSRP1 | 4.28E-05 | 0.51437 | 0.945 | 0.91 | 0.683871 | HC_vs_MS.Trt_F |
| MT-ND2 | 4.40E-05 | 0.36241 | 0.908 | 0.84 | 0.702733 | HC_vs_MS.Trt_F |
| KAT6A | 4.54E-05 | 0.3408 | 0.824 | 0.71 | 0.724671 | HC_vs_MS.Trt_F |
| PRRC2C | 4.57E-05 | 0.30669 | 0.847 | 0.69 | 0.730333 | HC_vs_MS.Trt_F |
| SH2D2A | 4.68E-05 | 0.91695 | 0.488 | 0.35 | 0.747459 | HC_vs_MS.Trt_F |

| | | | | | | |
|---|---|---|---|---|---|---|
| KRAS | 4.72E-05 | 0.34394 | 0.483 | 0.32 | 0.753406 | HC_vs_MS.Trt_F |
| BVES | 4.74E-05 | 0.47164 | 0.772 | 0.64 | 0.756661 | HC_vs_MS.Trt_F |
| CCS | 4.92E-05 | 0.99786 | 0.329 | 0.18 | 0.786236 | HC_vs_MS.Trt_F |
| HLA-DRB5 | 5.01E-05 | -0.8485 | 0.246 | 0.12 | 0.800537 | HC_vs_MS.Trt_F |
| MTM1 | 5.05E-05 | 0.25591 | 0.228 | 0.1 | 0.807393 | HC_vs_MS.Trt_F |
| NEBL | 5.10E-05 | 0.45564 | 0.37 | 0.22 | 0.814044 | HC_vs_MS.Trt_F |
| EIF5A | 5.13E-05 | -0.3259 | 0.575 | 0.71 | 0.819951 | HC_vs_MS.Trt_F |
| HIVEP2 | 5.19E-05 | 1.0312 | 0.382 | 0.23 | 0.828936 | HC_vs_MS.Trt_F |
| SYT17 | 5.28E-05 | 0.34664 | 0.439 | 0.28 | 0.842942 | HC_vs_MS.Trt_F |
| ATP10B | 5.29E-05 | 0.29962 | 0.824 | 0.69 | 0.845492 | HC_vs_MS.Trt_F |
| LFNG | 5.36E-05 | 0.69286 | 0.344 | 0.2 | 0.856561 | HC_vs_MS.Trt_F |
| NRP1 | 5.43E-05 | -0.4707 | 0.465 | 0.3 | 0.867556 | HC_vs_MS.Trt_F |
| CCDC171 | 5.46E-05 | 1.34007 | 0.569 | 0.41 | 0.872556 | HC_vs_MS.Trt_F |
| ITGB2 | 5.80E-05 | 0.7408 | 0.685 | 0.58 | 0.925855 | HC_vs_MS.Trt_F |
| TCHP | 5.86E-05 | 0.50703 | 0.613 | 0.45 | 0.93556 | HC_vs_MS.Trt_F |
| MOV10 | 5.89E-05 | 0.68654 | 0.584 | 0.42 | 0.940542 | HC_vs_MS.Trt_F |
| SFT2D2 | 5.95E-05 | 0.45022 | 0.991 | 0.97 | 0.9511 | HC_vs_MS.Trt_F |
| FAM49B | 6.05E-05 | 0.42869 | 0.986 | 0.96 | 0.96642 | HC_vs_MS.Trt_F |
| ODF2L | 6.05E-05 | 0.55823 | 0.986 | 0.94 | 0.966537 | HC_vs_MS.Trt_F |
| JMJD1C | 6.07E-05 | 0.79469 | 0.373 | 0.23 | 0.969039 | HC_vs_MS.Trt_F |
| EREG | 6.17E-05 | 0.767 | 0.853 | 0.76 | 0.985094 | HC_vs_MS.Trt_F |
| LRRTM4 | 6.30E-05 | 0.41959 | 0.633 | 0.48 | 1 | HC_vs_MS.Trt_F |
| PHKA2 | 6.38E-05 | 0.33623 | 0.734 | 0.6 | 1 | HC_vs_MS.Trt_F |
| SLC9A8 | 6.49E-05 | 0.44541 | 0.26 | 0.13 | 1 | HC_vs_MS.Trt_F |
| IFRD1 | 6.51E-05 | 0.46976 | 0.564 | 0.42 | 1 | HC_vs_MS.Trt_F |
| PDE4B | 6.53E-05 | 0.69965 | 0.445 | 0.3 | 1 | HC_vs_MS.Trt_F |
| EPHB4 | 6.60E-05 | -0.4298 | 0.751 | 0.81 | 1 | HC_vs_MS.Trt_F |
| BORCS5 | 6.60E-05 | 0.73933 | 0.74 | 0.6 | 1 | HC_vs_MS.Trt_F |
| ABCC9 | 6.75E-05 | 0.30473 | 0.98 | 0.92 | 1 | HC_vs_MS.Trt_F |
| TMEM182 | 7.01E-05 | 0.46933 | 0.78 | 0.66 | 1 | HC_vs_MS.Trt_F |
| KIAA1217 | 7.17E-05 | 0.90823 | 0.165 | 0.07 | 1 | HC_vs_MS.Trt_F |
| SETD1B | 7.23E-05 | 1.13046 | 0.249 | 0.13 | 1 | HC_vs_MS.Trt_F |
| GPR141 | 7.26E-05 | 0.26296 | 0.598 | 0.44 | 1 | HC_vs_MS.Trt_F |
| TBC1D1 | 7.50E-05 | 0.36381 | 0.688 | 0.52 | 1 | HC_vs_MS.Trt_F |
| ATN1 | 7.79E-05 | 0.61512 | 0.26 | 0.13 | 1 | HC_vs_MS.Trt_F |
| TAF7 | 7.83E-05 | 0.60274 | 0.448 | 0.29 | 1 | HC_vs_MS.Trt_F |
| C12orf50 | 7.90E-05 | 0.2969 | 0.922 | 0.89 | 1 | HC_vs_MS.Trt_F |
| RALGAPB | 8.09E-05 | 0.31082 | 0.332 | 0.18 | 1 | HC_vs_MS.Trt_F |
| RHOBTB3 | 8.09E-05 | 0.34547 | 0.749 | 0.57 | 1 | HC_vs_MS.Trt_F |
| KATNAL2 | 8.35E-05 | 0.85012 | 0.416 | 0.26 | 1 | HC_vs_MS.Trt_F |

| | | | | | | |
|---|---|---|---|---|---|---|
| SF3A1 | 8.42E-05 | 0.84613 | 0.425 | 0.28 | 1 | HC_vs_MS.Trt_F |
| CHTF8 | 8.54E-05 | 0.62168 | 0.301 | 0.16 | 1 | HC_vs_MS.Trt_F |
| PIAS3 | 8.60E-05 | 0.58624 | 0.249 | 0.12 | 1 | HC_vs_MS.Trt_F |
| EIF4G1 | 8.62E-05 | 0.46252 | 0.613 | 0.48 | 1 | HC_vs_MS.Trt_F |
| SLC11A2 | 8.87E-05 | 0.51421 | 0.801 | 0.68 | 1 | HC_vs_MS.Trt_F |
| JUND | 8.90E-05 | 0.43384 | 0.474 | 0.32 | 1 | HC_vs_MS.Trt_F |
| TMEM43 | 9.14E-05 | 0.72224 | 0.344 | 0.2 | 1 | HC_vs_MS.Trt_F |
| ETFDH | 9.21E-05 | -0.2561 | 0.74 | 0.62 | 1 | HC_vs_MS.Trt_F |
| ZDHHC13 | 9.25E-05 | -0.796 | 0.803 | 0.66 | 1 | HC_vs_MS.Trt_F |
| KMT2D | 1.01E-04 | 0.8306 | 0.87 | 0.84 | 1 | HC_vs_MS.Trt_F |
| HTRA4 | 1.01E-04 | 0.36015 | 0.535 | 0.37 | 1 | HC_vs_MS.Trt_F |
| AIF1L | 1.04E-04 | 0.4886 | 0.11 | 0.03 | 1 | HC_vs_MS.Trt_F |
| RCL1 | 1.06E-04 | 0.39199 | 0.546 | 0.4 | 1 | HC_vs_MS.Trt_F |
| TPP1 | 1.07E-04 | 0.39436 | 0.477 | 0.32 | 1 | HC_vs_MS.Trt_F |
| TMOD3 | 1.10E-04 | 0.2508 | 0.965 | 0.91 | 1 | HC_vs_MS.Trt_F |
| WFDC1 | 1.15E-04 | 0.2883 | 0.835 | 0.76 | 1 | HC_vs_MS.Trt_F |
| KLF7 | 1.15E-04 | -0.5127 | 0.266 | 0.13 | 1 | HC_vs_MS.Trt_F |
| SPATS2L | 1.17E-04 | 0.29571 | 0.968 | 0.94 | 1 | HC_vs_MS.Trt_F |
| FLVCR1 | 1.17E-04 | 0.86013 | 0.702 | 0.54 | 1 | HC_vs_MS.Trt_F |
| RP11-467J12 | 1.19E-04 | 0.34616 | 0.607 | 0.45 | 1 | HC_vs_MS.Trt_F |
| RPS6 | 1.20E-04 | -0.3134 | 0.919 | 0.93 | 1 | HC_vs_MS.Trt_F |
| HEATR5B | 1.21E-04 | 0.41448 | 0.523 | 0.35 | 1 | HC_vs_MS.Trt_F |
| NFKBIA | 1.22E-04 | 0.58816 | 0.468 | 0.31 | 1 | HC_vs_MS.Trt_F |
| ARF6 | 1.22E-04 | -0.8313 | 0.592 | 0.7 | 1 | HC_vs_MS.Trt_F |
| NUPL2 | 1.22E-04 | 0.3104 | 0.991 | 0.96 | 1 | HC_vs_MS.Trt_F |
| PLCB2 | 1.24E-04 | 0.91862 | 0.327 | 0.19 | 1 | HC_vs_MS.Trt_F |
| LIMS3 | 1.24E-04 | 0.26845 | 0.647 | 0.53 | 1 | HC_vs_MS.Trt_F |
| TXNRD1 | 1.24E-04 | 1.31928 | 0.376 | 0.23 | 1 | HC_vs_MS.Trt_F |
| CCR8 | 1.28E-04 | 1.0225 | 0.257 | 0.13 | 1 | HC_vs_MS.Trt_F |
| HSP90B1 | 1.33E-04 | 0.32217 | 0.405 | 0.26 | 1 | HC_vs_MS.Trt_F |
| BAZ2A | 0.00013 | 0.63117 | 0.731 | 0.59 | 1 | HC_vs_MS.Trt_F |
| TNIK | 1.33E-04 | 0.63187 | 0.451 | 0.31 | 1 | HC_vs_MS.Trt_F |
| GABARAPL1 | 1.33E-04 | 1.09232 | 0.251 | 0.13 | 1 | HC_vs_MS.Trt_F |
| IRF1 | 1.33E-04 | 0.45607 | 0.922 | 0.85 | 1 | HC_vs_MS.Trt_F |
| H2AFY | 1.35E-04 | 0.30087 | 0.321 | 0.18 | 1 | HC_vs_MS.Trt_F |
| COL28A1 | 1.36E-04 | 0.47694 | 0.488 | 0.34 | 1 | HC_vs_MS.Trt_F |
| TUBA1B | 1.37E-04 | 0.25627 | 0.442 | 0.28 | 1 | HC_vs_MS.Trt_F |
| CALD1 | 1.38E-04 | 1.18353 | 0.327 | 0.19 | 1 | HC_vs_MS.Trt_F |
| VNN3 | 1.39E-04 | 0.67349 | 0.477 | 0.32 | 1 | HC_vs_MS.Trt_F |
| GSTO2 | 1.40E-04 | 0.42074 | 0.896 | 0.77 | 1 | HC_vs_MS.Trt_F |

| | | | | | | |
|---|---|---|---|---|---|---|
| NOTCH2 | 1.42E-04 | 0.92162 | 0.309 | 0.17 | 1 | HC_vs_MS.Trt_F |
| GTPBP1 | 1.51E-04 | 0.65636 | 0.604 | 0.49 | 1 | HC_vs_MS.Trt_F |
| PFKP | 1.52E-04 | 0.30911 | 0.387 | 0.23 | 1 | HC_vs_MS.Trt_F |
| TTC39C | 1.55E-04 | -0.9098 | 0.824 | 0.82 | 1 | HC_vs_MS.Trt_F |
| PIP4K2A | 1.58E-04 | -0.6364 | 0.72 | 0.82 | 1 | HC_vs_MS.Trt_F |
| EHD2 | 1.60E-04 | 0.91774 | 0.884 | 0.8 | 1 | HC_vs_MS.Trt_F |
| RGR | 1.62E-04 | 0.64853 | 0.529 | 0.35 | 1 | HC_vs_MS.Trt_F |
| SLC30A3 | 1.63E-04 | 0.258 | 0.529 | 0.36 | 1 | HC_vs_MS.Trt_F |
| TUBA1A | 1.64E-04 | 1.06414 | 0.361 | 0.23 | 1 | HC_vs_MS.Trt_F |
| DDX21 | 1.70E-04 | 0.60735 | 0.393 | 0.25 | 1 | HC_vs_MS.Trt_F |
| NR4A3 | 1.77E-04 | 1.39177 | 0.246 | 0.13 | 1 | HC_vs_MS.Trt_F |
| PACS1 | 1.77E-04 | 0.43457 | 0.853 | 0.79 | 1 | HC_vs_MS.Trt_F |
| RPL6 | 1.82E-04 | -0.3644 | 0.945 | 0.95 | 1 | HC_vs_MS.Trt_F |
| ZSCAN30 | 1.89E-04 | 0.25346 | 0.835 | 0.67 | 1 | HC_vs_MS.Trt_F |
| CREM | 1.96E-04 | 0.66266 | 0.445 | 0.31 | 1 | HC_vs_MS.Trt_F |
| DAAM1 | 1.97E-04 | -0.3249 | 0.988 | 0.98 | 1 | HC_vs_MS.Trt_F |
| PPP2R1A | 1.99E-04 | 0.32979 | 0.775 | 0.64 | 1 | HC_vs_MS.Trt_F |
| JAK2 | 2.00E-04 | 0.9393 | 0.303 | 0.17 | 1 | HC_vs_MS.Trt_F |
| ARID1A | 2.01E-04 | 0.37927 | 0.965 | 0.91 | 1 | HC_vs_MS.Trt_F |
| AF011889.5 | 2.03E-04 | 0.51133 | 0.295 | 0.17 | 1 | HC_vs_MS.Trt_F |
| ACBD6 | 2.03E-04 | 0.54081 | 0.477 | 0.34 | 1 | HC_vs_MS.Trt_F |
| SETD1A | 2.04E-04 | 1.1601 | 0.168 | 0.07 | 1 | HC_vs_MS.Trt_F |
| VPS36 | 2.04E-04 | 0.31419 | 0.358 | 0.22 | 1 | HC_vs_MS.Trt_F |
| NEGR1 | 2.06E-04 | 0.4599 | 0.873 | 0.76 | 1 | HC_vs_MS.Trt_F |
| SYTL3 | 2.06E-04 | 0.39944 | 0.384 | 0.24 | 1 | HC_vs_MS.Trt_F |
| GDPD5 | 2.08E-04 | 0.91824 | 0.332 | 0.19 | 1 | HC_vs_MS.Trt_F |
| TMED3 | 2.12E-04 | 0.28149 | 0.699 | 0.56 | 1 | HC_vs_MS.Trt_F |
| FER | 2.16E-04 | 0.71533 | 0.575 | 0.42 | 1 | HC_vs_MS.Trt_F |
| CFAP74 | 2.28E-04 | 0.27812 | 0.882 | 0.8 | 1 | HC_vs_MS.Trt_F |
| MGAT4C | 2.33E-04 | 0.38551 | 0.835 | 0.72 | 1 | HC_vs_MS.Trt_F |
| CDH8 | 2.36E-04 | 0.67192 | 0.514 | 0.36 | 1 | HC_vs_MS.Trt_F |
| LTBP3 | 2.38E-04 | 0.5087 | 0.757 | 0.62 | 1 | HC_vs_MS.Trt_F |
| SLC38A2 | 2.44E-04 | 0.84139 | 0.425 | 0.28 | 1 | HC_vs_MS.Trt_F |
| EXOSC7 | 2.46E-04 | 0.61785 | 0.552 | 0.4 | 1 | HC_vs_MS.Trt_F |
| IGFBPL1 | 2.50E-04 | 0.36411 | 0.685 | 0.52 | 1 | HC_vs_MS.Trt_F |
| BAG6 | 2.61E-04 | 0.74754 | 0.691 | 0.55 | 1 | HC_vs_MS.Trt_F |
| ALPK1 | 2.61E-04 | 0.26221 | 0.806 | 0.7 | 1 | HC_vs_MS.Trt_F |
| RGPD1 | 0.00028 | 0.73241 | 0.364 | 0.22 | 1 | HC_vs_MS.Trt_F |
| EXOC3L4 | 2.77E-04 | 0.30917 | 0.254 | 0.14 | 1 | HC_vs_MS.Trt_F |
| IRS2 | 2.78E-04 | 1.5451 | 0.477 | 0.36 | 1 | HC_vs_MS.Trt_F |

| | | | | | | |
|---|---|---|---|---|---|---|
| ITGA11 | 2.80E-04 | 0.97585 | 0.708 | 0.56 | 1 | HC_vs_MS.Trt_F |
| SYNE2 | 2.82E-04 | 0.38456 | 0.962 | 0.94 | 1 | HC_vs_MS.Trt_F |
| SPDYE2 | 2.86E-04 | -0.4065 | 0.538 | 0.63 | 1 | HC_vs_MS.Trt_F |
| SNX9 | 2.92E-04 | 0.55735 | 0.254 | 0.14 | 1 | HC_vs_MS.Trt_F |
| ATP5B | 2.96E-04 | 0.41102 | 0.488 | 0.35 | 1 | HC_vs_MS.Trt_F |
| HNRNPL | 2.96E-04 | -0.7091 | 0.85 | 0.81 | 1 | HC_vs_MS.Trt_F |
| AP4E1 | 2.97E-04 | -0.2631 | 0.486 | 0.34 | 1 | HC_vs_MS.Trt_F |
| SETD5 | 3.01E-04 | 0.31544 | 0.665 | 0.52 | 1 | HC_vs_MS.Trt_F |
| SAFB2 | 3.03E-04 | 0.69369 | 0.908 | 0.83 | 1 | HC_vs_MS.Trt_F |
| TMEM91 | 0.00031 | 0.50532 | 0.598 | 0.45 | 1 | HC_vs_MS.Trt_F |
| RPL18A | 3.17E-04 | -0.3209 | 0.858 | 0.88 | 1 | HC_vs_MS.Trt_F |
| PRKCQ | 3.18E-04 | 0.29703 | 0.396 | 0.26 | 1 | HC_vs_MS.Trt_F |
| DIP2A | 3.25E-04 | 0.64238 | 0.884 | 0.81 | 1 | HC_vs_MS.Trt_F |
| JOSD1 | 3.25E-04 | 0.62833 | 0.639 | 0.5 | 1 | HC_vs_MS.Trt_F |
| MFAP5 | 3.34E-04 | 0.46946 | 0.454 | 0.32 | 1 | HC_vs_MS.Trt_F |
| HLA-F | 3.35E-04 | 0.69953 | 0.592 | 0.48 | 1 | HC_vs_MS.Trt_F |
| NMRK1 | 3.39E-04 | -0.9628 | 0.382 | 0.22 | 1 | HC_vs_MS.Trt_F |
| CHRDL1 | 3.46E-04 | 0.32997 | 0.182 | 0.09 | 1 | HC_vs_MS.Trt_F |
| KIAA0141 | 3.51E-04 | 1.12285 | 0.249 | 0.13 | 1 | HC_vs_MS.Trt_F |
| GPX3 | 3.57E-04 | 1.17362 | 0.205 | 0.1 | 1 | HC_vs_MS.Trt_F |
| ZNF445 | 3.60E-04 | 0.4428 | 0.988 | 0.98 | 1 | HC_vs_MS.Trt_F |
| MTBP | 3.68E-04 | 0.58671 | 0.676 | 0.55 | 1 | HC_vs_MS.Trt_F |
| ARHGEF1 | 3.68E-04 | 0.435 | 0.928 | 0.89 | 1 | HC_vs_MS.Trt_F |
| DLGAP4 | 3.68E-04 | 0.71896 | 0.257 | 0.14 | 1 | HC_vs_MS.Trt_F |
| SYCP1 | 3.76E-04 | 0.34961 | 0.384 | 0.25 | 1 | HC_vs_MS.Trt_F |
| BRD9 | 3.80E-04 | 0.91142 | 0.234 | 0.13 | 1 | HC_vs_MS.Trt_F |
| TGFBR2 | 3.80E-04 | 0.63603 | 0.277 | 0.16 | 1 | HC_vs_MS.Trt_F |
| CASP8 | 3.83E-04 | 0.42835 | 0.931 | 0.91 | 1 | HC_vs_MS.Trt_F |
| CD151 | 3.87E-04 | 0.44942 | 0.942 | 0.88 | 1 | HC_vs_MS.Trt_F |
| CAV1 | 3.92E-04 | 1.02687 | 0.171 | 0.08 | 1 | HC_vs_MS.Trt_F |
| GSG1 | 3.92E-04 | 0.36097 | 0.627 | 0.5 | 1 | HC_vs_MS.Trt_F |
| PRKACB | 3.95E-04 | 0.40677 | 0.35 | 0.22 | 1 | HC_vs_MS.Trt_F |
| ZNF83 | 3.99E-04 | 0.35915 | 0.87 | 0.82 | 1 | HC_vs_MS.Trt_F |
| NLK | 4.04E-04 | 0.83536 | 0.37 | 0.23 | 1 | HC_vs_MS.Trt_F |
| OPRM1 | 0.00041 | 0.44702 | 0.89 | 0.82 | 1 | HC_vs_MS.Trt_F |
| RAB8B | 4.09E-04 | 0.55788 | 0.399 | 0.27 | 1 | HC_vs_MS.Trt_F |
| ZNF287 | 4.14E-04 | 0.41345 | 0.517 | 0.38 | 1 | HC_vs_MS.Trt_F |
| LRCH3 | 4.16E-04 | 0.4938 | 0.988 | 0.97 | 1 | HC_vs_MS.Trt_F |
| GOLGA8A | 4.22E-04 | 0.41444 | 0.509 | 0.4 | 1 | HC_vs_MS.Trt_F |
| GABPB1 | 4.32E-04 | -0.5462 | 0.581 | 0.43 | 1 | HC_vs_MS.Trt_F |

| | | | | | | |
|---|---|---|---|---|---|---|
| TUBB4B | 4.34E-04 | 0.29971 | 0.283 | 0.16 | 1 | HC_vs_MS.Trt_F |
| 11-Sep | 4.39E-04 | 0.35477 | 0.538 | 0.39 | 1 | HC_vs_MS.Trt_F |
| APOD | 4.44E-04 | 0.66356 | 0.35 | 0.23 | 1 | HC_vs_MS.Trt_F |
| ATXN2 | 0.00045 | 0.32559 | 0.934 | 0.86 | 1 | HC_vs_MS.Trt_F |
| GTF2H2C | 4.54E-04 | -0.3618 | 0.477 | 0.34 | 1 | HC_vs_MS.Trt_F |
| DOCK2 | 4.57E-04 | 0.41038 | 0.509 | 0.36 | 1 | HC_vs_MS.Trt_F |
| PTPN1 | 4.60E-04 | 0.56723 | 0.41 | 0.26 | 1 | HC_vs_MS.Trt_F |
| KMT2A | 4.65E-04 | 0.25921 | 0.962 | 0.93 | 1 | HC_vs_MS.Trt_F |
| SRSF1 | 4.73E-04 | 0.30866 | 0.624 | 0.48 | 1 | HC_vs_MS.Trt_F |
| ADGRF5 | 4.73E-04 | -1.3388 | 0.133 | 0.05 | 1 | HC_vs_MS.Trt_F |
| STK4 | 4.74E-04 | 0.29067 | 0.925 | 0.86 | 1 | HC_vs_MS.Trt_F |
| FNBP4 | 4.75E-04 | 0.76209 | 0.546 | 0.42 | 1 | HC_vs_MS.Trt_F |
| MARK3 | 4.82E-04 | 0.63523 | 0.882 | 0.82 | 1 | HC_vs_MS.Trt_F |
| SERPING1 | 4.84E-04 | 0.90094 | 0.15 | 0.07 | 1 | HC_vs_MS.Trt_F |
| ANAPC1 | 4.95E-04 | 0.7405 | 0.65 | 0.5 | 1 | HC_vs_MS.Trt_F |
| ZNF850 | 5.06E-04 | 0.30117 | 0.957 | 0.9 | 1 | HC_vs_MS.Trt_F |
| MCMDC2 | 5.13E-04 | 1.10151 | 0.488 | 0.37 | 1 | HC_vs_MS.Trt_F |
| CTBP1 | 5.27E-04 | 0.38705 | 0.532 | 0.37 | 1 | HC_vs_MS.Trt_F |
| CAPN7 | 5.39E-04 | 0.55846 | 0.624 | 0.46 | 1 | HC_vs_MS.Trt_F |
| FBXO15 | 5.40E-04 | 0.41706 | 0.665 | 0.55 | 1 | HC_vs_MS.Trt_F |
| NIPBL | 5.40E-04 | 0.54722 | 0.624 | 0.48 | 1 | HC_vs_MS.Trt_F |
| LGMN | 5.40E-04 | 0.2581 | 0.971 | 0.94 | 1 | HC_vs_MS.Trt_F |
| PEX1 | 5.40E-04 | 0.4589 | 0.792 | 0.7 | 1 | HC_vs_MS.Trt_F |
| COL1A2 | 5.48E-04 | 1.46795 | 0.532 | 0.39 | 1 | HC_vs_MS.Trt_F |
| SLITRK2 | 5.54E-04 | 0.41256 | 0.353 | 0.23 | 1 | HC_vs_MS.Trt_F |
| PRDM2 | 5.55E-04 | 0.52207 | 0.52 | 0.39 | 1 | HC_vs_MS.Trt_F |
| PKM | 5.60E-04 | 0.41498 | 0.647 | 0.48 | 1 | HC_vs_MS.Trt_F |
| PHF1 | 5.63E-04 | 1.74344 | 0.234 | 0.13 | 1 | HC_vs_MS.Trt_F |
| JUNB | 5.63E-04 | 0.30779 | 0.676 | 0.57 | 1 | HC_vs_MS.Trt_F |
| FASTKD2 | 5.69E-04 | -0.5352 | 0.668 | 0.53 | 1 | HC_vs_MS.Trt_F |
| PICALM | 5.70E-04 | 0.40541 | 0.335 | 0.2 | 1 | HC_vs_MS.Trt_F |
| UNC80 | 5.70E-04 | 0.30303 | 0.864 | 0.79 | 1 | HC_vs_MS.Trt_F |
| RPS19 | 5.72E-04 | -0.3251 | 0.928 | 0.95 | 1 | HC_vs_MS.Trt_F |
| HGSNAT | 5.76E-04 | 0.71621 | 0.772 | 0.7 | 1 | HC_vs_MS.Trt_F |
| ZZEF1 | 5.83E-04 | 0.69261 | 0.592 | 0.46 | 1 | HC_vs_MS.Trt_F |
| RASA1 | 5.84E-04 | 0.40577 | 0.312 | 0.19 | 1 | HC_vs_MS.Trt_F |
| CYP3A5 | 5.92E-04 | 0.34766 | 0.884 | 0.8 | 1 | HC_vs_MS.Trt_F |
| FKBP14 | 6.06E-04 | 0.53844 | 0.728 | 0.62 | 1 | HC_vs_MS.Trt_F |
| CDC14A | 6.10E-04 | 0.6609 | 0.303 | 0.18 | 1 | HC_vs_MS.Trt_F |
| CD28 | 6.23E-04 | 0.34773 | 0.65 | 0.55 | 1 | HC_vs_MS.Trt_F |

| | | | | | | |
|---|---|---|---|---|---|---|
| FNTA | 6.38E-04 | 0.55826 | 0.896 | 0.82 | 1 | HC_vs_MS.Trt_F |
| RPL7 | 6.44E-04 | -0.3343 | 0.844 | 0.84 | 1 | HC_vs_MS.Trt_F |
| B3GAT1 | 6.55E-04 | 0.77716 | 0.312 | 0.19 | 1 | HC_vs_MS.Trt_F |
| SRSF3 | 6.57E-04 | -0.5961 | 0.902 | 0.9 | 1 | HC_vs_MS.Trt_F |
| CD40LG | 6.64E-04 | 1.13049 | 0.266 | 0.16 | 1 | HC_vs_MS.Trt_F |
| TEP1 | 6.81E-04 | 1.22067 | 0.488 | 0.37 | 1 | HC_vs_MS.Trt_F |
| SLC35C2 | 7.22E-04 | 0.48966 | 0.63 | 0.48 | 1 | HC_vs_MS.Trt_F |
| MFSD14C | 7.28E-04 | 0.64235 | 0.746 | 0.62 | 1 | HC_vs_MS.Trt_F |
| PPP1R15A | 7.29E-04 | 0.55867 | 0.702 | 0.65 | 1 | HC_vs_MS.Trt_F |
| CDK6 | 7.33E-04 | 0.98685 | 0.257 | 0.15 | 1 | HC_vs_MS.Trt_F |
| YTHDF3 | 7.48E-04 | 0.52174 | 0.754 | 0.64 | 1 | HC_vs_MS.Trt_F |
| ARMC9 | 7.58E-04 | 0.34721 | 0.757 | 0.69 | 1 | HC_vs_MS.Trt_F |
| RNF19A | 7.61E-04 | 0.65281 | 0.506 | 0.4 | 1 | HC_vs_MS.Trt_F |
| ULK4 | 7.86E-04 | 1.43013 | 0.431 | 0.29 | 1 | HC_vs_MS.Trt_F |
| EFCAB11 | 7.88E-04 | 0.27902 | 0.873 | 0.8 | 1 | HC_vs_MS.Trt_F |
| CDK5RAP1 | 7.92E-04 | 0.37903 | 0.48 | 0.34 | 1 | HC_vs_MS.Trt_F |
| DTX2 | 7.93E-04 | 0.64469 | 0.506 | 0.37 | 1 | HC_vs_MS.Trt_F |
| CLEC7A | 7.95E-04 | 0.8115 | 0.665 | 0.52 | 1 | HC_vs_MS.Trt_F |
| AKT2 | 7.95E-04 | 0.47745 | 0.968 | 0.95 | 1 | HC_vs_MS.Trt_F |
| CASC3 | 8.09E-04 | 0.27646 | 0.697 | 0.53 | 1 | HC_vs_MS.Trt_F |
| ATF7 | 8.10E-04 | 0.42115 | 0.538 | 0.4 | 1 | HC_vs_MS.Trt_F |
| PDLIM2 | 8.20E-04 | 0.40412 | 0.772 | 0.63 | 1 | HC_vs_MS.Trt_F |
| TFPI | 8.30E-04 | 0.28437 | 0.948 | 0.89 | 1 | HC_vs_MS.Trt_F |
| CBX6 | 8.38E-04 | -0.7356 | 0.532 | 0.6 | 1 | HC_vs_MS.Trt_F |
| RHBDD2 | 8.52E-04 | 0.39783 | 0.367 | 0.24 | 1 | HC_vs_MS.Trt_F |
| FLNB | 8.61E-04 | 0.26476 | 0.471 | 0.35 | 1 | HC_vs_MS.Trt_F |
| CAPS2 | 8.69E-04 | 1.17019 | 0.488 | 0.36 | 1 | HC_vs_MS.Trt_F |
| APOLD1 | 8.84E-04 | 0.59715 | 0.566 | 0.45 | 1 | HC_vs_MS.Trt_F |
| PHKG1 | 8.88E-04 | 0.48027 | 0.373 | 0.25 | 1 | HC_vs_MS.Trt_F |
| PLEC | 8.88E-04 | 0.32157 | 0.439 | 0.31 | 1 | HC_vs_MS.Trt_F |
| MBNL2 | 8.88E-04 | 0.70946 | 0.315 | 0.19 | 1 | HC_vs_MS.Trt_F |
| PPP6R2 | 8.94E-04 | 0.28569 | 0.428 | 0.29 | 1 | HC_vs_MS.Trt_F |
| THRA | 9.01E-04 | 0.79343 | 0.208 | 0.1 | 1 | HC_vs_MS.Trt_F |
| MYO9A | 9.10E-04 | 0.27355 | 0.402 | 0.27 | 1 | HC_vs_MS.Trt_F |
| C5orf45 | 9.10E-04 | 0.51534 | 0.913 | 0.85 | 1 | HC_vs_MS.Trt_F |
| SYVN1 | 9.17E-04 | 0.55543 | 0.298 | 0.18 | 1 | HC_vs_MS.Trt_F |
| MAPKBP1 | 9.26E-04 | 0.56006 | 0.552 | 0.41 | 1 | HC_vs_MS.Trt_F |
| SPARCL1 | 9.33E-04 | -0.3277 | 0.292 | 0.18 | 1 | HC_vs_MS.Trt_F |
| XRCC1 | 9.39E-04 | 0.562 | 0.228 | 0.12 | 1 | HC_vs_MS.Trt_F |
| CACNA1D | 9.41E-04 | 0.78523 | 0.728 | 0.63 | 1 | HC_vs_MS.Trt_F |

| | | | | | | |
|---|---|---|---|---|---|---|
| NCKAP1L | 9.54E-04 | 0.53494 | 0.815 | 0.7 | 1 | HC_vs_MS.Trt_F |
| IL1RAP | 9.55E-04 | 0.86808 | 0.659 | 0.55 | 1 | HC_vs_MS.Trt_F |
| RTN4 | 9.57E-04 | 0.50522 | 0.35 | 0.23 | 1 | HC_vs_MS.Trt_F |
| SEC23IP | 9.57E-04 | 0.30227 | 0.512 | 0.39 | 1 | HC_vs_MS.Trt_F |
| SRSF7 | 9.68E-04 | 0.28749 | 0.538 | 0.4 | 1 | HC_vs_MS.Trt_F |
| GPR174 | 9.80E-04 | 0.59135 | 0.263 | 0.16 | 1 | HC_vs_MS.Trt_F |
| WDR89 | 9.87E-04 | 0.73074 | 0.399 | 0.27 | 1 | HC_vs_MS.Trt_F |
| RASL10B | 9.91E-04 | 0.26454 | 0.977 | 0.97 | 1 | HC_vs_MS.Trt_F |
| SRSF6 | 9.98E-04 | -0.9168 | 0.627 | 0.72 | 1 | HC_vs_MS.Trt_F |
| PPIC | 9.98E-04 | 0.85472 | 0.173 | 0.09 | 1 | HC_vs_MS.Trt_F |
| DND1 | 0.001 | -0.4854 | 0.038 | 0.1 | 1 | HC_vs_MS.Trt_F |
| ABO | 0.00101 | 0.38498 | 0.462 | 0.33 | 1 | HC_vs_MS.Trt_F |
| AFDN | 0.00101 | 1.39012 | 0.185 | 0.09 | 1 | HC_vs_MS.Trt_F |
| BCO2 | 0.00101 | 0.60759 | 0.627 | 0.51 | 1 | HC_vs_MS.Trt_F |
| CMSS1 | 0.00101 | 0.46884 | 0.763 | 0.71 | 1 | HC_vs_MS.Trt_F |
| SP3 | 0.00102 | 0.76178 | 0.639 | 0.49 | 1 | HC_vs_MS.Trt_F |
| CSTF1 | 0.00103 | -1.0021 | 0.72 | 0.82 | 1 | HC_vs_MS.Trt_F |
| RIMS2 | 0.00105 | 0.31122 | 0.957 | 0.92 | 1 | HC_vs_MS.Trt_F |
| INO80C | 0.00105 | 0.3389 | 0.934 | 0.92 | 1 | HC_vs_MS.Trt_F |
| KIAA1328 | 0.00105 | 0.32626 | 0.986 | 0.94 | 1 | HC_vs_MS.Trt_F |
| PDGFRB | 0.00106 | 1.07898 | 0.116 | 0.04 | 1 | HC_vs_MS.Trt_F |
| CXorf38 | 0.00106 | 0.35104 | 0.847 | 0.82 | 1 | HC_vs_MS.Trt_F |
| CTNNB1 | 0.00108 | -0.5018 | 0.503 | 0.63 | 1 | HC_vs_MS.Trt_F |
| NDUFAF5 | 0.00111 | 0.33658 | 0.633 | 0.48 | 1 | HC_vs_MS.Trt_F |
| SLC17A5 | 0.00111 | 0.27962 | 0.569 | 0.42 | 1 | HC_vs_MS.Trt_F |
| SEC16A | 0.00111 | 0.49279 | 0.26 | 0.15 | 1 | HC_vs_MS.Trt_F |
| RHOC | 0.00112 | 0.47627 | 0.266 | 0.16 | 1 | HC_vs_MS.Trt_F |
| MT-ND3 | 0.00112 | 0.3804 | 0.532 | 0.42 | 1 | HC_vs_MS.Trt_F |
| DNAJB1 | 0.00114 | 0.39319 | 0.777 | 0.68 | 1 | HC_vs_MS.Trt_F |
| CCDC144A | 0.00114 | 0.61818 | 0.908 | 0.87 | 1 | HC_vs_MS.Trt_F |
| UBA1 | 0.00116 | 0.45713 | 0.844 | 0.76 | 1 | HC_vs_MS.Trt_F |
| MAP1LC3B | 0.00117 | 0.48261 | 0.442 | 0.31 | 1 | HC_vs_MS.Trt_F |
| XDH | 0.00117 | 0.39592 | 0.431 | 0.31 | 1 | HC_vs_MS.Trt_F |
| NRG4 | 0.00117 | 0.45343 | 0.72 | 0.59 | 1 | HC_vs_MS.Trt_F |
| FOSB | 0.00118 | 1.01596 | 0.474 | 0.36 | 1 | HC_vs_MS.Trt_F |
| NXF1 | 0.0012 | 0.41197 | 0.673 | 0.56 | 1 | HC_vs_MS.Trt_F |
| VPS13B | 0.0012 | 0.38962 | 0.76 | 0.63 | 1 | HC_vs_MS.Trt_F |
| KIAA1683 | 0.00121 | 0.67114 | 0.717 | 0.6 | 1 | HC_vs_MS.Trt_F |
| BAG1 | 0.00121 | 0.38603 | 0.954 | 0.87 | 1 | HC_vs_MS.Trt_F |
| CECR1 | 0.00122 | 0.73383 | 0.503 | 0.36 | 1 | HC_vs_MS.Trt_F |

| | | | | | | |
|---|---|---|---|---|---|---|
| CSF2RA | 0.00124 | 0.27408 | 0.988 | 0.97 | 1 | HC_vs_MS.Trt_F |
| EFTUD2 | 0.00125 | 0.44303 | 0.806 | 0.76 | 1 | HC_vs_MS.Trt_F |
| ZMAT5 | 0.00125 | 0.41597 | 0.777 | 0.68 | 1 | HC_vs_MS.Trt_F |
| APBB3 | 0.00126 | 0.94463 | 0.202 | 0.11 | 1 | HC_vs_MS.Trt_F |
| SV2B | 0.00127 | 0.27837 | 0.442 | 0.31 | 1 | HC_vs_MS.Trt_F |
| ARHGEF9 | 0.00128 | 0.80408 | 0.613 | 0.52 | 1 | HC_vs_MS.Trt_F |
| CXXC5 | 0.00129 | -0.682 | 0.569 | 0.45 | 1 | HC_vs_MS.Trt_F |
| BATF | 0.0013 | 0.63187 | 0.24 | 0.14 | 1 | HC_vs_MS.Trt_F |
| KDM4C | 0.00131 | 0.39711 | 0.332 | 0.22 | 1 | HC_vs_MS.Trt_F |
| ANKRD44 | 0.00131 | 0.30399 | 0.864 | 0.76 | 1 | HC_vs_MS.Trt_F |
| MT-ND6 | 0.00131 | 0.30395 | 0.777 | 0.65 | 1 | HC_vs_MS.Trt_F |
| SLC4A5 | 0.00132 | 0.59443 | 0.827 | 0.72 | 1 | HC_vs_MS.Trt_F |
| DNAJB11 | 0.00134 | 0.75289 | 0.269 | 0.16 | 1 | HC_vs_MS.Trt_F |
| CDH6 | 0.00135 | 0.26678 | 0.627 | 0.5 | 1 | HC_vs_MS.Trt_F |
| ABHD11 | 0.00135 | 0.57246 | 0.751 | 0.59 | 1 | HC_vs_MS.Trt_F |
| DUSP6 | 0.00135 | 1.44208 | 0.147 | 0.07 | 1 | HC_vs_MS.Trt_F |
| MYO10 | 0.00137 | 0.41577 | 0.74 | 0.62 | 1 | HC_vs_MS.Trt_F |
| GSN | 0.00146 | 0.71788 | 0.714 | 0.64 | 1 | HC_vs_MS.Trt_F |
| ZGPAT | 0.00146 | 1.23283 | 0.208 | 0.11 | 1 | HC_vs_MS.Trt_F |
| BTLA | 0.00148 | -0.3636 | 0.22 | 0.12 | 1 | HC_vs_MS.Trt_F |
| SDCBP | 0.00149 | 0.33103 | 0.309 | 0.2 | 1 | HC_vs_MS.Trt_F |
| PISD | 0.00149 | 0.51966 | 0.217 | 0.12 | 1 | HC_vs_MS.Trt_F |
| FRRS1L | 0.00151 | 0.25355 | 0.925 | 0.85 | 1 | HC_vs_MS.Trt_F |
| IRAK1BP1 | 0.00151 | 0.47069 | 0.734 | 0.61 | 1 | HC_vs_MS.Trt_F |
| PCDH10 | 0.00152 | 0.35199 | 0.214 | 0.12 | 1 | HC_vs_MS.Trt_F |
| INPP4B | 0.00154 | 0.51993 | 0.601 | 0.49 | 1 | HC_vs_MS.Trt_F |
| HMGA2 | 0.00155 | 0.2695 | 0.74 | 0.62 | 1 | HC_vs_MS.Trt_F |
| TBC1D5 | 0.00156 | 0.38266 | 0.647 | 0.54 | 1 | HC_vs_MS.Trt_F |
| FAM120C | 0.00156 | 0.53132 | 0.298 | 0.19 | 1 | HC_vs_MS.Trt_F |
| NUP35 | 0.00156 | -0.2705 | 0.662 | 0.5 | 1 | HC_vs_MS.Trt_F |
| TACC1 | 0.00158 | 0.45926 | 0.566 | 0.47 | 1 | HC_vs_MS.Trt_F |
| AP1G2 | 0.00158 | 0.28316 | 0.731 | 0.63 | 1 | HC_vs_MS.Trt_F |
| C10orf10 | 0.00158 | 0.57591 | 0.107 | 0.04 | 1 | HC_vs_MS.Trt_F |
| SPDYE1 | 0.00161 | -0.3825 | 0.483 | 0.59 | 1 | HC_vs_MS.Trt_F |
| BCLAF1 | 0.00162 | 0.54117 | 0.512 | 0.4 | 1 | HC_vs_MS.Trt_F |
| LRBA | 0.00164 | 0.85222 | 0.52 | 0.4 | 1 | HC_vs_MS.Trt_F |
| ARHGAP29 | 0.00166 | 0.31808 | 0.538 | 0.38 | 1 | HC_vs_MS.Trt_F |
| NR4A1 | 0.00168 | 0.31964 | 0.569 | 0.44 | 1 | HC_vs_MS.Trt_F |
| FREM2 | 0.00168 | -0.3411 | 0.598 | 0.67 | 1 | HC_vs_MS.Trt_F |
| RPS18 | 0.00169 | -0.2835 | 0.954 | 0.95 | 1 | HC_vs_MS.Trt_F |

| | | | | | | |
|---|---|---|---|---|---|---|
| FGF23 | 0.0017 | 0.42301 | 0.925 | 0.82 | 1 | HC_vs_MS.Trt_F |
| OPHN1 | 0.00172 | 0.27755 | 0.977 | 0.96 | 1 | HC_vs_MS.Trt_F |
| PTBP3 | 0.00173 | 0.41785 | 0.425 | 0.31 | 1 | HC_vs_MS.Trt_F |
| TKT | 0.00173 | 0.26282 | 0.705 | 0.58 | 1 | HC_vs_MS.Trt_F |
| ZNF212 | 0.00174 | -0.2991 | 0.506 | 0.37 | 1 | HC_vs_MS.Trt_F |
| CCDC7 | 0.00174 | 1.17193 | 0.191 | 0.1 | 1 | HC_vs_MS.Trt_F |
| CCDC57 | 0.00175 | 0.35762 | 0.997 | 0.99 | 1 | HC_vs_MS.Trt_F |
| GADD45B | 0.00175 | 1.1691 | 0.22 | 0.13 | 1 | HC_vs_MS.Trt_F |
| AIFM2 | 0.00175 | -0.402 | 0.514 | 0.37 | 1 | HC_vs_MS.Trt_F |
| GCNT4 | 0.00175 | -0.8987 | 0.344 | 0.44 | 1 | HC_vs_MS.Trt_F |
| LYRM2 | 0.00177 | 0.31734 | 0.876 | 0.81 | 1 | HC_vs_MS.Trt_F |
| CYB561D1 | 0.0018 | 0.52482 | 0.5 | 0.39 | 1 | HC_vs_MS.Trt_F |
| OSBPL10 | 0.00181 | 0.45361 | 0.468 | 0.36 | 1 | HC_vs_MS.Trt_F |
| SUCO | 0.00181 | 1.14667 | 0.194 | 0.1 | 1 | HC_vs_MS.Trt_F |
| KDM6B | 0.00183 | 0.88227 | 0.289 | 0.18 | 1 | HC_vs_MS.Trt_F |
| CPT2 | 0.00186 | 0.44765 | 0.818 | 0.73 | 1 | HC_vs_MS.Trt_F |
| KLC4 | 0.00186 | 0.59678 | 0.876 | 0.8 | 1 | HC_vs_MS.Trt_F |
| PSMB5 | 0.00186 | 1.05922 | 0.231 | 0.13 | 1 | HC_vs_MS.Trt_F |
| FOSL2 | 0.00187 | 0.94631 | 0.347 | 0.25 | 1 | HC_vs_MS.Trt_F |
| BAZ2B | 0.00187 | 0.66861 | 0.847 | 0.81 | 1 | HC_vs_MS.Trt_F |
| HNRNPDL | 0.00188 | 0.26565 | 0.636 | 0.51 | 1 | HC_vs_MS.Trt_F |
| KIF1B | 0.00191 | 1.02637 | 0.844 | 0.74 | 1 | HC_vs_MS.Trt_F |
| PTBP1 | 0.00192 | 0.76494 | 0.303 | 0.2 | 1 | HC_vs_MS.Trt_F |
| EIF2S3 | 0.00193 | -0.4325 | 0.957 | 0.96 | 1 | HC_vs_MS.Trt_F |
| GRIA4 | 0.00193 | 0.41043 | 0.191 | 0.1 | 1 | HC_vs_MS.Trt_F |
| GAPDH | 0.00193 | 0.2666 | 0.818 | 0.71 | 1 | HC_vs_MS.Trt_F |
| KDM5D | 0.00195 | 1.9068 | 0.39 | 0.27 | 1 | HC_vs_MS.Trt_F |
| MPRIP | 0.00197 | 0.77715 | 0.948 | 0.89 | 1 | HC_vs_MS.Trt_F |
| MRPS28 | 0.00197 | 0.29233 | 0.711 | 0.59 | 1 | HC_vs_MS.Trt_F |
| RASGEF1B | 0.00199 | 0.64483 | 0.812 | 0.7 | 1 | HC_vs_MS.Trt_F |
| TLDC1 | 0.002 | 0.53224 | 0.783 | 0.64 | 1 | HC_vs_MS.Trt_F |
| TOMM7 | 0.002 | 0.34423 | 0.564 | 0.44 | 1 | HC_vs_MS.Trt_F |
| DHX30 | 0.00201 | 0.37323 | 0.734 | 0.63 | 1 | HC_vs_MS.Trt_F |
| PIGG | 0.00201 | 0.56351 | 0.627 | 0.51 | 1 | HC_vs_MS.Trt_F |
| HOOK2 | 0.00201 | 0.34746 | 0.98 | 0.95 | 1 | HC_vs_MS.Trt_F |
| COL5A1 | 0.00202 | 1.02154 | 0.104 | 0.04 | 1 | HC_vs_MS.Trt_F |
| EMP2 | 0.00204 | 0.32614 | 0.957 | 0.91 | 1 | HC_vs_MS.Trt_F |
| POLR2G | 0.00205 | 0.60647 | 0.237 | 0.14 | 1 | HC_vs_MS.Trt_F |
| MFSD6 | 0.00206 | 0.86444 | 0.249 | 0.15 | 1 | HC_vs_MS.Trt_F |
| PHYH | 0.00207 | 0.68395 | 0.309 | 0.2 | 1 | HC_vs_MS.Trt_F |

| | | | | | | |
|---|---|---|---|---|---|---|
| CPB1 | 0.0021 | 0.44064 | 0.569 | 0.45 | 1 | HC_vs_MS.Trt_F |
| ELK1 | 0.00213 | 0.36672 | 0.171 | 0.08 | 1 | HC_vs_MS.Trt_F |
| ACAD9 | 0.00213 | 0.56924 | 0.338 | 0.22 | 1 | HC_vs_MS.Trt_F |
| MCF2L2 | 0.00215 | 0.68164 | 0.962 | 0.94 | 1 | HC_vs_MS.Trt_F |
| NPHP4 | 0.00218 | 0.89568 | 0.145 | 0.07 | 1 | HC_vs_MS.Trt_F |
| GRK4 | 0.00218 | 0.39016 | 0.653 | 0.53 | 1 | HC_vs_MS.Trt_F |
| CLCN7 | 0.0022 | 0.65062 | 0.387 | 0.26 | 1 | HC_vs_MS.Trt_F |
| SLC25A45 | 0.00222 | -0.4515 | 0.951 | 0.98 | 1 | HC_vs_MS.Trt_F |
| SPC25 | 0.00225 | 0.39152 | 0.763 | 0.65 | 1 | HC_vs_MS.Trt_F |
| GNB1L | 0.00226 | 0.88069 | 0.292 | 0.18 | 1 | HC_vs_MS.Trt_F |
| USP25 | 0.00226 | -0.8286 | 0.52 | 0.37 | 1 | HC_vs_MS.Trt_F |
| POU6F2 | 0.00226 | 0.29791 | 0.353 | 0.23 | 1 | HC_vs_MS.Trt_F |
| TMEM106A | 0.00227 | 0.45726 | 0.728 | 0.65 | 1 | HC_vs_MS.Trt_F |
| MYCBP2 | 0.00229 | 0.33341 | 0.465 | 0.34 | 1 | HC_vs_MS.Trt_F |
| HK1 | 0.00229 | 0.26933 | 0.647 | 0.52 | 1 | HC_vs_MS.Trt_F |
| GNL2 | 0.0023 | 0.32914 | 0.76 | 0.66 | 1 | HC_vs_MS.Trt_F |
| TRPM7 | 0.0023 | 0.88375 | 0.934 | 0.91 | 1 | HC_vs_MS.Trt_F |
| MNS1 | 0.0023 | 0.42342 | 0.234 | 0.14 | 1 | HC_vs_MS.Trt_F |
| ADGRE5 | 0.00232 | 0.47338 | 0.734 | 0.6 | 1 | HC_vs_MS.Trt_F |
| GCN1 | 0.00232 | 0.78365 | 0.217 | 0.12 | 1 | HC_vs_MS.Trt_F |
| BICDL1 | 0.00232 | 0.95955 | 0.662 | 0.58 | 1 | HC_vs_MS.Trt_F |
| GSTM3 | 0.00232 | 0.44027 | 0.598 | 0.47 | 1 | HC_vs_MS.Trt_F |
| ITGB1 | 0.00233 | -0.6736 | 0.592 | 0.64 | 1 | HC_vs_MS.Trt_F |
| ZNF398 | 0.00233 | 0.83542 | 0.269 | 0.17 | 1 | HC_vs_MS.Trt_F |
| STRN4 | 0.00234 | 0.40198 | 0.598 | 0.48 | 1 | HC_vs_MS.Trt_F |
| SUSD3 | 0.00236 | -0.7574 | 0.298 | 0.4 | 1 | HC_vs_MS.Trt_F |
| KIF27 | 0.00238 | -0.5144 | 0.474 | 0.35 | 1 | HC_vs_MS.Trt_F |
| KLHL33 | 0.00238 | 0.40268 | 0.59 | 0.48 | 1 | HC_vs_MS.Trt_F |
| KARS | 0.00239 | 0.30737 | 0.332 | 0.21 | 1 | HC_vs_MS.Trt_F |
| ATP2B4 | 0.00243 | 0.4398 | 0.578 | 0.44 | 1 | HC_vs_MS.Trt_F |
| RERE | 0.00244 | 0.55003 | 0.451 | 0.33 | 1 | HC_vs_MS.Trt_F |
| STXBP2 | 0.00245 | 0.97255 | 0.321 | 0.22 | 1 | HC_vs_MS.Trt_F |
| U2SURP | 0.00246 | 0.41687 | 0.364 | 0.25 | 1 | HC_vs_MS.Trt_F |
| GSTM4 | 0.00248 | 0.38874 | 0.211 | 0.12 | 1 | HC_vs_MS.Trt_F |
| PRDX4 | 0.00249 | 0.70764 | 0.176 | 0.09 | 1 | HC_vs_MS.Trt_F |
| ALDH3B1 | 0.0025 | 0.30348 | 0.809 | 0.75 | 1 | HC_vs_MS.Trt_F |
| PAX6 | 0.00251 | 0.43348 | 0.396 | 0.28 | 1 | HC_vs_MS.Trt_F |
| ULK3 | 0.00251 | 0.37375 | 0.251 | 0.14 | 1 | HC_vs_MS.Trt_F |
| EPB41L2 | 0.00253 | 0.34458 | 0.214 | 0.12 | 1 | HC_vs_MS.Trt_F |
| ZBTB40 | 0.00253 | 0.65066 | 0.526 | 0.4 | 1 | HC_vs_MS.Trt_F |

| | | | | | | |
|---|---|---|---|---|---|---|
| PRPF40A | 0.00255 | 0.38801 | 0.587 | 0.46 | 1 | HC_vs_MS.Trt_F |
| PCM1 | 0.00256 | 0.45999 | 0.887 | 0.78 | 1 | HC_vs_MS.Trt_F |
| YIPF6 | 0.00256 | 0.48175 | 0.908 | 0.82 | 1 | HC_vs_MS.Trt_F |
| TBC1D20 | 0.00257 | 0.50573 | 0.694 | 0.58 | 1 | HC_vs_MS.Trt_F |
| HSPA1B | 0.00257 | 0.33657 | 0.329 | 0.21 | 1 | HC_vs_MS.Trt_F |
| LRCH4 | 0.00258 | 0.45979 | 0.575 | 0.45 | 1 | HC_vs_MS.Trt_F |
| ARAF | 0.00259 | 0.30093 | 0.24 | 0.14 | 1 | HC_vs_MS.Trt_F |
| MSI2 | 0.00259 | 0.95831 | 0.942 | 0.86 | 1 | HC_vs_MS.Trt_F |
| ADSS | 0.0026 | 0.34284 | 0.231 | 0.14 | 1 | HC_vs_MS.Trt_F |
| JAZF1 | 0.0026 | 1.12812 | 0.194 | 0.11 | 1 | HC_vs_MS.Trt_F |
| BCL6 | 0.00263 | 0.92526 | 0.191 | 0.1 | 1 | HC_vs_MS.Trt_F |
| PHLDA1 | 0.00265 | 0.64092 | 0.624 | 0.53 | 1 | HC_vs_MS.Trt_F |
| TMX1 | 0.00266 | 0.49633 | 0.497 | 0.37 | 1 | HC_vs_MS.Trt_F |
| CNOT6 | 0.00267 | 1.03529 | 0.182 | 0.09 | 1 | HC_vs_MS.Trt_F |
| SYMPK | 0.00268 | 0.49576 | 0.922 | 0.86 | 1 | HC_vs_MS.Trt_F |
| POLR2I | 0.00271 | 0.5953 | 0.231 | 0.13 | 1 | HC_vs_MS.Trt_F |
| PRKAG2 | 0.00272 | 0.78639 | 0.422 | 0.31 | 1 | HC_vs_MS.Trt_F |
| IDE | 0.00272 | 0.63601 | 0.564 | 0.44 | 1 | HC_vs_MS.Trt_F |
| TTC39A | 0.00272 | 0.29805 | 0.376 | 0.27 | 1 | HC_vs_MS.Trt_F |
| REEP3 | 0.00273 | -2.0319 | 0.486 | 0.53 | 1 | HC_vs_MS.Trt_F |
| CTNND1 | 0.00273 | 0.68908 | 0.483 | 0.36 | 1 | HC_vs_MS.Trt_F |
| KHDRBS1 | 0.00276 | 0.42059 | 0.24 | 0.14 | 1 | HC_vs_MS.Trt_F |
| CTDSPL | 0.00276 | 0.55258 | 0.335 | 0.22 | 1 | HC_vs_MS.Trt_F |
| TXNRD2 | 0.00276 | 0.52433 | 0.925 | 0.83 | 1 | HC_vs_MS.Trt_F |
| MAPK8 | 0.00277 | 0.49498 | 0.497 | 0.37 | 1 | HC_vs_MS.Trt_F |
| PTTG1IP | 0.00277 | 0.60377 | 0.254 | 0.16 | 1 | HC_vs_MS.Trt_F |
| PSMB7 | 0.00278 | 0.37107 | 0.277 | 0.18 | 1 | HC_vs_MS.Trt_F |
| TTL | 0.00279 | 0.27011 | 0.939 | 0.88 | 1 | HC_vs_MS.Trt_F |
| SAMD15 | 0.00279 | 0.27306 | 0.361 | 0.25 | 1 | HC_vs_MS.Trt_F |
| LPCAT4 | 0.00279 | 0.26807 | 0.439 | 0.32 | 1 | HC_vs_MS.Trt_F |
| KIAA2026 | 0.0028 | 0.64122 | 0.269 | 0.17 | 1 | HC_vs_MS.Trt_F |
| DUSP10 | 0.00283 | 0.83671 | 0.335 | 0.22 | 1 | HC_vs_MS.Trt_F |
| HNF4G | 0.00284 | 0.44077 | 0.329 | 0.22 | 1 | HC_vs_MS.Trt_F |
| LYPLA2 | 0.00285 | 0.65052 | 0.208 | 0.12 | 1 | HC_vs_MS.Trt_F |
| USP9X | 0.00286 | 0.2719 | 0.645 | 0.53 | 1 | HC_vs_MS.Trt_F |
| ESYT2 | 0.00286 | 0.2524 | 0.668 | 0.56 | 1 | HC_vs_MS.Trt_F |
| MED26 | 0.00287 | 2.24686 | 0.124 | 0.06 | 1 | HC_vs_MS.Trt_F |
| SPTLC3 | 0.00293 | 0.61254 | 0.575 | 0.48 | 1 | HC_vs_MS.Trt_F |
| GON4L | 0.00294 | 0.52594 | 0.887 | 0.84 | 1 | HC_vs_MS.Trt_F |
| PARPBP | 0.00294 | -1.1594 | 0.465 | 0.34 | 1 | HC_vs_MS.Trt_F |

| | | | | | | |
|---|---|---|---|---|---|---|
| WSB1 | 0.00295 | 0.4369 | 0.899 | 0.83 | 1 | HC_vs_MS.Trt_F |
| SCAMP4 | 0.00295 | 0.41884 | 0.908 | 0.89 | 1 | HC_vs_MS.Trt_F |
| ZBTB16 | 0.00295 | 1.15229 | 0.442 | 0.33 | 1 | HC_vs_MS.Trt_F |
| ICAM1 | 0.00298 | 0.4424 | 0.555 | 0.42 | 1 | HC_vs_MS.Trt_F |
| PIEZO1 | 0.00298 | 0.38248 | 0.448 | 0.32 | 1 | HC_vs_MS.Trt_F |
| SZT2 | 0.00299 | 0.322 | 0.884 | 0.81 | 1 | HC_vs_MS.Trt_F |
| MAPK6 | 0.00303 | -1.0844 | 0.54 | 0.42 | 1 | HC_vs_MS.Trt_F |
| CSF3R | 0.00303 | 0.81987 | 0.711 | 0.58 | 1 | HC_vs_MS.Trt_F |
| PTGER4 | 0.00304 | 0.33699 | 0.277 | 0.17 | 1 | HC_vs_MS.Trt_F |
| UNKL | 0.00305 | -0.5808 | 0.379 | 0.48 | 1 | HC_vs_MS.Trt_F |
| PIK3R5 | 0.00306 | 0.37803 | 0.422 | 0.29 | 1 | HC_vs_MS.Trt_F |
| MAPRE2 | 0.00308 | 0.54657 | 0.338 | 0.24 | 1 | HC_vs_MS.Trt_F |
| DMAP1 | 0.00308 | 0.38046 | 0.572 | 0.43 | 1 | HC_vs_MS.Trt_F |
| CD55 | 0.00311 | 0.5458 | 0.633 | 0.54 | 1 | HC_vs_MS.Trt_F |
| ZNF623 | 0.00311 | 0.42782 | 0.72 | 0.63 | 1 | HC_vs_MS.Trt_F |
| PTRF | 0.00312 | 0.53193 | 0.474 | 0.35 | 1 | HC_vs_MS.Trt_F |
| EIF1B | 0.00312 | 0.49957 | 0.246 | 0.15 | 1 | HC_vs_MS.Trt_F |
| ZMYM2 | 0.00315 | 0.40711 | 0.454 | 0.34 | 1 | HC_vs_MS.Trt_F |
| TAGAP | 0.00316 | 0.54789 | 0.353 | 0.25 | 1 | HC_vs_MS.Trt_F |
| HACD2 | 0.00317 | 0.28862 | 0.968 | 0.91 | 1 | HC_vs_MS.Trt_F |
| DCN | 0.00321 | 0.82879 | 0.749 | 0.62 | 1 | HC_vs_MS.Trt_F |
| FAM118A | 0.00321 | 0.67733 | 0.621 | 0.52 | 1 | HC_vs_MS.Trt_F |
| SFXN3 | 0.00322 | 0.49947 | 0.419 | 0.31 | 1 | HC_vs_MS.Trt_F |
| PIGP | 0.00323 | 0.44198 | 0.835 | 0.78 | 1 | HC_vs_MS.Trt_F |
| AP006285.1 | 0.00327 | 0.29719 | 0.861 | 0.78 | 1 | HC_vs_MS.Trt_F |
| LYZ | 0.0033 | 0.43286 | 0.503 | 0.37 | 1 | HC_vs_MS.Trt_F |
| CARD16 | 0.00331 | 0.28514 | 0.361 | 0.24 | 1 | HC_vs_MS.Trt_F |
| SHANK2 | 0.00331 | 0.37653 | 0.384 | 0.28 | 1 | HC_vs_MS.Trt_F |
| RNF8 | 0.00332 | 0.77632 | 0.561 | 0.45 | 1 | HC_vs_MS.Trt_F |
| GPR137C | 0.00335 | 0.4032 | 0.63 | 0.55 | 1 | HC_vs_MS.Trt_F |
| PGD | 0.00336 | 0.84275 | 0.194 | 0.1 | 1 | HC_vs_MS.Trt_F |
| NPIPB6 | 0.00337 | 0.74295 | 0.289 | 0.19 | 1 | HC_vs_MS.Trt_F |
| TPCN2 | 0.00337 | 1.06197 | 0.682 | 0.59 | 1 | HC_vs_MS.Trt_F |
| PLOD2 | 0.00337 | 0.33856 | 0.439 | 0.34 | 1 | HC_vs_MS.Trt_F |
| TJP1 | 0.00338 | 1.00989 | 0.263 | 0.16 | 1 | HC_vs_MS.Trt_F |
| CCDC36 | 0.00341 | 0.26716 | 0.451 | 0.32 | 1 | HC_vs_MS.Trt_F |
| ARHGEF10 | 0.00345 | -0.7187 | 0.694 | 0.79 | 1 | HC_vs_MS.Trt_F |
| XPNPEP1 | 0.00346 | 0.3179 | 0.431 | 0.3 | 1 | HC_vs_MS.Trt_F |
| NFX1 | 0.00347 | 0.35277 | 0.899 | 0.78 | 1 | HC_vs_MS.Trt_F |
| SH2B1 | 0.00348 | 0.43947 | 0.642 | 0.51 | 1 | HC_vs_MS.Trt_F |

| | | | | | | |
|---|---|---|---|---|---|---|
| ADK | 0.00349 | 0.56643 | 0.289 | 0.19 | 1 | HC_vs_MS.Trt_F |
| CXCR4 | 0.0035 | 0.59373 | 0.37 | 0.27 | 1 | HC_vs_MS.Trt_F |
| ITPR3 | 0.0035 | 1.54602 | 0.205 | 0.12 | 1 | HC_vs_MS.Trt_F |
| 3-Mar | 0.00352 | 0.58582 | 0.396 | 0.27 | 1 | HC_vs_MS.Trt_F |
| PHF12 | 0.00355 | 0.39766 | 0.757 | 0.66 | 1 | HC_vs_MS.Trt_F |
| AP2A2 | 0.0036 | 0.32045 | 0.757 | 0.63 | 1 | HC_vs_MS.Trt_F |
| CITED2 | 0.00361 | 1.07456 | 0.249 | 0.16 | 1 | HC_vs_MS.Trt_F |
| NPIPB15 | 0.00363 | -0.9856 | 0.237 | 0.33 | 1 | HC_vs_MS.Trt_F |
| MAN1A2 | 0.00363 | 0.28327 | 0.367 | 0.25 | 1 | HC_vs_MS.Trt_F |
| CH507-42P1 | 0.00366 | 1.12486 | 0.257 | 0.17 | 1 | HC_vs_MS.Trt_F |
| SERPINB9 | 0.00367 | -0.3302 | 0.708 | 0.8 | 1 | HC_vs_MS.Trt_F |
| NUDCD1 | 0.00368 | 1.34908 | 0.162 | 0.08 | 1 | HC_vs_MS.Trt_F |
| NSMCE2 | 0.00371 | 0.28434 | 0.647 | 0.54 | 1 | HC_vs_MS.Trt_F |
| FSBP | 0.00371 | 0.70947 | 0.384 | 0.27 | 1 | HC_vs_MS.Trt_F |
| SGMS1 | 0.00375 | 0.631 | 0.286 | 0.18 | 1 | HC_vs_MS.Trt_F |
| KATNAL1 | 0.00376 | 0.93559 | 0.613 | 0.52 | 1 | HC_vs_MS.Trt_F |
| OGDH | 0.00378 | 0.27042 | 0.595 | 0.46 | 1 | HC_vs_MS.Trt_F |
| NUP214 | 0.0038 | 0.36654 | 0.694 | 0.62 | 1 | HC_vs_MS.Trt_F |
| TAOK2 | 0.00381 | -0.2887 | 0.179 | 0.09 | 1 | HC_vs_MS.Trt_F |
| RUNDC3B | 0.00383 | 0.4086 | 0.309 | 0.21 | 1 | HC_vs_MS.Trt_F |
| PLAUR | 0.00383 | 1.79111 | 0.647 | 0.56 | 1 | HC_vs_MS.Trt_F |
| STK17B | 0.00385 | -0.4904 | 0.948 | 0.94 | 1 | HC_vs_MS.Trt_F |
| CLEC2D | 0.00388 | -0.5661 | 0.803 | 0.83 | 1 | HC_vs_MS.Trt_F |
| ARFGEF1 | 0.00389 | 0.30342 | 0.697 | 0.57 | 1 | HC_vs_MS.Trt_F |
| SPON2 | 0.00389 | 0.94912 | 0.488 | 0.37 | 1 | HC_vs_MS.Trt_F |
| AKAP6 | 0.0039 | -0.4409 | 0.777 | 0.66 | 1 | HC_vs_MS.Trt_F |
| IL21R | 0.00393 | 0.60111 | 0.645 | 0.52 | 1 | HC_vs_MS.Trt_F |
| MTMR10 | 0.00395 | 1.15077 | 0.272 | 0.18 | 1 | HC_vs_MS.Trt_F |
| BRD1 | 0.00396 | 0.84111 | 0.199 | 0.12 | 1 | HC_vs_MS.Trt_F |
| ATXN2L | 0.00396 | 1.40931 | 0.298 | 0.21 | 1 | HC_vs_MS.Trt_F |
| AACS | 0.00397 | 0.39749 | 0.991 | 1 | 1 | HC_vs_MS.Trt_F |
| C1orf158 | 0.00398 | -0.5282 | 0.777 | 0.88 | 1 | HC_vs_MS.Trt_F |
| LHFP | 0.00399 | 0.29673 | 0.26 | 0.17 | 1 | HC_vs_MS.Trt_F |
| FGL2 | 0.004 | 0.83324 | 0.396 | 0.29 | 1 | HC_vs_MS.Trt_F |
| NBAS | 0.00401 | 0.99877 | 0.251 | 0.16 | 1 | HC_vs_MS.Trt_F |
| STK33 | 0.00401 | 0.31216 | 0.893 | 0.82 | 1 | HC_vs_MS.Trt_F |
| COPA | 0.00405 | -0.2614 | 0.35 | 0.23 | 1 | HC_vs_MS.Trt_F |
| MRPS5 | 0.00405 | 0.26995 | 0.832 | 0.74 | 1 | HC_vs_MS.Trt_F |
| MLANA | 0.00406 | 0.32816 | 0.506 | 0.39 | 1 | HC_vs_MS.Trt_F |
| ANKRD17 | 0.00406 | 0.98361 | 0.295 | 0.2 | 1 | HC_vs_MS.Trt_F |

| | | | | | | |
|---|---|---|---|---|---|---|
| STARD8 | 0.00413 | 0.45009 | 0.76 | 0.67 | 1 | HC_vs_MS.Trt_F |
| GALK1 | 0.00415 | 0.64077 | 0.402 | 0.28 | 1 | HC_vs_MS.Trt_F |
| TRAPPC3L | 0.0042 | 0.28212 | 0.616 | 0.48 | 1 | HC_vs_MS.Trt_F |
| STIL | 0.00423 | 0.31984 | 0.668 | 0.55 | 1 | HC_vs_MS.Trt_F |
| OTUD4 | 0.00425 | 0.67411 | 0.699 | 0.6 | 1 | HC_vs_MS.Trt_F |
| MED28 | 0.00429 | 0.47575 | 0.91 | 0.87 | 1 | HC_vs_MS.Trt_F |
| RALGDS | 0.00429 | 0.62441 | 0.373 | 0.27 | 1 | HC_vs_MS.Trt_F |
| MBD6 | 0.00432 | 0.78746 | 0.329 | 0.23 | 1 | HC_vs_MS.Trt_F |
| ANP32A | 0.00436 | 0.72776 | 0.312 | 0.21 | 1 | HC_vs_MS.Trt_F |
| SNU13 | 0.00438 | 0.25678 | 0.324 | 0.22 | 1 | HC_vs_MS.Trt_F |
| JRK | 0.00438 | 1.33191 | 0.483 | 0.35 | 1 | HC_vs_MS.Trt_F |
| RP11-449H3 | 0.00441 | 0.9944 | 0.341 | 0.23 | 1 | HC_vs_MS.Trt_F |
| ZNF789 | 0.00443 | -1.0495 | 0.734 | 0.8 | 1 | HC_vs_MS.Trt_F |
| DCTN4 | 0.00444 | 0.40768 | 0.399 | 0.29 | 1 | HC_vs_MS.Trt_F |
| NCALD | 0.00444 | -0.3224 | 0.78 | 0.65 | 1 | HC_vs_MS.Trt_F |
| CDHR3 | 0.00448 | 0.39549 | 0.676 | 0.51 | 1 | HC_vs_MS.Trt_F |
| HLA-DPA1 | 0.00453 | 0.28352 | 0.309 | 0.2 | 1 | HC_vs_MS.Trt_F |
| AKNA | 0.00454 | 0.3562 | 0.925 | 0.9 | 1 | HC_vs_MS.Trt_F |
| SECISBP2L | 0.00454 | 0.27523 | 0.792 | 0.68 | 1 | HC_vs_MS.Trt_F |
| LMCD1 | 0.00454 | 0.43904 | 0.37 | 0.27 | 1 | HC_vs_MS.Trt_F |
| TAB3 | 0.00454 | 1.09462 | 0.749 | 0.67 | 1 | HC_vs_MS.Trt_F |
| LETM1 | 0.00458 | -0.4693 | 0.838 | 0.89 | 1 | HC_vs_MS.Trt_F |
| PTP4A1 | 0.00465 | 0.41554 | 0.298 | 0.2 | 1 | HC_vs_MS.Trt_F |
| GIMAP1 | 0.00465 | 0.32034 | 0.509 | 0.38 | 1 | HC_vs_MS.Trt_F |
| MRPL43 | 0.0047 | 0.91448 | 0.185 | 0.1 | 1 | HC_vs_MS.Trt_F |
| NTRK1 | 0.0047 | 0.91525 | 0.249 | 0.16 | 1 | HC_vs_MS.Trt_F |
| PLP2 | 0.0048 | 0.3136 | 0.41 | 0.3 | 1 | HC_vs_MS.Trt_F |
| CUX1 | 0.0048 | 0.3686 | 0.358 | 0.25 | 1 | HC_vs_MS.Trt_F |
| MYL6 | 0.00481 | -0.3862 | 0.688 | 0.71 | 1 | HC_vs_MS.Trt_F |
| PPIP5K1 | 0.0049 | 0.41428 | 0.63 | 0.51 | 1 | HC_vs_MS.Trt_F |
| CIC | 0.0049 | 0.93965 | 0.15 | 0.08 | 1 | HC_vs_MS.Trt_F |
| ARHGAP33 | 0.00492 | 1.11964 | 0.197 | 0.12 | 1 | HC_vs_MS.Trt_F |
| NDE1 | 0.00492 | 0.44016 | 0.812 | 0.72 | 1 | HC_vs_MS.Trt_F |
| SP2 | 0.00493 | 1.36104 | 0.384 | 0.28 | 1 | HC_vs_MS.Trt_F |
| GIMAP7 | 0.00494 | -0.5381 | 0.41 | 0.49 | 1 | HC_vs_MS.Trt_F |
| CCT6A | 0.00495 | -0.2502 | 0.63 | 0.46 | 1 | HC_vs_MS.Trt_F |
| MAGI1 | 0.00497 | 0.29397 | 0.133 | 0.07 | 1 | HC_vs_MS.Trt_F |
| SPNS1 | 0.00498 | 0.59507 | 0.87 | 0.76 | 1 | HC_vs_MS.Trt_F |
| ZNF595 | 0.00499 | 0.43084 | 0.858 | 0.83 | 1 | HC_vs_MS.Trt_F |
| ADAMTSL3 | 0.005 | 0.35039 | 0.98 | 0.89 | 1 | HC_vs_MS.Trt_F |

| | | | | | | |
|---|---|---|---|---|---|---|
| MBOAT2 | 0.00508 | 0.71333 | 0.379 | 0.26 | 1 | HC_vs_MS.Trt_F |
| SUOX | 0.00511 | 0.46536 | 0.835 | 0.78 | 1 | HC_vs_MS.Trt_F |
| ZC3H12D | 0.00514 | 0.50447 | 0.876 | 0.82 | 1 | HC_vs_MS.Trt_F |
| DUSP28 | 0.00515 | 0.28986 | 0.754 | 0.64 | 1 | HC_vs_MS.Trt_F |
| LTA4H | 0.00521 | 0.40495 | 0.393 | 0.27 | 1 | HC_vs_MS.Trt_F |
| AP3B1 | 0.00522 | -0.3227 | 0.486 | 0.35 | 1 | HC_vs_MS.Trt_F |
| RFX7 | 0.00524 | 0.54746 | 0.234 | 0.15 | 1 | HC_vs_MS.Trt_F |
| HOMER2 | 0.00526 | 0.32705 | 0.616 | 0.51 | 1 | HC_vs_MS.Trt_F |
| RORA | 0.00527 | 0.35315 | 0.697 | 0.62 | 1 | HC_vs_MS.Trt_F |
| FAH | 0.0053 | 1.39295 | 0.205 | 0.12 | 1 | HC_vs_MS.Trt_F |
| ST8SIA3 | 0.00533 | 0.27588 | 0.191 | 0.11 | 1 | HC_vs_MS.Trt_F |
| CD47 | 0.00536 | -0.5299 | 0.887 | 0.88 | 1 | HC_vs_MS.Trt_F |
| PAM16 | 0.00536 | 1.08214 | 0.237 | 0.14 | 1 | HC_vs_MS.Trt_F |
| C2orf68 | 0.00538 | -0.8746 | 0.829 | 0.83 | 1 | HC_vs_MS.Trt_F |
| HN1 | 0.00541 | 0.33801 | 0.812 | 0.73 | 1 | HC_vs_MS.Trt_F |
| GBF1 | 0.00545 | 0.46813 | 0.243 | 0.15 | 1 | HC_vs_MS.Trt_F |
| PPP5C | 0.0055 | 0.47546 | 0.558 | 0.44 | 1 | HC_vs_MS.Trt_F |
| AMPD2 | 0.00551 | 1.35208 | 0.165 | 0.09 | 1 | HC_vs_MS.Trt_F |
| MRPS26 | 0.00553 | 0.53173 | 0.194 | 0.11 | 1 | HC_vs_MS.Trt_F |
| PINLYP | 0.00567 | 0.25216 | 0.621 | 0.51 | 1 | HC_vs_MS.Trt_F |
| HLA-DOA | 0.00569 | 0.52023 | 0.743 | 0.6 | 1 | HC_vs_MS.Trt_F |
| GSK3B | 0.00575 | 0.91059 | 0.766 | 0.68 | 1 | HC_vs_MS.Trt_F |
| FAM13B | 0.00582 | 0.6284 | 0.48 | 0.37 | 1 | HC_vs_MS.Trt_F |
| 9-Sep | 0.00585 | 0.56896 | 0.549 | 0.45 | 1 | HC_vs_MS.Trt_F |
| FAN1 | 0.00586 | -0.5175 | 0.309 | 0.21 | 1 | HC_vs_MS.Trt_F |
| ATAT1 | 0.00589 | 0.8652 | 0.121 | 0.06 | 1 | HC_vs_MS.Trt_F |
| KIF18A | 0.00594 | -0.2849 | 0.405 | 0.29 | 1 | HC_vs_MS.Trt_F |
| HSPA1A | 0.00598 | 0.6168 | 0.263 | 0.17 | 1 | HC_vs_MS.Trt_F |
| GCLM | 0.00598 | 0.39192 | 0.746 | 0.67 | 1 | HC_vs_MS.Trt_F |
| ACVR2B | 0.006 | 0.70618 | 0.491 | 0.4 | 1 | HC_vs_MS.Trt_F |
| NUDCD3 | 0.006 | 0.29719 | 0.344 | 0.24 | 1 | HC_vs_MS.Trt_F |
| RBBP9 | 0.00601 | 1.57793 | 0.486 | 0.38 | 1 | HC_vs_MS.Trt_F |
| AC024592.1 | 0.00607 | 1.00937 | 0.376 | 0.28 | 1 | HC_vs_MS.Trt_F |
| GBP4 | 0.00608 | 0.4344 | 0.838 | 0.72 | 1 | HC_vs_MS.Trt_F |
| ACSL4 | 0.00611 | 0.65887 | 0.217 | 0.13 | 1 | HC_vs_MS.Trt_F |
| TUBGCP2 | 0.00611 | 0.25066 | 0.801 | 0.72 | 1 | HC_vs_MS.Trt_F |
| DPY19L3 | 0.00611 | 0.25543 | 0.934 | 0.9 | 1 | HC_vs_MS.Trt_F |
| ARHGAP10 | 0.00615 | 0.68287 | 0.182 | 0.1 | 1 | HC_vs_MS.Trt_F |
| LGALS8 | 0.00615 | 0.43334 | 0.708 | 0.59 | 1 | HC_vs_MS.Trt_F |
| TNKS1BP1 | 0.00615 | 0.61469 | 0.118 | 0.06 | 1 | HC_vs_MS.Trt_F |

| | | | | | | |
|---|---|---|---|---|---|---|
| ATG2A | 0.00617 | 0.54006 | 0.801 | 0.71 | 1 | HC_vs_MS.Trt_F |
| EXOSC10 | 0.00617 | 0.41045 | 0.893 | 0.8 | 1 | HC_vs_MS.Trt_F |
| NCBP1 | 0.00618 | 0.78649 | 0.688 | 0.6 | 1 | HC_vs_MS.Trt_F |
| ECSIT | 0.00618 | 0.64907 | 0.327 | 0.23 | 1 | HC_vs_MS.Trt_F |
| THAP4 | 0.00618 | 0.56239 | 0.185 | 0.1 | 1 | HC_vs_MS.Trt_F |
| TANC2 | 0.00622 | -0.994 | 0.327 | 0.22 | 1 | HC_vs_MS.Trt_F |
| AP3M2 | 0.00623 | 0.55551 | 0.208 | 0.12 | 1 | HC_vs_MS.Trt_F |
| TRNT1 | 0.00623 | 0.50663 | 0.396 | 0.28 | 1 | HC_vs_MS.Trt_F |
| ADORA2A | 0.00635 | 0.79095 | 0.237 | 0.15 | 1 | HC_vs_MS.Trt_F |
| ILF3 | 0.00641 | 0.26759 | 0.803 | 0.72 | 1 | HC_vs_MS.Trt_F |
| ZNF544 | 0.00644 | 0.26251 | 0.954 | 0.92 | 1 | HC_vs_MS.Trt_F |
| YIPF5 | 0.00649 | 1.01818 | 0.569 | 0.5 | 1 | HC_vs_MS.Trt_F |
| NCK2 | 0.0065 | 0.91276 | 0.292 | 0.2 | 1 | HC_vs_MS.Trt_F |
| RPLP1 | 0.00657 | -0.2594 | 0.939 | 0.96 | 1 | HC_vs_MS.Trt_F |
| CALM1 | 0.00658 | -0.3845 | 0.803 | 0.8 | 1 | HC_vs_MS.Trt_F |
| TLR2 | 0.00659 | 0.33803 | 0.156 | 0.09 | 1 | HC_vs_MS.Trt_F |
| ZNF638 | 0.00663 | 0.30166 | 0.448 | 0.33 | 1 | HC_vs_MS.Trt_F |
| HSD17B8 | 0.00664 | -0.2901 | 0.153 | 0.08 | 1 | HC_vs_MS.Trt_F |
| METTL23 | 0.00667 | 0.38555 | 0.243 | 0.16 | 1 | HC_vs_MS.Trt_F |
| METTL22 | 0.00668 | 0.98136 | 0.705 | 0.62 | 1 | HC_vs_MS.Trt_F |
| CCT2 | 0.00668 | -0.2846 | 0.899 | 0.8 | 1 | HC_vs_MS.Trt_F |
| ATR | 0.00669 | 0.32687 | 0.561 | 0.44 | 1 | HC_vs_MS.Trt_F |
| ATIC | 0.00673 | 0.27763 | 0.451 | 0.34 | 1 | HC_vs_MS.Trt_F |
| SLC4A8 | 0.00674 | 0.39724 | 0.948 | 0.93 | 1 | HC_vs_MS.Trt_F |
| RGPD8 | 0.00675 | 0.79226 | 0.327 | 0.23 | 1 | HC_vs_MS.Trt_F |
| ALDH16A1 | 0.00676 | 0.3184 | 0.702 | 0.63 | 1 | HC_vs_MS.Trt_F |
| ITPK1 | 0.00679 | 0.31584 | 0.673 | 0.55 | 1 | HC_vs_MS.Trt_F |
| PTEN | 0.0068 | -1.2151 | 0.468 | 0.52 | 1 | HC_vs_MS.Trt_F |
| DMTF1 | 0.00682 | 0.50773 | 0.595 | 0.5 | 1 | HC_vs_MS.Trt_F |
| CPD | 0.00683 | 0.5737 | 0.743 | 0.65 | 1 | HC_vs_MS.Trt_F |
| CES2 | 0.00687 | -0.6997 | 0.422 | 0.31 | 1 | HC_vs_MS.Trt_F |
| FUS | 0.00699 | 0.56266 | 0.85 | 0.84 | 1 | HC_vs_MS.Trt_F |
| TLN1 | 0.00701 | 0.35938 | 0.685 | 0.57 | 1 | HC_vs_MS.Trt_F |
| NLE1 | 0.00703 | 1.3405 | 0.321 | 0.22 | 1 | HC_vs_MS.Trt_F |
| PI4KB | 0.00707 | 0.54932 | 0.251 | 0.16 | 1 | HC_vs_MS.Trt_F |
| EGR3 | 0.00709 | 0.74133 | 0.28 | 0.19 | 1 | HC_vs_MS.Trt_F |
| C2orf42 | 0.00713 | 0.40851 | 0.182 | 0.1 | 1 | HC_vs_MS.Trt_F |
| PML | 0.00714 | 0.2782 | 0.728 | 0.6 | 1 | HC_vs_MS.Trt_F |
| ZNF253 | 0.00717 | -0.5257 | 0.754 | 0.83 | 1 | HC_vs_MS.Trt_F |
| MCAM | 0.00718 | 1.47166 | 0.598 | 0.47 | 1 | HC_vs_MS.Trt_F |

| | | | | | | |
|---|---|---|---|---|---|---|
| PPIL3 | 0.0072 | -0.3479 | 0.841 | 0.73 | 1 | HC_vs_MS.Trt_F |
| LDHA | 0.00721 | 0.53646 | 0.754 | 0.66 | 1 | HC_vs_MS.Trt_F |
| KDM4B | 0.00726 | 0.75969 | 0.697 | 0.59 | 1 | HC_vs_MS.Trt_F |
| PUS7L | 0.00728 | 0.46053 | 0.91 | 0.87 | 1 | HC_vs_MS.Trt_F |
| IL7R | 0.00732 | 0.25139 | 0.905 | 0.9 | 1 | HC_vs_MS.Trt_F |
| PRR3 | 0.00733 | 0.52409 | 0.855 | 0.78 | 1 | HC_vs_MS.Trt_F |
| SKP2 | 0.00736 | -0.4024 | 0.616 | 0.49 | 1 | HC_vs_MS.Trt_F |
| CCDC80 | 0.00737 | 0.89157 | 0.783 | 0.7 | 1 | HC_vs_MS.Trt_F |
| ZCCHC2 | 0.00738 | 2.52059 | 0.139 | 0.07 | 1 | HC_vs_MS.Trt_F |
| CD300LG | 0.00741 | 0.47296 | 0.523 | 0.44 | 1 | HC_vs_MS.Trt_F |
| FPGS | 0.00747 | 0.31698 | 0.234 | 0.14 | 1 | HC_vs_MS.Trt_F |
| ZNF831 | 0.00747 | 0.33942 | 0.324 | 0.22 | 1 | HC_vs_MS.Trt_F |
| RBM14 | 0.00751 | 0.28358 | 0.251 | 0.16 | 1 | HC_vs_MS.Trt_F |
| MATK | 0.00753 | -0.2891 | 0.662 | 0.56 | 1 | HC_vs_MS.Trt_F |
| NEMP1 | 0.00754 | 1.34488 | 0.147 | 0.08 | 1 | HC_vs_MS.Trt_F |
| ARHGEF39 | 0.00754 | 0.84324 | 0.737 | 0.62 | 1 | HC_vs_MS.Trt_F |
| KCTD7 | 0.00755 | 0.49635 | 0.303 | 0.21 | 1 | HC_vs_MS.Trt_F |
| LMAN2L | 0.00759 | 0.5613 | 0.107 | 0.05 | 1 | HC_vs_MS.Trt_F |
| TBCK | 0.00759 | 0.60729 | 0.679 | 0.62 | 1 | HC_vs_MS.Trt_F |
| INIP | 0.0076 | -0.4375 | 0.757 | 0.6 | 1 | HC_vs_MS.Trt_F |
| RUNDC3A | 0.00761 | 0.46662 | 0.564 | 0.47 | 1 | HC_vs_MS.Trt_F |
| EIF4G2 | 0.00761 | -0.4994 | 0.552 | 0.6 | 1 | HC_vs_MS.Trt_F |
| ARL13B | 0.00762 | 0.62386 | 0.251 | 0.16 | 1 | HC_vs_MS.Trt_F |
| TARBP1 | 0.00763 | 1.5107 | 0.477 | 0.37 | 1 | HC_vs_MS.Trt_F |
| MADD | 0.00765 | 0.7939 | 0.283 | 0.19 | 1 | HC_vs_MS.Trt_F |
| AMZ2 | 0.0077 | 0.43999 | 0.656 | 0.53 | 1 | HC_vs_MS.Trt_F |
| PTK2B | 0.00773 | 0.51707 | 0.491 | 0.38 | 1 | HC_vs_MS.Trt_F |
| IQSEC1 | 0.00775 | 0.68363 | 0.286 | 0.19 | 1 | HC_vs_MS.Trt_F |
| KLF2 | 0.0078 | 0.2764 | 0.26 | 0.17 | 1 | HC_vs_MS.Trt_F |
| KRR1 | 0.00784 | 0.59108 | 0.717 | 0.64 | 1 | HC_vs_MS.Trt_F |
| CEP19 | 0.00785 | -0.5465 | 0.708 | 0.63 | 1 | HC_vs_MS.Trt_F |
| DXO | 0.00786 | 0.56034 | 0.171 | 0.1 | 1 | HC_vs_MS.Trt_F |
| ADRB2 | 0.00794 | -0.4667 | 0.309 | 0.21 | 1 | HC_vs_MS.Trt_F |
| CTTNBP2NL | 0.00795 | 0.5404 | 0.503 | 0.39 | 1 | HC_vs_MS.Trt_F |
| BCAS4 | 0.00797 | 0.30072 | 0.968 | 0.94 | 1 | HC_vs_MS.Trt_F |
| RUNX3 | 0.00797 | 0.29852 | 0.269 | 0.18 | 1 | HC_vs_MS.Trt_F |
| CEP350 | 0.00799 | 0.43458 | 0.405 | 0.3 | 1 | HC_vs_MS.Trt_F |
| PLA2G2C | 0.00801 | 0.37046 | 0.581 | 0.49 | 1 | HC_vs_MS.Trt_F |
| CLEC17A | 0.00802 | 0.62821 | 0.312 | 0.22 | 1 | HC_vs_MS.Trt_F |
| C2orf91 | 0.00803 | 0.4594 | 0.512 | 0.4 | 1 | HC_vs_MS.Trt_F |

| | | | | | | |
|---|---|---|---|---|---|---|
| RUFY1 | 0.00809 | 0.99044 | 0.35 | 0.25 | 1 | HC_vs_MS.Trt_F |
| SFSWAP | 0.0081 | 0.27976 | 0.61 | 0.52 | 1 | HC_vs_MS.Trt_F |
| PLEKHA3 | 0.0082 | 0.39446 | 0.835 | 0.76 | 1 | HC_vs_MS.Trt_F |
| ZFP36L2 | 0.00822 | 0.35028 | 0.595 | 0.49 | 1 | HC_vs_MS.Trt_F |
| HDAC4 | 0.00826 | 0.2707 | 0.601 | 0.53 | 1 | HC_vs_MS.Trt_F |
| DNAH1 | 0.0083 | 0.82132 | 0.211 | 0.13 | 1 | HC_vs_MS.Trt_F |
| NBPF3 | 0.00832 | 0.34726 | 0.488 | 0.4 | 1 | HC_vs_MS.Trt_F |
| PZP | 0.00834 | 0.82067 | 0.15 | 0.08 | 1 | HC_vs_MS.Trt_F |
| EP300 | 0.00838 | 0.40716 | 0.78 | 0.66 | 1 | HC_vs_MS.Trt_F |
| EYS | 0.0084 | 0.41486 | 0.801 | 0.75 | 1 | HC_vs_MS.Trt_F |
| PSMD7 | 0.00841 | 0.33873 | 0.37 | 0.27 | 1 | HC_vs_MS.Trt_F |
| IFIT1 | 0.00842 | 1.37184 | 0.312 | 0.21 | 1 | HC_vs_MS.Trt_F |
| CLIP4 | 0.00843 | 1.15549 | 0.162 | 0.09 | 1 | HC_vs_MS.Trt_F |
| STRN3 | 0.00846 | -0.6539 | 0.815 | 0.83 | 1 | HC_vs_MS.Trt_F |
| GSTZ1 | 0.00848 | 0.61719 | 0.595 | 0.48 | 1 | HC_vs_MS.Trt_F |
| CST3 | 0.00849 | 1.58054 | 0.153 | 0.09 | 1 | HC_vs_MS.Trt_F |
| CCDC141 | 0.0085 | 0.55311 | 0.702 | 0.65 | 1 | HC_vs_MS.Trt_F |
| MECR | 0.00858 | 0.45981 | 0.405 | 0.28 | 1 | HC_vs_MS.Trt_F |
| ZNF396 | 0.00858 | 0.29632 | 0.896 | 0.82 | 1 | HC_vs_MS.Trt_F |
| MMRN2 | 0.00858 | 0.70475 | 0.249 | 0.16 | 1 | HC_vs_MS.Trt_F |
| SEC14L6 | 0.0086 | 0.43149 | 0.798 | 0.7 | 1 | HC_vs_MS.Trt_F |
| ATAD5 | 0.00862 | 0.86507 | 0.618 | 0.5 | 1 | HC_vs_MS.Trt_F |
| PSMC1 | 0.00865 | 0.28733 | 0.454 | 0.34 | 1 | HC_vs_MS.Trt_F |
| CLK2 | 0.00872 | 0.86597 | 0.202 | 0.12 | 1 | HC_vs_MS.Trt_F |
| GNAQ | 0.00877 | 0.60639 | 0.257 | 0.17 | 1 | HC_vs_MS.Trt_F |
| MAP4K5 | 0.00879 | 0.70535 | 0.601 | 0.48 | 1 | HC_vs_MS.Trt_F |
| MTHFD1 | 0.0088 | 0.41207 | 0.896 | 0.87 | 1 | HC_vs_MS.Trt_F |
| CALCRL | 0.00882 | 0.36383 | 0.737 | 0.67 | 1 | HC_vs_MS.Trt_F |
| ZNF674 | 0.00882 | 0.55117 | 0.948 | 0.9 | 1 | HC_vs_MS.Trt_F |
| OBSCN | 0.00882 | 0.47208 | 0.269 | 0.18 | 1 | HC_vs_MS.Trt_F |
| CCNL1 | 0.00884 | 0.38859 | 0.445 | 0.34 | 1 | HC_vs_MS.Trt_F |
| COL6A1 | 0.00884 | 2.02508 | 0.107 | 0.05 | 1 | HC_vs_MS.Trt_F |
| CDC42 | 0.00889 | -0.4989 | 0.905 | 0.91 | 1 | HC_vs_MS.Trt_F |
| FAM46C | 0.00889 | 0.433 | 0.41 | 0.31 | 1 | HC_vs_MS.Trt_F |
| MAFF | 0.00892 | 0.48373 | 0.723 | 0.62 | 1 | HC_vs_MS.Trt_F |
| RGPD2 | 0.00897 | 0.43391 | 0.243 | 0.16 | 1 | HC_vs_MS.Trt_F |
| ZNF799 | 0.00898 | -0.9024 | 0.471 | 0.37 | 1 | HC_vs_MS.Trt_F |
| SMIM12 | 0.00899 | 0.39071 | 0.818 | 0.74 | 1 | HC_vs_MS.Trt_F |
| SLC41A1 | 0.00901 | 0.84951 | 0.179 | 0.1 | 1 | HC_vs_MS.Trt_F |
| SAMD9 | 0.00902 | 0.41249 | 0.234 | 0.15 | 1 | HC_vs_MS.Trt_F |

| | | | | | | |
|---|---|---|---|---|---|---|
| PPP2R2A | 0.00906 | 1.24298 | 0.208 | 0.13 | 1 | HC_vs_MS.Trt_F |
| RBM6 | 0.00906 | -0.5176 | 0.858 | 0.87 | 1 | HC_vs_MS.Trt_F |
| TKFC | 0.00909 | 0.29874 | 0.639 | 0.53 | 1 | HC_vs_MS.Trt_F |
| GOLM1 | 0.00909 | 1.06331 | 0.642 | 0.52 | 1 | HC_vs_MS.Trt_F |
| THG1L | 0.00911 | -0.8707 | 0.621 | 0.54 | 1 | HC_vs_MS.Trt_F |
| NME3 | 0.00922 | 0.46598 | 0.185 | 0.11 | 1 | HC_vs_MS.Trt_F |
| TUBG2 | 0.00923 | 0.49081 | 0.147 | 0.08 | 1 | HC_vs_MS.Trt_F |
| FAM114A1 | 0.00927 | 0.49917 | 0.618 | 0.51 | 1 | HC_vs_MS.Trt_F |
| PREX1 | 0.00933 | 0.4864 | 0.613 | 0.52 | 1 | HC_vs_MS.Trt_F |
| POLR3E | 0.00937 | 0.28847 | 0.474 | 0.4 | 1 | HC_vs_MS.Trt_F |
| CBS | 0.00941 | -0.4136 | 0.162 | 0.24 | 1 | HC_vs_MS.Trt_F |
| ZEB1 | 0.00942 | 0.53391 | 0.624 | 0.5 | 1 | HC_vs_MS.Trt_F |
| S1PR4 | 0.00943 | 0.46752 | 0.231 | 0.15 | 1 | HC_vs_MS.Trt_F |
| PLEKHM3 | 0.00947 | 0.75233 | 0.584 | 0.48 | 1 | HC_vs_MS.Trt_F |
| G6PC2 | 0.00947 | 0.32006 | 0.379 | 0.28 | 1 | HC_vs_MS.Trt_F |
| TUT1 | 0.00948 | -0.5564 | 0.191 | 0.11 | 1 | HC_vs_MS.Trt_F |
| TNFRSF18 | 0.0095 | 0.6208 | 0.289 | 0.2 | 1 | HC_vs_MS.Trt_F |
| KCNC2 | 0.00958 | 0.37498 | 0.442 | 0.35 | 1 | HC_vs_MS.Trt_F |
| MCL1 | 0.00965 | 0.38764 | 0.462 | 0.35 | 1 | HC_vs_MS.Trt_F |
| HYPK | 0.00965 | 0.58727 | 0.922 | 0.9 | 1 | HC_vs_MS.Trt_F |
| TLCD2 | 0.00967 | 0.37261 | 0.948 | 0.93 | 1 | HC_vs_MS.Trt_F |
| PARVB | 0.00975 | 0.55089 | 0.749 | 0.69 | 1 | HC_vs_MS.Trt_F |
| CNBD2 | 0.00976 | 0.45511 | 0.543 | 0.44 | 1 | HC_vs_MS.Trt_F |
| GADL1 | 0.0098 | 0.27778 | 0.731 | 0.66 | 1 | HC_vs_MS.Trt_F |
| AFF1 | 0.00981 | 0.41844 | 0.39 | 0.28 | 1 | HC_vs_MS.Trt_F |
| IL1R1 | 0.00983 | -0.4334 | 0.422 | 0.32 | 1 | HC_vs_MS.Trt_F |
| VKORC1 | 0.00986 | 0.27077 | 0.214 | 0.13 | 1 | HC_vs_MS.Trt_F |
| SCAMP3 | 0.00989 | 0.658 | 0.251 | 0.17 | 1 | HC_vs_MS.Trt_F |
| SPICE1 | 0.00991 | 0.92539 | 0.324 | 0.23 | 1 | HC_vs_MS.Trt_F |
| TRIM44 | 0.00996 | -0.2641 | 0.387 | 0.28 | 1 | HC_vs_MS.Trt_F |
| UBE4B | 0.00996 | 0.9159 | 0.659 | 0.55 | 1 | HC_vs_MS.Trt_F |
| SMAD1 | 0.00996 | -0.8336 | 0.442 | 0.33 | 1 | HC_vs_MS.Trt_F |
| GPM6A | 0.00998 | 0.74884 | 0.355 | 0.26 | 1 | HC_vs_MS.Trt_F |
| CASP9 | 0.01002 | 0.54708 | 0.471 | 0.37 | 1 | HC_vs_MS.Trt_F |
| WWOX | 0.01003 | 0.29257 | 0.286 | 0.19 | 1 | HC_vs_MS.Trt_F |
| COL4A2 | 0.01004 | 0.89625 | 0.121 | 0.06 | 1 | HC_vs_MS.Trt_F |
| RABL3 | 0.01004 | -0.4596 | 0.607 | 0.67 | 1 | HC_vs_MS.Trt_F |
| CCSER2 | 0.01005 | -0.3245 | 0.578 | 0.44 | 1 | HC_vs_MS.Trt_F |
| DHRS12 | 0.01009 | 0.75824 | 0.162 | 0.09 | 1 | HC_vs_MS.Trt_F |
| TMEM239 | 0.01012 | 0.34419 | 0.382 | 0.3 | 1 | HC_vs_MS.Trt_F |

| | | | | | | |
|---|---|---|---|---|---|---|
| SNX27 | 0.01014 | -0.4646 | 0.803 | 0.68 | 1 | HC_vs_MS.Trt_F |
| PLGLB1 | 0.01019 | 0.75785 | 0.185 | 0.11 | 1 | HC_vs_MS.Trt_F |
| RBMS1 | 0.01022 | 0.32505 | 0.731 | 0.61 | 1 | HC_vs_MS.Trt_F |
| FSD1L | 0.01022 | 0.8944 | 0.28 | 0.19 | 1 | HC_vs_MS.Trt_F |
| POLR3D | 0.01024 | 1.38467 | 0.445 | 0.35 | 1 | HC_vs_MS.Trt_F |
| HIPK1 | 0.01024 | 0.85672 | 0.263 | 0.18 | 1 | HC_vs_MS.Trt_F |
| TECRL | 0.01037 | 0.40882 | 0.286 | 0.2 | 1 | HC_vs_MS.Trt_F |
| CARD19 | 0.0104 | 0.73998 | 0.37 | 0.28 | 1 | HC_vs_MS.Trt_F |
| TBXA2R | 0.01041 | 0.29534 | 0.746 | 0.67 | 1 | HC_vs_MS.Trt_F |
| CACNB3 | 0.01042 | 0.43822 | 0.633 | 0.51 | 1 | HC_vs_MS.Trt_F |
| TPCN1 | 0.01045 | 0.31478 | 0.428 | 0.34 | 1 | HC_vs_MS.Trt_F |
| RAB13 | 0.01049 | 0.28736 | 0.572 | 0.46 | 1 | HC_vs_MS.Trt_F |
| ERICH3 | 0.0105 | 1.51711 | 0.118 | 0.06 | 1 | HC_vs_MS.Trt_F |
| NT5C | 0.01051 | 0.81042 | 0.205 | 0.13 | 1 | HC_vs_MS.Trt_F |
| TCERG1 | 0.01057 | 0.36834 | 0.341 | 0.25 | 1 | HC_vs_MS.Trt_F |
| KLC1 | 0.01057 | 0.42857 | 0.682 | 0.6 | 1 | HC_vs_MS.Trt_F |
| ELP2 | 0.01058 | 0.29785 | 0.688 | 0.57 | 1 | HC_vs_MS.Trt_F |
| GALNT2 | 0.01063 | -0.3434 | 0.286 | 0.19 | 1 | HC_vs_MS.Trt_F |
| CEP41 | 0.01068 | 0.34946 | 0.87 | 0.78 | 1 | HC_vs_MS.Trt_F |
| MAP7D1 | 0.01068 | 0.66336 | 0.59 | 0.46 | 1 | HC_vs_MS.Trt_F |
| MBD5 | 0.0107 | 0.69901 | 0.87 | 0.84 | 1 | HC_vs_MS.Trt_F |
| DOT1L | 0.01074 | 0.77819 | 0.818 | 0.75 | 1 | HC_vs_MS.Trt_F |
| ZWILCH | 0.01077 | -0.9158 | 0.694 | 0.59 | 1 | HC_vs_MS.Trt_F |
| TSC22D3 | 0.01081 | 0.31023 | 0.754 | 0.68 | 1 | HC_vs_MS.Trt_F |
| ULBP3 | 0.01081 | -0.4989 | 0.647 | 0.54 | 1 | HC_vs_MS.Trt_F |
| SMYD3 | 0.01088 | 0.35681 | 0.234 | 0.15 | 1 | HC_vs_MS.Trt_F |
| TTC5 | 0.01089 | 0.45512 | 0.624 | 0.51 | 1 | HC_vs_MS.Trt_F |
| C11orf24 | 0.01096 | 1.03574 | 0.275 | 0.19 | 1 | HC_vs_MS.Trt_F |
| MYO1G | 0.01102 | 0.34744 | 0.653 | 0.58 | 1 | HC_vs_MS.Trt_F |
| CYCS | 0.01107 | 0.45627 | 0.801 | 0.74 | 1 | HC_vs_MS.Trt_F |
| WDR74 | 0.01107 | 0.40976 | 0.653 | 0.53 | 1 | HC_vs_MS.Trt_F |
| C1orf123 | 0.01112 | 0.30853 | 0.22 | 0.14 | 1 | HC_vs_MS.Trt_F |
| RBM18 | 0.01125 | -0.5644 | 0.393 | 0.28 | 1 | HC_vs_MS.Trt_F |
| IGF2BP3 | 0.01126 | 0.34449 | 0.613 | 0.53 | 1 | HC_vs_MS.Trt_F |
| RGPD4 | 0.01132 | 0.62508 | 0.217 | 0.14 | 1 | HC_vs_MS.Trt_F |
| ZNF724 | 0.01133 | 0.47572 | 0.925 | 0.86 | 1 | HC_vs_MS.Trt_F |
| MET | 0.01136 | -0.3999 | 0.332 | 0.4 | 1 | HC_vs_MS.Trt_F |
| DMPK | 0.01139 | 0.61196 | 0.668 | 0.57 | 1 | HC_vs_MS.Trt_F |
| RNGTT | 0.0114 | 0.63659 | 0.439 | 0.34 | 1 | HC_vs_MS.Trt_F |
| LYST | 0.01144 | 0.62953 | 0.422 | 0.31 | 1 | HC_vs_MS.Trt_F |

| | | | | | | |
|---|---|---|---|---|---|---|
| NCAPG2 | 0.01152 | -1.5356 | 0.202 | 0.27 | 1 | HC_vs_MS.Trt_F |
| UTP18 | 0.01152 | 0.64715 | 0.202 | 0.13 | 1 | HC_vs_MS.Trt_F |
| ELMO1 | 0.01155 | 0.38039 | 0.841 | 0.79 | 1 | HC_vs_MS.Trt_F |
| SEL1L | 0.01156 | 0.48206 | 0.254 | 0.17 | 1 | HC_vs_MS.Trt_F |
| BMP1 | 0.01158 | 0.59483 | 0.639 | 0.54 | 1 | HC_vs_MS.Trt_F |
| TNKS2 | 0.01162 | 0.41012 | 0.347 | 0.26 | 1 | HC_vs_MS.Trt_F |
| CDC37 | 0.01162 | 0.36832 | 0.465 | 0.36 | 1 | HC_vs_MS.Trt_F |
| AKAP11 | 0.01169 | 0.55954 | 0.286 | 0.19 | 1 | HC_vs_MS.Trt_F |
| AC012314.1 | 0.01171 | -0.3221 | 0.676 | 0.69 | 1 | HC_vs_MS.Trt_F |
| AC012314.1 | 0.01171 | -0.3221 | 0.676 | 0.69 | 1 | HC_vs_MS.Trt_F |
| AC012314.1 | 0.01171 | -0.3221 | 0.676 | 0.69 | 1 | HC_vs_MS.Trt_F |
| AC012314.2 | 0.01171 | -0.3221 | 0.676 | 0.69 | 1 | HC_vs_MS.Trt_F |
| AC012314.4 | 0.01171 | -0.3221 | 0.676 | 0.69 | 1 | HC_vs_MS.Trt_F |
| AC012314.5 | 0.01171 | -0.3221 | 0.676 | 0.69 | 1 | HC_vs_MS.Trt_F |
| POLE4 | 0.01178 | 0.7483 | 0.188 | 0.11 | 1 | HC_vs_MS.Trt_F |
| R3HDM2 | 0.0118 | -0.5129 | 0.772 | 0.76 | 1 | HC_vs_MS.Trt_F |
| DNASE1L3 | 0.0118 | 0.38646 | 0.462 | 0.34 | 1 | HC_vs_MS.Trt_F |
| CRISPLD2 | 0.01186 | 0.39619 | 0.809 | 0.77 | 1 | HC_vs_MS.Trt_F |
| HNRNPU | 0.01197 | -0.4927 | 0.506 | 0.57 | 1 | HC_vs_MS.Trt_F |
| TMPRSS13 | 0.01203 | -0.8996 | 0.243 | 0.16 | 1 | HC_vs_MS.Trt_F |
| ASAP1 | 0.01204 | 1.31313 | 0.257 | 0.18 | 1 | HC_vs_MS.Trt_F |
| MNAT1 | 0.01207 | 0.86474 | 0.156 | 0.09 | 1 | HC_vs_MS.Trt_F |
| LIG1 | 0.01216 | 0.70579 | 0.194 | 0.12 | 1 | HC_vs_MS.Trt_F |
| PAK3 | 0.01219 | 0.44804 | 0.543 | 0.42 | 1 | HC_vs_MS.Trt_F |
| FAM131A | 0.01236 | -0.4738 | 0.882 | 0.8 | 1 | HC_vs_MS.Trt_F |
| NSUN4 | 0.01252 | 0.41502 | 0.806 | 0.73 | 1 | HC_vs_MS.Trt_F |
| RNPC3 | 0.01256 | 0.35153 | 0.405 | 0.3 | 1 | HC_vs_MS.Trt_F |
| ECE1 | 0.01257 | 0.33114 | 0.763 | 0.68 | 1 | HC_vs_MS.Trt_F |
| HCFC1 | 0.01258 | 0.44277 | 0.341 | 0.25 | 1 | HC_vs_MS.Trt_F |
| SULT2B1 | 0.01264 | 0.27496 | 0.569 | 0.46 | 1 | HC_vs_MS.Trt_F |
| LMF2 | 0.01265 | 0.5996 | 0.303 | 0.22 | 1 | HC_vs_MS.Trt_F |
| FADS6 | 0.01282 | 0.28071 | 0.639 | 0.53 | 1 | HC_vs_MS.Trt_F |
| PHC3 | 0.01289 | 0.31376 | 0.968 | 0.97 | 1 | HC_vs_MS.Trt_F |
| ARHGAP11B | 0.01291 | 0.25832 | 0.289 | 0.2 | 1 | HC_vs_MS.Trt_F |
| BMP8B | 0.01293 | 0.29052 | 0.514 | 0.4 | 1 | HC_vs_MS.Trt_F |
| EPG5 | 0.01301 | 0.56915 | 0.564 | 0.45 | 1 | HC_vs_MS.Trt_F |
| ZNF395 | 0.01303 | 0.61651 | 0.592 | 0.53 | 1 | HC_vs_MS.Trt_F |
| RPL35A | 0.01307 | -0.357 | 0.829 | 0.84 | 1 | HC_vs_MS.Trt_F |
| FXR2 | 0.0131 | 0.47254 | 0.714 | 0.63 | 1 | HC_vs_MS.Trt_F |
| SLC25A34 | 0.01311 | -0.5611 | 0.668 | 0.55 | 1 | HC_vs_MS.Trt_F |

| | | | | | | |
|---|---|---|---|---|---|---|
| RP3-461F17 | 0.01312 | 0.54795 | 0.283 | 0.19 | 1 | HC_vs_MS.Trt_F |
| TPH1 | 0.01319 | -0.6761 | 0.564 | 0.43 | 1 | HC_vs_MS.Trt_F |
| COL6A3 | 0.01322 | 1.64712 | 0.517 | 0.4 | 1 | HC_vs_MS.Trt_F |
| PIGH | 0.01327 | -0.4207 | 0.688 | 0.56 | 1 | HC_vs_MS.Trt_F |
| SIAH1 | 0.01329 | 0.66562 | 0.275 | 0.19 | 1 | HC_vs_MS.Trt_F |
| OFD1 | 0.0133 | 0.26845 | 0.298 | 0.2 | 1 | HC_vs_MS.Trt_F |
| CCNYL1 | 0.01332 | -0.7228 | 0.48 | 0.35 | 1 | HC_vs_MS.Trt_F |
| FIS1 | 0.01336 | 0.35076 | 0.367 | 0.26 | 1 | HC_vs_MS.Trt_F |
| ZNF343 | 0.01339 | 1.11947 | 0.52 | 0.43 | 1 | HC_vs_MS.Trt_F |
| PIM1 | 0.01339 | 0.25626 | 0.457 | 0.34 | 1 | HC_vs_MS.Trt_F |
| DCAF17 | 0.0134 | 0.30895 | 0.483 | 0.38 | 1 | HC_vs_MS.Trt_F |
| KCNJ6 | 0.01345 | 0.53937 | 0.873 | 0.83 | 1 | HC_vs_MS.Trt_F |
| YIF1A | 0.01347 | 0.51843 | 0.173 | 0.1 | 1 | HC_vs_MS.Trt_F |
| FAM45A | 0.01351 | 0.31121 | 0.754 | 0.67 | 1 | HC_vs_MS.Trt_F |
| KCNJ5 | 0.01352 | 0.36872 | 0.673 | 0.57 | 1 | HC_vs_MS.Trt_F |
| CUL1 | 0.01365 | 0.98644 | 0.237 | 0.16 | 1 | HC_vs_MS.Trt_F |
| FCHO1 | 0.01369 | 0.73607 | 0.818 | 0.73 | 1 | HC_vs_MS.Trt_F |
| CYP2C19 | 0.01374 | 0.40836 | 0.769 | 0.7 | 1 | HC_vs_MS.Trt_F |
| NFKBIZ | 0.01376 | 0.88589 | 0.445 | 0.36 | 1 | HC_vs_MS.Trt_F |
| RIPK1 | 0.01381 | 0.97254 | 0.237 | 0.16 | 1 | HC_vs_MS.Trt_F |
| LMTK2 | 0.01389 | 0.25881 | 0.303 | 0.22 | 1 | HC_vs_MS.Trt_F |
| DMD | 0.01394 | 0.30438 | 0.728 | 0.63 | 1 | HC_vs_MS.Trt_F |
| SLBP | 0.01397 | 0.2733 | 0.243 | 0.16 | 1 | HC_vs_MS.Trt_F |
| VPS25 | 0.01397 | 0.80787 | 0.338 | 0.25 | 1 | HC_vs_MS.Trt_F |
| ZC3H10 | 0.014 | 0.37529 | 0.613 | 0.51 | 1 | HC_vs_MS.Trt_F |
| USP38 | 0.01401 | 0.25753 | 0.384 | 0.29 | 1 | HC_vs_MS.Trt_F |
| PTPN3 | 0.01418 | 0.27175 | 0.835 | 0.74 | 1 | HC_vs_MS.Trt_F |
| KLF11 | 0.01419 | 1.06951 | 0.107 | 0.05 | 1 | HC_vs_MS.Trt_F |
| GBP2 | 0.01424 | 0.27705 | 0.46 | 0.36 | 1 | HC_vs_MS.Trt_F |
| CAPS | 0.01432 | 0.2638 | 0.971 | 0.95 | 1 | HC_vs_MS.Trt_F |
| CRY1 | 0.01448 | 0.49777 | 0.208 | 0.13 | 1 | HC_vs_MS.Trt_F |
| DUSP16 | 0.01449 | -0.4702 | 0.329 | 0.23 | 1 | HC_vs_MS.Trt_F |
| UQCC1 | 0.01457 | 0.48134 | 0.595 | 0.5 | 1 | HC_vs_MS.Trt_F |
| DYNLT3 | 0.01459 | 0.67448 | 0.691 | 0.62 | 1 | HC_vs_MS.Trt_F |
| TP53I13 | 0.01462 | 0.65 | 0.179 | 0.11 | 1 | HC_vs_MS.Trt_F |
| ERAL1 | 0.01466 | 0.80326 | 0.153 | 0.09 | 1 | HC_vs_MS.Trt_F |
| CFHR1 | 0.01468 | 0.34882 | 0.786 | 0.71 | 1 | HC_vs_MS.Trt_F |
| PBX2 | 0.01475 | 0.82427 | 0.173 | 0.11 | 1 | HC_vs_MS.Trt_F |
| ARHGDIA | 0.01476 | 1.09511 | 0.416 | 0.34 | 1 | HC_vs_MS.Trt_F |
| WTAP | 0.01483 | -0.2821 | 0.621 | 0.65 | 1 | HC_vs_MS.Trt_F |

| | | | | | | |
|---|---|---|---|---|---|---|
| GPAM | 0.01483 | 0.61129 | 0.113 | 0.06 | 1 | HC_vs_MS.Trt_F |
| CPPED1 | 0.01488 | 0.54076 | 0.65 | 0.58 | 1 | HC_vs_MS.Trt_F |
| DST | 0.01489 | 0.39267 | 0.757 | 0.67 | 1 | HC_vs_MS.Trt_F |
| CARMIL1 | 0.0149 | 0.54195 | 0.361 | 0.26 | 1 | HC_vs_MS.Trt_F |
| MAP4K1 | 0.01492 | 0.36523 | 0.91 | 0.88 | 1 | HC_vs_MS.Trt_F |
| KIAA0930 | 0.01507 | 1.05584 | 0.691 | 0.61 | 1 | HC_vs_MS.Trt_F |
| VPS37B | 0.0151 | 1.17041 | 0.694 | 0.62 | 1 | HC_vs_MS.Trt_F |
| PROSER3 | 0.01513 | 0.68197 | 0.916 | 0.84 | 1 | HC_vs_MS.Trt_F |
| GPCPD1 | 0.01514 | 0.33864 | 0.445 | 0.36 | 1 | HC_vs_MS.Trt_F |
| LSR | 0.01519 | 1.52591 | 0.237 | 0.17 | 1 | HC_vs_MS.Trt_F |
| LAT2 | 0.01525 | 0.70502 | 0.876 | 0.81 | 1 | HC_vs_MS.Trt_F |
| ERC1 | 0.0153 | 0.3081 | 0.931 | 0.89 | 1 | HC_vs_MS.Trt_F |
| RAN | 0.01535 | 0.36104 | 0.468 | 0.37 | 1 | HC_vs_MS.Trt_F |
| RANBP3 | 0.01538 | 0.31264 | 0.838 | 0.76 | 1 | HC_vs_MS.Trt_F |
| SIRT5 | 0.01545 | 0.45472 | 0.879 | 0.79 | 1 | HC_vs_MS.Trt_F |
| TRAM2 | 0.01548 | 0.37755 | 0.309 | 0.22 | 1 | HC_vs_MS.Trt_F |
| TRIP11 | 0.01565 | 0.31229 | 0.613 | 0.51 | 1 | HC_vs_MS.Trt_F |
| KLRG1 | 0.0157 | 0.50403 | 0.465 | 0.37 | 1 | HC_vs_MS.Trt_F |
| DENND4C | 0.01577 | 0.77506 | 0.442 | 0.34 | 1 | HC_vs_MS.Trt_F |
| CTSL | 0.01579 | -1.1868 | 0.078 | 0.14 | 1 | HC_vs_MS.Trt_F |
| ANKRD36B | 0.01586 | -0.2641 | 0.743 | 0.64 | 1 | HC_vs_MS.Trt_F |
| USP37 | 0.01591 | 0.53538 | 0.535 | 0.44 | 1 | HC_vs_MS.Trt_F |
| NUP210 | 0.01593 | 0.42031 | 0.578 | 0.5 | 1 | HC_vs_MS.Trt_F |
| TBC1D15 | 0.01601 | 0.79655 | 0.592 | 0.51 | 1 | HC_vs_MS.Trt_F |
| RABGAP1 | 0.01606 | 0.60266 | 0.468 | 0.36 | 1 | HC_vs_MS.Trt_F |
| ARPC4 | 0.01607 | -0.4879 | 0.645 | 0.68 | 1 | HC_vs_MS.Trt_F |
| GRSF1 | 0.01608 | -0.2528 | 0.76 | 0.68 | 1 | HC_vs_MS.Trt_F |
| ZNF106 | 0.01624 | 0.31351 | 0.905 | 0.85 | 1 | HC_vs_MS.Trt_F |
| ZNF592 | 0.0163 | 0.55131 | 0.246 | 0.16 | 1 | HC_vs_MS.Trt_F |
| EML6 | 0.01633 | 0.603 | 0.665 | 0.61 | 1 | HC_vs_MS.Trt_F |
| RABGGTB | 0.01639 | 0.85655 | 0.373 | 0.28 | 1 | HC_vs_MS.Trt_F |
| KPNA2 | 0.0164 | 0.44733 | 0.168 | 0.1 | 1 | HC_vs_MS.Trt_F |
| RP11-176H8 | 0.01641 | 0.77488 | 0.887 | 0.83 | 1 | HC_vs_MS.Trt_F |
| MTMR6 | 0.01647 | 1.27962 | 0.171 | 0.1 | 1 | HC_vs_MS.Trt_F |
| WBP2 | 0.01648 | 0.37105 | 0.723 | 0.6 | 1 | HC_vs_MS.Trt_F |
| RAB2B | 0.01654 | 0.26746 | 0.74 | 0.64 | 1 | HC_vs_MS.Trt_F |
| UGCG | 0.01655 | 0.36938 | 0.286 | 0.2 | 1 | HC_vs_MS.Trt_F |
| MSL2 | 0.01656 | 0.381 | 0.22 | 0.14 | 1 | HC_vs_MS.Trt_F |
| TGIF1 | 0.01657 | 0.37577 | 0.315 | 0.23 | 1 | HC_vs_MS.Trt_F |
| ICOSLG | 0.01663 | 0.6253 | 0.803 | 0.71 | 1 | HC_vs_MS.Trt_F |

| | | | | | | |
|---|---|---|---|---|---|---|
| SMC6 | 0.01665 | 0.83676 | 0.54 | 0.44 | 1 | HC_vs_MS.Trt_F |
| RLIM | 0.01666 | 0.70635 | 0.772 | 0.66 | 1 | HC_vs_MS.Trt_F |
| CTLA4 | 0.01673 | 0.37451 | 0.321 | 0.23 | 1 | HC_vs_MS.Trt_F |
| NAV1 | 0.01677 | 0.31694 | 0.908 | 0.87 | 1 | HC_vs_MS.Trt_F |
| WDFY2 | 0.01678 | 0.58202 | 0.645 | 0.55 | 1 | HC_vs_MS.Trt_F |
| UBE2B | 0.01679 | -0.2752 | 0.757 | 0.68 | 1 | HC_vs_MS.Trt_F |
| CCR4 | 0.01682 | -0.5934 | 0.922 | 0.91 | 1 | HC_vs_MS.Trt_F |
| TMEM219 | 0.0169 | 0.38729 | 0.327 | 0.24 | 1 | HC_vs_MS.Trt_F |
| ABR | 0.01694 | 0.46339 | 0.789 | 0.75 | 1 | HC_vs_MS.Trt_F |
| GINS3 | 0.01702 | 0.26922 | 0.89 | 0.84 | 1 | HC_vs_MS.Trt_F |
| PPP4R3B | 0.0171 | 0.43515 | 0.497 | 0.4 | 1 | HC_vs_MS.Trt_F |
| GALNT15 | 0.01713 | -0.2727 | 0.37 | 0.28 | 1 | HC_vs_MS.Trt_F |
| ACSS1 | 0.01714 | 0.83491 | 0.627 | 0.57 | 1 | HC_vs_MS.Trt_F |
| RIC1 | 0.0172 | 0.48811 | 0.202 | 0.13 | 1 | HC_vs_MS.Trt_F |
| ZNF726 | 0.0172 | 0.48174 | 0.902 | 0.86 | 1 | HC_vs_MS.Trt_F |
| PHPT1 | 0.01723 | -0.4228 | 0.228 | 0.14 | 1 | HC_vs_MS.Trt_F |
| NOX5 | 0.01724 | 0.38539 | 0.592 | 0.51 | 1 | HC_vs_MS.Trt_F |
| C4BPB | 0.01725 | 0.32899 | 0.85 | 0.81 | 1 | HC_vs_MS.Trt_F |
| ZNF345 | 0.01726 | 0.78287 | 0.85 | 0.8 | 1 | HC_vs_MS.Trt_F |
| OAS3 | 0.01733 | 0.65319 | 0.665 | 0.55 | 1 | HC_vs_MS.Trt_F |
| GINM1 | 0.01735 | 0.41825 | 0.598 | 0.54 | 1 | HC_vs_MS.Trt_F |
| FDFT1 | 0.01737 | 0.32183 | 0.364 | 0.28 | 1 | HC_vs_MS.Trt_F |
| UVSSA | 0.01738 | 0.75261 | 0.824 | 0.74 | 1 | HC_vs_MS.Trt_F |
| ABCA2 | 0.01743 | 0.84527 | 0.341 | 0.25 | 1 | HC_vs_MS.Trt_F |
| MAPK14 | 0.01745 | 0.5858 | 0.205 | 0.13 | 1 | HC_vs_MS.Trt_F |
| ZNF778 | 0.01746 | 0.37347 | 0.705 | 0.64 | 1 | HC_vs_MS.Trt_F |
| SMNDC1 | 0.01747 | 0.89739 | 0.197 | 0.13 | 1 | HC_vs_MS.Trt_F |
| FASTKD5 | 0.0175 | -0.7266 | 0.159 | 0.09 | 1 | HC_vs_MS.Trt_F |
| MR1 | 0.01751 | 0.51111 | 0.936 | 0.9 | 1 | HC_vs_MS.Trt_F |
| NREP | 0.01752 | 0.6366 | 0.205 | 0.14 | 1 | HC_vs_MS.Trt_F |
| ACAD10 | 0.01752 | -0.8256 | 0.506 | 0.38 | 1 | HC_vs_MS.Trt_F |
| CDKN1B | 0.01752 | 0.46921 | 0.355 | 0.27 | 1 | HC_vs_MS.Trt_F |
| RP4-635E18 | 0.01754 | -1.3274 | 0.173 | 0.23 | 1 | HC_vs_MS.Trt_F |
| TAF1 | 0.01757 | -0.393 | 0.815 | 0.75 | 1 | HC_vs_MS.Trt_F |
| RPS4Y1 | 0.0176 | 2.41091 | 0.136 | 0.08 | 1 | HC_vs_MS.Trt_F |
| MCM6 | 0.01762 | 0.89773 | 0.173 | 0.11 | 1 | HC_vs_MS.Trt_F |
| SOCS3 | 0.01764 | 0.32559 | 0.214 | 0.14 | 1 | HC_vs_MS.Trt_F |
| HOXB2 | 0.0177 | 0.33304 | 0.182 | 0.11 | 1 | HC_vs_MS.Trt_F |
| GMDS | 0.01774 | 0.43209 | 0.194 | 0.12 | 1 | HC_vs_MS.Trt_F |
| DENND5B | 0.01777 | 0.50897 | 0.642 | 0.53 | 1 | HC_vs_MS.Trt_F |

| | | | | | | |
|---|---|---|---|---|---|---|
| TMED4 | 0.01781 | -0.6157 | 0.951 | 0.95 | 1 | HC_vs_MS.Trt_F |
| CYP2C9 | 0.01794 | 0.28813 | 0.581 | 0.49 | 1 | HC_vs_MS.Trt_F |
| ENAH | 0.018 | 0.55282 | 0.723 | 0.62 | 1 | HC_vs_MS.Trt_F |
| CCR5 | 0.01804 | 0.99104 | 0.147 | 0.09 | 1 | HC_vs_MS.Trt_F |
| RAPGEF6 | 0.01806 | 0.5173 | 0.546 | 0.44 | 1 | HC_vs_MS.Trt_F |
| HAX1 | 0.0182 | 0.28787 | 0.269 | 0.18 | 1 | HC_vs_MS.Trt_F |
| TAF11 | 0.01826 | -0.2884 | 0.561 | 0.45 | 1 | HC_vs_MS.Trt_F |
| DHFR2 | 0.01826 | -0.4431 | 0.691 | 0.6 | 1 | HC_vs_MS.Trt_F |
| ADAM15 | 0.0183 | -0.3149 | 0.465 | 0.37 | 1 | HC_vs_MS.Trt_F |
| STK11 | 0.01831 | 0.60877 | 0.471 | 0.38 | 1 | HC_vs_MS.Trt_F |
| ATF2 | 0.01837 | 1.06959 | 0.251 | 0.18 | 1 | HC_vs_MS.Trt_F |
| BTG2 | 0.01838 | 0.2907 | 0.321 | 0.23 | 1 | HC_vs_MS.Trt_F |
| AKR1B1 | 0.0184 | -0.2952 | 0.803 | 0.69 | 1 | HC_vs_MS.Trt_F |
| USP33 | 0.01845 | -0.4959 | 0.965 | 0.87 | 1 | HC_vs_MS.Trt_F |
| MRPL16 | 0.01849 | 0.36517 | 0.211 | 0.14 | 1 | HC_vs_MS.Trt_F |
| ZNF879 | 0.01855 | 0.62946 | 0.367 | 0.27 | 1 | HC_vs_MS.Trt_F |
| SCN1A | 0.0186 | 0.39121 | 0.451 | 0.34 | 1 | HC_vs_MS.Trt_F |
| NMNAT1 | 0.01863 | 0.33503 | 0.986 | 0.99 | 1 | HC_vs_MS.Trt_F |
| TGFB1 | 0.01867 | 0.79412 | 0.393 | 0.3 | 1 | HC_vs_MS.Trt_F |
| CS | 0.0187 | 0.25062 | 0.864 | 0.78 | 1 | HC_vs_MS.Trt_F |
| PPCS | 0.01875 | -0.3289 | 0.367 | 0.26 | 1 | HC_vs_MS.Trt_F |
| ZFYVE28 | 0.01886 | 0.92065 | 0.231 | 0.16 | 1 | HC_vs_MS.Trt_F |
| ENOSF1 | 0.0189 | 0.72851 | 0.879 | 0.81 | 1 | HC_vs_MS.Trt_F |
| UTRN | 0.01893 | 0.29754 | 0.52 | 0.42 | 1 | HC_vs_MS.Trt_F |
| CIART | 0.01902 | 0.54657 | 0.558 | 0.48 | 1 | HC_vs_MS.Trt_F |
| LRFN1 | 0.01907 | 0.38814 | 0.173 | 0.11 | 1 | HC_vs_MS.Trt_F |
| TNXB | 0.01909 | 1.39763 | 0.168 | 0.1 | 1 | HC_vs_MS.Trt_F |
| SLC25A33 | 0.01912 | 0.54361 | 0.656 | 0.54 | 1 | HC_vs_MS.Trt_F |
| GABRE | 0.01919 | 0.39888 | 0.199 | 0.13 | 1 | HC_vs_MS.Trt_F |
| KCNJ15 | 0.01924 | 0.42372 | 0.705 | 0.6 | 1 | HC_vs_MS.Trt_F |
| ZFAND3 | 0.01928 | 0.65588 | 0.734 | 0.63 | 1 | HC_vs_MS.Trt_F |
| NMT1 | 0.01932 | -0.3632 | 0.367 | 0.26 | 1 | HC_vs_MS.Trt_F |
| CNOT1 | 0.01932 | 0.58433 | 0.835 | 0.75 | 1 | HC_vs_MS.Trt_F |
| ZCCHC14 | 0.01933 | 1.83092 | 0.139 | 0.08 | 1 | HC_vs_MS.Trt_F |
| INSR | 0.01942 | 0.478 | 0.517 | 0.44 | 1 | HC_vs_MS.Trt_F |
| TMEM98 | 0.01945 | 0.52402 | 0.61 | 0.51 | 1 | HC_vs_MS.Trt_F |
| IKBKAP | 0.01948 | 0.46607 | 0.338 | 0.26 | 1 | HC_vs_MS.Trt_F |
| KIAA1324L | 0.01949 | 0.6534 | 0.104 | 0.05 | 1 | HC_vs_MS.Trt_F |
| CLEC16A | 0.01952 | 0.40484 | 0.694 | 0.62 | 1 | HC_vs_MS.Trt_F |
| ZKSCAN3 | 0.01958 | -0.4922 | 0.896 | 0.84 | 1 | HC_vs_MS.Trt_F |

| | | | | | | |
|---|---|---|---|---|---|---|
| FOXM1 | 0.01967 | 0.35834 | 0.809 | 0.72 | 1 | HC_vs_MS.Trt_F |
| CNTNAP3B | 0.01975 | 0.25345 | 0.893 | 0.84 | 1 | HC_vs_MS.Trt_F |
| PLIN2 | 0.01986 | 0.32437 | 0.697 | 0.6 | 1 | HC_vs_MS.Trt_F |
| ZBTB3 | 0.01994 | 0.40567 | 0.85 | 0.78 | 1 | HC_vs_MS.Trt_F |
| CCNT1 | 0.01995 | 0.41313 | 0.249 | 0.17 | 1 | HC_vs_MS.Trt_F |
| GALNT6 | 0.01996 | 0.72506 | 0.78 | 0.66 | 1 | HC_vs_MS.Trt_F |
| WDR91 | 0.01999 | -0.3398 | 0.535 | 0.43 | 1 | HC_vs_MS.Trt_F |
| CFAP36 | 0.01999 | 0.60004 | 0.486 | 0.39 | 1 | HC_vs_MS.Trt_F |
| NSUN3 | 0.02009 | -0.4237 | 0.89 | 0.81 | 1 | HC_vs_MS.Trt_F |
| FAM120B | 0.02011 | 0.28192 | 0.364 | 0.27 | 1 | HC_vs_MS.Trt_F |
| ZNF385A | 0.02018 | 0.26673 | 0.717 | 0.64 | 1 | HC_vs_MS.Trt_F |
| TBC1D32 | 0.0202 | 0.50696 | 0.564 | 0.46 | 1 | HC_vs_MS.Trt_F |
| FAS | 0.02027 | 0.38992 | 0.442 | 0.34 | 1 | HC_vs_MS.Trt_F |
| MGAT5 | 0.02027 | 0.49941 | 0.214 | 0.14 | 1 | HC_vs_MS.Trt_F |
| MPP7 | 0.0203 | 1.63801 | 0.347 | 0.27 | 1 | HC_vs_MS.Trt_F |
| FRMD4B | 0.02033 | 1.20942 | 0.934 | 0.96 | 1 | HC_vs_MS.Trt_F |
| PLAG1 | 0.02035 | -0.8252 | 0.46 | 0.37 | 1 | HC_vs_MS.Trt_F |
| MED6 | 0.02045 | 0.53381 | 0.601 | 0.5 | 1 | HC_vs_MS.Trt_F |
| TEX9 | 0.02048 | 0.25011 | 0.841 | 0.75 | 1 | HC_vs_MS.Trt_F |
| MFSD11 | 0.02056 | 0.42954 | 0.497 | 0.42 | 1 | HC_vs_MS.Trt_F |
| PMAIP1 | 0.02056 | 0.32783 | 0.263 | 0.18 | 1 | HC_vs_MS.Trt_F |
| AP4B1 | 0.02058 | -0.2652 | 0.682 | 0.6 | 1 | HC_vs_MS.Trt_F |
| HEPACAM | 0.02063 | -0.261 | 0.376 | 0.3 | 1 | HC_vs_MS.Trt_F |
| EAF1 | 0.02063 | -0.4749 | 0.448 | 0.33 | 1 | HC_vs_MS.Trt_F |
| SYBU | 0.02073 | 1.18898 | 0.113 | 0.06 | 1 | HC_vs_MS.Trt_F |
| PDE4A | 0.02087 | 0.89638 | 0.176 | 0.12 | 1 | HC_vs_MS.Trt_F |
| SLC5A5 | 0.02095 | 0.577 | 0.517 | 0.43 | 1 | HC_vs_MS.Trt_F |
| TSSC4 | 0.02103 | -0.6432 | 0.127 | 0.19 | 1 | HC_vs_MS.Trt_F |
| POLR3GL | 0.02114 | 0.42394 | 0.243 | 0.17 | 1 | HC_vs_MS.Trt_F |
| USP11 | 0.02116 | 0.7068 | 0.266 | 0.19 | 1 | HC_vs_MS.Trt_F |
| CDKN1A | 0.02124 | 0.87596 | 0.182 | 0.12 | 1 | HC_vs_MS.Trt_F |
| COQ7 | 0.02126 | 0.32003 | 0.434 | 0.34 | 1 | HC_vs_MS.Trt_F |
| FAM19A2 | 0.0213 | 1.4632 | 0.289 | 0.21 | 1 | HC_vs_MS.Trt_F |
| NECAB3 | 0.0213 | -1.435 | 0.058 | 0.11 | 1 | HC_vs_MS.Trt_F |
| AMPD3 | 0.02135 | 0.49094 | 0.436 | 0.33 | 1 | HC_vs_MS.Trt_F |
| ZNF805 | 0.0214 | -0.3423 | 0.832 | 0.77 | 1 | HC_vs_MS.Trt_F |
| JAM2 | 0.02141 | 0.39439 | 0.607 | 0.53 | 1 | HC_vs_MS.Trt_F |
| ZC3H18 | 0.02143 | 0.38749 | 0.766 | 0.67 | 1 | HC_vs_MS.Trt_F |
| STEAP4 | 0.02143 | 0.26374 | 0.743 | 0.64 | 1 | HC_vs_MS.Trt_F |
| MACC1 | 0.0215 | 0.40686 | 0.786 | 0.72 | 1 | HC_vs_MS.Trt_F |

| | | | | | | |
|---|---|---|---|---|---|---|
| MCCC2 | 0.02157 | 0.64962 | 0.734 | 0.65 | 1 | HC_vs_MS.Trt_F |
| GLB1L3 | 0.02164 | 0.45339 | 0.763 | 0.69 | 1 | HC_vs_MS.Trt_F |
| SCARB1 | 0.02167 | 0.80337 | 0.587 | 0.48 | 1 | HC_vs_MS.Trt_F |
| CDKAL1 | 0.02168 | 0.63621 | 0.63 | 0.53 | 1 | HC_vs_MS.Trt_F |
| UBQLN2 | 0.02169 | 0.36484 | 0.292 | 0.21 | 1 | HC_vs_MS.Trt_F |
| AFMID | 0.02172 | 0.28349 | 0.957 | 0.91 | 1 | HC_vs_MS.Trt_F |
| ITPR2 | 0.02173 | 0.81714 | 0.283 | 0.21 | 1 | HC_vs_MS.Trt_F |
| SLC35E2 | 0.02178 | 0.4012 | 0.477 | 0.39 | 1 | HC_vs_MS.Trt_F |
| SNX19 | 0.0219 | 0.36876 | 0.269 | 0.19 | 1 | HC_vs_MS.Trt_F |
| MAPKAPK3 | 0.0219 | 0.67543 | 0.24 | 0.17 | 1 | HC_vs_MS.Trt_F |
| SBK1 | 0.02191 | 0.73338 | 0.246 | 0.17 | 1 | HC_vs_MS.Trt_F |
| KLHL20 | 0.02191 | 0.37539 | 0.197 | 0.13 | 1 | HC_vs_MS.Trt_F |
| WDR37 | 0.02196 | 0.55253 | 0.231 | 0.16 | 1 | HC_vs_MS.Trt_F |
| RNF152 | 0.02197 | 0.46976 | 0.509 | 0.4 | 1 | HC_vs_MS.Trt_F |
| DDX27 | 0.02203 | 0.25568 | 0.737 | 0.64 | 1 | HC_vs_MS.Trt_F |
| PHF3 | 0.02203 | 0.28484 | 0.494 | 0.4 | 1 | HC_vs_MS.Trt_F |
| WDTC1 | 0.02208 | 0.47411 | 0.399 | 0.29 | 1 | HC_vs_MS.Trt_F |
| GRHL2 | 0.02212 | 0.38545 | 0.28 | 0.2 | 1 | HC_vs_MS.Trt_F |
| HELQ | 0.02222 | 0.86091 | 0.197 | 0.13 | 1 | HC_vs_MS.Trt_F |
| PRR12 | 0.02224 | 1.17083 | 0.133 | 0.08 | 1 | HC_vs_MS.Trt_F |
| MYO9B | 0.02227 | 0.90364 | 0.275 | 0.2 | 1 | HC_vs_MS.Trt_F |
| SPATA5 | 0.02234 | 0.90012 | 0.798 | 0.7 | 1 | HC_vs_MS.Trt_F |
| TGFB1I1 | 0.02239 | 0.37621 | 0.332 | 0.25 | 1 | HC_vs_MS.Trt_F |
| ADCY4 | 0.02245 | 0.42026 | 0.15 | 0.09 | 1 | HC_vs_MS.Trt_F |
| PUS1 | 0.02247 | 0.41665 | 0.85 | 0.76 | 1 | HC_vs_MS.Trt_F |
| INO80D | 0.02248 | 0.48154 | 0.483 | 0.39 | 1 | HC_vs_MS.Trt_F |
| SPOCK2 | 0.0225 | 0.33433 | 0.613 | 0.55 | 1 | HC_vs_MS.Trt_F |
| PDE5A | 0.02255 | -0.2854 | 0.153 | 0.09 | 1 | HC_vs_MS.Trt_F |
| CALCOCO1 | 0.02271 | 0.25439 | 0.569 | 0.48 | 1 | HC_vs_MS.Trt_F |
| PRPF31 | 0.02272 | 0.33586 | 0.234 | 0.16 | 1 | HC_vs_MS.Trt_F |
| BBX | 0.02279 | 0.32945 | 0.367 | 0.27 | 1 | HC_vs_MS.Trt_F |
| FBXO25 | 0.02288 | 0.90399 | 0.454 | 0.36 | 1 | HC_vs_MS.Trt_F |
| RANBP6 | 0.02288 | -1.1289 | 0.168 | 0.1 | 1 | HC_vs_MS.Trt_F |
| CHTOP | 0.02317 | -0.4064 | 0.671 | 0.55 | 1 | HC_vs_MS.Trt_F |
| DPP6 | 0.02328 | -0.4898 | 0.601 | 0.51 | 1 | HC_vs_MS.Trt_F |
| ACADSB | 0.02328 | -0.6338 | 0.717 | 0.61 | 1 | HC_vs_MS.Trt_F |
| RPL22L1 | 0.02329 | -0.2619 | 0.332 | 0.24 | 1 | HC_vs_MS.Trt_F |
| FCAR | 0.02331 | 0.25289 | 0.882 | 0.84 | 1 | HC_vs_MS.Trt_F |
| EPHX2 | 0.02334 | 0.72298 | 0.353 | 0.27 | 1 | HC_vs_MS.Trt_F |
| CXorf40A | 0.02344 | 0.70345 | 0.188 | 0.12 | 1 | HC_vs_MS.Trt_F |

| | | | | | | |
|---|---|---|---|---|---|---|
| ZNF318 | 0.02352 | 1.25118 | 0.711 | 0.61 | 1 | HC_vs_MS.Trt_F |
| DNAH9 | 0.02356 | -0.3955 | 0.5 | 0.57 | 1 | HC_vs_MS.Trt_F |
| NMD3 | 0.02361 | -0.2998 | 0.792 | 0.71 | 1 | HC_vs_MS.Trt_F |
| SAYSD1 | 0.02367 | -0.4352 | 0.208 | 0.14 | 1 | HC_vs_MS.Trt_F |
| GATC | 0.02374 | 0.76366 | 0.604 | 0.53 | 1 | HC_vs_MS.Trt_F |
| ACSF2 | 0.02379 | -0.2783 | 0.561 | 0.45 | 1 | HC_vs_MS.Trt_F |
| CASZ1 | 0.0238 | 1.00851 | 0.341 | 0.26 | 1 | HC_vs_MS.Trt_F |
| UHRF1BP1 | 0.02399 | 0.34761 | 0.616 | 0.52 | 1 | HC_vs_MS.Trt_F |
| FAM227A | 0.02405 | 0.28771 | 0.942 | 0.87 | 1 | HC_vs_MS.Trt_F |
| C2CD2L | 0.02409 | 0.37677 | 0.546 | 0.43 | 1 | HC_vs_MS.Trt_F |
| BCL2L1 | 0.02414 | 0.30671 | 0.318 | 0.23 | 1 | HC_vs_MS.Trt_F |
| ABCA9 | 0.02417 | 0.55294 | 0.882 | 0.82 | 1 | HC_vs_MS.Trt_F |
| RCBTB1 | 0.02419 | 0.80027 | 0.168 | 0.1 | 1 | HC_vs_MS.Trt_F |
| KCNAB2 | 0.02432 | 0.3286 | 0.477 | 0.38 | 1 | HC_vs_MS.Trt_F |
| TMEM209 | 0.02435 | -0.3918 | 0.818 | 0.75 | 1 | HC_vs_MS.Trt_F |
| NBPF19 | 0.0245 | 0.71723 | 0.474 | 0.38 | 1 | HC_vs_MS.Trt_F |
| IKZF3 | 0.02459 | 0.27837 | 0.977 | 0.95 | 1 | HC_vs_MS.Trt_F |
| TBRG4 | 0.02464 | -0.4538 | 0.436 | 0.32 | 1 | HC_vs_MS.Trt_F |
| KCTD19 | 0.0248 | 0.26921 | 0.783 | 0.72 | 1 | HC_vs_MS.Trt_F |
| DIS3L2 | 0.0248 | 0.32485 | 0.569 | 0.49 | 1 | HC_vs_MS.Trt_F |
| KIF23 | 0.02495 | 0.2742 | 0.618 | 0.54 | 1 | HC_vs_MS.Trt_F |
| SASH1 | 0.02495 | 0.26235 | 0.399 | 0.31 | 1 | HC_vs_MS.Trt_F |
| NOS3 | 0.02511 | 0.3149 | 0.445 | 0.34 | 1 | HC_vs_MS.Trt_F |
| WAS | 0.0252 | 1.2611 | 0.332 | 0.26 | 1 | HC_vs_MS.Trt_F |
| TXNL4A | 0.02522 | 0.52275 | 0.572 | 0.48 | 1 | HC_vs_MS.Trt_F |
| IFT81 | 0.02531 | 0.36316 | 0.792 | 0.67 | 1 | HC_vs_MS.Trt_F |
| CASP8AP2 | 0.02532 | 0.44846 | 0.379 | 0.29 | 1 | HC_vs_MS.Trt_F |
| TMEM52B | 0.02532 | 0.25438 | 0.801 | 0.72 | 1 | HC_vs_MS.Trt_F |
| UBE2A | 0.02546 | -0.301 | 0.416 | 0.32 | 1 | HC_vs_MS.Trt_F |
| DDX1 | 0.0255 | 0.28072 | 0.205 | 0.13 | 1 | HC_vs_MS.Trt_F |
| EXOC6B | 0.02554 | 0.82114 | 0.182 | 0.12 | 1 | HC_vs_MS.Trt_F |
| RPUSD3 | 0.02567 | 0.36316 | 0.867 | 0.77 | 1 | HC_vs_MS.Trt_F |
| RHBDF2 | 0.02568 | -0.2949 | 0.939 | 0.91 | 1 | HC_vs_MS.Trt_F |
| TMEM41A | 0.02568 | 0.42998 | 0.858 | 0.8 | 1 | HC_vs_MS.Trt_F |
| CADM1 | 0.02576 | -0.7904 | 0.289 | 0.35 | 1 | HC_vs_MS.Trt_F |
| RGS2 | 0.02581 | 0.5901 | 0.202 | 0.14 | 1 | HC_vs_MS.Trt_F |
| ITGA5 | 0.02588 | -0.6187 | 0.168 | 0.1 | 1 | HC_vs_MS.Trt_F |
| FAM168A | 0.02595 | 0.33604 | 0.185 | 0.12 | 1 | HC_vs_MS.Trt_F |
| ADAM28 | 0.02601 | -0.7458 | 0.211 | 0.14 | 1 | HC_vs_MS.Trt_F |
| BCAR3 | 0.02602 | 0.63855 | 0.676 | 0.59 | 1 | HC_vs_MS.Trt_F |

| | | | | | | |
|---|---|---|---|---|---|---|
| UCKL1 | 0.02604 | 0.93016 | 0.188 | 0.12 | 1 | HC_vs_MS.Trt_F |
| EXOSC5 | 0.02614 | 0.32326 | 0.142 | 0.08 | 1 | HC_vs_MS.Trt_F |
| FAM65A | 0.02621 | 0.68303 | 0.197 | 0.13 | 1 | HC_vs_MS.Trt_F |
| POGK | 0.02625 | 0.50899 | 0.312 | 0.23 | 1 | HC_vs_MS.Trt_F |
| BIN3 | 0.02626 | -0.3453 | 0.457 | 0.35 | 1 | HC_vs_MS.Trt_F |
| MAPK8IP3 | 0.02634 | 0.55468 | 0.22 | 0.15 | 1 | HC_vs_MS.Trt_F |
| CENPF | 0.02645 | 0.72746 | 0.431 | 0.32 | 1 | HC_vs_MS.Trt_F |
| FAXC | 0.02647 | 0.28732 | 0.396 | 0.31 | 1 | HC_vs_MS.Trt_F |
| ZNF555 | 0.0265 | -0.3371 | 0.711 | 0.6 | 1 | HC_vs_MS.Trt_F |
| GTF3C1 | 0.02653 | 0.37018 | 0.182 | 0.12 | 1 | HC_vs_MS.Trt_F |
| UQCRH | 0.02659 | 0.31719 | 0.581 | 0.49 | 1 | HC_vs_MS.Trt_F |
| RBM38 | 0.02672 | 0.41269 | 0.202 | 0.13 | 1 | HC_vs_MS.Trt_F |
| HNRNPAB | 0.02674 | 0.41348 | 0.223 | 0.15 | 1 | HC_vs_MS.Trt_F |
| EVI5 | 0.02681 | 0.25021 | 0.864 | 0.79 | 1 | HC_vs_MS.Trt_F |
| DHX15 | 0.02693 | 0.41817 | 0.54 | 0.46 | 1 | HC_vs_MS.Trt_F |
| TM9SF3 | 0.02705 | -1.1866 | 0.139 | 0.2 | 1 | HC_vs_MS.Trt_F |
| TCEAL1 | 0.02705 | 0.64831 | 0.121 | 0.07 | 1 | HC_vs_MS.Trt_F |
| DDHD1 | 0.02707 | 0.30574 | 0.942 | 0.86 | 1 | HC_vs_MS.Trt_F |
| SCML4 | 0.02707 | 0.35469 | 0.662 | 0.57 | 1 | HC_vs_MS.Trt_F |
| ERLIN1 | 0.02711 | 1.4687 | 0.101 | 0.05 | 1 | HC_vs_MS.Trt_F |
| NABP1 | 0.02711 | -0.5662 | 0.766 | 0.8 | 1 | HC_vs_MS.Trt_F |
| HLA-DRB1 | 0.02723 | 0.2676 | 0.321 | 0.23 | 1 | HC_vs_MS.Trt_F |
| RBM48 | 0.02728 | -0.818 | 0.829 | 0.72 | 1 | HC_vs_MS.Trt_F |
| ZFP36L1 | 0.02734 | -0.3845 | 0.769 | 0.76 | 1 | HC_vs_MS.Trt_F |
| THOC7 | 0.02744 | -1.0507 | 0.159 | 0.22 | 1 | HC_vs_MS.Trt_F |
| ATG4B | 0.02745 | 0.4525 | 0.26 | 0.18 | 1 | HC_vs_MS.Trt_F |
| PITRM1 | 0.02753 | 0.33833 | 0.211 | 0.14 | 1 | HC_vs_MS.Trt_F |
| TTI2 | 0.02768 | 0.27405 | 0.512 | 0.4 | 1 | HC_vs_MS.Trt_F |
| PGAP1 | 0.0277 | 0.62215 | 0.457 | 0.37 | 1 | HC_vs_MS.Trt_F |
| NEK1 | 0.02775 | 0.70861 | 0.402 | 0.31 | 1 | HC_vs_MS.Trt_F |
| ACTN4 | 0.02777 | 0.26282 | 0.5 | 0.41 | 1 | HC_vs_MS.Trt_F |
| PPM1L | 0.02786 | -0.286 | 0.246 | 0.31 | 1 | HC_vs_MS.Trt_F |
| UBTF | 0.02789 | 0.57579 | 0.321 | 0.24 | 1 | HC_vs_MS.Trt_F |
| HEATR1 | 0.02792 | -1.0276 | 0.202 | 0.14 | 1 | HC_vs_MS.Trt_F |
| PBLD | 0.02794 | 0.34423 | 0.673 | 0.57 | 1 | HC_vs_MS.Trt_F |
| FOXJ2 | 0.02794 | 0.29964 | 0.503 | 0.39 | 1 | HC_vs_MS.Trt_F |
| CISD2 | 0.02802 | -0.4655 | 0.523 | 0.45 | 1 | HC_vs_MS.Trt_F |
| BBS5 | 0.02814 | 0.26716 | 0.361 | 0.28 | 1 | HC_vs_MS.Trt_F |
| TUBGCP5 | 0.02818 | 0.38604 | 0.697 | 0.64 | 1 | HC_vs_MS.Trt_F |
| BCAM | 0.02819 | 0.56906 | 0.786 | 0.7 | 1 | HC_vs_MS.Trt_F |

| | | | | | | |
|---|---|---|---|---|---|---|
| SMAD9 | 0.02834 | -0.6113 | 0.448 | 0.33 | 1 | HC_vs_MS.Trt_F |
| LIN52 | 0.02847 | -0.3698 | 0.777 | 0.68 | 1 | HC_vs_MS.Trt_F |
| FUT8 | 0.0285 | 0.47162 | 1 | 0.99 | 1 | HC_vs_MS.Trt_F |
| WHRN | 0.0285 | 0.78695 | 0.171 | 0.11 | 1 | HC_vs_MS.Trt_F |
| STYX | 0.02852 | -0.2813 | 0.624 | 0.5 | 1 | HC_vs_MS.Trt_F |
| MTG2 | 0.02859 | -0.6088 | 0.847 | 0.82 | 1 | HC_vs_MS.Trt_F |
| TRNAU1AP | 0.02864 | 0.55526 | 0.919 | 0.88 | 1 | HC_vs_MS.Trt_F |
| NUP93 | 0.02864 | -0.3905 | 0.737 | 0.66 | 1 | HC_vs_MS.Trt_F |
| SH3BP1 | 0.02873 | 0.56102 | 0.156 | 0.1 | 1 | HC_vs_MS.Trt_F |
| BOD1L1 | 0.02881 | 0.25723 | 0.757 | 0.7 | 1 | HC_vs_MS.Trt_F |
| TRABD | 0.02881 | 0.43738 | 0.382 | 0.31 | 1 | HC_vs_MS.Trt_F |
| DPH6 | 0.02914 | 0.25923 | 0.477 | 0.36 | 1 | HC_vs_MS.Trt_F |
| SLC10A3 | 0.02921 | -0.5032 | 0.159 | 0.1 | 1 | HC_vs_MS.Trt_F |
| CHD4 | 0.02922 | -0.3028 | 0.636 | 0.54 | 1 | HC_vs_MS.Trt_F |
| RAB17 | 0.02926 | 0.38617 | 0.246 | 0.18 | 1 | HC_vs_MS.Trt_F |
| DPYSL2 | 0.0293 | -0.7648 | 0.52 | 0.57 | 1 | HC_vs_MS.Trt_F |
| PLS3 | 0.02939 | -0.2664 | 0.46 | 0.35 | 1 | HC_vs_MS.Trt_F |
| PFDN1 | 0.02943 | 0.3541 | 0.237 | 0.16 | 1 | HC_vs_MS.Trt_F |
| ZDHHC15 | 0.02946 | 0.34819 | 0.647 | 0.55 | 1 | HC_vs_MS.Trt_F |
| KIF20B | 0.02948 | -0.4282 | 0.188 | 0.12 | 1 | HC_vs_MS.Trt_F |
| ATF6 | 0.02959 | 0.87866 | 0.283 | 0.2 | 1 | HC_vs_MS.Trt_F |
| PCYT1B | 0.0296 | 0.28213 | 0.526 | 0.44 | 1 | HC_vs_MS.Trt_F |
| ARFGAP1 | 0.0296 | -0.58 | 0.234 | 0.15 | 1 | HC_vs_MS.Trt_F |
| ADCK1 | 0.02972 | 1.14577 | 0.312 | 0.23 | 1 | HC_vs_MS.Trt_F |
| ELF4 | 0.02976 | 0.41251 | 0.153 | 0.09 | 1 | HC_vs_MS.Trt_F |
| CLDN11 | 0.02983 | -0.3551 | 0.699 | 0.57 | 1 | HC_vs_MS.Trt_F |
| CDK19 | 0.02988 | 0.50647 | 0.358 | 0.27 | 1 | HC_vs_MS.Trt_F |
| SF3A3 | 0.02993 | -0.4191 | 0.408 | 0.31 | 1 | HC_vs_MS.Trt_F |
| TAF15 | 0.02994 | 0.36413 | 0.254 | 0.18 | 1 | HC_vs_MS.Trt_F |
| C22orf29 | 0.02999 | -0.354 | 0.494 | 0.39 | 1 | HC_vs_MS.Trt_F |
| C12orf10 | 0.03002 | 0.57561 | 0.286 | 0.2 | 1 | HC_vs_MS.Trt_F |
| TCAF2 | 0.03006 | -0.3316 | 0.78 | 0.68 | 1 | HC_vs_MS.Trt_F |
| IFI35 | 0.03007 | 0.33863 | 0.873 | 0.8 | 1 | HC_vs_MS.Trt_F |
| RBMS2 | 0.03008 | 0.2586 | 0.939 | 0.87 | 1 | HC_vs_MS.Trt_F |
| BTBD9 | 0.03009 | 0.59668 | 0.621 | 0.57 | 1 | HC_vs_MS.Trt_F |
| WRNIP1 | 0.03009 | 0.59591 | 0.133 | 0.08 | 1 | HC_vs_MS.Trt_F |
| CCDC169 | 0.03024 | 0.47032 | 0.636 | 0.52 | 1 | HC_vs_MS.Trt_F |
| SMG7 | 0.03025 | 0.39542 | 0.24 | 0.16 | 1 | HC_vs_MS.Trt_F |
| MCM4 | 0.03026 | 0.29005 | 0.673 | 0.6 | 1 | HC_vs_MS.Trt_F |
| CWC22 | 0.03037 | 0.26726 | 0.179 | 0.12 | 1 | HC_vs_MS.Trt_F |

| | | | | | | |
|---|---|---|---|---|---|---|
| BCL3 | 0.03049 | 0.59406 | 0.147 | 0.09 | 1 | HC_vs_MS.Trt_F |
| RPS2 | 0.03059 | 0.37889 | 0.598 | 0.51 | 1 | HC_vs_MS.Trt_F |
| MRPL44 | 0.03064 | 0.46751 | 0.627 | 0.53 | 1 | HC_vs_MS.Trt_F |
| ZBTB4 | 0.03065 | 0.62919 | 0.194 | 0.13 | 1 | HC_vs_MS.Trt_F |
| CREBBP | 0.03073 | 0.65421 | 0.59 | 0.49 | 1 | HC_vs_MS.Trt_F |
| KDM2A | 0.03077 | 0.28549 | 0.919 | 0.92 | 1 | HC_vs_MS.Trt_F |
| ARL11 | 0.03085 | 0.32185 | 0.685 | 0.58 | 1 | HC_vs_MS.Trt_F |
| U2AF2 | 0.03085 | 1.10863 | 0.243 | 0.17 | 1 | HC_vs_MS.Trt_F |
| ARFGEF2 | 0.03086 | 0.78666 | 0.494 | 0.41 | 1 | HC_vs_MS.Trt_F |
| RASSF8 | 0.03094 | 0.32456 | 0.514 | 0.41 | 1 | HC_vs_MS.Trt_F |
| PCNX4 | 0.03107 | 0.25081 | 0.922 | 0.92 | 1 | HC_vs_MS.Trt_F |
| SPG21 | 0.03112 | 0.34733 | 0.913 | 0.88 | 1 | HC_vs_MS.Trt_F |
| SLC46A3 | 0.03113 | 0.25914 | 0.136 | 0.08 | 1 | HC_vs_MS.Trt_F |
| FDX1 | 0.0312 | -0.3023 | 0.384 | 0.28 | 1 | HC_vs_MS.Trt_F |
| JMY | 0.03142 | 0.96087 | 0.353 | 0.28 | 1 | HC_vs_MS.Trt_F |
| CHGB | 0.03142 | 1.03126 | 0.5 | 0.4 | 1 | HC_vs_MS.Trt_F |
| ERVK3-1 | 0.03144 | 0.57873 | 0.379 | 0.3 | 1 | HC_vs_MS.Trt_F |
| PLEKHM1 | 0.0316 | 0.3006 | 0.861 | 0.8 | 1 | HC_vs_MS.Trt_F |
| ATAD2B | 0.03161 | 0.8244 | 0.746 | 0.74 | 1 | HC_vs_MS.Trt_F |
| ATP1B3 | 0.03161 | 0.36612 | 0.211 | 0.14 | 1 | HC_vs_MS.Trt_F |
| PIGN | 0.03165 | 0.35489 | 0.653 | 0.58 | 1 | HC_vs_MS.Trt_F |
| MTHFD2L | 0.03168 | 0.71679 | 0.535 | 0.47 | 1 | HC_vs_MS.Trt_F |
| STK3 | 0.03169 | 0.40404 | 0.786 | 0.73 | 1 | HC_vs_MS.Trt_F |
| ZNF462 | 0.03173 | 0.73806 | 0.24 | 0.18 | 1 | HC_vs_MS.Trt_F |
| ZC3H12B | 0.03177 | -1.0783 | 0.63 | 0.5 | 1 | HC_vs_MS.Trt_F |
| HAUS6 | 0.03187 | 0.63774 | 0.39 | 0.31 | 1 | HC_vs_MS.Trt_F |
| HSF1 | 0.03188 | 0.47718 | 0.526 | 0.42 | 1 | HC_vs_MS.Trt_F |
| RP11-437B1 | 0.03195 | 0.27964 | 0.639 | 0.54 | 1 | HC_vs_MS.Trt_F |
| SF3B4 | 0.03199 | 0.43793 | 0.243 | 0.17 | 1 | HC_vs_MS.Trt_F |
| C1orf216 | 0.0321 | -0.2894 | 0.789 | 0.73 | 1 | HC_vs_MS.Trt_F |
| SYNRG | 0.03211 | 0.7254 | 0.971 | 0.93 | 1 | HC_vs_MS.Trt_F |
| KIAA1147 | 0.03214 | 0.68658 | 0.124 | 0.07 | 1 | HC_vs_MS.Trt_F |
| EIF1AX | 0.03224 | -0.4081 | 0.347 | 0.25 | 1 | HC_vs_MS.Trt_F |
| GOLGA8R | 0.03226 | 0.4949 | 0.558 | 0.48 | 1 | HC_vs_MS.Trt_F |
| TMEM39A | 0.03234 | 0.63902 | 0.211 | 0.14 | 1 | HC_vs_MS.Trt_F |
| TUBA1C | 0.03234 | 0.32819 | 0.679 | 0.58 | 1 | HC_vs_MS.Trt_F |
| CCNG1 | 0.03247 | -0.9085 | 0.211 | 0.27 | 1 | HC_vs_MS.Trt_F |
| TMEM50B | 0.03251 | 0.31073 | 0.734 | 0.67 | 1 | HC_vs_MS.Trt_F |
| ZNF91 | 0.03262 | 0.38958 | 0.806 | 0.73 | 1 | HC_vs_MS.Trt_F |
| KHNYN | 0.03263 | 2.01588 | 0.136 | 0.08 | 1 | HC_vs_MS.Trt_F |

| | | | | | | |
|---|---|---|---|---|---|---|
| GNAI2 | 0.03264 | 0.69981 | 0.451 | 0.37 | 1 | HC_vs_MS.Trt_F |
| EIF1AD | 0.03266 | -0.8189 | 0.694 | 0.73 | 1 | HC_vs_MS.Trt_F |
| NAPEPLD | 0.03266 | 0.59504 | 0.384 | 0.31 | 1 | HC_vs_MS.Trt_F |
| PKD1 | 0.03267 | 0.65282 | 0.454 | 0.35 | 1 | HC_vs_MS.Trt_F |
| BCL9L | 0.03268 | 1.22318 | 0.205 | 0.14 | 1 | HC_vs_MS.Trt_F |
| ZNF714 | 0.03268 | 0.45907 | 0.96 | 0.91 | 1 | HC_vs_MS.Trt_F |
| SLC9A9 | 0.03274 | 0.86408 | 0.171 | 0.11 | 1 | HC_vs_MS.Trt_F |
| IREB2 | 0.03274 | 0.37465 | 0.844 | 0.75 | 1 | HC_vs_MS.Trt_F |
| ZMYM3 | 0.03274 | 0.2864 | 0.408 | 0.34 | 1 | HC_vs_MS.Trt_F |
| PRKG2 | 0.03292 | 0.47849 | 0.387 | 0.3 | 1 | HC_vs_MS.Trt_F |
| IGSF9 | 0.03296 | 0.31354 | 0.344 | 0.26 | 1 | HC_vs_MS.Trt_F |
| SVIP | 0.03311 | 0.623 | 0.546 | 0.45 | 1 | HC_vs_MS.Trt_F |
| NCOA7 | 0.03334 | 0.5802 | 0.801 | 0.72 | 1 | HC_vs_MS.Trt_F |
| AZI2 | 0.03335 | 0.36001 | 0.197 | 0.13 | 1 | HC_vs_MS.Trt_F |
| MRPS35 | 0.03343 | 0.63396 | 0.173 | 0.11 | 1 | HC_vs_MS.Trt_F |
| P2RY11 | 0.03355 | -0.6866 | 0.092 | 0.15 | 1 | HC_vs_MS.Trt_F |
| DNAH12 | 0.03358 | -0.6954 | 0.107 | 0.16 | 1 | HC_vs_MS.Trt_F |
| RNF114 | 0.03362 | 0.33577 | 0.772 | 0.67 | 1 | HC_vs_MS.Trt_F |
| NOTCH4 | 0.03373 | 0.37058 | 0.85 | 0.78 | 1 | HC_vs_MS.Trt_F |
| OGG1 | 0.03378 | -0.38 | 0.431 | 0.34 | 1 | HC_vs_MS.Trt_F |
| RALGAPA2 | 0.0338 | 0.66262 | 0.205 | 0.14 | 1 | HC_vs_MS.Trt_F |
| TIAM2 | 0.03383 | 0.30327 | 0.815 | 0.75 | 1 | HC_vs_MS.Trt_F |
| OXSR1 | 0.03385 | -0.5393 | 0.124 | 0.18 | 1 | HC_vs_MS.Trt_F |
| CENPI | 0.03387 | 0.88179 | 0.471 | 0.39 | 1 | HC_vs_MS.Trt_F |
| OAS1 | 0.03397 | 0.27933 | 0.584 | 0.51 | 1 | HC_vs_MS.Trt_F |
| KAT8 | 0.03406 | 0.71112 | 0.165 | 0.1 | 1 | HC_vs_MS.Trt_F |
| SEMA4C | 0.03407 | 0.59476 | 0.801 | 0.73 | 1 | HC_vs_MS.Trt_F |
| FUT6 | 0.03421 | 0.36839 | 0.506 | 0.41 | 1 | HC_vs_MS.Trt_F |
| RBM43 | 0.03442 | 0.25153 | 0.682 | 0.6 | 1 | HC_vs_MS.Trt_F |
| PNRC1 | 0.03453 | 0.31171 | 0.35 | 0.27 | 1 | HC_vs_MS.Trt_F |
| FGFR1OP2 | 0.03454 | 0.30955 | 0.376 | 0.29 | 1 | HC_vs_MS.Trt_F |
| PDZD7 | 0.03482 | 0.42331 | 0.497 | 0.41 | 1 | HC_vs_MS.Trt_F |
| NT5C3B | 0.03486 | 0.71611 | 0.659 | 0.61 | 1 | HC_vs_MS.Trt_F |
| CYB561 | 0.0349 | 0.99713 | 0.39 | 0.31 | 1 | HC_vs_MS.Trt_F |
| SLX4 | 0.0349 | 0.43345 | 0.569 | 0.5 | 1 | HC_vs_MS.Trt_F |
| DNAJC1 | 0.03507 | 0.57224 | 0.199 | 0.14 | 1 | HC_vs_MS.Trt_F |
| MCC | 0.03509 | 0.96079 | 0.532 | 0.43 | 1 | HC_vs_MS.Trt_F |
| ZKSCAN8 | 0.03513 | 1.24495 | 0.303 | 0.24 | 1 | HC_vs_MS.Trt_F |
| LAX1 | 0.03515 | 0.36351 | 0.587 | 0.5 | 1 | HC_vs_MS.Trt_F |
| UNC79 | 0.03518 | 0.25302 | 0.408 | 0.32 | 1 | HC_vs_MS.Trt_F |

| | | | | | | |
|---|---|---|---|---|---|---|
| ACOXL | 0.03539 | 0.38221 | 0.72 | 0.66 | 1 | HC_vs_MS.Trt_F |
| NDRG1 | 0.03542 | 0.55649 | 0.191 | 0.13 | 1 | HC_vs_MS.Trt_F |
| DPP8 | 0.03545 | 0.39712 | 0.442 | 0.36 | 1 | HC_vs_MS.Trt_F |
| SLC35D1 | 0.03562 | 0.5483 | 0.35 | 0.26 | 1 | HC_vs_MS.Trt_F |
| ADAT2 | 0.03587 | -0.6966 | 0.486 | 0.38 | 1 | HC_vs_MS.Trt_F |
| PTPN9 | 0.0359 | 0.38384 | 0.355 | 0.27 | 1 | HC_vs_MS.Trt_F |
| RPS6KA1 | 0.03593 | 0.55681 | 0.237 | 0.17 | 1 | HC_vs_MS.Trt_F |
| ZNF277 | 0.03598 | -0.8557 | 0.616 | 0.5 | 1 | HC_vs_MS.Trt_F |
| FAM102B | 0.036 | 0.76273 | 0.315 | 0.22 | 1 | HC_vs_MS.Trt_F |
| CC2D2B | 0.036 | -0.6259 | 0.185 | 0.12 | 1 | HC_vs_MS.Trt_F |
| ZMYND8 | 0.03609 | 0.3037 | 0.419 | 0.33 | 1 | HC_vs_MS.Trt_F |
| MAF1 | 0.03616 | 0.36947 | 0.225 | 0.16 | 1 | HC_vs_MS.Trt_F |
| ICA1L | 0.0363 | 0.61535 | 0.812 | 0.77 | 1 | HC_vs_MS.Trt_F |
| CDH12 | 0.03636 | 0.31829 | 0.803 | 0.75 | 1 | HC_vs_MS.Trt_F |
| HPS4 | 0.03638 | 0.5165 | 0.818 | 0.74 | 1 | HC_vs_MS.Trt_F |
| UTP6 | 0.03645 | -0.5381 | 0.899 | 0.87 | 1 | HC_vs_MS.Trt_F |
| NUDT6 | 0.03647 | 0.83006 | 0.584 | 0.51 | 1 | HC_vs_MS.Trt_F |
| SLC1A5 | 0.03648 | 0.32549 | 0.251 | 0.18 | 1 | HC_vs_MS.Trt_F |
| SUZ12 | 0.03654 | 0.42373 | 0.717 | 0.62 | 1 | HC_vs_MS.Trt_F |
| KRT8 | 0.03664 | 0.25425 | 0.867 | 0.81 | 1 | HC_vs_MS.Trt_F |
| RIN3 | 0.0367 | 0.34857 | 0.249 | 0.18 | 1 | HC_vs_MS.Trt_F |
| DNHD1 | 0.03683 | 0.26399 | 0.39 | 0.3 | 1 | HC_vs_MS.Trt_F |
| MAP4 | 0.03683 | 0.38743 | 0.815 | 0.76 | 1 | HC_vs_MS.Trt_F |
| PLOD3 | 0.03686 | 1.14246 | 0.78 | 0.67 | 1 | HC_vs_MS.Trt_F |
| NFKBID | 0.03691 | -0.4567 | 0.642 | 0.53 | 1 | HC_vs_MS.Trt_F |
| SMAP2 | 0.03703 | -0.2523 | 0.37 | 0.28 | 1 | HC_vs_MS.Trt_F |
| BLOC1S2 | 0.03705 | -0.5017 | 0.598 | 0.5 | 1 | HC_vs_MS.Trt_F |
| SLC24A1 | 0.03725 | -0.9085 | 0.529 | 0.41 | 1 | HC_vs_MS.Trt_F |
| VPS4A | 0.03745 | 0.41766 | 0.688 | 0.6 | 1 | HC_vs_MS.Trt_F |
| RHNO1 | 0.03763 | 0.3843 | 0.74 | 0.64 | 1 | HC_vs_MS.Trt_F |
| METTL18 | 0.03765 | -0.7962 | 0.061 | 0.1 | 1 | HC_vs_MS.Trt_F |
| SLITRK5 | 0.03774 | 0.28741 | 0.801 | 0.73 | 1 | HC_vs_MS.Trt_F |
| GNA12 | 0.03779 | -0.2576 | 0.113 | 0.07 | 1 | HC_vs_MS.Trt_F |
| SLC25A19 | 0.03804 | 0.6422 | 0.465 | 0.38 | 1 | HC_vs_MS.Trt_F |
| ZNF142 | 0.03806 | 0.7909 | 0.136 | 0.08 | 1 | HC_vs_MS.Trt_F |
| TANGO6 | 0.03809 | 0.29504 | 0.87 | 0.76 | 1 | HC_vs_MS.Trt_F |
| AP2S1 | 0.03815 | -0.3523 | 0.22 | 0.14 | 1 | HC_vs_MS.Trt_F |
| SLIT3 | 0.03821 | 0.94036 | 0.616 | 0.53 | 1 | HC_vs_MS.Trt_F |
| CHDH | 0.03824 | -0.559 | 0.572 | 0.48 | 1 | HC_vs_MS.Trt_F |
| RSL24D1 | 0.03832 | -0.2779 | 0.309 | 0.22 | 1 | HC_vs_MS.Trt_F |

| | | | | | | |
|---|---|---|---|---|---|---|
| HPF1 | 0.03845 | 0.25396 | 0.145 | 0.09 | 1 | HC_vs_MS.Trt_F |
| FAM9C | 0.03851 | 0.30196 | 0.601 | 0.54 | 1 | HC_vs_MS.Trt_F |
| SFPQ | 0.03854 | -0.2646 | 0.639 | 0.54 | 1 | HC_vs_MS.Trt_F |
| ABCC1 | 0.03856 | 0.47891 | 0.402 | 0.32 | 1 | HC_vs_MS.Trt_F |
| AKR7A2 | 0.03868 | 0.25668 | 0.165 | 0.1 | 1 | HC_vs_MS.Trt_F |
| ZDHHC2 | 0.03873 | -1.2664 | 0.116 | 0.17 | 1 | HC_vs_MS.Trt_F |
| HSBP1L1 | 0.03874 | 0.37477 | 0.431 | 0.35 | 1 | HC_vs_MS.Trt_F |
| AP4M1 | 0.03884 | -0.3011 | 0.671 | 0.58 | 1 | HC_vs_MS.Trt_F |
| CREBL2 | 0.03886 | 0.6797 | 0.168 | 0.11 | 1 | HC_vs_MS.Trt_F |
| ARHGEF26 | 0.03891 | 0.52169 | 0.595 | 0.49 | 1 | HC_vs_MS.Trt_F |
| OAT | 0.03892 | 0.70044 | 0.156 | 0.1 | 1 | HC_vs_MS.Trt_F |
| SLC16A7 | 0.03893 | 0.82589 | 0.711 | 0.61 | 1 | HC_vs_MS.Trt_F |
| SLC26A11 | 0.03904 | 0.52007 | 0.697 | 0.6 | 1 | HC_vs_MS.Trt_F |
| RABGEF1 | 0.03906 | -0.2772 | 0.436 | 0.33 | 1 | HC_vs_MS.Trt_F |
| CHURC1 | 0.03907 | -0.3927 | 0.714 | 0.63 | 1 | HC_vs_MS.Trt_F |
| MED18 | 0.03915 | -0.4493 | 0.682 | 0.62 | 1 | HC_vs_MS.Trt_F |
| QSOX1 | 0.03915 | 0.54591 | 0.474 | 0.39 | 1 | HC_vs_MS.Trt_F |
| SERPINE1 | 0.03935 | 0.31911 | 0.266 | 0.19 | 1 | HC_vs_MS.Trt_F |
| SLC12A4 | 0.03954 | 0.68264 | 0.286 | 0.21 | 1 | HC_vs_MS.Trt_F |
| ARHGEF28 | 0.03955 | -0.2564 | 0.24 | 0.17 | 1 | HC_vs_MS.Trt_F |
| POLA1 | 0.03956 | 1.3048 | 0.136 | 0.08 | 1 | HC_vs_MS.Trt_F |
| CD84 | 0.03957 | 0.58486 | 0.76 | 0.72 | 1 | HC_vs_MS.Trt_F |
| TMEM241 | 0.03962 | 1.0071 | 0.694 | 0.61 | 1 | HC_vs_MS.Trt_F |
| SLC35D2 | 0.03966 | -1.222 | 0.064 | 0.11 | 1 | HC_vs_MS.Trt_F |
| INTS1 | 0.03986 | 1.00348 | 0.439 | 0.37 | 1 | HC_vs_MS.Trt_F |
| PTBP2 | 0.03991 | -0.992 | 0.682 | 0.55 | 1 | HC_vs_MS.Trt_F |
| NAA50 | 0.04 | -0.5186 | 0.512 | 0.43 | 1 | HC_vs_MS.Trt_F |
| ERVW-1 | 0.04003 | 0.40741 | 0.445 | 0.36 | 1 | HC_vs_MS.Trt_F |
| PLA2R1 | 0.04003 | 0.77219 | 0.592 | 0.64 | 1 | HC_vs_MS.Trt_F |
| TYW3 | 0.04005 | 0.50401 | 0.503 | 0.41 | 1 | HC_vs_MS.Trt_F |
| EXOC6 | 0.04007 | 1.02888 | 0.341 | 0.26 | 1 | HC_vs_MS.Trt_F |
| PELI1 | 0.04012 | 0.47586 | 0.237 | 0.17 | 1 | HC_vs_MS.Trt_F |
| UBA6 | 0.04012 | -0.2913 | 0.514 | 0.59 | 1 | HC_vs_MS.Trt_F |
| NUDT3 | 0.0402 | 0.33883 | 0.584 | 0.5 | 1 | HC_vs_MS.Trt_F |
| NUB1 | 0.04024 | 0.3106 | 0.876 | 0.85 | 1 | HC_vs_MS.Trt_F |
| CENPL | 0.04032 | 0.50236 | 0.789 | 0.72 | 1 | HC_vs_MS.Trt_F |
| ABCB9 | 0.04094 | 0.45627 | 0.85 | 0.81 | 1 | HC_vs_MS.Trt_F |
| MCOLN1 | 0.04097 | 0.41674 | 0.806 | 0.78 | 1 | HC_vs_MS.Trt_F |
| ZNF780A | 0.0412 | -0.4974 | 0.434 | 0.34 | 1 | HC_vs_MS.Trt_F |
| RCHY1 | 0.04139 | 0.31192 | 0.364 | 0.28 | 1 | HC_vs_MS.Trt_F |

| | | | | | | |
|---|---|---|---|---|---|---|
| HDAC8 | 0.0414 | 0.67401 | 0.243 | 0.18 | 1 | HC_vs_MS.Trt_F |
| FAM192A | 0.04146 | -0.6121 | 0.52 | 0.4 | 1 | HC_vs_MS.Trt_F |
| PHACTR3 | 0.04153 | 0.35824 | 0.176 | 0.12 | 1 | HC_vs_MS.Trt_F |
| THOP1 | 0.04168 | 0.34865 | 0.402 | 0.32 | 1 | HC_vs_MS.Trt_F |
| 6-Mar | 0.04168 | -0.5722 | 0.827 | 0.83 | 1 | HC_vs_MS.Trt_F |
| KIAA1586 | 0.04181 | -1.5312 | 0.104 | 0.16 | 1 | HC_vs_MS.Trt_F |
| TGS1 | 0.04182 | 0.32081 | 0.217 | 0.16 | 1 | HC_vs_MS.Trt_F |
| TOM1 | 0.04183 | 0.29308 | 0.335 | 0.26 | 1 | HC_vs_MS.Trt_F |
| MYH3 | 0.0419 | 0.76926 | 0.699 | 0.6 | 1 | HC_vs_MS.Trt_F |
| TMEM218 | 0.04191 | -1.9339 | 0.116 | 0.16 | 1 | HC_vs_MS.Trt_F |
| WDR26 | 0.04192 | 0.59134 | 0.717 | 0.63 | 1 | HC_vs_MS.Trt_F |
| EIF3I | 0.04202 | 0.33037 | 0.613 | 0.54 | 1 | HC_vs_MS.Trt_F |
| RNF2 | 0.04203 | 0.48657 | 0.494 | 0.4 | 1 | HC_vs_MS.Trt_F |
| KIAA0408 | 0.04204 | 0.56918 | 0.376 | 0.29 | 1 | HC_vs_MS.Trt_F |
| STMN3 | 0.04207 | 1.28737 | 0.249 | 0.18 | 1 | HC_vs_MS.Trt_F |
| CCDC149 | 0.04207 | 0.56734 | 0.468 | 0.37 | 1 | HC_vs_MS.Trt_F |
| SUDS3 | 0.04207 | 0.56589 | 0.841 | 0.81 | 1 | HC_vs_MS.Trt_F |
| NBPF26 | 0.04214 | 0.83199 | 0.35 | 0.27 | 1 | HC_vs_MS.Trt_F |
| LY6G5B | 0.04228 | 0.32682 | 0.864 | 0.82 | 1 | HC_vs_MS.Trt_F |
| PARP2 | 0.0423 | 1.67451 | 0.436 | 0.34 | 1 | HC_vs_MS.Trt_F |
| AHR | 0.0423 | 0.28347 | 0.688 | 0.65 | 1 | HC_vs_MS.Trt_F |
| GRID1 | 0.04235 | -0.5518 | 0.098 | 0.15 | 1 | HC_vs_MS.Trt_F |
| IGF1R | 0.04242 | 0.30233 | 0.387 | 0.29 | 1 | HC_vs_MS.Trt_F |
| AHCY | 0.04253 | -1.4884 | 0.098 | 0.15 | 1 | HC_vs_MS.Trt_F |
| PLEKHJ1 | 0.04257 | 0.31978 | 0.821 | 0.78 | 1 | HC_vs_MS.Trt_F |
| LSG1 | 0.043 | -0.2868 | 0.503 | 0.41 | 1 | HC_vs_MS.Trt_F |
| ARHGEF7 | 0.04309 | 0.31636 | 0.303 | 0.23 | 1 | HC_vs_MS.Trt_F |
| CDK16 | 0.04327 | 0.35799 | 0.254 | 0.18 | 1 | HC_vs_MS.Trt_F |
| MTMR9 | 0.04338 | 0.32325 | 0.361 | 0.28 | 1 | HC_vs_MS.Trt_F |
| STX7 | 0.04358 | 0.28182 | 0.795 | 0.72 | 1 | HC_vs_MS.Trt_F |
| LEMD3 | 0.04367 | -0.3732 | 0.714 | 0.59 | 1 | HC_vs_MS.Trt_F |
| CCNB1IP1 | 0.04375 | 0.35873 | 0.858 | 0.78 | 1 | HC_vs_MS.Trt_F |
| ACAT2 | 0.0439 | 1.45977 | 0.295 | 0.23 | 1 | HC_vs_MS.Trt_F |
| TWSG1 | 0.04391 | 0.41169 | 0.454 | 0.37 | 1 | HC_vs_MS.Trt_F |
| ATOX1 | 0.04392 | 1.28037 | 0.277 | 0.21 | 1 | HC_vs_MS.Trt_F |
| VPS13D | 0.04393 | -0.7216 | 0.344 | 0.25 | 1 | HC_vs_MS.Trt_F |
| CCDC50 | 0.04408 | 0.30071 | 0.503 | 0.42 | 1 | HC_vs_MS.Trt_F |
| PPP4R1 | 0.04419 | 0.44035 | 0.295 | 0.21 | 1 | HC_vs_MS.Trt_F |
| KMT2C | 0.04426 | 0.26707 | 0.451 | 0.37 | 1 | HC_vs_MS.Trt_F |
| RARA | 0.0446 | 0.71068 | 0.162 | 0.11 | 1 | HC_vs_MS.Trt_F |

| | | | | | | |
|---|---|---|---|---|---|---|
| BMPR2 | 0.04464 | 0.51762 | 0.604 | 0.54 | 1 | HC_vs_MS.Trt_F |
| C9orf3 | 0.04466 | 0.47005 | 0.78 | 0.7 | 1 | HC_vs_MS.Trt_F |
| CCDC180 | 0.04469 | 0.29668 | 0.818 | 0.71 | 1 | HC_vs_MS.Trt_F |
| ATP8B3 | 0.04471 | 0.35671 | 0.847 | 0.78 | 1 | HC_vs_MS.Trt_F |
| PIK3R4 | 0.04474 | 0.8634 | 0.182 | 0.12 | 1 | HC_vs_MS.Trt_F |
| GTF3C5 | 0.04483 | 0.35412 | 0.35 | 0.27 | 1 | HC_vs_MS.Trt_F |
| SYF2 | 0.04486 | 0.41864 | 0.5 | 0.41 | 1 | HC_vs_MS.Trt_F |
| TRIB3 | 0.04489 | -0.7632 | 0.121 | 0.07 | 1 | HC_vs_MS.Trt_F |
| ZNF627 | 0.0452 | 1.18873 | 0.633 | 0.55 | 1 | HC_vs_MS.Trt_F |
| SNRK | 0.0452 | 0.80608 | 0.52 | 0.45 | 1 | HC_vs_MS.Trt_F |
| HAUS7 | 0.04527 | 0.34053 | 0.147 | 0.09 | 1 | HC_vs_MS.Trt_F |
| PLG | 0.0453 | 0.59446 | 0.254 | 0.19 | 1 | HC_vs_MS.Trt_F |
| FBXL17 | 0.04533 | 1.62117 | 0.324 | 0.24 | 1 | HC_vs_MS.Trt_F |
| SLC25A35 | 0.04538 | 0.65232 | 0.202 | 0.14 | 1 | HC_vs_MS.Trt_F |
| ATP6V1H | 0.04539 | 0.7179 | 0.434 | 0.37 | 1 | HC_vs_MS.Trt_F |
| IFT74 | 0.04541 | 0.26536 | 0.465 | 0.37 | 1 | HC_vs_MS.Trt_F |
| G2E3 | 0.04547 | -0.3028 | 0.549 | 0.46 | 1 | HC_vs_MS.Trt_F |
| ZNF525 | 0.04547 | 0.8792 | 0.673 | 0.59 | 1 | HC_vs_MS.Trt_F |
| BAG3 | 0.04555 | 0.87396 | 0.118 | 0.07 | 1 | HC_vs_MS.Trt_F |
| NUCKS1 | 0.04558 | -0.5972 | 0.5 | 0.52 | 1 | HC_vs_MS.Trt_F |
| PHLDB1 | 0.04565 | 0.64508 | 0.986 | 0.98 | 1 | HC_vs_MS.Trt_F |
| ATF1 | 0.04577 | 0.89886 | 0.191 | 0.13 | 1 | HC_vs_MS.Trt_F |
| COLEC11 | 0.04584 | 0.43451 | 0.393 | 0.31 | 1 | HC_vs_MS.Trt_F |
| NGDN | 0.04603 | -0.3575 | 0.367 | 0.28 | 1 | HC_vs_MS.Trt_F |
| PCCB | 0.04613 | 0.40916 | 0.679 | 0.63 | 1 | HC_vs_MS.Trt_F |
| ATP13A4 | 0.04623 | 0.41885 | 0.503 | 0.42 | 1 | HC_vs_MS.Trt_F |
| SLC7A6 | 0.04633 | 0.38843 | 0.555 | 0.49 | 1 | HC_vs_MS.Trt_F |
| GPX1 | 0.04638 | 0.31038 | 0.205 | 0.14 | 1 | HC_vs_MS.Trt_F |
| MSRB1 | 0.0467 | -0.4992 | 0.13 | 0.08 | 1 | HC_vs_MS.Trt_F |
| MATN2 | 0.04687 | -0.4031 | 0.613 | 0.52 | 1 | HC_vs_MS.Trt_F |
| NOM1 | 0.04698 | 0.44769 | 0.714 | 0.62 | 1 | HC_vs_MS.Trt_F |
| HAT1 | 0.04706 | 0.35352 | 0.884 | 0.87 | 1 | HC_vs_MS.Trt_F |
| EPS15L1 | 0.04708 | 0.38266 | 0.827 | 0.8 | 1 | HC_vs_MS.Trt_F |
| EPM2AIP1 | 0.04709 | 0.34343 | 0.309 | 0.23 | 1 | HC_vs_MS.Trt_F |
| HERC5 | 0.04717 | 0.55324 | 0.214 | 0.15 | 1 | HC_vs_MS.Trt_F |
| VSTM4 | 0.04728 | 0.99433 | 0.777 | 0.68 | 1 | HC_vs_MS.Trt_F |
| TTC21B | 0.04729 | 0.56169 | 0.416 | 0.34 | 1 | HC_vs_MS.Trt_F |
| ZNF585B | 0.04746 | 0.58641 | 0.772 | 0.65 | 1 | HC_vs_MS.Trt_F |
| CLASP2 | 0.04752 | 0.74087 | 0.751 | 0.67 | 1 | HC_vs_MS.Trt_F |
| MAT2B | 0.0477 | -0.7348 | 0.746 | 0.76 | 1 | HC_vs_MS.Trt_F |

| | | | | | | |
|---|---|---|---|---|---|---|
| TAB2 | 0.04772 | 0.26483 | 0.578 | 0.49 | 1 | HC_vs_MS.Trt_F |
| HERC3 | 0.04774 | 0.57519 | 0.341 | 0.27 | 1 | HC_vs_MS.Trt_F |
| TDRD1 | 0.04781 | 0.77707 | 0.225 | 0.16 | 1 | HC_vs_MS.Trt_F |
| CORO7-PAM | 0.04793 | 0.96869 | 0.156 | 0.1 | 1 | HC_vs_MS.Trt_F |
| CEP290 | 0.04806 | -0.2986 | 0.173 | 0.11 | 1 | HC_vs_MS.Trt_F |
| STK25 | 0.04819 | -0.5209 | 0.76 | 0.63 | 1 | HC_vs_MS.Trt_F |
| CCDC24 | 0.04825 | -0.2762 | 0.714 | 0.75 | 1 | HC_vs_MS.Trt_F |
| SOD1 | 0.04832 | 0.25598 | 0.717 | 0.68 | 1 | HC_vs_MS.Trt_F |
| PDZK1 | 0.04842 | 0.47433 | 0.355 | 0.28 | 1 | HC_vs_MS.Trt_F |
| TSPYL4 | 0.0486 | 0.4017 | 0.168 | 0.11 | 1 | HC_vs_MS.Trt_F |
| CD109 | 0.04871 | 0.47433 | 0.477 | 0.4 | 1 | HC_vs_MS.Trt_F |
| ITSN1 | 0.04874 | 0.38603 | 0.936 | 0.85 | 1 | HC_vs_MS.Trt_F |
| ZBED1 | 0.04898 | 0.58962 | 0.546 | 0.47 | 1 | HC_vs_MS.Trt_F |
| SIK2 | 0.04905 | 0.28686 | 0.763 | 0.69 | 1 | HC_vs_MS.Trt_F |
| MAPRE1 | 0.04908 | 0.3186 | 0.28 | 0.21 | 1 | HC_vs_MS.Trt_F |
| SLC5A10 | 0.04909 | -1.0275 | 0.058 | 0.1 | 1 | HC_vs_MS.Trt_F |
| RUNX1T1 | 0.04912 | 0.28441 | 0.249 | 0.18 | 1 | HC_vs_MS.Trt_F |
| SCAF8 | 0.04937 | -0.2692 | 0.301 | 0.22 | 1 | HC_vs_MS.Trt_F |
| H2AFV | 0.04945 | 0.46844 | 0.691 | 0.64 | 1 | HC_vs_MS.Trt_F |
| IL12RB2 | 0.04965 | 0.48168 | 0.162 | 0.11 | 1 | HC_vs_MS.Trt_F |
| SPAG7 | 0.04967 | 0.46533 | 0.893 | 0.83 | 1 | HC_vs_MS.Trt_F |
| SLAIN2 | 0.05008 | -0.6744 | 0.162 | 0.21 | 1 | HC_vs_MS.Trt_F |
| PRPF4B | 0.05025 | -0.4886 | 0.364 | 0.28 | 1 | HC_vs_MS.Trt_F |
| NUP62 | 0.0503 | 0.28006 | 0.587 | 0.5 | 1 | HC_vs_MS.Trt_F |
| PFKM | 0.05031 | -0.3881 | 0.884 | 0.87 | 1 | HC_vs_MS.Trt_F |
| VRK2 | 0.05032 | -0.2628 | 0.295 | 0.22 | 1 | HC_vs_MS.Trt_F |
| BORCS7 | 0.0505 | 0.60909 | 0.419 | 0.33 | 1 | HC_vs_MS.Trt_F |
| TMEM55A | 0.05056 | 1.37408 | 0.113 | 0.07 | 1 | HC_vs_MS.Trt_F |
| ALKBH5 | 0.05068 | 0.29623 | 0.208 | 0.14 | 1 | HC_vs_MS.Trt_F |
| BET1L | 0.05089 | 0.26092 | 0.251 | 0.18 | 1 | HC_vs_MS.Trt_F |
| PRRG4 | 0.05104 | 0.28249 | 0.85 | 0.81 | 1 | HC_vs_MS.Trt_F |
| EMP1 | 0.05108 | 0.7027 | 0.225 | 0.16 | 1 | HC_vs_MS.Trt_F |
| ERN1 | 0.05126 | 0.27138 | 0.795 | 0.77 | 1 | HC_vs_MS.Trt_F |
| TRMT12 | 0.05137 | -0.3489 | 0.512 | 0.39 | 1 | HC_vs_MS.Trt_F |
| EMC2 | 0.0515 | 0.53451 | 0.517 | 0.43 | 1 | HC_vs_MS.Trt_F |
| RASGRF2 | 0.05154 | 0.92953 | 0.228 | 0.17 | 1 | HC_vs_MS.Trt_F |
| DCAF1 | 0.05157 | 0.28764 | 0.225 | 0.17 | 1 | HC_vs_MS.Trt_F |
| GAB3 | 0.05158 | -0.5666 | 0.451 | 0.36 | 1 | HC_vs_MS.Trt_F |
| PDE1B | 0.05193 | 0.42599 | 0.601 | 0.51 | 1 | HC_vs_MS.Trt_F |
| HDAC2 | 0.05194 | 0.42994 | 0.74 | 0.66 | 1 | HC_vs_MS.Trt_F |

| | | | | | | |
|---|---|---|---|---|---|---|
| CD7 | 0.05196 | 0.68115 | 0.289 | 0.22 | 1 | HC_vs_MS.Trt_F |
| IDH3B | 0.05241 | 0.29993 | 0.237 | 0.17 | 1 | HC_vs_MS.Trt_F |
| C6orf223 | 0.05245 | 0.39085 | 0.286 | 0.22 | 1 | HC_vs_MS.Trt_F |
| FAM193A | 0.0525 | 0.31972 | 0.341 | 0.27 | 1 | HC_vs_MS.Trt_F |
| C9orf131 | 0.05261 | 0.93682 | 0.182 | 0.13 | 1 | HC_vs_MS.Trt_F |
| COMTD1 | 0.05262 | -0.3413 | 0.173 | 0.11 | 1 | HC_vs_MS.Trt_F |
| DYNLT1 | 0.05289 | 0.62629 | 0.587 | 0.51 | 1 | HC_vs_MS.Trt_F |
| C14orf169 | 0.05291 | -1.1213 | 0.066 | 0.11 | 1 | HC_vs_MS.Trt_F |
| NAGK | 0.05309 | 0.29261 | 0.832 | 0.76 | 1 | HC_vs_MS.Trt_F |
| PLD2 | 0.05316 | 0.41825 | 0.142 | 0.09 | 1 | HC_vs_MS.Trt_F |
| BMT2 | 0.05329 | 1.4375 | 0.136 | 0.09 | 1 | HC_vs_MS.Trt_F |
| ZEB2 | 0.0535 | -0.7997 | 0.647 | 0.69 | 1 | HC_vs_MS.Trt_F |
| BRD2 | 0.05362 | 0.27037 | 0.679 | 0.6 | 1 | HC_vs_MS.Trt_F |
| EED | 0.05364 | -0.5833 | 0.231 | 0.16 | 1 | HC_vs_MS.Trt_F |
| HDAC7 | 0.05364 | 0.85003 | 0.321 | 0.26 | 1 | HC_vs_MS.Trt_F |
| PHACTR2 | 0.05388 | -0.7593 | 0.598 | 0.64 | 1 | HC_vs_MS.Trt_F |
| PURA | 0.05402 | -0.697 | 0.136 | 0.09 | 1 | HC_vs_MS.Trt_F |
| NXN | 0.05409 | 0.31275 | 0.578 | 0.5 | 1 | HC_vs_MS.Trt_F |
| DDX18 | 0.05418 | -0.3023 | 0.798 | 0.69 | 1 | HC_vs_MS.Trt_F |
| DENND4A | 0.05422 | 0.66347 | 0.439 | 0.38 | 1 | HC_vs_MS.Trt_F |
| CSNK1D | 0.05422 | 0.38088 | 0.954 | 0.92 | 1 | HC_vs_MS.Trt_F |
| DNM3 | 0.05427 | 1.12201 | 0.121 | 0.08 | 1 | HC_vs_MS.Trt_F |
| DYM | 0.05431 | 0.3492 | 0.74 | 0.69 | 1 | HC_vs_MS.Trt_F |
| FLT4 | 0.05438 | 0.50805 | 0.124 | 0.08 | 1 | HC_vs_MS.Trt_F |
| RPP30 | 0.05441 | 0.58607 | 0.679 | 0.64 | 1 | HC_vs_MS.Trt_F |
| NADK | 0.05446 | -0.3015 | 0.306 | 0.23 | 1 | HC_vs_MS.Trt_F |
| MRPS30 | 0.05449 | -0.4186 | 0.312 | 0.37 | 1 | HC_vs_MS.Trt_F |
| ASCC1 | 0.05454 | 0.66204 | 0.714 | 0.61 | 1 | HC_vs_MS.Trt_F |
| MMP11 | 0.05457 | 0.58609 | 0.431 | 0.36 | 1 | HC_vs_MS.Trt_F |
| SUN2 | 0.05467 | 0.25453 | 0.422 | 0.35 | 1 | HC_vs_MS.Trt_F |
| KRIT1 | 0.05487 | 0.72511 | 0.673 | 0.6 | 1 | HC_vs_MS.Trt_F |
| ANAPC15 | 0.05487 | -0.4415 | 0.815 | 0.76 | 1 | HC_vs_MS.Trt_F |
| VAMP1 | 0.05498 | 0.39691 | 0.185 | 0.13 | 1 | HC_vs_MS.Trt_F |
| ICE2 | 0.0551 | -0.2805 | 0.882 | 0.84 | 1 | HC_vs_MS.Trt_F |
| TBC1D12 | 0.05512 | 0.33549 | 0.445 | 0.38 | 1 | HC_vs_MS.Trt_F |
| AGFG2 | 0.05538 | 0.34612 | 0.355 | 0.28 | 1 | HC_vs_MS.Trt_F |
| MFSD12 | 0.05538 | 0.70692 | 0.295 | 0.22 | 1 | HC_vs_MS.Trt_F |
| RIOK2 | 0.05545 | -0.2795 | 0.39 | 0.31 | 1 | HC_vs_MS.Trt_F |
| FOXP3 | 0.05554 | 0.54602 | 0.503 | 0.42 | 1 | HC_vs_MS.Trt_F |
| SYNJ2 | 0.05568 | 0.29941 | 0.928 | 0.9 | 1 | HC_vs_MS.Trt_F |

| | | | | | | |
|---|---|---|---|---|---|---|
| YWHAE | 0.05572 | -0.6064 | 0.431 | 0.47 | 1 | HC_vs_MS.Trt_F |
| RYBP | 0.05579 | 0.44451 | 0.113 | 0.07 | 1 | HC_vs_MS.Trt_F |
| PIGT | 0.05587 | 1.17556 | 0.147 | 0.1 | 1 | HC_vs_MS.Trt_F |
| WDR70 | 0.05593 | 0.60766 | 0.315 | 0.23 | 1 | HC_vs_MS.Trt_F |
| FOXRED2 | 0.05597 | -0.254 | 0.59 | 0.5 | 1 | HC_vs_MS.Trt_F |
| UPK3BL | 0.05601 | -0.495 | 0.532 | 0.58 | 1 | HC_vs_MS.Trt_F |
| UMAD1 | 0.05609 | -0.3222 | 0.22 | 0.16 | 1 | HC_vs_MS.Trt_F |
| RTN3 | 0.05615 | 0.41873 | 0.211 | 0.16 | 1 | HC_vs_MS.Trt_F |
| ZSCAN29 | 0.05617 | -0.7417 | 0.445 | 0.48 | 1 | HC_vs_MS.Trt_F |
| FOXRED1 | 0.05621 | 1.52585 | 0.549 | 0.48 | 1 | HC_vs_MS.Trt_F |
| GET4 | 0.05628 | 0.29403 | 0.535 | 0.45 | 1 | HC_vs_MS.Trt_F |
| RAB3GAP1 | 0.05631 | -0.4292 | 0.402 | 0.31 | 1 | HC_vs_MS.Trt_F |
| RP11-613M1 | 0.0566 | 0.26374 | 0.22 | 0.16 | 1 | HC_vs_MS.Trt_F |
| RPL36AL | 0.05665 | -0.5674 | 0.509 | 0.53 | 1 | HC_vs_MS.Trt_F |
| LRRC8A | 0.05677 | 0.28525 | 0.173 | 0.12 | 1 | HC_vs_MS.Trt_F |
| TP63 | 0.0568 | 0.5372 | 0.465 | 0.39 | 1 | HC_vs_MS.Trt_F |
| KCNMB3 | 0.05705 | -0.5215 | 0.465 | 0.38 | 1 | HC_vs_MS.Trt_F |
| ALS2 | 0.05707 | -0.5762 | 0.497 | 0.4 | 1 | HC_vs_MS.Trt_F |
| CD83 | 0.0571 | 0.87436 | 0.179 | 0.13 | 1 | HC_vs_MS.Trt_F |
| LNPEP | 0.05716 | 0.34478 | 0.598 | 0.53 | 1 | HC_vs_MS.Trt_F |
| PMPCA | 0.05717 | 0.32018 | 0.775 | 0.71 | 1 | HC_vs_MS.Trt_F |
| OSBPL3 | 0.05724 | 0.26285 | 0.532 | 0.45 | 1 | HC_vs_MS.Trt_F |
| SMURF2 | 0.05724 | 0.93841 | 0.801 | 0.72 | 1 | HC_vs_MS.Trt_F |
| DHRS3 | 0.05735 | -0.6765 | 0.338 | 0.25 | 1 | HC_vs_MS.Trt_F |
| PHB | 0.05743 | 0.31992 | 0.199 | 0.14 | 1 | HC_vs_MS.Trt_F |
| XPA | 0.05758 | 0.56193 | 0.208 | 0.15 | 1 | HC_vs_MS.Trt_F |
| RBKS | 0.05759 | 0.82264 | 0.145 | 0.1 | 1 | HC_vs_MS.Trt_F |
| SART1 | 0.05759 | 0.70225 | 0.37 | 0.29 | 1 | HC_vs_MS.Trt_F |
| EVI2B | 0.05766 | 0.33334 | 0.289 | 0.22 | 1 | HC_vs_MS.Trt_F |
| PTPRG | 0.0577 | 1.11663 | 0.295 | 0.23 | 1 | HC_vs_MS.Trt_F |
| RRP15 | 0.05773 | -0.4339 | 0.636 | 0.53 | 1 | HC_vs_MS.Trt_F |
| SFT2D3 | 0.05775 | -0.5151 | 0.535 | 0.45 | 1 | HC_vs_MS.Trt_F |
| POLQ | 0.05777 | 0.44386 | 0.618 | 0.53 | 1 | HC_vs_MS.Trt_F |
| PECAM1 | 0.05799 | 0.26468 | 0.798 | 0.75 | 1 | HC_vs_MS.Trt_F |
| HNRNPF | 0.05805 | -0.6274 | 0.335 | 0.39 | 1 | HC_vs_MS.Trt_F |
| EEA1 | 0.05815 | -0.3307 | 0.277 | 0.21 | 1 | HC_vs_MS.Trt_F |
| SEC22B | 0.05821 | -0.7719 | 0.829 | 0.85 | 1 | HC_vs_MS.Trt_F |
| APOBEC3A | 0.05839 | 0.2654 | 0.462 | 0.38 | 1 | HC_vs_MS.Trt_F |
| ILVBL | 0.05841 | -0.7618 | 0.318 | 0.23 | 1 | HC_vs_MS.Trt_F |
| UBAP1L | 0.05843 | 0.79806 | 0.399 | 0.32 | 1 | HC_vs_MS.Trt_F |

| | | | | | | |
|---|---|---|---|---|---|---|
| RNF207 | 0.05844 | -0.4303 | 0.725 | 0.74 | 1 | HC_vs_MS.Trt_F |
| WDR48 | 0.05849 | -0.3995 | 0.211 | 0.14 | 1 | HC_vs_MS.Trt_F |
| ZNF460 | 0.0585 | 0.62569 | 0.22 | 0.16 | 1 | HC_vs_MS.Trt_F |
| STMN1 | 0.05856 | 0.30272 | 0.775 | 0.73 | 1 | HC_vs_MS.Trt_F |
| GATAD1 | 0.05866 | 0.27149 | 0.896 | 0.86 | 1 | HC_vs_MS.Trt_F |
| HSD3B7 | 0.05879 | 0.34288 | 0.506 | 0.42 | 1 | HC_vs_MS.Trt_F |
| CEP85L | 0.05879 | 0.30027 | 0.292 | 0.22 | 1 | HC_vs_MS.Trt_F |
| SLC27A1 | 0.05882 | 0.37517 | 0.87 | 0.78 | 1 | HC_vs_MS.Trt_F |
| MITF | 0.05894 | 0.28026 | 0.199 | 0.14 | 1 | HC_vs_MS.Trt_F |
| UGDH | 0.05905 | -0.4482 | 0.584 | 0.64 | 1 | HC_vs_MS.Trt_F |
| RGS9 | 0.05926 | 0.41796 | 0.633 | 0.57 | 1 | HC_vs_MS.Trt_F |
| VMAC | 0.05934 | 0.4322 | 0.691 | 0.63 | 1 | HC_vs_MS.Trt_F |
| SYNPO | 0.05942 | 0.29118 | 0.237 | 0.18 | 1 | HC_vs_MS.Trt_F |
| ZNF134 | 0.05947 | 1.26531 | 0.194 | 0.14 | 1 | HC_vs_MS.Trt_F |
| KCNMB4 | 0.05951 | 0.40077 | 0.37 | 0.31 | 1 | HC_vs_MS.Trt_F |
| NME1 | 0.05955 | 0.31068 | 0.639 | 0.73 | 1 | HC_vs_MS.Trt_F |
| ABI2 | 0.05956 | 0.39601 | 0.899 | 0.84 | 1 | HC_vs_MS.Trt_F |
| RECQL5 | 0.05976 | 1.42873 | 0.257 | 0.19 | 1 | HC_vs_MS.Trt_F |
| PRPF18 | 0.05993 | 0.69969 | 0.165 | 0.11 | 1 | HC_vs_MS.Trt_F |
| ANKMY1 | 0.05996 | 0.67241 | 0.462 | 0.37 | 1 | HC_vs_MS.Trt_F |
| SOCS1 | 0.05999 | 0.81013 | 0.139 | 0.09 | 1 | HC_vs_MS.Trt_F |
| TVP23B | 0.05999 | 0.33533 | 0.647 | 0.57 | 1 | HC_vs_MS.Trt_F |
| NDUFB6 | 0.06028 | 0.36321 | 0.561 | 0.49 | 1 | HC_vs_MS.Trt_F |
| CES3 | 0.0603 | 0.55121 | 0.327 | 0.26 | 1 | HC_vs_MS.Trt_F |
| HNRNPUL1 | 0.06035 | 0.6483 | 0.705 | 0.66 | 1 | HC_vs_MS.Trt_F |
| ZNF195 | 0.0604 | 0.52492 | 0.621 | 0.53 | 1 | HC_vs_MS.Trt_F |
| SMG9 | 0.06052 | -0.2813 | 0.688 | 0.59 | 1 | HC_vs_MS.Trt_F |
| BCCIP | 0.06056 | 0.64661 | 0.228 | 0.17 | 1 | HC_vs_MS.Trt_F |
| ESRP1 | 0.06061 | -0.7363 | 0.165 | 0.22 | 1 | HC_vs_MS.Trt_F |
| ZNF354B | 0.06062 | 0.37522 | 0.361 | 0.29 | 1 | HC_vs_MS.Trt_F |
| SLC20A2 | 0.06076 | 0.76822 | 0.179 | 0.13 | 1 | HC_vs_MS.Trt_F |
| RHOT1 | 0.06084 | 0.52234 | 0.254 | 0.18 | 1 | HC_vs_MS.Trt_F |
| CENPV | 0.06088 | 0.5012 | 0.801 | 0.72 | 1 | HC_vs_MS.Trt_F |
| MPP6 | 0.06101 | 1.17017 | 0.335 | 0.26 | 1 | HC_vs_MS.Trt_F |
| HIVEP1 | 0.06101 | 0.48395 | 0.665 | 0.63 | 1 | HC_vs_MS.Trt_F |
| HS3ST3B1 | 0.06104 | -0.4791 | 0.46 | 0.52 | 1 | HC_vs_MS.Trt_F |
| ANKRD6 | 0.06105 | 0.399 | 0.197 | 0.14 | 1 | HC_vs_MS.Trt_F |
| SERGEF | 0.06126 | 0.32085 | 0.185 | 0.13 | 1 | HC_vs_MS.Trt_F |
| CAMK2G | 0.06126 | 0.25205 | 0.676 | 0.58 | 1 | HC_vs_MS.Trt_F |
| LECT2 | 0.06127 | 0.39907 | 0.191 | 0.14 | 1 | HC_vs_MS.Trt_F |

| | | | | | | |
|---|---|---|---|---|---|---|
| TADA3 | 0.06139 | 0.4689 | 0.616 | 0.53 | 1 | HC_vs_MS.Trt_F |
| PTPRB | 0.06147 | 0.30717 | 0.624 | 0.54 | 1 | HC_vs_MS.Trt_F |
| NAA10 | 0.0615 | 0.55677 | 0.448 | 0.38 | 1 | HC_vs_MS.Trt_F |
| HSPA4 | 0.06152 | 0.27946 | 0.434 | 0.35 | 1 | HC_vs_MS.Trt_F |
| RNF41 | 0.06155 | -0.3452 | 0.801 | 0.72 | 1 | HC_vs_MS.Trt_F |
| ZNF816 | 0.06158 | -0.7622 | 0.506 | 0.43 | 1 | HC_vs_MS.Trt_F |
| YWHAH | 0.06168 | 0.48362 | 0.246 | 0.19 | 1 | HC_vs_MS.Trt_F |
| KIAA0556 | 0.06187 | 0.77131 | 0.358 | 0.28 | 1 | HC_vs_MS.Trt_F |
| CMTM3 | 0.06197 | 0.64394 | 0.243 | 0.18 | 1 | HC_vs_MS.Trt_F |
| FAM159A | 0.06203 | 0.58148 | 0.63 | 0.52 | 1 | HC_vs_MS.Trt_F |
| LGALS9 | 0.06209 | 0.53975 | 0.298 | 0.22 | 1 | HC_vs_MS.Trt_F |
| FSTL4 | 0.06221 | 0.38565 | 0.246 | 0.18 | 1 | HC_vs_MS.Trt_F |
| INO80E | 0.06229 | 0.68836 | 0.22 | 0.16 | 1 | HC_vs_MS.Trt_F |
| UHMK1 | 0.06246 | -0.3452 | 0.329 | 0.26 | 1 | HC_vs_MS.Trt_F |
| PIGF | 0.06279 | -0.2865 | 0.13 | 0.08 | 1 | HC_vs_MS.Trt_F |
| TNIP1 | 0.06286 | 1.02398 | 0.355 | 0.3 | 1 | HC_vs_MS.Trt_F |
| EZH1 | 0.063 | 0.43067 | 0.939 | 0.88 | 1 | HC_vs_MS.Trt_F |
| TRIM2 | 0.06306 | 0.60847 | 0.251 | 0.19 | 1 | HC_vs_MS.Trt_F |
| DDA1 | 0.06312 | 0.35843 | 0.598 | 0.52 | 1 | HC_vs_MS.Trt_F |
| CAPRIN1 | 0.06335 | -0.2569 | 0.347 | 0.26 | 1 | HC_vs_MS.Trt_F |
| ZNF720 | 0.06337 | 0.73689 | 0.584 | 0.5 | 1 | HC_vs_MS.Trt_F |
| TRAF2 | 0.0639 | -1.063 | 0.214 | 0.26 | 1 | HC_vs_MS.Trt_F |
| DENND2D | 0.06399 | -0.358 | 0.338 | 0.39 | 1 | HC_vs_MS.Trt_F |
| UBE3B | 0.06405 | 0.76281 | 0.393 | 0.33 | 1 | HC_vs_MS.Trt_F |
| PSMG2 | 0.0641 | 0.313 | 0.225 | 0.17 | 1 | HC_vs_MS.Trt_F |
| TMEM263 | 0.06418 | -0.5663 | 0.465 | 0.39 | 1 | HC_vs_MS.Trt_F |
| RPS26 | 0.06429 | -0.3061 | 0.702 | 0.69 | 1 | HC_vs_MS.Trt_F |
| FASN | 0.06439 | 1.6602 | 0.171 | 0.12 | 1 | HC_vs_MS.Trt_F |
| KCTD9 | 0.0647 | -1.1362 | 0.263 | 0.19 | 1 | HC_vs_MS.Trt_F |
| RPGR | 0.06471 | 1.08806 | 0.11 | 0.07 | 1 | HC_vs_MS.Trt_F |
| BMF | 0.06472 | -0.5035 | 0.15 | 0.1 | 1 | HC_vs_MS.Trt_F |
| TTC31 | 0.06477 | 0.85253 | 0.246 | 0.19 | 1 | HC_vs_MS.Trt_F |
| PLBD2 | 0.06479 | 0.72856 | 0.41 | 0.34 | 1 | HC_vs_MS.Trt_F |
| ANAPC5 | 0.06512 | 0.2987 | 0.962 | 0.94 | 1 | HC_vs_MS.Trt_F |
| CCDC58 | 0.06519 | 0.36676 | 0.35 | 0.27 | 1 | HC_vs_MS.Trt_F |
| CUBN | 0.06547 | 0.85105 | 0.477 | 0.39 | 1 | HC_vs_MS.Trt_F |
| ZBTB46 | 0.06549 | 0.40681 | 0.136 | 0.09 | 1 | HC_vs_MS.Trt_F |
| ZNF764 | 0.06551 | 0.69719 | 0.399 | 0.33 | 1 | HC_vs_MS.Trt_F |
| PHLDA3 | 0.06564 | 0.48725 | 0.425 | 0.35 | 1 | HC_vs_MS.Trt_F |
| DNAL1 | 0.06576 | 0.38658 | 0.951 | 0.9 | 1 | HC_vs_MS.Trt_F |

| | | | | | | |
|---|---|---|---|---|---|---|
| ROCK1 | 0.06576 | 0.28119 | 0.751 | 0.68 | 1 | HC_vs_MS.Trt_F |
| ZDHHC21 | 0.06584 | -0.5236 | 0.301 | 0.23 | 1 | HC_vs_MS.Trt_F |
| FOXN3 | 0.06589 | 0.34281 | 0.913 | 0.88 | 1 | HC_vs_MS.Trt_F |
| FEZ2 | 0.06598 | 1.17339 | 0.54 | 0.46 | 1 | HC_vs_MS.Trt_F |
| SERPINA10 | 0.06605 | 0.26543 | 0.627 | 0.53 | 1 | HC_vs_MS.Trt_F |
| ZNF664 | 0.06612 | 0.52174 | 0.301 | 0.23 | 1 | HC_vs_MS.Trt_F |
| FKBP10 | 0.06613 | 0.46281 | 0.428 | 0.35 | 1 | HC_vs_MS.Trt_F |
| ZNF780B | 0.06654 | -0.709 | 0.584 | 0.64 | 1 | HC_vs_MS.Trt_F |
| TPGS1 | 0.0666 | 0.99595 | 0.312 | 0.39 | 1 | HC_vs_MS.Trt_F |
| MEIS2 | 0.06661 | 0.39485 | 0.225 | 0.17 | 1 | HC_vs_MS.Trt_F |
| SWSAP1 | 0.06681 | 0.57207 | 0.783 | 0.7 | 1 | HC_vs_MS.Trt_F |
| TLE4 | 0.06704 | 0.28174 | 0.199 | 0.14 | 1 | HC_vs_MS.Trt_F |
| RABEP2 | 0.06743 | 0.52084 | 0.566 | 0.52 | 1 | HC_vs_MS.Trt_F |
| SDR42E1 | 0.06744 | 0.47605 | 0.471 | 0.38 | 1 | HC_vs_MS.Trt_F |
| PCBP3 | 0.06745 | 0.27502 | 0.506 | 0.44 | 1 | HC_vs_MS.Trt_F |
| FLNA | 0.06754 | 0.28053 | 0.592 | 0.51 | 1 | HC_vs_MS.Trt_F |
| RAI1 | 0.06768 | 1.20733 | 0.234 | 0.18 | 1 | HC_vs_MS.Trt_F |
| MLEC | 0.06777 | 0.67061 | 0.15 | 0.1 | 1 | HC_vs_MS.Trt_F |
| MELK | 0.06797 | -0.6281 | 0.584 | 0.59 | 1 | HC_vs_MS.Trt_F |
| NIT1 | 0.068 | 0.36903 | 0.676 | 0.62 | 1 | HC_vs_MS.Trt_F |
| KCTD11 | 0.06823 | 0.91165 | 0.173 | 0.12 | 1 | HC_vs_MS.Trt_F |
| SLC25A40 | 0.06827 | -0.6514 | 0.364 | 0.28 | 1 | HC_vs_MS.Trt_F |
| STARD6 | 0.06837 | -0.4051 | 0.176 | 0.13 | 1 | HC_vs_MS.Trt_F |
| DAG1 | 0.06841 | 1.49765 | 0.604 | 0.55 | 1 | HC_vs_MS.Trt_F |
| PRRC2B | 0.06842 | 0.65114 | 0.312 | 0.25 | 1 | HC_vs_MS.Trt_F |
| USO1 | 0.06851 | 0.51055 | 0.208 | 0.15 | 1 | HC_vs_MS.Trt_F |
| ESAM | 0.06851 | -1.2638 | 0.116 | 0.17 | 1 | HC_vs_MS.Trt_F |
| ZNF226 | 0.06853 | -0.4961 | 0.581 | 0.5 | 1 | HC_vs_MS.Trt_F |
| TBCEL | 0.06859 | 0.28149 | 0.113 | 0.16 | 1 | HC_vs_MS.Trt_F |
| SUPT20H | 0.06859 | 0.37655 | 0.491 | 0.41 | 1 | HC_vs_MS.Trt_F |
| CTC1 | 0.06876 | 0.35273 | 0.491 | 0.43 | 1 | HC_vs_MS.Trt_F |
| MYH10 | 0.06897 | 0.75924 | 0.147 | 0.1 | 1 | HC_vs_MS.Trt_F |
| NANOS1 | 0.069 | -0.2819 | 0.535 | 0.57 | 1 | HC_vs_MS.Trt_F |
| ARHGAP11A | 0.06906 | 0.47351 | 0.422 | 0.35 | 1 | HC_vs_MS.Trt_F |
| SUN3 | 0.06907 | 0.36674 | 0.413 | 0.34 | 1 | HC_vs_MS.Trt_F |
| C2CD5 | 0.06918 | -0.5128 | 0.211 | 0.15 | 1 | HC_vs_MS.Trt_F |
| SIRPB2 | 0.06927 | 0.43712 | 0.416 | 0.33 | 1 | HC_vs_MS.Trt_F |
| SPG20 | 0.06931 | 0.42102 | 0.165 | 0.11 | 1 | HC_vs_MS.Trt_F |
| SPHK2 | 0.06935 | 1.19198 | 0.393 | 0.32 | 1 | HC_vs_MS.Trt_F |
| DNAJB9 | 0.06942 | 0.79086 | 0.171 | 0.12 | 1 | HC_vs_MS.Trt_F |

| | | | | | | |
|---|---|---|---|---|---|---|
| ADRA1A | 0.06973 | 0.2835 | 0.618 | 0.58 | 1 | HC_vs_MS.Trt_F |
| TNS3 | 0.06978 | 0.65033 | 0.419 | 0.33 | 1 | HC_vs_MS.Trt_F |
| NOL11 | 0.06981 | -0.7517 | 0.578 | 0.48 | 1 | HC_vs_MS.Trt_F |
| THOC2 | 0.06984 | 0.26461 | 0.601 | 0.53 | 1 | HC_vs_MS.Trt_F |
| ADARB2 | 0.06992 | -0.6877 | 0.402 | 0.33 | 1 | HC_vs_MS.Trt_F |
| GANAB | 0.07 | 1.02252 | 0.503 | 0.45 | 1 | HC_vs_MS.Trt_F |
| AKT3 | 0.07014 | 0.31309 | 0.798 | 0.71 | 1 | HC_vs_MS.Trt_F |
| DCC | 0.07015 | 0.45464 | 0.116 | 0.17 | 1 | HC_vs_MS.Trt_F |
| MAST3 | 0.0702 | -0.4369 | 0.668 | 0.75 | 1 | HC_vs_MS.Trt_F |
| USP32 | 0.07036 | 0.66393 | 0.332 | 0.28 | 1 | HC_vs_MS.Trt_F |
| ALMS1 | 0.07042 | 1.58684 | 0.517 | 0.45 | 1 | HC_vs_MS.Trt_F |
| FRMD3 | 0.07043 | 0.36254 | 0.124 | 0.08 | 1 | HC_vs_MS.Trt_F |
| RP11-343C2 | 0.07048 | 0.56749 | 0.168 | 0.12 | 1 | HC_vs_MS.Trt_F |
| ABI3BP | 0.07052 | 0.26482 | 0.5 | 0.41 | 1 | HC_vs_MS.Trt_F |
| PPAT | 0.07062 | -0.7289 | 0.344 | 0.28 | 1 | HC_vs_MS.Trt_F |
| ACTR3B | 0.07095 | 0.49241 | 0.592 | 0.5 | 1 | HC_vs_MS.Trt_F |
| TTC14 | 0.07101 | 0.42349 | 0.442 | 0.35 | 1 | HC_vs_MS.Trt_F |
| ARHGAP17 | 0.0715 | 0.48325 | 0.63 | 0.55 | 1 | HC_vs_MS.Trt_F |
| C18orf21 | 0.07151 | 0.71972 | 0.118 | 0.08 | 1 | HC_vs_MS.Trt_F |
| COQ5 | 0.07156 | 0.29085 | 0.876 | 0.82 | 1 | HC_vs_MS.Trt_F |
| BNIP3L | 0.07162 | -0.3167 | 0.263 | 0.19 | 1 | HC_vs_MS.Trt_F |
| LAIR2 | 0.0719 | 0.34322 | 0.179 | 0.13 | 1 | HC_vs_MS.Trt_F |
| ARL17B | 0.07204 | -0.561 | 0.78 | 0.71 | 1 | HC_vs_MS.Trt_F |
| MIB1 | 0.07213 | 1.39005 | 0.171 | 0.12 | 1 | HC_vs_MS.Trt_F |
| GNAS | 0.07216 | -0.5177 | 0.497 | 0.51 | 1 | HC_vs_MS.Trt_F |
| CEP164 | 0.07221 | 0.34768 | 0.208 | 0.15 | 1 | HC_vs_MS.Trt_F |
| AASDH | 0.07226 | 0.40174 | 0.142 | 0.09 | 1 | HC_vs_MS.Trt_F |
| TNFRSF21 | 0.07228 | 0.98575 | 0.185 | 0.14 | 1 | HC_vs_MS.Trt_F |
| BAP1 | 0.0723 | -0.41 | 0.382 | 0.3 | 1 | HC_vs_MS.Trt_F |
| LZIC | 0.07244 | 0.40325 | 0.873 | 0.82 | 1 | HC_vs_MS.Trt_F |
| NCR1 | 0.07246 | 0.52106 | 0.266 | 0.21 | 1 | HC_vs_MS.Trt_F |
| NISCH | 0.07247 | 0.51321 | 0.454 | 0.4 | 1 | HC_vs_MS.Trt_F |
| KLHL3 | 0.07252 | 0.45575 | 0.277 | 0.21 | 1 | HC_vs_MS.Trt_F |
| GPR85 | 0.07264 | 0.28983 | 0.41 | 0.35 | 1 | HC_vs_MS.Trt_F |
| CDC42BPG | 0.07267 | 0.55495 | 0.616 | 0.54 | 1 | HC_vs_MS.Trt_F |
| ZNF782 | 0.07269 | 1.1148 | 0.147 | 0.1 | 1 | HC_vs_MS.Trt_F |
| GEMIN6 | 0.07272 | -0.2853 | 0.968 | 0.92 | 1 | HC_vs_MS.Trt_F |
| DET1 | 0.07275 | -0.5352 | 0.601 | 0.54 | 1 | HC_vs_MS.Trt_F |
| LRRC8C | 0.07278 | 0.51809 | 0.335 | 0.28 | 1 | HC_vs_MS.Trt_F |
| TTC28 | 0.07298 | -0.8112 | 0.46 | 0.38 | 1 | HC_vs_MS.Trt_F |

| | | | | | | |
|---|---|---|---|---|---|---|
| PRKDC | 0.0732 | 0.33979 | 0.63 | 0.58 | 1 | HC_vs_MS.Trt_F |
| NFAT5 | 0.07329 | 0.79651 | 0.893 | 0.84 | 1 | HC_vs_MS.Trt_F |
| NHLRC3 | 0.07358 | -0.3219 | 0.422 | 0.35 | 1 | HC_vs_MS.Trt_F |
| TTF1 | 0.0736 | 0.58358 | 0.512 | 0.45 | 1 | HC_vs_MS.Trt_F |
| ETHE1 | 0.07362 | 1.0021 | 0.136 | 0.09 | 1 | HC_vs_MS.Trt_F |
| SLAMF7 | 0.07363 | -0.8253 | 0.477 | 0.38 | 1 | HC_vs_MS.Trt_F |
| TXNDC11 | 0.07389 | 0.4168 | 0.39 | 0.32 | 1 | HC_vs_MS.Trt_F |
| FBXW5 | 0.07393 | 0.43623 | 0.257 | 0.19 | 1 | HC_vs_MS.Trt_F |
| EXOC4 | 0.07403 | 0.46403 | 0.575 | 0.49 | 1 | HC_vs_MS.Trt_F |
| LDLR | 0.07407 | 0.26194 | 0.98 | 0.92 | 1 | HC_vs_MS.Trt_F |
| RNF19B | 0.07416 | -0.5653 | 0.069 | 0.11 | 1 | HC_vs_MS.Trt_F |
| NBPF9 | 0.07419 | 0.573 | 0.702 | 0.65 | 1 | HC_vs_MS.Trt_F |
| BORCS6 | 0.07439 | 0.50907 | 0.133 | 0.09 | 1 | HC_vs_MS.Trt_F |
| FAM160B2 | 0.07448 | 0.60554 | 0.76 | 0.66 | 1 | HC_vs_MS.Trt_F |
| MGMT | 0.07453 | 0.59413 | 0.165 | 0.12 | 1 | HC_vs_MS.Trt_F |
| TMEM116 | 0.0746 | 0.29413 | 0.928 | 0.87 | 1 | HC_vs_MS.Trt_F |
| TSC2 | 0.07476 | 0.34623 | 0.731 | 0.66 | 1 | HC_vs_MS.Trt_F |
| ST8SIA6 | 0.07482 | -0.6997 | 0.46 | 0.37 | 1 | HC_vs_MS.Trt_F |
| ARNTL | 0.07483 | 0.54362 | 0.251 | 0.19 | 1 | HC_vs_MS.Trt_F |
| PIGM | 0.07485 | 0.27423 | 0.679 | 0.62 | 1 | HC_vs_MS.Trt_F |
| AFF4 | 0.07495 | 0.79025 | 0.532 | 0.48 | 1 | HC_vs_MS.Trt_F |
| PRPF38A | 0.07503 | -0.7136 | 0.673 | 0.59 | 1 | HC_vs_MS.Trt_F |
| FAM53B | 0.07525 | 0.61041 | 0.26 | 0.2 | 1 | HC_vs_MS.Trt_F |
| PQLC3 | 0.0753 | 0.449 | 0.191 | 0.13 | 1 | HC_vs_MS.Trt_F |
| PCDHGA6 | 0.0753 | 0.48727 | 0.85 | 0.81 | 1 | HC_vs_MS.Trt_F |
| RB1 | 0.0754 | 0.86532 | 0.179 | 0.13 | 1 | HC_vs_MS.Trt_F |
| SLC25A32 | 0.07545 | -0.5399 | 0.425 | 0.34 | 1 | HC_vs_MS.Trt_F |
| CYP4A11 | 0.07549 | 0.30337 | 0.13 | 0.09 | 1 | HC_vs_MS.Trt_F |
| VPS52 | 0.07563 | 0.29269 | 0.22 | 0.16 | 1 | HC_vs_MS.Trt_F |
| NBPF1 | 0.07579 | 0.30761 | 0.283 | 0.22 | 1 | HC_vs_MS.Trt_F |
| NOP16 | 0.07599 | 0.29194 | 0.228 | 0.17 | 1 | HC_vs_MS.Trt_F |
| IMPA1 | 0.07607 | -0.8289 | 0.422 | 0.34 | 1 | HC_vs_MS.Trt_F |
| RHOG | 0.0761 | 0.3911 | 0.428 | 0.36 | 1 | HC_vs_MS.Trt_F |
| RPS27 | 0.07621 | 0.27753 | 0.301 | 0.25 | 1 | HC_vs_MS.Trt_F |
| PRPF39 | 0.0764 | -0.3467 | 0.315 | 0.38 | 1 | HC_vs_MS.Trt_F |
| PPM1M | 0.07644 | 0.3333 | 0.197 | 0.14 | 1 | HC_vs_MS.Trt_F |
| GLTSCR1L | 0.07645 | 0.57613 | 0.231 | 0.17 | 1 | HC_vs_MS.Trt_F |
| ANP32E | 0.07656 | -1.1298 | 0.462 | 0.48 | 1 | HC_vs_MS.Trt_F |
| DPH1 | 0.07664 | 0.3131 | 0.91 | 0.86 | 1 | HC_vs_MS.Trt_F |
| KLRC1 | 0.07665 | -0.2868 | 0.338 | 0.27 | 1 | HC_vs_MS.Trt_F |

| | | | | | | |
|---|---|---|---|---|---|---|
| AKAP12 | 0.07673 | 0.68598 | 0.09 | 0.13 | 1 | HC_vs_MS.Trt_F |
| ADA | 0.07673 | -0.5735 | 0.107 | 0.15 | 1 | HC_vs_MS.Trt_F |
| NRDC | 0.07683 | 0.27901 | 0.419 | 0.34 | 1 | HC_vs_MS.Trt_F |
| ALAS1 | 0.07698 | 0.63214 | 0.159 | 0.11 | 1 | HC_vs_MS.Trt_F |
| ABCB6 | 0.07702 | 0.26188 | 0.194 | 0.14 | 1 | HC_vs_MS.Trt_F |
| FKBP4 | 0.07732 | 0.35455 | 0.159 | 0.11 | 1 | HC_vs_MS.Trt_F |
| NFIB | 0.07759 | -0.4493 | 0.191 | 0.14 | 1 | HC_vs_MS.Trt_F |
| CUL7 | 0.07806 | 0.88369 | 0.121 | 0.08 | 1 | HC_vs_MS.Trt_F |
| MAK | 0.0781 | 0.27223 | 0.249 | 0.19 | 1 | HC_vs_MS.Trt_F |
| WDR43 | 0.07811 | -0.2659 | 0.318 | 0.38 | 1 | HC_vs_MS.Trt_F |
| CEP162 | 0.07811 | -0.5206 | 0.217 | 0.16 | 1 | HC_vs_MS.Trt_F |
| PORCN | 0.07816 | 1.18292 | 0.121 | 0.08 | 1 | HC_vs_MS.Trt_F |
| TMEM135 | 0.07832 | 0.27892 | 0.803 | 0.76 | 1 | HC_vs_MS.Trt_F |
| CHSY1 | 0.07833 | -0.3384 | 0.324 | 0.26 | 1 | HC_vs_MS.Trt_F |
| NAA30 | 0.07837 | 0.42827 | 0.54 | 0.46 | 1 | HC_vs_MS.Trt_F |
| SVBP | 0.07843 | -0.8273 | 0.121 | 0.17 | 1 | HC_vs_MS.Trt_F |
| ACACB | 0.07845 | 0.29583 | 0.96 | 0.92 | 1 | HC_vs_MS.Trt_F |
| KLHL7 | 0.07879 | -0.3569 | 0.798 | 0.69 | 1 | HC_vs_MS.Trt_F |
| API5 | 0.07884 | -0.7962 | 0.327 | 0.23 | 1 | HC_vs_MS.Trt_F |
| SRPRB | 0.07901 | 0.3048 | 0.208 | 0.15 | 1 | HC_vs_MS.Trt_F |
| ADAL | 0.07947 | 0.73763 | 0.506 | 0.42 | 1 | HC_vs_MS.Trt_F |
| CXorf36 | 0.0796 | 0.5439 | 0.474 | 0.43 | 1 | HC_vs_MS.Trt_F |
| SOCS4 | 0.07979 | 1.14881 | 0.379 | 0.3 | 1 | HC_vs_MS.Trt_F |
| TMEM65 | 0.07982 | 0.48273 | 0.777 | 0.71 | 1 | HC_vs_MS.Trt_F |
| LTN1 | 0.07984 | 0.73266 | 0.197 | 0.14 | 1 | HC_vs_MS.Trt_F |
| UBR3 | 0.07991 | -0.4671 | 0.171 | 0.12 | 1 | HC_vs_MS.Trt_F |
| SH3PXD2A | 0.07994 | 0.39369 | 0.379 | 0.31 | 1 | HC_vs_MS.Trt_F |
| MMS19 | 0.07998 | 0.34976 | 0.225 | 0.17 | 1 | HC_vs_MS.Trt_F |
| HEMGN | 0.08019 | 0.49241 | 0.15 | 0.1 | 1 | HC_vs_MS.Trt_F |
| AIM2 | 0.08022 | 1.3939 | 0.341 | 0.28 | 1 | HC_vs_MS.Trt_F |
| SLC25A13 | 0.08033 | 0.57598 | 0.165 | 0.12 | 1 | HC_vs_MS.Trt_F |
| PAN3 | 0.08044 | -0.624 | 0.824 | 0.86 | 1 | HC_vs_MS.Trt_F |
| PKN2 | 0.08061 | 0.37923 | 0.379 | 0.31 | 1 | HC_vs_MS.Trt_F |
| PLAGL2 | 0.08092 | 1.16656 | 0.107 | 0.07 | 1 | HC_vs_MS.Trt_F |
| FAM206A | 0.081 | 1.69391 | 0.124 | 0.08 | 1 | HC_vs_MS.Trt_F |
| BCL9 | 0.08106 | 0.26218 | 0.136 | 0.09 | 1 | HC_vs_MS.Trt_F |
| MOG | 0.08115 | 0.41396 | 0.396 | 0.32 | 1 | HC_vs_MS.Trt_F |
| FBXL12 | 0.08125 | -0.3181 | 0.723 | 0.64 | 1 | HC_vs_MS.Trt_F |
| 9-Mar | 0.08128 | 0.94245 | 0.662 | 0.59 | 1 | HC_vs_MS.Trt_F |
| ERI2 | 0.08164 | 1.11866 | 0.197 | 0.15 | 1 | HC_vs_MS.Trt_F |

| | | | | | | |
|---|---|---|---|---|---|---|
| ANAPC2 | 0.08172 | 1.65081 | 0.124 | 0.08 | 1 | HC_vs_MS.Trt_F |
| BHLHB9 | 0.08183 | -0.7745 | 0.436 | 0.35 | 1 | HC_vs_MS.Trt_F |
| SPG11 | 0.08186 | 0.26082 | 0.853 | 0.82 | 1 | HC_vs_MS.Trt_F |
| JAML | 0.08211 | 0.43911 | 0.306 | 0.24 | 1 | HC_vs_MS.Trt_F |
| PEA15 | 0.08224 | -0.2912 | 0.194 | 0.13 | 1 | HC_vs_MS.Trt_F |
| TNRC6A | 0.08231 | 0.3964 | 0.887 | 0.84 | 1 | HC_vs_MS.Trt_F |
| ITGA6 | 0.08236 | 0.26028 | 0.199 | 0.14 | 1 | HC_vs_MS.Trt_F |
| MESDC2 | 0.0824 | -0.7998 | 0.613 | 0.5 | 1 | HC_vs_MS.Trt_F |
| ICMT | 0.08245 | 0.43759 | 0.74 | 0.65 | 1 | HC_vs_MS.Trt_F |
| CRYGS | 0.08276 | 1.0548 | 0.286 | 0.22 | 1 | HC_vs_MS.Trt_F |
| RAPGEF2 | 0.08294 | 0.63008 | 0.445 | 0.37 | 1 | HC_vs_MS.Trt_F |
| RRP36 | 0.08312 | 0.33451 | 0.514 | 0.44 | 1 | HC_vs_MS.Trt_F |
| ZNF606 | 0.08327 | 0.2757 | 0.815 | 0.81 | 1 | HC_vs_MS.Trt_F |
| SLFN13 | 0.0833 | -0.6025 | 0.731 | 0.63 | 1 | HC_vs_MS.Trt_F |
| FAM160B1 | 0.08332 | 0.75256 | 0.205 | 0.15 | 1 | HC_vs_MS.Trt_F |
| ZNF800 | 0.08345 | 0.73696 | 0.251 | 0.19 | 1 | HC_vs_MS.Trt_F |
| GZMM | 0.08351 | 0.27452 | 0.199 | 0.14 | 1 | HC_vs_MS.Trt_F |
| CSAD | 0.08353 | 0.46932 | 0.858 | 0.79 | 1 | HC_vs_MS.Trt_F |
| IL4R | 0.08361 | 1.11577 | 0.506 | 0.48 | 1 | HC_vs_MS.Trt_F |
| HOOK1 | 0.08379 | -0.5891 | 0.078 | 0.12 | 1 | HC_vs_MS.Trt_F |
| MFSD4B | 0.08392 | 0.35293 | 0.766 | 0.71 | 1 | HC_vs_MS.Trt_F |
| CLASRP | 0.08393 | 0.8257 | 0.202 | 0.15 | 1 | HC_vs_MS.Trt_F |
| NRROS | 0.08394 | 0.95165 | 0.173 | 0.13 | 1 | HC_vs_MS.Trt_F |
| SRSF11 | 0.08396 | -0.3583 | 0.803 | 0.72 | 1 | HC_vs_MS.Trt_F |
| TBCB | 0.08423 | -0.2754 | 0.714 | 0.74 | 1 | HC_vs_MS.Trt_F |
| AKAP1 | 0.08433 | 0.46642 | 0.15 | 0.1 | 1 | HC_vs_MS.Trt_F |
| PCED1B | 0.08452 | 0.4249 | 0.509 | 0.43 | 1 | HC_vs_MS.Trt_F |
| EIF3B | 0.0846 | -0.2753 | 0.422 | 0.35 | 1 | HC_vs_MS.Trt_F |
| SLC39A11 | 0.08475 | -0.6953 | 0.272 | 0.32 | 1 | HC_vs_MS.Trt_F |
| SKIV2L2 | 0.08496 | 0.30216 | 0.887 | 0.82 | 1 | HC_vs_MS.Trt_F |
| RAC1 | 0.08509 | -0.376 | 0.182 | 0.23 | 1 | HC_vs_MS.Trt_F |
| YPEL1 | 0.08514 | -1.2863 | 0.529 | 0.45 | 1 | HC_vs_MS.Trt_F |
| SLC2A8 | 0.08515 | 0.25065 | 0.327 | 0.27 | 1 | HC_vs_MS.Trt_F |
| LARS | 0.08527 | -0.6033 | 0.737 | 0.69 | 1 | HC_vs_MS.Trt_F |
| RIOK1 | 0.08528 | 0.29707 | 0.136 | 0.09 | 1 | HC_vs_MS.Trt_F |
| TRAPPC11 | 0.08529 | 0.75228 | 0.474 | 0.41 | 1 | HC_vs_MS.Trt_F |
| KLHL5 | 0.0855 | -0.3658 | 0.512 | 0.43 | 1 | HC_vs_MS.Trt_F |
| MIGA2 | 0.08552 | 0.9208 | 0.514 | 0.42 | 1 | HC_vs_MS.Trt_F |
| MAL | 0.08563 | 0.38241 | 0.237 | 0.18 | 1 | HC_vs_MS.Trt_F |
| SELENBP1 | 0.0858 | -1.0969 | 0.335 | 0.27 | 1 | HC_vs_MS.Trt_F |

| | | | | | | |
|---|---|---|---|---|---|---|
| GVQW1 | 0.08596 | -0.42 | 0.48 | 0.53 | 1 | HC_vs_MS.Trt_F |
| LARP4 | 0.08597 | 0.59818 | 0.408 | 0.35 | 1 | HC_vs_MS.Trt_F |
| UXT | 0.08632 | 0.25679 | 0.246 | 0.19 | 1 | HC_vs_MS.Trt_F |
| LCP1 | 0.08647 | -0.3732 | 0.621 | 0.63 | 1 | HC_vs_MS.Trt_F |
| ARL1 | 0.08651 | -0.8357 | 0.419 | 0.33 | 1 | HC_vs_MS.Trt_F |
| MRPL28 | 0.08657 | 0.76095 | 0.15 | 0.1 | 1 | HC_vs_MS.Trt_F |
| ITM2B | 0.0866 | -0.3879 | 0.584 | 0.6 | 1 | HC_vs_MS.Trt_F |
| ING1 | 0.08686 | -0.5417 | 0.656 | 0.6 | 1 | HC_vs_MS.Trt_F |
| LY6G5C | 0.08693 | 0.29445 | 0.613 | 0.55 | 1 | HC_vs_MS.Trt_F |
| ZNF583 | 0.08694 | -0.2776 | 0.882 | 0.78 | 1 | HC_vs_MS.Trt_F |
| TMEM261 | 0.08694 | -0.79 | 0.165 | 0.12 | 1 | HC_vs_MS.Trt_F |
| GOLPH3L | 0.08733 | 1.03223 | 0.263 | 0.2 | 1 | HC_vs_MS.Trt_F |
| AGPAT4 | 0.08751 | 0.48168 | 0.142 | 0.1 | 1 | HC_vs_MS.Trt_F |
| IRGQ | 0.08754 | 0.73666 | 0.991 | 0.98 | 1 | HC_vs_MS.Trt_F |
| ANKRD37 | 0.0876 | 1.03197 | 0.15 | 0.1 | 1 | HC_vs_MS.Trt_F |
| PGM5 | 0.08764 | 0.30153 | 0.104 | 0.07 | 1 | HC_vs_MS.Trt_F |
| ST6GAL1 | 0.08771 | 0.29125 | 0.434 | 0.37 | 1 | HC_vs_MS.Trt_F |
| VPS28 | 0.08772 | 0.31991 | 0.592 | 0.51 | 1 | HC_vs_MS.Trt_F |
| SUMF2 | 0.08787 | -0.3144 | 0.902 | 0.87 | 1 | HC_vs_MS.Trt_F |
| NUMBL | 0.08803 | 0.5502 | 0.627 | 0.55 | 1 | HC_vs_MS.Trt_F |
| PICK1 | 0.08823 | 1.12564 | 0.52 | 0.46 | 1 | HC_vs_MS.Trt_F |
| TSNARE1 | 0.08824 | 1.09535 | 0.133 | 0.09 | 1 | HC_vs_MS.Trt_F |
| MAT2A | 0.08828 | 0.38553 | 0.208 | 0.15 | 1 | HC_vs_MS.Trt_F |
| SEMA4F | 0.0883 | 0.52567 | 0.592 | 0.52 | 1 | HC_vs_MS.Trt_F |
| PSTK | 0.0884 | -0.3397 | 0.416 | 0.35 | 1 | HC_vs_MS.Trt_F |
| ZNF197 | 0.0884 | -0.3882 | 0.538 | 0.47 | 1 | HC_vs_MS.Trt_F |
| RNF165 | 0.08852 | 0.55988 | 0.54 | 0.46 | 1 | HC_vs_MS.Trt_F |
| KIAA0040 | 0.08854 | 0.54007 | 0.211 | 0.16 | 1 | HC_vs_MS.Trt_F |
| ZNF32 | 0.08855 | 1.59352 | 0.107 | 0.07 | 1 | HC_vs_MS.Trt_F |
| ANKEF1 | 0.08861 | -1.1013 | 0.269 | 0.2 | 1 | HC_vs_MS.Trt_F |
| MCM3AP | 0.08862 | 0.41532 | 0.546 | 0.45 | 1 | HC_vs_MS.Trt_F |
| COQ4 | 0.08887 | 0.39044 | 0.922 | 0.87 | 1 | HC_vs_MS.Trt_F |
| FARS2 | 0.08898 | 1.3701 | 0.127 | 0.09 | 1 | HC_vs_MS.Trt_F |
| 2-Mar | 0.08905 | -0.3399 | 0.636 | 0.56 | 1 | HC_vs_MS.Trt_F |
| TEFM | 0.08905 | 0.29927 | 0.763 | 0.7 | 1 | HC_vs_MS.Trt_F |
| MAP3K2 | 0.08927 | 0.5621 | 0.301 | 0.24 | 1 | HC_vs_MS.Trt_F |
| ANKUB1 | 0.08941 | 0.28703 | 0.491 | 0.41 | 1 | HC_vs_MS.Trt_F |
| ALDH8A1 | 0.08973 | 0.42743 | 0.59 | 0.53 | 1 | HC_vs_MS.Trt_F |
| CHCHD5 | 0.08975 | 0.79813 | 0.532 | 0.45 | 1 | HC_vs_MS.Trt_F |
| DFFB | 0.0898 | -0.4311 | 0.882 | 0.83 | 1 | HC_vs_MS.Trt_F |

| | | | | | | |
|---|---|---|---|---|---|---|
| TXLNG | 0.08985 | 0.3483 | 0.139 | 0.09 | 1 | HC_vs_MS.Trt_F |
| TRIM45 | 0.08992 | 0.92203 | 0.156 | 0.11 | 1 | HC_vs_MS.Trt_F |
| MMAB | 0.08993 | 0.33807 | 0.717 | 0.64 | 1 | HC_vs_MS.Trt_F |
| RNF24 | 0.08994 | 0.6328 | 0.581 | 0.53 | 1 | HC_vs_MS.Trt_F |
| ZADH2 | 0.09026 | -1.0501 | 0.861 | 0.82 | 1 | HC_vs_MS.Trt_F |
| CAPNS1 | 0.09036 | -0.2724 | 0.37 | 0.29 | 1 | HC_vs_MS.Trt_F |
| ANO9 | 0.09037 | 0.38585 | 0.289 | 0.23 | 1 | HC_vs_MS.Trt_F |
| TRIB2 | 0.09079 | -0.8881 | 0.367 | 0.39 | 1 | HC_vs_MS.Trt_F |
| ZNF28 | 0.09081 | -0.8195 | 0.829 | 0.78 | 1 | HC_vs_MS.Trt_F |
| IFI6 | 0.09099 | -0.4817 | 0.341 | 0.37 | 1 | HC_vs_MS.Trt_F |
| FANCG | 0.09099 | -0.8413 | 0.156 | 0.1 | 1 | HC_vs_MS.Trt_F |
| KDM5B | 0.09104 | 0.25436 | 0.327 | 0.37 | 1 | HC_vs_MS.Trt_F |
| TSPOAP1 | 0.09145 | 0.7149 | 0.321 | 0.26 | 1 | HC_vs_MS.Trt_F |
| AMT | 0.09149 | -0.7182 | 0.497 | 0.42 | 1 | HC_vs_MS.Trt_F |
| IPO9 | 0.09156 | -0.977 | 0.876 | 0.8 | 1 | HC_vs_MS.Trt_F |
| NFRKB | 0.09162 | 0.29811 | 0.185 | 0.13 | 1 | HC_vs_MS.Trt_F |
| WFDC11 | 0.09191 | 0.41414 | 0.289 | 0.23 | 1 | HC_vs_MS.Trt_F |
| WDR90 | 0.09204 | -0.4528 | 0.916 | 0.88 | 1 | HC_vs_MS.Trt_F |
| KAT14 | 0.0921 | -1.1246 | 0.072 | 0.11 | 1 | HC_vs_MS.Trt_F |
| LRRC14 | 0.09223 | 0.65681 | 0.121 | 0.08 | 1 | HC_vs_MS.Trt_F |
| CCT6B | 0.09253 | 0.65419 | 0.312 | 0.25 | 1 | HC_vs_MS.Trt_F |
| TMEM129 | 0.09269 | 0.44104 | 0.107 | 0.07 | 1 | HC_vs_MS.Trt_F |
| NIT2 | 0.09274 | 0.38974 | 0.676 | 0.58 | 1 | HC_vs_MS.Trt_F |
| CMTR1 | 0.09276 | 0.98664 | 0.208 | 0.16 | 1 | HC_vs_MS.Trt_F |
| MTO1 | 0.09303 | -0.2697 | 0.988 | 0.98 | 1 | HC_vs_MS.Trt_F |
| WBSCR22 | 0.09313 | 0.25873 | 0.789 | 0.75 | 1 | HC_vs_MS.Trt_F |
| MOB2 | 0.09335 | -0.5953 | 0.101 | 0.14 | 1 | HC_vs_MS.Trt_F |
| CSRNP2 | 0.09355 | -0.2999 | 0.139 | 0.09 | 1 | HC_vs_MS.Trt_F |
| NUP205 | 0.09367 | 0.86986 | 0.202 | 0.15 | 1 | HC_vs_MS.Trt_F |
| LIN54 | 0.09369 | -0.3917 | 0.653 | 0.57 | 1 | HC_vs_MS.Trt_F |
| TADA2A | 0.09374 | -0.3022 | 0.919 | 0.91 | 1 | HC_vs_MS.Trt_F |
| RAB26 | 0.09379 | 0.5126 | 0.179 | 0.13 | 1 | HC_vs_MS.Trt_F |
| SPINK9 | 0.09389 | 0.5936 | 0.688 | 0.59 | 1 | HC_vs_MS.Trt_F |
| TMX2 | 0.09423 | 0.28651 | 0.165 | 0.12 | 1 | HC_vs_MS.Trt_F |
| IQCK | 0.09546 | -0.3929 | 0.231 | 0.18 | 1 | HC_vs_MS.Trt_F |
| GGT1 | 0.09555 | 1.27037 | 0.156 | 0.11 | 1 | HC_vs_MS.Trt_F |
| RP11-111M1 | 0.09563 | 0.26353 | 0.908 | 0.86 | 1 | HC_vs_MS.Trt_F |
| ZNF7 | 0.09573 | 0.47207 | 0.934 | 0.95 | 1 | HC_vs_MS.Trt_F |
| RCOR1 | 0.0959 | 0.55957 | 0.225 | 0.17 | 1 | HC_vs_MS.Trt_F |
| MAP2K3 | 0.09593 | 0.55873 | 0.194 | 0.14 | 1 | HC_vs_MS.Trt_F |

| | | | | | | |
|---|---|---|---|---|---|---|
| LAMP1 | 0.09597 | 1.22074 | 0.41 | 0.35 | 1 | HC_vs_MS.Trt_F |
| AGTRAP | 0.09618 | 0.31396 | 0.179 | 0.13 | 1 | HC_vs_MS.Trt_F |
| EFL1 | 0.09622 | -0.9544 | 0.11 | 0.15 | 1 | HC_vs_MS.Trt_F |
| ANKRD10 | 0.09627 | -0.407 | 0.717 | 0.73 | 1 | HC_vs_MS.Trt_F |
| FCRL1 | 0.09634 | 0.73954 | 0.318 | 0.26 | 1 | HC_vs_MS.Trt_F |
| TTC9C | 0.09641 | 0.25825 | 0.387 | 0.31 | 1 | HC_vs_MS.Trt_F |
| ZNF43 | 0.09655 | -0.7394 | 0.772 | 0.77 | 1 | HC_vs_MS.Trt_F |
| AQP3 | 0.09685 | 0.27351 | 0.332 | 0.27 | 1 | HC_vs_MS.Trt_F |
| LSM6 | 0.09719 | 0.399 | 0.162 | 0.12 | 1 | HC_vs_MS.Trt_F |
| MRPS11 | 0.09743 | 0.55221 | 0.751 | 0.7 | 1 | HC_vs_MS.Trt_F |
| EPT1 | 0.09747 | 0.56864 | 0.431 | 0.36 | 1 | HC_vs_MS.Trt_F |
| GRAMD1A | 0.0975 | -0.4063 | 0.699 | 0.65 | 1 | HC_vs_MS.Trt_F |
| KCNRG | 0.09755 | 0.45455 | 0.425 | 0.35 | 1 | HC_vs_MS.Trt_F |
| RBFA | 0.09757 | 0.54418 | 0.364 | 0.3 | 1 | HC_vs_MS.Trt_F |
| NUDT22 | 0.0976 | -1.2245 | 0.676 | 0.72 | 1 | HC_vs_MS.Trt_F |
| MREG | 0.09761 | 0.49181 | 0.668 | 0.6 | 1 | HC_vs_MS.Trt_F |
| CD226 | 0.09771 | 0.67313 | 0.595 | 0.52 | 1 | HC_vs_MS.Trt_F |
| ZNF609 | 0.09773 | 0.43014 | 0.621 | 0.57 | 1 | HC_vs_MS.Trt_F |
| SGK3 | 0.09791 | 0.44911 | 0.951 | 0.87 | 1 | HC_vs_MS.Trt_F |
| GTPBP2 | 0.09808 | -0.5094 | 0.572 | 0.63 | 1 | HC_vs_MS.Trt_F |
| PCGF3 | 0.09824 | 0.27647 | 0.228 | 0.17 | 1 | HC_vs_MS.Trt_F |
| SACS | 0.09845 | 0.66826 | 0.153 | 0.11 | 1 | HC_vs_MS.Trt_F |
| NONO | 0.09847 | 0.28698 | 0.705 | 0.64 | 1 | HC_vs_MS.Trt_F |
| MED25 | 0.09882 | 0.46385 | 0.52 | 0.47 | 1 | HC_vs_MS.Trt_F |
| TBC1D19 | 0.09885 | 0.37965 | 0.153 | 0.11 | 1 | HC_vs_MS.Trt_F |
| KDELC2 | 0.09892 | -0.4761 | 0.65 | 0.59 | 1 | HC_vs_MS.Trt_F |
| MAN2A2 | 0.099 | 0.27137 | 0.364 | 0.31 | 1 | HC_vs_MS.Trt_F |
| SNED1 | 0.09906 | 0.56016 | 0.197 | 0.14 | 1 | HC_vs_MS.Trt_F |
| PRKCZ | 0.09909 | 0.74539 | 0.301 | 0.24 | 1 | HC_vs_MS.Trt_F |
| SORBS2 | 0.09911 | 0.29738 | 0.665 | 0.6 | 1 | HC_vs_MS.Trt_F |
| EHMT2 | 0.09923 | 0.45047 | 0.699 | 0.6 | 1 | HC_vs_MS.Trt_F |
| PGGHG | 0.09953 | 0.86391 | 0.176 | 0.13 | 1 | HC_vs_MS.Trt_F |
| PSD3 | 0.0996 | 0.5139 | 0.373 | 0.31 | 1 | HC_vs_MS.Trt_F |
| SRBD1 | 0.09964 | -1.8069 | 0.46 | 0.36 | 1 | HC_vs_MS.Trt_F |
| SF3B5 | 0.0997 | 0.80942 | 0.142 | 0.1 | 1 | HC_vs_MS.Trt_F |
| FGD4 | 0.09971 | 0.84166 | 0.861 | 0.76 | 1 | HC_vs_MS.Trt_F |
| RAD51 | 0.09976 | -0.5076 | 0.512 | 0.46 | 1 | HC_vs_MS.Trt_F |
| SLC25A36 | 0.09998 | 0.51226 | 0.176 | 0.13 | 1 | HC_vs_MS.Trt_F |
| PEX13 | 0.10005 | -1.0184 | 0.783 | 0.8 | 1 | HC_vs_MS.Trt_F |
| VPS8 | 0.10021 | -0.5586 | 0.691 | 0.6 | 1 | HC_vs_MS.Trt_F |

| | | | | | | |
|---|---|---|---|---|---|---|
| ZNF225 | 0.10029 | 0.27653 | 0.855 | 0.75 | 1 | HC_vs_MS.Trt_F |
| PEBP1 | 0.10034 | 0.27375 | 0.434 | 0.39 | 1 | HC_vs_MS.Trt_F |
| COPS7A | 0.10039 | 0.79122 | 0.286 | 0.23 | 1 | HC_vs_MS.Trt_F |
| CPNE1 | 0.1007 | -0.5824 | 0.523 | 0.52 | 1 | HC_vs_MS.Trt_F |
| SPTLC1 | 0.10089 | -0.2598 | 0.763 | 0.69 | 1 | HC_vs_MS.Trt_F |
| ARL5B | 0.10095 | 1.25745 | 0.153 | 0.11 | 1 | HC_vs_MS.Trt_F |
| AEBP2 | 0.10116 | -0.9181 | 0.754 | 0.69 | 1 | HC_vs_MS.Trt_F |
| FBXO11 | 0.10137 | 0.30463 | 0.526 | 0.45 | 1 | HC_vs_MS.Trt_F |
| IL17RA | 0.1015 | 0.3223 | 0.72 | 0.66 | 1 | HC_vs_MS.Trt_F |
| LRPAP1 | 0.10154 | -0.4557 | 0.887 | 0.84 | 1 | HC_vs_MS.Trt_F |
| PNPLA8 | 0.10156 | 0.46718 | 0.468 | 0.4 | 1 | HC_vs_MS.Trt_F |
| MCF2L | 0.1016 | 1.10453 | 0.37 | 0.3 | 1 | HC_vs_MS.Trt_F |
| ZNF84 | 0.10168 | -0.7633 | 0.312 | 0.25 | 1 | HC_vs_MS.Trt_F |
| GABBR1 | 0.10176 | 0.6812 | 0.208 | 0.16 | 1 | HC_vs_MS.Trt_F |
| TMEM167B | 0.10194 | 0.64411 | 0.792 | 0.77 | 1 | HC_vs_MS.Trt_F |
| PPP1R1C | 0.10214 | 0.34354 | 0.286 | 0.23 | 1 | HC_vs_MS.Trt_F |
| FANCI | 0.10218 | -0.3035 | 0.538 | 0.61 | 1 | HC_vs_MS.Trt_F |
| POLR2L | 0.10222 | 0.31411 | 0.321 | 0.26 | 1 | HC_vs_MS.Trt_F |
| CCDC94 | 0.10229 | 0.69393 | 0.194 | 0.14 | 1 | HC_vs_MS.Trt_F |
| SIKE1 | 0.10232 | -0.4246 | 0.523 | 0.45 | 1 | HC_vs_MS.Trt_F |
| GID8 | 0.10234 | 0.34128 | 0.191 | 0.14 | 1 | HC_vs_MS.Trt_F |
| FN1 | 0.10252 | -0.4803 | 0.5 | 0.4 | 1 | HC_vs_MS.Trt_F |
| TRAK1 | 0.10256 | 0.68043 | 0.136 | 0.09 | 1 | HC_vs_MS.Trt_F |
| TMSB15B | 0.1026 | 0.63365 | 0.361 | 0.3 | 1 | HC_vs_MS.Trt_F |
| SRSF8 | 0.10274 | -0.7997 | 0.671 | 0.69 | 1 | HC_vs_MS.Trt_F |
| ZNF528 | 0.10289 | 0.58421 | 0.65 | 0.58 | 1 | HC_vs_MS.Trt_F |
| PPRC1 | 0.10292 | 0.47363 | 0.165 | 0.12 | 1 | HC_vs_MS.Trt_F |
| OAZ1 | 0.10301 | -0.299 | 0.803 | 0.78 | 1 | HC_vs_MS.Trt_F |
| C12orf40 | 0.10314 | 0.63973 | 0.298 | 0.24 | 1 | HC_vs_MS.Trt_F |
| FAM117B | 0.10354 | 0.42253 | 0.474 | 0.41 | 1 | HC_vs_MS.Trt_F |
| BAG4 | 0.1037 | -0.4847 | 0.223 | 0.17 | 1 | HC_vs_MS.Trt_F |
| CCND2 | 0.10378 | -0.6634 | 0.465 | 0.52 | 1 | HC_vs_MS.Trt_F |
| FAM53C | 0.10394 | 0.64254 | 0.841 | 0.79 | 1 | HC_vs_MS.Trt_F |
| RNF170 | 0.10404 | 0.34778 | 0.642 | 0.55 | 1 | HC_vs_MS.Trt_F |
| CIR1 | 0.10404 | 0.31812 | 0.746 | 0.71 | 1 | HC_vs_MS.Trt_F |
| VPS37A | 0.10407 | 0.97184 | 0.191 | 0.14 | 1 | HC_vs_MS.Trt_F |
| SLC35B4 | 0.10465 | 0.61563 | 0.587 | 0.53 | 1 | HC_vs_MS.Trt_F |
| KIAA1468 | 0.10465 | 0.34468 | 0.361 | 0.3 | 1 | HC_vs_MS.Trt_F |
| KRTCAP2 | 0.10466 | -0.262 | 0.832 | 0.78 | 1 | HC_vs_MS.Trt_F |
| EEFSEC | 0.10497 | 1.13744 | 0.13 | 0.09 | 1 | HC_vs_MS.Trt_F |

| | | | | | | |
|---|---|---|---|---|---|---|
| EMCN | 0.10511 | 0.77811 | 0.509 | 0.45 | 1 | HC_vs_MS.Trt_F |
| P2RY8 | 0.10514 | 0.36496 | 0.434 | 0.37 | 1 | HC_vs_MS.Trt_F |
| EZH2 | 0.10516 | 0.36975 | 0.185 | 0.13 | 1 | HC_vs_MS.Trt_F |
| SNAP47 | 0.10519 | 0.36723 | 0.171 | 0.12 | 1 | HC_vs_MS.Trt_F |
| 8-Mar | 0.10521 | 0.55345 | 0.413 | 0.34 | 1 | HC_vs_MS.Trt_F |
| MYO1C | 0.10532 | -0.9374 | 0.269 | 0.2 | 1 | HC_vs_MS.Trt_F |
| HBG2 | 0.10538 | 0.31347 | 0.454 | 0.41 | 1 | HC_vs_MS.Trt_F |
| HEXA | 0.10544 | -0.4353 | 0.818 | 0.7 | 1 | HC_vs_MS.Trt_F |
| EMILIN2 | 0.10577 | -0.4958 | 0.48 | 0.42 | 1 | HC_vs_MS.Trt_F |
| PATL2 | 0.10671 | 0.38518 | 0.431 | 0.35 | 1 | HC_vs_MS.Trt_F |
| ABL2 | 0.10674 | 0.62105 | 0.803 | 0.73 | 1 | HC_vs_MS.Trt_F |
| NDUFB2 | 0.1068 | -0.4344 | 0.939 | 0.93 | 1 | HC_vs_MS.Trt_F |
| ORAI3 | 0.10696 | -1.0407 | 0.098 | 0.14 | 1 | HC_vs_MS.Trt_F |
| PEF1 | 0.10718 | 0.31768 | 0.211 | 0.16 | 1 | HC_vs_MS.Trt_F |
| FUK | 0.10721 | -0.551 | 0.616 | 0.53 | 1 | HC_vs_MS.Trt_F |
| ACSS3 | 0.1073 | 0.36232 | 0.298 | 0.25 | 1 | HC_vs_MS.Trt_F |
| DUS4L | 0.10755 | -0.6787 | 0.087 | 0.12 | 1 | HC_vs_MS.Trt_F |
| MCRS1 | 0.1079 | 0.31907 | 0.217 | 0.16 | 1 | HC_vs_MS.Trt_F |
| FAR2 | 0.10808 | 0.30403 | 0.237 | 0.19 | 1 | HC_vs_MS.Trt_F |
| RPTOR | 0.10816 | 0.29433 | 0.393 | 0.34 | 1 | HC_vs_MS.Trt_F |
| MROH1 | 0.10868 | 0.4937 | 0.604 | 0.55 | 1 | HC_vs_MS.Trt_F |
| MIEF2 | 0.10871 | 0.80042 | 0.54 | 0.46 | 1 | HC_vs_MS.Trt_F |
| CRLF2 | 0.10885 | 1.27766 | 0.37 | 0.3 | 1 | HC_vs_MS.Trt_F |
| SPIDR | 0.10912 | 0.57743 | 0.361 | 0.31 | 1 | HC_vs_MS.Trt_F |
| FOXO3 | 0.10936 | -0.7839 | 0.396 | 0.42 | 1 | HC_vs_MS.Trt_F |
| DRG2 | 0.10937 | 0.635 | 0.234 | 0.18 | 1 | HC_vs_MS.Trt_F |
| ERO1B | 0.10949 | 0.75407 | 0.133 | 0.09 | 1 | HC_vs_MS.Trt_F |
| RBBP8 | 0.1097 | 0.36471 | 0.142 | 0.1 | 1 | HC_vs_MS.Trt_F |
| CDAN1 | 0.10975 | 0.65952 | 0.529 | 0.55 | 1 | HC_vs_MS.Trt_F |
| PPP1R12B | 0.10991 | 1.12762 | 0.803 | 0.72 | 1 | HC_vs_MS.Trt_F |
| ZC3H7B | 0.11007 | 0.86285 | 0.853 | 0.77 | 1 | HC_vs_MS.Trt_F |
| PSMD13 | 0.1101 | -0.4017 | 0.54 | 0.6 | 1 | HC_vs_MS.Trt_F |
| ZNF324 | 0.11013 | -0.6356 | 0.066 | 0.1 | 1 | HC_vs_MS.Trt_F |
| MDH2 | 0.11014 | -0.306 | 0.269 | 0.2 | 1 | HC_vs_MS.Trt_F |
| HEATR6 | 0.11028 | 0.34062 | 0.607 | 0.53 | 1 | HC_vs_MS.Trt_F |
| ZAP70 | 0.11059 | 0.31161 | 0.494 | 0.44 | 1 | HC_vs_MS.Trt_F |
| SMARCA2 | 0.11069 | 0.45769 | 0.477 | 0.41 | 1 | HC_vs_MS.Trt_F |
| LANCL2 | 0.11074 | 0.2918 | 0.185 | 0.14 | 1 | HC_vs_MS.Trt_F |
| NVL | 0.11097 | -0.6615 | 0.835 | 0.74 | 1 | HC_vs_MS.Trt_F |
| PPP2R5E | 0.11111 | 0.36845 | 0.269 | 0.21 | 1 | HC_vs_MS.Trt_F |

| | | | | | | |
|---|---|---|---|---|---|---|
| JMJD6 | 0.11148 | 0.50117 | 0.494 | 0.42 | 1 | HC_vs_MS.Trt_F |
| SNX13 | 0.11149 | 0.55527 | 0.324 | 0.26 | 1 | HC_vs_MS.Trt_F |
| NET1 | 0.11157 | 0.39628 | 0.147 | 0.1 | 1 | HC_vs_MS.Trt_F |
| SLC8B1 | 0.11168 | 0.31097 | 0.538 | 0.45 | 1 | HC_vs_MS.Trt_F |
| C6orf106 | 0.1117 | 0.30825 | 0.165 | 0.12 | 1 | HC_vs_MS.Trt_F |
| TRAPPC2B | 0.11189 | 0.41881 | 0.436 | 0.37 | 1 | HC_vs_MS.Trt_F |
| PAIP2 | 0.11203 | -0.3697 | 0.607 | 0.67 | 1 | HC_vs_MS.Trt_F |
| PWP2 | 0.11216 | 0.37662 | 0.124 | 0.09 | 1 | HC_vs_MS.Trt_F |
| NXT1 | 0.1124 | 0.40917 | 0.118 | 0.08 | 1 | HC_vs_MS.Trt_F |
| ZNF596 | 0.11254 | -0.7699 | 0.512 | 0.45 | 1 | HC_vs_MS.Trt_F |
| ACSL3 | 0.11278 | 0.42555 | 0.162 | 0.12 | 1 | HC_vs_MS.Trt_F |
| MAP3K9 | 0.11291 | -0.4104 | 0.474 | 0.42 | 1 | HC_vs_MS.Trt_F |
| JKAMP | 0.11294 | 0.30808 | 0.847 | 0.8 | 1 | HC_vs_MS.Trt_F |
| AREL1 | 0.11318 | 0.51963 | 0.52 | 0.45 | 1 | HC_vs_MS.Trt_F |
| VASP | 0.11357 | 0.3053 | 0.645 | 0.57 | 1 | HC_vs_MS.Trt_F |
| NAV2 | 0.11365 | 1.07582 | 0.408 | 0.44 | 1 | HC_vs_MS.Trt_F |
| PXDN | 0.1138 | 1.00555 | 0.26 | 0.2 | 1 | HC_vs_MS.Trt_F |
| KLK2 | 0.11385 | 0.34461 | 0.379 | 0.3 | 1 | HC_vs_MS.Trt_F |
| APOL2 | 0.11395 | -0.6098 | 0.523 | 0.57 | 1 | HC_vs_MS.Trt_F |
| MAML1 | 0.11403 | 0.54189 | 0.208 | 0.16 | 1 | HC_vs_MS.Trt_F |
| BCAS1 | 0.11405 | 0.8609 | 0.425 | 0.35 | 1 | HC_vs_MS.Trt_F |
| ETNK2 | 0.11409 | 0.3578 | 0.249 | 0.2 | 1 | HC_vs_MS.Trt_F |
| TMEM9 | 0.11424 | 0.33487 | 0.133 | 0.09 | 1 | HC_vs_MS.Trt_F |
| RMND5B | 0.11427 | 0.658 | 0.682 | 0.61 | 1 | HC_vs_MS.Trt_F |
| USP49 | 0.11431 | 0.26157 | 0.694 | 0.75 | 1 | HC_vs_MS.Trt_F |
| BEST1 | 0.11512 | 0.30308 | 0.861 | 0.81 | 1 | HC_vs_MS.Trt_F |
| LARGE1 | 0.11514 | 0.35086 | 0.613 | 0.55 | 1 | HC_vs_MS.Trt_F |
| CCDC85C | 0.11527 | 0.43249 | 0.477 | 0.42 | 1 | HC_vs_MS.Trt_F |
| MSL1 | 0.11537 | -0.5398 | 0.624 | 0.53 | 1 | HC_vs_MS.Trt_F |
| SLC2A13 | 0.11547 | 0.42745 | 0.332 | 0.27 | 1 | HC_vs_MS.Trt_F |
| BAZ1B | 0.11559 | 0.30601 | 0.197 | 0.14 | 1 | HC_vs_MS.Trt_F |
| PPP1R16A | 0.11569 | 0.25589 | 0.477 | 0.41 | 1 | HC_vs_MS.Trt_F |
| MRM3 | 0.1157 | 0.62969 | 0.347 | 0.3 | 1 | HC_vs_MS.Trt_F |
| PLD6 | 0.11599 | -0.4698 | 0.477 | 0.41 | 1 | HC_vs_MS.Trt_F |
| TMC6 | 0.11607 | -0.4128 | 0.766 | 0.81 | 1 | HC_vs_MS.Trt_F |
| BUB3 | 0.11609 | -0.7213 | 0.668 | 0.66 | 1 | HC_vs_MS.Trt_F |
| CHMP7 | 0.11613 | -0.4379 | 0.465 | 0.51 | 1 | HC_vs_MS.Trt_F |
| CXCR3 | 0.11615 | 0.77422 | 0.228 | 0.18 | 1 | HC_vs_MS.Trt_F |
| THAP3 | 0.11624 | 0.37718 | 0.145 | 0.1 | 1 | HC_vs_MS.Trt_F |
| DNAJC21 | 0.11629 | 0.28582 | 0.428 | 0.38 | 1 | HC_vs_MS.Trt_F |

| | | | | | | |
|---|---|---|---|---|---|---|
| CDK13 | 0.11638 | -0.4545 | 0.908 | 0.84 | 1 | HC_vs_MS.Trt_F |
| ADARB1 | 0.11639 | 0.95219 | 0.263 | 0.21 | 1 | HC_vs_MS.Trt_F |
| BRPF3 | 0.11655 | -0.5067 | 0.066 | 0.1 | 1 | HC_vs_MS.Trt_F |
| HYAL4 | 0.11657 | 0.42147 | 0.549 | 0.46 | 1 | HC_vs_MS.Trt_F |
| HMGCS1 | 0.11658 | 0.32445 | 0.592 | 0.56 | 1 | HC_vs_MS.Trt_F |
| HDAC10 | 0.11683 | 1.4423 | 0.179 | 0.13 | 1 | HC_vs_MS.Trt_F |
| PRAF2 | 0.11691 | 0.35826 | 0.136 | 0.09 | 1 | HC_vs_MS.Trt_F |
| NUP153 | 0.11701 | 0.70996 | 0.249 | 0.19 | 1 | HC_vs_MS.Trt_F |
| CHST14 | 0.1171 | -0.6 | 0.303 | 0.25 | 1 | HC_vs_MS.Trt_F |
| ITGA2B | 0.1172 | 0.48754 | 0.471 | 0.41 | 1 | HC_vs_MS.Trt_F |
| ERBB3 | 0.11726 | -0.2681 | 0.803 | 0.72 | 1 | HC_vs_MS.Trt_F |
| METTL3 | 0.11727 | -0.2948 | 0.679 | 0.6 | 1 | HC_vs_MS.Trt_F |
| EXD2 | 0.11739 | -1.9529 | 0.275 | 0.21 | 1 | HC_vs_MS.Trt_F |
| NICN1 | 0.11747 | 0.37257 | 0.876 | 0.78 | 1 | HC_vs_MS.Trt_F |
| ST6GALNAC | 0.11748 | 0.70898 | 0.257 | 0.21 | 1 | HC_vs_MS.Trt_F |
| FAM174B | 0.11763 | 1.25098 | 0.173 | 0.13 | 1 | HC_vs_MS.Trt_F |
| ZNF497 | 0.11772 | 0.25205 | 0.353 | 0.29 | 1 | HC_vs_MS.Trt_F |
| MRPL34 | 0.11777 | -0.6396 | 0.329 | 0.26 | 1 | HC_vs_MS.Trt_F |
| ADIPOR2 | 0.11789 | -0.3945 | 0.159 | 0.2 | 1 | HC_vs_MS.Trt_F |
| AURKA | 0.11795 | -0.5479 | 0.358 | 0.31 | 1 | HC_vs_MS.Trt_F |
| METTL2B | 0.11797 | 0.32747 | 0.988 | 0.96 | 1 | HC_vs_MS.Trt_F |
| RNF220 | 0.11822 | 0.38965 | 0.699 | 0.6 | 1 | HC_vs_MS.Trt_F |
| SELL | 0.11825 | -0.4388 | 0.546 | 0.55 | 1 | HC_vs_MS.Trt_F |
| MBNL3 | 0.11827 | 0.35997 | 0.15 | 0.11 | 1 | HC_vs_MS.Trt_F |
| DTNB | 0.1183 | 0.46801 | 0.214 | 0.17 | 1 | HC_vs_MS.Trt_F |
| GATB | 0.11835 | 1.53621 | 0.292 | 0.24 | 1 | HC_vs_MS.Trt_F |
| LRRC32 | 0.1184 | 1.62954 | 0.101 | 0.07 | 1 | HC_vs_MS.Trt_F |
| TFAM | 0.11878 | 0.2672 | 0.321 | 0.26 | 1 | HC_vs_MS.Trt_F |
| RBM41 | 0.11878 | 0.30412 | 0.428 | 0.35 | 1 | HC_vs_MS.Trt_F |
| VWA7 | 0.11927 | 0.80883 | 0.387 | 0.32 | 1 | HC_vs_MS.Trt_F |
| ACTR5 | 0.1193 | 0.35053 | 0.168 | 0.12 | 1 | HC_vs_MS.Trt_F |
| IRAK1 | 0.11964 | 0.2867 | 0.292 | 0.23 | 1 | HC_vs_MS.Trt_F |
| ATRN | 0.1199 | 0.72321 | 0.165 | 0.12 | 1 | HC_vs_MS.Trt_F |
| C3orf33 | 0.11991 | -0.5365 | 0.532 | 0.5 | 1 | HC_vs_MS.Trt_F |
| TMEM53 | 0.12026 | -0.573 | 0.364 | 0.29 | 1 | HC_vs_MS.Trt_F |
| DNAJC19 | 0.12049 | -0.3838 | 0.298 | 0.23 | 1 | HC_vs_MS.Trt_F |
| WRAP73 | 0.12051 | -0.2738 | 0.37 | 0.3 | 1 | HC_vs_MS.Trt_F |
| FRG1 | 0.12071 | -0.6182 | 0.471 | 0.4 | 1 | HC_vs_MS.Trt_F |
| PIM3 | 0.12102 | 0.77875 | 0.176 | 0.13 | 1 | HC_vs_MS.Trt_F |
| METTL25 | 0.12114 | 0.38054 | 0.257 | 0.21 | 1 | HC_vs_MS.Trt_F |

| | | | | | | |
|---|---|---|---|---|---|---|
| ODC1 | 0.12121 | 0.38526 | 0.142 | 0.1 | 1 | HC_vs_MS.Trt_F |
| THAP9 | 0.12139 | 0.45002 | 0.639 | 0.58 | 1 | HC_vs_MS.Trt_F |
| HHLA3 | 0.12155 | 0.57979 | 0.355 | 0.3 | 1 | HC_vs_MS.Trt_F |
| HSD17B11 | 0.12161 | -0.438 | 0.532 | 0.45 | 1 | HC_vs_MS.Trt_F |
| HMG20A | 0.12168 | 0.28048 | 0.306 | 0.25 | 1 | HC_vs_MS.Trt_F |
| TMPRSS4-AS | 0.12177 | -0.3928 | 0.069 | 0.1 | 1 | HC_vs_MS.Trt_F |
| NUTF2 | 0.12182 | 0.36327 | 0.22 | 0.17 | 1 | HC_vs_MS.Trt_F |
| ZNF670 | 0.1222 | 1.18773 | 0.205 | 0.16 | 1 | HC_vs_MS.Trt_F |
| YAF2 | 0.12225 | -0.4794 | 0.769 | 0.69 | 1 | HC_vs_MS.Trt_F |
| SBF1 | 0.12298 | 0.80111 | 0.22 | 0.17 | 1 | HC_vs_MS.Trt_F |
| IFT43 | 0.12303 | -0.4173 | 0.069 | 0.1 | 1 | HC_vs_MS.Trt_F |
| FLYWCH1 | 0.12321 | 0.95219 | 0.832 | 0.76 | 1 | HC_vs_MS.Trt_F |
| ZNF468 | 0.12326 | -0.7027 | 0.529 | 0.46 | 1 | HC_vs_MS.Trt_F |
| ZNF335 | 0.12379 | 0.90258 | 0.376 | 0.33 | 1 | HC_vs_MS.Trt_F |
| ALG5 | 0.12384 | -0.4046 | 0.442 | 0.48 | 1 | HC_vs_MS.Trt_F |
| ATP6V0A2 | 0.12387 | 0.33889 | 0.971 | 0.95 | 1 | HC_vs_MS.Trt_F |
| MED7 | 0.12421 | -0.5608 | 0.081 | 0.12 | 1 | HC_vs_MS.Trt_F |
| GAMT | 0.1243 | 0.57235 | 0.63 | 0.54 | 1 | HC_vs_MS.Trt_F |
| KLF15 | 0.12439 | 0.31093 | 0.217 | 0.18 | 1 | HC_vs_MS.Trt_F |
| PRPF4 | 0.1246 | -0.3484 | 0.361 | 0.3 | 1 | HC_vs_MS.Trt_F |
| LMF1 | 0.12527 | 0.77831 | 0.156 | 0.12 | 1 | HC_vs_MS.Trt_F |
| HMGXB3 | 0.12566 | 0.84693 | 0.223 | 0.17 | 1 | HC_vs_MS.Trt_F |
| VAC14 | 0.12573 | 0.7045 | 0.133 | 0.09 | 1 | HC_vs_MS.Trt_F |
| HSD17B1 | 0.12587 | 0.46224 | 0.751 | 0.77 | 1 | HC_vs_MS.Trt_F |
| FAM208B | 0.1259 | -0.64 | 0.35 | 0.28 | 1 | HC_vs_MS.Trt_F |
| STOML2 | 0.12614 | 0.359 | 0.179 | 0.13 | 1 | HC_vs_MS.Trt_F |
| EML3 | 0.12614 | 0.66604 | 0.269 | 0.22 | 1 | HC_vs_MS.Trt_F |
| DDI2 | 0.12626 | 0.36933 | 0.812 | 0.77 | 1 | HC_vs_MS.Trt_F |
| PARD6G | 0.12641 | -0.4062 | 0.59 | 0.51 | 1 | HC_vs_MS.Trt_F |
| CHD6 | 0.12641 | -0.3542 | 0.251 | 0.19 | 1 | HC_vs_MS.Trt_F |
| RP11-307N1 | 0.12652 | 0.41358 | 0.116 | 0.08 | 1 | HC_vs_MS.Trt_F |
| FLYWCH2 | 0.1266 | 1.11421 | 0.107 | 0.07 | 1 | HC_vs_MS.Trt_F |
| SSH3 | 0.12694 | 1.1673 | 0.107 | 0.07 | 1 | HC_vs_MS.Trt_F |
| TRAK2 | 0.12694 | -0.3732 | 0.223 | 0.16 | 1 | HC_vs_MS.Trt_F |
| LCMT2 | 0.12711 | 0.51444 | 0.723 | 0.67 | 1 | HC_vs_MS.Trt_F |
| ZFPM2 | 0.12756 | 0.49968 | 0.442 | 0.41 | 1 | HC_vs_MS.Trt_F |
| FBXL3 | 0.12759 | 0.78067 | 0.176 | 0.13 | 1 | HC_vs_MS.Trt_F |
| DCTN3 | 0.1276 | -0.6162 | 0.402 | 0.45 | 1 | HC_vs_MS.Trt_F |
| CIAPIN1 | 0.12761 | -0.718 | 0.098 | 0.13 | 1 | HC_vs_MS.Trt_F |
| FAM76B | 0.12762 | 0.46955 | 0.142 | 0.1 | 1 | HC_vs_MS.Trt_F |

| | | | | | | |
|---|---|---|---|---|---|---|
| NUSAP1 | 0.12771 | 0.45753 | 0.783 | 0.73 | 1 | HC_vs_MS.Trt_F |
| UROS | 0.12808 | 0.948 | 0.509 | 0.45 | 1 | HC_vs_MS.Trt_F |
| HARS | 0.12824 | -0.5559 | 0.867 | 0.87 | 1 | HC_vs_MS.Trt_F |
| FCRL2 | 0.1283 | 0.51376 | 0.723 | 0.62 | 1 | HC_vs_MS.Trt_F |
| DDOST | 0.12831 | -0.2925 | 0.665 | 0.59 | 1 | HC_vs_MS.Trt_F |
| PHF20L1 | 0.12845 | -0.5453 | 0.662 | 0.67 | 1 | HC_vs_MS.Trt_F |
| ALKBH2 | 0.12869 | -0.6715 | 0.37 | 0.31 | 1 | HC_vs_MS.Trt_F |
| CEACAM21 | 0.12883 | -0.4895 | 0.275 | 0.31 | 1 | HC_vs_MS.Trt_F |
| SLC18B1 | 0.129 | 1.39238 | 0.116 | 0.08 | 1 | HC_vs_MS.Trt_F |
| PPP2R5C | 0.12929 | -0.4148 | 0.936 | 0.92 | 1 | HC_vs_MS.Trt_F |
| GIMAP5 | 0.12942 | -0.4705 | 0.292 | 0.34 | 1 | HC_vs_MS.Trt_F |
| ZNF683 | 0.12951 | 0.98131 | 0.075 | 0.11 | 1 | HC_vs_MS.Trt_F |
| KIAA0391 | 0.12953 | 0.32643 | 0.754 | 0.69 | 1 | HC_vs_MS.Trt_F |
| ADH5 | 0.12976 | -0.5865 | 0.555 | 0.45 | 1 | HC_vs_MS.Trt_F |
| OPRD1 | 0.12981 | 0.35593 | 0.855 | 0.8 | 1 | HC_vs_MS.Trt_F |
| ARFGAP3 | 0.13007 | 0.30052 | 0.376 | 0.33 | 1 | HC_vs_MS.Trt_F |
| CASP4 | 0.13031 | -0.6745 | 0.361 | 0.4 | 1 | HC_vs_MS.Trt_F |
| PDE12 | 0.13064 | -0.2515 | 0.919 | 0.86 | 1 | HC_vs_MS.Trt_F |
| PLEKHH2 | 0.13078 | 0.76645 | 0.697 | 0.67 | 1 | HC_vs_MS.Trt_F |
| TMEM138 | 0.1308 | 0.68946 | 0.867 | 0.81 | 1 | HC_vs_MS.Trt_F |
| TMEM242 | 0.13108 | 0.62538 | 0.171 | 0.13 | 1 | HC_vs_MS.Trt_F |
| ANGPTL4 | 0.13125 | 1.23884 | 0.35 | 0.28 | 1 | HC_vs_MS.Trt_F |
| AIMP2 | 0.13129 | -0.5772 | 0.358 | 0.29 | 1 | HC_vs_MS.Trt_F |
| NR1D1 | 0.13141 | 0.52984 | 0.153 | 0.11 | 1 | HC_vs_MS.Trt_F |
| ZKSCAN4 | 0.13153 | -0.7398 | 0.572 | 0.49 | 1 | HC_vs_MS.Trt_F |
| ABCF1 | 0.13162 | 0.38924 | 0.841 | 0.82 | 1 | HC_vs_MS.Trt_F |
| MRPL42 | 0.13173 | -0.5099 | 0.98 | 0.96 | 1 | HC_vs_MS.Trt_F |
| TMEM62 | 0.13176 | 0.49649 | 0.277 | 0.22 | 1 | HC_vs_MS.Trt_F |
| ADGRG2 | 0.1318 | 0.25454 | 0.665 | 0.59 | 1 | HC_vs_MS.Trt_F |
| SKIV2L | 0.13181 | 0.68451 | 0.185 | 0.14 | 1 | HC_vs_MS.Trt_F |
| TDP2 | 0.13184 | 0.29137 | 0.156 | 0.12 | 1 | HC_vs_MS.Trt_F |
| INPP5E | 0.13198 | -0.2866 | 0.312 | 0.25 | 1 | HC_vs_MS.Trt_F |
| C17orf75 | 0.13214 | -0.2829 | 0.864 | 0.83 | 1 | HC_vs_MS.Trt_F |
| IQCB1 | 0.13219 | 0.80785 | 0.142 | 0.1 | 1 | HC_vs_MS.Trt_F |
| DRC3 | 0.1322 | 0.45961 | 0.621 | 0.57 | 1 | HC_vs_MS.Trt_F |
| CENPH | 0.1322 | -0.5068 | 0.324 | 0.26 | 1 | HC_vs_MS.Trt_F |
| ZNF76 | 0.13232 | 0.45492 | 0.22 | 0.17 | 1 | HC_vs_MS.Trt_F |
| SRCAP | 0.13241 | 0.31731 | 0.705 | 0.62 | 1 | HC_vs_MS.Trt_F |
| SMIM8 | 0.13246 | -0.7579 | 0.685 | 0.6 | 1 | HC_vs_MS.Trt_F |
| AP4S1 | 0.13255 | 0.4061 | 0.965 | 0.95 | 1 | HC_vs_MS.Trt_F |

| | | | | | | |
|---|---|---|---|---|---|---|
| ZNF281 | 0.13265 | 0.85764 | 0.133 | 0.09 | 1 | HC_vs_MS.Trt_F |
| HMGXB4 | 0.13267 | 1.12609 | 0.156 | 0.12 | 1 | HC_vs_MS.Trt_F |
| CCDC85B | 0.13306 | 0.58405 | 0.153 | 0.11 | 1 | HC_vs_MS.Trt_F |
| DNAJC9 | 0.13348 | -0.8037 | 0.133 | 0.17 | 1 | HC_vs_MS.Trt_F |
| GLO1 | 0.13349 | 0.40797 | 0.124 | 0.09 | 1 | HC_vs_MS.Trt_F |
| RNF168 | 0.13367 | -0.9922 | 0.592 | 0.57 | 1 | HC_vs_MS.Trt_F |
| C1QTNF7 | 0.13386 | 0.44934 | 0.081 | 0.12 | 1 | HC_vs_MS.Trt_F |
| TADA2B | 0.13395 | 0.8373 | 0.361 | 0.3 | 1 | HC_vs_MS.Trt_F |
| ZNF552 | 0.13402 | 0.49292 | 0.486 | 0.43 | 1 | HC_vs_MS.Trt_F |
| GRAMD4 | 0.13404 | 1.1572 | 0.104 | 0.07 | 1 | HC_vs_MS.Trt_F |
| ARL5C | 0.13405 | 0.37703 | 0.451 | 0.38 | 1 | HC_vs_MS.Trt_F |
| CSNK1A1 | 0.13434 | -0.4345 | 0.841 | 0.89 | 1 | HC_vs_MS.Trt_F |
| CRYZL1 | 0.1344 | -0.3206 | 0.965 | 0.95 | 1 | HC_vs_MS.Trt_F |
| MEF2A | 0.13462 | -0.5089 | 0.572 | 0.5 | 1 | HC_vs_MS.Trt_F |
| NBR1 | 0.13472 | 0.31493 | 0.832 | 0.78 | 1 | HC_vs_MS.Trt_F |
| ZSWIM8 | 0.13474 | 0.37092 | 0.838 | 0.83 | 1 | HC_vs_MS.Trt_F |
| TMEM108 | 0.13515 | 1.11021 | 0.367 | 0.31 | 1 | HC_vs_MS.Trt_F |
| PRRG1 | 0.13528 | 0.44113 | 0.251 | 0.19 | 1 | HC_vs_MS.Trt_F |
| CMTR2 | 0.1353 | 0.32232 | 0.321 | 0.27 | 1 | HC_vs_MS.Trt_F |
| CHMP2A | 0.13569 | 0.25432 | 0.199 | 0.15 | 1 | HC_vs_MS.Trt_F |
| ARMC5 | 0.13576 | -0.4935 | 0.162 | 0.12 | 1 | HC_vs_MS.Trt_F |
| PRPS1 | 0.13581 | 0.47864 | 0.162 | 0.12 | 1 | HC_vs_MS.Trt_F |
| GNLY | 0.13584 | 0.50616 | 0.312 | 0.25 | 1 | HC_vs_MS.Trt_F |
| KIAA0825 | 0.13633 | -0.3873 | 0.361 | 0.3 | 1 | HC_vs_MS.Trt_F |
| TOM1L2 | 0.13643 | 0.26596 | 0.257 | 0.2 | 1 | HC_vs_MS.Trt_F |
| ATF5 | 0.13689 | -0.4088 | 0.341 | 0.26 | 1 | HC_vs_MS.Trt_F |
| TSC22D1 | 0.13708 | -0.261 | 0.116 | 0.08 | 1 | HC_vs_MS.Trt_F |
| RYK | 0.13722 | 1.00326 | 0.118 | 0.08 | 1 | HC_vs_MS.Trt_F |
| CAPN5 | 0.13734 | 0.35529 | 0.309 | 0.25 | 1 | HC_vs_MS.Trt_F |
| RP5-1022E2 | 0.1377 | -0.5549 | 0.113 | 0.15 | 1 | HC_vs_MS.Trt_F |
| SLAMF6 | 0.138 | -0.3394 | 0.549 | 0.47 | 1 | HC_vs_MS.Trt_F |
| AMY2B | 0.13815 | 0.31962 | 0.607 | 0.66 | 1 | HC_vs_MS.Trt_F |
| CYP46A1 | 0.13852 | -0.6417 | 0.63 | 0.55 | 1 | HC_vs_MS.Trt_F |
| RNF6 | 0.13904 | -0.8068 | 0.162 | 0.2 | 1 | HC_vs_MS.Trt_F |
| POLR3B | 0.13904 | 0.62687 | 0.182 | 0.14 | 1 | HC_vs_MS.Trt_F |
| ACSL1 | 0.13906 | -0.2981 | 0.28 | 0.23 | 1 | HC_vs_MS.Trt_F |
| ZNF316 | 0.1392 | -0.5109 | 0.627 | 0.53 | 1 | HC_vs_MS.Trt_F |
| WARS2 | 0.13931 | 0.29066 | 0.11 | 0.08 | 1 | HC_vs_MS.Trt_F |
| EXOSC3 | 0.14013 | 0.66057 | 0.465 | 0.39 | 1 | HC_vs_MS.Trt_F |
| ATG13 | 0.14035 | -0.3299 | 0.89 | 0.8 | 1 | HC_vs_MS.Trt_F |

| | | | | | | |
|---|---|---|---|---|---|---|
| WDR83OS | 0.14035 | 0.27727 | 0.231 | 0.18 | 1 | HC_vs_MS.Trt_F |
| TSEN34 | 0.14038 | 0.59007 | 0.159 | 0.21 | 1 | HC_vs_MS.Trt_F |
| C5orf22 | 0.14067 | -0.5596 | 0.355 | 0.29 | 1 | HC_vs_MS.Trt_F |
| PEX11B | 0.14096 | 0.60897 | 0.159 | 0.12 | 1 | HC_vs_MS.Trt_F |
| SAP130 | 0.14107 | 0.39977 | 0.194 | 0.15 | 1 | HC_vs_MS.Trt_F |
| MEI1 | 0.14114 | 0.36411 | 0.734 | 0.69 | 1 | HC_vs_MS.Trt_F |
| XRCC3 | 0.14123 | 0.32553 | 0.514 | 0.45 | 1 | HC_vs_MS.Trt_F |
| CD82 | 0.14126 | 0.27952 | 0.717 | 0.65 | 1 | HC_vs_MS.Trt_F |
| POFUT2 | 0.14168 | 0.94058 | 0.104 | 0.07 | 1 | HC_vs_MS.Trt_F |
| CHFR | 0.14172 | -0.4966 | 0.942 | 0.94 | 1 | HC_vs_MS.Trt_F |
| CCDC18 | 0.14175 | 0.42839 | 0.249 | 0.2 | 1 | HC_vs_MS.Trt_F |
| RCOR3 | 0.14175 | 0.29397 | 0.48 | 0.4 | 1 | HC_vs_MS.Trt_F |
| TM9SF1 | 0.14191 | -0.8204 | 0.318 | 0.26 | 1 | HC_vs_MS.Trt_F |
| ARF3 | 0.14195 | 0.28129 | 0.228 | 0.18 | 1 | HC_vs_MS.Trt_F |
| ZNF589 | 0.142 | -0.3439 | 0.538 | 0.45 | 1 | HC_vs_MS.Trt_F |
| TMA7 | 0.14219 | -0.6496 | 0.601 | 0.58 | 1 | HC_vs_MS.Trt_F |
| ERCC6L2 | 0.14226 | -0.438 | 0.234 | 0.18 | 1 | HC_vs_MS.Trt_F |
| RBPMS | 0.14234 | 0.36801 | 0.199 | 0.16 | 1 | HC_vs_MS.Trt_F |
| GCC2 | 0.14252 | -0.3603 | 0.803 | 0.75 | 1 | HC_vs_MS.Trt_F |
| SLC39A4 | 0.14276 | 0.40764 | 0.393 | 0.33 | 1 | HC_vs_MS.Trt_F |
| DENND3 | 0.14279 | 1.13818 | 0.564 | 0.49 | 1 | HC_vs_MS.Trt_F |
| ALG6 | 0.14324 | -0.7652 | 0.147 | 0.18 | 1 | HC_vs_MS.Trt_F |
| CNOT9 | 0.1436 | -0.4866 | 0.746 | 0.75 | 1 | HC_vs_MS.Trt_F |
| STK26 | 0.14383 | 0.63885 | 0.147 | 0.11 | 1 | HC_vs_MS.Trt_F |
| CEP68 | 0.14385 | -0.5731 | 0.598 | 0.51 | 1 | HC_vs_MS.Trt_F |
| 1-Mar | 0.14394 | 0.36193 | 0.344 | 0.28 | 1 | HC_vs_MS.Trt_F |
| SMARCD1 | 0.14432 | 0.98188 | 0.139 | 0.1 | 1 | HC_vs_MS.Trt_F |
| MAPT | 0.14435 | 0.50189 | 0.746 | 0.67 | 1 | HC_vs_MS.Trt_F |
| PFKFB2 | 0.14454 | -1.1358 | 0.225 | 0.28 | 1 | HC_vs_MS.Trt_F |
| TPD52L1 | 0.14459 | 0.47405 | 0.769 | 0.73 | 1 | HC_vs_MS.Trt_F |
| SPEF2 | 0.14481 | 0.88853 | 0.595 | 0.54 | 1 | HC_vs_MS.Trt_F |
| PLEK | 0.145 | -0.6334 | 0.084 | 0.12 | 1 | HC_vs_MS.Trt_F |
| CAPRIN2 | 0.14515 | 0.85403 | 0.757 | 0.67 | 1 | HC_vs_MS.Trt_F |
| RUFY2 | 0.14515 | -0.6496 | 0.592 | 0.48 | 1 | HC_vs_MS.Trt_F |
| ZFAND4 | 0.14533 | -0.6104 | 0.523 | 0.44 | 1 | HC_vs_MS.Trt_F |
| WASH1 | 0.1455 | 0.28497 | 0.266 | 0.22 | 1 | HC_vs_MS.Trt_F |
| IDNK | 0.14558 | -0.4331 | 0.454 | 0.37 | 1 | HC_vs_MS.Trt_F |
| MARVELD3 | 0.14564 | 0.28991 | 0.91 | 0.85 | 1 | HC_vs_MS.Trt_F |
| MED23 | 0.14577 | -0.3471 | 0.266 | 0.21 | 1 | HC_vs_MS.Trt_F |
| CC2D2A | 0.14587 | 0.71177 | 0.257 | 0.21 | 1 | HC_vs_MS.Trt_F |

| | | | | | | |
|---|---|---|---|---|---|---|
| C19orf12 | 0.14603 | 0.51769 | 0.434 | 0.36 | 1 | HC_vs_MS.Trt_F |
| ZC3H14 | 0.14606 | -0.3715 | 0.957 | 0.94 | 1 | HC_vs_MS.Trt_F |
| HID1 | 0.14619 | 0.25219 | 0.168 | 0.21 | 1 | HC_vs_MS.Trt_F |
| TFEB | 0.14622 | -0.5016 | 0.486 | 0.5 | 1 | HC_vs_MS.Trt_F |
| PRF1 | 0.14673 | -0.4491 | 0.223 | 0.26 | 1 | HC_vs_MS.Trt_F |
| MNT | 0.14676 | 0.38346 | 0.711 | 0.62 | 1 | HC_vs_MS.Trt_F |
| PTPA | 0.14683 | 0.28871 | 0.795 | 0.73 | 1 | HC_vs_MS.Trt_F |
| CHCHD3 | 0.14684 | 0.26823 | 0.173 | 0.13 | 1 | HC_vs_MS.Trt_F |
| BTD | 0.14707 | 0.60256 | 0.136 | 0.1 | 1 | HC_vs_MS.Trt_F |
| MTA1 | 0.14709 | 0.32494 | 0.445 | 0.39 | 1 | HC_vs_MS.Trt_F |
| FASTK | 0.14711 | 0.28558 | 0.188 | 0.14 | 1 | HC_vs_MS.Trt_F |
| KIF21A | 0.14726 | 0.50145 | 0.22 | 0.17 | 1 | HC_vs_MS.Trt_F |
| MRPS7 | 0.14741 | 0.27878 | 0.601 | 0.52 | 1 | HC_vs_MS.Trt_F |
| ZC3H4 | 0.14755 | -0.4478 | 0.341 | 0.37 | 1 | HC_vs_MS.Trt_F |
| ZNF551 | 0.14777 | 0.47763 | 0.543 | 0.49 | 1 | HC_vs_MS.Trt_F |
| FBXO18 | 0.1478 | -0.3786 | 0.448 | 0.38 | 1 | HC_vs_MS.Trt_F |
| WWC3 | 0.14781 | -0.3381 | 0.462 | 0.39 | 1 | HC_vs_MS.Trt_F |
| NPHP1 | 0.14793 | 0.30182 | 0.746 | 0.68 | 1 | HC_vs_MS.Trt_F |
| MAPK9 | 0.14891 | -1.0156 | 0.509 | 0.46 | 1 | HC_vs_MS.Trt_F |
| TMUB1 | 0.14893 | 0.56158 | 0.139 | 0.1 | 1 | HC_vs_MS.Trt_F |
| CCR2 | 0.14926 | 0.84059 | 0.153 | 0.12 | 1 | HC_vs_MS.Trt_F |
| SDF2L1 | 0.14938 | -0.6053 | 0.136 | 0.17 | 1 | HC_vs_MS.Trt_F |
| SGPP2 | 0.14943 | 0.41783 | 0.399 | 0.34 | 1 | HC_vs_MS.Trt_F |
| C3orf70 | 0.14972 | 0.32058 | 0.801 | 0.73 | 1 | HC_vs_MS.Trt_F |
| NEK9 | 0.15013 | 0.28 | 0.913 | 0.9 | 1 | HC_vs_MS.Trt_F |
| GRAMD3 | 0.15017 | 0.90146 | 0.757 | 0.68 | 1 | HC_vs_MS.Trt_F |
| NEK6 | 0.15034 | -0.3176 | 0.162 | 0.12 | 1 | HC_vs_MS.Trt_F |
| ZC3H12C | 0.15043 | 0.40637 | 0.133 | 0.1 | 1 | HC_vs_MS.Trt_F |
| PACRGL | 0.15057 | 0.83059 | 0.312 | 0.26 | 1 | HC_vs_MS.Trt_F |
| GNA11 | 0.15099 | 0.30733 | 0.572 | 0.5 | 1 | HC_vs_MS.Trt_F |
| KATNBL1 | 0.15112 | -0.4529 | 0.931 | 0.92 | 1 | HC_vs_MS.Trt_F |
| SUV39H2 | 0.15116 | 0.41451 | 0.488 | 0.4 | 1 | HC_vs_MS.Trt_F |
| PPP6R3 | 0.15119 | -0.3138 | 0.486 | 0.52 | 1 | HC_vs_MS.Trt_F |
| JMJD7 | 0.15132 | 0.36381 | 0.171 | 0.13 | 1 | HC_vs_MS.Trt_F |
| SLC4A7 | 0.15136 | -0.4475 | 0.295 | 0.34 | 1 | HC_vs_MS.Trt_F |
| LSM5 | 0.15137 | -0.3321 | 0.561 | 0.5 | 1 | HC_vs_MS.Trt_F |
| AMDHD2 | 0.15149 | 1.16764 | 0.121 | 0.09 | 1 | HC_vs_MS.Trt_F |
| CEP104 | 0.15164 | 0.69271 | 0.734 | 0.71 | 1 | HC_vs_MS.Trt_F |
| MXRA7 | 0.15171 | 0.34809 | 0.795 | 0.76 | 1 | HC_vs_MS.Trt_F |
| ERCC6L | 0.15213 | -1.5013 | 0.107 | 0.14 | 1 | HC_vs_MS.Trt_F |

| | | | | | | |
|---|---|---|---|---|---|---|
| TRMT5 | 0.15219 | -0.6227 | 0.165 | 0.21 | 1 | HC_vs_MS.Trt_F |
| PPIL2 | 0.15236 | 0.28265 | 0.454 | 0.39 | 1 | HC_vs_MS.Trt_F |
| ZNF260 | 0.15237 | -0.3617 | 0.627 | 0.53 | 1 | HC_vs_MS.Trt_F |
| CLSTN1 | 0.1524 | 0.26409 | 0.188 | 0.14 | 1 | HC_vs_MS.Trt_F |
| VCPIP1 | 0.15249 | -0.3246 | 0.439 | 0.37 | 1 | HC_vs_MS.Trt_F |
| AC013264.1 | 0.15252 | -0.5078 | 0.402 | 0.45 | 1 | HC_vs_MS.Trt_F |
| RGS12 | 0.15261 | 0.78399 | 0.887 | 0.86 | 1 | HC_vs_MS.Trt_F |
| CCDC120 | 0.15272 | 1.14162 | 0.587 | 0.51 | 1 | HC_vs_MS.Trt_F |
| CD200 | 0.15278 | -1.0287 | 0.662 | 0.7 | 1 | HC_vs_MS.Trt_F |
| SIRT3 | 0.15294 | 1.20462 | 0.318 | 0.37 | 1 | HC_vs_MS.Trt_F |
| SSFA2 | 0.15302 | 0.90916 | 0.191 | 0.14 | 1 | HC_vs_MS.Trt_F |
| FADS2 | 0.15305 | 0.46287 | 0.59 | 0.53 | 1 | HC_vs_MS.Trt_F |
| ZNF649 | 0.15319 | 0.42676 | 0.205 | 0.16 | 1 | HC_vs_MS.Trt_F |
| SETDB2 | 0.15329 | -0.414 | 0.896 | 0.85 | 1 | HC_vs_MS.Trt_F |
| MCCC1 | 0.15379 | 0.41246 | 0.555 | 0.49 | 1 | HC_vs_MS.Trt_F |
| ZDHHC20 | 0.15396 | 0.38716 | 0.526 | 0.46 | 1 | HC_vs_MS.Trt_F |
| NDOR1 | 0.15447 | 0.7662 | 0.113 | 0.08 | 1 | HC_vs_MS.Trt_F |
| NAB1 | 0.15472 | 0.66073 | 0.153 | 0.12 | 1 | HC_vs_MS.Trt_F |
| ITCH | 0.15494 | 0.38835 | 0.529 | 0.48 | 1 | HC_vs_MS.Trt_F |
| CDC42SE2 | 0.15507 | -0.413 | 0.847 | 0.82 | 1 | HC_vs_MS.Trt_F |
| VBP1 | 0.15537 | 0.53086 | 0.142 | 0.1 | 1 | HC_vs_MS.Trt_F |
| DDX51 | 0.15539 | -0.419 | 0.685 | 0.68 | 1 | HC_vs_MS.Trt_F |
| EFHD2 | 0.15547 | 0.57054 | 0.182 | 0.14 | 1 | HC_vs_MS.Trt_F |
| FAM86B1 | 0.15562 | -0.3001 | 0.081 | 0.12 | 1 | HC_vs_MS.Trt_F |
| ACACA | 0.15599 | 0.95329 | 0.645 | 0.59 | 1 | HC_vs_MS.Trt_F |
| SIPA1 | 0.15606 | -0.5665 | 0.662 | 0.53 | 1 | HC_vs_MS.Trt_F |
| CLK1 | 0.15628 | 0.28345 | 0.725 | 0.68 | 1 | HC_vs_MS.Trt_F |
| PHF2 | 0.15637 | 0.41255 | 0.104 | 0.07 | 1 | HC_vs_MS.Trt_F |
| ZCCHC6 | 0.15639 | -0.4115 | 0.318 | 0.24 | 1 | HC_vs_MS.Trt_F |
| SCFD1 | 0.15655 | 0.2568 | 0.798 | 0.74 | 1 | HC_vs_MS.Trt_F |
| TIE1 | 0.15656 | -0.5888 | 0.613 | 0.62 | 1 | HC_vs_MS.Trt_F |
| TNIP2 | 0.15662 | 0.45822 | 0.147 | 0.11 | 1 | HC_vs_MS.Trt_F |
| WBP1 | 0.15718 | 0.34947 | 0.147 | 0.11 | 1 | HC_vs_MS.Trt_F |
| CENPT | 0.15724 | -0.2927 | 0.269 | 0.21 | 1 | HC_vs_MS.Trt_F |
| EI24 | 0.1576 | -0.8194 | 0.786 | 0.72 | 1 | HC_vs_MS.Trt_F |
| PBDC1 | 0.15766 | 0.27641 | 0.139 | 0.1 | 1 | HC_vs_MS.Trt_F |
| ZNRF1 | 0.1578 | 0.30799 | 0.344 | 0.3 | 1 | HC_vs_MS.Trt_F |
| CCDC6 | 0.15791 | 1.1002 | 0.121 | 0.09 | 1 | HC_vs_MS.Trt_F |
| CES4A | 0.15796 | 0.68231 | 0.558 | 0.6 | 1 | HC_vs_MS.Trt_F |
| BRF2 | 0.15813 | 0.34688 | 0.194 | 0.15 | 1 | HC_vs_MS.Trt_F |

| | | | | | | |
|---|---|---|---|---|---|---|
| PKP2 | 0.15814 | 0.38541 | 0.676 | 0.66 | 1 | HC_vs_MS.Trt_F |
| MTRF1L | 0.15841 | -0.3261 | 0.133 | 0.09 | 1 | HC_vs_MS.Trt_F |
| HTATIP2 | 0.15842 | 0.51817 | 0.197 | 0.15 | 1 | HC_vs_MS.Trt_F |
| POLR3C | 0.15846 | 0.39587 | 0.616 | 0.54 | 1 | HC_vs_MS.Trt_F |
| CCDC14 | 0.15849 | 0.8813 | 0.228 | 0.19 | 1 | HC_vs_MS.Trt_F |
| MTHFD2 | 0.15856 | 0.29954 | 0.717 | 0.68 | 1 | HC_vs_MS.Trt_F |
| ZNF699 | 0.15871 | 1.20365 | 0.353 | 0.31 | 1 | HC_vs_MS.Trt_F |
| SLC35B1 | 0.15873 | 0.67503 | 0.708 | 0.66 | 1 | HC_vs_MS.Trt_F |
| SEC23B | 0.15881 | 0.36826 | 0.191 | 0.15 | 1 | HC_vs_MS.Trt_F |
| SYNC | 0.15893 | -0.6551 | 0.153 | 0.19 | 1 | HC_vs_MS.Trt_F |
| SLC25A14 | 0.15897 | 1.35657 | 0.324 | 0.26 | 1 | HC_vs_MS.Trt_F |
| CD2AP | 0.15931 | 0.44429 | 0.211 | 0.16 | 1 | HC_vs_MS.Trt_F |
| CDON | 0.15937 | 0.55032 | 0.526 | 0.46 | 1 | HC_vs_MS.Trt_F |
| VN1R1 | 0.15943 | 1.26701 | 0.211 | 0.16 | 1 | HC_vs_MS.Trt_F |
| TBC1D2B | 0.15947 | 0.2573 | 0.358 | 0.3 | 1 | HC_vs_MS.Trt_F |
| PDCD11 | 0.15973 | -0.5259 | 0.277 | 0.22 | 1 | HC_vs_MS.Trt_F |
| PKD2 | 0.15991 | -0.574 | 0.228 | 0.17 | 1 | HC_vs_MS.Trt_F |
| C11orf98 | 0.16019 | 0.25678 | 0.165 | 0.12 | 1 | HC_vs_MS.Trt_F |
| SBNO1 | 0.16019 | -0.4268 | 0.61 | 0.66 | 1 | HC_vs_MS.Trt_F |
| ACSL6 | 0.16038 | -0.287 | 0.618 | 0.66 | 1 | HC_vs_MS.Trt_F |
| TMEM59 | 0.1604 | -0.4806 | 0.882 | 0.9 | 1 | HC_vs_MS.Trt_F |
| CYLC2 | 0.16043 | 0.35855 | 0.118 | 0.09 | 1 | HC_vs_MS.Trt_F |
| CPSF1 | 0.16059 | 0.26011 | 0.214 | 0.17 | 1 | HC_vs_MS.Trt_F |
| ETV4 | 0.16074 | 0.61796 | 0.13 | 0.1 | 1 | HC_vs_MS.Trt_F |
| BTRC | 0.16075 | 0.94704 | 0.113 | 0.08 | 1 | HC_vs_MS.Trt_F |
| ALDH1L2 | 0.16085 | 0.93212 | 0.335 | 0.28 | 1 | HC_vs_MS.Trt_F |
| AKT1 | 0.16092 | 0.39487 | 0.254 | 0.21 | 1 | HC_vs_MS.Trt_F |
| ARPC2 | 0.16104 | -0.251 | 0.974 | 0.97 | 1 | HC_vs_MS.Trt_F |
| AK5 | 0.16115 | 0.48166 | 0.156 | 0.12 | 1 | HC_vs_MS.Trt_F |
| LRRC45 | 0.16141 | -0.3143 | 0.075 | 0.11 | 1 | HC_vs_MS.Trt_F |
| OSBP2 | 0.16189 | -0.3408 | 0.468 | 0.41 | 1 | HC_vs_MS.Trt_F |
| CPSF3L | 0.16199 | 0.27862 | 0.815 | 0.73 | 1 | HC_vs_MS.Trt_F |
| GOSR2 | 0.16206 | 0.346 | 0.171 | 0.13 | 1 | HC_vs_MS.Trt_F |
| PAQR3 | 0.16277 | 1.04314 | 0.159 | 0.12 | 1 | HC_vs_MS.Trt_F |
| NKD1 | 0.16294 | 0.30179 | 0.425 | 0.36 | 1 | HC_vs_MS.Trt_F |
| TRIP4 | 0.16309 | -0.4426 | 0.575 | 0.49 | 1 | HC_vs_MS.Trt_F |
| MON1A | 0.16318 | 1.23054 | 0.116 | 0.08 | 1 | HC_vs_MS.Trt_F |
| LGALS1 | 0.16321 | -0.3391 | 0.477 | 0.38 | 1 | HC_vs_MS.Trt_F |
| ZNF570 | 0.16328 | -0.4905 | 0.832 | 0.79 | 1 | HC_vs_MS.Trt_F |
| CCDC74A | 0.16338 | 1.05778 | 0.289 | 0.24 | 1 | HC_vs_MS.Trt_F |

| | | | | | | |
|---|---|---|---|---|---|---|
| RCC1 | 0.16341 | 0.53295 | 0.191 | 0.15 | 1 | HC_vs_MS.Trt_F |
| M1AP | 0.16425 | 0.3556 | 0.347 | 0.31 | 1 | HC_vs_MS.Trt_F |
| ANKAR | 0.16433 | 0.67464 | 0.179 | 0.14 | 1 | HC_vs_MS.Trt_F |
| NEDD4L | 0.16471 | 0.357 | 0.509 | 0.53 | 1 | HC_vs_MS.Trt_F |
| NUPR1 | 0.16519 | 0.28625 | 0.977 | 0.95 | 1 | HC_vs_MS.Trt_F |
| GRAP2 | 0.16526 | 0.25821 | 0.306 | 0.25 | 1 | HC_vs_MS.Trt_F |
| SSNA1 | 0.16563 | 0.31845 | 0.185 | 0.14 | 1 | HC_vs_MS.Trt_F |
| BFAR | 0.16606 | -0.469 | 0.526 | 0.54 | 1 | HC_vs_MS.Trt_F |
| TMEM186 | 0.1667 | 0.31333 | 0.526 | 0.48 | 1 | HC_vs_MS.Trt_F |
| ORC1 | 0.16685 | 0.58542 | 0.303 | 0.25 | 1 | HC_vs_MS.Trt_F |
| TSN | 0.16707 | -0.8876 | 0.428 | 0.36 | 1 | HC_vs_MS.Trt_F |
| ELP6 | 0.16758 | -1.1311 | 0.647 | 0.6 | 1 | HC_vs_MS.Trt_F |
| YIF1B | 0.1676 | -0.342 | 0.694 | 0.61 | 1 | HC_vs_MS.Trt_F |
| PPP1CA | 0.16763 | -0.2825 | 0.454 | 0.5 | 1 | HC_vs_MS.Trt_F |
| CNIH3 | 0.1679 | 0.78586 | 0.234 | 0.19 | 1 | HC_vs_MS.Trt_F |
| ATP6V0E2 | 0.16824 | 0.3993 | 0.234 | 0.19 | 1 | HC_vs_MS.Trt_F |
| PARP9 | 0.16836 | -0.5135 | 0.408 | 0.33 | 1 | HC_vs_MS.Trt_F |
| TRIM37 | 0.1684 | -0.2704 | 0.24 | 0.19 | 1 | HC_vs_MS.Trt_F |
| EME2 | 0.16843 | 0.51101 | 0.341 | 0.28 | 1 | HC_vs_MS.Trt_F |
| ESPL1 | 0.16852 | -0.7209 | 0.766 | 0.7 | 1 | HC_vs_MS.Trt_F |
| C12orf29 | 0.16865 | -0.4704 | 0.286 | 0.22 | 1 | HC_vs_MS.Trt_F |
| MYNN | 0.16887 | -0.2532 | 0.341 | 0.28 | 1 | HC_vs_MS.Trt_F |
| CCR7 | 0.1689 | -0.3918 | 0.379 | 0.42 | 1 | HC_vs_MS.Trt_F |
| C11orf21 | 0.16946 | 0.58229 | 0.173 | 0.13 | 1 | HC_vs_MS.Trt_F |
| MLYCD | 0.1695 | 1.00145 | 0.422 | 0.35 | 1 | HC_vs_MS.Trt_F |
| ITGA3 | 0.16966 | 0.45421 | 0.809 | 0.72 | 1 | HC_vs_MS.Trt_F |
| SIMC1 | 0.16991 | -0.3532 | 0.526 | 0.44 | 1 | HC_vs_MS.Trt_F |
| KIAA0232 | 0.16997 | 0.79465 | 0.156 | 0.12 | 1 | HC_vs_MS.Trt_F |
| POLR1A | 0.16998 | 0.55402 | 0.772 | 0.69 | 1 | HC_vs_MS.Trt_F |
| ENTPD6 | 0.17 | 0.50332 | 0.15 | 0.11 | 1 | HC_vs_MS.Trt_F |
| TIPARP | 0.17021 | 0.38881 | 0.165 | 0.13 | 1 | HC_vs_MS.Trt_F |
| COA6 | 0.17029 | -0.2503 | 0.156 | 0.12 | 1 | HC_vs_MS.Trt_F |
| FBXO5 | 0.17036 | 1.27819 | 0.431 | 0.38 | 1 | HC_vs_MS.Trt_F |
| ESYT3 | 0.17039 | 0.43707 | 0.584 | 0.5 | 1 | HC_vs_MS.Trt_F |
| FGFR2 | 0.17051 | 0.36788 | 0.509 | 0.45 | 1 | HC_vs_MS.Trt_F |
| PIAS2 | 0.17082 | 0.32378 | 0.815 | 0.73 | 1 | HC_vs_MS.Trt_F |
| GNAI3 | 0.17083 | -0.294 | 0.988 | 0.96 | 1 | HC_vs_MS.Trt_F |
| VCL | 0.17091 | 0.52377 | 0.188 | 0.14 | 1 | HC_vs_MS.Trt_F |
| SLC5A6 | 0.17094 | 0.47071 | 0.15 | 0.11 | 1 | HC_vs_MS.Trt_F |
| RBM17 | 0.17099 | -0.3047 | 0.425 | 0.34 | 1 | HC_vs_MS.Trt_F |

| | | | | | | |
|---|---|---|---|---|---|---|
| ZNF143 | 0.17105 | 0.39376 | 0.15 | 0.11 | 1 | HC_vs_MS.Trt_F |
| UACA | 0.17108 | 0.52675 | 0.532 | 0.47 | 1 | HC_vs_MS.Trt_F |
| DNM1 | 0.17119 | 0.39531 | 0.65 | 0.59 | 1 | HC_vs_MS.Trt_F |
| FEM1B | 0.17121 | 0.87806 | 0.124 | 0.09 | 1 | HC_vs_MS.Trt_F |
| ZNF74 | 0.17139 | 1.31165 | 0.494 | 0.45 | 1 | HC_vs_MS.Trt_F |
| CXorf56 | 0.17141 | -0.3198 | 0.78 | 0.7 | 1 | HC_vs_MS.Trt_F |
| ZNF286B | 0.17144 | -0.8736 | 0.558 | 0.47 | 1 | HC_vs_MS.Trt_F |
| NIPAL3 | 0.17169 | -0.4832 | 0.766 | 0.76 | 1 | HC_vs_MS.Trt_F |
| KIAA1109 | 0.17186 | -0.3688 | 0.373 | 0.31 | 1 | HC_vs_MS.Trt_F |
| ADGRD1 | 0.17193 | 0.65577 | 0.436 | 0.37 | 1 | HC_vs_MS.Trt_F |
| LRRC28 | 0.17209 | 0.30298 | 0.624 | 0.54 | 1 | HC_vs_MS.Trt_F |
| ATXN7L3B | 0.17214 | 0.39372 | 0.197 | 0.15 | 1 | HC_vs_MS.Trt_F |
| PRAMEF1 | 0.1725 | -0.2683 | 0.075 | 0.11 | 1 | HC_vs_MS.Trt_F |
| DAB2 | 0.17258 | -1.2725 | 0.191 | 0.23 | 1 | HC_vs_MS.Trt_F |
| C1QTNF2 | 0.1727 | 0.31304 | 0.344 | 0.28 | 1 | HC_vs_MS.Trt_F |
| RNF40 | 0.17272 | -0.811 | 0.098 | 0.13 | 1 | HC_vs_MS.Trt_F |
| BRAF | 0.17292 | 0.33058 | 0.191 | 0.15 | 1 | HC_vs_MS.Trt_F |
| WDR45 | 0.17293 | 0.34908 | 0.647 | 0.58 | 1 | HC_vs_MS.Trt_F |
| ZNF2 | 0.17315 | -1.6707 | 0.382 | 0.31 | 1 | HC_vs_MS.Trt_F |
| MKL2 | 0.17347 | 0.43026 | 0.769 | 0.71 | 1 | HC_vs_MS.Trt_F |
| BCAT2 | 0.17361 | -0.5299 | 0.243 | 0.19 | 1 | HC_vs_MS.Trt_F |
| CBX1 | 0.17374 | -0.2868 | 0.332 | 0.27 | 1 | HC_vs_MS.Trt_F |
| GGCT | 0.1739 | 1.04675 | 0.11 | 0.08 | 1 | HC_vs_MS.Trt_F |
| CAD | 0.17394 | 0.7442 | 0.777 | 0.73 | 1 | HC_vs_MS.Trt_F |
| GLUL | 0.17403 | 0.78359 | 0.263 | 0.21 | 1 | HC_vs_MS.Trt_F |
| CXADR | 0.17426 | -0.2947 | 0.289 | 0.33 | 1 | HC_vs_MS.Trt_F |
| ZC3H3 | 0.17428 | 0.42946 | 0.153 | 0.12 | 1 | HC_vs_MS.Trt_F |
| RAD21 | 0.17468 | -0.4357 | 0.277 | 0.31 | 1 | HC_vs_MS.Trt_F |
| POLR2A | 0.17504 | 0.68285 | 0.844 | 0.82 | 1 | HC_vs_MS.Trt_F |
| SERINC5 | 0.17521 | -0.3358 | 0.89 | 0.9 | 1 | HC_vs_MS.Trt_F |
| TBCA | 0.17566 | 0.26634 | 0.298 | 0.25 | 1 | HC_vs_MS.Trt_F |
| MIB2 | 0.17578 | -0.5551 | 0.202 | 0.25 | 1 | HC_vs_MS.Trt_F |
| RAB27A | 0.17579 | -0.2554 | 0.673 | 0.6 | 1 | HC_vs_MS.Trt_F |
| LMLN | 0.17593 | 0.3238 | 0.419 | 0.35 | 1 | HC_vs_MS.Trt_F |
| NLRP3 | 0.1761 | -0.7698 | 0.28 | 0.23 | 1 | HC_vs_MS.Trt_F |
| SEPHS2 | 0.17617 | 0.82433 | 0.145 | 0.11 | 1 | HC_vs_MS.Trt_F |
| RABEPK | 0.17621 | -0.3368 | 0.85 | 0.8 | 1 | HC_vs_MS.Trt_F |
| SETD4 | 0.17637 | -0.7803 | 0.564 | 0.51 | 1 | HC_vs_MS.Trt_F |
| SLC2A4RG | 0.1764 | 0.74913 | 0.11 | 0.08 | 1 | HC_vs_MS.Trt_F |
| HSD17B13 | 0.17641 | 0.5341 | 0.636 | 0.57 | 1 | HC_vs_MS.Trt_F |

| | | | | | | |
|---|---|---|---|---|---|---|
| TNS2 | 0.17649 | 1.04923 | 0.39 | 0.34 | 1 | HC_vs_MS.Trt_F |
| LARP7 | 0.17668 | -0.5476 | 0.257 | 0.2 | 1 | HC_vs_MS.Trt_F |
| CTR9 | 0.17671 | -0.4145 | 0.205 | 0.16 | 1 | HC_vs_MS.Trt_F |
| RUVBL1 | 0.17702 | -0.417 | 0.711 | 0.73 | 1 | HC_vs_MS.Trt_F |
| IPO11 | 0.17716 | 0.54332 | 0.173 | 0.13 | 1 | HC_vs_MS.Trt_F |
| DDAH2 | 0.17747 | -0.6416 | 0.118 | 0.08 | 1 | HC_vs_MS.Trt_F |
| ITFG1 | 0.17758 | 0.59875 | 0.211 | 0.17 | 1 | HC_vs_MS.Trt_F |
| RP11-577H5 | 0.1776 | -1.0644 | 0.104 | 0.13 | 1 | HC_vs_MS.Trt_F |
| PAPOLG | 0.17792 | -0.3545 | 0.827 | 0.73 | 1 | HC_vs_MS.Trt_F |
| BCDIN3D | 0.17804 | 0.68227 | 0.454 | 0.39 | 1 | HC_vs_MS.Trt_F |
| UBN1 | 0.17813 | -0.5354 | 0.194 | 0.15 | 1 | HC_vs_MS.Trt_F |
| VN1R2 | 0.17816 | 0.31656 | 0.11 | 0.08 | 1 | HC_vs_MS.Trt_F |
| MRPL52 | 0.17841 | -0.2733 | 0.682 | 0.59 | 1 | HC_vs_MS.Trt_F |
| SNUPN | 0.17854 | -0.845 | 0.887 | 0.79 | 1 | HC_vs_MS.Trt_F |
| RTCA | 0.1787 | -0.2684 | 0.645 | 0.58 | 1 | HC_vs_MS.Trt_F |
| SLC25A20 | 0.17917 | 0.66736 | 0.171 | 0.13 | 1 | HC_vs_MS.Trt_F |
| ANO8 | 0.17921 | 0.3425 | 0.136 | 0.1 | 1 | HC_vs_MS.Trt_F |
| VGLL4 | 0.17937 | 0.29182 | 0.951 | 0.94 | 1 | HC_vs_MS.Trt_F |
| CERS4 | 0.17938 | 0.57067 | 0.659 | 0.59 | 1 | HC_vs_MS.Trt_F |
| EFCAB5 | 0.17939 | 0.67046 | 0.503 | 0.44 | 1 | HC_vs_MS.Trt_F |
| TACR2 | 0.17945 | 0.59142 | 0.121 | 0.09 | 1 | HC_vs_MS.Trt_F |
| ATG9B | 0.17948 | -0.7729 | 0.124 | 0.16 | 1 | HC_vs_MS.Trt_F |
| RAD9B | 0.17979 | 0.28538 | 0.471 | 0.4 | 1 | HC_vs_MS.Trt_F |
| RSBN1 | 0.17979 | 0.30498 | 0.275 | 0.22 | 1 | HC_vs_MS.Trt_F |
| ZNF532 | 0.18016 | 0.26151 | 0.942 | 0.94 | 1 | HC_vs_MS.Trt_F |
| VPS54 | 0.18017 | -1.1392 | 0.419 | 0.36 | 1 | HC_vs_MS.Trt_F |
| PDE7B | 0.18027 | 0.30801 | 0.448 | 0.41 | 1 | HC_vs_MS.Trt_F |
| KLHL24 | 0.18074 | -0.38 | 0.882 | 0.85 | 1 | HC_vs_MS.Trt_F |
| USP1 | 0.18108 | -0.3629 | 0.197 | 0.15 | 1 | HC_vs_MS.Trt_F |
| RFX5 | 0.18119 | -0.3847 | 0.393 | 0.33 | 1 | HC_vs_MS.Trt_F |
| CYP51A1 | 0.18135 | -0.4182 | 0.532 | 0.46 | 1 | HC_vs_MS.Trt_F |
| ANKHD1 | 0.18142 | 0.28018 | 0.494 | 0.45 | 1 | HC_vs_MS.Trt_F |
| WDR76 | 0.18204 | -0.7836 | 0.656 | 0.69 | 1 | HC_vs_MS.Trt_F |
| LRRC59 | 0.18204 | 0.56457 | 0.176 | 0.14 | 1 | HC_vs_MS.Trt_F |
| UCHL5 | 0.18204 | -0.5312 | 0.223 | 0.17 | 1 | HC_vs_MS.Trt_F |
| AOAH | 0.18228 | -0.812 | 0.428 | 0.36 | 1 | HC_vs_MS.Trt_F |
| SFXN2 | 0.18249 | 0.84724 | 0.925 | 0.92 | 1 | HC_vs_MS.Trt_F |
| UBA2 | 0.18254 | 0.61269 | 0.624 | 0.57 | 1 | HC_vs_MS.Trt_F |
| GMEB1 | 0.18294 | 0.4761 | 0.821 | 0.79 | 1 | HC_vs_MS.Trt_F |
| PPME1 | 0.18302 | 0.57384 | 0.182 | 0.14 | 1 | HC_vs_MS.Trt_F |

| | | | | | | |
|---|---|---|---|---|---|---|
| C1orf112 | 0.18306 | -0.4042 | 0.546 | 0.47 | 1 | HC_vs_MS.Trt_F |
| ENPP2 | 0.18358 | 1.3484 | 0.156 | 0.12 | 1 | HC_vs_MS.Trt_F |
| WRN | 0.18379 | -0.6613 | 0.587 | 0.49 | 1 | HC_vs_MS.Trt_F |
| SURF2 | 0.18392 | -0.6338 | 0.13 | 0.16 | 1 | HC_vs_MS.Trt_F |
| XPO6 | 0.18436 | -0.577 | 0.286 | 0.32 | 1 | HC_vs_MS.Trt_F |
| SCAPER | 0.18453 | -0.3411 | 0.402 | 0.34 | 1 | HC_vs_MS.Trt_F |
| IBA57 | 0.18491 | 0.56531 | 0.829 | 0.81 | 1 | HC_vs_MS.Trt_F |
| PREPL | 0.18513 | 0.34559 | 0.708 | 0.65 | 1 | HC_vs_MS.Trt_F |
| PIGL | 0.18521 | 0.56876 | 0.714 | 0.67 | 1 | HC_vs_MS.Trt_F |
| RRP1 | 0.18529 | -0.5083 | 0.711 | 0.65 | 1 | HC_vs_MS.Trt_F |
| KBTBD3 | 0.18532 | -0.896 | 0.075 | 0.1 | 1 | HC_vs_MS.Trt_F |
| ARPC5 | 0.18577 | -0.4691 | 0.96 | 0.89 | 1 | HC_vs_MS.Trt_F |
| TPX2 | 0.18586 | -0.4032 | 0.286 | 0.23 | 1 | HC_vs_MS.Trt_F |
| AGBL3 | 0.18598 | -0.3685 | 0.28 | 0.23 | 1 | HC_vs_MS.Trt_F |
| MCOLN2 | 0.18643 | 0.25969 | 0.509 | 0.43 | 1 | HC_vs_MS.Trt_F |
| TMEM94 | 0.18677 | -0.4176 | 0.272 | 0.22 | 1 | HC_vs_MS.Trt_F |
| CLN6 | 0.18677 | 0.35356 | 0.824 | 0.79 | 1 | HC_vs_MS.Trt_F |
| TCP11L2 | 0.18702 | 0.42044 | 0.295 | 0.25 | 1 | HC_vs_MS.Trt_F |
| BTNL9 | 0.18716 | 0.39941 | 0.931 | 0.85 | 1 | HC_vs_MS.Trt_F |
| TSPYL1 | 0.1873 | -0.4979 | 0.639 | 0.62 | 1 | HC_vs_MS.Trt_F |
| DENND1A | 0.18752 | 0.43489 | 0.558 | 0.49 | 1 | HC_vs_MS.Trt_F |
| MKNK1 | 0.18754 | -0.3557 | 0.945 | 0.92 | 1 | HC_vs_MS.Trt_F |
| TRANK1 | 0.18755 | 0.66778 | 0.263 | 0.22 | 1 | HC_vs_MS.Trt_F |
| ZMAT1 | 0.1878 | 0.70822 | 0.39 | 0.34 | 1 | HC_vs_MS.Trt_F |
| DCAF13 | 0.18785 | -0.6036 | 0.766 | 0.66 | 1 | HC_vs_MS.Trt_F |
| TARDBP | 0.18787 | 0.351 | 0.636 | 0.59 | 1 | HC_vs_MS.Trt_F |
| HSPBAP1 | 0.18855 | 0.30576 | 0.584 | 0.53 | 1 | HC_vs_MS.Trt_F |
| SPAG9 | 0.18909 | 0.57243 | 0.913 | 0.88 | 1 | HC_vs_MS.Trt_F |
| ACVR2A | 0.18923 | -0.5916 | 0.315 | 0.25 | 1 | HC_vs_MS.Trt_F |
| HHIP | 0.18961 | 0.33146 | 0.405 | 0.35 | 1 | HC_vs_MS.Trt_F |
| SCUBE3 | 0.18967 | 0.39843 | 0.691 | 0.63 | 1 | HC_vs_MS.Trt_F |
| TLR6 | 0.18989 | -0.3417 | 0.309 | 0.26 | 1 | HC_vs_MS.Trt_F |
| CYB5R2 | 0.18991 | 0.54716 | 0.575 | 0.48 | 1 | HC_vs_MS.Trt_F |
| ENSA | 0.19 | -0.3088 | 0.821 | 0.82 | 1 | HC_vs_MS.Trt_F |
| NRAS | 0.1904 | 0.83184 | 0.361 | 0.3 | 1 | HC_vs_MS.Trt_F |
| CD101 | 0.19094 | -0.8835 | 0.075 | 0.1 | 1 | HC_vs_MS.Trt_F |
| NT5DC1 | 0.19129 | 0.77508 | 0.223 | 0.17 | 1 | HC_vs_MS.Trt_F |
| ANO10 | 0.19195 | -0.3913 | 0.671 | 0.62 | 1 | HC_vs_MS.Trt_F |
| WDR53 | 0.19203 | 0.32194 | 0.116 | 0.08 | 1 | HC_vs_MS.Trt_F |
| NCAPH2 | 0.19223 | 0.42274 | 0.757 | 0.71 | 1 | HC_vs_MS.Trt_F |

| | | | | | | |
|---|---|---|---|---|---|---|
| GPR155 | 0.19257 | 0.36163 | 0.939 | 0.9 | 1 | HC_vs_MS.Trt_F |
| ABCB10 | 0.19263 | -1.057 | 0.101 | 0.13 | 1 | HC_vs_MS.Trt_F |
| PDZD8 | 0.19267 | -0.6863 | 0.277 | 0.32 | 1 | HC_vs_MS.Trt_F |
| FOXK1 | 0.19273 | -0.4798 | 0.879 | 0.85 | 1 | HC_vs_MS.Trt_F |
| PPM1F | 0.19296 | 0.51914 | 0.832 | 0.76 | 1 | HC_vs_MS.Trt_F |
| ZNF845 | 0.19315 | 0.31153 | 0.711 | 0.65 | 1 | HC_vs_MS.Trt_F |
| HOMEZ | 0.19345 | -1.3012 | 0.26 | 0.3 | 1 | HC_vs_MS.Trt_F |
| MTR | 0.1937 | -0.6596 | 0.569 | 0.49 | 1 | HC_vs_MS.Trt_F |
| BLOC1S1 | 0.19372 | -0.3305 | 0.254 | 0.2 | 1 | HC_vs_MS.Trt_F |
| FMNL1 | 0.19372 | 0.42988 | 0.714 | 0.68 | 1 | HC_vs_MS.Trt_F |
| VTI1A | 0.19389 | -0.3972 | 0.902 | 0.87 | 1 | HC_vs_MS.Trt_F |
| USP46 | 0.1939 | 0.37755 | 0.731 | 0.68 | 1 | HC_vs_MS.Trt_F |
| SPRYD3 | 0.19394 | 0.58053 | 0.197 | 0.16 | 1 | HC_vs_MS.Trt_F |
| ACSL5 | 0.19437 | 0.37905 | 0.225 | 0.18 | 1 | HC_vs_MS.Trt_F |
| TAF10 | 0.19443 | 0.92189 | 0.133 | 0.1 | 1 | HC_vs_MS.Trt_F |
| GPX2 | 0.19446 | 0.8771 | 0.211 | 0.16 | 1 | HC_vs_MS.Trt_F |
| C2CD2 | 0.1948 | 0.69714 | 0.266 | 0.22 | 1 | HC_vs_MS.Trt_F |
| ANKRD39 | 0.19501 | -0.5235 | 0.214 | 0.25 | 1 | HC_vs_MS.Trt_F |
| FAM32A | 0.19509 | -0.3452 | 0.494 | 0.42 | 1 | HC_vs_MS.Trt_F |
| SPDYE3 | 0.19515 | 0.43002 | 0.506 | 0.59 | 1 | HC_vs_MS.Trt_F |
| RRAGA | 0.1953 | -0.4056 | 0.185 | 0.14 | 1 | HC_vs_MS.Trt_F |
| GGCX | 0.19537 | 0.27112 | 0.934 | 0.9 | 1 | HC_vs_MS.Trt_F |
| FGFR1 | 0.19542 | 0.36749 | 0.827 | 0.8 | 1 | HC_vs_MS.Trt_F |
| AHCYL2 | 0.19548 | 1.83426 | 0.194 | 0.16 | 1 | HC_vs_MS.Trt_F |
| IDH3G | 0.19556 | 0.26349 | 0.197 | 0.16 | 1 | HC_vs_MS.Trt_F |
| ZNF680 | 0.19577 | -0.7523 | 0.139 | 0.17 | 1 | HC_vs_MS.Trt_F |
| BBS1 | 0.19626 | -0.2728 | 0.656 | 0.61 | 1 | HC_vs_MS.Trt_F |
| RAB21 | 0.19628 | 0.40106 | 0.983 | 0.97 | 1 | HC_vs_MS.Trt_F |
| TXLNA | 0.1963 | -0.5423 | 0.373 | 0.39 | 1 | HC_vs_MS.Trt_F |
| LTB | 0.19635 | -0.365 | 0.584 | 0.59 | 1 | HC_vs_MS.Trt_F |
| BCOR | 0.19656 | -0.2768 | 0.217 | 0.17 | 1 | HC_vs_MS.Trt_F |
| MIER1 | 0.19675 | -0.2773 | 0.743 | 0.66 | 1 | HC_vs_MS.Trt_F |
| GPR137 | 0.19677 | -0.2875 | 0.292 | 0.24 | 1 | HC_vs_MS.Trt_F |
| SLC9B1 | 0.19689 | 0.39999 | 0.419 | 0.37 | 1 | HC_vs_MS.Trt_F |
| ESCO1 | 0.19691 | -0.3628 | 0.439 | 0.38 | 1 | HC_vs_MS.Trt_F |
| FSD2 | 0.19691 | 0.52221 | 0.488 | 0.46 | 1 | HC_vs_MS.Trt_F |
| SNRPD3 | 0.19697 | -0.4149 | 0.803 | 0.72 | 1 | HC_vs_MS.Trt_F |
| HBB | 0.19711 | -4.1882 | 0.075 | 0.1 | 1 | HC_vs_MS.Trt_F |
| DPM1 | 0.19719 | 0.72731 | 0.136 | 0.1 | 1 | HC_vs_MS.Trt_F |
| ABCC2 | 0.19801 | 0.25274 | 0.514 | 0.58 | 1 | HC_vs_MS.Trt_F |

| | | | | | | |
|---|---|---|---|---|---|---|
| ITGAX | 0.19827 | 1.255 | 0.558 | 0.48 | 1 | HC_vs_MS.Trt_F |
| MBD1 | 0.19847 | -0.3058 | 0.217 | 0.17 | 1 | HC_vs_MS.Trt_F |
| ARHGAP4 | 0.19855 | 0.33253 | 0.497 | 0.48 | 1 | HC_vs_MS.Trt_F |
| RNF38 | 0.19874 | 0.3263 | 0.237 | 0.19 | 1 | HC_vs_MS.Trt_F |
| VHL | 0.19905 | -0.2711 | 0.815 | 0.75 | 1 | HC_vs_MS.Trt_F |
| KDELR2 | 0.19908 | -0.318 | 0.208 | 0.16 | 1 | HC_vs_MS.Trt_F |
| PITPNA | 0.19919 | 1.13612 | 0.13 | 0.1 | 1 | HC_vs_MS.Trt_F |
| ZNF250 | 0.19931 | -1.2683 | 0.671 | 0.59 | 1 | HC_vs_MS.Trt_F |
| DPAGT1 | 0.19948 | -0.3877 | 0.084 | 0.11 | 1 | HC_vs_MS.Trt_F |
| CIAO1 | 0.19957 | -0.3878 | 0.355 | 0.29 | 1 | HC_vs_MS.Trt_F |
| RAP2C | 0.19962 | -0.2669 | 0.324 | 0.26 | 1 | HC_vs_MS.Trt_F |
| MAP3K13 | 0.19969 | 0.39595 | 0.85 | 0.82 | 1 | HC_vs_MS.Trt_F |
| RBM44 | 0.19989 | -0.724 | 0.231 | 0.27 | 1 | HC_vs_MS.Trt_F |
| CSGALNACT | 0.20042 | 0.39467 | 0.191 | 0.15 | 1 | HC_vs_MS.Trt_F |
| E2F6 | 0.20045 | 0.69012 | 0.272 | 0.31 | 1 | HC_vs_MS.Trt_F |
| FYTTD1 | 0.20064 | -0.5696 | 0.838 | 0.76 | 1 | HC_vs_MS.Trt_F |
| FCGRT | 0.20082 | 0.39189 | 0.237 | 0.19 | 1 | HC_vs_MS.Trt_F |
| BPNT1 | 0.20104 | -0.4903 | 0.775 | 0.68 | 1 | HC_vs_MS.Trt_F |
| NCAPD3 | 0.20106 | 1.7522 | 0.11 | 0.08 | 1 | HC_vs_MS.Trt_F |
| DCUN1D4 | 0.20111 | -0.3195 | 0.35 | 0.29 | 1 | HC_vs_MS.Trt_F |
| SLC35G2 | 0.20146 | -0.861 | 0.185 | 0.22 | 1 | HC_vs_MS.Trt_F |
| NAXD | 0.20154 | -1.13 | 0.087 | 0.12 | 1 | HC_vs_MS.Trt_F |
| CENPM | 0.20166 | -0.7078 | 0.382 | 0.31 | 1 | HC_vs_MS.Trt_F |
| SPAG4 | 0.20182 | 0.55498 | 0.28 | 0.25 | 1 | HC_vs_MS.Trt_F |
| TMEM256-P | 0.20186 | -0.3722 | 0.48 | 0.49 | 1 | HC_vs_MS.Trt_F |
| 5-Mar | 0.20193 | 0.38478 | 0.251 | 0.21 | 1 | HC_vs_MS.Trt_F |
| MYO1D | 0.2021 | 1.51714 | 0.202 | 0.17 | 1 | HC_vs_MS.Trt_F |
| ZNF85 | 0.20211 | -0.8036 | 0.72 | 0.65 | 1 | HC_vs_MS.Trt_F |
| MRPS2 | 0.20221 | 0.97845 | 0.101 | 0.07 | 1 | HC_vs_MS.Trt_F |
| BRF1 | 0.20244 | 0.63774 | 0.624 | 0.59 | 1 | HC_vs_MS.Trt_F |
| ANKZF1 | 0.20278 | 0.33789 | 0.835 | 0.82 | 1 | HC_vs_MS.Trt_F |
| HECTD1 | 0.20291 | 0.32687 | 0.587 | 0.52 | 1 | HC_vs_MS.Trt_F |
| KIF9 | 0.20301 | 0.51902 | 0.173 | 0.14 | 1 | HC_vs_MS.Trt_F |
| PLEKHG2 | 0.20312 | 0.52162 | 0.916 | 0.86 | 1 | HC_vs_MS.Trt_F |
| SPATA7 | 0.20317 | -0.2566 | 0.341 | 0.3 | 1 | HC_vs_MS.Trt_F |
| SS18 | 0.20324 | -0.3883 | 0.647 | 0.56 | 1 | HC_vs_MS.Trt_F |
| FNIP1 | 0.20325 | -0.443 | 0.587 | 0.63 | 1 | HC_vs_MS.Trt_F |
| WDR45B | 0.2033 | 0.2596 | 0.422 | 0.38 | 1 | HC_vs_MS.Trt_F |
| MYO15B | 0.20334 | 0.27803 | 0.246 | 0.2 | 1 | HC_vs_MS.Trt_F |
| VAPB | 0.20381 | -0.8187 | 0.217 | 0.17 | 1 | HC_vs_MS.Trt_F |

| | | | | | | |
|---|---|---|---|---|---|---|
| SUPT6H | 0.20394 | 0.37165 | 0.679 | 0.62 | 1 | HC_vs_MS.Trt_F |
| PSMG3 | 0.20416 | -0.9204 | 0.682 | 0.59 | 1 | HC_vs_MS.Trt_F |
| CEACAM4 | 0.20457 | 0.62037 | 0.136 | 0.1 | 1 | HC_vs_MS.Trt_F |
| LYRM1 | 0.20483 | -0.6316 | 0.616 | 0.53 | 1 | HC_vs_MS.Trt_F |
| MYOM3 | 0.20504 | 0.26071 | 0.633 | 0.6 | 1 | HC_vs_MS.Trt_F |
| TGFBR1 | 0.20513 | -0.3345 | 0.162 | 0.12 | 1 | HC_vs_MS.Trt_F |
| VNN2 | 0.20544 | -0.8484 | 0.188 | 0.22 | 1 | HC_vs_MS.Trt_F |
| ZBTB1 | 0.20568 | -0.2792 | 0.231 | 0.18 | 1 | HC_vs_MS.Trt_F |
| ERLEC1 | 0.20581 | 0.54881 | 0.413 | 0.36 | 1 | HC_vs_MS.Trt_F |
| GHDC | 0.20589 | -0.8408 | 0.335 | 0.28 | 1 | HC_vs_MS.Trt_F |
| NBPF12 | 0.2059 | -0.3142 | 0.347 | 0.37 | 1 | HC_vs_MS.Trt_F |
| SHANK1 | 0.20639 | 0.40824 | 0.329 | 0.3 | 1 | HC_vs_MS.Trt_F |
| AAMP | 0.20639 | -0.5575 | 0.234 | 0.18 | 1 | HC_vs_MS.Trt_F |
| KCNN3 | 0.20649 | -0.4841 | 0.474 | 0.5 | 1 | HC_vs_MS.Trt_F |
| PAQR8 | 0.20651 | -0.4664 | 0.607 | 0.51 | 1 | HC_vs_MS.Trt_F |
| LARP1B | 0.20669 | -1.5974 | 0.075 | 0.1 | 1 | HC_vs_MS.Trt_F |
| DCBLD1 | 0.20676 | 0.58022 | 0.61 | 0.54 | 1 | HC_vs_MS.Trt_F |
| LPAR2 | 0.20677 | -0.5352 | 0.165 | 0.13 | 1 | HC_vs_MS.Trt_F |
| NFXL1 | 0.20682 | 1.23986 | 0.145 | 0.11 | 1 | HC_vs_MS.Trt_F |
| TARS2 | 0.20688 | 0.27453 | 0.809 | 0.81 | 1 | HC_vs_MS.Trt_F |
| NEUROD1 | 0.20721 | 0.27832 | 0.113 | 0.08 | 1 | HC_vs_MS.Trt_F |
| TIMD4 | 0.20735 | 0.65753 | 0.101 | 0.07 | 1 | HC_vs_MS.Trt_F |
| DNAJB4 | 0.20736 | -0.2569 | 0.454 | 0.4 | 1 | HC_vs_MS.Trt_F |
| NFATC2 | 0.20751 | 0.32212 | 0.234 | 0.19 | 1 | HC_vs_MS.Trt_F |
| GFOD2 | 0.20782 | -0.2836 | 0.931 | 0.89 | 1 | HC_vs_MS.Trt_F |
| SMARCD2 | 0.20825 | -0.8397 | 0.118 | 0.15 | 1 | HC_vs_MS.Trt_F |
| ASRGL1 | 0.20864 | 0.40288 | 0.607 | 0.54 | 1 | HC_vs_MS.Trt_F |
| KLLN | 0.20867 | -0.4036 | 0.497 | 0.43 | 1 | HC_vs_MS.Trt_F |
| WDR5B | 0.20898 | 1.21871 | 0.24 | 0.19 | 1 | HC_vs_MS.Trt_F |
| TPRG1L | 0.20907 | 0.82613 | 0.298 | 0.26 | 1 | HC_vs_MS.Trt_F |
| PES1 | 0.20911 | 0.91136 | 0.142 | 0.11 | 1 | HC_vs_MS.Trt_F |
| GTF2H3 | 0.20913 | -0.6295 | 0.786 | 0.69 | 1 | HC_vs_MS.Trt_F |
| SRM | 0.20931 | -0.4227 | 0.199 | 0.16 | 1 | HC_vs_MS.Trt_F |
| NDC80 | 0.20954 | -0.7962 | 0.743 | 0.66 | 1 | HC_vs_MS.Trt_F |
| RPRD2 | 0.2096 | 0.29539 | 0.277 | 0.23 | 1 | HC_vs_MS.Trt_F |
| MAEL | 0.20979 | 0.29291 | 0.529 | 0.57 | 1 | HC_vs_MS.Trt_F |
| PDXK | 0.20992 | 0.2834 | 0.63 | 0.53 | 1 | HC_vs_MS.Trt_F |
| SP4 | 0.20996 | -0.5341 | 0.448 | 0.38 | 1 | HC_vs_MS.Trt_F |
| STK24 | 0.21013 | -0.59 | 0.347 | 0.29 | 1 | HC_vs_MS.Trt_F |
| ANKRD40 | 0.21028 | -0.7722 | 0.824 | 0.82 | 1 | HC_vs_MS.Trt_F |

| | | | | | | |
|---|---|---|---|---|---|---|
| MAP3K14 | 0.21043 | 0.28527 | 0.199 | 0.16 | 1 | HC_vs_MS.Trt_F |
| VAMP7 | 0.2105 | -0.5562 | 0.127 | 0.09 | 1 | HC_vs_MS.Trt_F |
| LRP2BP | 0.21058 | 0.99001 | 0.48 | 0.42 | 1 | HC_vs_MS.Trt_F |
| NOTCH1 | 0.21062 | 0.95001 | 0.156 | 0.12 | 1 | HC_vs_MS.Trt_F |
| GPA33 | 0.21136 | 0.93019 | 0.101 | 0.07 | 1 | HC_vs_MS.Trt_F |
| RPAP1 | 0.21136 | -0.4565 | 0.855 | 0.77 | 1 | HC_vs_MS.Trt_F |
| ZNF496 | 0.21152 | 0.66752 | 0.769 | 0.72 | 1 | HC_vs_MS.Trt_F |
| C3orf58 | 0.21153 | -1.0851 | 0.104 | 0.13 | 1 | HC_vs_MS.Trt_F |
| ZNF268 | 0.21155 | 0.37795 | 0.986 | 0.95 | 1 | HC_vs_MS.Trt_F |
| LANCL1 | 0.21155 | -0.2573 | 0.179 | 0.14 | 1 | HC_vs_MS.Trt_F |
| CREBRF | 0.21163 | -0.4991 | 0.777 | 0.69 | 1 | HC_vs_MS.Trt_F |
| ZNF783 | 0.2119 | 1.10113 | 0.442 | 0.38 | 1 | HC_vs_MS.Trt_F |
| PBX3 | 0.21202 | 0.61978 | 0.116 | 0.09 | 1 | HC_vs_MS.Trt_F |
| ATP6AP2 | 0.2124 | -0.3202 | 0.89 | 0.81 | 1 | HC_vs_MS.Trt_F |
| TCF3 | 0.21253 | 0.79419 | 0.11 | 0.08 | 1 | HC_vs_MS.Trt_F |
| CEP250 | 0.21282 | -0.7153 | 0.844 | 0.78 | 1 | HC_vs_MS.Trt_F |
| NAIP | 0.21284 | -0.3011 | 0.743 | 0.63 | 1 | HC_vs_MS.Trt_F |
| PTPRJ | 0.21292 | 0.32296 | 0.41 | 0.37 | 1 | HC_vs_MS.Trt_F |
| TEPSIN | 0.21297 | 0.67236 | 0.15 | 0.12 | 1 | HC_vs_MS.Trt_F |
| ZDHHC5 | 0.21321 | 0.75781 | 0.142 | 0.11 | 1 | HC_vs_MS.Trt_F |
| ESRRA | 0.21332 | 1.00579 | 0.127 | 0.1 | 1 | HC_vs_MS.Trt_F |
| INSL6 | 0.21343 | 0.37471 | 0.338 | 0.3 | 1 | HC_vs_MS.Trt_F |
| ZNF461 | 0.21376 | -0.7103 | 0.705 | 0.66 | 1 | HC_vs_MS.Trt_F |
| MCEE | 0.21388 | 0.37954 | 0.214 | 0.18 | 1 | HC_vs_MS.Trt_F |
| GPATCH2 | 0.21397 | -0.816 | 0.54 | 0.6 | 1 | HC_vs_MS.Trt_F |
| OLA1 | 0.21426 | -0.2707 | 0.494 | 0.42 | 1 | HC_vs_MS.Trt_F |
| SLC7A13 | 0.2143 | -0.436 | 0.121 | 0.16 | 1 | HC_vs_MS.Trt_F |
| NARF | 0.2145 | 0.46675 | 0.965 | 0.95 | 1 | HC_vs_MS.Trt_F |
| CTRL | 0.21486 | -0.3033 | 0.133 | 0.1 | 1 | HC_vs_MS.Trt_F |
| COPS5 | 0.21491 | 0.38708 | 0.893 | 0.82 | 1 | HC_vs_MS.Trt_F |
| WDR11 | 0.21509 | 0.6387 | 0.673 | 0.62 | 1 | HC_vs_MS.Trt_F |
| TTC22 | 0.21511 | 0.75598 | 0.494 | 0.43 | 1 | HC_vs_MS.Trt_F |
| DEPDC5 | 0.21517 | -0.4678 | 0.777 | 0.81 | 1 | HC_vs_MS.Trt_F |
| BICD1 | 0.21521 | 0.59385 | 0.118 | 0.09 | 1 | HC_vs_MS.Trt_F |
| CASP2 | 0.21562 | -0.4446 | 0.818 | 0.81 | 1 | HC_vs_MS.Trt_F |
| DMXL1 | 0.2157 | -0.3498 | 0.24 | 0.19 | 1 | HC_vs_MS.Trt_F |
| BRD3 | 0.21625 | -0.4599 | 0.237 | 0.18 | 1 | HC_vs_MS.Trt_F |
| AIDA | 0.21627 | 0.50385 | 0.139 | 0.1 | 1 | HC_vs_MS.Trt_F |
| ITGB8 | 0.21642 | 0.40849 | 0.78 | 0.81 | 1 | HC_vs_MS.Trt_F |
| ZFP91 | 0.21658 | 0.4313 | 0.104 | 0.08 | 1 | HC_vs_MS.Trt_F |

| | | | | | | |
|---|---|---|---|---|---|---|
| C1QTNF1 | 0.21761 | -0.5423 | 0.275 | 0.22 | 1 | HC_vs_MS.Trt_F |
| KIFAP3 | 0.21765 | -0.3988 | 0.188 | 0.15 | 1 | HC_vs_MS.Trt_F |
| GNB2 | 0.21781 | 0.70028 | 0.529 | 0.5 | 1 | HC_vs_MS.Trt_F |
| LGR4 | 0.21797 | 0.8129 | 0.075 | 0.1 | 1 | HC_vs_MS.Trt_F |
| BRI3BP | 0.21827 | 0.26045 | 0.902 | 0.87 | 1 | HC_vs_MS.Trt_F |
| MUM1 | 0.2186 | -0.4506 | 0.685 | 0.63 | 1 | HC_vs_MS.Trt_F |
| ARRDC3 | 0.21863 | -0.5458 | 0.454 | 0.48 | 1 | HC_vs_MS.Trt_F |
| LIMD1 | 0.21883 | 0.32269 | 0.879 | 0.83 | 1 | HC_vs_MS.Trt_F |
| FOXP4 | 0.21891 | 0.52882 | 0.422 | 0.36 | 1 | HC_vs_MS.Trt_F |
| RAD9A | 0.21892 | 0.29303 | 0.529 | 0.47 | 1 | HC_vs_MS.Trt_F |
| ENDOV | 0.21909 | 0.43493 | 0.659 | 0.6 | 1 | HC_vs_MS.Trt_F |
| B4GALT5 | 0.21919 | 0.53887 | 0.286 | 0.34 | 1 | HC_vs_MS.Trt_F |
| MPST | 0.21943 | -0.4782 | 0.887 | 0.82 | 1 | HC_vs_MS.Trt_F |
| PTCD3 | 0.2195 | -0.2922 | 0.922 | 0.9 | 1 | HC_vs_MS.Trt_F |
| BLOC1S4 | 0.21961 | 0.36013 | 0.171 | 0.21 | 1 | HC_vs_MS.Trt_F |
| CSF2RB | 0.21997 | 0.63337 | 0.124 | 0.09 | 1 | HC_vs_MS.Trt_F |
| CASC4 | 0.22009 | 0.38188 | 0.867 | 0.88 | 1 | HC_vs_MS.Trt_F |
| VPS33B | 0.22014 | 1.09327 | 0.416 | 0.48 | 1 | HC_vs_MS.Trt_F |
| SIK3 | 0.22044 | 0.34127 | 0.367 | 0.31 | 1 | HC_vs_MS.Trt_F |
| POP4 | 0.22104 | -0.332 | 0.168 | 0.13 | 1 | HC_vs_MS.Trt_F |
| ACPP | 0.22111 | 0.3301 | 0.225 | 0.19 | 1 | HC_vs_MS.Trt_F |
| FBLN1 | 0.22145 | 0.59058 | 0.188 | 0.15 | 1 | HC_vs_MS.Trt_F |
| AKR1C1 | 0.2216 | 0.41951 | 0.306 | 0.27 | 1 | HC_vs_MS.Trt_F |
| LAPTM4B | 0.22163 | 1.43827 | 0.225 | 0.18 | 1 | HC_vs_MS.Trt_F |
| TMEM120A | 0.22175 | -0.3144 | 0.179 | 0.14 | 1 | HC_vs_MS.Trt_F |
| MUTYH | 0.2218 | 0.55588 | 0.561 | 0.51 | 1 | HC_vs_MS.Trt_F |
| FZR1 | 0.22183 | 0.58276 | 0.168 | 0.13 | 1 | HC_vs_MS.Trt_F |
| CTB-133G6.: | 0.22184 | -0.6149 | 0.194 | 0.22 | 1 | HC_vs_MS.Trt_F |
| OTULIN | 0.22207 | 0.51065 | 0.497 | 0.44 | 1 | HC_vs_MS.Trt_F |
| STARD4 | 0.22231 | -0.9553 | 0.211 | 0.17 | 1 | HC_vs_MS.Trt_F |
| POLR2E | 0.22246 | -0.5782 | 0.564 | 0.58 | 1 | HC_vs_MS.Trt_F |
| ZNF584 | 0.22253 | 0.67282 | 0.746 | 0.69 | 1 | HC_vs_MS.Trt_F |
| SERTAD1 | 0.22261 | -0.2916 | 0.442 | 0.37 | 1 | HC_vs_MS.Trt_F |
| RNF141 | 0.22293 | 0.46091 | 0.445 | 0.4 | 1 | HC_vs_MS.Trt_F |
| LHPP | 0.22327 | 0.65509 | 0.15 | 0.12 | 1 | HC_vs_MS.Trt_F |
| GCA | 0.22342 | -0.8181 | 0.095 | 0.12 | 1 | HC_vs_MS.Trt_F |
| TAB1 | 0.22348 | 0.34719 | 0.121 | 0.09 | 1 | HC_vs_MS.Trt_F |
| THNSL1 | 0.2235 | -1.7367 | 0.153 | 0.12 | 1 | HC_vs_MS.Trt_F |
| HIPK2 | 0.22355 | -0.2906 | 0.13 | 0.09 | 1 | HC_vs_MS.Trt_F |
| TMEM164 | 0.22361 | 0.27037 | 0.107 | 0.08 | 1 | HC_vs_MS.Trt_F |

| | | | | | | |
|---|---|---|---|---|---|---|
| FCGR2B | 0.22401 | 0.915 | 0.075 | 0.1 | 1 | HC_vs_MS.Trt_F |
| COL15A1 | 0.22414 | 0.84246 | 0.425 | 0.36 | 1 | HC_vs_MS.Trt_F |
| TAPBP | 0.22424 | -0.2652 | 0.974 | 0.94 | 1 | HC_vs_MS.Trt_F |
| ALG12 | 0.2244 | 0.551 | 0.197 | 0.16 | 1 | HC_vs_MS.Trt_F |
| APPBP2 | 0.22441 | -0.4508 | 0.873 | 0.83 | 1 | HC_vs_MS.Trt_F |
| TMEM181 | 0.22444 | 0.30287 | 0.535 | 0.49 | 1 | HC_vs_MS.Trt_F |
| PLEKHG6 | 0.22524 | 0.35584 | 0.303 | 0.25 | 1 | HC_vs_MS.Trt_F |
| FBXO21 | 0.22541 | -0.3401 | 0.217 | 0.17 | 1 | HC_vs_MS.Trt_F |
| SIPA1L1 | 0.22549 | -0.4948 | 0.581 | 0.58 | 1 | HC_vs_MS.Trt_F |
| SPATA20 | 0.22573 | 0.96662 | 0.124 | 0.09 | 1 | HC_vs_MS.Trt_F |
| TRIM61 | 0.22576 | 0.50889 | 0.277 | 0.23 | 1 | HC_vs_MS.Trt_F |
| DLK2 | 0.22605 | -0.6518 | 0.332 | 0.36 | 1 | HC_vs_MS.Trt_F |
| PYCARD | 0.22613 | -0.9227 | 0.107 | 0.13 | 1 | HC_vs_MS.Trt_F |
| DCTN5 | 0.22637 | 0.36947 | 0.858 | 0.79 | 1 | HC_vs_MS.Trt_F |
| BRAT1 | 0.22641 | -0.274 | 0.338 | 0.29 | 1 | HC_vs_MS.Trt_F |
| TCAF1 | 0.2268 | 1.01273 | 0.387 | 0.34 | 1 | HC_vs_MS.Trt_F |
| GALE | 0.22689 | 0.32022 | 0.405 | 0.34 | 1 | HC_vs_MS.Trt_F |
| SFMBT2 | 0.22766 | 1.15415 | 0.379 | 0.34 | 1 | HC_vs_MS.Trt_F |
| CHML | 0.22771 | -1.1089 | 0.26 | 0.21 | 1 | HC_vs_MS.Trt_F |
| XPO4 | 0.22773 | -0.4326 | 0.147 | 0.11 | 1 | HC_vs_MS.Trt_F |
| AANAT | 0.22779 | 0.26379 | 0.101 | 0.13 | 1 | HC_vs_MS.Trt_F |
| ATP5S | 0.22782 | -0.6044 | 0.945 | 0.91 | 1 | HC_vs_MS.Trt_F |
| MMP25 | 0.22815 | 0.31526 | 0.197 | 0.16 | 1 | HC_vs_MS.Trt_F |
| UBQLN1 | 0.22848 | -0.4654 | 0.306 | 0.26 | 1 | HC_vs_MS.Trt_F |
| WEE1 | 0.22907 | -0.6788 | 0.769 | 0.69 | 1 | HC_vs_MS.Trt_F |
| CHST11 | 0.22925 | -0.3733 | 0.237 | 0.19 | 1 | HC_vs_MS.Trt_F |
| DIP2C | 0.22944 | 1.09754 | 0.116 | 0.09 | 1 | HC_vs_MS.Trt_F |
| ATXN1L | 0.22956 | 1.15495 | 0.127 | 0.1 | 1 | HC_vs_MS.Trt_F |
| SPPL2B | 0.22985 | 0.28143 | 0.416 | 0.37 | 1 | HC_vs_MS.Trt_F |
| SNIP1 | 0.23077 | -0.502 | 0.705 | 0.62 | 1 | HC_vs_MS.Trt_F |
| MDM2 | 0.23085 | -0.5127 | 0.988 | 0.99 | 1 | HC_vs_MS.Trt_F |
| SPATA33 | 0.23096 | 0.27815 | 0.668 | 0.59 | 1 | HC_vs_MS.Trt_F |
| SEC61A1 | 0.23104 | -0.6136 | 0.364 | 0.4 | 1 | HC_vs_MS.Trt_F |
| HCFC1R1 | 0.23144 | -0.3539 | 0.15 | 0.12 | 1 | HC_vs_MS.Trt_F |
| WDR20 | 0.23176 | -0.2959 | 0.517 | 0.43 | 1 | HC_vs_MS.Trt_F |
| MTCH2 | 0.23228 | 0.30322 | 0.124 | 0.09 | 1 | HC_vs_MS.Trt_F |
| C14orf93 | 0.23234 | -0.9888 | 0.335 | 0.28 | 1 | HC_vs_MS.Trt_F |
| NDUFAF6 | 0.23237 | -0.4208 | 0.662 | 0.59 | 1 | HC_vs_MS.Trt_F |
| ICK | 0.23266 | 0.723 | 0.246 | 0.2 | 1 | HC_vs_MS.Trt_F |
| ZNF511 | 0.2327 | -0.5104 | 0.092 | 0.12 | 1 | HC_vs_MS.Trt_F |

| | | | | | | |
|---|---|---|---|---|---|---|
| DLC1 | 0.23344 | 0.55162 | 0.202 | 0.17 | 1 | HC_vs_MS.Trt_F |
| TBC1D22A | 0.23345 | 0.29518 | 0.572 | 0.53 | 1 | HC_vs_MS.Trt_F |
| TMEM214 | 0.23356 | -0.7667 | 0.095 | 0.12 | 1 | HC_vs_MS.Trt_F |
| ZFP64 | 0.23398 | -0.4354 | 0.633 | 0.68 | 1 | HC_vs_MS.Trt_F |
| SEC14L4 | 0.23401 | 0.49154 | 0.613 | 0.56 | 1 | HC_vs_MS.Trt_F |
| GPC2 | 0.23403 | 0.32256 | 0.523 | 0.46 | 1 | HC_vs_MS.Trt_F |
| FAM63A | 0.2341 | 0.67167 | 0.581 | 0.55 | 1 | HC_vs_MS.Trt_F |
| CCDC28B | 0.23453 | -0.391 | 0.483 | 0.39 | 1 | HC_vs_MS.Trt_F |
| MYH11 | 0.23462 | 0.37842 | 0.942 | 0.94 | 1 | HC_vs_MS.Trt_F |
| GTSF1 | 0.23465 | -0.6163 | 0.324 | 0.27 | 1 | HC_vs_MS.Trt_F |
| SLC2A6 | 0.23475 | 1.50203 | 0.254 | 0.22 | 1 | HC_vs_MS.Trt_F |
| TMED8 | 0.23477 | 0.40533 | 0.642 | 0.6 | 1 | HC_vs_MS.Trt_F |
| HTN3 | 0.23507 | 0.28751 | 0.659 | 0.61 | 1 | HC_vs_MS.Trt_F |
| EPHB6 | 0.23541 | 0.36615 | 0.133 | 0.1 | 1 | HC_vs_MS.Trt_F |
| ASUN | 0.23558 | 0.32872 | 0.162 | 0.13 | 1 | HC_vs_MS.Trt_F |
| TSC22D4 | 0.23566 | 0.47491 | 0.303 | 0.27 | 1 | HC_vs_MS.Trt_F |
| PRELID2 | 0.23569 | 0.59949 | 0.572 | 0.49 | 1 | HC_vs_MS.Trt_F |
| RELN | 0.23588 | 0.27858 | 0.156 | 0.13 | 1 | HC_vs_MS.Trt_F |
| LMBR1L | 0.23626 | 0.32459 | 0.772 | 0.71 | 1 | HC_vs_MS.Trt_F |
| CCNK | 0.23631 | 0.43765 | 0.298 | 0.26 | 1 | HC_vs_MS.Trt_F |
| SMU1 | 0.23648 | -0.4424 | 0.786 | 0.77 | 1 | HC_vs_MS.Trt_F |
| SLA2 | 0.23688 | -0.3596 | 0.572 | 0.6 | 1 | HC_vs_MS.Trt_F |
| THAP5 | 0.23701 | -0.7621 | 0.934 | 0.9 | 1 | HC_vs_MS.Trt_F |
| UBN2 | 0.2371 | -0.665 | 0.708 | 0.63 | 1 | HC_vs_MS.Trt_F |
| AHRR | 0.23721 | 0.30604 | 0.136 | 0.17 | 1 | HC_vs_MS.Trt_F |
| C1orf52 | 0.23735 | 0.91123 | 0.35 | 0.3 | 1 | HC_vs_MS.Trt_F |
| OXCT1 | 0.23738 | 0.4388 | 0.408 | 0.36 | 1 | HC_vs_MS.Trt_F |
| MYLIP | 0.23742 | -0.6086 | 0.139 | 0.1 | 1 | HC_vs_MS.Trt_F |
| SOCS2 | 0.23767 | -0.2585 | 0.52 | 0.47 | 1 | HC_vs_MS.Trt_F |
| CISD1 | 0.23773 | -0.2812 | 0.124 | 0.09 | 1 | HC_vs_MS.Trt_F |
| TRA2A | 0.23824 | -0.4982 | 0.772 | 0.74 | 1 | HC_vs_MS.Trt_F |
| ZNF280B | 0.23831 | 0.62731 | 0.139 | 0.11 | 1 | HC_vs_MS.Trt_F |
| FBXO44 | 0.23839 | -0.3676 | 0.676 | 0.63 | 1 | HC_vs_MS.Trt_F |
| ID3 | 0.23844 | -0.5002 | 0.237 | 0.18 | 1 | HC_vs_MS.Trt_F |
| COX6A1 | 0.23906 | -0.5886 | 0.133 | 0.16 | 1 | HC_vs_MS.Trt_F |
| SLC44A2 | 0.23915 | -0.4122 | 0.728 | 0.74 | 1 | HC_vs_MS.Trt_F |
| BEND7 | 0.23916 | -0.3041 | 0.408 | 0.36 | 1 | HC_vs_MS.Trt_F |
| GMPS | 0.23998 | -0.3941 | 0.905 | 0.84 | 1 | HC_vs_MS.Trt_F |
| SLC35E2B | 0.24017 | 0.36786 | 0.616 | 0.55 | 1 | HC_vs_MS.Trt_F |
| PSMB3 | 0.2402 | -0.3418 | 0.269 | 0.3 | 1 | HC_vs_MS.Trt_F |

| | | | | | | |
|---|---|---|---|---|---|---|
| GOLGA8N | 0.24057 | -0.4234 | 0.101 | 0.08 | 1 | HC_vs_MS.Trt_F |
| PRMT3 | 0.24063 | -0.4657 | 0.358 | 0.3 | 1 | HC_vs_MS.Trt_F |
| TSTA3 | 0.24091 | -0.2938 | 0.205 | 0.16 | 1 | HC_vs_MS.Trt_F |
| DCK | 0.24115 | -0.2982 | 0.341 | 0.28 | 1 | HC_vs_MS.Trt_F |
| BMP8A | 0.24138 | 0.88787 | 0.668 | 0.6 | 1 | HC_vs_MS.Trt_F |
| RPS17 | 0.2414 | -0.4969 | 0.474 | 0.42 | 1 | HC_vs_MS.Trt_F |
| PAIP2B | 0.24169 | 0.76282 | 0.254 | 0.21 | 1 | HC_vs_MS.Trt_F |
| ZNF662 | 0.2417 | -1.0104 | 0.595 | 0.58 | 1 | HC_vs_MS.Trt_F |
| MAP7D3 | 0.24174 | -0.3175 | 0.662 | 0.6 | 1 | HC_vs_MS.Trt_F |
| PRKAB1 | 0.24174 | 0.34619 | 0.818 | 0.79 | 1 | HC_vs_MS.Trt_F |
| STT3A | 0.24192 | -0.8078 | 0.61 | 0.61 | 1 | HC_vs_MS.Trt_F |
| UBXN7 | 0.24225 | 0.73481 | 0.795 | 0.78 | 1 | HC_vs_MS.Trt_F |
| CD34 | 0.24226 | 0.34715 | 0.665 | 0.6 | 1 | HC_vs_MS.Trt_F |
| MRPS10 | 0.24232 | 0.53856 | 0.538 | 0.5 | 1 | HC_vs_MS.Trt_F |
| PHF14 | 0.24234 | 0.82099 | 0.15 | 0.12 | 1 | HC_vs_MS.Trt_F |
| PARL | 0.2424 | -0.2701 | 0.162 | 0.13 | 1 | HC_vs_MS.Trt_F |
| MDFIC | 0.24281 | 0.6338 | 0.408 | 0.36 | 1 | HC_vs_MS.Trt_F |
| ZBTB17 | 0.24286 | 0.33141 | 0.124 | 0.09 | 1 | HC_vs_MS.Trt_F |
| RRN3 | 0.2429 | -0.4403 | 0.306 | 0.26 | 1 | HC_vs_MS.Trt_F |
| CHD1L | 0.24307 | -0.3679 | 0.113 | 0.14 | 1 | HC_vs_MS.Trt_F |
| BCL2L11 | 0.24354 | -0.3279 | 0.549 | 0.5 | 1 | HC_vs_MS.Trt_F |
| VAMP8 | 0.24357 | -0.5023 | 0.298 | 0.32 | 1 | HC_vs_MS.Trt_F |
| MOCOS | 0.24432 | 0.46293 | 0.656 | 0.59 | 1 | HC_vs_MS.Trt_F |
| PANK4 | 0.24455 | 0.6964 | 0.107 | 0.08 | 1 | HC_vs_MS.Trt_F |
| CDIP1 | 0.24463 | -0.5934 | 0.223 | 0.26 | 1 | HC_vs_MS.Trt_F |
| INTS10 | 0.24464 | -0.256 | 0.613 | 0.54 | 1 | HC_vs_MS.Trt_F |
| SLC25A30 | 0.24473 | 0.33154 | 0.705 | 0.64 | 1 | HC_vs_MS.Trt_F |
| HEATR5A | 0.24491 | 0.27745 | 0.763 | 0.67 | 1 | HC_vs_MS.Trt_F |
| ZNF547 | 0.24492 | -0.6698 | 0.266 | 0.3 | 1 | HC_vs_MS.Trt_F |
| CCDC97 | 0.24516 | 0.3581 | 0.185 | 0.15 | 1 | HC_vs_MS.Trt_F |
| RYR3 | 0.24538 | -0.6612 | 0.532 | 0.48 | 1 | HC_vs_MS.Trt_F |
| FBXO36 | 0.2454 | 0.26189 | 0.691 | 0.69 | 1 | HC_vs_MS.Trt_F |
| CHI3L2 | 0.24547 | -1.0146 | 0.272 | 0.22 | 1 | HC_vs_MS.Trt_F |
| HSPH1 | 0.24593 | 0.27369 | 0.257 | 0.21 | 1 | HC_vs_MS.Trt_F |
| HMCES | 0.24598 | 0.59505 | 0.136 | 0.1 | 1 | HC_vs_MS.Trt_F |
| PCOLCE | 0.24623 | 1.25866 | 0.133 | 0.1 | 1 | HC_vs_MS.Trt_F |
| TOR4A | 0.24652 | 0.43493 | 0.254 | 0.22 | 1 | HC_vs_MS.Trt_F |
| FOXN2 | 0.24653 | -0.3201 | 0.272 | 0.22 | 1 | HC_vs_MS.Trt_F |
| C3orf18 | 0.24671 | 0.30434 | 0.315 | 0.27 | 1 | HC_vs_MS.Trt_F |
| PIK3CG | 0.24684 | -0.7906 | 0.355 | 0.3 | 1 | HC_vs_MS.Trt_F |

| | | | | | | |
|---|---|---|---|---|---|---|
| TYSND1 | 0.24712 | -0.3913 | 0.277 | 0.23 | 1 | HC_vs_MS.Trt_F |
| ZNF550 | 0.24731 | -0.7077 | 0.344 | 0.29 | 1 | HC_vs_MS.Trt_F |
| EIF5A2 | 0.24765 | -0.6159 | 0.419 | 0.35 | 1 | HC_vs_MS.Trt_F |
| OGFOD3 | 0.24767 | -0.4501 | 0.604 | 0.6 | 1 | HC_vs_MS.Trt_F |
| FBXO34 | 0.24788 | -0.7559 | 0.142 | 0.11 | 1 | HC_vs_MS.Trt_F |
| SMAD4 | 0.24793 | 0.65692 | 0.512 | 0.44 | 1 | HC_vs_MS.Trt_F |
| ANGPT4 | 0.24803 | -0.3184 | 0.448 | 0.49 | 1 | HC_vs_MS.Trt_F |
| MAP3K3 | 0.24846 | -0.4897 | 0.179 | 0.14 | 1 | HC_vs_MS.Trt_F |
| SGK494 | 0.24857 | -0.4937 | 0.777 | 0.7 | 1 | HC_vs_MS.Trt_F |
| ZBTB2 | 0.24887 | -1.2003 | 0.101 | 0.13 | 1 | HC_vs_MS.Trt_F |
| NPC1L1 | 0.249 | 0.33173 | 0.188 | 0.16 | 1 | HC_vs_MS.Trt_F |
| C5orf51 | 0.24907 | 0.26506 | 0.145 | 0.11 | 1 | HC_vs_MS.Trt_F |
| TUBG1 | 0.2494 | 0.79418 | 0.668 | 0.62 | 1 | HC_vs_MS.Trt_F |
| GPR35 | 0.24954 | 0.82542 | 0.321 | 0.27 | 1 | HC_vs_MS.Trt_F |
| TOB1 | 0.25 | -0.303 | 0.621 | 0.63 | 1 | HC_vs_MS.Trt_F |
| APIP | 0.25039 | -0.2536 | 0.509 | 0.48 | 1 | HC_vs_MS.Trt_F |
| ATF4 | 0.25048 | 0.51632 | 0.188 | 0.16 | 1 | HC_vs_MS.Trt_F |
| NPL | 0.25048 | 1.28698 | 0.223 | 0.27 | 1 | HC_vs_MS.Trt_F |
| ASB2 | 0.25049 | -1.2951 | 0.078 | 0.1 | 1 | HC_vs_MS.Trt_F |
| INTS3 | 0.25051 | 0.29416 | 0.769 | 0.73 | 1 | HC_vs_MS.Trt_F |
| UBE2Q2 | 0.25081 | 0.32994 | 0.15 | 0.12 | 1 | HC_vs_MS.Trt_F |
| WIZ | 0.25084 | 0.54269 | 0.355 | 0.31 | 1 | HC_vs_MS.Trt_F |
| LIFR | 0.25084 | -0.7781 | 0.142 | 0.18 | 1 | HC_vs_MS.Trt_F |
| NRBF2 | 0.25139 | -0.6155 | 0.139 | 0.17 | 1 | HC_vs_MS.Trt_F |
| GK | 0.25148 | 0.6359 | 0.228 | 0.19 | 1 | HC_vs_MS.Trt_F |
| DNAH6 | 0.25159 | -0.4943 | 0.191 | 0.22 | 1 | HC_vs_MS.Trt_F |
| USP20 | 0.25186 | -0.4348 | 0.249 | 0.28 | 1 | HC_vs_MS.Trt_F |
| CNDP1 | 0.25186 | 0.45685 | 0.671 | 0.62 | 1 | HC_vs_MS.Trt_F |
| ZBTB7A | 0.25187 | -0.6577 | 0.162 | 0.13 | 1 | HC_vs_MS.Trt_F |
| KIF13A | 0.25198 | 1.17076 | 0.364 | 0.31 | 1 | HC_vs_MS.Trt_F |
| MYD88 | 0.252 | 0.59945 | 0.176 | 0.14 | 1 | HC_vs_MS.Trt_F |
| CD300LD | 0.25235 | -1.1008 | 0.355 | 0.39 | 1 | HC_vs_MS.Trt_F |
| APAF1 | 0.25266 | -0.3154 | 0.676 | 0.6 | 1 | HC_vs_MS.Trt_F |
| DNPH1 | 0.25288 | -0.6712 | 0.185 | 0.21 | 1 | HC_vs_MS.Trt_F |
| USF1 | 0.25298 | -0.4665 | 0.223 | 0.18 | 1 | HC_vs_MS.Trt_F |
| NDST1 | 0.2532 | 0.86994 | 0.382 | 0.43 | 1 | HC_vs_MS.Trt_F |
| MRPL50 | 0.25335 | -0.5702 | 0.165 | 0.13 | 1 | HC_vs_MS.Trt_F |
| POLG | 0.25346 | -0.2659 | 0.179 | 0.14 | 1 | HC_vs_MS.Trt_F |
| PAG1 | 0.25365 | -0.421 | 0.757 | 0.69 | 1 | HC_vs_MS.Trt_F |
| ACTR2 | 0.25382 | -0.7345 | 0.413 | 0.42 | 1 | HC_vs_MS.Trt_F |

| | | | | | | |
|---|---|---|---|---|---|---|
| SIN3A | 0.25383 | -0.3572 | 0.832 | 0.78 | 1 | HC_vs_MS.Trt_F |
| NEIL1 | 0.25388 | 0.48864 | 0.824 | 0.74 | 1 | HC_vs_MS.Trt_F |
| GRN | 0.25393 | -0.6142 | 0.211 | 0.17 | 1 | HC_vs_MS.Trt_F |
| SLC30A4 | 0.25406 | -0.9816 | 0.905 | 0.85 | 1 | HC_vs_MS.Trt_F |
| RRP8 | 0.25433 | 0.25128 | 0.529 | 0.48 | 1 | HC_vs_MS.Trt_F |
| NDST2 | 0.25445 | 0.39022 | 0.124 | 0.09 | 1 | HC_vs_MS.Trt_F |
| EML2 | 0.25449 | 0.33525 | 0.714 | 0.69 | 1 | HC_vs_MS.Trt_F |
| CLMN | 0.2549 | -0.2569 | 0.711 | 0.63 | 1 | HC_vs_MS.Trt_F |
| FBXO7 | 0.25538 | 0.28589 | 0.315 | 0.28 | 1 | HC_vs_MS.Trt_F |
| C6orf163 | 0.25539 | -0.3932 | 0.474 | 0.42 | 1 | HC_vs_MS.Trt_F |
| MED19 | 0.25545 | 1.19328 | 0.113 | 0.09 | 1 | HC_vs_MS.Trt_F |
| P4HTM | 0.25557 | 0.39414 | 0.442 | 0.4 | 1 | HC_vs_MS.Trt_F |
| RGS11 | 0.25565 | 0.60699 | 0.217 | 0.18 | 1 | HC_vs_MS.Trt_F |
| PDHB | 0.25578 | -0.2746 | 0.844 | 0.82 | 1 | HC_vs_MS.Trt_F |
| CLSPN | 0.25666 | 0.27732 | 0.91 | 0.85 | 1 | HC_vs_MS.Trt_F |
| ADGRL1 | 0.25714 | 0.38834 | 0.101 | 0.08 | 1 | HC_vs_MS.Trt_F |
| PGM1 | 0.25724 | 0.37786 | 0.127 | 0.1 | 1 | HC_vs_MS.Trt_F |
| ZNF16 | 0.25773 | 0.28921 | 0.809 | 0.77 | 1 | HC_vs_MS.Trt_F |
| PDK3 | 0.25783 | -0.9838 | 0.41 | 0.33 | 1 | HC_vs_MS.Trt_F |
| ATG7 | 0.25784 | -0.8829 | 0.63 | 0.54 | 1 | HC_vs_MS.Trt_F |
| RUBCN | 0.25791 | -0.3771 | 0.873 | 0.87 | 1 | HC_vs_MS.Trt_F |
| SMIM20 | 0.25797 | -0.316 | 0.087 | 0.11 | 1 | HC_vs_MS.Trt_F |
| UBE2E1 | 0.25798 | 1.01161 | 0.11 | 0.08 | 1 | HC_vs_MS.Trt_F |
| TCTA | 0.25804 | 0.2507 | 0.633 | 0.58 | 1 | HC_vs_MS.Trt_F |
| TLE3 | 0.25805 | -0.3597 | 0.298 | 0.23 | 1 | HC_vs_MS.Trt_F |
| AES | 0.25852 | -0.2612 | 0.5 | 0.51 | 1 | HC_vs_MS.Trt_F |
| LGR6 | 0.25861 | -1.6737 | 0.168 | 0.14 | 1 | HC_vs_MS.Trt_F |
| SNX20 | 0.25869 | -0.2504 | 0.419 | 0.45 | 1 | HC_vs_MS.Trt_F |
| GRINA | 0.25914 | 0.36056 | 0.121 | 0.09 | 1 | HC_vs_MS.Trt_F |
| ANKRD27 | 0.25936 | -0.3434 | 0.26 | 0.21 | 1 | HC_vs_MS.Trt_F |
| OLR1 | 0.25945 | 0.41789 | 0.471 | 0.41 | 1 | HC_vs_MS.Trt_F |
| ADD1 | 0.25983 | -0.3417 | 0.777 | 0.78 | 1 | HC_vs_MS.Trt_F |
| FKBP11 | 0.26048 | 0.37625 | 0.72 | 0.7 | 1 | HC_vs_MS.Trt_F |
| USF3 | 0.26075 | 0.42809 | 0.737 | 0.73 | 1 | HC_vs_MS.Trt_F |
| DZIP1 | 0.26075 | 0.42272 | 0.393 | 0.36 | 1 | HC_vs_MS.Trt_F |
| RFT1 | 0.26098 | -0.2505 | 0.514 | 0.46 | 1 | HC_vs_MS.Trt_F |
| CXorf57 | 0.26113 | 1.48491 | 0.179 | 0.15 | 1 | HC_vs_MS.Trt_F |
| DEK | 0.26116 | -0.3516 | 0.454 | 0.41 | 1 | HC_vs_MS.Trt_F |
| MRM2 | 0.26124 | -0.7557 | 0.393 | 0.32 | 1 | HC_vs_MS.Trt_F |
| REPS2 | 0.26156 | 0.32026 | 0.647 | 0.63 | 1 | HC_vs_MS.Trt_F |

| | | | | | | |
|---|---|---|---|---|---|---|
| F2RL3 | 0.26168 | 0.49763 | 0.818 | 0.73 | 1 | HC_vs_MS.Trt_F |
| ANKS3 | 0.26171 | 0.38677 | 0.818 | 0.79 | 1 | HC_vs_MS.Trt_F |
| HEMK1 | 0.26181 | -0.4457 | 0.974 | 0.96 | 1 | HC_vs_MS.Trt_F |
| EIF3J | 0.26186 | 0.27256 | 0.202 | 0.17 | 1 | HC_vs_MS.Trt_F |
| PAK2 | 0.262 | 0.41695 | 0.74 | 0.68 | 1 | HC_vs_MS.Trt_F |
| ISYNA1 | 0.26273 | 1.04883 | 0.11 | 0.08 | 1 | HC_vs_MS.Trt_F |
| CNIH1 | 0.26284 | -0.6313 | 0.145 | 0.11 | 1 | HC_vs_MS.Trt_F |
| DEF8 | 0.26304 | -1.0521 | 0.512 | 0.46 | 1 | HC_vs_MS.Trt_F |
| BRMS1 | 0.26339 | -0.3797 | 0.185 | 0.14 | 1 | HC_vs_MS.Trt_F |
| ADCY9 | 0.26368 | -0.9498 | 0.335 | 0.29 | 1 | HC_vs_MS.Trt_F |
| KRBOX4 | 0.2637 | -0.6564 | 0.483 | 0.48 | 1 | HC_vs_MS.Trt_F |
| MRPL13 | 0.26374 | -0.2709 | 0.168 | 0.13 | 1 | HC_vs_MS.Trt_F |
| SNW1 | 0.26397 | 0.3532 | 0.275 | 0.23 | 1 | HC_vs_MS.Trt_F |
| LIPH | 0.26411 | 0.30087 | 0.697 | 0.67 | 1 | HC_vs_MS.Trt_F |
| CFH | 0.26418 | 0.44285 | 0.127 | 0.1 | 1 | HC_vs_MS.Trt_F |
| FAAP100 | 0.26422 | -0.4674 | 0.205 | 0.16 | 1 | HC_vs_MS.Trt_F |
| C18orf25 | 0.26442 | -0.2638 | 0.468 | 0.5 | 1 | HC_vs_MS.Trt_F |
| CCDC191 | 0.26447 | 0.55424 | 0.237 | 0.2 | 1 | HC_vs_MS.Trt_F |
| PXK | 0.26449 | -0.2722 | 0.153 | 0.12 | 1 | HC_vs_MS.Trt_F |
| POP7 | 0.26466 | -1.0252 | 0.081 | 0.1 | 1 | HC_vs_MS.Trt_F |
| TRIM8 | 0.26496 | -0.7828 | 0.087 | 0.11 | 1 | HC_vs_MS.Trt_F |
| FN3KRP | 0.26497 | -0.3156 | 0.145 | 0.11 | 1 | HC_vs_MS.Trt_F |
| ZNF230 | 0.26505 | 0.28981 | 0.633 | 0.57 | 1 | HC_vs_MS.Trt_F |
| EIF2AK4 | 0.26546 | -0.3325 | 0.705 | 0.62 | 1 | HC_vs_MS.Trt_F |
| DOPEY1 | 0.26583 | 0.73258 | 0.327 | 0.29 | 1 | HC_vs_MS.Trt_F |
| APLP1 | 0.26584 | 0.27262 | 0.676 | 0.61 | 1 | HC_vs_MS.Trt_F |
| MPP5 | 0.26612 | 0.37879 | 0.399 | 0.34 | 1 | HC_vs_MS.Trt_F |
| DUSP12 | 0.26615 | -0.6905 | 0.341 | 0.35 | 1 | HC_vs_MS.Trt_F |
| GGT7 | 0.26627 | 0.29877 | 0.194 | 0.16 | 1 | HC_vs_MS.Trt_F |
| NFYB | 0.26629 | -1.4496 | 0.303 | 0.33 | 1 | HC_vs_MS.Trt_F |
| AMACR | 0.26633 | -0.7467 | 0.075 | 0.1 | 1 | HC_vs_MS.Trt_F |
| SLC37A3 | 0.26645 | -0.4038 | 0.945 | 0.94 | 1 | HC_vs_MS.Trt_F |
| SMG8 | 0.26672 | -0.9889 | 0.107 | 0.13 | 1 | HC_vs_MS.Trt_F |
| MGST3 | 0.2674 | -0.3302 | 0.902 | 0.82 | 1 | HC_vs_MS.Trt_F |
| SCPEP1 | 0.26751 | 0.34086 | 0.772 | 0.79 | 1 | HC_vs_MS.Trt_F |
| RING1 | 0.26768 | 0.43746 | 0.104 | 0.08 | 1 | HC_vs_MS.Trt_F |
| SGSH | 0.26778 | 0.4618 | 0.558 | 0.53 | 1 | HC_vs_MS.Trt_F |
| THOC6 | 0.26787 | 0.65276 | 0.13 | 0.1 | 1 | HC_vs_MS.Trt_F |
| GPR183 | 0.26818 | 0.278 | 0.402 | 0.37 | 1 | HC_vs_MS.Trt_F |
| AGPAT1 | 0.26824 | -0.9802 | 0.118 | 0.14 | 1 | HC_vs_MS.Trt_F |

| | | | | | | |
|---|---|---|---|---|---|---|
| TOMM20 | 0.26825 | 0.28239 | 0.231 | 0.19 | 1 | HC_vs_MS.Trt_F |
| FYCO1 | 0.2684 | 0.48209 | 0.214 | 0.18 | 1 | HC_vs_MS.Trt_F |
| RFX3 | 0.2685 | 0.42997 | 0.228 | 0.19 | 1 | HC_vs_MS.Trt_F |
| TYMP | 0.26851 | 0.83281 | 0.367 | 0.32 | 1 | HC_vs_MS.Trt_F |
| ZNF415 | 0.26853 | -0.7326 | 0.332 | 0.3 | 1 | HC_vs_MS.Trt_F |
| POMT1 | 0.26866 | -0.271 | 0.584 | 0.55 | 1 | HC_vs_MS.Trt_F |
| IFT57 | 0.269 | 0.45109 | 0.142 | 0.11 | 1 | HC_vs_MS.Trt_F |
| CACUL1 | 0.26961 | -1.0654 | 0.783 | 0.77 | 1 | HC_vs_MS.Trt_F |
| ATG3 | 0.26966 | -0.2935 | 0.5 | 0.46 | 1 | HC_vs_MS.Trt_F |
| LRRC49 | 0.26972 | 1.16232 | 0.792 | 0.77 | 1 | HC_vs_MS.Trt_F |
| NKIRAS2 | 0.26985 | -0.8133 | 0.873 | 0.84 | 1 | HC_vs_MS.Trt_F |
| TOR2A | 0.27045 | -0.9262 | 0.133 | 0.1 | 1 | HC_vs_MS.Trt_F |
| ARHGEF40 | 0.27046 | -0.8513 | 0.217 | 0.18 | 1 | HC_vs_MS.Trt_F |
| HSD17B4 | 0.27053 | -0.667 | 0.656 | 0.59 | 1 | HC_vs_MS.Trt_F |
| BTBD7 | 0.27088 | -0.6955 | 0.818 | 0.79 | 1 | HC_vs_MS.Trt_F |
| HIPK3 | 0.27121 | 0.41637 | 0.179 | 0.15 | 1 | HC_vs_MS.Trt_F |
| PIK3R3 | 0.27145 | 0.42115 | 0.569 | 0.5 | 1 | HC_vs_MS.Trt_F |
| CDH26 | 0.27153 | 0.65872 | 0.173 | 0.14 | 1 | HC_vs_MS.Trt_F |
| MIF | 0.27157 | -0.6401 | 0.471 | 0.46 | 1 | HC_vs_MS.Trt_F |
| F3 | 0.27167 | 0.26551 | 0.535 | 0.5 | 1 | HC_vs_MS.Trt_F |
| RGP1 | 0.27193 | 0.58858 | 0.194 | 0.16 | 1 | HC_vs_MS.Trt_F |
| PIP5K1C | 0.27203 | 0.38026 | 0.13 | 0.1 | 1 | HC_vs_MS.Trt_F |
| KNG1 | 0.27222 | 0.73796 | 0.263 | 0.22 | 1 | HC_vs_MS.Trt_F |
| SFXN5 | 0.27251 | 0.47644 | 0.945 | 0.94 | 1 | HC_vs_MS.Trt_F |
| MTMR2 | 0.27257 | -0.9503 | 0.208 | 0.17 | 1 | HC_vs_MS.Trt_F |
| CAMSAP1 | 0.2729 | 0.4939 | 0.104 | 0.08 | 1 | HC_vs_MS.Trt_F |
| ZNF630 | 0.27358 | 0.77833 | 0.413 | 0.45 | 1 | HC_vs_MS.Trt_F |
| SKIL | 0.27409 | 0.2542 | 0.341 | 0.37 | 1 | HC_vs_MS.Trt_F |
| MATN1 | 0.27444 | 0.28252 | 0.116 | 0.09 | 1 | HC_vs_MS.Trt_F |
| C17orf89 | 0.27468 | -0.9225 | 0.095 | 0.12 | 1 | HC_vs_MS.Trt_F |
| UBE2L3 | 0.27476 | -0.4328 | 0.286 | 0.31 | 1 | HC_vs_MS.Trt_F |
| CAPN10 | 0.27493 | 0.33474 | 0.566 | 0.51 | 1 | HC_vs_MS.Trt_F |
| CH507-9B2.! | 0.27497 | 1.1669 | 0.142 | 0.11 | 1 | HC_vs_MS.Trt_F |
| PHF8 | 0.27523 | 0.66269 | 0.15 | 0.12 | 1 | HC_vs_MS.Trt_F |
| PARP8 | 0.27546 | -0.4249 | 0.514 | 0.45 | 1 | HC_vs_MS.Trt_F |
| ZNF569 | 0.27563 | 0.42727 | 0.228 | 0.19 | 1 | HC_vs_MS.Trt_F |
| SNX11 | 0.27628 | 0.47124 | 0.642 | 0.6 | 1 | HC_vs_MS.Trt_F |
| GTF2H1 | 0.27647 | 0.642 | 0.448 | 0.41 | 1 | HC_vs_MS.Trt_F |
| IKBKG | 0.27649 | 0.28479 | 0.199 | 0.17 | 1 | HC_vs_MS.Trt_F |
| POLI | 0.27651 | -0.9164 | 0.41 | 0.35 | 1 | HC_vs_MS.Trt_F |

| | | | | | | |
|---|---|---|---|---|---|---|
| ARV1 | 0.27656 | 0.83605 | 0.329 | 0.28 | 1 | HC_vs_MS.Trt_F |
| ITGB1BP1 | 0.27667 | -0.8019 | 0.191 | 0.22 | 1 | HC_vs_MS.Trt_F |
| RASSF3 | 0.27671 | -0.433 | 0.569 | 0.58 | 1 | HC_vs_MS.Trt_F |
| R3HCC1 | 0.27676 | 0.37367 | 0.139 | 0.11 | 1 | HC_vs_MS.Trt_F |
| PRKAR1B | 0.27684 | -0.5055 | 0.127 | 0.1 | 1 | HC_vs_MS.Trt_F |
| STAM2 | 0.27687 | -0.7653 | 0.552 | 0.58 | 1 | HC_vs_MS.Trt_F |
| CCR10 | 0.27724 | 0.37067 | 0.156 | 0.12 | 1 | HC_vs_MS.Trt_F |
| FAM189B | 0.27744 | 0.29286 | 0.746 | 0.71 | 1 | HC_vs_MS.Trt_F |
| KIZ | 0.27745 | 0.47919 | 0.147 | 0.12 | 1 | HC_vs_MS.Trt_F |
| DYDC2 | 0.27774 | 0.29321 | 0.101 | 0.08 | 1 | HC_vs_MS.Trt_F |
| CPSF3 | 0.27785 | 0.79859 | 0.136 | 0.11 | 1 | HC_vs_MS.Trt_F |
| CPSF4 | 0.27786 | -0.3807 | 0.497 | 0.42 | 1 | HC_vs_MS.Trt_F |
| ALG2 | 0.27808 | -0.8276 | 0.11 | 0.13 | 1 | HC_vs_MS.Trt_F |
| AC138969.4 | 0.27819 | 0.72659 | 0.448 | 0.4 | 1 | HC_vs_MS.Trt_F |
| ZNF30 | 0.27829 | -0.3176 | 0.078 | 0.1 | 1 | HC_vs_MS.Trt_F |
| DUS3L | 0.27843 | -0.3084 | 0.188 | 0.15 | 1 | HC_vs_MS.Trt_F |
| B3GAT2 | 0.27866 | -1.198 | 0.202 | 0.17 | 1 | HC_vs_MS.Trt_F |
| DGCR6L | 0.27879 | 0.261 | 0.171 | 0.14 | 1 | HC_vs_MS.Trt_F |
| LRWD1 | 0.27898 | 0.44758 | 0.61 | 0.63 | 1 | HC_vs_MS.Trt_F |
| ERMN | 0.27904 | -0.6189 | 0.197 | 0.23 | 1 | HC_vs_MS.Trt_F |
| C2orf50 | 0.27908 | -0.2691 | 0.41 | 0.41 | 1 | HC_vs_MS.Trt_F |
| TRAPPC10 | 0.27922 | 0.82362 | 0.295 | 0.26 | 1 | HC_vs_MS.Trt_F |
| SLC23A2 | 0.27922 | 0.54629 | 0.179 | 0.15 | 1 | HC_vs_MS.Trt_F |
| ENPP6 | 0.27926 | 0.34597 | 0.37 | 0.32 | 1 | HC_vs_MS.Trt_F |
| ZFX | 0.27978 | -0.6067 | 0.653 | 0.62 | 1 | HC_vs_MS.Trt_F |
| ZNF613 | 0.27979 | -1.414 | 0.422 | 0.36 | 1 | HC_vs_MS.Trt_F |
| C16orf58 | 0.27984 | -0.6398 | 0.179 | 0.14 | 1 | HC_vs_MS.Trt_F |
| SERPINH1 | 0.27988 | -0.3075 | 0.39 | 0.34 | 1 | HC_vs_MS.Trt_F |
| COX10 | 0.27999 | 0.83216 | 0.13 | 0.1 | 1 | HC_vs_MS.Trt_F |
| GPD1L | 0.28045 | -0.5904 | 0.136 | 0.16 | 1 | HC_vs_MS.Trt_F |
| PRIM1 | 0.28057 | 0.55757 | 0.176 | 0.14 | 1 | HC_vs_MS.Trt_F |
| SUFU | 0.28061 | 0.27547 | 0.121 | 0.09 | 1 | HC_vs_MS.Trt_F |
| CLPTM1 | 0.28137 | -0.4041 | 0.225 | 0.18 | 1 | HC_vs_MS.Trt_F |
| PRRC1 | 0.28139 | -0.9156 | 0.448 | 0.36 | 1 | HC_vs_MS.Trt_F |
| ZMAT2 | 0.28156 | -0.4765 | 0.136 | 0.1 | 1 | HC_vs_MS.Trt_F |
| QTRT1 | 0.28168 | 0.32668 | 0.844 | 0.82 | 1 | HC_vs_MS.Trt_F |
| ABHD17A | 0.28184 | 0.67244 | 0.136 | 0.11 | 1 | HC_vs_MS.Trt_F |
| PINK1 | 0.28187 | 0.50253 | 0.301 | 0.26 | 1 | HC_vs_MS.Trt_F |
| SMARCA4 | 0.28293 | 0.36463 | 0.829 | 0.81 | 1 | HC_vs_MS.Trt_F |
| EMD | 0.28295 | 0.30338 | 0.329 | 0.29 | 1 | HC_vs_MS.Trt_F |

| | | | | | | |
|---|---|---|---|---|---|---|
| ZNF706 | 0.28302 | -0.4006 | 0.373 | 0.4 | 1 | HC_vs_MS.Trt_F |
| ABCD4 | 0.28314 | -0.3477 | 0.855 | 0.8 | 1 | HC_vs_MS.Trt_F |
| HDAC3 | 0.2832 | -0.3283 | 0.156 | 0.12 | 1 | HC_vs_MS.Trt_F |
| ATP5G2 | 0.28325 | -0.3662 | 0.87 | 0.87 | 1 | HC_vs_MS.Trt_F |
| RECQL | 0.28341 | -1.0631 | 0.171 | 0.19 | 1 | HC_vs_MS.Trt_F |
| TOR1AIP2 | 0.28344 | -0.4885 | 0.994 | 0.98 | 1 | HC_vs_MS.Trt_F |
| MTHFR | 0.28357 | -0.3911 | 0.838 | 0.8 | 1 | HC_vs_MS.Trt_F |
| TNFAIP1 | 0.2842 | 0.80183 | 0.113 | 0.09 | 1 | HC_vs_MS.Trt_F |
| MKRN2 | 0.28491 | 0.38535 | 0.118 | 0.09 | 1 | HC_vs_MS.Trt_F |
| ARF4 | 0.28507 | -1.1549 | 0.182 | 0.21 | 1 | HC_vs_MS.Trt_F |
| PLCD4 | 0.28513 | 0.39573 | 0.344 | 0.3 | 1 | HC_vs_MS.Trt_F |
| SERTAD2 | 0.28514 | 0.61947 | 0.145 | 0.12 | 1 | HC_vs_MS.Trt_F |
| FAM76A | 0.28595 | -0.6411 | 0.855 | 0.86 | 1 | HC_vs_MS.Trt_F |
| ZNF524 | 0.286 | -0.6929 | 0.124 | 0.15 | 1 | HC_vs_MS.Trt_F |
| TRIM34 | 0.28605 | -0.4612 | 0.387 | 0.31 | 1 | HC_vs_MS.Trt_F |
| CTD-3088G3 | 0.28609 | 0.42244 | 0.561 | 0.5 | 1 | HC_vs_MS.Trt_F |
| TROVE2 | 0.28624 | -0.353 | 0.717 | 0.64 | 1 | HC_vs_MS.Trt_F |
| CDYL2 | 0.2866 | 0.26605 | 0.127 | 0.1 | 1 | HC_vs_MS.Trt_F |
| NAIF1 | 0.28675 | 0.27593 | 0.659 | 0.6 | 1 | HC_vs_MS.Trt_F |
| HARBI1 | 0.28687 | -0.6266 | 0.665 | 0.6 | 1 | HC_vs_MS.Trt_F |
| IARS2 | 0.28701 | -0.3387 | 0.171 | 0.14 | 1 | HC_vs_MS.Trt_F |
| DCAF6 | 0.28718 | 0.55433 | 0.269 | 0.22 | 1 | HC_vs_MS.Trt_F |
| LSM4 | 0.28748 | -0.2555 | 0.22 | 0.18 | 1 | HC_vs_MS.Trt_F |
| AP5B1 | 0.28774 | 0.58917 | 0.335 | 0.29 | 1 | HC_vs_MS.Trt_F |
| PDCL3 | 0.28808 | -0.5328 | 0.124 | 0.15 | 1 | HC_vs_MS.Trt_F |
| RNF111 | 0.28817 | -0.3216 | 0.335 | 0.28 | 1 | HC_vs_MS.Trt_F |
| ALG10B | 0.2882 | -0.2636 | 0.725 | 0.75 | 1 | HC_vs_MS.Trt_F |
| ATG5 | 0.28835 | -0.4913 | 0.185 | 0.15 | 1 | HC_vs_MS.Trt_F |
| ZNF33B | 0.2884 | -0.6987 | 0.488 | 0.51 | 1 | HC_vs_MS.Trt_F |
| DNAJC7 | 0.28861 | -0.4962 | 0.977 | 0.97 | 1 | HC_vs_MS.Trt_F |
| CDNF | 0.28882 | -0.8039 | 0.199 | 0.23 | 1 | HC_vs_MS.Trt_F |
| NCDN | 0.28922 | 0.32311 | 0.211 | 0.17 | 1 | HC_vs_MS.Trt_F |
| CELF1 | 0.28974 | -0.5872 | 0.477 | 0.39 | 1 | HC_vs_MS.Trt_F |
| SEMA5A | 0.2898 | 0.37135 | 0.39 | 0.41 | 1 | HC_vs_MS.Trt_F |
| TRIM35 | 0.28985 | -0.5598 | 0.202 | 0.22 | 1 | HC_vs_MS.Trt_F |
| COG8 | 0.29012 | -0.2858 | 0.679 | 0.67 | 1 | HC_vs_MS.Trt_F |
| DTWD1 | 0.29076 | -0.2855 | 0.659 | 0.62 | 1 | HC_vs_MS.Trt_F |
| SKI | 0.29083 | 0.6009 | 0.133 | 0.1 | 1 | HC_vs_MS.Trt_F |
| UTY | 0.29084 | 1.19011 | 0.497 | 0.43 | 1 | HC_vs_MS.Trt_F |
| FRMD8 | 0.29113 | -0.631 | 0.639 | 0.64 | 1 | HC_vs_MS.Trt_F |

| | | | | | | |
|---|---|---|---|---|---|---|
| MANEA | 0.29155 | 0.53409 | 0.145 | 0.12 | 1 | HC_vs_MS.Trt_F |
| TM2D2 | 0.29163 | -0.5754 | 0.454 | 0.39 | 1 | HC_vs_MS.Trt_F |
| PTRH2 | 0.29175 | -0.8219 | 0.217 | 0.18 | 1 | HC_vs_MS.Trt_F |
| PPP1R13B | 0.29197 | 0.36354 | 0.535 | 0.47 | 1 | HC_vs_MS.Trt_F |
| TET3 | 0.29248 | 0.67832 | 0.107 | 0.08 | 1 | HC_vs_MS.Trt_F |
| CEP63 | 0.29266 | -0.381 | 0.786 | 0.67 | 1 | HC_vs_MS.Trt_F |
| ZNF581 | 0.29266 | 1.05543 | 0.116 | 0.09 | 1 | HC_vs_MS.Trt_F |
| ARFRP1 | 0.29266 | 0.27359 | 0.509 | 0.48 | 1 | HC_vs_MS.Trt_F |
| ZNF737 | 0.29277 | 0.42416 | 0.899 | 0.85 | 1 | HC_vs_MS.Trt_F |
| RPUSD4 | 0.29309 | -0.4061 | 0.749 | 0.69 | 1 | HC_vs_MS.Trt_F |
| CDC26 | 0.29356 | -0.7615 | 0.231 | 0.25 | 1 | HC_vs_MS.Trt_F |
| AGAP2 | 0.29399 | 0.44435 | 0.225 | 0.19 | 1 | HC_vs_MS.Trt_F |
| PTPRF | 0.29401 | -0.2854 | 0.341 | 0.36 | 1 | HC_vs_MS.Trt_F |
| GALK2 | 0.29402 | 0.33622 | 0.91 | 0.89 | 1 | HC_vs_MS.Trt_F |
| DOCK4 | 0.29407 | 0.44896 | 0.318 | 0.28 | 1 | HC_vs_MS.Trt_F |
| MTMR3 | 0.29448 | 1.17976 | 0.272 | 0.24 | 1 | HC_vs_MS.Trt_F |
| GINS4 | 0.29452 | 0.29177 | 0.916 | 0.86 | 1 | HC_vs_MS.Trt_F |
| RIN1 | 0.29458 | 1.04029 | 0.199 | 0.23 | 1 | HC_vs_MS.Trt_F |
| CLDN12 | 0.29502 | 0.28464 | 0.471 | 0.44 | 1 | HC_vs_MS.Trt_F |
| CGREF1 | 0.29511 | 0.42259 | 0.101 | 0.13 | 1 | HC_vs_MS.Trt_F |
| ABCD3 | 0.29538 | -0.4902 | 0.176 | 0.14 | 1 | HC_vs_MS.Trt_F |
| TRMT44 | 0.2957 | 0.26304 | 0.28 | 0.23 | 1 | HC_vs_MS.Trt_F |
| SNRPD2 | 0.29579 | -0.331 | 0.509 | 0.53 | 1 | HC_vs_MS.Trt_F |
| ERP27 | 0.29604 | -0.2798 | 0.26 | 0.22 | 1 | HC_vs_MS.Trt_F |
| METTL9 | 0.29623 | -0.4869 | 0.656 | 0.59 | 1 | HC_vs_MS.Trt_F |
| ARL10 | 0.29631 | -0.453 | 0.922 | 0.9 | 1 | HC_vs_MS.Trt_F |
| IL24 | 0.29644 | 0.95384 | 0.136 | 0.11 | 1 | HC_vs_MS.Trt_F |
| NOMO1 | 0.29678 | -0.3856 | 0.199 | 0.17 | 1 | HC_vs_MS.Trt_F |
| TMEM230 | 0.29686 | -0.2889 | 0.46 | 0.5 | 1 | HC_vs_MS.Trt_F |
| SHISA2 | 0.29707 | 0.35693 | 0.298 | 0.26 | 1 | HC_vs_MS.Trt_F |
| NOVA1 | 0.29746 | 0.46126 | 0.113 | 0.09 | 1 | HC_vs_MS.Trt_F |
| LARP4B | 0.29777 | -1.0013 | 0.283 | 0.31 | 1 | HC_vs_MS.Trt_F |
| MIEF1 | 0.29786 | -0.5 | 0.477 | 0.43 | 1 | HC_vs_MS.Trt_F |
| CHMP2B | 0.29837 | -0.3233 | 0.176 | 0.14 | 1 | HC_vs_MS.Trt_F |
| AFAP1L1 | 0.29839 | 0.88146 | 0.569 | 0.54 | 1 | HC_vs_MS.Trt_F |
| TOMM5 | 0.29842 | -0.7926 | 0.194 | 0.22 | 1 | HC_vs_MS.Trt_F |
| PIK3AP1 | 0.29871 | 1.15142 | 0.147 | 0.12 | 1 | HC_vs_MS.Trt_F |
| SNAPC3 | 0.29881 | -0.3646 | 0.642 | 0.59 | 1 | HC_vs_MS.Trt_F |
| RAP1GAP2 | 0.29891 | -0.5949 | 0.246 | 0.27 | 1 | HC_vs_MS.Trt_F |
| TRAPPC5 | 0.29902 | -0.2769 | 0.942 | 0.91 | 1 | HC_vs_MS.Trt_F |

| | | | | | | |
|---|---|---|---|---|---|---|
| SLC35A2 | 0.29952 | 0.32022 | 0.104 | 0.08 | 1 | HC_vs_MS.Trt_F |
| PCNT | 0.29993 | -0.3133 | 0.199 | 0.23 | 1 | HC_vs_MS.Trt_F |
| TRAPPC2 | 0.30033 | -0.3781 | 0.948 | 0.89 | 1 | HC_vs_MS.Trt_F |
| YEATS4 | 0.30038 | -0.4939 | 0.225 | 0.19 | 1 | HC_vs_MS.Trt_F |
| GLTSCR2 | 0.30047 | 0.47994 | 0.876 | 0.87 | 1 | HC_vs_MS.Trt_F |
| WDR61 | 0.3005 | -0.2841 | 0.217 | 0.17 | 1 | HC_vs_MS.Trt_F |
| SIGLEC14 | 0.30084 | 0.55548 | 0.289 | 0.25 | 1 | HC_vs_MS.Trt_F |
| RELA | 0.30099 | 0.58221 | 0.214 | 0.18 | 1 | HC_vs_MS.Trt_F |
| FAM83G | 0.30106 | -0.3303 | 0.465 | 0.41 | 1 | HC_vs_MS.Trt_F |
| GRPEL1 | 0.30117 | -1.0039 | 0.156 | 0.18 | 1 | HC_vs_MS.Trt_F |
| EHBP1L1 | 0.30151 | -0.3099 | 0.246 | 0.2 | 1 | HC_vs_MS.Trt_F |
| PNO1 | 0.30176 | 0.26377 | 0.702 | 0.67 | 1 | HC_vs_MS.Trt_F |
| SIK1 | 0.30179 | 1.33338 | 0.159 | 0.13 | 1 | HC_vs_MS.Trt_F |
| POLM | 0.30206 | -0.3015 | 0.566 | 0.52 | 1 | HC_vs_MS.Trt_F |
| OCIAD1 | 0.30258 | -0.617 | 0.633 | 0.68 | 1 | HC_vs_MS.Trt_F |
| FAAH2 | 0.30294 | 0.78583 | 0.127 | 0.1 | 1 | HC_vs_MS.Trt_F |
| DCLK1 | 0.303 | 0.88284 | 0.361 | 0.31 | 1 | HC_vs_MS.Trt_F |
| NOP14 | 0.30357 | 0.31615 | 0.442 | 0.39 | 1 | HC_vs_MS.Trt_F |
| SLC14A1 | 0.30363 | -0.7495 | 0.552 | 0.5 | 1 | HC_vs_MS.Trt_F |
| RHOBTB2 | 0.3041 | 1.09926 | 0.104 | 0.08 | 1 | HC_vs_MS.Trt_F |
| MGRN1 | 0.30449 | 0.38667 | 0.864 | 0.84 | 1 | HC_vs_MS.Trt_F |
| COL1A1 | 0.30472 | 0.33397 | 0.558 | 0.53 | 1 | HC_vs_MS.Trt_F |
| SEC24B | 0.30473 | -0.2827 | 0.538 | 0.5 | 1 | HC_vs_MS.Trt_F |
| RALGPS2 | 0.30473 | 0.40566 | 0.37 | 0.42 | 1 | HC_vs_MS.Trt_F |
| KLHL2 | 0.30508 | -0.6335 | 0.367 | 0.34 | 1 | HC_vs_MS.Trt_F |
| MIER2 | 0.30528 | -0.6271 | 0.509 | 0.53 | 1 | HC_vs_MS.Trt_F |
| CD276 | 0.30532 | 0.43722 | 0.512 | 0.46 | 1 | HC_vs_MS.Trt_F |
| ELMSAN1 | 0.30553 | -0.9309 | 0.749 | 0.73 | 1 | HC_vs_MS.Trt_F |
| DENND6B | 0.3061 | 0.75642 | 0.399 | 0.35 | 1 | HC_vs_MS.Trt_F |
| NDUFS4 | 0.30669 | -0.5348 | 0.353 | 0.3 | 1 | HC_vs_MS.Trt_F |
| DOCK7 | 0.30744 | 0.29017 | 0.974 | 0.95 | 1 | HC_vs_MS.Trt_F |
| URB1 | 0.30751 | 1.25779 | 0.52 | 0.45 | 1 | HC_vs_MS.Trt_F |
| MBD3 | 0.30753 | 0.39455 | 0.775 | 0.71 | 1 | HC_vs_MS.Trt_F |
| SEL1L3 | 0.30754 | -0.4785 | 0.321 | 0.27 | 1 | HC_vs_MS.Trt_F |
| NEIL3 | 0.30765 | -0.5456 | 0.295 | 0.25 | 1 | HC_vs_MS.Trt_F |
| PRDX3 | 0.3078 | -0.3461 | 0.532 | 0.46 | 1 | HC_vs_MS.Trt_F |
| PABPC1L | 0.30789 | 0.80538 | 0.321 | 0.29 | 1 | HC_vs_MS.Trt_F |
| BECN1 | 0.30797 | -0.3099 | 0.775 | 0.73 | 1 | HC_vs_MS.Trt_F |
| SENP6 | 0.30802 | -0.4361 | 0.922 | 0.86 | 1 | HC_vs_MS.Trt_F |
| COX8A | 0.30878 | -0.3602 | 0.28 | 0.23 | 1 | HC_vs_MS.Trt_F |

| | | | | | | |
|---|---|---|---|---|---|---|
| CBX3 | 0.30887 | -0.3146 | 0.587 | 0.51 | 1 | HC_vs_MS.Trt_F |
| MAK16 | 0.30889 | -0.4373 | 0.107 | 0.13 | 1 | HC_vs_MS.Trt_F |
| SARDH | 0.30896 | 0.63098 | 0.486 | 0.44 | 1 | HC_vs_MS.Trt_F |
| ARHGAP5 | 0.30931 | 0.39711 | 0.462 | 0.41 | 1 | HC_vs_MS.Trt_F |
| IDH1 | 0.30951 | 1.10747 | 0.249 | 0.22 | 1 | HC_vs_MS.Trt_F |
| RP9 | 0.30958 | 0.28474 | 0.127 | 0.1 | 1 | HC_vs_MS.Trt_F |
| GRK3 | 0.30967 | -0.6717 | 0.408 | 0.35 | 1 | HC_vs_MS.Trt_F |
| ADAR | 0.31036 | 0.4523 | 0.841 | 0.81 | 1 | HC_vs_MS.Trt_F |
| UBLCP1 | 0.31073 | 0.3275 | 0.214 | 0.18 | 1 | HC_vs_MS.Trt_F |
| UBE3D | 0.31106 | -0.3483 | 0.671 | 0.64 | 1 | HC_vs_MS.Trt_F |
| C1orf131 | 0.31159 | -0.8503 | 0.636 | 0.59 | 1 | HC_vs_MS.Trt_F |
| CLIP2 | 0.3117 | 0.44458 | 0.405 | 0.36 | 1 | HC_vs_MS.Trt_F |
| HMGN3 | 0.31171 | -0.8223 | 0.211 | 0.23 | 1 | HC_vs_MS.Trt_F |
| LANCL3 | 0.31207 | -1.0397 | 0.358 | 0.31 | 1 | HC_vs_MS.Trt_F |
| RPE | 0.31213 | 0.43802 | 0.353 | 0.32 | 1 | HC_vs_MS.Trt_F |
| EMC1 | 0.31225 | -0.3675 | 0.876 | 0.83 | 1 | HC_vs_MS.Trt_F |
| AL354822.1 | 0.31229 | 0.48104 | 0.286 | 0.24 | 1 | HC_vs_MS.Trt_F |
| PSG2 | 0.3124 | 0.36754 | 0.124 | 0.1 | 1 | HC_vs_MS.Trt_F |
| SGPL1 | 0.31277 | -0.8523 | 0.471 | 0.51 | 1 | HC_vs_MS.Trt_F |
| AP1G1 | 0.31323 | 0.28377 | 0.737 | 0.68 | 1 | HC_vs_MS.Trt_F |
| PUS7 | 0.31353 | -0.5443 | 0.373 | 0.33 | 1 | HC_vs_MS.Trt_F |
| ZNF227 | 0.31358 | 0.35035 | 0.549 | 0.52 | 1 | HC_vs_MS.Trt_F |
| FAM21A | 0.3136 | -0.9136 | 0.225 | 0.18 | 1 | HC_vs_MS.Trt_F |
| ZNF678 | 0.31372 | 0.66233 | 0.228 | 0.19 | 1 | HC_vs_MS.Trt_F |
| C21orf91 | 0.31383 | -0.5239 | 0.22 | 0.18 | 1 | HC_vs_MS.Trt_F |
| ZNF407 | 0.31387 | 0.39483 | 0.124 | 0.1 | 1 | HC_vs_MS.Trt_F |
| INTS6 | 0.31429 | -0.4974 | 0.954 | 0.95 | 1 | HC_vs_MS.Trt_F |
| GGA3 | 0.31451 | -0.5135 | 0.191 | 0.21 | 1 | HC_vs_MS.Trt_F |
| OSBPL7 | 0.31469 | 0.47209 | 0.116 | 0.09 | 1 | HC_vs_MS.Trt_F |
| GRAMD1B | 0.3148 | -0.3629 | 0.312 | 0.33 | 1 | HC_vs_MS.Trt_F |
| CTSO | 0.31483 | 0.32611 | 0.124 | 0.1 | 1 | HC_vs_MS.Trt_F |
| CLN8 | 0.3152 | -0.3627 | 0.847 | 0.8 | 1 | HC_vs_MS.Trt_F |
| USP16 | 0.31533 | 0.36999 | 0.529 | 0.5 | 1 | HC_vs_MS.Trt_F |
| THAP7 | 0.31543 | 0.48027 | 0.139 | 0.11 | 1 | HC_vs_MS.Trt_F |
| FGF9 | 0.31545 | -0.3447 | 0.156 | 0.13 | 1 | HC_vs_MS.Trt_F |
| GRIN2C | 0.31559 | 0.59509 | 0.28 | 0.24 | 1 | HC_vs_MS.Trt_F |
| PAX5 | 0.31627 | -0.2515 | 0.159 | 0.13 | 1 | HC_vs_MS.Trt_F |
| SH3RF2 | 0.3164 | 0.55528 | 0.191 | 0.22 | 1 | HC_vs_MS.Trt_F |
| SP140 | 0.31647 | 0.26931 | 0.289 | 0.24 | 1 | HC_vs_MS.Trt_F |
| NRF1 | 0.31663 | -1.1625 | 0.358 | 0.31 | 1 | HC_vs_MS.Trt_F |

| | | | | | | |
|---|---|---|---|---|---|---|
| ARHGAP19 | 0.31664 | -0.9396 | 0.277 | 0.3 | 1 | HC_vs_MS.Trt_F |
| FBXO42 | 0.31688 | 0.50186 | 0.512 | 0.49 | 1 | HC_vs_MS.Trt_F |
| BLMH | 0.31688 | -0.7047 | 0.65 | 0.65 | 1 | HC_vs_MS.Trt_F |
| C1D | 0.31697 | -0.6432 | 0.139 | 0.16 | 1 | HC_vs_MS.Trt_F |
| GSE1 | 0.31698 | 0.56403 | 0.454 | 0.4 | 1 | HC_vs_MS.Trt_F |
| PDCD10 | 0.317 | -0.2814 | 0.243 | 0.2 | 1 | HC_vs_MS.Trt_F |
| UBE2L5P | 0.31707 | 0.55821 | 0.283 | 0.24 | 1 | HC_vs_MS.Trt_F |
| LINC00854 | 0.31797 | 0.72534 | 0.708 | 0.66 | 1 | HC_vs_MS.Trt_F |
| INTS6L | 0.31806 | -0.3879 | 0.142 | 0.11 | 1 | HC_vs_MS.Trt_F |
| WDYHV1 | 0.3182 | 0.52579 | 0.13 | 0.1 | 1 | HC_vs_MS.Trt_F |
| KLHDC2 | 0.31832 | -0.6611 | 0.246 | 0.27 | 1 | HC_vs_MS.Trt_F |
| SPTBN1 | 0.31835 | -0.3752 | 0.344 | 0.29 | 1 | HC_vs_MS.Trt_F |
| IMPDH1 | 0.31861 | -0.3644 | 0.139 | 0.17 | 1 | HC_vs_MS.Trt_F |
| SLC39A9 | 0.31862 | 0.31461 | 0.225 | 0.19 | 1 | HC_vs_MS.Trt_F |
| ZNF93 | 0.31875 | -0.5094 | 0.613 | 0.53 | 1 | HC_vs_MS.Trt_F |
| SPRED1 | 0.31938 | 0.27773 | 0.188 | 0.16 | 1 | HC_vs_MS.Trt_F |
| SLX1B | 0.31954 | -0.4239 | 0.5 | 0.5 | 1 | HC_vs_MS.Trt_F |
| CD40 | 0.31973 | 0.45481 | 0.185 | 0.15 | 1 | HC_vs_MS.Trt_F |
| STARD10 | 0.31977 | 0.2665 | 0.855 | 0.82 | 1 | HC_vs_MS.Trt_F |
| SNAPC5 | 0.31986 | -0.5266 | 0.616 | 0.6 | 1 | HC_vs_MS.Trt_F |
| ANGEL1 | 0.31996 | 0.34729 | 0.471 | 0.43 | 1 | HC_vs_MS.Trt_F |
| PPT1 | 0.32031 | -0.6546 | 0.214 | 0.23 | 1 | HC_vs_MS.Trt_F |
| MAGED2 | 0.32072 | -0.4791 | 0.24 | 0.19 | 1 | HC_vs_MS.Trt_F |
| CNOT7 | 0.32088 | -0.8124 | 0.546 | 0.54 | 1 | HC_vs_MS.Trt_F |
| POU6F1 | 0.32123 | -0.3487 | 0.558 | 0.49 | 1 | HC_vs_MS.Trt_F |
| SIGLEC8 | 0.32176 | 0.30014 | 0.578 | 0.52 | 1 | HC_vs_MS.Trt_F |
| MEIS3 | 0.3224 | -1.1802 | 0.127 | 0.1 | 1 | HC_vs_MS.Trt_F |
| COG1 | 0.32244 | 0.35632 | 0.902 | 0.85 | 1 | HC_vs_MS.Trt_F |
| DNAJC13 | 0.32264 | 0.36891 | 0.263 | 0.22 | 1 | HC_vs_MS.Trt_F |
| RND1 | 0.32268 | 0.28896 | 0.613 | 0.57 | 1 | HC_vs_MS.Trt_F |
| FAM210A | 0.32297 | -0.2803 | 0.39 | 0.33 | 1 | HC_vs_MS.Trt_F |
| ZCCHC11 | 0.32326 | 0.38618 | 0.945 | 0.92 | 1 | HC_vs_MS.Trt_F |
| ACSM6 | 0.32387 | 0.45274 | 0.147 | 0.12 | 1 | HC_vs_MS.Trt_F |
| IFIT3 | 0.32409 | -2.123 | 0.581 | 0.56 | 1 | HC_vs_MS.Trt_F |
| MS4A2 | 0.32416 | 0.51613 | 0.61 | 0.57 | 1 | HC_vs_MS.Trt_F |
| SLC22A23 | 0.32441 | 1.5538 | 0.101 | 0.08 | 1 | HC_vs_MS.Trt_F |
| CEP78 | 0.3253 | 0.79116 | 0.592 | 0.52 | 1 | HC_vs_MS.Trt_F |
| TRIM66 | 0.32538 | -0.3045 | 0.523 | 0.49 | 1 | HC_vs_MS.Trt_F |
| VPS29 | 0.32588 | -0.3339 | 0.535 | 0.48 | 1 | HC_vs_MS.Trt_F |
| TTC34 | 0.32695 | 0.44283 | 0.217 | 0.18 | 1 | HC_vs_MS.Trt_F |

| | | | | | | |
|---|---|---|---|---|---|---|
| AHSA2 | 0.32745 | 0.27373 | 0.283 | 0.24 | 1 | HC_vs_MS.Trt_F |
| ADPRH | 0.32753 | -0.3831 | 0.081 | 0.1 | 1 | HC_vs_MS.Trt_F |
| JARID2 | 0.32757 | -0.6718 | 0.26 | 0.21 | 1 | HC_vs_MS.Trt_F |
| GTF2H4 | 0.3278 | -0.2915 | 0.113 | 0.14 | 1 | HC_vs_MS.Trt_F |
| SEC22A | 0.32806 | -0.3934 | 0.702 | 0.64 | 1 | HC_vs_MS.Trt_F |
| TMF1 | 0.32852 | -0.2549 | 0.228 | 0.19 | 1 | HC_vs_MS.Trt_F |
| TBXAS1 | 0.32857 | 0.68399 | 0.434 | 0.39 | 1 | HC_vs_MS.Trt_F |
| ICE1 | 0.32861 | -0.3817 | 0.118 | 0.14 | 1 | HC_vs_MS.Trt_F |
| C1GALT1 | 0.32864 | -0.4397 | 0.613 | 0.6 | 1 | HC_vs_MS.Trt_F |
| NDUFS6 | 0.32868 | -0.4486 | 0.171 | 0.14 | 1 | HC_vs_MS.Trt_F |
| IL13RA1 | 0.32893 | 0.32397 | 0.329 | 0.28 | 1 | HC_vs_MS.Trt_F |
| B3GNT2 | 0.3292 | 0.25199 | 0.188 | 0.16 | 1 | HC_vs_MS.Trt_F |
| PARG | 0.32938 | -0.8664 | 0.188 | 0.16 | 1 | HC_vs_MS.Trt_F |
| PIP5KL1 | 0.32958 | 0.3167 | 0.503 | 0.5 | 1 | HC_vs_MS.Trt_F |
| DNAJC27 | 0.32958 | 0.41016 | 0.104 | 0.08 | 1 | HC_vs_MS.Trt_F |
| ABCF2 | 0.32984 | 0.53658 | 0.303 | 0.26 | 1 | HC_vs_MS.Trt_F |
| NR1H2 | 0.32985 | 0.28286 | 0.564 | 0.51 | 1 | HC_vs_MS.Trt_F |
| ADAMTS1 | 0.32992 | -0.2997 | 0.5 | 0.44 | 1 | HC_vs_MS.Trt_F |
| POGLUT1 | 0.3302 | -0.2588 | 0.269 | 0.3 | 1 | HC_vs_MS.Trt_F |
| FAM188B | 0.3303 | 0.70504 | 0.182 | 0.16 | 1 | HC_vs_MS.Trt_F |
| ATP5J | 0.33069 | -1.073 | 0.225 | 0.24 | 1 | HC_vs_MS.Trt_F |
| STARD9 | 0.33082 | 0.25653 | 0.931 | 0.9 | 1 | HC_vs_MS.Trt_F |
| STK16 | 0.33116 | -0.4792 | 0.092 | 0.12 | 1 | HC_vs_MS.Trt_F |
| LEMD2 | 0.33126 | 1.08625 | 0.118 | 0.09 | 1 | HC_vs_MS.Trt_F |
| ZSCAN2 | 0.33223 | 0.49022 | 0.829 | 0.76 | 1 | HC_vs_MS.Trt_F |
| RNFT1 | 0.33234 | -0.359 | 0.442 | 0.39 | 1 | HC_vs_MS.Trt_F |
| NOL10 | 0.33251 | 0.59246 | 0.202 | 0.17 | 1 | HC_vs_MS.Trt_F |
| TRIM33 | 0.33283 | 0.26355 | 0.396 | 0.36 | 1 | HC_vs_MS.Trt_F |
| CTD-2331H1 | 0.33285 | -0.748 | 0.217 | 0.25 | 1 | HC_vs_MS.Trt_F |
| TDG | 0.33314 | 0.58069 | 0.35 | 0.33 | 1 | HC_vs_MS.Trt_F |
| HSBP1 | 0.33336 | -0.6068 | 0.639 | 0.64 | 1 | HC_vs_MS.Trt_F |
| RBM47 | 0.33337 | 0.37089 | 0.494 | 0.44 | 1 | HC_vs_MS.Trt_F |
| TMEM55B | 0.33351 | 0.87583 | 0.113 | 0.09 | 1 | HC_vs_MS.Trt_F |
| VRK1 | 0.33414 | 0.74051 | 0.231 | 0.2 | 1 | HC_vs_MS.Trt_F |
| SRSF9 | 0.33466 | -0.6842 | 0.156 | 0.18 | 1 | HC_vs_MS.Trt_F |
| ZDHHC17 | 0.33477 | 0.82229 | 0.341 | 0.31 | 1 | HC_vs_MS.Trt_F |
| TMEM147 | 0.33492 | -0.403 | 0.165 | 0.13 | 1 | HC_vs_MS.Trt_F |
| ZNF22 | 0.33518 | 0.6641 | 0.118 | 0.09 | 1 | HC_vs_MS.Trt_F |
| TRPS1 | 0.33563 | 1.54946 | 0.127 | 0.1 | 1 | HC_vs_MS.Trt_F |
| BLVRA | 0.33597 | 0.34239 | 0.156 | 0.13 | 1 | HC_vs_MS.Trt_F |

| | | | | | | |
|---|---|---|---|---|---|---|
| PLCH2 | 0.33619 | 1.49775 | 0.104 | 0.08 | 1 | HC_vs_MS.Trt_F |
| MKKS | 0.33626 | -0.7546 | 0.104 | 0.13 | 1 | HC_vs_MS.Trt_F |
| MEGF8 | 0.33635 | 0.33127 | 0.532 | 0.51 | 1 | HC_vs_MS.Trt_F |
| TRIM38 | 0.3365 | 0.25311 | 0.905 | 0.85 | 1 | HC_vs_MS.Trt_F |
| CAB39 | 0.33659 | -0.5254 | 0.329 | 0.28 | 1 | HC_vs_MS.Trt_F |
| RAD17 | 0.33684 | -0.7338 | 0.471 | 0.42 | 1 | HC_vs_MS.Trt_F |
| IFITM1 | 0.33744 | -0.2511 | 0.546 | 0.55 | 1 | HC_vs_MS.Trt_F |
| RAB10 | 0.33814 | -0.7868 | 0.225 | 0.24 | 1 | HC_vs_MS.Trt_F |
| SLC16A6 | 0.33822 | -0.9454 | 0.275 | 0.23 | 1 | HC_vs_MS.Trt_F |
| TGDS | 0.33834 | -0.5368 | 0.098 | 0.12 | 1 | HC_vs_MS.Trt_F |
| CP | 0.33865 | -1.7015 | 0.15 | 0.12 | 1 | HC_vs_MS.Trt_F |
| FAM181B | 0.33922 | 0.25888 | 0.13 | 0.11 | 1 | HC_vs_MS.Trt_F |
| RNF180 | 0.33937 | -0.5564 | 0.35 | 0.31 | 1 | HC_vs_MS.Trt_F |
| MYB | 0.33943 | 0.3182 | 0.364 | 0.32 | 1 | HC_vs_MS.Trt_F |
| PFAS | 0.33962 | -0.893 | 0.662 | 0.6 | 1 | HC_vs_MS.Trt_F |
| DNAJC11 | 0.33972 | 0.61709 | 0.22 | 0.19 | 1 | HC_vs_MS.Trt_F |
| XXYLT1 | 0.33974 | -0.6644 | 0.217 | 0.18 | 1 | HC_vs_MS.Trt_F |
| HLTF | 0.33988 | 1.16363 | 0.494 | 0.44 | 1 | HC_vs_MS.Trt_F |
| NPAS3 | 0.3399 | 0.56781 | 0.448 | 0.4 | 1 | HC_vs_MS.Trt_F |
| L3HYPDH | 0.33995 | -0.5402 | 0.208 | 0.18 | 1 | HC_vs_MS.Trt_F |
| OPA3 | 0.34007 | 0.33925 | 0.977 | 0.96 | 1 | HC_vs_MS.Trt_F |
| DLGAP1 | 0.34011 | -0.4898 | 0.379 | 0.38 | 1 | HC_vs_MS.Trt_F |
| MAGEH1 | 0.34065 | 0.46261 | 0.11 | 0.09 | 1 | HC_vs_MS.Trt_F |
| RBM14-RBM | 0.34067 | 0.3436 | 0.139 | 0.11 | 1 | HC_vs_MS.Trt_F |
| TMEM159 | 0.34081 | 0.26625 | 0.61 | 0.62 | 1 | HC_vs_MS.Trt_F |
| B3GNT4 | 0.34083 | -0.7979 | 0.127 | 0.16 | 1 | HC_vs_MS.Trt_F |
| DCLRE1C | 0.34118 | -0.5804 | 0.717 | 0.69 | 1 | HC_vs_MS.Trt_F |
| ZNF177 | 0.34121 | 0.51412 | 0.107 | 0.13 | 1 | HC_vs_MS.Trt_F |
| SNRPE | 0.34125 | -0.4369 | 0.159 | 0.13 | 1 | HC_vs_MS.Trt_F |
| RPS6KC1 | 0.34127 | 0.87231 | 0.121 | 0.1 | 1 | HC_vs_MS.Trt_F |
| DERL2 | 0.34147 | -0.3664 | 0.341 | 0.3 | 1 | HC_vs_MS.Trt_F |
| UVRAG | 0.34168 | 0.38042 | 0.41 | 0.36 | 1 | HC_vs_MS.Trt_F |
| ID2 | 0.34171 | 0.5743 | 0.118 | 0.1 | 1 | HC_vs_MS.Trt_F |
| FKBP3 | 0.34182 | -0.4023 | 0.301 | 0.25 | 1 | HC_vs_MS.Trt_F |
| ERCC4 | 0.34211 | 0.38048 | 0.884 | 0.89 | 1 | HC_vs_MS.Trt_F |
| PRC1 | 0.34226 | -0.3906 | 0.61 | 0.55 | 1 | HC_vs_MS.Trt_F |
| MTMR4 | 0.34234 | 0.66063 | 0.298 | 0.26 | 1 | HC_vs_MS.Trt_F |
| CTNS | 0.34248 | 0.65076 | 0.182 | 0.16 | 1 | HC_vs_MS.Trt_F |
| DDX54 | 0.34274 | -0.6522 | 0.139 | 0.16 | 1 | HC_vs_MS.Trt_F |
| CCDC88B | 0.34309 | 0.55042 | 0.766 | 0.83 | 1 | HC_vs_MS.Trt_F |

| | | | | | | |
|---|---|---|---|---|---|---|
| DDRGK1 | 0.3435 | -0.4539 | 0.173 | 0.14 | 1 | HC_vs_MS.Trt_F |
| MRPS18A | 0.34362 | -0.5856 | 0.139 | 0.11 | 1 | HC_vs_MS.Trt_F |
| SLC26A2 | 0.34373 | 0.53089 | 0.775 | 0.76 | 1 | HC_vs_MS.Trt_F |
| AHCTF1 | 0.34381 | -0.5281 | 0.171 | 0.14 | 1 | HC_vs_MS.Trt_F |
| C4orf33 | 0.34433 | -0.9936 | 0.101 | 0.08 | 1 | HC_vs_MS.Trt_F |
| C11orf71 | 0.3445 | -0.3654 | 0.118 | 0.09 | 1 | HC_vs_MS.Trt_F |
| ANTXRL | 0.34456 | 0.28857 | 0.208 | 0.17 | 1 | HC_vs_MS.Trt_F |
| ADAMTS17 | 0.34464 | -0.2674 | 0.217 | 0.18 | 1 | HC_vs_MS.Trt_F |
| TCIRG1 | 0.34466 | -0.3104 | 0.315 | 0.33 | 1 | HC_vs_MS.Trt_F |
| COQ9 | 0.3448 | -0.9575 | 0.113 | 0.13 | 1 | HC_vs_MS.Trt_F |
| PRKD2 | 0.34483 | -0.3664 | 0.769 | 0.75 | 1 | HC_vs_MS.Trt_F |
| HNRNPUL2- | 0.34494 | -0.3603 | 0.127 | 0.1 | 1 | HC_vs_MS.Trt_F |
| PHF7 | 0.34505 | 0.76392 | 0.523 | 0.47 | 1 | HC_vs_MS.Trt_F |
| DHRS4L2 | 0.34523 | -0.7811 | 0.191 | 0.21 | 1 | HC_vs_MS.Trt_F |
| AASS | 0.34578 | -0.4284 | 0.422 | 0.35 | 1 | HC_vs_MS.Trt_F |
| DNAJB13 | 0.34583 | 0.29396 | 0.419 | 0.38 | 1 | HC_vs_MS.Trt_F |
| BCR | 0.34644 | -0.959 | 0.341 | 0.29 | 1 | HC_vs_MS.Trt_F |
| SUCLG1 | 0.34662 | -0.5997 | 0.425 | 0.44 | 1 | HC_vs_MS.Trt_F |
| CTD-2021K4 | 0.34692 | -0.6261 | 0.095 | 0.12 | 1 | HC_vs_MS.Trt_F |
| MS4A6A | 0.34703 | 1.17101 | 0.202 | 0.18 | 1 | HC_vs_MS.Trt_F |
| CD3E | 0.34712 | 0.44158 | 0.671 | 0.63 | 1 | HC_vs_MS.Trt_F |
| REXO2 | 0.3474 | -0.566 | 0.185 | 0.21 | 1 | HC_vs_MS.Trt_F |
| ANKFY1 | 0.34756 | 0.69729 | 0.697 | 0.65 | 1 | HC_vs_MS.Trt_F |
| IMMP2L | 0.34779 | 0.67641 | 0.127 | 0.1 | 1 | HC_vs_MS.Trt_F |
| MTMR12 | 0.34827 | 0.56006 | 0.488 | 0.45 | 1 | HC_vs_MS.Trt_F |
| CR1 | 0.34841 | 0.83555 | 0.618 | 0.54 | 1 | HC_vs_MS.Trt_F |
| ACOT13 | 0.34846 | 0.52417 | 0.408 | 0.37 | 1 | HC_vs_MS.Trt_F |
| CHD8 | 0.3485 | 0.80149 | 0.399 | 0.36 | 1 | HC_vs_MS.Trt_F |
| FUCA2 | 0.34891 | -0.6141 | 0.335 | 0.29 | 1 | HC_vs_MS.Trt_F |
| AGTPBP1 | 0.34894 | -0.5437 | 0.202 | 0.17 | 1 | HC_vs_MS.Trt_F |
| CA13 | 0.34904 | -0.4537 | 0.601 | 0.54 | 1 | HC_vs_MS.Trt_F |
| RNF20 | 0.34936 | 0.44647 | 0.15 | 0.12 | 1 | HC_vs_MS.Trt_F |
| FAM234B | 0.34944 | -0.3019 | 0.506 | 0.45 | 1 | HC_vs_MS.Trt_F |
| ABAT | 0.34999 | 0.25136 | 0.656 | 0.61 | 1 | HC_vs_MS.Trt_F |
| AP5M1 | 0.35003 | -0.4635 | 0.159 | 0.18 | 1 | HC_vs_MS.Trt_F |
| MBTPS1 | 0.35023 | -0.3668 | 0.555 | 0.57 | 1 | HC_vs_MS.Trt_F |
| COLGALT1 | 0.35024 | -0.2861 | 0.523 | 0.47 | 1 | HC_vs_MS.Trt_F |
| CPSF7 | 0.35027 | 0.38611 | 0.22 | 0.19 | 1 | HC_vs_MS.Trt_F |
| PPP1CC | 0.35038 | 0.34555 | 0.786 | 0.76 | 1 | HC_vs_MS.Trt_F |
| COQ10B | 0.3504 | -0.6224 | 0.688 | 0.6 | 1 | HC_vs_MS.Trt_F |

| | | | | | | |
|---|---|---|---|---|---|---|
| IL17RD | 0.35076 | -0.4864 | 0.636 | 0.59 | 1 | HC_vs_MS.Trt_F |
| GAREM1 | 0.35084 | 0.4851 | 0.078 | 0.1 | 1 | HC_vs_MS.Trt_F |
| DAD1 | 0.35119 | -0.4175 | 0.41 | 0.32 | 1 | HC_vs_MS.Trt_F |
| TMUB2 | 0.35161 | 0.30972 | 0.882 | 0.85 | 1 | HC_vs_MS.Trt_F |
| TMEM175 | 0.35168 | -0.3147 | 0.318 | 0.35 | 1 | HC_vs_MS.Trt_F |
| RIOK3 | 0.35191 | 0.2702 | 0.575 | 0.54 | 1 | HC_vs_MS.Trt_F |
| PLA2G4B | 0.35192 | -0.8016 | 0.52 | 0.47 | 1 | HC_vs_MS.Trt_F |
| MAN1B1 | 0.35193 | 0.43851 | 0.329 | 0.28 | 1 | HC_vs_MS.Trt_F |
| TXNL4B | 0.35249 | -0.7842 | 0.41 | 0.35 | 1 | HC_vs_MS.Trt_F |
| CBX7 | 0.35278 | -0.6062 | 0.361 | 0.38 | 1 | HC_vs_MS.Trt_F |
| DPH5 | 0.35281 | 0.32604 | 0.361 | 0.32 | 1 | HC_vs_MS.Trt_F |
| IDH2 | 0.35281 | -0.3242 | 0.587 | 0.51 | 1 | HC_vs_MS.Trt_F |
| TMEM168 | 0.35284 | -0.5539 | 0.592 | 0.54 | 1 | HC_vs_MS.Trt_F |
| WDR19 | 0.35335 | 1.00217 | 0.601 | 0.57 | 1 | HC_vs_MS.Trt_F |
| SMAGP | 0.35383 | -0.4536 | 0.636 | 0.62 | 1 | HC_vs_MS.Trt_F |
| TRIM13 | 0.35407 | -0.3546 | 0.682 | 0.61 | 1 | HC_vs_MS.Trt_F |
| KAZN | 0.3548 | 0.74675 | 0.558 | 0.49 | 1 | HC_vs_MS.Trt_F |
| FAM103A1 | 0.35487 | -1.0442 | 0.156 | 0.18 | 1 | HC_vs_MS.Trt_F |
| CUL2 | 0.3552 | 0.34719 | 0.272 | 0.23 | 1 | HC_vs_MS.Trt_F |
| STX5 | 0.35553 | 0.94498 | 0.159 | 0.13 | 1 | HC_vs_MS.Trt_F |
| ABHD10 | 0.35613 | -1.0118 | 0.116 | 0.14 | 1 | HC_vs_MS.Trt_F |
| YWHAG | 0.35628 | 0.30467 | 0.145 | 0.12 | 1 | HC_vs_MS.Trt_F |
| RIT1 | 0.35649 | -0.5966 | 0.477 | 0.42 | 1 | HC_vs_MS.Trt_F |
| CC2D1B | 0.3572 | 0.40855 | 0.65 | 0.64 | 1 | HC_vs_MS.Trt_F |
| TIMM8B | 0.35753 | -0.5985 | 0.118 | 0.09 | 1 | HC_vs_MS.Trt_F |
| PRKAA1 | 0.35901 | 0.56265 | 0.902 | 0.86 | 1 | HC_vs_MS.Trt_F |
| SARNP | 0.35929 | -0.5826 | 0.61 | 0.58 | 1 | HC_vs_MS.Trt_F |
| ANXA4 | 0.35939 | -0.3858 | 0.882 | 0.86 | 1 | HC_vs_MS.Trt_F |
| DNAJB5 | 0.3595 | -0.549 | 0.231 | 0.2 | 1 | HC_vs_MS.Trt_F |
| MCRIP2 | 0.35965 | -0.288 | 0.598 | 0.56 | 1 | HC_vs_MS.Trt_F |
| AKT1S1 | 0.35967 | -0.5018 | 0.22 | 0.18 | 1 | HC_vs_MS.Trt_F |
| PFDN4 | 0.35991 | -0.8159 | 0.098 | 0.12 | 1 | HC_vs_MS.Trt_F |
| SBNO2 | 0.36023 | 0.78389 | 0.228 | 0.19 | 1 | HC_vs_MS.Trt_F |
| MAP1B | 0.36034 | -1.3962 | 0.101 | 0.08 | 1 | HC_vs_MS.Trt_F |
| ERH | 0.36047 | -0.2691 | 0.223 | 0.18 | 1 | HC_vs_MS.Trt_F |
| CHIC1 | 0.36052 | 0.70407 | 0.107 | 0.09 | 1 | HC_vs_MS.Trt_F |
| ADI1 | 0.36114 | -0.2772 | 0.824 | 0.78 | 1 | HC_vs_MS.Trt_F |
| RTCB | 0.36125 | -0.6001 | 0.147 | 0.17 | 1 | HC_vs_MS.Trt_F |
| NETO2 | 0.36173 | -0.3809 | 0.587 | 0.6 | 1 | HC_vs_MS.Trt_F |
| PNPO | 0.36176 | 0.25547 | 0.723 | 0.66 | 1 | HC_vs_MS.Trt_F |

| | | | | | | |
|---|---|---|---|---|---|---|
| MED24 | 0.3618 | 0.60726 | 0.636 | 0.61 | 1 | HC_vs_MS.Trt_F |
| DYRK1A | 0.36186 | 0.68943 | 0.751 | 0.7 | 1 | HC_vs_MS.Trt_F |
| LEKR1 | 0.3625 | -0.3239 | 0.176 | 0.2 | 1 | HC_vs_MS.Trt_F |
| USP30 | 0.36251 | 0.41431 | 0.266 | 0.3 | 1 | HC_vs_MS.Trt_F |
| CEP112 | 0.36271 | 0.34165 | 0.948 | 0.91 | 1 | HC_vs_MS.Trt_F |
| PUS3 | 0.36294 | 0.56572 | 0.358 | 0.31 | 1 | HC_vs_MS.Trt_F |
| SV2A | 0.36322 | 0.4304 | 0.355 | 0.33 | 1 | HC_vs_MS.Trt_F |
| PTRHD1 | 0.36331 | -0.2672 | 0.277 | 0.24 | 1 | HC_vs_MS.Trt_F |
| SPA17 | 0.36393 | -0.3471 | 0.295 | 0.25 | 1 | HC_vs_MS.Trt_F |
| DCUN1D3 | 0.36406 | 0.87706 | 0.231 | 0.2 | 1 | HC_vs_MS.Trt_F |
| KAT5 | 0.36409 | -0.4916 | 0.191 | 0.16 | 1 | HC_vs_MS.Trt_F |
| VPS11 | 0.3644 | -0.4791 | 0.847 | 0.81 | 1 | HC_vs_MS.Trt_F |
| ATG14 | 0.36461 | 0.45997 | 0.72 | 0.71 | 1 | HC_vs_MS.Trt_F |
| CARNMT1 | 0.36497 | -0.4918 | 0.145 | 0.12 | 1 | HC_vs_MS.Trt_F |
| SDHB | 0.36499 | 0.46056 | 0.176 | 0.15 | 1 | HC_vs_MS.Trt_F |
| CNR2 | 0.36507 | 0.55674 | 0.844 | 0.82 | 1 | HC_vs_MS.Trt_F |
| NSA2 | 0.3651 | -0.4286 | 0.335 | 0.37 | 1 | HC_vs_MS.Trt_F |
| SLC25A29 | 0.36514 | 0.79624 | 0.387 | 0.35 | 1 | HC_vs_MS.Trt_F |
| VASH1 | 0.36543 | -0.6703 | 0.116 | 0.09 | 1 | HC_vs_MS.Trt_F |
| SLC13A4 | 0.36571 | 1.23861 | 0.373 | 0.35 | 1 | HC_vs_MS.Trt_F |
| UGT8 | 0.36601 | -0.3916 | 0.546 | 0.46 | 1 | HC_vs_MS.Trt_F |
| TENM1 | 0.36617 | 1.2659 | 0.139 | 0.12 | 1 | HC_vs_MS.Trt_F |
| UBAP2 | 0.36633 | 0.29813 | 0.749 | 0.68 | 1 | HC_vs_MS.Trt_F |
| C17orf51 | 0.36707 | -0.4767 | 0.408 | 0.36 | 1 | HC_vs_MS.Trt_F |
| LPAR6 | 0.36784 | -0.5934 | 0.416 | 0.42 | 1 | HC_vs_MS.Trt_F |
| DHDDS | 0.36792 | -0.4086 | 0.618 | 0.55 | 1 | HC_vs_MS.Trt_F |
| ZNF334 | 0.36807 | 0.76682 | 0.142 | 0.12 | 1 | HC_vs_MS.Trt_F |
| ZNF251 | 0.36832 | -0.3183 | 0.292 | 0.26 | 1 | HC_vs_MS.Trt_F |
| EPHA1 | 0.36844 | 0.43925 | 0.39 | 0.36 | 1 | HC_vs_MS.Trt_F |
| SLC15A2 | 0.36857 | -0.4058 | 0.543 | 0.49 | 1 | HC_vs_MS.Trt_F |
| XPO7 | 0.36867 | -0.4602 | 0.142 | 0.12 | 1 | HC_vs_MS.Trt_F |
| NME2 | 0.36893 | -0.4152 | 0.486 | 0.5 | 1 | HC_vs_MS.Trt_F |
| GPR157 | 0.36899 | 0.3301 | 0.847 | 0.8 | 1 | HC_vs_MS.Trt_F |
| ABC13-4748 | 0.36939 | 0.64602 | 0.101 | 0.08 | 1 | HC_vs_MS.Trt_F |
| MCM7 | 0.36951 | -0.752 | 0.153 | 0.17 | 1 | HC_vs_MS.Trt_F |
| MAP3K7CL | 0.36961 | 0.27609 | 0.465 | 0.42 | 1 | HC_vs_MS.Trt_F |
| GTF3C6 | 0.36964 | -0.489 | 0.199 | 0.16 | 1 | HC_vs_MS.Trt_F |
| CMIP | 0.3697 | 0.74205 | 0.11 | 0.09 | 1 | HC_vs_MS.Trt_F |
| GEMIN7 | 0.36982 | -0.6877 | 0.22 | 0.18 | 1 | HC_vs_MS.Trt_F |
| IFT172 | 0.37007 | 0.72789 | 0.65 | 0.59 | 1 | HC_vs_MS.Trt_F |

| | | | | | | |
|---|---|---|---|---|---|---|
| ZNF365 | 0.37033 | -0.7188 | 0.35 | 0.31 | 1 | HC_vs_MS.Trt_F |
| TMEM170A | 0.37093 | -0.262 | 0.861 | 0.83 | 1 | HC_vs_MS.Trt_F |
| SENP5 | 0.37106 | -0.5768 | 0.939 | 0.92 | 1 | HC_vs_MS.Trt_F |
| SETD2 | 0.37106 | -0.3415 | 0.399 | 0.34 | 1 | HC_vs_MS.Trt_F |
| ZNF701 | 0.37106 | -0.4371 | 0.477 | 0.44 | 1 | HC_vs_MS.Trt_F |
| PLPP5 | 0.37144 | -0.369 | 0.864 | 0.88 | 1 | HC_vs_MS.Trt_F |
| CCDC12 | 0.37153 | -0.3003 | 0.254 | 0.28 | 1 | HC_vs_MS.Trt_F |
| DGCR2 | 0.37159 | 0.27717 | 0.257 | 0.23 | 1 | HC_vs_MS.Trt_F |
| MASP2 | 0.3716 | -1.0593 | 0.335 | 0.3 | 1 | HC_vs_MS.Trt_F |
| DISP1 | 0.37167 | 1.26472 | 0.121 | 0.1 | 1 | HC_vs_MS.Trt_F |
| ZCCHC7 | 0.37188 | 0.40789 | 0.332 | 0.3 | 1 | HC_vs_MS.Trt_F |
| PIGS | 0.37268 | -0.5346 | 0.514 | 0.53 | 1 | HC_vs_MS.Trt_F |
| ARMCX3 | 0.37286 | 0.71571 | 0.116 | 0.09 | 1 | HC_vs_MS.Trt_F |
| FBL | 0.373 | -0.324 | 0.474 | 0.44 | 1 | HC_vs_MS.Trt_F |
| SLC39A7 | 0.37302 | -0.4968 | 0.113 | 0.13 | 1 | HC_vs_MS.Trt_F |
| METTL5 | 0.37311 | 0.48464 | 0.546 | 0.51 | 1 | HC_vs_MS.Trt_F |
| SAAL1 | 0.37316 | -0.4261 | 0.09 | 0.11 | 1 | HC_vs_MS.Trt_F |
| TCP11L1 | 0.37318 | 0.31269 | 0.199 | 0.23 | 1 | HC_vs_MS.Trt_F |
| CDCA3 | 0.37335 | -0.4607 | 0.575 | 0.57 | 1 | HC_vs_MS.Trt_F |
| JAGN1 | 0.37341 | -0.3025 | 0.11 | 0.13 | 1 | HC_vs_MS.Trt_F |
| ELP5 | 0.37345 | 1.01414 | 0.133 | 0.11 | 1 | HC_vs_MS.Trt_F |
| NFKB2 | 0.37351 | 1.18993 | 0.428 | 0.4 | 1 | HC_vs_MS.Trt_F |
| FBLN7 | 0.37357 | 0.66668 | 0.439 | 0.41 | 1 | HC_vs_MS.Trt_F |
| IRF2BPL | 0.3736 | -0.3286 | 0.474 | 0.49 | 1 | HC_vs_MS.Trt_F |
| CCDC138 | 0.37368 | 0.6433 | 0.217 | 0.19 | 1 | HC_vs_MS.Trt_F |
| SEMA6D | 0.37423 | 1.36796 | 0.147 | 0.13 | 1 | HC_vs_MS.Trt_F |
| RFX2 | 0.37444 | -0.3438 | 0.514 | 0.48 | 1 | HC_vs_MS.Trt_F |
| DRAM2 | 0.37453 | -0.2863 | 0.188 | 0.21 | 1 | HC_vs_MS.Trt_F |
| TSC22D2 | 0.37462 | -0.3816 | 0.488 | 0.44 | 1 | HC_vs_MS.Trt_F |
| GLRX3 | 0.37479 | -0.312 | 0.887 | 0.83 | 1 | HC_vs_MS.Trt_F |
| GNPTAB | 0.37532 | -0.3111 | 0.74 | 0.67 | 1 | HC_vs_MS.Trt_F |
| ALKBH6 | 0.37542 | 0.29375 | 0.538 | 0.57 | 1 | HC_vs_MS.Trt_F |
| PPOX | 0.37616 | -0.3883 | 0.263 | 0.23 | 1 | HC_vs_MS.Trt_F |
| DMWD | 0.37647 | 0.52382 | 0.879 | 0.84 | 1 | HC_vs_MS.Trt_F |
| CTD-3105H1 | 0.37651 | 0.36826 | 0.358 | 0.31 | 1 | HC_vs_MS.Trt_F |
| SMC1A | 0.37711 | 0.40707 | 0.263 | 0.23 | 1 | HC_vs_MS.Trt_F |
| STPG1 | 0.37767 | 0.45301 | 0.801 | 0.79 | 1 | HC_vs_MS.Trt_F |
| GDE1 | 0.37828 | 1.46721 | 0.315 | 0.28 | 1 | HC_vs_MS.Trt_F |
| PAN2 | 0.37856 | -0.5238 | 0.353 | 0.37 | 1 | HC_vs_MS.Trt_F |
| EBP | 0.37883 | -0.3073 | 0.234 | 0.19 | 1 | HC_vs_MS.Trt_F |

| | | | | | | |
|---|---|---|---|---|---|---|
| BGN | 0.37889 | 1.55507 | 0.084 | 0.11 | 1 | HC_vs_MS.Trt_F |
| SPCS2 | 0.37891 | -0.4149 | 0.324 | 0.35 | 1 | HC_vs_MS.Trt_F |
| MLH1 | 0.37976 | 0.27439 | 0.39 | 0.36 | 1 | HC_vs_MS.Trt_F |
| TSG101 | 0.37991 | -0.5151 | 0.506 | 0.44 | 1 | HC_vs_MS.Trt_F |
| C2orf69 | 0.38001 | -0.4002 | 0.303 | 0.26 | 1 | HC_vs_MS.Trt_F |
| AZIN1 | 0.38042 | -0.2717 | 0.399 | 0.34 | 1 | HC_vs_MS.Trt_F |
| ADNP | 0.38062 | -0.7614 | 0.483 | 0.43 | 1 | HC_vs_MS.Trt_F |
| PHF19 | 0.38072 | -0.4491 | 0.191 | 0.16 | 1 | HC_vs_MS.Trt_F |
| SRFBP1 | 0.38081 | -1.2217 | 0.471 | 0.46 | 1 | HC_vs_MS.Trt_F |
| ZNF696 | 0.38106 | -0.3647 | 0.344 | 0.36 | 1 | HC_vs_MS.Trt_F |
| GLRX5 | 0.38136 | -1.2549 | 0.101 | 0.12 | 1 | HC_vs_MS.Trt_F |
| TRIM28 | 0.38202 | 0.94737 | 0.113 | 0.09 | 1 | HC_vs_MS.Trt_F |
| CFAP20 | 0.38214 | -0.4783 | 0.162 | 0.18 | 1 | HC_vs_MS.Trt_F |
| ANKRD42 | 0.38243 | 1.09239 | 0.442 | 0.39 | 1 | HC_vs_MS.Trt_F |
| PGAM5 | 0.38262 | 0.33815 | 0.803 | 0.77 | 1 | HC_vs_MS.Trt_F |
| USP5 | 0.38264 | 0.69259 | 0.147 | 0.12 | 1 | HC_vs_MS.Trt_F |
| PNKP | 0.38276 | 0.68333 | 0.327 | 0.3 | 1 | HC_vs_MS.Trt_F |
| ASNSD1 | 0.38322 | -0.5663 | 0.165 | 0.13 | 1 | HC_vs_MS.Trt_F |
| ZNF140 | 0.38353 | -1.4524 | 0.223 | 0.19 | 1 | HC_vs_MS.Trt_F |
| C1orf186 | 0.38366 | 0.97009 | 0.587 | 0.55 | 1 | HC_vs_MS.Trt_F |
| BEX3 | 0.38372 | -0.3226 | 0.145 | 0.12 | 1 | HC_vs_MS.Trt_F |
| SLC37A2 | 0.38377 | -0.3163 | 0.514 | 0.51 | 1 | HC_vs_MS.Trt_F |
| SH2D1A | 0.3838 | -0.6895 | 0.292 | 0.24 | 1 | HC_vs_MS.Trt_F |
| VPS18 | 0.38395 | -0.644 | 0.251 | 0.23 | 1 | HC_vs_MS.Trt_F |
| SLC30A5 | 0.38396 | -0.4817 | 0.809 | 0.73 | 1 | HC_vs_MS.Trt_F |
| TRPM1 | 0.38397 | 0.77866 | 0.133 | 0.11 | 1 | HC_vs_MS.Trt_F |
| ATL3 | 0.38445 | -0.3491 | 0.838 | 0.81 | 1 | HC_vs_MS.Trt_F |
| TAL1 | 0.38446 | 0.43121 | 0.26 | 0.22 | 1 | HC_vs_MS.Trt_F |
| KIN | 0.38458 | -0.3283 | 0.772 | 0.74 | 1 | HC_vs_MS.Trt_F |
| NOL6 | 0.38478 | 0.38529 | 0.431 | 0.38 | 1 | HC_vs_MS.Trt_F |
| CCAR2 | 0.38499 | -0.7096 | 0.272 | 0.23 | 1 | HC_vs_MS.Trt_F |
| ZNF514 | 0.38545 | 0.74818 | 0.642 | 0.59 | 1 | HC_vs_MS.Trt_F |
| UNG | 0.38577 | -0.4657 | 0.24 | 0.21 | 1 | HC_vs_MS.Trt_F |
| RPF1 | 0.38603 | -0.3971 | 0.283 | 0.25 | 1 | HC_vs_MS.Trt_F |
| CHAMP1 | 0.38618 | 0.43868 | 0.673 | 0.64 | 1 | HC_vs_MS.Trt_F |
| PRKCI | 0.38626 | -0.3294 | 0.419 | 0.36 | 1 | HC_vs_MS.Trt_F |
| PCDH9 | 0.38666 | 0.63727 | 0.691 | 0.67 | 1 | HC_vs_MS.Trt_F |
| NME9 | 0.38706 | -0.709 | 0.275 | 0.24 | 1 | HC_vs_MS.Trt_F |
| JCHAIN | 0.38717 | -3.1749 | 0.11 | 0.13 | 1 | HC_vs_MS.Trt_F |
| HSF2 | 0.38728 | -0.5843 | 0.142 | 0.12 | 1 | HC_vs_MS.Trt_F |

| | | | | | | |
|---|---|---|---|---|---|---|
| DHRS11 | 0.38846 | 0.95057 | 0.344 | 0.29 | 1 | HC_vs_MS.Trt_F |
| BNIP1 | 0.38858 | -1.9509 | 0.087 | 0.1 | 1 | HC_vs_MS.Trt_F |
| ZNF235 | 0.38916 | -1.4359 | 0.159 | 0.13 | 1 | HC_vs_MS.Trt_F |
| LUC7L | 0.38935 | 0.58983 | 0.864 | 0.85 | 1 | HC_vs_MS.Trt_F |
| ZFYVE27 | 0.38957 | 0.36948 | 0.225 | 0.19 | 1 | HC_vs_MS.Trt_F |
| CSNK2A2 | 0.39024 | -0.4207 | 0.344 | 0.29 | 1 | HC_vs_MS.Trt_F |
| YES1 | 0.39034 | -0.368 | 0.347 | 0.37 | 1 | HC_vs_MS.Trt_F |
| NFS1 | 0.39052 | -0.9974 | 0.934 | 0.91 | 1 | HC_vs_MS.Trt_F |
| ZNF14 | 0.39056 | -0.6055 | 0.387 | 0.4 | 1 | HC_vs_MS.Trt_F |
| PDE8A | 0.39072 | 0.29125 | 0.526 | 0.46 | 1 | HC_vs_MS.Trt_F |
| HSDL2 | 0.39105 | -0.5903 | 0.185 | 0.16 | 1 | HC_vs_MS.Trt_F |
| RECK | 0.39147 | 0.27603 | 0.188 | 0.16 | 1 | HC_vs_MS.Trt_F |
| C19orf44 | 0.39195 | 0.59824 | 0.387 | 0.35 | 1 | HC_vs_MS.Trt_F |
| DLEC1 | 0.39232 | 1.4864 | 0.514 | 0.46 | 1 | HC_vs_MS.Trt_F |
| DEF6 | 0.39251 | -0.4752 | 0.428 | 0.43 | 1 | HC_vs_MS.Trt_F |
| EXO1 | 0.39302 | 0.42983 | 0.197 | 0.23 | 1 | HC_vs_MS.Trt_F |
| C5orf63 | 0.39368 | -0.4026 | 0.393 | 0.35 | 1 | HC_vs_MS.Trt_F |
| PIGW | 0.39388 | 1.03497 | 0.477 | 0.44 | 1 | HC_vs_MS.Trt_F |
| HELZ2 | 0.3942 | -0.4383 | 0.699 | 0.64 | 1 | HC_vs_MS.Trt_F |
| BCS1L | 0.39422 | -0.3337 | 0.09 | 0.11 | 1 | HC_vs_MS.Trt_F |
| USP22 | 0.39433 | -0.4591 | 0.847 | 0.82 | 1 | HC_vs_MS.Trt_F |
| FAM105A | 0.39444 | -0.287 | 0.28 | 0.25 | 1 | HC_vs_MS.Trt_F |
| SEPP1 | 0.39449 | 0.46743 | 0.188 | 0.16 | 1 | HC_vs_MS.Trt_F |
| ZNF600 | 0.39486 | 0.79763 | 0.231 | 0.2 | 1 | HC_vs_MS.Trt_F |
| DHX35 | 0.39498 | 0.4032 | 0.422 | 0.39 | 1 | HC_vs_MS.Trt_F |
| TC2N | 0.39501 | 0.30925 | 0.211 | 0.18 | 1 | HC_vs_MS.Trt_F |
| FKBP8 | 0.39511 | 1.35329 | 0.234 | 0.2 | 1 | HC_vs_MS.Trt_F |
| DHX40 | 0.39523 | -0.4454 | 0.147 | 0.12 | 1 | HC_vs_MS.Trt_F |
| NAALADL2 | 0.39527 | 0.31968 | 0.092 | 0.12 | 1 | HC_vs_MS.Trt_F |
| ASB8 | 0.3957 | 0.3952 | 0.74 | 0.75 | 1 | HC_vs_MS.Trt_F |
| ANKRD11 | 0.39572 | -0.4065 | 0.306 | 0.26 | 1 | HC_vs_MS.Trt_F |
| ZNF687 | 0.39593 | 0.54468 | 0.65 | 0.64 | 1 | HC_vs_MS.Trt_F |
| IFIT2 | 0.39595 | -0.8543 | 0.116 | 0.09 | 1 | HC_vs_MS.Trt_F |
| PHF10 | 0.39608 | 1.21532 | 0.113 | 0.09 | 1 | HC_vs_MS.Trt_F |
| TMEM220 | 0.39633 | 0.38223 | 0.572 | 0.57 | 1 | HC_vs_MS.Trt_F |
| MAVS | 0.39639 | 0.28495 | 0.962 | 0.95 | 1 | HC_vs_MS.Trt_F |
| SPOCD1 | 0.39659 | -0.4521 | 0.575 | 0.6 | 1 | HC_vs_MS.Trt_F |
| NDUFB3 | 0.39752 | -0.4759 | 0.199 | 0.22 | 1 | HC_vs_MS.Trt_F |
| PTP4A3 | 0.39772 | 0.50717 | 0.11 | 0.09 | 1 | HC_vs_MS.Trt_F |
| ZFYVE21 | 0.3978 | -0.7393 | 0.737 | 0.66 | 1 | HC_vs_MS.Trt_F |

| | | | | | | |
|---|---|---|---|---|---|---|
| VAV3 | 0.39841 | 0.40579 | 0.592 | 0.6 | 1 | HC_vs_MS.Trt_F |
| STK40 | 0.39878 | -0.3566 | 0.136 | 0.11 | 1 | HC_vs_MS.Trt_F |
| PLXDC2 | 0.39887 | 0.79678 | 0.74 | 0.7 | 1 | HC_vs_MS.Trt_F |
| MED8 | 0.39902 | 0.94137 | 0.116 | 0.09 | 1 | HC_vs_MS.Trt_F |
| MRE11A | 0.39913 | -0.8638 | 0.743 | 0.75 | 1 | HC_vs_MS.Trt_F |
| SWAP70 | 0.39914 | -1.8236 | 0.483 | 0.4 | 1 | HC_vs_MS.Trt_F |
| PRSS53 | 0.39954 | 1.41448 | 0.257 | 0.22 | 1 | HC_vs_MS.Trt_F |
| CCDC77 | 0.39971 | -0.411 | 0.714 | 0.63 | 1 | HC_vs_MS.Trt_F |
| SCAF4 | 0.3999 | -0.3214 | 0.28 | 0.24 | 1 | HC_vs_MS.Trt_F |
| TRMT6 | 0.40009 | -1.2812 | 0.118 | 0.14 | 1 | HC_vs_MS.Trt_F |
| HECA | 0.40013 | -0.5608 | 0.546 | 0.5 | 1 | HC_vs_MS.Trt_F |
| E2F3 | 0.40016 | 0.28299 | 0.225 | 0.19 | 1 | HC_vs_MS.Trt_F |
| CBLL1 | 0.40021 | -0.2678 | 0.243 | 0.21 | 1 | HC_vs_MS.Trt_F |
| KLRK1 | 0.40053 | -0.5955 | 0.711 | 0.66 | 1 | HC_vs_MS.Trt_F |
| ANKRD46 | 0.40056 | 0.72015 | 0.223 | 0.2 | 1 | HC_vs_MS.Trt_F |
| MORC3 | 0.4006 | -0.3381 | 0.503 | 0.48 | 1 | HC_vs_MS.Trt_F |
| CDC16 | 0.40089 | -0.4273 | 0.159 | 0.18 | 1 | HC_vs_MS.Trt_F |
| BAD | 0.40109 | 0.33378 | 0.147 | 0.12 | 1 | HC_vs_MS.Trt_F |
| ARRDC2 | 0.40139 | 0.32164 | 0.15 | 0.13 | 1 | HC_vs_MS.Trt_F |
| SAE1 | 0.40177 | -0.3065 | 0.22 | 0.19 | 1 | HC_vs_MS.Trt_F |
| RINL | 0.40211 | 0.36594 | 0.763 | 0.74 | 1 | HC_vs_MS.Trt_F |
| ZBTB21 | 0.40215 | -0.6268 | 0.127 | 0.15 | 1 | HC_vs_MS.Trt_F |
| SGO1 | 0.40237 | -0.3473 | 0.543 | 0.52 | 1 | HC_vs_MS.Trt_F |
| NDUFC2-KC | 0.40271 | -0.3677 | 0.092 | 0.11 | 1 | HC_vs_MS.Trt_F |
| PLEKHG4B | 0.40304 | 0.48363 | 0.509 | 0.44 | 1 | HC_vs_MS.Trt_F |
| UBXN8 | 0.40314 | 0.74216 | 0.22 | 0.2 | 1 | HC_vs_MS.Trt_F |
| KALRN | 0.40374 | 0.42165 | 0.329 | 0.3 | 1 | HC_vs_MS.Trt_F |
| LIPA | 0.40405 | 0.8143 | 0.318 | 0.28 | 1 | HC_vs_MS.Trt_F |
| OXSM | 0.40407 | -0.6782 | 0.153 | 0.13 | 1 | HC_vs_MS.Trt_F |
| SPHK1 | 0.40462 | -0.3425 | 0.223 | 0.19 | 1 | HC_vs_MS.Trt_F |
| AL078584.1 | 0.40478 | 0.346 | 0.471 | 0.42 | 1 | HC_vs_MS.Trt_F |
| SLF1 | 0.40495 | 0.61078 | 0.197 | 0.17 | 1 | HC_vs_MS.Trt_F |
| FKRP | 0.40565 | -0.3742 | 0.702 | 0.65 | 1 | HC_vs_MS.Trt_F |
| NIN | 0.40566 | -0.3792 | 0.199 | 0.22 | 1 | HC_vs_MS.Trt_F |
| CDK12 | 0.40576 | 0.39183 | 0.561 | 0.49 | 1 | HC_vs_MS.Trt_F |
| AQR | 0.40586 | -0.2987 | 0.671 | 0.63 | 1 | HC_vs_MS.Trt_F |
| ASAH1 | 0.40604 | -0.3267 | 0.76 | 0.73 | 1 | HC_vs_MS.Trt_F |
| ZNF585A | 0.4062 | -1.2435 | 0.353 | 0.36 | 1 | HC_vs_MS.Trt_F |
| GPR89A | 0.40629 | -0.3411 | 0.48 | 0.43 | 1 | HC_vs_MS.Trt_F |
| BBS4 | 0.40629 | 0.33778 | 0.107 | 0.13 | 1 | HC_vs_MS.Trt_F |

| | | | | | | |
|---|---|---|---|---|---|---|
| ABCC5 | 0.40634 | -0.6639 | 0.179 | 0.15 | 1 | HC_vs_MS.Trt_F |
| ARRDC1 | 0.40654 | -0.2855 | 0.306 | 0.27 | 1 | HC_vs_MS.Trt_F |
| CNN3 | 0.40696 | -0.362 | 0.289 | 0.26 | 1 | HC_vs_MS.Trt_F |
| LTBP4 | 0.40855 | 0.47252 | 0.561 | 0.5 | 1 | HC_vs_MS.Trt_F |
| TCEAL8 | 0.40871 | -0.4132 | 0.116 | 0.09 | 1 | HC_vs_MS.Trt_F |
| TRMT112 | 0.40879 | -0.4891 | 0.272 | 0.29 | 1 | HC_vs_MS.Trt_F |
| HRH2 | 0.40885 | 0.57601 | 0.136 | 0.12 | 1 | HC_vs_MS.Trt_F |
| ZNF433 | 0.4089 | -0.4583 | 0.185 | 0.16 | 1 | HC_vs_MS.Trt_F |
| ABHD6 | 0.40895 | -0.6853 | 0.532 | 0.49 | 1 | HC_vs_MS.Trt_F |
| IFIT5 | 0.40897 | -0.6682 | 0.104 | 0.08 | 1 | HC_vs_MS.Trt_F |
| H6PD | 0.40923 | 0.41467 | 0.87 | 0.85 | 1 | HC_vs_MS.Trt_F |
| PCYOX1L | 0.40974 | 1.29478 | 0.11 | 0.09 | 1 | HC_vs_MS.Trt_F |
| IL3RA | 0.40988 | 0.89519 | 0.121 | 0.1 | 1 | HC_vs_MS.Trt_F |
| ARHGEF15 | 0.41023 | 1.00266 | 0.454 | 0.42 | 1 | HC_vs_MS.Trt_F |
| KCNK10 | 0.41032 | 0.28128 | 0.173 | 0.16 | 1 | HC_vs_MS.Trt_F |
| MRPL18 | 0.41062 | -1.3672 | 0.145 | 0.16 | 1 | HC_vs_MS.Trt_F |
| AP1AR | 0.4108 | -0.5713 | 0.107 | 0.09 | 1 | HC_vs_MS.Trt_F |
| PCNA | 0.41081 | -0.3679 | 0.121 | 0.1 | 1 | HC_vs_MS.Trt_F |
| THOC3 | 0.41102 | -0.2884 | 0.532 | 0.53 | 1 | HC_vs_MS.Trt_F |
| STK38L | 0.41109 | -0.6831 | 0.702 | 0.66 | 1 | HC_vs_MS.Trt_F |
| AGO4 | 0.41137 | 0.99656 | 0.335 | 0.31 | 1 | HC_vs_MS.Trt_F |
| NAXE | 0.41179 | -0.5121 | 0.162 | 0.13 | 1 | HC_vs_MS.Trt_F |
| ZNF135 | 0.41195 | -1.2483 | 0.486 | 0.43 | 1 | HC_vs_MS.Trt_F |
| SNX12 | 0.41247 | -0.3184 | 0.116 | 0.09 | 1 | HC_vs_MS.Trt_F |
| C12orf49 | 0.41273 | -0.4082 | 0.74 | 0.74 | 1 | HC_vs_MS.Trt_F |
| ASF1A | 0.41277 | -0.8561 | 0.156 | 0.17 | 1 | HC_vs_MS.Trt_F |
| PCNX3 | 0.41335 | 0.40561 | 0.159 | 0.13 | 1 | HC_vs_MS.Trt_F |
| DCTPP1 | 0.41427 | -1.3482 | 0.095 | 0.11 | 1 | HC_vs_MS.Trt_F |
| PARVA | 0.41427 | 0.43901 | 0.749 | 0.72 | 1 | HC_vs_MS.Trt_F |
| SETDB1 | 0.41442 | -0.4965 | 0.63 | 0.6 | 1 | HC_vs_MS.Trt_F |
| SEMA4B | 0.41445 | -0.4352 | 0.376 | 0.35 | 1 | HC_vs_MS.Trt_F |
| NCR3LG1 | 0.41454 | 0.44308 | 0.084 | 0.1 | 1 | HC_vs_MS.Trt_F |
| TMEM39B | 0.41475 | 0.46975 | 0.176 | 0.16 | 1 | HC_vs_MS.Trt_F |
| CKAP5 | 0.41475 | -0.4795 | 0.168 | 0.14 | 1 | HC_vs_MS.Trt_F |
| ERCC8 | 0.41537 | -0.6079 | 0.344 | 0.36 | 1 | HC_vs_MS.Trt_F |
| ZSWIM1 | 0.41543 | 0.37997 | 0.734 | 0.74 | 1 | HC_vs_MS.Trt_F |
| LRP1 | 0.41612 | 1.16693 | 0.344 | 0.31 | 1 | HC_vs_MS.Trt_F |
| RALGPS1 | 0.41624 | 0.9575 | 0.477 | 0.41 | 1 | HC_vs_MS.Trt_F |
| RPS6KB2 | 0.41642 | 0.27406 | 0.861 | 0.85 | 1 | HC_vs_MS.Trt_F |
| DPY19L4 | 0.4168 | 0.87318 | 0.572 | 0.53 | 1 | HC_vs_MS.Trt_F |

| | | | | | | |
|---|---|---|---|---|---|---|
| PEX7 | 0.41703 | 0.91635 | 0.171 | 0.15 | 1 | HC_vs_MS.Trt_F |
| LINC00493 | 0.4172 | -0.3345 | 0.451 | 0.4 | 1 | HC_vs_MS.Trt_F |
| CCDC25 | 0.41721 | -1.3185 | 0.286 | 0.31 | 1 | HC_vs_MS.Trt_F |
| TCEB3 | 0.41725 | -0.5019 | 0.145 | 0.12 | 1 | HC_vs_MS.Trt_F |
| CFP | 0.41758 | -1.1781 | 0.422 | 0.36 | 1 | HC_vs_MS.Trt_F |
| CREG2 | 0.41773 | -0.3308 | 0.746 | 0.69 | 1 | HC_vs_MS.Trt_F |
| PAK1 | 0.4181 | -1.2097 | 0.202 | 0.17 | 1 | HC_vs_MS.Trt_F |
| CEP95 | 0.41838 | -0.2783 | 0.419 | 0.45 | 1 | HC_vs_MS.Trt_F |
| MED17 | 0.41855 | 0.27262 | 0.957 | 0.93 | 1 | HC_vs_MS.Trt_F |
| ATP6AP1 | 0.41869 | 0.45623 | 0.483 | 0.46 | 1 | HC_vs_MS.Trt_F |
| TFE3 | 0.41877 | 0.49589 | 0.11 | 0.09 | 1 | HC_vs_MS.Trt_F |
| RNF185 | 0.419 | -0.6343 | 0.306 | 0.27 | 1 | HC_vs_MS.Trt_F |
| TRMT1L | 0.41907 | 0.25587 | 0.124 | 0.1 | 1 | HC_vs_MS.Trt_F |
| PTGR1 | 0.41912 | 0.30087 | 0.639 | 0.6 | 1 | HC_vs_MS.Trt_F |
| NDUFA12 | 0.41921 | -0.6584 | 0.223 | 0.24 | 1 | HC_vs_MS.Trt_F |
| GPR132 | 0.42001 | 0.25096 | 0.419 | 0.38 | 1 | HC_vs_MS.Trt_F |
| TNFRSF25 | 0.42033 | -0.3643 | 0.717 | 0.68 | 1 | HC_vs_MS.Trt_F |
| CLASP1 | 0.4206 | 0.28423 | 0.312 | 0.27 | 1 | HC_vs_MS.Trt_F |
| ABCG8 | 0.42073 | 0.73532 | 0.633 | 0.59 | 1 | HC_vs_MS.Trt_F |
| FAM161B | 0.42105 | -0.6933 | 0.379 | 0.4 | 1 | HC_vs_MS.Trt_F |
| KCTD20 | 0.42144 | -0.3884 | 0.685 | 0.7 | 1 | HC_vs_MS.Trt_F |
| CLEC4E | 0.42151 | 0.64257 | 0.425 | 0.39 | 1 | HC_vs_MS.Trt_F |
| FTSJ1 | 0.42228 | -0.2527 | 0.168 | 0.14 | 1 | HC_vs_MS.Trt_F |
| SLIT1 | 0.42251 | 0.90024 | 0.277 | 0.25 | 1 | HC_vs_MS.Trt_F |
| LCORL | 0.42264 | -0.3557 | 0.136 | 0.11 | 1 | HC_vs_MS.Trt_F |
| CHKB | 0.42282 | 0.36537 | 0.277 | 0.25 | 1 | HC_vs_MS.Trt_F |
| AGPAT5 | 0.42316 | -1.1419 | 0.127 | 0.15 | 1 | HC_vs_MS.Trt_F |
| USP48 | 0.42333 | -0.4832 | 0.552 | 0.57 | 1 | HC_vs_MS.Trt_F |
| UBE2V1 | 0.42374 | -0.2562 | 0.483 | 0.42 | 1 | HC_vs_MS.Trt_F |
| FCER1G | 0.42387 | 0.30646 | 0.671 | 0.71 | 1 | HC_vs_MS.Trt_F |
| PIK3C2A | 0.42394 | 0.39453 | 0.431 | 0.45 | 1 | HC_vs_MS.Trt_F |
| BROX | 0.42396 | -0.9487 | 0.624 | 0.64 | 1 | HC_vs_MS.Trt_F |
| CD68 | 0.42409 | 0.42024 | 0.604 | 0.55 | 1 | HC_vs_MS.Trt_F |
| CHP1 | 0.42416 | -0.4635 | 0.806 | 0.78 | 1 | HC_vs_MS.Trt_F |
| ZNF557 | 0.42443 | -0.4033 | 0.879 | 0.87 | 1 | HC_vs_MS.Trt_F |
| UQCC3 | 0.42446 | -0.4057 | 0.48 | 0.43 | 1 | HC_vs_MS.Trt_F |
| RND2 | 0.42446 | 0.4931 | 0.425 | 0.39 | 1 | HC_vs_MS.Trt_F |
| ARHGEF4 | 0.42507 | 0.37064 | 0.627 | 0.59 | 1 | HC_vs_MS.Trt_F |
| LA16c-431H | 0.42522 | 0.62813 | 0.555 | 0.53 | 1 | HC_vs_MS.Trt_F |
| TIMM22 | 0.42542 | -0.6624 | 0.107 | 0.12 | 1 | HC_vs_MS.Trt_F |

| | | | | | | |
|---|---|---|---|---|---|---|
| A2ML1 | 0.42551 | 0.46115 | 0.685 | 0.63 | 1 | HC_vs_MS.Trt_F |
| FBXW4 | 0.42571 | -0.2697 | 0.506 | 0.45 | 1 | HC_vs_MS.Trt_F |
| ANK3 | 0.42578 | 0.47701 | 0.942 | 0.98 | 1 | HC_vs_MS.Trt_F |
| PLPP2 | 0.42663 | -0.2808 | 0.919 | 0.91 | 1 | HC_vs_MS.Trt_F |
| IRGM | 0.42695 | 0.43307 | 0.413 | 0.45 | 1 | HC_vs_MS.Trt_F |
| NEK8 | 0.42735 | -0.9169 | 0.621 | 0.57 | 1 | HC_vs_MS.Trt_F |
| UBE2K | 0.4276 | -0.5577 | 0.162 | 0.18 | 1 | HC_vs_MS.Trt_F |
| EIF2A | 0.42784 | -0.7369 | 0.621 | 0.61 | 1 | HC_vs_MS.Trt_F |
| VAMP5 | 0.42796 | -0.4901 | 0.182 | 0.15 | 1 | HC_vs_MS.Trt_F |
| HAUS5 | 0.42833 | -0.5777 | 0.523 | 0.53 | 1 | HC_vs_MS.Trt_F |
| RMND5A | 0.42915 | -0.5903 | 0.384 | 0.33 | 1 | HC_vs_MS.Trt_F |
| ZMYND11 | 0.42956 | -0.3598 | 0.13 | 0.11 | 1 | HC_vs_MS.Trt_F |
| HLA-DQB1 | 0.43031 | 0.99968 | 0.581 | 0.6 | 1 | HC_vs_MS.Trt_F |
| LBHD1 | 0.43071 | -1.0236 | 0.523 | 0.53 | 1 | HC_vs_MS.Trt_F |
| STAG3 | 0.4314 | 0.3385 | 0.234 | 0.21 | 1 | HC_vs_MS.Trt_F |
| ELOVL6 | 0.43156 | -0.2592 | 0.379 | 0.34 | 1 | HC_vs_MS.Trt_F |
| PLEKHG3 | 0.43161 | -0.3412 | 0.5 | 0.46 | 1 | HC_vs_MS.Trt_F |
| LOXL1 | 0.43198 | 0.58509 | 0.104 | 0.09 | 1 | HC_vs_MS.Trt_F |
| RPLP0 | 0.43226 | -0.2518 | 0.925 | 0.85 | 1 | HC_vs_MS.Trt_F |
| TAF6 | 0.43245 | -0.2949 | 0.584 | 0.53 | 1 | HC_vs_MS.Trt_F |
| KLHL18 | 0.43248 | 0.90583 | 0.399 | 0.38 | 1 | HC_vs_MS.Trt_F |
| SIRPB1 | 0.43277 | 0.82227 | 0.275 | 0.25 | 1 | HC_vs_MS.Trt_F |
| RELL1 | 0.43288 | -0.782 | 0.558 | 0.57 | 1 | HC_vs_MS.Trt_F |
| DPEP2 | 0.43314 | -0.6103 | 0.214 | 0.23 | 1 | HC_vs_MS.Trt_F |
| KIDINS220 | 0.43343 | -0.8265 | 0.298 | 0.26 | 1 | HC_vs_MS.Trt_F |
| STAT1 | 0.43349 | -0.3189 | 0.384 | 0.34 | 1 | HC_vs_MS.Trt_F |
| POM121C | 0.4336 | 0.34524 | 0.497 | 0.46 | 1 | HC_vs_MS.Trt_F |
| GPR137B | 0.43366 | 0.26323 | 0.124 | 0.1 | 1 | HC_vs_MS.Trt_F |
| RP11-849H4 | 0.43375 | 0.25151 | 0.587 | 0.56 | 1 | HC_vs_MS.Trt_F |
| FAM149A | 0.43397 | 0.82535 | 0.379 | 0.34 | 1 | HC_vs_MS.Trt_F |
| MCM8 | 0.43455 | -0.5224 | 0.78 | 0.73 | 1 | HC_vs_MS.Trt_F |
| NDUFS1 | 0.43473 | -0.3721 | 0.962 | 0.94 | 1 | HC_vs_MS.Trt_F |
| GTPBP8 | 0.43489 | -0.4496 | 0.702 | 0.64 | 1 | HC_vs_MS.Trt_F |
| MMGT1 | 0.43526 | 0.65014 | 0.118 | 0.1 | 1 | HC_vs_MS.Trt_F |
| C11orf57 | 0.43531 | 0.43796 | 0.769 | 0.75 | 1 | HC_vs_MS.Trt_F |
| CTC-454I21. | 0.43562 | -0.433 | 0.234 | 0.21 | 1 | HC_vs_MS.Trt_F |
| LONP2 | 0.43642 | -0.3473 | 0.798 | 0.73 | 1 | HC_vs_MS.Trt_F |
| CHMP6 | 0.43699 | 0.28251 | 0.145 | 0.12 | 1 | HC_vs_MS.Trt_F |
| TTC32 | 0.43727 | -0.7484 | 0.098 | 0.12 | 1 | HC_vs_MS.Trt_F |
| ACOT4 | 0.43727 | 0.39916 | 0.15 | 0.13 | 1 | HC_vs_MS.Trt_F |

| | | | | | | |
|---|---|---|---|---|---|---|
| FBXL8 | 0.43791 | 0.4403 | 0.618 | 0.55 | 1 | HC_vs_MS.Trt_F |
| RBM20 | 0.43796 | 0.31717 | 0.483 | 0.48 | 1 | HC_vs_MS.Trt_F |
| IL23R | 0.4384 | 0.32905 | 0.422 | 0.39 | 1 | HC_vs_MS.Trt_F |
| PRAM1 | 0.43853 | 0.27935 | 0.564 | 0.53 | 1 | HC_vs_MS.Trt_F |
| CRTAM | 0.43971 | -0.7786 | 0.442 | 0.38 | 1 | HC_vs_MS.Trt_F |
| TMEM56 | 0.44031 | -0.3417 | 0.428 | 0.48 | 1 | HC_vs_MS.Trt_F |
| LAS1L | 0.44035 | -0.3457 | 0.165 | 0.14 | 1 | HC_vs_MS.Trt_F |
| TNFSF9 | 0.44061 | 0.39417 | 0.11 | 0.09 | 1 | HC_vs_MS.Trt_F |
| LRIF1 | 0.44065 | 0.96085 | 0.121 | 0.1 | 1 | HC_vs_MS.Trt_F |
| PUS10 | 0.44085 | 0.43825 | 0.734 | 0.72 | 1 | HC_vs_MS.Trt_F |
| FAM134A | 0.44169 | -0.539 | 0.428 | 0.42 | 1 | HC_vs_MS.Trt_F |
| FAIM | 0.44201 | 0.41168 | 0.399 | 0.36 | 1 | HC_vs_MS.Trt_F |
| ZNF92 | 0.44212 | -0.5315 | 0.11 | 0.13 | 1 | HC_vs_MS.Trt_F |
| SCAMP2 | 0.44279 | -0.4501 | 0.329 | 0.34 | 1 | HC_vs_MS.Trt_F |
| ZNF624 | 0.44307 | 0.49355 | 0.434 | 0.38 | 1 | HC_vs_MS.Trt_F |
| SLC39A10 | 0.4432 | 0.41183 | 0.202 | 0.18 | 1 | HC_vs_MS.Trt_F |
| RBMXL1 | 0.44328 | 1.08546 | 0.153 | 0.13 | 1 | HC_vs_MS.Trt_F |
| PURB | 0.44392 | 0.60469 | 0.52 | 0.48 | 1 | HC_vs_MS.Trt_F |
| SLC2A1 | 0.44419 | -0.427 | 0.159 | 0.13 | 1 | HC_vs_MS.Trt_F |
| C15orf53 | 0.44429 | 0.56473 | 0.136 | 0.12 | 1 | HC_vs_MS.Trt_F |
| ZNF516 | 0.4444 | -0.5426 | 0.107 | 0.09 | 1 | HC_vs_MS.Trt_F |
| BIN2 | 0.44455 | -0.4481 | 0.543 | 0.53 | 1 | HC_vs_MS.Trt_F |
| STAT2 | 0.44481 | -0.338 | 0.931 | 0.89 | 1 | HC_vs_MS.Trt_F |
| KCNJ14 | 0.44496 | 0.41511 | 0.653 | 0.6 | 1 | HC_vs_MS.Trt_F |
| RFX1 | 0.44501 | 0.69408 | 0.379 | 0.35 | 1 | HC_vs_MS.Trt_F |
| ARHGEF5 | 0.44573 | 0.4642 | 0.121 | 0.1 | 1 | HC_vs_MS.Trt_F |
| GTF2F1 | 0.44576 | -0.3236 | 0.364 | 0.31 | 1 | HC_vs_MS.Trt_F |
| MOSPD3 | 0.44603 | 0.26642 | 0.124 | 0.1 | 1 | HC_vs_MS.Trt_F |
| FAM210B | 0.44632 | 0.35913 | 0.312 | 0.34 | 1 | HC_vs_MS.Trt_F |
| TRIM59 | 0.44701 | -0.63 | 0.853 | 0.82 | 1 | HC_vs_MS.Trt_F |
| UBE2N | 0.44706 | -0.3621 | 0.604 | 0.54 | 1 | HC_vs_MS.Trt_F |
| C12orf4 | 0.44722 | 0.25905 | 0.491 | 0.47 | 1 | HC_vs_MS.Trt_F |
| FBXO32 | 0.44745 | -0.5513 | 0.147 | 0.17 | 1 | HC_vs_MS.Trt_F |
| SPOUT1 | 0.44745 | -0.6563 | 0.188 | 0.16 | 1 | HC_vs_MS.Trt_F |
| CNOT8 | 0.44818 | -0.3817 | 0.249 | 0.22 | 1 | HC_vs_MS.Trt_F |
| GRAPL | 0.44879 | 0.9747 | 0.104 | 0.09 | 1 | HC_vs_MS.Trt_F |
| MRPS21 | 0.44961 | -0.3888 | 0.613 | 0.57 | 1 | HC_vs_MS.Trt_F |
| PAIP1 | 0.45021 | -0.6421 | 0.147 | 0.16 | 1 | HC_vs_MS.Trt_F |
| IST1 | 0.45033 | -0.4281 | 0.647 | 0.64 | 1 | HC_vs_MS.Trt_F |
| C21orf33 | 0.45035 | -0.2533 | 0.416 | 0.37 | 1 | HC_vs_MS.Trt_F |

| | | | | | | |
|---|---|---|---|---|---|---|
| CDV3 | 0.45084 | -0.951 | 0.251 | 0.26 | 1 | HC_vs_MS.Trt_F |
| OASL | 0.45117 | -0.6418 | 0.24 | 0.27 | 1 | HC_vs_MS.Trt_F |
| MRPL2 | 0.45154 | -0.5176 | 0.338 | 0.29 | 1 | HC_vs_MS.Trt_F |
| CSDE1 | 0.45193 | -0.3498 | 0.434 | 0.44 | 1 | HC_vs_MS.Trt_F |
| MAATS1 | 0.45261 | 0.28718 | 0.277 | 0.25 | 1 | HC_vs_MS.Trt_F |
| CYP2F1 | 0.45276 | 0.70438 | 0.185 | 0.17 | 1 | HC_vs_MS.Trt_F |
| SYT16 | 0.4528 | 0.50069 | 0.312 | 0.28 | 1 | HC_vs_MS.Trt_F |
| MEMO1 | 0.45311 | -0.4618 | 0.393 | 0.36 | 1 | HC_vs_MS.Trt_F |
| ROBO4 | 0.45324 | 1.1466 | 0.191 | 0.17 | 1 | HC_vs_MS.Trt_F |
| DNAJC8 | 0.45327 | -0.335 | 0.306 | 0.27 | 1 | HC_vs_MS.Trt_F |
| EXT2 | 0.45343 | -0.3626 | 0.191 | 0.17 | 1 | HC_vs_MS.Trt_F |
| RAD50 | 0.45401 | 0.306 | 0.225 | 0.2 | 1 | HC_vs_MS.Trt_F |
| GLCCI1 | 0.45417 | 0.69941 | 0.286 | 0.27 | 1 | HC_vs_MS.Trt_F |
| RANBP9 | 0.45438 | 0.30795 | 0.266 | 0.24 | 1 | HC_vs_MS.Trt_F |
| ZNF420 | 0.45438 | -0.8424 | 0.127 | 0.1 | 1 | HC_vs_MS.Trt_F |
| AKAP17A | 0.4544 | 0.51133 | 0.118 | 0.1 | 1 | HC_vs_MS.Trt_F |
| DAXX | 0.45448 | -0.4798 | 0.173 | 0.19 | 1 | HC_vs_MS.Trt_F |
| LRP12 | 0.45456 | -0.3033 | 0.682 | 0.6 | 1 | HC_vs_MS.Trt_F |
| PACSIN2 | 0.4546 | -0.5206 | 0.494 | 0.52 | 1 | HC_vs_MS.Trt_F |
| PRCC | 0.45521 | 0.52724 | 0.176 | 0.15 | 1 | HC_vs_MS.Trt_F |
| APOBEC3F | 0.45586 | -0.3547 | 0.928 | 0.92 | 1 | HC_vs_MS.Trt_F |
| CFL2 | 0.45591 | -0.4149 | 0.694 | 0.63 | 1 | HC_vs_MS.Trt_F |
| NCF1 | 0.45597 | 1.85892 | 0.543 | 0.5 | 1 | HC_vs_MS.Trt_F |
| HLX | 0.4561 | 0.28923 | 0.197 | 0.17 | 1 | HC_vs_MS.Trt_F |
| GFOD1 | 0.45629 | -1.0332 | 0.699 | 0.63 | 1 | HC_vs_MS.Trt_F |
| KDM5A | 0.45643 | -0.2529 | 0.673 | 0.65 | 1 | HC_vs_MS.Trt_F |
| MPHOSPH1( | 0.45757 | 0.31794 | 0.246 | 0.22 | 1 | HC_vs_MS.Trt_F |
| RASA3 | 0.45785 | -0.3576 | 0.442 | 0.46 | 1 | HC_vs_MS.Trt_F |
| PYHIN1 | 0.45852 | 0.27577 | 0.627 | 0.61 | 1 | HC_vs_MS.Trt_F |
| TIAL1 | 0.45863 | -0.2561 | 0.87 | 0.83 | 1 | HC_vs_MS.Trt_F |
| KPNA6 | 0.459 | -0.3616 | 0.592 | 0.59 | 1 | HC_vs_MS.Trt_F |
| CCDC93 | 0.45907 | -0.3525 | 0.179 | 0.15 | 1 | HC_vs_MS.Trt_F |
| DISC1 | 0.45915 | 0.25655 | 0.558 | 0.52 | 1 | HC_vs_MS.Trt_F |
| TECPR1 | 0.45915 | -0.4952 | 0.509 | 0.44 | 1 | HC_vs_MS.Trt_F |
| HNRNPM | 0.45941 | -0.298 | 0.266 | 0.23 | 1 | HC_vs_MS.Trt_F |
| ZNF561 | 0.45965 | -0.7429 | 0.601 | 0.51 | 1 | HC_vs_MS.Trt_F |
| EIF5B | 0.45965 | -0.2969 | 0.332 | 0.29 | 1 | HC_vs_MS.Trt_F |
| USP42 | 0.4597 | 0.43546 | 0.468 | 0.44 | 1 | HC_vs_MS.Trt_F |
| C8orf37 | 0.4613 | 0.45119 | 0.552 | 0.53 | 1 | HC_vs_MS.Trt_F |
| ZFP90 | 0.46173 | 0.61182 | 0.445 | 0.47 | 1 | HC_vs_MS.Trt_F |

| | | | | | | |
|---|---|---|---|---|---|---|
| DDHD2 | 0.46179 | 0.26645 | 0.864 | 0.82 | 1 | HC_vs_MS.Trt_F |
| PSMB1 | 0.46208 | -0.3028 | 0.358 | 0.31 | 1 | HC_vs_MS.Trt_F |
| NDUFB1 | 0.46219 | -0.304 | 0.78 | 0.72 | 1 | HC_vs_MS.Trt_F |
| CCDC153 | 0.46261 | -0.7481 | 0.269 | 0.25 | 1 | HC_vs_MS.Trt_F |
| CLPX | 0.46296 | -0.7427 | 0.731 | 0.67 | 1 | HC_vs_MS.Trt_F |
| RP11-505K9 | 0.463 | -0.3589 | 0.09 | 0.11 | 1 | HC_vs_MS.Trt_F |
| ISG20L2 | 0.46337 | 0.58908 | 0.85 | 0.82 | 1 | HC_vs_MS.Trt_F |
| TMEM222 | 0.46347 | 0.53754 | 0.153 | 0.13 | 1 | HC_vs_MS.Trt_F |
| ZNF484 | 0.46354 | 0.94593 | 0.107 | 0.09 | 1 | HC_vs_MS.Trt_F |
| RNF14 | 0.46412 | 0.64008 | 0.65 | 0.58 | 1 | HC_vs_MS.Trt_F |
| TMEM126B | 0.46448 | -0.49 | 0.234 | 0.21 | 1 | HC_vs_MS.Trt_F |
| ZNF518B | 0.4649 | -0.5743 | 0.116 | 0.09 | 1 | HC_vs_MS.Trt_F |
| ZDHHC14 | 0.46542 | 0.29851 | 0.338 | 0.36 | 1 | HC_vs_MS.Trt_F |
| RP11-1035H | 0.46561 | -0.8025 | 0.153 | 0.13 | 1 | HC_vs_MS.Trt_F |
| CSTF3 | 0.46584 | -1.3169 | 0.558 | 0.51 | 1 | HC_vs_MS.Trt_F |
| VPS51 | 0.46593 | 0.25438 | 0.283 | 0.26 | 1 | HC_vs_MS.Trt_F |
| GGPS1 | 0.46597 | -0.3216 | 0.454 | 0.47 | 1 | HC_vs_MS.Trt_F |
| PGM2 | 0.46622 | -0.7269 | 0.13 | 0.11 | 1 | HC_vs_MS.Trt_F |
| BTBD10 | 0.46656 | 0.30462 | 0.159 | 0.14 | 1 | HC_vs_MS.Trt_F |
| ZCRB1 | 0.4668 | -0.8468 | 0.165 | 0.18 | 1 | HC_vs_MS.Trt_F |
| SLC44A1 | 0.46697 | 0.86013 | 0.639 | 0.66 | 1 | HC_vs_MS.Trt_F |
| NNMT | 0.46706 | -0.416 | 0.139 | 0.12 | 1 | HC_vs_MS.Trt_F |
| CD24 | 0.46753 | -0.2593 | 0.373 | 0.41 | 1 | HC_vs_MS.Trt_F |
| ATG9A | 0.46791 | 0.70006 | 0.116 | 0.1 | 1 | HC_vs_MS.Trt_F |
| SNRPB2 | 0.46802 | -0.5964 | 0.208 | 0.22 | 1 | HC_vs_MS.Trt_F |
| NAAA | 0.46835 | -0.315 | 0.728 | 0.77 | 1 | HC_vs_MS.Trt_F |
| EGR1 | 0.46848 | -0.3476 | 0.121 | 0.14 | 1 | HC_vs_MS.Trt_F |
| HIVEP3 | 0.46852 | -0.293 | 0.092 | 0.11 | 1 | HC_vs_MS.Trt_F |
| DNAJB12 | 0.46877 | -0.5335 | 0.15 | 0.13 | 1 | HC_vs_MS.Trt_F |
| ZSCAN12 | 0.46877 | 0.53537 | 0.468 | 0.49 | 1 | HC_vs_MS.Trt_F |
| PTPRM | 0.4689 | 1.09781 | 0.225 | 0.2 | 1 | HC_vs_MS.Trt_F |
| TEX2 | 0.46924 | 0.56014 | 0.72 | 0.63 | 1 | HC_vs_MS.Trt_F |
| SAR1B | 0.46931 | -0.2565 | 0.977 | 0.97 | 1 | HC_vs_MS.Trt_F |
| LBR | 0.46952 | -0.6545 | 0.249 | 0.22 | 1 | HC_vs_MS.Trt_F |
| IRF7 | 0.46988 | -0.4393 | 0.145 | 0.16 | 1 | HC_vs_MS.Trt_F |
| CLCN6 | 0.47035 | 0.32256 | 0.59 | 0.53 | 1 | HC_vs_MS.Trt_F |
| CFD | 0.4711 | 0.74099 | 0.197 | 0.22 | 1 | HC_vs_MS.Trt_F |
| GDAP1 | 0.47173 | -0.3252 | 0.092 | 0.11 | 1 | HC_vs_MS.Trt_F |
| ANKDD1A | 0.47215 | 0.32398 | 0.581 | 0.56 | 1 | HC_vs_MS.Trt_F |
| MAP2K7 | 0.47267 | 0.39381 | 0.65 | 0.59 | 1 | HC_vs_MS.Trt_F |

| | | | | | | |
|---|---|---|---|---|---|---|
| FILIP1L | 0.47273 | 0.54841 | 0.442 | 0.39 | 1 | HC_vs_MS.Trt_F |
| ZZZ3 | 0.47295 | 0.27312 | 0.824 | 0.81 | 1 | HC_vs_MS.Trt_F |
| EHBP1 | 0.47303 | -0.3688 | 0.405 | 0.44 | 1 | HC_vs_MS.Trt_F |
| NAA20 | 0.47318 | -0.4525 | 0.156 | 0.13 | 1 | HC_vs_MS.Trt_F |
| DPM2 | 0.47326 | -0.6693 | 0.13 | 0.11 | 1 | HC_vs_MS.Trt_F |
| SLC25A17 | 0.47334 | -0.4718 | 0.312 | 0.32 | 1 | HC_vs_MS.Trt_F |
| DCAF4 | 0.47391 | -0.3622 | 0.587 | 0.54 | 1 | HC_vs_MS.Trt_F |
| SCLT1 | 0.47405 | 0.63699 | 0.673 | 0.62 | 1 | HC_vs_MS.Trt_F |
| COG3 | 0.47408 | -0.7273 | 0.168 | 0.18 | 1 | HC_vs_MS.Trt_F |
| CCHCR1 | 0.47415 | 1.66285 | 0.376 | 0.35 | 1 | HC_vs_MS.Trt_F |
| FRG2C | 0.47416 | 1.19696 | 0.194 | 0.17 | 1 | HC_vs_MS.Trt_F |
| PIP4K2B | 0.47437 | -0.443 | 0.153 | 0.13 | 1 | HC_vs_MS.Trt_F |
| ORC3 | 0.47507 | -0.3359 | 0.191 | 0.17 | 1 | HC_vs_MS.Trt_F |
| ZNF727 | 0.47514 | 1.25316 | 0.384 | 0.36 | 1 | HC_vs_MS.Trt_F |
| RNF113A | 0.47534 | -0.4695 | 0.139 | 0.12 | 1 | HC_vs_MS.Trt_F |
| COG6 | 0.47569 | -1.1474 | 0.159 | 0.18 | 1 | HC_vs_MS.Trt_F |
| C8orf59 | 0.47609 | -0.3185 | 0.442 | 0.45 | 1 | HC_vs_MS.Trt_F |
| POLD3 | 0.47612 | -1.4175 | 0.176 | 0.19 | 1 | HC_vs_MS.Trt_F |
| AAED1 | 0.47629 | -0.7417 | 0.829 | 0.78 | 1 | HC_vs_MS.Trt_F |
| H2AFJ | 0.47638 | 0.3977 | 0.15 | 0.13 | 1 | HC_vs_MS.Trt_F |
| LOXL2 | 0.47673 | 0.33318 | 0.74 | 0.73 | 1 | HC_vs_MS.Trt_F |
| FAM127B | 0.47679 | 0.34855 | 0.13 | 0.11 | 1 | HC_vs_MS.Trt_F |
| TARS | 0.47685 | -0.5781 | 0.257 | 0.22 | 1 | HC_vs_MS.Trt_F |
| INTS2 | 0.47738 | 0.28434 | 0.298 | 0.27 | 1 | HC_vs_MS.Trt_F |
| CAV2 | 0.4774 | 0.31259 | 0.61 | 0.59 | 1 | HC_vs_MS.Trt_F |
| PPP2CB | 0.47756 | 0.27577 | 0.564 | 0.59 | 1 | HC_vs_MS.Trt_F |
| RBMS3 | 0.47786 | 0.36894 | 0.697 | 0.67 | 1 | HC_vs_MS.Trt_F |
| HIGD2A | 0.47796 | -0.3535 | 0.335 | 0.29 | 1 | HC_vs_MS.Trt_F |
| CLEC2B | 0.47815 | -0.3992 | 0.251 | 0.22 | 1 | HC_vs_MS.Trt_F |
| LRP2 | 0.47861 | 0.38005 | 0.098 | 0.12 | 1 | HC_vs_MS.Trt_F |
| PCF11 | 0.47862 | -0.2754 | 0.512 | 0.46 | 1 | HC_vs_MS.Trt_F |
| LAMTOR3 | 0.47867 | -0.5691 | 0.353 | 0.36 | 1 | HC_vs_MS.Trt_F |
| PSMA4 | 0.47888 | -0.5413 | 0.402 | 0.36 | 1 | HC_vs_MS.Trt_F |
| ITGB4 | 0.47906 | -0.9719 | 0.237 | 0.25 | 1 | HC_vs_MS.Trt_F |
| FAM101B | 0.47943 | -0.3919 | 0.324 | 0.29 | 1 | HC_vs_MS.Trt_F |
| CEACAM1 | 0.47952 | 0.78468 | 0.882 | 0.92 | 1 | HC_vs_MS.Trt_F |
| GNG5 | 0.47982 | -0.3424 | 0.286 | 0.25 | 1 | HC_vs_MS.Trt_F |
| KYAT3 | 0.47983 | -0.3729 | 0.136 | 0.12 | 1 | HC_vs_MS.Trt_F |
| TREX1 | 0.48027 | 0.95092 | 0.121 | 0.1 | 1 | HC_vs_MS.Trt_F |
| EGLN2 | 0.48111 | -0.7079 | 0.251 | 0.26 | 1 | HC_vs_MS.Trt_F |

| | | | | | | |
|---|---|---|---|---|---|---|
| IFNAR1 | 0.48117 | -0.5176 | 0.561 | 0.5 | 1 | HC_vs_MS.Trt_F |
| PSMB8 | 0.48124 | -0.4969 | 0.382 | 0.39 | 1 | HC_vs_MS.Trt_F |
| KNTC1 | 0.48134 | -1.0291 | 0.838 | 0.81 | 1 | HC_vs_MS.Trt_F |
| TMX3 | 0.48145 | -0.388 | 0.462 | 0.43 | 1 | HC_vs_MS.Trt_F |
| SH2D1B | 0.48195 | 0.8628 | 0.263 | 0.23 | 1 | HC_vs_MS.Trt_F |
| PLEKHG5 | 0.48208 | -0.9761 | 0.182 | 0.16 | 1 | HC_vs_MS.Trt_F |
| ZNF254 | 0.48276 | -0.6804 | 0.416 | 0.45 | 1 | HC_vs_MS.Trt_F |
| PITPNM1 | 0.48317 | -0.6028 | 0.306 | 0.27 | 1 | HC_vs_MS.Trt_F |
| ZNF12 | 0.48387 | 0.94965 | 0.361 | 0.32 | 1 | HC_vs_MS.Trt_F |
| VCPKMT | 0.48414 | 0.59726 | 0.118 | 0.1 | 1 | HC_vs_MS.Trt_F |
| GPR161 | 0.48447 | -0.6279 | 0.436 | 0.41 | 1 | HC_vs_MS.Trt_F |
| NR2C2AP | 0.48454 | -0.7745 | 0.107 | 0.12 | 1 | HC_vs_MS.Trt_F |
| CH507-9B2.! | 0.48455 | -0.5462 | 0.517 | 0.46 | 1 | HC_vs_MS.Trt_F |
| SLC35B2 | 0.48462 | 0.64667 | 0.116 | 0.1 | 1 | HC_vs_MS.Trt_F |
| MORC2 | 0.48497 | -0.3822 | 0.434 | 0.37 | 1 | HC_vs_MS.Trt_F |
| JMJD7-PLA2 | 0.48548 | -0.7166 | 0.329 | 0.35 | 1 | HC_vs_MS.Trt_F |
| PLEKHO2 | 0.48566 | -0.4675 | 0.104 | 0.12 | 1 | HC_vs_MS.Trt_F |
| HYAL1 | 0.48636 | -0.4548 | 0.376 | 0.4 | 1 | HC_vs_MS.Trt_F |
| ZNF718 | 0.4872 | -0.4017 | 0.295 | 0.31 | 1 | HC_vs_MS.Trt_F |
| AIG1 | 0.48726 | -0.3634 | 0.642 | 0.58 | 1 | HC_vs_MS.Trt_F |
| TTC7A | 0.48736 | 0.28668 | 0.249 | 0.22 | 1 | HC_vs_MS.Trt_F |
| EGLN1 | 0.48765 | -1.1868 | 0.364 | 0.37 | 1 | HC_vs_MS.Trt_F |
| SEPSECS | 0.48771 | 0.3737 | 0.789 | 0.76 | 1 | HC_vs_MS.Trt_F |
| RCN1 | 0.48854 | -0.4236 | 0.116 | 0.1 | 1 | HC_vs_MS.Trt_F |
| ETNK1 | 0.48868 | -0.2763 | 0.503 | 0.51 | 1 | HC_vs_MS.Trt_F |
| EIF4EBP2 | 0.48896 | -0.3653 | 0.656 | 0.69 | 1 | HC_vs_MS.Trt_F |
| DUSP22 | 0.48933 | -0.4099 | 0.116 | 0.1 | 1 | HC_vs_MS.Trt_F |
| COMMD6 | 0.48936 | -0.3133 | 0.416 | 0.36 | 1 | HC_vs_MS.Trt_F |
| VPS50 | 0.48957 | 0.59287 | 0.665 | 0.6 | 1 | HC_vs_MS.Trt_F |
| C1orf159 | 0.49048 | 0.25454 | 0.208 | 0.18 | 1 | HC_vs_MS.Trt_F |
| ABRACL | 0.49049 | -0.3685 | 0.257 | 0.22 | 1 | HC_vs_MS.Trt_F |
| ZNF772 | 0.49089 | -0.6539 | 0.471 | 0.51 | 1 | HC_vs_MS.Trt_F |
| TRMU | 0.49091 | -0.409 | 0.199 | 0.22 | 1 | HC_vs_MS.Trt_F |
| IPPK | 0.49117 | -1.2443 | 0.101 | 0.08 | 1 | HC_vs_MS.Trt_F |
| RPA2 | 0.49222 | -0.5417 | 0.217 | 0.23 | 1 | HC_vs_MS.Trt_F |
| MOCS2 | 0.49405 | -0.3603 | 0.116 | 0.1 | 1 | HC_vs_MS.Trt_F |
| FANCA | 0.49424 | -0.2836 | 0.315 | 0.28 | 1 | HC_vs_MS.Trt_F |
| RNF7 | 0.49429 | 0.37565 | 0.682 | 0.64 | 1 | HC_vs_MS.Trt_F |
| COX18 | 0.49432 | -0.2612 | 0.887 | 0.85 | 1 | HC_vs_MS.Trt_F |
| CENPN | 0.49451 | 0.53859 | 0.217 | 0.2 | 1 | HC_vs_MS.Trt_F |

| | | | | | | |
|---|---|---|---|---|---|---|
| UBA5 | 0.49455 | -1.9633 | 0.136 | 0.15 | 1 | HC_vs_MS.Trt_F |
| EMC8 | 0.49523 | -0.8269 | 0.208 | 0.22 | 1 | HC_vs_MS.Trt_F |
| ZNF841 | 0.49546 | 0.80688 | 0.538 | 0.51 | 1 | HC_vs_MS.Trt_F |
| ARHGEF2 | 0.49561 | 0.41207 | 0.275 | 0.24 | 1 | HC_vs_MS.Trt_F |
| NDUFAF3 | 0.49562 | -0.43 | 0.185 | 0.16 | 1 | HC_vs_MS.Trt_F |
| HERC6 | 0.49583 | -0.4748 | 0.277 | 0.3 | 1 | HC_vs_MS.Trt_F |
| FCHO2 | 0.49602 | -0.3879 | 0.353 | 0.37 | 1 | HC_vs_MS.Trt_F |
| UBFD1 | 0.4965 | 1.31522 | 0.425 | 0.36 | 1 | HC_vs_MS.Trt_F |
| INSIG2 | 0.49719 | -1.0627 | 0.133 | 0.14 | 1 | HC_vs_MS.Trt_F |
| STX3 | 0.4972 | 0.73937 | 0.483 | 0.45 | 1 | HC_vs_MS.Trt_F |
| CARF | 0.49732 | -0.4737 | 0.546 | 0.51 | 1 | HC_vs_MS.Trt_F |
| AP003419.1 | 0.49765 | -0.4064 | 0.208 | 0.22 | 1 | HC_vs_MS.Trt_F |
| CDR2 | 0.49775 | 1.03487 | 0.168 | 0.15 | 1 | HC_vs_MS.Trt_F |
| CNEP1R1 | 0.49776 | -1.5601 | 0.555 | 0.58 | 1 | HC_vs_MS.Trt_F |
| PI16 | 0.49778 | -0.4314 | 0.142 | 0.16 | 1 | HC_vs_MS.Trt_F |
| APLF | 0.49782 | -0.8771 | 0.199 | 0.18 | 1 | HC_vs_MS.Trt_F |
| C14orf119 | 0.49817 | -0.8729 | 0.595 | 0.53 | 1 | HC_vs_MS.Trt_F |
| CDC37L1 | 0.49835 | 0.76893 | 0.118 | 0.1 | 1 | HC_vs_MS.Trt_F |
| AXL | 0.49842 | 0.47251 | 0.772 | 0.72 | 1 | HC_vs_MS.Trt_F |
| RNF121 | 0.49845 | 0.25336 | 0.361 | 0.32 | 1 | HC_vs_MS.Trt_F |
| ATP8A1 | 0.49864 | -0.2869 | 0.15 | 0.17 | 1 | HC_vs_MS.Trt_F |
| MDM1 | 0.49877 | 0.55308 | 0.11 | 0.09 | 1 | HC_vs_MS.Trt_F |
| VPS39 | 0.4993 | -0.8704 | 0.538 | 0.49 | 1 | HC_vs_MS.Trt_F |
| ARL8B | 0.5 | -0.4329 | 0.474 | 0.41 | 1 | HC_vs_MS.Trt_F |
| SURF1 | 0.50001 | -0.886 | 0.113 | 0.13 | 1 | HC_vs_MS.Trt_F |
| NUDT7 | 0.50038 | -0.3444 | 0.24 | 0.21 | 1 | HC_vs_MS.Trt_F |
| SLC35F2 | 0.50043 | 0.315 | 0.312 | 0.29 | 1 | HC_vs_MS.Trt_F |
| PCCA | 0.5006 | -0.4228 | 0.647 | 0.61 | 1 | HC_vs_MS.Trt_F |
| RXRA | 0.50063 | 0.58373 | 0.225 | 0.21 | 1 | HC_vs_MS.Trt_F |
| SEZ6 | 0.50123 | -0.3496 | 0.13 | 0.11 | 1 | HC_vs_MS.Trt_F |
| ZDHHC18 | 0.50147 | 0.34535 | 0.118 | 0.1 | 1 | HC_vs_MS.Trt_F |
| SUMF1 | 0.5019 | 0.76187 | 0.76 | 0.75 | 1 | HC_vs_MS.Trt_F |
| ATP5L2 | 0.50247 | 0.41836 | 0.353 | 0.32 | 1 | HC_vs_MS.Trt_F |
| GEMIN2 | 0.50256 | -0.4186 | 0.691 | 0.68 | 1 | HC_vs_MS.Trt_F |
| SLC22A5 | 0.5026 | 0.57094 | 0.428 | 0.39 | 1 | HC_vs_MS.Trt_F |
| IL12RB1 | 0.50297 | 0.25428 | 0.89 | 0.87 | 1 | HC_vs_MS.Trt_F |
| AGFG1 | 0.50309 | -0.6896 | 0.095 | 0.11 | 1 | HC_vs_MS.Trt_F |
| FURIN | 0.50332 | -0.4105 | 0.11 | 0.09 | 1 | HC_vs_MS.Trt_F |
| MBD2 | 0.50334 | -0.3468 | 0.095 | 0.11 | 1 | HC_vs_MS.Trt_F |
| RCC2 | 0.50351 | 0.26874 | 0.13 | 0.11 | 1 | HC_vs_MS.Trt_F |

| | | | | | | |
|---|---|---|---|---|---|---|
| NME6 | 0.50363 | 0.2764 | 0.931 | 0.87 | 1 | HC_vs_MS.Trt_F |
| SNAP23 | 0.50382 | -0.2626 | 0.682 | 0.66 | 1 | HC_vs_MS.Trt_F |
| TNFAIP8L1 | 0.5044 | -0.3366 | 0.882 | 0.85 | 1 | HC_vs_MS.Trt_F |
| PDGFA | 0.50442 | 0.26692 | 0.578 | 0.57 | 1 | HC_vs_MS.Trt_F |
| MRPL51 | 0.50464 | -0.359 | 0.156 | 0.17 | 1 | HC_vs_MS.Trt_F |
| IL18BP | 0.50504 | -0.3669 | 0.147 | 0.13 | 1 | HC_vs_MS.Trt_F |
| SYNE4 | 0.50515 | -0.4588 | 0.517 | 0.5 | 1 | HC_vs_MS.Trt_F |
| SVEP1 | 0.50557 | 2.15598 | 0.243 | 0.22 | 1 | HC_vs_MS.Trt_F |
| RNF146 | 0.50561 | -0.8142 | 0.113 | 0.13 | 1 | HC_vs_MS.Trt_F |
| LAMC3 | 0.50589 | -1.4061 | 0.283 | 0.3 | 1 | HC_vs_MS.Trt_F |
| PPP1R2 | 0.50595 | -0.7108 | 0.656 | 0.64 | 1 | HC_vs_MS.Trt_F |
| FAM173B | 0.50611 | 0.30573 | 0.197 | 0.18 | 1 | HC_vs_MS.Trt_F |
| WDR75 | 0.50631 | 0.5959 | 0.358 | 0.35 | 1 | HC_vs_MS.Trt_F |
| ZNF3 | 0.50637 | -0.7475 | 0.78 | 0.75 | 1 | HC_vs_MS.Trt_F |
| IMP4 | 0.50711 | -0.3902 | 0.263 | 0.28 | 1 | HC_vs_MS.Trt_F |
| ARAP2 | 0.50711 | -0.5192 | 0.335 | 0.29 | 1 | HC_vs_MS.Trt_F |
| PER3 | 0.50714 | 1.68394 | 0.101 | 0.09 | 1 | HC_vs_MS.Trt_F |
| MTIF2 | 0.50744 | 0.36685 | 0.408 | 0.42 | 1 | HC_vs_MS.Trt_F |
| PRICKLE3 | 0.50781 | 0.77055 | 0.555 | 0.5 | 1 | HC_vs_MS.Trt_F |
| ACP5 | 0.50808 | -0.4575 | 0.165 | 0.14 | 1 | HC_vs_MS.Trt_F |
| RARS | 0.5083 | -0.2782 | 0.532 | 0.49 | 1 | HC_vs_MS.Trt_F |
| XYLB | 0.50841 | 0.31378 | 0.488 | 0.47 | 1 | HC_vs_MS.Trt_F |
| TMIGD2 | 0.50894 | -0.6337 | 0.231 | 0.25 | 1 | HC_vs_MS.Trt_F |
| SLC25A23 | 0.50962 | -0.2607 | 0.462 | 0.49 | 1 | HC_vs_MS.Trt_F |
| BTF3 | 0.50981 | -0.254 | 0.555 | 0.55 | 1 | HC_vs_MS.Trt_F |
| TRRAP | 0.51013 | -0.3999 | 0.165 | 0.14 | 1 | HC_vs_MS.Trt_F |
| SMG6 | 0.51024 | 0.95722 | 0.251 | 0.24 | 1 | HC_vs_MS.Trt_F |
| ARMC1 | 0.51034 | -0.4962 | 0.127 | 0.14 | 1 | HC_vs_MS.Trt_F |
| RNPEP | 0.51118 | 0.2726 | 0.104 | 0.12 | 1 | HC_vs_MS.Trt_F |
| SLFN5 | 0.51129 | -0.3373 | 0.855 | 0.86 | 1 | HC_vs_MS.Trt_F |
| CLCN2 | 0.51137 | -0.2596 | 0.699 | 0.63 | 1 | HC_vs_MS.Trt_F |
| FOCAD | 0.5114 | -0.3377 | 0.561 | 0.52 | 1 | HC_vs_MS.Trt_F |
| C5orf15 | 0.5116 | -1.1298 | 0.156 | 0.17 | 1 | HC_vs_MS.Trt_F |
| TJP3 | 0.51213 | -0.5177 | 0.095 | 0.11 | 1 | HC_vs_MS.Trt_F |
| HABP4 | 0.51249 | -0.9964 | 0.182 | 0.17 | 1 | HC_vs_MS.Trt_F |
| FAR1 | 0.51268 | 0.98191 | 0.104 | 0.09 | 1 | HC_vs_MS.Trt_F |
| EIF5 | 0.51287 | -0.3534 | 0.598 | 0.58 | 1 | HC_vs_MS.Trt_F |
| B3GAT3 | 0.51314 | -0.4337 | 0.566 | 0.54 | 1 | HC_vs_MS.Trt_F |
| TMEM8B | 0.51321 | 0.34824 | 0.751 | 0.73 | 1 | HC_vs_MS.Trt_F |
| SPTSSA | 0.51326 | 0.34598 | 0.41 | 0.36 | 1 | HC_vs_MS.Trt_F |

| | | | | | | |
|---|---|---|---|---|---|---|
| USP9Y | 0.51358 | 0.90999 | 0.292 | 0.27 | 1 | HC_vs_MS.Trt_F |
| ALS2CL | 0.51375 | -0.3331 | 0.176 | 0.15 | 1 | HC_vs_MS.Trt_F |
| GPN1 | 0.51379 | -0.5309 | 0.147 | 0.17 | 1 | HC_vs_MS.Trt_F |
| RASAL2 | 0.5141 | 0.39646 | 0.555 | 0.51 | 1 | HC_vs_MS.Trt_F |
| NDUFC2 | 0.51447 | -0.571 | 0.613 | 0.57 | 1 | HC_vs_MS.Trt_F |
| ZC4H2 | 0.51501 | 1.73702 | 0.107 | 0.09 | 1 | HC_vs_MS.Trt_F |
| GNPAT | 0.51547 | -0.737 | 0.168 | 0.18 | 1 | HC_vs_MS.Trt_F |
| FPR1 | 0.51552 | 0.60779 | 0.506 | 0.48 | 1 | HC_vs_MS.Trt_F |
| MRPL32 | 0.51569 | -0.5751 | 0.168 | 0.18 | 1 | HC_vs_MS.Trt_F |
| TADA1 | 0.51571 | -0.5387 | 0.104 | 0.09 | 1 | HC_vs_MS.Trt_F |
| MED30 | 0.51573 | -0.6613 | 0.124 | 0.14 | 1 | HC_vs_MS.Trt_F |
| UAP1 | 0.51574 | -0.5797 | 0.092 | 0.11 | 1 | HC_vs_MS.Trt_F |
| VPS4B | 0.51588 | -0.4671 | 0.803 | 0.76 | 1 | HC_vs_MS.Trt_F |
| LYRM9 | 0.51616 | 0.52171 | 0.199 | 0.18 | 1 | HC_vs_MS.Trt_F |
| PAK6 | 0.51686 | 0.33675 | 0.179 | 0.16 | 1 | HC_vs_MS.Trt_F |
| NT5DC3 | 0.51704 | -0.4384 | 0.829 | 0.83 | 1 | HC_vs_MS.Trt_F |
| PTK6 | 0.51708 | 0.48274 | 0.405 | 0.39 | 1 | HC_vs_MS.Trt_F |
| WDR55 | 0.51709 | 0.25094 | 0.717 | 0.68 | 1 | HC_vs_MS.Trt_F |
| KRT10 | 0.51712 | -0.5944 | 0.113 | 0.13 | 1 | HC_vs_MS.Trt_F |
| AGA | 0.51745 | -0.4088 | 0.121 | 0.14 | 1 | HC_vs_MS.Trt_F |
| FASTKD3 | 0.51794 | -0.6524 | 0.09 | 0.1 | 1 | HC_vs_MS.Trt_F |
| STOM | 0.51804 | 0.27969 | 0.668 | 0.66 | 1 | HC_vs_MS.Trt_F |
| CELSR3 | 0.51811 | 0.52997 | 0.113 | 0.1 | 1 | HC_vs_MS.Trt_F |
| LRRC1 | 0.51852 | -1.031 | 0.306 | 0.32 | 1 | HC_vs_MS.Trt_F |
| OSTF1 | 0.5186 | -0.5888 | 0.327 | 0.29 | 1 | HC_vs_MS.Trt_F |
| MAP2K2 | 0.51894 | -0.8059 | 0.766 | 0.71 | 1 | HC_vs_MS.Trt_F |
| C2orf88 | 0.51926 | 0.55893 | 0.358 | 0.34 | 1 | HC_vs_MS.Trt_F |
| MYO7A | 0.51958 | 0.43948 | 0.361 | 0.39 | 1 | HC_vs_MS.Trt_F |
| SNRPB | 0.51977 | -0.4068 | 0.24 | 0.21 | 1 | HC_vs_MS.Trt_F |
| LRRC8D | 0.51984 | -0.7169 | 0.145 | 0.12 | 1 | HC_vs_MS.Trt_F |
| IDI1 | 0.52002 | -0.3665 | 0.243 | 0.21 | 1 | HC_vs_MS.Trt_F |
| ZNF37A | 0.52068 | -0.9318 | 0.916 | 0.86 | 1 | HC_vs_MS.Trt_F |
| SLC39A5 | 0.52072 | 0.63622 | 0.24 | 0.22 | 1 | HC_vs_MS.Trt_F |
| FAM188A | 0.52084 | -0.4153 | 0.133 | 0.12 | 1 | HC_vs_MS.Trt_F |
| HIST1H2AG | 0.52102 | 0.50453 | 0.37 | 0.34 | 1 | HC_vs_MS.Trt_F |
| CAMKMT | 0.52111 | -0.332 | 0.286 | 0.31 | 1 | HC_vs_MS.Trt_F |
| TECPR2 | 0.52133 | 1.65207 | 0.338 | 0.36 | 1 | HC_vs_MS.Trt_F |
| PRKCSH | 0.52185 | 0.25724 | 0.725 | 0.72 | 1 | HC_vs_MS.Trt_F |
| TP53BP1 | 0.52186 | -1.109 | 0.306 | 0.32 | 1 | HC_vs_MS.Trt_F |
| MRPL14 | 0.52192 | -0.5802 | 0.136 | 0.12 | 1 | HC_vs_MS.Trt_F |

| | | | | | | |
|---|---|---|---|---|---|---|
| DYNLL2 | 0.52194 | -0.6475 | 0.133 | 0.15 | 1 | HC_vs_MS.Trt_F |
| UTP23 | 0.52232 | -0.2606 | 0.59 | 0.57 | 1 | HC_vs_MS.Trt_F |
| COMMD5 | 0.5224 | -0.4734 | 0.098 | 0.11 | 1 | HC_vs_MS.Trt_F |
| LRRN3 | 0.52291 | -1.5035 | 0.121 | 0.14 | 1 | HC_vs_MS.Trt_F |
| GRAMD2 | 0.52335 | 0.45619 | 0.572 | 0.55 | 1 | HC_vs_MS.Trt_F |
| FNDC3B | 0.52344 | 0.50178 | 0.783 | 0.75 | 1 | HC_vs_MS.Trt_F |
| ZNF394 | 0.52444 | 0.28008 | 0.509 | 0.55 | 1 | HC_vs_MS.Trt_F |
| B3GALNT1 | 0.52446 | -0.4622 | 0.228 | 0.21 | 1 | HC_vs_MS.Trt_F |
| CASP7 | 0.52447 | 1.16993 | 0.191 | 0.17 | 1 | HC_vs_MS.Trt_F |
| SUCLA2 | 0.52448 | -0.3506 | 0.147 | 0.13 | 1 | HC_vs_MS.Trt_F |
| PADI2 | 0.52468 | 0.39496 | 0.315 | 0.28 | 1 | HC_vs_MS.Trt_F |
| HNRNPH3 | 0.52485 | -0.4037 | 0.251 | 0.22 | 1 | HC_vs_MS.Trt_F |
| LRTOMT | 0.5258 | -0.3879 | 0.87 | 0.85 | 1 | HC_vs_MS.Trt_F |
| HTATSF1 | 0.5266 | -1.1412 | 0.159 | 0.17 | 1 | HC_vs_MS.Trt_F |
| PPIL6 | 0.52664 | 0.36246 | 0.54 | 0.53 | 1 | HC_vs_MS.Trt_F |
| TMEM179B | 0.52679 | -0.5744 | 0.205 | 0.22 | 1 | HC_vs_MS.Trt_F |
| GCSAML | 0.52687 | 0.71556 | 0.338 | 0.32 | 1 | HC_vs_MS.Trt_F |
| IFI27L2 | 0.52717 | -0.8595 | 0.251 | 0.26 | 1 | HC_vs_MS.Trt_F |
| VPS26A | 0.52724 | -0.5903 | 0.408 | 0.38 | 1 | HC_vs_MS.Trt_F |
| TUBGCP3 | 0.52735 | -0.2828 | 0.165 | 0.14 | 1 | HC_vs_MS.Trt_F |
| STK38 | 0.52774 | -0.3947 | 0.46 | 0.45 | 1 | HC_vs_MS.Trt_F |
| APEX2 | 0.52776 | 0.64308 | 0.402 | 0.37 | 1 | HC_vs_MS.Trt_F |
| HIST2H2AA3 | 0.52788 | 0.28364 | 0.124 | 0.11 | 1 | HC_vs_MS.Trt_F |
| HINT1 | 0.52861 | -0.2759 | 0.772 | 0.72 | 1 | HC_vs_MS.Trt_F |
| SCARF1 | 0.52907 | 0.50371 | 0.486 | 0.46 | 1 | HC_vs_MS.Trt_F |
| ITGAV | 0.52917 | -0.2848 | 0.133 | 0.12 | 1 | HC_vs_MS.Trt_F |
| CCDC146 | 0.52952 | 0.37866 | 0.104 | 0.09 | 1 | HC_vs_MS.Trt_F |
| GNL3 | 0.52957 | -0.6482 | 0.162 | 0.14 | 1 | HC_vs_MS.Trt_F |
| TUNAR | 0.5297 | 1.31306 | 0.168 | 0.15 | 1 | HC_vs_MS.Trt_F |
| LEAP2 | 0.52994 | 0.25205 | 0.26 | 0.23 | 1 | HC_vs_MS.Trt_F |
| EPC2 | 0.53009 | 0.49992 | 0.118 | 0.1 | 1 | HC_vs_MS.Trt_F |
| DNAAF2 | 0.5301 | 0.25498 | 0.118 | 0.1 | 1 | HC_vs_MS.Trt_F |
| PSMC4 | 0.53015 | -0.4449 | 0.543 | 0.48 | 1 | HC_vs_MS.Trt_F |
| SLC35E1 | 0.53018 | 0.50083 | 0.991 | 0.97 | 1 | HC_vs_MS.Trt_F |
| FUT4 | 0.53048 | -0.3531 | 0.39 | 0.35 | 1 | HC_vs_MS.Trt_F |
| COPS4 | 0.5307 | -0.7863 | 0.491 | 0.45 | 1 | HC_vs_MS.Trt_F |
| RWDD2A | 0.53089 | -0.5461 | 0.52 | 0.5 | 1 | HC_vs_MS.Trt_F |
| CTD-2207O2 | 0.53089 | -0.6573 | 0.168 | 0.18 | 1 | HC_vs_MS.Trt_F |
| METTL12 | 0.53183 | 0.59391 | 0.725 | 0.66 | 1 | HC_vs_MS.Trt_F |
| DNAJA3 | 0.53282 | -0.4626 | 0.962 | 0.91 | 1 | HC_vs_MS.Trt_F |

| | | | | | | |
|---|---|---|---|---|---|---|
| DOCK9 | 0.53302 | 0.455 | 0.338 | 0.3 | 1 | HC_vs_MS.Trt_F |
| FAM111A | 0.53314 | -1.3078 | 0.552 | 0.53 | 1 | HC_vs_MS.Trt_F |
| SLC12A9 | 0.5344 | 0.34815 | 0.902 | 0.87 | 1 | HC_vs_MS.Trt_F |
| PAFAH1B3 | 0.53445 | 0.3338 | 0.11 | 0.09 | 1 | HC_vs_MS.Trt_F |
| ATAD3C | 0.53451 | -0.2608 | 0.824 | 0.86 | 1 | HC_vs_MS.Trt_F |
| RAP2B | 0.53619 | -0.347 | 0.697 | 0.64 | 1 | HC_vs_MS.Trt_F |
| ZNF70 | 0.53649 | -0.4046 | 0.893 | 0.87 | 1 | HC_vs_MS.Trt_F |
| ACTG2 | 0.53657 | 1.10628 | 0.277 | 0.26 | 1 | HC_vs_MS.Trt_F |
| CD27 | 0.53665 | -0.3518 | 0.517 | 0.5 | 1 | HC_vs_MS.Trt_F |
| IL5RA | 0.53685 | 1.64825 | 0.318 | 0.29 | 1 | HC_vs_MS.Trt_F |
| SYPL1 | 0.53698 | -0.35 | 0.439 | 0.41 | 1 | HC_vs_MS.Trt_F |
| DHX33 | 0.53699 | -0.3423 | 0.341 | 0.35 | 1 | HC_vs_MS.Trt_F |
| MED14 | 0.53707 | -0.4259 | 0.188 | 0.16 | 1 | HC_vs_MS.Trt_F |
| PPP1R9A | 0.53718 | 0.32472 | 0.225 | 0.24 | 1 | HC_vs_MS.Trt_F |
| MED13L | 0.53732 | -0.373 | 0.13 | 0.14 | 1 | HC_vs_MS.Trt_F |
| MROH8 | 0.53769 | -0.4695 | 0.373 | 0.39 | 1 | HC_vs_MS.Trt_F |
| ITIH5 | 0.53798 | 0.25107 | 0.818 | 0.81 | 1 | HC_vs_MS.Trt_F |
| LSM3 | 0.53856 | 0.46024 | 0.624 | 0.59 | 1 | HC_vs_MS.Trt_F |
| AFF3 | 0.53867 | 0.62421 | 0.806 | 0.75 | 1 | HC_vs_MS.Trt_F |
| SOS2 | 0.53909 | -0.8821 | 0.136 | 0.12 | 1 | HC_vs_MS.Trt_F |
| CHERP | 0.53914 | -0.285 | 0.191 | 0.17 | 1 | HC_vs_MS.Trt_F |
| PLRG1 | 0.53914 | 0.3183 | 0.145 | 0.13 | 1 | HC_vs_MS.Trt_F |
| BRD4 | 0.53956 | -0.3807 | 0.353 | 0.32 | 1 | HC_vs_MS.Trt_F |
| REXO1 | 0.53995 | 1.16748 | 0.118 | 0.1 | 1 | HC_vs_MS.Trt_F |
| TOR1B | 0.54003 | -0.5388 | 0.277 | 0.26 | 1 | HC_vs_MS.Trt_F |
| MAP3K10 | 0.54074 | 0.28918 | 0.39 | 0.34 | 1 | HC_vs_MS.Trt_F |
| ZNF836 | 0.54083 | -0.5936 | 0.251 | 0.22 | 1 | HC_vs_MS.Trt_F |
| ZNF80 | 0.54094 | -0.5134 | 0.246 | 0.22 | 1 | HC_vs_MS.Trt_F |
| SNURF | 0.54152 | -0.2654 | 0.199 | 0.18 | 1 | HC_vs_MS.Trt_F |
| C14orf80 | 0.54198 | 0.79627 | 0.324 | 0.28 | 1 | HC_vs_MS.Trt_F |
| KIAA0368 | 0.54206 | 0.42593 | 0.272 | 0.25 | 1 | HC_vs_MS.Trt_F |
| DDX39A | 0.54276 | -0.435 | 0.355 | 0.36 | 1 | HC_vs_MS.Trt_F |
| SPTB | 0.54308 | -0.2576 | 0.471 | 0.45 | 1 | HC_vs_MS.Trt_F |
| ZC3H15 | 0.54317 | -0.6876 | 0.188 | 0.2 | 1 | HC_vs_MS.Trt_F |
| KLHL21 | 0.5434 | 0.275 | 0.76 | 0.73 | 1 | HC_vs_MS.Trt_F |
| RRP9 | 0.54374 | 0.81416 | 0.118 | 0.1 | 1 | HC_vs_MS.Trt_F |
| NLRP2 | 0.54404 | -0.5067 | 0.85 | 0.82 | 1 | HC_vs_MS.Trt_F |
| KANK3 | 0.54405 | 0.34847 | 0.277 | 0.3 | 1 | HC_vs_MS.Trt_F |
| ATG12 | 0.5442 | -0.4634 | 0.789 | 0.77 | 1 | HC_vs_MS.Trt_F |
| CIITA | 0.54441 | -0.776 | 0.962 | 0.91 | 1 | HC_vs_MS.Trt_F |

| | | | | | | |
|---|---|---|---|---|---|---|
| ANKRD28 | 0.54489 | 0.71677 | 0.439 | 0.41 | 1 | HC_vs_MS.Trt_F |
| RNASET2 | 0.54545 | -0.4261 | 0.434 | 0.44 | 1 | HC_vs_MS.Trt_F |
| TMEM97 | 0.54583 | -1.1644 | 0.254 | 0.28 | 1 | HC_vs_MS.Trt_F |
| ERO1A | 0.54596 | -0.6957 | 0.905 | 0.9 | 1 | HC_vs_MS.Trt_F |
| AGMO | 0.54626 | 1.1944 | 0.272 | 0.24 | 1 | HC_vs_MS.Trt_F |
| PNPLA4 | 0.54631 | -0.6243 | 0.283 | 0.25 | 1 | HC_vs_MS.Trt_F |
| TMEM30A | 0.54648 | -1.1574 | 0.208 | 0.22 | 1 | HC_vs_MS.Trt_F |
| UBAP2L | 0.54661 | -0.3294 | 0.269 | 0.28 | 1 | HC_vs_MS.Trt_F |
| NTN5 | 0.54664 | -0.2751 | 0.159 | 0.14 | 1 | HC_vs_MS.Trt_F |
| SSB | 0.54711 | -0.5774 | 0.798 | 0.73 | 1 | HC_vs_MS.Trt_F |
| C8orf76 | 0.5472 | -0.2709 | 0.592 | 0.61 | 1 | HC_vs_MS.Trt_F |
| LAPTM4A | 0.54739 | -0.6372 | 0.228 | 0.2 | 1 | HC_vs_MS.Trt_F |
| TBPL1 | 0.54753 | -0.4911 | 0.277 | 0.25 | 1 | HC_vs_MS.Trt_F |
| RELT | 0.54791 | 0.55467 | 0.11 | 0.1 | 1 | HC_vs_MS.Trt_F |
| ZNF512 | 0.54793 | -0.8396 | 0.142 | 0.15 | 1 | HC_vs_MS.Trt_F |
| GMPPA | 0.54803 | -0.7785 | 0.292 | 0.3 | 1 | HC_vs_MS.Trt_F |
| COASY | 0.5484 | 0.31571 | 0.46 | 0.44 | 1 | HC_vs_MS.Trt_F |
| CDC27 | 0.5484 | -0.4149 | 0.474 | 0.5 | 1 | HC_vs_MS.Trt_F |
| GADD45GIP | 0.54873 | -0.5983 | 0.182 | 0.16 | 1 | HC_vs_MS.Trt_F |
| FCHSD2 | 0.54875 | -0.3639 | 0.162 | 0.18 | 1 | HC_vs_MS.Trt_F |
| UBE2W | 0.54887 | -0.4605 | 0.691 | 0.66 | 1 | HC_vs_MS.Trt_F |
| CDC25B | 0.54924 | 0.33869 | 0.312 | 0.29 | 1 | HC_vs_MS.Trt_F |
| PTDSS2 | 0.54963 | 0.92922 | 0.107 | 0.09 | 1 | HC_vs_MS.Trt_F |
| DIABLO | 0.54969 | -0.5719 | 0.749 | 0.74 | 1 | HC_vs_MS.Trt_F |
| LRRC37A | 0.54979 | 0.34821 | 0.434 | 0.4 | 1 | HC_vs_MS.Trt_F |
| NDUFB7 | 0.55043 | -0.6193 | 0.243 | 0.25 | 1 | HC_vs_MS.Trt_F |
| TTC33 | 0.5521 | -0.8756 | 0.246 | 0.26 | 1 | HC_vs_MS.Trt_F |
| MAN2B1 | 0.55214 | 0.27225 | 0.639 | 0.63 | 1 | HC_vs_MS.Trt_F |
| KLF3 | 0.55218 | -0.4789 | 0.618 | 0.62 | 1 | HC_vs_MS.Trt_F |
| SS18L1 | 0.55244 | -0.6131 | 0.465 | 0.46 | 1 | HC_vs_MS.Trt_F |
| C19orf48 | 0.55279 | 0.27522 | 0.405 | 0.37 | 1 | HC_vs_MS.Trt_F |
| RDX | 0.55333 | -0.871 | 0.286 | 0.25 | 1 | HC_vs_MS.Trt_F |
| YDJC | 0.55456 | -0.4079 | 0.15 | 0.16 | 1 | HC_vs_MS.Trt_F |
| FUT7 | 0.55491 | -0.5553 | 0.107 | 0.12 | 1 | HC_vs_MS.Trt_F |
| EXOSC8 | 0.55534 | -0.3079 | 0.512 | 0.52 | 1 | HC_vs_MS.Trt_F |
| TRMT10C | 0.55602 | -0.4871 | 0.121 | 0.1 | 1 | HC_vs_MS.Trt_F |
| ZSCAN20 | 0.55615 | -0.73 | 0.315 | 0.3 | 1 | HC_vs_MS.Trt_F |
| NAGA | 0.55635 | -0.6415 | 0.13 | 0.14 | 1 | HC_vs_MS.Trt_F |
| CCT3 | 0.5565 | 0.30215 | 0.581 | 0.55 | 1 | HC_vs_MS.Trt_F |
| OCIAD2 | 0.55715 | -0.3225 | 0.662 | 0.61 | 1 | HC_vs_MS.Trt_F |

| | | | | | | |
|---|---|---|---|---|---|---|
| UTP14C | 0.55746 | -1.3255 | 0.194 | 0.21 | 1 | HC_vs_MS.Trt_F |
| MCOLN3 | 0.55755 | 0.5354 | 0.517 | 0.51 | 1 | HC_vs_MS.Trt_F |
| ACOX3 | 0.55771 | -0.7854 | 0.121 | 0.13 | 1 | HC_vs_MS.Trt_F |
| MINK1 | 0.55781 | 0.35898 | 0.295 | 0.26 | 1 | HC_vs_MS.Trt_F |
| PLD1 | 0.55818 | 0.68509 | 0.549 | 0.58 | 1 | HC_vs_MS.Trt_F |
| LIN37 | 0.55833 | 0.5251 | 0.104 | 0.09 | 1 | HC_vs_MS.Trt_F |
| JADE1 | 0.55837 | -0.8352 | 0.364 | 0.38 | 1 | HC_vs_MS.Trt_F |
| CPEB4 | 0.55848 | -0.4161 | 0.197 | 0.17 | 1 | HC_vs_MS.Trt_F |
| C22orf46 | 0.55851 | -0.4418 | 0.39 | 0.36 | 1 | HC_vs_MS.Trt_F |
| IFI44 | 0.55921 | 0.28969 | 0.104 | 0.09 | 1 | HC_vs_MS.Trt_F |
| ERGIC2 | 0.55925 | 0.33538 | 0.526 | 0.47 | 1 | HC_vs_MS.Trt_F |
| MAP2K6 | 0.55937 | 0.39548 | 0.408 | 0.38 | 1 | HC_vs_MS.Trt_F |
| TRMT13 | 0.56012 | -0.4177 | 0.118 | 0.13 | 1 | HC_vs_MS.Trt_F |
| SMIM15 | 0.56087 | -0.3687 | 0.104 | 0.12 | 1 | HC_vs_MS.Trt_F |
| FHDC1 | 0.56135 | -0.5498 | 0.506 | 0.49 | 1 | HC_vs_MS.Trt_F |
| PODN | 0.56152 | 1.60664 | 0.436 | 0.42 | 1 | HC_vs_MS.Trt_F |
| IMMT | 0.56187 | -0.6796 | 0.486 | 0.44 | 1 | HC_vs_MS.Trt_F |
| KDM7A | 0.56189 | -0.2538 | 0.254 | 0.26 | 1 | HC_vs_MS.Trt_F |
| TFIP11 | 0.56223 | -1.0664 | 0.549 | 0.49 | 1 | HC_vs_MS.Trt_F |
| C15orf38-AI | 0.56245 | -0.5028 | 0.402 | 0.37 | 1 | HC_vs_MS.Trt_F |
| SETMAR | 0.56249 | -1.0434 | 0.332 | 0.33 | 1 | HC_vs_MS.Trt_F |
| COX6C | 0.56253 | -0.6048 | 0.503 | 0.49 | 1 | HC_vs_MS.Trt_F |
| MAGIX | 0.56268 | -0.3955 | 0.523 | 0.51 | 1 | HC_vs_MS.Trt_F |
| SULF2 | 0.563 | 0.27502 | 0.139 | 0.12 | 1 | HC_vs_MS.Trt_F |
| NFKB1 | 0.56306 | -1.5396 | 0.15 | 0.13 | 1 | HC_vs_MS.Trt_F |
| KTN1 | 0.56338 | -0.8273 | 0.604 | 0.57 | 1 | HC_vs_MS.Trt_F |
| OGFR | 0.56365 | 0.28825 | 0.223 | 0.2 | 1 | HC_vs_MS.Trt_F |
| P3H2 | 0.56402 | 0.5107 | 0.517 | 0.47 | 1 | HC_vs_MS.Trt_F |
| SERP1 | 0.56418 | -0.4921 | 0.584 | 0.59 | 1 | HC_vs_MS.Trt_F |
| AKAP8 | 0.56438 | 0.39726 | 0.422 | 0.39 | 1 | HC_vs_MS.Trt_F |
| AK1 | 0.56445 | -0.7825 | 0.121 | 0.1 | 1 | HC_vs_MS.Trt_F |
| SAXO2 | 0.56455 | 0.33233 | 0.269 | 0.24 | 1 | HC_vs_MS.Trt_F |
| CSMD1 | 0.56493 | 0.69077 | 0.361 | 0.33 | 1 | HC_vs_MS.Trt_F |
| ELMO2 | 0.56519 | -0.6409 | 0.246 | 0.21 | 1 | HC_vs_MS.Trt_F |
| STX2 | 0.5653 | -0.418 | 0.52 | 0.48 | 1 | HC_vs_MS.Trt_F |
| EID1 | 0.56581 | -0.3753 | 0.379 | 0.38 | 1 | HC_vs_MS.Trt_F |
| NLN | 0.56584 | -0.4423 | 0.754 | 0.73 | 1 | HC_vs_MS.Trt_F |
| MED1 | 0.56587 | -0.5271 | 0.396 | 0.36 | 1 | HC_vs_MS.Trt_F |
| TRIM68 | 0.5662 | 0.52166 | 0.618 | 0.56 | 1 | HC_vs_MS.Trt_F |
| TSEN15 | 0.56643 | -0.5102 | 0.237 | 0.21 | 1 | HC_vs_MS.Trt_F |

| | | | | | | |
|---|---|---|---|---|---|---|
| RAD52 | 0.56649 | 0.32966 | 0.254 | 0.27 | 1 | HC_vs_MS.Trt_F |
| FIGNL1 | 0.56666 | -0.3677 | 0.171 | 0.15 | 1 | HC_vs_MS.Trt_F |
| PIAS1 | 0.56781 | -0.2507 | 0.379 | 0.34 | 1 | HC_vs_MS.Trt_F |
| SPPL2A | 0.56799 | 0.44555 | 0.951 | 0.93 | 1 | HC_vs_MS.Trt_F |
| MAZ | 0.56805 | 0.31809 | 0.182 | 0.17 | 1 | HC_vs_MS.Trt_F |
| PRMT9 | 0.56841 | -0.2627 | 0.223 | 0.2 | 1 | HC_vs_MS.Trt_F |
| UBE2D4 | 0.56843 | -0.5714 | 0.699 | 0.72 | 1 | HC_vs_MS.Trt_F |
| NUDT19 | 0.56865 | -0.815 | 0.876 | 0.89 | 1 | HC_vs_MS.Trt_F |
| DIAPH2 | 0.56875 | 0.31181 | 0.341 | 0.32 | 1 | HC_vs_MS.Trt_F |
| CARD6 | 0.5689 | -0.4636 | 0.552 | 0.51 | 1 | HC_vs_MS.Trt_F |
| CLOCK | 0.56902 | -0.6497 | 0.171 | 0.18 | 1 | HC_vs_MS.Trt_F |
| RANGAP1 | 0.56927 | 0.40514 | 0.754 | 0.73 | 1 | HC_vs_MS.Trt_F |
| ZNF652 | 0.56957 | -1.0783 | 0.581 | 0.53 | 1 | HC_vs_MS.Trt_F |
| ATP5G1 | 0.56962 | -0.8036 | 0.306 | 0.32 | 1 | HC_vs_MS.Trt_F |
| FAM220A | 0.57051 | 0.85032 | 0.367 | 0.34 | 1 | HC_vs_MS.Trt_F |
| AKR1B10 | 0.57059 | 0.44907 | 0.488 | 0.48 | 1 | HC_vs_MS.Trt_F |
| FAM134C | 0.57066 | -0.39 | 0.694 | 0.63 | 1 | HC_vs_MS.Trt_F |
| LIG3 | 0.57122 | 1.16681 | 0.289 | 0.27 | 1 | HC_vs_MS.Trt_F |
| UNC119 | 0.5715 | -0.7565 | 0.101 | 0.11 | 1 | HC_vs_MS.Trt_F |
| GNB1 | 0.57195 | -0.3042 | 0.329 | 0.35 | 1 | HC_vs_MS.Trt_F |
| SHARPIN | 0.57212 | -0.327 | 0.139 | 0.15 | 1 | HC_vs_MS.Trt_F |
| ZNF587B | 0.57216 | -0.2719 | 0.35 | 0.31 | 1 | HC_vs_MS.Trt_F |
| POLE | 0.57246 | -0.3204 | 0.882 | 0.83 | 1 | HC_vs_MS.Trt_F |
| SNRNP48 | 0.57262 | 0.50558 | 0.208 | 0.22 | 1 | HC_vs_MS.Trt_F |
| BLM | 0.57263 | -0.3706 | 0.292 | 0.27 | 1 | HC_vs_MS.Trt_F |
| CCDC88A | 0.5727 | -0.2807 | 0.705 | 0.66 | 1 | HC_vs_MS.Trt_F |
| CUL3 | 0.57317 | -0.2622 | 0.347 | 0.31 | 1 | HC_vs_MS.Trt_F |
| ATP11A | 0.57321 | -0.2626 | 0.292 | 0.27 | 1 | HC_vs_MS.Trt_F |
| ZNF223 | 0.57391 | -1.0209 | 0.104 | 0.12 | 1 | HC_vs_MS.Trt_F |
| TMTC3 | 0.57392 | -0.3425 | 0.269 | 0.24 | 1 | HC_vs_MS.Trt_F |
| LSM12 | 0.57398 | -0.3804 | 0.656 | 0.67 | 1 | HC_vs_MS.Trt_F |
| MIS12 | 0.5741 | -0.3939 | 0.341 | 0.31 | 1 | HC_vs_MS.Trt_F |
| ARID5A | 0.57413 | -0.4509 | 0.225 | 0.23 | 1 | HC_vs_MS.Trt_F |
| PPT2 | 0.57416 | 0.63392 | 0.604 | 0.58 | 1 | HC_vs_MS.Trt_F |
| PDS5B | 0.57445 | -0.4446 | 0.344 | 0.36 | 1 | HC_vs_MS.Trt_F |
| STK39 | 0.57466 | 0.35114 | 0.162 | 0.14 | 1 | HC_vs_MS.Trt_F |
| C4orf3 | 0.57473 | -0.65 | 0.497 | 0.48 | 1 | HC_vs_MS.Trt_F |
| MTMR14 | 0.57478 | -0.3203 | 0.656 | 0.61 | 1 | HC_vs_MS.Trt_F |
| TNPO2 | 0.57587 | 1.12033 | 0.168 | 0.15 | 1 | HC_vs_MS.Trt_F |
| TXNDC16 | 0.57629 | -0.3481 | 0.133 | 0.12 | 1 | HC_vs_MS.Trt_F |

| | | | | | | |
|---|---|---|---|---|---|---|
| LST1 | 0.57663 | 0.44732 | 0.272 | 0.26 | 1 | HC_vs_MS.Trt_F |
| BTN2A1 | 0.57686 | -0.5326 | 0.156 | 0.17 | 1 | HC_vs_MS.Trt_F |
| NECAB1 | 0.57695 | 0.45463 | 0.156 | 0.14 | 1 | HC_vs_MS.Trt_F |
| UBAP1 | 0.5771 | -0.3013 | 0.153 | 0.13 | 1 | HC_vs_MS.Trt_F |
| CPSF6 | 0.57745 | 0.26554 | 0.396 | 0.42 | 1 | HC_vs_MS.Trt_F |
| ZC2HC1C | 0.57745 | -0.6451 | 0.124 | 0.14 | 1 | HC_vs_MS.Trt_F |
| SH2D3A | 0.57786 | 0.4445 | 0.945 | 0.95 | 1 | HC_vs_MS.Trt_F |
| SLC25A24 | 0.57787 | -0.4346 | 0.121 | 0.13 | 1 | HC_vs_MS.Trt_F |
| TNKS | 0.57792 | 0.68229 | 0.844 | 0.81 | 1 | HC_vs_MS.Trt_F |
| BEX4 | 0.57823 | -0.572 | 0.113 | 0.12 | 1 | HC_vs_MS.Trt_F |
| PYGB | 0.57828 | -0.2696 | 0.165 | 0.14 | 1 | HC_vs_MS.Trt_F |
| TACO1 | 0.57843 | 1.11676 | 0.393 | 0.39 | 1 | HC_vs_MS.Trt_F |
| ARID5B | 0.57862 | -0.5001 | 0.864 | 0.77 | 1 | HC_vs_MS.Trt_F |
| S1PR1 | 0.57865 | -0.3758 | 0.465 | 0.43 | 1 | HC_vs_MS.Trt_F |
| ANKRD13C | 0.57884 | -0.9607 | 0.145 | 0.12 | 1 | HC_vs_MS.Trt_F |
| CRAMP1 | 0.57912 | -0.616 | 0.104 | 0.09 | 1 | HC_vs_MS.Trt_F |
| RFC5 | 0.57944 | -0.5842 | 0.199 | 0.18 | 1 | HC_vs_MS.Trt_F |
| RGPD3 | 0.57962 | 0.32868 | 0.104 | 0.09 | 1 | HC_vs_MS.Trt_F |
| CECR5 | 0.57995 | -0.3118 | 0.214 | 0.19 | 1 | HC_vs_MS.Trt_F |
| RP11-15H20 | 0.57998 | 0.54066 | 0.136 | 0.15 | 1 | HC_vs_MS.Trt_F |
| SMS | 0.58001 | -0.3524 | 0.185 | 0.16 | 1 | HC_vs_MS.Trt_F |
| ZNF100 | 0.58034 | 0.27408 | 0.864 | 0.84 | 1 | HC_vs_MS.Trt_F |
| FBXO9 | 0.58039 | 0.25897 | 0.803 | 0.81 | 1 | HC_vs_MS.Trt_F |
| MGA | 0.58041 | -0.3231 | 0.908 | 0.88 | 1 | HC_vs_MS.Trt_F |
| AASDHPPT | 0.58064 | -0.2634 | 0.231 | 0.21 | 1 | HC_vs_MS.Trt_F |
| PYGO2 | 0.58114 | -0.4113 | 0.173 | 0.15 | 1 | HC_vs_MS.Trt_F |
| MRPL35 | 0.58193 | -1.0711 | 0.156 | 0.13 | 1 | HC_vs_MS.Trt_F |
| TOE1 | 0.58217 | -1.0977 | 0.09 | 0.1 | 1 | HC_vs_MS.Trt_F |
| COPS9 | 0.58233 | -0.5517 | 0.176 | 0.19 | 1 | HC_vs_MS.Trt_F |
| AEN | 0.58262 | 0.49814 | 0.153 | 0.14 | 1 | HC_vs_MS.Trt_F |
| PPP1R3F | 0.5827 | -0.3102 | 0.162 | 0.14 | 1 | HC_vs_MS.Trt_F |
| PPP6R1 | 0.5828 | -0.3079 | 0.355 | 0.31 | 1 | HC_vs_MS.Trt_F |
| ZNF576 | 0.58302 | -0.3641 | 0.546 | 0.54 | 1 | HC_vs_MS.Trt_F |
| NCBP2-AS2 | 0.58305 | -0.3236 | 0.139 | 0.12 | 1 | HC_vs_MS.Trt_F |
| FMR1 | 0.5831 | -0.3202 | 0.153 | 0.14 | 1 | HC_vs_MS.Trt_F |
| GALNT3 | 0.58319 | -0.6514 | 0.095 | 0.11 | 1 | HC_vs_MS.Trt_F |
| CWC27 | 0.58344 | 0.40178 | 0.168 | 0.15 | 1 | HC_vs_MS.Trt_F |
| XYLT1 | 0.58345 | 0.57558 | 0.162 | 0.14 | 1 | HC_vs_MS.Trt_F |
| TSPAN17 | 0.58376 | -0.4271 | 0.133 | 0.12 | 1 | HC_vs_MS.Trt_F |
| RAB9A | 0.58377 | -0.801 | 0.116 | 0.13 | 1 | HC_vs_MS.Trt_F |

| | | | | | | |
|---|---|---|---|---|---|---|
| GTF2E1 | 0.58389 | 0.94022 | 0.295 | 0.32 | 1 | HC_vs_MS.Trt_F |
| SQRDL | 0.58393 | -0.32 | 0.171 | 0.15 | 1 | HC_vs_MS.Trt_F |
| MRPS12 | 0.58418 | -0.3356 | 0.436 | 0.42 | 1 | HC_vs_MS.Trt_F |
| MRPS23 | 0.58422 | -0.4005 | 0.858 | 0.84 | 1 | HC_vs_MS.Trt_F |
| H2AFZ | 0.58433 | -0.2784 | 0.39 | 0.35 | 1 | HC_vs_MS.Trt_F |
| ZFP37 | 0.58447 | -0.8703 | 0.425 | 0.43 | 1 | HC_vs_MS.Trt_F |
| ARL6IP4 | 0.58475 | -0.3693 | 0.306 | 0.31 | 1 | HC_vs_MS.Trt_F |
| GART | 0.58521 | -0.2902 | 0.858 | 0.84 | 1 | HC_vs_MS.Trt_F |
| PYCR2 | 0.58531 | 0.42192 | 0.679 | 0.66 | 1 | HC_vs_MS.Trt_F |
| ABHD17B | 0.58578 | -0.4766 | 0.303 | 0.27 | 1 | HC_vs_MS.Trt_F |
| AKAP5 | 0.58614 | 0.3864 | 0.662 | 0.66 | 1 | HC_vs_MS.Trt_F |
| C14orf2 | 0.58619 | -0.5871 | 0.855 | 0.8 | 1 | HC_vs_MS.Trt_F |
| PMF1-BGLAI | 0.58667 | -0.4755 | 0.11 | 0.12 | 1 | HC_vs_MS.Trt_F |
| LEPR | 0.58696 | 0.42249 | 0.373 | 0.36 | 1 | HC_vs_MS.Trt_F |
| PGAP2 | 0.58699 | -0.3053 | 0.107 | 0.09 | 1 | HC_vs_MS.Trt_F |
| SEC61G | 0.58764 | -0.4963 | 0.312 | 0.32 | 1 | HC_vs_MS.Trt_F |
| CCDC117 | 0.58773 | -0.8679 | 0.416 | 0.37 | 1 | HC_vs_MS.Trt_F |
| SLC9A3R1 | 0.5883 | -0.4028 | 0.384 | 0.35 | 1 | HC_vs_MS.Trt_F |
| SLC48A1 | 0.58846 | -0.7931 | 0.633 | 0.59 | 1 | HC_vs_MS.Trt_F |
| ZNF574 | 0.5889 | 0.26495 | 0.373 | 0.39 | 1 | HC_vs_MS.Trt_F |
| VPS16 | 0.58915 | -0.6503 | 0.153 | 0.13 | 1 | HC_vs_MS.Trt_F |
| CACNG8 | 0.58931 | 0.28868 | 0.835 | 0.82 | 1 | HC_vs_MS.Trt_F |
| RBM19 | 0.58973 | -0.557 | 0.451 | 0.45 | 1 | HC_vs_MS.Trt_F |
| HNRNPK | 0.5907 | -0.263 | 0.708 | 0.67 | 1 | HC_vs_MS.Trt_F |
| CCDC126 | 0.59093 | -0.6202 | 0.118 | 0.1 | 1 | HC_vs_MS.Trt_F |
| MDC1 | 0.59127 | -0.4929 | 0.104 | 0.12 | 1 | HC_vs_MS.Trt_F |
| CNNM2 | 0.59166 | 1.38832 | 0.341 | 0.32 | 1 | HC_vs_MS.Trt_F |
| GABARAPL2 | 0.59204 | -0.2773 | 0.555 | 0.48 | 1 | HC_vs_MS.Trt_F |
| PDGFRA | 0.59253 | 0.38038 | 0.243 | 0.22 | 1 | HC_vs_MS.Trt_F |
| CTB-50L17.1 | 0.59287 | -0.3479 | 0.65 | 0.61 | 1 | HC_vs_MS.Trt_F |
| TANC1 | 0.59334 | 0.30358 | 0.376 | 0.38 | 1 | HC_vs_MS.Trt_F |
| CACNB2 | 0.5935 | -0.5339 | 0.471 | 0.47 | 1 | HC_vs_MS.Trt_F |
| UMPS | 0.59389 | -0.3372 | 0.688 | 0.68 | 1 | HC_vs_MS.Trt_F |
| TBC1D23 | 0.59397 | -0.2686 | 0.15 | 0.13 | 1 | HC_vs_MS.Trt_F |
| RNF13 | 0.59506 | -0.7893 | 0.199 | 0.17 | 1 | HC_vs_MS.Trt_F |
| SLC26A8 | 0.5954 | 0.32205 | 0.731 | 0.71 | 1 | HC_vs_MS.Trt_F |
| GPR22 | 0.59552 | -0.4437 | 0.127 | 0.11 | 1 | HC_vs_MS.Trt_F |
| MFSD14B | 0.59552 | -0.5552 | 0.139 | 0.12 | 1 | HC_vs_MS.Trt_F |
| MPG | 0.59562 | -0.4569 | 0.165 | 0.17 | 1 | HC_vs_MS.Trt_F |
| FEM1C | 0.5959 | -0.7043 | 0.147 | 0.16 | 1 | HC_vs_MS.Trt_F |

| | | | | | | |
|---|---|---|---|---|---|---|
| RAB5A | 0.59608 | -0.5438 | 0.176 | 0.16 | 1 | HC_vs_MS.Trt_F |
| ARPC5L | 0.59619 | -0.5477 | 0.197 | 0.21 | 1 | HC_vs_MS.Trt_F |
| C16orf45 | 0.59626 | 0.54087 | 0.22 | 0.2 | 1 | HC_vs_MS.Trt_F |
| ADPRHL2 | 0.59652 | -1.3096 | 0.095 | 0.1 | 1 | HC_vs_MS.Trt_F |
| C14orf159 | 0.59666 | 0.57914 | 0.592 | 0.62 | 1 | HC_vs_MS.Trt_F |
| KATNA1 | 0.59669 | -1.2952 | 0.113 | 0.12 | 1 | HC_vs_MS.Trt_F |
| ZNF432 | 0.59688 | 1.06183 | 0.642 | 0.62 | 1 | HC_vs_MS.Trt_F |
| FAM104A | 0.59705 | -0.7974 | 0.191 | 0.2 | 1 | HC_vs_MS.Trt_F |
| TTYH2 | 0.59732 | -0.3911 | 0.116 | 0.1 | 1 | HC_vs_MS.Trt_F |
| TNFAIP8 | 0.59759 | -0.5117 | 0.899 | 0.84 | 1 | HC_vs_MS.Trt_F |
| SAFB | 0.5977 | -0.6304 | 0.379 | 0.37 | 1 | HC_vs_MS.Trt_F |
| INPP5B | 0.59795 | -0.3183 | 0.96 | 0.95 | 1 | HC_vs_MS.Trt_F |
| NELFCD | 0.59807 | 0.40873 | 0.598 | 0.59 | 1 | HC_vs_MS.Trt_F |
| LRRC69 | 0.59872 | -0.2674 | 0.595 | 0.58 | 1 | HC_vs_MS.Trt_F |
| GIGYF2 | 0.59873 | -0.5194 | 0.925 | 0.89 | 1 | HC_vs_MS.Trt_F |
| ARHGAP44 | 0.59877 | -1.0486 | 0.329 | 0.31 | 1 | HC_vs_MS.Trt_F |
| TMEM19 | 0.599 | -0.4251 | 0.772 | 0.8 | 1 | HC_vs_MS.Trt_F |
| ZNF665 | 0.59909 | -0.3921 | 0.954 | 0.92 | 1 | HC_vs_MS.Trt_F |
| RAB36 | 0.59918 | 0.58427 | 0.39 | 0.36 | 1 | HC_vs_MS.Trt_F |
| AK6 | 0.59988 | -0.9605 | 0.234 | 0.24 | 1 | HC_vs_MS.Trt_F |
| LPCAT3 | 0.60037 | -0.4051 | 0.486 | 0.44 | 1 | HC_vs_MS.Trt_F |
| ADGRG1 | 0.6004 | -0.918 | 0.749 | 0.69 | 1 | HC_vs_MS.Trt_F |
| DDX20 | 0.60047 | -0.433 | 0.116 | 0.1 | 1 | HC_vs_MS.Trt_F |
| HSPE1-MOB | 0.60055 | -0.592 | 0.101 | 0.11 | 1 | HC_vs_MS.Trt_F |
| TBC1D9B | 0.60057 | -0.7356 | 0.503 | 0.45 | 1 | HC_vs_MS.Trt_F |
| DNMBP | 0.60093 | 0.56659 | 0.101 | 0.09 | 1 | HC_vs_MS.Trt_F |
| ZBTB25 | 0.6017 | -0.4568 | 0.983 | 0.98 | 1 | HC_vs_MS.Trt_F |
| SLC25A5 | 0.60206 | -0.3486 | 0.347 | 0.31 | 1 | HC_vs_MS.Trt_F |
| ARL2BP | 0.60209 | -0.4981 | 0.153 | 0.16 | 1 | HC_vs_MS.Trt_F |
| FAM65B | 0.60221 | -0.4239 | 0.673 | 0.65 | 1 | HC_vs_MS.Trt_F |
| ZC3H8 | 0.60232 | -0.3235 | 0.818 | 0.78 | 1 | HC_vs_MS.Trt_F |
| GNS | 0.60248 | 0.34408 | 0.775 | 0.8 | 1 | HC_vs_MS.Trt_F |
| PYM1 | 0.60255 | -0.5782 | 0.231 | 0.23 | 1 | HC_vs_MS.Trt_F |
| FAM185A | 0.60325 | 0.35321 | 0.113 | 0.13 | 1 | HC_vs_MS.Trt_F |
| SNX14 | 0.60354 | -0.4326 | 0.59 | 0.52 | 1 | HC_vs_MS.Trt_F |
| PEX11A | 0.60395 | -0.3166 | 0.578 | 0.57 | 1 | HC_vs_MS.Trt_F |
| BACE1 | 0.60403 | -0.594 | 0.613 | 0.54 | 1 | HC_vs_MS.Trt_F |
| SYT11 | 0.60404 | -0.3228 | 0.705 | 0.6 | 1 | HC_vs_MS.Trt_F |
| C1orf162 | 0.60407 | 0.37395 | 0.142 | 0.13 | 1 | HC_vs_MS.Trt_F |
| DENND5A | 0.60445 | 0.48527 | 0.275 | 0.26 | 1 | HC_vs_MS.Trt_F |

| | | | | | | |
|---|---|---|---|---|---|---|
| TIGD6 | 0.60452 | 0.86099 | 0.246 | 0.23 | 1 | HC_vs_MS.Trt_F |
| CNNM3 | 0.60517 | 0.27327 | 0.645 | 0.6 | 1 | HC_vs_MS.Trt_F |
| NEDD1 | 0.60572 | -0.292 | 0.15 | 0.16 | 1 | HC_vs_MS.Trt_F |
| ZNF721 | 0.60595 | -0.6804 | 0.376 | 0.37 | 1 | HC_vs_MS.Trt_F |
| MCM9 | 0.606 | -0.9695 | 0.341 | 0.35 | 1 | HC_vs_MS.Trt_F |
| DLD | 0.60609 | -0.4634 | 0.179 | 0.16 | 1 | HC_vs_MS.Trt_F |
| CMC1 | 0.60659 | -0.3231 | 0.217 | 0.19 | 1 | HC_vs_MS.Trt_F |
| C12orf43 | 0.60674 | 0.43805 | 0.922 | 0.9 | 1 | HC_vs_MS.Trt_F |
| PTAFR | 0.60678 | 0.48011 | 0.642 | 0.64 | 1 | HC_vs_MS.Trt_F |
| ZNF23 | 0.6073 | 0.56991 | 0.396 | 0.37 | 1 | HC_vs_MS.Trt_F |
| ZNF45 | 0.60744 | -0.5191 | 0.344 | 0.32 | 1 | HC_vs_MS.Trt_F |
| MRPL24 | 0.60824 | -0.2833 | 0.145 | 0.13 | 1 | HC_vs_MS.Trt_F |
| PUM2 | 0.60882 | -0.2769 | 0.338 | 0.35 | 1 | HC_vs_MS.Trt_F |
| ISY1 | 0.60931 | -0.3696 | 0.624 | 0.6 | 1 | HC_vs_MS.Trt_F |
| PIH1D1 | 0.60945 | 0.25207 | 0.338 | 0.31 | 1 | HC_vs_MS.Trt_F |
| BTN3A2 | 0.60956 | -0.3548 | 0.76 | 0.7 | 1 | HC_vs_MS.Trt_F |
| PAWR | 0.61043 | 0.25745 | 0.72 | 0.72 | 1 | HC_vs_MS.Trt_F |
| NOP10 | 0.61071 | -0.6447 | 0.228 | 0.23 | 1 | HC_vs_MS.Trt_F |
| ARMC2 | 0.61098 | 0.48938 | 0.11 | 0.1 | 1 | HC_vs_MS.Trt_F |
| SLC25A25 | 0.61157 | -0.6285 | 0.121 | 0.11 | 1 | HC_vs_MS.Trt_F |
| TMEM217 | 0.61184 | 0.70895 | 0.113 | 0.1 | 1 | HC_vs_MS.Trt_F |
| IP6K1 | 0.61297 | -0.5598 | 0.171 | 0.15 | 1 | HC_vs_MS.Trt_F |
| FAM78A | 0.61311 | 0.7045 | 0.364 | 0.34 | 1 | HC_vs_MS.Trt_F |
| HCST | 0.61342 | -0.3106 | 0.434 | 0.37 | 1 | HC_vs_MS.Trt_F |
| CREB1 | 0.61342 | -0.4564 | 0.665 | 0.63 | 1 | HC_vs_MS.Trt_F |
| CHRAC1 | 0.61437 | 0.5555 | 0.506 | 0.51 | 1 | HC_vs_MS.Trt_F |
| GSTT1 | 0.61448 | -0.3304 | 0.74 | 0.69 | 1 | HC_vs_MS.Trt_F |
| RAB28 | 0.61493 | 0.47821 | 0.136 | 0.12 | 1 | HC_vs_MS.Trt_F |
| RAB3D | 0.61536 | -0.5026 | 0.673 | 0.58 | 1 | HC_vs_MS.Trt_F |
| ZNF610 | 0.61544 | -0.4684 | 0.445 | 0.47 | 1 | HC_vs_MS.Trt_F |
| FAM208A | 0.61599 | -0.5822 | 0.711 | 0.74 | 1 | HC_vs_MS.Trt_F |
| CDC123 | 0.61643 | -0.3071 | 0.228 | 0.21 | 1 | HC_vs_MS.Trt_F |
| DBR1 | 0.61663 | -0.2578 | 0.786 | 0.79 | 1 | HC_vs_MS.Trt_F |
| ABCA6 | 0.61691 | 0.27885 | 0.685 | 0.69 | 1 | HC_vs_MS.Trt_F |
| HEXDC | 0.6172 | -0.646 | 0.405 | 0.37 | 1 | HC_vs_MS.Trt_F |
| TRMT2A | 0.61736 | -0.3677 | 0.185 | 0.16 | 1 | HC_vs_MS.Trt_F |
| QSER1 | 0.61746 | -1.107 | 0.11 | 0.1 | 1 | HC_vs_MS.Trt_F |
| RP11-277P1 | 0.61811 | 0.38943 | 0.09 | 0.1 | 1 | HC_vs_MS.Trt_F |
| PPARGC1B | 0.61815 | -0.3881 | 0.763 | 0.73 | 1 | HC_vs_MS.Trt_F |
| LRMP | 0.61824 | -0.3444 | 0.113 | 0.12 | 1 | HC_vs_MS.Trt_F |

| | | | | | | |
|---|---|---|---|---|---|---|
| KLHL26 | 0.61834 | 0.29864 | 0.552 | 0.56 | 1 | HC_vs_MS.Trt_F |
| ZHX1 | 0.61838 | -0.8404 | 0.598 | 0.57 | 1 | HC_vs_MS.Trt_F |
| ZNF181 | 0.61857 | -0.9635 | 0.194 | 0.17 | 1 | HC_vs_MS.Trt_F |
| EIF4E3 | 0.6187 | -0.8176 | 0.176 | 0.16 | 1 | HC_vs_MS.Trt_F |
| ZDHHC6 | 0.61872 | -0.6646 | 0.188 | 0.17 | 1 | HC_vs_MS.Trt_F |
| AK9 | 0.61929 | -0.6282 | 0.76 | 0.66 | 1 | HC_vs_MS.Trt_F |
| CCNT2 | 0.61999 | -0.4672 | 0.179 | 0.16 | 1 | HC_vs_MS.Trt_F |
| CTD-2192J1( | 0.62014 | 0.33607 | 0.512 | 0.48 | 1 | HC_vs_MS.Trt_F |
| SLC6A6 | 0.62017 | -0.4297 | 0.217 | 0.19 | 1 | HC_vs_MS.Trt_F |
| 7-Sep | 0.62053 | -0.3499 | 0.751 | 0.68 | 1 | HC_vs_MS.Trt_F |
| KLHL28 | 0.62094 | -0.3761 | 0.382 | 0.39 | 1 | HC_vs_MS.Trt_F |
| RRM2B | 0.6211 | 0.38219 | 0.182 | 0.17 | 1 | HC_vs_MS.Trt_F |
| SKA2 | 0.62148 | -0.6074 | 0.601 | 0.59 | 1 | HC_vs_MS.Trt_F |
| MOB1A | 0.62175 | -0.4118 | 0.85 | 0.82 | 1 | HC_vs_MS.Trt_F |
| COX5B | 0.6223 | -0.5669 | 0.832 | 0.79 | 1 | HC_vs_MS.Trt_F |
| CMC2 | 0.62256 | -0.5278 | 0.876 | 0.88 | 1 | HC_vs_MS.Trt_F |
| BOLA2B | 0.62259 | -0.3764 | 0.147 | 0.16 | 1 | HC_vs_MS.Trt_F |
| PDP2 | 0.62355 | 0.35929 | 0.835 | 0.8 | 1 | HC_vs_MS.Trt_F |
| CGGBP1 | 0.62376 | -0.7894 | 0.228 | 0.21 | 1 | HC_vs_MS.Trt_F |
| RPL24 | 0.62377 | -0.2908 | 0.777 | 0.75 | 1 | HC_vs_MS.Trt_F |
| SLC52A1 | 0.6239 | -0.2904 | 0.792 | 0.76 | 1 | HC_vs_MS.Trt_F |
| ZNF8 | 0.62427 | -0.3588 | 0.96 | 0.97 | 1 | HC_vs_MS.Trt_F |
| SCOC | 0.62433 | -0.3325 | 0.127 | 0.11 | 1 | HC_vs_MS.Trt_F |
| DBNDD1 | 0.62489 | 0.90115 | 0.182 | 0.19 | 1 | HC_vs_MS.Trt_F |
| SSSCA1 | 0.62496 | -0.3492 | 0.11 | 0.1 | 1 | HC_vs_MS.Trt_F |
| SSRP1 | 0.62523 | -0.3784 | 0.171 | 0.15 | 1 | HC_vs_MS.Trt_F |
| ANK1 | 0.62545 | -1.3238 | 0.217 | 0.23 | 1 | HC_vs_MS.Trt_F |
| S100B | 0.62569 | 0.64274 | 0.153 | 0.14 | 1 | HC_vs_MS.Trt_F |
| PLEKHG4 | 0.62571 | -0.4857 | 0.197 | 0.21 | 1 | HC_vs_MS.Trt_F |
| SDHA | 0.62579 | -0.3094 | 0.581 | 0.57 | 1 | HC_vs_MS.Trt_F |
| AARS2 | 0.62579 | -0.4495 | 0.457 | 0.42 | 1 | HC_vs_MS.Trt_F |
| TG | 0.62582 | 0.89558 | 0.098 | 0.11 | 1 | HC_vs_MS.Trt_F |
| PWWP2A | 0.62615 | -0.782 | 0.139 | 0.15 | 1 | HC_vs_MS.Trt_F |
| LYPLA1 | 0.62649 | -0.3143 | 0.561 | 0.52 | 1 | HC_vs_MS.Trt_F |
| PLBD1 | 0.62668 | 0.38677 | 0.5 | 0.48 | 1 | HC_vs_MS.Trt_F |
| ZSCAN22 | 0.62672 | -0.3357 | 0.48 | 0.49 | 1 | HC_vs_MS.Trt_F |
| APC2 | 0.62679 | 0.44389 | 0.104 | 0.12 | 1 | HC_vs_MS.Trt_F |
| LNPK | 0.6271 | -0.527 | 0.558 | 0.55 | 1 | HC_vs_MS.Trt_F |
| LPAR4 | 0.62715 | 0.34461 | 0.185 | 0.17 | 1 | HC_vs_MS.Trt_F |
| LEF1 | 0.62806 | -0.3082 | 0.379 | 0.38 | 1 | HC_vs_MS.Trt_F |

| | | | | | | |
|---|---|---|---|---|---|---|
| HIST1H1D | 0.62854 | 0.27744 | 0.095 | 0.1 | 1 | HC_vs_MS.Trt_F |
| FBRSL1 | 0.62917 | -0.2804 | 0.113 | 0.1 | 1 | HC_vs_MS.Trt_F |
| ACAA1 | 0.62924 | -0.4734 | 0.52 | 0.48 | 1 | HC_vs_MS.Trt_F |
| EIF3E | 0.62962 | -0.3978 | 0.538 | 0.46 | 1 | HC_vs_MS.Trt_F |
| HIP1R | 0.62973 | 0.89163 | 0.11 | 0.1 | 1 | HC_vs_MS.Trt_F |
| ZIK1 | 0.63009 | -1.2156 | 0.327 | 0.34 | 1 | HC_vs_MS.Trt_F |
| ZNF880 | 0.63019 | -0.6925 | 0.512 | 0.5 | 1 | HC_vs_MS.Trt_F |
| PLA2G6 | 0.63041 | 0.94369 | 0.428 | 0.43 | 1 | HC_vs_MS.Trt_F |
| MAPKAPK2 | 0.63093 | 0.81875 | 0.104 | 0.09 | 1 | HC_vs_MS.Trt_F |
| COX7A2 | 0.63113 | -0.4421 | 0.728 | 0.71 | 1 | HC_vs_MS.Trt_F |
| RAB35 | 0.63126 | 0.30352 | 0.168 | 0.15 | 1 | HC_vs_MS.Trt_F |
| TMEM115 | 0.63173 | -0.6683 | 0.107 | 0.12 | 1 | HC_vs_MS.Trt_F |
| CNOT11 | 0.63213 | 0.30429 | 0.133 | 0.12 | 1 | HC_vs_MS.Trt_F |
| FAM114A2 | 0.63245 | -0.4454 | 0.098 | 0.11 | 1 | HC_vs_MS.Trt_F |
| C6orf89 | 0.63312 | 0.3244 | 0.431 | 0.46 | 1 | HC_vs_MS.Trt_F |
| IFNLR1 | 0.63319 | 0.72661 | 0.13 | 0.14 | 1 | HC_vs_MS.Trt_F |
| TMEM136 | 0.63332 | -0.9385 | 0.512 | 0.54 | 1 | HC_vs_MS.Trt_F |
| SBDS | 0.63375 | -0.2842 | 0.309 | 0.28 | 1 | HC_vs_MS.Trt_F |
| EPHA4 | 0.63384 | -0.78 | 0.237 | 0.21 | 1 | HC_vs_MS.Trt_F |
| PIKFYVE | 0.63415 | 0.48702 | 0.41 | 0.4 | 1 | HC_vs_MS.Trt_F |
| GPN3 | 0.63441 | 0.99602 | 0.379 | 0.36 | 1 | HC_vs_MS.Trt_F |
| BSPRY | 0.63448 | 0.59969 | 0.11 | 0.1 | 1 | HC_vs_MS.Trt_F |
| ZC3H7A | 0.6345 | -0.6947 | 0.416 | 0.37 | 1 | HC_vs_MS.Trt_F |
| SMARCAL1 | 0.63516 | -0.3393 | 0.743 | 0.69 | 1 | HC_vs_MS.Trt_F |
| SAV1 | 0.63516 | -0.5561 | 0.329 | 0.34 | 1 | HC_vs_MS.Trt_F |
| PRPF6 | 0.63565 | -0.3217 | 0.231 | 0.21 | 1 | HC_vs_MS.Trt_F |
| PANK1 | 0.63584 | 0.43166 | 0.497 | 0.45 | 1 | HC_vs_MS.Trt_F |
| PNP | 0.63596 | 0.25771 | 0.416 | 0.39 | 1 | HC_vs_MS.Trt_F |
| DKC1 | 0.6362 | -0.294 | 0.373 | 0.39 | 1 | HC_vs_MS.Trt_F |
| RABL2B | 0.63659 | 0.43836 | 0.916 | 0.9 | 1 | HC_vs_MS.Trt_F |
| FAM227B | 0.63694 | -0.5517 | 0.384 | 0.39 | 1 | HC_vs_MS.Trt_F |
| MEAF6 | 0.63709 | -0.3337 | 0.277 | 0.25 | 1 | HC_vs_MS.Trt_F |
| TIGAR | 0.63713 | -0.5051 | 0.905 | 0.9 | 1 | HC_vs_MS.Trt_F |
| DDIAS | 0.63718 | 0.35118 | 0.46 | 0.47 | 1 | HC_vs_MS.Trt_F |
| COG2 | 0.63746 | -0.3736 | 0.37 | 0.35 | 1 | HC_vs_MS.Trt_F |
| ZNF761 | 0.63768 | -1.1535 | 0.737 | 0.73 | 1 | HC_vs_MS.Trt_F |
| RPRD1A | 0.6379 | -0.4358 | 0.179 | 0.16 | 1 | HC_vs_MS.Trt_F |
| PPP5D1 | 0.63802 | -0.3714 | 0.783 | 0.75 | 1 | HC_vs_MS.Trt_F |
| PDE6D | 0.63838 | -0.8824 | 0.147 | 0.16 | 1 | HC_vs_MS.Trt_F |
| RBX1 | 0.6385 | -0.9117 | 0.309 | 0.26 | 1 | HC_vs_MS.Trt_F |

| | | | | | | |
|---|---|---|---|---|---|---|
| COIL | 0.63871 | -0.6258 | 0.142 | 0.15 | 1 | HC_vs_MS.Trt_F |
| EID2B | 0.63877 | 0.46724 | 0.523 | 0.51 | 1 | HC_vs_MS.Trt_F |
| NAA38 | 0.63966 | -0.4374 | 0.147 | 0.13 | 1 | HC_vs_MS.Trt_F |
| PLAT | 0.64033 | -0.468 | 0.104 | 0.12 | 1 | HC_vs_MS.Trt_F |
| PLXDC1 | 0.64045 | -0.5905 | 0.731 | 0.69 | 1 | HC_vs_MS.Trt_F |
| ZMYM4 | 0.64056 | 0.77433 | 0.165 | 0.15 | 1 | HC_vs_MS.Trt_F |
| SCO1 | 0.64086 | -0.2733 | 0.85 | 0.79 | 1 | HC_vs_MS.Trt_F |
| ZNF808 | 0.64093 | -0.9814 | 0.526 | 0.54 | 1 | HC_vs_MS.Trt_F |
| PBX4 | 0.64105 | -0.4793 | 0.197 | 0.18 | 1 | HC_vs_MS.Trt_F |
| CTIF | 0.64118 | -0.8483 | 0.234 | 0.25 | 1 | HC_vs_MS.Trt_F |
| STXBP5 | 0.64174 | -0.4439 | 0.197 | 0.18 | 1 | HC_vs_MS.Trt_F |
| FLII | 0.64197 | -0.451 | 0.249 | 0.22 | 1 | HC_vs_MS.Trt_F |
| MTRF1 | 0.64231 | -0.2867 | 0.598 | 0.57 | 1 | HC_vs_MS.Trt_F |
| IL17RC | 0.64239 | 1.15471 | 0.439 | 0.43 | 1 | HC_vs_MS.Trt_F |
| TMPRSS11B | 0.64323 | -0.5077 | 0.373 | 0.37 | 1 | HC_vs_MS.Trt_F |
| CRB1 | 0.64351 | -0.4477 | 0.176 | 0.16 | 1 | HC_vs_MS.Trt_F |
| G6PD | 0.64372 | -0.4645 | 0.107 | 0.12 | 1 | HC_vs_MS.Trt_F |
| ADHFE1 | 0.64378 | -0.4328 | 0.179 | 0.16 | 1 | HC_vs_MS.Trt_F |
| CNST | 0.64388 | -0.4542 | 0.303 | 0.27 | 1 | HC_vs_MS.Trt_F |
| TMEM160 | 0.64407 | -0.3411 | 0.173 | 0.16 | 1 | HC_vs_MS.Trt_F |
| PMS2 | 0.64413 | 0.46911 | 0.685 | 0.62 | 1 | HC_vs_MS.Trt_F |
| MFSD13A | 0.64419 | -0.3846 | 0.777 | 0.75 | 1 | HC_vs_MS.Trt_F |
| HEATR4 | 0.64434 | -0.5057 | 0.413 | 0.4 | 1 | HC_vs_MS.Trt_F |
| MEGF6 | 0.64466 | -0.3412 | 0.162 | 0.14 | 1 | HC_vs_MS.Trt_F |
| NOD1 | 0.64533 | -0.2533 | 0.13 | 0.12 | 1 | HC_vs_MS.Trt_F |
| PRMT1 | 0.64576 | 0.26816 | 0.832 | 0.81 | 1 | HC_vs_MS.Trt_F |
| SPG7 | 0.64607 | -0.3906 | 0.986 | 0.96 | 1 | HC_vs_MS.Trt_F |
| CTD-2370N5 | 0.64625 | -0.2816 | 0.156 | 0.17 | 1 | HC_vs_MS.Trt_F |
| SULT1A1 | 0.6463 | 0.5273 | 0.618 | 0.57 | 1 | HC_vs_MS.Trt_F |
| ZNF749 | 0.64729 | -0.5864 | 0.457 | 0.46 | 1 | HC_vs_MS.Trt_F |
| SIRT1 | 0.64731 | -0.7997 | 0.153 | 0.14 | 1 | HC_vs_MS.Trt_F |
| MYO5C | 0.64742 | -0.8394 | 0.428 | 0.39 | 1 | HC_vs_MS.Trt_F |
| SH3GLB2 | 0.64785 | -0.3771 | 0.145 | 0.16 | 1 | HC_vs_MS.Trt_F |
| NPTN | 0.64786 | -0.4729 | 0.338 | 0.3 | 1 | HC_vs_MS.Trt_F |
| AKAP10 | 0.64806 | -0.5759 | 0.538 | 0.47 | 1 | HC_vs_MS.Trt_F |
| YPEL2 | 0.64906 | -0.3576 | 0.214 | 0.19 | 1 | HC_vs_MS.Trt_F |
| IDUA | 0.6498 | 0.26237 | 0.107 | 0.1 | 1 | HC_vs_MS.Trt_F |
| POLA2 | 0.65009 | -0.6296 | 0.662 | 0.66 | 1 | HC_vs_MS.Trt_F |
| ZNF763 | 0.65012 | -1.2508 | 0.202 | 0.18 | 1 | HC_vs_MS.Trt_F |
| MIOS | 0.65106 | 0.60599 | 0.306 | 0.29 | 1 | HC_vs_MS.Trt_F |

| | | | | | | |
|---|---|---|---|---|---|---|
| KMT5A | 0.65158 | -0.4388 | 0.28 | 0.25 | 1 | HC_vs_MS.Trt_F |
| OSBPL2 | 0.65206 | 0.31369 | 0.523 | 0.49 | 1 | HC_vs_MS.Trt_F |
| PLGRKT | 0.65207 | -0.8001 | 0.101 | 0.11 | 1 | HC_vs_MS.Trt_F |
| ZNF626 | 0.65219 | 0.40513 | 0.783 | 0.76 | 1 | HC_vs_MS.Trt_F |
| AP1S2 | 0.65242 | -0.4742 | 0.228 | 0.2 | 1 | HC_vs_MS.Trt_F |
| MED21 | 0.65263 | -0.4402 | 0.156 | 0.14 | 1 | HC_vs_MS.Trt_F |
| ZNF24 | 0.65276 | -0.4398 | 0.199 | 0.18 | 1 | HC_vs_MS.Trt_F |
| YIPF1 | 0.65294 | -0.3625 | 0.124 | 0.11 | 1 | HC_vs_MS.Trt_F |
| PTDSS1 | 0.65352 | -0.2747 | 0.419 | 0.38 | 1 | HC_vs_MS.Trt_F |
| TRIM11 | 0.65385 | -0.4747 | 0.604 | 0.55 | 1 | HC_vs_MS.Trt_F |
| COX16 | 0.6539 | -0.5571 | 0.552 | 0.56 | 1 | HC_vs_MS.Trt_F |
| PGBD2 | 0.65405 | -0.9261 | 0.373 | 0.33 | 1 | HC_vs_MS.Trt_F |
| PYCRL | 0.65411 | 0.81699 | 0.465 | 0.44 | 1 | HC_vs_MS.Trt_F |
| GARS | 0.65442 | -0.8379 | 0.214 | 0.19 | 1 | HC_vs_MS.Trt_F |
| SEC24D | 0.65508 | -0.7592 | 0.526 | 0.52 | 1 | HC_vs_MS.Trt_F |
| N6AMT1 | 0.65509 | -0.629 | 0.159 | 0.17 | 1 | HC_vs_MS.Trt_F |
| LAMA5 | 0.65526 | -1.0728 | 0.306 | 0.28 | 1 | HC_vs_MS.Trt_F |
| PARN | 0.65527 | -0.6326 | 0.746 | 0.76 | 1 | HC_vs_MS.Trt_F |
| CCZ1 | 0.65572 | 0.47092 | 0.72 | 0.67 | 1 | HC_vs_MS.Trt_F |
| RAB2A | 0.65687 | -0.2526 | 0.251 | 0.23 | 1 | HC_vs_MS.Trt_F |
| DCST1 | 0.65694 | -0.6101 | 0.145 | 0.16 | 1 | HC_vs_MS.Trt_F |
| RNF126 | 0.65712 | -0.4399 | 0.22 | 0.23 | 1 | HC_vs_MS.Trt_F |
| TGM2 | 0.65761 | -0.3365 | 0.613 | 0.61 | 1 | HC_vs_MS.Trt_F |
| HILPDA | 0.65826 | 0.40868 | 0.592 | 0.57 | 1 | HC_vs_MS.Trt_F |
| PIGK | 0.65883 | -0.6735 | 0.136 | 0.14 | 1 | HC_vs_MS.Trt_F |
| PDIA5 | 0.65926 | 0.43961 | 0.434 | 0.41 | 1 | HC_vs_MS.Trt_F |
| ZFAND1 | 0.66022 | -0.7908 | 0.156 | 0.14 | 1 | HC_vs_MS.Trt_F |
| USE1 | 0.6612 | -0.2944 | 0.887 | 0.92 | 1 | HC_vs_MS.Trt_F |
| GCLC | 0.66182 | 0.79 | 0.601 | 0.57 | 1 | HC_vs_MS.Trt_F |
| IFI44L | 0.66245 | -1.0445 | 0.228 | 0.21 | 1 | HC_vs_MS.Trt_F |
| FAM179A | 0.663 | -0.3258 | 0.838 | 0.85 | 1 | HC_vs_MS.Trt_F |
| DNASE2 | 0.66433 | -0.4839 | 0.48 | 0.45 | 1 | HC_vs_MS.Trt_F |
| ENTPD7 | 0.66482 | 1.43552 | 0.384 | 0.35 | 1 | HC_vs_MS.Trt_F |
| FNIP2 | 0.66503 | 0.37638 | 0.35 | 0.32 | 1 | HC_vs_MS.Trt_F |
| COPG1 | 0.66572 | -0.5987 | 0.173 | 0.18 | 1 | HC_vs_MS.Trt_F |
| CCT8 | 0.66584 | -0.3741 | 0.283 | 0.26 | 1 | HC_vs_MS.Trt_F |
| TMEM184B | 0.66603 | 0.40771 | 0.101 | 0.09 | 1 | HC_vs_MS.Trt_F |
| TOPBP1 | 0.66604 | 1.16695 | 0.133 | 0.12 | 1 | HC_vs_MS.Trt_F |
| KATNB1 | 0.66637 | -0.3063 | 0.098 | 0.11 | 1 | HC_vs_MS.Trt_F |
| CYB561D2 | 0.66638 | -0.3345 | 0.136 | 0.12 | 1 | HC_vs_MS.Trt_F |

| | | | | | | |
|---|---|---|---|---|---|---|
| GABPB2 | 0.66793 | -0.2656 | 0.977 | 0.98 | 1 | HC_vs_MS.Trt_F |
| FXR1 | 0.66928 | -0.4019 | 0.225 | 0.21 | 1 | HC_vs_MS.Trt_F |
| ITPKB | 0.66945 | -0.2616 | 0.63 | 0.6 | 1 | HC_vs_MS.Trt_F |
| TCF20 | 0.66975 | 0.47045 | 0.156 | 0.14 | 1 | HC_vs_MS.Trt_F |
| ARPC4-TTLL3 | 0.66986 | -1.0463 | 0.118 | 0.13 | 1 | HC_vs_MS.Trt_F |
| PCNP | 0.66996 | -0.5413 | 0.269 | 0.24 | 1 | HC_vs_MS.Trt_F |
| CUL4B | 0.67001 | -0.7151 | 0.145 | 0.13 | 1 | HC_vs_MS.Trt_F |
| C1QBP | 0.67012 | -0.35 | 0.301 | 0.27 | 1 | HC_vs_MS.Trt_F |
| B3GALNT2 | 0.6705 | 0.60253 | 0.668 | 0.63 | 1 | HC_vs_MS.Trt_F |
| MFN1 | 0.67197 | -0.6092 | 0.506 | 0.46 | 1 | HC_vs_MS.Trt_F |
| NCOA6 | 0.67202 | 0.26809 | 0.728 | 0.71 | 1 | HC_vs_MS.Trt_F |
| CA5B | 0.67278 | 0.277 | 0.879 | 0.84 | 1 | HC_vs_MS.Trt_F |
| COA7 | 0.6729 | 0.70183 | 0.893 | 0.91 | 1 | HC_vs_MS.Trt_F |
| KANK1 | 0.67305 | -0.6136 | 0.78 | 0.73 | 1 | HC_vs_MS.Trt_F |
| PELP1 | 0.67318 | 1.03204 | 0.277 | 0.26 | 1 | HC_vs_MS.Trt_F |
| TMEM9B | 0.67327 | -0.586 | 0.228 | 0.21 | 1 | HC_vs_MS.Trt_F |
| FAM184B | 0.67344 | 0.35085 | 0.142 | 0.16 | 1 | HC_vs_MS.Trt_F |
| PPP2R5A | 0.67354 | -1.0627 | 0.237 | 0.21 | 1 | HC_vs_MS.Trt_F |
| C3orf62 | 0.67391 | 0.48917 | 0.801 | 0.77 | 1 | HC_vs_MS.Trt_F |
| NFE2L1 | 0.67398 | 0.67645 | 0.101 | 0.11 | 1 | HC_vs_MS.Trt_F |
| EIF2B2 | 0.67419 | -0.466 | 0.616 | 0.54 | 1 | HC_vs_MS.Trt_F |
| TCTN3 | 0.67422 | -0.2771 | 0.133 | 0.12 | 1 | HC_vs_MS.Trt_F |
| C9orf72 | 0.67484 | -0.735 | 0.162 | 0.17 | 1 | HC_vs_MS.Trt_F |
| ATP5I | 0.67556 | -0.5346 | 0.283 | 0.26 | 1 | HC_vs_MS.Trt_F |
| CD180 | 0.67561 | -0.74 | 0.315 | 0.3 | 1 | HC_vs_MS.Trt_F |
| RDH11 | 0.67567 | -0.5188 | 0.558 | 0.5 | 1 | HC_vs_MS.Trt_F |
| C18orf8 | 0.67592 | 0.32193 | 0.353 | 0.34 | 1 | HC_vs_MS.Trt_F |
| PER2 | 0.67624 | -0.8882 | 0.54 | 0.51 | 1 | HC_vs_MS.Trt_F |
| PGGT1B | 0.67646 | -0.5487 | 0.202 | 0.18 | 1 | HC_vs_MS.Trt_F |
| EDEM3 | 0.67649 | -0.6213 | 0.309 | 0.28 | 1 | HC_vs_MS.Trt_F |
| MRI1 | 0.677 | -0.2826 | 0.749 | 0.72 | 1 | HC_vs_MS.Trt_F |
| ADIPOR1 | 0.67742 | -0.4316 | 0.289 | 0.27 | 1 | HC_vs_MS.Trt_F |
| STAU2 | 0.67803 | -0.2519 | 0.656 | 0.67 | 1 | HC_vs_MS.Trt_F |
| PPIE | 0.6781 | -0.4069 | 0.789 | 0.78 | 1 | HC_vs_MS.Trt_F |
| TSR2 | 0.6782 | -0.6458 | 0.46 | 0.45 | 1 | HC_vs_MS.Trt_F |
| SLC35E4 | 0.67823 | 0.94026 | 0.361 | 0.35 | 1 | HC_vs_MS.Trt_F |
| STRIP1 | 0.67868 | -0.5033 | 0.179 | 0.18 | 1 | HC_vs_MS.Trt_F |
| AP3D1 | 0.67922 | 0.32874 | 0.168 | 0.16 | 1 | HC_vs_MS.Trt_F |
| FAM213B | 0.67957 | -0.3706 | 0.116 | 0.12 | 1 | HC_vs_MS.Trt_F |
| MED22 | 0.68061 | 0.44091 | 0.613 | 0.63 | 1 | HC_vs_MS.Trt_F |

| | | | | | | |
|---|---|---|---|---|---|---|
| DYNC1LI1 | 0.68094 | -0.5304 | 0.46 | 0.48 | 1 | HC_vs_MS.Trt_F |
| CAMK4 | 0.68113 | 0.32564 | 0.578 | 0.59 | 1 | HC_vs_MS.Trt_F |
| PANK2 | 0.68115 | -0.5656 | 0.575 | 0.52 | 1 | HC_vs_MS.Trt_F |
| NCBP2 | 0.68151 | -0.3361 | 0.286 | 0.26 | 1 | HC_vs_MS.Trt_F |
| KLHDC1 | 0.6816 | 0.63608 | 0.332 | 0.31 | 1 | HC_vs_MS.Trt_F |
| SNX29 | 0.68161 | 0.60027 | 0.806 | 0.78 | 1 | HC_vs_MS.Trt_F |
| TRPM3 | 0.68172 | 0.25801 | 0.503 | 0.47 | 1 | HC_vs_MS.Trt_F |
| PCDHB16 | 0.68194 | 0.6498 | 0.358 | 0.35 | 1 | HC_vs_MS.Trt_F |
| HACE1 | 0.68198 | -0.299 | 0.688 | 0.64 | 1 | HC_vs_MS.Trt_F |
| ACBD5 | 0.68214 | 0.36303 | 0.514 | 0.5 | 1 | HC_vs_MS.Trt_F |
| ADSSL1 | 0.68297 | -0.2737 | 0.347 | 0.32 | 1 | HC_vs_MS.Trt_F |
| PLEKHA7 | 0.68303 | -0.5452 | 0.266 | 0.25 | 1 | HC_vs_MS.Trt_F |
| FUZ | 0.68329 | 0.6857 | 0.286 | 0.28 | 1 | HC_vs_MS.Trt_F |
| SCP2 | 0.68346 | -0.4825 | 0.306 | 0.28 | 1 | HC_vs_MS.Trt_F |
| ACSM3 | 0.68382 | 0.41568 | 0.679 | 0.61 | 1 | HC_vs_MS.Trt_F |
| STX18 | 0.68422 | -0.6062 | 0.159 | 0.14 | 1 | HC_vs_MS.Trt_F |
| TNFSF14 | 0.68474 | 0.27643 | 0.694 | 0.69 | 1 | HC_vs_MS.Trt_F |
| GOLGA1 | 0.68481 | -1.0314 | 0.127 | 0.13 | 1 | HC_vs_MS.Trt_F |
| GMCL1 | 0.68518 | -1.0767 | 0.384 | 0.36 | 1 | HC_vs_MS.Trt_F |
| ENOX2 | 0.68555 | -0.4038 | 0.145 | 0.13 | 1 | HC_vs_MS.Trt_F |
| TCEAL3 | 0.6858 | -0.3846 | 0.15 | 0.13 | 1 | HC_vs_MS.Trt_F |
| RPP38 | 0.68647 | -0.2828 | 0.465 | 0.43 | 1 | HC_vs_MS.Trt_F |
| NF2 | 0.68688 | -0.7368 | 0.821 | 0.77 | 1 | HC_vs_MS.Trt_F |
| ZNF527 | 0.68724 | 0.35659 | 0.934 | 0.9 | 1 | HC_vs_MS.Trt_F |
| NRIP2 | 0.68734 | -0.9837 | 0.335 | 0.33 | 1 | HC_vs_MS.Trt_F |
| RELL2 | 0.68767 | -0.2895 | 0.104 | 0.11 | 1 | HC_vs_MS.Trt_F |
| TMEM14A | 0.68767 | -0.4384 | 0.101 | 0.11 | 1 | HC_vs_MS.Trt_F |
| ZER1 | 0.68787 | -0.3784 | 0.145 | 0.15 | 1 | HC_vs_MS.Trt_F |
| WDR73 | 0.68787 | -0.3698 | 0.974 | 0.97 | 1 | HC_vs_MS.Trt_F |
| XKR8 | 0.68788 | -0.6299 | 0.185 | 0.19 | 1 | HC_vs_MS.Trt_F |
| HDHD2 | 0.6879 | -0.4105 | 0.147 | 0.16 | 1 | HC_vs_MS.Trt_F |
| EXOSC9 | 0.68795 | -0.6424 | 0.254 | 0.23 | 1 | HC_vs_MS.Trt_F |
| PQLC2 | 0.68838 | -0.6063 | 0.454 | 0.4 | 1 | HC_vs_MS.Trt_F |
| TIMM10 | 0.68859 | 0.26251 | 0.118 | 0.11 | 1 | HC_vs_MS.Trt_F |
| PSMD5 | 0.68876 | -0.2872 | 0.434 | 0.4 | 1 | HC_vs_MS.Trt_F |
| GRHPR | 0.68899 | -0.6782 | 0.549 | 0.51 | 1 | HC_vs_MS.Trt_F |
| SIGLEC6 | 0.68903 | 0.66873 | 0.535 | 0.5 | 1 | HC_vs_MS.Trt_F |
| SCARB2 | 0.68949 | -0.3643 | 0.341 | 0.32 | 1 | HC_vs_MS.Trt_F |
| F2RL1 | 0.68974 | -1.5436 | 0.214 | 0.22 | 1 | HC_vs_MS.Trt_F |
| FAM135A | 0.68998 | -0.6227 | 0.471 | 0.47 | 1 | HC_vs_MS.Trt_F |

| | | | | | | |
|---|---|---|---|---|---|---|
| IRS1 | 0.69115 | 0.33889 | 0.107 | 0.12 | 1 | HC_vs_MS.Trt_F |
| CYLD | 0.69144 | -0.2826 | 0.714 | 0.73 | 1 | HC_vs_MS.Trt_F |
| CD38 | 0.69158 | 0.62553 | 0.387 | 0.36 | 1 | HC_vs_MS.Trt_F |
| LINC00959 | 0.69161 | -0.7538 | 0.147 | 0.13 | 1 | HC_vs_MS.Trt_F |
| NF1 | 0.69176 | 0.38512 | 0.76 | 0.72 | 1 | HC_vs_MS.Trt_F |
| PON2 | 0.69181 | -0.7389 | 0.442 | 0.42 | 1 | HC_vs_MS.Trt_F |
| KLHL12 | 0.6919 | -0.2926 | 0.121 | 0.11 | 1 | HC_vs_MS.Trt_F |
| RCAN1 | 0.6919 | 0.43353 | 0.39 | 0.38 | 1 | HC_vs_MS.Trt_F |
| STRAP | 0.69249 | -0.3304 | 0.298 | 0.31 | 1 | HC_vs_MS.Trt_F |
| DYRK2 | 0.69312 | -0.4632 | 0.173 | 0.16 | 1 | HC_vs_MS.Trt_F |
| CROCC | 0.6934 | -0.3528 | 0.552 | 0.55 | 1 | HC_vs_MS.Trt_F |
| UBE2D2 | 0.69378 | -0.5045 | 0.41 | 0.39 | 1 | HC_vs_MS.Trt_F |
| PHTF1 | 0.69436 | -0.5669 | 0.569 | 0.54 | 1 | HC_vs_MS.Trt_F |
| TMCO6 | 0.69438 | -0.474 | 0.526 | 0.5 | 1 | HC_vs_MS.Trt_F |
| APC | 0.69455 | 0.46411 | 0.538 | 0.52 | 1 | HC_vs_MS.Trt_F |
| ZNF682 | 0.69461 | 0.86088 | 0.405 | 0.4 | 1 | HC_vs_MS.Trt_F |
| ZNF441 | 0.69462 | -1.3644 | 0.197 | 0.21 | 1 | HC_vs_MS.Trt_F |
| LARS2 | 0.69469 | 0.38719 | 0.546 | 0.54 | 1 | HC_vs_MS.Trt_F |
| USP6 | 0.69493 | 0.43814 | 0.139 | 0.15 | 1 | HC_vs_MS.Trt_F |
| TRIQK | 0.695 | -0.4123 | 0.558 | 0.55 | 1 | HC_vs_MS.Trt_F |
| SELM | 0.69514 | -0.3256 | 0.277 | 0.28 | 1 | HC_vs_MS.Trt_F |
| RUNDC1 | 0.69525 | -0.3337 | 0.775 | 0.75 | 1 | HC_vs_MS.Trt_F |
| BMPR1B | 0.69543 | 0.39911 | 0.176 | 0.17 | 1 | HC_vs_MS.Trt_F |
| PTGER2 | 0.6956 | -0.6358 | 0.217 | 0.22 | 1 | HC_vs_MS.Trt_F |
| PTRH1 | 0.69575 | -0.3882 | 0.777 | 0.71 | 1 | HC_vs_MS.Trt_F |
| GALNT10 | 0.69582 | 0.60547 | 0.578 | 0.55 | 1 | HC_vs_MS.Trt_F |
| VLDLR | 0.69582 | -1.4845 | 0.488 | 0.45 | 1 | HC_vs_MS.Trt_F |
| NCAPD2 | 0.69612 | -0.7319 | 0.454 | 0.45 | 1 | HC_vs_MS.Trt_F |
| TMEM5 | 0.69618 | -0.3335 | 0.523 | 0.5 | 1 | HC_vs_MS.Trt_F |
| HINT3 | 0.69632 | -0.3497 | 0.303 | 0.31 | 1 | HC_vs_MS.Trt_F |
| NUDC | 0.69664 | -0.2538 | 0.668 | 0.63 | 1 | HC_vs_MS.Trt_F |
| RP11-226E2 | 0.69681 | 0.35845 | 0.639 | 0.6 | 1 | HC_vs_MS.Trt_F |
| ZNF672 | 0.69696 | -0.6571 | 0.246 | 0.23 | 1 | HC_vs_MS.Trt_F |
| ZNF75A | 0.697 | -0.6833 | 0.783 | 0.77 | 1 | HC_vs_MS.Trt_F |
| GIMAP6 | 0.69716 | -0.473 | 0.202 | 0.21 | 1 | HC_vs_MS.Trt_F |
| TESPA1 | 0.69734 | -0.2955 | 0.451 | 0.41 | 1 | HC_vs_MS.Trt_F |
| FSD1 | 0.69745 | 0.25891 | 0.457 | 0.43 | 1 | HC_vs_MS.Trt_F |
| KAT6B | 0.69806 | 0.98764 | 0.136 | 0.12 | 1 | HC_vs_MS.Trt_F |
| RPL17-C18o | 0.69844 | 0.64278 | 0.09 | 0.1 | 1 | HC_vs_MS.Trt_F |
| CBX5 | 0.69866 | -0.305 | 0.934 | 0.93 | 1 | HC_vs_MS.Trt_F |

| | | | | | | |
|---|---|---|---|---|---|---|
| ZNF175 | 0.70057 | 0.48566 | 0.211 | 0.22 | 1 | HC_vs_MS.Trt_F |
| GBP3 | 0.70147 | -0.633 | 0.225 | 0.2 | 1 | HC_vs_MS.Trt_F |
| SLC35A5 | 0.7022 | -0.5573 | 0.468 | 0.49 | 1 | HC_vs_MS.Trt_F |
| GOPC | 0.70236 | -0.6908 | 0.145 | 0.13 | 1 | HC_vs_MS.Trt_F |
| CTSB | 0.70247 | -0.5815 | 0.974 | 0.92 | 1 | HC_vs_MS.Trt_F |
| ZKSCAN5 | 0.70344 | -0.5382 | 0.691 | 0.63 | 1 | HC_vs_MS.Trt_F |
| CYB5D1 | 0.70349 | 0.47583 | 0.205 | 0.19 | 1 | HC_vs_MS.Trt_F |
| EPHX1 | 0.70379 | 0.58396 | 0.107 | 0.1 | 1 | HC_vs_MS.Trt_F |
| TMEM11 | 0.70386 | -0.5781 | 0.124 | 0.13 | 1 | HC_vs_MS.Trt_F |
| RPS21 | 0.70407 | -0.2931 | 0.442 | 0.44 | 1 | HC_vs_MS.Trt_F |
| MTRR | 0.70449 | -0.4131 | 0.208 | 0.19 | 1 | HC_vs_MS.Trt_F |
| ANKRD26 | 0.70479 | -0.3643 | 0.159 | 0.14 | 1 | HC_vs_MS.Trt_F |
| AZIN2 | 0.70521 | 0.83447 | 0.379 | 0.36 | 1 | HC_vs_MS.Trt_F |
| HENMT1 | 0.70557 | -0.3486 | 0.327 | 0.32 | 1 | HC_vs_MS.Trt_F |
| POMK | 0.7056 | -0.3514 | 0.618 | 0.57 | 1 | HC_vs_MS.Trt_F |
| UFSP2 | 0.70721 | -0.3676 | 0.116 | 0.12 | 1 | HC_vs_MS.Trt_F |
| MBD4 | 0.70729 | -0.8766 | 0.263 | 0.26 | 1 | HC_vs_MS.Trt_F |
| IL20RB | 0.70747 | -0.5927 | 0.277 | 0.3 | 1 | HC_vs_MS.Trt_F |
| POT1 | 0.70776 | -1.1134 | 0.11 | 0.12 | 1 | HC_vs_MS.Trt_F |
| FBXO3 | 0.7078 | -0.4244 | 0.289 | 0.27 | 1 | HC_vs_MS.Trt_F |
| EBNA1BP2 | 0.70808 | -0.5334 | 0.552 | 0.54 | 1 | HC_vs_MS.Trt_F |
| JADE2 | 0.70817 | -0.4348 | 0.202 | 0.19 | 1 | HC_vs_MS.Trt_F |
| PITPNM2 | 0.7082 | 0.54101 | 0.445 | 0.41 | 1 | HC_vs_MS.Trt_F |
| BRCC3 | 0.70881 | -0.4559 | 0.272 | 0.25 | 1 | HC_vs_MS.Trt_F |
| BUD13 | 0.70919 | -0.6995 | 0.104 | 0.11 | 1 | HC_vs_MS.Trt_F |
| DSTYK | 0.70938 | -0.5421 | 0.838 | 0.8 | 1 | HC_vs_MS.Trt_F |
| FBXW7 | 0.7096 | -0.5113 | 0.182 | 0.17 | 1 | HC_vs_MS.Trt_F |
| WFDC8 | 0.70971 | -0.2707 | 0.329 | 0.3 | 1 | HC_vs_MS.Trt_F |
| FUBP1 | 0.70973 | -0.674 | 0.419 | 0.41 | 1 | HC_vs_MS.Trt_F |
| INO80 | 0.71061 | -0.4336 | 0.147 | 0.13 | 1 | HC_vs_MS.Trt_F |
| PLPP1 | 0.71072 | -0.3875 | 0.11 | 0.1 | 1 | HC_vs_MS.Trt_F |
| PSMD12 | 0.71075 | -0.6645 | 0.769 | 0.72 | 1 | HC_vs_MS.Trt_F |
| TM7SF3 | 0.71133 | -0.399 | 0.951 | 0.96 | 1 | HC_vs_MS.Trt_F |
| TRAPPC9 | 0.71145 | -1.0618 | 0.139 | 0.14 | 1 | HC_vs_MS.Trt_F |
| ZNF17 | 0.7116 | -1.123 | 0.312 | 0.32 | 1 | HC_vs_MS.Trt_F |
| BRDT | 0.71181 | -0.2505 | 0.598 | 0.55 | 1 | HC_vs_MS.Trt_F |
| LRRC40 | 0.71182 | -0.4424 | 0.156 | 0.14 | 1 | HC_vs_MS.Trt_F |
| C14orf1 | 0.71207 | -0.3641 | 0.306 | 0.29 | 1 | HC_vs_MS.Trt_F |
| TOB2 | 0.71217 | -0.2575 | 0.159 | 0.17 | 1 | HC_vs_MS.Trt_F |
| SLC31A1 | 0.71231 | 0.3709 | 0.78 | 0.78 | 1 | HC_vs_MS.Trt_F |

| | | | | | | |
|---|---|---|---|---|---|---|
| HIP1 | 0.71243 | -0.3889 | 0.665 | 0.66 | 1 | HC_vs_MS.Trt_F |
| TUBD1 | 0.71261 | 0.28659 | 0.775 | 0.76 | 1 | HC_vs_MS.Trt_F |
| DSE | 0.71304 | 0.51608 | 0.104 | 0.12 | 1 | HC_vs_MS.Trt_F |
| GTF2A2 | 0.71344 | -0.4861 | 0.633 | 0.6 | 1 | HC_vs_MS.Trt_F |
| ZNF418 | 0.71383 | 0.39064 | 0.194 | 0.21 | 1 | HC_vs_MS.Trt_F |
| ZNF350 | 0.714 | -1.1541 | 0.595 | 0.57 | 1 | HC_vs_MS.Trt_F |
| EIF2D | 0.71421 | -0.3412 | 0.723 | 0.64 | 1 | HC_vs_MS.Trt_F |
| SDK2 | 0.71422 | -0.3113 | 0.555 | 0.54 | 1 | HC_vs_MS.Trt_F |
| TMPO | 0.71442 | -0.4742 | 0.572 | 0.54 | 1 | HC_vs_MS.Trt_F |
| NBPF11 | 0.71454 | -0.3038 | 0.286 | 0.29 | 1 | HC_vs_MS.Trt_F |
| MRPL40 | 0.71516 | -0.5291 | 0.471 | 0.45 | 1 | HC_vs_MS.Trt_F |
| GORASP1 | 0.71528 | 0.442 | 0.092 | 0.1 | 1 | HC_vs_MS.Trt_F |
| ZNF211 | 0.7158 | -0.6462 | 0.202 | 0.21 | 1 | HC_vs_MS.Trt_F |
| ZBTB44 | 0.71612 | -0.4056 | 0.303 | 0.28 | 1 | HC_vs_MS.Trt_F |
| BTBD19 | 0.71614 | 0.49389 | 0.795 | 0.77 | 1 | HC_vs_MS.Trt_F |
| CRELD2 | 0.71616 | -0.8109 | 0.645 | 0.6 | 1 | HC_vs_MS.Trt_F |
| MINPP1 | 0.71633 | 1.31273 | 0.104 | 0.1 | 1 | HC_vs_MS.Trt_F |
| INCA1 | 0.71677 | -0.7727 | 0.211 | 0.22 | 1 | HC_vs_MS.Trt_F |
| GAL3ST4 | 0.71681 | -1.19 | 0.451 | 0.45 | 1 | HC_vs_MS.Trt_F |
| BCL2 | 0.71755 | -0.4203 | 0.277 | 0.26 | 1 | HC_vs_MS.Trt_F |
| GTPBP4 | 0.71766 | 0.81706 | 0.514 | 0.49 | 1 | HC_vs_MS.Trt_F |
| HLA-DMB | 0.71795 | 1.48397 | 0.107 | 0.1 | 1 | HC_vs_MS.Trt_F |
| PSME3 | 0.7181 | -0.2833 | 0.694 | 0.72 | 1 | HC_vs_MS.Trt_F |
| TRIP13 | 0.71835 | 0.30143 | 0.546 | 0.53 | 1 | HC_vs_MS.Trt_F |
| RASGRP2 | 0.71864 | 0.31105 | 0.301 | 0.31 | 1 | HC_vs_MS.Trt_F |
| ATL2 | 0.719 | 0.25499 | 0.879 | 0.84 | 1 | HC_vs_MS.Trt_F |
| LTV1 | 0.71945 | -0.373 | 0.139 | 0.14 | 1 | HC_vs_MS.Trt_F |
| LUZP1 | 0.71992 | 0.5339 | 0.561 | 0.55 | 1 | HC_vs_MS.Trt_F |
| PIGO | 0.72088 | 0.27144 | 0.384 | 0.38 | 1 | HC_vs_MS.Trt_F |
| SMARCB1 | 0.72147 | 0.45059 | 0.538 | 0.55 | 1 | HC_vs_MS.Trt_F |
| ADAM33 | 0.72156 | 0.74871 | 0.107 | 0.12 | 1 | HC_vs_MS.Trt_F |
| TAF3 | 0.72171 | -0.3667 | 0.179 | 0.17 | 1 | HC_vs_MS.Trt_F |
| ACTR3 | 0.72179 | -0.3905 | 0.775 | 0.74 | 1 | HC_vs_MS.Trt_F |
| RARS2 | 0.72184 | 0.27766 | 0.598 | 0.59 | 1 | HC_vs_MS.Trt_F |
| OST4 | 0.72187 | -0.4848 | 0.416 | 0.36 | 1 | HC_vs_MS.Trt_F |
| COMMD9 | 0.72202 | -0.9585 | 0.503 | 0.49 | 1 | HC_vs_MS.Trt_F |
| BTN3A3 | 0.72209 | 0.30435 | 0.243 | 0.23 | 1 | HC_vs_MS.Trt_F |
| PHLPP2 | 0.7222 | 0.27306 | 0.818 | 0.84 | 1 | HC_vs_MS.Trt_F |
| FKTN | 0.72239 | 0.60758 | 0.199 | 0.21 | 1 | HC_vs_MS.Trt_F |
| ZNF487 | 0.72261 | 0.55218 | 0.514 | 0.48 | 1 | HC_vs_MS.Trt_F |

| | | | | | | |
|---|---|---|---|---|---|---|
| TMEM14C | 0.72272 | -0.3597 | 0.142 | 0.13 | 1 | HC_vs_MS.Trt_F |
| MITD1 | 0.72285 | 0.33613 | 0.962 | 0.96 | 1 | HC_vs_MS.Trt_F |
| MKRN1 | 0.72307 | -0.5609 | 0.853 | 0.81 | 1 | HC_vs_MS.Trt_F |
| KRCC1 | 0.72317 | -0.4647 | 0.124 | 0.13 | 1 | HC_vs_MS.Trt_F |
| SCYL1 | 0.72338 | -0.4524 | 0.269 | 0.27 | 1 | HC_vs_MS.Trt_F |
| WDR97 | 0.7234 | -0.3321 | 0.546 | 0.5 | 1 | HC_vs_MS.Trt_F |
| AC240274.1 | 0.72367 | 0.59182 | 0.283 | 0.28 | 1 | HC_vs_MS.Trt_F |
| PHLDB3 | 0.72398 | 0.43595 | 0.665 | 0.64 | 1 | HC_vs_MS.Trt_F |
| PLA2G12A | 0.72418 | 0.45257 | 0.679 | 0.64 | 1 | HC_vs_MS.Trt_F |
| GOLGA7 | 0.72438 | -0.2856 | 0.199 | 0.18 | 1 | HC_vs_MS.Trt_F |
| CAMKK1 | 0.7245 | -0.3433 | 0.22 | 0.23 | 1 | HC_vs_MS.Trt_F |
| WBP1L | 0.72473 | -0.2741 | 0.124 | 0.11 | 1 | HC_vs_MS.Trt_F |
| ENOPH1 | 0.72476 | -0.3973 | 0.173 | 0.16 | 1 | HC_vs_MS.Trt_F |
| RNF31 | 0.72534 | 0.36793 | 0.312 | 0.32 | 1 | HC_vs_MS.Trt_F |
| RNF34 | 0.72538 | -0.5353 | 0.272 | 0.27 | 1 | HC_vs_MS.Trt_F |
| FAM91A1 | 0.72605 | 0.29757 | 0.344 | 0.34 | 1 | HC_vs_MS.Trt_F |
| PSMG1 | 0.7265 | -0.2973 | 0.188 | 0.17 | 1 | HC_vs_MS.Trt_F |
| TAF1C | 0.72658 | 0.40035 | 0.118 | 0.11 | 1 | HC_vs_MS.Trt_F |
| PDCD4 | 0.72664 | -0.686 | 0.656 | 0.63 | 1 | HC_vs_MS.Trt_F |
| AVIL | 0.72705 | 0.7057 | 0.408 | 0.39 | 1 | HC_vs_MS.Trt_F |
| SERPINB6 | 0.72722 | -0.5736 | 0.535 | 0.49 | 1 | HC_vs_MS.Trt_F |
| TMEM104 | 0.72725 | 0.52693 | 0.168 | 0.16 | 1 | HC_vs_MS.Trt_F |
| RP11-235E1 | 0.7273 | 0.63332 | 0.28 | 0.27 | 1 | HC_vs_MS.Trt_F |
| TOMM70 | 0.72771 | -0.8631 | 0.188 | 0.18 | 1 | HC_vs_MS.Trt_F |
| FAM234A | 0.72782 | 0.30057 | 0.272 | 0.26 | 1 | HC_vs_MS.Trt_F |
| TCOF1 | 0.72816 | -0.7429 | 0.809 | 0.77 | 1 | HC_vs_MS.Trt_F |
| UPF1 | 0.72848 | 0.41263 | 0.13 | 0.14 | 1 | HC_vs_MS.Trt_F |
| WDFY1 | 0.72876 | -0.4572 | 0.39 | 0.35 | 1 | HC_vs_MS.Trt_F |
| CHP2 | 0.7288 | -0.2932 | 0.471 | 0.41 | 1 | HC_vs_MS.Trt_F |
| FAM133B | 0.7289 | -0.4525 | 0.798 | 0.75 | 1 | HC_vs_MS.Trt_F |
| SMARCC1 | 0.72922 | 0.27704 | 0.462 | 0.44 | 1 | HC_vs_MS.Trt_F |
| COL4A3BP | 0.72939 | -0.2513 | 0.168 | 0.16 | 1 | HC_vs_MS.Trt_F |
| THUMPD3 | 0.72982 | -0.2697 | 0.434 | 0.41 | 1 | HC_vs_MS.Trt_F |
| POLR1E | 0.72989 | -0.566 | 0.142 | 0.13 | 1 | HC_vs_MS.Trt_F |
| MIEN1 | 0.72993 | -0.46 | 0.165 | 0.17 | 1 | HC_vs_MS.Trt_F |
| MGAT2 | 0.72999 | 0.25844 | 0.136 | 0.13 | 1 | HC_vs_MS.Trt_F |
| ATF6B | 0.73051 | -0.3486 | 0.942 | 0.92 | 1 | HC_vs_MS.Trt_F |
| VPS45 | 0.73088 | 0.25631 | 0.289 | 0.28 | 1 | HC_vs_MS.Trt_F |
| VWA5B1 | 0.7311 | -0.2881 | 0.627 | 0.59 | 1 | HC_vs_MS.Trt_F |
| ZNF565 | 0.73135 | -0.2822 | 0.113 | 0.1 | 1 | HC_vs_MS.Trt_F |

| | | | | | | |
|---|---|---|---|---|---|---|
| PHACTR1 | 0.73148 | 0.885 | 0.477 | 0.47 | 1 | HC_vs_MS.Trt_F |
| IDH3A | 0.73212 | -0.4484 | 0.754 | 0.73 | 1 | HC_vs_MS.Trt_F |
| SSU72 | 0.73334 | -0.272 | 0.928 | 0.91 | 1 | HC_vs_MS.Trt_F |
| ACAD11 | 0.73358 | -0.7651 | 0.182 | 0.17 | 1 | HC_vs_MS.Trt_F |
| 7-Mar | 0.7336 | -0.5855 | 0.263 | 0.27 | 1 | HC_vs_MS.Trt_F |
| TBC1D31 | 0.73361 | -0.2724 | 0.685 | 0.64 | 1 | HC_vs_MS.Trt_F |
| SMIM11A | 0.73473 | 0.54916 | 0.295 | 0.31 | 1 | HC_vs_MS.Trt_F |
| CAND1 | 0.73527 | -0.4866 | 0.867 | 0.88 | 1 | HC_vs_MS.Trt_F |
| NIPA1 | 0.73556 | 0.71317 | 0.191 | 0.2 | 1 | HC_vs_MS.Trt_F |
| AGRN | 0.73617 | 0.47907 | 0.416 | 0.41 | 1 | HC_vs_MS.Trt_F |
| NAGPA | 0.73629 | 0.46105 | 0.665 | 0.67 | 1 | HC_vs_MS.Trt_F |
| ZNF10 | 0.73654 | -1.0029 | 0.136 | 0.13 | 1 | HC_vs_MS.Trt_F |
| SUMO1 | 0.73656 | -0.3138 | 0.523 | 0.5 | 1 | HC_vs_MS.Trt_F |
| GNE | 0.73686 | -0.5916 | 0.225 | 0.23 | 1 | HC_vs_MS.Trt_F |
| COMMD2 | 0.73689 | -0.7697 | 0.376 | 0.35 | 1 | HC_vs_MS.Trt_F |
| RP11-644F5 | 0.73711 | -0.373 | 0.731 | 0.72 | 1 | HC_vs_MS.Trt_F |
| SOX4 | 0.73724 | -0.3822 | 0.104 | 0.1 | 1 | HC_vs_MS.Trt_F |
| C19orf54 | 0.73744 | -0.4017 | 0.486 | 0.49 | 1 | HC_vs_MS.Trt_F |
| ISCU | 0.73756 | -0.4232 | 0.321 | 0.29 | 1 | HC_vs_MS.Trt_F |
| PSMB4 | 0.73773 | -0.5773 | 0.387 | 0.37 | 1 | HC_vs_MS.Trt_F |
| POLR3H | 0.73787 | 1.15586 | 0.272 | 0.3 | 1 | HC_vs_MS.Trt_F |
| DECR2 | 0.73795 | -0.2982 | 0.847 | 0.78 | 1 | HC_vs_MS.Trt_F |
| FADS1 | 0.73816 | 0.40367 | 0.647 | 0.68 | 1 | HC_vs_MS.Trt_F |
| PODXL | 0.73905 | 0.39189 | 0.211 | 0.2 | 1 | HC_vs_MS.Trt_F |
| YY1AP1 | 0.73983 | -0.2642 | 0.543 | 0.55 | 1 | HC_vs_MS.Trt_F |
| ABHD5 | 0.73993 | 0.57041 | 0.136 | 0.13 | 1 | HC_vs_MS.Trt_F |
| SVIL | 0.74011 | 0.38941 | 0.185 | 0.19 | 1 | HC_vs_MS.Trt_F |
| DUSP23 | 0.74014 | -0.6678 | 0.136 | 0.12 | 1 | HC_vs_MS.Trt_F |
| ATP5L | 0.74014 | -0.3496 | 0.462 | 0.42 | 1 | HC_vs_MS.Trt_F |
| CD8B | 0.74025 | 1.29029 | 0.121 | 0.12 | 1 | HC_vs_MS.Trt_F |
| XYLT2 | 0.7408 | -0.5443 | 0.668 | 0.65 | 1 | HC_vs_MS.Trt_F |
| NOA1 | 0.74093 | 0.31067 | 0.121 | 0.11 | 1 | HC_vs_MS.Trt_F |
| CARS2 | 0.74129 | -0.265 | 0.78 | 0.75 | 1 | HC_vs_MS.Trt_F |
| C8orf33 | 0.74134 | -1.2721 | 0.249 | 0.25 | 1 | HC_vs_MS.Trt_F |
| RDH5 | 0.74136 | -0.3054 | 0.471 | 0.45 | 1 | HC_vs_MS.Trt_F |
| CRBN | 0.74141 | -0.5164 | 0.486 | 0.44 | 1 | HC_vs_MS.Trt_F |
| COQ6 | 0.74142 | -0.4684 | 0.841 | 0.76 | 1 | HC_vs_MS.Trt_F |
| SNRPG | 0.74175 | -0.3263 | 0.329 | 0.31 | 1 | HC_vs_MS.Trt_F |
| MGLL | 0.74212 | 0.28991 | 0.431 | 0.41 | 1 | HC_vs_MS.Trt_F |
| LXN | 0.74297 | -0.6012 | 0.202 | 0.19 | 1 | HC_vs_MS.Trt_F |

| | | | | | | |
|---|---|---|---|---|---|---|
| VCP | 0.74313 | -0.255 | 0.306 | 0.3 | 1 | HC_vs_MS.Trt_F |
| HLA-DRB4 | 0.74339 | 1.65898 | 0.124 | 0.12 | 1 | HC_vs_MS.Trt_F |
| SFT2D1 | 0.74345 | -0.6815 | 0.419 | 0.41 | 1 | HC_vs_MS.Trt_F |
| TRIM16 | 0.7437 | -1.0207 | 0.497 | 0.47 | 1 | HC_vs_MS.Trt_F |
| ZNF148 | 0.74376 | -0.5025 | 0.298 | 0.3 | 1 | HC_vs_MS.Trt_F |
| UBE3C | 0.74382 | -0.5655 | 0.613 | 0.58 | 1 | HC_vs_MS.Trt_F |
| LSS | 0.74394 | -0.4136 | 0.13 | 0.12 | 1 | HC_vs_MS.Trt_F |
| ZNF146 | 0.74404 | -0.4481 | 0.211 | 0.19 | 1 | HC_vs_MS.Trt_F |
| RP11-500M8 | 0.74422 | -0.8057 | 0.124 | 0.11 | 1 | HC_vs_MS.Trt_F |
| CCDC186 | 0.74469 | -0.3474 | 0.587 | 0.57 | 1 | HC_vs_MS.Trt_F |
| OSBPL11 | 0.74487 | -0.336 | 0.118 | 0.11 | 1 | HC_vs_MS.Trt_F |
| C5 | 0.74554 | -0.7066 | 0.465 | 0.46 | 1 | HC_vs_MS.Trt_F |
| UQCRQ | 0.7457 | -0.5642 | 0.598 | 0.57 | 1 | HC_vs_MS.Trt_F |
| MT1F | 0.7462 | -0.4497 | 0.101 | 0.11 | 1 | HC_vs_MS.Trt_F |
| COA3 | 0.74683 | -0.7868 | 0.145 | 0.15 | 1 | HC_vs_MS.Trt_F |
| C15orf52 | 0.74689 | 0.33683 | 0.425 | 0.39 | 1 | HC_vs_MS.Trt_F |
| OMA1 | 0.747 | -0.4732 | 0.627 | 0.59 | 1 | HC_vs_MS.Trt_F |
| DBT | 0.74714 | -0.5017 | 0.867 | 0.82 | 1 | HC_vs_MS.Trt_F |
| C2orf74 | 0.74762 | -0.3459 | 0.477 | 0.45 | 1 | HC_vs_MS.Trt_F |
| DNAJC3 | 0.74789 | -0.8359 | 0.592 | 0.53 | 1 | HC_vs_MS.Trt_F |
| TMEM128 | 0.74824 | -0.7886 | 0.171 | 0.16 | 1 | HC_vs_MS.Trt_F |
| CAMTA1 | 0.74832 | -0.654 | 0.159 | 0.17 | 1 | HC_vs_MS.Trt_F |
| FSCN1 | 0.74861 | 0.662 | 0.101 | 0.09 | 1 | HC_vs_MS.Trt_F |
| SRPX2 | 0.749 | 0.27206 | 0.251 | 0.26 | 1 | HC_vs_MS.Trt_F |
| IL18R1 | 0.74934 | 0.59986 | 0.142 | 0.13 | 1 | HC_vs_MS.Trt_F |
| FBXW8 | 0.7495 | 0.38846 | 0.321 | 0.33 | 1 | HC_vs_MS.Trt_F |
| DIDO1 | 0.74976 | -0.4684 | 0.199 | 0.2 | 1 | HC_vs_MS.Trt_F |
| TNIP3 | 0.75099 | 0.42624 | 0.512 | 0.48 | 1 | HC_vs_MS.Trt_F |
| ZNF549 | 0.75177 | -0.4007 | 0.688 | 0.67 | 1 | HC_vs_MS.Trt_F |
| ALDH1B1 | 0.75186 | -0.5971 | 0.662 | 0.63 | 1 | HC_vs_MS.Trt_F |
| MALSU1 | 0.75291 | -0.6812 | 0.121 | 0.13 | 1 | HC_vs_MS.Trt_F |
| PPP2R2D | 0.75353 | -0.4818 | 0.65 | 0.63 | 1 | HC_vs_MS.Trt_F |
| SIL1 | 0.75383 | -0.266 | 0.116 | 0.12 | 1 | HC_vs_MS.Trt_F |
| IRAK2 | 0.75387 | 0.41144 | 0.341 | 0.35 | 1 | HC_vs_MS.Trt_F |
| UCHL3 | 0.75422 | -0.7069 | 0.159 | 0.14 | 1 | HC_vs_MS.Trt_F |
| CAPG | 0.75454 | 0.41143 | 0.153 | 0.14 | 1 | HC_vs_MS.Trt_F |
| CASP3 | 0.75491 | -0.7454 | 0.173 | 0.16 | 1 | HC_vs_MS.Trt_F |
| TUBB3 | 0.75494 | 0.65781 | 0.292 | 0.3 | 1 | HC_vs_MS.Trt_F |
| CIPC | 0.75513 | -0.3751 | 0.488 | 0.45 | 1 | HC_vs_MS.Trt_F |
| TMED7 | 0.75572 | -0.4831 | 0.098 | 0.1 | 1 | HC_vs_MS.Trt_F |

| | | | | | | |
|---|---|---|---|---|---|---|
| FAM110C | 0.75622 | -0.3534 | 0.275 | 0.26 | 1 | HC_vs_MS.Trt_F |
| VAMP3 | 0.75754 | -0.2535 | 0.353 | 0.35 | 1 | HC_vs_MS.Trt_F |
| KIF21B | 0.75757 | 0.37248 | 0.434 | 0.43 | 1 | HC_vs_MS.Trt_F |
| GALNS | 0.75777 | -0.3987 | 0.566 | 0.53 | 1 | HC_vs_MS.Trt_F |
| XPOT | 0.75836 | -0.288 | 0.477 | 0.46 | 1 | HC_vs_MS.Trt_F |
| ZNF274 | 0.75854 | -0.3225 | 0.645 | 0.64 | 1 | HC_vs_MS.Trt_F |
| PPP3CA | 0.75893 | -0.609 | 0.566 | 0.56 | 1 | HC_vs_MS.Trt_F |
| FBXW11 | 0.75911 | -0.6704 | 0.303 | 0.28 | 1 | HC_vs_MS.Trt_F |
| RANGRF | 0.7592 | -0.3775 | 0.116 | 0.1 | 1 | HC_vs_MS.Trt_F |
| KLHL42 | 0.75944 | 0.51903 | 0.124 | 0.12 | 1 | HC_vs_MS.Trt_F |
| ASXL1 | 0.75946 | -0.681 | 0.214 | 0.19 | 1 | HC_vs_MS.Trt_F |
| UBXN6 | 0.75951 | -0.5589 | 0.095 | 0.1 | 1 | HC_vs_MS.Trt_F |
| TPSB2 | 0.75971 | 3.43554 | 0.286 | 0.28 | 1 | HC_vs_MS.Trt_F |
| RAD51B | 0.7601 | -0.496 | 0.327 | 0.31 | 1 | HC_vs_MS.Trt_F |
| MAGOHB | 0.76027 | 0.26353 | 0.098 | 0.1 | 1 | HC_vs_MS.Trt_F |
| SMPD4 | 0.76034 | -0.3315 | 0.318 | 0.3 | 1 | HC_vs_MS.Trt_F |
| BACE2 | 0.76052 | 0.66125 | 0.306 | 0.28 | 1 | HC_vs_MS.Trt_F |
| SRGAP1 | 0.76077 | 0.31849 | 0.954 | 0.96 | 1 | HC_vs_MS.Trt_F |
| NFU1 | 0.761 | 0.30535 | 0.162 | 0.17 | 1 | HC_vs_MS.Trt_F |
| PFDN5 | 0.76134 | -0.32 | 0.908 | 0.87 | 1 | HC_vs_MS.Trt_F |
| RPP21 | 0.76134 | -0.4291 | 0.197 | 0.18 | 1 | HC_vs_MS.Trt_F |
| DNPEP | 0.76151 | 0.61396 | 0.48 | 0.49 | 1 | HC_vs_MS.Trt_F |
| BLZF1 | 0.76164 | 0.47849 | 0.509 | 0.5 | 1 | HC_vs_MS.Trt_F |
| CNIH4 | 0.76189 | -0.2984 | 0.636 | 0.62 | 1 | HC_vs_MS.Trt_F |
| CCNY | 0.76214 | -0.5215 | 0.494 | 0.47 | 1 | HC_vs_MS.Trt_F |
| C16orf87 | 0.76246 | -0.351 | 0.812 | 0.79 | 1 | HC_vs_MS.Trt_F |
| SLC16A5 | 0.76306 | -0.2727 | 0.191 | 0.18 | 1 | HC_vs_MS.Trt_F |
| REC8 | 0.76323 | 0.54309 | 0.118 | 0.11 | 1 | HC_vs_MS.Trt_F |
| GRIN2B | 0.76328 | 0.5456 | 0.237 | 0.22 | 1 | HC_vs_MS.Trt_F |
| LYAR | 0.76341 | -0.6629 | 0.121 | 0.11 | 1 | HC_vs_MS.Trt_F |
| HYLS1 | 0.76345 | -0.6317 | 0.454 | 0.42 | 1 | HC_vs_MS.Trt_F |
| WBP11 | 0.76358 | -0.3339 | 0.327 | 0.3 | 1 | HC_vs_MS.Trt_F |
| RPUSD1 | 0.76409 | -0.4947 | 0.11 | 0.12 | 1 | HC_vs_MS.Trt_F |
| TAF4 | 0.76447 | 0.47933 | 0.133 | 0.13 | 1 | HC_vs_MS.Trt_F |
| ZNF781 | 0.76475 | -0.4454 | 0.301 | 0.28 | 1 | HC_vs_MS.Trt_F |
| YRDC | 0.76476 | 0.42191 | 0.113 | 0.11 | 1 | HC_vs_MS.Trt_F |
| RBMX | 0.76478 | -0.2512 | 0.815 | 0.77 | 1 | HC_vs_MS.Trt_F |
| MAD2L2 | 0.76485 | -0.3657 | 0.15 | 0.14 | 1 | HC_vs_MS.Trt_F |
| CDK2 | 0.76487 | 0.65252 | 0.113 | 0.1 | 1 | HC_vs_MS.Trt_F |
| GAA | 0.76512 | 0.63392 | 0.101 | 0.09 | 1 | HC_vs_MS.Trt_F |

| | | | | | | |
|---|---|---|---|---|---|---|
| FBN1 | 0.76524 | 0.2656 | 0.514 | 0.48 | 1 | HC_vs_MS.Trt_F |
| ZNF391 | 0.7654 | 0.56394 | 0.185 | 0.19 | 1 | HC_vs_MS.Trt_F |
| 15-Sep | 0.76554 | -0.359 | 0.237 | 0.24 | 1 | HC_vs_MS.Trt_F |
| DUSP3 | 0.76558 | 0.42315 | 0.772 | 0.77 | 1 | HC_vs_MS.Trt_F |
| CREB3 | 0.76571 | -0.3816 | 0.104 | 0.11 | 1 | HC_vs_MS.Trt_F |
| RNF26 | 0.76623 | -1.119 | 0.127 | 0.13 | 1 | HC_vs_MS.Trt_F |
| SLC20A1 | 0.76666 | -0.2978 | 0.26 | 0.24 | 1 | HC_vs_MS.Trt_F |
| CEP70 | 0.76724 | -0.7208 | 0.156 | 0.16 | 1 | HC_vs_MS.Trt_F |
| MXD1 | 0.76724 | -0.3947 | 0.171 | 0.17 | 1 | HC_vs_MS.Trt_F |
| HECTD3 | 0.76809 | -0.266 | 0.145 | 0.15 | 1 | HC_vs_MS.Trt_F |
| EXOSC6 | 0.76826 | -0.4182 | 0.928 | 0.9 | 1 | HC_vs_MS.Trt_F |
| TEKT3 | 0.76854 | 0.28008 | 0.179 | 0.17 | 1 | HC_vs_MS.Trt_F |
| TSPAN5 | 0.76861 | 0.25066 | 0.645 | 0.62 | 1 | HC_vs_MS.Trt_F |
| MORF4L1 | 0.76891 | -0.3907 | 0.812 | 0.81 | 1 | HC_vs_MS.Trt_F |
| ATP6V0C | 0.76923 | 0.6044 | 0.283 | 0.27 | 1 | HC_vs_MS.Trt_F |
| GLIPR1 | 0.7695 | -0.3179 | 0.309 | 0.31 | 1 | HC_vs_MS.Trt_F |
| WWP1 | 0.76953 | -0.2685 | 0.451 | 0.42 | 1 | HC_vs_MS.Trt_F |
| BCL11B | 0.76986 | -0.3629 | 0.465 | 0.45 | 1 | HC_vs_MS.Trt_F |
| CCP110 | 0.77099 | 0.44439 | 0.52 | 0.53 | 1 | HC_vs_MS.Trt_F |
| RIPK2 | 0.77223 | 0.25363 | 0.22 | 0.23 | 1 | HC_vs_MS.Trt_F |
| RGS5 | 0.77223 | 0.26151 | 0.11 | 0.1 | 1 | HC_vs_MS.Trt_F |
| CCDC90B | 0.77229 | -0.4331 | 0.951 | 0.91 | 1 | HC_vs_MS.Trt_F |
| ZNF284 | 0.77231 | -0.2779 | 0.613 | 0.58 | 1 | HC_vs_MS.Trt_F |
| PPM1A | 0.77248 | -0.3117 | 0.179 | 0.18 | 1 | HC_vs_MS.Trt_F |
| LSM11 | 0.7727 | 0.35947 | 0.607 | 0.56 | 1 | HC_vs_MS.Trt_F |
| PRDX6 | 0.77297 | -0.7922 | 0.173 | 0.17 | 1 | HC_vs_MS.Trt_F |
| KIF2A | 0.77301 | -0.5908 | 0.251 | 0.25 | 1 | HC_vs_MS.Trt_F |
| METTL2A | 0.77309 | -0.2849 | 0.74 | 0.68 | 1 | HC_vs_MS.Trt_F |
| MMP24 | 0.77344 | -0.2974 | 0.095 | 0.1 | 1 | HC_vs_MS.Trt_F |
| NUP54 | 0.77373 | -0.4012 | 0.208 | 0.19 | 1 | HC_vs_MS.Trt_F |
| ZNF491 | 0.77388 | 0.47683 | 0.488 | 0.45 | 1 | HC_vs_MS.Trt_F |
| SLCO4C1 | 0.7741 | -0.9179 | 0.188 | 0.19 | 1 | HC_vs_MS.Trt_F |
| TMEM25 | 0.77428 | -1.1996 | 0.402 | 0.4 | 1 | HC_vs_MS.Trt_F |
| MGAT4A | 0.77439 | -0.3129 | 0.445 | 0.41 | 1 | HC_vs_MS.Trt_F |
| TM2D3 | 0.77441 | 0.28216 | 0.815 | 0.78 | 1 | HC_vs_MS.Trt_F |
| TRIM52 | 0.77448 | -0.3988 | 0.908 | 0.86 | 1 | HC_vs_MS.Trt_F |
| PSMG4 | 0.77458 | -0.303 | 0.789 | 0.79 | 1 | HC_vs_MS.Trt_F |
| THBS1 | 0.77471 | 0.60863 | 0.263 | 0.25 | 1 | HC_vs_MS.Trt_F |
| CD302 | 0.77484 | -0.8366 | 0.425 | 0.4 | 1 | HC_vs_MS.Trt_F |
| LIPE | 0.77499 | -0.7804 | 0.396 | 0.34 | 1 | HC_vs_MS.Trt_F |

| | | | | | | |
|---|---|---|---|---|---|---|
| PLEKHB1 | 0.77521 | -0.7196 | 0.107 | 0.1 | 1 | HC_vs_MS.Trt_F |
| AGBL5 | 0.77555 | -0.9484 | 0.179 | 0.17 | 1 | HC_vs_MS.Trt_F |
| MGST2 | 0.77583 | -0.7303 | 0.214 | 0.22 | 1 | HC_vs_MS.Trt_F |
| KYAT1 | 0.77676 | -0.767 | 0.173 | 0.16 | 1 | HC_vs_MS.Trt_F |
| NPIPA5 | 0.77699 | 0.70364 | 0.104 | 0.11 | 1 | HC_vs_MS.Trt_F |
| SNX8 | 0.77775 | -0.7893 | 0.15 | 0.14 | 1 | HC_vs_MS.Trt_F |
| ZNF292 | 0.7779 | -0.3668 | 0.824 | 0.78 | 1 | HC_vs_MS.Trt_F |
| UBP1 | 0.7786 | 0.46721 | 0.13 | 0.12 | 1 | HC_vs_MS.Trt_F |
| RGL4 | 0.77926 | 0.35089 | 0.263 | 0.26 | 1 | HC_vs_MS.Trt_F |
| CSE1L | 0.77932 | -0.7904 | 0.225 | 0.22 | 1 | HC_vs_MS.Trt_F |
| MED31 | 0.77933 | -0.5252 | 0.133 | 0.14 | 1 | HC_vs_MS.Trt_F |
| AKR1C2 | 0.77976 | 1.09875 | 0.107 | 0.1 | 1 | HC_vs_MS.Trt_F |
| ALKBH3 | 0.77977 | 0.67316 | 0.113 | 0.1 | 1 | HC_vs_MS.Trt_F |
| ZXDC | 0.77997 | 0.59538 | 0.116 | 0.11 | 1 | HC_vs_MS.Trt_F |
| AP3S2 | 0.78046 | -0.4166 | 0.725 | 0.68 | 1 | HC_vs_MS.Trt_F |
| ZNF19 | 0.78065 | 0.30331 | 0.815 | 0.8 | 1 | HC_vs_MS.Trt_F |
| RPS6KA2 | 0.78102 | -0.4197 | 0.113 | 0.1 | 1 | HC_vs_MS.Trt_F |
| ABHD12 | 0.78155 | -0.2524 | 0.118 | 0.12 | 1 | HC_vs_MS.Trt_F |
| GANC | 0.78209 | -0.8154 | 0.665 | 0.65 | 1 | HC_vs_MS.Trt_F |
| LIMK2 | 0.7824 | -0.2881 | 0.182 | 0.19 | 1 | HC_vs_MS.Trt_F |
| THUMPD2 | 0.78351 | -0.5523 | 0.41 | 0.42 | 1 | HC_vs_MS.Trt_F |
| PTCD1 | 0.7844 | 0.46928 | 0.624 | 0.63 | 1 | HC_vs_MS.Trt_F |
| CECR2 | 0.78455 | 0.41214 | 0.558 | 0.6 | 1 | HC_vs_MS.Trt_F |
| RBBP5 | 0.78462 | -0.5989 | 0.59 | 0.57 | 1 | HC_vs_MS.Trt_F |
| MLPH | 0.78469 | 1.01299 | 0.514 | 0.48 | 1 | HC_vs_MS.Trt_F |
| WNT9A | 0.7854 | -0.5618 | 0.113 | 0.11 | 1 | HC_vs_MS.Trt_F |
| ZNF655 | 0.78615 | 0.25346 | 0.616 | 0.61 | 1 | HC_vs_MS.Trt_F |
| NDUFA13 | 0.78715 | -0.2687 | 0.506 | 0.48 | 1 | HC_vs_MS.Trt_F |
| TK2 | 0.78752 | 0.77386 | 0.168 | 0.17 | 1 | HC_vs_MS.Trt_F |
| BRWD3 | 0.78763 | -0.3155 | 0.153 | 0.14 | 1 | HC_vs_MS.Trt_F |
| ABLIM1 | 0.78853 | -0.3206 | 0.373 | 0.34 | 1 | HC_vs_MS.Trt_F |
| GBA | 0.78871 | -0.4675 | 0.506 | 0.48 | 1 | HC_vs_MS.Trt_F |
| FBXO28 | 0.78877 | 0.422 | 0.116 | 0.11 | 1 | HC_vs_MS.Trt_F |
| APOL1 | 0.78968 | -0.7509 | 0.702 | 0.7 | 1 | HC_vs_MS.Trt_F |
| BICD2 | 0.7904 | -0.5601 | 0.116 | 0.12 | 1 | HC_vs_MS.Trt_F |
| IL18 | 0.79053 | 0.78307 | 0.491 | 0.49 | 1 | HC_vs_MS.Trt_F |
| CCT5 | 0.79062 | -0.3287 | 0.223 | 0.21 | 1 | HC_vs_MS.Trt_F |
| SH3KBP1 | 0.79094 | -0.3366 | 0.497 | 0.47 | 1 | HC_vs_MS.Trt_F |
| TWISTNB | 0.79109 | -0.6602 | 0.136 | 0.14 | 1 | HC_vs_MS.Trt_F |
| KCMF1 | 0.79136 | -0.7844 | 0.54 | 0.52 | 1 | HC_vs_MS.Trt_F |

| | | | | | | |
|---|---|---|---|---|---|---|
| ZNF273 | 0.79232 | -0.4907 | 0.936 | 0.89 | 1 | HC_vs_MS.Trt_F |
| DBN1 | 0.79285 | -0.6115 | 0.249 | 0.23 | 1 | HC_vs_MS.Trt_F |
| PEX14 | 0.79336 | 0.56151 | 0.124 | 0.12 | 1 | HC_vs_MS.Trt_F |
| DNAJB7 | 0.7934 | -0.394 | 0.249 | 0.23 | 1 | HC_vs_MS.Trt_F |
| IL27RA | 0.79363 | -0.4721 | 0.402 | 0.39 | 1 | HC_vs_MS.Trt_F |
| SLC35A3 | 0.79399 | 0.30982 | 0.835 | 0.82 | 1 | HC_vs_MS.Trt_F |
| NUDT15 | 0.79445 | -1.0434 | 0.63 | 0.59 | 1 | HC_vs_MS.Trt_F |
| ATP5H | 0.79478 | -0.3887 | 0.806 | 0.75 | 1 | HC_vs_MS.Trt_F |
| BDH1 | 0.79509 | 0.33394 | 0.908 | 0.89 | 1 | HC_vs_MS.Trt_F |
| RABAC1 | 0.79531 | -0.5172 | 0.283 | 0.27 | 1 | HC_vs_MS.Trt_F |
| HIST1H2BE | 0.79556 | 0.47056 | 0.335 | 0.32 | 1 | HC_vs_MS.Trt_F |
| BOP1 | 0.79559 | 0.32389 | 0.249 | 0.24 | 1 | HC_vs_MS.Trt_F |
| RP11-701H2 | 0.79562 | -0.2876 | 0.136 | 0.12 | 1 | HC_vs_MS.Trt_F |
| TMEM205 | 0.79569 | -0.4403 | 0.118 | 0.11 | 1 | HC_vs_MS.Trt_F |
| MUS81 | 0.79601 | -0.2653 | 0.159 | 0.16 | 1 | HC_vs_MS.Trt_F |
| FRMD4A | 0.79609 | -0.281 | 0.656 | 0.65 | 1 | HC_vs_MS.Trt_F |
| ALG3 | 0.79647 | -0.7721 | 0.173 | 0.17 | 1 | HC_vs_MS.Trt_F |
| GTF2F2 | 0.7971 | -0.3512 | 0.211 | 0.2 | 1 | HC_vs_MS.Trt_F |
| TRAPPC6A | 0.79759 | -0.3724 | 0.185 | 0.19 | 1 | HC_vs_MS.Trt_F |
| FAM83H | 0.79782 | 0.49937 | 0.147 | 0.14 | 1 | HC_vs_MS.Trt_F |
| FLCN | 0.79788 | 0.25435 | 0.893 | 0.88 | 1 | HC_vs_MS.Trt_F |
| AGBL2 | 0.79811 | 0.64793 | 0.28 | 0.29 | 1 | HC_vs_MS.Trt_F |
| TOMM6 | 0.79815 | -0.5391 | 0.24 | 0.22 | 1 | HC_vs_MS.Trt_F |
| TNS4 | 0.79881 | 0.32042 | 0.572 | 0.6 | 1 | HC_vs_MS.Trt_F |
| FBXL22 | 0.79946 | 0.51028 | 0.13 | 0.14 | 1 | HC_vs_MS.Trt_F |
| SSR2 | 0.79949 | -0.3006 | 0.902 | 0.9 | 1 | HC_vs_MS.Trt_F |
| SHMT2 | 0.79961 | -0.3326 | 0.254 | 0.25 | 1 | HC_vs_MS.Trt_F |
| LRFN3 | 0.79964 | 0.27267 | 0.63 | 0.6 | 1 | HC_vs_MS.Trt_F |
| SLC39A14 | 0.80001 | 0.93137 | 0.436 | 0.46 | 1 | HC_vs_MS.Trt_F |
| TNPO3 | 0.80082 | 0.48603 | 0.173 | 0.18 | 1 | HC_vs_MS.Trt_F |
| CD33 | 0.80108 | 0.31815 | 0.618 | 0.6 | 1 | HC_vs_MS.Trt_F |
| ZNF773 | 0.80124 | 0.37398 | 0.662 | 0.65 | 1 | HC_vs_MS.Trt_F |
| CEP170 | 0.80157 | 0.586 | 0.572 | 0.6 | 1 | HC_vs_MS.Trt_F |
| IL10RA | 0.802 | -0.2577 | 0.578 | 0.55 | 1 | HC_vs_MS.Trt_F |
| DCTD | 0.80205 | -0.5592 | 0.153 | 0.14 | 1 | HC_vs_MS.Trt_F |
| AMMECR1 | 0.8022 | -0.2658 | 0.159 | 0.15 | 1 | HC_vs_MS.Trt_F |
| C19orf53 | 0.80224 | -0.5344 | 0.405 | 0.37 | 1 | HC_vs_MS.Trt_F |
| PSMC3 | 0.8025 | -0.7135 | 0.185 | 0.17 | 1 | HC_vs_MS.Trt_F |
| FADS3 | 0.80288 | -0.261 | 0.662 | 0.59 | 1 | HC_vs_MS.Trt_F |
| CTD-3148I1( | 0.80324 | -0.2946 | 0.11 | 0.12 | 1 | HC_vs_MS.Trt_F |

| | | | | | | |
|---|---|---|---|---|---|---|
| DOCK5 | 0.80341 | 0.34312 | 0.587 | 0.6 | 1 | HC_vs_MS.Trt_F |
| ZSCAN25 | 0.80409 | -0.4378 | 0.462 | 0.46 | 1 | HC_vs_MS.Trt_F |
| LETMD1 | 0.80431 | -0.3235 | 0.858 | 0.83 | 1 | HC_vs_MS.Trt_F |
| PPP1R3B | 0.80434 | -0.7261 | 0.286 | 0.28 | 1 | HC_vs_MS.Trt_F |
| NDUFS8 | 0.80444 | -0.2653 | 0.835 | 0.84 | 1 | HC_vs_MS.Trt_F |
| ACTR3C | 0.80514 | -0.4457 | 0.561 | 0.53 | 1 | HC_vs_MS.Trt_F |
| RALBP1 | 0.80534 | -0.3243 | 0.454 | 0.44 | 1 | HC_vs_MS.Trt_F |
| S1PR5 | 0.80545 | 0.25714 | 0.121 | 0.13 | 1 | HC_vs_MS.Trt_F |
| TMEM79 | 0.80578 | -0.4998 | 0.098 | 0.1 | 1 | HC_vs_MS.Trt_F |
| RNF103 | 0.80596 | -0.4503 | 0.639 | 0.63 | 1 | HC_vs_MS.Trt_F |
| EML5 | 0.80701 | 0.39504 | 0.243 | 0.23 | 1 | HC_vs_MS.Trt_F |
| PLXNA4 | 0.80718 | 0.26007 | 0.104 | 0.1 | 1 | HC_vs_MS.Trt_F |
| MRS2 | 0.8073 | 0.64321 | 0.812 | 0.86 | 1 | HC_vs_MS.Trt_F |
| NUP50 | 0.80736 | -0.5488 | 0.734 | 0.67 | 1 | HC_vs_MS.Trt_F |
| E2F5 | 0.80743 | -0.7842 | 0.393 | 0.4 | 1 | HC_vs_MS.Trt_F |
| BDH2 | 0.80753 | -0.5664 | 0.234 | 0.22 | 1 | HC_vs_MS.Trt_F |
| NAE1 | 0.80755 | -0.7512 | 0.737 | 0.73 | 1 | HC_vs_MS.Trt_F |
| ZCCHC17 | 0.80766 | -0.3523 | 0.142 | 0.13 | 1 | HC_vs_MS.Trt_F |
| UTP4 | 0.80787 | -0.3572 | 0.838 | 0.81 | 1 | HC_vs_MS.Trt_F |
| BPHL | 0.80854 | -0.2521 | 0.694 | 0.63 | 1 | HC_vs_MS.Trt_F |
| RSL1D1 | 0.80883 | -0.2904 | 0.884 | 0.85 | 1 | HC_vs_MS.Trt_F |
| CLDND1 | 0.8089 | -0.5796 | 0.379 | 0.37 | 1 | HC_vs_MS.Trt_F |
| REST | 0.80931 | -0.3702 | 0.188 | 0.19 | 1 | HC_vs_MS.Trt_F |
| DAZAP2 | 0.80945 | -0.289 | 0.861 | 0.81 | 1 | HC_vs_MS.Trt_F |
| RPUSD2 | 0.80954 | -0.8132 | 0.15 | 0.16 | 1 | HC_vs_MS.Trt_F |
| ZNF540 | 0.81003 | -0.8824 | 0.494 | 0.49 | 1 | HC_vs_MS.Trt_F |
| TAZ | 0.81016 | 0.30456 | 0.59 | 0.58 | 1 | HC_vs_MS.Trt_F |
| ZDHHC16 | 0.81041 | -0.4829 | 0.118 | 0.11 | 1 | HC_vs_MS.Trt_F |
| LCLAT1 | 0.81116 | -0.2566 | 0.835 | 0.8 | 1 | HC_vs_MS.Trt_F |
| VPS26B | 0.81117 | 0.25278 | 0.118 | 0.12 | 1 | HC_vs_MS.Trt_F |
| MUC20 | 0.81162 | 0.3292 | 0.558 | 0.54 | 1 | HC_vs_MS.Trt_F |
| AKAP8L | 0.81187 | 0.64955 | 0.104 | 0.1 | 1 | HC_vs_MS.Trt_F |
| IDO1 | 0.81212 | -0.2691 | 0.543 | 0.53 | 1 | HC_vs_MS.Trt_F |
| ZNF71 | 0.81246 | -0.516 | 0.292 | 0.27 | 1 | HC_vs_MS.Trt_F |
| LIPC | 0.81251 | 0.46347 | 0.332 | 0.34 | 1 | HC_vs_MS.Trt_F |
| TNFSF4 | 0.81252 | 0.95085 | 0.136 | 0.13 | 1 | HC_vs_MS.Trt_F |
| PLAA | 0.81305 | -0.3738 | 0.133 | 0.14 | 1 | HC_vs_MS.Trt_F |
| RAP1GDS1 | 0.81347 | -0.2579 | 0.379 | 0.36 | 1 | HC_vs_MS.Trt_F |
| EGFL8 | 0.81348 | 0.25633 | 0.569 | 0.55 | 1 | HC_vs_MS.Trt_F |
| TELO2 | 0.81357 | -0.5547 | 0.113 | 0.12 | 1 | HC_vs_MS.Trt_F |

| | | | | | | |
|---|---|---|---|---|---|---|
| KIAA1671 | 0.81513 | -0.3656 | 0.538 | 0.51 | 1 | HC_vs_MS.Trt_F |
| ACTR1A | 0.81533 | 0.30204 | 0.506 | 0.47 | 1 | HC_vs_MS.Trt_F |
| SF3B6 | 0.81578 | -0.4135 | 0.214 | 0.2 | 1 | HC_vs_MS.Trt_F |
| DTX3L | 0.8158 | -0.466 | 0.711 | 0.66 | 1 | HC_vs_MS.Trt_F |
| TXK | 0.81584 | -0.386 | 0.621 | 0.63 | 1 | HC_vs_MS.Trt_F |
| LRPPRC | 0.81623 | -0.5618 | 0.532 | 0.47 | 1 | HC_vs_MS.Trt_F |
| MSRB3 | 0.8164 | 0.40214 | 0.523 | 0.51 | 1 | HC_vs_MS.Trt_F |
| KIAA1429 | 0.81664 | -0.5157 | 0.855 | 0.84 | 1 | HC_vs_MS.Trt_F |
| PLCL1 | 0.8167 | -1.1975 | 0.13 | 0.12 | 1 | HC_vs_MS.Trt_F |
| CDRT4 | 0.8168 | -0.3998 | 0.139 | 0.13 | 1 | HC_vs_MS.Trt_F |
| MSH6 | 0.8171 | -0.264 | 0.841 | 0.85 | 1 | HC_vs_MS.Trt_F |
| PRKAR2A | 0.8179 | 0.33831 | 0.913 | 0.87 | 1 | HC_vs_MS.Trt_F |
| GYS1 | 0.81794 | -0.6043 | 0.165 | 0.16 | 1 | HC_vs_MS.Trt_F |
| TFB1M | 0.81888 | -0.4724 | 0.448 | 0.45 | 1 | HC_vs_MS.Trt_F |
| MTHFD1L | 0.81915 | -1.1539 | 0.139 | 0.14 | 1 | HC_vs_MS.Trt_F |
| CCDC43 | 0.81985 | -0.3454 | 0.113 | 0.12 | 1 | HC_vs_MS.Trt_F |
| TMEM185B | 0.81989 | 0.59876 | 0.335 | 0.34 | 1 | HC_vs_MS.Trt_F |
| ABHD2 | 0.81999 | -0.8705 | 0.659 | 0.65 | 1 | HC_vs_MS.Trt_F |
| CEP152 | 0.82001 | -0.5119 | 0.795 | 0.71 | 1 | HC_vs_MS.Trt_F |
| ATP6V1B2 | 0.82067 | -0.6023 | 0.711 | 0.67 | 1 | HC_vs_MS.Trt_F |
| TREML2 | 0.82168 | -1.0554 | 0.133 | 0.13 | 1 | HC_vs_MS.Trt_F |
| ZC3H11A | 0.82232 | -0.2576 | 0.231 | 0.21 | 1 | HC_vs_MS.Trt_F |
| HACD1 | 0.82236 | -0.6386 | 0.636 | 0.62 | 1 | HC_vs_MS.Trt_F |
| TCEAL4 | 0.82257 | -0.3738 | 0.159 | 0.16 | 1 | HC_vs_MS.Trt_F |
| MAP4K2 | 0.8226 | -0.3265 | 0.708 | 0.69 | 1 | HC_vs_MS.Trt_F |
| RALB | 0.82264 | -1.0798 | 0.13 | 0.13 | 1 | HC_vs_MS.Trt_F |
| LMO7 | 0.8229 | 0.37125 | 0.269 | 0.28 | 1 | HC_vs_MS.Trt_F |
| FBLN5 | 0.82291 | -1.0085 | 0.199 | 0.2 | 1 | HC_vs_MS.Trt_F |
| HBP1 | 0.82304 | 0.29528 | 0.812 | 0.8 | 1 | HC_vs_MS.Trt_F |
| UBL4A | 0.82314 | -0.4273 | 0.118 | 0.12 | 1 | HC_vs_MS.Trt_F |
| SLC6A16 | 0.82333 | -1.0021 | 0.266 | 0.25 | 1 | HC_vs_MS.Trt_F |
| DENND1B | 0.82392 | 0.42773 | 0.159 | 0.16 | 1 | HC_vs_MS.Trt_F |
| MRPL21 | 0.82459 | -0.401 | 0.124 | 0.13 | 1 | HC_vs_MS.Trt_F |
| RAB8A | 0.82483 | -0.2912 | 0.607 | 0.57 | 1 | HC_vs_MS.Trt_F |
| ZNF133 | 0.82535 | 0.83081 | 0.454 | 0.46 | 1 | HC_vs_MS.Trt_F |
| RASAL1 | 0.82555 | 0.94408 | 0.318 | 0.31 | 1 | HC_vs_MS.Trt_F |
| PLEKHF2 | 0.82602 | -0.9028 | 0.101 | 0.1 | 1 | HC_vs_MS.Trt_F |
| CACNA1F | 0.82608 | 0.55291 | 0.318 | 0.32 | 1 | HC_vs_MS.Trt_F |
| ZNF280C | 0.82627 | 0.69738 | 0.298 | 0.29 | 1 | HC_vs_MS.Trt_F |
| SRRD | 0.82651 | -0.2928 | 0.474 | 0.45 | 1 | HC_vs_MS.Trt_F |

| | | | | | | |
|---|---|---|---|---|---|---|
| RETSAT | 0.82747 | -0.5092 | 0.179 | 0.18 | 1 | HC_vs_MS.Trt_F |
| EIF3M | 0.82783 | -0.2778 | 0.815 | 0.79 | 1 | HC_vs_MS.Trt_F |
| SLC23A3 | 0.82824 | -0.7839 | 0.483 | 0.46 | 1 | HC_vs_MS.Trt_F |
| CHST3 | 0.82838 | 1.34398 | 0.188 | 0.2 | 1 | HC_vs_MS.Trt_F |
| CENPK | 0.82853 | -0.3422 | 0.824 | 0.79 | 1 | HC_vs_MS.Trt_F |
| C20orf194 | 0.82908 | -0.2782 | 0.344 | 0.33 | 1 | HC_vs_MS.Trt_F |
| HMGCL | 0.82982 | 0.63511 | 0.624 | 0.6 | 1 | HC_vs_MS.Trt_F |
| FDPS | 0.82993 | -0.3564 | 0.526 | 0.52 | 1 | HC_vs_MS.Trt_F |
| RP11-244H3 | 0.83045 | -1.7673 | 0.107 | 0.11 | 1 | HC_vs_MS.Trt_F |
| PHEX | 0.83055 | 0.91046 | 0.104 | 0.1 | 1 | HC_vs_MS.Trt_F |
| PRRT2 | 0.83162 | 1.19115 | 0.208 | 0.22 | 1 | HC_vs_MS.Trt_F |
| SEC23A | 0.83171 | -0.3607 | 0.96 | 0.95 | 1 | HC_vs_MS.Trt_F |
| ANXA2R | 0.83215 | -0.5456 | 0.182 | 0.18 | 1 | HC_vs_MS.Trt_F |
| MFGE8 | 0.83252 | -0.3925 | 0.604 | 0.55 | 1 | HC_vs_MS.Trt_F |
| UBE2V2 | 0.83394 | -0.7794 | 0.734 | 0.68 | 1 | HC_vs_MS.Trt_F |
| ATP6V1G1 | 0.83395 | -0.3973 | 0.425 | 0.41 | 1 | HC_vs_MS.Trt_F |
| RPL26L1 | 0.83407 | -0.6003 | 0.165 | 0.17 | 1 | HC_vs_MS.Trt_F |
| ZBTB33 | 0.83412 | -0.5524 | 0.101 | 0.09 | 1 | HC_vs_MS.Trt_F |
| RABEP1 | 0.83511 | -0.7142 | 0.225 | 0.21 | 1 | HC_vs_MS.Trt_F |
| ZNF830 | 0.83586 | -0.39 | 0.104 | 0.11 | 1 | HC_vs_MS.Trt_F |
| GOSR1 | 0.8359 | -0.3379 | 0.971 | 0.97 | 1 | HC_vs_MS.Trt_F |
| MED4 | 0.83593 | -0.3607 | 0.636 | 0.62 | 1 | HC_vs_MS.Trt_F |
| EAPP | 0.83614 | -0.2568 | 0.225 | 0.23 | 1 | HC_vs_MS.Trt_F |
| TRAT1 | 0.83621 | -0.3926 | 0.249 | 0.24 | 1 | HC_vs_MS.Trt_F |
| NEO1 | 0.83654 | -0.444 | 0.266 | 0.24 | 1 | HC_vs_MS.Trt_F |
| GIN1 | 0.8366 | -1.0796 | 0.379 | 0.37 | 1 | HC_vs_MS.Trt_F |
| NCR3 | 0.8369 | -0.7982 | 0.113 | 0.12 | 1 | HC_vs_MS.Trt_F |
| PRMT2 | 0.83695 | -0.4106 | 0.442 | 0.42 | 1 | HC_vs_MS.Trt_F |
| SETX | 0.83716 | -0.2738 | 0.338 | 0.31 | 1 | HC_vs_MS.Trt_F |
| CYTH2 | 0.83747 | 0.37273 | 0.676 | 0.64 | 1 | HC_vs_MS.Trt_F |
| LY75-CD302 | 0.83748 | 0.67754 | 0.104 | 0.1 | 1 | HC_vs_MS.Trt_F |
| FEZ1 | 0.83756 | 0.35059 | 0.621 | 0.61 | 1 | HC_vs_MS.Trt_F |
| C2orf61 | 0.8376 | -0.854 | 0.15 | 0.14 | 1 | HC_vs_MS.Trt_F |
| CRK | 0.83797 | 0.56077 | 0.639 | 0.63 | 1 | HC_vs_MS.Trt_F |
| PPAN | 0.83797 | 0.41363 | 0.419 | 0.42 | 1 | HC_vs_MS.Trt_F |
| ZFP3 | 0.83933 | 1.53273 | 0.11 | 0.12 | 1 | HC_vs_MS.Trt_F |
| NEMP2 | 0.83949 | -0.4877 | 0.118 | 0.12 | 1 | HC_vs_MS.Trt_F |
| ZFYVE19 | 0.83968 | 0.70536 | 0.113 | 0.11 | 1 | HC_vs_MS.Trt_F |
| SERHL2 | 0.83994 | -0.5948 | 0.367 | 0.34 | 1 | HC_vs_MS.Trt_F |
| ZNF563 | 0.84005 | -1.6277 | 0.555 | 0.55 | 1 | HC_vs_MS.Trt_F |

| | | | | | | |
|---|---|---|---|---|---|---|
| KLHL9 | 0.84055 | 0.49537 | 0.373 | 0.38 | 1 | HC_vs_MS.Trt_F |
| RNF145 | 0.84166 | -0.5804 | 0.191 | 0.19 | 1 | HC_vs_MS.Trt_F |
| DAAM2 | 0.84169 | -0.3871 | 0.468 | 0.42 | 1 | HC_vs_MS.Trt_F |
| ZNF507 | 0.84188 | -0.6719 | 0.197 | 0.18 | 1 | HC_vs_MS.Trt_F |
| CSNK2A1 | 0.8422 | -0.5084 | 0.76 | 0.71 | 1 | HC_vs_MS.Trt_F |
| NHLRC2 | 0.8424 | -0.2959 | 0.798 | 0.76 | 1 | HC_vs_MS.Trt_F |
| FAM110A | 0.84245 | -0.5306 | 0.197 | 0.19 | 1 | HC_vs_MS.Trt_F |
| EBLN2 | 0.84271 | -0.4909 | 0.11 | 0.1 | 1 | HC_vs_MS.Trt_F |
| C21orf59 | 0.84272 | -0.2603 | 0.46 | 0.44 | 1 | HC_vs_MS.Trt_F |
| PLPP3 | 0.84289 | 0.75727 | 0.315 | 0.3 | 1 | HC_vs_MS.Trt_F |
| NADK2 | 0.84291 | -0.9866 | 0.5 | 0.48 | 1 | HC_vs_MS.Trt_F |
| YKT6 | 0.84322 | -0.5829 | 0.124 | 0.12 | 1 | HC_vs_MS.Trt_F |
| HAVCR1 | 0.84381 | 0.71587 | 0.373 | 0.39 | 1 | HC_vs_MS.Trt_F |
| KIAA1549 | 0.84383 | 0.61111 | 0.396 | 0.38 | 1 | HC_vs_MS.Trt_F |
| NAA40 | 0.84416 | -0.433 | 0.561 | 0.56 | 1 | HC_vs_MS.Trt_F |
| TEX14 | 0.84416 | -0.4315 | 0.116 | 0.11 | 1 | HC_vs_MS.Trt_F |
| PLXNC1 | 0.8443 | 0.6926 | 0.095 | 0.1 | 1 | HC_vs_MS.Trt_F |
| MAN2A1 | 0.84487 | -0.4446 | 0.26 | 0.25 | 1 | HC_vs_MS.Trt_F |
| TUFT1 | 0.84496 | -0.2783 | 0.396 | 0.37 | 1 | HC_vs_MS.Trt_F |
| SLC7A1 | 0.84539 | -0.2823 | 0.098 | 0.1 | 1 | HC_vs_MS.Trt_F |
| TTBK2 | 0.84577 | 0.93613 | 0.555 | 0.53 | 1 | HC_vs_MS.Trt_F |
| MYO18A | 0.84663 | 0.29112 | 0.517 | 0.51 | 1 | HC_vs_MS.Trt_F |
| NR6A1 | 0.84692 | -0.6908 | 0.477 | 0.46 | 1 | HC_vs_MS.Trt_F |
| WDR92 | 0.84696 | 0.8012 | 0.743 | 0.71 | 1 | HC_vs_MS.Trt_F |
| TVP23A | 0.84703 | -0.3167 | 0.517 | 0.51 | 1 | HC_vs_MS.Trt_F |
| STAR | 0.84787 | 0.3932 | 0.286 | 0.29 | 1 | HC_vs_MS.Trt_F |
| ASPSCR1 | 0.84799 | -0.2558 | 0.668 | 0.64 | 1 | HC_vs_MS.Trt_F |
| EFCAB10 | 0.848 | 0.37019 | 0.205 | 0.2 | 1 | HC_vs_MS.Trt_F |
| ADPRM | 0.84811 | -0.4201 | 0.098 | 0.1 | 1 | HC_vs_MS.Trt_F |
| USP53 | 0.84824 | -0.6102 | 0.604 | 0.57 | 1 | HC_vs_MS.Trt_F |
| MRPS27 | 0.84843 | 0.73381 | 0.506 | 0.54 | 1 | HC_vs_MS.Trt_F |
| METTL10 | 0.84856 | -0.6752 | 0.324 | 0.32 | 1 | HC_vs_MS.Trt_F |
| TSTD2 | 0.84865 | -0.2544 | 0.121 | 0.13 | 1 | HC_vs_MS.Trt_F |
| PATZ1 | 0.84925 | -0.3844 | 0.118 | 0.11 | 1 | HC_vs_MS.Trt_F |
| ERF | 0.8493 | 0.3884 | 0.11 | 0.1 | 1 | HC_vs_MS.Trt_F |
| SUGP1 | 0.85019 | -0.4258 | 0.486 | 0.46 | 1 | HC_vs_MS.Trt_F |
| MYL6B | 0.85027 | -0.3636 | 0.604 | 0.57 | 1 | HC_vs_MS.Trt_F |
| RCCD1 | 0.85048 | 0.25714 | 0.202 | 0.2 | 1 | HC_vs_MS.Trt_F |
| CTPS1 | 0.85055 | 0.59325 | 0.676 | 0.66 | 1 | HC_vs_MS.Trt_F |
| PAXIP1 | 0.85067 | -0.5906 | 0.474 | 0.46 | 1 | HC_vs_MS.Trt_F |

| | | | | | | |
|---|---|---|---|---|---|---|
| SERPINE2 | 0.85079 | -1.2037 | 0.367 | 0.35 | 1 | HC_vs_MS.Trt_F |
| CYB5R1 | 0.85093 | 0.52562 | 0.104 | 0.11 | 1 | HC_vs_MS.Trt_F |
| TMEM150C | 0.85161 | 0.76598 | 0.569 | 0.59 | 1 | HC_vs_MS.Trt_F |
| MSMO1 | 0.85218 | -0.602 | 0.555 | 0.54 | 1 | HC_vs_MS.Trt_F |
| PIN4 | 0.85238 | -1.1508 | 0.133 | 0.13 | 1 | HC_vs_MS.Trt_F |
| NUDCD2 | 0.85292 | -0.8554 | 0.786 | 0.77 | 1 | HC_vs_MS.Trt_F |
| SNTN | 0.85355 | 0.33227 | 0.344 | 0.35 | 1 | HC_vs_MS.Trt_F |
| SLC25A53 | 0.85409 | 0.53273 | 0.483 | 0.46 | 1 | HC_vs_MS.Trt_F |
| SNX10 | 0.85446 | -0.8858 | 0.142 | 0.13 | 1 | HC_vs_MS.Trt_F |
| AVL9 | 0.85468 | -0.4755 | 0.922 | 0.9 | 1 | HC_vs_MS.Trt_F |
| TSC1 | 0.85469 | 0.46275 | 0.48 | 0.48 | 1 | HC_vs_MS.Trt_F |
| PGS1 | 0.85502 | 0.6926 | 0.633 | 0.65 | 1 | HC_vs_MS.Trt_F |
| SFMBT1 | 0.85532 | -0.6706 | 0.587 | 0.58 | 1 | HC_vs_MS.Trt_F |
| SARM1 | 0.85631 | -0.6307 | 0.422 | 0.42 | 1 | HC_vs_MS.Trt_F |
| TMEM45B | 0.85633 | 0.27155 | 0.604 | 0.59 | 1 | HC_vs_MS.Trt_F |
| TUSC2 | 0.85722 | -0.2615 | 0.113 | 0.12 | 1 | HC_vs_MS.Trt_F |
| MRFAP1 | 0.85744 | -0.4837 | 0.341 | 0.34 | 1 | HC_vs_MS.Trt_F |
| MGME1 | 0.85745 | 0.27803 | 0.118 | 0.11 | 1 | HC_vs_MS.Trt_F |
| NAF1 | 0.8578 | 0.59629 | 0.497 | 0.47 | 1 | HC_vs_MS.Trt_F |
| CHMP1A | 0.85797 | -0.3581 | 0.231 | 0.22 | 1 | HC_vs_MS.Trt_F |
| KNL1 | 0.85813 | 0.60693 | 0.451 | 0.43 | 1 | HC_vs_MS.Trt_F |
| PSMD1 | 0.85862 | -0.6195 | 0.217 | 0.21 | 1 | HC_vs_MS.Trt_F |
| RTKN2 | 0.85868 | -0.5034 | 0.532 | 0.5 | 1 | HC_vs_MS.Trt_F |
| WASF2 | 0.8598 | -0.2991 | 0.569 | 0.5 | 1 | HC_vs_MS.Trt_F |
| CAPZA2 | 0.8601 | -0.4085 | 0.908 | 0.9 | 1 | HC_vs_MS.Trt_F |
| COMMD8 | 0.86031 | -0.2631 | 0.118 | 0.11 | 1 | HC_vs_MS.Trt_F |
| MIS18BP1 | 0.86043 | -0.953 | 0.679 | 0.63 | 1 | HC_vs_MS.Trt_F |
| KSR1 | 0.86053 | 0.33557 | 0.28 | 0.29 | 1 | HC_vs_MS.Trt_F |
| RAD1 | 0.86057 | -0.2862 | 0.717 | 0.7 | 1 | HC_vs_MS.Trt_F |
| G6PC3 | 0.86085 | -0.4683 | 0.85 | 0.86 | 1 | HC_vs_MS.Trt_F |
| METRN | 0.86116 | -1.0646 | 0.121 | 0.12 | 1 | HC_vs_MS.Trt_F |
| PDIA4 | 0.86161 | 0.58674 | 0.101 | 0.1 | 1 | HC_vs_MS.Trt_F |
| CPNE8 | 0.86164 | -0.394 | 0.162 | 0.15 | 1 | HC_vs_MS.Trt_F |
| FMN1 | 0.86242 | 0.92793 | 0.269 | 0.27 | 1 | HC_vs_MS.Trt_F |
| UBE3A | 0.86266 | -0.5677 | 0.598 | 0.57 | 1 | HC_vs_MS.Trt_F |
| ATP1B1 | 0.8632 | 0.27349 | 0.306 | 0.31 | 1 | HC_vs_MS.Trt_F |
| CDK7 | 0.86366 | -0.8816 | 0.22 | 0.22 | 1 | HC_vs_MS.Trt_F |
| MCM3 | 0.86371 | -0.5394 | 0.225 | 0.22 | 1 | HC_vs_MS.Trt_F |
| NBEAL1 | 0.86374 | 0.3472 | 0.899 | 0.86 | 1 | HC_vs_MS.Trt_F |
| GOLGB1 | 0.86378 | -0.4261 | 0.616 | 0.59 | 1 | HC_vs_MS.Trt_F |

| | | | | | | |
|---|---|---|---|---|---|---|
| VIL1 | 0.86409 | 0.79039 | 0.217 | 0.21 | 1 | HC_vs_MS.Trt_F |
| CWC15 | 0.86449 | -0.57 | 0.208 | 0.2 | 1 | HC_vs_MS.Trt_F |
| SLC25A4 | 0.86478 | -0.5629 | 0.318 | 0.31 | 1 | HC_vs_MS.Trt_F |
| MTERF4 | 0.86491 | -0.3665 | 0.431 | 0.39 | 1 | HC_vs_MS.Trt_F |
| THYN1 | 0.86525 | -0.2886 | 0.156 | 0.16 | 1 | HC_vs_MS.Trt_F |
| GIT1 | 0.86538 | -0.3179 | 0.234 | 0.22 | 1 | HC_vs_MS.Trt_F |
| FAM120AOS | 0.86564 | 0.36079 | 0.688 | 0.67 | 1 | HC_vs_MS.Trt_F |
| GMEB2 | 0.8658 | 0.25008 | 0.13 | 0.12 | 1 | HC_vs_MS.Trt_F |
| RAB43 | 0.8666 | 0.7211 | 0.228 | 0.22 | 1 | HC_vs_MS.Trt_F |
| ATP2C1 | 0.86685 | 0.33564 | 0.191 | 0.18 | 1 | HC_vs_MS.Trt_F |
| CAAP1 | 0.86692 | -0.4185 | 0.165 | 0.16 | 1 | HC_vs_MS.Trt_F |
| ZSCAN32 | 0.86806 | -0.3283 | 0.246 | 0.23 | 1 | HC_vs_MS.Trt_F |
| TPRA1 | 0.86881 | 0.36181 | 0.182 | 0.18 | 1 | HC_vs_MS.Trt_F |
| VAMP4 | 0.87003 | -0.3093 | 0.231 | 0.22 | 1 | HC_vs_MS.Trt_F |
| IARS | 0.87122 | -0.5477 | 0.705 | 0.68 | 1 | HC_vs_MS.Trt_F |
| MAP3K7 | 0.87166 | -0.5528 | 0.159 | 0.15 | 1 | HC_vs_MS.Trt_F |
| SLC12A2 | 0.87173 | -0.782 | 0.572 | 0.57 | 1 | HC_vs_MS.Trt_F |
| NOP56 | 0.87229 | -0.5573 | 0.471 | 0.45 | 1 | HC_vs_MS.Trt_F |
| FANK1 | 0.87243 | -0.5774 | 0.211 | 0.19 | 1 | HC_vs_MS.Trt_F |
| APPL1 | 0.8728 | 0.90751 | 0.494 | 0.49 | 1 | HC_vs_MS.Trt_F |
| USP45 | 0.87342 | -0.9296 | 0.179 | 0.18 | 1 | HC_vs_MS.Trt_F |
| ARIH2 | 0.87344 | -0.681 | 0.994 | 0.97 | 1 | HC_vs_MS.Trt_F |
| SOCS7 | 0.87347 | 0.29669 | 0.439 | 0.45 | 1 | HC_vs_MS.Trt_F |
| GM2A | 0.87407 | 0.52598 | 0.552 | 0.57 | 1 | HC_vs_MS.Trt_F |
| SAMM50 | 0.87411 | -0.6616 | 0.465 | 0.45 | 1 | HC_vs_MS.Trt_F |
| ZNF573 | 0.87446 | 0.453 | 0.413 | 0.42 | 1 | HC_vs_MS.Trt_F |
| IMMP1L | 0.87479 | -0.6866 | 0.575 | 0.56 | 1 | HC_vs_MS.Trt_F |
| DYNC1I2 | 0.87593 | -0.732 | 0.231 | 0.21 | 1 | HC_vs_MS.Trt_F |
| COPS7B | 0.87597 | 0.39686 | 0.769 | 0.76 | 1 | HC_vs_MS.Trt_F |
| GALNT7 | 0.87676 | -0.9275 | 0.107 | 0.1 | 1 | HC_vs_MS.Trt_F |
| OSMR | 0.87722 | 0.88923 | 0.584 | 0.57 | 1 | HC_vs_MS.Trt_F |
| RNF43 | 0.87727 | -0.277 | 0.35 | 0.32 | 1 | HC_vs_MS.Trt_F |
| GABPA | 0.87735 | 0.77535 | 0.113 | 0.11 | 1 | HC_vs_MS.Trt_F |
| EIF3A | 0.87774 | -0.2674 | 0.448 | 0.44 | 1 | HC_vs_MS.Trt_F |
| ZNF425 | 0.87795 | -0.5635 | 0.445 | 0.44 | 1 | HC_vs_MS.Trt_F |
| CSGALNACT | 0.87838 | -0.2937 | 0.538 | 0.51 | 1 | HC_vs_MS.Trt_F |
| GCDH | 0.87867 | -0.3491 | 0.197 | 0.19 | 1 | HC_vs_MS.Trt_F |
| COA1 | 0.87888 | -0.3205 | 0.772 | 0.75 | 1 | HC_vs_MS.Trt_F |
| GTF2E2 | 0.87907 | -0.2726 | 0.639 | 0.6 | 1 | HC_vs_MS.Trt_F |
| C3orf67 | 0.87959 | -0.7954 | 0.439 | 0.44 | 1 | HC_vs_MS.Trt_F |

| | | | | | | |
|---|---|---|---|---|---|---|
| GLCE | 0.88042 | -0.6381 | 0.165 | 0.16 | 1 | HC_vs_MS.Trt_F |
| AD000671.6 | 0.88076 | -0.274 | 0.13 | 0.13 | 1 | HC_vs_MS.Trt_F |
| TMEM71 | 0.881 | 0.32081 | 0.116 | 0.12 | 1 | HC_vs_MS.Trt_F |
| ATAD1 | 0.88191 | -0.2963 | 0.173 | 0.18 | 1 | HC_vs_MS.Trt_F |
| MPHOSPH6 | 0.88202 | -0.3622 | 0.145 | 0.14 | 1 | HC_vs_MS.Trt_F |
| SUPT16H | 0.88203 | -0.3125 | 0.731 | 0.69 | 1 | HC_vs_MS.Trt_F |
| VPS41 | 0.88223 | -0.2845 | 0.517 | 0.5 | 1 | HC_vs_MS.Trt_F |
| SMARCE1 | 0.88301 | -0.6357 | 0.523 | 0.49 | 1 | HC_vs_MS.Trt_F |
| RIMKLB | 0.88326 | -0.844 | 0.118 | 0.12 | 1 | HC_vs_MS.Trt_F |
| NPTXR | 0.88403 | 0.90346 | 0.355 | 0.32 | 1 | HC_vs_MS.Trt_F |
| LIN7B | 0.88405 | -0.2932 | 0.338 | 0.33 | 1 | HC_vs_MS.Trt_F |
| ZNF213 | 0.88416 | -0.8806 | 0.523 | 0.5 | 1 | HC_vs_MS.Trt_F |
| TES | 0.88426 | -0.6078 | 0.87 | 0.81 | 1 | HC_vs_MS.Trt_F |
| CSNK1G2 | 0.88438 | -0.4555 | 0.133 | 0.12 | 1 | HC_vs_MS.Trt_F |
| TOP3B | 0.8844 | 0.36552 | 0.15 | 0.15 | 1 | HC_vs_MS.Trt_F |
| PDK2 | 0.88445 | -0.7302 | 0.292 | 0.28 | 1 | HC_vs_MS.Trt_F |
| HNRNPUL2 | 0.88466 | 0.57225 | 0.124 | 0.13 | 1 | HC_vs_MS.Trt_F |
| BRWD1 | 0.88497 | -0.4945 | 0.85 | 0.77 | 1 | HC_vs_MS.Trt_F |
| C8orf44-SGK | 0.88535 | 0.45844 | 0.101 | 0.1 | 1 | HC_vs_MS.Trt_F |
| DGUOK | 0.8863 | -0.4202 | 0.217 | 0.22 | 1 | HC_vs_MS.Trt_F |
| FAM134B | 0.88689 | 0.27259 | 0.113 | 0.12 | 1 | HC_vs_MS.Trt_F |
| UNK | 0.88774 | 0.55271 | 0.127 | 0.12 | 1 | HC_vs_MS.Trt_F |
| COPS6 | 0.88803 | -0.2793 | 0.708 | 0.72 | 1 | HC_vs_MS.Trt_F |
| NUAK2 | 0.88832 | 0.63562 | 0.347 | 0.33 | 1 | HC_vs_MS.Trt_F |
| IL1RL1 | 0.88833 | 1.46852 | 0.124 | 0.12 | 1 | HC_vs_MS.Trt_F |
| FAM65C | 0.88877 | 0.38043 | 0.41 | 0.39 | 1 | HC_vs_MS.Trt_F |
| CAMK2D | 0.88936 | -0.4365 | 0.225 | 0.23 | 1 | HC_vs_MS.Trt_F |
| MTPAP | 0.8896 | -0.5451 | 0.873 | 0.83 | 1 | HC_vs_MS.Trt_F |
| TIMELESS | 0.88966 | -0.4489 | 0.393 | 0.37 | 1 | HC_vs_MS.Trt_F |
| PCMTD1 | 0.8898 | -0.7694 | 0.312 | 0.3 | 1 | HC_vs_MS.Trt_F |
| ATMIN | 0.88986 | -0.6039 | 0.28 | 0.27 | 1 | HC_vs_MS.Trt_F |
| ANAPC11 | 0.89006 | -0.7519 | 0.422 | 0.4 | 1 | HC_vs_MS.Trt_F |
| BLOC1S3 | 0.89051 | -0.6989 | 0.65 | 0.6 | 1 | HC_vs_MS.Trt_F |
| KIAA1614 | 0.89115 | 0.46518 | 0.335 | 0.3 | 1 | HC_vs_MS.Trt_F |
| MAD2L1 | 0.89146 | -0.4221 | 0.477 | 0.45 | 1 | HC_vs_MS.Trt_F |
| TRAFD1 | 0.89153 | -0.3151 | 0.243 | 0.23 | 1 | HC_vs_MS.Trt_F |
| SAT2 | 0.89174 | 0.37889 | 0.127 | 0.12 | 1 | HC_vs_MS.Trt_F |
| THUMPD1 | 0.89194 | -0.393 | 0.806 | 0.79 | 1 | HC_vs_MS.Trt_F |
| RB1CC1 | 0.89219 | -0.3587 | 0.306 | 0.28 | 1 | HC_vs_MS.Trt_F |
| HDAC1 | 0.89223 | -0.2547 | 0.292 | 0.29 | 1 | HC_vs_MS.Trt_F |

| | | | | | | |
|---|---|---|---|---|---|---|
| NDUFB10 | 0.89229 | -0.6509 | 0.26 | 0.25 | 1 | HC_vs_MS.Trt_F |
| RGMB | 0.89295 | -0.5988 | 0.116 | 0.11 | 1 | HC_vs_MS.Trt_F |
| ZKSCAN1 | 0.89296 | -0.2547 | 0.974 | 0.96 | 1 | HC_vs_MS.Trt_F |
| KLHDC10 | 0.89299 | -0.3707 | 0.436 | 0.41 | 1 | HC_vs_MS.Trt_F |
| NAPB | 0.89344 | -1.0948 | 0.448 | 0.44 | 1 | HC_vs_MS.Trt_F |
| AHI1 | 0.89383 | 0.46805 | 0.428 | 0.43 | 1 | HC_vs_MS.Trt_F |
| PCIF1 | 0.89402 | -0.3599 | 0.347 | 0.32 | 1 | HC_vs_MS.Trt_F |
| MSS51 | 0.89457 | 0.59009 | 0.581 | 0.54 | 1 | HC_vs_MS.Trt_F |
| SPRYD7 | 0.89473 | -0.2747 | 0.728 | 0.69 | 1 | HC_vs_MS.Trt_F |
| TYMS | 0.89506 | -0.8174 | 0.251 | 0.24 | 1 | HC_vs_MS.Trt_F |
| HLA-DRB3 | 0.89537 | -0.5336 | 0.188 | 0.18 | 1 | HC_vs_MS.Trt_F |
| NEIL2 | 0.89548 | -0.3489 | 0.124 | 0.12 | 1 | HC_vs_MS.Trt_F |
| FAM229B | 0.89596 | 0.25419 | 0.376 | 0.35 | 1 | HC_vs_MS.Trt_F |
| CWC25 | 0.89596 | -0.5941 | 0.384 | 0.4 | 1 | HC_vs_MS.Trt_F |
| PTCD2 | 0.89607 | -0.8201 | 0.61 | 0.61 | 1 | HC_vs_MS.Trt_F |
| OSTC | 0.89621 | -0.3898 | 0.266 | 0.27 | 1 | HC_vs_MS.Trt_F |
| OVCA2 | 0.89631 | -0.8355 | 0.101 | 0.1 | 1 | HC_vs_MS.Trt_F |
| C10orf95 | 0.8965 | -0.4752 | 0.185 | 0.17 | 1 | HC_vs_MS.Trt_F |
| HINT2 | 0.89673 | -0.2506 | 0.107 | 0.1 | 1 | HC_vs_MS.Trt_F |
| SDC4 | 0.89701 | -0.7184 | 0.171 | 0.16 | 1 | HC_vs_MS.Trt_F |
| VSIG10 | 0.89743 | 0.25754 | 0.725 | 0.7 | 1 | HC_vs_MS.Trt_F |
| METTL15 | 0.89753 | 0.30611 | 0.396 | 0.41 | 1 | HC_vs_MS.Trt_F |
| RNF181 | 0.89768 | -0.5483 | 0.173 | 0.17 | 1 | HC_vs_MS.Trt_F |
| RSAD1 | 0.89812 | 0.38323 | 0.113 | 0.11 | 1 | HC_vs_MS.Trt_F |
| KLF10 | 0.89828 | -0.3284 | 0.098 | 0.1 | 1 | HC_vs_MS.Trt_F |
| UBA3 | 0.8984 | -0.3568 | 0.416 | 0.42 | 1 | HC_vs_MS.Trt_F |
| IGHMBP2 | 0.89873 | -0.7164 | 0.402 | 0.39 | 1 | HC_vs_MS.Trt_F |
| CD164 | 0.89885 | -0.3734 | 0.598 | 0.57 | 1 | HC_vs_MS.Trt_F |
| METTL17 | 0.89923 | -0.3883 | 0.685 | 0.63 | 1 | HC_vs_MS.Trt_F |
| GSTM1 | 0.8993 | -0.5568 | 0.194 | 0.18 | 1 | HC_vs_MS.Trt_F |
| PUF60 | 0.90006 | -0.2819 | 0.24 | 0.22 | 1 | HC_vs_MS.Trt_F |
| CDKN2AIP | 0.90059 | -0.5033 | 0.133 | 0.13 | 1 | HC_vs_MS.Trt_F |
| FBXL4 | 0.90074 | -0.4319 | 0.546 | 0.53 | 1 | HC_vs_MS.Trt_F |
| ADAMTSL4 | 0.9008 | -0.323 | 0.243 | 0.25 | 1 | HC_vs_MS.Trt_F |
| RFXANK | 0.901 | -0.9553 | 0.26 | 0.25 | 1 | HC_vs_MS.Trt_F |
| CCND3 | 0.90216 | -0.3096 | 0.575 | 0.54 | 1 | HC_vs_MS.Trt_F |
| PTPN18 | 0.90222 | -0.4192 | 0.434 | 0.41 | 1 | HC_vs_MS.Trt_F |
| UBE2O | 0.90243 | 0.54187 | 0.497 | 0.48 | 1 | HC_vs_MS.Trt_F |
| MTUS1 | 0.90264 | 0.45554 | 0.41 | 0.39 | 1 | HC_vs_MS.Trt_F |
| RALY | 0.90282 | -0.3013 | 0.37 | 0.37 | 1 | HC_vs_MS.Trt_F |

| | | | | | | |
|---|---|---|---|---|---|---|
| DNA2 | 0.90305 | -0.3356 | 0.231 | 0.23 | 1 | HC_vs_MS.Trt_F |
| SLCO2B1 | 0.90321 | 0.61676 | 0.379 | 0.36 | 1 | HC_vs_MS.Trt_F |
| NMI | 0.90416 | 0.28489 | 0.309 | 0.31 | 1 | HC_vs_MS.Trt_F |
| CHPT1 | 0.90418 | -0.4152 | 0.642 | 0.63 | 1 | HC_vs_MS.Trt_F |
| SPAG16 | 0.90444 | 0.5502 | 0.494 | 0.47 | 1 | HC_vs_MS.Trt_F |
| ZNF530 | 0.90554 | 0.64939 | 0.509 | 0.51 | 1 | HC_vs_MS.Trt_F |
| C3orf38 | 0.90575 | 0.63034 | 0.104 | 0.1 | 1 | HC_vs_MS.Trt_F |
| BTN3A1 | 0.90579 | -0.2985 | 0.61 | 0.64 | 1 | HC_vs_MS.Trt_F |
| TOLLIP | 0.90622 | -0.7314 | 0.142 | 0.13 | 1 | HC_vs_MS.Trt_F |
| CCDC59 | 0.90632 | -0.3644 | 0.162 | 0.16 | 1 | HC_vs_MS.Trt_F |
| NKIRAS1 | 0.90634 | -0.2543 | 0.298 | 0.29 | 1 | HC_vs_MS.Trt_F |
| PRIMPOL | 0.90722 | 0.48896 | 0.11 | 0.11 | 1 | HC_vs_MS.Trt_F |
| TRAPPC2L | 0.90749 | -0.5367 | 0.136 | 0.13 | 1 | HC_vs_MS.Trt_F |
| STX1B | 0.90776 | 0.52942 | 0.298 | 0.3 | 1 | HC_vs_MS.Trt_F |
| ZNF658 | 0.90802 | 0.62528 | 0.22 | 0.22 | 1 | HC_vs_MS.Trt_F |
| UXS1 | 0.90803 | -0.3313 | 0.254 | 0.25 | 1 | HC_vs_MS.Trt_F |
| MRPL19 | 0.90816 | -0.6763 | 0.283 | 0.27 | 1 | HC_vs_MS.Trt_F |
| VPS33A | 0.90944 | -0.7331 | 0.876 | 0.88 | 1 | HC_vs_MS.Trt_F |
| PSMD4 | 0.90971 | -0.5043 | 0.694 | 0.68 | 1 | HC_vs_MS.Trt_F |
| COQ10A | 0.90985 | 0.77704 | 0.549 | 0.54 | 1 | HC_vs_MS.Trt_F |
| TAF9 | 0.91012 | -0.2896 | 0.399 | 0.39 | 1 | HC_vs_MS.Trt_F |
| MRPL41 | 0.9104 | -0.5603 | 0.182 | 0.18 | 1 | HC_vs_MS.Trt_F |
| VDAC2 | 0.91045 | -0.2784 | 0.283 | 0.27 | 1 | HC_vs_MS.Trt_F |
| ZNRD1 | 0.91081 | -0.2722 | 0.153 | 0.15 | 1 | HC_vs_MS.Trt_F |
| MCU | 0.91197 | 0.68915 | 0.257 | 0.27 | 1 | HC_vs_MS.Trt_F |
| GOLGA3 | 0.91236 | -0.4842 | 0.798 | 0.8 | 1 | HC_vs_MS.Trt_F |
| MKRN3 | 0.91245 | 0.6996 | 0.587 | 0.6 | 1 | HC_vs_MS.Trt_F |
| CAPN15 | 0.91251 | 1.0075 | 0.205 | 0.21 | 1 | HC_vs_MS.Trt_F |
| RAB4A | 0.91255 | -0.7883 | 0.118 | 0.12 | 1 | HC_vs_MS.Trt_F |
| ZNF625 | 0.91298 | -0.307 | 0.289 | 0.28 | 1 | HC_vs_MS.Trt_F |
| TRPT1 | 0.91425 | 0.32927 | 0.118 | 0.12 | 1 | HC_vs_MS.Trt_F |
| MAD2L1BP | 0.91503 | -1.1248 | 0.191 | 0.18 | 1 | HC_vs_MS.Trt_F |
| MARK4 | 0.91696 | 0.25223 | 0.578 | 0.56 | 1 | HC_vs_MS.Trt_F |
| C15orf41 | 0.91722 | -0.4305 | 0.645 | 0.6 | 1 | HC_vs_MS.Trt_F |
| HTRA2 | 0.91734 | -0.5524 | 0.139 | 0.13 | 1 | HC_vs_MS.Trt_F |
| STRADB | 0.91738 | -0.5632 | 0.101 | 0.1 | 1 | HC_vs_MS.Trt_F |
| MAPK1 | 0.91745 | -0.4838 | 0.592 | 0.54 | 1 | HC_vs_MS.Trt_F |
| TCEB2 | 0.91774 | -0.5134 | 0.35 | 0.32 | 1 | HC_vs_MS.Trt_F |
| ZFYVE26 | 0.91804 | 0.54415 | 0.876 | 0.87 | 1 | HC_vs_MS.Trt_F |
| ZFAND2A | 0.91823 | -0.4004 | 0.121 | 0.12 | 1 | HC_vs_MS.Trt_F |

| | | | | | | |
|---|---|---|---|---|---|---|
| PHAX | 0.91875 | -0.2513 | 0.893 | 0.86 | 1 | HC_vs_MS.Trt_F |
| OPA1 | 0.91879 | -0.3545 | 0.156 | 0.16 | 1 | HC_vs_MS.Trt_F |
| ZNF862 | 0.91883 | -0.503 | 0.145 | 0.14 | 1 | HC_vs_MS.Trt_F |
| COA5 | 0.91929 | -0.5739 | 0.52 | 0.46 | 1 | HC_vs_MS.Trt_F |
| EMC7 | 0.92023 | -0.5908 | 0.22 | 0.21 | 1 | HC_vs_MS.Trt_F |
| RAB4B | 0.92035 | -0.4101 | 0.384 | 0.35 | 1 | HC_vs_MS.Trt_F |
| VEPH1 | 0.9207 | 0.36174 | 0.113 | 0.12 | 1 | HC_vs_MS.Trt_F |
| STK11IP | 0.9208 | -0.2612 | 0.101 | 0.1 | 1 | HC_vs_MS.Trt_F |
| IMPDH2 | 0.92105 | -0.3597 | 0.225 | 0.22 | 1 | HC_vs_MS.Trt_F |
| ADRM1 | 0.92116 | -0.3216 | 0.295 | 0.27 | 1 | HC_vs_MS.Trt_F |
| WDR7 | 0.92135 | -1.4352 | 0.373 | 0.36 | 1 | HC_vs_MS.Trt_F |
| ZNF224 | 0.92136 | -0.4418 | 0.451 | 0.45 | 1 | HC_vs_MS.Trt_F |
| SAMD3 | 0.92223 | -0.2509 | 0.295 | 0.28 | 1 | HC_vs_MS.Trt_F |
| TRAPPC4 | 0.92289 | -0.4509 | 0.936 | 0.9 | 1 | HC_vs_MS.Trt_F |
| ANKRD55 | 0.9233 | -0.3844 | 0.182 | 0.17 | 1 | HC_vs_MS.Trt_F |
| CEP128 | 0.92335 | -0.3219 | 0.202 | 0.2 | 1 | HC_vs_MS.Trt_F |
| HIST2H4B | 0.92357 | 0.64055 | 0.176 | 0.17 | 1 | HC_vs_MS.Trt_F |
| LINC00116 | 0.92385 | -0.6275 | 0.176 | 0.17 | 1 | HC_vs_MS.Trt_F |
| HELLS | 0.92419 | -0.5683 | 0.841 | 0.78 | 1 | HC_vs_MS.Trt_F |
| MYC | 0.92429 | -0.8642 | 0.168 | 0.16 | 1 | HC_vs_MS.Trt_F |
| ACTN1 | 0.92479 | -1.4149 | 0.653 | 0.6 | 1 | HC_vs_MS.Trt_F |
| PCGF1 | 0.92489 | -0.3308 | 0.402 | 0.4 | 1 | HC_vs_MS.Trt_F |
| OAZ2 | 0.92521 | -0.6006 | 0.815 | 0.78 | 1 | HC_vs_MS.Trt_F |
| RP11-458D2 | 0.92574 | -0.4551 | 0.127 | 0.13 | 1 | HC_vs_MS.Trt_F |
| XBP1 | 0.92656 | -0.3259 | 0.188 | 0.18 | 1 | HC_vs_MS.Trt_F |
| GLRA3 | 0.92778 | 0.27553 | 0.306 | 0.32 | 1 | HC_vs_MS.Trt_F |
| ACRC | 0.928 | -0.6628 | 0.454 | 0.44 | 1 | HC_vs_MS.Trt_F |
| TRIAP1 | 0.92811 | -0.3753 | 0.11 | 0.1 | 1 | HC_vs_MS.Trt_F |
| PDPN | 0.92868 | 0.53994 | 0.514 | 0.5 | 1 | HC_vs_MS.Trt_F |
| GID4 | 0.92942 | 0.30336 | 0.702 | 0.67 | 1 | HC_vs_MS.Trt_F |
| GLT8D1 | 0.93022 | -0.8604 | 0.312 | 0.3 | 1 | HC_vs_MS.Trt_F |
| NSUN6 | 0.93034 | -0.338 | 0.728 | 0.68 | 1 | HC_vs_MS.Trt_F |
| SRGAP3 | 0.93038 | 1.18726 | 0.879 | 0.84 | 1 | HC_vs_MS.Trt_F |
| SETD6 | 0.93045 | 0.31845 | 0.462 | 0.46 | 1 | HC_vs_MS.Trt_F |
| DCAF12 | 0.93057 | -0.4405 | 0.179 | 0.17 | 1 | HC_vs_MS.Trt_F |
| ZFAT | 0.93068 | 1.18924 | 0.338 | 0.33 | 1 | HC_vs_MS.Trt_F |
| CEPT1 | 0.93105 | -0.8201 | 0.361 | 0.34 | 1 | HC_vs_MS.Trt_F |
| BABAM1 | 0.93108 | -0.4669 | 0.63 | 0.62 | 1 | HC_vs_MS.Trt_F |
| SART3 | 0.93132 | -0.3797 | 0.185 | 0.18 | 1 | HC_vs_MS.Trt_F |
| EDEM1 | 0.93167 | -0.5364 | 0.335 | 0.32 | 1 | HC_vs_MS.Trt_F |

| | | | | | | |
|---|---|---|---|---|---|---|
| SYNJ2BP | 0.93175 | -0.2681 | 0.867 | 0.79 | 1 | HC_vs_MS.Trt_F |
| SFR1 | 0.93226 | 0.40526 | 0.292 | 0.3 | 1 | HC_vs_MS.Trt_F |
| TBC1D14 | 0.93239 | -0.3357 | 0.185 | 0.18 | 1 | HC_vs_MS.Trt_F |
| NUCB1 | 0.93289 | -0.3211 | 0.283 | 0.27 | 1 | HC_vs_MS.Trt_F |
| LLGL1 | 0.93334 | 0.71976 | 0.301 | 0.3 | 1 | HC_vs_MS.Trt_F |
| ROCK2 | 0.93378 | 0.88839 | 0.471 | 0.46 | 1 | HC_vs_MS.Trt_F |
| THAP11 | 0.93495 | -0.5228 | 0.101 | 0.1 | 1 | HC_vs_MS.Trt_F |
| EXT1 | 0.93499 | 0.68635 | 0.199 | 0.2 | 1 | HC_vs_MS.Trt_F |
| TIMP2 | 0.93512 | 1.03805 | 0.202 | 0.21 | 1 | HC_vs_MS.Trt_F |
| TP53INP1 | 0.93543 | -0.429 | 0.355 | 0.35 | 1 | HC_vs_MS.Trt_F |
| RRBP1 | 0.9356 | 0.52811 | 0.318 | 0.32 | 1 | HC_vs_MS.Trt_F |
| SNAPIN | 0.93561 | 0.30073 | 0.147 | 0.15 | 1 | HC_vs_MS.Trt_F |
| GPAT4 | 0.93576 | 0.31836 | 0.783 | 0.73 | 1 | HC_vs_MS.Trt_F |
| SSBP1 | 0.93636 | -0.3063 | 0.699 | 0.69 | 1 | HC_vs_MS.Trt_F |
| SNRPA | 0.93688 | -0.4835 | 0.298 | 0.28 | 1 | HC_vs_MS.Trt_F |
| ATP10D | 0.93712 | 0.5269 | 0.405 | 0.39 | 1 | HC_vs_MS.Trt_F |
| SGCB | 0.93727 | -0.5234 | 0.315 | 0.3 | 1 | HC_vs_MS.Trt_F |
| MCRIP1 | 0.93803 | -0.3984 | 0.41 | 0.41 | 1 | HC_vs_MS.Trt_F |
| ZC3H6 | 0.93831 | -0.5929 | 0.85 | 0.8 | 1 | HC_vs_MS.Trt_F |
| NDUFB4 | 0.93835 | -0.7513 | 0.254 | 0.24 | 1 | HC_vs_MS.Trt_F |
| CCT7 | 0.93841 | -0.2762 | 0.26 | 0.26 | 1 | HC_vs_MS.Trt_F |
| ZNF33A | 0.93886 | -0.504 | 0.584 | 0.55 | 1 | HC_vs_MS.Trt_F |
| N4BP2L1 | 0.93887 | -0.4829 | 0.653 | 0.65 | 1 | HC_vs_MS.Trt_F |
| COMMD10 | 0.93901 | -0.2662 | 0.341 | 0.32 | 1 | HC_vs_MS.Trt_F |
| BIRC2 | 0.93924 | -0.8417 | 0.202 | 0.19 | 1 | HC_vs_MS.Trt_F |
| NPIPB11 | 0.93943 | -0.4553 | 0.555 | 0.52 | 1 | HC_vs_MS.Trt_F |
| NECAP2 | 0.94073 | -0.5431 | 0.483 | 0.46 | 1 | HC_vs_MS.Trt_F |
| CNKSR1 | 0.94076 | 0.60786 | 0.176 | 0.17 | 1 | HC_vs_MS.Trt_F |
| NDUFB5 | 0.94102 | -0.3727 | 0.702 | 0.69 | 1 | HC_vs_MS.Trt_F |
| SCAF11 | 0.9413 | -0.6734 | 0.899 | 0.87 | 1 | HC_vs_MS.Trt_F |
| U2AF1 | 0.94143 | -0.3915 | 0.532 | 0.51 | 1 | HC_vs_MS.Trt_F |
| FCHSD1 | 0.94156 | -0.4155 | 0.327 | 0.3 | 1 | HC_vs_MS.Trt_F |
| COQ3 | 0.94222 | 0.29558 | 0.251 | 0.25 | 1 | HC_vs_MS.Trt_F |
| SRD5A3 | 0.9423 | -0.8327 | 0.413 | 0.39 | 1 | HC_vs_MS.Trt_F |
| CEBPZ | 0.9423 | -0.5816 | 0.246 | 0.24 | 1 | HC_vs_MS.Trt_F |
| CUTC | 0.94255 | -0.2761 | 0.162 | 0.16 | 1 | HC_vs_MS.Trt_F |
| TMBIM4 | 0.94281 | -0.3449 | 0.853 | 0.82 | 1 | HC_vs_MS.Trt_F |
| MTERF1 | 0.94315 | -1.1049 | 0.254 | 0.24 | 1 | HC_vs_MS.Trt_F |
| MZB1 | 0.94356 | -0.4661 | 0.656 | 0.63 | 1 | HC_vs_MS.Trt_F |
| CNGB1 | 0.94398 | 0.31554 | 0.338 | 0.33 | 1 | HC_vs_MS.Trt_F |

| | | | | | | |
|---|---|---|---|---|---|---|
| DAPP1 | 0.94458 | -0.5734 | 0.251 | 0.25 | 1 | HC_vs_MS.Trt_F |
| NR2C2 | 0.94461 | -0.6128 | 0.656 | 0.62 | 1 | HC_vs_MS.Trt_F |
| TMEM248 | 0.94503 | -0.338 | 0.22 | 0.21 | 1 | HC_vs_MS.Trt_F |
| ISOC1 | 0.94513 | -0.5953 | 0.124 | 0.12 | 1 | HC_vs_MS.Trt_F |
| OSTM1 | 0.94541 | -0.4156 | 0.795 | 0.82 | 1 | HC_vs_MS.Trt_F |
| PDE3B | 0.94561 | -0.6647 | 0.613 | 0.55 | 1 | HC_vs_MS.Trt_F |
| RGS13 | 0.94589 | -0.3415 | 0.442 | 0.4 | 1 | HC_vs_MS.Trt_F |
| PAXBP1 | 0.94594 | -0.9489 | 0.223 | 0.21 | 1 | HC_vs_MS.Trt_F |
| SSR1 | 0.94635 | -0.487 | 0.934 | 0.92 | 1 | HC_vs_MS.Trt_F |
| LGALS4 | 0.94695 | -0.5008 | 0.613 | 0.57 | 1 | HC_vs_MS.Trt_F |
| DHFR | 0.94704 | -0.2923 | 0.639 | 0.61 | 1 | HC_vs_MS.Trt_F |
| TPK1 | 0.94818 | 0.70634 | 0.396 | 0.39 | 1 | HC_vs_MS.Trt_F |
| ANKRD16 | 0.94842 | -0.8221 | 0.471 | 0.46 | 1 | HC_vs_MS.Trt_F |
| PLOD1 | 0.94861 | -1.2311 | 0.691 | 0.68 | 1 | HC_vs_MS.Trt_F |
| SLC31A2 | 0.94869 | -1.2542 | 0.13 | 0.13 | 1 | HC_vs_MS.Trt_F |
| MRPL53 | 0.94876 | -0.4378 | 0.133 | 0.13 | 1 | HC_vs_MS.Trt_F |
| SMYD2 | 0.94887 | -0.2785 | 0.156 | 0.16 | 1 | HC_vs_MS.Trt_F |
| HAUS4 | 0.94899 | -1.1871 | 0.156 | 0.15 | 1 | HC_vs_MS.Trt_F |
| BOLA2 | 0.94902 | -0.2539 | 0.145 | 0.14 | 1 | HC_vs_MS.Trt_F |
| KIF1BP | 0.94904 | -0.465 | 0.711 | 0.72 | 1 | HC_vs_MS.Trt_F |
| ASNS | 0.94947 | 0.70169 | 0.283 | 0.28 | 1 | HC_vs_MS.Trt_F |
| PAFAH1B2 | 0.9495 | -0.7172 | 0.91 | 0.82 | 1 | HC_vs_MS.Trt_F |
| HERPUD1 | 0.94981 | -0.394 | 0.746 | 0.75 | 1 | HC_vs_MS.Trt_F |
| GRPEL2 | 0.95081 | -0.6161 | 0.275 | 0.27 | 1 | HC_vs_MS.Trt_F |
| POC5 | 0.95123 | -0.582 | 0.113 | 0.11 | 1 | HC_vs_MS.Trt_F |
| COX17 | 0.95126 | -0.3089 | 0.335 | 0.32 | 1 | HC_vs_MS.Trt_F |
| RAE1 | 0.9522 | -0.3124 | 0.139 | 0.13 | 1 | HC_vs_MS.Trt_F |
| ACP1 | 0.95296 | -0.2982 | 0.266 | 0.26 | 1 | HC_vs_MS.Trt_F |
| TOPORS-AS1 | 0.95407 | -0.6938 | 0.116 | 0.12 | 1 | HC_vs_MS.Trt_F |
| DYX1C1 | 0.95482 | -0.307 | 0.529 | 0.52 | 1 | HC_vs_MS.Trt_F |
| ADTRP | 0.95526 | -0.2904 | 0.37 | 0.34 | 1 | HC_vs_MS.Trt_F |
| ABCA5 | 0.95559 | -0.6896 | 0.653 | 0.62 | 1 | HC_vs_MS.Trt_F |
| KDELR1 | 0.95585 | -0.5046 | 0.266 | 0.25 | 1 | HC_vs_MS.Trt_F |
| RANBP10 | 0.95602 | 0.2949 | 0.159 | 0.16 | 1 | HC_vs_MS.Trt_F |
| FOXP1 | 0.95613 | -0.269 | 0.89 | 0.86 | 1 | HC_vs_MS.Trt_F |
| MZF1 | 0.9566 | 0.40328 | 0.133 | 0.13 | 1 | HC_vs_MS.Trt_F |
| ZNF813 | 0.95687 | -0.3151 | 0.763 | 0.77 | 1 | HC_vs_MS.Trt_F |
| MAP2K1 | 0.95725 | -0.5165 | 0.153 | 0.15 | 1 | HC_vs_MS.Trt_F |
| PCID2 | 0.95733 | -0.4384 | 0.361 | 0.35 | 1 | HC_vs_MS.Trt_F |
| POLR2M | 0.95832 | -0.7379 | 0.165 | 0.16 | 1 | HC_vs_MS.Trt_F |

| | | | | | | |
|---|---|---|---|---|---|---|
| ZNF248 | 0.95902 | 0.29401 | 0.188 | 0.18 | 1 | HC_vs_MS.Trt_F |
| UBR2 | 0.95936 | 0.28142 | 0.549 | 0.54 | 1 | HC_vs_MS.Trt_F |
| ZNF654 | 0.95937 | -0.8138 | 0.145 | 0.14 | 1 | HC_vs_MS.Trt_F |
| SLC25A11 | 0.95957 | -0.4826 | 0.194 | 0.18 | 1 | HC_vs_MS.Trt_F |
| BOLA3 | 0.95987 | -0.472 | 0.54 | 0.52 | 1 | HC_vs_MS.Trt_F |
| MTFR1L | 0.96001 | -0.2964 | 0.578 | 0.57 | 1 | HC_vs_MS.Trt_F |
| IFI30 | 0.96006 | -0.3953 | 0.364 | 0.35 | 1 | HC_vs_MS.Trt_F |
| HS3ST1 | 0.96017 | -0.4363 | 0.338 | 0.33 | 1 | HC_vs_MS.Trt_F |
| DR1 | 0.96044 | -0.5253 | 0.827 | 0.8 | 1 | HC_vs_MS.Trt_F |
| PPP2R3B | 0.96099 | 0.40666 | 0.558 | 0.53 | 1 | HC_vs_MS.Trt_F |
| PAFAH2 | 0.96106 | -0.5723 | 0.566 | 0.54 | 1 | HC_vs_MS.Trt_F |
| GALNT1 | 0.96284 | -0.6477 | 0.101 | 0.1 | 1 | HC_vs_MS.Trt_F |
| BARD1 | 0.96327 | -0.501 | 0.159 | 0.15 | 1 | HC_vs_MS.Trt_F |
| SETD3 | 0.9634 | 0.33132 | 0.121 | 0.12 | 1 | HC_vs_MS.Trt_F |
| AGGF1 | 0.9636 | -0.4483 | 0.165 | 0.16 | 1 | HC_vs_MS.Trt_F |
| NR1H3 | 0.9639 | 1.12376 | 0.15 | 0.15 | 1 | HC_vs_MS.Trt_F |
| HSPA14 | 0.96399 | -0.4841 | 0.373 | 0.37 | 1 | HC_vs_MS.Trt_F |
| FARP2 | 0.96441 | -1.3568 | 0.376 | 0.35 | 1 | HC_vs_MS.Trt_F |
| DHTKD1 | 0.96441 | -0.5112 | 0.858 | 0.83 | 1 | HC_vs_MS.Trt_F |
| CTC-487M2[illegible] | 0.96469 | 0.51579 | 0.91 | 0.94 | 1 | HC_vs_MS.Trt_F |
| UQCR10 | 0.96498 | -0.537 | 0.364 | 0.34 | 1 | HC_vs_MS.Trt_F |
| SCRN2 | 0.96575 | -1.076 | 0.136 | 0.13 | 1 | HC_vs_MS.Trt_F |
| MOAP1 | 0.96577 | -0.5525 | 0.413 | 0.39 | 1 | HC_vs_MS.Trt_F |
| FAM124B | 0.96634 | -1.1365 | 0.118 | 0.12 | 1 | HC_vs_MS.Trt_F |
| SMYD4 | 0.96672 | -0.8193 | 0.855 | 0.81 | 1 | HC_vs_MS.Trt_F |
| ANKRD18A | 0.96699 | 0.55369 | 0.28 | 0.27 | 1 | HC_vs_MS.Trt_F |
| DNAH7 | 0.96779 | 0.3233 | 0.694 | 0.67 | 1 | HC_vs_MS.Trt_F |
| SEC61A2 | 0.96792 | -0.7279 | 0.113 | 0.11 | 1 | HC_vs_MS.Trt_F |
| GUCD1 | 0.96806 | -0.9516 | 0.512 | 0.48 | 1 | HC_vs_MS.Trt_F |
| ST20 | 0.96818 | 0.62886 | 0.254 | 0.25 | 1 | HC_vs_MS.Trt_F |
| ETAA1 | 0.96835 | -0.4956 | 0.15 | 0.15 | 1 | HC_vs_MS.Trt_F |
| DERA | 0.96877 | -0.9728 | 0.113 | 0.11 | 1 | HC_vs_MS.Trt_F |
| ATP2A3 | 0.96903 | -0.284 | 0.338 | 0.33 | 1 | HC_vs_MS.Trt_F |
| PIGA | 0.96912 | -0.3164 | 0.488 | 0.46 | 1 | HC_vs_MS.Trt_F |
| DOCK6 | 0.96916 | 0.89011 | 0.552 | 0.57 | 1 | HC_vs_MS.Trt_F |
| BTN2A2 | 0.97089 | 0.59185 | 0.101 | 0.1 | 1 | HC_vs_MS.Trt_F |
| CEP120 | 0.97112 | -0.6602 | 0.173 | 0.17 | 1 | HC_vs_MS.Trt_F |
| CLTB | 0.97148 | 0.42969 | 0.107 | 0.11 | 1 | HC_vs_MS.Trt_F |
| SP110 | 0.97213 | -0.3158 | 0.705 | 0.68 | 1 | HC_vs_MS.Trt_F |
| WDR47 | 0.97287 | -0.4957 | 0.231 | 0.23 | 1 | HC_vs_MS.Trt_F |

| | | | | | | |
|---|---|---|---|---|---|---|
| PRKRA | 0.97318 | -0.434 | 0.272 | 0.26 | 1 | HC_vs_MS.Trt_F |
| CBFB | 0.97326 | -0.2962 | 0.182 | 0.17 | 1 | HC_vs_MS.Trt_F |
| GORASP2 | 0.97357 | -0.4944 | 0.228 | 0.22 | 1 | HC_vs_MS.Trt_F |
| TXNDC9 | 0.97396 | -0.5191 | 0.142 | 0.14 | 1 | HC_vs_MS.Trt_F |
| RTEL1 | 0.97398 | 1.1277 | 0.116 | 0.12 | 1 | HC_vs_MS.Trt_F |
| WNT10A | 0.97431 | 0.72831 | 0.159 | 0.16 | 1 | HC_vs_MS.Trt_F |
| CRNKL1 | 0.97479 | -0.3888 | 0.116 | 0.11 | 1 | HC_vs_MS.Trt_F |
| WDR83 | 0.9749 | 0.50799 | 0.607 | 0.58 | 1 | HC_vs_MS.Trt_F |
| PALB2 | 0.97506 | -1.0918 | 0.254 | 0.24 | 1 | HC_vs_MS.Trt_F |
| ZNF622 | 0.97561 | -0.3708 | 0.147 | 0.14 | 1 | HC_vs_MS.Trt_F |
| RINT1 | 0.97686 | -0.8074 | 0.569 | 0.55 | 1 | HC_vs_MS.Trt_F |
| EXOC5 | 0.9769 | -0.2828 | 0.454 | 0.44 | 1 | HC_vs_MS.Trt_F |
| AGPS | 0.97693 | 0.35687 | 0.162 | 0.16 | 1 | HC_vs_MS.Trt_F |
| CEP135 | 0.97761 | 0.3288 | 0.824 | 0.8 | 1 | HC_vs_MS.Trt_F |
| SEH1L | 0.97787 | -0.3126 | 0.922 | 0.91 | 1 | HC_vs_MS.Trt_F |
| ZNF317 | 0.97788 | -0.5511 | 0.139 | 0.13 | 1 | HC_vs_MS.Trt_F |
| SPPL3 | 0.97803 | 0.62848 | 0.61 | 0.59 | 1 | HC_vs_MS.Trt_F |
| MRPL36 | 0.97829 | -0.9702 | 0.118 | 0.12 | 1 | HC_vs_MS.Trt_F |
| RP11-599B1 | 0.97895 | -0.2758 | 0.228 | 0.22 | 1 | HC_vs_MS.Trt_F |
| SLC30A7 | 0.97923 | -0.7219 | 0.734 | 0.68 | 1 | HC_vs_MS.Trt_F |
| JOSD2 | 0.97924 | -0.3098 | 0.11 | 0.11 | 1 | HC_vs_MS.Trt_F |
| F2R | 0.97952 | -0.5655 | 0.578 | 0.56 | 1 | HC_vs_MS.Trt_F |
| VTA1 | 0.98009 | -0.8802 | 0.425 | 0.41 | 1 | HC_vs_MS.Trt_F |
| RGS16 | 0.98015 | 0.39965 | 0.113 | 0.11 | 1 | HC_vs_MS.Trt_F |
| UQCC2 | 0.98028 | -0.6853 | 0.116 | 0.11 | 1 | HC_vs_MS.Trt_F |
| NDUFAF1 | 0.98063 | -0.271 | 0.454 | 0.45 | 1 | HC_vs_MS.Trt_F |
| ZHX2 | 0.98067 | -0.309 | 0.139 | 0.14 | 1 | HC_vs_MS.Trt_F |
| C9orf64 | 0.98072 | -0.2816 | 0.592 | 0.58 | 1 | HC_vs_MS.Trt_F |
| NOG | 0.98107 | -1.242 | 0.139 | 0.13 | 1 | HC_vs_MS.Trt_F |
| OTUD7A | 0.98149 | -0.5398 | 0.159 | 0.16 | 1 | HC_vs_MS.Trt_F |
| PSMA5 | 0.98158 | -0.3483 | 0.303 | 0.3 | 1 | HC_vs_MS.Trt_F |
| CLU | 0.98242 | -0.437 | 0.561 | 0.55 | 1 | HC_vs_MS.Trt_F |
| SNX3 | 0.98289 | -0.4142 | 0.182 | 0.17 | 1 | HC_vs_MS.Trt_F |
| PDCD6 | 0.98334 | -0.5088 | 0.725 | 0.67 | 1 | HC_vs_MS.Trt_F |
| MAMLD1 | 0.98361 | -0.4984 | 0.194 | 0.19 | 1 | HC_vs_MS.Trt_F |
| MRPS14 | 0.98405 | -0.5553 | 0.176 | 0.17 | 1 | HC_vs_MS.Trt_F |
| EPPK1 | 0.98432 | 0.2544 | 0.101 | 0.1 | 1 | HC_vs_MS.Trt_F |
| ATG16L1 | 0.98438 | -0.5101 | 0.179 | 0.17 | 1 | HC_vs_MS.Trt_F |
| SHTN1 | 0.98475 | -1.3596 | 0.118 | 0.12 | 1 | HC_vs_MS.Trt_F |
| PRKCE | 0.98479 | 0.98863 | 0.171 | 0.17 | 1 | HC_vs_MS.Trt_F |

| | | | | | | |
|---|---|---|---|---|---|---|
| MMACHC | 0.98536 | 0.38122 | 0.916 | 0.91 | 1 | HC_vs_MS.Trt_F |
| BSCL2 | 0.9855 | 0.466 | 0.202 | 0.2 | 1 | HC_vs_MS.Trt_F |
| ZNF180 | 0.98599 | -1.0478 | 0.355 | 0.35 | 1 | HC_vs_MS.Trt_F |
| MTIF3 | 0.98723 | -0.7356 | 0.24 | 0.22 | 1 | HC_vs_MS.Trt_F |
| TLK1 | 0.98726 | -0.4908 | 0.74 | 0.79 | 1 | HC_vs_MS.Trt_F |
| GCM1 | 0.9874 | -0.7443 | 0.616 | 0.58 | 1 | HC_vs_MS.Trt_F |
| ARSK | 0.98744 | 0.56991 | 0.604 | 0.59 | 1 | HC_vs_MS.Trt_F |
| ZNF829 | 0.98768 | -0.4211 | 0.896 | 0.88 | 1 | HC_vs_MS.Trt_F |
| CHMP4A | 0.98784 | -0.6019 | 0.636 | 0.57 | 1 | HC_vs_MS.Trt_F |
| EXOSC1 | 0.98804 | 0.37162 | 0.558 | 0.56 | 1 | HC_vs_MS.Trt_F |
| ZNF69 | 0.98827 | 0.68977 | 0.454 | 0.45 | 1 | HC_vs_MS.Trt_F |
| AP3S1 | 0.98913 | -0.4627 | 0.413 | 0.39 | 1 | HC_vs_MS.Trt_F |
| ZNF264 | 0.9893 | -0.2719 | 0.957 | 0.95 | 1 | HC_vs_MS.Trt_F |
| GFI1B | 0.98941 | 0.37285 | 0.581 | 0.58 | 1 | HC_vs_MS.Trt_F |
| THAP1 | 0.9895 | -1.0989 | 0.104 | 0.1 | 1 | HC_vs_MS.Trt_F |
| ZNF517 | 0.98955 | -0.3311 | 0.431 | 0.44 | 1 | HC_vs_MS.Trt_F |
| MLST8 | 0.9897 | 0.78385 | 0.434 | 0.42 | 1 | HC_vs_MS.Trt_F |
| EVI5L | 0.99023 | -0.3352 | 0.318 | 0.31 | 1 | HC_vs_MS.Trt_F |
| FAM217B | 0.99028 | -0.3282 | 0.399 | 0.39 | 1 | HC_vs_MS.Trt_F |
| UBE2Z | 0.9904 | -0.3441 | 0.286 | 0.29 | 1 | HC_vs_MS.Trt_F |
| FAM200A | 0.99052 | 1.29135 | 0.246 | 0.24 | 1 | HC_vs_MS.Trt_F |
| GPR107 | 0.9907 | -0.335 | 0.208 | 0.21 | 1 | HC_vs_MS.Trt_F |
| EIF4B | 0.99082 | -0.3106 | 0.743 | 0.73 | 1 | HC_vs_MS.Trt_F |
| TTI1 | 0.99122 | -0.6904 | 0.249 | 0.23 | 1 | HC_vs_MS.Trt_F |
| KLHDC3 | 0.99131 | -0.4104 | 0.173 | 0.17 | 1 | HC_vs_MS.Trt_F |
| DAPK3 | 0.99159 | -0.4332 | 0.116 | 0.11 | 1 | HC_vs_MS.Trt_F |
| ITIH4 | 0.99161 | -0.5705 | 0.26 | 0.25 | 1 | HC_vs_MS.Trt_F |
| TRMT61B | 0.9918 | -0.3226 | 0.133 | 0.13 | 1 | HC_vs_MS.Trt_F |
| MSL3 | 0.99183 | -0.9576 | 0.188 | 0.18 | 1 | HC_vs_MS.Trt_F |
| ZSWIM7 | 0.99265 | -0.5043 | 0.824 | 0.78 | 1 | HC_vs_MS.Trt_F |
| XRN2 | 0.9937 | -0.2962 | 0.231 | 0.23 | 1 | HC_vs_MS.Trt_F |
| RP4-583P15 | 0.99424 | 0.43177 | 0.15 | 0.15 | 1 | HC_vs_MS.Trt_F |
| TLR5 | 0.99486 | 0.66237 | 0.636 | 0.62 | 1 | HC_vs_MS.Trt_F |
| ULK2 | 0.99549 | -0.6516 | 0.725 | 0.73 | 1 | HC_vs_MS.Trt_F |
| TCP1 | 0.99583 | -0.5585 | 0.428 | 0.41 | 1 | HC_vs_MS.Trt_F |
| LILRB5 | 0.99589 | 0.51748 | 0.379 | 0.37 | 1 | HC_vs_MS.Trt_F |
| ZNF384 | 0.99593 | -0.2603 | 0.147 | 0.14 | 1 | HC_vs_MS.Trt_F |
| DEAF1 | 0.99744 | 0.60306 | 0.329 | 0.33 | 1 | HC_vs_MS.Trt_F |
| RP11-2C24.9 | 0.99785 | -0.6219 | 0.13 | 0.13 | 1 | HC_vs_MS.Trt_F |
| PPP1R3E | 0.99788 | 1.42894 | 0.246 | 0.25 | 1 | HC_vs_MS.Trt_F |

| | | | | | | |
|---|---|---|---|---|---|---|
| NQO1 | 0.99796 | -0.5486 | 0.598 | 0.58 | 1 | HC_vs_MS.Trt_F |
| USP28 | 0.99799 | 0.29461 | 0.41 | 0.39 | 1 | HC_vs_MS.Trt_F |
| ZBTB11 | 0.99906 | -0.5324 | 0.529 | 0.53 | 1 | HC_vs_MS.Trt_F |
| GMFB | 0.99938 | -0.2543 | 0.124 | 0.12 | 1 | HC_vs_MS.Trt_F |
| ZFP30 | 0.99946 | -0.5154 | 0.844 | 0.84 | 1 | HC_vs_MS.Trt_F |
| CEP126 | 0.99964 | 0.27453 | 0.783 | 0.73 | 1 | HC_vs_MS.Trt_F |
| TUBE1 | 0.99981 | -0.4368 | 0.514 | 0.51 | 1 | HC_vs_MS.Trt_F |
| TAC4 | 2.84E-23 | 0.70161 | 0.722 | 0.49 | 4.53E-19 | HC_vs_MS.Trt_B |
| MT-CO2 | 1.25E-22 | -0.8549 | 0.981 | 0.98 | 1.99E-18 | HC_vs_MS.Trt_B |
| SENP7 | 1.58E-21 | 0.42843 | 0.987 | 0.87 | 2.53E-17 | HC_vs_MS.Trt_B |
| PPDPF | 2.00E-21 | 1.01843 | 0.433 | 0.19 | 3.20E-17 | HC_vs_MS.Trt_B |
| ANKRD30BL | 2.19E-21 | 0.64247 | 0.584 | 0.31 | 3.50E-17 | HC_vs_MS.Trt_B |
| FAM227A | 1.17E-20 | 0.5069 | 0.909 | 0.74 | 1.87E-16 | HC_vs_MS.Trt_B |
| RPL3 | 1.38E-20 | -0.5348 | 0.977 | 1 | 2.21E-16 | HC_vs_MS.Trt_B |
| SLC35E3 | 1.91E-19 | 0.32053 | 0.999 | 0.98 | 3.05E-15 | HC_vs_MS.Trt_B |
| RASGRP3 | 3.24E-19 | 0.38254 | 0.991 | 0.96 | 5.18E-15 | HC_vs_MS.Trt_B |
| SFT2D2 | 3.17E-18 | 0.32297 | 0.98 | 0.88 | 5.07E-14 | HC_vs_MS.Trt_B |
| PTPN3 | 2.21E-17 | 0.29246 | 0.803 | 0.59 | 3.52E-13 | HC_vs_MS.Trt_B |
| C4orf36 | 3.01E-17 | 0.47991 | 0.838 | 0.64 | 4.80E-13 | HC_vs_MS.Trt_B |
| LRRC57 | 5.43E-17 | 0.26916 | 0.97 | 0.9 | 8.67E-13 | HC_vs_MS.Trt_B |
| CYSLTR1 | 6.28E-17 | 0.43061 | 0.946 | 0.81 | 1.00E-12 | HC_vs_MS.Trt_B |
| MGAT4C | 7.38E-17 | 0.4914 | 0.777 | 0.58 | 1.18E-12 | HC_vs_MS.Trt_B |
| MORC4 | 7.94E-17 | 0.3162 | 0.991 | 0.95 | 1.27E-12 | HC_vs_MS.Trt_B |
| ITGB3BP | 8.73E-17 | 0.51959 | 0.784 | 0.59 | 1.39E-12 | HC_vs_MS.Trt_B |
| ZNF283 | 8.77E-17 | 0.37171 | 0.984 | 0.95 | 1.40E-12 | HC_vs_MS.Trt_B |
| ACOT9 | 8.96E-17 | 1.04822 | 0.929 | 0.8 | 1.43E-12 | HC_vs_MS.Trt_B |
| TMSB4X | 1.59E-16 | -0.359 | 0.993 | 0.99 | 2.54E-12 | HC_vs_MS.Trt_B |
| SAMD12 | 2.74E-16 | 0.37605 | 0.972 | 0.87 | 4.37E-12 | HC_vs_MS.Trt_B |
| C5AR2 | 3.16E-16 | 0.33969 | 0.941 | 0.85 | 5.05E-12 | HC_vs_MS.Trt_B |
| CACNA1A | 5.76E-16 | 0.32439 | 1 | 0.99 | 9.20E-12 | HC_vs_MS.Trt_B |
| RGPD5 | 6.02E-16 | 0.68931 | 0.816 | 0.65 | 9.61E-12 | HC_vs_MS.Trt_B |
| PBX1 | 6.05E-16 | 0.56354 | 0.891 | 0.73 | 9.66E-12 | HC_vs_MS.Trt_B |
| PLPP2 | 6.09E-16 | 0.40308 | 0.891 | 0.75 | 9.72E-12 | HC_vs_MS.Trt_B |
| LPP | 7.05E-16 | 0.70811 | 0.971 | 0.87 | 1.13E-11 | HC_vs_MS.Trt_B |
| GIGYF2 | 8.28E-16 | 0.58848 | 0.91 | 0.8 | 1.32E-11 | HC_vs_MS.Trt_B |
| ZFP14 | 1.66E-15 | 0.77942 | 0.857 | 0.69 | 2.65E-11 | HC_vs_MS.Trt_B |
| TMEM45A | 1.71E-15 | 0.27303 | 0.996 | 0.95 | 2.72E-11 | HC_vs_MS.Trt_B |
| RPS14 | 1.93E-15 | -0.7435 | 0.959 | 0.95 | 3.08E-11 | HC_vs_MS.Trt_B |
| NDUFS1 | 2.55E-15 | 0.36875 | 0.952 | 0.83 | 4.07E-11 | HC_vs_MS.Trt_B |
| NUPL2 | 4.67E-15 | 0.42717 | 0.962 | 0.88 | 7.46E-11 | HC_vs_MS.Trt_B |

| | | | | | | |
|---|---|---|---|---|---|---|
| SRD5A1 | 5.37E-15 | 0.5226 | 0.701 | 0.47 | 8.58E-11 | HC_vs_MS.Trt_B |
| STARD8 | 5.38E-15 | 0.48656 | 0.69 | 0.49 | 8.60E-11 | HC_vs_MS.Trt_B |
| CHST12 | 5.79E-15 | 0.28461 | 0.99 | 0.96 | 9.25E-11 | HC_vs_MS.Trt_B |
| ZNF844 | 6.55E-15 | 0.41406 | 0.904 | 0.8 | 1.05E-10 | HC_vs_MS.Trt_B |
| ZNF445 | 1.09E-14 | 0.25168 | 0.978 | 0.91 | 1.74E-10 | HC_vs_MS.Trt_B |
| C12orf50 | 1.16E-14 | 0.40759 | 0.893 | 0.77 | 1.86E-10 | HC_vs_MS.Trt_B |
| C4orf32 | 1.55E-14 | 0.28303 | 0.986 | 0.94 | 2.48E-10 | HC_vs_MS.Trt_B |
| ZNF675 | 1.68E-14 | -0.6538 | 0.922 | 0.8 | 2.69E-10 | HC_vs_MS.Trt_B |
| ABI2 | 1.69E-14 | 0.38419 | 0.857 | 0.69 | 2.69E-10 | HC_vs_MS.Trt_B |
| CCDC125 | 2.12E-14 | 0.40569 | 0.936 | 0.8 | 3.39E-10 | HC_vs_MS.Trt_B |
| AKT2 | 2.69E-14 | 0.59721 | 0.946 | 0.88 | 4.29E-10 | HC_vs_MS.Trt_B |
| ZNF382 | 3.01E-14 | 0.48878 | 0.926 | 0.84 | 4.81E-10 | HC_vs_MS.Trt_B |
| FLAD1 | 3.34E-14 | 0.3782 | 0.67 | 0.46 | 5.33E-10 | HC_vs_MS.Trt_B |
| SLFN12L | 3.36E-14 | 0.34118 | 0.835 | 0.66 | 5.37E-10 | HC_vs_MS.Trt_B |
| EPN1 | 3.71E-14 | 0.42708 | 0.949 | 0.88 | 5.92E-10 | HC_vs_MS.Trt_B |
| TMEM106B | 4.04E-14 | 0.33969 | 0.888 | 0.73 | 6.45E-10 | HC_vs_MS.Trt_B |
| CYP1A2 | 4.06E-14 | 0.43178 | 0.864 | 0.71 | 6.48E-10 | HC_vs_MS.Trt_B |
| FTO | 4.49E-14 | 0.29945 | 0.99 | 0.95 | 7.17E-10 | HC_vs_MS.Trt_B |
| ITGA11 | 4.55E-14 | 0.47304 | 0.619 | 0.41 | 7.26E-10 | HC_vs_MS.Trt_B |
| EEF2 | 4.70E-14 | 0.43864 | 0.894 | 0.77 | 7.51E-10 | HC_vs_MS.Trt_B |
| WISP2 | 5.33E-14 | 0.44744 | 0.606 | 0.39 | 8.51E-10 | HC_vs_MS.Trt_B |
| CAMK2G | 5.42E-14 | 0.46275 | 0.641 | 0.4 | 8.65E-10 | HC_vs_MS.Trt_B |
| REL | 5.59E-14 | 0.45529 | 0.954 | 0.91 | 8.93E-10 | HC_vs_MS.Trt_B |
| SULT1B1 | 6.62E-14 | 0.40282 | 0.959 | 0.85 | 1.06E-09 | HC_vs_MS.Trt_B |
| NPIPB4 | 8.21E-14 | 0.39806 | 0.951 | 0.85 | 1.31E-09 | HC_vs_MS.Trt_B |
| NOS1 | 9.60E-14 | 0.36716 | 0.719 | 0.52 | 1.53E-09 | HC_vs_MS.Trt_B |
| CATSPERG | 1.02E-13 | 0.28272 | 0.893 | 0.77 | 1.63E-09 | HC_vs_MS.Trt_B |
| TBC1D8B | 1.45E-13 | 0.26694 | 0.957 | 0.81 | 2.32E-09 | HC_vs_MS.Trt_B |
| LILRA2 | 1.69E-13 | 0.3843 | 0.787 | 0.62 | 2.69E-09 | HC_vs_MS.Trt_B |
| CD151 | 1.79E-13 | 0.34866 | 0.91 | 0.77 | 2.86E-09 | HC_vs_MS.Trt_B |
| KDM2A | 2.02E-13 | 0.427 | 0.907 | 0.78 | 3.22E-09 | HC_vs_MS.Trt_B |
| RAB12 | 2.11E-13 | 2.45134 | 0.362 | 0.2 | 3.37E-09 | HC_vs_MS.Trt_B |
| UVSSA | 2.44E-13 | 1.00119 | 0.746 | 0.54 | 3.89E-09 | HC_vs_MS.Trt_B |
| GCNT3 | 2.60E-13 | 0.40893 | 0.877 | 0.74 | 4.15E-09 | HC_vs_MS.Trt_B |
| FBXL18 | 2.77E-13 | 0.41215 | 0.984 | 0.92 | 4.42E-09 | HC_vs_MS.Trt_B |
| IKZF3 | 2.92E-13 | 0.38345 | 0.972 | 0.86 | 4.66E-09 | HC_vs_MS.Trt_B |
| SGK3 | 3.29E-13 | 0.28196 | 0.906 | 0.77 | 5.26E-09 | HC_vs_MS.Trt_B |
| FAM131A | 3.51E-13 | 0.68253 | 0.806 | 0.6 | 5.60E-09 | HC_vs_MS.Trt_B |
| ZNF396 | 3.88E-13 | 0.4256 | 0.867 | 0.68 | 6.20E-09 | HC_vs_MS.Trt_B |
| P3H1 | 3.94E-13 | 0.65601 | 0.945 | 0.83 | 6.30E-09 | HC_vs_MS.Trt_B |

| | | | | | | |
|---|---|---|---|---|---|---|
| DTWD2 | 4.13E-13 | 0.59684 | 0.743 | 0.56 | 6.59E-09 | HC_vs_MS.Trt_B |
| ZNF785 | 4.13E-13 | 0.66234 | 0.906 | 0.77 | 6.59E-09 | HC_vs_MS.Trt_B |
| CLPB | 4.46E-13 | 0.33977 | 0.957 | 0.88 | 7.13E-09 | HC_vs_MS.Trt_B |
| PRPF3 | 4.80E-13 | 0.55896 | 0.859 | 0.74 | 7.66E-09 | HC_vs_MS.Trt_B |
| TTC5 | 5.35E-13 | 0.67542 | 0.601 | 0.41 | 8.55E-09 | HC_vs_MS.Trt_B |
| CHFR | 6.51E-13 | 0.32847 | 0.936 | 0.82 | 1.04E-08 | HC_vs_MS.Trt_B |
| LGMN | 6.85E-13 | 0.68621 | 0.958 | 0.9 | 1.09E-08 | HC_vs_MS.Trt_B |
| CFAP54 | 7.22E-13 | 0.30469 | 0.939 | 0.8 | 1.15E-08 | HC_vs_MS.Trt_B |
| C5AR1 | 7.41E-13 | 0.32733 | 0.959 | 0.85 | 1.18E-08 | HC_vs_MS.Trt_B |
| LETMD1 | 7.43E-13 | 0.32523 | 0.838 | 0.67 | 1.19E-08 | HC_vs_MS.Trt_B |
| AACS | 7.67E-13 | 0.3909 | 0.986 | 0.93 | 1.23E-08 | HC_vs_MS.Trt_B |
| FAM83F | 9.08E-13 | 0.377 | 0.87 | 0.73 | 1.45E-08 | HC_vs_MS.Trt_B |
| TMEM41B | 9.69E-13 | 0.672 | 0.806 | 0.66 | 1.55E-08 | HC_vs_MS.Trt_B |
| EP300 | 9.84E-13 | 0.31649 | 0.707 | 0.51 | 1.57E-08 | HC_vs_MS.Trt_B |
| ZNF567 | 9.93E-13 | 0.45958 | 0.868 | 0.69 | 1.59E-08 | HC_vs_MS.Trt_B |
| HELB | 1.03E-12 | 0.50959 | 0.899 | 0.77 | 1.64E-08 | HC_vs_MS.Trt_B |
| SLC25A16 | 1.14E-12 | 0.33514 | 0.957 | 0.89 | 1.82E-08 | HC_vs_MS.Trt_B |
| RPL37A | 1.18E-12 | 0.30932 | 0.817 | 0.6 | 1.89E-08 | HC_vs_MS.Trt_B |
| MRI1 | 1.30E-12 | 0.65951 | 0.719 | 0.57 | 2.07E-08 | HC_vs_MS.Trt_B |
| ASB18 | 1.73E-12 | 0.52574 | 0.567 | 0.38 | 2.77E-08 | HC_vs_MS.Trt_B |
| FLYWCH1 | 1.75E-12 | 0.44186 | 0.787 | 0.58 | 2.79E-08 | HC_vs_MS.Trt_B |
| PHLDB1 | 1.75E-12 | 0.34603 | 0.942 | 0.88 | 2.80E-08 | HC_vs_MS.Trt_B |
| TRIM22 | 1.83E-12 | 0.3106 | 0.951 | 0.8 | 2.92E-08 | HC_vs_MS.Trt_B |
| SMCHD1 | 1.85E-12 | 0.33481 | 0.778 | 0.58 | 2.95E-08 | HC_vs_MS.Trt_B |
| ZNF329 | 1.85E-12 | 0.26326 | 0.716 | 0.56 | 2.96E-08 | HC_vs_MS.Trt_B |
| TOR3A | 2.05E-12 | 0.51416 | 0.828 | 0.72 | 3.27E-08 | HC_vs_MS.Trt_B |
| MLLT6 | 2.81E-12 | 0.5214 | 0.91 | 0.82 | 4.48E-08 | HC_vs_MS.Trt_B |
| NPLOC4 | 2.85E-12 | 0.34253 | 0.952 | 0.85 | 4.55E-08 | HC_vs_MS.Trt_B |
| SHISA9 | 2.90E-12 | 0.31 | 0.925 | 0.83 | 4.64E-08 | HC_vs_MS.Trt_B |
| MARVELD3 | 3.17E-12 | 0.31661 | 0.867 | 0.72 | 5.07E-08 | HC_vs_MS.Trt_B |
| COQ4 | 3.18E-12 | 0.30155 | 0.894 | 0.76 | 5.07E-08 | HC_vs_MS.Trt_B |
| MPV17L | 3.35E-12 | 0.32396 | 0.975 | 0.93 | 5.36E-08 | HC_vs_MS.Trt_B |
| PAK2 | 3.38E-12 | 0.30091 | 0.745 | 0.54 | 5.40E-08 | HC_vs_MS.Trt_B |
| DSTYK | 3.40E-12 | 0.79459 | 0.816 | 0.66 | 5.42E-08 | HC_vs_MS.Trt_B |
| ZFP36L2 | 3.58E-12 | 0.50791 | 0.533 | 0.33 | 5.72E-08 | HC_vs_MS.Trt_B |
| TMEM212 | 3.76E-12 | 0.31425 | 0.942 | 0.83 | 6.01E-08 | HC_vs_MS.Trt_B |
| RNF115 | 3.86E-12 | 0.32852 | 0.964 | 0.87 | 6.17E-08 | HC_vs_MS.Trt_B |
| QPRT | 4.48E-12 | 0.33988 | 0.92 | 0.81 | 7.15E-08 | HC_vs_MS.Trt_B |
| DAGLB | 4.88E-12 | 0.37412 | 0.828 | 0.65 | 7.79E-08 | HC_vs_MS.Trt_B |
| FAM122C | 6.11E-12 | 0.33358 | 0.901 | 0.8 | 9.76E-08 | HC_vs_MS.Trt_B |

| | | | | | | |
|---|---|---|---|---|---|---|
| BAG6 | 6.25E-12 | 0.29383 | 0.622 | 0.44 | 9.99E-08 | HC_vs_MS.Trt_B |
| IQCC | 6.36E-12 | -0.4099 | 0.655 | 0.43 | 1.02E-07 | HC_vs_MS.Trt_B |
| CNKSR3 | 6.52E-12 | 0.29572 | 0.975 | 0.92 | 1.04E-07 | HC_vs_MS.Trt_B |
| FAXDC2 | 6.59E-12 | 0.5812 | 0.839 | 0.71 | 1.05E-07 | HC_vs_MS.Trt_B |
| NAE1 | 6.63E-12 | 0.27003 | 0.761 | 0.57 | 1.06E-07 | HC_vs_MS.Trt_B |
| ALDH7A1 | 7.29E-12 | 0.33143 | 0.991 | 0.94 | 1.16E-07 | HC_vs_MS.Trt_B |
| LRCH3 | 7.32E-12 | 0.51112 | 0.97 | 0.92 | 1.17E-07 | HC_vs_MS.Trt_B |
| CEP112 | 7.71E-12 | 0.31935 | 0.933 | 0.83 | 1.23E-07 | HC_vs_MS.Trt_B |
| SUPV3L1 | 8.04E-12 | 0.26465 | 0.688 | 0.49 | 1.28E-07 | HC_vs_MS.Trt_B |
| ALG9 | 8.04E-12 | 0.51026 | 0.875 | 0.71 | 1.28E-07 | HC_vs_MS.Trt_B |
| NUCB2 | 8.79E-12 | 0.28784 | 0.971 | 0.89 | 1.40E-07 | HC_vs_MS.Trt_B |
| NUPR1 | 8.89E-12 | 0.27365 | 0.941 | 0.86 | 1.42E-07 | HC_vs_MS.Trt_B |
| NARF | 8.98E-12 | 0.32901 | 0.959 | 0.87 | 1.43E-07 | HC_vs_MS.Trt_B |
| TXNDC15 | 8.98E-12 | 0.4881 | 0.687 | 0.51 | 1.43E-07 | HC_vs_MS.Trt_B |
| FLRT2 | 9.38E-12 | 0.35452 | 0.92 | 0.85 | 1.50E-07 | HC_vs_MS.Trt_B |
| ZNF26 | 1.00E-11 | 0.29471 | 0.996 | 0.95 | 1.60E-07 | HC_vs_MS.Trt_B |
| CRLF3 | 1.01E-11 | 0.4532 | 0.933 | 0.78 | 1.61E-07 | HC_vs_MS.Trt_B |
| FKBP14 | 1.01E-11 | 0.54696 | 0.629 | 0.44 | 1.62E-07 | HC_vs_MS.Trt_B |
| FUT2 | 1.05E-11 | 0.41232 | 0.741 | 0.56 | 1.68E-07 | HC_vs_MS.Trt_B |
| LYN | 1.15E-11 | 0.29119 | 0.587 | 0.39 | 1.84E-07 | HC_vs_MS.Trt_B |
| AP1S3 | 1.31E-11 | 0.36371 | 0.904 | 0.79 | 2.10E-07 | HC_vs_MS.Trt_B |
| PSMA6 | 1.44E-11 | 0.27605 | 0.93 | 0.78 | 2.29E-07 | HC_vs_MS.Trt_B |
| CORO7 | 1.49E-11 | 0.3442 | 0.858 | 0.7 | 2.38E-07 | HC_vs_MS.Trt_B |
| C1orf56 | 1.57E-11 | -0.8451 | 0.872 | 0.91 | 2.51E-07 | HC_vs_MS.Trt_B |
| NPIPB5 | 1.59E-11 | 0.30751 | 0.926 | 0.78 | 2.54E-07 | HC_vs_MS.Trt_B |
| NCBP3 | 1.73E-11 | 0.38908 | 0.877 | 0.73 | 2.77E-07 | HC_vs_MS.Trt_B |
| NDRG2 | 1.76E-11 | 0.27828 | 0.851 | 0.72 | 2.81E-07 | HC_vs_MS.Trt_B |
| ATP10B | 1.84E-11 | 0.41366 | 0.716 | 0.54 | 2.94E-07 | HC_vs_MS.Trt_B |
| RPL7 | 1.97E-11 | -0.4567 | 0.926 | 0.9 | 3.15E-07 | HC_vs_MS.Trt_B |
| GTSE1 | 1.99E-11 | 0.29916 | 0.568 | 0.36 | 3.18E-07 | HC_vs_MS.Trt_B |
| MYLK3 | 2.00E-11 | 0.27858 | 0.922 | 0.81 | 3.19E-07 | HC_vs_MS.Trt_B |
| FECH | 2.00E-11 | 0.5834 | 0.696 | 0.51 | 3.20E-07 | HC_vs_MS.Trt_B |
| ARL17A | 2.03E-11 | 0.76624 | 0.691 | 0.5 | 3.24E-07 | HC_vs_MS.Trt_B |
| TMEM181 | 2.15E-11 | 0.30231 | 0.504 | 0.29 | 3.44E-07 | HC_vs_MS.Trt_B |
| CNR2 | 2.22E-11 | 0.40606 | 0.799 | 0.65 | 3.55E-07 | HC_vs_MS.Trt_B |
| OPA3 | 2.22E-11 | 0.51739 | 0.978 | 0.9 | 3.55E-07 | HC_vs_MS.Trt_B |
| TMEM63A | 2.52E-11 | 0.42117 | 0.784 | 0.6 | 4.03E-07 | HC_vs_MS.Trt_B |
| SZT2 | 2.72E-11 | 0.5528 | 0.849 | 0.68 | 4.34E-07 | HC_vs_MS.Trt_B |
| SWSAP1 | 2.90E-11 | 0.28038 | 0.72 | 0.56 | 4.63E-07 | HC_vs_MS.Trt_B |
| FAM63B | 2.91E-11 | 0.90374 | 0.796 | 0.62 | 4.65E-07 | HC_vs_MS.Trt_B |

| | | | | | | |
|---|---|---|---|---|---|---|
| PRELP | 2.94E-11 | 0.25282 | 0.738 | 0.53 | 4.70E-07 | HC_vs_MS.Trt_B |
| PLA2G7 | 3.06E-11 | 0.82743 | 0.594 | 0.41 | 4.89E-07 | HC_vs_MS.Trt_B |
| EMC1 | 3.14E-11 | 0.5723 | 0.814 | 0.65 | 5.02E-07 | HC_vs_MS.Trt_B |
| TTYH1 | 3.20E-11 | -0.316 | 0.625 | 0.45 | 5.10E-07 | HC_vs_MS.Trt_B |
| RGPD6 | 3.73E-11 | 0.4908 | 0.588 | 0.38 | 5.97E-07 | HC_vs_MS.Trt_B |
| HOOK3 | 3.84E-11 | 0.55239 | 0.935 | 0.85 | 6.14E-07 | HC_vs_MS.Trt_B |
| CLN5 | 3.92E-11 | 0.25877 | 0.962 | 0.86 | 6.26E-07 | HC_vs_MS.Trt_B |
| CLSPN | 3.97E-11 | 0.29409 | 0.854 | 0.72 | 6.34E-07 | HC_vs_MS.Trt_B |
| MED28 | 4.10E-11 | 0.50825 | 0.855 | 0.74 | 6.55E-07 | HC_vs_MS.Trt_B |
| PKNOX1 | 4.24E-11 | 0.3171 | 0.77 | 0.63 | 6.78E-07 | HC_vs_MS.Trt_B |
| FFAR2 | 4.31E-11 | 0.28135 | 0.758 | 0.59 | 6.88E-07 | HC_vs_MS.Trt_B |
| WTIP | 4.35E-11 | 0.32256 | 0.964 | 0.9 | 6.95E-07 | HC_vs_MS.Trt_B |
| SIGLEC10 | 4.41E-11 | 0.46871 | 0.623 | 0.44 | 7.05E-07 | HC_vs_MS.Trt_B |
| C10orf11 | 4.56E-11 | 0.33189 | 0.959 | 0.9 | 7.29E-07 | HC_vs_MS.Trt_B |
| KIFC3 | 4.74E-11 | 0.3274 | 0.67 | 0.47 | 7.57E-07 | HC_vs_MS.Trt_B |
| RPL34 | 4.78E-11 | -0.6912 | 0.958 | 0.94 | 7.63E-07 | HC_vs_MS.Trt_B |
| HINFP | 4.82E-11 | 0.47535 | 0.99 | 0.95 | 7.71E-07 | HC_vs_MS.Trt_B |
| PCDHA4 | 4.93E-11 | 0.40003 | 0.904 | 0.79 | 7.87E-07 | HC_vs_MS.Trt_B |
| ZNF891 | 5.02E-11 | 0.4102 | 0.958 | 0.88 | 8.01E-07 | HC_vs_MS.Trt_B |
| TARSL2 | 5.05E-11 | 0.34928 | 0.97 | 0.89 | 8.06E-07 | HC_vs_MS.Trt_B |
| FAM213A | 5.05E-11 | 0.47431 | 0.807 | 0.64 | 8.06E-07 | HC_vs_MS.Trt_B |
| STRBP | 5.15E-11 | 0.53205 | 0.63 | 0.45 | 8.22E-07 | HC_vs_MS.Trt_B |
| IRF1 | 5.34E-11 | 0.45566 | 0.888 | 0.79 | 8.53E-07 | HC_vs_MS.Trt_B |
| P2RX5-TAX1 | 5.40E-11 | 0.40819 | 0.857 | 0.71 | 8.63E-07 | HC_vs_MS.Trt_B |
| SIK2 | 5.95E-11 | 0.38443 | 0.684 | 0.53 | 9.50E-07 | HC_vs_MS.Trt_B |
| CES3 | 6.15E-11 | 0.52121 | 0.309 | 0.14 | 9.82E-07 | HC_vs_MS.Trt_B |
| TRDMT1 | 6.41E-11 | 0.37371 | 0.726 | 0.54 | 1.02E-06 | HC_vs_MS.Trt_B |
| EIF1AY | 6.49E-11 | 0.50705 | 0.609 | 0.45 | 1.04E-06 | HC_vs_MS.Trt_B |
| LRRC37A3 | 6.68E-11 | 1.01862 | 0.528 | 0.36 | 1.07E-06 | HC_vs_MS.Trt_B |
| ZNF829 | 6.88E-11 | 0.37001 | 0.877 | 0.76 | 1.10E-06 | HC_vs_MS.Trt_B |
| ZNF37A | 7.02E-11 | 0.38727 | 0.874 | 0.75 | 1.12E-06 | HC_vs_MS.Trt_B |
| ATCAY | 7.40E-11 | 0.31277 | 0.897 | 0.8 | 1.18E-06 | HC_vs_MS.Trt_B |
| HFE | 7.42E-11 | 0.36275 | 0.683 | 0.5 | 1.19E-06 | HC_vs_MS.Trt_B |
| CHAF1B | 7.61E-11 | 0.35715 | 0.932 | 0.8 | 1.22E-06 | HC_vs_MS.Trt_B |
| PCDHB1 | 8.59E-11 | 0.34247 | 0.778 | 0.66 | 1.37E-06 | HC_vs_MS.Trt_B |
| ZNF496 | 9.27E-11 | 0.26686 | 0.681 | 0.48 | 1.48E-06 | HC_vs_MS.Trt_B |
| GTF2IRD2 | 9.31E-11 | 0.3442 | 0.954 | 0.86 | 1.49E-06 | HC_vs_MS.Trt_B |
| CPAMD8 | 9.48E-11 | 0.33672 | 0.699 | 0.51 | 1.51E-06 | HC_vs_MS.Trt_B |
| APBA3 | 9.68E-11 | 0.26877 | 0.746 | 0.56 | 1.55E-06 | HC_vs_MS.Trt_B |
| TAOK1 | 9.74E-11 | 0.79119 | 0.906 | 0.81 | 1.56E-06 | HC_vs_MS.Trt_B |

| | | | | | | |
|---|---|---|---|---|---|---|
| C16orf87 | 9.90E-11 | 0.36758 | 0.761 | 0.63 | 1.58E-06 | HC_vs_MS.Trt_B |
| SAR1A | 9.93E-11 | 0.30197 | 0.709 | 0.54 | 1.59E-06 | HC_vs_MS.Trt_B |
| PCSK1N | 9.94E-11 | 0.43862 | 0.372 | 0.2 | 1.59E-06 | HC_vs_MS.Trt_B |
| RPS19 | 1.00E-10 | -0.6042 | 0.933 | 0.91 | 1.60E-06 | HC_vs_MS.Trt_B |
| RHOH | 1.02E-10 | 0.40501 | 0.861 | 0.72 | 1.63E-06 | HC_vs_MS.Trt_B |
| SYT5 | 1.10E-10 | 0.3884 | 0.696 | 0.54 | 1.76E-06 | HC_vs_MS.Trt_B |
| ZNF431 | 1.16E-10 | 0.31136 | 0.987 | 0.94 | 1.85E-06 | HC_vs_MS.Trt_B |
| ANKLE2 | 1.18E-10 | 0.30513 | 0.984 | 0.95 | 1.88E-06 | HC_vs_MS.Trt_B |
| ARHGAP8 | 1.26E-10 | 0.3244 | 0.741 | 0.55 | 2.01E-06 | HC_vs_MS.Trt_B |
| TSPAN14 | 1.32E-10 | 0.27612 | 0.972 | 0.91 | 2.11E-06 | HC_vs_MS.Trt_B |
| KLF3 | 1.34E-10 | 0.32788 | 0.639 | 0.46 | 2.15E-06 | HC_vs_MS.Trt_B |
| DACT3 | 1.36E-10 | 0.26389 | 0.955 | 0.91 | 2.17E-06 | HC_vs_MS.Trt_B |
| RBMS2 | 1.36E-10 | 0.38947 | 0.861 | 0.73 | 2.17E-06 | HC_vs_MS.Trt_B |
| ZC3H12D | 1.39E-10 | 0.35725 | 0.812 | 0.65 | 2.22E-06 | HC_vs_MS.Trt_B |
| BCL11B | 1.40E-10 | 0.41986 | 0.493 | 0.29 | 2.24E-06 | HC_vs_MS.Trt_B |
| GGA2 | 1.43E-10 | 0.45851 | 0.658 | 0.48 | 2.28E-06 | HC_vs_MS.Trt_B |
| LCTL | 1.47E-10 | 0.3498 | 0.606 | 0.44 | 2.34E-06 | HC_vs_MS.Trt_B |
| AP006285.1 | 1.47E-10 | 0.26907 | 0.791 | 0.63 | 2.35E-06 | HC_vs_MS.Trt_B |
| EIF2AK2 | 1.52E-10 | 0.63583 | 0.825 | 0.68 | 2.43E-06 | HC_vs_MS.Trt_B |
| MOCS3 | 1.52E-10 | 0.27521 | 0.822 | 0.69 | 2.43E-06 | HC_vs_MS.Trt_B |
| ARPIN | 1.59E-10 | 0.34151 | 0.87 | 0.76 | 2.53E-06 | HC_vs_MS.Trt_B |
| ARID1B | 1.64E-10 | 0.34458 | 0.946 | 0.86 | 2.63E-06 | HC_vs_MS.Trt_B |
| EXPH5 | 1.65E-10 | 0.35482 | 0.7 | 0.56 | 2.64E-06 | HC_vs_MS.Trt_B |
| ZNF200 | 1.75E-10 | 0.38928 | 0.774 | 0.63 | 2.79E-06 | HC_vs_MS.Trt_B |
| SCAP | 1.78E-10 | 0.54301 | 0.617 | 0.43 | 2.84E-06 | HC_vs_MS.Trt_B |
| LRIG2 | 1.78E-10 | 0.28175 | 0.912 | 0.83 | 2.85E-06 | HC_vs_MS.Trt_B |
| SH3GL1 | 1.81E-10 | 0.56933 | 0.759 | 0.63 | 2.89E-06 | HC_vs_MS.Trt_B |
| CTC-487M2: | 1.82E-10 | 0.4206 | 0.917 | 0.85 | 2.91E-06 | HC_vs_MS.Trt_B |
| LAIR1 | 1.86E-10 | 1.14377 | 0.857 | 0.77 | 2.97E-06 | HC_vs_MS.Trt_B |
| FUT11 | 1.97E-10 | 1.11633 | 0.757 | 0.6 | 3.15E-06 | HC_vs_MS.Trt_B |
| CCZ1B | 2.00E-10 | 0.59833 | 0.712 | 0.53 | 3.19E-06 | HC_vs_MS.Trt_B |
| SLC13A5 | 2.19E-10 | 0.34419 | 0.746 | 0.56 | 3.50E-06 | HC_vs_MS.Trt_B |
| UBA1 | 2.25E-10 | 0.26727 | 0.762 | 0.61 | 3.60E-06 | HC_vs_MS.Trt_B |
| SEC22C | 2.40E-10 | 0.58876 | 0.691 | 0.52 | 3.84E-06 | HC_vs_MS.Trt_B |
| MPRIP | 2.44E-10 | 0.29134 | 0.936 | 0.83 | 3.90E-06 | HC_vs_MS.Trt_B |
| DNAL1 | 2.46E-10 | 0.48696 | 0.919 | 0.82 | 3.93E-06 | HC_vs_MS.Trt_B |
| ZKSCAN3 | 2.46E-10 | 0.69939 | 0.807 | 0.68 | 3.94E-06 | HC_vs_MS.Trt_B |
| ZNF90 | 2.50E-10 | 0.37929 | 0.88 | 0.71 | 4.00E-06 | HC_vs_MS.Trt_B |
| GPR141 | 2.56E-10 | 0.37569 | 0.509 | 0.31 | 4.09E-06 | HC_vs_MS.Trt_B |
| SPN | 2.66E-10 | 0.41583 | 0.98 | 0.93 | 4.25E-06 | HC_vs_MS.Trt_B |

| | | | | | | |
|---|---|---|---|---|---|---|
| EHMT1 | 2.70E-10 | 0.28046 | 0.941 | 0.83 | 4.31E-06 | HC_vs_MS.Trt_B |
| QPCTL | 2.74E-10 | 0.30275 | 0.742 | 0.6 | 4.38E-06 | HC_vs_MS.Trt_B |
| GLTSCR2 | 2.92E-10 | 0.40086 | 0.849 | 0.73 | 4.66E-06 | HC_vs_MS.Trt_B |
| GNL3L | 3.03E-10 | 0.33597 | 0.928 | 0.82 | 4.84E-06 | HC_vs_MS.Trt_B |
| EVI5 | 3.05E-10 | 0.39809 | 0.828 | 0.67 | 4.87E-06 | HC_vs_MS.Trt_B |
| SHOX | 3.24E-10 | 0.3164 | 0.839 | 0.72 | 5.17E-06 | HC_vs_MS.Trt_B |
| HNRNPH1 | 3.29E-10 | -0.7841 | 0.99 | 0.99 | 5.26E-06 | HC_vs_MS.Trt_B |
| INTS1 | 3.30E-10 | 1.27456 | 0.413 | 0.24 | 5.27E-06 | HC_vs_MS.Trt_B |
| GTPBP10 | 3.34E-10 | -0.3083 | 0.787 | 0.65 | 5.33E-06 | HC_vs_MS.Trt_B |
| MRPL3 | 3.37E-10 | 0.45245 | 0.548 | 0.38 | 5.38E-06 | HC_vs_MS.Trt_B |
| POLD4 | 3.40E-10 | 0.25241 | 0.797 | 0.65 | 5.43E-06 | HC_vs_MS.Trt_B |
| PHC3 | 3.42E-10 | 0.30144 | 0.958 | 0.88 | 5.46E-06 | HC_vs_MS.Trt_B |
| DIP2A | 3.49E-10 | 0.32394 | 0.826 | 0.66 | 5.58E-06 | HC_vs_MS.Trt_B |
| TLDC1 | 3.55E-10 | 0.60361 | 0.762 | 0.59 | 5.67E-06 | HC_vs_MS.Trt_B |
| TBC1D15 | 3.55E-10 | 0.67735 | 0.572 | 0.38 | 5.67E-06 | HC_vs_MS.Trt_B |
| HACD3 | 3.65E-10 | -0.261 | 0.784 | 0.58 | 5.84E-06 | HC_vs_MS.Trt_B |
| ADGRE2 | 3.67E-10 | 0.44147 | 0.757 | 0.59 | 5.86E-06 | HC_vs_MS.Trt_B |
| NDE1 | 3.69E-10 | 0.58033 | 0.717 | 0.53 | 5.89E-06 | HC_vs_MS.Trt_B |
| GFPT1 | 3.77E-10 | 0.96528 | 0.706 | 0.55 | 6.02E-06 | HC_vs_MS.Trt_B |
| KAZN | 3.79E-10 | 0.35208 | 0.47 | 0.29 | 6.05E-06 | HC_vs_MS.Trt_B |
| SCAI | 3.90E-10 | 0.25611 | 0.832 | 0.72 | 6.23E-06 | HC_vs_MS.Trt_B |
| TBC1D1 | 4.13E-10 | 0.35298 | 0.616 | 0.43 | 6.60E-06 | HC_vs_MS.Trt_B |
| NT5C2 | 4.31E-10 | 0.55275 | 0.735 | 0.59 | 6.88E-06 | HC_vs_MS.Trt_B |
| PPP2CA | 4.31E-10 | 0.62455 | 0.691 | 0.56 | 6.89E-06 | HC_vs_MS.Trt_B |
| CA5A | 4.38E-10 | 0.27361 | 0.977 | 0.92 | 6.99E-06 | HC_vs_MS.Trt_B |
| L3MBTL2 | 4.46E-10 | 0.47234 | 0.783 | 0.6 | 7.13E-06 | HC_vs_MS.Trt_B |
| CDK13 | 4.53E-10 | 0.57912 | 0.825 | 0.73 | 7.23E-06 | HC_vs_MS.Trt_B |
| ABL2 | 4.55E-10 | 0.43541 | 0.787 | 0.57 | 7.26E-06 | HC_vs_MS.Trt_B |
| MED9 | 4.64E-10 | 0.39654 | 0.59 | 0.4 | 7.41E-06 | HC_vs_MS.Trt_B |
| MED18 | 5.28E-10 | 0.52527 | 0.642 | 0.45 | 8.44E-06 | HC_vs_MS.Trt_B |
| CXorf23 | 5.39E-10 | 0.52312 | 0.741 | 0.56 | 8.61E-06 | HC_vs_MS.Trt_B |
| CTRC | 5.51E-10 | 0.34409 | 0.688 | 0.54 | 8.80E-06 | HC_vs_MS.Trt_B |
| SCRG1 | 5.68E-10 | 0.32593 | 0.714 | 0.57 | 9.07E-06 | HC_vs_MS.Trt_B |
| DDB1 | 5.82E-10 | 0.4127 | 0.901 | 0.76 | 9.29E-06 | HC_vs_MS.Trt_B |
| MT-ND4 | 6.41E-10 | -0.5335 | 0.99 | 0.99 | 1.02E-05 | HC_vs_MS.Trt_B |
| ACTA2 | 6.49E-10 | 0.58878 | 0.638 | 0.48 | 1.04E-05 | HC_vs_MS.Trt_B |
| RNF44 | 6.63E-10 | 0.34583 | 0.37 | 0.22 | 1.06E-05 | HC_vs_MS.Trt_B |
| GPR137C | 6.72E-10 | 0.29761 | 0.599 | 0.43 | 1.07E-05 | HC_vs_MS.Trt_B |
| TMEM138 | 6.87E-10 | 0.29431 | 0.855 | 0.73 | 1.10E-05 | HC_vs_MS.Trt_B |
| CFLAR | 6.91E-10 | 0.25078 | 1 | 0.99 | 1.10E-05 | HC_vs_MS.Trt_B |

| | | | | | | |
|---|---|---|---|---|---|---|
| PRDM7 | 6.97E-10 | 0.38277 | 0.509 | 0.34 | 1.11E-05 | HC_vs_MS.Trt_B |
| CPM | 7.03E-10 | 0.45952 | 0.803 | 0.64 | 1.12E-05 | HC_vs_MS.Trt_B |
| ARSD | 7.20E-10 | 0.45691 | 0.822 | 0.68 | 1.15E-05 | HC_vs_MS.Trt_B |
| SMARCC2 | 7.22E-10 | 0.28978 | 0.884 | 0.73 | 1.15E-05 | HC_vs_MS.Trt_B |
| MR1 | 7.29E-10 | 0.30098 | 0.913 | 0.78 | 1.16E-05 | HC_vs_MS.Trt_B |
| CACNB4 | 7.41E-10 | 0.37622 | 0.858 | 0.74 | 1.18E-05 | HC_vs_MS.Trt_B |
| ANKH | 7.50E-10 | 0.25769 | 0.619 | 0.44 | 1.20E-05 | HC_vs_MS.Trt_B |
| PAAF1 | 7.52E-10 | 0.27889 | 0.949 | 0.87 | 1.20E-05 | HC_vs_MS.Trt_B |
| LIN28A | 7.54E-10 | 0.26403 | 0.741 | 0.59 | 1.20E-05 | HC_vs_MS.Trt_B |
| TINF2 | 8.09E-10 | 0.50607 | 0.83 | 0.69 | 1.29E-05 | HC_vs_MS.Trt_B |
| NAIP | 8.22E-10 | 0.8733 | 0.675 | 0.51 | 1.31E-05 | HC_vs_MS.Trt_B |
| NFS1 | 8.32E-10 | 0.46009 | 0.886 | 0.78 | 1.33E-05 | HC_vs_MS.Trt_B |
| MACC1 | 8.35E-10 | 0.41896 | 0.672 | 0.52 | 1.33E-05 | HC_vs_MS.Trt_B |
| DDX6 | 8.40E-10 | 0.29901 | 0.972 | 0.93 | 1.34E-05 | HC_vs_MS.Trt_B |
| SLFN5 | 8.43E-10 | 0.44594 | 0.88 | 0.76 | 1.35E-05 | HC_vs_MS.Trt_B |
| SLC9A7 | 8.43E-10 | 0.41617 | 0.938 | 0.83 | 1.35E-05 | HC_vs_MS.Trt_B |
| MSI2 | 8.54E-10 | 0.45544 | 0.83 | 0.72 | 1.36E-05 | HC_vs_MS.Trt_B |
| IL6R | 8.84E-10 | 0.32543 | 0.6 | 0.43 | 1.41E-05 | HC_vs_MS.Trt_B |
| PCDHA10 | 8.90E-10 | 0.30418 | 0.538 | 0.36 | 1.42E-05 | HC_vs_MS.Trt_B |
| TEX9 | 9.04E-10 | 0.27422 | 0.752 | 0.57 | 1.44E-05 | HC_vs_MS.Trt_B |
| ZGRF1 | 9.06E-10 | 0.6413 | 0.755 | 0.59 | 1.45E-05 | HC_vs_MS.Trt_B |
| WEE1 | 9.10E-10 | 0.38613 | 0.684 | 0.51 | 1.45E-05 | HC_vs_MS.Trt_B |
| LRRC37B | 9.26E-10 | 0.58399 | 0.619 | 0.44 | 1.48E-05 | HC_vs_MS.Trt_B |
| GSDMD | 9.66E-10 | 0.2686 | 0.794 | 0.62 | 1.54E-05 | HC_vs_MS.Trt_B |
| C15orf59 | 9.75E-10 | 0.38553 | 0.654 | 0.5 | 1.56E-05 | HC_vs_MS.Trt_B |
| ADAT1 | 1.01E-09 | 0.45798 | 0.671 | 0.53 | 1.61E-05 | HC_vs_MS.Trt_B |
| KCNK6 | 1.02E-09 | 0.29898 | 0.959 | 0.88 | 1.64E-05 | HC_vs_MS.Trt_B |
| AP4S1 | 1.04E-09 | 0.39958 | 0.93 | 0.85 | 1.66E-05 | HC_vs_MS.Trt_B |
| CSF2RA | 1.08E-09 | 0.26181 | 0.984 | 0.92 | 1.72E-05 | HC_vs_MS.Trt_B |
| SUN3 | 1.10E-09 | 0.35144 | 0.387 | 0.22 | 1.76E-05 | HC_vs_MS.Trt_B |
| PPARA | 1.11E-09 | 0.47967 | 0.904 | 0.81 | 1.77E-05 | HC_vs_MS.Trt_B |
| TF | 1.11E-09 | 0.37677 | 0.888 | 0.8 | 1.78E-05 | HC_vs_MS.Trt_B |
| ADAMTS4 | 1.12E-09 | 0.57733 | 0.896 | 0.76 | 1.79E-05 | HC_vs_MS.Trt_B |
| TMEM170A | 1.12E-09 | 0.80794 | 0.81 | 0.71 | 1.79E-05 | HC_vs_MS.Trt_B |
| PLA2G15 | 1.13E-09 | 0.37665 | 0.78 | 0.63 | 1.81E-05 | HC_vs_MS.Trt_B |
| CXorf38 | 1.14E-09 | 0.63979 | 0.836 | 0.71 | 1.82E-05 | HC_vs_MS.Trt_B |
| ZNF169 | 1.14E-09 | 0.37762 | 0.832 | 0.7 | 1.82E-05 | HC_vs_MS.Trt_B |
| EDRF1 | 1.15E-09 | 0.65402 | 0.626 | 0.44 | 1.84E-05 | HC_vs_MS.Trt_B |
| TRAF3IP2 | 1.17E-09 | 0.78442 | 0.886 | 0.77 | 1.87E-05 | HC_vs_MS.Trt_B |
| HCAR1 | 1.19E-09 | 0.33276 | 0.868 | 0.74 | 1.90E-05 | HC_vs_MS.Trt_B |

| | | | | | | |
|---|---|---|---|---|---|---|
| SLITRK2 | 1.20E-09 | 0.42809 | 0.335 | 0.17 | 1.91E-05 | HC_vs_MS.Trt_B |
| FGD4 | 1.23E-09 | 0.32876 | 0.82 | 0.63 | 1.97E-05 | HC_vs_MS.Trt_B |
| EREG | 1.24E-09 | -0.7385 | 0.761 | 0.6 | 1.98E-05 | HC_vs_MS.Trt_B |
| RPS26 | 1.25E-09 | -0.412 | 0.661 | 0.68 | 1.99E-05 | HC_vs_MS.Trt_B |
| MYH11 | 1.25E-09 | 0.42805 | 0.917 | 0.85 | 2.00E-05 | HC_vs_MS.Trt_B |
| NICN1 | 1.42E-09 | 0.55216 | 0.77 | 0.6 | 2.26E-05 | HC_vs_MS.Trt_B |
| ATG7 | 1.56E-09 | 0.60292 | 0.561 | 0.36 | 2.50E-05 | HC_vs_MS.Trt_B |
| RAD51C | 1.57E-09 | 0.33665 | 0.909 | 0.79 | 2.51E-05 | HC_vs_MS.Trt_B |
| GCSAML | 1.59E-09 | 0.32593 | 0.328 | 0.17 | 2.54E-05 | HC_vs_MS.Trt_B |
| TIAM2 | 1.59E-09 | 0.30243 | 0.799 | 0.64 | 2.54E-05 | HC_vs_MS.Trt_B |
| IFT80 | 1.60E-09 | 0.49827 | 0.765 | 0.63 | 2.56E-05 | HC_vs_MS.Trt_B |
| AP3S2 | 1.63E-09 | 0.32462 | 0.694 | 0.55 | 2.61E-05 | HC_vs_MS.Trt_B |
| SUGT1 | 1.63E-09 | 0.49231 | 0.903 | 0.8 | 2.61E-05 | HC_vs_MS.Trt_B |
| FBXL4 | 1.64E-09 | 1.05214 | 0.554 | 0.39 | 2.63E-05 | HC_vs_MS.Trt_B |
| GADL1 | 1.66E-09 | 0.37092 | 0.622 | 0.46 | 2.65E-05 | HC_vs_MS.Trt_B |
| RP11-47I22. | 1.74E-09 | 0.37394 | 0.596 | 0.42 | 2.78E-05 | HC_vs_MS.Trt_B |
| PUS1 | 1.78E-09 | 0.53664 | 0.855 | 0.7 | 2.85E-05 | HC_vs_MS.Trt_B |
| CERCAM | 1.86E-09 | 0.28205 | 0.981 | 0.94 | 2.97E-05 | HC_vs_MS.Trt_B |
| SLC35E1 | 1.87E-09 | 0.41183 | 0.975 | 0.91 | 2.98E-05 | HC_vs_MS.Trt_B |
| SPATS2L | 1.87E-09 | 0.51432 | 0.941 | 0.86 | 2.98E-05 | HC_vs_MS.Trt_B |
| P2RX7 | 1.95E-09 | 0.56678 | 0.749 | 0.59 | 3.12E-05 | HC_vs_MS.Trt_B |
| CTCFL | 1.96E-09 | 0.27399 | 0.825 | 0.68 | 3.12E-05 | HC_vs_MS.Trt_B |
| SPATA6L | 2.02E-09 | 0.47499 | 0.781 | 0.63 | 3.22E-05 | HC_vs_MS.Trt_B |
| CNOT1 | 2.03E-09 | 0.2756 | 0.806 | 0.64 | 3.24E-05 | HC_vs_MS.Trt_B |
| PAX6 | 2.05E-09 | 0.71289 | 0.397 | 0.24 | 3.27E-05 | HC_vs_MS.Trt_B |
| IPCEF1 | 2.07E-09 | 0.34061 | 0.53 | 0.37 | 3.31E-05 | HC_vs_MS.Trt_B |
| ISG20L2 | 2.09E-09 | 0.26114 | 0.803 | 0.64 | 3.33E-05 | HC_vs_MS.Trt_B |
| BRI3BP | 2.09E-09 | 0.67615 | 0.846 | 0.69 | 3.34E-05 | HC_vs_MS.Trt_B |
| HSD17B4 | 2.10E-09 | 0.26741 | 0.642 | 0.48 | 3.35E-05 | HC_vs_MS.Trt_B |
| SSTR2 | 2.13E-09 | 0.27775 | 0.91 | 0.82 | 3.41E-05 | HC_vs_MS.Trt_B |
| PTCD3 | 2.14E-09 | 0.32314 | 0.912 | 0.77 | 3.42E-05 | HC_vs_MS.Trt_B |
| ZNF233 | 2.22E-09 | 0.51562 | 0.467 | 0.32 | 3.55E-05 | HC_vs_MS.Trt_B |
| AC004824.1 | 2.23E-09 | 0.31228 | 0.416 | 0.25 | 3.57E-05 | HC_vs_MS.Trt_B |
| TMEM213 | 2.32E-09 | 0.25626 | 0.816 | 0.67 | 3.71E-05 | HC_vs_MS.Trt_B |
| DNASE1 | 2.37E-09 | 0.34115 | 0.9 | 0.8 | 3.78E-05 | HC_vs_MS.Trt_B |
| RPS3A | 2.39E-09 | -0.3727 | 0.938 | 0.92 | 3.82E-05 | HC_vs_MS.Trt_B |
| TADA2A | 2.45E-09 | 0.7383 | 0.87 | 0.78 | 3.91E-05 | HC_vs_MS.Trt_B |
| EML1 | 2.50E-09 | 0.31905 | 0.719 | 0.55 | 3.99E-05 | HC_vs_MS.Trt_B |
| TCEANC2 | 2.70E-09 | 0.36426 | 0.881 | 0.75 | 4.32E-05 | HC_vs_MS.Trt_B |
| ACP7 | 2.71E-09 | 0.37095 | 0.472 | 0.31 | 4.33E-05 | HC_vs_MS.Trt_B |

| | | | | | | |
|---|---|---|---|---|---|---|
| HIF3A | 2.72E-09 | 0.31945 | 0.864 | 0.74 | 4.34E-05 | HC_vs_MS.Trt_B |
| APOL6 | 2.76E-09 | 0.77095 | 0.843 | 0.73 | 4.41E-05 | HC_vs_MS.Trt_B |
| LIMS1 | 2.79E-09 | 0.42171 | 0.884 | 0.78 | 4.45E-05 | HC_vs_MS.Trt_B |
| WFDC1 | 2.89E-09 | 0.2812 | 0.79 | 0.65 | 4.62E-05 | HC_vs_MS.Trt_B |
| DYNLT3 | 2.91E-09 | 0.5643 | 0.662 | 0.5 | 4.64E-05 | HC_vs_MS.Trt_B |
| PCNX4 | 2.93E-09 | 0.36218 | 0.877 | 0.75 | 4.67E-05 | HC_vs_MS.Trt_B |
| CDC5L | 2.95E-09 | 0.39914 | 0.59 | 0.4 | 4.71E-05 | HC_vs_MS.Trt_B |
| LPO | 2.97E-09 | 0.53247 | 0.61 | 0.46 | 4.74E-05 | HC_vs_MS.Trt_B |
| ANKS4B | 3.00E-09 | 0.28091 | 0.628 | 0.45 | 4.79E-05 | HC_vs_MS.Trt_B |
| PDS5A | 3.00E-09 | 0.28649 | 0.754 | 0.59 | 4.79E-05 | HC_vs_MS.Trt_B |
| PIGL | 3.06E-09 | 0.29817 | 0.674 | 0.5 | 4.89E-05 | HC_vs_MS.Trt_B |
| R3HDM4 | 3.18E-09 | 0.40631 | 0.555 | 0.36 | 5.08E-05 | HC_vs_MS.Trt_B |
| TRIM66 | 3.19E-09 | 0.48861 | 0.493 | 0.32 | 5.10E-05 | HC_vs_MS.Trt_B |
| FAM175A | 3.31E-09 | 0.28901 | 0.852 | 0.71 | 5.29E-05 | HC_vs_MS.Trt_B |
| MRPS33 | 3.33E-09 | 0.41266 | 0.916 | 0.83 | 5.32E-05 | HC_vs_MS.Trt_B |
| HAUS2 | 3.37E-09 | 0.47167 | 0.794 | 0.67 | 5.38E-05 | HC_vs_MS.Trt_B |
| PRR19 | 3.43E-09 | 0.33685 | 0.693 | 0.54 | 5.47E-05 | HC_vs_MS.Trt_B |
| CYP4F8 | 3.51E-09 | 0.35003 | 0.771 | 0.63 | 5.61E-05 | HC_vs_MS.Trt_B |
| IL17RA | 3.55E-09 | 0.44716 | 0.652 | 0.5 | 5.66E-05 | HC_vs_MS.Trt_B |
| ZNF385A | 3.67E-09 | 0.32156 | 0.696 | 0.52 | 5.85E-05 | HC_vs_MS.Trt_B |
| ATP5S | 3.70E-09 | 0.44066 | 0.888 | 0.79 | 5.91E-05 | HC_vs_MS.Trt_B |
| PCBD2 | 3.75E-09 | 0.47775 | 0.799 | 0.67 | 5.99E-05 | HC_vs_MS.Trt_B |
| HTRA4 | 3.76E-09 | 0.39633 | 0.455 | 0.28 | 6.00E-05 | HC_vs_MS.Trt_B |
| SYT12 | 3.80E-09 | 0.315 | 0.716 | 0.55 | 6.07E-05 | HC_vs_MS.Trt_B |
| MMAB | 3.83E-09 | -0.251 | 0.664 | 0.47 | 6.11E-05 | HC_vs_MS.Trt_B |
| FILIP1L | 3.87E-09 | 1.02535 | 0.39 | 0.23 | 6.18E-05 | HC_vs_MS.Trt_B |
| CYP2C19 | 4.08E-09 | 0.35651 | 0.719 | 0.56 | 6.52E-05 | HC_vs_MS.Trt_B |
| CCNI | 4.11E-09 | 0.28789 | 0.822 | 0.67 | 6.57E-05 | HC_vs_MS.Trt_B |
| LRRC74B | 4.15E-09 | 0.34298 | 0.748 | 0.59 | 6.63E-05 | HC_vs_MS.Trt_B |
| WDR66 | 4.25E-09 | 0.28946 | 0.948 | 0.89 | 6.78E-05 | HC_vs_MS.Trt_B |
| EFTUD2 | 4.44E-09 | 0.37188 | 0.836 | 0.7 | 7.09E-05 | HC_vs_MS.Trt_B |
| KANSL1 | 4.57E-09 | 0.90522 | 0.707 | 0.56 | 7.29E-05 | HC_vs_MS.Trt_B |
| KCNMB1 | 4.57E-09 | 0.28223 | 0.69 | 0.52 | 7.31E-05 | HC_vs_MS.Trt_B |
| EIF4EBP2 | 4.58E-09 | 0.43931 | 0.628 | 0.45 | 7.31E-05 | HC_vs_MS.Trt_B |
| CAPS2 | 4.60E-09 | 0.45366 | 0.452 | 0.29 | 7.34E-05 | HC_vs_MS.Trt_B |
| INPP5B | 4.62E-09 | 0.30383 | 0.942 | 0.86 | 7.38E-05 | HC_vs_MS.Trt_B |
| GINS3 | 4.73E-09 | 0.34621 | 0.849 | 0.75 | 7.55E-05 | HC_vs_MS.Trt_B |
| ZNF70 | 4.85E-09 | 0.3413 | 0.829 | 0.71 | 7.74E-05 | HC_vs_MS.Trt_B |
| KLHL6 | 4.85E-09 | 0.62987 | 0.729 | 0.59 | 7.75E-05 | HC_vs_MS.Trt_B |
| DHRSX | 4.94E-09 | 0.27842 | 0.725 | 0.58 | 7.90E-05 | HC_vs_MS.Trt_B |

| | | | | | | |
|---|---|---|---|---|---|---|
| RCL1 | 4.98E-09 | 0.6374 | 0.5 | 0.33 | 7.95E-05 | HC_vs_MS.Trt_B |
| SEC63 | 5.07E-09 | 0.26228 | 0.636 | 0.46 | 8.09E-05 | HC_vs_MS.Trt_B |
| DECR2 | 5.10E-09 | 0.58981 | 0.801 | 0.64 | 8.14E-05 | HC_vs_MS.Trt_B |
| ZNFX1 | 5.18E-09 | 0.33062 | 0.814 | 0.65 | 8.28E-05 | HC_vs_MS.Trt_B |
| SPC25 | 5.20E-09 | 0.47635 | 0.677 | 0.52 | 8.31E-05 | HC_vs_MS.Trt_B |
| ZNF724 | 5.26E-09 | 0.54981 | 0.899 | 0.78 | 8.41E-05 | HC_vs_MS.Trt_B |
| FMNL1 | 5.36E-09 | 0.61348 | 0.701 | 0.56 | 8.55E-05 | HC_vs_MS.Trt_B |
| MATN2 | 5.40E-09 | 0.30479 | 0.58 | 0.41 | 8.62E-05 | HC_vs_MS.Trt_B |
| PCDHGA8 | 5.41E-09 | 0.37904 | 0.596 | 0.46 | 8.65E-05 | HC_vs_MS.Trt_B |
| BPTF | 5.51E-09 | 0.29078 | 0.807 | 0.69 | 8.80E-05 | HC_vs_MS.Trt_B |
| KIF1B | 5.53E-09 | 0.3575 | 0.788 | 0.66 | 8.84E-05 | HC_vs_MS.Trt_B |
| ARHGEF9 | 5.55E-09 | 0.32144 | 0.571 | 0.42 | 8.86E-05 | HC_vs_MS.Trt_B |
| HKR1 | 5.61E-09 | 0.40313 | 0.967 | 0.9 | 8.96E-05 | HC_vs_MS.Trt_B |
| DFFB | 5.65E-09 | 0.33376 | 0.807 | 0.66 | 9.02E-05 | HC_vs_MS.Trt_B |
| RGMA | 5.94E-09 | 0.44222 | 0.652 | 0.49 | 9.48E-05 | HC_vs_MS.Trt_B |
| HTR3B | 5.95E-09 | 0.28338 | 0.632 | 0.48 | 9.51E-05 | HC_vs_MS.Trt_B |
| ZNF805 | 6.06E-09 | 0.3376 | 0.739 | 0.6 | 9.69E-05 | HC_vs_MS.Trt_B |
| OR52A1 | 6.09E-09 | 0.29754 | 0.749 | 0.63 | 9.73E-05 | HC_vs_MS.Trt_B |
| RPL18A | 6.47E-09 | -0.4209 | 0.914 | 0.88 | 1.03E-04 | HC_vs_MS.Trt_B |
| OLA1 | 6.51E-09 | 0.43077 | 0.455 | 0.28 | 1.04E-04 | HC_vs_MS.Trt_B |
| SHCBP1 | 6.52E-09 | 0.42873 | 0.707 | 0.55 | 1.04E-04 | HC_vs_MS.Trt_B |
| LYPD5 | 6.56E-09 | 0.33583 | 0.699 | 0.55 | 1.05E-04 | HC_vs_MS.Trt_B |
| METTL8 | 7.01E-09 | 0.49037 | 0.916 | 0.82 | 1.12E-04 | HC_vs_MS.Trt_B |
| SEMA3C | 7.08E-09 | 0.42225 | 0.662 | 0.5 | 1.13E-04 | HC_vs_MS.Trt_B |
| ZNF549 | 7.27E-09 | 0.51765 | 0.688 | 0.54 | 1.16E-04 | HC_vs_MS.Trt_B |
| MXRA7 | 7.32E-09 | 0.66135 | 0.78 | 0.65 | 1.17E-04 | HC_vs_MS.Trt_B |
| TSEN2 | 7.58E-09 | 0.38794 | 0.881 | 0.74 | 1.21E-04 | HC_vs_MS.Trt_B |
| HACD2 | 7.67E-09 | 0.30894 | 0.939 | 0.83 | 1.23E-04 | HC_vs_MS.Trt_B |
| AFF3 | 7.80E-09 | 0.6136 | 0.748 | 0.6 | 1.25E-04 | HC_vs_MS.Trt_B |
| ALDH3B1 | 8.13E-09 | 0.27524 | 0.735 | 0.57 | 1.30E-04 | HC_vs_MS.Trt_B |
| FBXO22 | 8.13E-09 | -0.4456 | 0.765 | 0.62 | 1.30E-04 | HC_vs_MS.Trt_B |
| DET1 | 8.31E-09 | 0.51787 | 0.536 | 0.36 | 1.33E-04 | HC_vs_MS.Trt_B |
| ZNF141 | 8.47E-09 | 0.5057 | 0.968 | 0.92 | 1.35E-04 | HC_vs_MS.Trt_B |
| ETFDH | 8.50E-09 | 0.39498 | 0.714 | 0.56 | 1.36E-04 | HC_vs_MS.Trt_B |
| POU5F2 | 8.54E-09 | 0.2562 | 0.678 | 0.51 | 1.36E-04 | HC_vs_MS.Trt_B |
| SKA1 | 8.66E-09 | 0.34993 | 0.677 | 0.52 | 1.38E-04 | HC_vs_MS.Trt_B |
| ADCY7 | 9.28E-09 | 0.41133 | 0.765 | 0.62 | 1.48E-04 | HC_vs_MS.Trt_B |
| GMEB1 | 9.40E-09 | 0.31362 | 0.803 | 0.64 | 1.50E-04 | HC_vs_MS.Trt_B |
| HIF1A | 9.41E-09 | 0.55308 | 0.914 | 0.8 | 1.50E-04 | HC_vs_MS.Trt_B |
| ZNF3 | 9.77E-09 | 0.41367 | 0.742 | 0.58 | 1.56E-04 | HC_vs_MS.Trt_B |

| | | | | | | |
|---|---|---|---|---|---|---|
| SS18 | 1.01E-08 | 0.63631 | 0.596 | 0.43 | 1.61E-04 | HC_vs_MS.Trt_B |
| POU6F1 | 1.03E-08 | 0.79532 | 0.522 | 0.37 | 1.65E-04 | HC_vs_MS.Trt_B |
| ZNF770 | 1.08E-08 | 1.0773 | 0.322 | 0.17 | 1.72E-04 | HC_vs_MS.Trt_B |
| MECOM | 1.11E-08 | 0.48736 | 0.378 | 0.21 | 0.000178 | HC_vs_MS.Trt_B |
| MAVS | 1.12E-08 | 0.53571 | 0.935 | 0.84 | 1.79E-04 | HC_vs_MS.Trt_B |
| METTL6 | 1.17E-08 | 0.32573 | 0.878 | 0.78 | 1.88E-04 | HC_vs_MS.Trt_B |
| ABCG8 | 1.18E-08 | 0.25993 | 0.614 | 0.46 | 1.88E-04 | HC_vs_MS.Trt_B |
| ARHGEF39 | 1.19E-08 | 0.91844 | 0.648 | 0.5 | 1.89E-04 | HC_vs_MS.Trt_B |
| PDDC1 | 1.20E-08 | 0.25851 | 0.941 | 0.86 | 1.91E-04 | HC_vs_MS.Trt_B |
| CSTF1 | 1.20E-08 | 0.2829 | 0.771 | 0.65 | 1.91E-04 | HC_vs_MS.Trt_B |
| ORAOV1 | 1.23E-08 | 0.83643 | 0.694 | 0.53 | 1.97E-04 | HC_vs_MS.Trt_B |
| HM13 | 1.28E-08 | 0.45292 | 0.83 | 0.72 | 2.04E-04 | HC_vs_MS.Trt_B |
| CYB561A3 | 1.31E-08 | 0.25447 | 0.981 | 0.93 | 2.09E-04 | HC_vs_MS.Trt_B |
| PCSK5 | 1.32E-08 | -0.3359 | 0.97 | 0.88 | 2.11E-04 | HC_vs_MS.Trt_B |
| AC013461.1 | 1.35E-08 | 0.35714 | 0.81 | 0.67 | 2.16E-04 | HC_vs_MS.Trt_B |
| MED24 | 1.36E-08 | -0.2899 | 0.622 | 0.46 | 2.17E-04 | HC_vs_MS.Trt_B |
| VASP | 1.37E-08 | 0.91296 | 0.562 | 0.41 | 2.18E-04 | HC_vs_MS.Trt_B |
| RAB3B | 1.38E-08 | 0.28748 | 0.745 | 0.57 | 2.20E-04 | HC_vs_MS.Trt_B |
| FRRS1L | 1.41E-08 | 0.26831 | 0.865 | 0.76 | 2.26E-04 | HC_vs_MS.Trt_B |
| FBXO40 | 1.44E-08 | 0.68769 | 0.328 | 0.18 | 2.30E-04 | HC_vs_MS.Trt_B |
| CDKL5 | 1.47E-08 | 0.28551 | 0.932 | 0.82 | 2.35E-04 | HC_vs_MS.Trt_B |
| WIPF2 | 1.48E-08 | 0.67098 | 0.772 | 0.62 | 2.37E-04 | HC_vs_MS.Trt_B |
| SLC4A8 | 1.54E-08 | -0.4494 | 0.88 | 0.78 | 2.46E-04 | HC_vs_MS.Trt_B |
| 6-Sep | 1.55E-08 | 0.28028 | 0.777 | 0.64 | 2.48E-04 | HC_vs_MS.Trt_B |
| ITGA3 | 1.57E-08 | 0.32482 | 0.754 | 0.6 | 2.50E-04 | HC_vs_MS.Trt_B |
| DISP2 | 1.57E-08 | 0.29721 | 0.913 | 0.83 | 2.50E-04 | HC_vs_MS.Trt_B |
| DIRAS2 | 1.62E-08 | 0.32571 | 0.787 | 0.66 | 2.58E-04 | HC_vs_MS.Trt_B |
| RFTN2 | 1.62E-08 | 0.39395 | 0.68 | 0.55 | 0.000258 | HC_vs_MS.Trt_B |
| PHF20L1 | 1.62E-08 | 0.67321 | 0.651 | 0.51 | 2.59E-04 | HC_vs_MS.Trt_B |
| ZBTB3 | 1.63E-08 | 0.47935 | 0.793 | 0.66 | 2.60E-04 | HC_vs_MS.Trt_B |
| DHFR2 | 1.72E-08 | 0.76331 | 0.59 | 0.42 | 2.75E-04 | HC_vs_MS.Trt_B |
| MTAP | 1.76E-08 | 0.39581 | 0.732 | 0.55 | 2.82E-04 | HC_vs_MS.Trt_B |
| POLR3A | 1.77E-08 | 0.30778 | 0.829 | 0.7 | 2.83E-04 | HC_vs_MS.Trt_B |
| BAZ2A | 1.78E-08 | 0.38516 | 0.665 | 0.5 | 2.84E-04 | HC_vs_MS.Trt_B |
| LSM3 | 1.89E-08 | -0.3179 | 0.588 | 0.42 | 3.02E-04 | HC_vs_MS.Trt_B |
| HSPBAP1 | 1.92E-08 | 0.52712 | 0.5 | 0.33 | 3.06E-04 | HC_vs_MS.Trt_B |
| PAPLN | 1.93E-08 | 0.67332 | 0.764 | 0.62 | 3.07E-04 | HC_vs_MS.Trt_B |
| RABGAP1L | 2.00E-08 | 0.63791 | 0.891 | 0.78 | 3.19E-04 | HC_vs_MS.Trt_B |
| C22orf39 | 2.01E-08 | 0.55479 | 0.688 | 0.5 | 3.20E-04 | HC_vs_MS.Trt_B |
| PXMP4 | 2.10E-08 | 0.56964 | 0.816 | 0.74 | 3.35E-04 | HC_vs_MS.Trt_B |

| | | | | | | |
|---|---|---|---|---|---|---|
| CREG2 | 2.16E-08 | 0.49628 | 0.677 | 0.55 | 3.46E-04 | HC_vs_MS.Trt_B |
| FKBP5 | 2.18E-08 | 0.47084 | 0.762 | 0.64 | 3.49E-04 | HC_vs_MS.Trt_B |
| P3H4 | 2.22E-08 | 0.31187 | 0.896 | 0.76 | 3.55E-04 | HC_vs_MS.Trt_B |
| C3orf62 | 2.23E-08 | 0.62686 | 0.758 | 0.6 | 3.56E-04 | HC_vs_MS.Trt_B |
| VPS13B | 2.23E-08 | 0.26382 | 0.735 | 0.57 | 3.56E-04 | HC_vs_MS.Trt_B |
| DCLRE1C | 2.23E-08 | -0.7008 | 0.681 | 0.51 | 3.57E-04 | HC_vs_MS.Trt_B |
| PTK2 | 2.24E-08 | 0.25168 | 0.948 | 0.83 | 3.57E-04 | HC_vs_MS.Trt_B |
| C15orf57 | 2.25E-08 | 0.97982 | 0.772 | 0.66 | 3.60E-04 | HC_vs_MS.Trt_B |
| HDAC4 | 2.27E-08 | 0.46162 | 0.559 | 0.38 | 3.63E-04 | HC_vs_MS.Trt_B |
| ATM | 2.29E-08 | 0.25042 | 0.952 | 0.9 | 3.66E-04 | HC_vs_MS.Trt_B |
| RBM28 | 2.36E-08 | 0.5899 | 0.877 | 0.78 | 3.76E-04 | HC_vs_MS.Trt_B |
| CBL | 2.44E-08 | 0.54444 | 0.609 | 0.46 | 3.90E-04 | HC_vs_MS.Trt_B |
| TMEM156 | 2.47E-08 | -0.5291 | 0.768 | 0.62 | 3.95E-04 | HC_vs_MS.Trt_B |
| MAGI2 | 2.58E-08 | 0.26046 | 0.787 | 0.61 | 4.12E-04 | HC_vs_MS.Trt_B |
| TNKS | 2.59E-08 | 0.60448 | 0.797 | 0.66 | 4.14E-04 | HC_vs_MS.Trt_B |
| PLPP5 | 2.61E-08 | 0.54888 | 0.848 | 0.71 | 4.16E-04 | HC_vs_MS.Trt_B |
| IMMP1L | 2.64E-08 | -0.4279 | 0.559 | 0.41 | 4.22E-04 | HC_vs_MS.Trt_B |
| AARSD1 | 2.67E-08 | 0.72042 | 0.678 | 0.54 | 4.26E-04 | HC_vs_MS.Trt_B |
| METTL9 | 2.71E-08 | 0.31273 | 0.601 | 0.43 | 4.32E-04 | HC_vs_MS.Trt_B |
| NAV1 | 2.72E-08 | 0.81139 | 0.845 | 0.75 | 4.34E-04 | HC_vs_MS.Trt_B |
| NHSL2 | 2.77E-08 | -0.3533 | 0.606 | 0.44 | 4.42E-04 | HC_vs_MS.Trt_B |
| TCTA | 2.79E-08 | 0.28743 | 0.61 | 0.43 | 4.46E-04 | HC_vs_MS.Trt_B |
| ZNF154 | 2.95E-08 | 0.43679 | 0.684 | 0.53 | 4.71E-04 | HC_vs_MS.Trt_B |
| HAUS1 | 2.97E-08 | 0.44035 | 0.635 | 0.46 | 4.75E-04 | HC_vs_MS.Trt_B |
| SNRNP70 | 3.00E-08 | 1.12505 | 0.299 | 0.16 | 4.79E-04 | HC_vs_MS.Trt_B |
| AHSA1 | 3.02E-08 | 0.56109 | 0.541 | 0.39 | 4.82E-04 | HC_vs_MS.Trt_B |
| KIAA2012 | 3.05E-08 | 0.36653 | 0.643 | 0.51 | 4.87E-04 | HC_vs_MS.Trt_B |
| PLEKHB2 | 3.14E-08 | 0.55637 | 0.843 | 0.73 | 5.02E-04 | HC_vs_MS.Trt_B |
| TRIM52 | 3.14E-08 | -0.374 | 0.799 | 0.66 | 5.02E-04 | HC_vs_MS.Trt_B |
| HACE1 | 3.24E-08 | 0.33236 | 0.655 | 0.48 | 5.17E-04 | HC_vs_MS.Trt_B |
| TIMM17B | 3.26E-08 | -0.4032 | 0.73 | 0.58 | 5.20E-04 | HC_vs_MS.Trt_B |
| SPACA6 | 3.36E-08 | 0.27311 | 0.87 | 0.8 | 5.37E-04 | HC_vs_MS.Trt_B |
| AAMDC | 3.37E-08 | 0.31123 | 0.899 | 0.83 | 5.38E-04 | HC_vs_MS.Trt_B |
| MCM3AP | 3.47E-08 | 0.46364 | 0.529 | 0.37 | 5.54E-04 | HC_vs_MS.Trt_B |
| SMYD4 | 3.66E-08 | 0.5426 | 0.833 | 0.67 | 5.85E-04 | HC_vs_MS.Trt_B |
| ZFP36 | 3.68E-08 | 0.68618 | 0.393 | 0.24 | 5.87E-04 | HC_vs_MS.Trt_B |
| TCAF1 | 3.75E-08 | 0.54335 | 0.336 | 0.19 | 6.00E-04 | HC_vs_MS.Trt_B |
| PNPT1 | 3.76E-08 | 0.60149 | 0.855 | 0.77 | 6.00E-04 | HC_vs_MS.Trt_B |
| EYS | 3.78E-08 | 0.37748 | 0.809 | 0.63 | 6.05E-04 | HC_vs_MS.Trt_B |
| LUZP2 | 3.85E-08 | 0.25486 | 0.393 | 0.24 | 6.15E-04 | HC_vs_MS.Trt_B |

| | | | | | | |
|---|---|---|---|---|---|---|
| KAT6A | 3.91E-08 | 0.66537 | 0.767 | 0.61 | 6.25E-04 | HC_vs_MS.Trt_B |
| ADGRF1 | 4.19E-08 | 0.27522 | 0.775 | 0.62 | 6.70E-04 | HC_vs_MS.Trt_B |
| KLHL36 | 4.31E-08 | 1.016 | 0.636 | 0.49 | 6.89E-04 | HC_vs_MS.Trt_B |
| ZKSCAN5 | 4.38E-08 | 0.33838 | 0.562 | 0.4 | 7.00E-04 | HC_vs_MS.Trt_B |
| TMEM67 | 4.38E-08 | 0.36601 | 0.884 | 0.8 | 7.00E-04 | HC_vs_MS.Trt_B |
| ZNF160 | 4.42E-08 | 0.41415 | 0.593 | 0.42 | 7.05E-04 | HC_vs_MS.Trt_B |
| LTBR | 4.57E-08 | 0.28701 | 0.929 | 0.82 | 7.30E-04 | HC_vs_MS.Trt_B |
| WBSCR22 | 4.67E-08 | 0.41066 | 0.726 | 0.58 | 7.45E-04 | HC_vs_MS.Trt_B |
| AKAP7 | 4.68E-08 | 0.66517 | 0.5 | 0.35 | 7.47E-04 | HC_vs_MS.Trt_B |
| MICAL2 | 4.75E-08 | 0.41752 | 0.822 | 0.68 | 7.59E-04 | HC_vs_MS.Trt_B |
| SCML4 | 4.97E-08 | 0.37051 | 0.629 | 0.46 | 7.93E-04 | HC_vs_MS.Trt_B |
| UBN2 | 4.98E-08 | 0.74142 | 0.606 | 0.46 | 7.96E-04 | HC_vs_MS.Trt_B |
| SMARCAL1 | 5.00E-08 | 0.45188 | 0.706 | 0.56 | 7.98E-04 | HC_vs_MS.Trt_B |
| ZNF623 | 5.02E-08 | -0.2981 | 0.629 | 0.48 | 8.02E-04 | HC_vs_MS.Trt_B |
| TGM2 | 5.02E-08 | 0.39763 | 0.552 | 0.4 | 8.02E-04 | HC_vs_MS.Trt_B |
| FAM189B | 5.17E-08 | 0.48923 | 0.69 | 0.56 | 8.26E-04 | HC_vs_MS.Trt_B |
| CDH3 | 5.19E-08 | 0.33703 | 0.846 | 0.75 | 8.29E-04 | HC_vs_MS.Trt_B |
| CYP2B6 | 5.26E-08 | 0.29904 | 0.607 | 0.46 | 8.40E-04 | HC_vs_MS.Trt_B |
| PLEKHG2 | 5.29E-08 | 0.34401 | 0.819 | 0.71 | 8.45E-04 | HC_vs_MS.Trt_B |
| TTC6 | 5.33E-08 | 0.34799 | 0.286 | 0.15 | 8.52E-04 | HC_vs_MS.Trt_B |
| ZFYVE26 | 5.36E-08 | 0.47372 | 0.836 | 0.71 | 8.56E-04 | HC_vs_MS.Trt_B |
| STARD10 | 5.43E-08 | 0.32332 | 0.825 | 0.67 | 8.68E-04 | HC_vs_MS.Trt_B |
| ZNF778 | 5.47E-08 | -1.0824 | 0.654 | 0.5 | 8.73E-04 | HC_vs_MS.Trt_B |
| PTCHD1 | 5.53E-08 | 0.49803 | 0.703 | 0.57 | 8.84E-04 | HC_vs_MS.Trt_B |
| PANK1 | 5.91E-08 | 0.27138 | 0.452 | 0.3 | 9.44E-04 | HC_vs_MS.Trt_B |
| ABHD15 | 5.92E-08 | 1.1551 | 0.329 | 0.18 | 9.46E-04 | HC_vs_MS.Trt_B |
| IRF9 | 5.96E-08 | 0.31762 | 0.891 | 0.76 | 9.52E-04 | HC_vs_MS.Trt_B |
| CYP2C9 | 6.00E-08 | 0.25468 | 0.588 | 0.43 | 9.58E-04 | HC_vs_MS.Trt_B |
| BET1 | 6.11E-08 | 0.43126 | 0.568 | 0.4 | 9.76E-04 | HC_vs_MS.Trt_B |
| SPTSSA | 6.18E-08 | 0.25419 | 0.401 | 0.24 | 9.88E-04 | HC_vs_MS.Trt_B |
| SH3PXD2A | 6.35E-08 | -0.2802 | 0.314 | 0.17 | 0.001014 | HC_vs_MS.Trt_B |
| ASB6 | 6.37E-08 | 0.8149 | 0.583 | 0.44 | 0.001018 | HC_vs_MS.Trt_B |
| ARHGAP22 | 6.41E-08 | 0.25391 | 0.643 | 0.46 | 0.001023 | HC_vs_MS.Trt_B |
| ZNF568 | 6.48E-08 | 0.42277 | 0.845 | 0.71 | 0.001035 | HC_vs_MS.Trt_B |
| PARP9 | 6.52E-08 | 0.68835 | 0.388 | 0.24 | 0.001041 | HC_vs_MS.Trt_B |
| SSH1 | 6.71E-08 | 0.55707 | 0.846 | 0.72 | 0.001071 | HC_vs_MS.Trt_B |
| PARVB | 6.72E-08 | 0.50655 | 0.677 | 0.55 | 0.001074 | HC_vs_MS.Trt_B |
| C4BPB | 6.93E-08 | 0.34072 | 0.78 | 0.65 | 0.001107 | HC_vs_MS.Trt_B |
| ZNF620 | 7.11E-08 | 0.30967 | 0.716 | 0.57 | 0.001135 | HC_vs_MS.Trt_B |
| SLC36A2 | 7.18E-08 | 0.38056 | 0.619 | 0.49 | 0.001147 | HC_vs_MS.Trt_B |

| | | | | | | |
|---|---|---|---|---|---|---|
| SLC37A3 | 7.24E-08 | 0.38864 | 0.936 | 0.85 | 0.001157 | HC_vs_MS.Trt_B |
| TNIK | 7.32E-08 | 1.00165 | 0.4 | 0.26 | 0.00117 | HC_vs_MS.Trt_B |
| PRRG4 | 7.36E-08 | 0.32123 | 0.757 | 0.64 | 0.001176 | HC_vs_MS.Trt_B |
| CALCOCO1 | 7.44E-08 | 0.57591 | 0.497 | 0.34 | 0.001188 | HC_vs_MS.Trt_B |
| ZNF611 | 7.75E-08 | 0.42073 | 0.951 | 0.85 | 0.001238 | HC_vs_MS.Trt_B |
| NWD1 | 7.78E-08 | 0.33957 | 0.487 | 0.34 | 0.001243 | HC_vs_MS.Trt_B |
| CREBBP | 7.81E-08 | 0.34538 | 0.564 | 0.4 | 0.001247 | HC_vs_MS.Trt_B |
| DENND2C | 7.96E-08 | 0.33355 | 0.751 | 0.61 | 0.001272 | HC_vs_MS.Trt_B |
| ASNS | 8.01E-08 | 0.73444 | 0.296 | 0.16 | 0.001279 | HC_vs_MS.Trt_B |
| ARL17B | 8.04E-08 | 0.26987 | 0.722 | 0.58 | 0.001284 | HC_vs_MS.Trt_B |
| SNUPN | 8.18E-08 | 0.31648 | 0.843 | 0.71 | 0.001306 | HC_vs_MS.Trt_B |
| MT-ND4L | 8.29E-08 | -0.5242 | 0.984 | 0.99 | 0.001324 | HC_vs_MS.Trt_B |
| IGSF9 | 8.41E-08 | 0.68689 | 0.304 | 0.17 | 0.001344 | HC_vs_MS.Trt_B |
| WDR12 | 8.44E-08 | -0.393 | 0.883 | 0.79 | 0.001348 | HC_vs_MS.Trt_B |
| STPG1 | 8.47E-08 | 0.34293 | 0.755 | 0.63 | 0.001352 | HC_vs_MS.Trt_B |
| POC1B | 8.49E-08 | 0.49341 | 0.742 | 0.59 | 0.001356 | HC_vs_MS.Trt_B |
| PPP5D1 | 8.91E-08 | 0.40403 | 0.729 | 0.6 | 0.001423 | HC_vs_MS.Trt_B |
| ST6GAL1 | 8.97E-08 | 0.40158 | 0.462 | 0.31 | 0.001432 | HC_vs_MS.Trt_B |
| TRIM25 | 9.02E-08 | 0.91851 | 0.33 | 0.19 | 0.001441 | HC_vs_MS.Trt_B |
| SMAD4 | 9.25E-08 | 0.26331 | 0.491 | 0.31 | 0.001477 | HC_vs_MS.Trt_B |
| DDX60L | 9.26E-08 | 0.53774 | 0.714 | 0.56 | 0.001479 | HC_vs_MS.Trt_B |
| ZNF28 | 9.27E-08 | 1.12834 | 0.777 | 0.63 | 0.001481 | HC_vs_MS.Trt_B |
| C10orf67 | 9.61E-08 | 0.67109 | 0.28 | 0.15 | 0.001535 | HC_vs_MS.Trt_B |
| PRKAG1 | 9.73E-08 | 0.33358 | 0.632 | 0.47 | 0.001554 | HC_vs_MS.Trt_B |
| SLC25A44 | 9.73E-08 | 0.29029 | 0.674 | 0.49 | 0.001554 | HC_vs_MS.Trt_B |
| RNF213 | 9.76E-08 | 0.27285 | 0.981 | 0.89 | 0.001559 | HC_vs_MS.Trt_B |
| SCO1 | 9.87E-08 | 0.48064 | 0.775 | 0.66 | 0.001576 | HC_vs_MS.Trt_B |
| PTPRK | 9.93E-08 | 0.35385 | 0.443 | 0.3 | 0.001587 | HC_vs_MS.Trt_B |
| PPP2R1B | 1.03E-07 | 0.41095 | 0.409 | 0.27 | 0.001637 | HC_vs_MS.Trt_B |
| CLN8 | 1.03E-07 | 0.52299 | 0.839 | 0.72 | 0.001638 | HC_vs_MS.Trt_B |
| MOSPD2 | 1.03E-07 | 0.28083 | 0.309 | 0.18 | 0.001641 | HC_vs_MS.Trt_B |
| LSM7 | 1.04E-07 | -0.7008 | 0.675 | 0.49 | 0.001656 | HC_vs_MS.Trt_B |
| PHLDB2 | 1.05E-07 | -1.0063 | 0.51 | 0.37 | 0.001685 | HC_vs_MS.Trt_B |
| RGS12 | 1.07E-07 | 0.90491 | 0.823 | 0.68 | 0.001708 | HC_vs_MS.Trt_B |
| SLC43A2 | 1.09E-07 | 0.36633 | 0.786 | 0.63 | 0.001738 | HC_vs_MS.Trt_B |
| ZNF483 | 1.10E-07 | 0.5361 | 0.826 | 0.7 | 0.00175 | HC_vs_MS.Trt_B |
| RNGTT | 1.10E-07 | 0.69841 | 0.394 | 0.26 | 0.001753 | HC_vs_MS.Trt_B |
| CDK5RAP1 | 1.11E-07 | 0.77434 | 0.384 | 0.25 | 0.001766 | HC_vs_MS.Trt_B |
| EIF4A2 | 1.11E-07 | 0.36201 | 0.554 | 0.4 | 0.001776 | HC_vs_MS.Trt_B |
| SPTLC2 | 1.16E-07 | 0.84392 | 0.697 | 0.59 | 0.001846 | HC_vs_MS.Trt_B |

| | | | | | | |
|---|---|---|---|---|---|---|
| HERC1 | 1.18E-07 | -0.2664 | 0.83 | 0.67 | 0.001888 | HC_vs_MS.Trt_B |
| ZNF124 | 1.18E-07 | 0.57091 | 0.665 | 0.51 | 0.00189 | HC_vs_MS.Trt_B |
| RICTOR | 1.19E-07 | 0.25743 | 0.796 | 0.66 | 0.001899 | HC_vs_MS.Trt_B |
| TSKU | 1.20E-07 | 0.29202 | 0.488 | 0.32 | 0.001917 | HC_vs_MS.Trt_B |
| PNLIPRP1 | 1.20E-07 | 0.25335 | 0.809 | 0.7 | 0.001921 | HC_vs_MS.Trt_B |
| GCNT2 | 1.21E-07 | 0.65615 | 0.732 | 0.58 | 0.001934 | HC_vs_MS.Trt_B |
| FBXO27 | 1.23E-07 | 0.27839 | 0.697 | 0.58 | 0.001968 | HC_vs_MS.Trt_B |
| MARVELD2 | 1.23E-07 | 0.58121 | 0.59 | 0.48 | 0.001972 | HC_vs_MS.Trt_B |
| SMARCA4 | 1.27E-07 | -0.5363 | 0.79 | 0.66 | 0.002036 | HC_vs_MS.Trt_B |
| TRIM65 | 1.29E-07 | 0.40207 | 0.684 | 0.53 | 0.002067 | HC_vs_MS.Trt_B |
| KCNJ5 | 1.30E-07 | 0.49971 | 0.597 | 0.47 | 0.002081 | HC_vs_MS.Trt_B |
| DDX46 | 1.31E-07 | 0.94756 | 0.52 | 0.39 | 0.002095 | HC_vs_MS.Trt_B |
| CDC14B | 1.32E-07 | 1.38676 | 0.322 | 0.18 | 0.002105 | HC_vs_MS.Trt_B |
| SIDT1 | 1.33E-07 | 0.55051 | 0.781 | 0.65 | 0.002117 | HC_vs_MS.Trt_B |
| ZCCHC8 | 1.35E-07 | 0.31168 | 0.813 | 0.69 | 0.002153 | HC_vs_MS.Trt_B |
| SCUBE2 | 1.36E-07 | 0.34863 | 0.304 | 0.17 | 0.002166 | HC_vs_MS.Trt_B |
| SRSF4 | 1.36E-07 | 0.37887 | 0.703 | 0.56 | 0.002179 | HC_vs_MS.Trt_B |
| WHSC1 | 1.37E-07 | 0.34814 | 0.707 | 0.54 | 0.002188 | HC_vs_MS.Trt_B |
| NRBP1 | 1.37E-07 | 0.39985 | 0.593 | 0.43 | 0.002191 | HC_vs_MS.Trt_B |
| ZNF345 | 1.38E-07 | 0.27195 | 0.814 | 0.68 | 0.002204 | HC_vs_MS.Trt_B |
| PCF11 | 1.39E-07 | 0.45774 | 0.519 | 0.36 | 0.002221 | HC_vs_MS.Trt_B |
| PNPLA3 | 1.49E-07 | 0.33137 | 0.53 | 0.39 | 0.002387 | HC_vs_MS.Trt_B |
| LIX1 | 1.50E-07 | 0.27872 | 0.387 | 0.24 | 0.002397 | HC_vs_MS.Trt_B |
| SLC35A3 | 1.51E-07 | 0.78269 | 0.799 | 0.68 | 0.002415 | HC_vs_MS.Trt_B |
| LRBA | 1.52E-07 | 0.38026 | 0.477 | 0.33 | 0.002426 | HC_vs_MS.Trt_B |
| SNX20 | 1.53E-07 | 0.37771 | 0.481 | 0.32 | 0.002447 | HC_vs_MS.Trt_B |
| CALCOCO2 | 1.53E-07 | -0.2812 | 0.988 | 0.95 | 0.002448 | HC_vs_MS.Trt_B |
| CHGB | 1.55E-07 | 0.68657 | 0.458 | 0.31 | 0.002474 | HC_vs_MS.Trt_B |
| NEK5 | 1.61E-07 | 0.31703 | 0.852 | 0.71 | 0.00257 | HC_vs_MS.Trt_B |
| INIP | 1.63E-07 | 0.38821 | 0.681 | 0.55 | 0.002607 | HC_vs_MS.Trt_B |
| OCLN | 1.65E-07 | 0.39257 | 0.796 | 0.63 | 0.002633 | HC_vs_MS.Trt_B |
| RASGRP1 | 1.68E-07 | 0.33527 | 0.354 | 0.22 | 0.002677 | HC_vs_MS.Trt_B |
| SORT1 | 1.70E-07 | 0.60689 | 0.528 | 0.38 | 0.002721 | HC_vs_MS.Trt_B |
| PIGB | 1.71E-07 | 0.42509 | 0.775 | 0.65 | 0.002733 | HC_vs_MS.Trt_B |
| MYADM | 1.73E-07 | 0.2528 | 0.728 | 0.6 | 0.002758 | HC_vs_MS.Trt_B |
| ATXN2 | 1.76E-07 | 0.61229 | 0.88 | 0.79 | 0.002808 | HC_vs_MS.Trt_B |
| HECTD4 | 1.76E-07 | 0.32851 | 0.662 | 0.52 | 0.002819 | HC_vs_MS.Trt_B |
| TAZ | 1.80E-07 | 0.48436 | 0.543 | 0.38 | 0.002876 | HC_vs_MS.Trt_B |
| ZNF546 | 1.82E-07 | 0.28468 | 0.893 | 0.8 | 0.002904 | HC_vs_MS.Trt_B |
| BANP | 1.84E-07 | 0.46475 | 0.658 | 0.52 | 0.002933 | HC_vs_MS.Trt_B |

| | | | | | | |
|---|---|---|---|---|---|---|
| PPARGC1B | 1.85E-07 | 0.8597 | 0.694 | 0.56 | 0.002961 | HC_vs_MS.Trt_B |
| NDUFAF5 | 1.86E-07 | 0.55188 | 0.538 | 0.38 | 0.00297 | HC_vs_MS.Trt_B |
| CCDC81 | 1.91E-07 | 0.35153 | 0.486 | 0.34 | 0.003055 | HC_vs_MS.Trt_B |
| HELLS | 1.92E-07 | 0.32876 | 0.774 | 0.65 | 0.003066 | HC_vs_MS.Trt_B |
| GNA11 | 1.92E-07 | 0.49262 | 0.53 | 0.39 | 0.003067 | HC_vs_MS.Trt_B |
| TPH1 | 1.95E-07 | -0.81 | 0.51 | 0.35 | 0.00311 | HC_vs_MS.Trt_B |
| SH2D6 | 2.00E-07 | 0.39224 | 0.417 | 0.27 | 0.003197 | HC_vs_MS.Trt_B |
| TTLL5 | 2.01E-07 | 0.75011 | 0.632 | 0.5 | 0.003205 | HC_vs_MS.Trt_B |
| AOAH | 2.01E-07 | -0.9112 | 0.386 | 0.24 | 0.00321 | HC_vs_MS.Trt_B |
| AP5S1 | 2.02E-07 | 0.54888 | 0.864 | 0.75 | 0.00322 | HC_vs_MS.Trt_B |
| NFE2L2 | 2.02E-07 | 0.39473 | 0.561 | 0.41 | 0.003227 | HC_vs_MS.Trt_B |
| TMEM135 | 2.03E-07 | -0.3079 | 0.768 | 0.62 | 0.003249 | HC_vs_MS.Trt_B |
| SNIP1 | 2.10E-07 | 0.36982 | 0.659 | 0.53 | 0.003353 | HC_vs_MS.Trt_B |
| COG8 | 2.16E-07 | 0.30405 | 0.629 | 0.48 | 0.003458 | HC_vs_MS.Trt_B |
| AZIN2 | 2.17E-07 | -0.8291 | 0.319 | 0.18 | 0.00346 | HC_vs_MS.Trt_B |
| LILRB4 | 2.18E-07 | 0.37272 | 0.336 | 0.2 | 0.003487 | HC_vs_MS.Trt_B |
| DDX10 | 2.20E-07 | 0.95906 | 0.612 | 0.48 | 0.003516 | HC_vs_MS.Trt_B |
| ZNF566 | 2.22E-07 | 0.39631 | 0.861 | 0.75 | 0.003538 | HC_vs_MS.Trt_B |
| TBC1D5 | 2.22E-07 | 0.25542 | 0.597 | 0.44 | 0.003546 | HC_vs_MS.Trt_B |
| TBX6 | 2.24E-07 | 0.28551 | 0.47 | 0.33 | 0.003583 | HC_vs_MS.Trt_B |
| UGGT2 | 2.26E-07 | 0.27119 | 0.803 | 0.65 | 0.003608 | HC_vs_MS.Trt_B |
| AGO3 | 2.28E-07 | 0.5198 | 0.965 | 0.92 | 0.003638 | HC_vs_MS.Trt_B |
| SIRT5 | 2.28E-07 | 0.43591 | 0.857 | 0.75 | 0.003641 | HC_vs_MS.Trt_B |
| NELFA | 2.29E-07 | 0.66665 | 0.786 | 0.66 | 0.003651 | HC_vs_MS.Trt_B |
| C16orf70 | 2.32E-07 | -0.355 | 0.661 | 0.51 | 0.003713 | HC_vs_MS.Trt_B |
| MYO1B | 2.36E-07 | 0.30237 | 0.393 | 0.25 | 0.003769 | HC_vs_MS.Trt_B |
| STX7 | 2.39E-07 | 0.28864 | 0.72 | 0.58 | 0.003813 | HC_vs_MS.Trt_B |
| AMN1 | 2.40E-07 | 0.25377 | 0.765 | 0.6 | 0.003828 | HC_vs_MS.Trt_B |
| MAPT | 2.41E-07 | 0.40988 | 0.677 | 0.51 | 0.003842 | HC_vs_MS.Trt_B |
| USP46 | 2.41E-07 | 0.25393 | 0.717 | 0.56 | 0.003851 | HC_vs_MS.Trt_B |
| TMEM189 | 2.43E-07 | 0.30287 | 0.829 | 0.69 | 0.003876 | HC_vs_MS.Trt_B |
| KLLN | 2.43E-07 | 0.36484 | 0.461 | 0.31 | 0.003886 | HC_vs_MS.Trt_B |
| PROCR | 2.46E-07 | 0.25764 | 0.839 | 0.73 | 0.003935 | HC_vs_MS.Trt_B |
| HSD17B2 | 2.48E-07 | 0.4511 | 0.749 | 0.62 | 0.003955 | HC_vs_MS.Trt_B |
| LARGE1 | 2.56E-07 | 0.31875 | 0.557 | 0.41 | 0.004094 | HC_vs_MS.Trt_B |
| ACAP2 | 2.60E-07 | 0.31821 | 0.687 | 0.55 | 0.004161 | HC_vs_MS.Trt_B |
| RNFT2 | 2.63E-07 | 0.34551 | 0.87 | 0.76 | 0.004194 | HC_vs_MS.Trt_B |
| NUP107 | 2.65E-07 | 0.62014 | 0.741 | 0.6 | 0.00424 | HC_vs_MS.Trt_B |
| ZNF347 | 2.66E-07 | 0.62125 | 0.91 | 0.82 | 0.00425 | HC_vs_MS.Trt_B |
| ESCO2 | 2.69E-07 | 0.33641 | 0.665 | 0.56 | 0.004289 | HC_vs_MS.Trt_B |

| | | | | | | |
|---|---|---|---|---|---|---|
| FGFR1OP | 2.73E-07 | -0.3305 | 0.835 | 0.7 | 0.004355 | HC_vs_MS.Trt_B |
| PIGX | 2.73E-07 | 0.53397 | 0.43 | 0.3 | 0.004364 | HC_vs_MS.Trt_B |
| ZFAND3 | 2.73E-07 | 0.83925 | 0.642 | 0.5 | 0.004368 | HC_vs_MS.Trt_B |
| ELK3 | 2.74E-07 | 0.33967 | 0.393 | 0.24 | 0.004375 | HC_vs_MS.Trt_B |
| SH3BP5 | 2.79E-07 | 0.37292 | 0.801 | 0.7 | 0.004454 | HC_vs_MS.Trt_B |
| ZNF320 | 2.85E-07 | 0.32923 | 0.977 | 0.92 | 0.00455 | HC_vs_MS.Trt_B |
| PNRC1 | 2.86E-07 | 0.77977 | 0.31 | 0.18 | 0.004561 | HC_vs_MS.Trt_B |
| CLNK | 2.88E-07 | 0.31398 | 0.626 | 0.49 | 0.004602 | HC_vs_MS.Trt_B |
| SEC14L1 | 2.91E-07 | 0.88474 | 0.665 | 0.53 | 0.004653 | HC_vs_MS.Trt_B |
| ASAP3 | 2.91E-07 | 0.29342 | 0.53 | 0.38 | 0.004655 | HC_vs_MS.Trt_B |
| SPIB | 2.92E-07 | 0.28821 | 0.822 | 0.75 | 0.004658 | HC_vs_MS.Trt_B |
| PDPK1 | 2.94E-07 | 0.4829 | 0.599 | 0.45 | 0.004699 | HC_vs_MS.Trt_B |
| STOML1 | 2.94E-07 | 0.31967 | 0.765 | 0.63 | 0.004701 | HC_vs_MS.Trt_B |
| COX20 | 2.95E-07 | -0.6146 | 0.652 | 0.48 | 0.004717 | HC_vs_MS.Trt_B |
| DDX58 | 2.96E-07 | 0.51825 | 0.338 | 0.2 | 0.004729 | HC_vs_MS.Trt_B |
| SLA2 | 2.98E-07 | 0.97076 | 0.522 | 0.38 | 0.004758 | HC_vs_MS.Trt_B |
| BICDL2 | 2.99E-07 | 0.30222 | 0.296 | 0.16 | 0.004769 | HC_vs_MS.Trt_B |
| HYOU1 | 3.02E-07 | 0.51586 | 0.475 | 0.31 | 0.004821 | HC_vs_MS.Trt_B |
| BLZF1 | 3.03E-07 | 0.7909 | 0.441 | 0.31 | 0.004837 | HC_vs_MS.Trt_B |
| GFOD2 | 3.06E-07 | 0.47498 | 0.886 | 0.81 | 0.004887 | HC_vs_MS.Trt_B |
| CPA4 | 3.09E-07 | 0.29805 | 0.525 | 0.38 | 0.004941 | HC_vs_MS.Trt_B |
| PTRF | 3.14E-07 | 0.31371 | 0.352 | 0.21 | 0.00502 | HC_vs_MS.Trt_B |
| USP37 | 3.15E-07 | -0.3453 | 0.467 | 0.31 | 0.005034 | HC_vs_MS.Trt_B |
| COQ5 | 3.15E-07 | 0.48026 | 0.864 | 0.69 | 0.005038 | HC_vs_MS.Trt_B |
| NNT | 3.19E-07 | 0.41562 | 0.638 | 0.5 | 0.005089 | HC_vs_MS.Trt_B |
| KRAS | 3.24E-07 | 0.42563 | 0.372 | 0.23 | 0.005173 | HC_vs_MS.Trt_B |
| DMBX1 | 3.28E-07 | 0.34337 | 0.394 | 0.25 | 0.005231 | HC_vs_MS.Trt_B |
| HEATR3 | 3.29E-07 | 1.07054 | 0.625 | 0.47 | 0.00526 | HC_vs_MS.Trt_B |
| CHD1 | 3.39E-07 | 0.39724 | 0.468 | 0.33 | 0.005422 | HC_vs_MS.Trt_B |
| ZNF525 | 3.40E-07 | -1.5593 | 0.597 | 0.45 | 0.005427 | HC_vs_MS.Trt_B |
| CAPZA2 | 3.44E-07 | -0.5154 | 0.891 | 0.82 | 0.005499 | HC_vs_MS.Trt_B |
| ARID3B | 3.51E-07 | 0.53658 | 0.712 | 0.6 | 0.005602 | HC_vs_MS.Trt_B |
| ZC3HAV1L | 3.53E-07 | 0.46136 | 0.433 | 0.29 | 0.005632 | HC_vs_MS.Trt_B |
| SRGAP3 | 3.57E-07 | -0.3258 | 0.862 | 0.75 | 0.005703 | HC_vs_MS.Trt_B |
| SRBD1 | 3.57E-07 | -0.4233 | 0.355 | 0.22 | 0.005704 | HC_vs_MS.Trt_B |
| RRM1 | 3.58E-07 | 0.39829 | 0.341 | 0.2 | 0.005724 | HC_vs_MS.Trt_B |
| GRAMD2 | 3.63E-07 | 0.27262 | 0.617 | 0.48 | 0.005793 | HC_vs_MS.Trt_B |
| SPNS1 | 3.69E-07 | 0.30294 | 0.787 | 0.67 | 0.005894 | HC_vs_MS.Trt_B |
| PARN | 3.75E-07 | 0.29025 | 0.714 | 0.61 | 0.005993 | HC_vs_MS.Trt_B |
| WDR90 | 3.75E-07 | 0.25378 | 0.878 | 0.78 | 0.005997 | HC_vs_MS.Trt_B |

| | | | | | | |
|---|---|---|---|---|---|---|
| GTF2H2 | 3.78E-07 | 0.32441 | 0.884 | 0.82 | 0.006033 | HC_vs_MS.Trt_B |
| HLA-DQB1 | 3.83E-07 | 0.5467 | 0.546 | 0.39 | 0.006113 | HC_vs_MS.Trt_B |
| NAMPT | 3.84E-07 | 0.9763 | 0.687 | 0.56 | 0.006138 | HC_vs_MS.Trt_B |
| SDE2 | 3.86E-07 | 0.40556 | 0.657 | 0.53 | 0.006159 | HC_vs_MS.Trt_B |
| DLST | 3.94E-07 | 0.42102 | 0.888 | 0.79 | 0.006298 | HC_vs_MS.Trt_B |
| SAG | 3.98E-07 | 0.25428 | 0.942 | 0.92 | 0.006349 | HC_vs_MS.Trt_B |
| SLC2A10 | 4.01E-07 | 0.36733 | 0.458 | 0.31 | 0.006398 | HC_vs_MS.Trt_B |
| SLC27A1 | 4.02E-07 | 0.50311 | 0.797 | 0.69 | 0.006426 | HC_vs_MS.Trt_B |
| SLC4A1 | 4.12E-07 | 0.33991 | 0.671 | 0.52 | 0.006579 | HC_vs_MS.Trt_B |
| BMS1 | 4.13E-07 | 0.31182 | 0.536 | 0.39 | 0.006597 | HC_vs_MS.Trt_B |
| PSPH | 4.16E-07 | 0.39742 | 0.83 | 0.71 | 0.006649 | HC_vs_MS.Trt_B |
| GNB4 | 4.23E-07 | 0.36305 | 0.791 | 0.68 | 0.006749 | HC_vs_MS.Trt_B |
| CHPT1 | 4.26E-07 | 0.37259 | 0.609 | 0.46 | 0.006803 | HC_vs_MS.Trt_B |
| GSTO2 | 4.32E-07 | 0.33038 | 0.858 | 0.72 | 0.006899 | HC_vs_MS.Trt_B |
| CEP41 | 4.34E-07 | 0.26911 | 0.751 | 0.62 | 0.006926 | HC_vs_MS.Trt_B |
| ZFPM2 | 4.36E-07 | 0.35834 | 0.417 | 0.28 | 0.006958 | HC_vs_MS.Trt_B |
| ULK2 | 4.36E-07 | 0.49073 | 0.701 | 0.57 | 0.006972 | HC_vs_MS.Trt_B |
| NFYC | 4.37E-07 | 0.27358 | 0.572 | 0.42 | 0.006976 | HC_vs_MS.Trt_B |
| GSK3B | 4.38E-07 | 0.5114 | 0.712 | 0.57 | 0.007002 | HC_vs_MS.Trt_B |
| KIAA0930 | 4.47E-07 | 0.44782 | 0.612 | 0.48 | 0.007134 | HC_vs_MS.Trt_B |
| PRPF38A | 4.56E-07 | -0.4081 | 0.612 | 0.46 | 0.007291 | HC_vs_MS.Trt_B |
| POM121C | 4.60E-07 | 0.552 | 0.494 | 0.35 | 0.007346 | HC_vs_MS.Trt_B |
| TAOK3 | 4.64E-07 | 0.35698 | 0.759 | 0.65 | 0.007408 | HC_vs_MS.Trt_B |
| SLC25A15 | 4.69E-07 | 0.28071 | 0.672 | 0.53 | 0.007493 | HC_vs_MS.Trt_B |
| TRAF6 | 4.70E-07 | 0.59887 | 0.712 | 0.56 | 0.007511 | HC_vs_MS.Trt_B |
| OSBPL1A | 4.76E-07 | 0.30992 | 0.736 | 0.59 | 0.007597 | HC_vs_MS.Trt_B |
| CCZ1 | 4.78E-07 | 0.34132 | 0.72 | 0.56 | 0.00764 | HC_vs_MS.Trt_B |
| SCHIP1 | 4.81E-07 | 0.39903 | 0.532 | 0.38 | 0.00769 | HC_vs_MS.Trt_B |
| ELMO1 | 4.91E-07 | 0.33262 | 0.754 | 0.64 | 0.007845 | HC_vs_MS.Trt_B |
| SLC25A34 | 4.93E-07 | 0.48056 | 0.622 | 0.47 | 0.007873 | HC_vs_MS.Trt_B |
| HIVEP2 | 5.05E-07 | 0.39396 | 0.396 | 0.25 | 0.00806 | HC_vs_MS.Trt_B |
| NABP1 | 5.09E-07 | 0.36646 | 0.757 | 0.64 | 0.008125 | HC_vs_MS.Trt_B |
| APOBEC3G | 5.17E-07 | 0.27481 | 0.77 | 0.65 | 0.008264 | HC_vs_MS.Trt_B |
| JMJD7-PLA2 | 5.19E-07 | -0.3091 | 0.386 | 0.25 | 0.008289 | HC_vs_MS.Trt_B |
| DPPA4 | 5.20E-07 | 0.30762 | 0.852 | 0.76 | 0.008306 | HC_vs_MS.Trt_B |
| TMEM107 | 5.20E-07 | -0.4287 | 0.583 | 0.42 | 0.008313 | HC_vs_MS.Trt_B |
| KIAA1841 | 5.21E-07 | 0.63814 | 0.846 | 0.75 | 0.008314 | HC_vs_MS.Trt_B |
| NXPE3 | 5.37E-07 | 0.60548 | 0.907 | 0.82 | 0.008577 | HC_vs_MS.Trt_B |
| UBXN7 | 5.38E-07 | 0.70498 | 0.787 | 0.65 | 0.008598 | HC_vs_MS.Trt_B |
| MCM8 | 5.45E-07 | 0.51012 | 0.696 | 0.57 | 0.008711 | HC_vs_MS.Trt_B |

| | | | | | | |
|---|---|---|---|---|---|---|
| MAP7D3 | 5.49E-07 | 1.02713 | 0.577 | 0.43 | 0.008769 | HC_vs_MS.Trt_B |
| SIN3B | 5.50E-07 | 0.61767 | 0.754 | 0.62 | 0.00878 | HC_vs_MS.Trt_B |
| LUZP1 | 5.56E-07 | 0.49686 | 0.478 | 0.34 | 0.008881 | HC_vs_MS.Trt_B |
| GAB1 | 5.61E-07 | 0.33647 | 0.532 | 0.4 | 0.008961 | HC_vs_MS.Trt_B |
| PRRC2A | 5.65E-07 | 0.50414 | 0.757 | 0.65 | 0.00903 | HC_vs_MS.Trt_B |
| USP22 | 5.67E-07 | 0.41813 | 0.83 | 0.7 | 0.009059 | HC_vs_MS.Trt_B |
| ZNF556 | 5.68E-07 | 0.27502 | 0.939 | 0.86 | 0.009076 | HC_vs_MS.Trt_B |
| OAZ2 | 5.73E-07 | 0.28475 | 0.755 | 0.63 | 0.009145 | HC_vs_MS.Trt_B |
| PGBD2 | 5.75E-07 | 0.49187 | 0.301 | 0.16 | 0.009182 | HC_vs_MS.Trt_B |
| PTRH1 | 6.02E-07 | 0.42355 | 0.736 | 0.6 | 0.009621 | HC_vs_MS.Trt_B |
| C17orf62 | 6.14E-07 | 0.44872 | 0.719 | 0.6 | 0.009808 | HC_vs_MS.Trt_B |
| DPH6 | 6.23E-07 | 0.59553 | 0.409 | 0.28 | 0.009949 | HC_vs_MS.Trt_B |
| HELZ2 | 6.26E-07 | 1.3043 | 0.633 | 0.5 | 0.009995 | HC_vs_MS.Trt_B |
| PDLIM5 | 6.33E-07 | 0.50591 | 0.872 | 0.77 | 0.010104 | HC_vs_MS.Trt_B |
| HAP1 | 6.33E-07 | 0.31482 | 0.746 | 0.6 | 0.010109 | HC_vs_MS.Trt_B |
| COPB2 | 6.37E-07 | 0.34404 | 0.896 | 0.82 | 0.010178 | HC_vs_MS.Trt_B |
| TMEM184A | 6.38E-07 | -0.5039 | 0.419 | 0.27 | 0.010184 | HC_vs_MS.Trt_B |
| FER | 6.45E-07 | 1.18601 | 0.504 | 0.36 | 0.010306 | HC_vs_MS.Trt_B |
| TAF1 | 6.49E-07 | 0.34806 | 0.799 | 0.68 | 0.010362 | HC_vs_MS.Trt_B |
| DENND4A | 6.61E-07 | 0.25493 | 0.388 | 0.25 | 0.010556 | HC_vs_MS.Trt_B |
| TMEM127 | 6.76E-07 | 0.44903 | 0.691 | 0.57 | 0.010791 | HC_vs_MS.Trt_B |
| SMPD3 | 6.77E-07 | 0.73022 | 0.696 | 0.53 | 0.010818 | HC_vs_MS.Trt_B |
| TRAPPC8 | 6.83E-07 | 0.34699 | 0.672 | 0.51 | 0.010909 | HC_vs_MS.Trt_B |
| SNTB2 | 6.83E-07 | 0.59282 | 0.774 | 0.6 | 0.010914 | HC_vs_MS.Trt_B |
| SCLT1 | 6.85E-07 | 0.39986 | 0.614 | 0.47 | 0.01094 | HC_vs_MS.Trt_B |
| OGFOD3 | 6.90E-07 | 0.97004 | 0.577 | 0.45 | 0.011013 | HC_vs_MS.Trt_B |
| URB1 | 6.95E-07 | 0.73611 | 0.471 | 0.34 | 0.011097 | HC_vs_MS.Trt_B |
| ZCCHC4 | 6.96E-07 | 0.5279 | 0.635 | 0.48 | 0.01112 | HC_vs_MS.Trt_B |
| ZNF561 | 6.99E-07 | 0.2635 | 0.532 | 0.4 | 0.011166 | HC_vs_MS.Trt_B |
| CCT6B | 7.08E-07 | 0.68002 | 0.255 | 0.14 | 0.011313 | HC_vs_MS.Trt_B |
| ARL10 | 7.14E-07 | 0.4902 | 0.849 | 0.78 | 0.011399 | HC_vs_MS.Trt_B |
| RP11-77H9.: | 7.27E-07 | 0.25875 | 0.497 | 0.36 | 0.011618 | HC_vs_MS.Trt_B |
| HSP90AA1 | 7.36E-07 | 0.29303 | 0.614 | 0.51 | 0.011757 | HC_vs_MS.Trt_B |
| ACSM2A | 7.43E-07 | 0.39774 | 0.467 | 0.34 | 0.011864 | HC_vs_MS.Trt_B |
| HBG2 | 7.55E-07 | -1.3504 | 0.457 | 0.32 | 0.012057 | HC_vs_MS.Trt_B |
| DNAJC18 | 7.55E-07 | 0.3442 | 0.826 | 0.74 | 0.012063 | HC_vs_MS.Trt_B |
| HSD17B13 | 7.60E-07 | 0.44193 | 0.57 | 0.43 | 0.012141 | HC_vs_MS.Trt_B |
| BICDL1 | 7.73E-07 | -0.381 | 0.581 | 0.46 | 0.012345 | HC_vs_MS.Trt_B |
| PANK3 | 7.79E-07 | 0.36143 | 0.429 | 0.29 | 0.012436 | HC_vs_MS.Trt_B |
| ZNF773 | 7.82E-07 | 0.95666 | 0.571 | 0.46 | 0.012495 | HC_vs_MS.Trt_B |

| | | | | | | |
|---|---|---|---|---|---|---|
| MZT2A | 7.84E-07 | 0.2835 | 0.786 | 0.66 | 0.012524 | HC_vs_MS.Trt_B |
| PPP1CB | 7.93E-07 | -0.9573 | 0.636 | 0.72 | 0.012662 | HC_vs_MS.Trt_B |
| TMEM44 | 7.93E-07 | 0.41836 | 0.59 | 0.45 | 0.01267 | HC_vs_MS.Trt_B |
| CACNA1D | 7.96E-07 | 0.68276 | 0.661 | 0.52 | 0.012713 | HC_vs_MS.Trt_B |
| RABL2B | 8.03E-07 | 0.37408 | 0.867 | 0.79 | 0.012827 | HC_vs_MS.Trt_B |
| MAPK6 | 8.11E-07 | 0.61852 | 0.446 | 0.31 | 0.012947 | HC_vs_MS.Trt_B |
| TNIP3 | 8.14E-07 | 0.31968 | 0.499 | 0.37 | 0.013008 | HC_vs_MS.Trt_B |
| RNASEL | 8.15E-07 | 0.6558 | 0.338 | 0.2 | 0.013014 | HC_vs_MS.Trt_B |
| BCL7A | 8.16E-07 | 0.42364 | 0.649 | 0.54 | 0.013037 | HC_vs_MS.Trt_B |
| NR4A1 | 8.19E-07 | 0.4032 | 0.435 | 0.3 | 0.013084 | HC_vs_MS.Trt_B |
| TRMT44 | 8.20E-07 | 0.51059 | 0.238 | 0.12 | 0.013091 | HC_vs_MS.Trt_B |
| CTLA4 | 8.43E-07 | 0.6062 | 0.248 | 0.13 | 0.013471 | HC_vs_MS.Trt_B |
| DUSP28 | 8.55E-07 | 0.4569 | 0.671 | 0.58 | 0.013663 | HC_vs_MS.Trt_B |
| TMEM98 | 8.59E-07 | 0.31072 | 0.528 | 0.38 | 0.013723 | HC_vs_MS.Trt_B |
| FAM57B | 8.61E-07 | 0.29475 | 0.672 | 0.55 | 0.013759 | HC_vs_MS.Trt_B |
| PAQR5 | 8.64E-07 | 0.25932 | 0.516 | 0.38 | 0.013794 | HC_vs_MS.Trt_B |
| CPPED1 | 8.75E-07 | -0.463 | 0.593 | 0.44 | 0.013984 | HC_vs_MS.Trt_B |
| TIMELESS | 8.90E-07 | -0.3413 | 0.301 | 0.17 | 0.014213 | HC_vs_MS.Trt_B |
| CABIN1 | 8.94E-07 | 0.95188 | 0.255 | 0.14 | 0.014285 | HC_vs_MS.Trt_B |
| NIPAL1 | 9.04E-07 | 0.26605 | 0.41 | 0.28 | 0.014438 | HC_vs_MS.Trt_B |
| MGA | 9.07E-07 | 0.50609 | 0.858 | 0.75 | 0.014483 | HC_vs_MS.Trt_B |
| TFIP11 | 9.12E-07 | 0.62171 | 0.507 | 0.35 | 0.014567 | HC_vs_MS.Trt_B |
| BRIP1 | 9.13E-07 | -0.4513 | 0.845 | 0.7 | 0.014585 | HC_vs_MS.Trt_B |
| PEX1 | 9.28E-07 | 0.46687 | 0.67 | 0.55 | 0.014825 | HC_vs_MS.Trt_B |
| JRK | 9.39E-07 | 1.29376 | 0.433 | 0.29 | 0.014994 | HC_vs_MS.Trt_B |
| RNF8 | 9.41E-07 | -0.2537 | 0.496 | 0.36 | 0.015034 | HC_vs_MS.Trt_B |
| TRPV3 | 9.48E-07 | 0.44745 | 0.596 | 0.46 | 0.015135 | HC_vs_MS.Trt_B |
| KCNE3 | 9.54E-07 | 0.25835 | 0.525 | 0.38 | 0.015242 | HC_vs_MS.Trt_B |
| GLYR1 | 9.62E-07 | 0.27296 | 0.391 | 0.24 | 0.01537 | HC_vs_MS.Trt_B |
| MNT | 9.64E-07 | 0.49971 | 0.623 | 0.5 | 0.015396 | HC_vs_MS.Trt_B |
| TLR6 | 9.67E-07 | 1.27863 | 0.303 | 0.18 | 0.015444 | HC_vs_MS.Trt_B |
| PNMA2 | 9.68E-07 | 0.35033 | 0.658 | 0.54 | 0.01547 | HC_vs_MS.Trt_B |
| CHP2 | 9.79E-07 | 0.27559 | 0.407 | 0.26 | 0.015632 | HC_vs_MS.Trt_B |
| NUB1 | 9.80E-07 | 0.36835 | 0.874 | 0.78 | 0.015649 | HC_vs_MS.Trt_B |
| C12orf40 | 9.86E-07 | 0.76528 | 0.288 | 0.16 | 0.015743 | HC_vs_MS.Trt_B |
| DENND1A | 9.93E-07 | 0.32941 | 0.435 | 0.31 | 0.015868 | HC_vs_MS.Trt_B |
| SCN3B | 9.99E-07 | 0.35629 | 0.68 | 0.59 | 0.01595 | HC_vs_MS.Trt_B |
| NRG4 | 1.01E-06 | 0.33089 | 0.667 | 0.54 | 0.016117 | HC_vs_MS.Trt_B |
| ATG10 | 1.02E-06 | 0.30831 | 0.936 | 0.85 | 0.016255 | HC_vs_MS.Trt_B |
| ACACB | 1.03E-06 | -0.5505 | 0.929 | 0.83 | 0.016532 | HC_vs_MS.Trt_B |

| | | | | | | |
|---|---|---|---|---|---|---|
| ANGEL2 | 1.04E-06 | -0.3165 | 0.696 | 0.57 | 0.016595 | HC_vs_MS.Trt_B |
| ZNF470 | 1.04E-06 | 0.41336 | 0.714 | 0.57 | 0.016616 | HC_vs_MS.Trt_B |
| LSAMP | 1.05E-06 | 0.29044 | 0.443 | 0.29 | 0.01672 | HC_vs_MS.Trt_B |
| HSD3B7 | 1.05E-06 | 0.76421 | 0.432 | 0.29 | 0.016819 | HC_vs_MS.Trt_B |
| ELAC2 | 1.10E-06 | 0.30039 | 0.674 | 0.52 | 0.017604 | HC_vs_MS.Trt_B |
| ZNF670-ZNF | 1.12E-06 | 0.29982 | 0.63 | 0.49 | 0.017867 | HC_vs_MS.Trt_B |
| PPFIA1 | 1.13E-06 | 0.44278 | 0.657 | 0.52 | 0.018007 | HC_vs_MS.Trt_B |
| KIAA1257 | 1.14E-06 | 0.37495 | 0.562 | 0.42 | 0.018137 | HC_vs_MS.Trt_B |
| MED25 | 1.14E-06 | 0.6949 | 0.486 | 0.34 | 0.018177 | HC_vs_MS.Trt_B |
| SPP2 | 1.14E-06 | 0.31458 | 0.691 | 0.57 | 0.018178 | HC_vs_MS.Trt_B |
| MRPS28 | 1.14E-06 | 0.48329 | 0.596 | 0.45 | 0.018186 | HC_vs_MS.Trt_B |
| PLEKHM1 | 1.15E-06 | 0.66863 | 0.816 | 0.66 | 0.018336 | HC_vs_MS.Trt_B |
| BAG1 | 1.15E-06 | 0.44316 | 0.926 | 0.83 | 0.01837 | HC_vs_MS.Trt_B |
| CRLS1 | 1.18E-06 | 0.50257 | 0.417 | 0.28 | 0.018828 | HC_vs_MS.Trt_B |
| ZFR2 | 1.19E-06 | 0.35774 | 0.48 | 0.36 | 0.019015 | HC_vs_MS.Trt_B |
| SOX6 | 1.21E-06 | 0.54912 | 0.429 | 0.29 | 0.019322 | HC_vs_MS.Trt_B |
| C9orf64 | 1.22E-06 | 0.70156 | 0.596 | 0.44 | 0.019491 | HC_vs_MS.Trt_B |
| WDTC1 | 1.22E-06 | 0.97991 | 0.359 | 0.23 | 0.01953 | HC_vs_MS.Trt_B |
| MSRA | 1.23E-06 | 0.48886 | 0.239 | 0.12 | 0.019655 | HC_vs_MS.Trt_B |
| BUD31 | 1.24E-06 | -0.4262 | 0.714 | 0.59 | 0.019853 | HC_vs_MS.Trt_B |
| PPIL2 | 1.25E-06 | 0.61146 | 0.394 | 0.26 | 0.019949 | HC_vs_MS.Trt_B |
| THG1L | 1.25E-06 | -0.4428 | 0.551 | 0.41 | 0.019954 | HC_vs_MS.Trt_B |
| MYOM3 | 1.25E-06 | 0.44399 | 0.561 | 0.44 | 0.019964 | HC_vs_MS.Trt_B |
| ACBD6 | 1.26E-06 | 0.44537 | 0.467 | 0.34 | 0.020166 | HC_vs_MS.Trt_B |
| NR2C1 | 1.29E-06 | 0.49609 | 0.648 | 0.48 | 0.020649 | HC_vs_MS.Trt_B |
| TMC8 | 1.30E-06 | 0.27853 | 0.748 | 0.65 | 0.020685 | HC_vs_MS.Trt_B |
| DHTKD1 | 1.30E-06 | 0.39066 | 0.806 | 0.69 | 0.020694 | HC_vs_MS.Trt_B |
| ZNF814 | 1.30E-06 | 0.41533 | 0.954 | 0.89 | 0.020808 | HC_vs_MS.Trt_B |
| PXYLP1 | 1.31E-06 | 1.04055 | 0.5 | 0.35 | 0.020936 | HC_vs_MS.Trt_B |
| ZNF7 | 1.31E-06 | -0.2756 | 0.928 | 0.85 | 0.02098 | HC_vs_MS.Trt_B |
| TEN1-CDK3 | 1.31E-06 | -0.2876 | 0.52 | 0.38 | 0.020998 | HC_vs_MS.Trt_B |
| SYNJ2 | 1.32E-06 | 0.2667 | 0.917 | 0.79 | 0.02103 | HC_vs_MS.Trt_B |
| BEST1 | 1.34E-06 | 0.56674 | 0.806 | 0.71 | 0.021327 | HC_vs_MS.Trt_B |
| TMEM199 | 1.35E-06 | 0.81292 | 0.578 | 0.44 | 0.02156 | HC_vs_MS.Trt_B |
| LRRC2 | 1.35E-06 | -0.4236 | 0.762 | 0.63 | 0.021581 | HC_vs_MS.Trt_B |
| TRAF3 | 1.36E-06 | 0.46489 | 0.342 | 0.21 | 0.021726 | HC_vs_MS.Trt_B |
| RAD17 | 1.39E-06 | 0.49874 | 0.496 | 0.36 | 0.022133 | HC_vs_MS.Trt_B |
| PAQR8 | 1.39E-06 | 1.01551 | 0.51 | 0.38 | 0.022196 | HC_vs_MS.Trt_B |
| ABC7-42404 | 1.40E-06 | 0.59044 | 0.707 | 0.59 | 0.022366 | HC_vs_MS.Trt_B |
| SLC35A5 | 1.40E-06 | -1.0132 | 0.436 | 0.3 | 0.022428 | HC_vs_MS.Trt_B |

| | | | | | | |
|---|---|---|---|---|---|---|
| NARFL | 1.44E-06 | 0.38508 | 0.762 | 0.62 | 0.022981 | HC_vs_MS.Trt_B |
| GALM | 1.44E-06 | -0.2849 | 0.561 | 0.42 | 0.023016 | HC_vs_MS.Trt_B |
| GGCX | 1.44E-06 | 0.80834 | 0.891 | 0.82 | 0.023059 | HC_vs_MS.Trt_B |
| ZSCAN2 | 1.45E-06 | 0.45123 | 0.761 | 0.61 | 0.02314 | HC_vs_MS.Trt_B |
| LETM2 | 1.45E-06 | 0.49068 | 0.667 | 0.54 | 0.023185 | HC_vs_MS.Trt_B |
| FGD6 | 1.45E-06 | 1.09566 | 0.43 | 0.29 | 0.023223 | HC_vs_MS.Trt_B |
| TARBP1 | 1.45E-06 | -0.4335 | 0.412 | 0.27 | 0.023228 | HC_vs_MS.Trt_B |
| ADD1 | 1.47E-06 | 0.28521 | 0.787 | 0.65 | 0.02354 | HC_vs_MS.Trt_B |
| PCCB | 1.48E-06 | 0.31887 | 0.628 | 0.48 | 0.0236 | HC_vs_MS.Trt_B |
| ZNF615 | 1.52E-06 | 0.67224 | 0.47 | 0.33 | 0.024303 | HC_vs_MS.Trt_B |
| PRDM2 | 1.52E-06 | 0.58015 | 0.426 | 0.29 | 0.024338 | HC_vs_MS.Trt_B |
| MLKL | 1.53E-06 | 0.30729 | 0.696 | 0.54 | 0.024461 | HC_vs_MS.Trt_B |
| AEBP2 | 1.53E-06 | -0.3895 | 0.696 | 0.53 | 0.024469 | HC_vs_MS.Trt_B |
| NAGPA | 1.53E-06 | 0.30988 | 0.638 | 0.51 | 0.02451 | HC_vs_MS.Trt_B |
| VWA5B1 | 1.56E-06 | 0.3225 | 0.574 | 0.43 | 0.024916 | HC_vs_MS.Trt_B |
| CAPRIN2 | 1.56E-06 | 0.53631 | 0.678 | 0.53 | 0.02493 | HC_vs_MS.Trt_B |
| RHBDF2 | 1.58E-06 | 0.44618 | 0.919 | 0.84 | 0.025162 | HC_vs_MS.Trt_B |
| RERE | 1.58E-06 | 0.59985 | 0.393 | 0.25 | 0.025165 | HC_vs_MS.Trt_B |
| RSPH10B | 1.58E-06 | 0.38963 | 0.603 | 0.46 | 0.025233 | HC_vs_MS.Trt_B |
| GALE | 1.59E-06 | -1.3843 | 0.323 | 0.19 | 0.025374 | HC_vs_MS.Trt_B |
| CATSPER2 | 1.59E-06 | 0.57892 | 0.771 | 0.65 | 0.025428 | HC_vs_MS.Trt_B |
| STMN1 | 1.59E-06 | 0.34728 | 0.691 | 0.6 | 0.025443 | HC_vs_MS.Trt_B |
| INSR | 1.62E-06 | 0.38239 | 0.462 | 0.31 | 0.025822 | HC_vs_MS.Trt_B |
| CBX7 | 1.62E-06 | 0.40002 | 0.413 | 0.28 | 0.025874 | HC_vs_MS.Trt_B |
| CSGALNACT | 1.65E-06 | 0.2832 | 0.514 | 0.37 | 0.026422 | HC_vs_MS.Trt_B |
| HYAL4 | 1.66E-06 | 0.36959 | 0.572 | 0.42 | 0.026573 | HC_vs_MS.Trt_B |
| ADAR | 1.68E-06 | 0.34156 | 0.796 | 0.74 | 0.026789 | HC_vs_MS.Trt_B |
| SLC38A3 | 1.68E-06 | 0.32177 | 0.281 | 0.16 | 0.026823 | HC_vs_MS.Trt_B |
| L2HGDH | 1.68E-06 | 0.5282 | 0.807 | 0.73 | 0.026829 | HC_vs_MS.Trt_B |
| RP11-435I1( | 1.68E-06 | 0.44612 | 0.362 | 0.24 | 0.026841 | HC_vs_MS.Trt_B |
| SIRT2 | 1.70E-06 | 0.31318 | 0.897 | 0.81 | 0.027186 | HC_vs_MS.Trt_B |
| IRGQ | 1.73E-06 | 0.43354 | 0.98 | 0.93 | 0.027639 | HC_vs_MS.Trt_B |
| RSRP1 | 1.76E-06 | 0.38406 | 0.929 | 0.84 | 0.028125 | HC_vs_MS.Trt_B |
| C1orf64 | 1.80E-06 | 0.27049 | 0.564 | 0.43 | 0.028734 | HC_vs_MS.Trt_B |
| RALY | 1.82E-06 | 0.45605 | 0.399 | 0.26 | 0.028999 | HC_vs_MS.Trt_B |
| MRPS16 | 1.83E-06 | 0.38334 | 0.764 | 0.65 | 0.029201 | HC_vs_MS.Trt_B |
| RAD51B | 1.85E-06 | 0.69634 | 0.287 | 0.17 | 0.029584 | HC_vs_MS.Trt_B |
| CCL16 | 1.85E-06 | 0.26531 | 0.607 | 0.47 | 0.029589 | HC_vs_MS.Trt_B |
| NFATC2IP | 1.86E-06 | -0.2913 | 0.657 | 0.53 | 0.02965 | HC_vs_MS.Trt_B |
| LIX1L | 1.86E-06 | 0.46242 | 0.775 | 0.7 | 0.029707 | HC_vs_MS.Trt_B |

| | | | | | | |
|---|---|---|---|---|---|---|
| MRRF | 1.87E-06 | 0.52841 | 0.625 | 0.51 | 0.029854 | HC_vs_MS.Trt_B |
| SLC35E2 | 1.87E-06 | 0.4416 | 0.464 | 0.33 | 0.02989 | HC_vs_MS.Trt_B |
| DIS3L2 | 1.87E-06 | 0.51608 | 0.478 | 0.33 | 0.029939 | HC_vs_MS.Trt_B |
| RUFY3 | 1.91E-06 | -0.3413 | 0.545 | 0.4 | 0.030533 | HC_vs_MS.Trt_B |
| ANKUB1 | 1.92E-06 | 0.32704 | 0.413 | 0.29 | 0.030632 | HC_vs_MS.Trt_B |
| DYM | 1.93E-06 | 0.45487 | 0.701 | 0.55 | 0.030832 | HC_vs_MS.Trt_B |
| TSC1 | 1.94E-06 | 0.2949 | 0.478 | 0.33 | 0.030985 | HC_vs_MS.Trt_B |
| ZDHHC3 | 1.95E-06 | 0.36838 | 0.467 | 0.34 | 0.031081 | HC_vs_MS.Trt_B |
| CLMN | 1.95E-06 | 0.3223 | 0.63 | 0.49 | 0.03109 | HC_vs_MS.Trt_B |
| HAGH | 1.96E-06 | 0.36461 | 0.759 | 0.61 | 0.03134 | HC_vs_MS.Trt_B |
| ACAA1 | 1.97E-06 | 0.28395 | 0.468 | 0.33 | 0.031388 | HC_vs_MS.Trt_B |
| KIAA0319L | 1.98E-06 | 0.51139 | 0.662 | 0.53 | 0.031634 | HC_vs_MS.Trt_B |
| AP4E1 | 1.99E-06 | -0.2778 | 0.397 | 0.26 | 0.03175 | HC_vs_MS.Trt_B |
| COA7 | 1.99E-06 | 0.36893 | 0.871 | 0.78 | 0.031857 | HC_vs_MS.Trt_B |
| PAK4 | 2.00E-06 | 0.28941 | 0.678 | 0.56 | 0.03192 | HC_vs_MS.Trt_B |
| PKIB | 2.01E-06 | 0.37018 | 0.7 | 0.6 | 0.032169 | HC_vs_MS.Trt_B |
| CMBL | 2.02E-06 | 0.3191 | 0.48 | 0.35 | 0.032302 | HC_vs_MS.Trt_B |
| DNAJC21 | 2.02E-06 | 0.74536 | 0.381 | 0.26 | 0.03232 | HC_vs_MS.Trt_B |
| FNBP4 | 2.03E-06 | 0.28175 | 0.481 | 0.33 | 0.03246 | HC_vs_MS.Trt_B |
| TUSC3 | 2.06E-06 | 0.3965 | 0.41 | 0.29 | 0.032953 | HC_vs_MS.Trt_B |
| PLK5 | 2.07E-06 | 0.53569 | 0.528 | 0.38 | 0.033059 | HC_vs_MS.Trt_B |
| PLEKHA2 | 2.10E-06 | 0.31203 | 0.793 | 0.68 | 0.033496 | HC_vs_MS.Trt_B |
| SLC3A1 | 2.11E-06 | 0.44845 | 0.684 | 0.54 | 0.033774 | HC_vs_MS.Trt_B |
| ESYT2 | 2.15E-06 | 0.29708 | 0.543 | 0.41 | 0.034267 | HC_vs_MS.Trt_B |
| COA1 | 2.16E-06 | 0.41942 | 0.759 | 0.64 | 0.034558 | HC_vs_MS.Trt_B |
| TPP2 | 2.17E-06 | 0.56636 | 0.53 | 0.41 | 0.034706 | HC_vs_MS.Trt_B |
| LAMP1 | 2.17E-06 | 0.63808 | 0.372 | 0.25 | 0.034731 | HC_vs_MS.Trt_B |
| SSX5 | 2.19E-06 | 0.37898 | 0.199 | 0.09 | 0.034926 | HC_vs_MS.Trt_B |
| EP400NL | 2.22E-06 | 0.67537 | 0.723 | 0.63 | 0.035522 | HC_vs_MS.Trt_B |
| H3F3B | 2.24E-06 | -0.55 | 0.781 | 0.79 | 0.035799 | HC_vs_MS.Trt_B |
| GLTSCR1 | 2.26E-06 | 0.47465 | 0.581 | 0.48 | 0.036043 | HC_vs_MS.Trt_B |
| ZNF225 | 2.28E-06 | 0.66985 | 0.768 | 0.68 | 0.036475 | HC_vs_MS.Trt_B |
| CCR4 | 2.29E-06 | 0.68774 | 0.861 | 0.79 | 0.036515 | HC_vs_MS.Trt_B |
| ZYG11B | 2.30E-06 | 1.06857 | 0.704 | 0.61 | 0.036771 | HC_vs_MS.Trt_B |
| CCDC13 | 2.32E-06 | 0.82472 | 0.32 | 0.19 | 0.037016 | HC_vs_MS.Trt_B |
| PTPRM | 2.33E-06 | 1.2688 | 0.203 | 0.1 | 0.037139 | HC_vs_MS.Trt_B |
| ZNF839 | 2.36E-06 | 0.25202 | 0.719 | 0.57 | 0.037722 | HC_vs_MS.Trt_B |
| OR6J1 | 2.37E-06 | 0.34343 | 0.68 | 0.55 | 0.037888 | HC_vs_MS.Trt_B |
| LCMT2 | 2.37E-06 | 1.05658 | 0.654 | 0.54 | 0.037919 | HC_vs_MS.Trt_B |
| CSAD | 2.41E-06 | 0.33002 | 0.806 | 0.68 | 0.038543 | HC_vs_MS.Trt_B |

| | | | | | | |
|---|---|---|---|---|---|---|
| RAB40B | 2.45E-06 | -0.3695 | 0.448 | 0.32 | 0.039146 | HC_vs_MS.Trt_B |
| RLIM | 2.46E-06 | 0.30128 | 0.707 | 0.56 | 0.039268 | HC_vs_MS.Trt_B |
| WDR11 | 2.46E-06 | 0.64031 | 0.62 | 0.5 | 0.039299 | HC_vs_MS.Trt_B |
| GP2 | 2.48E-06 | 0.27756 | 0.504 | 0.37 | 0.039544 | HC_vs_MS.Trt_B |
| LINC00854 | 2.48E-06 | 0.52268 | 0.635 | 0.51 | 0.039672 | HC_vs_MS.Trt_B |
| ACSF2 | 2.50E-06 | 0.74111 | 0.47 | 0.34 | 0.039995 | HC_vs_MS.Trt_B |
| ERAP1 | 2.51E-06 | 0.55016 | 0.584 | 0.45 | 0.040077 | HC_vs_MS.Trt_B |
| CD55 | 2.51E-06 | 0.2696 | 0.487 | 0.35 | 0.040106 | HC_vs_MS.Trt_B |
| CNOT4 | 2.52E-06 | 0.57556 | 0.686 | 0.55 | 0.040267 | HC_vs_MS.Trt_B |
| TMEM186 | 2.56E-06 | 0.42819 | 0.51 | 0.37 | 0.040882 | HC_vs_MS.Trt_B |
| LDHAL6A | 2.59E-06 | 0.25393 | 0.349 | 0.23 | 0.041299 | HC_vs_MS.Trt_B |
| FOPNL | 2.59E-06 | 0.4841 | 0.594 | 0.45 | 0.041445 | HC_vs_MS.Trt_B |
| RNF212 | 2.64E-06 | 0.65499 | 0.572 | 0.43 | 0.042186 | HC_vs_MS.Trt_B |
| ZNF841 | 2.64E-06 | 0.81705 | 0.504 | 0.37 | 0.042227 | HC_vs_MS.Trt_B |
| ONECUT3 | 2.67E-06 | 0.31562 | 0.436 | 0.31 | 0.042656 | HC_vs_MS.Trt_B |
| NEXN | 2.68E-06 | 0.43993 | 0.462 | 0.34 | 0.042873 | HC_vs_MS.Trt_B |
| LIPT1 | 2.70E-06 | 0.31374 | 0.671 | 0.52 | 0.04306 | HC_vs_MS.Trt_B |
| TCHP | 2.71E-06 | 0.80479 | 0.512 | 0.38 | 0.043303 | HC_vs_MS.Trt_B |
| TIGAR | 2.72E-06 | -0.3682 | 0.89 | 0.81 | 0.043443 | HC_vs_MS.Trt_B |
| IGFBP2 | 2.72E-06 | 0.26426 | 0.574 | 0.45 | 0.043467 | HC_vs_MS.Trt_B |
| LPIN3 | 2.72E-06 | 0.42092 | 0.432 | 0.3 | 0.043484 | HC_vs_MS.Trt_B |
| FLI1 | 2.73E-06 | 0.28837 | 0.5 | 0.36 | 0.043607 | HC_vs_MS.Trt_B |
| RPS6KL1 | 2.74E-06 | 0.27439 | 0.688 | 0.56 | 0.043728 | HC_vs_MS.Trt_B |
| CHD7 | 2.76E-06 | -0.3499 | 0.554 | 0.4 | 0.044055 | HC_vs_MS.Trt_B |
| CHTOP | 2.77E-06 | 0.29747 | 0.593 | 0.44 | 0.044185 | HC_vs_MS.Trt_B |
| TNFRSF10A | 2.78E-06 | 0.43661 | 0.567 | 0.46 | 0.04443 | HC_vs_MS.Trt_B |
| NFIC | 2.79E-06 | -0.6143 | 0.412 | 0.28 | 0.044621 | HC_vs_MS.Trt_B |
| PROM2 | 2.80E-06 | 0.30162 | 0.706 | 0.58 | 0.044713 | HC_vs_MS.Trt_B |
| ARNT | 2.80E-06 | 0.73389 | 0.757 | 0.63 | 0.044767 | HC_vs_MS.Trt_B |
| CSNK1D | 2.82E-06 | 0.366 | 0.936 | 0.86 | 0.045018 | HC_vs_MS.Trt_B |
| CEP135 | 2.85E-06 | 0.48155 | 0.757 | 0.63 | 0.045539 | HC_vs_MS.Trt_B |
| RPS2 | 2.92E-06 | 0.30312 | 0.606 | 0.47 | 0.04669 | HC_vs_MS.Trt_B |
| DDX39B | 2.93E-06 | 0.3627 | 0.691 | 0.56 | 0.046837 | HC_vs_MS.Trt_B |
| COQ8A | 2.96E-06 | 0.6901 | 0.633 | 0.5 | 0.047212 | HC_vs_MS.Trt_B |
| MAP2K2 | 3.00E-06 | -0.3426 | 0.704 | 0.55 | 0.047992 | HC_vs_MS.Trt_B |
| AKAP5 | 3.11E-06 | 0.75409 | 0.625 | 0.52 | 0.049686 | HC_vs_MS.Trt_B |
| PRSS27 | 3.11E-06 | 0.46356 | 0.755 | 0.64 | 0.049745 | HC_vs_MS.Trt_B |
| KCNJ6 | 3.18E-06 | 0.93959 | 0.799 | 0.72 | 0.050845 | HC_vs_MS.Trt_B |
| CPD | 3.21E-06 | 0.25901 | 0.693 | 0.54 | 0.051241 | HC_vs_MS.Trt_B |
| PNMAL1 | 3.22E-06 | 0.30746 | 0.428 | 0.3 | 0.051407 | HC_vs_MS.Trt_B |

| | | | | | | |
|---|---|---|---|---|---|---|
| IL21R | 3.22E-06 | 0.42967 | 0.523 | 0.39 | 0.05151 | HC_vs_MS.Trt_B |
| ZNF548 | 3.29E-06 | 0.43608 | 0.638 | 0.51 | 0.052517 | HC_vs_MS.Trt_B |
| LDLRAD4 | 3.29E-06 | 0.47926 | 0.343 | 0.22 | 0.052611 | HC_vs_MS.Trt_B |
| NUP50 | 3.32E-06 | 0.49157 | 0.649 | 0.54 | 0.052952 | HC_vs_MS.Trt_B |
| MKL2 | 3.32E-06 | 0.39127 | 0.696 | 0.57 | 0.053008 | HC_vs_MS.Trt_B |
| C21orf62 | 3.33E-06 | 0.33038 | 0.535 | 0.39 | 0.053215 | HC_vs_MS.Trt_B |
| TALDO1 | 3.38E-06 | 0.25808 | 0.662 | 0.54 | 0.05392 | HC_vs_MS.Trt_B |
| EFCAB5 | 3.38E-06 | -0.3088 | 0.436 | 0.31 | 0.053963 | HC_vs_MS.Trt_B |
| MAP3K9 | 3.45E-06 | 0.76129 | 0.448 | 0.33 | 0.055136 | HC_vs_MS.Trt_B |
| KCNN2 | 3.49E-06 | 0.29893 | 0.255 | 0.14 | 0.055795 | HC_vs_MS.Trt_B |
| GRSF1 | 3.58E-06 | 0.3773 | 0.73 | 0.62 | 0.05712 | HC_vs_MS.Trt_B |
| BMP1 | 3.59E-06 | -0.3526 | 0.535 | 0.38 | 0.057397 | HC_vs_MS.Trt_B |
| RNF10 | 3.60E-06 | 0.31563 | 0.807 | 0.7 | 0.057511 | HC_vs_MS.Trt_B |
| C2orf91 | 3.68E-06 | 0.27181 | 0.467 | 0.34 | 0.058842 | HC_vs_MS.Trt_B |
| RCHY1 | 3.73E-06 | 0.82176 | 0.309 | 0.19 | 0.059553 | HC_vs_MS.Trt_B |
| CERS4 | 3.73E-06 | 0.58254 | 0.613 | 0.48 | 0.059561 | HC_vs_MS.Trt_B |
| ZNF442 | 3.74E-06 | 0.32986 | 0.717 | 0.57 | 0.059688 | HC_vs_MS.Trt_B |
| SLC22A11 | 3.76E-06 | 0.25014 | 0.414 | 0.28 | 0.059979 | HC_vs_MS.Trt_B |
| ST3GAL3 | 3.83E-06 | 0.63151 | 0.793 | 0.65 | 0.061174 | HC_vs_MS.Trt_B |
| TSC2 | 3.86E-06 | 0.38923 | 0.684 | 0.53 | 0.061648 | HC_vs_MS.Trt_B |
| GLB1L3 | 3.86E-06 | 0.36848 | 0.7 | 0.59 | 0.061736 | HC_vs_MS.Trt_B |
| TMEM132A | 3.87E-06 | 0.43063 | 0.384 | 0.25 | 0.061843 | HC_vs_MS.Trt_B |
| HEMK1 | 3.90E-06 | 0.25214 | 0.951 | 0.87 | 0.062347 | HC_vs_MS.Trt_B |
| CYB561D1 | 3.91E-06 | 0.58586 | 0.461 | 0.33 | 0.062495 | HC_vs_MS.Trt_B |
| NEK4 | 3.96E-06 | 0.33402 | 0.767 | 0.61 | 0.063247 | HC_vs_MS.Trt_B |
| HERC2 | 3.98E-06 | 0.46341 | 0.851 | 0.76 | 0.063526 | HC_vs_MS.Trt_B |
| TMBIM1 | 3.99E-06 | 0.33441 | 0.588 | 0.44 | 0.063701 | HC_vs_MS.Trt_B |
| C14orf159 | 4.03E-06 | 0.7636 | 0.638 | 0.53 | 0.064351 | HC_vs_MS.Trt_B |
| CSNK1G1 | 4.03E-06 | 0.27812 | 0.564 | 0.42 | 0.064409 | HC_vs_MS.Trt_B |
| RNF141 | 4.05E-06 | 0.40301 | 0.425 | 0.29 | 0.06472 | HC_vs_MS.Trt_B |
| FITM2 | 4.06E-06 | -0.4342 | 0.69 | 0.57 | 0.064898 | HC_vs_MS.Trt_B |
| SMIM5 | 4.07E-06 | 0.33011 | 0.304 | 0.19 | 0.064949 | HC_vs_MS.Trt_B |
| ARHGAP26 | 4.13E-06 | 0.29208 | 0.626 | 0.49 | 0.065941 | HC_vs_MS.Trt_B |
| LRRC28 | 4.16E-06 | 0.8068 | 0.512 | 0.38 | 0.066395 | HC_vs_MS.Trt_B |
| BRF1 | 4.16E-06 | 0.46707 | 0.565 | 0.44 | 0.066474 | HC_vs_MS.Trt_B |
| CDK11A | 4.18E-06 | 0.27791 | 0.413 | 0.28 | 0.066737 | HC_vs_MS.Trt_B |
| TMEM25 | 4.21E-06 | 0.85362 | 0.394 | 0.25 | 0.067175 | HC_vs_MS.Trt_B |
| ARL16 | 4.29E-06 | 0.41688 | 0.745 | 0.61 | 0.06851 | HC_vs_MS.Trt_B |
| FGD5 | 4.41E-06 | 0.45145 | 0.525 | 0.38 | 0.07037 | HC_vs_MS.Trt_B |
| GDAP2 | 4.41E-06 | 0.51106 | 0.684 | 0.55 | 0.070424 | HC_vs_MS.Trt_B |

| | | | | | | |
|---|---|---|---|---|---|---|
| PPP1R12B | 4.43E-06 | 0.4081 | 0.696 | 0.57 | 0.070699 | HC_vs_MS.Trt_B |
| CCND2 | 4.45E-06 | 0.62046 | 0.381 | 0.26 | 0.071052 | HC_vs_MS.Trt_B |
| TFB1M | 4.47E-06 | 0.43591 | 0.422 | 0.3 | 0.071355 | HC_vs_MS.Trt_B |
| ACSS1 | 4.51E-06 | 1.19273 | 0.577 | 0.46 | 0.072097 | HC_vs_MS.Trt_B |
| NAA15 | 4.56E-06 | 0.35064 | 0.739 | 0.63 | 0.072858 | HC_vs_MS.Trt_B |
| GALNT10 | 4.57E-06 | 0.28159 | 0.53 | 0.4 | 0.072919 | HC_vs_MS.Trt_B |
| NOTCH2 | 4.66E-06 | 0.75117 | 0.204 | 0.1 | 0.074477 | HC_vs_MS.Trt_B |
| FBXW4 | 4.68E-06 | 0.47439 | 0.493 | 0.38 | 0.074785 | HC_vs_MS.Trt_B |
| C5orf42 | 4.69E-06 | -0.5987 | 0.51 | 0.39 | 0.074857 | HC_vs_MS.Trt_B |
| CCT6A | 4.80E-06 | 0.31282 | 0.47 | 0.33 | 0.076678 | HC_vs_MS.Trt_B |
| RNF130 | 4.81E-06 | -0.9169 | 0.633 | 0.49 | 0.076815 | HC_vs_MS.Trt_B |
| PRKX | 4.84E-06 | 0.35461 | 0.526 | 0.4 | 0.077295 | HC_vs_MS.Trt_B |
| PAXIP1 | 4.86E-06 | -0.8801 | 0.435 | 0.31 | 0.077578 | HC_vs_MS.Trt_B |
| GCM1 | 4.90E-06 | 0.3506 | 0.49 | 0.37 | 0.07823 | HC_vs_MS.Trt_B |
| PML | 4.98E-06 | 0.42702 | 0.596 | 0.46 | 0.079519 | HC_vs_MS.Trt_B |
| TRAPPC2 | 4.98E-06 | -0.2728 | 0.917 | 0.83 | 0.079577 | HC_vs_MS.Trt_B |
| PTGIS | 5.01E-06 | 0.26498 | 0.667 | 0.58 | 0.080066 | HC_vs_MS.Trt_B |
| CENPV | 5.15E-06 | 0.41814 | 0.71 | 0.58 | 0.082187 | HC_vs_MS.Trt_B |
| IQCG | 5.15E-06 | 0.51573 | 0.748 | 0.65 | 0.082319 | HC_vs_MS.Trt_B |
| EML6 | 5.21E-06 | 0.26064 | 0.578 | 0.46 | 0.083221 | HC_vs_MS.Trt_B |
| NVL | 5.22E-06 | 0.32408 | 0.8 | 0.66 | 0.083417 | HC_vs_MS.Trt_B |
| ZNF589 | 5.23E-06 | 0.33004 | 0.459 | 0.33 | 0.083495 | HC_vs_MS.Trt_B |
| EFCAB12 | 5.24E-06 | 0.30125 | 0.745 | 0.62 | 0.083728 | HC_vs_MS.Trt_B |
| AKAP6 | 5.42E-06 | 0.39239 | 0.697 | 0.57 | 0.086535 | HC_vs_MS.Trt_B |
| RNF19A | 5.42E-06 | 0.29247 | 0.412 | 0.28 | 0.08654 | HC_vs_MS.Trt_B |
| PABPC4 | 5.45E-06 | 0.33087 | 0.81 | 0.68 | 0.087112 | HC_vs_MS.Trt_B |
| UHRF1BP1 | 5.46E-06 | 0.84915 | 0.549 | 0.43 | 0.087236 | HC_vs_MS.Trt_B |
| KDELC2 | 5.64E-06 | 0.84532 | 0.554 | 0.44 | 0.090014 | HC_vs_MS.Trt_B |
| KIAA1958 | 5.66E-06 | 0.55873 | 0.699 | 0.57 | 0.090426 | HC_vs_MS.Trt_B |
| VEZT | 5.73E-06 | -0.6806 | 0.742 | 0.59 | 0.091485 | HC_vs_MS.Trt_B |
| RPH3A | 5.76E-06 | 0.25658 | 0.423 | 0.29 | 0.091945 | HC_vs_MS.Trt_B |
| PHLPP2 | 5.77E-06 | 0.67251 | 0.8 | 0.66 | 0.09218 | HC_vs_MS.Trt_B |
| SOX13 | 5.77E-06 | 0.2688 | 0.739 | 0.6 | 0.09218 | HC_vs_MS.Trt_B |
| QSOX1 | 5.83E-06 | 0.80665 | 0.37 | 0.25 | 0.093119 | HC_vs_MS.Trt_B |
| CTD-3088G3 | 5.88E-06 | -0.482 | 0.51 | 0.37 | 0.093921 | HC_vs_MS.Trt_B |
| BMP8B | 5.88E-06 | 0.41152 | 0.403 | 0.28 | 0.093976 | HC_vs_MS.Trt_B |
| SLC46A1 | 5.90E-06 | 0.48274 | 0.484 | 0.36 | 0.094213 | HC_vs_MS.Trt_B |
| MYL6 | 5.94E-06 | -0.6745 | 0.594 | 0.68 | 0.094886 | HC_vs_MS.Trt_B |
| ST8SIA6 | 5.95E-06 | 0.7294 | 0.443 | 0.32 | 0.095044 | HC_vs_MS.Trt_B |
| ZNF606 | 5.95E-06 | 0.40715 | 0.775 | 0.69 | 0.095106 | HC_vs_MS.Trt_B |

| | | | | | | |
|---|---|---|---|---|---|---|
| BORCS5 | 5.97E-06 | 0.6641 | 0.629 | 0.49 | 0.095337 | HC_vs_MS.Trt_B |
| PPP5C | 5.98E-06 | 0.40718 | 0.397 | 0.27 | 0.095589 | HC_vs_MS.Trt_B |
| HDAC2 | 5.99E-06 | -0.4334 | 0.713 | 0.56 | 0.095696 | HC_vs_MS.Trt_B |
| MICU1 | 5.99E-06 | 0.32812 | 0.642 | 0.53 | 0.095745 | HC_vs_MS.Trt_B |
| MOB1B | 6.07E-06 | 0.4521 | 0.61 | 0.47 | 0.096899 | HC_vs_MS.Trt_B |
| LRFN3 | 6.09E-06 | 0.34451 | 0.617 | 0.47 | 0.097268 | HC_vs_MS.Trt_B |
| KCNJ15 | 6.14E-06 | 0.46615 | 0.588 | 0.49 | 0.098009 | HC_vs_MS.Trt_B |
| RBM48 | 6.17E-06 | -0.4222 | 0.755 | 0.61 | 0.098515 | HC_vs_MS.Trt_B |
| SKIDA1 | 6.22E-06 | 0.26425 | 0.404 | 0.28 | 0.099417 | HC_vs_MS.Trt_B |
| UBE2O | 6.30E-06 | 0.38097 | 0.545 | 0.38 | 0.100556 | HC_vs_MS.Trt_B |
| PIAS2 | 6.32E-06 | 0.34925 | 0.777 | 0.68 | 0.100883 | HC_vs_MS.Trt_B |
| CCDC88B | 6.32E-06 | 0.41186 | 0.759 | 0.63 | 0.100893 | HC_vs_MS.Trt_B |
| NUP210 | 6.32E-06 | 0.255 | 0.52 | 0.38 | 0.100896 | HC_vs_MS.Trt_B |
| NCKAP5L | 6.32E-06 | 0.51936 | 0.593 | 0.46 | 0.1009 | HC_vs_MS.Trt_B |
| MRPL30 | 6.32E-06 | -0.3963 | 0.781 | 0.66 | 0.10094 | HC_vs_MS.Trt_B |
| RPL10A | 6.33E-06 | -0.3733 | 0.851 | 0.82 | 0.101156 | HC_vs_MS.Trt_B |
| ZNF557 | 6.34E-06 | -0.4972 | 0.857 | 0.76 | 0.101327 | HC_vs_MS.Trt_B |
| QRSL1 | 6.36E-06 | -0.5385 | 0.555 | 0.41 | 0.101627 | HC_vs_MS.Trt_B |
| ZC3H7A | 6.37E-06 | 0.52456 | 0.383 | 0.26 | 0.101687 | HC_vs_MS.Trt_B |
| AK7 | 6.44E-06 | 0.25714 | 0.793 | 0.69 | 0.102873 | HC_vs_MS.Trt_B |
| ZNF583 | 6.52E-06 | 0.48118 | 0.803 | 0.71 | 0.104107 | HC_vs_MS.Trt_B |
| TRIM24 | 6.52E-06 | -0.2805 | 0.458 | 0.33 | 0.10411 | HC_vs_MS.Trt_B |
| SLC38A6 | 6.52E-06 | 0.27065 | 0.643 | 0.5 | 0.104194 | HC_vs_MS.Trt_B |
| ABCA6 | 6.54E-06 | 0.25001 | 0.63 | 0.5 | 0.104515 | HC_vs_MS.Trt_B |
| SERINC3 | 6.55E-06 | 0.33069 | 0.661 | 0.54 | 0.104551 | HC_vs_MS.Trt_B |
| NEB | 6.58E-06 | 0.97854 | 0.532 | 0.4 | 0.105034 | HC_vs_MS.Trt_B |
| SLC4A1AP | 6.59E-06 | -0.3138 | 0.761 | 0.67 | 0.10533 | HC_vs_MS.Trt_B |
| SLC36A3 | 6.60E-06 | 0.29193 | 0.441 | 0.33 | 0.105438 | HC_vs_MS.Trt_B |
| ATP8B1 | 6.62E-06 | 0.50197 | 0.265 | 0.15 | 0.105809 | HC_vs_MS.Trt_B |
| SH2B1 | 6.63E-06 | 0.38192 | 0.57 | 0.46 | 0.105855 | HC_vs_MS.Trt_B |
| HNRNPUL1 | 6.65E-06 | 0.63046 | 0.688 | 0.58 | 0.106234 | HC_vs_MS.Trt_B |
| AGO2 | 6.69E-06 | 0.26057 | 0.413 | 0.29 | 0.106792 | HC_vs_MS.Trt_B |
| C9orf66 | 6.75E-06 | 0.37985 | 0.574 | 0.44 | 0.107808 | HC_vs_MS.Trt_B |
| LMLN | 6.83E-06 | -0.6926 | 0.435 | 0.3 | 0.109142 | HC_vs_MS.Trt_B |
| NOM1 | 6.87E-06 | 0.60772 | 0.614 | 0.53 | 0.109674 | HC_vs_MS.Trt_B |
| ATAD5 | 6.94E-06 | -0.6757 | 0.601 | 0.43 | 0.110778 | HC_vs_MS.Trt_B |
| TGFB1 | 7.00E-06 | 0.49643 | 0.316 | 0.2 | 0.111877 | HC_vs_MS.Trt_B |
| RPF2 | 7.01E-06 | 0.58731 | 0.304 | 0.19 | 0.11199 | HC_vs_MS.Trt_B |
| LEMD3 | 7.09E-06 | 0.49405 | 0.636 | 0.5 | 0.113224 | HC_vs_MS.Trt_B |
| PTMA | 7.16E-06 | -0.3753 | 0.941 | 0.91 | 0.114289 | HC_vs_MS.Trt_B |

| | | | | | | |
|---|---|---|---|---|---|---|
| VPS13C | 7.16E-06 | 0.29358 | 0.751 | 0.63 | 0.114435 | HC_vs_MS.Trt_B |
| SIMC1 | 7.17E-06 | 0.78937 | 0.42 | 0.3 | 0.114482 | HC_vs_MS.Trt_B |
| C2orf74 | 7.18E-06 | 0.30098 | 0.457 | 0.32 | 0.114659 | HC_vs_MS.Trt_B |
| SURF6 | 7.23E-06 | 0.34946 | 0.517 | 0.38 | 0.115503 | HC_vs_MS.Trt_B |
| GRAMD1A | 7.28E-06 | -0.352 | 0.716 | 0.54 | 0.116205 | HC_vs_MS.Trt_B |
| DYDC1 | 7.34E-06 | 0.26731 | 0.584 | 0.45 | 0.117294 | HC_vs_MS.Trt_B |
| SVIP | 7.38E-06 | 0.33971 | 0.535 | 0.39 | 0.117939 | HC_vs_MS.Trt_B |
| SLC2A3 | 7.40E-06 | 0.59648 | 0.732 | 0.63 | 0.118248 | HC_vs_MS.Trt_B |
| ARL5A | 7.42E-06 | 0.57554 | 0.187 | 0.09 | 0.118487 | HC_vs_MS.Trt_B |
| TERF2IP | 7.48E-06 | 0.66756 | 0.378 | 0.26 | 0.119425 | HC_vs_MS.Trt_B |
| EMCN | 7.52E-06 | 0.40711 | 0.438 | 0.31 | 0.12016 | HC_vs_MS.Trt_B |
| CTDSPL | 7.54E-06 | 0.3915 | 0.235 | 0.13 | 0.12038 | HC_vs_MS.Trt_B |
| BPIFB1 | 7.55E-06 | 0.2645 | 0.507 | 0.37 | 0.120607 | HC_vs_MS.Trt_B |
| RUVBL2 | 7.59E-06 | -0.3135 | 0.854 | 0.72 | 0.121198 | HC_vs_MS.Trt_B |
| EDC3 | 7.59E-06 | 0.38002 | 0.387 | 0.27 | 0.121201 | HC_vs_MS.Trt_B |
| ATXN10 | 7.65E-06 | 0.29128 | 0.48 | 0.34 | 0.122126 | HC_vs_MS.Trt_B |
| GPR85 | 7.80E-06 | 0.54682 | 0.388 | 0.27 | 0.124525 | HC_vs_MS.Trt_B |
| DLG1 | 7.81E-06 | 0.38391 | 0.581 | 0.46 | 0.124771 | HC_vs_MS.Trt_B |
| SIPA1L1 | 7.82E-06 | 0.8861 | 0.594 | 0.44 | 0.124873 | HC_vs_MS.Trt_B |
| CSNK2A2 | 7.84E-06 | 0.64683 | 0.27 | 0.16 | 0.125189 | HC_vs_MS.Trt_B |
| ZNF66 | 7.84E-06 | -0.8421 | 0.648 | 0.49 | 0.125217 | HC_vs_MS.Trt_B |
| POLQ | 7.94E-06 | 0.26721 | 0.545 | 0.41 | 0.126876 | HC_vs_MS.Trt_B |
| WDR91 | 7.95E-06 | 0.34286 | 0.477 | 0.36 | 0.126998 | HC_vs_MS.Trt_B |
| MCOLN2 | 8.00E-06 | -0.5478 | 0.446 | 0.32 | 0.127777 | HC_vs_MS.Trt_B |
| NBPF9 | 8.00E-06 | 0.25974 | 0.709 | 0.58 | 0.127838 | HC_vs_MS.Trt_B |
| XKR4 | 8.03E-06 | 0.32969 | 0.542 | 0.42 | 0.128266 | HC_vs_MS.Trt_B |
| PLXDC1 | 8.09E-06 | 0.54245 | 0.729 | 0.62 | 0.12926 | HC_vs_MS.Trt_B |
| REPS2 | 8.15E-06 | 0.26168 | 0.606 | 0.47 | 0.130238 | HC_vs_MS.Trt_B |
| SUGP2 | 8.19E-06 | -0.3746 | 0.709 | 0.59 | 0.130879 | HC_vs_MS.Trt_B |
| C15orf52 | 8.19E-06 | 1.09025 | 0.414 | 0.29 | 0.130898 | HC_vs_MS.Trt_B |
| LRRC58 | 8.21E-06 | 0.52646 | 0.716 | 0.6 | 0.131076 | HC_vs_MS.Trt_B |
| ACSS3 | 8.21E-06 | 0.42771 | 0.267 | 0.16 | 0.131194 | HC_vs_MS.Trt_B |
| HILPDA | 8.28E-06 | 0.28994 | 0.593 | 0.47 | 0.132327 | HC_vs_MS.Trt_B |
| SLC35B4 | 8.39E-06 | 0.27242 | 0.533 | 0.43 | 0.134089 | HC_vs_MS.Trt_B |
| STIL | 8.42E-06 | -0.6833 | 0.607 | 0.5 | 0.134537 | HC_vs_MS.Trt_B |
| SAMD4A | 8.46E-06 | 0.48988 | 0.584 | 0.44 | 0.135088 | HC_vs_MS.Trt_B |
| YTHDF3 | 8.51E-06 | 0.26964 | 0.642 | 0.51 | 0.135972 | HC_vs_MS.Trt_B |
| P4HB | 8.60E-06 | 0.35833 | 0.654 | 0.52 | 0.137307 | HC_vs_MS.Trt_B |
| MYLK | 8.61E-06 | -0.6971 | 0.562 | 0.41 | 0.137531 | HC_vs_MS.Trt_B |
| NME7 | 8.63E-06 | 0.45417 | 0.622 | 0.49 | 0.137846 | HC_vs_MS.Trt_B |

| | | | | | | |
|---|---|---|---|---|---|---|
| TCOF1 | 8.65E-06 | -0.4037 | 0.79 | 0.64 | 0.138171 | HC_vs_MS.Trt_B |
| IL1RAP | 8.76E-06 | 0.25772 | 0.658 | 0.53 | 0.139851 | HC_vs_MS.Trt_B |
| CHST5 | 8.76E-06 | 0.25472 | 0.457 | 0.34 | 0.139895 | HC_vs_MS.Trt_B |
| BRD2 | 8.78E-06 | 0.58357 | 0.546 | 0.43 | 0.140193 | HC_vs_MS.Trt_B |
| GANC | 8.98E-06 | 0.32006 | 0.607 | 0.48 | 0.143374 | HC_vs_MS.Trt_B |
| RPH3AL | 9.00E-06 | 0.93043 | 0.32 | 0.21 | 0.143813 | HC_vs_MS.Trt_B |
| POLM | 9.20E-06 | 0.39826 | 0.533 | 0.4 | 0.146986 | HC_vs_MS.Trt_B |
| KMT2E | 9.23E-06 | 0.39617 | 0.767 | 0.65 | 0.147495 | HC_vs_MS.Trt_B |
| PRICKLE3 | 9.24E-06 | 0.41542 | 0.51 | 0.38 | 0.147559 | HC_vs_MS.Trt_B |
| ARSK | 9.30E-06 | -0.301 | 0.538 | 0.42 | 0.148492 | HC_vs_MS.Trt_B |
| KATNAL1 | 9.39E-06 | -0.6377 | 0.523 | 0.41 | 0.150056 | HC_vs_MS.Trt_B |
| DDX52 | 9.41E-06 | -0.3561 | 0.871 | 0.78 | 0.150232 | HC_vs_MS.Trt_B |
| DHX30 | 9.44E-06 | 0.35774 | 0.639 | 0.52 | 0.150726 | HC_vs_MS.Trt_B |
| GNG4 | 9.49E-06 | 0.37724 | 0.677 | 0.55 | 0.151562 | HC_vs_MS.Trt_B |
| FBXO5 | 9.66E-06 | -0.8961 | 0.401 | 0.27 | 0.154234 | HC_vs_MS.Trt_B |
| DST | 9.66E-06 | 0.34838 | 0.755 | 0.61 | 0.154288 | HC_vs_MS.Trt_B |
| AP2A2 | 9.67E-06 | 0.32601 | 0.616 | 0.49 | 0.154466 | HC_vs_MS.Trt_B |
| DMD | 9.74E-06 | 0.34711 | 0.619 | 0.49 | 0.155609 | HC_vs_MS.Trt_B |
| ATP2A2 | 9.89E-06 | 0.9769 | 0.694 | 0.63 | 0.158022 | HC_vs_MS.Trt_B |
| WDR89 | 1.01E-05 | 0.55675 | 0.362 | 0.23 | 0.160774 | HC_vs_MS.Trt_B |
| SCGB3A2 | 1.01E-05 | 0.3445 | 0.28 | 0.17 | 0.160829 | HC_vs_MS.Trt_B |
| ANKRD45 | 1.01E-05 | 0.27094 | 0.543 | 0.43 | 0.161276 | HC_vs_MS.Trt_B |
| GPBP1L1 | 1.02E-05 | 0.75168 | 0.458 | 0.34 | 0.16258 | HC_vs_MS.Trt_B |
| TRAPPC12 | 1.02E-05 | 0.31578 | 0.599 | 0.46 | 0.162626 | HC_vs_MS.Trt_B |
| CD6 | 1.02E-05 | 0.37504 | 0.641 | 0.51 | 0.163664 | HC_vs_MS.Trt_B |
| NCAPH2 | 1.03E-05 | 0.70122 | 0.686 | 0.61 | 0.16411 | HC_vs_MS.Trt_B |
| EIF2D | 1.03E-05 | -0.5313 | 0.716 | 0.57 | 0.164413 | HC_vs_MS.Trt_B |
| CCDC25 | 1.04E-05 | -0.5801 | 0.286 | 0.17 | 0.165586 | HC_vs_MS.Trt_B |
| MOK | 1.04E-05 | 0.27935 | 0.79 | 0.66 | 0.166113 | HC_vs_MS.Trt_B |
| OAS1 | 1.06E-05 | 0.56401 | 0.503 | 0.41 | 0.168608 | HC_vs_MS.Trt_B |
| PSMB2 | 1.06E-05 | 0.28827 | 0.733 | 0.6 | 0.168672 | HC_vs_MS.Trt_B |
| MCOLN3 | 1.06E-05 | 0.25787 | 0.467 | 0.33 | 0.168723 | HC_vs_MS.Trt_B |
| TMPO | 1.06E-05 | 0.30317 | 0.535 | 0.41 | 0.16896 | HC_vs_MS.Trt_B |
| NSUN4 | 1.06E-05 | 0.32183 | 0.755 | 0.63 | 0.169274 | HC_vs_MS.Trt_B |
| ABHD12B | 1.07E-05 | 0.44673 | 0.614 | 0.48 | 0.170262 | HC_vs_MS.Trt_B |
| TRMT2B | 1.07E-05 | 0.53997 | 0.625 | 0.49 | 0.170466 | HC_vs_MS.Trt_B |
| FAM134C | 1.07E-05 | 0.36768 | 0.604 | 0.48 | 0.170843 | HC_vs_MS.Trt_B |
| TMEM231 | 1.08E-05 | 0.28014 | 0.709 | 0.58 | 0.172758 | HC_vs_MS.Trt_B |
| POLL | 1.09E-05 | 0.48324 | 0.404 | 0.28 | 0.17342 | HC_vs_MS.Trt_B |
| ZNF429 | 1.11E-05 | 0.3757 | 0.903 | 0.82 | 0.176757 | HC_vs_MS.Trt_B |

| | | | | | | |
|---|---|---|---|---|---|---|
| EIF1AD | 1.11E-05 | -0.3192 | 0.693 | 0.57 | 0.177626 | HC_vs_MS.Trt_B |
| WDR17 | 1.11E-05 | 0.37204 | 0.716 | 0.57 | 0.177733 | HC_vs_MS.Trt_B |
| SIN3A | 1.12E-05 | 0.26433 | 0.783 | 0.66 | 0.178738 | HC_vs_MS.Trt_B |
| RAB5B | 1.12E-05 | 0.40745 | 0.864 | 0.79 | 0.179079 | HC_vs_MS.Trt_B |
| ZNF585B | 1.12E-05 | 0.31127 | 0.723 | 0.6 | 0.179112 | HC_vs_MS.Trt_B |
| KIF3B | 1.14E-05 | 0.92048 | 0.438 | 0.33 | 0.181869 | HC_vs_MS.Trt_B |
| PTGIR | 1.15E-05 | -0.3621 | 0.671 | 0.54 | 0.183732 | HC_vs_MS.Trt_B |
| TRMT12 | 1.15E-05 | 0.36294 | 0.394 | 0.27 | 0.18398 | HC_vs_MS.Trt_B |
| POTEC | 1.15E-05 | 0.41331 | 0.406 | 0.28 | 0.183993 | HC_vs_MS.Trt_B |
| GCLM | 1.16E-05 | 0.49938 | 0.672 | 0.58 | 0.184637 | HC_vs_MS.Trt_B |
| ACBD5 | 1.17E-05 | -0.3168 | 0.467 | 0.34 | 0.186597 | HC_vs_MS.Trt_B |
| NEDD9 | 1.18E-05 | 0.74007 | 0.467 | 0.33 | 0.187825 | HC_vs_MS.Trt_B |
| IFRD1 | 1.18E-05 | 0.80514 | 0.387 | 0.26 | 0.188272 | HC_vs_MS.Trt_B |
| NUP188 | 1.18E-05 | 0.52844 | 0.526 | 0.41 | 0.188555 | HC_vs_MS.Trt_B |
| BCAS3 | 1.19E-05 | 0.40161 | 0.823 | 0.74 | 0.189769 | HC_vs_MS.Trt_B |
| ATP6V1E1 | 1.20E-05 | -0.2699 | 0.632 | 0.47 | 0.191423 | HC_vs_MS.Trt_B |
| ZZEF1 | 1.21E-05 | 0.42552 | 0.501 | 0.36 | 0.19333 | HC_vs_MS.Trt_B |
| SOAT1 | 1.21E-05 | 0.32511 | 0.7 | 0.61 | 0.193388 | HC_vs_MS.Trt_B |
| TESC | 1.22E-05 | 0.47758 | 0.232 | 0.13 | 0.194139 | HC_vs_MS.Trt_B |
| CH507-9B2.9 | 1.22E-05 | 0.3106 | 0.403 | 0.28 | 0.194426 | HC_vs_MS.Trt_B |
| FAM71D | 1.22E-05 | 0.54279 | 0.12 | 0.05 | 0.195298 | HC_vs_MS.Trt_B |
| ILF3 | 1.22E-05 | 0.28598 | 0.749 | 0.65 | 0.19531 | HC_vs_MS.Trt_B |
| SIGLEC11 | 1.22E-05 | 0.32556 | 0.548 | 0.42 | 0.195469 | HC_vs_MS.Trt_B |
| RFX3 | 1.23E-05 | 0.25691 | 0.194 | 0.1 | 0.196485 | HC_vs_MS.Trt_B |
| RP11-166B2 | 1.24E-05 | 0.38469 | 0.507 | 0.38 | 0.198027 | HC_vs_MS.Trt_B |
| CCDC171 | 1.25E-05 | 0.40193 | 0.462 | 0.34 | 0.199495 | HC_vs_MS.Trt_B |
| EIF2AK1 | 1.26E-05 | 0.5262 | 0.454 | 0.33 | 0.201014 | HC_vs_MS.Trt_B |
| EXOSC6 | 1.26E-05 | -0.2961 | 0.901 | 0.82 | 0.201648 | HC_vs_MS.Trt_B |
| TYK2 | 1.27E-05 | 0.29262 | 0.659 | 0.53 | 0.202563 | HC_vs_MS.Trt_B |
| FOXRED2 | 1.29E-05 | 0.29407 | 0.529 | 0.4 | 0.206344 | HC_vs_MS.Trt_B |
| FBXW12 | 1.29E-05 | 0.43211 | 0.323 | 0.21 | 0.206609 | HC_vs_MS.Trt_B |
| SULT1C4 | 1.29E-05 | 0.37063 | 0.554 | 0.46 | 0.206757 | HC_vs_MS.Trt_B |
| ANKRD54 | 1.30E-05 | 0.33709 | 0.693 | 0.56 | 0.207541 | HC_vs_MS.Trt_B |
| CYTH1 | 1.31E-05 | 0.26955 | 0.578 | 0.46 | 0.210022 | HC_vs_MS.Trt_B |
| PLCE1 | 1.32E-05 | -0.2655 | 0.916 | 0.8 | 0.210698 | HC_vs_MS.Trt_B |
| HEBP1 | 1.32E-05 | 1.00189 | 0.262 | 0.15 | 0.211564 | HC_vs_MS.Trt_B |
| IRAK4 | 1.34E-05 | -0.5861 | 0.594 | 0.47 | 0.214565 | HC_vs_MS.Trt_B |
| ZNF215 | 1.36E-05 | 0.36914 | 0.397 | 0.28 | 0.217578 | HC_vs_MS.Trt_B |
| GDF11 | 1.37E-05 | 0.26817 | 0.325 | 0.22 | 0.21921 | HC_vs_MS.Trt_B |
| RUNDC3B | 1.38E-05 | 0.43209 | 0.309 | 0.2 | 0.2197 | HC_vs_MS.Trt_B |

| | | | | | | |
|---|---|---|---|---|---|---|
| TFAP4 | 1.38E-05 | 0.31608 | 0.671 | 0.55 | 0.219887 | HC_vs_MS.Trt_B |
| WBSCR17 | 1.38E-05 | 0.30045 | 0.314 | 0.2 | 0.220674 | HC_vs_MS.Trt_B |
| KIAA0100 | 1.42E-05 | 0.57648 | 0.497 | 0.38 | 0.226584 | HC_vs_MS.Trt_B |
| ZNF626 | 1.42E-05 | -0.436 | 0.703 | 0.59 | 0.227088 | HC_vs_MS.Trt_B |
| BTG2 | 1.43E-05 | 0.82726 | 0.178 | 0.09 | 0.228024 | HC_vs_MS.Trt_B |
| XPO5 | 1.45E-05 | 0.38795 | 0.688 | 0.54 | 0.231183 | HC_vs_MS.Trt_B |
| SLC16A7 | 1.46E-05 | 0.80109 | 0.628 | 0.51 | 0.232784 | HC_vs_MS.Trt_B |
| GNAL | 1.47E-05 | -0.353 | 0.443 | 0.32 | 0.234383 | HC_vs_MS.Trt_B |
| HAPLN4 | 1.47E-05 | 0.27289 | 0.417 | 0.29 | 0.234468 | HC_vs_MS.Trt_B |
| RSPRY1 | 1.47E-05 | -0.3076 | 0.335 | 0.22 | 0.235331 | HC_vs_MS.Trt_B |
| PIK3C2A | 1.47E-05 | 1.03873 | 0.386 | 0.27 | 0.235389 | HC_vs_MS.Trt_B |
| CLUAP1 | 1.48E-05 | -0.2609 | 0.825 | 0.68 | 0.235686 | HC_vs_MS.Trt_B |
| HIBCH | 1.51E-05 | 0.63056 | 0.245 | 0.14 | 0.240596 | HC_vs_MS.Trt_B |
| RPTOR | 1.51E-05 | 0.28476 | 0.378 | 0.25 | 0.241522 | HC_vs_MS.Trt_B |
| CHD3 | 1.51E-05 | 0.38949 | 0.365 | 0.24 | 0.241979 | HC_vs_MS.Trt_B |
| TBC1D4 | 1.53E-05 | 0.4027 | 0.354 | 0.24 | 0.243641 | HC_vs_MS.Trt_B |
| DHX15 | 1.53E-05 | -0.2956 | 0.514 | 0.38 | 0.244856 | HC_vs_MS.Trt_B |
| DHX57 | 1.54E-05 | -0.6017 | 0.555 | 0.41 | 0.245883 | HC_vs_MS.Trt_B |
| ZNF543 | 1.54E-05 | -0.4708 | 0.477 | 0.34 | 0.246091 | HC_vs_MS.Trt_B |
| PPIL6 | 1.54E-05 | 0.29291 | 0.497 | 0.39 | 0.246682 | HC_vs_MS.Trt_B |
| CDH6 | 1.56E-05 | 0.2623 | 0.506 | 0.39 | 0.248596 | HC_vs_MS.Trt_B |
| PARVA | 1.56E-05 | 0.30594 | 0.719 | 0.6 | 0.248761 | HC_vs_MS.Trt_B |
| CECR1 | 1.57E-05 | 0.40667 | 0.486 | 0.36 | 0.250124 | HC_vs_MS.Trt_B |
| TAPT1 | 1.57E-05 | 0.65928 | 0.684 | 0.56 | 0.251281 | HC_vs_MS.Trt_B |
| DGAT2 | 1.58E-05 | 0.26616 | 0.306 | 0.2 | 0.252202 | HC_vs_MS.Trt_B |
| CREBRF | 1.58E-05 | 0.49213 | 0.768 | 0.63 | 0.252514 | HC_vs_MS.Trt_B |
| TRANK1 | 1.60E-05 | 0.80539 | 0.246 | 0.14 | 0.255427 | HC_vs_MS.Trt_B |
| SPATA4 | 1.63E-05 | 0.29071 | 0.587 | 0.49 | 0.261064 | HC_vs_MS.Trt_B |
| ST8SIA1 | 1.64E-05 | -0.3305 | 0.625 | 0.47 | 0.261425 | HC_vs_MS.Trt_B |
| SOWAHC | 1.64E-05 | 0.48965 | 0.42 | 0.31 | 0.261756 | HC_vs_MS.Trt_B |
| DTX2 | 1.66E-05 | 0.54674 | 0.394 | 0.28 | 0.264532 | HC_vs_MS.Trt_B |
| AGMAT | 1.66E-05 | 0.26557 | 0.848 | 0.77 | 0.265126 | HC_vs_MS.Trt_B |
| PTGES | 1.67E-05 | 0.30559 | 0.481 | 0.36 | 0.267157 | HC_vs_MS.Trt_B |
| CLEC2B | 1.68E-05 | 0.45601 | 0.209 | 0.11 | 0.26763 | HC_vs_MS.Trt_B |
| SMAGP | 1.68E-05 | 0.37454 | 0.639 | 0.51 | 0.267639 | HC_vs_MS.Trt_B |
| UVRAG | 1.68E-05 | 0.43889 | 0.423 | 0.3 | 0.267991 | HC_vs_MS.Trt_B |
| FNDC3A | 1.69E-05 | 0.35987 | 0.484 | 0.37 | 0.269538 | HC_vs_MS.Trt_B |
| MPI | 1.69E-05 | 0.3728 | 0.513 | 0.38 | 0.270638 | HC_vs_MS.Trt_B |
| MRPS17 | 1.72E-05 | 0.27152 | 0.845 | 0.77 | 0.274417 | HC_vs_MS.Trt_B |
| DHDDS | 1.73E-05 | 0.38508 | 0.543 | 0.41 | 0.276185 | HC_vs_MS.Trt_B |

| | | | | | | |
|---|---|---|---|---|---|---|
| GSDMB | 1.74E-05 | 0.29659 | 0.862 | 0.76 | 0.277275 | HC_vs_MS.Trt_B |
| RAB2B | 1.75E-05 | 0.303 | 0.652 | 0.53 | 0.278842 | HC_vs_MS.Trt_B |
| PLIN3 | 1.76E-05 | -0.2518 | 0.519 | 0.36 | 0.280442 | HC_vs_MS.Trt_B |
| KMT2D | 1.78E-05 | 0.29782 | 0.855 | 0.74 | 0.284807 | HC_vs_MS.Trt_B |
| AC013264.1 | 1.78E-05 | 0.52092 | 0.336 | 0.23 | 0.285014 | HC_vs_MS.Trt_B |
| FBXO24 | 1.79E-05 | 0.37097 | 0.423 | 0.29 | 0.286621 | HC_vs_MS.Trt_B |
| ALMS1 | 1.81E-05 | 1.13165 | 0.461 | 0.34 | 0.288474 | HC_vs_MS.Trt_B |
| ZNF343 | 1.81E-05 | 1.62168 | 0.433 | 0.31 | 0.288998 | HC_vs_MS.Trt_B |
| RGCC | 1.83E-05 | 0.74061 | 0.18 | 0.09 | 0.292004 | HC_vs_MS.Trt_B |
| RFC2 | 1.84E-05 | -0.3893 | 0.736 | 0.58 | 0.293781 | HC_vs_MS.Trt_B |
| PAPSS1 | 1.84E-05 | 0.4209 | 0.199 | 0.1 | 0.294077 | HC_vs_MS.Trt_B |
| SDHD | 1.85E-05 | -0.2795 | 0.654 | 0.51 | 0.296027 | HC_vs_MS.Trt_B |
| ZNF419 | 1.85E-05 | -0.3555 | 0.758 | 0.66 | 0.296058 | HC_vs_MS.Trt_B |
| MGP | 1.87E-05 | 0.28845 | 0.523 | 0.42 | 0.298724 | HC_vs_MS.Trt_B |
| AARS2 | 1.87E-05 | 1.4205 | 0.384 | 0.28 | 0.299368 | HC_vs_MS.Trt_B |
| PPFIBP1 | 1.89E-05 | 0.41085 | 0.735 | 0.6 | 0.301761 | HC_vs_MS.Trt_B |
| BEAN1 | 1.91E-05 | -0.5996 | 0.422 | 0.29 | 0.305035 | HC_vs_MS.Trt_B |
| ACTR3B | 1.92E-05 | 1.16226 | 0.493 | 0.38 | 0.307379 | HC_vs_MS.Trt_B |
| EGLN1 | 1.94E-05 | 1.05849 | 0.354 | 0.24 | 0.309803 | HC_vs_MS.Trt_B |
| MB21D1 | 1.95E-05 | 0.28594 | 0.72 | 0.59 | 0.310742 | HC_vs_MS.Trt_B |
| LRRC27 | 1.95E-05 | 0.25834 | 0.803 | 0.66 | 0.311278 | HC_vs_MS.Trt_B |
| ZC3H12B | 1.96E-05 | 0.57106 | 0.552 | 0.43 | 0.312436 | HC_vs_MS.Trt_B |
| PTPRA | 1.96E-05 | 0.98554 | 0.164 | 0.08 | 0.313459 | HC_vs_MS.Trt_B |
| UGDH | 1.97E-05 | 0.51565 | 0.528 | 0.42 | 0.314421 | HC_vs_MS.Trt_B |
| NEK2 | 1.97E-05 | 0.30092 | 0.629 | 0.53 | 0.314493 | HC_vs_MS.Trt_B |
| TBC1D7 | 1.98E-05 | 0.61347 | 0.513 | 0.39 | 0.315568 | HC_vs_MS.Trt_B |
| ZNF587B | 2.00E-05 | 0.29855 | 0.336 | 0.22 | 0.319056 | HC_vs_MS.Trt_B |
| C16orf89 | 2.00E-05 | 0.29001 | 0.27 | 0.17 | 0.319304 | HC_vs_MS.Trt_B |
| NOXO1 | 2.01E-05 | 0.35671 | 0.159 | 0.08 | 0.320917 | HC_vs_MS.Trt_B |
| SPAG16 | 2.01E-05 | -0.4353 | 0.441 | 0.31 | 0.321756 | HC_vs_MS.Trt_B |
| SLC4A5 | 2.02E-05 | 0.28902 | 0.7 | 0.56 | 0.321983 | HC_vs_MS.Trt_B |
| TRIM13 | 2.02E-05 | 0.35803 | 0.604 | 0.48 | 0.322039 | HC_vs_MS.Trt_B |
| KDM4B | 2.02E-05 | 0.60649 | 0.594 | 0.49 | 0.32218 | HC_vs_MS.Trt_B |
| ZNF44 | 2.02E-05 | -0.3971 | 0.535 | 0.4 | 0.322526 | HC_vs_MS.Trt_B |
| SNX1 | 2.02E-05 | 0.39592 | 0.733 | 0.64 | 0.322727 | HC_vs_MS.Trt_B |
| ERVW-1 | 2.05E-05 | 0.64506 | 0.365 | 0.25 | 0.327347 | HC_vs_MS.Trt_B |
| TIMM23B | 2.05E-05 | 0.6082 | 0.581 | 0.46 | 0.327742 | HC_vs_MS.Trt_B |
| C22orf46 | 2.10E-05 | 0.72002 | 0.364 | 0.25 | 0.334756 | HC_vs_MS.Trt_B |
| CXXC5 | 2.11E-05 | 0.81798 | 0.504 | 0.37 | 0.336471 | HC_vs_MS.Trt_B |
| ZNF519 | 2.11E-05 | 0.28465 | 0.867 | 0.72 | 0.337188 | HC_vs_MS.Trt_B |

| | | | | | | |
|---|---|---|---|---|---|---|
| UBE2V2 | 2.14E-05 | -0.5277 | 0.722 | 0.62 | 0.342154 | HC_vs_MS.Trt_B |
| GPR157 | 2.15E-05 | 0.27325 | 0.786 | 0.68 | 0.3427 | HC_vs_MS.Trt_B |
| NOL6 | 2.21E-05 | 0.83693 | 0.372 | 0.28 | 0.352963 | HC_vs_MS.Trt_B |
| SREK1IP1 | 2.21E-05 | 0.461 | 0.41 | 0.29 | 0.353621 | HC_vs_MS.Trt_B |
| PDZD7 | 2.22E-05 | 0.26144 | 0.483 | 0.34 | 0.354099 | HC_vs_MS.Trt_B |
| FUCA2 | 2.25E-05 | 0.35192 | 0.296 | 0.18 | 0.358611 | HC_vs_MS.Trt_B |
| PICK1 | 2.26E-05 | -0.7386 | 0.429 | 0.32 | 0.360801 | HC_vs_MS.Trt_B |
| IRAK3 | 2.27E-05 | 0.64534 | 0.413 | 0.28 | 0.361879 | HC_vs_MS.Trt_B |
| CNDP2 | 2.29E-05 | 0.32547 | 0.684 | 0.55 | 0.365248 | HC_vs_MS.Trt_B |
| APAF1 | 2.30E-05 | 0.30498 | 0.572 | 0.44 | 0.367131 | HC_vs_MS.Trt_B |
| RNF220 | 2.33E-05 | 0.28416 | 0.641 | 0.53 | 0.3721 | HC_vs_MS.Trt_B |
| GFI1B | 2.35E-05 | 0.3919 | 0.541 | 0.43 | 0.375307 | HC_vs_MS.Trt_B |
| SV2B | 2.35E-05 | 0.65918 | 0.383 | 0.26 | 0.376004 | HC_vs_MS.Trt_B |
| CLASP2 | 2.35E-05 | 0.70752 | 0.7 | 0.56 | 0.376148 | HC_vs_MS.Trt_B |
| APPBP2 | 2.38E-05 | 0.36362 | 0.832 | 0.76 | 0.380719 | HC_vs_MS.Trt_B |
| EID2B | 2.40E-05 | 0.51254 | 0.523 | 0.41 | 0.382679 | HC_vs_MS.Trt_B |
| UFD1L | 2.42E-05 | 0.25449 | 0.761 | 0.66 | 0.386939 | HC_vs_MS.Trt_B |
| ZNF843 | 2.44E-05 | 0.27618 | 0.693 | 0.59 | 0.388984 | HC_vs_MS.Trt_B |
| SCRN3 | 2.44E-05 | 0.35003 | 0.426 | 0.29 | 0.389486 | HC_vs_MS.Trt_B |
| CUL3 | 2.44E-05 | -0.3044 | 0.412 | 0.29 | 0.390001 | HC_vs_MS.Trt_B |
| AMPD3 | 2.44E-05 | 0.90253 | 0.43 | 0.32 | 0.390226 | HC_vs_MS.Trt_B |
| JOSD1 | 2.47E-05 | 0.30185 | 0.533 | 0.4 | 0.394016 | HC_vs_MS.Trt_B |
| FRMD4A | 2.47E-05 | 0.40132 | 0.629 | 0.5 | 0.394381 | HC_vs_MS.Trt_B |
| C21orf33 | 2.48E-05 | 0.39137 | 0.377 | 0.26 | 0.395399 | HC_vs_MS.Trt_B |
| LHFPL5 | 2.48E-05 | 0.26651 | 0.528 | 0.4 | 0.395717 | HC_vs_MS.Trt_B |
| OTUD3 | 2.49E-05 | 0.47513 | 0.5 | 0.38 | 0.398135 | HC_vs_MS.Trt_B |
| C11orf31 | 2.51E-05 | 0.42367 | 0.507 | 0.4 | 0.400713 | HC_vs_MS.Trt_B |
| OSBPL9 | 2.51E-05 | 0.28126 | 0.328 | 0.21 | 0.401516 | HC_vs_MS.Trt_B |
| ZFP64 | 2.52E-05 | 0.39633 | 0.58 | 0.46 | 0.402085 | HC_vs_MS.Trt_B |
| SREK1 | 2.52E-05 | -0.2561 | 0.523 | 0.39 | 0.402193 | HC_vs_MS.Trt_B |
| NFKBIZ | 2.52E-05 | 0.67593 | 0.336 | 0.23 | 0.40317 | HC_vs_MS.Trt_B |
| SRSF7 | 2.52E-05 | 0.324 | 0.384 | 0.28 | 0.403317 | HC_vs_MS.Trt_B |
| MINK1 | 2.54E-05 | 0.47711 | 0.252 | 0.15 | 0.405175 | HC_vs_MS.Trt_B |
| TNR | 2.58E-05 | 0.3527 | 0.29 | 0.19 | 0.412231 | HC_vs_MS.Trt_B |
| DDX11 | 2.59E-05 | 0.48218 | 0.414 | 0.3 | 0.41311 | HC_vs_MS.Trt_B |
| FAM83H-AS | 2.60E-05 | 0.2561 | 0.694 | 0.61 | 0.415985 | HC_vs_MS.Trt_B |
| SP140L | 2.62E-05 | 0.2611 | 0.757 | 0.62 | 0.418123 | HC_vs_MS.Trt_B |
| TRAM2 | 2.62E-05 | -0.338 | 0.248 | 0.14 | 0.419186 | HC_vs_MS.Trt_B |
| WWC3 | 2.65E-05 | -0.869 | 0.345 | 0.23 | 0.422539 | HC_vs_MS.Trt_B |
| PAN2 | 2.67E-05 | 0.90184 | 0.322 | 0.21 | 0.426375 | HC_vs_MS.Trt_B |

| | | | | | | |
|---|---|---|---|---|---|---|
| FOXP3 | 2.67E-05 | 0.64458 | 0.457 | 0.35 | 0.426737 | HC_vs_MS.Trt_B |
| NARS2 | 2.70E-05 | 0.69998 | 0.39 | 0.28 | 0.431023 | HC_vs_MS.Trt_B |
| PIGS | 2.70E-05 | 0.40648 | 0.48 | 0.37 | 0.43116 | HC_vs_MS.Trt_B |
| NUBPL | 2.71E-05 | 0.30021 | 0.752 | 0.6 | 0.433453 | HC_vs_MS.Trt_B |
| ARHGAP5 | 2.73E-05 | 0.34885 | 0.497 | 0.36 | 0.436374 | HC_vs_MS.Trt_B |
| COPS7B | 2.74E-05 | 0.31156 | 0.783 | 0.69 | 0.437132 | HC_vs_MS.Trt_B |
| TUBGCP2 | 2.75E-05 | -0.3654 | 0.761 | 0.62 | 0.438744 | HC_vs_MS.Trt_B |
| PDE3A | 2.77E-05 | 0.27004 | 0.22 | 0.12 | 0.442091 | HC_vs_MS.Trt_B |
| FILIP1 | 2.78E-05 | 0.35528 | 0.384 | 0.26 | 0.444563 | HC_vs_MS.Trt_B |
| STRN | 2.79E-05 | 0.32823 | 0.225 | 0.13 | 0.446015 | HC_vs_MS.Trt_B |
| CDK10 | 2.82E-05 | 0.5481 | 0.91 | 0.8 | 0.45116 | HC_vs_MS.Trt_B |
| FBXO45 | 2.83E-05 | -0.3579 | 0.423 | 0.3 | 0.451905 | HC_vs_MS.Trt_B |
| ZNF395 | 2.83E-05 | 0.26289 | 0.542 | 0.43 | 0.452744 | HC_vs_MS.Trt_B |
| SPEF2 | 2.85E-05 | 0.50995 | 0.546 | 0.42 | 0.454895 | HC_vs_MS.Trt_B |
| ARNTL2 | 2.85E-05 | 0.27705 | 0.614 | 0.53 | 0.455906 | HC_vs_MS.Trt_B |
| CPQ | 2.90E-05 | -0.3818 | 0.357 | 0.24 | 0.463803 | HC_vs_MS.Trt_B |
| SNX5 | 2.94E-05 | 0.43755 | 0.583 | 0.48 | 0.470024 | HC_vs_MS.Trt_B |
| ZNF362 | 2.95E-05 | 1.48954 | 0.117 | 0.05 | 0.471311 | HC_vs_MS.Trt_B |
| RASAL3 | 2.98E-05 | 0.97967 | 0.184 | 0.1 | 0.47543 | HC_vs_MS.Trt_B |
| RAD50 | 2.98E-05 | 0.27379 | 0.184 | 0.09 | 0.475459 | HC_vs_MS.Trt_B |
| AFDN | 3.00E-05 | 0.88495 | 0.113 | 0.04 | 0.479 | HC_vs_MS.Trt_B |
| ACSF3 | 3.00E-05 | 0.34825 | 0.799 | 0.68 | 0.479315 | HC_vs_MS.Trt_B |
| ZNF74 | 3.02E-05 | 0.33588 | 0.433 | 0.32 | 0.48215 | HC_vs_MS.Trt_B |
| SEMA5A | 3.03E-05 | 0.26609 | 0.39 | 0.28 | 0.483526 | HC_vs_MS.Trt_B |
| STRN4 | 3.05E-05 | 0.3143 | 0.483 | 0.37 | 0.486694 | HC_vs_MS.Trt_B |
| NCR1 | 3.08E-05 | 0.57966 | 0.223 | 0.12 | 0.492765 | HC_vs_MS.Trt_B |
| EFCAB13 | 3.11E-05 | 0.5059 | 0.32 | 0.21 | 0.496336 | HC_vs_MS.Trt_B |
| DHX8 | 3.12E-05 | 0.33249 | 0.652 | 0.52 | 0.497825 | HC_vs_MS.Trt_B |
| EIF4A1 | 3.16E-05 | 0.32813 | 0.491 | 0.38 | 0.504152 | HC_vs_MS.Trt_B |
| QRICH1 | 3.17E-05 | 0.26008 | 0.539 | 0.41 | 0.505768 | HC_vs_MS.Trt_B |
| MCL1 | 3.18E-05 | 0.36561 | 0.372 | 0.26 | 0.508633 | HC_vs_MS.Trt_B |
| GPN3 | 3.23E-05 | 1.20868 | 0.374 | 0.27 | 0.515415 | HC_vs_MS.Trt_B |
| PYROXD1 | 3.26E-05 | 0.83871 | 0.462 | 0.35 | 0.520735 | HC_vs_MS.Trt_B |
| 2-Mar | 3.27E-05 | 0.4884 | 0.516 | 0.4 | 0.522747 | HC_vs_MS.Trt_B |
| DPYD | 3.28E-05 | 0.63022 | 0.471 | 0.36 | 0.523757 | HC_vs_MS.Trt_B |
| QTRT2 | 3.28E-05 | 0.42272 | 0.787 | 0.66 | 0.524456 | HC_vs_MS.Trt_B |
| GRB2 | 3.31E-05 | 0.41014 | 0.645 | 0.53 | 0.528706 | HC_vs_MS.Trt_B |
| LILRA6 | 3.31E-05 | 0.27215 | 0.523 | 0.41 | 0.52945 | HC_vs_MS.Trt_B |
| ITGB8 | 3.35E-05 | -0.4504 | 0.758 | 0.64 | 0.535843 | HC_vs_MS.Trt_B |
| YARS | 3.38E-05 | 0.42089 | 0.803 | 0.75 | 0.539498 | HC_vs_MS.Trt_B |

| | | | | | | |
|---|---|---|---|---|---|---|
| AP4B1 | 3.42E-05 | -0.3095 | 0.622 | 0.51 | 0.545567 | HC_vs_MS.Trt_B |
| DCAF11 | 3.45E-05 | 0.90402 | 0.796 | 0.63 | 0.551327 | HC_vs_MS.Trt_B |
| MICU2 | 3.45E-05 | 0.8475 | 0.438 | 0.34 | 0.551596 | HC_vs_MS.Trt_B |
| RFT1 | 3.46E-05 | 0.42578 | 0.467 | 0.34 | 0.552464 | HC_vs_MS.Trt_B |
| FADS3 | 3.59E-05 | 0.2731 | 0.625 | 0.5 | 0.573261 | HC_vs_MS.Trt_B |
| MLH1 | 3.61E-05 | 1.32124 | 0.364 | 0.25 | 0.575873 | HC_vs_MS.Trt_B |
| CDHR3 | 3.65E-05 | 0.38295 | 0.617 | 0.51 | 0.582955 | HC_vs_MS.Trt_B |
| TSPAN32 | 3.67E-05 | 0.51976 | 0.274 | 0.17 | 0.586461 | HC_vs_MS.Trt_B |
| MCAM | 3.68E-05 | 0.27086 | 0.451 | 0.34 | 0.588358 | HC_vs_MS.Trt_B |
| ZNF831 | 3.69E-05 | 0.39212 | 0.236 | 0.14 | 0.589307 | HC_vs_MS.Trt_B |
| KRIT1 | 3.69E-05 | 0.3191 | 0.61 | 0.5 | 0.589441 | HC_vs_MS.Trt_B |
| DMPK | 3.73E-05 | -0.3272 | 0.588 | 0.46 | 0.595349 | HC_vs_MS.Trt_B |
| ZNF500 | 3.78E-05 | -0.4364 | 0.613 | 0.47 | 0.603036 | HC_vs_MS.Trt_B |
| FAM20B | 3.78E-05 | 0.39125 | 0.565 | 0.42 | 0.60384 | HC_vs_MS.Trt_B |
| MIER2 | 3.80E-05 | 0.50917 | 0.529 | 0.39 | 0.606472 | HC_vs_MS.Trt_B |
| ANKHD1 | 3.80E-05 | 0.33262 | 0.43 | 0.32 | 0.606809 | HC_vs_MS.Trt_B |
| ANKFY1 | 3.85E-05 | 0.47987 | 0.617 | 0.5 | 0.615079 | HC_vs_MS.Trt_B |
| ATPAF1 | 3.89E-05 | 1.03266 | 0.235 | 0.14 | 0.620748 | HC_vs_MS.Trt_B |
| FAM219B | 3.91E-05 | 0.39462 | 0.477 | 0.35 | 0.624264 | HC_vs_MS.Trt_B |
| ECHDC1 | 3.93E-05 | -0.6438 | 0.797 | 0.69 | 0.62811 | HC_vs_MS.Trt_B |
| MYEF2 | 3.95E-05 | 0.38462 | 0.451 | 0.33 | 0.630534 | HC_vs_MS.Trt_B |
| CEP76 | 3.97E-05 | 0.58353 | 0.787 | 0.7 | 0.63369 | HC_vs_MS.Trt_B |
| LIMK2 | 3.97E-05 | 0.67489 | 0.201 | 0.11 | 0.633985 | HC_vs_MS.Trt_B |
| IPP | 3.99E-05 | 0.2503 | 0.43 | 0.3 | 0.636972 | HC_vs_MS.Trt_B |
| PAM | 4.00E-05 | 0.87183 | 0.325 | 0.22 | 0.638815 | HC_vs_MS.Trt_B |
| BMF | 4.02E-05 | 0.35059 | 0.139 | 0.06 | 0.641532 | HC_vs_MS.Trt_B |
| TRIM6 | 4.06E-05 | 0.28243 | 0.522 | 0.39 | 0.648379 | HC_vs_MS.Trt_B |
| ADGRA3 | 4.08E-05 | 0.35974 | 0.393 | 0.29 | 0.651206 | HC_vs_MS.Trt_B |
| LIMD1 | 4.09E-05 | 0.35554 | 0.813 | 0.7 | 0.653198 | HC_vs_MS.Trt_B |
| HERPUD2 | 4.10E-05 | 0.37419 | 0.481 | 0.38 | 0.654177 | HC_vs_MS.Trt_B |
| ZNF486 | 4.13E-05 | 0.54977 | 0.649 | 0.56 | 0.659564 | HC_vs_MS.Trt_B |
| MTA3 | 4.16E-05 | -0.3501 | 0.641 | 0.52 | 0.664319 | HC_vs_MS.Trt_B |
| MED20 | 4.17E-05 | -0.5323 | 0.314 | 0.2 | 0.665459 | HC_vs_MS.Trt_B |
| CYBA | 4.18E-05 | 0.5205 | 0.255 | 0.16 | 0.667179 | HC_vs_MS.Trt_B |
| METTL16 | 4.20E-05 | 0.29672 | 0.759 | 0.66 | 0.671314 | HC_vs_MS.Trt_B |
| FOCAD | 4.24E-05 | 0.47534 | 0.529 | 0.4 | 0.677496 | HC_vs_MS.Trt_B |
| NDUFS4 | 4.25E-05 | 0.63184 | 0.267 | 0.17 | 0.678781 | HC_vs_MS.Trt_B |
| TBCK | 4.28E-05 | 0.30402 | 0.725 | 0.59 | 0.683371 | HC_vs_MS.Trt_B |
| DDX18 | 4.29E-05 | -0.2536 | 0.745 | 0.61 | 0.684997 | HC_vs_MS.Trt_B |
| PACS1 | 4.31E-05 | 0.34851 | 0.817 | 0.74 | 0.687648 | HC_vs_MS.Trt_B |

| | | | | | | |
|---|---|---|---|---|---|---|
| TAP1 | 4.31E-05 | 0.43229 | 0.617 | 0.51 | 0.688271 | HC_vs_MS.Trt_B |
| ITGB7 | 4.31E-05 | 0.52397 | 0.37 | 0.26 | 0.68875 | HC_vs_MS.Trt_B |
| GRAP | 4.32E-05 | 0.83025 | 0.204 | 0.12 | 0.690608 | HC_vs_MS.Trt_B |
| GTF2IRD1 | 4.33E-05 | 0.28155 | 0.645 | 0.53 | 0.691665 | HC_vs_MS.Trt_B |
| PGA3 | 4.35E-05 | 0.50892 | 0.291 | 0.19 | 0.695181 | HC_vs_MS.Trt_B |
| ACTN1 | 4.38E-05 | 0.321 | 0.599 | 0.48 | 0.699805 | HC_vs_MS.Trt_B |
| MAFF | 4.39E-05 | 0.37988 | 0.642 | 0.52 | 0.700902 | HC_vs_MS.Trt_B |
| EPT1 | 4.39E-05 | 0.28725 | 0.336 | 0.22 | 0.701602 | HC_vs_MS.Trt_B |
| ADCK1 | 4.40E-05 | 0.72471 | 0.274 | 0.18 | 0.703051 | HC_vs_MS.Trt_B |
| ZNF224 | 4.40E-05 | 0.75982 | 0.458 | 0.36 | 0.703504 | HC_vs_MS.Trt_B |
| CEACAM8 | 4.41E-05 | 0.30048 | 0.504 | 0.41 | 0.704683 | HC_vs_MS.Trt_B |
| B4GALT4 | 4.42E-05 | 0.56304 | 0.604 | 0.48 | 0.705826 | HC_vs_MS.Trt_B |
| CHI3L1 | 4.51E-05 | 0.34204 | 0.314 | 0.21 | 0.721045 | HC_vs_MS.Trt_B |
| FAM161A | 4.52E-05 | 0.26442 | 0.614 | 0.51 | 0.72273 | HC_vs_MS.Trt_B |
| CPSF4 | 4.56E-05 | 0.62855 | 0.357 | 0.24 | 0.728952 | HC_vs_MS.Trt_B |
| TMEM45B | 4.57E-05 | 0.32089 | 0.603 | 0.48 | 0.730123 | HC_vs_MS.Trt_B |
| PRPF40A | 4.59E-05 | -0.3068 | 0.465 | 0.32 | 0.733548 | HC_vs_MS.Trt_B |
| LRTOMT | 4.62E-05 | 0.25473 | 0.803 | 0.73 | 0.737654 | HC_vs_MS.Trt_B |
| HYPK | 4.65E-05 | 0.32054 | 0.903 | 0.8 | 0.742842 | HC_vs_MS.Trt_B |
| TRAF1 | 4.68E-05 | 0.46329 | 0.322 | 0.21 | 0.747334 | HC_vs_MS.Trt_B |
| RRAGC | 4.69E-05 | 0.69115 | 0.509 | 0.39 | 0.748769 | HC_vs_MS.Trt_B |
| PAGR1 | 4.69E-05 | 0.45714 | 0.183 | 0.1 | 0.749139 | HC_vs_MS.Trt_B |
| POM121 | 4.69E-05 | 0.36724 | 0.397 | 0.27 | 0.74921 | HC_vs_MS.Trt_B |
| THSD4 | 4.71E-05 | 0.41617 | 0.362 | 0.25 | 0.75227 | HC_vs_MS.Trt_B |
| NFATC2 | 4.75E-05 | 0.44249 | 0.178 | 0.09 | 0.758679 | HC_vs_MS.Trt_B |
| HOXB2 | 4.76E-05 | 0.87557 | 0.123 | 0.05 | 0.760874 | HC_vs_MS.Trt_B |
| PDCD6 | 4.77E-05 | -0.2842 | 0.667 | 0.53 | 0.761983 | HC_vs_MS.Trt_B |
| NUDCD2 | 4.77E-05 | -0.2522 | 0.767 | 0.65 | 0.762386 | HC_vs_MS.Trt_B |
| SLC50A1 | 4.83E-05 | 0.2616 | 0.745 | 0.66 | 0.771683 | HC_vs_MS.Trt_B |
| IKZF2 | 4.89E-05 | 0.28646 | 0.357 | 0.25 | 0.780567 | HC_vs_MS.Trt_B |
| BCAP29 | 4.90E-05 | 0.70964 | 0.374 | 0.25 | 0.782139 | HC_vs_MS.Trt_B |
| OTUB1 | 4.97E-05 | -0.3275 | 0.929 | 0.8 | 0.794286 | HC_vs_MS.Trt_B |
| NATD1 | 4.98E-05 | 0.63754 | 0.481 | 0.37 | 0.794913 | HC_vs_MS.Trt_B |
| MCUR1 | 5.03E-05 | 0.48193 | 0.786 | 0.69 | 0.804226 | HC_vs_MS.Trt_B |
| NUP43 | 5.04E-05 | -0.2997 | 0.843 | 0.72 | 0.80475 | HC_vs_MS.Trt_B |
| HPS1 | 5.07E-05 | 0.30405 | 0.6 | 0.45 | 0.809553 | HC_vs_MS.Trt_B |
| VENTX | 5.08E-05 | 0.26697 | 0.403 | 0.29 | 0.81122 | HC_vs_MS.Trt_B |
| GALNS | 5.15E-05 | 0.60222 | 0.542 | 0.42 | 0.822945 | HC_vs_MS.Trt_B |
| DDX25 | 5.16E-05 | 0.33732 | 0.654 | 0.54 | 0.823773 | HC_vs_MS.Trt_B |
| EVC | 5.21E-05 | 1.01548 | 0.242 | 0.15 | 0.83224 | HC_vs_MS.Trt_B |

| | | | | | | |
|---|---|---|---|---|---|---|
| PPARD | 5.23E-05 | 0.38068 | 0.529 | 0.41 | 0.834998 | HC_vs_MS.Trt_B |
| OPN3 | 5.25E-05 | 0.71067 | 0.294 | 0.19 | 0.838745 | HC_vs_MS.Trt_B |
| NSUN2 | 5.27E-05 | 0.47088 | 0.659 | 0.53 | 0.842331 | HC_vs_MS.Trt_B |
| ASGR1 | 5.28E-05 | 0.30399 | 0.455 | 0.35 | 0.843771 | HC_vs_MS.Trt_B |
| NXF1 | 5.29E-05 | 0.46159 | 0.559 | 0.43 | 0.844341 | HC_vs_MS.Trt_B |
| ZNF845 | 5.29E-05 | 0.41658 | 0.645 | 0.51 | 0.845146 | HC_vs_MS.Trt_B |
| HUNK | 5.36E-05 | 0.26184 | 0.419 | 0.31 | 0.856702 | HC_vs_MS.Trt_B |
| KLHL33 | 5.37E-05 | 0.25402 | 0.497 | 0.39 | 0.857134 | HC_vs_MS.Trt_B |
| PALM | 5.40E-05 | 0.26196 | 0.535 | 0.44 | 0.861987 | HC_vs_MS.Trt_B |
| STAM2 | 5.40E-05 | 0.52949 | 0.532 | 0.42 | 0.863075 | HC_vs_MS.Trt_B |
| FOXJ2 | 5.42E-05 | 0.53505 | 0.449 | 0.33 | 0.865046 | HC_vs_MS.Trt_B |
| COL1A1 | 5.43E-05 | 0.40026 | 0.497 | 0.39 | 0.868053 | HC_vs_MS.Trt_B |
| DCDC1 | 5.45E-05 | 0.38492 | 0.48 | 0.37 | 0.869902 | HC_vs_MS.Trt_B |
| UROS | 5.46E-05 | 0.6877 | 0.457 | 0.34 | 0.872107 | HC_vs_MS.Trt_B |
| HSD17B12 | 5.53E-05 | 0.34017 | 0.475 | 0.35 | 0.883746 | HC_vs_MS.Trt_B |
| GPA33 | 5.55E-05 | 0.73928 | 0.146 | 0.07 | 0.885773 | HC_vs_MS.Trt_B |
| GTF2E2 | 5.61E-05 | 0.61023 | 0.588 | 0.46 | 0.895506 | HC_vs_MS.Trt_B |
| PHGDH | 5.68E-05 | 0.28755 | 0.241 | 0.14 | 0.907843 | HC_vs_MS.Trt_B |
| PTPRS | 5.80E-05 | 0.32676 | 0.509 | 0.39 | 0.926203 | HC_vs_MS.Trt_B |
| SLC35G1 | 5.80E-05 | 0.61331 | 0.146 | 0.07 | 0.926975 | HC_vs_MS.Trt_B |
| TNFSF14 | 5.83E-05 | 0.57039 | 0.606 | 0.49 | 0.930761 | HC_vs_MS.Trt_B |
| ZNF430 | 5.84E-05 | -0.4071 | 0.923 | 0.83 | 0.933332 | HC_vs_MS.Trt_B |
| DNAJC7 | 5.86E-05 | 0.27423 | 0.977 | 0.95 | 0.935676 | HC_vs_MS.Trt_B |
| NT5C3B | 5.87E-05 | 0.46834 | 0.616 | 0.51 | 0.938155 | HC_vs_MS.Trt_B |
| CLU | 5.91E-05 | 0.4264 | 0.464 | 0.36 | 0.944626 | HC_vs_MS.Trt_B |
| TRIP13 | 5.92E-05 | 0.36244 | 0.543 | 0.44 | 0.945732 | HC_vs_MS.Trt_B |
| RBM33 | 5.97E-05 | 0.31938 | 0.29 | 0.19 | 0.954335 | HC_vs_MS.Trt_B |
| GNL2 | 5.99E-05 | 0.64818 | 0.63 | 0.53 | 0.956996 | HC_vs_MS.Trt_B |
| DDX19B | 6.02E-05 | -0.7254 | 0.432 | 0.31 | 0.961962 | HC_vs_MS.Trt_B |
| NDUFA9 | 6.03E-05 | -0.5052 | 0.868 | 0.73 | 0.962446 | HC_vs_MS.Trt_B |
| LSM8 | 6.04E-05 | -0.4742 | 0.861 | 0.75 | 0.965084 | HC_vs_MS.Trt_B |
| CCDC141 | 6.06E-05 | 0.26725 | 0.622 | 0.51 | 0.968007 | HC_vs_MS.Trt_B |
| PLK3 | 6.09E-05 | 0.45568 | 0.683 | 0.58 | 0.972745 | HC_vs_MS.Trt_B |
| C10orf76 | 6.12E-05 | 0.45839 | 0.333 | 0.22 | 0.978141 | HC_vs_MS.Trt_B |
| TAB2 | 6.23E-05 | -0.5006 | 0.584 | 0.44 | 0.994819 | HC_vs_MS.Trt_B |
| KIF23 | 6.26E-05 | 0.29657 | 0.523 | 0.39 | 1 | HC_vs_MS.Trt_B |
| PNPLA4 | 6.28E-05 | -0.8686 | 0.284 | 0.18 | 1 | HC_vs_MS.Trt_B |
| GTF3C2 | 6.34E-05 | 0.46115 | 0.532 | 0.41 | 1 | HC_vs_MS.Trt_B |
| FUK | 6.38E-05 | 0.32611 | 0.542 | 0.43 | 1 | HC_vs_MS.Trt_B |
| CCNY | 6.39E-05 | 0.31254 | 0.441 | 0.33 | 1 | HC_vs_MS.Trt_B |

| | | | | | | |
|---|---|---|---|---|---|---|
| HYLS1 | 6.39E-05 | 1.15696 | 0.39 | 0.28 | 1 | HC_vs_MS.Trt_B |
| FSD1 | 6.39E-05 | 0.25687 | 0.39 | 0.28 | 1 | HC_vs_MS.Trt_B |
| TUBGCP4 | 6.41E-05 | 0.62126 | 0.784 | 0.7 | 1 | HC_vs_MS.Trt_B |
| CREB1 | 6.44E-05 | 0.45945 | 0.643 | 0.55 | 1 | HC_vs_MS.Trt_B |
| LRFN1 | 6.44E-05 | 0.93387 | 0.142 | 0.07 | 1 | HC_vs_MS.Trt_B |
| SEMA4C | 6.48E-05 | -0.5898 | 0.762 | 0.64 | 1 | HC_vs_MS.Trt_B |
| CDKL1 | 6.51E-05 | -0.4399 | 0.704 | 0.59 | 1 | HC_vs_MS.Trt_B |
| FAM53C | 6.51E-05 | 0.42187 | 0.786 | 0.68 | 1 | HC_vs_MS.Trt_B |
| CYP4F11 | 6.58E-05 | 0.39161 | 0.367 | 0.28 | 1 | HC_vs_MS.Trt_B |
| MAP4K2 | 6.63E-05 | 0.57145 | 0.642 | 0.56 | 1 | HC_vs_MS.Trt_B |
| BEND6 | 6.67E-05 | 0.38501 | 0.197 | 0.11 | 1 | HC_vs_MS.Trt_B |
| PUM2 | 6.80E-05 | 0.76205 | 0.33 | 0.23 | 1 | HC_vs_MS.Trt_B |
| TATDN1 | 6.85E-05 | 0.32775 | 0.664 | 0.59 | 1 | HC_vs_MS.Trt_B |
| CDON | 6.87E-05 | 0.37382 | 0.458 | 0.35 | 1 | HC_vs_MS.Trt_B |
| UTRN | 6.88E-05 | 0.28867 | 0.546 | 0.44 | 1 | HC_vs_MS.Trt_B |
| GAN | 6.90E-05 | -0.3191 | 0.812 | 0.74 | 1 | HC_vs_MS.Trt_B |
| NEURL1 | 6.90E-05 | 0.25573 | 0.3 | 0.2 | 1 | HC_vs_MS.Trt_B |
| TBC1D24 | 6.94E-05 | -0.285 | 0.706 | 0.6 | 1 | HC_vs_MS.Trt_B |
| ATP8B2 | 6.95E-05 | 0.51765 | 0.807 | 0.71 | 1 | HC_vs_MS.Trt_B |
| KIAA1549 | 6.96E-05 | 0.25376 | 0.381 | 0.27 | 1 | HC_vs_MS.Trt_B |
| FAM78A | 6.96E-05 | 0.45176 | 0.365 | 0.26 | 1 | HC_vs_MS.Trt_B |
| TRIM5 | 6.97E-05 | 0.64156 | 0.468 | 0.36 | 1 | HC_vs_MS.Trt_B |
| LCLAT1 | 6.98E-05 | 0.25539 | 0.812 | 0.68 | 1 | HC_vs_MS.Trt_B |
| DRC3 | 7.00E-05 | 0.4416 | 0.591 | 0.49 | 1 | HC_vs_MS.Trt_B |
| ZMAT1 | 7.01E-05 | 0.91284 | 0.341 | 0.24 | 1 | HC_vs_MS.Trt_B |
| RBBP9 | 7.04E-05 | 0.96854 | 0.33 | 0.23 | 1 | HC_vs_MS.Trt_B |
| DMAP1 | 7.05E-05 | -0.3006 | 0.441 | 0.31 | 1 | HC_vs_MS.Trt_B |
| CCDC14 | 7.09E-05 | -0.3281 | 0.228 | 0.13 | 1 | HC_vs_MS.Trt_B |
| SCN1A | 7.22E-05 | 0.32565 | 0.372 | 0.26 | 1 | HC_vs_MS.Trt_B |
| SLC24A1 | 7.37E-05 | -0.3768 | 0.468 | 0.34 | 1 | HC_vs_MS.Trt_B |
| CCDC124 | 7.37E-05 | 0.26991 | 0.486 | 0.37 | 1 | HC_vs_MS.Trt_B |
| SPTLC1 | 7.41E-05 | -0.6268 | 0.723 | 0.6 | 1 | HC_vs_MS.Trt_B |
| PLEKHD1 | 7.42E-05 | 0.32499 | 0.706 | 0.61 | 1 | HC_vs_MS.Trt_B |
| MUC20 | 7.46E-05 | 0.61761 | 0.536 | 0.41 | 1 | HC_vs_MS.Trt_B |
| LCN12 | 7.50E-05 | 0.34728 | 0.274 | 0.18 | 1 | HC_vs_MS.Trt_B |
| MAP3K10 | 7.55E-05 | 0.54424 | 0.354 | 0.24 | 1 | HC_vs_MS.Trt_B |
| RC3H1 | 7.73E-05 | 0.26233 | 0.438 | 0.35 | 1 | HC_vs_MS.Trt_B |
| NAGK | 7.74E-05 | 0.55065 | 0.733 | 0.65 | 1 | HC_vs_MS.Trt_B |
| CD40LG | 7.75E-05 | 0.51766 | 0.22 | 0.13 | 1 | HC_vs_MS.Trt_B |
| APOL1 | 7.82E-05 | 0.63391 | 0.7 | 0.59 | 1 | HC_vs_MS.Trt_B |

| | | | | | | |
|---|---|---|---|---|---|---|
| ALS2 | 7.86E-05 | 0.34593 | 0.435 | 0.31 | 1 | HC_vs_MS.Trt_B |
| GPLD1 | 7.87E-05 | 0.93827 | 0.684 | 0.63 | 1 | HC_vs_MS.Trt_B |
| HOPX | 7.92E-05 | -0.2707 | 0.817 | 0.69 | 1 | HC_vs_MS.Trt_B |
| RAD9B | 7.92E-05 | -0.4373 | 0.391 | 0.29 | 1 | HC_vs_MS.Trt_B |
| AFAP1 | 7.97E-05 | -0.3149 | 0.391 | 0.28 | 1 | HC_vs_MS.Trt_B |
| STRA6 | 7.98E-05 | 0.29022 | 0.548 | 0.45 | 1 | HC_vs_MS.Trt_B |
| PKHD1L1 | 8.02E-05 | 0.36859 | 0.446 | 0.33 | 1 | HC_vs_MS.Trt_B |
| PHLPP1 | 8.05E-05 | 0.36154 | 0.719 | 0.6 | 1 | HC_vs_MS.Trt_B |
| ICE2 | 8.06E-05 | -0.3605 | 0.829 | 0.77 | 1 | HC_vs_MS.Trt_B |
| MPP7 | 8.09E-05 | 0.76991 | 0.255 | 0.16 | 1 | HC_vs_MS.Trt_B |
| CCNF | 8.09E-05 | 0.47726 | 0.439 | 0.34 | 1 | HC_vs_MS.Trt_B |
| TGFB1I1 | 8.16E-05 | 0.80854 | 0.258 | 0.17 | 1 | HC_vs_MS.Trt_B |
| BCKDHB | 8.31E-05 | -0.5422 | 0.584 | 0.48 | 1 | HC_vs_MS.Trt_B |
| GAB3 | 8.35E-05 | -0.449 | 0.346 | 0.23 | 1 | HC_vs_MS.Trt_B |
| CCDC9 | 8.40E-05 | 0.36477 | 0.265 | 0.17 | 1 | HC_vs_MS.Trt_B |
| ZDHHC13 | 8.46E-05 | -0.6456 | 0.739 | 0.6 | 1 | HC_vs_MS.Trt_B |
| ADSL | 8.49E-05 | 0.31422 | 0.872 | 0.77 | 1 | HC_vs_MS.Trt_B |
| ADAMTSL5 | 8.52E-05 | 0.28903 | 0.291 | 0.19 | 1 | HC_vs_MS.Trt_B |
| HAPLN3 | 8.57E-05 | 0.40678 | 0.196 | 0.11 | 1 | HC_vs_MS.Trt_B |
| C2orf88 | 8.59E-05 | 0.64549 | 0.345 | 0.24 | 1 | HC_vs_MS.Trt_B |
| NCALD | 8.60E-05 | 0.3436 | 0.699 | 0.61 | 1 | HC_vs_MS.Trt_B |
| RBM14 | 8.65E-05 | 0.55128 | 0.223 | 0.14 | 1 | HC_vs_MS.Trt_B |
| APOOL | 8.65E-05 | -0.3518 | 0.771 | 0.65 | 1 | HC_vs_MS.Trt_B |
| LDB1 | 8.67E-05 | 0.46755 | 0.142 | 0.07 | 1 | HC_vs_MS.Trt_B |
| HSP90AB1 | 8.83E-05 | 0.25847 | 0.532 | 0.41 | 1 | HC_vs_MS.Trt_B |
| KRBA2 | 8.90E-05 | 0.33162 | 0.37 | 0.26 | 1 | HC_vs_MS.Trt_B |
| PCBP3 | 8.95E-05 | 0.56219 | 0.459 | 0.34 | 1 | HC_vs_MS.Trt_B |
| IFT88 | 8.98E-05 | -0.9138 | 0.549 | 0.43 | 1 | HC_vs_MS.Trt_B |
| FUT10 | 9.07E-05 | -0.4448 | 0.358 | 0.28 | 1 | HC_vs_MS.Trt_B |
| CRTC2 | 9.12E-05 | 0.41238 | 0.387 | 0.27 | 1 | HC_vs_MS.Trt_B |
| KLK2 | 9.19E-05 | 0.35399 | 0.341 | 0.24 | 1 | HC_vs_MS.Trt_B |
| ATP1A3 | 9.22E-05 | 0.88479 | 0.361 | 0.26 | 1 | HC_vs_MS.Trt_B |
| WAC | 9.22E-05 | 0.38193 | 0.445 | 0.33 | 1 | HC_vs_MS.Trt_B |
| JUND | 9.29E-05 | 0.30834 | 0.162 | 0.09 | 1 | HC_vs_MS.Trt_B |
| DYNLL1 | 9.31E-05 | 0.504 | 0.31 | 0.22 | 1 | HC_vs_MS.Trt_B |
| CDK19 | 9.37E-05 | 0.68754 | 0.313 | 0.21 | 1 | HC_vs_MS.Trt_B |
| PPP3CC | 9.48E-05 | 0.58763 | 0.207 | 0.12 | 1 | HC_vs_MS.Trt_B |
| SHMT2 | 9.50E-05 | 0.30825 | 0.304 | 0.2 | 1 | HC_vs_MS.Trt_B |
| GRK4 | 9.58E-05 | 0.71586 | 0.565 | 0.49 | 1 | HC_vs_MS.Trt_B |
| ZNF726 | 9.61E-05 | -0.2899 | 0.858 | 0.74 | 1 | HC_vs_MS.Trt_B |

| | | | | | | |
|---|---|---|---|---|---|---|
| AGPAT5 | 9.61E-05 | 0.70019 | 0.141 | 0.07 | 1 | HC_vs_MS.Trt_B |
| RIF1 | 9.63E-05 | 0.4101 | 0.591 | 0.45 | 1 | HC_vs_MS.Trt_B |
| INHBE | 9.83E-05 | 0.30709 | 0.364 | 0.26 | 1 | HC_vs_MS.Trt_B |
| OCEL1 | 9.87E-05 | 0.62872 | 0.517 | 0.41 | 1 | HC_vs_MS.Trt_B |
| DBP | 9.88E-05 | 0.31604 | 0.661 | 0.54 | 1 | HC_vs_MS.Trt_B |
| SIAE | 9.89E-05 | 0.4557 | 0.599 | 0.49 | 1 | HC_vs_MS.Trt_B |
| SMC5 | 9.93E-05 | 0.35825 | 0.386 | 0.28 | 1 | HC_vs_MS.Trt_B |
| EXOSC2 | 9.93E-05 | -0.4844 | 0.751 | 0.63 | 1 | HC_vs_MS.Trt_B |
| TIAL1 | 1.00E-04 | -0.388 | 0.862 | 0.74 | 1 | HC_vs_MS.Trt_B |
| ROCK2 | 1.01E-04 | 0.56098 | 0.439 | 0.34 | 1 | HC_vs_MS.Trt_B |
| SLC44A1 | 1.01E-04 | 0.93812 | 0.628 | 0.57 | 1 | HC_vs_MS.Trt_B |
| DIAPH2 | 1.01E-04 | 0.40346 | 0.304 | 0.2 | 1 | HC_vs_MS.Trt_B |
| ATP5H | 1.01E-04 | -0.6159 | 0.738 | 0.64 | 1 | HC_vs_MS.Trt_B |
| BAZ1A | 1.01E-04 | -0.2695 | 0.374 | 0.26 | 1 | HC_vs_MS.Trt_B |
| PIGR | 1.02E-04 | 0.26678 | 0.557 | 0.45 | 1 | HC_vs_MS.Trt_B |
| RPAP1 | 1.03E-04 | 0.38833 | 0.803 | 0.69 | 1 | HC_vs_MS.Trt_B |
| LNPK | 1.03E-04 | -0.3782 | 0.519 | 0.41 | 1 | HC_vs_MS.Trt_B |
| JAM2 | 1.03E-04 | 0.2654 | 0.5 | 0.41 | 1 | HC_vs_MS.Trt_B |
| FBXO31 | 1.03E-04 | 0.91323 | 0.3 | 0.2 | 1 | HC_vs_MS.Trt_B |
| OR52I1 | 1.03E-04 | 0.48272 | 0.458 | 0.34 | 1 | HC_vs_MS.Trt_B |
| TRPV1 | 1.05E-04 | 1.05423 | 0.401 | 0.29 | 1 | HC_vs_MS.Trt_B |
| CEP95 | 1.05E-04 | -0.4368 | 0.426 | 0.29 | 1 | HC_vs_MS.Trt_B |
| PCDHGA1 | 1.05E-04 | -0.3645 | 0.317 | 0.21 | 1 | HC_vs_MS.Trt_B |
| HSPA1B | 1.06E-04 | 0.55935 | 0.201 | 0.12 | 1 | HC_vs_MS.Trt_B |
| MT-ND5 | 1.06E-04 | -0.5359 | 0.849 | 0.85 | 1 | HC_vs_MS.Trt_B |
| EFNA5 | 1.06E-04 | 0.44391 | 0.378 | 0.28 | 1 | HC_vs_MS.Trt_B |
| ATF7IP2 | 1.06E-04 | 0.29423 | 0.535 | 0.42 | 1 | HC_vs_MS.Trt_B |
| ZNF879 | 1.07E-04 | 0.4017 | 0.312 | 0.21 | 1 | HC_vs_MS.Trt_B |
| N4BP2L1 | 1.07E-04 | -0.3642 | 0.561 | 0.44 | 1 | HC_vs_MS.Trt_B |
| LGALS9 | 1.07E-04 | 0.31266 | 0.299 | 0.2 | 1 | HC_vs_MS.Trt_B |
| DCAF13 | 1.08E-04 | -0.3607 | 0.625 | 0.5 | 1 | HC_vs_MS.Trt_B |
| APBB2 | 1.09E-04 | 0.34662 | 0.317 | 0.22 | 1 | HC_vs_MS.Trt_B |
| DHX37 | 1.09E-04 | 0.30382 | 0.455 | 0.35 | 1 | HC_vs_MS.Trt_B |
| DHRS1 | 1.10E-04 | 0.30055 | 0.628 | 0.51 | 1 | HC_vs_MS.Trt_B |
| PANK2 | 1.12E-04 | 1.43898 | 0.517 | 0.42 | 1 | HC_vs_MS.Trt_B |
| NUP85 | 1.12E-04 | 0.63628 | 0.475 | 0.36 | 1 | HC_vs_MS.Trt_B |
| AC025335.1 | 1.12E-04 | 0.25858 | 0.613 | 0.49 | 1 | HC_vs_MS.Trt_B |
| TMEM184C | 1.12E-04 | 0.35422 | 0.533 | 0.41 | 1 | HC_vs_MS.Trt_B |
| CDCA3 | 1.14E-04 | 0.63184 | 0.57 | 0.46 | 1 | HC_vs_MS.Trt_B |
| ZNF619 | 1.15E-04 | 0.75967 | 0.716 | 0.61 | 1 | HC_vs_MS.Trt_B |

| | | | | | | |
|---|---|---|---|---|---|---|
| FUT6 | 1.15E-04 | 0.64527 | 0.404 | 0.29 | 1 | HC_vs_MS.Trt_B |
| CAPZA1 | 1.16E-04 | -0.668 | 0.757 | 0.81 | 1 | HC_vs_MS.Trt_B |
| PEX13 | 1.17E-04 | 0.45782 | 0.762 | 0.69 | 1 | HC_vs_MS.Trt_B |
| SNTG1 | 1.17E-04 | 0.2702 | 0.532 | 0.43 | 1 | HC_vs_MS.Trt_B |
| SIPA1L3 | 1.18E-04 | 0.28651 | 0.619 | 0.5 | 1 | HC_vs_MS.Trt_B |
| ALG13 | 1.19E-04 | 0.37069 | 0.284 | 0.19 | 1 | HC_vs_MS.Trt_B |
| TFR2 | 1.20E-04 | -0.3918 | 0.696 | 0.58 | 1 | HC_vs_MS.Trt_B |
| UBA3 | 1.21E-04 | 0.2737 | 0.446 | 0.33 | 1 | HC_vs_MS.Trt_B |
| PPAN | 1.21E-04 | 0.60249 | 0.368 | 0.27 | 1 | HC_vs_MS.Trt_B |
| CTH | 1.22E-04 | 0.41918 | 0.626 | 0.51 | 1 | HC_vs_MS.Trt_B |
| ENTPD7 | 1.23E-04 | 0.3774 | 0.345 | 0.24 | 1 | HC_vs_MS.Trt_B |
| NAA30 | 1.23E-04 | 0.33673 | 0.443 | 0.34 | 1 | HC_vs_MS.Trt_B |
| HUS1 | 1.23E-04 | -0.3123 | 0.475 | 0.37 | 1 | HC_vs_MS.Trt_B |
| ADRA1A | 1.23E-04 | 0.25376 | 0.593 | 0.49 | 1 | HC_vs_MS.Trt_B |
| PLCD4 | 1.24E-04 | 0.67747 | 0.286 | 0.19 | 1 | HC_vs_MS.Trt_B |
| GAL3ST4 | 1.25E-04 | 1.43192 | 0.449 | 0.35 | 1 | HC_vs_MS.Trt_B |
| FAH | 1.26E-04 | 0.34522 | 0.141 | 0.07 | 1 | HC_vs_MS.Trt_B |
| ITFG2 | 1.27E-04 | 0.25764 | 0.309 | 0.21 | 1 | HC_vs_MS.Trt_B |
| TRMT5 | 1.28E-04 | 1.09497 | 0.165 | 0.09 | 1 | HC_vs_MS.Trt_B |
| RENBP | 1.28E-04 | 0.31466 | 0.82 | 0.71 | 1 | HC_vs_MS.Trt_B |
| ZMYND8 | 1.28E-04 | -0.3919 | 0.33 | 0.22 | 1 | HC_vs_MS.Trt_B |
| CRTAP | 1.29E-04 | 0.41145 | 0.522 | 0.43 | 1 | HC_vs_MS.Trt_B |
| MMP24-AS1 | 1.29E-04 | 0.40168 | 0.725 | 0.61 | 1 | HC_vs_MS.Trt_B |
| MRPS21 | 1.29E-04 | -0.3254 | 0.554 | 0.42 | 1 | HC_vs_MS.Trt_B |
| ANKRD30B | 1.31E-04 | 0.40086 | 0.496 | 0.39 | 1 | HC_vs_MS.Trt_B |
| MAP2K7 | 1.31E-04 | 0.29445 | 0.609 | 0.51 | 1 | HC_vs_MS.Trt_B |
| GJB4 | 1.32E-04 | 0.2848 | 0.651 | 0.55 | 1 | HC_vs_MS.Trt_B |
| PLS1 | 1.32E-04 | 0.49848 | 0.283 | 0.19 | 1 | HC_vs_MS.Trt_B |
| TMX3 | 1.32E-04 | -0.2567 | 0.417 | 0.3 | 1 | HC_vs_MS.Trt_B |
| SH3BP1 | 1.33E-04 | 0.73658 | 0.109 | 0.05 | 1 | HC_vs_MS.Trt_B |
| RAPGEF6 | 1.33E-04 | 0.43336 | 0.536 | 0.44 | 1 | HC_vs_MS.Trt_B |
| CHRNA7 | 1.34E-04 | 0.29021 | 0.794 | 0.7 | 1 | HC_vs_MS.Trt_B |
| LARP4 | 1.34E-04 | 0.615 | 0.358 | 0.25 | 1 | HC_vs_MS.Trt_B |
| CHDH | 1.35E-04 | 0.34464 | 0.472 | 0.37 | 1 | HC_vs_MS.Trt_B |
| SF3A1 | 1.37E-04 | 0.53919 | 0.349 | 0.26 | 1 | HC_vs_MS.Trt_B |
| 1-Mar | 1.37E-04 | 0.42601 | 0.583 | 0.47 | 1 | HC_vs_MS.Trt_B |
| MSANTD1 | 1.37E-04 | -0.7334 | 0.441 | 0.35 | 1 | HC_vs_MS.Trt_B |
| SHQ1 | 1.38E-04 | -0.3398 | 0.69 | 0.58 | 1 | HC_vs_MS.Trt_B |
| GOT2 | 1.39E-04 | 0.29701 | 0.836 | 0.76 | 1 | HC_vs_MS.Trt_B |
| DMXL1 | 1.39E-04 | 0.43672 | 0.171 | 0.09 | 1 | HC_vs_MS.Trt_B |

| | | | | | | |
|---|---|---|---|---|---|---|
| PIP5KL1 | 1.40E-04 | 0.33664 | 0.454 | 0.35 | 1 | HC_vs_MS.Trt_B |
| GSTT1 | 1.40E-04 | -1.0212 | 0.723 | 0.58 | 1 | HC_vs_MS.Trt_B |
| BCAS1 | 1.41E-04 | 0.97715 | 0.345 | 0.25 | 1 | HC_vs_MS.Trt_B |
| BTBD1 | 1.41E-04 | 0.4005 | 0.572 | 0.47 | 1 | HC_vs_MS.Trt_B |
| PARPBP | 1.41E-04 | -0.6905 | 0.37 | 0.26 | 1 | HC_vs_MS.Trt_B |
| ABCD4 | 1.41E-04 | -0.2711 | 0.79 | 0.67 | 1 | HC_vs_MS.Trt_B |
| WDR59 | 1.42E-04 | -0.2822 | 0.954 | 0.85 | 1 | HC_vs_MS.Trt_B |
| CP | 1.42E-04 | 1.06034 | 0.148 | 0.08 | 1 | HC_vs_MS.Trt_B |
| APBB1 | 1.42E-04 | 0.32765 | 0.238 | 0.15 | 1 | HC_vs_MS.Trt_B |
| NRAS | 1.43E-04 | 0.91589 | 0.304 | 0.21 | 1 | HC_vs_MS.Trt_B |
| GID4 | 1.44E-04 | 0.47832 | 0.675 | 0.54 | 1 | HC_vs_MS.Trt_B |
| SLC26A11 | 1.45E-04 | 0.31149 | 0.654 | 0.54 | 1 | HC_vs_MS.Trt_B |
| KCNMB4 | 1.45E-04 | 0.64002 | 0.293 | 0.2 | 1 | HC_vs_MS.Trt_B |
| CCDC113 | 1.46E-04 | 0.29792 | 0.639 | 0.54 | 1 | HC_vs_MS.Trt_B |
| AL022397.1 | 1.46E-04 | 0.36418 | 0.238 | 0.15 | 1 | HC_vs_MS.Trt_B |
| ERGIC2 | 1.47E-04 | -0.4301 | 0.471 | 0.35 | 1 | HC_vs_MS.Trt_B |
| ZFYVE16 | 1.47E-04 | -0.4041 | 0.883 | 0.83 | 1 | HC_vs_MS.Trt_B |
| DDOST | 1.47E-04 | 0.29497 | 0.545 | 0.44 | 1 | HC_vs_MS.Trt_B |
| MPP6 | 1.48E-04 | 0.74644 | 0.278 | 0.19 | 1 | HC_vs_MS.Trt_B |
| PLA2G5 | 1.49E-04 | 0.28625 | 0.233 | 0.15 | 1 | HC_vs_MS.Trt_B |
| TMEM134 | 1.50E-04 | -0.3451 | 0.891 | 0.79 | 1 | HC_vs_MS.Trt_B |
| CFP | 1.50E-04 | -0.6045 | 0.357 | 0.26 | 1 | HC_vs_MS.Trt_B |
| KIAA2013 | 1.51E-04 | -0.3475 | 0.307 | 0.21 | 1 | HC_vs_MS.Trt_B |
| IKBIP | 1.53E-04 | 0.59574 | 0.439 | 0.33 | 1 | HC_vs_MS.Trt_B |
| TRAPPC11 | 1.54E-04 | 0.34361 | 0.413 | 0.31 | 1 | HC_vs_MS.Trt_B |
| SERTAD1 | 1.55E-04 | 0.82517 | 0.339 | 0.23 | 1 | HC_vs_MS.Trt_B |
| HSPA9 | 1.55E-04 | 0.54133 | 0.407 | 0.3 | 1 | HC_vs_MS.Trt_B |
| MTIF2 | 1.56E-04 | 1.95277 | 0.41 | 0.32 | 1 | HC_vs_MS.Trt_B |
| TPMT | 1.56E-04 | -0.428 | 0.726 | 0.61 | 1 | HC_vs_MS.Trt_B |
| SVEP1 | 1.57E-04 | 0.31505 | 0.21 | 0.12 | 1 | HC_vs_MS.Trt_B |
| ACTR3C | 1.57E-04 | 0.78115 | 0.523 | 0.42 | 1 | HC_vs_MS.Trt_B |
| VOPP1 | 1.58E-04 | 0.35341 | 0.326 | 0.23 | 1 | HC_vs_MS.Trt_B |
| PHTF2 | 1.59E-04 | 0.38708 | 0.345 | 0.25 | 1 | HC_vs_MS.Trt_B |
| ZNF781 | 1.59E-04 | 0.96068 | 0.291 | 0.19 | 1 | HC_vs_MS.Trt_B |
| DAAM2 | 1.59E-04 | 0.26377 | 0.381 | 0.28 | 1 | HC_vs_MS.Trt_B |
| METTL12 | 1.60E-04 | 0.6225 | 0.613 | 0.49 | 1 | HC_vs_MS.Trt_B |
| GYG1 | 1.60E-04 | 0.37027 | 0.542 | 0.43 | 1 | HC_vs_MS.Trt_B |
| TMEM2 | 1.62E-04 | -0.2622 | 0.584 | 0.47 | 1 | HC_vs_MS.Trt_B |
| NPIPB3 | 1.64E-04 | -0.266 | 0.668 | 0.52 | 1 | HC_vs_MS.Trt_B |
| ATP6V0A1 | 1.64E-04 | 0.97626 | 0.828 | 0.73 | 1 | HC_vs_MS.Trt_B |

| | | | | | | |
|---|---|---|---|---|---|---|
| PSMA2 | 1.65E-04 | -0.3342 | 0.554 | 0.44 | 1 | HC_vs_MS.Trt_B |
| GTF3C5 | 1.66E-04 | 0.98076 | 0.301 | 0.21 | 1 | HC_vs_MS.Trt_B |
| GIT1 | 1.66E-04 | 0.81717 | 0.209 | 0.13 | 1 | HC_vs_MS.Trt_B |
| MAD2L1 | 1.67E-04 | -0.2802 | 0.435 | 0.33 | 1 | HC_vs_MS.Trt_B |
| LAT2 | 1.67E-04 | 0.25811 | 0.854 | 0.77 | 1 | HC_vs_MS.Trt_B |
| C20orf196 | 1.68E-04 | 0.59145 | 0.355 | 0.24 | 1 | HC_vs_MS.Trt_B |
| CENPM | 1.68E-04 | -0.4817 | 0.3 | 0.19 | 1 | HC_vs_MS.Trt_B |
| SARM1 | 1.68E-04 | -0.2568 | 0.443 | 0.32 | 1 | HC_vs_MS.Trt_B |
| POGK | 1.68E-04 | 1.22936 | 0.265 | 0.18 | 1 | HC_vs_MS.Trt_B |
| POU5F1 | 1.71E-04 | 0.38017 | 0.478 | 0.39 | 1 | HC_vs_MS.Trt_B |
| EML2 | 1.72E-04 | 0.59294 | 0.7 | 0.58 | 1 | HC_vs_MS.Trt_B |
| ZBTB25 | 1.72E-04 | -0.3428 | 0.955 | 0.89 | 1 | HC_vs_MS.Trt_B |
| NFASC | 1.73E-04 | -0.3483 | 0.584 | 0.48 | 1 | HC_vs_MS.Trt_B |
| SETD1B | 1.75E-04 | 0.85747 | 0.201 | 0.12 | 1 | HC_vs_MS.Trt_B |
| MFSD13A | 1.77E-04 | 0.38859 | 0.712 | 0.58 | 1 | HC_vs_MS.Trt_B |
| TMEM68 | 1.77E-04 | 0.28864 | 0.871 | 0.8 | 1 | HC_vs_MS.Trt_B |
| CORIN | 1.77E-04 | 0.27358 | 0.574 | 0.47 | 1 | HC_vs_MS.Trt_B |
| SERPINB6 | 1.77E-04 | -0.3902 | 0.484 | 0.37 | 1 | HC_vs_MS.Trt_B |
| SLC25A43 | 1.77E-04 | 0.40665 | 0.241 | 0.15 | 1 | HC_vs_MS.Trt_B |
| PTCH1 | 1.78E-04 | 0.39908 | 0.606 | 0.47 | 1 | HC_vs_MS.Trt_B |
| GFOD1 | 1.79E-04 | 0.43018 | 0.593 | 0.5 | 1 | HC_vs_MS.Trt_B |
| ERO1A | 1.79E-04 | -0.251 | 0.867 | 0.79 | 1 | HC_vs_MS.Trt_B |
| KDM5D | 1.81E-04 | 1.91143 | 0.316 | 0.22 | 1 | HC_vs_MS.Trt_B |
| EIF2A | 1.82E-04 | -0.3036 | 0.597 | 0.47 | 1 | HC_vs_MS.Trt_B |
| MPHOSPH9 | 1.82E-04 | 0.28217 | 0.806 | 0.75 | 1 | HC_vs_MS.Trt_B |
| PCYT1B | 1.83E-04 | 0.29445 | 0.483 | 0.4 | 1 | HC_vs_MS.Trt_B |
| CBX1 | 1.85E-04 | 0.52267 | 0.272 | 0.18 | 1 | HC_vs_MS.Trt_B |
| LMBR1 | 1.86E-04 | 0.38261 | 0.507 | 0.39 | 1 | HC_vs_MS.Trt_B |
| NBPF19 | 1.87E-04 | 0.30757 | 0.446 | 0.34 | 1 | HC_vs_MS.Trt_B |
| PKD2L2 | 1.87E-04 | 0.26738 | 0.687 | 0.56 | 1 | HC_vs_MS.Trt_B |
| FBXL5 | 1.87E-04 | 0.32424 | 0.301 | 0.21 | 1 | HC_vs_MS.Trt_B |
| OR7D2 | 1.87E-04 | 0.45025 | 0.249 | 0.16 | 1 | HC_vs_MS.Trt_B |
| TMCO3 | 1.89E-04 | 0.43804 | 0.716 | 0.58 | 1 | HC_vs_MS.Trt_B |
| PSMD13 | 1.89E-04 | -0.3401 | 0.52 | 0.38 | 1 | HC_vs_MS.Trt_B |
| ENDOV | 1.90E-04 | 0.51477 | 0.577 | 0.48 | 1 | HC_vs_MS.Trt_B |
| NAT10 | 1.90E-04 | 0.25338 | 0.354 | 0.25 | 1 | HC_vs_MS.Trt_B |
| DPH1 | 1.90E-04 | 0.35623 | 0.855 | 0.8 | 1 | HC_vs_MS.Trt_B |
| LYL1 | 1.91E-04 | 0.43764 | 0.397 | 0.29 | 1 | HC_vs_MS.Trt_B |
| MAPK3 | 1.95E-04 | 0.26787 | 0.794 | 0.69 | 1 | HC_vs_MS.Trt_B |
| XPNPEP1 | 1.95E-04 | 0.38879 | 0.287 | 0.19 | 1 | HC_vs_MS.Trt_B |

| | | | | | | |
|---|---|---|---|---|---|---|
| CPT1A | 1.96E-04 | 0.40375 | 0.629 | 0.55 | 1 | HC_vs_MS.Trt_B |
| NIT1 | 1.96E-04 | 0.33887 | 0.604 | 0.47 | 1 | HC_vs_MS.Trt_B |
| TMEM33 | 1.98E-04 | -0.4806 | 0.87 | 0.8 | 1 | HC_vs_MS.Trt_B |
| SCYL3 | 1.99E-04 | 0.33435 | 0.659 | 0.58 | 1 | HC_vs_MS.Trt_B |
| SLK | 1.99E-04 | 0.36475 | 0.196 | 0.12 | 1 | HC_vs_MS.Trt_B |
| TLN1 | 2.01E-04 | 0.36142 | 0.575 | 0.48 | 1 | HC_vs_MS.Trt_B |
| BTF3L4 | 2.03E-04 | 0.56032 | 0.429 | 0.34 | 1 | HC_vs_MS.Trt_B |
| NUP133 | 2.03E-04 | 0.62729 | 0.499 | 0.39 | 1 | HC_vs_MS.Trt_B |
| PHLDA1 | 2.04E-04 | 0.5759 | 0.459 | 0.36 | 1 | HC_vs_MS.Trt_B |
| PMS2 | 2.07E-04 | 0.26109 | 0.662 | 0.56 | 1 | HC_vs_MS.Trt_B |
| SLC30A6 | 2.07E-04 | -0.4506 | 0.607 | 0.5 | 1 | HC_vs_MS.Trt_B |
| HOMER2 | 2.08E-04 | -0.9314 | 0.552 | 0.46 | 1 | HC_vs_MS.Trt_B |
| ATP7B | 2.09E-04 | 0.70655 | 0.2 | 0.12 | 1 | HC_vs_MS.Trt_B |
| PTK2B | 2.10E-04 | 0.4759 | 0.393 | 0.29 | 1 | HC_vs_MS.Trt_B |
| NRF1 | 2.10E-04 | 0.44663 | 0.29 | 0.2 | 1 | HC_vs_MS.Trt_B |
| IRGM | 2.13E-04 | 0.47808 | 0.396 | 0.29 | 1 | HC_vs_MS.Trt_B |
| FAM186A | 2.13E-04 | 0.31708 | 0.407 | 0.3 | 1 | HC_vs_MS.Trt_B |
| RP11-849H4 | 2.15E-04 | 0.27317 | 0.596 | 0.48 | 1 | HC_vs_MS.Trt_B |
| EME2 | 2.16E-04 | 0.68276 | 0.286 | 0.19 | 1 | HC_vs_MS.Trt_B |
| RPS6KA6 | 2.16E-04 | 0.32761 | 0.552 | 0.46 | 1 | HC_vs_MS.Trt_B |
| ZAP70 | 2.18E-04 | 0.36765 | 0.464 | 0.36 | 1 | HC_vs_MS.Trt_B |
| GP6 | 2.19E-04 | 0.64504 | 0.459 | 0.35 | 1 | HC_vs_MS.Trt_B |
| GNB2 | 2.19E-04 | 0.47199 | 0.491 | 0.38 | 1 | HC_vs_MS.Trt_B |
| BTBD11 | 2.19E-04 | 0.43013 | 0.457 | 0.34 | 1 | HC_vs_MS.Trt_B |
| RBM43 | 2.19E-04 | 0.66232 | 0.59 | 0.48 | 1 | HC_vs_MS.Trt_B |
| BFAR | 2.20E-04 | 0.73893 | 0.53 | 0.43 | 1 | HC_vs_MS.Trt_B |
| MME | 2.20E-04 | 0.33111 | 0.439 | 0.33 | 1 | HC_vs_MS.Trt_B |
| SIKE1 | 2.22E-04 | 0.56759 | 0.394 | 0.29 | 1 | HC_vs_MS.Trt_B |
| STAG3 | 2.24E-04 | 0.96943 | 0.18 | 0.1 | 1 | HC_vs_MS.Trt_B |
| FAM110C | 2.24E-04 | 0.86617 | 0.242 | 0.16 | 1 | HC_vs_MS.Trt_B |
| STIM1 | 2.25E-04 | 0.27984 | 0.523 | 0.41 | 1 | HC_vs_MS.Trt_B |
| WNT7A | 2.26E-04 | 1.58461 | 0.103 | 0.04 | 1 | HC_vs_MS.Trt_B |
| CNPPD1 | 2.26E-04 | 0.29809 | 0.293 | 0.2 | 1 | HC_vs_MS.Trt_B |
| ZBTB24 | 2.27E-04 | 0.32005 | 0.341 | 0.25 | 1 | HC_vs_MS.Trt_B |
| FEZ2 | 2.27E-04 | 0.75218 | 0.446 | 0.34 | 1 | HC_vs_MS.Trt_B |
| ZNF266 | 2.28E-04 | 0.4364 | 0.335 | 0.24 | 1 | HC_vs_MS.Trt_B |
| KCTD1 | 2.28E-04 | 0.27949 | 0.572 | 0.48 | 1 | HC_vs_MS.Trt_B |
| MAST4 | 2.28E-04 | 0.36171 | 0.591 | 0.48 | 1 | HC_vs_MS.Trt_B |
| TUBGCP3 | 2.28E-04 | 0.59841 | 0.167 | 0.09 | 1 | HC_vs_MS.Trt_B |
| TNRC6A | 2.29E-04 | 0.26126 | 0.764 | 0.7 | 1 | HC_vs_MS.Trt_B |

| | | | | | | |
|---|---|---|---|---|---|---|
| CATSPERB | 2.30E-04 | 0.4004 | 0.622 | 0.51 | 1 | HC_vs_MS.Trt_B |
| CHML | 2.31E-04 | 0.54617 | 0.235 | 0.15 | 1 | HC_vs_MS.Trt_B |
| USE1 | 2.32E-04 | 0.26003 | 0.852 | 0.77 | 1 | HC_vs_MS.Trt_B |
| NDUFA11 | 2.33E-04 | -0.3748 | 0.594 | 0.45 | 1 | HC_vs_MS.Trt_B |
| SNF8 | 2.34E-04 | -0.2592 | 0.826 | 0.73 | 1 | HC_vs_MS.Trt_B |
| KMT5B | 2.38E-04 | 1.11812 | 0.294 | 0.21 | 1 | HC_vs_MS.Trt_B |
| ELP4 | 2.40E-04 | -0.3542 | 0.529 | 0.44 | 1 | HC_vs_MS.Trt_B |
| TMEM219 | 2.42E-04 | 0.32254 | 0.236 | 0.15 | 1 | HC_vs_MS.Trt_B |
| NDUFS7 | 2.44E-04 | -0.3928 | 0.725 | 0.59 | 1 | HC_vs_MS.Trt_B |
| IFNGR2 | 2.44E-04 | 0.28503 | 0.31 | 0.23 | 1 | HC_vs_MS.Trt_B |
| SEC24D | 2.45E-04 | 0.45428 | 0.484 | 0.38 | 1 | HC_vs_MS.Trt_B |
| FCRL1 | 2.49E-04 | 1.17498 | 0.249 | 0.16 | 1 | HC_vs_MS.Trt_B |
| TRAPPC3 | 2.50E-04 | 0.35921 | 0.372 | 0.28 | 1 | HC_vs_MS.Trt_B |
| ZDHHC4 | 2.52E-04 | 0.36966 | 0.401 | 0.29 | 1 | HC_vs_MS.Trt_B |
| GOLGA2 | 2.52E-04 | 0.49828 | 0.636 | 0.5 | 1 | HC_vs_MS.Trt_B |
| ZNF835 | 2.54E-04 | 0.34258 | 0.558 | 0.45 | 1 | HC_vs_MS.Trt_B |
| SGTA | 2.55E-04 | 0.39393 | 0.506 | 0.4 | 1 | HC_vs_MS.Trt_B |
| NF1 | 2.55E-04 | 0.26683 | 0.723 | 0.62 | 1 | HC_vs_MS.Trt_B |
| MIEF1 | 2.59E-04 | -0.3013 | 0.465 | 0.35 | 1 | HC_vs_MS.Trt_B |
| USP10 | 2.61E-04 | 0.43343 | 0.243 | 0.16 | 1 | HC_vs_MS.Trt_B |
| ADAMTS17 | 2.61E-04 | -0.7978 | 0.194 | 0.12 | 1 | HC_vs_MS.Trt_B |
| ZNF681 | 2.63E-04 | -0.3481 | 0.649 | 0.55 | 1 | HC_vs_MS.Trt_B |
| FAS | 2.63E-04 | 0.35143 | 0.348 | 0.24 | 1 | HC_vs_MS.Trt_B |
| SLC47A2 | 2.63E-04 | -0.3159 | 0.233 | 0.15 | 1 | HC_vs_MS.Trt_B |
| MYOM1 | 2.64E-04 | 0.43624 | 0.264 | 0.17 | 1 | HC_vs_MS.Trt_B |
| WAS | 2.64E-04 | 0.82488 | 0.278 | 0.19 | 1 | HC_vs_MS.Trt_B |
| CD7 | 2.65E-04 | 0.96178 | 0.242 | 0.16 | 1 | HC_vs_MS.Trt_B |
| CWF19L2 | 2.71E-04 | 0.28675 | 0.316 | 0.22 | 1 | HC_vs_MS.Trt_B |
| CLCN5 | 2.72E-04 | 0.35658 | 0.28 | 0.18 | 1 | HC_vs_MS.Trt_B |
| SEL1L | 2.72E-04 | 0.9471 | 0.193 | 0.12 | 1 | HC_vs_MS.Trt_B |
| ZNF425 | 2.73E-04 | 0.35041 | 0.393 | 0.29 | 1 | HC_vs_MS.Trt_B |
| GPR132 | 2.73E-04 | 0.5492 | 0.372 | 0.27 | 1 | HC_vs_MS.Trt_B |
| HES2 | 2.73E-04 | 0.25181 | 0.587 | 0.51 | 1 | HC_vs_MS.Trt_B |
| SRSF1 | 2.74E-04 | 0.34103 | 0.496 | 0.41 | 1 | HC_vs_MS.Trt_B |
| PARP12 | 2.78E-04 | 0.3886 | 0.245 | 0.16 | 1 | HC_vs_MS.Trt_B |
| ATP6V1H | 2.78E-04 | 0.71292 | 0.371 | 0.27 | 1 | HC_vs_MS.Trt_B |
| NUCKS1 | 2.79E-04 | -0.2573 | 0.496 | 0.39 | 1 | HC_vs_MS.Trt_B |
| GPSM3 | 2.80E-04 | 0.26948 | 0.454 | 0.35 | 1 | HC_vs_MS.Trt_B |
| LMF2 | 2.81E-04 | 0.42034 | 0.213 | 0.13 | 1 | HC_vs_MS.Trt_B |
| PIM1 | 2.82E-04 | 0.37463 | 0.384 | 0.28 | 1 | HC_vs_MS.Trt_B |

| | | | | | | |
|---|---|---|---|---|---|---|
| ZBTB37 | 2.83E-04 | 0.49987 | 0.664 | 0.55 | 1 | HC_vs_MS.Trt_B |
| ABHD6 | 2.84E-04 | -0.4756 | 0.461 | 0.34 | 1 | HC_vs_MS.Trt_B |
| DNAJA1 | 2.85E-04 | 0.25295 | 0.286 | 0.19 | 1 | HC_vs_MS.Trt_B |
| TBRG4 | 2.85E-04 | 0.78673 | 0.391 | 0.29 | 1 | HC_vs_MS.Trt_B |
| SETD5 | 2.87E-04 | 0.39958 | 0.496 | 0.39 | 1 | HC_vs_MS.Trt_B |
| MDH1 | 2.90E-04 | -0.3559 | 0.609 | 0.49 | 1 | HC_vs_MS.Trt_B |
| SLC25A53 | 2.90E-04 | -0.5412 | 0.425 | 0.32 | 1 | HC_vs_MS.Trt_B |
| FUT9 | 2.90E-04 | 0.26692 | 0.368 | 0.28 | 1 | HC_vs_MS.Trt_B |
| RP11-226E2 | 2.90E-04 | -0.5873 | 0.635 | 0.52 | 1 | HC_vs_MS.Trt_B |
| ZNF213 | 2.91E-04 | 0.80713 | 0.455 | 0.33 | 1 | HC_vs_MS.Trt_B |
| ZNF720 | 2.92E-04 | -0.2667 | 0.497 | 0.38 | 1 | HC_vs_MS.Trt_B |
| C1orf27 | 2.94E-04 | 1.39708 | 0.155 | 0.08 | 1 | HC_vs_MS.Trt_B |
| RSL1D1 | 2.95E-04 | -0.3149 | 0.846 | 0.73 | 1 | HC_vs_MS.Trt_B |
| RALGAPA2 | 2.97E-04 | 0.81241 | 0.149 | 0.08 | 1 | HC_vs_MS.Trt_B |
| RSRC1 | 2.98E-04 | -0.555 | 0.739 | 0.6 | 1 | HC_vs_MS.Trt_B |
| DEPDC4 | 2.99E-04 | 0.40158 | 0.438 | 0.34 | 1 | HC_vs_MS.Trt_B |
| CTD-2192J1( | 3.01E-04 | 0.898 | 0.454 | 0.35 | 1 | HC_vs_MS.Trt_B |
| FBXL17 | 3.05E-04 | 0.47937 | 0.243 | 0.16 | 1 | HC_vs_MS.Trt_B |
| NSMCE3 | 3.05E-04 | 1.1357 | 0.319 | 0.22 | 1 | HC_vs_MS.Trt_B |
| MBP | 3.06E-04 | 0.40504 | 0.384 | 0.28 | 1 | HC_vs_MS.Trt_B |
| CES4A | 3.08E-04 | 0.36001 | 0.507 | 0.42 | 1 | HC_vs_MS.Trt_B |
| ZNF284 | 3.10E-04 | -0.8712 | 0.558 | 0.43 | 1 | HC_vs_MS.Trt_B |
| IL1R1 | 3.11E-04 | 0.72823 | 0.317 | 0.22 | 1 | HC_vs_MS.Trt_B |
| SPON2 | 3.13E-04 | 0.64771 | 0.383 | 0.28 | 1 | HC_vs_MS.Trt_B |
| LRRC69 | 3.13E-04 | 0.33512 | 0.575 | 0.47 | 1 | HC_vs_MS.Trt_B |
| USP38 | 3.15E-04 | 0.44587 | 0.312 | 0.22 | 1 | HC_vs_MS.Trt_B |
| RBM7 | 3.15E-04 | -0.3907 | 0.674 | 0.6 | 1 | HC_vs_MS.Trt_B |
| KIAA1755 | 3.15E-04 | 0.37909 | 0.194 | 0.12 | 1 | HC_vs_MS.Trt_B |
| MAPK1IP1L | 3.15E-04 | -0.2579 | 0.799 | 0.68 | 1 | HC_vs_MS.Trt_B |
| TBC1D2B | 3.16E-04 | 1.18352 | 0.294 | 0.21 | 1 | HC_vs_MS.Trt_B |
| SARDH | 3.17E-04 | 0.58806 | 0.388 | 0.29 | 1 | HC_vs_MS.Trt_B |
| BCL7B | 3.19E-04 | 0.41313 | 0.455 | 0.37 | 1 | HC_vs_MS.Trt_B |
| TECRL | 3.21E-04 | 0.27275 | 0.217 | 0.13 | 1 | HC_vs_MS.Trt_B |
| RABIF | 3.21E-04 | 0.87989 | 0.364 | 0.26 | 1 | HC_vs_MS.Trt_B |
| GUSB | 3.23E-04 | -0.2511 | 0.681 | 0.58 | 1 | HC_vs_MS.Trt_B |
| CDC42EP3 | 3.25E-04 | 0.26841 | 0.461 | 0.35 | 1 | HC_vs_MS.Trt_B |
| ELAVL1 | 3.27E-04 | 0.36976 | 0.961 | 0.92 | 1 | HC_vs_MS.Trt_B |
| RP11-9J18.1 | 3.27E-04 | 0.26928 | 0.262 | 0.17 | 1 | HC_vs_MS.Trt_B |
| ZNF696 | 3.27E-04 | -0.2829 | 0.316 | 0.22 | 1 | HC_vs_MS.Trt_B |
| GBP2 | 3.29E-04 | 0.39318 | 0.416 | 0.32 | 1 | HC_vs_MS.Trt_B |

| | | | | | | |
|---|---|---|---|---|---|---|
| CTB-133G6.1 | 3.35E-04 | 0.42841 | 0.268 | 0.18 | 1 | HC_vs_MS.Trt_B |
| C1GALT1 | 3.36E-04 | 0.27736 | 0.564 | 0.47 | 1 | HC_vs_MS.Trt_B |
| CD93 | 3.38E-04 | 0.50337 | 0.196 | 0.12 | 1 | HC_vs_MS.Trt_B |
| U2AF1L4 | 3.38E-04 | -0.2609 | 0.786 | 0.69 | 1 | HC_vs_MS.Trt_B |
| UBFD1 | 3.38E-04 | 0.59154 | 0.367 | 0.26 | 1 | HC_vs_MS.Trt_B |
| DNAJC14 | 3.40E-04 | 0.32047 | 0.599 | 0.5 | 1 | HC_vs_MS.Trt_B |
| FBXL8 | 3.41E-04 | 0.48978 | 0.539 | 0.45 | 1 | HC_vs_MS.Trt_B |
| XAB2 | 3.47E-04 | 0.69689 | 0.228 | 0.14 | 1 | HC_vs_MS.Trt_B |
| EPHX2 | 3.48E-04 | 0.66891 | 0.294 | 0.21 | 1 | HC_vs_MS.Trt_B |
| KCNK3 | 3.51E-04 | 0.35242 | 0.609 | 0.49 | 1 | HC_vs_MS.Trt_B |
| PTPN12 | 3.52E-04 | 0.43261 | 0.717 | 0.63 | 1 | HC_vs_MS.Trt_B |
| ANGPT2 | 3.53E-04 | 0.45954 | 0.726 | 0.65 | 1 | HC_vs_MS.Trt_B |
| WDR76 | 3.53E-04 | 0.26009 | 0.658 | 0.55 | 1 | HC_vs_MS.Trt_B |
| U2AF1L5 | 3.54E-04 | -0.2572 | 0.632 | 0.51 | 1 | HC_vs_MS.Trt_B |
| QSOX2 | 3.57E-04 | -0.7588 | 0.158 | 0.09 | 1 | HC_vs_MS.Trt_B |
| RCCD1 | 3.57E-04 | 0.26309 | 0.178 | 0.1 | 1 | HC_vs_MS.Trt_B |
| TAF11 | 3.57E-04 | 0.62468 | 0.483 | 0.35 | 1 | HC_vs_MS.Trt_B |
| MED11 | 3.58E-04 | -0.3163 | 0.691 | 0.6 | 1 | HC_vs_MS.Trt_B |
| FAM105A | 3.60E-04 | -0.2782 | 0.275 | 0.18 | 1 | HC_vs_MS.Trt_B |
| WDR41 | 3.60E-04 | -0.3354 | 0.422 | 0.32 | 1 | HC_vs_MS.Trt_B |
| PDE4D | 3.60E-04 | 0.49547 | 0.304 | 0.22 | 1 | HC_vs_MS.Trt_B |
| GSTM2 | 3.61E-04 | -0.3594 | 0.39 | 0.28 | 1 | HC_vs_MS.Trt_B |
| ZCCHC7 | 3.61E-04 | 0.28535 | 0.312 | 0.22 | 1 | HC_vs_MS.Trt_B |
| C8orf33 | 3.63E-04 | 0.96709 | 0.206 | 0.12 | 1 | HC_vs_MS.Trt_B |
| SUPT5H | 3.63E-04 | 0.34993 | 0.425 | 0.32 | 1 | HC_vs_MS.Trt_B |
| TNFRSF14 | 3.65E-04 | 0.28038 | 0.32 | 0.22 | 1 | HC_vs_MS.Trt_B |
| SPECC1L-AD | 3.69E-04 | 0.58515 | 0.187 | 0.11 | 1 | HC_vs_MS.Trt_B |
| CCDC120 | 3.70E-04 | 0.6161 | 0.464 | 0.36 | 1 | HC_vs_MS.Trt_B |
| HMGCR | 3.70E-04 | -0.4782 | 0.68 | 0.57 | 1 | HC_vs_MS.Trt_B |
| TIMM17A | 3.72E-04 | -0.3171 | 0.89 | 0.8 | 1 | HC_vs_MS.Trt_B |
| MFSD8 | 3.73E-04 | 0.37982 | 0.377 | 0.29 | 1 | HC_vs_MS.Trt_B |
| WTAP | 3.75E-04 | -0.6287 | 0.593 | 0.66 | 1 | HC_vs_MS.Trt_B |
| IGBP1 | 3.78E-04 | 0.58034 | 0.213 | 0.13 | 1 | HC_vs_MS.Trt_B |
| CCDC117 | 3.78E-04 | -0.7049 | 0.299 | 0.21 | 1 | HC_vs_MS.Trt_B |
| LIN54 | 3.78E-04 | 0.68618 | 0.548 | 0.44 | 1 | HC_vs_MS.Trt_B |
| ESR2 | 3.79E-04 | 0.40075 | 0.542 | 0.42 | 1 | HC_vs_MS.Trt_B |
| HERC5 | 3.79E-04 | 0.48473 | 0.21 | 0.13 | 1 | HC_vs_MS.Trt_B |
| PSMD3 | 3.82E-04 | 0.31464 | 0.51 | 0.39 | 1 | HC_vs_MS.Trt_B |
| PDZK1 | 3.84E-04 | 0.42261 | 0.299 | 0.21 | 1 | HC_vs_MS.Trt_B |
| NBEAL1 | 3.86E-04 | 0.46787 | 0.87 | 0.78 | 1 | HC_vs_MS.Trt_B |

| | | | | | | |
|---|---|---|---|---|---|---|
| C9orf85 | 3.87E-04 | -0.3396 | 0.865 | 0.8 | 1 | HC_vs_MS.Trt_B |
| NDRG3 | 3.87E-04 | 0.32513 | 0.374 | 0.28 | 1 | HC_vs_MS.Trt_B |
| ZNF783 | 3.88E-04 | 0.59785 | 0.442 | 0.34 | 1 | HC_vs_MS.Trt_B |
| MYO10 | 3.89E-04 | 0.41412 | 0.667 | 0.59 | 1 | HC_vs_MS.Trt_B |
| SEMA4F | 3.90E-04 | 0.65993 | 0.486 | 0.4 | 1 | HC_vs_MS.Trt_B |
| IL17RE | 3.90E-04 | 0.25198 | 0.259 | 0.17 | 1 | HC_vs_MS.Trt_B |
| TEX10 | 3.93E-04 | -0.4351 | 0.465 | 0.35 | 1 | HC_vs_MS.Trt_B |
| TMEM39B | 3.94E-04 | 0.50707 | 0.17 | 0.1 | 1 | HC_vs_MS.Trt_B |
| ABO | 3.94E-04 | 0.4201 | 0.362 | 0.27 | 1 | HC_vs_MS.Trt_B |
| RP11-1070B | 3.94E-04 | 0.26704 | 0.186 | 0.11 | 1 | HC_vs_MS.Trt_B |
| RRP1 | 3.94E-04 | -0.8036 | 0.638 | 0.54 | 1 | HC_vs_MS.Trt_B |
| PIP5K1A | 3.94E-04 | 0.47301 | 0.493 | 0.39 | 1 | HC_vs_MS.Trt_B |
| POLR1A | 3.96E-04 | -0.4075 | 0.672 | 0.56 | 1 | HC_vs_MS.Trt_B |
| SLC44A2 | 3.97E-04 | 0.27976 | 0.722 | 0.64 | 1 | HC_vs_MS.Trt_B |
| EPB41L4A | 3.98E-04 | -0.522 | 0.561 | 0.46 | 1 | HC_vs_MS.Trt_B |
| MARS | 3.99E-04 | 0.39568 | 0.381 | 0.29 | 1 | HC_vs_MS.Trt_B |
| ADAL | 4.01E-04 | -0.3713 | 0.414 | 0.3 | 1 | HC_vs_MS.Trt_B |
| HHLA3 | 4.01E-04 | 0.63139 | 0.343 | 0.25 | 1 | HC_vs_MS.Trt_B |
| PHF7 | 4.02E-04 | 0.46467 | 0.459 | 0.35 | 1 | HC_vs_MS.Trt_B |
| DDX3X | 4.03E-04 | -0.3104 | 0.329 | 0.22 | 1 | HC_vs_MS.Trt_B |
| CALM3 | 4.04E-04 | 0.32034 | 0.43 | 0.34 | 1 | HC_vs_MS.Trt_B |
| SNX22 | 4.04E-04 | 0.44616 | 0.803 | 0.66 | 1 | HC_vs_MS.Trt_B |
| ATF2 | 4.05E-04 | 0.35935 | 0.213 | 0.14 | 1 | HC_vs_MS.Trt_B |
| CRELD2 | 4.06E-04 | 0.28177 | 0.574 | 0.47 | 1 | HC_vs_MS.Trt_B |
| PRRG3 | 4.08E-04 | 1.0817 | 0.597 | 0.51 | 1 | HC_vs_MS.Trt_B |
| NTRK1 | 4.10E-04 | 0.54787 | 0.109 | 0.05 | 1 | HC_vs_MS.Trt_B |
| SPCS3 | 4.11E-04 | 0.25553 | 0.491 | 0.39 | 1 | HC_vs_MS.Trt_B |
| ADI1 | 4.14E-04 | -0.3804 | 0.735 | 0.65 | 1 | HC_vs_MS.Trt_B |
| POMK | 4.14E-04 | 0.52934 | 0.536 | 0.43 | 1 | HC_vs_MS.Trt_B |
| GPR83 | 4.16E-04 | -0.5469 | 0.458 | 0.36 | 1 | HC_vs_MS.Trt_B |
| ACACA | 4.17E-04 | 0.79793 | 0.59 | 0.46 | 1 | HC_vs_MS.Trt_B |
| FUBP1 | 4.19E-04 | 0.3522 | 0.362 | 0.27 | 1 | HC_vs_MS.Trt_B |
| NUDT15 | 4.21E-04 | 0.47381 | 0.545 | 0.44 | 1 | HC_vs_MS.Trt_B |
| UHRF2 | 4.22E-04 | 0.67008 | 0.251 | 0.17 | 1 | HC_vs_MS.Trt_B |
| NDUFS3 | 4.25E-04 | -0.5806 | 0.554 | 0.43 | 1 | HC_vs_MS.Trt_B |
| C2orf50 | 4.34E-04 | 0.39255 | 0.351 | 0.26 | 1 | HC_vs_MS.Trt_B |
| GNPNAT1 | 4.34E-04 | -0.3084 | 0.513 | 0.4 | 1 | HC_vs_MS.Trt_B |
| LZTS1 | 4.37E-04 | 0.32881 | 0.325 | 0.23 | 1 | HC_vs_MS.Trt_B |
| BPNT1 | 4.39E-04 | 0.38821 | 0.659 | 0.57 | 1 | HC_vs_MS.Trt_B |
| ARSG | 4.39E-04 | -0.2687 | 0.894 | 0.85 | 1 | HC_vs_MS.Trt_B |

| | | | | | | |
|---|---|---|---|---|---|---|
| CADPS | 4.40E-04 | 0.34294 | 0.33 | 0.24 | 1 | HC_vs_MS.Trt_B |
| AK5 | 4.40E-04 | 1.06369 | 0.142 | 0.07 | 1 | HC_vs_MS.Trt_B |
| GLDN | 4.42E-04 | 0.31217 | 0.206 | 0.13 | 1 | HC_vs_MS.Trt_B |
| MYO5A | 4.43E-04 | 0.3064 | 0.755 | 0.65 | 1 | HC_vs_MS.Trt_B |
| CTD-2207O2 | 4.44E-04 | 0.7868 | 0.167 | 0.1 | 1 | HC_vs_MS.Trt_B |
| TMEM5 | 4.44E-04 | -0.3758 | 0.497 | 0.39 | 1 | HC_vs_MS.Trt_B |
| MTHFR | 4.47E-04 | -0.2671 | 0.809 | 0.7 | 1 | HC_vs_MS.Trt_B |
| MTHFD2L | 4.47E-04 | 0.25776 | 0.497 | 0.39 | 1 | HC_vs_MS.Trt_B |
| WDFY2 | 4.49E-04 | 0.73236 | 0.558 | 0.47 | 1 | HC_vs_MS.Trt_B |
| CD38 | 4.50E-04 | 0.63956 | 0.335 | 0.25 | 1 | HC_vs_MS.Trt_B |
| PSEN1 | 4.51E-04 | -0.4502 | 0.68 | 0.58 | 1 | HC_vs_MS.Trt_B |
| C16orf54 | 4.52E-04 | -0.8031 | 0.761 | 0.73 | 1 | HC_vs_MS.Trt_B |
| ANAPC4 | 4.53E-04 | 0.56241 | 0.262 | 0.18 | 1 | HC_vs_MS.Trt_B |
| PCGF6 | 4.59E-04 | -0.3004 | 0.701 | 0.59 | 1 | HC_vs_MS.Trt_B |
| TXNL4A | 4.60E-04 | -0.5667 | 0.471 | 0.38 | 1 | HC_vs_MS.Trt_B |
| ETV3 | 4.63E-04 | 0.48286 | 0.12 | 0.06 | 1 | HC_vs_MS.Trt_B |
| METRN | 4.68E-04 | -0.4827 | 0.12 | 0.06 | 1 | HC_vs_MS.Trt_B |
| PQLC1 | 4.70E-04 | 0.28263 | 0.448 | 0.34 | 1 | HC_vs_MS.Trt_B |
| ABHD18 | 4.72E-04 | 0.26891 | 0.833 | 0.73 | 1 | HC_vs_MS.Trt_B |
| GABRE | 4.74E-04 | 0.3444 | 0.206 | 0.13 | 1 | HC_vs_MS.Trt_B |
| BTBD19 | 4.76E-04 | -0.2891 | 0.726 | 0.63 | 1 | HC_vs_MS.Trt_B |
| MCF2L | 4.79E-04 | 0.49735 | 0.303 | 0.21 | 1 | HC_vs_MS.Trt_B |
| AIFM2 | 4.80E-04 | -0.8014 | 0.354 | 0.26 | 1 | HC_vs_MS.Trt_B |
| BUB1B | 4.84E-04 | 0.32765 | 0.378 | 0.3 | 1 | HC_vs_MS.Trt_B |
| ARHGEF6 | 4.86E-04 | 0.37742 | 0.339 | 0.25 | 1 | HC_vs_MS.Trt_B |
| ERCC4 | 4.88E-04 | -0.2743 | 0.858 | 0.79 | 1 | HC_vs_MS.Trt_B |
| SLC7A11 | 4.88E-04 | 0.27549 | 0.449 | 0.37 | 1 | HC_vs_MS.Trt_B |
| HLTF | 4.90E-04 | -0.3526 | 0.513 | 0.4 | 1 | HC_vs_MS.Trt_B |
| RTF1 | 4.94E-04 | -0.3146 | 0.557 | 0.45 | 1 | HC_vs_MS.Trt_B |
| FAM179A | 4.95E-04 | -0.5843 | 0.772 | 0.68 | 1 | HC_vs_MS.Trt_B |
| CTTNBP2NL | 4.95E-04 | 0.43894 | 0.441 | 0.34 | 1 | HC_vs_MS.Trt_B |
| WDR1 | 4.96E-04 | 0.25196 | 0.377 | 0.28 | 1 | HC_vs_MS.Trt_B |
| CDC14A | 4.96E-04 | -0.3193 | 0.229 | 0.14 | 1 | HC_vs_MS.Trt_B |
| GTF2H2C | 4.97E-04 | 0.49608 | 0.439 | 0.35 | 1 | HC_vs_MS.Trt_B |
| NEIL1 | 4.97E-04 | 0.35157 | 0.757 | 0.65 | 1 | HC_vs_MS.Trt_B |
| UBE2W | 4.98E-04 | -0.258 | 0.626 | 0.5 | 1 | HC_vs_MS.Trt_B |
| GNG7 | 4.98E-04 | 0.29559 | 0.394 | 0.29 | 1 | HC_vs_MS.Trt_B |
| GLCE | 5.04E-04 | 0.96073 | 0.141 | 0.07 | 1 | HC_vs_MS.Trt_B |
| SGK1 | 5.07E-04 | 0.3745 | 0.371 | 0.27 | 1 | HC_vs_MS.Trt_B |
| C1orf216 | 5.10E-04 | -0.386 | 0.81 | 0.68 | 1 | HC_vs_MS.Trt_B |

| | | | | | | |
|---|---|---|---|---|---|---|
| GMIP | 5.11E-04 | 0.30763 | 0.254 | 0.17 | 1 | HC_vs_MS.Trt_B |
| RPL6 | 5.18E-04 | -0.262 | 0.962 | 0.92 | 1 | HC_vs_MS.Trt_B |
| PCSK2 | 5.18E-04 | 0.31005 | 0.28 | 0.19 | 1 | HC_vs_MS.Trt_B |
| FBXO16 | 5.18E-04 | 0.28833 | 0.262 | 0.18 | 1 | HC_vs_MS.Trt_B |
| NDUFA2 | 5.19E-04 | -0.4069 | 0.445 | 0.33 | 1 | HC_vs_MS.Trt_B |
| TAB3 | 5.19E-04 | -0.3839 | 0.674 | 0.56 | 1 | HC_vs_MS.Trt_B |
| CLASP1 | 5.20E-04 | 0.27425 | 0.241 | 0.16 | 1 | HC_vs_MS.Trt_B |
| TTC37 | 5.20E-04 | 0.47695 | 0.372 | 0.27 | 1 | HC_vs_MS.Trt_B |
| ZNF12 | 5.22E-04 | 1.50487 | 0.329 | 0.24 | 1 | HC_vs_MS.Trt_B |
| CARHSP1 | 5.25E-04 | 0.68741 | 0.632 | 0.54 | 1 | HC_vs_MS.Trt_B |
| ZEB2 | 5.26E-04 | -0.7558 | 0.6 | 0.51 | 1 | HC_vs_MS.Trt_B |
| SOX5 | 5.27E-04 | 0.28371 | 0.174 | 0.1 | 1 | HC_vs_MS.Trt_B |
| SMAP1 | 5.27E-04 | 0.38905 | 0.18 | 0.11 | 1 | HC_vs_MS.Trt_B |
| KCTD21 | 5.28E-04 | -0.4194 | 0.416 | 0.32 | 1 | HC_vs_MS.Trt_B |
| PEX11A | 5.29E-04 | -0.3751 | 0.552 | 0.45 | 1 | HC_vs_MS.Trt_B |
| AMBRA1 | 5.30E-04 | 0.7425 | 0.201 | 0.12 | 1 | HC_vs_MS.Trt_B |
| NEBL | 5.31E-04 | 0.352 | 0.287 | 0.2 | 1 | HC_vs_MS.Trt_B |
| NMNAT1 | 5.33E-04 | 0.30854 | 0.977 | 0.94 | 1 | HC_vs_MS.Trt_B |
| VPS36 | 5.35E-04 | 0.32769 | 0.28 | 0.19 | 1 | HC_vs_MS.Trt_B |
| SHMT1 | 5.36E-04 | -0.3143 | 0.367 | 0.27 | 1 | HC_vs_MS.Trt_B |
| TGIF2 | 5.36E-04 | 0.67902 | 0.343 | 0.26 | 1 | HC_vs_MS.Trt_B |
| SRSF12 | 5.38E-04 | 0.41587 | 0.297 | 0.21 | 1 | HC_vs_MS.Trt_B |
| IL23R | 5.40E-04 | 0.70066 | 0.343 | 0.25 | 1 | HC_vs_MS.Trt_B |
| ZNF627 | 5.40E-04 | -0.3144 | 0.568 | 0.45 | 1 | HC_vs_MS.Trt_B |
| TLR3 | 5.43E-04 | 0.31672 | 0.286 | 0.21 | 1 | HC_vs_MS.Trt_B |
| ERI2 | 5.47E-04 | 0.35991 | 0.148 | 0.08 | 1 | HC_vs_MS.Trt_B |
| MS4A2 | 5.49E-04 | 0.52006 | 0.555 | 0.49 | 1 | HC_vs_MS.Trt_B |
| RNF14 | 5.50E-04 | -0.6578 | 0.603 | 0.48 | 1 | HC_vs_MS.Trt_B |
| LMCD1 | 5.50E-04 | -0.2889 | 0.284 | 0.19 | 1 | HC_vs_MS.Trt_B |
| TRIOBP | 5.57E-04 | -0.4756 | 0.397 | 0.29 | 1 | HC_vs_MS.Trt_B |
| COPS4 | 5.61E-04 | -0.2836 | 0.468 | 0.35 | 1 | HC_vs_MS.Trt_B |
| ZNF45 | 5.63E-04 | -1.0582 | 0.284 | 0.19 | 1 | HC_vs_MS.Trt_B |
| ATP2C1 | 5.66E-04 | 0.47934 | 0.152 | 0.09 | 1 | HC_vs_MS.Trt_B |
| PDCD7 | 5.66E-04 | 0.4689 | 0.438 | 0.34 | 1 | HC_vs_MS.Trt_B |
| PITPNM2 | 5.67E-04 | 0.30982 | 0.422 | 0.32 | 1 | HC_vs_MS.Trt_B |
| ARHGAP11A | 5.67E-04 | 0.53227 | 0.37 | 0.28 | 1 | HC_vs_MS.Trt_B |
| SNX29 | 5.69E-04 | 0.29574 | 0.77 | 0.73 | 1 | HC_vs_MS.Trt_B |
| BLOC1S2 | 5.70E-04 | -0.3522 | 0.522 | 0.42 | 1 | HC_vs_MS.Trt_B |
| VEZF1 | 5.75E-04 | 0.34626 | 0.201 | 0.12 | 1 | HC_vs_MS.Trt_B |
| ZNRF1 | 5.75E-04 | 0.61561 | 0.274 | 0.19 | 1 | HC_vs_MS.Trt_B |

| | | | | | | |
|---|---|---|---|---|---|---|
| RP11-235E1 | 5.76E-04 | 0.44842 | 0.265 | 0.19 | 1 | HC_vs_MS.Trt_B |
| SHC1 | 5.86E-04 | 0.27599 | 0.63 | 0.55 | 1 | HC_vs_MS.Trt_B |
| SP140 | 5.89E-04 | 0.5377 | 0.212 | 0.13 | 1 | HC_vs_MS.Trt_B |
| C15orf61 | 5.93E-04 | -0.3288 | 0.58 | 0.45 | 1 | HC_vs_MS.Trt_B |
| POLR3D | 5.99E-04 | 0.49256 | 0.4 | 0.3 | 1 | HC_vs_MS.Trt_B |
| ATP6V1A | 6.00E-04 | 0.49226 | 0.472 | 0.38 | 1 | HC_vs_MS.Trt_B |
| ETV4 | 6.01E-04 | 0.47083 | 0.101 | 0.05 | 1 | HC_vs_MS.Trt_B |
| OSBPL2 | 6.04E-04 | -0.2915 | 0.475 | 0.38 | 1 | HC_vs_MS.Trt_B |
| TSC22D2 | 6.09E-04 | 0.44107 | 0.372 | 0.29 | 1 | HC_vs_MS.Trt_B |
| KIAA1468 | 6.15E-04 | -0.525 | 0.316 | 0.23 | 1 | HC_vs_MS.Trt_B |
| C2orf49 | 6.16E-04 | -0.4361 | 0.425 | 0.32 | 1 | HC_vs_MS.Trt_B |
| ZMYM1 | 6.18E-04 | 0.51984 | 0.549 | 0.45 | 1 | HC_vs_MS.Trt_B |
| TATDN3 | 6.19E-04 | 0.38567 | 0.775 | 0.7 | 1 | HC_vs_MS.Trt_B |
| TOP3A | 6.24E-04 | 0.45316 | 0.817 | 0.73 | 1 | HC_vs_MS.Trt_B |
| RALGAPB | 6.27E-04 | 0.54147 | 0.277 | 0.19 | 1 | HC_vs_MS.Trt_B |
| PYCR2 | 6.28E-04 | 0.31974 | 0.614 | 0.53 | 1 | HC_vs_MS.Trt_B |
| TTC39A | 6.31E-04 | 0.30888 | 0.333 | 0.24 | 1 | HC_vs_MS.Trt_B |
| ZNF514 | 6.34E-04 | 0.36486 | 0.597 | 0.48 | 1 | HC_vs_MS.Trt_B |
| NUP58 | 6.34E-04 | 0.55805 | 0.332 | 0.24 | 1 | HC_vs_MS.Trt_B |
| APCDD1 | 6.41E-04 | 0.26238 | 0.372 | 0.29 | 1 | HC_vs_MS.Trt_B |
| 9-Mar | 6.44E-04 | -0.3063 | 0.554 | 0.46 | 1 | HC_vs_MS.Trt_B |
| FANCL | 6.48E-04 | 0.69465 | 0.12 | 0.06 | 1 | HC_vs_MS.Trt_B |
| HNRNPL | 6.50E-04 | -0.3972 | 0.745 | 0.74 | 1 | HC_vs_MS.Trt_B |
| SERPINF1 | 6.51E-04 | 0.32022 | 0.752 | 0.65 | 1 | HC_vs_MS.Trt_B |
| TXNL4B | 6.54E-04 | 0.49292 | 0.354 | 0.26 | 1 | HC_vs_MS.Trt_B |
| TTC14 | 6.56E-04 | -0.2809 | 0.399 | 0.29 | 1 | HC_vs_MS.Trt_B |
| SCARB2 | 6.56E-04 | 0.26318 | 0.3 | 0.21 | 1 | HC_vs_MS.Trt_B |
| DGKD | 6.60E-04 | 0.5787 | 0.604 | 0.53 | 1 | HC_vs_MS.Trt_B |
| KIAA0586 | 6.62E-04 | -0.2662 | 0.796 | 0.71 | 1 | HC_vs_MS.Trt_B |
| ASZ1 | 6.67E-04 | 0.28695 | 0.454 | 0.36 | 1 | HC_vs_MS.Trt_B |
| TUBA1C | 6.69E-04 | 0.34796 | 0.588 | 0.48 | 1 | HC_vs_MS.Trt_B |
| FBXO25 | 6.70E-04 | 0.41789 | 0.358 | 0.26 | 1 | HC_vs_MS.Trt_B |
| CES2 | 6.77E-04 | 0.32805 | 0.316 | 0.22 | 1 | HC_vs_MS.Trt_B |
| WAPL | 6.83E-04 | 0.39876 | 0.452 | 0.36 | 1 | HC_vs_MS.Trt_B |
| ZNF880 | 6.86E-04 | -0.3769 | 0.47 | 0.37 | 1 | HC_vs_MS.Trt_B |
| TTC16 | 6.87E-04 | -0.5247 | 0.561 | 0.44 | 1 | HC_vs_MS.Trt_B |
| ZNF764 | 6.88E-04 | 0.68622 | 0.32 | 0.24 | 1 | HC_vs_MS.Trt_B |
| NOL12 | 6.93E-04 | -0.2928 | 0.483 | 0.37 | 1 | HC_vs_MS.Trt_B |
| MLANA | 7.14E-04 | 0.25103 | 0.394 | 0.3 | 1 | HC_vs_MS.Trt_B |
| ZNF260 | 7.24E-04 | 0.42604 | 0.506 | 0.4 | 1 | HC_vs_MS.Trt_B |

| | | | | | | |
|---|---|---|---|---|---|---|
| MORN4 | 7.25E-04 | 0.35741 | 0.532 | 0.43 | 1 | HC_vs_MS.Trt_B |
| RP11-449H3 | 7.26E-04 | 0.72122 | 0.255 | 0.17 | 1 | HC_vs_MS.Trt_B |
| FAM92A1 | 7.26E-04 | 0.5129 | 0.323 | 0.24 | 1 | HC_vs_MS.Trt_B |
| TMEM220 | 7.31E-04 | 0.396 | 0.532 | 0.45 | 1 | HC_vs_MS.Trt_B |
| SYT11 | 7.35E-04 | -0.5145 | 0.616 | 0.49 | 1 | HC_vs_MS.Trt_B |
| CEP164 | 7.40E-04 | 0.71578 | 0.157 | 0.09 | 1 | HC_vs_MS.Trt_B |
| ZSCAN10 | 7.44E-04 | 0.37945 | 0.352 | 0.26 | 1 | HC_vs_MS.Trt_B |
| KIAA1551 | 7.45E-04 | 0.2675 | 0.497 | 0.41 | 1 | HC_vs_MS.Trt_B |
| GMPPB | 7.49E-04 | 0.89475 | 0.41 | 0.32 | 1 | HC_vs_MS.Trt_B |
| HMBOX1 | 7.53E-04 | 0.29013 | 0.497 | 0.39 | 1 | HC_vs_MS.Trt_B |
| METTL15 | 7.55E-04 | 0.71944 | 0.377 | 0.29 | 1 | HC_vs_MS.Trt_B |
| ARHGAP17 | 7.56E-04 | 0.33482 | 0.558 | 0.47 | 1 | HC_vs_MS.Trt_B |
| RNF157 | 7.67E-04 | -0.6233 | 0.607 | 0.5 | 1 | HC_vs_MS.Trt_B |
| SP3 | 7.67E-04 | 0.32801 | 0.559 | 0.44 | 1 | HC_vs_MS.Trt_B |
| SLC7A9 | 7.67E-04 | 0.34278 | 0.403 | 0.32 | 1 | HC_vs_MS.Trt_B |
| ALG1 | 7.69E-04 | -0.5102 | 0.613 | 0.49 | 1 | HC_vs_MS.Trt_B |
| LAX1 | 7.70E-04 | 0.47963 | 0.467 | 0.38 | 1 | HC_vs_MS.Trt_B |
| PON2 | 7.71E-04 | 1.03736 | 0.42 | 0.33 | 1 | HC_vs_MS.Trt_B |
| PUS7 | 7.73E-04 | 0.82188 | 0.333 | 0.24 | 1 | HC_vs_MS.Trt_B |
| GREB1 | 7.74E-04 | 0.27978 | 0.836 | 0.74 | 1 | HC_vs_MS.Trt_B |
| TP63 | 7.75E-04 | 0.36508 | 0.39 | 0.3 | 1 | HC_vs_MS.Trt_B |
| SLC25A26 | 7.76E-04 | -0.3343 | 0.625 | 0.54 | 1 | HC_vs_MS.Trt_B |
| MARK4 | 7.82E-04 | 0.29754 | 0.58 | 0.48 | 1 | HC_vs_MS.Trt_B |
| RASA4B | 7.84E-04 | 0.38915 | 0.436 | 0.34 | 1 | HC_vs_MS.Trt_B |
| PRKCQ | 7.85E-04 | 0.32127 | 0.354 | 0.26 | 1 | HC_vs_MS.Trt_B |
| MTHFD2 | 7.87E-04 | -0.4882 | 0.625 | 0.53 | 1 | HC_vs_MS.Trt_B |
| CBLB | 7.87E-04 | 0.29697 | 0.355 | 0.26 | 1 | HC_vs_MS.Trt_B |
| SAXO2 | 7.90E-04 | -0.309 | 0.226 | 0.15 | 1 | HC_vs_MS.Trt_B |
| NMRAL1 | 7.91E-04 | -0.3852 | 0.496 | 0.37 | 1 | HC_vs_MS.Trt_B |
| NCOA4 | 7.92E-04 | 0.69193 | 0.204 | 0.13 | 1 | HC_vs_MS.Trt_B |
| GATA3 | 7.93E-04 | 0.42942 | 0.246 | 0.17 | 1 | HC_vs_MS.Trt_B |
| NPAP1 | 7.93E-04 | 0.27087 | 0.465 | 0.38 | 1 | HC_vs_MS.Trt_B |
| TTPAL | 7.94E-04 | -0.2763 | 0.791 | 0.69 | 1 | HC_vs_MS.Trt_B |
| SLC25A19 | 7.95E-04 | 0.52291 | 0.397 | 0.31 | 1 | HC_vs_MS.Trt_B |
| ZBED1 | 7.96E-04 | 0.51185 | 0.477 | 0.38 | 1 | HC_vs_MS.Trt_B |
| PLD1 | 7.96E-04 | 0.74796 | 0.491 | 0.39 | 1 | HC_vs_MS.Trt_B |
| VPS52 | 8.03E-04 | 0.63208 | 0.159 | 0.09 | 1 | HC_vs_MS.Trt_B |
| HSBP1 | 8.06E-04 | -0.6348 | 0.599 | 0.5 | 1 | HC_vs_MS.Trt_B |
| IMPAD1 | 8.08E-04 | 0.4412 | 0.472 | 0.37 | 1 | HC_vs_MS.Trt_B |
| ZDHHC17 | 8.14E-04 | 0.31066 | 0.345 | 0.26 | 1 | HC_vs_MS.Trt_B |

| | | | | | | |
|---|---|---|---|---|---|---|
| CLDN12 | 8.15E-04 | 0.28244 | 0.374 | 0.28 | 1 | HC_vs_MS.Trt_B |
| FLT4 | 8.19E-04 | 1.24513 | 0.104 | 0.05 | 1 | HC_vs_MS.Trt_B |
| FLCN | 8.20E-04 | 0.59763 | 0.859 | 0.8 | 1 | HC_vs_MS.Trt_B |
| KSR1 | 8.23E-04 | 0.44195 | 0.286 | 0.2 | 1 | HC_vs_MS.Trt_B |
| SMCR8 | 8.25E-04 | -0.5418 | 0.294 | 0.2 | 1 | HC_vs_MS.Trt_B |
| ZNF528 | 8.27E-04 | 0.67688 | 0.574 | 0.47 | 1 | HC_vs_MS.Trt_B |
| ILKAP | 8.30E-04 | -0.2574 | 0.588 | 0.46 | 1 | HC_vs_MS.Trt_B |
| ZNF687 | 8.35E-04 | 0.55409 | 0.593 | 0.5 | 1 | HC_vs_MS.Trt_B |
| SLC41A3 | 8.36E-04 | 0.40573 | 0.286 | 0.2 | 1 | HC_vs_MS.Trt_B |
| GALK1 | 8.39E-04 | 0.56634 | 0.297 | 0.22 | 1 | HC_vs_MS.Trt_B |
| NUDT19 | 8.44E-04 | -0.6489 | 0.87 | 0.77 | 1 | HC_vs_MS.Trt_B |
| DDX55 | 8.45E-04 | 0.49555 | 0.641 | 0.57 | 1 | HC_vs_MS.Trt_B |
| ZNF133 | 8.47E-04 | 0.86068 | 0.449 | 0.34 | 1 | HC_vs_MS.Trt_B |
| CCM2 | 8.53E-04 | 0.26321 | 0.649 | 0.56 | 1 | HC_vs_MS.Trt_B |
| ANKRD22 | 8.55E-04 | 0.40329 | 0.125 | 0.07 | 1 | HC_vs_MS.Trt_B |
| APOBEC3F | 8.56E-04 | -0.433 | 0.861 | 0.8 | 1 | HC_vs_MS.Trt_B |
| PLEKHG4 | 8.58E-04 | 0.6293 | 0.184 | 0.11 | 1 | HC_vs_MS.Trt_B |
| NLGN4Y | 8.60E-04 | 0.38549 | 0.355 | 0.26 | 1 | HC_vs_MS.Trt_B |
| VHL | 8.61E-04 | 0.33579 | 0.749 | 0.65 | 1 | HC_vs_MS.Trt_B |
| CARS2 | 8.62E-04 | 0.41537 | 0.758 | 0.65 | 1 | HC_vs_MS.Trt_B |
| ABLIM3 | 8.65E-04 | -0.6063 | 0.051 | 0.1 | 1 | HC_vs_MS.Trt_B |
| PARP4 | 8.66E-04 | 0.30783 | 0.271 | 0.19 | 1 | HC_vs_MS.Trt_B |
| HYAL1 | 8.66E-04 | 0.332 | 0.355 | 0.27 | 1 | HC_vs_MS.Trt_B |
| PAIP2B | 8.72E-04 | 0.27476 | 0.225 | 0.15 | 1 | HC_vs_MS.Trt_B |
| WDR19 | 8.79E-04 | 0.33433 | 0.512 | 0.42 | 1 | HC_vs_MS.Trt_B |
| ACSS2 | 8.81E-04 | 1.5672 | 0.549 | 0.47 | 1 | HC_vs_MS.Trt_B |
| ATAD2B | 8.83E-04 | 0.37739 | 0.628 | 0.55 | 1 | HC_vs_MS.Trt_B |
| SYT15 | 8.84E-04 | 0.41149 | 0.5 | 0.42 | 1 | HC_vs_MS.Trt_B |
| TMEM175 | 8.88E-04 | 0.41788 | 0.301 | 0.22 | 1 | HC_vs_MS.Trt_B |
| AREL1 | 8.88E-04 | 0.41919 | 0.446 | 0.35 | 1 | HC_vs_MS.Trt_B |
| SAMD15 | 8.89E-04 | 0.25339 | 0.323 | 0.23 | 1 | HC_vs_MS.Trt_B |
| TWSG1 | 8.90E-04 | 0.59031 | 0.371 | 0.28 | 1 | HC_vs_MS.Trt_B |
| IL5RA | 9.04E-04 | 1.334 | 0.252 | 0.17 | 1 | HC_vs_MS.Trt_B |
| TTBK2 | 9.06E-04 | -0.8377 | 0.493 | 0.4 | 1 | HC_vs_MS.Trt_B |
| CCL5 | 9.10E-04 | -0.4579 | 0.89 | 0.8 | 1 | HC_vs_MS.Trt_B |
| HDAC9 | 9.19E-04 | 0.73053 | 0.459 | 0.37 | 1 | HC_vs_MS.Trt_B |
| IRF2 | 9.26E-04 | 0.39566 | 0.332 | 0.24 | 1 | HC_vs_MS.Trt_B |
| PRKCI | 9.37E-04 | 0.49443 | 0.319 | 0.22 | 1 | HC_vs_MS.Trt_B |
| MBTPS1 | 9.38E-04 | 0.27057 | 0.517 | 0.41 | 1 | HC_vs_MS.Trt_B |
| NAPEPLD | 9.40E-04 | 0.63839 | 0.32 | 0.23 | 1 | HC_vs_MS.Trt_B |

| | | | | | | |
|---|---|---|---|---|---|---|
| AMMECR1L | 9.42E-04 | -0.2771 | 0.204 | 0.13 | 1 | HC_vs_MS.Trt_B |
| RNF214 | 9.43E-04 | 0.28318 | 0.223 | 0.15 | 1 | HC_vs_MS.Trt_B |
| TERB2 | 9.48E-04 | 0.25864 | 0.32 | 0.23 | 1 | HC_vs_MS.Trt_B |
| TAL1 | 9.49E-04 | 0.25949 | 0.212 | 0.14 | 1 | HC_vs_MS.Trt_B |
| WWC1 | 9.49E-04 | -0.2508 | 0.307 | 0.22 | 1 | HC_vs_MS.Trt_B |
| MESDC2 | 9.59E-04 | -0.2976 | 0.532 | 0.44 | 1 | HC_vs_MS.Trt_B |
| PABPC1L | 9.62E-04 | 1.48962 | 0.284 | 0.2 | 1 | HC_vs_MS.Trt_B |
| ZNF398 | 9.63E-04 | 0.81796 | 0.148 | 0.08 | 1 | HC_vs_MS.Trt_B |
| RABL6 | 9.68E-04 | -0.3833 | 0.509 | 0.42 | 1 | HC_vs_MS.Trt_B |
| TBC1D8 | 9.68E-04 | 0.33621 | 0.425 | 0.31 | 1 | HC_vs_MS.Trt_B |
| CPT2 | 9.76E-04 | -0.5631 | 0.746 | 0.63 | 1 | HC_vs_MS.Trt_B |
| ZNF674 | 9.78E-04 | 0.25742 | 0.904 | 0.8 | 1 | HC_vs_MS.Trt_B |
| MAPK13 | 9.79E-04 | -0.295 | 0.638 | 0.53 | 1 | HC_vs_MS.Trt_B |
| CSMD2 | 9.94E-04 | 0.31943 | 0.345 | 0.26 | 1 | HC_vs_MS.Trt_B |
| ANO10 | 9.98E-04 | 0.36193 | 0.568 | 0.45 | 1 | HC_vs_MS.Trt_B |
| ACLY | 0.001 | 0.4802 | 0.181 | 0.11 | 1 | HC_vs_MS.Trt_B |
| MTG2 | 0.00101 | 0.44395 | 0.775 | 0.71 | 1 | HC_vs_MS.Trt_B |
| KLHL5 | 0.00101 | -0.2648 | 0.419 | 0.33 | 1 | HC_vs_MS.Trt_B |
| ICMT | 0.00101 | -0.3059 | 0.609 | 0.53 | 1 | HC_vs_MS.Trt_B |
| FAM161B | 0.00102 | -0.3906 | 0.348 | 0.26 | 1 | HC_vs_MS.Trt_B |
| HECW2 | 0.00102 | 0.40079 | 0.403 | 0.32 | 1 | HC_vs_MS.Trt_B |
| ZNF551 | 0.00102 | 1.18384 | 0.513 | 0.4 | 1 | HC_vs_MS.Trt_B |
| SLX4IP | 0.00103 | -0.495 | 0.471 | 0.38 | 1 | HC_vs_MS.Trt_B |
| MRS2 | 0.00103 | 0.41073 | 0.764 | 0.71 | 1 | HC_vs_MS.Trt_B |
| RFX2 | 0.00104 | 0.53304 | 0.477 | 0.4 | 1 | HC_vs_MS.Trt_B |
| EXOSC7 | 0.00104 | -0.423 | 0.446 | 0.35 | 1 | HC_vs_MS.Trt_B |
| HDAC3 | 0.00105 | 0.98717 | 0.117 | 0.06 | 1 | HC_vs_MS.Trt_B |
| CAPN15 | 0.00105 | 0.5617 | 0.207 | 0.13 | 1 | HC_vs_MS.Trt_B |
| PPP2R2D | 0.00106 | -1.1857 | 0.622 | 0.53 | 1 | HC_vs_MS.Trt_B |
| ZNF540 | 0.00106 | -1.1032 | 0.428 | 0.34 | 1 | HC_vs_MS.Trt_B |
| C1orf127 | 0.00106 | 0.4099 | 0.272 | 0.2 | 1 | HC_vs_MS.Trt_B |
| PSMD4 | 0.00107 | -0.4244 | 0.642 | 0.54 | 1 | HC_vs_MS.Trt_B |
| NDUFC2 | 0.00107 | -0.3351 | 0.562 | 0.46 | 1 | HC_vs_MS.Trt_B |
| QKI | 0.00107 | 0.58757 | 0.386 | 0.29 | 1 | HC_vs_MS.Trt_B |
| CDC27 | 0.00108 | -0.2935 | 0.438 | 0.34 | 1 | HC_vs_MS.Trt_B |
| IVD | 0.00108 | 0.48617 | 0.9 | 0.85 | 1 | HC_vs_MS.Trt_B |
| NADK | 0.00109 | 0.27197 | 0.287 | 0.2 | 1 | HC_vs_MS.Trt_B |
| CCDC159 | 0.00109 | -0.4389 | 0.765 | 0.68 | 1 | HC_vs_MS.Trt_B |
| GBP3 | 0.00109 | 0.64012 | 0.194 | 0.12 | 1 | HC_vs_MS.Trt_B |
| PSPC1 | 0.00109 | 0.30933 | 0.309 | 0.22 | 1 | HC_vs_MS.Trt_B |

| | | | | | | |
|---|---|---|---|---|---|---|
| TVP23A | 0.0011 | 0.47435 | 0.454 | 0.36 | 1 | HC_vs_MS.Trt_B |
| SUV39H2 | 0.0011 | 0.53359 | 0.407 | 0.31 | 1 | HC_vs_MS.Trt_B |
| IQCE | 0.0011 | 0.54222 | 0.406 | 0.29 | 1 | HC_vs_MS.Trt_B |
| TMEM108 | 0.0011 | 0.83244 | 0.332 | 0.24 | 1 | HC_vs_MS.Trt_B |
| GALNT6 | 0.00111 | 0.2533 | 0.655 | 0.56 | 1 | HC_vs_MS.Trt_B |
| B3GALNT2 | 0.00111 | -0.4425 | 0.619 | 0.53 | 1 | HC_vs_MS.Trt_B |
| ANKDD1A | 0.00112 | -0.2784 | 0.522 | 0.43 | 1 | HC_vs_MS.Trt_B |
| SMG5 | 0.00112 | 0.42934 | 0.149 | 0.09 | 1 | HC_vs_MS.Trt_B |
| PCCA | 0.00112 | 0.32541 | 0.568 | 0.47 | 1 | HC_vs_MS.Trt_B |
| PTPN9 | 0.00113 | 0.75696 | 0.32 | 0.23 | 1 | HC_vs_MS.Trt_B |
| WWP1 | 0.00113 | -0.3665 | 0.432 | 0.32 | 1 | HC_vs_MS.Trt_B |
| ZNF564 | 0.00114 | -0.5681 | 0.629 | 0.52 | 1 | HC_vs_MS.Trt_B |
| TECPR2 | 0.00115 | 0.43107 | 0.345 | 0.25 | 1 | HC_vs_MS.Trt_B |
| TBC1D9B | 0.00115 | 0.48031 | 0.457 | 0.35 | 1 | HC_vs_MS.Trt_B |
| LAPTM4B | 0.00116 | -1.0218 | 0.194 | 0.12 | 1 | HC_vs_MS.Trt_B |
| VPS4B | 0.00116 | -0.4566 | 0.742 | 0.62 | 1 | HC_vs_MS.Trt_B |
| CA14 | 0.00116 | 0.26933 | 0.572 | 0.46 | 1 | HC_vs_MS.Trt_B |
| AP3M1 | 0.00118 | 0.76092 | 0.484 | 0.41 | 1 | HC_vs_MS.Trt_B |
| LAMTOR3 | 0.00118 | -0.4321 | 0.325 | 0.23 | 1 | HC_vs_MS.Trt_B |
| CDH26 | 0.00119 | 0.72459 | 0.159 | 0.1 | 1 | HC_vs_MS.Trt_B |
| NOP14 | 0.00119 | 0.5336 | 0.413 | 0.31 | 1 | HC_vs_MS.Trt_B |
| DNAJB12 | 0.0012 | 0.76403 | 0.12 | 0.06 | 1 | HC_vs_MS.Trt_B |
| HIKESHI | 0.0012 | -0.4608 | 0.43 | 0.31 | 1 | HC_vs_MS.Trt_B |
| ST7L | 0.00121 | 0.26498 | 0.468 | 0.38 | 1 | HC_vs_MS.Trt_B |
| TTC21B | 0.00121 | 0.97393 | 0.359 | 0.26 | 1 | HC_vs_MS.Trt_B |
| FBXO9 | 0.00122 | -0.485 | 0.752 | 0.67 | 1 | HC_vs_MS.Trt_B |
| KDM4C | 0.00123 | 0.35787 | 0.291 | 0.21 | 1 | HC_vs_MS.Trt_B |
| TBC1D32 | 0.00123 | -0.356 | 0.449 | 0.35 | 1 | HC_vs_MS.Trt_B |
| NFKB2 | 0.00123 | 0.3782 | 0.341 | 0.26 | 1 | HC_vs_MS.Trt_B |
| RHNO1 | 0.00123 | 0.42164 | 0.613 | 0.54 | 1 | HC_vs_MS.Trt_B |
| MTHFD1L | 0.00125 | -0.6757 | 0.125 | 0.07 | 1 | HC_vs_MS.Trt_B |
| HJURP | 0.00125 | 0.35976 | 0.37 | 0.28 | 1 | HC_vs_MS.Trt_B |
| DNHD1 | 0.00125 | 0.93434 | 0.316 | 0.24 | 1 | HC_vs_MS.Trt_B |
| UFM1 | 0.00126 | 0.34994 | 0.326 | 0.24 | 1 | HC_vs_MS.Trt_B |
| FKBP9 | 0.00127 | -0.866 | 0.445 | 0.34 | 1 | HC_vs_MS.Trt_B |
| DZIP3 | 0.00128 | -0.6008 | 0.257 | 0.17 | 1 | HC_vs_MS.Trt_B |
| CDK6 | 0.00128 | 0.61206 | 0.145 | 0.08 | 1 | HC_vs_MS.Trt_B |
| ZBP1 | 0.00129 | 0.32073 | 0.567 | 0.46 | 1 | HC_vs_MS.Trt_B |
| F2R | 0.00129 | -0.5523 | 0.525 | 0.44 | 1 | HC_vs_MS.Trt_B |
| IMMT | 0.0013 | -0.7424 | 0.443 | 0.34 | 1 | HC_vs_MS.Trt_B |

| | | | | | | |
|---|---|---|---|---|---|---|
| SNX30 | 0.0013 | -0.4755 | 0.435 | 0.34 | 1 | HC_vs_MS.Trt_B |
| SGSM2 | 0.00132 | 0.33114 | 0.264 | 0.19 | 1 | HC_vs_MS.Trt_B |
| PLA2G12A | 0.00133 | -0.5171 | 0.57 | 0.47 | 1 | HC_vs_MS.Trt_B |
| IER2 | 0.00134 | 0.25535 | 0.309 | 0.23 | 1 | HC_vs_MS.Trt_B |
| CHTF8 | 0.00135 | 0.25075 | 0.217 | 0.14 | 1 | HC_vs_MS.Trt_B |
| ALDH6A1 | 0.00135 | -0.5937 | 0.796 | 0.73 | 1 | HC_vs_MS.Trt_B |
| AMZ2 | 0.00137 | -0.3488 | 0.523 | 0.41 | 1 | HC_vs_MS.Trt_B |
| LGR6 | 0.00139 | 0.26243 | 0.178 | 0.11 | 1 | HC_vs_MS.Trt_B |
| USP14 | 0.0014 | 0.41862 | 0.575 | 0.47 | 1 | HC_vs_MS.Trt_B |
| CRCP | 0.0014 | -0.2702 | 0.862 | 0.76 | 1 | HC_vs_MS.Trt_B |
| PSMC1 | 0.00141 | -0.518 | 0.42 | 0.32 | 1 | HC_vs_MS.Trt_B |
| PARP10 | 0.00141 | 0.37694 | 0.157 | 0.09 | 1 | HC_vs_MS.Trt_B |
| FAT1 | 0.00141 | 0.3202 | 0.284 | 0.2 | 1 | HC_vs_MS.Trt_B |
| L3MBTL3 | 0.00141 | 1.02035 | 0.164 | 0.1 | 1 | HC_vs_MS.Trt_B |
| ALDH1L2 | 0.00142 | 0.45591 | 0.232 | 0.16 | 1 | HC_vs_MS.Trt_B |
| ARV1 | 0.00142 | -1.0771 | 0.232 | 0.16 | 1 | HC_vs_MS.Trt_B |
| ERI1 | 0.00143 | 0.25549 | 0.365 | 0.27 | 1 | HC_vs_MS.Trt_B |
| TJAP1 | 0.00143 | -0.7964 | 0.346 | 0.25 | 1 | HC_vs_MS.Trt_B |
| MED22 | 0.00144 | 0.29629 | 0.506 | 0.42 | 1 | HC_vs_MS.Trt_B |
| MAF1 | 0.00144 | 0.35236 | 0.193 | 0.12 | 1 | HC_vs_MS.Trt_B |
| ARRB2 | 0.00146 | 0.27661 | 0.771 | 0.66 | 1 | HC_vs_MS.Trt_B |
| DDX51 | 0.00147 | 0.6337 | 0.645 | 0.55 | 1 | HC_vs_MS.Trt_B |
| RABGEF1 | 0.00147 | 0.6689 | 0.297 | 0.21 | 1 | HC_vs_MS.Trt_B |
| FAM156A | 0.00148 | 0.42771 | 0.165 | 0.1 | 1 | HC_vs_MS.Trt_B |
| KLHL2 | 0.00148 | 0.3868 | 0.323 | 0.24 | 1 | HC_vs_MS.Trt_B |
| FAM149A | 0.00148 | 0.36446 | 0.306 | 0.22 | 1 | HC_vs_MS.Trt_B |
| SRRM1 | 0.00149 | -0.4148 | 0.868 | 0.75 | 1 | HC_vs_MS.Trt_B |
| CBWD2 | 0.00149 | 0.42695 | 0.414 | 0.33 | 1 | HC_vs_MS.Trt_B |
| VRTN | 0.0015 | -0.5578 | 0.155 | 0.09 | 1 | HC_vs_MS.Trt_B |
| XPR1 | 0.0015 | 0.2951 | 0.299 | 0.21 | 1 | HC_vs_MS.Trt_B |
| ATG14 | 0.00151 | 0.3479 | 0.654 | 0.57 | 1 | HC_vs_MS.Trt_B |
| SEC23IP | 0.00151 | -0.4018 | 0.412 | 0.32 | 1 | HC_vs_MS.Trt_B |
| PRR4 | 0.00152 | 0.28561 | 0.216 | 0.14 | 1 | HC_vs_MS.Trt_B |
| ARHGEF38 | 0.00152 | 0.31449 | 0.361 | 0.28 | 1 | HC_vs_MS.Trt_B |
| PRODH | 0.00153 | 0.74921 | 0.386 | 0.29 | 1 | HC_vs_MS.Trt_B |
| SLC39A3 | 0.00153 | -0.2713 | 0.581 | 0.5 | 1 | HC_vs_MS.Trt_B |
| PPM1B | 0.00154 | 0.47409 | 0.388 | 0.3 | 1 | HC_vs_MS.Trt_B |
| CDH8 | 0.00154 | 0.28669 | 0.477 | 0.39 | 1 | HC_vs_MS.Trt_B |
| SGPP2 | 0.00154 | -0.358 | 0.332 | 0.25 | 1 | HC_vs_MS.Trt_B |
| CLCN6 | 0.00154 | 0.90261 | 0.496 | 0.41 | 1 | HC_vs_MS.Trt_B |

| | | | | | | |
|---|---|---|---|---|---|---|
| ARRDC3 | 0.00154 | -0.6504 | 0.464 | 0.54 | 1 | HC_vs_MS.Trt_B |
| NBEAL2 | 0.00155 | 0.62762 | 0.206 | 0.14 | 1 | HC_vs_MS.Trt_B |
| NKD1 | 0.00156 | 0.50372 | 0.394 | 0.29 | 1 | HC_vs_MS.Trt_B |
| TROAP | 0.00156 | 0.25834 | 0.438 | 0.34 | 1 | HC_vs_MS.Trt_B |
| ACAN | 0.00157 | 0.97556 | 0.268 | 0.19 | 1 | HC_vs_MS.Trt_B |
| KDM6B | 0.00159 | 0.7127 | 0.161 | 0.1 | 1 | HC_vs_MS.Trt_B |
| GPCPD1 | 0.00159 | -0.2705 | 0.293 | 0.2 | 1 | HC_vs_MS.Trt_B |
| TRAF3IP1 | 0.00159 | 0.47256 | 0.371 | 0.29 | 1 | HC_vs_MS.Trt_B |
| DCBLD1 | 0.00159 | -0.3847 | 0.504 | 0.41 | 1 | HC_vs_MS.Trt_B |
| POLH | 0.00159 | -0.6554 | 0.541 | 0.44 | 1 | HC_vs_MS.Trt_B |
| FAM3A | 0.00159 | 0.27452 | 0.752 | 0.64 | 1 | HC_vs_MS.Trt_B |
| CCNC | 0.00159 | -0.3952 | 0.604 | 0.49 | 1 | HC_vs_MS.Trt_B |
| SPATA7 | 0.0016 | 0.27926 | 0.313 | 0.23 | 1 | HC_vs_MS.Trt_B |
| PRKRIP1 | 0.00161 | 0.88926 | 0.13 | 0.07 | 1 | HC_vs_MS.Trt_B |
| BRCA2 | 0.00161 | -0.6447 | 0.358 | 0.27 | 1 | HC_vs_MS.Trt_B |
| TUBA1B | 0.00162 | 0.31946 | 0.284 | 0.2 | 1 | HC_vs_MS.Trt_B |
| RABGAP1 | 0.00164 | 0.75757 | 0.381 | 0.29 | 1 | HC_vs_MS.Trt_B |
| ZNF677 | 0.00164 | 0.62395 | 0.794 | 0.74 | 1 | HC_vs_MS.Trt_B |
| ZNF69 | 0.00164 | 0.25272 | 0.452 | 0.37 | 1 | HC_vs_MS.Trt_B |
| MUM1 | 0.00165 | 0.32429 | 0.599 | 0.48 | 1 | HC_vs_MS.Trt_B |
| PHF20 | 0.00166 | -0.3827 | 0.478 | 0.38 | 1 | HC_vs_MS.Trt_B |
| XPO6 | 0.00166 | 0.48384 | 0.312 | 0.23 | 1 | HC_vs_MS.Trt_B |
| MED4 | 0.00166 | -0.4123 | 0.646 | 0.56 | 1 | HC_vs_MS.Trt_B |
| COMMD9 | 0.00166 | 0.26397 | 0.446 | 0.36 | 1 | HC_vs_MS.Trt_B |
| MTR | 0.00167 | 0.28098 | 0.478 | 0.41 | 1 | HC_vs_MS.Trt_B |
| FKBP8 | 0.00167 | 1.11191 | 0.196 | 0.13 | 1 | HC_vs_MS.Trt_B |
| TRIM2 | 0.00171 | 0.66869 | 0.23 | 0.16 | 1 | HC_vs_MS.Trt_B |
| UPF3A | 0.00171 | -0.3754 | 0.712 | 0.62 | 1 | HC_vs_MS.Trt_B |
| MAN1B1 | 0.00171 | 0.40338 | 0.354 | 0.26 | 1 | HC_vs_MS.Trt_B |
| BACH2 | 0.00173 | 0.53437 | 0.242 | 0.17 | 1 | HC_vs_MS.Trt_B |
| F2RL1 | 0.00173 | 1.13994 | 0.191 | 0.12 | 1 | HC_vs_MS.Trt_B |
| BRD8 | 0.00174 | -0.4562 | 0.735 | 0.65 | 1 | HC_vs_MS.Trt_B |
| TFAM | 0.00174 | 0.38894 | 0.312 | 0.24 | 1 | HC_vs_MS.Trt_B |
| RALGDS | 0.00175 | 0.25416 | 0.345 | 0.26 | 1 | HC_vs_MS.Trt_B |
| CITED2 | 0.00175 | 0.83945 | 0.139 | 0.08 | 1 | HC_vs_MS.Trt_B |
| FAM118A | 0.00177 | 0.64738 | 0.52 | 0.45 | 1 | HC_vs_MS.Trt_B |
| SCAMP3 | 0.00177 | 0.40872 | 0.209 | 0.14 | 1 | HC_vs_MS.Trt_B |
| CD3E | 0.00178 | 0.307 | 0.683 | 0.62 | 1 | HC_vs_MS.Trt_B |
| FHOD1 | 0.00178 | 0.58013 | 0.135 | 0.08 | 1 | HC_vs_MS.Trt_B |
| RHPN2 | 0.00178 | 0.35099 | 0.548 | 0.48 | 1 | HC_vs_MS.Trt_B |

| | | | | | | |
|---|---|---|---|---|---|---|
| ALDH1B1 | 0.00179 | 0.30726 | 0.591 | 0.5 | 1 | HC_vs_MS.Trt_B |
| RP11-80H18 | 0.00179 | 0.72194 | 0.267 | 0.19 | 1 | HC_vs_MS.Trt_B |
| EPS8 | 0.0018 | 0.30916 | 0.391 | 0.3 | 1 | HC_vs_MS.Trt_B |
| IFI44L | 0.00183 | 1.36453 | 0.2 | 0.13 | 1 | HC_vs_MS.Trt_B |
| SLX4 | 0.00184 | -0.4715 | 0.443 | 0.36 | 1 | HC_vs_MS.Trt_B |
| TEX35 | 0.00185 | 0.26485 | 0.312 | 0.24 | 1 | HC_vs_MS.Trt_B |
| GMPPA | 0.00188 | -0.3477 | 0.236 | 0.16 | 1 | HC_vs_MS.Trt_B |
| SMIM10L1 | 0.00192 | 0.76822 | 0.165 | 0.1 | 1 | HC_vs_MS.Trt_B |
| MIS18BP1 | 0.00194 | -0.4179 | 0.586 | 0.48 | 1 | HC_vs_MS.Trt_B |
| SLC23A3 | 0.00194 | 0.34227 | 0.41 | 0.31 | 1 | HC_vs_MS.Trt_B |
| MYO9A | 0.00194 | 0.58953 | 0.323 | 0.24 | 1 | HC_vs_MS.Trt_B |
| CENPC | 0.00194 | -0.3342 | 0.567 | 0.47 | 1 | HC_vs_MS.Trt_B |
| B4GALT5 | 0.00195 | 0.98765 | 0.251 | 0.18 | 1 | HC_vs_MS.Trt_B |
| ZNHIT6 | 0.00195 | 0.92236 | 0.393 | 0.29 | 1 | HC_vs_MS.Trt_B |
| CNTROB | 0.00195 | 0.78784 | 0.177 | 0.11 | 1 | HC_vs_MS.Trt_B |
| DPYSL5 | 0.00196 | 0.30664 | 0.591 | 0.53 | 1 | HC_vs_MS.Trt_B |
| OBFC1 | 0.00197 | 0.28845 | 0.255 | 0.18 | 1 | HC_vs_MS.Trt_B |
| NSD1 | 0.00197 | 0.28676 | 0.645 | 0.55 | 1 | HC_vs_MS.Trt_B |
| HPSE | 0.00198 | -0.5112 | 0.617 | 0.53 | 1 | HC_vs_MS.Trt_B |
| PAK3 | 0.00198 | 0.29144 | 0.384 | 0.3 | 1 | HC_vs_MS.Trt_B |
| KAT8 | 0.00199 | 0.59823 | 0.145 | 0.09 | 1 | HC_vs_MS.Trt_B |
| PBX2 | 0.00199 | 0.77981 | 0.12 | 0.07 | 1 | HC_vs_MS.Trt_B |
| PCTP | 0.00199 | -0.7087 | 0.391 | 0.31 | 1 | HC_vs_MS.Trt_B |
| RPS19BP1 | 0.002 | -0.2698 | 0.433 | 0.34 | 1 | HC_vs_MS.Trt_B |
| ERVK3-1 | 0.002 | 0.41059 | 0.274 | 0.2 | 1 | HC_vs_MS.Trt_B |
| ZNF280D | 0.002 | 0.85613 | 0.214 | 0.15 | 1 | HC_vs_MS.Trt_B |
| PWWP2A | 0.002 | 0.30505 | 0.165 | 0.1 | 1 | HC_vs_MS.Trt_B |
| SFN | 0.00201 | 0.28979 | 0.271 | 0.2 | 1 | HC_vs_MS.Trt_B |
| LBHD1 | 0.00201 | 1.0368 | 0.477 | 0.38 | 1 | HC_vs_MS.Trt_B |
| SFXN3 | 0.00201 | 0.7302 | 0.323 | 0.24 | 1 | HC_vs_MS.Trt_B |
| SLU7 | 0.00201 | -0.4932 | 0.368 | 0.27 | 1 | HC_vs_MS.Trt_B |
| MAGIX | 0.00202 | 0.67869 | 0.465 | 0.39 | 1 | HC_vs_MS.Trt_B |
| ELOVL7 | 0.00203 | 0.34896 | 0.319 | 0.24 | 1 | HC_vs_MS.Trt_B |
| CDNF | 0.00205 | -1.38 | 0.184 | 0.12 | 1 | HC_vs_MS.Trt_B |
| TRIM68 | 0.00205 | -0.5048 | 0.509 | 0.41 | 1 | HC_vs_MS.Trt_B |
| GPRC5A | 0.00205 | 0.31556 | 0.613 | 0.51 | 1 | HC_vs_MS.Trt_B |
| ZNF682 | 0.00205 | 0.51004 | 0.37 | 0.28 | 1 | HC_vs_MS.Trt_B |
| C2orf69 | 0.00205 | 0.29673 | 0.239 | 0.16 | 1 | HC_vs_MS.Trt_B |
| LRCH4 | 0.00206 | 0.63727 | 0.5 | 0.42 | 1 | HC_vs_MS.Trt_B |
| ATAD1 | 0.00206 | 0.25883 | 0.143 | 0.08 | 1 | HC_vs_MS.Trt_B |

| | | | | | | |
|---|---|---|---|---|---|---|
| ACVR2B | 0.00208 | 0.97107 | 0.383 | 0.3 | 1 | HC_vs_MS.Trt_B |
| TAF1A | 0.00209 | 0.87368 | 0.394 | 0.31 | 1 | HC_vs_MS.Trt_B |
| GCAT | 0.00209 | -0.6142 | 0.223 | 0.15 | 1 | HC_vs_MS.Trt_B |
| ADAP2 | 0.0021 | -0.3944 | 0.172 | 0.11 | 1 | HC_vs_MS.Trt_B |
| AMDHD1 | 0.00211 | 0.97733 | 0.196 | 0.13 | 1 | HC_vs_MS.Trt_B |
| MORC3 | 0.00211 | -0.2649 | 0.486 | 0.4 | 1 | HC_vs_MS.Trt_B |
| BUB3 | 0.00213 | -0.344 | 0.616 | 0.5 | 1 | HC_vs_MS.Trt_B |
| PBXIP1 | 0.00215 | 0.28479 | 0.539 | 0.47 | 1 | HC_vs_MS.Trt_B |
| RPUSD3 | 0.00215 | -0.4062 | 0.777 | 0.7 | 1 | HC_vs_MS.Trt_B |
| KALRN | 0.00215 | 0.85797 | 0.3 | 0.22 | 1 | HC_vs_MS.Trt_B |
| FBL | 0.00217 | 0.33696 | 0.426 | 0.34 | 1 | HC_vs_MS.Trt_B |
| SERPINH1 | 0.00218 | 1.23505 | 0.304 | 0.22 | 1 | HC_vs_MS.Trt_B |
| ZNF439 | 0.00218 | -0.8879 | 0.616 | 0.55 | 1 | HC_vs_MS.Trt_B |
| ZNF644 | 0.00218 | 0.55202 | 0.194 | 0.13 | 1 | HC_vs_MS.Trt_B |
| GPRIN3 | 0.00219 | 0.31016 | 0.428 | 0.35 | 1 | HC_vs_MS.Trt_B |
| NOC2L | 0.0022 | -0.3481 | 0.716 | 0.63 | 1 | HC_vs_MS.Trt_B |
| 8-Mar | 0.0022 | -0.4592 | 0.387 | 0.29 | 1 | HC_vs_MS.Trt_B |
| CCNYL1 | 0.0022 | 0.55276 | 0.359 | 0.27 | 1 | HC_vs_MS.Trt_B |
| IL6ST | 0.0022 | 0.45563 | 0.367 | 0.29 | 1 | HC_vs_MS.Trt_B |
| RSAD2 | 0.00221 | 0.88421 | 0.326 | 0.25 | 1 | HC_vs_MS.Trt_B |
| SLIT1 | 0.00221 | 0.32874 | 0.236 | 0.17 | 1 | HC_vs_MS.Trt_B |
| GSTM3 | 0.00222 | 0.62746 | 0.506 | 0.39 | 1 | HC_vs_MS.Trt_B |
| DBR1 | 0.00222 | -0.2793 | 0.749 | 0.65 | 1 | HC_vs_MS.Trt_B |
| XPO1 | 0.00223 | -0.2815 | 0.916 | 0.87 | 1 | HC_vs_MS.Trt_B |
| HIST1H2BG | 0.00223 | 0.87354 | 0.472 | 0.4 | 1 | HC_vs_MS.Trt_B |
| LRPAP1 | 0.00225 | -0.4312 | 0.868 | 0.77 | 1 | HC_vs_MS.Trt_B |
| ITPK1 | 0.00225 | -0.255 | 0.555 | 0.46 | 1 | HC_vs_MS.Trt_B |
| C5 | 0.00226 | 0.68754 | 0.4 | 0.32 | 1 | HC_vs_MS.Trt_B |
| URM1 | 0.00226 | 0.83569 | 0.138 | 0.08 | 1 | HC_vs_MS.Trt_B |
| VASH2 | 0.00227 | 0.27735 | 0.745 | 0.64 | 1 | HC_vs_MS.Trt_B |
| ZNF280C | 0.00227 | 0.64889 | 0.252 | 0.18 | 1 | HC_vs_MS.Trt_B |
| KCTD20 | 0.00227 | 0.27234 | 0.616 | 0.57 | 1 | HC_vs_MS.Trt_B |
| COL6A3 | 0.00228 | 0.76433 | 0.452 | 0.36 | 1 | HC_vs_MS.Trt_B |
| AC004381.6 | 0.00228 | 0.45687 | 0.196 | 0.12 | 1 | HC_vs_MS.Trt_B |
| ZNF485 | 0.00229 | 0.28878 | 0.186 | 0.12 | 1 | HC_vs_MS.Trt_B |
| ZKSCAN7 | 0.00229 | 0.2776 | 0.599 | 0.52 | 1 | HC_vs_MS.Trt_B |
| RAB37 | 0.0023 | 0.30473 | 0.38 | 0.3 | 1 | HC_vs_MS.Trt_B |
| C3orf70 | 0.0023 | 0.2549 | 0.699 | 0.66 | 1 | HC_vs_MS.Trt_B |
| SSB | 0.0023 | -0.3918 | 0.749 | 0.64 | 1 | HC_vs_MS.Trt_B |
| ZNF75A | 0.00231 | -0.5091 | 0.72 | 0.61 | 1 | HC_vs_MS.Trt_B |

| | | | | | | |
|---|---|---|---|---|---|---|
| PCED1A | 0.00231 | 0.37781 | 0.136 | 0.08 | 1 | HC_vs_MS.Trt_B |
| ANTXRL | 0.00232 | 0.32957 | 0.204 | 0.13 | 1 | HC_vs_MS.Trt_B |
| C11orf24 | 0.00233 | 1.13074 | 0.246 | 0.17 | 1 | HC_vs_MS.Trt_B |
| YIF1B | 0.00235 | -0.4494 | 0.583 | 0.5 | 1 | HC_vs_MS.Trt_B |
| GPR35 | 0.00235 | 0.566 | 0.28 | 0.2 | 1 | HC_vs_MS.Trt_B |
| VNN3 | 0.00235 | -0.2818 | 0.358 | 0.26 | 1 | HC_vs_MS.Trt_B |
| ZFAND4 | 0.00235 | 0.81749 | 0.396 | 0.3 | 1 | HC_vs_MS.Trt_B |
| GCLC | 0.00237 | 0.42241 | 0.497 | 0.41 | 1 | HC_vs_MS.Trt_B |
| CHSY1 | 0.00238 | 0.70207 | 0.257 | 0.18 | 1 | HC_vs_MS.Trt_B |
| ZNF763 | 0.00239 | 1.1379 | 0.172 | 0.11 | 1 | HC_vs_MS.Trt_B |
| SLC16A12 | 0.0024 | 0.28067 | 0.397 | 0.32 | 1 | HC_vs_MS.Trt_B |
| GPC2 | 0.00241 | 0.42568 | 0.397 | 0.33 | 1 | HC_vs_MS.Trt_B |
| SKA2 | 0.00242 | -0.4002 | 0.551 | 0.44 | 1 | HC_vs_MS.Trt_B |
| UTP23 | 0.00242 | 0.29414 | 0.499 | 0.42 | 1 | HC_vs_MS.Trt_B |
| VAC14 | 0.00242 | 0.5857 | 0.104 | 0.05 | 1 | HC_vs_MS.Trt_B |
| PYCRL | 0.00243 | -0.8431 | 0.406 | 0.31 | 1 | HC_vs_MS.Trt_B |
| D2HGDH | 0.00243 | 0.66791 | 0.439 | 0.35 | 1 | HC_vs_MS.Trt_B |
| LA16c-431H | 0.00245 | 0.35351 | 0.493 | 0.42 | 1 | HC_vs_MS.Trt_B |
| ACY1 | 0.00246 | -0.5041 | 0.58 | 0.47 | 1 | HC_vs_MS.Trt_B |
| RNF215 | 0.00247 | 0.4184 | 0.432 | 0.34 | 1 | HC_vs_MS.Trt_B |
| NPTN | 0.00249 | 0.432 | 0.248 | 0.17 | 1 | HC_vs_MS.Trt_B |
| ABHD2 | 0.0025 | -0.3182 | 0.622 | 0.52 | 1 | HC_vs_MS.Trt_B |
| ENTPD3 | 0.0025 | 0.27744 | 0.33 | 0.24 | 1 | HC_vs_MS.Trt_B |
| GPR137 | 0.00251 | 0.63888 | 0.207 | 0.14 | 1 | HC_vs_MS.Trt_B |
| RAD23B | 0.00251 | -0.3067 | 0.439 | 0.34 | 1 | HC_vs_MS.Trt_B |
| CHST14 | 0.00251 | 1.17577 | 0.225 | 0.16 | 1 | HC_vs_MS.Trt_B |
| TLK1 | 0.00253 | -0.2632 | 0.729 | 0.63 | 1 | HC_vs_MS.Trt_B |
| KLHL32 | 0.00255 | 0.78087 | 0.101 | 0.05 | 1 | HC_vs_MS.Trt_B |
| CA12 | 0.00255 | 0.67369 | 0.265 | 0.19 | 1 | HC_vs_MS.Trt_B |
| STARD4 | 0.00256 | 1.24726 | 0.143 | 0.09 | 1 | HC_vs_MS.Trt_B |
| CARF | 0.00257 | -0.4552 | 0.486 | 0.4 | 1 | HC_vs_MS.Trt_B |
| COX15 | 0.00257 | -0.4747 | 0.548 | 0.46 | 1 | HC_vs_MS.Trt_B |
| PROSC | 0.00259 | 0.29781 | 0.414 | 0.33 | 1 | HC_vs_MS.Trt_B |
| SUN1 | 0.0026 | 0.27974 | 0.593 | 0.52 | 1 | HC_vs_MS.Trt_B |
| TGFBR2 | 0.0026 | 1.09581 | 0.162 | 0.1 | 1 | HC_vs_MS.Trt_B |
| SARNP | 0.0026 | -0.2622 | 0.559 | 0.44 | 1 | HC_vs_MS.Trt_B |
| PTPN4 | 0.00261 | -0.3441 | 0.588 | 0.47 | 1 | HC_vs_MS.Trt_B |
| PPRC1 | 0.00262 | 0.5381 | 0.113 | 0.06 | 1 | HC_vs_MS.Trt_B |
| C11orf21 | 0.00262 | 0.71914 | 0.138 | 0.08 | 1 | HC_vs_MS.Trt_B |
| ELF3 | 0.00262 | 0.38867 | 0.545 | 0.44 | 1 | HC_vs_MS.Trt_B |

| | | | | | | |
|---|---|---|---|---|---|---|
| BCL9L | 0.00263 | 0.42471 | 0.17 | 0.11 | 1 | HC_vs_MS.Trt_B |
| TTC34 | 0.00263 | 0.45411 | 0.193 | 0.12 | 1 | HC_vs_MS.Trt_B |
| GRAMD1C | 0.00263 | 0.45943 | 0.184 | 0.12 | 1 | HC_vs_MS.Trt_B |
| SKAP2 | 0.00263 | 0.98704 | 0.184 | 0.12 | 1 | HC_vs_MS.Trt_B |
| STMN3 | 0.00264 | 0.33115 | 0.225 | 0.16 | 1 | HC_vs_MS.Trt_B |
| FAM101B | 0.00265 | -0.3754 | 0.246 | 0.17 | 1 | HC_vs_MS.Trt_B |
| C1orf131 | 0.00265 | -0.2882 | 0.575 | 0.49 | 1 | HC_vs_MS.Trt_B |
| MRPS26 | 0.00265 | 0.36465 | 0.122 | 0.07 | 1 | HC_vs_MS.Trt_B |
| CPSF7 | 0.00265 | 0.57665 | 0.212 | 0.15 | 1 | HC_vs_MS.Trt_B |
| MCM9 | 0.00265 | -0.2653 | 0.314 | 0.23 | 1 | HC_vs_MS.Trt_B |
| STAT5A | 0.00267 | 0.36359 | 0.17 | 0.11 | 1 | HC_vs_MS.Trt_B |
| CMTR2 | 0.00267 | 0.50392 | 0.257 | 0.18 | 1 | HC_vs_MS.Trt_B |
| AQP3 | 0.00267 | 0.38555 | 0.262 | 0.19 | 1 | HC_vs_MS.Trt_B |
| PSMA4 | 0.00269 | -0.319 | 0.349 | 0.26 | 1 | HC_vs_MS.Trt_B |
| BRK1 | 0.00269 | 0.26146 | 0.242 | 0.17 | 1 | HC_vs_MS.Trt_B |
| COQ6 | 0.0027 | 0.49476 | 0.761 | 0.67 | 1 | HC_vs_MS.Trt_B |
| GDPD5 | 0.0027 | 1.28278 | 0.267 | 0.2 | 1 | HC_vs_MS.Trt_B |
| SLC39A14 | 0.00271 | 0.58729 | 0.396 | 0.31 | 1 | HC_vs_MS.Trt_B |
| C19orf48 | 0.00273 | 1.00472 | 0.364 | 0.28 | 1 | HC_vs_MS.Trt_B |
| ZNF526 | 0.00273 | -0.3662 | 0.459 | 0.37 | 1 | HC_vs_MS.Trt_B |
| SEPN1 | 0.00275 | 0.28605 | 0.351 | 0.28 | 1 | HC_vs_MS.Trt_B |
| SULF1 | 0.00276 | 0.42961 | 0.174 | 0.11 | 1 | HC_vs_MS.Trt_B |
| RASGEF1B | 0.00278 | 0.2889 | 0.672 | 0.61 | 1 | HC_vs_MS.Trt_B |
| CDC40 | 0.00278 | 0.35721 | 0.532 | 0.45 | 1 | HC_vs_MS.Trt_B |
| STAT1 | 0.0028 | 0.40026 | 0.42 | 0.35 | 1 | HC_vs_MS.Trt_B |
| ACER1 | 0.00281 | 0.54641 | 0.438 | 0.33 | 1 | HC_vs_MS.Trt_B |
| PPP3R1 | 0.00281 | 0.75449 | 0.158 | 0.1 | 1 | HC_vs_MS.Trt_B |
| USP28 | 0.00282 | 0.55454 | 0.352 | 0.27 | 1 | HC_vs_MS.Trt_B |
| RABL2A | 0.00283 | 0.29745 | 0.488 | 0.42 | 1 | HC_vs_MS.Trt_B |
| ORC3 | 0.00284 | 0.50509 | 0.158 | 0.1 | 1 | HC_vs_MS.Trt_B |
| ERC1 | 0.00284 | 0.34149 | 0.861 | 0.82 | 1 | HC_vs_MS.Trt_B |
| ZNF699 | 0.00284 | -0.2774 | 0.251 | 0.18 | 1 | HC_vs_MS.Trt_B |
| LAPTM4A | 0.00285 | 0.39045 | 0.158 | 0.1 | 1 | HC_vs_MS.Trt_B |
| FARSB | 0.00285 | -0.4778 | 0.594 | 0.52 | 1 | HC_vs_MS.Trt_B |
| UBXN8 | 0.00286 | 1.3199 | 0.177 | 0.12 | 1 | HC_vs_MS.Trt_B |
| TMEM183A | 0.00287 | 0.53557 | 0.183 | 0.12 | 1 | HC_vs_MS.Trt_B |
| ZNF135 | 0.00288 | 0.42751 | 0.432 | 0.34 | 1 | HC_vs_MS.Trt_B |
| MTMR3 | 0.00289 | 0.39452 | 0.249 | 0.18 | 1 | HC_vs_MS.Trt_B |
| LARS2 | 0.0029 | 0.61235 | 0.478 | 0.4 | 1 | HC_vs_MS.Trt_B |
| COMMD1 | 0.00291 | -0.2596 | 0.536 | 0.42 | 1 | HC_vs_MS.Trt_B |

| | | | | | | |
|---|---|---|---|---|---|---|
| PROX2 | 0.00292 | 0.88347 | 0.123 | 0.07 | 1 | HC_vs_MS.Trt_B |
| AARS | 0.00292 | 0.38665 | 0.426 | 0.34 | 1 | HC_vs_MS.Trt_B |
| ZNF786 | 0.00294 | -0.4611 | 0.406 | 0.32 | 1 | HC_vs_MS.Trt_B |
| PARK2 | 0.00294 | 0.41348 | 0.604 | 0.53 | 1 | HC_vs_MS.Trt_B |
| FAM227B | 0.00296 | 0.53058 | 0.387 | 0.31 | 1 | HC_vs_MS.Trt_B |
| EXD1 | 0.00298 | 0.29714 | 0.472 | 0.4 | 1 | HC_vs_MS.Trt_B |
| PLEKHA7 | 0.00298 | -0.9154 | 0.184 | 0.12 | 1 | HC_vs_MS.Trt_B |
| SFMBT1 | 0.00302 | -0.4606 | 0.549 | 0.48 | 1 | HC_vs_MS.Trt_B |
| PDZD2 | 0.00303 | 0.4728 | 0.668 | 0.62 | 1 | HC_vs_MS.Trt_B |
| OGFOD1 | 0.00303 | -0.7335 | 0.596 | 0.53 | 1 | HC_vs_MS.Trt_B |
| PIEZO1 | 0.00303 | 0.33226 | 0.31 | 0.24 | 1 | HC_vs_MS.Trt_B |
| FCRL2 | 0.00306 | 0.2873 | 0.619 | 0.52 | 1 | HC_vs_MS.Trt_B |
| TADA3 | 0.00308 | 0.46967 | 0.52 | 0.44 | 1 | HC_vs_MS.Trt_B |
| MELK | 0.00309 | 0.27719 | 0.504 | 0.42 | 1 | HC_vs_MS.Trt_B |
| NPIPB8 | 0.00314 | 0.27525 | 0.123 | 0.07 | 1 | HC_vs_MS.Trt_B |
| PHRF1 | 0.00317 | 0.73962 | 0.142 | 0.09 | 1 | HC_vs_MS.Trt_B |
| RP11-619A1 | 0.00321 | 0.2539 | 0.545 | 0.45 | 1 | HC_vs_MS.Trt_B |
| LHFPL2 | 0.00324 | 0.92949 | 0.29 | 0.21 | 1 | HC_vs_MS.Trt_B |
| GSN | 0.00324 | 0.52179 | 0.554 | 0.45 | 1 | HC_vs_MS.Trt_B |
| TXLNA | 0.00325 | 0.25672 | 0.277 | 0.2 | 1 | HC_vs_MS.Trt_B |
| FGFR4 | 0.00329 | 0.29441 | 0.393 | 0.31 | 1 | HC_vs_MS.Trt_B |
| C14orf119 | 0.0033 | -0.2819 | 0.533 | 0.43 | 1 | HC_vs_MS.Trt_B |
| NPL | 0.00331 | -0.3868 | 0.223 | 0.15 | 1 | HC_vs_MS.Trt_B |
| MRPL10 | 0.00332 | -0.2517 | 0.4 | 0.3 | 1 | HC_vs_MS.Trt_B |
| PNPLA8 | 0.00332 | 0.4014 | 0.333 | 0.26 | 1 | HC_vs_MS.Trt_B |
| ICK | 0.00333 | -0.7141 | 0.207 | 0.14 | 1 | HC_vs_MS.Trt_B |
| MTMR14 | 0.00334 | -0.3075 | 0.571 | 0.45 | 1 | HC_vs_MS.Trt_B |
| NR4A2 | 0.00335 | 0.32072 | 0.146 | 0.09 | 1 | HC_vs_MS.Trt_B |
| PPAT | 0.00336 | 0.25203 | 0.271 | 0.2 | 1 | HC_vs_MS.Trt_B |
| SLC2A6 | 0.00337 | 0.33121 | 0.262 | 0.19 | 1 | HC_vs_MS.Trt_B |
| TANC1 | 0.00339 | 0.52623 | 0.339 | 0.26 | 1 | HC_vs_MS.Trt_B |
| RNF144A | 0.00342 | 0.40443 | 0.196 | 0.13 | 1 | HC_vs_MS.Trt_B |
| ZSCAN22 | 0.00342 | 0.31297 | 0.399 | 0.33 | 1 | HC_vs_MS.Trt_B |
| TAS2R4 | 0.00345 | 0.43844 | 0.341 | 0.26 | 1 | HC_vs_MS.Trt_B |
| DOCK6 | 0.00346 | -0.299 | 0.546 | 0.45 | 1 | HC_vs_MS.Trt_B |
| FGF9 | 0.00346 | 0.60467 | 0.13 | 0.08 | 1 | HC_vs_MS.Trt_B |
| MFSD4B | 0.00347 | 0.42685 | 0.626 | 0.56 | 1 | HC_vs_MS.Trt_B |
| POR | 0.00347 | 0.40191 | 0.301 | 0.23 | 1 | HC_vs_MS.Trt_B |
| FADS2 | 0.00348 | 0.54014 | 0.486 | 0.4 | 1 | HC_vs_MS.Trt_B |
| BRWD1 | 0.00349 | -0.467 | 0.813 | 0.68 | 1 | HC_vs_MS.Trt_B |

| | | | | | | |
|---|---|---|---|---|---|---|
| PKN1 | 0.00349 | 0.26251 | 0.803 | 0.69 | 1 | HC_vs_MS.Trt_B |
| CACUL1 | 0.00349 | -0.4925 | 0.664 | 0.56 | 1 | HC_vs_MS.Trt_B |
| FBXO34 | 0.0035 | 0.71484 | 0.113 | 0.06 | 1 | HC_vs_MS.Trt_B |
| LDHC | 0.00351 | 0.35699 | 0.184 | 0.12 | 1 | HC_vs_MS.Trt_B |
| STXBP6 | 0.00351 | 0.2873 | 0.161 | 0.1 | 1 | HC_vs_MS.Trt_B |
| AGO1 | 0.00353 | 0.5173 | 0.439 | 0.35 | 1 | HC_vs_MS.Trt_B |
| CSNK2A1 | 0.00355 | 0.26555 | 0.654 | 0.56 | 1 | HC_vs_MS.Trt_B |
| RABGGTB | 0.00356 | 0.34423 | 0.235 | 0.17 | 1 | HC_vs_MS.Trt_B |
| TMEM222 | 0.00356 | 0.82698 | 0.132 | 0.08 | 1 | HC_vs_MS.Trt_B |
| MAP3K4 | 0.00357 | 1.23007 | 0.183 | 0.12 | 1 | HC_vs_MS.Trt_B |
| TMEM132B | 0.00361 | 0.60187 | 0.267 | 0.19 | 1 | HC_vs_MS.Trt_B |
| ANKRD52 | 0.00362 | 0.43079 | 0.174 | 0.11 | 1 | HC_vs_MS.Trt_B |
| PFKFB3 | 0.00363 | 0.35724 | 0.23 | 0.16 | 1 | HC_vs_MS.Trt_B |
| MICALL1 | 0.00365 | 0.29522 | 0.462 | 0.37 | 1 | HC_vs_MS.Trt_B |
| HSDL2 | 0.00366 | 0.71522 | 0.142 | 0.09 | 1 | HC_vs_MS.Trt_B |
| MRPL2 | 0.00368 | -0.4901 | 0.262 | 0.18 | 1 | HC_vs_MS.Trt_B |
| GTSF1 | 0.00369 | 0.41941 | 0.219 | 0.15 | 1 | HC_vs_MS.Trt_B |
| MAB21L3 | 0.00369 | 0.35742 | 0.362 | 0.28 | 1 | HC_vs_MS.Trt_B |
| ZSCAN18 | 0.00369 | 0.29797 | 0.662 | 0.58 | 1 | HC_vs_MS.Trt_B |
| WWOX | 0.0037 | 0.60088 | 0.236 | 0.17 | 1 | HC_vs_MS.Trt_B |
| CHIC2 | 0.00374 | 0.54393 | 0.171 | 0.11 | 1 | HC_vs_MS.Trt_B |
| ZNF391 | 0.00376 | 0.34591 | 0.184 | 0.12 | 1 | HC_vs_MS.Trt_B |
| COPS6 | 0.00377 | -0.5858 | 0.671 | 0.54 | 1 | HC_vs_MS.Trt_B |
| GIN1 | 0.00378 | -0.8285 | 0.326 | 0.24 | 1 | HC_vs_MS.Trt_B |
| CWC25 | 0.00378 | 0.69933 | 0.333 | 0.26 | 1 | HC_vs_MS.Trt_B |
| ACVR1 | 0.00379 | -0.5653 | 0.193 | 0.13 | 1 | HC_vs_MS.Trt_B |
| KRT15 | 0.00384 | 0.36427 | 0.417 | 0.34 | 1 | HC_vs_MS.Trt_B |
| LEO1 | 0.00387 | 0.83897 | 0.548 | 0.49 | 1 | HC_vs_MS.Trt_B |
| CCDC163 | 0.0039 | 0.61342 | 0.538 | 0.45 | 1 | HC_vs_MS.Trt_B |
| C11orf57 | 0.00393 | -0.4416 | 0.768 | 0.68 | 1 | HC_vs_MS.Trt_B |
| JMJD7 | 0.00393 | 0.49094 | 0.145 | 0.09 | 1 | HC_vs_MS.Trt_B |
| SLC22A25 | 0.00393 | 0.30103 | 0.313 | 0.24 | 1 | HC_vs_MS.Trt_B |
| MTA1 | 0.00394 | 0.4635 | 0.403 | 0.32 | 1 | HC_vs_MS.Trt_B |
| GPHN | 0.00399 | 0.29559 | 0.387 | 0.31 | 1 | HC_vs_MS.Trt_B |
| GGNBP2 | 0.00403 | -0.35 | 0.681 | 0.6 | 1 | HC_vs_MS.Trt_B |
| PATL2 | 0.00403 | 0.46345 | 0.358 | 0.28 | 1 | HC_vs_MS.Trt_B |
| SLC39A4 | 0.00404 | -0.3332 | 0.386 | 0.3 | 1 | HC_vs_MS.Trt_B |
| MSS51 | 0.00404 | 0.2589 | 0.536 | 0.45 | 1 | HC_vs_MS.Trt_B |
| DPH5 | 0.00405 | -0.3617 | 0.343 | 0.26 | 1 | HC_vs_MS.Trt_B |
| COLEC11 | 0.00406 | 0.94947 | 0.367 | 0.28 | 1 | HC_vs_MS.Trt_B |

| | | | | | | |
|---|---|---|---|---|---|---|
| SFMBT2 | 0.00409 | 0.84699 | 0.293 | 0.22 | 1 | HC_vs_MS.Trt_B |
| MEF2A | 0.0041 | 0.90771 | 0.443 | 0.36 | 1 | HC_vs_MS.Trt_B |
| TACR2 | 0.0041 | -0.3579 | 0.129 | 0.08 | 1 | HC_vs_MS.Trt_B |
| TMEM260 | 0.00411 | 0.41767 | 0.4 | 0.33 | 1 | HC_vs_MS.Trt_B |
| PPP4R1 | 0.00412 | 0.29553 | 0.238 | 0.17 | 1 | HC_vs_MS.Trt_B |
| TMEM64 | 0.00413 | 0.37975 | 0.358 | 0.28 | 1 | HC_vs_MS.Trt_B |
| C2orf54 | 0.00413 | 0.3029 | 0.114 | 0.07 | 1 | HC_vs_MS.Trt_B |
| ASUN | 0.00413 | 0.69784 | 0.109 | 0.06 | 1 | HC_vs_MS.Trt_B |
| STXBP4 | 0.00413 | 0.61559 | 0.483 | 0.38 | 1 | HC_vs_MS.Trt_B |
| MAN2B1 | 0.00414 | 0.37744 | 0.57 | 0.48 | 1 | HC_vs_MS.Trt_B |
| SLC12A7 | 0.00418 | 0.76804 | 0.138 | 0.08 | 1 | HC_vs_MS.Trt_B |
| WDR13 | 0.00418 | -0.4298 | 0.814 | 0.73 | 1 | HC_vs_MS.Trt_B |
| IL4R | 0.00421 | 0.36308 | 0.425 | 0.34 | 1 | HC_vs_MS.Trt_B |
| NBPF12 | 0.00421 | 0.88939 | 0.312 | 0.24 | 1 | HC_vs_MS.Trt_B |
| ETF1 | 0.00421 | -0.4237 | 0.283 | 0.2 | 1 | HC_vs_MS.Trt_B |
| ABHD14A | 0.00421 | 0.33461 | 0.457 | 0.4 | 1 | HC_vs_MS.Trt_B |
| TMEM185A | 0.00422 | -0.4835 | 0.483 | 0.39 | 1 | HC_vs_MS.Trt_B |
| ARFGEF2 | 0.00427 | 0.35011 | 0.458 | 0.38 | 1 | HC_vs_MS.Trt_B |
| ACTR6 | 0.00427 | -0.574 | 0.513 | 0.41 | 1 | HC_vs_MS.Trt_B |
| SHF | 0.00428 | 0.32094 | 0.336 | 0.27 | 1 | HC_vs_MS.Trt_B |
| SLC30A8 | 0.00429 | 0.39751 | 0.401 | 0.32 | 1 | HC_vs_MS.Trt_B |
| G6PC2 | 0.0043 | 0.29369 | 0.272 | 0.2 | 1 | HC_vs_MS.Trt_B |
| ATG2B | 0.00431 | 0.69121 | 0.296 | 0.23 | 1 | HC_vs_MS.Trt_B |
| WDR82 | 0.00433 | 0.42357 | 0.384 | 0.31 | 1 | HC_vs_MS.Trt_B |
| POMZP3 | 0.00434 | -0.3719 | 0.435 | 0.34 | 1 | HC_vs_MS.Trt_B |
| CLTA | 0.00434 | 0.29392 | 0.575 | 0.5 | 1 | HC_vs_MS.Trt_B |
| SCAPER | 0.00435 | 0.49846 | 0.239 | 0.17 | 1 | HC_vs_MS.Trt_B |
| SPHK2 | 0.00435 | 0.26602 | 0.293 | 0.22 | 1 | HC_vs_MS.Trt_B |
| PHKB | 0.00436 | -0.3762 | 0.745 | 0.62 | 1 | HC_vs_MS.Trt_B |
| DPH3 | 0.00436 | 1.00564 | 0.238 | 0.17 | 1 | HC_vs_MS.Trt_B |
| VARS2 | 0.00438 | 1.43033 | 0.471 | 0.39 | 1 | HC_vs_MS.Trt_B |
| SRSF8 | 0.00441 | -0.2888 | 0.568 | 0.5 | 1 | HC_vs_MS.Trt_B |
| SKIL | 0.00442 | 0.70492 | 0.277 | 0.2 | 1 | HC_vs_MS.Trt_B |
| APPL2 | 0.00444 | 0.51038 | 0.348 | 0.27 | 1 | HC_vs_MS.Trt_B |
| HTT | 0.00449 | 0.30956 | 0.236 | 0.16 | 1 | HC_vs_MS.Trt_B |
| STK17A | 0.00449 | 0.32357 | 0.274 | 0.2 | 1 | HC_vs_MS.Trt_B |
| C1orf162 | 0.00449 | 0.5774 | 0.11 | 0.06 | 1 | HC_vs_MS.Trt_B |
| ZNF84 | 0.00449 | -0.3774 | 0.284 | 0.21 | 1 | HC_vs_MS.Trt_B |
| TTC33 | 0.00451 | -0.399 | 0.228 | 0.16 | 1 | HC_vs_MS.Trt_B |
| DDX47 | 0.00451 | -0.7227 | 0.507 | 0.41 | 1 | HC_vs_MS.Trt_B |

| | | | | | | |
|---|---|---|---|---|---|---|
| ST3GAL6 | 0.00452 | 0.38396 | 0.31 | 0.23 | 1 | HC_vs_MS.Trt_B |
| NECAP1 | 0.00454 | 0.2634 | 0.436 | 0.35 | 1 | HC_vs_MS.Trt_B |
| ASPSCR1 | 0.00456 | 0.86025 | 0.613 | 0.56 | 1 | HC_vs_MS.Trt_B |
| SET | 0.00456 | -0.5971 | 0.888 | 0.85 | 1 | HC_vs_MS.Trt_B |
| ADGRG1 | 0.00464 | -0.3078 | 0.654 | 0.58 | 1 | HC_vs_MS.Trt_B |
| HEBP2 | 0.00464 | -1.1199 | 0.377 | 0.29 | 1 | HC_vs_MS.Trt_B |
| HMGN4 | 0.00467 | 0.28822 | 0.241 | 0.18 | 1 | HC_vs_MS.Trt_B |
| BRE | 0.00468 | 0.30309 | 0.2 | 0.14 | 1 | HC_vs_MS.Trt_B |
| GULP1 | 0.00468 | 0.32843 | 0.42 | 0.35 | 1 | HC_vs_MS.Trt_B |
| SLC35G2 | 0.00469 | -1.0332 | 0.162 | 0.11 | 1 | HC_vs_MS.Trt_B |
| CTSA | 0.0047 | 0.34445 | 0.659 | 0.56 | 1 | HC_vs_MS.Trt_B |
| IGF2R | 0.00471 | -0.2868 | 0.401 | 0.34 | 1 | HC_vs_MS.Trt_B |
| HAUS5 | 0.00473 | -0.5582 | 0.422 | 0.32 | 1 | HC_vs_MS.Trt_B |
| HLX | 0.00475 | 0.39655 | 0.193 | 0.13 | 1 | HC_vs_MS.Trt_B |
| BTLA | 0.00476 | 1.31592 | 0.154 | 0.1 | 1 | HC_vs_MS.Trt_B |
| ADAM15 | 0.00477 | 0.51139 | 0.361 | 0.29 | 1 | HC_vs_MS.Trt_B |
| TRIM11 | 0.00479 | 0.43095 | 0.507 | 0.43 | 1 | HC_vs_MS.Trt_B |
| STAG2 | 0.00479 | -0.3922 | 0.765 | 0.67 | 1 | HC_vs_MS.Trt_B |
| WIZ | 0.00481 | 0.51126 | 0.287 | 0.22 | 1 | HC_vs_MS.Trt_B |
| RP11-722G7 | 0.00481 | 0.40291 | 0.438 | 0.36 | 1 | HC_vs_MS.Trt_B |
| DHRS11 | 0.00481 | -0.4635 | 0.283 | 0.21 | 1 | HC_vs_MS.Trt_B |
| DNAJB13 | 0.00482 | 0.42263 | 0.364 | 0.28 | 1 | HC_vs_MS.Trt_B |
| FDFT1 | 0.00484 | 0.26595 | 0.296 | 0.22 | 1 | HC_vs_MS.Trt_B |
| WDR36 | 0.00484 | 0.49214 | 0.142 | 0.09 | 1 | HC_vs_MS.Trt_B |
| TANC2 | 0.00484 | 0.59515 | 0.257 | 0.18 | 1 | HC_vs_MS.Trt_B |
| ARRDC1 | 0.00485 | 0.52535 | 0.235 | 0.17 | 1 | HC_vs_MS.Trt_B |
| ESPL1 | 0.00487 | 0.25897 | 0.68 | 0.58 | 1 | HC_vs_MS.Trt_B |
| FASTKD1 | 0.00488 | 0.38799 | 0.23 | 0.16 | 1 | HC_vs_MS.Trt_B |
| AR | 0.00493 | 0.46973 | 0.222 | 0.16 | 1 | HC_vs_MS.Trt_B |
| SDR42E1 | 0.00493 | 0.81509 | 0.386 | 0.31 | 1 | HC_vs_MS.Trt_B |
| COX6C | 0.00499 | -0.2833 | 0.467 | 0.37 | 1 | HC_vs_MS.Trt_B |
| RNF135 | 0.00502 | 0.54143 | 0.248 | 0.17 | 1 | HC_vs_MS.Trt_B |
| AP003419.1 | 0.00502 | 0.44797 | 0.155 | 0.1 | 1 | HC_vs_MS.Trt_B |
| CACNB2 | 0.00503 | 0.3411 | 0.449 | 0.37 | 1 | HC_vs_MS.Trt_B |
| ABCF2 | 0.00505 | 0.58309 | 0.226 | 0.16 | 1 | HC_vs_MS.Trt_B |
| ZNF441 | 0.00506 | 0.66693 | 0.174 | 0.11 | 1 | HC_vs_MS.Trt_B |
| THRA | 0.00507 | 0.47711 | 0.142 | 0.09 | 1 | HC_vs_MS.Trt_B |
| CHI3L2 | 0.00508 | 0.52207 | 0.201 | 0.14 | 1 | HC_vs_MS.Trt_B |
| KIAA0430 | 0.00509 | 0.27862 | 0.28 | 0.21 | 1 | HC_vs_MS.Trt_B |
| ZNF701 | 0.00509 | 0.32646 | 0.436 | 0.38 | 1 | HC_vs_MS.Trt_B |

| | | | | | | |
|---|---|---|---|---|---|---|
| NBPF20 | 0.00512 | 0.35368 | 0.317 | 0.24 | 1 | HC_vs_MS.Trt_B |
| C10orf54 | 0.00512 | 0.28794 | 0.355 | 0.28 | 1 | HC_vs_MS.Trt_B |
| CECR5 | 0.00514 | 0.41757 | 0.175 | 0.12 | 1 | HC_vs_MS.Trt_B |
| LOXL2 | 0.00515 | 0.31167 | 0.71 | 0.63 | 1 | HC_vs_MS.Trt_B |
| CYHR1 | 0.00518 | 0.42581 | 0.268 | 0.21 | 1 | HC_vs_MS.Trt_B |
| TRPC4AP | 0.00521 | 0.3564 | 0.264 | 0.19 | 1 | HC_vs_MS.Trt_B |
| WDR5B | 0.00521 | 0.52704 | 0.183 | 0.12 | 1 | HC_vs_MS.Trt_B |
| UNC45B | 0.00522 | 0.81314 | 0.12 | 0.07 | 1 | HC_vs_MS.Trt_B |
| NME4 | 0.00524 | 0.97277 | 0.116 | 0.07 | 1 | HC_vs_MS.Trt_B |
| CFTR | 0.00524 | 0.2595 | 0.264 | 0.19 | 1 | HC_vs_MS.Trt_B |
| CBY1 | 0.00524 | 0.53143 | 0.188 | 0.13 | 1 | HC_vs_MS.Trt_B |
| STARD7 | 0.00527 | 0.30918 | 0.499 | 0.43 | 1 | HC_vs_MS.Trt_B |
| HERC6 | 0.00528 | 0.83969 | 0.306 | 0.24 | 1 | HC_vs_MS.Trt_B |
| FAM111A | 0.00529 | 0.2525 | 0.474 | 0.4 | 1 | HC_vs_MS.Trt_B |
| CREB3L2 | 0.00529 | 0.30626 | 0.445 | 0.39 | 1 | HC_vs_MS.Trt_B |
| SLC25A42 | 0.00531 | -0.765 | 0.538 | 0.47 | 1 | HC_vs_MS.Trt_B |
| AAED1 | 0.00531 | -0.8303 | 0.78 | 0.69 | 1 | HC_vs_MS.Trt_B |
| SLC13A4 | 0.00534 | 0.37351 | 0.345 | 0.27 | 1 | HC_vs_MS.Trt_B |
| UTY | 0.00537 | 1.1482 | 0.422 | 0.35 | 1 | HC_vs_MS.Trt_B |
| PRNP | 0.00541 | 0.45674 | 0.187 | 0.13 | 1 | HC_vs_MS.Trt_B |
| GCNT1 | 0.00545 | 0.55138 | 0.249 | 0.18 | 1 | HC_vs_MS.Trt_B |
| GNA13 | 0.00548 | 0.63919 | 0.132 | 0.08 | 1 | HC_vs_MS.Trt_B |
| LETM1 | 0.00548 | -0.2697 | 0.828 | 0.76 | 1 | HC_vs_MS.Trt_B |
| ATP5B | 0.00548 | 0.27207 | 0.325 | 0.25 | 1 | HC_vs_MS.Trt_B |
| TNKS2 | 0.00549 | 0.33463 | 0.259 | 0.19 | 1 | HC_vs_MS.Trt_B |
| CHMP7 | 0.0055 | 0.35172 | 0.51 | 0.45 | 1 | HC_vs_MS.Trt_B |
| CASP3 | 0.00551 | 0.84741 | 0.12 | 0.07 | 1 | HC_vs_MS.Trt_B |
| DROSHA | 0.00553 | 0.28638 | 0.332 | 0.26 | 1 | HC_vs_MS.Trt_B |
| TIPIN | 0.00554 | 0.26259 | 0.562 | 0.48 | 1 | HC_vs_MS.Trt_B |
| UPK3B | 0.00554 | 0.41212 | 0.113 | 0.07 | 1 | HC_vs_MS.Trt_B |
| RPS15 | 0.00558 | -0.3247 | 0.928 | 0.86 | 1 | HC_vs_MS.Trt_B |
| IPO7 | 0.00558 | -0.501 | 0.4 | 0.32 | 1 | HC_vs_MS.Trt_B |
| KLHL11 | 0.00563 | 0.58388 | 0.475 | 0.41 | 1 | HC_vs_MS.Trt_B |
| TPO | 0.00565 | 0.61171 | 0.228 | 0.16 | 1 | HC_vs_MS.Trt_B |
| IMPG2 | 0.00567 | 0.25376 | 0.291 | 0.22 | 1 | HC_vs_MS.Trt_B |
| 3-Mar | 0.00568 | 0.92305 | 0.326 | 0.26 | 1 | HC_vs_MS.Trt_B |
| SREBF2 | 0.00568 | 0.55862 | 0.138 | 0.09 | 1 | HC_vs_MS.Trt_B |
| ANKRD40 | 0.00569 | -0.453 | 0.746 | 0.66 | 1 | HC_vs_MS.Trt_B |
| RBBP7 | 0.00571 | -0.2738 | 0.701 | 0.63 | 1 | HC_vs_MS.Trt_B |
| MOV10 | 0.00575 | 0.30433 | 0.449 | 0.36 | 1 | HC_vs_MS.Trt_B |

| | | | | | | |
|---|---|---|---|---|---|---|
| SMUG1 | 0.00579 | -0.4422 | 0.686 | 0.58 | 1 | HC_vs_MS.Trt_B |
| FAM208A | 0.00581 | -0.4357 | 0.736 | 0.65 | 1 | HC_vs_MS.Trt_B |
| THAP5 | 0.00583 | -0.6204 | 0.87 | 0.8 | 1 | HC_vs_MS.Trt_B |
| USP45 | 0.00586 | 0.77402 | 0.172 | 0.12 | 1 | HC_vs_MS.Trt_B |
| MAPKAP1 | 0.00586 | -0.2699 | 0.362 | 0.28 | 1 | HC_vs_MS.Trt_B |
| RUNDC1 | 0.00591 | 0.25253 | 0.701 | 0.63 | 1 | HC_vs_MS.Trt_B |
| PDGFA | 0.00592 | 0.6321 | 0.501 | 0.46 | 1 | HC_vs_MS.Trt_B |
| CHRNB1 | 0.00595 | -0.5542 | 0.584 | 0.48 | 1 | HC_vs_MS.Trt_B |
| TEX2 | 0.00596 | 0.62576 | 0.632 | 0.57 | 1 | HC_vs_MS.Trt_B |
| OLFML1 | 0.006 | 0.63196 | 0.225 | 0.17 | 1 | HC_vs_MS.Trt_B |
| APH1A | 0.006 | 0.34956 | 0.181 | 0.12 | 1 | HC_vs_MS.Trt_B |
| TPRG1L | 0.00601 | -0.9657 | 0.242 | 0.18 | 1 | HC_vs_MS.Trt_B |
| NOS3 | 0.00601 | 0.54569 | 0.362 | 0.29 | 1 | HC_vs_MS.Trt_B |
| ACRC | 0.00604 | 0.94959 | 0.423 | 0.34 | 1 | HC_vs_MS.Trt_B |
| USP25 | 0.00605 | -0.2992 | 0.449 | 0.36 | 1 | HC_vs_MS.Trt_B |
| C1orf52 | 0.00605 | 0.62979 | 0.283 | 0.22 | 1 | HC_vs_MS.Trt_B |
| UHRF1BP1L | 0.00615 | -0.4788 | 0.323 | 0.24 | 1 | HC_vs_MS.Trt_B |
| NQO1 | 0.00616 | 0.48478 | 0.517 | 0.46 | 1 | HC_vs_MS.Trt_B |
| IRF4 | 0.00617 | 0.30316 | 0.259 | 0.19 | 1 | HC_vs_MS.Trt_B |
| PHC1 | 0.00617 | 0.58585 | 0.12 | 0.07 | 1 | HC_vs_MS.Trt_B |
| PINK1 | 0.00619 | 0.41207 | 0.268 | 0.19 | 1 | HC_vs_MS.Trt_B |
| ZNF248 | 0.0062 | 0.27363 | 0.149 | 0.1 | 1 | HC_vs_MS.Trt_B |
| MTA2 | 0.00622 | -0.3908 | 0.249 | 0.32 | 1 | HC_vs_MS.Trt_B |
| GABARAPL2 | 0.00624 | -0.2781 | 0.443 | 0.37 | 1 | HC_vs_MS.Trt_B |
| FAM160A1 | 0.00624 | -0.8178 | 0.187 | 0.13 | 1 | HC_vs_MS.Trt_B |
| PPIAL4G | 0.00624 | -0.3425 | 0.619 | 0.56 | 1 | HC_vs_MS.Trt_B |
| IQCB1 | 0.0063 | 0.40038 | 0.11 | 0.06 | 1 | HC_vs_MS.Trt_B |
| ERGIC3 | 0.00633 | -0.5628 | 0.803 | 0.7 | 1 | HC_vs_MS.Trt_B |
| GRK7 | 0.00635 | -0.2936 | 0.329 | 0.26 | 1 | HC_vs_MS.Trt_B |
| MBD4 | 0.00635 | 0.40352 | 0.249 | 0.18 | 1 | HC_vs_MS.Trt_B |
| MTF1 | 0.00635 | -0.2602 | 0.267 | 0.2 | 1 | HC_vs_MS.Trt_B |
| ZBTB7A | 0.00635 | 0.49839 | 0.12 | 0.07 | 1 | HC_vs_MS.Trt_B |
| CD300LD | 0.00637 | 0.25887 | 0.362 | 0.28 | 1 | HC_vs_MS.Trt_B |
| SLC16A4 | 0.0064 | 0.30333 | 0.662 | 0.58 | 1 | HC_vs_MS.Trt_B |
| FGGY | 0.00643 | 0.64452 | 0.394 | 0.32 | 1 | HC_vs_MS.Trt_B |
| VWA7 | 0.00643 | 0.26275 | 0.313 | 0.24 | 1 | HC_vs_MS.Trt_B |
| SLC2A8 | 0.00645 | 0.38082 | 0.259 | 0.2 | 1 | HC_vs_MS.Trt_B |
| CCDC170 | 0.00646 | -0.2957 | 0.377 | 0.31 | 1 | HC_vs_MS.Trt_B |
| FAM91A1 | 0.00646 | 1.36738 | 0.299 | 0.23 | 1 | HC_vs_MS.Trt_B |
| PIM2 | 0.00647 | 0.26155 | 0.77 | 0.73 | 1 | HC_vs_MS.Trt_B |

| | | | | | | |
|---|---|---|---|---|---|---|
| EXO1 | 0.00652 | 0.27849 | 0.164 | 0.11 | 1 | HC_vs_MS.Trt_B |
| UST | 0.00653 | -0.7491 | 0.272 | 0.2 | 1 | HC_vs_MS.Trt_B |
| STRC | 0.00655 | -0.8177 | 0.503 | 0.43 | 1 | HC_vs_MS.Trt_B |
| DERL1 | 0.00656 | -0.2625 | 0.468 | 0.38 | 1 | HC_vs_MS.Trt_B |
| PLEKHG3 | 0.00659 | 0.31928 | 0.396 | 0.33 | 1 | HC_vs_MS.Trt_B |
| LGSN | 0.00662 | 0.28131 | 0.452 | 0.38 | 1 | HC_vs_MS.Trt_B |
| GNS | 0.00662 | 1.05374 | 0.739 | 0.7 | 1 | HC_vs_MS.Trt_B |
| PQLC2 | 0.00664 | -0.2911 | 0.396 | 0.32 | 1 | HC_vs_MS.Trt_B |
| CLDN4 | 0.00667 | 0.69224 | 0.543 | 0.45 | 1 | HC_vs_MS.Trt_B |
| CDH9 | 0.00667 | 0.28924 | 0.12 | 0.07 | 1 | HC_vs_MS.Trt_B |
| ALOX15 | 0.00669 | 0.28206 | 0.145 | 0.09 | 1 | HC_vs_MS.Trt_B |
| MRPS15 | 0.00671 | -0.3558 | 0.49 | 0.42 | 1 | HC_vs_MS.Trt_B |
| TUBA4A | 0.00674 | 0.2878 | 0.22 | 0.16 | 1 | HC_vs_MS.Trt_B |
| ARFGEF1 | 0.00687 | -0.5137 | 0.628 | 0.53 | 1 | HC_vs_MS.Trt_B |
| CCDC36 | 0.00691 | -0.7494 | 0.333 | 0.25 | 1 | HC_vs_MS.Trt_B |
| TRIM27 | 0.00692 | -0.316 | 0.384 | 0.31 | 1 | HC_vs_MS.Trt_B |
| DCAKD | 0.00692 | 0.44998 | 0.39 | 0.31 | 1 | HC_vs_MS.Trt_B |
| MYBBP1A | 0.00693 | 0.5253 | 0.143 | 0.09 | 1 | HC_vs_MS.Trt_B |
| PACS2 | 0.00694 | 0.48817 | 0.246 | 0.19 | 1 | HC_vs_MS.Trt_B |
| C6orf163 | 0.00699 | 0.41722 | 0.452 | 0.36 | 1 | HC_vs_MS.Trt_B |
| KIAA1033 | 0.00705 | 0.28707 | 0.281 | 0.21 | 1 | HC_vs_MS.Trt_B |
| ZNF197 | 0.00708 | 0.59631 | 0.467 | 0.39 | 1 | HC_vs_MS.Trt_B |
| NIN | 0.00708 | 0.33236 | 0.212 | 0.15 | 1 | HC_vs_MS.Trt_B |
| RRP1B | 0.00708 | 0.44375 | 0.199 | 0.14 | 1 | HC_vs_MS.Trt_B |
| WNT10A | 0.00711 | 0.74349 | 0.126 | 0.08 | 1 | HC_vs_MS.Trt_B |
| ACBD4 | 0.00718 | 0.38033 | 0.317 | 0.25 | 1 | HC_vs_MS.Trt_B |
| FAM50A | 0.00718 | 0.39085 | 0.114 | 0.07 | 1 | HC_vs_MS.Trt_B |
| HSD17B1 | 0.00719 | -0.3944 | 0.725 | 0.62 | 1 | HC_vs_MS.Trt_B |
| ETFA | 0.0072 | -0.2855 | 0.643 | 0.55 | 1 | HC_vs_MS.Trt_B |
| BZW2 | 0.00721 | 0.43676 | 0.133 | 0.08 | 1 | HC_vs_MS.Trt_B |
| USP49 | 0.00721 | 0.41123 | 0.694 | 0.6 | 1 | HC_vs_MS.Trt_B |
| TMX4 | 0.00725 | 0.27598 | 0.535 | 0.48 | 1 | HC_vs_MS.Trt_B |
| GALNT16 | 0.00726 | 0.34409 | 0.552 | 0.48 | 1 | HC_vs_MS.Trt_B |
| APBB3 | 0.00727 | 0.36591 | 0.123 | 0.07 | 1 | HC_vs_MS.Trt_B |
| ARF6 | 0.00728 | -0.6434 | 0.535 | 0.61 | 1 | HC_vs_MS.Trt_B |
| CTD-3105H1 | 0.00729 | 0.3482 | 0.293 | 0.22 | 1 | HC_vs_MS.Trt_B |
| KBTBD6 | 0.00737 | 1.02137 | 0.325 | 0.25 | 1 | HC_vs_MS.Trt_B |
| AC006449.1 | 0.00739 | 0.37737 | 0.116 | 0.07 | 1 | HC_vs_MS.Trt_B |
| C14orf80 | 0.00742 | -0.9389 | 0.262 | 0.2 | 1 | HC_vs_MS.Trt_B |
| H2AFV | 0.00745 | -0.4969 | 0.603 | 0.51 | 1 | HC_vs_MS.Trt_B |

| | | | | | | |
|---|---|---|---|---|---|---|
| KIAA1143 | 0.00745 | 0.52853 | 0.491 | 0.43 | 1 | HC_vs_MS.Trt_B |
| CNOT3 | 0.00747 | 0.99037 | 0.123 | 0.07 | 1 | HC_vs_MS.Trt_B |
| ZNF846 | 0.00749 | 0.52914 | 0.446 | 0.36 | 1 | HC_vs_MS.Trt_B |
| PLA2G4B | 0.00752 | -0.6222 | 0.448 | 0.38 | 1 | HC_vs_MS.Trt_B |
| ATE1 | 0.00756 | 0.5611 | 0.286 | 0.22 | 1 | HC_vs_MS.Trt_B |
| GM2A | 0.0076 | 0.70955 | 0.528 | 0.46 | 1 | HC_vs_MS.Trt_B |
| MAP1S | 0.00762 | 0.42155 | 0.581 | 0.52 | 1 | HC_vs_MS.Trt_B |
| MYSM1 | 0.00765 | 0.26777 | 0.206 | 0.15 | 1 | HC_vs_MS.Trt_B |
| PAPOLG | 0.00769 | -0.2775 | 0.733 | 0.65 | 1 | HC_vs_MS.Trt_B |
| SYNGR2 | 0.0077 | -0.4821 | 0.555 | 0.46 | 1 | HC_vs_MS.Trt_B |
| GTPBP3 | 0.00771 | -0.4746 | 0.632 | 0.57 | 1 | HC_vs_MS.Trt_B |
| SATB1 | 0.00772 | 0.38144 | 0.239 | 0.18 | 1 | HC_vs_MS.Trt_B |
| SERAC1 | 0.00775 | 0.38968 | 0.442 | 0.36 | 1 | HC_vs_MS.Trt_B |
| RP5-994D16 | 0.0078 | 0.91248 | 0.462 | 0.39 | 1 | HC_vs_MS.Trt_B |
| INPP5E | 0.00781 | 0.3949 | 0.27 | 0.2 | 1 | HC_vs_MS.Trt_B |
| CASC10 | 0.00787 | -0.4452 | 0.372 | 0.3 | 1 | HC_vs_MS.Trt_B |
| ZMYM4 | 0.00795 | -0.4098 | 0.126 | 0.08 | 1 | HC_vs_MS.Trt_B |
| TRIM34 | 0.00797 | -0.3104 | 0.314 | 0.25 | 1 | HC_vs_MS.Trt_B |
| CCDC90B | 0.00798 | -0.2537 | 0.957 | 0.87 | 1 | HC_vs_MS.Trt_B |
| CAMKK2 | 0.00798 | 0.84029 | 0.465 | 0.4 | 1 | HC_vs_MS.Trt_B |
| TMEM267 | 0.00799 | 0.77632 | 0.286 | 0.22 | 1 | HC_vs_MS.Trt_B |
| COX19 | 0.008 | 0.25599 | 0.601 | 0.5 | 1 | HC_vs_MS.Trt_B |
| COMMD2 | 0.00804 | 0.3257 | 0.329 | 0.26 | 1 | HC_vs_MS.Trt_B |
| RFTN1 | 0.00805 | 0.46971 | 0.346 | 0.28 | 1 | HC_vs_MS.Trt_B |
| GNLY | 0.00806 | 0.73852 | 0.226 | 0.17 | 1 | HC_vs_MS.Trt_B |
| SLC25A4 | 0.00808 | -0.79 | 0.272 | 0.21 | 1 | HC_vs_MS.Trt_B |
| KTN1 | 0.0081 | -0.5212 | 0.571 | 0.45 | 1 | HC_vs_MS.Trt_B |
| UBAP1L | 0.00811 | 0.65482 | 0.355 | 0.29 | 1 | HC_vs_MS.Trt_B |
| VN1R1 | 0.00815 | 0.26345 | 0.154 | 0.1 | 1 | HC_vs_MS.Trt_B |
| TLL2 | 0.00816 | 0.29915 | 0.316 | 0.25 | 1 | HC_vs_MS.Trt_B |
| KIAA0368 | 0.00816 | 0.36069 | 0.212 | 0.15 | 1 | HC_vs_MS.Trt_B |
| AURKA | 0.00818 | -1.1878 | 0.286 | 0.23 | 1 | HC_vs_MS.Trt_B |
| BLOC1S5 | 0.00821 | -0.2708 | 0.636 | 0.52 | 1 | HC_vs_MS.Trt_B |
| FIS1 | 0.00822 | 0.30275 | 0.235 | 0.17 | 1 | HC_vs_MS.Trt_B |
| YARS2 | 0.00823 | -0.3924 | 0.377 | 0.28 | 1 | HC_vs_MS.Trt_B |
| DPP9 | 0.00824 | 0.25185 | 0.649 | 0.58 | 1 | HC_vs_MS.Trt_B |
| OLAH | 0.00825 | 0.28126 | 0.432 | 0.37 | 1 | HC_vs_MS.Trt_B |
| USB1 | 0.00829 | 0.49937 | 0.119 | 0.07 | 1 | HC_vs_MS.Trt_B |
| ECHDC3 | 0.00837 | 0.77916 | 0.109 | 0.06 | 1 | HC_vs_MS.Trt_B |
| SENP8 | 0.00839 | -0.2573 | 0.445 | 0.39 | 1 | HC_vs_MS.Trt_B |

| | | | | | | |
|---|---|---|---|---|---|---|
| MYH14 | 0.00841 | 0.306 | 0.201 | 0.15 | 1 | HC_vs_MS.Trt_B |
| SLC6A6 | 0.00844 | 0.57752 | 0.148 | 0.1 | 1 | HC_vs_MS.Trt_B |
| LXN | 0.00846 | -0.4101 | 0.168 | 0.11 | 1 | HC_vs_MS.Trt_B |
| RAI1 | 0.00848 | 0.37647 | 0.209 | 0.15 | 1 | HC_vs_MS.Trt_B |
| RECQL5 | 0.00848 | 0.3197 | 0.196 | 0.14 | 1 | HC_vs_MS.Trt_B |
| METTL21B | 0.00851 | 0.63171 | 0.448 | 0.38 | 1 | HC_vs_MS.Trt_B |
| IQCK | 0.00854 | 0.55018 | 0.19 | 0.13 | 1 | HC_vs_MS.Trt_B |
| DEPDC1B | 0.00855 | -0.6965 | 0.206 | 0.15 | 1 | HC_vs_MS.Trt_B |
| SLC1A2 | 0.0086 | 0.56264 | 0.329 | 0.26 | 1 | HC_vs_MS.Trt_B |
| ANKRD16 | 0.00861 | 0.40245 | 0.429 | 0.35 | 1 | HC_vs_MS.Trt_B |
| ASB8 | 0.00865 | 0.28185 | 0.648 | 0.61 | 1 | HC_vs_MS.Trt_B |
| JADE2 | 0.0087 | 0.36405 | 0.155 | 0.1 | 1 | HC_vs_MS.Trt_B |
| ZHX1 | 0.00872 | 0.47659 | 0.548 | 0.49 | 1 | HC_vs_MS.Trt_B |
| DEAF1 | 0.00873 | 0.49946 | 0.296 | 0.22 | 1 | HC_vs_MS.Trt_B |
| SP1 | 0.00873 | 0.78397 | 0.168 | 0.12 | 1 | HC_vs_MS.Trt_B |
| SMN1 | 0.00876 | -0.2736 | 0.383 | 0.3 | 1 | HC_vs_MS.Trt_B |
| VAMP3 | 0.00878 | 0.27159 | 0.278 | 0.21 | 1 | HC_vs_MS.Trt_B |
| ELOVL6 | 0.00879 | 0.34276 | 0.274 | 0.21 | 1 | HC_vs_MS.Trt_B |
| CENPF | 0.00884 | 0.59376 | 0.343 | 0.26 | 1 | HC_vs_MS.Trt_B |
| UPP1 | 0.00886 | -0.3286 | 0.636 | 0.53 | 1 | HC_vs_MS.Trt_B |
| POLDIP3 | 0.00891 | 0.3671 | 0.119 | 0.07 | 1 | HC_vs_MS.Trt_B |
| CCDC102B | 0.00891 | 0.59157 | 0.157 | 0.1 | 1 | HC_vs_MS.Trt_B |
| TTC31 | 0.00891 | 0.26908 | 0.206 | 0.15 | 1 | HC_vs_MS.Trt_B |
| TRIM37 | 0.00891 | 0.73188 | 0.17 | 0.11 | 1 | HC_vs_MS.Trt_B |
| PAK6 | 0.00895 | 0.52447 | 0.155 | 0.1 | 1 | HC_vs_MS.Trt_B |
| MDN1 | 0.00902 | 0.25723 | 0.241 | 0.18 | 1 | HC_vs_MS.Trt_B |
| FOXM1 | 0.00914 | 0.67077 | 0.686 | 0.61 | 1 | HC_vs_MS.Trt_B |
| ZCCHC10 | 0.00914 | -0.6671 | 0.586 | 0.49 | 1 | HC_vs_MS.Trt_B |
| LIPC | 0.0092 | 0.95078 | 0.306 | 0.24 | 1 | HC_vs_MS.Trt_B |
| PSTPIP1 | 0.0093 | -0.3857 | 0.414 | 0.33 | 1 | HC_vs_MS.Trt_B |
| CISD2 | 0.00932 | 0.73043 | 0.439 | 0.35 | 1 | HC_vs_MS.Trt_B |
| CCDC66 | 0.00935 | -0.6155 | 0.601 | 0.51 | 1 | HC_vs_MS.Trt_B |
| P2RY10 | 0.00937 | 0.29188 | 0.309 | 0.23 | 1 | HC_vs_MS.Trt_B |
| OR5A2 | 0.00939 | 0.28174 | 0.219 | 0.16 | 1 | HC_vs_MS.Trt_B |
| N4BP2 | 0.00945 | -0.7469 | 0.332 | 0.26 | 1 | HC_vs_MS.Trt_B |
| TOR4A | 0.00947 | -0.6649 | 0.187 | 0.13 | 1 | HC_vs_MS.Trt_B |
| ATXN7 | 0.00953 | 0.41107 | 0.497 | 0.42 | 1 | HC_vs_MS.Trt_B |
| CDK17 | 0.00954 | 0.37879 | 0.172 | 0.12 | 1 | HC_vs_MS.Trt_B |
| DDX43 | 0.00955 | 0.90749 | 0.168 | 0.11 | 1 | HC_vs_MS.Trt_B |
| AC138645.2 | 0.00957 | 0.31988 | 0.112 | 0.07 | 1 | HC_vs_MS.Trt_B |

| | | | | | | |
|---|---|---|---|---|---|---|
| AKR1C1 | 0.00957 | 0.29497 | 0.229 | 0.17 | 1 | HC_vs_MS.Trt_B |
| GAREM2 | 0.00964 | 0.25336 | 0.499 | 0.4 | 1 | HC_vs_MS.Trt_B |
| KLRK1 | 0.00966 | -1.1062 | 0.643 | 0.58 | 1 | HC_vs_MS.Trt_B |
| STAT2 | 0.00969 | 0.25715 | 0.883 | 0.84 | 1 | HC_vs_MS.Trt_B |
| NHEJ1 | 0.0097 | 1.7771 | 0.161 | 0.11 | 1 | HC_vs_MS.Trt_B |
| CTSW | 0.0097 | -0.6966 | 0.622 | 0.53 | 1 | HC_vs_MS.Trt_B |
| ERCC3 | 0.00972 | -0.6929 | 0.361 | 0.29 | 1 | HC_vs_MS.Trt_B |
| AKT1 | 0.00975 | -0.4341 | 0.203 | 0.14 | 1 | HC_vs_MS.Trt_B |
| PROSER1 | 0.00976 | 0.59885 | 0.171 | 0.12 | 1 | HC_vs_MS.Trt_B |
| ALKBH2 | 0.00979 | 0.54979 | 0.307 | 0.24 | 1 | HC_vs_MS.Trt_B |
| ATN1 | 0.0098 | 0.41576 | 0.17 | 0.12 | 1 | HC_vs_MS.Trt_B |
| TRAPPC2B | 0.00985 | -0.4244 | 0.328 | 0.26 | 1 | HC_vs_MS.Trt_B |
| SART3 | 0.00987 | 0.74842 | 0.138 | 0.09 | 1 | HC_vs_MS.Trt_B |
| MYB | 0.00989 | 0.87035 | 0.28 | 0.21 | 1 | HC_vs_MS.Trt_B |
| TPRG1 | 0.0099 | 0.39193 | 0.18 | 0.12 | 1 | HC_vs_MS.Trt_B |
| DYNLRB1 | 0.0099 | 0.31005 | 0.177 | 0.12 | 1 | HC_vs_MS.Trt_B |
| SV2C | 0.00998 | 0.5144 | 0.53 | 0.44 | 1 | HC_vs_MS.Trt_B |
| ZSCAN32 | 0.01 | -0.6941 | 0.235 | 0.17 | 1 | HC_vs_MS.Trt_B |
| APEH | 0.01001 | 0.31256 | 0.19 | 0.13 | 1 | HC_vs_MS.Trt_B |
| NFKBIA | 0.01002 | 0.37102 | 0.243 | 0.18 | 1 | HC_vs_MS.Trt_B |
| NDST4 | 0.01002 | 0.43811 | 0.109 | 0.06 | 1 | HC_vs_MS.Trt_B |
| E2F8 | 0.01002 | 0.49351 | 0.133 | 0.09 | 1 | HC_vs_MS.Trt_B |
| ZBTB16 | 0.01004 | -0.7081 | 0.372 | 0.3 | 1 | HC_vs_MS.Trt_B |
| DENND6A | 0.01005 | 0.3284 | 0.559 | 0.48 | 1 | HC_vs_MS.Trt_B |
| MTX2 | 0.01017 | 0.35999 | 0.191 | 0.14 | 1 | HC_vs_MS.Trt_B |
| MRPS12 | 0.01019 | -0.4805 | 0.371 | 0.3 | 1 | HC_vs_MS.Trt_B |
| SHPK | 0.01033 | -0.7927 | 0.401 | 0.32 | 1 | HC_vs_MS.Trt_B |
| PKIG | 0.01033 | -0.2748 | 0.626 | 0.52 | 1 | HC_vs_MS.Trt_B |
| ZNF335 | 0.0104 | 0.30336 | 0.31 | 0.25 | 1 | HC_vs_MS.Trt_B |
| HEXIM1 | 0.01046 | 0.40842 | 0.193 | 0.14 | 1 | HC_vs_MS.Trt_B |
| SLC9A4 | 0.01047 | 0.26035 | 0.483 | 0.41 | 1 | HC_vs_MS.Trt_B |
| WNK3 | 0.0105 | -0.9881 | 0.288 | 0.22 | 1 | HC_vs_MS.Trt_B |
| QRICH2 | 0.01055 | 0.70222 | 0.174 | 0.12 | 1 | HC_vs_MS.Trt_B |
| RMI1 | 0.01057 | -0.3359 | 0.242 | 0.18 | 1 | HC_vs_MS.Trt_B |
| GTF2E1 | 0.01058 | -0.7507 | 0.254 | 0.19 | 1 | HC_vs_MS.Trt_B |
| AGBL2 | 0.01058 | -0.3441 | 0.299 | 0.22 | 1 | HC_vs_MS.Trt_B |
| SETMAR | 0.0106 | 0.59404 | 0.314 | 0.25 | 1 | HC_vs_MS.Trt_B |
| TCFL5 | 0.01061 | 0.69581 | 0.468 | 0.42 | 1 | HC_vs_MS.Trt_B |
| CDCP1 | 0.01062 | 0.91602 | 0.43 | 0.37 | 1 | HC_vs_MS.Trt_B |
| ANKRD6 | 0.01066 | 0.87542 | 0.165 | 0.11 | 1 | HC_vs_MS.Trt_B |

| | | | | | | |
|---|---|---|---|---|---|---|
| PIK3AP1 | 0.01067 | -0.9277 | 0.122 | 0.08 | 1 | HC_vs_MS.Trt_B |
| ATP8A1 | 0.01073 | 0.64028 | 0.161 | 0.11 | 1 | HC_vs_MS.Trt_B |
| AGFG2 | 0.01076 | 0.27058 | 0.336 | 0.27 | 1 | HC_vs_MS.Trt_B |
| SRL | 0.01077 | 0.2877 | 0.346 | 0.29 | 1 | HC_vs_MS.Trt_B |
| SPECC1L | 0.01077 | 0.3122 | 0.136 | 0.09 | 1 | HC_vs_MS.Trt_B |
| FOXO3 | 0.01079 | 0.40032 | 0.365 | 0.29 | 1 | HC_vs_MS.Trt_B |
| C19orf44 | 0.01083 | -0.4659 | 0.332 | 0.26 | 1 | HC_vs_MS.Trt_B |
| ARMC8 | 0.01087 | 0.26202 | 0.472 | 0.4 | 1 | HC_vs_MS.Trt_B |
| PPP4R3A | 0.01093 | 0.37955 | 0.204 | 0.15 | 1 | HC_vs_MS.Trt_B |
| NPRL3 | 0.01095 | 0.34721 | 0.299 | 0.23 | 1 | HC_vs_MS.Trt_B |
| SLC15A2 | 0.01097 | -0.4054 | 0.486 | 0.39 | 1 | HC_vs_MS.Trt_B |
| C2CD5 | 0.01103 | 0.38646 | 0.165 | 0.11 | 1 | HC_vs_MS.Trt_B |
| BMPR2 | 0.01103 | 0.41232 | 0.445 | 0.36 | 1 | HC_vs_MS.Trt_B |
| PLCG1 | 0.01109 | 0.27271 | 0.364 | 0.29 | 1 | HC_vs_MS.Trt_B |
| CPT1B | 0.01111 | 0.9326 | 0.171 | 0.12 | 1 | HC_vs_MS.Trt_B |
| CARNS1 | 0.01113 | 0.30424 | 0.197 | 0.14 | 1 | HC_vs_MS.Trt_B |
| ARFIP1 | 0.01116 | -0.472 | 0.212 | 0.15 | 1 | HC_vs_MS.Trt_B |
| CARD14 | 0.01123 | 0.31032 | 0.762 | 0.69 | 1 | HC_vs_MS.Trt_B |
| GRAMD3 | 0.01132 | -0.4588 | 0.69 | 0.62 | 1 | HC_vs_MS.Trt_B |
| IFT46 | 0.01134 | 0.67905 | 0.206 | 0.15 | 1 | HC_vs_MS.Trt_B |
| AIG1 | 0.01135 | 0.35794 | 0.545 | 0.45 | 1 | HC_vs_MS.Trt_B |
| SMC6 | 0.01157 | -0.4144 | 0.464 | 0.39 | 1 | HC_vs_MS.Trt_B |
| MRPL20 | 0.01162 | -0.2863 | 0.509 | 0.44 | 1 | HC_vs_MS.Trt_B |
| TRRAP | 0.01163 | 0.55018 | 0.158 | 0.11 | 1 | HC_vs_MS.Trt_B |
| RBM18 | 0.01163 | 0.3125 | 0.312 | 0.24 | 1 | HC_vs_MS.Trt_B |
| USP13 | 0.01168 | 0.69858 | 0.259 | 0.2 | 1 | HC_vs_MS.Trt_B |
| ARHGEF28 | 0.01173 | 0.27033 | 0.165 | 0.11 | 1 | HC_vs_MS.Trt_B |
| NBAS | 0.01173 | -0.3027 | 0.213 | 0.15 | 1 | HC_vs_MS.Trt_B |
| B3GAT3 | 0.01174 | -0.4544 | 0.532 | 0.42 | 1 | HC_vs_MS.Trt_B |
| TAF1B | 0.0118 | -0.288 | 0.464 | 0.38 | 1 | HC_vs_MS.Trt_B |
| C6orf25 | 0.01185 | 0.53519 | 0.436 | 0.37 | 1 | HC_vs_MS.Trt_B |
| PTK6 | 0.01188 | 0.66371 | 0.332 | 0.27 | 1 | HC_vs_MS.Trt_B |
| ZBTB40 | 0.01192 | 0.5399 | 0.365 | 0.29 | 1 | HC_vs_MS.Trt_B |
| SEPSECS | 0.01192 | 0.54649 | 0.758 | 0.68 | 1 | HC_vs_MS.Trt_B |
| NOP56 | 0.01193 | -0.5682 | 0.4 | 0.32 | 1 | HC_vs_MS.Trt_B |
| ZNF614 | 0.01195 | 0.95171 | 0.252 | 0.19 | 1 | HC_vs_MS.Trt_B |
| ZNF257 | 0.01195 | -0.4294 | 0.536 | 0.45 | 1 | HC_vs_MS.Trt_B |
| E2F5 | 0.01196 | 0.72704 | 0.332 | 0.28 | 1 | HC_vs_MS.Trt_B |
| LRWD1 | 0.01207 | -0.2504 | 0.541 | 0.48 | 1 | HC_vs_MS.Trt_B |
| PRIM2 | 0.01207 | 0.27183 | 0.436 | 0.36 | 1 | HC_vs_MS.Trt_B |

| | | | | | | |
|---|---|---|---|---|---|---|
| KCTD3 | 0.01212 | 0.32075 | 0.301 | 0.25 | 1 | HC_vs_MS.Trt_B |
| PCDHGB6 | 0.01213 | 0.38658 | 0.352 | 0.29 | 1 | HC_vs_MS.Trt_B |
| PLEKHM3 | 0.01223 | 0.67577 | 0.545 | 0.48 | 1 | HC_vs_MS.Trt_B |
| ZC3H11A | 0.01226 | 0.32213 | 0.204 | 0.15 | 1 | HC_vs_MS.Trt_B |
| RDH11 | 0.01226 | -0.2647 | 0.52 | 0.44 | 1 | HC_vs_MS.Trt_B |
| GRAP2 | 0.01236 | 0.36993 | 0.268 | 0.21 | 1 | HC_vs_MS.Trt_B |
| NDUFA12 | 0.01236 | -0.933 | 0.168 | 0.22 | 1 | HC_vs_MS.Trt_B |
| MPC2 | 0.01238 | -0.4738 | 0.428 | 0.36 | 1 | HC_vs_MS.Trt_B |
| ERLIN2 | 0.0124 | 0.29108 | 0.358 | 0.29 | 1 | HC_vs_MS.Trt_B |
| CERKL | 0.0124 | -0.4516 | 0.354 | 0.28 | 1 | HC_vs_MS.Trt_B |
| GYS1 | 0.01246 | 0.26822 | 0.139 | 0.09 | 1 | HC_vs_MS.Trt_B |
| PVR | 0.01259 | -0.4811 | 0.454 | 0.39 | 1 | HC_vs_MS.Trt_B |
| GCKR | 0.0126 | 0.44641 | 0.139 | 0.09 | 1 | HC_vs_MS.Trt_B |
| CD40 | 0.01261 | 0.2738 | 0.143 | 0.09 | 1 | HC_vs_MS.Trt_B |
| RASGRP2 | 0.01265 | 0.28236 | 0.384 | 0.32 | 1 | HC_vs_MS.Trt_B |
| LIN7C | 0.0127 | 0.38903 | 0.297 | 0.24 | 1 | HC_vs_MS.Trt_B |
| PSTK | 0.01271 | -0.4863 | 0.338 | 0.26 | 1 | HC_vs_MS.Trt_B |
| CMPK1 | 0.01279 | 0.33613 | 0.232 | 0.17 | 1 | HC_vs_MS.Trt_B |
| SDSL | 0.0128 | 0.48928 | 0.328 | 0.26 | 1 | HC_vs_MS.Trt_B |
| SPA17 | 0.01282 | 0.55061 | 0.212 | 0.15 | 1 | HC_vs_MS.Trt_B |
| ODF2 | 0.01284 | 0.45158 | 0.461 | 0.4 | 1 | HC_vs_MS.Trt_B |
| ANKRD20A3 | 0.01285 | 0.36845 | 0.112 | 0.07 | 1 | HC_vs_MS.Trt_B |
| VAPA | 0.01307 | -0.3315 | 0.732 | 0.64 | 1 | HC_vs_MS.Trt_B |
| CHAMP1 | 0.01308 | 0.52842 | 0.581 | 0.52 | 1 | HC_vs_MS.Trt_B |
| TANGO2 | 0.01321 | 0.33761 | 0.841 | 0.77 | 1 | HC_vs_MS.Trt_B |
| ZNF599 | 0.01325 | -0.3502 | 0.565 | 0.46 | 1 | HC_vs_MS.Trt_B |
| IFT74 | 0.01326 | -0.2781 | 0.375 | 0.31 | 1 | HC_vs_MS.Trt_B |
| AGGF1 | 0.0134 | 0.65834 | 0.123 | 0.08 | 1 | HC_vs_MS.Trt_B |
| DERL2 | 0.01343 | -0.3705 | 0.268 | 0.2 | 1 | HC_vs_MS.Trt_B |
| PIGBOS1 | 0.01347 | -0.4534 | 0.558 | 0.5 | 1 | HC_vs_MS.Trt_B |
| RGS14 | 0.01348 | 0.28629 | 0.194 | 0.14 | 1 | HC_vs_MS.Trt_B |
| SLC14A1 | 0.01349 | -0.2732 | 0.484 | 0.4 | 1 | HC_vs_MS.Trt_B |
| KIF9 | 0.01353 | 0.45005 | 0.106 | 0.06 | 1 | HC_vs_MS.Trt_B |
| ZNF563 | 0.01358 | 0.62388 | 0.516 | 0.42 | 1 | HC_vs_MS.Trt_B |
| MIS12 | 0.01363 | -0.3287 | 0.291 | 0.22 | 1 | HC_vs_MS.Trt_B |
| RUFY2 | 0.01363 | 0.35081 | 0.507 | 0.42 | 1 | HC_vs_MS.Trt_B |
| NRIP2 | 0.01365 | 0.74289 | 0.336 | 0.27 | 1 | HC_vs_MS.Trt_B |
| NEIL3 | 0.01369 | 0.57224 | 0.259 | 0.2 | 1 | HC_vs_MS.Trt_B |
| BAG2 | 0.01372 | 0.42784 | 0.23 | 0.17 | 1 | HC_vs_MS.Trt_B |
| RASSF1 | 0.01377 | 1.03209 | 0.155 | 0.11 | 1 | HC_vs_MS.Trt_B |

| | | | | | | |
|---|---|---|---|---|---|---|
| PET100 | 0.01379 | -0.4397 | 0.623 | 0.53 | 1 | HC_vs_MS.Trt_B |
| RAB26 | 0.0138 | -0.2618 | 0.132 | 0.09 | 1 | HC_vs_MS.Trt_B |
| PIGA | 0.01382 | -0.6467 | 0.39 | 0.32 | 1 | HC_vs_MS.Trt_B |
| AIRE | 0.01385 | 0.30542 | 0.217 | 0.16 | 1 | HC_vs_MS.Trt_B |
| SH3GLB1 | 0.01387 | 0.28043 | 0.439 | 0.39 | 1 | HC_vs_MS.Trt_B |
| HAVCR2 | 0.01393 | -0.3222 | 0.514 | 0.43 | 1 | HC_vs_MS.Trt_B |
| HSPB9 | 0.01398 | 0.59014 | 0.203 | 0.15 | 1 | HC_vs_MS.Trt_B |
| TMSB15B | 0.01398 | 0.6414 | 0.288 | 0.22 | 1 | HC_vs_MS.Trt_B |
| C1orf146 | 0.01407 | 0.35657 | 0.165 | 0.11 | 1 | HC_vs_MS.Trt_B |
| CNNM2 | 0.0141 | 0.47317 | 0.264 | 0.21 | 1 | HC_vs_MS.Trt_B |
| CC2D2A | 0.01415 | 0.31419 | 0.241 | 0.18 | 1 | HC_vs_MS.Trt_B |
| KLHL9 | 0.01418 | -0.6166 | 0.343 | 0.28 | 1 | HC_vs_MS.Trt_B |
| PES1 | 0.01432 | 0.57854 | 0.104 | 0.06 | 1 | HC_vs_MS.Trt_B |
| AGBL1 | 0.01433 | 0.28773 | 0.339 | 0.28 | 1 | HC_vs_MS.Trt_B |
| LST1 | 0.01434 | -0.3457 | 0.245 | 0.18 | 1 | HC_vs_MS.Trt_B |
| IGHMBP2 | 0.01442 | 1.09075 | 0.349 | 0.29 | 1 | HC_vs_MS.Trt_B |
| TGDS | 0.01454 | -0.3432 | 0.107 | 0.07 | 1 | HC_vs_MS.Trt_B |
| TMEM159 | 0.01456 | 0.38061 | 0.607 | 0.56 | 1 | HC_vs_MS.Trt_B |
| RNF170 | 0.0146 | -0.3046 | 0.546 | 0.47 | 1 | HC_vs_MS.Trt_B |
| TJP2 | 0.01464 | 0.52426 | 0.339 | 0.28 | 1 | HC_vs_MS.Trt_B |
| POLA2 | 0.01467 | 0.37293 | 0.597 | 0.53 | 1 | HC_vs_MS.Trt_B |
| RAC1 | 0.01468 | 0.25507 | 0.151 | 0.1 | 1 | HC_vs_MS.Trt_B |
| JUNB | 0.01468 | 0.28021 | 0.354 | 0.29 | 1 | HC_vs_MS.Trt_B |
| CNIH3 | 0.01469 | 0.40082 | 0.171 | 0.12 | 1 | HC_vs_MS.Trt_B |
| BBS1 | 0.01474 | 0.31215 | 0.577 | 0.5 | 1 | HC_vs_MS.Trt_B |
| PDE8A | 0.01481 | 0.72887 | 0.397 | 0.33 | 1 | HC_vs_MS.Trt_B |
| SORBS1 | 0.01483 | 0.43192 | 0.219 | 0.16 | 1 | HC_vs_MS.Trt_B |
| SNAP47 | 0.01486 | 0.41707 | 0.12 | 0.08 | 1 | HC_vs_MS.Trt_B |
| SLC39A9 | 0.01487 | 0.91007 | 0.174 | 0.12 | 1 | HC_vs_MS.Trt_B |
| GPR89B | 0.01492 | 0.2885 | 0.314 | 0.25 | 1 | HC_vs_MS.Trt_B |
| MRPL19 | 0.01497 | 0.58237 | 0.243 | 0.19 | 1 | HC_vs_MS.Trt_B |
| RAP1GAP2 | 0.01501 | 0.47885 | 0.191 | 0.14 | 1 | HC_vs_MS.Trt_B |
| FUNDC2 | 0.01529 | -0.6192 | 0.451 | 0.39 | 1 | HC_vs_MS.Trt_B |
| FMR1 | 0.01542 | 0.89578 | 0.103 | 0.06 | 1 | HC_vs_MS.Trt_B |
| VPS13A | 0.01544 | -0.3948 | 0.287 | 0.22 | 1 | HC_vs_MS.Trt_B |
| MITF | 0.01544 | 0.36338 | 0.139 | 0.09 | 1 | HC_vs_MS.Trt_B |
| C2CD2L | 0.01546 | 0.59598 | 0.409 | 0.36 | 1 | HC_vs_MS.Trt_B |
| TMUB2 | 0.01549 | -0.3264 | 0.813 | 0.74 | 1 | HC_vs_MS.Trt_B |
| AP3S1 | 0.0155 | -0.2538 | 0.319 | 0.26 | 1 | HC_vs_MS.Trt_B |
| CASP9 | 0.01551 | -0.6813 | 0.328 | 0.26 | 1 | HC_vs_MS.Trt_B |

| | | | | | | |
|---|---|---|---|---|---|---|
| TACO1 | 0.01552 | 0.59901 | 0.37 | 0.31 | 1 | HC_vs_MS.Trt_B |
| ZNF530 | 0.01556 | 0.69488 | 0.483 | 0.41 | 1 | HC_vs_MS.Trt_B |
| PNP | 0.01557 | 0.65822 | 0.316 | 0.24 | 1 | HC_vs_MS.Trt_B |
| MLYCD | 0.0156 | 0.32984 | 0.357 | 0.29 | 1 | HC_vs_MS.Trt_B |
| DOPEY1 | 0.01562 | 0.38927 | 0.288 | 0.22 | 1 | HC_vs_MS.Trt_B |
| FERMT2 | 0.01576 | 0.2515 | 0.452 | 0.39 | 1 | HC_vs_MS.Trt_B |
| YES1 | 0.01583 | -0.3987 | 0.358 | 0.29 | 1 | HC_vs_MS.Trt_B |
| FGF11 | 0.01589 | 0.39649 | 0.21 | 0.15 | 1 | HC_vs_MS.Trt_B |
| FBXO32 | 0.01601 | 0.55022 | 0.145 | 0.1 | 1 | HC_vs_MS.Trt_B |
| MFSD9 | 0.01604 | 0.4912 | 0.488 | 0.42 | 1 | HC_vs_MS.Trt_B |
| RPS6KA1 | 0.01604 | 0.37218 | 0.174 | 0.12 | 1 | HC_vs_MS.Trt_B |
| NUDT6 | 0.01609 | -0.6228 | 0.475 | 0.39 | 1 | HC_vs_MS.Trt_B |
| TTC38 | 0.01611 | 1.13657 | 0.429 | 0.36 | 1 | HC_vs_MS.Trt_B |
| SIK3 | 0.01613 | 1.07361 | 0.243 | 0.19 | 1 | HC_vs_MS.Trt_B |
| CCDC126 | 0.0162 | -0.6115 | 0.116 | 0.07 | 1 | HC_vs_MS.Trt_B |
| NBPF1 | 0.01628 | 0.80376 | 0.229 | 0.17 | 1 | HC_vs_MS.Trt_B |
| ZNF672 | 0.01632 | -0.2863 | 0.236 | 0.17 | 1 | HC_vs_MS.Trt_B |
| SSU72 | 0.01655 | -0.2915 | 0.89 | 0.82 | 1 | HC_vs_MS.Trt_B |
| THBS1 | 0.01661 | 0.45992 | 0.209 | 0.16 | 1 | HC_vs_MS.Trt_B |
| LCA5L | 0.01678 | 0.37106 | 0.342 | 0.28 | 1 | HC_vs_MS.Trt_B |
| GRWD1 | 0.0168 | 0.32247 | 0.457 | 0.38 | 1 | HC_vs_MS.Trt_B |
| EVI5L | 0.01685 | 0.42292 | 0.268 | 0.21 | 1 | HC_vs_MS.Trt_B |
| FAM188B | 0.01686 | 0.41451 | 0.165 | 0.11 | 1 | HC_vs_MS.Trt_B |
| ZNF287 | 0.01689 | 0.32445 | 0.365 | 0.31 | 1 | HC_vs_MS.Trt_B |
| UQCRB | 0.0169 | -0.562 | 0.677 | 0.58 | 1 | HC_vs_MS.Trt_B |
| OAS2 | 0.0169 | 0.33209 | 0.213 | 0.16 | 1 | HC_vs_MS.Trt_B |
| MASP1 | 0.017 | 0.28623 | 0.238 | 0.19 | 1 | HC_vs_MS.Trt_B |
| COX7A2 | 0.017 | -0.6436 | 0.719 | 0.63 | 1 | HC_vs_MS.Trt_B |
| ZNF789 | 0.01702 | -0.2751 | 0.728 | 0.65 | 1 | HC_vs_MS.Trt_B |
| FAM192A | 0.01714 | -0.3405 | 0.417 | 0.34 | 1 | HC_vs_MS.Trt_B |
| ARHGAP19 | 0.01715 | -0.8489 | 0.27 | 0.21 | 1 | HC_vs_MS.Trt_B |
| DENND2D | 0.01717 | 0.32379 | 0.384 | 0.32 | 1 | HC_vs_MS.Trt_B |
| LAMC3 | 0.0173 | -0.7488 | 0.264 | 0.2 | 1 | HC_vs_MS.Trt_B |
| ANKRD17 | 0.01732 | 0.48415 | 0.251 | 0.19 | 1 | HC_vs_MS.Trt_B |
| PKD2 | 0.01736 | 0.87098 | 0.183 | 0.13 | 1 | HC_vs_MS.Trt_B |
| FAM185A | 0.01737 | 0.3292 | 0.103 | 0.06 | 1 | HC_vs_MS.Trt_B |
| CCP110 | 0.0174 | 0.98975 | 0.481 | 0.43 | 1 | HC_vs_MS.Trt_B |
| LAMP3 | 0.01742 | 0.92481 | 0.394 | 0.34 | 1 | HC_vs_MS.Trt_B |
| UBE3D | 0.01745 | -0.8177 | 0.528 | 0.44 | 1 | HC_vs_MS.Trt_B |
| SIAH3 | 0.01749 | 0.25142 | 0.125 | 0.08 | 1 | HC_vs_MS.Trt_B |

| | | | | | | |
|---|---|---|---|---|---|---|
| MRPL35 | 0.01753 | 0.25404 | 0.106 | 0.07 | 1 | HC_vs_MS.Trt_B |
| KDM5C | 0.01767 | 0.50305 | 0.37 | 0.3 | 1 | HC_vs_MS.Trt_B |
| GPR174 | 0.01772 | 0.4024 | 0.154 | 0.11 | 1 | HC_vs_MS.Trt_B |
| KDELR1 | 0.01772 | 0.65546 | 0.243 | 0.19 | 1 | HC_vs_MS.Trt_B |
| GBP1 | 0.01773 | 0.41375 | 0.229 | 0.17 | 1 | HC_vs_MS.Trt_B |
| RCC1 | 0.01779 | 0.27924 | 0.159 | 0.11 | 1 | HC_vs_MS.Trt_B |
| MRPL34 | 0.01783 | -0.332 | 0.329 | 0.26 | 1 | HC_vs_MS.Trt_B |
| DAZAP1 | 0.01807 | 0.4611 | 0.17 | 0.12 | 1 | HC_vs_MS.Trt_B |
| EXD2 | 0.01811 | 1.0966 | 0.181 | 0.13 | 1 | HC_vs_MS.Trt_B |
| PLEKHG5 | 0.01811 | 0.43643 | 0.165 | 0.12 | 1 | HC_vs_MS.Trt_B |
| ZNF230 | 0.01813 | 0.86275 | 0.568 | 0.51 | 1 | HC_vs_MS.Trt_B |
| ZNF678 | 0.0182 | 1.33179 | 0.172 | 0.12 | 1 | HC_vs_MS.Trt_B |
| ATF6 | 0.0182 | 0.32543 | 0.209 | 0.16 | 1 | HC_vs_MS.Trt_B |
| SPPL2B | 0.01822 | 0.66945 | 0.368 | 0.31 | 1 | HC_vs_MS.Trt_B |
| ENTPD6 | 0.01828 | 0.74036 | 0.12 | 0.08 | 1 | HC_vs_MS.Trt_B |
| PRR13 | 0.01829 | -0.3991 | 0.781 | 0.68 | 1 | HC_vs_MS.Trt_B |
| NTPCR | 0.01833 | -0.3542 | 0.391 | 0.33 | 1 | HC_vs_MS.Trt_B |
| DPY30 | 0.01847 | -0.282 | 0.377 | 0.3 | 1 | HC_vs_MS.Trt_B |
| FBXL22 | 0.01847 | 0.38762 | 0.103 | 0.06 | 1 | HC_vs_MS.Trt_B |
| PGF | 0.01852 | -0.261 | 0.238 | 0.19 | 1 | HC_vs_MS.Trt_B |
| BAG5 | 0.01853 | -0.2689 | 0.323 | 0.26 | 1 | HC_vs_MS.Trt_B |
| CDKN2AIPNI | 0.01854 | -0.4472 | 0.51 | 0.41 | 1 | HC_vs_MS.Trt_B |
| GPM6A | 0.01857 | 0.31873 | 0.28 | 0.22 | 1 | HC_vs_MS.Trt_B |
| RAB8B | 0.01866 | 0.7026 | 0.178 | 0.13 | 1 | HC_vs_MS.Trt_B |
| GLOD4 | 0.01867 | -0.4189 | 0.612 | 0.53 | 1 | HC_vs_MS.Trt_B |
| IRF7 | 0.01878 | 0.90689 | 0.112 | 0.07 | 1 | HC_vs_MS.Trt_B |
| URGCP | 0.01883 | -0.5243 | 0.459 | 0.39 | 1 | HC_vs_MS.Trt_B |
| FANCC | 0.01889 | 0.95818 | 0.264 | 0.21 | 1 | HC_vs_MS.Trt_B |
| U2AF1 | 0.01894 | -0.6025 | 0.535 | 0.44 | 1 | HC_vs_MS.Trt_B |
| TCTN2 | 0.01898 | -0.3522 | 0.73 | 0.62 | 1 | HC_vs_MS.Trt_B |
| SMAD3 | 0.01917 | 0.53481 | 0.139 | 0.09 | 1 | HC_vs_MS.Trt_B |
| JMJD1C | 0.01919 | 0.27 | 0.22 | 0.17 | 1 | HC_vs_MS.Trt_B |
| ZNF107 | 0.01919 | 0.37642 | 0.41 | 0.36 | 1 | HC_vs_MS.Trt_B |
| ZNF594 | 0.0192 | -1.0913 | 0.419 | 0.35 | 1 | HC_vs_MS.Trt_B |
| DUSP4 | 0.01924 | 0.28943 | 0.164 | 0.11 | 1 | HC_vs_MS.Trt_B |
| MTUS1 | 0.01927 | 0.55302 | 0.339 | 0.27 | 1 | HC_vs_MS.Trt_B |
| ZNF592 | 0.01933 | 0.49346 | 0.167 | 0.12 | 1 | HC_vs_MS.Trt_B |
| GLB1L2 | 0.0194 | -0.5694 | 0.341 | 0.27 | 1 | HC_vs_MS.Trt_B |
| ANKRD7 | 0.01945 | 0.4615 | 0.171 | 0.12 | 1 | HC_vs_MS.Trt_B |
| BIRC2 | 0.01948 | 0.69614 | 0.133 | 0.09 | 1 | HC_vs_MS.Trt_B |

| | | | | | | |
|---|---|---|---|---|---|---|
| LRRC47 | 0.01965 | 0.3297 | 0.281 | 0.22 | 1 | HC_vs_MS.Trt_B |
| SMARCC1 | 0.01981 | 0.38349 | 0.423 | 0.36 | 1 | HC_vs_MS.Trt_B |
| USP42 | 0.01984 | 0.25837 | 0.449 | 0.39 | 1 | HC_vs_MS.Trt_B |
| NLE1 | 0.01994 | 0.9654 | 0.214 | 0.17 | 1 | HC_vs_MS.Trt_B |
| ANKRD9 | 0.01998 | 0.46602 | 0.47 | 0.41 | 1 | HC_vs_MS.Trt_B |
| SMARCD1 | 0.01999 | 0.50359 | 0.132 | 0.09 | 1 | HC_vs_MS.Trt_B |
| OAZ3 | 0.01999 | -0.7537 | 0.38 | 0.31 | 1 | HC_vs_MS.Trt_B |
| TOP3B | 0.02013 | 0.5208 | 0.133 | 0.09 | 1 | HC_vs_MS.Trt_B |
| SUB1 | 0.02026 | -0.3501 | 0.551 | 0.45 | 1 | HC_vs_MS.Trt_B |
| RALGPS2 | 0.02039 | -0.581 | 0.354 | 0.3 | 1 | HC_vs_MS.Trt_B |
| CCDC97 | 0.02043 | 0.34714 | 0.126 | 0.08 | 1 | HC_vs_MS.Trt_B |
| KLHL8 | 0.02044 | -0.3258 | 0.599 | 0.51 | 1 | HC_vs_MS.Trt_B |
| SNX3 | 0.02052 | -0.381 | 0.158 | 0.11 | 1 | HC_vs_MS.Trt_B |
| UBB | 0.02071 | -0.4754 | 0.503 | 0.54 | 1 | HC_vs_MS.Trt_B |
| POMGNT1 | 0.02081 | 0.67543 | 0.662 | 0.61 | 1 | HC_vs_MS.Trt_B |
| GABRA6 | 0.02087 | -0.2509 | 0.067 | 0.11 | 1 | HC_vs_MS.Trt_B |
| OCIAD2 | 0.02107 | -0.6436 | 0.626 | 0.54 | 1 | HC_vs_MS.Trt_B |
| USP7 | 0.02113 | -0.3298 | 0.304 | 0.24 | 1 | HC_vs_MS.Trt_B |
| TMEM106A | 0.02117 | -0.4499 | 0.677 | 0.6 | 1 | HC_vs_MS.Trt_B |
| ZBTB38 | 0.02125 | -0.2524 | 0.535 | 0.47 | 1 | HC_vs_MS.Trt_B |
| ATP10A | 0.02129 | 0.38782 | 0.101 | 0.06 | 1 | HC_vs_MS.Trt_B |
| ZNF688 | 0.0213 | -0.9712 | 0.529 | 0.48 | 1 | HC_vs_MS.Trt_B |
| KLHDC10 | 0.02146 | -0.3624 | 0.387 | 0.31 | 1 | HC_vs_MS.Trt_B |
| DDX56 | 0.02148 | 0.27967 | 0.217 | 0.17 | 1 | HC_vs_MS.Trt_B |
| APIP | 0.02148 | 0.53991 | 0.455 | 0.41 | 1 | HC_vs_MS.Trt_B |
| AHR | 0.02157 | 0.53955 | 0.607 | 0.55 | 1 | HC_vs_MS.Trt_B |
| CCDC137 | 0.02161 | -0.7638 | 0.225 | 0.17 | 1 | HC_vs_MS.Trt_B |
| FAM210B | 0.02164 | 0.31198 | 0.31 | 0.25 | 1 | HC_vs_MS.Trt_B |
| CCDC22 | 0.02165 | 0.42253 | 0.129 | 0.09 | 1 | HC_vs_MS.Trt_B |
| ARL13B | 0.02167 | 0.39209 | 0.187 | 0.14 | 1 | HC_vs_MS.Trt_B |
| GDI1 | 0.02193 | 0.57604 | 0.274 | 0.22 | 1 | HC_vs_MS.Trt_B |
| CTPS1 | 0.0221 | -0.6146 | 0.609 | 0.55 | 1 | HC_vs_MS.Trt_B |
| ZNF610 | 0.02211 | 0.40846 | 0.362 | 0.31 | 1 | HC_vs_MS.Trt_B |
| GSE1 | 0.02214 | 0.612 | 0.338 | 0.28 | 1 | HC_vs_MS.Trt_B |
| XKR8 | 0.02216 | 1.19785 | 0.17 | 0.12 | 1 | HC_vs_MS.Trt_B |
| TOR1A | 0.0222 | -0.5602 | 0.68 | 0.58 | 1 | HC_vs_MS.Trt_B |
| ZNF497 | 0.0223 | 1.48243 | 0.268 | 0.21 | 1 | HC_vs_MS.Trt_B |
| TBC1D31 | 0.02246 | 0.26629 | 0.597 | 0.52 | 1 | HC_vs_MS.Trt_B |
| TMEM42 | 0.02251 | -0.4048 | 0.113 | 0.07 | 1 | HC_vs_MS.Trt_B |
| POLR1C | 0.02254 | 0.52426 | 0.412 | 0.36 | 1 | HC_vs_MS.Trt_B |

| | | | | | | |
|---|---|---|---|---|---|---|
| PODXL | 0.02264 | 0.29783 | 0.193 | 0.15 | 1 | HC_vs_MS.Trt_B |
| CXorf21 | 0.02265 | 1.01641 | 0.286 | 0.23 | 1 | HC_vs_MS.Trt_B |
| PTER | 0.02267 | 0.54723 | 0.152 | 0.11 | 1 | HC_vs_MS.Trt_B |
| DUSP18 | 0.02273 | 0.28659 | 0.586 | 0.54 | 1 | HC_vs_MS.Trt_B |
| PGGHG | 0.02278 | 0.6005 | 0.146 | 0.1 | 1 | HC_vs_MS.Trt_B |
| FAR2 | 0.02287 | 0.5624 | 0.186 | 0.14 | 1 | HC_vs_MS.Trt_B |
| KLRC1 | 0.02288 | 0.36725 | 0.291 | 0.23 | 1 | HC_vs_MS.Trt_B |
| CPNE2 | 0.02296 | -0.7463 | 0.372 | 0.31 | 1 | HC_vs_MS.Trt_B |
| CLIP4 | 0.02303 | 1.02568 | 0.125 | 0.09 | 1 | HC_vs_MS.Trt_B |
| FSBP | 0.02306 | 0.89219 | 0.314 | 0.26 | 1 | HC_vs_MS.Trt_B |
| FAM160B1 | 0.02308 | 0.37751 | 0.191 | 0.14 | 1 | HC_vs_MS.Trt_B |
| CXXC1 | 0.02316 | 0.50735 | 0.149 | 0.11 | 1 | HC_vs_MS.Trt_B |
| INTS4 | 0.02322 | 0.45064 | 0.341 | 0.28 | 1 | HC_vs_MS.Trt_B |
| TNFRSF10B | 0.0233 | 0.72603 | 0.404 | 0.34 | 1 | HC_vs_MS.Trt_B |
| ZNF280B | 0.02332 | -0.5215 | 0.116 | 0.08 | 1 | HC_vs_MS.Trt_B |
| PTBP2 | 0.02334 | -0.5098 | 0.562 | 0.47 | 1 | HC_vs_MS.Trt_B |
| GTPBP2 | 0.02346 | -0.34 | 0.516 | 0.46 | 1 | HC_vs_MS.Trt_B |
| RIPK1 | 0.02349 | 0.74581 | 0.167 | 0.12 | 1 | HC_vs_MS.Trt_B |
| C12orf43 | 0.02355 | -0.2963 | 0.843 | 0.77 | 1 | HC_vs_MS.Trt_B |
| DCP1A | 0.02357 | 0.42535 | 0.355 | 0.29 | 1 | HC_vs_MS.Trt_B |
| C8orf76 | 0.02376 | -0.3068 | 0.572 | 0.5 | 1 | HC_vs_MS.Trt_B |
| JKAMP | 0.0238 | -0.4733 | 0.745 | 0.68 | 1 | HC_vs_MS.Trt_B |
| GSK3A | 0.0239 | 0.75551 | 0.328 | 0.27 | 1 | HC_vs_MS.Trt_B |
| ANKRD42 | 0.02398 | 0.28078 | 0.384 | 0.33 | 1 | HC_vs_MS.Trt_B |
| HEMGN | 0.02401 | -0.3607 | 0.117 | 0.08 | 1 | HC_vs_MS.Trt_B |
| SEC24C | 0.02402 | 0.56926 | 0.22 | 0.17 | 1 | HC_vs_MS.Trt_B |
| RTTN | 0.02412 | 0.70557 | 0.239 | 0.19 | 1 | HC_vs_MS.Trt_B |
| STK24 | 0.02419 | -0.3693 | 0.254 | 0.2 | 1 | HC_vs_MS.Trt_B |
| METTL10 | 0.02423 | -0.2588 | 0.317 | 0.25 | 1 | HC_vs_MS.Trt_B |
| PI4KB | 0.02424 | 0.60327 | 0.167 | 0.12 | 1 | HC_vs_MS.Trt_B |
| TMEM185B | 0.02426 | 1.01378 | 0.332 | 0.27 | 1 | HC_vs_MS.Trt_B |
| TK2 | 0.02427 | 0.25718 | 0.126 | 0.09 | 1 | HC_vs_MS.Trt_B |
| ECT2L | 0.02438 | 0.29119 | 0.277 | 0.22 | 1 | HC_vs_MS.Trt_B |
| COMMD10 | 0.02438 | -0.5509 | 0.275 | 0.22 | 1 | HC_vs_MS.Trt_B |
| MRPS18B | 0.02441 | -0.5023 | 0.661 | 0.58 | 1 | HC_vs_MS.Trt_B |
| TMEM248 | 0.02441 | 0.37447 | 0.168 | 0.12 | 1 | HC_vs_MS.Trt_B |
| ATP6V0C | 0.02445 | 0.35483 | 0.213 | 0.16 | 1 | HC_vs_MS.Trt_B |
| RAB11A | 0.02452 | -0.4974 | 0.626 | 0.56 | 1 | HC_vs_MS.Trt_B |
| NOLC1 | 0.02456 | 0.37581 | 0.103 | 0.07 | 1 | HC_vs_MS.Trt_B |
| NMI | 0.0246 | -0.5021 | 0.274 | 0.22 | 1 | HC_vs_MS.Trt_B |

| | | | | | | |
|---|---|---|---|---|---|---|
| ZNF33B | 0.02462 | 0.37669 | 0.497 | 0.4 | 1 | HC_vs_MS.Trt_B |
| LZTR1 | 0.02467 | 1.13482 | 0.248 | 0.2 | 1 | HC_vs_MS.Trt_B |
| PHYH | 0.02467 | 0.34679 | 0.165 | 0.12 | 1 | HC_vs_MS.Trt_B |
| BRMS1 | 0.0247 | 0.49925 | 0.141 | 0.1 | 1 | HC_vs_MS.Trt_B |
| TMA7 | 0.02473 | -0.7578 | 0.516 | 0.53 | 1 | HC_vs_MS.Trt_B |
| RADIL | 0.02477 | 0.33252 | 0.281 | 0.23 | 1 | HC_vs_MS.Trt_B |
| SSPO | 0.02497 | 0.97647 | 0.123 | 0.08 | 1 | HC_vs_MS.Trt_B |
| ZDHHC7 | 0.02501 | 0.72999 | 0.17 | 0.12 | 1 | HC_vs_MS.Trt_B |
| FRMD8 | 0.02503 | 0.31984 | 0.565 | 0.5 | 1 | HC_vs_MS.Trt_B |
| POTEI | 0.02504 | 0.3953 | 0.19 | 0.14 | 1 | HC_vs_MS.Trt_B |
| APEX2 | 0.02514 | 0.43452 | 0.307 | 0.25 | 1 | HC_vs_MS.Trt_B |
| MYO9B | 0.02517 | 0.46657 | 0.22 | 0.17 | 1 | HC_vs_MS.Trt_B |
| NUP54 | 0.02523 | 0.39952 | 0.18 | 0.13 | 1 | HC_vs_MS.Trt_B |
| VPS41 | 0.02527 | 0.59557 | 0.452 | 0.41 | 1 | HC_vs_MS.Trt_B |
| DNAH6 | 0.02532 | 0.49916 | 0.168 | 0.12 | 1 | HC_vs_MS.Trt_B |
| RASGRF2 | 0.02534 | 0.40889 | 0.165 | 0.12 | 1 | HC_vs_MS.Trt_B |
| USF1 | 0.02536 | 0.49659 | 0.146 | 0.1 | 1 | HC_vs_MS.Trt_B |
| YWHAH | 0.02544 | -0.279 | 0.125 | 0.08 | 1 | HC_vs_MS.Trt_B |
| SELENBP1 | 0.02545 | 0.32789 | 0.275 | 0.22 | 1 | HC_vs_MS.Trt_B |
| ING1 | 0.02548 | -0.6119 | 0.543 | 0.5 | 1 | HC_vs_MS.Trt_B |
| ESD | 0.02552 | 0.5191 | 0.114 | 0.08 | 1 | HC_vs_MS.Trt_B |
| AMMECR1 | 0.02552 | 0.62654 | 0.132 | 0.09 | 1 | HC_vs_MS.Trt_B |
| ARFGAP1 | 0.02562 | 0.47301 | 0.133 | 0.09 | 1 | HC_vs_MS.Trt_B |
| SEMA4B | 0.02566 | 0.27665 | 0.397 | 0.34 | 1 | HC_vs_MS.Trt_B |
| WDR70 | 0.02571 | 0.36792 | 0.264 | 0.21 | 1 | HC_vs_MS.Trt_B |
| SNTG2 | 0.02573 | 0.25057 | 0.219 | 0.17 | 1 | HC_vs_MS.Trt_B |
| SOCS7 | 0.02575 | 0.641 | 0.349 | 0.29 | 1 | HC_vs_MS.Trt_B |
| STX3 | 0.02577 | 0.35127 | 0.377 | 0.34 | 1 | HC_vs_MS.Trt_B |
| PSMB4 | 0.02592 | -0.4985 | 0.278 | 0.34 | 1 | HC_vs_MS.Trt_B |
| STON2 | 0.02595 | 0.45461 | 0.258 | 0.21 | 1 | HC_vs_MS.Trt_B |
| TXNL1 | 0.02601 | 0.36301 | 0.629 | 0.56 | 1 | HC_vs_MS.Trt_B |
| CDK7 | 0.02606 | 0.71245 | 0.135 | 0.09 | 1 | HC_vs_MS.Trt_B |
| C8orf59 | 0.02607 | -0.2916 | 0.348 | 0.28 | 1 | HC_vs_MS.Trt_B |
| CLDN15 | 0.02608 | 0.45641 | 0.394 | 0.33 | 1 | HC_vs_MS.Trt_B |
| ESF1 | 0.02609 | 0.2941 | 0.136 | 0.09 | 1 | HC_vs_MS.Trt_B |
| TRIP12 | 0.0262 | -0.3726 | 0.458 | 0.38 | 1 | HC_vs_MS.Trt_B |
| RELA | 0.02624 | 0.25746 | 0.154 | 0.11 | 1 | HC_vs_MS.Trt_B |
| RALGPS1 | 0.02629 | -0.5042 | 0.38 | 0.3 | 1 | HC_vs_MS.Trt_B |
| ZSCAN12 | 0.02635 | 0.31902 | 0.42 | 0.36 | 1 | HC_vs_MS.Trt_B |
| RP4-635E18 | 0.02641 | -0.8235 | 0.212 | 0.26 | 1 | HC_vs_MS.Trt_B |

| | | | | | | |
|---|---|---|---|---|---|---|
| NR1D1 | 0.02641 | 0.56473 | 0.11 | 0.07 | 1 | HC_vs_MS.Trt_B |
| SMC4 | 0.02644 | -0.61 | 0.449 | 0.36 | 1 | HC_vs_MS.Trt_B |
| NDUFS8 | 0.02667 | -0.2554 | 0.767 | 0.74 | 1 | HC_vs_MS.Trt_B |
| ARMC6 | 0.02667 | -0.4046 | 0.303 | 0.24 | 1 | HC_vs_MS.Trt_B |
| SBF1 | 0.02668 | 0.3404 | 0.2 | 0.15 | 1 | HC_vs_MS.Trt_B |
| JAKMIP2 | 0.02668 | 0.39469 | 0.352 | 0.31 | 1 | HC_vs_MS.Trt_B |
| FOXRED1 | 0.02669 | 0.5495 | 0.507 | 0.43 | 1 | HC_vs_MS.Trt_B |
| SLC12A4 | 0.02673 | 1.30485 | 0.19 | 0.14 | 1 | HC_vs_MS.Trt_B |
| TFRC | 0.02674 | -0.2899 | 0.574 | 0.49 | 1 | HC_vs_MS.Trt_B |
| BRD3 | 0.02681 | -0.2791 | 0.168 | 0.12 | 1 | HC_vs_MS.Trt_B |
| DTNB | 0.0269 | 0.61574 | 0.154 | 0.11 | 1 | HC_vs_MS.Trt_B |
| TRERF1 | 0.02692 | 0.28606 | 0.183 | 0.14 | 1 | HC_vs_MS.Trt_B |
| DIP2B | 0.02698 | 0.33294 | 0.359 | 0.31 | 1 | HC_vs_MS.Trt_B |
| RLF | 0.02743 | 0.27804 | 0.168 | 0.12 | 1 | HC_vs_MS.Trt_B |
| TRIM58 | 0.02746 | 0.3712 | 0.564 | 0.51 | 1 | HC_vs_MS.Trt_B |
| UNC13A | 0.02753 | 0.49648 | 0.261 | 0.21 | 1 | HC_vs_MS.Trt_B |
| LPAR2 | 0.02761 | 0.65836 | 0.11 | 0.07 | 1 | HC_vs_MS.Trt_B |
| MAPK8 | 0.02775 | -0.3479 | 0.393 | 0.33 | 1 | HC_vs_MS.Trt_B |
| SLC6A16 | 0.02775 | 0.71392 | 0.243 | 0.19 | 1 | HC_vs_MS.Trt_B |
| ZNF527 | 0.02785 | 0.25446 | 0.933 | 0.87 | 1 | HC_vs_MS.Trt_B |
| LIG3 | 0.02789 | 1.13738 | 0.33 | 0.28 | 1 | HC_vs_MS.Trt_B |
| ZNF664 | 0.02791 | -0.2881 | 0.233 | 0.18 | 1 | HC_vs_MS.Trt_B |
| EML3 | 0.02792 | 0.32084 | 0.216 | 0.16 | 1 | HC_vs_MS.Trt_B |
| ZNF134 | 0.02793 | -0.3984 | 0.174 | 0.13 | 1 | HC_vs_MS.Trt_B |
| COX5B | 0.02796 | -0.6009 | 0.759 | 0.68 | 1 | HC_vs_MS.Trt_B |
| PRKG2 | 0.0281 | 0.26922 | 0.391 | 0.31 | 1 | HC_vs_MS.Trt_B |
| DNA2 | 0.02816 | 0.29385 | 0.217 | 0.17 | 1 | HC_vs_MS.Trt_B |
| HACL1 | 0.02829 | 0.66844 | 0.125 | 0.09 | 1 | HC_vs_MS.Trt_B |
| IGF1R | 0.02831 | 0.50661 | 0.354 | 0.3 | 1 | HC_vs_MS.Trt_B |
| RAB17 | 0.02833 | 0.25411 | 0.23 | 0.18 | 1 | HC_vs_MS.Trt_B |
| DUSP10 | 0.02856 | 0.67168 | 0.141 | 0.1 | 1 | HC_vs_MS.Trt_B |
| PSIP1 | 0.0286 | -0.2532 | 0.758 | 0.69 | 1 | HC_vs_MS.Trt_B |
| RP4-583P15 | 0.02865 | -0.2562 | 0.283 | 0.22 | 1 | HC_vs_MS.Trt_B |
| LRRC8B | 0.02888 | 0.36803 | 0.409 | 0.35 | 1 | HC_vs_MS.Trt_B |
| MCMDC2 | 0.02904 | -0.4727 | 0.419 | 0.36 | 1 | HC_vs_MS.Trt_B |
| PREB | 0.02915 | 0.78699 | 0.116 | 0.08 | 1 | HC_vs_MS.Trt_B |
| ZNF221 | 0.0292 | 0.36516 | 0.245 | 0.19 | 1 | HC_vs_MS.Trt_B |
| RWDD2A | 0.02922 | 0.44723 | 0.416 | 0.35 | 1 | HC_vs_MS.Trt_B |
| USP36 | 0.02933 | 0.64322 | 0.239 | 0.19 | 1 | HC_vs_MS.Trt_B |
| B4GALT3 | 0.02943 | 0.46502 | 0.157 | 0.11 | 1 | HC_vs_MS.Trt_B |

| | | | | | | |
|---|---|---|---|---|---|---|
| NCAPG2 | 0.02944 | -2.4333 | 0.159 | 0.2 | 1 | HC_vs_MS.Trt_B |
| KNSTRN | 0.02947 | 0.37708 | 0.475 | 0.41 | 1 | HC_vs_MS.Trt_B |
| PIK3CG | 0.0295 | 0.47057 | 0.238 | 0.18 | 1 | HC_vs_MS.Trt_B |
| TONSL | 0.02962 | 0.33235 | 0.597 | 0.54 | 1 | HC_vs_MS.Trt_B |
| DUS3L | 0.02991 | 0.25109 | 0.123 | 0.08 | 1 | HC_vs_MS.Trt_B |
| SETDB1 | 0.02993 | 0.33983 | 0.517 | 0.47 | 1 | HC_vs_MS.Trt_B |
| ZFHX3 | 0.02994 | 0.85062 | 0.143 | 0.1 | 1 | HC_vs_MS.Trt_B |
| L3HYPDH | 0.02996 | -0.3225 | 0.194 | 0.14 | 1 | HC_vs_MS.Trt_B |
| NUMBL | 0.02999 | 0.43932 | 0.51 | 0.47 | 1 | HC_vs_MS.Trt_B |
| FBXO6 | 0.03027 | 0.42894 | 0.617 | 0.55 | 1 | HC_vs_MS.Trt_B |
| RAP1B | 0.0304 | -0.6194 | 0.993 | 0.96 | 1 | HC_vs_MS.Trt_B |
| CALD1 | 0.03046 | 0.27115 | 0.191 | 0.14 | 1 | HC_vs_MS.Trt_B |
| MMRN2 | 0.03064 | -0.8282 | 0.152 | 0.11 | 1 | HC_vs_MS.Trt_B |
| EXD3 | 0.03096 | 0.31487 | 0.43 | 0.37 | 1 | HC_vs_MS.Trt_B |
| GMFG | 0.03132 | -0.3482 | 0.772 | 0.63 | 1 | HC_vs_MS.Trt_B |
| EPHA4 | 0.03151 | -0.6652 | 0.184 | 0.13 | 1 | HC_vs_MS.Trt_B |
| ZNF428 | 0.03164 | 0.25067 | 0.157 | 0.11 | 1 | HC_vs_MS.Trt_B |
| DUSP3 | 0.03165 | 0.43088 | 0.732 | 0.72 | 1 | HC_vs_MS.Trt_B |
| FASTKD5 | 0.03181 | -0.5928 | 0.077 | 0.11 | 1 | HC_vs_MS.Trt_B |
| HABP4 | 0.03184 | -0.5442 | 0.104 | 0.07 | 1 | HC_vs_MS.Trt_B |
| ZNF71 | 0.03207 | 0.47206 | 0.204 | 0.16 | 1 | HC_vs_MS.Trt_B |
| SLC9A8 | 0.03214 | 0.41866 | 0.175 | 0.13 | 1 | HC_vs_MS.Trt_B |
| SSFA2 | 0.03219 | -0.2895 | 0.119 | 0.08 | 1 | HC_vs_MS.Trt_B |
| FAM173B | 0.0322 | -0.6204 | 0.172 | 0.13 | 1 | HC_vs_MS.Trt_B |
| LRRC1 | 0.0323 | 0.52219 | 0.258 | 0.21 | 1 | HC_vs_MS.Trt_B |
| SORD | 0.0323 | 0.73014 | 0.142 | 0.1 | 1 | HC_vs_MS.Trt_B |
| TXNDC12 | 0.03246 | 0.47457 | 0.152 | 0.11 | 1 | HC_vs_MS.Trt_B |
| ZNF700 | 0.03254 | 0.78726 | 0.148 | 0.11 | 1 | HC_vs_MS.Trt_B |
| ZNF749 | 0.03264 | 0.96865 | 0.38 | 0.34 | 1 | HC_vs_MS.Trt_B |
| PRKCZ | 0.03268 | 0.88696 | 0.239 | 0.19 | 1 | HC_vs_MS.Trt_B |
| ZNF736 | 0.03285 | 0.71729 | 0.47 | 0.42 | 1 | HC_vs_MS.Trt_B |
| ZNF432 | 0.03291 | 0.83517 | 0.552 | 0.48 | 1 | HC_vs_MS.Trt_B |
| ZNF721 | 0.03292 | 0.59931 | 0.296 | 0.25 | 1 | HC_vs_MS.Trt_B |
| ZMYM2 | 0.03294 | -0.3575 | 0.351 | 0.29 | 1 | HC_vs_MS.Trt_B |
| PPM1L | 0.03294 | 0.98791 | 0.236 | 0.19 | 1 | HC_vs_MS.Trt_B |
| FAM210A | 0.03304 | -0.361 | 0.257 | 0.21 | 1 | HC_vs_MS.Trt_B |
| KCTD10 | 0.03322 | 0.45777 | 0.371 | 0.31 | 1 | HC_vs_MS.Trt_B |
| PRRG1 | 0.03323 | 0.38765 | 0.174 | 0.13 | 1 | HC_vs_MS.Trt_B |
| MRPL49 | 0.03326 | -0.4199 | 0.455 | 0.39 | 1 | HC_vs_MS.Trt_B |
| AKR1B1 | 0.03336 | -0.3627 | 0.733 | 0.67 | 1 | HC_vs_MS.Trt_B |

| | | | | | | |
|---|---|---|---|---|---|---|
| ZNF354A | 0.03344 | 0.36918 | 0.18 | 0.14 | 1 | HC_vs_MS.Trt_B |
| CHST11 | 0.03349 | 0.29976 | 0.152 | 0.11 | 1 | HC_vs_MS.Trt_B |
| FAM214A | 0.03352 | -0.3581 | 0.193 | 0.14 | 1 | HC_vs_MS.Trt_B |
| WRN | 0.03363 | 0.62906 | 0.432 | 0.38 | 1 | HC_vs_MS.Trt_B |
| LPAR6 | 0.03363 | -0.2583 | 0.396 | 0.34 | 1 | HC_vs_MS.Trt_B |
| UBA5 | 0.03365 | -0.4735 | 0.142 | 0.1 | 1 | HC_vs_MS.Trt_B |
| EEF1E1 | 0.03384 | -0.3478 | 0.52 | 0.47 | 1 | HC_vs_MS.Trt_B |
| DNPEP | 0.03387 | 0.56374 | 0.448 | 0.4 | 1 | HC_vs_MS.Trt_B |
| CMC1 | 0.03399 | -0.5128 | 0.161 | 0.12 | 1 | HC_vs_MS.Trt_B |
| MAT2A | 0.03409 | 0.58647 | 0.126 | 0.09 | 1 | HC_vs_MS.Trt_B |
| ZNF552 | 0.03421 | -0.6077 | 0.399 | 0.33 | 1 | HC_vs_MS.Trt_B |
| PLD6 | 0.03442 | 0.45716 | 0.435 | 0.38 | 1 | HC_vs_MS.Trt_B |
| SIGMAR1 | 0.03448 | 0.36281 | 0.184 | 0.14 | 1 | HC_vs_MS.Trt_B |
| LRP1 | 0.03466 | 0.33152 | 0.216 | 0.17 | 1 | HC_vs_MS.Trt_B |
| PAM16 | 0.03482 | 0.59656 | 0.181 | 0.14 | 1 | HC_vs_MS.Trt_B |
| USP30 | 0.03482 | -0.4813 | 0.267 | 0.21 | 1 | HC_vs_MS.Trt_B |
| DIABLO | 0.03485 | -0.3481 | 0.674 | 0.59 | 1 | HC_vs_MS.Trt_B |
| HSPH1 | 0.03494 | 0.57719 | 0.165 | 0.12 | 1 | HC_vs_MS.Trt_B |
| ZNF517 | 0.035 | 0.56923 | 0.375 | 0.33 | 1 | HC_vs_MS.Trt_B |
| ANO8 | 0.03513 | 0.46771 | 0.133 | 0.09 | 1 | HC_vs_MS.Trt_B |
| CCDC138 | 0.03514 | -0.5912 | 0.157 | 0.11 | 1 | HC_vs_MS.Trt_B |
| STXBP2 | 0.03522 | 0.36115 | 0.252 | 0.21 | 1 | HC_vs_MS.Trt_B |
| EFR3A | 0.03528 | 0.78519 | 0.135 | 0.1 | 1 | HC_vs_MS.Trt_B |
| DDX60 | 0.03537 | 0.33893 | 0.123 | 0.09 | 1 | HC_vs_MS.Trt_B |
| ZNF740 | 0.0354 | 0.36028 | 0.142 | 0.1 | 1 | HC_vs_MS.Trt_B |
| AKAP2 | 0.03555 | 0.32875 | 0.254 | 0.2 | 1 | HC_vs_MS.Trt_B |
| FAM149B1 | 0.03558 | -0.7496 | 0.178 | 0.13 | 1 | HC_vs_MS.Trt_B |
| RP11-327P2 | 0.03565 | 0.40194 | 0.116 | 0.08 | 1 | HC_vs_MS.Trt_B |
| KIF27 | 0.03568 | -0.6177 | 0.303 | 0.24 | 1 | HC_vs_MS.Trt_B |
| RRN3 | 0.03599 | 0.4858 | 0.245 | 0.2 | 1 | HC_vs_MS.Trt_B |
| PGS1 | 0.0361 | 0.34112 | 0.57 | 0.51 | 1 | HC_vs_MS.Trt_B |
| STARD5 | 0.03611 | 0.34799 | 0.114 | 0.08 | 1 | HC_vs_MS.Trt_B |
| DLEC1 | 0.03615 | 0.43166 | 0.429 | 0.38 | 1 | HC_vs_MS.Trt_B |
| TAF4 | 0.03629 | 0.2633 | 0.123 | 0.08 | 1 | HC_vs_MS.Trt_B |
| SCYL2 | 0.03629 | 0.80789 | 0.11 | 0.07 | 1 | HC_vs_MS.Trt_B |
| CASP4 | 0.0365 | -0.5199 | 0.336 | 0.26 | 1 | HC_vs_MS.Trt_B |
| ANP32A | 0.03653 | 0.33395 | 0.274 | 0.23 | 1 | HC_vs_MS.Trt_B |
| TESMIN | 0.03672 | 0.30982 | 0.629 | 0.55 | 1 | HC_vs_MS.Trt_B |
| STX10 | 0.03685 | 0.58188 | 0.12 | 0.08 | 1 | HC_vs_MS.Trt_B |
| NDUFAF1 | 0.03701 | -0.2512 | 0.365 | 0.31 | 1 | HC_vs_MS.Trt_B |

| | | | | | | |
|---|---|---|---|---|---|---|
| SUPT16H | 0.03705 | -0.2714 | 0.678 | 0.63 | 1 | HC_vs_MS.Trt_B |
| EDA | 0.03706 | 0.62411 | 0.33 | 0.27 | 1 | HC_vs_MS.Trt_B |
| FO538757.1 | 0.03717 | 0.54259 | 0.162 | 0.12 | 1 | HC_vs_MS.Trt_B |
| TAPBPL | 0.03725 | 0.26228 | 0.2 | 0.16 | 1 | HC_vs_MS.Trt_B |
| PCGF3 | 0.03729 | 0.42739 | 0.229 | 0.18 | 1 | HC_vs_MS.Trt_B |
| KCNMB3 | 0.03735 | -0.6132 | 0.367 | 0.3 | 1 | HC_vs_MS.Trt_B |
| MYOZ3 | 0.03744 | 0.5592 | 0.243 | 0.2 | 1 | HC_vs_MS.Trt_B |
| EPB41L2 | 0.03753 | 0.53034 | 0.112 | 0.07 | 1 | HC_vs_MS.Trt_B |
| RGS1 | 0.03769 | 0.55768 | 0.129 | 0.09 | 1 | HC_vs_MS.Trt_B |
| ZNF812P | 0.03782 | 0.45242 | 0.109 | 0.07 | 1 | HC_vs_MS.Trt_B |
| MGLL | 0.03803 | 0.30693 | 0.319 | 0.27 | 1 | HC_vs_MS.Trt_B |
| BANK1 | 0.03858 | -0.8276 | 0.168 | 0.12 | 1 | HC_vs_MS.Trt_B |
| ARFIP2 | 0.03874 | 0.6607 | 0.432 | 0.37 | 1 | HC_vs_MS.Trt_B |
| GMDS | 0.03883 | 0.28293 | 0.112 | 0.08 | 1 | HC_vs_MS.Trt_B |
| ST3GAL5 | 0.03932 | -0.8643 | 0.284 | 0.23 | 1 | HC_vs_MS.Trt_B |
| CAMLG | 0.03944 | -0.4838 | 0.442 | 0.35 | 1 | HC_vs_MS.Trt_B |
| PTPN23 | 0.03959 | 0.43192 | 0.142 | 0.1 | 1 | HC_vs_MS.Trt_B |
| POU2F1 | 0.03995 | 0.40323 | 0.419 | 0.35 | 1 | HC_vs_MS.Trt_B |
| ZBTB4 | 0.03999 | 0.57077 | 0.126 | 0.09 | 1 | HC_vs_MS.Trt_B |
| EIF4E2 | 0.03999 | -0.5271 | 0.322 | 0.26 | 1 | HC_vs_MS.Trt_B |
| PNPLA1 | 0.04029 | 0.37374 | 0.309 | 0.25 | 1 | HC_vs_MS.Trt_B |
| PSMC4 | 0.04036 | -0.4518 | 0.457 | 0.39 | 1 | HC_vs_MS.Trt_B |
| C1orf159 | 0.04059 | 0.57172 | 0.114 | 0.08 | 1 | HC_vs_MS.Trt_B |
| MED1 | 0.0408 | 0.86476 | 0.307 | 0.27 | 1 | HC_vs_MS.Trt_B |
| SMAD5 | 0.04083 | 0.3954 | 0.441 | 0.37 | 1 | HC_vs_MS.Trt_B |
| RIPK2 | 0.04091 | 1.30303 | 0.161 | 0.12 | 1 | HC_vs_MS.Trt_B |
| NKIRAS1 | 0.04092 | 1.20775 | 0.239 | 0.2 | 1 | HC_vs_MS.Trt_B |
| ZFP2 | 0.04109 | 1.0484 | 0.19 | 0.14 | 1 | HC_vs_MS.Trt_B |
| SH2B3 | 0.04109 | 0.44301 | 0.142 | 0.1 | 1 | HC_vs_MS.Trt_B |
| C21orf2 | 0.04147 | 0.31664 | 0.123 | 0.09 | 1 | HC_vs_MS.Trt_B |
| DPP8 | 0.04148 | 0.28514 | 0.357 | 0.29 | 1 | HC_vs_MS.Trt_B |
| RNF138 | 0.04199 | 0.30963 | 0.145 | 0.11 | 1 | HC_vs_MS.Trt_B |
| EML5 | 0.0421 | 1.23494 | 0.223 | 0.17 | 1 | HC_vs_MS.Trt_B |
| LILRB3 | 0.04211 | 0.77811 | 0.123 | 0.09 | 1 | HC_vs_MS.Trt_B |
| IDH1 | 0.04211 | 0.48252 | 0.191 | 0.15 | 1 | HC_vs_MS.Trt_B |
| DSTN | 0.04212 | -0.7894 | 0.494 | 0.43 | 1 | HC_vs_MS.Trt_B |
| TNFRSF13C | 0.04225 | -0.2843 | 0.62 | 0.54 | 1 | HC_vs_MS.Trt_B |
| HIGD2A | 0.04249 | -0.2542 | 0.239 | 0.19 | 1 | HC_vs_MS.Trt_B |
| SH2D3C | 0.0425 | 0.45488 | 0.2 | 0.16 | 1 | HC_vs_MS.Trt_B |
| RBM10 | 0.04255 | 0.39261 | 0.175 | 0.13 | 1 | HC_vs_MS.Trt_B |

| | | | | | | |
|---|---|---|---|---|---|---|
| SPRYD3 | 0.04261 | 0.41661 | 0.136 | 0.1 | 1 | HC_vs_MS.Trt_B |
| DFNA5 | 0.04263 | 0.38125 | 0.12 | 0.08 | 1 | HC_vs_MS.Trt_B |
| TIRAP | 0.04268 | 1.17605 | 0.401 | 0.36 | 1 | HC_vs_MS.Trt_B |
| C3orf49 | 0.04277 | -0.3331 | 0.31 | 0.26 | 1 | HC_vs_MS.Trt_B |
| CCDC152 | 0.04281 | 0.73813 | 0.197 | 0.16 | 1 | HC_vs_MS.Trt_B |
| NCDN | 0.04291 | 0.46882 | 0.216 | 0.17 | 1 | HC_vs_MS.Trt_B |
| ABC13-4748 | 0.04296 | 0.30299 | 0.117 | 0.08 | 1 | HC_vs_MS.Trt_B |
| THEM4 | 0.04302 | -0.3175 | 0.796 | 0.72 | 1 | HC_vs_MS.Trt_B |
| EIF2S3 | 0.04304 | -0.3223 | 0.954 | 0.94 | 1 | HC_vs_MS.Trt_B |
| PTBP1 | 0.04307 | 0.42863 | 0.209 | 0.16 | 1 | HC_vs_MS.Trt_B |
| TMEM35B | 0.04311 | -0.307 | 0.152 | 0.11 | 1 | HC_vs_MS.Trt_B |
| DGCR8 | 0.04327 | 1.24055 | 0.116 | 0.08 | 1 | HC_vs_MS.Trt_B |
| USP31 | 0.04328 | -0.3509 | 0.135 | 0.09 | 1 | HC_vs_MS.Trt_B |
| ZNF292 | 0.04343 | -0.3509 | 0.814 | 0.73 | 1 | HC_vs_MS.Trt_B |
| MRE11A | 0.04356 | -0.6934 | 0.675 | 0.6 | 1 | HC_vs_MS.Trt_B |
| PRSS53 | 0.04363 | 0.33844 | 0.162 | 0.12 | 1 | HC_vs_MS.Trt_B |
| INPP5F | 0.04393 | 0.54646 | 0.307 | 0.25 | 1 | HC_vs_MS.Trt_B |
| NAF1 | 0.04404 | 0.40253 | 0.409 | 0.33 | 1 | HC_vs_MS.Trt_B |
| MBD6 | 0.04406 | 0.39327 | 0.199 | 0.15 | 1 | HC_vs_MS.Trt_B |
| MRPL57 | 0.04411 | -0.5535 | 0.672 | 0.63 | 1 | HC_vs_MS.Trt_B |
| RAB14 | 0.04417 | 0.44897 | 0.201 | 0.16 | 1 | HC_vs_MS.Trt_B |
| RASSF3 | 0.04419 | -0.6053 | 0.642 | 0.65 | 1 | HC_vs_MS.Trt_B |
| MIB2 | 0.04436 | 1.14817 | 0.177 | 0.14 | 1 | HC_vs_MS.Trt_B |
| TTC3 | 0.04437 | -0.2875 | 0.7 | 0.65 | 1 | HC_vs_MS.Trt_B |
| BRD7 | 0.04446 | -0.9518 | 0.522 | 0.45 | 1 | HC_vs_MS.Trt_B |
| KHDRBS1 | 0.04462 | 0.28673 | 0.183 | 0.14 | 1 | HC_vs_MS.Trt_B |
| RGP1 | 0.04467 | 0.58326 | 0.123 | 0.09 | 1 | HC_vs_MS.Trt_B |
| TRIM61 | 0.04471 | 0.39601 | 0.214 | 0.17 | 1 | HC_vs_MS.Trt_B |
| PDCD4 | 0.04494 | -0.536 | 0.6 | 0.5 | 1 | HC_vs_MS.Trt_B |
| NT5DC1 | 0.04546 | -0.3705 | 0.194 | 0.15 | 1 | HC_vs_MS.Trt_B |
| SEC61A1 | 0.04558 | 0.26418 | 0.299 | 0.25 | 1 | HC_vs_MS.Trt_B |
| PITRM1 | 0.04561 | 0.27622 | 0.172 | 0.13 | 1 | HC_vs_MS.Trt_B |
| FGF23 | 0.04583 | 1.65287 | 0.828 | 0.79 | 1 | HC_vs_MS.Trt_B |
| ARHGAP21 | 0.04604 | 0.73273 | 0.296 | 0.25 | 1 | HC_vs_MS.Trt_B |
| PTOV1 | 0.04612 | 0.49372 | 0.133 | 0.1 | 1 | HC_vs_MS.Trt_B |
| IKBKAP | 0.04614 | 0.50274 | 0.216 | 0.17 | 1 | HC_vs_MS.Trt_B |
| ITCH | 0.04651 | -0.2705 | 0.47 | 0.41 | 1 | HC_vs_MS.Trt_B |
| BDH2 | 0.04672 | 0.42907 | 0.183 | 0.14 | 1 | HC_vs_MS.Trt_B |
| GOLGB1 | 0.04672 | -0.4689 | 0.509 | 0.44 | 1 | HC_vs_MS.Trt_B |
| UBASH3B | 0.04691 | -0.2586 | 0.38 | 0.33 | 1 | HC_vs_MS.Trt_B |

| | | | | | | |
|---|---|---|---|---|---|---|
| DENND6B | 0.04692 | 0.28166 | 0.259 | 0.22 | 1 | HC_vs_MS.Trt_B |
| UTP18 | 0.04712 | 0.45196 | 0.157 | 0.12 | 1 | HC_vs_MS.Trt_B |
| ZDHHC2 | 0.04724 | -0.5517 | 0.106 | 0.14 | 1 | HC_vs_MS.Trt_B |
| SOS1 | 0.04738 | 0.64713 | 0.157 | 0.12 | 1 | HC_vs_MS.Trt_B |
| DCUN1D5 | 0.04765 | 0.57306 | 0.425 | 0.37 | 1 | HC_vs_MS.Trt_B |
| CMTR1 | 0.04778 | 0.8117 | 0.159 | 0.12 | 1 | HC_vs_MS.Trt_B |
| ITGB5 | 0.04787 | 0.33158 | 0.328 | 0.28 | 1 | HC_vs_MS.Trt_B |
| ZNF267 | 0.04815 | -0.3242 | 0.117 | 0.08 | 1 | HC_vs_MS.Trt_B |
| YPEL2 | 0.04819 | 0.28728 | 0.186 | 0.14 | 1 | HC_vs_MS.Trt_B |
| SBNO2 | 0.04827 | 0.56789 | 0.175 | 0.13 | 1 | HC_vs_MS.Trt_B |
| IL15 | 0.04886 | 0.26806 | 0.357 | 0.31 | 1 | HC_vs_MS.Trt_B |
| ZNF669 | 0.04896 | 0.77352 | 0.514 | 0.47 | 1 | HC_vs_MS.Trt_B |
| EMC8 | 0.04904 | 0.92553 | 0.132 | 0.1 | 1 | HC_vs_MS.Trt_B |
| MFSD4A | 0.0491 | -0.8292 | 0.475 | 0.42 | 1 | HC_vs_MS.Trt_B |
| HDAC10 | 0.04917 | 0.62339 | 0.123 | 0.09 | 1 | HC_vs_MS.Trt_B |
| ANXA2R | 0.04944 | -0.3765 | 0.178 | 0.13 | 1 | HC_vs_MS.Trt_B |
| HPCAL1 | 0.04958 | -0.4192 | 0.357 | 0.31 | 1 | HC_vs_MS.Trt_B |
| YDJC | 0.04973 | -0.3802 | 0.116 | 0.08 | 1 | HC_vs_MS.Trt_B |
| KDM3B | 0.04982 | 0.45549 | 0.381 | 0.32 | 1 | HC_vs_MS.Trt_B |
| FBXO21 | 0.05003 | 0.46214 | 0.152 | 0.11 | 1 | HC_vs_MS.Trt_B |
| RIT1 | 0.05009 | -0.5935 | 0.388 | 0.33 | 1 | HC_vs_MS.Trt_B |
| CIPC | 0.05011 | -0.5803 | 0.37 | 0.32 | 1 | HC_vs_MS.Trt_B |
| APPL1 | 0.05014 | -0.29 | 0.414 | 0.36 | 1 | HC_vs_MS.Trt_B |
| NEPRO | 0.05015 | 0.47156 | 0.194 | 0.15 | 1 | HC_vs_MS.Trt_B |
| RAB43 | 0.05022 | -0.2883 | 0.204 | 0.16 | 1 | HC_vs_MS.Trt_B |
| VAV3 | 0.05025 | -0.5259 | 0.551 | 0.49 | 1 | HC_vs_MS.Trt_B |
| LSM6 | 0.05048 | 0.46896 | 0.13 | 0.09 | 1 | HC_vs_MS.Trt_B |
| DHX33 | 0.05051 | 0.60472 | 0.317 | 0.27 | 1 | HC_vs_MS.Trt_B |
| SAP30BP | 0.05058 | 0.31309 | 0.206 | 0.16 | 1 | HC_vs_MS.Trt_B |
| ZNF417 | 0.05062 | 0.5587 | 0.693 | 0.65 | 1 | HC_vs_MS.Trt_B |
| TMPRSS3 | 0.05108 | -0.3294 | 0.342 | 0.29 | 1 | HC_vs_MS.Trt_B |
| OTUD4 | 0.05113 | -0.2543 | 0.597 | 0.56 | 1 | HC_vs_MS.Trt_B |
| RETSAT | 0.05114 | -0.5747 | 0.122 | 0.09 | 1 | HC_vs_MS.Trt_B |
| ERLEC1 | 0.05117 | -0.9591 | 0.358 | 0.29 | 1 | HC_vs_MS.Trt_B |
| MAATS1 | 0.05136 | 0.52013 | 0.2 | 0.16 | 1 | HC_vs_MS.Trt_B |
| PPCS | 0.05165 | -0.6309 | 0.322 | 0.26 | 1 | HC_vs_MS.Trt_B |
| FRS3 | 0.05165 | -1.2449 | 0.32 | 0.27 | 1 | HC_vs_MS.Trt_B |
| SHFM1 | 0.05201 | -0.333 | 0.881 | 0.85 | 1 | HC_vs_MS.Trt_B |
| FAM200B | 0.05229 | -0.2697 | 0.38 | 0.31 | 1 | HC_vs_MS.Trt_B |
| DBNDD1 | 0.05239 | 0.57574 | 0.165 | 0.12 | 1 | HC_vs_MS.Trt_B |

| | | | | | | |
|---|---|---|---|---|---|---|
| TEKT2 | 0.05258 | 0.49733 | 0.117 | 0.08 | 1 | HC_vs_MS.Trt_B |
| STK39 | 0.05258 | -0.402 | 0.103 | 0.07 | 1 | HC_vs_MS.Trt_B |
| SAYSD1 | 0.05276 | -0.6587 | 0.17 | 0.13 | 1 | HC_vs_MS.Trt_B |
| BTN3A3 | 0.05281 | 0.27421 | 0.223 | 0.18 | 1 | HC_vs_MS.Trt_B |
| GDI2 | 0.05301 | -0.2903 | 0.386 | 0.32 | 1 | HC_vs_MS.Trt_B |
| PIGW | 0.05318 | 1.28471 | 0.459 | 0.41 | 1 | HC_vs_MS.Trt_B |
| INVS | 0.05339 | 0.51133 | 0.135 | 0.1 | 1 | HC_vs_MS.Trt_B |
| ADTRP | 0.05346 | -0.3999 | 0.323 | 0.27 | 1 | HC_vs_MS.Trt_B |
| ARHGAP11B | 0.05346 | 0.39279 | 0.204 | 0.16 | 1 | HC_vs_MS.Trt_B |
| PIPOX | 0.0535 | -0.2624 | 0.507 | 0.45 | 1 | HC_vs_MS.Trt_B |
| MRPS22 | 0.05351 | 0.29535 | 0.47 | 0.43 | 1 | HC_vs_MS.Trt_B |
| ACSL4 | 0.05358 | 0.56524 | 0.145 | 0.11 | 1 | HC_vs_MS.Trt_B |
| MLF1 | 0.05359 | -0.526 | 0.293 | 0.25 | 1 | HC_vs_MS.Trt_B |
| FAM102B | 0.05362 | 0.57747 | 0.171 | 0.13 | 1 | HC_vs_MS.Trt_B |
| SEC22B | 0.05388 | -0.4151 | 0.801 | 0.74 | 1 | HC_vs_MS.Trt_B |
| ELAC1 | 0.05412 | 0.49159 | 0.439 | 0.39 | 1 | HC_vs_MS.Trt_B |
| COG3 | 0.05416 | 0.4024 | 0.168 | 0.13 | 1 | HC_vs_MS.Trt_B |
| BAK1 | 0.05426 | 0.89255 | 0.103 | 0.07 | 1 | HC_vs_MS.Trt_B |
| API5 | 0.05439 | -0.2858 | 0.235 | 0.19 | 1 | HC_vs_MS.Trt_B |
| COASY | 0.05455 | 0.83221 | 0.404 | 0.36 | 1 | HC_vs_MS.Trt_B |
| DLD | 0.05456 | -0.716 | 0.141 | 0.1 | 1 | HC_vs_MS.Trt_B |
| GPR37L1 | 0.05474 | 0.51681 | 0.32 | 0.27 | 1 | HC_vs_MS.Trt_B |
| ACTG2 | 0.05502 | 0.75663 | 0.241 | 0.2 | 1 | HC_vs_MS.Trt_B |
| LPIN2 | 0.0552 | -0.3948 | 0.125 | 0.09 | 1 | HC_vs_MS.Trt_B |
| NIFK | 0.05534 | -0.5381 | 0.522 | 0.45 | 1 | HC_vs_MS.Trt_B |
| IL13RA1 | 0.05544 | -0.3233 | 0.239 | 0.2 | 1 | HC_vs_MS.Trt_B |
| AP2A1 | 0.05557 | 0.82966 | 0.697 | 0.63 | 1 | HC_vs_MS.Trt_B |
| RNF167 | 0.05572 | 0.27381 | 0.27 | 0.22 | 1 | HC_vs_MS.Trt_B |
| SRM | 0.05579 | -0.4427 | 0.123 | 0.09 | 1 | HC_vs_MS.Trt_B |
| PAN3 | 0.05591 | -0.9229 | 0.757 | 0.77 | 1 | HC_vs_MS.Trt_B |
| GRTP1 | 0.05598 | 0.50633 | 0.506 | 0.46 | 1 | HC_vs_MS.Trt_B |
| PEX11G | 0.05605 | 0.35899 | 0.126 | 0.09 | 1 | HC_vs_MS.Trt_B |
| C6orf106 | 0.05642 | -0.3618 | 0.126 | 0.09 | 1 | HC_vs_MS.Trt_B |
| SIPA1 | 0.05652 | 0.46231 | 0.533 | 0.48 | 1 | HC_vs_MS.Trt_B |
| MAPRE1 | 0.05675 | 0.27418 | 0.155 | 0.12 | 1 | HC_vs_MS.Trt_B |
| NRROS | 0.05679 | 1.11379 | 0.101 | 0.07 | 1 | HC_vs_MS.Trt_B |
| COA4 | 0.05703 | -0.4446 | 0.275 | 0.22 | 1 | HC_vs_MS.Trt_B |
| PHACTR2 | 0.05727 | -0.6834 | 0.481 | 0.41 | 1 | HC_vs_MS.Trt_B |
| RNF38 | 0.05732 | 0.38405 | 0.181 | 0.14 | 1 | HC_vs_MS.Trt_B |
| ADCY9 | 0.05732 | -0.6571 | 0.261 | 0.21 | 1 | HC_vs_MS.Trt_B |

| | | | | | | |
|---|---|---|---|---|---|---|
| MVB12B | 0.05763 | 1.00989 | 0.109 | 0.08 | 1 | HC_vs_MS.Trt_B |
| MTMR12 | 0.05787 | 0.43308 | 0.401 | 0.35 | 1 | HC_vs_MS.Trt_B |
| DYNC1I2 | 0.05801 | 0.55478 | 0.174 | 0.14 | 1 | HC_vs_MS.Trt_B |
| RAB35 | 0.05802 | 0.38204 | 0.116 | 0.08 | 1 | HC_vs_MS.Trt_B |
| RBM42 | 0.05809 | 0.36097 | 0.172 | 0.13 | 1 | HC_vs_MS.Trt_B |
| C20orf194 | 0.05809 | -0.5137 | 0.225 | 0.18 | 1 | HC_vs_MS.Trt_B |
| ZNF142 | 0.05828 | 0.29607 | 0.101 | 0.07 | 1 | HC_vs_MS.Trt_B |
| TAF3 | 0.05829 | 0.28024 | 0.151 | 0.11 | 1 | HC_vs_MS.Trt_B |
| NR2C2 | 0.05841 | -0.3196 | 0.604 | 0.52 | 1 | HC_vs_MS.Trt_B |
| CXCR6 | 0.05848 | 0.60089 | 0.128 | 0.09 | 1 | HC_vs_MS.Trt_B |
| TAF12 | 0.05865 | -0.2976 | 0.128 | 0.09 | 1 | HC_vs_MS.Trt_B |
| CTD-2331H1 | 0.05888 | 0.7281 | 0.186 | 0.14 | 1 | HC_vs_MS.Trt_B |
| ACP6 | 0.05893 | 0.26117 | 0.549 | 0.46 | 1 | HC_vs_MS.Trt_B |
| TMEM259 | 0.05904 | 0.65393 | 0.113 | 0.08 | 1 | HC_vs_MS.Trt_B |
| STAM | 0.0592 | -0.3632 | 0.371 | 0.31 | 1 | HC_vs_MS.Trt_B |
| PFKP | 0.05923 | 0.37211 | 0.259 | 0.22 | 1 | HC_vs_MS.Trt_B |
| CTC-454I21. | 0.0593 | 0.31761 | 0.199 | 0.16 | 1 | HC_vs_MS.Trt_B |
| KNG1 | 0.05942 | -0.7735 | 0.187 | 0.14 | 1 | HC_vs_MS.Trt_B |
| LAMTOR4 | 0.05975 | -0.3782 | 0.441 | 0.38 | 1 | HC_vs_MS.Trt_B |
| ARPC2 | 0.05988 | -0.4086 | 0.965 | 0.93 | 1 | HC_vs_MS.Trt_B |
| SESN1 | 0.05995 | 0.3328 | 0.19 | 0.15 | 1 | HC_vs_MS.Trt_B |
| KCTD9 | 0.06025 | 1.08646 | 0.183 | 0.15 | 1 | HC_vs_MS.Trt_B |
| ACADM | 0.06049 | -0.2571 | 0.18 | 0.14 | 1 | HC_vs_MS.Trt_B |
| TAOK2 | 0.06067 | 0.4681 | 0.103 | 0.07 | 1 | HC_vs_MS.Trt_B |
| HPRT1 | 0.06105 | 0.47286 | 0.12 | 0.09 | 1 | HC_vs_MS.Trt_B |
| SMARCA5 | 0.06109 | -0.3169 | 0.328 | 0.27 | 1 | HC_vs_MS.Trt_B |
| TP53I13 | 0.06121 | 0.38891 | 0.116 | 0.08 | 1 | HC_vs_MS.Trt_B |
| ERMN | 0.06156 | -0.93 | 0.172 | 0.13 | 1 | HC_vs_MS.Trt_B |
| MGAT5 | 0.06158 | -0.4995 | 0.143 | 0.11 | 1 | HC_vs_MS.Trt_B |
| SLC20A1 | 0.0618 | 0.7722 | 0.142 | 0.11 | 1 | HC_vs_MS.Trt_B |
| ZNF341 | 0.06184 | -0.7213 | 0.371 | 0.31 | 1 | HC_vs_MS.Trt_B |
| FBXO3 | 0.06207 | 0.39094 | 0.291 | 0.25 | 1 | HC_vs_MS.Trt_B |
| PRR29 | 0.06281 | 0.51089 | 0.171 | 0.14 | 1 | HC_vs_MS.Trt_B |
| SULT1A3 | 0.06307 | -0.3142 | 0.606 | 0.55 | 1 | HC_vs_MS.Trt_B |
| NDUFAF7 | 0.06307 | 1.03714 | 0.37 | 0.33 | 1 | HC_vs_MS.Trt_B |
| ZFP41 | 0.06345 | -0.2767 | 0.104 | 0.07 | 1 | HC_vs_MS.Trt_B |
| PSMC3 | 0.06348 | -0.4728 | 0.103 | 0.14 | 1 | HC_vs_MS.Trt_B |
| PDPR | 0.06354 | 0.31514 | 0.364 | 0.31 | 1 | HC_vs_MS.Trt_B |
| RABAC1 | 0.06391 | -0.5195 | 0.17 | 0.21 | 1 | HC_vs_MS.Trt_B |
| PPM1A | 0.06402 | 0.59509 | 0.146 | 0.11 | 1 | HC_vs_MS.Trt_B |

| | | | | | | |
|---|---|---|---|---|---|---|
| SYTL2 | 0.06414 | -0.275 | 0.736 | 0.74 | 1 | HC_vs_MS.Trt_B |
| LRRC34 | 0.06466 | -0.4222 | 0.559 | 0.5 | 1 | HC_vs_MS.Trt_B |
| ARL1 | 0.06494 | 0.4096 | 0.346 | 0.29 | 1 | HC_vs_MS.Trt_B |
| TPRKB | 0.06498 | -0.3403 | 0.468 | 0.4 | 1 | HC_vs_MS.Trt_B |
| RBM3 | 0.06502 | -0.3417 | 0.803 | 0.72 | 1 | HC_vs_MS.Trt_B |
| TMPRSS7 | 0.06511 | 0.30136 | 0.257 | 0.21 | 1 | HC_vs_MS.Trt_B |
| E2F4 | 0.06522 | 0.78727 | 0.122 | 0.09 | 1 | HC_vs_MS.Trt_B |
| MOSPD1 | 0.0653 | -0.7611 | 0.178 | 0.14 | 1 | HC_vs_MS.Trt_B |
| SLC1A5 | 0.06534 | 0.80525 | 0.152 | 0.12 | 1 | HC_vs_MS.Trt_B |
| IWS1 | 0.06545 | 0.35882 | 0.128 | 0.09 | 1 | HC_vs_MS.Trt_B |
| SRPK1 | 0.06555 | -0.5566 | 0.132 | 0.17 | 1 | HC_vs_MS.Trt_B |
| ZSCAN20 | 0.06556 | 0.73378 | 0.238 | 0.2 | 1 | HC_vs_MS.Trt_B |
| ANKRD13C | 0.06573 | 0.2544 | 0.167 | 0.13 | 1 | HC_vs_MS.Trt_B |
| ZNF254 | 0.06578 | 0.4972 | 0.397 | 0.35 | 1 | HC_vs_MS.Trt_B |
| JMJD4 | 0.0658 | 0.72642 | 0.242 | 0.2 | 1 | HC_vs_MS.Trt_B |
| S1PR4 | 0.06601 | 0.32546 | 0.161 | 0.12 | 1 | HC_vs_MS.Trt_B |
| GGPS1 | 0.06617 | -0.3915 | 0.378 | 0.32 | 1 | HC_vs_MS.Trt_B |
| TCEAL3 | 0.06617 | -0.2888 | 0.11 | 0.08 | 1 | HC_vs_MS.Trt_B |
| FANCF | 0.06631 | 0.46077 | 0.236 | 0.2 | 1 | HC_vs_MS.Trt_B |
| TMEM97 | 0.06631 | 0.67391 | 0.23 | 0.19 | 1 | HC_vs_MS.Trt_B |
| CYB5B | 0.06677 | -0.5205 | 0.449 | 0.39 | 1 | HC_vs_MS.Trt_B |
| ZFP82 | 0.06684 | 0.46641 | 0.649 | 0.6 | 1 | HC_vs_MS.Trt_B |
| MYL6B | 0.06722 | -0.6198 | 0.506 | 0.46 | 1 | HC_vs_MS.Trt_B |
| GEMIN7 | 0.06748 | -0.4006 | 0.213 | 0.17 | 1 | HC_vs_MS.Trt_B |
| INPP4A | 0.06788 | 0.34776 | 0.33 | 0.28 | 1 | HC_vs_MS.Trt_B |
| KDM6A | 0.06811 | 0.3647 | 0.242 | 0.2 | 1 | HC_vs_MS.Trt_B |
| SMARCE1 | 0.06825 | -0.7051 | 0.435 | 0.37 | 1 | HC_vs_MS.Trt_B |
| NPIPB9 | 0.06828 | 0.40683 | 0.114 | 0.08 | 1 | HC_vs_MS.Trt_B |
| PPIG | 0.06832 | -0.3278 | 0.312 | 0.25 | 1 | HC_vs_MS.Trt_B |
| SSNA1 | 0.06834 | 0.35805 | 0.129 | 0.1 | 1 | HC_vs_MS.Trt_B |
| PLEKHG4B | 0.06839 | 0.25671 | 0.386 | 0.33 | 1 | HC_vs_MS.Trt_B |
| KIF13A | 0.06855 | -0.6172 | 0.27 | 0.22 | 1 | HC_vs_MS.Trt_B |
| NCAPD2 | 0.06877 | -0.4058 | 0.403 | 0.34 | 1 | HC_vs_MS.Trt_B |
| ERCC6L | 0.06891 | 1.51302 | 0.117 | 0.08 | 1 | HC_vs_MS.Trt_B |
| MAZ | 0.0692 | 0.32771 | 0.138 | 0.1 | 1 | HC_vs_MS.Trt_B |
| DNAH9 | 0.0692 | 0.28273 | 0.436 | 0.39 | 1 | HC_vs_MS.Trt_B |
| MYO7A | 0.0693 | 0.26056 | 0.278 | 0.23 | 1 | HC_vs_MS.Trt_B |
| IFNLR1 | 0.06973 | 0.35662 | 0.109 | 0.08 | 1 | HC_vs_MS.Trt_B |
| MTSS1 | 0.0699 | -0.9944 | 0.583 | 0.49 | 1 | HC_vs_MS.Trt_B |
| ZNF316 | 0.07021 | -0.5489 | 0.461 | 0.43 | 1 | HC_vs_MS.Trt_B |

| | | | | | | |
|---|---|---|---|---|---|---|
| GMPR2 | 0.07034 | 0.42588 | 0.107 | 0.08 | 1 | HC_vs_MS.Trt_B |
| LIPT2 | 0.07084 | 0.31141 | 0.123 | 0.09 | 1 | HC_vs_MS.Trt_B |
| CRIPAK | 0.07119 | 0.31691 | 0.21 | 0.17 | 1 | HC_vs_MS.Trt_B |
| OST4 | 0.0719 | -0.3081 | 0.388 | 0.32 | 1 | HC_vs_MS.Trt_B |
| SYNGAP1 | 0.07237 | 0.49779 | 0.136 | 0.1 | 1 | HC_vs_MS.Trt_B |
| CCR6 | 0.07262 | -0.7673 | 0.307 | 0.26 | 1 | HC_vs_MS.Trt_B |
| MFSD11 | 0.07277 | -0.3004 | 0.429 | 0.38 | 1 | HC_vs_MS.Trt_B |
| AXIN1 | 0.07302 | -0.5361 | 0.107 | 0.07 | 1 | HC_vs_MS.Trt_B |
| CPEB4 | 0.07324 | 1.0726 | 0.116 | 0.09 | 1 | HC_vs_MS.Trt_B |
| FARSA | 0.07337 | 0.29218 | 0.125 | 0.09 | 1 | HC_vs_MS.Trt_B |
| CHMP2A | 0.0738 | 0.3976 | 0.106 | 0.08 | 1 | HC_vs_MS.Trt_B |
| EIF3E | 0.07416 | -0.4906 | 0.539 | 0.46 | 1 | HC_vs_MS.Trt_B |
| MAML1 | 0.07423 | 0.44527 | 0.13 | 0.1 | 1 | HC_vs_MS.Trt_B |
| GPD1L | 0.07435 | -0.7851 | 0.112 | 0.08 | 1 | HC_vs_MS.Trt_B |
| TMEM87B | 0.07443 | 0.68005 | 0.109 | 0.08 | 1 | HC_vs_MS.Trt_B |
| EPHX1 | 0.07517 | -0.3085 | 0.107 | 0.07 | 1 | HC_vs_MS.Trt_B |
| ZNF76 | 0.07525 | 0.5702 | 0.143 | 0.11 | 1 | HC_vs_MS.Trt_B |
| PSMG4 | 0.07555 | -0.2869 | 0.728 | 0.69 | 1 | HC_vs_MS.Trt_B |
| HNRNPAB | 0.07589 | 0.61944 | 0.128 | 0.1 | 1 | HC_vs_MS.Trt_B |
| DNAJC28 | 0.07595 | 0.46129 | 0.374 | 0.33 | 1 | HC_vs_MS.Trt_B |
| IARS2 | 0.07643 | 0.46963 | 0.117 | 0.09 | 1 | HC_vs_MS.Trt_B |
| TP53BP2 | 0.0766 | 0.56001 | 0.135 | 0.1 | 1 | HC_vs_MS.Trt_B |
| HID1 | 0.07662 | -1.039 | 0.136 | 0.1 | 1 | HC_vs_MS.Trt_B |
| MCEE | 0.07668 | -0.4172 | 0.129 | 0.1 | 1 | HC_vs_MS.Trt_B |
| SLFN11 | 0.07703 | 0.67684 | 0.251 | 0.21 | 1 | HC_vs_MS.Trt_B |
| C19orf12 | 0.07727 | 0.62956 | 0.335 | 0.31 | 1 | HC_vs_MS.Trt_B |
| ZNF460 | 0.07742 | 0.75459 | 0.174 | 0.14 | 1 | HC_vs_MS.Trt_B |
| CHCHD3 | 0.07758 | 0.25961 | 0.148 | 0.11 | 1 | HC_vs_MS.Trt_B |
| TMEM140 | 0.07828 | 0.64357 | 0.178 | 0.14 | 1 | HC_vs_MS.Trt_B |
| CTD-2192J1( | 0.07872 | -1.1554 | 0.133 | 0.17 | 1 | HC_vs_MS.Trt_B |
| ANKRD23 | 0.07885 | -0.265 | 0.116 | 0.08 | 1 | HC_vs_MS.Trt_B |
| MCU | 0.07885 | -0.4971 | 0.246 | 0.2 | 1 | HC_vs_MS.Trt_B |
| FNIP2 | 0.07892 | 0.28808 | 0.296 | 0.25 | 1 | HC_vs_MS.Trt_B |
| CCNT1 | 0.07901 | 0.3584 | 0.116 | 0.09 | 1 | HC_vs_MS.Trt_B |
| TDRD3 | 0.07965 | -0.3825 | 0.519 | 0.47 | 1 | HC_vs_MS.Trt_B |
| CGRRF1 | 0.07966 | -0.7075 | 0.257 | 0.22 | 1 | HC_vs_MS.Trt_B |
| SH3BGRL | 0.07966 | -0.2543 | 0.328 | 0.27 | 1 | HC_vs_MS.Trt_B |
| ITGB1BP1 | 0.07985 | 0.48835 | 0.141 | 0.11 | 1 | HC_vs_MS.Trt_B |
| CNIH1 | 0.08018 | -0.5197 | 0.132 | 0.1 | 1 | HC_vs_MS.Trt_B |
| CASZ1 | 0.08036 | 0.56168 | 0.193 | 0.16 | 1 | HC_vs_MS.Trt_B |

| | | | | | | |
|---|---|---|---|---|---|---|
| CEP162 | 0.08045 | -0.7529 | 0.138 | 0.1 | 1 | HC_vs_MS.Trt_B |
| LLGL1 | 0.08048 | 1.06291 | 0.258 | 0.21 | 1 | HC_vs_MS.Trt_B |
| AASDHPPT | 0.08075 | 0.42903 | 0.123 | 0.09 | 1 | HC_vs_MS.Trt_B |
| ARID2 | 0.08081 | -0.3308 | 0.688 | 0.61 | 1 | HC_vs_MS.Trt_B |
| HINT3 | 0.08118 | 0.2821 | 0.296 | 0.25 | 1 | HC_vs_MS.Trt_B |
| BCDIN3D | 0.08121 | 0.40394 | 0.316 | 0.28 | 1 | HC_vs_MS.Trt_B |
| ATXN7L3B | 0.08138 | 0.47396 | 0.162 | 0.13 | 1 | HC_vs_MS.Trt_B |
| FAM32A | 0.08202 | -0.3189 | 0.436 | 0.38 | 1 | HC_vs_MS.Trt_B |
| PFKFB2 | 0.08229 | 0.36 | 0.2 | 0.16 | 1 | HC_vs_MS.Trt_B |
| SLC25A3 | 0.08261 | -0.3866 | 0.929 | 0.83 | 1 | HC_vs_MS.Trt_B |
| UBALD2 | 0.08268 | -0.5164 | 0.116 | 0.08 | 1 | HC_vs_MS.Trt_B |
| OASL | 0.08273 | 0.35717 | 0.177 | 0.14 | 1 | HC_vs_MS.Trt_B |
| SLC9A1 | 0.08282 | 0.74463 | 0.204 | 0.17 | 1 | HC_vs_MS.Trt_B |
| SRP54 | 0.08289 | -0.4394 | 0.168 | 0.13 | 1 | HC_vs_MS.Trt_B |
| C14orf1 | 0.08291 | -0.6905 | 0.259 | 0.21 | 1 | HC_vs_MS.Trt_B |
| LRRC8D | 0.08299 | -0.3508 | 0.148 | 0.11 | 1 | HC_vs_MS.Trt_B |
| BABAM1 | 0.08341 | -0.2614 | 0.607 | 0.54 | 1 | HC_vs_MS.Trt_B |
| ABCB8 | 0.08372 | 0.35678 | 0.533 | 0.51 | 1 | HC_vs_MS.Trt_B |
| RP11-793H1 | 0.08375 | 0.31684 | 0.257 | 0.22 | 1 | HC_vs_MS.Trt_B |
| TM9SF1 | 0.08467 | 0.3172 | 0.232 | 0.19 | 1 | HC_vs_MS.Trt_B |
| DNAJC19 | 0.08506 | -0.2756 | 0.228 | 0.19 | 1 | HC_vs_MS.Trt_B |
| AHRR | 0.08538 | -0.4809 | 0.113 | 0.08 | 1 | HC_vs_MS.Trt_B |
| MRPL40 | 0.08567 | -0.268 | 0.394 | 0.34 | 1 | HC_vs_MS.Trt_B |
| GANAB | 0.08587 | 0.56108 | 0.43 | 0.41 | 1 | HC_vs_MS.Trt_B |
| DCTN3 | 0.08623 | -0.591 | 0.361 | 0.31 | 1 | HC_vs_MS.Trt_B |
| CC2D2B | 0.08646 | 0.48123 | 0.125 | 0.09 | 1 | HC_vs_MS.Trt_B |
| MKRN2OS | 0.08652 | 0.32072 | 0.187 | 0.15 | 1 | HC_vs_MS.Trt_B |
| ENPP3 | 0.08658 | 0.38551 | 0.13 | 0.1 | 1 | HC_vs_MS.Trt_B |
| STT3A | 0.08662 | -0.4667 | 0.591 | 0.52 | 1 | HC_vs_MS.Trt_B |
| ACVR2A | 0.08669 | -0.5049 | 0.225 | 0.18 | 1 | HC_vs_MS.Trt_B |
| BCR | 0.08696 | -0.7568 | 0.261 | 0.22 | 1 | HC_vs_MS.Trt_B |
| PAQR7 | 0.08739 | 0.31874 | 0.486 | 0.46 | 1 | HC_vs_MS.Trt_B |
| USP40 | 0.0874 | 0.27022 | 0.404 | 0.34 | 1 | HC_vs_MS.Trt_B |
| ADAMTSL4 | 0.08755 | -0.3598 | 0.236 | 0.2 | 1 | HC_vs_MS.Trt_B |
| RHBDD1 | 0.08776 | 0.30385 | 0.317 | 0.28 | 1 | HC_vs_MS.Trt_B |
| ARF3 | 0.0881 | 0.31536 | 0.159 | 0.13 | 1 | HC_vs_MS.Trt_B |
| AP5Z1 | 0.08818 | 0.31127 | 0.143 | 0.11 | 1 | HC_vs_MS.Trt_B |
| EBLN2 | 0.08828 | 0.74831 | 0.114 | 0.09 | 1 | HC_vs_MS.Trt_B |
| TUFT1 | 0.08834 | 0.29599 | 0.349 | 0.31 | 1 | HC_vs_MS.Trt_B |
| TCEAL4 | 0.08857 | 0.28551 | 0.12 | 0.09 | 1 | HC_vs_MS.Trt_B |

| | | | | | | |
|---|---|---|---|---|---|---|
| RPS7 | 0.08863 | 0.45259 | 0.439 | 0.41 | 1 | HC_vs_MS.Trt_B |
| HMGA1 | 0.08882 | 0.26786 | 0.177 | 0.14 | 1 | HC_vs_MS.Trt_B |
| BRD9 | 0.0889 | -0.2713 | 0.146 | 0.11 | 1 | HC_vs_MS.Trt_B |
| HDDC2 | 0.08894 | 0.26239 | 0.116 | 0.09 | 1 | HC_vs_MS.Trt_B |
| BNC2 | 0.0892 | -0.4842 | 0.128 | 0.09 | 1 | HC_vs_MS.Trt_B |
| CRY2 | 0.08951 | -0.3992 | 0.443 | 0.41 | 1 | HC_vs_MS.Trt_B |
| IP6K2 | 0.08967 | -0.2899 | 0.314 | 0.26 | 1 | HC_vs_MS.Trt_B |
| MAK | 0.0897 | 0.26272 | 0.17 | 0.14 | 1 | HC_vs_MS.Trt_B |
| SNRPA1 | 0.08983 | -0.6104 | 0.652 | 0.57 | 1 | HC_vs_MS.Trt_B |
| NPTXR | 0.08998 | -0.2507 | 0.326 | 0.27 | 1 | HC_vs_MS.Trt_B |
| GSR | 0.08998 | 0.33391 | 0.203 | 0.17 | 1 | HC_vs_MS.Trt_B |
| USO1 | 0.0901 | 0.56382 | 0.133 | 0.1 | 1 | HC_vs_MS.Trt_B |
| MED16 | 0.09015 | -0.2615 | 0.146 | 0.19 | 1 | HC_vs_MS.Trt_B |
| MTERF1 | 0.0904 | 0.43679 | 0.196 | 0.16 | 1 | HC_vs_MS.Trt_B |
| PADI2 | 0.09067 | 0.48631 | 0.217 | 0.18 | 1 | HC_vs_MS.Trt_B |
| VPS11 | 0.09161 | -0.5319 | 0.767 | 0.71 | 1 | HC_vs_MS.Trt_B |
| TRADD | 0.09161 | -0.6596 | 0.133 | 0.17 | 1 | HC_vs_MS.Trt_B |
| SIRT1 | 0.09246 | -0.5615 | 0.103 | 0.07 | 1 | HC_vs_MS.Trt_B |
| SRPK3 | 0.09266 | -0.261 | 0.257 | 0.21 | 1 | HC_vs_MS.Trt_B |
| SQRDL | 0.09271 | -0.282 | 0.133 | 0.1 | 1 | HC_vs_MS.Trt_B |
| RDH10 | 0.0935 | -0.9093 | 0.167 | 0.13 | 1 | HC_vs_MS.Trt_B |
| PAEP | 0.09363 | -0.6981 | 0.393 | 0.35 | 1 | HC_vs_MS.Trt_B |
| GPR108 | 0.09381 | -0.5895 | 0.123 | 0.16 | 1 | HC_vs_MS.Trt_B |
| COA3 | 0.0939 | -1.0673 | 0.075 | 0.1 | 1 | HC_vs_MS.Trt_B |
| PLEC | 0.09398 | 0.33754 | 0.267 | 0.23 | 1 | HC_vs_MS.Trt_B |
| C12orf75 | 0.09409 | 0.32066 | 0.122 | 0.09 | 1 | HC_vs_MS.Trt_B |
| CORO7-PAM | 0.09416 | 0.29584 | 0.104 | 0.08 | 1 | HC_vs_MS.Trt_B |
| PHF6 | 0.09417 | -0.3293 | 0.667 | 0.57 | 1 | HC_vs_MS.Trt_B |
| MRPL37 | 0.09427 | -0.5612 | 0.126 | 0.16 | 1 | HC_vs_MS.Trt_B |
| RGPD2 | 0.09434 | -0.318 | 0.159 | 0.12 | 1 | HC_vs_MS.Trt_B |
| TPRA1 | 0.09439 | -0.3584 | 0.152 | 0.12 | 1 | HC_vs_MS.Trt_B |
| RP11-111H1 | 0.09496 | -0.4807 | 0.113 | 0.08 | 1 | HC_vs_MS.Trt_B |
| FAM63A | 0.09498 | -0.6033 | 0.509 | 0.43 | 1 | HC_vs_MS.Trt_B |
| IRF3 | 0.09515 | -0.2885 | 0.749 | 0.68 | 1 | HC_vs_MS.Trt_B |
| RRM2B | 0.09549 | -0.4017 | 0.117 | 0.09 | 1 | HC_vs_MS.Trt_B |
| PPP1R3F | 0.09553 | -0.9552 | 0.128 | 0.09 | 1 | HC_vs_MS.Trt_B |
| EFCAB10 | 0.09569 | -0.3433 | 0.168 | 0.13 | 1 | HC_vs_MS.Trt_B |
| ARL6IP5 | 0.0959 | -0.6119 | 0.578 | 0.6 | 1 | HC_vs_MS.Trt_B |
| FAIM | 0.096 | -0.5093 | 0.358 | 0.32 | 1 | HC_vs_MS.Trt_B |
| PIGO | 0.0963 | 0.45665 | 0.342 | 0.3 | 1 | HC_vs_MS.Trt_B |

| | | | | | | |
|---|---|---|---|---|---|---|
| HIST1H1D | 0.09638 | 0.50116 | 0.122 | 0.09 | 1 | HC_vs_MS.Trt_B |
| SNRPG | 0.09657 | -0.434 | 0.246 | 0.2 | 1 | HC_vs_MS.Trt_B |
| SAMD9L | 0.09663 | -0.2724 | 0.191 | 0.15 | 1 | HC_vs_MS.Trt_B |
| TGFBR3 | 0.09704 | 0.28607 | 0.388 | 0.34 | 1 | HC_vs_MS.Trt_B |
| ACP1 | 0.09749 | -0.7372 | 0.252 | 0.29 | 1 | HC_vs_MS.Trt_B |
| SCAMP1 | 0.09841 | -0.4454 | 0.442 | 0.4 | 1 | HC_vs_MS.Trt_B |
| TEPSIN | 0.09859 | 0.52317 | 0.104 | 0.08 | 1 | HC_vs_MS.Trt_B |
| 1-Mar | 0.09866 | -0.8398 | 0.31 | 0.26 | 1 | HC_vs_MS.Trt_B |
| PGGT1B | 0.09894 | -0.4437 | 0.158 | 0.12 | 1 | HC_vs_MS.Trt_B |
| SMURF1 | 0.09904 | -0.2932 | 0.103 | 0.07 | 1 | HC_vs_MS.Trt_B |
| CFD | 0.09935 | 0.84681 | 0.151 | 0.12 | 1 | HC_vs_MS.Trt_B |
| PNPLA2 | 0.09951 | -0.2955 | 0.17 | 0.13 | 1 | HC_vs_MS.Trt_B |
| CLHC1 | 0.10073 | -0.3933 | 0.467 | 0.41 | 1 | HC_vs_MS.Trt_B |
| TAP2 | 0.10081 | 0.49598 | 0.252 | 0.21 | 1 | HC_vs_MS.Trt_B |
| IFFO2 | 0.10148 | 0.51437 | 0.123 | 0.09 | 1 | HC_vs_MS.Trt_B |
| FTSJ1 | 0.10188 | 0.34379 | 0.123 | 0.09 | 1 | HC_vs_MS.Trt_B |
| HIPK1 | 0.102 | 0.44562 | 0.168 | 0.14 | 1 | HC_vs_MS.Trt_B |
| TNFRSF11A | 0.10233 | 0.68131 | 0.228 | 0.19 | 1 | HC_vs_MS.Trt_B |
| PSD3 | 0.10298 | 0.38887 | 0.31 | 0.26 | 1 | HC_vs_MS.Trt_B |
| ATP7A | 0.10313 | 0.63406 | 0.348 | 0.31 | 1 | HC_vs_MS.Trt_B |
| ALKBH5 | 0.10321 | 0.39464 | 0.123 | 0.09 | 1 | HC_vs_MS.Trt_B |
| GINS2 | 0.10345 | 0.69924 | 0.3 | 0.26 | 1 | HC_vs_MS.Trt_B |
| CDC26 | 0.10396 | -0.6958 | 0.162 | 0.2 | 1 | HC_vs_MS.Trt_B |
| NUP37 | 0.10417 | -1.0973 | 0.275 | 0.24 | 1 | HC_vs_MS.Trt_B |
| AGTPBP1 | 0.10431 | 0.79462 | 0.172 | 0.14 | 1 | HC_vs_MS.Trt_B |
| AGO4 | 0.10439 | -0.5934 | 0.28 | 0.24 | 1 | HC_vs_MS.Trt_B |
| IRAK2 | 0.10456 | -0.2889 | 0.288 | 0.24 | 1 | HC_vs_MS.Trt_B |
| HELQ | 0.1046 | 0.34464 | 0.132 | 0.1 | 1 | HC_vs_MS.Trt_B |
| TIGD6 | 0.10515 | 0.48168 | 0.188 | 0.15 | 1 | HC_vs_MS.Trt_B |
| RBMX | 0.1053 | -0.3245 | 0.762 | 0.66 | 1 | HC_vs_MS.Trt_B |
| ADAD2 | 0.1056 | -0.6384 | 0.225 | 0.19 | 1 | HC_vs_MS.Trt_B |
| TMEM39A | 0.10576 | -0.3392 | 0.117 | 0.09 | 1 | HC_vs_MS.Trt_B |
| CUTA | 0.10602 | -0.4828 | 0.27 | 0.3 | 1 | HC_vs_MS.Trt_B |
| PHKG1 | 0.10655 | 0.26178 | 0.267 | 0.22 | 1 | HC_vs_MS.Trt_B |
| MYO6 | 0.10674 | -1.2343 | 0.419 | 0.37 | 1 | HC_vs_MS.Trt_B |
| NDUFV2 | 0.10717 | -0.2574 | 0.406 | 0.35 | 1 | HC_vs_MS.Trt_B |
| CCDC174 | 0.10725 | -0.7095 | 0.12 | 0.15 | 1 | HC_vs_MS.Trt_B |
| FMNL3 | 0.10769 | 0.31666 | 0.235 | 0.2 | 1 | HC_vs_MS.Trt_B |
| NSF | 0.10809 | 0.26597 | 0.197 | 0.16 | 1 | HC_vs_MS.Trt_B |
| ARHGAP9 | 0.10837 | -0.2865 | 0.501 | 0.44 | 1 | HC_vs_MS.Trt_B |

| | | | | | | |
|---|---|---|---|---|---|---|
| PSENEN | 0.1084 | 0.30994 | 0.151 | 0.12 | 1 | HC_vs_MS.Trt_B |
| SYNE4 | 0.10848 | 0.40228 | 0.409 | 0.36 | 1 | HC_vs_MS.Trt_B |
| GRIA1 | 0.10851 | -0.3953 | 0.106 | 0.08 | 1 | HC_vs_MS.Trt_B |
| ARPC4 | 0.10852 | -0.4487 | 0.572 | 0.59 | 1 | HC_vs_MS.Trt_B |
| RTN3 | 0.10882 | 0.26534 | 0.128 | 0.1 | 1 | HC_vs_MS.Trt_B |
| GTPBP4 | 0.10926 | 0.47982 | 0.414 | 0.38 | 1 | HC_vs_MS.Trt_B |
| GLUL | 0.10949 | -0.4729 | 0.158 | 0.12 | 1 | HC_vs_MS.Trt_B |
| KIF2A | 0.10998 | -0.3374 | 0.306 | 0.26 | 1 | HC_vs_MS.Trt_B |
| SLF1 | 0.1101 | 0.57038 | 0.149 | 0.12 | 1 | HC_vs_MS.Trt_B |
| IL18BP | 0.11037 | 0.4907 | 0.106 | 0.08 | 1 | HC_vs_MS.Trt_B |
| RBM39 | 0.11052 | -0.3088 | 0.888 | 0.79 | 1 | HC_vs_MS.Trt_B |
| GLTSCR1L | 0.11064 | -0.6869 | 0.149 | 0.12 | 1 | HC_vs_MS.Trt_B |
| ZNF17 | 0.11075 | 0.4626 | 0.277 | 0.24 | 1 | HC_vs_MS.Trt_B |
| DNAJC11 | 0.11108 | 0.26325 | 0.168 | 0.14 | 1 | HC_vs_MS.Trt_B |
| POLR2H | 0.11121 | -0.3355 | 0.452 | 0.4 | 1 | HC_vs_MS.Trt_B |
| ICAM1 | 0.11123 | -0.5139 | 0.428 | 0.38 | 1 | HC_vs_MS.Trt_B |
| ZNF649 | 0.11147 | 0.26293 | 0.17 | 0.14 | 1 | HC_vs_MS.Trt_B |
| ORC5 | 0.11159 | 0.38294 | 0.239 | 0.2 | 1 | HC_vs_MS.Trt_B |
| CDAN1 | 0.112 | -0.3253 | 0.43 | 0.38 | 1 | HC_vs_MS.Trt_B |
| DISC1 | 0.11204 | 0.4044 | 0.486 | 0.42 | 1 | HC_vs_MS.Trt_B |
| AP5B1 | 0.11205 | 0.93517 | 0.235 | 0.2 | 1 | HC_vs_MS.Trt_B |
| PAK1 | 0.11209 | -0.3249 | 0.155 | 0.12 | 1 | HC_vs_MS.Trt_B |
| MSL2 | 0.11225 | -0.3701 | 0.141 | 0.11 | 1 | HC_vs_MS.Trt_B |
| DUSP1 | 0.11269 | 0.50025 | 0.168 | 0.14 | 1 | HC_vs_MS.Trt_B |
| MRPL16 | 0.1129 | -0.5173 | 0.142 | 0.11 | 1 | HC_vs_MS.Trt_B |
| ARHGAP12 | 0.11291 | -0.4126 | 0.139 | 0.11 | 1 | HC_vs_MS.Trt_B |
| ISCA2 | 0.11302 | -0.6621 | 0.467 | 0.41 | 1 | HC_vs_MS.Trt_B |
| NDUFB8 | 0.11311 | -0.8004 | 0.291 | 0.32 | 1 | HC_vs_MS.Trt_B |
| MRPL46 | 0.11317 | -0.3619 | 0.225 | 0.18 | 1 | HC_vs_MS.Trt_B |
| OTUD5 | 0.11357 | -0.3083 | 0.125 | 0.09 | 1 | HC_vs_MS.Trt_B |
| FBXW8 | 0.11369 | -0.5996 | 0.257 | 0.22 | 1 | HC_vs_MS.Trt_B |
| RNF144B | 0.11382 | -0.2503 | 0.293 | 0.25 | 1 | HC_vs_MS.Trt_B |
| POP5 | 0.11384 | -0.7751 | 0.623 | 0.55 | 1 | HC_vs_MS.Trt_B |
| PPP1R21 | 0.11414 | 0.5154 | 0.203 | 0.17 | 1 | HC_vs_MS.Trt_B |
| PDCD2 | 0.11437 | 0.32508 | 0.106 | 0.08 | 1 | HC_vs_MS.Trt_B |
| SLC22A18 | 0.11438 | 0.50144 | 0.503 | 0.45 | 1 | HC_vs_MS.Trt_B |
| ANKRD49 | 0.11445 | -0.6939 | 0.157 | 0.19 | 1 | HC_vs_MS.Trt_B |
| TNFRSF4 | 0.11446 | 0.38196 | 0.103 | 0.08 | 1 | HC_vs_MS.Trt_B |
| FANCI | 0.11447 | 0.92608 | 0.52 | 0.5 | 1 | HC_vs_MS.Trt_B |
| NRDE2 | 0.1145 | -0.3062 | 0.739 | 0.7 | 1 | HC_vs_MS.Trt_B |

| | | | | | | |
|---|---|---|---|---|---|---|
| USP16 | 0.11497 | -0.4554 | 0.449 | 0.41 | 1 | HC_vs_MS.Trt_B |
| NAA20 | 0.11501 | -0.4405 | 0.104 | 0.14 | 1 | HC_vs_MS.Trt_B |
| DIAPH3 | 0.11574 | 0.26017 | 0.329 | 0.29 | 1 | HC_vs_MS.Trt_B |
| TMEM167A | 0.11601 | -0.6186 | 0.329 | 0.28 | 1 | HC_vs_MS.Trt_B |
| CD2AP | 0.11609 | 0.35501 | 0.17 | 0.14 | 1 | HC_vs_MS.Trt_B |
| ACO2 | 0.11634 | 0.30486 | 0.12 | 0.09 | 1 | HC_vs_MS.Trt_B |
| C1orf109 | 0.11667 | 0.2708 | 0.868 | 0.82 | 1 | HC_vs_MS.Trt_B |
| TIMP2 | 0.11681 | -0.3346 | 0.165 | 0.13 | 1 | HC_vs_MS.Trt_B |
| CKS1B | 0.11744 | -0.6106 | 0.186 | 0.15 | 1 | HC_vs_MS.Trt_B |
| AGAP4 | 0.11767 | -0.6446 | 0.21 | 0.17 | 1 | HC_vs_MS.Trt_B |
| TAF13 | 0.11779 | -0.3566 | 0.116 | 0.09 | 1 | HC_vs_MS.Trt_B |
| AGPS | 0.1182 | 0.66528 | 0.119 | 0.09 | 1 | HC_vs_MS.Trt_B |
| SSRP1 | 0.11845 | 0.91611 | 0.113 | 0.09 | 1 | HC_vs_MS.Trt_B |
| CST7 | 0.11861 | -0.6031 | 0.084 | 0.11 | 1 | HC_vs_MS.Trt_B |
| RP11-894J1 | 0.11878 | 0.51418 | 0.112 | 0.08 | 1 | HC_vs_MS.Trt_B |
| SLC2A13 | 0.11902 | -0.3317 | 0.309 | 0.27 | 1 | HC_vs_MS.Trt_B |
| MT-CO3 | 0.11918 | -0.6362 | 0.597 | 0.59 | 1 | HC_vs_MS.Trt_B |
| VPS37A | 0.1193 | 0.65348 | 0.133 | 0.11 | 1 | HC_vs_MS.Trt_B |
| BROX | 0.11985 | -0.2801 | 0.567 | 0.52 | 1 | HC_vs_MS.Trt_B |
| CDK16 | 0.12177 | 0.72537 | 0.113 | 0.09 | 1 | HC_vs_MS.Trt_B |
| DNPH1 | 0.12227 | -0.7431 | 0.104 | 0.13 | 1 | HC_vs_MS.Trt_B |
| RAD23A | 0.12228 | -0.4601 | 0.719 | 0.68 | 1 | HC_vs_MS.Trt_B |
| TNFSF4 | 0.12229 | -0.7432 | 0.122 | 0.09 | 1 | HC_vs_MS.Trt_B |
| PRPF31 | 0.12324 | 0.48494 | 0.125 | 0.1 | 1 | HC_vs_MS.Trt_B |
| NEUROD2 | 0.1234 | 0.97527 | 0.188 | 0.23 | 1 | HC_vs_MS.Trt_B |
| HSPG2 | 0.12424 | 0.51893 | 0.335 | 0.31 | 1 | HC_vs_MS.Trt_B |
| EHBP1L1 | 0.1257 | 0.67895 | 0.184 | 0.16 | 1 | HC_vs_MS.Trt_B |
| IL24 | 0.12573 | -0.6412 | 0.106 | 0.14 | 1 | HC_vs_MS.Trt_B |
| FAM46C | 0.12588 | 0.38572 | 0.158 | 0.13 | 1 | HC_vs_MS.Trt_B |
| CXADR | 0.1259 | 0.44441 | 0.281 | 0.24 | 1 | HC_vs_MS.Trt_B |
| LEPR | 0.12629 | 0.26505 | 0.283 | 0.24 | 1 | HC_vs_MS.Trt_B |
| DIRC1 | 0.12669 | -0.3625 | 0.174 | 0.14 | 1 | HC_vs_MS.Trt_B |
| RPLP1 | 0.12729 | -0.254 | 0.948 | 0.9 | 1 | HC_vs_MS.Trt_B |
| TPP1 | 0.12757 | -0.4707 | 0.306 | 0.27 | 1 | HC_vs_MS.Trt_B |
| PATL1 | 0.12817 | 0.34581 | 0.133 | 0.11 | 1 | HC_vs_MS.Trt_B |
| GCN1 | 0.12922 | 0.34704 | 0.171 | 0.14 | 1 | HC_vs_MS.Trt_B |
| FAM135A | 0.12943 | -0.6101 | 0.404 | 0.35 | 1 | HC_vs_MS.Trt_B |
| HIGD1A | 0.13015 | -0.4105 | 0.148 | 0.12 | 1 | HC_vs_MS.Trt_B |
| ZNF181 | 0.13043 | 0.55531 | 0.157 | 0.13 | 1 | HC_vs_MS.Trt_B |
| TCTEX1D2 | 0.13103 | -0.4286 | 0.27 | 0.23 | 1 | HC_vs_MS.Trt_B |

| | | | | | | |
|---|---|---|---|---|---|---|
| MT-ND6 | 0.13163 | -0.3278 | 0.654 | 0.68 | 1 | HC_vs_MS.Trt_B |
| UBA2 | 0.1318 | -0.3701 | 0.513 | 0.46 | 1 | HC_vs_MS.Trt_B |
| PELI1 | 0.13181 | 0.49074 | 0.172 | 0.14 | 1 | HC_vs_MS.Trt_B |
| OFD1 | 0.13221 | 0.26505 | 0.246 | 0.22 | 1 | HC_vs_MS.Trt_B |
| SGF29 | 0.13365 | -0.696 | 0.271 | 0.22 | 1 | HC_vs_MS.Trt_B |
| NBPF11 | 0.134 | 0.39978 | 0.257 | 0.22 | 1 | HC_vs_MS.Trt_B |
| AK6 | 0.13635 | -0.3148 | 0.184 | 0.16 | 1 | HC_vs_MS.Trt_B |
| AAGAB | 0.13667 | 0.30416 | 0.113 | 0.09 | 1 | HC_vs_MS.Trt_B |
| MBD1 | 0.13693 | -0.6591 | 0.138 | 0.17 | 1 | HC_vs_MS.Trt_B |
| BCL2L1 | 0.13716 | 0.296 | 0.123 | 0.1 | 1 | HC_vs_MS.Trt_B |
| GNB1L | 0.13754 | -0.4671 | 0.183 | 0.15 | 1 | HC_vs_MS.Trt_B |
| MEMO1 | 0.138 | -0.2545 | 0.301 | 0.27 | 1 | HC_vs_MS.Trt_B |
| XPOT | 0.1388 | -0.3494 | 0.432 | 0.38 | 1 | HC_vs_MS.Trt_B |
| ITGAX | 0.13931 | -0.2513 | 0.471 | 0.41 | 1 | HC_vs_MS.Trt_B |
| FAM200A | 0.13952 | -0.291 | 0.207 | 0.17 | 1 | HC_vs_MS.Trt_B |
| REEP3 | 0.13964 | -1.248 | 0.429 | 0.38 | 1 | HC_vs_MS.Trt_B |
| SC5D | 0.13968 | -0.2856 | 0.517 | 0.47 | 1 | HC_vs_MS.Trt_B |
| PIN4 | 0.1397 | 0.43532 | 0.117 | 0.09 | 1 | HC_vs_MS.Trt_B |
| RFC4 | 0.14062 | -0.2947 | 0.228 | 0.19 | 1 | HC_vs_MS.Trt_B |
| DIEXF | 0.14079 | 0.66413 | 0.461 | 0.43 | 1 | HC_vs_MS.Trt_B |
| MAP2K3 | 0.1417 | 0.3594 | 0.116 | 0.09 | 1 | HC_vs_MS.Trt_B |
| NIPA2 | 0.14174 | -0.3836 | 0.091 | 0.12 | 1 | HC_vs_MS.Trt_B |
| CHD9 | 0.14175 | -0.3363 | 0.345 | 0.31 | 1 | HC_vs_MS.Trt_B |
| TTC7A | 0.14223 | 0.32785 | 0.164 | 0.13 | 1 | HC_vs_MS.Trt_B |
| NCLN | 0.14224 | 0.44812 | 0.122 | 0.1 | 1 | HC_vs_MS.Trt_B |
| OAZ1 | 0.14236 | -0.3319 | 0.796 | 0.76 | 1 | HC_vs_MS.Trt_B |
| DNAJB7 | 0.14262 | 0.3753 | 0.225 | 0.19 | 1 | HC_vs_MS.Trt_B |
| AFTPH | 0.14316 | 0.43307 | 0.162 | 0.13 | 1 | HC_vs_MS.Trt_B |
| TSN | 0.14342 | -0.7047 | 0.329 | 0.27 | 1 | HC_vs_MS.Trt_B |
| TMEM69 | 0.14344 | -0.2797 | 0.197 | 0.16 | 1 | HC_vs_MS.Trt_B |
| NOL10 | 0.14368 | 0.56572 | 0.133 | 0.11 | 1 | HC_vs_MS.Trt_B |
| VPS9D1 | 0.1448 | -0.7964 | 0.226 | 0.19 | 1 | HC_vs_MS.Trt_B |
| PPIL4 | 0.14498 | -0.3956 | 0.223 | 0.19 | 1 | HC_vs_MS.Trt_B |
| CDC42SE2 | 0.14595 | -0.2862 | 0.855 | 0.76 | 1 | HC_vs_MS.Trt_B |
| ZC3H3 | 0.14639 | 0.51306 | 0.125 | 0.1 | 1 | HC_vs_MS.Trt_B |
| CRIP2 | 0.1475 | 0.39432 | 0.12 | 0.1 | 1 | HC_vs_MS.Trt_B |
| PBX4 | 0.14922 | 0.27897 | 0.11 | 0.09 | 1 | HC_vs_MS.Trt_B |
| CYB5D1 | 0.1495 | 1.21177 | 0.138 | 0.11 | 1 | HC_vs_MS.Trt_B |
| UQCRC1 | 0.14967 | -0.6255 | 0.184 | 0.21 | 1 | HC_vs_MS.Trt_B |
| CPEB1 | 0.14987 | 0.28633 | 0.146 | 0.12 | 1 | HC_vs_MS.Trt_B |

| | | | | | | |
|---|---|---|---|---|---|---|
| TDRD1 | 0.15009 | 0.32151 | 0.168 | 0.14 | 1 | HC_vs_MS.Trt_B |
| SLC41A1 | 0.15048 | 0.33347 | 0.103 | 0.08 | 1 | HC_vs_MS.Trt_B |
| SULT1A2 | 0.15134 | 0.33182 | 0.159 | 0.13 | 1 | HC_vs_MS.Trt_B |
| PITHD1 | 0.15135 | 0.28057 | 0.145 | 0.12 | 1 | HC_vs_MS.Trt_B |
| ARID5A | 0.15205 | -0.3118 | 0.19 | 0.16 | 1 | HC_vs_MS.Trt_B |
| CBWD7 | 0.15237 | -0.3838 | 0.401 | 0.38 | 1 | HC_vs_MS.Trt_B |
| AGRN | 0.15323 | 0.34024 | 0.335 | 0.31 | 1 | HC_vs_MS.Trt_B |
| RFXANK | 0.15339 | -0.595 | 0.242 | 0.2 | 1 | HC_vs_MS.Trt_B |
| GBF1 | 0.15421 | 0.3879 | 0.148 | 0.12 | 1 | HC_vs_MS.Trt_B |
| EMC7 | 0.15422 | -0.6038 | 0.12 | 0.15 | 1 | HC_vs_MS.Trt_B |
| C1orf174 | 0.15465 | -0.2893 | 0.501 | 0.46 | 1 | HC_vs_MS.Trt_B |
| PSMD11 | 0.15585 | -0.4357 | 0.483 | 0.44 | 1 | HC_vs_MS.Trt_B |
| NELFCD | 0.15593 | 0.28863 | 0.519 | 0.49 | 1 | HC_vs_MS.Trt_B |
| ATP6V1B2 | 0.15641 | -0.7824 | 0.594 | 0.52 | 1 | HC_vs_MS.Trt_B |
| CHP1 | 0.15685 | -0.6738 | 0.754 | 0.68 | 1 | HC_vs_MS.Trt_B |
| NDUFB1 | 0.15764 | -0.397 | 0.73 | 0.65 | 1 | HC_vs_MS.Trt_B |
| PYROXD2 | 0.15779 | -0.2515 | 0.146 | 0.12 | 1 | HC_vs_MS.Trt_B |
| SLC48A1 | 0.15781 | -0.7915 | 0.583 | 0.54 | 1 | HC_vs_MS.Trt_B |
| HIST1H2BE | 0.15888 | 0.48144 | 0.265 | 0.22 | 1 | HC_vs_MS.Trt_B |
| LAMB1 | 0.15899 | -0.8514 | 0.239 | 0.2 | 1 | HC_vs_MS.Trt_B |
| RBM6 | 0.15923 | -0.3228 | 0.832 | 0.77 | 1 | HC_vs_MS.Trt_B |
| NUP153 | 0.15989 | -0.3836 | 0.159 | 0.13 | 1 | HC_vs_MS.Trt_B |
| TAF15 | 0.16013 | 0.39733 | 0.172 | 0.14 | 1 | HC_vs_MS.Trt_B |
| ERV3-1 | 0.16025 | 0.51228 | 0.484 | 0.44 | 1 | HC_vs_MS.Trt_B |
| ITIH4 | 0.1603 | 0.83895 | 0.201 | 0.17 | 1 | HC_vs_MS.Trt_B |
| NDUFB11 | 0.16037 | -0.3317 | 0.228 | 0.19 | 1 | HC_vs_MS.Trt_B |
| NME9 | 0.16044 | 0.79941 | 0.201 | 0.17 | 1 | HC_vs_MS.Trt_B |
| TCAIM | 0.16049 | -0.9322 | 0.184 | 0.15 | 1 | HC_vs_MS.Trt_B |
| VPS72 | 0.16056 | -0.2557 | 0.117 | 0.09 | 1 | HC_vs_MS.Trt_B |
| MAP3K14 | 0.16104 | 0.34296 | 0.141 | 0.11 | 1 | HC_vs_MS.Trt_B |
| DDX21 | 0.16197 | 0.35267 | 0.236 | 0.2 | 1 | HC_vs_MS.Trt_B |
| CD74 | 0.16199 | -0.4908 | 0.694 | 0.7 | 1 | HC_vs_MS.Trt_B |
| GNAS | 0.16203 | -0.2922 | 0.435 | 0.38 | 1 | HC_vs_MS.Trt_B |
| KIFAP3 | 0.16295 | 0.71531 | 0.141 | 0.12 | 1 | HC_vs_MS.Trt_B |
| SLC23A2 | 0.16336 | -0.2588 | 0.13 | 0.11 | 1 | HC_vs_MS.Trt_B |
| FAHD2A | 0.16366 | 0.52788 | 0.154 | 0.13 | 1 | HC_vs_MS.Trt_B |
| CAPNS1 | 0.16436 | -0.4998 | 0.259 | 0.22 | 1 | HC_vs_MS.Trt_B |
| RNF2 | 0.16473 | 0.26986 | 0.367 | 0.33 | 1 | HC_vs_MS.Trt_B |
| KLHL18 | 0.16489 | 0.29768 | 0.314 | 0.28 | 1 | HC_vs_MS.Trt_B |
| EXOG | 0.16525 | 0.3438 | 0.475 | 0.45 | 1 | HC_vs_MS.Trt_B |

| | | | | | | |
|---|---|---|---|---|---|---|
| YPEL5 | 0.16529 | -0.5554 | 0.175 | 0.14 | 1 | HC_vs_MS.Trt_B |
| CD47 | 0.16554 | -0.3648 | 0.912 | 0.85 | 1 | HC_vs_MS.Trt_B |
| PPP2R5D | 0.16582 | 0.31512 | 0.148 | 0.12 | 1 | HC_vs_MS.Trt_B |
| LRIG1 | 0.16631 | 0.40235 | 0.238 | 0.21 | 1 | HC_vs_MS.Trt_B |
| NAPB | 0.16729 | 0.44833 | 0.403 | 0.37 | 1 | HC_vs_MS.Trt_B |
| CNOT9 | 0.16729 | -0.35 | 0.72 | 0.66 | 1 | HC_vs_MS.Trt_B |
| ZFP37 | 0.16758 | -0.4693 | 0.328 | 0.3 | 1 | HC_vs_MS.Trt_B |
| NAPRT | 0.16792 | -0.2621 | 0.138 | 0.11 | 1 | HC_vs_MS.Trt_B |
| SLC39A11 | 0.16824 | -0.6436 | 0.251 | 0.21 | 1 | HC_vs_MS.Trt_B |
| PCMT1 | 0.16879 | 0.27109 | 0.174 | 0.15 | 1 | HC_vs_MS.Trt_B |
| FAM234A | 0.16888 | -0.2867 | 0.196 | 0.16 | 1 | HC_vs_MS.Trt_B |
| ATP5G2 | 0.16923 | -0.4405 | 0.852 | 0.76 | 1 | HC_vs_MS.Trt_B |
| SLC25A36 | 0.16944 | 0.53192 | 0.129 | 0.1 | 1 | HC_vs_MS.Trt_B |
| VAMP1 | 0.16974 | 0.26229 | 0.145 | 0.12 | 1 | HC_vs_MS.Trt_B |
| XRCC5 | 0.17035 | -0.2971 | 0.487 | 0.43 | 1 | HC_vs_MS.Trt_B |
| GALNT15 | 0.17092 | 0.83909 | 0.271 | 0.24 | 1 | HC_vs_MS.Trt_B |
| BRD4 | 0.17126 | 0.32813 | 0.351 | 0.32 | 1 | HC_vs_MS.Trt_B |
| MAN1A2 | 0.17184 | 0.25056 | 0.248 | 0.21 | 1 | HC_vs_MS.Trt_B |
| KCTD13 | 0.17193 | 0.38334 | 0.135 | 0.11 | 1 | HC_vs_MS.Trt_B |
| KIF21B | 0.17203 | -0.3183 | 0.378 | 0.33 | 1 | HC_vs_MS.Trt_B |
| PHF1 | 0.17288 | 0.31598 | 0.138 | 0.11 | 1 | HC_vs_MS.Trt_B |
| LRRN2 | 0.1738 | 1.18731 | 0.23 | 0.2 | 1 | HC_vs_MS.Trt_B |
| TMEM223 | 0.174 | 0.32533 | 0.133 | 0.11 | 1 | HC_vs_MS.Trt_B |
| ERCC2 | 0.17532 | -0.4944 | 0.465 | 0.43 | 1 | HC_vs_MS.Trt_B |
| USP6 | 0.17535 | -0.3048 | 0.128 | 0.1 | 1 | HC_vs_MS.Trt_B |
| EXOC1 | 0.17597 | -0.6379 | 0.164 | 0.14 | 1 | HC_vs_MS.Trt_B |
| CASP7 | 0.17655 | 0.63871 | 0.125 | 0.1 | 1 | HC_vs_MS.Trt_B |
| C7orf50 | 0.17724 | -0.3774 | 0.332 | 0.29 | 1 | HC_vs_MS.Trt_B |
| TAF2 | 0.17761 | -0.2531 | 0.43 | 0.38 | 1 | HC_vs_MS.Trt_B |
| LINC00094 | 0.17764 | -0.4667 | 0.457 | 0.4 | 1 | HC_vs_MS.Trt_B |
| SRRD | 0.17805 | -0.4784 | 0.37 | 0.32 | 1 | HC_vs_MS.Trt_B |
| DAZAP2 | 0.17873 | -0.4099 | 0.864 | 0.82 | 1 | HC_vs_MS.Trt_B |
| PRMT8 | 0.17892 | -0.6673 | 0.201 | 0.17 | 1 | HC_vs_MS.Trt_B |
| YPEL1 | 0.17934 | 0.35545 | 0.414 | 0.37 | 1 | HC_vs_MS.Trt_B |
| SLC38A10 | 0.18013 | 0.48527 | 0.304 | 0.28 | 1 | HC_vs_MS.Trt_B |
| PKIA | 0.1806 | -0.2758 | 0.104 | 0.08 | 1 | HC_vs_MS.Trt_B |
| MAP3K5 | 0.18089 | -0.3865 | 0.209 | 0.17 | 1 | HC_vs_MS.Trt_B |
| PRKAG2 | 0.18116 | -0.4792 | 0.299 | 0.26 | 1 | HC_vs_MS.Trt_B |
| MRPS9 | 0.1814 | -0.7678 | 0.087 | 0.11 | 1 | HC_vs_MS.Trt_B |
| TJP3 | 0.18193 | 0.39302 | 0.103 | 0.08 | 1 | HC_vs_MS.Trt_B |

| | | | | | | |
|---|---|---|---|---|---|---|
| IL20RB | 0.18212 | 0.33769 | 0.229 | 0.19 | 1 | HC_vs_MS.Trt_B |
| PIP5K1B | 0.18266 | 0.44316 | 0.116 | 0.09 | 1 | HC_vs_MS.Trt_B |
| CEACAM21 | 0.18288 | -0.4089 | 0.37 | 0.32 | 1 | HC_vs_MS.Trt_B |
| LANCL3 | 0.18291 | -0.3794 | 0.259 | 0.22 | 1 | HC_vs_MS.Trt_B |
| C5orf24 | 0.18311 | 0.51673 | 0.342 | 0.3 | 1 | HC_vs_MS.Trt_B |
| MCRIP1 | 0.18365 | -0.4167 | 0.381 | 0.35 | 1 | HC_vs_MS.Trt_B |
| GORASP2 | 0.18395 | 0.25572 | 0.155 | 0.13 | 1 | HC_vs_MS.Trt_B |
| ZFP36L1 | 0.18504 | -0.3609 | 0.701 | 0.7 | 1 | HC_vs_MS.Trt_B |
| CTNS | 0.1853 | 0.3126 | 0.106 | 0.08 | 1 | HC_vs_MS.Trt_B |
| CRIM1 | 0.18577 | -0.5894 | 0.139 | 0.12 | 1 | HC_vs_MS.Trt_B |
| TIPRL | 0.18581 | -0.3054 | 0.281 | 0.24 | 1 | HC_vs_MS.Trt_B |
| STEAP3 | 0.18634 | 0.37898 | 0.116 | 0.09 | 1 | HC_vs_MS.Trt_B |
| KCTD11 | 0.18697 | -0.4109 | 0.149 | 0.12 | 1 | HC_vs_MS.Trt_B |
| MRPL51 | 0.18757 | -0.6984 | 0.083 | 0.11 | 1 | HC_vs_MS.Trt_B |
| BCOR | 0.18765 | -0.3761 | 0.112 | 0.09 | 1 | HC_vs_MS.Trt_B |
| TTC28 | 0.18815 | 0.51301 | 0.325 | 0.29 | 1 | HC_vs_MS.Trt_B |
| SMN2 | 0.18824 | -0.3459 | 0.807 | 0.8 | 1 | HC_vs_MS.Trt_B |
| SAMD9 | 0.18838 | 0.30914 | 0.183 | 0.16 | 1 | HC_vs_MS.Trt_B |
| MIGA2 | 0.18843 | -0.5069 | 0.399 | 0.37 | 1 | HC_vs_MS.Trt_B |
| ATP2B1 | 0.18851 | -0.2593 | 0.368 | 0.33 | 1 | HC_vs_MS.Trt_B |
| ATG12 | 0.1887 | -0.452 | 0.696 | 0.65 | 1 | HC_vs_MS.Trt_B |
| SYTL3 | 0.18882 | -0.2794 | 0.122 | 0.1 | 1 | HC_vs_MS.Trt_B |
| PER2 | 0.18885 | -0.3931 | 0.452 | 0.4 | 1 | HC_vs_MS.Trt_B |
| A2M | 0.18888 | -0.6486 | 0.126 | 0.16 | 1 | HC_vs_MS.Trt_B |
| GNE | 0.19147 | 0.5269 | 0.193 | 0.16 | 1 | HC_vs_MS.Trt_B |
| MXD4 | 0.19155 | -0.2756 | 0.123 | 0.1 | 1 | HC_vs_MS.Trt_B |
| PYGB | 0.1919 | 0.67639 | 0.101 | 0.08 | 1 | HC_vs_MS.Trt_B |
| RUNX2 | 0.19259 | 0.34113 | 0.151 | 0.12 | 1 | HC_vs_MS.Trt_B |
| ZNF418 | 0.193 | 0.58284 | 0.172 | 0.14 | 1 | HC_vs_MS.Trt_B |
| F2RL2 | 0.19327 | 0.41194 | 0.229 | 0.2 | 1 | HC_vs_MS.Trt_B |
| SHH | 0.19378 | 1.42188 | 0.106 | 0.08 | 1 | HC_vs_MS.Trt_B |
| RFWD2 | 0.194 | -0.5648 | 0.291 | 0.27 | 1 | HC_vs_MS.Trt_B |
| C11orf98 | 0.19434 | -0.6362 | 0.103 | 0.13 | 1 | HC_vs_MS.Trt_B |
| ARF5 | 0.19482 | -0.4282 | 0.196 | 0.16 | 1 | HC_vs_MS.Trt_B |
| BAG4 | 0.19486 | -0.4074 | 0.141 | 0.11 | 1 | HC_vs_MS.Trt_B |
| TNFSF13B | 0.19605 | -0.7607 | 0.5 | 0.45 | 1 | HC_vs_MS.Trt_B |
| EIF3CL | 0.19614 | 0.30435 | 0.167 | 0.14 | 1 | HC_vs_MS.Trt_B |
| LPAR4 | 0.19697 | 0.65261 | 0.132 | 0.11 | 1 | HC_vs_MS.Trt_B |
| EEA1 | 0.19726 | 0.32459 | 0.181 | 0.16 | 1 | HC_vs_MS.Trt_B |
| APOBEC3C | 0.1984 | -0.4505 | 0.642 | 0.61 | 1 | HC_vs_MS.Trt_B |

| | | | | | | |
|---|---|---|---|---|---|---|
| JTB | 0.19846 | -0.4516 | 0.21 | 0.17 | 1 | HC_vs_MS.Trt_B |
| NPIPA7 | 0.19857 | -0.4156 | 0.154 | 0.12 | 1 | HC_vs_MS.Trt_B |
| TNS2 | 0.1988 | 0.27467 | 0.291 | 0.26 | 1 | HC_vs_MS.Trt_B |
| NFYB | 0.19906 | -0.9157 | 0.235 | 0.21 | 1 | HC_vs_MS.Trt_B |
| TRIM39 | 0.19915 | 0.65311 | 0.112 | 0.09 | 1 | HC_vs_MS.Trt_B |
| USMG5 | 0.19917 | -0.4601 | 0.188 | 0.16 | 1 | HC_vs_MS.Trt_B |
| C1orf228 | 0.20011 | -0.2552 | 0.242 | 0.21 | 1 | HC_vs_MS.Trt_B |
| BEX4 | 0.20079 | 0.48822 | 0.103 | 0.08 | 1 | HC_vs_MS.Trt_B |
| ANK1 | 0.20172 | -1.1388 | 0.225 | 0.19 | 1 | HC_vs_MS.Trt_B |
| RP11-500M8 | 0.20529 | -1.2132 | 0.083 | 0.1 | 1 | HC_vs_MS.Trt_B |
| KANK3 | 0.20563 | 0.39394 | 0.201 | 0.24 | 1 | HC_vs_MS.Trt_B |
| FOSB | 0.20582 | 0.5148 | 0.186 | 0.16 | 1 | HC_vs_MS.Trt_B |
| TFEB | 0.20642 | 0.60783 | 0.404 | 0.39 | 1 | HC_vs_MS.Trt_B |
| EIF2B1 | 0.20704 | -0.3126 | 0.329 | 0.29 | 1 | HC_vs_MS.Trt_B |
| MTERF3 | 0.20743 | -0.8428 | 0.18 | 0.15 | 1 | HC_vs_MS.Trt_B |
| SMAD1 | 0.20795 | 0.69436 | 0.32 | 0.29 | 1 | HC_vs_MS.Trt_B |
| VPS54 | 0.20798 | -0.6258 | 0.374 | 0.33 | 1 | HC_vs_MS.Trt_B |
| RP11-574F2 | 0.20834 | 0.2515 | 0.154 | 0.13 | 1 | HC_vs_MS.Trt_B |
| TPX2 | 0.20847 | 1.05916 | 0.181 | 0.15 | 1 | HC_vs_MS.Trt_B |
| SLC16A6 | 0.20848 | -0.6227 | 0.242 | 0.21 | 1 | HC_vs_MS.Trt_B |
| THYN1 | 0.20911 | -0.6697 | 0.13 | 0.16 | 1 | HC_vs_MS.Trt_B |
| MTMR2 | 0.20911 | 0.43556 | 0.158 | 0.13 | 1 | HC_vs_MS.Trt_B |
| TBC1D23 | 0.20922 | -0.5679 | 0.114 | 0.09 | 1 | HC_vs_MS.Trt_B |
| STX12 | 0.20939 | 0.36185 | 0.184 | 0.16 | 1 | HC_vs_MS.Trt_B |
| ZNF862 | 0.21086 | 0.2915 | 0.107 | 0.09 | 1 | HC_vs_MS.Trt_B |
| SRPRB | 0.2109 | 0.40987 | 0.139 | 0.12 | 1 | HC_vs_MS.Trt_B |
| P4HA1 | 0.21157 | 0.26748 | 0.161 | 0.13 | 1 | HC_vs_MS.Trt_B |
| ATOX1 | 0.2116 | 0.25502 | 0.174 | 0.15 | 1 | HC_vs_MS.Trt_B |
| PRKACA | 0.21162 | 0.33702 | 0.172 | 0.15 | 1 | HC_vs_MS.Trt_B |
| TAF5L | 0.21174 | 0.72364 | 0.104 | 0.08 | 1 | HC_vs_MS.Trt_B |
| TPI1 | 0.21192 | -0.2566 | 0.372 | 0.4 | 1 | HC_vs_MS.Trt_B |
| LCAT | 0.21228 | -0.5578 | 0.187 | 0.16 | 1 | HC_vs_MS.Trt_B |
| KPTN | 0.21256 | 0.79495 | 0.212 | 0.17 | 1 | HC_vs_MS.Trt_B |
| NDUFA13 | 0.21273 | -0.8168 | 0.409 | 0.41 | 1 | HC_vs_MS.Trt_B |
| KDM5B | 0.21393 | 0.30223 | 0.277 | 0.25 | 1 | HC_vs_MS.Trt_B |
| PPP1R9A | 0.2141 | 0.28141 | 0.158 | 0.13 | 1 | HC_vs_MS.Trt_B |
| FCGR2A | 0.21463 | -0.5941 | 0.162 | 0.14 | 1 | HC_vs_MS.Trt_B |
| GARNL3 | 0.21471 | 0.61721 | 0.155 | 0.13 | 1 | HC_vs_MS.Trt_B |
| ZNF622 | 0.21566 | -0.9801 | 0.081 | 0.1 | 1 | HC_vs_MS.Trt_B |
| COG6 | 0.21576 | -0.6622 | 0.12 | 0.1 | 1 | HC_vs_MS.Trt_B |

| | | | | | | |
|---|---|---|---|---|---|---|
| PTPRE | 0.21592 | -0.3036 | 0.355 | 0.32 | 1 | HC_vs_MS.Trt_B |
| ATG4B | 0.21628 | 0.33366 | 0.152 | 0.13 | 1 | HC_vs_MS.Trt_B |
| HSPD1 | 0.2163 | 0.27587 | 0.193 | 0.17 | 1 | HC_vs_MS.Trt_B |
| LSM12 | 0.21673 | -0.2906 | 0.622 | 0.58 | 1 | HC_vs_MS.Trt_B |
| BLMH | 0.21703 | 0.34665 | 0.581 | 0.55 | 1 | HC_vs_MS.Trt_B |
| DUSP7 | 0.21744 | -0.3164 | 0.109 | 0.09 | 1 | HC_vs_MS.Trt_B |
| TSPYL1 | 0.21768 | -0.2605 | 0.522 | 0.47 | 1 | HC_vs_MS.Trt_B |
| ZNF18 | 0.21768 | 0.31868 | 0.312 | 0.29 | 1 | HC_vs_MS.Trt_B |
| NSMAF | 0.21782 | 0.31749 | 0.162 | 0.14 | 1 | HC_vs_MS.Trt_B |
| HNF1B | 0.21795 | 0.30372 | 0.17 | 0.14 | 1 | HC_vs_MS.Trt_B |
| BCAT2 | 0.219 | -0.4282 | 0.142 | 0.12 | 1 | HC_vs_MS.Trt_B |
| HGS | 0.21928 | -0.3289 | 0.599 | 0.55 | 1 | HC_vs_MS.Trt_B |
| MPG | 0.21946 | -0.6407 | 0.094 | 0.12 | 1 | HC_vs_MS.Trt_B |
| ABCD3 | 0.21977 | 0.86169 | 0.125 | 0.1 | 1 | HC_vs_MS.Trt_B |
| VPS18 | 0.22118 | 0.44098 | 0.197 | 0.17 | 1 | HC_vs_MS.Trt_B |
| POLR2L | 0.22134 | -0.3126 | 0.212 | 0.18 | 1 | HC_vs_MS.Trt_B |
| CCDC115 | 0.22182 | -0.5625 | 0.086 | 0.11 | 1 | HC_vs_MS.Trt_B |
| STAU1 | 0.22188 | -0.5375 | 0.293 | 0.33 | 1 | HC_vs_MS.Trt_B |
| WDR37 | 0.22325 | -0.2734 | 0.161 | 0.13 | 1 | HC_vs_MS.Trt_B |
| PCNP | 0.22456 | -0.6344 | 0.235 | 0.2 | 1 | HC_vs_MS.Trt_B |
| XYLT1 | 0.22631 | 0.60044 | 0.116 | 0.1 | 1 | HC_vs_MS.Trt_B |
| SPEG | 0.22724 | 0.29541 | 0.23 | 0.21 | 1 | HC_vs_MS.Trt_B |
| CEP55 | 0.22858 | -0.661 | 0.126 | 0.1 | 1 | HC_vs_MS.Trt_B |
| UBE2K | 0.22977 | 0.5119 | 0.133 | 0.11 | 1 | HC_vs_MS.Trt_B |
| ENO2 | 0.22978 | 0.66923 | 0.199 | 0.17 | 1 | HC_vs_MS.Trt_B |
| RAPGEF2 | 0.23086 | -0.3418 | 0.348 | 0.32 | 1 | HC_vs_MS.Trt_B |
| CCDC15 | 0.23106 | 0.38817 | 0.216 | 0.19 | 1 | HC_vs_MS.Trt_B |
| MAP2K4 | 0.23239 | 1.06586 | 0.213 | 0.19 | 1 | HC_vs_MS.Trt_B |
| MRPL32 | 0.23409 | -0.4439 | 0.122 | 0.1 | 1 | HC_vs_MS.Trt_B |
| APOBR | 0.2353 | -0.6282 | 0.104 | 0.08 | 1 | HC_vs_MS.Trt_B |
| XPO7 | 0.23617 | -0.3884 | 0.1 | 0.12 | 1 | HC_vs_MS.Trt_B |
| RAD21 | 0.23662 | -0.3203 | 0.33 | 0.3 | 1 | HC_vs_MS.Trt_B |
| UPK3BL | 0.23752 | -0.5295 | 0.532 | 0.53 | 1 | HC_vs_MS.Trt_B |
| RHBDD3 | 0.23793 | 0.66924 | 0.467 | 0.44 | 1 | HC_vs_MS.Trt_B |
| GCNT4 | 0.23828 | -0.5822 | 0.475 | 0.5 | 1 | HC_vs_MS.Trt_B |
| PNISR | 0.23902 | -0.2902 | 0.554 | 0.48 | 1 | HC_vs_MS.Trt_B |
| C19orf52 | 0.23926 | 0.37392 | 0.241 | 0.22 | 1 | HC_vs_MS.Trt_B |
| AGBL3 | 0.24014 | -0.3384 | 0.248 | 0.21 | 1 | HC_vs_MS.Trt_B |
| MAP2K5 | 0.24212 | 0.30067 | 0.103 | 0.08 | 1 | HC_vs_MS.Trt_B |
| UBE2Z | 0.2423 | -0.2655 | 0.181 | 0.16 | 1 | HC_vs_MS.Trt_B |

| | | | | | | |
|---|---|---|---|---|---|---|
| C3orf18 | 0.24308 | -0.373 | 0.214 | 0.19 | 1 | HC_vs_MS.Trt_B |
| MED12 | 0.24336 | -0.3249 | 0.172 | 0.14 | 1 | HC_vs_MS.Trt_B |
| COIL | 0.24362 | -0.7521 | 0.106 | 0.08 | 1 | HC_vs_MS.Trt_B |
| TMED9 | 0.24392 | -0.4258 | 0.278 | 0.24 | 1 | HC_vs_MS.Trt_B |
| BLM | 0.24463 | 0.37913 | 0.239 | 0.22 | 1 | HC_vs_MS.Trt_B |
| MRPS31 | 0.24564 | -0.4695 | 0.387 | 0.35 | 1 | HC_vs_MS.Trt_B |
| ALG5 | 0.24606 | -0.4981 | 0.399 | 0.43 | 1 | HC_vs_MS.Trt_B |
| CTNNBL1 | 0.24648 | -0.3755 | 0.172 | 0.15 | 1 | HC_vs_MS.Trt_B |
| AGAP2 | 0.24663 | -0.3119 | 0.151 | 0.12 | 1 | HC_vs_MS.Trt_B |
| TRIP4 | 0.24673 | 0.32141 | 0.477 | 0.46 | 1 | HC_vs_MS.Trt_B |
| RCOR1 | 0.24685 | 0.3832 | 0.13 | 0.11 | 1 | HC_vs_MS.Trt_B |
| ALG6 | 0.24706 | 0.4571 | 0.101 | 0.08 | 1 | HC_vs_MS.Trt_B |
| ZFAND2B | 0.24762 | -0.411 | 0.172 | 0.14 | 1 | HC_vs_MS.Trt_B |
| CPNE8 | 0.24787 | -0.6966 | 0.088 | 0.11 | 1 | HC_vs_MS.Trt_B |
| TTC13 | 0.24822 | -0.3927 | 0.114 | 0.09 | 1 | HC_vs_MS.Trt_B |
| PRKD3 | 0.24847 | -0.3961 | 0.322 | 0.29 | 1 | HC_vs_MS.Trt_B |
| CAT | 0.24915 | -0.3013 | 0.133 | 0.11 | 1 | HC_vs_MS.Trt_B |
| COX16 | 0.24952 | -0.3394 | 0.487 | 0.53 | 1 | HC_vs_MS.Trt_B |
| CCDC18 | 0.25009 | -0.7573 | 0.159 | 0.13 | 1 | HC_vs_MS.Trt_B |
| DHX16 | 0.25231 | 0.29962 | 0.112 | 0.09 | 1 | HC_vs_MS.Trt_B |
| TMEM263 | 0.25322 | -0.4291 | 0.377 | 0.34 | 1 | HC_vs_MS.Trt_B |
| NSUN7 | 0.25433 | -1.353 | 0.229 | 0.2 | 1 | HC_vs_MS.Trt_B |
| ATP13A3 | 0.2546 | -0.3434 | 0.104 | 0.09 | 1 | HC_vs_MS.Trt_B |
| TNPO2 | 0.25491 | 0.49842 | 0.122 | 0.1 | 1 | HC_vs_MS.Trt_B |
| WDR83OS | 0.2557 | -0.2614 | 0.178 | 0.15 | 1 | HC_vs_MS.Trt_B |
| MUS81 | 0.25596 | 0.277 | 0.112 | 0.09 | 1 | HC_vs_MS.Trt_B |
| GADD45GIP | 0.25612 | -0.3206 | 0.126 | 0.1 | 1 | HC_vs_MS.Trt_B |
| FKBP3 | 0.25638 | -0.4328 | 0.268 | 0.24 | 1 | HC_vs_MS.Trt_B |
| MEGF9 | 0.25681 | -0.4357 | 0.436 | 0.39 | 1 | HC_vs_MS.Trt_B |
| ANKRD36C | 0.25705 | -0.2581 | 0.517 | 0.48 | 1 | HC_vs_MS.Trt_B |
| VAMP4 | 0.25728 | -1.0383 | 0.17 | 0.14 | 1 | HC_vs_MS.Trt_B |
| TRAM1 | 0.25894 | -0.278 | 0.383 | 0.34 | 1 | HC_vs_MS.Trt_B |
| HMGN1 | 0.25931 | -0.3514 | 0.414 | 0.38 | 1 | HC_vs_MS.Trt_B |
| TPD52L2 | 0.25944 | -0.3693 | 0.201 | 0.18 | 1 | HC_vs_MS.Trt_B |
| ARL14EP | 0.25961 | -0.5124 | 0.203 | 0.18 | 1 | HC_vs_MS.Trt_B |
| POLR3GL | 0.26402 | -0.2784 | 0.174 | 0.15 | 1 | HC_vs_MS.Trt_B |
| DHX29 | 0.2648 | 0.46248 | 0.128 | 0.11 | 1 | HC_vs_MS.Trt_B |
| SLCO4C1 | 0.26558 | 0.27535 | 0.157 | 0.13 | 1 | HC_vs_MS.Trt_B |
| ANAPC10 | 0.2663 | 0.34039 | 0.104 | 0.09 | 1 | HC_vs_MS.Trt_B |
| HLA-DRB5 | 0.26652 | -2.4197 | 0.087 | 0.11 | 1 | HC_vs_MS.Trt_B |

| | | | | | | |
|---|---|---|---|---|---|---|
| PHTF1 | 0.26726 | -0.8797 | 0.428 | 0.4 | 1 | HC_vs_MS.Trt_B |
| DSN1 | 0.26779 | 0.56122 | 0.225 | 0.2 | 1 | HC_vs_MS.Trt_B |
| LPCAT4 | 0.26791 | 0.56109 | 0.262 | 0.24 | 1 | HC_vs_MS.Trt_B |
| TOPBP1 | 0.26885 | 0.42125 | 0.106 | 0.09 | 1 | HC_vs_MS.Trt_B |
| LAYN | 0.26926 | -0.7926 | 0.193 | 0.17 | 1 | HC_vs_MS.Trt_B |
| TBCB | 0.26996 | -0.4584 | 0.645 | 0.6 | 1 | HC_vs_MS.Trt_B |
| BRAP | 0.27072 | -0.3407 | 0.28 | 0.25 | 1 | HC_vs_MS.Trt_B |
| DNAAF5 | 0.27107 | -1.5546 | 0.19 | 0.17 | 1 | HC_vs_MS.Trt_B |
| HDHD3 | 0.27109 | 0.37247 | 0.136 | 0.12 | 1 | HC_vs_MS.Trt_B |
| CDC42 | 0.2714 | -0.4711 | 0.868 | 0.81 | 1 | HC_vs_MS.Trt_B |
| CD96 | 0.27218 | -0.4104 | 0.549 | 0.51 | 1 | HC_vs_MS.Trt_B |
| TMIGD2 | 0.27398 | -0.3076 | 0.199 | 0.17 | 1 | HC_vs_MS.Trt_B |
| CDH15 | 0.27502 | 0.38169 | 0.403 | 0.38 | 1 | HC_vs_MS.Trt_B |
| IQGAP1 | 0.27637 | -0.3949 | 0.791 | 0.72 | 1 | HC_vs_MS.Trt_B |
| UBR3 | 0.27782 | -0.5947 | 0.116 | 0.1 | 1 | HC_vs_MS.Trt_B |
| CEBPZ | 0.27784 | -0.4123 | 0.2 | 0.17 | 1 | HC_vs_MS.Trt_B |
| TRIM46 | 0.27785 | -0.4202 | 0.417 | 0.38 | 1 | HC_vs_MS.Trt_B |
| HNRNPLL | 0.27796 | -0.2577 | 0.235 | 0.21 | 1 | HC_vs_MS.Trt_B |
| PPP2R2A | 0.27832 | -0.4762 | 0.126 | 0.11 | 1 | HC_vs_MS.Trt_B |
| CHST3 | 0.27845 | 0.33081 | 0.139 | 0.12 | 1 | HC_vs_MS.Trt_B |
| FN1 | 0.27911 | -0.6824 | 0.348 | 0.31 | 1 | HC_vs_MS.Trt_B |
| 6-Mar | 0.28003 | -0.4178 | 0.8 | 0.73 | 1 | HC_vs_MS.Trt_B |
| YPEL3 | 0.2807 | -0.296 | 0.235 | 0.2 | 1 | HC_vs_MS.Trt_B |
| LGALS1 | 0.28093 | -0.7321 | 0.175 | 0.2 | 1 | HC_vs_MS.Trt_B |
| BNIP3L | 0.28106 | -0.346 | 0.181 | 0.16 | 1 | HC_vs_MS.Trt_B |
| MTM1 | 0.28169 | -0.4423 | 0.135 | 0.11 | 1 | HC_vs_MS.Trt_B |
| CDRT4 | 0.28247 | 0.30178 | 0.101 | 0.08 | 1 | HC_vs_MS.Trt_B |
| SPAG4 | 0.28295 | -0.3729 | 0.188 | 0.16 | 1 | HC_vs_MS.Trt_B |
| CAGE1 | 0.28411 | -0.3566 | 0.291 | 0.25 | 1 | HC_vs_MS.Trt_B |
| UPF3B | 0.28436 | -0.2927 | 0.13 | 0.11 | 1 | HC_vs_MS.Trt_B |
| PRDX3 | 0.28536 | -0.8075 | 0.409 | 0.37 | 1 | HC_vs_MS.Trt_B |
| NPIPB6 | 0.28594 | 0.27122 | 0.186 | 0.17 | 1 | HC_vs_MS.Trt_B |
| MAD1L1 | 0.28684 | -0.5476 | 0.284 | 0.24 | 1 | HC_vs_MS.Trt_B |
| TRAFD1 | 0.28753 | 0.3371 | 0.201 | 0.18 | 1 | HC_vs_MS.Trt_B |
| SESN3 | 0.28796 | -0.2743 | 0.259 | 0.23 | 1 | HC_vs_MS.Trt_B |
| STX18 | 0.29023 | 0.6064 | 0.12 | 0.1 | 1 | HC_vs_MS.Trt_B |
| NCBP2-AS2 | 0.29047 | -0.359 | 0.116 | 0.14 | 1 | HC_vs_MS.Trt_B |
| MSL3 | 0.29135 | 0.2563 | 0.184 | 0.16 | 1 | HC_vs_MS.Trt_B |
| DAD1 | 0.29162 | -0.4323 | 0.307 | 0.27 | 1 | HC_vs_MS.Trt_B |
| HMGN3 | 0.29233 | -0.6676 | 0.17 | 0.19 | 1 | HC_vs_MS.Trt_B |

| | | | | | | |
|---|---|---|---|---|---|---|
| LARP1 | 0.29355 | -0.2997 | 0.18 | 0.15 | 1 | HC_vs_MS.Trt_B |
| TMTC3 | 0.2948 | -0.6117 | 0.181 | 0.16 | 1 | HC_vs_MS.Trt_B |
| GPR107 | 0.29522 | 0.65386 | 0.145 | 0.13 | 1 | HC_vs_MS.Trt_B |
| VRK1 | 0.2955 | -0.6219 | 0.206 | 0.18 | 1 | HC_vs_MS.Trt_B |
| TSEN34 | 0.29633 | -0.4284 | 0.112 | 0.09 | 1 | HC_vs_MS.Trt_B |
| CREM | 0.29919 | -1.085 | 0.135 | 0.16 | 1 | HC_vs_MS.Trt_B |
| BAP1 | 0.29931 | -0.5921 | 0.307 | 0.28 | 1 | HC_vs_MS.Trt_B |
| HAVCR1 | 0.29956 | -0.6856 | 0.335 | 0.3 | 1 | HC_vs_MS.Trt_B |
| SLC25A35 | 0.29978 | -0.6305 | 0.142 | 0.12 | 1 | HC_vs_MS.Trt_B |
| ISCU | 0.30008 | -0.3651 | 0.317 | 0.34 | 1 | HC_vs_MS.Trt_B |
| DCAF6 | 0.30095 | 0.67451 | 0.159 | 0.14 | 1 | HC_vs_MS.Trt_B |
| INCA1 | 0.30098 | -0.7482 | 0.18 | 0.16 | 1 | HC_vs_MS.Trt_B |
| TMEM216 | 0.3014 | -0.3815 | 0.322 | 0.29 | 1 | HC_vs_MS.Trt_B |
| G2E3 | 0.30181 | -0.2631 | 0.488 | 0.46 | 1 | HC_vs_MS.Trt_B |
| PTTG1IP | 0.30266 | 0.30063 | 0.139 | 0.12 | 1 | HC_vs_MS.Trt_B |
| KDM2B | 0.30292 | 0.26453 | 0.791 | 0.74 | 1 | HC_vs_MS.Trt_B |
| PSMB7 | 0.30447 | -0.2622 | 0.174 | 0.2 | 1 | HC_vs_MS.Trt_B |
| C5orf15 | 0.30671 | -0.4194 | 0.113 | 0.09 | 1 | HC_vs_MS.Trt_B |
| AGL | 0.3076 | 0.38236 | 0.13 | 0.11 | 1 | HC_vs_MS.Trt_B |
| AC240274.1 | 0.30835 | 1.13522 | 0.241 | 0.21 | 1 | HC_vs_MS.Trt_B |
| ATRX | 0.3084 | -0.3733 | 0.551 | 0.5 | 1 | HC_vs_MS.Trt_B |
| CCNT2 | 0.30898 | 0.33215 | 0.133 | 0.12 | 1 | HC_vs_MS.Trt_B |
| NDUFB3 | 0.30905 | -0.3097 | 0.143 | 0.12 | 1 | HC_vs_MS.Trt_B |
| PARP1 | 0.30986 | -0.3253 | 0.313 | 0.28 | 1 | HC_vs_MS.Trt_B |
| PRKCB | 0.30989 | -0.3623 | 0.291 | 0.26 | 1 | HC_vs_MS.Trt_B |
| IER3IP1 | 0.31137 | -0.7124 | 0.123 | 0.1 | 1 | HC_vs_MS.Trt_B |
| COMMD6 | 0.31202 | -0.7263 | 0.332 | 0.34 | 1 | HC_vs_MS.Trt_B |
| RXRA | 0.3128 | 1.1061 | 0.138 | 0.12 | 1 | HC_vs_MS.Trt_B |
| RNF168 | 0.31282 | -0.355 | 0.543 | 0.48 | 1 | HC_vs_MS.Trt_B |
| PPME1 | 0.31352 | 0.36494 | 0.116 | 0.1 | 1 | HC_vs_MS.Trt_B |
| LSR | 0.3136 | 0.34116 | 0.152 | 0.13 | 1 | HC_vs_MS.Trt_B |
| RAPH1 | 0.31402 | -0.2883 | 0.146 | 0.12 | 1 | HC_vs_MS.Trt_B |
| CBX3 | 0.31416 | -0.3445 | 0.535 | 0.49 | 1 | HC_vs_MS.Trt_B |
| MYO1D | 0.31552 | -0.7267 | 0.128 | 0.11 | 1 | HC_vs_MS.Trt_B |
| RANBP10 | 0.31782 | -0.2509 | 0.107 | 0.09 | 1 | HC_vs_MS.Trt_B |
| COX6B1 | 0.31845 | 0.41538 | 0.104 | 0.09 | 1 | HC_vs_MS.Trt_B |
| IL11RA | 0.31912 | -0.3456 | 0.145 | 0.12 | 1 | HC_vs_MS.Trt_B |
| MTPN | 0.31968 | 0.2589 | 0.162 | 0.14 | 1 | HC_vs_MS.Trt_B |
| EXOSC8 | 0.32204 | -0.6051 | 0.452 | 0.43 | 1 | HC_vs_MS.Trt_B |
| HMGXB3 | 0.3231 | 0.26741 | 0.128 | 0.11 | 1 | HC_vs_MS.Trt_B |

| | | | | | | |
|---|---|---|---|---|---|---|
| LYRM9 | 0.32358 | 0.79819 | 0.196 | 0.18 | 1 | HC_vs_MS.Trt_B |
| GNB1 | 0.32616 | -0.2884 | 0.29 | 0.26 | 1 | HC_vs_MS.Trt_B |
| ZNF223 | 0.32667 | -0.7218 | 0.101 | 0.08 | 1 | HC_vs_MS.Trt_B |
| TP53BP1 | 0.32669 | -0.3175 | 0.252 | 0.22 | 1 | HC_vs_MS.Trt_B |
| IKZF5 | 0.32758 | -0.3967 | 0.158 | 0.18 | 1 | HC_vs_MS.Trt_B |
| PEX7 | 0.32819 | 0.45612 | 0.141 | 0.12 | 1 | HC_vs_MS.Trt_B |
| ANKRD26 | 0.32874 | 0.72444 | 0.123 | 0.11 | 1 | HC_vs_MS.Trt_B |
| NPIPB15 | 0.33153 | -0.6609 | 0.193 | 0.21 | 1 | HC_vs_MS.Trt_B |
| CASP8AP2 | 0.33196 | -0.32 | 0.342 | 0.31 | 1 | HC_vs_MS.Trt_B |
| MYL12A | 0.33249 | -0.2611 | 0.894 | 0.79 | 1 | HC_vs_MS.Trt_B |
| PLCL1 | 0.3333 | -0.2523 | 0.129 | 0.11 | 1 | HC_vs_MS.Trt_B |
| NIPA1 | 0.3342 | 0.37562 | 0.101 | 0.12 | 1 | HC_vs_MS.Trt_B |
| GPS2 | 0.33688 | -0.4164 | 0.193 | 0.21 | 1 | HC_vs_MS.Trt_B |
| ZNF547 | 0.33704 | -1.2761 | 0.203 | 0.17 | 1 | HC_vs_MS.Trt_B |
| PRKRA | 0.33718 | -0.2625 | 0.216 | 0.2 | 1 | HC_vs_MS.Trt_B |
| GUCA1B | 0.33729 | -0.2795 | 0.29 | 0.26 | 1 | HC_vs_MS.Trt_B |
| ESAM | 0.33873 | 0.79273 | 0.109 | 0.09 | 1 | HC_vs_MS.Trt_B |
| PILRB | 0.3403 | -0.6446 | 0.419 | 0.39 | 1 | HC_vs_MS.Trt_B |
| XXYLT1 | 0.34095 | -0.7616 | 0.183 | 0.16 | 1 | HC_vs_MS.Trt_B |
| ZNF574 | 0.34131 | 0.38332 | 0.316 | 0.29 | 1 | HC_vs_MS.Trt_B |
| PACRGL | 0.34173 | 0.45621 | 0.271 | 0.25 | 1 | HC_vs_MS.Trt_B |
| NFXL1 | 0.3424 | 0.51708 | 0.103 | 0.09 | 1 | HC_vs_MS.Trt_B |
| ERP44 | 0.34252 | -0.5478 | 0.219 | 0.19 | 1 | HC_vs_MS.Trt_B |
| LSG1 | 0.34277 | -0.2766 | 0.361 | 0.34 | 1 | HC_vs_MS.Trt_B |
| ERICH1 | 0.34278 | -0.4509 | 0.178 | 0.16 | 1 | HC_vs_MS.Trt_B |
| HIST1H4C | 0.34295 | -0.4314 | 0.319 | 0.28 | 1 | HC_vs_MS.Trt_B |
| PCNT | 0.34527 | -0.2603 | 0.167 | 0.15 | 1 | HC_vs_MS.Trt_B |
| NDUFB10 | 0.34595 | -0.418 | 0.177 | 0.16 | 1 | HC_vs_MS.Trt_B |
| PAPD4 | 0.34669 | -0.577 | 0.226 | 0.2 | 1 | HC_vs_MS.Trt_B |
| BRICD5 | 0.34692 | 0.28637 | 0.251 | 0.23 | 1 | HC_vs_MS.Trt_B |
| ATRAID | 0.34744 | -0.4131 | 0.183 | 0.21 | 1 | HC_vs_MS.Trt_B |
| TMED10 | 0.34789 | -0.3965 | 0.641 | 0.6 | 1 | HC_vs_MS.Trt_B |
| RPA1 | 0.3482 | -0.4816 | 0.264 | 0.24 | 1 | HC_vs_MS.Trt_B |
| NME3 | 0.34967 | -0.3599 | 0.113 | 0.1 | 1 | HC_vs_MS.Trt_B |
| METTL5 | 0.35123 | -0.5683 | 0.464 | 0.43 | 1 | HC_vs_MS.Trt_B |
| RPS21 | 0.35139 | -0.4025 | 0.439 | 0.4 | 1 | HC_vs_MS.Trt_B |
| ZBTB1 | 0.35245 | -0.312 | 0.136 | 0.16 | 1 | HC_vs_MS.Trt_B |
| ADRM1 | 0.35475 | -0.4399 | 0.172 | 0.19 | 1 | HC_vs_MS.Trt_B |
| ZMYM5 | 0.3552 | -0.4066 | 0.2 | 0.18 | 1 | HC_vs_MS.Trt_B |
| FNDC3B | 0.35563 | -0.2723 | 0.667 | 0.66 | 1 | HC_vs_MS.Trt_B |

| | | | | | | |
|---|---|---|---|---|---|---|
| PATZ1 | 0.35799 | -0.2581 | 0.107 | 0.09 | 1 | HC_vs_MS.Trt_B |
| FAM168B | 0.35816 | -0.4252 | 0.106 | 0.09 | 1 | HC_vs_MS.Trt_B |
| NSUN5 | 0.35867 | -0.5344 | 0.312 | 0.28 | 1 | HC_vs_MS.Trt_B |
| SLC31A2 | 0.35918 | 0.39647 | 0.101 | 0.09 | 1 | HC_vs_MS.Trt_B |
| CD3G | 0.3595 | -0.4956 | 0.948 | 0.92 | 1 | HC_vs_MS.Trt_B |
| OARD1 | 0.35966 | -0.5666 | 0.197 | 0.17 | 1 | HC_vs_MS.Trt_B |
| LINC00116 | 0.35973 | -0.7292 | 0.106 | 0.12 | 1 | HC_vs_MS.Trt_B |
| RBM4B | 0.3604 | 0.26263 | 0.174 | 0.16 | 1 | HC_vs_MS.Trt_B |
| ADAM28 | 0.36052 | -0.3292 | 0.135 | 0.12 | 1 | HC_vs_MS.Trt_B |
| MAP3K7 | 0.36079 | -0.4505 | 0.133 | 0.12 | 1 | HC_vs_MS.Trt_B |
| HLA-DRB3 | 0.36127 | -1.3942 | 0.093 | 0.11 | 1 | HC_vs_MS.Trt_B |
| CPNE1 | 0.36239 | -0.3315 | 0.459 | 0.46 | 1 | HC_vs_MS.Trt_B |
| ARHGEF40 | 0.36441 | -0.7064 | 0.125 | 0.11 | 1 | HC_vs_MS.Trt_B |
| RPA3 | 0.36872 | -0.5213 | 0.178 | 0.16 | 1 | HC_vs_MS.Trt_B |
| SPIN1 | 0.36971 | -0.7114 | 0.155 | 0.14 | 1 | HC_vs_MS.Trt_B |
| AHCTF1 | 0.36999 | -0.5907 | 0.133 | 0.15 | 1 | HC_vs_MS.Trt_B |
| APOL2 | 0.37142 | -0.3679 | 0.561 | 0.5 | 1 | HC_vs_MS.Trt_B |
| COX17 | 0.37266 | -0.4423 | 0.245 | 0.22 | 1 | HC_vs_MS.Trt_B |
| UBE2I | 0.37293 | -0.2506 | 0.52 | 0.48 | 1 | HC_vs_MS.Trt_B |
| MRPL13 | 0.37355 | -0.5408 | 0.087 | 0.1 | 1 | HC_vs_MS.Trt_B |
| VAMP5 | 0.37461 | -0.3749 | 0.113 | 0.1 | 1 | HC_vs_MS.Trt_B |
| PSMA1 | 0.37687 | -0.5523 | 0.294 | 0.26 | 1 | HC_vs_MS.Trt_B |
| CYC1 | 0.37688 | -0.7518 | 0.099 | 0.11 | 1 | HC_vs_MS.Trt_B |
| DICER1 | 0.37919 | -0.5458 | 0.148 | 0.13 | 1 | HC_vs_MS.Trt_B |
| CARD16 | 0.37975 | -0.3606 | 0.252 | 0.22 | 1 | HC_vs_MS.Trt_B |
| EIF1AX | 0.37983 | -0.3503 | 0.242 | 0.22 | 1 | HC_vs_MS.Trt_B |
| CCDC93 | 0.38076 | 0.65387 | 0.114 | 0.1 | 1 | HC_vs_MS.Trt_B |
| ADRB2 | 0.38183 | 0.40254 | 0.187 | 0.17 | 1 | HC_vs_MS.Trt_B |
| GSTO1 | 0.38251 | -0.4122 | 0.11 | 0.13 | 1 | HC_vs_MS.Trt_B |
| CCT5 | 0.38259 | -0.5038 | 0.138 | 0.16 | 1 | HC_vs_MS.Trt_B |
| RRAGA | 0.38355 | 0.25591 | 0.11 | 0.1 | 1 | HC_vs_MS.Trt_B |
| COPE | 0.38364 | -0.2542 | 0.303 | 0.28 | 1 | HC_vs_MS.Trt_B |
| ADARB1 | 0.38446 | -0.4895 | 0.197 | 0.18 | 1 | HC_vs_MS.Trt_B |
| CDC123 | 0.38491 | -0.4655 | 0.188 | 0.17 | 1 | HC_vs_MS.Trt_B |
| UBN1 | 0.38521 | -0.412 | 0.117 | 0.1 | 1 | HC_vs_MS.Trt_B |
| FAM103A1 | 0.3858 | 0.45 | 0.101 | 0.09 | 1 | HC_vs_MS.Trt_B |
| MAMLD1 | 0.38697 | -0.7442 | 0.187 | 0.16 | 1 | HC_vs_MS.Trt_B |
| B3GNTL1 | 0.38708 | -0.3378 | 0.217 | 0.2 | 1 | HC_vs_MS.Trt_B |
| ALOX5AP | 0.38804 | -0.5575 | 0.213 | 0.23 | 1 | HC_vs_MS.Trt_B |
| ADCY4 | 0.38948 | -0.643 | 0.088 | 0.11 | 1 | HC_vs_MS.Trt_B |

| | | | | | | |
|---|---|---|---|---|---|---|
| PI4KA | 0.38997 | -0.4785 | 0.225 | 0.2 | 1 | HC_vs_MS.Trt_B |
| PIBF1 | 0.39191 | 0.37439 | 0.128 | 0.11 | 1 | HC_vs_MS.Trt_B |
| PSMB6 | 0.39338 | -0.7792 | 0.571 | 0.58 | 1 | HC_vs_MS.Trt_B |
| SGPL1 | 0.39367 | -0.4565 | 0.419 | 0.38 | 1 | HC_vs_MS.Trt_B |
| PSEN2 | 0.39367 | -1.2647 | 0.138 | 0.12 | 1 | HC_vs_MS.Trt_B |
| S100A6 | 0.39489 | -0.2972 | 0.496 | 0.45 | 1 | HC_vs_MS.Trt_B |
| DENND5A | 0.39499 | -0.5033 | 0.225 | 0.2 | 1 | HC_vs_MS.Trt_B |
| ZMIZ2 | 0.39599 | -0.3428 | 0.128 | 0.11 | 1 | HC_vs_MS.Trt_B |
| CUL1 | 0.39677 | -0.2629 | 0.158 | 0.14 | 1 | HC_vs_MS.Trt_B |
| SCP2 | 0.39684 | -0.441 | 0.184 | 0.16 | 1 | HC_vs_MS.Trt_B |
| PSME2 | 0.39779 | -0.4906 | 0.301 | 0.31 | 1 | HC_vs_MS.Trt_B |
| PAPD5 | 0.3984 | 0.55608 | 0.136 | 0.12 | 1 | HC_vs_MS.Trt_B |
| DOK2 | 0.3986 | -0.3588 | 0.333 | 0.34 | 1 | HC_vs_MS.Trt_B |
| FKBP4 | 0.39884 | -0.69 | 0.094 | 0.11 | 1 | HC_vs_MS.Trt_B |
| SLC16A1 | 0.39984 | 0.52889 | 0.103 | 0.09 | 1 | HC_vs_MS.Trt_B |
| HIST1H3E | 0.40039 | 1.16168 | 0.112 | 0.1 | 1 | HC_vs_MS.Trt_B |
| SLX1A | 0.40169 | -0.3464 | 0.459 | 0.42 | 1 | HC_vs_MS.Trt_B |
| NOG | 0.40223 | -0.6992 | 0.241 | 0.22 | 1 | HC_vs_MS.Trt_B |
| INF2 | 0.40342 | -0.6521 | 0.199 | 0.18 | 1 | HC_vs_MS.Trt_B |
| UBL3 | 0.40371 | -0.7145 | 0.165 | 0.18 | 1 | HC_vs_MS.Trt_B |
| PHACTR1 | 0.40507 | 0.84769 | 0.393 | 0.39 | 1 | HC_vs_MS.Trt_B |
| RB1CC1 | 0.40771 | -0.3495 | 0.293 | 0.27 | 1 | HC_vs_MS.Trt_B |
| TWF2 | 0.40823 | -0.4235 | 0.168 | 0.19 | 1 | HC_vs_MS.Trt_B |
| RNF181 | 0.40834 | -0.616 | 0.139 | 0.15 | 1 | HC_vs_MS.Trt_B |
| MAPRE2 | 0.4128 | -0.4139 | 0.184 | 0.21 | 1 | HC_vs_MS.Trt_B |
| MED14 | 0.41376 | 0.56434 | 0.141 | 0.13 | 1 | HC_vs_MS.Trt_B |
| GTF3C4 | 0.41429 | -0.2618 | 0.29 | 0.27 | 1 | HC_vs_MS.Trt_B |
| NEO1 | 0.41674 | -0.6985 | 0.245 | 0.22 | 1 | HC_vs_MS.Trt_B |
| ITM2B | 0.41757 | -0.2518 | 0.567 | 0.51 | 1 | HC_vs_MS.Trt_B |
| ATP5I | 0.41879 | -0.6412 | 0.219 | 0.2 | 1 | HC_vs_MS.Trt_B |
| CIB1 | 0.41909 | -0.5168 | 0.196 | 0.17 | 1 | HC_vs_MS.Trt_B |
| KLHDC3 | 0.42058 | 0.36771 | 0.117 | 0.11 | 1 | HC_vs_MS.Trt_B |
| ABCB6 | 0.42114 | 0.41688 | 0.113 | 0.1 | 1 | HC_vs_MS.Trt_B |
| SYF2 | 0.42144 | -0.4116 | 0.419 | 0.38 | 1 | HC_vs_MS.Trt_B |
| RANBP1 | 0.4226 | -0.8011 | 0.142 | 0.16 | 1 | HC_vs_MS.Trt_B |
| PAIP2 | 0.42305 | -0.5515 | 0.571 | 0.54 | 1 | HC_vs_MS.Trt_B |
| FBXW7 | 0.42389 | -0.3717 | 0.152 | 0.14 | 1 | HC_vs_MS.Trt_B |
| GTF3A | 0.42504 | -0.3058 | 0.394 | 0.34 | 1 | HC_vs_MS.Trt_B |
| BLOC1S1 | 0.42756 | -0.3826 | 0.165 | 0.14 | 1 | HC_vs_MS.Trt_B |
| ZNF600 | 0.42823 | 0.60377 | 0.142 | 0.13 | 1 | HC_vs_MS.Trt_B |

| | | | | | | |
|---|---|---|---|---|---|---|
| SEC61G | 0.43113 | -0.3293 | 0.216 | 0.2 | 1 | HC_vs_MS.Trt_B |
| PRR14 | 0.43137 | 0.30435 | 0.114 | 0.1 | 1 | HC_vs_MS.Trt_B |
| C7orf49 | 0.43157 | 0.32252 | 0.145 | 0.13 | 1 | HC_vs_MS.Trt_B |
| RNF31 | 0.43487 | -0.5054 | 0.28 | 0.25 | 1 | HC_vs_MS.Trt_B |
| GPX2 | 0.43512 | 1.19109 | 0.18 | 0.17 | 1 | HC_vs_MS.Trt_B |
| ATP11B | 0.43664 | -0.4978 | 0.204 | 0.19 | 1 | HC_vs_MS.Trt_B |
| ZNF573 | 0.43819 | 1.11465 | 0.368 | 0.36 | 1 | HC_vs_MS.Trt_B |
| CBLL1 | 0.43885 | -0.2843 | 0.272 | 0.25 | 1 | HC_vs_MS.Trt_B |
| C10orf95 | 0.44091 | -0.3548 | 0.162 | 0.18 | 1 | HC_vs_MS.Trt_B |
| ITM2A | 0.44263 | -0.301 | 0.219 | 0.24 | 1 | HC_vs_MS.Trt_B |
| KIAA0556 | 0.44476 | 0.63812 | 0.259 | 0.24 | 1 | HC_vs_MS.Trt_B |
| LSM4 | 0.44595 | -0.2996 | 0.165 | 0.15 | 1 | HC_vs_MS.Trt_B |
| DCUN1D4 | 0.446 | -0.3418 | 0.249 | 0.23 | 1 | HC_vs_MS.Trt_B |
| ENY2 | 0.45026 | -0.9656 | 0.525 | 0.48 | 1 | HC_vs_MS.Trt_B |
| DNAJC4 | 0.45059 | 0.2875 | 0.117 | 0.11 | 1 | HC_vs_MS.Trt_B |
| TSPYL2 | 0.45213 | -0.4253 | 0.152 | 0.17 | 1 | HC_vs_MS.Trt_B |
| SLC35F2 | 0.45386 | -0.7933 | 0.219 | 0.2 | 1 | HC_vs_MS.Trt_B |
| SNRPE | 0.45452 | -0.389 | 0.11 | 0.1 | 1 | HC_vs_MS.Trt_B |
| FLNB | 0.45609 | 0.39493 | 0.428 | 0.41 | 1 | HC_vs_MS.Trt_B |
| FAM21A | 0.45921 | -0.3731 | 0.206 | 0.19 | 1 | HC_vs_MS.Trt_B |
| RPP21 | 0.46351 | -0.5125 | 0.112 | 0.12 | 1 | HC_vs_MS.Trt_B |
| DNAJC3 | 0.46586 | -0.625 | 0.458 | 0.42 | 1 | HC_vs_MS.Trt_B |
| R3HDM2 | 0.467 | -0.4426 | 0.78 | 0.76 | 1 | HC_vs_MS.Trt_B |
| RBMXL1 | 0.4686 | 0.79527 | 0.12 | 0.11 | 1 | HC_vs_MS.Trt_B |
| BCAS2 | 0.46942 | -0.494 | 0.162 | 0.14 | 1 | HC_vs_MS.Trt_B |
| NSMCE4A | 0.47186 | -0.4191 | 0.32 | 0.3 | 1 | HC_vs_MS.Trt_B |
| DUT | 0.47282 | -0.4047 | 0.133 | 0.12 | 1 | HC_vs_MS.Trt_B |
| ENO1 | 0.47335 | -0.2618 | 0.377 | 0.4 | 1 | HC_vs_MS.Trt_B |
| FAM76A | 0.47351 | -0.71 | 0.761 | 0.73 | 1 | HC_vs_MS.Trt_B |
| IRF2BP2 | 0.47533 | -0.6314 | 0.197 | 0.17 | 1 | HC_vs_MS.Trt_B |
| RTCB | 0.47537 | 0.53919 | 0.12 | 0.11 | 1 | HC_vs_MS.Trt_B |
| WDR61 | 0.47541 | -0.5335 | 0.13 | 0.15 | 1 | HC_vs_MS.Trt_B |
| ELOVL1 | 0.47569 | -0.2813 | 0.117 | 0.13 | 1 | HC_vs_MS.Trt_B |
| ZC3H4 | 0.47655 | -0.4328 | 0.312 | 0.29 | 1 | HC_vs_MS.Trt_B |
| KYAT1 | 0.47749 | -0.5926 | 0.12 | 0.11 | 1 | HC_vs_MS.Trt_B |
| LIPA | 0.47931 | -0.3865 | 0.262 | 0.24 | 1 | HC_vs_MS.Trt_B |
| ROMO1 | 0.48097 | -0.4222 | 0.139 | 0.12 | 1 | HC_vs_MS.Trt_B |
| CWC27 | 0.48151 | -0.2727 | 0.101 | 0.09 | 1 | HC_vs_MS.Trt_B |
| SNRPA | 0.48204 | -0.3158 | 0.288 | 0.27 | 1 | HC_vs_MS.Trt_B |
| LPCAT3 | 0.48409 | 0.66225 | 0.409 | 0.39 | 1 | HC_vs_MS.Trt_B |

| | | | | | | |
|---|---|---|---|---|---|---|
| PPP2R5E | 0.48417 | -0.2558 | 0.188 | 0.17 | 1 | HC_vs_MS.Trt_B |
| PPID | 0.4844 | -0.6229 | 0.352 | 0.39 | 1 | HC_vs_MS.Trt_B |
| HSPE1 | 0.48494 | -0.6836 | 0.23 | 0.24 | 1 | HC_vs_MS.Trt_B |
| MAP1LC3B | 0.48534 | -0.4572 | 0.303 | 0.27 | 1 | HC_vs_MS.Trt_B |
| STRA13 | 0.48572 | -0.479 | 0.09 | 0.1 | 1 | HC_vs_MS.Trt_B |
| ZIK1 | 0.48608 | 0.38773 | 0.262 | 0.25 | 1 | HC_vs_MS.Trt_B |
| MCRS1 | 0.48673 | -0.3424 | 0.161 | 0.14 | 1 | HC_vs_MS.Trt_B |
| NIPAL2 | 0.48722 | -1.0594 | 0.281 | 0.26 | 1 | HC_vs_MS.Trt_B |
| TRIM26 | 0.48746 | -0.2635 | 0.157 | 0.14 | 1 | HC_vs_MS.Trt_B |
| PEPD | 0.48858 | 0.29322 | 0.104 | 0.09 | 1 | HC_vs_MS.Trt_B |
| EAPP | 0.48906 | -0.2959 | 0.203 | 0.19 | 1 | HC_vs_MS.Trt_B |
| MTRR | 0.49029 | -0.3629 | 0.133 | 0.15 | 1 | HC_vs_MS.Trt_B |
| MTHFS | 0.49156 | -0.489 | 0.138 | 0.12 | 1 | HC_vs_MS.Trt_B |
| SOCS2 | 0.49241 | 0.31562 | 0.346 | 0.33 | 1 | HC_vs_MS.Trt_B |
| URI1 | 0.49315 | -0.5816 | 0.187 | 0.17 | 1 | HC_vs_MS.Trt_B |
| TIA1 | 0.49365 | -0.3018 | 0.216 | 0.2 | 1 | HC_vs_MS.Trt_B |
| MCM5 | 0.4941 | -0.3719 | 0.164 | 0.15 | 1 | HC_vs_MS.Trt_B |
| TOB1 | 0.49531 | -0.2575 | 0.486 | 0.46 | 1 | HC_vs_MS.Trt_B |
| MYDGF | 0.49729 | -0.3147 | 0.113 | 0.1 | 1 | HC_vs_MS.Trt_B |
| PPIB | 0.499 | -0.3347 | 0.677 | 0.63 | 1 | HC_vs_MS.Trt_B |
| JCHAIN | 0.50069 | -0.424 | 0.114 | 0.1 | 1 | HC_vs_MS.Trt_B |
| MED10 | 0.50073 | -0.3468 | 0.217 | 0.2 | 1 | HC_vs_MS.Trt_B |
| NCOA2 | 0.5017 | -0.2595 | 0.201 | 0.18 | 1 | HC_vs_MS.Trt_B |
| PRPF39 | 0.50186 | -0.3575 | 0.258 | 0.24 | 1 | HC_vs_MS.Trt_B |
| E4F1 | 0.5028 | -0.4897 | 0.094 | 0.11 | 1 | HC_vs_MS.Trt_B |
| DDIT4 | 0.50372 | 0.32257 | 0.132 | 0.12 | 1 | HC_vs_MS.Trt_B |
| HNRNPD | 0.50388 | -0.2612 | 0.225 | 0.21 | 1 | HC_vs_MS.Trt_B |
| FAM69A | 0.50457 | -0.3541 | 0.106 | 0.09 | 1 | HC_vs_MS.Trt_B |
| FIGNL1 | 0.50499 | 1.36288 | 0.114 | 0.11 | 1 | HC_vs_MS.Trt_B |
| S1PR1 | 0.50523 | -0.2739 | 0.516 | 0.48 | 1 | HC_vs_MS.Trt_B |
| TSPO | 0.50744 | -0.2519 | 0.271 | 0.25 | 1 | HC_vs_MS.Trt_B |
| PPOX | 0.50746 | -0.3289 | 0.216 | 0.2 | 1 | HC_vs_MS.Trt_B |
| KIF22 | 0.50748 | -0.4238 | 0.188 | 0.17 | 1 | HC_vs_MS.Trt_B |
| TRMT61B | 0.50794 | -0.2574 | 0.107 | 0.1 | 1 | HC_vs_MS.Trt_B |
| DCP1B | 0.50986 | -0.4912 | 0.133 | 0.15 | 1 | HC_vs_MS.Trt_B |
| CHMP3 | 0.51036 | -0.4617 | 0.299 | 0.27 | 1 | HC_vs_MS.Trt_B |
| CTDSP1 | 0.51069 | -0.2923 | 0.165 | 0.15 | 1 | HC_vs_MS.Trt_B |
| FAHD2B | 0.51334 | -0.312 | 0.112 | 0.1 | 1 | HC_vs_MS.Trt_B |
| RIC8B | 0.51347 | -0.4734 | 0.123 | 0.11 | 1 | HC_vs_MS.Trt_B |
| PTEN | 0.51368 | -0.3141 | 0.478 | 0.45 | 1 | HC_vs_MS.Trt_B |

| | | | | | | |
|---|---|---|---|---|---|---|
| PEX19 | 0.51452 | -0.2716 | 0.093 | 0.11 | 1 | HC_vs_MS.Trt_B |
| RPL35 | 0.51475 | -0.3014 | 0.759 | 0.66 | 1 | HC_vs_MS.Trt_B |
| AGAP6 | 0.51545 | -0.5934 | 0.101 | 0.09 | 1 | HC_vs_MS.Trt_B |
| OSGEPL1 | 0.51641 | -0.4916 | 0.245 | 0.26 | 1 | HC_vs_MS.Trt_B |
| ST20 | 0.51675 | -0.5247 | 0.146 | 0.16 | 1 | HC_vs_MS.Trt_B |
| RANBP9 | 0.51893 | -0.5507 | 0.186 | 0.17 | 1 | HC_vs_MS.Trt_B |
| UBE2V1 | 0.5208 | -0.2785 | 0.391 | 0.37 | 1 | HC_vs_MS.Trt_B |
| PIP4K2A | 0.52098 | -0.4196 | 0.741 | 0.7 | 1 | HC_vs_MS.Trt_B |
| BMPR1B | 0.52169 | -1.3557 | 0.122 | 0.11 | 1 | HC_vs_MS.Trt_B |
| TMEM143 | 0.52269 | 0.69885 | 0.465 | 0.44 | 1 | HC_vs_MS.Trt_B |
| RAP1GDS1 | 0.523 | -0.4927 | 0.338 | 0.34 | 1 | HC_vs_MS.Trt_B |
| DNAJB4 | 0.5261 | 0.6764 | 0.284 | 0.27 | 1 | HC_vs_MS.Trt_B |
| ITGA5 | 0.52684 | -0.5776 | 0.103 | 0.11 | 1 | HC_vs_MS.Trt_B |
| PYURF | 0.52878 | -0.3217 | 0.171 | 0.16 | 1 | HC_vs_MS.Trt_B |
| PPIE | 0.5291 | -0.7029 | 0.752 | 0.67 | 1 | HC_vs_MS.Trt_B |
| RGSL1 | 0.52911 | -0.3216 | 0.158 | 0.17 | 1 | HC_vs_MS.Trt_B |
| BSCL2 | 0.52926 | 0.37087 | 0.188 | 0.18 | 1 | HC_vs_MS.Trt_B |
| AMFR | 0.53089 | 0.45533 | 0.514 | 0.5 | 1 | HC_vs_MS.Trt_B |
| GLIPR2 | 0.53304 | -0.5271 | 0.196 | 0.17 | 1 | HC_vs_MS.Trt_B |
| PSMB10 | 0.53629 | -0.3214 | 0.37 | 0.34 | 1 | HC_vs_MS.Trt_B |
| ANKRA2 | 0.53676 | -0.5576 | 0.11 | 0.1 | 1 | HC_vs_MS.Trt_B |
| LARP7 | 0.53743 | -0.5466 | 0.172 | 0.19 | 1 | HC_vs_MS.Trt_B |
| ARNTL | 0.53925 | -0.3099 | 0.191 | 0.18 | 1 | HC_vs_MS.Trt_B |
| RDX | 0.53925 | -0.5092 | 0.236 | 0.22 | 1 | HC_vs_MS.Trt_B |
| RP11-701H2 | 0.54011 | 0.27917 | 0.109 | 0.1 | 1 | HC_vs_MS.Trt_B |
| SEC61B | 0.54028 | -0.2853 | 0.159 | 0.17 | 1 | HC_vs_MS.Trt_B |
| CREBZF | 0.54117 | -0.4513 | 0.167 | 0.15 | 1 | HC_vs_MS.Trt_B |
| ARPC1A | 0.54395 | -0.312 | 0.23 | 0.22 | 1 | HC_vs_MS.Trt_B |
| LRRC56 | 0.54498 | 0.52293 | 0.38 | 0.38 | 1 | HC_vs_MS.Trt_B |
| IFI27L2 | 0.54984 | -0.8044 | 0.142 | 0.13 | 1 | HC_vs_MS.Trt_B |
| CEP120 | 0.5506 | -1.0442 | 0.094 | 0.11 | 1 | HC_vs_MS.Trt_B |
| RHBDD2 | 0.5519 | -0.3352 | 0.226 | 0.21 | 1 | HC_vs_MS.Trt_B |
| MFSD10 | 0.55195 | -0.4193 | 0.107 | 0.12 | 1 | HC_vs_MS.Trt_B |
| RP11-411B6 | 0.55447 | 0.58215 | 0.219 | 0.21 | 1 | HC_vs_MS.Trt_B |
| S100A11 | 0.55564 | -0.35 | 0.301 | 0.28 | 1 | HC_vs_MS.Trt_B |
| EMD | 0.55844 | 0.33454 | 0.194 | 0.19 | 1 | HC_vs_MS.Trt_B |
| VDAC2 | 0.55856 | -0.5062 | 0.235 | 0.25 | 1 | HC_vs_MS.Trt_B |
| IMPDH1 | 0.55889 | -0.6627 | 0.101 | 0.09 | 1 | HC_vs_MS.Trt_B |
| RBX1 | 0.56432 | -0.7399 | 0.23 | 0.24 | 1 | HC_vs_MS.Trt_B |
| ECSIT | 0.56548 | -0.325 | 0.365 | 0.35 | 1 | HC_vs_MS.Trt_B |

| | | | | | | |
|---|---|---|---|---|---|---|
| UBXN11 | 0.56587 | -0.4951 | 0.138 | 0.15 | 1 | HC_vs_MS.Trt_B |
| SCAND1 | 0.56715 | -0.3839 | 0.188 | 0.17 | 1 | HC_vs_MS.Trt_B |
| NDUFA1 | 0.56853 | -0.5768 | 0.209 | 0.22 | 1 | HC_vs_MS.Trt_B |
| BLVRA | 0.56895 | -0.3137 | 0.103 | 0.11 | 1 | HC_vs_MS.Trt_B |
| DCUN1D3 | 0.57021 | 0.7153 | 0.139 | 0.13 | 1 | HC_vs_MS.Trt_B |
| CWC15 | 0.57066 | -0.4784 | 0.19 | 0.2 | 1 | HC_vs_MS.Trt_B |
| RNF123 | 0.57118 | -0.3807 | 0.1 | 0.11 | 1 | HC_vs_MS.Trt_B |
| STON1 | 0.57353 | -0.9519 | 0.301 | 0.28 | 1 | HC_vs_MS.Trt_B |
| POLR2J | 0.57542 | -0.4244 | 0.152 | 0.17 | 1 | HC_vs_MS.Trt_B |
| DAXX | 0.57623 | -0.5466 | 0.18 | 0.19 | 1 | HC_vs_MS.Trt_B |
| AMPH | 0.57638 | -0.5164 | 0.136 | 0.13 | 1 | HC_vs_MS.Trt_B |
| UNG | 0.57817 | 0.55738 | 0.2 | 0.19 | 1 | HC_vs_MS.Trt_B |
| ZNF658 | 0.58071 | -0.8458 | 0.175 | 0.16 | 1 | HC_vs_MS.Trt_B |
| EHBP1 | 0.58075 | -0.6674 | 0.354 | 0.34 | 1 | HC_vs_MS.Trt_B |
| UCHL3 | 0.58375 | -0.5272 | 0.104 | 0.09 | 1 | HC_vs_MS.Trt_B |
| EWSR1 | 0.58424 | -0.2887 | 0.499 | 0.47 | 1 | HC_vs_MS.Trt_B |
| ROBO4 | 0.58496 | -0.2676 | 0.133 | 0.15 | 1 | HC_vs_MS.Trt_B |
| ZNF93 | 0.58807 | -0.7837 | 0.532 | 0.5 | 1 | HC_vs_MS.Trt_B |
| RGS10 | 0.58912 | -0.3126 | 0.177 | 0.17 | 1 | HC_vs_MS.Trt_B |
| COPS9 | 0.58968 | -0.5191 | 0.113 | 0.12 | 1 | HC_vs_MS.Trt_B |
| CTD-2370N5 | 0.59188 | -0.252 | 0.101 | 0.09 | 1 | HC_vs_MS.Trt_B |
| IFIT3 | 0.59505 | 0.45533 | 0.43 | 0.47 | 1 | HC_vs_MS.Trt_B |
| RIC1 | 0.59505 | 0.56818 | 0.103 | 0.1 | 1 | HC_vs_MS.Trt_B |
| MAN1C1 | 0.59525 | -0.2956 | 0.867 | 0.83 | 1 | HC_vs_MS.Trt_B |
| HERPUD1 | 0.59701 | -0.6369 | 0.643 | 0.63 | 1 | HC_vs_MS.Trt_B |
| CCNG2 | 0.59791 | -0.4841 | 0.265 | 0.25 | 1 | HC_vs_MS.Trt_B |
| CAMTA1 | 0.60116 | -0.6354 | 0.116 | 0.11 | 1 | HC_vs_MS.Trt_B |
| SELT | 0.602 | -0.3705 | 0.306 | 0.29 | 1 | HC_vs_MS.Trt_B |
| SRSF6 | 0.60201 | -0.456 | 0.607 | 0.6 | 1 | HC_vs_MS.Trt_B |
| PDE6D | 0.6022 | -0.3772 | 0.1 | 0.11 | 1 | HC_vs_MS.Trt_B |
| GNG5 | 0.6025 | -0.5272 | 0.18 | 0.19 | 1 | HC_vs_MS.Trt_B |
| FNTB | 0.60311 | 0.64082 | 0.126 | 0.12 | 1 | HC_vs_MS.Trt_B |
| LIAS | 0.60342 | 0.38445 | 0.094 | 0.11 | 1 | HC_vs_MS.Trt_B |
| ADSSL1 | 0.60727 | 0.26256 | 0.257 | 0.24 | 1 | HC_vs_MS.Trt_B |
| ATP6V0E1 | 0.60773 | -0.5091 | 0.27 | 0.24 | 1 | HC_vs_MS.Trt_B |
| THEMIS | 0.60784 | 0.42457 | 0.194 | 0.18 | 1 | HC_vs_MS.Trt_B |
| QARS | 0.60852 | -0.3279 | 0.177 | 0.19 | 1 | HC_vs_MS.Trt_B |
| MGST2 | 0.60868 | -0.4223 | 0.157 | 0.15 | 1 | HC_vs_MS.Trt_B |
| SDHB | 0.61354 | -0.6987 | 0.112 | 0.12 | 1 | HC_vs_MS.Trt_B |
| SRI | 0.61516 | -0.5997 | 0.367 | 0.34 | 1 | HC_vs_MS.Trt_B |

| | | | | | | |
|---|---|---|---|---|---|---|
| SH3RF2 | 0.6164 | 0.26231 | 0.165 | 0.16 | 1 | HC_vs_MS.Trt_B |
| TXN | 0.61641 | -0.3903 | 0.19 | 0.17 | 1 | HC_vs_MS.Trt_B |
| ATG5 | 0.61849 | -0.2872 | 0.116 | 0.13 | 1 | HC_vs_MS.Trt_B |
| PUS3 | 0.62098 | 0.5735 | 0.31 | 0.3 | 1 | HC_vs_MS.Trt_B |
| DUSP11 | 0.62158 | -0.3133 | 0.519 | 0.52 | 1 | HC_vs_MS.Trt_B |
| ATP5L | 0.62422 | -0.5071 | 0.375 | 0.37 | 1 | HC_vs_MS.Trt_B |
| HEXDC | 0.62423 | -0.4042 | 0.338 | 0.31 | 1 | HC_vs_MS.Trt_B |
| DNAJB11 | 0.62472 | -0.2804 | 0.142 | 0.13 | 1 | HC_vs_MS.Trt_B |
| CRYGS | 0.62683 | -1.1184 | 0.217 | 0.2 | 1 | HC_vs_MS.Trt_B |
| HMGN2 | 0.62744 | -0.3195 | 0.6 | 0.59 | 1 | HC_vs_MS.Trt_B |
| AC012314.1 | 0.6312 | -0.2953 | 0.117 | 0.12 | 1 | HC_vs_MS.Trt_B |
| AC012314.1 | 0.6312 | -0.2953 | 0.117 | 0.12 | 1 | HC_vs_MS.Trt_B |
| AC012314.1 | 0.6312 | -0.2953 | 0.117 | 0.12 | 1 | HC_vs_MS.Trt_B |
| AC012314.1 | 0.6312 | -0.2953 | 0.117 | 0.12 | 1 | HC_vs_MS.Trt_B |
| AC012314.3 | 0.6312 | -0.2953 | 0.117 | 0.12 | 1 | HC_vs_MS.Trt_B |
| AC012314.6 | 0.6312 | -0.2953 | 0.117 | 0.12 | 1 | HC_vs_MS.Trt_B |
| AC012314.7 | 0.6312 | -0.2953 | 0.117 | 0.12 | 1 | HC_vs_MS.Trt_B |
| AC012314.9 | 0.6312 | -0.2953 | 0.117 | 0.12 | 1 | HC_vs_MS.Trt_B |
| CASP2 | 0.63245 | -0.5194 | 0.793 | 0.73 | 1 | HC_vs_MS.Trt_B |
| ADCY1 | 0.63259 | 0.50803 | 0.135 | 0.13 | 1 | HC_vs_MS.Trt_B |
| MED23 | 0.63265 | -0.3339 | 0.213 | 0.2 | 1 | HC_vs_MS.Trt_B |
| RPF1 | 0.63274 | -0.904 | 0.232 | 0.21 | 1 | HC_vs_MS.Trt_B |
| SELK | 0.63334 | -0.8022 | 0.342 | 0.33 | 1 | HC_vs_MS.Trt_B |
| XPO4 | 0.63765 | -0.2862 | 0.112 | 0.1 | 1 | HC_vs_MS.Trt_B |
| CLOCK | 0.63822 | -0.4209 | 0.123 | 0.12 | 1 | HC_vs_MS.Trt_B |
| ACAT2 | 0.63977 | -0.5792 | 0.22 | 0.21 | 1 | HC_vs_MS.Trt_B |
| HDDC3 | 0.6409 | 0.44149 | 0.306 | 0.29 | 1 | HC_vs_MS.Trt_B |
| GK | 0.64103 | -1.3677 | 0.126 | 0.13 | 1 | HC_vs_MS.Trt_B |
| NCF4 | 0.6421 | -0.6234 | 0.161 | 0.15 | 1 | HC_vs_MS.Trt_B |
| EIF2S1 | 0.64303 | -0.6687 | 0.258 | 0.23 | 1 | HC_vs_MS.Trt_B |
| NHP2 | 0.64514 | -0.2771 | 0.172 | 0.16 | 1 | HC_vs_MS.Trt_B |
| VAPB | 0.64638 | 0.31215 | 0.157 | 0.15 | 1 | HC_vs_MS.Trt_B |
| ZNF586 | 0.6471 | -0.3549 | 0.149 | 0.14 | 1 | HC_vs_MS.Trt_B |
| ZNF146 | 0.6485 | -0.4303 | 0.164 | 0.15 | 1 | HC_vs_MS.Trt_B |
| SDAD1 | 0.64982 | -0.3056 | 0.138 | 0.13 | 1 | HC_vs_MS.Trt_B |
| FAM134A | 0.65143 | -0.5732 | 0.383 | 0.39 | 1 | HC_vs_MS.Trt_B |
| POLE4 | 0.65147 | -0.3219 | 0.097 | 0.11 | 1 | HC_vs_MS.Trt_B |
| JAKMIP1 | 0.65251 | -0.388 | 0.175 | 0.16 | 1 | HC_vs_MS.Trt_B |
| FBXL3 | 0.65286 | -0.5934 | 0.104 | 0.1 | 1 | HC_vs_MS.Trt_B |
| MIB1 | 0.65294 | 0.34029 | 0.101 | 0.09 | 1 | HC_vs_MS.Trt_B |

| | | | | | | |
|---|---|---|---|---|---|---|
| SCMH1 | 0.65511 | 0.29844 | 0.104 | 0.1 | 1 | HC_vs_MS.Trt_B |
| REXO2 | 0.65766 | -0.434 | 0.139 | 0.15 | 1 | HC_vs_MS.Trt_B |
| ASB13 | 0.65873 | -1.0372 | 0.097 | 0.11 | 1 | HC_vs_MS.Trt_B |
| IL17RC | 0.65914 | -0.5543 | 0.336 | 0.33 | 1 | HC_vs_MS.Trt_B |
| ERH | 0.66177 | -0.2662 | 0.201 | 0.19 | 1 | HC_vs_MS.Trt_B |
| ALDH9A1 | 0.66356 | -0.3726 | 0.11 | 0.12 | 1 | HC_vs_MS.Trt_B |
| UBQLN1 | 0.6654 | 0.28614 | 0.177 | 0.17 | 1 | HC_vs_MS.Trt_B |
| OSMR | 0.66723 | -0.5795 | 0.49 | 0.5 | 1 | HC_vs_MS.Trt_B |
| CTIF | 0.66894 | 0.90828 | 0.162 | 0.16 | 1 | HC_vs_MS.Trt_B |
| GPM6B | 0.66996 | -0.3438 | 0.114 | 0.11 | 1 | HC_vs_MS.Trt_B |
| SLC25A5 | 0.67055 | -0.5626 | 0.278 | 0.26 | 1 | HC_vs_MS.Trt_B |
| USP9Y | 0.67239 | 0.85575 | 0.235 | 0.22 | 1 | HC_vs_MS.Trt_B |
| HIST1H2AG | 0.67561 | 0.79214 | 0.278 | 0.28 | 1 | HC_vs_MS.Trt_B |
| PRDX2 | 0.67675 | -0.6377 | 0.212 | 0.22 | 1 | HC_vs_MS.Trt_B |
| ITPA | 0.67769 | -0.3388 | 0.113 | 0.11 | 1 | HC_vs_MS.Trt_B |
| GMPR | 0.67814 | -0.6053 | 0.348 | 0.32 | 1 | HC_vs_MS.Trt_B |
| APOL3 | 0.67865 | 0.37484 | 0.199 | 0.19 | 1 | HC_vs_MS.Trt_B |
| ATP1B1 | 0.67913 | 0.43011 | 0.161 | 0.17 | 1 | HC_vs_MS.Trt_B |
| UQCRFS1 | 0.67993 | -0.3498 | 0.229 | 0.21 | 1 | HC_vs_MS.Trt_B |
| DEDD2 | 0.68073 | -0.4394 | 0.312 | 0.3 | 1 | HC_vs_MS.Trt_B |
| IMP4 | 0.6832 | -0.3023 | 0.165 | 0.17 | 1 | HC_vs_MS.Trt_B |
| ARGLU1 | 0.687 | -0.4052 | 0.667 | 0.61 | 1 | HC_vs_MS.Trt_B |
| POMP | 0.69018 | -0.5956 | 0.167 | 0.17 | 1 | HC_vs_MS.Trt_B |
| UPK1B | 0.69085 | -0.2696 | 0.397 | 0.41 | 1 | HC_vs_MS.Trt_B |
| UBASH3A | 0.69181 | -0.5841 | 0.183 | 0.17 | 1 | HC_vs_MS.Trt_B |
| TTC39C | 0.69232 | -0.496 | 0.819 | 0.76 | 1 | HC_vs_MS.Trt_B |
| CLIC1 | 0.69249 | -0.4994 | 0.543 | 0.53 | 1 | HC_vs_MS.Trt_B |
| KCNAB3 | 0.69278 | -0.9851 | 0.343 | 0.33 | 1 | HC_vs_MS.Trt_B |
| SRRT | 0.69334 | 0.3753 | 0.145 | 0.14 | 1 | HC_vs_MS.Trt_B |
| NME1 | 0.69408 | -0.4318 | 0.542 | 0.58 | 1 | HC_vs_MS.Trt_B |
| CALHM2 | 0.69602 | -0.3172 | 0.122 | 0.13 | 1 | HC_vs_MS.Trt_B |
| ADNP | 0.69751 | -0.534 | 0.438 | 0.42 | 1 | HC_vs_MS.Trt_B |
| CYB5R3 | 0.70436 | -0.5001 | 0.474 | 0.45 | 1 | HC_vs_MS.Trt_B |
| PSMA7 | 0.70485 | -0.4787 | 0.367 | 0.36 | 1 | HC_vs_MS.Trt_B |
| HLA-DRA | 0.70609 | -0.6379 | 0.116 | 0.12 | 1 | HC_vs_MS.Trt_B |
| RPL36AL | 0.70713 | -0.3409 | 0.467 | 0.44 | 1 | HC_vs_MS.Trt_B |
| THOC7 | 0.7084 | -0.6083 | 0.145 | 0.13 | 1 | HC_vs_MS.Trt_B |
| PNN | 0.70905 | -0.4421 | 0.246 | 0.25 | 1 | HC_vs_MS.Trt_B |
| RPE | 0.71319 | -0.4398 | 0.288 | 0.28 | 1 | HC_vs_MS.Trt_B |
| CLEC18A | 0.71371 | 0.62635 | 0.281 | 0.27 | 1 | HC_vs_MS.Trt_B |

| | | | | | | |
|---|---|---|---|---|---|---|
| EIF6 | 0.71535 | -0.4229 | 0.125 | 0.13 | 1 | HC_vs_MS.Trt_B |
| KIF20B | 0.7161 | -0.5375 | 0.119 | 0.11 | 1 | HC_vs_MS.Trt_B |
| DBN1 | 0.71725 | 0.41175 | 0.217 | 0.21 | 1 | HC_vs_MS.Trt_B |
| PER1 | 0.71814 | 0.30326 | 0.332 | 0.34 | 1 | HC_vs_MS.Trt_B |
| RUVBL1 | 0.71867 | -0.3746 | 0.643 | 0.63 | 1 | HC_vs_MS.Trt_B |
| MDH2 | 0.71968 | -0.3656 | 0.18 | 0.19 | 1 | HC_vs_MS.Trt_B |
| ACAD9 | 0.71987 | -0.6002 | 0.248 | 0.24 | 1 | HC_vs_MS.Trt_B |
| HMGB2 | 0.72067 | -0.3162 | 0.22 | 0.21 | 1 | HC_vs_MS.Trt_B |
| BIN1 | 0.72095 | -0.443 | 0.262 | 0.26 | 1 | HC_vs_MS.Trt_B |
| GTF2F1 | 0.72243 | -1.0197 | 0.268 | 0.25 | 1 | HC_vs_MS.Trt_B |
| BAZ1B | 0.72318 | -0.2614 | 0.168 | 0.16 | 1 | HC_vs_MS.Trt_B |
| CEACAM1 | 0.72326 | 0.87368 | 0.79 | 0.82 | 1 | HC_vs_MS.Trt_B |
| RBBP6 | 0.72382 | -0.4584 | 0.22 | 0.21 | 1 | HC_vs_MS.Trt_B |
| EDEM3 | 0.72492 | -0.3301 | 0.274 | 0.27 | 1 | HC_vs_MS.Trt_B |
| CEP128 | 0.7274 | -0.4053 | 0.183 | 0.17 | 1 | HC_vs_MS.Trt_B |
| ADHFE1 | 0.72812 | -0.5579 | 0.129 | 0.12 | 1 | HC_vs_MS.Trt_B |
| TTC1 | 0.72867 | -0.6243 | 0.151 | 0.16 | 1 | HC_vs_MS.Trt_B |
| PTPN6 | 0.73243 | -0.3198 | 0.328 | 0.31 | 1 | HC_vs_MS.Trt_B |
| TMEM104 | 0.73263 | -0.2713 | 0.1 | 0.11 | 1 | HC_vs_MS.Trt_B |
| MIF | 0.73394 | -0.6112 | 0.361 | 0.35 | 1 | HC_vs_MS.Trt_B |
| FAM96B | 0.73537 | -0.4901 | 0.172 | 0.16 | 1 | HC_vs_MS.Trt_B |
| PVRIG | 0.73558 | -0.5279 | 0.267 | 0.25 | 1 | HC_vs_MS.Trt_B |
| AP1G2 | 0.73583 | -0.2926 | 0.661 | 0.61 | 1 | HC_vs_MS.Trt_B |
| ASF1A | 0.7368 | -0.6534 | 0.141 | 0.13 | 1 | HC_vs_MS.Trt_B |
| MTG1 | 0.73727 | -0.2694 | 0.148 | 0.14 | 1 | HC_vs_MS.Trt_B |
| ARL6IP6 | 0.73792 | -0.2833 | 0.103 | 0.1 | 1 | HC_vs_MS.Trt_B |
| RECQL | 0.73811 | 0.30821 | 0.128 | 0.12 | 1 | HC_vs_MS.Trt_B |
| IFITM1 | 0.73925 | -0.2578 | 0.583 | 0.54 | 1 | HC_vs_MS.Trt_B |
| CXCR3 | 0.73937 | -0.5017 | 0.103 | 0.1 | 1 | HC_vs_MS.Trt_B |
| WASF2 | 0.74242 | -0.2518 | 0.509 | 0.48 | 1 | HC_vs_MS.Trt_B |
| MRPS34 | 0.74249 | -0.3757 | 0.138 | 0.14 | 1 | HC_vs_MS.Trt_B |
| RSL24D1 | 0.74358 | -0.4274 | 0.249 | 0.24 | 1 | HC_vs_MS.Trt_B |
| ZNF211 | 0.74644 | -0.5984 | 0.152 | 0.16 | 1 | HC_vs_MS.Trt_B |
| MPV17L2 | 0.74823 | -0.8949 | 0.174 | 0.16 | 1 | HC_vs_MS.Trt_B |
| ZC2HC1C | 0.74984 | 0.45847 | 0.109 | 0.1 | 1 | HC_vs_MS.Trt_B |
| CCHCR1 | 0.7505 | -0.5615 | 0.293 | 0.29 | 1 | HC_vs_MS.Trt_B |
| MRPL41 | 0.75219 | -0.7185 | 0.128 | 0.13 | 1 | HC_vs_MS.Trt_B |
| ZNF236 | 0.75459 | -0.4044 | 0.106 | 0.11 | 1 | HC_vs_MS.Trt_B |
| FAM96A | 0.75478 | -0.3734 | 0.312 | 0.3 | 1 | HC_vs_MS.Trt_B |
| HOMEZ | 0.75478 | -0.6681 | 0.183 | 0.19 | 1 | HC_vs_MS.Trt_B |

| | | | | | | |
|---|---|---|---|---|---|---|
| AATF | 0.75597 | -0.3126 | 0.165 | 0.16 | 1 | HC_vs_MS.Trt_B |
| LSM2 | 0.757 | -0.2589 | 0.125 | 0.12 | 1 | HC_vs_MS.Trt_B |
| RRAGD | 0.75773 | 0.37302 | 0.119 | 0.11 | 1 | HC_vs_MS.Trt_B |
| ARID4A | 0.75869 | -0.3809 | 0.278 | 0.27 | 1 | HC_vs_MS.Trt_B |
| NOMO1 | 0.76268 | 0.51463 | 0.116 | 0.11 | 1 | HC_vs_MS.Trt_B |
| E2F6 | 0.76309 | 0.97152 | 0.217 | 0.22 | 1 | HC_vs_MS.Trt_B |
| MT1X | 0.76347 | -0.6662 | 0.101 | 0.1 | 1 | HC_vs_MS.Trt_B |
| ADAM19 | 0.7659 | -0.3198 | 0.319 | 0.31 | 1 | HC_vs_MS.Trt_B |
| SERF2 | 0.76628 | -0.4493 | 0.799 | 0.69 | 1 | HC_vs_MS.Trt_B |
| ZNF92 | 0.76777 | 0.5733 | 0.1 | 0.11 | 1 | HC_vs_MS.Trt_B |
| GLTP | 0.76912 | -0.2672 | 0.583 | 0.58 | 1 | HC_vs_MS.Trt_B |
| COX6A1 | 0.76981 | -0.3424 | 0.113 | 0.11 | 1 | HC_vs_MS.Trt_B |
| TRAF2 | 0.77014 | -0.2813 | 0.161 | 0.15 | 1 | HC_vs_MS.Trt_B |
| MLF2 | 0.77073 | -0.3094 | 0.152 | 0.14 | 1 | HC_vs_MS.Trt_B |
| TBC1D13 | 0.77093 | 0.47667 | 0.109 | 0.12 | 1 | HC_vs_MS.Trt_B |
| CTNNB1 | 0.77099 | -0.4572 | 0.532 | 0.5 | 1 | HC_vs_MS.Trt_B |
| RP11-599B1 | 0.7719 | -0.2861 | 0.259 | 0.25 | 1 | HC_vs_MS.Trt_B |
| STOM | 0.77639 | -0.5119 | 0.577 | 0.53 | 1 | HC_vs_MS.Trt_B |
| PPP2R5A | 0.77726 | -0.4097 | 0.187 | 0.19 | 1 | HC_vs_MS.Trt_B |
| TBPL1 | 0.77925 | -0.7016 | 0.188 | 0.18 | 1 | HC_vs_MS.Trt_B |
| TMED4 | 0.78052 | -0.3416 | 0.913 | 0.86 | 1 | HC_vs_MS.Trt_B |
| PDCD10 | 0.78083 | -0.532 | 0.174 | 0.17 | 1 | HC_vs_MS.Trt_B |
| MRPS14 | 0.78093 | -0.8647 | 0.116 | 0.11 | 1 | HC_vs_MS.Trt_B |
| MCM7 | 0.78375 | -0.4015 | 0.18 | 0.18 | 1 | HC_vs_MS.Trt_B |
| POLR2J2 | 0.78731 | -0.4108 | 0.261 | 0.27 | 1 | HC_vs_MS.Trt_B |
| ATG9B | 0.7875 | 0.29092 | 0.122 | 0.12 | 1 | HC_vs_MS.Trt_B |
| MAP3K3 | 0.78875 | -0.2536 | 0.142 | 0.15 | 1 | HC_vs_MS.Trt_B |
| ANP32E | 0.7892 | -0.4316 | 0.384 | 0.37 | 1 | HC_vs_MS.Trt_B |
| ARHGAP44 | 0.7918 | 0.54941 | 0.242 | 0.24 | 1 | HC_vs_MS.Trt_B |
| TOMM5 | 0.79573 | -0.4396 | 0.135 | 0.14 | 1 | HC_vs_MS.Trt_B |
| PLA2G4E | 0.79586 | -0.3421 | 0.114 | 0.11 | 1 | HC_vs_MS.Trt_B |
| GRAMD1B | 0.79586 | -0.4247 | 0.299 | 0.29 | 1 | HC_vs_MS.Trt_B |
| SAFB | 0.79625 | -0.3428 | 0.345 | 0.35 | 1 | HC_vs_MS.Trt_B |
| MLX | 0.79666 | -0.2716 | 0.399 | 0.41 | 1 | HC_vs_MS.Trt_B |
| DVL3 | 0.79793 | -0.5104 | 0.235 | 0.23 | 1 | HC_vs_MS.Trt_B |
| BTBD8 | 0.79793 | -0.6482 | 0.293 | 0.29 | 1 | HC_vs_MS.Trt_B |
| TRMT1 | 0.7992 | -0.2787 | 0.129 | 0.12 | 1 | HC_vs_MS.Trt_B |
| TCEB2 | 0.8029 | -0.4625 | 0.301 | 0.28 | 1 | HC_vs_MS.Trt_B |
| KCNJ3 | 0.80403 | 0.29926 | 0.214 | 0.22 | 1 | HC_vs_MS.Trt_B |
| RAB5A | 0.80627 | -0.5476 | 0.113 | 0.11 | 1 | HC_vs_MS.Trt_B |

| | | | | | | |
|---|---|---|---|---|---|---|
| THOC1 | 0.8075 | 0.37019 | 0.19 | 0.19 | 1 | HC_vs_MS.Trt_B |
| ATP6V1G1 | 0.80756 | -0.4783 | 0.426 | 0.4 | 1 | HC_vs_MS.Trt_B |
| NPM1 | 0.81214 | -0.35 | 0.648 | 0.58 | 1 | HC_vs_MS.Trt_B |
| PHPT1 | 0.81231 | -0.6846 | 0.106 | 0.11 | 1 | HC_vs_MS.Trt_B |
| ZNF836 | 0.81334 | -0.8247 | 0.161 | 0.16 | 1 | HC_vs_MS.Trt_B |
| ZNF641 | 0.81468 | -0.2796 | 0.383 | 0.37 | 1 | HC_vs_MS.Trt_B |
| UBE2L3 | 0.81591 | -0.4069 | 0.241 | 0.23 | 1 | HC_vs_MS.Trt_B |
| ZNF14 | 0.81595 | -0.3654 | 0.32 | 0.31 | 1 | HC_vs_MS.Trt_B |
| H2AFZ | 0.81712 | -0.3831 | 0.333 | 0.31 | 1 | HC_vs_MS.Trt_B |
| TCEAL8 | 0.8173 | -0.3705 | 0.109 | 0.11 | 1 | HC_vs_MS.Trt_B |
| GNG10 | 0.81877 | -1.0673 | 0.11 | 0.11 | 1 | HC_vs_MS.Trt_B |
| TMEM173 | 0.81939 | -0.4372 | 0.201 | 0.19 | 1 | HC_vs_MS.Trt_B |
| DHX35 | 0.81941 | -0.4381 | 0.339 | 0.33 | 1 | HC_vs_MS.Trt_B |
| PSMD7 | 0.82067 | -0.4314 | 0.275 | 0.27 | 1 | HC_vs_MS.Trt_B |
| CKLF | 0.82077 | -0.2701 | 0.148 | 0.14 | 1 | HC_vs_MS.Trt_B |
| CASP1 | 0.82417 | -0.4325 | 0.196 | 0.19 | 1 | HC_vs_MS.Trt_B |
| TM9SF3 | 0.82642 | -0.3434 | 0.11 | 0.11 | 1 | HC_vs_MS.Trt_B |
| DDIAS | 0.82912 | 0.41437 | 0.355 | 0.35 | 1 | HC_vs_MS.Trt_B |
| NDUFC1 | 0.83028 | -0.3279 | 0.2 | 0.21 | 1 | HC_vs_MS.Trt_B |
| SNRPD2 | 0.83171 | -0.6707 | 0.51 | 0.48 | 1 | HC_vs_MS.Trt_B |
| CNOT8 | 0.83205 | -0.3199 | 0.225 | 0.22 | 1 | HC_vs_MS.Trt_B |
| UQCR10 | 0.83261 | -0.8671 | 0.288 | 0.28 | 1 | HC_vs_MS.Trt_B |
| RPS20 | 0.83665 | -0.2957 | 0.791 | 0.76 | 1 | HC_vs_MS.Trt_B |
| BANF1 | 0.83677 | -0.5007 | 0.181 | 0.18 | 1 | HC_vs_MS.Trt_B |
| PITPNB | 0.83677 | -0.4113 | 0.161 | 0.17 | 1 | HC_vs_MS.Trt_B |
| NOL7 | 0.83696 | -0.3384 | 0.199 | 0.19 | 1 | HC_vs_MS.Trt_B |
| MAP1LC3B2 | 0.83705 | -0.6664 | 0.206 | 0.2 | 1 | HC_vs_MS.Trt_B |
| NPEPL1 | 0.83935 | -0.6453 | 0.11 | 0.11 | 1 | HC_vs_MS.Trt_B |
| MYL12B | 0.83958 | -0.2565 | 0.459 | 0.45 | 1 | HC_vs_MS.Trt_B |
| ACSL6 | 0.84192 | -0.385 | 0.509 | 0.55 | 1 | HC_vs_MS.Trt_B |
| ARL6IP4 | 0.84569 | -0.4064 | 0.284 | 0.28 | 1 | HC_vs_MS.Trt_B |
| DHX9 | 0.8479 | -0.4538 | 0.194 | 0.19 | 1 | HC_vs_MS.Trt_B |
| PNPLA6 | 0.84911 | -0.3858 | 0.126 | 0.13 | 1 | HC_vs_MS.Trt_B |
| FAM172A | 0.84937 | -0.3017 | 0.11 | 0.11 | 1 | HC_vs_MS.Trt_B |
| MKRN3 | 0.85232 | 0.97805 | 0.452 | 0.46 | 1 | HC_vs_MS.Trt_B |
| MVB12A | 0.85237 | -0.5054 | 0.514 | 0.5 | 1 | HC_vs_MS.Trt_B |
| ATP6V0B | 0.85724 | -0.2732 | 0.145 | 0.15 | 1 | HC_vs_MS.Trt_B |
| ATP5G1 | 0.85753 | -0.484 | 0.228 | 0.23 | 1 | HC_vs_MS.Trt_B |
| PSMB5 | 0.85768 | -0.2956 | 0.099 | 0.1 | 1 | HC_vs_MS.Trt_B |
| KLHL3 | 0.858 | -0.5751 | 0.184 | 0.19 | 1 | HC_vs_MS.Trt_B |

| | | | | | | |
|---|---|---|---|---|---|---|
| RNF13 | 0.85982 | -0.7176 | 0.135 | 0.13 | 1 | HC_vs_MS.Trt_B |
| PSMB8 | 0.86011 | -0.2844 | 0.346 | 0.34 | 1 | HC_vs_MS.Trt_B |
| SNRPC | 0.86389 | -0.3629 | 0.172 | 0.18 | 1 | HC_vs_MS.Trt_B |
| NUDT7 | 0.87024 | -0.4303 | 0.177 | 0.18 | 1 | HC_vs_MS.Trt_B |
| SFT2D1 | 0.8727 | -0.5488 | 0.307 | 0.31 | 1 | HC_vs_MS.Trt_B |
| PIP4K2B | 0.87291 | -1.1766 | 0.107 | 0.1 | 1 | HC_vs_MS.Trt_B |
| SRP9 | 0.87526 | -0.5843 | 0.345 | 0.34 | 1 | HC_vs_MS.Trt_B |
| MORF4L2 | 0.87924 | -0.356 | 0.155 | 0.16 | 1 | HC_vs_MS.Trt_B |
| AKIRIN2 | 0.8799 | -0.2867 | 0.132 | 0.14 | 1 | HC_vs_MS.Trt_B |
| RNF43 | 0.88037 | -0.7798 | 0.296 | 0.28 | 1 | HC_vs_MS.Trt_B |
| CHKB | 0.8811 | -0.7973 | 0.186 | 0.19 | 1 | HC_vs_MS.Trt_B |
| VAMP8 | 0.88772 | -0.5324 | 0.233 | 0.23 | 1 | HC_vs_MS.Trt_B |
| SH2D1A | 0.89053 | -0.4377 | 0.159 | 0.16 | 1 | HC_vs_MS.Trt_B |
| PDIA6 | 0.89096 | -0.3848 | 0.177 | 0.17 | 1 | HC_vs_MS.Trt_B |
| ZNF302 | 0.89558 | -0.4551 | 0.426 | 0.42 | 1 | HC_vs_MS.Trt_B |
| NDUFB7 | 0.89769 | -0.5474 | 0.155 | 0.15 | 1 | HC_vs_MS.Trt_B |
| TAF7 | 0.90261 | -0.4681 | 0.291 | 0.29 | 1 | HC_vs_MS.Trt_B |
| AIP | 0.90516 | -0.2938 | 0.228 | 0.22 | 1 | HC_vs_MS.Trt_B |
| CXorf57 | 0.90635 | -1.6091 | 0.11 | 0.11 | 1 | HC_vs_MS.Trt_B |
| CNN3 | 0.90808 | 0.39524 | 0.165 | 0.16 | 1 | HC_vs_MS.Trt_B |
| PPT1 | 0.90887 | -0.4009 | 0.168 | 0.17 | 1 | HC_vs_MS.Trt_B |
| HDHD2 | 0.9117 | -0.613 | 0.114 | 0.11 | 1 | HC_vs_MS.Trt_B |
| SMG7 | 0.91629 | -0.3804 | 0.162 | 0.16 | 1 | HC_vs_MS.Trt_B |
| CCT4 | 0.91699 | -0.3594 | 0.18 | 0.18 | 1 | HC_vs_MS.Trt_B |
| B3GAT2 | 0.91855 | 0.48944 | 0.151 | 0.15 | 1 | HC_vs_MS.Trt_B |
| ZCRB1 | 0.92007 | -0.4711 | 0.146 | 0.14 | 1 | HC_vs_MS.Trt_B |
| LBR | 0.92088 | -0.3165 | 0.206 | 0.2 | 1 | HC_vs_MS.Trt_B |
| TEX264 | 0.92183 | -0.321 | 0.154 | 0.15 | 1 | HC_vs_MS.Trt_B |
| TRAPPC13 | 0.92364 | 0.37685 | 0.145 | 0.15 | 1 | HC_vs_MS.Trt_B |
| ZNF180 | 0.92425 | -0.4197 | 0.332 | 0.34 | 1 | HC_vs_MS.Trt_B |
| BCAP31 | 0.9254 | 0.28206 | 0.122 | 0.12 | 1 | HC_vs_MS.Trt_B |
| TMEM256-P | 0.92984 | -0.2573 | 0.514 | 0.48 | 1 | HC_vs_MS.Trt_B |
| OPA1 | 0.92986 | 0.3454 | 0.122 | 0.12 | 1 | HC_vs_MS.Trt_B |
| CCDC59 | 0.93211 | -0.2786 | 0.114 | 0.11 | 1 | HC_vs_MS.Trt_B |
| EFHB | 0.93376 | -0.3147 | 0.249 | 0.25 | 1 | HC_vs_MS.Trt_B |
| SDC3 | 0.93381 | 0.38095 | 0.268 | 0.27 | 1 | HC_vs_MS.Trt_B |
| ANKLE1 | 0.93978 | -0.2948 | 0.154 | 0.16 | 1 | HC_vs_MS.Trt_B |
| OSER1 | 0.9444 | -0.747 | 0.157 | 0.16 | 1 | HC_vs_MS.Trt_B |
| LONP1 | 0.94535 | -0.4745 | 0.488 | 0.5 | 1 | HC_vs_MS.Trt_B |
| ASIC4 | 0.94542 | 0.34553 | 0.21 | 0.21 | 1 | HC_vs_MS.Trt_B |

| | | | | | | |
|---|---|---|---|---|---|---|
| LYPLA2 | 0.94638 | -0.3388 | 0.135 | 0.14 | 1 | HC_vs_MS.Trt_B |
| AP1S2 | 0.94871 | -0.4477 | 0.136 | 0.14 | 1 | HC_vs_MS.Trt_B |
| CSNK1A1 | 0.9493 | -0.2714 | 0.796 | 0.79 | 1 | HC_vs_MS.Trt_B |
| PARG | 0.95069 | 0.48191 | 0.104 | 0.11 | 1 | HC_vs_MS.Trt_B |
| CTBS | 0.95382 | -0.6598 | 0.125 | 0.13 | 1 | HC_vs_MS.Trt_B |
| WDR48 | 0.95399 | -0.2643 | 0.129 | 0.13 | 1 | HC_vs_MS.Trt_B |
| DCAF1 | 0.95415 | -0.2876 | 0.172 | 0.17 | 1 | HC_vs_MS.Trt_B |
| SPHK1 | 0.95464 | 0.33137 | 0.138 | 0.14 | 1 | HC_vs_MS.Trt_B |
| GID8 | 0.95529 | -0.4045 | 0.116 | 0.11 | 1 | HC_vs_MS.Trt_B |
| CALM1 | 0.95657 | -0.2662 | 0.816 | 0.73 | 1 | HC_vs_MS.Trt_B |
| SULT1A4 | 0.95693 | -0.4538 | 0.21 | 0.21 | 1 | HC_vs_MS.Trt_B |
| CNKSR1 | 0.95804 | 0.68801 | 0.138 | 0.14 | 1 | HC_vs_MS.Trt_B |
| NAXE | 0.959 | -0.2504 | 0.107 | 0.11 | 1 | HC_vs_MS.Trt_B |
| CEP70 | 0.9601 | -0.9363 | 0.123 | 0.13 | 1 | HC_vs_MS.Trt_B |
| CRIP1 | 0.96586 | -0.7464 | 0.296 | 0.28 | 1 | HC_vs_MS.Trt_B |
| IDI1 | 0.9661 | -0.2543 | 0.116 | 0.12 | 1 | HC_vs_MS.Trt_B |
| EBP | 0.96704 | -0.3806 | 0.129 | 0.13 | 1 | HC_vs_MS.Trt_B |
| NDUFA6 | 0.97005 | -0.4228 | 0.188 | 0.19 | 1 | HC_vs_MS.Trt_B |
| GOLGA7 | 0.97167 | -0.4891 | 0.216 | 0.22 | 1 | HC_vs_MS.Trt_B |
| NDUFS6 | 0.974 | -0.6071 | 0.107 | 0.11 | 1 | HC_vs_MS.Trt_B |
| ATP6V1F | 0.97441 | -0.4607 | 0.155 | 0.15 | 1 | HC_vs_MS.Trt_B |
| VPS29 | 0.97461 | -0.59 | 0.414 | 0.41 | 1 | HC_vs_MS.Trt_B |
| MRPL33 | 0.97464 | -0.5595 | 0.11 | 0.11 | 1 | HC_vs_MS.Trt_B |
| KIAA0408 | 0.97496 | -0.4025 | 0.27 | 0.28 | 1 | HC_vs_MS.Trt_B |
| STK17B | 0.97537 | -0.4578 | 0.933 | 0.87 | 1 | HC_vs_MS.Trt_B |
| SNRPB | 0.97563 | -0.4332 | 0.188 | 0.19 | 1 | HC_vs_MS.Trt_B |
| C12orf29 | 0.97738 | -0.4214 | 0.199 | 0.19 | 1 | HC_vs_MS.Trt_B |
| CDV3 | 0.97756 | -0.3511 | 0.188 | 0.19 | 1 | HC_vs_MS.Trt_B |
| TBP | 0.97798 | -0.2551 | 0.314 | 0.31 | 1 | HC_vs_MS.Trt_B |
| EMP3 | 0.97923 | -0.293 | 0.507 | 0.46 | 1 | HC_vs_MS.Trt_B |
| LTV1 | 0.98116 | -0.2648 | 0.101 | 0.1 | 1 | HC_vs_MS.Trt_B |
| PPP3CA | 0.98156 | -0.5552 | 0.425 | 0.42 | 1 | HC_vs_MS.Trt_B |
| SNRNP40 | 0.9822 | -0.2518 | 0.135 | 0.14 | 1 | HC_vs_MS.Trt_B |
| SF3B4 | 0.98886 | -0.3274 | 0.172 | 0.17 | 1 | HC_vs_MS.Trt_B |
| BOP1 | 0.99337 | 0.52599 | 0.186 | 0.19 | 1 | HC_vs_MS.Trt_B |
| SOD1 | 0.9937 | -0.3507 | 0.646 | 0.64 | 1 | HC_vs_MS.Trt_B |
| CCDC82 | 0.99403 | -0.4972 | 0.129 | 0.13 | 1 | HC_vs_MS.Trt_B |
| CHURC1-FN' | 0.99573 | -0.4869 | 0.165 | 0.16 | 1 | HC_vs_MS.Trt_B |
| PCSK5 | 2.08E-32 | 2.61124 | 0.994 | 0.93 | 3.32E-28 | HC_vs_MS.NoTrt_F |
| MT-ND4L | 1.15E-22 | 1.29296 | 0.974 | 1 | 1.84E-18 | HC_vs_MS.NoTrt_F |

| | | | | | | |
|---|---|---|---|---|---|---|
| ORC6 | 4.44E-21 | 1.1084 | 0.994 | 0.97 | 7.09E-17 | HC_vs_MS.NoTrt_F |
| METTL21A | 2.74E-17 | 1.0664 | 0.977 | 0.95 | 4.38E-13 | HC_vs_MS.NoTrt_F |
| MORC4 | 3.02E-16 | 0.60796 | 1 | 0.99 | 4.83E-12 | HC_vs_MS.NoTrt_F |
| RBPJ | 2.20E-15 | -0.3025 | 0.408 | 0.79 | 3.52E-11 | HC_vs_MS.NoTrt_F |
| RAD21 | 4.64E-12 | -0.9097 | 0.277 | 0.57 | 7.41E-08 | HC_vs_MS.NoTrt_F |
| HS3ST3B1 | 6.41E-12 | -0.7353 | 0.46 | 0.71 | 1.02E-07 | HC_vs_MS.NoTrt_F |
| TYR | 2.09E-11 | -0.6736 | 0.205 | 0.46 | 3.34E-07 | HC_vs_MS.NoTrt_F |
| C2CD3 | 5.02E-11 | 0.37303 | 1 | 0.99 | 8.02E-07 | HC_vs_MS.NoTrt_F |
| ARHGEF18 | 5.46E-11 | -0.3426 | 0.266 | 0.55 | 8.72E-07 | HC_vs_MS.NoTrt_F |
| TMEM45A | 1.90E-10 | 0.51473 | 1 | 0.99 | 3.04E-06 | HC_vs_MS.NoTrt_F |
| RPS3A | 2.75E-10 | -0.3521 | 0.882 | 0.99 | 4.39E-06 | HC_vs_MS.NoTrt_F |
| ARHGAP45 | 6.40E-10 | -0.5341 | 0.61 | 0.87 | 1.02E-05 | HC_vs_MS.NoTrt_F |
| MED15 | 7.14E-10 | -0.5498 | 0.54 | 0.76 | 1.14E-05 | HC_vs_MS.NoTrt_F |
| ITGB1 | 2.01E-09 | -0.5623 | 0.592 | 0.82 | 3.22E-05 | HC_vs_MS.NoTrt_F |
| RASA3 | 2.15E-09 | -0.5405 | 0.442 | 0.72 | 3.44E-05 | HC_vs_MS.NoTrt_F |
| RAD54L2 | 2.35E-09 | -0.7775 | 0.642 | 0.81 | 3.76E-05 | HC_vs_MS.NoTrt_F |
| EPB41 | 2.84E-09 | -0.4896 | 0.335 | 0.62 | 4.54E-05 | HC_vs_MS.NoTrt_F |
| ABCC9 | 2.99E-09 | 0.64826 | 0.98 | 0.96 | 4.78E-05 | HC_vs_MS.NoTrt_F |
| SLFN5 | 3.58E-09 | -0.9851 | 0.855 | 0.93 | 5.71E-05 | HC_vs_MS.NoTrt_F |
| AQP3 | 7.33E-09 | -0.6231 | 0.332 | 0.6 | 1.17E-04 | HC_vs_MS.NoTrt_F |
| BTN3A2 | 8.65E-09 | -0.6117 | 0.76 | 0.85 | 1.38E-04 | HC_vs_MS.NoTrt_F |
| PAN3 | 9.50E-09 | -0.3629 | 0.824 | 0.95 | 1.52E-04 | HC_vs_MS.NoTrt_F |
| SULT1B1 | 1.20E-08 | 0.65073 | 0.988 | 0.96 | 1.92E-04 | HC_vs_MS.NoTrt_F |
| LCP1 | 1.65E-08 | -0.4662 | 0.621 | 0.85 | 2.64E-04 | HC_vs_MS.NoTrt_F |
| ACTR2 | 1.71E-08 | -0.6056 | 0.413 | 0.66 | 2.73E-04 | HC_vs_MS.NoTrt_F |
| UBE2D2 | 1.73E-08 | -0.6844 | 0.41 | 0.65 | 2.76E-04 | HC_vs_MS.NoTrt_F |
| RPL26 | 1.88E-08 | -0.5077 | 0.899 | 0.97 | 3.00E-04 | HC_vs_MS.NoTrt_F |
| RPS4Y1 | 2.08E-08 | -1.4389 | 0.136 | 0.33 | 3.32E-04 | HC_vs_MS.NoTrt_F |
| FAM110A | 2.12E-08 | -0.4391 | 0.197 | 0.43 | 3.39E-04 | HC_vs_MS.NoTrt_F |
| RPS6 | 2.28E-08 | -0.3221 | 0.919 | 1 | 3.64E-04 | HC_vs_MS.NoTrt_F |
| NABP1 | 2.55E-08 | -0.5089 | 0.766 | 0.88 | 4.07E-04 | HC_vs_MS.NoTrt_F |
| HNRNPA0 | 2.90E-08 | -0.5292 | 0.338 | 0.59 | 4.63E-04 | HC_vs_MS.NoTrt_F |
| PIK3IP1 | 3.40E-08 | -0.2572 | 0.581 | 0.82 | 5.43E-04 | HC_vs_MS.NoTrt_F |
| CSDE1 | 6.24E-08 | -0.5824 | 0.434 | 0.67 | 9.96E-04 | HC_vs_MS.NoTrt_F |
| LSM6 | 6.48E-08 | -0.2843 | 0.162 | 0.38 | 0.001036 | HC_vs_MS.NoTrt_F |
| RNF6 | 6.54E-08 | -1.5414 | 0.162 | 0.35 | 0.001045 | HC_vs_MS.NoTrt_F |
| C1D | 7.61E-08 | -0.387 | 0.139 | 0.34 | 0.001216 | HC_vs_MS.NoTrt_F |
| GPSM3 | 8.08E-08 | -0.5089 | 0.451 | 0.67 | 0.00129 | HC_vs_MS.NoTrt_F |
| NRBF2 | 9.63E-08 | -0.6026 | 0.139 | 0.33 | 0.001537 | HC_vs_MS.NoTrt_F |
| ASF1A | 1.01E-07 | -0.4176 | 0.156 | 0.35 | 0.001617 | HC_vs_MS.NoTrt_F |

| | | | | | | |
|---|---|---|---|---|---|---|
| MT-CO2 | 1.03E-07 | 0.59896 | 0.983 | 1 | 0.001647 | HC_vs_MS.NoTrt_F |
| LDHB | 1.07E-07 | -0.3571 | 0.595 | 0.85 | 0.00171 | HC_vs_MS.NoTrt_F |
| LBH | 1.08E-07 | -0.4693 | 0.61 | 0.82 | 0.001723 | HC_vs_MS.NoTrt_F |
| CPA5 | 1.27E-07 | -1.6281 | 0.017 | 0.13 | 0.002023 | HC_vs_MS.NoTrt_F |
| PRRG3 | 1.34E-07 | 1.51696 | 0.737 | 0.55 | 0.002148 | HC_vs_MS.NoTrt_F |
| GNPAT | 1.35E-07 | -0.9723 | 0.168 | 0.37 | 0.002154 | HC_vs_MS.NoTrt_F |
| RIMS2 | 1.44E-07 | 0.58483 | 0.957 | 0.92 | 0.002298 | HC_vs_MS.NoTrt_F |
| HOXB2 | 1.56E-07 | -0.3358 | 0.182 | 0.39 | 0.002497 | HC_vs_MS.NoTrt_F |
| RBX1 | 1.60E-07 | -0.4543 | 0.309 | 0.54 | 0.002548 | HC_vs_MS.NoTrt_F |
| MOB2 | 1.68E-07 | -1.2395 | 0.101 | 0.27 | 0.00268 | HC_vs_MS.NoTrt_F |
| C4orf36 | 1.72E-07 | 0.56595 | 0.908 | 0.86 | 0.002744 | HC_vs_MS.NoTrt_F |
| RAB5A | 1.72E-07 | -0.6207 | 0.176 | 0.38 | 0.002747 | HC_vs_MS.NoTrt_F |
| HNRNPF | 1.75E-07 | -0.6694 | 0.335 | 0.55 | 0.002788 | HC_vs_MS.NoTrt_F |
| INTU | 1.77E-07 | 0.30378 | 0.991 | 0.99 | 0.002823 | HC_vs_MS.NoTrt_F |
| KIAA1551 | 1.79E-07 | -0.4495 | 0.48 | 0.71 | 0.002853 | HC_vs_MS.NoTrt_F |
| C12orf45 | 1.84E-07 | 0.35972 | 1 | 1 | 0.002932 | HC_vs_MS.NoTrt_F |
| MRPL18 | 1.85E-07 | -0.8618 | 0.145 | 0.33 | 0.002962 | HC_vs_MS.NoTrt_F |
| SOD2 | 1.90E-07 | 0.4163 | 1 | 1 | 0.003035 | HC_vs_MS.NoTrt_F |
| TMED7 | 1.98E-07 | -1.7154 | 0.098 | 0.27 | 0.003163 | HC_vs_MS.NoTrt_F |
| PTEN | 2.38E-07 | -0.4888 | 0.468 | 0.66 | 0.003807 | HC_vs_MS.NoTrt_F |
| AES | 2.57E-07 | -0.4076 | 0.5 | 0.74 | 0.004104 | HC_vs_MS.NoTrt_F |
| LIMD2 | 2.72E-07 | -0.3917 | 0.581 | 0.82 | 0.004352 | HC_vs_MS.NoTrt_F |
| TBC1D14 | 2.88E-07 | -0.6113 | 0.185 | 0.39 | 0.004601 | HC_vs_MS.NoTrt_F |
| COL8A1 | 2.97E-07 | 0.65658 | 0.916 | 0.81 | 0.00475 | HC_vs_MS.NoTrt_F |
| DAXX | 2.99E-07 | -0.7164 | 0.173 | 0.37 | 0.004783 | HC_vs_MS.NoTrt_F |
| BIN2 | 3.00E-07 | -0.6126 | 0.543 | 0.71 | 0.004795 | HC_vs_MS.NoTrt_F |
| EEF2 | 3.08E-07 | -0.4088 | 0.806 | 0.94 | 0.004916 | HC_vs_MS.NoTrt_F |
| ELF1 | 3.94E-07 | -0.5336 | 0.355 | 0.59 | 0.006292 | HC_vs_MS.NoTrt_F |
| TRMT13 | 4.10E-07 | -0.769 | 0.118 | 0.29 | 0.006544 | HC_vs_MS.NoTrt_F |
| TRMT112 | 4.58E-07 | -0.4567 | 0.272 | 0.49 | 0.007308 | HC_vs_MS.NoTrt_F |
| TCEAL8 | 4.75E-07 | -0.9028 | 0.116 | 0.29 | 0.007582 | HC_vs_MS.NoTrt_F |
| NME2 | 4.91E-07 | -0.5071 | 0.486 | 0.69 | 0.007848 | HC_vs_MS.NoTrt_F |
| A2M | 4.94E-07 | 1.47593 | 0.399 | 0.19 | 0.007894 | HC_vs_MS.NoTrt_F |
| MAT2B | 4.97E-07 | -0.5909 | 0.746 | 0.88 | 0.007947 | HC_vs_MS.NoTrt_F |
| CACYBP | 5.25E-07 | -0.4977 | 0.246 | 0.46 | 0.008394 | HC_vs_MS.NoTrt_F |
| ANXA2 | 5.26E-07 | -0.4174 | 0.743 | 0.86 | 0.008398 | HC_vs_MS.NoTrt_F |
| ANXA11 | 5.87E-07 | -0.5191 | 0.838 | 0.91 | 0.009375 | HC_vs_MS.NoTrt_F |
| ZNF408 | 6.02E-07 | -1.6608 | 0.026 | 0.14 | 0.009614 | HC_vs_MS.NoTrt_F |
| EIF4A2 | 6.53E-07 | -0.3829 | 0.621 | 0.83 | 0.010432 | HC_vs_MS.NoTrt_F |
| NEK5 | 6.68E-07 | 0.50679 | 0.902 | 0.86 | 0.010674 | HC_vs_MS.NoTrt_F |

| | | | | | | |
|---|---|---|---|---|---|---|
| CCDC14 | 6.71E-07 | -0.3148 | 0.228 | 0.45 | 0.010723 | HC_vs_MS.NoTrt_F |
| APOL2 | 7.34E-07 | -0.4549 | 0.523 | 0.71 | 0.011731 | HC_vs_MS.NoTrt_F |
| MCM7 | 7.87E-07 | -0.8088 | 0.153 | 0.33 | 0.012563 | HC_vs_MS.NoTrt_F |
| EMC4 | 7.87E-07 | 0.6273 | 0.997 | 0.99 | 0.012567 | HC_vs_MS.NoTrt_F |
| ANKRD13C | 8.09E-07 | -0.4346 | 0.145 | 0.33 | 0.012929 | HC_vs_MS.NoTrt_F |
| JAKMIP1 | 8.21E-07 | -0.4028 | 0.24 | 0.45 | 0.013119 | HC_vs_MS.NoTrt_F |
| LIMK2 | 8.29E-07 | -0.6358 | 0.182 | 0.38 | 0.013236 | HC_vs_MS.NoTrt_F |
| UPK3BL | 9.09E-07 | -0.4073 | 0.532 | 0.73 | 0.014513 | HC_vs_MS.NoTrt_F |
| FAM160B1 | 9.13E-07 | -0.6679 | 0.205 | 0.4 | 0.014579 | HC_vs_MS.NoTrt_F |
| UBL7 | 9.50E-07 | -0.7479 | 0.145 | 0.32 | 0.015167 | HC_vs_MS.NoTrt_F |
| GCNT4 | 9.52E-07 | -0.436 | 0.344 | 0.57 | 0.015201 | HC_vs_MS.NoTrt_F |
| FABP4 | 1.02E-06 | 1.25275 | 0.39 | 0.2 | 0.016304 | HC_vs_MS.NoTrt_F |
| CTB-133G6.: | 1.03E-06 | -0.7428 | 0.194 | 0.39 | 0.016521 | HC_vs_MS.NoTrt_F |
| KRT10 | 1.04E-06 | -0.6662 | 0.113 | 0.28 | 0.016593 | HC_vs_MS.NoTrt_F |
| THYN1 | 1.11E-06 | -0.532 | 0.156 | 0.34 | 0.017723 | HC_vs_MS.NoTrt_F |
| ASXL1 | 1.11E-06 | -0.4696 | 0.214 | 0.41 | 0.017751 | HC_vs_MS.NoTrt_F |
| DACT3 | 1.19E-06 | 0.43024 | 0.983 | 0.95 | 0.01901 | HC_vs_MS.NoTrt_F |
| DYNLL1 | 1.20E-06 | -0.5301 | 0.393 | 0.58 | 0.019178 | HC_vs_MS.NoTrt_F |
| WTAP | 1.20E-06 | -0.4041 | 0.621 | 0.81 | 0.019189 | HC_vs_MS.NoTrt_F |
| SPTBN5 | 1.23E-06 | -1.0813 | 0.02 | 0.12 | 0.01967 | HC_vs_MS.NoTrt_F |
| 6-Mar | 1.26E-06 | -0.3981 | 0.827 | 0.9 | 0.020173 | HC_vs_MS.NoTrt_F |
| MRPL34 | 1.31E-06 | -0.7576 | 0.329 | 0.51 | 0.02099 | HC_vs_MS.NoTrt_F |
| NARS | 1.37E-06 | -1.0132 | 0.176 | 0.35 | 0.021929 | HC_vs_MS.NoTrt_F |
| KDM8 | 1.38E-06 | -1.6443 | 0.052 | 0.18 | 0.022095 | HC_vs_MS.NoTrt_F |
| PDE9A | 1.45E-06 | -2.0845 | 0.043 | 0.17 | 0.023163 | HC_vs_MS.NoTrt_F |
| SPIN2B | 1.46E-06 | -0.7754 | 0.046 | 0.17 | 0.023357 | HC_vs_MS.NoTrt_F |
| EBP | 1.47E-06 | -0.5393 | 0.234 | 0.43 | 0.023455 | HC_vs_MS.NoTrt_F |
| IKBKE | 1.49E-06 | -0.4371 | 0.309 | 0.53 | 0.023734 | HC_vs_MS.NoTrt_F |
| ALKBH5 | 1.53E-06 | -0.5543 | 0.208 | 0.4 | 0.024506 | HC_vs_MS.NoTrt_F |
| SAMD12 | 1.54E-06 | 0.51898 | 0.994 | 0.98 | 0.024677 | HC_vs_MS.NoTrt_F |
| TNFAIP8L2 | 1.60E-06 | -1.0561 | 0.098 | 0.25 | 0.025503 | HC_vs_MS.NoTrt_F |
| YJEFN3 | 1.60E-06 | -1.3896 | 0.052 | 0.18 | 0.025543 | HC_vs_MS.NoTrt_F |
| CLDND1 | 1.73E-06 | -0.4249 | 0.379 | 0.6 | 0.027644 | HC_vs_MS.NoTrt_F |
| HSPA1A | 1.75E-06 | -0.295 | 0.263 | 0.47 | 0.027981 | HC_vs_MS.NoTrt_F |
| XXcos-LUCA | 1.80E-06 | -0.2551 | 0.029 | 0.14 | 0.02874 | HC_vs_MS.NoTrt_F |
| LTB | 1.80E-06 | -0.3207 | 0.584 | 0.81 | 0.028803 | HC_vs_MS.NoTrt_F |
| PWWP2A | 1.82E-06 | -0.6204 | 0.139 | 0.31 | 0.028996 | HC_vs_MS.NoTrt_F |
| IRF2 | 1.82E-06 | -0.5869 | 0.338 | 0.53 | 0.029014 | HC_vs_MS.NoTrt_F |
| ANK3 | 1.84E-06 | 0.63292 | 0.942 | 0.89 | 0.029409 | HC_vs_MS.NoTrt_F |
| TBC1D23 | 1.89E-06 | -0.8319 | 0.15 | 0.33 | 0.030114 | HC_vs_MS.NoTrt_F |

| | | | | | | |
|---|---|---|---|---|---|---|
| SDF2 | 1.89E-06 | -0.5069 | 0.243 | 0.44 | 0.030235 | HC_vs_MS.NoTrt_F |
| TTC39C | 1.93E-06 | -0.4305 | 0.824 | 0.9 | 0.030809 | HC_vs_MS.NoTrt_F |
| TRPV2 | 1.97E-06 | -0.4666 | 0.231 | 0.43 | 0.03147 | HC_vs_MS.NoTrt_F |
| PLCL1 | 2.17E-06 | -1.0972 | 0.13 | 0.29 | 0.034644 | HC_vs_MS.NoTrt_F |
| RNF40 | 2.33E-06 | -1.2612 | 0.098 | 0.25 | 0.037285 | HC_vs_MS.NoTrt_F |
| CCR4 | 2.40E-06 | -0.8511 | 0.922 | 0.94 | 0.038348 | HC_vs_MS.NoTrt_F |
| REXO4 | 2.53E-06 | -0.7443 | 0.061 | 0.19 | 0.040478 | HC_vs_MS.NoTrt_F |
| CBLL1 | 3.03E-06 | -0.575 | 0.243 | 0.44 | 0.048422 | HC_vs_MS.NoTrt_F |
| MT-ND2 | 3.18E-06 | 0.73669 | 0.908 | 0.89 | 0.050766 | HC_vs_MS.NoTrt_F |
| MEAF6 | 3.32E-06 | -0.4665 | 0.277 | 0.48 | 0.053009 | HC_vs_MS.NoTrt_F |
| GTF3A | 3.33E-06 | -0.2698 | 0.373 | 0.61 | 0.053121 | HC_vs_MS.NoTrt_F |
| ACLY | 3.38E-06 | -0.2779 | 0.211 | 0.4 | 0.053953 | HC_vs_MS.NoTrt_F |
| VAMP8 | 3.48E-06 | -0.3289 | 0.298 | 0.53 | 0.055642 | HC_vs_MS.NoTrt_F |
| SYNPO | 3.60E-06 | -0.3478 | 0.237 | 0.44 | 0.057479 | HC_vs_MS.NoTrt_F |
| NDUFB8 | 3.68E-06 | -0.3252 | 0.37 | 0.6 | 0.058808 | HC_vs_MS.NoTrt_F |
| LRRC8C | 3.80E-06 | -0.3104 | 0.335 | 0.54 | 0.060619 | HC_vs_MS.NoTrt_F |
| ARRDC5 | 3.85E-06 | -0.5466 | 0.101 | 0.25 | 0.06149 | HC_vs_MS.NoTrt_F |
| ALG10 | 3.87E-06 | -0.5897 | 0.069 | 0.2 | 0.061783 | HC_vs_MS.NoTrt_F |
| KRTAP3-1 | 3.95E-06 | -0.3405 | 0.075 | 0.22 | 0.063047 | HC_vs_MS.NoTrt_F |
| GIMAP7 | 3.96E-06 | -0.3189 | 0.41 | 0.65 | 0.063332 | HC_vs_MS.NoTrt_F |
| SASS6 | 4.06E-06 | -0.3611 | 0.055 | 0.18 | 0.06483 | HC_vs_MS.NoTrt_F |
| FAM104A | 4.11E-06 | -0.3888 | 0.191 | 0.37 | 0.065584 | HC_vs_MS.NoTrt_F |
| ARRDC3 | 4.16E-06 | -0.3335 | 0.454 | 0.69 | 0.066438 | HC_vs_MS.NoTrt_F |
| EGLN2 | 4.27E-06 | -0.3789 | 0.251 | 0.44 | 0.068173 | HC_vs_MS.NoTrt_F |
| ADA | 4.44E-06 | -0.7993 | 0.107 | 0.26 | 0.070914 | HC_vs_MS.NoTrt_F |
| RACGAP1 | 4.45E-06 | -0.6803 | 0.066 | 0.2 | 0.071002 | HC_vs_MS.NoTrt_F |
| SHISA9 | 4.54E-06 | 0.34247 | 0.971 | 0.92 | 0.072454 | HC_vs_MS.NoTrt_F |
| KYNU | 4.54E-06 | 0.4417 | 0.983 | 0.99 | 0.072568 | HC_vs_MS.NoTrt_F |
| CDPF1 | 4.62E-06 | -0.6029 | 0.058 | 0.18 | 0.073829 | HC_vs_MS.NoTrt_F |
| MSANTD4 | 4.65E-06 | -0.499 | 0.064 | 0.19 | 0.074221 | HC_vs_MS.NoTrt_F |
| STAT1 | 4.70E-06 | -0.583 | 0.384 | 0.58 | 0.075127 | HC_vs_MS.NoTrt_F |
| OCIAD2 | 4.72E-06 | -0.3778 | 0.662 | 0.82 | 0.075327 | HC_vs_MS.NoTrt_F |
| DDX28 | 4.91E-06 | -1.1938 | 0.061 | 0.19 | 0.078417 | HC_vs_MS.NoTrt_F |
| FRK | 5.17E-06 | 0.40249 | 0.997 | 1 | 0.082508 | HC_vs_MS.NoTrt_F |
| REST | 5.24E-06 | -0.3306 | 0.188 | 0.37 | 0.08366 | HC_vs_MS.NoTrt_F |
| ZBTB45 | 5.28E-06 | -0.4933 | 0.035 | 0.14 | 0.084396 | HC_vs_MS.NoTrt_F |
| MUS81 | 5.40E-06 | -0.5488 | 0.159 | 0.33 | 0.086262 | HC_vs_MS.NoTrt_F |
| TXNIP | 5.98E-06 | -0.3841 | 0.639 | 0.8 | 0.095471 | HC_vs_MS.NoTrt_F |
| ATAD3A | 6.07E-06 | -0.6255 | 0.061 | 0.19 | 0.096879 | HC_vs_MS.NoTrt_F |
| HSPA1B | 6.10E-06 | -0.9144 | 0.329 | 0.49 | 0.097406 | HC_vs_MS.NoTrt_F |

| | | | | | | |
|---|---|---|---|---|---|---|
| LGALS3 | 6.27E-06 | -0.4469 | 0.647 | 0.77 | 0.100186 | HC_vs_MS.NoTrt_F |
| TTC32 | 6.83E-06 | -0.6055 | 0.098 | 0.24 | 0.109142 | HC_vs_MS.NoTrt_F |
| RP11-343C2 | 7.28E-06 | -0.4607 | 0.168 | 0.34 | 0.116231 | HC_vs_MS.NoTrt_F |
| TSGA10 | 7.34E-06 | -0.7747 | 0.072 | 0.2 | 0.117256 | HC_vs_MS.NoTrt_F |
| ACAD11 | 7.45E-06 | -1.4462 | 0.182 | 0.35 | 0.118942 | HC_vs_MS.NoTrt_F |
| TRMU | 7.47E-06 | -0.7861 | 0.199 | 0.38 | 0.119309 | HC_vs_MS.NoTrt_F |
| TTC1 | 7.51E-06 | -0.5618 | 0.199 | 0.37 | 0.119923 | HC_vs_MS.NoTrt_F |
| NUDT2 | 7.55E-06 | -0.7298 | 0.055 | 0.18 | 0.120551 | HC_vs_MS.NoTrt_F |
| CYSLTR1 | 7.60E-06 | 0.57537 | 0.98 | 0.96 | 0.121371 | HC_vs_MS.NoTrt_F |
| CDRT1 | 7.69E-06 | -2.0846 | 0.02 | 0.11 | 0.122764 | HC_vs_MS.NoTrt_F |
| PUM2 | 8.08E-06 | -0.51 | 0.338 | 0.53 | 0.128997 | HC_vs_MS.NoTrt_F |
| TMBIM6 | 8.12E-06 | -0.3825 | 0.832 | 0.91 | 0.12972 | HC_vs_MS.NoTrt_F |
| RNF168 | 8.24E-06 | -0.3904 | 0.592 | 0.74 | 0.131644 | HC_vs_MS.NoTrt_F |
| MAPK7 | 8.25E-06 | -1.1955 | 0.046 | 0.16 | 0.131705 | HC_vs_MS.NoTrt_F |
| SUCLG2 | 8.28E-06 | -0.5066 | 0.199 | 0.38 | 0.13226 | HC_vs_MS.NoTrt_F |
| SLC35D2 | 8.31E-06 | -1.2036 | 0.064 | 0.19 | 0.132694 | HC_vs_MS.NoTrt_F |
| PLGRKT | 8.31E-06 | -0.7922 | 0.101 | 0.24 | 0.132791 | HC_vs_MS.NoTrt_F |
| RELL2 | 8.47E-06 | -0.7186 | 0.104 | 0.25 | 0.135287 | HC_vs_MS.NoTrt_F |
| MT-ND4 | 8.58E-06 | 0.45226 | 0.991 | 1 | 0.137127 | HC_vs_MS.NoTrt_F |
| CFAP54 | 8.63E-06 | 0.52647 | 0.965 | 0.97 | 0.137776 | HC_vs_MS.NoTrt_F |
| SLC25A22 | 8.69E-06 | -0.7032 | 0.066 | 0.19 | 0.138839 | HC_vs_MS.NoTrt_F |
| KCNA3 | 8.71E-06 | -0.607 | 0.234 | 0.41 | 0.139148 | HC_vs_MS.NoTrt_F |
| RAB11B | 8.73E-06 | -0.4381 | 0.419 | 0.62 | 0.139464 | HC_vs_MS.NoTrt_F |
| WDPCP | 8.88E-06 | 0.40949 | 0.988 | 0.96 | 0.141853 | HC_vs_MS.NoTrt_F |
| ARL5B | 8.98E-06 | 0.48491 | 0.153 | 0.33 | 0.143439 | HC_vs_MS.NoTrt_F |
| C11orf98 | 9.58E-06 | -0.5905 | 0.165 | 0.33 | 0.153013 | HC_vs_MS.NoTrt_F |
| EFCAB7 | 9.60E-06 | -1.6361 | 0.038 | 0.14 | 0.153397 | HC_vs_MS.NoTrt_F |
| HEBP2 | 9.66E-06 | -0.5849 | 0.35 | 0.53 | 0.154247 | HC_vs_MS.NoTrt_F |
| C5orf15 | 9.77E-06 | -0.6889 | 0.156 | 0.32 | 0.156112 | HC_vs_MS.NoTrt_F |
| NDUFA6 | 1.02E-05 | -0.4511 | 0.257 | 0.45 | 0.162238 | HC_vs_MS.NoTrt_F |
| METTL18 | 1.02E-05 | -0.9673 | 0.061 | 0.18 | 0.163178 | HC_vs_MS.NoTrt_F |
| UBAC1 | 1.02E-05 | -0.5836 | 0.04 | 0.15 | 0.163189 | HC_vs_MS.NoTrt_F |
| UNC119 | 1.07E-05 | -0.5559 | 0.101 | 0.24 | 0.170959 | HC_vs_MS.NoTrt_F |
| FUT7 | 1.07E-05 | -0.481 | 0.107 | 0.25 | 0.171257 | HC_vs_MS.NoTrt_F |
| SUV39H1 | 1.08E-05 | -0.4472 | 0.04 | 0.15 | 0.172217 | HC_vs_MS.NoTrt_F |
| TCEA1 | 1.08E-05 | -0.4965 | 0.341 | 0.54 | 0.172781 | HC_vs_MS.NoTrt_F |
| UGGT2 | 1.10E-05 | 0.57105 | 0.908 | 0.88 | 0.17499 | HC_vs_MS.NoTrt_F |
| CEP164 | 1.11E-05 | -0.2657 | 0.208 | 0.38 | 0.177451 | HC_vs_MS.NoTrt_F |
| CSGALNACT | 1.12E-05 | -0.6275 | 0.538 | 0.69 | 0.179547 | HC_vs_MS.NoTrt_F |
| PI4K2B | 1.13E-05 | -0.5268 | 0.032 | 0.13 | 0.179912 | HC_vs_MS.NoTrt_F |

| | | | | | | |
|---|---|---|---|---|---|---|
| ANXA2R | 1.14E-05 | -0.4757 | 0.182 | 0.35 | 0.182735 | HC_vs_MS.NoTrt_F |
| NOP9 | 1.21E-05 | -1.2464 | 0.038 | 0.14 | 0.192481 | HC_vs_MS.NoTrt_F |
| BAZ1B | 1.21E-05 | -0.5481 | 0.197 | 0.37 | 0.192864 | HC_vs_MS.NoTrt_F |
| HSPE1-MOB | 1.21E-05 | -1.0711 | 0.101 | 0.24 | 0.193681 | HC_vs_MS.NoTrt_F |
| ZNF706 | 1.22E-05 | -0.4244 | 0.373 | 0.57 | 0.194701 | HC_vs_MS.NoTrt_F |
| MT-ND5 | 1.23E-05 | 0.61895 | 0.899 | 0.88 | 0.195706 | HC_vs_MS.NoTrt_F |
| SRP54 | 1.25E-05 | -0.3772 | 0.251 | 0.43 | 0.199403 | HC_vs_MS.NoTrt_F |
| SLC9A9 | 1.25E-05 | -0.398 | 0.171 | 0.34 | 0.199741 | HC_vs_MS.NoTrt_F |
| ICAM3 | 1.26E-05 | -0.294 | 0.419 | 0.65 | 0.201596 | HC_vs_MS.NoTrt_F |
| APTX | 1.26E-05 | -0.5178 | 0.173 | 0.34 | 0.201663 | HC_vs_MS.NoTrt_F |
| SAFB | 1.27E-05 | -0.5149 | 0.379 | 0.55 | 0.202871 | HC_vs_MS.NoTrt_F |
| GBP5 | 1.27E-05 | -0.4749 | 0.691 | 0.81 | 0.203108 | HC_vs_MS.NoTrt_F |
| AMMECR1L | 1.31E-05 | -0.624 | 0.251 | 0.44 | 0.209736 | HC_vs_MS.NoTrt_F |
| CCDC167 | 1.32E-05 | -0.3152 | 0.173 | 0.34 | 0.210083 | HC_vs_MS.NoTrt_F |
| IL10RA | 1.41E-05 | -0.4756 | 0.578 | 0.74 | 0.225972 | HC_vs_MS.NoTrt_F |
| INTS9 | 1.43E-05 | -0.7533 | 0.13 | 0.28 | 0.228662 | HC_vs_MS.NoTrt_F |
| UFSP2 | 1.44E-05 | -0.9797 | 0.116 | 0.26 | 0.230403 | HC_vs_MS.NoTrt_F |
| C12orf4 | 1.45E-05 | -0.5147 | 0.491 | 0.72 | 0.231057 | HC_vs_MS.NoTrt_F |
| LPAR6 | 1.47E-05 | -0.6412 | 0.416 | 0.59 | 0.234771 | HC_vs_MS.NoTrt_F |
| IL6R | 1.47E-05 | -0.5982 | 0.682 | 0.79 | 0.23546 | HC_vs_MS.NoTrt_F |
| TMEM42 | 1.51E-05 | -0.4117 | 0.133 | 0.28 | 0.241279 | HC_vs_MS.NoTrt_F |
| TAF1C | 1.53E-05 | -0.6029 | 0.118 | 0.27 | 0.243843 | HC_vs_MS.NoTrt_F |
| FAM153A | 1.55E-05 | -0.3882 | 0.462 | 0.65 | 0.247074 | HC_vs_MS.NoTrt_F |
| ACTR1B | 1.56E-05 | -0.4975 | 0.188 | 0.36 | 0.249166 | HC_vs_MS.NoTrt_F |
| NEDD4 | 1.59E-05 | 0.51139 | 0.983 | 0.99 | 0.254673 | HC_vs_MS.NoTrt_F |
| C17orf80 | 1.61E-05 | -1.1524 | 0.052 | 0.17 | 0.256468 | HC_vs_MS.NoTrt_F |
| RASGRP1 | 1.62E-05 | -0.4424 | 0.384 | 0.6 | 0.258061 | HC_vs_MS.NoTrt_F |
| RPAP2 | 1.63E-05 | 0.3168 | 0.997 | 1 | 0.261016 | HC_vs_MS.NoTrt_F |
| RTCB | 1.66E-05 | -0.4686 | 0.147 | 0.3 | 0.26586 | HC_vs_MS.NoTrt_F |
| JTB | 1.67E-05 | -0.431 | 0.26 | 0.45 | 0.266214 | HC_vs_MS.NoTrt_F |
| RPA2 | 1.72E-05 | -0.6561 | 0.217 | 0.39 | 0.27512 | HC_vs_MS.NoTrt_F |
| RECQL | 1.72E-05 | -1.0729 | 0.171 | 0.33 | 0.275265 | HC_vs_MS.NoTrt_F |
| NUCB1 | 1.83E-05 | -0.577 | 0.283 | 0.47 | 0.292472 | HC_vs_MS.NoTrt_F |
| TRIM35 | 1.84E-05 | -0.2523 | 0.202 | 0.38 | 0.293514 | HC_vs_MS.NoTrt_F |
| UBAP1 | 1.84E-05 | -0.7628 | 0.153 | 0.3 | 0.294004 | HC_vs_MS.NoTrt_F |
| CCDC186 | 1.88E-05 | -0.6556 | 0.587 | 0.71 | 0.300837 | HC_vs_MS.NoTrt_F |
| SVBP | 1.90E-05 | -0.6061 | 0.121 | 0.27 | 0.303733 | HC_vs_MS.NoTrt_F |
| STARD5 | 1.91E-05 | -0.4013 | 0.116 | 0.25 | 0.304923 | HC_vs_MS.NoTrt_F |
| EIF6 | 1.92E-05 | -0.3092 | 0.168 | 0.33 | 0.307171 | HC_vs_MS.NoTrt_F |
| KRCC1 | 1.92E-05 | -0.4352 | 0.124 | 0.27 | 0.307331 | HC_vs_MS.NoTrt_F |

| | | | | | | |
|---|---|---|---|---|---|---|
| PPP1R2 | 1.93E-05 | -0.6476 | 0.656 | 0.75 | 0.30877 | HC_vs_MS.NoTrt_F |
| MTFP1 | 1.94E-05 | -0.407 | 0.142 | 0.3 | 0.310603 | HC_vs_MS.NoTrt_F |
| BTN3A1 | 2.00E-05 | -0.5688 | 0.61 | 0.74 | 0.319131 | HC_vs_MS.NoTrt_F |
| RRP12 | 2.00E-05 | 0.53571 | 0.934 | 0.87 | 0.320194 | HC_vs_MS.NoTrt_F |
| DNAJC9 | 2.02E-05 | -0.3625 | 0.133 | 0.28 | 0.32206 | HC_vs_MS.NoTrt_F |
| CD36 | 2.04E-05 | 1.58964 | 0.679 | 0.53 | 0.325069 | HC_vs_MS.NoTrt_F |
| KBTBD3 | 2.16E-05 | -0.7854 | 0.075 | 0.2 | 0.345331 | HC_vs_MS.NoTrt_F |
| ZNF775 | 2.17E-05 | -0.7928 | 0.043 | 0.15 | 0.346127 | HC_vs_MS.NoTrt_F |
| SAYSD1 | 2.18E-05 | -0.7077 | 0.208 | 0.38 | 0.348198 | HC_vs_MS.NoTrt_F |
| LACTB2 | 2.22E-05 | -0.7697 | 0.064 | 0.18 | 0.353986 | HC_vs_MS.NoTrt_F |
| OCM | 2.23E-05 | -0.6126 | 0.023 | 0.11 | 0.35658 | HC_vs_MS.NoTrt_F |
| RPS3 | 2.27E-05 | -0.2711 | 0.913 | 0.99 | 0.363255 | HC_vs_MS.NoTrt_F |
| POP7 | 2.30E-05 | -1.0723 | 0.081 | 0.2 | 0.368001 | HC_vs_MS.NoTrt_F |
| RPL9 | 2.34E-05 | -0.4283 | 0.884 | 0.95 | 0.37358 | HC_vs_MS.NoTrt_F |
| MRPL23 | 2.34E-05 | -0.3321 | 0.11 | 0.25 | 0.373773 | HC_vs_MS.NoTrt_F |
| METAP1D | 2.37E-05 | -0.5177 | 0.023 | 0.11 | 0.378108 | HC_vs_MS.NoTrt_F |
| PCYT2 | 2.39E-05 | -1.0196 | 0.035 | 0.13 | 0.381625 | HC_vs_MS.NoTrt_F |
| ZNF585A | 2.48E-05 | -1.907 | 0.353 | 0.53 | 0.396879 | HC_vs_MS.NoTrt_F |
| MIPOL1 | 2.51E-05 | 0.37756 | 1 | 1 | 0.401215 | HC_vs_MS.NoTrt_F |
| CCNE2 | 2.53E-05 | -0.7736 | 0.026 | 0.12 | 0.403703 | HC_vs_MS.NoTrt_F |
| EPS8L2 | 2.58E-05 | -0.4849 | 0.032 | 0.13 | 0.412786 | HC_vs_MS.NoTrt_F |
| POLR2K | 2.59E-05 | -0.3394 | 0.153 | 0.31 | 0.413818 | HC_vs_MS.NoTrt_F |
| IFIH1 | 2.60E-05 | -0.3584 | 0.026 | 0.12 | 0.415796 | HC_vs_MS.NoTrt_F |
| RP11-2C24.9 | 2.60E-05 | -0.5514 | 0.13 | 0.28 | 0.415914 | HC_vs_MS.NoTrt_F |
| PAGR1 | 2.74E-05 | -0.4657 | 0.173 | 0.34 | 0.437951 | HC_vs_MS.NoTrt_F |
| TPT1 | 2.77E-05 | -0.2651 | 0.879 | 0.96 | 0.44275 | HC_vs_MS.NoTrt_F |
| UBAP2L | 2.80E-05 | -0.5784 | 0.269 | 0.45 | 0.447358 | HC_vs_MS.NoTrt_F |
| THAP11 | 2.81E-05 | -0.7697 | 0.101 | 0.23 | 0.44922 | HC_vs_MS.NoTrt_F |
| DNAJB1 | 2.83E-05 | -0.4489 | 0.777 | 0.92 | 0.451941 | HC_vs_MS.NoTrt_F |
| ELF4 | 2.87E-05 | -0.6705 | 0.153 | 0.3 | 0.458721 | HC_vs_MS.NoTrt_F |
| HPCAL4 | 2.87E-05 | -1.4161 | 0.055 | 0.17 | 0.459126 | HC_vs_MS.NoTrt_F |
| ARL6IP1 | 2.94E-05 | -0.4898 | 0.506 | 0.66 | 0.470205 | HC_vs_MS.NoTrt_F |
| ZNF182 | 2.97E-05 | -0.477 | 0.061 | 0.18 | 0.474484 | HC_vs_MS.NoTrt_F |
| VAMP5 | 2.99E-05 | -0.6901 | 0.182 | 0.33 | 0.47839 | HC_vs_MS.NoTrt_F |
| BTN2A1 | 3.00E-05 | -0.4955 | 0.156 | 0.3 | 0.479002 | HC_vs_MS.NoTrt_F |
| MTX2 | 3.01E-05 | -0.2992 | 0.225 | 0.39 | 0.48072 | HC_vs_MS.NoTrt_F |
| GTF3C5 | 3.02E-05 | -0.4328 | 0.35 | 0.52 | 0.48275 | HC_vs_MS.NoTrt_F |
| PYCARD | 3.06E-05 | -0.3153 | 0.107 | 0.24 | 0.48943 | HC_vs_MS.NoTrt_F |
| SDHA | 3.12E-05 | -0.4753 | 0.581 | 0.74 | 0.498393 | HC_vs_MS.NoTrt_F |
| ANXA7 | 3.13E-05 | -0.3813 | 0.431 | 0.62 | 0.4997 | HC_vs_MS.NoTrt_F |

| | | | | | | |
|---|---|---|---|---|---|---|
| OXSR1 | 3.16E-05 | -0.2979 | 0.124 | 0.27 | 0.504559 | HC_vs_MS.NoTrt_F |
| CNKSR3 | 3.19E-05 | 0.34674 | 0.991 | 0.97 | 0.509889 | HC_vs_MS.NoTrt_F |
| GEMIN4 | 3.38E-05 | -0.8141 | 0.075 | 0.19 | 0.539318 | HC_vs_MS.NoTrt_F |
| PDCD10 | 3.48E-05 | -0.3999 | 0.243 | 0.41 | 0.555091 | HC_vs_MS.NoTrt_F |
| ZNF384 | 3.50E-05 | -0.5006 | 0.147 | 0.3 | 0.559129 | HC_vs_MS.NoTrt_F |
| UQCRFS1 | 3.51E-05 | -0.4785 | 0.225 | 0.39 | 0.561043 | HC_vs_MS.NoTrt_F |
| FGF13 | 3.52E-05 | 0.55798 | 0.968 | 0.98 | 0.56297 | HC_vs_MS.NoTrt_F |
| MED31 | 3.56E-05 | -0.5589 | 0.133 | 0.28 | 0.568915 | HC_vs_MS.NoTrt_F |
| VDAC3 | 3.65E-05 | -0.2645 | 0.194 | 0.37 | 0.583046 | HC_vs_MS.NoTrt_F |
| SNX20 | 3.68E-05 | -0.572 | 0.419 | 0.56 | 0.588095 | HC_vs_MS.NoTrt_F |
| GPBP1 | 3.71E-05 | -0.5649 | 0.35 | 0.52 | 0.592037 | HC_vs_MS.NoTrt_F |
| EBAG9 | 3.72E-05 | -0.4347 | 0.104 | 0.24 | 0.594282 | HC_vs_MS.NoTrt_F |
| TMEM206 | 3.81E-05 | -0.7939 | 0.066 | 0.18 | 0.608539 | HC_vs_MS.NoTrt_F |
| C15orf53 | 3.90E-05 | -0.3119 | 0.136 | 0.28 | 0.622886 | HC_vs_MS.NoTrt_F |
| BIRC2 | 3.90E-05 | -0.7013 | 0.202 | 0.36 | 0.623524 | HC_vs_MS.NoTrt_F |
| FANK1 | 3.92E-05 | -0.6884 | 0.211 | 0.35 | 0.626718 | HC_vs_MS.NoTrt_F |
| HNRNPH3 | 3.99E-05 | -0.5338 | 0.251 | 0.43 | 0.637723 | HC_vs_MS.NoTrt_F |
| ZDHHC2 | 3.99E-05 | -1.1698 | 0.116 | 0.25 | 0.637824 | HC_vs_MS.NoTrt_F |
| ZNF862 | 4.00E-05 | -0.8158 | 0.145 | 0.29 | 0.638691 | HC_vs_MS.NoTrt_F |
| CCT8 | 4.07E-05 | -0.2812 | 0.283 | 0.48 | 0.649537 | HC_vs_MS.NoTrt_F |
| MRPL51 | 4.07E-05 | -0.3749 | 0.156 | 0.31 | 0.650205 | HC_vs_MS.NoTrt_F |
| CDKN2C | 4.12E-05 | -1.2112 | 0.043 | 0.14 | 0.658579 | HC_vs_MS.NoTrt_F |
| ADGRD2 | 4.16E-05 | 0.44229 | 0.87 | 0.8 | 0.665273 | HC_vs_MS.NoTrt_F |
| CRBN | 4.24E-05 | -0.3081 | 0.486 | 0.65 | 0.67717 | HC_vs_MS.NoTrt_F |
| TMEM79 | 4.24E-05 | -1.0212 | 0.098 | 0.23 | 0.678031 | HC_vs_MS.NoTrt_F |
| HCCS | 4.25E-05 | -1.1132 | 0.046 | 0.15 | 0.679263 | HC_vs_MS.NoTrt_F |
| ITPA | 4.27E-05 | -0.5456 | 0.176 | 0.33 | 0.681939 | HC_vs_MS.NoTrt_F |
| PIGF | 4.38E-05 | -0.9824 | 0.13 | 0.27 | 0.699252 | HC_vs_MS.NoTrt_F |
| HMGN3 | 4.39E-05 | -0.6011 | 0.211 | 0.37 | 0.701811 | HC_vs_MS.NoTrt_F |
| ABLIM3 | 4.50E-05 | 1.85446 | 0.275 | 0.13 | 0.718236 | HC_vs_MS.NoTrt_F |
| FTO | 4.54E-05 | 0.40905 | 0.994 | 0.99 | 0.725035 | HC_vs_MS.NoTrt_F |
| USP8 | 4.58E-05 | 0.58847 | 1 | 0.99 | 0.731987 | HC_vs_MS.NoTrt_F |
| ARPC4-TTLL: | 4.61E-05 | -0.7262 | 0.118 | 0.25 | 0.737061 | HC_vs_MS.NoTrt_F |
| IL7 | 4.66E-05 | -0.3396 | 0.084 | 0.21 | 0.744283 | HC_vs_MS.NoTrt_F |
| C7orf55-LU( | 4.68E-05 | -0.7045 | 0.072 | 0.19 | 0.74724 | HC_vs_MS.NoTrt_F |
| SH3KBP1 | 4.70E-05 | -0.3447 | 0.497 | 0.67 | 0.751284 | HC_vs_MS.NoTrt_F |
| R3HCC1 | 4.77E-05 | -0.6254 | 0.139 | 0.28 | 0.76249 | HC_vs_MS.NoTrt_F |
| HNRNPK | 4.80E-05 | -0.3022 | 0.708 | 0.83 | 0.766239 | HC_vs_MS.NoTrt_F |
| ADAM12 | 4.82E-05 | -0.3738 | 0.205 | 0.37 | 0.769315 | HC_vs_MS.NoTrt_F |
| C7orf25 | 5.00E-05 | -1.6034 | 0.023 | 0.11 | 0.79939 | HC_vs_MS.NoTrt_F |

| | | | | | | |
|---|---|---|---|---|---|---|
| DUSP14 | 5.07E-05 | -1.1072 | 0.038 | 0.13 | 0.809715 | HC_vs_MS.NoTrt_F |
| ELK3 | 5.08E-05 | -0.6744 | 0.408 | 0.58 | 0.811163 | HC_vs_MS.NoTrt_F |
| RP5-1022E2 | 5.12E-05 | -0.7108 | 0.113 | 0.24 | 0.817065 | HC_vs_MS.NoTrt_F |
| VCP | 5.16E-05 | -0.4207 | 0.306 | 0.49 | 0.824361 | HC_vs_MS.NoTrt_F |
| CDK8 | 5.17E-05 | -0.5168 | 0.046 | 0.15 | 0.825133 | HC_vs_MS.NoTrt_F |
| LSS | 5.17E-05 | -1.4565 | 0.13 | 0.27 | 0.825405 | HC_vs_MS.NoTrt_F |
| ANKLE2 | 5.20E-05 | 0.40155 | 0.994 | 0.99 | 0.830955 | HC_vs_MS.NoTrt_F |
| RARRES3 | 5.22E-05 | -0.382 | 0.289 | 0.48 | 0.834262 | HC_vs_MS.NoTrt_F |
| PRPF6 | 5.27E-05 | -0.6212 | 0.231 | 0.38 | 0.841943 | HC_vs_MS.NoTrt_F |
| TRAPPC2L | 5.31E-05 | -0.8775 | 0.136 | 0.28 | 0.847755 | HC_vs_MS.NoTrt_F |
| SRSF3 | 5.32E-05 | -0.3474 | 0.902 | 0.92 | 0.849708 | HC_vs_MS.NoTrt_F |
| SUSD3 | 5.34E-05 | -0.4774 | 0.298 | 0.48 | 0.853366 | HC_vs_MS.NoTrt_F |
| VPS72 | 5.36E-05 | -0.4847 | 0.145 | 0.29 | 0.856537 | HC_vs_MS.NoTrt_F |
| DHX32 | 5.41E-05 | -1.5975 | 0.026 | 0.11 | 0.863551 | HC_vs_MS.NoTrt_F |
| CNOT11 | 5.41E-05 | -0.7394 | 0.133 | 0.27 | 0.863683 | HC_vs_MS.NoTrt_F |
| OSER1 | 5.44E-05 | -0.3042 | 0.225 | 0.4 | 0.869135 | HC_vs_MS.NoTrt_F |
| FAM50B | 5.46E-05 | -0.4647 | 0.04 | 0.14 | 0.872524 | HC_vs_MS.NoTrt_F |
| ARMCX5 | 5.47E-05 | -0.5752 | 0.075 | 0.19 | 0.873325 | HC_vs_MS.NoTrt_F |
| TIFA | 5.49E-05 | -0.4407 | 0.124 | 0.27 | 0.876879 | HC_vs_MS.NoTrt_F |
| MTA2 | 5.52E-05 | -1.0366 | 0.263 | 0.43 | 0.881771 | HC_vs_MS.NoTrt_F |
| ABRACL | 5.58E-05 | -0.3545 | 0.257 | 0.45 | 0.890624 | HC_vs_MS.NoTrt_F |
| C4orf3 | 5.62E-05 | -0.7851 | 0.497 | 0.61 | 0.898196 | HC_vs_MS.NoTrt_F |
| ATP13A1 | 5.63E-05 | -0.4019 | 0.283 | 0.44 | 0.899184 | HC_vs_MS.NoTrt_F |
| TBC1D2 | 5.64E-05 | -0.6357 | 0.064 | 0.18 | 0.900509 | HC_vs_MS.NoTrt_F |
| ZNF792 | 5.66E-05 | -0.397 | 0.032 | 0.12 | 0.904323 | HC_vs_MS.NoTrt_F |
| SDHAF3 | 5.77E-05 | -0.689 | 0.072 | 0.19 | 0.92193 | HC_vs_MS.NoTrt_F |
| CDKN1B | 5.81E-05 | -0.2544 | 0.355 | 0.55 | 0.928681 | HC_vs_MS.NoTrt_F |
| THAP1 | 5.82E-05 | -0.3003 | 0.104 | 0.23 | 0.929592 | HC_vs_MS.NoTrt_F |
| DYNLL2 | 5.83E-05 | -0.6026 | 0.133 | 0.27 | 0.930532 | HC_vs_MS.NoTrt_F |
| CETN2 | 5.85E-05 | -0.7071 | 0.035 | 0.13 | 0.934477 | HC_vs_MS.NoTrt_F |
| ENTPD6 | 5.90E-05 | -0.4905 | 0.15 | 0.29 | 0.943055 | HC_vs_MS.NoTrt_F |
| ABHD15 | 5.92E-05 | 1.40548 | 0.312 | 0.49 | 0.946336 | HC_vs_MS.NoTrt_F |
| CDKL5 | 5.93E-05 | 0.36602 | 0.962 | 0.93 | 0.947542 | HC_vs_MS.NoTrt_F |
| KCNJ6 | 6.14E-05 | 0.82418 | 0.873 | 0.83 | 0.981343 | HC_vs_MS.NoTrt_F |
| UBE2L6 | 6.14E-05 | -0.3134 | 0.257 | 0.44 | 0.981359 | HC_vs_MS.NoTrt_F |
| ACP1 | 6.29E-05 | -0.3138 | 0.266 | 0.44 | 1 | HC_vs_MS.NoTrt_F |
| PGM2 | 6.34E-05 | -1.1815 | 0.13 | 0.27 | 1 | HC_vs_MS.NoTrt_F |
| SAP18 | 6.40E-05 | -0.2959 | 0.367 | 0.56 | 1 | HC_vs_MS.NoTrt_F |
| ZBTB39 | 6.41E-05 | -0.7907 | 0.029 | 0.12 | 1 | HC_vs_MS.NoTrt_F |
| TRIM47 | 6.41E-05 | -0.9873 | 0.058 | 0.17 | 1 | HC_vs_MS.NoTrt_F |

| | | | | | | |
|---|---|---|---|---|---|---|
| FOXN2 | 6.42E-05 | -0.3471 | 0.272 | 0.45 | 1 | HC_vs_MS.NoTrt_F |
| ITFG1 | 6.46E-05 | -0.2641 | 0.211 | 0.37 | 1 | HC_vs_MS.NoTrt_F |
| GIMAP1-GIN | 6.46E-05 | -0.5778 | 0.072 | 0.19 | 1 | HC_vs_MS.NoTrt_F |
| CHKB-CPT1E | 6.57E-05 | -0.4876 | 0.029 | 0.12 | 1 | HC_vs_MS.NoTrt_F |
| POLR2M | 6.59E-05 | -1.1516 | 0.165 | 0.3 | 1 | HC_vs_MS.NoTrt_F |
| BBS7 | 7.02E-05 | -1.3894 | 0.055 | 0.16 | 1 | HC_vs_MS.NoTrt_F |
| OR52A1 | 7.03E-05 | 0.51766 | 0.829 | 0.75 | 1 | HC_vs_MS.NoTrt_F |
| PDIK1L | 7.04E-05 | -1.5659 | 0.066 | 0.18 | 1 | HC_vs_MS.NoTrt_F |
| ZNF263 | 7.12E-05 | -1.0849 | 0.064 | 0.17 | 1 | HC_vs_MS.NoTrt_F |
| TERF2IP | 7.20E-05 | -0.2977 | 0.396 | 0.56 | 1 | HC_vs_MS.NoTrt_F |
| ANTXR2 | 7.23E-05 | -0.3672 | 0.697 | 0.83 | 1 | HC_vs_MS.NoTrt_F |
| ARHGAP19 | 7.26E-05 | -0.5265 | 0.277 | 0.44 | 1 | HC_vs_MS.NoTrt_F |
| CUTC | 7.33E-05 | -0.7963 | 0.162 | 0.3 | 1 | HC_vs_MS.NoTrt_F |
| ZC3H3 | 7.51E-05 | -0.599 | 0.153 | 0.29 | 1 | HC_vs_MS.NoTrt_F |
| SNAP47 | 7.52E-05 | -0.4577 | 0.171 | 0.32 | 1 | HC_vs_MS.NoTrt_F |
| TRIM37 | 7.58E-05 | -0.9193 | 0.24 | 0.38 | 1 | HC_vs_MS.NoTrt_F |
| ARL3 | 7.86E-05 | -0.4097 | 0.069 | 0.18 | 1 | HC_vs_MS.NoTrt_F |
| CTD-3105H1 | 7.96E-05 | -0.2682 | 0.292 | 0.46 | 1 | HC_vs_MS.NoTrt_F |
| RNF20 | 8.02E-05 | -0.5681 | 0.15 | 0.29 | 1 | HC_vs_MS.NoTrt_F |
| SURF2 | 8.05E-05 | -0.6171 | 0.13 | 0.27 | 1 | HC_vs_MS.NoTrt_F |
| SLAIN2 | 8.13E-05 | -0.3606 | 0.162 | 0.3 | 1 | HC_vs_MS.NoTrt_F |
| EIF3D | 8.14E-05 | -0.2991 | 0.416 | 0.6 | 1 | HC_vs_MS.NoTrt_F |
| AATF | 8.21E-05 | -0.4706 | 0.208 | 0.36 | 1 | HC_vs_MS.NoTrt_F |
| RUNX2 | 8.27E-05 | -0.4514 | 0.202 | 0.37 | 1 | HC_vs_MS.NoTrt_F |
| STX18 | 8.39E-05 | -0.5086 | 0.159 | 0.3 | 1 | HC_vs_MS.NoTrt_F |
| C14orf169 | 8.49E-05 | -0.7189 | 0.066 | 0.18 | 1 | HC_vs_MS.NoTrt_F |
| EIF3G | 8.56E-05 | -0.3134 | 0.853 | 0.96 | 1 | HC_vs_MS.NoTrt_F |
| RPLP0 | 8.58E-05 | -0.2528 | 0.925 | 0.97 | 1 | HC_vs_MS.NoTrt_F |
| SEC24A | 8.61E-05 | -0.3783 | 0.066 | 0.18 | 1 | HC_vs_MS.NoTrt_F |
| SETD2 | 8.69E-05 | -0.5127 | 0.399 | 0.57 | 1 | HC_vs_MS.NoTrt_F |
| ORMDL1 | 8.76E-05 | -0.4043 | 0.318 | 0.5 | 1 | HC_vs_MS.NoTrt_F |
| YPEL2 | 8.81E-05 | -0.5315 | 0.214 | 0.36 | 1 | HC_vs_MS.NoTrt_F |
| CHPF2 | 9.02E-05 | -0.6226 | 0.095 | 0.22 | 1 | HC_vs_MS.NoTrt_F |
| ICAM2 | 9.03E-05 | -0.2529 | 0.743 | 0.84 | 1 | HC_vs_MS.NoTrt_F |
| GBP2 | 9.07E-05 | -0.3199 | 0.46 | 0.64 | 1 | HC_vs_MS.NoTrt_F |
| RPS29 | 9.13E-05 | 0.6643 | 0.948 | 0.94 | 1 | HC_vs_MS.NoTrt_F |
| SMC3 | 9.14E-05 | -0.3985 | 0.309 | 0.47 | 1 | HC_vs_MS.NoTrt_F |
| FADD | 9.24E-05 | -0.9011 | 0.092 | 0.21 | 1 | HC_vs_MS.NoTrt_F |
| USP20 | 9.36E-05 | -0.3533 | 0.249 | 0.41 | 1 | HC_vs_MS.NoTrt_F |
| EPAS1 | 9.48E-05 | 2.85906 | 0.272 | 0.14 | 1 | HC_vs_MS.NoTrt_F |

| | | | | | | |
|---|---|---|---|---|---|---|
| CYB5R1 | 9.60E-05 | -0.657 | 0.104 | 0.23 | 1 | HC_vs_MS.NoTrt_F |
| THOC7 | 9.76E-05 | -0.5195 | 0.159 | 0.3 | 1 | HC_vs_MS.NoTrt_F |
| CCL5 | 9.76E-05 | 1.05229 | 0.936 | 0.94 | 1 | HC_vs_MS.NoTrt_F |
| CDKN2A | 9.99E-05 | -0.4288 | 0.069 | 0.18 | 1 | HC_vs_MS.NoTrt_F |
| GTF2A1 | 1.00E-04 | -1.3779 | 0.061 | 0.17 | 1 | HC_vs_MS.NoTrt_F |
| CCDC13 | 1.02E-04 | -0.4982 | 0.289 | 0.47 | 1 | HC_vs_MS.NoTrt_F |
| TMEM115 | 1.02E-04 | -0.7923 | 0.107 | 0.23 | 1 | HC_vs_MS.NoTrt_F |
| CD5 | 1.03E-04 | -0.2896 | 0.457 | 0.66 | 1 | HC_vs_MS.NoTrt_F |
| GABARAPL2 | 1.04E-04 | -0.6787 | 0.555 | 0.66 | 1 | HC_vs_MS.NoTrt_F |
| RTKN2 | 1.05E-04 | -0.5306 | 0.532 | 0.66 | 1 | HC_vs_MS.NoTrt_F |
| ACAD10 | 1.06E-04 | -0.9702 | 0.506 | 0.64 | 1 | HC_vs_MS.NoTrt_F |
| SLC3A2 | 1.06E-04 | -0.3015 | 0.503 | 0.66 | 1 | HC_vs_MS.NoTrt_F |
| EID3 | 1.09E-04 | -0.6565 | 0.046 | 0.14 | 1 | HC_vs_MS.NoTrt_F |
| DPP7 | 1.09E-04 | -0.2505 | 0.341 | 0.53 | 1 | HC_vs_MS.NoTrt_F |
| MPC1 | 1.10E-04 | -0.5013 | 0.145 | 0.28 | 1 | HC_vs_MS.NoTrt_F |
| HDAC3 | 1.11E-04 | -0.7377 | 0.156 | 0.29 | 1 | HC_vs_MS.NoTrt_F |
| TAF9B | 1.14E-04 | -0.6602 | 0.118 | 0.25 | 1 | HC_vs_MS.NoTrt_F |
| ADCK2 | 1.14E-04 | -1.0924 | 0.038 | 0.13 | 1 | HC_vs_MS.NoTrt_F |
| CA6 | 1.14E-04 | 0.33417 | 0.821 | 0.69 | 1 | HC_vs_MS.NoTrt_F |
| STUB1 | 1.15E-04 | -0.6282 | 0.095 | 0.22 | 1 | HC_vs_MS.NoTrt_F |
| PPDPF | 1.16E-04 | 1.335 | 0.514 | 0.45 | 1 | HC_vs_MS.NoTrt_F |
| PIN4 | 1.16E-04 | -0.5328 | 0.133 | 0.26 | 1 | HC_vs_MS.NoTrt_F |
| YWHAZ | 1.16E-04 | 0.35317 | 0.968 | 0.98 | 1 | HC_vs_MS.NoTrt_F |
| CBX4 | 1.16E-04 | -0.3594 | 0.046 | 0.14 | 1 | HC_vs_MS.NoTrt_F |
| P2RY8 | 1.17E-04 | -0.2577 | 0.434 | 0.61 | 1 | HC_vs_MS.NoTrt_F |
| RAPGEF5 | 1.17E-04 | 1.22176 | 0.101 | 0.01 | 1 | HC_vs_MS.NoTrt_F |
| TJP3 | 1.18E-04 | -1.5444 | 0.095 | 0.21 | 1 | HC_vs_MS.NoTrt_F |
| POLDIP2 | 1.18E-04 | -0.4774 | 0.136 | 0.27 | 1 | HC_vs_MS.NoTrt_F |
| BUB3 | 1.19E-04 | -0.4186 | 0.668 | 0.8 | 1 | HC_vs_MS.NoTrt_F |
| PCMTD1 | 1.20E-04 | -0.5923 | 0.312 | 0.48 | 1 | HC_vs_MS.NoTrt_F |
| SCAMP2 | 1.21E-04 | -0.2591 | 0.329 | 0.52 | 1 | HC_vs_MS.NoTrt_F |
| RAB39B | 1.21E-04 | -0.898 | 0.035 | 0.12 | 1 | HC_vs_MS.NoTrt_F |
| GON7 | 1.21E-04 | -0.6046 | 0.035 | 0.12 | 1 | HC_vs_MS.NoTrt_F |
| OXLD1 | 1.22E-04 | -0.5586 | 0.127 | 0.26 | 1 | HC_vs_MS.NoTrt_F |
| SERINC4 | 1.22E-04 | -0.5781 | 0.058 | 0.16 | 1 | HC_vs_MS.NoTrt_F |
| TNFSF13B | 1.25E-04 | -0.2686 | 0.642 | 0.72 | 1 | HC_vs_MS.NoTrt_F |
| DOLK | 1.26E-04 | -0.6839 | 0.029 | 0.11 | 1 | HC_vs_MS.NoTrt_F |
| BLOC1S6 | 1.26E-04 | 0.44259 | 0.997 | 0.99 | 1 | HC_vs_MS.NoTrt_F |
| CLEC4A | 1.27E-04 | -0.2643 | 0.04 | 0.13 | 1 | HC_vs_MS.NoTrt_F |
| AGAP6 | 1.27E-04 | -0.4771 | 0.127 | 0.25 | 1 | HC_vs_MS.NoTrt_F |

| | | | | | | |
|---|---|---|---|---|---|---|
| MGAT4C | 1.27E-04 | 0.49108 | 0.835 | 0.7 | 1 | HC_vs_MS.NoTrt_F |
| POMT2 | 1.29E-04 | 0.47718 | 0.965 | 0.95 | 1 | HC_vs_MS.NoTrt_F |
| UTP3 | 1.30E-04 | -0.889 | 0.032 | 0.12 | 1 | HC_vs_MS.NoTrt_F |
| GPR18 | 1.30E-04 | -0.4549 | 0.072 | 0.18 | 1 | HC_vs_MS.NoTrt_F |
| PBX4 | 1.31E-04 | -0.4542 | 0.197 | 0.35 | 1 | HC_vs_MS.NoTrt_F |
| ZDHHC24 | 1.31E-04 | 0.30801 | 0.942 | 0.87 | 1 | HC_vs_MS.NoTrt_F |
| TSSC4 | 1.33E-04 | -0.3826 | 0.127 | 0.26 | 1 | HC_vs_MS.NoTrt_F |
| TMEM212 | 1.36E-04 | 0.38083 | 0.974 | 0.96 | 1 | HC_vs_MS.NoTrt_F |
| UBE2L3 | 1.37E-04 | -0.6191 | 0.286 | 0.45 | 1 | HC_vs_MS.NoTrt_F |
| ATP5J2-PTCI | 1.39E-04 | -0.5437 | 0.032 | 0.12 | 1 | HC_vs_MS.NoTrt_F |
| KPNA5 | 1.39E-04 | 0.29172 | 0.997 | 0.98 | 1 | HC_vs_MS.NoTrt_F |
| IFI27L1 | 1.40E-04 | 0.40532 | 0.035 | 0.12 | 1 | HC_vs_MS.NoTrt_F |
| SCAF11 | 1.40E-04 | -0.3537 | 0.899 | 0.93 | 1 | HC_vs_MS.NoTrt_F |
| ZNF92 | 1.40E-04 | -1.5066 | 0.11 | 0.23 | 1 | HC_vs_MS.NoTrt_F |
| MKRN2 | 1.40E-04 | -0.5509 | 0.118 | 0.24 | 1 | HC_vs_MS.NoTrt_F |
| PREP | 1.41E-04 | -0.3086 | 0.13 | 0.26 | 1 | HC_vs_MS.NoTrt_F |
| KCTD5 | 1.42E-04 | -1.1947 | 0.081 | 0.19 | 1 | HC_vs_MS.NoTrt_F |
| RAB10 | 1.43E-04 | -0.4532 | 0.225 | 0.37 | 1 | HC_vs_MS.NoTrt_F |
| CSK | 1.44E-04 | -0.298 | 0.283 | 0.46 | 1 | HC_vs_MS.NoTrt_F |
| NOC4L | 1.45E-04 | -0.4465 | 0.075 | 0.19 | 1 | HC_vs_MS.NoTrt_F |
| GNG11 | 1.45E-04 | 0.98684 | 0.136 | 0.03 | 1 | HC_vs_MS.NoTrt_F |
| TMEM230 | 1.47E-04 | -0.3458 | 0.46 | 0.63 | 1 | HC_vs_MS.NoTrt_F |
| TRIB2 | 1.47E-04 | -0.8776 | 0.367 | 0.5 | 1 | HC_vs_MS.NoTrt_F |
| CAP1 | 1.48E-04 | -0.3725 | 0.618 | 0.79 | 1 | HC_vs_MS.NoTrt_F |
| SMIM15 | 1.50E-04 | -0.9022 | 0.104 | 0.22 | 1 | HC_vs_MS.NoTrt_F |
| MARS2 | 1.50E-04 | -0.5101 | 0.055 | 0.16 | 1 | HC_vs_MS.NoTrt_F |
| CD164 | 1.52E-04 | -0.4038 | 0.598 | 0.73 | 1 | HC_vs_MS.NoTrt_F |
| COPG1 | 1.53E-04 | -0.8639 | 0.173 | 0.32 | 1 | HC_vs_MS.NoTrt_F |
| ZNF184 | 1.56E-04 | -0.4949 | 0.087 | 0.2 | 1 | HC_vs_MS.NoTrt_F |
| ZRANB2 | 1.59E-04 | -0.4769 | 0.353 | 0.51 | 1 | HC_vs_MS.NoTrt_F |
| MCM9 | 1.60E-04 | -1.1878 | 0.341 | 0.48 | 1 | HC_vs_MS.NoTrt_F |
| JUN | 1.60E-04 | -0.3529 | 0.491 | 0.64 | 1 | HC_vs_MS.NoTrt_F |
| RAB35 | 1.61E-04 | -0.748 | 0.168 | 0.3 | 1 | HC_vs_MS.NoTrt_F |
| UNC119B | 1.63E-04 | -0.5429 | 0.052 | 0.15 | 1 | HC_vs_MS.NoTrt_F |
| IFT43 | 1.64E-04 | -0.6274 | 0.069 | 0.18 | 1 | HC_vs_MS.NoTrt_F |
| BRPF1 | 1.64E-04 | -0.553 | 0.075 | 0.18 | 1 | HC_vs_MS.NoTrt_F |
| PRDX1 | 1.67E-04 | -0.2539 | 0.607 | 0.75 | 1 | HC_vs_MS.NoTrt_F |
| FAM173A | 1.67E-04 | 0.2604 | 0.069 | 0.18 | 1 | HC_vs_MS.NoTrt_F |
| NFYB | 1.68E-04 | -1.2086 | 0.303 | 0.43 | 1 | HC_vs_MS.NoTrt_F |
| SETD3 | 1.69E-04 | -0.5309 | 0.121 | 0.25 | 1 | HC_vs_MS.NoTrt_F |

| | | | | | | |
|---|---|---|---|---|---|---|
| CIAPIN1 | 1.69E-04 | -0.6661 | 0.098 | 0.22 | 1 | HC_vs_MS.NoTrt_F |
| TMEM9B | 1.70E-04 | -0.332 | 0.228 | 0.39 | 1 | HC_vs_MS.NoTrt_F |
| PRKAR1A | 1.71E-04 | -0.3249 | 0.353 | 0.52 | 1 | HC_vs_MS.NoTrt_F |
| UBE2K | 1.71E-04 | -0.5309 | 0.162 | 0.3 | 1 | HC_vs_MS.NoTrt_F |
| PCF11 | 1.71E-04 | -0.4111 | 0.512 | 0.65 | 1 | HC_vs_MS.NoTrt_F |
| CXXC1 | 1.74E-04 | -0.3966 | 0.211 | 0.37 | 1 | HC_vs_MS.NoTrt_F |
| KBTBD4 | 1.77E-04 | -0.7592 | 0.084 | 0.19 | 1 | HC_vs_MS.NoTrt_F |
| NPM1 | 1.78E-04 | -0.2795 | 0.636 | 0.78 | 1 | HC_vs_MS.NoTrt_F |
| DCAF15 | 1.79E-04 | -0.4178 | 0.052 | 0.15 | 1 | HC_vs_MS.NoTrt_F |
| CHMP7 | 1.86E-04 | -0.5478 | 0.465 | 0.6 | 1 | HC_vs_MS.NoTrt_F |
| PCDHGA6 | 1.87E-04 | 0.52171 | 0.85 | 0.75 | 1 | HC_vs_MS.NoTrt_F |
| YWHAG | 1.87E-04 | -0.3213 | 0.145 | 0.28 | 1 | HC_vs_MS.NoTrt_F |
| SGPP1 | 1.91E-04 | 0.26988 | 0.104 | 0.23 | 1 | HC_vs_MS.NoTrt_F |
| TCP1 | 1.96E-04 | -0.5431 | 0.428 | 0.58 | 1 | HC_vs_MS.NoTrt_F |
| NDC1 | 1.97E-04 | -0.5374 | 0.066 | 0.17 | 1 | HC_vs_MS.NoTrt_F |
| ZNF142 | 1.97E-04 | -0.6364 | 0.136 | 0.27 | 1 | HC_vs_MS.NoTrt_F |
| IK | 1.99E-04 | -0.3066 | 0.803 | 0.93 | 1 | HC_vs_MS.NoTrt_F |
| GOPC | 2.02E-04 | -0.661 | 0.145 | 0.28 | 1 | HC_vs_MS.NoTrt_F |
| ATP6V1G1 | 2.02E-04 | -0.2652 | 0.425 | 0.62 | 1 | HC_vs_MS.NoTrt_F |
| COMMD1 | 2.03E-04 | 0.53747 | 0.621 | 0.82 | 1 | HC_vs_MS.NoTrt_F |
| TMF1 | 2.03E-04 | -0.2707 | 0.228 | 0.39 | 1 | HC_vs_MS.NoTrt_F |
| EPSTI1 | 2.04E-04 | -0.7316 | 0.173 | 0.32 | 1 | HC_vs_MS.NoTrt_F |
| FH | 2.04E-04 | -0.2667 | 0.075 | 0.18 | 1 | HC_vs_MS.NoTrt_F |
| DNAJC10 | 2.05E-04 | 0.26479 | 0.974 | 0.97 | 1 | HC_vs_MS.NoTrt_F |
| PERP | 2.06E-04 | -0.5554 | 0.087 | 0.2 | 1 | HC_vs_MS.NoTrt_F |
| PDCD2L | 2.07E-04 | -1.3865 | 0.043 | 0.13 | 1 | HC_vs_MS.NoTrt_F |
| C3orf58 | 2.10E-04 | -0.6342 | 0.104 | 0.22 | 1 | HC_vs_MS.NoTrt_F |
| MED7 | 2.10E-04 | -0.4026 | 0.081 | 0.19 | 1 | HC_vs_MS.NoTrt_F |
| CFAP69 | 2.12E-04 | 0.53057 | 0.873 | 0.82 | 1 | HC_vs_MS.NoTrt_F |
| TIMP3 | 2.14E-04 | 1.86254 | 0.228 | 0.11 | 1 | HC_vs_MS.NoTrt_F |
| RB1CC1 | 2.20E-04 | -0.7781 | 0.306 | 0.44 | 1 | HC_vs_MS.NoTrt_F |
| HSPA1L | 2.20E-04 | -1.0201 | 0.04 | 0.13 | 1 | HC_vs_MS.NoTrt_F |
| DGUOK | 2.23E-04 | -0.3088 | 0.217 | 0.37 | 1 | HC_vs_MS.NoTrt_F |
| OPHN1 | 2.27E-04 | 0.3928 | 0.977 | 0.98 | 1 | HC_vs_MS.NoTrt_F |
| SHMT2 | 2.32E-04 | -0.6611 | 0.254 | 0.4 | 1 | HC_vs_MS.NoTrt_F |
| R3HCC1L | 2.35E-04 | -0.2531 | 0.298 | 0.45 | 1 | HC_vs_MS.NoTrt_F |
| ANKRD18A | 2.37E-04 | -0.6404 | 0.28 | 0.42 | 1 | HC_vs_MS.NoTrt_F |
| HKDC1 | 2.40E-04 | -0.7953 | 0.061 | 0.16 | 1 | HC_vs_MS.NoTrt_F |
| SEPHS1 | 2.40E-04 | -0.5041 | 0.075 | 0.18 | 1 | HC_vs_MS.NoTrt_F |
| MSN | 2.44E-04 | -0.3572 | 0.598 | 0.75 | 1 | HC_vs_MS.NoTrt_F |

| | | | | | | |
|---|---|---|---|---|---|---|
| NDUFAF3 | 2.46E-04 | -0.3815 | 0.185 | 0.33 | 1 | HC_vs_MS.NoTrt_F |
| EIF4E3 | 2.49E-04 | -0.7895 | 0.176 | 0.32 | 1 | HC_vs_MS.NoTrt_F |
| MT-ND1 | 2.50E-04 | 0.589 | 0.945 | 0.94 | 1 | HC_vs_MS.NoTrt_F |
| CLPTM1L | 2.52E-04 | 1.58615 | 0.078 | 0.19 | 1 | HC_vs_MS.NoTrt_F |
| STRIP1 | 2.54E-04 | -0.6823 | 0.179 | 0.31 | 1 | HC_vs_MS.NoTrt_F |
| MIF4GD | 2.55E-04 | -0.2631 | 0.142 | 0.28 | 1 | HC_vs_MS.NoTrt_F |
| CCDC65 | 2.57E-04 | -0.643 | 0.069 | 0.17 | 1 | HC_vs_MS.NoTrt_F |
| ZC3H7A | 2.60E-04 | -0.4297 | 0.416 | 0.56 | 1 | HC_vs_MS.NoTrt_F |
| METTL13 | 2.64E-04 | -0.5802 | 0.072 | 0.18 | 1 | HC_vs_MS.NoTrt_F |
| C9orf84 | 2.64E-04 | -0.8204 | 0.043 | 0.13 | 1 | HC_vs_MS.NoTrt_F |
| RP11-307N1 | 2.65E-04 | -0.3582 | 0.116 | 0.24 | 1 | HC_vs_MS.NoTrt_F |
| RUNDC3B | 2.70E-04 | -0.3196 | 0.309 | 0.44 | 1 | HC_vs_MS.NoTrt_F |
| GOLGA1 | 2.72E-04 | -1.0635 | 0.127 | 0.24 | 1 | HC_vs_MS.NoTrt_F |
| AHCTF1 | 2.72E-04 | -0.4295 | 0.171 | 0.3 | 1 | HC_vs_MS.NoTrt_F |
| CTTN | 2.74E-04 | 1.68121 | 0.295 | 0.16 | 1 | HC_vs_MS.NoTrt_F |
| ARHGAP9 | 2.78E-04 | -0.304 | 0.624 | 0.77 | 1 | HC_vs_MS.NoTrt_F |
| UBR7 | 2.79E-04 | -0.4768 | 0.13 | 0.25 | 1 | HC_vs_MS.NoTrt_F |
| RNF126 | 2.81E-04 | -0.3962 | 0.22 | 0.37 | 1 | HC_vs_MS.NoTrt_F |
| SPARC | 2.81E-04 | 2.20284 | 0.249 | 0.13 | 1 | HC_vs_MS.NoTrt_F |
| WBP1L | 2.81E-04 | -1.3442 | 0.124 | 0.24 | 1 | HC_vs_MS.NoTrt_F |
| IRF7 | 2.82E-04 | -0.268 | 0.145 | 0.28 | 1 | HC_vs_MS.NoTrt_F |
| TMEM205 | 2.84E-04 | -0.5513 | 0.118 | 0.24 | 1 | HC_vs_MS.NoTrt_F |
| SPATA4 | 2.85E-04 | 0.49935 | 0.72 | 0.54 | 1 | HC_vs_MS.NoTrt_F |
| RP11-500M8 | 2.87E-04 | -0.4272 | 0.124 | 0.25 | 1 | HC_vs_MS.NoTrt_F |
| DNTTIP2 | 2.87E-04 | 0.28743 | 0.96 | 0.95 | 1 | HC_vs_MS.NoTrt_F |
| STX16-NPEP | 2.89E-04 | -0.409 | 0.113 | 0.23 | 1 | HC_vs_MS.NoTrt_F |
| ZNF77 | 2.90E-04 | -1.4175 | 0.035 | 0.12 | 1 | HC_vs_MS.NoTrt_F |
| C6orf47 | 2.90E-04 | -0.7721 | 0.107 | 0.22 | 1 | HC_vs_MS.NoTrt_F |
| LEF1 | 2.90E-04 | -0.2701 | 0.379 | 0.55 | 1 | HC_vs_MS.NoTrt_F |
| ACOT8 | 2.92E-04 | -0.4936 | 0.13 | 0.25 | 1 | HC_vs_MS.NoTrt_F |
| LYPLA2 | 2.95E-04 | -0.2958 | 0.208 | 0.36 | 1 | HC_vs_MS.NoTrt_F |
| ZSWIM6 | 2.96E-04 | -0.2581 | 0.072 | 0.18 | 1 | HC_vs_MS.NoTrt_F |
| RGPD5 | 2.97E-04 | 0.65197 | 0.919 | 0.9 | 1 | HC_vs_MS.NoTrt_F |
| C2orf76 | 3.01E-04 | -0.7167 | 0.029 | 0.11 | 1 | HC_vs_MS.NoTrt_F |
| KCNN4 | 3.03E-04 | -0.3227 | 0.182 | 0.32 | 1 | HC_vs_MS.NoTrt_F |
| FAM21A | 3.05E-04 | -0.7007 | 0.225 | 0.38 | 1 | HC_vs_MS.NoTrt_F |
| RP11-108K1 | 3.09E-04 | -0.2665 | 0.029 | 0.11 | 1 | HC_vs_MS.NoTrt_F |
| LEKR1 | 3.10E-04 | -0.4909 | 0.176 | 0.32 | 1 | HC_vs_MS.NoTrt_F |
| BUB1 | 3.10E-04 | -1.3685 | 0.029 | 0.11 | 1 | HC_vs_MS.NoTrt_F |
| DUT | 3.11E-04 | -0.3037 | 0.165 | 0.3 | 1 | HC_vs_MS.NoTrt_F |

| | | | | | | |
|---|---|---|---|---|---|---|
| GOLGA5 | 3.13E-04 | -0.8264 | 0.142 | 0.27 | 1 | HC_vs_MS.NoTrt_F |
| SIRT6 | 3.13E-04 | -0.8016 | 0.066 | 0.17 | 1 | HC_vs_MS.NoTrt_F |
| SLC16A6 | 3.13E-04 | -0.6055 | 0.275 | 0.41 | 1 | HC_vs_MS.NoTrt_F |
| TMEM248 | 3.16E-04 | -0.5234 | 0.22 | 0.35 | 1 | HC_vs_MS.NoTrt_F |
| RNF121 | 3.18E-04 | -0.4548 | 0.361 | 0.51 | 1 | HC_vs_MS.NoTrt_F |
| RBM22 | 3.22E-04 | -0.2644 | 0.165 | 0.3 | 1 | HC_vs_MS.NoTrt_F |
| NPDC1 | 3.23E-04 | -0.2546 | 0.069 | 0.17 | 1 | HC_vs_MS.NoTrt_F |
| NKRF | 3.25E-04 | -0.3279 | 0.072 | 0.18 | 1 | HC_vs_MS.NoTrt_F |
| ALKBH7 | 3.27E-04 | -0.3707 | 0.133 | 0.26 | 1 | HC_vs_MS.NoTrt_F |
| RAB40C | 3.27E-04 | -0.3064 | 0.032 | 0.11 | 1 | HC_vs_MS.NoTrt_F |
| GOLGA7 | 3.37E-04 | -0.3517 | 0.199 | 0.34 | 1 | HC_vs_MS.NoTrt_F |
| GLYR1 | 3.39E-04 | -0.4138 | 0.419 | 0.54 | 1 | HC_vs_MS.NoTrt_F |
| ARHGEF11 | 3.42E-04 | 0.79 | 0.052 | 0.14 | 1 | HC_vs_MS.NoTrt_F |
| PRDM4 | 3.46E-04 | -0.4516 | 0.069 | 0.17 | 1 | HC_vs_MS.NoTrt_F |
| C7orf26 | 3.47E-04 | 0.31051 | 0.098 | 0.21 | 1 | HC_vs_MS.NoTrt_F |
| HCFC2 | 3.48E-04 | -0.9501 | 0.078 | 0.18 | 1 | HC_vs_MS.NoTrt_F |
| C7orf43 | 3.51E-04 | 0.421 | 0.052 | 0.14 | 1 | HC_vs_MS.NoTrt_F |
| HOXB3 | 3.52E-04 | 0.48124 | 0.052 | 0.14 | 1 | HC_vs_MS.NoTrt_F |
| PPCDC | 3.53E-04 | -0.5851 | 0.081 | 0.19 | 1 | HC_vs_MS.NoTrt_F |
| ZNF740 | 3.55E-04 | 0.30877 | 0.153 | 0.29 | 1 | HC_vs_MS.NoTrt_F |
| LTV1 | 3.56E-04 | -0.7082 | 0.139 | 0.26 | 1 | HC_vs_MS.NoTrt_F |
| CANT1 | 3.59E-04 | -0.8978 | 0.095 | 0.2 | 1 | HC_vs_MS.NoTrt_F |
| ZDHHC8 | 3.64E-04 | -0.3174 | 0.095 | 0.2 | 1 | HC_vs_MS.NoTrt_F |
| MPHOSPH6 | 3.67E-04 | -0.8846 | 0.145 | 0.27 | 1 | HC_vs_MS.NoTrt_F |
| AC127070.1 | 3.67E-04 | -0.3481 | 0.066 | 0.17 | 1 | HC_vs_MS.NoTrt_F |
| OPTN | 3.76E-04 | -0.4676 | 0.728 | 0.83 | 1 | HC_vs_MS.NoTrt_F |
| PRMT9 | 3.77E-04 | 0.70572 | 0.223 | 0.38 | 1 | HC_vs_MS.NoTrt_F |
| ACCS | 3.77E-04 | -1.1491 | 0.072 | 0.17 | 1 | HC_vs_MS.NoTrt_F |
| SS18L2 | 3.77E-04 | -0.4913 | 0.358 | 0.5 | 1 | HC_vs_MS.NoTrt_F |
| GIMAP6 | 3.81E-04 | -0.595 | 0.202 | 0.34 | 1 | HC_vs_MS.NoTrt_F |
| SLC12A7 | 3.82E-04 | -0.6987 | 0.147 | 0.28 | 1 | HC_vs_MS.NoTrt_F |
| MTRR | 3.84E-04 | -0.5363 | 0.208 | 0.35 | 1 | HC_vs_MS.NoTrt_F |
| FKBP8 | 3.84E-04 | 0.49944 | 0.234 | 0.38 | 1 | HC_vs_MS.NoTrt_F |
| C19orf43 | 3.86E-04 | -0.4185 | 0.182 | 0.32 | 1 | HC_vs_MS.NoTrt_F |
| FAM210B | 3.87E-04 | -0.8849 | 0.312 | 0.46 | 1 | HC_vs_MS.NoTrt_F |
| KIAA0196 | 3.90E-04 | -0.3746 | 0.153 | 0.28 | 1 | HC_vs_MS.NoTrt_F |
| GPR171 | 3.95E-04 | -1.6235 | 0.043 | 0.13 | 1 | HC_vs_MS.NoTrt_F |
| FBLN5 | 3.95E-04 | -0.6867 | 0.199 | 0.33 | 1 | HC_vs_MS.NoTrt_F |
| FAM98A | 3.96E-04 | -0.6619 | 0.064 | 0.16 | 1 | HC_vs_MS.NoTrt_F |
| STK32C | 3.96E-04 | -0.4956 | 0.046 | 0.13 | 1 | HC_vs_MS.NoTrt_F |

| | | | | | | |
|---|---|---|---|---|---|---|
| GPATCH3 | 3.96E-04 | -0.9311 | 0.072 | 0.17 | 1 | HC_vs_MS.NoTrt_F |
| RP11-159D1 | 3.97E-04 | -0.5334 | 0.139 | 0.26 | 1 | HC_vs_MS.NoTrt_F |
| LRRC61 | 3.98E-04 | -0.9762 | 0.072 | 0.17 | 1 | HC_vs_MS.NoTrt_F |
| PRMT5 | 3.98E-04 | -0.7445 | 0.061 | 0.16 | 1 | HC_vs_MS.NoTrt_F |
| TMEM14A | 3.99E-04 | -0.5399 | 0.101 | 0.21 | 1 | HC_vs_MS.NoTrt_F |
| GDI2 | 4.01E-04 | -0.2899 | 0.451 | 0.6 | 1 | HC_vs_MS.NoTrt_F |
| CA4 | 4.02E-04 | 1.91289 | 0.211 | 0.1 | 1 | HC_vs_MS.NoTrt_F |
| LBR | 4.03E-04 | -0.4897 | 0.249 | 0.39 | 1 | HC_vs_MS.NoTrt_F |
| GTF2F2 | 4.03E-04 | -0.6865 | 0.211 | 0.34 | 1 | HC_vs_MS.NoTrt_F |
| AP1S1 | 4.04E-04 | -0.42 | 0.064 | 0.16 | 1 | HC_vs_MS.NoTrt_F |
| HHIPL1 | 4.05E-04 | -0.3031 | 0.121 | 0.24 | 1 | HC_vs_MS.NoTrt_F |
| RNF123 | 4.09E-04 | -0.7403 | 0.156 | 0.29 | 1 | HC_vs_MS.NoTrt_F |
| EMC9 | 4.12E-04 | -0.529 | 0.049 | 0.14 | 1 | HC_vs_MS.NoTrt_F |
| VPS16 | 4.17E-04 | -0.2975 | 0.153 | 0.29 | 1 | HC_vs_MS.NoTrt_F |
| ZFP62 | 4.19E-04 | -1.0319 | 0.09 | 0.19 | 1 | HC_vs_MS.NoTrt_F |
| COG3 | 4.20E-04 | -0.7682 | 0.168 | 0.29 | 1 | HC_vs_MS.NoTrt_F |
| CEMP1 | 4.21E-04 | -0.2877 | 0.078 | 0.18 | 1 | HC_vs_MS.NoTrt_F |
| PASK | 4.25E-04 | -0.458 | 0.514 | 0.65 | 1 | HC_vs_MS.NoTrt_F |
| CP | 4.26E-04 | -0.5413 | 0.15 | 0.27 | 1 | HC_vs_MS.NoTrt_F |
| MCM2 | 4.26E-04 | -1.5081 | 0.043 | 0.13 | 1 | HC_vs_MS.NoTrt_F |
| AC009950.1 | 4.29E-04 | -0.4963 | 0.064 | 0.16 | 1 | HC_vs_MS.NoTrt_F |
| APH1B | 4.41E-04 | -0.8696 | 0.072 | 0.17 | 1 | HC_vs_MS.NoTrt_F |
| PDGFA | 4.41E-04 | -0.2654 | 0.578 | 0.68 | 1 | HC_vs_MS.NoTrt_F |
| AGO3 | 4.43E-04 | 0.32677 | 0.986 | 0.98 | 1 | HC_vs_MS.NoTrt_F |
| NUS1 | 4.45E-04 | -0.3349 | 0.066 | 0.17 | 1 | HC_vs_MS.NoTrt_F |
| CHST6 | 4.46E-04 | 0.38784 | 0.916 | 0.88 | 1 | HC_vs_MS.NoTrt_F |
| RAB2A | 4.47E-04 | -0.461 | 0.251 | 0.39 | 1 | HC_vs_MS.NoTrt_F |
| SOCS5 | 4.50E-04 | 0.5465 | 0.064 | 0.16 | 1 | HC_vs_MS.NoTrt_F |
| SESN2 | 4.52E-04 | -1.0065 | 0.043 | 0.13 | 1 | HC_vs_MS.NoTrt_F |
| PAPD7 | 4.54E-04 | -1.5087 | 0.087 | 0.19 | 1 | HC_vs_MS.NoTrt_F |
| CLEC2B | 4.58E-04 | -0.4592 | 0.251 | 0.38 | 1 | HC_vs_MS.NoTrt_F |
| CRIPT | 4.61E-04 | -0.3803 | 0.246 | 0.39 | 1 | HC_vs_MS.NoTrt_F |
| GEMIN7 | 4.62E-04 | -0.5448 | 0.22 | 0.35 | 1 | HC_vs_MS.NoTrt_F |
| ID3 | 4.63E-04 | -0.3312 | 0.237 | 0.38 | 1 | HC_vs_MS.NoTrt_F |
| THEM6 | 4.63E-04 | -0.5078 | 0.09 | 0.19 | 1 | HC_vs_MS.NoTrt_F |
| BLCAP | 4.65E-04 | -0.4811 | 0.353 | 0.5 | 1 | HC_vs_MS.NoTrt_F |
| PIK3CA | 4.65E-04 | 0.39591 | 0.046 | 0.13 | 1 | HC_vs_MS.NoTrt_F |
| ZNF232 | 4.67E-04 | -0.2886 | 0.043 | 0.13 | 1 | HC_vs_MS.NoTrt_F |
| TDRKH | 4.69E-04 | -0.4716 | 0.075 | 0.18 | 1 | HC_vs_MS.NoTrt_F |
| HNRNPM | 4.69E-04 | -0.4303 | 0.266 | 0.41 | 1 | HC_vs_MS.NoTrt_F |

| | | | | | | |
|---|---|---|---|---|---|---|
| UFM1 | 4.75E-04 | -0.2977 | 0.405 | 0.55 | 1 | HC_vs_MS.NoTrt_F |
| PEX6 | 4.75E-04 | -0.6084 | 0.043 | 0.13 | 1 | HC_vs_MS.NoTrt_F |
| C9orf69 | 4.76E-04 | -0.6061 | 0.043 | 0.13 | 1 | HC_vs_MS.NoTrt_F |
| CWC22 | 4.77E-04 | -0.294 | 0.179 | 0.31 | 1 | HC_vs_MS.NoTrt_F |
| SNX3 | 4.78E-04 | -0.5117 | 0.182 | 0.32 | 1 | HC_vs_MS.NoTrt_F |
| ADPRHL2 | 4.82E-04 | -1.1457 | 0.095 | 0.2 | 1 | HC_vs_MS.NoTrt_F |
| PHF14 | 4.84E-04 | -0.3129 | 0.15 | 0.28 | 1 | HC_vs_MS.NoTrt_F |
| RBM15B | 4.87E-04 | -1.2269 | 0.04 | 0.12 | 1 | HC_vs_MS.NoTrt_F |
| BYSL | 4.88E-04 | -1.2225 | 0.066 | 0.16 | 1 | HC_vs_MS.NoTrt_F |
| RRAGB | 4.89E-04 | -0.9331 | 0.084 | 0.18 | 1 | HC_vs_MS.NoTrt_F |
| PRDX6 | 4.89E-04 | -0.6748 | 0.173 | 0.29 | 1 | HC_vs_MS.NoTrt_F |
| AKIRIN1 | 4.93E-04 | -0.5788 | 0.535 | 0.68 | 1 | HC_vs_MS.NoTrt_F |
| DHRS4 | 4.94E-04 | -0.3841 | 0.116 | 0.23 | 1 | HC_vs_MS.NoTrt_F |
| OR2A7 | 4.95E-04 | -0.6911 | 0.043 | 0.13 | 1 | HC_vs_MS.NoTrt_F |
| CHCHD3 | 5.02E-04 | -0.4366 | 0.173 | 0.3 | 1 | HC_vs_MS.NoTrt_F |
| ZNF79 | 5.07E-04 | -0.4479 | 0.072 | 0.17 | 1 | HC_vs_MS.NoTrt_F |
| TRAPPC6A | 5.07E-04 | -0.3721 | 0.185 | 0.32 | 1 | HC_vs_MS.NoTrt_F |
| MTERF4 | 5.09E-04 | -0.3745 | 0.431 | 0.56 | 1 | HC_vs_MS.NoTrt_F |
| BEX3 | 5.12E-04 | -0.7163 | 0.145 | 0.26 | 1 | HC_vs_MS.NoTrt_F |
| IL2RA | 5.19E-04 | -0.3123 | 0.387 | 0.54 | 1 | HC_vs_MS.NoTrt_F |
| COA3 | 5.24E-04 | -0.4976 | 0.145 | 0.27 | 1 | HC_vs_MS.NoTrt_F |
| SMIM4 | 5.25E-04 | -0.7626 | 0.064 | 0.16 | 1 | HC_vs_MS.NoTrt_F |
| MORC2 | 5.26E-04 | -0.2659 | 0.434 | 0.54 | 1 | HC_vs_MS.NoTrt_F |
| PHC1 | 5.28E-04 | -0.6316 | 0.124 | 0.24 | 1 | HC_vs_MS.NoTrt_F |
| MS4A15 | 5.31E-04 | -0.4114 | 0.038 | 0.12 | 1 | HC_vs_MS.NoTrt_F |
| NAXD | 5.37E-04 | -0.9036 | 0.087 | 0.19 | 1 | HC_vs_MS.NoTrt_F |
| ATRIP | 5.38E-04 | -0.4116 | 0.127 | 0.24 | 1 | HC_vs_MS.NoTrt_F |
| ARPC5L | 5.40E-04 | -0.4211 | 0.197 | 0.33 | 1 | HC_vs_MS.NoTrt_F |
| METTL4 | 5.41E-04 | -0.8411 | 0.124 | 0.24 | 1 | HC_vs_MS.NoTrt_F |
| UBE2D3 | 5.45E-04 | -0.2788 | 0.717 | 0.82 | 1 | HC_vs_MS.NoTrt_F |
| CNN2 | 5.46E-04 | -0.3001 | 0.986 | 0.98 | 1 | HC_vs_MS.NoTrt_F |
| PAIP1 | 5.52E-04 | -0.2557 | 0.147 | 0.27 | 1 | HC_vs_MS.NoTrt_F |
| IRGQ | 5.57E-04 | 0.52871 | 0.991 | 0.98 | 1 | HC_vs_MS.NoTrt_F |
| TCEAL4 | 5.58E-04 | -0.8438 | 0.159 | 0.28 | 1 | HC_vs_MS.NoTrt_F |
| ZC3H11A | 5.60E-04 | -0.7571 | 0.231 | 0.35 | 1 | HC_vs_MS.NoTrt_F |
| TAF4 | 5.61E-04 | -0.4844 | 0.133 | 0.26 | 1 | HC_vs_MS.NoTrt_F |
| AKR1C3 | 5.64E-04 | -1.0339 | 0.035 | 0.11 | 1 | HC_vs_MS.NoTrt_F |
| CTD-2192J1( | 5.65E-04 | 0.5275 | 0.139 | 0.26 | 1 | HC_vs_MS.NoTrt_F |
| PLEKHF2 | 5.65E-04 | -1.159 | 0.101 | 0.2 | 1 | HC_vs_MS.NoTrt_F |
| RTFDC1 | 5.66E-04 | -0.3437 | 0.217 | 0.36 | 1 | HC_vs_MS.NoTrt_F |

| | | | | | | |
|---|---|---|---|---|---|---|
| HP1BP3 | 5.66E-04 | -0.3364 | 0.876 | 0.93 | 1 | HC_vs_MS.NoTrt_F |
| NDUFB3 | 5.67E-04 | 0.32817 | 0.199 | 0.34 | 1 | HC_vs_MS.NoTrt_F |
| C2orf47 | 5.68E-04 | -0.6741 | 0.035 | 0.11 | 1 | HC_vs_MS.NoTrt_F |
| PRADC1 | 5.71E-04 | -0.9033 | 0.035 | 0.11 | 1 | HC_vs_MS.NoTrt_F |
| ARL2BP | 5.75E-04 | -0.3958 | 0.153 | 0.28 | 1 | HC_vs_MS.NoTrt_F |
| CNST | 5.76E-04 | -0.6857 | 0.303 | 0.46 | 1 | HC_vs_MS.NoTrt_F |
| HSPG2 | 5.77E-04 | 1.60031 | 0.535 | 0.41 | 1 | HC_vs_MS.NoTrt_F |
| ERP44 | 5.79E-04 | -0.3435 | 0.289 | 0.43 | 1 | HC_vs_MS.NoTrt_F |
| CNOT3 | 5.80E-04 | -0.865 | 0.147 | 0.26 | 1 | HC_vs_MS.NoTrt_F |
| CCT4 | 5.82E-04 | -0.4772 | 0.286 | 0.43 | 1 | HC_vs_MS.NoTrt_F |
| BCAP31 | 5.87E-04 | -0.8368 | 0.185 | 0.31 | 1 | HC_vs_MS.NoTrt_F |
| SRA1 | 5.89E-04 | -0.302 | 0.165 | 0.29 | 1 | HC_vs_MS.NoTrt_F |
| PRG4 | 5.94E-04 | -0.5579 | 0.032 | 0.11 | 1 | HC_vs_MS.NoTrt_F |
| RBAK | 6.02E-04 | -2.0124 | 0.032 | 0.11 | 1 | HC_vs_MS.NoTrt_F |
| PPP1R7 | 6.13E-04 | -0.527 | 0.185 | 0.31 | 1 | HC_vs_MS.NoTrt_F |
| TTLL12 | 6.13E-04 | -0.599 | 0.058 | 0.15 | 1 | HC_vs_MS.NoTrt_F |
| RP11-371E8 | 6.20E-04 | -0.316 | 0.035 | 0.11 | 1 | HC_vs_MS.NoTrt_F |
| SNU13 | 6.23E-04 | -0.499 | 0.324 | 0.48 | 1 | HC_vs_MS.NoTrt_F |
| ACTR8 | 6.24E-04 | -0.3866 | 0.081 | 0.18 | 1 | HC_vs_MS.NoTrt_F |
| RTN4IP1 | 6.27E-04 | -0.8848 | 0.035 | 0.11 | 1 | HC_vs_MS.NoTrt_F |
| YIPF2 | 6.35E-04 | -0.5357 | 0.095 | 0.2 | 1 | HC_vs_MS.NoTrt_F |
| MICU2 | 6.36E-04 | -0.2914 | 0.474 | 0.62 | 1 | HC_vs_MS.NoTrt_F |
| U2AF1 | 6.40E-04 | -0.864 | 0.532 | 0.64 | 1 | HC_vs_MS.NoTrt_F |
| HDLBP | 6.42E-04 | -0.3758 | 0.486 | 0.6 | 1 | HC_vs_MS.NoTrt_F |
| PPM1A | 6.42E-04 | -0.7289 | 0.179 | 0.31 | 1 | HC_vs_MS.NoTrt_F |
| TCP11L2 | 6.45E-04 | -0.4297 | 0.295 | 0.42 | 1 | HC_vs_MS.NoTrt_F |
| KMT2D | 6.49E-04 | 1.03393 | 0.87 | 0.86 | 1 | HC_vs_MS.NoTrt_F |
| CLPTM1 | 6.50E-04 | -0.4463 | 0.225 | 0.35 | 1 | HC_vs_MS.NoTrt_F |
| FAM86C1 | 6.52E-04 | -0.7244 | 0.052 | 0.14 | 1 | HC_vs_MS.NoTrt_F |
| FUCA2 | 6.53E-04 | -0.3394 | 0.335 | 0.48 | 1 | HC_vs_MS.NoTrt_F |
| MBD2 | 6.53E-04 | -0.4711 | 0.095 | 0.2 | 1 | HC_vs_MS.NoTrt_F |
| ECH1 | 6.55E-04 | -0.3055 | 0.405 | 0.55 | 1 | HC_vs_MS.NoTrt_F |
| TCEA2 | 6.56E-04 | -0.5026 | 0.032 | 0.11 | 1 | HC_vs_MS.NoTrt_F |
| COPG2 | 6.61E-04 | 0.67173 | 0.092 | 0.2 | 1 | HC_vs_MS.NoTrt_F |
| HS3ST3A1 | 6.61E-04 | -0.306 | 0.084 | 0.19 | 1 | HC_vs_MS.NoTrt_F |
| SPARCL1 | 6.72E-04 | 1.47554 | 0.292 | 0.17 | 1 | HC_vs_MS.NoTrt_F |
| SH3BP5L | 6.78E-04 | -0.3555 | 0.052 | 0.14 | 1 | HC_vs_MS.NoTrt_F |
| DESI2 | 6.81E-04 | 0.25865 | 0.081 | 0.18 | 1 | HC_vs_MS.NoTrt_F |
| LARP1B | 6.83E-04 | -1.1848 | 0.075 | 0.17 | 1 | HC_vs_MS.NoTrt_F |
| ZNF511 | 6.83E-04 | -0.6889 | 0.092 | 0.19 | 1 | HC_vs_MS.NoTrt_F |

| | | | | | | |
|---|---|---|---|---|---|---|
| EIF4B | 6.87E-04 | -0.3117 | 0.743 | 0.8 | 1 | HC_vs_MS.NoTrt_F |
| TAGAP | 6.90E-04 | -0.2737 | 0.353 | 0.52 | 1 | HC_vs_MS.NoTrt_F |
| TAF5L | 6.95E-04 | -0.8243 | 0.171 | 0.3 | 1 | HC_vs_MS.NoTrt_F |
| LDLRAP1 | 6.96E-04 | -0.2979 | 0.431 | 0.58 | 1 | HC_vs_MS.NoTrt_F |
| PLEKHO2 | 6.98E-04 | -0.5035 | 0.104 | 0.21 | 1 | HC_vs_MS.NoTrt_F |
| JMJD7-PLA2 | 6.99E-04 | -0.5705 | 0.329 | 0.47 | 1 | HC_vs_MS.NoTrt_F |
| ALCAM | 7.00E-04 | -0.9368 | 0.066 | 0.16 | 1 | HC_vs_MS.NoTrt_F |
| GSPT2 | 7.07E-04 | 0.53433 | 0.052 | 0.14 | 1 | HC_vs_MS.NoTrt_F |
| OLFM2 | 7.16E-04 | -1.3431 | 0.098 | 0.2 | 1 | HC_vs_MS.NoTrt_F |
| LIG1 | 7.17E-04 | -0.5281 | 0.194 | 0.33 | 1 | HC_vs_MS.NoTrt_F |
| ELAVL1 | 7.21E-04 | 0.35753 | 0.983 | 0.97 | 1 | HC_vs_MS.NoTrt_F |
| SURF1 | 7.31E-04 | -0.506 | 0.113 | 0.22 | 1 | HC_vs_MS.NoTrt_F |
| PHLDB1 | 7.34E-04 | 0.74388 | 0.986 | 0.97 | 1 | HC_vs_MS.NoTrt_F |
| LINC00998 | 7.38E-04 | -0.2611 | 0.081 | 0.18 | 1 | HC_vs_MS.NoTrt_F |
| GCA | 7.39E-04 | -0.8813 | 0.095 | 0.19 | 1 | HC_vs_MS.NoTrt_F |
| ATP2C2 | 7.41E-04 | 0.43502 | 0.032 | 0.11 | 1 | HC_vs_MS.NoTrt_F |
| GPS1 | 7.48E-04 | -0.4691 | 0.162 | 0.28 | 1 | HC_vs_MS.NoTrt_F |
| DUSP22 | 7.48E-04 | -0.5295 | 0.116 | 0.23 | 1 | HC_vs_MS.NoTrt_F |
| MYH9 | 7.57E-04 | -0.313 | 0.829 | 0.95 | 1 | HC_vs_MS.NoTrt_F |
| KAT14 | 7.66E-04 | -0.6248 | 0.072 | 0.17 | 1 | HC_vs_MS.NoTrt_F |
| RNF113A | 7.70E-04 | -0.6093 | 0.139 | 0.25 | 1 | HC_vs_MS.NoTrt_F |
| PSAT1 | 7.71E-04 | -0.4678 | 0.064 | 0.16 | 1 | HC_vs_MS.NoTrt_F |
| LSP1 | 7.79E-04 | -0.2539 | 0.711 | 0.87 | 1 | HC_vs_MS.NoTrt_F |
| KPNA6 | 7.80E-04 | -0.3828 | 0.592 | 0.72 | 1 | HC_vs_MS.NoTrt_F |
| VAMP7 | 7.81E-04 | -1.0185 | 0.127 | 0.24 | 1 | HC_vs_MS.NoTrt_F |
| SYNJ2BP-CO | 7.93E-04 | -0.3068 | 0.168 | 0.3 | 1 | HC_vs_MS.NoTrt_F |
| VAMP2 | 8.00E-04 | -0.2613 | 0.445 | 0.61 | 1 | HC_vs_MS.NoTrt_F |
| YTHDF1 | 8.04E-04 | -0.2614 | 0.081 | 0.18 | 1 | HC_vs_MS.NoTrt_F |
| AGAP9 | 8.07E-04 | 0.58305 | 0.084 | 0.19 | 1 | HC_vs_MS.NoTrt_F |
| CAV1 | 8.10E-04 | 1.43021 | 0.171 | 0.07 | 1 | HC_vs_MS.NoTrt_F |
| LSM14A | 8.12E-04 | -0.3188 | 0.523 | 0.66 | 1 | HC_vs_MS.NoTrt_F |
| CUL4A | 8.19E-04 | -0.4877 | 0.723 | 0.83 | 1 | HC_vs_MS.NoTrt_F |
| MAN1B1 | 8.21E-04 | -0.4992 | 0.329 | 0.48 | 1 | HC_vs_MS.NoTrt_F |
| TLDC2 | 8.27E-04 | 1.17145 | 0.066 | 0.16 | 1 | HC_vs_MS.NoTrt_F |
| FBXO34 | 8.34E-04 | -1.155 | 0.142 | 0.25 | 1 | HC_vs_MS.NoTrt_F |
| GAR1 | 8.35E-04 | -0.8553 | 0.061 | 0.15 | 1 | HC_vs_MS.NoTrt_F |
| FIG4 | 8.39E-04 | -1.2956 | 0.061 | 0.15 | 1 | HC_vs_MS.NoTrt_F |
| FAM118B | 8.42E-04 | -0.373 | 0.072 | 0.17 | 1 | HC_vs_MS.NoTrt_F |
| SRSF9 | 8.45E-04 | -0.4945 | 0.156 | 0.28 | 1 | HC_vs_MS.NoTrt_F |
| ATG5 | 8.45E-04 | -0.3553 | 0.185 | 0.32 | 1 | HC_vs_MS.NoTrt_F |

| | | | | | | |
|---|---|---|---|---|---|---|
| ZFAND2B | 8.47E-04 | -0.8211 | 0.231 | 0.35 | 1 | HC_vs_MS.NoTrt_F |
| AGGF1 | 8.48E-04 | -0.6565 | 0.165 | 0.29 | 1 | HC_vs_MS.NoTrt_F |
| CEP85 | 8.59E-04 | -0.9338 | 0.139 | 0.25 | 1 | HC_vs_MS.NoTrt_F |
| PLAA | 8.66E-04 | -0.3041 | 0.133 | 0.25 | 1 | HC_vs_MS.NoTrt_F |
| HTATSF1 | 8.72E-04 | -0.5121 | 0.159 | 0.28 | 1 | HC_vs_MS.NoTrt_F |
| GIMAP8 | 8.81E-04 | -0.7362 | 0.087 | 0.18 | 1 | HC_vs_MS.NoTrt_F |
| TREX1 | 8.89E-04 | 0.30136 | 0.121 | 0.24 | 1 | HC_vs_MS.NoTrt_F |
| EMC7 | 8.92E-04 | -0.3281 | 0.22 | 0.34 | 1 | HC_vs_MS.NoTrt_F |
| ST7 | 8.95E-04 | -0.9224 | 0.084 | 0.18 | 1 | HC_vs_MS.NoTrt_F |
| SIGMAR1 | 9.15E-04 | -0.3444 | 0.202 | 0.33 | 1 | HC_vs_MS.NoTrt_F |
| SORBS2 | 9.18E-04 | 0.49471 | 0.665 | 0.52 | 1 | HC_vs_MS.NoTrt_F |
| STT3A | 9.18E-04 | -0.9181 | 0.61 | 0.73 | 1 | HC_vs_MS.NoTrt_F |
| ZNF155 | 9.34E-04 | -1.2845 | 0.043 | 0.12 | 1 | HC_vs_MS.NoTrt_F |
| TRIM23 | 9.37E-04 | -0.835 | 0.072 | 0.17 | 1 | HC_vs_MS.NoTrt_F |
| PAFAH1B3 | 9.42E-04 | -0.2587 | 0.11 | 0.22 | 1 | HC_vs_MS.NoTrt_F |
| KPNB1 | 9.45E-04 | -0.4361 | 0.679 | 0.74 | 1 | HC_vs_MS.NoTrt_F |
| PPP1R21 | 9.46E-04 | -0.3421 | 0.275 | 0.41 | 1 | HC_vs_MS.NoTrt_F |
| KLHDC7B | 9.46E-04 | 0.5899 | 0.046 | 0.13 | 1 | HC_vs_MS.NoTrt_F |
| MRPS2 | 9.49E-04 | -0.5006 | 0.101 | 0.2 | 1 | HC_vs_MS.NoTrt_F |
| RNF5 | 9.57E-04 | -0.4381 | 0.156 | 0.28 | 1 | HC_vs_MS.NoTrt_F |
| BRWD1 | 9.58E-04 | -0.619 | 0.85 | 0.84 | 1 | HC_vs_MS.NoTrt_F |
| TULP3 | 9.58E-04 | -0.35 | 0.046 | 0.13 | 1 | HC_vs_MS.NoTrt_F |
| ARHGAP19-! | 9.59E-04 | -0.7644 | 0.04 | 0.12 | 1 | HC_vs_MS.NoTrt_F |
| PHF1 | 9.60E-04 | 0.29806 | 0.234 | 0.38 | 1 | HC_vs_MS.NoTrt_F |
| ZNF691 | 9.61E-04 | 0.27367 | 0.046 | 0.13 | 1 | HC_vs_MS.NoTrt_F |
| RP4-635E18 | 9.65E-04 | -0.4712 | 0.173 | 0.29 | 1 | HC_vs_MS.NoTrt_F |
| DCTPP1 | 9.74E-04 | -0.9445 | 0.095 | 0.19 | 1 | HC_vs_MS.NoTrt_F |
| PCGF3 | 9.81E-04 | -0.3512 | 0.228 | 0.36 | 1 | HC_vs_MS.NoTrt_F |
| TECPR1 | 9.82E-04 | -0.6252 | 0.509 | 0.6 | 1 | HC_vs_MS.NoTrt_F |
| ATP5I | 9.83E-04 | 0.29425 | 0.283 | 0.47 | 1 | HC_vs_MS.NoTrt_F |
| CEACAM21 | 9.87E-04 | -0.6549 | 0.275 | 0.39 | 1 | HC_vs_MS.NoTrt_F |
| RP11-529K1 | 0.00101 | -0.3544 | 0.095 | 0.2 | 1 | HC_vs_MS.NoTrt_F |
| ZDHHC5 | 0.00101 | 0.3903 | 0.142 | 0.26 | 1 | HC_vs_MS.NoTrt_F |
| SMIM20 | 0.00102 | -0.2979 | 0.087 | 0.18 | 1 | HC_vs_MS.NoTrt_F |
| TOMM6 | 0.00102 | -0.2594 | 0.24 | 0.38 | 1 | HC_vs_MS.NoTrt_F |
| WRAP53 | 0.00104 | -0.4018 | 0.043 | 0.12 | 1 | HC_vs_MS.NoTrt_F |
| STK17A | 0.00105 | -0.4074 | 0.289 | 0.44 | 1 | HC_vs_MS.NoTrt_F |
| SLC38A1 | 0.00105 | -0.2934 | 0.783 | 0.87 | 1 | HC_vs_MS.NoTrt_F |
| ANKRD55 | 0.00106 | -0.3799 | 0.182 | 0.3 | 1 | HC_vs_MS.NoTrt_F |
| LAMP1 | 0.00106 | 0.26945 | 0.41 | 0.54 | 1 | HC_vs_MS.NoTrt_F |

| | | | | | | |
|---|---|---|---|---|---|---|
| NIPSNAP3A | 0.00106 | -0.4879 | 0.069 | 0.16 | 1 | HC_vs_MS.NoTrt_F |
| EIF3E | 0.00106 | -0.5107 | 0.538 | 0.64 | 1 | HC_vs_MS.NoTrt_F |
| FBXO28 | 0.00107 | -0.6791 | 0.116 | 0.22 | 1 | HC_vs_MS.NoTrt_F |
| C12orf75 | 0.00107 | -0.2958 | 0.176 | 0.3 | 1 | HC_vs_MS.NoTrt_F |
| TMEM41A | 0.00107 | 0.7468 | 0.858 | 0.79 | 1 | HC_vs_MS.NoTrt_F |
| TOPORS | 0.00107 | -0.3748 | 0.124 | 0.23 | 1 | HC_vs_MS.NoTrt_F |
| ZBTB33 | 0.00108 | -0.9978 | 0.101 | 0.2 | 1 | HC_vs_MS.NoTrt_F |
| SAV1 | 0.00108 | -0.3556 | 0.329 | 0.48 | 1 | HC_vs_MS.NoTrt_F |
| WAC | 0.00109 | -0.4515 | 0.448 | 0.59 | 1 | HC_vs_MS.NoTrt_F |
| ASNSD1 | 0.00109 | -0.2751 | 0.165 | 0.29 | 1 | HC_vs_MS.NoTrt_F |
| HSPA9 | 0.00109 | -0.261 | 0.54 | 0.66 | 1 | HC_vs_MS.NoTrt_F |
| C8orf44-SGK | 0.0011 | -0.2788 | 0.101 | 0.2 | 1 | HC_vs_MS.NoTrt_F |
| CELF1 | 0.0011 | -0.6609 | 0.477 | 0.58 | 1 | HC_vs_MS.NoTrt_F |
| RNF145 | 0.00111 | -0.6933 | 0.191 | 0.32 | 1 | HC_vs_MS.NoTrt_F |
| METTL10 | 0.00111 | -0.3656 | 0.324 | 0.46 | 1 | HC_vs_MS.NoTrt_F |
| TAX1BP3 | 0.00112 | -0.5903 | 0.081 | 0.18 | 1 | HC_vs_MS.NoTrt_F |
| TMPRSS4-AS | 0.00113 | -0.3901 | 0.069 | 0.16 | 1 | HC_vs_MS.NoTrt_F |
| ZNF606 | 0.00113 | 0.42577 | 0.815 | 0.78 | 1 | HC_vs_MS.NoTrt_F |
| PDGFRB | 0.00113 | 2.36476 | 0.116 | 0.03 | 1 | HC_vs_MS.NoTrt_F |
| RBM14-RBM | 0.00114 | -0.7545 | 0.139 | 0.25 | 1 | HC_vs_MS.NoTrt_F |
| PPP4C | 0.00114 | -0.3115 | 0.431 | 0.56 | 1 | HC_vs_MS.NoTrt_F |
| SPP2 | 0.00114 | 0.40905 | 0.809 | 0.75 | 1 | HC_vs_MS.NoTrt_F |
| NDUFA8 | 0.00114 | -0.3842 | 0.116 | 0.22 | 1 | HC_vs_MS.NoTrt_F |
| FNDC3B | 0.00115 | 1.07545 | 0.783 | 0.65 | 1 | HC_vs_MS.NoTrt_F |
| GNG10 | 0.00115 | -0.3016 | 0.136 | 0.25 | 1 | HC_vs_MS.NoTrt_F |
| MRPS14 | 0.00115 | -0.2856 | 0.176 | 0.3 | 1 | HC_vs_MS.NoTrt_F |
| MYC | 0.00115 | -0.8075 | 0.168 | 0.29 | 1 | HC_vs_MS.NoTrt_F |
| OGFOD2 | 0.00115 | -0.5283 | 0.121 | 0.23 | 1 | HC_vs_MS.NoTrt_F |
| SBDS | 0.00119 | -0.4082 | 0.309 | 0.45 | 1 | HC_vs_MS.NoTrt_F |
| RAB4B | 0.00119 | -0.485 | 0.384 | 0.5 | 1 | HC_vs_MS.NoTrt_F |
| LDOC1L | 0.00119 | -1.374 | 0.055 | 0.14 | 1 | HC_vs_MS.NoTrt_F |
| PRR7 | 0.00121 | 0.46611 | 0.043 | 0.12 | 1 | HC_vs_MS.NoTrt_F |
| FUNDC1 | 0.00121 | -0.4314 | 0.055 | 0.14 | 1 | HC_vs_MS.NoTrt_F |
| BTBD10 | 0.00121 | -0.4422 | 0.159 | 0.28 | 1 | HC_vs_MS.NoTrt_F |
| ZNF24 | 0.00123 | -0.3331 | 0.199 | 0.33 | 1 | HC_vs_MS.NoTrt_F |
| DUSP16 | 0.00123 | -0.4929 | 0.329 | 0.46 | 1 | HC_vs_MS.NoTrt_F |
| TIMM22 | 0.00124 | -0.6631 | 0.107 | 0.2 | 1 | HC_vs_MS.NoTrt_F |
| MKRN1 | 0.00124 | -0.8853 | 0.853 | 0.9 | 1 | HC_vs_MS.NoTrt_F |
| MYADM | 0.00125 | 0.93733 | 0.867 | 0.81 | 1 | HC_vs_MS.NoTrt_F |
| LAPTM5 | 0.00125 | -0.3272 | 0.639 | 0.76 | 1 | HC_vs_MS.NoTrt_F |

| | | | | | | |
|---|---|---|---|---|---|---|
| PPP1R11 | 0.00125 | -0.4144 | 0.211 | 0.34 | 1 | HC_vs_MS.NoTrt_F |
| SACM1L | 0.00125 | -0.5218 | 0.309 | 0.43 | 1 | HC_vs_MS.NoTrt_F |
| RP11-411B6 | 0.00126 | 0.40937 | 0.26 | 0.4 | 1 | HC_vs_MS.NoTrt_F |
| ZMYND11 | 0.00126 | -0.6933 | 0.13 | 0.24 | 1 | HC_vs_MS.NoTrt_F |
| APH1A | 0.00126 | -0.4996 | 0.202 | 0.33 | 1 | HC_vs_MS.NoTrt_F |
| MTCH2 | 0.00126 | -0.3493 | 0.124 | 0.23 | 1 | HC_vs_MS.NoTrt_F |
| PNMA1 | 0.00126 | -0.8746 | 0.09 | 0.18 | 1 | HC_vs_MS.NoTrt_F |
| SLC7A1 | 0.00127 | -1.091 | 0.098 | 0.19 | 1 | HC_vs_MS.NoTrt_F |
| SMG8 | 0.00127 | -0.642 | 0.107 | 0.21 | 1 | HC_vs_MS.NoTrt_F |
| CNIH1 | 0.00128 | -0.4123 | 0.145 | 0.25 | 1 | HC_vs_MS.NoTrt_F |
| OTUD6B | 0.00129 | -0.4833 | 0.084 | 0.18 | 1 | HC_vs_MS.NoTrt_F |
| KIAA1217 | 0.00129 | 1.41983 | 0.165 | 0.07 | 1 | HC_vs_MS.NoTrt_F |
| CDKN2AIP | 0.00131 | -0.4958 | 0.133 | 0.24 | 1 | HC_vs_MS.NoTrt_F |
| TRAPPC6B | 0.00133 | -0.5092 | 0.477 | 0.59 | 1 | HC_vs_MS.NoTrt_F |
| TMX2 | 0.00133 | -0.3095 | 0.165 | 0.28 | 1 | HC_vs_MS.NoTrt_F |
| MCUB | 0.00134 | -0.2638 | 0.306 | 0.45 | 1 | HC_vs_MS.NoTrt_F |
| TBC1D8B | 0.00134 | 0.42382 | 0.965 | 0.96 | 1 | HC_vs_MS.NoTrt_F |
| RPP14 | 0.00134 | 0.54493 | 0.951 | 0.93 | 1 | HC_vs_MS.NoTrt_F |
| DEF6 | 0.00135 | -0.2639 | 0.428 | 0.61 | 1 | HC_vs_MS.NoTrt_F |
| PLPP6 | 0.00135 | -0.2672 | 0.061 | 0.14 | 1 | HC_vs_MS.NoTrt_F |
| ADIPOR2 | 0.00135 | -0.3947 | 0.159 | 0.27 | 1 | HC_vs_MS.NoTrt_F |
| ATXN7L2 | 0.00136 | 0.29835 | 0.055 | 0.14 | 1 | HC_vs_MS.NoTrt_F |
| ICA1 | 0.00137 | 1.05645 | 0.676 | 0.56 | 1 | HC_vs_MS.NoTrt_F |
| ABHD4 | 0.00137 | 0.31812 | 0.069 | 0.16 | 1 | HC_vs_MS.NoTrt_F |
| FLT1 | 0.00138 | 0.45124 | 0.777 | 0.59 | 1 | HC_vs_MS.NoTrt_F |
| CCNT2 | 0.00138 | -0.3926 | 0.179 | 0.3 | 1 | HC_vs_MS.NoTrt_F |
| TRIM73 | 0.00138 | 0.35903 | 0.913 | 0.91 | 1 | HC_vs_MS.NoTrt_F |
| SYCP2 | 0.00138 | -0.9225 | 0.049 | 0.13 | 1 | HC_vs_MS.NoTrt_F |
| KLRD1 | 0.00139 | 0.43583 | 0.98 | 0.98 | 1 | HC_vs_MS.NoTrt_F |
| ERP29 | 0.0014 | -0.2932 | 0.858 | 0.91 | 1 | HC_vs_MS.NoTrt_F |
| ZFPL1 | 0.0014 | 0.9644 | 0.069 | 0.16 | 1 | HC_vs_MS.NoTrt_F |
| FAM8A1 | 0.0014 | 0.62529 | 0.058 | 0.14 | 1 | HC_vs_MS.NoTrt_F |
| XPO4 | 0.0014 | -0.7589 | 0.147 | 0.26 | 1 | HC_vs_MS.NoTrt_F |
| GRPEL2 | 0.00141 | -0.3427 | 0.275 | 0.4 | 1 | HC_vs_MS.NoTrt_F |
| OVCA2 | 0.00142 | -0.4097 | 0.101 | 0.2 | 1 | HC_vs_MS.NoTrt_F |
| CRNKL1 | 0.00142 | -0.3473 | 0.116 | 0.22 | 1 | HC_vs_MS.NoTrt_F |
| EPM2AIP1 | 0.00142 | -0.3723 | 0.309 | 0.42 | 1 | HC_vs_MS.NoTrt_F |
| COL4A2 | 0.00143 | 1.32199 | 0.121 | 0.04 | 1 | HC_vs_MS.NoTrt_F |
| TRPC4AP | 0.00143 | -0.4008 | 0.329 | 0.46 | 1 | HC_vs_MS.NoTrt_F |
| ALG6 | 0.00143 | -0.3821 | 0.147 | 0.26 | 1 | HC_vs_MS.NoTrt_F |

| | | | | | | |
|---|---|---|---|---|---|---|
| CTU2 | 0.00144 | -0.399 | 0.052 | 0.13 | 1 | HC_vs_MS.NoTrt_F |
| ARHGAP26 | 0.00145 | 0.72708 | 0.777 | 0.68 | 1 | HC_vs_MS.NoTrt_F |
| CCDC122 | 0.00145 | 0.31096 | 0.936 | 0.94 | 1 | HC_vs_MS.NoTrt_F |
| FAM156B | 0.00145 | 0.35543 | 0.095 | 0.19 | 1 | HC_vs_MS.NoTrt_F |
| RP11-507K1 | 0.00146 | 1.04662 | 0.052 | 0.13 | 1 | HC_vs_MS.NoTrt_F |
| RRM1 | 0.00147 | -0.2608 | 0.387 | 0.55 | 1 | HC_vs_MS.NoTrt_F |
| TRIM62 | 0.00147 | 0.4877 | 0.078 | 0.17 | 1 | HC_vs_MS.NoTrt_F |
| ZMYM6 | 0.00147 | 0.5027 | 0.884 | 0.79 | 1 | HC_vs_MS.NoTrt_F |
| TCF12 | 0.00148 | -0.79 | 0.306 | 0.43 | 1 | HC_vs_MS.NoTrt_F |
| FDXR | 0.00148 | -0.3215 | 0.09 | 0.18 | 1 | HC_vs_MS.NoTrt_F |
| ZNF507 | 0.00148 | -0.4791 | 0.197 | 0.32 | 1 | HC_vs_MS.NoTrt_F |
| L3MBTL3 | 0.00148 | -0.3399 | 0.118 | 0.22 | 1 | HC_vs_MS.NoTrt_F |
| RUNDC1 | 0.00149 | 0.51879 | 0.775 | 0.71 | 1 | HC_vs_MS.NoTrt_F |
| ARMC7 | 0.00149 | -1.0262 | 0.09 | 0.18 | 1 | HC_vs_MS.NoTrt_F |
| TANGO2 | 0.0015 | 0.6264 | 0.887 | 0.87 | 1 | HC_vs_MS.NoTrt_F |
| PFKL | 0.00151 | -0.6061 | 0.76 | 0.79 | 1 | HC_vs_MS.NoTrt_F |
| DNPH1 | 0.00151 | 0.32714 | 0.185 | 0.32 | 1 | HC_vs_MS.NoTrt_F |
| ZDHHC7 | 0.00152 | -0.3008 | 0.194 | 0.31 | 1 | HC_vs_MS.NoTrt_F |
| EIF3CL | 0.00154 | -0.4805 | 0.191 | 0.32 | 1 | HC_vs_MS.NoTrt_F |
| PEPD | 0.00154 | -0.2617 | 0.176 | 0.3 | 1 | HC_vs_MS.NoTrt_F |
| SFT2D2 | 0.00154 | 0.40661 | 0.991 | 0.98 | 1 | HC_vs_MS.NoTrt_F |
| AC138517.6 | 0.00154 | -0.9524 | 0.107 | 0.22 | 1 | HC_vs_MS.NoTrt_F |
| CHCHD1 | 0.00155 | -0.3503 | 0.116 | 0.22 | 1 | HC_vs_MS.NoTrt_F |
| POLR3D | 0.00156 | 0.6122 | 0.445 | 0.58 | 1 | HC_vs_MS.NoTrt_F |
| LTBR | 0.00156 | 0.2655 | 0.951 | 0.93 | 1 | HC_vs_MS.NoTrt_F |
| ATP5J2 | 0.00157 | 0.59534 | 0.353 | 0.53 | 1 | HC_vs_MS.NoTrt_F |
| ZNF407 | 0.00157 | -0.4554 | 0.124 | 0.23 | 1 | HC_vs_MS.NoTrt_F |
| RRAGA | 0.00157 | -0.7904 | 0.185 | 0.29 | 1 | HC_vs_MS.NoTrt_F |
| DUSP10 | 0.00157 | 0.44178 | 0.335 | 0.49 | 1 | HC_vs_MS.NoTrt_F |
| C5orf63 | 0.00158 | -0.5009 | 0.393 | 0.51 | 1 | HC_vs_MS.NoTrt_F |
| CRY1 | 0.00158 | -0.4483 | 0.208 | 0.34 | 1 | HC_vs_MS.NoTrt_F |
| PCBD1 | 0.00158 | -0.7758 | 0.104 | 0.2 | 1 | HC_vs_MS.NoTrt_F |
| SPDYE6 | 0.0016 | 0.4902 | 0.145 | 0.26 | 1 | HC_vs_MS.NoTrt_F |
| TFE3 | 0.00161 | -0.2659 | 0.11 | 0.21 | 1 | HC_vs_MS.NoTrt_F |
| POLR2C | 0.00161 | -0.5464 | 0.569 | 0.68 | 1 | HC_vs_MS.NoTrt_F |
| IDUA | 0.00161 | -0.3604 | 0.107 | 0.21 | 1 | HC_vs_MS.NoTrt_F |
| TMEM30A | 0.00161 | -0.7426 | 0.208 | 0.33 | 1 | HC_vs_MS.NoTrt_F |
| GNL3 | 0.00161 | -0.382 | 0.162 | 0.27 | 1 | HC_vs_MS.NoTrt_F |
| SQRDL | 0.00162 | -0.3467 | 0.171 | 0.29 | 1 | HC_vs_MS.NoTrt_F |
| ZNF253 | 0.00163 | 0.48291 | 0.754 | 0.59 | 1 | HC_vs_MS.NoTrt_F |

| | | | | | | |
|---|---|---|---|---|---|---|
| MED21 | 0.00163 | -0.6817 | 0.156 | 0.27 | 1 | HC_vs_MS.NoTrt_F |
| PYGL | 0.00165 | 0.41229 | 0.884 | 0.87 | 1 | HC_vs_MS.NoTrt_F |
| TTC14 | 0.00165 | -0.3712 | 0.442 | 0.54 | 1 | HC_vs_MS.NoTrt_F |
| DLAT | 0.00165 | 0.38505 | 0.075 | 0.17 | 1 | HC_vs_MS.NoTrt_F |
| INSL3 | 0.00166 | 0.27662 | 0.043 | 0.12 | 1 | HC_vs_MS.NoTrt_F |
| RNF219 | 0.00166 | -0.5598 | 0.072 | 0.16 | 1 | HC_vs_MS.NoTrt_F |
| PHF5A | 0.00166 | -0.4285 | 0.133 | 0.24 | 1 | HC_vs_MS.NoTrt_F |
| ARIH1 | 0.00166 | 0.31074 | 0.974 | 0.97 | 1 | HC_vs_MS.NoTrt_F |
| CRYZ | 0.00167 | -0.604 | 0.133 | 0.23 | 1 | HC_vs_MS.NoTrt_F |
| TOMM5 | 0.00168 | -0.2815 | 0.194 | 0.32 | 1 | HC_vs_MS.NoTrt_F |
| AIDA | 0.00168 | -0.7686 | 0.139 | 0.24 | 1 | HC_vs_MS.NoTrt_F |
| ABL1 | 0.00168 | 0.46946 | 0.075 | 0.17 | 1 | HC_vs_MS.NoTrt_F |
| MT-ND6 | 0.0017 | 0.80574 | 0.777 | 0.8 | 1 | HC_vs_MS.NoTrt_F |
| MAP3K4 | 0.0017 | -0.606 | 0.202 | 0.33 | 1 | HC_vs_MS.NoTrt_F |
| CAD | 0.0017 | 1.51577 | 0.777 | 0.7 | 1 | HC_vs_MS.NoTrt_F |
| NLGN3 | 0.00171 | -0.6354 | 0.064 | 0.15 | 1 | HC_vs_MS.NoTrt_F |
| INF2 | 0.00171 | -0.4941 | 0.249 | 0.38 | 1 | HC_vs_MS.NoTrt_F |
| LPCAT1 | 0.00173 | -0.3309 | 0.064 | 0.15 | 1 | HC_vs_MS.NoTrt_F |
| C10orf88 | 0.00173 | -0.3177 | 0.191 | 0.31 | 1 | HC_vs_MS.NoTrt_F |
| DOK1 | 0.00173 | -0.3757 | 0.107 | 0.2 | 1 | HC_vs_MS.NoTrt_F |
| SNRPE | 0.00175 | 0.39204 | 0.159 | 0.28 | 1 | HC_vs_MS.NoTrt_F |
| CRTC1 | 0.00175 | 0.41209 | 0.217 | 0.35 | 1 | HC_vs_MS.NoTrt_F |
| DGCR8 | 0.00175 | -0.3795 | 0.136 | 0.25 | 1 | HC_vs_MS.NoTrt_F |
| ZNF44 | 0.00175 | 0.58264 | 0.54 | 0.69 | 1 | HC_vs_MS.NoTrt_F |
| STMN3 | 0.00176 | 0.57137 | 0.249 | 0.39 | 1 | HC_vs_MS.NoTrt_F |
| RNF185 | 0.00176 | -0.5712 | 0.306 | 0.42 | 1 | HC_vs_MS.NoTrt_F |
| QARS | 0.00176 | -0.6036 | 0.246 | 0.36 | 1 | HC_vs_MS.NoTrt_F |
| CLASRP | 0.00177 | -0.5301 | 0.202 | 0.32 | 1 | HC_vs_MS.NoTrt_F |
| CDCP1 | 0.00177 | 0.6214 | 0.561 | 0.41 | 1 | HC_vs_MS.NoTrt_F |
| BEX2 | 0.00177 | -0.3204 | 0.092 | 0.19 | 1 | HC_vs_MS.NoTrt_F |
| HNRNPU | 0.00178 | -0.3831 | 0.506 | 0.61 | 1 | HC_vs_MS.NoTrt_F |
| TNIP1 | 0.00178 | 0.48718 | 0.355 | 0.52 | 1 | HC_vs_MS.NoTrt_F |
| IL15RA | 0.00178 | -0.3788 | 0.127 | 0.23 | 1 | HC_vs_MS.NoTrt_F |
| SMAD5 | 0.00179 | 0.67386 | 0.566 | 0.45 | 1 | HC_vs_MS.NoTrt_F |
| COPS3 | 0.0018 | -0.5743 | 0.286 | 0.41 | 1 | HC_vs_MS.NoTrt_F |
| ATG16L1 | 0.00181 | -0.7547 | 0.179 | 0.29 | 1 | HC_vs_MS.NoTrt_F |
| TMEM177 | 0.00183 | -1.5554 | 0.043 | 0.12 | 1 | HC_vs_MS.NoTrt_F |
| USMG5 | 0.00183 | 0.25431 | 0.266 | 0.43 | 1 | HC_vs_MS.NoTrt_F |
| ASB13 | 0.00184 | -1.6621 | 0.107 | 0.21 | 1 | HC_vs_MS.NoTrt_F |
| RAD17 | 0.00185 | -0.5987 | 0.471 | 0.6 | 1 | HC_vs_MS.NoTrt_F |

| | | | | | | |
|---|---|---|---|---|---|---|
| PCDHA4 | 0.00186 | 0.34639 | 0.957 | 0.91 | 1 | HC_vs_MS.NoTrt_F |
| CKAP2 | 0.00186 | -0.2861 | 0.147 | 0.26 | 1 | HC_vs_MS.NoTrt_F |
| LUC7L2 | 0.00186 | -0.3402 | 0.584 | 0.69 | 1 | HC_vs_MS.NoTrt_F |
| CIAO1 | 0.00187 | -0.2843 | 0.355 | 0.48 | 1 | HC_vs_MS.NoTrt_F |
| GEMIN6 | 0.00188 | 0.25176 | 0.968 | 0.93 | 1 | HC_vs_MS.NoTrt_F |
| KRBA1 | 0.00188 | -0.2503 | 0.043 | 0.12 | 1 | HC_vs_MS.NoTrt_F |
| KIAA1456 | 0.00188 | 0.33799 | 0.962 | 0.93 | 1 | HC_vs_MS.NoTrt_F |
| ZNF317 | 0.00188 | -0.4734 | 0.139 | 0.25 | 1 | HC_vs_MS.NoTrt_F |
| TRAF2 | 0.00189 | -0.7405 | 0.214 | 0.33 | 1 | HC_vs_MS.NoTrt_F |
| SNAPC4 | 0.0019 | -0.6884 | 0.058 | 0.14 | 1 | HC_vs_MS.NoTrt_F |
| NSMAF | 0.00191 | -0.3531 | 0.263 | 0.39 | 1 | HC_vs_MS.NoTrt_F |
| CLINT1 | 0.00191 | -0.3168 | 0.679 | 0.77 | 1 | HC_vs_MS.NoTrt_F |
| LAIR2 | 0.00191 | 0.28694 | 0.179 | 0.31 | 1 | HC_vs_MS.NoTrt_F |
| ESR2 | 0.00191 | 0.39359 | 0.555 | 0.7 | 1 | HC_vs_MS.NoTrt_F |
| NFKBIA | 0.00192 | 1.43023 | 0.468 | 0.38 | 1 | HC_vs_MS.NoTrt_F |
| ATG4A | 0.00196 | 0.35454 | 0.075 | 0.17 | 1 | HC_vs_MS.NoTrt_F |
| CPQ | 0.00196 | -0.7827 | 0.37 | 0.5 | 1 | HC_vs_MS.NoTrt_F |
| VWF | 0.00197 | 1.8118 | 0.237 | 0.13 | 1 | HC_vs_MS.NoTrt_F |
| LMF2 | 0.00198 | 0.28135 | 0.303 | 0.47 | 1 | HC_vs_MS.NoTrt_F |
| ARNTL | 0.00199 | 0.29911 | 0.251 | 0.39 | 1 | HC_vs_MS.NoTrt_F |
| CEP85L | 0.002 | -0.2737 | 0.292 | 0.41 | 1 | HC_vs_MS.NoTrt_F |
| HIBCH | 0.00201 | -0.2654 | 0.26 | 0.4 | 1 | HC_vs_MS.NoTrt_F |
| STAT5A | 0.00203 | -0.7132 | 0.211 | 0.33 | 1 | HC_vs_MS.NoTrt_F |
| THOP1 | 0.00205 | 0.26918 | 0.402 | 0.54 | 1 | HC_vs_MS.NoTrt_F |
| CDH5 | 0.00207 | 0.98698 | 0.101 | 0.03 | 1 | HC_vs_MS.NoTrt_F |
| TRIM28 | 0.00208 | -0.6471 | 0.113 | 0.21 | 1 | HC_vs_MS.NoTrt_F |
| PPP1R13L | 0.00208 | 0.2734 | 0.858 | 0.77 | 1 | HC_vs_MS.NoTrt_F |
| CDC16 | 0.00208 | 0.36095 | 0.159 | 0.27 | 1 | HC_vs_MS.NoTrt_F |
| MEGF11 | 0.00209 | -0.6474 | 0.043 | 0.12 | 1 | HC_vs_MS.NoTrt_F |
| ZNF322 | 0.00209 | 0.62407 | 0.043 | 0.12 | 1 | HC_vs_MS.NoTrt_F |
| GMIP | 0.0021 | -0.901 | 0.263 | 0.39 | 1 | HC_vs_MS.NoTrt_F |
| UBA3 | 0.00211 | -0.5167 | 0.416 | 0.54 | 1 | HC_vs_MS.NoTrt_F |
| RRS1 | 0.00213 | -0.5401 | 0.11 | 0.21 | 1 | HC_vs_MS.NoTrt_F |
| RP11-234B2 | 0.00213 | -0.3423 | 0.04 | 0.11 | 1 | HC_vs_MS.NoTrt_F |
| DCTD | 0.00213 | -0.308 | 0.153 | 0.26 | 1 | HC_vs_MS.NoTrt_F |
| ZBED6 | 0.00216 | -0.7615 | 0.058 | 0.14 | 1 | HC_vs_MS.NoTrt_F |
| C5orf58 | 0.00217 | -0.3098 | 0.04 | 0.11 | 1 | HC_vs_MS.NoTrt_F |
| SNAPIN | 0.00217 | 0.25541 | 0.147 | 0.26 | 1 | HC_vs_MS.NoTrt_F |
| FLII | 0.00218 | -0.3112 | 0.249 | 0.37 | 1 | HC_vs_MS.NoTrt_F |
| TMEM214 | 0.00219 | -0.5412 | 0.095 | 0.19 | 1 | HC_vs_MS.NoTrt_F |

| | | | | | | |
|---|---|---|---|---|---|---|
| VWA9 | 0.00221 | -1.3022 | 0.09 | 0.18 | 1 | HC_vs_MS.NoTrt_F |
| QDPR | 0.00222 | -0.3158 | 0.075 | 0.16 | 1 | HC_vs_MS.NoTrt_F |
| INPP1 | 0.00223 | 0.87257 | 0.058 | 0.14 | 1 | HC_vs_MS.NoTrt_F |
| KAT6B | 0.00224 | 0.59892 | 0.136 | 0.24 | 1 | HC_vs_MS.NoTrt_F |
| PLCXD2 | 0.00224 | -0.2724 | 0.081 | 0.17 | 1 | HC_vs_MS.NoTrt_F |
| LDOC1 | 0.00225 | -0.3238 | 0.078 | 0.17 | 1 | HC_vs_MS.NoTrt_F |
| RDX | 0.00225 | -0.6274 | 0.286 | 0.41 | 1 | HC_vs_MS.NoTrt_F |
| EDEM3 | 0.00226 | -0.3813 | 0.309 | 0.43 | 1 | HC_vs_MS.NoTrt_F |
| UCHL3 | 0.00226 | -0.2738 | 0.159 | 0.27 | 1 | HC_vs_MS.NoTrt_F |
| XAB2 | 0.00227 | 0.30888 | 0.24 | 0.37 | 1 | HC_vs_MS.NoTrt_F |
| DAPK3 | 0.00228 | -0.738 | 0.116 | 0.21 | 1 | HC_vs_MS.NoTrt_F |
| DDRGK1 | 0.00229 | -0.5559 | 0.173 | 0.28 | 1 | HC_vs_MS.NoTrt_F |
| L3HYPDH | 0.00232 | -0.7545 | 0.208 | 0.32 | 1 | HC_vs_MS.NoTrt_F |
| NMNAT1 | 0.00232 | 0.41129 | 0.986 | 1 | 1 | HC_vs_MS.NoTrt_F |
| SAAL1 | 0.00232 | -0.5952 | 0.09 | 0.18 | 1 | HC_vs_MS.NoTrt_F |
| IL17RB | 0.00233 | -0.2785 | 0.04 | 0.11 | 1 | HC_vs_MS.NoTrt_F |
| SLC9B2 | 0.00234 | 0.48797 | 0.066 | 0.15 | 1 | HC_vs_MS.NoTrt_F |
| STIM2 | 0.00235 | -0.3329 | 0.191 | 0.3 | 1 | HC_vs_MS.NoTrt_F |
| PYM1 | 0.00237 | -0.5118 | 0.231 | 0.35 | 1 | HC_vs_MS.NoTrt_F |
| KATNA1 | 0.00237 | -0.4103 | 0.113 | 0.21 | 1 | HC_vs_MS.NoTrt_F |
| HIVEP3 | 0.00238 | -0.4374 | 0.092 | 0.18 | 1 | HC_vs_MS.NoTrt_F |
| CASP3 | 0.00238 | -0.3852 | 0.173 | 0.29 | 1 | HC_vs_MS.NoTrt_F |
| YY1AP1 | 0.00238 | -0.3005 | 0.543 | 0.65 | 1 | HC_vs_MS.NoTrt_F |
| EIF4EBP1 | 0.00239 | 0.29833 | 0.055 | 0.13 | 1 | HC_vs_MS.NoTrt_F |
| SAT1 | 0.00239 | -0.3902 | 0.595 | 0.76 | 1 | HC_vs_MS.NoTrt_F |
| CNDP2 | 0.00242 | -0.5656 | 0.749 | 0.8 | 1 | HC_vs_MS.NoTrt_F |
| INTS5 | 0.00243 | -0.3974 | 0.09 | 0.18 | 1 | HC_vs_MS.NoTrt_F |
| HIST3H2A | 0.00245 | 0.32622 | 0.069 | 0.16 | 1 | HC_vs_MS.NoTrt_F |
| B4GALT7 | 0.00245 | 0.62128 | 0.078 | 0.17 | 1 | HC_vs_MS.NoTrt_F |
| PKIG | 0.00245 | 0.52398 | 0.685 | 0.71 | 1 | HC_vs_MS.NoTrt_F |
| ADAM8 | 0.00245 | 0.26391 | 0.205 | 0.34 | 1 | HC_vs_MS.NoTrt_F |
| SERBP1 | 0.00246 | -0.3555 | 0.642 | 0.74 | 1 | HC_vs_MS.NoTrt_F |
| NSMCE1 | 0.00248 | -0.4381 | 0.665 | 0.74 | 1 | HC_vs_MS.NoTrt_F |
| KAT5 | 0.00248 | -0.4302 | 0.191 | 0.3 | 1 | HC_vs_MS.NoTrt_F |
| HSPA4 | 0.0025 | -0.3488 | 0.434 | 0.53 | 1 | HC_vs_MS.NoTrt_F |
| BNIP1 | 0.00251 | -0.4869 | 0.087 | 0.18 | 1 | HC_vs_MS.NoTrt_F |
| TROVE2 | 0.00254 | -0.3526 | 0.717 | 0.78 | 1 | HC_vs_MS.NoTrt_F |
| UBE2E1 | 0.00254 | -0.388 | 0.11 | 0.2 | 1 | HC_vs_MS.NoTrt_F |
| HPCAL1 | 0.00254 | -0.3578 | 0.361 | 0.47 | 1 | HC_vs_MS.NoTrt_F |
| SRI | 0.00254 | -0.4467 | 0.471 | 0.59 | 1 | HC_vs_MS.NoTrt_F |

| | | | | | | |
|---|---|---|---|---|---|---|
| GOLT1B | 0.00255 | -0.2705 | 0.139 | 0.24 | 1 | HC_vs_MS.NoTrt_F |
| DUSP28 | 0.00255 | -0.3015 | 0.754 | 0.83 | 1 | HC_vs_MS.NoTrt_F |
| RBM12B | 0.00256 | -0.3932 | 0.165 | 0.28 | 1 | HC_vs_MS.NoTrt_F |
| C20orf24 | 0.00256 | -0.255 | 0.153 | 0.26 | 1 | HC_vs_MS.NoTrt_F |
| SUPT7L | 0.00258 | -0.4078 | 0.468 | 0.59 | 1 | HC_vs_MS.NoTrt_F |
| PJA1 | 0.00259 | -0.7379 | 0.075 | 0.16 | 1 | HC_vs_MS.NoTrt_F |
| RP11-286N2 | 0.00259 | 0.34571 | 0.977 | 0.97 | 1 | HC_vs_MS.NoTrt_F |
| C14orf159 | 0.0026 | 0.46359 | 0.592 | 0.75 | 1 | HC_vs_MS.NoTrt_F |
| CKAP5 | 0.0026 | -0.3594 | 0.168 | 0.29 | 1 | HC_vs_MS.NoTrt_F |
| UTP4 | 0.0026 | -0.5388 | 0.838 | 0.88 | 1 | HC_vs_MS.NoTrt_F |
| BPGM | 0.00261 | 0.50953 | 0.066 | 0.15 | 1 | HC_vs_MS.NoTrt_F |
| UCK2 | 0.00266 | 0.37392 | 0.09 | 0.18 | 1 | HC_vs_MS.NoTrt_F |
| HIST2H4A | 0.00266 | -0.3017 | 0.09 | 0.18 | 1 | HC_vs_MS.NoTrt_F |
| PPIH | 0.00267 | -0.5319 | 0.098 | 0.19 | 1 | HC_vs_MS.NoTrt_F |
| PSD | 0.00267 | -0.4491 | 0.098 | 0.19 | 1 | HC_vs_MS.NoTrt_F |
| TOMM22 | 0.00268 | -0.5723 | 0.214 | 0.33 | 1 | HC_vs_MS.NoTrt_F |
| GYS1 | 0.00269 | -0.2535 | 0.165 | 0.28 | 1 | HC_vs_MS.NoTrt_F |
| SCYL1 | 0.0027 | -0.4421 | 0.269 | 0.39 | 1 | HC_vs_MS.NoTrt_F |
| C4orf32 | 0.0027 | 0.28969 | 0.994 | 0.99 | 1 | HC_vs_MS.NoTrt_F |
| LGALS3BP | 0.00276 | 1.02146 | 0.711 | 0.6 | 1 | HC_vs_MS.NoTrt_F |
| LPP | 0.00276 | 0.40404 | 0.988 | 0.96 | 1 | HC_vs_MS.NoTrt_F |
| FLCN | 0.00277 | 0.54639 | 0.893 | 0.82 | 1 | HC_vs_MS.NoTrt_F |
| CLUL1 | 0.00277 | -0.3846 | 0.075 | 0.16 | 1 | HC_vs_MS.NoTrt_F |
| PRKG2 | 0.00279 | -0.3058 | 0.387 | 0.54 | 1 | HC_vs_MS.NoTrt_F |
| BFSP2 | 0.0028 | 0.67564 | 0.052 | 0.13 | 1 | HC_vs_MS.NoTrt_F |
| CTRL | 0.00281 | -1.1229 | 0.133 | 0.23 | 1 | HC_vs_MS.NoTrt_F |
| SLC2A6 | 0.00281 | 0.48871 | 0.254 | 0.38 | 1 | HC_vs_MS.NoTrt_F |
| ARF4 | 0.00283 | -0.6144 | 0.182 | 0.29 | 1 | HC_vs_MS.NoTrt_F |
| HOOK1 | 0.00283 | -0.3954 | 0.078 | 0.17 | 1 | HC_vs_MS.NoTrt_F |
| SNRPA | 0.00283 | -0.2728 | 0.298 | 0.41 | 1 | HC_vs_MS.NoTrt_F |
| LENG8 | 0.00284 | 0.78656 | 0.777 | 0.77 | 1 | HC_vs_MS.NoTrt_F |
| MRPL33 | 0.00286 | 0.28735 | 0.214 | 0.34 | 1 | HC_vs_MS.NoTrt_F |
| RSPH3 | 0.00287 | 0.29499 | 0.829 | 0.82 | 1 | HC_vs_MS.NoTrt_F |
| DIRAS2 | 0.00288 | 0.38849 | 0.873 | 0.83 | 1 | HC_vs_MS.NoTrt_F |
| CCDC71 | 0.00289 | 0.50207 | 0.072 | 0.16 | 1 | HC_vs_MS.NoTrt_F |
| CSTF2 | 0.00289 | -0.642 | 0.095 | 0.18 | 1 | HC_vs_MS.NoTrt_F |
| THAP2 | 0.0029 | -0.5341 | 0.107 | 0.2 | 1 | HC_vs_MS.NoTrt_F |
| DNAJB12 | 0.0029 | -0.4493 | 0.15 | 0.25 | 1 | HC_vs_MS.NoTrt_F |
| ALDH3A2 | 0.0029 | 0.2536 | 0.295 | 0.41 | 1 | HC_vs_MS.NoTrt_F |
| MSH2 | 0.00297 | -0.2706 | 0.13 | 0.23 | 1 | HC_vs_MS.NoTrt_F |

| | | | | | | |
|---|---|---|---|---|---|---|
| WDR77 | 0.00297 | -0.4727 | 0.136 | 0.24 | 1 | HC_vs_MS.NoTrt_F |
| POMGNT1 | 0.00298 | 0.30358 | 0.766 | 0.67 | 1 | HC_vs_MS.NoTrt_F |
| TNK2 | 0.00299 | -0.3143 | 0.182 | 0.29 | 1 | HC_vs_MS.NoTrt_F |
| VCPKMT | 0.003 | -0.4797 | 0.118 | 0.22 | 1 | HC_vs_MS.NoTrt_F |
| SRGAP2B | 0.00301 | -0.699 | 0.503 | 0.6 | 1 | HC_vs_MS.NoTrt_F |
| ZSCAN26 | 0.00305 | -0.2641 | 0.072 | 0.16 | 1 | HC_vs_MS.NoTrt_F |
| PIBF1 | 0.00306 | 0.41166 | 0.162 | 0.28 | 1 | HC_vs_MS.NoTrt_F |
| TRIM8 | 0.00307 | -0.9481 | 0.087 | 0.17 | 1 | HC_vs_MS.NoTrt_F |
| TBC1D22B | 0.00312 | -0.636 | 0.095 | 0.18 | 1 | HC_vs_MS.NoTrt_F |
| LRRC45 | 0.00315 | -0.3811 | 0.075 | 0.16 | 1 | HC_vs_MS.NoTrt_F |
| COX16 | 0.00315 | -0.5872 | 0.552 | 0.62 | 1 | HC_vs_MS.NoTrt_F |
| FAM86B1 | 0.00315 | -1.2341 | 0.081 | 0.17 | 1 | HC_vs_MS.NoTrt_F |
| ARHGEF12 | 0.00316 | 0.50931 | 0.988 | 0.98 | 1 | HC_vs_MS.NoTrt_F |
| MAN1A2 | 0.00317 | -0.3851 | 0.367 | 0.48 | 1 | HC_vs_MS.NoTrt_F |
| RAB11FIP5 | 0.00318 | 0.27827 | 0.049 | 0.12 | 1 | HC_vs_MS.NoTrt_F |
| POLDIP3 | 0.00318 | -0.575 | 0.191 | 0.3 | 1 | HC_vs_MS.NoTrt_F |
| DNTTIP1 | 0.00319 | 0.48699 | 0.066 | 0.15 | 1 | HC_vs_MS.NoTrt_F |
| TRAFD1 | 0.00319 | -0.608 | 0.243 | 0.35 | 1 | HC_vs_MS.NoTrt_F |
| ZNF26 | 0.0032 | 0.52682 | 0.994 | 0.99 | 1 | HC_vs_MS.NoTrt_F |
| KCNJ15 | 0.0032 | 0.5284 | 0.705 | 0.56 | 1 | HC_vs_MS.NoTrt_F |
| MRPL15 | 0.00322 | 0.56876 | 0.098 | 0.19 | 1 | HC_vs_MS.NoTrt_F |
| HMGB2 | 0.00323 | -0.3451 | 0.249 | 0.37 | 1 | HC_vs_MS.NoTrt_F |
| AL356053.1 | 0.00324 | 0.30045 | 0.049 | 0.12 | 1 | HC_vs_MS.NoTrt_F |
| MFSD3 | 0.00324 | 0.51302 | 0.064 | 0.14 | 1 | HC_vs_MS.NoTrt_F |
| ATP2C1 | 0.00326 | -0.3125 | 0.191 | 0.3 | 1 | HC_vs_MS.NoTrt_F |
| CTD-2021K4 | 0.0033 | -0.3418 | 0.095 | 0.18 | 1 | HC_vs_MS.NoTrt_F |
| CHIC2 | 0.0033 | -0.3171 | 0.214 | 0.33 | 1 | HC_vs_MS.NoTrt_F |
| TNRC6C | 0.00332 | 0.31876 | 0.263 | 0.39 | 1 | HC_vs_MS.NoTrt_F |
| YKT6 | 0.00334 | -0.3269 | 0.124 | 0.22 | 1 | HC_vs_MS.NoTrt_F |
| TCF19 | 0.00334 | -0.4549 | 0.072 | 0.16 | 1 | HC_vs_MS.NoTrt_F |
| SLC52A2 | 0.00334 | -0.7743 | 0.043 | 0.11 | 1 | HC_vs_MS.NoTrt_F |
| TTC33 | 0.00335 | -0.5947 | 0.246 | 0.36 | 1 | HC_vs_MS.NoTrt_F |
| B4GALT3 | 0.00335 | -0.2626 | 0.214 | 0.34 | 1 | HC_vs_MS.NoTrt_F |
| MCM3 | 0.00335 | -0.307 | 0.225 | 0.34 | 1 | HC_vs_MS.NoTrt_F |
| TMEM218 | 0.00338 | -1.5066 | 0.116 | 0.2 | 1 | HC_vs_MS.NoTrt_F |
| FAM213A | 0.00339 | 0.81118 | 0.893 | 0.87 | 1 | HC_vs_MS.NoTrt_F |
| TARBP2 | 0.00343 | -0.5456 | 0.095 | 0.18 | 1 | HC_vs_MS.NoTrt_F |
| UBR2 | 0.00344 | 0.46447 | 0.549 | 0.67 | 1 | HC_vs_MS.NoTrt_F |
| CPEB4 | 0.00345 | -0.2969 | 0.197 | 0.3 | 1 | HC_vs_MS.NoTrt_F |
| TRMT6 | 0.00347 | -0.3586 | 0.118 | 0.22 | 1 | HC_vs_MS.NoTrt_F |

| | | | | | | |
|---|---|---|---|---|---|---|
| RC3H2 | 0.00348 | -0.5411 | 0.503 | 0.61 | 1 | HC_vs_MS.NoTrt_F |
| PIGV | 0.00348 | -0.8175 | 0.072 | 0.16 | 1 | HC_vs_MS.NoTrt_F |
| C3orf38 | 0.00351 | -0.6243 | 0.104 | 0.19 | 1 | HC_vs_MS.NoTrt_F |
| RNF34 | 0.00352 | -0.3593 | 0.272 | 0.4 | 1 | HC_vs_MS.NoTrt_F |
| AGPAT1 | 0.00352 | -0.9705 | 0.118 | 0.21 | 1 | HC_vs_MS.NoTrt_F |
| TMPPE | 0.00352 | -0.4599 | 0.061 | 0.14 | 1 | HC_vs_MS.NoTrt_F |
| DENND4B | 0.00353 | -0.5224 | 0.22 | 0.34 | 1 | HC_vs_MS.NoTrt_F |
| COMMD5 | 0.00355 | -0.4764 | 0.098 | 0.19 | 1 | HC_vs_MS.NoTrt_F |
| AMACR | 0.00356 | -1.2004 | 0.075 | 0.16 | 1 | HC_vs_MS.NoTrt_F |
| ASTN2 | 0.00357 | 0.3847 | 0.977 | 0.96 | 1 | HC_vs_MS.NoTrt_F |
| ZNF689 | 0.00358 | 0.92273 | 0.046 | 0.12 | 1 | HC_vs_MS.NoTrt_F |
| OSBPL11 | 0.00361 | -0.6015 | 0.118 | 0.21 | 1 | HC_vs_MS.NoTrt_F |
| ARL17B | 0.00362 | -0.2726 | 0.78 | 0.85 | 1 | HC_vs_MS.NoTrt_F |
| ZNF224 | 0.00364 | -1.2469 | 0.451 | 0.56 | 1 | HC_vs_MS.NoTrt_F |
| SLC35A2 | 0.00364 | -0.5624 | 0.104 | 0.19 | 1 | HC_vs_MS.NoTrt_F |
| NOG | 0.00365 | 0.30807 | 0.139 | 0.24 | 1 | HC_vs_MS.NoTrt_F |
| PES1 | 0.00366 | -0.2727 | 0.142 | 0.24 | 1 | HC_vs_MS.NoTrt_F |
| HSPA13 | 0.00369 | -1.2074 | 0.043 | 0.11 | 1 | HC_vs_MS.NoTrt_F |
| C7 | 0.0037 | -0.4205 | 0.387 | 0.51 | 1 | HC_vs_MS.NoTrt_F |
| SNX24 | 0.00371 | 0.32816 | 0.043 | 0.11 | 1 | HC_vs_MS.NoTrt_F |
| NUDT16 | 0.00371 | -0.9116 | 0.087 | 0.17 | 1 | HC_vs_MS.NoTrt_F |
| NBN | 0.00372 | -0.3023 | 0.136 | 0.23 | 1 | HC_vs_MS.NoTrt_F |
| PDHA1 | 0.00374 | -0.2553 | 0.249 | 0.37 | 1 | HC_vs_MS.NoTrt_F |
| PIP4K2C | 0.00374 | -0.4996 | 0.243 | 0.34 | 1 | HC_vs_MS.NoTrt_F |
| CCR6 | 0.00377 | -0.5568 | 0.419 | 0.53 | 1 | HC_vs_MS.NoTrt_F |
| UMAD1 | 0.00377 | -0.3189 | 0.22 | 0.34 | 1 | HC_vs_MS.NoTrt_F |
| UBE2Q2 | 0.00378 | -0.2787 | 0.15 | 0.26 | 1 | HC_vs_MS.NoTrt_F |
| CENPJ | 0.00379 | -0.6745 | 0.055 | 0.13 | 1 | HC_vs_MS.NoTrt_F |
| CRAMP1 | 0.00379 | -1.4929 | 0.104 | 0.19 | 1 | HC_vs_MS.NoTrt_F |
| DLG3 | 0.00382 | 0.38383 | 0.046 | 0.12 | 1 | HC_vs_MS.NoTrt_F |
| VPS36 | 0.00383 | -0.5047 | 0.358 | 0.48 | 1 | HC_vs_MS.NoTrt_F |
| FKBP9 | 0.00386 | 0.53353 | 0.59 | 0.48 | 1 | HC_vs_MS.NoTrt_F |
| IL33 | 0.00387 | 0.49433 | 0.046 | 0.12 | 1 | HC_vs_MS.NoTrt_F |
| EEPD1 | 0.00388 | 0.29542 | 0.092 | 0.18 | 1 | HC_vs_MS.NoTrt_F |
| FBXO41 | 0.00389 | 0.64854 | 0.061 | 0.14 | 1 | HC_vs_MS.NoTrt_F |
| AL592183.1 | 0.0039 | -0.3973 | 0.118 | 0.21 | 1 | HC_vs_MS.NoTrt_F |
| UBALD2 | 0.0039 | -0.3975 | 0.118 | 0.21 | 1 | HC_vs_MS.NoTrt_F |
| NUP205 | 0.00392 | -0.3925 | 0.202 | 0.31 | 1 | HC_vs_MS.NoTrt_F |
| CD7 | 0.00393 | 0.28746 | 0.289 | 0.41 | 1 | HC_vs_MS.NoTrt_F |
| MMGT1 | 0.00397 | -0.7081 | 0.118 | 0.21 | 1 | HC_vs_MS.NoTrt_F |

| | | | | | | |
|---|---|---|---|---|---|---|
| ERCC6L2 | 0.00397 | -0.5777 | 0.234 | 0.35 | 1 | HC_vs_MS.NoTrt_F |
| AP5S1 | 0.00397 | 0.78529 | 0.876 | 0.8 | 1 | HC_vs_MS.NoTrt_F |
| CCNI2 | 0.00398 | -0.2872 | 0.043 | 0.11 | 1 | HC_vs_MS.NoTrt_F |
| PLEKHB1 | 0.004 | -0.7488 | 0.107 | 0.19 | 1 | HC_vs_MS.NoTrt_F |
| FAM122B | 0.00402 | -0.2861 | 0.13 | 0.23 | 1 | HC_vs_MS.NoTrt_F |
| TRIAP1 | 0.00402 | -0.3362 | 0.11 | 0.2 | 1 | HC_vs_MS.NoTrt_F |
| FAM221A | 0.00406 | 0.30568 | 0.983 | 0.98 | 1 | HC_vs_MS.NoTrt_F |
| WDR53 | 0.00406 | -0.2861 | 0.116 | 0.21 | 1 | HC_vs_MS.NoTrt_F |
| ANAPC11 | 0.00406 | 0.33127 | 0.422 | 0.54 | 1 | HC_vs_MS.NoTrt_F |
| RBL2 | 0.00407 | -0.3098 | 0.621 | 0.71 | 1 | HC_vs_MS.NoTrt_F |
| PHF10 | 0.00409 | 0.95188 | 0.113 | 0.2 | 1 | HC_vs_MS.NoTrt_F |
| TRAPPC10 | 0.0041 | 0.80618 | 0.295 | 0.42 | 1 | HC_vs_MS.NoTrt_F |
| WIPF1 | 0.00411 | -0.3606 | 0.711 | 0.8 | 1 | HC_vs_MS.NoTrt_F |
| LTA | 0.00415 | -0.2828 | 0.09 | 0.18 | 1 | HC_vs_MS.NoTrt_F |
| PRPS1 | 0.00415 | -0.2596 | 0.162 | 0.27 | 1 | HC_vs_MS.NoTrt_F |
| MMADHC | 0.00416 | -0.4388 | 0.422 | 0.54 | 1 | HC_vs_MS.NoTrt_F |
| TMED1 | 0.00417 | 0.27503 | 0.069 | 0.15 | 1 | HC_vs_MS.NoTrt_F |
| HLA-DQB1 | 0.00417 | 0.79311 | 0.581 | 0.7 | 1 | HC_vs_MS.NoTrt_F |
| GPLD1 | 0.00418 | 0.59499 | 0.78 | 0.71 | 1 | HC_vs_MS.NoTrt_F |
| RGMB | 0.00418 | -0.8065 | 0.116 | 0.2 | 1 | HC_vs_MS.NoTrt_F |
| SCRIB | 0.0042 | 0.40834 | 0.043 | 0.11 | 1 | HC_vs_MS.NoTrt_F |
| ZFYVE1 | 0.0042 | -0.259 | 0.095 | 0.18 | 1 | HC_vs_MS.NoTrt_F |
| ADAM17 | 0.00422 | -0.5608 | 0.078 | 0.16 | 1 | HC_vs_MS.NoTrt_F |
| TXNDC16 | 0.00422 | 0.45165 | 0.133 | 0.23 | 1 | HC_vs_MS.NoTrt_F |
| POLR2E | 0.00422 | -0.3005 | 0.564 | 0.66 | 1 | HC_vs_MS.NoTrt_F |
| CDH13 | 0.00424 | 0.34759 | 0.841 | 0.74 | 1 | HC_vs_MS.NoTrt_F |
| KIAA0922 | 0.00426 | -0.2572 | 0.468 | 0.58 | 1 | HC_vs_MS.NoTrt_F |
| MAP3K7 | 0.00427 | -0.374 | 0.159 | 0.26 | 1 | HC_vs_MS.NoTrt_F |
| INTS6L | 0.00428 | -0.6202 | 0.142 | 0.24 | 1 | HC_vs_MS.NoTrt_F |
| RGS1 | 0.0043 | -0.3324 | 0.575 | 0.7 | 1 | HC_vs_MS.NoTrt_F |
| MRM1 | 0.0043 | -0.4163 | 0.04 | 0.11 | 1 | HC_vs_MS.NoTrt_F |
| FBF1 | 0.00433 | 1.02341 | 0.058 | 0.13 | 1 | HC_vs_MS.NoTrt_F |
| FXN | 0.00435 | -0.3047 | 0.22 | 0.33 | 1 | HC_vs_MS.NoTrt_F |
| ANKMY2 | 0.00436 | -0.287 | 0.078 | 0.16 | 1 | HC_vs_MS.NoTrt_F |
| RAB11FIP2 | 0.00437 | -0.5086 | 0.066 | 0.14 | 1 | HC_vs_MS.NoTrt_F |
| FAM185A | 0.00437 | -0.4838 | 0.113 | 0.2 | 1 | HC_vs_MS.NoTrt_F |
| RPTOR | 0.00438 | 0.61479 | 0.393 | 0.53 | 1 | HC_vs_MS.NoTrt_F |
| ALDH9A1 | 0.00439 | -0.469 | 0.205 | 0.32 | 1 | HC_vs_MS.NoTrt_F |
| DUSP7 | 0.0044 | 0.39351 | 0.118 | 0.21 | 1 | HC_vs_MS.NoTrt_F |
| ZNF324 | 0.0044 | 0.37694 | 0.066 | 0.14 | 1 | HC_vs_MS.NoTrt_F |

| | | | | | | |
|---|---|---|---|---|---|---|
| MDM1 | 0.00441 | -0.2854 | 0.11 | 0.2 | 1 | HC_vs_MS.NoTrt_F |
| MS4A10 | 0.00442 | 0.30697 | 0.746 | 0.64 | 1 | HC_vs_MS.NoTrt_F |
| E4F1 | 0.00442 | -0.4127 | 0.159 | 0.25 | 1 | HC_vs_MS.NoTrt_F |
| DHX58 | 0.00445 | 1.19148 | 0.04 | 0.11 | 1 | HC_vs_MS.NoTrt_F |
| EDEM2 | 0.00449 | -0.3141 | 0.171 | 0.27 | 1 | HC_vs_MS.NoTrt_F |
| FAM103A1 | 0.00449 | -0.3407 | 0.156 | 0.26 | 1 | HC_vs_MS.NoTrt_F |
| CCAR2 | 0.0045 | -0.8662 | 0.272 | 0.39 | 1 | HC_vs_MS.NoTrt_F |
| RDH14 | 0.0045 | 0.8466 | 0.058 | 0.13 | 1 | HC_vs_MS.NoTrt_F |
| RIOK1 | 0.00452 | -0.4098 | 0.136 | 0.23 | 1 | HC_vs_MS.NoTrt_F |
| ATP6V1C1 | 0.00452 | 0.36847 | 0.494 | 0.6 | 1 | HC_vs_MS.NoTrt_F |
| CDIP1 | 0.00452 | -0.4094 | 0.223 | 0.33 | 1 | HC_vs_MS.NoTrt_F |
| APEH | 0.00458 | -0.3894 | 0.211 | 0.32 | 1 | HC_vs_MS.NoTrt_F |
| RBM25 | 0.0046 | -0.3198 | 0.587 | 0.67 | 1 | HC_vs_MS.NoTrt_F |
| PARL | 0.00461 | -0.3858 | 0.162 | 0.27 | 1 | HC_vs_MS.NoTrt_F |
| RAB9A | 0.00464 | -0.6091 | 0.116 | 0.2 | 1 | HC_vs_MS.NoTrt_F |
| BORCS8 | 0.00469 | 0.32891 | 0.962 | 0.95 | 1 | HC_vs_MS.NoTrt_F |
| TAS2R4 | 0.00469 | -0.7233 | 0.355 | 0.47 | 1 | HC_vs_MS.NoTrt_F |
| DDX39A | 0.00469 | -0.3272 | 0.355 | 0.48 | 1 | HC_vs_MS.NoTrt_F |
| BBS4 | 0.0047 | -0.3064 | 0.107 | 0.19 | 1 | HC_vs_MS.NoTrt_F |
| CTCF | 0.00471 | 0.94282 | 0.133 | 0.23 | 1 | HC_vs_MS.NoTrt_F |
| CARNMT1 | 0.00471 | -0.3769 | 0.145 | 0.24 | 1 | HC_vs_MS.NoTrt_F |
| MZF1 | 0.00474 | 0.35321 | 0.133 | 0.23 | 1 | HC_vs_MS.NoTrt_F |
| LRRC37B | 0.00474 | 0.2612 | 0.61 | 0.8 | 1 | HC_vs_MS.NoTrt_F |
| DDX42 | 0.00476 | 0.28405 | 0.373 | 0.49 | 1 | HC_vs_MS.NoTrt_F |
| NUP155 | 0.0048 | 0.64625 | 0.942 | 0.91 | 1 | HC_vs_MS.NoTrt_F |
| UBA5 | 0.00481 | -0.8927 | 0.136 | 0.23 | 1 | HC_vs_MS.NoTrt_F |
| CHERP | 0.00484 | -0.4843 | 0.191 | 0.29 | 1 | HC_vs_MS.NoTrt_F |
| IGSF8 | 0.00485 | -0.4311 | 0.179 | 0.28 | 1 | HC_vs_MS.NoTrt_F |
| RWDD3 | 0.00485 | -0.45 | 0.124 | 0.22 | 1 | HC_vs_MS.NoTrt_F |
| OGDH | 0.00489 | -0.3533 | 0.595 | 0.69 | 1 | HC_vs_MS.NoTrt_F |
| RAB14 | 0.00492 | -0.3201 | 0.269 | 0.38 | 1 | HC_vs_MS.NoTrt_F |
| TCEAL3 | 0.00492 | -0.7259 | 0.15 | 0.24 | 1 | HC_vs_MS.NoTrt_F |
| COMMD8 | 0.00494 | -0.3657 | 0.118 | 0.21 | 1 | HC_vs_MS.NoTrt_F |
| GPR108 | 0.00494 | -0.5429 | 0.214 | 0.32 | 1 | HC_vs_MS.NoTrt_F |
| GMPPA | 0.00495 | -0.5404 | 0.292 | 0.41 | 1 | HC_vs_MS.NoTrt_F |
| RBM27 | 0.00497 | -0.3727 | 0.145 | 0.24 | 1 | HC_vs_MS.NoTrt_F |
| CHD1L | 0.00497 | 0.5427 | 0.113 | 0.2 | 1 | HC_vs_MS.NoTrt_F |
| ZNF595 | 0.00497 | 0.47042 | 0.858 | 0.81 | 1 | HC_vs_MS.NoTrt_F |
| WAS | 0.00498 | 0.85635 | 0.332 | 0.46 | 1 | HC_vs_MS.NoTrt_F |
| RALA | 0.00498 | -0.47 | 0.246 | 0.35 | 1 | HC_vs_MS.NoTrt_F |

| | | | | | | |
|---|---|---|---|---|---|---|
| PRR4 | 0.00499 | 0.45574 | 0.26 | 0.4 | 1 | HC_vs_MS.NoTrt_F |
| RAB28 | 0.005 | 0.25722 | 0.136 | 0.24 | 1 | HC_vs_MS.NoTrt_F |
| TMEM173 | 0.00503 | -0.326 | 0.327 | 0.45 | 1 | HC_vs_MS.NoTrt_F |
| SERPING1 | 0.00504 | 1.80111 | 0.15 | 0.07 | 1 | HC_vs_MS.NoTrt_F |
| HOPX | 0.00504 | 0.4944 | 0.893 | 0.85 | 1 | HC_vs_MS.NoTrt_F |
| RAB21 | 0.00505 | 0.68082 | 0.983 | 0.98 | 1 | HC_vs_MS.NoTrt_F |
| RITA1 | 0.00508 | -0.2952 | 0.064 | 0.14 | 1 | HC_vs_MS.NoTrt_F |
| ECHS1 | 0.00508 | 0.33968 | 0.153 | 0.25 | 1 | HC_vs_MS.NoTrt_F |
| ARHGAP28 | 0.00511 | 0.32948 | 0.887 | 0.9 | 1 | HC_vs_MS.NoTrt_F |
| PET117 | 0.00511 | -0.2904 | 0.095 | 0.18 | 1 | HC_vs_MS.NoTrt_F |
| GNAL | 0.00512 | -0.2617 | 0.526 | 0.63 | 1 | HC_vs_MS.NoTrt_F |
| ZNF544 | 0.00516 | 0.33193 | 0.954 | 0.93 | 1 | HC_vs_MS.NoTrt_F |
| XKR9 | 0.00518 | 0.33298 | 0.699 | 0.62 | 1 | HC_vs_MS.NoTrt_F |
| TFPT | 0.00519 | 0.30001 | 0.101 | 0.19 | 1 | HC_vs_MS.NoTrt_F |
| ERBIN | 0.00519 | -0.5334 | 0.286 | 0.41 | 1 | HC_vs_MS.NoTrt_F |
| DPAGT1 | 0.0052 | -0.984 | 0.084 | 0.16 | 1 | HC_vs_MS.NoTrt_F |
| CSNK2A2 | 0.00523 | -0.9278 | 0.344 | 0.44 | 1 | HC_vs_MS.NoTrt_F |
| GOLGB1 | 0.00525 | -0.4006 | 0.616 | 0.7 | 1 | HC_vs_MS.NoTrt_F |
| SDHAF1 | 0.00527 | -0.5896 | 0.087 | 0.17 | 1 | HC_vs_MS.NoTrt_F |
| GBP3 | 0.00531 | -0.3598 | 0.225 | 0.33 | 1 | HC_vs_MS.NoTrt_F |
| PRPS2 | 0.00533 | -0.5226 | 0.136 | 0.23 | 1 | HC_vs_MS.NoTrt_F |
| ISOC1 | 0.00534 | -0.2522 | 0.124 | 0.22 | 1 | HC_vs_MS.NoTrt_F |
| TMEM117 | 0.00535 | -0.6363 | 0.075 | 0.16 | 1 | HC_vs_MS.NoTrt_F |
| PRPF38A | 0.00536 | -0.7749 | 0.673 | 0.73 | 1 | HC_vs_MS.NoTrt_F |
| ARFIP1 | 0.00536 | 0.53453 | 0.251 | 0.37 | 1 | HC_vs_MS.NoTrt_F |
| ARRB2 | 0.00537 | -0.6509 | 0.829 | 0.88 | 1 | HC_vs_MS.NoTrt_F |
| CDC42SE1 | 0.0054 | -0.568 | 0.491 | 0.57 | 1 | HC_vs_MS.NoTrt_F |
| STRA13 | 0.00541 | 0.28265 | 0.168 | 0.28 | 1 | HC_vs_MS.NoTrt_F |
| ABHD10 | 0.00542 | -0.7367 | 0.116 | 0.2 | 1 | HC_vs_MS.NoTrt_F |
| GSN | 0.00546 | 1.49689 | 0.714 | 0.64 | 1 | HC_vs_MS.NoTrt_F |
| NELFB | 0.00547 | -0.7939 | 0.072 | 0.15 | 1 | HC_vs_MS.NoTrt_F |
| RAVER1 | 0.00548 | -0.3839 | 0.173 | 0.28 | 1 | HC_vs_MS.NoTrt_F |
| ERAP2 | 0.00548 | -0.715 | 0.679 | 0.77 | 1 | HC_vs_MS.NoTrt_F |
| IL23A | 0.00549 | 0.32962 | 0.052 | 0.12 | 1 | HC_vs_MS.NoTrt_F |
| RASGEF1B | 0.00553 | 1.13851 | 0.812 | 0.76 | 1 | HC_vs_MS.NoTrt_F |
| BTN2A2 | 0.00555 | 0.37993 | 0.101 | 0.19 | 1 | HC_vs_MS.NoTrt_F |
| GGA3 | 0.00557 | -0.2538 | 0.191 | 0.29 | 1 | HC_vs_MS.NoTrt_F |
| SYAP1 | 0.00557 | 0.48267 | 0.98 | 0.98 | 1 | HC_vs_MS.NoTrt_F |
| TMEM229B | 0.00558 | -0.5909 | 0.231 | 0.35 | 1 | HC_vs_MS.NoTrt_F |
| ZNF671 | 0.00558 | -0.6499 | 0.098 | 0.18 | 1 | HC_vs_MS.NoTrt_F |

| | | | | | | |
|---|---|---|---|---|---|---|
| APOBEC2 | 0.0056 | 0.40013 | 0.064 | 0.14 | 1 | HC_vs_MS.NoTrt_F |
| RIC8B | 0.00562 | -0.3354 | 0.165 | 0.26 | 1 | HC_vs_MS.NoTrt_F |
| SEMA3C | 0.00563 | -0.2834 | 0.656 | 0.81 | 1 | HC_vs_MS.NoTrt_F |
| THAP4 | 0.00565 | 0.3094 | 0.185 | 0.29 | 1 | HC_vs_MS.NoTrt_F |
| OPA1 | 0.00566 | -0.2691 | 0.156 | 0.26 | 1 | HC_vs_MS.NoTrt_F |
| LANCL1 | 0.00567 | -0.5248 | 0.179 | 0.28 | 1 | HC_vs_MS.NoTrt_F |
| FOXK2 | 0.00567 | -0.3934 | 0.318 | 0.42 | 1 | HC_vs_MS.NoTrt_F |
| TLCD2 | 0.0057 | 0.37908 | 0.948 | 0.96 | 1 | HC_vs_MS.NoTrt_F |
| GORASP2 | 0.00571 | -0.3425 | 0.228 | 0.33 | 1 | HC_vs_MS.NoTrt_F |
| IARS | 0.00573 | -0.2907 | 0.705 | 0.79 | 1 | HC_vs_MS.NoTrt_F |
| SKIL | 0.00574 | 0.53526 | 0.341 | 0.47 | 1 | HC_vs_MS.NoTrt_F |
| CDK12 | 0.00575 | -0.3138 | 0.561 | 0.65 | 1 | HC_vs_MS.NoTrt_F |
| ZNF646 | 0.00575 | -0.4132 | 0.092 | 0.18 | 1 | HC_vs_MS.NoTrt_F |
| SLC39A6 | 0.00576 | -0.6038 | 0.095 | 0.18 | 1 | HC_vs_MS.NoTrt_F |
| PRKAR1B | 0.00577 | -0.3899 | 0.127 | 0.22 | 1 | HC_vs_MS.NoTrt_F |
| ATP10B | 0.00578 | 0.39152 | 0.824 | 0.76 | 1 | HC_vs_MS.NoTrt_F |
| FRMD5 | 0.00578 | 0.36981 | 0.049 | 0.12 | 1 | HC_vs_MS.NoTrt_F |
| FIP1L1 | 0.00584 | -0.2511 | 0.295 | 0.41 | 1 | HC_vs_MS.NoTrt_F |
| ATAD3B | 0.00585 | -0.5589 | 0.812 | 0.89 | 1 | HC_vs_MS.NoTrt_F |
| ARHGAP22 | 0.00586 | 0.25777 | 0.734 | 0.65 | 1 | HC_vs_MS.NoTrt_F |
| KIF27 | 0.00588 | 0.35214 | 0.474 | 0.36 | 1 | HC_vs_MS.NoTrt_F |
| KDELR2 | 0.00592 | -0.4783 | 0.208 | 0.32 | 1 | HC_vs_MS.NoTrt_F |
| EXOSC8 | 0.00596 | -0.9148 | 0.512 | 0.56 | 1 | HC_vs_MS.NoTrt_F |
| KIFC1 | 0.00597 | 0.52872 | 0.081 | 0.16 | 1 | HC_vs_MS.NoTrt_F |
| SNX12 | 0.00602 | -1.1809 | 0.116 | 0.2 | 1 | HC_vs_MS.NoTrt_F |
| GPD1L | 0.00602 | -0.725 | 0.136 | 0.23 | 1 | HC_vs_MS.NoTrt_F |
| YBEY | 0.00606 | -0.5457 | 0.072 | 0.15 | 1 | HC_vs_MS.NoTrt_F |
| NUDT5 | 0.00612 | 0.25293 | 0.24 | 0.37 | 1 | HC_vs_MS.NoTrt_F |
| LCMT2 | 0.00613 | -0.2891 | 0.723 | 0.77 | 1 | HC_vs_MS.NoTrt_F |
| RCN1 | 0.00614 | -0.5923 | 0.116 | 0.2 | 1 | HC_vs_MS.NoTrt_F |
| MAF | 0.00619 | 1.0016 | 0.538 | 0.46 | 1 | HC_vs_MS.NoTrt_F |
| MVK | 0.00622 | -0.4755 | 0.058 | 0.13 | 1 | HC_vs_MS.NoTrt_F |
| PPP1R10 | 0.00623 | -0.681 | 0.564 | 0.65 | 1 | HC_vs_MS.NoTrt_F |
| OSGEPL1 | 0.00624 | 0.25981 | 0.257 | 0.39 | 1 | HC_vs_MS.NoTrt_F |
| MAP2K3 | 0.00626 | -0.2927 | 0.194 | 0.3 | 1 | HC_vs_MS.NoTrt_F |
| C8orf33 | 0.00626 | -0.3986 | 0.249 | 0.37 | 1 | HC_vs_MS.NoTrt_F |
| SLU7 | 0.00627 | -0.4421 | 0.442 | 0.55 | 1 | HC_vs_MS.NoTrt_F |
| AC138645.2 | 0.00628 | -0.3926 | 0.124 | 0.22 | 1 | HC_vs_MS.NoTrt_F |
| KIAA0513 | 0.00628 | 0.41516 | 0.743 | 0.64 | 1 | HC_vs_MS.NoTrt_F |
| C5orf51 | 0.00634 | -0.45 | 0.145 | 0.24 | 1 | HC_vs_MS.NoTrt_F |

| | | | | | | |
|---|---|---|---|---|---|---|
| EPN1 | 0.00635 | 0.45148 | 0.957 | 0.96 | 1 | HC_vs_MS.NoTrt_F |
| FAM196B | 0.00635 | 0.29643 | 0.092 | 0.18 | 1 | HC_vs_MS.NoTrt_F |
| UBE2G1 | 0.00638 | -0.3137 | 0.225 | 0.34 | 1 | HC_vs_MS.NoTrt_F |
| UTRN | 0.00639 | -0.3101 | 0.52 | 0.62 | 1 | HC_vs_MS.NoTrt_F |
| SUMO2 | 0.00642 | -0.2554 | 0.893 | 0.9 | 1 | HC_vs_MS.NoTrt_F |
| EIF2AK3 | 0.00642 | 0.30757 | 0.072 | 0.15 | 1 | HC_vs_MS.NoTrt_F |
| AHCYL1 | 0.00643 | 0.69207 | 0.072 | 0.15 | 1 | HC_vs_MS.NoTrt_F |
| GPR25 | 0.00648 | 0.7446 | 0.046 | 0.11 | 1 | HC_vs_MS.NoTrt_F |
| 2-Sep | 0.00648 | -0.4153 | 0.925 | 0.93 | 1 | HC_vs_MS.NoTrt_F |
| FKBP14 | 0.00655 | 0.29512 | 0.728 | 0.65 | 1 | HC_vs_MS.NoTrt_F |
| PAK1IP1 | 0.00659 | 0.28223 | 0.061 | 0.13 | 1 | HC_vs_MS.NoTrt_F |
| UCK1 | 0.00659 | 0.97905 | 0.061 | 0.13 | 1 | HC_vs_MS.NoTrt_F |
| ATG4C | 0.0066 | 0.49556 | 0.061 | 0.13 | 1 | HC_vs_MS.NoTrt_F |
| LXN | 0.0066 | -0.3545 | 0.202 | 0.3 | 1 | HC_vs_MS.NoTrt_F |
| STOML1 | 0.00661 | 0.35395 | 0.847 | 0.77 | 1 | HC_vs_MS.NoTrt_F |
| PNPLA6 | 0.00662 | 0.28064 | 0.185 | 0.29 | 1 | HC_vs_MS.NoTrt_F |
| AP2M1 | 0.00662 | -0.261 | 0.361 | 0.48 | 1 | HC_vs_MS.NoTrt_F |
| TSPAN18 | 0.00664 | 0.25097 | 0.084 | 0.17 | 1 | HC_vs_MS.NoTrt_F |
| FMC1 | 0.00665 | -0.2842 | 0.662 | 0.78 | 1 | HC_vs_MS.NoTrt_F |
| IPO4 | 0.00666 | -0.4649 | 0.043 | 0.11 | 1 | HC_vs_MS.NoTrt_F |
| ZNF7 | 0.00667 | 0.32795 | 0.934 | 0.96 | 1 | HC_vs_MS.NoTrt_F |
| FAM114A1 | 0.00673 | 0.58688 | 0.618 | 0.5 | 1 | HC_vs_MS.NoTrt_F |
| DYNC1LI1 | 0.00674 | -0.4085 | 0.46 | 0.57 | 1 | HC_vs_MS.NoTrt_F |
| TFB2M | 0.00676 | 0.42218 | 0.069 | 0.14 | 1 | HC_vs_MS.NoTrt_F |
| SLC5A10 | 0.00678 | -0.5213 | 0.058 | 0.13 | 1 | HC_vs_MS.NoTrt_F |
| SPATS2L | 0.0068 | 0.35023 | 0.968 | 0.94 | 1 | HC_vs_MS.NoTrt_F |
| MIS18A | 0.00681 | 0.96769 | 0.075 | 0.16 | 1 | HC_vs_MS.NoTrt_F |
| SLC37A1 | 0.00682 | -0.3871 | 0.087 | 0.17 | 1 | HC_vs_MS.NoTrt_F |
| CXorf65 | 0.00685 | 0.63414 | 0.092 | 0.18 | 1 | HC_vs_MS.NoTrt_F |
| ARMT1 | 0.00687 | -0.2681 | 0.09 | 0.17 | 1 | HC_vs_MS.NoTrt_F |
| SAMM50 | 0.00689 | -0.3304 | 0.465 | 0.56 | 1 | HC_vs_MS.NoTrt_F |
| WDR44 | 0.0069 | -0.6308 | 0.133 | 0.22 | 1 | HC_vs_MS.NoTrt_F |
| ANKRD39 | 0.0069 | 0.37292 | 0.214 | 0.33 | 1 | HC_vs_MS.NoTrt_F |
| AC013461.1 | 0.00692 | 0.48449 | 0.844 | 0.77 | 1 | HC_vs_MS.NoTrt_F |
| ZNF211 | 0.00694 | 0.30935 | 0.202 | 0.3 | 1 | HC_vs_MS.NoTrt_F |
| WDR31 | 0.00695 | 0.28088 | 0.882 | 0.84 | 1 | HC_vs_MS.NoTrt_F |
| SPC25 | 0.00696 | 0.43323 | 0.763 | 0.71 | 1 | HC_vs_MS.NoTrt_F |
| RAB4A | 0.00697 | -0.5483 | 0.118 | 0.2 | 1 | HC_vs_MS.NoTrt_F |
| DHRS13 | 0.00698 | 0.5666 | 0.061 | 0.13 | 1 | HC_vs_MS.NoTrt_F |
| RCC2 | 0.00699 | -0.65 | 0.13 | 0.22 | 1 | HC_vs_MS.NoTrt_F |

| | | | | | | |
|---|---|---|---|---|---|---|
| GPATCH1 | 0.00701 | -0.3206 | 0.413 | 0.53 | 1 | HC_vs_MS.NoTrt_F |
| UPRT | 0.00706 | 0.3501 | 0.069 | 0.14 | 1 | HC_vs_MS.NoTrt_F |
| MRPL55 | 0.00706 | 0.66316 | 0.121 | 0.21 | 1 | HC_vs_MS.NoTrt_F |
| NUPR1 | 0.0071 | 0.37894 | 0.977 | 0.97 | 1 | HC_vs_MS.NoTrt_F |
| RPGRIP1L | 0.00715 | 0.27903 | 0.827 | 0.74 | 1 | HC_vs_MS.NoTrt_F |
| SNX16 | 0.00718 | -0.7033 | 0.043 | 0.11 | 1 | HC_vs_MS.NoTrt_F |
| PDCD6IP | 0.00718 | -0.258 | 0.338 | 0.46 | 1 | HC_vs_MS.NoTrt_F |
| ATP8B2 | 0.0072 | -0.3996 | 0.838 | 0.86 | 1 | HC_vs_MS.NoTrt_F |
| AQP11 | 0.00725 | 0.29135 | 0.046 | 0.11 | 1 | HC_vs_MS.NoTrt_F |
| TMEM101 | 0.00728 | 0.26 | 0.095 | 0.18 | 1 | HC_vs_MS.NoTrt_F |
| KIAA0368 | 0.00729 | -0.4256 | 0.272 | 0.38 | 1 | HC_vs_MS.NoTrt_F |
| CNNM4 | 0.00733 | 1.52591 | 0.046 | 0.11 | 1 | HC_vs_MS.NoTrt_F |
| B3GAT2 | 0.00734 | -0.4678 | 0.202 | 0.31 | 1 | HC_vs_MS.NoTrt_F |
| MBD1 | 0.00739 | -0.7075 | 0.217 | 0.32 | 1 | HC_vs_MS.NoTrt_F |
| PTAR1 | 0.00742 | -0.8192 | 0.22 | 0.32 | 1 | HC_vs_MS.NoTrt_F |
| BROX | 0.00742 | -0.297 | 0.624 | 0.71 | 1 | HC_vs_MS.NoTrt_F |
| ZFAND6 | 0.00742 | -0.4103 | 0.26 | 0.37 | 1 | HC_vs_MS.NoTrt_F |
| FAM175B | 0.00743 | -0.4864 | 0.078 | 0.16 | 1 | HC_vs_MS.NoTrt_F |
| NOP14 | 0.00744 | -1.1447 | 0.442 | 0.55 | 1 | HC_vs_MS.NoTrt_F |
| MUL1 | 0.00746 | -0.5159 | 0.09 | 0.17 | 1 | HC_vs_MS.NoTrt_F |
| CNEP1R1 | 0.0075 | -0.8249 | 0.555 | 0.66 | 1 | HC_vs_MS.NoTrt_F |
| MRPL35 | 0.00751 | -1.2272 | 0.156 | 0.24 | 1 | HC_vs_MS.NoTrt_F |
| PPP1R12C | 0.00753 | -0.3329 | 0.081 | 0.16 | 1 | HC_vs_MS.NoTrt_F |
| NT5C | 0.00756 | 0.26334 | 0.205 | 0.32 | 1 | HC_vs_MS.NoTrt_F |
| HIPK2 | 0.00759 | -0.6141 | 0.13 | 0.21 | 1 | HC_vs_MS.NoTrt_F |
| NOC3L | 0.00762 | -0.3195 | 0.078 | 0.16 | 1 | HC_vs_MS.NoTrt_F |
| UTY | 0.00768 | -0.6945 | 0.497 | 0.57 | 1 | HC_vs_MS.NoTrt_F |
| TCEB3 | 0.00771 | -0.505 | 0.145 | 0.24 | 1 | HC_vs_MS.NoTrt_F |
| GATSL3 | 0.00773 | 0.42496 | 0.043 | 0.11 | 1 | HC_vs_MS.NoTrt_F |
| TSSC1 | 0.00776 | -0.3281 | 0.136 | 0.23 | 1 | HC_vs_MS.NoTrt_F |
| SNRNP48 | 0.00776 | -0.442 | 0.208 | 0.3 | 1 | HC_vs_MS.NoTrt_F |
| CLCN5 | 0.00779 | -0.4119 | 0.263 | 0.37 | 1 | HC_vs_MS.NoTrt_F |
| TMEM254 | 0.00781 | -0.6821 | 0.121 | 0.21 | 1 | HC_vs_MS.NoTrt_F |
| PIAS1 | 0.00781 | -0.3174 | 0.379 | 0.48 | 1 | HC_vs_MS.NoTrt_F |
| EBLN2 | 0.00782 | -1.2801 | 0.11 | 0.19 | 1 | HC_vs_MS.NoTrt_F |
| STX1B | 0.00782 | 0.29509 | 0.298 | 0.4 | 1 | HC_vs_MS.NoTrt_F |
| XPO7 | 0.00783 | -0.5281 | 0.142 | 0.23 | 1 | HC_vs_MS.NoTrt_F |
| TTC31 | 0.00787 | -0.3339 | 0.246 | 0.35 | 1 | HC_vs_MS.NoTrt_F |
| VIPAS39 | 0.00787 | -0.9369 | 0.107 | 0.19 | 1 | HC_vs_MS.NoTrt_F |
| PRR3 | 0.00788 | 0.55456 | 0.855 | 0.8 | 1 | HC_vs_MS.NoTrt_F |

| | | | | | | |
|---|---|---|---|---|---|---|
| IFITM3 | 0.00788 | 1.08263 | 0.688 | 0.66 | 1 | HC_vs_MS.NoTrt_F |
| POC5 | 0.00789 | -0.8753 | 0.113 | 0.2 | 1 | HC_vs_MS.NoTrt_F |
| TRMT61A | 0.0079 | 0.58382 | 0.058 | 0.13 | 1 | HC_vs_MS.NoTrt_F |
| GCC1 | 0.00792 | -1.4967 | 0.361 | 0.45 | 1 | HC_vs_MS.NoTrt_F |
| TOMM34 | 0.00793 | 0.4268 | 0.078 | 0.16 | 1 | HC_vs_MS.NoTrt_F |
| ZNF654 | 0.00794 | -1.0323 | 0.145 | 0.23 | 1 | HC_vs_MS.NoTrt_F |
| MANBA | 0.00801 | -0.2907 | 0.22 | 0.32 | 1 | HC_vs_MS.NoTrt_F |
| EFHC1 | 0.00801 | 0.51658 | 0.922 | 0.87 | 1 | HC_vs_MS.NoTrt_F |
| RP11-49K24 | 0.00805 | -0.3947 | 0.165 | 0.27 | 1 | HC_vs_MS.NoTrt_F |
| CAMK2D | 0.00806 | 0.63233 | 0.225 | 0.35 | 1 | HC_vs_MS.NoTrt_F |
| PUM3 | 0.00806 | -0.6787 | 0.191 | 0.29 | 1 | HC_vs_MS.NoTrt_F |
| EFCAB11 | 0.0081 | 0.39744 | 0.873 | 0.83 | 1 | HC_vs_MS.NoTrt_F |
| UBXN6 | 0.00811 | 0.38085 | 0.095 | 0.18 | 1 | HC_vs_MS.NoTrt_F |
| C15orf61 | 0.00812 | 0.38506 | 0.746 | 0.63 | 1 | HC_vs_MS.NoTrt_F |
| UBFD1 | 0.00814 | 0.94441 | 0.425 | 0.54 | 1 | HC_vs_MS.NoTrt_F |
| PSMC3 | 0.00815 | -0.5308 | 0.185 | 0.28 | 1 | HC_vs_MS.NoTrt_F |
| BCAS2 | 0.00816 | 0.31433 | 0.329 | 0.47 | 1 | HC_vs_MS.NoTrt_F |
| GPA33 | 0.00822 | 0.83829 | 0.101 | 0.18 | 1 | HC_vs_MS.NoTrt_F |
| ZNF554 | 0.00822 | 0.36123 | 0.968 | 0.96 | 1 | HC_vs_MS.NoTrt_F |
| CASP8 | 0.00828 | -0.3403 | 0.931 | 0.97 | 1 | HC_vs_MS.NoTrt_F |
| COQ10B | 0.00828 | -0.5987 | 0.688 | 0.79 | 1 | HC_vs_MS.NoTrt_F |
| PEX11B | 0.0083 | 0.85465 | 0.159 | 0.27 | 1 | HC_vs_MS.NoTrt_F |
| NF2 | 0.00832 | -0.51 | 0.821 | 0.85 | 1 | HC_vs_MS.NoTrt_F |
| RHOQ | 0.00833 | 0.33981 | 0.09 | 0.17 | 1 | HC_vs_MS.NoTrt_F |
| BRAP | 0.00834 | -0.6027 | 0.37 | 0.47 | 1 | HC_vs_MS.NoTrt_F |
| SPCS2 | 0.00834 | -0.3716 | 0.324 | 0.44 | 1 | HC_vs_MS.NoTrt_F |
| DYNC1I2 | 0.00835 | -0.6433 | 0.231 | 0.33 | 1 | HC_vs_MS.NoTrt_F |
| ACSL4 | 0.00838 | -0.6943 | 0.217 | 0.33 | 1 | HC_vs_MS.NoTrt_F |
| SDAD1 | 0.00842 | -0.3742 | 0.199 | 0.3 | 1 | HC_vs_MS.NoTrt_F |
| RANGRF | 0.00844 | -0.3354 | 0.116 | 0.2 | 1 | HC_vs_MS.NoTrt_F |
| CSE1L | 0.00844 | -0.4749 | 0.225 | 0.33 | 1 | HC_vs_MS.NoTrt_F |
| ZNF721 | 0.00846 | -0.3324 | 0.376 | 0.46 | 1 | HC_vs_MS.NoTrt_F |
| FAM134C | 0.00848 | -0.6923 | 0.694 | 0.74 | 1 | HC_vs_MS.NoTrt_F |
| RNF187 | 0.00855 | -0.2849 | 0.095 | 0.18 | 1 | HC_vs_MS.NoTrt_F |
| LIPH | 0.0086 | 0.36391 | 0.697 | 0.59 | 1 | HC_vs_MS.NoTrt_F |
| MRPL27 | 0.00863 | 0.29329 | 0.176 | 0.27 | 1 | HC_vs_MS.NoTrt_F |
| GALNT7 | 0.00863 | -0.4586 | 0.107 | 0.19 | 1 | HC_vs_MS.NoTrt_F |
| PQLC3 | 0.00864 | -0.6073 | 0.191 | 0.28 | 1 | HC_vs_MS.NoTrt_F |
| LYRM5 | 0.00867 | -0.7047 | 0.092 | 0.17 | 1 | HC_vs_MS.NoTrt_F |
| KIF20B | 0.00867 | -0.2564 | 0.188 | 0.29 | 1 | HC_vs_MS.NoTrt_F |

| | | | | | | |
|---|---|---|---|---|---|---|
| KIAA1462 | 0.00871 | 0.47134 | 0.39 | 0.28 | 1 | HC_vs_MS.NoTrt_F |
| CD3EAP | 0.00871 | 0.52013 | 0.89 | 0.77 | 1 | HC_vs_MS.NoTrt_F |
| WTIP | 0.00874 | 0.3012 | 0.986 | 1 | 1 | HC_vs_MS.NoTrt_F |
| ELMOD2 | 0.00875 | 0.29563 | 0.095 | 0.18 | 1 | HC_vs_MS.NoTrt_F |
| NANOG | 0.00878 | 0.31869 | 0.936 | 0.88 | 1 | HC_vs_MS.NoTrt_F |
| SLC25A25 | 0.00879 | -0.3616 | 0.121 | 0.21 | 1 | HC_vs_MS.NoTrt_F |
| GCLM | 0.00879 | 0.26183 | 0.746 | 0.7 | 1 | HC_vs_MS.NoTrt_F |
| HNRNPUL2 | 0.00879 | 0.60188 | 0.124 | 0.21 | 1 | HC_vs_MS.NoTrt_F |
| RPS6KA4 | 0.0088 | 1.74679 | 0.043 | 0.11 | 1 | HC_vs_MS.NoTrt_F |
| GLYCTK | 0.00884 | -0.4931 | 0.136 | 0.23 | 1 | HC_vs_MS.NoTrt_F |
| SLC25A1 | 0.00885 | 0.50378 | 0.11 | 0.19 | 1 | HC_vs_MS.NoTrt_F |
| FAM65B | 0.00887 | -0.3783 | 0.673 | 0.74 | 1 | HC_vs_MS.NoTrt_F |
| FAM105A | 0.00899 | -0.6742 | 0.28 | 0.38 | 1 | HC_vs_MS.NoTrt_F |
| MGMT | 0.00899 | 0.41645 | 0.165 | 0.27 | 1 | HC_vs_MS.NoTrt_F |
| GGA2 | 0.00901 | -0.4478 | 0.72 | 0.8 | 1 | HC_vs_MS.NoTrt_F |
| SLC25A38 | 0.00904 | -0.4913 | 0.118 | 0.2 | 1 | HC_vs_MS.NoTrt_F |
| GPX7 | 0.00904 | 0.40716 | 0.081 | 0.16 | 1 | HC_vs_MS.NoTrt_F |
| CEPT1 | 0.00905 | -0.3958 | 0.361 | 0.45 | 1 | HC_vs_MS.NoTrt_F |
| SURF6 | 0.00906 | -0.4312 | 0.598 | 0.67 | 1 | HC_vs_MS.NoTrt_F |
| CAMKMT | 0.00911 | 0.4078 | 0.286 | 0.4 | 1 | HC_vs_MS.NoTrt_F |
| TSC22D4 | 0.00919 | 0.36031 | 0.303 | 0.45 | 1 | HC_vs_MS.NoTrt_F |
| PEX7 | 0.00922 | 0.29857 | 0.171 | 0.27 | 1 | HC_vs_MS.NoTrt_F |
| ZNF30 | 0.00923 | 1.34315 | 0.078 | 0.15 | 1 | HC_vs_MS.NoTrt_F |
| ZDHHC3 | 0.00931 | -0.4094 | 0.532 | 0.63 | 1 | HC_vs_MS.NoTrt_F |
| ARHGEF15 | 0.00932 | 1.22181 | 0.454 | 0.34 | 1 | HC_vs_MS.NoTrt_F |
| TRADD | 0.00935 | -0.2796 | 0.225 | 0.32 | 1 | HC_vs_MS.NoTrt_F |
| KDM7A | 0.00935 | -0.5984 | 0.254 | 0.35 | 1 | HC_vs_MS.NoTrt_F |
| ING4 | 0.00937 | -0.3787 | 0.303 | 0.4 | 1 | HC_vs_MS.NoTrt_F |
| MGAT2 | 0.00939 | -0.2945 | 0.136 | 0.23 | 1 | HC_vs_MS.NoTrt_F |
| TAOK1 | 0.00939 | 0.59961 | 0.945 | 0.95 | 1 | HC_vs_MS.NoTrt_F |
| SIRT7 | 0.00943 | -0.367 | 0.419 | 0.51 | 1 | HC_vs_MS.NoTrt_F |
| BMI1 | 0.00944 | 0.56535 | 0.072 | 0.14 | 1 | HC_vs_MS.NoTrt_F |
| WDR81 | 0.00944 | 0.77053 | 0.084 | 0.16 | 1 | HC_vs_MS.NoTrt_F |
| IDH3B | 0.00947 | 0.27659 | 0.237 | 0.35 | 1 | HC_vs_MS.NoTrt_F |
| FAM114A2 | 0.00949 | 0.25528 | 0.098 | 0.18 | 1 | HC_vs_MS.NoTrt_F |
| TIMM23 | 0.00951 | 0.27679 | 0.269 | 0.4 | 1 | HC_vs_MS.NoTrt_F |
| ZNF850 | 0.00952 | 0.28776 | 0.957 | 0.91 | 1 | HC_vs_MS.NoTrt_F |
| PTK2 | 0.00953 | 0.28005 | 0.951 | 0.96 | 1 | HC_vs_MS.NoTrt_F |
| ARF3 | 0.00953 | 0.26904 | 0.228 | 0.34 | 1 | HC_vs_MS.NoTrt_F |
| BICD1 | 0.00953 | -0.3893 | 0.118 | 0.2 | 1 | HC_vs_MS.NoTrt_F |

| | | | | | | | |
|---|---|---|---|---|---|---|---|
| MAST2 | 0.00955 | 1.18038 | 0.055 | 0.12 | | 1 | HC_vs_MS.NoTrt_F |
| CYLD | 0.00955 | -0.2806 | 0.714 | 0.8 | | 1 | HC_vs_MS.NoTrt_F |
| SWI5 | 0.00958 | -0.2883 | 0.084 | 0.16 | | 1 | HC_vs_MS.NoTrt_F |
| TMEM208 | 0.0096 | 0.44697 | 0.179 | 0.29 | | 1 | HC_vs_MS.NoTrt_F |
| MRPS36 | 0.00961 | 0.43996 | 0.171 | 0.28 | | 1 | HC_vs_MS.NoTrt_F |
| PDZD11 | 0.00962 | -0.3408 | 0.121 | 0.2 | | 1 | HC_vs_MS.NoTrt_F |
| ZFP36 | 0.00965 | 0.82832 | 0.673 | 0.68 | | 1 | HC_vs_MS.NoTrt_F |
| TEFM | 0.00965 | 0.42549 | 0.763 | 0.67 | | 1 | HC_vs_MS.NoTrt_F |
| ACP2 | 0.0097 | 0.28661 | 0.084 | 0.16 | | 1 | HC_vs_MS.NoTrt_F |
| RHNO1 | 0.0097 | 0.2859 | 0.74 | 0.64 | | 1 | HC_vs_MS.NoTrt_F |
| CNTROB | 0.00972 | -0.5562 | 0.197 | 0.29 | | 1 | HC_vs_MS.NoTrt_F |
| AP006285.1 | 0.00977 | 0.44206 | 0.861 | 0.81 | | 1 | HC_vs_MS.NoTrt_F |
| CCDC85B | 0.00982 | 0.58096 | 0.153 | 0.25 | | 1 | HC_vs_MS.NoTrt_F |
| KCNAB1 | 0.00985 | 0.44845 | 0.064 | 0.13 | | 1 | HC_vs_MS.NoTrt_F |
| GBP1 | 0.00988 | -0.2536 | 0.289 | 0.4 | | 1 | HC_vs_MS.NoTrt_F |
| TEX30 | 0.00998 | 0.51521 | 0.075 | 0.15 | | 1 | HC_vs_MS.NoTrt_F |
| RPRD1A | 0.01001 | -0.3628 | 0.179 | 0.27 | | 1 | HC_vs_MS.NoTrt_F |
| MAP2K1 | 0.01003 | -0.5112 | 0.153 | 0.24 | | 1 | HC_vs_MS.NoTrt_F |
| ZFYVE16 | 0.01006 | 0.34791 | 0.942 | 0.9 | | 1 | HC_vs_MS.NoTrt_F |
| CTD-2369P2 | 0.01007 | 0.66496 | 0.084 | 0.16 | | 1 | HC_vs_MS.NoTrt_F |
| C10orf67 | 0.01009 | 0.26215 | 0.269 | 0.38 | | 1 | HC_vs_MS.NoTrt_F |
| PFDN4 | 0.0101 | 0.3146 | 0.098 | 0.18 | | 1 | HC_vs_MS.NoTrt_F |
| CEP72 | 0.0101 | 0.91302 | 0.064 | 0.13 | | 1 | HC_vs_MS.NoTrt_F |
| MTERF2 | 0.0101 | -0.6187 | 0.095 | 0.17 | | 1 | HC_vs_MS.NoTrt_F |
| SEC14L2 | 0.01011 | -0.3324 | 0.165 | 0.25 | | 1 | HC_vs_MS.NoTrt_F |
| RGL2 | 0.01014 | -0.7217 | 0.179 | 0.27 | | 1 | HC_vs_MS.NoTrt_F |
| GLB1 | 0.01015 | -0.5287 | 0.272 | 0.37 | | 1 | HC_vs_MS.NoTrt_F |
| ANAPC13 | 0.01018 | 0.27306 | 0.124 | 0.21 | | 1 | HC_vs_MS.NoTrt_F |
| KNG1 | 0.01018 | 0.55814 | 0.263 | 0.37 | | 1 | HC_vs_MS.NoTrt_F |
| PRKAB2 | 0.01029 | 0.83285 | 0.084 | 0.16 | | 1 | HC_vs_MS.NoTrt_F |
| EEFSEC | 0.01029 | -0.711 | 0.13 | 0.22 | | 1 | HC_vs_MS.NoTrt_F |
| FOSB | 0.0103 | 1.87414 | 0.474 | 0.4 | | 1 | HC_vs_MS.NoTrt_F |
| SLC7A6OS | 0.01033 | 0.37574 | 0.197 | 0.3 | | 1 | HC_vs_MS.NoTrt_F |
| SREBF1 | 0.0104 | -1.2108 | 0.061 | 0.13 | | 1 | HC_vs_MS.NoTrt_F |
| ZBTB3 | 0.01041 | 0.68462 | 0.85 | 0.84 | | 1 | HC_vs_MS.NoTrt_F |
| ZNF845 | 0.01042 | 1.13705 | 0.711 | 0.59 | | 1 | HC_vs_MS.NoTrt_F |
| CDC7 | 0.01044 | 0.3878 | 0.052 | 0.12 | | 1 | HC_vs_MS.NoTrt_F |
| USP21 | 0.01047 | -0.4825 | 0.139 | 0.23 | | 1 | HC_vs_MS.NoTrt_F |
| QTRT2 | 0.01048 | 0.46281 | 0.821 | 0.75 | | 1 | HC_vs_MS.NoTrt_F |
| CCDC125 | 0.0105 | 0.3236 | 0.96 | 0.96 | | 1 | HC_vs_MS.NoTrt_F |

| | | | | | | |
|---|---|---|---|---|---|---|
| TTC4 | 0.01053 | -0.7978 | 0.118 | 0.2 | 1 | HC_vs_MS.NoTrt_F |
| ZBTB44 | 0.01055 | -0.2789 | 0.303 | 0.4 | 1 | HC_vs_MS.NoTrt_F |
| SCAP | 0.01056 | -0.3839 | 0.653 | 0.77 | 1 | HC_vs_MS.NoTrt_F |
| PPP2R3C | 0.01057 | -0.2788 | 0.205 | 0.3 | 1 | HC_vs_MS.NoTrt_F |
| ZNF726 | 0.01068 | 0.4533 | 0.902 | 0.86 | 1 | HC_vs_MS.NoTrt_F |
| TWF1 | 0.01068 | -0.6562 | 0.11 | 0.19 | 1 | HC_vs_MS.NoTrt_F |
| ABHD14A-A( | 0.01071 | -0.3661 | 0.049 | 0.11 | 1 | HC_vs_MS.NoTrt_F |
| CRAT | 0.01074 | 0.78539 | 0.061 | 0.13 | 1 | HC_vs_MS.NoTrt_F |
| SLC9A7 | 0.01081 | 0.48796 | 0.977 | 0.92 | 1 | HC_vs_MS.NoTrt_F |
| MARK2 | 0.01081 | -0.3151 | 0.188 | 0.29 | 1 | HC_vs_MS.NoTrt_F |
| LPAR5 | 0.01081 | 0.89496 | 0.072 | 0.14 | 1 | HC_vs_MS.NoTrt_F |
| AMPH | 0.01084 | -0.2821 | 0.168 | 0.26 | 1 | HC_vs_MS.NoTrt_F |
| RYBP | 0.01085 | -0.5958 | 0.113 | 0.19 | 1 | HC_vs_MS.NoTrt_F |
| RNF13 | 0.01091 | -0.2666 | 0.199 | 0.3 | 1 | HC_vs_MS.NoTrt_F |
| PHLDA1 | 0.01096 | 1.5286 | 0.624 | 0.56 | 1 | HC_vs_MS.NoTrt_F |
| LMBRD2 | 0.01096 | -0.4243 | 0.13 | 0.22 | 1 | HC_vs_MS.NoTrt_F |
| KIF21B | 0.01096 | 0.7914 | 0.434 | 0.55 | 1 | HC_vs_MS.NoTrt_F |
| FHOD1 | 0.01101 | 1.24147 | 0.194 | 0.3 | 1 | HC_vs_MS.NoTrt_F |
| KLRK1 | 0.01101 | 1.18846 | 0.711 | 0.64 | 1 | HC_vs_MS.NoTrt_F |
| TMEM126A | 0.01102 | 0.37324 | 0.084 | 0.16 | 1 | HC_vs_MS.NoTrt_F |
| LTBP1 | 0.01102 | 0.7071 | 0.107 | 0.04 | 1 | HC_vs_MS.NoTrt_F |
| NCKIPSD | 0.01102 | -0.9948 | 0.046 | 0.11 | 1 | HC_vs_MS.NoTrt_F |
| PBDC1 | 0.01103 | 0.5398 | 0.139 | 0.23 | 1 | HC_vs_MS.NoTrt_F |
| ZNF83 | 0.01103 | 0.32407 | 0.87 | 0.84 | 1 | HC_vs_MS.NoTrt_F |
| SIRT1 | 0.01104 | -0.4584 | 0.153 | 0.24 | 1 | HC_vs_MS.NoTrt_F |
| MRPL49 | 0.01105 | -0.4005 | 0.436 | 0.53 | 1 | HC_vs_MS.NoTrt_F |
| SPATA20 | 0.01105 | 0.42932 | 0.124 | 0.21 | 1 | HC_vs_MS.NoTrt_F |
| UBE2O | 0.01105 | 0.53139 | 0.497 | 0.6 | 1 | HC_vs_MS.NoTrt_F |
| PHACTR4 | 0.01106 | 0.57585 | 0.948 | 0.98 | 1 | HC_vs_MS.NoTrt_F |
| AR | 0.01111 | -0.5765 | 0.234 | 0.33 | 1 | HC_vs_MS.NoTrt_F |
| POLR3GL | 0.01111 | 0.35626 | 0.243 | 0.37 | 1 | HC_vs_MS.NoTrt_F |
| TBC1D3L | 0.01112 | 1.00836 | 0.095 | 0.17 | 1 | HC_vs_MS.NoTrt_F |
| P3H1 | 0.01113 | 0.41567 | 0.96 | 0.98 | 1 | HC_vs_MS.NoTrt_F |
| SH2D3C | 0.01117 | -0.3452 | 0.257 | 0.35 | 1 | HC_vs_MS.NoTrt_F |
| CSGALNACT | 0.01118 | -0.4122 | 0.191 | 0.29 | 1 | HC_vs_MS.NoTrt_F |
| C6orf106 | 0.01119 | -0.6786 | 0.165 | 0.25 | 1 | HC_vs_MS.NoTrt_F |
| CSTF3 | 0.01124 | -1.2198 | 0.558 | 0.63 | 1 | HC_vs_MS.NoTrt_F |
| CRYBG3 | 0.01126 | -0.3105 | 0.09 | 0.17 | 1 | HC_vs_MS.NoTrt_F |
| EGLN1 | 0.01127 | -0.8734 | 0.364 | 0.45 | 1 | HC_vs_MS.NoTrt_F |
| NAAA | 0.0113 | -0.4327 | 0.728 | 0.77 | 1 | HC_vs_MS.NoTrt_F |

| | | | | | | |
|---|---|---|---|---|---|---|
| PTPRA | 0.01132 | 0.31422 | 0.188 | 0.29 | 1 | HC_vs_MS.NoTrt_F |
| CTSK | 0.01138 | 0.84399 | 0.049 | 0.11 | 1 | HC_vs_MS.NoTrt_F |
| C6orf226 | 0.0114 | 0.34589 | 0.061 | 0.13 | 1 | HC_vs_MS.NoTrt_F |
| EHD3 | 0.01143 | 0.52032 | 0.098 | 0.18 | 1 | HC_vs_MS.NoTrt_F |
| GDAP1 | 0.01145 | 0.33885 | 0.092 | 0.17 | 1 | HC_vs_MS.NoTrt_F |
| MC1R | 0.01146 | 0.40188 | 0.072 | 0.14 | 1 | HC_vs_MS.NoTrt_F |
| SLC19A1 | 0.01149 | -0.3403 | 0.087 | 0.16 | 1 | HC_vs_MS.NoTrt_F |
| CLMP | 0.01155 | -0.8206 | 0.419 | 0.53 | 1 | HC_vs_MS.NoTrt_F |
| HADHB | 0.01157 | -0.3958 | 0.555 | 0.62 | 1 | HC_vs_MS.NoTrt_F |
| ZNF202 | 0.01162 | 0.78965 | 0.049 | 0.11 | 1 | HC_vs_MS.NoTrt_F |
| SPOPL | 0.01166 | 0.25516 | 0.084 | 0.16 | 1 | HC_vs_MS.NoTrt_F |
| CTR9 | 0.0117 | -0.3689 | 0.205 | 0.3 | 1 | HC_vs_MS.NoTrt_F |
| ACOX3 | 0.01171 | -0.3794 | 0.121 | 0.2 | 1 | HC_vs_MS.NoTrt_F |
| TMPO | 0.01172 | -0.6064 | 0.572 | 0.65 | 1 | HC_vs_MS.NoTrt_F |
| SBF1 | 0.01174 | 0.60997 | 0.22 | 0.33 | 1 | HC_vs_MS.NoTrt_F |
| CENPH | 0.01193 | -0.8103 | 0.324 | 0.41 | 1 | HC_vs_MS.NoTrt_F |
| HIST1H4I | 0.01199 | -0.4297 | 0.049 | 0.11 | 1 | HC_vs_MS.NoTrt_F |
| FBL | 0.012 | -0.6602 | 0.474 | 0.56 | 1 | HC_vs_MS.NoTrt_F |
| CTD-2207O2 | 0.01205 | -0.5701 | 0.168 | 0.25 | 1 | HC_vs_MS.NoTrt_F |
| CELSR3 | 0.01208 | 0.32408 | 0.113 | 0.19 | 1 | HC_vs_MS.NoTrt_F |
| TMCO6 | 0.01208 | -0.3381 | 0.526 | 0.62 | 1 | HC_vs_MS.NoTrt_F |
| SYNC | 0.0121 | 0.31098 | 0.153 | 0.25 | 1 | HC_vs_MS.NoTrt_F |
| ZSCAN16 | 0.01212 | 0.39857 | 0.058 | 0.12 | 1 | HC_vs_MS.NoTrt_F |
| ZNF174 | 0.01213 | 1.42415 | 0.046 | 0.11 | 1 | HC_vs_MS.NoTrt_F |
| MED20 | 0.0122 | -0.9299 | 0.376 | 0.48 | 1 | HC_vs_MS.NoTrt_F |
| MIER3 | 0.01221 | 0.96044 | 0.061 | 0.13 | 1 | HC_vs_MS.NoTrt_F |
| HMGXB4 | 0.01223 | 0.31809 | 0.156 | 0.25 | 1 | HC_vs_MS.NoTrt_F |
| HEMK1 | 0.01226 | 0.28174 | 0.974 | 0.97 | 1 | HC_vs_MS.NoTrt_F |
| MEOX1 | 0.01227 | -0.4544 | 0.087 | 0.16 | 1 | HC_vs_MS.NoTrt_F |
| KDM6A | 0.01241 | -0.2656 | 0.263 | 0.36 | 1 | HC_vs_MS.NoTrt_F |
| SLC38A5 | 0.01245 | -0.7822 | 0.092 | 0.17 | 1 | HC_vs_MS.NoTrt_F |
| TRAPPC13 | 0.01247 | 0.57985 | 0.156 | 0.24 | 1 | HC_vs_MS.NoTrt_F |
| WDR47 | 0.01248 | -0.7068 | 0.231 | 0.33 | 1 | HC_vs_MS.NoTrt_F |
| SPECC1 | 0.01253 | 0.55324 | 0.954 | 0.93 | 1 | HC_vs_MS.NoTrt_F |
| SLC23A2 | 0.01256 | 0.66816 | 0.179 | 0.28 | 1 | HC_vs_MS.NoTrt_F |
| MED26 | 0.01261 | 0.92319 | 0.124 | 0.22 | 1 | HC_vs_MS.NoTrt_F |
| HIST2H4B | 0.01261 | 0.42419 | 0.176 | 0.27 | 1 | HC_vs_MS.NoTrt_F |
| GNL1 | 0.01264 | -0.492 | 0.118 | 0.2 | 1 | HC_vs_MS.NoTrt_F |
| ZNF672 | 0.01266 | 0.3246 | 0.246 | 0.35 | 1 | HC_vs_MS.NoTrt_F |
| LYPLAL1 | 0.01271 | 0.2793 | 0.092 | 0.17 | 1 | HC_vs_MS.NoTrt_F |

| | | | | | | |
|---|---|---|---|---|---|---|
| ALDH2 | 0.01272 | 0.36678 | 0.986 | 0.97 | 1 | HC_vs_MS.NoTrt_F |
| ESRRA | 0.01273 | -0.2872 | 0.127 | 0.21 | 1 | HC_vs_MS.NoTrt_F |
| PGAM1 | 0.01281 | -0.2787 | 0.355 | 0.45 | 1 | HC_vs_MS.NoTrt_F |
| ALKBH3 | 0.01285 | 0.82147 | 0.113 | 0.19 | 1 | HC_vs_MS.NoTrt_F |
| VTI1B | 0.01287 | 0.29245 | 0.28 | 0.38 | 1 | HC_vs_MS.NoTrt_F |
| AP3D1 | 0.01296 | 0.59949 | 0.168 | 0.27 | 1 | HC_vs_MS.NoTrt_F |
| ZBTB2 | 0.01297 | -0.5271 | 0.101 | 0.18 | 1 | HC_vs_MS.NoTrt_F |
| FBXO38 | 0.01299 | -0.7363 | 0.523 | 0.6 | 1 | HC_vs_MS.NoTrt_F |
| FUOM | 0.01299 | 0.33359 | 0.046 | 0.11 | 1 | HC_vs_MS.NoTrt_F |
| DTNBP1 | 0.01301 | -0.5235 | 0.142 | 0.23 | 1 | HC_vs_MS.NoTrt_F |
| CENPB | 0.01312 | 0.49332 | 0.046 | 0.11 | 1 | HC_vs_MS.NoTrt_F |
| PDAP1 | 0.0133 | -0.2501 | 0.428 | 0.53 | 1 | HC_vs_MS.NoTrt_F |
| SLMAP | 0.01332 | -0.4521 | 0.772 | 0.83 | 1 | HC_vs_MS.NoTrt_F |
| ADGRV1 | 0.01333 | -0.4718 | 0.211 | 0.3 | 1 | HC_vs_MS.NoTrt_F |
| ATOX1 | 0.01333 | 1.01938 | 0.277 | 0.39 | 1 | HC_vs_MS.NoTrt_F |
| LYRM7 | 0.01333 | 0.41787 | 0.96 | 0.9 | 1 | HC_vs_MS.NoTrt_F |
| BIVM | 0.01335 | 0.48512 | 0.332 | 0.44 | 1 | HC_vs_MS.NoTrt_F |
| LMNA | 0.01338 | 0.75543 | 0.801 | 0.76 | 1 | HC_vs_MS.NoTrt_F |
| VAPB | 0.01339 | -0.8404 | 0.217 | 0.3 | 1 | HC_vs_MS.NoTrt_F |
| C3 | 0.01345 | 0.36765 | 0.968 | 0.93 | 1 | HC_vs_MS.NoTrt_F |
| EPHA4 | 0.01353 | -1.3896 | 0.237 | 0.34 | 1 | HC_vs_MS.NoTrt_F |
| FAM227B | 0.01361 | -0.5412 | 0.384 | 0.49 | 1 | HC_vs_MS.NoTrt_F |
| NUDT22 | 0.01361 | -0.4033 | 0.676 | 0.71 | 1 | HC_vs_MS.NoTrt_F |
| INVS | 0.01364 | -0.9113 | 0.173 | 0.26 | 1 | HC_vs_MS.NoTrt_F |
| CASP4 | 0.01367 | -0.3583 | 0.361 | 0.46 | 1 | HC_vs_MS.NoTrt_F |
| BCS1L | 0.01368 | 0.41917 | 0.09 | 0.16 | 1 | HC_vs_MS.NoTrt_F |
| PPCS | 0.0137 | -0.2795 | 0.367 | 0.48 | 1 | HC_vs_MS.NoTrt_F |
| HTT | 0.01372 | -0.2885 | 0.266 | 0.37 | 1 | HC_vs_MS.NoTrt_F |
| TNIP3 | 0.01387 | 0.8967 | 0.512 | 0.64 | 1 | HC_vs_MS.NoTrt_F |
| DLGAP4 | 0.0139 | -0.2879 | 0.257 | 0.36 | 1 | HC_vs_MS.NoTrt_F |
| MIPEP | 0.01392 | 0.34368 | 0.078 | 0.15 | 1 | HC_vs_MS.NoTrt_F |
| SLFN13 | 0.01396 | 0.99262 | 0.731 | 0.67 | 1 | HC_vs_MS.NoTrt_F |
| PDE6D | 0.01396 | -0.4991 | 0.147 | 0.23 | 1 | HC_vs_MS.NoTrt_F |
| DRAXIN | 0.01397 | 0.29286 | 0.896 | 0.85 | 1 | HC_vs_MS.NoTrt_F |
| COL1A2 | 0.01398 | 1.72866 | 0.532 | 0.44 | 1 | HC_vs_MS.NoTrt_F |
| CCDC43 | 0.014 | -0.5473 | 0.113 | 0.19 | 1 | HC_vs_MS.NoTrt_F |
| COL15A1 | 0.01407 | 1.5417 | 0.425 | 0.32 | 1 | HC_vs_MS.NoTrt_F |
| TRMT61B | 0.01408 | 0.79104 | 0.133 | 0.22 | 1 | HC_vs_MS.NoTrt_F |
| AGAP2 | 0.01411 | 0.26688 | 0.225 | 0.33 | 1 | HC_vs_MS.NoTrt_F |
| MEFV | 0.01421 | 0.27724 | 0.841 | 0.79 | 1 | HC_vs_MS.NoTrt_F |

| | | | | | | |
|---|---|---|---|---|---|---|
| STK35 | 0.01422 | 1.39934 | 0.081 | 0.16 | 1 | HC_vs_MS.NoTrt_F |
| ADAMTSL3 | 0.01423 | 0.52164 | 0.98 | 0.98 | 1 | HC_vs_MS.NoTrt_F |
| NBPF11 | 0.01425 | 0.3406 | 0.286 | 0.4 | 1 | HC_vs_MS.NoTrt_F |
| LGR4 | 0.01439 | 0.46972 | 0.075 | 0.14 | 1 | HC_vs_MS.NoTrt_F |
| VPS37A | 0.01445 | 0.27736 | 0.191 | 0.28 | 1 | HC_vs_MS.NoTrt_F |
| PAFAH2 | 0.01449 | -0.4275 | 0.566 | 0.64 | 1 | HC_vs_MS.NoTrt_F |
| VCPIP1 | 0.01453 | -0.7136 | 0.439 | 0.53 | 1 | HC_vs_MS.NoTrt_F |
| GBE1 | 0.01453 | 0.44399 | 0.066 | 0.13 | 1 | HC_vs_MS.NoTrt_F |
| PUDP | 0.01454 | -0.4454 | 0.113 | 0.19 | 1 | HC_vs_MS.NoTrt_F |
| PAFAH1B2 | 0.01455 | -0.2872 | 0.91 | 0.95 | 1 | HC_vs_MS.NoTrt_F |
| PCYOX1L | 0.01462 | 0.46459 | 0.11 | 0.19 | 1 | HC_vs_MS.NoTrt_F |
| ENGASE | 0.01466 | 0.87879 | 0.084 | 0.16 | 1 | HC_vs_MS.NoTrt_F |
| TRAF7 | 0.01469 | 0.77002 | 0.15 | 0.24 | 1 | HC_vs_MS.NoTrt_F |
| NOD1 | 0.01473 | -0.4895 | 0.13 | 0.21 | 1 | HC_vs_MS.NoTrt_F |
| MARS | 0.01477 | -0.5013 | 0.419 | 0.5 | 1 | HC_vs_MS.NoTrt_F |
| GPRC5A | 0.0148 | 0.28322 | 0.723 | 0.66 | 1 | HC_vs_MS.NoTrt_F |
| IKZF4 | 0.01481 | 0.94649 | 0.211 | 0.32 | 1 | HC_vs_MS.NoTrt_F |
| ACAD9 | 0.01484 | -0.4829 | 0.338 | 0.42 | 1 | HC_vs_MS.NoTrt_F |
| ABC13-4748 | 0.01484 | 0.56155 | 0.101 | 0.18 | 1 | HC_vs_MS.NoTrt_F |
| SRRM1 | 0.01487 | -0.3339 | 0.919 | 0.9 | 1 | HC_vs_MS.NoTrt_F |
| POLR3F | 0.0149 | -0.3374 | 0.087 | 0.16 | 1 | HC_vs_MS.NoTrt_F |
| TARS | 0.0149 | -0.3705 | 0.257 | 0.34 | 1 | HC_vs_MS.NoTrt_F |
| GXYLT1 | 0.01498 | -0.4102 | 0.434 | 0.53 | 1 | HC_vs_MS.NoTrt_F |
| LRRFIP2 | 0.01498 | -0.4611 | 0.766 | 0.81 | 1 | HC_vs_MS.NoTrt_F |
| SGSM3 | 0.01499 | 0.28163 | 0.223 | 0.33 | 1 | HC_vs_MS.NoTrt_F |
| FUCA1 | 0.01499 | 0.68302 | 0.078 | 0.15 | 1 | HC_vs_MS.NoTrt_F |
| JADE1 | 0.015 | -0.602 | 0.364 | 0.46 | 1 | HC_vs_MS.NoTrt_F |
| MAMLD1 | 0.01502 | 0.4963 | 0.194 | 0.29 | 1 | HC_vs_MS.NoTrt_F |
| DGCR2 | 0.01503 | -0.2693 | 0.257 | 0.36 | 1 | HC_vs_MS.NoTrt_F |
| CDC27 | 0.01505 | -0.3716 | 0.474 | 0.57 | 1 | HC_vs_MS.NoTrt_F |
| CDC23 | 0.01505 | -0.2741 | 0.116 | 0.19 | 1 | HC_vs_MS.NoTrt_F |
| GK | 0.01508 | -0.3646 | 0.228 | 0.34 | 1 | HC_vs_MS.NoTrt_F |
| DISP2 | 0.01508 | 0.34538 | 0.934 | 0.93 | 1 | HC_vs_MS.NoTrt_F |
| PAPD5 | 0.01508 | -0.7321 | 0.185 | 0.28 | 1 | HC_vs_MS.NoTrt_F |
| DHRS3 | 0.01511 | 1.34215 | 0.338 | 0.25 | 1 | HC_vs_MS.NoTrt_F |
| RETSAT | 0.01514 | -0.4203 | 0.179 | 0.28 | 1 | HC_vs_MS.NoTrt_F |
| ADD3 | 0.01518 | -0.45 | 0.827 | 0.82 | 1 | HC_vs_MS.NoTrt_F |
| ZSCAN32 | 0.01521 | -0.3065 | 0.246 | 0.33 | 1 | HC_vs_MS.NoTrt_F |
| LSAMP | 0.01525 | 0.47644 | 0.566 | 0.5 | 1 | HC_vs_MS.NoTrt_F |
| NPTN | 0.01528 | -0.2612 | 0.338 | 0.43 | 1 | HC_vs_MS.NoTrt_F |

| | | | | | | |
|---|---|---|---|---|---|---|
| GMFB | 0.01529 | -0.362 | 0.124 | 0.2 | 1 | HC_vs_MS.NoTrt_F |
| ZNF32 | 0.01531 | 0.334 | 0.107 | 0.18 | 1 | HC_vs_MS.NoTrt_F |
| ABCC3 | 0.01531 | 0.32388 | 0.861 | 0.83 | 1 | HC_vs_MS.NoTrt_F |
| PCMT1 | 0.01532 | -0.5008 | 0.228 | 0.32 | 1 | HC_vs_MS.NoTrt_F |
| SLC44A1 | 0.01537 | -0.5671 | 0.639 | 0.71 | 1 | HC_vs_MS.NoTrt_F |
| VWA5A | 0.01538 | -0.2645 | 0.09 | 0.16 | 1 | HC_vs_MS.NoTrt_F |
| TPH2 | 0.01541 | 0.38602 | 0.711 | 0.68 | 1 | HC_vs_MS.NoTrt_F |
| C19orf54 | 0.01549 | -0.608 | 0.486 | 0.58 | 1 | HC_vs_MS.NoTrt_F |
| ZNF680 | 0.01549 | 0.46401 | 0.139 | 0.22 | 1 | HC_vs_MS.NoTrt_F |
| EIF3J | 0.01553 | 0.6733 | 0.202 | 0.32 | 1 | HC_vs_MS.NoTrt_F |
| RELA | 0.01554 | -0.3731 | 0.214 | 0.31 | 1 | HC_vs_MS.NoTrt_F |
| GPR160 | 0.01556 | 0.30311 | 0.055 | 0.12 | 1 | HC_vs_MS.NoTrt_F |
| SCO2 | 0.01565 | 0.75722 | 0.078 | 0.15 | 1 | HC_vs_MS.NoTrt_F |
| ZNF275 | 0.01565 | -0.3197 | 0.072 | 0.14 | 1 | HC_vs_MS.NoTrt_F |
| SETMAR | 0.01565 | -0.4405 | 0.332 | 0.45 | 1 | HC_vs_MS.NoTrt_F |
| PEMT | 0.0157 | 0.49659 | 0.084 | 0.16 | 1 | HC_vs_MS.NoTrt_F |
| PSMC4 | 0.01571 | -0.4623 | 0.543 | 0.6 | 1 | HC_vs_MS.NoTrt_F |
| ABCC2 | 0.01571 | 0.31547 | 0.514 | 0.66 | 1 | HC_vs_MS.NoTrt_F |
| BAG4 | 0.01573 | 0.29759 | 0.223 | 0.33 | 1 | HC_vs_MS.NoTrt_F |
| EMC8 | 0.01582 | -0.9105 | 0.208 | 0.29 | 1 | HC_vs_MS.NoTrt_F |
| SNRPC | 0.01586 | 0.38227 | 0.272 | 0.39 | 1 | HC_vs_MS.NoTrt_F |
| SAMD9L | 0.01589 | -0.3013 | 0.246 | 0.34 | 1 | HC_vs_MS.NoTrt_F |
| MKKS | 0.0159 | -0.6266 | 0.104 | 0.18 | 1 | HC_vs_MS.NoTrt_F |
| UPF1 | 0.016 | 0.78999 | 0.13 | 0.21 | 1 | HC_vs_MS.NoTrt_F |
| NAV1 | 0.01601 | 0.56542 | 0.908 | 0.89 | 1 | HC_vs_MS.NoTrt_F |
| SUMO4 | 0.01603 | 1.4947 | 0.049 | 0.11 | 1 | HC_vs_MS.NoTrt_F |
| KIR3DX1 | 0.01608 | 0.31334 | 0.931 | 0.92 | 1 | HC_vs_MS.NoTrt_F |
| KCNE4 | 0.01618 | 0.28472 | 0.919 | 0.87 | 1 | HC_vs_MS.NoTrt_F |
| LGMN | 0.01624 | 0.3168 | 0.971 | 0.96 | 1 | HC_vs_MS.NoTrt_F |
| ROCK2 | 0.01627 | 0.44045 | 0.471 | 0.59 | 1 | HC_vs_MS.NoTrt_F |
| SURF4 | 0.01628 | 0.33403 | 0.306 | 0.44 | 1 | HC_vs_MS.NoTrt_F |
| NPC2 | 0.01629 | 0.3233 | 0.15 | 0.24 | 1 | HC_vs_MS.NoTrt_F |
| ANKHD1-EIF | 0.01631 | 0.96491 | 0.064 | 0.13 | 1 | HC_vs_MS.NoTrt_F |
| RAB3GAP1 | 0.01631 | -0.3789 | 0.402 | 0.49 | 1 | HC_vs_MS.NoTrt_F |
| ZNF821 | 0.01632 | 0.98896 | 0.084 | 0.16 | 1 | HC_vs_MS.NoTrt_F |
| KLF12 | 0.01633 | -0.3142 | 0.376 | 0.5 | 1 | HC_vs_MS.NoTrt_F |
| PPM1L | 0.01636 | -1.1112 | 0.246 | 0.33 | 1 | HC_vs_MS.NoTrt_F |
| ZNF783 | 0.0164 | 0.34666 | 0.442 | 0.52 | 1 | HC_vs_MS.NoTrt_F |
| MS4A7 | 0.01644 | 0.36502 | 0.829 | 0.8 | 1 | HC_vs_MS.NoTrt_F |
| UBLCP1 | 0.01644 | 0.64802 | 0.214 | 0.31 | 1 | HC_vs_MS.NoTrt_F |

| | | | | | | |
|---|---|---|---|---|---|---|
| SEMA3G | 0.01654 | -0.54 | 0.049 | 0.11 | 1 | HC_vs_MS.NoTrt_F |
| CENPT | 0.01658 | -0.564 | 0.269 | 0.37 | 1 | HC_vs_MS.NoTrt_F |
| ATG9A | 0.01659 | 0.34127 | 0.116 | 0.19 | 1 | HC_vs_MS.NoTrt_F |
| FAM91A1 | 0.01659 | -0.6854 | 0.344 | 0.45 | 1 | HC_vs_MS.NoTrt_F |
| EPHX1 | 0.01664 | 0.25407 | 0.107 | 0.18 | 1 | HC_vs_MS.NoTrt_F |
| ADH7 | 0.01671 | -0.369 | 0.188 | 0.28 | 1 | HC_vs_MS.NoTrt_F |
| SCGB2B2 | 0.01672 | 1.33156 | 0.066 | 0.13 | 1 | HC_vs_MS.NoTrt_F |
| RCSD1 | 0.01677 | -0.3264 | 0.936 | 0.96 | 1 | HC_vs_MS.NoTrt_F |
| RPL7L1 | 0.01679 | 0.26575 | 1 | 1 | 1 | HC_vs_MS.NoTrt_F |
| CCDC22 | 0.01679 | -0.3411 | 0.162 | 0.25 | 1 | HC_vs_MS.NoTrt_F |
| PARVG | 0.01683 | 0.25008 | 0.315 | 0.44 | 1 | HC_vs_MS.NoTrt_F |
| BDH2 | 0.01693 | -0.5139 | 0.234 | 0.32 | 1 | HC_vs_MS.NoTrt_F |
| RCE1 | 0.01699 | -0.6472 | 0.136 | 0.21 | 1 | HC_vs_MS.NoTrt_F |
| TIMM8B | 0.01706 | -0.4281 | 0.118 | 0.19 | 1 | HC_vs_MS.NoTrt_F |
| BOLA1 | 0.01708 | 0.6415 | 0.072 | 0.14 | 1 | HC_vs_MS.NoTrt_F |
| STRADB | 0.01708 | -0.4406 | 0.101 | 0.17 | 1 | HC_vs_MS.NoTrt_F |
| EEF2KMT | 0.01709 | 0.39699 | 0.104 | 0.18 | 1 | HC_vs_MS.NoTrt_F |
| ZNF143 | 0.0171 | -0.2732 | 0.15 | 0.23 | 1 | HC_vs_MS.NoTrt_F |
| AGFG2 | 0.01711 | 0.26337 | 0.355 | 0.46 | 1 | HC_vs_MS.NoTrt_F |
| FRG2C | 0.01713 | 0.89533 | 0.194 | 0.29 | 1 | HC_vs_MS.NoTrt_F |
| MMP24 | 0.01714 | 0.40263 | 0.095 | 0.17 | 1 | HC_vs_MS.NoTrt_F |
| TOP3B | 0.01714 | 0.6917 | 0.15 | 0.24 | 1 | HC_vs_MS.NoTrt_F |
| SLC35G1 | 0.01718 | -0.3776 | 0.176 | 0.25 | 1 | HC_vs_MS.NoTrt_F |
| ANP32A | 0.01724 | -0.2559 | 0.312 | 0.4 | 1 | HC_vs_MS.NoTrt_F |
| PSMD4 | 0.01727 | -0.3967 | 0.694 | 0.75 | 1 | HC_vs_MS.NoTrt_F |
| PEX19 | 0.01731 | 0.25472 | 0.156 | 0.24 | 1 | HC_vs_MS.NoTrt_F |
| RP11-949J7. | 0.01731 | -1.3165 | 0.087 | 0.16 | 1 | HC_vs_MS.NoTrt_F |
| ATF6 | 0.01734 | 0.44149 | 0.283 | 0.38 | 1 | HC_vs_MS.NoTrt_F |
| FUT8 | 0.01737 | 0.43061 | 1 | 0.99 | 1 | HC_vs_MS.NoTrt_F |
| HEATR1 | 0.0175 | -0.5564 | 0.202 | 0.3 | 1 | HC_vs_MS.NoTrt_F |
| SMYD3 | 0.0175 | -0.406 | 0.234 | 0.33 | 1 | HC_vs_MS.NoTrt_F |
| LIN7C | 0.01752 | -0.253 | 0.347 | 0.45 | 1 | HC_vs_MS.NoTrt_F |
| NCF2 | 0.01759 | 0.4456 | 0.064 | 0.13 | 1 | HC_vs_MS.NoTrt_F |
| MT-CYB | 0.01775 | 0.58309 | 0.792 | 0.79 | 1 | HC_vs_MS.NoTrt_F |
| FARP2 | 0.01776 | -1.3256 | 0.376 | 0.46 | 1 | HC_vs_MS.NoTrt_F |
| FAR1 | 0.01776 | 0.29609 | 0.104 | 0.18 | 1 | HC_vs_MS.NoTrt_F |
| LRRFIP1 | 0.0178 | 0.3876 | 0.997 | 1 | 1 | HC_vs_MS.NoTrt_F |
| DPM3 | 0.01782 | 0.42765 | 0.066 | 0.13 | 1 | HC_vs_MS.NoTrt_F |
| GALNT8 | 0.01786 | 0.38987 | 0.052 | 0.11 | 1 | HC_vs_MS.NoTrt_F |
| ALG3 | 0.01787 | 0.35786 | 0.173 | 0.27 | 1 | HC_vs_MS.NoTrt_F |

| | | | | | | |
|---|---|---|---|---|---|---|
| PLEKHA4 | 0.01788 | -0.3144 | 0.081 | 0.15 | 1 | HC_vs_MS.NoTrt_F |
| AL078584.1 | 0.01789 | -0.5562 | 0.471 | 0.53 | 1 | HC_vs_MS.NoTrt_F |
| FAM50A | 0.01795 | 0.43136 | 0.142 | 0.22 | 1 | HC_vs_MS.NoTrt_F |
| NUDT16L1 | 0.01797 | 0.27801 | 0.153 | 0.24 | 1 | HC_vs_MS.NoTrt_F |
| RPS6KB1 | 0.01798 | -0.3861 | 0.87 | 0.95 | 1 | HC_vs_MS.NoTrt_F |
| LLNLF-187D8 | 0.01805 | 0.31058 | 0.659 | 0.54 | 1 | HC_vs_MS.NoTrt_F |
| REL | 0.0181 | 0.62002 | 0.983 | 0.97 | 1 | HC_vs_MS.NoTrt_F |
| PLEKHM2 | 0.01817 | -0.6047 | 0.049 | 0.11 | 1 | HC_vs_MS.NoTrt_F |
| SLC25A53 | 0.01817 | 0.31868 | 0.483 | 0.59 | 1 | HC_vs_MS.NoTrt_F |
| KCTD10 | 0.01818 | 1.15361 | 0.483 | 0.37 | 1 | HC_vs_MS.NoTrt_F |
| PGM3 | 0.01823 | 0.7831 | 0.263 | 0.37 | 1 | HC_vs_MS.NoTrt_F |
| AC068533.7 | 0.01825 | 0.38477 | 0.052 | 0.11 | 1 | HC_vs_MS.NoTrt_F |
| PLCG2 | 0.01826 | 0.9376 | 0.113 | 0.19 | 1 | HC_vs_MS.NoTrt_F |
| ZNF223 | 0.01826 | 0.43716 | 0.104 | 0.18 | 1 | HC_vs_MS.NoTrt_F |
| DCAF12 | 0.0183 | -0.5425 | 0.179 | 0.27 | 1 | HC_vs_MS.NoTrt_F |
| NDUFB5 | 0.01833 | -0.6783 | 0.702 | 0.75 | 1 | HC_vs_MS.NoTrt_F |
| HDHD3 | 0.01833 | -0.2558 | 0.194 | 0.28 | 1 | HC_vs_MS.NoTrt_F |
| PKP2 | 0.01836 | 0.42461 | 0.676 | 0.58 | 1 | HC_vs_MS.NoTrt_F |
| VAMP3 | 0.01846 | -0.3594 | 0.353 | 0.44 | 1 | HC_vs_MS.NoTrt_F |
| ZDHHC6 | 0.01853 | -0.2616 | 0.188 | 0.28 | 1 | HC_vs_MS.NoTrt_F |
| PDK2 | 0.01855 | -0.4097 | 0.292 | 0.39 | 1 | HC_vs_MS.NoTrt_F |
| DERL1 | 0.01864 | -0.291 | 0.529 | 0.58 | 1 | HC_vs_MS.NoTrt_F |
| MTMR2 | 0.01878 | -0.3533 | 0.208 | 0.29 | 1 | HC_vs_MS.NoTrt_F |
| FNTB | 0.01878 | -0.5717 | 0.162 | 0.24 | 1 | HC_vs_MS.NoTrt_F |
| AGFG1 | 0.01879 | -0.4818 | 0.095 | 0.17 | 1 | HC_vs_MS.NoTrt_F |
| ISYNA1 | 0.01892 | 0.43067 | 0.11 | 0.18 | 1 | HC_vs_MS.NoTrt_F |
| ALDOC | 0.01892 | 0.42806 | 0.095 | 0.17 | 1 | HC_vs_MS.NoTrt_F |
| CLSTN1 | 0.01894 | 0.25395 | 0.188 | 0.28 | 1 | HC_vs_MS.NoTrt_F |
| UTP14C | 0.01897 | -0.9396 | 0.194 | 0.28 | 1 | HC_vs_MS.NoTrt_F |
| LRG1 | 0.019 | 0.32195 | 0.835 | 0.76 | 1 | HC_vs_MS.NoTrt_F |
| PHLPP2 | 0.019 | 0.30465 | 0.818 | 0.75 | 1 | HC_vs_MS.NoTrt_F |
| BMT2 | 0.019 | 0.76215 | 0.136 | 0.22 | 1 | HC_vs_MS.NoTrt_F |
| ETV7 | 0.01902 | -0.5968 | 0.058 | 0.12 | 1 | HC_vs_MS.NoTrt_F |
| CXCR5 | 0.01906 | 0.29259 | 0.153 | 0.24 | 1 | HC_vs_MS.NoTrt_F |
| NCKAP1 | 0.01924 | 0.77787 | 0.743 | 0.65 | 1 | HC_vs_MS.NoTrt_F |
| ZNF512 | 0.01926 | -0.2943 | 0.142 | 0.22 | 1 | HC_vs_MS.NoTrt_F |
| SEC24D | 0.01931 | -0.3417 | 0.526 | 0.6 | 1 | HC_vs_MS.NoTrt_F |
| RSAD1 | 0.01934 | 0.45258 | 0.113 | 0.19 | 1 | HC_vs_MS.NoTrt_F |
| SMARCA5 | 0.01937 | -0.2826 | 0.384 | 0.49 | 1 | HC_vs_MS.NoTrt_F |
| ABCA3 | 0.01946 | 0.27836 | 0.078 | 0.14 | 1 | HC_vs_MS.NoTrt_F |

| | | | | | | |
|---|---|---|---|---|---|---|
| TMCC1 | 0.01948 | 0.33687 | 0.118 | 0.19 | 1 | HC_vs_MS.NoTrt_F |
| APOD | 0.01949 | 2.30074 | 0.35 | 0.26 | 1 | HC_vs_MS.NoTrt_F |
| SP4 | 0.01951 | -0.6904 | 0.448 | 0.5 | 1 | HC_vs_MS.NoTrt_F |
| CAPN12 | 0.01952 | 0.93599 | 0.613 | 0.54 | 1 | HC_vs_MS.NoTrt_F |
| MFSD13A | 0.01955 | -0.5909 | 0.777 | 0.85 | 1 | HC_vs_MS.NoTrt_F |
| DDX10 | 0.01956 | -0.8294 | 0.694 | 0.75 | 1 | HC_vs_MS.NoTrt_F |
| PIGK | 0.01958 | -0.5249 | 0.136 | 0.21 | 1 | HC_vs_MS.NoTrt_F |
| STEAP2 | 0.01958 | 0.29282 | 0.78 | 0.73 | 1 | HC_vs_MS.NoTrt_F |
| VPS54 | 0.01963 | -0.6261 | 0.419 | 0.53 | 1 | HC_vs_MS.NoTrt_F |
| C21orf2 | 0.01966 | -0.2673 | 0.185 | 0.27 | 1 | HC_vs_MS.NoTrt_F |
| SLC39A11 | 0.0197 | 0.39539 | 0.272 | 0.38 | 1 | HC_vs_MS.NoTrt_F |
| LCMT1 | 0.01972 | 0.29064 | 0.202 | 0.3 | 1 | HC_vs_MS.NoTrt_F |
| ZMYM5 | 0.01976 | -0.3956 | 0.292 | 0.38 | 1 | HC_vs_MS.NoTrt_F |
| CYTH2 | 0.01981 | 0.43795 | 0.676 | 0.77 | 1 | HC_vs_MS.NoTrt_F |
| IMPA2 | 0.01989 | 0.28782 | 0.87 | 0.82 | 1 | HC_vs_MS.NoTrt_F |
| CSNK1G3 | 0.0199 | -0.2693 | 0.133 | 0.22 | 1 | HC_vs_MS.NoTrt_F |
| BLK | 0.01993 | 0.74156 | 0.069 | 0.13 | 1 | HC_vs_MS.NoTrt_F |
| RPE | 0.01994 | -0.3849 | 0.353 | 0.45 | 1 | HC_vs_MS.NoTrt_F |
| NCAPD3 | 0.01995 | 0.38038 | 0.11 | 0.18 | 1 | HC_vs_MS.NoTrt_F |
| AQP7 | 0.01995 | 1.05078 | 0.147 | 0.08 | 1 | HC_vs_MS.NoTrt_F |
| ZNF692 | 0.02007 | 0.31518 | 0.13 | 0.21 | 1 | HC_vs_MS.NoTrt_F |
| PTBP2 | 0.02015 | -0.4525 | 0.682 | 0.72 | 1 | HC_vs_MS.NoTrt_F |
| CTC-432M1! | 0.02027 | 1.27432 | 0.061 | 0.12 | 1 | HC_vs_MS.NoTrt_F |
| FAM199X | 0.02044 | 1.23167 | 0.072 | 0.14 | 1 | HC_vs_MS.NoTrt_F |
| SLC25A37 | 0.02045 | 0.64431 | 0.65 | 0.8 | 1 | HC_vs_MS.NoTrt_F |
| CTC-479C5.1 | 0.02045 | -0.547 | 0.601 | 0.63 | 1 | HC_vs_MS.NoTrt_F |
| PYGO2 | 0.02053 | -0.6085 | 0.173 | 0.25 | 1 | HC_vs_MS.NoTrt_F |
| VPS4A | 0.02055 | -0.3256 | 0.688 | 0.75 | 1 | HC_vs_MS.NoTrt_F |
| ORC3 | 0.02057 | 0.51305 | 0.191 | 0.28 | 1 | HC_vs_MS.NoTrt_F |
| ABCD2 | 0.02059 | 0.32709 | 0.118 | 0.19 | 1 | HC_vs_MS.NoTrt_F |
| WNT10A | 0.0206 | 1.08854 | 0.159 | 0.25 | 1 | HC_vs_MS.NoTrt_F |
| KIAA1147 | 0.02061 | -0.2686 | 0.124 | 0.2 | 1 | HC_vs_MS.NoTrt_F |
| HINT2 | 0.02065 | 0.46784 | 0.107 | 0.18 | 1 | HC_vs_MS.NoTrt_F |
| SLC16A4 | 0.02067 | 0.49692 | 0.749 | 0.69 | 1 | HC_vs_MS.NoTrt_F |
| MRPL24 | 0.02068 | -0.3324 | 0.145 | 0.22 | 1 | HC_vs_MS.NoTrt_F |
| HAUS4 | 0.0207 | -0.2697 | 0.156 | 0.24 | 1 | HC_vs_MS.NoTrt_F |
| LINC00959 | 0.02074 | -0.8013 | 0.147 | 0.23 | 1 | HC_vs_MS.NoTrt_F |
| CARMIL2 | 0.02074 | 1.58046 | 0.095 | 0.17 | 1 | HC_vs_MS.NoTrt_F |
| GJB7 | 0.02074 | 0.66073 | 0.09 | 0.16 | 1 | HC_vs_MS.NoTrt_F |
| LSM1 | 0.02079 | 0.29321 | 0.162 | 0.25 | 1 | HC_vs_MS.NoTrt_F |

| | | | | | | |
|---|---|---|---|---|---|---|
| CIDEB | 0.02082 | 0.61495 | 0.072 | 0.14 | 1 | HC_vs_MS.NoTrt_F |
| ZC2HC1A | 0.0209 | 0.29994 | 0.22 | 0.32 | 1 | HC_vs_MS.NoTrt_F |
| INO80E | 0.02092 | -0.3564 | 0.22 | 0.3 | 1 | HC_vs_MS.NoTrt_F |
| ZNF394 | 0.02096 | -0.5022 | 0.509 | 0.61 | 1 | HC_vs_MS.NoTrt_F |
| CCDC7 | 0.02096 | 0.78983 | 0.191 | 0.29 | 1 | HC_vs_MS.NoTrt_F |
| TTC8 | 0.02101 | 0.32932 | 0.757 | 0.69 | 1 | HC_vs_MS.NoTrt_F |
| MED1 | 0.02102 | -0.6677 | 0.396 | 0.48 | 1 | HC_vs_MS.NoTrt_F |
| PLEKHG4 | 0.02103 | -0.5872 | 0.197 | 0.28 | 1 | HC_vs_MS.NoTrt_F |
| ISPD | 0.02109 | 0.45715 | 0.882 | 0.82 | 1 | HC_vs_MS.NoTrt_F |
| FBXO7 | 0.02117 | -0.3758 | 0.315 | 0.41 | 1 | HC_vs_MS.NoTrt_F |
| RNPEPL1 | 0.02127 | 0.77274 | 0.171 | 0.26 | 1 | HC_vs_MS.NoTrt_F |
| NANS | 0.02127 | 0.69777 | 0.812 | 0.81 | 1 | HC_vs_MS.NoTrt_F |
| MYCBP | 0.02131 | 0.48331 | 0.145 | 0.23 | 1 | HC_vs_MS.NoTrt_F |
| CEMIP | 0.02132 | -0.2802 | 0.069 | 0.13 | 1 | HC_vs_MS.NoTrt_F |
| AHRR | 0.02138 | 0.38533 | 0.136 | 0.22 | 1 | HC_vs_MS.NoTrt_F |
| XBP1 | 0.0215 | -0.2561 | 0.188 | 0.28 | 1 | HC_vs_MS.NoTrt_F |
| SMARCE1 | 0.02152 | -0.4938 | 0.523 | 0.6 | 1 | HC_vs_MS.NoTrt_F |
| EML6 | 0.02162 | 0.59769 | 0.665 | 0.62 | 1 | HC_vs_MS.NoTrt_F |
| TTYH2 | 0.02166 | -0.4124 | 0.116 | 0.19 | 1 | HC_vs_MS.NoTrt_F |
| SEMA6A | 0.0217 | 0.31546 | 0.066 | 0.13 | 1 | HC_vs_MS.NoTrt_F |
| CNOT7 | 0.02174 | -0.32 | 0.546 | 0.62 | 1 | HC_vs_MS.NoTrt_F |
| TEPSIN | 0.02174 | 0.85323 | 0.15 | 0.23 | 1 | HC_vs_MS.NoTrt_F |
| IFI30 | 0.02176 | -0.4829 | 0.364 | 0.48 | 1 | HC_vs_MS.NoTrt_F |
| BTBD19 | 0.02179 | 0.79658 | 0.795 | 0.72 | 1 | HC_vs_MS.NoTrt_F |
| ATP8A1 | 0.0218 | 0.45804 | 0.15 | 0.23 | 1 | HC_vs_MS.NoTrt_F |
| RBMS2 | 0.02183 | 0.38714 | 0.939 | 0.88 | 1 | HC_vs_MS.NoTrt_F |
| BUB1B | 0.02184 | -0.5556 | 0.448 | 0.34 | 1 | HC_vs_MS.NoTrt_F |
| MIEN1 | 0.02185 | 0.33209 | 0.165 | 0.25 | 1 | HC_vs_MS.NoTrt_F |
| ROMO1 | 0.02192 | 0.44447 | 0.234 | 0.34 | 1 | HC_vs_MS.NoTrt_F |
| PLIN2 | 0.02193 | 0.25659 | 0.697 | 0.76 | 1 | HC_vs_MS.NoTrt_F |
| TEN1 | 0.02194 | -0.7796 | 0.329 | 0.41 | 1 | HC_vs_MS.NoTrt_F |
| MYCBP2 | 0.02197 | 0.4651 | 0.465 | 0.64 | 1 | HC_vs_MS.NoTrt_F |
| PTGES3L | 0.022 | 0.32166 | 0.844 | 0.82 | 1 | HC_vs_MS.NoTrt_F |
| CCDC144A | 0.02201 | 0.78938 | 0.908 | 0.91 | 1 | HC_vs_MS.NoTrt_F |
| C10orf76 | 0.02211 | -0.3447 | 0.376 | 0.47 | 1 | HC_vs_MS.NoTrt_F |
| ADNP2 | 0.02212 | 0.27821 | 0.078 | 0.14 | 1 | HC_vs_MS.NoTrt_F |
| PGK1 | 0.02215 | -0.258 | 0.717 | 0.79 | 1 | HC_vs_MS.NoTrt_F |
| EXOC7 | 0.02219 | -0.3006 | 0.465 | 0.53 | 1 | HC_vs_MS.NoTrt_F |
| SMTN | 0.02219 | 0.79054 | 0.442 | 0.32 | 1 | HC_vs_MS.NoTrt_F |
| ZNF362 | 0.02223 | 0.41195 | 0.15 | 0.23 | 1 | HC_vs_MS.NoTrt_F |

| | | | | | | |
|---|---|---|---|---|---|---|
| TAOK3 | 0.02229 | 0.34126 | 0.792 | 0.85 | 1 | HC_vs_MS.NoTrt_F |
| DNAJB9 | 0.02232 | 0.25575 | 0.171 | 0.26 | 1 | HC_vs_MS.NoTrt_F |
| ZBED6CL | 0.02239 | 1.60425 | 0.069 | 0.13 | 1 | HC_vs_MS.NoTrt_F |
| TPP2 | 0.02248 | -0.3114 | 0.517 | 0.64 | 1 | HC_vs_MS.NoTrt_F |
| MRPL14 | 0.02251 | 0.28889 | 0.136 | 0.22 | 1 | HC_vs_MS.NoTrt_F |
| CD300LD | 0.02265 | 0.25859 | 0.355 | 0.46 | 1 | HC_vs_MS.NoTrt_F |
| MRPL45 | 0.0227 | 0.57157 | 0.078 | 0.14 | 1 | HC_vs_MS.NoTrt_F |
| DNAJC13 | 0.02272 | 0.53034 | 0.263 | 0.36 | 1 | HC_vs_MS.NoTrt_F |
| NPIPB5 | 0.02272 | 0.41306 | 0.971 | 0.99 | 1 | HC_vs_MS.NoTrt_F |
| CDC14A | 0.0228 | 0.2562 | 0.303 | 0.42 | 1 | HC_vs_MS.NoTrt_F |
| ZCWPW1 | 0.0228 | -0.7977 | 0.095 | 0.17 | 1 | HC_vs_MS.NoTrt_F |
| EIF2A | 0.02287 | -0.3985 | 0.621 | 0.68 | 1 | HC_vs_MS.NoTrt_F |
| XXbac-BPG2 | 0.02291 | 0.55697 | 0.081 | 0.15 | 1 | HC_vs_MS.NoTrt_F |
| TMEM145 | 0.02294 | 0.26317 | 0.942 | 0.97 | 1 | HC_vs_MS.NoTrt_F |
| POLD2 | 0.02299 | 0.42164 | 0.124 | 0.2 | 1 | HC_vs_MS.NoTrt_F |
| ZBTB32 | 0.02302 | 0.48209 | 0.069 | 0.13 | 1 | HC_vs_MS.NoTrt_F |
| EXTL2 | 0.02302 | 0.46881 | 0.066 | 0.13 | 1 | HC_vs_MS.NoTrt_F |
| SPATS2 | 0.02302 | 0.4452 | 0.977 | 0.95 | 1 | HC_vs_MS.NoTrt_F |
| RUSC1 | 0.02315 | 0.50486 | 0.084 | 0.15 | 1 | HC_vs_MS.NoTrt_F |
| DOCK11 | 0.02316 | 0.3037 | 0.555 | 0.73 | 1 | HC_vs_MS.NoTrt_F |
| ELP5 | 0.02329 | 0.65822 | 0.133 | 0.22 | 1 | HC_vs_MS.NoTrt_F |
| MTHFSD | 0.02342 | -0.9803 | 0.087 | 0.16 | 1 | HC_vs_MS.NoTrt_F |
| API5 | 0.0235 | -0.6263 | 0.327 | 0.4 | 1 | HC_vs_MS.NoTrt_F |
| MCMBP | 0.02352 | 0.55641 | 0.523 | 0.6 | 1 | HC_vs_MS.NoTrt_F |
| ZMYM3 | 0.02354 | -0.5319 | 0.408 | 0.51 | 1 | HC_vs_MS.NoTrt_F |
| C6orf1 | 0.02356 | 0.51961 | 0.159 | 0.25 | 1 | HC_vs_MS.NoTrt_F |
| HBB | 0.02357 | -3.8156 | 0.075 | 0.14 | 1 | HC_vs_MS.NoTrt_F |
| SDHD | 0.02358 | -0.4135 | 0.685 | 0.75 | 1 | HC_vs_MS.NoTrt_F |
| ZNF626 | 0.02361 | 0.71867 | 0.783 | 0.72 | 1 | HC_vs_MS.NoTrt_F |
| LYAR | 0.02367 | 0.27202 | 0.121 | 0.2 | 1 | HC_vs_MS.NoTrt_F |
| SLA2 | 0.02369 | -0.5128 | 0.572 | 0.66 | 1 | HC_vs_MS.NoTrt_F |
| ACTR3B | 0.02377 | 0.79032 | 0.592 | 0.5 | 1 | HC_vs_MS.NoTrt_F |
| ICOS | 0.02387 | 0.31787 | 0.405 | 0.54 | 1 | HC_vs_MS.NoTrt_F |
| PPARD | 0.02392 | -0.4949 | 0.561 | 0.65 | 1 | HC_vs_MS.NoTrt_F |
| C5orf45 | 0.02394 | 0.68159 | 0.913 | 0.92 | 1 | HC_vs_MS.NoTrt_F |
| ARPP19 | 0.02402 | 0.30195 | 0.486 | 0.61 | 1 | HC_vs_MS.NoTrt_F |
| SSSCA1 | 0.02407 | 0.29881 | 0.11 | 0.18 | 1 | HC_vs_MS.NoTrt_F |
| RP11-458D2 | 0.02407 | -0.7876 | 0.127 | 0.2 | 1 | HC_vs_MS.NoTrt_F |
| ANXA1 | 0.02409 | 0.58322 | 0.422 | 0.36 | 1 | HC_vs_MS.NoTrt_F |
| RP11-192H2 | 0.0241 | 0.41063 | 0.737 | 0.75 | 1 | HC_vs_MS.NoTrt_F |

| | | | | | | |
|---|---|---|---|---|---|---|
| DDX58 | 0.02417 | -0.5949 | 0.364 | 0.45 | 1 | HC_vs_MS.NoTrt_F |
| ATP1A1 | 0.0242 | -0.4021 | 0.749 | 0.81 | 1 | HC_vs_MS.NoTrt_F |
| KIAA0825 | 0.02421 | 0.9537 | 0.361 | 0.47 | 1 | HC_vs_MS.NoTrt_F |
| DBT | 0.02423 | -0.535 | 0.867 | 0.92 | 1 | HC_vs_MS.NoTrt_F |
| CREB1 | 0.02432 | -0.5218 | 0.665 | 0.71 | 1 | HC_vs_MS.NoTrt_F |
| NMRK1 | 0.02438 | -0.6412 | 0.382 | 0.44 | 1 | HC_vs_MS.NoTrt_F |
| LRFN1 | 0.02439 | -0.618 | 0.173 | 0.25 | 1 | HC_vs_MS.NoTrt_F |
| AMIGO2 | 0.0244 | -0.4029 | 0.058 | 0.12 | 1 | HC_vs_MS.NoTrt_F |
| ZNF624 | 0.02447 | -0.306 | 0.434 | 0.52 | 1 | HC_vs_MS.NoTrt_F |
| NSA2 | 0.02455 | 0.28959 | 0.335 | 0.44 | 1 | HC_vs_MS.NoTrt_F |
| ZNF420 | 0.02459 | -0.4557 | 0.127 | 0.2 | 1 | HC_vs_MS.NoTrt_F |
| IFI35 | 0.0246 | 0.64612 | 0.873 | 0.83 | 1 | HC_vs_MS.NoTrt_F |
| POT1 | 0.02467 | -0.5996 | 0.11 | 0.18 | 1 | HC_vs_MS.NoTrt_F |
| TNPO1 | 0.02468 | -0.297 | 0.549 | 0.63 | 1 | HC_vs_MS.NoTrt_F |
| ZYG11B | 0.02468 | -0.2537 | 0.763 | 0.82 | 1 | HC_vs_MS.NoTrt_F |
| KIF5C | 0.02468 | -0.4608 | 0.058 | 0.12 | 1 | HC_vs_MS.NoTrt_F |
| MGST2 | 0.02469 | -0.5834 | 0.214 | 0.29 | 1 | HC_vs_MS.NoTrt_F |
| TTC27 | 0.02471 | 1.04718 | 0.145 | 0.23 | 1 | HC_vs_MS.NoTrt_F |
| CRB3 | 0.02476 | 0.2909 | 0.064 | 0.12 | 1 | HC_vs_MS.NoTrt_F |
| ARL8B | 0.02477 | -0.4521 | 0.474 | 0.55 | 1 | HC_vs_MS.NoTrt_F |
| HTRA2 | 0.0248 | -0.3245 | 0.139 | 0.22 | 1 | HC_vs_MS.NoTrt_F |
| DOT1L | 0.02486 | 0.67271 | 0.818 | 0.79 | 1 | HC_vs_MS.NoTrt_F |
| ATR | 0.02488 | 0.25921 | 0.561 | 0.66 | 1 | HC_vs_MS.NoTrt_F |
| DDX52 | 0.02488 | 0.41339 | 0.916 | 0.89 | 1 | HC_vs_MS.NoTrt_F |
| CLYBL | 0.02489 | -0.2891 | 0.124 | 0.2 | 1 | HC_vs_MS.NoTrt_F |
| SEMA4B | 0.02489 | -1.0754 | 0.376 | 0.48 | 1 | HC_vs_MS.NoTrt_F |
| ACTR6 | 0.02496 | -0.2957 | 0.564 | 0.65 | 1 | HC_vs_MS.NoTrt_F |
| GFM2 | 0.025 | -1.0105 | 0.254 | 0.34 | 1 | HC_vs_MS.NoTrt_F |
| SLF1 | 0.02513 | 0.4522 | 0.197 | 0.29 | 1 | HC_vs_MS.NoTrt_F |
| PYGM | 0.02514 | 0.33902 | 0.801 | 0.77 | 1 | HC_vs_MS.NoTrt_F |
| ZER1 | 0.02517 | -0.3386 | 0.145 | 0.22 | 1 | HC_vs_MS.NoTrt_F |
| EVI2A | 0.02521 | 0.4573 | 0.225 | 0.33 | 1 | HC_vs_MS.NoTrt_F |
| ZNF649 | 0.02521 | 0.38378 | 0.205 | 0.29 | 1 | HC_vs_MS.NoTrt_F |
| GRPEL1 | 0.02531 | -0.2855 | 0.156 | 0.24 | 1 | HC_vs_MS.NoTrt_F |
| NUTM2D | 0.02535 | 0.27501 | 0.095 | 0.17 | 1 | HC_vs_MS.NoTrt_F |
| KRBOX4 | 0.02543 | -0.4259 | 0.483 | 0.58 | 1 | HC_vs_MS.NoTrt_F |
| TMEM261 | 0.02544 | 0.29106 | 0.165 | 0.25 | 1 | HC_vs_MS.NoTrt_F |
| RFXANK | 0.0255 | -0.7303 | 0.26 | 0.35 | 1 | HC_vs_MS.NoTrt_F |
| GPR22 | 0.02554 | -0.5446 | 0.127 | 0.2 | 1 | HC_vs_MS.NoTrt_F |
| PGBD2 | 0.02567 | -0.6783 | 0.373 | 0.48 | 1 | HC_vs_MS.NoTrt_F |

| | | | | | | |
|---|---|---|---|---|---|---|
| CCR3 | 0.02568 | 0.55481 | 0.078 | 0.14 | 1 | HC_vs_MS.NoTrt_F |
| AKT3 | 0.02569 | 0.67487 | 0.798 | 0.76 | 1 | HC_vs_MS.NoTrt_F |
| CKS2 | 0.02571 | 0.38027 | 0.069 | 0.13 | 1 | HC_vs_MS.NoTrt_F |
| ZBTB41 | 0.02573 | 1.44545 | 0.066 | 0.13 | 1 | HC_vs_MS.NoTrt_F |
| CCDC142 | 0.02575 | 0.29073 | 0.676 | 0.61 | 1 | HC_vs_MS.NoTrt_F |
| SLC41A3 | 0.02578 | -0.4494 | 0.393 | 0.49 | 1 | HC_vs_MS.NoTrt_F |
| NDUFC1 | 0.02578 | 0.48581 | 0.289 | 0.4 | 1 | HC_vs_MS.NoTrt_F |
| SIRT3 | 0.02588 | 0.81748 | 0.318 | 0.41 | 1 | HC_vs_MS.NoTrt_F |
| GPRASP1 | 0.02595 | 0.92884 | 0.075 | 0.14 | 1 | HC_vs_MS.NoTrt_F |
| PAIP2B | 0.02602 | -0.521 | 0.254 | 0.34 | 1 | HC_vs_MS.NoTrt_F |
| PLIN3 | 0.02604 | -0.5368 | 0.561 | 0.6 | 1 | HC_vs_MS.NoTrt_F |
| ZFP37 | 0.02606 | -1.0024 | 0.425 | 0.51 | 1 | HC_vs_MS.NoTrt_F |
| DDX19B | 0.02623 | -0.3826 | 0.509 | 0.57 | 1 | HC_vs_MS.NoTrt_F |
| TMEM98 | 0.02623 | 0.35597 | 0.61 | 0.53 | 1 | HC_vs_MS.NoTrt_F |
| GRHL1 | 0.02631 | -0.3787 | 0.509 | 0.59 | 1 | HC_vs_MS.NoTrt_F |
| BLVRB | 0.02631 | 0.35098 | 0.413 | 0.31 | 1 | HC_vs_MS.NoTrt_F |
| RARG | 0.02653 | 0.85628 | 0.055 | 0.11 | 1 | HC_vs_MS.NoTrt_F |
| MKS1 | 0.02655 | 0.57608 | 0.081 | 0.14 | 1 | HC_vs_MS.NoTrt_F |
| NGDN | 0.02658 | -0.3714 | 0.367 | 0.45 | 1 | HC_vs_MS.NoTrt_F |
| QSER1 | 0.02669 | -0.6178 | 0.11 | 0.18 | 1 | HC_vs_MS.NoTrt_F |
| LILRA2 | 0.0267 | 0.34871 | 0.838 | 0.75 | 1 | HC_vs_MS.NoTrt_F |
| MRPS30 | 0.02672 | -0.2804 | 0.312 | 0.39 | 1 | HC_vs_MS.NoTrt_F |
| TLE4 | 0.02675 | 0.65428 | 0.199 | 0.29 | 1 | HC_vs_MS.NoTrt_F |
| DOCK7 | 0.02678 | 0.41506 | 0.974 | 0.96 | 1 | HC_vs_MS.NoTrt_F |
| KBTBD6 | 0.02678 | -0.2727 | 0.399 | 0.46 | 1 | HC_vs_MS.NoTrt_F |
| KIAA1586 | 0.02683 | -0.3012 | 0.104 | 0.17 | 1 | HC_vs_MS.NoTrt_F |
| IL1RL1 | 0.02693 | 1.65097 | 0.124 | 0.2 | 1 | HC_vs_MS.NoTrt_F |
| KIZ | 0.02693 | 0.85964 | 0.147 | 0.23 | 1 | HC_vs_MS.NoTrt_F |
| GGPS1 | 0.02702 | -0.4101 | 0.454 | 0.55 | 1 | HC_vs_MS.NoTrt_F |
| SLC35C1 | 0.02707 | 0.40367 | 0.072 | 0.13 | 1 | HC_vs_MS.NoTrt_F |
| CASD1 | 0.02709 | 1.1443 | 0.075 | 0.14 | 1 | HC_vs_MS.NoTrt_F |
| UCKL1 | 0.02715 | 0.29813 | 0.188 | 0.28 | 1 | HC_vs_MS.NoTrt_F |
| IFFO2 | 0.02717 | -0.4123 | 0.176 | 0.25 | 1 | HC_vs_MS.NoTrt_F |
| NLRP2 | 0.02742 | 0.2755 | 0.85 | 0.8 | 1 | HC_vs_MS.NoTrt_F |
| NPEPL1 | 0.02745 | 0.85174 | 0.176 | 0.27 | 1 | HC_vs_MS.NoTrt_F |
| ARRDC1 | 0.02763 | -0.2994 | 0.306 | 0.38 | 1 | HC_vs_MS.NoTrt_F |
| REV1 | 0.02768 | -0.8029 | 0.211 | 0.29 | 1 | HC_vs_MS.NoTrt_F |
| ZNF627 | 0.02775 | 1.29517 | 0.633 | 0.54 | 1 | HC_vs_MS.NoTrt_F |
| JCHAIN | 0.02779 | 1.2011 | 0.11 | 0.18 | 1 | HC_vs_MS.NoTrt_F |
| MLYCD | 0.02782 | -0.3175 | 0.422 | 0.51 | 1 | HC_vs_MS.NoTrt_F |

| | | | | | | |
|---|---|---|---|---|---|---|
| GSTP1 | 0.02782 | -0.7314 | 0.075 | 0.13 | 1 | HC_vs_MS.NoTrt_F |
| KLF10 | 0.02786 | -0.4701 | 0.098 | 0.17 | 1 | HC_vs_MS.NoTrt_F |
| DPP4 | 0.0279 | 0.32843 | 0.124 | 0.2 | 1 | HC_vs_MS.NoTrt_F |
| UBR3 | 0.02792 | -0.4905 | 0.171 | 0.24 | 1 | HC_vs_MS.NoTrt_F |
| CFAP70 | 0.02804 | -0.7922 | 0.269 | 0.35 | 1 | HC_vs_MS.NoTrt_F |
| ABHD16A | 0.02805 | -0.3987 | 0.208 | 0.29 | 1 | HC_vs_MS.NoTrt_F |
| BUD13 | 0.02807 | 0.34257 | 0.104 | 0.17 | 1 | HC_vs_MS.NoTrt_F |
| GPR68 | 0.02816 | 0.53036 | 0.139 | 0.22 | 1 | HC_vs_MS.NoTrt_F |
| FAAH2 | 0.02818 | 0.80573 | 0.127 | 0.2 | 1 | HC_vs_MS.NoTrt_F |
| KDELR1 | 0.0282 | -0.8008 | 0.266 | 0.35 | 1 | HC_vs_MS.NoTrt_F |
| ALAD | 0.02825 | -0.3243 | 0.431 | 0.52 | 1 | HC_vs_MS.NoTrt_F |
| CENPI | 0.02826 | 0.86492 | 0.471 | 0.39 | 1 | HC_vs_MS.NoTrt_F |
| RAD50 | 0.02827 | 0.5948 | 0.225 | 0.32 | 1 | HC_vs_MS.NoTrt_F |
| RP11-244H3 | 0.02837 | -0.8998 | 0.107 | 0.17 | 1 | HC_vs_MS.NoTrt_F |
| LILRA5 | 0.0284 | 0.32205 | 0.754 | 0.76 | 1 | HC_vs_MS.NoTrt_F |
| GEMIN5 | 0.02843 | -0.6496 | 0.052 | 0.11 | 1 | HC_vs_MS.NoTrt_F |
| GID8 | 0.02849 | 0.3463 | 0.191 | 0.28 | 1 | HC_vs_MS.NoTrt_F |
| ST20-MTHFS | 0.02859 | 0.29932 | 0.145 | 0.22 | 1 | HC_vs_MS.NoTrt_F |
| CEP55 | 0.02865 | -0.5437 | 0.15 | 0.23 | 1 | HC_vs_MS.NoTrt_F |
| CMIP | 0.02867 | -0.3542 | 0.11 | 0.18 | 1 | HC_vs_MS.NoTrt_F |
| IMPACT | 0.02869 | -0.3314 | 0.061 | 0.12 | 1 | HC_vs_MS.NoTrt_F |
| GDF5 | 0.0287 | 0.27179 | 0.081 | 0.14 | 1 | HC_vs_MS.NoTrt_F |
| NXT1 | 0.02871 | -0.3873 | 0.118 | 0.19 | 1 | HC_vs_MS.NoTrt_F |
| RSBN1 | 0.02873 | 0.41995 | 0.275 | 0.38 | 1 | HC_vs_MS.NoTrt_F |
| KIAA0556 | 0.02875 | -0.5659 | 0.358 | 0.45 | 1 | HC_vs_MS.NoTrt_F |
| LDLR | 0.02881 | 0.54216 | 0.98 | 0.96 | 1 | HC_vs_MS.NoTrt_F |
| VPS18 | 0.02889 | -0.7827 | 0.251 | 0.34 | 1 | HC_vs_MS.NoTrt_F |
| SCFD2 | 0.0289 | 0.7387 | 0.087 | 0.16 | 1 | HC_vs_MS.NoTrt_F |
| WDR60 | 0.02898 | 0.29163 | 0.15 | 0.23 | 1 | HC_vs_MS.NoTrt_F |
| CCDC88C | 0.02905 | 0.25637 | 0.569 | 0.67 | 1 | HC_vs_MS.NoTrt_F |
| TMEM187 | 0.02906 | 0.91531 | 0.064 | 0.12 | 1 | HC_vs_MS.NoTrt_F |
| KDM1A | 0.02917 | 0.35345 | 0.084 | 0.15 | 1 | HC_vs_MS.NoTrt_F |
| IPPK | 0.02924 | -0.8342 | 0.101 | 0.17 | 1 | HC_vs_MS.NoTrt_F |
| AC009133.2 | 0.02928 | 0.29179 | 0.124 | 0.2 | 1 | HC_vs_MS.NoTrt_F |
| PECAM1 | 0.02939 | 0.60858 | 0.798 | 0.76 | 1 | HC_vs_MS.NoTrt_F |
| LAMC3 | 0.02945 | 0.25718 | 0.283 | 0.39 | 1 | HC_vs_MS.NoTrt_F |
| HMCES | 0.02963 | 0.35356 | 0.136 | 0.21 | 1 | HC_vs_MS.NoTrt_F |
| BAHD1 | 0.02973 | 1.2215 | 0.075 | 0.14 | 1 | HC_vs_MS.NoTrt_F |
| KXD1 | 0.02974 | -0.361 | 0.858 | 0.87 | 1 | HC_vs_MS.NoTrt_F |
| PNPLA7 | 0.02975 | 0.7539 | 0.098 | 0.17 | 1 | HC_vs_MS.NoTrt_F |

| | | | | | | |
|---|---|---|---|---|---|---|
| H1FX | 0.02981 | 0.80075 | 0.064 | 0.12 | 1 | HC_vs_MS.NoTrt_F |
| MOB3C | 0.02993 | 1.38282 | 0.072 | 0.13 | 1 | HC_vs_MS.NoTrt_F |
| GPAT2 | 0.02998 | 0.26311 | 0.052 | 0.11 | 1 | HC_vs_MS.NoTrt_F |
| PPIC | 0.03 | 0.34336 | 0.173 | 0.11 | 1 | HC_vs_MS.NoTrt_F |
| ZNF124 | 0.03006 | 0.98197 | 0.769 | 0.67 | 1 | HC_vs_MS.NoTrt_F |
| C1GALT1 | 0.03023 | -0.298 | 0.613 | 0.67 | 1 | HC_vs_MS.NoTrt_F |
| KLRB1 | 0.03032 | 0.35352 | 0.199 | 0.29 | 1 | HC_vs_MS.NoTrt_F |
| CTSA | 0.03039 | -0.6678 | 0.668 | 0.74 | 1 | HC_vs_MS.NoTrt_F |
| GPX3 | 0.03041 | 0.89896 | 0.205 | 0.14 | 1 | HC_vs_MS.NoTrt_F |
| IDH3G | 0.03042 | 0.29183 | 0.197 | 0.29 | 1 | HC_vs_MS.NoTrt_F |
| RASAL3 | 0.03049 | 0.36001 | 0.22 | 0.32 | 1 | HC_vs_MS.NoTrt_F |
| GGT1 | 0.03049 | 0.83345 | 0.156 | 0.24 | 1 | HC_vs_MS.NoTrt_F |
| FLYWCH1 | 0.03052 | 0.27267 | 0.832 | 0.9 | 1 | HC_vs_MS.NoTrt_F |
| ACAN | 0.03053 | 0.30336 | 0.269 | 0.37 | 1 | HC_vs_MS.NoTrt_F |
| TRMT10C | 0.03057 | 0.44571 | 0.121 | 0.19 | 1 | HC_vs_MS.NoTrt_F |
| SLC25A51 | 0.03065 | -0.6107 | 0.575 | 0.64 | 1 | HC_vs_MS.NoTrt_F |
| CISD3 | 0.03071 | 0.43972 | 0.118 | 0.19 | 1 | HC_vs_MS.NoTrt_F |
| ICE1 | 0.03071 | 0.86411 | 0.118 | 0.19 | 1 | HC_vs_MS.NoTrt_F |
| ENSA | 0.03075 | -0.5501 | 0.821 | 0.88 | 1 | HC_vs_MS.NoTrt_F |
| GRK3 | 0.03079 | -0.581 | 0.408 | 0.48 | 1 | HC_vs_MS.NoTrt_F |
| SLC10A7 | 0.03086 | -0.6162 | 0.107 | 0.17 | 1 | HC_vs_MS.NoTrt_F |
| XXYLT1 | 0.03093 | -0.788 | 0.217 | 0.3 | 1 | HC_vs_MS.NoTrt_F |
| AGTRAP | 0.03117 | 0.35026 | 0.179 | 0.27 | 1 | HC_vs_MS.NoTrt_F |
| C15orf39 | 0.0312 | 0.51911 | 0.098 | 0.17 | 1 | HC_vs_MS.NoTrt_F |
| SFXN2 | 0.03136 | 0.44914 | 0.925 | 0.93 | 1 | HC_vs_MS.NoTrt_F |
| EIF4A3 | 0.03138 | -1.2689 | 0.786 | 0.8 | 1 | HC_vs_MS.NoTrt_F |
| GRK6 | 0.0314 | 0.34304 | 0.223 | 0.32 | 1 | HC_vs_MS.NoTrt_F |
| TOR2A | 0.03143 | -0.4392 | 0.133 | 0.2 | 1 | HC_vs_MS.NoTrt_F |
| C12orf50 | 0.03144 | 0.3163 | 0.922 | 0.95 | 1 | HC_vs_MS.NoTrt_F |
| STAMBP | 0.03148 | 0.29057 | 0.162 | 0.24 | 1 | HC_vs_MS.NoTrt_F |
| EIF4E | 0.03166 | 0.25847 | 0.988 | 0.99 | 1 | HC_vs_MS.NoTrt_F |
| MEPCE | 0.03166 | 1.15419 | 0.084 | 0.15 | 1 | HC_vs_MS.NoTrt_F |
| XKR6 | 0.03175 | 0.59348 | 0.084 | 0.15 | 1 | HC_vs_MS.NoTrt_F |
| SUN1 | 0.0318 | 0.48208 | 0.717 | 0.7 | 1 | HC_vs_MS.NoTrt_F |
| GRIN2C | 0.03188 | 0.35821 | 0.28 | 0.38 | 1 | HC_vs_MS.NoTrt_F |
| FMR1 | 0.03189 | -0.3894 | 0.153 | 0.23 | 1 | HC_vs_MS.NoTrt_F |
| SCMH1 | 0.03193 | 0.46262 | 0.176 | 0.26 | 1 | HC_vs_MS.NoTrt_F |
| CHRNA6 | 0.03195 | 0.76589 | 0.367 | 0.28 | 1 | HC_vs_MS.NoTrt_F |
| C4orf48 | 0.03198 | 0.39541 | 0.162 | 0.25 | 1 | HC_vs_MS.NoTrt_F |
| GLCE | 0.03199 | -1.2587 | 0.165 | 0.23 | 1 | HC_vs_MS.NoTrt_F |

| | | | | | | |
|---|---|---|---|---|---|---|
| TXNDC12 | 0.03203 | 0.30283 | 0.199 | 0.28 | 1 | HC_vs_MS.NoTrt_F |
| GSAP | 0.03204 | 0.71762 | 0.09 | 0.16 | 1 | HC_vs_MS.NoTrt_F |
| ZNF586 | 0.03206 | 0.29522 | 0.211 | 0.3 | 1 | HC_vs_MS.NoTrt_F |
| ABCC5 | 0.03209 | -0.4578 | 0.179 | 0.25 | 1 | HC_vs_MS.NoTrt_F |
| PGGT1B | 0.03209 | 0.3298 | 0.202 | 0.29 | 1 | HC_vs_MS.NoTrt_F |
| TRIM58 | 0.03222 | 0.45768 | 0.691 | 0.61 | 1 | HC_vs_MS.NoTrt_F |
| VPS51 | 0.03225 | 0.32856 | 0.283 | 0.39 | 1 | HC_vs_MS.NoTrt_F |
| CAND1 | 0.03226 | -0.4314 | 0.867 | 0.91 | 1 | HC_vs_MS.NoTrt_F |
| MFSD5 | 0.03232 | -0.5348 | 0.092 | 0.16 | 1 | HC_vs_MS.NoTrt_F |
| OSBP | 0.03234 | 0.81173 | 0.095 | 0.16 | 1 | HC_vs_MS.NoTrt_F |
| USP13 | 0.03234 | -0.3427 | 0.347 | 0.43 | 1 | HC_vs_MS.NoTrt_F |
| RBM26 | 0.03235 | 0.2687 | 0.506 | 0.6 | 1 | HC_vs_MS.NoTrt_F |
| MED17 | 0.03243 | 0.32588 | 0.957 | 0.95 | 1 | HC_vs_MS.NoTrt_F |
| NFE2L3 | 0.0325 | -0.8618 | 0.116 | 0.18 | 1 | HC_vs_MS.NoTrt_F |
| EIF4ENIF1 | 0.03257 | 0.43386 | 0.653 | 0.71 | 1 | HC_vs_MS.NoTrt_F |
| SSX5 | 0.03264 | -0.3889 | 0.188 | 0.27 | 1 | HC_vs_MS.NoTrt_F |
| SARM1 | 0.03264 | 0.26929 | 0.422 | 0.51 | 1 | HC_vs_MS.NoTrt_F |
| RORC | 0.03266 | 0.95205 | 0.121 | 0.19 | 1 | HC_vs_MS.NoTrt_F |
| EIF2B3 | 0.03267 | -0.5251 | 0.477 | 0.53 | 1 | HC_vs_MS.NoTrt_F |
| CRYL1 | 0.03267 | 0.42197 | 0.11 | 0.18 | 1 | HC_vs_MS.NoTrt_F |
| KIFC3 | 0.03274 | 0.53703 | 0.751 | 0.67 | 1 | HC_vs_MS.NoTrt_F |
| PIK3R5 | 0.03281 | 0.2763 | 0.422 | 0.55 | 1 | HC_vs_MS.NoTrt_F |
| NAXE | 0.03289 | 0.417 | 0.162 | 0.24 | 1 | HC_vs_MS.NoTrt_F |
| GNPDA2 | 0.03292 | 0.47601 | 0.092 | 0.16 | 1 | HC_vs_MS.NoTrt_F |
| C17orf89 | 0.03293 | 0.59181 | 0.095 | 0.16 | 1 | HC_vs_MS.NoTrt_F |
| HDAC6 | 0.03298 | -0.3415 | 0.896 | 0.92 | 1 | HC_vs_MS.NoTrt_F |
| FAM53C | 0.03302 | 0.62401 | 0.841 | 0.82 | 1 | HC_vs_MS.NoTrt_F |
| RAB7A | 0.0331 | -0.3646 | 0.474 | 0.55 | 1 | HC_vs_MS.NoTrt_F |
| FBXL15 | 0.03312 | 0.3826 | 0.162 | 0.24 | 1 | HC_vs_MS.NoTrt_F |
| CCDC174 | 0.03314 | 0.34669 | 0.211 | 0.3 | 1 | HC_vs_MS.NoTrt_F |
| NOL9 | 0.03333 | 0.35286 | 0.968 | 0.96 | 1 | HC_vs_MS.NoTrt_F |
| ENG | 0.03336 | 1.33939 | 0.205 | 0.14 | 1 | HC_vs_MS.NoTrt_F |
| CTD-3148I1( | 0.03342 | 0.26487 | 0.11 | 0.18 | 1 | HC_vs_MS.NoTrt_F |
| TNFAIP8 | 0.03342 | -0.3033 | 0.899 | 0.91 | 1 | HC_vs_MS.NoTrt_F |
| CYP2E1 | 0.03351 | 0.75691 | 0.087 | 0.15 | 1 | HC_vs_MS.NoTrt_F |
| ZFP36L1 | 0.03353 | 0.61538 | 0.769 | 0.76 | 1 | HC_vs_MS.NoTrt_F |
| LAMTOR1 | 0.03354 | 0.31383 | 0.254 | 0.37 | 1 | HC_vs_MS.NoTrt_F |
| BPIFB1 | 0.03362 | -0.3407 | 0.486 | 0.59 | 1 | HC_vs_MS.NoTrt_F |
| ODF2L | 0.03368 | 0.44658 | 0.986 | 0.97 | 1 | HC_vs_MS.NoTrt_F |
| PTPN22 | 0.03372 | -0.3723 | 0.416 | 0.51 | 1 | HC_vs_MS.NoTrt_F |

| | | | | | | |
|---|---|---|---|---|---|---|
| TXLNG | 0.03372 | 0.30384 | 0.139 | 0.22 | 1 | HC_vs_MS.NoTrt_F |
| SULT1A2 | 0.03378 | 0.26127 | 0.197 | 0.28 | 1 | HC_vs_MS.NoTrt_F |
| HEXB | 0.03385 | -0.2913 | 0.197 | 0.28 | 1 | HC_vs_MS.NoTrt_F |
| ZNF780B | 0.03387 | -1.1558 | 0.584 | 0.65 | 1 | HC_vs_MS.NoTrt_F |
| ZNF236 | 0.03391 | 0.43655 | 0.182 | 0.27 | 1 | HC_vs_MS.NoTrt_F |
| AKAP17A | 0.03405 | -0.3813 | 0.118 | 0.19 | 1 | HC_vs_MS.NoTrt_F |
| SHTN1 | 0.03412 | -1.0076 | 0.118 | 0.19 | 1 | HC_vs_MS.NoTrt_F |
| LY6G5B | 0.03413 | 0.28473 | 0.864 | 0.82 | 1 | HC_vs_MS.NoTrt_F |
| ATPAF1 | 0.03423 | -0.3834 | 0.263 | 0.34 | 1 | HC_vs_MS.NoTrt_F |
| CDKL1 | 0.03438 | 0.72585 | 0.803 | 0.73 | 1 | HC_vs_MS.NoTrt_F |
| PHACTR1 | 0.03439 | 0.90943 | 0.477 | 0.38 | 1 | HC_vs_MS.NoTrt_F |
| KRT8 | 0.03445 | 0.30109 | 0.867 | 0.83 | 1 | HC_vs_MS.NoTrt_F |
| EIF4E2 | 0.03449 | -0.293 | 0.393 | 0.47 | 1 | HC_vs_MS.NoTrt_F |
| MINA | 0.03455 | 0.5129 | 0.127 | 0.2 | 1 | HC_vs_MS.NoTrt_F |
| KYAT1 | 0.03464 | -0.259 | 0.173 | 0.25 | 1 | HC_vs_MS.NoTrt_F |
| ZNF333 | 0.03466 | 0.73961 | 0.942 | 0.94 | 1 | HC_vs_MS.NoTrt_F |
| PIAS4 | 0.03485 | 0.57454 | 0.231 | 0.32 | 1 | HC_vs_MS.NoTrt_F |
| TP63 | 0.03486 | 0.32298 | 0.465 | 0.57 | 1 | HC_vs_MS.NoTrt_F |
| STX8 | 0.03486 | 0.53668 | 0.884 | 0.87 | 1 | HC_vs_MS.NoTrt_F |
| IGSF9 | 0.03487 | -0.3181 | 0.344 | 0.41 | 1 | HC_vs_MS.NoTrt_F |
| ZDHHC14 | 0.03493 | 0.56253 | 0.338 | 0.44 | 1 | HC_vs_MS.NoTrt_F |
| ZNF428 | 0.03495 | -0.2599 | 0.234 | 0.32 | 1 | HC_vs_MS.NoTrt_F |
| CCDC82 | 0.03504 | 0.62292 | 0.194 | 0.28 | 1 | HC_vs_MS.NoTrt_F |
| HSF5 | 0.03507 | 0.47012 | 0.098 | 0.16 | 1 | HC_vs_MS.NoTrt_F |
| CREBBP | 0.03518 | 0.5893 | 0.59 | 0.66 | 1 | HC_vs_MS.NoTrt_F |
| RNF125 | 0.03524 | 0.3778 | 0.951 | 0.95 | 1 | HC_vs_MS.NoTrt_F |
| LHPP | 0.03524 | 0.52188 | 0.15 | 0.23 | 1 | HC_vs_MS.NoTrt_F |
| GRHPR | 0.03525 | -0.6422 | 0.549 | 0.6 | 1 | HC_vs_MS.NoTrt_F |
| ZDHHC16 | 0.03541 | -0.5693 | 0.118 | 0.18 | 1 | HC_vs_MS.NoTrt_F |
| CYB5D2 | 0.03564 | 0.31185 | 0.431 | 0.53 | 1 | HC_vs_MS.NoTrt_F |
| PPP1R3E | 0.03565 | 0.41893 | 0.246 | 0.33 | 1 | HC_vs_MS.NoTrt_F |
| CCDC170 | 0.03568 | 0.29263 | 0.506 | 0.4 | 1 | HC_vs_MS.NoTrt_F |
| SLFN12 | 0.03568 | 0.61289 | 0.078 | 0.14 | 1 | HC_vs_MS.NoTrt_F |
| N4BP1 | 0.03573 | 0.25897 | 0.168 | 0.25 | 1 | HC_vs_MS.NoTrt_F |
| FPR2 | 0.03577 | -0.3385 | 0.546 | 0.43 | 1 | HC_vs_MS.NoTrt_F |
| ZNF287 | 0.03587 | 0.48535 | 0.517 | 0.44 | 1 | HC_vs_MS.NoTrt_F |
| PLSCR1 | 0.03593 | 0.26069 | 0.058 | 0.11 | 1 | HC_vs_MS.NoTrt_F |
| C10orf10 | 0.036 | 1.73517 | 0.107 | 0.06 | 1 | HC_vs_MS.NoTrt_F |
| PPIF | 0.03611 | -0.5693 | 0.116 | 0.18 | 1 | HC_vs_MS.NoTrt_F |
| EXO1 | 0.03616 | 0.37791 | 0.197 | 0.28 | 1 | HC_vs_MS.NoTrt_F |

| | | | | | | |
|---|---|---|---|---|---|---|
| ZNF445 | 0.03627 | 0.36001 | 0.988 | 0.98 | 1 | HC_vs_MS.NoTrt_F |
| ADGRE2 | 0.03637 | 0.7809 | 0.855 | 0.86 | 1 | HC_vs_MS.NoTrt_F |
| SCRG1 | 0.03641 | 0.33203 | 0.806 | 0.75 | 1 | HC_vs_MS.NoTrt_F |
| SORL1 | 0.0365 | -0.2954 | 0.723 | 0.77 | 1 | HC_vs_MS.NoTrt_F |
| CDH15 | 0.03655 | 0.25505 | 0.535 | 0.44 | 1 | HC_vs_MS.NoTrt_F |
| TRIM11 | 0.03663 | -1.0853 | 0.604 | 0.64 | 1 | HC_vs_MS.NoTrt_F |
| BLVRA | 0.03665 | 0.47324 | 0.156 | 0.23 | 1 | HC_vs_MS.NoTrt_F |
| RAC1 | 0.03674 | 0.68899 | 0.182 | 0.27 | 1 | HC_vs_MS.NoTrt_F |
| MT-ATP8 | 0.03679 | 0.4555 | 0.783 | 0.8 | 1 | HC_vs_MS.NoTrt_F |
| DNAH1 | 0.03681 | 0.32148 | 0.211 | 0.3 | 1 | HC_vs_MS.NoTrt_F |
| TET1 | 0.03684 | 0.44603 | 0.095 | 0.16 | 1 | HC_vs_MS.NoTrt_F |
| NT5DC1 | 0.03689 | 0.4443 | 0.223 | 0.3 | 1 | HC_vs_MS.NoTrt_F |
| LEMD2 | 0.03699 | 0.59781 | 0.118 | 0.19 | 1 | HC_vs_MS.NoTrt_F |
| NUP153 | 0.037 | 0.27634 | 0.249 | 0.34 | 1 | HC_vs_MS.NoTrt_F |
| CHRNB1 | 0.03717 | 0.78097 | 0.659 | 0.6 | 1 | HC_vs_MS.NoTrt_F |
| PSEN2 | 0.03719 | -1.0237 | 0.15 | 0.23 | 1 | HC_vs_MS.NoTrt_F |
| CD151 | 0.03721 | 0.37281 | 0.942 | 0.92 | 1 | HC_vs_MS.NoTrt_F |
| ALKBH6 | 0.03725 | -0.5775 | 0.538 | 0.6 | 1 | HC_vs_MS.NoTrt_F |
| TK2 | 0.0373 | 0.77038 | 0.168 | 0.24 | 1 | HC_vs_MS.NoTrt_F |
| ITPK1 | 0.03734 | -0.4934 | 0.673 | 0.58 | 1 | HC_vs_MS.NoTrt_F |
| NRIP2 | 0.03737 | -0.474 | 0.335 | 0.4 | 1 | HC_vs_MS.NoTrt_F |
| ZNF514 | 0.03743 | -0.5481 | 0.642 | 0.69 | 1 | HC_vs_MS.NoTrt_F |
| C1orf123 | 0.03761 | -0.4288 | 0.22 | 0.29 | 1 | HC_vs_MS.NoTrt_F |
| SLC4A2 | 0.03761 | 0.33055 | 0.09 | 0.16 | 1 | HC_vs_MS.NoTrt_F |
| ADAM32 | 0.03765 | 0.32103 | 0.896 | 0.93 | 1 | HC_vs_MS.NoTrt_F |
| TNRC18 | 0.03767 | 0.672 | 0.847 | 0.81 | 1 | HC_vs_MS.NoTrt_F |
| ADTRP | 0.03768 | -0.7724 | 0.37 | 0.43 | 1 | HC_vs_MS.NoTrt_F |
| ZNF496 | 0.03779 | 0.80959 | 0.769 | 0.74 | 1 | HC_vs_MS.NoTrt_F |
| DBNDD2 | 0.03783 | 0.28011 | 0.066 | 0.12 | 1 | HC_vs_MS.NoTrt_F |
| C2CD5 | 0.0379 | -0.2929 | 0.211 | 0.29 | 1 | HC_vs_MS.NoTrt_F |
| NAALADL2 | 0.03799 | 0.27534 | 0.092 | 0.16 | 1 | HC_vs_MS.NoTrt_F |
| VIL1 | 0.03812 | 0.62297 | 0.217 | 0.29 | 1 | HC_vs_MS.NoTrt_F |
| TRMT5 | 0.03819 | -1.3333 | 0.165 | 0.23 | 1 | HC_vs_MS.NoTrt_F |
| TCP11L1 | 0.03823 | -0.3457 | 0.199 | 0.28 | 1 | HC_vs_MS.NoTrt_F |
| MAGEH1 | 0.0383 | 0.48735 | 0.11 | 0.18 | 1 | HC_vs_MS.NoTrt_F |
| TSPAN32 | 0.03831 | 0.40431 | 0.251 | 0.36 | 1 | HC_vs_MS.NoTrt_F |
| EHMT2 | 0.03842 | -0.4708 | 0.699 | 0.77 | 1 | HC_vs_MS.NoTrt_F |
| RBM15 | 0.03853 | 0.60324 | 0.078 | 0.14 | 1 | HC_vs_MS.NoTrt_F |
| RAB3D | 0.03862 | -0.6986 | 0.673 | 0.72 | 1 | HC_vs_MS.NoTrt_F |
| LRPPRC | 0.03871 | -0.5625 | 0.532 | 0.58 | 1 | HC_vs_MS.NoTrt_F |

| | | | | | | |
|---|---|---|---|---|---|---|
| PPP2R5A | 0.03892 | -1.2995 | 0.237 | 0.3 | 1 | HC_vs_MS.NoTrt_F |
| CAMSAP1 | 0.03895 | 0.35301 | 0.104 | 0.17 | 1 | HC_vs_MS.NoTrt_F |
| CEBPZ | 0.039 | -0.6783 | 0.246 | 0.33 | 1 | HC_vs_MS.NoTrt_F |
| KTN1 | 0.03905 | -0.4239 | 0.604 | 0.71 | 1 | HC_vs_MS.NoTrt_F |
| MAPK6 | 0.03909 | -0.3295 | 0.54 | 0.59 | 1 | HC_vs_MS.NoTrt_F |
| CARD11 | 0.03909 | 0.62512 | 0.202 | 0.29 | 1 | HC_vs_MS.NoTrt_F |
| ZNF134 | 0.03909 | -0.3203 | 0.194 | 0.27 | 1 | HC_vs_MS.NoTrt_F |
| BCAT1 | 0.03914 | 0.31663 | 0.483 | 0.55 | 1 | HC_vs_MS.NoTrt_F |
| LRIF1 | 0.03916 | 0.3134 | 0.121 | 0.19 | 1 | HC_vs_MS.NoTrt_F |
| FEM1C | 0.03917 | -0.5817 | 0.147 | 0.22 | 1 | HC_vs_MS.NoTrt_F |
| LPIN2 | 0.03937 | 0.31711 | 0.211 | 0.3 | 1 | HC_vs_MS.NoTrt_F |
| GIMAP2 | 0.0394 | -0.4055 | 0.887 | 0.91 | 1 | HC_vs_MS.NoTrt_F |
| BRPF3 | 0.03947 | 1.73573 | 0.066 | 0.12 | 1 | HC_vs_MS.NoTrt_F |
| ABCB10 | 0.03949 | -0.702 | 0.101 | 0.16 | 1 | HC_vs_MS.NoTrt_F |
| CWC27 | 0.03958 | 0.565 | 0.168 | 0.24 | 1 | HC_vs_MS.NoTrt_F |
| NRM | 0.03959 | 0.52682 | 0.058 | 0.11 | 1 | HC_vs_MS.NoTrt_F |
| UBASH3B | 0.03959 | -0.3865 | 0.439 | 0.51 | 1 | HC_vs_MS.NoTrt_F |
| SULT1A1 | 0.03967 | 0.29405 | 0.618 | 0.69 | 1 | HC_vs_MS.NoTrt_F |
| MACC1 | 0.0397 | 0.51734 | 0.786 | 0.69 | 1 | HC_vs_MS.NoTrt_F |
| SUPT16H | 0.03973 | -0.4977 | 0.731 | 0.77 | 1 | HC_vs_MS.NoTrt_F |
| LRRC59 | 0.03993 | 0.29972 | 0.176 | 0.26 | 1 | HC_vs_MS.NoTrt_F |
| FBXO8 | 0.04006 | 1.13892 | 0.101 | 0.17 | 1 | HC_vs_MS.NoTrt_F |
| HIST1H1D | 0.04035 | 0.82844 | 0.095 | 0.16 | 1 | HC_vs_MS.NoTrt_F |
| DPM2 | 0.04037 | -0.6307 | 0.13 | 0.19 | 1 | HC_vs_MS.NoTrt_F |
| SIGIRR | 0.04041 | 0.39281 | 0.358 | 0.48 | 1 | HC_vs_MS.NoTrt_F |
| TNFAIP1 | 0.04046 | 0.25488 | 0.113 | 0.18 | 1 | HC_vs_MS.NoTrt_F |
| KLHL28 | 0.04058 | -0.3949 | 0.382 | 0.43 | 1 | HC_vs_MS.NoTrt_F |
| LRRC14 | 0.0406 | 0.96104 | 0.121 | 0.19 | 1 | HC_vs_MS.NoTrt_F |
| BTRC | 0.04061 | 0.37371 | 0.113 | 0.18 | 1 | HC_vs_MS.NoTrt_F |
| BAG1 | 0.04063 | 0.27057 | 0.954 | 0.97 | 1 | HC_vs_MS.NoTrt_F |
| APOE | 0.04064 | 0.34319 | 0.792 | 0.81 | 1 | HC_vs_MS.NoTrt_F |
| AFMID | 0.04065 | 0.30815 | 0.957 | 0.95 | 1 | HC_vs_MS.NoTrt_F |
| ZNF674 | 0.04085 | 0.59741 | 0.948 | 0.92 | 1 | HC_vs_MS.NoTrt_F |
| TINF2 | 0.04093 | -0.3554 | 0.818 | 0.88 | 1 | HC_vs_MS.NoTrt_F |
| BATF | 0.04104 | 0.36924 | 0.24 | 0.34 | 1 | HC_vs_MS.NoTrt_F |
| TRA2A | 0.04104 | -0.3029 | 0.772 | 0.81 | 1 | HC_vs_MS.NoTrt_F |
| FLYWCH2 | 0.04106 | 0.50321 | 0.107 | 0.17 | 1 | HC_vs_MS.NoTrt_F |
| NAPRT | 0.04108 | 0.50815 | 0.214 | 0.3 | 1 | HC_vs_MS.NoTrt_F |
| GRK4 | 0.04118 | 0.27449 | 0.653 | 0.54 | 1 | HC_vs_MS.NoTrt_F |
| PRIM1 | 0.04122 | 0.39064 | 0.176 | 0.25 | 1 | HC_vs_MS.NoTrt_F |

| | | | | | | |
|---|---|---|---|---|---|---|
| CHST7 | 0.0413 | 0.74884 | 0.092 | 0.16 | 1 | HC_vs_MS.NoTrt_F |
| SEC61A2 | 0.0413 | 0.32324 | 0.113 | 0.18 | 1 | HC_vs_MS.NoTrt_F |
| CSTF2T | 0.04137 | -0.3111 | 0.13 | 0.2 | 1 | HC_vs_MS.NoTrt_F |
| TASP1 | 0.0415 | -0.2868 | 0.15 | 0.22 | 1 | HC_vs_MS.NoTrt_F |
| SELP | 0.04152 | 1.42467 | 0.095 | 0.16 | 1 | HC_vs_MS.NoTrt_F |
| KSR1 | 0.04154 | 1.17404 | 0.28 | 0.38 | 1 | HC_vs_MS.NoTrt_F |
| PCBP4 | 0.04156 | 0.43583 | 0.113 | 0.18 | 1 | HC_vs_MS.NoTrt_F |
| RCCD1 | 0.04173 | -0.7161 | 0.202 | 0.27 | 1 | HC_vs_MS.NoTrt_F |
| IMMP2L | 0.04174 | -0.3838 | 0.127 | 0.19 | 1 | HC_vs_MS.NoTrt_F |
| DYRK2 | 0.04178 | 0.4482 | 0.173 | 0.25 | 1 | HC_vs_MS.NoTrt_F |
| FER1L5 | 0.04179 | 0.35689 | 0.824 | 0.83 | 1 | HC_vs_MS.NoTrt_F |
| SERPINB8 | 0.04193 | -0.9651 | 0.064 | 0.12 | 1 | HC_vs_MS.NoTrt_F |
| MST1 | 0.042 | 0.26307 | 0.055 | 0.11 | 1 | HC_vs_MS.NoTrt_F |
| TMTC2 | 0.04207 | 0.49028 | 0.095 | 0.16 | 1 | HC_vs_MS.NoTrt_F |
| SCFD1 | 0.0421 | 0.65266 | 0.798 | 0.76 | 1 | HC_vs_MS.NoTrt_F |
| TNF | 0.0421 | 0.54519 | 0.156 | 0.23 | 1 | HC_vs_MS.NoTrt_F |
| HDAC8 | 0.04236 | 0.85705 | 0.243 | 0.34 | 1 | HC_vs_MS.NoTrt_F |
| USPL1 | 0.04246 | 0.41323 | 0.142 | 0.21 | 1 | HC_vs_MS.NoTrt_F |
| SLC35E2B | 0.04255 | 0.54689 | 0.616 | 0.69 | 1 | HC_vs_MS.NoTrt_F |
| SNX8 | 0.04278 | -1.1827 | 0.15 | 0.22 | 1 | HC_vs_MS.NoTrt_F |
| KLHL33 | 0.04283 | 0.59024 | 0.59 | 0.49 | 1 | HC_vs_MS.NoTrt_F |
| HBA2 | 0.04286 | -2.932 | 0.055 | 0.11 | 1 | HC_vs_MS.NoTrt_F |
| VPS13D | 0.04287 | -0.4918 | 0.344 | 0.41 | 1 | HC_vs_MS.NoTrt_F |
| CTDP1 | 0.04293 | 1.49232 | 0.078 | 0.14 | 1 | HC_vs_MS.NoTrt_F |
| GATA3 | 0.04305 | 0.27648 | 0.355 | 0.46 | 1 | HC_vs_MS.NoTrt_F |
| STAT4 | 0.04305 | 0.88156 | 0.439 | 0.39 | 1 | HC_vs_MS.NoTrt_F |
| CLIC2 | 0.04311 | -0.2937 | 0.448 | 0.53 | 1 | HC_vs_MS.NoTrt_F |
| UNG | 0.04315 | -0.6189 | 0.24 | 0.32 | 1 | HC_vs_MS.NoTrt_F |
| CBFB | 0.04315 | 0.76214 | 0.182 | 0.26 | 1 | HC_vs_MS.NoTrt_F |
| RAB43 | 0.04322 | 1.05731 | 0.228 | 0.31 | 1 | HC_vs_MS.NoTrt_F |
| MTSS1 | 0.04326 | 1.31712 | 0.708 | 0.67 | 1 | HC_vs_MS.NoTrt_F |
| SMARCD1 | 0.04334 | 0.48282 | 0.139 | 0.21 | 1 | HC_vs_MS.NoTrt_F |
| RNF26 | 0.04341 | 0.47558 | 0.127 | 0.19 | 1 | HC_vs_MS.NoTrt_F |
| AHNAK | 0.04345 | 0.43548 | 0.89 | 0.9 | 1 | HC_vs_MS.NoTrt_F |
| MROH7 | 0.04349 | 0.28847 | 0.613 | 0.53 | 1 | HC_vs_MS.NoTrt_F |
| WDR5 | 0.04354 | 0.32609 | 0.139 | 0.21 | 1 | HC_vs_MS.NoTrt_F |
| SAP130 | 0.04375 | 0.33159 | 0.194 | 0.28 | 1 | HC_vs_MS.NoTrt_F |
| MOGS | 0.04376 | 0.29598 | 0.121 | 0.19 | 1 | HC_vs_MS.NoTrt_F |
| GREB1 | 0.04379 | 0.35042 | 0.85 | 0.82 | 1 | HC_vs_MS.NoTrt_F |
| CCNJ | 0.04383 | 0.41305 | 0.075 | 0.13 | 1 | HC_vs_MS.NoTrt_F |

| | | | | | | |
|---|---|---|---|---|---|---|
| LYG2 | 0.04401 | 0.64167 | 0.075 | 0.13 | 1 | HC_vs_MS.NoTrt_F |
| NMI | 0.04402 | -0.6038 | 0.309 | 0.38 | 1 | HC_vs_MS.NoTrt_F |
| ME2 | 0.04416 | 0.39321 | 0.939 | 0.92 | 1 | HC_vs_MS.NoTrt_F |
| SLC8B1 | 0.04426 | -0.4027 | 0.538 | 0.59 | 1 | HC_vs_MS.NoTrt_F |
| PHKA2 | 0.04429 | 0.33791 | 0.734 | 0.82 | 1 | HC_vs_MS.NoTrt_F |
| RBM45 | 0.04432 | 0.91995 | 0.095 | 0.16 | 1 | HC_vs_MS.NoTrt_F |
| CHD6 | 0.04436 | -0.3353 | 0.251 | 0.33 | 1 | HC_vs_MS.NoTrt_F |
| HAUS8 | 0.04441 | -0.3656 | 0.751 | 0.79 | 1 | HC_vs_MS.NoTrt_F |
| ADCY2 | 0.04452 | 0.26884 | 0.815 | 0.8 | 1 | HC_vs_MS.NoTrt_F |
| CHN1 | 0.04456 | 0.3436 | 0.072 | 0.13 | 1 | HC_vs_MS.NoTrt_F |
| GPR63 | 0.04456 | -0.28 | 0.39 | 0.47 | 1 | HC_vs_MS.NoTrt_F |
| NEO1 | 0.04466 | 0.30946 | 0.266 | 0.36 | 1 | HC_vs_MS.NoTrt_F |
| MBIP | 0.04476 | 0.88785 | 0.098 | 0.16 | 1 | HC_vs_MS.NoTrt_F |
| SYNGAP1 | 0.04476 | 0.42414 | 0.185 | 0.27 | 1 | HC_vs_MS.NoTrt_F |
| CXorf40A | 0.04485 | 0.25419 | 0.188 | 0.27 | 1 | HC_vs_MS.NoTrt_F |
| HGSNAT | 0.04492 | 0.85697 | 0.772 | 0.73 | 1 | HC_vs_MS.NoTrt_F |
| KIRREL3 | 0.04493 | 0.33166 | 0.618 | 0.55 | 1 | HC_vs_MS.NoTrt_F |
| MCTP2 | 0.04494 | 0.88734 | 0.564 | 0.46 | 1 | HC_vs_MS.NoTrt_F |
| IFI27L2 | 0.04495 | -0.4843 | 0.251 | 0.32 | 1 | HC_vs_MS.NoTrt_F |
| PIGM | 0.04499 | 0.76237 | 0.679 | 0.59 | 1 | HC_vs_MS.NoTrt_F |
| GLRX5 | 0.04499 | -0.6591 | 0.101 | 0.16 | 1 | HC_vs_MS.NoTrt_F |
| STX5 | 0.04503 | 0.74289 | 0.159 | 0.23 | 1 | HC_vs_MS.NoTrt_F |
| XRCC4 | 0.04505 | 1.22598 | 0.084 | 0.14 | 1 | HC_vs_MS.NoTrt_F |
| TBCC | 0.04506 | -0.2565 | 0.341 | 0.42 | 1 | HC_vs_MS.NoTrt_F |
| ZNF827 | 0.04508 | 1.10126 | 0.055 | 0.11 | 1 | HC_vs_MS.NoTrt_F |
| ZNF365 | 0.04528 | -0.557 | 0.35 | 0.46 | 1 | HC_vs_MS.NoTrt_F |
| TRAK2 | 0.04529 | -0.3333 | 0.223 | 0.29 | 1 | HC_vs_MS.NoTrt_F |
| BLOC1S1 | 0.0453 | 0.25113 | 0.254 | 0.35 | 1 | HC_vs_MS.NoTrt_F |
| OXSM | 0.04534 | -0.8924 | 0.153 | 0.22 | 1 | HC_vs_MS.NoTrt_F |
| NWD1 | 0.04537 | -0.3588 | 0.627 | 0.53 | 1 | HC_vs_MS.NoTrt_F |
| RAP1GDS1 | 0.04554 | 0.34918 | 0.379 | 0.5 | 1 | HC_vs_MS.NoTrt_F |
| ALG2 | 0.04573 | 0.30205 | 0.11 | 0.17 | 1 | HC_vs_MS.NoTrt_F |
| MBD4 | 0.04586 | -0.2605 | 0.263 | 0.33 | 1 | HC_vs_MS.NoTrt_F |
| STAU2 | 0.04608 | -0.4027 | 0.656 | 0.71 | 1 | HC_vs_MS.NoTrt_F |
| LYL1 | 0.04615 | 0.26091 | 0.486 | 0.57 | 1 | HC_vs_MS.NoTrt_F |
| TRMT44 | 0.04632 | -0.3068 | 0.28 | 0.37 | 1 | HC_vs_MS.NoTrt_F |
| TSPAN5 | 0.0464 | -0.3805 | 0.645 | 0.67 | 1 | HC_vs_MS.NoTrt_F |
| PCCB | 0.04647 | 0.78131 | 0.679 | 0.66 | 1 | HC_vs_MS.NoTrt_F |
| CD81 | 0.04652 | 1.35189 | 0.055 | 0.11 | 1 | HC_vs_MS.NoTrt_F |
| PELP1 | 0.04657 | 0.66775 | 0.277 | 0.36 | 1 | HC_vs_MS.NoTrt_F |

| | | | | | | |
|---|---|---|---|---|---|---|
| KANSL2 | 0.04665 | 0.52591 | 0.188 | 0.27 | 1 | HC_vs_MS.NoTrt_F |
| PPP3R1 | 0.04666 | -0.6413 | 0.173 | 0.24 | 1 | HC_vs_MS.NoTrt_F |
| HACD1 | 0.04681 | -0.6411 | 0.636 | 0.71 | 1 | HC_vs_MS.NoTrt_F |
| FASTKD1 | 0.04689 | -0.3815 | 0.306 | 0.39 | 1 | HC_vs_MS.NoTrt_F |
| ZNF747 | 0.04693 | 0.30458 | 0.751 | 0.66 | 1 | HC_vs_MS.NoTrt_F |
| FBXO48 | 0.04706 | -0.5914 | 0.694 | 0.8 | 1 | HC_vs_MS.NoTrt_F |
| GATAD2B | 0.04709 | 0.36442 | 0.5 | 0.65 | 1 | HC_vs_MS.NoTrt_F |
| TFIP11 | 0.04709 | -1.0429 | 0.549 | 0.61 | 1 | HC_vs_MS.NoTrt_F |
| CTBS | 0.04717 | 0.39761 | 0.147 | 0.22 | 1 | HC_vs_MS.NoTrt_F |
| PLIN5 | 0.04733 | 0.3001 | 0.864 | 0.8 | 1 | HC_vs_MS.NoTrt_F |
| CD8A | 0.04738 | 1.65464 | 0.081 | 0.14 | 1 | HC_vs_MS.NoTrt_F |
| IGFLR1 | 0.0474 | 0.51971 | 0.578 | 0.71 | 1 | HC_vs_MS.NoTrt_F |
| GP1BA | 0.04751 | 0.92711 | 0.055 | 0.11 | 1 | HC_vs_MS.NoTrt_F |
| VPS4B | 0.04757 | -0.2847 | 0.803 | 0.8 | 1 | HC_vs_MS.NoTrt_F |
| MYSM1 | 0.04758 | 0.31333 | 0.269 | 0.35 | 1 | HC_vs_MS.NoTrt_F |
| DPH2 | 0.04768 | 0.67388 | 0.098 | 0.16 | 1 | HC_vs_MS.NoTrt_F |
| ZNF41 | 0.0477 | 1.30716 | 0.055 | 0.11 | 1 | HC_vs_MS.NoTrt_F |
| LTB4R | 0.04774 | -0.3779 | 0.376 | 0.45 | 1 | HC_vs_MS.NoTrt_F |
| PCNX3 | 0.0478 | 0.76669 | 0.159 | 0.23 | 1 | HC_vs_MS.NoTrt_F |
| GFRAL | 0.04809 | 0.32603 | 0.465 | 0.39 | 1 | HC_vs_MS.NoTrt_F |
| UQCRC1 | 0.04826 | 0.29227 | 0.263 | 0.37 | 1 | HC_vs_MS.NoTrt_F |
| N6AMT1 | 0.04827 | -0.5673 | 0.159 | 0.23 | 1 | HC_vs_MS.NoTrt_F |
| NEMP2 | 0.04842 | 0.28917 | 0.118 | 0.18 | 1 | HC_vs_MS.NoTrt_F |
| SCOC | 0.04844 | 0.25063 | 0.127 | 0.19 | 1 | HC_vs_MS.NoTrt_F |
| ATL3 | 0.04858 | -0.5557 | 0.838 | 0.87 | 1 | HC_vs_MS.NoTrt_F |
| C17orf51 | 0.04862 | -0.5251 | 0.408 | 0.48 | 1 | HC_vs_MS.NoTrt_F |
| ZNF22 | 0.04863 | 0.51073 | 0.118 | 0.18 | 1 | HC_vs_MS.NoTrt_F |
| SYNE1 | 0.04878 | 0.41482 | 0.965 | 0.96 | 1 | HC_vs_MS.NoTrt_F |
| LMLN | 0.0488 | -1.363 | 0.419 | 0.5 | 1 | HC_vs_MS.NoTrt_F |
| YTHDC1 | 0.04881 | -0.2732 | 0.512 | 0.6 | 1 | HC_vs_MS.NoTrt_F |
| SPACA6 | 0.04898 | 0.28832 | 0.893 | 0.91 | 1 | HC_vs_MS.NoTrt_F |
| NAIP | 0.04908 | -0.4847 | 0.743 | 0.76 | 1 | HC_vs_MS.NoTrt_F |
| FAM234A | 0.0491 | 0.28628 | 0.272 | 0.36 | 1 | HC_vs_MS.NoTrt_F |
| ACVR2A | 0.04911 | -0.4794 | 0.315 | 0.4 | 1 | HC_vs_MS.NoTrt_F |
| MCU | 0.04917 | 0.47329 | 0.257 | 0.35 | 1 | HC_vs_MS.NoTrt_F |
| MRPS9 | 0.04922 | 0.30224 | 0.168 | 0.24 | 1 | HC_vs_MS.NoTrt_F |
| PLRG1 | 0.04939 | 0.3074 | 0.145 | 0.22 | 1 | HC_vs_MS.NoTrt_F |
| GRIN2B | 0.04939 | 0.48207 | 0.237 | 0.32 | 1 | HC_vs_MS.NoTrt_F |
| CCDC25 | 0.04946 | -0.2976 | 0.286 | 0.37 | 1 | HC_vs_MS.NoTrt_F |
| SS18 | 0.0495 | -0.3058 | 0.647 | 0.7 | 1 | HC_vs_MS.NoTrt_F |

| | | | | | | |
|---|---|---|---|---|---|---|
| UBE2E3 | 0.04966 | 0.98848 | 0.072 | 0.13 | 1 | HC_vs_MS.NoTrt_F |
| NOB1 | 0.04972 | -0.4109 | 0.199 | 0.27 | 1 | HC_vs_MS.NoTrt_F |
| NEK8 | 0.04979 | -0.8507 | 0.621 | 0.67 | 1 | HC_vs_MS.NoTrt_F |
| ISOC2 | 0.04993 | 1.28075 | 0.072 | 0.13 | 1 | HC_vs_MS.NoTrt_F |
| AL109923.1 | 0.04994 | 0.36862 | 0.09 | 0.15 | 1 | HC_vs_MS.NoTrt_F |
| S100B | 0.05011 | 0.61637 | 0.153 | 0.23 | 1 | HC_vs_MS.NoTrt_F |
| TNS1 | 0.05016 | 0.972 | 0.39 | 0.31 | 1 | HC_vs_MS.NoTrt_F |
| NEIL2 | 0.05019 | 0.61478 | 0.124 | 0.19 | 1 | HC_vs_MS.NoTrt_F |
| SMG1 | 0.05024 | 0.33578 | 0.986 | 0.98 | 1 | HC_vs_MS.NoTrt_F |
| SH3BP1 | 0.05041 | -0.3777 | 0.156 | 0.23 | 1 | HC_vs_MS.NoTrt_F |
| NSFL1C | 0.05042 | 0.30006 | 0.173 | 0.25 | 1 | HC_vs_MS.NoTrt_F |
| SF3B4 | 0.05046 | -0.4734 | 0.243 | 0.32 | 1 | HC_vs_MS.NoTrt_F |
| CNNM2 | 0.05053 | 1.86139 | 0.341 | 0.43 | 1 | HC_vs_MS.NoTrt_F |
| ERF | 0.05065 | 0.99212 | 0.11 | 0.18 | 1 | HC_vs_MS.NoTrt_F |
| MFSD14C | 0.05069 | 0.30099 | 0.746 | 0.73 | 1 | HC_vs_MS.NoTrt_F |
| STMND1 | 0.05076 | 0.25786 | 0.867 | 0.83 | 1 | HC_vs_MS.NoTrt_F |
| PRKACA | 0.05085 | -0.3647 | 0.249 | 0.33 | 1 | HC_vs_MS.NoTrt_F |
| EPN3 | 0.05094 | 0.35034 | 0.081 | 0.14 | 1 | HC_vs_MS.NoTrt_F |
| ZNF569 | 0.05096 | 0.52441 | 0.228 | 0.3 | 1 | HC_vs_MS.NoTrt_F |
| AFF3 | 0.05099 | 0.75563 | 0.806 | 0.74 | 1 | HC_vs_MS.NoTrt_F |
| WDR5B | 0.05102 | -0.7684 | 0.24 | 0.3 | 1 | HC_vs_MS.NoTrt_F |
| CAPN15 | 0.05112 | 0.9615 | 0.205 | 0.29 | 1 | HC_vs_MS.NoTrt_F |
| TRIP12 | 0.05118 | -0.3096 | 0.543 | 0.6 | 1 | HC_vs_MS.NoTrt_F |
| ATMIN | 0.05124 | -0.3348 | 0.28 | 0.35 | 1 | HC_vs_MS.NoTrt_F |
| GEMIN8 | 0.05125 | 0.32013 | 0.078 | 0.13 | 1 | HC_vs_MS.NoTrt_F |
| RANBP9 | 0.0513 | 0.37249 | 0.266 | 0.37 | 1 | HC_vs_MS.NoTrt_F |
| CABYR | 0.05135 | 0.44016 | 0.584 | 0.51 | 1 | HC_vs_MS.NoTrt_F |
| HLA-DMB | 0.05136 | 0.33319 | 0.107 | 0.17 | 1 | HC_vs_MS.NoTrt_F |
| BAG3 | 0.05144 | 0.49279 | 0.118 | 0.19 | 1 | HC_vs_MS.NoTrt_F |
| ESD | 0.05154 | 0.31245 | 0.191 | 0.27 | 1 | HC_vs_MS.NoTrt_F |
| STARD9 | 0.05161 | 0.61589 | 0.931 | 0.91 | 1 | HC_vs_MS.NoTrt_F |
| OCLN | 0.05175 | 0.5049 | 0.812 | 0.82 | 1 | HC_vs_MS.NoTrt_F |
| ZNF770 | 0.05203 | -0.529 | 0.35 | 0.41 | 1 | HC_vs_MS.NoTrt_F |
| AMOTL1 | 0.05204 | -0.9341 | 0.075 | 0.13 | 1 | HC_vs_MS.NoTrt_F |
| ZNF181 | 0.0521 | -1.4208 | 0.194 | 0.27 | 1 | HC_vs_MS.NoTrt_F |
| DCTN3 | 0.0522 | -0.2943 | 0.402 | 0.49 | 1 | HC_vs_MS.NoTrt_F |
| RCAN1 | 0.05238 | 0.65579 | 0.39 | 0.48 | 1 | HC_vs_MS.NoTrt_F |
| ADCY3 | 0.05242 | 0.31732 | 0.11 | 0.17 | 1 | HC_vs_MS.NoTrt_F |
| HAUS7 | 0.05244 | -0.9888 | 0.147 | 0.21 | 1 | HC_vs_MS.NoTrt_F |
| GSG1 | 0.05266 | 0.54104 | 0.627 | 0.59 | 1 | HC_vs_MS.NoTrt_F |

| | | | | | | |
|---|---|---|---|---|---|---|
| RNF213 | 0.05266 | 0.26904 | 0.994 | 0.99 | 1 | HC_vs_MS.NoTrt_F |
| UNC45B | 0.05299 | -0.7097 | 0.142 | 0.21 | 1 | HC_vs_MS.NoTrt_F |
| NCOR2 | 0.0531 | 1.20536 | 0.552 | 0.5 | 1 | HC_vs_MS.NoTrt_F |
| TRAK1 | 0.05313 | 0.40952 | 0.136 | 0.19 | 1 | HC_vs_MS.NoTrt_F |
| SLC13A4 | 0.05337 | 0.55235 | 0.373 | 0.46 | 1 | HC_vs_MS.NoTrt_F |
| CLASP1 | 0.05357 | 0.43383 | 0.312 | 0.4 | 1 | HC_vs_MS.NoTrt_F |
| PRAF2 | 0.0536 | 0.51614 | 0.136 | 0.2 | 1 | HC_vs_MS.NoTrt_F |
| AEN | 0.05363 | 0.89404 | 0.153 | 0.23 | 1 | HC_vs_MS.NoTrt_F |
| PIP5KL1 | 0.05371 | 0.40795 | 0.503 | 0.58 | 1 | HC_vs_MS.NoTrt_F |
| CGNL1 | 0.05383 | -0.5894 | 0.347 | 0.41 | 1 | HC_vs_MS.NoTrt_F |
| PAXBP1 | 0.05384 | -0.5848 | 0.223 | 0.29 | 1 | HC_vs_MS.NoTrt_F |
| C4BPA | 0.05384 | -0.3165 | 0.361 | 0.45 | 1 | HC_vs_MS.NoTrt_F |
| SHPRH | 0.05385 | -0.3894 | 0.751 | 0.77 | 1 | HC_vs_MS.NoTrt_F |
| ZNF254 | 0.05389 | -0.4182 | 0.416 | 0.48 | 1 | HC_vs_MS.NoTrt_F |
| GIN1 | 0.05399 | -0.582 | 0.379 | 0.46 | 1 | HC_vs_MS.NoTrt_F |
| XRCC1 | 0.05408 | 0.48151 | 0.228 | 0.31 | 1 | HC_vs_MS.NoTrt_F |
| ZNF195 | 0.0541 | 0.54142 | 0.621 | 0.67 | 1 | HC_vs_MS.NoTrt_F |
| PFAS | 0.05428 | -0.5493 | 0.662 | 0.58 | 1 | HC_vs_MS.NoTrt_F |
| GPX1 | 0.05428 | 0.35926 | 0.205 | 0.29 | 1 | HC_vs_MS.NoTrt_F |
| LSM2 | 0.05428 | 0.256 | 0.197 | 0.28 | 1 | HC_vs_MS.NoTrt_F |
| ZC4H2 | 0.05437 | 1.06374 | 0.107 | 0.17 | 1 | HC_vs_MS.NoTrt_F |
| SLC11A1 | 0.05445 | 0.30653 | 0.954 | 0.93 | 1 | HC_vs_MS.NoTrt_F |
| WDR48 | 0.05459 | -0.3809 | 0.211 | 0.28 | 1 | HC_vs_MS.NoTrt_F |
| MIS18BP1 | 0.0546 | -0.9071 | 0.679 | 0.71 | 1 | HC_vs_MS.NoTrt_F |
| FASTK | 0.05461 | 0.422 | 0.188 | 0.27 | 1 | HC_vs_MS.NoTrt_F |
| UQCR10 | 0.05462 | 0.30309 | 0.364 | 0.48 | 1 | HC_vs_MS.NoTrt_F |
| TAB1 | 0.05465 | 0.28221 | 0.121 | 0.18 | 1 | HC_vs_MS.NoTrt_F |
| INPPL1 | 0.05476 | 0.8225 | 0.121 | 0.19 | 1 | HC_vs_MS.NoTrt_F |
| TOM1 | 0.05487 | 0.28151 | 0.335 | 0.43 | 1 | HC_vs_MS.NoTrt_F |
| TMUB1 | 0.0549 | 0.53994 | 0.139 | 0.21 | 1 | HC_vs_MS.NoTrt_F |
| CDC34 | 0.05495 | 0.54298 | 0.061 | 0.11 | 1 | HC_vs_MS.NoTrt_F |
| BACH1 | 0.05499 | -0.378 | 0.303 | 0.39 | 1 | HC_vs_MS.NoTrt_F |
| OSTM1 | 0.05501 | -0.8822 | 0.795 | 0.85 | 1 | HC_vs_MS.NoTrt_F |
| TRMT1 | 0.05516 | 0.26964 | 0.162 | 0.23 | 1 | HC_vs_MS.NoTrt_F |
| SNAP29 | 0.05521 | -0.5893 | 0.72 | 0.77 | 1 | HC_vs_MS.NoTrt_F |
| PSMA2 | 0.05533 | 0.34976 | 0.595 | 0.66 | 1 | HC_vs_MS.NoTrt_F |
| PARP15 | 0.05538 | 0.53411 | 0.942 | 0.93 | 1 | HC_vs_MS.NoTrt_F |
| ASB7 | 0.05549 | 0.76987 | 0.11 | 0.17 | 1 | HC_vs_MS.NoTrt_F |
| NR1H3 | 0.0555 | 1.82807 | 0.15 | 0.22 | 1 | HC_vs_MS.NoTrt_F |
| CASS4 | 0.05572 | 0.68571 | 0.566 | 0.49 | 1 | HC_vs_MS.NoTrt_F |

| | | | | | | |
|---|---|---|---|---|---|---|
| NSG1 | 0.05581 | -0.2653 | 0.668 | 0.74 | 1 | HC_vs_MS.NoTrt_F |
| RP11-701H2 | 0.05584 | -0.4837 | 0.136 | 0.19 | 1 | HC_vs_MS.NoTrt_F |
| C16orf54 | 0.0559 | 0.2529 | 0.737 | 0.87 | 1 | HC_vs_MS.NoTrt_F |
| EPHA1 | 0.05591 | 0.44058 | 0.39 | 0.48 | 1 | HC_vs_MS.NoTrt_F |
| CASP7 | 0.05595 | -0.567 | 0.191 | 0.26 | 1 | HC_vs_MS.NoTrt_F |
| APOPT1 | 0.05605 | 0.26966 | 0.931 | 0.9 | 1 | HC_vs_MS.NoTrt_F |
| TRPS1 | 0.05621 | 0.3587 | 0.127 | 0.19 | 1 | HC_vs_MS.NoTrt_F |
| TDRD1 | 0.05623 | 0.70821 | 0.225 | 0.3 | 1 | HC_vs_MS.NoTrt_F |
| SORBS3 | 0.05623 | 1.06598 | 0.078 | 0.13 | 1 | HC_vs_MS.NoTrt_F |
| PNKP | 0.05624 | 0.8187 | 0.327 | 0.41 | 1 | HC_vs_MS.NoTrt_F |
| ZNF571 | 0.0563 | -0.6923 | 0.434 | 0.54 | 1 | HC_vs_MS.NoTrt_F |
| KIAA1143 | 0.05638 | -0.6561 | 0.523 | 0.61 | 1 | HC_vs_MS.NoTrt_F |
| ATG101 | 0.05646 | -0.4185 | 0.113 | 0.17 | 1 | HC_vs_MS.NoTrt_F |
| SLC25A6 | 0.05661 | 0.26453 | 0.523 | 0.7 | 1 | HC_vs_MS.NoTrt_F |
| STRBP | 0.05665 | -0.5709 | 0.65 | 0.72 | 1 | HC_vs_MS.NoTrt_F |
| NCAPG2 | 0.05669 | 1.04254 | 0.202 | 0.28 | 1 | HC_vs_MS.NoTrt_F |
| AP1AR | 0.05675 | 0.32588 | 0.107 | 0.17 | 1 | HC_vs_MS.NoTrt_F |
| SMC4 | 0.05694 | -0.3944 | 0.483 | 0.55 | 1 | HC_vs_MS.NoTrt_F |
| SH2D2A | 0.057 | 1.08367 | 0.488 | 0.47 | 1 | HC_vs_MS.NoTrt_F |
| SGSM1 | 0.057 | -0.3033 | 0.543 | 0.58 | 1 | HC_vs_MS.NoTrt_F |
| ERO1B | 0.05702 | 0.61603 | 0.133 | 0.2 | 1 | HC_vs_MS.NoTrt_F |
| ETS1 | 0.05708 | -0.3027 | 0.986 | 0.98 | 1 | HC_vs_MS.NoTrt_F |
| ARHGEF2 | 0.05709 | 0.48022 | 0.275 | 0.35 | 1 | HC_vs_MS.NoTrt_F |
| CRADD | 0.05714 | 0.57948 | 0.107 | 0.17 | 1 | HC_vs_MS.NoTrt_F |
| YY1 | 0.05718 | 0.39838 | 0.35 | 0.43 | 1 | HC_vs_MS.NoTrt_F |
| TMEM185A | 0.05721 | 0.26761 | 0.587 | 0.48 | 1 | HC_vs_MS.NoTrt_F |
| HIST1H3E | 0.05733 | -0.6568 | 0.136 | 0.2 | 1 | HC_vs_MS.NoTrt_F |
| UBL3 | 0.05737 | 0.30621 | 0.197 | 0.27 | 1 | HC_vs_MS.NoTrt_F |
| AHCY | 0.05737 | 0.49948 | 0.098 | 0.16 | 1 | HC_vs_MS.NoTrt_F |
| IDI1 | 0.05747 | 0.25691 | 0.243 | 0.33 | 1 | HC_vs_MS.NoTrt_F |
| RBM10 | 0.05762 | 0.29276 | 0.205 | 0.28 | 1 | HC_vs_MS.NoTrt_F |
| SNAPC1 | 0.05762 | -0.3628 | 0.705 | 0.82 | 1 | HC_vs_MS.NoTrt_F |
| NOL10 | 0.05769 | -0.4085 | 0.202 | 0.27 | 1 | HC_vs_MS.NoTrt_F |
| KIAA1210 | 0.05773 | -0.2657 | 0.275 | 0.34 | 1 | HC_vs_MS.NoTrt_F |
| CLIP4 | 0.05774 | -0.336 | 0.162 | 0.23 | 1 | HC_vs_MS.NoTrt_F |
| SMAGP | 0.05776 | -0.4584 | 0.636 | 0.72 | 1 | HC_vs_MS.NoTrt_F |
| GSDMB | 0.0578 | 0.52789 | 0.908 | 0.91 | 1 | HC_vs_MS.NoTrt_F |
| TMEM5 | 0.05786 | 0.91697 | 0.523 | 0.6 | 1 | HC_vs_MS.NoTrt_F |
| RBPMS | 0.05789 | 1.81681 | 0.199 | 0.14 | 1 | HC_vs_MS.NoTrt_F |
| LIAS | 0.0579 | -0.4368 | 0.133 | 0.19 | 1 | HC_vs_MS.NoTrt_F |

| | | | | | | |
|---|---|---|---|---|---|---|
| INTS10 | 0.05797 | -0.5481 | 0.613 | 0.69 | 1 | HC_vs_MS.NoTrt_F |
| FAM162A | 0.05798 | -0.3513 | 0.439 | 0.5 | 1 | HC_vs_MS.NoTrt_F |
| CNOT10 | 0.05805 | 0.42898 | 0.936 | 0.92 | 1 | HC_vs_MS.NoTrt_F |
| CCDC34 | 0.05806 | 0.36492 | 0.118 | 0.18 | 1 | HC_vs_MS.NoTrt_F |
| UTS2 | 0.0581 | 0.32894 | 0.124 | 0.19 | 1 | HC_vs_MS.NoTrt_F |
| COQ9 | 0.0582 | 0.2692 | 0.113 | 0.17 | 1 | HC_vs_MS.NoTrt_F |
| FUNDC2 | 0.05821 | -0.3399 | 0.561 | 0.62 | 1 | HC_vs_MS.NoTrt_F |
| STK25 | 0.05823 | -0.97 | 0.76 | 0.78 | 1 | HC_vs_MS.NoTrt_F |
| TRMT2B | 0.05839 | -0.4771 | 0.731 | 0.81 | 1 | HC_vs_MS.NoTrt_F |
| PHF19 | 0.05847 | 0.27591 | 0.191 | 0.27 | 1 | HC_vs_MS.NoTrt_F |
| SNX22 | 0.05847 | -0.3185 | 0.838 | 0.81 | 1 | HC_vs_MS.NoTrt_F |
| NFIA | 0.05849 | 0.90107 | 0.156 | 0.1 | 1 | HC_vs_MS.NoTrt_F |
| TICAM1 | 0.05851 | 0.5537 | 0.095 | 0.16 | 1 | HC_vs_MS.NoTrt_F |
| TSC1 | 0.05853 | 0.66732 | 0.48 | 0.56 | 1 | HC_vs_MS.NoTrt_F |
| SNAPC2 | 0.05868 | 0.50908 | 0.087 | 0.14 | 1 | HC_vs_MS.NoTrt_F |
| RNF144B | 0.05879 | 0.37215 | 0.312 | 0.41 | 1 | HC_vs_MS.NoTrt_F |
| AKAP10 | 0.05889 | -0.2714 | 0.538 | 0.57 | 1 | HC_vs_MS.NoTrt_F |
| PRR14 | 0.05892 | 0.58377 | 0.153 | 0.22 | 1 | HC_vs_MS.NoTrt_F |
| ZSCAN22 | 0.05895 | -1.0739 | 0.48 | 0.57 | 1 | HC_vs_MS.NoTrt_F |
| ACADM | 0.05901 | 0.35487 | 0.231 | 0.32 | 1 | HC_vs_MS.NoTrt_F |
| VPS41 | 0.05903 | -0.2974 | 0.517 | 0.58 | 1 | HC_vs_MS.NoTrt_F |
| TRIP13 | 0.05905 | 0.32317 | 0.546 | 0.61 | 1 | HC_vs_MS.NoTrt_F |
| FABP7 | 0.0591 | -0.2698 | 0.197 | 0.26 | 1 | HC_vs_MS.NoTrt_F |
| IFI44L | 0.05914 | 1.07098 | 0.228 | 0.32 | 1 | HC_vs_MS.NoTrt_F |
| PDXK | 0.05931 | 1.08199 | 0.63 | 0.56 | 1 | HC_vs_MS.NoTrt_F |
| FGF23 | 0.05932 | 0.69161 | 0.925 | 0.85 | 1 | HC_vs_MS.NoTrt_F |
| STRN4 | 0.05935 | 1.03331 | 0.598 | 0.55 | 1 | HC_vs_MS.NoTrt_F |
| ACTG2 | 0.05942 | 0.73351 | 0.277 | 0.35 | 1 | HC_vs_MS.NoTrt_F |
| CYP4F8 | 0.05951 | 0.26835 | 0.847 | 0.8 | 1 | HC_vs_MS.NoTrt_F |
| RUNDC3A | 0.05954 | 0.41021 | 0.564 | 0.51 | 1 | HC_vs_MS.NoTrt_F |
| PCID2 | 0.05959 | -0.3563 | 0.361 | 0.43 | 1 | HC_vs_MS.NoTrt_F |
| LTN1 | 0.05972 | 0.67311 | 0.197 | 0.28 | 1 | HC_vs_MS.NoTrt_F |
| MRPL39 | 0.05987 | 0.42767 | 0.159 | 0.23 | 1 | HC_vs_MS.NoTrt_F |
| NOL7 | 0.05989 | 0.58888 | 0.257 | 0.36 | 1 | HC_vs_MS.NoTrt_F |
| NUDT21 | 0.05991 | -0.3772 | 0.725 | 0.8 | 1 | HC_vs_MS.NoTrt_F |
| CD300LG | 0.05998 | 0.812 | 0.523 | 0.46 | 1 | HC_vs_MS.NoTrt_F |
| TWISTNB | 0.06 | -0.4766 | 0.136 | 0.2 | 1 | HC_vs_MS.NoTrt_F |
| TATDN2 | 0.06002 | 0.25054 | 0.13 | 0.19 | 1 | HC_vs_MS.NoTrt_F |
| PALB2 | 0.06004 | -0.9604 | 0.254 | 0.32 | 1 | HC_vs_MS.NoTrt_F |
| EIF5A2 | 0.06018 | -0.4367 | 0.419 | 0.46 | 1 | HC_vs_MS.NoTrt_F |

| | | | | | | |
|---|---|---|---|---|---|---|
| CDC42BPB | 0.06021 | 0.65315 | 0.081 | 0.13 | 1 | HC_vs_MS.NoTrt_F |
| C16orf72 | 0.06022 | 0.529 | 0.887 | 0.85 | 1 | HC_vs_MS.NoTrt_F |
| PARP6 | 0.06039 | -0.2741 | 0.697 | 0.8 | 1 | HC_vs_MS.NoTrt_F |
| SLC25A33 | 0.06049 | 0.65324 | 0.656 | 0.56 | 1 | HC_vs_MS.NoTrt_F |
| DCAF16 | 0.06052 | -0.4181 | 0.812 | 0.82 | 1 | HC_vs_MS.NoTrt_F |
| RASL10B | 0.06056 | 0.2625 | 0.977 | 0.98 | 1 | HC_vs_MS.NoTrt_F |
| PLAC8 | 0.06058 | 0.82378 | 0.809 | 0.79 | 1 | HC_vs_MS.NoTrt_F |
| RING1 | 0.06071 | 0.7473 | 0.104 | 0.17 | 1 | HC_vs_MS.NoTrt_F |
| HSPD1 | 0.06076 | 0.2744 | 0.303 | 0.4 | 1 | HC_vs_MS.NoTrt_F |
| DPPA4 | 0.06079 | 0.30316 | 0.864 | 0.84 | 1 | HC_vs_MS.NoTrt_F |
| WAPL | 0.06099 | -0.5783 | 0.523 | 0.58 | 1 | HC_vs_MS.NoTrt_F |
| COASY | 0.0611 | -0.3381 | 0.46 | 0.53 | 1 | HC_vs_MS.NoTrt_F |
| SGTB | 0.06125 | -0.3105 | 0.694 | 0.74 | 1 | HC_vs_MS.NoTrt_F |
| FANCL | 0.06126 | -0.7175 | 0.173 | 0.24 | 1 | HC_vs_MS.NoTrt_F |
| AGAP4 | 0.06127 | 0.35417 | 0.254 | 0.34 | 1 | HC_vs_MS.NoTrt_F |
| PDIA4 | 0.06148 | 1.51041 | 0.101 | 0.16 | 1 | HC_vs_MS.NoTrt_F |
| RAB27A | 0.06151 | -0.282 | 0.673 | 0.77 | 1 | HC_vs_MS.NoTrt_F |
| NDRG1 | 0.06158 | 0.62699 | 0.191 | 0.28 | 1 | HC_vs_MS.NoTrt_F |
| TP53I13 | 0.0616 | 0.37031 | 0.179 | 0.25 | 1 | HC_vs_MS.NoTrt_F |
| MVB12B | 0.06162 | 0.26417 | 0.159 | 0.22 | 1 | HC_vs_MS.NoTrt_F |
| SEC11C | 0.0618 | 0.26061 | 0.434 | 0.51 | 1 | HC_vs_MS.NoTrt_F |
| RP11-226E2 | 0.06187 | 0.36738 | 0.639 | 0.74 | 1 | HC_vs_MS.NoTrt_F |
| COQ3 | 0.06192 | 0.89513 | 0.251 | 0.33 | 1 | HC_vs_MS.NoTrt_F |
| AMMECR1 | 0.06198 | -0.4534 | 0.159 | 0.23 | 1 | HC_vs_MS.NoTrt_F |
| SLC22A23 | 0.06199 | 0.65292 | 0.101 | 0.16 | 1 | HC_vs_MS.NoTrt_F |
| TAF10 | 0.06231 | 0.27176 | 0.133 | 0.2 | 1 | HC_vs_MS.NoTrt_F |
| FAM104B | 0.06232 | 0.4364 | 0.087 | 0.14 | 1 | HC_vs_MS.NoTrt_F |
| THUMPD2 | 0.06242 | 0.46025 | 0.41 | 0.5 | 1 | HC_vs_MS.NoTrt_F |
| UGDH | 0.06243 | -0.7922 | 0.584 | 0.67 | 1 | HC_vs_MS.NoTrt_F |
| CACNB3 | 0.06247 | 0.27501 | 0.633 | 0.7 | 1 | HC_vs_MS.NoTrt_F |
| ETV3 | 0.06252 | -0.2601 | 0.243 | 0.32 | 1 | HC_vs_MS.NoTrt_F |
| EFHD2 | 0.06269 | 0.33797 | 0.182 | 0.26 | 1 | HC_vs_MS.NoTrt_F |
| DEGS1 | 0.06283 | -0.4483 | 0.841 | 0.88 | 1 | HC_vs_MS.NoTrt_F |
| SERPINI1 | 0.06289 | -0.4826 | 0.095 | 0.15 | 1 | HC_vs_MS.NoTrt_F |
| PAG1 | 0.06305 | -0.3882 | 0.757 | 0.8 | 1 | HC_vs_MS.NoTrt_F |
| OPRM1 | 0.06306 | 0.30284 | 0.89 | 0.87 | 1 | HC_vs_MS.NoTrt_F |
| TCF4 | 0.06309 | 0.70212 | 0.818 | 0.76 | 1 | HC_vs_MS.NoTrt_F |
| MYO9A | 0.06312 | -0.3162 | 0.402 | 0.49 | 1 | HC_vs_MS.NoTrt_F |
| MRPL46 | 0.06346 | -0.5983 | 0.26 | 0.33 | 1 | HC_vs_MS.NoTrt_F |
| NRF1 | 0.06346 | -0.8899 | 0.358 | 0.44 | 1 | HC_vs_MS.NoTrt_F |

| | | | | | | |
|---|---|---|---|---|---|---|
| GHRL | 0.06354 | 0.72689 | 0.107 | 0.17 | 1 | HC_vs_MS.NoTrt_F |
| TBC1D20 | 0.06359 | 0.35481 | 0.694 | 0.61 | 1 | HC_vs_MS.NoTrt_F |
| INIP | 0.06367 | -0.7128 | 0.757 | 0.81 | 1 | HC_vs_MS.NoTrt_F |
| ERCC6 | 0.06372 | -0.3497 | 0.275 | 0.34 | 1 | HC_vs_MS.NoTrt_F |
| GALK1 | 0.06403 | 1.52264 | 0.402 | 0.33 | 1 | HC_vs_MS.NoTrt_F |
| FAAP20 | 0.06406 | 0.29108 | 0.91 | 0.87 | 1 | HC_vs_MS.NoTrt_F |
| NRAS | 0.06417 | 0.6194 | 0.361 | 0.46 | 1 | HC_vs_MS.NoTrt_F |
| FBRS | 0.06424 | 0.36579 | 0.185 | 0.27 | 1 | HC_vs_MS.NoTrt_F |
| UBQLN1 | 0.06428 | -0.2757 | 0.306 | 0.39 | 1 | HC_vs_MS.NoTrt_F |
| FAM134A | 0.06447 | 0.26986 | 0.428 | 0.5 | 1 | HC_vs_MS.NoTrt_F |
| CAPRIN2 | 0.06478 | -0.3625 | 0.757 | 0.8 | 1 | HC_vs_MS.NoTrt_F |
| REXO1 | 0.06487 | 0.99149 | 0.118 | 0.18 | 1 | HC_vs_MS.NoTrt_F |
| NOP16 | 0.06495 | 0.4359 | 0.228 | 0.3 | 1 | HC_vs_MS.NoTrt_F |
| TOB2 | 0.06502 | 0.43408 | 0.159 | 0.23 | 1 | HC_vs_MS.NoTrt_F |
| SIN3B | 0.06514 | 0.76619 | 0.812 | 0.78 | 1 | HC_vs_MS.NoTrt_F |
| HS2ST1 | 0.06543 | -0.5357 | 0.127 | 0.18 | 1 | HC_vs_MS.NoTrt_F |
| CMTM3 | 0.06551 | 0.57308 | 0.243 | 0.33 | 1 | HC_vs_MS.NoTrt_F |
| ZMIZ2 | 0.06551 | 0.35137 | 0.199 | 0.28 | 1 | HC_vs_MS.NoTrt_F |
| PHF20L1 | 0.06557 | -0.7009 | 0.662 | 0.69 | 1 | HC_vs_MS.NoTrt_F |
| EPPIN | 0.06557 | 0.37432 | 0.627 | 0.56 | 1 | HC_vs_MS.NoTrt_F |
| CYP7B1 | 0.06563 | 0.49018 | 0.457 | 0.54 | 1 | HC_vs_MS.NoTrt_F |
| PSMG1 | 0.06566 | -0.2642 | 0.188 | 0.26 | 1 | HC_vs_MS.NoTrt_F |
| RNF157 | 0.06567 | -0.251 | 0.653 | 0.74 | 1 | HC_vs_MS.NoTrt_F |
| XAF1 | 0.06569 | 0.37835 | 0.844 | 0.84 | 1 | HC_vs_MS.NoTrt_F |
| ERI3 | 0.06582 | -0.4715 | 0.142 | 0.2 | 1 | HC_vs_MS.NoTrt_F |
| GIPR | 0.06587 | 0.29968 | 0.627 | 0.71 | 1 | HC_vs_MS.NoTrt_F |
| HCFC1R1 | 0.06588 | -0.2782 | 0.15 | 0.22 | 1 | HC_vs_MS.NoTrt_F |
| LIN37 | 0.06618 | 0.30181 | 0.104 | 0.16 | 1 | HC_vs_MS.NoTrt_F |
| CHD9 | 0.06645 | -0.5263 | 0.477 | 0.54 | 1 | HC_vs_MS.NoTrt_F |
| ADAM19 | 0.06665 | 0.33538 | 0.419 | 0.55 | 1 | HC_vs_MS.NoTrt_F |
| WARS2 | 0.06665 | 0.44447 | 0.11 | 0.17 | 1 | HC_vs_MS.NoTrt_F |
| CLTB | 0.06698 | 0.5445 | 0.107 | 0.17 | 1 | HC_vs_MS.NoTrt_F |
| FAM20A | 0.06709 | -0.8292 | 0.171 | 0.24 | 1 | HC_vs_MS.NoTrt_F |
| GSTM1 | 0.06722 | 0.60077 | 0.194 | 0.27 | 1 | HC_vs_MS.NoTrt_F |
| IER3IP1 | 0.06728 | 0.31285 | 0.182 | 0.25 | 1 | HC_vs_MS.NoTrt_F |
| FAM188B | 0.06767 | 0.52146 | 0.182 | 0.25 | 1 | HC_vs_MS.NoTrt_F |
| MRPL30 | 0.06771 | -0.6222 | 0.827 | 0.88 | 1 | HC_vs_MS.NoTrt_F |
| KLRC1 | 0.06778 | 0.5838 | 0.338 | 0.44 | 1 | HC_vs_MS.NoTrt_F |
| IVD | 0.06804 | 0.42429 | 0.954 | 0.93 | 1 | HC_vs_MS.NoTrt_F |
| ZC3H18 | 0.06821 | 0.30614 | 0.766 | 0.83 | 1 | HC_vs_MS.NoTrt_F |

| | | | | | | |
|---|---|---|---|---|---|---|
| CCDC6 | 0.0683 | 0.97061 | 0.121 | 0.18 | 1 | HC_vs_MS.NoTrt_F |
| DHX8 | 0.06833 | -0.4796 | 0.691 | 0.73 | 1 | HC_vs_MS.NoTrt_F |
| NDUFAB1 | 0.06838 | 0.26127 | 0.275 | 0.36 | 1 | HC_vs_MS.NoTrt_F |
| PAPSS2 | 0.06839 | 0.50766 | 0.072 | 0.12 | 1 | HC_vs_MS.NoTrt_F |
| TMED3 | 0.06844 | 0.63417 | 0.699 | 0.65 | 1 | HC_vs_MS.NoTrt_F |
| RNF25 | 0.06845 | 0.31406 | 0.113 | 0.17 | 1 | HC_vs_MS.NoTrt_F |
| NPIPA5 | 0.06847 | 0.34942 | 0.104 | 0.16 | 1 | HC_vs_MS.NoTrt_F |
| DPH1 | 0.06848 | 0.35772 | 0.91 | 0.88 | 1 | HC_vs_MS.NoTrt_F |
| DAZAP1 | 0.0685 | -0.2666 | 0.194 | 0.27 | 1 | HC_vs_MS.NoTrt_F |
| MTERF3 | 0.06888 | -1.1351 | 0.283 | 0.34 | 1 | HC_vs_MS.NoTrt_F |
| SCO1 | 0.06889 | -0.5371 | 0.85 | 0.85 | 1 | HC_vs_MS.NoTrt_F |
| APOLD1 | 0.06897 | 0.63349 | 0.566 | 0.52 | 1 | HC_vs_MS.NoTrt_F |
| HTRA4 | 0.06897 | 0.30339 | 0.535 | 0.46 | 1 | HC_vs_MS.NoTrt_F |
| ZGRF1 | 0.06898 | 0.55885 | 0.85 | 0.83 | 1 | HC_vs_MS.NoTrt_F |
| CDR2 | 0.06898 | 1.34938 | 0.168 | 0.24 | 1 | HC_vs_MS.NoTrt_F |
| SPCS3 | 0.06907 | 0.27489 | 0.512 | 0.61 | 1 | HC_vs_MS.NoTrt_F |
| DNAL1 | 0.06913 | 0.48262 | 0.951 | 0.94 | 1 | HC_vs_MS.NoTrt_F |
| CIITA | 0.06918 | -0.4382 | 0.962 | 0.9 | 1 | HC_vs_MS.NoTrt_F |
| VBP1 | 0.06919 | 0.66572 | 0.142 | 0.21 | 1 | HC_vs_MS.NoTrt_F |
| GGCX | 0.06922 | 0.80139 | 0.934 | 0.95 | 1 | HC_vs_MS.NoTrt_F |
| FAM46A | 0.06938 | -0.3846 | 0.113 | 0.17 | 1 | HC_vs_MS.NoTrt_F |
| UNK | 0.06946 | 0.36122 | 0.127 | 0.19 | 1 | HC_vs_MS.NoTrt_F |
| MIIP | 0.06946 | 0.27034 | 0.199 | 0.28 | 1 | HC_vs_MS.NoTrt_F |
| ZCCHC9 | 0.06955 | 0.27605 | 0.775 | 0.68 | 1 | HC_vs_MS.NoTrt_F |
| POU2AF1 | 0.06967 | 0.54925 | 0.766 | 0.73 | 1 | HC_vs_MS.NoTrt_F |
| SNRPG | 0.06996 | 0.37894 | 0.329 | 0.45 | 1 | HC_vs_MS.NoTrt_F |
| HS1BP3 | 0.07007 | -0.3639 | 0.188 | 0.25 | 1 | HC_vs_MS.NoTrt_F |
| SVIL | 0.07048 | 1.21552 | 0.185 | 0.26 | 1 | HC_vs_MS.NoTrt_F |
| VAT1 | 0.07052 | 0.41775 | 0.272 | 0.19 | 1 | HC_vs_MS.NoTrt_F |
| UAP1 | 0.07092 | 0.76553 | 0.092 | 0.15 | 1 | HC_vs_MS.NoTrt_F |
| UBL4A | 0.07123 | 0.32537 | 0.118 | 0.18 | 1 | HC_vs_MS.NoTrt_F |
| UBP1 | 0.07135 | 0.69815 | 0.13 | 0.19 | 1 | HC_vs_MS.NoTrt_F |
| UNC50 | 0.07139 | 0.42307 | 0.142 | 0.2 | 1 | HC_vs_MS.NoTrt_F |
| PDE3B | 0.07139 | -0.6067 | 0.613 | 0.63 | 1 | HC_vs_MS.NoTrt_F |
| SLC25A46 | 0.07159 | 0.52013 | 0.162 | 0.23 | 1 | HC_vs_MS.NoTrt_F |
| BTBD11 | 0.07161 | 0.97063 | 0.48 | 0.59 | 1 | HC_vs_MS.NoTrt_F |
| TMX3 | 0.07182 | -0.3261 | 0.462 | 0.56 | 1 | HC_vs_MS.NoTrt_F |
| AHSA1 | 0.07214 | -0.6292 | 0.587 | 0.61 | 1 | HC_vs_MS.NoTrt_F |
| DENND6A | 0.07238 | -0.2806 | 0.595 | 0.66 | 1 | HC_vs_MS.NoTrt_F |
| SPOP | 0.07253 | -0.3745 | 0.98 | 0.98 | 1 | HC_vs_MS.NoTrt_F |

| | | | | | | |
|---|---|---|---|---|---|---|
| E2F6 | 0.07254 | -0.3853 | 0.272 | 0.34 | 1 | HC_vs_MS.NoTrt_F |
| CECR1 | 0.07269 | -0.3226 | 0.503 | 0.59 | 1 | HC_vs_MS.NoTrt_F |
| FBXO15 | 0.07287 | 0.42441 | 0.665 | 0.61 | 1 | HC_vs_MS.NoTrt_F |
| EXOC1 | 0.07294 | 0.35861 | 0.22 | 0.29 | 1 | HC_vs_MS.NoTrt_F |
| UBXN7 | 0.07295 | 0.35812 | 0.795 | 0.88 | 1 | HC_vs_MS.NoTrt_F |
| MRE11A | 0.07299 | -0.4659 | 0.743 | 0.82 | 1 | HC_vs_MS.NoTrt_F |
| FAM219A | 0.07301 | 0.67496 | 0.075 | 0.13 | 1 | HC_vs_MS.NoTrt_F |
| PARVB | 0.07313 | 0.42841 | 0.749 | 0.73 | 1 | HC_vs_MS.NoTrt_F |
| CSMD1 | 0.07314 | 0.91582 | 0.361 | 0.29 | 1 | HC_vs_MS.NoTrt_F |
| JMY | 0.07319 | 0.78857 | 0.353 | 0.45 | 1 | HC_vs_MS.NoTrt_F |
| NDUFS2 | 0.07323 | -0.255 | 0.454 | 0.5 | 1 | HC_vs_MS.NoTrt_F |
| MANF | 0.07333 | 0.56831 | 0.064 | 0.11 | 1 | HC_vs_MS.NoTrt_F |
| ANAPC7 | 0.07333 | 0.39962 | 0.59 | 0.67 | 1 | HC_vs_MS.NoTrt_F |
| MINPP1 | 0.07354 | 0.37599 | 0.104 | 0.16 | 1 | HC_vs_MS.NoTrt_F |
| CLCN6 | 0.07382 | 1.26025 | 0.59 | 0.53 | 1 | HC_vs_MS.NoTrt_F |
| PARN | 0.07413 | -0.7027 | 0.746 | 0.79 | 1 | HC_vs_MS.NoTrt_F |
| SLC25A16 | 0.07448 | 0.26506 | 0.974 | 0.98 | 1 | HC_vs_MS.NoTrt_F |
| HSD11B1L | 0.07451 | 0.36218 | 0.731 | 0.7 | 1 | HC_vs_MS.NoTrt_F |
| AP3S1 | 0.07456 | -0.3032 | 0.413 | 0.45 | 1 | HC_vs_MS.NoTrt_F |
| BTLA | 0.07464 | 0.74129 | 0.22 | 0.28 | 1 | HC_vs_MS.NoTrt_F |
| CFH | 0.07466 | 0.37646 | 0.127 | 0.19 | 1 | HC_vs_MS.NoTrt_F |
| LINC01125 | 0.0747 | 0.41316 | 0.957 | 0.96 | 1 | HC_vs_MS.NoTrt_F |
| HIP1R | 0.07483 | 1.11255 | 0.11 | 0.17 | 1 | HC_vs_MS.NoTrt_F |
| RGL4 | 0.07491 | 0.73108 | 0.263 | 0.36 | 1 | HC_vs_MS.NoTrt_F |
| SOAT1 | 0.07535 | -0.3765 | 0.743 | 0.79 | 1 | HC_vs_MS.NoTrt_F |
| IFITM2 | 0.07536 | 0.53938 | 0.824 | 0.81 | 1 | HC_vs_MS.NoTrt_F |
| SLC35E4 | 0.07546 | 1.03973 | 0.361 | 0.45 | 1 | HC_vs_MS.NoTrt_F |
| TMEM135 | 0.0755 | 0.4286 | 0.803 | 0.83 | 1 | HC_vs_MS.NoTrt_F |
| DUS1L | 0.07558 | 1.16849 | 0.118 | 0.18 | 1 | HC_vs_MS.NoTrt_F |
| FAM156A | 0.07589 | -0.6307 | 0.208 | 0.27 | 1 | HC_vs_MS.NoTrt_F |
| PGM1 | 0.07593 | 0.43701 | 0.127 | 0.19 | 1 | HC_vs_MS.NoTrt_F |
| SBNO2 | 0.07625 | 0.3962 | 0.228 | 0.3 | 1 | HC_vs_MS.NoTrt_F |
| TOPBP1 | 0.07629 | 1.19191 | 0.133 | 0.19 | 1 | HC_vs_MS.NoTrt_F |
| CCDC47 | 0.07643 | -0.3612 | 0.691 | 0.79 | 1 | HC_vs_MS.NoTrt_F |
| DXO | 0.07661 | 0.372 | 0.171 | 0.24 | 1 | HC_vs_MS.NoTrt_F |
| HIST1H2AC | 0.07666 | 1.01405 | 0.081 | 0.13 | 1 | HC_vs_MS.NoTrt_F |
| ZBTB1 | 0.07666 | -0.4833 | 0.231 | 0.3 | 1 | HC_vs_MS.NoTrt_F |
| CCDC146 | 0.0767 | -0.3375 | 0.104 | 0.16 | 1 | HC_vs_MS.NoTrt_F |
| ACTR5 | 0.07693 | 0.25114 | 0.168 | 0.23 | 1 | HC_vs_MS.NoTrt_F |
| SLC36A1 | 0.07693 | 0.27421 | 0.098 | 0.16 | 1 | HC_vs_MS.NoTrt_F |

| | | | | | | |
|---|---|---|---|---|---|---|
| PLA2G6 | 0.07697 | 0.51984 | 0.428 | 0.52 | 1 | HC_vs_MS.NoTrt_F |
| FCRL5 | 0.07716 | 0.38062 | 0.569 | 0.52 | 1 | HC_vs_MS.NoTrt_F |
| PCDHGA8 | 0.07725 | 0.34405 | 0.662 | 0.56 | 1 | HC_vs_MS.NoTrt_F |
| AKT1S1 | 0.07733 | -1.0134 | 0.22 | 0.28 | 1 | HC_vs_MS.NoTrt_F |
| ZUFSP | 0.07749 | 0.4012 | 0.098 | 0.16 | 1 | HC_vs_MS.NoTrt_F |
| YLPM1 | 0.07772 | 0.65574 | 0.749 | 0.67 | 1 | HC_vs_MS.NoTrt_F |
| ENPP3 | 0.07773 | 1.19951 | 0.139 | 0.2 | 1 | HC_vs_MS.NoTrt_F |
| IL5RA | 0.07793 | 1.58305 | 0.318 | 0.4 | 1 | HC_vs_MS.NoTrt_F |
| RPL17-C18o | 0.07821 | 0.42244 | 0.09 | 0.14 | 1 | HC_vs_MS.NoTrt_F |
| PPP2R2D | 0.07825 | -0.4284 | 0.65 | 0.73 | 1 | HC_vs_MS.NoTrt_F |
| RNF169 | 0.07837 | 0.89712 | 0.092 | 0.14 | 1 | HC_vs_MS.NoTrt_F |
| SFMBT1 | 0.07841 | -0.6554 | 0.587 | 0.68 | 1 | HC_vs_MS.NoTrt_F |
| TMEM14C | 0.07842 | 0.25281 | 0.142 | 0.2 | 1 | HC_vs_MS.NoTrt_F |
| TNS3 | 0.07846 | 0.78672 | 0.419 | 0.35 | 1 | HC_vs_MS.NoTrt_F |
| CHIC1 | 0.07848 | 0.55747 | 0.107 | 0.17 | 1 | HC_vs_MS.NoTrt_F |
| ARL5A | 0.0788 | 0.45787 | 0.205 | 0.28 | 1 | HC_vs_MS.NoTrt_F |
| USO1 | 0.07886 | 0.28492 | 0.208 | 0.28 | 1 | HC_vs_MS.NoTrt_F |
| TMCO4 | 0.079 | 0.25178 | 0.136 | 0.19 | 1 | HC_vs_MS.NoTrt_F |
| HOMEZ | 0.07924 | -0.7393 | 0.26 | 0.32 | 1 | HC_vs_MS.NoTrt_F |
| POLR2H | 0.0794 | 0.40908 | 0.448 | 0.54 | 1 | HC_vs_MS.NoTrt_F |
| ELAC1 | 0.07952 | -0.5642 | 0.512 | 0.58 | 1 | HC_vs_MS.NoTrt_F |
| PHF2 | 0.07957 | 0.9088 | 0.104 | 0.16 | 1 | HC_vs_MS.NoTrt_F |
| RMI1 | 0.07957 | 0.77225 | 0.286 | 0.38 | 1 | HC_vs_MS.NoTrt_F |
| GLTSCR2 | 0.07967 | 0.28155 | 0.876 | 0.92 | 1 | HC_vs_MS.NoTrt_F |
| ZNF670-ZNF | 0.07971 | 0.45067 | 0.671 | 0.67 | 1 | HC_vs_MS.NoTrt_F |
| RAD51 | 0.07981 | -0.5346 | 0.512 | 0.61 | 1 | HC_vs_MS.NoTrt_F |
| ICAM4 | 0.0799 | 0.29062 | 0.668 | 0.61 | 1 | HC_vs_MS.NoTrt_F |
| DHCR7 | 0.08004 | 0.40712 | 0.081 | 0.13 | 1 | HC_vs_MS.NoTrt_F |
| GAA | 0.08012 | 0.97096 | 0.101 | 0.16 | 1 | HC_vs_MS.NoTrt_F |
| SRRD | 0.08013 | 0.34208 | 0.474 | 0.54 | 1 | HC_vs_MS.NoTrt_F |
| POLG2 | 0.08025 | -0.4875 | 0.538 | 0.56 | 1 | HC_vs_MS.NoTrt_F |
| DDX20 | 0.08029 | 0.64218 | 0.116 | 0.17 | 1 | HC_vs_MS.NoTrt_F |
| UBE3A | 0.08041 | -0.6308 | 0.598 | 0.64 | 1 | HC_vs_MS.NoTrt_F |
| TUNAR | 0.08053 | 1.47386 | 0.168 | 0.24 | 1 | HC_vs_MS.NoTrt_F |
| TREML2 | 0.08066 | 0.45067 | 0.133 | 0.2 | 1 | HC_vs_MS.NoTrt_F |
| TYMP | 0.08077 | 0.4026 | 0.367 | 0.45 | 1 | HC_vs_MS.NoTrt_F |
| BRWD3 | 0.08079 | -0.292 | 0.153 | 0.22 | 1 | HC_vs_MS.NoTrt_F |
| HMGCR | 0.08083 | 0.38222 | 0.711 | 0.81 | 1 | HC_vs_MS.NoTrt_F |
| BLOC1S4 | 0.08089 | 0.76551 | 0.171 | 0.23 | 1 | HC_vs_MS.NoTrt_F |
| ERI1 | 0.08094 | 0.33271 | 0.451 | 0.54 | 1 | HC_vs_MS.NoTrt_F |

| | | | | | | |
|---|---|---|---|---|---|---|
| WDR83OS | 0.08102 | 0.30225 | 0.231 | 0.31 | 1 | HC_vs_MS.NoTrt_F |
| MSH3 | 0.08128 | -0.6663 | 0.474 | 0.51 | 1 | HC_vs_MS.NoTrt_F |
| PACSIN2 | 0.0813 | -0.4956 | 0.494 | 0.58 | 1 | HC_vs_MS.NoTrt_F |
| CHMP6 | 0.0815 | 0.67523 | 0.145 | 0.21 | 1 | HC_vs_MS.NoTrt_F |
| KCTD11 | 0.08182 | 1.47795 | 0.173 | 0.24 | 1 | HC_vs_MS.NoTrt_F |
| INTS7 | 0.082 | -0.2993 | 0.457 | 0.53 | 1 | HC_vs_MS.NoTrt_F |
| PKP4 | 0.08211 | 0.3314 | 0.116 | 0.17 | 1 | HC_vs_MS.NoTrt_F |
| ZNF282 | 0.08235 | 0.91211 | 0.081 | 0.13 | 1 | HC_vs_MS.NoTrt_F |
| CORO2A | 0.08241 | 0.49775 | 0.861 | 0.78 | 1 | HC_vs_MS.NoTrt_F |
| TTLL1 | 0.08244 | 1.07335 | 0.064 | 0.11 | 1 | HC_vs_MS.NoTrt_F |
| LPGAT1 | 0.0825 | 0.71727 | 0.61 | 0.51 | 1 | HC_vs_MS.NoTrt_F |
| C3orf70 | 0.08257 | 0.56628 | 0.801 | 0.74 | 1 | HC_vs_MS.NoTrt_F |
| PODXL | 0.08277 | 0.38442 | 0.211 | 0.28 | 1 | HC_vs_MS.NoTrt_F |
| SKI | 0.08289 | 0.51865 | 0.133 | 0.19 | 1 | HC_vs_MS.NoTrt_F |
| C1orf162 | 0.08305 | 0.45928 | 0.142 | 0.2 | 1 | HC_vs_MS.NoTrt_F |
| ING3 | 0.08308 | 0.39217 | 0.176 | 0.24 | 1 | HC_vs_MS.NoTrt_F |
| FAM117A | 0.08308 | 0.3362 | 0.717 | 0.79 | 1 | HC_vs_MS.NoTrt_F |
| PANK4 | 0.08323 | 0.30322 | 0.107 | 0.16 | 1 | HC_vs_MS.NoTrt_F |
| PTPRB | 0.08338 | 0.4742 | 0.624 | 0.55 | 1 | HC_vs_MS.NoTrt_F |
| SNAPC5 | 0.08351 | -0.4947 | 0.616 | 0.66 | 1 | HC_vs_MS.NoTrt_F |
| FADS3 | 0.08377 | -0.3385 | 0.662 | 0.71 | 1 | HC_vs_MS.NoTrt_F |
| STX16 | 0.08395 | -0.3501 | 0.506 | 0.57 | 1 | HC_vs_MS.NoTrt_F |
| MIS12 | 0.08434 | -0.505 | 0.341 | 0.42 | 1 | HC_vs_MS.NoTrt_F |
| IP6K2 | 0.08447 | -0.2847 | 0.373 | 0.44 | 1 | HC_vs_MS.NoTrt_F |
| URB1 | 0.0848 | 1.07415 | 0.52 | 0.58 | 1 | HC_vs_MS.NoTrt_F |
| ZNF280B | 0.08525 | 0.9712 | 0.139 | 0.2 | 1 | HC_vs_MS.NoTrt_F |
| RBMS1 | 0.08535 | 1.30502 | 0.731 | 0.71 | 1 | HC_vs_MS.NoTrt_F |
| COL4A3BP | 0.08541 | 0.36891 | 0.168 | 0.24 | 1 | HC_vs_MS.NoTrt_F |
| GZMA | 0.08561 | 0.56417 | 0.133 | 0.19 | 1 | HC_vs_MS.NoTrt_F |
| RECK | 0.08562 | 0.47057 | 0.188 | 0.25 | 1 | HC_vs_MS.NoTrt_F |
| ARHGEF6 | 0.08565 | 0.31875 | 0.379 | 0.49 | 1 | HC_vs_MS.NoTrt_F |
| ACAA1 | 0.08569 | -0.3269 | 0.52 | 0.58 | 1 | HC_vs_MS.NoTrt_F |
| APIP | 0.08583 | 0.36618 | 0.509 | 0.61 | 1 | HC_vs_MS.NoTrt_F |
| CAPNS1 | 0.0859 | -0.3095 | 0.37 | 0.44 | 1 | HC_vs_MS.NoTrt_F |
| MTPN | 0.08603 | 0.46724 | 0.257 | 0.33 | 1 | HC_vs_MS.NoTrt_F |
| GLUD1 | 0.08637 | 0.34799 | 0.275 | 0.35 | 1 | HC_vs_MS.NoTrt_F |
| RGCC | 0.08658 | 0.37255 | 0.41 | 0.53 | 1 | HC_vs_MS.NoTrt_F |
| PLD1 | 0.08667 | 0.65923 | 0.549 | 0.65 | 1 | HC_vs_MS.NoTrt_F |
| PRDM10 | 0.08676 | 1.41966 | 0.09 | 0.14 | 1 | HC_vs_MS.NoTrt_F |
| PRPF4B | 0.08677 | -0.2661 | 0.364 | 0.45 | 1 | HC_vs_MS.NoTrt_F |

| | | | | | | |
|---|---|---|---|---|---|---|
| AP1S2 | 0.08686 | 0.30767 | 0.228 | 0.3 | 1 | HC_vs_MS.NoTrt_F |
| MITD1 | 0.08723 | 0.34861 | 0.962 | 0.97 | 1 | HC_vs_MS.NoTrt_F |
| MAP4K2 | 0.08727 | -0.2873 | 0.708 | 0.76 | 1 | HC_vs_MS.NoTrt_F |
| TRIM16L | 0.08739 | 0.39273 | 0.783 | 0.8 | 1 | HC_vs_MS.NoTrt_F |
| EMC6 | 0.08748 | 0.63515 | 0.116 | 0.17 | 1 | HC_vs_MS.NoTrt_F |
| NBAS | 0.08754 | 0.58195 | 0.251 | 0.33 | 1 | HC_vs_MS.NoTrt_F |
| KLHDC1 | 0.08777 | 0.38158 | 0.332 | 0.4 | 1 | HC_vs_MS.NoTrt_F |
| SCYL3 | 0.08789 | 0.48909 | 0.818 | 0.72 | 1 | HC_vs_MS.NoTrt_F |
| ALG1 | 0.088 | -0.3186 | 0.601 | 0.66 | 1 | HC_vs_MS.NoTrt_F |
| LAMTOR2 | 0.08803 | 0.27916 | 0.191 | 0.26 | 1 | HC_vs_MS.NoTrt_F |
| KPTN | 0.08863 | -0.832 | 0.22 | 0.28 | 1 | HC_vs_MS.NoTrt_F |
| EIF4G3 | 0.08873 | 0.32293 | 0.301 | 0.37 | 1 | HC_vs_MS.NoTrt_F |
| SESN1 | 0.08874 | 0.49044 | 0.214 | 0.29 | 1 | HC_vs_MS.NoTrt_F |
| CCDC183 | 0.0888 | 0.62416 | 0.078 | 0.13 | 1 | HC_vs_MS.NoTrt_F |
| NOMO1 | 0.0889 | 0.46623 | 0.199 | 0.27 | 1 | HC_vs_MS.NoTrt_F |
| GNPNAT1 | 0.08905 | 1.03094 | 0.581 | 0.52 | 1 | HC_vs_MS.NoTrt_F |
| DFNA5 | 0.08911 | -0.5866 | 0.15 | 0.22 | 1 | HC_vs_MS.NoTrt_F |
| SLC26A6 | 0.08916 | 0.80656 | 0.113 | 0.17 | 1 | HC_vs_MS.NoTrt_F |
| KIAA1328 | 0.08928 | 0.2814 | 0.986 | 0.99 | 1 | HC_vs_MS.NoTrt_F |
| PCDH9 | 0.09016 | 0.32306 | 0.691 | 0.75 | 1 | HC_vs_MS.NoTrt_F |
| KYAT3 | 0.09023 | -0.3031 | 0.136 | 0.19 | 1 | HC_vs_MS.NoTrt_F |
| ZNF468 | 0.09027 | -0.5151 | 0.529 | 0.62 | 1 | HC_vs_MS.NoTrt_F |
| CTD-2192J1( | 0.09028 | 0.39541 | 0.512 | 0.61 | 1 | HC_vs_MS.NoTrt_F |
| MAZ | 0.0903 | 0.42085 | 0.182 | 0.25 | 1 | HC_vs_MS.NoTrt_F |
| PTP4A3 | 0.09047 | 0.82373 | 0.11 | 0.17 | 1 | HC_vs_MS.NoTrt_F |
| FCGR2B | 0.09077 | 0.75049 | 0.075 | 0.12 | 1 | HC_vs_MS.NoTrt_F |
| ZNF425 | 0.09083 | -1.3679 | 0.445 | 0.51 | 1 | HC_vs_MS.NoTrt_F |
| C5AR1 | 0.09094 | 0.36908 | 0.98 | 0.97 | 1 | HC_vs_MS.NoTrt_F |
| NIPA1 | 0.09125 | 0.86557 | 0.191 | 0.25 | 1 | HC_vs_MS.NoTrt_F |
| AP3B1 | 0.09137 | -0.4294 | 0.486 | 0.54 | 1 | HC_vs_MS.NoTrt_F |
| PPP2CB | 0.09147 | 0.48704 | 0.564 | 0.6 | 1 | HC_vs_MS.NoTrt_F |
| SRGAP1 | 0.0916 | 0.55536 | 0.954 | 0.94 | 1 | HC_vs_MS.NoTrt_F |
| TNC | 0.09164 | 0.27861 | 0.344 | 0.44 | 1 | HC_vs_MS.NoTrt_F |
| MGME1 | 0.09178 | 0.58197 | 0.118 | 0.17 | 1 | HC_vs_MS.NoTrt_F |
| CCZ1 | 0.09188 | -0.4012 | 0.72 | 0.77 | 1 | HC_vs_MS.NoTrt_F |
| MFGE8 | 0.09212 | -1.247 | 0.604 | 0.62 | 1 | HC_vs_MS.NoTrt_F |
| PPP3CB | 0.09213 | 0.68303 | 0.136 | 0.19 | 1 | HC_vs_MS.NoTrt_F |
| LIMS3 | 0.09216 | 0.29399 | 0.647 | 0.56 | 1 | HC_vs_MS.NoTrt_F |
| MAGED2 | 0.09217 | -0.2845 | 0.24 | 0.29 | 1 | HC_vs_MS.NoTrt_F |
| SNX21 | 0.09218 | 0.82915 | 0.468 | 0.56 | 1 | HC_vs_MS.NoTrt_F |

| | | | | | | |
|---|---|---|---|---|---|---|
| L3MBTL4 | 0.09237 | -0.7244 | 0.494 | 0.59 | 1 | HC_vs_MS.NoTrt_F |
| WDR46 | 0.09266 | 0.90786 | 0.133 | 0.19 | 1 | HC_vs_MS.NoTrt_F |
| CASP9 | 0.09272 | 0.8365 | 0.471 | 0.4 | 1 | HC_vs_MS.NoTrt_F |
| NFE2L1 | 0.09273 | 0.35127 | 0.101 | 0.16 | 1 | HC_vs_MS.NoTrt_F |
| IFIT5 | 0.09277 | 0.32816 | 0.104 | 0.16 | 1 | HC_vs_MS.NoTrt_F |
| PRKCZ | 0.0928 | 0.53665 | 0.301 | 0.38 | 1 | HC_vs_MS.NoTrt_F |
| POLR3H | 0.09289 | 0.69014 | 0.272 | 0.35 | 1 | HC_vs_MS.NoTrt_F |
| MON1A | 0.0929 | 0.27101 | 0.116 | 0.17 | 1 | HC_vs_MS.NoTrt_F |
| MAD2L1BP | 0.09295 | -0.8362 | 0.191 | 0.25 | 1 | HC_vs_MS.NoTrt_F |
| ZNF701 | 0.093 | 0.47264 | 0.477 | 0.57 | 1 | HC_vs_MS.NoTrt_F |
| E2F5 | 0.09309 | -0.3264 | 0.393 | 0.46 | 1 | HC_vs_MS.NoTrt_F |
| MRPL50 | 0.09336 | -0.3407 | 0.165 | 0.22 | 1 | HC_vs_MS.NoTrt_F |
| CDH8 | 0.09345 | 0.2922 | 0.514 | 0.59 | 1 | HC_vs_MS.NoTrt_F |
| MAP4K3 | 0.0935 | 0.43919 | 0.084 | 0.13 | 1 | HC_vs_MS.NoTrt_F |
| GZMM | 0.09357 | 0.53537 | 0.199 | 0.28 | 1 | HC_vs_MS.NoTrt_F |
| ZFAND3 | 0.09382 | 0.4939 | 0.734 | 0.79 | 1 | HC_vs_MS.NoTrt_F |
| LOXL1 | 0.09404 | 0.86594 | 0.104 | 0.16 | 1 | HC_vs_MS.NoTrt_F |
| NAGA | 0.0941 | 0.6279 | 0.13 | 0.19 | 1 | HC_vs_MS.NoTrt_F |
| PAPLN | 0.09418 | 0.43471 | 0.847 | 0.82 | 1 | HC_vs_MS.NoTrt_F |
| DAP3 | 0.0942 | -0.569 | 0.913 | 0.93 | 1 | HC_vs_MS.NoTrt_F |
| SRPK2 | 0.09425 | 0.4299 | 0.341 | 0.44 | 1 | HC_vs_MS.NoTrt_F |
| CFHR1 | 0.09428 | 0.42723 | 0.786 | 0.77 | 1 | HC_vs_MS.NoTrt_F |
| RAD51B | 0.0943 | -0.405 | 0.327 | 0.4 | 1 | HC_vs_MS.NoTrt_F |
| MTM1 | 0.09435 | -0.2541 | 0.228 | 0.29 | 1 | HC_vs_MS.NoTrt_F |
| MRPS15 | 0.09442 | -0.3066 | 0.584 | 0.62 | 1 | HC_vs_MS.NoTrt_F |
| COG6 | 0.09443 | -0.2547 | 0.159 | 0.22 | 1 | HC_vs_MS.NoTrt_F |
| INTS12 | 0.09449 | 0.28658 | 0.136 | 0.19 | 1 | HC_vs_MS.NoTrt_F |
| UHRF1BP1L | 0.09452 | -0.8899 | 0.393 | 0.47 | 1 | HC_vs_MS.NoTrt_F |
| CD58 | 0.09456 | 0.48336 | 0.303 | 0.39 | 1 | HC_vs_MS.NoTrt_F |
| CCDC137 | 0.09464 | 0.36724 | 0.298 | 0.37 | 1 | HC_vs_MS.NoTrt_F |
| BST1 | 0.0947 | 0.25639 | 0.555 | 0.63 | 1 | HC_vs_MS.NoTrt_F |
| TMUB2 | 0.09473 | 0.32362 | 0.882 | 0.86 | 1 | HC_vs_MS.NoTrt_F |
| GS1-114I9.3 | 0.09488 | 1.14849 | 0.078 | 0.13 | 1 | HC_vs_MS.NoTrt_F |
| ARFGAP1 | 0.09497 | -0.646 | 0.234 | 0.28 | 1 | HC_vs_MS.NoTrt_F |
| CHMP1B | 0.09499 | 0.49955 | 0.855 | 0.83 | 1 | HC_vs_MS.NoTrt_F |
| SLAMF6 | 0.09501 | -0.8046 | 0.549 | 0.56 | 1 | HC_vs_MS.NoTrt_F |
| PIM3 | 0.09507 | 0.42834 | 0.176 | 0.24 | 1 | HC_vs_MS.NoTrt_F |
| GOLM1 | 0.09517 | 0.94322 | 0.642 | 0.62 | 1 | HC_vs_MS.NoTrt_F |
| NSUN5 | 0.09521 | 0.26416 | 0.35 | 0.41 | 1 | HC_vs_MS.NoTrt_F |
| SLC11A2 | 0.09522 | 0.26541 | 0.801 | 0.74 | 1 | HC_vs_MS.NoTrt_F |

| | | | | | | |
|---|---|---|---|---|---|---|
| BMP1 | 0.09523 | 0.80593 | 0.639 | 0.6 | 1 | HC_vs_MS.NoTrt_F |
| PRKRIP1 | 0.09534 | -0.3457 | 0.211 | 0.26 | 1 | HC_vs_MS.NoTrt_F |
| AAMP | 0.09555 | -0.4899 | 0.234 | 0.28 | 1 | HC_vs_MS.NoTrt_F |
| ZNF800 | 0.09569 | 0.72477 | 0.251 | 0.33 | 1 | HC_vs_MS.NoTrt_F |
| POU2F1 | 0.09585 | 0.34906 | 0.422 | 0.48 | 1 | HC_vs_MS.NoTrt_F |
| COG5 | 0.09588 | 1.45336 | 0.263 | 0.34 | 1 | HC_vs_MS.NoTrt_F |
| PNLIPRP1 | 0.09593 | 0.30818 | 0.853 | 0.86 | 1 | HC_vs_MS.NoTrt_F |
| TMEM161B | 0.09596 | -0.3722 | 0.642 | 0.71 | 1 | HC_vs_MS.NoTrt_F |
| ZNF484 | 0.09616 | 0.92722 | 0.107 | 0.16 | 1 | HC_vs_MS.NoTrt_F |
| ANKRD26 | 0.09616 | 0.97296 | 0.159 | 0.22 | 1 | HC_vs_MS.NoTrt_F |
| ESRP2 | 0.09658 | 0.50378 | 0.081 | 0.13 | 1 | HC_vs_MS.NoTrt_F |
| CCNG2 | 0.09661 | 0.51096 | 0.35 | 0.41 | 1 | HC_vs_MS.NoTrt_F |
| TMEM185B | 0.09675 | -0.9183 | 0.335 | 0.39 | 1 | HC_vs_MS.NoTrt_F |
| TUBGCP5 | 0.09679 | 0.41286 | 0.697 | 0.79 | 1 | HC_vs_MS.NoTrt_F |
| HARS2 | 0.09684 | 0.93973 | 0.153 | 0.22 | 1 | HC_vs_MS.NoTrt_F |
| ARHGAP11A | 0.0971 | -0.947 | 0.422 | 0.5 | 1 | HC_vs_MS.NoTrt_F |
| ABCA9 | 0.0973 | 0.72222 | 0.882 | 0.88 | 1 | HC_vs_MS.NoTrt_F |
| AKAP6 | 0.09738 | 0.27194 | 0.777 | 0.76 | 1 | HC_vs_MS.NoTrt_F |
| ARHGAP10 | 0.09744 | 0.49603 | 0.182 | 0.25 | 1 | HC_vs_MS.NoTrt_F |
| CSF1R | 0.0975 | -0.8237 | 0.197 | 0.27 | 1 | HC_vs_MS.NoTrt_F |
| RYK | 0.09752 | 0.38026 | 0.118 | 0.17 | 1 | HC_vs_MS.NoTrt_F |
| POLQ | 0.09754 | 0.53313 | 0.618 | 0.55 | 1 | HC_vs_MS.NoTrt_F |
| SYNJ2BP | 0.09779 | -0.2778 | 0.867 | 0.91 | 1 | HC_vs_MS.NoTrt_F |
| PRDM15 | 0.09798 | -0.4884 | 0.636 | 0.58 | 1 | HC_vs_MS.NoTrt_F |
| ATP5G1 | 0.09806 | -0.2765 | 0.306 | 0.38 | 1 | HC_vs_MS.NoTrt_F |
| RAB12 | 0.09836 | 1.75106 | 0.361 | 0.35 | 1 | HC_vs_MS.NoTrt_F |
| TCF7 | 0.09836 | 1.04052 | 0.78 | 0.76 | 1 | HC_vs_MS.NoTrt_F |
| OSCAR | 0.09856 | 0.27191 | 0.066 | 0.11 | 1 | HC_vs_MS.NoTrt_F |
| FAM58A | 0.09858 | 0.62345 | 0.107 | 0.16 | 1 | HC_vs_MS.NoTrt_F |
| SLC25A39 | 0.09875 | 0.25467 | 0.72 | 0.77 | 1 | HC_vs_MS.NoTrt_F |
| ZNF429 | 0.09882 | 0.76096 | 0.928 | 0.91 | 1 | HC_vs_MS.NoTrt_F |
| RMND5A | 0.09926 | 0.94436 | 0.384 | 0.32 | 1 | HC_vs_MS.NoTrt_F |
| USP34 | 0.09927 | 0.28979 | 0.671 | 0.72 | 1 | HC_vs_MS.NoTrt_F |
| EDARADD | 0.0995 | -0.323 | 0.107 | 0.16 | 1 | HC_vs_MS.NoTrt_F |
| APP | 0.09971 | 0.88519 | 0.731 | 0.72 | 1 | HC_vs_MS.NoTrt_F |
| NEPRO | 0.09983 | -0.4008 | 0.197 | 0.25 | 1 | HC_vs_MS.NoTrt_F |
| CEP83 | 0.09983 | 0.38384 | 0.754 | 0.69 | 1 | HC_vs_MS.NoTrt_F |
| USP6NL | 0.09998 | 0.26591 | 0.827 | 0.8 | 1 | HC_vs_MS.NoTrt_F |
| ADPRHL1 | 0.1002 | -0.5225 | 0.315 | 0.38 | 1 | HC_vs_MS.NoTrt_F |
| C14orf93 | 0.10036 | -0.5197 | 0.335 | 0.39 | 1 | HC_vs_MS.NoTrt_F |

| | | | | | | |
|---|---|---|---|---|---|---|
| ORC5 | 0.10103 | 0.33274 | 0.263 | 0.34 | 1 | HC_vs_MS.NoTrt_F |
| NUDT18 | 0.10103 | 0.41193 | 0.066 | 0.11 | 1 | HC_vs_MS.NoTrt_F |
| TNKS2 | 0.10111 | 0.54544 | 0.347 | 0.45 | 1 | HC_vs_MS.NoTrt_F |
| CHMP3 | 0.1014 | -0.3272 | 0.367 | 0.42 | 1 | HC_vs_MS.NoTrt_F |
| MLKL | 0.102 | -0.3705 | 0.829 | 0.72 | 1 | HC_vs_MS.NoTrt_F |
| TACR2 | 0.10205 | 0.53417 | 0.121 | 0.18 | 1 | HC_vs_MS.NoTrt_F |
| SLC7A6 | 0.10216 | 0.45323 | 0.555 | 0.66 | 1 | HC_vs_MS.NoTrt_F |
| PSTPIP2 | 0.10237 | -0.3407 | 0.408 | 0.48 | 1 | HC_vs_MS.NoTrt_F |
| ANAPC10 | 0.1025 | 0.33982 | 0.168 | 0.23 | 1 | HC_vs_MS.NoTrt_F |
| RBL1 | 0.10262 | -0.2973 | 0.642 | 0.66 | 1 | HC_vs_MS.NoTrt_F |
| TBX19 | 0.10282 | 0.30904 | 0.11 | 0.16 | 1 | HC_vs_MS.NoTrt_F |
| TMEM234 | 0.10286 | 0.62204 | 0.127 | 0.18 | 1 | HC_vs_MS.NoTrt_F |
| NDUFS5 | 0.10333 | 0.26958 | 0.277 | 0.37 | 1 | HC_vs_MS.NoTrt_F |
| LAPTM4A | 0.1034 | -0.3026 | 0.228 | 0.28 | 1 | HC_vs_MS.NoTrt_F |
| MAPT | 0.10358 | 0.65183 | 0.746 | 0.72 | 1 | HC_vs_MS.NoTrt_F |
| VTI1A | 0.10361 | -0.4154 | 0.902 | 0.92 | 1 | HC_vs_MS.NoTrt_F |
| MTOR | 0.1037 | -0.6029 | 0.243 | 0.29 | 1 | HC_vs_MS.NoTrt_F |
| TMEM38A | 0.10372 | 0.37167 | 0.581 | 0.49 | 1 | HC_vs_MS.NoTrt_F |
| FXR2 | 0.10375 | 0.61496 | 0.714 | 0.69 | 1 | HC_vs_MS.NoTrt_F |
| DTX3L | 0.10377 | 0.34099 | 0.711 | 0.79 | 1 | HC_vs_MS.NoTrt_F |
| TSG101 | 0.1038 | -0.3713 | 0.506 | 0.54 | 1 | HC_vs_MS.NoTrt_F |
| DIMT1 | 0.10452 | 0.75441 | 0.162 | 0.23 | 1 | HC_vs_MS.NoTrt_F |
| LRCH3 | 0.10459 | 0.27527 | 0.988 | 0.98 | 1 | HC_vs_MS.NoTrt_F |
| MIB2 | 0.1046 | 0.4545 | 0.202 | 0.26 | 1 | HC_vs_MS.NoTrt_F |
| PMF1-BGLAI | 0.10463 | 1.01511 | 0.11 | 0.17 | 1 | HC_vs_MS.NoTrt_F |
| TUT1 | 0.10479 | -1.2889 | 0.191 | 0.24 | 1 | HC_vs_MS.NoTrt_F |
| FAM227A | 0.10509 | 0.28929 | 0.942 | 0.94 | 1 | HC_vs_MS.NoTrt_F |
| XKR8 | 0.10514 | -0.98 | 0.185 | 0.24 | 1 | HC_vs_MS.NoTrt_F |
| KIDINS220 | 0.1052 | -0.5699 | 0.298 | 0.35 | 1 | HC_vs_MS.NoTrt_F |
| CENPP | 0.1052 | -0.2823 | 0.064 | 0.11 | 1 | HC_vs_MS.NoTrt_F |
| PCED1A | 0.10525 | 0.40746 | 0.185 | 0.25 | 1 | HC_vs_MS.NoTrt_F |
| ZNF169 | 0.10534 | 0.44995 | 0.908 | 0.87 | 1 | HC_vs_MS.NoTrt_F |
| MIXL1 | 0.10544 | 0.28637 | 0.725 | 0.6 | 1 | HC_vs_MS.NoTrt_F |
| DNAJC5 | 0.10579 | 1.04448 | 0.11 | 0.16 | 1 | HC_vs_MS.NoTrt_F |
| EML3 | 0.10585 | 0.46222 | 0.269 | 0.34 | 1 | HC_vs_MS.NoTrt_F |
| L3MBTL1 | 0.10589 | 0.58575 | 0.685 | 0.65 | 1 | HC_vs_MS.NoTrt_F |
| TBRG4 | 0.106 | -0.4509 | 0.436 | 0.48 | 1 | HC_vs_MS.NoTrt_F |
| STXBP3 | 0.10602 | -0.5071 | 0.636 | 0.65 | 1 | HC_vs_MS.NoTrt_F |
| RP11-619A1 | 0.10642 | -0.4902 | 0.63 | 0.7 | 1 | HC_vs_MS.NoTrt_F |
| ALKBH2 | 0.10649 | -0.4367 | 0.37 | 0.44 | 1 | HC_vs_MS.NoTrt_F |

| | | | | | | |
|---|---|---|---|---|---|---|
| MAP3K7CL | 0.10658 | -0.3195 | 0.465 | 0.53 | 1 | HC_vs_MS.NoTrt_F |
| TPCN1 | 0.10673 | -0.3596 | 0.428 | 0.51 | 1 | HC_vs_MS.NoTrt_F |
| CRELD2 | 0.10688 | -0.4291 | 0.645 | 0.67 | 1 | HC_vs_MS.NoTrt_F |
| NFXL1 | 0.10692 | 0.48928 | 0.145 | 0.2 | 1 | HC_vs_MS.NoTrt_F |
| MARVELD2 | 0.10717 | 0.48842 | 0.725 | 0.72 | 1 | HC_vs_MS.NoTrt_F |
| CSTF1 | 0.10723 | -0.82 | 0.72 | 0.76 | 1 | HC_vs_MS.NoTrt_F |
| CXCL16 | 0.10731 | 0.50329 | 0.662 | 0.61 | 1 | HC_vs_MS.NoTrt_F |
| CYP2C19 | 0.10739 | 0.54536 | 0.769 | 0.76 | 1 | HC_vs_MS.NoTrt_F |
| BPHL | 0.10742 | 0.48194 | 0.694 | 0.75 | 1 | HC_vs_MS.NoTrt_F |
| ZACN | 0.10745 | 0.49531 | 0.084 | 0.13 | 1 | HC_vs_MS.NoTrt_F |
| DMXL1 | 0.1077 | 0.68519 | 0.24 | 0.31 | 1 | HC_vs_MS.NoTrt_F |
| BICD2 | 0.10771 | 0.57706 | 0.116 | 0.17 | 1 | HC_vs_MS.NoTrt_F |
| CNBD2 | 0.10774 | 0.84345 | 0.543 | 0.49 | 1 | HC_vs_MS.NoTrt_F |
| KRBA2 | 0.10777 | -1.2259 | 0.387 | 0.44 | 1 | HC_vs_MS.NoTrt_F |
| ERGIC1 | 0.1078 | 0.29072 | 0.882 | 0.86 | 1 | HC_vs_MS.NoTrt_F |
| TMEM164 | 0.1079 | 0.79788 | 0.107 | 0.16 | 1 | HC_vs_MS.NoTrt_F |
| C18orf25 | 0.10797 | -0.349 | 0.468 | 0.5 | 1 | HC_vs_MS.NoTrt_F |
| EXOC3L4 | 0.10803 | -0.5343 | 0.254 | 0.31 | 1 | HC_vs_MS.NoTrt_F |
| CSF2RB | 0.10809 | 0.47763 | 0.124 | 0.18 | 1 | HC_vs_MS.NoTrt_F |
| TMED2 | 0.10817 | -0.3332 | 0.78 | 0.77 | 1 | HC_vs_MS.NoTrt_F |
| C12orf60 | 0.10831 | 0.52321 | 0.676 | 0.62 | 1 | HC_vs_MS.NoTrt_F |
| GOLGA6L1 | 0.10833 | 0.43551 | 0.471 | 0.43 | 1 | HC_vs_MS.NoTrt_F |
| PTPN7 | 0.10863 | 0.75035 | 0.775 | 0.75 | 1 | HC_vs_MS.NoTrt_F |
| SMG6 | 0.10869 | 0.3556 | 0.251 | 0.33 | 1 | HC_vs_MS.NoTrt_F |
| PTTG1IP | 0.10877 | 0.32178 | 0.254 | 0.34 | 1 | HC_vs_MS.NoTrt_F |
| MAP3K12 | 0.10881 | 0.39211 | 0.731 | 0.79 | 1 | HC_vs_MS.NoTrt_F |
| STX4 | 0.10887 | 0.28122 | 0.223 | 0.28 | 1 | HC_vs_MS.NoTrt_F |
| TSNARE1 | 0.10902 | 0.54603 | 0.133 | 0.19 | 1 | HC_vs_MS.NoTrt_F |
| NFYA | 0.10908 | 0.72455 | 0.159 | 0.22 | 1 | HC_vs_MS.NoTrt_F |
| SERPINA10 | 0.1091 | 0.3855 | 0.627 | 0.58 | 1 | HC_vs_MS.NoTrt_F |
| ZBTB46 | 0.10911 | -0.5737 | 0.136 | 0.19 | 1 | HC_vs_MS.NoTrt_F |
| VPS8 | 0.10912 | -0.7169 | 0.691 | 0.75 | 1 | HC_vs_MS.NoTrt_F |
| AK5 | 0.10913 | 0.68038 | 0.156 | 0.22 | 1 | HC_vs_MS.NoTrt_F |
| RP11-864I4. | 0.10914 | -0.4836 | 0.084 | 0.13 | 1 | HC_vs_MS.NoTrt_F |
| NRP1 | 0.10919 | 0.43353 | 0.465 | 0.41 | 1 | HC_vs_MS.NoTrt_F |
| SSH1 | 0.10925 | 0.40721 | 0.887 | 0.89 | 1 | HC_vs_MS.NoTrt_F |
| ZFP36L2 | 0.10939 | 1.00349 | 0.595 | 0.64 | 1 | HC_vs_MS.NoTrt_F |
| CNOT6 | 0.10952 | 0.27927 | 0.182 | 0.23 | 1 | HC_vs_MS.NoTrt_F |
| SZT2 | 0.10958 | 0.31557 | 0.884 | 0.9 | 1 | HC_vs_MS.NoTrt_F |
| HERPUD2 | 0.10981 | 0.34328 | 0.561 | 0.63 | 1 | HC_vs_MS.NoTrt_F |

| | | | | | | |
|---|---|---|---|---|---|---|
| KHNYN | 0.10995 | 0.87201 | 0.136 | 0.19 | 1 | HC_vs_MS.NoTrt_F |
| CPM | 0.11 | 0.58262 | 0.832 | 0.77 | 1 | HC_vs_MS.NoTrt_F |
| ERP27 | 0.1102 | 0.28636 | 0.26 | 0.34 | 1 | HC_vs_MS.NoTrt_F |
| NOL6 | 0.11022 | 1.29405 | 0.431 | 0.5 | 1 | HC_vs_MS.NoTrt_F |
| MRPL16 | 0.11022 | 0.4555 | 0.211 | 0.28 | 1 | HC_vs_MS.NoTrt_F |
| CDK7 | 0.11052 | 0.36046 | 0.22 | 0.29 | 1 | HC_vs_MS.NoTrt_F |
| AIM2 | 0.11092 | 1.18655 | 0.341 | 0.27 | 1 | HC_vs_MS.NoTrt_F |
| SLC26A2 | 0.11111 | 0.47065 | 0.775 | 0.71 | 1 | HC_vs_MS.NoTrt_F |
| TNFSF8 | 0.11116 | 0.58528 | 0.159 | 0.22 | 1 | HC_vs_MS.NoTrt_F |
| WBP4 | 0.11133 | 0.66444 | 0.116 | 0.17 | 1 | HC_vs_MS.NoTrt_F |
| CENPO | 0.11137 | 0.89932 | 0.072 | 0.12 | 1 | HC_vs_MS.NoTrt_F |
| DISP1 | 0.11145 | 0.55953 | 0.121 | 0.17 | 1 | HC_vs_MS.NoTrt_F |
| SUN2 | 0.11158 | 0.36516 | 0.422 | 0.54 | 1 | HC_vs_MS.NoTrt_F |
| OMA1 | 0.11167 | -0.4374 | 0.627 | 0.69 | 1 | HC_vs_MS.NoTrt_F |
| CCDC80 | 0.11168 | 0.88295 | 0.783 | 0.75 | 1 | HC_vs_MS.NoTrt_F |
| USP31 | 0.11181 | 0.61598 | 0.133 | 0.19 | 1 | HC_vs_MS.NoTrt_F |
| MAEA | 0.11186 | -0.2818 | 0.653 | 0.71 | 1 | HC_vs_MS.NoTrt_F |
| CLSTN3 | 0.11193 | 0.39078 | 0.199 | 0.27 | 1 | HC_vs_MS.NoTrt_F |
| PI4KA | 0.11221 | 0.59409 | 0.373 | 0.32 | 1 | HC_vs_MS.NoTrt_F |
| C15orf41 | 0.11268 | 0.3356 | 0.645 | 0.71 | 1 | HC_vs_MS.NoTrt_F |
| PLXDC2 | 0.11278 | 1.19338 | 0.74 | 0.69 | 1 | HC_vs_MS.NoTrt_F |
| RPL37A | 0.11359 | 0.38302 | 0.867 | 0.91 | 1 | HC_vs_MS.NoTrt_F |
| PCGF1 | 0.11361 | -0.3515 | 0.402 | 0.46 | 1 | HC_vs_MS.NoTrt_F |
| LNPK | 0.11389 | -0.2769 | 0.558 | 0.62 | 1 | HC_vs_MS.NoTrt_F |
| THAP9 | 0.11401 | 0.42207 | 0.639 | 0.57 | 1 | HC_vs_MS.NoTrt_F |
| ANAPC2 | 0.11422 | 0.92242 | 0.124 | 0.18 | 1 | HC_vs_MS.NoTrt_F |
| ZNF3 | 0.11427 | -0.7099 | 0.78 | 0.86 | 1 | HC_vs_MS.NoTrt_F |
| B4GALNT2 | 0.11453 | 0.41541 | 0.341 | 0.44 | 1 | HC_vs_MS.NoTrt_F |
| EHBP1L1 | 0.11461 | 0.56315 | 0.246 | 0.31 | 1 | HC_vs_MS.NoTrt_F |
| INPP4A | 0.11478 | 0.34149 | 0.405 | 0.46 | 1 | HC_vs_MS.NoTrt_F |
| KIAA1549 | 0.11491 | 0.47293 | 0.396 | 0.47 | 1 | HC_vs_MS.NoTrt_F |
| TP53BP2 | 0.11511 | 1.30739 | 0.179 | 0.24 | 1 | HC_vs_MS.NoTrt_F |
| PLK1 | 0.11512 | 0.5312 | 0.465 | 0.53 | 1 | HC_vs_MS.NoTrt_F |
| FAM63B | 0.11532 | -0.4399 | 0.769 | 0.79 | 1 | HC_vs_MS.NoTrt_F |
| PITHD1 | 0.11598 | 0.36613 | 0.191 | 0.25 | 1 | HC_vs_MS.NoTrt_F |
| ZNF581 | 0.11599 | 0.75411 | 0.116 | 0.17 | 1 | HC_vs_MS.NoTrt_F |
| SRSF8 | 0.11613 | -0.8861 | 0.671 | 0.7 | 1 | HC_vs_MS.NoTrt_F |
| C6orf136 | 0.11622 | 0.31684 | 0.121 | 0.17 | 1 | HC_vs_MS.NoTrt_F |
| AKAP2 | 0.11625 | -0.3311 | 0.312 | 0.38 | 1 | HC_vs_MS.NoTrt_F |
| EIF1AX | 0.11639 | 0.58692 | 0.347 | 0.29 | 1 | HC_vs_MS.NoTrt_F |

| | | | | | | |
|---|---|---|---|---|---|---|
| RAB8A | 0.11645 | -0.3303 | 0.607 | 0.63 | 1 | HC_vs_MS.NoTrt_F |
| CPEB3 | 0.11649 | -0.9173 | 0.075 | 0.12 | 1 | HC_vs_MS.NoTrt_F |
| SCP2 | 0.11653 | 0.30901 | 0.306 | 0.39 | 1 | HC_vs_MS.NoTrt_F |
| TMEM241 | 0.11665 | 1.09743 | 0.694 | 0.65 | 1 | HC_vs_MS.NoTrt_F |
| TMEM39A | 0.11673 | 0.56142 | 0.211 | 0.28 | 1 | HC_vs_MS.NoTrt_F |
| AMDHD1 | 0.11681 | 0.40565 | 0.214 | 0.28 | 1 | HC_vs_MS.NoTrt_F |
| TAF9 | 0.11685 | -0.3327 | 0.399 | 0.46 | 1 | HC_vs_MS.NoTrt_F |
| TBC1D2B | 0.11694 | 0.59444 | 0.358 | 0.41 | 1 | HC_vs_MS.NoTrt_F |
| ZNF778 | 0.11712 | 0.73911 | 0.705 | 0.66 | 1 | HC_vs_MS.NoTrt_F |
| C1QTNF7 | 0.11717 | 0.48453 | 0.081 | 0.13 | 1 | HC_vs_MS.NoTrt_F |
| ARMCX4 | 0.11725 | 0.68255 | 0.223 | 0.29 | 1 | HC_vs_MS.NoTrt_F |
| ZNF100 | 0.11736 | 0.90313 | 0.864 | 0.83 | 1 | HC_vs_MS.NoTrt_F |
| MTPAP | 0.11737 | -0.4266 | 0.873 | 0.91 | 1 | HC_vs_MS.NoTrt_F |
| SUGP1 | 0.11739 | -0.3651 | 0.486 | 0.53 | 1 | HC_vs_MS.NoTrt_F |
| RPS7 | 0.11745 | -0.2923 | 0.538 | 0.59 | 1 | HC_vs_MS.NoTrt_F |
| CCDC28A | 0.11745 | -0.4882 | 0.251 | 0.31 | 1 | HC_vs_MS.NoTrt_F |
| TLK2 | 0.11754 | -0.2713 | 0.431 | 0.49 | 1 | HC_vs_MS.NoTrt_F |
| POLD4 | 0.11763 | 0.36298 | 0.85 | 0.87 | 1 | HC_vs_MS.NoTrt_F |
| SELO | 0.11787 | 1.11056 | 0.081 | 0.13 | 1 | HC_vs_MS.NoTrt_F |
| C2orf42 | 0.11804 | 0.25276 | 0.182 | 0.24 | 1 | HC_vs_MS.NoTrt_F |
| NCLN | 0.11805 | 0.32413 | 0.185 | 0.24 | 1 | HC_vs_MS.NoTrt_F |
| PDDC1 | 0.1181 | 0.31297 | 0.977 | 0.98 | 1 | HC_vs_MS.NoTrt_F |
| DISC1 | 0.11815 | 0.32328 | 0.558 | 0.48 | 1 | HC_vs_MS.NoTrt_F |
| ARL17A | 0.1182 | 0.26675 | 0.679 | 0.77 | 1 | HC_vs_MS.NoTrt_F |
| RP11-894J1 | 0.11828 | -0.4211 | 0.139 | 0.19 | 1 | HC_vs_MS.NoTrt_F |
| FARP1 | 0.11867 | 0.31239 | 0.968 | 0.95 | 1 | HC_vs_MS.NoTrt_F |
| TIPARP | 0.11869 | 0.39117 | 0.165 | 0.23 | 1 | HC_vs_MS.NoTrt_F |
| ZNF33A | 0.11869 | -0.702 | 0.584 | 0.61 | 1 | HC_vs_MS.NoTrt_F |
| PPP1R9A | 0.11872 | 0.33465 | 0.225 | 0.28 | 1 | HC_vs_MS.NoTrt_F |
| C12orf66 | 0.11905 | -0.5212 | 0.382 | 0.44 | 1 | HC_vs_MS.NoTrt_F |
| FUK | 0.1191 | -1.1421 | 0.616 | 0.65 | 1 | HC_vs_MS.NoTrt_F |
| ZHX3 | 0.11911 | 0.67498 | 0.344 | 0.41 | 1 | HC_vs_MS.NoTrt_F |
| CMAS | 0.11917 | 0.32356 | 0.13 | 0.18 | 1 | HC_vs_MS.NoTrt_F |
| CYP4B1 | 0.11921 | 0.35892 | 0.699 | 0.63 | 1 | HC_vs_MS.NoTrt_F |
| PDHB | 0.11941 | -0.3025 | 0.844 | 0.9 | 1 | HC_vs_MS.NoTrt_F |
| LUC7L3 | 0.11955 | 0.2859 | 0.572 | 0.7 | 1 | HC_vs_MS.NoTrt_F |
| DNAJC7 | 0.11958 | -0.3308 | 0.977 | 0.97 | 1 | HC_vs_MS.NoTrt_F |
| MED13 | 0.11983 | 0.59202 | 0.595 | 0.56 | 1 | HC_vs_MS.NoTrt_F |
| HACL1 | 0.11995 | 0.40798 | 0.162 | 0.22 | 1 | HC_vs_MS.NoTrt_F |
| BMPR1A | 0.12003 | 0.38912 | 0.803 | 0.79 | 1 | HC_vs_MS.NoTrt_F |

| | | | | | | |
|---|---|---|---|---|---|---|
| USE1 | 0.12031 | 0.39663 | 0.887 | 0.86 | 1 | HC_vs_MS.NoTrt_F |
| USP49 | 0.12046 | 0.64553 | 0.694 | 0.73 | 1 | HC_vs_MS.NoTrt_F |
| ZWILCH | 0.12048 | -0.7484 | 0.694 | 0.73 | 1 | HC_vs_MS.NoTrt_F |
| DGKQ | 0.12048 | 1.5097 | 0.09 | 0.14 | 1 | HC_vs_MS.NoTrt_F |
| MAPK8IP3 | 0.12049 | 0.43131 | 0.22 | 0.29 | 1 | HC_vs_MS.NoTrt_F |
| FAM53B | 0.1205 | 0.47979 | 0.26 | 0.34 | 1 | HC_vs_MS.NoTrt_F |
| ZNF880 | 0.12066 | 0.28196 | 0.512 | 0.56 | 1 | HC_vs_MS.NoTrt_F |
| DHX40 | 0.12081 | 0.47976 | 0.147 | 0.2 | 1 | HC_vs_MS.NoTrt_F |
| ZNF623 | 0.12083 | 0.89588 | 0.72 | 0.72 | 1 | HC_vs_MS.NoTrt_F |
| PIP4K2B | 0.12144 | -0.5431 | 0.153 | 0.2 | 1 | HC_vs_MS.NoTrt_F |
| SLC39A7 | 0.12148 | 0.25511 | 0.113 | 0.16 | 1 | HC_vs_MS.NoTrt_F |
| POLR2J3 | 0.12155 | 0.342 | 0.997 | 0.99 | 1 | HC_vs_MS.NoTrt_F |
| QPRT | 0.1216 | 0.33593 | 0.936 | 0.88 | 1 | HC_vs_MS.NoTrt_F |
| TOE1 | 0.12162 | -0.5688 | 0.09 | 0.13 | 1 | HC_vs_MS.NoTrt_F |
| C2orf81 | 0.12198 | 0.31913 | 0.864 | 0.87 | 1 | HC_vs_MS.NoTrt_F |
| ABI2 | 0.12202 | 0.50536 | 0.899 | 0.88 | 1 | HC_vs_MS.NoTrt_F |
| POLR2J | 0.12254 | 0.40582 | 0.231 | 0.3 | 1 | HC_vs_MS.NoTrt_F |
| CSF3R | 0.12275 | 1.03795 | 0.711 | 0.71 | 1 | HC_vs_MS.NoTrt_F |
| NCOA7 | 0.12276 | 0.7109 | 0.801 | 0.77 | 1 | HC_vs_MS.NoTrt_F |
| AP2A1 | 0.12313 | -0.397 | 0.734 | 0.82 | 1 | HC_vs_MS.NoTrt_F |
| FUBP3 | 0.12314 | 0.35557 | 0.104 | 0.16 | 1 | HC_vs_MS.NoTrt_F |
| CLEC16A | 0.1233 | 0.5887 | 0.694 | 0.67 | 1 | HC_vs_MS.NoTrt_F |
| AIMP2 | 0.12366 | -1.1267 | 0.358 | 0.42 | 1 | HC_vs_MS.NoTrt_F |
| NPTXR | 0.12374 | 0.96982 | 0.355 | 0.42 | 1 | HC_vs_MS.NoTrt_F |
| TMEM156 | 0.12381 | -0.301 | 0.832 | 0.9 | 1 | HC_vs_MS.NoTrt_F |
| IL18BP | 0.12397 | 0.39114 | 0.147 | 0.2 | 1 | HC_vs_MS.NoTrt_F |
| ATP1B1 | 0.12397 | 1.12401 | 0.306 | 0.39 | 1 | HC_vs_MS.NoTrt_F |
| SYS1 | 0.12418 | 0.55576 | 0.529 | 0.46 | 1 | HC_vs_MS.NoTrt_F |
| NEMP1 | 0.1245 | 0.57482 | 0.147 | 0.2 | 1 | HC_vs_MS.NoTrt_F |
| COG4 | 0.12466 | -0.4707 | 0.751 | 0.79 | 1 | HC_vs_MS.NoTrt_F |
| ST8SIA4 | 0.1249 | -0.5014 | 0.769 | 0.81 | 1 | HC_vs_MS.NoTrt_F |
| TBC1D3I | 0.12498 | 0.36578 | 0.223 | 0.29 | 1 | HC_vs_MS.NoTrt_F |
| ZNF235 | 0.12507 | -1.1208 | 0.159 | 0.22 | 1 | HC_vs_MS.NoTrt_F |
| ZSCAN29 | 0.12517 | 0.59183 | 0.445 | 0.48 | 1 | HC_vs_MS.NoTrt_F |
| AMY2B | 0.12577 | -0.4156 | 0.607 | 0.64 | 1 | HC_vs_MS.NoTrt_F |
| NBPF20 | 0.1258 | 0.91165 | 0.347 | 0.41 | 1 | HC_vs_MS.NoTrt_F |
| USP46 | 0.12582 | 0.46247 | 0.731 | 0.87 | 1 | HC_vs_MS.NoTrt_F |
| MARK4 | 0.126 | 0.27073 | 0.578 | 0.66 | 1 | HC_vs_MS.NoTrt_F |
| TIAM1 | 0.1261 | -0.3301 | 0.54 | 0.6 | 1 | HC_vs_MS.NoTrt_F |
| RP4-583P15 | 0.12623 | 0.29033 | 0.15 | 0.2 | 1 | HC_vs_MS.NoTrt_F |

| | | | | | | |
|---|---|---|---|---|---|---|
| KDM5D | 0.12634 | -1.018 | 0.39 | 0.43 | 1 | HC_vs_MS.NoTrt_F |
| SOX6 | 0.12637 | -0.3284 | 0.448 | 0.5 | 1 | HC_vs_MS.NoTrt_F |
| PHAX | 0.12644 | -0.4686 | 0.893 | 0.9 | 1 | HC_vs_MS.NoTrt_F |
| CAMKK1 | 0.12664 | -0.2983 | 0.22 | 0.28 | 1 | HC_vs_MS.NoTrt_F |
| ATAD2B | 0.1267 | 0.51947 | 0.746 | 0.74 | 1 | HC_vs_MS.NoTrt_F |
| NUP35 | 0.12673 | 0.37249 | 0.662 | 0.61 | 1 | HC_vs_MS.NoTrt_F |
| FOXP2 | 0.12699 | 0.27851 | 0.257 | 0.32 | 1 | HC_vs_MS.NoTrt_F |
| TMEM186 | 0.12748 | -0.4805 | 0.526 | 0.61 | 1 | HC_vs_MS.NoTrt_F |
| KLHDC2 | 0.1279 | 0.45853 | 0.246 | 0.3 | 1 | HC_vs_MS.NoTrt_F |
| PLCD1 | 0.12822 | 1.24741 | 0.116 | 0.17 | 1 | HC_vs_MS.NoTrt_F |
| TMX4 | 0.12827 | -0.4095 | 0.636 | 0.62 | 1 | HC_vs_MS.NoTrt_F |
| UTP11 | 0.12884 | 0.69962 | 0.549 | 0.63 | 1 | HC_vs_MS.NoTrt_F |
| SERPINF1 | 0.12896 | 0.25038 | 0.827 | 0.88 | 1 | HC_vs_MS.NoTrt_F |
| AC013264.1 | 0.12901 | 0.3999 | 0.402 | 0.53 | 1 | HC_vs_MS.NoTrt_F |
| THTPA | 0.12908 | 0.43716 | 0.139 | 0.19 | 1 | HC_vs_MS.NoTrt_F |
| CCNY | 0.1292 | -0.3245 | 0.494 | 0.43 | 1 | HC_vs_MS.NoTrt_F |
| TMEM258 | 0.12927 | 0.52 | 0.361 | 0.45 | 1 | HC_vs_MS.NoTrt_F |
| KSR2 | 0.12931 | -0.5024 | 0.249 | 0.3 | 1 | HC_vs_MS.NoTrt_F |
| AKR1C2 | 0.12933 | 0.41071 | 0.107 | 0.16 | 1 | HC_vs_MS.NoTrt_F |
| ATP10A | 0.12936 | 2.05817 | 0.116 | 0.17 | 1 | HC_vs_MS.NoTrt_F |
| DNPEP | 0.12941 | -0.3689 | 0.48 | 0.52 | 1 | HC_vs_MS.NoTrt_F |
| ZNF234 | 0.12949 | 0.59627 | 0.835 | 0.79 | 1 | HC_vs_MS.NoTrt_F |
| EZH2 | 0.1296 | -0.3061 | 0.185 | 0.24 | 1 | HC_vs_MS.NoTrt_F |
| TFAM | 0.12985 | 0.8184 | 0.321 | 0.39 | 1 | HC_vs_MS.NoTrt_F |
| PROSER1 | 0.12997 | 0.29845 | 0.208 | 0.27 | 1 | HC_vs_MS.NoTrt_F |
| SOS1 | 0.13012 | 0.37411 | 0.199 | 0.26 | 1 | HC_vs_MS.NoTrt_F |
| ZNF665 | 0.13016 | 0.2905 | 0.954 | 0.95 | 1 | HC_vs_MS.NoTrt_F |
| CEP162 | 0.13052 | -0.5307 | 0.217 | 0.28 | 1 | HC_vs_MS.NoTrt_F |
| ZXDB | 0.1306 | 1.43786 | 0.087 | 0.13 | 1 | HC_vs_MS.NoTrt_F |
| C8orf59 | 0.13088 | -0.269 | 0.442 | 0.5 | 1 | HC_vs_MS.NoTrt_F |
| ALDH3B1 | 0.13098 | 0.2779 | 0.809 | 0.77 | 1 | HC_vs_MS.NoTrt_F |
| ELP6 | 0.13111 | -1.1237 | 0.647 | 0.69 | 1 | HC_vs_MS.NoTrt_F |
| TLR5 | 0.1313 | 0.59711 | 0.636 | 0.71 | 1 | HC_vs_MS.NoTrt_F |
| DHDDS | 0.13133 | -0.6436 | 0.618 | 0.62 | 1 | HC_vs_MS.NoTrt_F |
| ZNF563 | 0.13138 | -0.2716 | 0.555 | 0.62 | 1 | HC_vs_MS.NoTrt_F |
| NAB1 | 0.13151 | 0.57876 | 0.153 | 0.21 | 1 | HC_vs_MS.NoTrt_F |
| TDP1 | 0.13154 | -0.2856 | 0.382 | 0.42 | 1 | HC_vs_MS.NoTrt_F |
| TRMT1L | 0.13157 | 0.63432 | 0.124 | 0.17 | 1 | HC_vs_MS.NoTrt_F |
| RRM2B | 0.13168 | -0.5607 | 0.182 | 0.24 | 1 | HC_vs_MS.NoTrt_F |
| ADK | 0.13171 | 0.36126 | 0.289 | 0.36 | 1 | HC_vs_MS.NoTrt_F |

| | | | | | | |
|---|---|---|---|---|---|---|
| SWSAP1 | 0.13219 | 0.27072 | 0.783 | 0.71 | 1 | HC_vs_MS.NoTrt_F |
| WNT2B | 0.13228 | 0.51913 | 0.509 | 0.56 | 1 | HC_vs_MS.NoTrt_F |
| FSD2 | 0.13242 | 0.31623 | 0.488 | 0.56 | 1 | HC_vs_MS.NoTrt_F |
| NUCKS1 | 0.13252 | 0.29778 | 0.5 | 0.6 | 1 | HC_vs_MS.NoTrt_F |
| PSENEN | 0.13262 | 0.31641 | 0.246 | 0.3 | 1 | HC_vs_MS.NoTrt_F |
| CLHC1 | 0.13299 | -0.4373 | 0.526 | 0.59 | 1 | HC_vs_MS.NoTrt_F |
| RCHY1 | 0.13307 | -0.2751 | 0.364 | 0.43 | 1 | HC_vs_MS.NoTrt_F |
| PRH1 | 0.13313 | 0.46291 | 0.133 | 0.19 | 1 | HC_vs_MS.NoTrt_F |
| UBA6 | 0.1332 | -0.292 | 0.514 | 0.56 | 1 | HC_vs_MS.NoTrt_F |
| PRRC1 | 0.13321 | -0.7356 | 0.448 | 0.49 | 1 | HC_vs_MS.NoTrt_F |
| SLC35D1 | 0.1333 | 0.3837 | 0.35 | 0.43 | 1 | HC_vs_MS.NoTrt_F |
| NR4A2 | 0.13343 | 0.54969 | 0.607 | 0.61 | 1 | HC_vs_MS.NoTrt_F |
| FSIP2 | 0.13346 | 0.34461 | 0.847 | 0.82 | 1 | HC_vs_MS.NoTrt_F |
| ODC1 | 0.13375 | 0.91185 | 0.142 | 0.2 | 1 | HC_vs_MS.NoTrt_F |
| LAT2 | 0.13395 | 1.05921 | 0.876 | 0.88 | 1 | HC_vs_MS.NoTrt_F |
| MTMR12 | 0.1346 | 0.43272 | 0.488 | 0.53 | 1 | HC_vs_MS.NoTrt_F |
| LIX1L | 0.13473 | 0.4374 | 0.847 | 0.79 | 1 | HC_vs_MS.NoTrt_F |
| RAE1 | 0.13483 | 0.7524 | 0.139 | 0.19 | 1 | HC_vs_MS.NoTrt_F |
| CTNNA1 | 0.1349 | 0.26573 | 0.662 | 0.65 | 1 | HC_vs_MS.NoTrt_F |
| TFDP2 | 0.13495 | 0.40429 | 0.945 | 0.94 | 1 | HC_vs_MS.NoTrt_F |
| TP53BP1 | 0.13502 | -0.4589 | 0.306 | 0.37 | 1 | HC_vs_MS.NoTrt_F |
| MMACHC | 0.13547 | 0.65276 | 0.916 | 0.89 | 1 | HC_vs_MS.NoTrt_F |
| RNF19B | 0.13548 | 0.89706 | 0.069 | 0.11 | 1 | HC_vs_MS.NoTrt_F |
| SOX4 | 0.13565 | 1.12009 | 0.104 | 0.16 | 1 | HC_vs_MS.NoTrt_F |
| SGPL1 | 0.13581 | -0.4907 | 0.471 | 0.52 | 1 | HC_vs_MS.NoTrt_F |
| TRPM7 | 0.13587 | 1.1885 | 0.934 | 0.94 | 1 | HC_vs_MS.NoTrt_F |
| DDX1 | 0.13592 | 0.78156 | 0.205 | 0.27 | 1 | HC_vs_MS.NoTrt_F |
| ZNF99 | 0.13601 | 0.25821 | 0.627 | 0.55 | 1 | HC_vs_MS.NoTrt_F |
| MCFD2 | 0.13624 | 0.40381 | 0.934 | 0.95 | 1 | HC_vs_MS.NoTrt_F |
| ICK | 0.13624 | -0.719 | 0.246 | 0.29 | 1 | HC_vs_MS.NoTrt_F |
| LY75-CD302 | 0.13642 | 2.44632 | 0.104 | 0.15 | 1 | HC_vs_MS.NoTrt_F |
| PFKFB2 | 0.13655 | -0.4211 | 0.225 | 0.28 | 1 | HC_vs_MS.NoTrt_F |
| P2RX5 | 0.13676 | 0.62987 | 0.113 | 0.16 | 1 | HC_vs_MS.NoTrt_F |
| BTNL9 | 0.13686 | 0.8184 | 0.931 | 0.88 | 1 | HC_vs_MS.NoTrt_F |
| C1orf27 | 0.13701 | -0.3578 | 0.24 | 0.29 | 1 | HC_vs_MS.NoTrt_F |
| FDX1 | 0.13709 | -0.3233 | 0.384 | 0.42 | 1 | HC_vs_MS.NoTrt_F |
| REEP3 | 0.13709 | -0.6827 | 0.486 | 0.53 | 1 | HC_vs_MS.NoTrt_F |
| PPFIBP2 | 0.13718 | -0.9262 | 0.286 | 0.34 | 1 | HC_vs_MS.NoTrt_F |
| TBL2 | 0.13729 | -0.3586 | 0.746 | 0.67 | 1 | HC_vs_MS.NoTrt_F |
| RBM38 | 0.13747 | 0.51969 | 0.202 | 0.27 | 1 | HC_vs_MS.NoTrt_F |

| | | | | | | |
|---|---|---|---|---|---|---|
| FASN | 0.13756 | 0.73449 | 0.171 | 0.23 | 1 | HC_vs_MS.NoTrt_F |
| CDYL2 | 0.13764 | 0.37353 | 0.127 | 0.18 | 1 | HC_vs_MS.NoTrt_F |
| RNF31 | 0.13837 | 0.42299 | 0.312 | 0.38 | 1 | HC_vs_MS.NoTrt_F |
| SPDYE5 | 0.1386 | 0.25961 | 0.494 | 0.57 | 1 | HC_vs_MS.NoTrt_F |
| WRN | 0.13869 | -0.4787 | 0.587 | 0.58 | 1 | HC_vs_MS.NoTrt_F |
| GRIA4 | 0.13875 | 0.32223 | 0.191 | 0.14 | 1 | HC_vs_MS.NoTrt_F |
| PNPT1 | 0.13884 | 0.38699 | 0.882 | 0.92 | 1 | HC_vs_MS.NoTrt_F |
| GLCCI1 | 0.13905 | 0.3145 | 0.286 | 0.36 | 1 | HC_vs_MS.NoTrt_F |
| TFEB | 0.13927 | -0.7281 | 0.486 | 0.52 | 1 | HC_vs_MS.NoTrt_F |
| ZBTB7A | 0.13958 | 0.37178 | 0.162 | 0.22 | 1 | HC_vs_MS.NoTrt_F |
| SLCO2B1 | 0.13964 | 0.75084 | 0.379 | 0.45 | 1 | HC_vs_MS.NoTrt_F |
| ITGB3BP | 0.13964 | 0.36104 | 0.78 | 0.83 | 1 | HC_vs_MS.NoTrt_F |
| DMPK | 0.1399 | 0.26931 | 0.668 | 0.74 | 1 | HC_vs_MS.NoTrt_F |
| PDCD2 | 0.14011 | 0.70028 | 0.15 | 0.2 | 1 | HC_vs_MS.NoTrt_F |
| PER1 | 0.14018 | 1.01816 | 0.587 | 0.56 | 1 | HC_vs_MS.NoTrt_F |
| HEATR6 | 0.14018 | 0.35834 | 0.607 | 0.66 | 1 | HC_vs_MS.NoTrt_F |
| FAIM | 0.14025 | 0.50484 | 0.399 | 0.47 | 1 | HC_vs_MS.NoTrt_F |
| TBC1D15 | 0.14026 | -0.8507 | 0.592 | 0.66 | 1 | HC_vs_MS.NoTrt_F |
| SRL | 0.14053 | 0.27192 | 0.483 | 0.43 | 1 | HC_vs_MS.NoTrt_F |
| B4GALT5 | 0.1407 | 0.40026 | 0.286 | 0.35 | 1 | HC_vs_MS.NoTrt_F |
| TMPRSS4 | 0.14092 | 0.28073 | 0.659 | 0.61 | 1 | HC_vs_MS.NoTrt_F |
| KIAA1683 | 0.14096 | 0.35828 | 0.717 | 0.67 | 1 | HC_vs_MS.NoTrt_F |
| CMSS1 | 0.14115 | 0.4678 | 0.763 | 0.79 | 1 | HC_vs_MS.NoTrt_F |
| DUSP23 | 0.14119 | 0.50171 | 0.136 | 0.19 | 1 | HC_vs_MS.NoTrt_F |
| GPR146 | 0.14136 | 0.97255 | 0.237 | 0.18 | 1 | HC_vs_MS.NoTrt_F |
| ANKRD30BL | 0.14139 | 0.28472 | 0.621 | 0.61 | 1 | HC_vs_MS.NoTrt_F |
| AGBL5 | 0.14141 | -0.7153 | 0.179 | 0.23 | 1 | HC_vs_MS.NoTrt_F |
| SART1 | 0.14153 | -0.5647 | 0.37 | 0.42 | 1 | HC_vs_MS.NoTrt_F |
| LMTK2 | 0.14169 | 0.62725 | 0.303 | 0.37 | 1 | HC_vs_MS.NoTrt_F |
| FAM188A | 0.14223 | 0.62097 | 0.133 | 0.18 | 1 | HC_vs_MS.NoTrt_F |
| C18orf21 | 0.14243 | 0.31677 | 0.118 | 0.17 | 1 | HC_vs_MS.NoTrt_F |
| RCBTB1 | 0.14257 | 0.75623 | 0.168 | 0.23 | 1 | HC_vs_MS.NoTrt_F |
| ARHGAP31 | 0.14259 | 0.322 | 0.373 | 0.41 | 1 | HC_vs_MS.NoTrt_F |
| MSRB2 | 0.14311 | -0.2664 | 0.092 | 0.14 | 1 | HC_vs_MS.NoTrt_F |
| ENY2 | 0.14315 | 0.4786 | 0.63 | 0.55 | 1 | HC_vs_MS.NoTrt_F |
| MAFF | 0.14315 | 0.33337 | 0.723 | 0.67 | 1 | HC_vs_MS.NoTrt_F |
| MAP4 | 0.14359 | 0.58594 | 0.815 | 0.82 | 1 | HC_vs_MS.NoTrt_F |
| CDC73 | 0.14398 | 0.49738 | 0.642 | 0.55 | 1 | HC_vs_MS.NoTrt_F |
| BCAT2 | 0.14398 | -0.4417 | 0.243 | 0.29 | 1 | HC_vs_MS.NoTrt_F |
| MAATS1 | 0.14422 | 0.27869 | 0.277 | 0.35 | 1 | HC_vs_MS.NoTrt_F |

| | | | | | | |
|---|---|---|---|---|---|---|
| TRAM2 | 0.14453 | 0.28042 | 0.309 | 0.36 | 1 | HC_vs_MS.NoTrt_F |
| C4BPB | 0.1446 | 0.32893 | 0.85 | 0.85 | 1 | HC_vs_MS.NoTrt_F |
| ELL | 0.14474 | 0.84075 | 0.197 | 0.27 | 1 | HC_vs_MS.NoTrt_F |
| SETD5 | 0.14503 | 0.50211 | 0.665 | 0.7 | 1 | HC_vs_MS.NoTrt_F |
| CCDC153 | 0.14507 | 0.5415 | 0.269 | 0.35 | 1 | HC_vs_MS.NoTrt_F |
| SIK1 | 0.14514 | 1.27007 | 0.159 | 0.22 | 1 | HC_vs_MS.NoTrt_F |
| GALNT1 | 0.14544 | 1.8548 | 0.101 | 0.15 | 1 | HC_vs_MS.NoTrt_F |
| ZNF462 | 0.1455 | 0.73382 | 0.24 | 0.19 | 1 | HC_vs_MS.NoTrt_F |
| TM2D2 | 0.14578 | 0.80221 | 0.454 | 0.4 | 1 | HC_vs_MS.NoTrt_F |
| DLG5 | 0.14609 | 0.42396 | 0.517 | 0.45 | 1 | HC_vs_MS.NoTrt_F |
| SIAH1 | 0.14672 | 0.57158 | 0.275 | 0.34 | 1 | HC_vs_MS.NoTrt_F |
| KCTD9 | 0.14681 | -1.0421 | 0.263 | 0.32 | 1 | HC_vs_MS.NoTrt_F |
| RBM41 | 0.14702 | 0.35009 | 0.428 | 0.5 | 1 | HC_vs_MS.NoTrt_F |
| THG1L | 0.14722 | -0.5976 | 0.621 | 0.71 | 1 | HC_vs_MS.NoTrt_F |
| SIGLEC8 | 0.14776 | 0.54618 | 0.578 | 0.54 | 1 | HC_vs_MS.NoTrt_F |
| MRRF | 0.14795 | -1.1172 | 0.685 | 0.67 | 1 | HC_vs_MS.NoTrt_F |
| TKFC | 0.14812 | 0.84759 | 0.639 | 0.61 | 1 | HC_vs_MS.NoTrt_F |
| MTERF1 | 0.14815 | -0.959 | 0.254 | 0.3 | 1 | HC_vs_MS.NoTrt_F |
| PIGU | 0.14823 | 0.97449 | 0.136 | 0.19 | 1 | HC_vs_MS.NoTrt_F |
| ACOT4 | 0.14838 | 0.33165 | 0.15 | 0.21 | 1 | HC_vs_MS.NoTrt_F |
| CDYL | 0.14838 | -0.4064 | 0.107 | 0.16 | 1 | HC_vs_MS.NoTrt_F |
| DNAH7 | 0.14858 | 0.47966 | 0.694 | 0.67 | 1 | HC_vs_MS.NoTrt_F |
| C11orf21 | 0.14879 | 0.54414 | 0.173 | 0.23 | 1 | HC_vs_MS.NoTrt_F |
| DUSP2 | 0.14881 | 0.48332 | 0.332 | 0.43 | 1 | HC_vs_MS.NoTrt_F |
| ATPIF1 | 0.1489 | 0.68726 | 0.298 | 0.4 | 1 | HC_vs_MS.NoTrt_F |
| COL6A1 | 0.149 | 3.10279 | 0.107 | 0.07 | 1 | HC_vs_MS.NoTrt_F |
| FDPS | 0.14925 | -0.2521 | 0.526 | 0.55 | 1 | HC_vs_MS.NoTrt_F |
| SNX19 | 0.14931 | 0.25145 | 0.269 | 0.34 | 1 | HC_vs_MS.NoTrt_F |
| ERGIC2 | 0.14937 | -0.3919 | 0.526 | 0.55 | 1 | HC_vs_MS.NoTrt_F |
| RXRA | 0.14941 | 0.36461 | 0.225 | 0.29 | 1 | HC_vs_MS.NoTrt_F |
| ZNF549 | 0.14944 | 0.54241 | 0.688 | 0.73 | 1 | HC_vs_MS.NoTrt_F |
| ZDHHC17 | 0.14948 | 0.70341 | 0.341 | 0.42 | 1 | HC_vs_MS.NoTrt_F |
| NUDCD3 | 0.14948 | 0.51328 | 0.344 | 0.41 | 1 | HC_vs_MS.NoTrt_F |
| GRAPL | 0.14955 | 0.93587 | 0.104 | 0.15 | 1 | HC_vs_MS.NoTrt_F |
| G2E3 | 0.1499 | -0.8927 | 0.549 | 0.58 | 1 | HC_vs_MS.NoTrt_F |
| SULT1A3 | 0.14994 | 0.29498 | 0.679 | 0.73 | 1 | HC_vs_MS.NoTrt_F |
| ZNF76 | 0.15011 | 0.42226 | 0.22 | 0.28 | 1 | HC_vs_MS.NoTrt_F |
| RIPK1 | 0.15035 | 0.33446 | 0.237 | 0.29 | 1 | HC_vs_MS.NoTrt_F |
| ATXN2 | 0.15056 | 0.26402 | 0.934 | 0.92 | 1 | HC_vs_MS.NoTrt_F |
| LRRC28 | 0.15072 | -0.4645 | 0.624 | 0.67 | 1 | HC_vs_MS.NoTrt_F |

| | | | | | | |
|---|---|---|---|---|---|---|
| MGRN1 | 0.15079 | 0.31648 | 0.864 | 0.79 | 1 | HC_vs_MS.NoTrt_F |
| NAMPT | 0.1509 | 0.85291 | 0.824 | 0.8 | 1 | HC_vs_MS.NoTrt_F |
| VWA7 | 0.15112 | 0.57718 | 0.387 | 0.46 | 1 | HC_vs_MS.NoTrt_F |
| LCA5L | 0.15122 | 0.58693 | 0.416 | 0.5 | 1 | HC_vs_MS.NoTrt_F |
| TRPM1 | 0.15136 | 0.34384 | 0.133 | 0.18 | 1 | HC_vs_MS.NoTrt_F |
| RHOC | 0.15137 | 0.90251 | 0.266 | 0.22 | 1 | HC_vs_MS.NoTrt_F |
| AARS | 0.15166 | -0.432 | 0.428 | 0.5 | 1 | HC_vs_MS.NoTrt_F |
| TLR6 | 0.15169 | -0.4803 | 0.309 | 0.38 | 1 | HC_vs_MS.NoTrt_F |
| PRKCSH | 0.15175 | 0.81194 | 0.725 | 0.72 | 1 | HC_vs_MS.NoTrt_F |
| ATP11C | 0.15202 | 1.19588 | 0.09 | 0.13 | 1 | HC_vs_MS.NoTrt_F |
| HNRNPAB | 0.15225 | 0.40186 | 0.223 | 0.29 | 1 | HC_vs_MS.NoTrt_F |
| LAX1 | 0.15248 | 0.28508 | 0.587 | 0.62 | 1 | HC_vs_MS.NoTrt_F |
| TNKS1BP1 | 0.15286 | 0.5442 | 0.118 | 0.08 | 1 | HC_vs_MS.NoTrt_F |
| WRAP73 | 0.15303 | -0.502 | 0.37 | 0.42 | 1 | HC_vs_MS.NoTrt_F |
| ADAT1 | 0.15304 | -0.6067 | 0.717 | 0.73 | 1 | HC_vs_MS.NoTrt_F |
| MCAM | 0.1531 | 0.94133 | 0.598 | 0.53 | 1 | HC_vs_MS.NoTrt_F |
| PRPSAP2 | 0.1531 | -0.5903 | 0.653 | 0.71 | 1 | HC_vs_MS.NoTrt_F |
| RABGGTA | 0.15313 | 0.2678 | 0.153 | 0.21 | 1 | HC_vs_MS.NoTrt_F |
| PLA2G5 | 0.15348 | 0.29261 | 0.309 | 0.26 | 1 | HC_vs_MS.NoTrt_F |
| BCAM | 0.15392 | 0.54922 | 0.786 | 0.75 | 1 | HC_vs_MS.NoTrt_F |
| HIF3A | 0.15508 | 0.37245 | 0.908 | 0.88 | 1 | HC_vs_MS.NoTrt_F |
| ALG12 | 0.15526 | 0.29616 | 0.197 | 0.25 | 1 | HC_vs_MS.NoTrt_F |
| SMN2 | 0.15547 | 0.27314 | 0.844 | 0.73 | 1 | HC_vs_MS.NoTrt_F |
| ST8SIA5 | 0.15582 | 0.27874 | 0.723 | 0.69 | 1 | HC_vs_MS.NoTrt_F |
| NEIL3 | 0.15592 | -0.7101 | 0.295 | 0.34 | 1 | HC_vs_MS.NoTrt_F |
| BIRC5 | 0.15599 | 0.50049 | 0.725 | 0.72 | 1 | HC_vs_MS.NoTrt_F |
| MPP7 | 0.15601 | 0.96308 | 0.347 | 0.41 | 1 | HC_vs_MS.NoTrt_F |
| DDX43 | 0.15605 | -0.6255 | 0.185 | 0.24 | 1 | HC_vs_MS.NoTrt_F |
| MT1X | 0.15634 | 0.39782 | 0.165 | 0.22 | 1 | HC_vs_MS.NoTrt_F |
| DESI1 | 0.15646 | -0.3953 | 0.633 | 0.68 | 1 | HC_vs_MS.NoTrt_F |
| HELZ | 0.15657 | 0.29729 | 0.699 | 0.76 | 1 | HC_vs_MS.NoTrt_F |
| B3GNT5 | 0.15663 | -0.8232 | 0.587 | 0.62 | 1 | HC_vs_MS.NoTrt_F |
| HIST1H2BG | 0.15673 | 0.25481 | 0.497 | 0.43 | 1 | HC_vs_MS.NoTrt_F |
| TMEM154 | 0.15701 | 0.2849 | 0.945 | 0.93 | 1 | HC_vs_MS.NoTrt_F |
| DNMBP | 0.15706 | 1.66475 | 0.101 | 0.14 | 1 | HC_vs_MS.NoTrt_F |
| MPRIP | 0.15708 | 0.36257 | 0.948 | 0.92 | 1 | HC_vs_MS.NoTrt_F |
| GOLGA8N | 0.15722 | 0.6332 | 0.101 | 0.15 | 1 | HC_vs_MS.NoTrt_F |
| PCED1B | 0.15761 | 0.62124 | 0.509 | 0.62 | 1 | HC_vs_MS.NoTrt_F |
| PLXNA4 | 0.15777 | 0.30286 | 0.104 | 0.15 | 1 | HC_vs_MS.NoTrt_F |
| TPCN2 | 0.15786 | -0.5477 | 0.682 | 0.75 | 1 | HC_vs_MS.NoTrt_F |

| | | | | | | |
|---|---|---|---|---|---|---|
| NAV3 | 0.15841 | 0.3023 | 0.754 | 0.72 | 1 | HC_vs_MS.NoTrt_F |
| TLL2 | 0.1585 | 0.32792 | 0.462 | 0.42 | 1 | HC_vs_MS.NoTrt_F |
| RNF114 | 0.15946 | -0.287 | 0.772 | 0.8 | 1 | HC_vs_MS.NoTrt_F |
| PDP2 | 0.15951 | -0.2905 | 0.835 | 0.8 | 1 | HC_vs_MS.NoTrt_F |
| SUFU | 0.15953 | 0.26708 | 0.121 | 0.17 | 1 | HC_vs_MS.NoTrt_F |
| CYP20A1 | 0.15957 | -0.4028 | 0.919 | 0.92 | 1 | HC_vs_MS.NoTrt_F |
| WDR7 | 0.15971 | -1.5682 | 0.373 | 0.42 | 1 | HC_vs_MS.NoTrt_F |
| FANCM | 0.15971 | -0.6884 | 0.133 | 0.18 | 1 | HC_vs_MS.NoTrt_F |
| ABCD3 | 0.15978 | -0.9316 | 0.176 | 0.23 | 1 | HC_vs_MS.NoTrt_F |
| NUDT7 | 0.15984 | 0.26667 | 0.24 | 0.29 | 1 | HC_vs_MS.NoTrt_F |
| SGF29 | 0.15989 | 0.4323 | 0.306 | 0.37 | 1 | HC_vs_MS.NoTrt_F |
| ZNF227 | 0.16041 | 0.48206 | 0.549 | 0.51 | 1 | HC_vs_MS.NoTrt_F |
| COQ5 | 0.16045 | 0.61952 | 0.876 | 0.9 | 1 | HC_vs_MS.NoTrt_F |
| C11orf80 | 0.16081 | 0.49684 | 0.494 | 0.56 | 1 | HC_vs_MS.NoTrt_F |
| RABL3 | 0.16135 | -0.7066 | 0.607 | 0.64 | 1 | HC_vs_MS.NoTrt_F |
| GPN3 | 0.1615 | 0.74643 | 0.379 | 0.45 | 1 | HC_vs_MS.NoTrt_F |
| RCC1L | 0.16164 | -0.5515 | 0.301 | 0.35 | 1 | HC_vs_MS.NoTrt_F |
| SYTL2 | 0.16196 | 0.37566 | 0.754 | 0.74 | 1 | HC_vs_MS.NoTrt_F |
| CARHSP1 | 0.16273 | -0.634 | 0.688 | 0.72 | 1 | HC_vs_MS.NoTrt_F |
| 7-Mar | 0.16295 | 0.50107 | 0.263 | 0.33 | 1 | HC_vs_MS.NoTrt_F |
| WRNIP1 | 0.16308 | 0.85866 | 0.133 | 0.18 | 1 | HC_vs_MS.NoTrt_F |
| NCKAP5L | 0.16334 | 0.55261 | 0.714 | 0.67 | 1 | HC_vs_MS.NoTrt_F |
| PSKH1 | 0.1634 | 0.86853 | 0.13 | 0.18 | 1 | HC_vs_MS.NoTrt_F |
| NCOA1 | 0.16342 | 0.82282 | 0.223 | 0.28 | 1 | HC_vs_MS.NoTrt_F |
| TNFRSF19 | 0.16352 | 0.36542 | 0.182 | 0.24 | 1 | HC_vs_MS.NoTrt_F |
| ART4 | 0.1638 | 0.40161 | 0.572 | 0.5 | 1 | HC_vs_MS.NoTrt_F |
| SORBS1 | 0.1639 | -0.3483 | 0.269 | 0.33 | 1 | HC_vs_MS.NoTrt_F |
| ZNF66 | 0.16436 | -0.5092 | 0.671 | 0.62 | 1 | HC_vs_MS.NoTrt_F |
| KHSRP | 0.16441 | 0.26601 | 0.153 | 0.2 | 1 | HC_vs_MS.NoTrt_F |
| GTSF1 | 0.16442 | -0.9734 | 0.324 | 0.27 | 1 | HC_vs_MS.NoTrt_F |
| CHMP2A | 0.16445 | 0.4299 | 0.199 | 0.26 | 1 | HC_vs_MS.NoTrt_F |
| MLLT1 | 0.16451 | 1.29797 | 0.098 | 0.14 | 1 | HC_vs_MS.NoTrt_F |
| ZC2HC1C | 0.1647 | -0.3479 | 0.124 | 0.17 | 1 | HC_vs_MS.NoTrt_F |
| CECR2 | 0.16483 | 0.61333 | 0.558 | 0.53 | 1 | HC_vs_MS.NoTrt_F |
| USP4 | 0.16499 | 0.28926 | 0.977 | 0.94 | 1 | HC_vs_MS.NoTrt_F |
| PAN2 | 0.16512 | 0.25689 | 0.353 | 0.4 | 1 | HC_vs_MS.NoTrt_F |
| MORN4 | 0.16517 | -0.4747 | 0.61 | 0.65 | 1 | HC_vs_MS.NoTrt_F |
| WIPI1 | 0.16545 | 0.34684 | 0.893 | 0.89 | 1 | HC_vs_MS.NoTrt_F |
| GPAA1 | 0.16577 | 0.6171 | 0.182 | 0.24 | 1 | HC_vs_MS.NoTrt_F |
| CNTLN | 0.16612 | 0.35923 | 0.436 | 0.36 | 1 | HC_vs_MS.NoTrt_F |

| | | | | | | |
|---|---|---|---|---|---|---|
| PAK1 | 0.16658 | -0.5927 | 0.202 | 0.26 | 1 | HC_vs_MS.NoTrt_F |
| SCARB1 | 0.16677 | 0.51796 | 0.587 | 0.51 | 1 | HC_vs_MS.NoTrt_F |
| CDAN1 | 0.16682 | 0.60993 | 0.529 | 0.56 | 1 | HC_vs_MS.NoTrt_F |
| ZNF460 | 0.16689 | 0.58331 | 0.22 | 0.28 | 1 | HC_vs_MS.NoTrt_F |
| TUBD1 | 0.16699 | -0.5026 | 0.775 | 0.82 | 1 | HC_vs_MS.NoTrt_F |
| POLD3 | 0.16707 | -1.2502 | 0.176 | 0.22 | 1 | HC_vs_MS.NoTrt_F |
| ENOX2 | 0.16711 | 0.68845 | 0.145 | 0.19 | 1 | HC_vs_MS.NoTrt_F |
| MALT1 | 0.16786 | 0.49404 | 0.153 | 0.2 | 1 | HC_vs_MS.NoTrt_F |
| SENP1 | 0.16813 | 0.74014 | 0.231 | 0.3 | 1 | HC_vs_MS.NoTrt_F |
| IPO11 | 0.16817 | 0.2956 | 0.173 | 0.23 | 1 | HC_vs_MS.NoTrt_F |
| ZNF385D | 0.16853 | 0.34725 | 0.327 | 0.4 | 1 | HC_vs_MS.NoTrt_F |
| SGTA | 0.16868 | -0.3223 | 0.59 | 0.64 | 1 | HC_vs_MS.NoTrt_F |
| ANKS1B | 0.16885 | 0.29819 | 0.743 | 0.62 | 1 | HC_vs_MS.NoTrt_F |
| CEP63 | 0.16934 | -0.5262 | 0.786 | 0.8 | 1 | HC_vs_MS.NoTrt_F |
| ST13 | 0.16939 | 0.31241 | 0.355 | 0.44 | 1 | HC_vs_MS.NoTrt_F |
| SEC14L6 | 0.16943 | 0.44821 | 0.798 | 0.76 | 1 | HC_vs_MS.NoTrt_F |
| FSD1 | 0.16945 | 0.73595 | 0.457 | 0.42 | 1 | HC_vs_MS.NoTrt_F |
| STEAP4 | 0.16965 | 0.40369 | 0.743 | 0.76 | 1 | HC_vs_MS.NoTrt_F |
| MT-ATP6 | 0.16965 | 0.32731 | 0.89 | 0.92 | 1 | HC_vs_MS.NoTrt_F |
| CADM1 | 0.17014 | -0.9639 | 0.289 | 0.33 | 1 | HC_vs_MS.NoTrt_F |
| EHD2 | 0.17046 | 0.26547 | 0.884 | 0.91 | 1 | HC_vs_MS.NoTrt_F |
| AGAP3 | 0.1705 | 0.83934 | 0.072 | 0.11 | 1 | HC_vs_MS.NoTrt_F |
| TRABD | 0.17063 | 0.27456 | 0.382 | 0.48 | 1 | HC_vs_MS.NoTrt_F |
| ABCA10 | 0.17078 | 1.30191 | 0.095 | 0.14 | 1 | HC_vs_MS.NoTrt_F |
| KIAA1614 | 0.17093 | 0.63313 | 0.335 | 0.38 | 1 | HC_vs_MS.NoTrt_F |
| ZNRF2 | 0.17094 | 1.31511 | 0.098 | 0.14 | 1 | HC_vs_MS.NoTrt_F |
| RPS6KA2 | 0.17094 | 0.46781 | 0.113 | 0.16 | 1 | HC_vs_MS.NoTrt_F |
| RIMKLB | 0.17112 | 0.42754 | 0.118 | 0.16 | 1 | HC_vs_MS.NoTrt_F |
| RP11-111H1 | 0.17155 | -0.2915 | 0.136 | 0.18 | 1 | HC_vs_MS.NoTrt_F |
| SPECC1L-AD | 0.17255 | -0.9572 | 0.231 | 0.28 | 1 | HC_vs_MS.NoTrt_F |
| GOLGA6L9 | 0.17266 | 1.08479 | 0.072 | 0.11 | 1 | HC_vs_MS.NoTrt_F |
| UBE2Z | 0.17286 | 0.29687 | 0.286 | 0.34 | 1 | HC_vs_MS.NoTrt_F |
| SLC25A29 | 0.17289 | 0.3386 | 0.387 | 0.45 | 1 | HC_vs_MS.NoTrt_F |
| HSDL2 | 0.17294 | -0.3759 | 0.185 | 0.24 | 1 | HC_vs_MS.NoTrt_F |
| GSTM4 | 0.1734 | 2.5792 | 0.211 | 0.17 | 1 | HC_vs_MS.NoTrt_F |
| GRAMD1B | 0.17341 | -0.4083 | 0.312 | 0.37 | 1 | HC_vs_MS.NoTrt_F |
| MCOLN1 | 0.17416 | 0.37107 | 0.806 | 0.82 | 1 | HC_vs_MS.NoTrt_F |
| DUSP1 | 0.17421 | 0.28455 | 0.624 | 0.72 | 1 | HC_vs_MS.NoTrt_F |
| RPS27 | 0.17438 | 0.277 | 0.301 | 0.39 | 1 | HC_vs_MS.NoTrt_F |
| MLEC | 0.1746 | 0.32869 | 0.15 | 0.2 | 1 | HC_vs_MS.NoTrt_F |

| | | | | | | |
|---|---|---|---|---|---|---|
| ZFP64 | 0.17464 | -0.5411 | 0.633 | 0.64 | 1 | HC_vs_MS.NoTrt_F |
| RELN | 0.17503 | 0.2675 | 0.156 | 0.21 | 1 | HC_vs_MS.NoTrt_F |
| MAPK8 | 0.17506 | 0.33036 | 0.497 | 0.56 | 1 | HC_vs_MS.NoTrt_F |
| SPDYA | 0.17514 | 0.27379 | 0.118 | 0.17 | 1 | HC_vs_MS.NoTrt_F |
| TESPA1 | 0.17518 | 0.26832 | 0.451 | 0.52 | 1 | HC_vs_MS.NoTrt_F |
| TRIM5 | 0.17561 | 0.41183 | 0.503 | 0.56 | 1 | HC_vs_MS.NoTrt_F |
| CAMK2N1 | 0.17569 | -0.3923 | 0.113 | 0.16 | 1 | HC_vs_MS.NoTrt_F |
| VNN3 | 0.17576 | 1.31362 | 0.477 | 0.47 | 1 | HC_vs_MS.NoTrt_F |
| LAMA4 | 0.176 | 0.78124 | 0.355 | 0.3 | 1 | HC_vs_MS.NoTrt_F |
| MOSPD1 | 0.17607 | -0.8129 | 0.217 | 0.27 | 1 | HC_vs_MS.NoTrt_F |
| ZSCAN31 | 0.17622 | 0.4762 | 0.139 | 0.1 | 1 | HC_vs_MS.NoTrt_F |
| CCDC86 | 0.17635 | 1.13947 | 0.09 | 0.13 | 1 | HC_vs_MS.NoTrt_F |
| PHF6 | 0.17649 | -0.957 | 0.714 | 0.74 | 1 | HC_vs_MS.NoTrt_F |
| TSC22D1 | 0.17668 | 1.7624 | 0.116 | 0.08 | 1 | HC_vs_MS.NoTrt_F |
| PABPC1L | 0.17684 | 0.43717 | 0.321 | 0.39 | 1 | HC_vs_MS.NoTrt_F |
| SLC25A15 | 0.177 | -0.3245 | 0.754 | 0.7 | 1 | HC_vs_MS.NoTrt_F |
| PTPN20 | 0.17765 | 1.08868 | 0.081 | 0.12 | 1 | HC_vs_MS.NoTrt_F |
| DDX39B | 0.17771 | 0.5121 | 0.783 | 0.8 | 1 | HC_vs_MS.NoTrt_F |
| GDAP2 | 0.17789 | 0.63287 | 0.766 | 0.69 | 1 | HC_vs_MS.NoTrt_F |
| SPATA5 | 0.17806 | 0.86897 | 0.798 | 0.73 | 1 | HC_vs_MS.NoTrt_F |
| ABCB8 | 0.17811 | 0.43427 | 0.627 | 0.56 | 1 | HC_vs_MS.NoTrt_F |
| SKIV2L | 0.17819 | 0.69694 | 0.185 | 0.24 | 1 | HC_vs_MS.NoTrt_F |
| FGD4 | 0.17837 | 0.93402 | 0.861 | 0.84 | 1 | HC_vs_MS.NoTrt_F |
| ATM | 0.17839 | 0.51293 | 0.977 | 0.95 | 1 | HC_vs_MS.NoTrt_F |
| ORMDL2 | 0.17842 | -0.2819 | 0.549 | 0.56 | 1 | HC_vs_MS.NoTrt_F |
| TSTD2 | 0.17849 | 0.54748 | 0.121 | 0.17 | 1 | HC_vs_MS.NoTrt_F |
| C2orf49 | 0.17874 | -0.3607 | 0.462 | 0.52 | 1 | HC_vs_MS.NoTrt_F |
| TCOF1 | 0.1788 | -0.3502 | 0.809 | 0.88 | 1 | HC_vs_MS.NoTrt_F |
| OSGEP | 0.17883 | -0.5989 | 0.671 | 0.68 | 1 | HC_vs_MS.NoTrt_F |
| SEC11A | 0.17884 | -0.3462 | 0.853 | 0.88 | 1 | HC_vs_MS.NoTrt_F |
| AGO2 | 0.17886 | 0.78262 | 0.468 | 0.45 | 1 | HC_vs_MS.NoTrt_F |
| FBXO18 | 0.17905 | -0.4667 | 0.448 | 0.5 | 1 | HC_vs_MS.NoTrt_F |
| ATG7 | 0.17917 | -0.621 | 0.63 | 0.69 | 1 | HC_vs_MS.NoTrt_F |
| UBXN8 | 0.17928 | 1.47479 | 0.22 | 0.28 | 1 | HC_vs_MS.NoTrt_F |
| ARID5A | 0.17939 | 0.39062 | 0.225 | 0.29 | 1 | HC_vs_MS.NoTrt_F |
| NETO2 | 0.17943 | -0.4947 | 0.587 | 0.61 | 1 | HC_vs_MS.NoTrt_F |
| IFNAR1 | 0.17943 | 0.51377 | 0.561 | 0.51 | 1 | HC_vs_MS.NoTrt_F |
| SLC10A1 | 0.17955 | 1.6032 | 0.072 | 0.11 | 1 | HC_vs_MS.NoTrt_F |
| CENPM | 0.18027 | -0.4916 | 0.382 | 0.43 | 1 | HC_vs_MS.NoTrt_F |
| GLA | 0.18035 | 1.92113 | 0.084 | 0.13 | 1 | HC_vs_MS.NoTrt_F |

| | | | | | | |
|---|---|---|---|---|---|---|
| AAR2 | 0.18036 | 0.66064 | 0.266 | 0.32 | 1 | HC_vs_MS.NoTrt_F |
| 1-Dec | 0.18052 | 0.33106 | 0.575 | 0.54 | 1 | HC_vs_MS.NoTrt_F |
| DAG1 | 0.18062 | 1.02417 | 0.604 | 0.59 | 1 | HC_vs_MS.NoTrt_F |
| IQCG | 0.18092 | 0.37832 | 0.827 | 0.75 | 1 | HC_vs_MS.NoTrt_F |
| COPS6 | 0.18094 | 0.3283 | 0.708 | 0.66 | 1 | HC_vs_MS.NoTrt_F |
| ATXN2L | 0.18102 | 0.95147 | 0.298 | 0.38 | 1 | HC_vs_MS.NoTrt_F |
| NFKB2 | 0.1814 | -0.3638 | 0.428 | 0.49 | 1 | HC_vs_MS.NoTrt_F |
| USF2 | 0.18155 | 0.31536 | 0.627 | 0.65 | 1 | HC_vs_MS.NoTrt_F |
| PHYH | 0.18197 | 0.37674 | 0.309 | 0.35 | 1 | HC_vs_MS.NoTrt_F |
| EDEM1 | 0.18201 | 0.48101 | 0.335 | 0.43 | 1 | HC_vs_MS.NoTrt_F |
| NPHP1 | 0.1821 | 0.38909 | 0.746 | 0.72 | 1 | HC_vs_MS.NoTrt_F |
| RP9 | 0.1822 | 0.32669 | 0.127 | 0.17 | 1 | HC_vs_MS.NoTrt_F |
| NTHL1 | 0.18224 | -0.5076 | 0.075 | 0.11 | 1 | HC_vs_MS.NoTrt_F |
| TRDMT1 | 0.18244 | -0.2544 | 0.78 | 0.77 | 1 | HC_vs_MS.NoTrt_F |
| CPPED1 | 0.1826 | -0.2804 | 0.65 | 0.7 | 1 | HC_vs_MS.NoTrt_F |
| TGFBR2 | 0.18291 | 0.85064 | 0.277 | 0.23 | 1 | HC_vs_MS.NoTrt_F |
| GALNT9 | 0.18309 | -0.2693 | 0.223 | 0.27 | 1 | HC_vs_MS.NoTrt_F |
| CKAP4 | 0.18312 | 1.58346 | 0.11 | 0.16 | 1 | HC_vs_MS.NoTrt_F |
| TIMM44 | 0.18323 | 0.68332 | 0.673 | 0.77 | 1 | HC_vs_MS.NoTrt_F |
| EXOC6 | 0.18327 | 0.45261 | 0.341 | 0.38 | 1 | HC_vs_MS.NoTrt_F |
| HSD17B13 | 0.18331 | 0.65951 | 0.636 | 0.58 | 1 | HC_vs_MS.NoTrt_F |
| WHSC1 | 0.18342 | 0.5212 | 0.74 | 0.74 | 1 | HC_vs_MS.NoTrt_F |
| JOSD1 | 0.1835 | 0.4183 | 0.639 | 0.68 | 1 | HC_vs_MS.NoTrt_F |
| CTIF | 0.18352 | 0.3701 | 0.234 | 0.29 | 1 | HC_vs_MS.NoTrt_F |
| CLEC7A | 0.18368 | 0.88226 | 0.665 | 0.66 | 1 | HC_vs_MS.NoTrt_F |
| PPIL4 | 0.1838 | 0.28646 | 0.277 | 0.34 | 1 | HC_vs_MS.NoTrt_F |
| WDR59 | 0.18393 | -0.3836 | 0.951 | 0.98 | 1 | HC_vs_MS.NoTrt_F |
| CBLB | 0.18398 | 1.12503 | 0.431 | 0.4 | 1 | HC_vs_MS.NoTrt_F |
| C1orf186 | 0.18406 | 1.12905 | 0.587 | 0.69 | 1 | HC_vs_MS.NoTrt_F |
| OSBPL7 | 0.18432 | 0.81391 | 0.116 | 0.16 | 1 | HC_vs_MS.NoTrt_F |
| FLT4 | 0.18439 | 0.65109 | 0.124 | 0.17 | 1 | HC_vs_MS.NoTrt_F |
| RP11-1212A | 0.18441 | 0.25014 | 0.546 | 0.49 | 1 | HC_vs_MS.NoTrt_F |
| ADGRE5 | 0.18448 | 0.54126 | 0.734 | 0.69 | 1 | HC_vs_MS.NoTrt_F |
| HPF1 | 0.18473 | 0.93886 | 0.145 | 0.19 | 1 | HC_vs_MS.NoTrt_F |
| PTPRM | 0.18489 | 0.99146 | 0.225 | 0.29 | 1 | HC_vs_MS.NoTrt_F |
| TMEM52B | 0.18509 | 0.38474 | 0.801 | 0.79 | 1 | HC_vs_MS.NoTrt_F |
| LMNB1 | 0.1851 | 0.30634 | 0.147 | 0.19 | 1 | HC_vs_MS.NoTrt_F |
| UBC | 0.18528 | 0.70412 | 0.578 | 0.55 | 1 | HC_vs_MS.NoTrt_F |
| ATG14 | 0.18604 | 0.54629 | 0.72 | 0.66 | 1 | HC_vs_MS.NoTrt_F |
| WDR33 | 0.18618 | 0.66263 | 0.266 | 0.33 | 1 | HC_vs_MS.NoTrt_F |

| | | | | | | |
|---|---|---|---|---|---|---|
| GSTZ1 | 0.18624 | 0.80268 | 0.595 | 0.54 | 1 | HC_vs_MS.NoTrt_F |
| CCDC28B | 0.18652 | -0.4802 | 0.483 | 0.51 | 1 | HC_vs_MS.NoTrt_F |
| C11orf71 | 0.1868 | 0.5999 | 0.118 | 0.16 | 1 | HC_vs_MS.NoTrt_F |
| ZNF559-ZNF | 0.18682 | 1.18968 | 0.069 | 0.11 | 1 | HC_vs_MS.NoTrt_F |
| TUBG2 | 0.18703 | -0.5329 | 0.147 | 0.19 | 1 | HC_vs_MS.NoTrt_F |
| ZNF566 | 0.18704 | -0.2986 | 0.934 | 0.87 | 1 | HC_vs_MS.NoTrt_F |
| CHRNA3 | 0.18745 | -0.3417 | 0.194 | 0.24 | 1 | HC_vs_MS.NoTrt_F |
| LRRC58 | 0.18747 | 0.81852 | 0.731 | 0.74 | 1 | HC_vs_MS.NoTrt_F |
| DDX25 | 0.18763 | 0.272 | 0.728 | 0.71 | 1 | HC_vs_MS.NoTrt_F |
| GPAT4 | 0.18773 | -0.2927 | 0.783 | 0.85 | 1 | HC_vs_MS.NoTrt_F |
| TPD52L1 | 0.18832 | 0.57888 | 0.769 | 0.79 | 1 | HC_vs_MS.NoTrt_F |
| PCBP1 | 0.1886 | 0.8223 | 0.561 | 0.56 | 1 | HC_vs_MS.NoTrt_F |
| IFT80 | 0.18942 | 0.67436 | 0.792 | 0.79 | 1 | HC_vs_MS.NoTrt_F |
| CYP2F1 | 0.18957 | 0.68876 | 0.185 | 0.24 | 1 | HC_vs_MS.NoTrt_F |
| DERA | 0.18958 | 0.66124 | 0.113 | 0.16 | 1 | HC_vs_MS.NoTrt_F |
| PILRB | 0.18967 | 0.4961 | 0.48 | 0.54 | 1 | HC_vs_MS.NoTrt_F |
| HDDC3 | 0.18971 | -1.0737 | 0.396 | 0.43 | 1 | HC_vs_MS.NoTrt_F |
| MIOS | 0.18975 | 0.63142 | 0.306 | 0.37 | 1 | HC_vs_MS.NoTrt_F |
| TOM1L2 | 0.18975 | 0.47992 | 0.257 | 0.32 | 1 | HC_vs_MS.NoTrt_F |
| STYXL1 | 0.18977 | 0.43621 | 0.306 | 0.35 | 1 | HC_vs_MS.NoTrt_F |
| RAD23A | 0.18988 | -0.2802 | 0.766 | 0.77 | 1 | HC_vs_MS.NoTrt_F |
| GMDS | 0.19066 | 0.53673 | 0.194 | 0.25 | 1 | HC_vs_MS.NoTrt_F |
| RPS6KB2 | 0.19078 | -0.285 | 0.861 | 0.87 | 1 | HC_vs_MS.NoTrt_F |
| PTRH1 | 0.19098 | 0.36729 | 0.777 | 0.74 | 1 | HC_vs_MS.NoTrt_F |
| ATXN1L | 0.19113 | 0.57372 | 0.127 | 0.17 | 1 | HC_vs_MS.NoTrt_F |
| ITGAE | 0.19135 | -0.3427 | 0.601 | 0.67 | 1 | HC_vs_MS.NoTrt_F |
| PDZK1 | 0.19155 | 0.48017 | 0.355 | 0.29 | 1 | HC_vs_MS.NoTrt_F |
| EML2 | 0.19157 | 0.59603 | 0.714 | 0.68 | 1 | HC_vs_MS.NoTrt_F |
| ACRC | 0.1916 | -0.3929 | 0.454 | 0.49 | 1 | HC_vs_MS.NoTrt_F |
| LAP3 | 0.19185 | 0.71998 | 0.197 | 0.25 | 1 | HC_vs_MS.NoTrt_F |
| GFI1B | 0.19198 | 0.38262 | 0.581 | 0.64 | 1 | HC_vs_MS.NoTrt_F |
| THBS4 | 0.19201 | 1.63067 | 0.078 | 0.12 | 1 | HC_vs_MS.NoTrt_F |
| CXCR3 | 0.19234 | 0.43348 | 0.228 | 0.3 | 1 | HC_vs_MS.NoTrt_F |
| BMP8A | 0.19245 | 0.35244 | 0.668 | 0.71 | 1 | HC_vs_MS.NoTrt_F |
| ZNF395 | 0.19251 | 0.92662 | 0.592 | 0.67 | 1 | HC_vs_MS.NoTrt_F |
| ATAD1 | 0.19273 | 0.26779 | 0.173 | 0.23 | 1 | HC_vs_MS.NoTrt_F |
| METTL8 | 0.19309 | 0.39252 | 0.948 | 0.93 | 1 | HC_vs_MS.NoTrt_F |
| ZSCAN12 | 0.19312 | 1.44593 | 0.468 | 0.4 | 1 | HC_vs_MS.NoTrt_F |
| GMEB2 | 0.19312 | 0.93773 | 0.13 | 0.17 | 1 | HC_vs_MS.NoTrt_F |
| PLCH2 | 0.19334 | 2.73371 | 0.104 | 0.15 | 1 | HC_vs_MS.NoTrt_F |

| | | | | | | |
|---|---|---|---|---|---|---|
| BCAS1 | 0.19392 | -1.0458 | 0.425 | 0.46 | 1 | HC_vs_MS.NoTrt_F |
| EXD1 | 0.19416 | 0.30791 | 0.581 | 0.55 | 1 | HC_vs_MS.NoTrt_F |
| NDUFV1 | 0.19426 | 0.25299 | 0.338 | 0.4 | 1 | HC_vs_MS.NoTrt_F |
| AC025335.1 | 0.19437 | -0.2838 | 0.685 | 0.58 | 1 | HC_vs_MS.NoTrt_F |
| TMEM38B | 0.19454 | -0.5615 | 0.272 | 0.32 | 1 | HC_vs_MS.NoTrt_F |
| U2AF2 | 0.19463 | 0.53809 | 0.243 | 0.3 | 1 | HC_vs_MS.NoTrt_F |
| KMT2C | 0.1947 | 0.3464 | 0.451 | 0.56 | 1 | HC_vs_MS.NoTrt_F |
| PLPP2 | 0.19493 | 0.33243 | 0.919 | 0.87 | 1 | HC_vs_MS.NoTrt_F |
| ABCF2 | 0.1952 | 0.43667 | 0.303 | 0.35 | 1 | HC_vs_MS.NoTrt_F |
| ABHD17A | 0.19524 | 0.46368 | 0.136 | 0.18 | 1 | HC_vs_MS.NoTrt_F |
| LARP1 | 0.1956 | -0.3668 | 0.275 | 0.33 | 1 | HC_vs_MS.NoTrt_F |
| STX11 | 0.19569 | 0.83941 | 0.095 | 0.14 | 1 | HC_vs_MS.NoTrt_F |
| TCEA3 | 0.1957 | 1.56798 | 0.084 | 0.12 | 1 | HC_vs_MS.NoTrt_F |
| TMEM131 | 0.1957 | 0.34456 | 0.197 | 0.25 | 1 | HC_vs_MS.NoTrt_F |
| THAP3 | 0.19573 | 0.29955 | 0.145 | 0.19 | 1 | HC_vs_MS.NoTrt_F |
| MELTF | 0.19581 | 0.42711 | 0.121 | 0.17 | 1 | HC_vs_MS.NoTrt_F |
| LARP4B | 0.19584 | 0.68645 | 0.283 | 0.36 | 1 | HC_vs_MS.NoTrt_F |
| ATPAF2 | 0.19615 | 0.40788 | 0.503 | 0.56 | 1 | HC_vs_MS.NoTrt_F |
| MED19 | 0.19632 | 1.45044 | 0.113 | 0.16 | 1 | HC_vs_MS.NoTrt_F |
| AHSA2 | 0.1966 | 0.72582 | 0.283 | 0.34 | 1 | HC_vs_MS.NoTrt_F |
| P2RX5-TAX1 | 0.19661 | 0.3296 | 0.887 | 0.87 | 1 | HC_vs_MS.NoTrt_F |
| TRERF1 | 0.19681 | 0.57852 | 0.251 | 0.32 | 1 | HC_vs_MS.NoTrt_F |
| COQ4 | 0.19722 | 0.62514 | 0.922 | 0.9 | 1 | HC_vs_MS.NoTrt_F |
| ATP6V1H | 0.1976 | 0.31378 | 0.434 | 0.5 | 1 | HC_vs_MS.NoTrt_F |
| ASH2L | 0.19766 | -0.4041 | 0.633 | 0.67 | 1 | HC_vs_MS.NoTrt_F |
| HMGXB3 | 0.19795 | 0.88296 | 0.223 | 0.27 | 1 | HC_vs_MS.NoTrt_F |
| COG2 | 0.19797 | 0.93461 | 0.37 | 0.44 | 1 | HC_vs_MS.NoTrt_F |
| PITPNB | 0.19804 | 0.28923 | 0.228 | 0.29 | 1 | HC_vs_MS.NoTrt_F |
| SPATA17 | 0.19812 | 0.25721 | 0.717 | 0.64 | 1 | HC_vs_MS.NoTrt_F |
| NLK | 0.19821 | 0.77986 | 0.37 | 0.31 | 1 | HC_vs_MS.NoTrt_F |
| KLHL20 | 0.19842 | 0.2645 | 0.197 | 0.25 | 1 | HC_vs_MS.NoTrt_F |
| CTD-2331H1 | 0.19854 | -0.9121 | 0.217 | 0.27 | 1 | HC_vs_MS.NoTrt_F |
| GCHFR | 0.19863 | 0.66891 | 0.136 | 0.18 | 1 | HC_vs_MS.NoTrt_F |
| COX8A | 0.19866 | 0.27424 | 0.28 | 0.36 | 1 | HC_vs_MS.NoTrt_F |
| FAM129C | 0.19871 | 0.29392 | 0.89 | 0.85 | 1 | HC_vs_MS.NoTrt_F |
| ENTPD7 | 0.19908 | 0.94204 | 0.384 | 0.44 | 1 | HC_vs_MS.NoTrt_F |
| MFSD12 | 0.1992 | 0.28561 | 0.295 | 0.34 | 1 | HC_vs_MS.NoTrt_F |
| PPAN | 0.19945 | 0.36227 | 0.419 | 0.48 | 1 | HC_vs_MS.NoTrt_F |
| PCDHA9 | 0.19951 | 0.36191 | 0.673 | 0.62 | 1 | HC_vs_MS.NoTrt_F |
| GGCT | 0.19993 | 1.06821 | 0.11 | 0.16 | 1 | HC_vs_MS.NoTrt_F |

| | | | | | | |
|---|---|---|---|---|---|---|
| PHB | 0.2001 | 0.33018 | 0.199 | 0.25 | 1 | HC_vs_MS.NoTrt_F |
| IFI44 | 0.20018 | 1.08417 | 0.104 | 0.14 | 1 | HC_vs_MS.NoTrt_F |
| GRAMD3 | 0.20033 | 0.43464 | 0.757 | 0.86 | 1 | HC_vs_MS.NoTrt_F |
| HINFP | 0.20036 | -0.5896 | 0.991 | 0.99 | 1 | HC_vs_MS.NoTrt_F |
| PDCD1 | 0.20048 | 0.27752 | 0.127 | 0.17 | 1 | HC_vs_MS.NoTrt_F |
| ITGAV | 0.20067 | 0.60571 | 0.133 | 0.18 | 1 | HC_vs_MS.NoTrt_F |
| SLC25A13 | 0.20077 | 0.70617 | 0.165 | 0.22 | 1 | HC_vs_MS.NoTrt_F |
| WDR37 | 0.20085 | 0.8105 | 0.231 | 0.29 | 1 | HC_vs_MS.NoTrt_F |
| ANKHD1 | 0.20104 | 0.32702 | 0.494 | 0.6 | 1 | HC_vs_MS.NoTrt_F |
| CYBA | 0.20127 | 0.56129 | 0.324 | 0.39 | 1 | HC_vs_MS.NoTrt_F |
| LINC01420 | 0.20146 | 0.61497 | 0.223 | 0.28 | 1 | HC_vs_MS.NoTrt_F |
| RAB30 | 0.20155 | 0.38849 | 0.382 | 0.43 | 1 | HC_vs_MS.NoTrt_F |
| STOM | 0.20164 | 1.09729 | 0.668 | 0.64 | 1 | HC_vs_MS.NoTrt_F |
| FBXL8 | 0.20188 | 0.60592 | 0.618 | 0.64 | 1 | HC_vs_MS.NoTrt_F |
| ZCCHC14 | 0.20191 | 0.99721 | 0.139 | 0.18 | 1 | HC_vs_MS.NoTrt_F |
| OTUB1 | 0.20197 | 0.33764 | 0.931 | 0.98 | 1 | HC_vs_MS.NoTrt_F |
| C12orf40 | 0.20236 | -0.2853 | 0.298 | 0.36 | 1 | HC_vs_MS.NoTrt_F |
| HEMGN | 0.20249 | -0.2532 | 0.15 | 0.2 | 1 | HC_vs_MS.NoTrt_F |
| FBXW4 | 0.2025 | 0.25719 | 0.506 | 0.56 | 1 | HC_vs_MS.NoTrt_F |
| AASDH | 0.20265 | 0.72713 | 0.142 | 0.19 | 1 | HC_vs_MS.NoTrt_F |
| TTC39B | 0.20282 | 0.90496 | 0.838 | 0.77 | 1 | HC_vs_MS.NoTrt_F |
| ZSWIM4 | 0.20289 | 1.19198 | 0.087 | 0.13 | 1 | HC_vs_MS.NoTrt_F |
| THBS1 | 0.20302 | -0.4925 | 0.263 | 0.3 | 1 | HC_vs_MS.NoTrt_F |
| CCND2 | 0.20302 | 0.38618 | 0.465 | 0.59 | 1 | HC_vs_MS.NoTrt_F |
| TBK1 | 0.20335 | 0.84029 | 0.127 | 0.17 | 1 | HC_vs_MS.NoTrt_F |
| NKAP | 0.20381 | 0.31467 | 0.951 | 0.94 | 1 | HC_vs_MS.NoTrt_F |
| PANK2 | 0.20386 | -0.8444 | 0.575 | 0.56 | 1 | HC_vs_MS.NoTrt_F |
| IFT57 | 0.20405 | 0.83774 | 0.142 | 0.19 | 1 | HC_vs_MS.NoTrt_F |
| AL354822.1 | 0.20421 | 0.6512 | 0.286 | 0.33 | 1 | HC_vs_MS.NoTrt_F |
| PIK3C2A | 0.20428 | 0.64338 | 0.431 | 0.48 | 1 | HC_vs_MS.NoTrt_F |
| FKBP2 | 0.20456 | 0.93411 | 0.13 | 0.18 | 1 | HC_vs_MS.NoTrt_F |
| TRANK1 | 0.20466 | 0.36731 | 0.263 | 0.32 | 1 | HC_vs_MS.NoTrt_F |
| LIPC | 0.20476 | 0.40152 | 0.332 | 0.39 | 1 | HC_vs_MS.NoTrt_F |
| STXBP1 | 0.20483 | 0.83435 | 0.069 | 0.11 | 1 | HC_vs_MS.NoTrt_F |
| OFD1 | 0.2051 | 0.81242 | 0.298 | 0.37 | 1 | HC_vs_MS.NoTrt_F |
| TRIP4 | 0.20513 | -0.4674 | 0.575 | 0.6 | 1 | HC_vs_MS.NoTrt_F |
| SPAG4 | 0.20515 | 0.56435 | 0.28 | 0.35 | 1 | HC_vs_MS.NoTrt_F |
| CCNC | 0.20532 | 0.28663 | 0.624 | 0.69 | 1 | HC_vs_MS.NoTrt_F |
| FAM193B | 0.20533 | 0.55206 | 0.512 | 0.56 | 1 | HC_vs_MS.NoTrt_F |
| CCNB1IP1 | 0.20589 | 0.28453 | 0.858 | 0.83 | 1 | HC_vs_MS.NoTrt_F |

| | | | | | | |
|---|---|---|---|---|---|---|
| HELLS | 0.20594 | -0.4419 | 0.841 | 0.85 | 1 | HC_vs_MS.NoTrt_F |
| DIABLO | 0.20599 | -0.543 | 0.749 | 0.8 | 1 | HC_vs_MS.NoTrt_F |
| WWTR1 | 0.20619 | 0.44567 | 0.474 | 0.52 | 1 | HC_vs_MS.NoTrt_F |
| BICDL1 | 0.20626 | 0.44884 | 0.662 | 0.66 | 1 | HC_vs_MS.NoTrt_F |
| NEUROD2 | 0.2064 | 0.40557 | 0.277 | 0.22 | 1 | HC_vs_MS.NoTrt_F |
| C20orf194 | 0.2066 | 0.62827 | 0.344 | 0.41 | 1 | HC_vs_MS.NoTrt_F |
| MLLT11 | 0.2066 | 0.28586 | 0.194 | 0.25 | 1 | HC_vs_MS.NoTrt_F |
| AP1G2 | 0.20661 | 0.43218 | 0.731 | 0.74 | 1 | HC_vs_MS.NoTrt_F |
| LBHD1 | 0.207 | -0.2985 | 0.523 | 0.56 | 1 | HC_vs_MS.NoTrt_F |
| SLC39A13 | 0.20708 | 0.76505 | 0.179 | 0.23 | 1 | HC_vs_MS.NoTrt_F |
| TES | 0.20708 | -0.5288 | 0.87 | 0.89 | 1 | HC_vs_MS.NoTrt_F |
| LIPE | 0.20722 | 0.81772 | 0.396 | 0.44 | 1 | HC_vs_MS.NoTrt_F |
| NR1D2 | 0.20739 | 0.29217 | 0.185 | 0.24 | 1 | HC_vs_MS.NoTrt_F |
| LETM2 | 0.20751 | 0.35683 | 0.798 | 0.81 | 1 | HC_vs_MS.NoTrt_F |
| ANGEL1 | 0.20768 | 0.58985 | 0.471 | 0.51 | 1 | HC_vs_MS.NoTrt_F |
| LIN54 | 0.20776 | -0.705 | 0.653 | 0.66 | 1 | HC_vs_MS.NoTrt_F |
| TEX14 | 0.20795 | 1.13284 | 0.116 | 0.16 | 1 | HC_vs_MS.NoTrt_F |
| HIST1H4C | 0.20796 | 0.5924 | 0.387 | 0.51 | 1 | HC_vs_MS.NoTrt_F |
| TAPT1 | 0.20803 | 0.39294 | 0.78 | 0.76 | 1 | HC_vs_MS.NoTrt_F |
| AFAP1L1 | 0.20812 | 1.16873 | 0.569 | 0.48 | 1 | HC_vs_MS.NoTrt_F |
| BANK1 | 0.20824 | 0.79856 | 0.197 | 0.25 | 1 | HC_vs_MS.NoTrt_F |
| KBTBD2 | 0.20843 | 0.35215 | 0.72 | 0.69 | 1 | HC_vs_MS.NoTrt_F |
| BRD7 | 0.20845 | 0.69609 | 0.673 | 0.63 | 1 | HC_vs_MS.NoTrt_F |
| FAM76A | 0.20875 | -0.2741 | 0.855 | 0.85 | 1 | HC_vs_MS.NoTrt_F |
| WSB2 | 0.20879 | 0.58618 | 0.832 | 0.85 | 1 | HC_vs_MS.NoTrt_F |
| HIST1H2AG | 0.20886 | 1.06074 | 0.37 | 0.42 | 1 | HC_vs_MS.NoTrt_F |
| SLC5A3 | 0.20927 | 0.6312 | 0.162 | 0.21 | 1 | HC_vs_MS.NoTrt_F |
| TFDP1 | 0.21039 | 0.43428 | 0.075 | 0.11 | 1 | HC_vs_MS.NoTrt_F |
| HIPK3 | 0.21041 | 0.65917 | 0.179 | 0.23 | 1 | HC_vs_MS.NoTrt_F |
| FAM101B | 0.21042 | 0.80178 | 0.324 | 0.4 | 1 | HC_vs_MS.NoTrt_F |
| BCDIN3D | 0.21053 | -0.8659 | 0.454 | 0.46 | 1 | HC_vs_MS.NoTrt_F |
| NDUFS4 | 0.21055 | 0.34906 | 0.353 | 0.4 | 1 | HC_vs_MS.NoTrt_F |
| BVES | 0.21079 | 0.25518 | 0.772 | 0.75 | 1 | HC_vs_MS.NoTrt_F |
| SLC41A1 | 0.2112 | 0.51254 | 0.179 | 0.23 | 1 | HC_vs_MS.NoTrt_F |
| ZNF526 | 0.21126 | -0.3235 | 0.532 | 0.56 | 1 | HC_vs_MS.NoTrt_F |
| FBXO9 | 0.21158 | -0.2558 | 0.803 | 0.84 | 1 | HC_vs_MS.NoTrt_F |
| SMCR8 | 0.21162 | 0.68087 | 0.309 | 0.36 | 1 | HC_vs_MS.NoTrt_F |
| FBXO5 | 0.2118 | 1.28927 | 0.431 | 0.49 | 1 | HC_vs_MS.NoTrt_F |
| PLA2G2C | 0.2119 | 0.34814 | 0.581 | 0.53 | 1 | HC_vs_MS.NoTrt_F |
| MBNL3 | 0.21199 | 0.68784 | 0.15 | 0.2 | 1 | HC_vs_MS.NoTrt_F |

| | | | | | | |
|---|---|---|---|---|---|---|
| SRP72 | 0.21212 | 0.35198 | 0.295 | 0.35 | 1 | HC_vs_MS.NoTrt_F |
| SUCLG1 | 0.21217 | -0.3119 | 0.425 | 0.46 | 1 | HC_vs_MS.NoTrt_F |
| C9orf64 | 0.21255 | -0.6484 | 0.592 | 0.62 | 1 | HC_vs_MS.NoTrt_F |
| ANKRD28 | 0.21256 | 0.91503 | 0.439 | 0.5 | 1 | HC_vs_MS.NoTrt_F |
| GTF3C4 | 0.21275 | 0.5341 | 0.35 | 0.4 | 1 | HC_vs_MS.NoTrt_F |
| LEMD3 | 0.21279 | -0.6567 | 0.714 | 0.74 | 1 | HC_vs_MS.NoTrt_F |
| SMN1 | 0.21284 | 0.25233 | 0.402 | 0.48 | 1 | HC_vs_MS.NoTrt_F |
| UBR4 | 0.21309 | 0.25919 | 0.775 | 0.82 | 1 | HC_vs_MS.NoTrt_F |
| UVRAG | 0.2131 | 0.55042 | 0.41 | 0.46 | 1 | HC_vs_MS.NoTrt_F |
| ACSM3 | 0.2133 | 0.32333 | 0.679 | 0.69 | 1 | HC_vs_MS.NoTrt_F |
| PTGIS | 0.21331 | 0.76981 | 0.746 | 0.75 | 1 | HC_vs_MS.NoTrt_F |
| ARHGAP18 | 0.21336 | 0.56745 | 0.15 | 0.19 | 1 | HC_vs_MS.NoTrt_F |
| SLC7A5 | 0.21344 | 0.25862 | 0.147 | 0.19 | 1 | HC_vs_MS.NoTrt_F |
| MCRS1 | 0.21344 | 0.49637 | 0.217 | 0.28 | 1 | HC_vs_MS.NoTrt_F |
| ZNF274 | 0.2136 | -0.584 | 0.645 | 0.7 | 1 | HC_vs_MS.NoTrt_F |
| PGAP3 | 0.21363 | -0.7616 | 0.153 | 0.2 | 1 | HC_vs_MS.NoTrt_F |
| ZDHHC20 | 0.21415 | 0.35729 | 0.526 | 0.56 | 1 | HC_vs_MS.NoTrt_F |
| EPS8 | 0.2144 | 0.3322 | 0.488 | 0.54 | 1 | HC_vs_MS.NoTrt_F |
| CCDC120 | 0.21458 | 1.40704 | 0.587 | 0.65 | 1 | HC_vs_MS.NoTrt_F |
| CCDC163 | 0.21473 | 0.6506 | 0.616 | 0.55 | 1 | HC_vs_MS.NoTrt_F |
| KDM4A | 0.21481 | 0.38689 | 0.188 | 0.24 | 1 | HC_vs_MS.NoTrt_F |
| ARSB | 0.21529 | 0.62214 | 0.685 | 0.59 | 1 | HC_vs_MS.NoTrt_F |
| MYBBP1A | 0.21529 | -0.3278 | 0.176 | 0.22 | 1 | HC_vs_MS.NoTrt_F |
| IQCK | 0.21557 | -0.4906 | 0.231 | 0.28 | 1 | HC_vs_MS.NoTrt_F |
| SARNP | 0.21558 | -0.3008 | 0.61 | 0.65 | 1 | HC_vs_MS.NoTrt_F |
| ARMCX3 | 0.21657 | 0.59213 | 0.116 | 0.16 | 1 | HC_vs_MS.NoTrt_F |
| TNFRSF10B | 0.21677 | 0.41169 | 0.46 | 0.51 | 1 | HC_vs_MS.NoTrt_F |
| ZNF318 | 0.2168 | 1.19097 | 0.711 | 0.7 | 1 | HC_vs_MS.NoTrt_F |
| QRICH2 | 0.21682 | 0.5063 | 0.217 | 0.27 | 1 | HC_vs_MS.NoTrt_F |
| PIPOX | 0.2171 | -0.3613 | 0.584 | 0.6 | 1 | HC_vs_MS.NoTrt_F |
| ESF1 | 0.21714 | -0.4482 | 0.205 | 0.25 | 1 | HC_vs_MS.NoTrt_F |
| ELOVL1 | 0.21742 | 0.42656 | 0.205 | 0.26 | 1 | HC_vs_MS.NoTrt_F |
| TMEM217 | 0.2175 | 0.54219 | 0.113 | 0.16 | 1 | HC_vs_MS.NoTrt_F |
| LGALS9 | 0.21755 | 0.5838 | 0.298 | 0.36 | 1 | HC_vs_MS.NoTrt_F |
| HOOK2 | 0.21792 | 0.25547 | 0.98 | 0.97 | 1 | HC_vs_MS.NoTrt_F |
| CFD | 0.21821 | 1.27397 | 0.197 | 0.24 | 1 | HC_vs_MS.NoTrt_F |
| TFG | 0.21843 | 0.26743 | 0.191 | 0.24 | 1 | HC_vs_MS.NoTrt_F |
| WDTC1 | 0.21848 | 0.28972 | 0.399 | 0.44 | 1 | HC_vs_MS.NoTrt_F |
| YAF2 | 0.21873 | -0.4647 | 0.769 | 0.81 | 1 | HC_vs_MS.NoTrt_F |
| MCM8 | 0.21879 | -0.3541 | 0.78 | 0.72 | 1 | HC_vs_MS.NoTrt_F |

| | | | | | | |
|---|---|---|---|---|---|---|
| PIK3R4 | 0.21884 | 0.96774 | 0.182 | 0.23 | 1 | HC_vs_MS.NoTrt_F |
| LRRC1 | 0.21887 | -0.4565 | 0.306 | 0.35 | 1 | HC_vs_MS.NoTrt_F |
| EIF2B4 | 0.21895 | 0.48848 | 0.54 | 0.61 | 1 | HC_vs_MS.NoTrt_F |
| CC2D2A | 0.21898 | 0.67843 | 0.257 | 0.31 | 1 | HC_vs_MS.NoTrt_F |
| NEDD1 | 0.21926 | 0.68223 | 0.15 | 0.2 | 1 | HC_vs_MS.NoTrt_F |
| GSDMA | 0.21945 | 0.28034 | 0.734 | 0.68 | 1 | HC_vs_MS.NoTrt_F |
| CINP | 0.2196 | 0.40715 | 0.899 | 0.9 | 1 | HC_vs_MS.NoTrt_F |
| LMF1 | 0.21971 | 0.78239 | 0.156 | 0.2 | 1 | HC_vs_MS.NoTrt_F |
| RNASEL | 0.21995 | 0.32725 | 0.39 | 0.45 | 1 | HC_vs_MS.NoTrt_F |
| LIFR | 0.22002 | 0.464 | 0.142 | 0.19 | 1 | HC_vs_MS.NoTrt_F |
| POLH | 0.22022 | -0.3938 | 0.598 | 0.61 | 1 | HC_vs_MS.NoTrt_F |
| TOP1 | 0.22044 | 0.40462 | 0.225 | 0.28 | 1 | HC_vs_MS.NoTrt_F |
| CDKN1A | 0.22073 | 0.41647 | 0.182 | 0.24 | 1 | HC_vs_MS.NoTrt_F |
| FOPNL | 0.22079 | -0.2808 | 0.65 | 0.64 | 1 | HC_vs_MS.NoTrt_F |
| RPP21 | 0.22117 | 0.34606 | 0.197 | 0.25 | 1 | HC_vs_MS.NoTrt_F |
| PIK3AP1 | 0.22121 | 1.13514 | 0.147 | 0.19 | 1 | HC_vs_MS.NoTrt_F |
| PEX13 | 0.22125 | -0.7493 | 0.783 | 0.77 | 1 | HC_vs_MS.NoTrt_F |
| MOB4 | 0.22143 | -0.5469 | 0.419 | 0.44 | 1 | HC_vs_MS.NoTrt_F |
| HPS3 | 0.22149 | 0.54663 | 0.341 | 0.37 | 1 | HC_vs_MS.NoTrt_F |
| CXorf21 | 0.22157 | -1.4175 | 0.355 | 0.41 | 1 | HC_vs_MS.NoTrt_F |
| PXK | 0.22205 | 0.57196 | 0.153 | 0.2 | 1 | HC_vs_MS.NoTrt_F |
| RFC2 | 0.22252 | 0.37369 | 0.751 | 0.77 | 1 | HC_vs_MS.NoTrt_F |
| RELT | 0.22292 | 0.69577 | 0.11 | 0.16 | 1 | HC_vs_MS.NoTrt_F |
| PIKFYVE | 0.22303 | 0.55394 | 0.41 | 0.46 | 1 | HC_vs_MS.NoTrt_F |
| INTS6 | 0.22321 | -0.3973 | 0.954 | 0.96 | 1 | HC_vs_MS.NoTrt_F |
| CAPN3 | 0.22352 | -0.5628 | 0.098 | 0.13 | 1 | HC_vs_MS.NoTrt_F |
| SLC48A1 | 0.22406 | -0.5457 | 0.633 | 0.72 | 1 | HC_vs_MS.NoTrt_F |
| GALNT11 | 0.22425 | 0.29526 | 0.512 | 0.44 | 1 | HC_vs_MS.NoTrt_F |
| ATP7A | 0.22439 | 0.82863 | 0.399 | 0.45 | 1 | HC_vs_MS.NoTrt_F |
| GCDH | 0.22452 | 0.27423 | 0.197 | 0.24 | 1 | HC_vs_MS.NoTrt_F |
| DDAH2 | 0.22477 | -0.5969 | 0.118 | 0.16 | 1 | HC_vs_MS.NoTrt_F |
| ST3GAL5 | 0.22508 | 0.27517 | 0.295 | 0.35 | 1 | HC_vs_MS.NoTrt_F |
| CTRC | 0.2252 | 0.33487 | 0.702 | 0.7 | 1 | HC_vs_MS.NoTrt_F |
| C2orf61 | 0.2259 | -0.5727 | 0.15 | 0.19 | 1 | HC_vs_MS.NoTrt_F |
| DSN1 | 0.2259 | -0.6698 | 0.289 | 0.34 | 1 | HC_vs_MS.NoTrt_F |
| RAB11FIP3 | 0.22595 | 0.69172 | 0.211 | 0.26 | 1 | HC_vs_MS.NoTrt_F |
| CEP76 | 0.2261 | 0.42047 | 0.827 | 0.83 | 1 | HC_vs_MS.NoTrt_F |
| SNX14 | 0.22613 | -0.3687 | 0.59 | 0.62 | 1 | HC_vs_MS.NoTrt_F |
| ATP5E | 0.22632 | 0.47564 | 0.491 | 0.67 | 1 | HC_vs_MS.NoTrt_F |
| PLEK | 0.22646 | 2.31614 | 0.084 | 0.12 | 1 | HC_vs_MS.NoTrt_F |

| | | | | | | |
|---|---|---|---|---|---|---|
| ATP6V1B2 | 0.22663 | -0.4327 | 0.711 | 0.75 | 1 | HC_vs_MS.NoTrt_F |
| ALPK1 | 0.22681 | 0.60768 | 0.806 | 0.77 | 1 | HC_vs_MS.NoTrt_F |
| TBL1X | 0.22682 | -0.4569 | 0.65 | 0.6 | 1 | HC_vs_MS.NoTrt_F |
| AKT1 | 0.22716 | 0.806 | 0.254 | 0.31 | 1 | HC_vs_MS.NoTrt_F |
| CLPX | 0.22744 | -0.9346 | 0.731 | 0.76 | 1 | HC_vs_MS.NoTrt_F |
| PRCP | 0.22745 | 0.65576 | 0.858 | 0.79 | 1 | HC_vs_MS.NoTrt_F |
| RP11-6L6.2 | 0.22763 | -0.3628 | 0.078 | 0.11 | 1 | HC_vs_MS.NoTrt_F |
| STEAP3 | 0.22764 | -0.7493 | 0.185 | 0.23 | 1 | HC_vs_MS.NoTrt_F |
| ZNF714 | 0.2277 | 0.36899 | 0.96 | 0.93 | 1 | HC_vs_MS.NoTrt_F |
| AP2A2 | 0.228 | 0.30441 | 0.757 | 0.7 | 1 | HC_vs_MS.NoTrt_F |
| FAM133B | 0.22811 | -0.4982 | 0.798 | 0.81 | 1 | HC_vs_MS.NoTrt_F |
| DNAH3 | 0.22816 | -0.2715 | 0.538 | 0.54 | 1 | HC_vs_MS.NoTrt_F |
| FAM96B | 0.2284 | 0.27344 | 0.257 | 0.31 | 1 | HC_vs_MS.NoTrt_F |
| DICER1 | 0.22885 | -0.2869 | 0.237 | 0.28 | 1 | HC_vs_MS.NoTrt_F |
| DROSHA | 0.22916 | 0.43063 | 0.37 | 0.43 | 1 | HC_vs_MS.NoTrt_F |
| TNRC6A | 0.22946 | 0.34181 | 0.887 | 0.85 | 1 | HC_vs_MS.NoTrt_F |
| GGT7 | 0.23022 | 0.47717 | 0.194 | 0.24 | 1 | HC_vs_MS.NoTrt_F |
| LARGE1 | 0.23056 | 0.53974 | 0.613 | 0.6 | 1 | HC_vs_MS.NoTrt_F |
| PZP | 0.23095 | 1.34678 | 0.15 | 0.12 | 1 | HC_vs_MS.NoTrt_F |
| PAK6 | 0.23099 | 0.39284 | 0.179 | 0.23 | 1 | HC_vs_MS.NoTrt_F |
| URM1 | 0.23169 | 0.406 | 0.194 | 0.24 | 1 | HC_vs_MS.NoTrt_F |
| GRAP | 0.23179 | 0.39212 | 0.225 | 0.28 | 1 | HC_vs_MS.NoTrt_F |
| PLEKHG3 | 0.23179 | -0.5011 | 0.5 | 0.53 | 1 | HC_vs_MS.NoTrt_F |
| IBA57 | 0.23183 | 1.13087 | 0.829 | 0.86 | 1 | HC_vs_MS.NoTrt_F |
| HSBP1 | 0.23193 | 0.42828 | 0.639 | 0.7 | 1 | HC_vs_MS.NoTrt_F |
| NARS2 | 0.232 | -0.8835 | 0.448 | 0.49 | 1 | HC_vs_MS.NoTrt_F |
| TBC1D12 | 0.23239 | 0.28895 | 0.445 | 0.4 | 1 | HC_vs_MS.NoTrt_F |
| EDA | 0.23276 | -0.7599 | 0.428 | 0.48 | 1 | HC_vs_MS.NoTrt_F |
| DHX35 | 0.2328 | -0.4675 | 0.422 | 0.45 | 1 | HC_vs_MS.NoTrt_F |
| FUT3 | 0.23282 | -0.4461 | 0.228 | 0.27 | 1 | HC_vs_MS.NoTrt_F |
| ZNF550 | 0.23339 | -0.2822 | 0.344 | 0.38 | 1 | HC_vs_MS.NoTrt_F |
| ASCC3 | 0.23391 | 0.38453 | 0.338 | 0.41 | 1 | HC_vs_MS.NoTrt_F |
| SLC43A1 | 0.23395 | 0.26698 | 0.156 | 0.2 | 1 | HC_vs_MS.NoTrt_F |
| RNF135 | 0.23444 | 0.81668 | 0.309 | 0.37 | 1 | HC_vs_MS.NoTrt_F |
| ZNF543 | 0.2347 | 0.93005 | 0.607 | 0.69 | 1 | HC_vs_MS.NoTrt_F |
| PPP4R3A | 0.23484 | 0.2534 | 0.306 | 0.37 | 1 | HC_vs_MS.NoTrt_F |
| RTN2 | 0.23493 | 0.53879 | 0.309 | 0.26 | 1 | HC_vs_MS.NoTrt_F |
| METAP2 | 0.23495 | -0.2864 | 0.477 | 0.55 | 1 | HC_vs_MS.NoTrt_F |
| CKS1B | 0.23517 | 0.50499 | 0.272 | 0.32 | 1 | HC_vs_MS.NoTrt_F |
| INTS2 | 0.23518 | -0.2585 | 0.298 | 0.34 | 1 | HC_vs_MS.NoTrt_F |

| | | | | | | |
|---|---|---|---|---|---|---|
| ACSBG1 | 0.23525 | 0.42693 | 0.428 | 0.5 | 1 | HC_vs_MS.NoTrt_F |
| IL17RE | 0.23525 | 0.66991 | 0.373 | 0.33 | 1 | HC_vs_MS.NoTrt_F |
| EPHB1 | 0.2354 | 1.29058 | 0.087 | 0.12 | 1 | HC_vs_MS.NoTrt_F |
| INTS4 | 0.2358 | -0.344 | 0.413 | 0.44 | 1 | HC_vs_MS.NoTrt_F |
| ZNF257 | 0.23586 | 0.40664 | 0.569 | 0.51 | 1 | HC_vs_MS.NoTrt_F |
| PSMD11 | 0.2359 | 0.25364 | 0.558 | 0.59 | 1 | HC_vs_MS.NoTrt_F |
| RINL | 0.23591 | -0.3286 | 0.763 | 0.77 | 1 | HC_vs_MS.NoTrt_F |
| ALOX12 | 0.23609 | 0.49118 | 0.104 | 0.14 | 1 | HC_vs_MS.NoTrt_F |
| MRPL43 | 0.23624 | 0.60541 | 0.185 | 0.24 | 1 | HC_vs_MS.NoTrt_F |
| CD276 | 0.23671 | 0.50975 | 0.512 | 0.46 | 1 | HC_vs_MS.NoTrt_F |
| GSTO2 | 0.23711 | 0.48074 | 0.896 | 0.92 | 1 | HC_vs_MS.NoTrt_F |
| RTF1 | 0.23716 | -0.4042 | 0.645 | 0.66 | 1 | HC_vs_MS.NoTrt_F |
| FNIP1 | 0.2373 | 0.39186 | 0.587 | 0.67 | 1 | HC_vs_MS.NoTrt_F |
| ASB6 | 0.23739 | -0.548 | 0.691 | 0.69 | 1 | HC_vs_MS.NoTrt_F |
| COMTD1 | 0.2375 | 0.34919 | 0.173 | 0.21 | 1 | HC_vs_MS.NoTrt_F |
| HEG1 | 0.23763 | 1.20094 | 0.182 | 0.14 | 1 | HC_vs_MS.NoTrt_F |
| FAM149A | 0.23794 | 0.79212 | 0.379 | 0.42 | 1 | HC_vs_MS.NoTrt_F |
| KHDRBS1 | 0.23795 | 0.67566 | 0.24 | 0.3 | 1 | HC_vs_MS.NoTrt_F |
| DENND6B | 0.23796 | 0.79169 | 0.399 | 0.34 | 1 | HC_vs_MS.NoTrt_F |
| HLA-DRB3 | 0.23811 | 0.3068 | 0.188 | 0.24 | 1 | HC_vs_MS.NoTrt_F |
| MIER2 | 0.23822 | 0.29037 | 0.509 | 0.54 | 1 | HC_vs_MS.NoTrt_F |
| IKBKB | 0.23843 | -0.3018 | 0.864 | 0.9 | 1 | HC_vs_MS.NoTrt_F |
| SLC20A2 | 0.23849 | 0.72875 | 0.179 | 0.23 | 1 | HC_vs_MS.NoTrt_F |
| B3GALT4 | 0.2385 | -0.8615 | 0.165 | 0.2 | 1 | HC_vs_MS.NoTrt_F |
| ABCA2 | 0.23851 | 0.55221 | 0.341 | 0.39 | 1 | HC_vs_MS.NoTrt_F |
| C3orf18 | 0.23855 | 0.72049 | 0.315 | 0.35 | 1 | HC_vs_MS.NoTrt_F |
| CCSAP | 0.23874 | 0.77752 | 0.188 | 0.24 | 1 | HC_vs_MS.NoTrt_F |
| MRPS12 | 0.23908 | -0.3358 | 0.436 | 0.48 | 1 | HC_vs_MS.NoTrt_F |
| KMT5B | 0.23919 | 0.45128 | 0.353 | 0.4 | 1 | HC_vs_MS.NoTrt_F |
| ABCC4 | 0.23932 | 0.39709 | 0.789 | 0.81 | 1 | HC_vs_MS.NoTrt_F |
| WHAMM | 0.23934 | 0.25916 | 0.873 | 0.87 | 1 | HC_vs_MS.NoTrt_F |
| CUL1 | 0.23997 | 0.87741 | 0.237 | 0.3 | 1 | HC_vs_MS.NoTrt_F |
| LST1 | 0.24018 | 0.88307 | 0.272 | 0.33 | 1 | HC_vs_MS.NoTrt_F |
| KIAA0141 | 0.24059 | 0.50347 | 0.249 | 0.3 | 1 | HC_vs_MS.NoTrt_F |
| C16orf87 | 0.24133 | -0.2752 | 0.812 | 0.85 | 1 | HC_vs_MS.NoTrt_F |
| DHX15 | 0.24144 | 0.59194 | 0.54 | 0.59 | 1 | HC_vs_MS.NoTrt_F |
| SCARB2 | 0.2416 | 0.25736 | 0.341 | 0.4 | 1 | HC_vs_MS.NoTrt_F |
| PACS1 | 0.2416 | 0.5502 | 0.853 | 0.85 | 1 | HC_vs_MS.NoTrt_F |
| LRPAP1 | 0.24165 | -0.2994 | 0.887 | 0.9 | 1 | HC_vs_MS.NoTrt_F |
| HLA-F | 0.24169 | 0.50132 | 0.592 | 0.67 | 1 | HC_vs_MS.NoTrt_F |

| | | | | | | |
|---|---|---|---|---|---|---|
| CTSL | 0.24204 | 0.44096 | 0.078 | 0.11 | 1 | HC_vs_MS.NoTrt_F |
| NMT1 | 0.24225 | -0.4643 | 0.367 | 0.42 | 1 | HC_vs_MS.NoTrt_F |
| PPP5C | 0.24233 | 0.65895 | 0.558 | 0.51 | 1 | HC_vs_MS.NoTrt_F |
| GTF3C3 | 0.2424 | 0.4539 | 0.699 | 0.72 | 1 | HC_vs_MS.NoTrt_F |
| APOBR | 0.24254 | 0.39827 | 0.176 | 0.23 | 1 | HC_vs_MS.NoTrt_F |
| BAZ2A | 0.24311 | 1.0718 | 0.731 | 0.71 | 1 | HC_vs_MS.NoTrt_F |
| H2AFJ | 0.24312 | 0.79874 | 0.15 | 0.19 | 1 | HC_vs_MS.NoTrt_F |
| GPATCH11 | 0.24315 | -0.361 | 0.535 | 0.59 | 1 | HC_vs_MS.NoTrt_F |
| FDXACB1 | 0.2432 | -0.3418 | 0.445 | 0.51 | 1 | HC_vs_MS.NoTrt_F |
| C21orf33 | 0.24327 | 0.30873 | 0.416 | 0.46 | 1 | HC_vs_MS.NoTrt_F |
| ORAI1 | 0.24328 | 0.5456 | 0.121 | 0.16 | 1 | HC_vs_MS.NoTrt_F |
| IDNK | 0.24341 | -0.8123 | 0.454 | 0.48 | 1 | HC_vs_MS.NoTrt_F |
| ARL16 | 0.24349 | -0.4002 | 0.783 | 0.8 | 1 | HC_vs_MS.NoTrt_F |
| PNN | 0.24351 | 0.36572 | 0.301 | 0.37 | 1 | HC_vs_MS.NoTrt_F |
| MEX3C | 0.24362 | 0.61176 | 0.124 | 0.17 | 1 | HC_vs_MS.NoTrt_F |
| ZNF350 | 0.24374 | -1.1764 | 0.595 | 0.62 | 1 | HC_vs_MS.NoTrt_F |
| RRP8 | 0.24376 | 0.27865 | 0.529 | 0.45 | 1 | HC_vs_MS.NoTrt_F |
| SPTLC3 | 0.2439 | 0.53811 | 0.575 | 0.56 | 1 | HC_vs_MS.NoTrt_F |
| LILRB3 | 0.24397 | -0.682 | 0.162 | 0.2 | 1 | HC_vs_MS.NoTrt_F |
| APBB3 | 0.24398 | 0.71772 | 0.202 | 0.25 | 1 | HC_vs_MS.NoTrt_F |
| TEKT3 | 0.24398 | 0.26289 | 0.179 | 0.23 | 1 | HC_vs_MS.NoTrt_F |
| CPA4 | 0.24415 | 0.25529 | 0.584 | 0.51 | 1 | HC_vs_MS.NoTrt_F |
| PSMD14 | 0.24418 | 0.26674 | 0.179 | 0.23 | 1 | HC_vs_MS.NoTrt_F |
| CXCR2 | 0.24464 | 0.72785 | 0.341 | 0.38 | 1 | HC_vs_MS.NoTrt_F |
| IMMT | 0.24465 | -0.5423 | 0.486 | 0.54 | 1 | HC_vs_MS.NoTrt_F |
| MICAL2 | 0.24471 | 0.86781 | 0.89 | 0.86 | 1 | HC_vs_MS.NoTrt_F |
| SLC39A5 | 0.24474 | 0.68921 | 0.24 | 0.29 | 1 | HC_vs_MS.NoTrt_F |
| NHP2 | 0.24475 | 0.33153 | 0.266 | 0.31 | 1 | HC_vs_MS.NoTrt_F |
| SHROOM3 | 0.2448 | 0.25242 | 0.327 | 0.29 | 1 | HC_vs_MS.NoTrt_F |
| EPPK1 | 0.24491 | 1.41059 | 0.101 | 0.14 | 1 | HC_vs_MS.NoTrt_F |
| SETD1A | 0.24503 | 2.19993 | 0.168 | 0.13 | 1 | HC_vs_MS.NoTrt_F |
| ORC1 | 0.24503 | 0.90074 | 0.303 | 0.27 | 1 | HC_vs_MS.NoTrt_F |
| AP1S3 | 0.24583 | 0.27535 | 0.951 | 0.95 | 1 | HC_vs_MS.NoTrt_F |
| GDPGP1 | 0.24593 | 0.27711 | 0.948 | 0.96 | 1 | HC_vs_MS.NoTrt_F |
| CDH26 | 0.24618 | 0.74118 | 0.173 | 0.22 | 1 | HC_vs_MS.NoTrt_F |
| UTP6 | 0.24622 | -0.2796 | 0.899 | 0.91 | 1 | HC_vs_MS.NoTrt_F |
| DUS3L | 0.24653 | 0.59642 | 0.188 | 0.24 | 1 | HC_vs_MS.NoTrt_F |
| PBRM1 | 0.24662 | 0.40505 | 0.725 | 0.79 | 1 | HC_vs_MS.NoTrt_F |
| DTD1 | 0.24689 | -0.7973 | 0.382 | 0.4 | 1 | HC_vs_MS.NoTrt_F |
| ATP13A3 | 0.24699 | 0.51615 | 0.173 | 0.22 | 1 | HC_vs_MS.NoTrt_F |

| | | | | | | |
|---|---|---|---|---|---|---|
| APAF1 | 0.24702 | -0.6323 | 0.676 | 0.68 | 1 | HC_vs_MS.NoTrt_F |
| AXL | 0.24749 | -0.4122 | 0.772 | 0.79 | 1 | HC_vs_MS.NoTrt_F |
| CAPN7 | 0.24752 | -0.2645 | 0.624 | 0.63 | 1 | HC_vs_MS.NoTrt_F |
| S100A4 | 0.24816 | 0.38442 | 0.757 | 0.9 | 1 | HC_vs_MS.NoTrt_F |
| WNK1 | 0.24831 | 0.26378 | 0.613 | 0.72 | 1 | HC_vs_MS.NoTrt_F |
| AL022397.1 | 0.24853 | 0.29001 | 0.292 | 0.37 | 1 | HC_vs_MS.NoTrt_F |
| SOCS1 | 0.24855 | 0.54519 | 0.139 | 0.19 | 1 | HC_vs_MS.NoTrt_F |
| UBE2V2 | 0.24895 | -0.3987 | 0.734 | 0.77 | 1 | HC_vs_MS.NoTrt_F |
| NEK1 | 0.24896 | 0.35321 | 0.402 | 0.45 | 1 | HC_vs_MS.NoTrt_F |
| OPN3 | 0.24899 | -0.7347 | 0.263 | 0.3 | 1 | HC_vs_MS.NoTrt_F |
| SLC8A1 | 0.24924 | 0.80699 | 0.347 | 0.29 | 1 | HC_vs_MS.NoTrt_F |
| RAI1 | 0.24929 | 0.55287 | 0.234 | 0.29 | 1 | HC_vs_MS.NoTrt_F |
| AVPR1A | 0.24953 | 0.6625 | 0.246 | 0.2 | 1 | HC_vs_MS.NoTrt_F |
| BCAP29 | 0.24955 | -0.6533 | 0.41 | 0.48 | 1 | HC_vs_MS.NoTrt_F |
| WASL | 0.24967 | 0.37386 | 0.136 | 0.18 | 1 | HC_vs_MS.NoTrt_F |
| KAT2B | 0.25104 | 0.44893 | 0.315 | 0.39 | 1 | HC_vs_MS.NoTrt_F |
| JAZF1 | 0.25133 | 0.93519 | 0.194 | 0.24 | 1 | HC_vs_MS.NoTrt_F |
| TBC1D22A | 0.25167 | 0.26027 | 0.572 | 0.62 | 1 | HC_vs_MS.NoTrt_F |
| ATP7B | 0.25204 | 1.29209 | 0.197 | 0.24 | 1 | HC_vs_MS.NoTrt_F |
| PGM2L1 | 0.2522 | -0.3242 | 0.546 | 0.6 | 1 | HC_vs_MS.NoTrt_F |
| IL13RA1 | 0.25238 | 0.29033 | 0.329 | 0.39 | 1 | HC_vs_MS.NoTrt_F |
| LAMP3 | 0.25276 | 0.48435 | 0.494 | 0.55 | 1 | HC_vs_MS.NoTrt_F |
| MADD | 0.25325 | 0.48952 | 0.283 | 0.35 | 1 | HC_vs_MS.NoTrt_F |
| TUFT1 | 0.25341 | -0.6534 | 0.396 | 0.44 | 1 | HC_vs_MS.NoTrt_F |
| WNK3 | 0.25352 | -0.3297 | 0.405 | 0.34 | 1 | HC_vs_MS.NoTrt_F |
| MYH3 | 0.25398 | 0.59154 | 0.699 | 0.68 | 1 | HC_vs_MS.NoTrt_F |
| FAHD2A | 0.2543 | 0.6988 | 0.199 | 0.24 | 1 | HC_vs_MS.NoTrt_F |
| KDELC2 | 0.25466 | -0.5018 | 0.65 | 0.58 | 1 | HC_vs_MS.NoTrt_F |
| ARL10 | 0.25505 | -0.2502 | 0.922 | 0.85 | 1 | HC_vs_MS.NoTrt_F |
| C9orf131 | 0.25511 | -0.3026 | 0.182 | 0.22 | 1 | HC_vs_MS.NoTrt_F |
| HRH2 | 0.25547 | -0.4567 | 0.136 | 0.17 | 1 | HC_vs_MS.NoTrt_F |
| MXD4 | 0.25601 | 0.49601 | 0.254 | 0.3 | 1 | HC_vs_MS.NoTrt_F |
| MCEE | 0.25631 | 0.5413 | 0.214 | 0.26 | 1 | HC_vs_MS.NoTrt_F |
| GNA15 | 0.25665 | 0.75544 | 0.081 | 0.12 | 1 | HC_vs_MS.NoTrt_F |
| GPNMB | 0.25676 | 0.26146 | 0.734 | 0.69 | 1 | HC_vs_MS.NoTrt_F |
| MSRB1 | 0.25689 | -0.3596 | 0.13 | 0.17 | 1 | HC_vs_MS.NoTrt_F |
| HIST4H4 | 0.25701 | -0.3102 | 0.561 | 0.61 | 1 | HC_vs_MS.NoTrt_F |
| C15orf38-AF | 0.25733 | -0.8334 | 0.402 | 0.43 | 1 | HC_vs_MS.NoTrt_F |
| PPIL2 | 0.25747 | 0.58115 | 0.454 | 0.49 | 1 | HC_vs_MS.NoTrt_F |
| C14orf1 | 0.25792 | -0.3023 | 0.306 | 0.34 | 1 | HC_vs_MS.NoTrt_F |

| | | | | | | |
|---|---|---|---|---|---|---|
| NCOA6 | 0.25799 | 0.39718 | 0.728 | 0.81 | 1 | HC_vs_MS.NoTrt_F |
| TMEM168 | 0.2581 | 0.49513 | 0.592 | 0.51 | 1 | HC_vs_MS.NoTrt_F |
| TMEM65 | 0.25825 | 0.34481 | 0.777 | 0.7 | 1 | HC_vs_MS.NoTrt_F |
| NSMCE4A | 0.25844 | 0.36918 | 0.379 | 0.42 | 1 | HC_vs_MS.NoTrt_F |
| TMED8 | 0.25896 | 0.58234 | 0.642 | 0.66 | 1 | HC_vs_MS.NoTrt_F |
| CDC42BPG | 0.25915 | 0.80987 | 0.616 | 0.56 | 1 | HC_vs_MS.NoTrt_F |
| BRCA2 | 0.25958 | -0.298 | 0.393 | 0.43 | 1 | HC_vs_MS.NoTrt_F |
| LGALS1 | 0.25974 | 0.6353 | 0.477 | 0.47 | 1 | HC_vs_MS.NoTrt_F |
| ERCC3 | 0.26013 | -0.2991 | 0.434 | 0.48 | 1 | HC_vs_MS.NoTrt_F |
| ARFGAP2 | 0.26043 | -0.2671 | 0.566 | 0.64 | 1 | HC_vs_MS.NoTrt_F |
| RGPD3 | 0.26066 | 0.38005 | 0.104 | 0.14 | 1 | HC_vs_MS.NoTrt_F |
| LCLAT1 | 0.26078 | 0.36263 | 0.835 | 0.85 | 1 | HC_vs_MS.NoTrt_F |
| WISP2 | 0.26105 | 0.5245 | 0.685 | 0.61 | 1 | HC_vs_MS.NoTrt_F |
| LRRC8A | 0.26125 | -0.354 | 0.173 | 0.21 | 1 | HC_vs_MS.NoTrt_F |
| STT3B | 0.26148 | 0.77545 | 0.618 | 0.58 | 1 | HC_vs_MS.NoTrt_F |
| TMEM91 | 0.26168 | 0.58631 | 0.598 | 0.56 | 1 | HC_vs_MS.NoTrt_F |
| LILRB5 | 0.26177 | 0.4804 | 0.379 | 0.43 | 1 | HC_vs_MS.NoTrt_F |
| DOPEY1 | 0.26186 | 0.6304 | 0.327 | 0.38 | 1 | HC_vs_MS.NoTrt_F |
| ALDH1B1 | 0.26186 | -0.3113 | 0.662 | 0.67 | 1 | HC_vs_MS.NoTrt_F |
| ZSWIM1 | 0.26208 | 0.74562 | 0.734 | 0.72 | 1 | HC_vs_MS.NoTrt_F |
| ITSN1 | 0.26224 | 0.50119 | 0.936 | 0.92 | 1 | HC_vs_MS.NoTrt_F |
| FMNL3 | 0.26228 | 0.52341 | 0.431 | 0.39 | 1 | HC_vs_MS.NoTrt_F |
| GPM6B | 0.26277 | 0.35059 | 0.171 | 0.21 | 1 | HC_vs_MS.NoTrt_F |
| TXNL4A | 0.26297 | 0.55041 | 0.572 | 0.53 | 1 | HC_vs_MS.NoTrt_F |
| MOBP | 0.26304 | 0.25387 | 0.656 | 0.6 | 1 | HC_vs_MS.NoTrt_F |
| KIF3A | 0.26382 | 0.33914 | 0.786 | 0.72 | 1 | HC_vs_MS.NoTrt_F |
| GOSR2 | 0.26385 | 1.10779 | 0.171 | 0.22 | 1 | HC_vs_MS.NoTrt_F |
| RBP7 | 0.26395 | 1.12912 | 0.376 | 0.32 | 1 | HC_vs_MS.NoTrt_F |
| LHFP | 0.26396 | 1.05346 | 0.26 | 0.23 | 1 | HC_vs_MS.NoTrt_F |
| PLEKHA5 | 0.26432 | -0.4276 | 0.945 | 0.95 | 1 | HC_vs_MS.NoTrt_F |
| GNB1L | 0.26447 | 0.42971 | 0.292 | 0.33 | 1 | HC_vs_MS.NoTrt_F |
| HINT3 | 0.26453 | -0.6104 | 0.303 | 0.35 | 1 | HC_vs_MS.NoTrt_F |
| AL928654.7 | 0.26455 | 0.40825 | 0.078 | 0.11 | 1 | HC_vs_MS.NoTrt_F |
| IL3RA | 0.26468 | 1.41453 | 0.121 | 0.16 | 1 | HC_vs_MS.NoTrt_F |
| PNISR | 0.26473 | 0.34326 | 0.555 | 0.67 | 1 | HC_vs_MS.NoTrt_F |
| BRAF | 0.26476 | 0.70995 | 0.191 | 0.24 | 1 | HC_vs_MS.NoTrt_F |
| NDC80 | 0.26481 | -0.5356 | 0.743 | 0.77 | 1 | HC_vs_MS.NoTrt_F |
| PCTP | 0.26483 | 0.40693 | 0.442 | 0.49 | 1 | HC_vs_MS.NoTrt_F |
| RP11-1035H | 0.26486 | 0.43367 | 0.153 | 0.19 | 1 | HC_vs_MS.NoTrt_F |
| RMND5B | 0.26489 | 0.58824 | 0.682 | 0.64 | 1 | HC_vs_MS.NoTrt_F |

| | | | | | | |
|---|---|---|---|---|---|---|
| KCTD20 | 0.26504 | 0.4584 | 0.685 | 0.72 | 1 | HC_vs_MS.NoTrt_F |
| EBPL | 0.26508 | 0.32828 | 0.717 | 0.71 | 1 | HC_vs_MS.NoTrt_F |
| ZNF609 | 0.26511 | -0.6406 | 0.621 | 0.66 | 1 | HC_vs_MS.NoTrt_F |
| TCTA | 0.26526 | 0.68343 | 0.633 | 0.64 | 1 | HC_vs_MS.NoTrt_F |
| ZNF331 | 0.26534 | 0.56077 | 0.757 | 0.78 | 1 | HC_vs_MS.NoTrt_F |
| KLHL24 | 0.26535 | -0.4769 | 0.882 | 0.88 | 1 | HC_vs_MS.NoTrt_F |
| MBOAT7 | 0.2654 | 0.74676 | 0.139 | 0.18 | 1 | HC_vs_MS.NoTrt_F |
| INPP5K | 0.26544 | -0.2765 | 0.621 | 0.69 | 1 | HC_vs_MS.NoTrt_F |
| FASTKD5 | 0.26607 | -0.312 | 0.159 | 0.2 | 1 | HC_vs_MS.NoTrt_F |
| EMC10 | 0.26629 | -0.2518 | 0.864 | 0.87 | 1 | HC_vs_MS.NoTrt_F |
| TMEM216 | 0.26638 | -0.4521 | 0.471 | 0.52 | 1 | HC_vs_MS.NoTrt_F |
| AGPAT5 | 0.26694 | 0.59298 | 0.127 | 0.17 | 1 | HC_vs_MS.NoTrt_F |
| DEPDC5 | 0.26718 | -0.4779 | 0.777 | 0.81 | 1 | HC_vs_MS.NoTrt_F |
| SLC16A3 | 0.26724 | 0.77779 | 0.595 | 0.55 | 1 | HC_vs_MS.NoTrt_F |
| NKPD1 | 0.26724 | 0.3368 | 0.098 | 0.13 | 1 | HC_vs_MS.NoTrt_F |
| MGST3 | 0.26748 | -0.3185 | 0.902 | 0.89 | 1 | HC_vs_MS.NoTrt_F |
| CXADR | 0.26768 | -0.8831 | 0.289 | 0.34 | 1 | HC_vs_MS.NoTrt_F |
| ANKRD44 | 0.26836 | 0.29825 | 0.864 | 0.91 | 1 | HC_vs_MS.NoTrt_F |
| SEC14L4 | 0.26845 | 0.61124 | 0.613 | 0.56 | 1 | HC_vs_MS.NoTrt_F |
| QSOX2 | 0.26865 | 0.39878 | 0.194 | 0.24 | 1 | HC_vs_MS.NoTrt_F |
| CAMK1D | 0.26866 | -0.2841 | 0.855 | 0.84 | 1 | HC_vs_MS.NoTrt_F |
| ZNF655 | 0.26869 | 0.2652 | 0.616 | 0.69 | 1 | HC_vs_MS.NoTrt_F |
| SLC38A6 | 0.26919 | 0.50998 | 0.671 | 0.67 | 1 | HC_vs_MS.NoTrt_F |
| HYKK | 0.26929 | 0.54815 | 0.89 | 0.89 | 1 | HC_vs_MS.NoTrt_F |
| SLC9A6 | 0.26948 | -0.3949 | 0.462 | 0.54 | 1 | HC_vs_MS.NoTrt_F |
| ZNF782 | 0.26974 | 1.3323 | 0.147 | 0.19 | 1 | HC_vs_MS.NoTrt_F |
| NUDT9 | 0.26976 | -0.2879 | 0.662 | 0.59 | 1 | HC_vs_MS.NoTrt_F |
| C6orf25 | 0.26989 | 0.55523 | 0.506 | 0.55 | 1 | HC_vs_MS.NoTrt_F |
| SCG3 | 0.26991 | 0.28182 | 0.26 | 0.32 | 1 | HC_vs_MS.NoTrt_F |
| CCDC169 | 0.27013 | 0.44108 | 0.636 | 0.61 | 1 | HC_vs_MS.NoTrt_F |
| SNRNP27 | 0.2704 | 0.59114 | 0.277 | 0.34 | 1 | HC_vs_MS.NoTrt_F |
| FEM1B | 0.27042 | 0.28392 | 0.124 | 0.16 | 1 | HC_vs_MS.NoTrt_F |
| GSTT1 | 0.27053 | 0.33504 | 0.74 | 0.81 | 1 | HC_vs_MS.NoTrt_F |
| CLCN3 | 0.2709 | -0.3755 | 0.399 | 0.44 | 1 | HC_vs_MS.NoTrt_F |
| NOCT | 0.27104 | 0.40008 | 0.728 | 0.71 | 1 | HC_vs_MS.NoTrt_F |
| C15orf52 | 0.27115 | 0.51038 | 0.425 | 0.48 | 1 | HC_vs_MS.NoTrt_F |
| JARID2 | 0.27131 | -0.6591 | 0.26 | 0.3 | 1 | HC_vs_MS.NoTrt_F |
| MAP1LC3A | 0.27131 | 0.41637 | 0.098 | 0.13 | 1 | HC_vs_MS.NoTrt_F |
| SASH1 | 0.27161 | 1.0606 | 0.399 | 0.34 | 1 | HC_vs_MS.NoTrt_F |
| RPP38 | 0.27165 | -0.7318 | 0.465 | 0.5 | 1 | HC_vs_MS.NoTrt_F |

| | | | | | | |
|---|---|---|---|---|---|---|
| SERPINB1 | 0.27173 | -0.2585 | 0.723 | 0.72 | 1 | HC_vs_MS.NoTrt_F |
| FCRL2 | 0.27177 | 0.73155 | 0.723 | 0.67 | 1 | HC_vs_MS.NoTrt_F |
| ZNF652 | 0.27207 | -0.5635 | 0.581 | 0.61 | 1 | HC_vs_MS.NoTrt_F |
| NUP43 | 0.2722 | 0.31327 | 0.905 | 0.85 | 1 | HC_vs_MS.NoTrt_F |
| 3-Sep | 0.27222 | 0.7253 | 0.075 | 0.11 | 1 | HC_vs_MS.NoTrt_F |
| PDCD7 | 0.27242 | 0.57006 | 0.549 | 0.48 | 1 | HC_vs_MS.NoTrt_F |
| ATG13 | 0.27278 | 0.3831 | 0.89 | 0.84 | 1 | HC_vs_MS.NoTrt_F |
| CLVS1 | 0.27334 | 0.25302 | 0.685 | 0.66 | 1 | HC_vs_MS.NoTrt_F |
| KIAA0930 | 0.27374 | 0.83863 | 0.691 | 0.64 | 1 | HC_vs_MS.NoTrt_F |
| ETFDH | 0.27385 | 0.34352 | 0.74 | 0.74 | 1 | HC_vs_MS.NoTrt_F |
| RALGPS2 | 0.27407 | 0.48624 | 0.37 | 0.41 | 1 | HC_vs_MS.NoTrt_F |
| USF1 | 0.27436 | -1.2134 | 0.223 | 0.25 | 1 | HC_vs_MS.NoTrt_F |
| PRDX5 | 0.27457 | 0.55105 | 0.286 | 0.36 | 1 | HC_vs_MS.NoTrt_F |
| NPIPA7 | 0.27495 | 0.31091 | 0.182 | 0.23 | 1 | HC_vs_MS.NoTrt_F |
| IMPDH2 | 0.27574 | 0.27642 | 0.225 | 0.28 | 1 | HC_vs_MS.NoTrt_F |
| SLC2A5 | 0.2758 | 0.28644 | 0.74 | 0.72 | 1 | HC_vs_MS.NoTrt_F |
| VAV1 | 0.27673 | 0.29807 | 0.876 | 0.91 | 1 | HC_vs_MS.NoTrt_F |
| CST3 | 0.27683 | 1.55964 | 0.153 | 0.12 | 1 | HC_vs_MS.NoTrt_F |
| DLEC1 | 0.2772 | 0.49126 | 0.514 | 0.54 | 1 | HC_vs_MS.NoTrt_F |
| STPG1 | 0.27759 | -0.2772 | 0.801 | 0.81 | 1 | HC_vs_MS.NoTrt_F |
| KIAA1109 | 0.27786 | -0.2576 | 0.373 | 0.43 | 1 | HC_vs_MS.NoTrt_F |
| COPA | 0.2779 | -0.3065 | 0.35 | 0.39 | 1 | HC_vs_MS.NoTrt_F |
| ZNF316 | 0.27835 | -1.1066 | 0.627 | 0.53 | 1 | HC_vs_MS.NoTrt_F |
| SLC9A8 | 0.2784 | 0.31839 | 0.26 | 0.31 | 1 | HC_vs_MS.NoTrt_F |
| VWA2 | 0.27846 | -0.3834 | 0.78 | 0.79 | 1 | HC_vs_MS.NoTrt_F |
| CES3 | 0.27864 | 0.25938 | 0.327 | 0.39 | 1 | HC_vs_MS.NoTrt_F |
| SERPINB10 | 0.27886 | 0.27319 | 0.448 | 0.43 | 1 | HC_vs_MS.NoTrt_F |
| MAN1C1 | 0.27897 | 0.45242 | 0.905 | 0.85 | 1 | HC_vs_MS.NoTrt_F |
| F11R | 0.27913 | 0.80696 | 0.691 | 0.66 | 1 | HC_vs_MS.NoTrt_F |
| C17orf105 | 0.27934 | 0.2549 | 0.74 | 0.73 | 1 | HC_vs_MS.NoTrt_F |
| NDST2 | 0.27945 | 1.1322 | 0.124 | 0.16 | 1 | HC_vs_MS.NoTrt_F |
| BSPRY | 0.27968 | 0.76799 | 0.11 | 0.14 | 1 | HC_vs_MS.NoTrt_F |
| APEX1 | 0.27974 | 0.69125 | 0.211 | 0.26 | 1 | HC_vs_MS.NoTrt_F |
| RINT1 | 0.27995 | -0.4322 | 0.569 | 0.62 | 1 | HC_vs_MS.NoTrt_F |
| MRPS26 | 0.28007 | 0.44271 | 0.194 | 0.23 | 1 | HC_vs_MS.NoTrt_F |
| ATG2B | 0.28018 | 0.35579 | 0.39 | 0.43 | 1 | HC_vs_MS.NoTrt_F |
| HLA-DPA1 | 0.28032 | 0.28484 | 0.309 | 0.36 | 1 | HC_vs_MS.NoTrt_F |
| COLEC11 | 0.28046 | 0.26738 | 0.393 | 0.45 | 1 | HC_vs_MS.NoTrt_F |
| GTF2H2C.2 | 0.28065 | 0.60047 | 0.289 | 0.34 | 1 | HC_vs_MS.NoTrt_F |
| RP11-644F5 | 0.28139 | -0.358 | 0.731 | 0.67 | 1 | HC_vs_MS.NoTrt_F |

| | | | | | | |
|---|---|---|---|---|---|---|
| ZKSCAN8 | 0.28142 | 1.42268 | 0.303 | 0.38 | 1 | HC_vs_MS.NoTrt_F |
| KCMF1 | 0.28165 | 0.56442 | 0.54 | 0.55 | 1 | HC_vs_MS.NoTrt_F |
| CLK3 | 0.28194 | -0.2998 | 0.818 | 0.83 | 1 | HC_vs_MS.NoTrt_F |
| BCL2L1 | 0.28206 | 0.7242 | 0.318 | 0.28 | 1 | HC_vs_MS.NoTrt_F |
| SMAD4 | 0.28209 | 0.28521 | 0.512 | 0.53 | 1 | HC_vs_MS.NoTrt_F |
| DCLK1 | 0.28244 | 0.89555 | 0.361 | 0.4 | 1 | HC_vs_MS.NoTrt_F |
| KIAA0040 | 0.28247 | 0.49238 | 0.211 | 0.26 | 1 | HC_vs_MS.NoTrt_F |
| CSNK1G2 | 0.28247 | -0.2793 | 0.133 | 0.17 | 1 | HC_vs_MS.NoTrt_F |
| NUMA1 | 0.28258 | 0.38051 | 0.893 | 0.87 | 1 | HC_vs_MS.NoTrt_F |
| SREBF2 | 0.28298 | 0.44701 | 0.22 | 0.27 | 1 | HC_vs_MS.NoTrt_F |
| CLK2 | 0.28379 | 1.19946 | 0.202 | 0.17 | 1 | HC_vs_MS.NoTrt_F |
| ADSL | 0.28389 | -0.3607 | 0.873 | 0.89 | 1 | HC_vs_MS.NoTrt_F |
| CD59 | 0.28394 | -0.5092 | 0.757 | 0.79 | 1 | HC_vs_MS.NoTrt_F |
| AGBL2 | 0.28403 | 0.48753 | 0.28 | 0.32 | 1 | HC_vs_MS.NoTrt_F |
| LSM3 | 0.28442 | 0.40836 | 0.624 | 0.65 | 1 | HC_vs_MS.NoTrt_F |
| CD302 | 0.28462 | 0.26998 | 0.425 | 0.35 | 1 | HC_vs_MS.NoTrt_F |
| TENM1 | 0.28474 | 1.03102 | 0.139 | 0.18 | 1 | HC_vs_MS.NoTrt_F |
| NUDCD2 | 0.28555 | -0.5676 | 0.786 | 0.76 | 1 | HC_vs_MS.NoTrt_F |
| SLAMF7 | 0.28557 | 0.47121 | 0.477 | 0.46 | 1 | HC_vs_MS.NoTrt_F |
| CRTC3 | 0.28564 | 0.25029 | 0.575 | 0.63 | 1 | HC_vs_MS.NoTrt_F |
| RPS6KA3 | 0.28564 | -0.3244 | 0.535 | 0.6 | 1 | HC_vs_MS.NoTrt_F |
| PHF7 | 0.28588 | 1.02849 | 0.523 | 0.49 | 1 | HC_vs_MS.NoTrt_F |
| SOCS7 | 0.28606 | 0.96618 | 0.439 | 0.51 | 1 | HC_vs_MS.NoTrt_F |
| CCNF | 0.28614 | -0.8946 | 0.503 | 0.53 | 1 | HC_vs_MS.NoTrt_F |
| AKR1B10 | 0.28638 | 0.39841 | 0.488 | 0.51 | 1 | HC_vs_MS.NoTrt_F |
| RPN1 | 0.28647 | 0.39953 | 0.76 | 0.78 | 1 | HC_vs_MS.NoTrt_F |
| NME4 | 0.28654 | 0.2937 | 0.168 | 0.2 | 1 | HC_vs_MS.NoTrt_F |
| YIPF1 | 0.28667 | 1.4102 | 0.124 | 0.16 | 1 | HC_vs_MS.NoTrt_F |
| VPS39 | 0.28702 | -0.5271 | 0.538 | 0.56 | 1 | HC_vs_MS.NoTrt_F |
| ARPC1A | 0.28727 | 0.49098 | 0.332 | 0.38 | 1 | HC_vs_MS.NoTrt_F |
| STK26 | 0.28745 | 0.6765 | 0.147 | 0.19 | 1 | HC_vs_MS.NoTrt_F |
| GCSAML | 0.28752 | 0.65158 | 0.338 | 0.39 | 1 | HC_vs_MS.NoTrt_F |
| GTF2E2 | 0.28859 | 0.26542 | 0.639 | 0.7 | 1 | HC_vs_MS.NoTrt_F |
| MTFR1L | 0.28874 | -0.288 | 0.578 | 0.59 | 1 | HC_vs_MS.NoTrt_F |
| TBC1D19 | 0.28946 | 0.6068 | 0.153 | 0.19 | 1 | HC_vs_MS.NoTrt_F |
| SFXN5 | 0.28951 | 0.41838 | 0.945 | 0.95 | 1 | HC_vs_MS.NoTrt_F |
| ZNF620 | 0.2899 | -0.2559 | 0.754 | 0.81 | 1 | HC_vs_MS.NoTrt_F |
| RRP9 | 0.28994 | 0.94766 | 0.118 | 0.16 | 1 | HC_vs_MS.NoTrt_F |
| DCAF11 | 0.29004 | -0.2611 | 0.847 | 0.85 | 1 | HC_vs_MS.NoTrt_F |
| ACSF2 | 0.29028 | 0.4208 | 0.561 | 0.54 | 1 | HC_vs_MS.NoTrt_F |

| | | | | | | |
|---|---|---|---|---|---|---|
| IGHMBP2 | 0.29079 | 0.54424 | 0.402 | 0.43 | 1 | HC_vs_MS.NoTrt_F |
| OSMR | 0.29121 | 0.75962 | 0.584 | 0.63 | 1 | HC_vs_MS.NoTrt_F |
| APOBEC3C | 0.29151 | 0.57558 | 0.777 | 0.85 | 1 | HC_vs_MS.NoTrt_F |
| UNC45A | 0.29181 | 0.2798 | 0.737 | 0.72 | 1 | HC_vs_MS.NoTrt_F |
| TSPYL4 | 0.29181 | 0.83418 | 0.168 | 0.21 | 1 | HC_vs_MS.NoTrt_F |
| EPG5 | 0.2921 | 0.42937 | 0.564 | 0.6 | 1 | HC_vs_MS.NoTrt_F |
| ZNF805 | 0.29214 | 0.8201 | 0.832 | 0.79 | 1 | HC_vs_MS.NoTrt_F |
| WDFY1 | 0.29306 | -0.2994 | 0.39 | 0.43 | 1 | HC_vs_MS.NoTrt_F |
| ZNF781 | 0.29332 | -0.428 | 0.301 | 0.35 | 1 | HC_vs_MS.NoTrt_F |
| PIGW | 0.29388 | 0.34783 | 0.477 | 0.43 | 1 | HC_vs_MS.NoTrt_F |
| IL18R1 | 0.29404 | 0.40965 | 0.142 | 0.18 | 1 | HC_vs_MS.NoTrt_F |
| TPK1 | 0.29413 | 0.49535 | 0.396 | 0.44 | 1 | HC_vs_MS.NoTrt_F |
| ZKSCAN7 | 0.29458 | 0.34109 | 0.647 | 0.58 | 1 | HC_vs_MS.NoTrt_F |
| CLSPN | 0.29466 | 0.26871 | 0.91 | 0.91 | 1 | HC_vs_MS.NoTrt_F |
| CTNND1 | 0.29468 | 0.98182 | 0.483 | 0.45 | 1 | HC_vs_MS.NoTrt_F |
| GPR89B | 0.29483 | 0.32561 | 0.402 | 0.44 | 1 | HC_vs_MS.NoTrt_F |
| OAS1 | 0.29532 | -0.301 | 0.584 | 0.58 | 1 | HC_vs_MS.NoTrt_F |
| LA16c-431H | 0.2956 | 0.72073 | 0.555 | 0.51 | 1 | HC_vs_MS.NoTrt_F |
| CFP | 0.29617 | 1.14544 | 0.422 | 0.38 | 1 | HC_vs_MS.NoTrt_F |
| CCDC88B | 0.29641 | 0.71183 | 0.766 | 0.84 | 1 | HC_vs_MS.NoTrt_F |
| CAV2 | 0.29663 | 0.44242 | 0.61 | 0.58 | 1 | HC_vs_MS.NoTrt_F |
| MON2 | 0.29703 | 0.55788 | 0.431 | 0.49 | 1 | HC_vs_MS.NoTrt_F |
| MSI2 | 0.29709 | 0.40352 | 0.942 | 0.91 | 1 | HC_vs_MS.NoTrt_F |
| TP53RK | 0.29722 | 0.3065 | 0.266 | 0.31 | 1 | HC_vs_MS.NoTrt_F |
| C10orf128 | 0.29726 | -0.2657 | 0.142 | 0.18 | 1 | HC_vs_MS.NoTrt_F |
| OSBP2 | 0.2973 | -0.3826 | 0.468 | 0.52 | 1 | HC_vs_MS.NoTrt_F |
| HIST1H2BE | 0.29795 | 0.39306 | 0.335 | 0.39 | 1 | HC_vs_MS.NoTrt_F |
| SCAMP3 | 0.29806 | 0.27503 | 0.251 | 0.31 | 1 | HC_vs_MS.NoTrt_F |
| AHR | 0.29849 | 0.52003 | 0.688 | 0.66 | 1 | HC_vs_MS.NoTrt_F |
| CRACR2A | 0.29894 | -0.3377 | 0.694 | 0.71 | 1 | HC_vs_MS.NoTrt_F |
| MTHFD2 | 0.29928 | -0.4152 | 0.717 | 0.75 | 1 | HC_vs_MS.NoTrt_F |
| TIMM23B | 0.2994 | 0.48151 | 0.647 | 0.66 | 1 | HC_vs_MS.NoTrt_F |
| ZNF683 | 0.2995 | 2.07889 | 0.075 | 0.11 | 1 | HC_vs_MS.NoTrt_F |
| ZNF264 | 0.30016 | 0.59955 | 0.957 | 0.95 | 1 | HC_vs_MS.NoTrt_F |
| PAWR | 0.3004 | 0.45437 | 0.72 | 0.73 | 1 | HC_vs_MS.NoTrt_F |
| OGFOD1 | 0.30043 | -0.5021 | 0.679 | 0.69 | 1 | HC_vs_MS.NoTrt_F |
| RBM12 | 0.30065 | 0.8202 | 0.289 | 0.27 | 1 | HC_vs_MS.NoTrt_F |
| TTC7A | 0.30104 | 0.70048 | 0.249 | 0.3 | 1 | HC_vs_MS.NoTrt_F |
| ZNRF1 | 0.30121 | 0.83947 | 0.344 | 0.31 | 1 | HC_vs_MS.NoTrt_F |
| FAM175A | 0.3017 | 0.26033 | 0.867 | 0.88 | 1 | HC_vs_MS.NoTrt_F |

| | | | | | | |
|---|---|---|---|---|---|---|
| GTPBP8 | 0.30201 | -0.4907 | 0.702 | 0.74 | 1 | HC_vs_MS.NoTrt_F |
| GOLGA6L22 | 0.30207 | 0.29119 | 0.434 | 0.41 | 1 | HC_vs_MS.NoTrt_F |
| UQCC3 | 0.30207 | 0.43607 | 0.48 | 0.5 | 1 | HC_vs_MS.NoTrt_F |
| TNFAIP3 | 0.3023 | 0.72698 | 0.633 | 0.65 | 1 | HC_vs_MS.NoTrt_F |
| TMEM182 | 0.30243 | 0.29419 | 0.78 | 0.82 | 1 | HC_vs_MS.NoTrt_F |
| FAM234B | 0.30248 | -0.5471 | 0.506 | 0.54 | 1 | HC_vs_MS.NoTrt_F |
| TGFB1 | 0.30259 | 0.73198 | 0.393 | 0.45 | 1 | HC_vs_MS.NoTrt_F |
| ACADL | 0.30263 | -0.3134 | 0.332 | 0.37 | 1 | HC_vs_MS.NoTrt_F |
| SPECC1L | 0.30266 | 0.25296 | 0.188 | 0.23 | 1 | HC_vs_MS.NoTrt_F |
| SDC4 | 0.30309 | 0.48559 | 0.171 | 0.21 | 1 | HC_vs_MS.NoTrt_F |
| RNF144A | 0.30342 | 1.76714 | 0.208 | 0.25 | 1 | HC_vs_MS.NoTrt_F |
| ZNF574 | 0.3036 | 0.51558 | 0.373 | 0.39 | 1 | HC_vs_MS.NoTrt_F |
| ANKAR | 0.3041 | 0.3203 | 0.179 | 0.22 | 1 | HC_vs_MS.NoTrt_F |
| ZNF267 | 0.30419 | 0.62049 | 0.225 | 0.28 | 1 | HC_vs_MS.NoTrt_F |
| IL1R1 | 0.30501 | 1.78254 | 0.422 | 0.39 | 1 | HC_vs_MS.NoTrt_F |
| SMARCA2 | 0.30528 | 0.42186 | 0.477 | 0.55 | 1 | HC_vs_MS.NoTrt_F |
| SCAPER | 0.30529 | -0.3284 | 0.402 | 0.43 | 1 | HC_vs_MS.NoTrt_F |
| KDM5B | 0.30533 | 0.89049 | 0.327 | 0.35 | 1 | HC_vs_MS.NoTrt_F |
| ZC3H7B | 0.30598 | 0.25644 | 0.853 | 0.83 | 1 | HC_vs_MS.NoTrt_F |
| NBPF9 | 0.30635 | 0.55427 | 0.702 | 0.8 | 1 | HC_vs_MS.NoTrt_F |
| MRTO4 | 0.30649 | 0.45115 | 0.538 | 0.61 | 1 | HC_vs_MS.NoTrt_F |
| ST20 | 0.30739 | 0.89956 | 0.254 | 0.22 | 1 | HC_vs_MS.NoTrt_F |
| WDR74 | 0.30742 | 0.34196 | 0.653 | 0.62 | 1 | HC_vs_MS.NoTrt_F |
| ARL1 | 0.30747 | -0.5372 | 0.419 | 0.45 | 1 | HC_vs_MS.NoTrt_F |
| TBXA2R | 0.30758 | 0.4086 | 0.746 | 0.77 | 1 | HC_vs_MS.NoTrt_F |
| CACNB1 | 0.30763 | 0.91025 | 0.145 | 0.18 | 1 | HC_vs_MS.NoTrt_F |
| SMIM8 | 0.30782 | 0.53257 | 0.685 | 0.63 | 1 | HC_vs_MS.NoTrt_F |
| NOTCH1 | 0.30787 | 0.71326 | 0.156 | 0.19 | 1 | HC_vs_MS.NoTrt_F |
| RLIM | 0.30787 | 0.63317 | 0.772 | 0.8 | 1 | HC_vs_MS.NoTrt_F |
| NAF1 | 0.30828 | 0.62609 | 0.497 | 0.53 | 1 | HC_vs_MS.NoTrt_F |
| PHLDB3 | 0.30845 | 0.46026 | 0.665 | 0.66 | 1 | HC_vs_MS.NoTrt_F |
| MOCS3 | 0.30876 | -0.3731 | 0.879 | 0.88 | 1 | HC_vs_MS.NoTrt_F |
| STX12 | 0.30943 | 0.534 | 0.306 | 0.35 | 1 | HC_vs_MS.NoTrt_F |
| FAM206A | 0.30946 | 1.32877 | 0.124 | 0.16 | 1 | HC_vs_MS.NoTrt_F |
| ERLEC1 | 0.3096 | 0.51469 | 0.413 | 0.48 | 1 | HC_vs_MS.NoTrt_F |
| NSMCE3 | 0.3097 | -0.3715 | 0.373 | 0.41 | 1 | HC_vs_MS.NoTrt_F |
| ATL2 | 0.30971 | 0.40895 | 0.879 | 0.88 | 1 | HC_vs_MS.NoTrt_F |
| ARHGEF26 | 0.30992 | 1.15981 | 0.595 | 0.53 | 1 | HC_vs_MS.NoTrt_F |
| ZBED5 | 0.31041 | -0.3477 | 0.705 | 0.73 | 1 | HC_vs_MS.NoTrt_F |
| GYG1 | 0.31059 | -0.5674 | 0.618 | 0.64 | 1 | HC_vs_MS.NoTrt_F |

| | | | | | | |
|---|---|---|---|---|---|---|
| RAB11FIP1 | 0.31064 | 0.57642 | 0.832 | 0.83 | 1 | HC_vs_MS.NoTrt_F |
| FN3KRP | 0.31081 | 0.54488 | 0.145 | 0.18 | 1 | HC_vs_MS.NoTrt_F |
| NUP50 | 0.31105 | -0.2889 | 0.734 | 0.75 | 1 | HC_vs_MS.NoTrt_F |
| NEU1 | 0.31123 | 1.14648 | 0.682 | 0.62 | 1 | HC_vs_MS.NoTrt_F |
| FO538757.1 | 0.31152 | 0.41492 | 0.225 | 0.27 | 1 | HC_vs_MS.NoTrt_F |
| CUL7 | 0.312 | 0.91826 | 0.121 | 0.16 | 1 | HC_vs_MS.NoTrt_F |
| ADH4 | 0.31215 | 0.30167 | 0.347 | 0.33 | 1 | HC_vs_MS.NoTrt_F |
| TSKU | 0.31215 | 0.60482 | 0.587 | 0.6 | 1 | HC_vs_MS.NoTrt_F |
| OGG1 | 0.31251 | -0.2576 | 0.431 | 0.46 | 1 | HC_vs_MS.NoTrt_F |
| RBM34 | 0.31266 | 0.52278 | 0.717 | 0.72 | 1 | HC_vs_MS.NoTrt_F |
| PHRF1 | 0.31285 | 0.72451 | 0.202 | 0.25 | 1 | HC_vs_MS.NoTrt_F |
| SOGA1 | 0.31359 | 0.51049 | 0.711 | 0.65 | 1 | HC_vs_MS.NoTrt_F |
| DDX59 | 0.31371 | 0.32946 | 0.491 | 0.45 | 1 | HC_vs_MS.NoTrt_F |
| CNOT2 | 0.31392 | -0.2801 | 0.936 | 0.96 | 1 | HC_vs_MS.NoTrt_F |
| ERCC2 | 0.31405 | -0.8081 | 0.561 | 0.59 | 1 | HC_vs_MS.NoTrt_F |
| QRSL1 | 0.31446 | 0.29155 | 0.592 | 0.66 | 1 | HC_vs_MS.NoTrt_F |
| STON1 | 0.31503 | -0.7962 | 0.384 | 0.43 | 1 | HC_vs_MS.NoTrt_F |
| SERPINE2 | 0.31623 | -1.1395 | 0.367 | 0.4 | 1 | HC_vs_MS.NoTrt_F |
| NEBL | 0.3163 | -0.5149 | 0.37 | 0.42 | 1 | HC_vs_MS.NoTrt_F |
| CARF | 0.31631 | -0.5947 | 0.546 | 0.57 | 1 | HC_vs_MS.NoTrt_F |
| ITFG2 | 0.31633 | 0.32441 | 0.376 | 0.42 | 1 | HC_vs_MS.NoTrt_F |
| GSTO1 | 0.31675 | 0.29604 | 0.217 | 0.26 | 1 | HC_vs_MS.NoTrt_F |
| NUDT6 | 0.31676 | -0.2783 | 0.584 | 0.61 | 1 | HC_vs_MS.NoTrt_F |
| FAM117B | 0.31681 | 0.27707 | 0.474 | 0.51 | 1 | HC_vs_MS.NoTrt_F |
| KATNAL1 | 0.3169 | 0.30773 | 0.613 | 0.64 | 1 | HC_vs_MS.NoTrt_F |
| PPP1R15A | 0.31701 | 0.36303 | 0.702 | 0.71 | 1 | HC_vs_MS.NoTrt_F |
| MUTYH | 0.31726 | -0.2642 | 0.561 | 0.57 | 1 | HC_vs_MS.NoTrt_F |
| WDR3 | 0.31801 | -0.9401 | 0.783 | 0.79 | 1 | HC_vs_MS.NoTrt_F |
| PIGN | 0.31815 | 0.52949 | 0.653 | 0.64 | 1 | HC_vs_MS.NoTrt_F |
| PPP4R2 | 0.31829 | -0.2516 | 0.523 | 0.56 | 1 | HC_vs_MS.NoTrt_F |
| UBAP1L | 0.31837 | 0.48259 | 0.399 | 0.45 | 1 | HC_vs_MS.NoTrt_F |
| HSPA14 | 0.31843 | -0.3805 | 0.373 | 0.41 | 1 | HC_vs_MS.NoTrt_F |
| DOCK9 | 0.31852 | 0.8925 | 0.338 | 0.37 | 1 | HC_vs_MS.NoTrt_F |
| LENG1 | 0.31856 | 0.73891 | 0.147 | 0.19 | 1 | HC_vs_MS.NoTrt_F |
| ARHGAP4 | 0.31862 | 0.30778 | 0.497 | 0.6 | 1 | HC_vs_MS.NoTrt_F |
| GGT5 | 0.31867 | 0.3806 | 0.769 | 0.75 | 1 | HC_vs_MS.NoTrt_F |
| A2ML1 | 0.31878 | 0.28105 | 0.685 | 0.71 | 1 | HC_vs_MS.NoTrt_F |
| GOLGA8R | 0.31898 | 0.54345 | 0.558 | 0.54 | 1 | HC_vs_MS.NoTrt_F |
| RP11-544M2 | 0.31908 | 0.59403 | 0.272 | 0.31 | 1 | HC_vs_MS.NoTrt_F |
| ATAT1 | 0.31916 | 0.73072 | 0.121 | 0.16 | 1 | HC_vs_MS.NoTrt_F |

| | | | | | | |
|---|---|---|---|---|---|---|
| MFAP1 | 0.31926 | 0.62818 | 0.139 | 0.18 | 1 | HC_vs_MS.NoTrt_F |
| ILF3 | 0.31944 | 0.51779 | 0.803 | 0.83 | 1 | HC_vs_MS.NoTrt_F |
| KATNAL2 | 0.31977 | 0.66295 | 0.416 | 0.38 | 1 | HC_vs_MS.NoTrt_F |
| WIPF2 | 0.31982 | 0.40588 | 0.841 | 0.84 | 1 | HC_vs_MS.NoTrt_F |
| BRF1 | 0.32021 | 0.69407 | 0.624 | 0.58 | 1 | HC_vs_MS.NoTrt_F |
| NME1 | 0.32023 | 0.6361 | 0.639 | 0.56 | 1 | HC_vs_MS.NoTrt_F |
| ZMYND8 | 0.32024 | 0.32625 | 0.419 | 0.47 | 1 | HC_vs_MS.NoTrt_F |
| ZC3H14 | 0.32037 | -0.3334 | 0.957 | 0.97 | 1 | HC_vs_MS.NoTrt_F |
| WDR91 | 0.32051 | -0.2668 | 0.535 | 0.58 | 1 | HC_vs_MS.NoTrt_F |
| EMP2 | 0.32064 | 0.28064 | 0.957 | 0.95 | 1 | HC_vs_MS.NoTrt_F |
| HSD17B12 | 0.32086 | -0.4139 | 0.474 | 0.52 | 1 | HC_vs_MS.NoTrt_F |
| WDR75 | 0.32145 | 0.30134 | 0.358 | 0.43 | 1 | HC_vs_MS.NoTrt_F |
| ZNF19 | 0.32149 | 0.40876 | 0.815 | 0.84 | 1 | HC_vs_MS.NoTrt_F |
| CALU | 0.32152 | 0.4132 | 0.425 | 0.49 | 1 | HC_vs_MS.NoTrt_F |
| KCTD6 | 0.32186 | -0.2661 | 0.656 | 0.65 | 1 | HC_vs_MS.NoTrt_F |
| ZNF528 | 0.32243 | 0.518 | 0.65 | 0.59 | 1 | HC_vs_MS.NoTrt_F |
| PLGLB1 | 0.32245 | 0.37708 | 0.185 | 0.23 | 1 | HC_vs_MS.NoTrt_F |
| RUNX1T1 | 0.32245 | 0.49408 | 0.249 | 0.28 | 1 | HC_vs_MS.NoTrt_F |
| GPT2 | 0.3227 | -0.34 | 0.549 | 0.5 | 1 | HC_vs_MS.NoTrt_F |
| ZFYVE26 | 0.32273 | 0.8853 | 0.876 | 0.86 | 1 | HC_vs_MS.NoTrt_F |
| CALCOCO2 | 0.32293 | -0.4132 | 0.994 | 1 | 1 | HC_vs_MS.NoTrt_F |
| ARSD | 0.32329 | 0.51357 | 0.838 | 0.83 | 1 | HC_vs_MS.NoTrt_F |
| MGP | 0.32334 | 0.8542 | 0.61 | 0.58 | 1 | HC_vs_MS.NoTrt_F |
| EMILIN2 | 0.32335 | 0.5033 | 0.48 | 0.53 | 1 | HC_vs_MS.NoTrt_F |
| UBE2G2 | 0.32371 | 0.50229 | 0.754 | 0.72 | 1 | HC_vs_MS.NoTrt_F |
| SLC16A5 | 0.32378 | 0.42232 | 0.191 | 0.23 | 1 | HC_vs_MS.NoTrt_F |
| LUC7L | 0.32391 | 0.66167 | 0.864 | 0.85 | 1 | HC_vs_MS.NoTrt_F |
| VKORC1 | 0.3244 | 0.60374 | 0.214 | 0.25 | 1 | HC_vs_MS.NoTrt_F |
| ENPP2 | 0.32442 | 1.53059 | 0.156 | 0.13 | 1 | HC_vs_MS.NoTrt_F |
| IFIT2 | 0.32454 | -1.3089 | 0.116 | 0.15 | 1 | HC_vs_MS.NoTrt_F |
| YRDC | 0.32465 | 0.96956 | 0.113 | 0.15 | 1 | HC_vs_MS.NoTrt_F |
| CCDC92 | 0.32471 | 0.33747 | 0.095 | 0.13 | 1 | HC_vs_MS.NoTrt_F |
| SLC9B1 | 0.32474 | 0.39921 | 0.419 | 0.47 | 1 | HC_vs_MS.NoTrt_F |
| AP1B1 | 0.32474 | 0.73902 | 0.28 | 0.25 | 1 | HC_vs_MS.NoTrt_F |
| TGFB1I1 | 0.3253 | 0.90669 | 0.332 | 0.38 | 1 | HC_vs_MS.NoTrt_F |
| ZNF215 | 0.32538 | 0.51817 | 0.422 | 0.5 | 1 | HC_vs_MS.NoTrt_F |
| HARS | 0.32579 | -0.2675 | 0.867 | 0.87 | 1 | HC_vs_MS.NoTrt_F |
| TRAPPC2B | 0.32582 | 0.25869 | 0.436 | 0.39 | 1 | HC_vs_MS.NoTrt_F |
| DIP2C | 0.32584 | 0.64759 | 0.116 | 0.15 | 1 | HC_vs_MS.NoTrt_F |
| SEC14L1 | 0.32586 | -0.314 | 0.731 | 0.79 | 1 | HC_vs_MS.NoTrt_F |

| | | | | | | |
|---|---|---|---|---|---|---|
| CCDC149 | 0.32598 | 0.52768 | 0.468 | 0.49 | 1 | HC_vs_MS.NoTrt_F |
| CCDC107 | 0.32653 | 0.39731 | 0.266 | 0.32 | 1 | HC_vs_MS.NoTrt_F |
| U2SURP | 0.32679 | 0.55113 | 0.364 | 0.43 | 1 | HC_vs_MS.NoTrt_F |
| DGKD | 0.32679 | 0.26977 | 0.691 | 0.64 | 1 | HC_vs_MS.NoTrt_F |
| IDH2 | 0.3268 | 0.49045 | 0.587 | 0.56 | 1 | HC_vs_MS.NoTrt_F |
| ZGPAT | 0.32692 | 0.72054 | 0.208 | 0.25 | 1 | HC_vs_MS.NoTrt_F |
| CHPT1 | 0.32694 | 0.38657 | 0.642 | 0.61 | 1 | HC_vs_MS.NoTrt_F |
| UBE3D | 0.327 | -0.3109 | 0.671 | 0.69 | 1 | HC_vs_MS.NoTrt_F |
| PTPN9 | 0.3274 | 0.85885 | 0.355 | 0.39 | 1 | HC_vs_MS.NoTrt_F |
| PRR5-ARHG/ | 0.32745 | -0.6246 | 0.107 | 0.14 | 1 | HC_vs_MS.NoTrt_F |
| FURIN | 0.32754 | -0.4407 | 0.11 | 0.14 | 1 | HC_vs_MS.NoTrt_F |
| TMEM126B | 0.3281 | 0.31322 | 0.234 | 0.27 | 1 | HC_vs_MS.NoTrt_F |
| NXN | 0.32835 | 0.379 | 0.578 | 0.55 | 1 | HC_vs_MS.NoTrt_F |
| EDF1 | 0.32859 | 0.28603 | 0.477 | 0.58 | 1 | HC_vs_MS.NoTrt_F |
| IFNLR1 | 0.32863 | -1.1648 | 0.13 | 0.17 | 1 | HC_vs_MS.NoTrt_F |
| RGS5 | 0.32883 | 1.68935 | 0.11 | 0.14 | 1 | HC_vs_MS.NoTrt_F |
| BHLHE40 | 0.32937 | 0.65463 | 0.156 | 0.2 | 1 | HC_vs_MS.NoTrt_F |
| AL589743.1 | 0.32956 | 0.25175 | 0.642 | 0.59 | 1 | HC_vs_MS.NoTrt_F |
| SYF2 | 0.32972 | 0.79538 | 0.5 | 0.48 | 1 | HC_vs_MS.NoTrt_F |
| WNT7A | 0.33028 | 0.81202 | 0.098 | 0.13 | 1 | HC_vs_MS.NoTrt_F |
| VPS11 | 0.3304 | -0.7219 | 0.847 | 0.81 | 1 | HC_vs_MS.NoTrt_F |
| CDC14B | 0.33043 | 0.4482 | 0.347 | 0.37 | 1 | HC_vs_MS.NoTrt_F |
| PBX2 | 0.33044 | 0.49162 | 0.173 | 0.21 | 1 | HC_vs_MS.NoTrt_F |
| PTPN1 | 0.33063 | 0.7811 | 0.41 | 0.39 | 1 | HC_vs_MS.NoTrt_F |
| CYP4V2 | 0.33087 | -0.3893 | 0.763 | 0.82 | 1 | HC_vs_MS.NoTrt_F |
| FCHO1 | 0.33113 | -0.5374 | 0.818 | 0.82 | 1 | HC_vs_MS.NoTrt_F |
| KCNC4 | 0.33196 | 0.78433 | 0.786 | 0.82 | 1 | HC_vs_MS.NoTrt_F |
| ZNF354B | 0.33199 | 0.35699 | 0.361 | 0.41 | 1 | HC_vs_MS.NoTrt_F |
| NR3C1 | 0.33206 | 0.57725 | 0.338 | 0.3 | 1 | HC_vs_MS.NoTrt_F |
| QTRT1 | 0.33239 | 0.40282 | 0.844 | 0.83 | 1 | HC_vs_MS.NoTrt_F |
| USP12 | 0.33248 | 0.91378 | 0.124 | 0.16 | 1 | HC_vs_MS.NoTrt_F |
| SREK1 | 0.33248 | 0.56658 | 0.572 | 0.62 | 1 | HC_vs_MS.NoTrt_F |
| SF3A1 | 0.33261 | 0.35786 | 0.425 | 0.48 | 1 | HC_vs_MS.NoTrt_F |
| MCM6 | 0.33272 | 0.64069 | 0.173 | 0.22 | 1 | HC_vs_MS.NoTrt_F |
| CS | 0.33367 | 0.26784 | 0.864 | 0.83 | 1 | HC_vs_MS.NoTrt_F |
| FAM120A | 0.33476 | 0.46011 | 0.211 | 0.25 | 1 | HC_vs_MS.NoTrt_F |
| ABAT | 0.33479 | -0.3939 | 0.656 | 0.67 | 1 | HC_vs_MS.NoTrt_F |
| FSCN1 | 0.33494 | 1.278 | 0.101 | 0.13 | 1 | HC_vs_MS.NoTrt_F |
| BACH2 | 0.33509 | 0.78089 | 0.292 | 0.26 | 1 | HC_vs_MS.NoTrt_F |
| CDCA7L | 0.33521 | 0.69955 | 0.127 | 0.16 | 1 | HC_vs_MS.NoTrt_F |

| | | | | | | |
|---|---|---|---|---|---|---|
| USP28 | 0.33537 | 0.89152 | 0.41 | 0.44 | 1 | HC_vs_MS.NoTrt_F |
| NTRK1 | 0.33549 | 0.72318 | 0.249 | 0.3 | 1 | HC_vs_MS.NoTrt_F |
| BCL3 | 0.33564 | 0.42564 | 0.147 | 0.19 | 1 | HC_vs_MS.NoTrt_F |
| SRGAP2C | 0.33577 | -0.2585 | 0.549 | 0.58 | 1 | HC_vs_MS.NoTrt_F |
| DOCK6 | 0.33589 | 1.00247 | 0.552 | 0.6 | 1 | HC_vs_MS.NoTrt_F |
| CFDP1 | 0.33593 | -0.2925 | 0.676 | 0.7 | 1 | HC_vs_MS.NoTrt_F |
| CORO1C | 0.3362 | 1.46861 | 0.081 | 0.11 | 1 | HC_vs_MS.NoTrt_F |
| DZANK1 | 0.33654 | 0.27633 | 0.763 | 0.7 | 1 | HC_vs_MS.NoTrt_F |
| STK40 | 0.33665 | 0.67775 | 0.136 | 0.17 | 1 | HC_vs_MS.NoTrt_F |
| TPST2 | 0.33688 | 0.61333 | 0.717 | 0.65 | 1 | HC_vs_MS.NoTrt_F |
| POGK | 0.33695 | 0.28343 | 0.312 | 0.35 | 1 | HC_vs_MS.NoTrt_F |
| UNC13D | 0.3371 | 0.28952 | 0.642 | 0.71 | 1 | HC_vs_MS.NoTrt_F |
| DUSP19 | 0.33774 | 0.26477 | 0.737 | 0.71 | 1 | HC_vs_MS.NoTrt_F |
| EIF1AY | 0.33844 | -0.6474 | 0.671 | 0.67 | 1 | HC_vs_MS.NoTrt_F |
| KRI1 | 0.33867 | -0.2808 | 0.549 | 0.56 | 1 | HC_vs_MS.NoTrt_F |
| FGD2 | 0.33935 | 0.52186 | 0.783 | 0.78 | 1 | HC_vs_MS.NoTrt_F |
| CHD1 | 0.33972 | 0.25024 | 0.558 | 0.6 | 1 | HC_vs_MS.NoTrt_F |
| CLEC4E | 0.34018 | 0.64572 | 0.425 | 0.49 | 1 | HC_vs_MS.NoTrt_F |
| DIP2B | 0.34021 | 0.42557 | 0.436 | 0.51 | 1 | HC_vs_MS.NoTrt_F |
| MORN1 | 0.34038 | 0.772 | 0.598 | 0.55 | 1 | HC_vs_MS.NoTrt_F |
| CLEC18A | 0.34069 | 0.58233 | 0.405 | 0.43 | 1 | HC_vs_MS.NoTrt_F |
| NPIPA1 | 0.34073 | 0.29564 | 0.702 | 0.77 | 1 | HC_vs_MS.NoTrt_F |
| TAF1B | 0.34086 | -0.5136 | 0.61 | 0.63 | 1 | HC_vs_MS.NoTrt_F |
| GPATCH4 | 0.34146 | 0.3241 | 0.315 | 0.35 | 1 | HC_vs_MS.NoTrt_F |
| SH2D1B | 0.34191 | 1.78847 | 0.263 | 0.24 | 1 | HC_vs_MS.NoTrt_F |
| ZNF14 | 0.34191 | 1.06757 | 0.387 | 0.41 | 1 | HC_vs_MS.NoTrt_F |
| MAP1B | 0.34192 | 0.40162 | 0.101 | 0.08 | 1 | HC_vs_MS.NoTrt_F |
| TNFRSF4 | 0.34195 | 0.53841 | 0.24 | 0.29 | 1 | HC_vs_MS.NoTrt_F |
| BCL2L13 | 0.34209 | -0.6292 | 0.59 | 0.6 | 1 | HC_vs_MS.NoTrt_F |
| AZIN2 | 0.34232 | 1.06111 | 0.379 | 0.36 | 1 | HC_vs_MS.NoTrt_F |
| FOXRED1 | 0.34291 | 0.98183 | 0.549 | 0.52 | 1 | HC_vs_MS.NoTrt_F |
| PMS2 | 0.34314 | 0.81167 | 0.685 | 0.73 | 1 | HC_vs_MS.NoTrt_F |
| SIN3A | 0.34327 | -0.2871 | 0.832 | 0.84 | 1 | HC_vs_MS.NoTrt_F |
| FRMD3 | 0.34329 | 0.29652 | 0.124 | 0.1 | 1 | HC_vs_MS.NoTrt_F |
| BTD | 0.34353 | 1.09606 | 0.136 | 0.17 | 1 | HC_vs_MS.NoTrt_F |
| SPON2 | 0.34354 | 0.61108 | 0.488 | 0.53 | 1 | HC_vs_MS.NoTrt_F |
| CHMP5 | 0.34358 | 0.38356 | 0.168 | 0.2 | 1 | HC_vs_MS.NoTrt_F |
| PAM16 | 0.34366 | 0.74364 | 0.237 | 0.28 | 1 | HC_vs_MS.NoTrt_F |
| PCYOX1 | 0.34371 | -0.3371 | 0.841 | 0.83 | 1 | HC_vs_MS.NoTrt_F |
| PGAP2 | 0.34374 | 0.61454 | 0.107 | 0.14 | 1 | HC_vs_MS.NoTrt_F |

| | | | | | | |
|---|---|---|---|---|---|---|
| DCC | 0.34376 | 0.37545 | 0.116 | 0.15 | 1 | HC_vs_MS.NoTrt_F |
| REV3L | 0.34378 | 0.30166 | 0.74 | 0.79 | 1 | HC_vs_MS.NoTrt_F |
| NCSTN | 0.34395 | 0.27259 | 0.24 | 0.29 | 1 | HC_vs_MS.NoTrt_F |
| SNX30 | 0.34416 | 0.47597 | 0.543 | 0.5 | 1 | HC_vs_MS.NoTrt_F |
| SMC1A | 0.34426 | 0.62393 | 0.263 | 0.32 | 1 | HC_vs_MS.NoTrt_F |
| HPS4 | 0.3443 | 0.68515 | 0.818 | 0.85 | 1 | HC_vs_MS.NoTrt_F |
| PDE1B | 0.34447 | 0.6482 | 0.601 | 0.56 | 1 | HC_vs_MS.NoTrt_F |
| FMNL1 | 0.34461 | 0.76576 | 0.714 | 0.85 | 1 | HC_vs_MS.NoTrt_F |
| DHRS11 | 0.34479 | 0.54498 | 0.344 | 0.37 | 1 | HC_vs_MS.NoTrt_F |
| ANKRD49 | 0.34496 | 0.88422 | 0.22 | 0.27 | 1 | HC_vs_MS.NoTrt_F |
| KCTD18 | 0.34499 | 0.25352 | 0.832 | 0.86 | 1 | HC_vs_MS.NoTrt_F |
| ZNF491 | 0.34514 | 0.50651 | 0.488 | 0.53 | 1 | HC_vs_MS.NoTrt_F |
| CACNA1F | 0.34538 | 0.73469 | 0.318 | 0.37 | 1 | HC_vs_MS.NoTrt_F |
| ADGRL1 | 0.3455 | 1.29421 | 0.101 | 0.13 | 1 | HC_vs_MS.NoTrt_F |
| PDE6B | 0.34565 | 0.50587 | 0.711 | 0.66 | 1 | HC_vs_MS.NoTrt_F |
| DDX21 | 0.34574 | 0.42973 | 0.393 | 0.45 | 1 | HC_vs_MS.NoTrt_F |
| AK3 | 0.34581 | -0.4547 | 0.497 | 0.51 | 1 | HC_vs_MS.NoTrt_F |
| RBBP5 | 0.34592 | 0.27567 | 0.59 | 0.63 | 1 | HC_vs_MS.NoTrt_F |
| APBA2 | 0.34593 | 0.51167 | 0.549 | 0.61 | 1 | HC_vs_MS.NoTrt_F |
| POLR3E | 0.34599 | 0.32389 | 0.474 | 0.56 | 1 | HC_vs_MS.NoTrt_F |
| CHSY1 | 0.34734 | -0.7648 | 0.324 | 0.36 | 1 | HC_vs_MS.NoTrt_F |
| ZNF587B | 0.34747 | 0.52905 | 0.35 | 0.38 | 1 | HC_vs_MS.NoTrt_F |
| N4BP2L1 | 0.3475 | -0.5297 | 0.653 | 0.7 | 1 | HC_vs_MS.NoTrt_F |
| CCS | 0.34873 | 0.35398 | 0.329 | 0.29 | 1 | HC_vs_MS.NoTrt_F |
| ZNF419 | 0.34879 | 0.40638 | 0.876 | 0.85 | 1 | HC_vs_MS.NoTrt_F |
| MPI | 0.34896 | 0.65213 | 0.517 | 0.56 | 1 | HC_vs_MS.NoTrt_F |
| ADPRH | 0.34903 | 2.88411 | 0.081 | 0.11 | 1 | HC_vs_MS.NoTrt_F |
| LRP1 | 0.34925 | 1.24574 | 0.344 | 0.29 | 1 | HC_vs_MS.NoTrt_F |
| CSNK2A1 | 0.34964 | -0.6011 | 0.76 | 0.78 | 1 | HC_vs_MS.NoTrt_F |
| RHOT1 | 0.34979 | 0.69507 | 0.254 | 0.29 | 1 | HC_vs_MS.NoTrt_F |
| NECAB1 | 0.35037 | 0.72071 | 0.156 | 0.19 | 1 | HC_vs_MS.NoTrt_F |
| FBXO24 | 0.35042 | -0.3135 | 0.517 | 0.51 | 1 | HC_vs_MS.NoTrt_F |
| GTF2IRD2B | 0.35078 | -0.2553 | 0.879 | 0.9 | 1 | HC_vs_MS.NoTrt_F |
| FOXP3 | 0.3515 | 0.27633 | 0.503 | 0.56 | 1 | HC_vs_MS.NoTrt_F |
| PLAUR | 0.3516 | 1.92813 | 0.647 | 0.61 | 1 | HC_vs_MS.NoTrt_F |
| DIEXF | 0.35185 | 0.46709 | 0.555 | 0.49 | 1 | HC_vs_MS.NoTrt_F |
| ZBTB4 | 0.35217 | 0.61048 | 0.194 | 0.23 | 1 | HC_vs_MS.NoTrt_F |
| CTSW | 0.35226 | 1.15458 | 0.688 | 0.61 | 1 | HC_vs_MS.NoTrt_F |
| ATN1 | 0.3523 | 0.78761 | 0.26 | 0.23 | 1 | HC_vs_MS.NoTrt_F |
| SLC12A2 | 0.35274 | 0.64997 | 0.572 | 0.59 | 1 | HC_vs_MS.NoTrt_F |

| | | | | | | |
|---|---|---|---|---|---|---|
| TPP1 | 0.353 | 0.847 | 0.477 | 0.5 | 1 | HC_vs_MS.NoTrt_F |
| HACE1 | 0.35346 | -0.2952 | 0.688 | 0.68 | 1 | HC_vs_MS.NoTrt_F |
| ELP3 | 0.35365 | -0.3711 | 0.497 | 0.5 | 1 | HC_vs_MS.NoTrt_F |
| ZNF160 | 0.35375 | 0.72851 | 0.621 | 0.59 | 1 | HC_vs_MS.NoTrt_F |
| FAM222B | 0.3539 | 0.67165 | 0.22 | 0.26 | 1 | HC_vs_MS.NoTrt_F |
| KDM2A | 0.35411 | 0.35147 | 0.919 | 0.93 | 1 | HC_vs_MS.NoTrt_F |
| VPS33B | 0.35425 | 1.12924 | 0.416 | 0.46 | 1 | HC_vs_MS.NoTrt_F |
| PFKFB3 | 0.35428 | -0.2648 | 0.344 | 0.4 | 1 | HC_vs_MS.NoTrt_F |
| MSTO1 | 0.3544 | 1.27652 | 0.139 | 0.18 | 1 | HC_vs_MS.NoTrt_F |
| NCR3LG1 | 0.35571 | -0.8276 | 0.084 | 0.11 | 1 | HC_vs_MS.NoTrt_F |
| PPM1N | 0.35605 | 0.72117 | 0.162 | 0.19 | 1 | HC_vs_MS.NoTrt_F |
| RABIF | 0.35605 | 0.40618 | 0.422 | 0.45 | 1 | HC_vs_MS.NoTrt_F |
| TM2D1 | 0.35627 | 0.345 | 0.916 | 0.92 | 1 | HC_vs_MS.NoTrt_F |
| BAG6 | 0.35679 | 1.06272 | 0.691 | 0.59 | 1 | HC_vs_MS.NoTrt_F |
| FIBP | 0.35711 | 0.59636 | 0.561 | 0.53 | 1 | HC_vs_MS.NoTrt_F |
| MTR | 0.35795 | -0.6453 | 0.569 | 0.55 | 1 | HC_vs_MS.NoTrt_F |
| SMAD3 | 0.35814 | 0.6829 | 0.214 | 0.25 | 1 | HC_vs_MS.NoTrt_F |
| WDR92 | 0.35837 | 0.35787 | 0.743 | 0.73 | 1 | HC_vs_MS.NoTrt_F |
| MCC | 0.35842 | 0.43485 | 0.532 | 0.53 | 1 | HC_vs_MS.NoTrt_F |
| CC2D1B | 0.35881 | 0.29688 | 0.65 | 0.63 | 1 | HC_vs_MS.NoTrt_F |
| CTSH | 0.35917 | 0.57618 | 0.116 | 0.14 | 1 | HC_vs_MS.NoTrt_F |
| TXNL1 | 0.3592 | 0.46842 | 0.668 | 0.71 | 1 | HC_vs_MS.NoTrt_F |
| ZNF45 | 0.35925 | -0.4997 | 0.344 | 0.38 | 1 | HC_vs_MS.NoTrt_F |
| KCNMB4 | 0.35925 | 0.40735 | 0.37 | 0.41 | 1 | HC_vs_MS.NoTrt_F |
| CEP126 | 0.35947 | 0.41468 | 0.783 | 0.81 | 1 | HC_vs_MS.NoTrt_F |
| SH2B3 | 0.35976 | 0.36636 | 0.208 | 0.24 | 1 | HC_vs_MS.NoTrt_F |
| NR4A3 | 0.35997 | 0.70076 | 0.246 | 0.22 | 1 | HC_vs_MS.NoTrt_F |
| APLP1 | 0.36018 | 0.38797 | 0.676 | 0.63 | 1 | HC_vs_MS.NoTrt_F |
| ZNF75D | 0.36062 | 0.86063 | 0.176 | 0.15 | 1 | HC_vs_MS.NoTrt_F |
| CBWD1 | 0.36065 | 0.56627 | 0.708 | 0.74 | 1 | HC_vs_MS.NoTrt_F |
| AOAH | 0.3609 | 0.33887 | 0.428 | 0.47 | 1 | HC_vs_MS.NoTrt_F |
| CTSC | 0.36108 | 0.32872 | 0.928 | 0.91 | 1 | HC_vs_MS.NoTrt_F |
| RND2 | 0.36109 | 0.38346 | 0.425 | 0.44 | 1 | HC_vs_MS.NoTrt_F |
| RFK | 0.36123 | 0.60498 | 0.5 | 0.53 | 1 | HC_vs_MS.NoTrt_F |
| KIAA0319L | 0.36128 | -0.2916 | 0.725 | 0.75 | 1 | HC_vs_MS.NoTrt_F |
| BCL9L | 0.36175 | 0.97601 | 0.205 | 0.25 | 1 | HC_vs_MS.NoTrt_F |
| KIAA1161 | 0.36201 | 0.26801 | 0.876 | 0.89 | 1 | HC_vs_MS.NoTrt_F |
| PLA2G15 | 0.36277 | -0.6937 | 0.841 | 0.82 | 1 | HC_vs_MS.NoTrt_F |
| PEAK1 | 0.36285 | 0.34083 | 0.957 | 0.96 | 1 | HC_vs_MS.NoTrt_F |
| UQCRH | 0.36299 | 0.79857 | 0.581 | 0.59 | 1 | HC_vs_MS.NoTrt_F |

| | | | | | | |
|---|---|---|---|---|---|---|
| ANKRD10 | 0.36336 | 0.30254 | 0.717 | 0.8 | 1 | HC_vs_MS.NoTrt_F |
| LRRC49 | 0.36342 | 1.11468 | 0.792 | 0.87 | 1 | HC_vs_MS.NoTrt_F |
| MPP6 | 0.36388 | 0.91906 | 0.335 | 0.38 | 1 | HC_vs_MS.NoTrt_F |
| RELB | 0.36446 | 0.55797 | 0.159 | 0.19 | 1 | HC_vs_MS.NoTrt_F |
| CD99L2 | 0.36449 | 0.7122 | 0.168 | 0.2 | 1 | HC_vs_MS.NoTrt_F |
| MIB1 | 0.3648 | 0.50237 | 0.171 | 0.2 | 1 | HC_vs_MS.NoTrt_F |
| TMEM144 | 0.36648 | 0.29359 | 0.205 | 0.24 | 1 | HC_vs_MS.NoTrt_F |
| C14orf80 | 0.36757 | 1.21002 | 0.324 | 0.35 | 1 | HC_vs_MS.NoTrt_F |
| SYT2 | 0.36767 | 1.01382 | 0.081 | 0.11 | 1 | HC_vs_MS.NoTrt_F |
| RANBP2 | 0.36777 | 0.28638 | 0.592 | 0.6 | 1 | HC_vs_MS.NoTrt_F |
| FAR2 | 0.3682 | 0.26344 | 0.237 | 0.28 | 1 | HC_vs_MS.NoTrt_F |
| ATP6V0D1 | 0.3684 | -0.2711 | 0.717 | 0.74 | 1 | HC_vs_MS.NoTrt_F |
| PLEKHG2 | 0.36843 | 0.71549 | 0.916 | 0.88 | 1 | HC_vs_MS.NoTrt_F |
| SPPL3 | 0.36875 | 0.47961 | 0.61 | 0.65 | 1 | HC_vs_MS.NoTrt_F |
| DNHD1 | 0.36884 | 0.4473 | 0.39 | 0.35 | 1 | HC_vs_MS.NoTrt_F |
| METTL21B | 0.36899 | 0.78441 | 0.535 | 0.53 | 1 | HC_vs_MS.NoTrt_F |
| HAT1 | 0.36931 | 0.55197 | 0.884 | 0.91 | 1 | HC_vs_MS.NoTrt_F |
| ZNF839 | 0.36941 | -0.2934 | 0.751 | 0.8 | 1 | HC_vs_MS.NoTrt_F |
| LDAH | 0.36956 | -0.2798 | 0.153 | 0.18 | 1 | HC_vs_MS.NoTrt_F |
| SEZ6 | 0.36973 | 0.7851 | 0.13 | 0.16 | 1 | HC_vs_MS.NoTrt_F |
| PIN1 | 0.37014 | 0.76039 | 0.795 | 0.79 | 1 | HC_vs_MS.NoTrt_F |
| HIST1H1C | 0.37042 | 0.43236 | 0.182 | 0.22 | 1 | HC_vs_MS.NoTrt_F |
| PDE4A | 0.37089 | 0.27661 | 0.176 | 0.22 | 1 | HC_vs_MS.NoTrt_F |
| GARNL3 | 0.37142 | 0.73654 | 0.223 | 0.25 | 1 | HC_vs_MS.NoTrt_F |
| MIEF1 | 0.37152 | -0.6255 | 0.477 | 0.5 | 1 | HC_vs_MS.NoTrt_F |
| ATP13A2 | 0.37255 | 0.33695 | 0.5 | 0.53 | 1 | HC_vs_MS.NoTrt_F |
| SORD | 0.37258 | 0.42055 | 0.214 | 0.25 | 1 | HC_vs_MS.NoTrt_F |
| ZNF71 | 0.37268 | -0.3213 | 0.292 | 0.32 | 1 | HC_vs_MS.NoTrt_F |
| KDM3B | 0.37306 | -0.284 | 0.425 | 0.43 | 1 | HC_vs_MS.NoTrt_F |
| TONSL | 0.37337 | 0.52283 | 0.668 | 0.66 | 1 | HC_vs_MS.NoTrt_F |
| RIN2 | 0.37339 | 0.64663 | 0.361 | 0.3 | 1 | HC_vs_MS.NoTrt_F |
| ADAMTS17 | 0.37363 | 0.90188 | 0.217 | 0.25 | 1 | HC_vs_MS.NoTrt_F |
| CDK4 | 0.37394 | 0.82184 | 0.636 | 0.64 | 1 | HC_vs_MS.NoTrt_F |
| ZNF396 | 0.37422 | 0.27284 | 0.896 | 0.92 | 1 | HC_vs_MS.NoTrt_F |
| ZFAND4 | 0.37463 | -0.5546 | 0.523 | 0.47 | 1 | HC_vs_MS.NoTrt_F |
| LCAT | 0.37504 | 0.49564 | 0.263 | 0.3 | 1 | HC_vs_MS.NoTrt_F |
| IL17RC | 0.3751 | 0.5039 | 0.439 | 0.37 | 1 | HC_vs_MS.NoTrt_F |
| SMAD1 | 0.37518 | 0.81431 | 0.442 | 0.4 | 1 | HC_vs_MS.NoTrt_F |
| PPP1R16B | 0.37555 | 0.97301 | 0.355 | 0.43 | 1 | HC_vs_MS.NoTrt_F |
| C15orf57 | 0.37607 | -0.269 | 0.841 | 0.84 | 1 | HC_vs_MS.NoTrt_F |

| | | | | | | |
|---|---|---|---|---|---|---|
| EEF1E1 | 0.37609 | 0.45719 | 0.621 | 0.56 | 1 | HC_vs_MS.NoTrt_F |
| ALDH5A1 | 0.37615 | -0.428 | 0.529 | 0.57 | 1 | HC_vs_MS.NoTrt_F |
| DIS3 | 0.37624 | -0.3105 | 0.731 | 0.76 | 1 | HC_vs_MS.NoTrt_F |
| SMURF2 | 0.37668 | 0.78318 | 0.801 | 0.86 | 1 | HC_vs_MS.NoTrt_F |
| ERCC5 | 0.37677 | 0.40359 | 0.436 | 0.48 | 1 | HC_vs_MS.NoTrt_F |
| ARHGEF40 | 0.37686 | -0.5311 | 0.217 | 0.25 | 1 | HC_vs_MS.NoTrt_F |
| FAM174B | 0.37701 | 1.11019 | 0.173 | 0.21 | 1 | HC_vs_MS.NoTrt_F |
| ZNF418 | 0.37708 | 0.90439 | 0.194 | 0.23 | 1 | HC_vs_MS.NoTrt_F |
| QSOX1 | 0.37743 | 0.97541 | 0.474 | 0.43 | 1 | HC_vs_MS.NoTrt_F |
| USP36 | 0.37785 | 0.32341 | 0.535 | 0.62 | 1 | HC_vs_MS.NoTrt_F |
| RNF165 | 0.37789 | 0.50451 | 0.54 | 0.51 | 1 | HC_vs_MS.NoTrt_F |
| UBA1 | 0.37821 | 0.56352 | 0.844 | 0.85 | 1 | HC_vs_MS.NoTrt_F |
| SOCS4 | 0.37825 | 0.72207 | 0.379 | 0.41 | 1 | HC_vs_MS.NoTrt_F |
| PANK3 | 0.37847 | 0.79538 | 0.471 | 0.44 | 1 | HC_vs_MS.NoTrt_F |
| PPP2R1B | 0.37872 | 0.3485 | 0.514 | 0.47 | 1 | HC_vs_MS.NoTrt_F |
| OSBPL2 | 0.37906 | -0.304 | 0.523 | 0.55 | 1 | HC_vs_MS.NoTrt_F |
| BBIP1 | 0.37937 | 0.26296 | 0.665 | 0.65 | 1 | HC_vs_MS.NoTrt_F |
| RP11-80H18 | 0.37944 | 0.48401 | 0.318 | 0.37 | 1 | HC_vs_MS.NoTrt_F |
| DENND4C | 0.37958 | 0.51846 | 0.442 | 0.48 | 1 | HC_vs_MS.NoTrt_F |
| MPV17L2 | 0.37992 | 0.57165 | 0.243 | 0.28 | 1 | HC_vs_MS.NoTrt_F |
| ADGRD1 | 0.38048 | 0.61474 | 0.436 | 0.52 | 1 | HC_vs_MS.NoTrt_F |
| VEPH1 | 0.3818 | 0.35207 | 0.113 | 0.14 | 1 | HC_vs_MS.NoTrt_F |
| CYHR1 | 0.38185 | 0.27826 | 0.306 | 0.35 | 1 | HC_vs_MS.NoTrt_F |
| NAV2 | 0.38228 | 1.23072 | 0.408 | 0.42 | 1 | HC_vs_MS.NoTrt_F |
| GRINA | 0.38231 | 0.85156 | 0.121 | 0.16 | 1 | HC_vs_MS.NoTrt_F |
| UBE2L5P | 0.38255 | 0.42285 | 0.283 | 0.31 | 1 | HC_vs_MS.NoTrt_F |
| C14orf2 | 0.38288 | -0.2603 | 0.855 | 0.9 | 1 | HC_vs_MS.NoTrt_F |
| CERS5 | 0.38306 | -0.3481 | 0.988 | 0.98 | 1 | HC_vs_MS.NoTrt_F |
| RAB26 | 0.38334 | 0.50178 | 0.179 | 0.22 | 1 | HC_vs_MS.NoTrt_F |
| SS18L1 | 0.38349 | -0.3694 | 0.465 | 0.5 | 1 | HC_vs_MS.NoTrt_F |
| CH507-9B2.! | 0.3839 | 0.60634 | 0.142 | 0.17 | 1 | HC_vs_MS.NoTrt_F |
| ZNF175 | 0.38392 | 0.39351 | 0.211 | 0.24 | 1 | HC_vs_MS.NoTrt_F |
| AP4M1 | 0.38418 | -0.3833 | 0.671 | 0.69 | 1 | HC_vs_MS.NoTrt_F |
| NKD1 | 0.38421 | 0.50832 | 0.425 | 0.49 | 1 | HC_vs_MS.NoTrt_F |
| RNF141 | 0.38421 | 0.59979 | 0.445 | 0.46 | 1 | HC_vs_MS.NoTrt_F |
| B3GNT2 | 0.38477 | 0.37956 | 0.188 | 0.23 | 1 | HC_vs_MS.NoTrt_F |
| RGS3 | 0.38539 | 0.36542 | 0.558 | 0.49 | 1 | HC_vs_MS.NoTrt_F |
| PPME1 | 0.38623 | 0.8948 | 0.182 | 0.22 | 1 | HC_vs_MS.NoTrt_F |
| GBA | 0.3865 | -0.8355 | 0.506 | 0.5 | 1 | HC_vs_MS.NoTrt_F |
| AKAP12 | 0.38666 | 0.97899 | 0.09 | 0.12 | 1 | HC_vs_MS.NoTrt_F |

| | | | | | | |
|---|---|---|---|---|---|---|
| NSMF | 0.38687 | 2.12296 | 0.11 | 0.14 | 1 | HC_vs_MS.NoTrt_F |
| THEMIS | 0.38699 | 0.85145 | 0.272 | 0.31 | 1 | HC_vs_MS.NoTrt_F |
| HOMER2 | 0.38704 | -0.5737 | 0.616 | 0.64 | 1 | HC_vs_MS.NoTrt_F |
| MYBL1 | 0.38761 | 0.78161 | 0.078 | 0.11 | 1 | HC_vs_MS.NoTrt_F |
| SLC2A8 | 0.38773 | -1.1174 | 0.327 | 0.38 | 1 | HC_vs_MS.NoTrt_F |
| PRRT2 | 0.38788 | 1.06041 | 0.208 | 0.25 | 1 | HC_vs_MS.NoTrt_F |
| PDZD2 | 0.38802 | 0.63418 | 0.697 | 0.66 | 1 | HC_vs_MS.NoTrt_F |
| CAPRIN1 | 0.38834 | -0.4189 | 0.347 | 0.37 | 1 | HC_vs_MS.NoTrt_F |
| C1orf52 | 0.38853 | 0.27341 | 0.35 | 0.38 | 1 | HC_vs_MS.NoTrt_F |
| ATP1B3 | 0.38882 | 0.68956 | 0.211 | 0.26 | 1 | HC_vs_MS.NoTrt_F |
| CHKA | 0.38947 | 1.76078 | 0.092 | 0.12 | 1 | HC_vs_MS.NoTrt_F |
| C1QTNF6 | 0.38958 | 0.37526 | 0.627 | 0.67 | 1 | HC_vs_MS.NoTrt_F |
| FIS1 | 0.38994 | 0.64362 | 0.367 | 0.41 | 1 | HC_vs_MS.NoTrt_F |
| RECQL5 | 0.39008 | 0.49306 | 0.257 | 0.29 | 1 | HC_vs_MS.NoTrt_F |
| TRIM39 | 0.39082 | 0.4285 | 0.191 | 0.23 | 1 | HC_vs_MS.NoTrt_F |
| GATAD2A | 0.39082 | 0.58484 | 0.749 | 0.77 | 1 | HC_vs_MS.NoTrt_F |
| F2R | 0.39101 | 0.5837 | 0.578 | 0.56 | 1 | HC_vs_MS.NoTrt_F |
| HAUS6 | 0.39195 | 0.39528 | 0.39 | 0.43 | 1 | HC_vs_MS.NoTrt_F |
| NIFK | 0.39242 | -0.4197 | 0.572 | 0.59 | 1 | HC_vs_MS.NoTrt_F |
| MVD | 0.39267 | 0.6523 | 0.552 | 0.58 | 1 | HC_vs_MS.NoTrt_F |
| POU5F1 | 0.39285 | 0.28362 | 0.564 | 0.55 | 1 | HC_vs_MS.NoTrt_F |
| MAGI1 | 0.39366 | 0.25571 | 0.133 | 0.11 | 1 | HC_vs_MS.NoTrt_F |
| SPDYE3 | 0.39368 | 0.74258 | 0.506 | 0.56 | 1 | HC_vs_MS.NoTrt_F |
| BGN | 0.3941 | 1.59323 | 0.084 | 0.11 | 1 | HC_vs_MS.NoTrt_F |
| COL5A1 | 0.39414 | 1.10833 | 0.104 | 0.08 | 1 | HC_vs_MS.NoTrt_F |
| SHKBP1 | 0.39426 | 0.27478 | 0.775 | 0.83 | 1 | HC_vs_MS.NoTrt_F |
| PANK1 | 0.39446 | 0.63756 | 0.497 | 0.47 | 1 | HC_vs_MS.NoTrt_F |
| APOBEC3D | 0.39449 | 0.34316 | 0.723 | 0.75 | 1 | HC_vs_MS.NoTrt_F |
| PTK6 | 0.39451 | 0.46191 | 0.405 | 0.38 | 1 | HC_vs_MS.NoTrt_F |
| IGF2R | 0.3947 | 0.33273 | 0.486 | 0.51 | 1 | HC_vs_MS.NoTrt_F |
| RYR3 | 0.3948 | -0.3803 | 0.532 | 0.5 | 1 | HC_vs_MS.NoTrt_F |
| MS4A2 | 0.39487 | 0.87302 | 0.61 | 0.59 | 1 | HC_vs_MS.NoTrt_F |
| COX7C | 0.39499 | 0.26433 | 0.48 | 0.6 | 1 | HC_vs_MS.NoTrt_F |
| CD8B | 0.3952 | 2.67337 | 0.121 | 0.16 | 1 | HC_vs_MS.NoTrt_F |
| LSM11 | 0.39532 | 0.47027 | 0.607 | 0.64 | 1 | HC_vs_MS.NoTrt_F |
| FSBP | 0.39596 | 0.59333 | 0.384 | 0.43 | 1 | HC_vs_MS.NoTrt_F |
| APOL1 | 0.39606 | -0.6384 | 0.702 | 0.72 | 1 | HC_vs_MS.NoTrt_F |
| ZNF197 | 0.39756 | -0.691 | 0.538 | 0.59 | 1 | HC_vs_MS.NoTrt_F |
| SFR1 | 0.39773 | 0.31994 | 0.292 | 0.32 | 1 | HC_vs_MS.NoTrt_F |
| PPP1R3G | 0.39789 | 0.27009 | 0.269 | 0.32 | 1 | HC_vs_MS.NoTrt_F |

| | | | | | | |
|---|---|---|---|---|---|---|
| DNAJC1 | 0.39792 | 2.13834 | 0.199 | 0.18 | 1 | HC_vs_MS.NoTrt_F |
| DCAF6 | 0.39833 | 0.82921 | 0.269 | 0.31 | 1 | HC_vs_MS.NoTrt_F |
| IFT81 | 0.39871 | 0.27872 | 0.792 | 0.79 | 1 | HC_vs_MS.NoTrt_F |
| FAM111A | 0.39895 | -0.8234 | 0.552 | 0.54 | 1 | HC_vs_MS.NoTrt_F |
| RAI14 | 0.3993 | 0.36288 | 0.121 | 0.15 | 1 | HC_vs_MS.NoTrt_F |
| GLTSCR1 | 0.39967 | 0.40377 | 0.688 | 0.64 | 1 | HC_vs_MS.NoTrt_F |
| STXBP5 | 0.40032 | 0.45946 | 0.197 | 0.23 | 1 | HC_vs_MS.NoTrt_F |
| INPP4B | 0.4004 | 0.57896 | 0.601 | 0.6 | 1 | HC_vs_MS.NoTrt_F |
| SEC13 | 0.40046 | 0.50604 | 0.22 | 0.26 | 1 | HC_vs_MS.NoTrt_F |
| ZNF662 | 0.4005 | 0.32807 | 0.595 | 0.58 | 1 | HC_vs_MS.NoTrt_F |
| FABP2 | 0.4006 | 0.34593 | 0.442 | 0.44 | 1 | HC_vs_MS.NoTrt_F |
| POLR2L | 0.40071 | 0.33372 | 0.321 | 0.37 | 1 | HC_vs_MS.NoTrt_F |
| ETHE1 | 0.40139 | 0.66161 | 0.136 | 0.17 | 1 | HC_vs_MS.NoTrt_F |
| RP11-435I1( | 0.40164 | -0.2846 | 0.399 | 0.45 | 1 | HC_vs_MS.NoTrt_F |
| USP1 | 0.4018 | 0.78617 | 0.197 | 0.23 | 1 | HC_vs_MS.NoTrt_F |
| GALNT2 | 0.40184 | -0.3472 | 0.286 | 0.32 | 1 | HC_vs_MS.NoTrt_F |
| MBTPS2 | 0.40194 | -0.4701 | 0.353 | 0.38 | 1 | HC_vs_MS.NoTrt_F |
| DDX11 | 0.40207 | -0.2731 | 0.486 | 0.5 | 1 | HC_vs_MS.NoTrt_F |
| MX2 | 0.4027 | 0.37085 | 0.913 | 0.9 | 1 | HC_vs_MS.NoTrt_F |
| ORC2 | 0.40282 | 0.30824 | 0.694 | 0.73 | 1 | HC_vs_MS.NoTrt_F |
| TM7SF3 | 0.40343 | -0.5773 | 0.951 | 0.93 | 1 | HC_vs_MS.NoTrt_F |
| ATP6V0B | 0.40364 | 0.612 | 0.26 | 0.32 | 1 | HC_vs_MS.NoTrt_F |
| TIPIN | 0.4043 | 0.51172 | 0.639 | 0.62 | 1 | HC_vs_MS.NoTrt_F |
| FNIP2 | 0.40448 | 0.43734 | 0.35 | 0.4 | 1 | HC_vs_MS.NoTrt_F |
| PRKCE | 0.40454 | 0.68166 | 0.171 | 0.2 | 1 | HC_vs_MS.NoTrt_F |
| FCHO2 | 0.40497 | -0.2802 | 0.353 | 0.39 | 1 | HC_vs_MS.NoTrt_F |
| KIF19 | 0.40587 | -0.5643 | 0.425 | 0.45 | 1 | HC_vs_MS.NoTrt_F |
| ITGA11 | 0.40594 | 0.69783 | 0.708 | 0.7 | 1 | HC_vs_MS.NoTrt_F |
| PRKAG1 | 0.40594 | 0.5254 | 0.63 | 0.68 | 1 | HC_vs_MS.NoTrt_F |
| PTDSS2 | 0.40655 | 2.39343 | 0.107 | 0.14 | 1 | HC_vs_MS.NoTrt_F |
| OBSCN | 0.40668 | 0.28756 | 0.269 | 0.3 | 1 | HC_vs_MS.NoTrt_F |
| RP11-793H1 | 0.40675 | 0.36847 | 0.329 | 0.37 | 1 | HC_vs_MS.NoTrt_F |
| BBS2 | 0.40683 | 0.31059 | 0.737 | 0.77 | 1 | HC_vs_MS.NoTrt_F |
| INO80D | 0.40701 | 0.35756 | 0.483 | 0.54 | 1 | HC_vs_MS.NoTrt_F |
| ZNF165 | 0.40703 | 0.91898 | 0.09 | 0.12 | 1 | HC_vs_MS.NoTrt_F |
| TBC1D8 | 0.40709 | 0.29652 | 0.445 | 0.5 | 1 | HC_vs_MS.NoTrt_F |
| FBLN1 | 0.40743 | 0.58333 | 0.188 | 0.23 | 1 | HC_vs_MS.NoTrt_F |
| FOXP4 | 0.40745 | 0.4137 | 0.422 | 0.44 | 1 | HC_vs_MS.NoTrt_F |
| GRK5 | 0.40786 | 0.46373 | 0.22 | 0.26 | 1 | HC_vs_MS.NoTrt_F |
| PRPF40B | 0.40811 | 0.60806 | 0.387 | 0.36 | 1 | HC_vs_MS.NoTrt_F |

| | | | | | | |
|---|---|---|---|---|---|---|
| NOL11 | 0.4096 | -0.2812 | 0.578 | 0.56 | 1 | HC_vs_MS.NoTrt_F |
| RSAD2 | 0.40982 | -0.6274 | 0.425 | 0.39 | 1 | HC_vs_MS.NoTrt_F |
| PICK1 | 0.40999 | 1.40396 | 0.52 | 0.5 | 1 | HC_vs_MS.NoTrt_F |
| LRRC2 | 0.41 | -0.3587 | 0.795 | 0.79 | 1 | HC_vs_MS.NoTrt_F |
| HSD17B4 | 0.4102 | -0.3166 | 0.656 | 0.7 | 1 | HC_vs_MS.NoTrt_F |
| COMMD9 | 0.41047 | -0.4989 | 0.503 | 0.46 | 1 | HC_vs_MS.NoTrt_F |
| CENPF | 0.41052 | 0.35585 | 0.431 | 0.49 | 1 | HC_vs_MS.NoTrt_F |
| UVSSA | 0.41108 | 0.2623 | 0.824 | 0.81 | 1 | HC_vs_MS.NoTrt_F |
| MDGA1 | 0.41143 | 0.56339 | 0.087 | 0.11 | 1 | HC_vs_MS.NoTrt_F |
| MAML1 | 0.41173 | 0.71037 | 0.208 | 0.25 | 1 | HC_vs_MS.NoTrt_F |
| CYB5R2 | 0.41176 | 0.62375 | 0.575 | 0.53 | 1 | HC_vs_MS.NoTrt_F |
| PWP2 | 0.41206 | 0.34238 | 0.124 | 0.16 | 1 | HC_vs_MS.NoTrt_F |
| ELL2 | 0.41306 | 0.37408 | 0.09 | 0.12 | 1 | HC_vs_MS.NoTrt_F |
| EID2B | 0.41347 | 1.09578 | 0.523 | 0.57 | 1 | HC_vs_MS.NoTrt_F |
| SSC5D | 0.41359 | -0.3635 | 0.092 | 0.12 | 1 | HC_vs_MS.NoTrt_F |
| PIK3R3 | 0.41369 | 0.37276 | 0.569 | 0.65 | 1 | HC_vs_MS.NoTrt_F |
| HIPK1 | 0.41374 | 0.80451 | 0.263 | 0.31 | 1 | HC_vs_MS.NoTrt_F |
| CCDC124 | 0.41388 | 0.37641 | 0.488 | 0.54 | 1 | HC_vs_MS.NoTrt_F |
| ING5 | 0.41457 | -0.2627 | 0.948 | 0.94 | 1 | HC_vs_MS.NoTrt_F |
| DNAH11 | 0.41481 | 0.32739 | 0.329 | 0.4 | 1 | HC_vs_MS.NoTrt_F |
| PLOD2 | 0.41483 | 0.42662 | 0.439 | 0.45 | 1 | HC_vs_MS.NoTrt_F |
| AC240274.1 | 0.4153 | 0.65781 | 0.283 | 0.33 | 1 | HC_vs_MS.NoTrt_F |
| BHMT | 0.41552 | 0.3391 | 0.595 | 0.57 | 1 | HC_vs_MS.NoTrt_F |
| SOWAHC | 0.41572 | -0.3156 | 0.514 | 0.55 | 1 | HC_vs_MS.NoTrt_F |
| TMEM44 | 0.41591 | -0.3984 | 0.679 | 0.64 | 1 | HC_vs_MS.NoTrt_F |
| OAZ2 | 0.4162 | -0.5297 | 0.815 | 0.85 | 1 | HC_vs_MS.NoTrt_F |
| TMEM71 | 0.41641 | 1.32261 | 0.116 | 0.14 | 1 | HC_vs_MS.NoTrt_F |
| NTAN1 | 0.41668 | 0.27078 | 0.772 | 0.77 | 1 | HC_vs_MS.NoTrt_F |
| TMEM80 | 0.41713 | 0.50516 | 0.552 | 0.6 | 1 | HC_vs_MS.NoTrt_F |
| CARD16 | 0.41732 | 0.46533 | 0.361 | 0.41 | 1 | HC_vs_MS.NoTrt_F |
| MRPL22 | 0.41734 | 0.56071 | 0.28 | 0.32 | 1 | HC_vs_MS.NoTrt_F |
| GON4L | 0.41757 | 0.41235 | 0.887 | 0.88 | 1 | HC_vs_MS.NoTrt_F |
| CYP3A5 | 0.41767 | 0.26553 | 0.884 | 0.93 | 1 | HC_vs_MS.NoTrt_F |
| CHST3 | 0.41798 | 1.18331 | 0.188 | 0.22 | 1 | HC_vs_MS.NoTrt_F |
| ZNF335 | 0.41826 | 0.47853 | 0.376 | 0.41 | 1 | HC_vs_MS.NoTrt_F |
| RBMXL1 | 0.41861 | 0.38054 | 0.153 | 0.18 | 1 | HC_vs_MS.NoTrt_F |
| PLXNA3 | 0.41872 | 0.72061 | 0.861 | 0.85 | 1 | HC_vs_MS.NoTrt_F |
| CPNE2 | 0.41877 | -0.4716 | 0.477 | 0.47 | 1 | HC_vs_MS.NoTrt_F |
| PQLC1 | 0.41888 | 0.29741 | 0.569 | 0.54 | 1 | HC_vs_MS.NoTrt_F |
| LINC00116 | 0.41894 | 0.61731 | 0.176 | 0.21 | 1 | HC_vs_MS.NoTrt_F |

| | | | | | | |
|---|---|---|---|---|---|---|
| TOMM70 | 0.419 | 0.65197 | 0.188 | 0.22 | 1 | HC_vs_MS.NoTrt_F |
| CA5B | 0.41919 | 0.67097 | 0.879 | 0.9 | 1 | HC_vs_MS.NoTrt_F |
| RBM33 | 0.41922 | 0.9 | 0.277 | 0.33 | 1 | HC_vs_MS.NoTrt_F |
| BACE2 | 0.41956 | 0.73508 | 0.306 | 0.33 | 1 | HC_vs_MS.NoTrt_F |
| PCDHB16 | 0.42031 | 0.66723 | 0.358 | 0.39 | 1 | HC_vs_MS.NoTrt_F |
| LSM5 | 0.42105 | 0.64742 | 0.561 | 0.5 | 1 | HC_vs_MS.NoTrt_F |
| SLC35E2 | 0.42114 | 0.5373 | 0.477 | 0.52 | 1 | HC_vs_MS.NoTrt_F |
| FAHD1 | 0.42115 | -0.4921 | 0.358 | 0.39 | 1 | HC_vs_MS.NoTrt_F |
| RP3-461F17 | 0.42135 | 0.80294 | 0.283 | 0.32 | 1 | HC_vs_MS.NoTrt_F |
| STARD10 | 0.42142 | 0.33245 | 0.855 | 0.88 | 1 | HC_vs_MS.NoTrt_F |
| ABHD2 | 0.42142 | -0.252 | 0.659 | 0.69 | 1 | HC_vs_MS.NoTrt_F |
| CHST14 | 0.42146 | 0.31997 | 0.303 | 0.28 | 1 | HC_vs_MS.NoTrt_F |
| NTPCR | 0.42169 | 0.44148 | 0.468 | 0.5 | 1 | HC_vs_MS.NoTrt_F |
| C7orf50 | 0.42171 | 0.57471 | 0.355 | 0.39 | 1 | HC_vs_MS.NoTrt_F |
| CKAP2L | 0.42226 | 0.31618 | 0.512 | 0.57 | 1 | HC_vs_MS.NoTrt_F |
| HN1L | 0.42254 | 0.68678 | 0.416 | 0.39 | 1 | HC_vs_MS.NoTrt_F |
| TXNL4B | 0.42265 | -0.4468 | 0.41 | 0.43 | 1 | HC_vs_MS.NoTrt_F |
| SUPT6H | 0.42315 | 0.60359 | 0.679 | 0.7 | 1 | HC_vs_MS.NoTrt_F |
| GPR35 | 0.42409 | -1.0018 | 0.321 | 0.33 | 1 | HC_vs_MS.NoTrt_F |
| AHCYL2 | 0.42474 | 1.68089 | 0.194 | 0.23 | 1 | HC_vs_MS.NoTrt_F |
| CACNG8 | 0.42479 | 0.38462 | 0.835 | 0.74 | 1 | HC_vs_MS.NoTrt_F |
| TAOK2 | 0.42525 | -0.6502 | 0.179 | 0.2 | 1 | HC_vs_MS.NoTrt_F |
| ZADH2 | 0.42529 | 0.26771 | 0.861 | 0.83 | 1 | HC_vs_MS.NoTrt_F |
| MME | 0.4254 | 0.45374 | 0.48 | 0.52 | 1 | HC_vs_MS.NoTrt_F |
| PRR29 | 0.4254 | -0.3329 | 0.26 | 0.28 | 1 | HC_vs_MS.NoTrt_F |
| RBM5 | 0.42572 | 0.36926 | 0.861 | 0.87 | 1 | HC_vs_MS.NoTrt_F |
| ZNF670 | 0.42577 | 1.14007 | 0.205 | 0.24 | 1 | HC_vs_MS.NoTrt_F |
| TAL1 | 0.42614 | 0.5617 | 0.26 | 0.23 | 1 | HC_vs_MS.NoTrt_F |
| SNTN | 0.42692 | 0.31287 | 0.344 | 0.38 | 1 | HC_vs_MS.NoTrt_F |
| ZNF773 | 0.42708 | 0.39929 | 0.662 | 0.6 | 1 | HC_vs_MS.NoTrt_F |
| CSAD | 0.42734 | 0.2976 | 0.858 | 0.86 | 1 | HC_vs_MS.NoTrt_F |
| BBS5 | 0.42785 | 0.2826 | 0.361 | 0.33 | 1 | HC_vs_MS.NoTrt_F |
| MAPKBP1 | 0.42786 | 0.98174 | 0.552 | 0.51 | 1 | HC_vs_MS.NoTrt_F |
| FBXL22 | 0.4279 | 1.12964 | 0.13 | 0.16 | 1 | HC_vs_MS.NoTrt_F |
| ZZZ3 | 0.42865 | 0.38926 | 0.824 | 0.82 | 1 | HC_vs_MS.NoTrt_F |
| KLF9 | 0.42874 | 1.50547 | 0.09 | 0.12 | 1 | HC_vs_MS.NoTrt_F |
| THNSL1 | 0.42893 | -0.3086 | 0.153 | 0.19 | 1 | HC_vs_MS.NoTrt_F |
| SACS | 0.4293 | 0.38756 | 0.153 | 0.18 | 1 | HC_vs_MS.NoTrt_F |
| HCST | 0.42959 | 0.35157 | 0.434 | 0.54 | 1 | HC_vs_MS.NoTrt_F |
| TRIB3 | 0.42983 | -0.416 | 0.121 | 0.15 | 1 | HC_vs_MS.NoTrt_F |

| | | | | | | |
|---|---|---|---|---|---|---|
| LIN7B | 0.43098 | -0.5914 | 0.338 | 0.37 | 1 | HC_vs_MS.NoTrt_F |
| RLF | 0.43132 | 0.46712 | 0.275 | 0.32 | 1 | HC_vs_MS.NoTrt_F |
| ZNF441 | 0.43143 | 0.33062 | 0.197 | 0.23 | 1 | HC_vs_MS.NoTrt_F |
| PEX1 | 0.43144 | 0.48379 | 0.792 | 0.79 | 1 | HC_vs_MS.NoTrt_F |
| GPR65 | 0.4319 | 0.70479 | 0.335 | 0.37 | 1 | HC_vs_MS.NoTrt_F |
| ZNF675 | 0.43194 | 0.51779 | 0.954 | 0.96 | 1 | HC_vs_MS.NoTrt_F |
| SEC31B | 0.43213 | 0.27344 | 0.451 | 0.49 | 1 | HC_vs_MS.NoTrt_F |
| DENND5A | 0.43224 | 0.42559 | 0.275 | 0.32 | 1 | HC_vs_MS.NoTrt_F |
| NBPF12 | 0.43317 | 0.64003 | 0.347 | 0.38 | 1 | HC_vs_MS.NoTrt_F |
| PALM2 | 0.43352 | 0.26248 | 0.497 | 0.48 | 1 | HC_vs_MS.NoTrt_F |
| RALGAPA2 | 0.43396 | 1.20069 | 0.205 | 0.25 | 1 | HC_vs_MS.NoTrt_F |
| ANKRD42 | 0.43424 | 0.5436 | 0.442 | 0.48 | 1 | HC_vs_MS.NoTrt_F |
| AKAP8 | 0.4349 | 0.30895 | 0.422 | 0.45 | 1 | HC_vs_MS.NoTrt_F |
| NSUN4 | 0.43508 | 0.68611 | 0.806 | 0.81 | 1 | HC_vs_MS.NoTrt_F |
| ZC3HC1 | 0.43517 | 0.40957 | 0.517 | 0.56 | 1 | HC_vs_MS.NoTrt_F |
| FERMT2 | 0.43543 | 0.26392 | 0.569 | 0.53 | 1 | HC_vs_MS.NoTrt_F |
| TPRG1 | 0.4355 | 0.63568 | 0.263 | 0.3 | 1 | HC_vs_MS.NoTrt_F |
| SF1 | 0.43593 | 0.49243 | 0.604 | 0.72 | 1 | HC_vs_MS.NoTrt_F |
| RIN1 | 0.43671 | 1.11661 | 0.199 | 0.23 | 1 | HC_vs_MS.NoTrt_F |
| TECPR2 | 0.43676 | 1.81942 | 0.338 | 0.37 | 1 | HC_vs_MS.NoTrt_F |
| SPACA9 | 0.43736 | -0.3444 | 0.685 | 0.64 | 1 | HC_vs_MS.NoTrt_F |
| SWAP70 | 0.43762 | -1.4823 | 0.483 | 0.49 | 1 | HC_vs_MS.NoTrt_F |
| NCBP1 | 0.43804 | 0.54006 | 0.688 | 0.7 | 1 | HC_vs_MS.NoTrt_F |
| CARD6 | 0.43861 | -0.6439 | 0.552 | 0.5 | 1 | HC_vs_MS.NoTrt_F |
| TM4SF19 | 0.43937 | 0.28502 | 0.616 | 0.62 | 1 | HC_vs_MS.NoTrt_F |
| PPT2 | 0.44012 | 0.29619 | 0.604 | 0.56 | 1 | HC_vs_MS.NoTrt_F |
| HNRNPA1L2 | 0.44015 | -0.5219 | 0.341 | 0.38 | 1 | HC_vs_MS.NoTrt_F |
| SIRT5 | 0.44061 | 0.76966 | 0.879 | 0.86 | 1 | HC_vs_MS.NoTrt_F |
| ITGB2 | 0.44127 | 0.57819 | 0.685 | 0.74 | 1 | HC_vs_MS.NoTrt_F |
| TNS4 | 0.44131 | 0.51172 | 0.572 | 0.57 | 1 | HC_vs_MS.NoTrt_F |
| NBPF19 | 0.44141 | 1.15965 | 0.474 | 0.46 | 1 | HC_vs_MS.NoTrt_F |
| RHBDD1 | 0.44158 | 0.301 | 0.422 | 0.44 | 1 | HC_vs_MS.NoTrt_F |
| CITED2 | 0.44211 | 1.28519 | 0.249 | 0.3 | 1 | HC_vs_MS.NoTrt_F |
| MLF1 | 0.44266 | 0.91724 | 0.367 | 0.41 | 1 | HC_vs_MS.NoTrt_F |
| TRHDE | 0.44274 | 0.36543 | 0.283 | 0.26 | 1 | HC_vs_MS.NoTrt_F |
| RP11-437B1 | 0.44312 | 0.44247 | 0.639 | 0.68 | 1 | HC_vs_MS.NoTrt_F |
| LZTR1 | 0.44343 | -0.4505 | 0.332 | 0.34 | 1 | HC_vs_MS.NoTrt_F |
| PRKCI | 0.4442 | -0.4023 | 0.419 | 0.42 | 1 | HC_vs_MS.NoTrt_F |
| FEZ1 | 0.44422 | 0.25138 | 0.621 | 0.6 | 1 | HC_vs_MS.NoTrt_F |
| ABHD14A | 0.44428 | 0.28729 | 0.564 | 0.59 | 1 | HC_vs_MS.NoTrt_F |

| | | | | | | |
|---|---|---|---|---|---|---|
| ZMIZ1 | 0.44433 | 0.92277 | 0.162 | 0.19 | 1 | HC_vs_MS.NoTrt_F |
| MCOLN3 | 0.4444 | 0.51413 | 0.517 | 0.56 | 1 | HC_vs_MS.NoTrt_F |
| CHP1 | 0.44445 | 0.44916 | 0.806 | 0.82 | 1 | HC_vs_MS.NoTrt_F |
| NFKBIZ | 0.4452 | 0.51919 | 0.445 | 0.5 | 1 | HC_vs_MS.NoTrt_F |
| CTTNBP2NL | 0.44522 | 0.61822 | 0.503 | 0.54 | 1 | HC_vs_MS.NoTrt_F |
| MRPL12 | 0.44529 | 1.91308 | 0.095 | 0.12 | 1 | HC_vs_MS.NoTrt_F |
| NCAPD2 | 0.44569 | -0.3514 | 0.454 | 0.46 | 1 | HC_vs_MS.NoTrt_F |
| ZNF846 | 0.44582 | 0.37839 | 0.494 | 0.53 | 1 | HC_vs_MS.NoTrt_F |
| ELN | 0.44679 | 0.40462 | 0.72 | 0.74 | 1 | HC_vs_MS.NoTrt_F |
| NUMBL | 0.44736 | 0.37756 | 0.627 | 0.58 | 1 | HC_vs_MS.NoTrt_F |
| POGZ | 0.44778 | 0.41534 | 0.876 | 0.86 | 1 | HC_vs_MS.NoTrt_F |
| MKRN3 | 0.4478 | 0.89173 | 0.587 | 0.54 | 1 | HC_vs_MS.NoTrt_F |
| YARS | 0.44788 | -0.2593 | 0.89 | 0.9 | 1 | HC_vs_MS.NoTrt_F |
| GLT8D1 | 0.44789 | -0.5477 | 0.312 | 0.33 | 1 | HC_vs_MS.NoTrt_F |
| TIMM50 | 0.44856 | 0.32022 | 0.98 | 0.99 | 1 | HC_vs_MS.NoTrt_F |
| NBPF1 | 0.44857 | 0.56478 | 0.283 | 0.33 | 1 | HC_vs_MS.NoTrt_F |
| SSPO | 0.44934 | 0.28795 | 0.199 | 0.23 | 1 | HC_vs_MS.NoTrt_F |
| LYZ | 0.44935 | 1.4033 | 0.503 | 0.48 | 1 | HC_vs_MS.NoTrt_F |
| MCM3AP | 0.44948 | 0.36467 | 0.546 | 0.56 | 1 | HC_vs_MS.NoTrt_F |
| MKLN1 | 0.44981 | 0.63759 | 0.665 | 0.66 | 1 | HC_vs_MS.NoTrt_F |
| SEPN1 | 0.44987 | 0.35359 | 0.442 | 0.43 | 1 | HC_vs_MS.NoTrt_F |
| MTUS1 | 0.44988 | 0.51799 | 0.41 | 0.44 | 1 | HC_vs_MS.NoTrt_F |
| RBKS | 0.45003 | 0.66139 | 0.145 | 0.18 | 1 | HC_vs_MS.NoTrt_F |
| SRRM2 | 0.45009 | 0.34511 | 0.754 | 0.79 | 1 | HC_vs_MS.NoTrt_F |
| CH507-42P1 | 0.45051 | 0.99735 | 0.257 | 0.3 | 1 | HC_vs_MS.NoTrt_F |
| ICMT | 0.45057 | -0.6872 | 0.74 | 0.66 | 1 | HC_vs_MS.NoTrt_F |
| EIF2AK2 | 0.45066 | 0.28417 | 0.829 | 0.85 | 1 | HC_vs_MS.NoTrt_F |
| IL18 | 0.45133 | 0.84965 | 0.491 | 0.53 | 1 | HC_vs_MS.NoTrt_F |
| MMAB | 0.4516 | -0.351 | 0.717 | 0.71 | 1 | HC_vs_MS.NoTrt_F |
| RSPRY1 | 0.45166 | 0.37184 | 0.416 | 0.46 | 1 | HC_vs_MS.NoTrt_F |
| CXCR4 | 0.45171 | 0.31411 | 0.37 | 0.44 | 1 | HC_vs_MS.NoTrt_F |
| LPIN3 | 0.45209 | -0.26 | 0.552 | 0.49 | 1 | HC_vs_MS.NoTrt_F |
| ESCO1 | 0.45239 | 0.33446 | 0.439 | 0.51 | 1 | HC_vs_MS.NoTrt_F |
| C8orf44 | 0.45259 | -0.3604 | 0.526 | 0.58 | 1 | HC_vs_MS.NoTrt_F |
| HBS1L | 0.45293 | 0.2847 | 0.618 | 0.68 | 1 | HC_vs_MS.NoTrt_F |
| RAD1 | 0.45331 | 0.26568 | 0.717 | 0.74 | 1 | HC_vs_MS.NoTrt_F |
| MYH10 | 0.45411 | 0.67354 | 0.147 | 0.17 | 1 | HC_vs_MS.NoTrt_F |
| BHLHB9 | 0.45416 | -0.2797 | 0.436 | 0.48 | 1 | HC_vs_MS.NoTrt_F |
| EML4 | 0.45427 | 0.26177 | 0.777 | 0.82 | 1 | HC_vs_MS.NoTrt_F |
| ZBTB40 | 0.45532 | 0.29499 | 0.526 | 0.48 | 1 | HC_vs_MS.NoTrt_F |

| | | | | | | |
|---|---|---|---|---|---|---|
| FUS | 0.45565 | 0.2757 | 0.85 | 0.9 | 1 | HC_vs_MS.NoTrt_F |
| ST3GAL2 | 0.45664 | 0.37423 | 0.962 | 0.99 | 1 | HC_vs_MS.NoTrt_F |
| PDRG1 | 0.45721 | -0.2707 | 0.162 | 0.18 | 1 | HC_vs_MS.NoTrt_F |
| CCNK | 0.45809 | 0.92668 | 0.298 | 0.35 | 1 | HC_vs_MS.NoTrt_F |
| COL24A1 | 0.45854 | 1.28442 | 0.092 | 0.12 | 1 | HC_vs_MS.NoTrt_F |
| ZCCHC7 | 0.45857 | 0.68871 | 0.332 | 0.39 | 1 | HC_vs_MS.NoTrt_F |
| RND1 | 0.45863 | 0.28871 | 0.613 | 0.62 | 1 | HC_vs_MS.NoTrt_F |
| PLA2G12A | 0.45884 | 0.37309 | 0.679 | 0.72 | 1 | HC_vs_MS.NoTrt_F |
| GTF2E1 | 0.45937 | 0.85951 | 0.295 | 0.33 | 1 | HC_vs_MS.NoTrt_F |
| TNS2 | 0.4599 | 1.31201 | 0.39 | 0.41 | 1 | HC_vs_MS.NoTrt_F |
| HLA-DOA | 0.45996 | -0.468 | 0.743 | 0.67 | 1 | HC_vs_MS.NoTrt_F |
| DHRS1 | 0.46021 | -0.3789 | 0.671 | 0.66 | 1 | HC_vs_MS.NoTrt_F |
| NCF1 | 0.46064 | 2.31063 | 0.543 | 0.6 | 1 | HC_vs_MS.NoTrt_F |
| VPS50 | 0.46102 | -0.4194 | 0.665 | 0.64 | 1 | HC_vs_MS.NoTrt_F |
| QKI | 0.46111 | 0.34855 | 0.448 | 0.42 | 1 | HC_vs_MS.NoTrt_F |
| SEMA5A | 0.4615 | 0.60307 | 0.39 | 0.42 | 1 | HC_vs_MS.NoTrt_F |
| RGPD6 | 0.46229 | 0.39279 | 0.685 | 0.69 | 1 | HC_vs_MS.NoTrt_F |
| C19orf52 | 0.46239 | 0.26956 | 0.353 | 0.38 | 1 | HC_vs_MS.NoTrt_F |
| RGS2 | 0.4624 | 1.47429 | 0.202 | 0.24 | 1 | HC_vs_MS.NoTrt_F |
| HLA-DRA | 0.46305 | 0.29981 | 0.298 | 0.26 | 1 | HC_vs_MS.NoTrt_F |
| POMK | 0.46318 | -0.4797 | 0.618 | 0.66 | 1 | HC_vs_MS.NoTrt_F |
| PLOD3 | 0.46342 | 0.68387 | 0.78 | 0.78 | 1 | HC_vs_MS.NoTrt_F |
| USP16 | 0.46547 | 0.55233 | 0.529 | 0.58 | 1 | HC_vs_MS.NoTrt_F |
| LRRK1 | 0.46569 | 0.30602 | 0.85 | 0.86 | 1 | HC_vs_MS.NoTrt_F |
| IFT122 | 0.46624 | -0.3255 | 0.916 | 0.93 | 1 | HC_vs_MS.NoTrt_F |
| RDH11 | 0.4663 | -0.2631 | 0.558 | 0.55 | 1 | HC_vs_MS.NoTrt_F |
| HAGH | 0.46674 | -0.2933 | 0.783 | 0.79 | 1 | HC_vs_MS.NoTrt_F |
| ADRA1A | 0.46692 | 0.34032 | 0.618 | 0.62 | 1 | HC_vs_MS.NoTrt_F |
| CD93 | 0.46734 | 0.65183 | 0.361 | 0.33 | 1 | HC_vs_MS.NoTrt_F |
| CCNT1 | 0.4674 | 1.14591 | 0.249 | 0.29 | 1 | HC_vs_MS.NoTrt_F |
| FBXO6 | 0.46744 | 0.68988 | 0.697 | 0.65 | 1 | HC_vs_MS.NoTrt_F |
| TRIM22 | 0.46757 | 0.27948 | 0.934 | 0.91 | 1 | HC_vs_MS.NoTrt_F |
| MCM5 | 0.46763 | 0.53979 | 0.263 | 0.29 | 1 | HC_vs_MS.NoTrt_F |
| CDK16 | 0.46822 | 0.35162 | 0.254 | 0.29 | 1 | HC_vs_MS.NoTrt_F |
| ACTR10 | 0.46872 | -0.5364 | 0.783 | 0.82 | 1 | HC_vs_MS.NoTrt_F |
| TMEM204 | 0.46886 | 0.299 | 0.234 | 0.25 | 1 | HC_vs_MS.NoTrt_F |
| COPS7B | 0.46921 | 0.5565 | 0.769 | 0.85 | 1 | HC_vs_MS.NoTrt_F |
| SRGN | 0.46978 | 0.36217 | 0.642 | 0.7 | 1 | HC_vs_MS.NoTrt_F |
| APEX2 | 0.47006 | 0.61702 | 0.402 | 0.37 | 1 | HC_vs_MS.NoTrt_F |
| PXMP4 | 0.47019 | 0.34702 | 0.876 | 0.87 | 1 | HC_vs_MS.NoTrt_F |

| | | | | | | |
|---|---|---|---|---|---|---|
| LEPR | 0.47035 | 0.93985 | 0.373 | 0.33 | 1 | HC_vs_MS.NoTrt_F |
| RBMS3 | 0.47083 | 0.59825 | 0.697 | 0.78 | 1 | HC_vs_MS.NoTrt_F |
| TEX2 | 0.47097 | 0.98453 | 0.72 | 0.73 | 1 | HC_vs_MS.NoTrt_F |
| SH3YL1 | 0.47098 | -0.7612 | 0.757 | 0.71 | 1 | HC_vs_MS.NoTrt_F |
| RNF24 | 0.47107 | 0.44932 | 0.581 | 0.61 | 1 | HC_vs_MS.NoTrt_F |
| ZFP41 | 0.47175 | 1.11945 | 0.168 | 0.19 | 1 | HC_vs_MS.NoTrt_F |
| ANKMY1 | 0.47196 | 0.9392 | 0.462 | 0.44 | 1 | HC_vs_MS.NoTrt_F |
| TMEM231 | 0.47264 | 0.34441 | 0.766 | 0.82 | 1 | HC_vs_MS.NoTrt_F |
| LEO1 | 0.47305 | 0.34939 | 0.61 | 0.56 | 1 | HC_vs_MS.NoTrt_F |
| GNLY | 0.47355 | 2.88652 | 0.312 | 0.34 | 1 | HC_vs_MS.NoTrt_F |
| B9D2 | 0.47382 | 0.47701 | 0.393 | 0.43 | 1 | HC_vs_MS.NoTrt_F |
| RFX2 | 0.47391 | 0.28949 | 0.514 | 0.53 | 1 | HC_vs_MS.NoTrt_F |
| KLHL36 | 0.47399 | 0.35642 | 0.688 | 0.67 | 1 | HC_vs_MS.NoTrt_F |
| TYW3 | 0.47464 | 0.69823 | 0.503 | 0.49 | 1 | HC_vs_MS.NoTrt_F |
| YIPF6 | 0.4754 | 0.5233 | 0.908 | 0.89 | 1 | HC_vs_MS.NoTrt_F |
| ZNF354A | 0.47542 | 0.45837 | 0.251 | 0.28 | 1 | HC_vs_MS.NoTrt_F |
| GNB1 | 0.47559 | 0.75376 | 0.329 | 0.35 | 1 | HC_vs_MS.NoTrt_F |
| UTP15 | 0.47597 | -0.9136 | 0.367 | 0.39 | 1 | HC_vs_MS.NoTrt_F |
| ERI2 | 0.476 | 0.90099 | 0.197 | 0.23 | 1 | HC_vs_MS.NoTrt_F |
| ZC3H12D | 0.47674 | 0.31502 | 0.876 | 0.91 | 1 | HC_vs_MS.NoTrt_F |
| CLIP2 | 0.47708 | -0.2579 | 0.405 | 0.37 | 1 | HC_vs_MS.NoTrt_F |
| SLCO4C1 | 0.47802 | 0.26358 | 0.188 | 0.22 | 1 | HC_vs_MS.NoTrt_F |
| CENPN | 0.47812 | 0.35509 | 0.217 | 0.25 | 1 | HC_vs_MS.NoTrt_F |
| HIST2H2AA3 | 0.47814 | 0.98273 | 0.124 | 0.16 | 1 | HC_vs_MS.NoTrt_F |
| GABBR1 | 0.47839 | 0.90327 | 0.208 | 0.24 | 1 | HC_vs_MS.NoTrt_F |
| NINJ1 | 0.47852 | 0.46686 | 0.613 | 0.65 | 1 | HC_vs_MS.NoTrt_F |
| MAPKAPK3 | 0.47871 | 0.91418 | 0.24 | 0.28 | 1 | HC_vs_MS.NoTrt_F |
| ISG20L2 | 0.47873 | 0.35687 | 0.85 | 0.87 | 1 | HC_vs_MS.NoTrt_F |
| RAB6A | 0.4791 | 0.29201 | 0.436 | 0.49 | 1 | HC_vs_MS.NoTrt_F |
| MTF1 | 0.47916 | -0.5392 | 0.327 | 0.35 | 1 | HC_vs_MS.NoTrt_F |
| ANKRD12 | 0.4792 | 0.26828 | 0.867 | 0.87 | 1 | HC_vs_MS.NoTrt_F |
| GRB10 | 0.47957 | 1.00248 | 0.341 | 0.32 | 1 | HC_vs_MS.NoTrt_F |
| GDE1 | 0.47966 | 0.52942 | 0.315 | 0.34 | 1 | HC_vs_MS.NoTrt_F |
| ST7L | 0.47989 | 0.31237 | 0.529 | 0.48 | 1 | HC_vs_MS.NoTrt_F |
| SERPINH1 | 0.47994 | -0.2612 | 0.39 | 0.41 | 1 | HC_vs_MS.NoTrt_F |
| HLA-DRB5 | 0.48004 | -1.9688 | 0.246 | 0.21 | 1 | HC_vs_MS.NoTrt_F |
| GIGYF2 | 0.48046 | -0.4166 | 0.925 | 0.92 | 1 | HC_vs_MS.NoTrt_F |
| PLPP3 | 0.48057 | 0.74996 | 0.315 | 0.33 | 1 | HC_vs_MS.NoTrt_F |
| CYTH4 | 0.48129 | -0.2766 | 0.844 | 0.82 | 1 | HC_vs_MS.NoTrt_F |
| ANKRD17 | 0.48136 | 0.45338 | 0.295 | 0.34 | 1 | HC_vs_MS.NoTrt_F |

| | | | | | | |
|---|---|---|---|---|---|---|
| SLC4A5 | 0.48148 | 0.38411 | 0.827 | 0.85 | 1 | HC_vs_MS.NoTrt_F |
| ZNF551 | 0.48177 | 1.18321 | 0.543 | 0.56 | 1 | HC_vs_MS.NoTrt_F |
| SC5D | 0.48264 | 0.44323 | 0.679 | 0.63 | 1 | HC_vs_MS.NoTrt_F |
| FAF2 | 0.48264 | 0.6293 | 0.191 | 0.22 | 1 | HC_vs_MS.NoTrt_F |
| FAM98B | 0.48303 | 0.28702 | 0.523 | 0.5 | 1 | HC_vs_MS.NoTrt_F |
| ZNF260 | 0.48304 | -0.2741 | 0.627 | 0.65 | 1 | HC_vs_MS.NoTrt_F |
| RILPL1 | 0.48331 | 0.3287 | 0.954 | 0.98 | 1 | HC_vs_MS.NoTrt_F |
| RER1 | 0.48371 | 0.72322 | 0.257 | 0.3 | 1 | HC_vs_MS.NoTrt_F |
| RNFT1 | 0.48399 | 0.38131 | 0.442 | 0.45 | 1 | HC_vs_MS.NoTrt_F |
| BRAT1 | 0.48419 | 0.47527 | 0.338 | 0.37 | 1 | HC_vs_MS.NoTrt_F |
| ZRANB3 | 0.48421 | 0.3103 | 0.737 | 0.76 | 1 | HC_vs_MS.NoTrt_F |
| UST | 0.48461 | 0.36905 | 0.37 | 0.33 | 1 | HC_vs_MS.NoTrt_F |
| SAA2 | 0.48505 | 0.25794 | 0.457 | 0.41 | 1 | HC_vs_MS.NoTrt_F |
| HERC2 | 0.4852 | 0.25821 | 0.91 | 0.91 | 1 | HC_vs_MS.NoTrt_F |
| TRPV1 | 0.48635 | 0.49158 | 0.477 | 0.54 | 1 | HC_vs_MS.NoTrt_F |
| FBXL4 | 0.48672 | 0.41672 | 0.546 | 0.58 | 1 | HC_vs_MS.NoTrt_F |
| RABAC1 | 0.48676 | 0.54408 | 0.283 | 0.33 | 1 | HC_vs_MS.NoTrt_F |
| RPP30 | 0.48756 | 0.47052 | 0.679 | 0.66 | 1 | HC_vs_MS.NoTrt_F |
| TOR4A | 0.4878 | 0.409 | 0.254 | 0.28 | 1 | HC_vs_MS.NoTrt_F |
| SUDS3 | 0.48826 | -0.3129 | 0.841 | 0.81 | 1 | HC_vs_MS.NoTrt_F |
| USP33 | 0.48828 | 0.2686 | 0.965 | 0.95 | 1 | HC_vs_MS.NoTrt_F |
| ABCB9 | 0.48834 | 0.42904 | 0.85 | 0.9 | 1 | HC_vs_MS.NoTrt_F |
| ARRB1 | 0.48852 | 0.67118 | 0.763 | 0.81 | 1 | HC_vs_MS.NoTrt_F |
| NPIPB3 | 0.48887 | 0.62245 | 0.777 | 0.83 | 1 | HC_vs_MS.NoTrt_F |
| MICU1 | 0.48887 | 0.36403 | 0.74 | 0.73 | 1 | HC_vs_MS.NoTrt_F |
| EXOG | 0.489 | 0.25839 | 0.561 | 0.58 | 1 | HC_vs_MS.NoTrt_F |
| MTAP | 0.48907 | 0.53855 | 0.789 | 0.87 | 1 | HC_vs_MS.NoTrt_F |
| TUBG1 | 0.48926 | 1.16756 | 0.668 | 0.61 | 1 | HC_vs_MS.NoTrt_F |
| TSR2 | 0.48943 | -0.3452 | 0.46 | 0.46 | 1 | HC_vs_MS.NoTrt_F |
| CR1 | 0.48979 | 0.93842 | 0.618 | 0.57 | 1 | HC_vs_MS.NoTrt_F |
| SLC25A14 | 0.49045 | 0.86658 | 0.324 | 0.35 | 1 | HC_vs_MS.NoTrt_F |
| ANKEF1 | 0.49069 | -1.4581 | 0.269 | 0.29 | 1 | HC_vs_MS.NoTrt_F |
| OTUD4 | 0.49074 | 0.52941 | 0.699 | 0.71 | 1 | HC_vs_MS.NoTrt_F |
| CHCHD5 | 0.49148 | 0.65142 | 0.532 | 0.53 | 1 | HC_vs_MS.NoTrt_F |
| MAGED1 | 0.49165 | -0.3658 | 0.775 | 0.79 | 1 | HC_vs_MS.NoTrt_F |
| TRIM33 | 0.49178 | 0.40211 | 0.396 | 0.35 | 1 | HC_vs_MS.NoTrt_F |
| GOLPH3L | 0.49192 | 0.65606 | 0.263 | 0.28 | 1 | HC_vs_MS.NoTrt_F |
| ZNF772 | 0.49195 | 0.37249 | 0.471 | 0.52 | 1 | HC_vs_MS.NoTrt_F |
| CRLF2 | 0.49219 | 1.03394 | 0.37 | 0.41 | 1 | HC_vs_MS.NoTrt_F |
| CLU | 0.49284 | 0.85762 | 0.561 | 0.6 | 1 | HC_vs_MS.NoTrt_F |

| | | | | | | |
|---|---|---|---|---|---|---|
| FAM161A | 0.49313 | 0.33594 | 0.725 | 0.73 | 1 | HC_vs_MS.NoTrt_F |
| SPIB | 0.4933 | 0.28035 | 0.818 | 0.86 | 1 | HC_vs_MS.NoTrt_F |
| ACSS1 | 0.49339 | 0.61474 | 0.627 | 0.67 | 1 | HC_vs_MS.NoTrt_F |
| PTK2B | 0.49354 | 0.56783 | 0.491 | 0.49 | 1 | HC_vs_MS.NoTrt_F |
| CCDC85C | 0.49385 | 0.59366 | 0.477 | 0.47 | 1 | HC_vs_MS.NoTrt_F |
| DNAJB5 | 0.49434 | 0.39351 | 0.231 | 0.21 | 1 | HC_vs_MS.NoTrt_F |
| KLHDC10 | 0.4947 | -0.7439 | 0.436 | 0.4 | 1 | HC_vs_MS.NoTrt_F |
| PDE3A | 0.49479 | -0.4139 | 0.246 | 0.28 | 1 | HC_vs_MS.NoTrt_F |
| MECR | 0.49498 | -0.318 | 0.405 | 0.44 | 1 | HC_vs_MS.NoTrt_F |
| C22orf39 | 0.49499 | 0.37368 | 0.763 | 0.75 | 1 | HC_vs_MS.NoTrt_F |
| HIKESHI | 0.4951 | 0.28597 | 0.532 | 0.57 | 1 | HC_vs_MS.NoTrt_F |
| RHOF | 0.49524 | 0.27796 | 0.757 | 0.78 | 1 | HC_vs_MS.NoTrt_F |
| IFT74 | 0.49587 | 0.32032 | 0.465 | 0.49 | 1 | HC_vs_MS.NoTrt_F |
| MYO7A | 0.49621 | 0.91252 | 0.361 | 0.41 | 1 | HC_vs_MS.NoTrt_F |
| NAA40 | 0.49651 | -0.3647 | 0.561 | 0.55 | 1 | HC_vs_MS.NoTrt_F |
| HACD3 | 0.4968 | -0.3551 | 0.821 | 0.81 | 1 | HC_vs_MS.NoTrt_F |
| MEGF8 | 0.49689 | 1.48569 | 0.532 | 0.54 | 1 | HC_vs_MS.NoTrt_F |
| PRKX | 0.49722 | 0.30631 | 0.636 | 0.68 | 1 | HC_vs_MS.NoTrt_F |
| GSTM3 | 0.49726 | 0.74971 | 0.598 | 0.6 | 1 | HC_vs_MS.NoTrt_F |
| MAPRE2 | 0.49736 | 0.33988 | 0.338 | 0.39 | 1 | HC_vs_MS.NoTrt_F |
| TRAT1 | 0.49756 | 0.74881 | 0.249 | 0.24 | 1 | HC_vs_MS.NoTrt_F |
| SLIT1 | 0.49835 | 0.65344 | 0.277 | 0.32 | 1 | HC_vs_MS.NoTrt_F |
| CDT1 | 0.49853 | 0.3474 | 0.5 | 0.49 | 1 | HC_vs_MS.NoTrt_F |
| KCNRG | 0.499 | 0.35161 | 0.425 | 0.46 | 1 | HC_vs_MS.NoTrt_F |
| GATS | 0.49938 | -0.3737 | 0.393 | 0.41 | 1 | HC_vs_MS.NoTrt_F |
| MT-CO3 | 0.49944 | 0.29206 | 0.679 | 0.67 | 1 | HC_vs_MS.NoTrt_F |
| REPS1 | 0.4995 | 0.25754 | 0.662 | 0.72 | 1 | HC_vs_MS.NoTrt_F |
| TAB3 | 0.49952 | 0.68674 | 0.749 | 0.81 | 1 | HC_vs_MS.NoTrt_F |
| EMC2 | 0.4996 | 0.3939 | 0.517 | 0.54 | 1 | HC_vs_MS.NoTrt_F |
| DZIP1 | 0.5003 | 0.39705 | 0.393 | 0.35 | 1 | HC_vs_MS.NoTrt_F |
| TSPOAP1 | 0.50046 | 0.97045 | 0.321 | 0.29 | 1 | HC_vs_MS.NoTrt_F |
| IFITM1 | 0.50047 | 0.32851 | 0.546 | 0.66 | 1 | HC_vs_MS.NoTrt_F |
| KIF13A | 0.50052 | 1.32119 | 0.364 | 0.35 | 1 | HC_vs_MS.NoTrt_F |
| CLNK | 0.50084 | 0.29603 | 0.743 | 0.69 | 1 | HC_vs_MS.NoTrt_F |
| CERS4 | 0.50152 | 0.73517 | 0.659 | 0.64 | 1 | HC_vs_MS.NoTrt_F |
| GNG7 | 0.50196 | -0.5571 | 0.413 | 0.46 | 1 | HC_vs_MS.NoTrt_F |
| ADGRG1 | 0.50269 | 0.53656 | 0.749 | 0.72 | 1 | HC_vs_MS.NoTrt_F |
| GATC | 0.50279 | 0.35916 | 0.604 | 0.6 | 1 | HC_vs_MS.NoTrt_F |
| FLNB | 0.50285 | 0.77068 | 0.471 | 0.43 | 1 | HC_vs_MS.NoTrt_F |
| P3H2 | 0.50289 | 0.59593 | 0.517 | 0.56 | 1 | HC_vs_MS.NoTrt_F |

| | | | | | | |
|---|---|---|---|---|---|---|
| VCL | 0.50294 | 0.97418 | 0.188 | 0.22 | 1 | HC_vs_MS.NoTrt_F |
| THEM4 | 0.50336 | 0.46968 | 0.838 | 0.79 | 1 | HC_vs_MS.NoTrt_F |
| FOS | 0.50343 | 1.03496 | 0.451 | 0.5 | 1 | HC_vs_MS.NoTrt_F |
| TERF2 | 0.50372 | 0.28312 | 0.91 | 0.87 | 1 | HC_vs_MS.NoTrt_F |
| EYS | 0.50388 | 0.34538 | 0.801 | 0.83 | 1 | HC_vs_MS.NoTrt_F |
| B4GALT1 | 0.50391 | 0.54103 | 0.702 | 0.74 | 1 | HC_vs_MS.NoTrt_F |
| ZSCAN21 | 0.50406 | -0.7195 | 0.28 | 0.31 | 1 | HC_vs_MS.NoTrt_F |
| PHACTR3 | 0.50432 | 0.46741 | 0.176 | 0.16 | 1 | HC_vs_MS.NoTrt_F |
| ASAP1 | 0.50435 | 1.2913 | 0.257 | 0.24 | 1 | HC_vs_MS.NoTrt_F |
| ZNF639 | 0.50454 | -0.4117 | 0.942 | 0.93 | 1 | HC_vs_MS.NoTrt_F |
| MRPS6 | 0.50564 | 0.25915 | 0.283 | 0.32 | 1 | HC_vs_MS.NoTrt_F |
| DCTN6 | 0.50623 | -0.2693 | 0.613 | 0.62 | 1 | HC_vs_MS.NoTrt_F |
| MED24 | 0.50646 | -0.4096 | 0.636 | 0.64 | 1 | HC_vs_MS.NoTrt_F |
| DBF4 | 0.50658 | 0.53764 | 0.488 | 0.45 | 1 | HC_vs_MS.NoTrt_F |
| CCDC152 | 0.50713 | -1.1744 | 0.251 | 0.27 | 1 | HC_vs_MS.NoTrt_F |
| ZNF131 | 0.50762 | 0.31453 | 0.861 | 0.87 | 1 | HC_vs_MS.NoTrt_F |
| RNF215 | 0.50793 | -0.3855 | 0.523 | 0.54 | 1 | HC_vs_MS.NoTrt_F |
| ERVW-1 | 0.50808 | 0.29284 | 0.445 | 0.41 | 1 | HC_vs_MS.NoTrt_F |
| FRRS1 | 0.50823 | 0.31587 | 0.552 | 0.55 | 1 | HC_vs_MS.NoTrt_F |
| USF3 | 0.50857 | 0.85432 | 0.737 | 0.77 | 1 | HC_vs_MS.NoTrt_F |
| ZC3H12B | 0.50897 | -0.4226 | 0.63 | 0.62 | 1 | HC_vs_MS.NoTrt_F |
| KCNB1 | 0.50908 | 0.40059 | 0.925 | 0.91 | 1 | HC_vs_MS.NoTrt_F |
| NISCH | 0.50965 | 0.77154 | 0.454 | 0.49 | 1 | HC_vs_MS.NoTrt_F |
| PUS7 | 0.51 | -0.7111 | 0.373 | 0.4 | 1 | HC_vs_MS.NoTrt_F |
| PADI2 | 0.51039 | 0.49626 | 0.315 | 0.29 | 1 | HC_vs_MS.NoTrt_F |
| NCAPH2 | 0.51065 | 0.76538 | 0.757 | 0.81 | 1 | HC_vs_MS.NoTrt_F |
| APELA | 0.51079 | 0.27257 | 0.431 | 0.45 | 1 | HC_vs_MS.NoTrt_F |
| ITPR3 | 0.51081 | 0.75933 | 0.205 | 0.24 | 1 | HC_vs_MS.NoTrt_F |
| EVA1C | 0.51098 | -0.335 | 0.202 | 0.23 | 1 | HC_vs_MS.NoTrt_F |
| CHTF8 | 0.51104 | 0.44933 | 0.301 | 0.33 | 1 | HC_vs_MS.NoTrt_F |
| SMG9 | 0.51117 | 0.41634 | 0.688 | 0.66 | 1 | HC_vs_MS.NoTrt_F |
| HDAC9 | 0.51163 | 0.35958 | 0.465 | 0.49 | 1 | HC_vs_MS.NoTrt_F |
| ZNF397 | 0.51195 | -0.265 | 0.962 | 0.96 | 1 | HC_vs_MS.NoTrt_F |
| RNASE4 | 0.51215 | 0.32596 | 0.084 | 0.11 | 1 | HC_vs_MS.NoTrt_F |
| CATIP | 0.51264 | 0.27124 | 0.882 | 0.91 | 1 | HC_vs_MS.NoTrt_F |
| CLASP2 | 0.51281 | 0.6492 | 0.751 | 0.82 | 1 | HC_vs_MS.NoTrt_F |
| PLEKHA1 | 0.51291 | -0.2689 | 0.96 | 0.95 | 1 | HC_vs_MS.NoTrt_F |
| MBTD1 | 0.51296 | 0.60971 | 0.509 | 0.53 | 1 | HC_vs_MS.NoTrt_F |
| ZBTB8OS | 0.51303 | 0.34671 | 0.948 | 0.98 | 1 | HC_vs_MS.NoTrt_F |
| ANKRD36B | 0.51343 | 0.29663 | 0.743 | 0.77 | 1 | HC_vs_MS.NoTrt_F |

| | | | | | | |
|---|---|---|---|---|---|---|
| KIAA1324L | 0.5135 | 1.17697 | 0.104 | 0.13 | 1 | HC_vs_MS.NoTrt_F |
| PACRGL | 0.51369 | 0.71574 | 0.312 | 0.29 | 1 | HC_vs_MS.NoTrt_F |
| SRPK3 | 0.51421 | -0.2871 | 0.315 | 0.34 | 1 | HC_vs_MS.NoTrt_F |
| PBX1 | 0.51427 | 0.27593 | 0.934 | 0.95 | 1 | HC_vs_MS.NoTrt_F |
| ANKRD16 | 0.51467 | 0.39211 | 0.471 | 0.49 | 1 | HC_vs_MS.NoTrt_F |
| HIVEP2 | 0.51495 | 0.38463 | 0.382 | 0.4 | 1 | HC_vs_MS.NoTrt_F |
| PSMF1 | 0.51513 | 0.31813 | 0.315 | 0.36 | 1 | HC_vs_MS.NoTrt_F |
| EPN2 | 0.51565 | 0.45145 | 0.858 | 0.9 | 1 | HC_vs_MS.NoTrt_F |
| STARD8 | 0.51572 | 0.37966 | 0.76 | 0.76 | 1 | HC_vs_MS.NoTrt_F |
| F5 | 0.51635 | 0.59383 | 0.257 | 0.3 | 1 | HC_vs_MS.NoTrt_F |
| HEATR4 | 0.51646 | 0.4911 | 0.413 | 0.44 | 1 | HC_vs_MS.NoTrt_F |
| MYO1C | 0.51663 | 0.92731 | 0.269 | 0.25 | 1 | HC_vs_MS.NoTrt_F |
| DCP1A | 0.51708 | 0.43393 | 0.451 | 0.4 | 1 | HC_vs_MS.NoTrt_F |
| CD40 | 0.51726 | -0.3196 | 0.185 | 0.21 | 1 | HC_vs_MS.NoTrt_F |
| GATM | 0.51755 | 0.31732 | 0.237 | 0.26 | 1 | HC_vs_MS.NoTrt_F |
| MBOAT2 | 0.51812 | 0.564 | 0.379 | 0.33 | 1 | HC_vs_MS.NoTrt_F |
| MAP3K13 | 0.51829 | 0.50697 | 0.85 | 0.83 | 1 | HC_vs_MS.NoTrt_F |
| HFE | 0.51898 | -0.4065 | 0.665 | 0.67 | 1 | HC_vs_MS.NoTrt_F |
| GAN | 0.51959 | 0.44503 | 0.876 | 0.85 | 1 | HC_vs_MS.NoTrt_F |
| SPAG7 | 0.51965 | 0.37039 | 0.893 | 0.87 | 1 | HC_vs_MS.NoTrt_F |
| MAPKAPK2 | 0.52019 | 1.13818 | 0.104 | 0.13 | 1 | HC_vs_MS.NoTrt_F |
| CC2D2B | 0.5202 | 0.40174 | 0.185 | 0.21 | 1 | HC_vs_MS.NoTrt_F |
| BCLAF1 | 0.52069 | 0.40968 | 0.512 | 0.59 | 1 | HC_vs_MS.NoTrt_F |
| MED29 | 0.52071 | 0.25779 | 0.627 | 0.67 | 1 | HC_vs_MS.NoTrt_F |
| MTMR4 | 0.52119 | 0.69965 | 0.298 | 0.33 | 1 | HC_vs_MS.NoTrt_F |
| ZHX1 | 0.52188 | -0.5545 | 0.598 | 0.61 | 1 | HC_vs_MS.NoTrt_F |
| UHRF1BP1 | 0.5219 | 0.41583 | 0.616 | 0.64 | 1 | HC_vs_MS.NoTrt_F |
| F3 | 0.52289 | 0.30945 | 0.535 | 0.5 | 1 | HC_vs_MS.NoTrt_F |
| TNKS | 0.52294 | 0.75808 | 0.844 | 0.84 | 1 | HC_vs_MS.NoTrt_F |
| ABCD4 | 0.52329 | 0.36103 | 0.855 | 0.83 | 1 | HC_vs_MS.NoTrt_F |
| DHX37 | 0.52406 | 0.40524 | 0.561 | 0.57 | 1 | HC_vs_MS.NoTrt_F |
| TSN | 0.52434 | -0.6345 | 0.428 | 0.44 | 1 | HC_vs_MS.NoTrt_F |
| AQR | 0.52521 | -0.363 | 0.671 | 0.67 | 1 | HC_vs_MS.NoTrt_F |
| PIGA | 0.52538 | -0.5708 | 0.488 | 0.51 | 1 | HC_vs_MS.NoTrt_F |
| 11-Sep | 0.52539 | 0.72621 | 0.538 | 0.51 | 1 | HC_vs_MS.NoTrt_F |
| PDIA5 | 0.52541 | 0.45356 | 0.434 | 0.46 | 1 | HC_vs_MS.NoTrt_F |
| SEPP1 | 0.52558 | 1.30706 | 0.188 | 0.22 | 1 | HC_vs_MS.NoTrt_F |
| GATB | 0.52628 | 1.0095 | 0.292 | 0.27 | 1 | HC_vs_MS.NoTrt_F |
| ABCG8 | 0.52643 | 0.69402 | 0.633 | 0.67 | 1 | HC_vs_MS.NoTrt_F |
| SLC41A2 | 0.52676 | 0.28298 | 0.28 | 0.29 | 1 | HC_vs_MS.NoTrt_F |

| | | | | | | |
|---|---|---|---|---|---|---|
| MED28 | 0.52697 | -0.4094 | 0.91 | 0.92 | 1 | HC_vs_MS.NoTrt_F |
| MTRF1L | 0.52703 | 0.69871 | 0.133 | 0.16 | 1 | HC_vs_MS.NoTrt_F |
| ADAM33 | 0.52705 | 0.56805 | 0.107 | 0.13 | 1 | HC_vs_MS.NoTrt_F |
| ZNF284 | 0.52713 | 0.26021 | 0.613 | 0.61 | 1 | HC_vs_MS.NoTrt_F |
| LPAR4 | 0.5274 | -0.481 | 0.185 | 0.2 | 1 | HC_vs_MS.NoTrt_F |
| TKT | 0.52744 | 0.41178 | 0.705 | 0.66 | 1 | HC_vs_MS.NoTrt_F |
| SLC36A4 | 0.52768 | -0.405 | 0.104 | 0.13 | 1 | HC_vs_MS.NoTrt_F |
| SLC31A2 | 0.52795 | 0.48508 | 0.13 | 0.16 | 1 | HC_vs_MS.NoTrt_F |
| COX19 | 0.52882 | 0.632 | 0.685 | 0.66 | 1 | HC_vs_MS.NoTrt_F |
| PUS3 | 0.52888 | 1.44949 | 0.358 | 0.39 | 1 | HC_vs_MS.NoTrt_F |
| HHIP | 0.5292 | 0.43207 | 0.405 | 0.38 | 1 | HC_vs_MS.NoTrt_F |
| PLD6 | 0.52945 | -0.6199 | 0.477 | 0.48 | 1 | HC_vs_MS.NoTrt_F |
| FUT10 | 0.52969 | 0.38474 | 0.428 | 0.44 | 1 | HC_vs_MS.NoTrt_F |
| HSPA2 | 0.53012 | -0.4696 | 0.624 | 0.59 | 1 | HC_vs_MS.NoTrt_F |
| VAPA | 0.53021 | 0.31312 | 0.789 | 0.77 | 1 | HC_vs_MS.NoTrt_F |
| TLE3 | 0.53074 | 0.7707 | 0.298 | 0.34 | 1 | HC_vs_MS.NoTrt_F |
| SPDYE1 | 0.53081 | 0.6151 | 0.483 | 0.47 | 1 | HC_vs_MS.NoTrt_F |
| TIMP2 | 0.53124 | 1.21164 | 0.202 | 0.23 | 1 | HC_vs_MS.NoTrt_F |
| FUT2 | 0.53184 | 0.3392 | 0.754 | 0.77 | 1 | HC_vs_MS.NoTrt_F |
| C12orf57 | 0.53191 | 0.56107 | 0.451 | 0.41 | 1 | HC_vs_MS.NoTrt_F |
| ATXN1 | 0.53205 | 0.35985 | 0.39 | 0.42 | 1 | HC_vs_MS.NoTrt_F |
| PRAM1 | 0.53271 | 0.34211 | 0.564 | 0.6 | 1 | HC_vs_MS.NoTrt_F |
| LINC00672 | 0.53301 | -0.3904 | 0.572 | 0.59 | 1 | HC_vs_MS.NoTrt_F |
| SFMBT2 | 0.53362 | 2.82159 | 0.379 | 0.38 | 1 | HC_vs_MS.NoTrt_F |
| STK3 | 0.5339 | 0.61489 | 0.786 | 0.82 | 1 | HC_vs_MS.NoTrt_F |
| DYNC1H1 | 0.53408 | 0.53741 | 0.957 | 0.93 | 1 | HC_vs_MS.NoTrt_F |
| ZFPM2 | 0.5342 | 0.36547 | 0.442 | 0.5 | 1 | HC_vs_MS.NoTrt_F |
| PLEKHG6 | 0.53454 | 0.39672 | 0.303 | 0.33 | 1 | HC_vs_MS.NoTrt_F |
| EIF3B | 0.53462 | 0.56493 | 0.422 | 0.4 | 1 | HC_vs_MS.NoTrt_F |
| ZNF75A | 0.53483 | -0.5199 | 0.783 | 0.83 | 1 | HC_vs_MS.NoTrt_F |
| CGREF1 | 0.53497 | 0.49131 | 0.101 | 0.12 | 1 | HC_vs_MS.NoTrt_F |
| MGLL | 0.53499 | 0.34251 | 0.431 | 0.4 | 1 | HC_vs_MS.NoTrt_F |
| PINK1 | 0.53505 | 0.75474 | 0.301 | 0.33 | 1 | HC_vs_MS.NoTrt_F |
| LIMS2 | 0.5355 | 1.2844 | 0.142 | 0.13 | 1 | HC_vs_MS.NoTrt_F |
| MAB21L3 | 0.53557 | 0.30793 | 0.399 | 0.4 | 1 | HC_vs_MS.NoTrt_F |
| MAP3K11 | 0.53558 | 0.3808 | 0.717 | 0.7 | 1 | HC_vs_MS.NoTrt_F |
| MYO1D | 0.53592 | 1.90377 | 0.202 | 0.23 | 1 | HC_vs_MS.NoTrt_F |
| CX3CL1 | 0.53596 | 0.31547 | 0.604 | 0.59 | 1 | HC_vs_MS.NoTrt_F |
| WARS | 0.53634 | 0.74629 | 0.315 | 0.34 | 1 | HC_vs_MS.NoTrt_F |
| JAM2 | 0.53646 | 0.32735 | 0.607 | 0.58 | 1 | HC_vs_MS.NoTrt_F |

| | | | | | | |
|---|---|---|---|---|---|---|
| OLFML1 | 0.53669 | 0.4118 | 0.263 | 0.28 | 1 | HC_vs_MS.NoTrt_F |
| FBXO45 | 0.537 | 0.71375 | 0.509 | 0.46 | 1 | HC_vs_MS.NoTrt_F |
| INPP5B | 0.53723 | -0.2538 | 0.96 | 0.95 | 1 | HC_vs_MS.NoTrt_F |
| FBXL12 | 0.53734 | -0.4608 | 0.723 | 0.71 | 1 | HC_vs_MS.NoTrt_F |
| MRM2 | 0.53736 | -0.3956 | 0.393 | 0.39 | 1 | HC_vs_MS.NoTrt_F |
| MAD2L1 | 0.53744 | -0.3648 | 0.477 | 0.49 | 1 | HC_vs_MS.NoTrt_F |
| DTWD1 | 0.53772 | 0.32475 | 0.659 | 0.64 | 1 | HC_vs_MS.NoTrt_F |
| UBN2 | 0.53779 | -0.7725 | 0.708 | 0.64 | 1 | HC_vs_MS.NoTrt_F |
| ZNF177 | 0.5382 | 1.01897 | 0.107 | 0.13 | 1 | HC_vs_MS.NoTrt_F |
| LAPTM4B | 0.53833 | 0.64982 | 0.225 | 0.24 | 1 | HC_vs_MS.NoTrt_F |
| 8-Mar | 0.53838 | 1.03484 | 0.413 | 0.46 | 1 | HC_vs_MS.NoTrt_F |
| PROSER3 | 0.53844 | 0.6415 | 0.916 | 0.9 | 1 | HC_vs_MS.NoTrt_F |
| TMED9 | 0.5391 | 0.27575 | 0.431 | 0.47 | 1 | HC_vs_MS.NoTrt_F |
| GMEB1 | 0.53926 | -0.4422 | 0.821 | 0.83 | 1 | HC_vs_MS.NoTrt_F |
| DCTN2 | 0.53931 | -0.2537 | 0.662 | 0.65 | 1 | HC_vs_MS.NoTrt_F |
| DNM3 | 0.53943 | 0.95334 | 0.121 | 0.14 | 1 | HC_vs_MS.NoTrt_F |
| KLHL11 | 0.53951 | 0.60988 | 0.584 | 0.59 | 1 | HC_vs_MS.NoTrt_F |
| DNM1 | 0.53981 | 0.4316 | 0.65 | 0.65 | 1 | HC_vs_MS.NoTrt_F |
| C9orf3 | 0.53987 | 0.86283 | 0.78 | 0.79 | 1 | HC_vs_MS.NoTrt_F |
| AMPD3 | 0.54021 | -0.8584 | 0.436 | 0.47 | 1 | HC_vs_MS.NoTrt_F |
| GRAMD4 | 0.54053 | 1.02795 | 0.104 | 0.13 | 1 | HC_vs_MS.NoTrt_F |
| ZBTB20 | 0.54059 | 0.32019 | 0.801 | 0.8 | 1 | HC_vs_MS.NoTrt_F |
| ASCC1 | 0.54099 | 0.29344 | 0.714 | 0.73 | 1 | HC_vs_MS.NoTrt_F |
| DSCR3 | 0.54125 | 0.27284 | 0.734 | 0.76 | 1 | HC_vs_MS.NoTrt_F |
| KIAA1958 | 0.54159 | -0.317 | 0.728 | 0.75 | 1 | HC_vs_MS.NoTrt_F |
| ZNF764 | 0.54301 | 1.46572 | 0.399 | 0.4 | 1 | HC_vs_MS.NoTrt_F |
| UBE2J2 | 0.54325 | 0.65532 | 0.361 | 0.39 | 1 | HC_vs_MS.NoTrt_F |
| MBD6 | 0.544 | 0.77238 | 0.329 | 0.32 | 1 | HC_vs_MS.NoTrt_F |
| SYVN1 | 0.54416 | 0.30648 | 0.298 | 0.33 | 1 | HC_vs_MS.NoTrt_F |
| CARD19 | 0.54424 | 0.62757 | 0.37 | 0.41 | 1 | HC_vs_MS.NoTrt_F |
| SLC4A8 | 0.5443 | 0.38472 | 0.948 | 0.92 | 1 | HC_vs_MS.NoTrt_F |
| MAP3K2 | 0.54443 | 0.39815 | 0.301 | 0.34 | 1 | HC_vs_MS.NoTrt_F |
| CRIM1 | 0.54469 | 1.06465 | 0.289 | 0.27 | 1 | HC_vs_MS.NoTrt_F |
| SERTAD1 | 0.54493 | 0.36686 | 0.442 | 0.45 | 1 | HC_vs_MS.NoTrt_F |
| C16orf45 | 0.54626 | 0.7138 | 0.22 | 0.25 | 1 | HC_vs_MS.NoTrt_F |
| ECI2 | 0.54675 | 0.38187 | 0.743 | 0.7 | 1 | HC_vs_MS.NoTrt_F |
| CXorf57 | 0.54697 | 0.79527 | 0.179 | 0.16 | 1 | HC_vs_MS.NoTrt_F |
| HIST1H1E | 0.54711 | 1.41641 | 0.136 | 0.16 | 1 | HC_vs_MS.NoTrt_F |
| COX7B | 0.54721 | 0.27425 | 0.552 | 0.59 | 1 | HC_vs_MS.NoTrt_F |
| NDUFS1 | 0.54752 | -0.6178 | 0.962 | 0.94 | 1 | HC_vs_MS.NoTrt_F |

| | | | | | | |
|---|---|---|---|---|---|---|
| PTGER4 | 0.5478 | 0.76039 | 0.277 | 0.32 | 1 | HC_vs_MS.NoTrt_F |
| SYMPK | 0.54784 | 0.53058 | 0.922 | 0.92 | 1 | HC_vs_MS.NoTrt_F |
| TIMM21 | 0.54791 | -0.3955 | 0.688 | 0.64 | 1 | HC_vs_MS.NoTrt_F |
| AP3M2 | 0.54826 | 0.65243 | 0.208 | 0.24 | 1 | HC_vs_MS.NoTrt_F |
| PAPOLG | 0.54859 | 0.7023 | 0.827 | 0.8 | 1 | HC_vs_MS.NoTrt_F |
| DLC1 | 0.54888 | 0.47699 | 0.202 | 0.23 | 1 | HC_vs_MS.NoTrt_F |
| PNP | 0.54896 | 0.80722 | 0.416 | 0.4 | 1 | HC_vs_MS.NoTrt_F |
| BCO2 | 0.549 | 0.29665 | 0.627 | 0.58 | 1 | HC_vs_MS.NoTrt_F |
| PHPT1 | 0.54933 | 0.3948 | 0.228 | 0.25 | 1 | HC_vs_MS.NoTrt_F |
| GAL3ST4 | 0.54956 | -0.3299 | 0.451 | 0.45 | 1 | HC_vs_MS.NoTrt_F |
| CRYGS | 0.54957 | 1.09897 | 0.286 | 0.32 | 1 | HC_vs_MS.NoTrt_F |
| FBXO16 | 0.54961 | 0.28134 | 0.327 | 0.3 | 1 | HC_vs_MS.NoTrt_F |
| MRI1 | 0.54963 | -0.3247 | 0.749 | 0.77 | 1 | HC_vs_MS.NoTrt_F |
| KIN | 0.55038 | -0.5741 | 0.772 | 0.82 | 1 | HC_vs_MS.NoTrt_F |
| TCAF2 | 0.55072 | -0.3228 | 0.78 | 0.81 | 1 | HC_vs_MS.NoTrt_F |
| ZNF589 | 0.55142 | 0.35474 | 0.538 | 0.55 | 1 | HC_vs_MS.NoTrt_F |
| PHLDA3 | 0.5517 | 0.76326 | 0.425 | 0.4 | 1 | HC_vs_MS.NoTrt_F |
| ASPH | 0.55183 | 0.72868 | 0.569 | 0.56 | 1 | HC_vs_MS.NoTrt_F |
| CERKL | 0.55252 | 0.25912 | 0.451 | 0.45 | 1 | HC_vs_MS.NoTrt_F |
| PAQR8 | 0.55326 | 0.29083 | 0.607 | 0.61 | 1 | HC_vs_MS.NoTrt_F |
| MYDGF | 0.55358 | 0.5258 | 0.231 | 0.26 | 1 | HC_vs_MS.NoTrt_F |
| PPIL6 | 0.55367 | 0.43922 | 0.54 | 0.54 | 1 | HC_vs_MS.NoTrt_F |
| FOCAD | 0.55453 | 0.35812 | 0.561 | 0.52 | 1 | HC_vs_MS.NoTrt_F |
| SVEP1 | 0.55486 | 2.16173 | 0.243 | 0.22 | 1 | HC_vs_MS.NoTrt_F |
| MS4A6A | 0.55508 | 1.55636 | 0.202 | 0.23 | 1 | HC_vs_MS.NoTrt_F |
| IL15 | 0.55564 | -0.8237 | 0.387 | 0.41 | 1 | HC_vs_MS.NoTrt_F |
| CAPN10 | 0.55587 | 0.57547 | 0.566 | 0.58 | 1 | HC_vs_MS.NoTrt_F |
| PLEKHG4B | 0.55589 | 0.51204 | 0.509 | 0.53 | 1 | HC_vs_MS.NoTrt_F |
| CARMIL1 | 0.55606 | 1.24672 | 0.361 | 0.32 | 1 | HC_vs_MS.NoTrt_F |
| KLRG1 | 0.55606 | 1.3109 | 0.465 | 0.48 | 1 | HC_vs_MS.NoTrt_F |
| ZNF615 | 0.5563 | 0.30203 | 0.549 | 0.62 | 1 | HC_vs_MS.NoTrt_F |
| HDAC4 | 0.55636 | 0.46581 | 0.601 | 0.65 | 1 | HC_vs_MS.NoTrt_F |
| METTL25 | 0.55642 | 1.25334 | 0.257 | 0.24 | 1 | HC_vs_MS.NoTrt_F |
| ATXN10 | 0.55738 | 0.48977 | 0.587 | 0.62 | 1 | HC_vs_MS.NoTrt_F |
| POMT1 | 0.5575 | 0.32155 | 0.584 | 0.58 | 1 | HC_vs_MS.NoTrt_F |
| PARP12 | 0.55775 | 1.2341 | 0.269 | 0.29 | 1 | HC_vs_MS.NoTrt_F |
| ZNF746 | 0.55825 | 0.47169 | 0.324 | 0.35 | 1 | HC_vs_MS.NoTrt_F |
| CNPY4 | 0.55836 | -0.288 | 0.725 | 0.72 | 1 | HC_vs_MS.NoTrt_F |
| PPARGC1B | 0.5585 | 0.47466 | 0.763 | 0.76 | 1 | HC_vs_MS.NoTrt_F |
| TAF1 | 0.55883 | -0.6712 | 0.815 | 0.85 | 1 | HC_vs_MS.NoTrt_F |

| | | | | | | |
|---|---|---|---|---|---|---|
| ZFYVE27 | 0.55915 | 0.92868 | 0.225 | 0.26 | 1 | HC_vs_MS.NoTrt_F |
| SLC5A12 | 0.56012 | 0.4281 | 0.124 | 0.15 | 1 | HC_vs_MS.NoTrt_F |
| FAM168A | 0.56013 | 0.44296 | 0.185 | 0.17 | 1 | HC_vs_MS.NoTrt_F |
| KLK2 | 0.5604 | 0.28453 | 0.379 | 0.41 | 1 | HC_vs_MS.NoTrt_F |
| MMP11 | 0.56106 | 0.48603 | 0.431 | 0.39 | 1 | HC_vs_MS.NoTrt_F |
| PCBP2 | 0.56112 | 0.27605 | 0.653 | 0.68 | 1 | HC_vs_MS.NoTrt_F |
| TTC34 | 0.56134 | 0.35445 | 0.217 | 0.24 | 1 | HC_vs_MS.NoTrt_F |
| BCL7B | 0.56134 | 1.24803 | 0.52 | 0.48 | 1 | HC_vs_MS.NoTrt_F |
| TUBB3 | 0.56146 | 0.53237 | 0.292 | 0.32 | 1 | HC_vs_MS.NoTrt_F |
| GSK3A | 0.56148 | -0.6261 | 0.353 | 0.37 | 1 | HC_vs_MS.NoTrt_F |
| SELK | 0.562 | 0.33205 | 0.54 | 0.52 | 1 | HC_vs_MS.NoTrt_F |
| RREB1 | 0.56213 | 0.50627 | 0.812 | 0.8 | 1 | HC_vs_MS.NoTrt_F |
| FBXO46 | 0.56236 | 0.38884 | 0.315 | 0.34 | 1 | HC_vs_MS.NoTrt_F |
| NFATC2IP | 0.56254 | 0.46938 | 0.749 | 0.73 | 1 | HC_vs_MS.NoTrt_F |
| POMZP3 | 0.56274 | 0.50604 | 0.538 | 0.54 | 1 | HC_vs_MS.NoTrt_F |
| NAA30 | 0.56278 | -0.7133 | 0.54 | 0.54 | 1 | HC_vs_MS.NoTrt_F |
| COL1A1 | 0.56286 | 0.92228 | 0.558 | 0.55 | 1 | HC_vs_MS.NoTrt_F |
| SCML4 | 0.56289 | 0.7982 | 0.662 | 0.75 | 1 | HC_vs_MS.NoTrt_F |
| SNRNP200 | 0.56333 | 0.35017 | 0.815 | 0.83 | 1 | HC_vs_MS.NoTrt_F |
| TPSB2 | 0.56354 | 3.56604 | 0.286 | 0.27 | 1 | HC_vs_MS.NoTrt_F |
| NUDCD1 | 0.56441 | 1.79166 | 0.162 | 0.14 | 1 | HC_vs_MS.NoTrt_F |
| IL4R | 0.56521 | 0.36855 | 0.506 | 0.49 | 1 | HC_vs_MS.NoTrt_F |
| ALDH1L2 | 0.56636 | 0.8532 | 0.335 | 0.31 | 1 | HC_vs_MS.NoTrt_F |
| CEP57L1 | 0.56688 | -0.4122 | 0.601 | 0.61 | 1 | HC_vs_MS.NoTrt_F |
| ZNF573 | 0.56735 | -0.2707 | 0.413 | 0.43 | 1 | HC_vs_MS.NoTrt_F |
| GPR174 | 0.56736 | 1.02708 | 0.263 | 0.26 | 1 | HC_vs_MS.NoTrt_F |
| RGS11 | 0.56744 | 0.89106 | 0.217 | 0.2 | 1 | HC_vs_MS.NoTrt_F |
| ZFYVE21 | 0.56758 | -0.29 | 0.737 | 0.73 | 1 | HC_vs_MS.NoTrt_F |
| ALS2 | 0.5678 | -0.9981 | 0.497 | 0.52 | 1 | HC_vs_MS.NoTrt_F |
| MAPK1IP1L | 0.56785 | 0.33688 | 0.896 | 0.9 | 1 | HC_vs_MS.NoTrt_F |
| NLE1 | 0.56915 | 0.33293 | 0.321 | 0.35 | 1 | HC_vs_MS.NoTrt_F |
| STX7 | 0.56944 | 0.87951 | 0.795 | 0.76 | 1 | HC_vs_MS.NoTrt_F |
| NSUN6 | 0.56972 | -0.2697 | 0.728 | 0.76 | 1 | HC_vs_MS.NoTrt_F |
| PKN2 | 0.57012 | 1.60886 | 0.379 | 0.43 | 1 | HC_vs_MS.NoTrt_F |
| SRD5A1 | 0.57024 | -0.5118 | 0.749 | 0.75 | 1 | HC_vs_MS.NoTrt_F |
| RNGTT | 0.57048 | 0.79512 | 0.439 | 0.46 | 1 | HC_vs_MS.NoTrt_F |
| GJB4 | 0.57055 | 0.32094 | 0.74 | 0.73 | 1 | HC_vs_MS.NoTrt_F |
| DDX27 | 0.57102 | 0.54887 | 0.737 | 0.72 | 1 | HC_vs_MS.NoTrt_F |
| DENR | 0.57103 | 0.26951 | 0.751 | 0.77 | 1 | HC_vs_MS.NoTrt_F |
| VHL | 0.57105 | -0.2897 | 0.815 | 0.82 | 1 | HC_vs_MS.NoTrt_F |

| | | | | | | |
|---|---|---|---|---|---|---|
| TTC22 | 0.57136 | -0.3479 | 0.494 | 0.52 | 1 | HC_vs_MS.NoTrt_F |
| GET4 | 0.57213 | 0.40219 | 0.535 | 0.56 | 1 | HC_vs_MS.NoTrt_F |
| ID2 | 0.57217 | 0.63986 | 0.118 | 0.14 | 1 | HC_vs_MS.NoTrt_F |
| CTPS2 | 0.57232 | 0.60942 | 0.142 | 0.17 | 1 | HC_vs_MS.NoTrt_F |
| MYO9B | 0.57238 | 0.80465 | 0.275 | 0.32 | 1 | HC_vs_MS.NoTrt_F |
| MRS2 | 0.57246 | 0.27826 | 0.812 | 0.82 | 1 | HC_vs_MS.NoTrt_F |
| FAM124B | 0.57296 | -0.9682 | 0.118 | 0.14 | 1 | HC_vs_MS.NoTrt_F |
| ZNF813 | 0.57304 | 0.66061 | 0.763 | 0.77 | 1 | HC_vs_MS.NoTrt_F |
| TMEM107 | 0.57318 | 1.0017 | 0.581 | 0.58 | 1 | HC_vs_MS.NoTrt_F |
| GNAQ | 0.57346 | 0.59689 | 0.257 | 0.29 | 1 | HC_vs_MS.NoTrt_F |
| RNPC3 | 0.57386 | 0.98265 | 0.405 | 0.4 | 1 | HC_vs_MS.NoTrt_F |
| PDE12 | 0.57389 | -0.3324 | 0.919 | 0.91 | 1 | HC_vs_MS.NoTrt_F |
| HIF1AN | 0.57398 | 0.49672 | 0.89 | 0.87 | 1 | HC_vs_MS.NoTrt_F |
| IQCF3 | 0.57489 | 0.27712 | 0.327 | 0.37 | 1 | HC_vs_MS.NoTrt_F |
| RP11-552F3 | 0.57572 | -0.4257 | 0.22 | 0.24 | 1 | HC_vs_MS.NoTrt_F |
| TMPRSS3 | 0.57634 | -0.7085 | 0.379 | 0.38 | 1 | HC_vs_MS.NoTrt_F |
| BCCIP | 0.57638 | 0.56748 | 0.228 | 0.25 | 1 | HC_vs_MS.NoTrt_F |
| ZMAT1 | 0.57643 | 0.28452 | 0.39 | 0.42 | 1 | HC_vs_MS.NoTrt_F |
| SPHK2 | 0.57662 | 0.88765 | 0.393 | 0.37 | 1 | HC_vs_MS.NoTrt_F |
| TBP | 0.57667 | 0.70849 | 0.431 | 0.48 | 1 | HC_vs_MS.NoTrt_F |
| ELOF1 | 0.57699 | 0.3504 | 0.61 | 0.64 | 1 | HC_vs_MS.NoTrt_F |
| FAM219B | 0.57757 | 0.90905 | 0.566 | 0.53 | 1 | HC_vs_MS.NoTrt_F |
| IRS1 | 0.5776 | 0.46606 | 0.107 | 0.13 | 1 | HC_vs_MS.NoTrt_F |
| PSMB6 | 0.57775 | 0.51581 | 0.645 | 0.62 | 1 | HC_vs_MS.NoTrt_F |
| PSTPIP1 | 0.57778 | 0.47771 | 0.535 | 0.51 | 1 | HC_vs_MS.NoTrt_F |
| FAM19A2 | 0.57809 | 1.17738 | 0.289 | 0.32 | 1 | HC_vs_MS.NoTrt_F |
| RGS17 | 0.57832 | 0.43471 | 0.488 | 0.51 | 1 | HC_vs_MS.NoTrt_F |
| MAK | 0.57833 | 0.65159 | 0.249 | 0.23 | 1 | HC_vs_MS.NoTrt_F |
| AEBP2 | 0.57947 | 0.36932 | 0.754 | 0.77 | 1 | HC_vs_MS.NoTrt_F |
| CCDC69 | 0.57964 | -0.5592 | 0.902 | 0.88 | 1 | HC_vs_MS.NoTrt_F |
| BOP1 | 0.58001 | 0.69978 | 0.249 | 0.26 | 1 | HC_vs_MS.NoTrt_F |
| KIAA1324 | 0.58027 | -0.5967 | 0.798 | 0.81 | 1 | HC_vs_MS.NoTrt_F |
| PDE8A | 0.58044 | 0.40089 | 0.526 | 0.48 | 1 | HC_vs_MS.NoTrt_F |
| SYBU | 0.5807 | 1.12276 | 0.113 | 0.1 | 1 | HC_vs_MS.NoTrt_F |
| FYTTD1 | 0.58079 | -0.3629 | 0.838 | 0.86 | 1 | HC_vs_MS.NoTrt_F |
| GIT2 | 0.58116 | 0.33927 | 0.751 | 0.76 | 1 | HC_vs_MS.NoTrt_F |
| UIMC1 | 0.5817 | -0.4577 | 0.879 | 0.86 | 1 | HC_vs_MS.NoTrt_F |
| SHFM1 | 0.58266 | 0.343 | 0.922 | 0.92 | 1 | HC_vs_MS.NoTrt_F |
| WIPI2 | 0.58276 | 0.34361 | 0.549 | 0.55 | 1 | HC_vs_MS.NoTrt_F |
| AREL1 | 0.58319 | 0.46849 | 0.52 | 0.54 | 1 | HC_vs_MS.NoTrt_F |

| | | | | | | |
|---|---|---|---|---|---|---|
| CABIN1 | 0.58366 | 0.50342 | 0.37 | 0.41 | 1 | HC_vs_MS.NoTrt_F |
| MRPL48 | 0.58366 | 0.27429 | 0.942 | 0.95 | 1 | HC_vs_MS.NoTrt_F |
| PDE4D | 0.58398 | 0.90765 | 0.405 | 0.4 | 1 | HC_vs_MS.NoTrt_F |
| ACTL6A | 0.58465 | 0.27199 | 0.327 | 0.34 | 1 | HC_vs_MS.NoTrt_F |
| RAB13 | 0.58489 | 0.2531 | 0.572 | 0.51 | 1 | HC_vs_MS.NoTrt_F |
| MXD1 | 0.58514 | 0.9312 | 0.171 | 0.19 | 1 | HC_vs_MS.NoTrt_F |
| LFNG | 0.58546 | 0.53451 | 0.344 | 0.32 | 1 | HC_vs_MS.NoTrt_F |
| SLTM | 0.58549 | 0.52431 | 0.471 | 0.5 | 1 | HC_vs_MS.NoTrt_F |
| MIEF2 | 0.58558 | 0.39552 | 0.54 | 0.56 | 1 | HC_vs_MS.NoTrt_F |
| SERPINB6 | 0.58575 | 0.75413 | 0.535 | 0.56 | 1 | HC_vs_MS.NoTrt_F |
| KLHL23 | 0.58578 | 0.40051 | 0.223 | 0.24 | 1 | HC_vs_MS.NoTrt_F |
| EP400NL | 0.58603 | -0.323 | 0.801 | 0.79 | 1 | HC_vs_MS.NoTrt_F |
| PAM | 0.5866 | 0.45101 | 0.445 | 0.45 | 1 | HC_vs_MS.NoTrt_F |
| CARNS1 | 0.58699 | 0.72441 | 0.243 | 0.26 | 1 | HC_vs_MS.NoTrt_F |
| ZNF585B | 0.58728 | 0.38495 | 0.772 | 0.75 | 1 | HC_vs_MS.NoTrt_F |
| COL16A1 | 0.5873 | 1.20293 | 0.092 | 0.11 | 1 | HC_vs_MS.NoTrt_F |
| SPTY2D1 | 0.58733 | 0.42503 | 0.422 | 0.44 | 1 | HC_vs_MS.NoTrt_F |
| FBXO25 | 0.58777 | 1.66756 | 0.454 | 0.41 | 1 | HC_vs_MS.NoTrt_F |
| LRP2 | 0.58788 | 0.43939 | 0.098 | 0.12 | 1 | HC_vs_MS.NoTrt_F |
| SETD6 | 0.58794 | 0.47951 | 0.462 | 0.49 | 1 | HC_vs_MS.NoTrt_F |
| RNF7 | 0.58828 | 0.31461 | 0.682 | 0.68 | 1 | HC_vs_MS.NoTrt_F |
| SH3BP5 | 0.58834 | -0.2849 | 0.841 | 0.83 | 1 | HC_vs_MS.NoTrt_F |
| MSRB3 | 0.58851 | 0.27615 | 0.523 | 0.52 | 1 | HC_vs_MS.NoTrt_F |
| SGK1 | 0.58887 | 1.53417 | 0.416 | 0.43 | 1 | HC_vs_MS.NoTrt_F |
| POLI | 0.58952 | -0.8614 | 0.41 | 0.41 | 1 | HC_vs_MS.NoTrt_F |
| CLEC17A | 0.58952 | 0.73588 | 0.312 | 0.3 | 1 | HC_vs_MS.NoTrt_F |
| FYN | 0.58953 | 0.27669 | 0.543 | 0.61 | 1 | HC_vs_MS.NoTrt_F |
| PPIE | 0.58979 | 0.329 | 0.789 | 0.82 | 1 | HC_vs_MS.NoTrt_F |
| RABL6 | 0.59097 | 0.34976 | 0.578 | 0.6 | 1 | HC_vs_MS.NoTrt_F |
| MEGF6 | 0.591 | 1.05626 | 0.162 | 0.18 | 1 | HC_vs_MS.NoTrt_F |
| NBPF26 | 0.59197 | 0.94714 | 0.35 | 0.37 | 1 | HC_vs_MS.NoTrt_F |
| HENMT1 | 0.59231 | -0.9526 | 0.327 | 0.35 | 1 | HC_vs_MS.NoTrt_F |
| UBE2D4 | 0.59264 | 0.56788 | 0.699 | 0.64 | 1 | HC_vs_MS.NoTrt_F |
| TMEM150C | 0.59278 | 0.58113 | 0.569 | 0.56 | 1 | HC_vs_MS.NoTrt_F |
| ARHGAP27 | 0.59326 | 0.44147 | 0.645 | 0.61 | 1 | HC_vs_MS.NoTrt_F |
| KIAA0232 | 0.59328 | 1.49015 | 0.156 | 0.18 | 1 | HC_vs_MS.NoTrt_F |
| PLCB2 | 0.59351 | 0.90225 | 0.327 | 0.34 | 1 | HC_vs_MS.NoTrt_F |
| FBLN7 | 0.59473 | 0.77655 | 0.439 | 0.43 | 1 | HC_vs_MS.NoTrt_F |
| ATRN | 0.59532 | 1.15489 | 0.165 | 0.19 | 1 | HC_vs_MS.NoTrt_F |
| SETD1B | 0.59547 | 0.65925 | 0.249 | 0.28 | 1 | HC_vs_MS.NoTrt_F |

| | | | | | | |
|---|---|---|---|---|---|---|
| DCAF17 | 0.59556 | -0.2522 | 0.483 | 0.45 | 1 | HC_vs_MS.NoTrt_F |
| RP11-235E1 | 0.59569 | 0.8588 | 0.28 | 0.27 | 1 | HC_vs_MS.NoTrt_F |
| C8orf37 | 0.59571 | 0.61575 | 0.552 | 0.56 | 1 | HC_vs_MS.NoTrt_F |
| MICAL3 | 0.59572 | 0.45894 | 0.855 | 0.83 | 1 | HC_vs_MS.NoTrt_F |
| ASPA | 0.59592 | 0.28655 | 0.801 | 0.81 | 1 | HC_vs_MS.NoTrt_F |
| IRAK3 | 0.59603 | -0.4385 | 0.474 | 0.48 | 1 | HC_vs_MS.NoTrt_F |
| CDC5L | 0.59604 | -0.3343 | 0.697 | 0.74 | 1 | HC_vs_MS.NoTrt_F |
| EXD2 | 0.59611 | -1.51 | 0.275 | 0.25 | 1 | HC_vs_MS.NoTrt_F |
| ZNF346 | 0.59689 | -0.3618 | 0.809 | 0.8 | 1 | HC_vs_MS.NoTrt_F |
| C11orf1 | 0.5973 | 0.31021 | 0.835 | 0.82 | 1 | HC_vs_MS.NoTrt_F |
| STX17 | 0.59743 | 0.45922 | 0.558 | 0.55 | 1 | HC_vs_MS.NoTrt_F |
| CXorf36 | 0.59756 | 0.5468 | 0.474 | 0.46 | 1 | HC_vs_MS.NoTrt_F |
| NHSL2 | 0.59765 | 0.42839 | 0.749 | 0.74 | 1 | HC_vs_MS.NoTrt_F |
| ITGA2B | 0.59857 | 0.33192 | 0.471 | 0.48 | 1 | HC_vs_MS.NoTrt_F |
| TDRD3 | 0.59874 | 0.93244 | 0.535 | 0.55 | 1 | HC_vs_MS.NoTrt_F |
| FAH | 0.59897 | 1.69521 | 0.205 | 0.19 | 1 | HC_vs_MS.NoTrt_F |
| AIFM1 | 0.59958 | -0.5426 | 0.636 | 0.63 | 1 | HC_vs_MS.NoTrt_F |
| RNF207 | 0.6004 | 0.33299 | 0.725 | 0.7 | 1 | HC_vs_MS.NoTrt_F |
| ZC3H12C | 0.60075 | -0.5181 | 0.133 | 0.15 | 1 | HC_vs_MS.NoTrt_F |
| PRSS53 | 0.60101 | 1.24872 | 0.257 | 0.27 | 1 | HC_vs_MS.NoTrt_F |
| ZNF678 | 0.60104 | 1.63828 | 0.228 | 0.25 | 1 | HC_vs_MS.NoTrt_F |
| NUDT14 | 0.60168 | 0.61679 | 0.509 | 0.54 | 1 | HC_vs_MS.NoTrt_F |
| PODN | 0.60224 | 1.54156 | 0.436 | 0.46 | 1 | HC_vs_MS.NoTrt_F |
| RAD9A | 0.60249 | 0.40863 | 0.529 | 0.54 | 1 | HC_vs_MS.NoTrt_F |
| AIFM2 | 0.60274 | 0.63329 | 0.514 | 0.49 | 1 | HC_vs_MS.NoTrt_F |
| CD177 | 0.60281 | 1.70046 | 0.095 | 0.12 | 1 | HC_vs_MS.NoTrt_F |
| AP5B1 | 0.60301 | -0.4141 | 0.335 | 0.37 | 1 | HC_vs_MS.NoTrt_F |
| TRAP1 | 0.6031 | 0.33358 | 0.766 | 0.83 | 1 | HC_vs_MS.NoTrt_F |
| KLHL2 | 0.60349 | 0.57656 | 0.367 | 0.39 | 1 | HC_vs_MS.NoTrt_F |
| UBE2R2 | 0.60357 | 0.90962 | 0.156 | 0.14 | 1 | HC_vs_MS.NoTrt_F |
| LDHC | 0.60363 | -0.48 | 0.225 | 0.24 | 1 | HC_vs_MS.NoTrt_F |
| CD79B | 0.60424 | 1.03464 | 0.173 | 0.2 | 1 | HC_vs_MS.NoTrt_F |
| TCTN2 | 0.60435 | 0.3328 | 0.795 | 0.78 | 1 | HC_vs_MS.NoTrt_F |
| EIF5A | 0.60481 | 0.77614 | 0.575 | 0.65 | 1 | HC_vs_MS.NoTrt_F |
| CD68 | 0.60529 | 0.43676 | 0.604 | 0.62 | 1 | HC_vs_MS.NoTrt_F |
| RP11-1070B | 0.60613 | 0.92709 | 0.243 | 0.23 | 1 | HC_vs_MS.NoTrt_F |
| RHBDD3 | 0.60632 | -0.6673 | 0.613 | 0.58 | 1 | HC_vs_MS.NoTrt_F |
| NPIPB9 | 0.60665 | 0.39068 | 0.205 | 0.23 | 1 | HC_vs_MS.NoTrt_F |
| NF1 | 0.60703 | 0.29554 | 0.76 | 0.75 | 1 | HC_vs_MS.NoTrt_F |
| BRD1 | 0.60864 | 1.68649 | 0.199 | 0.19 | 1 | HC_vs_MS.NoTrt_F |

| | | | | | | |
|---|---|---|---|---|---|---|
| SLC28A2 | 0.6087 | -0.3279 | 0.809 | 0.8 | 1 | HC_vs_MS.NoTrt_F |
| FOXM1 | 0.60872 | 0.49285 | 0.809 | 0.8 | 1 | HC_vs_MS.NoTrt_F |
| B3GNT7 | 0.60926 | 0.70698 | 0.231 | 0.21 | 1 | HC_vs_MS.NoTrt_F |
| TANC1 | 0.61004 | 0.59539 | 0.376 | 0.39 | 1 | HC_vs_MS.NoTrt_F |
| MAP3K3 | 0.61029 | 0.35806 | 0.179 | 0.2 | 1 | HC_vs_MS.NoTrt_F |
| RBM18 | 0.61029 | -0.5695 | 0.393 | 0.4 | 1 | HC_vs_MS.NoTrt_F |
| CREM | 0.61072 | 0.75951 | 0.445 | 0.45 | 1 | HC_vs_MS.NoTrt_F |
| MFSD4A | 0.61078 | -0.3238 | 0.535 | 0.5 | 1 | HC_vs_MS.NoTrt_F |
| SF3B3 | 0.61099 | 0.30374 | 0.971 | 0.97 | 1 | HC_vs_MS.NoTrt_F |
| MTHFD1L | 0.61111 | 0.8892 | 0.139 | 0.16 | 1 | HC_vs_MS.NoTrt_F |
| PKNOX1 | 0.61155 | -0.4313 | 0.824 | 0.8 | 1 | HC_vs_MS.NoTrt_F |
| FAM92A1 | 0.61167 | -0.357 | 0.358 | 0.39 | 1 | HC_vs_MS.NoTrt_F |
| TAZ | 0.61197 | 0.28743 | 0.59 | 0.62 | 1 | HC_vs_MS.NoTrt_F |
| VNN2 | 0.61213 | 0.95698 | 0.188 | 0.2 | 1 | HC_vs_MS.NoTrt_F |
| PMPCB | 0.61252 | 0.6897 | 0.676 | 0.65 | 1 | HC_vs_MS.NoTrt_F |
| USP10 | 0.61313 | 0.39728 | 0.321 | 0.35 | 1 | HC_vs_MS.NoTrt_F |
| GLB1L3 | 0.61366 | 0.29974 | 0.763 | 0.75 | 1 | HC_vs_MS.NoTrt_F |
| NBPF3 | 0.61373 | 1.30898 | 0.488 | 0.54 | 1 | HC_vs_MS.NoTrt_F |
| THUMPD3 | 0.61374 | 0.38022 | 0.434 | 0.45 | 1 | HC_vs_MS.NoTrt_F |
| PTPRJ | 0.61378 | 0.66526 | 0.41 | 0.45 | 1 | HC_vs_MS.NoTrt_F |
| ZNF74 | 0.61422 | 0.59454 | 0.494 | 0.51 | 1 | HC_vs_MS.NoTrt_F |
| NLRP12 | 0.61526 | 0.35181 | 0.775 | 0.76 | 1 | HC_vs_MS.NoTrt_F |
| GLUL | 0.61552 | 0.77849 | 0.263 | 0.24 | 1 | HC_vs_MS.NoTrt_F |
| CENPV | 0.61568 | 0.39941 | 0.801 | 0.77 | 1 | HC_vs_MS.NoTrt_F |
| HELZ2 | 0.61573 | -0.2579 | 0.699 | 0.71 | 1 | HC_vs_MS.NoTrt_F |
| LRCH4 | 0.61588 | 0.75272 | 0.575 | 0.61 | 1 | HC_vs_MS.NoTrt_F |
| C16orf74 | 0.61621 | -0.4325 | 0.277 | 0.29 | 1 | HC_vs_MS.NoTrt_F |
| MRPL44 | 0.61625 | -0.2695 | 0.627 | 0.64 | 1 | HC_vs_MS.NoTrt_F |
| SLC15A4 | 0.61665 | 0.26361 | 0.775 | 0.8 | 1 | HC_vs_MS.NoTrt_F |
| ABCG1 | 0.61674 | -0.2985 | 0.523 | 0.5 | 1 | HC_vs_MS.NoTrt_F |
| SFT2D3 | 0.61699 | 0.26047 | 0.535 | 0.53 | 1 | HC_vs_MS.NoTrt_F |
| NHLRC2 | 0.61724 | 0.26953 | 0.798 | 0.79 | 1 | HC_vs_MS.NoTrt_F |
| FASLG | 0.61754 | 1.49597 | 0.124 | 0.14 | 1 | HC_vs_MS.NoTrt_F |
| NEXN | 0.61778 | 0.37832 | 0.535 | 0.51 | 1 | HC_vs_MS.NoTrt_F |
| MFN1 | 0.61779 | -0.3988 | 0.506 | 0.5 | 1 | HC_vs_MS.NoTrt_F |
| PNPLA8 | 0.61797 | 0.56856 | 0.468 | 0.45 | 1 | HC_vs_MS.NoTrt_F |
| PSG2 | 0.61828 | 0.33587 | 0.124 | 0.11 | 1 | HC_vs_MS.NoTrt_F |
| ZNF527 | 0.61835 | 0.47227 | 0.934 | 0.95 | 1 | HC_vs_MS.NoTrt_F |
| GSPT1 | 0.61856 | 0.30834 | 0.601 | 0.61 | 1 | HC_vs_MS.NoTrt_F |
| CCT6B | 0.61866 | 0.66497 | 0.312 | 0.33 | 1 | HC_vs_MS.NoTrt_F |

| | | | | | | |
|---|---|---|---|---|---|---|
| CUL2 | 0.61903 | 0.56511 | 0.272 | 0.29 | 1 | HC_vs_MS.NoTrt_F |
| HYAL1 | 0.61925 | -0.3631 | 0.376 | 0.4 | 1 | HC_vs_MS.NoTrt_F |
| TTC28 | 0.61999 | -0.3451 | 0.46 | 0.44 | 1 | HC_vs_MS.NoTrt_F |
| NADK2 | 0.62022 | -0.9597 | 0.5 | 0.48 | 1 | HC_vs_MS.NoTrt_F |
| CREG1 | 0.62044 | 0.67116 | 0.355 | 0.38 | 1 | HC_vs_MS.NoTrt_F |
| NR4A1 | 0.62053 | 1.20789 | 0.569 | 0.57 | 1 | HC_vs_MS.NoTrt_F |
| WDR19 | 0.62058 | 0.77448 | 0.601 | 0.61 | 1 | HC_vs_MS.NoTrt_F |
| ANKRD54 | 0.62094 | 0.56941 | 0.754 | 0.8 | 1 | HC_vs_MS.NoTrt_F |
| PON2 | 0.62135 | 1.02763 | 0.442 | 0.46 | 1 | HC_vs_MS.NoTrt_F |
| ACE | 0.62141 | 0.31781 | 0.647 | 0.7 | 1 | HC_vs_MS.NoTrt_F |
| SLC35B1 | 0.62173 | 0.3895 | 0.708 | 0.72 | 1 | HC_vs_MS.NoTrt_F |
| MCCC2 | 0.62218 | 0.6442 | 0.734 | 0.75 | 1 | HC_vs_MS.NoTrt_F |
| HAVCR1 | 0.62226 | 0.66535 | 0.373 | 0.39 | 1 | HC_vs_MS.NoTrt_F |
| PLEKHH3 | 0.62257 | 0.2882 | 0.633 | 0.6 | 1 | HC_vs_MS.NoTrt_F |
| ST3GAL3 | 0.62258 | 0.39147 | 0.832 | 0.89 | 1 | HC_vs_MS.NoTrt_F |
| SCLT1 | 0.6232 | 0.87841 | 0.673 | 0.67 | 1 | HC_vs_MS.NoTrt_F |
| LIPT2 | 0.62341 | 0.80099 | 0.173 | 0.2 | 1 | HC_vs_MS.NoTrt_F |
| MRPS10 | 0.62342 | 0.50113 | 0.538 | 0.5 | 1 | HC_vs_MS.NoTrt_F |
| FAM200A | 0.62374 | 0.84587 | 0.246 | 0.26 | 1 | HC_vs_MS.NoTrt_F |
| GADD45B | 0.62387 | 0.58869 | 0.22 | 0.25 | 1 | HC_vs_MS.NoTrt_F |
| CREB3L4 | 0.62443 | 0.41632 | 0.405 | 0.44 | 1 | HC_vs_MS.NoTrt_F |
| RIC3 | 0.62448 | 0.27564 | 0.488 | 0.52 | 1 | HC_vs_MS.NoTrt_F |
| BCR | 0.62449 | 0.99402 | 0.341 | 0.32 | 1 | HC_vs_MS.NoTrt_F |
| ACSL1 | 0.62455 | 0.29539 | 0.28 | 0.29 | 1 | HC_vs_MS.NoTrt_F |
| PPP6C | 0.6247 | 0.33181 | 0.272 | 0.3 | 1 | HC_vs_MS.NoTrt_F |
| PLXDC1 | 0.62481 | 0.49771 | 0.731 | 0.75 | 1 | HC_vs_MS.NoTrt_F |
| VPS37B | 0.62524 | 0.59521 | 0.694 | 0.74 | 1 | HC_vs_MS.NoTrt_F |
| ETV6 | 0.62532 | 0.31429 | 0.246 | 0.27 | 1 | HC_vs_MS.NoTrt_F |
| RBM43 | 0.62541 | -0.3346 | 0.682 | 0.66 | 1 | HC_vs_MS.NoTrt_F |
| TGIF1 | 0.62558 | 0.53589 | 0.315 | 0.35 | 1 | HC_vs_MS.NoTrt_F |
| LAMP2 | 0.62575 | -0.2556 | 0.564 | 0.55 | 1 | HC_vs_MS.NoTrt_F |
| ZNF707 | 0.6262 | 0.61675 | 0.821 | 0.79 | 1 | HC_vs_MS.NoTrt_F |
| B3GAT1 | 0.62691 | 0.47212 | 0.312 | 0.35 | 1 | HC_vs_MS.NoTrt_F |
| NR1H2 | 0.62708 | 0.69891 | 0.564 | 0.61 | 1 | HC_vs_MS.NoTrt_F |
| TPX2 | 0.62737 | 0.39361 | 0.286 | 0.27 | 1 | HC_vs_MS.NoTrt_F |
| IRF2BPL | 0.62808 | 0.45237 | 0.474 | 0.51 | 1 | HC_vs_MS.NoTrt_F |
| PI4KB | 0.62842 | 0.58169 | 0.251 | 0.28 | 1 | HC_vs_MS.NoTrt_F |
| PLEKHH2 | 0.62849 | 0.79101 | 0.697 | 0.69 | 1 | HC_vs_MS.NoTrt_F |
| MLPH | 0.62882 | 1.17654 | 0.514 | 0.52 | 1 | HC_vs_MS.NoTrt_F |
| CNR2 | 0.62887 | 0.43265 | 0.844 | 0.9 | 1 | HC_vs_MS.NoTrt_F |

| | | | | | | |
|---|---|---|---|---|---|---|
| CDC6 | 0.6295 | -0.3201 | 0.301 | 0.32 | 1 | HC_vs_MS.NoTrt_F |
| THAP12 | 0.62973 | -0.7355 | 0.679 | 0.66 | 1 | HC_vs_MS.NoTrt_F |
| SEMA4C | 0.63054 | 0.79138 | 0.801 | 0.84 | 1 | HC_vs_MS.NoTrt_F |
| MBNL2 | 0.63073 | 0.8293 | 0.315 | 0.3 | 1 | HC_vs_MS.NoTrt_F |
| BLMH | 0.63125 | -0.7635 | 0.65 | 0.7 | 1 | HC_vs_MS.NoTrt_F |
| LRP2BP | 0.6316 | 0.38177 | 0.48 | 0.49 | 1 | HC_vs_MS.NoTrt_F |
| CHGB | 0.6324 | 0.46928 | 0.5 | 0.49 | 1 | HC_vs_MS.NoTrt_F |
| ERICH3 | 0.63269 | 1.35229 | 0.118 | 0.14 | 1 | HC_vs_MS.NoTrt_F |
| SPHK1 | 0.63278 | -0.4895 | 0.223 | 0.2 | 1 | HC_vs_MS.NoTrt_F |
| ST6GALNAC | 0.63323 | 0.89359 | 0.257 | 0.26 | 1 | HC_vs_MS.NoTrt_F |
| SIK3 | 0.63337 | 0.76073 | 0.367 | 0.39 | 1 | HC_vs_MS.NoTrt_F |
| CD40LG | 0.63342 | 1.02457 | 0.266 | 0.29 | 1 | HC_vs_MS.NoTrt_F |
| COA5 | 0.63499 | -0.5164 | 0.52 | 0.51 | 1 | HC_vs_MS.NoTrt_F |
| YIF1A | 0.63676 | 0.88158 | 0.173 | 0.2 | 1 | HC_vs_MS.NoTrt_F |
| IL10RB | 0.63682 | 1.11029 | 0.523 | 0.51 | 1 | HC_vs_MS.NoTrt_F |
| HLA-DRB4 | 0.63725 | 0.97987 | 0.124 | 0.14 | 1 | HC_vs_MS.NoTrt_F |
| KLHL18 | 0.63727 | 0.81255 | 0.399 | 0.42 | 1 | HC_vs_MS.NoTrt_F |
| ENTPD5 | 0.63747 | 0.82234 | 0.497 | 0.53 | 1 | HC_vs_MS.NoTrt_F |
| TNFRSF25 | 0.63836 | 0.33068 | 0.717 | 0.75 | 1 | HC_vs_MS.NoTrt_F |
| CD34 | 0.63841 | 0.29949 | 0.665 | 0.69 | 1 | HC_vs_MS.NoTrt_F |
| RBM47 | 0.63857 | 0.4452 | 0.494 | 0.47 | 1 | HC_vs_MS.NoTrt_F |
| IGF1R | 0.63908 | 1.78012 | 0.387 | 0.37 | 1 | HC_vs_MS.NoTrt_F |
| SIRPB2 | 0.63947 | 0.44359 | 0.416 | 0.44 | 1 | HC_vs_MS.NoTrt_F |
| TNFSF4 | 0.63965 | 0.7134 | 0.136 | 0.16 | 1 | HC_vs_MS.NoTrt_F |
| ST5 | 0.64003 | 0.42837 | 0.393 | 0.38 | 1 | HC_vs_MS.NoTrt_F |
| TSC22D2 | 0.6413 | 0.44453 | 0.488 | 0.48 | 1 | HC_vs_MS.NoTrt_F |
| TYW1 | 0.64166 | 0.26509 | 0.312 | 0.34 | 1 | HC_vs_MS.NoTrt_F |
| TOMM40 | 0.64171 | -0.3178 | 0.653 | 0.65 | 1 | HC_vs_MS.NoTrt_F |
| MED25 | 0.64175 | 0.5787 | 0.52 | 0.5 | 1 | HC_vs_MS.NoTrt_F |
| DDX3X | 0.64238 | 0.50946 | 0.367 | 0.4 | 1 | HC_vs_MS.NoTrt_F |
| FILIP1L | 0.64457 | 0.66496 | 0.442 | 0.42 | 1 | HC_vs_MS.NoTrt_F |
| TRAPPC11 | 0.64492 | 0.51368 | 0.474 | 0.5 | 1 | HC_vs_MS.NoTrt_F |
| ADCK1 | 0.64525 | -0.321 | 0.312 | 0.33 | 1 | HC_vs_MS.NoTrt_F |
| TGFBR1 | 0.64529 | 1.1925 | 0.162 | 0.18 | 1 | HC_vs_MS.NoTrt_F |
| SCN11A | 0.64562 | 0.26746 | 0.419 | 0.46 | 1 | HC_vs_MS.NoTrt_F |
| WDR89 | 0.64587 | 0.96687 | 0.399 | 0.43 | 1 | HC_vs_MS.NoTrt_F |
| GTPBP3 | 0.64591 | 0.29044 | 0.711 | 0.75 | 1 | HC_vs_MS.NoTrt_F |
| PCNX1 | 0.64619 | 0.27576 | 0.861 | 0.86 | 1 | HC_vs_MS.NoTrt_F |
| DNAJB11 | 0.64629 | 0.38963 | 0.269 | 0.3 | 1 | HC_vs_MS.NoTrt_F |
| PRKDC | 0.64643 | 0.73962 | 0.63 | 0.65 | 1 | HC_vs_MS.NoTrt_F |

| | | | | | | |
|---|---|---|---|---|---|---|
| TBRG1 | 0.64649 | 0.3143 | 0.737 | 0.74 | 1 | HC_vs_MS.NoTrt_F |
| SLX4IP | 0.64707 | 0.2756 | 0.506 | 0.49 | 1 | HC_vs_MS.NoTrt_F |
| SEC16A | 0.64737 | 0.60385 | 0.26 | 0.29 | 1 | HC_vs_MS.NoTrt_F |
| ZNF547 | 0.64811 | 0.80416 | 0.266 | 0.29 | 1 | HC_vs_MS.NoTrt_F |
| ZNF808 | 0.64853 | 0.42417 | 0.526 | 0.54 | 1 | HC_vs_MS.NoTrt_F |
| FADS2 | 0.64858 | 0.9546 | 0.59 | 0.58 | 1 | HC_vs_MS.NoTrt_F |
| RGPD4 | 0.64904 | 0.72676 | 0.217 | 0.21 | 1 | HC_vs_MS.NoTrt_F |
| TMEM53 | 0.64913 | -0.4552 | 0.364 | 0.34 | 1 | HC_vs_MS.NoTrt_F |
| LCOR | 0.6496 | 0.6146 | 0.393 | 0.44 | 1 | HC_vs_MS.NoTrt_F |
| CPSF6 | 0.65056 | 0.4152 | 0.396 | 0.41 | 1 | HC_vs_MS.NoTrt_F |
| GRN | 0.65121 | 0.9183 | 0.211 | 0.19 | 1 | HC_vs_MS.NoTrt_F |
| SUMF2 | 0.65151 | 0.28294 | 0.902 | 0.91 | 1 | HC_vs_MS.NoTrt_F |
| ZNF230 | 0.65165 | 0.73553 | 0.633 | 0.62 | 1 | HC_vs_MS.NoTrt_F |
| TMEM116 | 0.65213 | -0.2715 | 0.928 | 0.91 | 1 | HC_vs_MS.NoTrt_F |
| HMGCL | 0.65231 | -0.3816 | 0.624 | 0.6 | 1 | HC_vs_MS.NoTrt_F |
| ADORA2A | 0.6525 | 0.53521 | 0.237 | 0.27 | 1 | HC_vs_MS.NoTrt_F |
| POLN | 0.65279 | -0.3114 | 0.691 | 0.72 | 1 | HC_vs_MS.NoTrt_F |
| TTLL5 | 0.65326 | 0.49243 | 0.749 | 0.76 | 1 | HC_vs_MS.NoTrt_F |
| ZNF835 | 0.6549 | 0.40413 | 0.65 | 0.64 | 1 | HC_vs_MS.NoTrt_F |
| SAMSN1 | 0.6552 | 0.25532 | 0.555 | 0.56 | 1 | HC_vs_MS.NoTrt_F |
| NFIB | 0.65521 | 1.51037 | 0.191 | 0.18 | 1 | HC_vs_MS.NoTrt_F |
| TGOLN2 | 0.65543 | 0.45966 | 0.653 | 0.72 | 1 | HC_vs_MS.NoTrt_F |
| NFS1 | 0.6556 | -0.4605 | 0.934 | 0.92 | 1 | HC_vs_MS.NoTrt_F |
| ARHGAP17 | 0.65567 | 0.64576 | 0.63 | 0.65 | 1 | HC_vs_MS.NoTrt_F |
| C16orf52 | 0.65571 | 0.85719 | 0.098 | 0.12 | 1 | HC_vs_MS.NoTrt_F |
| SLC39A8 | 0.65597 | -0.5621 | 0.384 | 0.39 | 1 | HC_vs_MS.NoTrt_F |
| KLHL3 | 0.65599 | 0.796 | 0.277 | 0.29 | 1 | HC_vs_MS.NoTrt_F |
| OSBPL10 | 0.65671 | 0.83979 | 0.468 | 0.5 | 1 | HC_vs_MS.NoTrt_F |
| PAAF1 | 0.65727 | -0.3252 | 0.98 | 0.99 | 1 | HC_vs_MS.NoTrt_F |
| ULK4 | 0.65787 | 0.80851 | 0.431 | 0.4 | 1 | HC_vs_MS.NoTrt_F |
| CIRBP | 0.65896 | 0.36672 | 0.604 | 0.67 | 1 | HC_vs_MS.NoTrt_F |
| GOLGA8B | 0.659 | 0.30244 | 0.821 | 0.82 | 1 | HC_vs_MS.NoTrt_F |
| WDFY2 | 0.65949 | 0.4822 | 0.645 | 0.65 | 1 | HC_vs_MS.NoTrt_F |
| ZNF727 | 0.66047 | 1.175 | 0.384 | 0.41 | 1 | HC_vs_MS.NoTrt_F |
| FAM102B | 0.66126 | 1.10766 | 0.315 | 0.32 | 1 | HC_vs_MS.NoTrt_F |
| ZNF85 | 0.66127 | -0.674 | 0.72 | 0.74 | 1 | HC_vs_MS.NoTrt_F |
| TRAF3IP1 | 0.66186 | 0.32078 | 0.431 | 0.4 | 1 | HC_vs_MS.NoTrt_F |
| KCTD3 | 0.66197 | 0.32937 | 0.367 | 0.38 | 1 | HC_vs_MS.NoTrt_F |
| ZKSCAN4 | 0.66211 | -0.9402 | 0.572 | 0.54 | 1 | HC_vs_MS.NoTrt_F |
| DYRK1A | 0.66239 | 0.80196 | 0.751 | 0.77 | 1 | HC_vs_MS.NoTrt_F |

| | | | | | | |
|---|---|---|---|---|---|---|
| STK38L | 0.66277 | -0.6179 | 0.702 | 0.78 | 1 | HC_vs_MS.NoTrt_F |
| TMEM184C | 0.66297 | 0.28903 | 0.592 | 0.6 | 1 | HC_vs_MS.NoTrt_F |
| C4orf22 | 0.66426 | 0.35567 | 0.451 | 0.45 | 1 | HC_vs_MS.NoTrt_F |
| ZFP14 | 0.66476 | 0.34471 | 0.887 | 0.9 | 1 | HC_vs_MS.NoTrt_F |
| OPRD1 | 0.6649 | 0.39088 | 0.855 | 0.88 | 1 | HC_vs_MS.NoTrt_F |
| ANGPTL4 | 0.66491 | 1.34338 | 0.35 | 0.35 | 1 | HC_vs_MS.NoTrt_F |
| NKIRAS1 | 0.66522 | -0.3278 | 0.298 | 0.28 | 1 | HC_vs_MS.NoTrt_F |
| PLOD1 | 0.66525 | -0.2795 | 0.691 | 0.66 | 1 | HC_vs_MS.NoTrt_F |
| ATP6V1D | 0.66537 | 0.42043 | 0.607 | 0.66 | 1 | HC_vs_MS.NoTrt_F |
| NEDD4L | 0.66558 | -0.5815 | 0.509 | 0.46 | 1 | HC_vs_MS.NoTrt_F |
| SLIT3 | 0.66561 | 0.97118 | 0.616 | 0.62 | 1 | HC_vs_MS.NoTrt_F |
| WDR54 | 0.66572 | 0.40332 | 0.592 | 0.58 | 1 | HC_vs_MS.NoTrt_F |
| SAP30BP | 0.66582 | 0.37627 | 0.318 | 0.31 | 1 | HC_vs_MS.NoTrt_F |
| SMC6 | 0.66606 | 0.44698 | 0.54 | 0.56 | 1 | HC_vs_MS.NoTrt_F |
| TBCCD1 | 0.66661 | 0.32451 | 0.439 | 0.41 | 1 | HC_vs_MS.NoTrt_F |
| FAM120AOS | 0.6672 | 0.56942 | 0.688 | 0.67 | 1 | HC_vs_MS.NoTrt_F |
| CEP152 | 0.66733 | -0.83 | 0.795 | 0.7 | 1 | HC_vs_MS.NoTrt_F |
| FAM65C | 0.66735 | 0.48123 | 0.41 | 0.4 | 1 | HC_vs_MS.NoTrt_F |
| PLEKHM1 | 0.66758 | 0.32084 | 0.861 | 0.92 | 1 | HC_vs_MS.NoTrt_F |
| CEP170 | 0.66764 | 0.37609 | 0.572 | 0.59 | 1 | HC_vs_MS.NoTrt_F |
| SIRPB1 | 0.66822 | 1.55536 | 0.275 | 0.31 | 1 | HC_vs_MS.NoTrt_F |
| RP11-231C1 | 0.66825 | 0.60261 | 0.705 | 0.79 | 1 | HC_vs_MS.NoTrt_F |
| PHF21A | 0.66837 | 0.35496 | 0.318 | 0.34 | 1 | HC_vs_MS.NoTrt_F |
| SLC14A1 | 0.66866 | 0.31832 | 0.552 | 0.54 | 1 | HC_vs_MS.NoTrt_F |
| RNF170 | 0.6693 | -0.3406 | 0.642 | 0.64 | 1 | HC_vs_MS.NoTrt_F |
| PRPF8 | 0.6699 | 0.36221 | 0.893 | 0.87 | 1 | HC_vs_MS.NoTrt_F |
| SLX1B | 0.67001 | 0.28354 | 0.5 | 0.49 | 1 | HC_vs_MS.NoTrt_F |
| GAMT | 0.67038 | 0.54219 | 0.63 | 0.61 | 1 | HC_vs_MS.NoTrt_F |
| NELFCD | 0.67051 | 1.07915 | 0.598 | 0.59 | 1 | HC_vs_MS.NoTrt_F |
| TPMT | 0.67057 | 0.49198 | 0.809 | 0.8 | 1 | HC_vs_MS.NoTrt_F |
| SLC25A40 | 0.67088 | -0.7105 | 0.364 | 0.38 | 1 | HC_vs_MS.NoTrt_F |
| CELF2 | 0.67097 | 0.35718 | 0.592 | 0.69 | 1 | HC_vs_MS.NoTrt_F |
| ZNF286B | 0.67185 | 0.44637 | 0.558 | 0.53 | 1 | HC_vs_MS.NoTrt_F |
| ANKS3 | 0.67199 | 0.29244 | 0.818 | 0.79 | 1 | HC_vs_MS.NoTrt_F |
| JMJD6 | 0.67293 | 0.65956 | 0.494 | 0.46 | 1 | HC_vs_MS.NoTrt_F |
| LIN52 | 0.67295 | 0.32596 | 0.777 | 0.78 | 1 | HC_vs_MS.NoTrt_F |
| TBXAS1 | 0.67305 | 1.49245 | 0.434 | 0.46 | 1 | HC_vs_MS.NoTrt_F |
| IDE | 0.67378 | 0.68991 | 0.564 | 0.55 | 1 | HC_vs_MS.NoTrt_F |
| KIF13B | 0.6739 | 0.29425 | 0.159 | 0.18 | 1 | HC_vs_MS.NoTrt_F |
| INTS1 | 0.67394 | 0.42779 | 0.439 | 0.46 | 1 | HC_vs_MS.NoTrt_F |

| | | | | | | |
|---|---|---|---|---|---|---|
| MRPL20 | 0.67494 | 0.32984 | 0.595 | 0.56 | 1 | HC_vs_MS.NoTrt_F |
| NRROS | 0.67511 | 1.21567 | 0.173 | 0.19 | 1 | HC_vs_MS.NoTrt_F |
| SMARCAL1 | 0.67561 | 0.32394 | 0.743 | 0.74 | 1 | HC_vs_MS.NoTrt_F |
| SUZ12 | 0.67574 | 0.65123 | 0.717 | 0.71 | 1 | HC_vs_MS.NoTrt_F |
| NABP2 | 0.67615 | 0.56024 | 0.494 | 0.48 | 1 | HC_vs_MS.NoTrt_F |
| CERS2 | 0.67636 | 0.25074 | 0.298 | 0.27 | 1 | HC_vs_MS.NoTrt_F |
| HSPB1 | 0.67686 | 0.2761 | 0.436 | 0.44 | 1 | HC_vs_MS.NoTrt_F |
| CYB561 | 0.67699 | 0.88835 | 0.39 | 0.37 | 1 | HC_vs_MS.NoTrt_F |
| CCDC74A | 0.67705 | 1.11928 | 0.289 | 0.32 | 1 | HC_vs_MS.NoTrt_F |
| NIT2 | 0.67744 | 0.30404 | 0.676 | 0.68 | 1 | HC_vs_MS.NoTrt_F |
| IGFL2 | 0.67797 | 0.2503 | 0.422 | 0.47 | 1 | HC_vs_MS.NoTrt_F |
| MAPK9 | 0.67824 | 0.31663 | 0.509 | 0.5 | 1 | HC_vs_MS.NoTrt_F |
| SLX4 | 0.67864 | -0.302 | 0.569 | 0.56 | 1 | HC_vs_MS.NoTrt_F |
| MNS1 | 0.68015 | 0.27795 | 0.234 | 0.23 | 1 | HC_vs_MS.NoTrt_F |
| TUBB4A | 0.6805 | -0.4905 | 0.197 | 0.22 | 1 | HC_vs_MS.NoTrt_F |
| CCDC15 | 0.6807 | 0.89658 | 0.28 | 0.29 | 1 | HC_vs_MS.NoTrt_F |
| RCL1 | 0.6807 | 0.78381 | 0.546 | 0.52 | 1 | HC_vs_MS.NoTrt_F |
| SPPL2B | 0.68071 | 0.29054 | 0.416 | 0.4 | 1 | HC_vs_MS.NoTrt_F |
| CARM1 | 0.68114 | -0.3096 | 0.194 | 0.22 | 1 | HC_vs_MS.NoTrt_F |
| SPDYE16 | 0.68148 | 0.52645 | 0.821 | 0.89 | 1 | HC_vs_MS.NoTrt_F |
| A1BG | 0.68178 | 0.29652 | 0.902 | 0.87 | 1 | HC_vs_MS.NoTrt_F |
| MTIF2 | 0.68185 | 0.5453 | 0.408 | 0.41 | 1 | HC_vs_MS.NoTrt_F |
| PACS2 | 0.6819 | 0.47697 | 0.338 | 0.35 | 1 | HC_vs_MS.NoTrt_F |
| SCARF1 | 0.68302 | 0.58128 | 0.486 | 0.5 | 1 | HC_vs_MS.NoTrt_F |
| EGR3 | 0.68318 | 0.53146 | 0.28 | 0.3 | 1 | HC_vs_MS.NoTrt_F |
| RFC5 | 0.6834 | -0.2731 | 0.199 | 0.21 | 1 | HC_vs_MS.NoTrt_F |
| PSMB9 | 0.68354 | 0.43803 | 0.775 | 0.8 | 1 | HC_vs_MS.NoTrt_F |
| AHI1 | 0.68402 | 0.29147 | 0.428 | 0.41 | 1 | HC_vs_MS.NoTrt_F |
| DNAJB13 | 0.68404 | 0.38452 | 0.419 | 0.45 | 1 | HC_vs_MS.NoTrt_F |
| ACY1 | 0.68435 | 0.73663 | 0.627 | 0.62 | 1 | HC_vs_MS.NoTrt_F |
| FGGY | 0.68515 | 0.39749 | 0.506 | 0.53 | 1 | HC_vs_MS.NoTrt_F |
| PRKAA1 | 0.68606 | 0.39297 | 0.902 | 0.95 | 1 | HC_vs_MS.NoTrt_F |
| C14orf119 | 0.68622 | -0.5538 | 0.595 | 0.56 | 1 | HC_vs_MS.NoTrt_F |
| TNFAIP8L1 | 0.6866 | -0.5137 | 0.882 | 0.87 | 1 | HC_vs_MS.NoTrt_F |
| SECISBP2 | 0.68682 | 0.32422 | 0.676 | 0.66 | 1 | HC_vs_MS.NoTrt_F |
| NIPAL1 | 0.68703 | -0.315 | 0.454 | 0.49 | 1 | HC_vs_MS.NoTrt_F |
| DDHD2 | 0.68765 | 0.50403 | 0.864 | 0.88 | 1 | HC_vs_MS.NoTrt_F |
| SPATA7 | 0.68789 | 0.26101 | 0.341 | 0.37 | 1 | HC_vs_MS.NoTrt_F |
| PSMB7 | 0.68822 | 0.2672 | 0.277 | 0.3 | 1 | HC_vs_MS.NoTrt_F |
| PITPNM2 | 0.68829 | 1.3165 | 0.445 | 0.48 | 1 | HC_vs_MS.NoTrt_F |

| | | | | | | |
|---|---|---|---|---|---|---|
| ASRGL1 | 0.68862 | 0.88878 | 0.607 | 0.63 | 1 | HC_vs_MS.NoTrt_F |
| CRK | 0.6887 | 1.02123 | 0.639 | 0.69 | 1 | HC_vs_MS.NoTrt_F |
| SLX1A | 0.68881 | 0.31439 | 0.546 | 0.56 | 1 | HC_vs_MS.NoTrt_F |
| 2-Mar | 0.68882 | -0.3432 | 0.636 | 0.65 | 1 | HC_vs_MS.NoTrt_F |
| HDAC7 | 0.68903 | 0.70601 | 0.321 | 0.35 | 1 | HC_vs_MS.NoTrt_F |
| EFCAB13 | 0.68906 | -0.5912 | 0.327 | 0.35 | 1 | HC_vs_MS.NoTrt_F |
| SLC22A18 | 0.68945 | 0.4104 | 0.601 | 0.6 | 1 | HC_vs_MS.NoTrt_F |
| FAM3A | 0.68977 | 0.44513 | 0.775 | 0.8 | 1 | HC_vs_MS.NoTrt_F |
| ZNF16 | 0.68995 | 0.57561 | 0.809 | 0.83 | 1 | HC_vs_MS.NoTrt_F |
| TRIM16 | 0.69001 | -0.7549 | 0.497 | 0.49 | 1 | HC_vs_MS.NoTrt_F |
| VPS13A | 0.6902 | 0.52786 | 0.367 | 0.38 | 1 | HC_vs_MS.NoTrt_F |
| IL12RB2 | 0.69088 | 1.47038 | 0.162 | 0.16 | 1 | HC_vs_MS.NoTrt_F |
| ZNF664 | 0.69088 | 0.73821 | 0.301 | 0.34 | 1 | HC_vs_MS.NoTrt_F |
| MAN2C1 | 0.69119 | 0.31008 | 0.847 | 0.85 | 1 | HC_vs_MS.NoTrt_F |
| DST | 0.69143 | 0.35812 | 0.757 | 0.8 | 1 | HC_vs_MS.NoTrt_F |
| PCGF6 | 0.69265 | -0.399 | 0.792 | 0.79 | 1 | HC_vs_MS.NoTrt_F |
| TBC1D32 | 0.6927 | 1.20839 | 0.564 | 0.53 | 1 | HC_vs_MS.NoTrt_F |
| ZNF157 | 0.69277 | -0.3326 | 0.803 | 0.83 | 1 | HC_vs_MS.NoTrt_F |
| C11orf31 | 0.6932 | 0.26897 | 0.575 | 0.57 | 1 | HC_vs_MS.NoTrt_F |
| EXOSC6 | 0.69563 | 0.30157 | 0.928 | 0.96 | 1 | HC_vs_MS.NoTrt_F |
| AKR1B15 | 0.69743 | 0.26291 | 0.618 | 0.67 | 1 | HC_vs_MS.NoTrt_F |
| ZNF43 | 0.69808 | 0.34667 | 0.772 | 0.79 | 1 | HC_vs_MS.NoTrt_F |
| SLC24A1 | 0.69829 | -0.7334 | 0.529 | 0.51 | 1 | HC_vs_MS.NoTrt_F |
| MBD3 | 0.69861 | 1.09661 | 0.775 | 0.77 | 1 | HC_vs_MS.NoTrt_F |
| MTMR9 | 0.6987 | 0.57062 | 0.361 | 0.39 | 1 | HC_vs_MS.NoTrt_F |
| TBC1D17 | 0.69891 | -0.2849 | 0.893 | 0.89 | 1 | HC_vs_MS.NoTrt_F |
| ZNF559 | 0.6992 | 0.36099 | 0.685 | 0.71 | 1 | HC_vs_MS.NoTrt_F |
| URI1 | 0.69925 | 0.5167 | 0.269 | 0.29 | 1 | HC_vs_MS.NoTrt_F |
| GNL2 | 0.69962 | 0.31701 | 0.76 | 0.73 | 1 | HC_vs_MS.NoTrt_F |
| HSD17B7 | 0.69964 | -0.4546 | 0.598 | 0.64 | 1 | HC_vs_MS.NoTrt_F |
| ZDHHC21 | 0.69968 | -0.3345 | 0.301 | 0.32 | 1 | HC_vs_MS.NoTrt_F |
| SGSH | 0.69985 | 0.60329 | 0.558 | 0.61 | 1 | HC_vs_MS.NoTrt_F |
| MMS19 | 0.70004 | 0.4842 | 0.225 | 0.25 | 1 | HC_vs_MS.NoTrt_F |
| TUBA1A | 0.70004 | 0.5093 | 0.361 | 0.36 | 1 | HC_vs_MS.NoTrt_F |
| GPCPD1 | 0.70008 | 0.64518 | 0.445 | 0.48 | 1 | HC_vs_MS.NoTrt_F |
| STAR | 0.70105 | 0.35664 | 0.286 | 0.3 | 1 | HC_vs_MS.NoTrt_F |
| TTC37 | 0.70195 | 0.3048 | 0.413 | 0.43 | 1 | HC_vs_MS.NoTrt_F |
| RRP36 | 0.70203 | 0.25488 | 0.514 | 0.48 | 1 | HC_vs_MS.NoTrt_F |
| CIC | 0.70206 | 0.53422 | 0.15 | 0.17 | 1 | HC_vs_MS.NoTrt_F |
| ZNF516 | 0.70232 | 1.44996 | 0.107 | 0.12 | 1 | HC_vs_MS.NoTrt_F |

| | | | | | | |
|---|---|---|---|---|---|---|
| MAP2K6 | 0.70245 | 0.87178 | 0.408 | 0.44 | 1 | HC_vs_MS.NoTrt_F |
| IFIT3 | 0.70261 | -1.047 | 0.581 | 0.55 | 1 | HC_vs_MS.NoTrt_F |
| DYNLRB1 | 0.70312 | 0.51276 | 0.269 | 0.31 | 1 | HC_vs_MS.NoTrt_F |
| AP3S2 | 0.7032 | -0.3675 | 0.725 | 0.75 | 1 | HC_vs_MS.NoTrt_F |
| IDS | 0.70357 | 0.27763 | 0.879 | 0.88 | 1 | HC_vs_MS.NoTrt_F |
| IBTK | 0.70388 | -0.533 | 0.613 | 0.62 | 1 | HC_vs_MS.NoTrt_F |
| LRRC56 | 0.70389 | -0.2856 | 0.436 | 0.38 | 1 | HC_vs_MS.NoTrt_F |
| MORN3 | 0.70393 | -0.2506 | 0.882 | 0.87 | 1 | HC_vs_MS.NoTrt_F |
| RIPK2 | 0.70421 | 0.66469 | 0.22 | 0.23 | 1 | HC_vs_MS.NoTrt_F |
| BRIP1 | 0.70425 | -0.4465 | 0.876 | 0.87 | 1 | HC_vs_MS.NoTrt_F |
| SMYD4 | 0.7049 | -0.2785 | 0.855 | 0.87 | 1 | HC_vs_MS.NoTrt_F |
| ZNF493 | 0.70496 | 0.6658 | 0.821 | 0.8 | 1 | HC_vs_MS.NoTrt_F |
| GALNT6 | 0.70503 | 0.70288 | 0.78 | 0.81 | 1 | HC_vs_MS.NoTrt_F |
| LASP1 | 0.70511 | -0.3643 | 0.78 | 0.82 | 1 | HC_vs_MS.NoTrt_F |
| P4HA2 | 0.70552 | 0.87217 | 0.514 | 0.53 | 1 | HC_vs_MS.NoTrt_F |
| 1-Mar | 0.70578 | 0.36166 | 0.639 | 0.66 | 1 | HC_vs_MS.NoTrt_F |
| PRKCA | 0.70579 | 0.28837 | 0.775 | 0.76 | 1 | HC_vs_MS.NoTrt_F |
| ARSK | 0.70585 | 0.59875 | 0.604 | 0.61 | 1 | HC_vs_MS.NoTrt_F |
| COBLL1 | 0.70612 | 0.25167 | 0.598 | 0.58 | 1 | HC_vs_MS.NoTrt_F |
| SPICE1 | 0.70635 | 0.57015 | 0.324 | 0.34 | 1 | HC_vs_MS.NoTrt_F |
| ZBED1 | 0.70652 | -0.2575 | 0.546 | 0.55 | 1 | HC_vs_MS.NoTrt_F |
| GDPD5 | 0.70661 | 1.13426 | 0.332 | 0.34 | 1 | HC_vs_MS.NoTrt_F |
| PLBD2 | 0.70746 | 0.58211 | 0.41 | 0.43 | 1 | HC_vs_MS.NoTrt_F |
| EPT1 | 0.70753 | 0.35437 | 0.431 | 0.4 | 1 | HC_vs_MS.NoTrt_F |
| MKL2 | 0.70766 | -0.4002 | 0.769 | 0.8 | 1 | HC_vs_MS.NoTrt_F |
| PBX3 | 0.70806 | 0.89361 | 0.116 | 0.13 | 1 | HC_vs_MS.NoTrt_F |
| GSE1 | 0.70825 | 1.42186 | 0.454 | 0.46 | 1 | HC_vs_MS.NoTrt_F |
| TRIM26 | 0.70853 | 0.61314 | 0.249 | 0.28 | 1 | HC_vs_MS.NoTrt_F |
| C11orf63 | 0.70993 | 0.70815 | 0.523 | 0.51 | 1 | HC_vs_MS.NoTrt_F |
| HPS5 | 0.7101 | -0.5475 | 0.795 | 0.82 | 1 | HC_vs_MS.NoTrt_F |
| LMCD1 | 0.71043 | 1.47032 | 0.37 | 0.39 | 1 | HC_vs_MS.NoTrt_F |
| RAB8B | 0.71051 | 0.29556 | 0.399 | 0.39 | 1 | HC_vs_MS.NoTrt_F |
| SNUPN | 0.71071 | -0.3795 | 0.887 | 0.92 | 1 | HC_vs_MS.NoTrt_F |
| XPNPEP1 | 0.71091 | 0.42073 | 0.431 | 0.43 | 1 | HC_vs_MS.NoTrt_F |
| PM20D2 | 0.71119 | 0.44563 | 0.78 | 0.85 | 1 | HC_vs_MS.NoTrt_F |
| ZZEF1 | 0.71139 | 0.51497 | 0.592 | 0.58 | 1 | HC_vs_MS.NoTrt_F |
| GAB1 | 0.71157 | 0.60084 | 0.587 | 0.6 | 1 | HC_vs_MS.NoTrt_F |
| ZNF708 | 0.71168 | 0.45351 | 0.827 | 0.85 | 1 | HC_vs_MS.NoTrt_F |
| POC1B | 0.71181 | 0.4103 | 0.812 | 0.83 | 1 | HC_vs_MS.NoTrt_F |
| GAPVD1 | 0.71249 | 0.26182 | 0.855 | 0.85 | 1 | HC_vs_MS.NoTrt_F |

| | | | | | | |
|---|---|---|---|---|---|---|
| PRR12 | 0.71312 | 1.25445 | 0.133 | 0.16 | 1 | HC_vs_MS.NoTrt_F |
| RALBP1 | 0.71325 | 0.53679 | 0.454 | 0.44 | 1 | HC_vs_MS.NoTrt_F |
| DRG1 | 0.71413 | 0.96938 | 0.228 | 0.25 | 1 | HC_vs_MS.NoTrt_F |
| ZNF343 | 0.71444 | 0.30001 | 0.52 | 0.54 | 1 | HC_vs_MS.NoTrt_F |
| CTBP1 | 0.71445 | 0.64048 | 0.532 | 0.56 | 1 | HC_vs_MS.NoTrt_F |
| AKR1C1 | 0.71476 | 0.43641 | 0.306 | 0.31 | 1 | HC_vs_MS.NoTrt_F |
| PRKD2 | 0.71476 | 0.45196 | 0.769 | 0.79 | 1 | HC_vs_MS.NoTrt_F |
| KIF21A | 0.71524 | 0.72299 | 0.22 | 0.25 | 1 | HC_vs_MS.NoTrt_F |
| DCN | 0.71591 | 0.84425 | 0.749 | 0.77 | 1 | HC_vs_MS.NoTrt_F |
| LRRN4CL | 0.71594 | 0.268 | 0.598 | 0.59 | 1 | HC_vs_MS.NoTrt_F |
| SPIN1 | 0.71595 | 0.72324 | 0.228 | 0.24 | 1 | HC_vs_MS.NoTrt_F |
| GP6 | 0.71602 | 0.6306 | 0.486 | 0.53 | 1 | HC_vs_MS.NoTrt_F |
| GSR | 0.71767 | 0.48416 | 0.329 | 0.32 | 1 | HC_vs_MS.NoTrt_F |
| STXBP4 | 0.71771 | 0.57598 | 0.584 | 0.59 | 1 | HC_vs_MS.NoTrt_F |
| TMEM256-P | 0.7178 | 0.42662 | 0.48 | 0.53 | 1 | HC_vs_MS.NoTrt_F |
| POU6F1 | 0.71786 | -0.3644 | 0.558 | 0.54 | 1 | HC_vs_MS.NoTrt_F |
| SHISA2 | 0.71798 | 0.35796 | 0.298 | 0.33 | 1 | HC_vs_MS.NoTrt_F |
| PATL2 | 0.71879 | 0.45966 | 0.431 | 0.43 | 1 | HC_vs_MS.NoTrt_F |
| RGS16 | 0.71896 | 1.11171 | 0.113 | 0.13 | 1 | HC_vs_MS.NoTrt_F |
| MGA | 0.7195 | -0.639 | 0.908 | 0.89 | 1 | HC_vs_MS.NoTrt_F |
| CD83 | 0.71968 | 1.07642 | 0.179 | 0.18 | 1 | HC_vs_MS.NoTrt_F |
| MLLT6 | 0.71978 | 0.3486 | 0.913 | 0.91 | 1 | HC_vs_MS.NoTrt_F |
| PARP2 | 0.71984 | 1.18116 | 0.436 | 0.42 | 1 | HC_vs_MS.NoTrt_F |
| FRYL | 0.72037 | 0.44922 | 0.792 | 0.89 | 1 | HC_vs_MS.NoTrt_F |
| SEMA6D | 0.7204 | 1.35698 | 0.147 | 0.17 | 1 | HC_vs_MS.NoTrt_F |
| LSR | 0.72048 | 0.78932 | 0.237 | 0.26 | 1 | HC_vs_MS.NoTrt_F |
| PHTF2 | 0.72052 | 0.76269 | 0.552 | 0.57 | 1 | HC_vs_MS.NoTrt_F |
| TALDO1 | 0.72068 | 0.25172 | 0.743 | 0.76 | 1 | HC_vs_MS.NoTrt_F |
| ZNF341 | 0.72072 | -0.3189 | 0.486 | 0.49 | 1 | HC_vs_MS.NoTrt_F |
| C1orf159 | 0.7208 | 1.50723 | 0.208 | 0.23 | 1 | HC_vs_MS.NoTrt_F |
| GALNT15 | 0.72118 | 0.42782 | 0.37 | 0.38 | 1 | HC_vs_MS.NoTrt_F |
| DCPS | 0.72127 | 0.28029 | 0.509 | 0.51 | 1 | HC_vs_MS.NoTrt_F |
| PITPNM1 | 0.72197 | 0.47576 | 0.306 | 0.32 | 1 | HC_vs_MS.NoTrt_F |
| MTFMT | 0.72239 | 0.29617 | 0.867 | 0.92 | 1 | HC_vs_MS.NoTrt_F |
| ERAL1 | 0.72256 | 0.82659 | 0.153 | 0.17 | 1 | HC_vs_MS.NoTrt_F |
| EMP1 | 0.72257 | 1.31693 | 0.225 | 0.25 | 1 | HC_vs_MS.NoTrt_F |
| KAZN | 0.72263 | 0.74446 | 0.558 | 0.57 | 1 | HC_vs_MS.NoTrt_F |
| MOCOS | 0.72271 | 0.43565 | 0.656 | 0.68 | 1 | HC_vs_MS.NoTrt_F |
| FLG2 | 0.72283 | -0.5711 | 0.532 | 0.5 | 1 | HC_vs_MS.NoTrt_F |
| ERBB3 | 0.72317 | 0.35341 | 0.803 | 0.82 | 1 | HC_vs_MS.NoTrt_F |

| | | | | | | |
|---|---|---|---|---|---|---|
| CTDSPL2 | 0.72317 | -0.4889 | 0.723 | 0.76 | 1 | HC_vs_MS.NoTrt_F |
| TYK2 | 0.72319 | 0.67608 | 0.665 | 0.74 | 1 | HC_vs_MS.NoTrt_F |
| PRKAG2 | 0.72361 | 0.72861 | 0.422 | 0.44 | 1 | HC_vs_MS.NoTrt_F |
| ZNRF3 | 0.72373 | -0.2634 | 0.367 | 0.34 | 1 | HC_vs_MS.NoTrt_F |
| CPT1A | 0.72389 | 0.62894 | 0.746 | 0.7 | 1 | HC_vs_MS.NoTrt_F |
| CEP97 | 0.72433 | 0.59114 | 0.734 | 0.78 | 1 | HC_vs_MS.NoTrt_F |
| MOAP1 | 0.72449 | 0.53976 | 0.413 | 0.42 | 1 | HC_vs_MS.NoTrt_F |
| DIS3L2 | 0.72494 | 0.4389 | 0.569 | 0.57 | 1 | HC_vs_MS.NoTrt_F |
| HCFC1 | 0.72519 | 0.51493 | 0.341 | 0.38 | 1 | HC_vs_MS.NoTrt_F |
| WDR78 | 0.7254 | -0.2791 | 0.529 | 0.54 | 1 | HC_vs_MS.NoTrt_F |
| SSH2 | 0.72548 | 0.34969 | 0.801 | 0.86 | 1 | HC_vs_MS.NoTrt_F |
| CD320 | 0.72577 | 0.44748 | 0.766 | 0.8 | 1 | HC_vs_MS.NoTrt_F |
| FSTL4 | 0.72581 | 0.29134 | 0.246 | 0.27 | 1 | HC_vs_MS.NoTrt_F |
| TRIM45 | 0.72607 | 0.31326 | 0.156 | 0.17 | 1 | HC_vs_MS.NoTrt_F |
| GPR89A | 0.72637 | 0.54998 | 0.48 | 0.52 | 1 | HC_vs_MS.NoTrt_F |
| COMMD4 | 0.72644 | 0.36316 | 0.945 | 0.95 | 1 | HC_vs_MS.NoTrt_F |
| UAP1L1 | 0.72765 | 0.86184 | 0.355 | 0.36 | 1 | HC_vs_MS.NoTrt_F |
| MNT | 0.72786 | 0.34671 | 0.711 | 0.7 | 1 | HC_vs_MS.NoTrt_F |
| TADA2A | 0.72787 | -0.4798 | 0.919 | 0.92 | 1 | HC_vs_MS.NoTrt_F |
| OXCT1 | 0.72846 | 1.25642 | 0.408 | 0.46 | 1 | HC_vs_MS.NoTrt_F |
| CCDC114 | 0.72934 | -0.7025 | 0.494 | 0.49 | 1 | HC_vs_MS.NoTrt_F |
| C9orf85 | 0.72941 | 0.2603 | 0.919 | 0.93 | 1 | HC_vs_MS.NoTrt_F |
| UBB | 0.72945 | 0.30648 | 0.621 | 0.73 | 1 | HC_vs_MS.NoTrt_F |
| SLC35E1 | 0.72963 | 0.30881 | 0.991 | 0.97 | 1 | HC_vs_MS.NoTrt_F |
| MREG | 0.7304 | 0.39681 | 0.668 | 0.66 | 1 | HC_vs_MS.NoTrt_F |
| ZNF417 | 0.73074 | 0.74031 | 0.769 | 0.75 | 1 | HC_vs_MS.NoTrt_F |
| ZNF684 | 0.73239 | -0.2836 | 0.751 | 0.74 | 1 | HC_vs_MS.NoTrt_F |
| CHDH | 0.73253 | -0.6863 | 0.572 | 0.55 | 1 | HC_vs_MS.NoTrt_F |
| TMEM220 | 0.7329 | 0.30124 | 0.572 | 0.56 | 1 | HC_vs_MS.NoTrt_F |
| HTN3 | 0.73359 | 0.37791 | 0.659 | 0.69 | 1 | HC_vs_MS.NoTrt_F |
| LRWD1 | 0.73377 | 0.49832 | 0.61 | 0.59 | 1 | HC_vs_MS.NoTrt_F |
| SLC35B4 | 0.73449 | 0.70388 | 0.587 | 0.59 | 1 | HC_vs_MS.NoTrt_F |
| BTAF1 | 0.735 | 0.31813 | 0.347 | 0.37 | 1 | HC_vs_MS.NoTrt_F |
| TARSL2 | 0.73535 | 0.34703 | 0.974 | 0.98 | 1 | HC_vs_MS.NoTrt_F |
| CPD | 0.7359 | 0.52629 | 0.743 | 0.77 | 1 | HC_vs_MS.NoTrt_F |
| MED22 | 0.73669 | 0.47051 | 0.613 | 0.6 | 1 | HC_vs_MS.NoTrt_F |
| CDK5RAP1 | 0.737 | 0.32628 | 0.48 | 0.45 | 1 | HC_vs_MS.NoTrt_F |
| UBOX5 | 0.73738 | -0.2566 | 0.682 | 0.66 | 1 | HC_vs_MS.NoTrt_F |
| ZNF879 | 0.7378 | 0.54907 | 0.367 | 0.35 | 1 | HC_vs_MS.NoTrt_F |
| STARD4 | 0.7381 | 0.41601 | 0.211 | 0.19 | 1 | HC_vs_MS.NoTrt_F |

| | | | | | | |
|---|---|---|---|---|---|---|
| FGFR2 | 0.73812 | 0.28368 | 0.509 | 0.51 | 1 | HC_vs_MS.NoTrt_F |
| XPA | 0.73822 | 1.20798 | 0.208 | 0.23 | 1 | HC_vs_MS.NoTrt_F |
| RNF130 | 0.73835 | 0.40654 | 0.723 | 0.69 | 1 | HC_vs_MS.NoTrt_F |
| CNP | 0.73837 | 0.30412 | 0.783 | 0.74 | 1 | HC_vs_MS.NoTrt_F |
| NUDC | 0.73902 | -0.2671 | 0.668 | 0.62 | 1 | HC_vs_MS.NoTrt_F |
| KIF22 | 0.73904 | 0.60473 | 0.246 | 0.24 | 1 | HC_vs_MS.NoTrt_F |
| NFKBID | 0.73942 | 0.39676 | 0.642 | 0.64 | 1 | HC_vs_MS.NoTrt_F |
| AGPAT3 | 0.73963 | 0.53751 | 0.694 | 0.68 | 1 | HC_vs_MS.NoTrt_F |
| ALG10B | 0.74077 | -0.6263 | 0.725 | 0.67 | 1 | HC_vs_MS.NoTrt_F |
| PIP5K1C | 0.74102 | 1.15707 | 0.13 | 0.14 | 1 | HC_vs_MS.NoTrt_F |
| VSIG1 | 0.74167 | 0.49771 | 0.633 | 0.65 | 1 | HC_vs_MS.NoTrt_F |
| AASS | 0.74177 | 0.78219 | 0.422 | 0.44 | 1 | HC_vs_MS.NoTrt_F |
| OCEL1 | 0.74206 | 0.4043 | 0.514 | 0.51 | 1 | HC_vs_MS.NoTrt_F |
| CCNL1 | 0.74215 | 0.60644 | 0.445 | 0.5 | 1 | HC_vs_MS.NoTrt_F |
| PPP1R14D | 0.7425 | 0.27212 | 0.11 | 0.12 | 1 | HC_vs_MS.NoTrt_F |
| YTHDC2 | 0.74332 | 0.38475 | 0.506 | 0.51 | 1 | HC_vs_MS.NoTrt_F |
| TMEM199 | 0.74431 | -0.6151 | 0.702 | 0.7 | 1 | HC_vs_MS.NoTrt_F |
| SSR3 | 0.74457 | 0.38681 | 0.676 | 0.66 | 1 | HC_vs_MS.NoTrt_F |
| PRELID2 | 0.74546 | 0.72181 | 0.572 | 0.54 | 1 | HC_vs_MS.NoTrt_F |
| E2F3 | 0.74556 | 0.44115 | 0.225 | 0.24 | 1 | HC_vs_MS.NoTrt_F |
| SLC7A11 | 0.74571 | 0.26714 | 0.532 | 0.51 | 1 | HC_vs_MS.NoTrt_F |
| CSF1 | 0.74654 | 0.66164 | 0.176 | 0.19 | 1 | HC_vs_MS.NoTrt_F |
| NUP214 | 0.747 | 0.28358 | 0.694 | 0.74 | 1 | HC_vs_MS.NoTrt_F |
| HAUS5 | 0.74732 | -0.5064 | 0.523 | 0.51 | 1 | HC_vs_MS.NoTrt_F |
| LRCH1 | 0.74819 | 0.69192 | 0.159 | 0.15 | 1 | HC_vs_MS.NoTrt_F |
| MROH8 | 0.74835 | -0.5185 | 0.373 | 0.39 | 1 | HC_vs_MS.NoTrt_F |
| C1orf112 | 0.74895 | 0.30547 | 0.546 | 0.55 | 1 | HC_vs_MS.NoTrt_F |
| ADAM28 | 0.7496 | 0.52478 | 0.211 | 0.23 | 1 | HC_vs_MS.NoTrt_F |
| CCDC159 | 0.74973 | -0.5375 | 0.795 | 0.79 | 1 | HC_vs_MS.NoTrt_F |
| LRRC32 | 0.75026 | 1.58056 | 0.101 | 0.12 | 1 | HC_vs_MS.NoTrt_F |
| RERE | 0.75039 | 0.66328 | 0.451 | 0.47 | 1 | HC_vs_MS.NoTrt_F |
| GPM6A | 0.7508 | 0.70504 | 0.355 | 0.36 | 1 | HC_vs_MS.NoTrt_F |
| IRS2 | 0.75097 | 1.09472 | 0.477 | 0.49 | 1 | HC_vs_MS.NoTrt_F |
| BCKDHB | 0.75133 | -0.537 | 0.656 | 0.65 | 1 | HC_vs_MS.NoTrt_F |
| PPP2R3A | 0.75182 | 0.26597 | 0.798 | 0.81 | 1 | HC_vs_MS.NoTrt_F |
| CAMK1 | 0.75202 | 0.35441 | 0.561 | 0.59 | 1 | HC_vs_MS.NoTrt_F |
| GTPBP1 | 0.75233 | 0.46922 | 0.604 | 0.65 | 1 | HC_vs_MS.NoTrt_F |
| RANBP6 | 0.75256 | -0.7377 | 0.168 | 0.16 | 1 | HC_vs_MS.NoTrt_F |
| NIT1 | 0.75265 | 0.60717 | 0.676 | 0.69 | 1 | HC_vs_MS.NoTrt_F |
| LMOD3 | 0.75268 | -0.419 | 0.514 | 0.51 | 1 | HC_vs_MS.NoTrt_F |

| | | | | | | |
|---|---|---|---|---|---|---|
| GALE | 0.75271 | 0.66966 | 0.405 | 0.4 | 1 | HC_vs_MS.NoTrt_F |
| DNAJC22 | 0.75383 | -0.3633 | 0.402 | 0.4 | 1 | HC_vs_MS.NoTrt_F |
| PRKCD | 0.75426 | 0.25229 | 0.566 | 0.55 | 1 | HC_vs_MS.NoTrt_F |
| KCNJ5 | 0.75458 | 0.30367 | 0.673 | 0.65 | 1 | HC_vs_MS.NoTrt_F |
| ICA1L | 0.75478 | 0.42816 | 0.812 | 0.81 | 1 | HC_vs_MS.NoTrt_F |
| ITPKB | 0.75488 | 0.50533 | 0.63 | 0.62 | 1 | HC_vs_MS.NoTrt_F |
| ACSS3 | 0.75523 | 0.28459 | 0.298 | 0.32 | 1 | HC_vs_MS.NoTrt_F |
| RASEF | 0.7556 | 0.3086 | 0.899 | 0.95 | 1 | HC_vs_MS.NoTrt_F |
| HES2 | 0.75635 | 0.27705 | 0.688 | 0.66 | 1 | HC_vs_MS.NoTrt_F |
| PLA2R1 | 0.75654 | 0.67505 | 0.592 | 0.54 | 1 | HC_vs_MS.NoTrt_F |
| CD37 | 0.75675 | 0.30085 | 0.798 | 0.86 | 1 | HC_vs_MS.NoTrt_F |
| CDC40 | 0.75679 | 0.34135 | 0.546 | 0.56 | 1 | HC_vs_MS.NoTrt_F |
| TNFSF9 | 0.75684 | 0.53145 | 0.11 | 0.11 | 1 | HC_vs_MS.NoTrt_F |
| UMPS | 0.75706 | -0.6716 | 0.688 | 0.7 | 1 | HC_vs_MS.NoTrt_F |
| KIAA2013 | 0.7574 | 0.59508 | 0.361 | 0.37 | 1 | HC_vs_MS.NoTrt_F |
| SMPD3 | 0.75757 | 0.46841 | 0.728 | 0.74 | 1 | HC_vs_MS.NoTrt_F |
| BAIAP2L1 | 0.75794 | 0.29322 | 0.587 | 0.57 | 1 | HC_vs_MS.NoTrt_F |
| CMTR2 | 0.75807 | 0.35254 | 0.321 | 0.31 | 1 | HC_vs_MS.NoTrt_F |
| ECE1 | 0.75811 | 0.3584 | 0.763 | 0.72 | 1 | HC_vs_MS.NoTrt_F |
| RASAL1 | 0.75846 | 0.84746 | 0.318 | 0.33 | 1 | HC_vs_MS.NoTrt_F |
| ABCB6 | 0.75875 | 0.25694 | 0.194 | 0.21 | 1 | HC_vs_MS.NoTrt_F |
| TGFBRAP1 | 0.75887 | 0.46534 | 0.535 | 0.55 | 1 | HC_vs_MS.NoTrt_F |
| DENND3 | 0.75901 | 1.08944 | 0.564 | 0.59 | 1 | HC_vs_MS.NoTrt_F |
| EI24 | 0.75928 | -0.3906 | 0.786 | 0.84 | 1 | HC_vs_MS.NoTrt_F |
| UBE4B | 0.75949 | 1.22105 | 0.659 | 0.68 | 1 | HC_vs_MS.NoTrt_F |
| ENDOV | 0.75997 | -0.3736 | 0.659 | 0.66 | 1 | HC_vs_MS.NoTrt_F |
| ASIC4 | 0.76019 | 0.28982 | 0.266 | 0.25 | 1 | HC_vs_MS.NoTrt_F |
| ZNF433 | 0.76049 | 0.632 | 0.185 | 0.2 | 1 | HC_vs_MS.NoTrt_F |
| APPL1 | 0.76153 | 0.37493 | 0.494 | 0.48 | 1 | HC_vs_MS.NoTrt_F |
| PCOLCE | 0.76158 | 0.97095 | 0.133 | 0.13 | 1 | HC_vs_MS.NoTrt_F |
| CCDC77 | 0.76204 | -0.3885 | 0.714 | 0.7 | 1 | HC_vs_MS.NoTrt_F |
| RGPD8 | 0.76249 | 0.95713 | 0.327 | 0.37 | 1 | HC_vs_MS.NoTrt_F |
| RCOR1 | 0.76259 | 0.92041 | 0.225 | 0.22 | 1 | HC_vs_MS.NoTrt_F |
| ERLIN2 | 0.76417 | 0.49572 | 0.422 | 0.42 | 1 | HC_vs_MS.NoTrt_F |
| HILPDA | 0.76434 | 0.5442 | 0.592 | 0.62 | 1 | HC_vs_MS.NoTrt_F |
| BRI3BP | 0.76449 | 0.30467 | 0.902 | 0.95 | 1 | HC_vs_MS.NoTrt_F |
| S1PR5 | 0.76509 | 0.54129 | 0.121 | 0.13 | 1 | HC_vs_MS.NoTrt_F |
| TAF6 | 0.76512 | -0.6147 | 0.584 | 0.58 | 1 | HC_vs_MS.NoTrt_F |
| FBXO36 | 0.76543 | 0.55392 | 0.691 | 0.66 | 1 | HC_vs_MS.NoTrt_F |
| PTBP3 | 0.76548 | 0.31719 | 0.425 | 0.46 | 1 | HC_vs_MS.NoTrt_F |

| | | | | | | |
|---|---|---|---|---|---|---|
| RSRC1 | 0.76558 | 0.34426 | 0.757 | 0.76 | 1 | HC_vs_MS.NoTrt_F |
| TAF8 | 0.76591 | -0.5106 | 0.971 | 0.97 | 1 | HC_vs_MS.NoTrt_F |
| PTBP1 | 0.76664 | 0.44787 | 0.303 | 0.33 | 1 | HC_vs_MS.NoTrt_F |
| CLEC4C | 0.76736 | 0.28673 | 0.442 | 0.45 | 1 | HC_vs_MS.NoTrt_F |
| EGR1 | 0.76777 | 2.22425 | 0.121 | 0.12 | 1 | HC_vs_MS.NoTrt_F |
| RGR | 0.76846 | 0.61744 | 0.529 | 0.51 | 1 | HC_vs_MS.NoTrt_F |
| ALMS1 | 0.76894 | 1.04252 | 0.517 | 0.48 | 1 | HC_vs_MS.NoTrt_F |
| FMN1 | 0.76917 | 0.70153 | 0.269 | 0.25 | 1 | HC_vs_MS.NoTrt_F |
| HSPBAP1 | 0.7696 | 0.80512 | 0.584 | 0.57 | 1 | HC_vs_MS.NoTrt_F |
| CRKL | 0.76966 | 0.5012 | 0.607 | 0.64 | 1 | HC_vs_MS.NoTrt_F |
| TNFAIP2 | 0.76977 | 0.70061 | 0.39 | 0.39 | 1 | HC_vs_MS.NoTrt_F |
| KLC4 | 0.77049 | 0.31885 | 0.876 | 0.86 | 1 | HC_vs_MS.NoTrt_F |
| FKTN | 0.77075 | 1.22807 | 0.199 | 0.22 | 1 | HC_vs_MS.NoTrt_F |
| DENND5B | 0.77164 | 0.33578 | 0.642 | 0.63 | 1 | HC_vs_MS.NoTrt_F |
| KRR1 | 0.77238 | 0.35695 | 0.717 | 0.71 | 1 | HC_vs_MS.NoTrt_F |
| KDM6B | 0.77373 | 0.5771 | 0.289 | 0.29 | 1 | HC_vs_MS.NoTrt_F |
| PPP1R16A | 0.77436 | 0.377 | 0.477 | 0.5 | 1 | HC_vs_MS.NoTrt_F |
| COL6A3 | 0.77449 | 2.01699 | 0.517 | 0.54 | 1 | HC_vs_MS.NoTrt_F |
| RASAL2 | 0.77484 | 0.75374 | 0.555 | 0.56 | 1 | HC_vs_MS.NoTrt_F |
| LMBR1 | 0.77527 | 0.74603 | 0.532 | 0.54 | 1 | HC_vs_MS.NoTrt_F |
| IFRD1 | 0.77535 | 0.72868 | 0.564 | 0.58 | 1 | HC_vs_MS.NoTrt_F |
| ZNF700 | 0.7754 | 0.42396 | 0.243 | 0.26 | 1 | HC_vs_MS.NoTrt_F |
| DPH6 | 0.77571 | 0.73156 | 0.477 | 0.45 | 1 | HC_vs_MS.NoTrt_F |
| MRPS34 | 0.77602 | 0.88956 | 0.254 | 0.29 | 1 | HC_vs_MS.NoTrt_F |
| NAPEPLD | 0.77631 | 0.53739 | 0.384 | 0.38 | 1 | HC_vs_MS.NoTrt_F |
| ERLIN1 | 0.77643 | -0.3846 | 0.101 | 0.11 | 1 | HC_vs_MS.NoTrt_F |
| RPUSD4 | 0.77659 | -0.5812 | 0.749 | 0.8 | 1 | HC_vs_MS.NoTrt_F |
| SRSF10 | 0.7767 | 0.26587 | 0.806 | 0.83 | 1 | HC_vs_MS.NoTrt_F |
| ANO10 | 0.77676 | -0.8692 | 0.671 | 0.66 | 1 | HC_vs_MS.NoTrt_F |
| DCAKD | 0.77677 | 0.69224 | 0.471 | 0.44 | 1 | HC_vs_MS.NoTrt_F |
| PHF20 | 0.77682 | 0.43735 | 0.598 | 0.61 | 1 | HC_vs_MS.NoTrt_F |
| PPP1CB | 0.77706 | 0.32935 | 0.749 | 0.84 | 1 | HC_vs_MS.NoTrt_F |
| ATP5L2 | 0.77763 | 0.34171 | 0.353 | 0.36 | 1 | HC_vs_MS.NoTrt_F |
| ZCCHC4 | 0.77771 | -0.5356 | 0.691 | 0.7 | 1 | HC_vs_MS.NoTrt_F |
| TRMT12 | 0.77834 | -0.2656 | 0.512 | 0.5 | 1 | HC_vs_MS.NoTrt_F |
| TTC21B | 0.77867 | 0.32214 | 0.416 | 0.41 | 1 | HC_vs_MS.NoTrt_F |
| FAM126A | 0.77891 | 0.34655 | 0.728 | 0.71 | 1 | HC_vs_MS.NoTrt_F |
| ALG11 | 0.77921 | 0.28316 | 0.867 | 0.89 | 1 | HC_vs_MS.NoTrt_F |
| FCRL3 | 0.7795 | 0.74113 | 0.347 | 0.38 | 1 | HC_vs_MS.NoTrt_F |
| HGS | 0.77969 | 0.35688 | 0.691 | 0.7 | 1 | HC_vs_MS.NoTrt_F |

| | | | | | | |
|---|---|---|---|---|---|---|
| USP11 | 0.78024 | 0.36023 | 0.266 | 0.29 | 1 | HC_vs_MS.NoTrt_F |
| ZNF681 | 0.78106 | -0.4178 | 0.708 | 0.69 | 1 | HC_vs_MS.NoTrt_F |
| DIAPH3 | 0.7821 | 0.29161 | 0.364 | 0.38 | 1 | HC_vs_MS.NoTrt_F |
| DMWD | 0.78253 | 0.41255 | 0.879 | 0.92 | 1 | HC_vs_MS.NoTrt_F |
| DCUN1D3 | 0.78264 | 0.96424 | 0.231 | 0.24 | 1 | HC_vs_MS.NoTrt_F |
| SULT2B1 | 0.78277 | -0.3218 | 0.569 | 0.6 | 1 | HC_vs_MS.NoTrt_F |
| FGFR4 | 0.78302 | 0.25842 | 0.445 | 0.44 | 1 | HC_vs_MS.NoTrt_F |
| LRBA | 0.78307 | 0.6443 | 0.52 | 0.53 | 1 | HC_vs_MS.NoTrt_F |
| SAXO2 | 0.78337 | 0.52469 | 0.269 | 0.28 | 1 | HC_vs_MS.NoTrt_F |
| MT-ND3 | 0.7837 | 0.80666 | 0.532 | 0.67 | 1 | HC_vs_MS.NoTrt_F |
| SLC23A3 | 0.78388 | 0.3338 | 0.483 | 0.49 | 1 | HC_vs_MS.NoTrt_F |
| MASP2 | 0.7839 | -0.7525 | 0.335 | 0.34 | 1 | HC_vs_MS.NoTrt_F |
| PARPBP | 0.784 | -0.5684 | 0.465 | 0.45 | 1 | HC_vs_MS.NoTrt_F |
| TWSG1 | 0.78457 | 0.56789 | 0.454 | 0.46 | 1 | HC_vs_MS.NoTrt_F |
| SLC9A1 | 0.78509 | 1.17351 | 0.292 | 0.32 | 1 | HC_vs_MS.NoTrt_F |
| MCPH1 | 0.78512 | -0.3843 | 0.743 | 0.75 | 1 | HC_vs_MS.NoTrt_F |
| POLR1C | 0.78515 | 0.83821 | 0.48 | 0.49 | 1 | HC_vs_MS.NoTrt_F |
| BCL6 | 0.78549 | 0.75463 | 0.191 | 0.19 | 1 | HC_vs_MS.NoTrt_F |
| PPP2R5C | 0.7858 | 0.2995 | 0.936 | 0.96 | 1 | HC_vs_MS.NoTrt_F |
| PTPRG | 0.78635 | 1.13828 | 0.295 | 0.32 | 1 | HC_vs_MS.NoTrt_F |
| CAPS2 | 0.78664 | 0.94371 | 0.488 | 0.51 | 1 | HC_vs_MS.NoTrt_F |
| DDX19A | 0.78667 | -0.3286 | 0.974 | 0.98 | 1 | HC_vs_MS.NoTrt_F |
| CTPS1 | 0.7868 | 0.26851 | 0.676 | 0.66 | 1 | HC_vs_MS.NoTrt_F |
| RPAP1 | 0.78685 | -0.6738 | 0.855 | 0.86 | 1 | HC_vs_MS.NoTrt_F |
| CCNYL1 | 0.78701 | -0.3049 | 0.48 | 0.5 | 1 | HC_vs_MS.NoTrt_F |
| PIGL | 0.78719 | 0.74995 | 0.714 | 0.75 | 1 | HC_vs_MS.NoTrt_F |
| TTF1 | 0.7875 | 0.25224 | 0.512 | 0.51 | 1 | HC_vs_MS.NoTrt_F |
| DRAM2 | 0.78752 | 0.69036 | 0.188 | 0.2 | 1 | HC_vs_MS.NoTrt_F |
| HSF2 | 0.78774 | 0.71893 | 0.142 | 0.16 | 1 | HC_vs_MS.NoTrt_F |
| DNA2 | 0.78791 | -0.3032 | 0.231 | 0.24 | 1 | HC_vs_MS.NoTrt_F |
| INSR | 0.78823 | 0.90281 | 0.517 | 0.51 | 1 | HC_vs_MS.NoTrt_F |
| GRHL2 | 0.78834 | 0.36647 | 0.28 | 0.31 | 1 | HC_vs_MS.NoTrt_F |
| IL12RB1 | 0.78851 | 0.29042 | 0.89 | 0.88 | 1 | HC_vs_MS.NoTrt_F |
| NARFL | 0.78854 | 0.74337 | 0.809 | 0.83 | 1 | HC_vs_MS.NoTrt_F |
| CYB5D1 | 0.78864 | -0.822 | 0.205 | 0.21 | 1 | HC_vs_MS.NoTrt_F |
| ZEB2 | 0.78881 | 1.59743 | 0.647 | 0.61 | 1 | HC_vs_MS.NoTrt_F |
| LIMA1 | 0.78883 | 0.63679 | 0.514 | 0.48 | 1 | HC_vs_MS.NoTrt_F |
| IFI27 | 0.78966 | 0.73835 | 0.471 | 0.46 | 1 | HC_vs_MS.NoTrt_F |
| NOSTRIN | 0.79069 | 0.49521 | 0.335 | 0.35 | 1 | HC_vs_MS.NoTrt_F |
| RAB36 | 0.79077 | 1.10618 | 0.39 | 0.38 | 1 | HC_vs_MS.NoTrt_F |

| | | | | | | |
|---|---|---|---|---|---|---|
| SPRYD4 | 0.79111 | 0.54904 | 0.89 | 0.93 | 1 | HC_vs_MS.NoTrt_F |
| FAM13A | 0.79125 | 0.63641 | 0.277 | 0.28 | 1 | HC_vs_MS.NoTrt_F |
| CD55 | 0.79186 | 0.7942 | 0.633 | 0.67 | 1 | HC_vs_MS.NoTrt_F |
| IL1RAP | 0.79208 | 1.02003 | 0.659 | 0.71 | 1 | HC_vs_MS.NoTrt_F |
| EDC3 | 0.79248 | 0.35372 | 0.448 | 0.46 | 1 | HC_vs_MS.NoTrt_F |
| TRIM66 | 0.79315 | 0.3217 | 0.523 | 0.55 | 1 | HC_vs_MS.NoTrt_F |
| GPAM | 0.79324 | -0.6385 | 0.113 | 0.12 | 1 | HC_vs_MS.NoTrt_F |
| ULK2 | 0.7935 | -0.9846 | 0.725 | 0.72 | 1 | HC_vs_MS.NoTrt_F |
| NSMCE2 | 0.79352 | 0.44807 | 0.647 | 0.67 | 1 | HC_vs_MS.NoTrt_F |
| OSBPL3 | 0.79363 | 0.5179 | 0.532 | 0.57 | 1 | HC_vs_MS.NoTrt_F |
| RDH10 | 0.79373 | -0.4297 | 0.246 | 0.25 | 1 | HC_vs_MS.NoTrt_F |
| NAPG | 0.79396 | 0.34645 | 0.659 | 0.67 | 1 | HC_vs_MS.NoTrt_F |
| AC016577.1 | 0.79414 | 0.32102 | 0.436 | 0.5 | 1 | HC_vs_MS.NoTrt_F |
| DAAM2 | 0.79553 | 0.32889 | 0.468 | 0.44 | 1 | HC_vs_MS.NoTrt_F |
| TYMS | 0.79632 | 0.30804 | 0.251 | 0.26 | 1 | HC_vs_MS.NoTrt_F |
| RRM2 | 0.79673 | 0.47005 | 0.526 | 0.56 | 1 | HC_vs_MS.NoTrt_F |
| CPB1 | 0.79673 | 0.28083 | 0.569 | 0.61 | 1 | HC_vs_MS.NoTrt_F |
| APMAP | 0.79713 | 0.65234 | 0.249 | 0.26 | 1 | HC_vs_MS.NoTrt_F |
| G6PC2 | 0.79761 | 0.2732 | 0.379 | 0.41 | 1 | HC_vs_MS.NoTrt_F |
| DGKE | 0.79881 | 0.3856 | 0.74 | 0.75 | 1 | HC_vs_MS.NoTrt_F |
| GEMIN2 | 0.79982 | 0.38566 | 0.691 | 0.71 | 1 | HC_vs_MS.NoTrt_F |
| ZC3H6 | 0.80036 | -0.4717 | 0.85 | 0.79 | 1 | HC_vs_MS.NoTrt_F |
| SPEG | 0.80068 | 0.70294 | 0.303 | 0.32 | 1 | HC_vs_MS.NoTrt_F |
| C11orf70 | 0.80078 | 0.25264 | 0.266 | 0.29 | 1 | HC_vs_MS.NoTrt_F |
| ZSWIM8 | 0.8008 | 0.83497 | 0.838 | 0.86 | 1 | HC_vs_MS.NoTrt_F |
| PARP11 | 0.80101 | 0.30222 | 0.543 | 0.51 | 1 | HC_vs_MS.NoTrt_F |
| CCDC171 | 0.80214 | 0.96672 | 0.569 | 0.6 | 1 | HC_vs_MS.NoTrt_F |
| POLK | 0.80221 | 0.37304 | 0.835 | 0.87 | 1 | HC_vs_MS.NoTrt_F |
| POLR2D | 0.8027 | -0.289 | 0.789 | 0.77 | 1 | HC_vs_MS.NoTrt_F |
| CDK6 | 0.80277 | 1.44989 | 0.257 | 0.27 | 1 | HC_vs_MS.NoTrt_F |
| MFAP3 | 0.80308 | 0.2961 | 0.682 | 0.69 | 1 | HC_vs_MS.NoTrt_F |
| ARHGAP33 | 0.80329 | 0.96279 | 0.197 | 0.22 | 1 | HC_vs_MS.NoTrt_F |
| NLN | 0.80353 | -0.4906 | 0.754 | 0.71 | 1 | HC_vs_MS.NoTrt_F |
| C22orf46 | 0.80393 | 0.63235 | 0.39 | 0.38 | 1 | HC_vs_MS.NoTrt_F |
| ZNF616 | 0.8041 | 0.51127 | 0.766 | 0.83 | 1 | HC_vs_MS.NoTrt_F |
| FAM63A | 0.80441 | 0.89762 | 0.581 | 0.59 | 1 | HC_vs_MS.NoTrt_F |
| WBP2NL | 0.80488 | 0.51911 | 0.419 | 0.43 | 1 | HC_vs_MS.NoTrt_F |
| TPGS1 | 0.8052 | 1.51317 | 0.312 | 0.29 | 1 | HC_vs_MS.NoTrt_F |
| MPDU1 | 0.80626 | -0.3253 | 0.861 | 0.89 | 1 | HC_vs_MS.NoTrt_F |
| PIEZO1 | 0.80651 | 0.2793 | 0.448 | 0.43 | 1 | HC_vs_MS.NoTrt_F |

| | | | | | | |
|---|---|---|---|---|---|---|
| NBPF10 | 0.80684 | -0.4204 | 0.303 | 0.3 | 1 | HC_vs_MS.NoTrt_F |
| ASAH2B | 0.80699 | 0.32938 | 0.422 | 0.41 | 1 | HC_vs_MS.NoTrt_F |
| ZNF2 | 0.80714 | -0.764 | 0.382 | 0.38 | 1 | HC_vs_MS.NoTrt_F |
| C1QTNF2 | 0.80763 | 0.27058 | 0.344 | 0.35 | 1 | HC_vs_MS.NoTrt_F |
| PGM5 | 0.80794 | 0.41393 | 0.104 | 0.12 | 1 | HC_vs_MS.NoTrt_F |
| ANAPC5 | 0.80883 | 0.26661 | 0.962 | 0.96 | 1 | HC_vs_MS.NoTrt_F |
| RP11-327P2 | 0.80926 | 0.38556 | 0.191 | 0.2 | 1 | HC_vs_MS.NoTrt_F |
| H2AFV | 0.8094 | 0.32688 | 0.691 | 0.73 | 1 | HC_vs_MS.NoTrt_F |
| PXDN | 0.80943 | 1.1111 | 0.26 | 0.25 | 1 | HC_vs_MS.NoTrt_F |
| NR6A1 | 0.81113 | -0.9328 | 0.477 | 0.47 | 1 | HC_vs_MS.NoTrt_F |
| SRGAP3 | 0.81144 | 0.87047 | 0.879 | 0.88 | 1 | HC_vs_MS.NoTrt_F |
| GPATCH2L | 0.81161 | 0.52312 | 0.769 | 0.72 | 1 | HC_vs_MS.NoTrt_F |
| HLTF | 0.81225 | 2.33788 | 0.494 | 0.51 | 1 | HC_vs_MS.NoTrt_F |
| ARHGEF4 | 0.81228 | 0.55031 | 0.627 | 0.64 | 1 | HC_vs_MS.NoTrt_F |
| ZNF720 | 0.81263 | 0.80409 | 0.584 | 0.59 | 1 | HC_vs_MS.NoTrt_F |
| ESYT3 | 0.81321 | 0.46639 | 0.584 | 0.57 | 1 | HC_vs_MS.NoTrt_F |
| MTG2 | 0.81448 | -0.4226 | 0.847 | 0.83 | 1 | HC_vs_MS.NoTrt_F |
| LARS2 | 0.81522 | 0.62289 | 0.546 | 0.54 | 1 | HC_vs_MS.NoTrt_F |
| ATAD5 | 0.81527 | 0.28319 | 0.618 | 0.64 | 1 | HC_vs_MS.NoTrt_F |
| LTBP4 | 0.81554 | 0.53459 | 0.561 | 0.57 | 1 | HC_vs_MS.NoTrt_F |
| BRD9 | 0.81605 | 0.79591 | 0.234 | 0.26 | 1 | HC_vs_MS.NoTrt_F |
| POM121C | 0.81645 | 0.42395 | 0.497 | 0.5 | 1 | HC_vs_MS.NoTrt_F |
| C21orf58 | 0.81666 | 0.35382 | 0.532 | 0.55 | 1 | HC_vs_MS.NoTrt_F |
| ZMAT5 | 0.81751 | 0.38029 | 0.777 | 0.78 | 1 | HC_vs_MS.NoTrt_F |
| CYP51A1 | 0.81756 | 0.49122 | 0.532 | 0.54 | 1 | HC_vs_MS.NoTrt_F |
| NOX5 | 0.81768 | 0.38423 | 0.592 | 0.62 | 1 | HC_vs_MS.NoTrt_F |
| EFHD1 | 0.81781 | -0.3898 | 0.315 | 0.34 | 1 | HC_vs_MS.NoTrt_F |
| MAP3K10 | 0.81844 | 0.78453 | 0.39 | 0.38 | 1 | HC_vs_MS.NoTrt_F |
| CLTC | 0.81884 | -0.4379 | 0.483 | 0.44 | 1 | HC_vs_MS.NoTrt_F |
| ZNF530 | 0.81889 | -0.5322 | 0.509 | 0.52 | 1 | HC_vs_MS.NoTrt_F |
| ATP10D | 0.81917 | 1.29709 | 0.405 | 0.4 | 1 | HC_vs_MS.NoTrt_F |
| NAA50 | 0.81965 | -0.4099 | 0.512 | 0.53 | 1 | HC_vs_MS.NoTrt_F |
| NLRP3 | 0.8206 | 1.01756 | 0.28 | 0.29 | 1 | HC_vs_MS.NoTrt_F |
| EIF2B2 | 0.82082 | -0.4999 | 0.616 | 0.6 | 1 | HC_vs_MS.NoTrt_F |
| NOTCH2 | 0.82124 | 0.81009 | 0.309 | 0.33 | 1 | HC_vs_MS.NoTrt_F |
| NEK6 | 0.82131 | 0.64339 | 0.162 | 0.18 | 1 | HC_vs_MS.NoTrt_F |
| NCR1 | 0.82172 | 1.28906 | 0.266 | 0.28 | 1 | HC_vs_MS.NoTrt_F |
| TRIM34 | 0.82198 | 1.2162 | 0.387 | 0.37 | 1 | HC_vs_MS.NoTrt_F |
| CD109 | 0.82233 | 0.38235 | 0.477 | 0.46 | 1 | HC_vs_MS.NoTrt_F |
| CLCN7 | 0.82233 | 0.27103 | 0.387 | 0.4 | 1 | HC_vs_MS.NoTrt_F |

| | | | | | | |
|---|---|---|---|---|---|---|
| LANCL2 | 0.82261 | 0.6775 | 0.185 | 0.2 | 1 | HC_vs_MS.NoTrt_F |
| SLC25A20 | 0.82266 | 0.49075 | 0.171 | 0.18 | 1 | HC_vs_MS.NoTrt_F |
| STIL | 0.82322 | -0.4499 | 0.668 | 0.63 | 1 | HC_vs_MS.NoTrt_F |
| PRDX4 | 0.82327 | 0.3731 | 0.176 | 0.17 | 1 | HC_vs_MS.NoTrt_F |
| GINS4 | 0.82333 | 0.39203 | 0.916 | 0.93 | 1 | HC_vs_MS.NoTrt_F |
| TNXB | 0.82366 | 1.70903 | 0.168 | 0.18 | 1 | HC_vs_MS.NoTrt_F |
| TNFRSF21 | 0.82383 | 0.92969 | 0.185 | 0.19 | 1 | HC_vs_MS.NoTrt_F |
| UPF3A | 0.82403 | -0.291 | 0.775 | 0.79 | 1 | HC_vs_MS.NoTrt_F |
| KCNK10 | 0.82425 | 0.33346 | 0.173 | 0.18 | 1 | HC_vs_MS.NoTrt_F |
| SMIM11A | 0.82427 | 0.71712 | 0.295 | 0.32 | 1 | HC_vs_MS.NoTrt_F |
| CTDSPL | 0.82445 | 0.4868 | 0.335 | 0.32 | 1 | HC_vs_MS.NoTrt_F |
| FGF11 | 0.8249 | 0.31838 | 0.306 | 0.31 | 1 | HC_vs_MS.NoTrt_F |
| SIGLEC6 | 0.82493 | 0.66175 | 0.535 | 0.49 | 1 | HC_vs_MS.NoTrt_F |
| PQLC2 | 0.82531 | -0.974 | 0.454 | 0.45 | 1 | HC_vs_MS.NoTrt_F |
| C17orf62 | 0.82556 | 0.38061 | 0.731 | 0.76 | 1 | HC_vs_MS.NoTrt_F |
| GUCD1 | 0.82571 | -0.7311 | 0.512 | 0.48 | 1 | HC_vs_MS.NoTrt_F |
| OLR1 | 0.82589 | 0.31419 | 0.471 | 0.48 | 1 | HC_vs_MS.NoTrt_F |
| ARHGEF9 | 0.82596 | 0.56039 | 0.613 | 0.62 | 1 | HC_vs_MS.NoTrt_F |
| SRBD1 | 0.82637 | -0.6972 | 0.46 | 0.45 | 1 | HC_vs_MS.NoTrt_F |
| KIF3B | 0.82641 | 0.40887 | 0.5 | 0.49 | 1 | HC_vs_MS.NoTrt_F |
| ITGA2 | 0.82656 | -0.3376 | 0.477 | 0.45 | 1 | HC_vs_MS.NoTrt_F |
| SCAND1 | 0.82659 | 0.4776 | 0.272 | 0.29 | 1 | HC_vs_MS.NoTrt_F |
| ADCY4 | 0.82674 | 0.31167 | 0.15 | 0.17 | 1 | HC_vs_MS.NoTrt_F |
| TEP1 | 0.8269 | 0.77494 | 0.488 | 0.52 | 1 | HC_vs_MS.NoTrt_F |
| CNOT6L | 0.82704 | 0.49515 | 0.329 | 0.35 | 1 | HC_vs_MS.NoTrt_F |
| METTL14 | 0.82784 | 0.35426 | 0.543 | 0.51 | 1 | HC_vs_MS.NoTrt_F |
| SEMA3D | 0.82832 | 0.28405 | 0.595 | 0.56 | 1 | HC_vs_MS.NoTrt_F |
| SARDH | 0.82834 | 0.54162 | 0.486 | 0.49 | 1 | HC_vs_MS.NoTrt_F |
| MATN1 | 0.82892 | 1.28377 | 0.116 | 0.13 | 1 | HC_vs_MS.NoTrt_F |
| MYOM1 | 0.82925 | -0.3515 | 0.402 | 0.39 | 1 | HC_vs_MS.NoTrt_F |
| PHB2 | 0.82927 | 0.36337 | 0.275 | 0.26 | 1 | HC_vs_MS.NoTrt_F |
| TJP1 | 0.82933 | 1.18971 | 0.263 | 0.27 | 1 | HC_vs_MS.NoTrt_F |
| ZFPM1 | 0.82953 | 0.83602 | 0.11 | 0.12 | 1 | HC_vs_MS.NoTrt_F |
| ZNF578 | 0.82971 | 0.29196 | 0.962 | 0.97 | 1 | HC_vs_MS.NoTrt_F |
| IGSF11 | 0.83034 | 0.25033 | 0.095 | 0.11 | 1 | HC_vs_MS.NoTrt_F |
| FOXRED2 | 0.83061 | 0.36039 | 0.59 | 0.6 | 1 | HC_vs_MS.NoTrt_F |
| TMEM2 | 0.83107 | 0.39477 | 0.708 | 0.71 | 1 | HC_vs_MS.NoTrt_F |
| TP53I11 | 0.83155 | 1.29312 | 0.162 | 0.16 | 1 | HC_vs_MS.NoTrt_F |
| PRMT3 | 0.83159 | -0.3622 | 0.358 | 0.36 | 1 | HC_vs_MS.NoTrt_F |
| ZNF669 | 0.83189 | -0.3202 | 0.624 | 0.66 | 1 | HC_vs_MS.NoTrt_F |

| | | | | | | |
|---|---|---|---|---|---|---|
| LSG1 | 0.8323 | -0.4328 | 0.503 | 0.48 | 1 | HC_vs_MS.NoTrt_F |
| LTA4H | 0.83321 | 0.49792 | 0.393 | 0.4 | 1 | HC_vs_MS.NoTrt_F |
| ZNF277 | 0.83377 | -0.4056 | 0.616 | 0.61 | 1 | HC_vs_MS.NoTrt_F |
| COPB2 | 0.83377 | -0.5128 | 0.908 | 0.96 | 1 | HC_vs_MS.NoTrt_F |
| TMEM129 | 0.83385 | 0.43751 | 0.107 | 0.12 | 1 | HC_vs_MS.NoTrt_F |
| MAEL | 0.83401 | 0.56148 | 0.529 | 0.54 | 1 | HC_vs_MS.NoTrt_F |
| TMEM260 | 0.8346 | -0.5829 | 0.46 | 0.45 | 1 | HC_vs_MS.NoTrt_F |
| NQO2 | 0.83468 | -0.3926 | 0.829 | 0.8 | 1 | HC_vs_MS.NoTrt_F |
| ITGAX | 0.83534 | 1.60816 | 0.558 | 0.56 | 1 | HC_vs_MS.NoTrt_F |
| LRRC37A | 0.83569 | 0.73191 | 0.434 | 0.45 | 1 | HC_vs_MS.NoTrt_F |
| IQGAP1 | 0.83608 | 0.29314 | 0.847 | 0.87 | 1 | HC_vs_MS.NoTrt_F |
| CNDP1 | 0.8364 | 0.44424 | 0.671 | 0.69 | 1 | HC_vs_MS.NoTrt_F |
| METTL22 | 0.83727 | 0.61946 | 0.705 | 0.7 | 1 | HC_vs_MS.NoTrt_F |
| IPO9 | 0.83731 | -0.9641 | 0.876 | 0.86 | 1 | HC_vs_MS.NoTrt_F |
| IFIT1 | 0.83734 | 0.63804 | 0.312 | 0.32 | 1 | HC_vs_MS.NoTrt_F |
| IAPP | 0.83735 | 0.25106 | 0.46 | 0.45 | 1 | HC_vs_MS.NoTrt_F |
| PAX5 | 0.83748 | 0.49739 | 0.159 | 0.16 | 1 | HC_vs_MS.NoTrt_F |
| LYRM1 | 0.83793 | -0.8703 | 0.616 | 0.57 | 1 | HC_vs_MS.NoTrt_F |
| DNAJC14 | 0.83799 | -0.4747 | 0.65 | 0.62 | 1 | HC_vs_MS.NoTrt_F |
| VN1R1 | 0.83806 | 1.24335 | 0.211 | 0.2 | 1 | HC_vs_MS.NoTrt_F |
| FAM20B | 0.83832 | 0.37617 | 0.63 | 0.62 | 1 | HC_vs_MS.NoTrt_F |
| PHEX | 0.83837 | -0.2588 | 0.104 | 0.11 | 1 | HC_vs_MS.NoTrt_F |
| TMPRSS7 | 0.83847 | 0.32807 | 0.306 | 0.3 | 1 | HC_vs_MS.NoTrt_F |
| F2RL3 | 0.8388 | 0.52833 | 0.818 | 0.78 | 1 | HC_vs_MS.NoTrt_F |
| FBXL5 | 0.83909 | 0.62208 | 0.376 | 0.42 | 1 | HC_vs_MS.NoTrt_F |
| N4BP2 | 0.84025 | -0.31 | 0.451 | 0.44 | 1 | HC_vs_MS.NoTrt_F |
| RAD51C | 0.84084 | -0.2706 | 0.968 | 0.97 | 1 | HC_vs_MS.NoTrt_F |
| DENND4A | 0.84113 | 0.61167 | 0.439 | 0.46 | 1 | HC_vs_MS.NoTrt_F |
| CHD7 | 0.84118 | -0.3335 | 0.636 | 0.6 | 1 | HC_vs_MS.NoTrt_F |
| APBA1 | 0.84133 | -1.1415 | 0.451 | 0.45 | 1 | HC_vs_MS.NoTrt_F |
| CDK19 | 0.84199 | 0.3614 | 0.358 | 0.37 | 1 | HC_vs_MS.NoTrt_F |
| VPS13B | 0.84318 | 0.32291 | 0.76 | 0.77 | 1 | HC_vs_MS.NoTrt_F |
| C1orf109 | 0.84505 | -0.4178 | 0.922 | 0.86 | 1 | HC_vs_MS.NoTrt_F |
| WFDC11 | 0.84512 | 0.3437 | 0.289 | 0.3 | 1 | HC_vs_MS.NoTrt_F |
| ICAM1 | 0.84582 | 0.70379 | 0.555 | 0.61 | 1 | HC_vs_MS.NoTrt_F |
| PPWD1 | 0.84627 | 0.33636 | 0.301 | 0.33 | 1 | HC_vs_MS.NoTrt_F |
| POLA2 | 0.84695 | -0.3733 | 0.662 | 0.66 | 1 | HC_vs_MS.NoTrt_F |
| FAM210A | 0.84724 | -0.5388 | 0.39 | 0.4 | 1 | HC_vs_MS.NoTrt_F |
| DENND1A | 0.84735 | 0.49348 | 0.558 | 0.56 | 1 | HC_vs_MS.NoTrt_F |
| COX20 | 0.84737 | 0.53712 | 0.671 | 0.65 | 1 | HC_vs_MS.NoTrt_F |

| | | | | | | |
|---|---|---|---|---|---|---|
| DCAF13 | 0.84745 | 0.36596 | 0.766 | 0.72 | 1 | HC_vs_MS.NoTrt_F |
| ITGA4 | 0.84754 | 0.54484 | 0.272 | 0.3 | 1 | HC_vs_MS.NoTrt_F |
| NAA15 | 0.84822 | 0.27718 | 0.801 | 0.79 | 1 | HC_vs_MS.NoTrt_F |
| TARS2 | 0.84824 | 0.49508 | 0.809 | 0.87 | 1 | HC_vs_MS.NoTrt_F |
| RNF220 | 0.8484 | 0.74795 | 0.699 | 0.73 | 1 | HC_vs_MS.NoTrt_F |
| SPATA6 | 0.84853 | 0.57455 | 0.173 | 0.18 | 1 | HC_vs_MS.NoTrt_F |
| TGIF2 | 0.84882 | -0.5834 | 0.413 | 0.41 | 1 | HC_vs_MS.NoTrt_F |
| PLD3 | 0.84992 | 0.42015 | 0.772 | 0.82 | 1 | HC_vs_MS.NoTrt_F |
| MX1 | 0.85001 | 0.2826 | 0.786 | 0.82 | 1 | HC_vs_MS.NoTrt_F |
| MTHFR | 0.85045 | 0.32979 | 0.838 | 0.9 | 1 | HC_vs_MS.NoTrt_F |
| RBP1 | 0.85072 | 0.32207 | 0.468 | 0.48 | 1 | HC_vs_MS.NoTrt_F |
| MTF2 | 0.85091 | 0.52384 | 0.402 | 0.38 | 1 | HC_vs_MS.NoTrt_F |
| SIMC1 | 0.8521 | 0.67942 | 0.526 | 0.53 | 1 | HC_vs_MS.NoTrt_F |
| DPH5 | 0.85218 | 0.40538 | 0.361 | 0.36 | 1 | HC_vs_MS.NoTrt_F |
| CHD8 | 0.85223 | 0.71136 | 0.399 | 0.41 | 1 | HC_vs_MS.NoTrt_F |
| SLC35F6 | 0.85274 | 0.39098 | 0.85 | 0.9 | 1 | HC_vs_MS.NoTrt_F |
| SYT16 | 0.85286 | 0.45798 | 0.312 | 0.33 | 1 | HC_vs_MS.NoTrt_F |
| EEA1 | 0.85305 | -0.4912 | 0.277 | 0.28 | 1 | HC_vs_MS.NoTrt_F |
| NOTCH2NL | 0.85339 | 0.91233 | 0.292 | 0.3 | 1 | HC_vs_MS.NoTrt_F |
| RP11-176H8 | 0.854 | 0.543 | 0.887 | 0.87 | 1 | HC_vs_MS.NoTrt_F |
| HSD17B8 | 0.85402 | 0.50805 | 0.153 | 0.16 | 1 | HC_vs_MS.NoTrt_F |
| ACSM6 | 0.85479 | 0.48861 | 0.147 | 0.15 | 1 | HC_vs_MS.NoTrt_F |
| AFDN | 0.85488 | 1.47015 | 0.185 | 0.18 | 1 | HC_vs_MS.NoTrt_F |
| MTRF1 | 0.85534 | -0.4205 | 0.598 | 0.56 | 1 | HC_vs_MS.NoTrt_F |
| FBXO11 | 0.85556 | 0.42352 | 0.526 | 0.56 | 1 | HC_vs_MS.NoTrt_F |
| PLEC | 0.85587 | 0.70239 | 0.439 | 0.48 | 1 | HC_vs_MS.NoTrt_F |
| RGPD2 | 0.8561 | 1.23052 | 0.243 | 0.27 | 1 | HC_vs_MS.NoTrt_F |
| UROS | 0.85682 | 0.54929 | 0.509 | 0.5 | 1 | HC_vs_MS.NoTrt_F |
| MAST4 | 0.85685 | -0.54 | 0.676 | 0.65 | 1 | HC_vs_MS.NoTrt_F |
| NBEAL1 | 0.85688 | 0.83808 | 0.899 | 0.91 | 1 | HC_vs_MS.NoTrt_F |
| BRD8 | 0.85724 | -0.4775 | 0.786 | 0.77 | 1 | HC_vs_MS.NoTrt_F |
| NDUFAF6 | 0.85777 | -0.608 | 0.662 | 0.66 | 1 | HC_vs_MS.NoTrt_F |
| EXT1 | 0.85832 | 1.17707 | 0.199 | 0.21 | 1 | HC_vs_MS.NoTrt_F |
| CNN3 | 0.85872 | 0.45322 | 0.289 | 0.3 | 1 | HC_vs_MS.NoTrt_F |
| CSRNP1 | 0.85974 | 0.92763 | 0.327 | 0.37 | 1 | HC_vs_MS.NoTrt_F |
| RP11-347C1 | 0.86022 | 0.28374 | 0.809 | 0.83 | 1 | HC_vs_MS.NoTrt_F |
| EREG | 0.86053 | 0.54277 | 0.853 | 0.88 | 1 | HC_vs_MS.NoTrt_F |
| LIG3 | 0.86056 | 0.54692 | 0.289 | 0.28 | 1 | HC_vs_MS.NoTrt_F |
| CCR8 | 0.86133 | 0.89908 | 0.257 | 0.26 | 1 | HC_vs_MS.NoTrt_F |
| CEP68 | 0.86158 | -0.8498 | 0.598 | 0.55 | 1 | HC_vs_MS.NoTrt_F |

| | | | | | | |
|---|---|---|---|---|---|---|
| ADRB2 | 0.8619 | 0.41708 | 0.309 | 0.31 | 1 | HC_vs_MS.NoTrt_F |
| VPS33A | 0.86216 | -0.4146 | 0.876 | 0.86 | 1 | HC_vs_MS.NoTrt_F |
| ABCA6 | 0.86285 | 0.47239 | 0.685 | 0.67 | 1 | HC_vs_MS.NoTrt_F |
| TCF25 | 0.86315 | 0.3075 | 0.936 | 0.96 | 1 | HC_vs_MS.NoTrt_F |
| FPR1 | 0.86337 | 0.6673 | 0.506 | 0.53 | 1 | HC_vs_MS.NoTrt_F |
| XRCC3 | 0.8638 | -0.3355 | 0.514 | 0.5 | 1 | HC_vs_MS.NoTrt_F |
| EXOSC7 | 0.86543 | 0.30398 | 0.552 | 0.56 | 1 | HC_vs_MS.NoTrt_F |
| ZDHHC4 | 0.86571 | -0.3152 | 0.52 | 0.52 | 1 | HC_vs_MS.NoTrt_F |
| PGAP1 | 0.8658 | 0.9182 | 0.457 | 0.46 | 1 | HC_vs_MS.NoTrt_F |
| TIMM10B | 0.86601 | 0.43683 | 0.616 | 0.58 | 1 | HC_vs_MS.NoTrt_F |
| DOCK4 | 0.86602 | 0.36553 | 0.318 | 0.32 | 1 | HC_vs_MS.NoTrt_F |
| FOSL2 | 0.86658 | 0.65381 | 0.347 | 0.39 | 1 | HC_vs_MS.NoTrt_F |
| FLOT1 | 0.86666 | 0.644 | 0.223 | 0.22 | 1 | HC_vs_MS.NoTrt_F |
| ZNF836 | 0.86702 | -1.2043 | 0.251 | 0.24 | 1 | HC_vs_MS.NoTrt_F |
| USP32 | 0.8677 | 2.47248 | 0.332 | 0.37 | 1 | HC_vs_MS.NoTrt_F |
| PRKCB | 0.8683 | 0.41131 | 0.298 | 0.3 | 1 | HC_vs_MS.NoTrt_F |
| ZFYVE28 | 0.86845 | 0.83707 | 0.231 | 0.25 | 1 | HC_vs_MS.NoTrt_F |
| RIT1 | 0.86944 | 0.48892 | 0.477 | 0.5 | 1 | HC_vs_MS.NoTrt_F |
| PAQR3 | 0.86945 | 1.10743 | 0.159 | 0.17 | 1 | HC_vs_MS.NoTrt_F |
| RRP1 | 0.86985 | 0.37295 | 0.711 | 0.71 | 1 | HC_vs_MS.NoTrt_F |
| CCDC138 | 0.8701 | 0.35232 | 0.217 | 0.22 | 1 | HC_vs_MS.NoTrt_F |
| SLC22A5 | 0.87047 | 0.45778 | 0.428 | 0.42 | 1 | HC_vs_MS.NoTrt_F |
| AC138969.4 | 0.87068 | 0.65083 | 0.448 | 0.45 | 1 | HC_vs_MS.NoTrt_F |
| VMP1 | 0.87085 | 0.27502 | 0.858 | 0.88 | 1 | HC_vs_MS.NoTrt_F |
| HERC3 | 0.87142 | 0.75002 | 0.341 | 0.37 | 1 | HC_vs_MS.NoTrt_F |
| CD160 | 0.87199 | 1.28843 | 0.318 | 0.32 | 1 | HC_vs_MS.NoTrt_F |
| POU5F1B | 0.87247 | 0.2583 | 0.451 | 0.45 | 1 | HC_vs_MS.NoTrt_F |
| MCF2L | 0.87266 | 0.53572 | 0.37 | 0.35 | 1 | HC_vs_MS.NoTrt_F |
| ZNF555 | 0.87322 | -0.5513 | 0.711 | 0.71 | 1 | HC_vs_MS.NoTrt_F |
| COLGALT1 | 0.8734 | 0.31286 | 0.523 | 0.54 | 1 | HC_vs_MS.NoTrt_F |
| TXN2 | 0.87342 | 0.47501 | 0.382 | 0.38 | 1 | HC_vs_MS.NoTrt_F |
| PMM2 | 0.87393 | -0.3352 | 0.777 | 0.79 | 1 | HC_vs_MS.NoTrt_F |
| PIGH | 0.87449 | 0.35488 | 0.688 | 0.68 | 1 | HC_vs_MS.NoTrt_F |
| CIART | 0.87545 | 0.36231 | 0.558 | 0.57 | 1 | HC_vs_MS.NoTrt_F |
| PRRC2A | 0.87579 | 0.36236 | 0.809 | 0.85 | 1 | HC_vs_MS.NoTrt_F |
| ALDH8A1 | 0.87584 | 0.37848 | 0.59 | 0.61 | 1 | HC_vs_MS.NoTrt_F |
| MRM3 | 0.87601 | 0.85875 | 0.347 | 0.36 | 1 | HC_vs_MS.NoTrt_F |
| RAB40B | 0.87619 | 0.3528 | 0.601 | 0.62 | 1 | HC_vs_MS.NoTrt_F |
| MAGT1 | 0.87723 | 0.42748 | 0.867 | 0.84 | 1 | HC_vs_MS.NoTrt_F |
| SNX11 | 0.87802 | 0.82823 | 0.642 | 0.67 | 1 | HC_vs_MS.NoTrt_F |

| | | | | | | |
|---|---|---|---|---|---|---|
| PRRC2C | 0.8784 | 0.26535 | 0.847 | 0.86 | 1 | HC_vs_MS.NoTrt_F |
| PHKG1 | 0.87858 | 0.65184 | 0.373 | 0.4 | 1 | HC_vs_MS.NoTrt_F |
| C3orf33 | 0.87956 | -0.4546 | 0.532 | 0.58 | 1 | HC_vs_MS.NoTrt_F |
| PIAS3 | 0.88035 | 0.85972 | 0.249 | 0.25 | 1 | HC_vs_MS.NoTrt_F |
| NFIC | 0.88051 | 0.41771 | 0.483 | 0.49 | 1 | HC_vs_MS.NoTrt_F |
| AGRN | 0.88064 | 0.53368 | 0.416 | 0.42 | 1 | HC_vs_MS.NoTrt_F |
| ANKFY1 | 0.88067 | 0.5894 | 0.697 | 0.71 | 1 | HC_vs_MS.NoTrt_F |
| EMG1 | 0.88091 | 0.42796 | 0.225 | 0.23 | 1 | HC_vs_MS.NoTrt_F |
| AGMO | 0.88115 | 1.36365 | 0.272 | 0.27 | 1 | HC_vs_MS.NoTrt_F |
| ZNF69 | 0.88136 | 0.77832 | 0.454 | 0.44 | 1 | HC_vs_MS.NoTrt_F |
| NEGR1 | 0.88152 | 0.32251 | 0.873 | 0.89 | 1 | HC_vs_MS.NoTrt_F |
| ZCCHC11 | 0.88186 | 0.38522 | 0.945 | 0.97 | 1 | HC_vs_MS.NoTrt_F |
| ACAT1 | 0.88196 | 0.28055 | 0.688 | 0.72 | 1 | HC_vs_MS.NoTrt_F |
| SIGLEC14 | 0.88197 | 0.60008 | 0.289 | 0.3 | 1 | HC_vs_MS.NoTrt_F |
| DGAT2 | 0.88283 | 0.33009 | 0.353 | 0.37 | 1 | HC_vs_MS.NoTrt_F |
| CNOT1 | 0.88342 | 0.77016 | 0.835 | 0.85 | 1 | HC_vs_MS.NoTrt_F |
| CACNA1D | 0.88349 | 0.50732 | 0.728 | 0.74 | 1 | HC_vs_MS.NoTrt_F |
| BAG2 | 0.88364 | 0.27149 | 0.306 | 0.29 | 1 | HC_vs_MS.NoTrt_F |
| LGR6 | 0.88365 | -0.5132 | 0.168 | 0.18 | 1 | HC_vs_MS.NoTrt_F |
| TMEM219 | 0.88367 | 0.5705 | 0.327 | 0.34 | 1 | HC_vs_MS.NoTrt_F |
| MLXIPL | 0.88375 | 0.34582 | 0.494 | 0.52 | 1 | HC_vs_MS.NoTrt_F |
| ANKRD13D | 0.88393 | 0.74514 | 0.734 | 0.74 | 1 | HC_vs_MS.NoTrt_F |
| CNIH3 | 0.88394 | 0.41249 | 0.234 | 0.23 | 1 | HC_vs_MS.NoTrt_F |
| PSMB5 | 0.88451 | 0.50928 | 0.231 | 0.22 | 1 | HC_vs_MS.NoTrt_F |
| WDR83 | 0.88463 | 0.34911 | 0.607 | 0.6 | 1 | HC_vs_MS.NoTrt_F |
| PDE4B | 0.88519 | 0.77127 | 0.445 | 0.54 | 1 | HC_vs_MS.NoTrt_F |
| CCDC157 | 0.88547 | 0.31129 | 0.694 | 0.7 | 1 | HC_vs_MS.NoTrt_F |
| IFT88 | 0.88594 | 0.73366 | 0.581 | 0.58 | 1 | HC_vs_MS.NoTrt_F |
| HYAL4 | 0.88595 | 0.4286 | 0.549 | 0.56 | 1 | HC_vs_MS.NoTrt_F |
| AC024592.1 | 0.88659 | 0.90962 | 0.376 | 0.39 | 1 | HC_vs_MS.NoTrt_F |
| PEA15 | 0.8866 | 0.62047 | 0.194 | 0.19 | 1 | HC_vs_MS.NoTrt_F |
| SLC25A42 | 0.88666 | 1.03741 | 0.607 | 0.61 | 1 | HC_vs_MS.NoTrt_F |
| COX5A | 0.8874 | 0.38731 | 0.35 | 0.35 | 1 | HC_vs_MS.NoTrt_F |
| TMEM143 | 0.88743 | 0.83026 | 0.543 | 0.55 | 1 | HC_vs_MS.NoTrt_F |
| DHTKD1 | 0.8899 | 0.37579 | 0.858 | 0.85 | 1 | HC_vs_MS.NoTrt_F |
| GPR155 | 0.88994 | -0.2836 | 0.939 | 0.91 | 1 | HC_vs_MS.NoTrt_F |
| DARS | 0.88999 | 0.54918 | 0.428 | 0.44 | 1 | HC_vs_MS.NoTrt_F |
| KCNJ14 | 0.89002 | 0.3556 | 0.653 | 0.64 | 1 | HC_vs_MS.NoTrt_F |
| TMEM127 | 0.89046 | 0.45753 | 0.754 | 0.71 | 1 | HC_vs_MS.NoTrt_F |
| NR2C1 | 0.89124 | 0.56343 | 0.746 | 0.76 | 1 | HC_vs_MS.NoTrt_F |

| | | | | | | |
|---|---|---|---|---|---|---|
| PRMT1 | 0.89159 | 0.39336 | 0.832 | 0.85 | 1 | HC_vs_MS.NoTrt_F |
| C14orf79 | 0.89197 | 0.35957 | 0.639 | 0.63 | 1 | HC_vs_MS.NoTrt_F |
| POLRMT | 0.89224 | 1.2547 | 0.176 | 0.18 | 1 | HC_vs_MS.NoTrt_F |
| EMCN | 0.8928 | 0.55674 | 0.509 | 0.5 | 1 | HC_vs_MS.NoTrt_F |
| TBC1D10B | 0.89303 | 0.28026 | 0.697 | 0.73 | 1 | HC_vs_MS.NoTrt_F |
| BAG5 | 0.89323 | -0.7064 | 0.416 | 0.4 | 1 | HC_vs_MS.NoTrt_F |
| VPS25 | 0.89326 | 0.75677 | 0.338 | 0.34 | 1 | HC_vs_MS.NoTrt_F |
| FBXL20 | 0.89331 | 0.43547 | 0.951 | 0.94 | 1 | HC_vs_MS.NoTrt_F |
| PAXIP1 | 0.89419 | -0.5291 | 0.474 | 0.45 | 1 | HC_vs_MS.NoTrt_F |
| CEP135 | 0.89428 | -0.2754 | 0.824 | 0.8 | 1 | HC_vs_MS.NoTrt_F |
| MAP7D1 | 0.89444 | 0.27903 | 0.59 | 0.58 | 1 | HC_vs_MS.NoTrt_F |
| PRDM2 | 0.89519 | 0.66869 | 0.52 | 0.57 | 1 | HC_vs_MS.NoTrt_F |
| OSBPL9 | 0.89542 | -0.2974 | 0.425 | 0.41 | 1 | HC_vs_MS.NoTrt_F |
| GNAI2 | 0.89542 | 0.59897 | 0.451 | 0.45 | 1 | HC_vs_MS.NoTrt_F |
| INTS3 | 0.89559 | 0.34998 | 0.769 | 0.79 | 1 | HC_vs_MS.NoTrt_F |
| MCOLN2 | 0.89578 | 0.27641 | 0.509 | 0.51 | 1 | HC_vs_MS.NoTrt_F |
| ERV3-1 | 0.89629 | -0.3396 | 0.581 | 0.54 | 1 | HC_vs_MS.NoTrt_F |
| YTHDF3 | 0.89632 | 0.29198 | 0.754 | 0.75 | 1 | HC_vs_MS.NoTrt_F |
| STX3 | 0.89685 | 1.20927 | 0.483 | 0.49 | 1 | HC_vs_MS.NoTrt_F |
| FCRL1 | 0.89863 | 0.90713 | 0.318 | 0.34 | 1 | HC_vs_MS.NoTrt_F |
| SLC26A8 | 0.89911 | 0.35763 | 0.731 | 0.77 | 1 | HC_vs_MS.NoTrt_F |
| SATB1 | 0.8992 | 1.30702 | 0.249 | 0.25 | 1 | HC_vs_MS.NoTrt_F |
| GMPR | 0.89929 | 0.35492 | 0.39 | 0.4 | 1 | HC_vs_MS.NoTrt_F |
| C1orf216 | 0.90005 | 0.49245 | 0.789 | 0.84 | 1 | HC_vs_MS.NoTrt_F |
| MAN2A2 | 0.90032 | 1.39889 | 0.364 | 0.37 | 1 | HC_vs_MS.NoTrt_F |
| MTA3 | 0.90103 | 0.4891 | 0.697 | 0.7 | 1 | HC_vs_MS.NoTrt_F |
| DPYSL2 | 0.90108 | -0.396 | 0.52 | 0.49 | 1 | HC_vs_MS.NoTrt_F |
| DUSP6 | 0.90129 | 1.19166 | 0.147 | 0.16 | 1 | HC_vs_MS.NoTrt_F |
| UACA | 0.90141 | 0.73123 | 0.532 | 0.5 | 1 | HC_vs_MS.NoTrt_F |
| KIF9 | 0.9018 | 0.29836 | 0.173 | 0.19 | 1 | HC_vs_MS.NoTrt_F |
| IRGM | 0.90199 | 0.64854 | 0.413 | 0.43 | 1 | HC_vs_MS.NoTrt_F |
| FAM179A | 0.90206 | -0.379 | 0.838 | 0.86 | 1 | HC_vs_MS.NoTrt_F |
| POLR1A | 0.9027 | 0.56311 | 0.772 | 0.76 | 1 | HC_vs_MS.NoTrt_F |
| ZNF326 | 0.9032 | -0.2928 | 0.639 | 0.62 | 1 | HC_vs_MS.NoTrt_F |
| DEF8 | 0.90335 | -0.6076 | 0.512 | 0.53 | 1 | HC_vs_MS.NoTrt_F |
| NDST1 | 0.9038 | 0.66642 | 0.382 | 0.36 | 1 | HC_vs_MS.NoTrt_F |
| CCDC18 | 0.90383 | 0.69329 | 0.249 | 0.26 | 1 | HC_vs_MS.NoTrt_F |
| ERCC1 | 0.9039 | 0.47258 | 0.861 | 0.81 | 1 | HC_vs_MS.NoTrt_F |
| PIK3CB | 0.90433 | -0.616 | 0.517 | 0.49 | 1 | HC_vs_MS.NoTrt_F |
| RTCA | 0.90456 | 0.41661 | 0.645 | 0.64 | 1 | HC_vs_MS.NoTrt_F |

| | | | | | | |
|---|---|---|---|---|---|---|
| SAMD4A | 0.9046 | 0.52475 | 0.63 | 0.65 | 1 | HC_vs_MS.NoTrt_F |
| TTC38 | 0.90464 | 0.80032 | 0.538 | 0.52 | 1 | HC_vs_MS.NoTrt_F |
| ADM2 | 0.90467 | -0.4346 | 0.358 | 0.35 | 1 | HC_vs_MS.NoTrt_F |
| FAM60A | 0.90504 | 0.27344 | 0.535 | 0.56 | 1 | HC_vs_MS.NoTrt_F |
| SSR4 | 0.90556 | 0.55465 | 0.448 | 0.45 | 1 | HC_vs_MS.NoTrt_F |
| MTMR1 | 0.90671 | 0.90516 | 0.254 | 0.27 | 1 | HC_vs_MS.NoTrt_F |
| PTPN23 | 0.90749 | 0.45422 | 0.205 | 0.2 | 1 | HC_vs_MS.NoTrt_F |
| CXCL14 | 0.90757 | 1.64757 | 0.194 | 0.2 | 1 | HC_vs_MS.NoTrt_F |
| MECP2 | 0.90814 | 0.32012 | 0.962 | 0.97 | 1 | HC_vs_MS.NoTrt_F |
| EXOSC2 | 0.90874 | 0.32557 | 0.792 | 0.78 | 1 | HC_vs_MS.NoTrt_F |
| LINS1 | 0.90912 | 0.30766 | 0.627 | 0.69 | 1 | HC_vs_MS.NoTrt_F |
| CDS2 | 0.90955 | 0.36162 | 0.827 | 0.86 | 1 | HC_vs_MS.NoTrt_F |
| LRRC8B | 0.90959 | 0.74294 | 0.488 | 0.48 | 1 | HC_vs_MS.NoTrt_F |
| SFN | 0.90964 | 0.46742 | 0.353 | 0.35 | 1 | HC_vs_MS.NoTrt_F |
| TMEM159 | 0.9098 | 0.35615 | 0.61 | 0.59 | 1 | HC_vs_MS.NoTrt_F |
| SHPK | 0.91024 | -0.373 | 0.512 | 0.49 | 1 | HC_vs_MS.NoTrt_F |
| PURA | 0.91086 | -0.5563 | 0.136 | 0.14 | 1 | HC_vs_MS.NoTrt_F |
| SUN3 | 0.91171 | 0.33724 | 0.413 | 0.4 | 1 | HC_vs_MS.NoTrt_F |
| EPHX2 | 0.91177 | 0.91176 | 0.353 | 0.38 | 1 | HC_vs_MS.NoTrt_F |
| IPO8 | 0.91195 | 0.271 | 0.575 | 0.62 | 1 | HC_vs_MS.NoTrt_F |
| ARHGEF39 | 0.91207 | 0.30326 | 0.737 | 0.76 | 1 | HC_vs_MS.NoTrt_F |
| MMP25 | 0.91263 | 0.90739 | 0.197 | 0.21 | 1 | HC_vs_MS.NoTrt_F |
| MEIS2 | 0.9136 | 0.4025 | 0.225 | 0.23 | 1 | HC_vs_MS.NoTrt_F |
| SLC13A5 | 0.91381 | 0.33982 | 0.789 | 0.82 | 1 | HC_vs_MS.NoTrt_F |
| NDUFA5 | 0.91408 | 0.32812 | 0.659 | 0.69 | 1 | HC_vs_MS.NoTrt_F |
| STRADA | 0.9143 | 0.3354 | 0.656 | 0.62 | 1 | HC_vs_MS.NoTrt_F |
| MACF1 | 0.9145 | 0.35898 | 0.91 | 0.92 | 1 | HC_vs_MS.NoTrt_F |
| MAPK10 | 0.9146 | 0.52763 | 0.332 | 0.33 | 1 | HC_vs_MS.NoTrt_F |
| USP9X | 0.91483 | 0.27251 | 0.645 | 0.65 | 1 | HC_vs_MS.NoTrt_F |
| KIF18A | 0.91494 | 0.35272 | 0.405 | 0.42 | 1 | HC_vs_MS.NoTrt_F |
| ZKSCAN5 | 0.91543 | -0.5571 | 0.691 | 0.67 | 1 | HC_vs_MS.NoTrt_F |
| NOS3 | 0.91548 | -0.4746 | 0.445 | 0.43 | 1 | HC_vs_MS.NoTrt_F |
| ETF1 | 0.91597 | 0.30757 | 0.448 | 0.48 | 1 | HC_vs_MS.NoTrt_F |
| SLC38A2 | 0.9163 | 0.59551 | 0.425 | 0.44 | 1 | HC_vs_MS.NoTrt_F |
| ZNF141 | 0.91769 | -0.2651 | 0.971 | 0.97 | 1 | HC_vs_MS.NoTrt_F |
| UGT1A1 | 0.91802 | 0.32764 | 0.569 | 0.61 | 1 | HC_vs_MS.NoTrt_F |
| LATS1 | 0.91815 | 0.37189 | 0.717 | 0.72 | 1 | HC_vs_MS.NoTrt_F |
| ZBTB16 | 0.91848 | 1.78046 | 0.442 | 0.44 | 1 | HC_vs_MS.NoTrt_F |
| USP37 | 0.91916 | -0.3809 | 0.535 | 0.51 | 1 | HC_vs_MS.NoTrt_F |
| ICE2 | 0.91987 | -0.7083 | 0.882 | 0.85 | 1 | HC_vs_MS.NoTrt_F |

| | | | | | | |
|---|---|---|---|---|---|---|
| SKIV2L2 | 0.92072 | 0.36992 | 0.887 | 0.9 | 1 | HC_vs_MS.NoTrt_F |
| CMPK1 | 0.92092 | 0.35295 | 0.361 | 0.37 | 1 | HC_vs_MS.NoTrt_F |
| BMP8B | 0.92168 | 0.2658 | 0.514 | 0.5 | 1 | HC_vs_MS.NoTrt_F |
| ZNF699 | 0.92192 | 0.68382 | 0.353 | 0.36 | 1 | HC_vs_MS.NoTrt_F |
| APPBP2 | 0.92203 | -0.6992 | 0.873 | 0.85 | 1 | HC_vs_MS.NoTrt_F |
| AKR1A1 | 0.92232 | 0.39656 | 0.757 | 0.86 | 1 | HC_vs_MS.NoTrt_F |
| ALG13 | 0.92256 | 0.29711 | 0.41 | 0.43 | 1 | HC_vs_MS.NoTrt_F |
| PDE5A | 0.92383 | -0.3328 | 0.153 | 0.15 | 1 | HC_vs_MS.NoTrt_F |
| INPP5F | 0.92444 | 0.79454 | 0.35 | 0.38 | 1 | HC_vs_MS.NoTrt_F |
| FUZ | 0.92568 | 0.58944 | 0.286 | 0.3 | 1 | HC_vs_MS.NoTrt_F |
| ENAH | 0.92591 | 0.50145 | 0.723 | 0.76 | 1 | HC_vs_MS.NoTrt_F |
| CCHCR1 | 0.92631 | 0.57004 | 0.376 | 0.37 | 1 | HC_vs_MS.NoTrt_F |
| ERBB2 | 0.92632 | 0.26823 | 0.847 | 0.86 | 1 | HC_vs_MS.NoTrt_F |
| DAGLB | 0.92657 | -0.2852 | 0.861 | 0.88 | 1 | HC_vs_MS.NoTrt_F |
| TMTC3 | 0.9269 | 0.82857 | 0.269 | 0.28 | 1 | HC_vs_MS.NoTrt_F |
| SLC38A9 | 0.92699 | 1.07859 | 0.434 | 0.45 | 1 | HC_vs_MS.NoTrt_F |
| TXK | 0.9272 | 0.7495 | 0.621 | 0.64 | 1 | HC_vs_MS.NoTrt_F |
| NIPBL | 0.92774 | 0.5624 | 0.624 | 0.63 | 1 | HC_vs_MS.NoTrt_F |
| GUF1 | 0.92784 | 0.8968 | 0.448 | 0.45 | 1 | HC_vs_MS.NoTrt_F |
| WEE1 | 0.92857 | -0.4456 | 0.769 | 0.76 | 1 | HC_vs_MS.NoTrt_F |
| CTB-50L17.1 | 0.92857 | -0.2566 | 0.65 | 0.64 | 1 | HC_vs_MS.NoTrt_F |
| ZFP30 | 0.92896 | -0.4991 | 0.844 | 0.91 | 1 | HC_vs_MS.NoTrt_F |
| VSTM4 | 0.92987 | 0.95633 | 0.777 | 0.76 | 1 | HC_vs_MS.NoTrt_F |
| TNFRSF18 | 0.93024 | 1.3483 | 0.289 | 0.33 | 1 | HC_vs_MS.NoTrt_F |
| BAZ1A | 0.9307 | 0.37251 | 0.483 | 0.49 | 1 | HC_vs_MS.NoTrt_F |
| SKP1 | 0.93123 | 0.34967 | 0.853 | 0.86 | 1 | HC_vs_MS.NoTrt_F |
| TCEAL1 | 0.93142 | 0.86919 | 0.121 | 0.13 | 1 | HC_vs_MS.NoTrt_F |
| KLF7 | 0.93193 | 0.44521 | 0.266 | 0.25 | 1 | HC_vs_MS.NoTrt_F |
| RPH3AL | 0.93205 | 0.66122 | 0.393 | 0.38 | 1 | HC_vs_MS.NoTrt_F |
| BPTF | 0.9325 | 0.34712 | 0.832 | 0.83 | 1 | HC_vs_MS.NoTrt_F |
| ZNF658 | 0.93266 | 1.22326 | 0.22 | 0.22 | 1 | HC_vs_MS.NoTrt_F |
| NEIL1 | 0.93292 | 0.53024 | 0.824 | 0.81 | 1 | HC_vs_MS.NoTrt_F |
| TEX10 | 0.93309 | 0.2915 | 0.526 | 0.52 | 1 | HC_vs_MS.NoTrt_F |
| PYCRL | 0.93407 | 0.89421 | 0.465 | 0.46 | 1 | HC_vs_MS.NoTrt_F |
| DBNDD1 | 0.93457 | 0.99572 | 0.182 | 0.19 | 1 | HC_vs_MS.NoTrt_F |
| BCL2L11 | 0.93462 | 0.34929 | 0.549 | 0.55 | 1 | HC_vs_MS.NoTrt_F |
| FCER1G | 0.93475 | 1.17028 | 0.671 | 0.72 | 1 | HC_vs_MS.NoTrt_F |
| TINAGL1 | 0.93548 | 0.32323 | 0.54 | 0.55 | 1 | HC_vs_MS.NoTrt_F |
| SPDYE2B | 0.93587 | 0.857 | 0.37 | 0.4 | 1 | HC_vs_MS.NoTrt_F |
| MTFR1 | 0.93644 | 0.41109 | 0.208 | 0.21 | 1 | HC_vs_MS.NoTrt_F |

| | | | | | | |
|---|---|---|---|---|---|---|
| CTSB | 0.93682 | -0.2718 | 0.974 | 0.96 | 1 | HC_vs_MS.NoTrt_F |
| EML5 | 0.93709 | 0.46651 | 0.243 | 0.24 | 1 | HC_vs_MS.NoTrt_F |
| SYNE4 | 0.93749 | 0.32609 | 0.517 | 0.53 | 1 | HC_vs_MS.NoTrt_F |
| PLAGL1 | 0.93768 | 0.4247 | 0.13 | 0.14 | 1 | HC_vs_MS.NoTrt_F |
| UPB1 | 0.93789 | 0.32208 | 0.624 | 0.66 | 1 | HC_vs_MS.NoTrt_F |
| FLVCR1 | 0.93806 | 0.78166 | 0.702 | 0.7 | 1 | HC_vs_MS.NoTrt_F |
| ULBP3 | 0.9383 | 0.27599 | 0.647 | 0.6 | 1 | HC_vs_MS.NoTrt_F |
| HSD3B7 | 0.93902 | 0.33911 | 0.506 | 0.51 | 1 | HC_vs_MS.NoTrt_F |
| CALD1 | 0.93903 | 1.90197 | 0.327 | 0.32 | 1 | HC_vs_MS.NoTrt_F |
| SCNM1 | 0.94033 | -0.3169 | 0.824 | 0.78 | 1 | HC_vs_MS.NoTrt_F |
| ZSCAN18 | 0.94034 | 0.44274 | 0.731 | 0.69 | 1 | HC_vs_MS.NoTrt_F |
| PPAT | 0.94056 | 0.91526 | 0.344 | 0.34 | 1 | HC_vs_MS.NoTrt_F |
| TADA2B | 0.94071 | 1.91342 | 0.361 | 0.36 | 1 | HC_vs_MS.NoTrt_F |
| EPB41L2 | 0.94102 | -0.7496 | 0.214 | 0.22 | 1 | HC_vs_MS.NoTrt_F |
| FSD1L | 0.94145 | 0.41917 | 0.28 | 0.27 | 1 | HC_vs_MS.NoTrt_F |
| ZNF785 | 0.94276 | -0.2738 | 0.916 | 0.93 | 1 | HC_vs_MS.NoTrt_F |
| LTBP3 | 0.9438 | 0.70163 | 0.757 | 0.76 | 1 | HC_vs_MS.NoTrt_F |
| NDUFA10 | 0.94394 | 0.43048 | 0.39 | 0.39 | 1 | HC_vs_MS.NoTrt_F |
| EMC3 | 0.94396 | -0.2601 | 0.928 | 0.91 | 1 | HC_vs_MS.NoTrt_F |
| PECR | 0.94485 | 0.46491 | 0.751 | 0.72 | 1 | HC_vs_MS.NoTrt_F |
| TBL3 | 0.94517 | 0.55996 | 0.344 | 0.34 | 1 | HC_vs_MS.NoTrt_F |
| MMRN2 | 0.9452 | 0.29218 | 0.249 | 0.26 | 1 | HC_vs_MS.NoTrt_F |
| ESYT2 | 0.9454 | 0.56722 | 0.668 | 0.71 | 1 | HC_vs_MS.NoTrt_F |
| GOLGA8A | 0.9456 | 0.74203 | 0.509 | 0.59 | 1 | HC_vs_MS.NoTrt_F |
| BTBD8 | 0.94626 | 0.3061 | 0.361 | 0.36 | 1 | HC_vs_MS.NoTrt_F |
| EXOC5 | 0.94704 | 0.68885 | 0.454 | 0.47 | 1 | HC_vs_MS.NoTrt_F |
| TMEM255B | 0.94711 | 0.87811 | 0.243 | 0.25 | 1 | HC_vs_MS.NoTrt_F |
| CARS2 | 0.94775 | 0.29956 | 0.78 | 0.77 | 1 | HC_vs_MS.NoTrt_F |
| CYB561D1 | 0.94837 | 0.55418 | 0.5 | 0.48 | 1 | HC_vs_MS.NoTrt_F |
| PCBP3 | 0.9488 | 0.3249 | 0.506 | 0.53 | 1 | HC_vs_MS.NoTrt_F |
| EXD3 | 0.94926 | 0.43275 | 0.569 | 0.56 | 1 | HC_vs_MS.NoTrt_F |
| RFX5 | 0.95044 | 0.40136 | 0.393 | 0.38 | 1 | HC_vs_MS.NoTrt_F |
| OTULIN | 0.95059 | 0.42763 | 0.497 | 0.47 | 1 | HC_vs_MS.NoTrt_F |
| MFSD11 | 0.95059 | 0.54732 | 0.497 | 0.5 | 1 | HC_vs_MS.NoTrt_F |
| MCMDC2 | 0.95071 | 0.83692 | 0.488 | 0.5 | 1 | HC_vs_MS.NoTrt_F |
| NEURL1 | 0.9509 | -0.4772 | 0.303 | 0.29 | 1 | HC_vs_MS.NoTrt_F |
| FER | 0.95149 | -0.2501 | 0.575 | 0.59 | 1 | HC_vs_MS.NoTrt_F |
| PLEKHG5 | 0.9516 | 0.51007 | 0.182 | 0.19 | 1 | HC_vs_MS.NoTrt_F |
| APBB2 | 0.95168 | 0.87622 | 0.376 | 0.39 | 1 | HC_vs_MS.NoTrt_F |
| CGRRF1 | 0.95179 | 1.22639 | 0.332 | 0.33 | 1 | HC_vs_MS.NoTrt_F |

| | | | | | | |
|---|---|---|---|---|---|---|
| ZNF841 | 0.95234 | 1.05151 | 0.538 | 0.54 | 1 | HC_vs_MS.NoTrt_F |
| GBA2 | 0.95269 | 0.29334 | 0.656 | 0.61 | 1 | HC_vs_MS.NoTrt_F |
| TMEM108 | 0.9529 | 0.70323 | 0.367 | 0.37 | 1 | HC_vs_MS.NoTrt_F |
| SLC38A10 | 0.95293 | 0.37948 | 0.448 | 0.43 | 1 | HC_vs_MS.NoTrt_F |
| PRRG1 | 0.953 | 0.40072 | 0.251 | 0.25 | 1 | HC_vs_MS.NoTrt_F |
| ZNF799 | 0.95366 | 0.57886 | 0.471 | 0.48 | 1 | HC_vs_MS.NoTrt_F |
| XPR1 | 0.95388 | 0.26369 | 0.445 | 0.45 | 1 | HC_vs_MS.NoTrt_F |
| PHKB | 0.95478 | -0.5855 | 0.777 | 0.77 | 1 | HC_vs_MS.NoTrt_F |
| ATP1A3 | 0.95479 | -0.4624 | 0.405 | 0.4 | 1 | HC_vs_MS.NoTrt_F |
| ZNF562 | 0.95483 | -0.7213 | 0.977 | 0.97 | 1 | HC_vs_MS.NoTrt_F |
| RP11-849H4 | 0.95497 | 0.47895 | 0.587 | 0.58 | 1 | HC_vs_MS.NoTrt_F |
| NOM1 | 0.95536 | -0.5249 | 0.714 | 0.69 | 1 | HC_vs_MS.NoTrt_F |
| PTRF | 0.95539 | 1.15998 | 0.474 | 0.46 | 1 | HC_vs_MS.NoTrt_F |
| ACSS2 | 0.95569 | 1.12944 | 0.572 | 0.55 | 1 | HC_vs_MS.NoTrt_F |
| SH3GLB1 | 0.95592 | 0.36579 | 0.555 | 0.56 | 1 | HC_vs_MS.NoTrt_F |
| MRPS23 | 0.95684 | -0.5601 | 0.858 | 0.84 | 1 | HC_vs_MS.NoTrt_F |
| NUDT3 | 0.95741 | 0.35561 | 0.584 | 0.6 | 1 | HC_vs_MS.NoTrt_F |
| NFKB1 | 0.95767 | 1.51975 | 0.15 | 0.16 | 1 | HC_vs_MS.NoTrt_F |
| ATP2A2 | 0.95822 | 0.69441 | 0.749 | 0.72 | 1 | HC_vs_MS.NoTrt_F |
| RNF19A | 0.95856 | 0.57728 | 0.506 | 0.56 | 1 | HC_vs_MS.NoTrt_F |
| RASGRF2 | 0.95889 | 1.32876 | 0.228 | 0.23 | 1 | HC_vs_MS.NoTrt_F |
| SPINK9 | 0.9589 | 0.66753 | 0.688 | 0.76 | 1 | HC_vs_MS.NoTrt_F |
| ZNF140 | 0.95896 | -0.6102 | 0.223 | 0.23 | 1 | HC_vs_MS.NoTrt_F |
| NOP2 | 0.95897 | -0.4124 | 0.474 | 0.45 | 1 | HC_vs_MS.NoTrt_F |
| SHQ1 | 0.95929 | -0.5941 | 0.754 | 0.76 | 1 | HC_vs_MS.NoTrt_F |
| KLLN | 0.96123 | 0.45362 | 0.497 | 0.52 | 1 | HC_vs_MS.NoTrt_F |
| NSL1 | 0.96128 | 0.34118 | 0.821 | 0.78 | 1 | HC_vs_MS.NoTrt_F |
| NOTCH3 | 0.96137 | 0.30932 | 0.616 | 0.63 | 1 | HC_vs_MS.NoTrt_F |
| KCNN2 | 0.96139 | 0.36688 | 0.309 | 0.32 | 1 | HC_vs_MS.NoTrt_F |
| DDB1 | 0.962 | -0.4132 | 0.922 | 0.88 | 1 | HC_vs_MS.NoTrt_F |
| KANK3 | 0.9628 | 0.51756 | 0.277 | 0.28 | 1 | HC_vs_MS.NoTrt_F |
| RALGPS1 | 0.96289 | 0.87281 | 0.477 | 0.45 | 1 | HC_vs_MS.NoTrt_F |
| C5 | 0.96321 | -0.9637 | 0.465 | 0.48 | 1 | HC_vs_MS.NoTrt_F |
| 4-Sep | 0.96347 | 0.8641 | 0.529 | 0.5 | 1 | HC_vs_MS.NoTrt_F |
| NAE1 | 0.96526 | -0.5167 | 0.737 | 0.73 | 1 | HC_vs_MS.NoTrt_F |
| ST8SIA1 | 0.96547 | 1.00792 | 0.647 | 0.64 | 1 | HC_vs_MS.NoTrt_F |
| THOC2 | 0.96547 | 0.52884 | 0.601 | 0.61 | 1 | HC_vs_MS.NoTrt_F |
| ARFRP1 | 0.96573 | 0.41155 | 0.509 | 0.53 | 1 | HC_vs_MS.NoTrt_F |
| IFFO1 | 0.96629 | 0.76782 | 0.474 | 0.46 | 1 | HC_vs_MS.NoTrt_F |
| CTD-2521M1 | 0.96675 | 0.3167 | 0.37 | 0.39 | 1 | HC_vs_MS.NoTrt_F |

| | | | | | | |
|---|---|---|---|---|---|---|
| NME9 | 0.96712 | 0.77475 | 0.275 | 0.29 | 1 | HC_vs_MS.NoTrt_F |
| RNF152 | 0.96733 | 0.50397 | 0.509 | 0.51 | 1 | HC_vs_MS.NoTrt_F |
| SOS2 | 0.96738 | 0.64529 | 0.136 | 0.14 | 1 | HC_vs_MS.NoTrt_F |
| CLDN12 | 0.9675 | 0.30554 | 0.471 | 0.5 | 1 | HC_vs_MS.NoTrt_F |
| SLC12A4 | 0.96764 | 0.89912 | 0.286 | 0.28 | 1 | HC_vs_MS.NoTrt_F |
| WDR13 | 0.96799 | 0.47552 | 0.884 | 0.87 | 1 | HC_vs_MS.NoTrt_F |
| NATD1 | 0.9687 | 0.25917 | 0.561 | 0.54 | 1 | HC_vs_MS.NoTrt_F |
| TMEM64 | 0.96872 | 0.50672 | 0.428 | 0.46 | 1 | HC_vs_MS.NoTrt_F |
| HIVEP1 | 0.96923 | 0.61402 | 0.665 | 0.74 | 1 | HC_vs_MS.NoTrt_F |
| SLC10A3 | 0.96944 | 0.38161 | 0.159 | 0.16 | 1 | HC_vs_MS.NoTrt_F |
| ZNF577 | 0.96945 | 0.6613 | 0.428 | 0.42 | 1 | HC_vs_MS.NoTrt_F |
| ZNF532 | 0.96946 | -0.3061 | 0.942 | 0.97 | 1 | HC_vs_MS.NoTrt_F |
| NDUFA9 | 0.97042 | -0.3492 | 0.905 | 0.9 | 1 | HC_vs_MS.NoTrt_F |
| CDK17 | 0.97111 | 0.83819 | 0.289 | 0.31 | 1 | HC_vs_MS.NoTrt_F |
| CHD3 | 0.97123 | 0.44072 | 0.373 | 0.38 | 1 | HC_vs_MS.NoTrt_F |
| GIPC1 | 0.97133 | 0.47175 | 0.552 | 0.59 | 1 | HC_vs_MS.NoTrt_F |
| KNL1 | 0.97166 | 0.92303 | 0.451 | 0.43 | 1 | HC_vs_MS.NoTrt_F |
| FOXJ3 | 0.97179 | 0.56358 | 0.627 | 0.61 | 1 | HC_vs_MS.NoTrt_F |
| SYNGR2 | 0.97196 | 0.37664 | 0.601 | 0.62 | 1 | HC_vs_MS.NoTrt_F |
| ALG8 | 0.97203 | -0.5613 | 0.543 | 0.51 | 1 | HC_vs_MS.NoTrt_F |
| EXOC8 | 0.97271 | 0.36898 | 0.46 | 0.46 | 1 | HC_vs_MS.NoTrt_F |
| DNAJA1 | 0.97349 | 0.34928 | 0.494 | 0.5 | 1 | HC_vs_MS.NoTrt_F |
| CTC-487M23 | 0.97355 | 0.32049 | 0.91 | 0.95 | 1 | HC_vs_MS.NoTrt_F |
| BCAR3 | 0.974 | 0.5842 | 0.676 | 0.66 | 1 | HC_vs_MS.NoTrt_F |
| TRAPPC3L | 0.97427 | 0.25846 | 0.616 | 0.64 | 1 | HC_vs_MS.NoTrt_F |
| HPS1 | 0.97474 | 0.32894 | 0.668 | 0.67 | 1 | HC_vs_MS.NoTrt_F |
| RABGGTB | 0.97508 | 0.32277 | 0.373 | 0.38 | 1 | HC_vs_MS.NoTrt_F |
| DDIAS | 0.97519 | 0.33439 | 0.46 | 0.44 | 1 | HC_vs_MS.NoTrt_F |
| BTBD2 | 0.97576 | 0.91485 | 0.431 | 0.44 | 1 | HC_vs_MS.NoTrt_F |
| MTHFD2L | 0.9761 | 0.62642 | 0.535 | 0.54 | 1 | HC_vs_MS.NoTrt_F |
| LYST | 0.97622 | 0.66868 | 0.422 | 0.43 | 1 | HC_vs_MS.NoTrt_F |
| COQ7 | 0.97758 | 0.41102 | 0.434 | 0.43 | 1 | HC_vs_MS.NoTrt_F |
| SPCS1 | 0.97763 | 0.25041 | 0.899 | 0.93 | 1 | HC_vs_MS.NoTrt_F |
| SLC39A3 | 0.97766 | 0.35366 | 0.65 | 0.68 | 1 | HC_vs_MS.NoTrt_F |
| FAM120C | 0.97784 | 0.4203 | 0.298 | 0.32 | 1 | HC_vs_MS.NoTrt_F |
| WDR26 | 0.9781 | 0.98386 | 0.717 | 0.72 | 1 | HC_vs_MS.NoTrt_F |
| PTPRCAP | 0.9791 | 0.26818 | 0.668 | 0.82 | 1 | HC_vs_MS.NoTrt_F |
| ZNF212 | 0.97936 | -1.1865 | 0.506 | 0.48 | 1 | HC_vs_MS.NoTrt_F |
| ZNF483 | 0.97953 | -0.324 | 0.882 | 0.87 | 1 | HC_vs_MS.NoTrt_F |
| IL23R | 0.97953 | -1.1608 | 0.422 | 0.4 | 1 | HC_vs_MS.NoTrt_F |

| | | | | | | |
|---|---|---|---|---|---|---|
| GTF2A2 | 0.98121 | -0.7015 | 0.633 | 0.62 | 1 | HC_vs_MS.NoTrt_F |
| PGD | 0.98145 | 0.32409 | 0.194 | 0.19 | 1 | HC_vs_MS.NoTrt_F |
| CEP95 | 0.98233 | 0.33782 | 0.419 | 0.43 | 1 | HC_vs_MS.NoTrt_F |
| FAM159A | 0.98245 | 0.38555 | 0.63 | 0.63 | 1 | HC_vs_MS.NoTrt_F |
| POFUT1 | 0.98311 | 0.49701 | 0.616 | 0.6 | 1 | HC_vs_MS.NoTrt_F |
| NFAT5 | 0.98339 | 0.25976 | 0.893 | 0.89 | 1 | HC_vs_MS.NoTrt_F |
| ITGA10 | 0.9838 | -0.2812 | 0.543 | 0.55 | 1 | HC_vs_MS.NoTrt_F |
| ENOSF1 | 0.98411 | 1.09111 | 0.879 | 0.9 | 1 | HC_vs_MS.NoTrt_F |
| BPNT1 | 0.98469 | -0.3082 | 0.775 | 0.7 | 1 | HC_vs_MS.NoTrt_F |
| MEIOC | 0.98481 | -0.3349 | 0.699 | 0.69 | 1 | HC_vs_MS.NoTrt_F |
| TMEM191C | 0.98524 | 0.30522 | 0.772 | 0.83 | 1 | HC_vs_MS.NoTrt_F |
| SGPP2 | 0.9856 | 0.71099 | 0.399 | 0.41 | 1 | HC_vs_MS.NoTrt_F |
| ITM2A | 0.98623 | 0.5776 | 0.347 | 0.37 | 1 | HC_vs_MS.NoTrt_F |
| RABL2A | 0.98653 | -0.2568 | 0.601 | 0.56 | 1 | HC_vs_MS.NoTrt_F |
| SYTL3 | 0.98687 | 0.33331 | 0.384 | 0.4 | 1 | HC_vs_MS.NoTrt_F |
| PRR27 | 0.98698 | 0.26038 | 0.517 | 0.53 | 1 | HC_vs_MS.NoTrt_F |
| MAP7D3 | 0.98722 | 0.48133 | 0.662 | 0.65 | 1 | HC_vs_MS.NoTrt_F |
| NPAS3 | 0.98759 | 0.32646 | 0.448 | 0.44 | 1 | HC_vs_MS.NoTrt_F |
| SLC27A1 | 0.98796 | 0.58081 | 0.87 | 0.87 | 1 | HC_vs_MS.NoTrt_F |
| WASH1 | 0.98864 | 0.75029 | 0.266 | 0.28 | 1 | HC_vs_MS.NoTrt_F |
| TBC1D31 | 0.98948 | -0.4718 | 0.685 | 0.68 | 1 | HC_vs_MS.NoTrt_F |
| SLC50A1 | 0.98986 | 0.32068 | 0.803 | 0.8 | 1 | HC_vs_MS.NoTrt_F |
| HPGD | 0.98997 | 0.74699 | 0.275 | 0.29 | 1 | HC_vs_MS.NoTrt_F |
| SYK | 0.99017 | 0.47526 | 0.431 | 0.45 | 1 | HC_vs_MS.NoTrt_F |
| CAMTA2 | 0.99041 | 0.26437 | 0.549 | 0.55 | 1 | HC_vs_MS.NoTrt_F |
| ZNF334 | 0.99055 | 0.76415 | 0.142 | 0.15 | 1 | HC_vs_MS.NoTrt_F |
| KLF11 | 0.99063 | 0.70645 | 0.107 | 0.11 | 1 | HC_vs_MS.NoTrt_F |
| MOB3B | 0.99097 | 0.34668 | 0.575 | 0.57 | 1 | HC_vs_MS.NoTrt_F |
| RNF41 | 0.99106 | -0.5978 | 0.801 | 0.82 | 1 | HC_vs_MS.NoTrt_F |
| ITIH5 | 0.99106 | 0.37022 | 0.818 | 0.85 | 1 | HC_vs_MS.NoTrt_F |
| ZNF91 | 0.99203 | 0.7552 | 0.806 | 0.82 | 1 | HC_vs_MS.NoTrt_F |
| POLR2A | 0.99205 | 0.45379 | 0.844 | 0.85 | 1 | HC_vs_MS.NoTrt_F |
| DDI2 | 0.99226 | 0.38669 | 0.812 | 0.81 | 1 | HC_vs_MS.NoTrt_F |
| SLC5A5 | 0.99232 | -0.3016 | 0.517 | 0.5 | 1 | HC_vs_MS.NoTrt_F |
| COX15 | 0.99234 | -0.5208 | 0.639 | 0.62 | 1 | HC_vs_MS.NoTrt_F |
| JAKMIP2 | 0.99307 | 0.73338 | 0.512 | 0.51 | 1 | HC_vs_MS.NoTrt_F |
| C20orf196 | 0.99336 | 0.55655 | 0.373 | 0.38 | 1 | HC_vs_MS.NoTrt_F |
| YEATS2 | 0.9939 | 0.30609 | 0.728 | 0.7 | 1 | HC_vs_MS.NoTrt_F |
| SPPL2A | 0.99399 | 0.28183 | 0.951 | 0.95 | 1 | HC_vs_MS.NoTrt_F |
| TARBP1 | 0.99403 | 0.58204 | 0.477 | 0.49 | 1 | HC_vs_MS.NoTrt_F |

| | | | | | | |
|---|---|---|---|---|---|---|
| DHX29 | 0.99463 | 0.49575 | 0.249 | 0.25 | 1 | HC_vs_MS.NoTrt_F |
| AP4E1 | 0.99505 | -0.4915 | 0.486 | 0.46 | 1 | HC_vs_MS.NoTrt_F |
| BTBD9 | 0.99528 | 1.22913 | 0.621 | 0.63 | 1 | HC_vs_MS.NoTrt_F |
| DCUN1D5 | 0.99582 | -0.3448 | 0.483 | 0.45 | 1 | HC_vs_MS.NoTrt_F |
| KPNA4 | 0.99583 | 0.52989 | 0.659 | 0.69 | 1 | HC_vs_MS.NoTrt_F |
| WSB1 | 0.99639 | 0.25443 | 0.899 | 0.92 | 1 | HC_vs_MS.NoTrt_F |
| SCAF4 | 0.99667 | 0.3048 | 0.28 | 0.29 | 1 | HC_vs_MS.NoTrt_F |
| PRICKLE3 | 0.99721 | 1.03219 | 0.555 | 0.53 | 1 | HC_vs_MS.NoTrt_F |
| HYLS1 | 0.99737 | 0.70296 | 0.454 | 0.45 | 1 | HC_vs_MS.NoTrt_F |
| CAGE1 | 0.99751 | 0.2503 | 0.373 | 0.38 | 1 | HC_vs_MS.NoTrt_F |
| SH3RF2 | 0.99754 | 1.26477 | 0.191 | 0.19 | 1 | HC_vs_MS.NoTrt_F |
| ZNF233 | 0.99768 | 0.30856 | 0.491 | 0.5 | 1 | HC_vs_MS.NoTrt_F |
| COQ8A | 0.9978 | 0.30176 | 0.705 | 0.7 | 1 | HC_vs_MS.NoTrt_F |
| OAS3 | 0.99804 | 0.48545 | 0.665 | 0.69 | 1 | HC_vs_MS.NoTrt_F |
| PPP1R12B | 0.99855 | 1.21116 | 0.803 | 0.8 | 1 | HC_vs_MS.NoTrt_F |
| RFX1 | 0.9989 | 0.92149 | 0.379 | 0.38 | 1 | HC_vs_MS.NoTrt_F |
| FBN1 | 0.99898 | 0.45433 | 0.514 | 0.53 | 1 | HC_vs_MS.NoTrt_F |
| DOCK5 | 0.99901 | 0.47928 | 0.587 | 0.63 | 1 | HC_vs_MS.NoTrt_F |
| BRD4 | 1 | 0.69096 | 0.353 | 0.37 | 1 | HC_vs_MS.NoTrt_F |
| FADS1 | 1 | 0.57275 | 0.647 | 0.69 | 1 | HC_vs_MS.NoTrt_F |
| RBBP9 | 1 | 0.46167 | 0.486 | 0.5 | 1 | HC_vs_MS.NoTrt_F |
| FAM135A | 1 | -0.9954 | 0.471 | 0.46 | 1 | HC_vs_MS.NoTrt_F |
| HLA-B | 1.14E-36 | -0.8265 | 0.906 | 0.98 | 1.82E-32 | HC_vs_MS.NoTrt_B |
| RPS4Y1 | 2.18E-25 | -2.5784 | 0.072 | 0.35 | 3.48E-21 | HC_vs_MS.NoTrt_B |
| FGF23 | 5.33E-24 | 2.1637 | 0.828 | 0.54 | 8.52E-20 | HC_vs_MS.NoTrt_B |
| TMSB10 | 3.97E-19 | -0.6517 | 0.913 | 1 | 6.34E-15 | HC_vs_MS.NoTrt_B |
| RPL13A | 1.02E-18 | -0.5325 | 0.904 | 1 | 1.63E-14 | HC_vs_MS.NoTrt_B |
| ARRDC3 | 1.17E-18 | -0.5935 | 0.464 | 0.84 | 1.87E-14 | HC_vs_MS.NoTrt_B |
| IL32 | 7.12E-16 | -0.5172 | 0.851 | 0.97 | 1.14E-11 | HC_vs_MS.NoTrt_B |
| TMSB4X | 1.23E-15 | -0.4266 | 0.993 | 1 | 1.97E-11 | HC_vs_MS.NoTrt_B |
| S100A4 | 1.40E-15 | -0.7662 | 0.491 | 0.8 | 2.24E-11 | HC_vs_MS.NoTrt_B |
| MIPOL1 | 2.42E-15 | 0.57393 | 0.997 | 0.98 | 3.87E-11 | HC_vs_MS.NoTrt_B |
| CD99 | 5.40E-15 | -0.6729 | 0.377 | 0.71 | 8.63E-11 | HC_vs_MS.NoTrt_B |
| TMEM45A | 7.51E-15 | 0.53795 | 0.996 | 1 | 1.20E-10 | HC_vs_MS.NoTrt_B |
| RPS3 | 2.09E-14 | -0.4008 | 0.92 | 1 | 3.34E-10 | HC_vs_MS.NoTrt_B |
| FRK | 3.01E-14 | 0.47036 | 0.991 | 1 | 4.81E-10 | HC_vs_MS.NoTrt_B |
| B2M | 3.14E-14 | -0.257 | 1 | 1 | 5.01E-10 | HC_vs_MS.NoTrt_B |
| CRIP1 | 5.53E-14 | -0.8146 | 0.296 | 0.57 | 8.83E-10 | HC_vs_MS.NoTrt_B |
| CD52 | 9.12E-14 | -0.5357 | 0.742 | 0.94 | 1.46E-09 | HC_vs_MS.NoTrt_B |
| ANXA2 | 1.18E-13 | -0.7037 | 0.607 | 0.82 | 1.89E-09 | HC_vs_MS.NoTrt_B |

| | | | | | | |
|---|---|---|---|---|---|---|
| RPL27 | 2.25E-13 | -0.4687 | 0.606 | 0.89 | 3.59E-09 | HC_vs_MS.NoTrt_B |
| MT-ND4L | 5.17E-13 | 0.9432 | 0.984 | 0.99 | 8.26E-09 | HC_vs_MS.NoTrt_B |
| MORC4 | 5.21E-13 | 0.49767 | 0.991 | 1 | 8.33E-09 | HC_vs_MS.NoTrt_B |
| LSP1 | 5.92E-13 | -0.5472 | 0.601 | 0.87 | 9.45E-09 | HC_vs_MS.NoTrt_B |
| RPS27A | 7.25E-13 | -0.5093 | 0.788 | 0.97 | 1.16E-08 | HC_vs_MS.NoTrt_B |
| RPL36 | 1.18E-12 | -0.5285 | 0.593 | 0.87 | 1.89E-08 | HC_vs_MS.NoTrt_B |
| RPL7A | 1.63E-12 | -0.4636 | 0.764 | 0.97 | 2.61E-08 | HC_vs_MS.NoTrt_B |
| UQCR11 | 2.34E-12 | -0.5315 | 0.504 | 0.77 | 3.74E-08 | HC_vs_MS.NoTrt_B |
| RPL27A | 2.47E-12 | -0.4915 | 0.928 | 0.98 | 3.95E-08 | HC_vs_MS.NoTrt_B |
| RPL28 | 4.43E-12 | -0.4177 | 0.925 | 1 | 7.07E-08 | HC_vs_MS.NoTrt_B |
| ARHGDIB | 7.27E-12 | -0.352 | 0.9 | 1 | 1.16E-07 | HC_vs_MS.NoTrt_B |
| HLA-A | 7.55E-12 | -0.5364 | 0.83 | 0.94 | 1.21E-07 | HC_vs_MS.NoTrt_B |
| RAN | 1.12E-11 | -0.5964 | 0.39 | 0.68 | 1.79E-07 | HC_vs_MS.NoTrt_B |
| RPL19 | 1.15E-11 | -0.4556 | 0.797 | 0.94 | 1.84E-07 | HC_vs_MS.NoTrt_B |
| TMEM50A | 1.48E-11 | -0.7002 | 0.233 | 0.48 | 2.36E-07 | HC_vs_MS.NoTrt_B |
| C12orf45 | 1.86E-11 | 0.38453 | 0.997 | 1 | 2.97E-07 | HC_vs_MS.NoTrt_B |
| RPL24 | 2.57E-11 | -0.5451 | 0.728 | 0.91 | 4.11E-07 | HC_vs_MS.NoTrt_B |
| INTU | 3.77E-11 | 0.50744 | 0.987 | 0.97 | 6.02E-07 | HC_vs_MS.NoTrt_B |
| RPAP2 | 5.94E-11 | 0.54234 | 0.997 | 1 | 9.48E-07 | HC_vs_MS.NoTrt_B |
| MTRNR2L12 | 6.15E-11 | 0.38999 | 1 | 1 | 9.82E-07 | HC_vs_MS.NoTrt_B |
| GAPDH | 6.44E-11 | -0.5155 | 0.703 | 0.89 | 1.03E-06 | HC_vs_MS.NoTrt_B |
| RPLP1 | 6.81E-11 | -0.4273 | 0.948 | 1 | 1.09E-06 | HC_vs_MS.NoTrt_B |
| RPS21 | 8.26E-11 | -0.5301 | 0.439 | 0.76 | 1.32E-06 | HC_vs_MS.NoTrt_B |
| COMMD1 | 9.12E-11 | -0.5678 | 0.536 | 0.8 | 1.46E-06 | HC_vs_MS.NoTrt_B |
| SPATA4 | 1.07E-10 | 0.56432 | 0.587 | 0.36 | 1.71E-06 | HC_vs_MS.NoTrt_B |
| PSME1 | 1.16E-10 | -0.5041 | 0.522 | 0.76 | 1.85E-06 | HC_vs_MS.NoTrt_B |
| S100A6 | 1.36E-10 | -0.4649 | 0.496 | 0.76 | 2.18E-06 | HC_vs_MS.NoTrt_B |
| JAKMIP1 | 1.75E-10 | -1.0861 | 0.175 | 0.38 | 2.79E-06 | HC_vs_MS.NoTrt_B |
| FXYD5 | 2.06E-10 | -0.4349 | 0.713 | 0.89 | 3.29E-06 | HC_vs_MS.NoTrt_B |
| LRRC57 | 2.33E-10 | 0.2855 | 0.97 | 0.95 | 3.73E-06 | HC_vs_MS.NoTrt_B |
| RPS8 | 2.75E-10 | -0.35 | 0.949 | 1 | 4.40E-06 | HC_vs_MS.NoTrt_B |
| FTH1 | 2.99E-10 | -0.3891 | 0.759 | 0.93 | 4.78E-06 | HC_vs_MS.NoTrt_B |
| NACA | 3.24E-10 | -0.4678 | 0.89 | 0.98 | 5.18E-06 | HC_vs_MS.NoTrt_B |
| ANK3 | 3.40E-10 | 0.80186 | 0.904 | 0.82 | 5.43E-06 | HC_vs_MS.NoTrt_B |
| RPS20 | 5.33E-10 | -0.5579 | 0.791 | 0.92 | 8.52E-06 | HC_vs_MS.NoTrt_B |
| NDUFA6 | 5.93E-10 | -0.5316 | 0.188 | 0.41 | 9.47E-06 | HC_vs_MS.NoTrt_B |
| H3F3A | 7.92E-10 | -0.3484 | 0.61 | 0.86 | 1.27E-05 | HC_vs_MS.NoTrt_B |
| MYL6 | 9.89E-10 | -0.3875 | 0.594 | 0.85 | 1.58E-05 | HC_vs_MS.NoTrt_B |
| MIF | 1.21E-09 | -0.4506 | 0.361 | 0.64 | 1.94E-05 | HC_vs_MS.NoTrt_B |
| EPN1 | 1.31E-09 | 0.41406 | 0.949 | 0.93 | 2.09E-05 | HC_vs_MS.NoTrt_B |

| | | | | | | |
|---|---|---|---|---|---|---|
| RPSA | 1.39E-09 | -0.4874 | 0.652 | 0.83 | 2.23E-05 | HC_vs_MS.NoTrt_B |
| ORC6 | 1.40E-09 | 0.57032 | 0.977 | 0.97 | 2.23E-05 | HC_vs_MS.NoTrt_B |
| RPS18 | 1.68E-09 | -0.3579 | 0.968 | 0.99 | 2.68E-05 | HC_vs_MS.NoTrt_B |
| CKLF | 1.77E-09 | -0.5965 | 0.148 | 0.34 | 2.83E-05 | HC_vs_MS.NoTrt_B |
| CCR6 | 1.81E-09 | -1.2676 | 0.307 | 0.52 | 2.90E-05 | HC_vs_MS.NoTrt_B |
| PRDM1 | 1.97E-09 | -0.3164 | 0.414 | 0.71 | 3.15E-05 | HC_vs_MS.NoTrt_B |
| EHD1 | 2.01E-09 | -0.6957 | 0.264 | 0.49 | 3.20E-05 | HC_vs_MS.NoTrt_B |
| RPS16 | 2.04E-09 | -0.4112 | 0.775 | 0.92 | 3.27E-05 | HC_vs_MS.NoTrt_B |
| S100A10 | 2.25E-09 | -0.4668 | 0.491 | 0.76 | 3.59E-05 | HC_vs_MS.NoTrt_B |
| COX7C | 2.37E-09 | -0.5624 | 0.31 | 0.56 | 3.78E-05 | HC_vs_MS.NoTrt_B |
| NEDD4 | 2.46E-09 | 0.63483 | 0.964 | 0.97 | 3.92E-05 | HC_vs_MS.NoTrt_B |
| PKM | 2.52E-09 | -0.3758 | 0.487 | 0.76 | 4.02E-05 | HC_vs_MS.NoTrt_B |
| FTL | 2.68E-09 | -0.3195 | 0.917 | 1 | 4.29E-05 | HC_vs_MS.NoTrt_B |
| RPS23 | 2.72E-09 | -0.3447 | 0.838 | 0.97 | 4.35E-05 | HC_vs_MS.NoTrt_B |
| SULT1B1 | 2.78E-09 | 0.70301 | 0.959 | 0.96 | 4.44E-05 | HC_vs_MS.NoTrt_B |
| HNRNPA1 | 3.02E-09 | -0.3682 | 0.748 | 0.93 | 4.82E-05 | HC_vs_MS.NoTrt_B |
| EMP3 | 3.30E-09 | -0.4538 | 0.507 | 0.75 | 5.26E-05 | HC_vs_MS.NoTrt_B |
| COL8A1 | 3.54E-09 | 0.55947 | 0.826 | 0.74 | 5.66E-05 | HC_vs_MS.NoTrt_B |
| ALOX5AP | 4.17E-09 | -0.6587 | 0.213 | 0.43 | 6.66E-05 | HC_vs_MS.NoTrt_B |
| GLIPR2 | 4.43E-09 | -0.8907 | 0.196 | 0.38 | 7.08E-05 | HC_vs_MS.NoTrt_B |
| RPL37 | 4.67E-09 | -0.3383 | 0.942 | 0.99 | 7.46E-05 | HC_vs_MS.NoTrt_B |
| CSTB | 4.95E-09 | -0.3421 | 0.406 | 0.65 | 7.91E-05 | HC_vs_MS.NoTrt_B |
| XRCC6 | 5.85E-09 | -0.4805 | 0.546 | 0.76 | 9.35E-05 | HC_vs_MS.NoTrt_B |
| CFAP54 | 6.91E-09 | 0.59023 | 0.939 | 0.96 | 1.10E-04 | HC_vs_MS.NoTrt_B |
| RPL32 | 9.31E-09 | -0.3824 | 0.919 | 0.99 | 1.49E-04 | HC_vs_MS.NoTrt_B |
| RPL12 | 1.03E-08 | -0.3786 | 0.832 | 0.95 | 1.65E-04 | HC_vs_MS.NoTrt_B |
| METTL21A | 1.06E-08 | 0.57804 | 0.942 | 0.97 | 1.69E-04 | HC_vs_MS.NoTrt_B |
| ACTG1 | 1.16E-08 | -0.4013 | 0.878 | 0.96 | 1.85E-04 | HC_vs_MS.NoTrt_B |
| NDUFA1 | 1.23E-08 | -0.4469 | 0.209 | 0.43 | 1.97E-04 | HC_vs_MS.NoTrt_B |
| RPL35A | 1.32E-08 | -0.398 | 0.886 | 0.97 | 2.11E-04 | HC_vs_MS.NoTrt_B |
| LIME1 | 1.44E-08 | -0.4768 | 0.365 | 0.62 | 2.30E-04 | HC_vs_MS.NoTrt_B |
| H3F3B | 1.66E-08 | -0.2565 | 0.781 | 0.96 | 2.64E-04 | HC_vs_MS.NoTrt_B |
| RPS13 | 1.68E-08 | -0.3543 | 0.796 | 0.96 | 2.68E-04 | HC_vs_MS.NoTrt_B |
| ENO1 | 1.69E-08 | -0.5089 | 0.377 | 0.63 | 2.71E-04 | HC_vs_MS.NoTrt_B |
| NDUFA4 | 1.82E-08 | -0.4976 | 0.558 | 0.75 | 2.91E-04 | HC_vs_MS.NoTrt_B |
| ATP5E | 2.00E-08 | -0.3266 | 0.372 | 0.61 | 3.20E-04 | HC_vs_MS.NoTrt_B |
| CDC37 | 2.34E-08 | -0.476 | 0.323 | 0.56 | 3.74E-04 | HC_vs_MS.NoTrt_B |
| TPI1 | 2.37E-08 | -0.5483 | 0.372 | 0.6 | 3.78E-04 | HC_vs_MS.NoTrt_B |
| RPL18 | 2.51E-08 | -0.3465 | 0.694 | 0.92 | 4.02E-04 | HC_vs_MS.NoTrt_B |
| CCDC167 | 2.71E-08 | -0.9461 | 0.099 | 0.25 | 4.33E-04 | HC_vs_MS.NoTrt_B |

| | | | | | | |
|---|---|---|---|---|---|---|
| DBI | 2.87E-08 | -0.806 | 0.149 | 0.33 | 4.59E-04 | HC_vs_MS.NoTrt_B |
| RPL30 | 2.90E-08 | -0.361 | 0.852 | 0.97 | 4.63E-04 | HC_vs_MS.NoTrt_B |
| RPL13 | 2.93E-08 | -0.2993 | 0.929 | 0.99 | 4.67E-04 | HC_vs_MS.NoTrt_B |
| SLC25A6 | 2.95E-08 | -0.4534 | 0.554 | 0.76 | 4.71E-04 | HC_vs_MS.NoTrt_B |
| TXNIP | 3.29E-08 | -0.3614 | 0.813 | 0.94 | 5.26E-04 | HC_vs_MS.NoTrt_B |
| COX7A2L | 3.30E-08 | -0.4434 | 0.258 | 0.48 | 5.27E-04 | HC_vs_MS.NoTrt_B |
| COPE | 3.77E-08 | -0.405 | 0.303 | 0.55 | 6.03E-04 | HC_vs_MS.NoTrt_B |
| TOMM7 | 4.03E-08 | -0.4244 | 0.554 | 0.75 | 6.44E-04 | HC_vs_MS.NoTrt_B |
| IRGQ | 4.16E-08 | 0.73794 | 0.98 | 0.99 | 6.64E-04 | HC_vs_MS.NoTrt_B |
| PRDX1 | 4.39E-08 | -0.4907 | 0.465 | 0.67 | 7.01E-04 | HC_vs_MS.NoTrt_B |
| VAMP5 | 4.68E-08 | -0.7381 | 0.113 | 0.28 | 7.47E-04 | HC_vs_MS.NoTrt_B |
| RPL29 | 5.21E-08 | -0.3332 | 0.836 | 0.98 | 8.32E-04 | HC_vs_MS.NoTrt_B |
| RAB12 | 5.75E-08 | 2.66349 | 0.362 | 0.19 | 9.18E-04 | HC_vs_MS.NoTrt_B |
| MYL12A | 5.77E-08 | -0.2756 | 0.894 | 1 | 9.22E-04 | HC_vs_MS.NoTrt_B |
| BIN1 | 5.84E-08 | -0.5106 | 0.262 | 0.49 | 9.33E-04 | HC_vs_MS.NoTrt_B |
| TMEM173 | 6.22E-08 | -0.9196 | 0.201 | 0.38 | 9.94E-04 | HC_vs_MS.NoTrt_B |
| ANKRD30BL | 6.33E-08 | 0.45776 | 0.584 | 0.41 | 0.001012 | HC_vs_MS.NoTrt_B |
| NDUFA13 | 6.92E-08 | -0.5236 | 0.409 | 0.62 | 0.001105 | HC_vs_MS.NoTrt_B |
| RPL23A | 7.10E-08 | -0.3456 | 0.875 | 0.98 | 0.001134 | HC_vs_MS.NoTrt_B |
| RPS11 | 8.46E-08 | -0.3518 | 0.649 | 0.87 | 0.001351 | HC_vs_MS.NoTrt_B |
| ASTN2 | 8.86E-08 | 0.52638 | 0.943 | 0.95 | 0.001416 | HC_vs_MS.NoTrt_B |
| RPL41 | 9.36E-08 | -0.3609 | 0.957 | 1 | 0.001496 | HC_vs_MS.NoTrt_B |
| RASL10B | 1.01E-07 | 0.34068 | 0.972 | 0.97 | 0.001615 | HC_vs_MS.NoTrt_B |
| ANXA5 | 1.08E-07 | -0.6797 | 0.23 | 0.42 | 0.001729 | HC_vs_MS.NoTrt_B |
| CEACAM21 | 1.13E-07 | -0.3918 | 0.37 | 0.57 | 0.001805 | HC_vs_MS.NoTrt_B |
| RPS19 | 1.23E-07 | -0.3454 | 0.933 | 1 | 0.001969 | HC_vs_MS.NoTrt_B |
| RPS25 | 1.36E-07 | -0.3122 | 0.771 | 0.95 | 0.002176 | HC_vs_MS.NoTrt_B |
| PSMB7 | 1.38E-07 | -0.5979 | 0.174 | 0.35 | 0.002211 | HC_vs_MS.NoTrt_B |
| ADAM32 | 1.45E-07 | 0.70725 | 0.854 | 0.82 | 0.002318 | HC_vs_MS.NoTrt_B |
| GMFG | 1.51E-07 | -0.3106 | 0.772 | 0.91 | 0.002414 | HC_vs_MS.NoTrt_B |
| NSRP1 | 1.61E-07 | 0.309 | 0.999 | 1 | 0.002567 | HC_vs_MS.NoTrt_B |
| TAGLN2 | 1.61E-07 | -0.4302 | 0.496 | 0.71 | 0.002571 | HC_vs_MS.NoTrt_B |
| RPS15A | 1.72E-07 | -0.3336 | 0.991 | 1 | 0.002739 | HC_vs_MS.NoTrt_B |
| C9orf16 | 1.99E-07 | -0.3637 | 0.206 | 0.4 | 0.003185 | HC_vs_MS.NoTrt_B |
| RP11-49K24 | 2.00E-07 | -0.4531 | 0.117 | 0.28 | 0.003189 | HC_vs_MS.NoTrt_B |
| RPL22 | 2.04E-07 | -0.3 | 0.584 | 0.8 | 0.003253 | HC_vs_MS.NoTrt_B |
| VIM | 2.16E-07 | -0.3319 | 0.788 | 0.92 | 0.003447 | HC_vs_MS.NoTrt_B |
| TXN | 2.17E-07 | -0.3877 | 0.19 | 0.38 | 0.003461 | HC_vs_MS.NoTrt_B |
| FGF13 | 2.24E-07 | 0.46722 | 0.943 | 0.95 | 0.00358 | HC_vs_MS.NoTrt_B |
| MSN | 2.24E-07 | -0.3983 | 0.558 | 0.74 | 0.003582 | HC_vs_MS.NoTrt_B |

| | | | | | | |
|---|---|---|---|---|---|---|
| EBP | 2.33E-07 | -0.8695 | 0.129 | 0.28 | 0.00372 | HC_vs_MS.NoTrt_B |
| ASB3 | 2.40E-07 | 0.44495 | 0.936 | 0.94 | 0.003838 | HC_vs_MS.NoTrt_B |
| THYN1 | 2.68E-07 | -0.6641 | 0.13 | 0.29 | 0.004273 | HC_vs_MS.NoTrt_B |
| COX8A | 2.75E-07 | -0.3553 | 0.18 | 0.37 | 0.004393 | HC_vs_MS.NoTrt_B |
| VCP | 2.78E-07 | -0.4796 | 0.249 | 0.45 | 0.004439 | HC_vs_MS.NoTrt_B |
| RPL18A | 2.79E-07 | -0.2931 | 0.914 | 1 | 0.004457 | HC_vs_MS.NoTrt_B |
| ITM2B | 2.80E-07 | -0.3014 | 0.567 | 0.83 | 0.004476 | HC_vs_MS.NoTrt_B |
| RPL17 | 2.86E-07 | -0.2942 | 0.903 | 1 | 0.004562 | HC_vs_MS.NoTrt_B |
| RPS12 | 2.95E-07 | -0.3346 | 0.887 | 0.98 | 0.004713 | HC_vs_MS.NoTrt_B |
| SOD2 | 2.99E-07 | 0.30232 | 1 | 1 | 0.004773 | HC_vs_MS.NoTrt_B |
| PRRG3 | 3.07E-07 | 1.6224 | 0.597 | 0.42 | 0.004904 | HC_vs_MS.NoTrt_B |
| UBE2D2 | 3.15E-07 | -0.3075 | 0.377 | 0.6 | 0.005032 | HC_vs_MS.NoTrt_B |
| UPK3BL | 3.29E-07 | -0.5215 | 0.532 | 0.67 | 0.005252 | HC_vs_MS.NoTrt_B |
| RPL11 | 3.30E-07 | -0.257 | 0.957 | 1 | 0.005268 | HC_vs_MS.NoTrt_B |
| TSPO | 3.35E-07 | -0.3553 | 0.271 | 0.5 | 0.005357 | HC_vs_MS.NoTrt_B |
| LDHB | 3.66E-07 | -0.3454 | 0.726 | 0.91 | 0.00585 | HC_vs_MS.NoTrt_B |
| HLA-C | 4.10E-07 | -0.6103 | 0.886 | 0.92 | 0.006549 | HC_vs_MS.NoTrt_B |
| RPL35 | 4.14E-07 | -0.3868 | 0.759 | 0.9 | 0.006612 | HC_vs_MS.NoTrt_B |
| RPL10A | 4.16E-07 | -0.3521 | 0.851 | 0.97 | 0.006647 | HC_vs_MS.NoTrt_B |
| ATP6V1F | 4.16E-07 | -0.5866 | 0.155 | 0.32 | 0.006649 | HC_vs_MS.NoTrt_B |
| SRP9 | 4.81E-07 | -0.2694 | 0.345 | 0.56 | 0.007682 | HC_vs_MS.NoTrt_B |
| PIK3IP1 | 5.01E-07 | -0.294 | 0.664 | 0.84 | 0.008 | HC_vs_MS.NoTrt_B |
| MYL12B | 5.03E-07 | -0.3895 | 0.459 | 0.69 | 0.008034 | HC_vs_MS.NoTrt_B |
| SLC25A3 | 5.06E-07 | -0.479 | 0.929 | 0.96 | 0.008089 | HC_vs_MS.NoTrt_B |
| ANXA6 | 5.23E-07 | -0.3758 | 0.426 | 0.64 | 0.008357 | HC_vs_MS.NoTrt_B |
| ANXA11 | 5.38E-07 | -0.3935 | 0.797 | 0.88 | 0.008598 | HC_vs_MS.NoTrt_B |
| C2CD3 | 5.39E-07 | 0.2629 | 0.991 | 0.97 | 0.008615 | HC_vs_MS.NoTrt_B |
| SERF2 | 6.80E-07 | -0.2658 | 0.799 | 0.95 | 0.010864 | HC_vs_MS.NoTrt_B |
| RP11-286N2 | 6.86E-07 | 0.41814 | 0.958 | 0.93 | 0.01095 | HC_vs_MS.NoTrt_B |
| ARHGAP30 | 6.95E-07 | -0.4634 | 0.355 | 0.56 | 0.011102 | HC_vs_MS.NoTrt_B |
| CSGALNACT | 6.95E-07 | -0.4909 | 0.514 | 0.69 | 0.011105 | HC_vs_MS.NoTrt_B |
| RPS28 | 7.01E-07 | -0.3932 | 0.826 | 0.94 | 0.011192 | HC_vs_MS.NoTrt_B |
| CHCHD2 | 7.15E-07 | -0.4594 | 0.346 | 0.55 | 0.011418 | HC_vs_MS.NoTrt_B |
| SOD1 | 7.81E-07 | -0.3128 | 0.646 | 0.8 | 0.012483 | HC_vs_MS.NoTrt_B |
| SH3BGRL3 | 7.86E-07 | -0.3022 | 0.787 | 0.91 | 0.012548 | HC_vs_MS.NoTrt_B |
| CCDC107 | 7.86E-07 | -0.7354 | 0.107 | 0.25 | 0.012549 | HC_vs_MS.NoTrt_B |
| SNRPD2 | 8.06E-07 | -0.3902 | 0.51 | 0.7 | 0.012881 | HC_vs_MS.NoTrt_B |
| PSMC5 | 8.95E-07 | -0.6312 | 0.235 | 0.42 | 0.014295 | HC_vs_MS.NoTrt_B |
| CACNA1A | 9.96E-07 | 0.29596 | 1 | 1 | 0.015911 | HC_vs_MS.NoTrt_B |
| COPS9 | 1.00E-06 | -0.482 | 0.113 | 0.26 | 0.016014 | HC_vs_MS.NoTrt_B |

| | | | | | | |
|---|---|---|---|---|---|---|
| CD3D | 1.01E-06 | -0.278 | 0.528 | 0.77 | 0.016132 | HC_vs_MS.NoTrt_B |
| SLC15A1 | 1.02E-06 | 0.4266 | 0.838 | 0.74 | 0.016295 | HC_vs_MS.NoTrt_B |
| GARS | 1.11E-06 | -0.5445 | 0.136 | 0.29 | 0.017683 | HC_vs_MS.NoTrt_B |
| NASP | 1.16E-06 | 0.70284 | 0.92 | 0.88 | 0.01855 | HC_vs_MS.NoTrt_B |
| TMEM14C | 1.17E-06 | -0.6725 | 0.081 | 0.21 | 0.018755 | HC_vs_MS.NoTrt_B |
| FBXO7 | 1.19E-06 | -0.4638 | 0.264 | 0.44 | 0.019011 | HC_vs_MS.NoTrt_B |
| OAZ1 | 1.22E-06 | -0.3217 | 0.796 | 0.91 | 0.019514 | HC_vs_MS.NoTrt_B |
| ATP6V1G1 | 1.24E-06 | -0.2977 | 0.426 | 0.64 | 0.019857 | HC_vs_MS.NoTrt_B |
| PFN1 | 1.34E-06 | -0.2605 | 0.952 | 1 | 0.021423 | HC_vs_MS.NoTrt_B |
| IL2RA | 1.35E-06 | -0.5582 | 0.335 | 0.51 | 0.02153 | HC_vs_MS.NoTrt_B |
| ABCC9 | 1.41E-06 | 0.50711 | 0.959 | 0.95 | 0.022593 | HC_vs_MS.NoTrt_B |
| 1-Sep | 1.44E-06 | -0.3154 | 0.665 | 0.84 | 0.022991 | HC_vs_MS.NoTrt_B |
| UBE2I | 1.47E-06 | -0.3164 | 0.52 | 0.72 | 0.023517 | HC_vs_MS.NoTrt_B |
| RPL36A | 1.49E-06 | -0.3908 | 0.713 | 0.84 | 0.023866 | HC_vs_MS.NoTrt_B |
| PSMD8 | 1.55E-06 | -0.6675 | 0.568 | 0.73 | 0.024735 | HC_vs_MS.NoTrt_B |
| GTF3A | 1.57E-06 | -0.3125 | 0.394 | 0.63 | 0.025029 | HC_vs_MS.NoTrt_B |
| PKIG | 1.66E-06 | 0.50481 | 0.626 | 0.54 | 0.026445 | HC_vs_MS.NoTrt_B |
| HOXB2 | 1.73E-06 | -0.3884 | 0.123 | 0.27 | 0.027703 | HC_vs_MS.NoTrt_B |
| FAM98C | 1.89E-06 | 0.3392 | 0.968 | 0.99 | 0.030203 | HC_vs_MS.NoTrt_B |
| RP11-529K1 | 1.90E-06 | -0.7084 | 0.054 | 0.16 | 0.030278 | HC_vs_MS.NoTrt_B |
| TCEB2 | 1.94E-06 | -0.3423 | 0.301 | 0.5 | 0.030962 | HC_vs_MS.NoTrt_B |
| PSMB3 | 2.01E-06 | -0.3812 | 0.216 | 0.4 | 0.032131 | HC_vs_MS.NoTrt_B |
| CDKL5 | 2.08E-06 | 0.34957 | 0.932 | 0.94 | 0.033273 | HC_vs_MS.NoTrt_B |
| COPZ1 | 2.12E-06 | -0.4187 | 0.152 | 0.31 | 0.03379 | HC_vs_MS.NoTrt_B |
| MRPL28 | 2.13E-06 | -0.8472 | 0.083 | 0.21 | 0.03407 | HC_vs_MS.NoTrt_B |
| CCDC144A | 2.22E-06 | 0.93531 | 0.878 | 0.84 | 0.035434 | HC_vs_MS.NoTrt_B |
| RACK1 | 2.26E-06 | -0.255 | 0.999 | 1 | 0.036172 | HC_vs_MS.NoTrt_B |
| ORC4 | 2.30E-06 | 0.52122 | 0.98 | 1 | 0.036768 | HC_vs_MS.NoTrt_B |
| TSTA3 | 2.36E-06 | -0.6448 | 0.114 | 0.25 | 0.037675 | HC_vs_MS.NoTrt_B |
| EIF4H | 2.42E-06 | -0.371 | 0.346 | 0.55 | 0.038632 | HC_vs_MS.NoTrt_B |
| NUPR1 | 2.43E-06 | 0.31926 | 0.941 | 0.96 | 0.03875 | HC_vs_MS.NoTrt_B |
| C4orf32 | 2.57E-06 | 0.38995 | 0.986 | 1 | 0.041046 | HC_vs_MS.NoTrt_B |
| APOL2 | 2.70E-06 | -0.3693 | 0.561 | 0.7 | 0.043125 | HC_vs_MS.NoTrt_B |
| GPS2 | 2.71E-06 | -0.4784 | 0.193 | 0.36 | 0.043213 | HC_vs_MS.NoTrt_B |
| CTLA4 | 2.73E-06 | -0.5181 | 0.248 | 0.41 | 0.043649 | HC_vs_MS.NoTrt_B |
| KPNA5 | 2.78E-06 | 0.40835 | 0.983 | 0.98 | 0.044472 | HC_vs_MS.NoTrt_B |
| SELT | 2.80E-06 | -0.5634 | 0.306 | 0.47 | 0.044757 | HC_vs_MS.NoTrt_B |
| EEF1B2 | 2.81E-06 | -0.329 | 0.467 | 0.7 | 0.044925 | HC_vs_MS.NoTrt_B |
| ARPC3 | 2.83E-06 | -0.3577 | 0.428 | 0.64 | 0.04521 | HC_vs_MS.NoTrt_B |
| RASGRP3 | 2.86E-06 | 0.26265 | 0.991 | 1 | 0.045678 | HC_vs_MS.NoTrt_B |

| | | | | | | |
|---|---|---|---|---|---|---|
| WDR31 | 2.99E-06 | 0.48237 | 0.836 | 0.81 | 0.047783 | HC_vs_MS.NoTrt_B |
| RNF26 | 2.99E-06 | -0.5553 | 0.077 | 0.2 | 0.047796 | HC_vs_MS.NoTrt_B |
| DGCR2 | 3.35E-06 | -0.4568 | 0.18 | 0.34 | 0.053589 | HC_vs_MS.NoTrt_B |
| DIRAS2 | 3.47E-06 | 0.5663 | 0.787 | 0.75 | 0.055449 | HC_vs_MS.NoTrt_B |
| TUBA1A | 3.54E-06 | -0.6566 | 0.186 | 0.34 | 0.056505 | HC_vs_MS.NoTrt_B |
| MCM7 | 3.54E-06 | -0.5932 | 0.18 | 0.33 | 0.056572 | HC_vs_MS.NoTrt_B |
| TWF2 | 3.62E-06 | -0.4893 | 0.168 | 0.32 | 0.057859 | HC_vs_MS.NoTrt_B |
| HMGB2 | 3.74E-06 | -0.5999 | 0.22 | 0.39 | 0.059663 | HC_vs_MS.NoTrt_B |
| PYM1 | 3.92E-06 | -0.5613 | 0.245 | 0.41 | 0.062644 | HC_vs_MS.NoTrt_B |
| TMEM182 | 3.95E-06 | 0.73827 | 0.762 | 0.79 | 0.063057 | HC_vs_MS.NoTrt_B |
| SHISA9 | 4.04E-06 | 0.4045 | 0.925 | 0.95 | 0.064499 | HC_vs_MS.NoTrt_B |
| CBR4 | 4.20E-06 | -0.2929 | 0.135 | 0.28 | 0.067161 | HC_vs_MS.NoTrt_B |
| UCP2 | 4.32E-06 | -0.4115 | 0.738 | 0.86 | 0.068942 | HC_vs_MS.NoTrt_B |
| TRPC3 | 4.36E-06 | 0.42327 | 0.362 | 0.18 | 0.069699 | HC_vs_MS.NoTrt_B |
| FLII | 4.58E-06 | -0.4288 | 0.172 | 0.33 | 0.07312 | HC_vs_MS.NoTrt_B |
| ELP5 | 4.63E-06 | -1.1769 | 0.054 | 0.15 | 0.074016 | HC_vs_MS.NoTrt_B |
| SEC61G | 4.64E-06 | -0.6709 | 0.216 | 0.37 | 0.074192 | HC_vs_MS.NoTrt_B |
| DEF6 | 4.82E-06 | -0.3319 | 0.399 | 0.6 | 0.077007 | HC_vs_MS.NoTrt_B |
| GBP5 | 5.05E-06 | -0.3242 | 0.641 | 0.78 | 0.080731 | HC_vs_MS.NoTrt_B |
| WDPCP | 5.24E-06 | 0.46984 | 0.971 | 0.98 | 0.083654 | HC_vs_MS.NoTrt_B |
| TMC2 | 5.24E-06 | 0.54454 | 0.665 | 0.57 | 0.083678 | HC_vs_MS.NoTrt_B |
| RARRES3 | 5.24E-06 | -0.4886 | 0.277 | 0.46 | 0.083768 | HC_vs_MS.NoTrt_B |
| RPS10 | 5.25E-06 | -0.3881 | 0.041 | 0.13 | 0.083809 | HC_vs_MS.NoTrt_B |
| TMBIM6 | 5.26E-06 | -0.3163 | 0.809 | 0.92 | 0.084 | HC_vs_MS.NoTrt_B |
| RPLP2 | 5.34E-06 | -0.2999 | 0.777 | 0.96 | 0.085306 | HC_vs_MS.NoTrt_B |
| RABAC1 | 5.59E-06 | -0.5568 | 0.17 | 0.32 | 0.089273 | HC_vs_MS.NoTrt_B |
| HCST | 5.62E-06 | -0.4341 | 0.287 | 0.48 | 0.089835 | HC_vs_MS.NoTrt_B |
| ITGB1 | 5.81E-06 | -0.351 | 0.617 | 0.76 | 0.092734 | HC_vs_MS.NoTrt_B |
| RPS6KA1 | 6.11E-06 | -0.6263 | 0.174 | 0.32 | 0.097613 | HC_vs_MS.NoTrt_B |
| MED23 | 6.19E-06 | -0.4416 | 0.213 | 0.37 | 0.098888 | HC_vs_MS.NoTrt_B |
| HINFP | 6.36E-06 | 0.34703 | 0.99 | 0.99 | 0.101605 | HC_vs_MS.NoTrt_B |
| DNPH1 | 6.50E-06 | -0.495 | 0.104 | 0.23 | 0.103902 | HC_vs_MS.NoTrt_B |
| P2RY8 | 6.57E-06 | -0.5427 | 0.491 | 0.64 | 0.104891 | HC_vs_MS.NoTrt_B |
| TRIM56 | 6.64E-06 | 0.5329 | 0.964 | 0.98 | 0.10604 | HC_vs_MS.NoTrt_B |
| POLR2G | 6.64E-06 | -0.4514 | 0.132 | 0.28 | 0.106097 | HC_vs_MS.NoTrt_B |
| NDUFB4 | 6.68E-06 | -0.3817 | 0.199 | 0.36 | 0.106691 | HC_vs_MS.NoTrt_B |
| DNAJC9 | 7.11E-06 | -0.6069 | 0.11 | 0.24 | 0.113636 | HC_vs_MS.NoTrt_B |
| LGALS3 | 7.19E-06 | -0.6508 | 0.528 | 0.67 | 0.114866 | HC_vs_MS.NoTrt_B |
| PYCARD | 7.44E-06 | -0.6008 | 0.072 | 0.18 | 0.118804 | HC_vs_MS.NoTrt_B |
| IL10RA | 7.75E-06 | -0.3278 | 0.613 | 0.75 | 0.123787 | HC_vs_MS.NoTrt_B |

| | | | | | | |
|---|---|---|---|---|---|---|
| WDR45 | 7.81E-06 | -0.6302 | 0.549 | 0.72 | 0.124738 | HC_vs_MS.NoTrt_B |
| NUDT2 | 8.52E-06 | -1.0782 | 0.03 | 0.11 | 0.13614 | HC_vs_MS.NoTrt_B |
| CTB-133G6.: | 8.52E-06 | -0.2622 | 0.268 | 0.45 | 0.136161 | HC_vs_MS.NoTrt_B |
| BLOC1S1 | 8.90E-06 | -0.5281 | 0.165 | 0.31 | 0.142188 | HC_vs_MS.NoTrt_B |
| ARHGAP25 | 9.01E-06 | -0.4358 | 0.425 | 0.59 | 0.143901 | HC_vs_MS.NoTrt_B |
| CD5 | 9.14E-06 | -0.3286 | 0.439 | 0.65 | 0.146002 | HC_vs_MS.NoTrt_B |
| ERAP2 | 9.27E-06 | -0.7805 | 0.619 | 0.72 | 0.147991 | HC_vs_MS.NoTrt_B |
| EZR | 9.47E-06 | -0.449 | 0.63 | 0.74 | 0.151307 | HC_vs_MS.NoTrt_B |
| SAMD12 | 9.70E-06 | 0.45979 | 0.972 | 1 | 0.154947 | HC_vs_MS.NoTrt_B |
| MTRR | 9.71E-06 | -0.3679 | 0.133 | 0.27 | 0.155143 | HC_vs_MS.NoTrt_B |
| COX6C | 1.06E-05 | -0.3047 | 0.467 | 0.62 | 0.169422 | HC_vs_MS.NoTrt_B |
| DDRGK1 | 1.06E-05 | -0.7178 | 0.087 | 0.2 | 0.169886 | HC_vs_MS.NoTrt_B |
| CD74 | 1.08E-05 | -0.3457 | 0.694 | 0.81 | 0.172118 | HC_vs_MS.NoTrt_B |
| GLYCTK | 1.08E-05 | -1.4398 | 0.067 | 0.17 | 0.173084 | HC_vs_MS.NoTrt_B |
| PDCD5 | 1.09E-05 | -0.4908 | 0.149 | 0.29 | 0.173414 | HC_vs_MS.NoTrt_B |
| KCNJ6 | 1.09E-05 | 1.15561 | 0.799 | 0.75 | 0.173895 | HC_vs_MS.NoTrt_B |
| ERP44 | 1.12E-05 | -0.4071 | 0.219 | 0.37 | 0.178714 | HC_vs_MS.NoTrt_B |
| CNN2 | 1.14E-05 | -0.4399 | 0.983 | 0.99 | 0.182119 | HC_vs_MS.NoTrt_B |
| MTRNR2L8 | 1.16E-05 | 0.27565 | 0.997 | 1 | 0.185735 | HC_vs_MS.NoTrt_B |
| PSMA7 | 1.17E-05 | -0.2862 | 0.367 | 0.55 | 0.186366 | HC_vs_MS.NoTrt_B |
| EID1 | 1.19E-05 | -0.3336 | 0.336 | 0.54 | 0.189764 | HC_vs_MS.NoTrt_B |
| FAM122C | 1.21E-05 | 0.32808 | 0.901 | 0.91 | 0.193232 | HC_vs_MS.NoTrt_B |
| CASP8 | 1.26E-05 | -0.4933 | 0.914 | 0.95 | 0.201109 | HC_vs_MS.NoTrt_B |
| KYNU | 1.26E-05 | 0.33024 | 0.968 | 0.99 | 0.201307 | HC_vs_MS.NoTrt_B |
| RPLP0 | 1.27E-05 | -0.2548 | 0.916 | 0.97 | 0.202907 | HC_vs_MS.NoTrt_B |
| MS4A10 | 1.30E-05 | 0.40763 | 0.683 | 0.56 | 0.207625 | HC_vs_MS.NoTrt_B |
| CST7 | 1.31E-05 | -0.7721 | 0.084 | 0.2 | 0.20875 | HC_vs_MS.NoTrt_B |
| RPS5 | 1.31E-05 | -0.2553 | 0.833 | 0.95 | 0.209403 | HC_vs_MS.NoTrt_B |
| CA5A | 1.33E-05 | 0.3194 | 0.977 | 0.96 | 0.212702 | HC_vs_MS.NoTrt_B |
| MSH5 | 1.34E-05 | 0.54832 | 0.929 | 0.93 | 0.213976 | HC_vs_MS.NoTrt_B |
| REEP5 | 1.43E-05 | -0.265 | 0.293 | 0.45 | 0.227799 | HC_vs_MS.NoTrt_B |
| COX7A2 | 1.43E-05 | -0.363 | 0.719 | 0.82 | 0.229111 | HC_vs_MS.NoTrt_B |
| GJC1 | 1.46E-05 | 0.3455 | 0.938 | 0.94 | 0.232635 | HC_vs_MS.NoTrt_B |
| MAD1L1 | 1.48E-05 | -0.4397 | 0.284 | 0.45 | 0.236212 | HC_vs_MS.NoTrt_B |
| LDOC1 | 1.49E-05 | -0.9691 | 0.035 | 0.11 | 0.237621 | HC_vs_MS.NoTrt_B |
| MRPL33 | 1.60E-05 | -0.4951 | 0.11 | 0.23 | 0.254872 | HC_vs_MS.NoTrt_B |
| PEBP1 | 1.62E-05 | -0.391 | 0.341 | 0.52 | 0.259004 | HC_vs_MS.NoTrt_B |
| DOK2 | 1.66E-05 | -0.3901 | 0.333 | 0.51 | 0.265016 | HC_vs_MS.NoTrt_B |
| RNF181 | 1.73E-05 | -0.8344 | 0.139 | 0.27 | 0.276687 | HC_vs_MS.NoTrt_B |
| UQCR10 | 1.82E-05 | -0.37 | 0.288 | 0.45 | 0.290138 | HC_vs_MS.NoTrt_B |

| | | | | | | |
|---|---|---|---|---|---|---|
| TSC22D3 | 1.83E-05 | -0.3222 | 0.555 | 0.75 | 0.291629 | HC_vs_MS.NoTrt_B |
| CLIC1 | 1.85E-05 | -0.4707 | 0.543 | 0.69 | 0.294785 | HC_vs_MS.NoTrt_B |
| SPATA20 | 1.91E-05 | -1.1206 | 0.048 | 0.14 | 0.304351 | HC_vs_MS.NoTrt_B |
| SNRPC | 1.91E-05 | -0.4005 | 0.172 | 0.32 | 0.305522 | HC_vs_MS.NoTrt_B |
| C12orf75 | 1.91E-05 | -0.2835 | 0.122 | 0.25 | 0.305607 | HC_vs_MS.NoTrt_B |
| ALDH2 | 1.96E-05 | 0.30954 | 0.965 | 0.97 | 0.313747 | HC_vs_MS.NoTrt_B |
| ARID4A | 2.03E-05 | -0.3914 | 0.278 | 0.44 | 0.32398 | HC_vs_MS.NoTrt_B |
| METTL23 | 2.03E-05 | -0.2731 | 0.138 | 0.28 | 0.324737 | HC_vs_MS.NoTrt_B |
| MTRNR2L10 | 2.03E-05 | 0.35719 | 0.968 | 1 | 0.324881 | HC_vs_MS.NoTrt_B |
| MPG | 2.07E-05 | -0.6021 | 0.094 | 0.21 | 0.330452 | HC_vs_MS.NoTrt_B |
| CD53 | 2.15E-05 | -0.2944 | 0.617 | 0.82 | 0.343098 | HC_vs_MS.NoTrt_B |
| YBEY | 2.18E-05 | -1.0772 | 0.042 | 0.12 | 0.347484 | HC_vs_MS.NoTrt_B |
| MYO1G | 2.19E-05 | -0.4263 | 0.529 | 0.68 | 0.350322 | HC_vs_MS.NoTrt_B |
| NAT6 | 2.34E-05 | -1.4199 | 0.03 | 0.1 | 0.373252 | HC_vs_MS.NoTrt_B |
| PSMB4 | 2.34E-05 | -0.2649 | 0.278 | 0.45 | 0.373263 | HC_vs_MS.NoTrt_B |
| EYS | 2.37E-05 | 0.78462 | 0.809 | 0.77 | 0.378546 | HC_vs_MS.NoTrt_B |
| ATP5B | 2.40E-05 | -0.3721 | 0.325 | 0.5 | 0.382977 | HC_vs_MS.NoTrt_B |
| TSSC1 | 2.44E-05 | -0.8106 | 0.065 | 0.16 | 0.390213 | HC_vs_MS.NoTrt_B |
| BLOC1S6 | 2.46E-05 | 0.47649 | 0.986 | 1 | 0.39329 | HC_vs_MS.NoTrt_B |
| KARS | 2.47E-05 | -0.4918 | 0.236 | 0.39 | 0.394737 | HC_vs_MS.NoTrt_B |
| RPL31 | 2.50E-05 | -0.2514 | 0.8 | 0.94 | 0.399127 | HC_vs_MS.NoTrt_B |
| MTRNR2L1 | 2.51E-05 | 0.25549 | 0.997 | 1 | 0.400557 | HC_vs_MS.NoTrt_B |
| PTK2 | 2.52E-05 | 0.39265 | 0.948 | 0.91 | 0.402544 | HC_vs_MS.NoTrt_B |
| ARHGEF12 | 2.58E-05 | 0.34935 | 0.97 | 0.98 | 0.412876 | HC_vs_MS.NoTrt_B |
| MRPL55 | 2.60E-05 | -1.1705 | 0.062 | 0.16 | 0.415882 | HC_vs_MS.NoTrt_B |
| HDHD2 | 2.64E-05 | -0.6624 | 0.114 | 0.23 | 0.421838 | HC_vs_MS.NoTrt_B |
| NUP155 | 2.65E-05 | 0.87391 | 0.906 | 0.89 | 0.422877 | HC_vs_MS.NoTrt_B |
| MRPL51 | 2.67E-05 | -0.5367 | 0.083 | 0.19 | 0.426384 | HC_vs_MS.NoTrt_B |
| RASA4 | 2.69E-05 | -0.3307 | 0.664 | 0.77 | 0.429252 | HC_vs_MS.NoTrt_B |
| NDUFB7 | 2.75E-05 | -0.6213 | 0.155 | 0.29 | 0.439108 | HC_vs_MS.NoTrt_B |
| NDUFB8 | 2.76E-05 | -0.2862 | 0.291 | 0.47 | 0.441112 | HC_vs_MS.NoTrt_B |
| MRPL18 | 2.87E-05 | -0.3347 | 0.088 | 0.2 | 0.457988 | HC_vs_MS.NoTrt_B |
| PSME2 | 2.93E-05 | -0.4228 | 0.301 | 0.47 | 0.467226 | HC_vs_MS.NoTrt_B |
| APRT | 3.03E-05 | -0.3558 | 0.33 | 0.51 | 0.483372 | HC_vs_MS.NoTrt_B |
| PPP1CA | 3.15E-05 | -0.3742 | 0.414 | 0.6 | 0.503188 | HC_vs_MS.NoTrt_B |
| TTC7A | 3.15E-05 | -0.4026 | 0.164 | 0.3 | 0.503502 | HC_vs_MS.NoTrt_B |
| ELAVL1 | 3.20E-05 | 0.47894 | 0.961 | 0.95 | 0.511223 | HC_vs_MS.NoTrt_B |
| PSMB10 | 3.46E-05 | -0.344 | 0.37 | 0.55 | 0.552633 | HC_vs_MS.NoTrt_B |
| HLA-F | 3.52E-05 | -0.3569 | 0.517 | 0.65 | 0.561557 | HC_vs_MS.NoTrt_B |
| DRAP1 | 3.55E-05 | -0.3702 | 0.158 | 0.3 | 0.567369 | HC_vs_MS.NoTrt_B |

| | | | | | | |
|---|---|---|---|---|---|---|
| TEX264 | 3.62E-05 | -0.5648 | 0.154 | 0.29 | 0.578396 | HC_vs_MS.NoTrt_B |
| CD247 | 3.68E-05 | -0.2733 | 0.438 | 0.63 | 0.587024 | HC_vs_MS.NoTrt_B |
| PLCXD1 | 3.70E-05 | 0.50296 | 0.919 | 0.91 | 0.590391 | HC_vs_MS.NoTrt_B |
| SBDS | 3.80E-05 | -0.2517 | 0.259 | 0.42 | 0.606288 | HC_vs_MS.NoTrt_B |
| AQP3 | 3.81E-05 | -0.5818 | 0.262 | 0.41 | 0.6079 | HC_vs_MS.NoTrt_B |
| ZFAND2B | 3.81E-05 | -0.6026 | 0.172 | 0.3 | 0.608199 | HC_vs_MS.NoTrt_B |
| NME2 | 3.83E-05 | -0.269 | 0.464 | 0.65 | 0.611262 | HC_vs_MS.NoTrt_B |
| RGS10 | 3.86E-05 | -0.6589 | 0.177 | 0.31 | 0.61652 | HC_vs_MS.NoTrt_B |
| FBXW5 | 3.90E-05 | -0.2954 | 0.197 | 0.35 | 0.622429 | HC_vs_MS.NoTrt_B |
| CEACAM1 | 3.91E-05 | 0.70584 | 0.79 | 0.69 | 0.625106 | HC_vs_MS.NoTrt_B |
| ZNF37A | 3.97E-05 | 0.7882 | 0.874 | 0.85 | 0.634915 | HC_vs_MS.NoTrt_B |
| C11orf54 | 4.11E-05 | 0.56099 | 0.807 | 0.76 | 0.656223 | HC_vs_MS.NoTrt_B |
| TERB1 | 4.23E-05 | 0.40656 | 0.675 | 0.6 | 0.676155 | HC_vs_MS.NoTrt_B |
| RPS9 | 4.44E-05 | -0.2724 | 0.923 | 0.97 | 0.709773 | HC_vs_MS.NoTrt_B |
| FAM84B | 4.50E-05 | -0.8517 | 0.041 | 0.12 | 0.718311 | HC_vs_MS.NoTrt_B |
| DHRS4 | 4.53E-05 | -0.8384 | 0.072 | 0.17 | 0.723555 | HC_vs_MS.NoTrt_B |
| TFPI | 4.62E-05 | 0.41637 | 0.939 | 0.98 | 0.738444 | HC_vs_MS.NoTrt_B |
| ACTN4 | 4.64E-05 | -0.4394 | 0.309 | 0.46 | 0.741665 | HC_vs_MS.NoTrt_B |
| SSBP2 | 4.87E-05 | 0.34001 | 0.925 | 0.92 | 0.777907 | HC_vs_MS.NoTrt_B |
| HCLS1 | 4.93E-05 | -0.2542 | 0.562 | 0.75 | 0.787707 | HC_vs_MS.NoTrt_B |
| POMP | 5.07E-05 | -0.5604 | 0.167 | 0.3 | 0.809839 | HC_vs_MS.NoTrt_B |
| TMEM256 | 5.11E-05 | -0.3989 | 0.046 | 0.13 | 0.816249 | HC_vs_MS.NoTrt_B |
| LGALS1 | 5.27E-05 | -0.6499 | 0.175 | 0.31 | 0.841631 | HC_vs_MS.NoTrt_B |
| PBXIP1 | 5.47E-05 | -0.3025 | 0.539 | 0.71 | 0.874037 | HC_vs_MS.NoTrt_B |
| CCDC122 | 5.63E-05 | 0.36203 | 0.909 | 0.91 | 0.899086 | HC_vs_MS.NoTrt_B |
| PERP | 5.80E-05 | -1.1 | 0.042 | 0.12 | 0.926762 | HC_vs_MS.NoTrt_B |
| COA3 | 5.82E-05 | -0.6812 | 0.075 | 0.17 | 0.929853 | HC_vs_MS.NoTrt_B |
| VDAC3 | 6.19E-05 | -0.4573 | 0.196 | 0.34 | 0.988743 | HC_vs_MS.NoTrt_B |
| FAM118A | 6.20E-05 | -0.5913 | 0.52 | 0.68 | 0.990775 | HC_vs_MS.NoTrt_B |
| SNRPE | 6.32E-05 | -0.2943 | 0.11 | 0.23 | 1 | HC_vs_MS.NoTrt_B |
| AFMID | 6.52E-05 | 0.37121 | 0.93 | 0.9 | 1 | HC_vs_MS.NoTrt_B |
| ZSCAN5A | 6.59E-05 | 0.43962 | 0.788 | 0.7 | 1 | HC_vs_MS.NoTrt_B |
| SLAMF1 | 6.70E-05 | -0.2776 | 0.177 | 0.32 | 1 | HC_vs_MS.NoTrt_B |
| CCR8 | 6.73E-05 | -0.9765 | 0.087 | 0.19 | 1 | HC_vs_MS.NoTrt_B |
| FGD3 | 6.80E-05 | -0.3037 | 0.516 | 0.68 | 1 | HC_vs_MS.NoTrt_B |
| ZNF431 | 6.81E-05 | 0.51735 | 0.987 | 1 | 1 | HC_vs_MS.NoTrt_B |
| C1D | 6.92E-05 | -0.4802 | 0.093 | 0.2 | 1 | HC_vs_MS.NoTrt_B |
| SLC19A3 | 7.08E-05 | 0.32829 | 0.8 | 0.73 | 1 | HC_vs_MS.NoTrt_B |
| GLB1 | 7.10E-05 | -0.7105 | 0.201 | 0.34 | 1 | HC_vs_MS.NoTrt_B |
| MRPS34 | 7.32E-05 | -0.3859 | 0.138 | 0.27 | 1 | HC_vs_MS.NoTrt_B |

| | | | | | | |
|---|---|---|---|---|---|---|
| ZNF268 | 7.33E-05 | 0.2577 | 0.958 | 0.92 | 1 | HC_vs_MS.NoTrt_B |
| CRYZ | 7.33E-05 | -0.4644 | 0.055 | 0.14 | 1 | HC_vs_MS.NoTrt_B |
| MTRNR2L6 | 7.63E-05 | 0.30619 | 0.974 | 1 | 1 | HC_vs_MS.NoTrt_B |
| CHTF8 | 7.66E-05 | -0.4641 | 0.217 | 0.35 | 1 | HC_vs_MS.NoTrt_B |
| HARS2 | 7.87E-05 | -0.5061 | 0.106 | 0.22 | 1 | HC_vs_MS.NoTrt_B |
| AP1S1 | 7.90E-05 | -0.6218 | 0.039 | 0.11 | 1 | HC_vs_MS.NoTrt_B |
| RPS6KA5 | 7.94E-05 | 0.37181 | 0.999 | 1 | 1 | HC_vs_MS.NoTrt_B |
| TMEM212 | 8.19E-05 | 0.45266 | 0.942 | 0.95 | 1 | HC_vs_MS.NoTrt_B |
| FAHD2B | 8.40E-05 | -1.006 | 0.112 | 0.22 | 1 | HC_vs_MS.NoTrt_B |
| CCDC125 | 8.57E-05 | 0.52229 | 0.936 | 0.96 | 1 | HC_vs_MS.NoTrt_B |
| R3HDM1 | 8.61E-05 | 0.78652 | 0.838 | 0.84 | 1 | HC_vs_MS.NoTrt_B |
| KDELR2 | 8.67E-05 | -0.451 | 0.117 | 0.23 | 1 | HC_vs_MS.NoTrt_B |
| PARP1 | 8.91E-05 | -0.4342 | 0.313 | 0.45 | 1 | HC_vs_MS.NoTrt_B |
| FDXR | 9.05E-05 | -0.8098 | 0.033 | 0.1 | 1 | HC_vs_MS.NoTrt_B |
| INSIG1 | 9.26E-05 | -0.3335 | 0.084 | 0.18 | 1 | HC_vs_MS.NoTrt_B |
| ARSD | 9.34E-05 | 0.59573 | 0.822 | 0.82 | 1 | HC_vs_MS.NoTrt_B |
| EGLN2 | 9.44E-05 | -0.2969 | 0.232 | 0.38 | 1 | HC_vs_MS.NoTrt_B |
| CNKSR3 | 9.71E-05 | 0.31191 | 0.975 | 1 | 1 | HC_vs_MS.NoTrt_B |
| MPC1 | 9.73E-05 | -0.4726 | 0.077 | 0.17 | 1 | HC_vs_MS.NoTrt_B |
| NDUFA2 | 9.83E-05 | -0.3252 | 0.445 | 0.58 | 1 | HC_vs_MS.NoTrt_B |
| IMPDH1 | 9.84E-05 | -0.8533 | 0.101 | 0.21 | 1 | HC_vs_MS.NoTrt_B |
| PPM1M | 9.92E-05 | -0.405 | 0.133 | 0.25 | 1 | HC_vs_MS.NoTrt_B |
| TRMT1L | 9.95E-05 | -1.0943 | 0.054 | 0.14 | 1 | HC_vs_MS.NoTrt_B |
| ODF2L | 1.02E-04 | 0.25046 | 0.951 | 0.92 | 1 | HC_vs_MS.NoTrt_B |
| SCD5 | 1.06E-04 | 0.44731 | 0.803 | 0.75 | 1 | HC_vs_MS.NoTrt_B |
| PKP2 | 1.07E-04 | 0.34455 | 0.594 | 0.47 | 1 | HC_vs_MS.NoTrt_B |
| RBX1 | 1.08E-04 | -0.4273 | 0.23 | 0.37 | 1 | HC_vs_MS.NoTrt_B |
| MYLK3 | 1.09E-04 | 0.38757 | 0.922 | 0.94 | 1 | HC_vs_MS.NoTrt_B |
| MGAT4C | 1.14E-04 | 0.44529 | 0.777 | 0.68 | 1 | HC_vs_MS.NoTrt_B |
| TMEM107 | 1.15E-04 | 0.71357 | 0.583 | 0.45 | 1 | HC_vs_MS.NoTrt_B |
| LITAF | 1.16E-04 | -0.3191 | 0.533 | 0.65 | 1 | HC_vs_MS.NoTrt_B |
| PQLC3 | 1.17E-04 | -0.4519 | 0.12 | 0.23 | 1 | HC_vs_MS.NoTrt_B |
| S100A11 | 1.18E-04 | -0.2507 | 0.301 | 0.47 | 1 | HC_vs_MS.NoTrt_B |
| CYSLTR1 | 1.19E-04 | 0.51576 | 0.946 | 0.92 | 1 | HC_vs_MS.NoTrt_B |
| CYP1A2 | 1.20E-04 | 0.45337 | 0.864 | 0.8 | 1 | HC_vs_MS.NoTrt_B |
| PPDPF | 1.21E-04 | 1.04247 | 0.433 | 0.35 | 1 | HC_vs_MS.NoTrt_B |
| UBE2L3 | 1.22E-04 | -0.2933 | 0.241 | 0.38 | 1 | HC_vs_MS.NoTrt_B |
| TMEM213 | 1.22E-04 | 0.4293 | 0.816 | 0.75 | 1 | HC_vs_MS.NoTrt_B |
| ADAMTSL3 | 1.23E-04 | 0.35941 | 0.945 | 0.98 | 1 | HC_vs_MS.NoTrt_B |
| PPIB | 1.24E-04 | -0.4423 | 0.677 | 0.74 | 1 | HC_vs_MS.NoTrt_B |

| | | | | | | |
|---|---|---|---|---|---|---|
| ATP5J2 | 1.24E-04 | -0.3041 | 0.278 | 0.43 | 1 | HC_vs_MS.NoTrt_B |
| CCDC115 | 1.26E-04 | -0.6916 | 0.086 | 0.18 | 1 | HC_vs_MS.NoTrt_B |
| CACNB3 | 1.26E-04 | 0.86457 | 0.633 | 0.56 | 1 | HC_vs_MS.NoTrt_B |
| RP4-583P15 | 1.31E-04 | 0.35066 | 0.099 | 0.21 | 1 | HC_vs_MS.NoTrt_B |
| NCL | 1.33E-04 | -0.3239 | 0.499 | 0.64 | 1 | HC_vs_MS.NoTrt_B |
| HINT2 | 1.36E-04 | -0.7462 | 0.071 | 0.16 | 1 | HC_vs_MS.NoTrt_B |
| LSM4 | 1.37E-04 | -0.3599 | 0.165 | 0.3 | 1 | HC_vs_MS.NoTrt_B |
| SLC1A6 | 1.39E-04 | 0.39501 | 0.799 | 0.7 | 1 | HC_vs_MS.NoTrt_B |
| MIXL1 | 1.44E-04 | 0.37252 | 0.577 | 0.42 | 1 | HC_vs_MS.NoTrt_B |
| AKT1S1 | 1.46E-04 | -0.4101 | 0.136 | 0.25 | 1 | HC_vs_MS.NoTrt_B |
| STARD9 | 1.49E-04 | 0.5068 | 0.903 | 0.93 | 1 | HC_vs_MS.NoTrt_B |
| MGAT1 | 1.50E-04 | -0.442 | 0.487 | 0.63 | 1 | HC_vs_MS.NoTrt_B |
| KCNE4 | 1.50E-04 | 0.45638 | 0.864 | 0.87 | 1 | HC_vs_MS.NoTrt_B |
| SLC3A2 | 1.50E-04 | -0.6202 | 0.441 | 0.56 | 1 | HC_vs_MS.NoTrt_B |
| NONO | 1.51E-04 | -0.2726 | 0.651 | 0.78 | 1 | HC_vs_MS.NoTrt_B |
| USP8 | 1.53E-04 | 0.26586 | 0.99 | 0.99 | 1 | HC_vs_MS.NoTrt_B |
| TRABD | 1.53E-04 | -0.3159 | 0.32 | 0.46 | 1 | HC_vs_MS.NoTrt_B |
| FARP1 | 1.53E-04 | 0.44881 | 0.954 | 0.95 | 1 | HC_vs_MS.NoTrt_B |
| SLBP | 1.54E-04 | -0.4153 | 0.097 | 0.2 | 1 | HC_vs_MS.NoTrt_B |
| THEMIS2 | 1.55E-04 | -0.5207 | 0.094 | 0.2 | 1 | HC_vs_MS.NoTrt_B |
| PA2G4 | 1.58E-04 | -0.3404 | 0.339 | 0.49 | 1 | HC_vs_MS.NoTrt_B |
| ATP5O | 1.59E-04 | -0.3792 | 0.367 | 0.52 | 1 | HC_vs_MS.NoTrt_B |
| WTIP | 1.59E-04 | 0.36716 | 0.964 | 0.97 | 1 | HC_vs_MS.NoTrt_B |
| GLOD4 | 1.64E-04 | -0.4774 | 0.612 | 0.72 | 1 | HC_vs_MS.NoTrt_B |
| MRPS6 | 1.66E-04 | -0.4711 | 0.148 | 0.27 | 1 | HC_vs_MS.NoTrt_B |
| TUFM | 1.66E-04 | -0.2665 | 0.243 | 0.4 | 1 | HC_vs_MS.NoTrt_B |
| OPHN1 | 1.68E-04 | 0.38957 | 0.949 | 0.97 | 1 | HC_vs_MS.NoTrt_B |
| HSPA1B | 1.68E-04 | 0.66864 | 0.201 | 0.08 | 1 | HC_vs_MS.NoTrt_B |
| MRPL11 | 1.69E-04 | -0.6995 | 0.08 | 0.17 | 1 | HC_vs_MS.NoTrt_B |
| HLA-DRB1 | 1.69E-04 | -0.8313 | 0.174 | 0.29 | 1 | HC_vs_MS.NoTrt_B |
| CAPN2 | 1.77E-04 | -0.4268 | 0.742 | 0.85 | 1 | HC_vs_MS.NoTrt_B |
| IK | 1.78E-04 | -0.3941 | 0.79 | 0.87 | 1 | HC_vs_MS.NoTrt_B |
| RPL39 | 1.82E-04 | -0.3173 | 0.736 | 0.92 | 1 | HC_vs_MS.NoTrt_B |
| SIT1 | 1.87E-04 | -0.3054 | 0.219 | 0.36 | 1 | HC_vs_MS.NoTrt_B |
| IRF3 | 1.87E-04 | -0.5153 | 0.749 | 0.82 | 1 | HC_vs_MS.NoTrt_B |
| TACC2 | 1.88E-04 | 0.371 | 0.9 | 0.91 | 1 | HC_vs_MS.NoTrt_B |
| PSMC3 | 1.90E-04 | -0.491 | 0.103 | 0.21 | 1 | HC_vs_MS.NoTrt_B |
| BEX3 | 1.90E-04 | -0.3641 | 0.104 | 0.21 | 1 | HC_vs_MS.NoTrt_B |
| ARHGAP4 | 1.92E-04 | -0.305 | 0.429 | 0.58 | 1 | HC_vs_MS.NoTrt_B |
| SAG | 1.92E-04 | 0.28359 | 0.942 | 0.98 | 1 | HC_vs_MS.NoTrt_B |

| | | | | | | |
|---|---|---|---|---|---|---|
| PPP6C | 1.93E-04 | -0.3887 | 0.175 | 0.3 | 1 | HC_vs_MS.NoTrt_B |
| LAMTOR2 | 1.94E-04 | -0.4693 | 0.077 | 0.17 | 1 | HC_vs_MS.NoTrt_B |
| UBXN1 | 2.01E-04 | -0.352 | 0.241 | 0.39 | 1 | HC_vs_MS.NoTrt_B |
| GRK4 | 2.01E-04 | 0.52392 | 0.565 | 0.45 | 1 | HC_vs_MS.NoTrt_B |
| ZFP36L2 | 2.03E-04 | 1.19553 | 0.533 | 0.45 | 1 | HC_vs_MS.NoTrt_B |
| PTPMT1 | 2.09E-04 | -0.4932 | 0.08 | 0.17 | 1 | HC_vs_MS.NoTrt_B |
| NDUFC1 | 2.13E-04 | -0.3512 | 0.2 | 0.34 | 1 | HC_vs_MS.NoTrt_B |
| ASNA1 | 2.17E-04 | -0.7398 | 0.042 | 0.11 | 1 | HC_vs_MS.NoTrt_B |
| TDP2 | 2.19E-04 | -0.5417 | 0.094 | 0.19 | 1 | HC_vs_MS.NoTrt_B |
| TNFRSF1B | 2.23E-04 | -0.4005 | 0.723 | 0.76 | 1 | HC_vs_MS.NoTrt_B |
| SF3B4 | 2.24E-04 | -0.5172 | 0.172 | 0.29 | 1 | HC_vs_MS.NoTrt_B |
| FCRL3 | 2.26E-04 | 0.75895 | 0.397 | 0.25 | 1 | HC_vs_MS.NoTrt_B |
| MMACHC | 2.27E-04 | 0.53234 | 0.846 | 0.82 | 1 | HC_vs_MS.NoTrt_B |
| NDUFS2 | 2.28E-04 | -0.5457 | 0.384 | 0.5 | 1 | HC_vs_MS.NoTrt_B |
| LRR1 | 2.31E-04 | -1.6411 | 0.072 | 0.16 | 1 | HC_vs_MS.NoTrt_B |
| CPAMD8 | 2.33E-04 | 0.46267 | 0.699 | 0.61 | 1 | HC_vs_MS.NoTrt_B |
| DNAJB11 | 2.34E-04 | -0.3552 | 0.142 | 0.25 | 1 | HC_vs_MS.NoTrt_B |
| MYDGF | 2.36E-04 | -0.591 | 0.113 | 0.22 | 1 | HC_vs_MS.NoTrt_B |
| NPEPL1 | 2.41E-04 | -0.3205 | 0.11 | 0.22 | 1 | HC_vs_MS.NoTrt_B |
| NCBP2-AS2 | 2.44E-04 | -0.3379 | 0.116 | 0.23 | 1 | HC_vs_MS.NoTrt_B |
| RNF7 | 2.45E-04 | -0.4929 | 0.609 | 0.72 | 1 | HC_vs_MS.NoTrt_B |
| ITGB1BP1 | 2.46E-04 | -0.3393 | 0.141 | 0.25 | 1 | HC_vs_MS.NoTrt_B |
| C10orf95 | 2.49E-04 | -0.7197 | 0.162 | 0.27 | 1 | HC_vs_MS.NoTrt_B |
| ZNF124 | 2.50E-04 | 1.33847 | 0.665 | 0.63 | 1 | HC_vs_MS.NoTrt_B |
| JTB | 2.52E-04 | -0.3868 | 0.21 | 0.35 | 1 | HC_vs_MS.NoTrt_B |
| CTSD | 2.53E-04 | -0.5466 | 0.158 | 0.28 | 1 | HC_vs_MS.NoTrt_B |
| TNFRSF25 | 2.57E-04 | -0.2774 | 0.664 | 0.76 | 1 | HC_vs_MS.NoTrt_B |
| SLC36A3 | 2.62E-04 | 0.40005 | 0.441 | 0.31 | 1 | HC_vs_MS.NoTrt_B |
| HAMP | 2.63E-04 | 0.38365 | 0.823 | 0.8 | 1 | HC_vs_MS.NoTrt_B |
| ZNF776 | 2.68E-04 | 0.49753 | 0.751 | 0.68 | 1 | HC_vs_MS.NoTrt_B |
| ZNF891 | 2.69E-04 | 0.58085 | 0.958 | 0.97 | 1 | HC_vs_MS.NoTrt_B |
| AP5M1 | 2.69E-04 | -0.2594 | 0.109 | 0.21 | 1 | HC_vs_MS.NoTrt_B |
| KIAA1328 | 2.70E-04 | 0.42073 | 0.968 | 1 | 1 | HC_vs_MS.NoTrt_B |
| PTPRCAP | 2.71E-04 | -0.3518 | 0.67 | 0.78 | 1 | HC_vs_MS.NoTrt_B |
| LRRC2 | 2.71E-04 | 0.62787 | 0.762 | 0.68 | 1 | HC_vs_MS.NoTrt_B |
| C19orf53 | 2.72E-04 | -0.2882 | 0.371 | 0.51 | 1 | HC_vs_MS.NoTrt_B |
| LIPG | 2.77E-04 | 0.32813 | 0.858 | 0.81 | 1 | HC_vs_MS.NoTrt_B |
| ACP5 | 2.79E-04 | -0.6998 | 0.094 | 0.19 | 1 | HC_vs_MS.NoTrt_B |
| ZNF785 | 2.80E-04 | 0.32746 | 0.906 | 0.9 | 1 | HC_vs_MS.NoTrt_B |
| GPATCH2 | 2.81E-04 | 0.47938 | 0.504 | 0.36 | 1 | HC_vs_MS.NoTrt_B |

| | | | | | | |
|---|---|---|---|---|---|---|
| ZYX | 2.81E-04 | -0.321 | 0.239 | 0.38 | 1 | HC_vs_MS.NoTrt_B |
| CBR1 | 2.81E-04 | -0.4333 | 0.057 | 0.14 | 1 | HC_vs_MS.NoTrt_B |
| ATP1B3 | 2.83E-04 | -0.5193 | 0.086 | 0.18 | 1 | HC_vs_MS.NoTrt_B |
| GLRX | 2.86E-04 | -0.4179 | 0.683 | 0.78 | 1 | HC_vs_MS.NoTrt_B |
| FAM104A | 2.87E-04 | -0.4539 | 0.159 | 0.28 | 1 | HC_vs_MS.NoTrt_B |
| HLA-DPA1 | 2.94E-04 | -0.5441 | 0.155 | 0.27 | 1 | HC_vs_MS.NoTrt_B |
| OPA3 | 2.97E-04 | 0.25528 | 0.978 | 0.97 | 1 | HC_vs_MS.NoTrt_B |
| CASP1 | 3.00E-04 | -0.2695 | 0.196 | 0.32 | 1 | HC_vs_MS.NoTrt_B |
| RNF166 | 3.01E-04 | -0.3258 | 0.187 | 0.31 | 1 | HC_vs_MS.NoTrt_B |
| MRPS9 | 3.05E-04 | -0.6337 | 0.087 | 0.18 | 1 | HC_vs_MS.NoTrt_B |
| NUCB2 | 3.06E-04 | 0.738 | 0.971 | 0.94 | 1 | HC_vs_MS.NoTrt_B |
| ZBTB43 | 3.07E-04 | 0.6641 | 0.717 | 0.62 | 1 | HC_vs_MS.NoTrt_B |
| CASP10 | 3.12E-04 | 0.34973 | 0.91 | 0.88 | 1 | HC_vs_MS.NoTrt_B |
| ZNF546 | 3.12E-04 | 0.50803 | 0.893 | 0.91 | 1 | HC_vs_MS.NoTrt_B |
| RCN1 | 3.13E-04 | -0.5772 | 0.068 | 0.15 | 1 | HC_vs_MS.NoTrt_B |
| ZNF814 | 3.18E-04 | 0.45422 | 0.954 | 0.94 | 1 | HC_vs_MS.NoTrt_B |
| SVBP | 3.18E-04 | -0.9483 | 0.097 | 0.19 | 1 | HC_vs_MS.NoTrt_B |
| MDH2 | 3.33E-04 | -0.4652 | 0.18 | 0.3 | 1 | HC_vs_MS.NoTrt_B |
| ZNF253 | 3.33E-04 | 0.40047 | 0.672 | 0.53 | 1 | HC_vs_MS.NoTrt_B |
| DNASE1 | 3.34E-04 | 0.52734 | 0.9 | 0.86 | 1 | HC_vs_MS.NoTrt_B |
| SGSM3 | 3.36E-04 | -0.3399 | 0.146 | 0.26 | 1 | HC_vs_MS.NoTrt_B |
| CWC15 | 3.37E-04 | -0.3072 | 0.19 | 0.31 | 1 | HC_vs_MS.NoTrt_B |
| HMGN3 | 3.42E-04 | -0.2789 | 0.17 | 0.29 | 1 | HC_vs_MS.NoTrt_B |
| ZBTB8A | 3.46E-04 | 0.27837 | 0.939 | 0.94 | 1 | HC_vs_MS.NoTrt_B |
| TBC1D8 | 3.48E-04 | 0.34285 | 0.425 | 0.61 | 1 | HC_vs_MS.NoTrt_B |
| RNF114 | 3.51E-04 | -0.3717 | 0.694 | 0.83 | 1 | HC_vs_MS.NoTrt_B |
| MDM2 | 3.53E-04 | 0.49365 | 0.991 | 1 | 1 | HC_vs_MS.NoTrt_B |
| TMEM179B | 3.57E-04 | -0.4205 | 0.175 | 0.3 | 1 | HC_vs_MS.NoTrt_B |
| CD3EAP | 3.58E-04 | 0.27588 | 0.825 | 0.71 | 1 | HC_vs_MS.NoTrt_B |
| ATP10B | 3.65E-04 | 0.74245 | 0.716 | 0.69 | 1 | HC_vs_MS.NoTrt_B |
| ADRM1 | 3.69E-04 | -0.4489 | 0.172 | 0.29 | 1 | HC_vs_MS.NoTrt_B |
| SAFB | 3.79E-04 | -0.4221 | 0.345 | 0.46 | 1 | HC_vs_MS.NoTrt_B |
| TMED9 | 3.79E-04 | -0.4208 | 0.278 | 0.41 | 1 | HC_vs_MS.NoTrt_B |
| PTTG1 | 3.85E-04 | -0.4672 | 0.075 | 0.16 | 1 | HC_vs_MS.NoTrt_B |
| PSKH1 | 3.94E-04 | -1.3191 | 0.059 | 0.14 | 1 | HC_vs_MS.NoTrt_B |
| PAPD4 | 4.01E-04 | -0.4686 | 0.226 | 0.35 | 1 | HC_vs_MS.NoTrt_B |
| PIAS2 | 4.01E-04 | 1.03595 | 0.777 | 0.7 | 1 | HC_vs_MS.NoTrt_B |
| ROMO1 | 4.05E-04 | -0.4124 | 0.139 | 0.25 | 1 | HC_vs_MS.NoTrt_B |
| SMIM10L1 | 4.07E-04 | -0.447 | 0.165 | 0.28 | 1 | HC_vs_MS.NoTrt_B |
| OPN3 | 4.20E-04 | 0.87054 | 0.294 | 0.17 | 1 | HC_vs_MS.NoTrt_B |

| | | | | | | |
|---|---|---|---|---|---|---|
| CYBRD1 | 4.24E-04 | 0.3917 | 0.833 | 0.82 | 1 | HC_vs_MS.NoTrt_B |
| SLC44A1 | 4.26E-04 | 0.86474 | 0.628 | 0.54 | 1 | HC_vs_MS.NoTrt_B |
| CD48 | 4.35E-04 | -0.2509 | 0.241 | 0.38 | 1 | HC_vs_MS.NoTrt_B |
| RNASEK | 4.44E-04 | -0.4254 | 0.696 | 0.73 | 1 | HC_vs_MS.NoTrt_B |
| CSNK2B | 4.45E-04 | -0.3472 | 0.625 | 0.74 | 1 | HC_vs_MS.NoTrt_B |
| ACLY | 4.49E-04 | -0.2673 | 0.181 | 0.3 | 1 | HC_vs_MS.NoTrt_B |
| CCM2 | 4.50E-04 | -0.2617 | 0.649 | 0.74 | 1 | HC_vs_MS.NoTrt_B |
| SYNPO2L | 4.51E-04 | 0.38495 | 0.526 | 0.38 | 1 | HC_vs_MS.NoTrt_B |
| SDAD1 | 4.59E-04 | -0.3845 | 0.138 | 0.24 | 1 | HC_vs_MS.NoTrt_B |
| CHMP6 | 4.66E-04 | -0.8188 | 0.052 | 0.12 | 1 | HC_vs_MS.NoTrt_B |
| ST3GAL2 | 4.70E-04 | 0.58797 | 0.938 | 0.97 | 1 | HC_vs_MS.NoTrt_B |
| MIEN1 | 4.72E-04 | -0.3724 | 0.087 | 0.18 | 1 | HC_vs_MS.NoTrt_B |
| CCDC57 | 4.73E-04 | 0.27912 | 0.991 | 1 | 1 | HC_vs_MS.NoTrt_B |
| ARHGAP22 | 4.75E-04 | 0.47364 | 0.643 | 0.57 | 1 | HC_vs_MS.NoTrt_B |
| MOSPD2 | 4.79E-04 | 0.80026 | 0.309 | 0.19 | 1 | HC_vs_MS.NoTrt_B |
| KLHL7 | 4.80E-04 | 0.52545 | 0.712 | 0.67 | 1 | HC_vs_MS.NoTrt_B |
| HNRNPH1 | 4.81E-04 | 0.31748 | 0.99 | 1 | 1 | HC_vs_MS.NoTrt_B |
| AC013461.1 | 4.82E-04 | 0.32198 | 0.81 | 0.76 | 1 | HC_vs_MS.NoTrt_B |
| PDIA3 | 4.90E-04 | -0.2513 | 0.309 | 0.45 | 1 | HC_vs_MS.NoTrt_B |
| SNRPB2 | 4.93E-04 | -0.3903 | 0.164 | 0.28 | 1 | HC_vs_MS.NoTrt_B |
| GLIPR1 | 4.95E-04 | -0.3269 | 0.216 | 0.34 | 1 | HC_vs_MS.NoTrt_B |
| SPG7 | 4.96E-04 | 0.41906 | 0.978 | 0.98 | 1 | HC_vs_MS.NoTrt_B |
| RP11-111M: | 5.06E-04 | 0.43281 | 0.862 | 0.88 | 1 | HC_vs_MS.NoTrt_B |
| MRPL9 | 5.08E-04 | -0.3648 | 0.168 | 0.28 | 1 | HC_vs_MS.NoTrt_B |
| ZNF445 | 5.09E-04 | 0.47645 | 0.978 | 0.98 | 1 | HC_vs_MS.NoTrt_B |
| ACAP1 | 5.18E-04 | -0.2693 | 0.893 | 0.93 | 1 | HC_vs_MS.NoTrt_B |
| C14orf166 | 5.28E-04 | -0.4144 | 0.526 | 0.64 | 1 | HC_vs_MS.NoTrt_B |
| ACP2 | 5.30E-04 | -1.0301 | 0.039 | 0.1 | 1 | HC_vs_MS.NoTrt_B |
| DPF2 | 5.32E-04 | 0.48899 | 0.901 | 0.87 | 1 | HC_vs_MS.NoTrt_B |
| HOOK3 | 5.37E-04 | 0.57806 | 0.935 | 0.95 | 1 | HC_vs_MS.NoTrt_B |
| DHX16 | 5.46E-04 | -0.3504 | 0.112 | 0.21 | 1 | HC_vs_MS.NoTrt_B |
| RHOG | 5.46E-04 | -0.2751 | 0.314 | 0.46 | 1 | HC_vs_MS.NoTrt_B |
| SLC4A8 | 5.49E-04 | 0.51634 | 0.88 | 0.9 | 1 | HC_vs_MS.NoTrt_B |
| KLRD1 | 5.55E-04 | 0.31807 | 0.965 | 0.98 | 1 | HC_vs_MS.NoTrt_B |
| NDUFB10 | 5.64E-04 | -0.282 | 0.177 | 0.3 | 1 | HC_vs_MS.NoTrt_B |
| MARVELD2 | 5.67E-04 | 0.69802 | 0.59 | 0.53 | 1 | HC_vs_MS.NoTrt_B |
| CDC26 | 5.81E-04 | -0.2779 | 0.162 | 0.28 | 1 | HC_vs_MS.NoTrt_B |
| CRY1 | 5.81E-04 | -0.7073 | 0.091 | 0.18 | 1 | HC_vs_MS.NoTrt_B |
| ERICH1 | 5.82E-04 | -0.402 | 0.178 | 0.29 | 1 | HC_vs_MS.NoTrt_B |
| ZNF157 | 5.85E-04 | 0.44902 | 0.758 | 0.72 | 1 | HC_vs_MS.NoTrt_B |

| | | | | | | |
|---|---|---|---|---|---|---|
| GCNT2 | 5.89E-04 | 0.97514 | 0.732 | 0.67 | 1 | HC_vs_MS.NoTrt_B |
| SIVA1 | 5.90E-04 | -0.431 | 0.343 | 0.47 | 1 | HC_vs_MS.NoTrt_B |
| SMIM14 | 5.91E-04 | 0.3254 | 0.983 | 0.99 | 1 | HC_vs_MS.NoTrt_B |
| AURKAIP1 | 5.95E-04 | -0.3766 | 0.141 | 0.25 | 1 | HC_vs_MS.NoTrt_B |
| CHAF1B | 5.97E-04 | 0.33144 | 0.932 | 0.94 | 1 | HC_vs_MS.NoTrt_B |
| ZNF737 | 5.97E-04 | 0.3091 | 0.833 | 0.82 | 1 | HC_vs_MS.NoTrt_B |
| TARSL2 | 6.04E-04 | 0.28486 | 0.97 | 0.98 | 1 | HC_vs_MS.NoTrt_B |
| RHBG | 6.07E-04 | 0.33588 | 0.768 | 0.75 | 1 | HC_vs_MS.NoTrt_B |
| KCNJ3 | 6.08E-04 | 0.37788 | 0.214 | 0.1 | 1 | HC_vs_MS.NoTrt_B |
| H2AFJ | 6.23E-04 | -0.4682 | 0.086 | 0.17 | 1 | HC_vs_MS.NoTrt_B |
| SPOCK2 | 6.28E-04 | -0.2532 | 0.594 | 0.71 | 1 | HC_vs_MS.NoTrt_B |
| TAOK1 | 6.36E-04 | 0.90824 | 0.906 | 0.9 | 1 | HC_vs_MS.NoTrt_B |
| COX6A1 | 6.36E-04 | -0.3444 | 0.113 | 0.21 | 1 | HC_vs_MS.NoTrt_B |
| NPIPB4 | 6.38E-04 | 0.34719 | 0.951 | 0.98 | 1 | HC_vs_MS.NoTrt_B |
| POLR3GL | 6.44E-04 | -0.3359 | 0.174 | 0.29 | 1 | HC_vs_MS.NoTrt_B |
| LDLRAP1 | 6.48E-04 | -0.3952 | 0.449 | 0.55 | 1 | HC_vs_MS.NoTrt_B |
| GDPGP1 | 6.50E-04 | 0.47405 | 0.926 | 0.94 | 1 | HC_vs_MS.NoTrt_B |
| HTN1 | 6.50E-04 | 0.56621 | 0.768 | 0.75 | 1 | HC_vs_MS.NoTrt_B |
| CFDP1 | 6.52E-04 | -0.7905 | 0.641 | 0.71 | 1 | HC_vs_MS.NoTrt_B |
| UFSP2 | 6.54E-04 | -0.3249 | 0.074 | 0.16 | 1 | HC_vs_MS.NoTrt_B |
| KAT2B | 6.56E-04 | 0.40495 | 0.139 | 0.25 | 1 | HC_vs_MS.NoTrt_B |
| ATP1A4 | 6.58E-04 | 0.46628 | 0.783 | 0.75 | 1 | HC_vs_MS.NoTrt_B |
| ZNF524 | 6.58E-04 | -0.6607 | 0.057 | 0.13 | 1 | HC_vs_MS.NoTrt_B |
| TRIB1 | 6.61E-04 | -0.7557 | 0.046 | 0.11 | 1 | HC_vs_MS.NoTrt_B |
| PHLDB2 | 6.75E-04 | 0.44079 | 0.51 | 0.41 | 1 | HC_vs_MS.NoTrt_B |
| KANK2 | 6.79E-04 | 0.371 | 0.894 | 0.89 | 1 | HC_vs_MS.NoTrt_B |
| FAAP100 | 6.95E-04 | -0.5534 | 0.119 | 0.22 | 1 | HC_vs_MS.NoTrt_B |
| EIF3G | 7.02E-04 | -0.3572 | 0.816 | 0.89 | 1 | HC_vs_MS.NoTrt_B |
| STK16 | 7.03E-04 | -0.7969 | 0.043 | 0.11 | 1 | HC_vs_MS.NoTrt_B |
| POLE | 7.06E-04 | 0.5495 | 0.848 | 0.83 | 1 | HC_vs_MS.NoTrt_B |
| STXBP2 | 7.07E-04 | -0.2695 | 0.252 | 0.38 | 1 | HC_vs_MS.NoTrt_B |
| TAF1C | 7.12E-04 | -0.3549 | 0.075 | 0.16 | 1 | HC_vs_MS.NoTrt_B |
| MKKS | 7.23E-04 | -0.3945 | 0.046 | 0.11 | 1 | HC_vs_MS.NoTrt_B |
| PSMB6 | 7.37E-04 | -0.4143 | 0.571 | 0.7 | 1 | HC_vs_MS.NoTrt_B |
| GOSR2 | 7.52E-04 | -0.8144 | 0.086 | 0.17 | 1 | HC_vs_MS.NoTrt_B |
| ERLEC1 | 7.60E-04 | -0.6649 | 0.358 | 0.5 | 1 | HC_vs_MS.NoTrt_B |
| TBC1D8B | 7.67E-04 | 0.51127 | 0.957 | 0.96 | 1 | HC_vs_MS.NoTrt_B |
| DUSP16 | 7.77E-04 | -0.3516 | 0.206 | 0.32 | 1 | HC_vs_MS.NoTrt_B |
| UTP6 | 7.80E-04 | 0.35293 | 0.87 | 0.88 | 1 | HC_vs_MS.NoTrt_B |
| MACC1 | 7.95E-04 | 0.46971 | 0.672 | 0.59 | 1 | HC_vs_MS.NoTrt_B |

| | | | | | | |
|---|---|---|---|---|---|---|
| NUP37 | 8.08E-04 | -1.0157 | 0.275 | 0.4 | 1 | HC_vs_MS.NoTrt_B |
| PYGL | 8.08E-04 | 0.37074 | 0.888 | 0.87 | 1 | HC_vs_MS.NoTrt_B |
| EIF6 | 8.15E-04 | -0.5152 | 0.125 | 0.22 | 1 | HC_vs_MS.NoTrt_B |
| NT5C | 8.18E-04 | -0.3366 | 0.104 | 0.2 | 1 | HC_vs_MS.NoTrt_B |
| PCDHA4 | 8.23E-04 | 0.41007 | 0.904 | 0.86 | 1 | HC_vs_MS.NoTrt_B |
| MILR1 | 8.43E-04 | 0.36806 | 0.854 | 0.86 | 1 | HC_vs_MS.NoTrt_B |
| TAF1 | 8.54E-04 | 0.9269 | 0.799 | 0.8 | 1 | HC_vs_MS.NoTrt_B |
| ZNF660 | 8.58E-04 | 0.47395 | 0.564 | 0.51 | 1 | HC_vs_MS.NoTrt_B |
| RAP2C | 8.63E-04 | -0.3749 | 0.245 | 0.36 | 1 | HC_vs_MS.NoTrt_B |
| PCSK5 | 8.65E-04 | -0.426 | 0.97 | 0.95 | 1 | HC_vs_MS.NoTrt_B |
| PRMT2 | 8.79E-04 | -0.2734 | 0.343 | 0.47 | 1 | HC_vs_MS.NoTrt_B |
| VCPKMT | 8.85E-04 | -0.38 | 0.07 | 0.15 | 1 | HC_vs_MS.NoTrt_B |
| NIF3L1 | 8.88E-04 | -0.9727 | 0.074 | 0.15 | 1 | HC_vs_MS.NoTrt_B |
| DPPA4 | 9.01E-04 | 0.4884 | 0.852 | 0.87 | 1 | HC_vs_MS.NoTrt_B |
| IL1RAP | 9.07E-04 | 0.29398 | 0.658 | 0.59 | 1 | HC_vs_MS.NoTrt_B |
| RP11-77H9.: | 9.20E-04 | 0.36756 | 0.497 | 0.37 | 1 | HC_vs_MS.NoTrt_B |
| AP000304.1 | 9.35E-04 | -0.3867 | 0.197 | 0.31 | 1 | HC_vs_MS.NoTrt_B |
| SATB1 | 9.47E-04 | 0.89026 | 0.239 | 0.12 | 1 | HC_vs_MS.NoTrt_B |
| POLR2L | 9.58E-04 | -0.6095 | 0.212 | 0.32 | 1 | HC_vs_MS.NoTrt_B |
| NEK4 | 9.61E-04 | 0.51919 | 0.767 | 0.72 | 1 | HC_vs_MS.NoTrt_B |
| TMEM159 | 9.62E-04 | 0.30172 | 0.607 | 0.49 | 1 | HC_vs_MS.NoTrt_B |
| MUL1 | 9.67E-04 | -0.6609 | 0.041 | 0.1 | 1 | HC_vs_MS.NoTrt_B |
| SCIN | 9.71E-04 | 0.28229 | 0.88 | 0.89 | 1 | HC_vs_MS.NoTrt_B |
| CCR10 | 9.74E-04 | -0.6725 | 0.081 | 0.16 | 1 | HC_vs_MS.NoTrt_B |
| CCDC191 | 9.89E-04 | -0.4642 | 0.181 | 0.3 | 1 | HC_vs_MS.NoTrt_B |
| PHLDB1 | 0.00101 | 0.37983 | 0.942 | 0.95 | 1 | HC_vs_MS.NoTrt_B |
| RELL2 | 0.00101 | -0.4349 | 0.077 | 0.16 | 1 | HC_vs_MS.NoTrt_B |
| ZBTB8B | 0.00102 | 0.26411 | 0.954 | 0.94 | 1 | HC_vs_MS.NoTrt_B |
| RALB | 0.00103 | -0.274 | 0.048 | 0.11 | 1 | HC_vs_MS.NoTrt_B |
| OPTN | 0.00103 | -0.4555 | 0.696 | 0.78 | 1 | HC_vs_MS.NoTrt_B |
| EIF2S1 | 0.00104 | -0.3727 | 0.258 | 0.36 | 1 | HC_vs_MS.NoTrt_B |
| CPT1C | 0.00105 | 0.33963 | 0.883 | 0.92 | 1 | HC_vs_MS.NoTrt_B |
| MPV17L | 0.00105 | 0.33476 | 0.975 | 0.98 | 1 | HC_vs_MS.NoTrt_B |
| LRRK1 | 0.00106 | 0.48286 | 0.843 | 0.81 | 1 | HC_vs_MS.NoTrt_B |
| NPFFR1 | 0.00107 | 0.27608 | 0.891 | 0.85 | 1 | HC_vs_MS.NoTrt_B |
| EIF3I | 0.00108 | -0.7166 | 0.468 | 0.56 | 1 | HC_vs_MS.NoTrt_B |
| MALSU1 | 0.00109 | -0.4954 | 0.081 | 0.16 | 1 | HC_vs_MS.NoTrt_B |
| SHOX | 0.0011 | 0.37302 | 0.839 | 0.78 | 1 | HC_vs_MS.NoTrt_B |
| TCF7 | 0.0011 | 0.6601 | 0.787 | 0.73 | 1 | HC_vs_MS.NoTrt_B |
| MRPL53 | 0.00111 | -0.2821 | 0.087 | 0.17 | 1 | HC_vs_MS.NoTrt_B |

| | | | | | | |
|---|---|---|---|---|---|---|
| LYPLA2 | 0.00112 | -0.4417 | 0.135 | 0.23 | 1 | HC_vs_MS.NoTrt_B |
| ZNF511 | 0.00112 | -0.8128 | 0.052 | 0.12 | 1 | HC_vs_MS.NoTrt_B |
| ABC7-42404 | 0.00113 | 0.26791 | 0.707 | 0.63 | 1 | HC_vs_MS.NoTrt_B |
| PEPD | 0.00114 | -0.7777 | 0.104 | 0.19 | 1 | HC_vs_MS.NoTrt_B |
| NELFA | 0.00114 | 0.43885 | 0.786 | 0.76 | 1 | HC_vs_MS.NoTrt_B |
| TBCA | 0.00115 | -0.3375 | 0.213 | 0.32 | 1 | HC_vs_MS.NoTrt_B |
| SKA1 | 0.00115 | 0.33357 | 0.677 | 0.6 | 1 | HC_vs_MS.NoTrt_B |
| PLAA | 0.00115 | -0.6719 | 0.084 | 0.16 | 1 | HC_vs_MS.NoTrt_B |
| ITPA | 0.00116 | -0.4093 | 0.113 | 0.21 | 1 | HC_vs_MS.NoTrt_B |
| TSPYL4 | 0.00116 | -0.4855 | 0.104 | 0.2 | 1 | HC_vs_MS.NoTrt_B |
| ZNF347 | 0.00117 | 0.29006 | 0.91 | 0.9 | 1 | HC_vs_MS.NoTrt_B |
| CUTA | 0.00117 | -0.3235 | 0.27 | 0.4 | 1 | HC_vs_MS.NoTrt_B |
| DIDO1 | 0.00117 | -0.3881 | 0.213 | 0.33 | 1 | HC_vs_MS.NoTrt_B |
| UFD1L | 0.00119 | -0.304 | 0.761 | 0.83 | 1 | HC_vs_MS.NoTrt_B |
| NOSIP | 0.00119 | -0.2894 | 0.484 | 0.61 | 1 | HC_vs_MS.NoTrt_B |
| IBA57 | 0.00119 | 0.88504 | 0.772 | 0.73 | 1 | HC_vs_MS.NoTrt_B |
| AFAP1L1 | 0.00119 | 0.30122 | 0.461 | 0.32 | 1 | HC_vs_MS.NoTrt_B |
| ZNF829 | 0.00121 | 0.5972 | 0.877 | 0.89 | 1 | HC_vs_MS.NoTrt_B |
| EFHC1 | 0.00121 | 0.31877 | 0.903 | 0.9 | 1 | HC_vs_MS.NoTrt_B |
| B3GAT3 | 0.00121 | -0.3989 | 0.532 | 0.62 | 1 | HC_vs_MS.NoTrt_B |
| TRIOBP | 0.00122 | 0.48697 | 0.397 | 0.27 | 1 | HC_vs_MS.NoTrt_B |
| MTG1 | 0.00123 | -0.4097 | 0.148 | 0.25 | 1 | HC_vs_MS.NoTrt_B |
| AXL | 0.00125 | 0.27751 | 0.668 | 0.56 | 1 | HC_vs_MS.NoTrt_B |
| RP2 | 0.00125 | 0.58734 | 0.646 | 0.62 | 1 | HC_vs_MS.NoTrt_B |
| CAMLG | 0.00126 | -0.4122 | 0.442 | 0.56 | 1 | HC_vs_MS.NoTrt_B |
| SENP7 | 0.00129 | 0.40502 | 0.987 | 0.97 | 1 | HC_vs_MS.NoTrt_B |
| COX5A | 0.0013 | -0.2991 | 0.206 | 0.33 | 1 | HC_vs_MS.NoTrt_B |
| ARIH1 | 0.00131 | 0.36837 | 0.964 | 0.95 | 1 | HC_vs_MS.NoTrt_B |
| GBP3 | 0.00131 | -0.5928 | 0.194 | 0.29 | 1 | HC_vs_MS.NoTrt_B |
| CD79B | 0.00133 | -0.5335 | 0.059 | 0.13 | 1 | HC_vs_MS.NoTrt_B |
| TERF1 | 0.00133 | 0.45197 | 0.72 | 0.66 | 1 | HC_vs_MS.NoTrt_B |
| MAZ | 0.00134 | -0.5384 | 0.138 | 0.23 | 1 | HC_vs_MS.NoTrt_B |
| PPIL3 | 0.00135 | 0.2988 | 0.796 | 0.78 | 1 | HC_vs_MS.NoTrt_B |
| BTD | 0.00136 | -0.5562 | 0.075 | 0.15 | 1 | HC_vs_MS.NoTrt_B |
| NCF4 | 0.00136 | -0.3878 | 0.161 | 0.26 | 1 | HC_vs_MS.NoTrt_B |
| AKR7A2 | 0.00136 | -0.3663 | 0.113 | 0.21 | 1 | HC_vs_MS.NoTrt_B |
| G6PC3 | 0.00136 | 0.35804 | 0.819 | 0.77 | 1 | HC_vs_MS.NoTrt_B |
| RPA1 | 0.00137 | -0.6561 | 0.264 | 0.37 | 1 | HC_vs_MS.NoTrt_B |
| WDR61 | 0.00137 | -0.3097 | 0.13 | 0.23 | 1 | HC_vs_MS.NoTrt_B |
| HSPA4L | 0.00139 | 0.33369 | 0.874 | 0.86 | 1 | HC_vs_MS.NoTrt_B |

| | | | | | | |
|---|---|---|---|---|---|---|
| ARRDC5 | 0.00139 | -0.7419 | 0.087 | 0.17 | 1 | HC_vs_MS.NoTrt_B |
| ZNF708 | 0.00141 | 0.60589 | 0.784 | 0.75 | 1 | HC_vs_MS.NoTrt_B |
| TTC12 | 0.00141 | -0.6196 | 0.097 | 0.18 | 1 | HC_vs_MS.NoTrt_B |
| STAG1 | 0.00142 | 0.33508 | 0.206 | 0.33 | 1 | HC_vs_MS.NoTrt_B |
| PDCD10 | 0.00142 | -0.2799 | 0.174 | 0.28 | 1 | HC_vs_MS.NoTrt_B |
| XPR1 | 0.00143 | -0.3805 | 0.299 | 0.41 | 1 | HC_vs_MS.NoTrt_B |
| ZNF100 | 0.00143 | 0.63291 | 0.835 | 0.82 | 1 | HC_vs_MS.NoTrt_B |
| DENND2D | 0.00143 | 0.81997 | 0.384 | 0.28 | 1 | HC_vs_MS.NoTrt_B |
| NFS1 | 0.00144 | -0.5151 | 0.886 | 0.87 | 1 | HC_vs_MS.NoTrt_B |
| TRIM27 | 0.00144 | -0.5364 | 0.384 | 0.48 | 1 | HC_vs_MS.NoTrt_B |
| SPACA6 | 0.00146 | 0.26458 | 0.87 | 0.86 | 1 | HC_vs_MS.NoTrt_B |
| COX5B | 0.00147 | -0.2773 | 0.759 | 0.83 | 1 | HC_vs_MS.NoTrt_B |
| SFT2D2 | 0.00148 | 0.30923 | 0.98 | 0.98 | 1 | HC_vs_MS.NoTrt_B |
| SF3B5 | 0.00151 | -0.4119 | 0.086 | 0.17 | 1 | HC_vs_MS.NoTrt_B |
| FBXL2 | 0.00151 | 0.57749 | 0.636 | 0.58 | 1 | HC_vs_MS.NoTrt_B |
| ANKLE2 | 0.00152 | 0.3595 | 0.984 | 0.99 | 1 | HC_vs_MS.NoTrt_B |
| VAV3 | 0.00153 | 0.54898 | 0.551 | 0.43 | 1 | HC_vs_MS.NoTrt_B |
| TMEM145 | 0.00153 | 0.2932 | 0.893 | 0.94 | 1 | HC_vs_MS.NoTrt_B |
| DDX24 | 0.00153 | -0.3289 | 0.51 | 0.64 | 1 | HC_vs_MS.NoTrt_B |
| SLITRK5 | 0.00153 | 0.34446 | 0.733 | 0.65 | 1 | HC_vs_MS.NoTrt_B |
| LIG1 | 0.00154 | -0.4343 | 0.13 | 0.22 | 1 | HC_vs_MS.NoTrt_B |
| AACS | 0.00154 | 0.56014 | 0.986 | 0.98 | 1 | HC_vs_MS.NoTrt_B |
| FN3K | 0.00155 | 0.43254 | 0.783 | 0.76 | 1 | HC_vs_MS.NoTrt_B |
| BAD | 0.00155 | -0.4021 | 0.061 | 0.13 | 1 | HC_vs_MS.NoTrt_B |
| LYRM7 | 0.00155 | 0.54568 | 0.919 | 0.91 | 1 | HC_vs_MS.NoTrt_B |
| EMC6 | 0.00156 | -0.5078 | 0.057 | 0.12 | 1 | HC_vs_MS.NoTrt_B |
| TMEM126A | 0.00156 | -0.7606 | 0.054 | 0.12 | 1 | HC_vs_MS.NoTrt_B |
| NIM1K | 0.00158 | 0.26014 | 0.703 | 0.62 | 1 | HC_vs_MS.NoTrt_B |
| MOCS3 | 0.00158 | 0.80225 | 0.822 | 0.81 | 1 | HC_vs_MS.NoTrt_B |
| MAP7D3 | 0.00159 | 0.62186 | 0.577 | 0.49 | 1 | HC_vs_MS.NoTrt_B |
| ERP29 | 0.00159 | -0.3414 | 0.848 | 0.88 | 1 | HC_vs_MS.NoTrt_B |
| GPI | 0.00161 | -0.4142 | 0.886 | 0.92 | 1 | HC_vs_MS.NoTrt_B |
| TCF23 | 0.00161 | 0.2753 | 0.875 | 0.88 | 1 | HC_vs_MS.NoTrt_B |
| RCE1 | 0.00161 | -0.4445 | 0.061 | 0.13 | 1 | HC_vs_MS.NoTrt_B |
| ZNF26 | 0.00165 | 0.4159 | 0.996 | 0.99 | 1 | HC_vs_MS.NoTrt_B |
| PRF1 | 0.00167 | -0.3794 | 0.155 | 0.25 | 1 | HC_vs_MS.NoTrt_B |
| EIF3K | 0.00169 | -0.3188 | 0.664 | 0.78 | 1 | HC_vs_MS.NoTrt_B |
| GATAD1 | 0.0017 | 0.46783 | 0.877 | 0.88 | 1 | HC_vs_MS.NoTrt_B |
| SDHA | 0.0017 | -0.411 | 0.613 | 0.7 | 1 | HC_vs_MS.NoTrt_B |
| DBN1 | 0.00173 | 0.68761 | 0.217 | 0.12 | 1 | HC_vs_MS.NoTrt_B |

| | | | | | | |
|---|---|---|---|---|---|---|
| ZNF426 | 0.00174 | 1.02474 | 0.912 | 0.9 | 1 | HC_vs_MS.NoTrt_B |
| VPS16 | 0.00178 | -0.7214 | 0.086 | 0.16 | 1 | HC_vs_MS.NoTrt_B |
| LINC01125 | 0.00179 | 0.30813 | 0.949 | 0.96 | 1 | HC_vs_MS.NoTrt_B |
| MED11 | 0.0018 | 0.45822 | 0.691 | 0.62 | 1 | HC_vs_MS.NoTrt_B |
| CBWD2 | 0.00181 | -0.2678 | 0.414 | 0.54 | 1 | HC_vs_MS.NoTrt_B |
| STRN3 | 0.00182 | 0.78988 | 0.78 | 0.71 | 1 | HC_vs_MS.NoTrt_B |
| ZNF284 | 0.00183 | -0.9565 | 0.558 | 0.67 | 1 | HC_vs_MS.NoTrt_B |
| SPAG4 | 0.00183 | -0.6323 | 0.188 | 0.3 | 1 | HC_vs_MS.NoTrt_B |
| LGMN | 0.00184 | 0.62657 | 0.958 | 0.98 | 1 | HC_vs_MS.NoTrt_B |
| MPC2 | 0.00185 | -0.2737 | 0.428 | 0.53 | 1 | HC_vs_MS.NoTrt_B |
| PGM1 | 0.00186 | -0.9474 | 0.062 | 0.13 | 1 | HC_vs_MS.NoTrt_B |
| HERC3 | 0.00188 | -0.3949 | 0.283 | 0.38 | 1 | HC_vs_MS.NoTrt_B |
| GMPR2 | 0.00191 | -0.7047 | 0.107 | 0.19 | 1 | HC_vs_MS.NoTrt_B |
| YPEL5 | 0.00193 | -0.2745 | 0.175 | 0.28 | 1 | HC_vs_MS.NoTrt_B |
| LPXN | 0.00196 | -0.2577 | 0.3 | 0.41 | 1 | HC_vs_MS.NoTrt_B |
| SSNA1 | 0.00202 | -0.2964 | 0.129 | 0.23 | 1 | HC_vs_MS.NoTrt_B |
| NMNAT1 | 0.00203 | 0.44078 | 0.977 | 1 | 1 | HC_vs_MS.NoTrt_B |
| SPTLC2 | 0.00205 | 0.25124 | 0.697 | 0.6 | 1 | HC_vs_MS.NoTrt_B |
| SLC8B1 | 0.00207 | -0.292 | 0.433 | 0.55 | 1 | HC_vs_MS.NoTrt_B |
| ABCA8 | 0.00208 | 0.40947 | 0.441 | 0.32 | 1 | HC_vs_MS.NoTrt_B |
| NBPF10 | 0.0021 | 1.01818 | 0.278 | 0.17 | 1 | HC_vs_MS.NoTrt_B |
| FAM172A | 0.00211 | -0.3485 | 0.11 | 0.2 | 1 | HC_vs_MS.NoTrt_B |
| FMO3 | 0.00211 | 0.45764 | 0.567 | 0.51 | 1 | HC_vs_MS.NoTrt_B |
| FTO | 0.00211 | 0.35961 | 0.99 | 1 | 1 | HC_vs_MS.NoTrt_B |
| NEK5 | 0.00211 | 0.38294 | 0.852 | 0.83 | 1 | HC_vs_MS.NoTrt_B |
| CACNA1D | 0.00215 | 0.63228 | 0.661 | 0.58 | 1 | HC_vs_MS.NoTrt_B |
| PTGES3L | 0.00216 | 0.43775 | 0.82 | 0.82 | 1 | HC_vs_MS.NoTrt_B |
| CTTNBP2NL | 0.00217 | 0.6919 | 0.441 | 0.34 | 1 | HC_vs_MS.NoTrt_B |
| SMPD1 | 0.00217 | -0.9052 | 0.055 | 0.12 | 1 | HC_vs_MS.NoTrt_B |
| ATRAID | 0.00219 | -0.3075 | 0.183 | 0.29 | 1 | HC_vs_MS.NoTrt_B |
| VDAC1 | 0.00222 | -0.3941 | 0.719 | 0.84 | 1 | HC_vs_MS.NoTrt_B |
| TALDO1 | 0.00223 | -0.488 | 0.662 | 0.78 | 1 | HC_vs_MS.NoTrt_B |
| VTI1B | 0.00228 | -0.3706 | 0.217 | 0.32 | 1 | HC_vs_MS.NoTrt_B |
| STMND1 | 0.00229 | 0.32297 | 0.826 | 0.78 | 1 | HC_vs_MS.NoTrt_B |
| GSTO1 | 0.0023 | -0.3983 | 0.11 | 0.2 | 1 | HC_vs_MS.NoTrt_B |
| ST7 | 0.0023 | -1.5295 | 0.045 | 0.1 | 1 | HC_vs_MS.NoTrt_B |
| ERMARD | 0.00233 | -0.6232 | 0.071 | 0.14 | 1 | HC_vs_MS.NoTrt_B |
| SPP2 | 0.00236 | 0.62904 | 0.691 | 0.68 | 1 | HC_vs_MS.NoTrt_B |
| HADHA | 0.00236 | -0.3257 | 0.509 | 0.63 | 1 | HC_vs_MS.NoTrt_B |
| ZNF506 | 0.00237 | 0.29145 | 0.987 | 1 | 1 | HC_vs_MS.NoTrt_B |

| | | | | | | |
|---|---|---|---|---|---|---|
| EIF4E | 0.00243 | 0.29837 | 0.988 | 1 | 1 | HC_vs_MS.NoTrt_B |
| TBC1D30 | 0.00244 | 0.25197 | 0.887 | 0.85 | 1 | HC_vs_MS.NoTrt_B |
| TOMM6 | 0.00246 | -0.2917 | 0.199 | 0.31 | 1 | HC_vs_MS.NoTrt_B |
| PCBP4 | 0.00249 | -1.1149 | 0.049 | 0.11 | 1 | HC_vs_MS.NoTrt_B |
| CASP4 | 0.00249 | -0.3833 | 0.336 | 0.45 | 1 | HC_vs_MS.NoTrt_B |
| APH1A | 0.0025 | -0.4582 | 0.181 | 0.28 | 1 | HC_vs_MS.NoTrt_B |
| DOK1 | 0.0025 | -0.6994 | 0.045 | 0.1 | 1 | HC_vs_MS.NoTrt_B |
| OS9 | 0.00252 | -0.3433 | 0.226 | 0.34 | 1 | HC_vs_MS.NoTrt_B |
| PHF6 | 0.00253 | 0.83269 | 0.667 | 0.59 | 1 | HC_vs_MS.NoTrt_B |
| FAM32A | 0.00254 | -0.5253 | 0.436 | 0.53 | 1 | HC_vs_MS.NoTrt_B |
| HNRNPH2 | 0.00254 | -0.5902 | 0.155 | 0.24 | 1 | HC_vs_MS.NoTrt_B |
| PHLPP2 | 0.00256 | 0.86704 | 0.8 | 0.76 | 1 | HC_vs_MS.NoTrt_B |
| HPF1 | 0.00257 | -0.4136 | 0.078 | 0.15 | 1 | HC_vs_MS.NoTrt_B |
| CTNS | 0.00257 | -1.1094 | 0.106 | 0.18 | 1 | HC_vs_MS.NoTrt_B |
| CREB3 | 0.00258 | -0.8515 | 0.052 | 0.11 | 1 | HC_vs_MS.NoTrt_B |
| KDM4A | 0.00259 | -0.3158 | 0.088 | 0.17 | 1 | HC_vs_MS.NoTrt_B |
| MOCOS | 0.00261 | 0.36521 | 0.625 | 0.53 | 1 | HC_vs_MS.NoTrt_B |
| LY6G5C | 0.00261 | -0.2976 | 0.594 | 0.5 | 1 | HC_vs_MS.NoTrt_B |
| ALKBH7 | 0.00262 | -0.4838 | 0.106 | 0.19 | 1 | HC_vs_MS.NoTrt_B |
| DAGLB | 0.00262 | 0.63929 | 0.828 | 0.84 | 1 | HC_vs_MS.NoTrt_B |
| TM4SF18 | 0.00262 | 0.33289 | 0.267 | 0.16 | 1 | HC_vs_MS.NoTrt_B |
| CLN5 | 0.00263 | 0.31667 | 0.962 | 0.97 | 1 | HC_vs_MS.NoTrt_B |
| TRIP12 | 0.00267 | -0.2854 | 0.458 | 0.58 | 1 | HC_vs_MS.NoTrt_B |
| ZNF329 | 0.00267 | 0.37947 | 0.716 | 0.68 | 1 | HC_vs_MS.NoTrt_B |
| C11orf98 | 0.00268 | -0.7188 | 0.103 | 0.18 | 1 | HC_vs_MS.NoTrt_B |
| TM7SF3 | 0.0027 | 0.42201 | 0.935 | 0.96 | 1 | HC_vs_MS.NoTrt_B |
| MRPL21 | 0.00271 | -0.491 | 0.087 | 0.16 | 1 | HC_vs_MS.NoTrt_B |
| EEF2KMT | 0.00273 | -0.6001 | 0.057 | 0.12 | 1 | HC_vs_MS.NoTrt_B |
| ATP6V0D2 | 0.00273 | 0.30954 | 0.778 | 0.74 | 1 | HC_vs_MS.NoTrt_B |
| ZKSCAN1 | 0.00276 | 0.30469 | 0.939 | 0.92 | 1 | HC_vs_MS.NoTrt_B |
| SULT1A4 | 0.00277 | -0.7241 | 0.21 | 0.3 | 1 | HC_vs_MS.NoTrt_B |
| BCL2L1 | 0.00277 | -0.3053 | 0.123 | 0.21 | 1 | HC_vs_MS.NoTrt_B |
| ZNF554 | 0.00277 | 0.2662 | 0.951 | 0.96 | 1 | HC_vs_MS.NoTrt_B |
| ZNF844 | 0.00279 | 0.30405 | 0.904 | 0.89 | 1 | HC_vs_MS.NoTrt_B |
| GSTT1 | 0.00281 | 0.38841 | 0.723 | 0.64 | 1 | HC_vs_MS.NoTrt_B |
| FSIP2 | 0.00282 | 0.40775 | 0.793 | 0.76 | 1 | HC_vs_MS.NoTrt_B |
| C15orf59 | 0.00282 | 0.3703 | 0.654 | 0.6 | 1 | HC_vs_MS.NoTrt_B |
| PTRHD1 | 0.00284 | -0.5672 | 0.22 | 0.32 | 1 | HC_vs_MS.NoTrt_B |
| HSD17B10 | 0.00284 | -0.4644 | 0.09 | 0.17 | 1 | HC_vs_MS.NoTrt_B |
| DPP8 | 0.0029 | -0.2557 | 0.357 | 0.47 | 1 | HC_vs_MS.NoTrt_B |

| | | | | | | |
|---|---|---|---|---|---|---|
| C1orf186 | 0.00295 | 0.4829 | 0.614 | 0.57 | 1 | HC_vs_MS.NoTrt_B |
| LAMTOR4 | 0.00295 | -0.4185 | 0.441 | 0.54 | 1 | HC_vs_MS.NoTrt_B |
| HLA-DPB1 | 0.00296 | -0.591 | 0.443 | 0.53 | 1 | HC_vs_MS.NoTrt_B |
| HACD2 | 0.00299 | 0.35055 | 0.939 | 0.97 | 1 | HC_vs_MS.NoTrt_B |
| MAP4K3 | 0.003 | -0.7823 | 0.049 | 0.11 | 1 | HC_vs_MS.NoTrt_B |
| POLR2J | 0.00302 | -0.379 | 0.152 | 0.24 | 1 | HC_vs_MS.NoTrt_B |
| FEM1A | 0.00303 | 0.2686 | 0.933 | 0.92 | 1 | HC_vs_MS.NoTrt_B |
| SRI | 0.00303 | -0.3002 | 0.367 | 0.47 | 1 | HC_vs_MS.NoTrt_B |
| BCAS2 | 0.00305 | -0.3769 | 0.162 | 0.25 | 1 | HC_vs_MS.NoTrt_B |
| THOC7 | 0.00306 | -0.3506 | 0.145 | 0.24 | 1 | HC_vs_MS.NoTrt_B |
| RAB8A | 0.00306 | -0.4518 | 0.507 | 0.6 | 1 | HC_vs_MS.NoTrt_B |
| CCL5 | 0.00307 | 0.29928 | 0.89 | 0.88 | 1 | HC_vs_MS.NoTrt_B |
| PPP5D1 | 0.00309 | 0.58477 | 0.729 | 0.7 | 1 | HC_vs_MS.NoTrt_B |
| FAM127A | 0.00312 | -0.7011 | 0.065 | 0.13 | 1 | HC_vs_MS.NoTrt_B |
| CDK7 | 0.00312 | -0.3417 | 0.135 | 0.22 | 1 | HC_vs_MS.NoTrt_B |
| C1orf109 | 0.00321 | 0.26901 | 0.868 | 0.83 | 1 | HC_vs_MS.NoTrt_B |
| PLPP2 | 0.00321 | 0.28002 | 0.891 | 0.87 | 1 | HC_vs_MS.NoTrt_B |
| PROM2 | 0.00324 | 0.50063 | 0.706 | 0.67 | 1 | HC_vs_MS.NoTrt_B |
| SLC5A5 | 0.00326 | 1.20816 | 0.455 | 0.36 | 1 | HC_vs_MS.NoTrt_B |
| EXOC4 | 0.00326 | -0.2871 | 0.564 | 0.67 | 1 | HC_vs_MS.NoTrt_B |
| SDHAF3 | 0.00326 | 0.29117 | 0.045 | 0.1 | 1 | HC_vs_MS.NoTrt_B |
| SLC38A6 | 0.00333 | 0.31152 | 0.643 | 0.61 | 1 | HC_vs_MS.NoTrt_B |
| FYCO1 | 0.00333 | -0.4126 | 0.1 | 0.18 | 1 | HC_vs_MS.NoTrt_B |
| WDR62 | 0.00334 | 0.38711 | 0.884 | 0.88 | 1 | HC_vs_MS.NoTrt_B |
| LILRA2 | 0.00334 | 0.27367 | 0.787 | 0.73 | 1 | HC_vs_MS.NoTrt_B |
| COG5 | 0.00335 | 0.60595 | 0.199 | 0.3 | 1 | HC_vs_MS.NoTrt_B |
| SCGB3A2 | 0.00335 | 0.53742 | 0.28 | 0.17 | 1 | HC_vs_MS.NoTrt_B |
| NHP2 | 0.00335 | -0.3395 | 0.172 | 0.27 | 1 | HC_vs_MS.NoTrt_B |
| SNRPA | 0.00336 | -0.2523 | 0.288 | 0.4 | 1 | HC_vs_MS.NoTrt_B |
| SSX5 | 0.00337 | 0.5115 | 0.199 | 0.11 | 1 | HC_vs_MS.NoTrt_B |
| KCNN3 | 0.00337 | 0.25336 | 0.412 | 0.32 | 1 | HC_vs_MS.NoTrt_B |
| COL1A2 | 0.00337 | 0.35764 | 0.417 | 0.32 | 1 | HC_vs_MS.NoTrt_B |
| AP5Z1 | 0.00341 | -0.7006 | 0.143 | 0.23 | 1 | HC_vs_MS.NoTrt_B |
| NDUFAB1 | 0.00343 | -0.2996 | 0.17 | 0.27 | 1 | HC_vs_MS.NoTrt_B |
| KCNAB3 | 0.00343 | -0.383 | 0.343 | 0.47 | 1 | HC_vs_MS.NoTrt_B |
| KCNJ15 | 0.00346 | 0.59095 | 0.588 | 0.52 | 1 | HC_vs_MS.NoTrt_B |
| ANKS3 | 0.00347 | -0.288 | 0.746 | 0.7 | 1 | HC_vs_MS.NoTrt_B |
| NAPRT | 0.00347 | -0.391 | 0.138 | 0.23 | 1 | HC_vs_MS.NoTrt_B |
| CCDC12 | 0.00347 | -0.4047 | 0.281 | 0.4 | 1 | HC_vs_MS.NoTrt_B |
| POP4 | 0.00351 | -0.5625 | 0.101 | 0.18 | 1 | HC_vs_MS.NoTrt_B |

| | | | | | | |
|---|---|---|---|---|---|---|
| PMVK | 0.00354 | -0.3604 | 0.096 | 0.17 | 1 | HC_vs_MS.NoTrt_B |
| DDX39A | 0.00354 | -0.3116 | 0.268 | 0.38 | 1 | HC_vs_MS.NoTrt_B |
| SYMPK | 0.00355 | 0.2833 | 0.887 | 0.89 | 1 | HC_vs_MS.NoTrt_B |
| MANSC1 | 0.00358 | 0.26192 | 0.916 | 0.92 | 1 | HC_vs_MS.NoTrt_B |
| CAMTA1 | 0.0036 | -0.4694 | 0.116 | 0.2 | 1 | HC_vs_MS.NoTrt_B |
| PLAG1 | 0.0036 | 1.06119 | 0.358 | 0.25 | 1 | HC_vs_MS.NoTrt_B |
| TERF2 | 0.00364 | 0.45044 | 0.864 | 0.84 | 1 | HC_vs_MS.NoTrt_B |
| COMTD1 | 0.00364 | -0.6531 | 0.062 | 0.12 | 1 | HC_vs_MS.NoTrt_B |
| HECW2 | 0.00365 | 0.4915 | 0.403 | 0.3 | 1 | HC_vs_MS.NoTrt_B |
| HTRA4 | 0.00367 | 0.47611 | 0.455 | 0.35 | 1 | HC_vs_MS.NoTrt_B |
| STIM2 | 0.00369 | -0.2946 | 0.093 | 0.17 | 1 | HC_vs_MS.NoTrt_B |
| CTRL | 0.00373 | -0.679 | 0.062 | 0.12 | 1 | HC_vs_MS.NoTrt_B |
| YPEL3 | 0.00373 | -0.5075 | 0.235 | 0.32 | 1 | HC_vs_MS.NoTrt_B |
| SLC35A1 | 0.00374 | -0.255 | 0.081 | 0.15 | 1 | HC_vs_MS.NoTrt_B |
| PGK1 | 0.00377 | -0.3059 | 0.577 | 0.66 | 1 | HC_vs_MS.NoTrt_B |
| LILRA5 | 0.00379 | 0.40421 | 0.728 | 0.67 | 1 | HC_vs_MS.NoTrt_B |
| C3 | 0.00379 | 0.30361 | 0.933 | 0.96 | 1 | HC_vs_MS.NoTrt_B |
| ZFYVE16 | 0.0038 | 0.65072 | 0.883 | 0.88 | 1 | HC_vs_MS.NoTrt_B |
| FLT1 | 0.0038 | 0.26988 | 0.683 | 0.59 | 1 | HC_vs_MS.NoTrt_B |
| MITD1 | 0.00381 | 0.45628 | 0.943 | 0.96 | 1 | HC_vs_MS.NoTrt_B |
| ATP2A2 | 0.00386 | 1.26448 | 0.694 | 0.67 | 1 | HC_vs_MS.NoTrt_B |
| USF1 | 0.0039 | -2.0201 | 0.146 | 0.22 | 1 | HC_vs_MS.NoTrt_B |
| SRRM2 | 0.00391 | 0.67376 | 0.742 | 0.72 | 1 | HC_vs_MS.NoTrt_B |
| NEUROD2 | 0.00393 | 1.05532 | 0.188 | 0.1 | 1 | HC_vs_MS.NoTrt_B |
| PHB | 0.00399 | -0.5129 | 0.123 | 0.21 | 1 | HC_vs_MS.NoTrt_B |
| HOOK2 | 0.00399 | 0.5358 | 0.952 | 0.96 | 1 | HC_vs_MS.NoTrt_B |
| NELFE | 0.00403 | -0.3979 | 0.114 | 0.2 | 1 | HC_vs_MS.NoTrt_B |
| ACAD9 | 0.00404 | -1.0218 | 0.248 | 0.34 | 1 | HC_vs_MS.NoTrt_B |
| ZNF492 | 0.00404 | 0.50093 | 0.49 | 0.4 | 1 | HC_vs_MS.NoTrt_B |
| ITGB3BP | 0.00405 | 0.47102 | 0.784 | 0.79 | 1 | HC_vs_MS.NoTrt_B |
| CHRNA5 | 0.00407 | 0.39011 | 0.52 | 0.46 | 1 | HC_vs_MS.NoTrt_B |
| MAGED2 | 0.00409 | -0.4235 | 0.138 | 0.22 | 1 | HC_vs_MS.NoTrt_B |
| RANBP1 | 0.0041 | -0.5548 | 0.142 | 0.23 | 1 | HC_vs_MS.NoTrt_B |
| EZH2 | 0.00414 | -0.5369 | 0.099 | 0.17 | 1 | HC_vs_MS.NoTrt_B |
| PNKD | 0.00418 | -0.4883 | 0.099 | 0.17 | 1 | HC_vs_MS.NoTrt_B |
| KBTBD4 | 0.0042 | -0.6442 | 0.051 | 0.11 | 1 | HC_vs_MS.NoTrt_B |
| SZT2 | 0.00426 | 0.30632 | 0.849 | 0.82 | 1 | HC_vs_MS.NoTrt_B |
| BLVRA | 0.00428 | -0.6134 | 0.103 | 0.18 | 1 | HC_vs_MS.NoTrt_B |
| FKBP3 | 0.00428 | 0.29045 | 0.268 | 0.38 | 1 | HC_vs_MS.NoTrt_B |
| ARMC1 | 0.00431 | -0.5833 | 0.083 | 0.15 | 1 | HC_vs_MS.NoTrt_B |

| | | | | | | |
|---|---|---|---|---|---|---|
| ARF5 | 0.00434 | -0.3772 | 0.196 | 0.29 | 1 | HC_vs_MS.NoTrt_B |
| SLC25A11 | 0.00436 | -0.2792 | 0.104 | 0.18 | 1 | HC_vs_MS.NoTrt_B |
| LBHD1 | 0.00436 | 1.95219 | 0.477 | 0.39 | 1 | HC_vs_MS.NoTrt_B |
| PARK7 | 0.00437 | -0.2876 | 0.714 | 0.78 | 1 | HC_vs_MS.NoTrt_B |
| PCDHB1 | 0.0044 | 0.33225 | 0.778 | 0.79 | 1 | HC_vs_MS.NoTrt_B |
| ARPC1B | 0.00444 | -0.2924 | 0.928 | 0.98 | 1 | HC_vs_MS.NoTrt_B |
| PPP1R11 | 0.00445 | -0.3516 | 0.13 | 0.22 | 1 | HC_vs_MS.NoTrt_B |
| DERL2 | 0.00445 | 0.27103 | 0.268 | 0.37 | 1 | HC_vs_MS.NoTrt_B |
| RP11-577H5 | 0.00447 | -0.8579 | 0.075 | 0.14 | 1 | HC_vs_MS.NoTrt_B |
| UGGT2 | 0.00448 | 0.47809 | 0.803 | 0.78 | 1 | HC_vs_MS.NoTrt_B |
| FKBP15 | 0.00449 | -0.4966 | 0.07 | 0.14 | 1 | HC_vs_MS.NoTrt_B |
| SRD5A1 | 0.0045 | -0.2527 | 0.701 | 0.68 | 1 | HC_vs_MS.NoTrt_B |
| GABRP | 0.00452 | 0.64436 | 0.323 | 0.22 | 1 | HC_vs_MS.NoTrt_B |
| QPRT | 0.00454 | 0.35387 | 0.92 | 0.92 | 1 | HC_vs_MS.NoTrt_B |
| GPR85 | 0.00459 | 0.51441 | 0.388 | 0.28 | 1 | HC_vs_MS.NoTrt_B |
| AHSA2 | 0.00462 | 0.33148 | 0.219 | 0.32 | 1 | HC_vs_MS.NoTrt_B |
| SPICE1 | 0.00462 | -0.5376 | 0.235 | 0.35 | 1 | HC_vs_MS.NoTrt_B |
| MAP3K1 | 0.00476 | 0.34287 | 0.464 | 0.64 | 1 | HC_vs_MS.NoTrt_B |
| FKBP8 | 0.00477 | 0.53215 | 0.196 | 0.31 | 1 | HC_vs_MS.NoTrt_B |
| ETHE1 | 0.00477 | -0.4172 | 0.055 | 0.11 | 1 | HC_vs_MS.NoTrt_B |
| GABARAPL2 | 0.00478 | -0.4537 | 0.443 | 0.54 | 1 | HC_vs_MS.NoTrt_B |
| H6PD | 0.0048 | 0.86177 | 0.793 | 0.83 | 1 | HC_vs_MS.NoTrt_B |
| SCN7A | 0.00481 | 0.46695 | 0.522 | 0.44 | 1 | HC_vs_MS.NoTrt_B |
| TMEM147 | 0.00482 | -0.4338 | 0.11 | 0.19 | 1 | HC_vs_MS.NoTrt_B |
| NIPA2 | 0.00485 | -0.5552 | 0.091 | 0.16 | 1 | HC_vs_MS.NoTrt_B |
| UQCRH | 0.0049 | -0.2848 | 0.514 | 0.62 | 1 | HC_vs_MS.NoTrt_B |
| SLC16A12 | 0.00491 | 0.48321 | 0.397 | 0.31 | 1 | HC_vs_MS.NoTrt_B |
| RGL4 | 0.00492 | -0.3814 | 0.194 | 0.29 | 1 | HC_vs_MS.NoTrt_B |
| KATNA1 | 0.00492 | -0.8265 | 0.064 | 0.12 | 1 | HC_vs_MS.NoTrt_B |
| BRF1 | 0.00494 | 0.58827 | 0.565 | 0.47 | 1 | HC_vs_MS.NoTrt_B |
| SQSTM1 | 0.005 | -0.3303 | 0.81 | 0.84 | 1 | HC_vs_MS.NoTrt_B |
| PTPRC | 0.005 | 0.37744 | 0.962 | 0.98 | 1 | HC_vs_MS.NoTrt_B |
| DYNC1H1 | 0.0051 | 0.37969 | 0.938 | 0.96 | 1 | HC_vs_MS.NoTrt_B |
| BEX2 | 0.00516 | -0.5285 | 0.067 | 0.13 | 1 | HC_vs_MS.NoTrt_B |
| ST3GAL6 | 0.00519 | 0.70621 | 0.31 | 0.23 | 1 | HC_vs_MS.NoTrt_B |
| CEP19 | 0.00521 | 0.9677 | 0.643 | 0.58 | 1 | HC_vs_MS.NoTrt_B |
| MTMR9 | 0.00525 | -0.3529 | 0.274 | 0.38 | 1 | HC_vs_MS.NoTrt_B |
| EEFSEC | 0.00527 | -0.4033 | 0.083 | 0.15 | 1 | HC_vs_MS.NoTrt_B |
| SLC43A1 | 0.00531 | -0.7066 | 0.075 | 0.14 | 1 | HC_vs_MS.NoTrt_B |
| DPP6 | 0.00535 | 0.34768 | 0.477 | 0.39 | 1 | HC_vs_MS.NoTrt_B |

| | | | | | | |
|---|---|---|---|---|---|---|
| ASNSD1 | 0.00537 | -0.5806 | 0.109 | 0.18 | 1 | HC_vs_MS.NoTrt_B |
| TMEM109 | 0.00541 | -0.2913 | 0.141 | 0.23 | 1 | HC_vs_MS.NoTrt_B |
| UCHL3 | 0.00542 | -0.2815 | 0.104 | 0.18 | 1 | HC_vs_MS.NoTrt_B |
| PGAM1 | 0.00544 | -0.2654 | 0.206 | 0.3 | 1 | HC_vs_MS.NoTrt_B |
| LRRC36 | 0.00545 | 0.30066 | 0.836 | 0.82 | 1 | HC_vs_MS.NoTrt_B |
| KDM2A | 0.00548 | 0.72291 | 0.907 | 0.88 | 1 | HC_vs_MS.NoTrt_B |
| NLRP12 | 0.00549 | 0.37316 | 0.738 | 0.71 | 1 | HC_vs_MS.NoTrt_B |
| PIGR | 0.00553 | 0.37085 | 0.557 | 0.47 | 1 | HC_vs_MS.NoTrt_B |
| DACT3 | 0.00559 | 0.25139 | 0.955 | 0.96 | 1 | HC_vs_MS.NoTrt_B |
| TMEM161A | 0.00564 | -0.6754 | 0.135 | 0.22 | 1 | HC_vs_MS.NoTrt_B |
| FAM131A | 0.0057 | 0.7976 | 0.806 | 0.76 | 1 | HC_vs_MS.NoTrt_B |
| MRPL39 | 0.00572 | -0.5623 | 0.084 | 0.15 | 1 | HC_vs_MS.NoTrt_B |
| AC025335.1 | 0.00574 | 0.65455 | 0.613 | 0.51 | 1 | HC_vs_MS.NoTrt_B |
| PIP4K2C | 0.00576 | -0.8039 | 0.107 | 0.18 | 1 | HC_vs_MS.NoTrt_B |
| SGO2 | 0.00577 | 0.44873 | 0.791 | 0.8 | 1 | HC_vs_MS.NoTrt_B |
| ASMTL | 0.00586 | -0.52 | 0.068 | 0.13 | 1 | HC_vs_MS.NoTrt_B |
| C14orf2 | 0.00586 | -0.2511 | 0.833 | 0.88 | 1 | HC_vs_MS.NoTrt_B |
| C17orf89 | 0.00588 | -0.2844 | 0.071 | 0.14 | 1 | HC_vs_MS.NoTrt_B |
| TMEM120A | 0.00593 | -0.2826 | 0.096 | 0.17 | 1 | HC_vs_MS.NoTrt_B |
| RP11-437B1 | 0.00596 | 0.65109 | 0.574 | 0.5 | 1 | HC_vs_MS.NoTrt_B |
| SHMT1 | 0.00598 | -0.8469 | 0.367 | 0.48 | 1 | HC_vs_MS.NoTrt_B |
| AATF | 0.00601 | -0.5244 | 0.165 | 0.25 | 1 | HC_vs_MS.NoTrt_B |
| TECRL | 0.00601 | 0.48128 | 0.217 | 0.13 | 1 | HC_vs_MS.NoTrt_B |
| RUSC1 | 0.00603 | -1.0978 | 0.049 | 0.1 | 1 | HC_vs_MS.NoTrt_B |
| CD40LG | 0.00603 | 1.18053 | 0.22 | 0.14 | 1 | HC_vs_MS.NoTrt_B |
| TBCB | 0.00603 | -0.3506 | 0.645 | 0.71 | 1 | HC_vs_MS.NoTrt_B |
| MRPL54 | 0.00604 | -0.4188 | 0.087 | 0.16 | 1 | HC_vs_MS.NoTrt_B |
| TOP3B | 0.00607 | -0.2881 | 0.133 | 0.22 | 1 | HC_vs_MS.NoTrt_B |
| ICA1 | 0.00607 | 1.03903 | 0.596 | 0.54 | 1 | HC_vs_MS.NoTrt_B |
| TSEN2 | 0.00609 | 0.52654 | 0.881 | 0.88 | 1 | HC_vs_MS.NoTrt_B |
| TTC31 | 0.00614 | -0.6069 | 0.206 | 0.29 | 1 | HC_vs_MS.NoTrt_B |
| SLC30A5 | 0.00615 | 0.40913 | 0.777 | 0.71 | 1 | HC_vs_MS.NoTrt_B |
| ADCY2 | 0.00615 | 0.35312 | 0.717 | 0.68 | 1 | HC_vs_MS.NoTrt_B |
| METTL8 | 0.00616 | 0.30703 | 0.916 | 0.94 | 1 | HC_vs_MS.NoTrt_B |
| RAB21 | 0.00623 | 0.49039 | 0.964 | 0.97 | 1 | HC_vs_MS.NoTrt_B |
| POGZ | 0.00625 | 0.67181 | 0.822 | 0.8 | 1 | HC_vs_MS.NoTrt_B |
| C1orf216 | 0.00627 | -0.4039 | 0.81 | 0.8 | 1 | HC_vs_MS.NoTrt_B |
| METTL26 | 0.00628 | -0.2644 | 0.187 | 0.28 | 1 | HC_vs_MS.NoTrt_B |
| GFOD2 | 0.00629 | 0.66633 | 0.886 | 0.9 | 1 | HC_vs_MS.NoTrt_B |
| HSPA1A | 0.00631 | -0.3184 | 0.122 | 0.2 | 1 | HC_vs_MS.NoTrt_B |

| | | | | | | |
|---|---|---|---|---|---|---|
| TMEM223 | 0.00632 | -0.472 | 0.133 | 0.21 | 1 | HC_vs_MS.NoTrt_B |
| CHD7 | 0.00636 | 2.01048 | 0.554 | 0.51 | 1 | HC_vs_MS.NoTrt_B |
| SNRPB | 0.00643 | -0.2524 | 0.188 | 0.28 | 1 | HC_vs_MS.NoTrt_B |
| STARD10 | 0.00644 | 0.34172 | 0.825 | 0.81 | 1 | HC_vs_MS.NoTrt_B |
| ZNF821 | 0.00645 | -0.6753 | 0.049 | 0.1 | 1 | HC_vs_MS.NoTrt_B |
| CKAP2L | 0.00648 | 0.50712 | 0.49 | 0.41 | 1 | HC_vs_MS.NoTrt_B |
| TNFAIP8L2 | 0.00662 | -0.4233 | 0.084 | 0.15 | 1 | HC_vs_MS.NoTrt_B |
| C1orf159 | 0.00668 | -0.4078 | 0.114 | 0.19 | 1 | HC_vs_MS.NoTrt_B |
| SKA2 | 0.00674 | 0.25042 | 0.551 | 0.65 | 1 | HC_vs_MS.NoTrt_B |
| ANXA3 | 0.00678 | 0.30808 | 0.486 | 0.38 | 1 | HC_vs_MS.NoTrt_B |
| CACNB4 | 0.00678 | 0.47433 | 0.858 | 0.9 | 1 | HC_vs_MS.NoTrt_B |
| PRDX6 | 0.0068 | -0.2835 | 0.157 | 0.24 | 1 | HC_vs_MS.NoTrt_B |
| FAM135B | 0.00687 | 0.30778 | 0.291 | 0.2 | 1 | HC_vs_MS.NoTrt_B |
| ATXN2 | 0.00687 | 0.42737 | 0.88 | 0.85 | 1 | HC_vs_MS.NoTrt_B |
| DNAJA2 | 0.00689 | -0.4122 | 0.367 | 0.45 | 1 | HC_vs_MS.NoTrt_B |
| FAM83F | 0.0069 | 0.32123 | 0.87 | 0.85 | 1 | HC_vs_MS.NoTrt_B |
| HAUS3 | 0.00693 | 0.50475 | 0.822 | 0.85 | 1 | HC_vs_MS.NoTrt_B |
| DNAL1 | 0.00695 | 0.37931 | 0.919 | 0.94 | 1 | HC_vs_MS.NoTrt_B |
| CPQ | 0.00703 | -0.95 | 0.357 | 0.47 | 1 | HC_vs_MS.NoTrt_B |
| MRPL24 | 0.00704 | -0.5267 | 0.081 | 0.15 | 1 | HC_vs_MS.NoTrt_B |
| EPB41L4A | 0.00712 | -0.3803 | 0.561 | 0.49 | 1 | HC_vs_MS.NoTrt_B |
| C12orf57 | 0.00712 | -0.516 | 0.349 | 0.45 | 1 | HC_vs_MS.NoTrt_B |
| LRRC74B | 0.00713 | 0.36778 | 0.748 | 0.71 | 1 | HC_vs_MS.NoTrt_B |
| RASGRP1 | 0.00718 | -0.2722 | 0.354 | 0.45 | 1 | HC_vs_MS.NoTrt_B |
| POLDIP3 | 0.00728 | -0.3626 | 0.119 | 0.2 | 1 | HC_vs_MS.NoTrt_B |
| CCT6B | 0.00729 | 0.66192 | 0.255 | 0.16 | 1 | HC_vs_MS.NoTrt_B |
| IGFBPL1 | 0.00737 | 0.35129 | 0.562 | 0.5 | 1 | HC_vs_MS.NoTrt_B |
| BTBD9 | 0.00738 | 0.57877 | 0.552 | 0.46 | 1 | HC_vs_MS.NoTrt_B |
| TRAT1 | 0.0074 | 1.10139 | 0.249 | 0.17 | 1 | HC_vs_MS.NoTrt_B |
| AGBL3 | 0.00742 | -0.5023 | 0.248 | 0.34 | 1 | HC_vs_MS.NoTrt_B |
| ZYG11B | 0.00749 | 0.76995 | 0.704 | 0.62 | 1 | HC_vs_MS.NoTrt_B |
| PPP1R7 | 0.0075 | -0.3385 | 0.152 | 0.23 | 1 | HC_vs_MS.NoTrt_B |
| BRPF1 | 0.0075 | 0.76125 | 0.057 | 0.11 | 1 | HC_vs_MS.NoTrt_B |
| CMTR1 | 0.00751 | -0.3204 | 0.159 | 0.24 | 1 | HC_vs_MS.NoTrt_B |
| STRA13 | 0.00761 | -0.5848 | 0.09 | 0.16 | 1 | HC_vs_MS.NoTrt_B |
| SPDYE2B | 0.00764 | 1.15135 | 0.335 | 0.27 | 1 | HC_vs_MS.NoTrt_B |
| WDR74 | 0.00765 | 0.37801 | 0.596 | 0.48 | 1 | HC_vs_MS.NoTrt_B |
| R3HCC1 | 0.00765 | -0.2514 | 0.088 | 0.16 | 1 | HC_vs_MS.NoTrt_B |
| POM121 | 0.00767 | 0.25148 | 0.397 | 0.52 | 1 | HC_vs_MS.NoTrt_B |
| FGF22 | 0.00773 | 0.30205 | 0.37 | 0.29 | 1 | HC_vs_MS.NoTrt_B |

| | | | | | | |
|---|---|---|---|---|---|---|
| RBMS1 | 0.00777 | 0.58038 | 0.665 | 0.61 | 1 | HC_vs_MS.NoTrt_B |
| SPIRE2 | 0.00778 | 0.37401 | 0.917 | 0.92 | 1 | HC_vs_MS.NoTrt_B |
| CHEK1 | 0.00778 | 0.74355 | 0.767 | 0.77 | 1 | HC_vs_MS.NoTrt_B |
| TLR10 | 0.0078 | 0.26379 | 0.404 | 0.3 | 1 | HC_vs_MS.NoTrt_B |
| TOR1B | 0.00781 | 1.4084 | 0.2 | 0.11 | 1 | HC_vs_MS.NoTrt_B |
| DAPK3 | 0.00782 | -0.2724 | 0.061 | 0.12 | 1 | HC_vs_MS.NoTrt_B |
| ADA | 0.00784 | -0.4997 | 0.065 | 0.12 | 1 | HC_vs_MS.NoTrt_B |
| TNRC6B | 0.00787 | 0.50179 | 0.951 | 1 | 1 | HC_vs_MS.NoTrt_B |
| ZNF316 | 0.00791 | 0.53659 | 0.461 | 0.38 | 1 | HC_vs_MS.NoTrt_B |
| SNTB2 | 0.00792 | 0.50339 | 0.774 | 0.78 | 1 | HC_vs_MS.NoTrt_B |
| MIS18A | 0.00794 | -0.3355 | 0.051 | 0.1 | 1 | HC_vs_MS.NoTrt_B |
| PLPP5 | 0.00796 | 0.69604 | 0.848 | 0.84 | 1 | HC_vs_MS.NoTrt_B |
| CBFA2T2 | 0.008 | 0.33636 | 0.92 | 0.95 | 1 | HC_vs_MS.NoTrt_B |
| CLDN11 | 0.00802 | 0.59103 | 0.559 | 0.47 | 1 | HC_vs_MS.NoTrt_B |
| SLITRK2 | 0.00806 | 0.42236 | 0.335 | 0.25 | 1 | HC_vs_MS.NoTrt_B |
| SNIP1 | 0.0081 | 0.41407 | 0.659 | 0.6 | 1 | HC_vs_MS.NoTrt_B |
| NSA2 | 0.0081 | -0.4256 | 0.317 | 0.41 | 1 | HC_vs_MS.NoTrt_B |
| APOBEC2 | 0.0081 | -0.2917 | 0.049 | 0.1 | 1 | HC_vs_MS.NoTrt_B |
| ZNF428 | 0.00813 | -0.3192 | 0.157 | 0.23 | 1 | HC_vs_MS.NoTrt_B |
| SYT12 | 0.00813 | 0.29188 | 0.716 | 0.72 | 1 | HC_vs_MS.NoTrt_B |
| GGA3 | 0.00819 | -0.332 | 0.139 | 0.22 | 1 | HC_vs_MS.NoTrt_B |
| PRPF6 | 0.0082 | -0.5455 | 0.18 | 0.26 | 1 | HC_vs_MS.NoTrt_B |
| C9orf72 | 0.00823 | -0.3974 | 0.094 | 0.16 | 1 | HC_vs_MS.NoTrt_B |
| AICDA | 0.00824 | 0.39034 | 0.604 | 0.54 | 1 | HC_vs_MS.NoTrt_B |
| JOSD2 | 0.00825 | -0.411 | 0.065 | 0.12 | 1 | HC_vs_MS.NoTrt_B |
| SAT2 | 0.00825 | -0.4229 | 0.086 | 0.15 | 1 | HC_vs_MS.NoTrt_B |
| LSS | 0.00828 | -0.3049 | 0.078 | 0.14 | 1 | HC_vs_MS.NoTrt_B |
| DSEL | 0.00831 | 0.46028 | 0.657 | 0.6 | 1 | HC_vs_MS.NoTrt_B |
| FCAR | 0.00836 | 0.30641 | 0.87 | 0.87 | 1 | HC_vs_MS.NoTrt_B |
| PAX5 | 0.00841 | 0.49237 | 0.119 | 0.05 | 1 | HC_vs_MS.NoTrt_B |
| ABCF3 | 0.00845 | -0.5987 | 0.129 | 0.2 | 1 | HC_vs_MS.NoTrt_B |
| IZUMO2 | 0.00845 | 0.28362 | 0.897 | 0.93 | 1 | HC_vs_MS.NoTrt_B |
| KLHL33 | 0.00848 | 0.41378 | 0.497 | 0.42 | 1 | HC_vs_MS.NoTrt_B |
| MAB21L3 | 0.00856 | 0.75314 | 0.362 | 0.28 | 1 | HC_vs_MS.NoTrt_B |
| EI24 | 0.00859 | 0.39026 | 0.736 | 0.72 | 1 | HC_vs_MS.NoTrt_B |
| SCLY | 0.00859 | 0.49441 | 0.958 | 0.96 | 1 | HC_vs_MS.NoTrt_B |
| MRPS33 | 0.00866 | 0.29641 | 0.916 | 0.93 | 1 | HC_vs_MS.NoTrt_B |
| BEST1 | 0.00872 | 0.89933 | 0.806 | 0.8 | 1 | HC_vs_MS.NoTrt_B |
| DENND2C | 0.00881 | 0.34851 | 0.751 | 0.73 | 1 | HC_vs_MS.NoTrt_B |
| ZEB2 | 0.00881 | 0.36516 | 0.6 | 0.53 | 1 | HC_vs_MS.NoTrt_B |

| | | | | | | |
|---|---|---|---|---|---|---|
| NIPA1 | 0.00884 | -0.4735 | 0.101 | 0.17 | 1 | HC_vs_MS.NoTrt_B |
| OCLN | 0.0089 | 0.73303 | 0.796 | 0.77 | 1 | HC_vs_MS.NoTrt_B |
| ZNF619 | 0.00894 | 0.53285 | 0.716 | 0.7 | 1 | HC_vs_MS.NoTrt_B |
| ADCY1 | 0.00896 | 0.38258 | 0.135 | 0.22 | 1 | HC_vs_MS.NoTrt_B |
| OSBPL11 | 0.00904 | -0.3151 | 0.054 | 0.11 | 1 | HC_vs_MS.NoTrt_B |
| TMEM106B | 0.00907 | 0.56203 | 0.888 | 0.89 | 1 | HC_vs_MS.NoTrt_B |
| DENR | 0.00907 | 0.27469 | 0.752 | 0.68 | 1 | HC_vs_MS.NoTrt_B |
| MMP25 | 0.00908 | 1.0808 | 0.112 | 0.05 | 1 | HC_vs_MS.NoTrt_B |
| IGF2R | 0.00908 | 0.5916 | 0.401 | 0.32 | 1 | HC_vs_MS.NoTrt_B |
| PDGFA | 0.00909 | 0.66913 | 0.501 | 0.42 | 1 | HC_vs_MS.NoTrt_B |
| FUT11 | 0.0091 | 0.52784 | 0.757 | 0.72 | 1 | HC_vs_MS.NoTrt_B |
| OR52A1 | 0.00914 | 0.36632 | 0.749 | 0.72 | 1 | HC_vs_MS.NoTrt_B |
| MOK | 0.00923 | 0.27328 | 0.79 | 0.8 | 1 | HC_vs_MS.NoTrt_B |
| GCNT4 | 0.00929 | 0.87284 | 0.475 | 0.43 | 1 | HC_vs_MS.NoTrt_B |
| ZDHHC12 | 0.00931 | -0.5727 | 0.096 | 0.16 | 1 | HC_vs_MS.NoTrt_B |
| SLC25A25 | 0.0094 | -0.3209 | 0.086 | 0.15 | 1 | HC_vs_MS.NoTrt_B |
| CDK5RAP3 | 0.00943 | -0.2832 | 0.581 | 0.69 | 1 | HC_vs_MS.NoTrt_B |
| PSMA4 | 0.00944 | -0.5267 | 0.349 | 0.43 | 1 | HC_vs_MS.NoTrt_B |
| CHRNA7 | 0.00946 | 0.26603 | 0.794 | 0.78 | 1 | HC_vs_MS.NoTrt_B |
| TUSC2 | 0.00946 | -0.3958 | 0.062 | 0.12 | 1 | HC_vs_MS.NoTrt_B |
| FGFR1 | 0.00949 | 0.5007 | 0.783 | 0.78 | 1 | HC_vs_MS.NoTrt_B |
| POLK | 0.00951 | 0.56084 | 0.842 | 0.79 | 1 | HC_vs_MS.NoTrt_B |
| FAM234A | 0.00956 | -0.8955 | 0.196 | 0.28 | 1 | HC_vs_MS.NoTrt_B |
| TLR6 | 0.00961 | 1.22441 | 0.303 | 0.22 | 1 | HC_vs_MS.NoTrt_B |
| SLFN12L | 0.00963 | 0.25096 | 0.835 | 0.81 | 1 | HC_vs_MS.NoTrt_B |
| TUBGCP4 | 0.00966 | 0.53704 | 0.784 | 0.8 | 1 | HC_vs_MS.NoTrt_B |
| ARMC2 | 0.00966 | -0.3553 | 0.051 | 0.1 | 1 | HC_vs_MS.NoTrt_B |
| REPS1 | 0.00967 | 0.75001 | 0.632 | 0.62 | 1 | HC_vs_MS.NoTrt_B |
| DHRS4L2 | 0.00975 | -0.2933 | 0.142 | 0.22 | 1 | HC_vs_MS.NoTrt_B |
| IKZF5 | 0.00981 | 0.52621 | 0.158 | 0.25 | 1 | HC_vs_MS.NoTrt_B |
| ISG20L2 | 0.00982 | 0.49235 | 0.803 | 0.78 | 1 | HC_vs_MS.NoTrt_B |
| FRMPD1 | 0.00983 | 0.28763 | 0.83 | 0.85 | 1 | HC_vs_MS.NoTrt_B |
| LYPD5 | 0.00986 | 0.40483 | 0.699 | 0.71 | 1 | HC_vs_MS.NoTrt_B |
| RGPD5 | 0.00988 | 0.76576 | 0.816 | 0.83 | 1 | HC_vs_MS.NoTrt_B |
| TMEM236 | 0.0099 | 0.4642 | 0.73 | 0.71 | 1 | HC_vs_MS.NoTrt_B |
| CYTH2 | 0.00993 | -0.2504 | 0.591 | 0.75 | 1 | HC_vs_MS.NoTrt_B |
| ARPIN | 0.00993 | 0.29685 | 0.87 | 0.85 | 1 | HC_vs_MS.NoTrt_B |
| ACSS2 | 0.00998 | 1.7802 | 0.549 | 0.5 | 1 | HC_vs_MS.NoTrt_B |
| TUBA4A | 0.01001 | -0.3214 | 0.22 | 0.31 | 1 | HC_vs_MS.NoTrt_B |
| SLC18B1 | 0.01002 | -0.6767 | 0.055 | 0.11 | 1 | HC_vs_MS.NoTrt_B |

| | | | | | | |
|---|---|---|---|---|---|---|
| AKAP8L | 0.01003 | -0.4326 | 0.059 | 0.11 | 1 | HC_vs_MS.NoTrt_B |
| HIST1H2BN | 0.01008 | 0.27795 | 0.652 | 0.59 | 1 | HC_vs_MS.NoTrt_B |
| FIS1 | 0.01008 | -0.398 | 0.235 | 0.32 | 1 | HC_vs_MS.NoTrt_B |
| C16orf72 | 0.0101 | 0.70576 | 0.826 | 0.82 | 1 | HC_vs_MS.NoTrt_B |
| ZNF578 | 0.01011 | 0.39501 | 0.939 | 0.94 | 1 | HC_vs_MS.NoTrt_B |
| MIA2 | 0.01014 | 0.34119 | 0.38 | 0.3 | 1 | HC_vs_MS.NoTrt_B |
| ANAPC5 | 0.01026 | -0.2869 | 0.946 | 0.98 | 1 | HC_vs_MS.NoTrt_B |
| HIF3A | 0.01027 | 0.30915 | 0.864 | 0.84 | 1 | HC_vs_MS.NoTrt_B |
| ZNF480 | 0.01028 | 1.23772 | 0.722 | 0.72 | 1 | HC_vs_MS.NoTrt_B |
| REL | 0.01039 | 0.29628 | 0.954 | 0.95 | 1 | HC_vs_MS.NoTrt_B |
| KIAA1456 | 0.01043 | 0.26345 | 0.903 | 0.92 | 1 | HC_vs_MS.NoTrt_B |
| CSNK2A2 | 0.01045 | -0.4035 | 0.27 | 0.35 | 1 | HC_vs_MS.NoTrt_B |
| TRPM7 | 0.01048 | 0.26945 | 0.92 | 0.9 | 1 | HC_vs_MS.NoTrt_B |
| RP11-411B6 | 0.01049 | 0.59922 | 0.219 | 0.32 | 1 | HC_vs_MS.NoTrt_B |
| ADH1B | 0.01049 | 0.38225 | 0.322 | 0.25 | 1 | HC_vs_MS.NoTrt_B |
| AGO3 | 0.01052 | 0.64197 | 0.965 | 0.96 | 1 | HC_vs_MS.NoTrt_B |
| FAM111B | 0.01052 | 0.35158 | 0.572 | 0.55 | 1 | HC_vs_MS.NoTrt_B |
| RPGRIP1L | 0.01053 | 0.2587 | 0.722 | 0.7 | 1 | HC_vs_MS.NoTrt_B |
| ORMDL3 | 0.01062 | -0.393 | 0.091 | 0.16 | 1 | HC_vs_MS.NoTrt_B |
| RUVBL1 | 0.01072 | 0.57655 | 0.643 | 0.58 | 1 | HC_vs_MS.NoTrt_B |
| HLA-DQB1 | 0.01072 | -0.5362 | 0.546 | 0.65 | 1 | HC_vs_MS.NoTrt_B |
| LIX1L | 0.01075 | 0.59315 | 0.775 | 0.76 | 1 | HC_vs_MS.NoTrt_B |
| LIPA | 0.01076 | -0.3621 | 0.262 | 0.35 | 1 | HC_vs_MS.NoTrt_B |
| A2M | 0.01078 | 0.73998 | 0.126 | 0.06 | 1 | HC_vs_MS.NoTrt_B |
| PTPRB | 0.01085 | 0.38281 | 0.514 | 0.46 | 1 | HC_vs_MS.NoTrt_B |
| PIH1D1 | 0.01086 | -0.4144 | 0.245 | 0.32 | 1 | HC_vs_MS.NoTrt_B |
| DDIT4 | 0.01087 | -0.3991 | 0.132 | 0.21 | 1 | HC_vs_MS.NoTrt_B |
| TOR2A | 0.01095 | -0.5647 | 0.052 | 0.1 | 1 | HC_vs_MS.NoTrt_B |
| KCNK6 | 0.01098 | 0.51298 | 0.959 | 0.96 | 1 | HC_vs_MS.NoTrt_B |
| CA5B | 0.01098 | 0.60786 | 0.872 | 0.89 | 1 | HC_vs_MS.NoTrt_B |
| PLA2G4C | 0.01107 | 0.27238 | 0.938 | 0.95 | 1 | HC_vs_MS.NoTrt_B |
| GNG10 | 0.01109 | -0.2956 | 0.11 | 0.18 | 1 | HC_vs_MS.NoTrt_B |
| SGK3 | 0.01109 | 0.38429 | 0.906 | 0.89 | 1 | HC_vs_MS.NoTrt_B |
| IQGAP1 | 0.01112 | -0.3118 | 0.791 | 0.91 | 1 | HC_vs_MS.NoTrt_B |
| GNB4 | 0.01113 | 0.46274 | 0.791 | 0.77 | 1 | HC_vs_MS.NoTrt_B |
| PIGW | 0.01114 | 0.73937 | 0.459 | 0.38 | 1 | HC_vs_MS.NoTrt_B |
| C4BPB | 0.01115 | 0.44509 | 0.78 | 0.82 | 1 | HC_vs_MS.NoTrt_B |
| AFF3 | 0.01118 | 1.25058 | 0.748 | 0.73 | 1 | HC_vs_MS.NoTrt_B |
| AP1B1 | 0.01129 | -0.5555 | 0.145 | 0.22 | 1 | HC_vs_MS.NoTrt_B |
| CLEC18A | 0.01129 | 0.72485 | 0.281 | 0.38 | 1 | HC_vs_MS.NoTrt_B |

| | | | | | | |
|---|---|---|---|---|---|---|
| PPA2 | 0.01133 | -0.3172 | 0.633 | 0.74 | 1 | HC_vs_MS.NoTrt_B |
| KIR3DX1 | 0.01133 | 0.27268 | 0.861 | 0.91 | 1 | HC_vs_MS.NoTrt_B |
| PRDX3 | 0.01134 | -0.2639 | 0.409 | 0.51 | 1 | HC_vs_MS.NoTrt_B |
| FOXL2NB | 0.01134 | 0.43764 | 0.726 | 0.7 | 1 | HC_vs_MS.NoTrt_B |
| IFT122 | 0.01135 | 0.61539 | 0.862 | 0.88 | 1 | HC_vs_MS.NoTrt_B |
| MCRIP1 | 0.01143 | -0.3217 | 0.381 | 0.48 | 1 | HC_vs_MS.NoTrt_B |
| MED6 | 0.01144 | 0.33549 | 0.571 | 0.48 | 1 | HC_vs_MS.NoTrt_B |
| SPIB | 0.01145 | 0.32299 | 0.822 | 0.85 | 1 | HC_vs_MS.NoTrt_B |
| IL15RA | 0.01149 | -0.6522 | 0.072 | 0.13 | 1 | HC_vs_MS.NoTrt_B |
| RNF115 | 0.01152 | 0.31776 | 0.964 | 0.98 | 1 | HC_vs_MS.NoTrt_B |
| ZNF333 | 0.01152 | 0.31508 | 0.9 | 0.89 | 1 | HC_vs_MS.NoTrt_B |
| RTEL1 | 0.01157 | -0.4043 | 0.072 | 0.13 | 1 | HC_vs_MS.NoTrt_B |
| STIL | 0.01157 | 0.42101 | 0.607 | 0.55 | 1 | HC_vs_MS.NoTrt_B |
| CYB561D1 | 0.01158 | 0.32649 | 0.461 | 0.37 | 1 | HC_vs_MS.NoTrt_B |
| ERGIC3 | 0.01164 | -0.5607 | 0.803 | 0.87 | 1 | HC_vs_MS.NoTrt_B |
| GLTSCR1L | 0.01165 | -0.3245 | 0.149 | 0.23 | 1 | HC_vs_MS.NoTrt_B |
| KPNA2 | 0.01169 | -0.7324 | 0.072 | 0.13 | 1 | HC_vs_MS.NoTrt_B |
| ARRB2 | 0.01169 | 0.36641 | 0.771 | 0.71 | 1 | HC_vs_MS.NoTrt_B |
| AGTRAP | 0.01169 | -0.6174 | 0.077 | 0.14 | 1 | HC_vs_MS.NoTrt_B |
| ZC3H6 | 0.01172 | 0.57037 | 0.794 | 0.83 | 1 | HC_vs_MS.NoTrt_B |
| FAHD2A | 0.01174 | 0.45438 | 0.154 | 0.24 | 1 | HC_vs_MS.NoTrt_B |
| IRF2BP2 | 0.01176 | 0.83208 | 0.197 | 0.12 | 1 | HC_vs_MS.NoTrt_B |
| ANXA2R | 0.01177 | -0.3672 | 0.178 | 0.26 | 1 | HC_vs_MS.NoTrt_B |
| COG2 | 0.01195 | -0.2619 | 0.286 | 0.38 | 1 | HC_vs_MS.NoTrt_B |
| AGRN | 0.01199 | 0.50723 | 0.335 | 0.26 | 1 | HC_vs_MS.NoTrt_B |
| SNRPA1 | 0.01205 | -0.2697 | 0.652 | 0.73 | 1 | HC_vs_MS.NoTrt_B |
| ADGRD2 | 0.0121 | 0.32866 | 0.754 | 0.76 | 1 | HC_vs_MS.NoTrt_B |
| ZNF841 | 0.01213 | 0.87687 | 0.504 | 0.45 | 1 | HC_vs_MS.NoTrt_B |
| KLRK1 | 0.01214 | 0.56239 | 0.643 | 0.62 | 1 | HC_vs_MS.NoTrt_B |
| SMYD2 | 0.01214 | -0.4583 | 0.139 | 0.21 | 1 | HC_vs_MS.NoTrt_B |
| NALCN | 0.01224 | 0.28328 | 0.558 | 0.45 | 1 | HC_vs_MS.NoTrt_B |
| C1orf43 | 0.01224 | -0.3501 | 0.512 | 0.62 | 1 | HC_vs_MS.NoTrt_B |
| VWA2 | 0.01226 | 0.30818 | 0.725 | 0.68 | 1 | HC_vs_MS.NoTrt_B |
| RNF126 | 0.01231 | -0.2551 | 0.18 | 0.26 | 1 | HC_vs_MS.NoTrt_B |
| ECHS1 | 0.01231 | -0.4764 | 0.093 | 0.16 | 1 | HC_vs_MS.NoTrt_B |
| INPP5K | 0.01235 | -0.6957 | 0.458 | 0.55 | 1 | HC_vs_MS.NoTrt_B |
| UBR5 | 0.01237 | 0.57179 | 0.816 | 0.81 | 1 | HC_vs_MS.NoTrt_B |
| USMG5 | 0.01245 | -0.3461 | 0.188 | 0.28 | 1 | HC_vs_MS.NoTrt_B |
| CLDN19 | 0.01246 | -0.3924 | 0.183 | 0.28 | 1 | HC_vs_MS.NoTrt_B |
| HTRA2 | 0.01248 | -0.286 | 0.088 | 0.15 | 1 | HC_vs_MS.NoTrt_B |

| | | | | | | |
|---|---|---|---|---|---|---|
| EPHA1 | 0.01251 | -0.3369 | 0.328 | 0.43 | 1 | HC_vs_MS.NoTrt_B |
| PPM1F | 0.01254 | 0.73666 | 0.796 | 0.81 | 1 | HC_vs_MS.NoTrt_B |
| TRAPPC4 | 0.01256 | 0.50064 | 0.903 | 0.92 | 1 | HC_vs_MS.NoTrt_B |
| U2AF1L5 | 0.0126 | -0.316 | 0.632 | 0.7 | 1 | HC_vs_MS.NoTrt_B |
| RGS12 | 0.01261 | 0.95106 | 0.823 | 0.84 | 1 | HC_vs_MS.NoTrt_B |
| BOLA2B | 0.01266 | 0.32009 | 0.094 | 0.16 | 1 | HC_vs_MS.NoTrt_B |
| DDX27 | 0.01267 | -0.2654 | 0.58 | 0.68 | 1 | HC_vs_MS.NoTrt_B |
| IFT22 | 0.01268 | 0.33877 | 0.964 | 0.94 | 1 | HC_vs_MS.NoTrt_B |
| TCF7L1 | 0.01269 | 0.39815 | 0.655 | 0.61 | 1 | HC_vs_MS.NoTrt_B |
| LAS1L | 0.01272 | -0.4267 | 0.125 | 0.2 | 1 | HC_vs_MS.NoTrt_B |
| CEP112 | 0.01272 | 0.27826 | 0.933 | 0.91 | 1 | HC_vs_MS.NoTrt_B |
| BIRC2 | 0.01275 | 0.32392 | 0.133 | 0.21 | 1 | HC_vs_MS.NoTrt_B |
| CXCR4 | 0.01286 | 0.97128 | 0.188 | 0.11 | 1 | HC_vs_MS.NoTrt_B |
| GGT5 | 0.01288 | 0.3147 | 0.659 | 0.62 | 1 | HC_vs_MS.NoTrt_B |
| HNRNPH3 | 0.0129 | -0.2872 | 0.291 | 0.38 | 1 | HC_vs_MS.NoTrt_B |
| CREG2 | 0.01294 | 0.50122 | 0.677 | 0.63 | 1 | HC_vs_MS.NoTrt_B |
| PCDHB11 | 0.01296 | 0.3189 | 0.642 | 0.6 | 1 | HC_vs_MS.NoTrt_B |
| FAXDC2 | 0.01306 | 0.53619 | 0.839 | 0.85 | 1 | HC_vs_MS.NoTrt_B |
| LIMK2 | 0.01308 | -0.2849 | 0.201 | 0.29 | 1 | HC_vs_MS.NoTrt_B |
| ARF6 | 0.01312 | 0.33887 | 0.535 | 0.64 | 1 | HC_vs_MS.NoTrt_B |
| CYP3A43 | 0.01313 | 0.33928 | 0.696 | 0.71 | 1 | HC_vs_MS.NoTrt_B |
| MXRA7 | 0.01318 | 0.79826 | 0.78 | 0.79 | 1 | HC_vs_MS.NoTrt_B |
| CGRRF1 | 0.01325 | -0.3885 | 0.257 | 0.35 | 1 | HC_vs_MS.NoTrt_B |
| AMACR | 0.0133 | -1.1905 | 0.052 | 0.1 | 1 | HC_vs_MS.NoTrt_B |
| EVI5 | 0.01346 | 0.47732 | 0.828 | 0.81 | 1 | HC_vs_MS.NoTrt_B |
| C12orf76 | 0.01347 | 0.27792 | 0.952 | 0.96 | 1 | HC_vs_MS.NoTrt_B |
| ADAMTS4 | 0.01349 | 0.51984 | 0.896 | 0.91 | 1 | HC_vs_MS.NoTrt_B |
| BANF1 | 0.01363 | -0.3347 | 0.181 | 0.27 | 1 | HC_vs_MS.NoTrt_B |
| SRCIN1 | 0.01367 | 0.35724 | 0.516 | 0.45 | 1 | HC_vs_MS.NoTrt_B |
| POLA2 | 0.01368 | -0.4216 | 0.597 | 0.7 | 1 | HC_vs_MS.NoTrt_B |
| PHLDA3 | 0.01373 | -0.3187 | 0.303 | 0.41 | 1 | HC_vs_MS.NoTrt_B |
| RABGGTA | 0.01374 | -0.4476 | 0.099 | 0.16 | 1 | HC_vs_MS.NoTrt_B |
| CCR4 | 0.01375 | -0.3374 | 0.861 | 0.91 | 1 | HC_vs_MS.NoTrt_B |
| USP49 | 0.0138 | 0.56559 | 0.694 | 0.65 | 1 | HC_vs_MS.NoTrt_B |
| CHMP4A | 0.01385 | -0.4054 | 0.629 | 0.69 | 1 | HC_vs_MS.NoTrt_B |
| NDUFV3 | 0.0139 | 0.36916 | 0.954 | 0.97 | 1 | HC_vs_MS.NoTrt_B |
| UBE2F | 0.01392 | -0.3813 | 0.661 | 0.76 | 1 | HC_vs_MS.NoTrt_B |
| TPCN2 | 0.01396 | 0.55847 | 0.619 | 0.57 | 1 | HC_vs_MS.NoTrt_B |
| VGLL4 | 0.01402 | 0.45024 | 0.941 | 0.95 | 1 | HC_vs_MS.NoTrt_B |
| GPR1 | 0.01422 | 0.35065 | 0.243 | 0.16 | 1 | HC_vs_MS.NoTrt_B |

| | | | | | | |
|---|---|---|---|---|---|---|
| ZNF611 | 0.01429 | 0.48286 | 0.951 | 0.96 | 1 | HC_vs_MS.NoTrt_B |
| ABCF2 | 0.01433 | -0.2505 | 0.226 | 0.32 | 1 | HC_vs_MS.NoTrt_B |
| PRKAA1 | 0.01434 | 0.48109 | 0.891 | 0.9 | 1 | HC_vs_MS.NoTrt_B |
| NCOR2 | 0.01436 | 0.52094 | 0.404 | 0.31 | 1 | HC_vs_MS.NoTrt_B |
| SPATA17 | 0.01443 | 0.32706 | 0.613 | 0.56 | 1 | HC_vs_MS.NoTrt_B |
| SART3 | 0.01461 | -0.2626 | 0.138 | 0.21 | 1 | HC_vs_MS.NoTrt_B |
| MINPP1 | 0.01461 | -0.4172 | 0.072 | 0.13 | 1 | HC_vs_MS.NoTrt_B |
| LILRA6 | 0.01463 | 0.29613 | 0.523 | 0.44 | 1 | HC_vs_MS.NoTrt_B |
| PSEN1 | 0.01464 | -0.4018 | 0.68 | 0.77 | 1 | HC_vs_MS.NoTrt_B |
| PTGIS | 0.01464 | 0.30814 | 0.667 | 0.64 | 1 | HC_vs_MS.NoTrt_B |
| ZNF483 | 0.01469 | 0.65352 | 0.826 | 0.84 | 1 | HC_vs_MS.NoTrt_B |
| COL1A1 | 0.0147 | 0.49846 | 0.497 | 0.45 | 1 | HC_vs_MS.NoTrt_B |
| AKAP6 | 0.01473 | 0.4557 | 0.697 | 0.63 | 1 | HC_vs_MS.NoTrt_B |
| CHD6 | 0.01475 | -0.3726 | 0.183 | 0.27 | 1 | HC_vs_MS.NoTrt_B |
| CHMP1A | 0.01477 | -0.4324 | 0.172 | 0.25 | 1 | HC_vs_MS.NoTrt_B |
| PSMD13 | 0.01481 | -0.3803 | 0.52 | 0.62 | 1 | HC_vs_MS.NoTrt_B |
| KRCC1 | 0.01481 | -0.4837 | 0.099 | 0.16 | 1 | HC_vs_MS.NoTrt_B |
| CRISPLD2 | 0.01484 | 0.3229 | 0.772 | 0.8 | 1 | HC_vs_MS.NoTrt_B |
| ALDH1L1 | 0.01488 | 0.31072 | 0.583 | 0.54 | 1 | HC_vs_MS.NoTrt_B |
| FER | 0.0149 | 1.46334 | 0.504 | 0.46 | 1 | HC_vs_MS.NoTrt_B |
| ZNF724 | 0.01493 | 0.59521 | 0.899 | 0.92 | 1 | HC_vs_MS.NoTrt_B |
| TCF4 | 0.01494 | 0.62374 | 0.697 | 0.67 | 1 | HC_vs_MS.NoTrt_B |
| STAMBP | 0.01499 | -0.2585 | 0.112 | 0.18 | 1 | HC_vs_MS.NoTrt_B |
| ZNF385C | 0.01501 | 0.27154 | 0.581 | 0.52 | 1 | HC_vs_MS.NoTrt_B |
| TLR5 | 0.01528 | 1.14705 | 0.636 | 0.61 | 1 | HC_vs_MS.NoTrt_B |
| TRABD2A | 0.01544 | 0.49307 | 0.833 | 0.79 | 1 | HC_vs_MS.NoTrt_B |
| MIF4GD | 0.01547 | -0.3582 | 0.09 | 0.15 | 1 | HC_vs_MS.NoTrt_B |
| CXCR3 | 0.0155 | -0.3751 | 0.103 | 0.17 | 1 | HC_vs_MS.NoTrt_B |
| SEC24B | 0.01554 | 0.29573 | 0.494 | 0.63 | 1 | HC_vs_MS.NoTrt_B |
| SIM1 | 0.0156 | 0.52697 | 0.261 | 0.18 | 1 | HC_vs_MS.NoTrt_B |
| TPD52L2 | 0.01561 | -0.5328 | 0.201 | 0.28 | 1 | HC_vs_MS.NoTrt_B |
| MCM5 | 0.01574 | -0.6709 | 0.164 | 0.23 | 1 | HC_vs_MS.NoTrt_B |
| NICN1 | 0.01576 | 0.5794 | 0.77 | 0.73 | 1 | HC_vs_MS.NoTrt_B |
| SLC16A6 | 0.01578 | 0.29996 | 0.242 | 0.34 | 1 | HC_vs_MS.NoTrt_B |
| QDPR | 0.01592 | -0.7512 | 0.058 | 0.11 | 1 | HC_vs_MS.NoTrt_B |
| TMEM127 | 0.01593 | 0.70143 | 0.691 | 0.63 | 1 | HC_vs_MS.NoTrt_B |
| NPIPB5 | 0.01594 | 0.54502 | 0.926 | 0.95 | 1 | HC_vs_MS.NoTrt_B |
| C4orf36 | 0.01596 | 0.45106 | 0.838 | 0.89 | 1 | HC_vs_MS.NoTrt_B |
| DSN1 | 0.016 | -0.4962 | 0.225 | 0.31 | 1 | HC_vs_MS.NoTrt_B |
| HVCN1 | 0.01609 | 0.70045 | 0.501 | 0.42 | 1 | HC_vs_MS.NoTrt_B |

| | | | | | | |
|---|---|---|---|---|---|---|
| TMEM167A | 0.01609 | -0.3367 | 0.329 | 0.43 | 1 | HC_vs_MS.NoTrt_B |
| DAND5 | 0.01611 | 0.36145 | 0.749 | 0.69 | 1 | HC_vs_MS.NoTrt_B |
| COA7 | 0.01616 | 0.28643 | 0.871 | 0.86 | 1 | HC_vs_MS.NoTrt_B |
| RP11-343C2 | 0.01618 | -0.3868 | 0.112 | 0.18 | 1 | HC_vs_MS.NoTrt_B |
| TMEM35B | 0.0162 | -0.5814 | 0.152 | 0.23 | 1 | HC_vs_MS.NoTrt_B |
| TIPARP | 0.01622 | -0.3723 | 0.062 | 0.11 | 1 | HC_vs_MS.NoTrt_B |
| SLC25A5 | 0.01623 | -0.2981 | 0.278 | 0.37 | 1 | HC_vs_MS.NoTrt_B |
| FLRT2 | 0.01626 | 0.26023 | 0.92 | 0.9 | 1 | HC_vs_MS.NoTrt_B |
| PNPLA1 | 0.01637 | 0.62558 | 0.309 | 0.23 | 1 | HC_vs_MS.NoTrt_B |
| PPCDC | 0.01641 | -0.2965 | 0.058 | 0.11 | 1 | HC_vs_MS.NoTrt_B |
| BAIAP2L1 | 0.01647 | 0.47042 | 0.533 | 0.46 | 1 | HC_vs_MS.NoTrt_B |
| FBN1 | 0.01652 | 0.39578 | 0.416 | 0.35 | 1 | HC_vs_MS.NoTrt_B |
| ZNF570 | 0.01652 | 0.47346 | 0.78 | 0.75 | 1 | HC_vs_MS.NoTrt_B |
| FAM117B | 0.01662 | 0.90262 | 0.449 | 0.37 | 1 | HC_vs_MS.NoTrt_B |
| FAM221A | 0.01672 | 0.3037 | 0.961 | 0.96 | 1 | HC_vs_MS.NoTrt_B |
| PEF1 | 0.01683 | -0.405 | 0.128 | 0.2 | 1 | HC_vs_MS.NoTrt_B |
| AK3 | 0.01684 | 0.40028 | 0.464 | 0.38 | 1 | HC_vs_MS.NoTrt_B |
| RTF1 | 0.01687 | -0.5348 | 0.557 | 0.64 | 1 | HC_vs_MS.NoTrt_B |
| LRRC56 | 0.01691 | 0.39307 | 0.38 | 0.29 | 1 | HC_vs_MS.NoTrt_B |
| SCRN3 | 0.01697 | -0.4021 | 0.426 | 0.51 | 1 | HC_vs_MS.NoTrt_B |
| CISD2 | 0.01709 | 0.59593 | 0.439 | 0.53 | 1 | HC_vs_MS.NoTrt_B |
| VPS13C | 0.0171 | 0.26561 | 0.751 | 0.85 | 1 | HC_vs_MS.NoTrt_B |
| MICAL1 | 0.01711 | -0.3861 | 0.219 | 0.3 | 1 | HC_vs_MS.NoTrt_B |
| DCTN3 | 0.01725 | -0.4993 | 0.361 | 0.45 | 1 | HC_vs_MS.NoTrt_B |
| RP11-722G7 | 0.01729 | 0.55867 | 0.438 | 0.33 | 1 | HC_vs_MS.NoTrt_B |
| XXYLT1 | 0.01733 | -1.4796 | 0.183 | 0.25 | 1 | HC_vs_MS.NoTrt_B |
| EIF5A2 | 0.01735 | 0.44178 | 0.296 | 0.39 | 1 | HC_vs_MS.NoTrt_B |
| ATM | 0.01739 | 0.75435 | 0.952 | 0.98 | 1 | HC_vs_MS.NoTrt_B |
| TF | 0.01742 | 0.41069 | 0.888 | 0.91 | 1 | HC_vs_MS.NoTrt_B |
| NRF1 | 0.01742 | -0.4535 | 0.29 | 0.38 | 1 | HC_vs_MS.NoTrt_B |
| SAMD4B | 0.01747 | 0.79822 | 0.842 | 0.85 | 1 | HC_vs_MS.NoTrt_B |
| DDT | 0.01755 | -0.4827 | 0.132 | 0.2 | 1 | HC_vs_MS.NoTrt_B |
| ITGA3 | 0.01755 | 0.42148 | 0.754 | 0.7 | 1 | HC_vs_MS.NoTrt_B |
| ERGIC2 | 0.01757 | 0.33996 | 0.471 | 0.58 | 1 | HC_vs_MS.NoTrt_B |
| CCR7 | 0.01761 | 0.5033 | 0.613 | 0.57 | 1 | HC_vs_MS.NoTrt_B |
| PRKAG2 | 0.01763 | -0.5258 | 0.299 | 0.39 | 1 | HC_vs_MS.NoTrt_B |
| GCNT3 | 0.01785 | 0.33041 | 0.877 | 0.86 | 1 | HC_vs_MS.NoTrt_B |
| DDX17 | 0.01786 | 0.45022 | 0.907 | 0.95 | 1 | HC_vs_MS.NoTrt_B |
| KMT2D | 0.01806 | 0.71514 | 0.855 | 0.83 | 1 | HC_vs_MS.NoTrt_B |
| CLCN7 | 0.01813 | 0.32075 | 0.284 | 0.4 | 1 | HC_vs_MS.NoTrt_B |

| | | | | | | |
|---|---|---|---|---|---|---|
| SERTAD1 | 0.01815 | 0.91301 | 0.339 | 0.25 | 1 | HC_vs_MS.NoTrt_B |
| XAB2 | 0.01816 | 0.41709 | 0.228 | 0.32 | 1 | HC_vs_MS.NoTrt_B |
| CEACAM8 | 0.01816 | 0.37794 | 0.504 | 0.44 | 1 | HC_vs_MS.NoTrt_B |
| SULT1A3 | 0.01821 | -0.5439 | 0.606 | 0.63 | 1 | HC_vs_MS.NoTrt_B |
| RFTN2 | 0.01827 | 0.36776 | 0.68 | 0.62 | 1 | HC_vs_MS.NoTrt_B |
| KCNA3 | 0.01827 | -0.3848 | 0.168 | 0.24 | 1 | HC_vs_MS.NoTrt_B |
| CDC123 | 0.01833 | -0.2717 | 0.188 | 0.27 | 1 | HC_vs_MS.NoTrt_B |
| CDK10 | 0.01848 | 0.58225 | 0.91 | 0.92 | 1 | HC_vs_MS.NoTrt_B |
| PCBD1 | 0.0186 | -0.3935 | 0.062 | 0.11 | 1 | HC_vs_MS.NoTrt_B |
| TNFRSF18 | 0.01865 | -0.6705 | 0.067 | 0.12 | 1 | HC_vs_MS.NoTrt_B |
| LDHA | 0.01871 | -0.3985 | 0.732 | 0.76 | 1 | HC_vs_MS.NoTrt_B |
| MPLKIP | 0.01876 | 0.39628 | 0.941 | 0.96 | 1 | HC_vs_MS.NoTrt_B |
| LGR6 | 0.01878 | 0.31948 | 0.178 | 0.11 | 1 | HC_vs_MS.NoTrt_B |
| LAT2 | 0.01883 | 0.48144 | 0.854 | 0.84 | 1 | HC_vs_MS.NoTrt_B |
| DPH1 | 0.01893 | 0.25935 | 0.855 | 0.81 | 1 | HC_vs_MS.NoTrt_B |
| VPS53 | 0.01894 | 0.25865 | 0.941 | 0.95 | 1 | HC_vs_MS.NoTrt_B |
| PCDHA10 | 0.01897 | 0.25793 | 0.538 | 0.45 | 1 | HC_vs_MS.NoTrt_B |
| PTPN23 | 0.01909 | 1.53678 | 0.142 | 0.08 | 1 | HC_vs_MS.NoTrt_B |
| AHR | 0.01914 | 0.44484 | 0.607 | 0.51 | 1 | HC_vs_MS.NoTrt_B |
| INO80 | 0.01927 | -0.5133 | 0.075 | 0.13 | 1 | HC_vs_MS.NoTrt_B |
| FAM153A | 0.0194 | -0.2973 | 0.452 | 0.54 | 1 | HC_vs_MS.NoTrt_B |
| SEMA5A | 0.01942 | 0.33672 | 0.39 | 0.31 | 1 | HC_vs_MS.NoTrt_B |
| ZBTB40 | 0.01948 | 0.30486 | 0.365 | 0.45 | 1 | HC_vs_MS.NoTrt_B |
| CDH1 | 0.01957 | 0.27737 | 0.728 | 0.7 | 1 | HC_vs_MS.NoTrt_B |
| ZNF638 | 0.01957 | 0.86454 | 0.388 | 0.31 | 1 | HC_vs_MS.NoTrt_B |
| IFT88 | 0.0196 | -0.3873 | 0.549 | 0.47 | 1 | HC_vs_MS.NoTrt_B |
| RAET1E | 0.01961 | 0.41944 | 0.307 | 0.23 | 1 | HC_vs_MS.NoTrt_B |
| CPM | 0.01968 | 0.52184 | 0.803 | 0.8 | 1 | HC_vs_MS.NoTrt_B |
| FAM120A | 0.01971 | 0.89573 | 0.158 | 0.09 | 1 | HC_vs_MS.NoTrt_B |
| RPS6KB1 | 0.01974 | 0.35194 | 0.839 | 0.82 | 1 | HC_vs_MS.NoTrt_B |
| GK | 0.0198 | -0.8257 | 0.126 | 0.19 | 1 | HC_vs_MS.NoTrt_B |
| ENAH | 0.01981 | 0.31129 | 0.687 | 0.66 | 1 | HC_vs_MS.NoTrt_B |
| CCT8 | 0.01988 | -0.2636 | 0.21 | 0.29 | 1 | HC_vs_MS.NoTrt_B |
| GNL3 | 0.01991 | -0.394 | 0.093 | 0.15 | 1 | HC_vs_MS.NoTrt_B |
| ZNF317 | 0.01991 | -0.323 | 0.088 | 0.15 | 1 | HC_vs_MS.NoTrt_B |
| RNF222 | 0.01991 | 0.39363 | 0.401 | 0.33 | 1 | HC_vs_MS.NoTrt_B |
| RHBDD3 | 0.01997 | 0.74832 | 0.467 | 0.39 | 1 | HC_vs_MS.NoTrt_B |
| NEURL1 | 0.01998 | 0.68229 | 0.3 | 0.22 | 1 | HC_vs_MS.NoTrt_B |
| PMS1 | 0.02001 | 0.37803 | 0.564 | 0.67 | 1 | HC_vs_MS.NoTrt_B |
| ALG10 | 0.02004 | 0.36818 | 0.055 | 0.1 | 1 | HC_vs_MS.NoTrt_B |

| | | | | | | |
|---|---|---|---|---|---|---|
| STEAP2 | 0.02009 | 0.34584 | 0.68 | 0.69 | 1 | HC_vs_MS.NoTrt_B |
| IFRD2 | 0.02014 | -0.4762 | 0.084 | 0.14 | 1 | HC_vs_MS.NoTrt_B |
| POU2AF1 | 0.02015 | 0.59235 | 0.694 | 0.7 | 1 | HC_vs_MS.NoTrt_B |
| ATP1B1 | 0.02017 | -0.6339 | 0.161 | 0.23 | 1 | HC_vs_MS.NoTrt_B |
| UBL3 | 0.0202 | -0.2965 | 0.165 | 0.24 | 1 | HC_vs_MS.NoTrt_B |
| CDC27 | 0.0202 | -0.5272 | 0.438 | 0.51 | 1 | HC_vs_MS.NoTrt_B |
| FBXO4 | 0.02031 | -0.2906 | 0.201 | 0.28 | 1 | HC_vs_MS.NoTrt_B |
| VEZT | 0.02048 | -0.326 | 0.742 | 0.7 | 1 | HC_vs_MS.NoTrt_B |
| LENG8 | 0.0205 | 0.63542 | 0.741 | 0.75 | 1 | HC_vs_MS.NoTrt_B |
| PANK1 | 0.02064 | 0.37429 | 0.452 | 0.41 | 1 | HC_vs_MS.NoTrt_B |
| CATSPERB | 0.02097 | 0.58202 | 0.622 | 0.58 | 1 | HC_vs_MS.NoTrt_B |
| MTMR14 | 0.02098 | -0.3807 | 0.571 | 0.69 | 1 | HC_vs_MS.NoTrt_B |
| DDX42 | 0.02123 | 0.50943 | 0.339 | 0.45 | 1 | HC_vs_MS.NoTrt_B |
| PDE4DIP | 0.02134 | 0.27741 | 0.864 | 0.89 | 1 | HC_vs_MS.NoTrt_B |
| MAGI2 | 0.02135 | 0.28673 | 0.787 | 0.78 | 1 | HC_vs_MS.NoTrt_B |
| ARHGAP19 | 0.02136 | -0.4785 | 0.27 | 0.36 | 1 | HC_vs_MS.NoTrt_B |
| TDRD12 | 0.02143 | 1.00847 | 0.48 | 0.42 | 1 | HC_vs_MS.NoTrt_B |
| FOPNL | 0.02146 | 0.55785 | 0.594 | 0.52 | 1 | HC_vs_MS.NoTrt_B |
| TPH2 | 0.02147 | 0.40227 | 0.632 | 0.61 | 1 | HC_vs_MS.NoTrt_B |
| PLK5 | 0.02156 | 0.65115 | 0.528 | 0.47 | 1 | HC_vs_MS.NoTrt_B |
| ZNF512 | 0.02157 | -0.4177 | 0.117 | 0.18 | 1 | HC_vs_MS.NoTrt_B |
| TC2N | 0.02161 | -0.4019 | 0.174 | 0.24 | 1 | HC_vs_MS.NoTrt_B |
| EFCAB11 | 0.02165 | 0.37819 | 0.813 | 0.8 | 1 | HC_vs_MS.NoTrt_B |
| ZNF584 | 0.02165 | 0.52571 | 0.665 | 0.65 | 1 | HC_vs_MS.NoTrt_B |
| CNTNAP3 | 0.02169 | 0.3975 | 0.593 | 0.54 | 1 | HC_vs_MS.NoTrt_B |
| ARFGAP3 | 0.02175 | -0.3476 | 0.322 | 0.42 | 1 | HC_vs_MS.NoTrt_B |
| TRUB2 | 0.02177 | -0.282 | 0.955 | 0.98 | 1 | HC_vs_MS.NoTrt_B |
| TEX9 | 0.02182 | 0.41732 | 0.752 | 0.77 | 1 | HC_vs_MS.NoTrt_B |
| TSPAN17 | 0.02182 | -0.3232 | 0.064 | 0.11 | 1 | HC_vs_MS.NoTrt_B |
| PGD | 0.02183 | -1.0513 | 0.078 | 0.13 | 1 | HC_vs_MS.NoTrt_B |
| RNF6 | 0.02189 | -0.5149 | 0.155 | 0.22 | 1 | HC_vs_MS.NoTrt_B |
| MLLT6 | 0.02191 | 0.50404 | 0.91 | 0.89 | 1 | HC_vs_MS.NoTrt_B |
| GZMM | 0.02193 | -0.3558 | 0.11 | 0.17 | 1 | HC_vs_MS.NoTrt_B |
| ACVR2B | 0.022 | 0.40771 | 0.383 | 0.49 | 1 | HC_vs_MS.NoTrt_B |
| NIPAL1 | 0.02204 | 0.44661 | 0.41 | 0.36 | 1 | HC_vs_MS.NoTrt_B |
| NQO2 | 0.0221 | 0.26002 | 0.78 | 0.77 | 1 | HC_vs_MS.NoTrt_B |
| IGSF1 | 0.02225 | 0.39138 | 0.388 | 0.31 | 1 | HC_vs_MS.NoTrt_B |
| ZNF99 | 0.02232 | 0.27633 | 0.529 | 0.46 | 1 | HC_vs_MS.NoTrt_B |
| ASB7 | 0.02233 | -0.4111 | 0.064 | 0.11 | 1 | HC_vs_MS.NoTrt_B |
| FAM227A | 0.0224 | 0.2933 | 0.909 | 0.92 | 1 | HC_vs_MS.NoTrt_B |

| | | | | | | |
|---|---|---|---|---|---|---|
| TRADD | 0.02248 | -0.4026 | 0.133 | 0.2 | 1 | HC_vs_MS.NoTrt_B |
| SLC2A8 | 0.02272 | -0.5582 | 0.259 | 0.35 | 1 | HC_vs_MS.NoTrt_B |
| FCER1G | 0.02273 | 0.42889 | 0.674 | 0.62 | 1 | HC_vs_MS.NoTrt_B |
| TMEM120B | 0.02274 | 0.28044 | 0.972 | 0.97 | 1 | HC_vs_MS.NoTrt_B |
| ANKRD16 | 0.02285 | 0.39364 | 0.429 | 0.36 | 1 | HC_vs_MS.NoTrt_B |
| FAM83A | 0.02286 | 0.27883 | 0.199 | 0.13 | 1 | HC_vs_MS.NoTrt_B |
| ATP9B | 0.02292 | 0.44028 | 0.909 | 0.93 | 1 | HC_vs_MS.NoTrt_B |
| HNMT | 0.02293 | 0.30682 | 0.796 | 0.81 | 1 | HC_vs_MS.NoTrt_B |
| DOC2A | 0.02294 | 0.3511 | 0.475 | 0.43 | 1 | HC_vs_MS.NoTrt_B |
| TRIP13 | 0.02302 | 0.40523 | 0.543 | 0.49 | 1 | HC_vs_MS.NoTrt_B |
| MSRA | 0.02321 | 0.48785 | 0.239 | 0.17 | 1 | HC_vs_MS.NoTrt_B |
| RSPH10B | 0.02326 | 0.3604 | 0.603 | 0.52 | 1 | HC_vs_MS.NoTrt_B |
| UBLCP1 | 0.02336 | -0.5076 | 0.148 | 0.21 | 1 | HC_vs_MS.NoTrt_B |
| NBPF19 | 0.02337 | 0.71218 | 0.446 | 0.36 | 1 | HC_vs_MS.NoTrt_B |
| YIPF1 | 0.02341 | -0.6199 | 0.07 | 0.12 | 1 | HC_vs_MS.NoTrt_B |
| MRPS12 | 0.02342 | -0.4176 | 0.371 | 0.47 | 1 | HC_vs_MS.NoTrt_B |
| COPS3 | 0.02343 | -0.3566 | 0.22 | 0.29 | 1 | HC_vs_MS.NoTrt_B |
| SH3BP5 | 0.0235 | 0.38253 | 0.801 | 0.75 | 1 | HC_vs_MS.NoTrt_B |
| PPIL6 | 0.02351 | 0.34568 | 0.497 | 0.45 | 1 | HC_vs_MS.NoTrt_B |
| CATSPER2 | 0.02353 | 0.47503 | 0.771 | 0.71 | 1 | HC_vs_MS.NoTrt_B |
| NCBP3 | 0.02356 | 0.43521 | 0.877 | 0.9 | 1 | HC_vs_MS.NoTrt_B |
| ARNTL | 0.02363 | -0.4416 | 0.191 | 0.26 | 1 | HC_vs_MS.NoTrt_B |
| NCKAP1L | 0.02379 | -0.314 | 0.761 | 0.8 | 1 | HC_vs_MS.NoTrt_B |
| BPIFB1 | 0.02381 | 0.39813 | 0.507 | 0.44 | 1 | HC_vs_MS.NoTrt_B |
| NOTCH3 | 0.02388 | 0.33292 | 0.549 | 0.47 | 1 | HC_vs_MS.NoTrt_B |
| HSPG2 | 0.02389 | -0.266 | 0.335 | 0.26 | 1 | HC_vs_MS.NoTrt_B |
| SMAD9 | 0.02395 | -0.4045 | 0.29 | 0.37 | 1 | HC_vs_MS.NoTrt_B |
| PQBP1 | 0.02398 | -0.3775 | 0.135 | 0.2 | 1 | HC_vs_MS.NoTrt_B |
| AS3MT | 0.02401 | 0.41178 | 0.772 | 0.71 | 1 | HC_vs_MS.NoTrt_B |
| MIB1 | 0.02402 | 0.84573 | 0.101 | 0.16 | 1 | HC_vs_MS.NoTrt_B |
| E2F3 | 0.02405 | 0.38927 | 0.125 | 0.2 | 1 | HC_vs_MS.NoTrt_B |
| AL592183.1 | 0.02413 | -0.4574 | 0.078 | 0.13 | 1 | HC_vs_MS.NoTrt_B |
| SMG8 | 0.02415 | -0.4043 | 0.078 | 0.13 | 1 | HC_vs_MS.NoTrt_B |
| TEX261 | 0.02416 | -0.3147 | 0.057 | 0.1 | 1 | HC_vs_MS.NoTrt_B |
| STARD5 | 0.02417 | 1.49153 | 0.114 | 0.06 | 1 | HC_vs_MS.NoTrt_B |
| CDH13 | 0.02417 | 0.31211 | 0.732 | 0.73 | 1 | HC_vs_MS.NoTrt_B |
| TNFSF15 | 0.02419 | 0.275 | 0.583 | 0.49 | 1 | HC_vs_MS.NoTrt_B |
| ASAH1 | 0.02421 | -0.3231 | 0.678 | 0.76 | 1 | HC_vs_MS.NoTrt_B |
| ABCC3 | 0.02421 | 0.42319 | 0.804 | 0.79 | 1 | HC_vs_MS.NoTrt_B |
| PROCR | 0.02422 | 0.34031 | 0.839 | 0.84 | 1 | HC_vs_MS.NoTrt_B |

| | | | | | | |
|---|---|---|---|---|---|---|
| ZNF677 | 0.02426 | 0.33807 | 0.794 | 0.76 | 1 | HC_vs_MS.NoTrt_B |
| LMNA | 0.02428 | -0.6111 | 0.486 | 0.56 | 1 | HC_vs_MS.NoTrt_B |
| CCDC180 | 0.0244 | 0.39322 | 0.757 | 0.7 | 1 | HC_vs_MS.NoTrt_B |
| ATP5G3 | 0.02441 | -0.2613 | 0.343 | 0.42 | 1 | HC_vs_MS.NoTrt_B |
| KCTD19 | 0.0246 | 0.33841 | 0.743 | 0.69 | 1 | HC_vs_MS.NoTrt_B |
| KMT2B | 0.02462 | 0.6801 | 0.29 | 0.22 | 1 | HC_vs_MS.NoTrt_B |
| ARPC1A | 0.02476 | -0.7879 | 0.23 | 0.3 | 1 | HC_vs_MS.NoTrt_B |
| C2 | 0.02489 | 0.53669 | 0.696 | 0.68 | 1 | HC_vs_MS.NoTrt_B |
| C6orf203 | 0.0249 | 0.64969 | 0.599 | 0.54 | 1 | HC_vs_MS.NoTrt_B |
| TMEM69 | 0.0249 | -0.4481 | 0.197 | 0.28 | 1 | HC_vs_MS.NoTrt_B |
| ZNF451 | 0.02505 | 0.44665 | 0.758 | 0.71 | 1 | HC_vs_MS.NoTrt_B |
| CDK13 | 0.02506 | 0.2767 | 0.825 | 0.83 | 1 | HC_vs_MS.NoTrt_B |
| PSMA5 | 0.02507 | -0.2517 | 0.232 | 0.31 | 1 | HC_vs_MS.NoTrt_B |
| ARHGAP28 | 0.02509 | 0.29076 | 0.845 | 0.85 | 1 | HC_vs_MS.NoTrt_B |
| RRBP1 | 0.0251 | 0.68974 | 0.213 | 0.3 | 1 | HC_vs_MS.NoTrt_B |
| ITGA11 | 0.02511 | 0.41049 | 0.619 | 0.63 | 1 | HC_vs_MS.NoTrt_B |
| DNAAF5 | 0.02515 | 0.98781 | 0.19 | 0.12 | 1 | HC_vs_MS.NoTrt_B |
| SMIM8 | 0.02528 | 0.51333 | 0.599 | 0.52 | 1 | HC_vs_MS.NoTrt_B |
| SPDYA | 0.02532 | 0.56868 | 0.059 | 0.11 | 1 | HC_vs_MS.NoTrt_B |
| IFIT1 | 0.02536 | 0.5596 | 0.257 | 0.18 | 1 | HC_vs_MS.NoTrt_B |
| SPINT2 | 0.0254 | -0.3903 | 0.526 | 0.44 | 1 | HC_vs_MS.NoTrt_B |
| CHMP5 | 0.02544 | -0.3249 | 0.094 | 0.15 | 1 | HC_vs_MS.NoTrt_B |
| GADD45B | 0.02551 | -0.8596 | 0.075 | 0.12 | 1 | HC_vs_MS.NoTrt_B |
| TMC7 | 0.02552 | 0.34933 | 0.72 | 0.72 | 1 | HC_vs_MS.NoTrt_B |
| ACBD7 | 0.02553 | 0.27207 | 0.836 | 0.84 | 1 | HC_vs_MS.NoTrt_B |
| SDC3 | 0.0256 | 0.50248 | 0.268 | 0.18 | 1 | HC_vs_MS.NoTrt_B |
| VIMP | 0.02573 | -0.5957 | 0.091 | 0.15 | 1 | HC_vs_MS.NoTrt_B |
| ZNF843 | 0.02574 | 0.27098 | 0.693 | 0.62 | 1 | HC_vs_MS.NoTrt_B |
| C4orf19 | 0.02581 | 0.33299 | 0.742 | 0.76 | 1 | HC_vs_MS.NoTrt_B |
| ARHGDIA | 0.02597 | 0.29029 | 0.357 | 0.49 | 1 | HC_vs_MS.NoTrt_B |
| METTL16 | 0.02602 | 0.73861 | 0.759 | 0.69 | 1 | HC_vs_MS.NoTrt_B |
| ZC3H12A | 0.02603 | -0.4067 | 0.096 | 0.15 | 1 | HC_vs_MS.NoTrt_B |
| HTN3 | 0.02617 | 0.3979 | 0.614 | 0.58 | 1 | HC_vs_MS.NoTrt_B |
| ATP7A | 0.02624 | 1.39849 | 0.348 | 0.28 | 1 | HC_vs_MS.NoTrt_B |
| ZNF581 | 0.02634 | -0.3148 | 0.061 | 0.11 | 1 | HC_vs_MS.NoTrt_B |
| DEDD2 | 0.0264 | -0.4035 | 0.312 | 0.4 | 1 | HC_vs_MS.NoTrt_B |
| TARDBP | 0.02647 | 0.25701 | 0.594 | 0.7 | 1 | HC_vs_MS.NoTrt_B |
| ILK | 0.02647 | -0.2742 | 0.133 | 0.2 | 1 | HC_vs_MS.NoTrt_B |
| LRRC59 | 0.02654 | -0.3807 | 0.1 | 0.16 | 1 | HC_vs_MS.NoTrt_B |
| NUTF2 | 0.02682 | -0.7142 | 0.129 | 0.19 | 1 | HC_vs_MS.NoTrt_B |

| | | | | | | |
|---|---|---|---|---|---|---|
| ZNF782 | 0.02688 | 0.66385 | 0.081 | 0.14 | 1 | HC_vs_MS.NoTrt_B |
| ZNF726 | 0.02713 | 0.27761 | 0.858 | 0.84 | 1 | HC_vs_MS.NoTrt_B |
| SIN3B | 0.02724 | 0.54694 | 0.754 | 0.74 | 1 | HC_vs_MS.NoTrt_B |
| NUDCD3 | 0.02725 | -0.4478 | 0.313 | 0.37 | 1 | HC_vs_MS.NoTrt_B |
| TCAIM | 0.02726 | 0.47417 | 0.184 | 0.27 | 1 | HC_vs_MS.NoTrt_B |
| SFPQ | 0.02734 | 0.36671 | 0.513 | 0.42 | 1 | HC_vs_MS.NoTrt_B |
| ABHD12B | 0.02742 | 0.66934 | 0.614 | 0.6 | 1 | HC_vs_MS.NoTrt_B |
| TBC1D24 | 0.0275 | 0.67749 | 0.706 | 0.68 | 1 | HC_vs_MS.NoTrt_B |
| RETSAT | 0.02761 | -0.5754 | 0.122 | 0.18 | 1 | HC_vs_MS.NoTrt_B |
| BAG6 | 0.02765 | 0.72389 | 0.622 | 0.56 | 1 | HC_vs_MS.NoTrt_B |
| RSPH10B2 | 0.02767 | 0.27623 | 0.6 | 0.52 | 1 | HC_vs_MS.NoTrt_B |
| ZRANB2 | 0.02778 | 0.25075 | 0.37 | 0.49 | 1 | HC_vs_MS.NoTrt_B |
| MNS1 | 0.0278 | 0.32739 | 0.174 | 0.11 | 1 | HC_vs_MS.NoTrt_B |
| ZFP82 | 0.02796 | 0.69631 | 0.649 | 0.6 | 1 | HC_vs_MS.NoTrt_B |
| PRAF2 | 0.02799 | -0.6578 | 0.083 | 0.14 | 1 | HC_vs_MS.NoTrt_B |
| MAP4 | 0.02804 | -0.4396 | 0.801 | 0.81 | 1 | HC_vs_MS.NoTrt_B |
| SLC25A34 | 0.02813 | 0.48477 | 0.622 | 0.56 | 1 | HC_vs_MS.NoTrt_B |
| PRDM5 | 0.02815 | 0.35147 | 0.672 | 0.67 | 1 | HC_vs_MS.NoTrt_B |
| TMEM41A | 0.02835 | 0.3433 | 0.835 | 0.81 | 1 | HC_vs_MS.NoTrt_B |
| PAEP | 0.02848 | 0.40903 | 0.393 | 0.34 | 1 | HC_vs_MS.NoTrt_B |
| MTO1 | 0.02856 | -0.2758 | 0.975 | 0.99 | 1 | HC_vs_MS.NoTrt_B |
| CMTM3 | 0.02873 | -0.3851 | 0.142 | 0.21 | 1 | HC_vs_MS.NoTrt_B |
| CD276 | 0.02874 | 0.28943 | 0.459 | 0.37 | 1 | HC_vs_MS.NoTrt_B |
| WIPF1 | 0.02901 | 0.26143 | 0.691 | 0.83 | 1 | HC_vs_MS.NoTrt_B |
| LHFPL5 | 0.02907 | 0.34898 | 0.528 | 0.47 | 1 | HC_vs_MS.NoTrt_B |
| ARHGEF4 | 0.02908 | 0.39488 | 0.546 | 0.47 | 1 | HC_vs_MS.NoTrt_B |
| ZNF528 | 0.02909 | 0.94002 | 0.574 | 0.51 | 1 | HC_vs_MS.NoTrt_B |
| RBM28 | 0.02913 | 0.4164 | 0.877 | 0.85 | 1 | HC_vs_MS.NoTrt_B |
| LRG1 | 0.02923 | 0.27586 | 0.755 | 0.74 | 1 | HC_vs_MS.NoTrt_B |
| TMCO3 | 0.02927 | 0.49987 | 0.716 | 0.68 | 1 | HC_vs_MS.NoTrt_B |
| DRG1 | 0.02933 | -0.415 | 0.142 | 0.21 | 1 | HC_vs_MS.NoTrt_B |
| CEP68 | 0.0294 | -0.596 | 0.516 | 0.58 | 1 | HC_vs_MS.NoTrt_B |
| SYS1 | 0.02958 | -0.368 | 0.423 | 0.52 | 1 | HC_vs_MS.NoTrt_B |
| PPFIBP2 | 0.02961 | 0.55374 | 0.251 | 0.18 | 1 | HC_vs_MS.NoTrt_B |
| ZSCAN21 | 0.02967 | 1.48351 | 0.22 | 0.15 | 1 | HC_vs_MS.NoTrt_B |
| LCLAT1 | 0.02971 | 0.34389 | 0.812 | 0.86 | 1 | HC_vs_MS.NoTrt_B |
| CHMP1B | 0.02978 | 0.49079 | 0.775 | 0.78 | 1 | HC_vs_MS.NoTrt_B |
| ZNF468 | 0.02987 | 0.26167 | 0.494 | 0.62 | 1 | HC_vs_MS.NoTrt_B |
| ALDH6A1 | 0.0299 | -0.687 | 0.796 | 0.87 | 1 | HC_vs_MS.NoTrt_B |
| PLCB1 | 0.02992 | 0.31852 | 0.323 | 0.25 | 1 | HC_vs_MS.NoTrt_B |

| | | | | | | |
|---|---|---|---|---|---|---|
| FAM228B | 0.02993 | 0.26746 | 0.945 | 0.95 | 1 | HC_vs_MS.NoTrt_B |
| ZNF83 | 0.03 | 0.54496 | 0.849 | 0.83 | 1 | HC_vs_MS.NoTrt_B |
| MUT | 0.03006 | -0.3471 | 0.058 | 0.1 | 1 | HC_vs_MS.NoTrt_B |
| ZBTB3 | 0.03008 | 0.42412 | 0.793 | 0.74 | 1 | HC_vs_MS.NoTrt_B |
| APMAP | 0.03013 | -0.3437 | 0.119 | 0.18 | 1 | HC_vs_MS.NoTrt_B |
| ITSN1 | 0.03021 | 0.57511 | 0.859 | 0.85 | 1 | HC_vs_MS.NoTrt_B |
| HCFC1R1 | 0.03023 | -0.3937 | 0.058 | 0.1 | 1 | HC_vs_MS.NoTrt_B |
| CCL22 | 0.03023 | 0.2642 | 0.741 | 0.75 | 1 | HC_vs_MS.NoTrt_B |
| WDR18 | 0.03024 | -0.2515 | 0.078 | 0.13 | 1 | HC_vs_MS.NoTrt_B |
| FAM161A | 0.03032 | -0.6511 | 0.614 | 0.76 | 1 | HC_vs_MS.NoTrt_B |
| SEC14L6 | 0.03035 | 0.37638 | 0.723 | 0.71 | 1 | HC_vs_MS.NoTrt_B |
| ATP5L2 | 0.03046 | 0.37947 | 0.303 | 0.23 | 1 | HC_vs_MS.NoTrt_B |
| PTS | 0.03082 | -0.3269 | 0.062 | 0.11 | 1 | HC_vs_MS.NoTrt_B |
| ASAP1 | 0.03082 | 0.46871 | 0.146 | 0.09 | 1 | HC_vs_MS.NoTrt_B |
| GPLD1 | 0.0309 | 0.67885 | 0.684 | 0.63 | 1 | HC_vs_MS.NoTrt_B |
| NFIC | 0.03096 | 0.8456 | 0.412 | 0.34 | 1 | HC_vs_MS.NoTrt_B |
| MBTD1 | 0.03116 | 0.84035 | 0.429 | 0.35 | 1 | HC_vs_MS.NoTrt_B |
| ZNF154 | 0.03122 | 1.23936 | 0.684 | 0.69 | 1 | HC_vs_MS.NoTrt_B |
| CSNK1G2 | 0.03127 | -0.7562 | 0.062 | 0.11 | 1 | HC_vs_MS.NoTrt_B |
| COQ6 | 0.03153 | 0.31778 | 0.761 | 0.71 | 1 | HC_vs_MS.NoTrt_B |
| HDAC7 | 0.03155 | 0.47956 | 0.286 | 0.21 | 1 | HC_vs_MS.NoTrt_B |
| ABCC2 | 0.03174 | 0.6727 | 0.536 | 0.51 | 1 | HC_vs_MS.NoTrt_B |
| ANGPT4 | 0.03175 | 0.36621 | 0.396 | 0.33 | 1 | HC_vs_MS.NoTrt_B |
| TSSC4 | 0.03178 | -0.2603 | 0.104 | 0.16 | 1 | HC_vs_MS.NoTrt_B |
| PIGT | 0.03185 | -0.4234 | 0.08 | 0.13 | 1 | HC_vs_MS.NoTrt_B |
| C9orf142 | 0.03196 | -0.2757 | 0.207 | 0.28 | 1 | HC_vs_MS.NoTrt_B |
| PCYOX1L | 0.032 | -0.3966 | 0.062 | 0.11 | 1 | HC_vs_MS.NoTrt_B |
| SLC15A2 | 0.032 | -0.5065 | 0.486 | 0.61 | 1 | HC_vs_MS.NoTrt_B |
| TNFRSF13C | 0.03203 | 0.67178 | 0.62 | 0.58 | 1 | HC_vs_MS.NoTrt_B |
| ARID1B | 0.03221 | 0.53572 | 0.946 | 0.94 | 1 | HC_vs_MS.NoTrt_B |
| SLC12A4 | 0.0323 | 0.74732 | 0.19 | 0.12 | 1 | HC_vs_MS.NoTrt_B |
| DCAF12 | 0.03232 | -0.634 | 0.09 | 0.14 | 1 | HC_vs_MS.NoTrt_B |
| GTF2F1 | 0.0325 | -0.7387 | 0.268 | 0.34 | 1 | HC_vs_MS.NoTrt_B |
| FAM53C | 0.03258 | 0.35364 | 0.786 | 0.78 | 1 | HC_vs_MS.NoTrt_B |
| SFMBT2 | 0.0326 | 1.09799 | 0.293 | 0.23 | 1 | HC_vs_MS.NoTrt_B |
| C11orf49 | 0.03266 | 0.33483 | 0.755 | 0.75 | 1 | HC_vs_MS.NoTrt_B |
| USP32 | 0.03283 | 0.45684 | 0.219 | 0.29 | 1 | HC_vs_MS.NoTrt_B |
| LHPP | 0.03284 | -0.4116 | 0.11 | 0.17 | 1 | HC_vs_MS.NoTrt_B |
| ETV3 | 0.0329 | 1.0659 | 0.12 | 0.07 | 1 | HC_vs_MS.NoTrt_B |
| POLR2C | 0.03293 | -0.4607 | 0.552 | 0.63 | 1 | HC_vs_MS.NoTrt_B |

| | | | | | | |
|---|---|---|---|---|---|---|
| NPRL3 | 0.03302 | 0.47677 | 0.299 | 0.39 | 1 | HC_vs_MS.NoTrt_B |
| CKAP2 | 0.03309 | -0.4979 | 0.103 | 0.16 | 1 | HC_vs_MS.NoTrt_B |
| FOXO3 | 0.03309 | -0.3579 | 0.365 | 0.43 | 1 | HC_vs_MS.NoTrt_B |
| RFX5 | 0.03315 | -0.3924 | 0.281 | 0.35 | 1 | HC_vs_MS.NoTrt_B |
| TMBIM1 | 0.0332 | -0.4719 | 0.588 | 0.64 | 1 | HC_vs_MS.NoTrt_B |
| ACAT2 | 0.03328 | -0.2932 | 0.22 | 0.29 | 1 | HC_vs_MS.NoTrt_B |
| THAP4 | 0.03331 | -0.3308 | 0.106 | 0.16 | 1 | HC_vs_MS.NoTrt_B |
| ROBO1 | 0.0334 | 0.41988 | 0.48 | 0.42 | 1 | HC_vs_MS.NoTrt_B |
| SIM2 | 0.03348 | 0.34693 | 0.617 | 0.59 | 1 | HC_vs_MS.NoTrt_B |
| MRPS18A | 0.03356 | -0.3062 | 0.097 | 0.15 | 1 | HC_vs_MS.NoTrt_B |
| GRIK2 | 0.03357 | 0.27086 | 0.532 | 0.47 | 1 | HC_vs_MS.NoTrt_B |
| TRMT1 | 0.03375 | -0.3752 | 0.129 | 0.19 | 1 | HC_vs_MS.NoTrt_B |
| HIF1AN | 0.03396 | -0.3427 | 0.78 | 0.88 | 1 | HC_vs_MS.NoTrt_B |
| WWC3 | 0.03398 | 0.30084 | 0.345 | 0.44 | 1 | HC_vs_MS.NoTrt_B |
| ZFP36L1 | 0.03412 | 0.49292 | 0.701 | 0.69 | 1 | HC_vs_MS.NoTrt_B |
| LTBR | 0.03417 | 0.2701 | 0.929 | 0.95 | 1 | HC_vs_MS.NoTrt_B |
| NFATC3 | 0.03429 | 0.55409 | 0.668 | 0.58 | 1 | HC_vs_MS.NoTrt_B |
| DOK6 | 0.03449 | -0.4131 | 0.258 | 0.19 | 1 | HC_vs_MS.NoTrt_B |
| NPC2 | 0.03455 | -0.4022 | 0.084 | 0.14 | 1 | HC_vs_MS.NoTrt_B |
| TMEM243 | 0.0346 | 0.28133 | 0.255 | 0.36 | 1 | HC_vs_MS.NoTrt_B |
| POLD1 | 0.0346 | -0.4824 | 0.141 | 0.2 | 1 | HC_vs_MS.NoTrt_B |
| ZNF696 | 0.03462 | 0.4671 | 0.316 | 0.25 | 1 | HC_vs_MS.NoTrt_B |
| TMPRSS7 | 0.03467 | 0.36692 | 0.257 | 0.19 | 1 | HC_vs_MS.NoTrt_B |
| CCNT2 | 0.0347 | 0.33171 | 0.133 | 0.2 | 1 | HC_vs_MS.NoTrt_B |
| BCL11B | 0.03483 | 0.63791 | 0.493 | 0.43 | 1 | HC_vs_MS.NoTrt_B |
| CHKB | 0.03484 | -0.3808 | 0.186 | 0.25 | 1 | HC_vs_MS.NoTrt_B |
| NCKAP5L | 0.03485 | 0.75997 | 0.593 | 0.53 | 1 | HC_vs_MS.NoTrt_B |
| PLAUR | 0.03489 | 0.29128 | 0.574 | 0.52 | 1 | HC_vs_MS.NoTrt_B |
| CHID1 | 0.03491 | 0.67311 | 0.314 | 0.24 | 1 | HC_vs_MS.NoTrt_B |
| DPY19L3 | 0.03492 | 0.35067 | 0.913 | 0.92 | 1 | HC_vs_MS.NoTrt_B |
| TIPRL | 0.03502 | -0.5403 | 0.281 | 0.34 | 1 | HC_vs_MS.NoTrt_B |
| EPN2 | 0.03502 | 0.37628 | 0.771 | 0.81 | 1 | HC_vs_MS.NoTrt_B |
| GIGYF2 | 0.03505 | 0.3339 | 0.91 | 0.92 | 1 | HC_vs_MS.NoTrt_B |
| RBMX2 | 0.03527 | -0.2693 | 0.091 | 0.15 | 1 | HC_vs_MS.NoTrt_B |
| FILIP1L | 0.03534 | 0.54828 | 0.39 | 0.33 | 1 | HC_vs_MS.NoTrt_B |
| CYP2R1 | 0.03539 | -0.6039 | 0.081 | 0.13 | 1 | HC_vs_MS.NoTrt_B |
| PXMP4 | 0.03547 | 0.3231 | 0.816 | 0.82 | 1 | HC_vs_MS.NoTrt_B |
| GPR157 | 0.03559 | 0.31565 | 0.786 | 0.81 | 1 | HC_vs_MS.NoTrt_B |
| CCDC69 | 0.03567 | 0.33604 | 0.859 | 0.83 | 1 | HC_vs_MS.NoTrt_B |
| POC5 | 0.03567 | -0.4895 | 0.081 | 0.13 | 1 | HC_vs_MS.NoTrt_B |

| | | | | | | |
|---|---|---|---|---|---|---|
| PAWR | 0.03574 | 0.3487 | 0.699 | 0.66 | 1 | HC_vs_MS.NoTrt_B |
| S1PR3 | 0.03581 | 0.29355 | 0.757 | 0.7 | 1 | HC_vs_MS.NoTrt_B |
| ORAOV1 | 0.03584 | 0.27222 | 0.694 | 0.64 | 1 | HC_vs_MS.NoTrt_B |
| SOX5 | 0.03584 | 0.30121 | 0.174 | 0.11 | 1 | HC_vs_MS.NoTrt_B |
| PI16 | 0.0359 | -0.6128 | 0.113 | 0.17 | 1 | HC_vs_MS.NoTrt_B |
| NPC1L1 | 0.03593 | 0.28863 | 0.175 | 0.12 | 1 | HC_vs_MS.NoTrt_B |
| SLMAP | 0.03626 | 0.338 | 0.791 | 0.77 | 1 | HC_vs_MS.NoTrt_B |
| KDM5B | 0.03629 | 0.94349 | 0.277 | 0.37 | 1 | HC_vs_MS.NoTrt_B |
| MBD5 | 0.03632 | 0.563 | 0.843 | 0.88 | 1 | HC_vs_MS.NoTrt_B |
| TMC8 | 0.03639 | 0.49188 | 0.748 | 0.72 | 1 | HC_vs_MS.NoTrt_B |
| DHX34 | 0.03649 | -0.6027 | 0.059 | 0.1 | 1 | HC_vs_MS.NoTrt_B |
| POLE2 | 0.03659 | 0.51401 | 0.322 | 0.25 | 1 | HC_vs_MS.NoTrt_B |
| TOP1 | 0.03661 | -0.2891 | 0.129 | 0.19 | 1 | HC_vs_MS.NoTrt_B |
| SGSM2 | 0.03666 | 1.54937 | 0.264 | 0.2 | 1 | HC_vs_MS.NoTrt_B |
| CDC37L1 | 0.03686 | -0.4523 | 0.099 | 0.15 | 1 | HC_vs_MS.NoTrt_B |
| CEP152 | 0.03696 | 0.79932 | 0.683 | 0.67 | 1 | HC_vs_MS.NoTrt_B |
| DCBLD2 | 0.03697 | 0.52756 | 0.67 | 0.7 | 1 | HC_vs_MS.NoTrt_B |
| PPP2R5D | 0.03704 | -0.2628 | 0.148 | 0.21 | 1 | HC_vs_MS.NoTrt_B |
| ARV1 | 0.03742 | -0.6698 | 0.232 | 0.31 | 1 | HC_vs_MS.NoTrt_B |
| GPR68 | 0.03743 | -0.5185 | 0.068 | 0.11 | 1 | HC_vs_MS.NoTrt_B |
| GAB1 | 0.03749 | 0.3331 | 0.532 | 0.47 | 1 | HC_vs_MS.NoTrt_B |
| YLPM1 | 0.03763 | 0.79515 | 0.6 | 0.54 | 1 | HC_vs_MS.NoTrt_B |
| LAIR2 | 0.03769 | -0.5152 | 0.103 | 0.16 | 1 | HC_vs_MS.NoTrt_B |
| KLHL18 | 0.03779 | -0.572 | 0.314 | 0.39 | 1 | HC_vs_MS.NoTrt_B |
| RHOC | 0.0378 | -0.8816 | 0.074 | 0.12 | 1 | HC_vs_MS.NoTrt_B |
| GLO1 | 0.03784 | -0.3095 | 0.064 | 0.11 | 1 | HC_vs_MS.NoTrt_B |
| POLN | 0.0379 | 0.26886 | 0.636 | 0.6 | 1 | HC_vs_MS.NoTrt_B |
| DNAJC28 | 0.03799 | 0.50874 | 0.374 | 0.29 | 1 | HC_vs_MS.NoTrt_B |
| CRADD | 0.03805 | -0.5043 | 0.077 | 0.12 | 1 | HC_vs_MS.NoTrt_B |
| SLC35F2 | 0.03807 | -0.313 | 0.219 | 0.28 | 1 | HC_vs_MS.NoTrt_B |
| FMR1 | 0.03813 | 0.44391 | 0.103 | 0.16 | 1 | HC_vs_MS.NoTrt_B |
| ZNF816 | 0.03846 | 0.59624 | 0.407 | 0.5 | 1 | HC_vs_MS.NoTrt_B |
| RBMS2 | 0.03861 | 0.38904 | 0.861 | 0.85 | 1 | HC_vs_MS.NoTrt_B |
| GP2 | 0.03863 | 0.36133 | 0.504 | 0.47 | 1 | HC_vs_MS.NoTrt_B |
| ELMO2 | 0.03864 | -0.4919 | 0.203 | 0.27 | 1 | HC_vs_MS.NoTrt_B |
| PTH2R | 0.03868 | 0.34208 | 0.194 | 0.14 | 1 | HC_vs_MS.NoTrt_B |
| KCNE3 | 0.03873 | 0.34717 | 0.525 | 0.47 | 1 | HC_vs_MS.NoTrt_B |
| MYH7B | 0.03888 | -0.4034 | 0.786 | 0.78 | 1 | HC_vs_MS.NoTrt_B |
| LRRC27 | 0.03888 | 0.45576 | 0.803 | 0.81 | 1 | HC_vs_MS.NoTrt_B |
| FAM216A | 0.03889 | 0.73165 | 0.558 | 0.51 | 1 | HC_vs_MS.NoTrt_B |

| | | | | | | |
|---|---|---|---|---|---|---|
| TOLLIP | 0.0389 | -0.3611 | 0.077 | 0.12 | 1 | HC_vs_MS.NoTrt_B |
| ERCC3 | 0.03902 | -0.9704 | 0.361 | 0.43 | 1 | HC_vs_MS.NoTrt_B |
| FAM153C | 0.03906 | 0.28617 | 0.455 | 0.39 | 1 | HC_vs_MS.NoTrt_B |
| C1S | 0.03917 | 0.30201 | 0.63 | 0.6 | 1 | HC_vs_MS.NoTrt_B |
| GRAMD2 | 0.03927 | 0.30271 | 0.617 | 0.56 | 1 | HC_vs_MS.NoTrt_B |
| SSSCA1 | 0.03932 | -0.401 | 0.059 | 0.1 | 1 | HC_vs_MS.NoTrt_B |
| PRDM15 | 0.03939 | -0.2632 | 0.523 | 0.64 | 1 | HC_vs_MS.NoTrt_B |
| NPHP1 | 0.03942 | 0.33942 | 0.719 | 0.67 | 1 | HC_vs_MS.NoTrt_B |
| ARID2 | 0.03946 | 0.29627 | 0.688 | 0.74 | 1 | HC_vs_MS.NoTrt_B |
| RP11-235E1 | 0.03957 | 0.42846 | 0.265 | 0.2 | 1 | HC_vs_MS.NoTrt_B |
| ANO10 | 0.03972 | -0.4533 | 0.568 | 0.51 | 1 | HC_vs_MS.NoTrt_B |
| PORCN | 0.03983 | -0.806 | 0.061 | 0.1 | 1 | HC_vs_MS.NoTrt_B |
| STARD3NL | 0.03989 | 0.2828 | 0.775 | 0.76 | 1 | HC_vs_MS.NoTrt_B |
| XAF1 | 0.0401 | 0.28104 | 0.848 | 0.82 | 1 | HC_vs_MS.NoTrt_B |
| TBC1D13 | 0.04044 | 0.41005 | 0.109 | 0.16 | 1 | HC_vs_MS.NoTrt_B |
| RCC1 | 0.04048 | -0.3869 | 0.159 | 0.23 | 1 | HC_vs_MS.NoTrt_B |
| PDIA5 | 0.04054 | 0.25115 | 0.367 | 0.29 | 1 | HC_vs_MS.NoTrt_B |
| TRAF6 | 0.04055 | 0.89361 | 0.712 | 0.65 | 1 | HC_vs_MS.NoTrt_B |
| HDAC3 | 0.04055 | -0.4518 | 0.117 | 0.17 | 1 | HC_vs_MS.NoTrt_B |
| RP1L1 | 0.04068 | 0.32006 | 0.6 | 0.52 | 1 | HC_vs_MS.NoTrt_B |
| WNT7A | 0.0407 | 1.02631 | 0.103 | 0.05 | 1 | HC_vs_MS.NoTrt_B |
| RP11-47I22. | 0.04077 | 0.47278 | 0.596 | 0.58 | 1 | HC_vs_MS.NoTrt_B |
| IRAK3 | 0.04081 | 0.8775 | 0.413 | 0.37 | 1 | HC_vs_MS.NoTrt_B |
| ISPD | 0.04084 | 0.25063 | 0.787 | 0.78 | 1 | HC_vs_MS.NoTrt_B |
| AP5B1 | 0.04114 | 0.73405 | 0.235 | 0.17 | 1 | HC_vs_MS.NoTrt_B |
| C17orf99 | 0.04119 | 0.2931 | 0.403 | 0.35 | 1 | HC_vs_MS.NoTrt_B |
| NDUFA9 | 0.04132 | -0.2752 | 0.868 | 0.92 | 1 | HC_vs_MS.NoTrt_B |
| TKT | 0.04132 | -0.297 | 0.557 | 0.61 | 1 | HC_vs_MS.NoTrt_B |
| SNAP47 | 0.04137 | -0.6654 | 0.12 | 0.17 | 1 | HC_vs_MS.NoTrt_B |
| SCRN2 | 0.04172 | -0.3687 | 0.064 | 0.11 | 1 | HC_vs_MS.NoTrt_B |
| NOG | 0.04173 | 1.41208 | 0.241 | 0.19 | 1 | HC_vs_MS.NoTrt_B |
| KLRB1 | 0.04178 | -0.302 | 0.081 | 0.13 | 1 | HC_vs_MS.NoTrt_B |
| EIF4E3 | 0.04181 | 0.84132 | 0.18 | 0.26 | 1 | HC_vs_MS.NoTrt_B |
| FAM92A1 | 0.04194 | 0.48161 | 0.323 | 0.26 | 1 | HC_vs_MS.NoTrt_B |
| PLA2G2D | 0.04195 | 0.31973 | 0.426 | 0.37 | 1 | HC_vs_MS.NoTrt_B |
| WNT4 | 0.04199 | 0.58098 | 0.075 | 0.12 | 1 | HC_vs_MS.NoTrt_B |
| AGPAT5 | 0.04206 | 0.77352 | 0.141 | 0.09 | 1 | HC_vs_MS.NoTrt_B |
| STT3B | 0.04207 | 0.37892 | 0.561 | 0.51 | 1 | HC_vs_MS.NoTrt_B |
| TMEM5 | 0.04212 | 0.94359 | 0.497 | 0.45 | 1 | HC_vs_MS.NoTrt_B |
| CCDC150 | 0.04216 | 0.36215 | 0.672 | 0.7 | 1 | HC_vs_MS.NoTrt_B |

| | | | | | | |
|---|---|---|---|---|---|---|
| CASS4 | 0.04217 | 0.46171 | 0.454 | 0.39 | 1 | HC_vs_MS.NoTrt_B |
| ID3 | 0.0422 | 0.71943 | 0.222 | 0.16 | 1 | HC_vs_MS.NoTrt_B |
| TBC1D5 | 0.04226 | -0.2861 | 0.597 | 0.68 | 1 | HC_vs_MS.NoTrt_B |
| ALKBH2 | 0.04239 | -0.3026 | 0.307 | 0.39 | 1 | HC_vs_MS.NoTrt_B |
| SMN2 | 0.04251 | 0.38645 | 0.807 | 0.7 | 1 | HC_vs_MS.NoTrt_B |
| SPAG9 | 0.04261 | 0.46411 | 0.862 | 0.87 | 1 | HC_vs_MS.NoTrt_B |
| GIMAP5 | 0.04263 | 0.47114 | 0.478 | 0.41 | 1 | HC_vs_MS.NoTrt_B |
| ZNF365 | 0.04266 | -0.6074 | 0.27 | 0.35 | 1 | HC_vs_MS.NoTrt_B |
| ING4 | 0.04276 | -0.4062 | 0.241 | 0.31 | 1 | HC_vs_MS.NoTrt_B |
| CITED2 | 0.04276 | 1.26055 | 0.139 | 0.09 | 1 | HC_vs_MS.NoTrt_B |
| MTRNR2L3 | 0.04281 | -0.3825 | 0.229 | 0.17 | 1 | HC_vs_MS.NoTrt_B |
| DDX51 | 0.04293 | 0.33899 | 0.645 | 0.6 | 1 | HC_vs_MS.NoTrt_B |
| SLC24A2 | 0.04294 | 0.29538 | 0.636 | 0.6 | 1 | HC_vs_MS.NoTrt_B |
| PSME3 | 0.04313 | -0.44 | 0.601 | 0.69 | 1 | HC_vs_MS.NoTrt_B |
| ZIM3 | 0.0432 | 0.31824 | 0.529 | 0.5 | 1 | HC_vs_MS.NoTrt_B |
| SPATA31A7 | 0.04352 | 0.34873 | 0.533 | 0.45 | 1 | HC_vs_MS.NoTrt_B |
| NBPF14 | 0.04355 | 0.38517 | 0.41 | 0.33 | 1 | HC_vs_MS.NoTrt_B |
| XKR9 | 0.04356 | 0.50441 | 0.559 | 0.56 | 1 | HC_vs_MS.NoTrt_B |
| GSDMA | 0.04359 | 0.67905 | 0.601 | 0.58 | 1 | HC_vs_MS.NoTrt_B |
| ZNF614 | 0.04368 | 1.56508 | 0.252 | 0.18 | 1 | HC_vs_MS.NoTrt_B |
| PITPNM2 | 0.04389 | 1.11626 | 0.422 | 0.37 | 1 | HC_vs_MS.NoTrt_B |
| PRICKLE3 | 0.04391 | 0.43717 | 0.51 | 0.45 | 1 | HC_vs_MS.NoTrt_B |
| GTF2I | 0.04405 | 0.7216 | 0.887 | 0.88 | 1 | HC_vs_MS.NoTrt_B |
| PADI4 | 0.04408 | 0.52941 | 0.512 | 0.49 | 1 | HC_vs_MS.NoTrt_B |
| PICALM | 0.04411 | 0.32138 | 0.22 | 0.31 | 1 | HC_vs_MS.NoTrt_B |
| WDR70 | 0.04421 | -0.4166 | 0.264 | 0.35 | 1 | HC_vs_MS.NoTrt_B |
| FRMD4A | 0.04432 | 0.5779 | 0.629 | 0.58 | 1 | HC_vs_MS.NoTrt_B |
| NAIF1 | 0.04466 | 0.38218 | 0.588 | 0.54 | 1 | HC_vs_MS.NoTrt_B |
| GEMIN4 | 0.04473 | -0.2919 | 0.061 | 0.1 | 1 | HC_vs_MS.NoTrt_B |
| PSMD2 | 0.04483 | -0.2957 | 0.149 | 0.21 | 1 | HC_vs_MS.NoTrt_B |
| CRIPT | 0.04498 | 0.54772 | 0.18 | 0.25 | 1 | HC_vs_MS.NoTrt_B |
| RADIL | 0.04507 | 0.38944 | 0.281 | 0.21 | 1 | HC_vs_MS.NoTrt_B |
| RARA | 0.04509 | -0.5778 | 0.074 | 0.12 | 1 | HC_vs_MS.NoTrt_B |
| DDX19B | 0.04547 | -1.0849 | 0.432 | 0.48 | 1 | HC_vs_MS.NoTrt_B |
| MRPL42 | 0.0455 | 0.28689 | 0.964 | 0.98 | 1 | HC_vs_MS.NoTrt_B |
| PPP1R21 | 0.04557 | 0.25854 | 0.203 | 0.27 | 1 | HC_vs_MS.NoTrt_B |
| GNG7 | 0.04559 | 0.70644 | 0.394 | 0.35 | 1 | HC_vs_MS.NoTrt_B |
| RCOR1 | 0.0456 | 0.41658 | 0.13 | 0.19 | 1 | HC_vs_MS.NoTrt_B |
| CALM3 | 0.04574 | 0.34781 | 0.43 | 0.57 | 1 | HC_vs_MS.NoTrt_B |
| PISD | 0.04587 | -0.3115 | 0.119 | 0.17 | 1 | HC_vs_MS.NoTrt_B |

| | | | | | | |
|---|---|---|---|---|---|---|
| PIK3R1 | 0.04593 | 0.78081 | 0.31 | 0.24 | 1 | HC_vs_MS.NoTrt_B |
| RAB11FIP4 | 0.0461 | -0.571 | 0.687 | 0.75 | 1 | HC_vs_MS.NoTrt_B |
| TBCC | 0.04614 | -0.3878 | 0.364 | 0.43 | 1 | HC_vs_MS.NoTrt_B |
| UBE2O | 0.04624 | 0.45106 | 0.545 | 0.49 | 1 | HC_vs_MS.NoTrt_B |
| GALR1 | 0.04626 | 0.35232 | 0.616 | 0.58 | 1 | HC_vs_MS.NoTrt_B |
| NF2 | 0.04628 | -0.2502 | 0.762 | 0.81 | 1 | HC_vs_MS.NoTrt_B |
| VASH2 | 0.04633 | 0.46463 | 0.745 | 0.71 | 1 | HC_vs_MS.NoTrt_B |
| RAD54L2 | 0.04646 | -0.3151 | 0.61 | 0.68 | 1 | HC_vs_MS.NoTrt_B |
| CCDC59 | 0.04667 | -0.2799 | 0.114 | 0.17 | 1 | HC_vs_MS.NoTrt_B |
| GATC | 0.0467 | 0.42064 | 0.559 | 0.67 | 1 | HC_vs_MS.NoTrt_B |
| CSMD3 | 0.04675 | 0.33719 | 0.458 | 0.39 | 1 | HC_vs_MS.NoTrt_B |
| DYNLL2 | 0.0469 | 0.36895 | 0.091 | 0.14 | 1 | HC_vs_MS.NoTrt_B |
| PNPLA3 | 0.04697 | 0.32466 | 0.53 | 0.49 | 1 | HC_vs_MS.NoTrt_B |
| RASA4B | 0.04703 | 0.38081 | 0.436 | 0.52 | 1 | HC_vs_MS.NoTrt_B |
| INO80E | 0.04717 | 0.54631 | 0.158 | 0.23 | 1 | HC_vs_MS.NoTrt_B |
| PPARA | 0.04719 | 0.52236 | 0.904 | 0.91 | 1 | HC_vs_MS.NoTrt_B |
| RCBTB2 | 0.04721 | -0.8283 | 0.1 | 0.15 | 1 | HC_vs_MS.NoTrt_B |
| SH3GLB2 | 0.04734 | 0.75392 | 0.152 | 0.1 | 1 | HC_vs_MS.NoTrt_B |
| TRAPPC3L | 0.04767 | 0.41675 | 0.572 | 0.58 | 1 | HC_vs_MS.NoTrt_B |
| MACF1 | 0.04777 | 0.53954 | 0.871 | 0.84 | 1 | HC_vs_MS.NoTrt_B |
| BBIP1 | 0.0478 | 0.28088 | 0.529 | 0.61 | 1 | HC_vs_MS.NoTrt_B |
| FAM179B | 0.04795 | 0.27741 | 0.757 | 0.77 | 1 | HC_vs_MS.NoTrt_B |
| HCN1 | 0.04799 | 0.25344 | 0.361 | 0.29 | 1 | HC_vs_MS.NoTrt_B |
| CD82 | 0.04808 | -0.374 | 0.614 | 0.68 | 1 | HC_vs_MS.NoTrt_B |
| VSNL1 | 0.04822 | 0.29847 | 0.593 | 0.57 | 1 | HC_vs_MS.NoTrt_B |
| SCARF1 | 0.04827 | 0.26548 | 0.361 | 0.29 | 1 | HC_vs_MS.NoTrt_B |
| ZNF674 | 0.04847 | 0.31403 | 0.904 | 0.9 | 1 | HC_vs_MS.NoTrt_B |
| ABCG5 | 0.04849 | -0.2744 | 0.171 | 0.24 | 1 | HC_vs_MS.NoTrt_B |
| URI1 | 0.04851 | 0.58107 | 0.187 | 0.13 | 1 | HC_vs_MS.NoTrt_B |
| UPF2 | 0.04863 | -0.539 | 0.246 | 0.32 | 1 | HC_vs_MS.NoTrt_B |
| RASEF | 0.04877 | 0.34084 | 0.868 | 0.96 | 1 | HC_vs_MS.NoTrt_B |
| LARP4 | 0.04886 | 0.45218 | 0.358 | 0.43 | 1 | HC_vs_MS.NoTrt_B |
| TRIM6 | 0.04909 | 0.38489 | 0.522 | 0.47 | 1 | HC_vs_MS.NoTrt_B |
| C19orf60 | 0.0491 | -0.2576 | 0.865 | 0.91 | 1 | HC_vs_MS.NoTrt_B |
| SNAPC3 | 0.04928 | 0.57218 | 0.586 | 0.55 | 1 | HC_vs_MS.NoTrt_B |
| ZNF490 | 0.04933 | -0.3834 | 0.865 | 0.87 | 1 | HC_vs_MS.NoTrt_B |
| ZC3H10 | 0.04935 | 0.56695 | 0.507 | 0.48 | 1 | HC_vs_MS.NoTrt_B |
| TMX2 | 0.04953 | -0.2808 | 0.104 | 0.16 | 1 | HC_vs_MS.NoTrt_B |
| ZNF577 | 0.04962 | 0.49613 | 0.332 | 0.43 | 1 | HC_vs_MS.NoTrt_B |
| FGD4 | 0.04971 | 0.28479 | 0.82 | 0.83 | 1 | HC_vs_MS.NoTrt_B |

| | | | | | | |
|---|---|---|---|---|---|---|
| ICMT | 0.04974 | 0.67605 | 0.609 | 0.55 | 1 | HC_vs_MS.NoTrt_B |
| OSBPL1A | 0.0498 | 0.27156 | 0.736 | 0.71 | 1 | HC_vs_MS.NoTrt_B |
| PPOX | 0.04983 | -0.4488 | 0.216 | 0.28 | 1 | HC_vs_MS.NoTrt_B |
| MYH3 | 0.05001 | 0.38708 | 0.645 | 0.63 | 1 | HC_vs_MS.NoTrt_B |
| NCK1 | 0.05003 | 0.3519 | 0.364 | 0.48 | 1 | HC_vs_MS.NoTrt_B |
| PPP1R3B | 0.05003 | -0.2695 | 0.291 | 0.22 | 1 | HC_vs_MS.NoTrt_B |
| C19orf66 | 0.05026 | -0.3513 | 0.616 | 0.67 | 1 | HC_vs_MS.NoTrt_B |
| EXPH5 | 0.05027 | 0.7144 | 0.7 | 0.71 | 1 | HC_vs_MS.NoTrt_B |
| PPP2R5E | 0.05034 | 0.54109 | 0.188 | 0.27 | 1 | HC_vs_MS.NoTrt_B |
| ZNF114 | 0.05041 | 0.36893 | 0.471 | 0.43 | 1 | HC_vs_MS.NoTrt_B |
| CELF1 | 0.05042 | -0.2618 | 0.367 | 0.44 | 1 | HC_vs_MS.NoTrt_B |
| SURF1 | 0.0505 | -0.2819 | 0.07 | 0.11 | 1 | HC_vs_MS.NoTrt_B |
| SAMD15 | 0.05066 | 0.41514 | 0.323 | 0.25 | 1 | HC_vs_MS.NoTrt_B |
| RBM19 | 0.05079 | -0.2567 | 0.467 | 0.54 | 1 | HC_vs_MS.NoTrt_B |
| TIA1 | 0.05083 | 0.2511 | 0.216 | 0.29 | 1 | HC_vs_MS.NoTrt_B |
| ELF3 | 0.05088 | 0.60179 | 0.545 | 0.5 | 1 | HC_vs_MS.NoTrt_B |
| SLC27A1 | 0.05096 | 0.57409 | 0.797 | 0.84 | 1 | HC_vs_MS.NoTrt_B |
| BCL9L | 0.05097 | 0.81201 | 0.17 | 0.11 | 1 | HC_vs_MS.NoTrt_B |
| PGAM5 | 0.051 | 0.30238 | 0.714 | 0.65 | 1 | HC_vs_MS.NoTrt_B |
| HLA-DRB5 | 0.05101 | -2.5348 | 0.087 | 0.13 | 1 | HC_vs_MS.NoTrt_B |
| ZSCAN25 | 0.05105 | 0.57523 | 0.433 | 0.36 | 1 | HC_vs_MS.NoTrt_B |
| DNAH7 | 0.0516 | 0.53577 | 0.657 | 0.61 | 1 | HC_vs_MS.NoTrt_B |
| POU2F3 | 0.05171 | 0.35683 | 0.426 | 0.37 | 1 | HC_vs_MS.NoTrt_B |
| GMPPA | 0.05179 | -0.6151 | 0.236 | 0.31 | 1 | HC_vs_MS.NoTrt_B |
| THTPA | 0.05182 | -0.2845 | 0.08 | 0.12 | 1 | HC_vs_MS.NoTrt_B |
| DHRS7B | 0.05187 | 0.44488 | 0.516 | 0.47 | 1 | HC_vs_MS.NoTrt_B |
| RP11-1070B | 0.05188 | 0.92993 | 0.186 | 0.13 | 1 | HC_vs_MS.NoTrt_B |
| MATN2 | 0.05194 | 0.48424 | 0.58 | 0.56 | 1 | HC_vs_MS.NoTrt_B |
| MED8 | 0.05204 | -0.4678 | 0.067 | 0.11 | 1 | HC_vs_MS.NoTrt_B |
| KBTBD6 | 0.05213 | -0.42 | 0.325 | 0.43 | 1 | HC_vs_MS.NoTrt_B |
| DGKD | 0.05224 | 1.02367 | 0.604 | 0.57 | 1 | HC_vs_MS.NoTrt_B |
| EIF2S2 | 0.0523 | 0.26489 | 0.257 | 0.35 | 1 | HC_vs_MS.NoTrt_B |
| C12orf4 | 0.05239 | 0.299 | 0.588 | 0.72 | 1 | HC_vs_MS.NoTrt_B |
| IVD | 0.05268 | 0.35149 | 0.9 | 0.93 | 1 | HC_vs_MS.NoTrt_B |
| BAG1 | 0.05269 | 0.46938 | 0.926 | 0.93 | 1 | HC_vs_MS.NoTrt_B |
| GPR89B | 0.05273 | -0.361 | 0.314 | 0.38 | 1 | HC_vs_MS.NoTrt_B |
| FNTA | 0.05279 | 0.78922 | 0.854 | 0.84 | 1 | HC_vs_MS.NoTrt_B |
| RGPD6 | 0.05281 | 0.95241 | 0.588 | 0.56 | 1 | HC_vs_MS.NoTrt_B |
| SMAP1 | 0.05283 | -0.5284 | 0.18 | 0.24 | 1 | HC_vs_MS.NoTrt_B |
| ABHD1 | 0.05306 | 0.36142 | 0.758 | 0.74 | 1 | HC_vs_MS.NoTrt_B |

| | | | | | | |
|---|---|---|---|---|---|---|
| RBFOX3 | 0.05312 | 0.27622 | 0.783 | 0.78 | 1 | HC_vs_MS.NoTrt_B |
| PYGM | 0.05315 | 0.73136 | 0.667 | 0.69 | 1 | HC_vs_MS.NoTrt_B |
| CLTA | 0.05321 | -0.3632 | 0.575 | 0.64 | 1 | HC_vs_MS.NoTrt_B |
| ADGRE2 | 0.05322 | 0.55084 | 0.757 | 0.75 | 1 | HC_vs_MS.NoTrt_B |
| MALL | 0.05333 | 0.33913 | 0.638 | 0.65 | 1 | HC_vs_MS.NoTrt_B |
| SLC35A3 | 0.05344 | 0.31667 | 0.799 | 0.76 | 1 | HC_vs_MS.NoTrt_B |
| TMEM170A | 0.05352 | 0.273 | 0.81 | 0.77 | 1 | HC_vs_MS.NoTrt_B |
| SHKBP1 | 0.05357 | -0.2783 | 0.723 | 0.76 | 1 | HC_vs_MS.NoTrt_B |
| SORBS2 | 0.05364 | 0.28589 | 0.545 | 0.5 | 1 | HC_vs_MS.NoTrt_B |
| LIN28A | 0.05365 | 0.3118 | 0.741 | 0.76 | 1 | HC_vs_MS.NoTrt_B |
| MORC2 | 0.05368 | -0.3117 | 0.388 | 0.44 | 1 | HC_vs_MS.NoTrt_B |
| ZBTB20 | 0.0538 | 0.58243 | 0.796 | 0.74 | 1 | HC_vs_MS.NoTrt_B |
| PLAC8 | 0.05391 | 0.55769 | 0.755 | 0.77 | 1 | HC_vs_MS.NoTrt_B |
| TFAP4 | 0.05392 | 0.62883 | 0.671 | 0.67 | 1 | HC_vs_MS.NoTrt_B |
| ABCB9 | 0.05405 | 0.28641 | 0.833 | 0.84 | 1 | HC_vs_MS.NoTrt_B |
| FAM159A | 0.05417 | 0.93766 | 0.587 | 0.56 | 1 | HC_vs_MS.NoTrt_B |
| TRAPPC8 | 0.0542 | 0.49548 | 0.672 | 0.64 | 1 | HC_vs_MS.NoTrt_B |
| HLTF | 0.05441 | 0.77012 | 0.513 | 0.44 | 1 | HC_vs_MS.NoTrt_B |
| ZNF101 | 0.05446 | -0.2786 | 0.951 | 0.96 | 1 | HC_vs_MS.NoTrt_B |
| COQ4 | 0.05466 | 0.26917 | 0.894 | 0.94 | 1 | HC_vs_MS.NoTrt_B |
| ZNF75A | 0.05466 | -0.6377 | 0.72 | 0.8 | 1 | HC_vs_MS.NoTrt_B |
| CLIP2 | 0.05485 | 0.63369 | 0.362 | 0.32 | 1 | HC_vs_MS.NoTrt_B |
| KCTD1 | 0.05489 | 0.49719 | 0.572 | 0.55 | 1 | HC_vs_MS.NoTrt_B |
| NPHP3 | 0.05492 | 0.51371 | 0.893 | 0.91 | 1 | HC_vs_MS.NoTrt_B |
| SLX4IP | 0.05497 | -0.5043 | 0.471 | 0.52 | 1 | HC_vs_MS.NoTrt_B |
| FBXW8 | 0.05522 | -0.6705 | 0.257 | 0.33 | 1 | HC_vs_MS.NoTrt_B |
| SHH | 0.05527 | 1.16983 | 0.106 | 0.06 | 1 | HC_vs_MS.NoTrt_B |
| PLOD1 | 0.05528 | 0.39089 | 0.594 | 0.52 | 1 | HC_vs_MS.NoTrt_B |
| SCYL1 | 0.05531 | -0.3641 | 0.226 | 0.29 | 1 | HC_vs_MS.NoTrt_B |
| DNAJB4 | 0.05534 | -0.5921 | 0.284 | 0.36 | 1 | HC_vs_MS.NoTrt_B |
| AF011889.5 | 0.05552 | -0.2688 | 0.186 | 0.25 | 1 | HC_vs_MS.NoTrt_B |
| DMAP1 | 0.05561 | -0.4025 | 0.441 | 0.51 | 1 | HC_vs_MS.NoTrt_B |
| 1-Mar | 0.05562 | 0.5852 | 0.31 | 0.25 | 1 | HC_vs_MS.NoTrt_B |
| OLFM2 | 0.05579 | 0.63439 | 0.065 | 0.11 | 1 | HC_vs_MS.NoTrt_B |
| COG7 | 0.05592 | 0.30423 | 0.528 | 0.48 | 1 | HC_vs_MS.NoTrt_B |
| C6orf1 | 0.05604 | -0.306 | 0.097 | 0.15 | 1 | HC_vs_MS.NoTrt_B |
| HCAR1 | 0.05625 | 0.2856 | 0.868 | 0.91 | 1 | HC_vs_MS.NoTrt_B |
| ZNFX1 | 0.0563 | 0.52837 | 0.814 | 0.77 | 1 | HC_vs_MS.NoTrt_B |
| TBX6 | 0.05657 | 0.3147 | 0.47 | 0.4 | 1 | HC_vs_MS.NoTrt_B |
| ANKRD6 | 0.05661 | 0.364 | 0.165 | 0.11 | 1 | HC_vs_MS.NoTrt_B |

| | | | | | | |
|---|---|---|---|---|---|---|
| NLN | 0.05685 | 0.31285 | 0.742 | 0.72 | 1 | HC_vs_MS.NoTrt_B |
| EIF3B | 0.05695 | 0.52183 | 0.33 | 0.43 | 1 | HC_vs_MS.NoTrt_B |
| ANAPC1 | 0.05707 | 0.80491 | 0.614 | 0.56 | 1 | HC_vs_MS.NoTrt_B |
| ULK4 | 0.05746 | -0.4912 | 0.277 | 0.35 | 1 | HC_vs_MS.NoTrt_B |
| HAAO | 0.0575 | -0.5431 | 0.075 | 0.12 | 1 | HC_vs_MS.NoTrt_B |
| RBMXL1 | 0.05758 | 0.63363 | 0.12 | 0.07 | 1 | HC_vs_MS.NoTrt_B |
| PSMA2 | 0.05772 | -0.2622 | 0.554 | 0.61 | 1 | HC_vs_MS.NoTrt_B |
| PARP15 | 0.05786 | 0.3181 | 0.899 | 0.88 | 1 | HC_vs_MS.NoTrt_B |
| FBXO27 | 0.058 | 0.32229 | 0.697 | 0.68 | 1 | HC_vs_MS.NoTrt_B |
| XPNPEP1 | 0.05802 | -0.6008 | 0.287 | 0.35 | 1 | HC_vs_MS.NoTrt_B |
| CARS2 | 0.05811 | 0.74686 | 0.758 | 0.77 | 1 | HC_vs_MS.NoTrt_B |
| FASLG | 0.05819 | 0.2783 | 0.071 | 0.11 | 1 | HC_vs_MS.NoTrt_B |
| CCDC40 | 0.05823 | 0.27912 | 0.772 | 0.75 | 1 | HC_vs_MS.NoTrt_B |
| TRIM36 | 0.05824 | 0.29112 | 0.371 | 0.31 | 1 | HC_vs_MS.NoTrt_B |
| DUSP19 | 0.05844 | 0.3502 | 0.668 | 0.64 | 1 | HC_vs_MS.NoTrt_B |
| SNX22 | 0.05864 | 0.36828 | 0.803 | 0.8 | 1 | HC_vs_MS.NoTrt_B |
| DAP | 0.05873 | -0.2601 | 0.11 | 0.16 | 1 | HC_vs_MS.NoTrt_B |
| BTBD2 | 0.05905 | 0.98507 | 0.378 | 0.34 | 1 | HC_vs_MS.NoTrt_B |
| C15orf57 | 0.0595 | 0.65052 | 0.772 | 0.74 | 1 | HC_vs_MS.NoTrt_B |
| QTRT1 | 0.05968 | 0.67015 | 0.814 | 0.82 | 1 | HC_vs_MS.NoTrt_B |
| SOWAHC | 0.05969 | 0.48992 | 0.42 | 0.36 | 1 | HC_vs_MS.NoTrt_B |
| SPTLC3 | 0.05987 | 0.61399 | 0.513 | 0.51 | 1 | HC_vs_MS.NoTrt_B |
| ELK1 | 0.05997 | -1.0511 | 0.07 | 0.11 | 1 | HC_vs_MS.NoTrt_B |
| CHMP3 | 0.06009 | -0.3102 | 0.299 | 0.36 | 1 | HC_vs_MS.NoTrt_B |
| TMEM38A | 0.06026 | 0.53074 | 0.503 | 0.45 | 1 | HC_vs_MS.NoTrt_B |
| KCNE1 | 0.0603 | 0.32484 | 0.546 | 0.52 | 1 | HC_vs_MS.NoTrt_B |
| IGF1R | 0.06038 | 0.72136 | 0.354 | 0.29 | 1 | HC_vs_MS.NoTrt_B |
| CDK9 | 0.06052 | -0.662 | 0.072 | 0.11 | 1 | HC_vs_MS.NoTrt_B |
| PLXDC1 | 0.06071 | 0.34564 | 0.729 | 0.72 | 1 | HC_vs_MS.NoTrt_B |
| PNPLA6 | 0.0608 | -0.3735 | 0.126 | 0.18 | 1 | HC_vs_MS.NoTrt_B |
| HSD17B4 | 0.06082 | 0.71276 | 0.642 | 0.63 | 1 | HC_vs_MS.NoTrt_B |
| OSBPL6 | 0.06093 | 0.27152 | 0.562 | 0.55 | 1 | HC_vs_MS.NoTrt_B |
| LMCD1 | 0.06093 | -0.4401 | 0.284 | 0.22 | 1 | HC_vs_MS.NoTrt_B |
| SLC30A4 | 0.06095 | 0.34514 | 0.871 | 0.87 | 1 | HC_vs_MS.NoTrt_B |
| LRIG1 | 0.0611 | 0.30306 | 0.238 | 0.31 | 1 | HC_vs_MS.NoTrt_B |
| AGXT | 0.0616 | 0.29166 | 0.643 | 0.64 | 1 | HC_vs_MS.NoTrt_B |
| ERI1 | 0.06177 | 0.61462 | 0.365 | 0.45 | 1 | HC_vs_MS.NoTrt_B |
| DMXL2 | 0.06178 | 0.37864 | 0.462 | 0.43 | 1 | HC_vs_MS.NoTrt_B |
| EXTL3 | 0.0618 | 0.27727 | 0.923 | 0.92 | 1 | HC_vs_MS.NoTrt_B |
| SH2D1A | 0.06183 | 0.33608 | 0.159 | 0.22 | 1 | HC_vs_MS.NoTrt_B |

| | | | | | | |
|---|---|---|---|---|---|---|
| GLTSCR1 | 0.06229 | 0.47218 | 0.581 | 0.51 | 1 | HC_vs_MS.NoTrt_B |
| CD200 | 0.06237 | 0.74521 | 0.532 | 0.51 | 1 | HC_vs_MS.NoTrt_B |
| FRS3 | 0.06247 | 0.52135 | 0.32 | 0.27 | 1 | HC_vs_MS.NoTrt_B |
| CYC1 | 0.06256 | -0.2525 | 0.099 | 0.15 | 1 | HC_vs_MS.NoTrt_B |
| TMEM263 | 0.0627 | -0.2571 | 0.377 | 0.44 | 1 | HC_vs_MS.NoTrt_B |
| TTLL10 | 0.06291 | 0.40462 | 0.617 | 0.6 | 1 | HC_vs_MS.NoTrt_B |
| KSR1 | 0.06295 | 0.45342 | 0.286 | 0.22 | 1 | HC_vs_MS.NoTrt_B |
| E2F5 | 0.06299 | 0.83202 | 0.332 | 0.28 | 1 | HC_vs_MS.NoTrt_B |
| ATP6AP1 | 0.06307 | -0.3855 | 0.365 | 0.42 | 1 | HC_vs_MS.NoTrt_B |
| CXCR6 | 0.06308 | -0.8518 | 0.128 | 0.18 | 1 | HC_vs_MS.NoTrt_B |
| TRAPPC12 | 0.06308 | 0.60613 | 0.599 | 0.57 | 1 | HC_vs_MS.NoTrt_B |
| TTYH2 | 0.06315 | -0.4652 | 0.077 | 0.12 | 1 | HC_vs_MS.NoTrt_B |
| RNPC3 | 0.06321 | 0.43698 | 0.362 | 0.48 | 1 | HC_vs_MS.NoTrt_B |
| STK11 | 0.06335 | -0.2963 | 0.417 | 0.36 | 1 | HC_vs_MS.NoTrt_B |
| CALR | 0.06344 | -0.4044 | 0.752 | 0.82 | 1 | HC_vs_MS.NoTrt_B |
| AKAP5 | 0.06357 | 0.66501 | 0.625 | 0.58 | 1 | HC_vs_MS.NoTrt_B |
| TOPORS | 0.0637 | 0.31675 | 0.083 | 0.13 | 1 | HC_vs_MS.NoTrt_B |
| MRPL19 | 0.06404 | 0.49929 | 0.243 | 0.32 | 1 | HC_vs_MS.NoTrt_B |
| TEP1 | 0.06406 | 0.41891 | 0.399 | 0.49 | 1 | HC_vs_MS.NoTrt_B |
| SGPL1 | 0.06427 | -0.5418 | 0.419 | 0.48 | 1 | HC_vs_MS.NoTrt_B |
| SLC5A3 | 0.06434 | 0.77727 | 0.067 | 0.11 | 1 | HC_vs_MS.NoTrt_B |
| TTLL7 | 0.0644 | 0.25749 | 0.78 | 0.77 | 1 | HC_vs_MS.NoTrt_B |
| ZNF845 | 0.06446 | 0.87626 | 0.645 | 0.6 | 1 | HC_vs_MS.NoTrt_B |
| QPCTL | 0.0645 | 0.33143 | 0.742 | 0.76 | 1 | HC_vs_MS.NoTrt_B |
| DDX41 | 0.06463 | -0.3854 | 0.086 | 0.13 | 1 | HC_vs_MS.NoTrt_B |
| CTD-2369P2 | 0.06491 | 0.6383 | 0.284 | 0.24 | 1 | HC_vs_MS.NoTrt_B |
| RP11-166B2 | 0.06497 | 0.77676 | 0.507 | 0.48 | 1 | HC_vs_MS.NoTrt_B |
| STK26 | 0.06508 | 0.52149 | 0.078 | 0.12 | 1 | HC_vs_MS.NoTrt_B |
| CTH | 0.06517 | 0.36256 | 0.626 | 0.62 | 1 | HC_vs_MS.NoTrt_B |
| NFE2L2 | 0.06541 | 0.68199 | 0.561 | 0.5 | 1 | HC_vs_MS.NoTrt_B |
| RAD51 | 0.06556 | -0.2791 | 0.478 | 0.43 | 1 | HC_vs_MS.NoTrt_B |
| LOXL2 | 0.0657 | 0.6463 | 0.71 | 0.73 | 1 | HC_vs_MS.NoTrt_B |
| DMD | 0.06595 | 0.49404 | 0.619 | 0.62 | 1 | HC_vs_MS.NoTrt_B |
| B3GNTL1 | 0.06599 | -0.6043 | 0.217 | 0.29 | 1 | HC_vs_MS.NoTrt_B |
| NME3 | 0.0663 | -0.2816 | 0.113 | 0.16 | 1 | HC_vs_MS.NoTrt_B |
| RASGRF2 | 0.0663 | 0.65328 | 0.165 | 0.11 | 1 | HC_vs_MS.NoTrt_B |
| YKT6 | 0.06652 | -0.2961 | 0.077 | 0.12 | 1 | HC_vs_MS.NoTrt_B |
| PLCE1 | 0.06667 | 0.31825 | 0.916 | 0.88 | 1 | HC_vs_MS.NoTrt_B |
| DCUN1D1 | 0.06691 | 0.29626 | 0.372 | 0.44 | 1 | HC_vs_MS.NoTrt_B |
| CASK | 0.06699 | 0.27762 | 0.12 | 0.17 | 1 | HC_vs_MS.NoTrt_B |

| | | | | | | |
|---|---|---|---|---|---|---|
| RASSF8 | 0.06716 | 0.27971 | 0.461 | 0.41 | 1 | HC_vs_MS.NoTrt_B |
| ZNF264 | 0.06717 | 0.45532 | 0.917 | 0.91 | 1 | HC_vs_MS.NoTrt_B |
| BRWD3 | 0.0673 | 0.39886 | 0.117 | 0.07 | 1 | HC_vs_MS.NoTrt_B |
| ART4 | 0.06753 | 0.28553 | 0.5 | 0.45 | 1 | HC_vs_MS.NoTrt_B |
| ERCC1 | 0.06755 | -0.5448 | 0.784 | 0.88 | 1 | HC_vs_MS.NoTrt_B |
| APOL1 | 0.06758 | 0.92413 | 0.7 | 0.65 | 1 | HC_vs_MS.NoTrt_B |
| TMEM106A | 0.06767 | 0.28024 | 0.677 | 0.66 | 1 | HC_vs_MS.NoTrt_B |
| XPO7 | 0.06791 | -0.3681 | 0.1 | 0.15 | 1 | HC_vs_MS.NoTrt_B |
| PRSS27 | 0.06799 | 0.29709 | 0.755 | 0.74 | 1 | HC_vs_MS.NoTrt_B |
| RP11-849H4 | 0.06801 | 0.75464 | 0.596 | 0.51 | 1 | HC_vs_MS.NoTrt_B |
| SCYL3 | 0.06838 | 0.90711 | 0.659 | 0.64 | 1 | HC_vs_MS.NoTrt_B |
| ADGRG1 | 0.06848 | 0.67104 | 0.654 | 0.65 | 1 | HC_vs_MS.NoTrt_B |
| PTPRZ1 | 0.06883 | 0.28749 | 0.21 | 0.16 | 1 | HC_vs_MS.NoTrt_B |
| NOP56 | 0.06905 | -0.2619 | 0.4 | 0.47 | 1 | HC_vs_MS.NoTrt_B |
| GGPS1 | 0.06914 | -0.2652 | 0.378 | 0.47 | 1 | HC_vs_MS.NoTrt_B |
| DCAF10 | 0.06914 | 0.445 | 0.793 | 0.79 | 1 | HC_vs_MS.NoTrt_B |
| CWC27 | 0.06918 | -0.5302 | 0.101 | 0.15 | 1 | HC_vs_MS.NoTrt_B |
| ZC3H4 | 0.06955 | 0.46356 | 0.312 | 0.42 | 1 | HC_vs_MS.NoTrt_B |
| NAV2 | 0.0696 | 0.7217 | 0.372 | 0.32 | 1 | HC_vs_MS.NoTrt_B |
| ACOX3 | 0.06984 | -0.5258 | 0.077 | 0.12 | 1 | HC_vs_MS.NoTrt_B |
| UNC13C | 0.06995 | 0.32777 | 0.607 | 0.62 | 1 | HC_vs_MS.NoTrt_B |
| AGAP2 | 0.06998 | -0.5967 | 0.151 | 0.2 | 1 | HC_vs_MS.NoTrt_B |
| IPO9 | 0.07 | 0.30194 | 0.825 | 0.83 | 1 | HC_vs_MS.NoTrt_B |
| SYDE2 | 0.07023 | 0.4547 | 0.48 | 0.44 | 1 | HC_vs_MS.NoTrt_B |
| DCAF6 | 0.07023 | 0.53868 | 0.159 | 0.22 | 1 | HC_vs_MS.NoTrt_B |
| MELK | 0.0703 | 0.64573 | 0.504 | 0.48 | 1 | HC_vs_MS.NoTrt_B |
| REEP4 | 0.07031 | -0.5585 | 0.065 | 0.1 | 1 | HC_vs_MS.NoTrt_B |
| PMF1 | 0.07031 | -0.3355 | 0.462 | 0.51 | 1 | HC_vs_MS.NoTrt_B |
| UGT2B4 | 0.07036 | 0.28115 | 0.367 | 0.3 | 1 | HC_vs_MS.NoTrt_B |
| SLU7 | 0.07041 | -0.2933 | 0.368 | 0.43 | 1 | HC_vs_MS.NoTrt_B |
| IFRD1 | 0.07045 | 0.4726 | 0.387 | 0.33 | 1 | HC_vs_MS.NoTrt_B |
| CLNK | 0.07046 | 0.68878 | 0.626 | 0.64 | 1 | HC_vs_MS.NoTrt_B |
| TTC17 | 0.07058 | -0.284 | 0.41 | 0.46 | 1 | HC_vs_MS.NoTrt_B |
| GAS7 | 0.07088 | 0.37163 | 0.586 | 0.56 | 1 | HC_vs_MS.NoTrt_B |
| RBBP8 | 0.0711 | 0.62876 | 0.072 | 0.11 | 1 | HC_vs_MS.NoTrt_B |
| PSMD14 | 0.0713 | -0.536 | 0.087 | 0.13 | 1 | HC_vs_MS.NoTrt_B |
| CXorf36 | 0.07134 | 0.42226 | 0.383 | 0.34 | 1 | HC_vs_MS.NoTrt_B |
| ARSA | 0.07137 | -0.2758 | 0.68 | 0.73 | 1 | HC_vs_MS.NoTrt_B |
| SOS2 | 0.07141 | -0.4173 | 0.083 | 0.12 | 1 | HC_vs_MS.NoTrt_B |
| MEGF6 | 0.07201 | -0.3782 | 0.11 | 0.16 | 1 | HC_vs_MS.NoTrt_B |

| | | | | | | |
|---|---|---|---|---|---|---|
| TANC2 | 0.07202 | 1.63645 | 0.257 | 0.2 | 1 | HC_vs_MS.NoTrt_B |
| NUMB | 0.07217 | -0.2643 | 0.709 | 0.76 | 1 | HC_vs_MS.NoTrt_B |
| ARHGAP18 | 0.0724 | -1.0381 | 0.074 | 0.11 | 1 | HC_vs_MS.NoTrt_B |
| RAD23B | 0.07245 | -0.7109 | 0.439 | 0.53 | 1 | HC_vs_MS.NoTrt_B |
| PDCL | 0.07257 | -0.3673 | 0.375 | 0.44 | 1 | HC_vs_MS.NoTrt_B |
| COPS6 | 0.07257 | -0.4798 | 0.671 | 0.69 | 1 | HC_vs_MS.NoTrt_B |
| FPGT | 0.07267 | -0.5466 | 0.087 | 0.13 | 1 | HC_vs_MS.NoTrt_B |
| ST6GALNAC | 0.07268 | 0.27772 | 0.187 | 0.14 | 1 | HC_vs_MS.NoTrt_B |
| TGFBR3 | 0.07288 | 0.53691 | 0.388 | 0.47 | 1 | HC_vs_MS.NoTrt_B |
| UBE4A | 0.07289 | 0.40078 | 0.367 | 0.46 | 1 | HC_vs_MS.NoTrt_B |
| ATP6V1B2 | 0.0729 | -0.3643 | 0.594 | 0.67 | 1 | HC_vs_MS.NoTrt_B |
| OSER1 | 0.07295 | -0.4154 | 0.157 | 0.21 | 1 | HC_vs_MS.NoTrt_B |
| EIF2AK2 | 0.07303 | 0.57323 | 0.825 | 0.84 | 1 | HC_vs_MS.NoTrt_B |
| REXO1 | 0.07337 | -0.4244 | 0.068 | 0.11 | 1 | HC_vs_MS.NoTrt_B |
| SLC3A1 | 0.07345 | 1.10578 | 0.684 | 0.7 | 1 | HC_vs_MS.NoTrt_B |
| MAGEH1 | 0.07361 | -0.3955 | 0.074 | 0.11 | 1 | HC_vs_MS.NoTrt_B |
| UBASH3A | 0.07392 | -0.3821 | 0.183 | 0.24 | 1 | HC_vs_MS.NoTrt_B |
| TLE2 | 0.07395 | 0.87301 | 0.487 | 0.44 | 1 | HC_vs_MS.NoTrt_B |
| GLB1L3 | 0.07396 | 0.63113 | 0.7 | 0.69 | 1 | HC_vs_MS.NoTrt_B |
| EGR3 | 0.07406 | 0.25981 | 0.193 | 0.14 | 1 | HC_vs_MS.NoTrt_B |
| ARRB1 | 0.07418 | -0.3669 | 0.716 | 0.69 | 1 | HC_vs_MS.NoTrt_B |
| RNF44 | 0.07472 | 0.97334 | 0.37 | 0.32 | 1 | HC_vs_MS.NoTrt_B |
| TNFSF4 | 0.07488 | 0.69032 | 0.122 | 0.08 | 1 | HC_vs_MS.NoTrt_B |
| GBP6 | 0.07496 | 0.27802 | 0.445 | 0.38 | 1 | HC_vs_MS.NoTrt_B |
| LEO1 | 0.07514 | 0.87684 | 0.548 | 0.49 | 1 | HC_vs_MS.NoTrt_B |
| LLGL2 | 0.07535 | 0.4057 | 0.386 | 0.34 | 1 | HC_vs_MS.NoTrt_B |
| PEX16 | 0.07536 | -0.4648 | 0.712 | 0.8 | 1 | HC_vs_MS.NoTrt_B |
| BECN1 | 0.07543 | -0.3382 | 0.704 | 0.74 | 1 | HC_vs_MS.NoTrt_B |
| KBTBD2 | 0.07546 | 0.4193 | 0.632 | 0.6 | 1 | HC_vs_MS.NoTrt_B |
| BCAM | 0.07549 | 0.28263 | 0.741 | 0.75 | 1 | HC_vs_MS.NoTrt_B |
| MRTO4 | 0.07569 | -0.3479 | 0.449 | 0.55 | 1 | HC_vs_MS.NoTrt_B |
| MAN1C1 | 0.07573 | 0.55283 | 0.867 | 0.85 | 1 | HC_vs_MS.NoTrt_B |
| STRN | 0.07608 | 1.19538 | 0.225 | 0.17 | 1 | HC_vs_MS.NoTrt_B |
| FCRL2 | 0.07615 | 0.52967 | 0.619 | 0.58 | 1 | HC_vs_MS.NoTrt_B |
| TMEM132B | 0.07625 | 0.64248 | 0.267 | 0.21 | 1 | HC_vs_MS.NoTrt_B |
| DDI2 | 0.07628 | 0.288 | 0.751 | 0.7 | 1 | HC_vs_MS.NoTrt_B |
| PRKAR2A | 0.07633 | 0.36074 | 0.883 | 0.88 | 1 | HC_vs_MS.NoTrt_B |
| SERTAD2 | 0.07637 | 1.29 | 0.119 | 0.08 | 1 | HC_vs_MS.NoTrt_B |
| RAP1B | 0.07642 | 0.29309 | 0.993 | 0.99 | 1 | HC_vs_MS.NoTrt_B |
| RPP14 | 0.07695 | 0.25965 | 0.907 | 0.91 | 1 | HC_vs_MS.NoTrt_B |

| | | | | | | |
|---|---|---|---|---|---|---|
| PSPN | 0.07696 | 0.32103 | 0.688 | 0.66 | 1 | HC_vs_MS.NoTrt_B |
| C1orf35 | 0.07706 | -0.5645 | 0.2 | 0.25 | 1 | HC_vs_MS.NoTrt_B |
| TRAPPC6B | 0.07713 | 0.39794 | 0.503 | 0.47 | 1 | HC_vs_MS.NoTrt_B |
| GPATCH11 | 0.07717 | 0.65791 | 0.507 | 0.47 | 1 | HC_vs_MS.NoTrt_B |
| AGAP4 | 0.07719 | 0.6209 | 0.21 | 0.29 | 1 | HC_vs_MS.NoTrt_B |
| AMPH | 0.07729 | -0.5241 | 0.136 | 0.19 | 1 | HC_vs_MS.NoTrt_B |
| CRIP2 | 0.07794 | -0.3183 | 0.12 | 0.17 | 1 | HC_vs_MS.NoTrt_B |
| PPFIBP1 | 0.07795 | 0.43574 | 0.735 | 0.72 | 1 | HC_vs_MS.NoTrt_B |
| XRN1 | 0.07798 | 0.48827 | 0.909 | 0.92 | 1 | HC_vs_MS.NoTrt_B |
| DHFR2 | 0.07826 | -0.4014 | 0.59 | 0.63 | 1 | HC_vs_MS.NoTrt_B |
| PCGF5 | 0.07858 | -0.3771 | 0.329 | 0.38 | 1 | HC_vs_MS.NoTrt_B |
| ATP11B | 0.07869 | 0.38318 | 0.204 | 0.27 | 1 | HC_vs_MS.NoTrt_B |
| PDZD2 | 0.07888 | 0.51308 | 0.668 | 0.64 | 1 | HC_vs_MS.NoTrt_B |
| SMAD1 | 0.07935 | 1.17196 | 0.32 | 0.27 | 1 | HC_vs_MS.NoTrt_B |
| ADAR | 0.07965 | 0.48323 | 0.796 | 0.84 | 1 | HC_vs_MS.NoTrt_B |
| CACNB2 | 0.07983 | 0.9678 | 0.449 | 0.4 | 1 | HC_vs_MS.NoTrt_B |
| PRR5 | 0.08005 | 0.57868 | 0.51 | 0.48 | 1 | HC_vs_MS.NoTrt_B |
| SIGLEC10 | 0.08031 | 0.33979 | 0.623 | 0.61 | 1 | HC_vs_MS.NoTrt_B |
| C5orf42 | 0.08041 | 0.66163 | 0.51 | 0.5 | 1 | HC_vs_MS.NoTrt_B |
| C15orf53 | 0.08048 | -0.3253 | 0.065 | 0.1 | 1 | HC_vs_MS.NoTrt_B |
| MARK4 | 0.08063 | 0.48393 | 0.58 | 0.55 | 1 | HC_vs_MS.NoTrt_B |
| 11-Sep | 0.08065 | 0.5326 | 0.387 | 0.32 | 1 | HC_vs_MS.NoTrt_B |
| DEK | 0.08072 | 0.29872 | 0.414 | 0.52 | 1 | HC_vs_MS.NoTrt_B |
| WASH1 | 0.08103 | 0.71567 | 0.2 | 0.28 | 1 | HC_vs_MS.NoTrt_B |
| MAPKBP1 | 0.08104 | 0.38435 | 0.387 | 0.48 | 1 | HC_vs_MS.NoTrt_B |
| CWF19L2 | 0.08109 | 0.312 | 0.316 | 0.4 | 1 | HC_vs_MS.NoTrt_B |
| BRI3BP | 0.08118 | 0.60402 | 0.846 | 0.86 | 1 | HC_vs_MS.NoTrt_B |
| UHRF1BP1 | 0.08121 | -0.5553 | 0.549 | 0.65 | 1 | HC_vs_MS.NoTrt_B |
| LENG1 | 0.08157 | -0.6202 | 0.07 | 0.11 | 1 | HC_vs_MS.NoTrt_B |
| SLC9A9 | 0.08174 | -0.356 | 0.129 | 0.18 | 1 | HC_vs_MS.NoTrt_B |
| CCT7 | 0.08187 | -0.271 | 0.204 | 0.26 | 1 | HC_vs_MS.NoTrt_B |
| CCDC91 | 0.0819 | 0.50858 | 0.486 | 0.43 | 1 | HC_vs_MS.NoTrt_B |
| LHFPL2 | 0.08202 | 0.78664 | 0.29 | 0.36 | 1 | HC_vs_MS.NoTrt_B |
| SBF1 | 0.08214 | 0.88451 | 0.2 | 0.15 | 1 | HC_vs_MS.NoTrt_B |
| MRPS17 | 0.08221 | 0.32881 | 0.845 | 0.83 | 1 | HC_vs_MS.NoTrt_B |
| AMFR | 0.08256 | -0.3857 | 0.514 | 0.42 | 1 | HC_vs_MS.NoTrt_B |
| MROH8 | 0.0827 | 0.25333 | 0.329 | 0.27 | 1 | HC_vs_MS.NoTrt_B |
| FCHSD1 | 0.08278 | -0.2817 | 0.281 | 0.35 | 1 | HC_vs_MS.NoTrt_B |
| PPIH | 0.08279 | -0.5583 | 0.075 | 0.11 | 1 | HC_vs_MS.NoTrt_B |
| FAM13A | 0.08325 | 0.37551 | 0.207 | 0.15 | 1 | HC_vs_MS.NoTrt_B |

| | | | | | | |
|---|---|---|---|---|---|---|
| MAFF | 0.08336 | 0.41441 | 0.642 | 0.6 | 1 | HC_vs_MS.NoTrt_B |
| ITGA2B | 0.08348 | 0.2716 | 0.325 | 0.27 | 1 | HC_vs_MS.NoTrt_B |
| LIN7B | 0.08355 | 0.35635 | 0.309 | 0.25 | 1 | HC_vs_MS.NoTrt_B |
| ARRDC2 | 0.08372 | -0.318 | 0.084 | 0.12 | 1 | HC_vs_MS.NoTrt_B |
| GANC | 0.08372 | 0.38334 | 0.607 | 0.54 | 1 | HC_vs_MS.NoTrt_B |
| KLHL5 | 0.08373 | -0.6455 | 0.419 | 0.53 | 1 | HC_vs_MS.NoTrt_B |
| LYAR | 0.08376 | -0.2998 | 0.07 | 0.11 | 1 | HC_vs_MS.NoTrt_B |
| STK17B | 0.08377 | 0.29553 | 0.933 | 0.96 | 1 | HC_vs_MS.NoTrt_B |
| IGFL2 | 0.08378 | 0.4107 | 0.329 | 0.28 | 1 | HC_vs_MS.NoTrt_B |
| DDOST | 0.08381 | -0.2868 | 0.545 | 0.57 | 1 | HC_vs_MS.NoTrt_B |
| RNF113A | 0.08396 | -0.2516 | 0.096 | 0.14 | 1 | HC_vs_MS.NoTrt_B |
| RBMS3 | 0.08431 | 0.74365 | 0.672 | 0.67 | 1 | HC_vs_MS.NoTrt_B |
| EMCN | 0.08444 | 0.36274 | 0.438 | 0.37 | 1 | HC_vs_MS.NoTrt_B |
| SCAI | 0.08458 | 0.75139 | 0.832 | 0.83 | 1 | HC_vs_MS.NoTrt_B |
| PDK1 | 0.08462 | 0.254 | 0.797 | 0.86 | 1 | HC_vs_MS.NoTrt_B |
| TTC32 | 0.08468 | 0.47461 | 0.081 | 0.12 | 1 | HC_vs_MS.NoTrt_B |
| CXorf23 | 0.08477 | 0.49166 | 0.741 | 0.72 | 1 | HC_vs_MS.NoTrt_B |
| SYT14 | 0.08491 | 0.32474 | 0.316 | 0.27 | 1 | HC_vs_MS.NoTrt_B |
| PECAM1 | 0.08525 | 0.93905 | 0.748 | 0.74 | 1 | HC_vs_MS.NoTrt_B |
| MAP2K1 | 0.08528 | 1.19391 | 0.113 | 0.07 | 1 | HC_vs_MS.NoTrt_B |
| ZNF583 | 0.08531 | 0.4206 | 0.803 | 0.85 | 1 | HC_vs_MS.NoTrt_B |
| PDS5B | 0.08534 | 0.34713 | 0.312 | 0.38 | 1 | HC_vs_MS.NoTrt_B |
| ZDHHC18 | 0.08538 | -0.4544 | 0.065 | 0.1 | 1 | HC_vs_MS.NoTrt_B |
| SLC9A6 | 0.08549 | 0.33778 | 0.407 | 0.34 | 1 | HC_vs_MS.NoTrt_B |
| POLDIP2 | 0.08551 | -0.3577 | 0.093 | 0.14 | 1 | HC_vs_MS.NoTrt_B |
| LRRD1 | 0.08583 | 0.2874 | 0.522 | 0.48 | 1 | HC_vs_MS.NoTrt_B |
| ITGA1 | 0.08613 | 0.26698 | 0.661 | 0.67 | 1 | HC_vs_MS.NoTrt_B |
| TMEM199 | 0.08617 | -0.4097 | 0.578 | 0.67 | 1 | HC_vs_MS.NoTrt_B |
| HPSE | 0.0862 | -0.2653 | 0.617 | 0.57 | 1 | HC_vs_MS.NoTrt_B |
| RNF220 | 0.08623 | 0.75038 | 0.641 | 0.65 | 1 | HC_vs_MS.NoTrt_B |
| RHBDD2 | 0.08636 | -0.3146 | 0.226 | 0.28 | 1 | HC_vs_MS.NoTrt_B |
| SEMA4C | 0.08645 | 0.46075 | 0.762 | 0.76 | 1 | HC_vs_MS.NoTrt_B |
| CDH18 | 0.08648 | 0.31207 | 0.73 | 0.76 | 1 | HC_vs_MS.NoTrt_B |
| LINS1 | 0.08652 | 0.75505 | 0.572 | 0.55 | 1 | HC_vs_MS.NoTrt_B |
| TACR2 | 0.08662 | 0.47089 | 0.129 | 0.09 | 1 | HC_vs_MS.NoTrt_B |
| ZNF496 | 0.08665 | 0.71233 | 0.681 | 0.67 | 1 | HC_vs_MS.NoTrt_B |
| DPY19L4 | 0.08669 | 1.09219 | 0.488 | 0.45 | 1 | HC_vs_MS.NoTrt_B |
| PVR | 0.08746 | 0.28221 | 0.454 | 0.4 | 1 | HC_vs_MS.NoTrt_B |
| ARMC9 | 0.08769 | 0.53867 | 0.668 | 0.68 | 1 | HC_vs_MS.NoTrt_B |
| STXBP6 | 0.08788 | 0.28773 | 0.161 | 0.11 | 1 | HC_vs_MS.NoTrt_B |

| | | | | | | |
|---|---|---|---|---|---|---|
| RP11-176H8 | 0.08789 | 0.28179 | 0.823 | 0.82 | 1 | HC_vs_MS.NoTrt_B |
| ARL10 | 0.08807 | 0.66896 | 0.849 | 0.83 | 1 | HC_vs_MS.NoTrt_B |
| ABCC4 | 0.08809 | 0.52881 | 0.703 | 0.75 | 1 | HC_vs_MS.NoTrt_B |
| SCN3B | 0.08811 | 0.28634 | 0.68 | 0.69 | 1 | HC_vs_MS.NoTrt_B |
| AARSD1 | 0.08854 | 0.5044 | 0.678 | 0.68 | 1 | HC_vs_MS.NoTrt_B |
| SHROOM4 | 0.08855 | 0.2918 | 0.643 | 0.61 | 1 | HC_vs_MS.NoTrt_B |
| GRAMD3 | 0.08874 | -0.3 | 0.69 | 0.78 | 1 | HC_vs_MS.NoTrt_B |
| APPL2 | 0.08882 | 0.39851 | 0.348 | 0.29 | 1 | HC_vs_MS.NoTrt_B |
| ZNF225 | 0.08895 | 0.39493 | 0.768 | 0.74 | 1 | HC_vs_MS.NoTrt_B |
| LRP8 | 0.08899 | 0.29887 | 0.503 | 0.47 | 1 | HC_vs_MS.NoTrt_B |
| STPG1 | 0.08902 | 0.34093 | 0.755 | 0.77 | 1 | HC_vs_MS.NoTrt_B |
| BCAS1 | 0.08909 | -0.8132 | 0.345 | 0.42 | 1 | HC_vs_MS.NoTrt_B |
| NEO1 | 0.08914 | 0.27658 | 0.245 | 0.33 | 1 | HC_vs_MS.NoTrt_B |
| TGFBR2 | 0.08953 | 0.97734 | 0.162 | 0.11 | 1 | HC_vs_MS.NoTrt_B |
| SLC25A20 | 0.08955 | -0.8326 | 0.093 | 0.14 | 1 | HC_vs_MS.NoTrt_B |
| LLNLF-187D | 0.08961 | 0.32024 | 0.546 | 0.54 | 1 | HC_vs_MS.NoTrt_B |
| CAPG | 0.08989 | -0.6558 | 0.071 | 0.11 | 1 | HC_vs_MS.NoTrt_B |
| INTS9 | 0.08992 | -0.5286 | 0.094 | 0.14 | 1 | HC_vs_MS.NoTrt_B |
| RPF1 | 0.08997 | -0.4791 | 0.232 | 0.29 | 1 | HC_vs_MS.NoTrt_B |
| APOE | 0.09008 | 0.30233 | 0.743 | 0.74 | 1 | HC_vs_MS.NoTrt_B |
| DNAH9 | 0.09017 | 0.50787 | 0.436 | 0.38 | 1 | HC_vs_MS.NoTrt_B |
| ADCK1 | 0.09031 | 1.03046 | 0.274 | 0.22 | 1 | HC_vs_MS.NoTrt_B |
| ITGA4 | 0.09038 | 0.75839 | 0.223 | 0.17 | 1 | HC_vs_MS.NoTrt_B |
| SERF1B | 0.09042 | 0.27632 | 0.454 | 0.39 | 1 | HC_vs_MS.NoTrt_B |
| GREB1 | 0.09044 | 0.375 | 0.836 | 0.85 | 1 | HC_vs_MS.NoTrt_B |
| ZNF684 | 0.09058 | 0.61193 | 0.694 | 0.69 | 1 | HC_vs_MS.NoTrt_B |
| CD300LG | 0.0907 | 0.36666 | 0.4 | 0.36 | 1 | HC_vs_MS.NoTrt_B |
| C7orf73 | 0.09076 | -0.2857 | 0.61 | 0.67 | 1 | HC_vs_MS.NoTrt_B |
| RNF157 | 0.0909 | 0.3148 | 0.607 | 0.56 | 1 | HC_vs_MS.NoTrt_B |
| CEP89 | 0.09091 | 0.35932 | 0.868 | 0.92 | 1 | HC_vs_MS.NoTrt_B |
| SSR4 | 0.09106 | -0.4671 | 0.333 | 0.38 | 1 | HC_vs_MS.NoTrt_B |
| C3orf70 | 0.09138 | 0.37547 | 0.699 | 0.65 | 1 | HC_vs_MS.NoTrt_B |
| PNO1 | 0.09165 | 0.25294 | 0.686 | 0.67 | 1 | HC_vs_MS.NoTrt_B |
| MFSD10 | 0.09168 | -0.3869 | 0.107 | 0.15 | 1 | HC_vs_MS.NoTrt_B |
| AKR1D1 | 0.09177 | 0.2702 | 0.564 | 0.55 | 1 | HC_vs_MS.NoTrt_B |
| FOXP3 | 0.09211 | 0.4061 | 0.457 | 0.57 | 1 | HC_vs_MS.NoTrt_B |
| PPARGC1B | 0.09216 | 0.47511 | 0.694 | 0.67 | 1 | HC_vs_MS.NoTrt_B |
| GFM1 | 0.09224 | 0.27035 | 0.87 | 0.9 | 1 | HC_vs_MS.NoTrt_B |
| FBXO48 | 0.09228 | 0.75891 | 0.691 | 0.68 | 1 | HC_vs_MS.NoTrt_B |
| JRK | 0.09229 | 1.43347 | 0.433 | 0.4 | 1 | HC_vs_MS.NoTrt_B |

| | | | | | | |
|---|---|---|---|---|---|---|
| KIAA2012 | 0.09239 | 0.32356 | 0.643 | 0.63 | 1 | HC_vs_MS.NoTrt_B |
| RSL1D1 | 0.09257 | 0.26154 | 0.846 | 0.86 | 1 | HC_vs_MS.NoTrt_B |
| ZNF595 | 0.09263 | 0.96153 | 0.793 | 0.81 | 1 | HC_vs_MS.NoTrt_B |
| SEC14L4 | 0.09296 | 0.2953 | 0.516 | 0.48 | 1 | HC_vs_MS.NoTrt_B |
| PPP2R3B | 0.09301 | -0.8895 | 0.445 | 0.53 | 1 | HC_vs_MS.NoTrt_B |
| CLCC1 | 0.09305 | 0.46087 | 0.793 | 0.83 | 1 | HC_vs_MS.NoTrt_B |
| PIP4K2B | 0.09403 | -0.3213 | 0.107 | 0.15 | 1 | HC_vs_MS.NoTrt_B |
| POGLUT1 | 0.0942 | 0.64678 | 0.303 | 0.25 | 1 | HC_vs_MS.NoTrt_B |
| EFCAB12 | 0.09447 | 0.36535 | 0.745 | 0.74 | 1 | HC_vs_MS.NoTrt_B |
| COPS4 | 0.09502 | 0.25612 | 0.468 | 0.55 | 1 | HC_vs_MS.NoTrt_B |
| PIGQ | 0.09504 | -0.6979 | 0.403 | 0.44 | 1 | HC_vs_MS.NoTrt_B |
| HLA-DRB3 | 0.09523 | -0.4742 | 0.093 | 0.14 | 1 | HC_vs_MS.NoTrt_B |
| ZNF396 | 0.09527 | 0.26912 | 0.867 | 0.9 | 1 | HC_vs_MS.NoTrt_B |
| ZNF180 | 0.09535 | -0.2736 | 0.332 | 0.27 | 1 | HC_vs_MS.NoTrt_B |
| TMTC4 | 0.09547 | -0.472 | 0.075 | 0.11 | 1 | HC_vs_MS.NoTrt_B |
| F2R | 0.09556 | -0.4436 | 0.525 | 0.49 | 1 | HC_vs_MS.NoTrt_B |
| TANGO2 | 0.09559 | -0.6114 | 0.841 | 0.85 | 1 | HC_vs_MS.NoTrt_B |
| ACYP1 | 0.09566 | 0.33972 | 0.204 | 0.27 | 1 | HC_vs_MS.NoTrt_B |
| MYO15B | 0.09571 | -0.2657 | 0.217 | 0.27 | 1 | HC_vs_MS.NoTrt_B |
| C22orf29 | 0.09591 | -0.2842 | 0.367 | 0.42 | 1 | HC_vs_MS.NoTrt_B |
| APLP1 | 0.09598 | 0.27878 | 0.62 | 0.64 | 1 | HC_vs_MS.NoTrt_B |
| DNAJC12 | 0.09635 | 0.26007 | 0.367 | 0.33 | 1 | HC_vs_MS.NoTrt_B |
| MUTYH | 0.09637 | -0.4343 | 0.514 | 0.58 | 1 | HC_vs_MS.NoTrt_B |
| C20orf27 | 0.09639 | -0.3667 | 0.067 | 0.1 | 1 | HC_vs_MS.NoTrt_B |
| ALDH1B1 | 0.09663 | 1.08471 | 0.591 | 0.59 | 1 | HC_vs_MS.NoTrt_B |
| IKBKE | 0.09675 | -0.4285 | 0.293 | 0.35 | 1 | HC_vs_MS.NoTrt_B |
| ANXA1 | 0.09698 | 0.6778 | 0.178 | 0.13 | 1 | HC_vs_MS.NoTrt_B |
| FLCN | 0.09717 | 0.73206 | 0.859 | 0.86 | 1 | HC_vs_MS.NoTrt_B |
| MIER2 | 0.0972 | -0.372 | 0.529 | 0.45 | 1 | HC_vs_MS.NoTrt_B |
| FAXC | 0.09746 | 0.32746 | 0.325 | 0.28 | 1 | HC_vs_MS.NoTrt_B |
| ERGIC1 | 0.09756 | 0.41482 | 0.828 | 0.86 | 1 | HC_vs_MS.NoTrt_B |
| SEMA6B | 0.09757 | 0.25245 | 0.886 | 0.89 | 1 | HC_vs_MS.NoTrt_B |
| C2orf81 | 0.09761 | 0.26727 | 0.835 | 0.87 | 1 | HC_vs_MS.NoTrt_B |
| MAPK6 | 0.09792 | 0.35946 | 0.446 | 0.54 | 1 | HC_vs_MS.NoTrt_B |
| SPATA21 | 0.09819 | -0.2766 | 0.433 | 0.5 | 1 | HC_vs_MS.NoTrt_B |
| SMARCD1 | 0.09825 | 0.46127 | 0.132 | 0.18 | 1 | HC_vs_MS.NoTrt_B |
| NHLRC3 | 0.09827 | 0.26658 | 0.37 | 0.44 | 1 | HC_vs_MS.NoTrt_B |
| PIBF1 | 0.0984 | 0.2865 | 0.128 | 0.18 | 1 | HC_vs_MS.NoTrt_B |
| PARP12 | 0.09857 | 0.31431 | 0.245 | 0.18 | 1 | HC_vs_MS.NoTrt_B |
| MROH7 | 0.09877 | 0.31538 | 0.464 | 0.43 | 1 | HC_vs_MS.NoTrt_B |

| | | | | | | |
|---|---|---|---|---|---|---|
| DPYSL5 | 0.09882 | 0.37816 | 0.591 | 0.59 | 1 | HC_vs_MS.NoTrt_B |
| KIFC3 | 0.09907 | 0.42328 | 0.67 | 0.67 | 1 | HC_vs_MS.NoTrt_B |
| LYRM2 | 0.09918 | -0.3444 | 0.83 | 0.81 | 1 | HC_vs_MS.NoTrt_B |
| GJB4 | 0.09923 | 0.4029 | 0.651 | 0.63 | 1 | HC_vs_MS.NoTrt_B |
| FAM65B | 0.09959 | 0.49624 | 0.7 | 0.67 | 1 | HC_vs_MS.NoTrt_B |
| ITFG1 | 0.09979 | 0.42495 | 0.174 | 0.23 | 1 | HC_vs_MS.NoTrt_B |
| AIPL1 | 0.09982 | 0.29671 | 0.365 | 0.32 | 1 | HC_vs_MS.NoTrt_B |
| LRRN2 | 0.10007 | 1.10474 | 0.23 | 0.18 | 1 | HC_vs_MS.NoTrt_B |
| FUT2 | 0.1001 | 0.408 | 0.741 | 0.76 | 1 | HC_vs_MS.NoTrt_B |
| EVL | 0.10027 | 0.33705 | 0.836 | 0.89 | 1 | HC_vs_MS.NoTrt_B |
| RB1 | 0.10028 | 0.26704 | 0.125 | 0.17 | 1 | HC_vs_MS.NoTrt_B |
| ZNF613 | 0.10028 | -0.2983 | 0.413 | 0.49 | 1 | HC_vs_MS.NoTrt_B |
| GPA33 | 0.1004 | 0.93459 | 0.146 | 0.1 | 1 | HC_vs_MS.NoTrt_B |
| UTY | 0.10043 | -1.2223 | 0.422 | 0.47 | 1 | HC_vs_MS.NoTrt_B |
| ARHGAP11A | 0.10082 | -0.481 | 0.37 | 0.44 | 1 | HC_vs_MS.NoTrt_B |
| PGBD2 | 0.1011 | 0.61391 | 0.301 | 0.25 | 1 | HC_vs_MS.NoTrt_B |
| BMPR2 | 0.10129 | 0.28051 | 0.445 | 0.52 | 1 | HC_vs_MS.NoTrt_B |
| MTHFSD | 0.10133 | -0.2871 | 0.075 | 0.11 | 1 | HC_vs_MS.NoTrt_B |
| ATP1A3 | 0.10135 | 0.4371 | 0.361 | 0.31 | 1 | HC_vs_MS.NoTrt_B |
| ATP13A3 | 0.10138 | 0.56008 | 0.104 | 0.15 | 1 | HC_vs_MS.NoTrt_B |
| AGO2 | 0.10141 | 0.69763 | 0.413 | 0.37 | 1 | HC_vs_MS.NoTrt_B |
| EIF2S3 | 0.10148 | 0.35501 | 0.954 | 0.95 | 1 | HC_vs_MS.NoTrt_B |
| KRT8 | 0.10173 | 0.28215 | 0.822 | 0.79 | 1 | HC_vs_MS.NoTrt_B |
| PUS1 | 0.10175 | 0.51663 | 0.855 | 0.88 | 1 | HC_vs_MS.NoTrt_B |
| RORC | 0.10202 | -1.3053 | 0.077 | 0.11 | 1 | HC_vs_MS.NoTrt_B |
| TRIM16 | 0.10205 | -1.1063 | 0.438 | 0.48 | 1 | HC_vs_MS.NoTrt_B |
| CYP2C9 | 0.10212 | 0.27148 | 0.588 | 0.58 | 1 | HC_vs_MS.NoTrt_B |
| LARP7 | 0.10212 | 0.3069 | 0.172 | 0.23 | 1 | HC_vs_MS.NoTrt_B |
| TCTEX1D2 | 0.10238 | -0.2528 | 0.27 | 0.33 | 1 | HC_vs_MS.NoTrt_B |
| HYDIN | 0.10251 | 0.81444 | 0.299 | 0.24 | 1 | HC_vs_MS.NoTrt_B |
| ZNF497 | 0.10283 | 0.29534 | 0.268 | 0.33 | 1 | HC_vs_MS.NoTrt_B |
| SSH1 | 0.10317 | 0.44 | 0.846 | 0.8 | 1 | HC_vs_MS.NoTrt_B |
| RRAGD | 0.10321 | 0.74887 | 0.119 | 0.17 | 1 | HC_vs_MS.NoTrt_B |
| AP4B1 | 0.10325 | 0.33046 | 0.622 | 0.61 | 1 | HC_vs_MS.NoTrt_B |
| SRSF12 | 0.10327 | 0.43941 | 0.297 | 0.24 | 1 | HC_vs_MS.NoTrt_B |
| LINC00961 | 0.10331 | 0.49932 | 0.297 | 0.25 | 1 | HC_vs_MS.NoTrt_B |
| NABP2 | 0.1035 | -0.7732 | 0.412 | 0.49 | 1 | HC_vs_MS.NoTrt_B |
| AKAP7 | 0.10364 | 0.69239 | 0.5 | 0.47 | 1 | HC_vs_MS.NoTrt_B |
| CCNK | 0.10364 | 0.32601 | 0.222 | 0.29 | 1 | HC_vs_MS.NoTrt_B |
| NDUFA3 | 0.10392 | 0.34404 | 0.946 | 0.92 | 1 | HC_vs_MS.NoTrt_B |

| | | | | | | |
|---|---|---|---|---|---|---|
| MAVS | 0.10398 | 0.6079 | 0.935 | 0.96 | 1 | HC_vs_MS.NoTrt_B |
| SMARCA5 | 0.10408 | 0.32079 | 0.328 | 0.4 | 1 | HC_vs_MS.NoTrt_B |
| HIST1H1E | 0.10424 | 0.67556 | 0.11 | 0.07 | 1 | HC_vs_MS.NoTrt_B |
| RAB3B | 0.10431 | 0.31812 | 0.745 | 0.72 | 1 | HC_vs_MS.NoTrt_B |
| TTC6 | 0.105 | 0.37552 | 0.286 | 0.24 | 1 | HC_vs_MS.NoTrt_B |
| DNASE1L3 | 0.10513 | 0.34737 | 0.397 | 0.35 | 1 | HC_vs_MS.NoTrt_B |
| ANKIB1 | 0.10536 | 0.50165 | 0.304 | 0.37 | 1 | HC_vs_MS.NoTrt_B |
| PPFIA1 | 0.10551 | 0.28913 | 0.657 | 0.64 | 1 | HC_vs_MS.NoTrt_B |
| SLC38A10 | 0.10585 | 0.30145 | 0.304 | 0.38 | 1 | HC_vs_MS.NoTrt_B |
| HSD17B13 | 0.10591 | 0.47621 | 0.57 | 0.54 | 1 | HC_vs_MS.NoTrt_B |
| C1orf131 | 0.10592 | -0.5764 | 0.575 | 0.67 | 1 | HC_vs_MS.NoTrt_B |
| ICA1L | 0.10596 | 0.4346 | 0.739 | 0.71 | 1 | HC_vs_MS.NoTrt_B |
| SLC25A4 | 0.10601 | -0.4469 | 0.272 | 0.35 | 1 | HC_vs_MS.NoTrt_B |
| MCRIP2 | 0.10624 | 0.49122 | 0.561 | 0.51 | 1 | HC_vs_MS.NoTrt_B |
| ZNF345 | 0.1063 | 0.33474 | 0.814 | 0.85 | 1 | HC_vs_MS.NoTrt_B |
| UIMC1 | 0.10646 | 0.57756 | 0.813 | 0.82 | 1 | HC_vs_MS.NoTrt_B |
| USP47 | 0.10649 | 0.29392 | 0.206 | 0.27 | 1 | HC_vs_MS.NoTrt_B |
| DEAF1 | 0.10651 | 0.58383 | 0.296 | 0.36 | 1 | HC_vs_MS.NoTrt_B |
| U2AF1 | 0.10661 | -0.3799 | 0.535 | 0.56 | 1 | HC_vs_MS.NoTrt_B |
| PNMAL1 | 0.10706 | 0.26456 | 0.428 | 0.38 | 1 | HC_vs_MS.NoTrt_B |
| RNF2 | 0.10737 | -0.3018 | 0.367 | 0.45 | 1 | HC_vs_MS.NoTrt_B |
| HNRNPUL1 | 0.10768 | 0.8828 | 0.688 | 0.68 | 1 | HC_vs_MS.NoTrt_B |
| ARFGAP2 | 0.10773 | -0.372 | 0.525 | 0.6 | 1 | HC_vs_MS.NoTrt_B |
| PFKL | 0.10784 | -0.3313 | 0.729 | 0.73 | 1 | HC_vs_MS.NoTrt_B |
| RUFY3 | 0.10796 | 0.43588 | 0.545 | 0.51 | 1 | HC_vs_MS.NoTrt_B |
| KCNMB1 | 0.10828 | 0.27876 | 0.69 | 0.7 | 1 | HC_vs_MS.NoTrt_B |
| MKRN3 | 0.10905 | 1.08034 | 0.452 | 0.4 | 1 | HC_vs_MS.NoTrt_B |
| ACADS | 0.10943 | -0.3134 | 0.077 | 0.11 | 1 | HC_vs_MS.NoTrt_B |
| TMEM205 | 0.10949 | -0.3037 | 0.068 | 0.1 | 1 | HC_vs_MS.NoTrt_B |
| DYX1C1 | 0.10973 | 0.37078 | 0.455 | 0.43 | 1 | HC_vs_MS.NoTrt_B |
| CSAD | 0.1098 | 0.62516 | 0.806 | 0.81 | 1 | HC_vs_MS.NoTrt_B |
| ZNF280D | 0.10982 | 0.7452 | 0.214 | 0.17 | 1 | HC_vs_MS.NoTrt_B |
| MEIOC | 0.11026 | 0.3828 | 0.645 | 0.61 | 1 | HC_vs_MS.NoTrt_B |
| DCUN1D4 | 0.11028 | 0.65476 | 0.249 | 0.31 | 1 | HC_vs_MS.NoTrt_B |
| AIMP2 | 0.11048 | -0.8221 | 0.304 | 0.36 | 1 | HC_vs_MS.NoTrt_B |
| ANGPT2 | 0.11055 | 0.40045 | 0.726 | 0.71 | 1 | HC_vs_MS.NoTrt_B |
| DFFB | 0.11066 | 0.33186 | 0.807 | 0.79 | 1 | HC_vs_MS.NoTrt_B |
| USP25 | 0.11069 | 0.52383 | 0.449 | 0.54 | 1 | HC_vs_MS.NoTrt_B |
| ZNF622 | 0.11072 | -0.2527 | 0.081 | 0.12 | 1 | HC_vs_MS.NoTrt_B |
| CCHCR1 | 0.11093 | -0.9863 | 0.293 | 0.36 | 1 | HC_vs_MS.NoTrt_B |

| | | | | | | |
|---|---|---|---|---|---|---|
| DHRS3 | 0.11093 | 0.54036 | 0.229 | 0.18 | 1 | HC_vs_MS.NoTrt_B |
| TUFT1 | 0.11139 | 0.36595 | 0.349 | 0.44 | 1 | HC_vs_MS.NoTrt_B |
| CEP70 | 0.11145 | 0.32712 | 0.123 | 0.17 | 1 | HC_vs_MS.NoTrt_B |
| CLCN2 | 0.11158 | 0.38855 | 0.643 | 0.64 | 1 | HC_vs_MS.NoTrt_B |
| MICAL3 | 0.11205 | 0.3687 | 0.791 | 0.8 | 1 | HC_vs_MS.NoTrt_B |
| CORO7-PAM | 0.11215 | -0.4498 | 0.104 | 0.15 | 1 | HC_vs_MS.NoTrt_B |
| THAP9 | 0.11226 | 0.47455 | 0.587 | 0.6 | 1 | HC_vs_MS.NoTrt_B |
| PHKA2 | 0.11248 | 0.36273 | 0.599 | 0.67 | 1 | HC_vs_MS.NoTrt_B |
| RDH5 | 0.11254 | -0.4448 | 0.442 | 0.51 | 1 | HC_vs_MS.NoTrt_B |
| PCGF1 | 0.11256 | -0.3376 | 0.372 | 0.44 | 1 | HC_vs_MS.NoTrt_B |
| FGD2 | 0.11331 | 0.25568 | 0.703 | 0.71 | 1 | HC_vs_MS.NoTrt_B |
| SEPSECS | 0.11332 | 0.82252 | 0.758 | 0.8 | 1 | HC_vs_MS.NoTrt_B |
| PAFAH1B3 | 0.11347 | -0.3163 | 0.072 | 0.11 | 1 | HC_vs_MS.NoTrt_B |
| NPIPB15 | 0.11358 | -0.3079 | 0.193 | 0.24 | 1 | HC_vs_MS.NoTrt_B |
| KMT5C | 0.11374 | 0.44366 | 0.533 | 0.55 | 1 | HC_vs_MS.NoTrt_B |
| MCMDC2 | 0.11381 | -0.3319 | 0.419 | 0.48 | 1 | HC_vs_MS.NoTrt_B |
| CAMK2G | 0.11384 | 0.25776 | 0.641 | 0.61 | 1 | HC_vs_MS.NoTrt_B |
| ZMYM2 | 0.1141 | -0.369 | 0.351 | 0.39 | 1 | HC_vs_MS.NoTrt_B |
| NTRK1 | 0.11417 | 0.5459 | 0.109 | 0.07 | 1 | HC_vs_MS.NoTrt_B |
| KCTD11 | 0.11422 | -0.2589 | 0.149 | 0.2 | 1 | HC_vs_MS.NoTrt_B |
| NIT1 | 0.11446 | -0.34 | 0.604 | 0.56 | 1 | HC_vs_MS.NoTrt_B |
| SOX13 | 0.11462 | 0.36861 | 0.739 | 0.75 | 1 | HC_vs_MS.NoTrt_B |
| IRAK1BP1 | 0.11465 | 0.27532 | 0.674 | 0.67 | 1 | HC_vs_MS.NoTrt_B |
| FKBP10 | 0.11466 | 0.28514 | 0.345 | 0.29 | 1 | HC_vs_MS.NoTrt_B |
| SNX1 | 0.11479 | 0.3986 | 0.733 | 0.69 | 1 | HC_vs_MS.NoTrt_B |
| CRLS1 | 0.11488 | 0.74968 | 0.417 | 0.36 | 1 | HC_vs_MS.NoTrt_B |
| C1orf127 | 0.11525 | 0.43057 | 0.272 | 0.22 | 1 | HC_vs_MS.NoTrt_B |
| TMEM138 | 0.11528 | 0.37303 | 0.855 | 0.85 | 1 | HC_vs_MS.NoTrt_B |
| ATN1 | 0.1153 | 0.79684 | 0.17 | 0.12 | 1 | HC_vs_MS.NoTrt_B |
| POLR2M | 0.11532 | 0.41464 | 0.167 | 0.12 | 1 | HC_vs_MS.NoTrt_B |
| OLR1 | 0.11537 | 0.35244 | 0.414 | 0.37 | 1 | HC_vs_MS.NoTrt_B |
| UGDH | 0.11544 | 0.40476 | 0.528 | 0.64 | 1 | HC_vs_MS.NoTrt_B |
| NKTR | 0.11564 | 0.45288 | 0.849 | 0.83 | 1 | HC_vs_MS.NoTrt_B |
| BACH2 | 0.11592 | 0.68617 | 0.242 | 0.19 | 1 | HC_vs_MS.NoTrt_B |
| NET1 | 0.11597 | 0.51218 | 0.081 | 0.12 | 1 | HC_vs_MS.NoTrt_B |
| LTBP3 | 0.11653 | 0.4279 | 0.581 | 0.6 | 1 | HC_vs_MS.NoTrt_B |
| SPATA6L | 0.11656 | 0.35667 | 0.781 | 0.78 | 1 | HC_vs_MS.NoTrt_B |
| BEST3 | 0.11668 | 0.28811 | 0.161 | 0.12 | 1 | HC_vs_MS.NoTrt_B |
| PRDX2 | 0.11687 | -0.2767 | 0.212 | 0.26 | 1 | HC_vs_MS.NoTrt_B |
| RNPS1 | 0.11704 | -0.2539 | 0.896 | 0.94 | 1 | HC_vs_MS.NoTrt_B |

| | | | | | | |
|---|---|---|---|---|---|---|
| ZNF574 | 0.11764 | 1.04961 | 0.316 | 0.27 | 1 | HC_vs_MS.NoTrt_B |
| SQRDL | 0.11814 | -0.3251 | 0.133 | 0.18 | 1 | HC_vs_MS.NoTrt_B |
| DENND5A | 0.11835 | 1.33583 | 0.225 | 0.17 | 1 | HC_vs_MS.NoTrt_B |
| TGFB1 | 0.11842 | 1.13585 | 0.316 | 0.27 | 1 | HC_vs_MS.NoTrt_B |
| LRP12 | 0.11845 | 0.29555 | 0.609 | 0.61 | 1 | HC_vs_MS.NoTrt_B |
| TYW5 | 0.1186 | 1.01036 | 0.872 | 0.91 | 1 | HC_vs_MS.NoTrt_B |
| GPAA1 | 0.11868 | 0.26088 | 0.119 | 0.17 | 1 | HC_vs_MS.NoTrt_B |
| ZNF605 | 0.11871 | 0.2599 | 0.884 | 0.86 | 1 | HC_vs_MS.NoTrt_B |
| TMEM204 | 0.11873 | 0.86817 | 0.174 | 0.13 | 1 | HC_vs_MS.NoTrt_B |
| PKD1 | 0.11899 | 1.20872 | 0.436 | 0.38 | 1 | HC_vs_MS.NoTrt_B |
| C1orf106 | 0.11911 | 0.25752 | 0.738 | 0.72 | 1 | HC_vs_MS.NoTrt_B |
| LAMC2 | 0.11922 | 0.26999 | 0.555 | 0.54 | 1 | HC_vs_MS.NoTrt_B |
| RBM7 | 0.11929 | 0.66577 | 0.674 | 0.67 | 1 | HC_vs_MS.NoTrt_B |
| ADRA1A | 0.11951 | 0.33917 | 0.593 | 0.56 | 1 | HC_vs_MS.NoTrt_B |
| CDH3 | 0.1197 | 0.25264 | 0.846 | 0.87 | 1 | HC_vs_MS.NoTrt_B |
| VENTX | 0.12016 | 0.29258 | 0.403 | 0.37 | 1 | HC_vs_MS.NoTrt_B |
| TRMU | 0.12065 | 0.41418 | 0.188 | 0.25 | 1 | HC_vs_MS.NoTrt_B |
| ENDOV | 0.12096 | 1.00245 | 0.577 | 0.54 | 1 | HC_vs_MS.NoTrt_B |
| GAPVD1 | 0.12097 | 0.45535 | 0.819 | 0.88 | 1 | HC_vs_MS.NoTrt_B |
| ZRSR2 | 0.12121 | 0.56164 | 0.739 | 0.76 | 1 | HC_vs_MS.NoTrt_B |
| TBL2 | 0.12125 | 0.30914 | 0.69 | 0.71 | 1 | HC_vs_MS.NoTrt_B |
| BLM | 0.12133 | 0.70401 | 0.239 | 0.2 | 1 | HC_vs_MS.NoTrt_B |
| SERAC1 | 0.1214 | 0.47168 | 0.442 | 0.38 | 1 | HC_vs_MS.NoTrt_B |
| SUGP1 | 0.12142 | -0.3234 | 0.412 | 0.48 | 1 | HC_vs_MS.NoTrt_B |
| SLC25A14 | 0.12143 | -0.3089 | 0.213 | 0.27 | 1 | HC_vs_MS.NoTrt_B |
| GNG4 | 0.12157 | 0.3795 | 0.677 | 0.64 | 1 | HC_vs_MS.NoTrt_B |
| ZNF620 | 0.12187 | 0.49436 | 0.716 | 0.72 | 1 | HC_vs_MS.NoTrt_B |
| MTIF3 | 0.12215 | -0.3882 | 0.183 | 0.23 | 1 | HC_vs_MS.NoTrt_B |
| HNF1B | 0.12262 | 0.37943 | 0.17 | 0.12 | 1 | HC_vs_MS.NoTrt_B |
| TPCN1 | 0.12262 | 0.3391 | 0.42 | 0.5 | 1 | HC_vs_MS.NoTrt_B |
| JAZF1 | 0.12281 | 0.88381 | 0.086 | 0.12 | 1 | HC_vs_MS.NoTrt_B |
| PTCHD4 | 0.12281 | 0.37632 | 0.168 | 0.12 | 1 | HC_vs_MS.NoTrt_B |
| IQSEC1 | 0.12284 | 1.36014 | 0.254 | 0.21 | 1 | HC_vs_MS.NoTrt_B |
| USP30 | 0.12306 | -0.2689 | 0.267 | 0.33 | 1 | HC_vs_MS.NoTrt_B |
| MYT1L | 0.12317 | 0.26657 | 0.623 | 0.58 | 1 | HC_vs_MS.NoTrt_B |
| SLC13A5 | 0.12342 | 0.29422 | 0.746 | 0.77 | 1 | HC_vs_MS.NoTrt_B |
| ACAD11 | 0.12364 | -0.3676 | 0.184 | 0.23 | 1 | HC_vs_MS.NoTrt_B |
| PPRC1 | 0.12366 | 1.25316 | 0.113 | 0.08 | 1 | HC_vs_MS.NoTrt_B |
| GGCX | 0.12388 | 0.78852 | 0.891 | 0.94 | 1 | HC_vs_MS.NoTrt_B |
| CFAP69 | 0.12396 | 0.34124 | 0.801 | 0.86 | 1 | HC_vs_MS.NoTrt_B |

| | | | | | | |
|---|---|---|---|---|---|---|
| SERINC5 | 0.12433 | 0.43447 | 0.899 | 0.89 | 1 | HC_vs_MS.NoTrt_B |
| RPL26L1 | 0.1244 | -0.356 | 0.112 | 0.15 | 1 | HC_vs_MS.NoTrt_B |
| PSPH | 0.12452 | 0.36114 | 0.83 | 0.82 | 1 | HC_vs_MS.NoTrt_B |
| PEX13 | 0.12472 | 0.43304 | 0.762 | 0.76 | 1 | HC_vs_MS.NoTrt_B |
| ZNF470 | 0.125 | 0.81894 | 0.714 | 0.74 | 1 | HC_vs_MS.NoTrt_B |
| TDGF1 | 0.1251 | 0.35173 | 0.516 | 0.5 | 1 | HC_vs_MS.NoTrt_B |
| TEX10 | 0.12518 | -0.4742 | 0.465 | 0.52 | 1 | HC_vs_MS.NoTrt_B |
| SLK | 0.12534 | 0.8597 | 0.196 | 0.15 | 1 | HC_vs_MS.NoTrt_B |
| ISCA2 | 0.12564 | -0.2584 | 0.467 | 0.54 | 1 | HC_vs_MS.NoTrt_B |
| PDLIM5 | 0.12566 | 0.69722 | 0.872 | 0.88 | 1 | HC_vs_MS.NoTrt_B |
| HSPE1 | 0.12572 | 0.28488 | 0.23 | 0.29 | 1 | HC_vs_MS.NoTrt_B |
| NCOA7 | 0.12576 | 0.49713 | 0.745 | 0.74 | 1 | HC_vs_MS.NoTrt_B |
| TIGD6 | 0.12584 | 1.65883 | 0.188 | 0.14 | 1 | HC_vs_MS.NoTrt_B |
| RTN4 | 0.12623 | 0.69711 | 0.264 | 0.22 | 1 | HC_vs_MS.NoTrt_B |
| STX3 | 0.12653 | 0.39344 | 0.377 | 0.34 | 1 | HC_vs_MS.NoTrt_B |
| PHF7 | 0.12692 | -0.68 | 0.459 | 0.54 | 1 | HC_vs_MS.NoTrt_B |
| ADGRF1 | 0.12706 | 0.35474 | 0.775 | 0.79 | 1 | HC_vs_MS.NoTrt_B |
| AMMECR1 | 0.12712 | 0.28148 | 0.132 | 0.18 | 1 | HC_vs_MS.NoTrt_B |
| MMS22L | 0.12718 | 0.3698 | 0.909 | 0.92 | 1 | HC_vs_MS.NoTrt_B |
| MED17 | 0.12721 | 0.26665 | 0.945 | 0.97 | 1 | HC_vs_MS.NoTrt_B |
| DUSP12 | 0.12723 | -0.357 | 0.291 | 0.35 | 1 | HC_vs_MS.NoTrt_B |
| KIAA0408 | 0.12729 | 0.34496 | 0.27 | 0.22 | 1 | HC_vs_MS.NoTrt_B |
| MON1B | 0.12746 | 0.32743 | 0.667 | 0.61 | 1 | HC_vs_MS.NoTrt_B |
| FN3KRP | 0.12749 | -0.3525 | 0.07 | 0.1 | 1 | HC_vs_MS.NoTrt_B |
| MPI | 0.1275 | -0.4876 | 0.513 | 0.54 | 1 | HC_vs_MS.NoTrt_B |
| PHF20 | 0.12756 | 0.31084 | 0.478 | 0.56 | 1 | HC_vs_MS.NoTrt_B |
| CD164 | 0.12767 | 0.3211 | 0.63 | 0.69 | 1 | HC_vs_MS.NoTrt_B |
| TMEM169 | 0.12772 | -0.3286 | 0.488 | 0.6 | 1 | HC_vs_MS.NoTrt_B |
| MVB12A | 0.12801 | -0.3879 | 0.514 | 0.6 | 1 | HC_vs_MS.NoTrt_B |
| ABCB7 | 0.12806 | 0.66995 | 0.613 | 0.7 | 1 | HC_vs_MS.NoTrt_B |
| SLC23A2 | 0.12834 | -0.2851 | 0.13 | 0.17 | 1 | HC_vs_MS.NoTrt_B |
| PXYLP1 | 0.12839 | 1.32187 | 0.5 | 0.47 | 1 | HC_vs_MS.NoTrt_B |
| GTF2H4 | 0.12844 | -1.032 | 0.07 | 0.1 | 1 | HC_vs_MS.NoTrt_B |
| TBC1D9B | 0.12847 | -0.3589 | 0.457 | 0.51 | 1 | HC_vs_MS.NoTrt_B |
| ZBTB38 | 0.12849 | 0.52149 | 0.535 | 0.61 | 1 | HC_vs_MS.NoTrt_B |
| MPPE1 | 0.1286 | -0.3003 | 0.849 | 0.87 | 1 | HC_vs_MS.NoTrt_B |
| PPIP5K2 | 0.12868 | 0.68727 | 0.809 | 0.86 | 1 | HC_vs_MS.NoTrt_B |
| GPRC5A | 0.12911 | 0.42101 | 0.613 | 0.6 | 1 | HC_vs_MS.NoTrt_B |
| KIF6 | 0.12929 | 0.32141 | 0.849 | 0.9 | 1 | HC_vs_MS.NoTrt_B |
| LPAR3 | 0.12938 | 0.26472 | 0.393 | 0.35 | 1 | HC_vs_MS.NoTrt_B |

| | | | | | | |
|---|---|---|---|---|---|---|
| SLC9A4 | 0.12986 | 0.44562 | 0.483 | 0.47 | 1 | HC_vs_MS.NoTrt_B |
| LRRC37B | 0.12989 | 0.33369 | 0.619 | 0.59 | 1 | HC_vs_MS.NoTrt_B |
| SDSL | 0.1299 | 0.74129 | 0.328 | 0.29 | 1 | HC_vs_MS.NoTrt_B |
| TNFRSF10A | 0.12998 | 0.39392 | 0.567 | 0.51 | 1 | HC_vs_MS.NoTrt_B |
| TM9SF1 | 0.13008 | -0.2748 | 0.232 | 0.29 | 1 | HC_vs_MS.NoTrt_B |
| SLC4A1 | 0.13012 | 0.32668 | 0.671 | 0.67 | 1 | HC_vs_MS.NoTrt_B |
| KLHL21 | 0.13039 | -0.6143 | 0.646 | 0.7 | 1 | HC_vs_MS.NoTrt_B |
| TXLNG | 0.1304 | 1.34226 | 0.119 | 0.08 | 1 | HC_vs_MS.NoTrt_B |
| ARID3B | 0.13047 | 0.26767 | 0.712 | 0.69 | 1 | HC_vs_MS.NoTrt_B |
| MROH1 | 0.13058 | 0.26195 | 0.499 | 0.43 | 1 | HC_vs_MS.NoTrt_B |
| ZNF121 | 0.13074 | 0.25619 | 0.878 | 0.9 | 1 | HC_vs_MS.NoTrt_B |
| CLASP2 | 0.13092 | 0.88919 | 0.7 | 0.71 | 1 | HC_vs_MS.NoTrt_B |
| RIN1 | 0.13092 | -0.4424 | 0.197 | 0.25 | 1 | HC_vs_MS.NoTrt_B |
| KIN | 0.13121 | 0.31301 | 0.72 | 0.74 | 1 | HC_vs_MS.NoTrt_B |
| WNT2B | 0.13124 | 0.83225 | 0.455 | 0.41 | 1 | HC_vs_MS.NoTrt_B |
| PLEKHD1 | 0.13128 | 0.38255 | 0.706 | 0.7 | 1 | HC_vs_MS.NoTrt_B |
| BCL7A | 0.13137 | 0.39502 | 0.649 | 0.64 | 1 | HC_vs_MS.NoTrt_B |
| TRPM8 | 0.13155 | 0.25669 | 0.659 | 0.67 | 1 | HC_vs_MS.NoTrt_B |
| SELK | 0.13159 | -0.3247 | 0.342 | 0.38 | 1 | HC_vs_MS.NoTrt_B |
| RYK | 0.13161 | 1.16785 | 0.071 | 0.11 | 1 | HC_vs_MS.NoTrt_B |
| FAH | 0.13174 | 0.39612 | 0.141 | 0.19 | 1 | HC_vs_MS.NoTrt_B |
| SLC36A2 | 0.13178 | 0.3551 | 0.619 | 0.67 | 1 | HC_vs_MS.NoTrt_B |
| INTS6 | 0.13203 | 0.32651 | 0.928 | 0.94 | 1 | HC_vs_MS.NoTrt_B |
| FCRL5 | 0.13206 | 0.53365 | 0.472 | 0.46 | 1 | HC_vs_MS.NoTrt_B |
| DEDD | 0.13211 | 0.28961 | 0.125 | 0.17 | 1 | HC_vs_MS.NoTrt_B |
| TNFRSF11A | 0.13213 | 0.88842 | 0.228 | 0.18 | 1 | HC_vs_MS.NoTrt_B |
| PAXIP1 | 0.13237 | -0.5899 | 0.435 | 0.5 | 1 | HC_vs_MS.NoTrt_B |
| DECR2 | 0.13253 | 0.52514 | 0.801 | 0.8 | 1 | HC_vs_MS.NoTrt_B |
| DRC3 | 0.13262 | 0.40144 | 0.591 | 0.54 | 1 | HC_vs_MS.NoTrt_B |
| OSMR | 0.13298 | 0.39469 | 0.49 | 0.46 | 1 | HC_vs_MS.NoTrt_B |
| RNF144A | 0.13315 | 0.89957 | 0.196 | 0.15 | 1 | HC_vs_MS.NoTrt_B |
| FOSB | 0.13328 | 0.4701 | 0.186 | 0.24 | 1 | HC_vs_MS.NoTrt_B |
| CTD-3105H1 | 0.13336 | 0.48399 | 0.293 | 0.24 | 1 | HC_vs_MS.NoTrt_B |
| TRMT10B | 0.1334 | 0.55296 | 0.851 | 0.87 | 1 | HC_vs_MS.NoTrt_B |
| ZCCHC10 | 0.1335 | 0.35432 | 0.586 | 0.56 | 1 | HC_vs_MS.NoTrt_B |
| EPB41 | 0.1335 | 0.57068 | 0.454 | 0.59 | 1 | HC_vs_MS.NoTrt_B |
| GATA3 | 0.1339 | 0.4762 | 0.246 | 0.32 | 1 | HC_vs_MS.NoTrt_B |
| AC138517.6 | 0.1347 | 0.97104 | 0.081 | 0.12 | 1 | HC_vs_MS.NoTrt_B |
| ATP5A1 | 0.13473 | -0.3069 | 0.996 | 1 | 1 | HC_vs_MS.NoTrt_B |
| DENND1A | 0.13482 | 0.55098 | 0.435 | 0.38 | 1 | HC_vs_MS.NoTrt_B |

| | | | | | | |
|---|---|---|---|---|---|---|
| MGAT5 | 0.13489 | -0.4794 | 0.143 | 0.18 | 1 | HC_vs_MS.NoTrt_B |
| MS4A2 | 0.1349 | 0.2635 | 0.555 | 0.62 | 1 | HC_vs_MS.NoTrt_B |
| CACNG8 | 0.13498 | 0.2597 | 0.751 | 0.76 | 1 | HC_vs_MS.NoTrt_B |
| FAM193A | 0.13499 | 0.99963 | 0.332 | 0.29 | 1 | HC_vs_MS.NoTrt_B |
| ATRNL1 | 0.13513 | 0.31156 | 0.354 | 0.33 | 1 | HC_vs_MS.NoTrt_B |
| TTPAL | 0.13517 | 0.35723 | 0.791 | 0.8 | 1 | HC_vs_MS.NoTrt_B |
| CD24 | 0.13529 | 0.35441 | 0.352 | 0.3 | 1 | HC_vs_MS.NoTrt_B |
| SH3PXD2A | 0.13535 | 0.34975 | 0.314 | 0.27 | 1 | HC_vs_MS.NoTrt_B |
| TMEM132A | 0.13539 | 0.50486 | 0.384 | 0.37 | 1 | HC_vs_MS.NoTrt_B |
| MRRF | 0.13546 | 0.74483 | 0.625 | 0.6 | 1 | HC_vs_MS.NoTrt_B |
| TACO1 | 0.13558 | 1.08993 | 0.37 | 0.32 | 1 | HC_vs_MS.NoTrt_B |
| GOLGA5 | 0.13615 | -0.2527 | 0.084 | 0.12 | 1 | HC_vs_MS.NoTrt_B |
| SAA2 | 0.13637 | 0.3181 | 0.396 | 0.35 | 1 | HC_vs_MS.NoTrt_B |
| UBAP2 | 0.13669 | 0.63851 | 0.686 | 0.69 | 1 | HC_vs_MS.NoTrt_B |
| AIM1 | 0.13682 | -0.2775 | 0.486 | 0.55 | 1 | HC_vs_MS.NoTrt_B |
| ZBTB33 | 0.13685 | 0.3886 | 0.083 | 0.12 | 1 | HC_vs_MS.NoTrt_B |
| SLC8A1 | 0.137 | -0.3837 | 0.304 | 0.24 | 1 | HC_vs_MS.NoTrt_B |
| NSMCE4A | 0.13712 | -0.4346 | 0.32 | 0.37 | 1 | HC_vs_MS.NoTrt_B |
| SARNP | 0.13756 | -0.277 | 0.559 | 0.61 | 1 | HC_vs_MS.NoTrt_B |
| PPP1CC | 0.13769 | 0.38871 | 0.764 | 0.75 | 1 | HC_vs_MS.NoTrt_B |
| NRIP3 | 0.13773 | 0.34316 | 0.483 | 0.45 | 1 | HC_vs_MS.NoTrt_B |
| DOCK10 | 0.1379 | 0.31805 | 0.599 | 0.68 | 1 | HC_vs_MS.NoTrt_B |
| NOM1 | 0.13872 | 0.37595 | 0.614 | 0.58 | 1 | HC_vs_MS.NoTrt_B |
| KLHL12 | 0.13878 | -0.2803 | 0.075 | 0.11 | 1 | HC_vs_MS.NoTrt_B |
| MRPS2 | 0.13882 | -0.2519 | 0.074 | 0.11 | 1 | HC_vs_MS.NoTrt_B |
| DHX38 | 0.13928 | -0.2518 | 0.08 | 0.11 | 1 | HC_vs_MS.NoTrt_B |
| FAM186A | 0.13929 | 0.40679 | 0.407 | 0.37 | 1 | HC_vs_MS.NoTrt_B |
| SLC35D1 | 0.13934 | 0.81142 | 0.27 | 0.35 | 1 | HC_vs_MS.NoTrt_B |
| TMEM150C | 0.13944 | 0.28838 | 0.541 | 0.5 | 1 | HC_vs_MS.NoTrt_B |
| BRDT | 0.13964 | 0.28568 | 0.546 | 0.52 | 1 | HC_vs_MS.NoTrt_B |
| WBP2NL | 0.14009 | 0.56909 | 0.384 | 0.34 | 1 | HC_vs_MS.NoTrt_B |
| MCTP2 | 0.1401 | 0.62349 | 0.426 | 0.41 | 1 | HC_vs_MS.NoTrt_B |
| GSTM3 | 0.14023 | 0.6664 | 0.506 | 0.46 | 1 | HC_vs_MS.NoTrt_B |
| NOL12 | 0.14032 | -0.2805 | 0.483 | 0.42 | 1 | HC_vs_MS.NoTrt_B |
| SP2 | 0.14046 | 0.40724 | 0.3 | 0.37 | 1 | HC_vs_MS.NoTrt_B |
| MT-CYB | 0.14058 | 0.32931 | 0.738 | 0.76 | 1 | HC_vs_MS.NoTrt_B |
| SMARCB1 | 0.14074 | -0.2654 | 0.436 | 0.47 | 1 | HC_vs_MS.NoTrt_B |
| FAM136A | 0.14146 | -0.3518 | 0.288 | 0.35 | 1 | HC_vs_MS.NoTrt_B |
| STAMBPL1 | 0.14168 | 0.4541 | 0.507 | 0.48 | 1 | HC_vs_MS.NoTrt_B |
| CTCFL | 0.14184 | 0.2698 | 0.825 | 0.87 | 1 | HC_vs_MS.NoTrt_B |

| | | | | | | |
|---|---|---|---|---|---|---|
| HLA-DRA | 0.14209 | -0.8158 | 0.116 | 0.15 | 1 | HC_vs_MS.NoTrt_B |
| FMNL2 | 0.14222 | -0.2585 | 0.248 | 0.3 | 1 | HC_vs_MS.NoTrt_B |
| SMDT1 | 0.14251 | 0.31262 | 0.21 | 0.27 | 1 | HC_vs_MS.NoTrt_B |
| CATSPERG | 0.14265 | 0.28215 | 0.893 | 0.91 | 1 | HC_vs_MS.NoTrt_B |
| NME9 | 0.14271 | 0.90198 | 0.201 | 0.16 | 1 | HC_vs_MS.NoTrt_B |
| PARK2 | 0.14272 | 0.67599 | 0.604 | 0.57 | 1 | HC_vs_MS.NoTrt_B |
| FAAP20 | 0.14298 | 0.63827 | 0.865 | 0.86 | 1 | HC_vs_MS.NoTrt_B |
| AL078584.1 | 0.14312 | -0.4255 | 0.409 | 0.47 | 1 | HC_vs_MS.NoTrt_B |
| LDHC | 0.14314 | 0.35526 | 0.184 | 0.14 | 1 | HC_vs_MS.NoTrt_B |
| THOP1 | 0.14353 | -0.5165 | 0.332 | 0.41 | 1 | HC_vs_MS.NoTrt_B |
| AURKB | 0.14359 | -0.3822 | 0.423 | 0.38 | 1 | HC_vs_MS.NoTrt_B |
| PSTPIP2 | 0.14383 | -0.283 | 0.342 | 0.44 | 1 | HC_vs_MS.NoTrt_B |
| PSMG3 | 0.14391 | 0.27887 | 0.593 | 0.69 | 1 | HC_vs_MS.NoTrt_B |
| PCDHGA6 | 0.14457 | 0.26064 | 0.729 | 0.78 | 1 | HC_vs_MS.NoTrt_B |
| DDX54 | 0.14457 | -0.2509 | 0.099 | 0.14 | 1 | HC_vs_MS.NoTrt_B |
| GRAMD1C | 0.14564 | 0.44523 | 0.184 | 0.14 | 1 | HC_vs_MS.NoTrt_B |
| CEP126 | 0.14566 | 0.66726 | 0.726 | 0.73 | 1 | HC_vs_MS.NoTrt_B |
| HNF4G | 0.14578 | 0.35405 | 0.333 | 0.3 | 1 | HC_vs_MS.NoTrt_B |
| PHGDH | 0.14599 | 0.34479 | 0.241 | 0.2 | 1 | HC_vs_MS.NoTrt_B |
| KMT2A | 0.14632 | 0.41219 | 0.943 | 0.93 | 1 | HC_vs_MS.NoTrt_B |
| PRR14 | 0.14641 | 0.73818 | 0.114 | 0.16 | 1 | HC_vs_MS.NoTrt_B |
| EML1 | 0.1465 | 0.31984 | 0.719 | 0.75 | 1 | HC_vs_MS.NoTrt_B |
| PILRB | 0.14672 | -0.6662 | 0.419 | 0.44 | 1 | HC_vs_MS.NoTrt_B |
| ZMYM1 | 0.14685 | 1.06021 | 0.549 | 0.51 | 1 | HC_vs_MS.NoTrt_B |
| LMBR1 | 0.14696 | 0.56427 | 0.507 | 0.48 | 1 | HC_vs_MS.NoTrt_B |
| P2RX7 | 0.14703 | 0.30807 | 0.749 | 0.75 | 1 | HC_vs_MS.NoTrt_B |
| ZGRF1 | 0.14703 | 0.75681 | 0.755 | 0.78 | 1 | HC_vs_MS.NoTrt_B |
| PJA2 | 0.14788 | 0.47116 | 0.219 | 0.28 | 1 | HC_vs_MS.NoTrt_B |
| C3orf62 | 0.1482 | 0.7283 | 0.758 | 0.77 | 1 | HC_vs_MS.NoTrt_B |
| SFXN2 | 0.14822 | 0.63162 | 0.935 | 0.9 | 1 | HC_vs_MS.NoTrt_B |
| NIPAL3 | 0.14824 | 0.2991 | 0.765 | 0.77 | 1 | HC_vs_MS.NoTrt_B |
| PMPCA | 0.14842 | -0.2829 | 0.675 | 0.76 | 1 | HC_vs_MS.NoTrt_B |
| DTWD2 | 0.14858 | 0.37699 | 0.743 | 0.76 | 1 | HC_vs_MS.NoTrt_B |
| NR1H3 | 0.14875 | 0.88006 | 0.112 | 0.15 | 1 | HC_vs_MS.NoTrt_B |
| MAGOHB | 0.14886 | 0.76436 | 0.078 | 0.11 | 1 | HC_vs_MS.NoTrt_B |
| DMWD | 0.14905 | 0.34961 | 0.836 | 0.85 | 1 | HC_vs_MS.NoTrt_B |
| MTCH2 | 0.14913 | -0.4648 | 0.1 | 0.14 | 1 | HC_vs_MS.NoTrt_B |
| NFKB2 | 0.14971 | 0.37178 | 0.341 | 0.43 | 1 | HC_vs_MS.NoTrt_B |
| ACSF2 | 0.14982 | -0.3476 | 0.47 | 0.44 | 1 | HC_vs_MS.NoTrt_B |
| AFAP1 | 0.14991 | 0.54864 | 0.391 | 0.35 | 1 | HC_vs_MS.NoTrt_B |

| | | | | | | |
|---|---|---|---|---|---|---|
| CALB1 | 0.14996 | 0.44443 | 0.151 | 0.11 | 1 | HC_vs_MS.NoTrt_B |
| IKZF1 | 0.15009 | 0.27161 | 0.348 | 0.43 | 1 | HC_vs_MS.NoTrt_B |
| ZNF592 | 0.1501 | 0.34089 | 0.167 | 0.22 | 1 | HC_vs_MS.NoTrt_B |
| RIOK2 | 0.15042 | 0.79101 | 0.319 | 0.27 | 1 | HC_vs_MS.NoTrt_B |
| GOLGA8R | 0.15058 | 0.4096 | 0.439 | 0.53 | 1 | HC_vs_MS.NoTrt_B |
| SPECC1L | 0.1506 | 0.87681 | 0.136 | 0.1 | 1 | HC_vs_MS.NoTrt_B |
| IZUMO1 | 0.15071 | 0.26006 | 0.628 | 0.61 | 1 | HC_vs_MS.NoTrt_B |
| MRPL57 | 0.15076 | -0.2576 | 0.672 | 0.71 | 1 | HC_vs_MS.NoTrt_B |
| GALNT4 | 0.15103 | -0.3051 | 0.386 | 0.44 | 1 | HC_vs_MS.NoTrt_B |
| GGA1 | 0.15134 | 0.42988 | 0.968 | 0.97 | 1 | HC_vs_MS.NoTrt_B |
| KMT5B | 0.15139 | 0.63144 | 0.294 | 0.24 | 1 | HC_vs_MS.NoTrt_B |
| RAB14 | 0.15203 | 0.50379 | 0.201 | 0.26 | 1 | HC_vs_MS.NoTrt_B |
| DPH3 | 0.15205 | 0.71034 | 0.238 | 0.19 | 1 | HC_vs_MS.NoTrt_B |
| SLC9A7 | 0.15245 | 0.39646 | 0.938 | 0.94 | 1 | HC_vs_MS.NoTrt_B |
| TXNL1 | 0.15275 | 0.3442 | 0.629 | 0.71 | 1 | HC_vs_MS.NoTrt_B |
| NKD1 | 0.15299 | 0.25577 | 0.394 | 0.49 | 1 | HC_vs_MS.NoTrt_B |
| TMEM130 | 0.15309 | 0.31416 | 0.661 | 0.67 | 1 | HC_vs_MS.NoTrt_B |
| DST | 0.15313 | 0.49404 | 0.755 | 0.79 | 1 | HC_vs_MS.NoTrt_B |
| TMEM64 | 0.15319 | 0.71936 | 0.358 | 0.32 | 1 | HC_vs_MS.NoTrt_B |
| MAPK13 | 0.15321 | -0.3601 | 0.638 | 0.71 | 1 | HC_vs_MS.NoTrt_B |
| TEX35 | 0.15373 | 0.28458 | 0.312 | 0.27 | 1 | HC_vs_MS.NoTrt_B |
| BMP1 | 0.15383 | 0.35604 | 0.535 | 0.51 | 1 | HC_vs_MS.NoTrt_B |
| RP11-9J18.1 | 0.15387 | 0.54045 | 0.262 | 0.22 | 1 | HC_vs_MS.NoTrt_B |
| ACSM2A | 0.1539 | 0.47639 | 0.467 | 0.49 | 1 | HC_vs_MS.NoTrt_B |
| EPT1 | 0.15399 | 0.27037 | 0.336 | 0.4 | 1 | HC_vs_MS.NoTrt_B |
| MGMT | 0.15402 | -0.3011 | 0.071 | 0.1 | 1 | HC_vs_MS.NoTrt_B |
| KLHL26 | 0.15416 | 0.92262 | 0.574 | 0.57 | 1 | HC_vs_MS.NoTrt_B |
| MAATS1 | 0.15452 | 0.40674 | 0.2 | 0.25 | 1 | HC_vs_MS.NoTrt_B |
| TXNL4A | 0.15466 | -0.6112 | 0.471 | 0.52 | 1 | HC_vs_MS.NoTrt_B |
| USP33 | 0.15536 | 0.2911 | 0.945 | 0.94 | 1 | HC_vs_MS.NoTrt_B |
| TAF4 | 0.15539 | 0.36476 | 0.123 | 0.16 | 1 | HC_vs_MS.NoTrt_B |
| ZNF672 | 0.15604 | -0.4826 | 0.236 | 0.28 | 1 | HC_vs_MS.NoTrt_B |
| CHD8 | 0.1562 | 0.41777 | 0.365 | 0.31 | 1 | HC_vs_MS.NoTrt_B |
| NFASC | 0.15647 | 0.42038 | 0.584 | 0.59 | 1 | HC_vs_MS.NoTrt_B |
| ZNF133 | 0.15679 | -0.2831 | 0.449 | 0.41 | 1 | HC_vs_MS.NoTrt_B |
| NDUFAF5 | 0.15684 | 0.62797 | 0.538 | 0.54 | 1 | HC_vs_MS.NoTrt_B |
| AUH | 0.15801 | -0.2517 | 0.09 | 0.12 | 1 | HC_vs_MS.NoTrt_B |
| CDKL1 | 0.15802 | 0.42444 | 0.704 | 0.68 | 1 | HC_vs_MS.NoTrt_B |
| GMCL1 | 0.15821 | 0.55972 | 0.32 | 0.38 | 1 | HC_vs_MS.NoTrt_B |
| WDR82 | 0.15824 | -0.3892 | 0.384 | 0.44 | 1 | HC_vs_MS.NoTrt_B |

| | | | | | | |
|---|---|---|---|---|---|---|
| ABCB5 | 0.15846 | 0.28348 | 0.583 | 0.55 | 1 | HC_vs_MS.NoTrt_B |
| TRDMT1 | 0.1591 | 0.71832 | 0.726 | 0.74 | 1 | HC_vs_MS.NoTrt_B |
| CEP170 | 0.15925 | 0.4836 | 0.549 | 0.63 | 1 | HC_vs_MS.NoTrt_B |
| SIL1 | 0.15937 | -0.6804 | 0.077 | 0.11 | 1 | HC_vs_MS.NoTrt_B |
| RDX | 0.15942 | -0.3105 | 0.236 | 0.29 | 1 | HC_vs_MS.NoTrt_B |
| THRA | 0.15943 | 1.17669 | 0.142 | 0.1 | 1 | HC_vs_MS.NoTrt_B |
| TMEM267 | 0.15951 | 0.53243 | 0.286 | 0.23 | 1 | HC_vs_MS.NoTrt_B |
| LSG1 | 0.15984 | -0.5265 | 0.361 | 0.42 | 1 | HC_vs_MS.NoTrt_B |
| CCDC27 | 0.1599 | 0.29889 | 0.564 | 0.54 | 1 | HC_vs_MS.NoTrt_B |
| MTM1 | 0.16062 | -0.3775 | 0.135 | 0.17 | 1 | HC_vs_MS.NoTrt_B |
| KLF3 | 0.16073 | 0.2871 | 0.639 | 0.74 | 1 | HC_vs_MS.NoTrt_B |
| NIP7 | 0.16085 | -0.5556 | 0.606 | 0.66 | 1 | HC_vs_MS.NoTrt_B |
| MYO15A | 0.161 | 0.28321 | 0.781 | 0.82 | 1 | HC_vs_MS.NoTrt_B |
| OLFML1 | 0.16108 | 0.4545 | 0.225 | 0.28 | 1 | HC_vs_MS.NoTrt_B |
| USP42 | 0.16135 | 0.44839 | 0.449 | 0.41 | 1 | HC_vs_MS.NoTrt_B |
| MZF1 | 0.16145 | 0.42334 | 0.084 | 0.12 | 1 | HC_vs_MS.NoTrt_B |
| ADAMTS16 | 0.16151 | 0.55923 | 0.323 | 0.29 | 1 | HC_vs_MS.NoTrt_B |
| CCDC155 | 0.16162 | -0.4621 | 0.117 | 0.16 | 1 | HC_vs_MS.NoTrt_B |
| NBPF11 | 0.16192 | 0.42472 | 0.257 | 0.32 | 1 | HC_vs_MS.NoTrt_B |
| GRPEL2 | 0.16197 | 0.33686 | 0.294 | 0.34 | 1 | HC_vs_MS.NoTrt_B |
| CRBN | 0.16204 | 0.44642 | 0.467 | 0.55 | 1 | HC_vs_MS.NoTrt_B |
| SLA2 | 0.16225 | -0.3711 | 0.522 | 0.58 | 1 | HC_vs_MS.NoTrt_B |
| SUGP2 | 0.16249 | -0.3902 | 0.709 | 0.78 | 1 | HC_vs_MS.NoTrt_B |
| GNS | 0.16283 | 0.82762 | 0.739 | 0.72 | 1 | HC_vs_MS.NoTrt_B |
| EPHA4 | 0.16295 | -0.3299 | 0.184 | 0.23 | 1 | HC_vs_MS.NoTrt_B |
| DNHD1 | 0.16339 | 0.90922 | 0.316 | 0.27 | 1 | HC_vs_MS.NoTrt_B |
| FLOT1 | 0.16341 | 0.70843 | 0.123 | 0.09 | 1 | HC_vs_MS.NoTrt_B |
| COQ9 | 0.1642 | -0.4364 | 0.072 | 0.1 | 1 | HC_vs_MS.NoTrt_B |
| KIDINS220 | 0.16428 | 0.30974 | 0.272 | 0.22 | 1 | HC_vs_MS.NoTrt_B |
| WDFY3 | 0.16434 | 0.25606 | 0.261 | 0.22 | 1 | HC_vs_MS.NoTrt_B |
| BRIX1 | 0.16453 | 0.3431 | 0.665 | 0.61 | 1 | HC_vs_MS.NoTrt_B |
| KCNN2 | 0.16455 | 0.3781 | 0.255 | 0.22 | 1 | HC_vs_MS.NoTrt_B |
| BMP8B | 0.16472 | 0.43277 | 0.403 | 0.37 | 1 | HC_vs_MS.NoTrt_B |
| CFAP36 | 0.16474 | 0.35868 | 0.412 | 0.38 | 1 | HC_vs_MS.NoTrt_B |
| ZNF543 | 0.1648 | 0.5549 | 0.477 | 0.45 | 1 | HC_vs_MS.NoTrt_B |
| TSC1 | 0.16483 | -0.3599 | 0.478 | 0.55 | 1 | HC_vs_MS.NoTrt_B |
| PDE7B | 0.16488 | 0.47923 | 0.381 | 0.34 | 1 | HC_vs_MS.NoTrt_B |
| LATS1 | 0.16491 | 0.42347 | 0.654 | 0.74 | 1 | HC_vs_MS.NoTrt_B |
| PAK6 | 0.1655 | 0.55456 | 0.155 | 0.11 | 1 | HC_vs_MS.NoTrt_B |
| CRCP | 0.16558 | -0.4253 | 0.862 | 0.89 | 1 | HC_vs_MS.NoTrt_B |

| | | | | | | |
|---|---|---|---|---|---|---|
| DONSON | 0.16564 | -0.7036 | 0.154 | 0.2 | 1 | HC_vs_MS.NoTrt_B |
| ZNF330 | 0.16569 | -0.2946 | 0.072 | 0.1 | 1 | HC_vs_MS.NoTrt_B |
| IGSF9 | 0.16577 | 0.42872 | 0.304 | 0.27 | 1 | HC_vs_MS.NoTrt_B |
| RP5-994D16 | 0.16601 | 0.35183 | 0.462 | 0.57 | 1 | HC_vs_MS.NoTrt_B |
| DDX55 | 0.16605 | 0.50224 | 0.641 | 0.71 | 1 | HC_vs_MS.NoTrt_B |
| TRIM33 | 0.16607 | 0.32576 | 0.288 | 0.34 | 1 | HC_vs_MS.NoTrt_B |
| TOB1 | 0.16617 | 0.43017 | 0.486 | 0.48 | 1 | HC_vs_MS.NoTrt_B |
| MSH3 | 0.16643 | 0.48567 | 0.42 | 0.4 | 1 | HC_vs_MS.NoTrt_B |
| DEPDC5 | 0.16674 | -0.4896 | 0.783 | 0.77 | 1 | HC_vs_MS.NoTrt_B |
| MEI1 | 0.1675 | -0.5947 | 0.725 | 0.76 | 1 | HC_vs_MS.NoTrt_B |
| SCHIP1 | 0.16758 | 0.50134 | 0.532 | 0.53 | 1 | HC_vs_MS.NoTrt_B |
| RP11-544M: | 0.16804 | -0.5713 | 0.219 | 0.18 | 1 | HC_vs_MS.NoTrt_B |
| CNN1 | 0.16808 | 0.3363 | 0.543 | 0.54 | 1 | HC_vs_MS.NoTrt_B |
| DNAH1 | 0.16861 | 1.01269 | 0.143 | 0.11 | 1 | HC_vs_MS.NoTrt_B |
| NBPF3 | 0.16908 | 1.06006 | 0.451 | 0.44 | 1 | HC_vs_MS.NoTrt_B |
| MLH3 | 0.1691 | 0.48592 | 0.086 | 0.12 | 1 | HC_vs_MS.NoTrt_B |
| HYAL1 | 0.1692 | 0.50841 | 0.355 | 0.32 | 1 | HC_vs_MS.NoTrt_B |
| GSK3B | 0.16924 | 0.28647 | 0.712 | 0.72 | 1 | HC_vs_MS.NoTrt_B |
| VIPR2 | 0.16925 | 0.28928 | 0.241 | 0.2 | 1 | HC_vs_MS.NoTrt_B |
| OPRM1 | 0.16945 | 0.303 | 0.836 | 0.89 | 1 | HC_vs_MS.NoTrt_B |
| AGBL2 | 0.16945 | 0.75102 | 0.299 | 0.25 | 1 | HC_vs_MS.NoTrt_B |
| AAR2 | 0.17006 | 0.86824 | 0.249 | 0.21 | 1 | HC_vs_MS.NoTrt_B |
| RNF20 | 0.17014 | -0.3771 | 0.091 | 0.12 | 1 | HC_vs_MS.NoTrt_B |
| ZNF589 | 0.17021 | -0.4921 | 0.459 | 0.52 | 1 | HC_vs_MS.NoTrt_B |
| CDK19 | 0.17022 | 1.53721 | 0.313 | 0.28 | 1 | HC_vs_MS.NoTrt_B |
| BRCC3 | 0.17024 | 0.30814 | 0.248 | 0.2 | 1 | HC_vs_MS.NoTrt_B |
| ANGEL1 | 0.17064 | -0.8915 | 0.374 | 0.42 | 1 | HC_vs_MS.NoTrt_B |
| ABCG8 | 0.17072 | 0.26584 | 0.614 | 0.62 | 1 | HC_vs_MS.NoTrt_B |
| FAM103A1 | 0.17087 | 0.30276 | 0.101 | 0.14 | 1 | HC_vs_MS.NoTrt_B |
| BRD7 | 0.17134 | 0.85991 | 0.522 | 0.52 | 1 | HC_vs_MS.NoTrt_B |
| SPINK9 | 0.17136 | 0.44546 | 0.672 | 0.75 | 1 | HC_vs_MS.NoTrt_B |
| HEXIM1 | 0.17148 | 0.41178 | 0.193 | 0.25 | 1 | HC_vs_MS.NoTrt_B |
| PIGM | 0.17164 | -0.3613 | 0.587 | 0.54 | 1 | HC_vs_MS.NoTrt_B |
| SPAG7 | 0.17186 | -0.2931 | 0.851 | 0.88 | 1 | HC_vs_MS.NoTrt_B |
| AZU1 | 0.17238 | 0.30016 | 0.201 | 0.16 | 1 | HC_vs_MS.NoTrt_B |
| GLMP | 0.17248 | 0.26999 | 0.755 | 0.76 | 1 | HC_vs_MS.NoTrt_B |
| GJB7 | 0.17249 | 0.78185 | 0.1 | 0.14 | 1 | HC_vs_MS.NoTrt_B |
| SNX9 | 0.17249 | 0.26201 | 0.087 | 0.12 | 1 | HC_vs_MS.NoTrt_B |
| HABP4 | 0.17264 | 0.28812 | 0.104 | 0.14 | 1 | HC_vs_MS.NoTrt_B |
| TP53BP2 | 0.1727 | 0.959 | 0.135 | 0.18 | 1 | HC_vs_MS.NoTrt_B |

| | | | | | | |
|---|---|---|---|---|---|---|
| ZNF138 | 0.17275 | 1.11256 | 0.072 | 0.1 | 1 | HC_vs_MS.NoTrt_B |
| ADAMTS17 | 0.17279 | 0.45847 | 0.194 | 0.16 | 1 | HC_vs_MS.NoTrt_B |
| ATPAF2 | 0.1729 | 0.48508 | 0.465 | 0.45 | 1 | HC_vs_MS.NoTrt_B |
| IGSF6 | 0.17309 | 0.26062 | 0.626 | 0.62 | 1 | HC_vs_MS.NoTrt_B |
| THUMPD2 | 0.1732 | -0.3101 | 0.345 | 0.4 | 1 | HC_vs_MS.NoTrt_B |
| MFSD6 | 0.17329 | 0.55677 | 0.149 | 0.11 | 1 | HC_vs_MS.NoTrt_B |
| HSPB9 | 0.17336 | 0.44291 | 0.203 | 0.25 | 1 | HC_vs_MS.NoTrt_B |
| KIAA2013 | 0.17344 | -0.7327 | 0.307 | 0.36 | 1 | HC_vs_MS.NoTrt_B |
| SDF2 | 0.17357 | -0.3444 | 0.162 | 0.2 | 1 | HC_vs_MS.NoTrt_B |
| WDR97 | 0.17384 | 0.29263 | 0.474 | 0.44 | 1 | HC_vs_MS.NoTrt_B |
| LPCAT2 | 0.17389 | 0.36093 | 0.325 | 0.29 | 1 | HC_vs_MS.NoTrt_B |
| CDC20B | 0.17393 | 0.28151 | 0.601 | 0.63 | 1 | HC_vs_MS.NoTrt_B |
| NBPF20 | 0.17406 | 0.56817 | 0.317 | 0.27 | 1 | HC_vs_MS.NoTrt_B |
| UVSSA | 0.17419 | 0.7793 | 0.746 | 0.72 | 1 | HC_vs_MS.NoTrt_B |
| AKTIP | 0.1744 | -0.2844 | 0.562 | 0.63 | 1 | HC_vs_MS.NoTrt_B |
| ILVBL | 0.17443 | -0.4917 | 0.259 | 0.21 | 1 | HC_vs_MS.NoTrt_B |
| TRAF3IP1 | 0.17489 | 0.87378 | 0.371 | 0.35 | 1 | HC_vs_MS.NoTrt_B |
| CYLD | 0.17509 | 0.42059 | 0.728 | 0.72 | 1 | HC_vs_MS.NoTrt_B |
| FOXM1 | 0.17547 | 0.41498 | 0.686 | 0.68 | 1 | HC_vs_MS.NoTrt_B |
| RAB11FIP3 | 0.17593 | 0.33924 | 0.213 | 0.17 | 1 | HC_vs_MS.NoTrt_B |
| PHLPP1 | 0.17617 | 0.28615 | 0.719 | 0.74 | 1 | HC_vs_MS.NoTrt_B |
| MAT2A | 0.17641 | 0.30181 | 0.126 | 0.17 | 1 | HC_vs_MS.NoTrt_B |
| HEXA | 0.17651 | -0.4042 | 0.72 | 0.76 | 1 | HC_vs_MS.NoTrt_B |
| PAQR8 | 0.17742 | 0.69033 | 0.51 | 0.47 | 1 | HC_vs_MS.NoTrt_B |
| STXBP5 | 0.17818 | 0.58483 | 0.199 | 0.16 | 1 | HC_vs_MS.NoTrt_B |
| MVB12B | 0.17831 | 1.33183 | 0.109 | 0.08 | 1 | HC_vs_MS.NoTrt_B |
| SETD1B | 0.17837 | 0.66269 | 0.201 | 0.16 | 1 | HC_vs_MS.NoTrt_B |
| ATP6V0D1 | 0.17914 | -0.6893 | 0.601 | 0.62 | 1 | HC_vs_MS.NoTrt_B |
| RHNO1 | 0.17918 | -0.4351 | 0.613 | 0.68 | 1 | HC_vs_MS.NoTrt_B |
| GCSAML | 0.17927 | 0.3453 | 0.328 | 0.3 | 1 | HC_vs_MS.NoTrt_B |
| BSDC1 | 0.17927 | -0.4024 | 0.484 | 0.51 | 1 | HC_vs_MS.NoTrt_B |
| WRN | 0.1796 | 0.45081 | 0.432 | 0.4 | 1 | HC_vs_MS.NoTrt_B |
| COPS7A | 0.17971 | 0.25903 | 0.284 | 0.23 | 1 | HC_vs_MS.NoTrt_B |
| CPEB4 | 0.17994 | 0.51137 | 0.116 | 0.08 | 1 | HC_vs_MS.NoTrt_B |
| INTS7 | 0.18029 | 0.96999 | 0.412 | 0.36 | 1 | HC_vs_MS.NoTrt_B |
| FADS3 | 0.18041 | 0.43865 | 0.625 | 0.58 | 1 | HC_vs_MS.NoTrt_B |
| ZCCHC9 | 0.18044 | -0.4448 | 0.628 | 0.73 | 1 | HC_vs_MS.NoTrt_B |
| IKBIP | 0.18083 | 0.4705 | 0.439 | 0.4 | 1 | HC_vs_MS.NoTrt_B |
| PLA2G4D | 0.18101 | 0.26956 | 0.346 | 0.3 | 1 | HC_vs_MS.NoTrt_B |
| IPP | 0.18134 | 0.46772 | 0.43 | 0.38 | 1 | HC_vs_MS.NoTrt_B |

| | | | | | | |
|---|---|---|---|---|---|---|
| PCMTD1 | 0.1814 | 0.33029 | 0.275 | 0.35 | 1 | HC_vs_MS.NoTrt_B |
| YBX3 | 0.18145 | 0.32772 | 0.706 | 0.74 | 1 | HC_vs_MS.NoTrt_B |
| SHPRH | 0.18163 | 0.59031 | 0.696 | 0.67 | 1 | HC_vs_MS.NoTrt_B |
| PLEKHA3 | 0.18187 | 0.60775 | 0.783 | 0.75 | 1 | HC_vs_MS.NoTrt_B |
| ARHGAP17 | 0.18194 | 0.25821 | 0.558 | 0.64 | 1 | HC_vs_MS.NoTrt_B |
| TIMMDC1 | 0.18219 | 0.38183 | 0.086 | 0.12 | 1 | HC_vs_MS.NoTrt_B |
| MIER1 | 0.1824 | 0.83015 | 0.707 | 0.75 | 1 | HC_vs_MS.NoTrt_B |
| GTF3C4 | 0.18262 | 0.39268 | 0.29 | 0.35 | 1 | HC_vs_MS.NoTrt_B |
| DENND5B | 0.18264 | 0.27819 | 0.555 | 0.54 | 1 | HC_vs_MS.NoTrt_B |
| LILRB3 | 0.1827 | -0.8082 | 0.123 | 0.16 | 1 | HC_vs_MS.NoTrt_B |
| NRG4 | 0.18273 | 0.33378 | 0.667 | 0.66 | 1 | HC_vs_MS.NoTrt_B |
| C14orf1 | 0.18276 | -0.6209 | 0.259 | 0.3 | 1 | HC_vs_MS.NoTrt_B |
| C20orf144 | 0.18291 | 0.3486 | 0.362 | 0.33 | 1 | HC_vs_MS.NoTrt_B |
| KIAA0556 | 0.1832 | 0.30609 | 0.259 | 0.31 | 1 | HC_vs_MS.NoTrt_B |
| RIN3 | 0.18321 | 0.41602 | 0.401 | 0.36 | 1 | HC_vs_MS.NoTrt_B |
| UBE2S | 0.18346 | 0.31758 | 0.652 | 0.66 | 1 | HC_vs_MS.NoTrt_B |
| PIKFYVE | 0.18392 | 0.26627 | 0.365 | 0.42 | 1 | HC_vs_MS.NoTrt_B |
| HECTD1 | 0.18418 | 0.31732 | 0.519 | 0.62 | 1 | HC_vs_MS.NoTrt_B |
| XRCC3 | 0.18424 | 0.74558 | 0.422 | 0.38 | 1 | HC_vs_MS.NoTrt_B |
| NR4A2 | 0.18452 | 1.70569 | 0.146 | 0.11 | 1 | HC_vs_MS.NoTrt_B |
| CAMK2D | 0.18474 | 0.3853 | 0.236 | 0.3 | 1 | HC_vs_MS.NoTrt_B |
| ADSL | 0.18477 | 0.51279 | 0.872 | 0.82 | 1 | HC_vs_MS.NoTrt_B |
| IRF4 | 0.18513 | 0.52808 | 0.259 | 0.21 | 1 | HC_vs_MS.NoTrt_B |
| DZANK1 | 0.18538 | 0.40123 | 0.707 | 0.72 | 1 | HC_vs_MS.NoTrt_B |
| VTA1 | 0.1855 | 0.57341 | 0.33 | 0.4 | 1 | HC_vs_MS.NoTrt_B |
| CHM | 0.18569 | 0.62473 | 0.49 | 0.44 | 1 | HC_vs_MS.NoTrt_B |
| ABHD14A | 0.18578 | -0.3736 | 0.457 | 0.51 | 1 | HC_vs_MS.NoTrt_B |
| BORCS5 | 0.18632 | 0.68095 | 0.629 | 0.62 | 1 | HC_vs_MS.NoTrt_B |
| CCDC58 | 0.18642 | 0.4481 | 0.299 | 0.36 | 1 | HC_vs_MS.NoTrt_B |
| CD163 | 0.18645 | 0.37155 | 0.683 | 0.74 | 1 | HC_vs_MS.NoTrt_B |
| APCDD1 | 0.18674 | 0.2592 | 0.372 | 0.34 | 1 | HC_vs_MS.NoTrt_B |
| AMY2B | 0.18679 | 0.34926 | 0.545 | 0.63 | 1 | HC_vs_MS.NoTrt_B |
| CEP120 | 0.18715 | -0.9462 | 0.094 | 0.12 | 1 | HC_vs_MS.NoTrt_B |
| PEX11G | 0.1874 | 0.3429 | 0.126 | 0.17 | 1 | HC_vs_MS.NoTrt_B |
| INSR | 0.18742 | 0.36699 | 0.462 | 0.44 | 1 | HC_vs_MS.NoTrt_B |
| SUZ12 | 0.1876 | 0.41455 | 0.622 | 0.61 | 1 | HC_vs_MS.NoTrt_B |
| PABPN1 | 0.18798 | 0.89793 | 0.196 | 0.16 | 1 | HC_vs_MS.NoTrt_B |
| ZC3H18 | 0.18859 | 0.41528 | 0.725 | 0.71 | 1 | HC_vs_MS.NoTrt_B |
| BRD9 | 0.18987 | 1.19297 | 0.146 | 0.11 | 1 | HC_vs_MS.NoTrt_B |
| TTC39B | 0.19001 | 0.27357 | 0.733 | 0.71 | 1 | HC_vs_MS.NoTrt_B |

| | | | | | | |
|---|---|---|---|---|---|---|
| FRG2C | 0.19006 | 0.2716 | 0.162 | 0.13 | 1 | HC_vs_MS.NoTrt_B |
| IKZF4 | 0.19011 | 0.5971 | 0.174 | 0.21 | 1 | HC_vs_MS.NoTrt_B |
| SUN3 | 0.19015 | 0.25563 | 0.387 | 0.35 | 1 | HC_vs_MS.NoTrt_B |
| ADIPOR2 | 0.19034 | -0.4818 | 0.128 | 0.16 | 1 | HC_vs_MS.NoTrt_B |
| BOD1L1 | 0.19038 | 0.8101 | 0.681 | 0.73 | 1 | HC_vs_MS.NoTrt_B |
| TEX14 | 0.19042 | 0.83681 | 0.075 | 0.11 | 1 | HC_vs_MS.NoTrt_B |
| DIEXF | 0.19061 | 0.34803 | 0.461 | 0.41 | 1 | HC_vs_MS.NoTrt_B |
| HMGXB4 | 0.19087 | 0.31395 | 0.091 | 0.12 | 1 | HC_vs_MS.NoTrt_B |
| ZNF558 | 0.1911 | 0.31952 | 0.684 | 0.69 | 1 | HC_vs_MS.NoTrt_B |
| DARS | 0.19117 | 0.40603 | 0.296 | 0.37 | 1 | HC_vs_MS.NoTrt_B |
| SLC38A9 | 0.19133 | 0.50828 | 0.371 | 0.45 | 1 | HC_vs_MS.NoTrt_B |
| ITGA2 | 0.19136 | 0.39188 | 0.367 | 0.33 | 1 | HC_vs_MS.NoTrt_B |
| KIAA1109 | 0.19217 | 0.40908 | 0.323 | 0.38 | 1 | HC_vs_MS.NoTrt_B |
| MAPK14 | 0.19227 | 1.27977 | 0.142 | 0.11 | 1 | HC_vs_MS.NoTrt_B |
| ZFP14 | 0.19228 | 0.54477 | 0.857 | 0.84 | 1 | HC_vs_MS.NoTrt_B |
| TEPSIN | 0.19262 | 0.33274 | 0.104 | 0.14 | 1 | HC_vs_MS.NoTrt_B |
| RAB42 | 0.19292 | 0.3005 | 0.567 | 0.57 | 1 | HC_vs_MS.NoTrt_B |
| ZC3H12C | 0.19308 | 0.3933 | 0.097 | 0.13 | 1 | HC_vs_MS.NoTrt_B |
| BRD4 | 0.19312 | 0.39536 | 0.351 | 0.41 | 1 | HC_vs_MS.NoTrt_B |
| SLC2A6 | 0.19316 | -0.3351 | 0.262 | 0.22 | 1 | HC_vs_MS.NoTrt_B |
| CP | 0.1932 | 1.16136 | 0.148 | 0.11 | 1 | HC_vs_MS.NoTrt_B |
| LIPC | 0.1937 | 1.02786 | 0.306 | 0.26 | 1 | HC_vs_MS.NoTrt_B |
| BEND6 | 0.19398 | 0.27069 | 0.197 | 0.16 | 1 | HC_vs_MS.NoTrt_B |
| BEST2 | 0.19409 | 0.2764 | 0.306 | 0.27 | 1 | HC_vs_MS.NoTrt_B |
| NDUFAF7 | 0.19422 | 0.40031 | 0.37 | 0.33 | 1 | HC_vs_MS.NoTrt_B |
| TRIM21 | 0.19427 | -0.3108 | 0.116 | 0.15 | 1 | HC_vs_MS.NoTrt_B |
| VCPIP1 | 0.19448 | 0.51994 | 0.449 | 0.41 | 1 | HC_vs_MS.NoTrt_B |
| IPO7 | 0.19482 | 0.46805 | 0.4 | 0.36 | 1 | HC_vs_MS.NoTrt_B |
| USP1 | 0.19488 | 0.40769 | 0.119 | 0.16 | 1 | HC_vs_MS.NoTrt_B |
| JUND | 0.19539 | 0.60431 | 0.162 | 0.13 | 1 | HC_vs_MS.NoTrt_B |
| DUSP10 | 0.19544 | 0.26593 | 0.141 | 0.18 | 1 | HC_vs_MS.NoTrt_B |
| PNPLA4 | 0.19554 | -0.3253 | 0.284 | 0.34 | 1 | HC_vs_MS.NoTrt_B |
| UQCC2 | 0.19573 | -0.323 | 0.074 | 0.1 | 1 | HC_vs_MS.NoTrt_B |
| DHFR | 0.1959 | 0.30762 | 0.546 | 0.52 | 1 | HC_vs_MS.NoTrt_B |
| HYLS1 | 0.19628 | 0.90595 | 0.39 | 0.37 | 1 | HC_vs_MS.NoTrt_B |
| PSG5 | 0.19631 | 0.25163 | 0.317 | 0.28 | 1 | HC_vs_MS.NoTrt_B |
| ACADM | 0.19636 | 0.25064 | 0.18 | 0.14 | 1 | HC_vs_MS.NoTrt_B |
| TIMD4 | 0.1964 | 0.37628 | 0.072 | 0.1 | 1 | HC_vs_MS.NoTrt_B |
| KLK2 | 0.19641 | 0.48971 | 0.341 | 0.31 | 1 | HC_vs_MS.NoTrt_B |
| DCP1B | 0.19653 | -0.3463 | 0.133 | 0.17 | 1 | HC_vs_MS.NoTrt_B |

| | | | | | | |
|---|---|---|---|---|---|---|
| CEP85 | 0.1969 | -0.3777 | 0.083 | 0.11 | 1 | HC_vs_MS.NoTrt_B |
| UBE2H | 0.19714 | 0.73718 | 0.117 | 0.16 | 1 | HC_vs_MS.NoTrt_B |
| IPO8 | 0.19722 | 0.77134 | 0.523 | 0.5 | 1 | HC_vs_MS.NoTrt_B |
| ZNF248 | 0.19728 | 0.91071 | 0.149 | 0.11 | 1 | HC_vs_MS.NoTrt_B |
| PHF14 | 0.19783 | 1.34346 | 0.117 | 0.09 | 1 | HC_vs_MS.NoTrt_B |
| AKR1C1 | 0.19841 | 0.28872 | 0.229 | 0.28 | 1 | HC_vs_MS.NoTrt_B |
| MRPL14 | 0.19847 | 0.32903 | 0.077 | 0.11 | 1 | HC_vs_MS.NoTrt_B |
| NFKBIZ | 0.19908 | 0.25456 | 0.336 | 0.41 | 1 | HC_vs_MS.NoTrt_B |
| PSMC1 | 0.19924 | -0.2532 | 0.42 | 0.44 | 1 | HC_vs_MS.NoTrt_B |
| HSD17B7 | 0.19957 | 0.35783 | 0.548 | 0.51 | 1 | HC_vs_MS.NoTrt_B |
| ZNF682 | 0.19978 | -0.2542 | 0.37 | 0.42 | 1 | HC_vs_MS.NoTrt_B |
| TMEM181 | 0.19985 | -0.3297 | 0.504 | 0.55 | 1 | HC_vs_MS.NoTrt_B |
| BAX | 0.20005 | -0.2541 | 0.367 | 0.4 | 1 | HC_vs_MS.NoTrt_B |
| NAPB | 0.20007 | 0.45823 | 0.403 | 0.36 | 1 | HC_vs_MS.NoTrt_B |
| ZNF17 | 0.20012 | 1.36183 | 0.277 | 0.24 | 1 | HC_vs_MS.NoTrt_B |
| GIPC1 | 0.20017 | 0.29536 | 0.474 | 0.43 | 1 | HC_vs_MS.NoTrt_B |
| KIAA0513 | 0.20098 | -0.7666 | 0.593 | 0.56 | 1 | HC_vs_MS.NoTrt_B |
| HERC2 | 0.20123 | 0.48912 | 0.851 | 0.85 | 1 | HC_vs_MS.NoTrt_B |
| ATF1 | 0.20127 | 0.47572 | 0.133 | 0.17 | 1 | HC_vs_MS.NoTrt_B |
| TMPRSS4 | 0.20137 | 0.28376 | 0.53 | 0.52 | 1 | HC_vs_MS.NoTrt_B |
| MECOM | 0.2014 | 0.44588 | 0.378 | 0.36 | 1 | HC_vs_MS.NoTrt_B |
| RIC8B | 0.2014 | 0.3175 | 0.123 | 0.09 | 1 | HC_vs_MS.NoTrt_B |
| TCP11L1 | 0.20143 | 0.71711 | 0.161 | 0.2 | 1 | HC_vs_MS.NoTrt_B |
| C20orf196 | 0.20151 | -0.542 | 0.355 | 0.41 | 1 | HC_vs_MS.NoTrt_B |
| WDR75 | 0.20163 | 0.83428 | 0.32 | 0.27 | 1 | HC_vs_MS.NoTrt_B |
| CAMKK1 | 0.20186 | 0.82137 | 0.148 | 0.2 | 1 | HC_vs_MS.NoTrt_B |
| NMI | 0.20227 | -0.3282 | 0.274 | 0.31 | 1 | HC_vs_MS.NoTrt_B |
| ABHD11 | 0.20229 | 0.35845 | 0.629 | 0.62 | 1 | HC_vs_MS.NoTrt_B |
| ACAN | 0.20274 | 0.45348 | 0.268 | 0.34 | 1 | HC_vs_MS.NoTrt_B |
| RREB1 | 0.20275 | 0.81985 | 0.729 | 0.74 | 1 | HC_vs_MS.NoTrt_B |
| NAA60 | 0.2028 | 0.25064 | 0.925 | 0.94 | 1 | HC_vs_MS.NoTrt_B |
| ZNF559 | 0.20292 | 0.27059 | 0.67 | 0.69 | 1 | HC_vs_MS.NoTrt_B |
| MECP2 | 0.2032 | 0.45899 | 0.907 | 0.92 | 1 | HC_vs_MS.NoTrt_B |
| PIWIL3 | 0.20333 | 0.42699 | 0.345 | 0.31 | 1 | HC_vs_MS.NoTrt_B |
| CYP2B6 | 0.20337 | 0.33802 | 0.607 | 0.62 | 1 | HC_vs_MS.NoTrt_B |
| FADS6 | 0.20355 | 0.2702 | 0.539 | 0.52 | 1 | HC_vs_MS.NoTrt_B |
| SVEP1 | 0.20377 | 0.34463 | 0.21 | 0.17 | 1 | HC_vs_MS.NoTrt_B |
| CD36 | 0.2038 | 0.38178 | 0.555 | 0.55 | 1 | HC_vs_MS.NoTrt_B |
| NAA15 | 0.20402 | 0.64187 | 0.739 | 0.74 | 1 | HC_vs_MS.NoTrt_B |
| SEMA3D | 0.20428 | 0.33635 | 0.514 | 0.5 | 1 | HC_vs_MS.NoTrt_B |

| | | | | | | |
|---|---|---|---|---|---|---|
| UTP20 | 0.20444 | 0.59979 | 0.486 | 0.57 | 1 | HC_vs_MS.NoTrt_B |
| CLCN6 | 0.20447 | 0.92975 | 0.496 | 0.47 | 1 | HC_vs_MS.NoTrt_B |
| ZNF585A | 0.20448 | 0.58276 | 0.364 | 0.31 | 1 | HC_vs_MS.NoTrt_B |
| STARD8 | 0.20465 | 0.39106 | 0.69 | 0.75 | 1 | HC_vs_MS.NoTrt_B |
| FBXO25 | 0.20481 | 0.27898 | 0.358 | 0.32 | 1 | HC_vs_MS.NoTrt_B |
| HIST1H1D | 0.20483 | 1.37375 | 0.122 | 0.09 | 1 | HC_vs_MS.NoTrt_B |
| ULBP3 | 0.20544 | -0.483 | 0.491 | 0.59 | 1 | HC_vs_MS.NoTrt_B |
| RP11-458D2 | 0.20588 | 0.52411 | 0.106 | 0.08 | 1 | HC_vs_MS.NoTrt_B |
| ZNF28 | 0.20596 | 1.22298 | 0.777 | 0.77 | 1 | HC_vs_MS.NoTrt_B |
| UBFD1 | 0.20597 | 0.35089 | 0.367 | 0.33 | 1 | HC_vs_MS.NoTrt_B |
| NOTCH2NL | 0.2063 | 0.69309 | 0.217 | 0.18 | 1 | HC_vs_MS.NoTrt_B |
| BPHL | 0.20656 | 0.43216 | 0.612 | 0.6 | 1 | HC_vs_MS.NoTrt_B |
| ACSS3 | 0.20667 | 0.30354 | 0.267 | 0.23 | 1 | HC_vs_MS.NoTrt_B |
| TUBGCP6 | 0.20745 | 0.31273 | 0.583 | 0.63 | 1 | HC_vs_MS.NoTrt_B |
| MOB3B | 0.20759 | 0.26778 | 0.577 | 0.56 | 1 | HC_vs_MS.NoTrt_B |
| ZFYVE21 | 0.20778 | 0.6222 | 0.655 | 0.66 | 1 | HC_vs_MS.NoTrt_B |
| AGK | 0.20788 | 0.34964 | 0.18 | 0.23 | 1 | HC_vs_MS.NoTrt_B |
| EPHX2 | 0.20795 | 0.56655 | 0.294 | 0.25 | 1 | HC_vs_MS.NoTrt_B |
| RENBP | 0.20857 | 0.43219 | 0.82 | 0.84 | 1 | HC_vs_MS.NoTrt_B |
| SLC5A6 | 0.20873 | -0.3873 | 0.075 | 0.1 | 1 | HC_vs_MS.NoTrt_B |
| MKRN2OS | 0.2088 | 0.5916 | 0.187 | 0.16 | 1 | HC_vs_MS.NoTrt_B |
| IFNGR2 | 0.20881 | 0.48751 | 0.31 | 0.38 | 1 | HC_vs_MS.NoTrt_B |
| MRPS28 | 0.20893 | 0.31539 | 0.596 | 0.59 | 1 | HC_vs_MS.NoTrt_B |
| MYH10 | 0.20901 | 0.27026 | 0.083 | 0.11 | 1 | HC_vs_MS.NoTrt_B |
| EFCAB14 | 0.20938 | 0.49877 | 0.28 | 0.24 | 1 | HC_vs_MS.NoTrt_B |
| LAMB4 | 0.20993 | 0.4245 | 0.422 | 0.42 | 1 | HC_vs_MS.NoTrt_B |
| ZNF197 | 0.2104 | 0.58474 | 0.467 | 0.44 | 1 | HC_vs_MS.NoTrt_B |
| DNAJC19 | 0.2107 | 0.42959 | 0.228 | 0.28 | 1 | HC_vs_MS.NoTrt_B |
| MYOZ3 | 0.21077 | 0.40652 | 0.243 | 0.3 | 1 | HC_vs_MS.NoTrt_B |
| LUC7L | 0.21082 | 0.25194 | 0.839 | 0.86 | 1 | HC_vs_MS.NoTrt_B |
| ZNF12 | 0.21132 | 0.51446 | 0.329 | 0.28 | 1 | HC_vs_MS.NoTrt_B |
| TCERG1 | 0.21156 | 0.63503 | 0.309 | 0.28 | 1 | HC_vs_MS.NoTrt_B |
| TFAM | 0.21166 | 0.44413 | 0.312 | 0.27 | 1 | HC_vs_MS.NoTrt_B |
| ELK4 | 0.2119 | 0.72345 | 0.542 | 0.52 | 1 | HC_vs_MS.NoTrt_B |
| FAM178B | 0.21206 | 0.32082 | 0.613 | 0.6 | 1 | HC_vs_MS.NoTrt_B |
| TFDP2 | 0.21209 | 0.31578 | 0.903 | 0.91 | 1 | HC_vs_MS.NoTrt_B |
| ATF7 | 0.21254 | 0.34168 | 0.42 | 0.5 | 1 | HC_vs_MS.NoTrt_B |
| DYDC1 | 0.21256 | 0.26842 | 0.584 | 0.58 | 1 | HC_vs_MS.NoTrt_B |
| OR11A1 | 0.21285 | 0.37126 | 0.267 | 0.24 | 1 | HC_vs_MS.NoTrt_B |
| PTPRM | 0.21289 | 1.1977 | 0.203 | 0.17 | 1 | HC_vs_MS.NoTrt_B |

| | | | | | | |
|---|---|---|---|---|---|---|
| NABP1 | 0.21289 | 0.28645 | 0.757 | 0.82 | 1 | HC_vs_MS.NoTrt_B |
| CYP4V2 | 0.21308 | 0.79036 | 0.707 | 0.69 | 1 | HC_vs_MS.NoTrt_B |
| ABHD2 | 0.21312 | -0.3495 | 0.622 | 0.71 | 1 | HC_vs_MS.NoTrt_B |
| ZNF131 | 0.21367 | -0.3224 | 0.812 | 0.84 | 1 | HC_vs_MS.NoTrt_B |
| ARPC4-TTLL: | 0.21381 | -0.3149 | 0.084 | 0.11 | 1 | HC_vs_MS.NoTrt_B |
| JMJD1C | 0.21417 | 0.47793 | 0.22 | 0.28 | 1 | HC_vs_MS.NoTrt_B |
| ZNF678 | 0.21433 | 0.42838 | 0.172 | 0.14 | 1 | HC_vs_MS.NoTrt_B |
| PLGLB1 | 0.21491 | 0.39108 | 0.119 | 0.16 | 1 | HC_vs_MS.NoTrt_B |
| ERLIN2 | 0.21501 | 0.51365 | 0.358 | 0.32 | 1 | HC_vs_MS.NoTrt_B |
| NUFIP2 | 0.21522 | 0.54797 | 0.471 | 0.46 | 1 | HC_vs_MS.NoTrt_B |
| COTL1 | 0.21554 | -0.3122 | 0.88 | 0.91 | 1 | HC_vs_MS.NoTrt_B |
| SORT1 | 0.21571 | 0.58768 | 0.528 | 0.52 | 1 | HC_vs_MS.NoTrt_B |
| TRAM2 | 0.21579 | -0.3668 | 0.248 | 0.29 | 1 | HC_vs_MS.NoTrt_B |
| MGST2 | 0.2158 | -0.2589 | 0.157 | 0.2 | 1 | HC_vs_MS.NoTrt_B |
| LYZ | 0.21581 | 0.81195 | 0.406 | 0.38 | 1 | HC_vs_MS.NoTrt_B |
| PINLYP | 0.21593 | 0.29278 | 0.597 | 0.61 | 1 | HC_vs_MS.NoTrt_B |
| RCL1 | 0.21599 | 1.30934 | 0.5 | 0.5 | 1 | HC_vs_MS.NoTrt_B |
| IKBKAP | 0.21678 | 0.71749 | 0.216 | 0.18 | 1 | HC_vs_MS.NoTrt_B |
| SRPRA | 0.21691 | 0.33413 | 0.238 | 0.29 | 1 | HC_vs_MS.NoTrt_B |
| FANK1 | 0.21715 | -0.4481 | 0.158 | 0.19 | 1 | HC_vs_MS.NoTrt_B |
| TOPBP1 | 0.2178 | 0.57527 | 0.106 | 0.14 | 1 | HC_vs_MS.NoTrt_B |
| ZNF286B | 0.21806 | 1.00028 | 0.481 | 0.48 | 1 | HC_vs_MS.NoTrt_B |
| PRKAB1 | 0.21818 | 0.28433 | 0.784 | 0.76 | 1 | HC_vs_MS.NoTrt_B |
| THEMIS | 0.21819 | 1.06151 | 0.194 | 0.16 | 1 | HC_vs_MS.NoTrt_B |
| MAP1LC3C | 0.21867 | 0.4297 | 0.374 | 0.36 | 1 | HC_vs_MS.NoTrt_B |
| NDUFB5 | 0.21891 | -0.3474 | 0.68 | 0.74 | 1 | HC_vs_MS.NoTrt_B |
| FAN1 | 0.21911 | 0.76449 | 0.288 | 0.34 | 1 | HC_vs_MS.NoTrt_B |
| TMEM168 | 0.21944 | 0.62944 | 0.51 | 0.48 | 1 | HC_vs_MS.NoTrt_B |
| DLGAP4 | 0.21955 | 0.26639 | 0.149 | 0.19 | 1 | HC_vs_MS.NoTrt_B |
| ZNF302 | 0.21957 | 0.30696 | 0.426 | 0.35 | 1 | HC_vs_MS.NoTrt_B |
| NOTCH2 | 0.21968 | 0.88112 | 0.204 | 0.17 | 1 | HC_vs_MS.NoTrt_B |
| MAP2K3 | 0.21991 | -0.3741 | 0.116 | 0.15 | 1 | HC_vs_MS.NoTrt_B |
| ADRB2 | 0.22054 | 1.51838 | 0.187 | 0.16 | 1 | HC_vs_MS.NoTrt_B |
| UBN2 | 0.22079 | 0.7307 | 0.606 | 0.6 | 1 | HC_vs_MS.NoTrt_B |
| SEMA4B | 0.22115 | 0.37922 | 0.397 | 0.36 | 1 | HC_vs_MS.NoTrt_B |
| PIGA | 0.22157 | -0.2672 | 0.39 | 0.43 | 1 | HC_vs_MS.NoTrt_B |
| MFHAS1 | 0.22179 | -0.2681 | 0.075 | 0.1 | 1 | HC_vs_MS.NoTrt_B |
| GLYR1 | 0.22195 | -0.267 | 0.391 | 0.43 | 1 | HC_vs_MS.NoTrt_B |
| NOS3 | 0.22214 | 0.50074 | 0.362 | 0.44 | 1 | HC_vs_MS.NoTrt_B |
| SLC12A2 | 0.22217 | 0.60533 | 0.526 | 0.63 | 1 | HC_vs_MS.NoTrt_B |

| | | | | | | |
|---|---|---|---|---|---|---|
| HOMER3 | 0.22234 | -0.2588 | 0.32 | 0.28 | 1 | HC_vs_MS.NoTrt_B |
| FAM46C | 0.22285 | 1.01279 | 0.158 | 0.12 | 1 | HC_vs_MS.NoTrt_B |
| VSTM1 | 0.2233 | 0.27166 | 0.296 | 0.25 | 1 | HC_vs_MS.NoTrt_B |
| TUBGCP2 | 0.22333 | -0.2783 | 0.761 | 0.78 | 1 | HC_vs_MS.NoTrt_B |
| RP11-295P9 | 0.22339 | -0.4955 | 0.08 | 0.11 | 1 | HC_vs_MS.NoTrt_B |
| PHPT1 | 0.22361 | -0.2729 | 0.106 | 0.14 | 1 | HC_vs_MS.NoTrt_B |
| RAB1A | 0.22402 | 0.64276 | 0.135 | 0.17 | 1 | HC_vs_MS.NoTrt_B |
| PCBD2 | 0.22469 | 0.53632 | 0.799 | 0.83 | 1 | HC_vs_MS.NoTrt_B |
| RP11-644F5 | 0.2252 | 0.46687 | 0.625 | 0.64 | 1 | HC_vs_MS.NoTrt_B |
| ZFX | 0.22561 | 0.25996 | 0.651 | 0.6 | 1 | HC_vs_MS.NoTrt_B |
| MFN1 | 0.22569 | 0.3911 | 0.439 | 0.38 | 1 | HC_vs_MS.NoTrt_B |
| LZTS1 | 0.22575 | 0.28017 | 0.325 | 0.4 | 1 | HC_vs_MS.NoTrt_B |
| GCLM | 0.22579 | 0.54619 | 0.672 | 0.68 | 1 | HC_vs_MS.NoTrt_B |
| EFL1 | 0.2259 | -0.3897 | 0.091 | 0.12 | 1 | HC_vs_MS.NoTrt_B |
| HAP1 | 0.22649 | 0.37888 | 0.746 | 0.76 | 1 | HC_vs_MS.NoTrt_B |
| SAAL1 | 0.22658 | 0.62561 | 0.074 | 0.1 | 1 | HC_vs_MS.NoTrt_B |
| CALCRL | 0.22697 | 0.31483 | 0.646 | 0.68 | 1 | HC_vs_MS.NoTrt_B |
| KRT10 | 0.2273 | 0.41816 | 0.081 | 0.11 | 1 | HC_vs_MS.NoTrt_B |
| ZNF160 | 0.22733 | 0.47107 | 0.593 | 0.56 | 1 | HC_vs_MS.NoTrt_B |
| PNPLA8 | 0.22783 | 0.46909 | 0.333 | 0.4 | 1 | HC_vs_MS.NoTrt_B |
| PTK6 | 0.22791 | 0.54117 | 0.332 | 0.4 | 1 | HC_vs_MS.NoTrt_B |
| RFX3 | 0.22795 | 1.03389 | 0.194 | 0.16 | 1 | HC_vs_MS.NoTrt_B |
| CTDSPL | 0.22797 | 0.36808 | 0.235 | 0.2 | 1 | HC_vs_MS.NoTrt_B |
| SOCS4 | 0.22802 | 0.3248 | 0.246 | 0.3 | 1 | HC_vs_MS.NoTrt_B |
| HNRNPU | 0.22855 | 0.3051 | 0.48 | 0.56 | 1 | HC_vs_MS.NoTrt_B |
| BRAT1 | 0.2286 | 0.32953 | 0.317 | 0.27 | 1 | HC_vs_MS.NoTrt_B |
| FXN | 0.2286 | 0.34705 | 0.157 | 0.2 | 1 | HC_vs_MS.NoTrt_B |
| NCR1 | 0.22868 | 0.72159 | 0.223 | 0.18 | 1 | HC_vs_MS.NoTrt_B |
| PPP4R2 | 0.22876 | 0.48708 | 0.457 | 0.44 | 1 | HC_vs_MS.NoTrt_B |
| APBB2 | 0.22889 | 0.44265 | 0.317 | 0.29 | 1 | HC_vs_MS.NoTrt_B |
| SGPP2 | 0.22904 | 0.31782 | 0.332 | 0.28 | 1 | HC_vs_MS.NoTrt_B |
| EPB41L2 | 0.2295 | -0.4465 | 0.112 | 0.15 | 1 | HC_vs_MS.NoTrt_B |
| UST | 0.22961 | -0.3521 | 0.272 | 0.31 | 1 | HC_vs_MS.NoTrt_B |
| MTF1 | 0.22999 | 0.58297 | 0.267 | 0.23 | 1 | HC_vs_MS.NoTrt_B |
| EIF4E2 | 0.2301 | -0.3256 | 0.322 | 0.37 | 1 | HC_vs_MS.NoTrt_B |
| ZNF257 | 0.23034 | 0.572 | 0.536 | 0.5 | 1 | HC_vs_MS.NoTrt_B |
| BMP8A | 0.23065 | -0.3225 | 0.565 | 0.62 | 1 | HC_vs_MS.NoTrt_B |
| ZNF700 | 0.2307 | 0.8493 | 0.148 | 0.2 | 1 | HC_vs_MS.NoTrt_B |
| ST3GAL5 | 0.23086 | -0.532 | 0.284 | 0.33 | 1 | HC_vs_MS.NoTrt_B |
| HARS | 0.23087 | -0.2977 | 0.823 | 0.88 | 1 | HC_vs_MS.NoTrt_B |

| | | | | | | |
|---|---|---|---|---|---|---|
| TTC9C | 0.23117 | -0.3161 | 0.357 | 0.3 | 1 | HC_vs_MS.NoTrt_B |
| BTG3 | 0.23143 | -0.2853 | 0.232 | 0.27 | 1 | HC_vs_MS.NoTrt_B |
| LAMA4 | 0.23166 | 0.3428 | 0.229 | 0.2 | 1 | HC_vs_MS.NoTrt_B |
| JMY | 0.23172 | 0.3159 | 0.259 | 0.22 | 1 | HC_vs_MS.NoTrt_B |
| SULT2A1 | 0.23204 | 0.30159 | 0.7 | 0.72 | 1 | HC_vs_MS.NoTrt_B |
| RTTN | 0.23292 | 0.30529 | 0.239 | 0.22 | 1 | HC_vs_MS.NoTrt_B |
| C6orf25 | 0.23345 | 0.7871 | 0.436 | 0.42 | 1 | HC_vs_MS.NoTrt_B |
| PSD3 | 0.23348 | 0.8521 | 0.31 | 0.27 | 1 | HC_vs_MS.NoTrt_B |
| FAM35A | 0.23365 | 0.26294 | 0.103 | 0.14 | 1 | HC_vs_MS.NoTrt_B |
| POLR3A | 0.2338 | 0.33135 | 0.829 | 0.84 | 1 | HC_vs_MS.NoTrt_B |
| RASGRP4 | 0.23387 | -0.5839 | 0.472 | 0.5 | 1 | HC_vs_MS.NoTrt_B |
| PBX2 | 0.23396 | 1.17527 | 0.12 | 0.09 | 1 | HC_vs_MS.NoTrt_B |
| MRPL40 | 0.23417 | -0.3348 | 0.394 | 0.45 | 1 | HC_vs_MS.NoTrt_B |
| MITF | 0.2345 | 0.4327 | 0.139 | 0.18 | 1 | HC_vs_MS.NoTrt_B |
| FLT4 | 0.2346 | 0.33643 | 0.104 | 0.14 | 1 | HC_vs_MS.NoTrt_B |
| IFI44L | 0.23469 | 0.83009 | 0.2 | 0.16 | 1 | HC_vs_MS.NoTrt_B |
| SCN2B | 0.23481 | -0.6572 | 0.312 | 0.27 | 1 | HC_vs_MS.NoTrt_B |
| TTC14 | 0.23507 | 0.43746 | 0.399 | 0.46 | 1 | HC_vs_MS.NoTrt_B |
| SETX | 0.23534 | 0.40732 | 0.283 | 0.35 | 1 | HC_vs_MS.NoTrt_B |
| NREP | 0.23543 | 0.67901 | 0.191 | 0.16 | 1 | HC_vs_MS.NoTrt_B |
| GEN1 | 0.23578 | 0.27099 | 0.741 | 0.77 | 1 | HC_vs_MS.NoTrt_B |
| APOOL | 0.23625 | 0.3879 | 0.771 | 0.78 | 1 | HC_vs_MS.NoTrt_B |
| MIOS | 0.23639 | 0.54082 | 0.271 | 0.32 | 1 | HC_vs_MS.NoTrt_B |
| FAM222B | 0.23656 | 0.29042 | 0.136 | 0.17 | 1 | HC_vs_MS.NoTrt_B |
| FSD1L | 0.23657 | 0.32772 | 0.229 | 0.2 | 1 | HC_vs_MS.NoTrt_B |
| FOXK2 | 0.23675 | -0.2934 | 0.252 | 0.29 | 1 | HC_vs_MS.NoTrt_B |
| OVCA2 | 0.23678 | 0.32932 | 0.091 | 0.12 | 1 | HC_vs_MS.NoTrt_B |
| TRAPPC6A | 0.23738 | -0.2506 | 0.178 | 0.22 | 1 | HC_vs_MS.NoTrt_B |
| ZCCHC11 | 0.23747 | 0.37095 | 0.93 | 0.9 | 1 | HC_vs_MS.NoTrt_B |
| BCAP29 | 0.23797 | 0.32559 | 0.374 | 0.34 | 1 | HC_vs_MS.NoTrt_B |
| FAM107B | 0.2385 | 0.47054 | 0.348 | 0.32 | 1 | HC_vs_MS.NoTrt_B |
| PPARG | 0.23865 | 0.28581 | 0.377 | 0.35 | 1 | HC_vs_MS.NoTrt_B |
| SORBS1 | 0.23885 | 0.44021 | 0.219 | 0.18 | 1 | HC_vs_MS.NoTrt_B |
| LETM2 | 0.23907 | 0.2626 | 0.667 | 0.67 | 1 | HC_vs_MS.NoTrt_B |
| ATOX1 | 0.23918 | 0.69308 | 0.174 | 0.14 | 1 | HC_vs_MS.NoTrt_B |
| PALLD | 0.23948 | -0.3246 | 0.338 | 0.3 | 1 | HC_vs_MS.NoTrt_B |
| SCMH1 | 0.23962 | 0.41168 | 0.104 | 0.14 | 1 | HC_vs_MS.NoTrt_B |
| MTPN | 0.23964 | 0.28474 | 0.162 | 0.21 | 1 | HC_vs_MS.NoTrt_B |
| F5 | 0.23967 | 0.69634 | 0.272 | 0.24 | 1 | HC_vs_MS.NoTrt_B |
| ZC3H11A | 0.2402 | 0.4749 | 0.204 | 0.17 | 1 | HC_vs_MS.NoTrt_B |

| | | | | | | |
|---|---|---|---|---|---|---|
| ZMYM3 | 0.241 | 0.54198 | 0.419 | 0.4 | 1 | HC_vs_MS.NoTrt_B |
| ELF4 | 0.24101 | 0.29897 | 0.112 | 0.15 | 1 | HC_vs_MS.NoTrt_B |
| SBNO1 | 0.24139 | 0.33023 | 0.645 | 0.6 | 1 | HC_vs_MS.NoTrt_B |
| ANKRD49 | 0.24143 | 0.7782 | 0.157 | 0.2 | 1 | HC_vs_MS.NoTrt_B |
| EDEM1 | 0.2419 | 0.33039 | 0.314 | 0.37 | 1 | HC_vs_MS.NoTrt_B |
| KLHL6 | 0.24193 | 0.47187 | 0.729 | 0.73 | 1 | HC_vs_MS.NoTrt_B |
| DISC1 | 0.24219 | 1.15268 | 0.486 | 0.44 | 1 | HC_vs_MS.NoTrt_B |
| WAC | 0.24268 | 0.35998 | 0.445 | 0.51 | 1 | HC_vs_MS.NoTrt_B |
| WIPF2 | 0.24278 | 0.56526 | 0.772 | 0.82 | 1 | HC_vs_MS.NoTrt_B |
| CYP7B1 | 0.24285 | 0.47445 | 0.428 | 0.42 | 1 | HC_vs_MS.NoTrt_B |
| CDC14B | 0.24296 | 0.44214 | 0.322 | 0.29 | 1 | HC_vs_MS.NoTrt_B |
| GABRG3 | 0.24311 | 0.25758 | 0.348 | 0.31 | 1 | HC_vs_MS.NoTrt_B |
| DMKN | 0.24328 | 0.27028 | 0.471 | 0.43 | 1 | HC_vs_MS.NoTrt_B |
| HSD17B2 | 0.24354 | 0.25247 | 0.749 | 0.77 | 1 | HC_vs_MS.NoTrt_B |
| PELI1 | 0.24355 | 0.54771 | 0.172 | 0.22 | 1 | HC_vs_MS.NoTrt_B |
| WNT10A | 0.2438 | 0.72518 | 0.126 | 0.16 | 1 | HC_vs_MS.NoTrt_B |
| CA12 | 0.24381 | 0.6508 | 0.265 | 0.22 | 1 | HC_vs_MS.NoTrt_B |
| WDR60 | 0.24383 | 0.7474 | 0.133 | 0.1 | 1 | HC_vs_MS.NoTrt_B |
| SMAD4 | 0.24425 | 0.38092 | 0.491 | 0.44 | 1 | HC_vs_MS.NoTrt_B |
| EARS2 | 0.24443 | 0.29079 | 0.604 | 0.65 | 1 | HC_vs_MS.NoTrt_B |
| HKR1 | 0.24448 | 0.26229 | 0.967 | 0.98 | 1 | HC_vs_MS.NoTrt_B |
| THSD4 | 0.24481 | 0.49948 | 0.362 | 0.32 | 1 | HC_vs_MS.NoTrt_B |
| TADA2B | 0.24514 | 1.73335 | 0.317 | 0.3 | 1 | HC_vs_MS.NoTrt_B |
| HSBP1L1 | 0.24535 | -0.2561 | 0.365 | 0.41 | 1 | HC_vs_MS.NoTrt_B |
| SMURF2 | 0.24541 | 0.49212 | 0.794 | 0.77 | 1 | HC_vs_MS.NoTrt_B |
| ZFPM2 | 0.24576 | 0.43493 | 0.417 | 0.4 | 1 | HC_vs_MS.NoTrt_B |
| FOXJ2 | 0.24592 | 0.50996 | 0.449 | 0.51 | 1 | HC_vs_MS.NoTrt_B |
| TMEM40 | 0.24604 | 0.26829 | 0.772 | 0.79 | 1 | HC_vs_MS.NoTrt_B |
| MYB | 0.2469 | 0.55689 | 0.28 | 0.33 | 1 | HC_vs_MS.NoTrt_B |
| BBS4 | 0.2473 | -0.3594 | 0.077 | 0.1 | 1 | HC_vs_MS.NoTrt_B |
| CACNA1G | 0.24754 | 0.25174 | 0.184 | 0.16 | 1 | HC_vs_MS.NoTrt_B |
| ABCD4 | 0.24825 | 0.26683 | 0.79 | 0.79 | 1 | HC_vs_MS.NoTrt_B |
| PLEKHG3 | 0.24838 | -0.6858 | 0.396 | 0.47 | 1 | HC_vs_MS.NoTrt_B |
| SPDYE3 | 0.24881 | 0.52955 | 0.588 | 0.55 | 1 | HC_vs_MS.NoTrt_B |
| TAOK2 | 0.24892 | 0.73822 | 0.103 | 0.08 | 1 | HC_vs_MS.NoTrt_B |
| BID | 0.24894 | -0.4515 | 0.478 | 0.51 | 1 | HC_vs_MS.NoTrt_B |
| FBXL17 | 0.24914 | 0.54128 | 0.243 | 0.21 | 1 | HC_vs_MS.NoTrt_B |
| ZNF587B | 0.24922 | 0.42022 | 0.336 | 0.3 | 1 | HC_vs_MS.NoTrt_B |
| BBS9 | 0.24923 | -0.3597 | 0.096 | 0.12 | 1 | HC_vs_MS.NoTrt_B |
| AC240274.1 | 0.24935 | 1.10199 | 0.241 | 0.21 | 1 | HC_vs_MS.NoTrt_B |

| | | | | | | |
|---|---|---|---|---|---|---|
| VRK2 | 0.24969 | -0.3362 | 0.212 | 0.25 | 1 | HC_vs_MS.NoTrt_B |
| NATD1 | 0.24975 | 0.75876 | 0.481 | 0.46 | 1 | HC_vs_MS.NoTrt_B |
| AFTPH | 0.25026 | 0.41605 | 0.162 | 0.21 | 1 | HC_vs_MS.NoTrt_B |
| DPP3 | 0.25048 | 0.39616 | 0.187 | 0.15 | 1 | HC_vs_MS.NoTrt_B |
| GPR107 | 0.25094 | 0.74361 | 0.145 | 0.11 | 1 | HC_vs_MS.NoTrt_B |
| KIF5B | 0.25113 | 0.43871 | 0.262 | 0.34 | 1 | HC_vs_MS.NoTrt_B |
| GDE1 | 0.25113 | -1.1311 | 0.245 | 0.29 | 1 | HC_vs_MS.NoTrt_B |
| TRPV1 | 0.25125 | -0.2652 | 0.401 | 0.39 | 1 | HC_vs_MS.NoTrt_B |
| CRLF2 | 0.25126 | 0.31376 | 0.3 | 0.28 | 1 | HC_vs_MS.NoTrt_B |
| SYCP1 | 0.25163 | 0.30172 | 0.312 | 0.29 | 1 | HC_vs_MS.NoTrt_B |
| PLA2G12A | 0.25169 | -0.2558 | 0.57 | 0.65 | 1 | HC_vs_MS.NoTrt_B |
| PRKRIP1 | 0.25175 | 1.64814 | 0.13 | 0.1 | 1 | HC_vs_MS.NoTrt_B |
| ADAT2 | 0.25184 | 0.5942 | 0.404 | 0.38 | 1 | HC_vs_MS.NoTrt_B |
| CERKL | 0.25218 | 0.27024 | 0.354 | 0.43 | 1 | HC_vs_MS.NoTrt_B |
| CCDC13 | 0.25244 | 0.76892 | 0.32 | 0.3 | 1 | HC_vs_MS.NoTrt_B |
| VSX1 | 0.25262 | 0.26493 | 0.17 | 0.21 | 1 | HC_vs_MS.NoTrt_B |
| FUCA2 | 0.25337 | 0.36323 | 0.296 | 0.27 | 1 | HC_vs_MS.NoTrt_B |
| DPH5 | 0.25339 | -0.4792 | 0.343 | 0.38 | 1 | HC_vs_MS.NoTrt_B |
| CCDC130 | 0.25359 | 0.28894 | 0.429 | 0.49 | 1 | HC_vs_MS.NoTrt_B |
| USP13 | 0.25385 | 0.79554 | 0.259 | 0.23 | 1 | HC_vs_MS.NoTrt_B |
| KIF1B | 0.25394 | 0.47498 | 0.788 | 0.81 | 1 | HC_vs_MS.NoTrt_B |
| BLOC1S4 | 0.25407 | 0.46283 | 0.149 | 0.18 | 1 | HC_vs_MS.NoTrt_B |
| RFC4 | 0.25412 | -0.5512 | 0.228 | 0.27 | 1 | HC_vs_MS.NoTrt_B |
| UACA | 0.25452 | 0.35716 | 0.404 | 0.38 | 1 | HC_vs_MS.NoTrt_B |
| GFI1B | 0.25461 | 0.41866 | 0.541 | 0.54 | 1 | HC_vs_MS.NoTrt_B |
| ABCA5 | 0.25501 | 0.37475 | 0.6 | 0.68 | 1 | HC_vs_MS.NoTrt_B |
| P3H4 | 0.25533 | 0.36294 | 0.896 | 0.9 | 1 | HC_vs_MS.NoTrt_B |
| SCUBE2 | 0.25718 | 0.25153 | 0.304 | 0.28 | 1 | HC_vs_MS.NoTrt_B |
| CEP295 | 0.25792 | 0.95381 | 0.565 | 0.56 | 1 | HC_vs_MS.NoTrt_B |
| NPAP1 | 0.25802 | 0.38139 | 0.465 | 0.46 | 1 | HC_vs_MS.NoTrt_B |
| FSTL4 | 0.25891 | 0.5457 | 0.155 | 0.13 | 1 | HC_vs_MS.NoTrt_B |
| RANGAP1 | 0.25904 | -0.442 | 0.659 | 0.71 | 1 | HC_vs_MS.NoTrt_B |
| ZNF514 | 0.25905 | 0.38413 | 0.597 | 0.72 | 1 | HC_vs_MS.NoTrt_B |
| GNLY | 0.25909 | 1.27653 | 0.226 | 0.2 | 1 | HC_vs_MS.NoTrt_B |
| C14orf80 | 0.25926 | -1.0775 | 0.262 | 0.32 | 1 | HC_vs_MS.NoTrt_B |
| ATRIP | 0.25941 | -0.3963 | 0.078 | 0.1 | 1 | HC_vs_MS.NoTrt_B |
| ZNF346 | 0.25969 | -0.2697 | 0.743 | 0.77 | 1 | HC_vs_MS.NoTrt_B |
| TNFRSF9 | 0.25978 | -0.4545 | 0.899 | 0.91 | 1 | HC_vs_MS.NoTrt_B |
| JARID2 | 0.25981 | 0.26499 | 0.18 | 0.22 | 1 | HC_vs_MS.NoTrt_B |
| DLG1 | 0.25983 | 0.62155 | 0.581 | 0.56 | 1 | HC_vs_MS.NoTrt_B |

| | | | | | | |
|---|---|---|---|---|---|---|
| SVIL | 0.26048 | 0.53411 | 0.148 | 0.12 | 1 | HC_vs_MS.NoTrt_B |
| OXNAD1 | 0.26058 | 0.6846 | 0.31 | 0.28 | 1 | HC_vs_MS.NoTrt_B |
| PSEN2 | 0.26122 | -1.3555 | 0.138 | 0.17 | 1 | HC_vs_MS.NoTrt_B |
| NCAPH | 0.26153 | -0.4839 | 0.455 | 0.42 | 1 | HC_vs_MS.NoTrt_B |
| ALG3 | 0.26161 | 0.48571 | 0.12 | 0.09 | 1 | HC_vs_MS.NoTrt_B |
| CCDC85C | 0.26195 | 0.84942 | 0.407 | 0.37 | 1 | HC_vs_MS.NoTrt_B |
| SRFBP1 | 0.26216 | 0.66209 | 0.429 | 0.4 | 1 | HC_vs_MS.NoTrt_B |
| MME | 0.26227 | 0.45039 | 0.439 | 0.41 | 1 | HC_vs_MS.NoTrt_B |
| SLC33A1 | 0.2624 | -0.4354 | 0.893 | 0.89 | 1 | HC_vs_MS.NoTrt_B |
| GPM6A | 0.2626 | 0.26235 | 0.28 | 0.24 | 1 | HC_vs_MS.NoTrt_B |
| ARHGAP35 | 0.26278 | -0.5228 | 0.161 | 0.19 | 1 | HC_vs_MS.NoTrt_B |
| GSDMD | 0.26299 | -0.375 | 0.794 | 0.82 | 1 | HC_vs_MS.NoTrt_B |
| RGMA | 0.26302 | 0.35164 | 0.652 | 0.64 | 1 | HC_vs_MS.NoTrt_B |
| RP11-111H1 | 0.26328 | 0.30183 | 0.113 | 0.15 | 1 | HC_vs_MS.NoTrt_B |
| METTL5 | 0.26359 | -0.3984 | 0.464 | 0.48 | 1 | HC_vs_MS.NoTrt_B |
| ACSM2B | 0.2639 | 0.32701 | 0.522 | 0.53 | 1 | HC_vs_MS.NoTrt_B |
| SULT1C4 | 0.26408 | 0.35271 | 0.554 | 0.57 | 1 | HC_vs_MS.NoTrt_B |
| KLHL35 | 0.26443 | -0.2895 | 0.364 | 0.42 | 1 | HC_vs_MS.NoTrt_B |
| SCNN1A | 0.2645 | 0.30874 | 0.729 | 0.78 | 1 | HC_vs_MS.NoTrt_B |
| CCND2 | 0.26464 | 0.33504 | 0.381 | 0.44 | 1 | HC_vs_MS.NoTrt_B |
| JAM2 | 0.26477 | 0.28784 | 0.5 | 0.51 | 1 | HC_vs_MS.NoTrt_B |
| PTGES | 0.26487 | 0.31685 | 0.481 | 0.48 | 1 | HC_vs_MS.NoTrt_B |
| SHQ1 | 0.26525 | 0.67022 | 0.69 | 0.67 | 1 | HC_vs_MS.NoTrt_B |
| ATG2B | 0.26578 | 0.67929 | 0.296 | 0.36 | 1 | HC_vs_MS.NoTrt_B |
| ZNF780B | 0.26581 | 0.87412 | 0.588 | 0.6 | 1 | HC_vs_MS.NoTrt_B |
| PALB2 | 0.26616 | -0.3248 | 0.186 | 0.22 | 1 | HC_vs_MS.NoTrt_B |
| UBXN7 | 0.26645 | 0.68921 | 0.787 | 0.78 | 1 | HC_vs_MS.NoTrt_B |
| ZNF573 | 0.26647 | 0.28243 | 0.368 | 0.42 | 1 | HC_vs_MS.NoTrt_B |
| DGKE | 0.26665 | 0.2841 | 0.661 | 0.74 | 1 | HC_vs_MS.NoTrt_B |
| C1orf27 | 0.26667 | 0.45056 | 0.155 | 0.19 | 1 | HC_vs_MS.NoTrt_B |
| CHD9 | 0.26674 | 0.30717 | 0.345 | 0.4 | 1 | HC_vs_MS.NoTrt_B |
| ZFP30 | 0.26684 | 0.3477 | 0.828 | 0.81 | 1 | HC_vs_MS.NoTrt_B |
| ORAI3 | 0.26705 | -0.5096 | 0.093 | 0.12 | 1 | HC_vs_MS.NoTrt_B |
| PCSK6 | 0.26705 | 0.34542 | 0.633 | 0.63 | 1 | HC_vs_MS.NoTrt_B |
| PREPL | 0.26715 | 0.53751 | 0.683 | 0.67 | 1 | HC_vs_MS.NoTrt_B |
| ELMO1 | 0.26718 | 0.35771 | 0.754 | 0.75 | 1 | HC_vs_MS.NoTrt_B |
| PNPT1 | 0.26721 | 0.43406 | 0.855 | 0.88 | 1 | HC_vs_MS.NoTrt_B |
| PTPRK | 0.2675 | 0.34293 | 0.443 | 0.43 | 1 | HC_vs_MS.NoTrt_B |
| DNAJC15 | 0.26777 | -0.3661 | 0.175 | 0.21 | 1 | HC_vs_MS.NoTrt_B |
| TPGS1 | 0.26833 | 0.77004 | 0.254 | 0.22 | 1 | HC_vs_MS.NoTrt_B |

| | | | | | | |
|---|---|---|---|---|---|---|
| THRAP3 | 0.2684 | 0.30435 | 0.349 | 0.42 | 1 | HC_vs_MS.NoTrt_B |
| ZNF226 | 0.26858 | -0.4618 | 0.523 | 0.47 | 1 | HC_vs_MS.NoTrt_B |
| DHTKD1 | 0.26867 | 0.45139 | 0.806 | 0.83 | 1 | HC_vs_MS.NoTrt_B |
| EBPL | 0.26878 | 0.37858 | 0.706 | 0.73 | 1 | HC_vs_MS.NoTrt_B |
| ELOF1 | 0.26905 | -0.4669 | 0.581 | 0.62 | 1 | HC_vs_MS.NoTrt_B |
| MED14 | 0.26966 | 1.34503 | 0.141 | 0.11 | 1 | HC_vs_MS.NoTrt_B |
| OR6J1 | 0.26991 | 0.43605 | 0.68 | 0.69 | 1 | HC_vs_MS.NoTrt_B |
| HIST1H2BE | 0.26992 | 0.65719 | 0.265 | 0.31 | 1 | HC_vs_MS.NoTrt_B |
| NLK | 0.27021 | 1.25558 | 0.307 | 0.27 | 1 | HC_vs_MS.NoTrt_B |
| OR52I1 | 0.27051 | 0.57014 | 0.458 | 0.44 | 1 | HC_vs_MS.NoTrt_B |
| NRBP1 | 0.27057 | -0.2904 | 0.593 | 0.63 | 1 | HC_vs_MS.NoTrt_B |
| NAPEPLD | 0.27082 | 0.64394 | 0.32 | 0.29 | 1 | HC_vs_MS.NoTrt_B |
| PTPN2 | 0.27136 | -0.3528 | 0.965 | 0.98 | 1 | HC_vs_MS.NoTrt_B |
| UFL1 | 0.27143 | 0.4189 | 0.152 | 0.19 | 1 | HC_vs_MS.NoTrt_B |
| KIAA1755 | 0.27147 | 0.39782 | 0.194 | 0.17 | 1 | HC_vs_MS.NoTrt_B |
| THAP12 | 0.27152 | 0.50549 | 0.596 | 0.59 | 1 | HC_vs_MS.NoTrt_B |
| RIC1 | 0.27159 | 0.68979 | 0.103 | 0.08 | 1 | HC_vs_MS.NoTrt_B |
| CABYR | 0.27185 | 0.33505 | 0.486 | 0.49 | 1 | HC_vs_MS.NoTrt_B |
| ZSWIM1 | 0.27255 | 0.35224 | 0.681 | 0.76 | 1 | HC_vs_MS.NoTrt_B |
| CACNA1F | 0.27287 | 0.29838 | 0.251 | 0.22 | 1 | HC_vs_MS.NoTrt_B |
| ACBD4 | 0.27295 | -0.5623 | 0.317 | 0.35 | 1 | HC_vs_MS.NoTrt_B |
| REST | 0.27313 | 0.33428 | 0.17 | 0.21 | 1 | HC_vs_MS.NoTrt_B |
| LYN | 0.27322 | 0.27252 | 0.587 | 0.61 | 1 | HC_vs_MS.NoTrt_B |
| FANCF | 0.2735 | 0.37317 | 0.236 | 0.2 | 1 | HC_vs_MS.NoTrt_B |
| B3GALT5 | 0.27362 | 0.30911 | 0.342 | 0.32 | 1 | HC_vs_MS.NoTrt_B |
| RTCA | 0.27364 | -0.4008 | 0.526 | 0.57 | 1 | HC_vs_MS.NoTrt_B |
| CDKN2AIP | 0.27427 | -0.2591 | 0.078 | 0.1 | 1 | HC_vs_MS.NoTrt_B |
| PCNX2 | 0.27428 | 0.29389 | 0.925 | 0.95 | 1 | HC_vs_MS.NoTrt_B |
| MLST8 | 0.27503 | -0.2801 | 0.355 | 0.4 | 1 | HC_vs_MS.NoTrt_B |
| NXF1 | 0.27536 | 0.26707 | 0.559 | 0.64 | 1 | HC_vs_MS.NoTrt_B |
| GDF11 | 0.27569 | 0.45345 | 0.325 | 0.29 | 1 | HC_vs_MS.NoTrt_B |
| SLC1A4 | 0.27587 | 0.73995 | 0.574 | 0.61 | 1 | HC_vs_MS.NoTrt_B |
| ZDHHC17 | 0.27591 | 0.53459 | 0.345 | 0.3 | 1 | HC_vs_MS.NoTrt_B |
| SLC27A5 | 0.27604 | 0.29677 | 0.864 | 0.89 | 1 | HC_vs_MS.NoTrt_B |
| MYO1F | 0.27623 | -0.262 | 0.848 | 0.85 | 1 | HC_vs_MS.NoTrt_B |
| SLC13A3 | 0.27626 | 0.34877 | 0.299 | 0.28 | 1 | HC_vs_MS.NoTrt_B |
| PON2 | 0.27684 | 1.45831 | 0.42 | 0.41 | 1 | HC_vs_MS.NoTrt_B |
| DIAPH3 | 0.27701 | 0.45816 | 0.329 | 0.3 | 1 | HC_vs_MS.NoTrt_B |
| FAM120C | 0.2771 | 0.69403 | 0.274 | 0.25 | 1 | HC_vs_MS.NoTrt_B |
| CTSC | 0.27732 | -0.321 | 0.894 | 0.92 | 1 | HC_vs_MS.NoTrt_B |

| | | | | | | |
|---|---|---|---|---|---|---|
| STEAP3 | 0.27758 | 0.5027 | 0.116 | 0.15 | 1 | HC_vs_MS.NoTrt_B |
| SLC22A18 | 0.27876 | 1.07824 | 0.503 | 0.6 | 1 | HC_vs_MS.NoTrt_B |
| AP1G1 | 0.27899 | 0.3536 | 0.667 | 0.74 | 1 | HC_vs_MS.NoTrt_B |
| ERCC6L | 0.27938 | 0.87446 | 0.117 | 0.15 | 1 | HC_vs_MS.NoTrt_B |
| GP6 | 0.27955 | 0.41558 | 0.459 | 0.44 | 1 | HC_vs_MS.NoTrt_B |
| ANKRD13D | 0.28008 | 0.26532 | 0.668 | 0.67 | 1 | HC_vs_MS.NoTrt_B |
| FASTKD1 | 0.28095 | -0.2912 | 0.23 | 0.27 | 1 | HC_vs_MS.NoTrt_B |
| ZNF71 | 0.28105 | 0.97759 | 0.204 | 0.25 | 1 | HC_vs_MS.NoTrt_B |
| MED25 | 0.28151 | 0.85002 | 0.486 | 0.45 | 1 | HC_vs_MS.NoTrt_B |
| GSKIP | 0.28211 | -0.419 | 0.096 | 0.12 | 1 | HC_vs_MS.NoTrt_B |
| HECTD3 | 0.28249 | 1.15612 | 0.129 | 0.1 | 1 | HC_vs_MS.NoTrt_B |
| FBXL20 | 0.28266 | 0.26151 | 0.912 | 0.9 | 1 | HC_vs_MS.NoTrt_B |
| NOL6 | 0.28291 | -0.5676 | 0.372 | 0.42 | 1 | HC_vs_MS.NoTrt_B |
| ZNF224 | 0.28302 | 0.90378 | 0.458 | 0.47 | 1 | HC_vs_MS.NoTrt_B |
| EID2B | 0.28312 | -0.3047 | 0.523 | 0.5 | 1 | HC_vs_MS.NoTrt_B |
| OSBPL10 | 0.28366 | 0.498 | 0.391 | 0.35 | 1 | HC_vs_MS.NoTrt_B |
| PABPC1 | 0.28378 | 0.26387 | 0.93 | 0.95 | 1 | HC_vs_MS.NoTrt_B |
| MDM4 | 0.28414 | 0.3096 | 0.991 | 1 | 1 | HC_vs_MS.NoTrt_B |
| PATL1 | 0.28423 | 0.38989 | 0.133 | 0.17 | 1 | HC_vs_MS.NoTrt_B |
| FGFR4 | 0.2844 | 0.54976 | 0.393 | 0.36 | 1 | HC_vs_MS.NoTrt_B |
| NRP2 | 0.28455 | 0.32696 | 0.372 | 0.35 | 1 | HC_vs_MS.NoTrt_B |
| IFFO2 | 0.28466 | 0.54671 | 0.123 | 0.1 | 1 | HC_vs_MS.NoTrt_B |
| MDFIC | 0.28485 | 0.52466 | 0.296 | 0.35 | 1 | HC_vs_MS.NoTrt_B |
| ITGAX | 0.28486 | 0.30033 | 0.471 | 0.54 | 1 | HC_vs_MS.NoTrt_B |
| DHRS2 | 0.28513 | 0.35427 | 0.329 | 0.3 | 1 | HC_vs_MS.NoTrt_B |
| PPM1L | 0.28558 | 0.41542 | 0.236 | 0.2 | 1 | HC_vs_MS.NoTrt_B |
| CCDC88B | 0.28561 | 0.56702 | 0.759 | 0.82 | 1 | HC_vs_MS.NoTrt_B |
| CRX | 0.28584 | 0.30795 | 0.697 | 0.76 | 1 | HC_vs_MS.NoTrt_B |
| HACE1 | 0.28592 | 0.30423 | 0.655 | 0.65 | 1 | HC_vs_MS.NoTrt_B |
| BAZ2A | 0.28608 | 0.38609 | 0.665 | 0.68 | 1 | HC_vs_MS.NoTrt_B |
| SFXN3 | 0.28622 | -0.408 | 0.323 | 0.35 | 1 | HC_vs_MS.NoTrt_B |
| VPS13B | 0.28623 | 0.40746 | 0.735 | 0.71 | 1 | HC_vs_MS.NoTrt_B |
| NUP210 | 0.28629 | 0.67755 | 0.52 | 0.5 | 1 | HC_vs_MS.NoTrt_B |
| CD302 | 0.28643 | 0.98825 | 0.326 | 0.29 | 1 | HC_vs_MS.NoTrt_B |
| SCYL2 | 0.28651 | 0.34551 | 0.11 | 0.14 | 1 | HC_vs_MS.NoTrt_B |
| GSE1 | 0.28665 | 1.23496 | 0.338 | 0.32 | 1 | HC_vs_MS.NoTrt_B |
| PIP5K1A | 0.28678 | -0.2901 | 0.493 | 0.54 | 1 | HC_vs_MS.NoTrt_B |
| BNIP3L | 0.28736 | -0.2551 | 0.181 | 0.21 | 1 | HC_vs_MS.NoTrt_B |
| SGK494 | 0.28757 | 0.25841 | 0.738 | 0.71 | 1 | HC_vs_MS.NoTrt_B |
| TNFSF14 | 0.28764 | 0.48369 | 0.606 | 0.64 | 1 | HC_vs_MS.NoTrt_B |

| | | | | | | |
|---|---|---|---|---|---|---|
| PTRF | 0.28782 | 0.29199 | 0.352 | 0.33 | 1 | HC_vs_MS.NoTrt_B |
| CGGBP1 | 0.28789 | 0.48381 | 0.18 | 0.22 | 1 | HC_vs_MS.NoTrt_B |
| TRIM25 | 0.28811 | 1.14202 | 0.33 | 0.3 | 1 | HC_vs_MS.NoTrt_B |
| KCNG4 | 0.28817 | 0.25064 | 0.548 | 0.52 | 1 | HC_vs_MS.NoTrt_B |
| DNAJB14 | 0.28826 | 0.44155 | 0.48 | 0.52 | 1 | HC_vs_MS.NoTrt_B |
| ALDH3B1 | 0.28859 | 0.25894 | 0.735 | 0.76 | 1 | HC_vs_MS.NoTrt_B |
| APC | 0.28873 | 0.26921 | 0.501 | 0.46 | 1 | HC_vs_MS.NoTrt_B |
| EIF1AY | 0.28989 | -0.7459 | 0.609 | 0.65 | 1 | HC_vs_MS.NoTrt_B |
| C9orf64 | 0.29016 | 0.54492 | 0.596 | 0.59 | 1 | HC_vs_MS.NoTrt_B |
| NARF | 0.29022 | -0.2617 | 0.959 | 0.97 | 1 | HC_vs_MS.NoTrt_B |
| TBC1D7 | 0.29043 | 0.93066 | 0.513 | 0.48 | 1 | HC_vs_MS.NoTrt_B |
| SERINC1 | 0.29082 | 0.38964 | 0.645 | 0.63 | 1 | HC_vs_MS.NoTrt_B |
| SNX11 | 0.29095 | 0.42818 | 0.645 | 0.65 | 1 | HC_vs_MS.NoTrt_B |
| GMEB1 | 0.29116 | 0.42569 | 0.803 | 0.84 | 1 | HC_vs_MS.NoTrt_B |
| CDCP1 | 0.2914 | 1.42267 | 0.43 | 0.38 | 1 | HC_vs_MS.NoTrt_B |
| TMEM87B | 0.29179 | 0.73678 | 0.109 | 0.14 | 1 | HC_vs_MS.NoTrt_B |
| CDAN1 | 0.29188 | 0.2931 | 0.43 | 0.41 | 1 | HC_vs_MS.NoTrt_B |
| TSEN15 | 0.29206 | -0.5745 | 0.201 | 0.23 | 1 | HC_vs_MS.NoTrt_B |
| OTOA | 0.2921 | 0.32909 | 0.358 | 0.33 | 1 | HC_vs_MS.NoTrt_B |
| CPPED1 | 0.29228 | -0.6363 | 0.593 | 0.57 | 1 | HC_vs_MS.NoTrt_B |
| CCDC163 | 0.29238 | -0.2661 | 0.538 | 0.58 | 1 | HC_vs_MS.NoTrt_B |
| AP3M1 | 0.29282 | 0.40473 | 0.484 | 0.44 | 1 | HC_vs_MS.NoTrt_B |
| NEDD1 | 0.29351 | -0.3015 | 0.094 | 0.12 | 1 | HC_vs_MS.NoTrt_B |
| TPM1 | 0.29367 | 0.42287 | 0.493 | 0.45 | 1 | HC_vs_MS.NoTrt_B |
| PSTK | 0.29367 | -0.5618 | 0.338 | 0.29 | 1 | HC_vs_MS.NoTrt_B |
| RP11-347C1 | 0.29399 | 0.42922 | 0.664 | 0.7 | 1 | HC_vs_MS.NoTrt_B |
| C2CD2L | 0.29415 | 0.52138 | 0.409 | 0.47 | 1 | HC_vs_MS.NoTrt_B |
| CTD-3105H1 | 0.29467 | 0.29061 | 0.27 | 0.24 | 1 | HC_vs_MS.NoTrt_B |
| SLC16A7 | 0.29486 | 0.28728 | 0.628 | 0.6 | 1 | HC_vs_MS.NoTrt_B |
| ARMC10 | 0.29504 | 0.69999 | 0.632 | 0.64 | 1 | HC_vs_MS.NoTrt_B |
| ANKDD1A | 0.29559 | 0.47328 | 0.522 | 0.52 | 1 | HC_vs_MS.NoTrt_B |
| LAP3 | 0.29588 | -0.2801 | 0.119 | 0.15 | 1 | HC_vs_MS.NoTrt_B |
| TMEM183A | 0.29596 | 0.68677 | 0.183 | 0.15 | 1 | HC_vs_MS.NoTrt_B |
| TLN1 | 0.29618 | 0.85261 | 0.575 | 0.57 | 1 | HC_vs_MS.NoTrt_B |
| TDRD1 | 0.29638 | 0.49513 | 0.168 | 0.14 | 1 | HC_vs_MS.NoTrt_B |
| CRKL | 0.29709 | 0.67046 | 0.574 | 0.57 | 1 | HC_vs_MS.NoTrt_B |
| C5orf63 | 0.2975 | -0.2918 | 0.335 | 0.37 | 1 | HC_vs_MS.NoTrt_B |
| TMOD2 | 0.29759 | 0.26363 | 0.454 | 0.47 | 1 | HC_vs_MS.NoTrt_B |
| UVRAG | 0.29782 | 0.2633 | 0.423 | 0.5 | 1 | HC_vs_MS.NoTrt_B |
| SH3RF2 | 0.29785 | -0.5469 | 0.165 | 0.14 | 1 | HC_vs_MS.NoTrt_B |

| | | | | | | |
|---|---|---|---|---|---|---|
| CLDN4 | 0.29791 | 0.72266 | 0.543 | 0.64 | 1 | HC_vs_MS.NoTrt_B |
| MARK2 | 0.29826 | 0.45617 | 0.13 | 0.1 | 1 | HC_vs_MS.NoTrt_B |
| CTSW | 0.29837 | -0.4575 | 0.622 | 0.56 | 1 | HC_vs_MS.NoTrt_B |
| ZNF699 | 0.29854 | 0.36061 | 0.251 | 0.29 | 1 | HC_vs_MS.NoTrt_B |
| MRPS25 | 0.29914 | 0.26216 | 0.487 | 0.44 | 1 | HC_vs_MS.NoTrt_B |
| TRA2B | 0.29928 | 0.43452 | 0.543 | 0.51 | 1 | HC_vs_MS.NoTrt_B |
| ZNF85 | 0.29968 | -0.3419 | 0.652 | 0.72 | 1 | HC_vs_MS.NoTrt_B |
| CD38 | 0.30112 | 1.23157 | 0.335 | 0.31 | 1 | HC_vs_MS.NoTrt_B |
| KANK3 | 0.30134 | 0.47132 | 0.201 | 0.24 | 1 | HC_vs_MS.NoTrt_B |
| SLC35E4 | 0.30178 | 0.48775 | 0.333 | 0.3 | 1 | HC_vs_MS.NoTrt_B |
| NUP43 | 0.30195 | 0.28692 | 0.843 | 0.8 | 1 | HC_vs_MS.NoTrt_B |
| JMJD7-PLA2 | 0.30223 | 0.27154 | 0.386 | 0.43 | 1 | HC_vs_MS.NoTrt_B |
| ITGB5 | 0.30248 | 0.25207 | 0.328 | 0.38 | 1 | HC_vs_MS.NoTrt_B |
| ATXN1L | 0.3028 | 0.5807 | 0.078 | 0.1 | 1 | HC_vs_MS.NoTrt_B |
| GADL1 | 0.30284 | 0.47124 | 0.622 | 0.7 | 1 | HC_vs_MS.NoTrt_B |
| FDPS | 0.30295 | -0.4389 | 0.445 | 0.48 | 1 | HC_vs_MS.NoTrt_B |
| GGT6 | 0.3035 | 0.2923 | 0.454 | 0.43 | 1 | HC_vs_MS.NoTrt_B |
| CCDC159 | 0.3043 | -0.4492 | 0.765 | 0.84 | 1 | HC_vs_MS.NoTrt_B |
| SRSF10 | 0.30436 | 0.3748 | 0.722 | 0.79 | 1 | HC_vs_MS.NoTrt_B |
| NME7 | 0.30457 | 0.39627 | 0.622 | 0.64 | 1 | HC_vs_MS.NoTrt_B |
| TSC2 | 0.30469 | 0.45529 | 0.684 | 0.68 | 1 | HC_vs_MS.NoTrt_B |
| USP31 | 0.30542 | 0.33017 | 0.135 | 0.11 | 1 | HC_vs_MS.NoTrt_B |
| ZNF233 | 0.30561 | 0.40608 | 0.467 | 0.48 | 1 | HC_vs_MS.NoTrt_B |
| ZC3H13 | 0.30577 | 0.3552 | 0.326 | 0.29 | 1 | HC_vs_MS.NoTrt_B |
| DET1 | 0.30612 | 1.11749 | 0.536 | 0.5 | 1 | HC_vs_MS.NoTrt_B |
| MBTPS2 | 0.30618 | 0.49811 | 0.312 | 0.28 | 1 | HC_vs_MS.NoTrt_B |
| MIB2 | 0.30626 | 1.27913 | 0.177 | 0.22 | 1 | HC_vs_MS.NoTrt_B |
| PLS3 | 0.30651 | 0.25172 | 0.358 | 0.43 | 1 | HC_vs_MS.NoTrt_B |
| TESMIN | 0.30658 | 0.3651 | 0.629 | 0.7 | 1 | HC_vs_MS.NoTrt_B |
| ZNF680 | 0.30673 | 0.79944 | 0.096 | 0.12 | 1 | HC_vs_MS.NoTrt_B |
| C10orf82 | 0.30677 | 0.29001 | 0.535 | 0.54 | 1 | HC_vs_MS.NoTrt_B |
| YTHDC2 | 0.30682 | 0.84429 | 0.416 | 0.39 | 1 | HC_vs_MS.NoTrt_B |
| BNIP3 | 0.30697 | 0.2685 | 0.583 | 0.68 | 1 | HC_vs_MS.NoTrt_B |
| EPPIN | 0.30701 | 0.41051 | 0.462 | 0.47 | 1 | HC_vs_MS.NoTrt_B |
| ERAP1 | 0.30725 | 0.98371 | 0.584 | 0.62 | 1 | HC_vs_MS.NoTrt_B |
| NDUFAF1 | 0.30748 | 0.45126 | 0.365 | 0.42 | 1 | HC_vs_MS.NoTrt_B |
| ZC3H3 | 0.30813 | 0.53738 | 0.125 | 0.1 | 1 | HC_vs_MS.NoTrt_B |
| MAP3K10 | 0.30832 | 0.4401 | 0.354 | 0.34 | 1 | HC_vs_MS.NoTrt_B |
| CENPV | 0.30844 | 0.33045 | 0.71 | 0.71 | 1 | HC_vs_MS.NoTrt_B |
| CD160 | 0.30846 | 0.60138 | 0.294 | 0.27 | 1 | HC_vs_MS.NoTrt_B |

| | | | | | | |
|---|---|---|---|---|---|---|
| TRRAP | 0.30872 | 0.3536 | 0.158 | 0.13 | 1 | HC_vs_MS.NoTrt_B |
| LSM6 | 0.30888 | 0.31715 | 0.13 | 0.16 | 1 | HC_vs_MS.NoTrt_B |
| ARHGAP11B | 0.30901 | -0.2671 | 0.204 | 0.23 | 1 | HC_vs_MS.NoTrt_B |
| SLC16A5 | 0.30925 | 0.71463 | 0.154 | 0.12 | 1 | HC_vs_MS.NoTrt_B |
| NELFCD | 0.30955 | 0.25079 | 0.519 | 0.57 | 1 | HC_vs_MS.NoTrt_B |
| LRPPRC | 0.3096 | 0.51883 | 0.519 | 0.54 | 1 | HC_vs_MS.NoTrt_B |
| C1orf167 | 0.30987 | 0.26003 | 0.632 | 0.65 | 1 | HC_vs_MS.NoTrt_B |
| C2orf68 | 0.30995 | 0.28097 | 0.822 | 0.77 | 1 | HC_vs_MS.NoTrt_B |
| FUT6 | 0.31014 | 0.4751 | 0.404 | 0.47 | 1 | HC_vs_MS.NoTrt_B |
| WDTC1 | 0.31057 | 0.40228 | 0.359 | 0.35 | 1 | HC_vs_MS.NoTrt_B |
| SF3A1 | 0.31065 | 0.312 | 0.349 | 0.43 | 1 | HC_vs_MS.NoTrt_B |
| BAZ1A | 0.31152 | 0.27692 | 0.374 | 0.34 | 1 | HC_vs_MS.NoTrt_B |
| RNF130 | 0.31155 | -0.275 | 0.633 | 0.67 | 1 | HC_vs_MS.NoTrt_B |
| FAM234B | 0.31178 | -0.7043 | 0.414 | 0.38 | 1 | HC_vs_MS.NoTrt_B |
| PMS2 | 0.31213 | 0.76357 | 0.662 | 0.7 | 1 | HC_vs_MS.NoTrt_B |
| BEND2 | 0.31216 | 0.26862 | 0.555 | 0.58 | 1 | HC_vs_MS.NoTrt_B |
| SERPINB9 | 0.31231 | 0.5656 | 0.699 | 0.8 | 1 | HC_vs_MS.NoTrt_B |
| COL28A1 | 0.3125 | 0.33371 | 0.428 | 0.41 | 1 | HC_vs_MS.NoTrt_B |
| KIAA1210 | 0.31288 | -0.2533 | 0.203 | 0.23 | 1 | HC_vs_MS.NoTrt_B |
| SKIL | 0.31308 | 0.46891 | 0.277 | 0.24 | 1 | HC_vs_MS.NoTrt_B |
| OSBPL9 | 0.31349 | 0.51912 | 0.328 | 0.3 | 1 | HC_vs_MS.NoTrt_B |
| ACOT4 | 0.31367 | -0.4845 | 0.1 | 0.12 | 1 | HC_vs_MS.NoTrt_B |
| SYNRG | 0.31406 | 0.31764 | 0.929 | 0.96 | 1 | HC_vs_MS.NoTrt_B |
| PIAS3 | 0.31425 | 0.42204 | 0.113 | 0.14 | 1 | HC_vs_MS.NoTrt_B |
| OTUD5 | 0.31449 | -0.3206 | 0.125 | 0.15 | 1 | HC_vs_MS.NoTrt_B |
| EML6 | 0.31461 | 0.35687 | 0.578 | 0.57 | 1 | HC_vs_MS.NoTrt_B |
| HAUS2 | 0.31494 | 0.25858 | 0.794 | 0.83 | 1 | HC_vs_MS.NoTrt_B |
| NRIP2 | 0.31502 | 0.71073 | 0.336 | 0.32 | 1 | HC_vs_MS.NoTrt_B |
| ADAD2 | 0.31508 | 0.29107 | 0.225 | 0.28 | 1 | HC_vs_MS.NoTrt_B |
| CD300LD | 0.31557 | 0.27803 | 0.362 | 0.32 | 1 | HC_vs_MS.NoTrt_B |
| IFI27 | 0.31587 | 0.35528 | 0.355 | 0.34 | 1 | HC_vs_MS.NoTrt_B |
| HPS1 | 0.31625 | 0.34827 | 0.6 | 0.65 | 1 | HC_vs_MS.NoTrt_B |
| KDELR1 | 0.31626 | 0.46176 | 0.243 | 0.21 | 1 | HC_vs_MS.NoTrt_B |
| CAMTA2 | 0.31628 | 0.32364 | 0.501 | 0.45 | 1 | HC_vs_MS.NoTrt_B |
| EIF4A3 | 0.31715 | -0.4176 | 0.649 | 0.71 | 1 | HC_vs_MS.NoTrt_B |
| TRIM2 | 0.3174 | 0.9182 | 0.23 | 0.21 | 1 | HC_vs_MS.NoTrt_B |
| MKL2 | 0.31746 | 0.74961 | 0.696 | 0.72 | 1 | HC_vs_MS.NoTrt_B |
| STRBP | 0.31762 | -0.5416 | 0.63 | 0.61 | 1 | HC_vs_MS.NoTrt_B |
| CES2 | 0.31771 | -0.3477 | 0.316 | 0.27 | 1 | HC_vs_MS.NoTrt_B |
| FAM63A | 0.31827 | 0.53947 | 0.509 | 0.58 | 1 | HC_vs_MS.NoTrt_B |

| | | | | | | |
|---|---|---|---|---|---|---|
| LEKR1 | 0.31837 | -0.3904 | 0.174 | 0.21 | 1 | HC_vs_MS.NoTrt_B |
| ACSM3 | 0.31856 | 0.3389 | 0.594 | 0.6 | 1 | HC_vs_MS.NoTrt_B |
| LARP1 | 0.3187 | 0.50487 | 0.18 | 0.15 | 1 | HC_vs_MS.NoTrt_B |
| SCML4 | 0.31916 | 0.72169 | 0.629 | 0.64 | 1 | HC_vs_MS.NoTrt_B |
| SYT11 | 0.31932 | -0.3233 | 0.616 | 0.63 | 1 | HC_vs_MS.NoTrt_B |
| WAPL | 0.31947 | 0.65888 | 0.452 | 0.45 | 1 | HC_vs_MS.NoTrt_B |
| RBM10 | 0.3196 | 0.53047 | 0.175 | 0.15 | 1 | HC_vs_MS.NoTrt_B |
| UBE2G1 | 0.32003 | 0.29132 | 0.22 | 0.26 | 1 | HC_vs_MS.NoTrt_B |
| OAS1 | 0.32012 | 0.28578 | 0.503 | 0.49 | 1 | HC_vs_MS.NoTrt_B |
| SMC5 | 0.32027 | 0.34143 | 0.386 | 0.35 | 1 | HC_vs_MS.NoTrt_B |
| ALG10B | 0.32042 | 0.25948 | 0.639 | 0.62 | 1 | HC_vs_MS.NoTrt_B |
| MSMO1 | 0.32046 | 0.68061 | 0.525 | 0.52 | 1 | HC_vs_MS.NoTrt_B |
| PARP10 | 0.3207 | 0.67146 | 0.157 | 0.13 | 1 | HC_vs_MS.NoTrt_B |
| RUNDC1 | 0.3213 | 0.64156 | 0.701 | 0.69 | 1 | HC_vs_MS.NoTrt_B |
| RIOK3 | 0.32138 | 0.34926 | 0.514 | 0.48 | 1 | HC_vs_MS.NoTrt_B |
| ZNF879 | 0.32148 | 0.40545 | 0.312 | 0.36 | 1 | HC_vs_MS.NoTrt_B |
| NPIPA7 | 0.32151 | 0.26334 | 0.154 | 0.19 | 1 | HC_vs_MS.NoTrt_B |
| ATG16L1 | 0.32207 | 0.53674 | 0.112 | 0.09 | 1 | HC_vs_MS.NoTrt_B |
| GIPR | 0.32207 | 0.28686 | 0.619 | 0.69 | 1 | HC_vs_MS.NoTrt_B |
| HYAL4 | 0.32222 | 0.40977 | 0.572 | 0.56 | 1 | HC_vs_MS.NoTrt_B |
| POM121C | 0.32223 | 0.28179 | 0.494 | 0.52 | 1 | HC_vs_MS.NoTrt_B |
| GDNF | 0.32225 | 0.27508 | 0.294 | 0.27 | 1 | HC_vs_MS.NoTrt_B |
| RFX7 | 0.32235 | 0.40898 | 0.13 | 0.16 | 1 | HC_vs_MS.NoTrt_B |
| NAA40 | 0.3226 | -0.3563 | 0.548 | 0.53 | 1 | HC_vs_MS.NoTrt_B |
| AL354822.1 | 0.32277 | 0.53208 | 0.242 | 0.21 | 1 | HC_vs_MS.NoTrt_B |
| CCDC18 | 0.32314 | -0.2868 | 0.159 | 0.19 | 1 | HC_vs_MS.NoTrt_B |
| KHDRBS1 | 0.3234 | 0.91491 | 0.183 | 0.16 | 1 | HC_vs_MS.NoTrt_B |
| SIDT1 | 0.32352 | 0.4317 | 0.781 | 0.74 | 1 | HC_vs_MS.NoTrt_B |
| FAM110C | 0.32355 | 0.69367 | 0.242 | 0.29 | 1 | HC_vs_MS.NoTrt_B |
| PRPF8 | 0.3239 | -0.2641 | 0.868 | 0.87 | 1 | HC_vs_MS.NoTrt_B |
| HNRNPUL2 | 0.32484 | -0.3162 | 0.084 | 0.11 | 1 | HC_vs_MS.NoTrt_B |
| FGF9 | 0.32549 | 0.47025 | 0.13 | 0.1 | 1 | HC_vs_MS.NoTrt_B |
| MCEE | 0.32597 | 0.70836 | 0.129 | 0.16 | 1 | HC_vs_MS.NoTrt_B |
| DHX36 | 0.32604 | 0.48622 | 0.713 | 0.77 | 1 | HC_vs_MS.NoTrt_B |
| RALGPS2 | 0.32614 | 0.34833 | 0.354 | 0.32 | 1 | HC_vs_MS.NoTrt_B |
| ALG6 | 0.32632 | 0.46033 | 0.101 | 0.13 | 1 | HC_vs_MS.NoTrt_B |
| SERPINH1 | 0.32656 | 0.94967 | 0.304 | 0.36 | 1 | HC_vs_MS.NoTrt_B |
| BRAF | 0.32676 | 0.97361 | 0.106 | 0.14 | 1 | HC_vs_MS.NoTrt_B |
| ZDHHC24 | 0.32712 | 0.25306 | 0.864 | 0.91 | 1 | HC_vs_MS.NoTrt_B |
| CEP85L | 0.32726 | 0.49667 | 0.258 | 0.3 | 1 | HC_vs_MS.NoTrt_B |

| | | | | | | |
|---|---|---|---|---|---|---|
| CCDC126 | 0.3274 | 0.59673 | 0.116 | 0.15 | 1 | HC_vs_MS.NoTrt_B |
| RAB8B | 0.32746 | 0.76841 | 0.178 | 0.23 | 1 | HC_vs_MS.NoTrt_B |
| SLC25A46 | 0.32751 | 0.43118 | 0.099 | 0.12 | 1 | HC_vs_MS.NoTrt_B |
| CRELD2 | 0.32754 | -0.6468 | 0.574 | 0.61 | 1 | HC_vs_MS.NoTrt_B |
| L3MBTL1 | 0.32755 | 0.47454 | 0.616 | 0.6 | 1 | HC_vs_MS.NoTrt_B |
| GINM1 | 0.32805 | -0.4949 | 0.529 | 0.48 | 1 | HC_vs_MS.NoTrt_B |
| NDUFB6 | 0.32815 | 0.25041 | 0.5 | 0.56 | 1 | HC_vs_MS.NoTrt_B |
| DEPDC1B | 0.32827 | -0.9796 | 0.206 | 0.24 | 1 | HC_vs_MS.NoTrt_B |
| DUSP1 | 0.32885 | 1.09858 | 0.168 | 0.15 | 1 | HC_vs_MS.NoTrt_B |
| EBF2 | 0.32959 | 0.26121 | 0.288 | 0.26 | 1 | HC_vs_MS.NoTrt_B |
| TPO | 0.3298 | 0.58384 | 0.228 | 0.27 | 1 | HC_vs_MS.NoTrt_B |
| HES2 | 0.3299 | 0.27463 | 0.587 | 0.6 | 1 | HC_vs_MS.NoTrt_B |
| KIF21B | 0.33021 | 0.74877 | 0.378 | 0.37 | 1 | HC_vs_MS.NoTrt_B |
| CLSTN1 | 0.33027 | 0.36358 | 0.204 | 0.24 | 1 | HC_vs_MS.NoTrt_B |
| WDR59 | 0.33139 | 0.31948 | 0.954 | 0.94 | 1 | HC_vs_MS.NoTrt_B |
| SV2B | 0.3322 | 0.75008 | 0.383 | 0.37 | 1 | HC_vs_MS.NoTrt_B |
| ADGRA3 | 0.33237 | 0.34696 | 0.393 | 0.39 | 1 | HC_vs_MS.NoTrt_B |
| CTIF | 0.33261 | 0.92988 | 0.162 | 0.2 | 1 | HC_vs_MS.NoTrt_B |
| SAMD9L | 0.33274 | 0.62995 | 0.191 | 0.16 | 1 | HC_vs_MS.NoTrt_B |
| ACTN1 | 0.33307 | 0.54178 | 0.599 | 0.63 | 1 | HC_vs_MS.NoTrt_B |
| SLC38A2 | 0.33359 | 0.55432 | 0.239 | 0.21 | 1 | HC_vs_MS.NoTrt_B |
| NT5C3B | 0.33404 | 0.837 | 0.616 | 0.64 | 1 | HC_vs_MS.NoTrt_B |
| NOL11 | 0.33442 | -0.3502 | 0.53 | 0.49 | 1 | HC_vs_MS.NoTrt_B |
| SF3A2 | 0.33443 | 0.63209 | 0.365 | 0.34 | 1 | HC_vs_MS.NoTrt_B |
| TUBD1 | 0.3346 | 0.43135 | 0.822 | 0.8 | 1 | HC_vs_MS.NoTrt_B |
| TMPO | 0.33464 | 0.29798 | 0.535 | 0.5 | 1 | HC_vs_MS.NoTrt_B |
| PRPF38B | 0.33465 | 0.26403 | 0.47 | 0.52 | 1 | HC_vs_MS.NoTrt_B |
| COQ10A | 0.33471 | 0.85075 | 0.475 | 0.47 | 1 | HC_vs_MS.NoTrt_B |
| RAB5A | 0.33501 | 0.31436 | 0.113 | 0.14 | 1 | HC_vs_MS.NoTrt_B |
| KANK1 | 0.33629 | -0.2899 | 0.701 | 0.71 | 1 | HC_vs_MS.NoTrt_B |
| TCF20 | 0.33671 | 0.45925 | 0.109 | 0.14 | 1 | HC_vs_MS.NoTrt_B |
| CABIN1 | 0.33724 | 0.6065 | 0.255 | 0.23 | 1 | HC_vs_MS.NoTrt_B |
| PDCD6 | 0.33781 | -0.485 | 0.667 | 0.71 | 1 | HC_vs_MS.NoTrt_B |
| F3 | 0.33791 | 0.41747 | 0.497 | 0.52 | 1 | HC_vs_MS.NoTrt_B |
| SLC46A3 | 0.33804 | 0.6139 | 0.078 | 0.1 | 1 | HC_vs_MS.NoTrt_B |
| IMMT | 0.33807 | -0.3601 | 0.443 | 0.48 | 1 | HC_vs_MS.NoTrt_B |
| CDH15 | 0.33901 | 0.50209 | 0.403 | 0.37 | 1 | HC_vs_MS.NoTrt_B |
| STYX | 0.33919 | 0.35247 | 0.496 | 0.57 | 1 | HC_vs_MS.NoTrt_B |
| BTLA | 0.33936 | 0.99686 | 0.154 | 0.18 | 1 | HC_vs_MS.NoTrt_B |
| FUT10 | 0.33942 | 0.46087 | 0.358 | 0.35 | 1 | HC_vs_MS.NoTrt_B |

| | | | | | | |
|---|---|---|---|---|---|---|
| TMEM17 | 0.33972 | 0.26144 | 0.497 | 0.49 | 1 | HC_vs_MS.NoTrt_B |
| VAC14 | 0.33996 | 0.35695 | 0.104 | 0.08 | 1 | HC_vs_MS.NoTrt_B |
| KDM6B | 0.34027 | 0.47328 | 0.161 | 0.14 | 1 | HC_vs_MS.NoTrt_B |
| LMNB1 | 0.34076 | -0.4244 | 0.083 | 0.1 | 1 | HC_vs_MS.NoTrt_B |
| ZNF587 | 0.3413 | 0.33946 | 0.945 | 0.95 | 1 | HC_vs_MS.NoTrt_B |
| XBP1 | 0.34146 | 0.53108 | 0.093 | 0.12 | 1 | HC_vs_MS.NoTrt_B |
| TMEM185A | 0.3417 | 0.38405 | 0.483 | 0.46 | 1 | HC_vs_MS.NoTrt_B |
| PRR19 | 0.34177 | 0.32011 | 0.693 | 0.75 | 1 | HC_vs_MS.NoTrt_B |
| NUP98 | 0.3419 | 0.32214 | 0.758 | 0.78 | 1 | HC_vs_MS.NoTrt_B |
| SEC23B | 0.34212 | -0.3682 | 0.112 | 0.14 | 1 | HC_vs_MS.NoTrt_B |
| ZNF615 | 0.34214 | 1.18535 | 0.47 | 0.46 | 1 | HC_vs_MS.NoTrt_B |
| KLF13 | 0.34267 | 0.57792 | 0.297 | 0.35 | 1 | HC_vs_MS.NoTrt_B |
| EFTUD2 | 0.34269 | -0.3158 | 0.836 | 0.86 | 1 | HC_vs_MS.NoTrt_B |
| SLC35B4 | 0.34287 | 0.46102 | 0.533 | 0.54 | 1 | HC_vs_MS.NoTrt_B |
| TNKS2 | 0.34308 | 0.48788 | 0.259 | 0.3 | 1 | HC_vs_MS.NoTrt_B |
| ERN2 | 0.34317 | 0.25832 | 0.51 | 0.5 | 1 | HC_vs_MS.NoTrt_B |
| NAPG | 0.34326 | 0.60694 | 0.571 | 0.58 | 1 | HC_vs_MS.NoTrt_B |
| HYPK | 0.34351 | 0.38609 | 0.903 | 0.92 | 1 | HC_vs_MS.NoTrt_B |
| MANBA | 0.3442 | 0.51447 | 0.181 | 0.15 | 1 | HC_vs_MS.NoTrt_B |
| ABRA | 0.34456 | 0.52784 | 0.084 | 0.11 | 1 | HC_vs_MS.NoTrt_B |
| CHIT1 | 0.34458 | -1.072 | 0.09 | 0.11 | 1 | HC_vs_MS.NoTrt_B |
| RASAL3 | 0.34471 | 1.2962 | 0.184 | 0.16 | 1 | HC_vs_MS.NoTrt_B |
| OBSCN | 0.34507 | 0.43763 | 0.213 | 0.18 | 1 | HC_vs_MS.NoTrt_B |
| TNFAIP3 | 0.34513 | 0.34947 | 0.355 | 0.42 | 1 | HC_vs_MS.NoTrt_B |
| PGGT1B | 0.34558 | 0.66835 | 0.158 | 0.13 | 1 | HC_vs_MS.NoTrt_B |
| CTD-3088G3 | 0.3456 | 0.48625 | 0.51 | 0.5 | 1 | HC_vs_MS.NoTrt_B |
| PPP2R2D | 0.34578 | 0.26763 | 0.622 | 0.64 | 1 | HC_vs_MS.NoTrt_B |
| CYSTM1 | 0.34591 | -0.486 | 0.265 | 0.3 | 1 | HC_vs_MS.NoTrt_B |
| PPP2R3C | 0.34598 | -0.3145 | 0.138 | 0.16 | 1 | HC_vs_MS.NoTrt_B |
| LINC00371 | 0.34612 | 0.35624 | 0.136 | 0.11 | 1 | HC_vs_MS.NoTrt_B |
| WISP2 | 0.3463 | 0.28979 | 0.606 | 0.59 | 1 | HC_vs_MS.NoTrt_B |
| KIAA0825 | 0.3464 | 0.52233 | 0.294 | 0.27 | 1 | HC_vs_MS.NoTrt_B |
| RFWD2 | 0.34671 | 0.43854 | 0.291 | 0.34 | 1 | HC_vs_MS.NoTrt_B |
| WHSC1L1 | 0.34728 | 0.26961 | 0.596 | 0.69 | 1 | HC_vs_MS.NoTrt_B |
| NPAS2 | 0.34763 | 0.25349 | 0.426 | 0.4 | 1 | HC_vs_MS.NoTrt_B |
| DEPDC4 | 0.34785 | 0.60088 | 0.438 | 0.45 | 1 | HC_vs_MS.NoTrt_B |
| ZNF441 | 0.34787 | -0.5515 | 0.174 | 0.21 | 1 | HC_vs_MS.NoTrt_B |
| HIST2H4B | 0.348 | 0.53729 | 0.123 | 0.15 | 1 | HC_vs_MS.NoTrt_B |
| FBXO3 | 0.3482 | 0.39924 | 0.291 | 0.33 | 1 | HC_vs_MS.NoTrt_B |
| MT-ATP8 | 0.34889 | 0.47481 | 0.71 | 0.84 | 1 | HC_vs_MS.NoTrt_B |

| | | | | | | |
|---|---|---|---|---|---|---|
| CNOT3 | 0.34895 | 0.56319 | 0.123 | 0.1 | 1 | HC_vs_MS.NoTrt_B |
| PCDHGB6 | 0.34933 | 0.42648 | 0.352 | 0.33 | 1 | HC_vs_MS.NoTrt_B |
| DNAJC11 | 0.34942 | -0.2779 | 0.168 | 0.2 | 1 | HC_vs_MS.NoTrt_B |
| CTNNA1 | 0.34945 | -0.5312 | 0.562 | 0.63 | 1 | HC_vs_MS.NoTrt_B |
| CSRP1 | 0.35021 | -0.5896 | 0.497 | 0.52 | 1 | HC_vs_MS.NoTrt_B |
| TAF13 | 0.35026 | 0.25468 | 0.116 | 0.09 | 1 | HC_vs_MS.NoTrt_B |
| HSPB11 | 0.35058 | 0.26006 | 0.764 | 0.74 | 1 | HC_vs_MS.NoTrt_B |
| ATXN2L | 0.35108 | 0.55486 | 0.259 | 0.23 | 1 | HC_vs_MS.NoTrt_B |
| MORC3 | 0.35127 | 0.37256 | 0.486 | 0.44 | 1 | HC_vs_MS.NoTrt_B |
| ZBTB11 | 0.35129 | 1.03111 | 0.533 | 0.51 | 1 | HC_vs_MS.NoTrt_B |
| TMEM41B | 0.35168 | 0.48718 | 0.806 | 0.87 | 1 | HC_vs_MS.NoTrt_B |
| GSN | 0.35173 | -0.2938 | 0.554 | 0.56 | 1 | HC_vs_MS.NoTrt_B |
| GRK6 | 0.3518 | 0.48243 | 0.191 | 0.23 | 1 | HC_vs_MS.NoTrt_B |
| SPG21 | 0.35214 | -0.2583 | 0.872 | 0.88 | 1 | HC_vs_MS.NoTrt_B |
| CYB561 | 0.35219 | 0.69179 | 0.272 | 0.26 | 1 | HC_vs_MS.NoTrt_B |
| SULT1A1 | 0.35226 | -0.3352 | 0.535 | 0.6 | 1 | HC_vs_MS.NoTrt_B |
| MTA2 | 0.35241 | -0.3854 | 0.249 | 0.29 | 1 | HC_vs_MS.NoTrt_B |
| PRKCZ | 0.35247 | 0.5409 | 0.239 | 0.21 | 1 | HC_vs_MS.NoTrt_B |
| TNK2 | 0.35267 | 0.76347 | 0.132 | 0.11 | 1 | HC_vs_MS.NoTrt_B |
| CKAP5 | 0.3531 | 0.68538 | 0.151 | 0.12 | 1 | HC_vs_MS.NoTrt_B |
| BOLL | 0.3533 | 0.40776 | 0.154 | 0.13 | 1 | HC_vs_MS.NoTrt_B |
| ZNF212 | 0.35356 | -0.7444 | 0.371 | 0.33 | 1 | HC_vs_MS.NoTrt_B |
| GM2A | 0.35452 | -0.3143 | 0.528 | 0.57 | 1 | HC_vs_MS.NoTrt_B |
| TJP1 | 0.35515 | 0.42733 | 0.165 | 0.15 | 1 | HC_vs_MS.NoTrt_B |
| PPP4R3A | 0.35549 | 0.29631 | 0.204 | 0.24 | 1 | HC_vs_MS.NoTrt_B |
| TVP23A | 0.35557 | 0.31394 | 0.454 | 0.52 | 1 | HC_vs_MS.NoTrt_B |
| IRS2 | 0.35559 | 0.41244 | 0.322 | 0.3 | 1 | HC_vs_MS.NoTrt_B |
| ING1 | 0.3557 | 0.26224 | 0.543 | 0.61 | 1 | HC_vs_MS.NoTrt_B |
| MGA | 0.35573 | 0.39637 | 0.858 | 0.9 | 1 | HC_vs_MS.NoTrt_B |
| KIAA0430 | 0.35663 | 0.43858 | 0.28 | 0.32 | 1 | HC_vs_MS.NoTrt_B |
| HECA | 0.35674 | 0.38561 | 0.548 | 0.61 | 1 | HC_vs_MS.NoTrt_B |
| FAM174B | 0.35712 | -0.2999 | 0.12 | 0.1 | 1 | HC_vs_MS.NoTrt_B |
| MTRF1 | 0.35714 | -0.3381 | 0.528 | 0.57 | 1 | HC_vs_MS.NoTrt_B |
| MTA1 | 0.35751 | 0.36131 | 0.403 | 0.38 | 1 | HC_vs_MS.NoTrt_B |
| IFIT3 | 0.35774 | 0.81862 | 0.43 | 0.48 | 1 | HC_vs_MS.NoTrt_B |
| VPS26A | 0.35815 | 0.41366 | 0.361 | 0.42 | 1 | HC_vs_MS.NoTrt_B |
| EIF1B | 0.35881 | 0.37926 | 0.159 | 0.2 | 1 | HC_vs_MS.NoTrt_B |
| FAM189B | 0.3589 | 0.42679 | 0.69 | 0.74 | 1 | HC_vs_MS.NoTrt_B |
| LSM14A | 0.35912 | 0.32585 | 0.497 | 0.44 | 1 | HC_vs_MS.NoTrt_B |
| NOL10 | 0.3592 | 0.76055 | 0.133 | 0.16 | 1 | HC_vs_MS.NoTrt_B |

| | | | | | | |
|---|---|---|---|---|---|---|
| PPP1R10 | 0.35932 | -0.322 | 0.486 | 0.5 | 1 | HC_vs_MS.NoTrt_B |
| FBXL8 | 0.35963 | 0.71689 | 0.539 | 0.55 | 1 | HC_vs_MS.NoTrt_B |
| SLX4 | 0.35996 | 1.13403 | 0.443 | 0.45 | 1 | HC_vs_MS.NoTrt_B |
| NAXE | 0.35997 | -0.3195 | 0.107 | 0.13 | 1 | HC_vs_MS.NoTrt_B |
| EXOSC7 | 0.35998 | -0.3989 | 0.446 | 0.49 | 1 | HC_vs_MS.NoTrt_B |
| C7orf49 | 0.36007 | 0.32516 | 0.145 | 0.12 | 1 | HC_vs_MS.NoTrt_B |
| ZC3H12B | 0.3604 | 1.29246 | 0.552 | 0.52 | 1 | HC_vs_MS.NoTrt_B |
| PLGRKT | 0.3605 | -0.3663 | 0.09 | 0.11 | 1 | HC_vs_MS.NoTrt_B |
| ARHGAP5 | 0.36093 | 0.91201 | 0.497 | 0.49 | 1 | HC_vs_MS.NoTrt_B |
| ITGA5 | 0.36109 | -0.4011 | 0.103 | 0.12 | 1 | HC_vs_MS.NoTrt_B |
| LRP2BP | 0.36109 | -0.4253 | 0.435 | 0.47 | 1 | HC_vs_MS.NoTrt_B |
| FAM229B | 0.36143 | 0.39699 | 0.303 | 0.28 | 1 | HC_vs_MS.NoTrt_B |
| SELENBP1 | 0.36163 | 0.38229 | 0.275 | 0.24 | 1 | HC_vs_MS.NoTrt_B |
| HUWE1 | 0.36267 | 0.51155 | 0.452 | 0.42 | 1 | HC_vs_MS.NoTrt_B |
| MTHFS | 0.36268 | 0.79634 | 0.138 | 0.11 | 1 | HC_vs_MS.NoTrt_B |
| COLEC11 | 0.36282 | 0.97606 | 0.367 | 0.34 | 1 | HC_vs_MS.NoTrt_B |
| ZNF207 | 0.36288 | 0.31179 | 0.913 | 0.9 | 1 | HC_vs_MS.NoTrt_B |
| AGL | 0.36324 | 1.08662 | 0.13 | 0.11 | 1 | HC_vs_MS.NoTrt_B |
| ARL8B | 0.36355 | 0.53662 | 0.378 | 0.35 | 1 | HC_vs_MS.NoTrt_B |
| PAPOLA | 0.36419 | 0.45978 | 0.874 | 0.85 | 1 | HC_vs_MS.NoTrt_B |
| HIPK1 | 0.36438 | 0.56796 | 0.168 | 0.14 | 1 | HC_vs_MS.NoTrt_B |
| NSMCE2 | 0.36483 | -0.3766 | 0.538 | 0.61 | 1 | HC_vs_MS.NoTrt_B |
| ZNF391 | 0.36548 | -0.3712 | 0.184 | 0.22 | 1 | HC_vs_MS.NoTrt_B |
| SAMD4A | 0.36592 | 0.62223 | 0.584 | 0.63 | 1 | HC_vs_MS.NoTrt_B |
| C2orf50 | 0.36608 | 0.49652 | 0.351 | 0.33 | 1 | HC_vs_MS.NoTrt_B |
| OR7A5 | 0.36649 | 0.29275 | 0.42 | 0.42 | 1 | HC_vs_MS.NoTrt_B |
| SH2D2A | 0.36665 | 0.82151 | 0.171 | 0.15 | 1 | HC_vs_MS.NoTrt_B |
| RAPGEF6 | 0.36713 | 0.89433 | 0.536 | 0.54 | 1 | HC_vs_MS.NoTrt_B |
| TOR3A | 0.36721 | -0.3798 | 0.828 | 0.85 | 1 | HC_vs_MS.NoTrt_B |
| NIPBL | 0.36724 | 0.49582 | 0.584 | 0.57 | 1 | HC_vs_MS.NoTrt_B |
| CTC-454I21. | 0.3677 | 0.28163 | 0.199 | 0.17 | 1 | HC_vs_MS.NoTrt_B |
| TOP2B | 0.36781 | 0.55326 | 0.261 | 0.23 | 1 | HC_vs_MS.NoTrt_B |
| ERI2 | 0.36801 | 0.4447 | 0.148 | 0.18 | 1 | HC_vs_MS.NoTrt_B |
| TSPAN6 | 0.36809 | 0.51744 | 0.53 | 0.55 | 1 | HC_vs_MS.NoTrt_B |
| MICALL1 | 0.3684 | 0.34772 | 0.462 | 0.55 | 1 | HC_vs_MS.NoTrt_B |
| ZNF337 | 0.36844 | 0.25468 | 0.081 | 0.1 | 1 | HC_vs_MS.NoTrt_B |
| PRPF4B | 0.36859 | 0.30332 | 0.343 | 0.4 | 1 | HC_vs_MS.NoTrt_B |
| IL6ST | 0.36875 | 0.60933 | 0.367 | 0.34 | 1 | HC_vs_MS.NoTrt_B |
| IRS1 | 0.36898 | -0.2609 | 0.11 | 0.09 | 1 | HC_vs_MS.NoTrt_B |
| TIMM23 | 0.3694 | 0.68774 | 0.239 | 0.22 | 1 | HC_vs_MS.NoTrt_B |

| | | | | | | |
|---|---|---|---|---|---|---|
| TRAK2 | 0.36973 | 0.5501 | 0.197 | 0.24 | 1 | HC_vs_MS.NoTrt_B |
| CD209 | 0.36989 | 0.28043 | 0.433 | 0.43 | 1 | HC_vs_MS.NoTrt_B |
| MED20 | 0.36998 | -0.2874 | 0.314 | 0.36 | 1 | HC_vs_MS.NoTrt_B |
| AP2A2 | 0.36998 | 0.76239 | 0.616 | 0.64 | 1 | HC_vs_MS.NoTrt_B |
| TRIM69 | 0.37025 | -0.2789 | 0.829 | 0.82 | 1 | HC_vs_MS.NoTrt_B |
| ITPR3 | 0.37027 | 0.40788 | 0.119 | 0.15 | 1 | HC_vs_MS.NoTrt_B |
| TEN1-CDK3 | 0.37051 | 0.52 | 0.52 | 0.49 | 1 | HC_vs_MS.NoTrt_B |
| ATP10D | 0.37076 | 0.90055 | 0.3 | 0.35 | 1 | HC_vs_MS.NoTrt_B |
| B3GAT2 | 0.37082 | 0.76329 | 0.151 | 0.12 | 1 | HC_vs_MS.NoTrt_B |
| GPATCH2L | 0.37123 | 0.33528 | 0.67 | 0.78 | 1 | HC_vs_MS.NoTrt_B |
| FZD3 | 0.37123 | 0.57109 | 0.614 | 0.63 | 1 | HC_vs_MS.NoTrt_B |
| IL1R1 | 0.37147 | 0.28567 | 0.317 | 0.29 | 1 | HC_vs_MS.NoTrt_B |
| CTBP1 | 0.37163 | 0.25704 | 0.506 | 0.46 | 1 | HC_vs_MS.NoTrt_B |
| PTCHD1 | 0.37179 | 0.43302 | 0.703 | 0.75 | 1 | HC_vs_MS.NoTrt_B |
| ZZEF1 | 0.37185 | 0.42423 | 0.501 | 0.55 | 1 | HC_vs_MS.NoTrt_B |
| ACVR2A | 0.3719 | -0.354 | 0.225 | 0.26 | 1 | HC_vs_MS.NoTrt_B |
| PLEKHG5 | 0.37232 | 1.19664 | 0.165 | 0.14 | 1 | HC_vs_MS.NoTrt_B |
| ZNF763 | 0.37278 | 0.45313 | 0.172 | 0.15 | 1 | HC_vs_MS.NoTrt_B |
| MYOM3 | 0.37284 | 0.35634 | 0.561 | 0.57 | 1 | HC_vs_MS.NoTrt_B |
| PRKG2 | 0.37286 | 0.35198 | 0.391 | 0.46 | 1 | HC_vs_MS.NoTrt_B |
| ETFA | 0.37286 | -0.3665 | 0.643 | 0.65 | 1 | HC_vs_MS.NoTrt_B |
| GTF2E1 | 0.37303 | -0.3923 | 0.254 | 0.22 | 1 | HC_vs_MS.NoTrt_B |
| TRANK1 | 0.37317 | 0.73642 | 0.246 | 0.22 | 1 | HC_vs_MS.NoTrt_B |
| ADAT1 | 0.37332 | -0.5966 | 0.671 | 0.74 | 1 | HC_vs_MS.NoTrt_B |
| CNDP1 | 0.37334 | 0.26778 | 0.577 | 0.58 | 1 | HC_vs_MS.NoTrt_B |
| BTBD8 | 0.37349 | 0.62094 | 0.293 | 0.27 | 1 | HC_vs_MS.NoTrt_B |
| ZNF569 | 0.37367 | 0.88799 | 0.183 | 0.22 | 1 | HC_vs_MS.NoTrt_B |
| EPHB4 | 0.37407 | 0.25578 | 0.79 | 0.89 | 1 | HC_vs_MS.NoTrt_B |
| NRBF2 | 0.37421 | 0.57669 | 0.086 | 0.11 | 1 | HC_vs_MS.NoTrt_B |
| TSC22D4 | 0.37436 | 0.3042 | 0.283 | 0.32 | 1 | HC_vs_MS.NoTrt_B |
| POU5F1B | 0.37436 | 0.32071 | 0.352 | 0.34 | 1 | HC_vs_MS.NoTrt_B |
| WWP1 | 0.37439 | 0.27597 | 0.432 | 0.51 | 1 | HC_vs_MS.NoTrt_B |
| POU2F1 | 0.37471 | 0.4973 | 0.419 | 0.46 | 1 | HC_vs_MS.NoTrt_B |
| CBY1 | 0.37476 | 0.53191 | 0.188 | 0.16 | 1 | HC_vs_MS.NoTrt_B |
| UGT1A1 | 0.37513 | 0.34776 | 0.551 | 0.57 | 1 | HC_vs_MS.NoTrt_B |
| CEP350 | 0.37527 | 0.30033 | 0.323 | 0.37 | 1 | HC_vs_MS.NoTrt_B |
| KCNJ5 | 0.3753 | 0.27659 | 0.597 | 0.6 | 1 | HC_vs_MS.NoTrt_B |
| MAPK1 | 0.37575 | 0.50821 | 0.483 | 0.55 | 1 | HC_vs_MS.NoTrt_B |
| TAF2 | 0.37611 | 0.98154 | 0.43 | 0.42 | 1 | HC_vs_MS.NoTrt_B |
| POLL | 0.37634 | -0.6397 | 0.404 | 0.42 | 1 | HC_vs_MS.NoTrt_B |

| | | | | | | |
|---|---|---|---|---|---|---|
| PYROXD1 | 0.3764 | 0.632 | 0.462 | 0.51 | 1 | HC_vs_MS.NoTrt_B |
| MFSD12 | 0.3769 | 1.06422 | 0.261 | 0.23 | 1 | HC_vs_MS.NoTrt_B |
| PNMA2 | 0.37771 | 0.32523 | 0.658 | 0.71 | 1 | HC_vs_MS.NoTrt_B |
| CLVS1 | 0.37795 | 0.26019 | 0.568 | 0.59 | 1 | HC_vs_MS.NoTrt_B |
| NCDN | 0.37931 | 0.90054 | 0.216 | 0.19 | 1 | HC_vs_MS.NoTrt_B |
| ANKRD36 | 0.37934 | 0.442 | 0.714 | 0.66 | 1 | HC_vs_MS.NoTrt_B |
| TDRD3 | 0.38026 | -0.682 | 0.519 | 0.55 | 1 | HC_vs_MS.NoTrt_B |
| PLXNA4 | 0.3805 | 0.2629 | 0.083 | 0.1 | 1 | HC_vs_MS.NoTrt_B |
| DNAJC24 | 0.3814 | 0.33913 | 0.629 | 0.68 | 1 | HC_vs_MS.NoTrt_B |
| HIST1H2AG | 0.38146 | 1.32028 | 0.278 | 0.34 | 1 | HC_vs_MS.NoTrt_B |
| SOS1 | 0.38157 | 0.76086 | 0.157 | 0.14 | 1 | HC_vs_MS.NoTrt_B |
| NECTIN4 | 0.38278 | -0.3061 | 0.178 | 0.21 | 1 | HC_vs_MS.NoTrt_B |
| PLCG1 | 0.38279 | 0.37357 | 0.364 | 0.42 | 1 | HC_vs_MS.NoTrt_B |
| KLHDC1 | 0.38305 | 0.46034 | 0.251 | 0.3 | 1 | HC_vs_MS.NoTrt_B |
| XKR8 | 0.38331 | 0.44256 | 0.17 | 0.14 | 1 | HC_vs_MS.NoTrt_B |
| ATMIN | 0.38392 | 0.74245 | 0.287 | 0.27 | 1 | HC_vs_MS.NoTrt_B |
| ITGA10 | 0.38414 | 0.37196 | 0.449 | 0.54 | 1 | HC_vs_MS.NoTrt_B |
| WDR6 | 0.38425 | 0.32777 | 0.41 | 0.38 | 1 | HC_vs_MS.NoTrt_B |
| MTG2 | 0.38437 | 0.25777 | 0.775 | 0.78 | 1 | HC_vs_MS.NoTrt_B |
| EML5 | 0.38443 | 1.01818 | 0.223 | 0.25 | 1 | HC_vs_MS.NoTrt_B |
| LEMD2 | 0.38486 | 0.37812 | 0.09 | 0.11 | 1 | HC_vs_MS.NoTrt_B |
| ZNF418 | 0.38526 | 0.57834 | 0.172 | 0.15 | 1 | HC_vs_MS.NoTrt_B |
| SNRNP70 | 0.38536 | 0.71238 | 0.299 | 0.28 | 1 | HC_vs_MS.NoTrt_B |
| GTF2H2C.2 | 0.38539 | 0.32162 | 0.293 | 0.27 | 1 | HC_vs_MS.NoTrt_B |
| KDM6A | 0.38576 | 0.5247 | 0.242 | 0.22 | 1 | HC_vs_MS.NoTrt_B |
| CYP3A5 | 0.38633 | 0.28186 | 0.89 | 0.95 | 1 | HC_vs_MS.NoTrt_B |
| RARS2 | 0.38644 | -0.277 | 0.586 | 0.62 | 1 | HC_vs_MS.NoTrt_B |
| SCRG1 | 0.38688 | 0.3172 | 0.714 | 0.74 | 1 | HC_vs_MS.NoTrt_B |
| ZNF551 | 0.38712 | 1.02567 | 0.513 | 0.6 | 1 | HC_vs_MS.NoTrt_B |
| DYNLT3 | 0.38759 | 0.34048 | 0.662 | 0.69 | 1 | HC_vs_MS.NoTrt_B |
| ZBTB44 | 0.38777 | 0.56978 | 0.309 | 0.28 | 1 | HC_vs_MS.NoTrt_B |
| AKT2 | 0.38836 | 0.43984 | 0.946 | 0.98 | 1 | HC_vs_MS.NoTrt_B |
| PPME1 | 0.38918 | -0.5211 | 0.116 | 0.14 | 1 | HC_vs_MS.NoTrt_B |
| CCDC50 | 0.38963 | 1.29292 | 0.443 | 0.42 | 1 | HC_vs_MS.NoTrt_B |
| RP11-506B6 | 0.38991 | 0.26993 | 0.364 | 0.35 | 1 | HC_vs_MS.NoTrt_B |
| KAT8 | 0.39023 | 0.41259 | 0.145 | 0.12 | 1 | HC_vs_MS.NoTrt_B |
| CEP83 | 0.391 | 0.3595 | 0.658 | 0.63 | 1 | HC_vs_MS.NoTrt_B |
| RUFY2 | 0.39178 | 0.50971 | 0.507 | 0.54 | 1 | HC_vs_MS.NoTrt_B |
| CARD14 | 0.39184 | 0.45355 | 0.762 | 0.8 | 1 | HC_vs_MS.NoTrt_B |
| TMEM242 | 0.39315 | 0.38076 | 0.117 | 0.1 | 1 | HC_vs_MS.NoTrt_B |

| | | | | | | |
|---|---|---|---|---|---|---|
| TMX3 | 0.39318 | 0.65614 | 0.417 | 0.41 | 1 | HC_vs_MS.NoTrt_B |
| VPS39 | 0.39374 | 0.45607 | 0.496 | 0.55 | 1 | HC_vs_MS.NoTrt_B |
| DDHD1 | 0.39387 | 0.42868 | 0.919 | 0.95 | 1 | HC_vs_MS.NoTrt_B |
| PQLC2 | 0.39461 | -0.9951 | 0.396 | 0.45 | 1 | HC_vs_MS.NoTrt_B |
| PAGR1 | 0.39464 | 0.6715 | 0.183 | 0.22 | 1 | HC_vs_MS.NoTrt_B |
| DHX33 | 0.39472 | -0.3185 | 0.317 | 0.36 | 1 | HC_vs_MS.NoTrt_B |
| EFCAB10 | 0.39524 | 0.27691 | 0.168 | 0.21 | 1 | HC_vs_MS.NoTrt_B |
| MYEF2 | 0.39528 | 0.35119 | 0.451 | 0.44 | 1 | HC_vs_MS.NoTrt_B |
| CAPN1 | 0.3953 | -0.4652 | 0.572 | 0.59 | 1 | HC_vs_MS.NoTrt_B |
| OSBP2 | 0.39554 | -1.0204 | 0.37 | 0.35 | 1 | HC_vs_MS.NoTrt_B |
| PIGN | 0.39572 | 0.39642 | 0.586 | 0.6 | 1 | HC_vs_MS.NoTrt_B |
| BRCA1 | 0.39579 | 0.25925 | 0.813 | 0.83 | 1 | HC_vs_MS.NoTrt_B |
| MSI2 | 0.39597 | 0.7417 | 0.83 | 0.85 | 1 | HC_vs_MS.NoTrt_B |
| STX5 | 0.39604 | 0.25401 | 0.083 | 0.1 | 1 | HC_vs_MS.NoTrt_B |
| VSIG1 | 0.39605 | -0.2894 | 0.593 | 0.58 | 1 | HC_vs_MS.NoTrt_B |
| HGSNAT | 0.39616 | -0.2985 | 0.67 | 0.67 | 1 | HC_vs_MS.NoTrt_B |
| SLC2A3 | 0.39622 | 0.30662 | 0.732 | 0.81 | 1 | HC_vs_MS.NoTrt_B |
| POU5F1 | 0.39625 | 0.36267 | 0.478 | 0.49 | 1 | HC_vs_MS.NoTrt_B |
| NUP188 | 0.39653 | 0.28383 | 0.526 | 0.52 | 1 | HC_vs_MS.NoTrt_B |
| PDE10A | 0.39659 | 0.26214 | 0.3 | 0.33 | 1 | HC_vs_MS.NoTrt_B |
| NEB | 0.39725 | 0.4719 | 0.532 | 0.62 | 1 | HC_vs_MS.NoTrt_B |
| IVNS1ABP | 0.39744 | 0.54617 | 0.33 | 0.3 | 1 | HC_vs_MS.NoTrt_B |
| CFAP70 | 0.39768 | -0.2925 | 0.22 | 0.2 | 1 | HC_vs_MS.NoTrt_B |
| TNFAIP8 | 0.39816 | 0.3485 | 0.864 | 0.88 | 1 | HC_vs_MS.NoTrt_B |
| LUC7L2 | 0.39852 | 0.62491 | 0.519 | 0.56 | 1 | HC_vs_MS.NoTrt_B |
| POMK | 0.39872 | 0.52791 | 0.536 | 0.57 | 1 | HC_vs_MS.NoTrt_B |
| RNASE10 | 0.39884 | 0.27666 | 0.152 | 0.13 | 1 | HC_vs_MS.NoTrt_B |
| CDC40 | 0.39905 | 0.51509 | 0.532 | 0.57 | 1 | HC_vs_MS.NoTrt_B |
| ELOVL6 | 0.39944 | -1.1521 | 0.274 | 0.3 | 1 | HC_vs_MS.NoTrt_B |
| WDR93 | 0.39984 | 0.26019 | 0.412 | 0.41 | 1 | HC_vs_MS.NoTrt_B |
| QSOX2 | 0.40011 | -0.3734 | 0.158 | 0.18 | 1 | HC_vs_MS.NoTrt_B |
| KALRN | 0.40014 | 0.57053 | 0.3 | 0.36 | 1 | HC_vs_MS.NoTrt_B |
| NIFK | 0.40027 | -0.3269 | 0.522 | 0.57 | 1 | HC_vs_MS.NoTrt_B |
| LRCH4 | 0.40042 | 0.54431 | 0.5 | 0.51 | 1 | HC_vs_MS.NoTrt_B |
| ITIH1 | 0.40048 | 0.28874 | 0.172 | 0.16 | 1 | HC_vs_MS.NoTrt_B |
| TIMELESS | 0.40074 | -0.6666 | 0.301 | 0.27 | 1 | HC_vs_MS.NoTrt_B |
| NADK | 0.40112 | 0.55338 | 0.287 | 0.27 | 1 | HC_vs_MS.NoTrt_B |
| NRP1 | 0.40123 | 0.40635 | 0.386 | 0.37 | 1 | HC_vs_MS.NoTrt_B |
| KLHL11 | 0.40127 | 0.3007 | 0.475 | 0.46 | 1 | HC_vs_MS.NoTrt_B |
| ADARB2 | 0.40279 | 0.43741 | 0.348 | 0.35 | 1 | HC_vs_MS.NoTrt_B |

| | | | | | | |
|---|---|---|---|---|---|---|
| UROS | 0.40291 | -0.3275 | 0.457 | 0.49 | 1 | HC_vs_MS.NoTrt_B |
| MICA | 0.40321 | 0.40099 | 0.716 | 0.82 | 1 | HC_vs_MS.NoTrt_B |
| ETV4 | 0.40355 | 0.46056 | 0.101 | 0.08 | 1 | HC_vs_MS.NoTrt_B |
| YWHAH | 0.40364 | -0.3191 | 0.125 | 0.15 | 1 | HC_vs_MS.NoTrt_B |
| SCARB2 | 0.40369 | 0.66581 | 0.3 | 0.29 | 1 | HC_vs_MS.NoTrt_B |
| UBR2 | 0.40373 | 0.31834 | 0.538 | 0.58 | 1 | HC_vs_MS.NoTrt_B |
| TIAF1 | 0.40401 | -0.4579 | 0.536 | 0.49 | 1 | HC_vs_MS.NoTrt_B |
| PWWP2A | 0.40433 | 0.67462 | 0.165 | 0.2 | 1 | HC_vs_MS.NoTrt_B |
| CSNK1G3 | 0.40497 | 0.40668 | 0.087 | 0.11 | 1 | HC_vs_MS.NoTrt_B |
| SH3BP2 | 0.40508 | 0.28373 | 0.771 | 0.77 | 1 | HC_vs_MS.NoTrt_B |
| C2orf88 | 0.40511 | 0.4242 | 0.345 | 0.32 | 1 | HC_vs_MS.NoTrt_B |
| WDR5B | 0.40534 | 0.38182 | 0.183 | 0.22 | 1 | HC_vs_MS.NoTrt_B |
| UTS2B | 0.40562 | 0.32182 | 0.23 | 0.27 | 1 | HC_vs_MS.NoTrt_B |
| ADARB1 | 0.40582 | 0.67587 | 0.197 | 0.17 | 1 | HC_vs_MS.NoTrt_B |
| TM9SF3 | 0.40712 | 0.62533 | 0.11 | 0.14 | 1 | HC_vs_MS.NoTrt_B |
| LACE1 | 0.40712 | -0.3106 | 0.542 | 0.6 | 1 | HC_vs_MS.NoTrt_B |
| BCO2 | 0.40721 | 0.36291 | 0.526 | 0.56 | 1 | HC_vs_MS.NoTrt_B |
| RNF14 | 0.40734 | -0.3533 | 0.603 | 0.63 | 1 | HC_vs_MS.NoTrt_B |
| ZBTB1 | 0.40759 | 0.43638 | 0.136 | 0.16 | 1 | HC_vs_MS.NoTrt_B |
| C16orf70 | 0.40769 | 0.27035 | 0.661 | 0.68 | 1 | HC_vs_MS.NoTrt_B |
| CASP3 | 0.40775 | 0.36924 | 0.12 | 0.15 | 1 | HC_vs_MS.NoTrt_B |
| CDKN2AIPNI | 0.40782 | -0.4 | 0.51 | 0.55 | 1 | HC_vs_MS.NoTrt_B |
| LUC7L3 | 0.40793 | 0.28685 | 0.604 | 0.72 | 1 | HC_vs_MS.NoTrt_B |
| NSUN6 | 0.40811 | 0.48053 | 0.675 | 0.69 | 1 | HC_vs_MS.NoTrt_B |
| USP10 | 0.40845 | 0.37765 | 0.243 | 0.28 | 1 | HC_vs_MS.NoTrt_B |
| FANCL | 0.40861 | 0.25454 | 0.12 | 0.1 | 1 | HC_vs_MS.NoTrt_B |
| EGLN1 | 0.40897 | 0.79306 | 0.354 | 0.4 | 1 | HC_vs_MS.NoTrt_B |
| ABC13-4748 | 0.41007 | 0.42577 | 0.117 | 0.1 | 1 | HC_vs_MS.NoTrt_B |
| MYO7A | 0.4103 | 0.45609 | 0.278 | 0.27 | 1 | HC_vs_MS.NoTrt_B |
| RINL | 0.41057 | -0.4419 | 0.725 | 0.76 | 1 | HC_vs_MS.NoTrt_B |
| ZNF649 | 0.41079 | -0.3744 | 0.17 | 0.2 | 1 | HC_vs_MS.NoTrt_B |
| PCIF1 | 0.4121 | 0.31003 | 0.303 | 0.35 | 1 | HC_vs_MS.NoTrt_B |
| PEX11A | 0.41215 | 0.44485 | 0.552 | 0.63 | 1 | HC_vs_MS.NoTrt_B |
| RRP15 | 0.41226 | 0.56121 | 0.567 | 0.57 | 1 | HC_vs_MS.NoTrt_B |
| LPCAT3 | 0.41269 | -0.2977 | 0.409 | 0.41 | 1 | HC_vs_MS.NoTrt_B |
| CCSAP | 0.41286 | 0.8338 | 0.139 | 0.12 | 1 | HC_vs_MS.NoTrt_B |
| TRAF3 | 0.413 | 0.3809 | 0.342 | 0.38 | 1 | HC_vs_MS.NoTrt_B |
| AGAP6 | 0.41345 | 0.94777 | 0.101 | 0.12 | 1 | HC_vs_MS.NoTrt_B |
| BIRC6 | 0.41349 | 0.38172 | 0.422 | 0.48 | 1 | HC_vs_MS.NoTrt_B |
| DNAH17 | 0.41474 | 0.42838 | 0.309 | 0.28 | 1 | HC_vs_MS.NoTrt_B |

| | | | | | | |
|---|---|---|---|---|---|---|
| PDZD8 | 0.41494 | 0.68056 | 0.249 | 0.22 | 1 | HC_vs_MS.NoTrt_B |
| WDR7 | 0.41494 | -0.3507 | 0.388 | 0.41 | 1 | HC_vs_MS.NoTrt_B |
| PANK3 | 0.41507 | 0.29945 | 0.429 | 0.4 | 1 | HC_vs_MS.NoTrt_B |
| RP11-345J4. | 0.41516 | 0.29847 | 0.757 | 0.75 | 1 | HC_vs_MS.NoTrt_B |
| SPTBN1 | 0.4156 | 0.28445 | 0.294 | 0.27 | 1 | HC_vs_MS.NoTrt_B |
| TLK2 | 0.4156 | 0.50678 | 0.371 | 0.35 | 1 | HC_vs_MS.NoTrt_B |
| FBXO15 | 0.41568 | 0.4532 | 0.604 | 0.58 | 1 | HC_vs_MS.NoTrt_B |
| POMGNT1 | 0.41572 | 0.50788 | 0.662 | 0.69 | 1 | HC_vs_MS.NoTrt_B |
| EDRF1 | 0.41572 | 0.41302 | 0.626 | 0.62 | 1 | HC_vs_MS.NoTrt_B |
| NUDT6 | 0.41622 | -0.3424 | 0.475 | 0.55 | 1 | HC_vs_MS.NoTrt_B |
| UBE2V2 | 0.4169 | 0.51967 | 0.722 | 0.72 | 1 | HC_vs_MS.NoTrt_B |
| FBXO11 | 0.41701 | 0.8664 | 0.441 | 0.49 | 1 | HC_vs_MS.NoTrt_B |
| NHLRC2 | 0.41718 | 0.65714 | 0.707 | 0.72 | 1 | HC_vs_MS.NoTrt_B |
| ZNF862 | 0.41785 | 0.46114 | 0.107 | 0.13 | 1 | HC_vs_MS.NoTrt_B |
| GGT1 | 0.418 | -0.4478 | 0.088 | 0.11 | 1 | HC_vs_MS.NoTrt_B |
| KLHL36 | 0.41831 | 0.68909 | 0.636 | 0.65 | 1 | HC_vs_MS.NoTrt_B |
| NEDD9 | 0.41861 | 0.35244 | 0.467 | 0.42 | 1 | HC_vs_MS.NoTrt_B |
| CC2D2B | 0.41867 | 0.96456 | 0.125 | 0.11 | 1 | HC_vs_MS.NoTrt_B |
| MYLK | 0.41869 | -0.9015 | 0.562 | 0.55 | 1 | HC_vs_MS.NoTrt_B |
| FMNL1 | 0.41869 | 0.44397 | 0.701 | 0.71 | 1 | HC_vs_MS.NoTrt_B |
| TMEM98 | 0.41903 | 0.25417 | 0.528 | 0.54 | 1 | HC_vs_MS.NoTrt_B |
| G6PC2 | 0.41925 | 0.26893 | 0.272 | 0.25 | 1 | HC_vs_MS.NoTrt_B |
| LSM10 | 0.41944 | 0.31591 | 0.119 | 0.1 | 1 | HC_vs_MS.NoTrt_B |
| ZNF81 | 0.41961 | 0.48735 | 0.29 | 0.27 | 1 | HC_vs_MS.NoTrt_B |
| ZHX3 | 0.41967 | 0.38105 | 0.401 | 0.4 | 1 | HC_vs_MS.NoTrt_B |
| MUC20 | 0.41992 | 0.97978 | 0.536 | 0.56 | 1 | HC_vs_MS.NoTrt_B |
| SLC24A1 | 0.42017 | 0.79319 | 0.468 | 0.45 | 1 | HC_vs_MS.NoTrt_B |
| ALG12 | 0.4204 | 0.51131 | 0.146 | 0.12 | 1 | HC_vs_MS.NoTrt_B |
| PTGR1 | 0.42053 | 0.25676 | 0.6 | 0.69 | 1 | HC_vs_MS.NoTrt_B |
| GMEB2 | 0.42083 | 0.33825 | 0.088 | 0.11 | 1 | HC_vs_MS.NoTrt_B |
| MSH6 | 0.42148 | 0.33971 | 0.865 | 0.86 | 1 | HC_vs_MS.NoTrt_B |
| C12orf29 | 0.42149 | -0.3271 | 0.199 | 0.22 | 1 | HC_vs_MS.NoTrt_B |
| CNTLN | 0.42152 | 0.38466 | 0.328 | 0.31 | 1 | HC_vs_MS.NoTrt_B |
| SLC39A9 | 0.42187 | 0.62176 | 0.174 | 0.15 | 1 | HC_vs_MS.NoTrt_B |
| WAS | 0.42195 | 0.72355 | 0.278 | 0.32 | 1 | HC_vs_MS.NoTrt_B |
| CPEB1 | 0.422 | -0.2773 | 0.146 | 0.12 | 1 | HC_vs_MS.NoTrt_B |
| TIGD5 | 0.42204 | 0.86206 | 0.199 | 0.17 | 1 | HC_vs_MS.NoTrt_B |
| MBNL1 | 0.42205 | 0.35659 | 0.859 | 0.91 | 1 | HC_vs_MS.NoTrt_B |
| ALG1 | 0.42232 | -0.2509 | 0.613 | 0.65 | 1 | HC_vs_MS.NoTrt_B |
| DENND6B | 0.42252 | -0.3485 | 0.259 | 0.3 | 1 | HC_vs_MS.NoTrt_B |

| | | | | | | |
|---|---|---|---|---|---|---|
| RND1 | 0.42265 | 0.33581 | 0.551 | 0.56 | 1 | HC_vs_MS.NoTrt_B |
| PRKCB | 0.42325 | 0.28048 | 0.291 | 0.32 | 1 | HC_vs_MS.NoTrt_B |
| PHACTR2 | 0.42354 | 0.31326 | 0.481 | 0.5 | 1 | HC_vs_MS.NoTrt_B |
| KLHL9 | 0.42363 | 0.31129 | 0.343 | 0.34 | 1 | HC_vs_MS.NoTrt_B |
| IL18BP | 0.42371 | 0.70221 | 0.106 | 0.09 | 1 | HC_vs_MS.NoTrt_B |
| SLC25A36 | 0.42372 | 0.47406 | 0.129 | 0.15 | 1 | HC_vs_MS.NoTrt_B |
| TOR1AIP1 | 0.42387 | -0.343 | 0.53 | 0.49 | 1 | HC_vs_MS.NoTrt_B |
| ASRGL1 | 0.42401 | 0.38658 | 0.548 | 0.54 | 1 | HC_vs_MS.NoTrt_B |
| DMXL1 | 0.42482 | 0.58673 | 0.171 | 0.15 | 1 | HC_vs_MS.NoTrt_B |
| UMAD1 | 0.42495 | 0.4054 | 0.138 | 0.16 | 1 | HC_vs_MS.NoTrt_B |
| ZNF208 | 0.42531 | 0.47152 | 0.816 | 0.85 | 1 | HC_vs_MS.NoTrt_B |
| SIRPB1 | 0.42543 | 1.70955 | 0.216 | 0.21 | 1 | HC_vs_MS.NoTrt_B |
| EIF4G3 | 0.42556 | 0.90872 | 0.21 | 0.19 | 1 | HC_vs_MS.NoTrt_B |
| GTF3C6 | 0.42619 | 0.43661 | 0.151 | 0.13 | 1 | HC_vs_MS.NoTrt_B |
| ACTR3B | 0.42656 | 0.58212 | 0.493 | 0.55 | 1 | HC_vs_MS.NoTrt_B |
| TBXAS1 | 0.42705 | -0.3638 | 0.371 | 0.42 | 1 | HC_vs_MS.NoTrt_B |
| EIF2B1 | 0.42723 | -0.3939 | 0.329 | 0.34 | 1 | HC_vs_MS.NoTrt_B |
| FAM208B | 0.42781 | 0.36688 | 0.294 | 0.28 | 1 | HC_vs_MS.NoTrt_B |
| ZNF384 | 0.42901 | 0.54519 | 0.145 | 0.12 | 1 | HC_vs_MS.NoTrt_B |
| ERVK3-1 | 0.42902 | 0.32176 | 0.274 | 0.25 | 1 | HC_vs_MS.NoTrt_B |
| C5orf51 | 0.4291 | 0.2671 | 0.099 | 0.12 | 1 | HC_vs_MS.NoTrt_B |
| EXO1 | 0.42929 | -0.8971 | 0.164 | 0.2 | 1 | HC_vs_MS.NoTrt_B |
| PRR27 | 0.42946 | 0.26292 | 0.477 | 0.49 | 1 | HC_vs_MS.NoTrt_B |
| NLGN4Y | 0.42953 | 0.39826 | 0.355 | 0.35 | 1 | HC_vs_MS.NoTrt_B |
| TAF3 | 0.4296 | 0.47467 | 0.151 | 0.13 | 1 | HC_vs_MS.NoTrt_B |
| PAFAH2 | 0.42998 | 0.38779 | 0.47 | 0.52 | 1 | HC_vs_MS.NoTrt_B |
| KDM3B | 0.43019 | 0.54493 | 0.381 | 0.35 | 1 | HC_vs_MS.NoTrt_B |
| PBRM1 | 0.43065 | 0.43542 | 0.662 | 0.7 | 1 | HC_vs_MS.NoTrt_B |
| ZSCAN22 | 0.43069 | -0.8336 | 0.399 | 0.41 | 1 | HC_vs_MS.NoTrt_B |
| CD59 | 0.43116 | -0.2731 | 0.675 | 0.7 | 1 | HC_vs_MS.NoTrt_B |
| ESRRG | 0.43118 | 0.26146 | 0.368 | 0.35 | 1 | HC_vs_MS.NoTrt_B |
| DDX43 | 0.43135 | 0.85005 | 0.168 | 0.15 | 1 | HC_vs_MS.NoTrt_B |
| ERI3 | 0.43172 | 0.52529 | 0.087 | 0.11 | 1 | HC_vs_MS.NoTrt_B |
| HS1BP3 | 0.43234 | -0.2575 | 0.154 | 0.18 | 1 | HC_vs_MS.NoTrt_B |
| ACADSB | 0.43257 | 0.34595 | 0.628 | 0.72 | 1 | HC_vs_MS.NoTrt_B |
| EDA | 0.43274 | 0.84701 | 0.33 | 0.37 | 1 | HC_vs_MS.NoTrt_B |
| ABCB8 | 0.43291 | -0.3843 | 0.533 | 0.5 | 1 | HC_vs_MS.NoTrt_B |
| GALNT10 | 0.43316 | -0.3413 | 0.53 | 0.57 | 1 | HC_vs_MS.NoTrt_B |
| NECAP1 | 0.43334 | 0.50227 | 0.436 | 0.41 | 1 | HC_vs_MS.NoTrt_B |
| DOT1L | 0.43336 | 0.68016 | 0.729 | 0.77 | 1 | HC_vs_MS.NoTrt_B |

| | | | | | | |
|---|---|---|---|---|---|---|
| PIP5K1B | 0.43364 | 0.82816 | 0.116 | 0.1 | 1 | HC_vs_MS.NoTrt_B |
| HSH2D | 0.43457 | -0.6167 | 0.601 | 0.63 | 1 | HC_vs_MS.NoTrt_B |
| ABHD15 | 0.43494 | 0.35639 | 0.329 | 0.37 | 1 | HC_vs_MS.NoTrt_B |
| CMC1 | 0.4354 | 0.81381 | 0.161 | 0.19 | 1 | HC_vs_MS.NoTrt_B |
| BCKDHA | 0.43556 | -0.3114 | 0.175 | 0.2 | 1 | HC_vs_MS.NoTrt_B |
| EMSY | 0.43564 | 0.41397 | 0.49 | 0.48 | 1 | HC_vs_MS.NoTrt_B |
| DALRD3 | 0.43617 | -0.3801 | 0.116 | 0.14 | 1 | HC_vs_MS.NoTrt_B |
| ATP6V1A | 0.43646 | -0.3996 | 0.472 | 0.5 | 1 | HC_vs_MS.NoTrt_B |
| TUBB4A | 0.43655 | -0.3186 | 0.19 | 0.17 | 1 | HC_vs_MS.NoTrt_B |
| ZKSCAN4 | 0.43663 | -0.2911 | 0.435 | 0.42 | 1 | HC_vs_MS.NoTrt_B |
| SMC1A | 0.43664 | 0.65775 | 0.249 | 0.23 | 1 | HC_vs_MS.NoTrt_B |
| SLC25A23 | 0.43668 | 0.83951 | 0.507 | 0.52 | 1 | HC_vs_MS.NoTrt_B |
| ALDH3A2 | 0.43729 | -0.5874 | 0.262 | 0.29 | 1 | HC_vs_MS.NoTrt_B |
| BICDL1 | 0.4374 | 0.25604 | 0.581 | 0.57 | 1 | HC_vs_MS.NoTrt_B |
| TMEM52B | 0.43783 | 0.44563 | 0.658 | 0.7 | 1 | HC_vs_MS.NoTrt_B |
| CHRNA6 | 0.43792 | -0.4292 | 0.425 | 0.5 | 1 | HC_vs_MS.NoTrt_B |
| FAM20B | 0.43822 | -0.8039 | 0.565 | 0.62 | 1 | HC_vs_MS.NoTrt_B |
| ZFAND6 | 0.43879 | 0.3989 | 0.203 | 0.18 | 1 | HC_vs_MS.NoTrt_B |
| PKIB | 0.43919 | 0.28454 | 0.7 | 0.74 | 1 | HC_vs_MS.NoTrt_B |
| HIST1H2BG | 0.43931 | 1.2591 | 0.472 | 0.47 | 1 | HC_vs_MS.NoTrt_B |
| LZTFL1 | 0.43952 | -0.3502 | 0.532 | 0.57 | 1 | HC_vs_MS.NoTrt_B |
| VDR | 0.43969 | -0.5619 | 0.288 | 0.25 | 1 | HC_vs_MS.NoTrt_B |
| GRIPAP1 | 0.43978 | -0.2895 | 0.729 | 0.77 | 1 | HC_vs_MS.NoTrt_B |
| THADA | 0.43982 | 0.40249 | 0.854 | 0.85 | 1 | HC_vs_MS.NoTrt_B |
| SLFN13 | 0.43993 | 0.45322 | 0.574 | 0.57 | 1 | HC_vs_MS.NoTrt_B |
| IL21R | 0.44004 | 0.25062 | 0.523 | 0.57 | 1 | HC_vs_MS.NoTrt_B |
| ATP8B2 | 0.44039 | 0.25104 | 0.807 | 0.83 | 1 | HC_vs_MS.NoTrt_B |
| CCDC192 | 0.44062 | 0.3227 | 0.093 | 0.11 | 1 | HC_vs_MS.NoTrt_B |
| MAP3K4 | 0.4412 | 0.82391 | 0.183 | 0.16 | 1 | HC_vs_MS.NoTrt_B |
| PLIN3 | 0.44123 | -0.4675 | 0.519 | 0.48 | 1 | HC_vs_MS.NoTrt_B |
| NLE1 | 0.44156 | 1.12252 | 0.214 | 0.2 | 1 | HC_vs_MS.NoTrt_B |
| ACSBG1 | 0.44164 | 0.26348 | 0.361 | 0.35 | 1 | HC_vs_MS.NoTrt_B |
| NCAPD2 | 0.44208 | -0.325 | 0.403 | 0.44 | 1 | HC_vs_MS.NoTrt_B |
| DSC2 | 0.44247 | -1.0585 | 0.291 | 0.27 | 1 | HC_vs_MS.NoTrt_B |
| MED12 | 0.44254 | 0.31944 | 0.172 | 0.2 | 1 | HC_vs_MS.NoTrt_B |
| HELB | 0.44264 | 0.37032 | 0.899 | 0.91 | 1 | HC_vs_MS.NoTrt_B |
| PCED1A | 0.44303 | 0.27343 | 0.136 | 0.16 | 1 | HC_vs_MS.NoTrt_B |
| HMGA2 | 0.44311 | 0.25277 | 0.612 | 0.62 | 1 | HC_vs_MS.NoTrt_B |
| ZNF254 | 0.44328 | 0.61499 | 0.397 | 0.41 | 1 | HC_vs_MS.NoTrt_B |
| FSD1 | 0.44457 | 0.25422 | 0.39 | 0.38 | 1 | HC_vs_MS.NoTrt_B |

| | | | | | | |
|---|---|---|---|---|---|---|
| CLN6 | 0.44458 | 0.32776 | 0.759 | 0.82 | 1 | HC_vs_MS.NoTrt_B |
| NBPF12 | 0.44485 | 0.84204 | 0.312 | 0.35 | 1 | HC_vs_MS.NoTrt_B |
| ARL17A | 0.44513 | 0.26192 | 0.691 | 0.75 | 1 | HC_vs_MS.NoTrt_B |
| INTS1 | 0.44543 | 0.527 | 0.413 | 0.37 | 1 | HC_vs_MS.NoTrt_B |
| FAM217B | 0.44559 | 0.58982 | 0.352 | 0.35 | 1 | HC_vs_MS.NoTrt_B |
| NCLN | 0.44588 | 0.36631 | 0.122 | 0.15 | 1 | HC_vs_MS.NoTrt_B |
| SETD2 | 0.44629 | 0.36037 | 0.342 | 0.37 | 1 | HC_vs_MS.NoTrt_B |
| PARVA | 0.44635 | 0.343 | 0.719 | 0.72 | 1 | HC_vs_MS.NoTrt_B |
| ZNF644 | 0.44669 | 0.63618 | 0.194 | 0.17 | 1 | HC_vs_MS.NoTrt_B |
| IGF2BP3 | 0.44675 | 0.26493 | 0.545 | 0.55 | 1 | HC_vs_MS.NoTrt_B |
| NDUFS8 | 0.44689 | -0.2902 | 0.767 | 0.84 | 1 | HC_vs_MS.NoTrt_B |
| NUP85 | 0.44736 | 0.87072 | 0.475 | 0.49 | 1 | HC_vs_MS.NoTrt_B |
| SCLT1 | 0.44762 | 0.29656 | 0.614 | 0.63 | 1 | HC_vs_MS.NoTrt_B |
| ZNF706 | 0.44774 | 0.25308 | 0.372 | 0.42 | 1 | HC_vs_MS.NoTrt_B |
| APBB1 | 0.44786 | 0.34482 | 0.238 | 0.28 | 1 | HC_vs_MS.NoTrt_B |
| GALNT16 | 0.44798 | 0.56369 | 0.552 | 0.58 | 1 | HC_vs_MS.NoTrt_B |
| PCDHGA8 | 0.44818 | -0.3388 | 0.596 | 0.58 | 1 | HC_vs_MS.NoTrt_B |
| ANKRD17 | 0.44829 | 0.32785 | 0.251 | 0.28 | 1 | HC_vs_MS.NoTrt_B |
| EIF5A | 0.44849 | 1.21909 | 0.577 | 0.67 | 1 | HC_vs_MS.NoTrt_B |
| NETO2 | 0.44858 | 0.42018 | 0.532 | 0.55 | 1 | HC_vs_MS.NoTrt_B |
| ACSL1 | 0.44898 | 0.36175 | 0.239 | 0.21 | 1 | HC_vs_MS.NoTrt_B |
| SYNE4 | 0.44928 | 0.60967 | 0.409 | 0.41 | 1 | HC_vs_MS.NoTrt_B |
| ENOX2 | 0.44938 | 0.58457 | 0.088 | 0.11 | 1 | HC_vs_MS.NoTrt_B |
| ORMDL2 | 0.44958 | 0.2741 | 0.452 | 0.49 | 1 | HC_vs_MS.NoTrt_B |
| MATK | 0.44961 | 0.36755 | 0.559 | 0.61 | 1 | HC_vs_MS.NoTrt_B |
| PTAR1 | 0.44981 | 0.27509 | 0.168 | 0.2 | 1 | HC_vs_MS.NoTrt_B |
| C11orf24 | 0.44986 | 1.95302 | 0.246 | 0.23 | 1 | HC_vs_MS.NoTrt_B |
| GRHL1 | 0.45016 | 0.37721 | 0.484 | 0.49 | 1 | HC_vs_MS.NoTrt_B |
| MIS12 | 0.45017 | -0.4792 | 0.291 | 0.34 | 1 | HC_vs_MS.NoTrt_B |
| SNX3 | 0.45094 | 0.56773 | 0.158 | 0.14 | 1 | HC_vs_MS.NoTrt_B |
| FNIP1 | 0.45097 | 0.62115 | 0.575 | 0.55 | 1 | HC_vs_MS.NoTrt_B |
| ARHGEF10 | 0.45098 | 0.34669 | 0.607 | 0.58 | 1 | HC_vs_MS.NoTrt_B |
| RIT1 | 0.45107 | 0.26676 | 0.388 | 0.42 | 1 | HC_vs_MS.NoTrt_B |
| TNS1 | 0.4513 | 0.2931 | 0.245 | 0.23 | 1 | HC_vs_MS.NoTrt_B |
| ACTR6 | 0.45161 | -0.2664 | 0.513 | 0.54 | 1 | HC_vs_MS.NoTrt_B |
| KPNB1 | 0.4525 | 0.51037 | 0.697 | 0.69 | 1 | HC_vs_MS.NoTrt_B |
| BBX | 0.45262 | 0.26628 | 0.325 | 0.37 | 1 | HC_vs_MS.NoTrt_B |
| MASP2 | 0.4529 | -0.2589 | 0.236 | 0.27 | 1 | HC_vs_MS.NoTrt_B |
| ZNF91 | 0.45328 | 0.71442 | 0.787 | 0.79 | 1 | HC_vs_MS.NoTrt_B |
| ANKRD27 | 0.45343 | 0.41729 | 0.193 | 0.22 | 1 | HC_vs_MS.NoTrt_B |

| | | | | | | |
|---|---|---|---|---|---|---|
| COL6A3 | 0.45351 | 0.28699 | 0.452 | 0.45 | 1 | HC_vs_MS.NoTrt_B |
| DCBLD1 | 0.45378 | 0.80916 | 0.504 | 0.52 | 1 | HC_vs_MS.NoTrt_B |
| GSK3A | 0.45381 | 0.83017 | 0.328 | 0.32 | 1 | HC_vs_MS.NoTrt_B |
| GDI1 | 0.45417 | 0.69055 | 0.274 | 0.32 | 1 | HC_vs_MS.NoTrt_B |
| NR1H2 | 0.45443 | 0.30978 | 0.507 | 0.55 | 1 | HC_vs_MS.NoTrt_B |
| FAM210A | 0.45471 | 0.69832 | 0.257 | 0.3 | 1 | HC_vs_MS.NoTrt_B |
| ZNF485 | 0.45477 | 0.58658 | 0.186 | 0.17 | 1 | HC_vs_MS.NoTrt_B |
| TRIM34 | 0.45509 | -0.5758 | 0.314 | 0.3 | 1 | HC_vs_MS.NoTrt_B |
| ERCC8 | 0.45587 | 0.35772 | 0.341 | 0.37 | 1 | HC_vs_MS.NoTrt_B |
| CBX6 | 0.45605 | 0.36072 | 0.455 | 0.53 | 1 | HC_vs_MS.NoTrt_B |
| MTHFD1L | 0.45643 | 0.77747 | 0.125 | 0.11 | 1 | HC_vs_MS.NoTrt_B |
| TIPIN | 0.45713 | 0.51341 | 0.562 | 0.59 | 1 | HC_vs_MS.NoTrt_B |
| SLAIN2 | 0.4574 | 0.2781 | 0.148 | 0.17 | 1 | HC_vs_MS.NoTrt_B |
| EXOC3 | 0.45744 | -0.354 | 0.214 | 0.24 | 1 | HC_vs_MS.NoTrt_B |
| CNOT6 | 0.45755 | 0.3358 | 0.09 | 0.11 | 1 | HC_vs_MS.NoTrt_B |
| TANGO6 | 0.45788 | 0.31097 | 0.797 | 0.85 | 1 | HC_vs_MS.NoTrt_B |
| ZNF2 | 0.45789 | 1.23586 | 0.316 | 0.36 | 1 | HC_vs_MS.NoTrt_B |
| GOPC | 0.45812 | 0.90277 | 0.103 | 0.09 | 1 | HC_vs_MS.NoTrt_B |
| KTN1 | 0.45828 | 0.32264 | 0.571 | 0.67 | 1 | HC_vs_MS.NoTrt_B |
| SLC2A1 | 0.4595 | -0.2885 | 0.086 | 0.1 | 1 | HC_vs_MS.NoTrt_B |
| HDDC2 | 0.45958 | 0.4698 | 0.116 | 0.1 | 1 | HC_vs_MS.NoTrt_B |
| DGKZ | 0.45977 | 0.45732 | 0.422 | 0.46 | 1 | HC_vs_MS.NoTrt_B |
| CCNT1 | 0.46013 | 0.50254 | 0.116 | 0.1 | 1 | HC_vs_MS.NoTrt_B |
| STK39 | 0.46046 | 0.42374 | 0.103 | 0.12 | 1 | HC_vs_MS.NoTrt_B |
| ZNF624 | 0.46118 | 0.47564 | 0.436 | 0.43 | 1 | HC_vs_MS.NoTrt_B |
| WDR89 | 0.46122 | 0.30037 | 0.362 | 0.41 | 1 | HC_vs_MS.NoTrt_B |
| UFC1 | 0.46179 | -0.2572 | 0.778 | 0.76 | 1 | HC_vs_MS.NoTrt_B |
| PNRC1 | 0.46227 | 0.82826 | 0.31 | 0.3 | 1 | HC_vs_MS.NoTrt_B |
| UBE3A | 0.46274 | 0.59822 | 0.568 | 0.55 | 1 | HC_vs_MS.NoTrt_B |
| CYP3A4 | 0.46345 | 0.28952 | 0.157 | 0.14 | 1 | HC_vs_MS.NoTrt_B |
| DZIP3 | 0.46378 | 0.8417 | 0.257 | 0.24 | 1 | HC_vs_MS.NoTrt_B |
| CARF | 0.464 | -0.5537 | 0.486 | 0.51 | 1 | HC_vs_MS.NoTrt_B |
| EIF2AK4 | 0.46405 | 0.29335 | 0.642 | 0.67 | 1 | HC_vs_MS.NoTrt_B |
| PRKCE | 0.46406 | 0.63055 | 0.136 | 0.12 | 1 | HC_vs_MS.NoTrt_B |
| LTA4H | 0.46408 | 0.4162 | 0.33 | 0.3 | 1 | HC_vs_MS.NoTrt_B |
| CLDN15 | 0.46455 | 0.47454 | 0.394 | 0.37 | 1 | HC_vs_MS.NoTrt_B |
| AK6 | 0.46527 | -0.259 | 0.184 | 0.21 | 1 | HC_vs_MS.NoTrt_B |
| ZNF44 | 0.46618 | -0.5002 | 0.535 | 0.52 | 1 | HC_vs_MS.NoTrt_B |
| ZNF461 | 0.46637 | -0.3746 | 0.664 | 0.64 | 1 | HC_vs_MS.NoTrt_B |
| FAM47E-STE | 0.46691 | 0.4321 | 0.174 | 0.21 | 1 | HC_vs_MS.NoTrt_B |

| | | | | | | |
|---|---|---|---|---|---|---|
| C22orf46 | 0.46814 | 0.41222 | 0.364 | 0.35 | 1 | HC_vs_MS.NoTrt_B |
| NEMF | 0.46877 | 0.63174 | 0.777 | 0.8 | 1 | HC_vs_MS.NoTrt_B |
| RRM2B | 0.46939 | 0.55336 | 0.117 | 0.14 | 1 | HC_vs_MS.NoTrt_B |
| PPP4C | 0.46942 | 0.25749 | 0.455 | 0.42 | 1 | HC_vs_MS.NoTrt_B |
| ZNF548 | 0.46961 | 0.46024 | 0.638 | 0.67 | 1 | HC_vs_MS.NoTrt_B |
| ZFHX3 | 0.46978 | 1.1166 | 0.143 | 0.12 | 1 | HC_vs_MS.NoTrt_B |
| RIPK2 | 0.47013 | 0.52264 | 0.161 | 0.18 | 1 | HC_vs_MS.NoTrt_B |
| BRD2 | 0.47044 | 0.34273 | 0.546 | 0.6 | 1 | HC_vs_MS.NoTrt_B |
| KCNV1 | 0.4724 | 0.27507 | 0.581 | 0.6 | 1 | HC_vs_MS.NoTrt_B |
| PRH1 | 0.47247 | 0.25697 | 0.086 | 0.1 | 1 | HC_vs_MS.NoTrt_B |
| ATR | 0.47273 | 0.3771 | 0.468 | 0.52 | 1 | HC_vs_MS.NoTrt_B |
| RRP8 | 0.47277 | 0.56595 | 0.486 | 0.47 | 1 | HC_vs_MS.NoTrt_B |
| GBA | 0.47285 | -0.8293 | 0.458 | 0.5 | 1 | HC_vs_MS.NoTrt_B |
| ST7L | 0.473 | 0.27679 | 0.468 | 0.47 | 1 | HC_vs_MS.NoTrt_B |
| FAM71D | 0.47313 | 0.31731 | 0.12 | 0.1 | 1 | HC_vs_MS.NoTrt_B |
| NPIPB9 | 0.47314 | 0.51336 | 0.114 | 0.1 | 1 | HC_vs_MS.NoTrt_B |
| CXorf57 | 0.47328 | -0.4623 | 0.11 | 0.09 | 1 | HC_vs_MS.NoTrt_B |
| VPS25 | 0.47328 | -0.3821 | 0.255 | 0.28 | 1 | HC_vs_MS.NoTrt_B |
| PRKCD | 0.47365 | 0.52606 | 0.477 | 0.48 | 1 | HC_vs_MS.NoTrt_B |
| FRMPD2 | 0.4738 | 0.32859 | 0.13 | 0.11 | 1 | HC_vs_MS.NoTrt_B |
| RBM20 | 0.47397 | 0.32569 | 0.438 | 0.42 | 1 | HC_vs_MS.NoTrt_B |
| MAP2K6 | 0.47434 | 1.15654 | 0.333 | 0.32 | 1 | HC_vs_MS.NoTrt_B |
| WDR17 | 0.4745 | 0.30338 | 0.716 | 0.76 | 1 | HC_vs_MS.NoTrt_B |
| OTUD3 | 0.47491 | 1.45506 | 0.5 | 0.49 | 1 | HC_vs_MS.NoTrt_B |
| TP53I13 | 0.47512 | 0.77309 | 0.116 | 0.14 | 1 | HC_vs_MS.NoTrt_B |
| PPA1 | 0.47546 | -0.3828 | 0.797 | 0.84 | 1 | HC_vs_MS.NoTrt_B |
| COQ3 | 0.47562 | -0.2523 | 0.23 | 0.26 | 1 | HC_vs_MS.NoTrt_B |
| FKBP4 | 0.47638 | -0.7941 | 0.094 | 0.11 | 1 | HC_vs_MS.NoTrt_B |
| RUVBL2 | 0.4769 | -0.6884 | 0.854 | 0.92 | 1 | HC_vs_MS.NoTrt_B |
| MCMBP | 0.47708 | 1.24821 | 0.423 | 0.41 | 1 | HC_vs_MS.NoTrt_B |
| SSRP1 | 0.47714 | 0.59417 | 0.113 | 0.14 | 1 | HC_vs_MS.NoTrt_B |
| ZNF250 | 0.47717 | -0.2568 | 0.552 | 0.6 | 1 | HC_vs_MS.NoTrt_B |
| TMEM184C | 0.47788 | 0.64508 | 0.533 | 0.58 | 1 | HC_vs_MS.NoTrt_B |
| SLC46A1 | 0.47798 | 0.39762 | 0.484 | 0.45 | 1 | HC_vs_MS.NoTrt_B |
| SFT2D3 | 0.47799 | 0.41847 | 0.419 | 0.47 | 1 | HC_vs_MS.NoTrt_B |
| INTS2 | 0.47874 | 0.66745 | 0.235 | 0.22 | 1 | HC_vs_MS.NoTrt_B |
| BMF | 0.47907 | 0.25283 | 0.139 | 0.12 | 1 | HC_vs_MS.NoTrt_B |
| C6orf163 | 0.4792 | 0.43453 | 0.452 | 0.49 | 1 | HC_vs_MS.NoTrt_B |
| PARP14 | 0.47986 | 0.56981 | 0.409 | 0.39 | 1 | HC_vs_MS.NoTrt_B |
| RBM45 | 0.47986 | 0.40039 | 0.086 | 0.1 | 1 | HC_vs_MS.NoTrt_B |

| | | | | | | |
|---|---|---|---|---|---|---|
| MYO9B | 0.48006 | 0.53927 | 0.22 | 0.26 | 1 | HC_vs_MS.NoTrt_B |
| CHTOP | 0.4801 | 0.62069 | 0.593 | 0.59 | 1 | HC_vs_MS.NoTrt_B |
| ZNF549 | 0.48026 | 1.24669 | 0.688 | 0.7 | 1 | HC_vs_MS.NoTrt_B |
| AKAP11 | 0.48028 | 0.53535 | 0.226 | 0.21 | 1 | HC_vs_MS.NoTrt_B |
| IFNLR1 | 0.48037 | 0.26085 | 0.109 | 0.13 | 1 | HC_vs_MS.NoTrt_B |
| TEKT2 | 0.48055 | 0.46229 | 0.117 | 0.14 | 1 | HC_vs_MS.NoTrt_B |
| UBXN8 | 0.48107 | 0.67516 | 0.177 | 0.21 | 1 | HC_vs_MS.NoTrt_B |
| SLC41A1 | 0.48138 | 0.48906 | 0.103 | 0.09 | 1 | HC_vs_MS.NoTrt_B |
| SLC18A1 | 0.4825 | 0.26881 | 0.143 | 0.17 | 1 | HC_vs_MS.NoTrt_B |
| CTNND1 | 0.48255 | 0.52487 | 0.304 | 0.3 | 1 | HC_vs_MS.NoTrt_B |
| RFC5 | 0.48311 | -0.5294 | 0.119 | 0.14 | 1 | HC_vs_MS.NoTrt_B |
| EME2 | 0.48318 | 0.95579 | 0.286 | 0.32 | 1 | HC_vs_MS.NoTrt_B |
| MLLT10 | 0.48325 | 0.69222 | 0.3 | 0.33 | 1 | HC_vs_MS.NoTrt_B |
| SMIM15 | 0.48326 | 0.82883 | 0.088 | 0.11 | 1 | HC_vs_MS.NoTrt_B |
| SYNC | 0.48405 | 0.78115 | 0.123 | 0.15 | 1 | HC_vs_MS.NoTrt_B |
| DENND4B | 0.48415 | 0.38123 | 0.207 | 0.24 | 1 | HC_vs_MS.NoTrt_B |
| MRPS27 | 0.48433 | -0.6554 | 0.514 | 0.51 | 1 | HC_vs_MS.NoTrt_B |
| CHP2 | 0.48451 | 0.32382 | 0.407 | 0.41 | 1 | HC_vs_MS.NoTrt_B |
| PPID | 0.48456 | -0.2994 | 0.352 | 0.37 | 1 | HC_vs_MS.NoTrt_B |
| FADS2 | 0.48495 | 0.46751 | 0.486 | 0.54 | 1 | HC_vs_MS.NoTrt_B |
| MOCS2 | 0.48517 | 0.58865 | 0.084 | 0.1 | 1 | HC_vs_MS.NoTrt_B |
| U2AF2 | 0.48567 | 0.32536 | 0.186 | 0.17 | 1 | HC_vs_MS.NoTrt_B |
| SIDT2 | 0.48589 | 0.50346 | 0.751 | 0.79 | 1 | HC_vs_MS.NoTrt_B |
| RAB4B | 0.48665 | 0.28116 | 0.377 | 0.35 | 1 | HC_vs_MS.NoTrt_B |
| REV3L | 0.48702 | 0.38727 | 0.675 | 0.72 | 1 | HC_vs_MS.NoTrt_B |
| FSBP | 0.48713 | 0.87182 | 0.314 | 0.36 | 1 | HC_vs_MS.NoTrt_B |
| ELAC1 | 0.48717 | 0.29548 | 0.439 | 0.44 | 1 | HC_vs_MS.NoTrt_B |
| SLC30A8 | 0.48721 | 0.29605 | 0.401 | 0.38 | 1 | HC_vs_MS.NoTrt_B |
| XPOT | 0.48748 | 0.39241 | 0.432 | 0.41 | 1 | HC_vs_MS.NoTrt_B |
| HYKK | 0.48762 | 0.41152 | 0.813 | 0.86 | 1 | HC_vs_MS.NoTrt_B |
| GOLPH3L | 0.48783 | 0.44043 | 0.21 | 0.24 | 1 | HC_vs_MS.NoTrt_B |
| TONSL | 0.48813 | 0.78766 | 0.597 | 0.62 | 1 | HC_vs_MS.NoTrt_B |
| CASC3 | 0.48829 | 0.32466 | 0.533 | 0.52 | 1 | HC_vs_MS.NoTrt_B |
| ATP13A1 | 0.48858 | 0.65139 | 0.322 | 0.3 | 1 | HC_vs_MS.NoTrt_B |
| FAM214A | 0.48874 | 0.62962 | 0.193 | 0.22 | 1 | HC_vs_MS.NoTrt_B |
| AMDHD1 | 0.48966 | 0.55997 | 0.196 | 0.17 | 1 | HC_vs_MS.NoTrt_B |
| CC2D2A | 0.4897 | 0.28028 | 0.241 | 0.27 | 1 | HC_vs_MS.NoTrt_B |
| OPRD1 | 0.48984 | 0.29233 | 0.746 | 0.82 | 1 | HC_vs_MS.NoTrt_B |
| AASS | 0.49002 | 0.25455 | 0.312 | 0.29 | 1 | HC_vs_MS.NoTrt_B |
| PIGBOS1 | 0.49015 | -0.4166 | 0.558 | 0.62 | 1 | HC_vs_MS.NoTrt_B |

| | | | | | | |
|---|---|---|---|---|---|---|
| TTF1 | 0.49025 | -0.4172 | 0.396 | 0.42 | 1 | HC_vs_MS.NoTrt_B |
| ZNF781 | 0.49059 | 0.83047 | 0.291 | 0.27 | 1 | HC_vs_MS.NoTrt_B |
| CSNK1D | 0.49087 | 0.30102 | 0.936 | 0.96 | 1 | HC_vs_MS.NoTrt_B |
| BCAT1 | 0.49104 | -0.6079 | 0.446 | 0.45 | 1 | HC_vs_MS.NoTrt_B |
| TK2 | 0.49106 | 0.27005 | 0.126 | 0.15 | 1 | HC_vs_MS.NoTrt_B |
| MYADM | 0.49152 | 0.31517 | 0.728 | 0.77 | 1 | HC_vs_MS.NoTrt_B |
| DDX25 | 0.49242 | 0.30228 | 0.654 | 0.65 | 1 | HC_vs_MS.NoTrt_B |
| ENO3 | 0.49278 | 0.28193 | 0.467 | 0.47 | 1 | HC_vs_MS.NoTrt_B |
| CPSF4 | 0.49331 | 0.55898 | 0.357 | 0.36 | 1 | HC_vs_MS.NoTrt_B |
| AP4E1 | 0.49399 | 0.45485 | 0.397 | 0.38 | 1 | HC_vs_MS.NoTrt_B |
| TRAF5 | 0.49403 | 0.40505 | 0.716 | 0.72 | 1 | HC_vs_MS.NoTrt_B |
| NCALD | 0.49446 | 0.38033 | 0.699 | 0.73 | 1 | HC_vs_MS.NoTrt_B |
| GNE | 0.4948 | 0.86828 | 0.193 | 0.17 | 1 | HC_vs_MS.NoTrt_B |
| RNFT1 | 0.49531 | 0.39065 | 0.401 | 0.45 | 1 | HC_vs_MS.NoTrt_B |
| SIMC1 | 0.49627 | 0.52898 | 0.42 | 0.48 | 1 | HC_vs_MS.NoTrt_B |
| LCP2 | 0.49651 | 0.34007 | 0.719 | 0.77 | 1 | HC_vs_MS.NoTrt_B |
| ARL14EP | 0.49655 | -0.6574 | 0.203 | 0.22 | 1 | HC_vs_MS.NoTrt_B |
| LPCAT4 | 0.4966 | 0.39267 | 0.262 | 0.3 | 1 | HC_vs_MS.NoTrt_B |
| ZSCAN20 | 0.49715 | 0.78366 | 0.238 | 0.22 | 1 | HC_vs_MS.NoTrt_B |
| USF3 | 0.4975 | 0.30325 | 0.699 | 0.74 | 1 | HC_vs_MS.NoTrt_B |
| CAMK4 | 0.4978 | 0.36131 | 0.648 | 0.7 | 1 | HC_vs_MS.NoTrt_B |
| CHIC2 | 0.49784 | 0.51312 | 0.171 | 0.2 | 1 | HC_vs_MS.NoTrt_B |
| SEMA3B | 0.49893 | 0.55083 | 0.226 | 0.26 | 1 | HC_vs_MS.NoTrt_B |
| KIAA1143 | 0.49933 | 0.29706 | 0.491 | 0.47 | 1 | HC_vs_MS.NoTrt_B |
| SLC16A10 | 0.49937 | 0.30939 | 0.823 | 0.81 | 1 | HC_vs_MS.NoTrt_B |
| MRPS23 | 0.4994 | -0.4629 | 0.78 | 0.82 | 1 | HC_vs_MS.NoTrt_B |
| HSD3B7 | 0.49953 | 0.67077 | 0.432 | 0.44 | 1 | HC_vs_MS.NoTrt_B |
| PTOV1 | 0.49976 | 0.91716 | 0.133 | 0.12 | 1 | HC_vs_MS.NoTrt_B |
| UNKL | 0.50005 | 0.58301 | 0.43 | 0.4 | 1 | HC_vs_MS.NoTrt_B |
| ZNF181 | 0.50006 | 0.57083 | 0.157 | 0.18 | 1 | HC_vs_MS.NoTrt_B |
| CFP | 0.5002 | -0.613 | 0.357 | 0.41 | 1 | HC_vs_MS.NoTrt_B |
| C9orf3 | 0.5004 | -0.4882 | 0.704 | 0.73 | 1 | HC_vs_MS.NoTrt_B |
| CADM1 | 0.50044 | 1.38883 | 0.19 | 0.17 | 1 | HC_vs_MS.NoTrt_B |
| DYRK1A | 0.50132 | 0.35739 | 0.681 | 0.68 | 1 | HC_vs_MS.NoTrt_B |
| GNA13 | 0.50147 | 0.74213 | 0.132 | 0.16 | 1 | HC_vs_MS.NoTrt_B |
| MRPS15 | 0.50194 | -0.5048 | 0.49 | 0.44 | 1 | HC_vs_MS.NoTrt_B |
| RP11-231C1 | 0.50321 | 0.34804 | 0.655 | 0.69 | 1 | HC_vs_MS.NoTrt_B |
| RASGRP2 | 0.5035 | 0.43987 | 0.384 | 0.43 | 1 | HC_vs_MS.NoTrt_B |
| JUNB | 0.50423 | 0.87297 | 0.354 | 0.38 | 1 | HC_vs_MS.NoTrt_B |
| ZBED5 | 0.50508 | 0.28 | 0.678 | 0.7 | 1 | HC_vs_MS.NoTrt_B |

| | | | | | | |
|---|---|---|---|---|---|---|
| CADPS | 0.50522 | 0.34992 | 0.33 | 0.32 | 1 | HC_vs_MS.NoTrt_B |
| MRFAP1 | 0.50559 | -0.2553 | 0.268 | 0.29 | 1 | HC_vs_MS.NoTrt_B |
| GALNT6 | 0.5058 | 0.7158 | 0.655 | 0.71 | 1 | HC_vs_MS.NoTrt_B |
| MFGE8 | 0.50623 | 0.51979 | 0.504 | 0.54 | 1 | HC_vs_MS.NoTrt_B |
| TGDS | 0.50663 | 0.62304 | 0.107 | 0.09 | 1 | HC_vs_MS.NoTrt_B |
| F2RL1 | 0.50687 | 0.96207 | 0.191 | 0.17 | 1 | HC_vs_MS.NoTrt_B |
| ZNF836 | 0.50689 | 0.42013 | 0.161 | 0.18 | 1 | HC_vs_MS.NoTrt_B |
| OPA1 | 0.50702 | 0.31682 | 0.122 | 0.14 | 1 | HC_vs_MS.NoTrt_B |
| DNAJC3 | 0.50714 | 0.29335 | 0.458 | 0.5 | 1 | HC_vs_MS.NoTrt_B |
| NLRP2 | 0.50735 | 0.38675 | 0.794 | 0.81 | 1 | HC_vs_MS.NoTrt_B |
| TAPT1 | 0.50773 | 0.58308 | 0.684 | 0.7 | 1 | HC_vs_MS.NoTrt_B |
| CDK12 | 0.50774 | 0.29226 | 0.512 | 0.56 | 1 | HC_vs_MS.NoTrt_B |
| NOL4L | 0.50824 | 0.38818 | 0.597 | 0.69 | 1 | HC_vs_MS.NoTrt_B |
| RBM18 | 0.50829 | 0.39523 | 0.312 | 0.29 | 1 | HC_vs_MS.NoTrt_B |
| RNF212 | 0.50829 | 0.26263 | 0.572 | 0.58 | 1 | HC_vs_MS.NoTrt_B |
| SF1 | 0.5089 | 0.44556 | 0.512 | 0.63 | 1 | HC_vs_MS.NoTrt_B |
| RAB2B | 0.509 | 0.38627 | 0.652 | 0.65 | 1 | HC_vs_MS.NoTrt_B |
| SLC41A3 | 0.50909 | -0.6546 | 0.286 | 0.31 | 1 | HC_vs_MS.NoTrt_B |
| ZNF84 | 0.50917 | 0.62915 | 0.284 | 0.26 | 1 | HC_vs_MS.NoTrt_B |
| ZNF530 | 0.50953 | 0.7984 | 0.483 | 0.45 | 1 | HC_vs_MS.NoTrt_B |
| PIK3C2A | 0.50979 | 1.19235 | 0.386 | 0.37 | 1 | HC_vs_MS.NoTrt_B |
| KCNMB4 | 0.51048 | 0.4031 | 0.293 | 0.33 | 1 | HC_vs_MS.NoTrt_B |
| CCDC124 | 0.51107 | 0.33935 | 0.486 | 0.49 | 1 | HC_vs_MS.NoTrt_B |
| LNPEP | 0.51167 | 0.30968 | 0.583 | 0.65 | 1 | HC_vs_MS.NoTrt_B |
| SIN3A | 0.51174 | 0.36179 | 0.783 | 0.78 | 1 | HC_vs_MS.NoTrt_B |
| FANCA | 0.51179 | -0.4688 | 0.259 | 0.28 | 1 | HC_vs_MS.NoTrt_B |
| XPO1 | 0.51197 | 0.25944 | 0.916 | 0.93 | 1 | HC_vs_MS.NoTrt_B |
| CDC5L | 0.51199 | 0.47795 | 0.59 | 0.6 | 1 | HC_vs_MS.NoTrt_B |
| HAUS7 | 0.51202 | 0.33532 | 0.091 | 0.11 | 1 | HC_vs_MS.NoTrt_B |
| NPAT | 0.51242 | 0.56137 | 0.71 | 0.79 | 1 | HC_vs_MS.NoTrt_B |
| MCUR1 | 0.51258 | 0.531 | 0.786 | 0.82 | 1 | HC_vs_MS.NoTrt_B |
| ZNF835 | 0.51266 | 0.29381 | 0.558 | 0.57 | 1 | HC_vs_MS.NoTrt_B |
| ZC3HC1 | 0.51299 | -0.4696 | 0.458 | 0.5 | 1 | HC_vs_MS.NoTrt_B |
| KCTD13 | 0.51317 | 0.83539 | 0.135 | 0.12 | 1 | HC_vs_MS.NoTrt_B |
| EPS8 | 0.51318 | -0.4496 | 0.391 | 0.38 | 1 | HC_vs_MS.NoTrt_B |
| DTD1 | 0.51368 | 0.26153 | 0.307 | 0.28 | 1 | HC_vs_MS.NoTrt_B |
| PIGP | 0.5137 | 0.27031 | 0.775 | 0.82 | 1 | HC_vs_MS.NoTrt_B |
| RBM38 | 0.51377 | 0.3866 | 0.125 | 0.15 | 1 | HC_vs_MS.NoTrt_B |
| TTN | 0.5138 | 0.51849 | 0.628 | 0.69 | 1 | HC_vs_MS.NoTrt_B |
| TTC21B | 0.51427 | 0.37916 | 0.359 | 0.35 | 1 | HC_vs_MS.NoTrt_B |

| | | | | | | |
|---|---|---|---|---|---|---|
| ANO8 | 0.51429 | 0.64309 | 0.133 | 0.12 | 1 | HC_vs_MS.NoTrt_B |
| XRCC1 | 0.51458 | -0.3572 | 0.161 | 0.18 | 1 | HC_vs_MS.NoTrt_B |
| PADI2 | 0.51467 | 0.44736 | 0.217 | 0.2 | 1 | HC_vs_MS.NoTrt_B |
| ESF1 | 0.51474 | 0.78616 | 0.136 | 0.12 | 1 | HC_vs_MS.NoTrt_B |
| CCDC152 | 0.51492 | 0.52492 | 0.197 | 0.23 | 1 | HC_vs_MS.NoTrt_B |
| SPC24 | 0.51504 | -0.2884 | 0.859 | 0.82 | 1 | HC_vs_MS.NoTrt_B |
| PRRC1 | 0.51524 | 0.35042 | 0.352 | 0.38 | 1 | HC_vs_MS.NoTrt_B |
| SNCAIP | 0.51563 | 0.31371 | 0.384 | 0.4 | 1 | HC_vs_MS.NoTrt_B |
| DIABLO | 0.51578 | -0.3673 | 0.674 | 0.72 | 1 | HC_vs_MS.NoTrt_B |
| SEC31B | 0.51626 | 0.34579 | 0.446 | 0.47 | 1 | HC_vs_MS.NoTrt_B |
| C3orf49 | 0.51633 | 0.36536 | 0.31 | 0.3 | 1 | HC_vs_MS.NoTrt_B |
| RAB39A | 0.5165 | 0.28222 | 0.349 | 0.36 | 1 | HC_vs_MS.NoTrt_B |
| ZNF69 | 0.51658 | -0.3653 | 0.452 | 0.43 | 1 | HC_vs_MS.NoTrt_B |
| ARHGAP27 | 0.51664 | 0.35887 | 0.526 | 0.59 | 1 | HC_vs_MS.NoTrt_B |
| FAM69A | 0.51667 | 0.62342 | 0.106 | 0.12 | 1 | HC_vs_MS.NoTrt_B |
| PKHD1L1 | 0.51676 | 0.38679 | 0.446 | 0.47 | 1 | HC_vs_MS.NoTrt_B |
| RGS1 | 0.5169 | 0.54309 | 0.129 | 0.11 | 1 | HC_vs_MS.NoTrt_B |
| ADGB | 0.51738 | 0.27261 | 0.158 | 0.15 | 1 | HC_vs_MS.NoTrt_B |
| CDC6 | 0.51756 | 0.28593 | 0.27 | 0.25 | 1 | HC_vs_MS.NoTrt_B |
| MRPL49 | 0.51766 | -0.3354 | 0.455 | 0.46 | 1 | HC_vs_MS.NoTrt_B |
| NUDT14 | 0.5179 | 0.54537 | 0.438 | 0.48 | 1 | HC_vs_MS.NoTrt_B |
| ITGAE | 0.51796 | 0.40903 | 0.574 | 0.6 | 1 | HC_vs_MS.NoTrt_B |
| CC2D1A | 0.51896 | 0.34402 | 0.681 | 0.74 | 1 | HC_vs_MS.NoTrt_B |
| PROX2 | 0.51932 | 0.346 | 0.123 | 0.11 | 1 | HC_vs_MS.NoTrt_B |
| PSMG1 | 0.51948 | 0.39145 | 0.126 | 0.15 | 1 | HC_vs_MS.NoTrt_B |
| RP11-526A4 | 0.52104 | 0.2725 | 0.338 | 0.34 | 1 | HC_vs_MS.NoTrt_B |
| CDC14A | 0.52171 | 0.51193 | 0.229 | 0.26 | 1 | HC_vs_MS.NoTrt_B |
| NAV3 | 0.52188 | 0.30157 | 0.704 | 0.75 | 1 | HC_vs_MS.NoTrt_B |
| PAQR7 | 0.52204 | 0.46365 | 0.486 | 0.49 | 1 | HC_vs_MS.NoTrt_B |
| RPS17 | 0.52206 | 0.2637 | 0.394 | 0.42 | 1 | HC_vs_MS.NoTrt_B |
| LRRC37A | 0.5222 | 0.69015 | 0.399 | 0.44 | 1 | HC_vs_MS.NoTrt_B |
| LGALS9 | 0.52239 | 0.63541 | 0.299 | 0.29 | 1 | HC_vs_MS.NoTrt_B |
| KDM5C | 0.52377 | 0.83974 | 0.37 | 0.42 | 1 | HC_vs_MS.NoTrt_B |
| SELM | 0.52409 | 0.28497 | 0.259 | 0.29 | 1 | HC_vs_MS.NoTrt_B |
| PTPRS | 0.52448 | 0.25887 | 0.509 | 0.57 | 1 | HC_vs_MS.NoTrt_B |
| ZNF140 | 0.52615 | -0.4394 | 0.149 | 0.17 | 1 | HC_vs_MS.NoTrt_B |
| ITPKB | 0.52618 | 0.58724 | 0.562 | 0.59 | 1 | HC_vs_MS.NoTrt_B |
| MAP3K11 | 0.52625 | 0.87721 | 0.612 | 0.64 | 1 | HC_vs_MS.NoTrt_B |
| WDFY2 | 0.52643 | -0.3337 | 0.558 | 0.58 | 1 | HC_vs_MS.NoTrt_B |
| WDR77 | 0.52648 | 0.29676 | 0.106 | 0.12 | 1 | HC_vs_MS.NoTrt_B |

| | | | | | | |
|---|---|---|---|---|---|---|
| HENMT1 | 0.52688 | -0.4032 | 0.332 | 0.36 | 1 | HC_vs_MS.NoTrt_B |
| PITPNM1 | 0.52718 | 0.5369 | 0.246 | 0.23 | 1 | HC_vs_MS.NoTrt_B |
| SLC30A7 | 0.52723 | 0.28614 | 0.599 | 0.64 | 1 | HC_vs_MS.NoTrt_B |
| MTFR1 | 0.52743 | 0.57791 | 0.125 | 0.15 | 1 | HC_vs_MS.NoTrt_B |
| MRPS30 | 0.5278 | 0.54643 | 0.328 | 0.32 | 1 | HC_vs_MS.NoTrt_B |
| ZFP36 | 0.52808 | 0.32884 | 0.393 | 0.45 | 1 | HC_vs_MS.NoTrt_B |
| WDR36 | 0.52817 | 0.57702 | 0.142 | 0.12 | 1 | HC_vs_MS.NoTrt_B |
| PGGHG | 0.52842 | 0.49569 | 0.146 | 0.13 | 1 | HC_vs_MS.NoTrt_B |
| ST20-MTHFS | 0.52856 | 0.68867 | 0.116 | 0.1 | 1 | HC_vs_MS.NoTrt_B |
| SLC25A40 | 0.5294 | -0.2758 | 0.297 | 0.29 | 1 | HC_vs_MS.NoTrt_B |
| FBXW12 | 0.52944 | 0.26035 | 0.323 | 0.36 | 1 | HC_vs_MS.NoTrt_B |
| USP21 | 0.52983 | 0.53706 | 0.113 | 0.1 | 1 | HC_vs_MS.NoTrt_B |
| ZNF223 | 0.52985 | 0.97938 | 0.101 | 0.09 | 1 | HC_vs_MS.NoTrt_B |
| PDS5A | 0.5302 | 0.37109 | 0.754 | 0.8 | 1 | HC_vs_MS.NoTrt_B |
| MEGF8 | 0.5306 | 0.71103 | 0.393 | 0.38 | 1 | HC_vs_MS.NoTrt_B |
| RNF103 | 0.53081 | 0.44304 | 0.574 | 0.58 | 1 | HC_vs_MS.NoTrt_B |
| ZNRF1 | 0.53163 | 0.4272 | 0.274 | 0.26 | 1 | HC_vs_MS.NoTrt_B |
| CDK16 | 0.53181 | 0.53599 | 0.113 | 0.1 | 1 | HC_vs_MS.NoTrt_B |
| ZP3 | 0.53197 | 0.47663 | 0.601 | 0.61 | 1 | HC_vs_MS.NoTrt_B |
| ALDH5A1 | 0.53206 | 0.92253 | 0.468 | 0.51 | 1 | HC_vs_MS.NoTrt_B |
| ADNP | 0.53207 | 0.8264 | 0.438 | 0.47 | 1 | HC_vs_MS.NoTrt_B |
| BOLA2 | 0.5323 | 0.43484 | 0.123 | 0.15 | 1 | HC_vs_MS.NoTrt_B |
| NIPSNAP1 | 0.53234 | -0.5037 | 0.393 | 0.35 | 1 | HC_vs_MS.NoTrt_B |
| TGOLN2 | 0.53264 | 0.50824 | 0.619 | 0.65 | 1 | HC_vs_MS.NoTrt_B |
| PPP2CA | 0.53268 | 0.38495 | 0.691 | 0.78 | 1 | HC_vs_MS.NoTrt_B |
| RSBN1 | 0.53269 | 0.30897 | 0.249 | 0.23 | 1 | HC_vs_MS.NoTrt_B |
| ILDR2 | 0.53273 | 0.26563 | 0.304 | 0.29 | 1 | HC_vs_MS.NoTrt_B |
| WHAMM | 0.53282 | 0.38886 | 0.814 | 0.84 | 1 | HC_vs_MS.NoTrt_B |
| C1GALT1 | 0.53326 | 0.57017 | 0.564 | 0.58 | 1 | HC_vs_MS.NoTrt_B |
| MUM1 | 0.53372 | -0.2648 | 0.599 | 0.58 | 1 | HC_vs_MS.NoTrt_B |
| OGG1 | 0.5339 | 0.31712 | 0.364 | 0.34 | 1 | HC_vs_MS.NoTrt_B |
| AP3M2 | 0.53427 | 0.36237 | 0.101 | 0.09 | 1 | HC_vs_MS.NoTrt_B |
| PCGF3 | 0.53451 | 0.28189 | 0.229 | 0.25 | 1 | HC_vs_MS.NoTrt_B |
| CREB1 | 0.53463 | 0.37051 | 0.643 | 0.71 | 1 | HC_vs_MS.NoTrt_B |
| TLR3 | 0.53466 | 0.28799 | 0.286 | 0.28 | 1 | HC_vs_MS.NoTrt_B |
| NDC80 | 0.53497 | -0.8181 | 0.684 | 0.7 | 1 | HC_vs_MS.NoTrt_B |
| TXK | 0.53499 | 0.54844 | 0.599 | 0.63 | 1 | HC_vs_MS.NoTrt_B |
| STOML1 | 0.53504 | -0.4654 | 0.765 | 0.82 | 1 | HC_vs_MS.NoTrt_B |
| NMT2 | 0.53509 | -0.4577 | 0.925 | 0.92 | 1 | HC_vs_MS.NoTrt_B |
| TBC1D2B | 0.5352 | 0.97333 | 0.294 | 0.33 | 1 | HC_vs_MS.NoTrt_B |

| | | | | | | |
|---|---|---|---|---|---|---|
| CCDC38 | 0.53528 | 0.29132 | 0.378 | 0.38 | 1 | HC_vs_MS.NoTrt_B |
| NR1I3 | 0.5353 | 0.30228 | 0.399 | 0.4 | 1 | HC_vs_MS.NoTrt_B |
| NR6A1 | 0.53548 | 0.27936 | 0.378 | 0.39 | 1 | HC_vs_MS.NoTrt_B |
| DCDC1 | 0.53553 | 0.36257 | 0.48 | 0.47 | 1 | HC_vs_MS.NoTrt_B |
| C3orf33 | 0.53578 | 0.25406 | 0.475 | 0.48 | 1 | HC_vs_MS.NoTrt_B |
| CTDSP2 | 0.53604 | 0.52992 | 0.216 | 0.25 | 1 | HC_vs_MS.NoTrt_B |
| GMIP | 0.53605 | 0.99454 | 0.254 | 0.3 | 1 | HC_vs_MS.NoTrt_B |
| RBCK1 | 0.53695 | 0.32094 | 0.201 | 0.23 | 1 | HC_vs_MS.NoTrt_B |
| FIGNL1 | 0.53721 | 0.54579 | 0.114 | 0.14 | 1 | HC_vs_MS.NoTrt_B |
| FOXRED2 | 0.53724 | 0.35396 | 0.529 | 0.55 | 1 | HC_vs_MS.NoTrt_B |
| OFD1 | 0.53726 | 0.53375 | 0.246 | 0.24 | 1 | HC_vs_MS.NoTrt_B |
| PAM16 | 0.53741 | 0.4562 | 0.181 | 0.2 | 1 | HC_vs_MS.NoTrt_B |
| ASIC4 | 0.53757 | 0.44771 | 0.21 | 0.2 | 1 | HC_vs_MS.NoTrt_B |
| CDK6 | 0.53811 | 0.51167 | 0.145 | 0.17 | 1 | HC_vs_MS.NoTrt_B |
| SLC39A4 | 0.53886 | 0.38626 | 0.386 | 0.37 | 1 | HC_vs_MS.NoTrt_B |
| ZNF410 | 0.53932 | -0.4941 | 0.535 | 0.5 | 1 | HC_vs_MS.NoTrt_B |
| USP6NL | 0.53936 | 0.6505 | 0.768 | 0.82 | 1 | HC_vs_MS.NoTrt_B |
| MRPL52 | 0.53975 | 0.52627 | 0.626 | 0.67 | 1 | HC_vs_MS.NoTrt_B |
| ARSK | 0.54003 | -0.26 | 0.538 | 0.6 | 1 | HC_vs_MS.NoTrt_B |
| GALE | 0.54073 | -1.1128 | 0.323 | 0.36 | 1 | HC_vs_MS.NoTrt_B |
| RELA | 0.54078 | 0.58918 | 0.154 | 0.18 | 1 | HC_vs_MS.NoTrt_B |
| JAKMIP2 | 0.54155 | -0.8136 | 0.352 | 0.38 | 1 | HC_vs_MS.NoTrt_B |
| STON2 | 0.54224 | 0.36666 | 0.258 | 0.29 | 1 | HC_vs_MS.NoTrt_B |
| ZNF260 | 0.54268 | 0.33217 | 0.506 | 0.55 | 1 | HC_vs_MS.NoTrt_B |
| OGFOD1 | 0.54309 | -0.2837 | 0.596 | 0.68 | 1 | HC_vs_MS.NoTrt_B |
| GPATCH4 | 0.54309 | 0.92404 | 0.277 | 0.31 | 1 | HC_vs_MS.NoTrt_B |
| HYOU1 | 0.5432 | 0.33328 | 0.475 | 0.47 | 1 | HC_vs_MS.NoTrt_B |
| WDFY1 | 0.54367 | 0.57006 | 0.339 | 0.33 | 1 | HC_vs_MS.NoTrt_B |
| PROSER1 | 0.54478 | 0.28855 | 0.171 | 0.2 | 1 | HC_vs_MS.NoTrt_B |
| PDE8A | 0.54482 | 0.99782 | 0.397 | 0.44 | 1 | HC_vs_MS.NoTrt_B |
| NCOA3 | 0.54507 | 0.32575 | 0.739 | 0.78 | 1 | HC_vs_MS.NoTrt_B |
| HDAC9 | 0.54508 | 0.95118 | 0.459 | 0.51 | 1 | HC_vs_MS.NoTrt_B |
| FAM83G | 0.54512 | 0.70084 | 0.432 | 0.43 | 1 | HC_vs_MS.NoTrt_B |
| ERCC6L2 | 0.54586 | 0.33918 | 0.203 | 0.18 | 1 | HC_vs_MS.NoTrt_B |
| CCNG2 | 0.54615 | 0.29276 | 0.265 | 0.29 | 1 | HC_vs_MS.NoTrt_B |
| TBRG1 | 0.5462 | 0.42253 | 0.632 | 0.68 | 1 | HC_vs_MS.NoTrt_B |
| KIAA1841 | 0.54624 | 0.41216 | 0.846 | 0.88 | 1 | HC_vs_MS.NoTrt_B |
| PCF11 | 0.54636 | 0.35902 | 0.519 | 0.53 | 1 | HC_vs_MS.NoTrt_B |
| USP48 | 0.54647 | 0.44347 | 0.499 | 0.54 | 1 | HC_vs_MS.NoTrt_B |
| DMGDH | 0.5468 | 0.2827 | 0.397 | 0.45 | 1 | HC_vs_MS.NoTrt_B |

| | | | | | | |
|---|---|---|---|---|---|---|
| TIMM21 | 0.5469 | 0.5045 | 0.633 | 0.63 | 1 | HC_vs_MS.NoTrt_B |
| SRSF11 | 0.54691 | 0.25922 | 0.771 | 0.82 | 1 | HC_vs_MS.NoTrt_B |
| UBR3 | 0.54703 | 0.71417 | 0.116 | 0.14 | 1 | HC_vs_MS.NoTrt_B |
| GLUL | 0.54745 | 0.37374 | 0.158 | 0.18 | 1 | HC_vs_MS.NoTrt_B |
| CEP76 | 0.54752 | 0.50365 | 0.787 | 0.82 | 1 | HC_vs_MS.NoTrt_B |
| SC5D | 0.54762 | 0.32426 | 0.517 | 0.51 | 1 | HC_vs_MS.NoTrt_B |
| ZNF565 | 0.54788 | 0.82941 | 0.093 | 0.11 | 1 | HC_vs_MS.NoTrt_B |
| GOLGA8A | 0.54833 | 0.61235 | 0.412 | 0.47 | 1 | HC_vs_MS.NoTrt_B |
| C2CD2 | 0.54834 | 0.71936 | 0.22 | 0.21 | 1 | HC_vs_MS.NoTrt_B |
| C16orf58 | 0.54844 | 0.28228 | 0.107 | 0.09 | 1 | HC_vs_MS.NoTrt_B |
| ANKRD18A | 0.54867 | 0.42564 | 0.232 | 0.22 | 1 | HC_vs_MS.NoTrt_B |
| UNG | 0.54894 | 0.79614 | 0.2 | 0.18 | 1 | HC_vs_MS.NoTrt_B |
| FOCAD | 0.54904 | 0.77713 | 0.529 | 0.55 | 1 | HC_vs_MS.NoTrt_B |
| SUDS3 | 0.54925 | 0.38783 | 0.775 | 0.81 | 1 | HC_vs_MS.NoTrt_B |
| VIL1 | 0.54959 | 0.42713 | 0.155 | 0.17 | 1 | HC_vs_MS.NoTrt_B |
| ANKRD13C | 0.54989 | 0.3054 | 0.167 | 0.19 | 1 | HC_vs_MS.NoTrt_B |
| NBEAL1 | 0.5502 | 0.74932 | 0.87 | 0.89 | 1 | HC_vs_MS.NoTrt_B |
| PPP4R3B | 0.55031 | 0.30498 | 0.425 | 0.45 | 1 | HC_vs_MS.NoTrt_B |
| PYCRL | 0.55053 | -0.6606 | 0.406 | 0.45 | 1 | HC_vs_MS.NoTrt_B |
| TMEM65 | 0.55066 | 0.46751 | 0.683 | 0.72 | 1 | HC_vs_MS.NoTrt_B |
| COA1 | 0.55105 | 0.3784 | 0.759 | 0.82 | 1 | HC_vs_MS.NoTrt_B |
| PLEKHS1 | 0.5517 | 0.25707 | 0.152 | 0.14 | 1 | HC_vs_MS.NoTrt_B |
| SFR1 | 0.55171 | 0.29141 | 0.242 | 0.27 | 1 | HC_vs_MS.NoTrt_B |
| KIAA0100 | 0.55181 | 1.1156 | 0.497 | 0.5 | 1 | HC_vs_MS.NoTrt_B |
| VNN3 | 0.55188 | 0.32597 | 0.358 | 0.37 | 1 | HC_vs_MS.NoTrt_B |
| TPX2 | 0.55193 | 0.69425 | 0.181 | 0.2 | 1 | HC_vs_MS.NoTrt_B |
| PDLIM2 | 0.55228 | 0.25303 | 0.729 | 0.78 | 1 | HC_vs_MS.NoTrt_B |
| ATG3 | 0.55232 | 0.2962 | 0.409 | 0.36 | 1 | HC_vs_MS.NoTrt_B |
| FBXL4 | 0.55243 | 0.47816 | 0.554 | 0.55 | 1 | HC_vs_MS.NoTrt_B |
| VAMP7 | 0.55341 | 0.25201 | 0.099 | 0.11 | 1 | HC_vs_MS.NoTrt_B |
| MMAB | 0.55368 | 0.41311 | 0.664 | 0.67 | 1 | HC_vs_MS.NoTrt_B |
| ZNF805 | 0.554 | 0.4463 | 0.739 | 0.74 | 1 | HC_vs_MS.NoTrt_B |
| CCDC7 | 0.55416 | 0.2994 | 0.122 | 0.14 | 1 | HC_vs_MS.NoTrt_B |
| DENND3 | 0.55419 | 0.46187 | 0.454 | 0.45 | 1 | HC_vs_MS.NoTrt_B |
| ELOVL7 | 0.55447 | 0.29996 | 0.319 | 0.31 | 1 | HC_vs_MS.NoTrt_B |
| PCDHB9 | 0.55468 | 0.26596 | 0.348 | 0.4 | 1 | HC_vs_MS.NoTrt_B |
| MAP3K9 | 0.55476 | 0.6477 | 0.448 | 0.52 | 1 | HC_vs_MS.NoTrt_B |
| PTCD1 | 0.55501 | 0.38332 | 0.59 | 0.61 | 1 | HC_vs_MS.NoTrt_B |
| RCAN3 | 0.55514 | 0.2539 | 0.781 | 0.81 | 1 | HC_vs_MS.NoTrt_B |
| PDZD7 | 0.55526 | 0.34143 | 0.483 | 0.51 | 1 | HC_vs_MS.NoTrt_B |

| | | | | | | |
|---|---|---|---|---|---|---|
| LRRC8B | 0.55602 | 1.20692 | 0.409 | 0.47 | 1 | HC_vs_MS.NoTrt_B |
| METTL25 | 0.55609 | 0.67581 | 0.18 | 0.16 | 1 | HC_vs_MS.NoTrt_B |
| ATP6V1E1 | 0.55616 | -0.7225 | 0.632 | 0.66 | 1 | HC_vs_MS.NoTrt_B |
| SGCB | 0.55693 | 0.44389 | 0.267 | 0.24 | 1 | HC_vs_MS.NoTrt_B |
| HELZ2 | 0.55734 | 0.83472 | 0.633 | 0.7 | 1 | HC_vs_MS.NoTrt_B |
| TUBGCP3 | 0.5574 | 0.55262 | 0.167 | 0.19 | 1 | HC_vs_MS.NoTrt_B |
| TTC19 | 0.55825 | 0.29856 | 0.762 | 0.75 | 1 | HC_vs_MS.NoTrt_B |
| HECTD4 | 0.55852 | 0.87907 | 0.662 | 0.69 | 1 | HC_vs_MS.NoTrt_B |
| RSAD1 | 0.55933 | 0.60001 | 0.087 | 0.1 | 1 | HC_vs_MS.NoTrt_B |
| AP3S2 | 0.55977 | -0.2924 | 0.694 | 0.68 | 1 | HC_vs_MS.NoTrt_B |
| CNOT4 | 0.55978 | 0.61042 | 0.686 | 0.74 | 1 | HC_vs_MS.NoTrt_B |
| ULK2 | 0.56046 | 0.37757 | 0.701 | 0.74 | 1 | HC_vs_MS.NoTrt_B |
| KDM3A | 0.56167 | 0.2655 | 0.146 | 0.13 | 1 | HC_vs_MS.NoTrt_B |
| ACAD10 | 0.56174 | 0.35018 | 0.446 | 0.49 | 1 | HC_vs_MS.NoTrt_B |
| MKLN1 | 0.56224 | 0.37165 | 0.572 | 0.6 | 1 | HC_vs_MS.NoTrt_B |
| ATP7B | 0.56226 | 0.74119 | 0.2 | 0.18 | 1 | HC_vs_MS.NoTrt_B |
| SORD | 0.56242 | 0.83578 | 0.142 | 0.16 | 1 | HC_vs_MS.NoTrt_B |
| POLR2B | 0.56243 | 0.50198 | 0.446 | 0.48 | 1 | HC_vs_MS.NoTrt_B |
| ICOSLG | 0.56244 | 0.25701 | 0.722 | 0.75 | 1 | HC_vs_MS.NoTrt_B |
| HERC5 | 0.56259 | 0.35205 | 0.21 | 0.23 | 1 | HC_vs_MS.NoTrt_B |
| FNBP4 | 0.56276 | 0.41546 | 0.481 | 0.48 | 1 | HC_vs_MS.NoTrt_B |
| SOCS7 | 0.56278 | 0.55938 | 0.349 | 0.4 | 1 | HC_vs_MS.NoTrt_B |
| CMBL | 0.5631 | 0.2577 | 0.48 | 0.5 | 1 | HC_vs_MS.NoTrt_B |
| LSR | 0.56324 | -0.3384 | 0.152 | 0.17 | 1 | HC_vs_MS.NoTrt_B |
| GBA2 | 0.56328 | 0.51606 | 0.567 | 0.59 | 1 | HC_vs_MS.NoTrt_B |
| FKBP5 | 0.56347 | 0.3229 | 0.762 | 0.88 | 1 | HC_vs_MS.NoTrt_B |
| TMIGD2 | 0.56358 | 0.61304 | 0.199 | 0.19 | 1 | HC_vs_MS.NoTrt_B |
| ST8SIA6 | 0.56361 | 0.86325 | 0.443 | 0.51 | 1 | HC_vs_MS.NoTrt_B |
| RWDD3 | 0.56423 | 0.389 | 0.126 | 0.15 | 1 | HC_vs_MS.NoTrt_B |
| SEC62 | 0.56465 | 0.38835 | 0.414 | 0.44 | 1 | HC_vs_MS.NoTrt_B |
| ATG16L2 | 0.56475 | 0.28276 | 0.816 | 0.87 | 1 | HC_vs_MS.NoTrt_B |
| MCL1 | 0.5649 | 0.63179 | 0.372 | 0.38 | 1 | HC_vs_MS.NoTrt_B |
| SLC37A2 | 0.56508 | 0.36725 | 0.525 | 0.59 | 1 | HC_vs_MS.NoTrt_B |
| RHPN2 | 0.56526 | 0.39009 | 0.548 | 0.57 | 1 | HC_vs_MS.NoTrt_B |
| ZFYVE28 | 0.5657 | 0.48151 | 0.133 | 0.15 | 1 | HC_vs_MS.NoTrt_B |
| TDG | 0.56577 | 0.52164 | 0.339 | 0.32 | 1 | HC_vs_MS.NoTrt_B |
| UTP14A | 0.5662 | -0.5297 | 0.18 | 0.16 | 1 | HC_vs_MS.NoTrt_B |
| ZNF652 | 0.56666 | 0.46368 | 0.493 | 0.53 | 1 | HC_vs_MS.NoTrt_B |
| RAD1 | 0.56708 | 0.27566 | 0.72 | 0.73 | 1 | HC_vs_MS.NoTrt_B |
| SPTLC1 | 0.56724 | -0.2547 | 0.723 | 0.74 | 1 | HC_vs_MS.NoTrt_B |

| | | | | | | |
|---|---|---|---|---|---|---|
| COIL | 0.56725 | 0.78633 | 0.106 | 0.12 | 1 | HC_vs_MS.NoTrt_B |
| ZNF790 | 0.56821 | -0.6342 | 0.488 | 0.52 | 1 | HC_vs_MS.NoTrt_B |
| HIST1H3E | 0.56845 | -0.2716 | 0.112 | 0.1 | 1 | HC_vs_MS.NoTrt_B |
| ZNF285 | 0.5687 | 0.25435 | 0.297 | 0.29 | 1 | HC_vs_MS.NoTrt_B |
| ABO | 0.5689 | -0.459 | 0.362 | 0.36 | 1 | HC_vs_MS.NoTrt_B |
| RSRC1 | 0.56965 | -0.2519 | 0.739 | 0.71 | 1 | HC_vs_MS.NoTrt_B |
| UBA2 | 0.57006 | 0.40777 | 0.513 | 0.54 | 1 | HC_vs_MS.NoTrt_B |
| KIF3A | 0.57017 | 0.44737 | 0.675 | 0.72 | 1 | HC_vs_MS.NoTrt_B |
| DDHD2 | 0.5703 | 0.45335 | 0.793 | 0.81 | 1 | HC_vs_MS.NoTrt_B |
| PSTPIP1 | 0.57048 | -0.381 | 0.414 | 0.43 | 1 | HC_vs_MS.NoTrt_B |
| PSMD10 | 0.57136 | 0.5281 | 0.316 | 0.3 | 1 | HC_vs_MS.NoTrt_B |
| ZNF471 | 0.5715 | 0.26253 | 0.819 | 0.89 | 1 | HC_vs_MS.NoTrt_B |
| UBTF | 0.57155 | 0.76263 | 0.212 | 0.2 | 1 | HC_vs_MS.NoTrt_B |
| SHCBP1 | 0.57162 | 0.25611 | 0.707 | 0.76 | 1 | HC_vs_MS.NoTrt_B |
| CAMK1 | 0.57189 | 0.41843 | 0.516 | 0.56 | 1 | HC_vs_MS.NoTrt_B |
| PITHD1 | 0.572 | 0.34434 | 0.145 | 0.13 | 1 | HC_vs_MS.NoTrt_B |
| NBPF9 | 0.57204 | 0.51128 | 0.709 | 0.72 | 1 | HC_vs_MS.NoTrt_B |
| GAB3 | 0.5722 | 1.08466 | 0.346 | 0.39 | 1 | HC_vs_MS.NoTrt_B |
| POLR1C | 0.57226 | 0.35689 | 0.412 | 0.38 | 1 | HC_vs_MS.NoTrt_B |
| RPF2 | 0.5724 | 0.25294 | 0.304 | 0.29 | 1 | HC_vs_MS.NoTrt_B |
| ANTXRL | 0.57323 | 0.32596 | 0.204 | 0.24 | 1 | HC_vs_MS.NoTrt_B |
| HSPD1 | 0.57355 | 0.48404 | 0.193 | 0.22 | 1 | HC_vs_MS.NoTrt_B |
| PATL2 | 0.57379 | 1.009 | 0.358 | 0.37 | 1 | HC_vs_MS.NoTrt_B |
| PPP2R3A | 0.57402 | 0.26068 | 0.701 | 0.76 | 1 | HC_vs_MS.NoTrt_B |
| FHIT | 0.57421 | 0.42628 | 0.657 | 0.63 | 1 | HC_vs_MS.NoTrt_B |
| PDE4B | 0.57427 | 0.3909 | 0.28 | 0.31 | 1 | HC_vs_MS.NoTrt_B |
| PRKRA | 0.57432 | 0.37713 | 0.216 | 0.24 | 1 | HC_vs_MS.NoTrt_B |
| SPC25 | 0.57459 | 0.41347 | 0.677 | 0.72 | 1 | HC_vs_MS.NoTrt_B |
| GPATCH1 | 0.57464 | -0.6043 | 0.361 | 0.38 | 1 | HC_vs_MS.NoTrt_B |
| APPL1 | 0.57468 | 0.89083 | 0.414 | 0.41 | 1 | HC_vs_MS.NoTrt_B |
| DSP | 0.57484 | 0.27508 | 0.096 | 0.11 | 1 | HC_vs_MS.NoTrt_B |
| RASAL2 | 0.5749 | 0.26485 | 0.481 | 0.47 | 1 | HC_vs_MS.NoTrt_B |
| KIAA1551 | 0.57518 | 0.32942 | 0.497 | 0.58 | 1 | HC_vs_MS.NoTrt_B |
| NUP205 | 0.57598 | 0.79733 | 0.174 | 0.2 | 1 | HC_vs_MS.NoTrt_B |
| SEMA4F | 0.57648 | -0.2637 | 0.486 | 0.52 | 1 | HC_vs_MS.NoTrt_B |
| TAB2 | 0.57655 | 0.32321 | 0.584 | 0.57 | 1 | HC_vs_MS.NoTrt_B |
| FAF1 | 0.57657 | 0.29497 | 0.436 | 0.47 | 1 | HC_vs_MS.NoTrt_B |
| C2orf49 | 0.5773 | 0.30873 | 0.425 | 0.42 | 1 | HC_vs_MS.NoTrt_B |
| MFSD9 | 0.57749 | -0.5223 | 0.488 | 0.47 | 1 | HC_vs_MS.NoTrt_B |
| COPA | 0.57753 | 0.25341 | 0.223 | 0.25 | 1 | HC_vs_MS.NoTrt_B |

| | | | | | | |
|---|---|---|---|---|---|---|
| TMEM161B | 0.57762 | 0.36678 | 0.641 | 0.68 | 1 | HC_vs_MS.NoTrt_B |
| CDH26 | 0.57783 | 0.39493 | 0.159 | 0.15 | 1 | HC_vs_MS.NoTrt_B |
| SARM1 | 0.57784 | -0.4109 | 0.443 | 0.43 | 1 | HC_vs_MS.NoTrt_B |
| EP300 | 0.57827 | 0.44062 | 0.707 | 0.72 | 1 | HC_vs_MS.NoTrt_B |
| CROCC | 0.57836 | 0.3783 | 0.48 | 0.47 | 1 | HC_vs_MS.NoTrt_B |
| KLF7 | 0.57855 | -1.0472 | 0.188 | 0.16 | 1 | HC_vs_MS.NoTrt_B |
| TJP3 | 0.57923 | 0.4531 | 0.103 | 0.12 | 1 | HC_vs_MS.NoTrt_B |
| ZNF517 | 0.57934 | 0.3625 | 0.375 | 0.37 | 1 | HC_vs_MS.NoTrt_B |
| ACER1 | 0.57954 | 0.42495 | 0.438 | 0.46 | 1 | HC_vs_MS.NoTrt_B |
| UBE2G2 | 0.58049 | -0.3027 | 0.713 | 0.74 | 1 | HC_vs_MS.NoTrt_B |
| DCUN1D3 | 0.58051 | 0.33259 | 0.139 | 0.12 | 1 | HC_vs_MS.NoTrt_B |
| ANKRD40 | 0.58128 | 0.26526 | 0.746 | 0.78 | 1 | HC_vs_MS.NoTrt_B |
| MAML2 | 0.58156 | 0.53559 | 0.272 | 0.26 | 1 | HC_vs_MS.NoTrt_B |
| PPP1R3E | 0.58158 | -0.3094 | 0.209 | 0.23 | 1 | HC_vs_MS.NoTrt_B |
| SMG6 | 0.58192 | -0.3877 | 0.188 | 0.2 | 1 | HC_vs_MS.NoTrt_B |
| YARS2 | 0.58224 | 0.39891 | 0.377 | 0.36 | 1 | HC_vs_MS.NoTrt_B |
| CUBN | 0.58247 | 0.53767 | 0.397 | 0.41 | 1 | HC_vs_MS.NoTrt_B |
| FAR2 | 0.58295 | 0.95768 | 0.186 | 0.21 | 1 | HC_vs_MS.NoTrt_B |
| MAP2K7 | 0.58343 | 0.43115 | 0.609 | 0.64 | 1 | HC_vs_MS.NoTrt_B |
| SLC26A11 | 0.58364 | -0.3498 | 0.654 | 0.71 | 1 | HC_vs_MS.NoTrt_B |
| NDFIP2 | 0.58367 | 0.60267 | 0.132 | 0.12 | 1 | HC_vs_MS.NoTrt_B |
| GRK5 | 0.58374 | -0.3142 | 0.155 | 0.17 | 1 | HC_vs_MS.NoTrt_B |
| PLA2G7 | 0.58403 | 0.38841 | 0.594 | 0.62 | 1 | HC_vs_MS.NoTrt_B |
| CCDC22 | 0.58426 | 0.56649 | 0.129 | 0.11 | 1 | HC_vs_MS.NoTrt_B |
| SFXN1 | 0.58436 | 0.37974 | 0.775 | 0.82 | 1 | HC_vs_MS.NoTrt_B |
| IL20RB | 0.58453 | 0.77741 | 0.229 | 0.22 | 1 | HC_vs_MS.NoTrt_B |
| SETD4 | 0.58537 | 0.53698 | 0.483 | 0.48 | 1 | HC_vs_MS.NoTrt_B |
| LARGE1 | 0.58566 | 0.25936 | 0.557 | 0.55 | 1 | HC_vs_MS.NoTrt_B |
| 3-Mar | 0.58586 | 0.50043 | 0.326 | 0.36 | 1 | HC_vs_MS.NoTrt_B |
| CMTR2 | 0.58631 | 0.57392 | 0.257 | 0.24 | 1 | HC_vs_MS.NoTrt_B |
| TMEM39B | 0.58645 | 0.49188 | 0.17 | 0.19 | 1 | HC_vs_MS.NoTrt_B |
| SESN3 | 0.58662 | 0.83803 | 0.259 | 0.25 | 1 | HC_vs_MS.NoTrt_B |
| CNNM3 | 0.58662 | 0.61325 | 0.609 | 0.63 | 1 | HC_vs_MS.NoTrt_B |
| SUN1 | 0.58684 | 0.88044 | 0.593 | 0.61 | 1 | HC_vs_MS.NoTrt_B |
| SESN1 | 0.58695 | 0.66383 | 0.19 | 0.18 | 1 | HC_vs_MS.NoTrt_B |
| HEMGN | 0.58723 | 0.30006 | 0.117 | 0.14 | 1 | HC_vs_MS.NoTrt_B |
| CGN | 0.58793 | 0.287 | 0.201 | 0.2 | 1 | HC_vs_MS.NoTrt_B |
| NFATC1 | 0.588 | 0.57749 | 0.164 | 0.15 | 1 | HC_vs_MS.NoTrt_B |
| ZNF770 | 0.5883 | 0.2951 | 0.322 | 0.34 | 1 | HC_vs_MS.NoTrt_B |
| SP1 | 0.58897 | 0.84551 | 0.168 | 0.16 | 1 | HC_vs_MS.NoTrt_B |

| | | | | | | |
|---|---|---|---|---|---|---|
| NBN | 0.58899 | 0.3671 | 0.104 | 0.12 | 1 | HC_vs_MS.NoTrt_B |
| TP53 | 0.58953 | 0.41599 | 0.772 | 0.84 | 1 | HC_vs_MS.NoTrt_B |
| SNX19 | 0.58968 | 0.26664 | 0.21 | 0.23 | 1 | HC_vs_MS.NoTrt_B |
| PHTF1 | 0.58996 | -0.4894 | 0.428 | 0.45 | 1 | HC_vs_MS.NoTrt_B |
| TFR2 | 0.59004 | 0.28036 | 0.696 | 0.71 | 1 | HC_vs_MS.NoTrt_B |
| CTBS | 0.5901 | 0.43222 | 0.125 | 0.14 | 1 | HC_vs_MS.NoTrt_B |
| MTMR10 | 0.59012 | 0.59932 | 0.159 | 0.18 | 1 | HC_vs_MS.NoTrt_B |
| LPAR4 | 0.59016 | 0.53489 | 0.132 | 0.12 | 1 | HC_vs_MS.NoTrt_B |
| TBC1D20 | 0.59102 | 0.75943 | 0.568 | 0.63 | 1 | HC_vs_MS.NoTrt_B |
| PPP5C | 0.59141 | 0.30984 | 0.397 | 0.39 | 1 | HC_vs_MS.NoTrt_B |
| PUM3 | 0.59241 | 0.34257 | 0.135 | 0.12 | 1 | HC_vs_MS.NoTrt_B |
| BBS2 | 0.59254 | 0.26655 | 0.661 | 0.72 | 1 | HC_vs_MS.NoTrt_B |
| RHEB | 0.5928 | 0.4365 | 0.128 | 0.15 | 1 | HC_vs_MS.NoTrt_B |
| WRAP73 | 0.59281 | 0.37992 | 0.33 | 0.31 | 1 | HC_vs_MS.NoTrt_B |
| SGK1 | 0.59285 | -0.436 | 0.371 | 0.37 | 1 | HC_vs_MS.NoTrt_B |
| SKIDA1 | 0.5931 | -0.2708 | 0.404 | 0.41 | 1 | HC_vs_MS.NoTrt_B |
| TRIP11 | 0.59336 | 0.41956 | 0.52 | 0.5 | 1 | HC_vs_MS.NoTrt_B |
| DDX60L | 0.59342 | 0.55615 | 0.714 | 0.76 | 1 | HC_vs_MS.NoTrt_B |
| P4HB | 0.59351 | -0.2889 | 0.654 | 0.65 | 1 | HC_vs_MS.NoTrt_B |
| MBD6 | 0.5936 | 0.73152 | 0.199 | 0.18 | 1 | HC_vs_MS.NoTrt_B |
| SNX5 | 0.59383 | 0.26703 | 0.583 | 0.63 | 1 | HC_vs_MS.NoTrt_B |
| RBM26 | 0.59394 | 0.78335 | 0.468 | 0.5 | 1 | HC_vs_MS.NoTrt_B |
| C18orf32 | 0.59434 | 0.27222 | 0.835 | 0.87 | 1 | HC_vs_MS.NoTrt_B |
| FBXO6 | 0.59539 | 0.73462 | 0.617 | 0.71 | 1 | HC_vs_MS.NoTrt_B |
| KCNK3 | 0.59644 | 0.37123 | 0.609 | 0.68 | 1 | HC_vs_MS.NoTrt_B |
| PHLDA1 | 0.59674 | -0.3754 | 0.459 | 0.44 | 1 | HC_vs_MS.NoTrt_B |
| GLT8D1 | 0.59711 | -0.4195 | 0.229 | 0.24 | 1 | HC_vs_MS.NoTrt_B |
| AC138645.2 | 0.59725 | 0.39149 | 0.112 | 0.1 | 1 | HC_vs_MS.NoTrt_B |
| PDP2 | 0.59748 | -0.4121 | 0.778 | 0.87 | 1 | HC_vs_MS.NoTrt_B |
| JOSD1 | 0.59818 | 0.72105 | 0.533 | 0.5 | 1 | HC_vs_MS.NoTrt_B |
| POLR3H | 0.59829 | 0.46249 | 0.238 | 0.22 | 1 | HC_vs_MS.NoTrt_B |
| QRSL1 | 0.59891 | 0.45391 | 0.555 | 0.6 | 1 | HC_vs_MS.NoTrt_B |
| CEP63 | 0.59903 | -0.4151 | 0.699 | 0.76 | 1 | HC_vs_MS.NoTrt_B |
| POU5F2 | 0.60107 | 0.64302 | 0.678 | 0.68 | 1 | HC_vs_MS.NoTrt_B |
| IWS1 | 0.60131 | 0.6096 | 0.128 | 0.15 | 1 | HC_vs_MS.NoTrt_B |
| CUL9 | 0.60162 | 0.47362 | 0.154 | 0.17 | 1 | HC_vs_MS.NoTrt_B |
| BET1 | 0.60168 | 0.27131 | 0.568 | 0.56 | 1 | HC_vs_MS.NoTrt_B |
| CLEC4E | 0.60198 | 0.29501 | 0.343 | 0.4 | 1 | HC_vs_MS.NoTrt_B |
| ZFP2 | 0.60319 | 1.18095 | 0.19 | 0.18 | 1 | HC_vs_MS.NoTrt_B |
| CTD-2192J1( | 0.60325 | 0.71261 | 0.133 | 0.12 | 1 | HC_vs_MS.NoTrt_B |

| | | | | | | |
|---|---|---|---|---|---|---|
| ZNF235 | 0.60408 | 0.79173 | 0.103 | 0.09 | 1 | HC_vs_MS.NoTrt_B |
| ANKEF1 | 0.60427 | 0.68701 | 0.214 | 0.25 | 1 | HC_vs_MS.NoTrt_B |
| ZNF280B | 0.60441 | 0.58291 | 0.116 | 0.13 | 1 | HC_vs_MS.NoTrt_B |
| GSTO2 | 0.60449 | 0.39328 | 0.858 | 0.91 | 1 | HC_vs_MS.NoTrt_B |
| PCYT1A | 0.60466 | -0.3964 | 0.507 | 0.5 | 1 | HC_vs_MS.NoTrt_B |
| PPP4R1 | 0.60476 | -0.637 | 0.238 | 0.25 | 1 | HC_vs_MS.NoTrt_B |
| WDR33 | 0.60499 | 0.74397 | 0.254 | 0.25 | 1 | HC_vs_MS.NoTrt_B |
| GTF2E2 | 0.60512 | 0.7937 | 0.588 | 0.62 | 1 | HC_vs_MS.NoTrt_B |
| PAK1 | 0.60539 | -1.137 | 0.155 | 0.17 | 1 | HC_vs_MS.NoTrt_B |
| UBE2A | 0.60582 | 0.55767 | 0.346 | 0.32 | 1 | HC_vs_MS.NoTrt_B |
| SLC25A51 | 0.6059 | 0.51059 | 0.538 | 0.57 | 1 | HC_vs_MS.NoTrt_B |
| ZNF555 | 0.60664 | 0.85512 | 0.587 | 0.61 | 1 | HC_vs_MS.NoTrt_B |
| VARS2 | 0.60728 | 0.5324 | 0.471 | 0.45 | 1 | HC_vs_MS.NoTrt_B |
| TUT1 | 0.60734 | -0.6244 | 0.097 | 0.11 | 1 | HC_vs_MS.NoTrt_B |
| EGFL8 | 0.60737 | 0.32735 | 0.483 | 0.52 | 1 | HC_vs_MS.NoTrt_B |
| KIF9 | 0.60747 | 0.34358 | 0.106 | 0.09 | 1 | HC_vs_MS.NoTrt_B |
| CFD | 0.6076 | 0.79785 | 0.151 | 0.14 | 1 | HC_vs_MS.NoTrt_B |
| TECPR2 | 0.60778 | 1.09013 | 0.345 | 0.34 | 1 | HC_vs_MS.NoTrt_B |
| NAA30 | 0.6078 | 0.84427 | 0.443 | 0.46 | 1 | HC_vs_MS.NoTrt_B |
| RAB3IP | 0.60808 | 0.71358 | 0.674 | 0.67 | 1 | HC_vs_MS.NoTrt_B |
| PHC3 | 0.60848 | 0.36866 | 0.958 | 0.98 | 1 | HC_vs_MS.NoTrt_B |
| GPN2 | 0.60856 | -0.259 | 0.774 | 0.76 | 1 | HC_vs_MS.NoTrt_B |
| TMPRSS3 | 0.60887 | 0.50856 | 0.342 | 0.33 | 1 | HC_vs_MS.NoTrt_B |
| ANGEL2 | 0.60952 | 0.36485 | 0.696 | 0.69 | 1 | HC_vs_MS.NoTrt_B |
| TASP1 | 0.60953 | 0.37507 | 0.11 | 0.1 | 1 | HC_vs_MS.NoTrt_B |
| CLDND1 | 0.6096 | 0.32657 | 0.355 | 0.4 | 1 | HC_vs_MS.NoTrt_B |
| PPM1K | 0.61011 | -0.3715 | 0.875 | 0.89 | 1 | HC_vs_MS.NoTrt_B |
| PRR23C | 0.61013 | 0.34255 | 0.4 | 0.42 | 1 | HC_vs_MS.NoTrt_B |
| ASUN | 0.61063 | 0.54994 | 0.109 | 0.12 | 1 | HC_vs_MS.NoTrt_B |
| ROCK2 | 0.6107 | 0.9213 | 0.439 | 0.48 | 1 | HC_vs_MS.NoTrt_B |
| PRRC2B | 0.61081 | 0.38926 | 0.268 | 0.25 | 1 | HC_vs_MS.NoTrt_B |
| ALS2 | 0.61121 | 0.89365 | 0.435 | 0.45 | 1 | HC_vs_MS.NoTrt_B |
| MAN2C1 | 0.61137 | 0.31972 | 0.8 | 0.78 | 1 | HC_vs_MS.NoTrt_B |
| KIAA0922 | 0.61165 | 0.32998 | 0.49 | 0.53 | 1 | HC_vs_MS.NoTrt_B |
| D2HGDH | 0.61165 | 0.86984 | 0.439 | 0.43 | 1 | HC_vs_MS.NoTrt_B |
| IL17RE | 0.61299 | -0.6155 | 0.259 | 0.24 | 1 | HC_vs_MS.NoTrt_B |
| NBAS | 0.61356 | 0.53701 | 0.213 | 0.2 | 1 | HC_vs_MS.NoTrt_B |
| ABCE1 | 0.6137 | 0.4406 | 0.346 | 0.33 | 1 | HC_vs_MS.NoTrt_B |
| BRD8 | 0.6139 | -0.332 | 0.735 | 0.77 | 1 | HC_vs_MS.NoTrt_B |
| NFATC2 | 0.61413 | 0.74409 | 0.178 | 0.17 | 1 | HC_vs_MS.NoTrt_B |

| | | | | | | |
|---|---|---|---|---|---|---|
| RP11-894J14 | 0.61452 | 0.27765 | 0.112 | 0.12 | 1 | HC_vs_MS.NoTrt_B |
| PLEKHG4B | 0.61519 | 0.38236 | 0.386 | 0.44 | 1 | HC_vs_MS.NoTrt_B |
| HSD11B1L | 0.61722 | 0.26496 | 0.649 | 0.7 | 1 | HC_vs_MS.NoTrt_B |
| FBXO5 | 0.6176 | -0.3253 | 0.401 | 0.4 | 1 | HC_vs_MS.NoTrt_B |
| DYNAP | 0.61767 | 0.26572 | 0.187 | 0.18 | 1 | HC_vs_MS.NoTrt_B |
| TMEM191C | 0.6178 | 0.50039 | 0.662 | 0.72 | 1 | HC_vs_MS.NoTrt_B |
| PPP2R2A | 0.61861 | 0.50828 | 0.126 | 0.14 | 1 | HC_vs_MS.NoTrt_B |
| 1-Mar | 0.619 | 0.48766 | 0.583 | 0.61 | 1 | HC_vs_MS.NoTrt_B |
| MARK3 | 0.6191 | 0.29286 | 0.875 | 0.92 | 1 | HC_vs_MS.NoTrt_B |
| ABLIM1 | 0.61961 | 0.57555 | 0.413 | 0.43 | 1 | HC_vs_MS.NoTrt_B |
| PRMT1 | 0.6199 | -0.3192 | 0.81 | 0.82 | 1 | HC_vs_MS.NoTrt_B |
| MSL1 | 0.61991 | 0.41222 | 0.554 | 0.64 | 1 | HC_vs_MS.NoTrt_B |
| PLK1 | 0.61999 | -0.2911 | 0.383 | 0.42 | 1 | HC_vs_MS.NoTrt_B |
| SCUBE3 | 0.62012 | 0.34861 | 0.601 | 0.67 | 1 | HC_vs_MS.NoTrt_B |
| ANTXR2 | 0.62023 | -0.3507 | 0.672 | 0.67 | 1 | HC_vs_MS.NoTrt_B |
| TGIF2 | 0.62028 | 0.32999 | 0.343 | 0.32 | 1 | HC_vs_MS.NoTrt_B |
| RDH16 | 0.62037 | 0.26599 | 0.684 | 0.73 | 1 | HC_vs_MS.NoTrt_B |
| FGD6 | 0.62047 | 0.97519 | 0.43 | 0.43 | 1 | HC_vs_MS.NoTrt_B |
| HEATR3 | 0.62076 | 0.53278 | 0.625 | 0.68 | 1 | HC_vs_MS.NoTrt_B |
| PCNX3 | 0.62087 | 0.61104 | 0.103 | 0.09 | 1 | HC_vs_MS.NoTrt_B |
| PNPLA2 | 0.62122 | 0.35586 | 0.17 | 0.16 | 1 | HC_vs_MS.NoTrt_B |
| RNF43 | 0.62189 | 0.25557 | 0.296 | 0.27 | 1 | HC_vs_MS.NoTrt_B |
| SNRK | 0.62204 | 0.55529 | 0.545 | 0.55 | 1 | HC_vs_MS.NoTrt_B |
| RAB36 | 0.6221 | 0.34828 | 0.304 | 0.3 | 1 | HC_vs_MS.NoTrt_B |
| CD28 | 0.62224 | 0.31339 | 0.597 | 0.65 | 1 | HC_vs_MS.NoTrt_B |
| SENP1 | 0.62281 | 0.59932 | 0.167 | 0.15 | 1 | HC_vs_MS.NoTrt_B |
| CAMKMT | 0.62302 | 0.84442 | 0.291 | 0.28 | 1 | HC_vs_MS.NoTrt_B |
| GPCPD1 | 0.62343 | 0.70321 | 0.293 | 0.29 | 1 | HC_vs_MS.NoTrt_B |
| PDXK | 0.62363 | 0.81061 | 0.49 | 0.51 | 1 | HC_vs_MS.NoTrt_B |
| EMC2 | 0.62374 | -0.2997 | 0.465 | 0.5 | 1 | HC_vs_MS.NoTrt_B |
| LURAP1 | 0.62457 | 0.26654 | 0.339 | 0.35 | 1 | HC_vs_MS.NoTrt_B |
| PHC1 | 0.62501 | 0.44893 | 0.12 | 0.11 | 1 | HC_vs_MS.NoTrt_B |
| SGSM1 | 0.62504 | 0.36453 | 0.477 | 0.48 | 1 | HC_vs_MS.NoTrt_B |
| PES1 | 0.62573 | 0.28055 | 0.104 | 0.09 | 1 | HC_vs_MS.NoTrt_B |
| STX7 | 0.62621 | 0.78442 | 0.72 | 0.74 | 1 | HC_vs_MS.NoTrt_B |
| TMEM186 | 0.62628 | 0.48893 | 0.51 | 0.56 | 1 | HC_vs_MS.NoTrt_B |
| LYRM1 | 0.62631 | 0.84812 | 0.548 | 0.57 | 1 | HC_vs_MS.NoTrt_B |
| SLCO2A1 | 0.62643 | 0.33566 | 0.357 | 0.37 | 1 | HC_vs_MS.NoTrt_B |
| BET1L | 0.62644 | 0.35252 | 0.213 | 0.2 | 1 | HC_vs_MS.NoTrt_B |
| C9orf66 | 0.62697 | 0.25499 | 0.574 | 0.58 | 1 | HC_vs_MS.NoTrt_B |

| | | | | | | |
|---|---|---|---|---|---|---|
| PARP9 | 0.62743 | 0.73509 | 0.388 | 0.4 | 1 | HC_vs_MS.NoTrt_B |
| PIGO | 0.62746 | 0.31539 | 0.342 | 0.32 | 1 | HC_vs_MS.NoTrt_B |
| NUDC | 0.62759 | 0.2586 | 0.546 | 0.62 | 1 | HC_vs_MS.NoTrt_B |
| SNAPC5 | 0.62772 | 0.50836 | 0.536 | 0.57 | 1 | HC_vs_MS.NoTrt_B |
| SLC17A5 | 0.62797 | -0.5338 | 0.5 | 0.51 | 1 | HC_vs_MS.NoTrt_B |
| ZFAT | 0.62832 | -0.8355 | 0.319 | 0.31 | 1 | HC_vs_MS.NoTrt_B |
| ABCA2 | 0.62961 | 1.0538 | 0.238 | 0.23 | 1 | HC_vs_MS.NoTrt_B |
| ZC2HC1C | 0.62969 | 0.48988 | 0.109 | 0.12 | 1 | HC_vs_MS.NoTrt_B |
| FBLN1 | 0.62974 | 0.27855 | 0.146 | 0.14 | 1 | HC_vs_MS.NoTrt_B |
| UQCC1 | 0.62989 | 0.29014 | 0.517 | 0.52 | 1 | HC_vs_MS.NoTrt_B |
| LEAP2 | 0.63089 | 0.27541 | 0.191 | 0.21 | 1 | HC_vs_MS.NoTrt_B |
| ABCC5 | 0.63092 | -0.3206 | 0.136 | 0.15 | 1 | HC_vs_MS.NoTrt_B |
| GPX2 | 0.63097 | 0.49231 | 0.18 | 0.2 | 1 | HC_vs_MS.NoTrt_B |
| PTCD2 | 0.63154 | 0.44959 | 0.541 | 0.61 | 1 | HC_vs_MS.NoTrt_B |
| FAM185A | 0.63177 | 0.37645 | 0.103 | 0.09 | 1 | HC_vs_MS.NoTrt_B |
| LRRC47 | 0.6318 | 0.34631 | 0.281 | 0.3 | 1 | HC_vs_MS.NoTrt_B |
| TRA2A | 0.63203 | 0.31318 | 0.759 | 0.81 | 1 | HC_vs_MS.NoTrt_B |
| RNGTT | 0.63255 | 0.35905 | 0.394 | 0.39 | 1 | HC_vs_MS.NoTrt_B |
| KIAA1614 | 0.63294 | 0.34643 | 0.303 | 0.34 | 1 | HC_vs_MS.NoTrt_B |
| FMNL3 | 0.63318 | 0.47191 | 0.235 | 0.22 | 1 | HC_vs_MS.NoTrt_B |
| ZNF486 | 0.63349 | 0.25133 | 0.649 | 0.65 | 1 | HC_vs_MS.NoTrt_B |
| NOMO2 | 0.6344 | 0.81811 | 0.184 | 0.18 | 1 | HC_vs_MS.NoTrt_B |
| MAPK9 | 0.63441 | 0.28757 | 0.442 | 0.48 | 1 | HC_vs_MS.NoTrt_B |
| IPCEF1 | 0.63508 | 0.31917 | 0.53 | 0.52 | 1 | HC_vs_MS.NoTrt_B |
| ZNF267 | 0.63513 | 0.32337 | 0.117 | 0.13 | 1 | HC_vs_MS.NoTrt_B |
| CCDC102B | 0.63553 | 0.34234 | 0.157 | 0.17 | 1 | HC_vs_MS.NoTrt_B |
| VHL | 0.63578 | 0.26286 | 0.749 | 0.77 | 1 | HC_vs_MS.NoTrt_B |
| CWF19L1 | 0.63605 | -0.5522 | 0.784 | 0.83 | 1 | HC_vs_MS.NoTrt_B |
| CXCR2 | 0.63624 | 0.28089 | 0.28 | 0.28 | 1 | HC_vs_MS.NoTrt_B |
| INCA1 | 0.63636 | -0.4682 | 0.18 | 0.21 | 1 | HC_vs_MS.NoTrt_B |
| GRTP1 | 0.63649 | 0.32489 | 0.506 | 0.51 | 1 | HC_vs_MS.NoTrt_B |
| CNKSR1 | 0.63654 | -0.6097 | 0.138 | 0.15 | 1 | HC_vs_MS.NoTrt_B |
| ENTPD7 | 0.63669 | 0.51312 | 0.345 | 0.34 | 1 | HC_vs_MS.NoTrt_B |
| NAT9 | 0.63781 | 0.34768 | 0.18 | 0.2 | 1 | HC_vs_MS.NoTrt_B |
| ADAM15 | 0.63812 | -0.4213 | 0.361 | 0.34 | 1 | HC_vs_MS.NoTrt_B |
| DIAPH2 | 0.63824 | 0.66993 | 0.304 | 0.33 | 1 | HC_vs_MS.NoTrt_B |
| QSOX1 | 0.63869 | 0.33359 | 0.37 | 0.36 | 1 | HC_vs_MS.NoTrt_B |
| MED13 | 0.63872 | 0.35642 | 0.588 | 0.58 | 1 | HC_vs_MS.NoTrt_B |
| UCKL1 | 0.6388 | -0.269 | 0.113 | 0.12 | 1 | HC_vs_MS.NoTrt_B |
| PINK1 | 0.63906 | -0.2789 | 0.268 | 0.28 | 1 | HC_vs_MS.NoTrt_B |

| | | | | | | |
|---|---|---|---|---|---|---|
| IQCK | 0.63918 | 0.57614 | 0.19 | 0.22 | 1 | HC_vs_MS.NoTrt_B |
| VPS9D1 | 0.63961 | 0.26185 | 0.226 | 0.24 | 1 | HC_vs_MS.NoTrt_B |
| MTUS1 | 0.6397 | 1.07571 | 0.339 | 0.37 | 1 | HC_vs_MS.NoTrt_B |
| NUP50 | 0.63974 | 0.31945 | 0.649 | 0.69 | 1 | HC_vs_MS.NoTrt_B |
| FANCI | 0.64106 | 0.29718 | 0.52 | 0.51 | 1 | HC_vs_MS.NoTrt_B |
| ARFIP1 | 0.64112 | -0.3127 | 0.212 | 0.22 | 1 | HC_vs_MS.NoTrt_B |
| CRIM1 | 0.6412 | 0.37542 | 0.139 | 0.13 | 1 | HC_vs_MS.NoTrt_B |
| GPR137B | 0.64131 | 0.80855 | 0.088 | 0.1 | 1 | HC_vs_MS.NoTrt_B |
| SNTG2 | 0.64132 | 0.26751 | 0.219 | 0.21 | 1 | HC_vs_MS.NoTrt_B |
| SIK3 | 0.6426 | 0.54107 | 0.243 | 0.27 | 1 | HC_vs_MS.NoTrt_B |
| ZNF43 | 0.64286 | -0.3541 | 0.726 | 0.72 | 1 | HC_vs_MS.NoTrt_B |
| VPS18 | 0.64467 | -0.3727 | 0.197 | 0.21 | 1 | HC_vs_MS.NoTrt_B |
| CALD1 | 0.64503 | -0.6657 | 0.191 | 0.18 | 1 | HC_vs_MS.NoTrt_B |
| GNB2 | 0.64531 | 0.43034 | 0.491 | 0.5 | 1 | HC_vs_MS.NoTrt_B |
| ZNF831 | 0.6456 | 0.67001 | 0.236 | 0.23 | 1 | HC_vs_MS.NoTrt_B |
| MAPK8IP3 | 0.64565 | 0.35716 | 0.148 | 0.16 | 1 | HC_vs_MS.NoTrt_B |
| TEFM | 0.6464 | 0.43718 | 0.652 | 0.7 | 1 | HC_vs_MS.NoTrt_B |
| C16orf45 | 0.64656 | 0.33631 | 0.203 | 0.18 | 1 | HC_vs_MS.NoTrt_B |
| RNASEH1 | 0.64711 | -0.2553 | 0.729 | 0.74 | 1 | HC_vs_MS.NoTrt_B |
| MEF2A | 0.64784 | 0.4606 | 0.443 | 0.48 | 1 | HC_vs_MS.NoTrt_B |
| EHBP1 | 0.64784 | 0.4653 | 0.354 | 0.38 | 1 | HC_vs_MS.NoTrt_B |
| PCNX1 | 0.64791 | 0.2695 | 0.857 | 0.88 | 1 | HC_vs_MS.NoTrt_B |
| FCHO2 | 0.64829 | 0.60263 | 0.309 | 0.3 | 1 | HC_vs_MS.NoTrt_B |
| CNOT6L | 0.64883 | 0.69162 | 0.274 | 0.27 | 1 | HC_vs_MS.NoTrt_B |
| EIF2B4 | 0.64889 | 0.70239 | 0.486 | 0.52 | 1 | HC_vs_MS.NoTrt_B |
| CH507-9B2.! | 0.64936 | 0.52994 | 0.403 | 0.43 | 1 | HC_vs_MS.NoTrt_B |
| ACVR1 | 0.64958 | -0.394 | 0.193 | 0.21 | 1 | HC_vs_MS.NoTrt_B |
| UFM1 | 0.65004 | 0.40404 | 0.326 | 0.34 | 1 | HC_vs_MS.NoTrt_B |
| CIPC | 0.65036 | -0.4269 | 0.37 | 0.4 | 1 | HC_vs_MS.NoTrt_B |
| GCN1 | 0.65036 | 0.4142 | 0.171 | 0.16 | 1 | HC_vs_MS.NoTrt_B |
| SLC12A7 | 0.65062 | 0.36098 | 0.138 | 0.15 | 1 | HC_vs_MS.NoTrt_B |
| ZNF720 | 0.65065 | 0.25003 | 0.497 | 0.51 | 1 | HC_vs_MS.NoTrt_B |
| CREBL2 | 0.65161 | 0.75949 | 0.09 | 0.1 | 1 | HC_vs_MS.NoTrt_B |
| NAF1 | 0.65166 | 0.53705 | 0.409 | 0.4 | 1 | HC_vs_MS.NoTrt_B |
| RFX2 | 0.65174 | 1.00337 | 0.477 | 0.49 | 1 | HC_vs_MS.NoTrt_B |
| STX8 | 0.65177 | -0.3175 | 0.854 | 0.91 | 1 | HC_vs_MS.NoTrt_B |
| ATP6V1C1 | 0.65184 | -0.3072 | 0.47 | 0.45 | 1 | HC_vs_MS.NoTrt_B |
| RFXANK | 0.65206 | -0.7873 | 0.242 | 0.24 | 1 | HC_vs_MS.NoTrt_B |
| NUCKS1 | 0.65243 | 0.32455 | 0.496 | 0.55 | 1 | HC_vs_MS.NoTrt_B |
| JMJD6 | 0.65272 | -0.4988 | 0.455 | 0.44 | 1 | HC_vs_MS.NoTrt_B |

| | | | | | | |
|---|---|---|---|---|---|---|
| DDIAS | 0.65282 | 0.59857 | 0.355 | 0.35 | 1 | HC_vs_MS.NoTrt_B |
| GHRL | 0.65285 | 0.35941 | 0.088 | 0.1 | 1 | HC_vs_MS.NoTrt_B |
| PML | 0.65302 | 0.4629 | 0.596 | 0.6 | 1 | HC_vs_MS.NoTrt_B |
| PHF19 | 0.65307 | 0.46629 | 0.12 | 0.14 | 1 | HC_vs_MS.NoTrt_B |
| PRR29 | 0.65328 | 0.44862 | 0.171 | 0.2 | 1 | HC_vs_MS.NoTrt_B |
| LEF1 | 0.65329 | 0.34787 | 0.604 | 0.63 | 1 | HC_vs_MS.NoTrt_B |
| FBXW7 | 0.65332 | 0.49759 | 0.152 | 0.14 | 1 | HC_vs_MS.NoTrt_B |
| PAPD5 | 0.65357 | 0.49016 | 0.136 | 0.15 | 1 | HC_vs_MS.NoTrt_B |
| COL4A3BP | 0.65393 | 0.59041 | 0.129 | 0.15 | 1 | HC_vs_MS.NoTrt_B |
| RASGEF1B | 0.65416 | 0.56961 | 0.672 | 0.72 | 1 | HC_vs_MS.NoTrt_B |
| SMIM19 | 0.65428 | 0.54801 | 0.472 | 0.47 | 1 | HC_vs_MS.NoTrt_B |
| TRIM44 | 0.6543 | 0.67841 | 0.275 | 0.27 | 1 | HC_vs_MS.NoTrt_B |
| ARHGAP26 | 0.65478 | 0.40411 | 0.626 | 0.68 | 1 | HC_vs_MS.NoTrt_B |
| CSPP1 | 0.65481 | 1.13986 | 0.397 | 0.4 | 1 | HC_vs_MS.NoTrt_B |
| FMN1 | 0.65518 | 0.38646 | 0.206 | 0.2 | 1 | HC_vs_MS.NoTrt_B |
| FAM21C | 0.6553 | 0.29067 | 0.203 | 0.22 | 1 | HC_vs_MS.NoTrt_B |
| HCFC1 | 0.65548 | 0.43387 | 0.296 | 0.29 | 1 | HC_vs_MS.NoTrt_B |
| PTK7 | 0.65555 | 0.25675 | 0.564 | 0.58 | 1 | HC_vs_MS.NoTrt_B |
| RBM14-RBM | 0.65608 | 0.60487 | 0.09 | 0.1 | 1 | HC_vs_MS.NoTrt_B |
| DYRK2 | 0.65693 | 0.52379 | 0.181 | 0.2 | 1 | HC_vs_MS.NoTrt_B |
| OAZ2 | 0.65719 | -0.2576 | 0.755 | 0.83 | 1 | HC_vs_MS.NoTrt_B |
| RHOT1 | 0.65753 | 0.27635 | 0.178 | 0.19 | 1 | HC_vs_MS.NoTrt_B |
| CUL4B | 0.65762 | 0.60571 | 0.107 | 0.12 | 1 | HC_vs_MS.NoTrt_B |
| HEATR5B | 0.65769 | 0.32824 | 0.438 | 0.46 | 1 | HC_vs_MS.NoTrt_B |
| HACL1 | 0.65816 | 0.40536 | 0.125 | 0.14 | 1 | HC_vs_MS.NoTrt_B |
| FCGRT | 0.65824 | -0.4129 | 0.217 | 0.23 | 1 | HC_vs_MS.NoTrt_B |
| GNA11 | 0.65892 | 0.25863 | 0.53 | 0.51 | 1 | HC_vs_MS.NoTrt_B |
| RGCC | 0.65894 | 0.50165 | 0.18 | 0.2 | 1 | HC_vs_MS.NoTrt_B |
| ATIC | 0.65916 | 0.26439 | 0.377 | 0.41 | 1 | HC_vs_MS.NoTrt_B |
| SSBP3 | 0.66037 | -0.3106 | 0.394 | 0.43 | 1 | HC_vs_MS.NoTrt_B |
| ERC1 | 0.66038 | 0.42622 | 0.861 | 0.92 | 1 | HC_vs_MS.NoTrt_B |
| ARHGAP31 | 0.66091 | 0.3416 | 0.293 | 0.29 | 1 | HC_vs_MS.NoTrt_B |
| NANOS1 | 0.66132 | 0.26296 | 0.459 | 0.48 | 1 | HC_vs_MS.NoTrt_B |
| TMEM229B | 0.6617 | 0.86264 | 0.172 | 0.19 | 1 | HC_vs_MS.NoTrt_B |
| PTBP3 | 0.66196 | 0.69387 | 0.325 | 0.36 | 1 | HC_vs_MS.NoTrt_B |
| PARP11 | 0.66216 | 0.66522 | 0.454 | 0.49 | 1 | HC_vs_MS.NoTrt_B |
| NANP | 0.66229 | -0.7841 | 0.275 | 0.3 | 1 | HC_vs_MS.NoTrt_B |
| WDR55 | 0.6624 | -0.2648 | 0.613 | 0.67 | 1 | HC_vs_MS.NoTrt_B |
| MAP4K4 | 0.66244 | 0.27341 | 0.538 | 0.57 | 1 | HC_vs_MS.NoTrt_B |
| ZNF746 | 0.66282 | 0.25161 | 0.194 | 0.21 | 1 | HC_vs_MS.NoTrt_B |

| | | | | | | |
|---|---|---|---|---|---|---|
| AC138969.4 | 0.66292 | 0.51139 | 0.436 | 0.47 | 1 | HC_vs_MS.NoTrt_B |
| DBT | 0.66299 | -0.5405 | 0.851 | 0.85 | 1 | HC_vs_MS.NoTrt_B |
| C2orf54 | 0.66303 | 0.29742 | 0.114 | 0.11 | 1 | HC_vs_MS.NoTrt_B |
| MLF1 | 0.66312 | 0.29795 | 0.293 | 0.29 | 1 | HC_vs_MS.NoTrt_B |
| FKRP | 0.66421 | 0.34536 | 0.633 | 0.67 | 1 | HC_vs_MS.NoTrt_B |
| GYG1 | 0.66446 | -0.4402 | 0.542 | 0.52 | 1 | HC_vs_MS.NoTrt_B |
| PPP2R5A | 0.66522 | 0.53803 | 0.187 | 0.2 | 1 | HC_vs_MS.NoTrt_B |
| TSG101 | 0.66534 | 0.25524 | 0.432 | 0.45 | 1 | HC_vs_MS.NoTrt_B |
| VWA7 | 0.66542 | 0.75625 | 0.313 | 0.31 | 1 | HC_vs_MS.NoTrt_B |
| BEND7 | 0.66643 | 0.28857 | 0.316 | 0.34 | 1 | HC_vs_MS.NoTrt_B |
| AHCTF1 | 0.66657 | 0.49181 | 0.133 | 0.15 | 1 | HC_vs_MS.NoTrt_B |
| LRRC8D | 0.66704 | 0.40831 | 0.148 | 0.14 | 1 | HC_vs_MS.NoTrt_B |
| EHD3 | 0.66794 | 0.46012 | 0.091 | 0.1 | 1 | HC_vs_MS.NoTrt_B |
| PACSIN2 | 0.66814 | 0.26862 | 0.5 | 0.5 | 1 | HC_vs_MS.NoTrt_B |
| C19orf12 | 0.66827 | 0.44208 | 0.335 | 0.32 | 1 | HC_vs_MS.NoTrt_B |
| ZNF417 | 0.66864 | 0.80927 | 0.693 | 0.75 | 1 | HC_vs_MS.NoTrt_B |
| GCC2 | 0.66902 | 0.26145 | 0.806 | 0.83 | 1 | HC_vs_MS.NoTrt_B |
| PACS2 | 0.66916 | 0.51232 | 0.246 | 0.24 | 1 | HC_vs_MS.NoTrt_B |
| CCDC127 | 0.67012 | -0.4013 | 0.716 | 0.79 | 1 | HC_vs_MS.NoTrt_B |
| CLK4 | 0.6705 | 0.37098 | 0.657 | 0.69 | 1 | HC_vs_MS.NoTrt_B |
| YPEL1 | 0.67067 | 0.65833 | 0.414 | 0.41 | 1 | HC_vs_MS.NoTrt_B |
| STYXL1 | 0.67092 | -0.5666 | 0.268 | 0.25 | 1 | HC_vs_MS.NoTrt_B |
| TMEM2 | 0.67094 | 0.42338 | 0.584 | 0.57 | 1 | HC_vs_MS.NoTrt_B |
| TMEM19 | 0.67104 | -0.4421 | 0.743 | 0.78 | 1 | HC_vs_MS.NoTrt_B |
| KIF21A | 0.67156 | 0.90207 | 0.142 | 0.16 | 1 | HC_vs_MS.NoTrt_B |
| TEX11 | 0.67189 | 0.27607 | 0.239 | 0.23 | 1 | HC_vs_MS.NoTrt_B |
| HBS1L | 0.67194 | 0.36359 | 0.568 | 0.58 | 1 | HC_vs_MS.NoTrt_B |
| ZNF487 | 0.6724 | -0.8022 | 0.433 | 0.43 | 1 | HC_vs_MS.NoTrt_B |
| ZNF518A | 0.67399 | 0.35848 | 0.113 | 0.12 | 1 | HC_vs_MS.NoTrt_B |
| SHC1 | 0.67406 | 0.30639 | 0.63 | 0.65 | 1 | HC_vs_MS.NoTrt_B |
| IL27RA | 0.67488 | 0.35711 | 0.359 | 0.36 | 1 | HC_vs_MS.NoTrt_B |
| AHRR | 0.67502 | 0.50282 | 0.113 | 0.13 | 1 | HC_vs_MS.NoTrt_B |
| TMCO6 | 0.67512 | -0.5575 | 0.475 | 0.51 | 1 | HC_vs_MS.NoTrt_B |
| SRPK2 | 0.6756 | 0.4609 | 0.328 | 0.36 | 1 | HC_vs_MS.NoTrt_B |
| NSUN5 | 0.67642 | 0.79759 | 0.312 | 0.35 | 1 | HC_vs_MS.NoTrt_B |
| SLC28A2 | 0.6767 | 0.28478 | 0.746 | 0.81 | 1 | HC_vs_MS.NoTrt_B |
| SLC12A6 | 0.67696 | 0.33936 | 0.759 | 0.8 | 1 | HC_vs_MS.NoTrt_B |
| CENPN | 0.6772 | 0.4237 | 0.155 | 0.15 | 1 | HC_vs_MS.NoTrt_B |
| LRSAM1 | 0.67794 | 0.48043 | 0.132 | 0.12 | 1 | HC_vs_MS.NoTrt_B |
| MYO9A | 0.67857 | 0.68748 | 0.323 | 0.35 | 1 | HC_vs_MS.NoTrt_B |

| | | | | | | |
|---|---|---|---|---|---|---|
| NSD1 | 0.67861 | 0.44284 | 0.645 | 0.65 | 1 | HC_vs_MS.NoTrt_B |
| ERCC4 | 0.67909 | 0.39636 | 0.858 | 0.89 | 1 | HC_vs_MS.NoTrt_B |
| FAIM | 0.67921 | 0.55148 | 0.358 | 0.37 | 1 | HC_vs_MS.NoTrt_B |
| APLF | 0.68015 | -0.259 | 0.128 | 0.14 | 1 | HC_vs_MS.NoTrt_B |
| RABEP1 | 0.68019 | 0.38824 | 0.201 | 0.2 | 1 | HC_vs_MS.NoTrt_B |
| RBM44 | 0.68053 | 0.38059 | 0.174 | 0.2 | 1 | HC_vs_MS.NoTrt_B |
| ZNF286A | 0.68056 | 0.73577 | 0.354 | 0.38 | 1 | HC_vs_MS.NoTrt_B |
| AGPS | 0.68064 | 0.50478 | 0.119 | 0.13 | 1 | HC_vs_MS.NoTrt_B |
| RSAD2 | 0.68075 | 0.99017 | 0.326 | 0.37 | 1 | HC_vs_MS.NoTrt_B |
| RC3H1 | 0.68085 | 0.59397 | 0.438 | 0.5 | 1 | HC_vs_MS.NoTrt_B |
| WDR76 | 0.68116 | -0.3395 | 0.658 | 0.69 | 1 | HC_vs_MS.NoTrt_B |
| STON1 | 0.68168 | -0.5956 | 0.301 | 0.31 | 1 | HC_vs_MS.NoTrt_B |
| TBRG4 | 0.68372 | 0.25222 | 0.391 | 0.38 | 1 | HC_vs_MS.NoTrt_B |
| RNF207 | 0.68421 | -0.6164 | 0.661 | 0.73 | 1 | HC_vs_MS.NoTrt_B |
| ITCH | 0.68457 | 0.46338 | 0.47 | 0.5 | 1 | HC_vs_MS.NoTrt_B |
| LAX1 | 0.68485 | -0.5344 | 0.467 | 0.42 | 1 | HC_vs_MS.NoTrt_B |
| NOP16 | 0.68526 | 0.29707 | 0.201 | 0.2 | 1 | HC_vs_MS.NoTrt_B |
| POC1B | 0.68559 | 0.26062 | 0.742 | 0.74 | 1 | HC_vs_MS.NoTrt_B |
| SLC16A4 | 0.68562 | 0.50099 | 0.662 | 0.68 | 1 | HC_vs_MS.NoTrt_B |
| MS4A6A | 0.68626 | 0.32647 | 0.162 | 0.16 | 1 | HC_vs_MS.NoTrt_B |
| KIAA1033 | 0.68725 | 0.3931 | 0.281 | 0.3 | 1 | HC_vs_MS.NoTrt_B |
| HDAC10 | 0.68737 | 0.94556 | 0.123 | 0.14 | 1 | HC_vs_MS.NoTrt_B |
| L2HGDH | 0.6884 | 0.54344 | 0.807 | 0.89 | 1 | HC_vs_MS.NoTrt_B |
| GTF2B | 0.68845 | 0.38613 | 0.167 | 0.18 | 1 | HC_vs_MS.NoTrt_B |
| GMPS | 0.68955 | -0.5611 | 0.861 | 0.87 | 1 | HC_vs_MS.NoTrt_B |
| RUFY1 | 0.68976 | 0.42378 | 0.309 | 0.35 | 1 | HC_vs_MS.NoTrt_B |
| RRAGC | 0.68992 | 0.41911 | 0.509 | 0.58 | 1 | HC_vs_MS.NoTrt_B |
| ASXL1 | 0.68994 | 0.4162 | 0.199 | 0.22 | 1 | HC_vs_MS.NoTrt_B |
| MRPS11 | 0.69012 | -0.3123 | 0.713 | 0.7 | 1 | HC_vs_MS.NoTrt_B |
| P4HA1 | 0.69014 | 0.28904 | 0.161 | 0.17 | 1 | HC_vs_MS.NoTrt_B |
| MFSD11 | 0.69098 | -0.6164 | 0.429 | 0.42 | 1 | HC_vs_MS.NoTrt_B |
| NSMCE3 | 0.69138 | 1.18544 | 0.319 | 0.32 | 1 | HC_vs_MS.NoTrt_B |
| SLCO2B1 | 0.6914 | 0.25996 | 0.309 | 0.31 | 1 | HC_vs_MS.NoTrt_B |
| SH3BP1 | 0.69153 | 0.92649 | 0.109 | 0.1 | 1 | HC_vs_MS.NoTrt_B |
| LRRC37A2 | 0.69223 | 0.33442 | 0.688 | 0.76 | 1 | HC_vs_MS.NoTrt_B |
| SRCAP | 0.69234 | 0.32213 | 0.662 | 0.69 | 1 | HC_vs_MS.NoTrt_B |
| TP53INP1 | 0.69249 | 0.54691 | 0.351 | 0.37 | 1 | HC_vs_MS.NoTrt_B |
| NUDT16L1 | 0.69291 | 0.56276 | 0.11 | 0.1 | 1 | HC_vs_MS.NoTrt_B |
| MRPL32 | 0.69314 | 0.41443 | 0.122 | 0.14 | 1 | HC_vs_MS.NoTrt_B |
| N4BP1 | 0.69336 | 0.69345 | 0.11 | 0.1 | 1 | HC_vs_MS.NoTrt_B |

| | | | | | | |
|---|---|---|---|---|---|---|
| ZNF772 | 0.69357 | 0.45374 | 0.42 | 0.45 | 1 | HC_vs_MS.NoTrt_B |
| PIGS | 0.69362 | 0.32364 | 0.48 | 0.51 | 1 | HC_vs_MS.NoTrt_B |
| PODXL | 0.69408 | 0.28384 | 0.193 | 0.22 | 1 | HC_vs_MS.NoTrt_B |
| ZNF343 | 0.69446 | 1.66803 | 0.433 | 0.43 | 1 | HC_vs_MS.NoTrt_B |
| ZNF681 | 0.69522 | 0.29003 | 0.649 | 0.68 | 1 | HC_vs_MS.NoTrt_B |
| LANCL3 | 0.69549 | -0.4066 | 0.259 | 0.28 | 1 | HC_vs_MS.NoTrt_B |
| CDHR3 | 0.69568 | -0.272 | 0.617 | 0.62 | 1 | HC_vs_MS.NoTrt_B |
| CELA2A | 0.69569 | 0.25104 | 0.519 | 0.53 | 1 | HC_vs_MS.NoTrt_B |
| AKT3 | 0.69584 | 0.45814 | 0.649 | 0.7 | 1 | HC_vs_MS.NoTrt_B |
| ADSSL1 | 0.69621 | 0.50408 | 0.257 | 0.25 | 1 | HC_vs_MS.NoTrt_B |
| RPTOR | 0.69658 | 0.70897 | 0.378 | 0.4 | 1 | HC_vs_MS.NoTrt_B |
| CYB5R3 | 0.69743 | -0.272 | 0.474 | 0.49 | 1 | HC_vs_MS.NoTrt_B |
| DVL3 | 0.69746 | 0.79064 | 0.235 | 0.26 | 1 | HC_vs_MS.NoTrt_B |
| VPS13D | 0.69781 | 0.60253 | 0.317 | 0.34 | 1 | HC_vs_MS.NoTrt_B |
| GPRC5C | 0.69815 | 0.25118 | 0.264 | 0.26 | 1 | HC_vs_MS.NoTrt_B |
| GON4L | 0.69858 | 0.41361 | 0.833 | 0.89 | 1 | HC_vs_MS.NoTrt_B |
| URB1 | 0.69896 | 0.57671 | 0.471 | 0.5 | 1 | HC_vs_MS.NoTrt_B |
| DCTN4 | 0.69897 | -0.2795 | 0.332 | 0.34 | 1 | HC_vs_MS.NoTrt_B |
| UBR4 | 0.69909 | 0.31438 | 0.752 | 0.78 | 1 | HC_vs_MS.NoTrt_B |
| NFXL1 | 0.69918 | 0.37345 | 0.103 | 0.11 | 1 | HC_vs_MS.NoTrt_B |
| CCS | 0.70013 | -0.4166 | 0.204 | 0.22 | 1 | HC_vs_MS.NoTrt_B |
| GPBP1L1 | 0.70023 | 0.75756 | 0.458 | 0.44 | 1 | HC_vs_MS.NoTrt_B |
| HSPBAP1 | 0.70055 | 0.31742 | 0.5 | 0.51 | 1 | HC_vs_MS.NoTrt_B |
| CYP51A1 | 0.70091 | -0.3069 | 0.454 | 0.5 | 1 | HC_vs_MS.NoTrt_B |
| ZNF764 | 0.70135 | 0.27141 | 0.32 | 0.32 | 1 | HC_vs_MS.NoTrt_B |
| C1orf210 | 0.70221 | 0.33387 | 0.394 | 0.39 | 1 | HC_vs_MS.NoTrt_B |
| CBLB | 0.70232 | 0.26958 | 0.355 | 0.36 | 1 | HC_vs_MS.NoTrt_B |
| ISCA1 | 0.7031 | -0.294 | 0.659 | 0.64 | 1 | HC_vs_MS.NoTrt_B |
| MAP3K2 | 0.7032 | 0.40179 | 0.171 | 0.16 | 1 | HC_vs_MS.NoTrt_B |
| SIRT3 | 0.70345 | 0.57944 | 0.294 | 0.32 | 1 | HC_vs_MS.NoTrt_B |
| ABHD6 | 0.70352 | -0.5569 | 0.461 | 0.46 | 1 | HC_vs_MS.NoTrt_B |
| IQCE | 0.70382 | 0.31421 | 0.406 | 0.44 | 1 | HC_vs_MS.NoTrt_B |
| TMEM39A | 0.70465 | 0.56345 | 0.117 | 0.11 | 1 | HC_vs_MS.NoTrt_B |
| AARS | 0.70482 | 0.36424 | 0.426 | 0.42 | 1 | HC_vs_MS.NoTrt_B |
| TIMP2 | 0.70488 | 0.33172 | 0.165 | 0.16 | 1 | HC_vs_MS.NoTrt_B |
| RPS6KA3 | 0.70501 | 0.42218 | 0.491 | 0.55 | 1 | HC_vs_MS.NoTrt_B |
| LZTR1 | 0.7052 | 1.26526 | 0.248 | 0.28 | 1 | HC_vs_MS.NoTrt_B |
| HEMK1 | 0.70521 | -0.3731 | 0.951 | 0.98 | 1 | HC_vs_MS.NoTrt_B |
| FRA10AC1 | 0.70537 | 0.92074 | 0.532 | 0.61 | 1 | HC_vs_MS.NoTrt_B |
| STX17 | 0.7057 | 0.80089 | 0.452 | 0.5 | 1 | HC_vs_MS.NoTrt_B |

| | | | | | | |
|---|---|---|---|---|---|---|
| RCC1L | 0.70594 | 0.44821 | 0.265 | 0.27 | 1 | HC_vs_MS.NoTrt_B |
| GPRIN3 | 0.70662 | 0.957 | 0.428 | 0.49 | 1 | HC_vs_MS.NoTrt_B |
| EML3 | 0.70673 | 0.33174 | 0.216 | 0.2 | 1 | HC_vs_MS.NoTrt_B |
| C22orf39 | 0.70689 | 0.61275 | 0.688 | 0.69 | 1 | HC_vs_MS.NoTrt_B |
| C8orf44 | 0.70705 | 0.26659 | 0.535 | 0.57 | 1 | HC_vs_MS.NoTrt_B |
| ISY1 | 0.70785 | -0.4581 | 0.612 | 0.64 | 1 | HC_vs_MS.NoTrt_B |
| MGAT4A | 0.70839 | 0.566 | 0.406 | 0.41 | 1 | HC_vs_MS.NoTrt_B |
| LANCL1 | 0.70859 | 1.03386 | 0.122 | 0.14 | 1 | HC_vs_MS.NoTrt_B |
| SF3B3 | 0.70859 | -0.4088 | 0.954 | 0.96 | 1 | HC_vs_MS.NoTrt_B |
| NARS2 | 0.70903 | 0.30732 | 0.39 | 0.42 | 1 | HC_vs_MS.NoTrt_B |
| BCOR | 0.70971 | 0.38488 | 0.112 | 0.1 | 1 | HC_vs_MS.NoTrt_B |
| SAMD9 | 0.70972 | 0.71545 | 0.183 | 0.18 | 1 | HC_vs_MS.NoTrt_B |
| SARDH | 0.7098 | 0.46372 | 0.388 | 0.38 | 1 | HC_vs_MS.NoTrt_B |
| BLOC1S3 | 0.70996 | 0.27704 | 0.586 | 0.61 | 1 | HC_vs_MS.NoTrt_B |
| NUDT1 | 0.71021 | -0.4536 | 0.391 | 0.38 | 1 | HC_vs_MS.NoTrt_B |
| SPPL3 | 0.71025 | -0.4583 | 0.583 | 0.62 | 1 | HC_vs_MS.NoTrt_B |
| SGTB | 0.71038 | 0.38005 | 0.654 | 0.65 | 1 | HC_vs_MS.NoTrt_B |
| KRAS | 0.71055 | 0.58538 | 0.372 | 0.37 | 1 | HC_vs_MS.NoTrt_B |
| ACBD5 | 0.71124 | -0.513 | 0.467 | 0.45 | 1 | HC_vs_MS.NoTrt_B |
| LPAR6 | 0.71129 | 0.36217 | 0.396 | 0.38 | 1 | HC_vs_MS.NoTrt_B |
| CCNH | 0.71138 | 0.2743 | 0.178 | 0.19 | 1 | HC_vs_MS.NoTrt_B |
| ZNF407 | 0.7116 | 0.71165 | 0.097 | 0.11 | 1 | HC_vs_MS.NoTrt_B |
| BIVM | 0.71203 | 0.73896 | 0.328 | 0.32 | 1 | HC_vs_MS.NoTrt_B |
| SIPA1L1 | 0.71216 | 0.65908 | 0.594 | 0.64 | 1 | HC_vs_MS.NoTrt_B |
| CCDC174 | 0.71216 | 0.63642 | 0.12 | 0.11 | 1 | HC_vs_MS.NoTrt_B |
| FAM208A | 0.71218 | 0.30283 | 0.736 | 0.72 | 1 | HC_vs_MS.NoTrt_B |
| CD22 | 0.71244 | 0.25379 | 0.778 | 0.83 | 1 | HC_vs_MS.NoTrt_B |
| NCKAP1 | 0.7135 | 0.33184 | 0.636 | 0.7 | 1 | HC_vs_MS.NoTrt_B |
| RRNAD1 | 0.71367 | 0.38966 | 0.643 | 0.67 | 1 | HC_vs_MS.NoTrt_B |
| PRKDC | 0.71407 | 0.58605 | 0.622 | 0.65 | 1 | HC_vs_MS.NoTrt_B |
| FCRL1 | 0.71412 | 0.4638 | 0.249 | 0.24 | 1 | HC_vs_MS.NoTrt_B |
| TRNT1 | 0.71417 | -0.4613 | 0.319 | 0.28 | 1 | HC_vs_MS.NoTrt_B |
| UBIAD1 | 0.7142 | -0.2893 | 0.684 | 0.7 | 1 | HC_vs_MS.NoTrt_B |
| PDCD2 | 0.71455 | 0.50958 | 0.106 | 0.1 | 1 | HC_vs_MS.NoTrt_B |
| PKIA | 0.71483 | 0.74828 | 0.104 | 0.1 | 1 | HC_vs_MS.NoTrt_B |
| LAMC3 | 0.71498 | -0.8617 | 0.264 | 0.27 | 1 | HC_vs_MS.NoTrt_B |
| DUS4L | 0.7151 | -0.319 | 0.096 | 0.1 | 1 | HC_vs_MS.NoTrt_B |
| RCC2 | 0.71531 | 0.55482 | 0.093 | 0.1 | 1 | HC_vs_MS.NoTrt_B |
| CDC73 | 0.71539 | 0.66009 | 0.516 | 0.54 | 1 | HC_vs_MS.NoTrt_B |
| KLHDC10 | 0.71542 | 0.77859 | 0.387 | 0.4 | 1 | HC_vs_MS.NoTrt_B |

| | | | | | | |
|---|---|---|---|---|---|---|
| ZFC3H1 | 0.71559 | 0.39653 | 0.367 | 0.4 | 1 | HC_vs_MS.NoTrt_B |
| CARNMT1 | 0.71559 | 1.05517 | 0.097 | 0.11 | 1 | HC_vs_MS.NoTrt_B |
| ZNF66 | 0.71563 | -0.3135 | 0.648 | 0.69 | 1 | HC_vs_MS.NoTrt_B |
| SDR42E1 | 0.71625 | 1.19334 | 0.386 | 0.43 | 1 | HC_vs_MS.NoTrt_B |
| TIAL1 | 0.71657 | 0.6212 | 0.862 | 0.89 | 1 | HC_vs_MS.NoTrt_B |
| FAM3A | 0.7167 | 0.68607 | 0.752 | 0.84 | 1 | HC_vs_MS.NoTrt_B |
| PFKFB2 | 0.71671 | 0.88501 | 0.2 | 0.2 | 1 | HC_vs_MS.NoTrt_B |
| PHF3 | 0.71674 | 0.45534 | 0.483 | 0.52 | 1 | HC_vs_MS.NoTrt_B |
| DDX23 | 0.71687 | -0.4158 | 0.662 | 0.68 | 1 | HC_vs_MS.NoTrt_B |
| ESYT3 | 0.71713 | 0.59372 | 0.462 | 0.51 | 1 | HC_vs_MS.NoTrt_B |
| NIT2 | 0.71722 | 0.38584 | 0.655 | 0.69 | 1 | HC_vs_MS.NoTrt_B |
| TRIM24 | 0.71723 | 0.29688 | 0.458 | 0.47 | 1 | HC_vs_MS.NoTrt_B |
| CPB1 | 0.71789 | 0.32407 | 0.483 | 0.51 | 1 | HC_vs_MS.NoTrt_B |
| PCTP | 0.71817 | -0.4809 | 0.391 | 0.43 | 1 | HC_vs_MS.NoTrt_B |
| CCNF | 0.71855 | 1.08134 | 0.439 | 0.46 | 1 | HC_vs_MS.NoTrt_B |
| C5 | 0.71905 | -0.2826 | 0.4 | 0.39 | 1 | HC_vs_MS.NoTrt_B |
| ATP8A1 | 0.71981 | 0.3264 | 0.161 | 0.17 | 1 | HC_vs_MS.NoTrt_B |
| GCLC | 0.72018 | 0.56733 | 0.497 | 0.55 | 1 | HC_vs_MS.NoTrt_B |
| RGL2 | 0.72034 | 0.45838 | 0.135 | 0.15 | 1 | HC_vs_MS.NoTrt_B |
| GTF2IRD1 | 0.72089 | -0.299 | 0.645 | 0.64 | 1 | HC_vs_MS.NoTrt_B |
| AK5 | 0.72109 | 0.70915 | 0.142 | 0.14 | 1 | HC_vs_MS.NoTrt_B |
| IGF2BP1 | 0.72154 | 0.29917 | 0.533 | 0.55 | 1 | HC_vs_MS.NoTrt_B |
| CBX1 | 0.72216 | 0.26046 | 0.272 | 0.29 | 1 | HC_vs_MS.NoTrt_B |
| DOPEY2 | 0.72241 | -0.3317 | 0.768 | 0.8 | 1 | HC_vs_MS.NoTrt_B |
| TBC1D1 | 0.72254 | 0.55903 | 0.616 | 0.69 | 1 | HC_vs_MS.NoTrt_B |
| SPDYE6 | 0.72266 | 0.7258 | 0.122 | 0.14 | 1 | HC_vs_MS.NoTrt_B |
| CAND1 | 0.7231 | 0.34764 | 0.886 | 0.91 | 1 | HC_vs_MS.NoTrt_B |
| TCEAL8 | 0.72423 | 0.295 | 0.109 | 0.12 | 1 | HC_vs_MS.NoTrt_B |
| KLC1 | 0.72495 | 0.32165 | 0.586 | 0.67 | 1 | HC_vs_MS.NoTrt_B |
| NEBL | 0.72496 | 0.61869 | 0.287 | 0.32 | 1 | HC_vs_MS.NoTrt_B |
| UNC13A | 0.72551 | 0.54613 | 0.261 | 0.29 | 1 | HC_vs_MS.NoTrt_B |
| GNPNAT1 | 0.72569 | 0.33577 | 0.513 | 0.58 | 1 | HC_vs_MS.NoTrt_B |
| NBEAL2 | 0.72686 | 0.3596 | 0.206 | 0.22 | 1 | HC_vs_MS.NoTrt_B |
| PARD6G | 0.7277 | 0.27082 | 0.41 | 0.41 | 1 | HC_vs_MS.NoTrt_B |
| POTEI | 0.72777 | 0.32204 | 0.19 | 0.18 | 1 | HC_vs_MS.NoTrt_B |
| GFAP | 0.72779 | 0.25036 | 0.733 | 0.82 | 1 | HC_vs_MS.NoTrt_B |
| IGFBP2 | 0.72815 | 0.25284 | 0.574 | 0.63 | 1 | HC_vs_MS.NoTrt_B |
| LMAN1 | 0.72816 | 0.60597 | 0.162 | 0.18 | 1 | HC_vs_MS.NoTrt_B |
| E2F8 | 0.72856 | 0.47069 | 0.133 | 0.15 | 1 | HC_vs_MS.NoTrt_B |
| ZCCHC14 | 0.72879 | 0.65428 | 0.104 | 0.1 | 1 | HC_vs_MS.NoTrt_B |

| | | | | | | |
|---|---|---|---|---|---|---|
| TOMM70 | 0.72939 | 0.53739 | 0.129 | 0.14 | 1 | HC_vs_MS.NoTrt_B |
| SLC35F6 | 0.72974 | -0.2607 | 0.759 | 0.81 | 1 | HC_vs_MS.NoTrt_B |
| LACTB | 0.73068 | -0.3044 | 0.558 | 0.58 | 1 | HC_vs_MS.NoTrt_B |
| MTMR4 | 0.73076 | 0.52222 | 0.236 | 0.23 | 1 | HC_vs_MS.NoTrt_B |
| TRMT112 | 0.73148 | 0.27696 | 0.254 | 0.29 | 1 | HC_vs_MS.NoTrt_B |
| DUSP18 | 0.73154 | 0.26959 | 0.586 | 0.58 | 1 | HC_vs_MS.NoTrt_B |
| LTB4R | 0.73226 | -0.7259 | 0.339 | 0.31 | 1 | HC_vs_MS.NoTrt_B |
| CCDC82 | 0.73234 | 0.45181 | 0.129 | 0.14 | 1 | HC_vs_MS.NoTrt_B |
| ATAD2B | 0.73236 | 0.36775 | 0.628 | 0.69 | 1 | HC_vs_MS.NoTrt_B |
| POTEC | 0.73241 | 0.50513 | 0.406 | 0.44 | 1 | HC_vs_MS.NoTrt_B |
| SNRNP27 | 0.73333 | 0.35712 | 0.278 | 0.31 | 1 | HC_vs_MS.NoTrt_B |
| HINT3 | 0.73356 | 0.49019 | 0.296 | 0.28 | 1 | HC_vs_MS.NoTrt_B |
| EVC | 0.73367 | 0.91448 | 0.242 | 0.27 | 1 | HC_vs_MS.NoTrt_B |
| TNPO2 | 0.73375 | 0.47833 | 0.122 | 0.11 | 1 | HC_vs_MS.NoTrt_B |
| MCC | 0.73376 | 0.36537 | 0.438 | 0.48 | 1 | HC_vs_MS.NoTrt_B |
| RECQL5 | 0.73409 | 0.50333 | 0.196 | 0.21 | 1 | HC_vs_MS.NoTrt_B |
| APOBR | 0.7344 | 0.73501 | 0.104 | 0.11 | 1 | HC_vs_MS.NoTrt_B |
| FAM179A | 0.73447 | -0.4375 | 0.772 | 0.78 | 1 | HC_vs_MS.NoTrt_B |
| MAD2L1 | 0.73475 | -0.3863 | 0.435 | 0.46 | 1 | HC_vs_MS.NoTrt_B |
| CYB5D1 | 0.7348 | 0.43973 | 0.138 | 0.13 | 1 | HC_vs_MS.NoTrt_B |
| GAREM2 | 0.73499 | 0.40568 | 0.499 | 0.52 | 1 | HC_vs_MS.NoTrt_B |
| UBE2W | 0.73502 | 0.37034 | 0.626 | 0.64 | 1 | HC_vs_MS.NoTrt_B |
| PMM2 | 0.73666 | -0.3693 | 0.742 | 0.73 | 1 | HC_vs_MS.NoTrt_B |
| KIAA0907 | 0.73697 | 0.55832 | 0.762 | 0.78 | 1 | HC_vs_MS.NoTrt_B |
| TENM1 | 0.73707 | 0.90592 | 0.093 | 0.1 | 1 | HC_vs_MS.NoTrt_B |
| MINK1 | 0.73786 | 0.89223 | 0.252 | 0.26 | 1 | HC_vs_MS.NoTrt_B |
| BTBD11 | 0.738 | 1.37083 | 0.457 | 0.53 | 1 | HC_vs_MS.NoTrt_B |
| AGGF1 | 0.73803 | 1.04381 | 0.123 | 0.12 | 1 | HC_vs_MS.NoTrt_B |
| OGFR | 0.73824 | 0.29463 | 0.187 | 0.18 | 1 | HC_vs_MS.NoTrt_B |
| FAM120B | 0.73844 | 0.54171 | 0.346 | 0.38 | 1 | HC_vs_MS.NoTrt_B |
| PLOD2 | 0.73855 | 0.34552 | 0.387 | 0.42 | 1 | HC_vs_MS.NoTrt_B |
| SUGT1 | 0.73859 | 0.42047 | 0.903 | 0.94 | 1 | HC_vs_MS.NoTrt_B |
| ZNF846 | 0.73901 | 1.11268 | 0.446 | 0.44 | 1 | HC_vs_MS.NoTrt_B |
| HHLA3 | 0.7391 | 0.64816 | 0.343 | 0.38 | 1 | HC_vs_MS.NoTrt_B |
| XPO4 | 0.73938 | 1.17453 | 0.112 | 0.12 | 1 | HC_vs_MS.NoTrt_B |
| HDAC4 | 0.73954 | 0.30379 | 0.559 | 0.58 | 1 | HC_vs_MS.NoTrt_B |
| TMEM230 | 0.74063 | 0.28184 | 0.497 | 0.52 | 1 | HC_vs_MS.NoTrt_B |
| FGF11 | 0.74087 | 0.41328 | 0.21 | 0.21 | 1 | HC_vs_MS.NoTrt_B |
| USP54 | 0.74136 | 0.6115 | 0.58 | 0.63 | 1 | HC_vs_MS.NoTrt_B |
| CS | 0.74226 | 0.35607 | 0.842 | 0.82 | 1 | HC_vs_MS.NoTrt_B |

| | | | | | | |
|---|---|---|---|---|---|---|
| CCDC15 | 0.74267 | -0.8345 | 0.216 | 0.21 | 1 | HC_vs_MS.NoTrt_B |
| ANKRD46 | 0.74305 | 0.47912 | 0.171 | 0.18 | 1 | HC_vs_MS.NoTrt_B |
| FBXO31 | 0.7432 | 1.32485 | 0.3 | 0.31 | 1 | HC_vs_MS.NoTrt_B |
| DCAKD | 0.74373 | 0.69157 | 0.39 | 0.37 | 1 | HC_vs_MS.NoTrt_B |
| CIAO1 | 0.74393 | 0.3047 | 0.286 | 0.3 | 1 | HC_vs_MS.NoTrt_B |
| ETAA1 | 0.74411 | 0.31243 | 0.096 | 0.1 | 1 | HC_vs_MS.NoTrt_B |
| MAP2K4 | 0.74536 | 1.29971 | 0.213 | 0.21 | 1 | HC_vs_MS.NoTrt_B |
| MAGIX | 0.74588 | 0.69113 | 0.465 | 0.48 | 1 | HC_vs_MS.NoTrt_B |
| TTLL4 | 0.74598 | 0.5484 | 0.29 | 0.32 | 1 | HC_vs_MS.NoTrt_B |
| KCTD9 | 0.7461 | 0.6073 | 0.183 | 0.18 | 1 | HC_vs_MS.NoTrt_B |
| IDH2 | 0.74643 | -0.3666 | 0.513 | 0.51 | 1 | HC_vs_MS.NoTrt_B |
| MOB1A | 0.74839 | 0.37724 | 0.825 | 0.88 | 1 | HC_vs_MS.NoTrt_B |
| RALGAPB | 0.74883 | 0.25739 | 0.277 | 0.29 | 1 | HC_vs_MS.NoTrt_B |
| PLCB2 | 0.7492 | 0.29464 | 0.223 | 0.22 | 1 | HC_vs_MS.NoTrt_B |
| TNIP3 | 0.74927 | -0.2904 | 0.499 | 0.56 | 1 | HC_vs_MS.NoTrt_B |
| AEBP2 | 0.74974 | 0.38105 | 0.696 | 0.7 | 1 | HC_vs_MS.NoTrt_B |
| EBLN2 | 0.75065 | 0.46377 | 0.114 | 0.12 | 1 | HC_vs_MS.NoTrt_B |
| B4GALT5 | 0.75067 | 0.92545 | 0.251 | 0.28 | 1 | HC_vs_MS.NoTrt_B |
| ARHGAP44 | 0.75102 | 1.27795 | 0.242 | 0.24 | 1 | HC_vs_MS.NoTrt_B |
| C8orf76 | 0.75107 | -0.7425 | 0.572 | 0.58 | 1 | HC_vs_MS.NoTrt_B |
| OLAH | 0.75113 | 0.25842 | 0.432 | 0.47 | 1 | HC_vs_MS.NoTrt_B |
| ADORA2A | 0.75184 | -0.2702 | 0.117 | 0.12 | 1 | HC_vs_MS.NoTrt_B |
| AP2B1 | 0.752 | 0.37716 | 0.274 | 0.29 | 1 | HC_vs_MS.NoTrt_B |
| C21orf58 | 0.75224 | -0.272 | 0.464 | 0.49 | 1 | HC_vs_MS.NoTrt_B |
| FAF2 | 0.75273 | 0.73708 | 0.12 | 0.11 | 1 | HC_vs_MS.NoTrt_B |
| DGCR8 | 0.75352 | 0.46534 | 0.116 | 0.11 | 1 | HC_vs_MS.NoTrt_B |
| POLR2A | 0.75383 | 0.39337 | 0.781 | 0.81 | 1 | HC_vs_MS.NoTrt_B |
| POMT1 | 0.75484 | -0.6152 | 0.538 | 0.58 | 1 | HC_vs_MS.NoTrt_B |
| SLC25A43 | 0.75491 | 0.47345 | 0.241 | 0.22 | 1 | HC_vs_MS.NoTrt_B |
| OR8H2 | 0.75503 | -0.3266 | 0.186 | 0.2 | 1 | HC_vs_MS.NoTrt_B |
| TNFSF13B | 0.75539 | -0.2945 | 0.5 | 0.52 | 1 | HC_vs_MS.NoTrt_B |
| AR | 0.7557 | 0.31644 | 0.222 | 0.24 | 1 | HC_vs_MS.NoTrt_B |
| U2SURP | 0.75644 | 0.27782 | 0.333 | 0.34 | 1 | HC_vs_MS.NoTrt_B |
| TCEAL3 | 0.7566 | -0.2589 | 0.11 | 0.12 | 1 | HC_vs_MS.NoTrt_B |
| CAPN15 | 0.75702 | 0.50701 | 0.207 | 0.2 | 1 | HC_vs_MS.NoTrt_B |
| NUPL2 | 0.75747 | 0.26713 | 0.962 | 0.98 | 1 | HC_vs_MS.NoTrt_B |
| RPRD1A | 0.75821 | 0.29927 | 0.158 | 0.15 | 1 | HC_vs_MS.NoTrt_B |
| ZDHHC20 | 0.75873 | 0.29751 | 0.448 | 0.44 | 1 | HC_vs_MS.NoTrt_B |
| IL4R | 0.75944 | 0.38148 | 0.425 | 0.43 | 1 | HC_vs_MS.NoTrt_B |
| SNX30 | 0.75962 | 0.58353 | 0.435 | 0.45 | 1 | HC_vs_MS.NoTrt_B |

| | | | | | | |
|---|---|---|---|---|---|---|
| LRFN1 | 0.75969 | 0.52933 | 0.142 | 0.15 | 1 | HC_vs_MS.NoTrt_B |
| HIVEP1 | 0.7604 | 0.93709 | 0.654 | 0.73 | 1 | HC_vs_MS.NoTrt_B |
| KLRC1 | 0.76062 | 0.53224 | 0.291 | 0.3 | 1 | HC_vs_MS.NoTrt_B |
| RANBP9 | 0.76098 | 0.37068 | 0.186 | 0.18 | 1 | HC_vs_MS.NoTrt_B |
| PANK2 | 0.76194 | 0.64923 | 0.517 | 0.54 | 1 | HC_vs_MS.NoTrt_B |
| SH2B3 | 0.76196 | 0.43291 | 0.142 | 0.15 | 1 | HC_vs_MS.NoTrt_B |
| LPIN3 | 0.76228 | 0.45873 | 0.432 | 0.43 | 1 | HC_vs_MS.NoTrt_B |
| IRGM | 0.76243 | 0.32684 | 0.396 | 0.42 | 1 | HC_vs_MS.NoTrt_B |
| PPIF | 0.76249 | 0.32108 | 0.094 | 0.1 | 1 | HC_vs_MS.NoTrt_B |
| DICER1 | 0.76327 | 0.74748 | 0.148 | 0.16 | 1 | HC_vs_MS.NoTrt_B |
| TP53BP1 | 0.76347 | 0.30027 | 0.252 | 0.24 | 1 | HC_vs_MS.NoTrt_B |
| VPS41 | 0.76367 | 0.34924 | 0.452 | 0.44 | 1 | HC_vs_MS.NoTrt_B |
| EHBP1L1 | 0.76369 | 0.66408 | 0.184 | 0.18 | 1 | HC_vs_MS.NoTrt_B |
| UNC45B | 0.76374 | 0.33277 | 0.12 | 0.11 | 1 | HC_vs_MS.NoTrt_B |
| JADE1 | 0.76474 | 0.59817 | 0.326 | 0.32 | 1 | HC_vs_MS.NoTrt_B |
| ZNF395 | 0.76503 | 0.93578 | 0.542 | 0.61 | 1 | HC_vs_MS.NoTrt_B |
| FBXO42 | 0.76521 | 0.54843 | 0.449 | 0.49 | 1 | HC_vs_MS.NoTrt_B |
| RP11-1212A | 0.76531 | 0.3319 | 0.481 | 0.5 | 1 | HC_vs_MS.NoTrt_B |
| PPP1R16B | 0.76561 | 0.56318 | 0.184 | 0.19 | 1 | HC_vs_MS.NoTrt_B |
| MRM2 | 0.76622 | 0.41685 | 0.31 | 0.32 | 1 | HC_vs_MS.NoTrt_B |
| PSMC4 | 0.76685 | -0.5503 | 0.457 | 0.45 | 1 | HC_vs_MS.NoTrt_B |
| RAC1 | 0.76701 | 0.76581 | 0.151 | 0.17 | 1 | HC_vs_MS.NoTrt_B |
| ZDHHC7 | 0.76713 | 0.35492 | 0.17 | 0.18 | 1 | HC_vs_MS.NoTrt_B |
| ZNF230 | 0.76746 | 0.76429 | 0.568 | 0.56 | 1 | HC_vs_MS.NoTrt_B |
| MRS2 | 0.76761 | -0.3366 | 0.764 | 0.77 | 1 | HC_vs_MS.NoTrt_B |
| CRIPAK | 0.76793 | 0.35296 | 0.21 | 0.23 | 1 | HC_vs_MS.NoTrt_B |
| KIF1BP | 0.76803 | 0.48967 | 0.617 | 0.63 | 1 | HC_vs_MS.NoTrt_B |
| TRIM39 | 0.76843 | 0.32756 | 0.112 | 0.12 | 1 | HC_vs_MS.NoTrt_B |
| RSPRY1 | 0.76857 | -0.3502 | 0.335 | 0.36 | 1 | HC_vs_MS.NoTrt_B |
| LYST | 0.76882 | 1.25836 | 0.291 | 0.3 | 1 | HC_vs_MS.NoTrt_B |
| CLCN3 | 0.76922 | 1.0692 | 0.364 | 0.39 | 1 | HC_vs_MS.NoTrt_B |
| DNAJA4 | 0.76951 | 1.77675 | 0.112 | 0.11 | 1 | HC_vs_MS.NoTrt_B |
| MGLL | 0.76952 | 0.38841 | 0.319 | 0.36 | 1 | HC_vs_MS.NoTrt_B |
| IMMP1L | 0.7696 | 0.44107 | 0.559 | 0.58 | 1 | HC_vs_MS.NoTrt_B |
| TBC1D31 | 0.76977 | 0.45834 | 0.597 | 0.65 | 1 | HC_vs_MS.NoTrt_B |
| TJP2 | 0.76988 | 0.50051 | 0.339 | 0.37 | 1 | HC_vs_MS.NoTrt_B |
| MAN2A2 | 0.77171 | 0.59289 | 0.252 | 0.25 | 1 | HC_vs_MS.NoTrt_B |
| CTF1 | 0.77203 | -0.3236 | 0.248 | 0.24 | 1 | HC_vs_MS.NoTrt_B |
| MTERF3 | 0.77222 | -0.678 | 0.18 | 0.18 | 1 | HC_vs_MS.NoTrt_B |
| SMARCAD1 | 0.77303 | 0.8914 | 0.123 | 0.12 | 1 | HC_vs_MS.NoTrt_B |

| | | | | | | |
|---|---|---|---|---|---|---|
| RUNX3 | 0.77353 | 0.3711 | 0.136 | 0.13 | 1 | HC_vs_MS.NoTrt_B |
| EXOG | 0.77377 | 0.45634 | 0.475 | 0.5 | 1 | HC_vs_MS.NoTrt_B |
| PTPN4 | 0.77427 | 0.25013 | 0.588 | 0.58 | 1 | HC_vs_MS.NoTrt_B |
| BAG2 | 0.7743 | 0.76572 | 0.23 | 0.25 | 1 | HC_vs_MS.NoTrt_B |
| TLE3 | 0.77473 | 0.66891 | 0.193 | 0.19 | 1 | HC_vs_MS.NoTrt_B |
| SYNGAP1 | 0.77478 | 1.03308 | 0.136 | 0.15 | 1 | HC_vs_MS.NoTrt_B |
| DDX11 | 0.77491 | -0.2761 | 0.414 | 0.38 | 1 | HC_vs_MS.NoTrt_B |
| POLR1A | 0.77622 | -0.4391 | 0.672 | 0.69 | 1 | HC_vs_MS.NoTrt_B |
| ZMAT1 | 0.77652 | 0.93182 | 0.341 | 0.34 | 1 | HC_vs_MS.NoTrt_B |
| TMEM259 | 0.77671 | 1.21702 | 0.113 | 0.11 | 1 | HC_vs_MS.NoTrt_B |
| MAP3K3 | 0.7772 | 0.68065 | 0.142 | 0.14 | 1 | HC_vs_MS.NoTrt_B |
| ADCY9 | 0.77761 | 0.37111 | 0.261 | 0.28 | 1 | HC_vs_MS.NoTrt_B |
| CREB3L4 | 0.77821 | 1.282 | 0.399 | 0.4 | 1 | HC_vs_MS.NoTrt_B |
| EXT2 | 0.77859 | -0.4842 | 0.146 | 0.14 | 1 | HC_vs_MS.NoTrt_B |
| CEACAM4 | 0.77924 | -0.2674 | 0.097 | 0.1 | 1 | HC_vs_MS.NoTrt_B |
| TCEB3 | 0.77966 | 0.55162 | 0.094 | 0.1 | 1 | HC_vs_MS.NoTrt_B |
| NAA16 | 0.77975 | 0.26098 | 0.406 | 0.43 | 1 | HC_vs_MS.NoTrt_B |
| TNS2 | 0.77995 | 0.52858 | 0.291 | 0.31 | 1 | HC_vs_MS.NoTrt_B |
| ZNF625 | 0.78006 | -0.4276 | 0.243 | 0.25 | 1 | HC_vs_MS.NoTrt_B |
| TMEM119 | 0.78052 | -0.3904 | 0.555 | 0.57 | 1 | HC_vs_MS.NoTrt_B |
| APEX2 | 0.78109 | 0.53592 | 0.307 | 0.32 | 1 | HC_vs_MS.NoTrt_B |
| TNKS | 0.78154 | 0.26188 | 0.797 | 0.82 | 1 | HC_vs_MS.NoTrt_B |
| CENPH | 0.78231 | -0.2659 | 0.249 | 0.25 | 1 | HC_vs_MS.NoTrt_B |
| SLC22A5 | 0.78272 | 0.85893 | 0.375 | 0.41 | 1 | HC_vs_MS.NoTrt_B |
| HIGD1A | 0.78273 | 0.31138 | 0.148 | 0.16 | 1 | HC_vs_MS.NoTrt_B |
| MYO1B | 0.78273 | 0.25416 | 0.393 | 0.41 | 1 | HC_vs_MS.NoTrt_B |
| POGK | 0.78437 | 0.60192 | 0.265 | 0.26 | 1 | HC_vs_MS.NoTrt_B |
| CDK2AP2 | 0.78441 | 0.35369 | 0.199 | 0.22 | 1 | HC_vs_MS.NoTrt_B |
| DDX52 | 0.78516 | -0.4938 | 0.871 | 0.87 | 1 | HC_vs_MS.NoTrt_B |
| HJURP | 0.78614 | -0.5954 | 0.37 | 0.39 | 1 | HC_vs_MS.NoTrt_B |
| RNASEL | 0.78654 | 0.74181 | 0.338 | 0.37 | 1 | HC_vs_MS.NoTrt_B |
| NEK7 | 0.78726 | 0.59013 | 0.12 | 0.11 | 1 | HC_vs_MS.NoTrt_B |
| KANSL1 | 0.78776 | 0.43002 | 0.707 | 0.73 | 1 | HC_vs_MS.NoTrt_B |
| ZNF107 | 0.78851 | 0.89062 | 0.41 | 0.42 | 1 | HC_vs_MS.NoTrt_B |
| LYL1 | 0.78917 | 0.31277 | 0.397 | 0.43 | 1 | HC_vs_MS.NoTrt_B |
| ZMYM6 | 0.79008 | 0.34348 | 0.814 | 0.89 | 1 | HC_vs_MS.NoTrt_B |
| FOXRED1 | 0.79018 | 0.29566 | 0.507 | 0.51 | 1 | HC_vs_MS.NoTrt_B |
| CAMKK2 | 0.79068 | 0.94105 | 0.465 | 0.51 | 1 | HC_vs_MS.NoTrt_B |
| PHYKPL | 0.79141 | 0.32476 | 0.425 | 0.44 | 1 | HC_vs_MS.NoTrt_B |
| SIKE1 | 0.79204 | 0.44473 | 0.394 | 0.39 | 1 | HC_vs_MS.NoTrt_B |

| | | | | | | |
|---|---|---|---|---|---|---|
| COX10 | 0.79235 | 0.49328 | 0.11 | 0.12 | 1 | HC_vs_MS.NoTrt_B |
| SYVN1 | 0.79424 | 0.31534 | 0.229 | 0.23 | 1 | HC_vs_MS.NoTrt_B |
| PTRH2 | 0.79452 | 0.98649 | 0.132 | 0.12 | 1 | HC_vs_MS.NoTrt_B |
| GBAS | 0.79545 | 0.45788 | 0.691 | 0.71 | 1 | HC_vs_MS.NoTrt_B |
| DIMT1 | 0.79566 | 0.38666 | 0.174 | 0.17 | 1 | HC_vs_MS.NoTrt_B |
| PEX1 | 0.79623 | 0.32533 | 0.67 | 0.71 | 1 | HC_vs_MS.NoTrt_B |
| MPP6 | 0.79637 | 0.52007 | 0.278 | 0.3 | 1 | HC_vs_MS.NoTrt_B |
| HLX | 0.79661 | 0.2946 | 0.193 | 0.21 | 1 | HC_vs_MS.NoTrt_B |
| CPD | 0.79762 | 0.60545 | 0.693 | 0.74 | 1 | HC_vs_MS.NoTrt_B |
| EXD1 | 0.79791 | 0.3615 | 0.472 | 0.51 | 1 | HC_vs_MS.NoTrt_B |
| TNFAIP8L1 | 0.79821 | -0.4159 | 0.797 | 0.8 | 1 | HC_vs_MS.NoTrt_B |
| NARFL | 0.79848 | 0.40092 | 0.762 | 0.78 | 1 | HC_vs_MS.NoTrt_B |
| MIGA1 | 0.79863 | 0.42686 | 0.413 | 0.45 | 1 | HC_vs_MS.NoTrt_B |
| TTC33 | 0.79926 | 0.29729 | 0.228 | 0.22 | 1 | HC_vs_MS.NoTrt_B |
| PITPNC1 | 0.79972 | 0.47554 | 0.212 | 0.2 | 1 | HC_vs_MS.NoTrt_B |
| TG | 0.80006 | 0.42237 | 0.093 | 0.1 | 1 | HC_vs_MS.NoTrt_B |
| NOP2 | 0.8001 | 0.62898 | 0.381 | 0.38 | 1 | HC_vs_MS.NoTrt_B |
| SPG11 | 0.80033 | 0.36587 | 0.807 | 0.84 | 1 | HC_vs_MS.NoTrt_B |
| UHRF2 | 0.80039 | 0.77254 | 0.251 | 0.25 | 1 | HC_vs_MS.NoTrt_B |
| UBA6 | 0.80045 | 0.44035 | 0.491 | 0.49 | 1 | HC_vs_MS.NoTrt_B |
| PIK3CG | 0.80068 | 1.12233 | 0.238 | 0.23 | 1 | HC_vs_MS.NoTrt_B |
| PRIM1 | 0.80092 | 0.2931 | 0.143 | 0.15 | 1 | HC_vs_MS.NoTrt_B |
| CD69 | 0.80107 | 0.37391 | 0.174 | 0.18 | 1 | HC_vs_MS.NoTrt_B |
| ARHGEF38 | 0.80115 | 0.414 | 0.361 | 0.39 | 1 | HC_vs_MS.NoTrt_B |
| SEPN1 | 0.80116 | 0.42952 | 0.351 | 0.36 | 1 | HC_vs_MS.NoTrt_B |
| TPRA1 | 0.80143 | 0.50936 | 0.152 | 0.16 | 1 | HC_vs_MS.NoTrt_B |
| CA14 | 0.80184 | 0.27619 | 0.572 | 0.59 | 1 | HC_vs_MS.NoTrt_B |
| TPRG1 | 0.80199 | -0.2895 | 0.18 | 0.19 | 1 | HC_vs_MS.NoTrt_B |
| GMPPB | 0.80203 | 0.26332 | 0.41 | 0.41 | 1 | HC_vs_MS.NoTrt_B |
| HNRNPM | 0.80204 | 0.42423 | 0.274 | 0.3 | 1 | HC_vs_MS.NoTrt_B |
| ADAM28 | 0.80209 | 0.25961 | 0.135 | 0.13 | 1 | HC_vs_MS.NoTrt_B |
| MCOLN3 | 0.80211 | 0.36533 | 0.467 | 0.5 | 1 | HC_vs_MS.NoTrt_B |
| COG3 | 0.80287 | 0.61002 | 0.168 | 0.18 | 1 | HC_vs_MS.NoTrt_B |
| THAP5 | 0.80303 | 0.34583 | 0.87 | 0.89 | 1 | HC_vs_MS.NoTrt_B |
| IL5RA | 0.80346 | 1.29479 | 0.252 | 0.27 | 1 | HC_vs_MS.NoTrt_B |
| ECHDC3 | 0.80403 | 0.72035 | 0.109 | 0.1 | 1 | HC_vs_MS.NoTrt_B |
| IMPAD1 | 0.80438 | 0.47894 | 0.472 | 0.49 | 1 | HC_vs_MS.NoTrt_B |
| FBXW4 | 0.80482 | 0.44094 | 0.493 | 0.55 | 1 | HC_vs_MS.NoTrt_B |
| TBCEL | 0.8055 | 0.68415 | 0.116 | 0.11 | 1 | HC_vs_MS.NoTrt_B |
| CR1 | 0.80663 | 0.45159 | 0.51 | 0.55 | 1 | HC_vs_MS.NoTrt_B |

| | | | | | | |
|---|---|---|---|---|---|---|
| HEBP1 | 0.80665 | 0.40994 | 0.262 | 0.27 | 1 | HC_vs_MS.NoTrt_B |
| SH2D6 | 0.80674 | 0.33371 | 0.417 | 0.43 | 1 | HC_vs_MS.NoTrt_B |
| ANKRD20A3 | 0.80727 | 0.36184 | 0.112 | 0.12 | 1 | HC_vs_MS.NoTrt_B |
| WSB2 | 0.80757 | 0.56597 | 0.767 | 0.81 | 1 | HC_vs_MS.NoTrt_B |
| IFI27L2 | 0.8077 | 0.35466 | 0.142 | 0.15 | 1 | HC_vs_MS.NoTrt_B |
| IL23R | 0.80781 | 0.66969 | 0.343 | 0.36 | 1 | HC_vs_MS.NoTrt_B |
| ZBTB16 | 0.80801 | 0.46895 | 0.372 | 0.4 | 1 | HC_vs_MS.NoTrt_B |
| MRPL27 | 0.80841 | 0.66377 | 0.155 | 0.17 | 1 | HC_vs_MS.NoTrt_B |
| FAM47E | 0.80852 | 0.31788 | 0.642 | 0.69 | 1 | HC_vs_MS.NoTrt_B |
| ENTPD1 | 0.8088 | -0.3258 | 0.961 | 0.97 | 1 | HC_vs_MS.NoTrt_B |
| GTF2F2 | 0.80901 | 0.51726 | 0.204 | 0.2 | 1 | HC_vs_MS.NoTrt_B |
| GALNT2 | 0.80946 | 0.3956 | 0.187 | 0.18 | 1 | HC_vs_MS.NoTrt_B |
| PRODH | 0.80954 | 0.85993 | 0.386 | 0.42 | 1 | HC_vs_MS.NoTrt_B |
| GSPT1 | 0.81035 | 0.28471 | 0.522 | 0.56 | 1 | HC_vs_MS.NoTrt_B |
| VPS50 | 0.81079 | 0.3859 | 0.526 | 0.54 | 1 | HC_vs_MS.NoTrt_B |
| LRBA | 0.811 | 0.34379 | 0.477 | 0.49 | 1 | HC_vs_MS.NoTrt_B |
| TIAM1 | 0.81154 | 0.26069 | 0.539 | 0.56 | 1 | HC_vs_MS.NoTrt_B |
| THBS1 | 0.81168 | 0.54743 | 0.209 | 0.22 | 1 | HC_vs_MS.NoTrt_B |
| NDUFA8 | 0.81169 | 0.50434 | 0.1 | 0.11 | 1 | HC_vs_MS.NoTrt_B |
| NPL | 0.81186 | 0.64273 | 0.223 | 0.24 | 1 | HC_vs_MS.NoTrt_B |
| C1orf123 | 0.8126 | 0.37646 | 0.119 | 0.11 | 1 | HC_vs_MS.NoTrt_B |
| MOB1B | 0.81264 | 0.3143 | 0.61 | 0.62 | 1 | HC_vs_MS.NoTrt_B |
| SETD6 | 0.81329 | -0.3024 | 0.436 | 0.45 | 1 | HC_vs_MS.NoTrt_B |
| COA5 | 0.81344 | 0.49195 | 0.465 | 0.49 | 1 | HC_vs_MS.NoTrt_B |
| RABL6 | 0.81441 | 0.71043 | 0.509 | 0.52 | 1 | HC_vs_MS.NoTrt_B |
| CYHR1 | 0.81477 | 0.36886 | 0.268 | 0.27 | 1 | HC_vs_MS.NoTrt_B |
| TNFSF10 | 0.81549 | 0.41875 | 0.401 | 0.41 | 1 | HC_vs_MS.NoTrt_B |
| SLC25A44 | 0.81614 | 0.30995 | 0.674 | 0.7 | 1 | HC_vs_MS.NoTrt_B |
| DOCK11 | 0.81623 | 0.41615 | 0.514 | 0.57 | 1 | HC_vs_MS.NoTrt_B |
| BIN3 | 0.8166 | 0.50102 | 0.364 | 0.37 | 1 | HC_vs_MS.NoTrt_B |
| TLCD1 | 0.81662 | 0.30343 | 0.203 | 0.2 | 1 | HC_vs_MS.NoTrt_B |
| GANAB | 0.81689 | 0.35703 | 0.43 | 0.44 | 1 | HC_vs_MS.NoTrt_B |
| TET2 | 0.81712 | 0.82699 | 0.454 | 0.51 | 1 | HC_vs_MS.NoTrt_B |
| LEPR | 0.81769 | 0.76228 | 0.283 | 0.3 | 1 | HC_vs_MS.NoTrt_B |
| PRKCA | 0.81812 | -0.4714 | 0.739 | 0.78 | 1 | HC_vs_MS.NoTrt_B |
| BTG1 | 0.81842 | 0.33796 | 0.599 | 0.63 | 1 | HC_vs_MS.NoTrt_B |
| THG1L | 0.81891 | -0.4798 | 0.551 | 0.57 | 1 | HC_vs_MS.NoTrt_B |
| MRPL35 | 0.81916 | 0.68518 | 0.106 | 0.1 | 1 | HC_vs_MS.NoTrt_B |
| DDX3X | 0.81932 | 0.25004 | 0.329 | 0.32 | 1 | HC_vs_MS.NoTrt_B |
| PTPN12 | 0.81948 | 0.43175 | 0.717 | 0.82 | 1 | HC_vs_MS.NoTrt_B |

| | | | | | | |
|---|---|---|---|---|---|---|
| MECR | 0.8204 | -1.2073 | 0.306 | 0.3 | 1 | HC_vs_MS.NoTrt_B |
| INPP5F | 0.82047 | 1.13999 | 0.307 | 0.32 | 1 | HC_vs_MS.NoTrt_B |
| HEXDC | 0.82068 | -0.2609 | 0.338 | 0.34 | 1 | HC_vs_MS.NoTrt_B |
| NME4 | 0.821 | 0.60871 | 0.116 | 0.12 | 1 | HC_vs_MS.NoTrt_B |
| LPGAT1 | 0.82113 | -0.5016 | 0.446 | 0.47 | 1 | HC_vs_MS.NoTrt_B |
| STK24 | 0.8212 | 0.38775 | 0.254 | 0.27 | 1 | HC_vs_MS.NoTrt_B |
| MKNK2 | 0.82121 | 0.28878 | 0.199 | 0.21 | 1 | HC_vs_MS.NoTrt_B |
| DDX49 | 0.82208 | -0.5636 | 0.135 | 0.14 | 1 | HC_vs_MS.NoTrt_B |
| KLF8 | 0.82216 | -0.5017 | 0.726 | 0.72 | 1 | HC_vs_MS.NoTrt_B |
| NUP133 | 0.82241 | 0.54557 | 0.499 | 0.54 | 1 | HC_vs_MS.NoTrt_B |
| ARFGEF2 | 0.82265 | 0.78181 | 0.458 | 0.48 | 1 | HC_vs_MS.NoTrt_B |
| NCOA6 | 0.82275 | 0.59392 | 0.671 | 0.69 | 1 | HC_vs_MS.NoTrt_B |
| NHLRC1 | 0.82275 | -0.6518 | 0.233 | 0.23 | 1 | HC_vs_MS.NoTrt_B |
| RNF215 | 0.82278 | 0.36121 | 0.432 | 0.44 | 1 | HC_vs_MS.NoTrt_B |
| FDX1 | 0.82328 | 0.57806 | 0.287 | 0.29 | 1 | HC_vs_MS.NoTrt_B |
| SYNJ2BP | 0.82388 | 0.27178 | 0.823 | 0.82 | 1 | HC_vs_MS.NoTrt_B |
| KRBOX4 | 0.82431 | 0.38511 | 0.525 | 0.55 | 1 | HC_vs_MS.NoTrt_B |
| PEX7 | 0.82462 | -0.3257 | 0.141 | 0.15 | 1 | HC_vs_MS.NoTrt_B |
| PRR14L | 0.82481 | 0.43763 | 0.401 | 0.42 | 1 | HC_vs_MS.NoTrt_B |
| FSD2 | 0.82548 | 0.40433 | 0.452 | 0.49 | 1 | HC_vs_MS.NoTrt_B |
| ELK3 | 0.82576 | 0.39042 | 0.393 | 0.4 | 1 | HC_vs_MS.NoTrt_B |
| ABHD18 | 0.82593 | 0.27016 | 0.833 | 0.87 | 1 | HC_vs_MS.NoTrt_B |
| TTBK2 | 0.82604 | 0.27043 | 0.493 | 0.5 | 1 | HC_vs_MS.NoTrt_B |
| DNPEP | 0.82626 | 0.28441 | 0.448 | 0.47 | 1 | HC_vs_MS.NoTrt_B |
| CLEC2B | 0.8278 | 0.6257 | 0.209 | 0.22 | 1 | HC_vs_MS.NoTrt_B |
| WFDC11 | 0.82783 | 0.25426 | 0.196 | 0.21 | 1 | HC_vs_MS.NoTrt_B |
| PABPC1L | 0.82788 | 0.49833 | 0.284 | 0.31 | 1 | HC_vs_MS.NoTrt_B |
| AMPD3 | 0.82799 | -0.2639 | 0.43 | 0.45 | 1 | HC_vs_MS.NoTrt_B |
| ANKRD9 | 0.82868 | 0.42792 | 0.47 | 0.51 | 1 | HC_vs_MS.NoTrt_B |
| ARHGAP21 | 0.82885 | 0.63835 | 0.296 | 0.3 | 1 | HC_vs_MS.NoTrt_B |
| ADPRHL1 | 0.82935 | 0.27648 | 0.235 | 0.25 | 1 | HC_vs_MS.NoTrt_B |
| STK38L | 0.82937 | -0.2631 | 0.684 | 0.7 | 1 | HC_vs_MS.NoTrt_B |
| DNAH6 | 0.82943 | 0.61101 | 0.168 | 0.16 | 1 | HC_vs_MS.NoTrt_B |
| FAM126A | 0.83038 | 0.83418 | 0.661 | 0.73 | 1 | HC_vs_MS.NoTrt_B |
| CEP162 | 0.83041 | -0.84 | 0.138 | 0.13 | 1 | HC_vs_MS.NoTrt_B |
| TRPT1 | 0.83113 | 0.43739 | 0.096 | 0.1 | 1 | HC_vs_MS.NoTrt_B |
| SAMM50 | 0.83155 | -0.3919 | 0.483 | 0.48 | 1 | HC_vs_MS.NoTrt_B |
| RLF | 0.83201 | 0.55028 | 0.168 | 0.16 | 1 | HC_vs_MS.NoTrt_B |
| SLC25A35 | 0.83254 | -0.3971 | 0.142 | 0.14 | 1 | HC_vs_MS.NoTrt_B |
| LCAT | 0.83306 | -0.2561 | 0.187 | 0.2 | 1 | HC_vs_MS.NoTrt_B |

| | | | | | | |
|---|---|---|---|---|---|---|
| DNAJC21 | 0.83334 | 0.75382 | 0.381 | 0.4 | 1 | HC_vs_MS.NoTrt_B |
| TRIM61 | 0.83458 | -0.2564 | 0.214 | 0.21 | 1 | HC_vs_MS.NoTrt_B |
| GALNT15 | 0.83479 | 0.90424 | 0.271 | 0.28 | 1 | HC_vs_MS.NoTrt_B |
| ZKSCAN3 | 0.83479 | 0.39357 | 0.807 | 0.84 | 1 | HC_vs_MS.NoTrt_B |
| APOBEC3D | 0.83486 | -0.3726 | 0.612 | 0.6 | 1 | HC_vs_MS.NoTrt_B |
| KRBA2 | 0.83511 | 1.1211 | 0.37 | 0.41 | 1 | HC_vs_MS.NoTrt_B |
| RASA1 | 0.83532 | 0.67511 | 0.209 | 0.22 | 1 | HC_vs_MS.NoTrt_B |
| PTPRH | 0.83533 | -0.3533 | 0.125 | 0.12 | 1 | HC_vs_MS.NoTrt_B |
| PUM2 | 0.83552 | 0.43505 | 0.33 | 0.36 | 1 | HC_vs_MS.NoTrt_B |
| SDHAF2 | 0.8365 | 0.30642 | 0.223 | 0.24 | 1 | HC_vs_MS.NoTrt_B |
| ZNHIT6 | 0.8367 | 0.37655 | 0.393 | 0.38 | 1 | HC_vs_MS.NoTrt_B |
| DNAJC4 | 0.83675 | 0.57893 | 0.117 | 0.11 | 1 | HC_vs_MS.NoTrt_B |
| GLS | 0.83694 | 0.64292 | 0.628 | 0.64 | 1 | HC_vs_MS.NoTrt_B |
| EFR3A | 0.83764 | 0.38895 | 0.135 | 0.13 | 1 | HC_vs_MS.NoTrt_B |
| GRK3 | 0.83816 | -0.2609 | 0.333 | 0.34 | 1 | HC_vs_MS.NoTrt_B |
| RBM43 | 0.83839 | 0.28355 | 0.59 | 0.68 | 1 | HC_vs_MS.NoTrt_B |
| STMN1 | 0.8388 | -0.3528 | 0.691 | 0.7 | 1 | HC_vs_MS.NoTrt_B |
| LPP | 0.83887 | 0.29221 | 0.971 | 0.95 | 1 | HC_vs_MS.NoTrt_B |
| PPAN | 0.83902 | 1.03863 | 0.368 | 0.39 | 1 | HC_vs_MS.NoTrt_B |
| SAP130 | 0.83918 | 0.64915 | 0.158 | 0.16 | 1 | HC_vs_MS.NoTrt_B |
| TTC34 | 0.83946 | 0.97058 | 0.193 | 0.21 | 1 | HC_vs_MS.NoTrt_B |
| ETNK1 | 0.83989 | 0.45688 | 0.494 | 0.57 | 1 | HC_vs_MS.NoTrt_B |
| UHMK1 | 0.84036 | 0.48559 | 0.238 | 0.25 | 1 | HC_vs_MS.NoTrt_B |
| RAD51B | 0.84098 | 0.93871 | 0.287 | 0.29 | 1 | HC_vs_MS.NoTrt_B |
| TSC22D2 | 0.84099 | 0.82023 | 0.372 | 0.38 | 1 | HC_vs_MS.NoTrt_B |
| TMEM53 | 0.84134 | 1.02397 | 0.314 | 0.34 | 1 | HC_vs_MS.NoTrt_B |
| APBB3 | 0.84136 | 0.26079 | 0.123 | 0.13 | 1 | HC_vs_MS.NoTrt_B |
| LRRC8A | 0.84165 | -0.3027 | 0.104 | 0.1 | 1 | HC_vs_MS.NoTrt_B |
| KATNAL1 | 0.84189 | -0.8776 | 0.523 | 0.5 | 1 | HC_vs_MS.NoTrt_B |
| PRPSAP1 | 0.84228 | -0.4713 | 0.714 | 0.71 | 1 | HC_vs_MS.NoTrt_B |
| CEP97 | 0.8423 | 0.72644 | 0.628 | 0.65 | 1 | HC_vs_MS.NoTrt_B |
| SLC35E2 | 0.84236 | 0.28035 | 0.464 | 0.45 | 1 | HC_vs_MS.NoTrt_B |
| CXADR | 0.84306 | 0.33175 | 0.281 | 0.27 | 1 | HC_vs_MS.NoTrt_B |
| RAB5B | 0.84369 | 0.33293 | 0.864 | 0.92 | 1 | HC_vs_MS.NoTrt_B |
| MAN1A2 | 0.84404 | 0.47228 | 0.248 | 0.24 | 1 | HC_vs_MS.NoTrt_B |
| PRSS53 | 0.84423 | 0.26058 | 0.162 | 0.17 | 1 | HC_vs_MS.NoTrt_B |
| PNPO | 0.84446 | -0.4321 | 0.584 | 0.61 | 1 | HC_vs_MS.NoTrt_B |
| KAT6B | 0.84465 | 0.5096 | 0.106 | 0.11 | 1 | HC_vs_MS.NoTrt_B |
| MBD3 | 0.84476 | 0.55197 | 0.733 | 0.8 | 1 | HC_vs_MS.NoTrt_B |
| SPEN | 0.84526 | 0.3853 | 0.283 | 0.28 | 1 | HC_vs_MS.NoTrt_B |

| | | | | | | |
|---|---|---|---|---|---|---|
| RRP1B | 0.84556 | 0.39471 | 0.199 | 0.2 | 1 | HC_vs_MS.NoTrt_B |
| OAS2 | 0.8456 | 0.3475 | 0.213 | 0.22 | 1 | HC_vs_MS.NoTrt_B |
| PPIL4 | 0.84561 | 0.55927 | 0.223 | 0.22 | 1 | HC_vs_MS.NoTrt_B |
| KEAP1 | 0.84576 | -0.4156 | 0.448 | 0.43 | 1 | HC_vs_MS.NoTrt_B |
| IRF7 | 0.8458 | 0.44462 | 0.112 | 0.12 | 1 | HC_vs_MS.NoTrt_B |
| SH2D3C | 0.84601 | 0.42882 | 0.2 | 0.21 | 1 | HC_vs_MS.NoTrt_B |
| CREBBP | 0.84601 | 0.31253 | 0.564 | 0.61 | 1 | HC_vs_MS.NoTrt_B |
| ZNF213 | 0.84603 | 0.59952 | 0.455 | 0.48 | 1 | HC_vs_MS.NoTrt_B |
| PPP1R12B | 0.84703 | 0.47192 | 0.696 | 0.68 | 1 | HC_vs_MS.NoTrt_B |
| CAPN7 | 0.84727 | 0.26897 | 0.552 | 0.58 | 1 | HC_vs_MS.NoTrt_B |
| NPIPA5 | 0.84742 | 0.37793 | 0.101 | 0.1 | 1 | HC_vs_MS.NoTrt_B |
| C9orf131 | 0.84749 | 0.79798 | 0.148 | 0.15 | 1 | HC_vs_MS.NoTrt_B |
| RPP30 | 0.84813 | 0.42456 | 0.604 | 0.65 | 1 | HC_vs_MS.NoTrt_B |
| SLC35G1 | 0.84891 | 0.25824 | 0.146 | 0.15 | 1 | HC_vs_MS.NoTrt_B |
| AGO1 | 0.84893 | 0.27313 | 0.439 | 0.43 | 1 | HC_vs_MS.NoTrt_B |
| QKI | 0.84914 | -0.3711 | 0.386 | 0.39 | 1 | HC_vs_MS.NoTrt_B |
| ARAP2 | 0.84932 | 0.28222 | 0.265 | 0.27 | 1 | HC_vs_MS.NoTrt_B |
| KLHDC4 | 0.84988 | 0.44657 | 0.706 | 0.7 | 1 | HC_vs_MS.NoTrt_B |
| ANKFY1 | 0.85042 | 0.69001 | 0.617 | 0.71 | 1 | HC_vs_MS.NoTrt_B |
| UGCG | 0.85064 | 0.55419 | 0.178 | 0.17 | 1 | HC_vs_MS.NoTrt_B |
| DLG5 | 0.85075 | 0.35254 | 0.429 | 0.45 | 1 | HC_vs_MS.NoTrt_B |
| ALMS1 | 0.85139 | 0.43765 | 0.461 | 0.46 | 1 | HC_vs_MS.NoTrt_B |
| CCDC138 | 0.8524 | 0.34811 | 0.157 | 0.15 | 1 | HC_vs_MS.NoTrt_B |
| TMEM260 | 0.85244 | 0.44757 | 0.4 | 0.43 | 1 | HC_vs_MS.NoTrt_B |
| ALG13 | 0.85253 | 0.68042 | 0.284 | 0.3 | 1 | HC_vs_MS.NoTrt_B |
| WDR48 | 0.85271 | 0.48887 | 0.129 | 0.14 | 1 | HC_vs_MS.NoTrt_B |
| CA13 | 0.85302 | 0.34262 | 0.513 | 0.54 | 1 | HC_vs_MS.NoTrt_B |
| CHST14 | 0.85306 | 0.5255 | 0.225 | 0.23 | 1 | HC_vs_MS.NoTrt_B |
| CLPX | 0.85319 | 0.50179 | 0.683 | 0.73 | 1 | HC_vs_MS.NoTrt_B |
| PLEKHM1 | 0.85323 | 0.54786 | 0.816 | 0.87 | 1 | HC_vs_MS.NoTrt_B |
| HPCAL1 | 0.85324 | 0.27551 | 0.357 | 0.36 | 1 | HC_vs_MS.NoTrt_B |
| PLD6 | 0.85325 | 0.6304 | 0.435 | 0.46 | 1 | HC_vs_MS.NoTrt_B |
| NKIRAS1 | 0.85333 | 0.3986 | 0.239 | 0.25 | 1 | HC_vs_MS.NoTrt_B |
| TRIM58 | 0.85383 | 0.52125 | 0.564 | 0.58 | 1 | HC_vs_MS.NoTrt_B |
| OLA1 | 0.85481 | 0.26025 | 0.455 | 0.44 | 1 | HC_vs_MS.NoTrt_B |
| RAD50 | 0.85591 | 0.40121 | 0.184 | 0.19 | 1 | HC_vs_MS.NoTrt_B |
| MLH1 | 0.85603 | 0.48145 | 0.364 | 0.38 | 1 | HC_vs_MS.NoTrt_B |
| RABIF | 0.85605 | 0.5981 | 0.364 | 0.37 | 1 | HC_vs_MS.NoTrt_B |
| C11orf63 | 0.85611 | 0.34 | 0.42 | 0.47 | 1 | HC_vs_MS.NoTrt_B |
| TBCCD1 | 0.85749 | 0.48686 | 0.275 | 0.28 | 1 | HC_vs_MS.NoTrt_B |

| | | | | | | |
|---|---|---|---|---|---|---|
| THNSL1 | 0.85756 | 0.6479 | 0.096 | 0.1 | 1 | HC_vs_MS.NoTrt_B |
| ZBTB4 | 0.85758 | 0.78209 | 0.126 | 0.12 | 1 | HC_vs_MS.NoTrt_B |
| DDX56 | 0.8578 | 0.46652 | 0.217 | 0.23 | 1 | HC_vs_MS.NoTrt_B |
| FAM91A1 | 0.85809 | 0.49832 | 0.299 | 0.32 | 1 | HC_vs_MS.NoTrt_B |
| PPP3R1 | 0.85839 | 1.09571 | 0.158 | 0.16 | 1 | HC_vs_MS.NoTrt_B |
| CEP55 | 0.85856 | -0.702 | 0.126 | 0.13 | 1 | HC_vs_MS.NoTrt_B |
| FAM65C | 0.85895 | 0.37381 | 0.326 | 0.34 | 1 | HC_vs_MS.NoTrt_B |
| MAP2K5 | 0.85914 | 0.41346 | 0.103 | 0.11 | 1 | HC_vs_MS.NoTrt_B |
| METTL21B | 0.85917 | 0.73303 | 0.448 | 0.47 | 1 | HC_vs_MS.NoTrt_B |
| AL589743.1 | 0.85981 | -0.2959 | 0.519 | 0.55 | 1 | HC_vs_MS.NoTrt_B |
| SNRNP48 | 0.86 | 0.43034 | 0.207 | 0.21 | 1 | HC_vs_MS.NoTrt_B |
| SHPK | 0.86049 | -0.916 | 0.401 | 0.42 | 1 | HC_vs_MS.NoTrt_B |
| ZNF142 | 0.86065 | 0.26476 | 0.101 | 0.11 | 1 | HC_vs_MS.NoTrt_B |
| FBLN5 | 0.86068 | 0.45844 | 0.148 | 0.16 | 1 | HC_vs_MS.NoTrt_B |
| RAB2A | 0.86083 | 0.40829 | 0.214 | 0.23 | 1 | HC_vs_MS.NoTrt_B |
| ZNF749 | 0.86126 | 1.02026 | 0.38 | 0.41 | 1 | HC_vs_MS.NoTrt_B |
| STARD4 | 0.86154 | 0.29685 | 0.143 | 0.15 | 1 | HC_vs_MS.NoTrt_B |
| LIN54 | 0.8616 | 1.31616 | 0.548 | 0.63 | 1 | HC_vs_MS.NoTrt_B |
| CDC42EP3 | 0.86178 | 0.45722 | 0.461 | 0.49 | 1 | HC_vs_MS.NoTrt_B |
| GRM4 | 0.86244 | 0.27876 | 0.22 | 0.22 | 1 | HC_vs_MS.NoTrt_B |
| ARL1 | 0.86267 | 0.46703 | 0.346 | 0.35 | 1 | HC_vs_MS.NoTrt_B |
| ATXN7L3B | 0.86285 | 0.77787 | 0.162 | 0.17 | 1 | HC_vs_MS.NoTrt_B |
| GINS2 | 0.86286 | 0.86052 | 0.3 | 0.31 | 1 | HC_vs_MS.NoTrt_B |
| PGA3 | 0.86312 | 0.42397 | 0.291 | 0.3 | 1 | HC_vs_MS.NoTrt_B |
| AC013264.1 | 0.86342 | 0.7503 | 0.336 | 0.35 | 1 | HC_vs_MS.NoTrt_B |
| MNT | 0.86388 | 0.55077 | 0.623 | 0.7 | 1 | HC_vs_MS.NoTrt_B |
| LZIC | 0.86419 | 0.30152 | 0.788 | 0.85 | 1 | HC_vs_MS.NoTrt_B |
| INHBE | 0.86484 | 0.27088 | 0.364 | 0.39 | 1 | HC_vs_MS.NoTrt_B |
| BRCA2 | 0.86513 | 0.4995 | 0.358 | 0.38 | 1 | HC_vs_MS.NoTrt_B |
| TRIM68 | 0.86517 | -0.8748 | 0.509 | 0.54 | 1 | HC_vs_MS.NoTrt_B |
| GCKR | 0.86527 | 0.45401 | 0.139 | 0.15 | 1 | HC_vs_MS.NoTrt_B |
| AP4M1 | 0.86569 | -0.4988 | 0.538 | 0.61 | 1 | HC_vs_MS.NoTrt_B |
| SDCBP | 0.86598 | 0.43327 | 0.203 | 0.2 | 1 | HC_vs_MS.NoTrt_B |
| DLK2 | 0.86634 | 0.27711 | 0.312 | 0.32 | 1 | HC_vs_MS.NoTrt_B |
| MYCBP2 | 0.86666 | 0.69105 | 0.413 | 0.43 | 1 | HC_vs_MS.NoTrt_B |
| PAPOLG | 0.86677 | 0.35842 | 0.733 | 0.78 | 1 | HC_vs_MS.NoTrt_B |
| FAM149A | 0.86794 | -0.2562 | 0.306 | 0.3 | 1 | HC_vs_MS.NoTrt_B |
| MRPL10 | 0.86795 | -0.7189 | 0.4 | 0.39 | 1 | HC_vs_MS.NoTrt_B |
| GCNT1 | 0.86866 | 1.41795 | 0.249 | 0.27 | 1 | HC_vs_MS.NoTrt_B |
| CTSA | 0.86905 | -0.3317 | 0.659 | 0.65 | 1 | HC_vs_MS.NoTrt_B |

| | | | | | | |
|---|---|---|---|---|---|---|
| SPECC1L-AD | 0.86912 | 0.39749 | 0.187 | 0.18 | 1 | HC_vs_MS.NoTrt_B |
| KRT15 | 0.86919 | 0.29971 | 0.417 | 0.43 | 1 | HC_vs_MS.NoTrt_B |
| SGF29 | 0.86922 | -0.3481 | 0.271 | 0.26 | 1 | HC_vs_MS.NoTrt_B |
| RNF145 | 0.86937 | 0.70622 | 0.157 | 0.16 | 1 | HC_vs_MS.NoTrt_B |
| FAM168B | 0.86998 | 0.63277 | 0.106 | 0.1 | 1 | HC_vs_MS.NoTrt_B |
| AVL9 | 0.87007 | 0.38251 | 0.894 | 0.92 | 1 | HC_vs_MS.NoTrt_B |
| EFCAB3 | 0.87026 | 0.38765 | 0.522 | 0.61 | 1 | HC_vs_MS.NoTrt_B |
| DCTN6 | 0.8707 | -0.3365 | 0.519 | 0.55 | 1 | HC_vs_MS.NoTrt_B |
| SULF2 | 0.87087 | 0.31527 | 0.149 | 0.15 | 1 | HC_vs_MS.NoTrt_B |
| SLC23A3 | 0.8714 | -0.2526 | 0.41 | 0.42 | 1 | HC_vs_MS.NoTrt_B |
| NBPF1 | 0.87145 | 1.10302 | 0.229 | 0.25 | 1 | HC_vs_MS.NoTrt_B |
| KLHL2 | 0.87161 | 0.63796 | 0.323 | 0.34 | 1 | HC_vs_MS.NoTrt_B |
| RNF24 | 0.87165 | -0.281 | 0.545 | 0.56 | 1 | HC_vs_MS.NoTrt_B |
| VNN2 | 0.87179 | 0.34852 | 0.158 | 0.15 | 1 | HC_vs_MS.NoTrt_B |
| CTPS1 | 0.87189 | -0.4762 | 0.609 | 0.62 | 1 | HC_vs_MS.NoTrt_B |
| CHERP | 0.87213 | 0.32892 | 0.145 | 0.15 | 1 | HC_vs_MS.NoTrt_B |
| ESAM | 0.87214 | 0.64592 | 0.109 | 0.11 | 1 | HC_vs_MS.NoTrt_B |
| GDPD5 | 0.87232 | 0.30943 | 0.267 | 0.27 | 1 | HC_vs_MS.NoTrt_B |
| BTG2 | 0.8725 | 0.25041 | 0.178 | 0.17 | 1 | HC_vs_MS.NoTrt_B |
| RRP1 | 0.87355 | -0.2658 | 0.638 | 0.7 | 1 | HC_vs_MS.NoTrt_B |
| HTT | 0.87357 | 0.65478 | 0.236 | 0.24 | 1 | HC_vs_MS.NoTrt_B |
| NCAPG2 | 0.87377 | -0.9993 | 0.159 | 0.16 | 1 | HC_vs_MS.NoTrt_B |
| ENTPD6 | 0.87386 | 0.44181 | 0.12 | 0.12 | 1 | HC_vs_MS.NoTrt_B |
| CCDC186 | 0.87448 | 0.32472 | 0.575 | 0.57 | 1 | HC_vs_MS.NoTrt_B |
| ZNF670 | 0.87522 | 0.38167 | 0.161 | 0.17 | 1 | HC_vs_MS.NoTrt_B |
| GMDS | 0.87558 | 0.3711 | 0.112 | 0.11 | 1 | HC_vs_MS.NoTrt_B |
| GRAP | 0.87571 | 0.40583 | 0.204 | 0.21 | 1 | HC_vs_MS.NoTrt_B |
| PDE7A | 0.87603 | 0.61768 | 0.309 | 0.32 | 1 | HC_vs_MS.NoTrt_B |
| AP1M1 | 0.8771 | -0.2932 | 0.99 | 0.97 | 1 | HC_vs_MS.NoTrt_B |
| SP3 | 0.87722 | 0.70292 | 0.559 | 0.58 | 1 | HC_vs_MS.NoTrt_B |
| LAPTM4B | 0.87741 | -0.5564 | 0.194 | 0.2 | 1 | HC_vs_MS.NoTrt_B |
| AGTPBP1 | 0.87748 | 0.6073 | 0.172 | 0.17 | 1 | HC_vs_MS.NoTrt_B |
| GFPT1 | 0.87758 | 0.40401 | 0.706 | 0.74 | 1 | HC_vs_MS.NoTrt_B |
| ZNF227 | 0.87763 | 0.56805 | 0.465 | 0.5 | 1 | HC_vs_MS.NoTrt_B |
| CEP104 | 0.87806 | 0.29676 | 0.652 | 0.65 | 1 | HC_vs_MS.NoTrt_B |
| DPH6 | 0.87811 | 0.5897 | 0.409 | 0.44 | 1 | HC_vs_MS.NoTrt_B |
| INPP4B | 0.87814 | 0.27342 | 0.504 | 0.5 | 1 | HC_vs_MS.NoTrt_B |
| NAT10 | 0.87844 | -0.408 | 0.354 | 0.36 | 1 | HC_vs_MS.NoTrt_B |
| NTMT1 | 0.87855 | 0.6832 | 0.214 | 0.22 | 1 | HC_vs_MS.NoTrt_B |
| LIG3 | 0.87856 | 0.39043 | 0.33 | 0.33 | 1 | HC_vs_MS.NoTrt_B |

| | | | | | | |
|---|---|---|---|---|---|---|
| FIBP | 0.87876 | -0.2866 | 0.509 | 0.51 | 1 | HC_vs_MS.NoTrt_B |
| GLUD1 | 0.87888 | 0.39697 | 0.158 | 0.16 | 1 | HC_vs_MS.NoTrt_B |
| SCAF8 | 0.87906 | 0.45486 | 0.239 | 0.25 | 1 | HC_vs_MS.NoTrt_B |
| TBC1D32 | 0.87929 | -0.3348 | 0.449 | 0.46 | 1 | HC_vs_MS.NoTrt_B |
| FAM124B | 0.87966 | -0.3316 | 0.106 | 0.1 | 1 | HC_vs_MS.NoTrt_B |
| USP38 | 0.8804 | 0.27565 | 0.312 | 0.32 | 1 | HC_vs_MS.NoTrt_B |
| NOLC1 | 0.88081 | 0.61976 | 0.103 | 0.11 | 1 | HC_vs_MS.NoTrt_B |
| NDRG4 | 0.88083 | 0.28477 | 0.13 | 0.14 | 1 | HC_vs_MS.NoTrt_B |
| CCND3 | 0.88092 | 0.38933 | 0.516 | 0.55 | 1 | HC_vs_MS.NoTrt_B |
| NFATC2IP | 0.88093 | -0.2712 | 0.657 | 0.7 | 1 | HC_vs_MS.NoTrt_B |
| FBXO45 | 0.88151 | 0.5326 | 0.423 | 0.44 | 1 | HC_vs_MS.NoTrt_B |
| LMLN | 0.8819 | -0.4853 | 0.435 | 0.45 | 1 | HC_vs_MS.NoTrt_B |
| SLC15A4 | 0.88193 | -0.2592 | 0.722 | 0.73 | 1 | HC_vs_MS.NoTrt_B |
| DHRSX | 0.8824 | -0.3557 | 0.725 | 0.77 | 1 | HC_vs_MS.NoTrt_B |
| TMEM241 | 0.88243 | -0.5595 | 0.632 | 0.67 | 1 | HC_vs_MS.NoTrt_B |
| SULF1 | 0.88263 | 0.25786 | 0.174 | 0.17 | 1 | HC_vs_MS.NoTrt_B |
| C12orf40 | 0.88284 | 0.35908 | 0.288 | 0.3 | 1 | HC_vs_MS.NoTrt_B |
| MMRN2 | 0.88308 | -0.9349 | 0.152 | 0.15 | 1 | HC_vs_MS.NoTrt_B |
| IMPA1 | 0.88311 | 0.30769 | 0.336 | 0.34 | 1 | HC_vs_MS.NoTrt_B |
| CLTC | 0.88361 | -0.3531 | 0.355 | 0.35 | 1 | HC_vs_MS.NoTrt_B |
| ATG10 | 0.88371 | 0.31408 | 0.936 | 0.94 | 1 | HC_vs_MS.NoTrt_B |
| KDM4B | 0.88378 | 0.32663 | 0.594 | 0.6 | 1 | HC_vs_MS.NoTrt_B |
| PSMD9 | 0.8838 | -0.2938 | 0.954 | 0.98 | 1 | HC_vs_MS.NoTrt_B |
| CCDC117 | 0.88381 | 0.37639 | 0.299 | 0.31 | 1 | HC_vs_MS.NoTrt_B |
| FILIP1 | 0.88404 | 0.28471 | 0.384 | 0.41 | 1 | HC_vs_MS.NoTrt_B |
| CHI3L2 | 0.8842 | 0.4582 | 0.201 | 0.21 | 1 | HC_vs_MS.NoTrt_B |
| ATXN7L1 | 0.88438 | 0.51914 | 0.332 | 0.37 | 1 | HC_vs_MS.NoTrt_B |
| PLD2 | 0.88481 | -0.5916 | 0.107 | 0.1 | 1 | HC_vs_MS.NoTrt_B |
| SYPL1 | 0.88581 | 0.44136 | 0.383 | 0.39 | 1 | HC_vs_MS.NoTrt_B |
| MSS51 | 0.88612 | 0.39634 | 0.536 | 0.56 | 1 | HC_vs_MS.NoTrt_B |
| NPIPA1 | 0.88729 | 0.30635 | 0.683 | 0.72 | 1 | HC_vs_MS.NoTrt_B |
| TSPYL1 | 0.88732 | 0.37978 | 0.522 | 0.56 | 1 | HC_vs_MS.NoTrt_B |
| LST1 | 0.88779 | -0.3505 | 0.245 | 0.23 | 1 | HC_vs_MS.NoTrt_B |
| TCAF1 | 0.8879 | -0.4282 | 0.336 | 0.34 | 1 | HC_vs_MS.NoTrt_B |
| IQCB1 | 0.88836 | 0.88298 | 0.11 | 0.11 | 1 | HC_vs_MS.NoTrt_B |
| UBAP1L | 0.88874 | 1.16421 | 0.355 | 0.38 | 1 | HC_vs_MS.NoTrt_B |
| G2E3 | 0.88882 | 0.42136 | 0.488 | 0.5 | 1 | HC_vs_MS.NoTrt_B |
| CPSF6 | 0.88941 | 0.83321 | 0.409 | 0.42 | 1 | HC_vs_MS.NoTrt_B |
| HERPUD2 | 0.89002 | 0.65679 | 0.481 | 0.49 | 1 | HC_vs_MS.NoTrt_B |
| MAK | 0.89005 | 0.26048 | 0.17 | 0.17 | 1 | HC_vs_MS.NoTrt_B |

| | | | | | | |
|---|---|---|---|---|---|---|
| TSPOAP1 | 0.89082 | -0.4423 | 0.223 | 0.22 | 1 | HC_vs_MS.NoTrt_B |
| ZFAND3 | 0.8909 | 0.30596 | 0.642 | 0.67 | 1 | HC_vs_MS.NoTrt_B |
| PLEKHH2 | 0.89122 | 0.46338 | 0.619 | 0.68 | 1 | HC_vs_MS.NoTrt_B |
| PDPK1 | 0.89127 | 0.69393 | 0.599 | 0.62 | 1 | HC_vs_MS.NoTrt_B |
| DNAJB7 | 0.89168 | 0.45394 | 0.225 | 0.23 | 1 | HC_vs_MS.NoTrt_B |
| JUN | 0.89174 | 0.98309 | 0.119 | 0.12 | 1 | HC_vs_MS.NoTrt_B |
| ABL2 | 0.89214 | 0.8766 | 0.787 | 0.83 | 1 | HC_vs_MS.NoTrt_B |
| PDE3B | 0.89264 | 0.27758 | 0.591 | 0.61 | 1 | HC_vs_MS.NoTrt_B |
| GPR37L1 | 0.89305 | 0.64156 | 0.32 | 0.34 | 1 | HC_vs_MS.NoTrt_B |
| EHMT1 | 0.89403 | 0.29523 | 0.941 | 0.96 | 1 | HC_vs_MS.NoTrt_B |
| C12orf66 | 0.89408 | -0.3842 | 0.33 | 0.32 | 1 | HC_vs_MS.NoTrt_B |
| TYSND1 | 0.89461 | -0.535 | 0.257 | 0.26 | 1 | HC_vs_MS.NoTrt_B |
| FAM156A | 0.89584 | -0.5016 | 0.165 | 0.16 | 1 | HC_vs_MS.NoTrt_B |
| AFDN | 0.89595 | 0.3238 | 0.113 | 0.12 | 1 | HC_vs_MS.NoTrt_B |
| FYN | 0.89622 | 0.31991 | 0.523 | 0.54 | 1 | HC_vs_MS.NoTrt_B |
| ZNF799 | 0.8966 | 0.35437 | 0.39 | 0.42 | 1 | HC_vs_MS.NoTrt_B |
| PELP1 | 0.89721 | 0.42826 | 0.226 | 0.24 | 1 | HC_vs_MS.NoTrt_B |
| CARS | 0.89724 | -0.4324 | 0.757 | 0.75 | 1 | HC_vs_MS.NoTrt_B |
| ARPC5 | 0.89725 | 0.31192 | 0.901 | 0.94 | 1 | HC_vs_MS.NoTrt_B |
| LAMTOR3 | 0.89733 | 0.93441 | 0.325 | 0.34 | 1 | HC_vs_MS.NoTrt_B |
| FAM126B | 0.89738 | 0.28647 | 0.87 | 0.92 | 1 | HC_vs_MS.NoTrt_B |
| ZNF600 | 0.89825 | 0.80557 | 0.142 | 0.15 | 1 | HC_vs_MS.NoTrt_B |
| NAMPT | 0.89849 | 0.52477 | 0.687 | 0.71 | 1 | HC_vs_MS.NoTrt_B |
| NOTCH1 | 0.89966 | 0.27578 | 0.117 | 0.11 | 1 | HC_vs_MS.NoTrt_B |
| ANKRD13A | 0.89987 | 0.25853 | 0.787 | 0.81 | 1 | HC_vs_MS.NoTrt_B |
| TRIM4 | 0.89991 | 0.48339 | 0.583 | 0.62 | 1 | HC_vs_MS.NoTrt_B |
| ASXL2 | 0.90015 | 0.44185 | 0.193 | 0.2 | 1 | HC_vs_MS.NoTrt_B |
| PAXBP1 | 0.90032 | 0.59393 | 0.168 | 0.17 | 1 | HC_vs_MS.NoTrt_B |
| CHRM3 | 0.90074 | 0.43817 | 0.191 | 0.2 | 1 | HC_vs_MS.NoTrt_B |
| MCM3AP | 0.90101 | 0.35256 | 0.529 | 0.54 | 1 | HC_vs_MS.NoTrt_B |
| ASCC3 | 0.90109 | 0.6002 | 0.275 | 0.29 | 1 | HC_vs_MS.NoTrt_B |
| ZNF135 | 0.90131 | 0.64792 | 0.432 | 0.45 | 1 | HC_vs_MS.NoTrt_B |
| MREG | 0.90157 | 0.48844 | 0.614 | 0.69 | 1 | HC_vs_MS.NoTrt_B |
| USP15 | 0.90159 | 0.35693 | 0.822 | 0.87 | 1 | HC_vs_MS.NoTrt_B |
| CAPN10 | 0.90161 | 0.85664 | 0.462 | 0.48 | 1 | HC_vs_MS.NoTrt_B |
| CD7 | 0.90165 | 0.84553 | 0.242 | 0.25 | 1 | HC_vs_MS.NoTrt_B |
| ARF3 | 0.90227 | 0.58984 | 0.159 | 0.16 | 1 | HC_vs_MS.NoTrt_B |
| DTWD1 | 0.90258 | 0.28451 | 0.597 | 0.65 | 1 | HC_vs_MS.NoTrt_B |
| UAP1L1 | 0.90313 | -0.3303 | 0.301 | 0.3 | 1 | HC_vs_MS.NoTrt_B |
| ZNF14 | 0.90333 | 2.37971 | 0.32 | 0.34 | 1 | HC_vs_MS.NoTrt_B |

| | | | | | | |
|---|---|---|---|---|---|---|
| GTPBP3 | 0.90369 | -0.2726 | 0.632 | 0.65 | 1 | HC_vs_MS.NoTrt_B |
| ANKRA2 | 0.90388 | 0.467 | 0.11 | 0.11 | 1 | HC_vs_MS.NoTrt_B |
| TPP2 | 0.9042 | 0.26429 | 0.53 | 0.52 | 1 | HC_vs_MS.NoTrt_B |
| CAPN8 | 0.90443 | -0.3903 | 0.261 | 0.25 | 1 | HC_vs_MS.NoTrt_B |
| SMARCA2 | 0.90447 | 0.4388 | 0.413 | 0.43 | 1 | HC_vs_MS.NoTrt_B |
| RBBP6 | 0.90532 | 0.33331 | 0.22 | 0.23 | 1 | HC_vs_MS.NoTrt_B |
| MIEF1 | 0.90552 | -0.3898 | 0.465 | 0.44 | 1 | HC_vs_MS.NoTrt_B |
| AIFM1 | 0.90603 | -0.563 | 0.546 | 0.56 | 1 | HC_vs_MS.NoTrt_B |
| ZNF432 | 0.9061 | 0.30233 | 0.552 | 0.6 | 1 | HC_vs_MS.NoTrt_B |
| IRF5 | 0.90648 | 0.41789 | 0.377 | 0.38 | 1 | HC_vs_MS.NoTrt_B |
| ZC3HAV1L | 0.90671 | 0.27576 | 0.433 | 0.47 | 1 | HC_vs_MS.NoTrt_B |
| L3HYPDH | 0.90691 | 0.83152 | 0.194 | 0.2 | 1 | HC_vs_MS.NoTrt_B |
| WWOX | 0.90703 | 0.78087 | 0.236 | 0.25 | 1 | HC_vs_MS.NoTrt_B |
| SLAMF7 | 0.90719 | 0.51956 | 0.407 | 0.44 | 1 | HC_vs_MS.NoTrt_B |
| DCUN1D5 | 0.90722 | 0.36704 | 0.425 | 0.43 | 1 | HC_vs_MS.NoTrt_B |
| ENPP3 | 0.90746 | 0.82017 | 0.13 | 0.13 | 1 | HC_vs_MS.NoTrt_B |
| HIPK3 | 0.90834 | 0.29618 | 0.122 | 0.12 | 1 | HC_vs_MS.NoTrt_B |
| NR4A1 | 0.90853 | 0.26132 | 0.435 | 0.45 | 1 | HC_vs_MS.NoTrt_B |
| HIVEP2 | 0.90912 | 0.51805 | 0.396 | 0.43 | 1 | HC_vs_MS.NoTrt_B |
| IRAK2 | 0.90913 | 0.71498 | 0.288 | 0.3 | 1 | HC_vs_MS.NoTrt_B |
| GPR35 | 0.90917 | 0.6236 | 0.28 | 0.29 | 1 | HC_vs_MS.NoTrt_B |
| OTUD4 | 0.90931 | 0.48483 | 0.597 | 0.63 | 1 | HC_vs_MS.NoTrt_B |
| FARP2 | 0.90961 | -0.8412 | 0.381 | 0.39 | 1 | HC_vs_MS.NoTrt_B |
| KLHL23 | 0.90985 | 0.31463 | 0.157 | 0.16 | 1 | HC_vs_MS.NoTrt_B |
| UPB1 | 0.91017 | 0.2691 | 0.552 | 0.58 | 1 | HC_vs_MS.NoTrt_B |
| CD40 | 0.91026 | -0.2931 | 0.143 | 0.15 | 1 | HC_vs_MS.NoTrt_B |
| TTC39A | 0.91103 | 0.37221 | 0.333 | 0.36 | 1 | HC_vs_MS.NoTrt_B |
| TCOF1 | 0.91144 | -0.3496 | 0.79 | 0.79 | 1 | HC_vs_MS.NoTrt_B |
| RAB26 | 0.9118 | -0.3272 | 0.132 | 0.13 | 1 | HC_vs_MS.NoTrt_B |
| ZMAT3 | 0.91182 | 0.7215 | 0.564 | 0.6 | 1 | HC_vs_MS.NoTrt_B |
| FAM110A | 0.91243 | 0.38789 | 0.165 | 0.17 | 1 | HC_vs_MS.NoTrt_B |
| SLC6A16 | 0.91251 | 0.68249 | 0.243 | 0.24 | 1 | HC_vs_MS.NoTrt_B |
| MDN1 | 0.91274 | 0.59191 | 0.241 | 0.24 | 1 | HC_vs_MS.NoTrt_B |
| SLC43A3 | 0.91323 | -0.2606 | 0.423 | 0.4 | 1 | HC_vs_MS.NoTrt_B |
| PHACTR1 | 0.91329 | 0.62378 | 0.393 | 0.41 | 1 | HC_vs_MS.NoTrt_B |
| CKS1B | 0.91335 | 0.25647 | 0.186 | 0.19 | 1 | HC_vs_MS.NoTrt_B |
| ZNF175 | 0.91342 | -1.3163 | 0.167 | 0.17 | 1 | HC_vs_MS.NoTrt_B |
| ADH4 | 0.9139 | 0.28521 | 0.328 | 0.35 | 1 | HC_vs_MS.NoTrt_B |
| WDR3 | 0.9141 | 0.39921 | 0.712 | 0.73 | 1 | HC_vs_MS.NoTrt_B |
| GULP1 | 0.91444 | 0.49306 | 0.42 | 0.47 | 1 | HC_vs_MS.NoTrt_B |

| | | | | | | |
|---|---|---|---|---|---|---|
| APOL4 | 0.91535 | 0.31776 | 0.43 | 0.46 | 1 | HC_vs_MS.NoTrt_B |
| SEMA4D | 0.91553 | 0.3852 | 0.512 | 0.52 | 1 | HC_vs_MS.NoTrt_B |
| C21orf59 | 0.91617 | -0.3264 | 0.446 | 0.42 | 1 | HC_vs_MS.NoTrt_B |
| SCAMP1 | 0.9164 | 0.40741 | 0.442 | 0.44 | 1 | HC_vs_MS.NoTrt_B |
| FAM102B | 0.9171 | 0.72145 | 0.171 | 0.17 | 1 | HC_vs_MS.NoTrt_B |
| CAAP1 | 0.91738 | 0.79263 | 0.109 | 0.11 | 1 | HC_vs_MS.NoTrt_B |
| SRL | 0.91751 | 0.25601 | 0.346 | 0.37 | 1 | HC_vs_MS.NoTrt_B |
| KLRG1 | 0.91777 | 0.587 | 0.374 | 0.42 | 1 | HC_vs_MS.NoTrt_B |
| CLIP4 | 0.918 | 0.64603 | 0.125 | 0.13 | 1 | HC_vs_MS.NoTrt_B |
| C2orf42 | 0.91801 | 0.50374 | 0.1 | 0.1 | 1 | HC_vs_MS.NoTrt_B |
| FAM210B | 0.91821 | 0.38728 | 0.31 | 0.31 | 1 | HC_vs_MS.NoTrt_B |
| GID4 | 0.91834 | 0.65834 | 0.675 | 0.71 | 1 | HC_vs_MS.NoTrt_B |
| SORL1 | 0.91888 | 0.31829 | 0.799 | 0.78 | 1 | HC_vs_MS.NoTrt_B |
| RABEPK | 0.91952 | -0.3649 | 0.749 | 0.75 | 1 | HC_vs_MS.NoTrt_B |
| SLC1A2 | 0.91962 | 0.54492 | 0.329 | 0.36 | 1 | HC_vs_MS.NoTrt_B |
| LXN | 0.91986 | 0.32634 | 0.168 | 0.17 | 1 | HC_vs_MS.NoTrt_B |
| C1orf52 | 0.92065 | 0.35622 | 0.283 | 0.29 | 1 | HC_vs_MS.NoTrt_B |
| PPP2CB | 0.9208 | 0.52233 | 0.57 | 0.6 | 1 | HC_vs_MS.NoTrt_B |
| ZKSCAN8 | 0.92171 | 1.77357 | 0.281 | 0.29 | 1 | HC_vs_MS.NoTrt_B |
| FKBP11 | 0.92187 | -0.3053 | 0.684 | 0.7 | 1 | HC_vs_MS.NoTrt_B |
| FKTN | 0.92191 | 1.04143 | 0.143 | 0.14 | 1 | HC_vs_MS.NoTrt_B |
| AKT1 | 0.92227 | -0.3267 | 0.203 | 0.2 | 1 | HC_vs_MS.NoTrt_B |
| CREBRF | 0.92299 | 0.30579 | 0.768 | 0.81 | 1 | HC_vs_MS.NoTrt_B |
| RGS13 | 0.92305 | 0.367 | 0.338 | 0.37 | 1 | HC_vs_MS.NoTrt_B |
| IKZF2 | 0.92335 | 0.52845 | 0.357 | 0.39 | 1 | HC_vs_MS.NoTrt_B |
| RBL1 | 0.92428 | 0.62393 | 0.561 | 0.57 | 1 | HC_vs_MS.NoTrt_B |
| PIM1 | 0.92467 | 0.2863 | 0.384 | 0.41 | 1 | HC_vs_MS.NoTrt_B |
| PAK2 | 0.92483 | 0.31487 | 0.745 | 0.78 | 1 | HC_vs_MS.NoTrt_B |
| CARM1 | 0.92492 | -0.2541 | 0.132 | 0.14 | 1 | HC_vs_MS.NoTrt_B |
| METTL15 | 0.92531 | 0.63833 | 0.377 | 0.39 | 1 | HC_vs_MS.NoTrt_B |
| PPP1R15B | 0.92545 | -0.4205 | 0.281 | 0.28 | 1 | HC_vs_MS.NoTrt_B |
| C19orf54 | 0.92562 | 0.4258 | 0.43 | 0.46 | 1 | HC_vs_MS.NoTrt_B |
| KIAA1324 | 0.9261 | -0.3468 | 0.719 | 0.78 | 1 | HC_vs_MS.NoTrt_B |
| 2-Mar | 0.92634 | -0.6628 | 0.516 | 0.54 | 1 | HC_vs_MS.NoTrt_B |
| DOPEY1 | 0.92682 | 1.60979 | 0.288 | 0.31 | 1 | HC_vs_MS.NoTrt_B |
| RNF138 | 0.92693 | 0.54843 | 0.145 | 0.15 | 1 | HC_vs_MS.NoTrt_B |
| CNIH4 | 0.92699 | 0.9373 | 0.559 | 0.62 | 1 | HC_vs_MS.NoTrt_B |
| SYT15 | 0.92701 | 0.26229 | 0.5 | 0.54 | 1 | HC_vs_MS.NoTrt_B |
| CDT1 | 0.92746 | 0.43802 | 0.416 | 0.42 | 1 | HC_vs_MS.NoTrt_B |
| ITM2A | 0.92755 | 0.25331 | 0.219 | 0.22 | 1 | HC_vs_MS.NoTrt_B |

| | | | | | | |
|---|---|---|---|---|---|---|
| SLC4A5 | 0.9277 | 0.28437 | 0.7 | 0.77 | 1 | HC_vs_MS.NoTrt_B |
| CELF5 | 0.92777 | -0.3105 | 0.264 | 0.28 | 1 | HC_vs_MS.NoTrt_B |
| CARD6 | 0.92876 | -0.7033 | 0.503 | 0.55 | 1 | HC_vs_MS.NoTrt_B |
| AKAP8 | 0.92959 | -0.3306 | 0.37 | 0.37 | 1 | HC_vs_MS.NoTrt_B |
| PACS1 | 0.9298 | 0.29595 | 0.817 | 0.83 | 1 | HC_vs_MS.NoTrt_B |
| CNN3 | 0.93063 | 1.21231 | 0.165 | 0.17 | 1 | HC_vs_MS.NoTrt_B |
| MAF | 0.93148 | 0.89823 | 0.27 | 0.29 | 1 | HC_vs_MS.NoTrt_B |
| PRKAR1A | 0.93164 | 0.36056 | 0.254 | 0.26 | 1 | HC_vs_MS.NoTrt_B |
| NNT | 0.93185 | -0.3054 | 0.638 | 0.63 | 1 | HC_vs_MS.NoTrt_B |
| MAD2L1BP | 0.93207 | -0.5966 | 0.164 | 0.16 | 1 | HC_vs_MS.NoTrt_B |
| SMAD5 | 0.93286 | 0.81079 | 0.441 | 0.47 | 1 | HC_vs_MS.NoTrt_B |
| LRRC37A3 | 0.93306 | 0.52126 | 0.528 | 0.56 | 1 | HC_vs_MS.NoTrt_B |
| FAM19A2 | 0.93352 | 0.91051 | 0.217 | 0.22 | 1 | HC_vs_MS.NoTrt_B |
| TPRG1L | 0.93419 | 0.9177 | 0.242 | 0.24 | 1 | HC_vs_MS.NoTrt_B |
| UBAP1 | 0.93439 | 0.27079 | 0.122 | 0.12 | 1 | HC_vs_MS.NoTrt_B |
| ELMSAN1 | 0.93499 | 0.31564 | 0.674 | 0.68 | 1 | HC_vs_MS.NoTrt_B |
| RBPMS | 0.93546 | 0.27602 | 0.1 | 0.1 | 1 | HC_vs_MS.NoTrt_B |
| TAF11 | 0.93655 | 0.29571 | 0.483 | 0.51 | 1 | HC_vs_MS.NoTrt_B |
| FGD5 | 0.93685 | 0.51176 | 0.525 | 0.57 | 1 | HC_vs_MS.NoTrt_B |
| UBXN2B | 0.93712 | 0.59894 | 0.343 | 0.36 | 1 | HC_vs_MS.NoTrt_B |
| AC016577.1 | 0.93713 | 0.27265 | 0.391 | 0.42 | 1 | HC_vs_MS.NoTrt_B |
| CXXC5 | 0.93744 | 0.86587 | 0.504 | 0.55 | 1 | HC_vs_MS.NoTrt_B |
| ZNF354A | 0.93766 | 0.87152 | 0.18 | 0.19 | 1 | HC_vs_MS.NoTrt_B |
| KMT2C | 0.93839 | 0.46141 | 0.422 | 0.44 | 1 | HC_vs_MS.NoTrt_B |
| IKBKB | 0.93872 | 0.458 | 0.83 | 0.87 | 1 | HC_vs_MS.NoTrt_B |
| ZNF552 | 0.93931 | -0.3506 | 0.399 | 0.42 | 1 | HC_vs_MS.NoTrt_B |
| WNK1 | 0.93958 | 0.34308 | 0.526 | 0.57 | 1 | HC_vs_MS.NoTrt_B |
| TTC16 | 0.93967 | 0.33108 | 0.561 | 0.62 | 1 | HC_vs_MS.NoTrt_B |
| MVD | 0.93988 | 0.29118 | 0.454 | 0.5 | 1 | HC_vs_MS.NoTrt_B |
| MKL1 | 0.93999 | 0.45858 | 0.387 | 0.4 | 1 | HC_vs_MS.NoTrt_B |
| MIGA2 | 0.94046 | -0.3334 | 0.399 | 0.38 | 1 | HC_vs_MS.NoTrt_B |
| TBCK | 0.94058 | 0.47655 | 0.725 | 0.74 | 1 | HC_vs_MS.NoTrt_B |
| SWAP70 | 0.94139 | -0.4067 | 0.458 | 0.48 | 1 | HC_vs_MS.NoTrt_B |
| MBNL2 | 0.94159 | 0.26226 | 0.18 | 0.18 | 1 | HC_vs_MS.NoTrt_B |
| CCDC113 | 0.94205 | 0.31151 | 0.639 | 0.68 | 1 | HC_vs_MS.NoTrt_B |
| YEATS2 | 0.94218 | 0.5477 | 0.623 | 0.66 | 1 | HC_vs_MS.NoTrt_B |
| NEDD4L | 0.94255 | -0.3198 | 0.42 | 0.43 | 1 | HC_vs_MS.NoTrt_B |
| NUP35 | 0.94264 | -0.484 | 0.462 | 0.5 | 1 | HC_vs_MS.NoTrt_B |
| ZNF23 | 0.94294 | 0.38663 | 0.326 | 0.35 | 1 | HC_vs_MS.NoTrt_B |
| SHF | 0.94298 | 0.27822 | 0.336 | 0.35 | 1 | HC_vs_MS.NoTrt_B |

| | | | | | | |
|---|---|---|---|---|---|---|
| XYLT2 | 0.94343 | 0.33762 | 0.586 | 0.62 | 1 | HC_vs_MS.NoTrt_B |
| RNF4 | 0.94344 | 0.2726 | 0.765 | 0.77 | 1 | HC_vs_MS.NoTrt_B |
| SZRD1 | 0.94347 | 0.40978 | 0.193 | 0.2 | 1 | HC_vs_MS.NoTrt_B |
| 9-Sep | 0.94377 | 0.64781 | 0.587 | 0.67 | 1 | HC_vs_MS.NoTrt_B |
| TANC1 | 0.94388 | 0.65635 | 0.339 | 0.35 | 1 | HC_vs_MS.NoTrt_B |
| SLC25A37 | 0.94404 | 0.49144 | 0.649 | 0.68 | 1 | HC_vs_MS.NoTrt_B |
| RBFA | 0.94468 | 0.91501 | 0.288 | 0.29 | 1 | HC_vs_MS.NoTrt_B |
| MSL3 | 0.94481 | 0.38102 | 0.184 | 0.2 | 1 | HC_vs_MS.NoTrt_B |
| DHRS11 | 0.94574 | -0.6868 | 0.283 | 0.29 | 1 | HC_vs_MS.NoTrt_B |
| MEMO1 | 0.94577 | 0.28308 | 0.301 | 0.3 | 1 | HC_vs_MS.NoTrt_B |
| PPP1R16A | 0.94618 | 0.9898 | 0.436 | 0.45 | 1 | HC_vs_MS.NoTrt_B |
| ITPR1 | 0.94686 | 0.2734 | 0.154 | 0.15 | 1 | HC_vs_MS.NoTrt_B |
| ATPAF1 | 0.9472 | -0.5697 | 0.235 | 0.23 | 1 | HC_vs_MS.NoTrt_B |
| CENPM | 0.94734 | -0.7481 | 0.3 | 0.29 | 1 | HC_vs_MS.NoTrt_B |
| RGPD8 | 0.9475 | 0.96503 | 0.188 | 0.2 | 1 | HC_vs_MS.NoTrt_B |
| RAI1 | 0.94761 | 0.69547 | 0.209 | 0.21 | 1 | HC_vs_MS.NoTrt_B |
| RXRA | 0.94775 | 1.32092 | 0.138 | 0.14 | 1 | HC_vs_MS.NoTrt_B |
| UBP1 | 0.94819 | 0.71373 | 0.099 | 0.1 | 1 | HC_vs_MS.NoTrt_B |
| RC3H2 | 0.94823 | 0.43972 | 0.496 | 0.5 | 1 | HC_vs_MS.NoTrt_B |
| HEATR1 | 0.94842 | 0.41643 | 0.138 | 0.14 | 1 | HC_vs_MS.NoTrt_B |
| USP36 | 0.94874 | 0.52788 | 0.239 | 0.24 | 1 | HC_vs_MS.NoTrt_B |
| PTAFR | 0.94878 | 0.39232 | 0.567 | 0.61 | 1 | HC_vs_MS.NoTrt_B |
| LINC00094 | 0.94951 | 0.68923 | 0.457 | 0.46 | 1 | HC_vs_MS.NoTrt_B |
| TMEM140 | 0.94969 | 0.4227 | 0.178 | 0.18 | 1 | HC_vs_MS.NoTrt_B |
| SERPINF1 | 0.95028 | 0.36437 | 0.752 | 0.84 | 1 | HC_vs_MS.NoTrt_B |
| PTPN18 | 0.95051 | 1.03071 | 0.37 | 0.4 | 1 | HC_vs_MS.NoTrt_B |
| TMEM221 | 0.95053 | 0.49408 | 0.17 | 0.17 | 1 | HC_vs_MS.NoTrt_B |
| DHX30 | 0.95074 | -0.3588 | 0.639 | 0.64 | 1 | HC_vs_MS.NoTrt_B |
| DCTN5 | 0.95114 | -0.3157 | 0.781 | 0.82 | 1 | HC_vs_MS.NoTrt_B |
| ZEB1 | 0.95123 | 0.90818 | 0.565 | 0.56 | 1 | HC_vs_MS.NoTrt_B |
| STK17A | 0.9521 | 1.04039 | 0.274 | 0.29 | 1 | HC_vs_MS.NoTrt_B |
| NCBP1 | 0.95253 | 0.46896 | 0.649 | 0.71 | 1 | HC_vs_MS.NoTrt_B |
| TRIM46 | 0.95314 | -0.7854 | 0.417 | 0.44 | 1 | HC_vs_MS.NoTrt_B |
| ZNF397 | 0.95335 | -0.4223 | 0.946 | 0.96 | 1 | HC_vs_MS.NoTrt_B |
| ANK1 | 0.95337 | -1.2155 | 0.225 | 0.22 | 1 | HC_vs_MS.NoTrt_B |
| HMG20A | 0.95337 | 0.7713 | 0.264 | 0.28 | 1 | HC_vs_MS.NoTrt_B |
| XYLT1 | 0.95339 | 0.31194 | 0.116 | 0.11 | 1 | HC_vs_MS.NoTrt_B |
| PGM3 | 0.95418 | 1.0872 | 0.245 | 0.25 | 1 | HC_vs_MS.NoTrt_B |
| P2RY10 | 0.95428 | 0.41014 | 0.309 | 0.3 | 1 | HC_vs_MS.NoTrt_B |
| KDM4C | 0.95433 | 0.35212 | 0.291 | 0.3 | 1 | HC_vs_MS.NoTrt_B |

| | | | | | | |
|---|---|---|---|---|---|---|
| NIPAL2 | 0.95435 | -0.4003 | 0.281 | 0.28 | 1 | HC_vs_MS.NoTrt_B |
| PUS3 | 0.9545 | 0.30183 | 0.31 | 0.32 | 1 | HC_vs_MS.NoTrt_B |
| NDUFAF6 | 0.95457 | 0.33648 | 0.622 | 0.63 | 1 | HC_vs_MS.NoTrt_B |
| ZNF786 | 0.95502 | -0.3793 | 0.406 | 0.41 | 1 | HC_vs_MS.NoTrt_B |
| TBL3 | 0.95611 | 0.35919 | 0.268 | 0.27 | 1 | HC_vs_MS.NoTrt_B |
| PER1 | 0.95647 | 0.50674 | 0.332 | 0.34 | 1 | HC_vs_MS.NoTrt_B |
| TBC1D15 | 0.95649 | 0.50404 | 0.572 | 0.6 | 1 | HC_vs_MS.NoTrt_B |
| GUSB | 0.95682 | -0.3899 | 0.681 | 0.72 | 1 | HC_vs_MS.NoTrt_B |
| ZNF740 | 0.95735 | 0.37201 | 0.142 | 0.14 | 1 | HC_vs_MS.NoTrt_B |
| SSPO | 0.95815 | 0.91102 | 0.123 | 0.12 | 1 | HC_vs_MS.NoTrt_B |
| ALG14 | 0.95828 | -0.3641 | 0.899 | 0.93 | 1 | HC_vs_MS.NoTrt_B |
| AGO4 | 0.95953 | -1.1417 | 0.28 | 0.28 | 1 | HC_vs_MS.NoTrt_B |
| CC2D1B | 0.95962 | 0.25195 | 0.59 | 0.6 | 1 | HC_vs_MS.NoTrt_B |
| JAK2 | 0.95972 | 0.34888 | 0.223 | 0.22 | 1 | HC_vs_MS.NoTrt_B |
| LIPT2 | 0.96021 | -0.4154 | 0.123 | 0.12 | 1 | HC_vs_MS.NoTrt_B |
| KATNAL2 | 0.96041 | 0.6897 | 0.31 | 0.32 | 1 | HC_vs_MS.NoTrt_B |
| DHDDS | 0.96044 | 0.575 | 0.543 | 0.57 | 1 | HC_vs_MS.NoTrt_B |
| WDR44 | 0.96045 | 0.44103 | 0.1 | 0.1 | 1 | HC_vs_MS.NoTrt_B |
| GPC2 | 0.96093 | 0.57023 | 0.397 | 0.43 | 1 | HC_vs_MS.NoTrt_B |
| MTPAP | 0.9615 | 0.69441 | 0.832 | 0.89 | 1 | HC_vs_MS.NoTrt_B |
| SPEF2 | 0.9621 | 0.43267 | 0.546 | 0.56 | 1 | HC_vs_MS.NoTrt_B |
| ZNF658 | 0.96228 | 0.46582 | 0.175 | 0.18 | 1 | HC_vs_MS.NoTrt_B |
| YY1 | 0.96288 | 0.66524 | 0.314 | 0.32 | 1 | HC_vs_MS.NoTrt_B |
| PRPF40B | 0.96319 | 0.33848 | 0.204 | 0.21 | 1 | HC_vs_MS.NoTrt_B |
| EXOSC3 | 0.96327 | 0.33146 | 0.429 | 0.44 | 1 | HC_vs_MS.NoTrt_B |
| ZNF812P | 0.96347 | 0.36805 | 0.109 | 0.11 | 1 | HC_vs_MS.NoTrt_B |
| PRRC2A | 0.96409 | 0.46833 | 0.757 | 0.82 | 1 | HC_vs_MS.NoTrt_B |
| AMD1 | 0.9652 | -0.4456 | 0.713 | 0.74 | 1 | HC_vs_MS.NoTrt_B |
| RALGAPA2 | 0.96527 | 0.82817 | 0.149 | 0.15 | 1 | HC_vs_MS.NoTrt_B |
| ABCD3 | 0.96606 | 0.43388 | 0.125 | 0.12 | 1 | HC_vs_MS.NoTrt_B |
| SETD5 | 0.96608 | 0.77967 | 0.496 | 0.53 | 1 | HC_vs_MS.NoTrt_B |
| SYK | 0.9663 | 0.28353 | 0.333 | 0.35 | 1 | HC_vs_MS.NoTrt_B |
| MAP4K5 | 0.96661 | 0.54615 | 0.494 | 0.53 | 1 | HC_vs_MS.NoTrt_B |
| CLMN | 0.96725 | 0.29752 | 0.63 | 0.66 | 1 | HC_vs_MS.NoTrt_B |
| TAF1B | 0.96806 | 0.32272 | 0.464 | 0.49 | 1 | HC_vs_MS.NoTrt_B |
| KCTD21 | 0.96814 | -1.0453 | 0.416 | 0.42 | 1 | HC_vs_MS.NoTrt_B |
| LCOR | 0.96838 | 0.32914 | 0.367 | 0.37 | 1 | HC_vs_MS.NoTrt_B |
| 6-Mar | 0.96839 | 0.33677 | 0.8 | 0.85 | 1 | HC_vs_MS.NoTrt_B |
| MAGEA10 | 0.96907 | 0.27874 | 0.543 | 0.56 | 1 | HC_vs_MS.NoTrt_B |
| COMMD2 | 0.96949 | 0.38757 | 0.329 | 0.33 | 1 | HC_vs_MS.NoTrt_B |

| | | | | | | |
|---|---|---|---|---|---|---|
| PFKFB3 | 0.9702 | 0.78718 | 0.23 | 0.23 | 1 | HC_vs_MS.NoTrt_B |
| C1R | 0.97106 | 0.65506 | 0.346 | 0.38 | 1 | HC_vs_MS.NoTrt_B |
| PPAT | 0.97131 | 0.40008 | 0.271 | 0.28 | 1 | HC_vs_MS.NoTrt_B |
| DNAJB5 | 0.9715 | -1.38 | 0.146 | 0.15 | 1 | HC_vs_MS.NoTrt_B |
| COLGALT1 | 0.97382 | 0.29257 | 0.436 | 0.47 | 1 | HC_vs_MS.NoTrt_B |
| TNRC6A | 0.97383 | 0.3434 | 0.764 | 0.81 | 1 | HC_vs_MS.NoTrt_B |
| NEPRO | 0.97395 | 0.42993 | 0.194 | 0.2 | 1 | HC_vs_MS.NoTrt_B |
| C21orf2 | 0.97439 | 0.33997 | 0.123 | 0.12 | 1 | HC_vs_MS.NoTrt_B |
| CNOT1 | 0.97532 | 0.34149 | 0.806 | 0.84 | 1 | HC_vs_MS.NoTrt_B |
| RCAN1 | 0.976 | 0.37401 | 0.303 | 0.32 | 1 | HC_vs_MS.NoTrt_B |
| C2CD5 | 0.97619 | 0.62717 | 0.165 | 0.17 | 1 | HC_vs_MS.NoTrt_B |
| HSPH1 | 0.97659 | 0.70527 | 0.165 | 0.17 | 1 | HC_vs_MS.NoTrt_B |
| ZCCHC17 | 0.97665 | 0.36023 | 0.117 | 0.12 | 1 | HC_vs_MS.NoTrt_B |
| VPS4B | 0.97667 | -0.2508 | 0.742 | 0.78 | 1 | HC_vs_MS.NoTrt_B |
| BCR | 0.977 | -0.2746 | 0.261 | 0.27 | 1 | HC_vs_MS.NoTrt_B |
| KPTN | 0.97825 | 0.41266 | 0.212 | 0.22 | 1 | HC_vs_MS.NoTrt_B |
| MYSM1 | 0.9784 | 0.82348 | 0.206 | 0.21 | 1 | HC_vs_MS.NoTrt_B |
| ACTG2 | 0.97854 | 0.56918 | 0.241 | 0.24 | 1 | HC_vs_MS.NoTrt_B |
| HHLA1 | 0.97869 | 0.34889 | 0.381 | 0.41 | 1 | HC_vs_MS.NoTrt_B |
| NPIPB11 | 0.97915 | 0.6605 | 0.446 | 0.49 | 1 | HC_vs_MS.NoTrt_B |
| ZNF561 | 0.97944 | -0.4657 | 0.532 | 0.55 | 1 | HC_vs_MS.NoTrt_B |
| STAT2 | 0.98011 | 0.62061 | 0.883 | 0.92 | 1 | HC_vs_MS.NoTrt_B |
| NKIRAS2 | 0.98056 | -0.2904 | 0.806 | 0.85 | 1 | HC_vs_MS.NoTrt_B |
| ABCB6 | 0.9817 | 0.30791 | 0.113 | 0.11 | 1 | HC_vs_MS.NoTrt_B |
| CTD-2192J1( | 0.98217 | 0.56049 | 0.454 | 0.48 | 1 | HC_vs_MS.NoTrt_B |
| HAUS1 | 0.98241 | -0.359 | 0.635 | 0.62 | 1 | HC_vs_MS.NoTrt_B |
| ZNF664 | 0.98257 | 0.36638 | 0.233 | 0.24 | 1 | HC_vs_MS.NoTrt_B |
| RNF19A | 0.98261 | 0.36603 | 0.412 | 0.43 | 1 | HC_vs_MS.NoTrt_B |
| MYO6 | 0.98283 | 0.41294 | 0.419 | 0.45 | 1 | HC_vs_MS.NoTrt_B |
| AC004381.6 | 0.98285 | 0.57317 | 0.196 | 0.2 | 1 | HC_vs_MS.NoTrt_B |
| PCBP1 | 0.98322 | 0.37019 | 0.514 | 0.53 | 1 | HC_vs_MS.NoTrt_B |
| ANKRD22 | 0.98337 | 0.27063 | 0.125 | 0.12 | 1 | HC_vs_MS.NoTrt_B |
| CEP135 | 0.9834 | 0.44364 | 0.757 | 0.8 | 1 | HC_vs_MS.NoTrt_B |
| DCAF7 | 0.98394 | -0.2818 | 0.348 | 0.32 | 1 | HC_vs_MS.NoTrt_B |
| MORN4 | 0.98403 | 0.41882 | 0.532 | 0.58 | 1 | HC_vs_MS.NoTrt_B |
| ZNF33A | 0.98422 | 0.2608 | 0.574 | 0.58 | 1 | HC_vs_MS.NoTrt_B |
| CNPY4 | 0.98436 | 0.47607 | 0.69 | 0.75 | 1 | HC_vs_MS.NoTrt_B |
| LDHAL6A | 0.98439 | 0.50884 | 0.349 | 0.37 | 1 | HC_vs_MS.NoTrt_B |
| ERMN | 0.98452 | 0.26158 | 0.172 | 0.17 | 1 | HC_vs_MS.NoTrt_B |
| ATP13A4 | 0.98509 | 0.70079 | 0.397 | 0.42 | 1 | HC_vs_MS.NoTrt_B |

| | | | | | | |
|---|---|---|---|---|---|---|
| GPR174 | 0.98524 | 0.31224 | 0.154 | 0.16 | 1 | HC_vs_MS.NoTrt_B |
| PDE4D | 0.98577 | 0.35438 | 0.304 | 0.32 | 1 | HC_vs_MS.NoTrt_B |
| SIRT2 | 0.98613 | -0.4039 | 0.897 | 0.91 | 1 | HC_vs_MS.NoTrt_B |
| ETFB | 0.98639 | -0.2832 | 0.868 | 0.9 | 1 | HC_vs_MS.NoTrt_B |
| METTL12 | 0.98695 | 0.5879 | 0.613 | 0.67 | 1 | HC_vs_MS.NoTrt_B |
| SIRT1 | 0.98761 | 0.44903 | 0.103 | 0.1 | 1 | HC_vs_MS.NoTrt_B |
| CHD2 | 0.98811 | 0.30586 | 0.987 | 0.99 | 1 | HC_vs_MS.NoTrt_B |
| SSFA2 | 0.98837 | 0.49553 | 0.119 | 0.12 | 1 | HC_vs_MS.NoTrt_B |
| INPP5E | 0.98928 | 0.36167 | 0.27 | 0.27 | 1 | HC_vs_MS.NoTrt_B |
| HAUS8 | 0.98955 | -0.3307 | 0.726 | 0.75 | 1 | HC_vs_MS.NoTrt_B |
| CNIH3 | 0.99124 | -0.5556 | 0.171 | 0.17 | 1 | HC_vs_MS.NoTrt_B |
| FAM60A | 0.99142 | 0.53348 | 0.522 | 0.55 | 1 | HC_vs_MS.NoTrt_B |
| TLL2 | 0.99162 | 0.32112 | 0.316 | 0.33 | 1 | HC_vs_MS.NoTrt_B |
| XYLB | 0.99163 | 0.27943 | 0.433 | 0.45 | 1 | HC_vs_MS.NoTrt_B |
| ZNF287 | 0.99166 | -0.4026 | 0.365 | 0.37 | 1 | HC_vs_MS.NoTrt_B |
| MTBP | 0.99189 | 0.40515 | 0.597 | 0.62 | 1 | HC_vs_MS.NoTrt_B |
| MLANA | 0.99204 | 0.3098 | 0.394 | 0.44 | 1 | HC_vs_MS.NoTrt_B |
| PATJ | 0.99213 | 0.33512 | 0.662 | 0.68 | 1 | HC_vs_MS.NoTrt_B |
| USO1 | 0.99227 | 1.07089 | 0.133 | 0.14 | 1 | HC_vs_MS.NoTrt_B |
| NDST4 | 0.9925 | 0.35532 | 0.109 | 0.11 | 1 | HC_vs_MS.NoTrt_B |
| APBA2 | 0.99308 | 0.56246 | 0.522 | 0.54 | 1 | HC_vs_MS.NoTrt_B |
| USP3 | 0.99367 | 0.4486 | 0.743 | 0.74 | 1 | HC_vs_MS.NoTrt_B |
| CHAMP1 | 0.99376 | 0.35159 | 0.581 | 0.62 | 1 | HC_vs_MS.NoTrt_B |
| MED22 | 0.9943 | 0.60565 | 0.506 | 0.55 | 1 | HC_vs_MS.NoTrt_B |
| CCDC120 | 0.99433 | 0.43668 | 0.464 | 0.5 | 1 | HC_vs_MS.NoTrt_B |
| AURKA | 0.99445 | 0.53858 | 0.286 | 0.3 | 1 | HC_vs_MS.NoTrt_B |
| KNSTRN | 0.99562 | 0.46094 | 0.475 | 0.49 | 1 | HC_vs_MS.NoTrt_B |
| SLC7A6 | 0.99582 | 0.29393 | 0.51 | 0.5 | 1 | HC_vs_MS.NoTrt_B |
| KCNMB3 | 0.99622 | 0.48283 | 0.367 | 0.38 | 1 | HC_vs_MS.NoTrt_B |
| GARNL3 | 0.9969 | 1.36895 | 0.155 | 0.16 | 1 | HC_vs_MS.NoTrt_B |
| ANKRD7 | 0.99702 | 0.53258 | 0.171 | 0.18 | 1 | HC_vs_MS.NoTrt_B |
| EXD2 | 0.99747 | 1.04718 | 0.181 | 0.18 | 1 | HC_vs_MS.NoTrt_B |
| ATP10A | 0.99751 | 0.33151 | 0.101 | 0.1 | 1 | HC_vs_MS.NoTrt_B |
| CLMP | 0.99775 | 0.50767 | 0.378 | 0.4 | 1 | HC_vs_MS.NoTrt_B |
| KDM5D | 0.99921 | -0.9367 | 0.316 | 0.31 | 1 | HC_vs_MS.NoTrt_B |
| MMP24-AS1 | 0.99947 | 0.59597 | 0.725 | 0.76 | 1 | HC_vs_MS.NoTrt_B |
| CHD3 | 0.9997 | 0.37066 | 0.365 | 0.38 | 1 | HC_vs_MS.NoTrt_B |
| FBXO40 | 1 | 1.06782 | 0.328 | 0.34 | 1 | HC_vs_MS.NoTrt_B |
| FBXL22 | 1 | 0.86429 | 0.103 | 0.1 | 1 | HC_vs_MS.NoTrt_B |
| MT-ND4L | 2.06E-22 | 1.36987 | 0.989 | 1 | 3.29E-18 | MS.Trt_vs_MS.NoTrt_F |

| | | | | | | |
|---|---|---|---|---|---|---|
| MT-CO2 | 1.18E-18 | 1.11448 | 0.996 | 1 | 1.89E-14 | MS.Trt_vs_MS.NoTrt_F |
| RBPJ | 1.41E-18 | -0.3463 | 0.36 | 0.79 | 2.25E-14 | MS.Trt_vs_MS.NoTrt_F |
| HLA-B | 1.32E-17 | -0.6715 | 0.881 | 0.99 | 2.10E-13 | MS.Trt_vs_MS.NoTrt_F |
| TAC4 | 3.62E-14 | -0.9884 | 0.561 | 0.8 | 5.78E-10 | MS.Trt_vs_MS.NoTrt_F |
| CD69 | 4.93E-14 | -0.9566 | 0.277 | 0.66 | 7.88E-10 | MS.Trt_vs_MS.NoTrt_F |
| RGS1 | 8.40E-14 | -1.0624 | 0.356 | 0.7 | 1.34E-09 | MS.Trt_vs_MS.NoTrt_F |
| PCSK5 | 9.46E-14 | 1.55027 | 0.928 | 0.93 | 1.51E-09 | MS.Trt_vs_MS.NoTrt_F |
| ARHGAP9 | 1.08E-13 | -0.6735 | 0.46 | 0.77 | 1.73E-09 | MS.Trt_vs_MS.NoTrt_F |
| EEF1A1 | 2.56E-13 | -0.4483 | 0.964 | 1 | 4.09E-09 | MS.Trt_vs_MS.NoTrt_F |
| RPS4Y1 | 2.75E-13 | -3.8498 | 0.079 | 0.33 | 4.39E-09 | MS.Trt_vs_MS.NoTrt_F |
| RPS3 | 3.19E-13 | -0.4772 | 0.853 | 0.99 | 5.09E-09 | MS.Trt_vs_MS.NoTrt_F |
| HOXB2 | 2.36E-12 | -0.6688 | 0.112 | 0.39 | 3.77E-08 | MS.Trt_vs_MS.NoTrt_F |
| HSPA1A | 3.95E-12 | -0.9118 | 0.169 | 0.47 | 6.31E-08 | MS.Trt_vs_MS.NoTrt_F |
| EIF4A2 | 4.27E-12 | -0.611 | 0.46 | 0.83 | 6.83E-08 | MS.Trt_vs_MS.NoTrt_F |
| AQP3 | 4.29E-12 | -0.8966 | 0.27 | 0.6 | 6.85E-08 | MS.Trt_vs_MS.NoTrt_F |
| SNRNP70 | 4.34E-12 | -1.0312 | 0.183 | 0.48 | 6.92E-08 | MS.Trt_vs_MS.NoTrt_F |
| SRSF7 | 5.15E-12 | -0.4925 | 0.396 | 0.76 | 8.23E-08 | MS.Trt_vs_MS.NoTrt_F |
| ECSIT | 5.90E-12 | -0.8493 | 0.227 | 0.54 | 9.42E-08 | MS.Trt_vs_MS.NoTrt_F |
| RPL9 | 9.16E-12 | -0.6516 | 0.809 | 0.95 | 1.46E-07 | MS.Trt_vs_MS.NoTrt_F |
| HNRNPH1 | 9.83E-12 | 0.98686 | 0.993 | 1 | 1.57E-07 | MS.Trt_vs_MS.NoTrt_F |
| DNAJB1 | 2.04E-11 | -0.8421 | 0.683 | 0.92 | 3.26E-07 | MS.Trt_vs_MS.NoTrt_F |
| GBP5 | 3.13E-11 | -0.5737 | 0.54 | 0.81 | 4.99E-07 | MS.Trt_vs_MS.NoTrt_F |
| LSM6 | 4.34E-11 | -0.6833 | 0.115 | 0.38 | 6.93E-07 | MS.Trt_vs_MS.NoTrt_F |
| PBX1 | 6.33E-11 | -0.5771 | 0.878 | 0.95 | 1.01E-06 | MS.Trt_vs_MS.NoTrt_F |
| HSPA1B | 7.57E-11 | -1.2509 | 0.212 | 0.49 | 1.21E-06 | MS.Trt_vs_MS.NoTrt_F |
| METTL21A | 1.09E-10 | 0.91762 | 0.953 | 0.95 | 1.74E-06 | MS.Trt_vs_MS.NoTrt_F |
| MTRNR2L4 | 1.88E-10 | -0.4194 | 0.924 | 0.99 | 3.01E-06 | MS.Trt_vs_MS.NoTrt_F |
| TYR | 2.58E-10 | -0.6029 | 0.194 | 0.46 | 4.12E-06 | MS.Trt_vs_MS.NoTrt_F |
| JUN | 2.59E-10 | -0.4896 | 0.327 | 0.64 | 4.13E-06 | MS.Trt_vs_MS.NoTrt_F |
| PHF1 | 2.93E-10 | -1.4454 | 0.129 | 0.38 | 4.68E-06 | MS.Trt_vs_MS.NoTrt_F |
| GPSM3 | 5.21E-10 | -0.7046 | 0.374 | 0.67 | 8.31E-06 | MS.Trt_vs_MS.NoTrt_F |
| ELF1 | 5.25E-10 | -0.6808 | 0.295 | 0.59 | 8.38E-06 | MS.Trt_vs_MS.NoTrt_F |
| RPL26 | 5.79E-10 | -0.4218 | 0.835 | 0.97 | 9.25E-06 | MS.Trt_vs_MS.NoTrt_F |
| ALKBH5 | 6.86E-10 | -0.8505 | 0.144 | 0.4 | 1.10E-05 | MS.Trt_vs_MS.NoTrt_F |
| MAML2 | 7.19E-10 | -1.3733 | 0.14 | 0.39 | 1.15E-05 | MS.Trt_vs_MS.NoTrt_F |
| SYNPO | 7.97E-10 | -0.639 | 0.18 | 0.44 | 1.27E-05 | MS.Trt_vs_MS.NoTrt_F |
| ATP2B1 | 9.19E-10 | -0.2781 | 0.378 | 0.71 | 1.47E-05 | MS.Trt_vs_MS.NoTrt_F |
| LYPLA2 | 9.68E-10 | -0.9463 | 0.119 | 0.36 | 1.55E-05 | MS.Trt_vs_MS.NoTrt_F |
| LRRC8C | 1.02E-09 | -0.8285 | 0.277 | 0.54 | 1.63E-05 | MS.Trt_vs_MS.NoTrt_F |
| JUNB | 1.10E-09 | -0.5103 | 0.565 | 0.85 | 1.76E-05 | MS.Trt_vs_MS.NoTrt_F |

| | | | | | | |
|---|---|---|---|---|---|---|
| ANXA2 | 1.18E-09 | -0.502 | 0.637 | 0.86 | 1.89E-05 | MS.Trt_vs_MS.NoTrt_F |
| S100A4 | 1.23E-09 | -0.3053 | 0.55 | 0.9 | 1.97E-05 | MS.Trt_vs_MS.NoTrt_F |
| FAM160B1 | 1.33E-09 | -1.4204 | 0.147 | 0.4 | 2.12E-05 | MS.Trt_vs_MS.NoTrt_F |
| C1orf56 | 1.45E-09 | 0.93396 | 0.842 | 0.8 | 2.31E-05 | MS.Trt_vs_MS.NoTrt_F |
| POLR2G | 1.57E-09 | -0.755 | 0.137 | 0.39 | 2.51E-05 | MS.Trt_vs_MS.NoTrt_F |
| TNIK | 1.62E-09 | -0.4818 | 0.306 | 0.61 | 2.59E-05 | MS.Trt_vs_MS.NoTrt_F |
| CCDC14 | 1.66E-09 | -1.1961 | 0.187 | 0.45 | 2.65E-05 | MS.Trt_vs_MS.NoTrt_F |
| ANP32B | 2.21E-09 | -0.405 | 0.237 | 0.51 | 3.52E-05 | MS.Trt_vs_MS.NoTrt_F |
| VPS13C | 2.40E-09 | -0.6122 | 0.716 | 0.91 | 3.83E-05 | MS.Trt_vs_MS.NoTrt_F |
| TCEA1 | 2.50E-09 | -0.3421 | 0.252 | 0.54 | 4.00E-05 | MS.Trt_vs_MS.NoTrt_F |
| KIAA1551 | 2.65E-09 | -0.5045 | 0.403 | 0.71 | 4.23E-05 | MS.Trt_vs_MS.NoTrt_F |
| RUNDC3B | 2.74E-09 | -0.7282 | 0.212 | 0.44 | 4.38E-05 | MS.Trt_vs_MS.NoTrt_F |
| SLC9A9 | 3.07E-09 | -1.2621 | 0.108 | 0.34 | 4.90E-05 | MS.Trt_vs_MS.NoTrt_F |
| VPS36 | 3.11E-09 | -0.8189 | 0.219 | 0.48 | 4.96E-05 | MS.Trt_vs_MS.NoTrt_F |
| SAYSD1 | 3.36E-09 | -0.2725 | 0.137 | 0.38 | 5.36E-05 | MS.Trt_vs_MS.NoTrt_F |
| PKM | 3.63E-09 | -0.4899 | 0.478 | 0.79 | 5.80E-05 | MS.Trt_vs_MS.NoTrt_F |
| SF1 | 3.76E-09 | -0.4497 | 0.424 | 0.72 | 6.01E-05 | MS.Trt_vs_MS.NoTrt_F |
| CDKN1B | 4.04E-09 | -0.7236 | 0.266 | 0.55 | 6.46E-05 | MS.Trt_vs_MS.NoTrt_F |
| TAGAP | 4.06E-09 | -0.8216 | 0.245 | 0.52 | 6.48E-05 | MS.Trt_vs_MS.NoTrt_F |
| SMCHD1 | 5.31E-09 | -0.5247 | 0.608 | 0.85 | 8.48E-05 | MS.Trt_vs_MS.NoTrt_F |
| TAGLN2 | 5.36E-09 | -0.3632 | 0.471 | 0.82 | 8.56E-05 | MS.Trt_vs_MS.NoTrt_F |
| RASA2 | 5.54E-09 | -0.4398 | 0.327 | 0.61 | 8.84E-05 | MS.Trt_vs_MS.NoTrt_F |
| SMC3 | 6.35E-09 | -0.4404 | 0.205 | 0.47 | 1.01E-04 | MS.Trt_vs_MS.NoTrt_F |
| FLNA | 6.51E-09 | -0.423 | 0.511 | 0.8 | 1.04E-04 | MS.Trt_vs_MS.NoTrt_F |
| ARL5B | 6.96E-09 | -0.7725 | 0.112 | 0.33 | 1.11E-04 | MS.Trt_vs_MS.NoTrt_F |
| GBP2 | 7.31E-09 | -0.597 | 0.356 | 0.64 | 1.17E-04 | MS.Trt_vs_MS.NoTrt_F |
| TXNIP | 7.35E-09 | -0.4972 | 0.504 | 0.8 | 1.17E-04 | MS.Trt_vs_MS.NoTrt_F |
| ELF4 | 7.71E-09 | -1.083 | 0.094 | 0.3 | 1.23E-04 | MS.Trt_vs_MS.NoTrt_F |
| STXBP2 | 8.89E-09 | -1.1342 | 0.216 | 0.46 | 1.42E-04 | MS.Trt_vs_MS.NoTrt_F |
| RP11-343C2 | 9.30E-09 | -1.0282 | 0.115 | 0.34 | 1.49E-04 | MS.Trt_vs_MS.NoTrt_F |
| PRKCQ | 9.38E-09 | -0.5091 | 0.259 | 0.54 | 1.50E-04 | MS.Trt_vs_MS.NoTrt_F |
| ABCC1 | 9.61E-09 | -0.4485 | 0.32 | 0.6 | 1.53E-04 | MS.Trt_vs_MS.NoTrt_F |
| ORC6 | 1.06E-08 | 1.01191 | 0.964 | 0.97 | 1.69E-04 | MS.Trt_vs_MS.NoTrt_F |
| EFNA5 | 1.14E-08 | -0.325 | 0.32 | 0.57 | 1.82E-04 | MS.Trt_vs_MS.NoTrt_F |
| CEACAM1 | 1.22E-08 | -0.5398 | 0.917 | 0.75 | 1.94E-04 | MS.Trt_vs_MS.NoTrt_F |
| MAP1LC3B | 1.22E-08 | -0.325 | 0.313 | 0.59 | 1.94E-04 | MS.Trt_vs_MS.NoTrt_F |
| SNU13 | 1.33E-08 | -0.7558 | 0.216 | 0.48 | 2.12E-04 | MS.Trt_vs_MS.NoTrt_F |
| CD28 | 1.37E-08 | -0.2834 | 0.547 | 0.77 | 2.19E-04 | MS.Trt_vs_MS.NoTrt_F |
| 9-Sep | 1.62E-08 | -0.3375 | 0.453 | 0.71 | 2.58E-04 | MS.Trt_vs_MS.NoTrt_F |
| C10orf11 | 1.66E-08 | -0.3516 | 0.968 | 0.98 | 2.66E-04 | MS.Trt_vs_MS.NoTrt_F |

| | | | | | | |
|---|---|---|---|---|---|---|
| DUSP10 | 1.72E-08 | -0.3949 | 0.223 | 0.49 | 2.74E-04 | MS.Trt_vs_MS.NoTrt_F |
| SDF2 | 1.72E-08 | -0.5896 | 0.183 | 0.44 | 2.74E-04 | MS.Trt_vs_MS.NoTrt_F |
| ARHGEF18 | 1.73E-08 | -0.4355 | 0.277 | 0.55 | 2.76E-04 | MS.Trt_vs_MS.NoTrt_F |
| CEP164 | 1.75E-08 | -0.6134 | 0.147 | 0.38 | 2.80E-04 | MS.Trt_vs_MS.NoTrt_F |
| CCDC167 | 1.80E-08 | -0.2586 | 0.122 | 0.34 | 2.88E-04 | MS.Trt_vs_MS.NoTrt_F |
| GTF3C5 | 1.99E-08 | -0.7869 | 0.273 | 0.52 | 3.17E-04 | MS.Trt_vs_MS.NoTrt_F |
| EPB41 | 2.00E-08 | -0.291 | 0.309 | 0.62 | 3.19E-04 | MS.Trt_vs_MS.NoTrt_F |
| MED15 | 2.32E-08 | -0.465 | 0.55 | 0.76 | 3.71E-04 | MS.Trt_vs_MS.NoTrt_F |
| CACYBP | 2.33E-08 | -0.4309 | 0.201 | 0.46 | 3.72E-04 | MS.Trt_vs_MS.NoTrt_F |
| GAPDH | 2.53E-08 | -0.3973 | 0.705 | 0.93 | 4.04E-04 | MS.Trt_vs_MS.NoTrt_F |
| HNRNPA2B1 | 2.62E-08 | -0.3396 | 0.612 | 0.9 | 4.19E-04 | MS.Trt_vs_MS.NoTrt_F |
| HNRNPA0 | 2.73E-08 | -0.5673 | 0.309 | 0.59 | 4.35E-04 | MS.Trt_vs_MS.NoTrt_F |
| JMJD1C | 2.73E-08 | -0.6993 | 0.23 | 0.5 | 4.36E-04 | MS.Trt_vs_MS.NoTrt_F |
| IL6R | 3.19E-08 | -0.5899 | 0.576 | 0.79 | 5.09E-04 | MS.Trt_vs_MS.NoTrt_F |
| ACTR1B | 3.62E-08 | -0.6398 | 0.133 | 0.36 | 5.79E-04 | MS.Trt_vs_MS.NoTrt_F |
| MDN1 | 3.89E-08 | -0.6499 | 0.147 | 0.37 | 6.21E-04 | MS.Trt_vs_MS.NoTrt_F |
| KDM4C | 4.84E-08 | -0.6033 | 0.216 | 0.46 | 7.73E-04 | MS.Trt_vs_MS.NoTrt_F |
| DLGAP4 | 5.21E-08 | -1.0069 | 0.14 | 0.36 | 8.33E-04 | MS.Trt_vs_MS.NoTrt_F |
| ARHGDIA | 5.25E-08 | -0.9319 | 0.342 | 0.58 | 8.38E-04 | MS.Trt_vs_MS.NoTrt_F |
| PIGF | 5.51E-08 | -0.6959 | 0.083 | 0.27 | 8.80E-04 | MS.Trt_vs_MS.NoTrt_F |
| CDC25B | 5.55E-08 | -0.5379 | 0.291 | 0.55 | 8.86E-04 | MS.Trt_vs_MS.NoTrt_F |
| KIAA2026 | 5.77E-08 | -0.6218 | 0.165 | 0.39 | 9.22E-04 | MS.Trt_vs_MS.NoTrt_F |
| AMMECR1L | 6.79E-08 | -0.4761 | 0.191 | 0.44 | 0.001084 | MS.Trt_vs_MS.NoTrt_F |
| PRDX1 | 7.06E-08 | -0.4345 | 0.5 | 0.75 | 0.001127 | MS.Trt_vs_MS.NoTrt_F |
| NCKAP1L | 7.14E-08 | -0.7131 | 0.701 | 0.86 | 0.00114 | MS.Trt_vs_MS.NoTrt_F |
| BAZ1B | 7.17E-08 | -0.8542 | 0.144 | 0.37 | 0.001145 | MS.Trt_vs_MS.NoTrt_F |
| USMG5 | 7.43E-08 | 0.41199 | 0.18 | 0.43 | 0.001186 | MS.Trt_vs_MS.NoTrt_F |
| TCEAL8 | 7.48E-08 | -0.4896 | 0.094 | 0.29 | 0.001195 | MS.Trt_vs_MS.NoTrt_F |
| LIG1 | 7.48E-08 | -1.2339 | 0.119 | 0.33 | 0.001196 | MS.Trt_vs_MS.NoTrt_F |
| NARS | 7.76E-08 | -0.7966 | 0.137 | 0.35 | 0.00124 | MS.Trt_vs_MS.NoTrt_F |
| COMMD1 | 8.26E-08 | 0.42387 | 0.493 | 0.82 | 0.00132 | MS.Trt_vs_MS.NoTrt_F |
| NDUFB8 | 8.74E-08 | -0.251 | 0.309 | 0.6 | 0.001396 | MS.Trt_vs_MS.NoTrt_F |
| C11orf98 | 9.03E-08 | -0.8473 | 0.122 | 0.33 | 0.001442 | MS.Trt_vs_MS.NoTrt_F |
| FAM118A | 9.06E-08 | -0.9157 | 0.522 | 0.71 | 0.001446 | MS.Trt_vs_MS.NoTrt_F |
| LMF2 | 9.68E-08 | -0.3183 | 0.219 | 0.47 | 0.001546 | MS.Trt_vs_MS.NoTrt_F |
| IL2RA | 9.88E-08 | -0.4478 | 0.288 | 0.54 | 0.001578 | MS.Trt_vs_MS.NoTrt_F |
| PDLIM2 | 1.02E-07 | -0.6394 | 0.629 | 0.85 | 0.001634 | MS.Trt_vs_MS.NoTrt_F |
| LDHB | 1.02E-07 | -0.4051 | 0.579 | 0.85 | 0.001637 | MS.Trt_vs_MS.NoTrt_F |
| WWOX | 1.03E-07 | -0.3895 | 0.194 | 0.43 | 0.001653 | MS.Trt_vs_MS.NoTrt_F |
| SYNE2 | 1.05E-07 | -0.4536 | 0.942 | 0.98 | 0.001681 | MS.Trt_vs_MS.NoTrt_F |

| | | | | | | |
|---|---|---|---|---|---|---|
| SERBP1 | 1.09E-07 | -0.461 | 0.507 | 0.74 | 0.001743 | MS.Trt_vs_MS.NoTrt_F |
| ZNF142 | 1.10E-07 | -1.4273 | 0.083 | 0.27 | 0.001763 | MS.Trt_vs_MS.NoTrt_F |
| FAM46C | 1.11E-07 | -0.5427 | 0.309 | 0.57 | 0.00178 | MS.Trt_vs_MS.NoTrt_F |
| NCK2 | 1.15E-07 | -0.7539 | 0.198 | 0.43 | 0.001834 | MS.Trt_vs_MS.NoTrt_F |
| COL8A1 | 1.23E-07 | 0.51738 | 0.957 | 0.81 | 0.00196 | MS.Trt_vs_MS.NoTrt_F |
| JAKMIP1 | 1.25E-07 | -0.276 | 0.205 | 0.45 | 0.001998 | MS.Trt_vs_MS.NoTrt_F |
| ATF2 | 1.30E-07 | -1.0264 | 0.176 | 0.41 | 0.002073 | MS.Trt_vs_MS.NoTrt_F |
| CD3E | 1.37E-07 | -0.6649 | 0.626 | 0.85 | 0.002187 | MS.Trt_vs_MS.NoTrt_F |
| GABARAPL1 | 1.40E-07 | -1.0894 | 0.129 | 0.33 | 0.00223 | MS.Trt_vs_MS.NoTrt_F |
| CDC14A | 1.40E-07 | -0.4047 | 0.183 | 0.42 | 0.002231 | MS.Trt_vs_MS.NoTrt_F |
| CASP8 | 1.46E-07 | -0.7687 | 0.91 | 0.97 | 0.002335 | MS.Trt_vs_MS.NoTrt_F |
| SUPT5H | 1.53E-07 | -1.2279 | 0.36 | 0.59 | 0.002449 | MS.Trt_vs_MS.NoTrt_F |
| BTN3A2 | 1.56E-07 | -0.2569 | 0.701 | 0.85 | 0.002492 | MS.Trt_vs_MS.NoTrt_F |
| CNN2 | 1.59E-07 | -0.4948 | 0.964 | 0.98 | 0.002545 | MS.Trt_vs_MS.NoTrt_F |
| RFTN1 | 1.60E-07 | -0.3442 | 0.299 | 0.55 | 0.00256 | MS.Trt_vs_MS.NoTrt_F |
| ANTXR2 | 1.63E-07 | -0.3866 | 0.644 | 0.83 | 0.002606 | MS.Trt_vs_MS.NoTrt_F |
| TRERF1 | 1.63E-07 | -0.5838 | 0.112 | 0.32 | 0.002612 | MS.Trt_vs_MS.NoTrt_F |
| FAN1 | 1.71E-07 | 0.51251 | 0.209 | 0.44 | 0.002731 | MS.Trt_vs_MS.NoTrt_F |
| LGALS3 | 1.84E-07 | -0.4614 | 0.579 | 0.77 | 0.002937 | MS.Trt_vs_MS.NoTrt_F |
| DYNLL1 | 1.92E-07 | -0.3501 | 0.335 | 0.58 | 0.003068 | MS.Trt_vs_MS.NoTrt_F |
| MTMR6 | 2.06E-07 | -1.3148 | 0.104 | 0.29 | 0.003294 | MS.Trt_vs_MS.NoTrt_F |
| MED26 | 2.10E-07 | -1.3237 | 0.058 | 0.22 | 0.00336 | MS.Trt_vs_MS.NoTrt_F |
| GABRE | 2.16E-07 | -0.5648 | 0.129 | 0.32 | 0.003447 | MS.Trt_vs_MS.NoTrt_F |
| USP36 | 2.17E-07 | -0.314 | 0.353 | 0.62 | 0.00347 | MS.Trt_vs_MS.NoTrt_F |
| POLR3D | 2.30E-07 | -0.7725 | 0.345 | 0.58 | 0.003672 | MS.Trt_vs_MS.NoTrt_F |
| BBX | 2.32E-07 | -0.2961 | 0.273 | 0.54 | 0.003704 | MS.Trt_vs_MS.NoTrt_F |
| PAX6 | 2.33E-07 | -0.5186 | 0.281 | 0.5 | 0.003717 | MS.Trt_vs_MS.NoTrt_F |
| TMBIM6 | 2.35E-07 | -0.3487 | 0.745 | 0.91 | 0.003758 | MS.Trt_vs_MS.NoTrt_F |
| ATP5B | 2.40E-07 | -0.4615 | 0.345 | 0.61 | 0.003832 | MS.Trt_vs_MS.NoTrt_F |
| TAF7 | 2.42E-07 | -0.5259 | 0.288 | 0.53 | 0.003873 | MS.Trt_vs_MS.NoTrt_F |
| CWC22 | 2.50E-07 | -0.5612 | 0.115 | 0.31 | 0.003992 | MS.Trt_vs_MS.NoTrt_F |
| CRY1 | 2.51E-07 | -0.946 | 0.133 | 0.34 | 0.004006 | MS.Trt_vs_MS.NoTrt_F |
| MTM1 | 2.56E-07 | -0.51 | 0.101 | 0.29 | 0.004081 | MS.Trt_vs_MS.NoTrt_F |
| PLA2G7 | 2.63E-07 | -0.7442 | 0.482 | 0.71 | 0.004208 | MS.Trt_vs_MS.NoTrt_F |
| CYP4F11 | 2.67E-07 | -0.5358 | 0.302 | 0.52 | 0.004271 | MS.Trt_vs_MS.NoTrt_F |
| RALGDS | 2.78E-07 | -0.6182 | 0.27 | 0.5 | 0.004439 | MS.Trt_vs_MS.NoTrt_F |
| SPOCK2 | 2.88E-07 | -0.3813 | 0.55 | 0.76 | 0.004594 | MS.Trt_vs_MS.NoTrt_F |
| TNFRSF1B | 2.88E-07 | -0.3266 | 0.759 | 0.91 | 0.004603 | MS.Trt_vs_MS.NoTrt_F |
| ANXA11 | 2.89E-07 | -0.422 | 0.799 | 0.91 | 0.004615 | MS.Trt_vs_MS.NoTrt_F |
| RTN4 | 2.90E-07 | -0.6895 | 0.23 | 0.45 | 0.004633 | MS.Trt_vs_MS.NoTrt_F |

| | | | | | | |
|---|---|---|---|---|---|---|
| UBE2D3 | 2.91E-07 | -0.3038 | 0.612 | 0.82 | 0.004648 | MS.Trt_vs_MS.NoTrt_F |
| CSRNP1 | 2.96E-07 | -0.6089 | 0.147 | 0.37 | 0.004724 | MS.Trt_vs_MS.NoTrt_F |
| STK4 | 2.97E-07 | -0.3783 | 0.863 | 0.96 | 0.00475 | MS.Trt_vs_MS.NoTrt_F |
| PRKAR1A | 2.98E-07 | -0.483 | 0.281 | 0.52 | 0.004754 | MS.Trt_vs_MS.NoTrt_F |
| P2RY8 | 3.08E-07 | -0.6227 | 0.374 | 0.61 | 0.004925 | MS.Trt_vs_MS.NoTrt_F |
| SNAP47 | 3.16E-07 | -0.8249 | 0.122 | 0.32 | 0.005041 | MS.Trt_vs_MS.NoTrt_F |
| EEF2 | 3.17E-07 | -0.4488 | 0.817 | 0.94 | 0.005061 | MS.Trt_vs_MS.NoTrt_F |
| HTATIP2 | 3.19E-07 | -0.467 | 0.151 | 0.37 | 0.005103 | MS.Trt_vs_MS.NoTrt_F |
| CCNH | 3.21E-07 | -0.3592 | 0.223 | 0.46 | 0.005133 | MS.Trt_vs_MS.NoTrt_F |
| PREX1 | 3.26E-07 | -0.6786 | 0.518 | 0.73 | 0.00521 | MS.Trt_vs_MS.NoTrt_F |
| MT2A | 3.27E-07 | -0.3064 | 0.198 | 0.43 | 0.00522 | MS.Trt_vs_MS.NoTrt_F |
| HNRNPA1 | 3.27E-07 | -0.3309 | 0.629 | 0.92 | 0.00522 | MS.Trt_vs_MS.NoTrt_F |
| DUSP28 | 3.28E-07 | -0.5914 | 0.637 | 0.83 | 0.005232 | MS.Trt_vs_MS.NoTrt_F |
| RAD21 | 3.33E-07 | -0.4741 | 0.309 | 0.57 | 0.005318 | MS.Trt_vs_MS.NoTrt_F |
| ZNF253 | 3.36E-07 | 1.0086 | 0.831 | 0.59 | 0.00536 | MS.Trt_vs_MS.NoTrt_F |
| MAN1A2 | 3.60E-07 | -0.6684 | 0.245 | 0.48 | 0.005758 | MS.Trt_vs_MS.NoTrt_F |
| ZFP36 | 3.62E-07 | -0.5591 | 0.41 | 0.68 | 0.005781 | MS.Trt_vs_MS.NoTrt_F |
| LITAF | 3.64E-07 | -0.4191 | 0.46 | 0.7 | 0.005819 | MS.Trt_vs_MS.NoTrt_F |
| MYH9 | 3.84E-07 | -0.3555 | 0.719 | 0.95 | 0.006131 | MS.Trt_vs_MS.NoTrt_F |
| WNK1 | 3.88E-07 | -0.3963 | 0.464 | 0.72 | 0.006193 | MS.Trt_vs_MS.NoTrt_F |
| RP11-717K1 | 3.91E-07 | -0.6735 | 0.079 | 0.25 | 0.006252 | MS.Trt_vs_MS.NoTrt_F |
| SCAP | 4.26E-07 | -0.347 | 0.554 | 0.77 | 0.00681 | MS.Trt_vs_MS.NoTrt_F |
| SLAMF1 | 4.29E-07 | -0.5191 | 0.219 | 0.45 | 0.006858 | MS.Trt_vs_MS.NoTrt_F |
| CTLA4 | 4.41E-07 | -0.3525 | 0.234 | 0.45 | 0.007041 | MS.Trt_vs_MS.NoTrt_F |
| OSER1 | 4.60E-07 | -0.4581 | 0.18 | 0.4 | 0.007354 | MS.Trt_vs_MS.NoTrt_F |
| TIGIT | 4.64E-07 | -0.3498 | 0.313 | 0.55 | 0.007419 | MS.Trt_vs_MS.NoTrt_F |
| CCDC7 | 4.68E-07 | -0.3821 | 0.101 | 0.29 | 0.007474 | MS.Trt_vs_MS.NoTrt_F |
| ABHD14B | 4.82E-07 | -0.3434 | 0.165 | 0.39 | 0.0077 | MS.Trt_vs_MS.NoTrt_F |
| SEL1L | 4.86E-07 | -0.2864 | 0.169 | 0.39 | 0.007758 | MS.Trt_vs_MS.NoTrt_F |
| THAP4 | 4.89E-07 | -0.253 | 0.104 | 0.29 | 0.007805 | MS.Trt_vs_MS.NoTrt_F |
| CBLL1 | 5.01E-07 | -0.3072 | 0.209 | 0.44 | 0.007998 | MS.Trt_vs_MS.NoTrt_F |
| DOCK2 | 5.36E-07 | -0.3123 | 0.356 | 0.63 | 0.008566 | MS.Trt_vs_MS.NoTrt_F |
| C2CD3 | 5.51E-07 | 0.48066 | 0.986 | 0.99 | 0.008797 | MS.Trt_vs_MS.NoTrt_F |
| SGMS1 | 5.86E-07 | -0.8108 | 0.183 | 0.4 | 0.009368 | MS.Trt_vs_MS.NoTrt_F |
| ZRANB2 | 6.17E-07 | -0.3782 | 0.273 | 0.51 | 0.009849 | MS.Trt_vs_MS.NoTrt_F |
| PIP4K2C | 6.34E-07 | -0.6719 | 0.14 | 0.34 | 0.010132 | MS.Trt_vs_MS.NoTrt_F |
| CCR5 | 6.46E-07 | -1.0556 | 0.086 | 0.25 | 0.010318 | MS.Trt_vs_MS.NoTrt_F |
| HSP90AA1 | 6.56E-07 | -0.2632 | 0.673 | 0.85 | 0.010477 | MS.Trt_vs_MS.NoTrt_F |
| MT-ND4 | 6.61E-07 | 0.54094 | 1 | 1 | 0.010556 | MS.Trt_vs_MS.NoTrt_F |
| C12orf4 | 6.66E-07 | -0.7737 | 0.471 | 0.72 | 0.010631 | MS.Trt_vs_MS.NoTrt_F |

| | | | | | | |
|---|---|---|---|---|---|---|
| HSPA9 | 6.93E-07 | -0.5103 | 0.439 | 0.66 | 0.011074 | MS.Trt_vs_MS.NoTrt_F |
| RUNX2 | 7.05E-07 | -0.6034 | 0.158 | 0.37 | 0.011255 | MS.Trt_vs_MS.NoTrt_F |
| ZCCHC2 | 7.10E-07 | -2.4033 | 0.072 | 0.23 | 0.011346 | MS.Trt_vs_MS.NoTrt_F |
| ENTPD6 | 7.33E-07 | -0.9938 | 0.112 | 0.29 | 0.011707 | MS.Trt_vs_MS.NoTrt_F |
| GPBP1 | 7.47E-07 | -0.338 | 0.273 | 0.52 | 0.011936 | MS.Trt_vs_MS.NoTrt_F |
| PHC3 | 7.94E-07 | -0.4104 | 0.968 | 0.98 | 0.012677 | MS.Trt_vs_MS.NoTrt_F |
| NT5C | 8.15E-07 | -0.5471 | 0.129 | 0.32 | 0.013024 | MS.Trt_vs_MS.NoTrt_F |
| OGDH | 8.31E-07 | -0.6237 | 0.464 | 0.69 | 0.013279 | MS.Trt_vs_MS.NoTrt_F |
| ACSL4 | 8.75E-07 | -1.3532 | 0.133 | 0.33 | 0.013978 | MS.Trt_vs_MS.NoTrt_F |
| RASGRP1 | 8.90E-07 | -0.3072 | 0.345 | 0.6 | 0.014217 | MS.Trt_vs_MS.NoTrt_F |
| ITFG1 | 9.79E-07 | -0.8629 | 0.165 | 0.37 | 0.015635 | MS.Trt_vs_MS.NoTrt_F |
| ANKRD13C | 9.86E-07 | 0.52609 | 0.122 | 0.33 | 0.015753 | MS.Trt_vs_MS.NoTrt_F |
| UBE2L6 | 9.86E-07 | -0.3732 | 0.209 | 0.44 | 0.015756 | MS.Trt_vs_MS.NoTrt_F |
| ZC3H3 | 9.87E-07 | -1.0285 | 0.115 | 0.29 | 0.015758 | MS.Trt_vs_MS.NoTrt_F |
| TBC1D23 | 1.01E-06 | -0.5633 | 0.133 | 0.33 | 0.016101 | MS.Trt_vs_MS.NoTrt_F |
| SLFN5 | 1.02E-06 | -0.6477 | 0.856 | 0.93 | 0.016327 | MS.Trt_vs_MS.NoTrt_F |
| C6orf62 | 1.06E-06 | 0.95396 | 0.838 | 0.85 | 0.016962 | MS.Trt_vs_MS.NoTrt_F |
| BATF | 1.09E-06 | -0.2626 | 0.137 | 0.34 | 0.017422 | MS.Trt_vs_MS.NoTrt_F |
| RBX1 | 1.10E-06 | 0.45747 | 0.263 | 0.54 | 0.017552 | MS.Trt_vs_MS.NoTrt_F |
| AFF1 | 1.15E-06 | -0.5322 | 0.284 | 0.51 | 0.018335 | MS.Trt_vs_MS.NoTrt_F |
| CATIP | 1.19E-06 | 0.59942 | 0.892 | 0.91 | 0.019028 | MS.Trt_vs_MS.NoTrt_F |
| ARHGEF10 | 1.23E-06 | 0.64779 | 0.791 | 0.54 | 0.019686 | MS.Trt_vs_MS.NoTrt_F |
| COPS7A | 1.25E-06 | -0.7612 | 0.23 | 0.44 | 0.019952 | MS.Trt_vs_MS.NoTrt_F |
| SLC51A | 1.26E-06 | 0.28623 | 0.888 | 0.73 | 0.020131 | MS.Trt_vs_MS.NoTrt_F |
| STMN3 | 1.27E-06 | -0.716 | 0.183 | 0.39 | 0.020285 | MS.Trt_vs_MS.NoTrt_F |
| IDS | 1.27E-06 | -0.3303 | 0.748 | 0.88 | 0.020297 | MS.Trt_vs_MS.NoTrt_F |
| HS3ST3B1 | 1.28E-06 | -0.2562 | 0.518 | 0.71 | 0.020464 | MS.Trt_vs_MS.NoTrt_F |
| RPSA | 1.28E-06 | -0.3387 | 0.59 | 0.82 | 0.020524 | MS.Trt_vs_MS.NoTrt_F |
| STAT1 | 1.30E-06 | -0.2641 | 0.335 | 0.58 | 0.020723 | MS.Trt_vs_MS.NoTrt_F |
| RAD54L2 | 1.34E-06 | -0.5621 | 0.665 | 0.81 | 0.021445 | MS.Trt_vs_MS.NoTrt_F |
| NMRK1 | 1.37E-06 | 0.32155 | 0.223 | 0.44 | 0.021826 | MS.Trt_vs_MS.NoTrt_F |
| TRIM37 | 1.39E-06 | -0.6489 | 0.187 | 0.38 | 0.022177 | MS.Trt_vs_MS.NoTrt_F |
| FAM153A | 1.42E-06 | -0.4392 | 0.403 | 0.65 | 0.022664 | MS.Trt_vs_MS.NoTrt_F |
| TMEM63A | 1.44E-06 | -0.3068 | 0.665 | 0.88 | 0.022984 | MS.Trt_vs_MS.NoTrt_F |
| TNIP1 | 1.46E-06 | -0.5368 | 0.295 | 0.52 | 0.023311 | MS.Trt_vs_MS.NoTrt_F |
| PEBP1 | 1.47E-06 | -0.5052 | 0.385 | 0.6 | 0.023538 | MS.Trt_vs_MS.NoTrt_F |
| HEBP2 | 1.51E-06 | -0.5608 | 0.324 | 0.53 | 0.024113 | MS.Trt_vs_MS.NoTrt_F |
| STAT5A | 1.54E-06 | -0.7462 | 0.133 | 0.33 | 0.024566 | MS.Trt_vs_MS.NoTrt_F |
| ACAD9 | 1.55E-06 | -1.0522 | 0.216 | 0.42 | 0.024718 | MS.Trt_vs_MS.NoTrt_F |
| HDAC10 | 1.56E-06 | -1.5756 | 0.133 | 0.32 | 0.024971 | MS.Trt_vs_MS.NoTrt_F |

| | | | | | | |
|---|---|---|---|---|---|---|
| RP11-307N1 | 1.59E-06 | -0.7718 | 0.079 | 0.24 | 0.025328 | MS.Trt_vs_MS.NoTrt_F |
| CXXC1 | 1.68E-06 | -0.5942 | 0.158 | 0.37 | 0.02683 | MS.Trt_vs_MS.NoTrt_F |
| CRIP2 | 1.70E-06 | -0.2554 | 0.126 | 0.32 | 0.02717 | MS.Trt_vs_MS.NoTrt_F |
| WAS | 1.72E-06 | -0.4048 | 0.259 | 0.46 | 0.027518 | MS.Trt_vs_MS.NoTrt_F |
| RPS6KA1 | 1.75E-06 | -0.7667 | 0.169 | 0.37 | 0.028019 | MS.Trt_vs_MS.NoTrt_F |
| AATF | 1.80E-06 | -0.5152 | 0.158 | 0.36 | 0.028713 | MS.Trt_vs_MS.NoTrt_F |
| RALA | 1.82E-06 | -0.6273 | 0.155 | 0.35 | 0.029024 | MS.Trt_vs_MS.NoTrt_F |
| TBC1D14 | 1.89E-06 | -0.2756 | 0.176 | 0.39 | 0.030199 | MS.Trt_vs_MS.NoTrt_F |
| SLC3A2 | 1.89E-06 | -0.3793 | 0.453 | 0.66 | 0.030252 | MS.Trt_vs_MS.NoTrt_F |
| RARRES3 | 1.90E-06 | -0.3253 | 0.237 | 0.48 | 0.030277 | MS.Trt_vs_MS.NoTrt_F |
| UBQLN2 | 1.94E-06 | -0.5888 | 0.209 | 0.42 | 0.030942 | MS.Trt_vs_MS.NoTrt_F |
| HNRNPDL | 1.98E-06 | -0.2682 | 0.514 | 0.77 | 0.031631 | MS.Trt_vs_MS.NoTrt_F |
| RFX7 | 2.00E-06 | -0.3954 | 0.147 | 0.34 | 0.031898 | MS.Trt_vs_MS.NoTrt_F |
| CDH8 | 2.07E-06 | -0.3797 | 0.36 | 0.59 | 0.033041 | MS.Trt_vs_MS.NoTrt_F |
| AF011889.5 | 2.13E-06 | -0.5481 | 0.173 | 0.37 | 0.034006 | MS.Trt_vs_MS.NoTrt_F |
| PGK1 | 2.15E-06 | -0.3042 | 0.568 | 0.79 | 0.034265 | MS.Trt_vs_MS.NoTrt_F |
| SARS | 2.17E-06 | -0.2928 | 0.223 | 0.46 | 0.034614 | MS.Trt_vs_MS.NoTrt_F |
| ACOT8 | 2.17E-06 | -0.7102 | 0.09 | 0.25 | 0.034715 | MS.Trt_vs_MS.NoTrt_F |
| AKIRIN1 | 2.27E-06 | -0.3715 | 0.45 | 0.68 | 0.03622 | MS.Trt_vs_MS.NoTrt_F |
| TNFAIP8L2 | 2.32E-06 | -1.1085 | 0.086 | 0.25 | 0.036984 | MS.Trt_vs_MS.NoTrt_F |
| WHRN | 2.33E-06 | -0.5712 | 0.108 | 0.28 | 0.03717 | MS.Trt_vs_MS.NoTrt_F |
| R3HCC1 | 2.33E-06 | -0.9991 | 0.108 | 0.28 | 0.037226 | MS.Trt_vs_MS.NoTrt_F |
| ARF5 | 2.39E-06 | -0.2726 | 0.194 | 0.41 | 0.038127 | MS.Trt_vs_MS.NoTrt_F |
| MAF1 | 2.51E-06 | -0.2733 | 0.158 | 0.36 | 0.040164 | MS.Trt_vs_MS.NoTrt_F |
| CD48 | 2.64E-06 | -0.3275 | 0.252 | 0.49 | 0.042095 | MS.Trt_vs_MS.NoTrt_F |
| RIMS1 | 2.72E-06 | -0.4521 | 0.212 | 0.4 | 0.043438 | MS.Trt_vs_MS.NoTrt_F |
| PFKP | 2.84E-06 | -0.5478 | 0.23 | 0.44 | 0.0453 | MS.Trt_vs_MS.NoTrt_F |
| PPP1R1C | 2.84E-06 | -0.4195 | 0.23 | 0.43 | 0.045408 | MS.Trt_vs_MS.NoTrt_F |
| SNRNP40 | 2.95E-06 | -0.3081 | 0.165 | 0.36 | 0.047082 | MS.Trt_vs_MS.NoTrt_F |
| HDLBP | 3.11E-06 | -0.477 | 0.432 | 0.6 | 0.049653 | MS.Trt_vs_MS.NoTrt_F |
| EIF4A3 | 3.15E-06 | -1.2425 | 0.665 | 0.8 | 0.050276 | MS.Trt_vs_MS.NoTrt_F |
| FMNL2 | 3.22E-06 | -0.615 | 0.173 | 0.35 | 0.051485 | MS.Trt_vs_MS.NoTrt_F |
| FNBP4 | 3.23E-06 | -0.5601 | 0.424 | 0.64 | 0.05158 | MS.Trt_vs_MS.NoTrt_F |
| EXOC3L4 | 3.31E-06 | -0.8435 | 0.137 | 0.31 | 0.05285 | MS.Trt_vs_MS.NoTrt_F |
| ANP32A | 3.36E-06 | -0.9836 | 0.205 | 0.4 | 0.053653 | MS.Trt_vs_MS.NoTrt_F |
| ANO9 | 3.46E-06 | -0.4724 | 0.227 | 0.44 | 0.055336 | MS.Trt_vs_MS.NoTrt_F |
| TRAF1 | 3.48E-06 | 0.26531 | 0.273 | 0.51 | 0.055526 | MS.Trt_vs_MS.NoTrt_F |
| MYO9A | 3.62E-06 | -0.5897 | 0.27 | 0.49 | 0.057817 | MS.Trt_vs_MS.NoTrt_F |
| EPM2AIP1 | 3.62E-06 | -0.7157 | 0.23 | 0.42 | 0.057893 | MS.Trt_vs_MS.NoTrt_F |
| PRKG2 | 3.69E-06 | -0.7843 | 0.295 | 0.54 | 0.058979 | MS.Trt_vs_MS.NoTrt_F |

| | | | | | | |
|---|---|---|---|---|---|---|
| PSMD7 | 3.99E-06 | -0.5322 | 0.266 | 0.48 | 0.06381 | MS.Trt_vs_MS.NoTrt_F |
| SLC41A3 | 4.03E-06 | -0.5993 | 0.273 | 0.49 | 0.064435 | MS.Trt_vs_MS.NoTrt_F |
| GMIP | 4.15E-06 | -1.0644 | 0.191 | 0.39 | 0.066316 | MS.Trt_vs_MS.NoTrt_F |
| UBL7 | 4.23E-06 | -0.5152 | 0.137 | 0.32 | 0.067592 | MS.Trt_vs_MS.NoTrt_F |
| SYT17 | 4.32E-06 | -0.37 | 0.277 | 0.48 | 0.069051 | MS.Trt_vs_MS.NoTrt_F |
| PTAR1 | 4.39E-06 | -0.8272 | 0.137 | 0.32 | 0.07019 | MS.Trt_vs_MS.NoTrt_F |
| C15orf53 | 4.46E-06 | -0.8766 | 0.115 | 0.28 | 0.071282 | MS.Trt_vs_MS.NoTrt_F |
| EIF3D | 4.63E-06 | -0.3157 | 0.371 | 0.6 | 0.073948 | MS.Trt_vs_MS.NoTrt_F |
| TMED7 | 4.68E-06 | -1.2323 | 0.104 | 0.27 | 0.074803 | MS.Trt_vs_MS.NoTrt_F |
| ADSS | 4.70E-06 | -0.2727 | 0.137 | 0.32 | 0.075063 | MS.Trt_vs_MS.NoTrt_F |
| RRM1 | 4.72E-06 | -0.2633 | 0.331 | 0.55 | 0.075439 | MS.Trt_vs_MS.NoTrt_F |
| SP2 | 4.76E-06 | -1.3482 | 0.277 | 0.48 | 0.076039 | MS.Trt_vs_MS.NoTrt_F |
| MSANTD4 | 4.79E-06 | -1.6116 | 0.058 | 0.19 | 0.076548 | MS.Trt_vs_MS.NoTrt_F |
| IKBKE | 4.80E-06 | -0.2959 | 0.291 | 0.53 | 0.076641 | MS.Trt_vs_MS.NoTrt_F |
| ULK3 | 4.86E-06 | -0.5578 | 0.144 | 0.33 | 0.07768 | MS.Trt_vs_MS.NoTrt_F |
| LAMP1 | 4.86E-06 | -0.9513 | 0.349 | 0.54 | 0.077687 | MS.Trt_vs_MS.NoTrt_F |
| BORCS5 | 4.94E-06 | -0.5977 | 0.597 | 0.79 | 0.07893 | MS.Trt_vs_MS.NoTrt_F |
| RPL7A | 5.01E-06 | -0.3246 | 0.68 | 0.92 | 0.080023 | MS.Trt_vs_MS.NoTrt_F |
| ASUN | 5.10E-06 | -0.5755 | 0.126 | 0.3 | 0.081495 | MS.Trt_vs_MS.NoTrt_F |
| EHD2 | 5.24E-06 | -0.6523 | 0.799 | 0.91 | 0.083676 | MS.Trt_vs_MS.NoTrt_F |
| GIMAP2 | 5.31E-06 | -0.5109 | 0.853 | 0.91 | 0.084757 | MS.Trt_vs_MS.NoTrt_F |
| BTG1 | 5.32E-06 | -0.3241 | 0.619 | 0.85 | 0.084955 | MS.Trt_vs_MS.NoTrt_F |
| NDUFA4 | 5.56E-06 | 0.26861 | 0.554 | 0.77 | 0.08876 | MS.Trt_vs_MS.NoTrt_F |
| NDUFA6 | 5.61E-06 | -0.284 | 0.23 | 0.45 | 0.089588 | MS.Trt_vs_MS.NoTrt_F |
| RALGAPB | 5.61E-06 | -0.2993 | 0.18 | 0.38 | 0.089659 | MS.Trt_vs_MS.NoTrt_F |
| SLC44A5 | 5.64E-06 | 0.49139 | 0.817 | 0.83 | 0.090063 | MS.Trt_vs_MS.NoTrt_F |
| INTS9 | 5.65E-06 | -0.8819 | 0.112 | 0.28 | 0.090264 | MS.Trt_vs_MS.NoTrt_F |
| LBH | 5.65E-06 | -0.3713 | 0.651 | 0.82 | 0.090301 | MS.Trt_vs_MS.NoTrt_F |
| RPL13A | 5.66E-06 | -0.2924 | 0.813 | 0.99 | 0.090343 | MS.Trt_vs_MS.NoTrt_F |
| POLR2I | 5.82E-06 | -0.4556 | 0.133 | 0.31 | 0.092904 | MS.Trt_vs_MS.NoTrt_F |
| CHCHD3 | 5.97E-06 | -0.7049 | 0.129 | 0.3 | 0.09528 | MS.Trt_vs_MS.NoTrt_F |
| NFATC3 | 6.13E-06 | -0.2793 | 0.629 | 0.77 | 0.097876 | MS.Trt_vs_MS.NoTrt_F |
| PCGF3 | 6.13E-06 | -0.6277 | 0.173 | 0.36 | 0.097973 | MS.Trt_vs_MS.NoTrt_F |
| FBXW5 | 6.24E-06 | -0.3098 | 0.194 | 0.4 | 0.099735 | MS.Trt_vs_MS.NoTrt_F |
| TCERG1 | 6.30E-06 | -0.345 | 0.248 | 0.46 | 0.100566 | MS.Trt_vs_MS.NoTrt_F |
| BTLA | 6.35E-06 | 1.10494 | 0.122 | 0.28 | 0.101419 | MS.Trt_vs_MS.NoTrt_F |
| NEBL | 6.40E-06 | -0.9706 | 0.216 | 0.42 | 0.102189 | MS.Trt_vs_MS.NoTrt_F |
| ACLY | 6.61E-06 | -0.4883 | 0.201 | 0.4 | 0.105529 | MS.Trt_vs_MS.NoTrt_F |
| UTRN | 6.70E-06 | -0.6076 | 0.421 | 0.62 | 0.106951 | MS.Trt_vs_MS.NoTrt_F |
| BST2 | 6.72E-06 | -0.4297 | 0.155 | 0.34 | 0.107345 | MS.Trt_vs_MS.NoTrt_F |

| | | | | | | |
|---|---|---|---|---|---|---|
| PRRC2B | 6.79E-06 | -0.5413 | 0.245 | 0.46 | 0.108385 | MS.Trt_vs_MS.NoTrt_F |
| EPHB4 | 6.85E-06 | 0.66496 | 0.813 | 0.8 | 0.109483 | MS.Trt_vs_MS.NoTrt_F |
| RNF20 | 6.89E-06 | -1.0145 | 0.122 | 0.29 | 0.10999 | MS.Trt_vs_MS.NoTrt_F |
| UBXN11 | 6.98E-06 | -0.2939 | 0.173 | 0.37 | 0.111416 | MS.Trt_vs_MS.NoTrt_F |
| PTP4A1 | 7.02E-06 | -0.3228 | 0.198 | 0.4 | 0.1122 | MS.Trt_vs_MS.NoTrt_F |
| COX10 | 7.24E-06 | -0.9099 | 0.101 | 0.26 | 0.115716 | MS.Trt_vs_MS.NoTrt_F |
| NDUFC1 | 7.30E-06 | 0.33226 | 0.191 | 0.4 | 0.116546 | MS.Trt_vs_MS.NoTrt_F |
| RASA1 | 7.30E-06 | -0.6371 | 0.191 | 0.38 | 0.116581 | MS.Trt_vs_MS.NoTrt_F |
| SF3A1 | 7.36E-06 | -0.4883 | 0.277 | 0.48 | 0.117534 | MS.Trt_vs_MS.NoTrt_F |
| CEP85L | 7.42E-06 | -0.574 | 0.219 | 0.41 | 0.118447 | MS.Trt_vs_MS.NoTrt_F |
| C2orf91 | 7.65E-06 | -0.3872 | 0.403 | 0.59 | 0.122131 | MS.Trt_vs_MS.NoTrt_F |
| HDAC3 | 7.70E-06 | -0.4094 | 0.122 | 0.29 | 0.122918 | MS.Trt_vs_MS.NoTrt_F |
| CD7 | 7.70E-06 | -0.3937 | 0.219 | 0.41 | 0.122991 | MS.Trt_vs_MS.NoTrt_F |
| TADA3 | 7.87E-06 | -0.7092 | 0.529 | 0.74 | 0.125733 | MS.Trt_vs_MS.NoTrt_F |
| GIMAP1 | 7.88E-06 | -0.5206 | 0.378 | 0.58 | 0.125801 | MS.Trt_vs_MS.NoTrt_F |
| AKAP8L | 7.89E-06 | -0.7831 | 0.097 | 0.25 | 0.126069 | MS.Trt_vs_MS.NoTrt_F |
| UQCRFS1 | 7.89E-06 | -0.4252 | 0.194 | 0.39 | 0.126101 | MS.Trt_vs_MS.NoTrt_F |
| YJEFN3 | 7.90E-06 | -2.4733 | 0.054 | 0.18 | 0.126217 | MS.Trt_vs_MS.NoTrt_F |
| SRA1 | 8.00E-06 | -0.5322 | 0.126 | 0.29 | 0.127713 | MS.Trt_vs_MS.NoTrt_F |
| RIN3 | 8.01E-06 | -0.5432 | 0.176 | 0.37 | 0.127893 | MS.Trt_vs_MS.NoTrt_F |
| SNX9 | 8.23E-06 | -0.47 | 0.14 | 0.32 | 0.131455 | MS.Trt_vs_MS.NoTrt_F |
| MKRN2 | 8.32E-06 | -0.9362 | 0.09 | 0.24 | 0.13292 | MS.Trt_vs_MS.NoTrt_F |
| CECR1 | 8.44E-06 | -1.0564 | 0.363 | 0.59 | 0.134874 | MS.Trt_vs_MS.NoTrt_F |
| HAX1 | 8.86E-06 | -0.2697 | 0.18 | 0.38 | 0.141596 | MS.Trt_vs_MS.NoTrt_F |
| SAMD9 | 8.94E-06 | -0.5043 | 0.147 | 0.33 | 0.142743 | MS.Trt_vs_MS.NoTrt_F |
| IL21R | 8.94E-06 | -0.6075 | 0.522 | 0.75 | 0.142877 | MS.Trt_vs_MS.NoTrt_F |
| TMX2 | 9.02E-06 | -0.596 | 0.115 | 0.28 | 0.144028 | MS.Trt_vs_MS.NoTrt_F |
| TTC14 | 9.18E-06 | -0.7947 | 0.353 | 0.54 | 0.1466 | MS.Trt_vs_MS.NoTrt_F |
| MPC1 | 9.27E-06 | -0.508 | 0.119 | 0.28 | 0.148081 | MS.Trt_vs_MS.NoTrt_F |
| ACBD6 | 9.40E-06 | -0.3147 | 0.335 | 0.55 | 0.150128 | MS.Trt_vs_MS.NoTrt_F |
| OTUD5 | 9.47E-06 | -0.2926 | 0.137 | 0.32 | 0.151303 | MS.Trt_vs_MS.NoTrt_F |
| NUTF2 | 9.53E-06 | -0.325 | 0.169 | 0.35 | 0.152187 | MS.Trt_vs_MS.NoTrt_F |
| ST8SIA3 | 9.56E-06 | -0.3519 | 0.112 | 0.27 | 0.15267 | MS.Trt_vs_MS.NoTrt_F |
| SOCS3 | 9.88E-06 | -0.4605 | 0.14 | 0.32 | 0.157867 | MS.Trt_vs_MS.NoTrt_F |
| ZNF331 | 1.00E-05 | -0.8128 | 0.615 | 0.78 | 0.160022 | MS.Trt_vs_MS.NoTrt_F |
| CLASRP | 1.00E-05 | -1.3558 | 0.147 | 0.32 | 0.160156 | MS.Trt_vs_MS.NoTrt_F |
| HSPE1 | 1.01E-05 | -0.2634 | 0.259 | 0.46 | 0.160722 | MS.Trt_vs_MS.NoTrt_F |
| RANBP2 | 1.01E-05 | -0.3854 | 0.349 | 0.6 | 0.162061 | MS.Trt_vs_MS.NoTrt_F |
| MYL12B | 1.02E-05 | -0.288 | 0.489 | 0.75 | 0.163259 | MS.Trt_vs_MS.NoTrt_F |
| SMYD3 | 1.04E-05 | -0.7628 | 0.147 | 0.33 | 0.165681 | MS.Trt_vs_MS.NoTrt_F |

| | | | | | | |
|---|---|---|---|---|---|---|
| JRK | 1.04E-05 | -1.4194 | 0.353 | 0.55 | 0.165867 | MS.Trt_vs_MS.NoTrt_F |
| GLTSCR1L | 1.04E-05 | -0.428 | 0.173 | 0.37 | 0.166405 | MS.Trt_vs_MS.NoTrt_F |
| ATG4B | 1.04E-05 | -0.5422 | 0.183 | 0.38 | 0.166529 | MS.Trt_vs_MS.NoTrt_F |
| ACAD11 | 1.05E-05 | -0.6811 | 0.165 | 0.35 | 0.167052 | MS.Trt_vs_MS.NoTrt_F |
| ITPA | 1.11E-05 | -0.3754 | 0.151 | 0.33 | 0.177926 | MS.Trt_vs_MS.NoTrt_F |
| PSMG2 | 1.14E-05 | -0.2719 | 0.165 | 0.35 | 0.181879 | MS.Trt_vs_MS.NoTrt_F |
| IRF2 | 1.15E-05 | -0.6034 | 0.32 | 0.53 | 0.184467 | MS.Trt_vs_MS.NoTrt_F |
| MAPK10 | 1.16E-05 | -0.2941 | 0.155 | 0.33 | 0.184931 | MS.Trt_vs_MS.NoTrt_F |
| LRRC37B | 1.16E-05 | 0.39094 | 0.583 | 0.8 | 0.185224 | MS.Trt_vs_MS.NoTrt_F |
| UBAP1 | 1.17E-05 | -0.4615 | 0.133 | 0.3 | 0.186689 | MS.Trt_vs_MS.NoTrt_F |
| CAMK4 | 1.20E-05 | -0.3033 | 0.586 | 0.76 | 0.191814 | MS.Trt_vs_MS.NoTrt_F |
| NSMAF | 1.20E-05 | -0.4508 | 0.198 | 0.39 | 0.191995 | MS.Trt_vs_MS.NoTrt_F |
| TOMM20 | 1.21E-05 | -0.3899 | 0.194 | 0.38 | 0.193445 | MS.Trt_vs_MS.NoTrt_F |
| TCP11L2 | 1.25E-05 | -0.8501 | 0.248 | 0.42 | 0.19992 | MS.Trt_vs_MS.NoTrt_F |
| KLF6 | 1.29E-05 | -0.319 | 0.68 | 0.87 | 0.205321 | MS.Trt_vs_MS.NoTrt_F |
| MUC15 | 1.31E-05 | -0.3522 | 0.104 | 0.25 | 0.209651 | MS.Trt_vs_MS.NoTrt_F |
| CHKB | 1.35E-05 | -0.5026 | 0.245 | 0.44 | 0.216175 | MS.Trt_vs_MS.NoTrt_F |
| LIMK2 | 1.35E-05 | -0.3477 | 0.187 | 0.38 | 0.216263 | MS.Trt_vs_MS.NoTrt_F |
| ZNF281 | 1.37E-05 | -0.9774 | 0.094 | 0.24 | 0.218283 | MS.Trt_vs_MS.NoTrt_F |
| PHF14 | 1.37E-05 | -1.1339 | 0.119 | 0.28 | 0.218961 | MS.Trt_vs_MS.NoTrt_F |
| PPP6R2 | 1.38E-05 | -0.3936 | 0.288 | 0.5 | 0.220395 | MS.Trt_vs_MS.NoTrt_F |
| ARRDC5 | 1.39E-05 | -0.3387 | 0.097 | 0.25 | 0.221865 | MS.Trt_vs_MS.NoTrt_F |
| TAF1C | 1.46E-05 | -1.0033 | 0.108 | 0.27 | 0.232549 | MS.Trt_vs_MS.NoTrt_F |
| TIFA | 1.49E-05 | -0.3176 | 0.104 | 0.27 | 0.237448 | MS.Trt_vs_MS.NoTrt_F |
| NIM1K | 1.49E-05 | 0.41583 | 0.799 | 0.62 | 0.238309 | MS.Trt_vs_MS.NoTrt_F |
| PISD | 1.51E-05 | -0.3669 | 0.119 | 0.28 | 0.240679 | MS.Trt_vs_MS.NoTrt_F |
| KIAA0141 | 1.51E-05 | -0.6194 | 0.133 | 0.3 | 0.241818 | MS.Trt_vs_MS.NoTrt_F |
| YWHAG | 1.54E-05 | -0.626 | 0.119 | 0.28 | 0.245747 | MS.Trt_vs_MS.NoTrt_F |
| CD3G | 1.54E-05 | 0.88709 | 0.953 | 0.98 | 0.246445 | MS.Trt_vs_MS.NoTrt_F |
| HNF4G | 1.55E-05 | -0.3148 | 0.223 | 0.41 | 0.246809 | MS.Trt_vs_MS.NoTrt_F |
| RARA | 1.56E-05 | -0.8803 | 0.108 | 0.26 | 0.248551 | MS.Trt_vs_MS.NoTrt_F |
| PNPLA6 | 1.59E-05 | 0.25931 | 0.122 | 0.29 | 0.25372 | MS.Trt_vs_MS.NoTrt_F |
| PTTG1IP | 1.61E-05 | -0.282 | 0.158 | 0.34 | 0.257901 | MS.Trt_vs_MS.NoTrt_F |
| GDF5OS | 1.62E-05 | -0.3288 | 0.299 | 0.49 | 0.258425 | MS.Trt_vs_MS.NoTrt_F |
| DNAH1 | 1.62E-05 | -0.4998 | 0.129 | 0.3 | 0.259221 | MS.Trt_vs_MS.NoTrt_F |
| EPSTI1 | 1.63E-05 | -0.5271 | 0.14 | 0.32 | 0.259684 | MS.Trt_vs_MS.NoTrt_F |
| SAMD9L | 1.64E-05 | -0.3602 | 0.162 | 0.34 | 0.261585 | MS.Trt_vs_MS.NoTrt_F |
| TMEM55A | 1.64E-05 | -1.459 | 0.068 | 0.2 | 0.262234 | MS.Trt_vs_MS.NoTrt_F |
| RASA4 | 1.67E-05 | 0.25124 | 0.586 | 0.81 | 0.26604 | MS.Trt_vs_MS.NoTrt_F |
| MAPK6 | 1.67E-05 | 0.75499 | 0.421 | 0.59 | 0.267054 | MS.Trt_vs_MS.NoTrt_F |

| | | | | | | |
|---|---|---|---|---|---|---|
| STIM2 | 1.69E-05 | -0.3947 | 0.137 | 0.3 | 0.269167 | MS.Trt_vs_MS.NoTrt_F |
| GCN1 | 1.70E-05 | -1.0178 | 0.119 | 0.28 | 0.272288 | MS.Trt_vs_MS.NoTrt_F |
| PGM2 | 1.72E-05 | -0.4546 | 0.108 | 0.27 | 0.275292 | MS.Trt_vs_MS.NoTrt_F |
| MRPL28 | 1.72E-05 | -0.8364 | 0.104 | 0.25 | 0.275319 | MS.Trt_vs_MS.NoTrt_F |
| 4-Sep | 1.73E-05 | 0.32054 | 0.331 | 0.5 | 0.276835 | MS.Trt_vs_MS.NoTrt_F |
| PRKACB | 1.74E-05 | -0.4112 | 0.216 | 0.41 | 0.277414 | MS.Trt_vs_MS.NoTrt_F |
| HSPA4 | 1.79E-05 | -0.6283 | 0.349 | 0.53 | 0.286693 | MS.Trt_vs_MS.NoTrt_F |
| LDLRAD4 | 1.87E-05 | -0.6048 | 0.353 | 0.55 | 0.298788 | MS.Trt_vs_MS.NoTrt_F |
| VAMP2 | 1.92E-05 | -0.3803 | 0.367 | 0.61 | 0.306709 | MS.Trt_vs_MS.NoTrt_F |
| HP1BP3 | 1.94E-05 | -0.3043 | 0.82 | 0.93 | 0.309763 | MS.Trt_vs_MS.NoTrt_F |
| PREP | 1.95E-05 | -0.285 | 0.104 | 0.26 | 0.311993 | MS.Trt_vs_MS.NoTrt_F |
| 2-Sep | 1.98E-05 | -0.2586 | 0.874 | 0.93 | 0.316462 | MS.Trt_vs_MS.NoTrt_F |
| RRN3 | 2.04E-05 | 0.38972 | 0.263 | 0.48 | 0.325839 | MS.Trt_vs_MS.NoTrt_F |
| MAP1LC3B2 | 2.06E-05 | 0.43451 | 0.237 | 0.45 | 0.32893 | MS.Trt_vs_MS.NoTrt_F |
| WDR70 | 2.08E-05 | -0.6303 | 0.234 | 0.43 | 0.331777 | MS.Trt_vs_MS.NoTrt_F |
| ELP2 | 2.09E-05 | -0.4969 | 0.572 | 0.74 | 0.333569 | MS.Trt_vs_MS.NoTrt_F |
| JUND | 2.12E-05 | -0.3849 | 0.32 | 0.53 | 0.338499 | MS.Trt_vs_MS.NoTrt_F |
| SUCLG2 | 2.13E-05 | -0.367 | 0.194 | 0.38 | 0.339621 | MS.Trt_vs_MS.NoTrt_F |
| SUCO | 2.13E-05 | -0.9239 | 0.104 | 0.25 | 0.340773 | MS.Trt_vs_MS.NoTrt_F |
| AMDHD2 | 2.15E-05 | -1.319 | 0.086 | 0.23 | 0.34399 | MS.Trt_vs_MS.NoTrt_F |
| PRPF18 | 2.17E-05 | -0.7354 | 0.112 | 0.27 | 0.346144 | MS.Trt_vs_MS.NoTrt_F |
| NPIPB8 | 2.18E-05 | -0.374 | 0.108 | 0.26 | 0.347646 | MS.Trt_vs_MS.NoTrt_F |
| MSN | 2.19E-05 | -0.3399 | 0.55 | 0.75 | 0.349096 | MS.Trt_vs_MS.NoTrt_F |
| PPFIBP2 | 2.19E-05 | -0.9859 | 0.169 | 0.34 | 0.349867 | MS.Trt_vs_MS.NoTrt_F |
| NBEAL2 | 2.19E-05 | -0.3401 | 0.216 | 0.41 | 0.350212 | MS.Trt_vs_MS.NoTrt_F |
| CCDC13 | 2.22E-05 | -0.6833 | 0.259 | 0.47 | 0.354247 | MS.Trt_vs_MS.NoTrt_F |
| EXOC4 | 2.22E-05 | -0.5148 | 0.486 | 0.69 | 0.354764 | MS.Trt_vs_MS.NoTrt_F |
| NFRKB | 2.23E-05 | -0.4839 | 0.133 | 0.3 | 0.355449 | MS.Trt_vs_MS.NoTrt_F |
| CABIN1 | 2.26E-05 | -0.4768 | 0.223 | 0.41 | 0.361639 | MS.Trt_vs_MS.NoTrt_F |
| EFCAB6 | 2.29E-05 | -0.2511 | 0.349 | 0.56 | 0.365047 | MS.Trt_vs_MS.NoTrt_F |
| MRPS2 | 2.29E-05 | -1.479 | 0.072 | 0.2 | 0.365325 | MS.Trt_vs_MS.NoTrt_F |
| PPP4R3B | 2.29E-05 | -0.3438 | 0.403 | 0.58 | 0.365617 | MS.Trt_vs_MS.NoTrt_F |
| FAM84B | 2.34E-05 | -0.8869 | 0.054 | 0.18 | 0.373479 | MS.Trt_vs_MS.NoTrt_F |
| FXN | 2.37E-05 | -0.4542 | 0.158 | 0.33 | 0.37864 | MS.Trt_vs_MS.NoTrt_F |
| NME3 | 2.38E-05 | -0.433 | 0.108 | 0.26 | 0.379843 | MS.Trt_vs_MS.NoTrt_F |
| SMNDC1 | 2.39E-05 | -0.6791 | 0.126 | 0.29 | 0.382303 | MS.Trt_vs_MS.NoTrt_F |
| ARHGEF5 | 2.41E-05 | -0.4162 | 0.101 | 0.25 | 0.38457 | MS.Trt_vs_MS.NoTrt_F |
| RPTOR | 2.41E-05 | 0.32046 | 0.338 | 0.53 | 0.385357 | MS.Trt_vs_MS.NoTrt_F |
| GPR183 | 2.45E-05 | -0.3655 | 0.371 | 0.59 | 0.39073 | MS.Trt_vs_MS.NoTrt_F |
| HEATR1 | 2.47E-05 | 0.47126 | 0.137 | 0.3 | 0.395035 | MS.Trt_vs_MS.NoTrt_F |

| | | | | | | |
|---|---|---|---|---|---|---|
| PPM1M | 2.48E-05 | -0.4465 | 0.14 | 0.31 | 0.396015 | MS.Trt_vs_MS.NoTrt_F |
| DDX28 | 2.49E-05 | -1.3213 | 0.061 | 0.19 | 0.39729 | MS.Trt_vs_MS.NoTrt_F |
| SEPHS2 | 2.57E-05 | -0.6205 | 0.108 | 0.26 | 0.410888 | MS.Trt_vs_MS.NoTrt_F |
| WNT7B | 2.57E-05 | -0.3412 | 0.259 | 0.45 | 0.411304 | MS.Trt_vs_MS.NoTrt_F |
| CNOT11 | 2.58E-05 | -1.0437 | 0.119 | 0.27 | 0.411926 | MS.Trt_vs_MS.NoTrt_F |
| SLC39A10 | 2.59E-05 | -0.4648 | 0.176 | 0.36 | 0.4143 | MS.Trt_vs_MS.NoTrt_F |
| TRPC4AP | 2.59E-05 | -0.2949 | 0.259 | 0.46 | 0.414404 | MS.Trt_vs_MS.NoTrt_F |
| ZNF592 | 2.64E-05 | -0.3833 | 0.158 | 0.34 | 0.421216 | MS.Trt_vs_MS.NoTrt_F |
| ZDHHC5 | 2.66E-05 | -0.3675 | 0.108 | 0.26 | 0.424948 | MS.Trt_vs_MS.NoTrt_F |
| FTSJ3 | 2.67E-05 | -0.2957 | 0.09 | 0.23 | 0.425729 | MS.Trt_vs_MS.NoTrt_F |
| FIP1L1 | 2.67E-05 | -0.5 | 0.223 | 0.41 | 0.425772 | MS.Trt_vs_MS.NoTrt_F |
| OXLD1 | 2.67E-05 | -0.4688 | 0.108 | 0.26 | 0.426083 | MS.Trt_vs_MS.NoTrt_F |
| NBAS | 2.69E-05 | -0.4168 | 0.158 | 0.33 | 0.429602 | MS.Trt_vs_MS.NoTrt_F |
| EBAG9 | 2.70E-05 | -0.5069 | 0.094 | 0.24 | 0.431817 | MS.Trt_vs_MS.NoTrt_F |
| CCT4 | 2.75E-05 | -0.3916 | 0.241 | 0.43 | 0.438738 | MS.Trt_vs_MS.NoTrt_F |
| MAP3K4 | 2.76E-05 | -0.4802 | 0.158 | 0.33 | 0.440863 | MS.Trt_vs_MS.NoTrt_F |
| PCDH10 | 2.77E-05 | -0.3984 | 0.122 | 0.27 | 0.442086 | MS.Trt_vs_MS.NoTrt_F |
| MPZL3 | 2.77E-05 | -1.031 | 0.633 | 0.76 | 0.442551 | MS.Trt_vs_MS.NoTrt_F |
| LDHA | 2.78E-05 | -0.3678 | 0.655 | 0.81 | 0.444611 | MS.Trt_vs_MS.NoTrt_F |
| CHURC1 | 2.79E-05 | 0.38271 | 0.629 | 0.82 | 0.445407 | MS.Trt_vs_MS.NoTrt_F |
| LRFN1 | 2.79E-05 | -1.0062 | 0.108 | 0.25 | 0.446401 | MS.Trt_vs_MS.NoTrt_F |
| KCNA3 | 2.83E-05 | -0.3673 | 0.216 | 0.41 | 0.451614 | MS.Trt_vs_MS.NoTrt_F |
| TMEM35B | 2.83E-05 | -0.4044 | 0.162 | 0.34 | 0.451981 | MS.Trt_vs_MS.NoTrt_F |
| MMADHC | 2.85E-05 | -0.2826 | 0.356 | 0.54 | 0.455708 | MS.Trt_vs_MS.NoTrt_F |
| ATOX1 | 2.85E-05 | -0.261 | 0.212 | 0.39 | 0.455883 | MS.Trt_vs_MS.NoTrt_F |
| TMEM147 | 2.87E-05 | 0.25593 | 0.133 | 0.3 | 0.458314 | MS.Trt_vs_MS.NoTrt_F |
| HEATR5B | 2.87E-05 | -0.2545 | 0.349 | 0.58 | 0.459021 | MS.Trt_vs_MS.NoTrt_F |
| DCTN1 | 2.91E-05 | -0.3892 | 0.216 | 0.4 | 0.464282 | MS.Trt_vs_MS.NoTrt_F |
| VAMP7 | 2.95E-05 | -0.4623 | 0.094 | 0.24 | 0.470785 | MS.Trt_vs_MS.NoTrt_F |
| CCDC186 | 2.95E-05 | -0.3083 | 0.568 | 0.71 | 0.470887 | MS.Trt_vs_MS.NoTrt_F |
| CORO7-PAM | 2.95E-05 | -0.7254 | 0.101 | 0.25 | 0.470892 | MS.Trt_vs_MS.NoTrt_F |
| IL7 | 2.99E-05 | -0.9054 | 0.076 | 0.21 | 0.4783 | MS.Trt_vs_MS.NoTrt_F |
| PPP2R1A | 3.01E-05 | -0.4788 | 0.64 | 0.8 | 0.480816 | MS.Trt_vs_MS.NoTrt_F |
| TCHP | 3.07E-05 | -0.5687 | 0.446 | 0.62 | 0.489758 | MS.Trt_vs_MS.NoTrt_F |
| HACD3 | 3.07E-05 | -0.7375 | 0.665 | 0.81 | 0.489822 | MS.Trt_vs_MS.NoTrt_F |
| PINLYP | 3.10E-05 | -0.2866 | 0.511 | 0.69 | 0.494454 | MS.Trt_vs_MS.NoTrt_F |
| RIOK1 | 3.11E-05 | -0.7069 | 0.09 | 0.23 | 0.496573 | MS.Trt_vs_MS.NoTrt_F |
| SSNA1 | 3.11E-05 | -0.3889 | 0.14 | 0.31 | 0.496763 | MS.Trt_vs_MS.NoTrt_F |
| SGPP1 | 3.15E-05 | 0.43412 | 0.086 | 0.23 | 0.503202 | MS.Trt_vs_MS.NoTrt_F |
| KPNB1 | 3.15E-05 | -0.6843 | 0.644 | 0.74 | 0.503335 | MS.Trt_vs_MS.NoTrt_F |

| | | | | | | |
|---|---|---|---|---|---|---|
| FAM13B | 3.16E-05 | -0.8054 | 0.367 | 0.54 | 0.505002 | MS.Trt_vs_MS.NoTrt_F |
| SNRK | 3.19E-05 | -0.7091 | 0.45 | 0.63 | 0.508822 | MS.Trt_vs_MS.NoTrt_F |
| TTC31 | 3.22E-05 | -1.1864 | 0.187 | 0.35 | 0.515051 | MS.Trt_vs_MS.NoTrt_F |
| RAN | 3.25E-05 | -0.3387 | 0.371 | 0.58 | 0.518658 | MS.Trt_vs_MS.NoTrt_F |
| DTNBP1 | 3.26E-05 | -0.7353 | 0.086 | 0.23 | 0.520053 | MS.Trt_vs_MS.NoTrt_F |
| NUP205 | 3.29E-05 | -1.2624 | 0.147 | 0.31 | 0.525707 | MS.Trt_vs_MS.NoTrt_F |
| PHF3 | 3.30E-05 | -0.2711 | 0.403 | 0.6 | 0.52638 | MS.Trt_vs_MS.NoTrt_F |
| MARVELD2 | 3.31E-05 | -0.2627 | 0.536 | 0.72 | 0.528751 | MS.Trt_vs_MS.NoTrt_F |
| ARL17B | 3.38E-05 | 0.28835 | 0.712 | 0.85 | 0.54019 | MS.Trt_vs_MS.NoTrt_F |
| PICALM | 3.44E-05 | -0.4733 | 0.201 | 0.39 | 0.549592 | MS.Trt_vs_MS.NoTrt_F |
| RCBTB2 | 3.47E-05 | -0.329 | 0.115 | 0.27 | 0.554825 | MS.Trt_vs_MS.NoTrt_F |
| CERK | 3.50E-05 | -0.4504 | 0.439 | 0.63 | 0.55931 | MS.Trt_vs_MS.NoTrt_F |
| THAP7 | 3.55E-05 | -0.6809 | 0.112 | 0.26 | 0.566992 | MS.Trt_vs_MS.NoTrt_F |
| HNRNPL | 3.57E-05 | 0.50651 | 0.809 | 0.75 | 0.569774 | MS.Trt_vs_MS.NoTrt_F |
| ZDHHC3 | 3.60E-05 | -0.5475 | 0.435 | 0.63 | 0.575376 | MS.Trt_vs_MS.NoTrt_F |
| TUSC3 | 3.61E-05 | -0.3477 | 0.367 | 0.54 | 0.576161 | MS.Trt_vs_MS.NoTrt_F |
| HIVEP2 | 3.66E-05 | -0.6466 | 0.23 | 0.4 | 0.58535 | MS.Trt_vs_MS.NoTrt_F |
| OST4 | 3.67E-05 | 0.28355 | 0.36 | 0.66 | 0.586368 | MS.Trt_vs_MS.NoTrt_F |
| CP | 3.82E-05 | 1.16025 | 0.122 | 0.27 | 0.610095 | MS.Trt_vs_MS.NoTrt_F |
| DDX50 | 3.87E-05 | -0.2948 | 0.183 | 0.37 | 0.618282 | MS.Trt_vs_MS.NoTrt_F |
| PAN3 | 3.91E-05 | 0.26111 | 0.856 | 0.95 | 0.624506 | MS.Trt_vs_MS.NoTrt_F |
| TRMT13 | 3.91E-05 | -0.3513 | 0.129 | 0.29 | 0.624928 | MS.Trt_vs_MS.NoTrt_F |
| HLA-A | 4.01E-05 | -0.367 | 0.748 | 0.92 | 0.639955 | MS.Trt_vs_MS.NoTrt_F |
| ARRDC2 | 4.03E-05 | -0.3576 | 0.126 | 0.28 | 0.643673 | MS.Trt_vs_MS.NoTrt_F |
| RB1 | 4.03E-05 | -0.8223 | 0.129 | 0.29 | 0.644079 | MS.Trt_vs_MS.NoTrt_F |
| C1orf123 | 4.07E-05 | -0.7373 | 0.137 | 0.29 | 0.650355 | MS.Trt_vs_MS.NoTrt_F |
| ATF6 | 4.10E-05 | -0.4372 | 0.201 | 0.38 | 0.654154 | MS.Trt_vs_MS.NoTrt_F |
| INF2 | 4.12E-05 | -0.3306 | 0.198 | 0.38 | 0.658567 | MS.Trt_vs_MS.NoTrt_F |
| REXO4 | 4.14E-05 | -1.2415 | 0.068 | 0.19 | 0.661992 | MS.Trt_vs_MS.NoTrt_F |
| ZFP36L1 | 4.15E-05 | 0.99988 | 0.759 | 0.76 | 0.662708 | MS.Trt_vs_MS.NoTrt_F |
| ATP6V0E2 | 4.15E-05 | -0.4941 | 0.187 | 0.37 | 0.662854 | MS.Trt_vs_MS.NoTrt_F |
| SETD1B | 4.15E-05 | -0.4712 | 0.126 | 0.28 | 0.663252 | MS.Trt_vs_MS.NoTrt_F |
| CLIP4 | 4.17E-05 | -1.4915 | 0.09 | 0.23 | 0.665403 | MS.Trt_vs_MS.NoTrt_F |
| SLC16A6 | 4.17E-05 | 0.33995 | 0.23 | 0.41 | 0.666723 | MS.Trt_vs_MS.NoTrt_F |
| RTFDC1 | 4.18E-05 | -0.4222 | 0.183 | 0.36 | 0.667985 | MS.Trt_vs_MS.NoTrt_F |
| SF3B5 | 4.19E-05 | -0.7851 | 0.097 | 0.24 | 0.668605 | MS.Trt_vs_MS.NoTrt_F |
| SUMO2 | 4.21E-05 | -0.397 | 0.831 | 0.9 | 0.672878 | MS.Trt_vs_MS.NoTrt_F |
| CSGALNACT | 4.25E-05 | -0.3338 | 0.511 | 0.69 | 0.679136 | MS.Trt_vs_MS.NoTrt_F |
| MAP2K3 | 4.30E-05 | -0.8514 | 0.144 | 0.3 | 0.687112 | MS.Trt_vs_MS.NoTrt_F |
| PUM3 | 4.31E-05 | -0.8221 | 0.129 | 0.29 | 0.689024 | MS.Trt_vs_MS.NoTrt_F |

| | | | | | | |
|---|---|---|---|---|---|---|
| KIAA1147 | 4.32E-05 | -0.9551 | 0.072 | 0.2 | 0.689479 | MS.Trt_vs_MS.NoTrt_F |
| TRMT2B | 4.37E-05 | -0.5874 | 0.594 | 0.81 | 0.698009 | MS.Trt_vs_MS.NoTrt_F |
| HSPH1 | 4.38E-05 | -0.3899 | 0.209 | 0.4 | 0.700096 | MS.Trt_vs_MS.NoTrt_F |
| MNAT1 | 4.42E-05 | -0.9016 | 0.09 | 0.23 | 0.706545 | MS.Trt_vs_MS.NoTrt_F |
| RYBP | 4.45E-05 | -1.0403 | 0.068 | 0.19 | 0.711349 | MS.Trt_vs_MS.NoTrt_F |
| FBXO34 | 4.46E-05 | -0.3991 | 0.108 | 0.25 | 0.71275 | MS.Trt_vs_MS.NoTrt_F |
| ARL13B | 4.48E-05 | -0.5775 | 0.158 | 0.32 | 0.715322 | MS.Trt_vs_MS.NoTrt_F |
| MTCH2 | 4.50E-05 | -0.6525 | 0.094 | 0.23 | 0.718388 | MS.Trt_vs_MS.NoTrt_F |
| CNOT6 | 4.53E-05 | -0.756 | 0.094 | 0.23 | 0.723598 | MS.Trt_vs_MS.NoTrt_F |
| TRIP12 | 4.58E-05 | -0.4562 | 0.439 | 0.6 | 0.731328 | MS.Trt_vs_MS.NoTrt_F |
| UCKL1 | 4.60E-05 | -0.632 | 0.122 | 0.28 | 0.734812 | MS.Trt_vs_MS.NoTrt_F |
| FAM192A | 4.65E-05 | 0.59268 | 0.403 | 0.63 | 0.742885 | MS.Trt_vs_MS.NoTrt_F |
| MRPS35 | 4.71E-05 | -0.4385 | 0.112 | 0.27 | 0.752127 | MS.Trt_vs_MS.NoTrt_F |
| JAK2 | 4.74E-05 | -0.8099 | 0.165 | 0.33 | 0.756367 | MS.Trt_vs_MS.NoTrt_F |
| DICER1 | 4.74E-05 | -0.3917 | 0.119 | 0.28 | 0.756567 | MS.Trt_vs_MS.NoTrt_F |
| EFR3A | 4.75E-05 | -0.2812 | 0.14 | 0.31 | 0.759176 | MS.Trt_vs_MS.NoTrt_F |
| MRPL47 | 4.79E-05 | -0.4109 | 0.079 | 0.21 | 0.765223 | MS.Trt_vs_MS.NoTrt_F |
| KPNA2 | 4.79E-05 | -0.5038 | 0.101 | 0.24 | 0.765543 | MS.Trt_vs_MS.NoTrt_F |
| ROMO1 | 4.82E-05 | 0.47105 | 0.162 | 0.34 | 0.769717 | MS.Trt_vs_MS.NoTrt_F |
| PDGFA | 4.84E-05 | -0.5324 | 0.565 | 0.68 | 0.772341 | MS.Trt_vs_MS.NoTrt_F |
| PRPF6 | 4.87E-05 | -0.2995 | 0.205 | 0.38 | 0.77755 | MS.Trt_vs_MS.NoTrt_F |
| ZMYM3 | 4.87E-05 | -0.8183 | 0.335 | 0.51 | 0.777683 | MS.Trt_vs_MS.NoTrt_F |
| RBBP6 | 4.90E-05 | -0.4487 | 0.151 | 0.32 | 0.781994 | MS.Trt_vs_MS.NoTrt_F |
| PRPF31 | 4.96E-05 | -0.2534 | 0.155 | 0.33 | 0.791657 | MS.Trt_vs_MS.NoTrt_F |
| TRPV2 | 5.11E-05 | -0.5664 | 0.252 | 0.43 | 0.816122 | MS.Trt_vs_MS.NoTrt_F |
| ZDHHC13 | 5.14E-05 | 0.66805 | 0.662 | 0.86 | 0.820752 | MS.Trt_vs_MS.NoTrt_F |
| SLC38A1 | 5.15E-05 | -0.2651 | 0.73 | 0.87 | 0.823392 | MS.Trt_vs_MS.NoTrt_F |
| TBC1D5 | 5.16E-05 | -0.558 | 0.536 | 0.7 | 0.824143 | MS.Trt_vs_MS.NoTrt_F |
| RSRP1 | 5.19E-05 | -0.3998 | 0.91 | 0.97 | 0.829501 | MS.Trt_vs_MS.NoTrt_F |
| PPP1R11 | 5.26E-05 | -0.299 | 0.165 | 0.34 | 0.840137 | MS.Trt_vs_MS.NoTrt_F |
| MTFP1 | 5.28E-05 | -0.646 | 0.14 | 0.3 | 0.843973 | MS.Trt_vs_MS.NoTrt_F |
| DENND4B | 5.30E-05 | -0.7485 | 0.173 | 0.34 | 0.846568 | MS.Trt_vs_MS.NoTrt_F |
| SLC2A4RG | 5.39E-05 | -0.597 | 0.079 | 0.21 | 0.860858 | MS.Trt_vs_MS.NoTrt_F |
| NUCB1 | 5.41E-05 | -0.2559 | 0.273 | 0.47 | 0.863671 | MS.Trt_vs_MS.NoTrt_F |
| RPL14 | 5.43E-05 | -0.2898 | 0.968 | 0.99 | 0.867658 | MS.Trt_vs_MS.NoTrt_F |
| KAT6A | 5.44E-05 | -0.4856 | 0.709 | 0.85 | 0.868232 | MS.Trt_vs_MS.NoTrt_F |
| FKBP8 | 5.46E-05 | -0.8539 | 0.201 | 0.38 | 0.872607 | MS.Trt_vs_MS.NoTrt_F |
| CCDC141 | 5.53E-05 | -0.4481 | 0.647 | 0.79 | 0.883353 | MS.Trt_vs_MS.NoTrt_F |
| CDC5L | 5.59E-05 | -0.3264 | 0.536 | 0.74 | 0.892383 | MS.Trt_vs_MS.NoTrt_F |
| PELI1 | 5.59E-05 | -0.3855 | 0.169 | 0.34 | 0.893515 | MS.Trt_vs_MS.NoTrt_F |

| | | | | | | |
|---|---|---|---|---|---|---|
| LSS | 5.62E-05 | -1.0429 | 0.119 | 0.27 | 0.897169 | MS.Trt_vs_MS.NoTrt_F |
| TTC9C | 5.70E-05 | -0.293 | 0.313 | 0.49 | 0.909726 | MS.Trt_vs_MS.NoTrt_F |
| FBXL3 | 5.79E-05 | -0.9076 | 0.129 | 0.28 | 0.92457 | MS.Trt_vs_MS.NoTrt_F |
| METTL26 | 5.79E-05 | 0.33863 | 0.176 | 0.36 | 0.924924 | MS.Trt_vs_MS.NoTrt_F |
| BTN3A1 | 5.82E-05 | -0.2703 | 0.64 | 0.74 | 0.929046 | MS.Trt_vs_MS.NoTrt_F |
| C1D | 5.82E-05 | 0.25623 | 0.162 | 0.34 | 0.930182 | MS.Trt_vs_MS.NoTrt_F |
| ZYX | 5.99E-05 | -0.3361 | 0.291 | 0.49 | 0.95636 | MS.Trt_vs_MS.NoTrt_F |
| XPO4 | 5.99E-05 | -0.3263 | 0.112 | 0.26 | 0.956381 | MS.Trt_vs_MS.NoTrt_F |
| NOB1 | 6.07E-05 | -0.5119 | 0.115 | 0.27 | 0.970097 | MS.Trt_vs_MS.NoTrt_F |
| FAM160B2 | 6.09E-05 | -0.4526 | 0.662 | 0.83 | 0.973311 | MS.Trt_vs_MS.NoTrt_F |
| IL24 | 6.21E-05 | -0.9527 | 0.108 | 0.25 | 0.992656 | MS.Trt_vs_MS.NoTrt_F |
| ENTPD4 | 6.24E-05 | -0.4699 | 0.529 | 0.72 | 0.996639 | MS.Trt_vs_MS.NoTrt_F |
| RNF5 | 6.24E-05 | -0.3816 | 0.122 | 0.28 | 0.996656 | MS.Trt_vs_MS.NoTrt_F |
| ASCC3 | 6.26E-05 | 0.25402 | 0.216 | 0.41 | 0.999487 | MS.Trt_vs_MS.NoTrt_F |
| PRPS1 | 6.27E-05 | -0.7382 | 0.119 | 0.27 | 1 | MS.Trt_vs_MS.NoTrt_F |
| WBP2 | 6.31E-05 | -0.4215 | 0.601 | 0.77 | 1 | MS.Trt_vs_MS.NoTrt_F |
| DMTF1 | 6.35E-05 | -0.543 | 0.504 | 0.67 | 1 | MS.Trt_vs_MS.NoTrt_F |
| TNIP2 | 6.51E-05 | -0.4371 | 0.108 | 0.25 | 1 | MS.Trt_vs_MS.NoTrt_F |
| AIDA | 6.51E-05 | -1.2725 | 0.104 | 0.24 | 1 | MS.Trt_vs_MS.NoTrt_F |
| SH2B1 | 6.55E-05 | -0.2717 | 0.511 | 0.72 | 1 | MS.Trt_vs_MS.NoTrt_F |
| ATXN2L | 6.56E-05 | -0.4578 | 0.205 | 0.38 | 1 | MS.Trt_vs_MS.NoTrt_F |
| ELK3 | 6.63E-05 | -0.7933 | 0.396 | 0.58 | 1 | MS.Trt_vs_MS.NoTrt_F |
| CXXC5 | 6.65E-05 | 0.60059 | 0.446 | 0.64 | 1 | MS.Trt_vs_MS.NoTrt_F |
| PPP1CB | 6.67E-05 | 1.2403 | 0.849 | 0.84 | 1 | MS.Trt_vs_MS.NoTrt_F |
| KDM5D | 6.67E-05 | -2.9248 | 0.273 | 0.43 | 1 | MS.Trt_vs_MS.NoTrt_F |
| TXNRD1 | 6.70E-05 | -1.3035 | 0.23 | 0.39 | 1 | MS.Trt_vs_MS.NoTrt_F |
| PMAIP1 | 6.79E-05 | -0.5268 | 0.183 | 0.36 | 1 | MS.Trt_vs_MS.NoTrt_F |
| UTP14A | 6.82E-05 | -0.2511 | 0.158 | 0.32 | 1 | MS.Trt_vs_MS.NoTrt_F |
| NR3C2 | 6.90E-05 | -1.2241 | 0.058 | 0.17 | 1 | MS.Trt_vs_MS.NoTrt_F |
| IGSF8 | 6.96E-05 | -0.3034 | 0.122 | 0.28 | 1 | MS.Trt_vs_MS.NoTrt_F |
| PNPLA2 | 6.97E-05 | -0.4456 | 0.151 | 0.32 | 1 | MS.Trt_vs_MS.NoTrt_F |
| MRPL23 | 7.02E-05 | -0.3522 | 0.108 | 0.25 | 1 | MS.Trt_vs_MS.NoTrt_F |
| PQLC3 | 7.06E-05 | -1.0563 | 0.133 | 0.28 | 1 | MS.Trt_vs_MS.NoTrt_F |
| PDE9A | 7.25E-05 | -0.6903 | 0.054 | 0.17 | 1 | MS.Trt_vs_MS.NoTrt_F |
| MARK3 | 7.26E-05 | -0.4642 | 0.82 | 0.95 | 1 | MS.Trt_vs_MS.NoTrt_F |
| GABRA6 | 7.31E-05 | -0.2502 | 0.104 | 0.24 | 1 | MS.Trt_vs_MS.NoTrt_F |
| CRIPT | 7.37E-05 | -0.349 | 0.216 | 0.39 | 1 | MS.Trt_vs_MS.NoTrt_F |
| RAB35 | 7.38E-05 | -1.0516 | 0.151 | 0.3 | 1 | MS.Trt_vs_MS.NoTrt_F |
| FBXW7 | 7.38E-05 | 0.56184 | 0.165 | 0.34 | 1 | MS.Trt_vs_MS.NoTrt_F |
| CLMP | 7.44E-05 | -0.9771 | 0.356 | 0.53 | 1 | MS.Trt_vs_MS.NoTrt_F |

| | | | | | | |
|---|---|---|---|---|---|---|
| CNDP2 | 7.45E-05 | -0.5777 | 0.669 | 0.8 | 1 | MS.Trt_vs_MS.NoTrt_F |
| ZFAND2B | 7.51E-05 | -1.054 | 0.198 | 0.35 | 1 | MS.Trt_vs_MS.NoTrt_F |
| GPR63 | 7.51E-05 | -0.3962 | 0.27 | 0.47 | 1 | MS.Trt_vs_MS.NoTrt_F |
| TRIB1 | 7.54E-05 | -1.934 | 0.068 | 0.19 | 1 | MS.Trt_vs_MS.NoTrt_F |
| SRSF1 | 7.59E-05 | -0.2812 | 0.482 | 0.66 | 1 | MS.Trt_vs_MS.NoTrt_F |
| CPQ | 7.64E-05 | -0.7747 | 0.313 | 0.5 | 1 | MS.Trt_vs_MS.NoTrt_F |
| NUDT2 | 7.65E-05 | -1.5324 | 0.061 | 0.18 | 1 | MS.Trt_vs_MS.NoTrt_F |
| CXorf40A | 7.69E-05 | -0.4493 | 0.122 | 0.27 | 1 | MS.Trt_vs_MS.NoTrt_F |
| TP53I13 | 7.85E-05 | -0.2797 | 0.108 | 0.25 | 1 | MS.Trt_vs_MS.NoTrt_F |
| BMT2 | 7.90E-05 | -0.6753 | 0.086 | 0.22 | 1 | MS.Trt_vs_MS.NoTrt_F |
| ATP1A1 | 7.92E-05 | -0.2501 | 0.651 | 0.81 | 1 | MS.Trt_vs_MS.NoTrt_F |
| SLITRK2 | 8.00E-05 | -0.2705 | 0.23 | 0.39 | 1 | MS.Trt_vs_MS.NoTrt_F |
| GEMIN4 | 8.15E-05 | -0.6923 | 0.072 | 0.19 | 1 | MS.Trt_vs_MS.NoTrt_F |
| DDX3Y | 8.17E-05 | -2.0643 | 0.04 | 0.14 | 1 | MS.Trt_vs_MS.NoTrt_F |
| UBR7 | 8.28E-05 | -0.4676 | 0.112 | 0.25 | 1 | MS.Trt_vs_MS.NoTrt_F |
| RBM22 | 8.29E-05 | -0.3215 | 0.137 | 0.3 | 1 | MS.Trt_vs_MS.NoTrt_F |
| MRPL1 | 8.41E-05 | -0.3542 | 0.129 | 0.28 | 1 | MS.Trt_vs_MS.NoTrt_F |
| DBI | 8.45E-05 | -0.2654 | 0.194 | 0.37 | 1 | MS.Trt_vs_MS.NoTrt_F |
| RRAGA | 8.49E-05 | -0.3848 | 0.137 | 0.29 | 1 | MS.Trt_vs_MS.NoTrt_F |
| PLG | 8.56E-05 | -0.6604 | 0.191 | 0.34 | 1 | MS.Trt_vs_MS.NoTrt_F |
| CEP85 | 8.74E-05 | -0.7251 | 0.112 | 0.25 | 1 | MS.Trt_vs_MS.NoTrt_F |
| ATRIP | 8.85E-05 | -0.511 | 0.104 | 0.24 | 1 | MS.Trt_vs_MS.NoTrt_F |
| MUS81 | 8.93E-05 | -0.2834 | 0.162 | 0.33 | 1 | MS.Trt_vs_MS.NoTrt_F |
| TNFRSF13B | 8.98E-05 | -1.1769 | 0.036 | 0.13 | 1 | MS.Trt_vs_MS.NoTrt_F |
| PREB | 9.02E-05 | -0.3354 | 0.14 | 0.3 | 1 | MS.Trt_vs_MS.NoTrt_F |
| UCHL5 | 9.05E-05 | 0.46785 | 0.173 | 0.34 | 1 | MS.Trt_vs_MS.NoTrt_F |
| SRP54 | 9.08E-05 | -0.4143 | 0.252 | 0.43 | 1 | MS.Trt_vs_MS.NoTrt_F |
| TLN1 | 9.08E-05 | -0.3449 | 0.572 | 0.71 | 1 | MS.Trt_vs_MS.NoTrt_F |
| C6orf47 | 9.16E-05 | -0.8976 | 0.09 | 0.22 | 1 | MS.Trt_vs_MS.NoTrt_F |
| TTC1 | 9.16E-05 | -0.584 | 0.201 | 0.37 | 1 | MS.Trt_vs_MS.NoTrt_F |
| ASF1A | 9.22E-05 | 0.43851 | 0.169 | 0.35 | 1 | MS.Trt_vs_MS.NoTrt_F |
| REV1 | 9.31E-05 | -0.643 | 0.137 | 0.29 | 1 | MS.Trt_vs_MS.NoTrt_F |
| NOA1 | 9.37E-05 | -0.3196 | 0.112 | 0.25 | 1 | MS.Trt_vs_MS.NoTrt_F |
| CTC-479C5.1 | 9.38E-05 | -0.3874 | 0.464 | 0.63 | 1 | MS.Trt_vs_MS.NoTrt_F |
| HAAO | 9.39E-05 | 0.53897 | 0.086 | 0.22 | 1 | MS.Trt_vs_MS.NoTrt_F |
| SLC2A6 | 9.59E-05 | -1.0133 | 0.216 | 0.38 | 1 | MS.Trt_vs_MS.NoTrt_F |
| SYNRG | 9.69E-05 | -0.6104 | 0.928 | 0.97 | 1 | MS.Trt_vs_MS.NoTrt_F |
| RPL34 | 9.74E-05 | 0.61861 | 0.95 | 0.98 | 1 | MS.Trt_vs_MS.NoTrt_F |
| GABARAPL2 | 9.77E-05 | -0.4014 | 0.478 | 0.66 | 1 | MS.Trt_vs_MS.NoTrt_F |
| MANBA | 9.80E-05 | -0.5038 | 0.169 | 0.32 | 1 | MS.Trt_vs_MS.NoTrt_F |

| | | | | | | |
|---|---|---|---|---|---|---|
| EXOSC10 | 9.83E-05 | -0.3055 | 0.802 | 0.9 | 1 | MS.Trt_vs_MS.NoTrt_F |
| FAM78A | 9.87E-05 | -0.8199 | 0.338 | 0.51 | 1 | MS.Trt_vs_MS.NoTrt_F |
| BEX3 | 9.93E-05 | -0.3937 | 0.115 | 0.26 | 1 | MS.Trt_vs_MS.NoTrt_F |
| KCTD7 | 9.95E-05 | -0.6468 | 0.205 | 0.37 | 1 | MS.Trt_vs_MS.NoTrt_F |
| NOL8 | 1.00E-04 | -0.2586 | 0.191 | 0.38 | 1 | MS.Trt_vs_MS.NoTrt_F |
| GABPB1 | 1.00E-04 | 0.42667 | 0.428 | 0.61 | 1 | MS.Trt_vs_MS.NoTrt_F |
| FAM193A | 1.00E-04 | -0.3647 | 0.266 | 0.45 | 1 | MS.Trt_vs_MS.NoTrt_F |
| SASS6 | 1.01E-04 | -0.3681 | 0.065 | 0.18 | 1 | MS.Trt_vs_MS.NoTrt_F |
| KHDRBS1 | 1.01E-04 | 0.25506 | 0.144 | 0.3 | 1 | MS.Trt_vs_MS.NoTrt_F |
| SFN | 1.01E-04 | -0.3873 | 0.205 | 0.35 | 1 | MS.Trt_vs_MS.NoTrt_F |
| NRBP1 | 1.01E-04 | -0.429 | 0.5 | 0.67 | 1 | MS.Trt_vs_MS.NoTrt_F |
| C19orf43 | 1.02E-04 | -0.2782 | 0.155 | 0.32 | 1 | MS.Trt_vs_MS.NoTrt_F |
| TMEM9B | 1.02E-04 | 0.25396 | 0.209 | 0.39 | 1 | MS.Trt_vs_MS.NoTrt_F |
| PHYH | 1.03E-04 | -0.3072 | 0.198 | 0.35 | 1 | MS.Trt_vs_MS.NoTrt_F |
| RAB7A | 1.03E-04 | -0.3948 | 0.345 | 0.55 | 1 | MS.Trt_vs_MS.NoTrt_F |
| RNF121 | 1.03E-04 | -0.7082 | 0.32 | 0.51 | 1 | MS.Trt_vs_MS.NoTrt_F |
| GBP1 | 1.04E-04 | -0.3104 | 0.219 | 0.4 | 1 | MS.Trt_vs_MS.NoTrt_F |
| KIAA0196 | 1.04E-04 | -0.4472 | 0.133 | 0.28 | 1 | MS.Trt_vs_MS.NoTrt_F |
| ZNF862 | 1.05E-04 | -0.3128 | 0.137 | 0.29 | 1 | MS.Trt_vs_MS.NoTrt_F |
| TERF2IP | 1.06E-04 | -0.3628 | 0.374 | 0.56 | 1 | MS.Trt_vs_MS.NoTrt_F |
| SAT1 | 1.06E-04 | -0.5205 | 0.54 | 0.76 | 1 | MS.Trt_vs_MS.NoTrt_F |
| DTX2 | 1.06E-04 | -0.5892 | 0.371 | 0.54 | 1 | MS.Trt_vs_MS.NoTrt_F |
| TPCN2 | 1.06E-04 | -1.6097 | 0.586 | 0.75 | 1 | MS.Trt_vs_MS.NoTrt_F |
| TPP2 | 1.06E-04 | -0.2591 | 0.439 | 0.64 | 1 | MS.Trt_vs_MS.NoTrt_F |
| NEMP1 | 1.06E-04 | -0.7701 | 0.079 | 0.2 | 1 | MS.Trt_vs_MS.NoTrt_F |
| NEK5 | 1.09E-04 | 0.44207 | 0.899 | 0.86 | 1 | MS.Trt_vs_MS.NoTrt_F |
| OAT | 1.09E-04 | -0.7426 | 0.101 | 0.23 | 1 | MS.Trt_vs_MS.NoTrt_F |
| SPNS3 | 1.09E-04 | 0.42838 | 0.068 | 0.19 | 1 | MS.Trt_vs_MS.NoTrt_F |
| AP2M1 | 1.09E-04 | -0.4213 | 0.288 | 0.48 | 1 | MS.Trt_vs_MS.NoTrt_F |
| VPS4A | 1.09E-04 | -0.7433 | 0.604 | 0.75 | 1 | MS.Trt_vs_MS.NoTrt_F |
| MAN1B1 | 1.10E-04 | -0.9378 | 0.284 | 0.48 | 1 | MS.Trt_vs_MS.NoTrt_F |
| COPS3 | 1.11E-04 | -0.6085 | 0.237 | 0.41 | 1 | MS.Trt_vs_MS.NoTrt_F |
| PRPF40A | 1.11E-04 | -0.3533 | 0.457 | 0.63 | 1 | MS.Trt_vs_MS.NoTrt_F |
| INIP | 1.11E-04 | -0.2753 | 0.601 | 0.81 | 1 | MS.Trt_vs_MS.NoTrt_F |
| ACIN1 | 1.12E-04 | -0.5745 | 0.46 | 0.63 | 1 | MS.Trt_vs_MS.NoTrt_F |
| MXD4 | 1.13E-04 | 0.35095 | 0.14 | 0.3 | 1 | MS.Trt_vs_MS.NoTrt_F |
| TTPA | 1.15E-04 | -0.3756 | 0.356 | 0.54 | 1 | MS.Trt_vs_MS.NoTrt_F |
| GGA2 | 1.15E-04 | -0.4517 | 0.629 | 0.8 | 1 | MS.Trt_vs_MS.NoTrt_F |
| SIK2 | 1.16E-04 | -0.5142 | 0.691 | 0.82 | 1 | MS.Trt_vs_MS.NoTrt_F |
| TAP2 | 1.16E-04 | -0.5461 | 0.183 | 0.34 | 1 | MS.Trt_vs_MS.NoTrt_F |

| | | | | | | |
|---|---|---|---|---|---|---|
| FUCA2 | 1.16E-04 | 0.27471 | 0.288 | 0.48 | 1 | MS.Trt_vs_MS.NoTrt_F |
| RBM14-RBM | 1.16E-04 | -1.098 | 0.112 | 0.25 | 1 | MS.Trt_vs_MS.NoTrt_F |
| ZAP70 | 1.17E-04 | -0.3167 | 0.442 | 0.66 | 1 | MS.Trt_vs_MS.NoTrt_F |
| DIP2A | 1.18E-04 | -0.5669 | 0.813 | 0.87 | 1 | MS.Trt_vs_MS.NoTrt_F |
| DIP2B | 1.18E-04 | 0.37994 | 0.309 | 0.51 | 1 | MS.Trt_vs_MS.NoTrt_F |
| MRPL18 | 1.19E-04 | 0.50547 | 0.158 | 0.33 | 1 | MS.Trt_vs_MS.NoTrt_F |
| EEFSEC | 1.19E-04 | -1.8484 | 0.09 | 0.22 | 1 | MS.Trt_vs_MS.NoTrt_F |
| RGPD6 | 1.19E-04 | -0.4014 | 0.522 | 0.69 | 1 | MS.Trt_vs_MS.NoTrt_F |
| CNOT2 | 1.19E-04 | -0.3875 | 0.921 | 0.96 | 1 | MS.Trt_vs_MS.NoTrt_F |
| EHD1 | 1.19E-04 | -0.2992 | 0.291 | 0.49 | 1 | MS.Trt_vs_MS.NoTrt_F |
| WDR53 | 1.19E-04 | -0.6081 | 0.083 | 0.21 | 1 | MS.Trt_vs_MS.NoTrt_F |
| UTP18 | 1.20E-04 | -0.6772 | 0.126 | 0.27 | 1 | MS.Trt_vs_MS.NoTrt_F |
| SERINC4 | 1.21E-04 | -1.2327 | 0.054 | 0.16 | 1 | MS.Trt_vs_MS.NoTrt_F |
| TPCN1 | 1.22E-04 | -0.6744 | 0.335 | 0.51 | 1 | MS.Trt_vs_MS.NoTrt_F |
| ERAP2 | 1.22E-04 | -0.7304 | 0.619 | 0.77 | 1 | MS.Trt_vs_MS.NoTrt_F |
| PES1 | 1.22E-04 | -1.1841 | 0.108 | 0.24 | 1 | MS.Trt_vs_MS.NoTrt_F |
| UBE2E1 | 1.23E-04 | -1.3996 | 0.083 | 0.2 | 1 | MS.Trt_vs_MS.NoTrt_F |
| TRAPPC2L | 1.23E-04 | -0.3408 | 0.129 | 0.28 | 1 | MS.Trt_vs_MS.NoTrt_F |
| STRA13 | 1.23E-04 | 0.27851 | 0.126 | 0.28 | 1 | MS.Trt_vs_MS.NoTrt_F |
| PPP2R5D | 1.24E-04 | -0.2532 | 0.155 | 0.32 | 1 | MS.Trt_vs_MS.NoTrt_F |
| CRYZ | 1.27E-04 | -0.8424 | 0.101 | 0.23 | 1 | MS.Trt_vs_MS.NoTrt_F |
| TMEM62 | 1.27E-04 | -0.6598 | 0.223 | 0.39 | 1 | MS.Trt_vs_MS.NoTrt_F |
| TMBIM1 | 1.27E-04 | -0.2892 | 0.529 | 0.76 | 1 | MS.Trt_vs_MS.NoTrt_F |
| CDC37L1 | 1.28E-04 | -1.0099 | 0.101 | 0.23 | 1 | MS.Trt_vs_MS.NoTrt_F |
| DEDD2 | 1.28E-04 | -0.2674 | 0.349 | 0.53 | 1 | MS.Trt_vs_MS.NoTrt_F |
| PTTG1 | 1.28E-04 | 0.36216 | 0.126 | 0.28 | 1 | MS.Trt_vs_MS.NoTrt_F |
| RAB14 | 1.29E-04 | -0.4468 | 0.209 | 0.38 | 1 | MS.Trt_vs_MS.NoTrt_F |
| AP5S1 | 1.29E-04 | 0.94212 | 0.906 | 0.8 | 1 | MS.Trt_vs_MS.NoTrt_F |
| CAPN7 | 1.30E-04 | -0.823 | 0.46 | 0.63 | 1 | MS.Trt_vs_MS.NoTrt_F |
| CTRL | 1.31E-04 | -0.8196 | 0.097 | 0.23 | 1 | MS.Trt_vs_MS.NoTrt_F |
| PDCD6IP | 1.32E-04 | -0.2704 | 0.284 | 0.46 | 1 | MS.Trt_vs_MS.NoTrt_F |
| CCDC47 | 1.33E-04 | -0.3486 | 0.608 | 0.79 | 1 | MS.Trt_vs_MS.NoTrt_F |
| ZNF32 | 1.33E-04 | -1.2595 | 0.068 | 0.18 | 1 | MS.Trt_vs_MS.NoTrt_F |
| ZNF79 | 1.33E-04 | -1.298 | 0.061 | 0.17 | 1 | MS.Trt_vs_MS.NoTrt_F |
| TBCA | 1.36E-04 | -0.3166 | 0.248 | 0.41 | 1 | MS.Trt_vs_MS.NoTrt_F |
| ERVK3-1 | 1.37E-04 | -0.5774 | 0.295 | 0.47 | 1 | MS.Trt_vs_MS.NoTrt_F |
| CMTR1 | 1.38E-04 | -1.1049 | 0.155 | 0.3 | 1 | MS.Trt_vs_MS.NoTrt_F |
| ZNF407 | 1.39E-04 | -0.8502 | 0.097 | 0.23 | 1 | MS.Trt_vs_MS.NoTrt_F |
| SNRPE | 1.39E-04 | 0.82893 | 0.129 | 0.28 | 1 | MS.Trt_vs_MS.NoTrt_F |
| SH3BP1 | 1.39E-04 | -0.9387 | 0.097 | 0.23 | 1 | MS.Trt_vs_MS.NoTrt_F |

| | | | | | | |
|---|---|---|---|---|---|---|
| MAL | 1.40E-04 | -0.341 | 0.176 | 0.34 | 1 | MS.Trt_vs_MS.NoTrt_F |
| NT5C3A | 1.40E-04 | 0.37577 | 0.212 | 0.4 | 1 | MS.Trt_vs_MS.NoTrt_F |
| KBTBD4 | 1.40E-04 | -0.5745 | 0.076 | 0.19 | 1 | MS.Trt_vs_MS.NoTrt_F |
| ARGLU1 | 1.41E-04 | -0.2954 | 0.655 | 0.84 | 1 | MS.Trt_vs_MS.NoTrt_F |
| TREX1 | 1.42E-04 | -0.6496 | 0.104 | 0.24 | 1 | MS.Trt_vs_MS.NoTrt_F |
| HADHB | 1.43E-04 | -0.3307 | 0.471 | 0.62 | 1 | MS.Trt_vs_MS.NoTrt_F |
| MIF4GD | 1.43E-04 | -0.3562 | 0.129 | 0.28 | 1 | MS.Trt_vs_MS.NoTrt_F |
| ABHD4 | 1.44E-04 | -0.9121 | 0.054 | 0.16 | 1 | MS.Trt_vs_MS.NoTrt_F |
| CCDC171 | 1.44E-04 | -0.3734 | 0.414 | 0.6 | 1 | MS.Trt_vs_MS.NoTrt_F |
| ITPR2 | 1.45E-04 | -0.6461 | 0.205 | 0.37 | 1 | MS.Trt_vs_MS.NoTrt_F |
| VAC14 | 1.45E-04 | -0.7744 | 0.094 | 0.22 | 1 | MS.Trt_vs_MS.NoTrt_F |
| BAG4 | 1.45E-04 | 0.78229 | 0.169 | 0.33 | 1 | MS.Trt_vs_MS.NoTrt_F |
| UFM1 | 1.46E-04 | -0.3483 | 0.392 | 0.55 | 1 | MS.Trt_vs_MS.NoTrt_F |
| POLA1 | 1.46E-04 | -1.3122 | 0.083 | 0.2 | 1 | MS.Trt_vs_MS.NoTrt_F |
| CD84 | 1.46E-04 | -0.4855 | 0.716 | 0.8 | 1 | MS.Trt_vs_MS.NoTrt_F |
| LCMT2 | 1.46E-04 | -0.8036 | 0.673 | 0.77 | 1 | MS.Trt_vs_MS.NoTrt_F |
| TUBGCP2 | 1.46E-04 | -0.4661 | 0.716 | 0.83 | 1 | MS.Trt_vs_MS.NoTrt_F |
| CD5 | 1.46E-04 | -0.2501 | 0.446 | 0.66 | 1 | MS.Trt_vs_MS.NoTrt_F |
| PRMT9 | 1.47E-04 | 0.96838 | 0.198 | 0.38 | 1 | MS.Trt_vs_MS.NoTrt_F |
| PKIA | 1.49E-04 | -0.2854 | 0.112 | 0.25 | 1 | MS.Trt_vs_MS.NoTrt_F |
| DRG2 | 1.49E-04 | -0.6774 | 0.176 | 0.34 | 1 | MS.Trt_vs_MS.NoTrt_F |
| C21orf2 | 1.49E-04 | -0.266 | 0.126 | 0.27 | 1 | MS.Trt_vs_MS.NoTrt_F |
| EDEM2 | 1.49E-04 | -0.3079 | 0.126 | 0.27 | 1 | MS.Trt_vs_MS.NoTrt_F |
| R3HCC1L | 1.52E-04 | -0.3574 | 0.281 | 0.45 | 1 | MS.Trt_vs_MS.NoTrt_F |
| RNF38 | 1.52E-04 | -0.2768 | 0.194 | 0.36 | 1 | MS.Trt_vs_MS.NoTrt_F |
| BRAP | 1.54E-04 | -0.3826 | 0.291 | 0.47 | 1 | MS.Trt_vs_MS.NoTrt_F |
| GNB2 | 1.54E-04 | -0.5766 | 0.496 | 0.65 | 1 | MS.Trt_vs_MS.NoTrt_F |
| MED8 | 1.56E-04 | -0.9666 | 0.094 | 0.22 | 1 | MS.Trt_vs_MS.NoTrt_F |
| AHNAK | 1.58E-04 | -0.281 | 0.759 | 0.9 | 1 | MS.Trt_vs_MS.NoTrt_F |
| C6orf106 | 1.59E-04 | -0.9868 | 0.119 | 0.25 | 1 | MS.Trt_vs_MS.NoTrt_F |
| NOTCH2NL | 1.59E-04 | -0.3695 | 0.151 | 0.3 | 1 | MS.Trt_vs_MS.NoTrt_F |
| INO80E | 1.59E-04 | -1.0448 | 0.158 | 0.3 | 1 | MS.Trt_vs_MS.NoTrt_F |
| ERP29 | 1.60E-04 | -0.3031 | 0.806 | 0.91 | 1 | MS.Trt_vs_MS.NoTrt_F |
| TRIM2 | 1.63E-04 | -0.472 | 0.187 | 0.34 | 1 | MS.Trt_vs_MS.NoTrt_F |
| VAMP1 | 1.64E-04 | -0.4686 | 0.126 | 0.27 | 1 | MS.Trt_vs_MS.NoTrt_F |
| PAK2 | 1.65E-04 | -0.3365 | 0.68 | 0.86 | 1 | MS.Trt_vs_MS.NoTrt_F |
| MECP2 | 1.65E-04 | -0.3269 | 0.917 | 0.97 | 1 | MS.Trt_vs_MS.NoTrt_F |
| RUFY1 | 1.67E-04 | -0.8171 | 0.252 | 0.41 | 1 | MS.Trt_vs_MS.NoTrt_F |
| AGAP5 | 1.67E-04 | -1.173 | 0.076 | 0.19 | 1 | MS.Trt_vs_MS.NoTrt_F |
| NDUFB4 | 1.68E-04 | 0.50332 | 0.237 | 0.44 | 1 | MS.Trt_vs_MS.NoTrt_F |

| | | | | | | |
|---|---|---|---|---|---|---|
| DNAJB9 | 1.68E-04 | -0.5351 | 0.119 | 0.26 | 1 | MS.Trt_vs_MS.NoTrt_F |
| TBC1D2 | 1.68E-04 | -1.2694 | 0.065 | 0.18 | 1 | MS.Trt_vs_MS.NoTrt_F |
| C7orf26 | 1.68E-04 | -0.5739 | 0.086 | 0.21 | 1 | MS.Trt_vs_MS.NoTrt_F |
| ARL6IP1 | 1.69E-04 | -0.299 | 0.482 | 0.66 | 1 | MS.Trt_vs_MS.NoTrt_F |
| RBM14 | 1.69E-04 | -0.2624 | 0.162 | 0.32 | 1 | MS.Trt_vs_MS.NoTrt_F |
| VIM | 1.69E-04 | -0.3282 | 0.766 | 0.9 | 1 | MS.Trt_vs_MS.NoTrt_F |
| TAF15 | 1.69E-04 | -0.3244 | 0.18 | 0.34 | 1 | MS.Trt_vs_MS.NoTrt_F |
| FAM156B | 1.71E-04 | 0.51903 | 0.076 | 0.19 | 1 | MS.Trt_vs_MS.NoTrt_F |
| RELL2 | 1.71E-04 | -0.4291 | 0.112 | 0.25 | 1 | MS.Trt_vs_MS.NoTrt_F |
| THAP1 | 1.72E-04 | 0.79864 | 0.101 | 0.23 | 1 | MS.Trt_vs_MS.NoTrt_F |
| UBE2Q2 | 1.72E-04 | -0.6086 | 0.119 | 0.26 | 1 | MS.Trt_vs_MS.NoTrt_F |
| NPC2 | 1.73E-04 | 0.4517 | 0.101 | 0.24 | 1 | MS.Trt_vs_MS.NoTrt_F |
| IGSF9 | 1.73E-04 | -0.6316 | 0.259 | 0.41 | 1 | MS.Trt_vs_MS.NoTrt_F |
| PEPD | 1.74E-04 | -0.3343 | 0.144 | 0.3 | 1 | MS.Trt_vs_MS.NoTrt_F |
| THG1L | 1.77E-04 | 0.27313 | 0.54 | 0.71 | 1 | MS.Trt_vs_MS.NoTrt_F |
| ZNF831 | 1.79E-04 | -0.3007 | 0.223 | 0.4 | 1 | MS.Trt_vs_MS.NoTrt_F |
| MPHOSPH1( | 1.80E-04 | -0.4535 | 0.216 | 0.38 | 1 | MS.Trt_vs_MS.NoTrt_F |
| CD93 | 1.82E-04 | 0.66113 | 0.176 | 0.33 | 1 | MS.Trt_vs_MS.NoTrt_F |
| CCDC28A | 1.82E-04 | -0.4828 | 0.155 | 0.31 | 1 | MS.Trt_vs_MS.NoTrt_F |
| ABCA7 | 1.83E-04 | 0.28198 | 0.194 | 0.37 | 1 | MS.Trt_vs_MS.NoTrt_F |
| TSC22D3 | 1.84E-04 | -0.2743 | 0.683 | 0.89 | 1 | MS.Trt_vs_MS.NoTrt_F |
| HMGXB4 | 0.00018 | -0.808 | 0.115 | 0.25 | 1 | MS.Trt_vs_MS.NoTrt_F |
| LARP4 | 1.84E-04 | -0.4053 | 0.353 | 0.51 | 1 | MS.Trt_vs_MS.NoTrt_F |
| ERP44 | 1.86E-04 | -0.4596 | 0.27 | 0.43 | 1 | MS.Trt_vs_MS.NoTrt_F |
| RAB30 | 1.87E-04 | 0.3248 | 0.263 | 0.43 | 1 | MS.Trt_vs_MS.NoTrt_F |
| PDHA1 | 1.89E-04 | -0.3356 | 0.198 | 0.37 | 1 | MS.Trt_vs_MS.NoTrt_F |
| ASNSD1 | 1.89E-04 | 0.29119 | 0.133 | 0.29 | 1 | MS.Trt_vs_MS.NoTrt_F |
| MRPL43 | 1.91E-04 | -0.3091 | 0.104 | 0.24 | 1 | MS.Trt_vs_MS.NoTrt_F |
| VPS16 | 1.93E-04 | 0.35277 | 0.133 | 0.29 | 1 | MS.Trt_vs_MS.NoTrt_F |
| VPS37A | 1.94E-04 | -0.6945 | 0.14 | 0.28 | 1 | MS.Trt_vs_MS.NoTrt_F |
| RABGAP1L | 1.95E-04 | -0.4128 | 0.835 | 0.9 | 1 | MS.Trt_vs_MS.NoTrt_F |
| GOLGA5 | 1.95E-04 | -1.0109 | 0.126 | 0.27 | 1 | MS.Trt_vs_MS.NoTrt_F |
| ZGPAT | 1.95E-04 | -0.5123 | 0.112 | 0.25 | 1 | MS.Trt_vs_MS.NoTrt_F |
| ZNRD1 | 1.98E-04 | 0.36912 | 0.147 | 0.3 | 1 | MS.Trt_vs_MS.NoTrt_F |
| ACSL5 | 1.99E-04 | -0.5883 | 0.183 | 0.34 | 1 | MS.Trt_vs_MS.NoTrt_F |
| TRIP11 | 2.00E-04 | -0.4442 | 0.511 | 0.66 | 1 | MS.Trt_vs_MS.NoTrt_F |
| C16orf54 | 2.01E-04 | 1.33361 | 0.802 | 0.87 | 1 | MS.Trt_vs_MS.NoTrt_F |
| CKAP2 | 2.01E-04 | -0.4415 | 0.122 | 0.26 | 1 | MS.Trt_vs_MS.NoTrt_F |
| FH | 2.02E-04 | -0.5228 | 0.068 | 0.18 | 1 | MS.Trt_vs_MS.NoTrt_F |
| ACTN4 | 2.02E-04 | -0.4298 | 0.414 | 0.59 | 1 | MS.Trt_vs_MS.NoTrt_F |

| | | | | | | |
|---|---|---|---|---|---|---|
| LMBRD2 | 2.02E-04 | -0.3295 | 0.09 | 0.22 | 1 | MS.Trt_vs_MS.NoTrt_F |
| DCTN4 | 2.03E-04 | -0.5844 | 0.288 | 0.48 | 1 | MS.Trt_vs_MS.NoTrt_F |
| SH2D3C | 2.03E-04 | -0.2531 | 0.183 | 0.35 | 1 | MS.Trt_vs_MS.NoTrt_F |
| GBF1 | 2.03E-04 | -0.3919 | 0.151 | 0.3 | 1 | MS.Trt_vs_MS.NoTrt_F |
| RNF6 | 2.04E-04 | -0.7346 | 0.198 | 0.35 | 1 | MS.Trt_vs_MS.NoTrt_F |
| SLC25A44 | 2.04E-04 | 0.45875 | 0.586 | 0.77 | 1 | MS.Trt_vs_MS.NoTrt_F |
| DUSP4 | 2.05E-04 | -0.3105 | 0.277 | 0.45 | 1 | MS.Trt_vs_MS.NoTrt_F |
| TATDN1 | 2.05E-04 | -1.3556 | 0.644 | 0.76 | 1 | MS.Trt_vs_MS.NoTrt_F |
| SLC30A3 | 2.06E-04 | -0.315 | 0.356 | 0.54 | 1 | MS.Trt_vs_MS.NoTrt_F |
| LUC7L2 | 2.07E-04 | -0.3826 | 0.518 | 0.69 | 1 | MS.Trt_vs_MS.NoTrt_F |
| MTOR | 2.08E-04 | -0.5647 | 0.14 | 0.29 | 1 | MS.Trt_vs_MS.NoTrt_F |
| ATXN7L3B | 2.08E-04 | -0.3884 | 0.151 | 0.3 | 1 | MS.Trt_vs_MS.NoTrt_F |
| ABHD16A | 2.10E-04 | -0.3471 | 0.14 | 0.29 | 1 | MS.Trt_vs_MS.NoTrt_F |
| LPIN2 | 2.10E-04 | 0.25589 | 0.147 | 0.3 | 1 | MS.Trt_vs_MS.NoTrt_F |
| XPC | 2.11E-04 | -0.2949 | 0.223 | 0.39 | 1 | MS.Trt_vs_MS.NoTrt_F |
| EIF1B | 2.11E-04 | -0.3977 | 0.147 | 0.3 | 1 | MS.Trt_vs_MS.NoTrt_F |
| DAZAP1 | 2.12E-04 | -0.338 | 0.126 | 0.27 | 1 | MS.Trt_vs_MS.NoTrt_F |
| HS1BP3 | 2.12E-04 | -0.4624 | 0.119 | 0.25 | 1 | MS.Trt_vs_MS.NoTrt_F |
| RCE1 | 2.17E-04 | -0.507 | 0.086 | 0.21 | 1 | MS.Trt_vs_MS.NoTrt_F |
| GADD45GIP | 2.19E-04 | 0.60928 | 0.155 | 0.32 | 1 | MS.Trt_vs_MS.NoTrt_F |
| ILK | 2.19E-04 | -0.3087 | 0.187 | 0.35 | 1 | MS.Trt_vs_MS.NoTrt_F |
| ZMYND11 | 2.20E-04 | -0.3334 | 0.108 | 0.24 | 1 | MS.Trt_vs_MS.NoTrt_F |
| SLC39A4 | 2.22E-04 | -0.3768 | 0.331 | 0.5 | 1 | MS.Trt_vs_MS.NoTrt_F |
| TUT1 | 2.23E-04 | -0.7325 | 0.108 | 0.24 | 1 | MS.Trt_vs_MS.NoTrt_F |
| SHANK2 | 2.24E-04 | -0.318 | 0.281 | 0.45 | 1 | MS.Trt_vs_MS.NoTrt_F |
| SLC25A36 | 2.25E-04 | -0.5324 | 0.129 | 0.27 | 1 | MS.Trt_vs_MS.NoTrt_F |
| ATP6V1A | 2.28E-04 | -0.3991 | 0.518 | 0.67 | 1 | MS.Trt_vs_MS.NoTrt_F |
| TLE4 | 2.29E-04 | 0.37254 | 0.14 | 0.29 | 1 | MS.Trt_vs_MS.NoTrt_F |
| TFE3 | 2.29E-04 | -0.7618 | 0.09 | 0.21 | 1 | MS.Trt_vs_MS.NoTrt_F |
| UFSP2 | 2.33E-04 | -0.6122 | 0.122 | 0.26 | 1 | MS.Trt_vs_MS.NoTrt_F |
| BAG3 | 2.34E-04 | -0.3812 | 0.072 | 0.19 | 1 | MS.Trt_vs_MS.NoTrt_F |
| SLC46A3 | 2.39E-04 | -0.3185 | 0.083 | 0.2 | 1 | MS.Trt_vs_MS.NoTrt_F |
| PPP1R10 | 2.39E-04 | -0.653 | 0.478 | 0.65 | 1 | MS.Trt_vs_MS.NoTrt_F |
| FADD | 2.39E-04 | -0.7885 | 0.09 | 0.21 | 1 | MS.Trt_vs_MS.NoTrt_F |
| PBDC1 | 2.41E-04 | 0.26339 | 0.101 | 0.23 | 1 | MS.Trt_vs_MS.NoTrt_F |
| C11orf24 | 2.41E-04 | -1.0649 | 0.194 | 0.34 | 1 | MS.Trt_vs_MS.NoTrt_F |
| FAM104A | 2.43E-04 | 0.40855 | 0.198 | 0.37 | 1 | MS.Trt_vs_MS.NoTrt_F |
| BTBD10 | 2.44E-04 | -0.7469 | 0.137 | 0.28 | 1 | MS.Trt_vs_MS.NoTrt_F |
| SUPT7L | 2.46E-04 | -0.4081 | 0.403 | 0.59 | 1 | MS.Trt_vs_MS.NoTrt_F |
| ALG10 | 2.49E-04 | -1.7512 | 0.086 | 0.2 | 1 | MS.Trt_vs_MS.NoTrt_F |

| | | | | | | |
|---|---|---|---|---|---|---|
| DUT | 2.49E-04 | -0.2519 | 0.151 | 0.3 | 1 | MS.Trt_vs_MS.NoTrt_F |
| NUP210 | 2.51E-04 | -0.4069 | 0.5 | 0.67 | 1 | MS.Trt_vs_MS.NoTrt_F |
| CYP7B1 | 2.51E-04 | 0.39627 | 0.374 | 0.54 | 1 | MS.Trt_vs_MS.NoTrt_F |
| CHST6 | 2.54E-04 | 0.31791 | 0.932 | 0.88 | 1 | MS.Trt_vs_MS.NoTrt_F |
| INSIG1 | 2.57E-04 | -0.3236 | 0.165 | 0.32 | 1 | MS.Trt_vs_MS.NoTrt_F |
| GOLGA8A | 2.58E-04 | 0.32759 | 0.399 | 0.59 | 1 | MS.Trt_vs_MS.NoTrt_F |
| PPP1R7 | 2.58E-04 | -0.3919 | 0.162 | 0.31 | 1 | MS.Trt_vs_MS.NoTrt_F |
| VPS72 | 2.59E-04 | -0.3554 | 0.14 | 0.29 | 1 | MS.Trt_vs_MS.NoTrt_F |
| SF3B4 | 2.60E-04 | -0.9113 | 0.169 | 0.32 | 1 | MS.Trt_vs_MS.NoTrt_F |
| GNB1L | 2.63E-04 | -0.451 | 0.183 | 0.33 | 1 | MS.Trt_vs_MS.NoTrt_F |
| TMEM261 | 2.63E-04 | 1.0811 | 0.115 | 0.25 | 1 | MS.Trt_vs_MS.NoTrt_F |
| RBM12B | 2.66E-04 | -0.3046 | 0.133 | 0.28 | 1 | MS.Trt_vs_MS.NoTrt_F |
| LINC01420 | 2.66E-04 | 0.62093 | 0.129 | 0.28 | 1 | MS.Trt_vs_MS.NoTrt_F |
| KIAA0556 | 2.67E-04 | -1.3372 | 0.284 | 0.45 | 1 | MS.Trt_vs_MS.NoTrt_F |
| WBP1L | 2.67E-04 | -1.0701 | 0.112 | 0.24 | 1 | MS.Trt_vs_MS.NoTrt_F |
| SLC35A2 | 2.68E-04 | -0.8826 | 0.079 | 0.19 | 1 | MS.Trt_vs_MS.NoTrt_F |
| DNAJC30 | 2.68E-04 | -0.442 | 0.234 | 0.41 | 1 | MS.Trt_vs_MS.NoTrt_F |
| TMEM79 | 2.70E-04 | -0.5215 | 0.104 | 0.23 | 1 | MS.Trt_vs_MS.NoTrt_F |
| SACM1L | 2.70E-04 | -0.3629 | 0.241 | 0.43 | 1 | MS.Trt_vs_MS.NoTrt_F |
| THOC6 | 2.72E-04 | -0.5869 | 0.101 | 0.23 | 1 | MS.Trt_vs_MS.NoTrt_F |
| TSNAX | 2.74E-04 | -0.3683 | 0.662 | 0.8 | 1 | MS.Trt_vs_MS.NoTrt_F |
| TBC1D1 | 2.76E-04 | -0.4634 | 0.522 | 0.74 | 1 | MS.Trt_vs_MS.NoTrt_F |
| PAFAH1B3 | 2.79E-04 | -0.5925 | 0.094 | 0.22 | 1 | MS.Trt_vs_MS.NoTrt_F |
| NREP | 2.79E-04 | -0.6554 | 0.137 | 0.27 | 1 | MS.Trt_vs_MS.NoTrt_F |
| TAF9B | 2.80E-04 | -0.5352 | 0.119 | 0.25 | 1 | MS.Trt_vs_MS.NoTrt_F |
| LRIG1 | 2.80E-04 | 0.36398 | 0.191 | 0.35 | 1 | MS.Trt_vs_MS.NoTrt_F |
| ESR2 | 2.80E-04 | 0.31321 | 0.493 | 0.7 | 1 | MS.Trt_vs_MS.NoTrt_F |
| LGALS8 | 2.81E-04 | -0.5963 | 0.594 | 0.75 | 1 | MS.Trt_vs_MS.NoTrt_F |
| TOPORS | 2.84E-04 | -0.3443 | 0.104 | 0.23 | 1 | MS.Trt_vs_MS.NoTrt_F |
| TRIM28 | 2.85E-04 | -1.5944 | 0.09 | 0.21 | 1 | MS.Trt_vs_MS.NoTrt_F |
| MFSD6 | 2.90E-04 | -0.9289 | 0.147 | 0.29 | 1 | MS.Trt_vs_MS.NoTrt_F |
| SYCP1 | 2.90E-04 | -0.3708 | 0.248 | 0.43 | 1 | MS.Trt_vs_MS.NoTrt_F |
| TNPO2 | 2.92E-04 | -0.9052 | 0.151 | 0.29 | 1 | MS.Trt_vs_MS.NoTrt_F |
| SERGEF | 2.93E-04 | -0.3262 | 0.129 | 0.27 | 1 | MS.Trt_vs_MS.NoTrt_F |
| TRIM47 | 2.94E-04 | -0.7628 | 0.061 | 0.17 | 1 | MS.Trt_vs_MS.NoTrt_F |
| DTNB | 2.94E-04 | -0.292 | 0.165 | 0.32 | 1 | MS.Trt_vs_MS.NoTrt_F |
| GPR108 | 2.96E-04 | -0.4018 | 0.169 | 0.32 | 1 | MS.Trt_vs_MS.NoTrt_F |
| ZNF134 | 2.96E-04 | -1.5856 | 0.137 | 0.27 | 1 | MS.Trt_vs_MS.NoTrt_F |
| NOC4L | 2.98E-04 | -0.7591 | 0.076 | 0.19 | 1 | MS.Trt_vs_MS.NoTrt_F |
| STAT3 | 2.98E-04 | -0.277 | 0.795 | 0.87 | 1 | MS.Trt_vs_MS.NoTrt_F |

| | | | | | | |
|---|---|---|---|---|---|---|
| APOA2 | 2.99E-04 | -0.3131 | 0.205 | 0.35 | 1 | MS.Trt_vs_MS.NoTrt_F |
| RP11-500M8 | 2.99E-04 | 0.37853 | 0.112 | 0.25 | 1 | MS.Trt_vs_MS.NoTrt_F |
| BARD1 | 3.01E-04 | 0.41214 | 0.151 | 0.31 | 1 | MS.Trt_vs_MS.NoTrt_F |
| NDC1 | 3.02E-04 | -1.1019 | 0.065 | 0.17 | 1 | MS.Trt_vs_MS.NoTrt_F |
| TMEM208 | 3.09E-04 | 0.41598 | 0.137 | 0.29 | 1 | MS.Trt_vs_MS.NoTrt_F |
| ZC3H7A | 3.13E-04 | 0.265 | 0.367 | 0.56 | 1 | MS.Trt_vs_MS.NoTrt_F |
| LANCL1 | 3.13E-04 | -0.2675 | 0.137 | 0.28 | 1 | MS.Trt_vs_MS.NoTrt_F |
| ANKMY2 | 3.14E-04 | -0.6401 | 0.058 | 0.16 | 1 | MS.Trt_vs_MS.NoTrt_F |
| ATF7 | 3.16E-04 | -0.3224 | 0.403 | 0.58 | 1 | MS.Trt_vs_MS.NoTrt_F |
| RNF123 | 3.16E-04 | -0.689 | 0.144 | 0.29 | 1 | MS.Trt_vs_MS.NoTrt_F |
| CRLS1 | 3.17E-04 | -0.2549 | 0.288 | 0.45 | 1 | MS.Trt_vs_MS.NoTrt_F |
| HSPE1-MOB | 3.19E-04 | -0.4791 | 0.112 | 0.24 | 1 | MS.Trt_vs_MS.NoTrt_F |
| PPM1G | 3.21E-04 | -0.255 | 0.665 | 0.82 | 1 | MS.Trt_vs_MS.NoTrt_F |
| SNED1 | 3.21E-04 | -0.6271 | 0.144 | 0.29 | 1 | MS.Trt_vs_MS.NoTrt_F |
| NOL7 | 3.21E-04 | 0.45676 | 0.191 | 0.36 | 1 | MS.Trt_vs_MS.NoTrt_F |
| GLYCTK | 3.21E-04 | -0.4224 | 0.101 | 0.23 | 1 | MS.Trt_vs_MS.NoTrt_F |
| TXN | 3.21E-04 | 0.29536 | 0.212 | 0.39 | 1 | MS.Trt_vs_MS.NoTrt_F |
| ACAT2 | 3.25E-04 | -1.4281 | 0.234 | 0.37 | 1 | MS.Trt_vs_MS.NoTrt_F |
| OGFOD2 | 3.25E-04 | -0.6314 | 0.101 | 0.23 | 1 | MS.Trt_vs_MS.NoTrt_F |
| HS3ST3A1 | 3.26E-04 | -0.2985 | 0.076 | 0.19 | 1 | MS.Trt_vs_MS.NoTrt_F |
| RGCC | 3.26E-04 | 0.29413 | 0.317 | 0.53 | 1 | MS.Trt_vs_MS.NoTrt_F |
| PARP6 | 3.32E-04 | -0.5042 | 0.626 | 0.8 | 1 | MS.Trt_vs_MS.NoTrt_F |
| POLR2M | 3.34E-04 | -0.4138 | 0.158 | 0.3 | 1 | MS.Trt_vs_MS.NoTrt_F |
| HNRNPA3 | 3.35E-04 | -0.3012 | 0.399 | 0.55 | 1 | MS.Trt_vs_MS.NoTrt_F |
| CARS | 3.36E-04 | -0.3425 | 0.781 | 0.83 | 1 | MS.Trt_vs_MS.NoTrt_F |
| CUTC | 3.38E-04 | -0.5202 | 0.158 | 0.3 | 1 | MS.Trt_vs_MS.NoTrt_F |
| PDIK1L | 3.41E-04 | -1.056 | 0.068 | 0.18 | 1 | MS.Trt_vs_MS.NoTrt_F |
| MAT2A | 3.43E-04 | -0.4273 | 0.151 | 0.3 | 1 | MS.Trt_vs_MS.NoTrt_F |
| BCAT1 | 3.45E-04 | 0.33971 | 0.41 | 0.55 | 1 | MS.Trt_vs_MS.NoTrt_F |
| AP2S1 | 3.46E-04 | 0.48271 | 0.144 | 0.3 | 1 | MS.Trt_vs_MS.NoTrt_F |
| GCC2 | 3.46E-04 | 0.26682 | 0.748 | 0.9 | 1 | MS.Trt_vs_MS.NoTrt_F |
| MAP4K1 | 3.48E-04 | -0.4148 | 0.878 | 0.94 | 1 | MS.Trt_vs_MS.NoTrt_F |
| PPP2R2A | 3.49E-04 | -1.2659 | 0.129 | 0.27 | 1 | MS.Trt_vs_MS.NoTrt_F |
| PTBP2 | 3.51E-04 | 0.5395 | 0.554 | 0.72 | 1 | MS.Trt_vs_MS.NoTrt_F |
| S1PR4 | 3.51E-04 | -0.3971 | 0.147 | 0.29 | 1 | MS.Trt_vs_MS.NoTrt_F |
| GGT1 | 3.51E-04 | -0.4369 | 0.112 | 0.24 | 1 | MS.Trt_vs_MS.NoTrt_F |
| HELQ | 3.51E-04 | -0.6303 | 0.126 | 0.27 | 1 | MS.Trt_vs_MS.NoTrt_F |
| NPIPB6 | 3.52E-04 | -0.5432 | 0.194 | 0.33 | 1 | MS.Trt_vs_MS.NoTrt_F |
| SPATA13 | 3.53E-04 | -0.4398 | 0.46 | 0.6 | 1 | MS.Trt_vs_MS.NoTrt_F |
| SLK | 3.53E-04 | -0.4187 | 0.176 | 0.32 | 1 | MS.Trt_vs_MS.NoTrt_F |

| | | | | | | |
|---|---|---|---|---|---|---|
| LMTK2 | 3.56E-04 | 0.36844 | 0.216 | 0.37 | 1 | MS.Trt_vs_MS.NoTrt_F |
| GIMAP1-GIM | 3.57E-04 | -0.4469 | 0.076 | 0.19 | 1 | MS.Trt_vs_MS.NoTrt_F |
| ATAD3B | 3.59E-04 | -0.5234 | 0.745 | 0.89 | 1 | MS.Trt_vs_MS.NoTrt_F |
| ELF2 | 3.63E-04 | -0.3533 | 0.626 | 0.75 | 1 | MS.Trt_vs_MS.NoTrt_F |
| SDHAF3 | 3.65E-04 | -0.3396 | 0.076 | 0.19 | 1 | MS.Trt_vs_MS.NoTrt_F |
| C5orf51 | 3.66E-04 | -0.715 | 0.112 | 0.24 | 1 | MS.Trt_vs_MS.NoTrt_F |
| GTF2F2 | 3.69E-04 | -0.3352 | 0.201 | 0.34 | 1 | MS.Trt_vs_MS.NoTrt_F |
| FAS | 3.69E-04 | -0.5074 | 0.342 | 0.51 | 1 | MS.Trt_vs_MS.NoTrt_F |
| PTPN22 | 3.70E-04 | -0.3241 | 0.356 | 0.51 | 1 | MS.Trt_vs_MS.NoTrt_F |
| BACH1 | 3.72E-04 | -0.4391 | 0.223 | 0.39 | 1 | MS.Trt_vs_MS.NoTrt_F |
| EIF3CL | 3.77E-04 | -0.2897 | 0.162 | 0.32 | 1 | MS.Trt_vs_MS.NoTrt_F |
| CYB5R1 | 3.78E-04 | -1.1826 | 0.108 | 0.23 | 1 | MS.Trt_vs_MS.NoTrt_F |
| WDR77 | 3.80E-04 | -0.6554 | 0.112 | 0.24 | 1 | MS.Trt_vs_MS.NoTrt_F |
| METTL4 | 3.80E-04 | -0.6468 | 0.112 | 0.24 | 1 | MS.Trt_vs_MS.NoTrt_F |
| ACP5 | 3.82E-04 | 0.28723 | 0.14 | 0.29 | 1 | MS.Trt_vs_MS.NoTrt_F |
| SRD5A1 | 3.84E-04 | -0.7183 | 0.59 | 0.75 | 1 | MS.Trt_vs_MS.NoTrt_F |
| MT-ND3 | 3.85E-04 | 0.42627 | 0.421 | 0.67 | 1 | MS.Trt_vs_MS.NoTrt_F |
| HAUS7 | 3.85E-04 | -1.3293 | 0.094 | 0.21 | 1 | MS.Trt_vs_MS.NoTrt_F |
| TAS2R4 | 3.85E-04 | -0.8202 | 0.317 | 0.47 | 1 | MS.Trt_vs_MS.NoTrt_F |
| CTNNBL1 | 3.86E-04 | -0.3002 | 0.144 | 0.29 | 1 | MS.Trt_vs_MS.NoTrt_F |
| SLC41A1 | 3.87E-04 | -0.337 | 0.104 | 0.23 | 1 | MS.Trt_vs_MS.NoTrt_F |
| YTHDF1 | 3.87E-04 | 0.26019 | 0.068 | 0.18 | 1 | MS.Trt_vs_MS.NoTrt_F |
| LAPTM5 | 3.88E-04 | -0.2885 | 0.586 | 0.76 | 1 | MS.Trt_vs_MS.NoTrt_F |
| TAF4 | 3.89E-04 | -0.9637 | 0.126 | 0.26 | 1 | MS.Trt_vs_MS.NoTrt_F |
| DHFR2 | 3.89E-04 | 0.38928 | 0.597 | 0.69 | 1 | MS.Trt_vs_MS.NoTrt_F |
| MBD1 | 3.90E-04 | -0.4017 | 0.169 | 0.32 | 1 | MS.Trt_vs_MS.NoTrt_F |
| CENPH | 3.91E-04 | -0.3035 | 0.255 | 0.41 | 1 | MS.Trt_vs_MS.NoTrt_F |
| NDUFA1 | 3.93E-04 | 0.26343 | 0.259 | 0.45 | 1 | MS.Trt_vs_MS.NoTrt_F |
| B3GAT2 | 3.93E-04 | 0.73021 | 0.169 | 0.31 | 1 | MS.Trt_vs_MS.NoTrt_F |
| RIPK1 | 3.95E-04 | -0.6381 | 0.158 | 0.29 | 1 | MS.Trt_vs_MS.NoTrt_F |
| DMPK | 3.97E-04 | -0.3426 | 0.565 | 0.74 | 1 | MS.Trt_vs_MS.NoTrt_F |
| MADD | 3.98E-04 | -0.3044 | 0.194 | 0.35 | 1 | MS.Trt_vs_MS.NoTrt_F |
| CPSF7 | 4.00E-04 | -0.5184 | 0.187 | 0.34 | 1 | MS.Trt_vs_MS.NoTrt_F |
| B3GAT1 | 4.02E-04 | -0.305 | 0.191 | 0.35 | 1 | MS.Trt_vs_MS.NoTrt_F |
| CLPTM1L | 4.02E-04 | 0.72209 | 0.076 | 0.19 | 1 | MS.Trt_vs_MS.NoTrt_F |
| FABP7 | 4.03E-04 | -0.4291 | 0.133 | 0.26 | 1 | MS.Trt_vs_MS.NoTrt_F |
| SETD3 | 4.05E-04 | -0.8622 | 0.119 | 0.25 | 1 | MS.Trt_vs_MS.NoTrt_F |
| KRAS | 4.05E-04 | -0.3367 | 0.317 | 0.48 | 1 | MS.Trt_vs_MS.NoTrt_F |
| RIPK3 | 4.08E-04 | -0.2973 | 0.076 | 0.19 | 1 | MS.Trt_vs_MS.NoTrt_F |
| AKAP11 | 4.09E-04 | -0.7975 | 0.194 | 0.34 | 1 | MS.Trt_vs_MS.NoTrt_F |

| | | | | | | |
|---|---|---|---|---|---|---|
| FARS2 | 4.14E-04 | -1.5259 | 0.086 | 0.2 | 1 | MS.Trt_vs_MS.NoTrt_F |
| EPHB6 | 4.16E-04 | -0.4318 | 0.101 | 0.22 | 1 | MS.Trt_vs_MS.NoTrt_F |
| TMEM38B | 4.18E-04 | -0.4016 | 0.176 | 0.32 | 1 | MS.Trt_vs_MS.NoTrt_F |
| GPS1 | 4.20E-04 | -0.314 | 0.14 | 0.28 | 1 | MS.Trt_vs_MS.NoTrt_F |
| WAC | 4.20E-04 | -0.4006 | 0.417 | 0.59 | 1 | MS.Trt_vs_MS.NoTrt_F |
| DOK1 | 4.21E-04 | -0.3584 | 0.09 | 0.2 | 1 | MS.Trt_vs_MS.NoTrt_F |
| BET1L | 4.21E-04 | -0.3225 | 0.183 | 0.33 | 1 | MS.Trt_vs_MS.NoTrt_F |
| NDE1 | 4.23E-04 | -0.4187 | 0.723 | 0.85 | 1 | MS.Trt_vs_MS.NoTrt_F |
| CKS1B | 4.25E-04 | 0.51081 | 0.173 | 0.32 | 1 | MS.Trt_vs_MS.NoTrt_F |
| MAGED1 | 4.25E-04 | -0.5422 | 0.633 | 0.79 | 1 | MS.Trt_vs_MS.NoTrt_F |
| THRA | 4.25E-04 | -0.8313 | 0.104 | 0.23 | 1 | MS.Trt_vs_MS.NoTrt_F |
| SNX13 | 4.26E-04 | -0.4012 | 0.255 | 0.43 | 1 | MS.Trt_vs_MS.NoTrt_F |
| SMG5 | 4.30E-04 | -0.3133 | 0.187 | 0.34 | 1 | MS.Trt_vs_MS.NoTrt_F |
| SPSB3 | 4.32E-04 | 0.39728 | 0.266 | 0.46 | 1 | MS.Trt_vs_MS.NoTrt_F |
| TNFSF12 | 4.34E-04 | -0.3053 | 0.09 | 0.2 | 1 | MS.Trt_vs_MS.NoTrt_F |
| SPRYD3 | 4.35E-04 | -0.3955 | 0.155 | 0.3 | 1 | MS.Trt_vs_MS.NoTrt_F |
| PHC1 | 4.36E-04 | -0.5897 | 0.112 | 0.24 | 1 | MS.Trt_vs_MS.NoTrt_F |
| APH1A | 4.37E-04 | -0.3327 | 0.18 | 0.33 | 1 | MS.Trt_vs_MS.NoTrt_F |
| BYSL | 4.43E-04 | -2.5246 | 0.061 | 0.16 | 1 | MS.Trt_vs_MS.NoTrt_F |
| TUBG2 | 4.45E-04 | -1.0237 | 0.079 | 0.19 | 1 | MS.Trt_vs_MS.NoTrt_F |
| SERPINI1 | 4.46E-04 | -1.6192 | 0.054 | 0.15 | 1 | MS.Trt_vs_MS.NoTrt_F |
| SGTB | 4.47E-04 | -0.3876 | 0.633 | 0.74 | 1 | MS.Trt_vs_MS.NoTrt_F |
| CD200 | 4.48E-04 | 0.94851 | 0.698 | 0.53 | 1 | MS.Trt_vs_MS.NoTrt_F |
| TRMU | 4.48E-04 | -0.3771 | 0.216 | 0.38 | 1 | MS.Trt_vs_MS.NoTrt_F |
| FAM76B | 0.00045 | -0.3828 | 0.101 | 0.22 | 1 | MS.Trt_vs_MS.NoTrt_F |
| TCF12 | 4.52E-04 | -0.7002 | 0.273 | 0.43 | 1 | MS.Trt_vs_MS.NoTrt_F |
| ZNF395 | 4.53E-04 | 0.31011 | 0.529 | 0.67 | 1 | MS.Trt_vs_MS.NoTrt_F |
| BLCAP | 4.57E-04 | -0.3873 | 0.342 | 0.5 | 1 | MS.Trt_vs_MS.NoTrt_F |
| DNAJC27 | 4.61E-04 | -0.4223 | 0.083 | 0.19 | 1 | MS.Trt_vs_MS.NoTrt_F |
| PIAS3 | 4.61E-04 | 0.27348 | 0.115 | 0.25 | 1 | MS.Trt_vs_MS.NoTrt_F |
| MS4A10 | 4.62E-04 | 0.37402 | 0.773 | 0.64 | 1 | MS.Trt_vs_MS.NoTrt_F |
| TMEM42 | 4.65E-04 | -0.5052 | 0.147 | 0.28 | 1 | MS.Trt_vs_MS.NoTrt_F |
| BICD1 | 4.66E-04 | -0.9831 | 0.086 | 0.2 | 1 | MS.Trt_vs_MS.NoTrt_F |
| CSGALNACT | 4.66E-04 | -0.8068 | 0.151 | 0.29 | 1 | MS.Trt_vs_MS.NoTrt_F |
| PRPSAP2 | 4.69E-04 | -0.697 | 0.568 | 0.71 | 1 | MS.Trt_vs_MS.NoTrt_F |
| USP5 | 4.73E-04 | -0.6044 | 0.122 | 0.25 | 1 | MS.Trt_vs_MS.NoTrt_F |
| SLU7 | 4.73E-04 | -0.5711 | 0.396 | 0.55 | 1 | MS.Trt_vs_MS.NoTrt_F |
| ATP6V1C1 | 4.75E-04 | 0.4959 | 0.439 | 0.6 | 1 | MS.Trt_vs_MS.NoTrt_F |
| AMPD2 | 4.78E-04 | -1.1047 | 0.09 | 0.2 | 1 | MS.Trt_vs_MS.NoTrt_F |
| S100A11 | 4.79E-04 | 0.40684 | 0.374 | 0.62 | 1 | MS.Trt_vs_MS.NoTrt_F |

| | | | | | | |
|---|---|---|---|---|---|---|
| MTRNR2L3 | 4.81E-04 | 0.26993 | 0.41 | 0.25 | 1 | MS.Trt_vs_MS.NoTrt_F |
| FOXK2 | 4.85E-04 | -0.5384 | 0.281 | 0.42 | 1 | MS.Trt_vs_MS.NoTrt_F |
| MRPS18A | 4.86E-04 | 0.33598 | 0.112 | 0.24 | 1 | MS.Trt_vs_MS.NoTrt_F |
| ERBIN | 4.86E-04 | -0.391 | 0.248 | 0.41 | 1 | MS.Trt_vs_MS.NoTrt_F |
| PABPC4 | 4.89E-04 | -0.3945 | 0.766 | 0.87 | 1 | MS.Trt_vs_MS.NoTrt_F |
| ZMYM2 | 4.90E-04 | -0.2989 | 0.342 | 0.5 | 1 | MS.Trt_vs_MS.NoTrt_F |
| EIF2S3 | 4.91E-04 | 0.57767 | 0.957 | 0.95 | 1 | MS.Trt_vs_MS.NoTrt_F |
| KIAA0825 | 4.91E-04 | 1.34103 | 0.302 | 0.47 | 1 | MS.Trt_vs_MS.NoTrt_F |
| DPM1 | 4.95E-04 | -0.7646 | 0.101 | 0.22 | 1 | MS.Trt_vs_MS.NoTrt_F |
| NCDN | 4.99E-04 | -0.3609 | 0.173 | 0.32 | 1 | MS.Trt_vs_MS.NoTrt_F |
| ESRRA | 5.03E-04 | -1.293 | 0.097 | 0.21 | 1 | MS.Trt_vs_MS.NoTrt_F |
| YIPF2 | 5.06E-04 | -0.4512 | 0.09 | 0.2 | 1 | MS.Trt_vs_MS.NoTrt_F |
| SPATA20 | 5.07E-04 | -0.5373 | 0.094 | 0.21 | 1 | MS.Trt_vs_MS.NoTrt_F |
| C5orf15 | 5.07E-04 | 0.44091 | 0.169 | 0.32 | 1 | MS.Trt_vs_MS.NoTrt_F |
| SLC35F2 | 5.08E-04 | -0.3549 | 0.288 | 0.43 | 1 | MS.Trt_vs_MS.NoTrt_F |
| ZNF646 | 5.10E-04 | -0.8961 | 0.072 | 0.18 | 1 | MS.Trt_vs_MS.NoTrt_F |
| UBTF | 5.11E-04 | -0.5336 | 0.237 | 0.38 | 1 | MS.Trt_vs_MS.NoTrt_F |
| NARS2 | 5.13E-04 | -0.7722 | 0.317 | 0.49 | 1 | MS.Trt_vs_MS.NoTrt_F |
| ALDH16A1 | 5.17E-04 | -0.3975 | 0.626 | 0.77 | 1 | MS.Trt_vs_MS.NoTrt_F |
| C15orf39 | 5.18E-04 | -0.6847 | 0.065 | 0.17 | 1 | MS.Trt_vs_MS.NoTrt_F |
| ALKBH7 | 5.18E-04 | -0.2944 | 0.126 | 0.26 | 1 | MS.Trt_vs_MS.NoTrt_F |
| DNAJC11 | 5.18E-04 | -0.5964 | 0.187 | 0.33 | 1 | MS.Trt_vs_MS.NoTrt_F |
| STUB1 | 5.19E-04 | -0.2513 | 0.097 | 0.22 | 1 | MS.Trt_vs_MS.NoTrt_F |
| ZNF746 | 5.20E-04 | 0.32618 | 0.194 | 0.35 | 1 | MS.Trt_vs_MS.NoTrt_F |
| FAAP100 | 5.26E-04 | 0.50869 | 0.162 | 0.32 | 1 | MS.Trt_vs_MS.NoTrt_F |
| NPHP4 | 5.31E-04 | -0.7115 | 0.068 | 0.17 | 1 | MS.Trt_vs_MS.NoTrt_F |
| MZT1 | 5.33E-04 | -0.5644 | 0.058 | 0.16 | 1 | MS.Trt_vs_MS.NoTrt_F |
| HIPK2 | 5.40E-04 | -0.3235 | 0.094 | 0.21 | 1 | MS.Trt_vs_MS.NoTrt_F |
| SS18L2 | 5.41E-04 | -0.3431 | 0.345 | 0.5 | 1 | MS.Trt_vs_MS.NoTrt_F |
| AC138517.6 | 5.45E-04 | -1.0463 | 0.097 | 0.22 | 1 | MS.Trt_vs_MS.NoTrt_F |
| ZSCAN26 | 5.45E-04 | -1.2114 | 0.058 | 0.16 | 1 | MS.Trt_vs_MS.NoTrt_F |
| ZBTB7B | 5.46E-04 | -0.5089 | 0.097 | 0.21 | 1 | MS.Trt_vs_MS.NoTrt_F |
| PITPNA | 5.47E-04 | -1.0272 | 0.097 | 0.22 | 1 | MS.Trt_vs_MS.NoTrt_F |
| SDHAF4 | 5.47E-04 | -0.2762 | 0.655 | 0.79 | 1 | MS.Trt_vs_MS.NoTrt_F |
| NXT1 | 5.47E-04 | -0.7965 | 0.079 | 0.19 | 1 | MS.Trt_vs_MS.NoTrt_F |
| SSX5 | 5.49E-04 | -0.5262 | 0.137 | 0.27 | 1 | MS.Trt_vs_MS.NoTrt_F |
| ARFGEF2 | 5.51E-04 | -0.7787 | 0.406 | 0.57 | 1 | MS.Trt_vs_MS.NoTrt_F |
| CXCR4 | 5.51E-04 | -0.2796 | 0.27 | 0.44 | 1 | MS.Trt_vs_MS.NoTrt_F |
| SLC2A13 | 5.60E-04 | -0.5427 | 0.266 | 0.42 | 1 | MS.Trt_vs_MS.NoTrt_F |
| PEF1 | 5.60E-04 | -0.4294 | 0.158 | 0.3 | 1 | MS.Trt_vs_MS.NoTrt_F |

| | | | | | | |
|---|---|---|---|---|---|---|
| KHNYN | 5.66E-04 | -1.1439 | 0.083 | 0.19 | 1 | MS.Trt_vs_MS.NoTrt_F |
| AGPAT4 | 5.66E-04 | -0.2867 | 0.097 | 0.21 | 1 | MS.Trt_vs_MS.NoTrt_F |
| RPL18A | 5.67E-04 | 0.44979 | 0.881 | 0.97 | 1 | MS.Trt_vs_MS.NoTrt_F |
| ZNF143 | 5.72E-04 | -0.667 | 0.112 | 0.23 | 1 | MS.Trt_vs_MS.NoTrt_F |
| ATP5I | 5.73E-04 | 0.82885 | 0.255 | 0.47 | 1 | MS.Trt_vs_MS.NoTrt_F |
| SCAF11 | 5.75E-04 | 0.31973 | 0.871 | 0.93 | 1 | MS.Trt_vs_MS.NoTrt_F |
| RB1CC1 | 5.78E-04 | -0.4194 | 0.277 | 0.44 | 1 | MS.Trt_vs_MS.NoTrt_F |
| CAB39 | 5.79E-04 | 0.28517 | 0.277 | 0.44 | 1 | MS.Trt_vs_MS.NoTrt_F |
| SEC14L2 | 5.82E-04 | -0.372 | 0.122 | 0.25 | 1 | MS.Trt_vs_MS.NoTrt_F |
| RACGAP1 | 5.85E-04 | -1.0072 | 0.086 | 0.2 | 1 | MS.Trt_vs_MS.NoTrt_F |
| C12orf10 | 5.87E-04 | -0.4013 | 0.201 | 0.35 | 1 | MS.Trt_vs_MS.NoTrt_F |
| ZEB1 | 5.89E-04 | -0.6342 | 0.504 | 0.69 | 1 | MS.Trt_vs_MS.NoTrt_F |
| NAPRT | 5.90E-04 | 0.34419 | 0.155 | 0.3 | 1 | MS.Trt_vs_MS.NoTrt_F |
| GNPDA1 | 5.91E-04 | -0.3221 | 0.076 | 0.18 | 1 | MS.Trt_vs_MS.NoTrt_F |
| PIBF1 | 5.91E-04 | 0.31034 | 0.14 | 0.28 | 1 | MS.Trt_vs_MS.NoTrt_F |
| NUDT16 | 5.91E-04 | -0.6456 | 0.068 | 0.17 | 1 | MS.Trt_vs_MS.NoTrt_F |
| TBC1D15 | 5.95E-04 | -1.6472 | 0.507 | 0.66 | 1 | MS.Trt_vs_MS.NoTrt_F |
| RP11-977G1 | 5.97E-04 | -0.2581 | 0.065 | 0.17 | 1 | MS.Trt_vs_MS.NoTrt_F |
| TG | 5.98E-04 | -0.7932 | 0.112 | 0.23 | 1 | MS.Trt_vs_MS.NoTrt_F |
| MPHOSPH6 | 6.00E-04 | -0.5224 | 0.137 | 0.27 | 1 | MS.Trt_vs_MS.NoTrt_F |
| EIF2S1 | 6.01E-04 | 0.30752 | 0.252 | 0.43 | 1 | MS.Trt_vs_MS.NoTrt_F |
| ATAD3A | 6.04E-04 | 0.36597 | 0.079 | 0.19 | 1 | MS.Trt_vs_MS.NoTrt_F |
| THEM6 | 6.12E-04 | -0.984 | 0.086 | 0.19 | 1 | MS.Trt_vs_MS.NoTrt_F |
| SLC2A3 | 6.13E-04 | -0.2738 | 0.845 | 0.93 | 1 | MS.Trt_vs_MS.NoTrt_F |
| KDM8 | 6.17E-04 | -0.4327 | 0.076 | 0.18 | 1 | MS.Trt_vs_MS.NoTrt_F |
| BCL9 | 6.18E-04 | -0.4016 | 0.09 | 0.2 | 1 | MS.Trt_vs_MS.NoTrt_F |
| AR | 6.18E-04 | -0.6518 | 0.187 | 0.33 | 1 | MS.Trt_vs_MS.NoTrt_F |
| GPATCH3 | 6.19E-04 | -0.8843 | 0.068 | 0.17 | 1 | MS.Trt_vs_MS.NoTrt_F |
| MAP2K7 | 6.22E-04 | -0.2737 | 0.594 | 0.77 | 1 | MS.Trt_vs_MS.NoTrt_F |
| KIAA1468 | 6.27E-04 | -0.3223 | 0.295 | 0.45 | 1 | MS.Trt_vs_MS.NoTrt_F |
| ATF1 | 6.33E-04 | -1.0417 | 0.129 | 0.26 | 1 | MS.Trt_vs_MS.NoTrt_F |
| ATP13A1 | 6.33E-04 | -0.615 | 0.32 | 0.44 | 1 | MS.Trt_vs_MS.NoTrt_F |
| PSTPIP2 | 6.35E-04 | -0.5775 | 0.317 | 0.48 | 1 | MS.Trt_vs_MS.NoTrt_F |
| KATNAL1 | 6.35E-04 | -0.6279 | 0.522 | 0.64 | 1 | MS.Trt_vs_MS.NoTrt_F |
| FRA10AC1 | 6.38E-04 | 0.30736 | 0.489 | 0.69 | 1 | MS.Trt_vs_MS.NoTrt_F |
| LRRC14 | 6.49E-04 | 0.30423 | 0.079 | 0.19 | 1 | MS.Trt_vs_MS.NoTrt_F |
| CCDC58 | 6.58E-04 | -0.4048 | 0.27 | 0.43 | 1 | MS.Trt_vs_MS.NoTrt_F |
| B3GALT4 | 6.60E-04 | -1.0302 | 0.09 | 0.2 | 1 | MS.Trt_vs_MS.NoTrt_F |
| CDKN2A | 6.60E-04 | -0.4086 | 0.076 | 0.18 | 1 | MS.Trt_vs_MS.NoTrt_F |
| VCPKMT | 6.66E-04 | -1.077 | 0.101 | 0.22 | 1 | MS.Trt_vs_MS.NoTrt_F |

| | | | | | | |
|---|---|---|---|---|---|---|
| LNPEP | 6.67E-04 | -0.3477 | 0.529 | 0.67 | 1 | MS.Trt_vs_MS.NoTrt_F |
| PHF10 | 6.71E-04 | -0.2634 | 0.09 | 0.2 | 1 | MS.Trt_vs_MS.NoTrt_F |
| PTPN2 | 6.75E-04 | -0.6282 | 0.96 | 0.98 | 1 | MS.Trt_vs_MS.NoTrt_F |
| PITRM1 | 6.75E-04 | -0.2892 | 0.14 | 0.28 | 1 | MS.Trt_vs_MS.NoTrt_F |
| MOB2 | 6.77E-04 | -0.6442 | 0.14 | 0.27 | 1 | MS.Trt_vs_MS.NoTrt_F |
| VCPIP1 | 6.81E-04 | -0.389 | 0.371 | 0.53 | 1 | MS.Trt_vs_MS.NoTrt_F |
| CENPT | 6.82E-04 | -0.2713 | 0.205 | 0.37 | 1 | MS.Trt_vs_MS.NoTrt_F |
| PIGT | 6.85E-04 | -1.3005 | 0.097 | 0.21 | 1 | MS.Trt_vs_MS.NoTrt_F |
| IDUA | 0.00069 | -0.6228 | 0.097 | 0.21 | 1 | MS.Trt_vs_MS.NoTrt_F |
| MPP7 | 6.88E-04 | -0.6749 | 0.266 | 0.41 | 1 | MS.Trt_vs_MS.NoTrt_F |
| VIPAS39 | 6.88E-04 | -1.0356 | 0.083 | 0.19 | 1 | MS.Trt_vs_MS.NoTrt_F |
| GLS | 6.98E-04 | -0.3372 | 0.637 | 0.75 | 1 | MS.Trt_vs_MS.NoTrt_F |
| GNL3 | 7.08E-04 | 0.26616 | 0.137 | 0.27 | 1 | MS.Trt_vs_MS.NoTrt_F |
| DDX1 | 7.11E-04 | 0.50084 | 0.133 | 0.27 | 1 | MS.Trt_vs_MS.NoTrt_F |
| NHLRC3 | 7.20E-04 | 0.25246 | 0.345 | 0.5 | 1 | MS.Trt_vs_MS.NoTrt_F |
| ARPC4-TTLL3 | 7.22E-04 | 0.32018 | 0.126 | 0.25 | 1 | MS.Trt_vs_MS.NoTrt_F |
| ATF4 | 7.22E-04 | -0.3638 | 0.155 | 0.29 | 1 | MS.Trt_vs_MS.NoTrt_F |
| NOP14 | 7.22E-04 | -1.4609 | 0.392 | 0.55 | 1 | MS.Trt_vs_MS.NoTrt_F |
| CHKB-CPT1B | 7.23E-04 | -1.1268 | 0.036 | 0.12 | 1 | MS.Trt_vs_MS.NoTrt_F |
| MMGT1 | 7.25E-04 | -1.3583 | 0.097 | 0.21 | 1 | MS.Trt_vs_MS.NoTrt_F |
| GRAMD1A | 7.27E-04 | 0.25371 | 0.651 | 0.79 | 1 | MS.Trt_vs_MS.NoTrt_F |
| CASP8AP2 | 7.27E-04 | -0.5643 | 0.291 | 0.44 | 1 | MS.Trt_vs_MS.NoTrt_F |
| ROBO1 | 7.28E-04 | 0.30348 | 0.651 | 0.48 | 1 | MS.Trt_vs_MS.NoTrt_F |
| ZC2HC1A | 7.31E-04 | 0.34623 | 0.169 | 0.32 | 1 | MS.Trt_vs_MS.NoTrt_F |
| C1QTNF1 | 7.31E-04 | 0.5803 | 0.219 | 0.39 | 1 | MS.Trt_vs_MS.NoTrt_F |
| EMC4 | 7.32E-04 | 0.6978 | 1 | 0.99 | 1 | MS.Trt_vs_MS.NoTrt_F |
| ARHGEF6 | 7.35E-04 | 0.27908 | 0.309 | 0.49 | 1 | MS.Trt_vs_MS.NoTrt_F |
| OLFM2 | 7.39E-04 | -0.7547 | 0.09 | 0.2 | 1 | MS.Trt_vs_MS.NoTrt_F |
| RAB11B | 7.40E-04 | -0.4568 | 0.453 | 0.62 | 1 | MS.Trt_vs_MS.NoTrt_F |
| OFD1 | 7.49E-04 | 0.54397 | 0.201 | 0.37 | 1 | MS.Trt_vs_MS.NoTrt_F |
| UTY | 7.50E-04 | -1.8846 | 0.428 | 0.57 | 1 | MS.Trt_vs_MS.NoTrt_F |
| CPPED1 | 7.51E-04 | -0.8211 | 0.579 | 0.7 | 1 | MS.Trt_vs_MS.NoTrt_F |
| FAM122B | 7.52E-04 | -0.4033 | 0.115 | 0.23 | 1 | MS.Trt_vs_MS.NoTrt_F |
| SPON2 | 7.58E-04 | -0.338 | 0.367 | 0.53 | 1 | MS.Trt_vs_MS.NoTrt_F |
| CHID1 | 7.65E-04 | -0.4639 | 0.248 | 0.4 | 1 | MS.Trt_vs_MS.NoTrt_F |
| ANKRD18A | 7.66E-04 | -1.194 | 0.273 | 0.42 | 1 | MS.Trt_vs_MS.NoTrt_F |
| PAM16 | 7.75E-04 | -0.3385 | 0.144 | 0.28 | 1 | MS.Trt_vs_MS.NoTrt_F |
| SPDYE16 | 7.79E-04 | 0.72943 | 0.838 | 0.89 | 1 | MS.Trt_vs_MS.NoTrt_F |
| IQSEC1 | 7.82E-04 | -0.4836 | 0.191 | 0.34 | 1 | MS.Trt_vs_MS.NoTrt_F |
| SPDYE1 | 7.82E-04 | 0.99755 | 0.586 | 0.47 | 1 | MS.Trt_vs_MS.NoTrt_F |

| | | | | | | |
|---|---|---|---|---|---|---|
| KARS | 7.82E-04 | -0.469 | 0.212 | 0.36 | 1 | MS.Trt_vs_MS.NoTrt_F |
| NPRL2 | 7.82E-04 | -0.3032 | 0.151 | 0.29 | 1 | MS.Trt_vs_MS.NoTrt_F |
| INTS5 | 7.83E-04 | -1.123 | 0.076 | 0.18 | 1 | MS.Trt_vs_MS.NoTrt_F |
| CDK8 | 7.84E-04 | -1.7238 | 0.058 | 0.15 | 1 | MS.Trt_vs_MS.NoTrt_F |
| ZNF408 | 7.84E-04 | -1.7789 | 0.05 | 0.14 | 1 | MS.Trt_vs_MS.NoTrt_F |
| CANT1 | 7.87E-04 | -0.82 | 0.094 | 0.2 | 1 | MS.Trt_vs_MS.NoTrt_F |
| B4GALT3 | 7.88E-04 | -0.3068 | 0.191 | 0.34 | 1 | MS.Trt_vs_MS.NoTrt_F |
| RWDD4 | 7.98E-04 | 0.33041 | 0.094 | 0.21 | 1 | MS.Trt_vs_MS.NoTrt_F |
| RNF8 | 8.00E-04 | -0.7738 | 0.453 | 0.64 | 1 | MS.Trt_vs_MS.NoTrt_F |
| GRB10 | 8.02E-04 | -0.2556 | 0.183 | 0.32 | 1 | MS.Trt_vs_MS.NoTrt_F |
| IFRD1 | 8.05E-04 | 0.25892 | 0.417 | 0.58 | 1 | MS.Trt_vs_MS.NoTrt_F |
| TMEM120A | 8.07E-04 | 0.48782 | 0.137 | 0.28 | 1 | MS.Trt_vs_MS.NoTrt_F |
| NADK | 8.07E-04 | 0.48808 | 0.23 | 0.38 | 1 | MS.Trt_vs_MS.NoTrt_F |
| SERTAD2 | 8.13E-04 | -0.474 | 0.115 | 0.24 | 1 | MS.Trt_vs_MS.NoTrt_F |
| ZNF398 | 8.14E-04 | -0.6843 | 0.165 | 0.3 | 1 | MS.Trt_vs_MS.NoTrt_F |
| PRCC | 8.15E-04 | -0.432 | 0.151 | 0.29 | 1 | MS.Trt_vs_MS.NoTrt_F |
| KCTD5 | 8.25E-04 | -1.0012 | 0.086 | 0.19 | 1 | MS.Trt_vs_MS.NoTrt_F |
| CCDC97 | 8.30E-04 | -0.5336 | 0.147 | 0.28 | 1 | MS.Trt_vs_MS.NoTrt_F |
| NUP153 | 8.32E-04 | -0.4336 | 0.194 | 0.34 | 1 | MS.Trt_vs_MS.NoTrt_F |
| ERI3 | 8.35E-04 | -0.6879 | 0.094 | 0.2 | 1 | MS.Trt_vs_MS.NoTrt_F |
| FLYWCH1 | 8.36E-04 | -0.6795 | 0.759 | 0.9 | 1 | MS.Trt_vs_MS.NoTrt_F |
| SOAT1 | 8.39E-04 | -0.4026 | 0.676 | 0.79 | 1 | MS.Trt_vs_MS.NoTrt_F |
| ZNF717 | 8.41E-04 | -0.3225 | 0.05 | 0.14 | 1 | MS.Trt_vs_MS.NoTrt_F |
| ITGB3BP | 8.46E-04 | -0.3326 | 0.665 | 0.83 | 1 | MS.Trt_vs_MS.NoTrt_F |
| DYNLT1 | 8.55E-04 | -0.3962 | 0.511 | 0.72 | 1 | MS.Trt_vs_MS.NoTrt_F |
| SFXN3 | 8.56E-04 | -0.2713 | 0.306 | 0.46 | 1 | MS.Trt_vs_MS.NoTrt_F |
| ZNF740 | 8.58E-04 | 0.30987 | 0.151 | 0.29 | 1 | MS.Trt_vs_MS.NoTrt_F |
| IKZF3 | 8.68E-04 | -0.272 | 0.953 | 1 | 1 | MS.Trt_vs_MS.NoTrt_F |
| CETN2 | 8.69E-04 | -0.741 | 0.043 | 0.13 | 1 | MS.Trt_vs_MS.NoTrt_F |
| C6orf226 | 8.71E-04 | -0.2539 | 0.043 | 0.13 | 1 | MS.Trt_vs_MS.NoTrt_F |
| MGAT5 | 8.75E-04 | -0.3223 | 0.14 | 0.27 | 1 | MS.Trt_vs_MS.NoTrt_F |
| ARAP2 | 8.75E-04 | 0.35792 | 0.291 | 0.45 | 1 | MS.Trt_vs_MS.NoTrt_F |
| FYCO1 | 8.76E-04 | -0.2586 | 0.176 | 0.32 | 1 | MS.Trt_vs_MS.NoTrt_F |
| IL15RA | 8.76E-04 | -0.2598 | 0.112 | 0.23 | 1 | MS.Trt_vs_MS.NoTrt_F |
| POLDIP3 | 8.81E-04 | -0.5577 | 0.165 | 0.3 | 1 | MS.Trt_vs_MS.NoTrt_F |
| SCAMP3 | 8.81E-04 | -0.383 | 0.169 | 0.31 | 1 | MS.Trt_vs_MS.NoTrt_F |
| CISD1 | 8.85E-04 | 0.31667 | 0.094 | 0.2 | 1 | MS.Trt_vs_MS.NoTrt_F |
| ADAM12 | 8.86E-04 | -0.2915 | 0.212 | 0.37 | 1 | MS.Trt_vs_MS.NoTrt_F |
| ZC3H11A | 8.86E-04 | -0.4995 | 0.212 | 0.35 | 1 | MS.Trt_vs_MS.NoTrt_F |
| GADD45B | 8.89E-04 | -0.5804 | 0.129 | 0.25 | 1 | MS.Trt_vs_MS.NoTrt_F |

| | | | | | | |
|---|---|---|---|---|---|---|
| RAVER1 | 8.92E-04 | -0.5209 | 0.147 | 0.28 | 1 | MS.Trt_vs_MS.NoTrt_F |
| HCFC2 | 8.94E-04 | -1.1554 | 0.079 | 0.18 | 1 | MS.Trt_vs_MS.NoTrt_F |
| DNAJB11 | 8.97E-04 | -0.3633 | 0.162 | 0.3 | 1 | MS.Trt_vs_MS.NoTrt_F |
| SMPD1 | 9.02E-04 | 0.25969 | 0.112 | 0.23 | 1 | MS.Trt_vs_MS.NoTrt_F |
| EGLN2 | 9.04E-04 | 0.32895 | 0.255 | 0.44 | 1 | MS.Trt_vs_MS.NoTrt_F |
| POLR2C | 9.04E-04 | -0.4826 | 0.547 | 0.68 | 1 | MS.Trt_vs_MS.NoTrt_F |
| GLYR1 | 9.04E-04 | -0.5595 | 0.414 | 0.54 | 1 | MS.Trt_vs_MS.NoTrt_F |
| NCAPD3 | 9.04E-04 | -1.3718 | 0.079 | 0.18 | 1 | MS.Trt_vs_MS.NoTrt_F |
| ATG101 | 9.07E-04 | -0.625 | 0.072 | 0.17 | 1 | MS.Trt_vs_MS.NoTrt_F |
| FEZ2 | 9.07E-04 | -1.0302 | 0.46 | 0.61 | 1 | MS.Trt_vs_MS.NoTrt_F |
| MED20 | 9.07E-04 | -0.7926 | 0.32 | 0.48 | 1 | MS.Trt_vs_MS.NoTrt_F |
| EXOSC7 | 9.07E-04 | -0.3139 | 0.403 | 0.56 | 1 | MS.Trt_vs_MS.NoTrt_F |
| INSL6 | 9.09E-04 | -0.3059 | 0.295 | 0.46 | 1 | MS.Trt_vs_MS.NoTrt_F |
| VN1R2 | 9.11E-04 | -0.3416 | 0.079 | 0.18 | 1 | MS.Trt_vs_MS.NoTrt_F |
| SIGMAR1 | 9.13E-04 | -0.2687 | 0.183 | 0.33 | 1 | MS.Trt_vs_MS.NoTrt_F |
| SLC18B1 | 9.15E-04 | -1.2069 | 0.079 | 0.18 | 1 | MS.Trt_vs_MS.NoTrt_F |
| TLDC1 | 9.20E-04 | -0.5625 | 0.64 | 0.85 | 1 | MS.Trt_vs_MS.NoTrt_F |
| STARD5 | 9.22E-04 | -0.4592 | 0.133 | 0.25 | 1 | MS.Trt_vs_MS.NoTrt_F |
| HLX | 9.22E-04 | -0.2703 | 0.173 | 0.3 | 1 | MS.Trt_vs_MS.NoTrt_F |
| HIGD1A | 9.22E-04 | 0.46527 | 0.155 | 0.29 | 1 | MS.Trt_vs_MS.NoTrt_F |
| LCORL | 9.28E-04 | 0.40455 | 0.112 | 0.23 | 1 | MS.Trt_vs_MS.NoTrt_F |
| DDX19B | 9.28E-04 | -0.5033 | 0.414 | 0.57 | 1 | MS.Trt_vs_MS.NoTrt_F |
| ATP23 | 9.29E-04 | -0.2819 | 0.119 | 0.24 | 1 | MS.Trt_vs_MS.NoTrt_F |
| COA6 | 9.29E-04 | 0.32542 | 0.115 | 0.24 | 1 | MS.Trt_vs_MS.NoTrt_F |
| WRNIP1 | 9.34E-04 | 0.26276 | 0.079 | 0.18 | 1 | MS.Trt_vs_MS.NoTrt_F |
| NELFB | 9.39E-04 | -0.5396 | 0.058 | 0.15 | 1 | MS.Trt_vs_MS.NoTrt_F |
| TIMD4 | 9.42E-04 | -0.4887 | 0.072 | 0.17 | 1 | MS.Trt_vs_MS.NoTrt_F |
| CUEDC2 | 9.45E-04 | 0.39459 | 0.115 | 0.24 | 1 | MS.Trt_vs_MS.NoTrt_F |
| TENM4 | 9.45E-04 | -0.2855 | 0.234 | 0.38 | 1 | MS.Trt_vs_MS.NoTrt_F |
| C19orf53 | 9.48E-04 | 0.29635 | 0.374 | 0.55 | 1 | MS.Trt_vs_MS.NoTrt_F |
| CTD-2583A1 | 9.48E-04 | -0.4162 | 0.076 | 0.18 | 1 | MS.Trt_vs_MS.NoTrt_F |
| PTGER4 | 9.49E-04 | 0.4234 | 0.173 | 0.32 | 1 | MS.Trt_vs_MS.NoTrt_F |
| FLYWCH2 | 9.50E-04 | -0.611 | 0.072 | 0.17 | 1 | MS.Trt_vs_MS.NoTrt_F |
| KPNA5 | 9.50E-04 | -0.3605 | 0.928 | 0.98 | 1 | MS.Trt_vs_MS.NoTrt_F |
| ZBTB45 | 9.53E-04 | -0.7242 | 0.054 | 0.14 | 1 | MS.Trt_vs_MS.NoTrt_F |
| TCF20 | 9.54E-04 | -0.6751 | 0.144 | 0.27 | 1 | MS.Trt_vs_MS.NoTrt_F |
| RIF1 | 9.54E-04 | -0.2969 | 0.522 | 0.71 | 1 | MS.Trt_vs_MS.NoTrt_F |
| ORMDL2 | 9.54E-04 | -0.4784 | 0.41 | 0.56 | 1 | MS.Trt_vs_MS.NoTrt_F |
| CTBP1 | 9.55E-04 | 0.25343 | 0.367 | 0.56 | 1 | MS.Trt_vs_MS.NoTrt_F |
| C1orf158 | 9.57E-04 | 0.63972 | 0.878 | 0.79 | 1 | MS.Trt_vs_MS.NoTrt_F |

| | | | | | | |
|---|---|---|---|---|---|---|
| AIM1L | 9.60E-04 | -0.5889 | 0.061 | 0.16 | 1 | MS.Trt_vs_MS.NoTrt_F |
| HK1 | 9.60E-04 | -0.5192 | 0.522 | 0.66 | 1 | MS.Trt_vs_MS.NoTrt_F |
| VPS54 | 9.62E-04 | 0.51317 | 0.356 | 0.53 | 1 | MS.Trt_vs_MS.NoTrt_F |
| GSKIP | 9.62E-04 | -0.3555 | 0.112 | 0.23 | 1 | MS.Trt_vs_MS.NoTrt_F |
| MYCBP | 9.64E-04 | 0.39543 | 0.112 | 0.23 | 1 | MS.Trt_vs_MS.NoTrt_F |
| GNG5 | 9.64E-04 | 0.27041 | 0.252 | 0.43 | 1 | MS.Trt_vs_MS.NoTrt_F |
| KLRC1 | 9.64E-04 | 0.87059 | 0.266 | 0.44 | 1 | MS.Trt_vs_MS.NoTrt_F |
| TDP2 | 9.66E-04 | -0.2985 | 0.115 | 0.23 | 1 | MS.Trt_vs_MS.NoTrt_F |
| WAPL | 9.67E-04 | -0.4565 | 0.406 | 0.58 | 1 | MS.Trt_vs_MS.NoTrt_F |
| FAM96A | 9.70E-04 | -0.2506 | 0.295 | 0.45 | 1 | MS.Trt_vs_MS.NoTrt_F |
| FBXO28 | 9.72E-04 | -1.1011 | 0.108 | 0.22 | 1 | MS.Trt_vs_MS.NoTrt_F |
| ZCCHC14 | 9.73E-04 | -0.8337 | 0.079 | 0.18 | 1 | MS.Trt_vs_MS.NoTrt_F |
| TYSND1 | 9.76E-04 | 0.54694 | 0.23 | 0.38 | 1 | MS.Trt_vs_MS.NoTrt_F |
| TMTC2 | 9.77E-04 | -0.7839 | 0.065 | 0.16 | 1 | MS.Trt_vs_MS.NoTrt_F |
| CNOT3 | 9.84E-04 | -0.7312 | 0.137 | 0.26 | 1 | MS.Trt_vs_MS.NoTrt_F |
| CDKN1A | 9.87E-04 | -0.4595 | 0.119 | 0.24 | 1 | MS.Trt_vs_MS.NoTrt_F |
| PKD2 | 9.88E-04 | 0.44085 | 0.173 | 0.31 | 1 | MS.Trt_vs_MS.NoTrt_F |
| LPAR2 | 9.89E-04 | 0.74236 | 0.129 | 0.25 | 1 | MS.Trt_vs_MS.NoTrt_F |
| COX6A1 | 9.90E-04 | 0.37728 | 0.162 | 0.3 | 1 | MS.Trt_vs_MS.NoTrt_F |
| ARFIP1 | 9.90E-04 | 0.30766 | 0.23 | 0.37 | 1 | MS.Trt_vs_MS.NoTrt_F |
| C1GALT1C1 | 9.92E-04 | -0.5373 | 0.068 | 0.17 | 1 | MS.Trt_vs_MS.NoTrt_F |
| MDM1 | 9.99E-04 | -0.8385 | 0.094 | 0.2 | 1 | MS.Trt_vs_MS.NoTrt_F |
| UHMK1 | 0.001 | 0.27704 | 0.255 | 0.41 | 1 | MS.Trt_vs_MS.NoTrt_F |
| SCFD2 | 0.001 | -0.2711 | 0.061 | 0.16 | 1 | MS.Trt_vs_MS.NoTrt_F |
| FUS | 0.00101 | -0.287 | 0.842 | 0.9 | 1 | MS.Trt_vs_MS.NoTrt_F |
| ZSWIM6 | 0.00103 | -0.3073 | 0.076 | 0.18 | 1 | MS.Trt_vs_MS.NoTrt_F |
| FAM118B | 0.00103 | -0.5598 | 0.068 | 0.17 | 1 | MS.Trt_vs_MS.NoTrt_F |
| GINM1 | 0.00103 | -0.3022 | 0.536 | 0.67 | 1 | MS.Trt_vs_MS.NoTrt_F |
| PPRC1 | 0.00104 | -0.5127 | 0.119 | 0.24 | 1 | MS.Trt_vs_MS.NoTrt_F |
| ALDH9A1 | 0.00104 | -0.4249 | 0.176 | 0.32 | 1 | MS.Trt_vs_MS.NoTrt_F |
| TMEM140 | 0.00104 | -0.2636 | 0.155 | 0.28 | 1 | MS.Trt_vs_MS.NoTrt_F |
| CACNA2D4 | 0.00104 | -0.3526 | 0.068 | 0.17 | 1 | MS.Trt_vs_MS.NoTrt_F |
| KCTD11 | 0.00104 | 0.5663 | 0.122 | 0.24 | 1 | MS.Trt_vs_MS.NoTrt_F |
| MYO5C | 0.00104 | 0.70357 | 0.392 | 0.55 | 1 | MS.Trt_vs_MS.NoTrt_F |
| MASTL | 0.00105 | 0.4201 | 0.076 | 0.18 | 1 | MS.Trt_vs_MS.NoTrt_F |
| ESF1 | 0.00105 | -0.3182 | 0.129 | 0.25 | 1 | MS.Trt_vs_MS.NoTrt_F |
| SP3 | 0.00105 | -1.0105 | 0.489 | 0.64 | 1 | MS.Trt_vs_MS.NoTrt_F |
| CWC15 | 0.00105 | 0.36146 | 0.201 | 0.37 | 1 | MS.Trt_vs_MS.NoTrt_F |
| PHF8 | 0.00106 | -0.5198 | 0.119 | 0.24 | 1 | MS.Trt_vs_MS.NoTrt_F |
| TCF19 | 0.00106 | -0.4161 | 0.061 | 0.16 | 1 | MS.Trt_vs_MS.NoTrt_F |

| | | | | | | |
|---|---|---|---|---|---|---|
| EZH1 | 0.00106 | -0.6423 | 0.881 | 0.95 | 1 | MS.Trt_vs_MS.NoTrt_F |
| N4BP1 | 0.00106 | 0.25328 | 0.122 | 0.25 | 1 | MS.Trt_vs_MS.NoTrt_F |
| TAOK2 | 0.00106 | -0.3615 | 0.094 | 0.2 | 1 | MS.Trt_vs_MS.NoTrt_F |
| TAX1BP3 | 0.00107 | -0.6869 | 0.076 | 0.18 | 1 | MS.Trt_vs_MS.NoTrt_F |
| SLC25A35 | 0.00107 | -0.5763 | 0.137 | 0.27 | 1 | MS.Trt_vs_MS.NoTrt_F |
| PARP10 | 0.00107 | -0.2863 | 0.194 | 0.33 | 1 | MS.Trt_vs_MS.NoTrt_F |
| ZNF182 | 0.00107 | -0.9829 | 0.076 | 0.18 | 1 | MS.Trt_vs_MS.NoTrt_F |
| C4orf48 | 0.00107 | 0.62083 | 0.122 | 0.25 | 1 | MS.Trt_vs_MS.NoTrt_F |
| GK | 0.00108 | -1.0005 | 0.187 | 0.34 | 1 | MS.Trt_vs_MS.NoTrt_F |
| DHRS12 | 0.00108 | -0.6946 | 0.09 | 0.2 | 1 | MS.Trt_vs_MS.NoTrt_F |
| STK25 | 0.0011 | -0.4491 | 0.633 | 0.78 | 1 | MS.Trt_vs_MS.NoTrt_F |
| CDK12 | 0.00111 | -0.7056 | 0.489 | 0.65 | 1 | MS.Trt_vs_MS.NoTrt_F |
| APTX | 0.00111 | -0.3238 | 0.191 | 0.34 | 1 | MS.Trt_vs_MS.NoTrt_F |
| EHMT2 | 0.00111 | -0.9213 | 0.604 | 0.77 | 1 | MS.Trt_vs_MS.NoTrt_F |
| PIN4 | 0.00112 | 0.61806 | 0.129 | 0.26 | 1 | MS.Trt_vs_MS.NoTrt_F |
| SFSWAP | 0.00112 | -0.4934 | 0.522 | 0.66 | 1 | MS.Trt_vs_MS.NoTrt_F |
| UBE2J1 | 0.00112 | 0.31439 | 0.18 | 0.32 | 1 | MS.Trt_vs_MS.NoTrt_F |
| COPG2 | 0.00112 | 1.2396 | 0.09 | 0.2 | 1 | MS.Trt_vs_MS.NoTrt_F |
| PDE4D | 0.00112 | 0.43058 | 0.252 | 0.4 | 1 | MS.Trt_vs_MS.NoTrt_F |
| FUNDC2 | 0.00113 | -0.501 | 0.478 | 0.62 | 1 | MS.Trt_vs_MS.NoTrt_F |
| MRPS36 | 0.00114 | 0.43435 | 0.14 | 0.28 | 1 | MS.Trt_vs_MS.NoTrt_F |
| SNX12 | 0.00115 | -0.8625 | 0.094 | 0.2 | 1 | MS.Trt_vs_MS.NoTrt_F |
| EIF3J | 0.00115 | 0.40074 | 0.165 | 0.32 | 1 | MS.Trt_vs_MS.NoTrt_F |
| PLGLB1 | 0.00115 | -0.3808 | 0.112 | 0.23 | 1 | MS.Trt_vs_MS.NoTrt_F |
| DHX38 | 0.00115 | -0.2949 | 0.119 | 0.24 | 1 | MS.Trt_vs_MS.NoTrt_F |
| MAPK14 | 0.00117 | -0.6249 | 0.133 | 0.25 | 1 | MS.Trt_vs_MS.NoTrt_F |
| TMEM50B | 0.00117 | -0.3526 | 0.669 | 0.81 | 1 | MS.Trt_vs_MS.NoTrt_F |
| PHTF2 | 0.00117 | 0.32042 | 0.374 | 0.57 | 1 | MS.Trt_vs_MS.NoTrt_F |
| CALD1 | 0.00118 | 0.71844 | 0.191 | 0.32 | 1 | MS.Trt_vs_MS.NoTrt_F |
| TIAM2 | 0.00119 | -0.3191 | 0.748 | 0.86 | 1 | MS.Trt_vs_MS.NoTrt_F |
| YBX1 | 0.00119 | 0.35636 | 0.496 | 0.7 | 1 | MS.Trt_vs_MS.NoTrt_F |
| ZNF783 | 0.00119 | -0.7545 | 0.381 | 0.52 | 1 | MS.Trt_vs_MS.NoTrt_F |
| RTN3 | 0.0012 | -0.3032 | 0.155 | 0.28 | 1 | MS.Trt_vs_MS.NoTrt_F |
| SAFB2 | 0.0012 | -0.5552 | 0.831 | 0.9 | 1 | MS.Trt_vs_MS.NoTrt_F |
| SIM1 | 0.0012 | 0.90772 | 0.158 | 0.28 | 1 | MS.Trt_vs_MS.NoTrt_F |
| RP11-411B6 | 0.0012 | 0.25763 | 0.234 | 0.4 | 1 | MS.Trt_vs_MS.NoTrt_F |
| NCOA5 | 0.00122 | -0.293 | 0.129 | 0.25 | 1 | MS.Trt_vs_MS.NoTrt_F |
| WDR37 | 0.00123 | 0.25797 | 0.155 | 0.29 | 1 | MS.Trt_vs_MS.NoTrt_F |
| PSENEN | 0.00123 | 0.41817 | 0.169 | 0.3 | 1 | MS.Trt_vs_MS.NoTrt_F |
| FAM214A | 0.00123 | -0.2822 | 0.194 | 0.33 | 1 | MS.Trt_vs_MS.NoTrt_F |

| | | | | | | |
|---|---|---|---|---|---|---|
| SMG7 | 0.00125 | -0.3253 | 0.162 | 0.3 | 1 | MS.Trt_vs_MS.NoTrt_F |
| NPDC1 | 0.00125 | -0.3307 | 0.072 | 0.17 | 1 | MS.Trt_vs_MS.NoTrt_F |
| SLFN12 | 0.00125 | -0.5597 | 0.054 | 0.14 | 1 | MS.Trt_vs_MS.NoTrt_F |
| HEXB | 0.00125 | -0.3883 | 0.151 | 0.28 | 1 | MS.Trt_vs_MS.NoTrt_F |
| PPP2R3C | 0.00125 | -0.5144 | 0.176 | 0.3 | 1 | MS.Trt_vs_MS.NoTrt_F |
| CHMP2B | 0.00126 | 0.28747 | 0.144 | 0.27 | 1 | MS.Trt_vs_MS.NoTrt_F |
| LMNA | 0.00126 | -0.3833 | 0.651 | 0.76 | 1 | MS.Trt_vs_MS.NoTrt_F |
| TOMM6 | 0.00128 | 0.27968 | 0.219 | 0.38 | 1 | MS.Trt_vs_MS.NoTrt_F |
| IGFBPL1 | 0.00129 | -0.2613 | 0.518 | 0.65 | 1 | MS.Trt_vs_MS.NoTrt_F |
| ATP5J2 | 0.00129 | 0.78572 | 0.331 | 0.53 | 1 | MS.Trt_vs_MS.NoTrt_F |
| SLMAP | 0.00129 | -0.5722 | 0.759 | 0.83 | 1 | MS.Trt_vs_MS.NoTrt_F |
| SNW1 | 0.00129 | -0.264 | 0.234 | 0.38 | 1 | MS.Trt_vs_MS.NoTrt_F |
| ARHGEF1 | 0.0013 | -0.3023 | 0.885 | 0.93 | 1 | MS.Trt_vs_MS.NoTrt_F |
| DDX60 | 0.0013 | -0.3627 | 0.126 | 0.25 | 1 | MS.Trt_vs_MS.NoTrt_F |
| FBXL5 | 0.00131 | 0.70129 | 0.234 | 0.42 | 1 | MS.Trt_vs_MS.NoTrt_F |
| NFE2L3 | 0.00131 | -0.8955 | 0.083 | 0.18 | 1 | MS.Trt_vs_MS.NoTrt_F |
| SMARCD1 | 0.00131 | -0.4991 | 0.101 | 0.21 | 1 | MS.Trt_vs_MS.NoTrt_F |
| PDCD5 | 0.00131 | 0.27741 | 0.18 | 0.33 | 1 | MS.Trt_vs_MS.NoTrt_F |
| TMEM9 | 0.00131 | -0.3051 | 0.09 | 0.19 | 1 | MS.Trt_vs_MS.NoTrt_F |
| ADD3 | 0.00131 | -0.3588 | 0.716 | 0.82 | 1 | MS.Trt_vs_MS.NoTrt_F |
| SSBP4 | 0.00132 | -0.6776 | 0.083 | 0.18 | 1 | MS.Trt_vs_MS.NoTrt_F |
| PLAG1 | 0.00132 | 0.76961 | 0.367 | 0.52 | 1 | MS.Trt_vs_MS.NoTrt_F |
| CAMTA1 | 0.00133 | 0.6125 | 0.165 | 0.3 | 1 | MS.Trt_vs_MS.NoTrt_F |
| KLF7 | 0.00133 | 0.9579 | 0.133 | 0.25 | 1 | MS.Trt_vs_MS.NoTrt_F |
| SUPT20H | 0.00133 | -0.4569 | 0.414 | 0.56 | 1 | MS.Trt_vs_MS.NoTrt_F |
| FBXW11 | 0.00133 | 0.52074 | 0.277 | 0.45 | 1 | MS.Trt_vs_MS.NoTrt_F |
| ZBTB33 | 0.00133 | -0.4454 | 0.094 | 0.2 | 1 | MS.Trt_vs_MS.NoTrt_F |
| PLAT | 0.00135 | 0.27225 | 0.115 | 0.23 | 1 | MS.Trt_vs_MS.NoTrt_F |
| CSNK2A2 | 0.00135 | -0.507 | 0.288 | 0.44 | 1 | MS.Trt_vs_MS.NoTrt_F |
| TEP1 | 0.00136 | -0.4457 | 0.371 | 0.52 | 1 | MS.Trt_vs_MS.NoTrt_F |
| SLC12A7 | 0.00136 | -0.4508 | 0.147 | 0.28 | 1 | MS.Trt_vs_MS.NoTrt_F |
| CRAMP1 | 0.00136 | -0.8769 | 0.09 | 0.19 | 1 | MS.Trt_vs_MS.NoTrt_F |
| PEX7 | 0.00136 | -0.6178 | 0.147 | 0.27 | 1 | MS.Trt_vs_MS.NoTrt_F |
| ISYNA1 | 0.00136 | -0.6182 | 0.083 | 0.18 | 1 | MS.Trt_vs_MS.NoTrt_F |
| SLC35C2 | 0.00136 | -0.4652 | 0.475 | 0.64 | 1 | MS.Trt_vs_MS.NoTrt_F |
| HNRNPA1L2 | 0.00137 | -0.7001 | 0.227 | 0.38 | 1 | MS.Trt_vs_MS.NoTrt_F |
| HDHD2 | 0.00137 | 0.44248 | 0.155 | 0.29 | 1 | MS.Trt_vs_MS.NoTrt_F |
| GPR18 | 0.00137 | -0.6659 | 0.083 | 0.18 | 1 | MS.Trt_vs_MS.NoTrt_F |
| METAP1D | 0.00138 | -1.5767 | 0.036 | 0.11 | 1 | MS.Trt_vs_MS.NoTrt_F |
| HRAS | 0.00138 | -1.4819 | 0.04 | 0.12 | 1 | MS.Trt_vs_MS.NoTrt_F |

| | | | | | | |
|---|---|---|---|---|---|---|
| FAM156A | 0.00139 | -0.5424 | 0.155 | 0.27 | 1 | MS.Trt_vs_MS.NoTrt_F |
| POLDIP2 | 0.0014 | -0.3783 | 0.144 | 0.27 | 1 | MS.Trt_vs_MS.NoTrt_F |
| SF3A3 | 0.0014 | 0.28132 | 0.306 | 0.48 | 1 | MS.Trt_vs_MS.NoTrt_F |
| NFATC2 | 0.0014 | -0.2695 | 0.191 | 0.33 | 1 | MS.Trt_vs_MS.NoTrt_F |
| XAB2 | 0.00141 | 0.49881 | 0.219 | 0.37 | 1 | MS.Trt_vs_MS.NoTrt_F |
| NAA10 | 0.00141 | -0.5423 | 0.378 | 0.52 | 1 | MS.Trt_vs_MS.NoTrt_F |
| ATP9A | 0.00141 | -0.277 | 0.219 | 0.35 | 1 | MS.Trt_vs_MS.NoTrt_F |
| CDC42 | 0.00141 | 0.68497 | 0.91 | 0.97 | 1 | MS.Trt_vs_MS.NoTrt_F |
| WDR4 | 0.00142 | -0.792 | 0.086 | 0.19 | 1 | MS.Trt_vs_MS.NoTrt_F |
| TWF1 | 0.00142 | -0.816 | 0.086 | 0.19 | 1 | MS.Trt_vs_MS.NoTrt_F |
| GLO1 | 0.00143 | -0.541 | 0.086 | 0.19 | 1 | MS.Trt_vs_MS.NoTrt_F |
| CDRT1 | 0.00143 | -1.77 | 0.036 | 0.11 | 1 | MS.Trt_vs_MS.NoTrt_F |
| ITPR3 | 0.00143 | -0.7867 | 0.119 | 0.24 | 1 | MS.Trt_vs_MS.NoTrt_F |
| ETV3 | 0.00144 | -0.342 | 0.18 | 0.32 | 1 | MS.Trt_vs_MS.NoTrt_F |
| C7orf55-LU( | 0.00144 | -0.5892 | 0.086 | 0.19 | 1 | MS.Trt_vs_MS.NoTrt_F |
| ADCK2 | 0.00144 | -2.0453 | 0.047 | 0.13 | 1 | MS.Trt_vs_MS.NoTrt_F |
| AGAP6 | 0.00145 | -0.3997 | 0.14 | 0.25 | 1 | MS.Trt_vs_MS.NoTrt_F |
| BEX2 | 0.00145 | -0.6151 | 0.086 | 0.19 | 1 | MS.Trt_vs_MS.NoTrt_F |
| FBXO7 | 0.00145 | -0.6616 | 0.277 | 0.41 | 1 | MS.Trt_vs_MS.NoTrt_F |
| C8orf44-SGk | 0.00145 | -0.7373 | 0.097 | 0.2 | 1 | MS.Trt_vs_MS.NoTrt_F |
| TAB3 | 0.00146 | -0.4079 | 0.665 | 0.81 | 1 | MS.Trt_vs_MS.NoTrt_F |
| LRRC61 | 0.00148 | -1.432 | 0.076 | 0.17 | 1 | MS.Trt_vs_MS.NoTrt_F |
| BCAP31 | 0.00148 | -0.6384 | 0.176 | 0.31 | 1 | MS.Trt_vs_MS.NoTrt_F |
| OSBPL10 | 0.00148 | 0.38619 | 0.356 | 0.5 | 1 | MS.Trt_vs_MS.NoTrt_F |
| SPATC1L | 0.00148 | -1.1392 | 0.04 | 0.12 | 1 | MS.Trt_vs_MS.NoTrt_F |
| MKRN1 | 0.00148 | -0.3244 | 0.813 | 0.9 | 1 | MS.Trt_vs_MS.NoTrt_F |
| PAM | 0.00148 | 0.28533 | 0.302 | 0.45 | 1 | MS.Trt_vs_MS.NoTrt_F |
| MLYCD | 0.00148 | -1.3189 | 0.345 | 0.51 | 1 | MS.Trt_vs_MS.NoTrt_F |
| KAT6B | 0.00148 | -0.3887 | 0.122 | 0.24 | 1 | MS.Trt_vs_MS.NoTrt_F |
| ANKRD17 | 0.00149 | -0.5302 | 0.198 | 0.34 | 1 | MS.Trt_vs_MS.NoTrt_F |
| SRGAP2B | 0.0015 | -0.6419 | 0.439 | 0.6 | 1 | MS.Trt_vs_MS.NoTrt_F |
| TRAK1 | 0.0015 | -0.2709 | 0.09 | 0.19 | 1 | MS.Trt_vs_MS.NoTrt_F |
| MTERF3 | 0.00151 | -1.1101 | 0.216 | 0.34 | 1 | MS.Trt_vs_MS.NoTrt_F |
| TTC27 | 0.00151 | 0.88332 | 0.112 | 0.23 | 1 | MS.Trt_vs_MS.NoTrt_F |
| VKORC1 | 0.00151 | 0.33297 | 0.129 | 0.25 | 1 | MS.Trt_vs_MS.NoTrt_F |
| HOMER2 | 0.00151 | -0.9007 | 0.507 | 0.64 | 1 | MS.Trt_vs_MS.NoTrt_F |
| TTC4 | 0.00152 | -0.7962 | 0.094 | 0.2 | 1 | MS.Trt_vs_MS.NoTrt_F |
| RXRB | 0.00152 | -0.3127 | 0.183 | 0.32 | 1 | MS.Trt_vs_MS.NoTrt_F |
| PFDN1 | 0.00153 | -0.2873 | 0.162 | 0.29 | 1 | MS.Trt_vs_MS.NoTrt_F |
| BTRC | 0.00153 | -0.5733 | 0.079 | 0.18 | 1 | MS.Trt_vs_MS.NoTrt_F |

| | | | | | | |
|---|---|---|---|---|---|---|
| DCAF15 | 0.00153 | -1.2338 | 0.061 | 0.15 | 1 | MS.Trt_vs_MS.NoTrt_F |
| GNL1 | 0.00154 | -0.6679 | 0.094 | 0.2 | 1 | MS.Trt_vs_MS.NoTrt_F |
| RNF40 | 0.00155 | -0.4502 | 0.129 | 0.25 | 1 | MS.Trt_vs_MS.NoTrt_F |
| TMEM206 | 0.00155 | -0.9222 | 0.083 | 0.18 | 1 | MS.Trt_vs_MS.NoTrt_F |
| ABCA2 | 0.00155 | -0.2931 | 0.245 | 0.39 | 1 | MS.Trt_vs_MS.NoTrt_F |
| PI4K2A | 0.00156 | -0.918 | 0.072 | 0.17 | 1 | MS.Trt_vs_MS.NoTrt_F |
| SHMT2 | 0.00157 | -0.3285 | 0.252 | 0.4 | 1 | MS.Trt_vs_MS.NoTrt_F |
| SURF6 | 0.00157 | -0.2589 | 0.5 | 0.67 | 1 | MS.Trt_vs_MS.NoTrt_F |
| BCAP29 | 0.00157 | -0.61 | 0.327 | 0.48 | 1 | MS.Trt_vs_MS.NoTrt_F |
| CBX7 | 0.00158 | 0.35784 | 0.381 | 0.54 | 1 | MS.Trt_vs_MS.NoTrt_F |
| RTF1 | 0.00158 | -0.4312 | 0.529 | 0.66 | 1 | MS.Trt_vs_MS.NoTrt_F |
| TXNDC16 | 0.00158 | 0.79972 | 0.115 | 0.23 | 1 | MS.Trt_vs_MS.NoTrt_F |
| C10orf128 | 0.00158 | -0.2941 | 0.079 | 0.18 | 1 | MS.Trt_vs_MS.NoTrt_F |
| ZBTB46 | 0.00159 | -0.9805 | 0.09 | 0.19 | 1 | MS.Trt_vs_MS.NoTrt_F |
| PI4K2B | 0.00159 | -1.0513 | 0.05 | 0.13 | 1 | MS.Trt_vs_MS.NoTrt_F |
| RP5-864K19 | 0.00159 | -0.4914 | 0.047 | 0.13 | 1 | MS.Trt_vs_MS.NoTrt_F |
| TMEM43 | 0.00159 | -0.5474 | 0.198 | 0.33 | 1 | MS.Trt_vs_MS.NoTrt_F |
| ISCU | 0.0016 | 0.35716 | 0.288 | 0.48 | 1 | MS.Trt_vs_MS.NoTrt_F |
| RELA | 0.0016 | -0.9553 | 0.18 | 0.31 | 1 | MS.Trt_vs_MS.NoTrt_F |
| PPIH | 0.0016 | -0.4625 | 0.09 | 0.19 | 1 | MS.Trt_vs_MS.NoTrt_F |
| ITGA5 | 0.0016 | 0.82321 | 0.104 | 0.22 | 1 | MS.Trt_vs_MS.NoTrt_F |
| MTMR2 | 0.00161 | 0.59696 | 0.169 | 0.29 | 1 | MS.Trt_vs_MS.NoTrt_F |
| FAM105A | 0.00161 | -0.3872 | 0.245 | 0.38 | 1 | MS.Trt_vs_MS.NoTrt_F |
| MIGA2 | 0.00161 | -0.8413 | 0.424 | 0.55 | 1 | MS.Trt_vs_MS.NoTrt_F |
| DHX8 | 0.00161 | -0.4565 | 0.576 | 0.73 | 1 | MS.Trt_vs_MS.NoTrt_F |
| YWHAH | 0.00161 | -0.3792 | 0.187 | 0.32 | 1 | MS.Trt_vs_MS.NoTrt_F |
| FGL2 | 0.00162 | -0.97 | 0.291 | 0.42 | 1 | MS.Trt_vs_MS.NoTrt_F |
| TMEM229B | 0.00162 | -0.8012 | 0.205 | 0.35 | 1 | MS.Trt_vs_MS.NoTrt_F |
| RCHY1 | 0.00162 | -0.587 | 0.284 | 0.43 | 1 | MS.Trt_vs_MS.NoTrt_F |
| MTERF2 | 0.00162 | -0.511 | 0.076 | 0.17 | 1 | MS.Trt_vs_MS.NoTrt_F |
| ARHGAP11A | 0.00163 | -1.4205 | 0.353 | 0.5 | 1 | MS.Trt_vs_MS.NoTrt_F |
| EIF5A | 0.00163 | 1.10204 | 0.709 | 0.65 | 1 | MS.Trt_vs_MS.NoTrt_F |
| TXNDC11 | 0.00163 | -0.2621 | 0.324 | 0.49 | 1 | MS.Trt_vs_MS.NoTrt_F |
| CCR10 | 0.00164 | -0.3052 | 0.122 | 0.24 | 1 | MS.Trt_vs_MS.NoTrt_F |
| MAPK7 | 0.00164 | -0.8184 | 0.068 | 0.16 | 1 | MS.Trt_vs_MS.NoTrt_F |
| SLC8B1 | 0.00164 | -0.7136 | 0.446 | 0.59 | 1 | MS.Trt_vs_MS.NoTrt_F |
| RCC1 | 0.00164 | -0.6308 | 0.147 | 0.28 | 1 | MS.Trt_vs_MS.NoTrt_F |
| MARS2 | 0.00164 | -0.4498 | 0.065 | 0.16 | 1 | MS.Trt_vs_MS.NoTrt_F |
| CDCP1 | 0.00164 | 0.55602 | 0.554 | 0.41 | 1 | MS.Trt_vs_MS.NoTrt_F |
| BPIFB1 | 0.00165 | -0.5165 | 0.428 | 0.59 | 1 | MS.Trt_vs_MS.NoTrt_F |

| | | | | | | |
|---|---|---|---|---|---|---|
| SCMH1 | 0.00165 | 0.63874 | 0.137 | 0.26 | 1 | MS.Trt_vs_MS.NoTrt_F |
| MTX2 | 0.00166 | -0.4882 | 0.245 | 0.39 | 1 | MS.Trt_vs_MS.NoTrt_F |
| FASTKD5 | 0.00166 | 0.41459 | 0.094 | 0.2 | 1 | MS.Trt_vs_MS.NoTrt_F |
| COPG1 | 0.00166 | -0.2652 | 0.183 | 0.32 | 1 | MS.Trt_vs_MS.NoTrt_F |
| ARFGEF1 | 0.00166 | -0.3406 | 0.572 | 0.74 | 1 | MS.Trt_vs_MS.NoTrt_F |
| PBX2 | 0.00166 | -0.3326 | 0.108 | 0.21 | 1 | MS.Trt_vs_MS.NoTrt_F |
| SSRP1 | 0.00166 | 0.3405 | 0.151 | 0.28 | 1 | MS.Trt_vs_MS.NoTrt_F |
| PDE3A | 0.00166 | -0.6145 | 0.158 | 0.28 | 1 | MS.Trt_vs_MS.NoTrt_F |
| PGGHG | 0.00167 | -0.8064 | 0.129 | 0.24 | 1 | MS.Trt_vs_MS.NoTrt_F |
| ELK1 | 0.00167 | -0.5342 | 0.083 | 0.18 | 1 | MS.Trt_vs_MS.NoTrt_F |
| PPCDC | 0.00167 | 0.35818 | 0.086 | 0.19 | 1 | MS.Trt_vs_MS.NoTrt_F |
| EED | 0.00167 | 0.55504 | 0.162 | 0.3 | 1 | MS.Trt_vs_MS.NoTrt_F |
| HIRA | 0.00167 | 0.26623 | 0.061 | 0.15 | 1 | MS.Trt_vs_MS.NoTrt_F |
| MITF | 0.00167 | -0.2963 | 0.144 | 0.26 | 1 | MS.Trt_vs_MS.NoTrt_F |
| TCF3 | 0.00167 | -0.6086 | 0.079 | 0.18 | 1 | MS.Trt_vs_MS.NoTrt_F |
| ITGA6 | 0.00168 | -0.3134 | 0.14 | 0.27 | 1 | MS.Trt_vs_MS.NoTrt_F |
| TRIM25 | 0.00168 | -0.4538 | 0.324 | 0.46 | 1 | MS.Trt_vs_MS.NoTrt_F |
| UBALD2 | 0.00169 | -0.5386 | 0.104 | 0.21 | 1 | MS.Trt_vs_MS.NoTrt_F |
| CCZ1B | 0.0017 | -0.2513 | 0.662 | 0.77 | 1 | MS.Trt_vs_MS.NoTrt_F |
| ACTR8 | 0.00171 | -0.2701 | 0.083 | 0.18 | 1 | MS.Trt_vs_MS.NoTrt_F |
| ZW10 | 0.00171 | -0.7126 | 0.068 | 0.16 | 1 | MS.Trt_vs_MS.NoTrt_F |
| BORCS6 | 0.00171 | -0.7489 | 0.086 | 0.19 | 1 | MS.Trt_vs_MS.NoTrt_F |
| RBM17 | 0.00172 | 0.2877 | 0.342 | 0.53 | 1 | MS.Trt_vs_MS.NoTrt_F |
| ARMC5 | 0.00172 | 0.31919 | 0.115 | 0.23 | 1 | MS.Trt_vs_MS.NoTrt_F |
| PRKAB2 | 0.00172 | 0.97699 | 0.068 | 0.16 | 1 | MS.Trt_vs_MS.NoTrt_F |
| RPL41 | 0.00172 | 0.33822 | 0.942 | 1 | 1 | MS.Trt_vs_MS.NoTrt_F |
| HPF1 | 0.00173 | 0.68491 | 0.09 | 0.19 | 1 | MS.Trt_vs_MS.NoTrt_F |
| MANEA | 0.00174 | -0.493 | 0.115 | 0.23 | 1 | MS.Trt_vs_MS.NoTrt_F |
| TMEM222 | 0.00174 | -0.5334 | 0.133 | 0.25 | 1 | MS.Trt_vs_MS.NoTrt_F |
| ACP2 | 0.00175 | -1.4832 | 0.068 | 0.16 | 1 | MS.Trt_vs_MS.NoTrt_F |
| TEX264 | 0.00175 | 0.26753 | 0.198 | 0.34 | 1 | MS.Trt_vs_MS.NoTrt_F |
| FLVCR2 | 0.00175 | -0.5761 | 0.058 | 0.14 | 1 | MS.Trt_vs_MS.NoTrt_F |
| EID3 | 0.00175 | -1.0648 | 0.058 | 0.14 | 1 | MS.Trt_vs_MS.NoTrt_F |
| CEP128 | 0.00176 | 0.29106 | 0.198 | 0.33 | 1 | MS.Trt_vs_MS.NoTrt_F |
| FASN | 0.00176 | -0.9257 | 0.122 | 0.23 | 1 | MS.Trt_vs_MS.NoTrt_F |
| H2AFZ | 0.00177 | 0.35439 | 0.345 | 0.56 | 1 | MS.Trt_vs_MS.NoTrt_F |
| E4F1 | 0.00177 | -0.6194 | 0.14 | 0.25 | 1 | MS.Trt_vs_MS.NoTrt_F |
| ABHD15 | 0.00177 | 1.41825 | 0.324 | 0.49 | 1 | MS.Trt_vs_MS.NoTrt_F |
| LRRC59 | 0.00177 | -0.2648 | 0.137 | 0.26 | 1 | MS.Trt_vs_MS.NoTrt_F |
| SVIP | 0.00178 | -0.5771 | 0.45 | 0.62 | 1 | MS.Trt_vs_MS.NoTrt_F |

| | | | | | | |
|---|---|---|---|---|---|---|
| REV3L | 0.00179 | -0.2786 | 0.597 | 0.79 | 1 | MS.Trt_vs_MS.NoTrt_F |
| RPE | 0.00179 | -0.823 | 0.32 | 0.45 | 1 | MS.Trt_vs_MS.NoTrt_F |
| DDX58 | 0.00179 | -0.7539 | 0.306 | 0.45 | 1 | MS.Trt_vs_MS.NoTrt_F |
| AMZ2 | 0.00179 | -0.2841 | 0.529 | 0.71 | 1 | MS.Trt_vs_MS.NoTrt_F |
| RCC2 | 0.00179 | -0.9188 | 0.112 | 0.22 | 1 | MS.Trt_vs_MS.NoTrt_F |
| NUTM2D | 0.0018 | -0.7434 | 0.072 | 0.17 | 1 | MS.Trt_vs_MS.NoTrt_F |
| ZFAND6 | 0.0018 | -0.5086 | 0.237 | 0.37 | 1 | MS.Trt_vs_MS.NoTrt_F |
| FAM122A | 0.0018 | -0.5825 | 0.065 | 0.16 | 1 | MS.Trt_vs_MS.NoTrt_F |
| GZMM | 0.00181 | 0.26086 | 0.144 | 0.28 | 1 | MS.Trt_vs_MS.NoTrt_F |
| TACC1 | 0.00181 | -0.4131 | 0.468 | 0.6 | 1 | MS.Trt_vs_MS.NoTrt_F |
| U2AF2 | 0.00181 | -0.5705 | 0.173 | 0.3 | 1 | MS.Trt_vs_MS.NoTrt_F |
| RABEP1 | 0.00183 | 0.67338 | 0.209 | 0.36 | 1 | MS.Trt_vs_MS.NoTrt_F |
| OLAH | 0.00184 | -0.3635 | 0.396 | 0.54 | 1 | MS.Trt_vs_MS.NoTrt_F |
| MRPL53 | 0.00184 | 0.40926 | 0.126 | 0.25 | 1 | MS.Trt_vs_MS.NoTrt_F |
| SPOUT1 | 0.00185 | 0.75878 | 0.158 | 0.29 | 1 | MS.Trt_vs_MS.NoTrt_F |
| FASTKD1 | 0.00186 | -0.3414 | 0.248 | 0.39 | 1 | MS.Trt_vs_MS.NoTrt_F |
| SLC19A1 | 0.00186 | 0.45916 | 0.068 | 0.16 | 1 | MS.Trt_vs_MS.NoTrt_F |
| EXOC6B | 0.00186 | -0.7136 | 0.115 | 0.23 | 1 | MS.Trt_vs_MS.NoTrt_F |
| ZCCHC17 | 0.00186 | 0.43947 | 0.133 | 0.25 | 1 | MS.Trt_vs_MS.NoTrt_F |
| ACCS | 0.00187 | -0.9284 | 0.076 | 0.17 | 1 | MS.Trt_vs_MS.NoTrt_F |
| HKDC1 | 0.00187 | 0.34004 | 0.068 | 0.16 | 1 | MS.Trt_vs_MS.NoTrt_F |
| ILVBL | 0.00187 | 0.6941 | 0.234 | 0.39 | 1 | MS.Trt_vs_MS.NoTrt_F |
| GOLGA8B | 0.00187 | 0.27967 | 0.719 | 0.82 | 1 | MS.Trt_vs_MS.NoTrt_F |
| KLRB1 | 0.00187 | 0.52558 | 0.158 | 0.29 | 1 | MS.Trt_vs_MS.NoTrt_F |
| IKZF4 | 0.00188 | 0.79175 | 0.18 | 0.32 | 1 | MS.Trt_vs_MS.NoTrt_F |
| YY1 | 0.00188 | 0.43112 | 0.288 | 0.43 | 1 | MS.Trt_vs_MS.NoTrt_F |
| ICOS | 0.00188 | 0.40037 | 0.338 | 0.54 | 1 | MS.Trt_vs_MS.NoTrt_F |
| KIFAP3 | 0.00188 | 0.49197 | 0.147 | 0.27 | 1 | MS.Trt_vs_MS.NoTrt_F |
| U2AF1 | 0.00188 | -0.4724 | 0.507 | 0.64 | 1 | MS.Trt_vs_MS.NoTrt_F |
| MCMDC2 | 0.00188 | -0.2646 | 0.367 | 0.5 | 1 | MS.Trt_vs_MS.NoTrt_F |
| TMEM55B | 0.00189 | -0.9852 | 0.09 | 0.19 | 1 | MS.Trt_vs_MS.NoTrt_F |
| RC3H1 | 0.00189 | 0.30978 | 0.464 | 0.66 | 1 | MS.Trt_vs_MS.NoTrt_F |
| HTT | 0.00191 | -0.4501 | 0.23 | 0.37 | 1 | MS.Trt_vs_MS.NoTrt_F |
| KIAA0368 | 0.00191 | -0.8515 | 0.252 | 0.38 | 1 | MS.Trt_vs_MS.NoTrt_F |
| LRRC8D | 0.00193 | 0.82499 | 0.122 | 0.24 | 1 | MS.Trt_vs_MS.NoTrt_F |
| CASP3 | 0.00194 | 0.36023 | 0.158 | 0.29 | 1 | MS.Trt_vs_MS.NoTrt_F |
| PTBP1 | 0.00194 | -0.3171 | 0.198 | 0.33 | 1 | MS.Trt_vs_MS.NoTrt_F |
| MTHFD1 | 0.00194 | -0.5731 | 0.867 | 0.92 | 1 | MS.Trt_vs_MS.NoTrt_F |
| IARS2 | 0.00194 | 0.25959 | 0.137 | 0.25 | 1 | MS.Trt_vs_MS.NoTrt_F |
| FER | 0.00195 | -0.9654 | 0.424 | 0.59 | 1 | MS.Trt_vs_MS.NoTrt_F |

| | | | | | | |
|---|---|---|---|---|---|---|
| WDYHV1 | 0.00197 | -0.4115 | 0.104 | 0.21 | 1 | MS.Trt_vs_MS.NoTrt_F |
| MCPH1 | 0.00197 | -0.354 | 0.601 | 0.75 | 1 | MS.Trt_vs_MS.NoTrt_F |
| ZNF415 | 0.00197 | 0.61621 | 0.295 | 0.44 | 1 | MS.Trt_vs_MS.NoTrt_F |
| MLH1 | 0.00198 | -0.4798 | 0.356 | 0.49 | 1 | MS.Trt_vs_MS.NoTrt_F |
| FBLN5 | 0.00198 | 0.3218 | 0.201 | 0.33 | 1 | MS.Trt_vs_MS.NoTrt_F |
| MRPL37 | 0.00199 | -0.4041 | 0.155 | 0.28 | 1 | MS.Trt_vs_MS.NoTrt_F |
| EFTUD2 | 0.00199 | -0.5326 | 0.759 | 0.83 | 1 | MS.Trt_vs_MS.NoTrt_F |
| DGCR2 | 0.002 | -0.5464 | 0.227 | 0.36 | 1 | MS.Trt_vs_MS.NoTrt_F |
| BRPF1 | 0.00202 | -0.3714 | 0.083 | 0.18 | 1 | MS.Trt_vs_MS.NoTrt_F |
| SSH3 | 0.00203 | -0.9243 | 0.072 | 0.17 | 1 | MS.Trt_vs_MS.NoTrt_F |
| 3-Mar | 0.00203 | -0.6418 | 0.273 | 0.41 | 1 | MS.Trt_vs_MS.NoTrt_F |
| SIGIRR | 0.00204 | 0.28344 | 0.302 | 0.48 | 1 | MS.Trt_vs_MS.NoTrt_F |
| ANAPC2 | 0.00205 | -0.7284 | 0.083 | 0.18 | 1 | MS.Trt_vs_MS.NoTrt_F |
| TMX1 | 0.00205 | -0.3087 | 0.374 | 0.54 | 1 | MS.Trt_vs_MS.NoTrt_F |
| ARHGAP29 | 0.00206 | -0.3251 | 0.381 | 0.53 | 1 | MS.Trt_vs_MS.NoTrt_F |
| ZNF585A | 0.00207 | -0.6636 | 0.356 | 0.53 | 1 | MS.Trt_vs_MS.NoTrt_F |
| LRRFIP2 | 0.00207 | -0.3457 | 0.68 | 0.81 | 1 | MS.Trt_vs_MS.NoTrt_F |
| HERC5 | 0.00207 | -0.5388 | 0.147 | 0.27 | 1 | MS.Trt_vs_MS.NoTrt_F |
| ARHGEF28 | 0.00208 | 0.34056 | 0.173 | 0.3 | 1 | MS.Trt_vs_MS.NoTrt_F |
| B3GNT4 | 0.00208 | 0.5663 | 0.155 | 0.27 | 1 | MS.Trt_vs_MS.NoTrt_F |
| SLC25A45 | 0.00209 | 0.36983 | 0.978 | 0.98 | 1 | MS.Trt_vs_MS.NoTrt_F |
| NDOR1 | 0.00209 | -0.7868 | 0.079 | 0.17 | 1 | MS.Trt_vs_MS.NoTrt_F |
| KIZ | 0.00211 | 0.38045 | 0.119 | 0.23 | 1 | MS.Trt_vs_MS.NoTrt_F |
| TMEM181 | 0.00211 | -0.3095 | 0.489 | 0.63 | 1 | MS.Trt_vs_MS.NoTrt_F |
| GPX2 | 0.00212 | -0.8826 | 0.162 | 0.29 | 1 | MS.Trt_vs_MS.NoTrt_F |
| DUSP6 | 0.00213 | -0.2504 | 0.068 | 0.16 | 1 | MS.Trt_vs_MS.NoTrt_F |
| ZDHHC8 | 0.00213 | -0.3193 | 0.101 | 0.2 | 1 | MS.Trt_vs_MS.NoTrt_F |
| PAIP2B | 0.00214 | -1.2838 | 0.209 | 0.34 | 1 | MS.Trt_vs_MS.NoTrt_F |
| BLOC1S1 | 0.00214 | 0.58166 | 0.198 | 0.35 | 1 | MS.Trt_vs_MS.NoTrt_F |
| TMEM216 | 0.00215 | -0.4286 | 0.385 | 0.52 | 1 | MS.Trt_vs_MS.NoTrt_F |
| FBL | 0.00216 | -0.3362 | 0.435 | 0.56 | 1 | MS.Trt_vs_MS.NoTrt_F |
| RNF114 | 0.00216 | -0.6227 | 0.665 | 0.8 | 1 | MS.Trt_vs_MS.NoTrt_F |
| MCM6 | 0.00218 | -0.257 | 0.108 | 0.22 | 1 | MS.Trt_vs_MS.NoTrt_F |
| PTGES3L | 0.00218 | 0.32848 | 0.845 | 0.82 | 1 | MS.Trt_vs_MS.NoTrt_F |
| IQCB1 | 0.00218 | -0.8233 | 0.101 | 0.2 | 1 | MS.Trt_vs_MS.NoTrt_F |
| TUBA1A | 0.00218 | -0.5548 | 0.234 | 0.36 | 1 | MS.Trt_vs_MS.NoTrt_F |
| UCHL3 | 0.00219 | 0.43312 | 0.144 | 0.27 | 1 | MS.Trt_vs_MS.NoTrt_F |
| AGTPBP1 | 0.00219 | 0.5015 | 0.165 | 0.3 | 1 | MS.Trt_vs_MS.NoTrt_F |
| TMEM128 | 0.00221 | 0.62881 | 0.155 | 0.28 | 1 | MS.Trt_vs_MS.NoTrt_F |
| RC3H2 | 0.00222 | -0.3285 | 0.482 | 0.61 | 1 | MS.Trt_vs_MS.NoTrt_F |

| | | | | | | |
|---|---|---|---|---|---|---|
| RHEB | 0.00223 | -0.2924 | 0.158 | 0.28 | 1 | MS.Trt_vs_MS.NoTrt_F |
| NDUFS6 | 0.00225 | 0.40729 | 0.137 | 0.26 | 1 | MS.Trt_vs_MS.NoTrt_F |
| LRCH4 | 0.00226 | 0.29292 | 0.446 | 0.61 | 1 | MS.Trt_vs_MS.NoTrt_F |
| FANCL | 0.00227 | -0.4817 | 0.126 | 0.24 | 1 | MS.Trt_vs_MS.NoTrt_F |
| RP11-529K1 | 0.00227 | -0.4252 | 0.097 | 0.2 | 1 | MS.Trt_vs_MS.NoTrt_F |
| SLC35B2 | 0.00227 | -0.852 | 0.097 | 0.2 | 1 | MS.Trt_vs_MS.NoTrt_F |
| GRPEL2 | 0.00228 | 0.2734 | 0.266 | 0.4 | 1 | MS.Trt_vs_MS.NoTrt_F |
| SPIDR | 0.00229 | -0.725 | 0.306 | 0.42 | 1 | MS.Trt_vs_MS.NoTrt_F |
| TSNARE1 | 0.00231 | -0.5493 | 0.09 | 0.19 | 1 | MS.Trt_vs_MS.NoTrt_F |
| SNRPC | 0.00231 | 0.52066 | 0.227 | 0.39 | 1 | MS.Trt_vs_MS.NoTrt_F |
| FKBP15 | 0.00232 | -0.2975 | 0.133 | 0.25 | 1 | MS.Trt_vs_MS.NoTrt_F |
| VRK2 | 0.00233 | 0.488 | 0.223 | 0.36 | 1 | MS.Trt_vs_MS.NoTrt_F |
| SEPHS1 | 0.00233 | -0.7488 | 0.086 | 0.18 | 1 | MS.Trt_vs_MS.NoTrt_F |
| TSGA10 | 0.00234 | -0.629 | 0.101 | 0.2 | 1 | MS.Trt_vs_MS.NoTrt_F |
| BTN3A3 | 0.00236 | -0.4085 | 0.227 | 0.35 | 1 | MS.Trt_vs_MS.NoTrt_F |
| UBLCP1 | 0.0024 | 0.32051 | 0.176 | 0.31 | 1 | MS.Trt_vs_MS.NoTrt_F |
| CACNA1D | 0.0024 | -0.2779 | 0.633 | 0.74 | 1 | MS.Trt_vs_MS.NoTrt_F |
| PCYOX1L | 0.00241 | -0.8302 | 0.09 | 0.19 | 1 | MS.Trt_vs_MS.NoTrt_F |
| ATP6V1H | 0.00241 | -0.4041 | 0.371 | 0.5 | 1 | MS.Trt_vs_MS.NoTrt_F |
| POLR3C | 0.00243 | -0.3607 | 0.543 | 0.74 | 1 | MS.Trt_vs_MS.NoTrt_F |
| ST3GAL3 | 0.00245 | 0.44523 | 0.766 | 0.89 | 1 | MS.Trt_vs_MS.NoTrt_F |
| FAM49B | 0.00245 | -0.5988 | 0.96 | 0.98 | 1 | MS.Trt_vs_MS.NoTrt_F |
| METTL5 | 0.00245 | -0.5257 | 0.507 | 0.62 | 1 | MS.Trt_vs_MS.NoTrt_F |
| RPS19 | 0.00245 | 0.43318 | 0.953 | 0.98 | 1 | MS.Trt_vs_MS.NoTrt_F |
| MED4 | 0.00245 | 0.31515 | 0.622 | 0.79 | 1 | MS.Trt_vs_MS.NoTrt_F |
| GTF3C6 | 0.00246 | 0.51562 | 0.162 | 0.3 | 1 | MS.Trt_vs_MS.NoTrt_F |
| PSD4 | 0.00248 | -0.3157 | 0.896 | 0.94 | 1 | MS.Trt_vs_MS.NoTrt_F |
| ATP6V0C | 0.00249 | -0.4084 | 0.27 | 0.41 | 1 | MS.Trt_vs_MS.NoTrt_F |
| CDKAL1 | 0.00252 | -0.6446 | 0.529 | 0.67 | 1 | MS.Trt_vs_MS.NoTrt_F |
| P2RY10 | 0.00252 | -0.2509 | 0.338 | 0.48 | 1 | MS.Trt_vs_MS.NoTrt_F |
| SMIM15 | 0.00253 | -0.5335 | 0.115 | 0.22 | 1 | MS.Trt_vs_MS.NoTrt_F |
| GYS1 | 0.00255 | 0.35077 | 0.155 | 0.28 | 1 | MS.Trt_vs_MS.NoTrt_F |
| SUV39H1 | 0.00256 | -1.2977 | 0.065 | 0.15 | 1 | MS.Trt_vs_MS.NoTrt_F |
| DCTD | 0.0026 | 0.25124 | 0.14 | 0.26 | 1 | MS.Trt_vs_MS.NoTrt_F |
| PLEKHF2 | 0.0026 | -0.2562 | 0.104 | 0.2 | 1 | MS.Trt_vs_MS.NoTrt_F |
| NPB | 0.0026 | -0.2627 | 0.05 | 0.13 | 1 | MS.Trt_vs_MS.NoTrt_F |
| TCEAL4 | 0.00261 | -0.47 | 0.162 | 0.28 | 1 | MS.Trt_vs_MS.NoTrt_F |
| ACYP1 | 0.00261 | -0.2523 | 0.216 | 0.35 | 1 | MS.Trt_vs_MS.NoTrt_F |
| ZNF391 | 0.00262 | -0.3439 | 0.191 | 0.32 | 1 | MS.Trt_vs_MS.NoTrt_F |
| C18orf21 | 0.00263 | -0.4029 | 0.076 | 0.17 | 1 | MS.Trt_vs_MS.NoTrt_F |

| | | | | | | |
|---|---|---|---|---|---|---|
| TJP3 | 0.00263 | -1.0267 | 0.108 | 0.21 | 1 | MS.Trt_vs_MS.NoTrt_F |
| CCR6 | 0.00264 | -0.4546 | 0.399 | 0.53 | 1 | MS.Trt_vs_MS.NoTrt_F |
| FAM21C | 0.00265 | -0.2897 | 0.212 | 0.35 | 1 | MS.Trt_vs_MS.NoTrt_F |
| ZMYM4 | 0.00266 | -0.9075 | 0.151 | 0.26 | 1 | MS.Trt_vs_MS.NoTrt_F |
| NIPSNAP3A | 0.00267 | -0.7539 | 0.072 | 0.16 | 1 | MS.Trt_vs_MS.NoTrt_F |
| UNC119B | 0.00268 | -1.6639 | 0.065 | 0.15 | 1 | MS.Trt_vs_MS.NoTrt_F |
| AL078584.1 | 0.00269 | -0.9022 | 0.424 | 0.53 | 1 | MS.Trt_vs_MS.NoTrt_F |
| NOP9 | 0.00271 | -1.0439 | 0.061 | 0.14 | 1 | MS.Trt_vs_MS.NoTrt_F |
| CGGBP1 | 0.00271 | 0.58754 | 0.205 | 0.34 | 1 | MS.Trt_vs_MS.NoTrt_F |
| APEH | 0.00271 | -0.2559 | 0.191 | 0.32 | 1 | MS.Trt_vs_MS.NoTrt_F |
| NEK1 | 0.00271 | -0.3554 | 0.313 | 0.45 | 1 | MS.Trt_vs_MS.NoTrt_F |
| TRIO | 0.00271 | -0.7488 | 0.065 | 0.15 | 1 | MS.Trt_vs_MS.NoTrt_F |
| TXNDC12 | 0.00271 | 0.52033 | 0.158 | 0.28 | 1 | MS.Trt_vs_MS.NoTrt_F |
| NOP10 | 0.00272 | 0.54621 | 0.234 | 0.4 | 1 | MS.Trt_vs_MS.NoTrt_F |
| PFKL | 0.00272 | -0.4787 | 0.709 | 0.79 | 1 | MS.Trt_vs_MS.NoTrt_F |
| USP25 | 0.00273 | 0.9334 | 0.374 | 0.56 | 1 | MS.Trt_vs_MS.NoTrt_F |
| ZNHIT6 | 0.00273 | -0.3613 | 0.392 | 0.51 | 1 | MS.Trt_vs_MS.NoTrt_F |
| RHOBTB2 | 0.00273 | -0.9747 | 0.079 | 0.17 | 1 | MS.Trt_vs_MS.NoTrt_F |
| RGL2 | 0.00274 | -0.4777 | 0.151 | 0.27 | 1 | MS.Trt_vs_MS.NoTrt_F |
| FTSJ1 | 0.00275 | 0.36845 | 0.14 | 0.26 | 1 | MS.Trt_vs_MS.NoTrt_F |
| BLOC1S2 | 0.00277 | 0.27139 | 0.5 | 0.66 | 1 | MS.Trt_vs_MS.NoTrt_F |
| ARRB2 | 0.00277 | -0.5162 | 0.766 | 0.88 | 1 | MS.Trt_vs_MS.NoTrt_F |
| OSBPL11 | 0.00278 | -0.2655 | 0.108 | 0.21 | 1 | MS.Trt_vs_MS.NoTrt_F |
| ETAA1 | 0.00278 | 0.35125 | 0.147 | 0.27 | 1 | MS.Trt_vs_MS.NoTrt_F |
| FOSL2 | 0.00278 | -0.2925 | 0.245 | 0.39 | 1 | MS.Trt_vs_MS.NoTrt_F |
| HMGA1 | 0.00278 | -0.2541 | 0.176 | 0.3 | 1 | MS.Trt_vs_MS.NoTrt_F |
| IRS2 | 0.00279 | -0.4504 | 0.363 | 0.49 | 1 | MS.Trt_vs_MS.NoTrt_F |
| PIGV | 0.00279 | -0.7876 | 0.068 | 0.16 | 1 | MS.Trt_vs_MS.NoTrt_F |
| KCTD13 | 0.00279 | 0.41836 | 0.158 | 0.28 | 1 | MS.Trt_vs_MS.NoTrt_F |
| IL17RB | 0.0028 | -2.2636 | 0.04 | 0.11 | 1 | MS.Trt_vs_MS.NoTrt_F |
| MRPS14 | 0.0028 | 0.26966 | 0.169 | 0.3 | 1 | MS.Trt_vs_MS.NoTrt_F |
| TOMM22 | 0.00283 | -0.4443 | 0.201 | 0.33 | 1 | MS.Trt_vs_MS.NoTrt_F |
| C2CD2 | 0.00283 | -0.9367 | 0.219 | 0.33 | 1 | MS.Trt_vs_MS.NoTrt_F |
| OCM | 0.00286 | -0.6639 | 0.04 | 0.11 | 1 | MS.Trt_vs_MS.NoTrt_F |
| ZNF12 | 0.00286 | -1.0712 | 0.317 | 0.45 | 1 | MS.Trt_vs_MS.NoTrt_F |
| MRPL55 | 0.00286 | 0.89556 | 0.108 | 0.21 | 1 | MS.Trt_vs_MS.NoTrt_F |
| ELP5 | 0.00287 | -0.3559 | 0.112 | 0.22 | 1 | MS.Trt_vs_MS.NoTrt_F |
| FHOD1 | 0.00287 | 1.12701 | 0.169 | 0.3 | 1 | MS.Trt_vs_MS.NoTrt_F |
| SLC14A2 | 0.00288 | -0.2554 | 0.633 | 0.79 | 1 | MS.Trt_vs_MS.NoTrt_F |
| NUDT5 | 0.00288 | 0.40666 | 0.209 | 0.37 | 1 | MS.Trt_vs_MS.NoTrt_F |

| | | | | | | |
|---|---|---|---|---|---|---|
| BBS7 | 0.00289 | -0.8809 | 0.072 | 0.16 | 1 | MS.Trt_vs_MS.NoTrt_F |
| ALCAM | 0.00289 | -0.929 | 0.072 | 0.16 | 1 | MS.Trt_vs_MS.NoTrt_F |
| FBXO21 | 0.0029 | 0.25641 | 0.169 | 0.3 | 1 | MS.Trt_vs_MS.NoTrt_F |
| G2E3 | 0.00291 | -0.5898 | 0.457 | 0.58 | 1 | MS.Trt_vs_MS.NoTrt_F |
| DHX34 | 0.00291 | 0.38725 | 0.094 | 0.19 | 1 | MS.Trt_vs_MS.NoTrt_F |
| ZBTB39 | 0.00292 | -1.5226 | 0.043 | 0.12 | 1 | MS.Trt_vs_MS.NoTrt_F |
| PDE4A | 0.00293 | -0.6198 | 0.115 | 0.22 | 1 | MS.Trt_vs_MS.NoTrt_F |
| NFX1 | 0.00293 | -0.5521 | 0.784 | 0.9 | 1 | MS.Trt_vs_MS.NoTrt_F |
| PHF5A | 0.00293 | -0.3253 | 0.129 | 0.24 | 1 | MS.Trt_vs_MS.NoTrt_F |
| COG5 | 0.00293 | 1.39773 | 0.198 | 0.34 | 1 | MS.Trt_vs_MS.NoTrt_F |
| ADGRE1 | 0.00293 | -0.3666 | 0.047 | 0.12 | 1 | MS.Trt_vs_MS.NoTrt_F |
| PSMA4 | 0.00294 | 0.37355 | 0.356 | 0.5 | 1 | MS.Trt_vs_MS.NoTrt_F |
| PLEC | 0.00294 | 0.38083 | 0.309 | 0.48 | 1 | MS.Trt_vs_MS.NoTrt_F |
| ZNF275 | 0.00295 | -1.6346 | 0.058 | 0.14 | 1 | MS.Trt_vs_MS.NoTrt_F |
| POFUT2 | 0.00295 | -0.9693 | 0.072 | 0.16 | 1 | MS.Trt_vs_MS.NoTrt_F |
| MAP3K14 | 0.00296 | -0.3013 | 0.158 | 0.28 | 1 | MS.Trt_vs_MS.NoTrt_F |
| NKRF | 0.00296 | -0.5851 | 0.083 | 0.18 | 1 | MS.Trt_vs_MS.NoTrt_F |
| DUS2 | 0.00296 | -0.9697 | 0.076 | 0.17 | 1 | MS.Trt_vs_MS.NoTrt_F |
| NR1D1 | 0.00296 | -0.5956 | 0.112 | 0.22 | 1 | MS.Trt_vs_MS.NoTrt_F |
| NFKBIZ | 0.00296 | -0.3667 | 0.36 | 0.5 | 1 | MS.Trt_vs_MS.NoTrt_F |
| DDX24 | 0.00296 | -0.2797 | 0.532 | 0.67 | 1 | MS.Trt_vs_MS.NoTrt_F |
| LMLN | 0.00296 | -1.6868 | 0.353 | 0.5 | 1 | MS.Trt_vs_MS.NoTrt_F |
| UNC79 | 0.00297 | -0.3878 | 0.317 | 0.45 | 1 | MS.Trt_vs_MS.NoTrt_F |
| SPNS1 | 0.00299 | -0.5545 | 0.755 | 0.92 | 1 | MS.Trt_vs_MS.NoTrt_F |
| LEAP2 | 0.00299 | -0.4845 | 0.23 | 0.38 | 1 | MS.Trt_vs_MS.NoTrt_F |
| SNAPC4 | 0.00299 | -0.2663 | 0.058 | 0.14 | 1 | MS.Trt_vs_MS.NoTrt_F |
| INVS | 0.00299 | -0.9219 | 0.147 | 0.26 | 1 | MS.Trt_vs_MS.NoTrt_F |
| EXOC6 | 0.003 | -0.5763 | 0.259 | 0.38 | 1 | MS.Trt_vs_MS.NoTrt_F |
| AC009950.1 | 0.00301 | -0.4503 | 0.072 | 0.16 | 1 | MS.Trt_vs_MS.NoTrt_F |
| SWT1 | 0.00301 | -0.4859 | 0.076 | 0.17 | 1 | MS.Trt_vs_MS.NoTrt_F |
| NT5DC1 | 0.00301 | -0.3308 | 0.173 | 0.3 | 1 | MS.Trt_vs_MS.NoTrt_F |
| PAGR1 | 0.00302 | -0.2583 | 0.205 | 0.34 | 1 | MS.Trt_vs_MS.NoTrt_F |
| IFIH1 | 0.00304 | -1.2839 | 0.043 | 0.12 | 1 | MS.Trt_vs_MS.NoTrt_F |
| FAM175B | 0.00304 | -0.69 | 0.068 | 0.16 | 1 | MS.Trt_vs_MS.NoTrt_F |
| TDP1 | 0.00305 | -0.3221 | 0.299 | 0.42 | 1 | MS.Trt_vs_MS.NoTrt_F |
| GLOD4 | 0.00306 | -0.3111 | 0.594 | 0.72 | 1 | MS.Trt_vs_MS.NoTrt_F |
| CCT6A | 0.00309 | 0.30839 | 0.46 | 0.67 | 1 | MS.Trt_vs_MS.NoTrt_F |
| TMEM242 | 0.0031 | -0.452 | 0.129 | 0.24 | 1 | MS.Trt_vs_MS.NoTrt_F |
| NDUFA13 | 0.0031 | 0.32206 | 0.475 | 0.65 | 1 | MS.Trt_vs_MS.NoTrt_F |
| ATIC | 0.0031 | -0.3846 | 0.338 | 0.46 | 1 | MS.Trt_vs_MS.NoTrt_F |

| | | | | | | |
|---|---|---|---|---|---|---|
| TAOK3 | 0.00312 | 0.41092 | 0.745 | 0.85 | 1 | MS.Trt_vs_MS.NoTrt_F |
| TNPO3 | 0.00312 | -0.5592 | 0.18 | 0.3 | 1 | MS.Trt_vs_MS.NoTrt_F |
| STRBP | 0.00312 | -0.6763 | 0.572 | 0.72 | 1 | MS.Trt_vs_MS.NoTrt_F |
| TGIF2 | 0.00312 | -0.5618 | 0.27 | 0.41 | 1 | MS.Trt_vs_MS.NoTrt_F |
| ARMC2 | 0.00313 | -0.5293 | 0.097 | 0.19 | 1 | MS.Trt_vs_MS.NoTrt_F |
| CMPK1 | 0.00314 | 0.26081 | 0.23 | 0.37 | 1 | MS.Trt_vs_MS.NoTrt_F |
| CPNE2 | 0.00314 | -0.5647 | 0.349 | 0.47 | 1 | MS.Trt_vs_MS.NoTrt_F |
| TRMO | 0.00314 | -0.2823 | 0.144 | 0.27 | 1 | MS.Trt_vs_MS.NoTrt_F |
| LTV1 | 0.00315 | -0.3352 | 0.144 | 0.26 | 1 | MS.Trt_vs_MS.NoTrt_F |
| NAT6 | 0.00316 | 0.38463 | 0.076 | 0.17 | 1 | MS.Trt_vs_MS.NoTrt_F |
| MMP13 | 0.00316 | -0.3166 | 0.101 | 0.2 | 1 | MS.Trt_vs_MS.NoTrt_F |
| ZNF600 | 0.00319 | -0.7043 | 0.201 | 0.32 | 1 | MS.Trt_vs_MS.NoTrt_F |
| LYRM5 | 0.0032 | -0.5879 | 0.079 | 0.17 | 1 | MS.Trt_vs_MS.NoTrt_F |
| SNX20 | 0.0032 | -0.3216 | 0.446 | 0.56 | 1 | MS.Trt_vs_MS.NoTrt_F |
| MLANA | 0.00323 | -0.3215 | 0.385 | 0.55 | 1 | MS.Trt_vs_MS.NoTrt_F |
| PHF2 | 0.00323 | 0.49625 | 0.072 | 0.16 | 1 | MS.Trt_vs_MS.NoTrt_F |
| FAM98A | 0.00324 | 0.37517 | 0.072 | 0.16 | 1 | MS.Trt_vs_MS.NoTrt_F |
| CTD-2116N1 | 0.00324 | -0.4207 | 0.047 | 0.12 | 1 | MS.Trt_vs_MS.NoTrt_F |
| C10orf76 | 0.00324 | -0.4974 | 0.331 | 0.47 | 1 | MS.Trt_vs_MS.NoTrt_F |
| RP11-619A1 | 0.00325 | -0.4157 | 0.565 | 0.7 | 1 | MS.Trt_vs_MS.NoTrt_F |
| LZTS2 | 0.00326 | -0.6518 | 0.058 | 0.14 | 1 | MS.Trt_vs_MS.NoTrt_F |
| EIF3C | 0.00327 | -0.3336 | 0.748 | 0.83 | 1 | MS.Trt_vs_MS.NoTrt_F |
| SLC25A25 | 0.00329 | 0.26691 | 0.108 | 0.21 | 1 | MS.Trt_vs_MS.NoTrt_F |
| SART1 | 0.0033 | -1.2669 | 0.288 | 0.42 | 1 | MS.Trt_vs_MS.NoTrt_F |
| TTLL12 | 0.00331 | -0.7772 | 0.065 | 0.15 | 1 | MS.Trt_vs_MS.NoTrt_F |
| FAM8A1 | 0.00331 | 0.34721 | 0.061 | 0.14 | 1 | MS.Trt_vs_MS.NoTrt_F |
| TGFB1I1 | 0.00331 | 0.53048 | 0.245 | 0.38 | 1 | MS.Trt_vs_MS.NoTrt_F |
| LSM1 | 0.00334 | 0.26942 | 0.133 | 0.25 | 1 | MS.Trt_vs_MS.NoTrt_F |
| DEK | 0.00336 | 0.36797 | 0.41 | 0.55 | 1 | MS.Trt_vs_MS.NoTrt_F |
| INPPL1 | 0.00338 | 0.66991 | 0.09 | 0.19 | 1 | MS.Trt_vs_MS.NoTrt_F |
| USPL1 | 0.00338 | 0.53422 | 0.108 | 0.21 | 1 | MS.Trt_vs_MS.NoTrt_F |
| ZDHHC18 | 0.00339 | -0.3082 | 0.101 | 0.2 | 1 | MS.Trt_vs_MS.NoTrt_F |
| FCHO1 | 0.0034 | -1.2734 | 0.73 | 0.82 | 1 | MS.Trt_vs_MS.NoTrt_F |
| TAF10 | 0.0034 | -0.6501 | 0.101 | 0.2 | 1 | MS.Trt_vs_MS.NoTrt_F |
| KLC4 | 0.00341 | -0.2779 | 0.795 | 0.86 | 1 | MS.Trt_vs_MS.NoTrt_F |
| NUTM2B | 0.00342 | -0.4166 | 0.065 | 0.15 | 1 | MS.Trt_vs_MS.NoTrt_F |
| ZNF224 | 0.00346 | -0.8051 | 0.45 | 0.56 | 1 | MS.Trt_vs_MS.NoTrt_F |
| CREBL2 | 0.00348 | -0.4668 | 0.112 | 0.21 | 1 | MS.Trt_vs_MS.NoTrt_F |
| TMEM258 | 0.00348 | 0.58709 | 0.309 | 0.45 | 1 | MS.Trt_vs_MS.NoTrt_F |
| DIDO1 | 0.0035 | 0.30603 | 0.201 | 0.34 | 1 | MS.Trt_vs_MS.NoTrt_F |

| | | | | | | |
|---|---|---|---|---|---|---|
| BEX4 | 0.0035 | 0.33623 | 0.122 | 0.24 | 1 | MS.Trt_vs_MS.NoTrt_F |
| TCEAL3 | 0.00351 | -0.3413 | 0.133 | 0.24 | 1 | MS.Trt_vs_MS.NoTrt_F |
| ARL15 | 0.00354 | 0.47588 | 0.083 | 0.18 | 1 | MS.Trt_vs_MS.NoTrt_F |
| NET1 | 0.00355 | -0.4668 | 0.104 | 0.2 | 1 | MS.Trt_vs_MS.NoTrt_F |
| PLD2 | 0.00356 | -0.503 | 0.09 | 0.18 | 1 | MS.Trt_vs_MS.NoTrt_F |
| PIM3 | 0.00356 | -0.3504 | 0.129 | 0.24 | 1 | MS.Trt_vs_MS.NoTrt_F |
| TBC1D22B | 0.00356 | -0.8381 | 0.09 | 0.18 | 1 | MS.Trt_vs_MS.NoTrt_F |
| TOR2A | 0.00357 | 0.48698 | 0.101 | 0.2 | 1 | MS.Trt_vs_MS.NoTrt_F |
| UBE2B | 0.00358 | 0.29944 | 0.676 | 0.79 | 1 | MS.Trt_vs_MS.NoTrt_F |
| SEC23B | 0.0036 | -0.4197 | 0.147 | 0.26 | 1 | MS.Trt_vs_MS.NoTrt_F |
| ESD | 0.0036 | 0.36067 | 0.151 | 0.27 | 1 | MS.Trt_vs_MS.NoTrt_F |
| DNAJC8 | 0.00361 | 0.278 | 0.266 | 0.43 | 1 | MS.Trt_vs_MS.NoTrt_F |
| ZNF92 | 0.00361 | -0.9751 | 0.126 | 0.23 | 1 | MS.Trt_vs_MS.NoTrt_F |
| DAGLB | 0.00361 | -0.3525 | 0.741 | 0.88 | 1 | MS.Trt_vs_MS.NoTrt_F |
| TLK2 | 0.00361 | -0.4096 | 0.345 | 0.49 | 1 | MS.Trt_vs_MS.NoTrt_F |
| RP11-49K24 | 0.00363 | -0.4736 | 0.147 | 0.27 | 1 | MS.Trt_vs_MS.NoTrt_F |
| PIK3R1 | 0.00364 | -0.4046 | 0.327 | 0.45 | 1 | MS.Trt_vs_MS.NoTrt_F |
| WDR44 | 0.00364 | -0.5433 | 0.119 | 0.22 | 1 | MS.Trt_vs_MS.NoTrt_F |
| THAP2 | 0.00365 | -0.4626 | 0.104 | 0.2 | 1 | MS.Trt_vs_MS.NoTrt_F |
| PSMC1 | 0.00368 | -0.3415 | 0.342 | 0.46 | 1 | MS.Trt_vs_MS.NoTrt_F |
| IMPA1 | 0.00368 | 0.58343 | 0.338 | 0.48 | 1 | MS.Trt_vs_MS.NoTrt_F |
| LAMTOR1 | 0.0037 | 0.45802 | 0.219 | 0.37 | 1 | MS.Trt_vs_MS.NoTrt_F |
| RP3-461F17 | 0.00371 | 0.25499 | 0.194 | 0.32 | 1 | MS.Trt_vs_MS.NoTrt_F |
| PPM1A | 0.00371 | -0.4171 | 0.183 | 0.31 | 1 | MS.Trt_vs_MS.NoTrt_F |
| PLEKHM3 | 0.00373 | -0.7622 | 0.478 | 0.62 | 1 | MS.Trt_vs_MS.NoTrt_F |
| PRPS2 | 0.00373 | -0.6151 | 0.126 | 0.23 | 1 | MS.Trt_vs_MS.NoTrt_F |
| CIC | 0.00375 | -0.4054 | 0.079 | 0.17 | 1 | MS.Trt_vs_MS.NoTrt_F |
| DUS1L | 0.00376 | 1.09009 | 0.086 | 0.18 | 1 | MS.Trt_vs_MS.NoTrt_F |
| ATP2C1 | 0.00377 | -0.6482 | 0.183 | 0.3 | 1 | MS.Trt_vs_MS.NoTrt_F |
| RYK | 0.00377 | -0.623 | 0.083 | 0.17 | 1 | MS.Trt_vs_MS.NoTrt_F |
| ATG9A | 0.00377 | -0.3588 | 0.097 | 0.19 | 1 | MS.Trt_vs_MS.NoTrt_F |
| RPS6KC1 | 0.00378 | -0.68 | 0.097 | 0.19 | 1 | MS.Trt_vs_MS.NoTrt_F |
| MLEC | 0.0038 | -0.3419 | 0.101 | 0.2 | 1 | MS.Trt_vs_MS.NoTrt_F |
| CMIP | 0.0038 | -1.0963 | 0.09 | 0.18 | 1 | MS.Trt_vs_MS.NoTrt_F |
| PRDM8 | 0.00381 | -0.5484 | 0.086 | 0.18 | 1 | MS.Trt_vs_MS.NoTrt_F |
| RRS1 | 0.00381 | -0.519 | 0.108 | 0.21 | 1 | MS.Trt_vs_MS.NoTrt_F |
| ADAM8 | 0.00382 | 0.26937 | 0.201 | 0.34 | 1 | MS.Trt_vs_MS.NoTrt_F |
| NLE1 | 0.00385 | -1.0076 | 0.223 | 0.35 | 1 | MS.Trt_vs_MS.NoTrt_F |
| CUL4A | 0.00386 | -0.3837 | 0.759 | 0.83 | 1 | MS.Trt_vs_MS.NoTrt_F |
| MT1F | 0.00386 | 0.3705 | 0.108 | 0.21 | 1 | MS.Trt_vs_MS.NoTrt_F |

| | | | | | | |
|---|---|---|---|---|---|---|
| TNFAIP1 | 0.00387 | -0.547 | 0.086 | 0.18 | 1 | MS.Trt_vs_MS.NoTrt_F |
| PSD3 | 0.00387 | -0.5998 | 0.306 | 0.45 | 1 | MS.Trt_vs_MS.NoTrt_F |
| NBPF3 | 0.00387 | 0.96172 | 0.403 | 0.54 | 1 | MS.Trt_vs_MS.NoTrt_F |
| ZRANB1 | 0.00387 | -0.3666 | 0.122 | 0.22 | 1 | MS.Trt_vs_MS.NoTrt_F |
| FAM196B | 0.00388 | -0.5295 | 0.086 | 0.18 | 1 | MS.Trt_vs_MS.NoTrt_F |
| SLC25A37 | 0.00388 | 0.62514 | 0.622 | 0.8 | 1 | MS.Trt_vs_MS.NoTrt_F |
| C6orf223 | 0.00388 | -0.362 | 0.219 | 0.34 | 1 | MS.Trt_vs_MS.NoTrt_F |
| SEMA4B | 0.00389 | -0.6403 | 0.349 | 0.48 | 1 | MS.Trt_vs_MS.NoTrt_F |
| CAPG | 0.0039 | -0.3483 | 0.144 | 0.25 | 1 | MS.Trt_vs_MS.NoTrt_F |
| POLE4 | 0.00392 | -0.5307 | 0.112 | 0.21 | 1 | MS.Trt_vs_MS.NoTrt_F |
| METTL10 | 0.00392 | 0.30961 | 0.324 | 0.46 | 1 | MS.Trt_vs_MS.NoTrt_F |
| CAPRIN2 | 0.00392 | -1.2165 | 0.673 | 0.8 | 1 | MS.Trt_vs_MS.NoTrt_F |
| DCAF4 | 0.00394 | 0.46598 | 0.536 | 0.66 | 1 | MS.Trt_vs_MS.NoTrt_F |
| RBM27 | 0.00395 | -0.4295 | 0.133 | 0.24 | 1 | MS.Trt_vs_MS.NoTrt_F |
| ZNF44 | 0.00395 | 0.39606 | 0.579 | 0.69 | 1 | MS.Trt_vs_MS.NoTrt_F |
| EVI2A | 0.00396 | 0.60927 | 0.191 | 0.33 | 1 | MS.Trt_vs_MS.NoTrt_F |
| HPS3 | 0.00397 | 0.38062 | 0.241 | 0.37 | 1 | MS.Trt_vs_MS.NoTrt_F |
| SDHD | 0.00398 | -0.4118 | 0.633 | 0.75 | 1 | MS.Trt_vs_MS.NoTrt_F |
| TNFRSF4 | 0.00398 | 0.52142 | 0.165 | 0.29 | 1 | MS.Trt_vs_MS.NoTrt_F |
| ASNA1 | 0.00399 | -0.327 | 0.086 | 0.18 | 1 | MS.Trt_vs_MS.NoTrt_F |
| HOXB3 | 0.00401 | -0.3121 | 0.065 | 0.14 | 1 | MS.Trt_vs_MS.NoTrt_F |
| TRIM23 | 0.00401 | -1.555 | 0.079 | 0.17 | 1 | MS.Trt_vs_MS.NoTrt_F |
| C8orf44 | 0.00402 | -0.6029 | 0.439 | 0.58 | 1 | MS.Trt_vs_MS.NoTrt_F |
| RCL1 | 0.00403 | 0.39182 | 0.396 | 0.52 | 1 | MS.Trt_vs_MS.NoTrt_F |
| VPS52 | 0.00404 | -0.4973 | 0.158 | 0.27 | 1 | MS.Trt_vs_MS.NoTrt_F |
| SNX10 | 0.00405 | 0.65226 | 0.133 | 0.24 | 1 | MS.Trt_vs_MS.NoTrt_F |
| RAB37 | 0.00406 | -0.442 | 0.32 | 0.46 | 1 | MS.Trt_vs_MS.NoTrt_F |
| HIGD2A | 0.00408 | 0.44408 | 0.288 | 0.46 | 1 | MS.Trt_vs_MS.NoTrt_F |
| GRB2 | 0.0041 | 0.36806 | 0.647 | 0.79 | 1 | MS.Trt_vs_MS.NoTrt_F |
| FIS1 | 0.0041 | 0.29286 | 0.259 | 0.41 | 1 | MS.Trt_vs_MS.NoTrt_F |
| TRPT1 | 0.0041 | -0.4605 | 0.115 | 0.22 | 1 | MS.Trt_vs_MS.NoTrt_F |
| CYCS | 0.00414 | -0.6067 | 0.741 | 0.81 | 1 | MS.Trt_vs_MS.NoTrt_F |
| ERMARD | 0.00414 | -0.292 | 0.104 | 0.2 | 1 | MS.Trt_vs_MS.NoTrt_F |
| ADORA2A | 0.00415 | -0.2557 | 0.151 | 0.27 | 1 | MS.Trt_vs_MS.NoTrt_F |
| MED28 | 0.00416 | -0.8852 | 0.867 | 0.92 | 1 | MS.Trt_vs_MS.NoTrt_F |
| YKT6 | 0.00417 | 0.25601 | 0.115 | 0.22 | 1 | MS.Trt_vs_MS.NoTrt_F |
| KLF12 | 0.00417 | -0.3948 | 0.353 | 0.5 | 1 | MS.Trt_vs_MS.NoTrt_F |
| SLC9A1 | 0.00417 | 0.94512 | 0.187 | 0.32 | 1 | MS.Trt_vs_MS.NoTrt_F |
| MAN1C1 | 0.00418 | 0.63112 | 0.899 | 0.85 | 1 | MS.Trt_vs_MS.NoTrt_F |
| ZFYVE19 | 0.00418 | -0.6919 | 0.108 | 0.2 | 1 | MS.Trt_vs_MS.NoTrt_F |

| | | | | | | |
|---|---|---|---|---|---|---|
| FRG2C | 0.00418 | -0.3016 | 0.169 | 0.29 | 1 | MS.Trt_vs_MS.NoTrt_F |
| SOCS1 | 0.00419 | -0.2649 | 0.094 | 0.19 | 1 | MS.Trt_vs_MS.NoTrt_F |
| ADARB1 | 0.00419 | -0.9446 | 0.209 | 0.34 | 1 | MS.Trt_vs_MS.NoTrt_F |
| XPR1 | 0.00419 | 0.30261 | 0.302 | 0.45 | 1 | MS.Trt_vs_MS.NoTrt_F |
| C6orf1 | 0.0042 | 0.28327 | 0.133 | 0.25 | 1 | MS.Trt_vs_MS.NoTrt_F |
| PAPD7 | 0.00421 | -1.4353 | 0.097 | 0.19 | 1 | MS.Trt_vs_MS.NoTrt_F |
| MECR | 0.00421 | -0.7778 | 0.284 | 0.44 | 1 | MS.Trt_vs_MS.NoTrt_F |
| PORCN | 0.00421 | -1.0586 | 0.079 | 0.17 | 1 | MS.Trt_vs_MS.NoTrt_F |
| LAMTOR3 | 0.00422 | 0.32121 | 0.356 | 0.53 | 1 | MS.Trt_vs_MS.NoTrt_F |
| ICAM1 | 0.00424 | 0.26139 | 0.424 | 0.61 | 1 | MS.Trt_vs_MS.NoTrt_F |
| RABEP2 | 0.00425 | -0.3942 | 0.518 | 0.63 | 1 | MS.Trt_vs_MS.NoTrt_F |
| TAF1B | 0.00425 | -0.3065 | 0.468 | 0.63 | 1 | MS.Trt_vs_MS.NoTrt_F |
| HECA | 0.00426 | 0.47034 | 0.496 | 0.65 | 1 | MS.Trt_vs_MS.NoTrt_F |
| CASC3 | 0.00426 | -0.3904 | 0.529 | 0.65 | 1 | MS.Trt_vs_MS.NoTrt_F |
| AKAP2 | 0.00427 | -0.3052 | 0.234 | 0.38 | 1 | MS.Trt_vs_MS.NoTrt_F |
| IMPAD1 | 0.00427 | -0.312 | 0.475 | 0.62 | 1 | MS.Trt_vs_MS.NoTrt_F |
| C3orf38 | 0.00429 | -1.2547 | 0.101 | 0.19 | 1 | MS.Trt_vs_MS.NoTrt_F |
| CLCN7 | 0.00429 | -0.3796 | 0.263 | 0.4 | 1 | MS.Trt_vs_MS.NoTrt_F |
| AAAS | 0.0043 | -0.3282 | 0.119 | 0.22 | 1 | MS.Trt_vs_MS.NoTrt_F |
| WDR5B | 0.0043 | -1.9872 | 0.191 | 0.3 | 1 | MS.Trt_vs_MS.NoTrt_F |
| DIAPH2 | 0.00431 | -0.5394 | 0.317 | 0.46 | 1 | MS.Trt_vs_MS.NoTrt_F |
| DENND4C | 0.00431 | -0.2566 | 0.338 | 0.48 | 1 | MS.Trt_vs_MS.NoTrt_F |
| 5-Mar | 0.00432 | -0.3318 | 0.209 | 0.33 | 1 | MS.Trt_vs_MS.NoTrt_F |
| RIC1 | 0.00432 | -0.3765 | 0.126 | 0.23 | 1 | MS.Trt_vs_MS.NoTrt_F |
| CDK6 | 0.00432 | 0.46304 | 0.147 | 0.27 | 1 | MS.Trt_vs_MS.NoTrt_F |
| ABT1 | 0.00434 | 0.34965 | 0.194 | 0.33 | 1 | MS.Trt_vs_MS.NoTrt_F |
| SLC6A6 | 0.00435 | 0.48731 | 0.194 | 0.33 | 1 | MS.Trt_vs_MS.NoTrt_F |
| MRRF | 0.00435 | -0.9689 | 0.583 | 0.67 | 1 | MS.Trt_vs_MS.NoTrt_F |
| CHPF2 | 0.00435 | -0.4101 | 0.115 | 0.22 | 1 | MS.Trt_vs_MS.NoTrt_F |
| CPSF3 | 0.00435 | -0.6558 | 0.108 | 0.2 | 1 | MS.Trt_vs_MS.NoTrt_F |
| RPGRIP1 | 0.00438 | 0.38378 | 0.248 | 0.38 | 1 | MS.Trt_vs_MS.NoTrt_F |
| FAM46A | 0.00438 | -0.3704 | 0.083 | 0.17 | 1 | MS.Trt_vs_MS.NoTrt_F |
| REC8 | 0.00438 | -0.6153 | 0.112 | 0.21 | 1 | MS.Trt_vs_MS.NoTrt_F |
| GAR1 | 0.00438 | -0.4605 | 0.068 | 0.15 | 1 | MS.Trt_vs_MS.NoTrt_F |
| SUB1 | 0.00439 | 0.29926 | 0.554 | 0.74 | 1 | MS.Trt_vs_MS.NoTrt_F |
| COA3 | 0.00441 | 0.28921 | 0.147 | 0.27 | 1 | MS.Trt_vs_MS.NoTrt_F |
| AP1S1 | 0.00441 | -0.4326 | 0.076 | 0.16 | 1 | MS.Trt_vs_MS.NoTrt_F |
| SPG20 | 0.00442 | -0.3711 | 0.112 | 0.21 | 1 | MS.Trt_vs_MS.NoTrt_F |
| SLC25A1 | 0.0045 | 0.28339 | 0.097 | 0.19 | 1 | MS.Trt_vs_MS.NoTrt_F |
| NPEPL1 | 0.00451 | 0.7314 | 0.147 | 0.27 | 1 | MS.Trt_vs_MS.NoTrt_F |

| | | | | | | |
|---|---|---|---|---|---|---|
| LMCD1 | 0.00452 | 1.03128 | 0.266 | 0.39 | 1 | MS.Trt_vs_MS.NoTrt_F |
| RP11-159D1 | 0.00452 | -0.3031 | 0.147 | 0.26 | 1 | MS.Trt_vs_MS.NoTrt_F |
| CPA5 | 0.00453 | -1.2044 | 0.054 | 0.13 | 1 | MS.Trt_vs_MS.NoTrt_F |
| KIAA1143 | 0.00453 | -0.8576 | 0.475 | 0.61 | 1 | MS.Trt_vs_MS.NoTrt_F |
| KCNC2 | 0.00454 | -0.2516 | 0.349 | 0.48 | 1 | MS.Trt_vs_MS.NoTrt_F |
| EZH2 | 0.00454 | -0.6758 | 0.133 | 0.24 | 1 | MS.Trt_vs_MS.NoTrt_F |
| KLHL42 | 0.00455 | -0.4751 | 0.115 | 0.21 | 1 | MS.Trt_vs_MS.NoTrt_F |
| FAR1 | 0.00456 | -0.6858 | 0.09 | 0.18 | 1 | MS.Trt_vs_MS.NoTrt_F |
| MCMBP | 0.00458 | 0.48902 | 0.464 | 0.6 | 1 | MS.Trt_vs_MS.NoTrt_F |
| TRIM35 | 0.00459 | 0.30755 | 0.223 | 0.38 | 1 | MS.Trt_vs_MS.NoTrt_F |
| MON1A | 0.00459 | -0.9595 | 0.083 | 0.17 | 1 | MS.Trt_vs_MS.NoTrt_F |
| GIMAP8 | 0.00461 | 0.5001 | 0.09 | 0.18 | 1 | MS.Trt_vs_MS.NoTrt_F |
| MRPL38 | 0.00462 | -0.2919 | 0.406 | 0.53 | 1 | MS.Trt_vs_MS.NoTrt_F |
| SECISBP2L | 0.00465 | -0.2634 | 0.683 | 0.83 | 1 | MS.Trt_vs_MS.NoTrt_F |
| PRR4 | 0.00466 | 0.33659 | 0.259 | 0.4 | 1 | MS.Trt_vs_MS.NoTrt_F |
| SLC25A38 | 0.00467 | -0.6696 | 0.104 | 0.2 | 1 | MS.Trt_vs_MS.NoTrt_F |
| ODC1 | 0.00468 | 0.52659 | 0.101 | 0.2 | 1 | MS.Trt_vs_MS.NoTrt_F |
| SLC7A1 | 0.00468 | -0.8087 | 0.101 | 0.19 | 1 | MS.Trt_vs_MS.NoTrt_F |
| RASSF1 | 0.00469 | -0.2781 | 0.223 | 0.35 | 1 | MS.Trt_vs_MS.NoTrt_F |
| RITA1 | 0.0047 | -0.7143 | 0.061 | 0.14 | 1 | MS.Trt_vs_MS.NoTrt_F |
| TDRKH | 0.0047 | -0.3195 | 0.086 | 0.18 | 1 | MS.Trt_vs_MS.NoTrt_F |
| WWP1 | 0.00472 | 0.29466 | 0.424 | 0.59 | 1 | MS.Trt_vs_MS.NoTrt_F |
| PRKAG1 | 0.00473 | 0.51102 | 0.525 | 0.68 | 1 | MS.Trt_vs_MS.NoTrt_F |
| PPP1R12C | 0.00475 | -0.3627 | 0.076 | 0.16 | 1 | MS.Trt_vs_MS.NoTrt_F |
| FAM50B | 0.00475 | -1.3568 | 0.061 | 0.14 | 1 | MS.Trt_vs_MS.NoTrt_F |
| SCN1A | 0.00476 | -0.2752 | 0.342 | 0.48 | 1 | MS.Trt_vs_MS.NoTrt_F |
| CHTOP | 0.00476 | 0.31842 | 0.55 | 0.7 | 1 | MS.Trt_vs_MS.NoTrt_F |
| GRIP1 | 0.00476 | -0.507 | 0.061 | 0.14 | 1 | MS.Trt_vs_MS.NoTrt_F |
| CCDC65 | 0.00478 | -0.3406 | 0.083 | 0.17 | 1 | MS.Trt_vs_MS.NoTrt_F |
| RGPD1 | 0.00478 | -0.4906 | 0.223 | 0.35 | 1 | MS.Trt_vs_MS.NoTrt_F |
| CRIP1 | 0.00478 | 0.35446 | 0.432 | 0.64 | 1 | MS.Trt_vs_MS.NoTrt_F |
| CFAP36 | 0.00479 | -0.8438 | 0.392 | 0.5 | 1 | MS.Trt_vs_MS.NoTrt_F |
| KAT2B | 0.0048 | 0.37517 | 0.237 | 0.39 | 1 | MS.Trt_vs_MS.NoTrt_F |
| ADAM17 | 0.00481 | -0.2836 | 0.076 | 0.16 | 1 | MS.Trt_vs_MS.NoTrt_F |
| GBP3 | 0.00481 | 0.27325 | 0.201 | 0.33 | 1 | MS.Trt_vs_MS.NoTrt_F |
| TRAPPC13 | 0.00481 | 0.46194 | 0.14 | 0.24 | 1 | MS.Trt_vs_MS.NoTrt_F |
| TET3 | 0.00483 | -0.6413 | 0.083 | 0.17 | 1 | MS.Trt_vs_MS.NoTrt_F |
| AP4B1 | 0.00484 | 0.36239 | 0.604 | 0.74 | 1 | MS.Trt_vs_MS.NoTrt_F |
| PDCD11 | 0.00484 | 0.58147 | 0.219 | 0.35 | 1 | MS.Trt_vs_MS.NoTrt_F |
| DDX11 | 0.00484 | -0.258 | 0.385 | 0.5 | 1 | MS.Trt_vs_MS.NoTrt_F |

| | | | | | | |
|---|---|---|---|---|---|---|
| SNX2 | 0.00485 | -0.2953 | 0.633 | 0.74 | 1 | MS.Trt_vs_MS.NoTrt_F |
| ZNF770 | 0.00488 | -0.5107 | 0.291 | 0.41 | 1 | MS.Trt_vs_MS.NoTrt_F |
| LEMD2 | 0.00488 | -0.4884 | 0.094 | 0.19 | 1 | MS.Trt_vs_MS.NoTrt_F |
| SLC25A28 | 0.00488 | -0.8105 | 0.061 | 0.14 | 1 | MS.Trt_vs_MS.NoTrt_F |
| NAA20 | 0.00488 | 0.42713 | 0.133 | 0.24 | 1 | MS.Trt_vs_MS.NoTrt_F |
| C16orf58 | 0.00489 | 0.52286 | 0.14 | 0.25 | 1 | MS.Trt_vs_MS.NoTrt_F |
| DAPK3 | 0.0049 | -0.3049 | 0.112 | 0.21 | 1 | MS.Trt_vs_MS.NoTrt_F |
| ASB13 | 0.00491 | -1.6754 | 0.112 | 0.21 | 1 | MS.Trt_vs_MS.NoTrt_F |
| ZYG11B | 0.00492 | -0.3787 | 0.687 | 0.82 | 1 | MS.Trt_vs_MS.NoTrt_F |
| GTF2H2C | 0.00492 | 0.52116 | 0.342 | 0.49 | 1 | MS.Trt_vs_MS.NoTrt_F |
| WDR81 | 0.00494 | 0.62584 | 0.076 | 0.16 | 1 | MS.Trt_vs_MS.NoTrt_F |
| PHF12 | 0.00495 | -0.387 | 0.662 | 0.75 | 1 | MS.Trt_vs_MS.NoTrt_F |
| TAF5L | 0.00495 | -0.7645 | 0.173 | 0.3 | 1 | MS.Trt_vs_MS.NoTrt_F |
| MFSD12 | 0.00497 | -0.4213 | 0.219 | 0.34 | 1 | MS.Trt_vs_MS.NoTrt_F |
| C1orf131 | 0.00497 | 0.86801 | 0.586 | 0.71 | 1 | MS.Trt_vs_MS.NoTrt_F |
| TCEB2 | 0.00498 | 0.64579 | 0.32 | 0.52 | 1 | MS.Trt_vs_MS.NoTrt_F |
| DDX31 | 0.005 | -0.9862 | 0.068 | 0.15 | 1 | MS.Trt_vs_MS.NoTrt_F |
| HBS1L | 0.005 | 0.45766 | 0.522 | 0.68 | 1 | MS.Trt_vs_MS.NoTrt_F |
| MYO18A | 0.00501 | -0.5371 | 0.514 | 0.65 | 1 | MS.Trt_vs_MS.NoTrt_F |
| LIPA | 0.00502 | -0.572 | 0.284 | 0.41 | 1 | MS.Trt_vs_MS.NoTrt_F |
| C9orf69 | 0.00502 | -1.0173 | 0.054 | 0.13 | 1 | MS.Trt_vs_MS.NoTrt_F |
| GPRASP1 | 0.00503 | -0.7913 | 0.061 | 0.14 | 1 | MS.Trt_vs_MS.NoTrt_F |
| MATR3 | 0.00503 | -0.3613 | 0.932 | 0.96 | 1 | MS.Trt_vs_MS.NoTrt_F |
| C4orf33 | 0.00503 | 0.8848 | 0.079 | 0.17 | 1 | MS.Trt_vs_MS.NoTrt_F |
| UBXN7 | 0.00503 | -0.3767 | 0.777 | 0.88 | 1 | MS.Trt_vs_MS.NoTrt_F |
| HHLA3 | 0.00504 | -0.4798 | 0.302 | 0.43 | 1 | MS.Trt_vs_MS.NoTrt_F |
| ZNF446 | 0.00505 | -0.914 | 0.054 | 0.13 | 1 | MS.Trt_vs_MS.NoTrt_F |
| ZNF644 | 0.00506 | 0.38966 | 0.209 | 0.34 | 1 | MS.Trt_vs_MS.NoTrt_F |
| MARK2 | 0.00507 | -0.4798 | 0.173 | 0.29 | 1 | MS.Trt_vs_MS.NoTrt_F |
| DLG3 | 0.00507 | -1.0683 | 0.047 | 0.12 | 1 | MS.Trt_vs_MS.NoTrt_F |
| GPR34 | 0.00508 | -0.6162 | 0.068 | 0.15 | 1 | MS.Trt_vs_MS.NoTrt_F |
| ST3GAL1 | 0.00509 | -0.2991 | 0.234 | 0.34 | 1 | MS.Trt_vs_MS.NoTrt_F |
| ATP5E | 0.00511 | 0.65283 | 0.417 | 0.67 | 1 | MS.Trt_vs_MS.NoTrt_F |
| C21orf91 | 0.00512 | 0.53481 | 0.18 | 0.3 | 1 | MS.Trt_vs_MS.NoTrt_F |
| RING1 | 0.00512 | 0.30983 | 0.079 | 0.17 | 1 | MS.Trt_vs_MS.NoTrt_F |
| CHML | 0.00514 | 1.25672 | 0.209 | 0.34 | 1 | MS.Trt_vs_MS.NoTrt_F |
| DMAP1 | 0.00515 | -0.5633 | 0.432 | 0.55 | 1 | MS.Trt_vs_MS.NoTrt_F |
| C7orf25 | 0.00515 | -2.0323 | 0.04 | 0.11 | 1 | MS.Trt_vs_MS.NoTrt_F |
| TADA1 | 0.00516 | 0.32313 | 0.086 | 0.18 | 1 | MS.Trt_vs_MS.NoTrt_F |
| ARHGAP19 | 0.00518 | 0.41317 | 0.299 | 0.44 | 1 | MS.Trt_vs_MS.NoTrt_F |

| | | | | | | |
|---|---|---|---|---|---|---|
| ST8SIA4 | 0.00518 | -0.4626 | 0.737 | 0.81 | 1 | MS.Trt_vs_MS.NoTrt_F |
| RBM12 | 0.00518 | 0.26738 | 0.147 | 0.27 | 1 | MS.Trt_vs_MS.NoTrt_F |
| AFF4 | 0.00518 | -0.6844 | 0.482 | 0.61 | 1 | MS.Trt_vs_MS.NoTrt_F |
| TPP1 | 0.00519 | 0.45264 | 0.317 | 0.5 | 1 | MS.Trt_vs_MS.NoTrt_F |
| TRAFD1 | 0.00524 | -0.2929 | 0.227 | 0.35 | 1 | MS.Trt_vs_MS.NoTrt_F |
| OVCA2 | 0.00524 | 0.42589 | 0.101 | 0.2 | 1 | MS.Trt_vs_MS.NoTrt_F |
| RAB2B | 0.00524 | -0.3764 | 0.637 | 0.74 | 1 | MS.Trt_vs_MS.NoTrt_F |
| GSK3B | 0.00526 | -0.8361 | 0.683 | 0.81 | 1 | MS.Trt_vs_MS.NoTrt_F |
| GPR146 | 0.00526 | 0.33952 | 0.09 | 0.18 | 1 | MS.Trt_vs_MS.NoTrt_F |
| FBF1 | 0.00527 | 0.60712 | 0.058 | 0.13 | 1 | MS.Trt_vs_MS.NoTrt_F |
| HEMGN | 0.00528 | -0.7456 | 0.104 | 0.2 | 1 | MS.Trt_vs_MS.NoTrt_F |
| STK17A | 0.00528 | -0.3136 | 0.291 | 0.44 | 1 | MS.Trt_vs_MS.NoTrt_F |
| DHX33 | 0.0053 | 0.26885 | 0.349 | 0.49 | 1 | MS.Trt_vs_MS.NoTrt_F |
| IFT52 | 0.00531 | -0.3071 | 0.086 | 0.18 | 1 | MS.Trt_vs_MS.NoTrt_F |
| GABPA | 0.00531 | -0.8383 | 0.108 | 0.2 | 1 | MS.Trt_vs_MS.NoTrt_F |
| ATP1B3 | 0.00532 | 0.32345 | 0.144 | 0.26 | 1 | MS.Trt_vs_MS.NoTrt_F |
| PRR7 | 0.00533 | -0.3432 | 0.05 | 0.12 | 1 | MS.Trt_vs_MS.NoTrt_F |
| PDHX | 0.00534 | -0.2792 | 0.637 | 0.76 | 1 | MS.Trt_vs_MS.NoTrt_F |
| SNX29 | 0.00536 | -0.7403 | 0.781 | 0.86 | 1 | MS.Trt_vs_MS.NoTrt_F |
| GAB3 | 0.0054 | 0.7765 | 0.363 | 0.51 | 1 | MS.Trt_vs_MS.NoTrt_F |
| DNAH8 | 0.0054 | 0.88362 | 0.065 | 0.14 | 1 | MS.Trt_vs_MS.NoTrt_F |
| EIF2B3 | 0.00541 | -0.6144 | 0.424 | 0.53 | 1 | MS.Trt_vs_MS.NoTrt_F |
| CEACAM4 | 0.00541 | -0.3749 | 0.101 | 0.19 | 1 | MS.Trt_vs_MS.NoTrt_F |
| RPP25L | 0.00541 | 0.45506 | 0.072 | 0.16 | 1 | MS.Trt_vs_MS.NoTrt_F |
| PLS3 | 0.00545 | 0.38427 | 0.353 | 0.52 | 1 | MS.Trt_vs_MS.NoTrt_F |
| EPC2 | 0.00547 | -0.6496 | 0.101 | 0.19 | 1 | MS.Trt_vs_MS.NoTrt_F |
| AASDH | 0.00547 | 0.3254 | 0.094 | 0.19 | 1 | MS.Trt_vs_MS.NoTrt_F |
| RNF13 | 0.00548 | 0.52271 | 0.173 | 0.3 | 1 | MS.Trt_vs_MS.NoTrt_F |
| ADAL | 0.00552 | -0.7735 | 0.421 | 0.58 | 1 | MS.Trt_vs_MS.NoTrt_F |
| KIAA1324L | 0.00553 | 0.52357 | 0.054 | 0.13 | 1 | MS.Trt_vs_MS.NoTrt_F |
| ARPC1A | 0.00554 | 0.51129 | 0.245 | 0.38 | 1 | MS.Trt_vs_MS.NoTrt_F |
| GCHFR | 0.00556 | 0.69776 | 0.09 | 0.18 | 1 | MS.Trt_vs_MS.NoTrt_F |
| LEMD3 | 0.00556 | -0.2835 | 0.594 | 0.74 | 1 | MS.Trt_vs_MS.NoTrt_F |
| COX7A2 | 0.00558 | 0.29256 | 0.705 | 0.8 | 1 | MS.Trt_vs_MS.NoTrt_F |
| DFNA5 | 0.00559 | -0.3485 | 0.119 | 0.22 | 1 | MS.Trt_vs_MS.NoTrt_F |
| ATP6V0B | 0.00562 | 0.37231 | 0.187 | 0.32 | 1 | MS.Trt_vs_MS.NoTrt_F |
| C9orf131 | 0.00563 | -1.2394 | 0.126 | 0.22 | 1 | MS.Trt_vs_MS.NoTrt_F |
| NAA38 | 0.00563 | 0.55823 | 0.129 | 0.24 | 1 | MS.Trt_vs_MS.NoTrt_F |
| C2orf68 | 0.00564 | 0.69774 | 0.827 | 0.8 | 1 | MS.Trt_vs_MS.NoTrt_F |
| ADGRV1 | 0.00565 | -0.2575 | 0.183 | 0.3 | 1 | MS.Trt_vs_MS.NoTrt_F |

| | | | | | | |
|---|---|---|---|---|---|---|
| HLA-DRB5 | 0.00567 | -1.1204 | 0.115 | 0.21 | 1 | MS.Trt_vs_MS.NoTrt_F |
| MGAT2 | 0.00568 | -0.5529 | 0.126 | 0.23 | 1 | MS.Trt_vs_MS.NoTrt_F |
| RAB40C | 0.00568 | -1.0094 | 0.043 | 0.11 | 1 | MS.Trt_vs_MS.NoTrt_F |
| DDB1 | 0.0057 | -0.2736 | 0.881 | 0.88 | 1 | MS.Trt_vs_MS.NoTrt_F |
| INTS10 | 0.00573 | -0.2921 | 0.543 | 0.69 | 1 | MS.Trt_vs_MS.NoTrt_F |
| CTD-2369P2 | 0.00573 | 0.94027 | 0.076 | 0.16 | 1 | MS.Trt_vs_MS.NoTrt_F |
| ZNF146 | 0.00573 | 0.35756 | 0.194 | 0.32 | 1 | MS.Trt_vs_MS.NoTrt_F |
| RAI1 | 0.00574 | -0.6545 | 0.176 | 0.29 | 1 | MS.Trt_vs_MS.NoTrt_F |
| PEMT | 0.00576 | 0.69877 | 0.072 | 0.16 | 1 | MS.Trt_vs_MS.NoTrt_F |
| PGM3 | 0.00577 | 1.02161 | 0.23 | 0.37 | 1 | MS.Trt_vs_MS.NoTrt_F |
| COG4 | 0.00577 | -0.4312 | 0.68 | 0.79 | 1 | MS.Trt_vs_MS.NoTrt_F |
| FAM134C | 0.00577 | -0.3024 | 0.629 | 0.74 | 1 | MS.Trt_vs_MS.NoTrt_F |
| NCOR2 | 0.00582 | 0.27901 | 0.363 | 0.5 | 1 | MS.Trt_vs_MS.NoTrt_F |
| PPIF | 0.00588 | -0.6984 | 0.094 | 0.18 | 1 | MS.Trt_vs_MS.NoTrt_F |
| MPG | 0.00589 | 0.25714 | 0.173 | 0.3 | 1 | MS.Trt_vs_MS.NoTrt_F |
| GLB1 | 0.0059 | -0.6687 | 0.248 | 0.37 | 1 | MS.Trt_vs_MS.NoTrt_F |
| WDR45B | 0.0059 | -0.3624 | 0.381 | 0.53 | 1 | MS.Trt_vs_MS.NoTrt_F |
| SPECC1L | 0.00593 | 0.27609 | 0.122 | 0.23 | 1 | MS.Trt_vs_MS.NoTrt_F |
| FBXO41 | 0.00594 | 0.33106 | 0.061 | 0.14 | 1 | MS.Trt_vs_MS.NoTrt_F |
| SMIM4 | 0.00598 | -0.8831 | 0.079 | 0.16 | 1 | MS.Trt_vs_MS.NoTrt_F |
| COX6B1 | 0.00599 | 0.27956 | 0.18 | 0.29 | 1 | MS.Trt_vs_MS.NoTrt_F |
| ORC3 | 0.00599 | 0.849 | 0.165 | 0.28 | 1 | MS.Trt_vs_MS.NoTrt_F |
| S1PR3 | 0.00601 | 0.33909 | 0.802 | 0.71 | 1 | MS.Trt_vs_MS.NoTrt_F |
| PER3 | 0.00602 | -1.4539 | 0.086 | 0.17 | 1 | MS.Trt_vs_MS.NoTrt_F |
| RHBDD2 | 0.00604 | -0.3427 | 0.241 | 0.37 | 1 | MS.Trt_vs_MS.NoTrt_F |
| FAM210B | 0.00604 | -1.2441 | 0.338 | 0.46 | 1 | MS.Trt_vs_MS.NoTrt_F |
| HIST2H4A | 0.00605 | -0.5052 | 0.094 | 0.18 | 1 | MS.Trt_vs_MS.NoTrt_F |
| SAT2 | 0.00605 | -0.3261 | 0.122 | 0.22 | 1 | MS.Trt_vs_MS.NoTrt_F |
| TMEM254 | 0.00608 | -0.5406 | 0.112 | 0.21 | 1 | MS.Trt_vs_MS.NoTrt_F |
| FKBP10 | 0.00608 | -0.4151 | 0.349 | 0.5 | 1 | MS.Trt_vs_MS.NoTrt_F |
| TRMT44 | 0.00609 | -0.5698 | 0.23 | 0.37 | 1 | MS.Trt_vs_MS.NoTrt_F |
| DSN1 | 0.00609 | -0.6251 | 0.219 | 0.34 | 1 | MS.Trt_vs_MS.NoTrt_F |
| MRPL41 | 0.00611 | 0.34617 | 0.176 | 0.3 | 1 | MS.Trt_vs_MS.NoTrt_F |
| SIRT1 | 0.00611 | 0.34125 | 0.137 | 0.24 | 1 | MS.Trt_vs_MS.NoTrt_F |
| EMC7 | 0.00611 | 0.26273 | 0.205 | 0.34 | 1 | MS.Trt_vs_MS.NoTrt_F |
| DNASE1L3 | 0.00612 | -0.2768 | 0.338 | 0.5 | 1 | MS.Trt_vs_MS.NoTrt_F |
| KDM4B | 0.00613 | -0.8109 | 0.59 | 0.71 | 1 | MS.Trt_vs_MS.NoTrt_F |
| AIMP2 | 0.00614 | -0.5495 | 0.288 | 0.42 | 1 | MS.Trt_vs_MS.NoTrt_F |
| SUV39H2 | 0.00619 | -0.5528 | 0.399 | 0.56 | 1 | MS.Trt_vs_MS.NoTrt_F |
| TFG | 0.00622 | 0.25524 | 0.133 | 0.24 | 1 | MS.Trt_vs_MS.NoTrt_F |

| | | | | | | |
|---|---|---|---|---|---|---|
| COMTD1 | 0.00622 | 0.69049 | 0.112 | 0.21 | 1 | MS.Trt_vs_MS.NoTrt_F |
| DGCR8 | 0.00625 | -0.2805 | 0.137 | 0.25 | 1 | MS.Trt_vs_MS.NoTrt_F |
| LARP7 | 0.00626 | 0.53989 | 0.201 | 0.32 | 1 | MS.Trt_vs_MS.NoTrt_F |
| MYD88 | 0.00627 | -0.6176 | 0.144 | 0.25 | 1 | MS.Trt_vs_MS.NoTrt_F |
| EID1 | 0.00627 | 0.2557 | 0.381 | 0.56 | 1 | MS.Trt_vs_MS.NoTrt_F |
| PPP1R16B | 0.0063 | 0.81886 | 0.291 | 0.43 | 1 | MS.Trt_vs_MS.NoTrt_F |
| ZNF672 | 0.00632 | 0.98172 | 0.234 | 0.35 | 1 | MS.Trt_vs_MS.NoTrt_F |
| FBXO32 | 0.00634 | 0.40178 | 0.165 | 0.28 | 1 | MS.Trt_vs_MS.NoTrt_F |
| TMEM156 | 0.00634 | -0.5229 | 0.781 | 0.9 | 1 | MS.Trt_vs_MS.NoTrt_F |
| HPCAL4 | 0.00634 | -0.5991 | 0.083 | 0.17 | 1 | MS.Trt_vs_MS.NoTrt_F |
| RAB8B | 0.00634 | -0.2623 | 0.266 | 0.39 | 1 | MS.Trt_vs_MS.NoTrt_F |
| CCDC94 | 0.00635 | -0.6946 | 0.14 | 0.24 | 1 | MS.Trt_vs_MS.NoTrt_F |
| QDPR | 0.00636 | 0.33008 | 0.076 | 0.16 | 1 | MS.Trt_vs_MS.NoTrt_F |
| KRIT1 | 0.00636 | -0.8838 | 0.597 | 0.73 | 1 | MS.Trt_vs_MS.NoTrt_F |
| ARL3 | 0.0064 | 0.30389 | 0.094 | 0.18 | 1 | MS.Trt_vs_MS.NoTrt_F |
| IMMP1L | 0.00641 | 0.92871 | 0.561 | 0.69 | 1 | MS.Trt_vs_MS.NoTrt_F |
| CHMP4A | 0.00642 | 0.37852 | 0.572 | 0.75 | 1 | MS.Trt_vs_MS.NoTrt_F |
| OAS1 | 0.00642 | -0.5803 | 0.507 | 0.58 | 1 | MS.Trt_vs_MS.NoTrt_F |
| PYCR2 | 0.00643 | -0.2508 | 0.662 | 0.78 | 1 | MS.Trt_vs_MS.NoTrt_F |
| APIP | 0.00643 | 0.61977 | 0.482 | 0.61 | 1 | MS.Trt_vs_MS.NoTrt_F |
| YEATS4 | 0.00646 | 0.25127 | 0.191 | 0.3 | 1 | MS.Trt_vs_MS.NoTrt_F |
| TACO1 | 0.00646 | -1.3595 | 0.385 | 0.48 | 1 | MS.Trt_vs_MS.NoTrt_F |
| HSPB9 | 0.00648 | -0.4366 | 0.187 | 0.3 | 1 | MS.Trt_vs_MS.NoTrt_F |
| ZNF420 | 0.00651 | 0.38672 | 0.104 | 0.2 | 1 | MS.Trt_vs_MS.NoTrt_F |
| ASNS | 0.00652 | -0.519 | 0.281 | 0.4 | 1 | MS.Trt_vs_MS.NoTrt_F |
| SLC13A4 | 0.00653 | -0.6863 | 0.349 | 0.46 | 1 | MS.Trt_vs_MS.NoTrt_F |
| ASB6 | 0.00654 | -0.5892 | 0.586 | 0.69 | 1 | MS.Trt_vs_MS.NoTrt_F |
| SEMA3C | 0.00655 | -0.3847 | 0.673 | 0.81 | 1 | MS.Trt_vs_MS.NoTrt_F |
| ZNF543 | 0.00656 | 0.89994 | 0.532 | 0.69 | 1 | MS.Trt_vs_MS.NoTrt_F |
| TSPYL4 | 0.00657 | 0.43248 | 0.108 | 0.21 | 1 | MS.Trt_vs_MS.NoTrt_F |
| TPM4 | 0.00658 | -0.3387 | 0.917 | 0.95 | 1 | MS.Trt_vs_MS.NoTrt_F |
| NAXE | 0.00658 | 0.92907 | 0.133 | 0.24 | 1 | MS.Trt_vs_MS.NoTrt_F |
| LARS | 0.0066 | 0.50947 | 0.691 | 0.79 | 1 | MS.Trt_vs_MS.NoTrt_F |
| EPC1 | 0.00661 | -0.3717 | 0.716 | 0.82 | 1 | MS.Trt_vs_MS.NoTrt_F |
| POP4 | 0.00662 | 0.44221 | 0.129 | 0.24 | 1 | MS.Trt_vs_MS.NoTrt_F |
| ZNF337 | 0.00663 | -0.5966 | 0.097 | 0.19 | 1 | MS.Trt_vs_MS.NoTrt_F |
| ANAPC13 | 0.00663 | 0.51733 | 0.112 | 0.21 | 1 | MS.Trt_vs_MS.NoTrt_F |
| CCNT1 | 0.00663 | 0.73278 | 0.165 | 0.29 | 1 | MS.Trt_vs_MS.NoTrt_F |
| EPHA4 | 0.00665 | -0.6096 | 0.212 | 0.34 | 1 | MS.Trt_vs_MS.NoTrt_F |
| SDHAF1 | 0.00665 | 0.41477 | 0.083 | 0.17 | 1 | MS.Trt_vs_MS.NoTrt_F |

| | | | | | | |
|---|---|---|---|---|---|---|
| BEST1 | 0.00668 | -0.2507 | 0.806 | 0.91 | 1 | MS.Trt_vs_MS.NoTrt_F |
| TRAPPC9 | 0.00668 | 0.89392 | 0.144 | 0.25 | 1 | MS.Trt_vs_MS.NoTrt_F |
| PCBD1 | 0.00668 | -0.5579 | 0.108 | 0.2 | 1 | MS.Trt_vs_MS.NoTrt_F |
| AP3D1 | 0.00669 | 0.27075 | 0.155 | 0.27 | 1 | MS.Trt_vs_MS.NoTrt_F |
| PTPN9 | 0.0067 | 0.47501 | 0.273 | 0.39 | 1 | MS.Trt_vs_MS.NoTrt_F |
| LYPLAL1 | 0.0067 | -0.5297 | 0.083 | 0.17 | 1 | MS.Trt_vs_MS.NoTrt_F |
| NTMT1 | 0.00674 | -0.2552 | 0.212 | 0.31 | 1 | MS.Trt_vs_MS.NoTrt_F |
| PCMT1 | 0.00676 | -0.2871 | 0.212 | 0.32 | 1 | MS.Trt_vs_MS.NoTrt_F |
| DNAJC2 | 0.00676 | -0.3748 | 0.752 | 0.82 | 1 | MS.Trt_vs_MS.NoTrt_F |
| AGAP4 | 0.00677 | 0.46367 | 0.209 | 0.34 | 1 | MS.Trt_vs_MS.NoTrt_F |
| IARS | 0.00677 | 0.25694 | 0.683 | 0.79 | 1 | MS.Trt_vs_MS.NoTrt_F |
| WBP1 | 0.0068 | -0.3729 | 0.108 | 0.2 | 1 | MS.Trt_vs_MS.NoTrt_F |
| BCORL1 | 0.00681 | -0.5537 | 0.083 | 0.17 | 1 | MS.Trt_vs_MS.NoTrt_F |
| ALG1 | 0.00684 | -0.2758 | 0.558 | 0.66 | 1 | MS.Trt_vs_MS.NoTrt_F |
| TFAM | 0.00684 | 0.5512 | 0.255 | 0.39 | 1 | MS.Trt_vs_MS.NoTrt_F |
| FUK | 0.00685 | -0.5911 | 0.532 | 0.65 | 1 | MS.Trt_vs_MS.NoTrt_F |
| SP1 | 0.00685 | 0.25327 | 0.169 | 0.29 | 1 | MS.Trt_vs_MS.NoTrt_F |
| ZNF280B | 0.00688 | 0.3439 | 0.108 | 0.2 | 1 | MS.Trt_vs_MS.NoTrt_F |
| DCUN1D4 | 0.00689 | 0.30688 | 0.288 | 0.43 | 1 | MS.Trt_vs_MS.NoTrt_F |
| FUT10 | 0.00693 | 0.32601 | 0.313 | 0.44 | 1 | MS.Trt_vs_MS.NoTrt_F |
| TRIM62 | 0.00694 | -1.2868 | 0.086 | 0.17 | 1 | MS.Trt_vs_MS.NoTrt_F |
| PYCARD | 0.00695 | 0.60742 | 0.133 | 0.24 | 1 | MS.Trt_vs_MS.NoTrt_F |
| IMMP2L | 0.00697 | -1.0602 | 0.101 | 0.19 | 1 | MS.Trt_vs_MS.NoTrt_F |
| YIF1A | 0.00697 | 0.36316 | 0.104 | 0.2 | 1 | MS.Trt_vs_MS.NoTrt_F |
| UBFD1 | 0.00697 | -0.3708 | 0.363 | 0.54 | 1 | MS.Trt_vs_MS.NoTrt_F |
| RAD50 | 0.00698 | 0.2888 | 0.198 | 0.32 | 1 | MS.Trt_vs_MS.NoTrt_F |
| CENPF | 0.00701 | -0.3716 | 0.324 | 0.49 | 1 | MS.Trt_vs_MS.NoTrt_F |
| MRPL14 | 0.00704 | 0.8691 | 0.115 | 0.22 | 1 | MS.Trt_vs_MS.NoTrt_F |
| NUDT6 | 0.00706 | -1.1084 | 0.507 | 0.61 | 1 | MS.Trt_vs_MS.NoTrt_F |
| GRHL1 | 0.00709 | -0.5227 | 0.471 | 0.59 | 1 | MS.Trt_vs_MS.NoTrt_F |
| HBG2 | 0.00709 | -0.2929 | 0.406 | 0.55 | 1 | MS.Trt_vs_MS.NoTrt_F |
| MRPL36 | 0.00709 | 0.97038 | 0.115 | 0.22 | 1 | MS.Trt_vs_MS.NoTrt_F |
| HNRNPH2 | 0.00711 | 0.25057 | 0.219 | 0.34 | 1 | MS.Trt_vs_MS.NoTrt_F |
| ORAOV1 | 0.00713 | 0.44568 | 0.633 | 0.75 | 1 | MS.Trt_vs_MS.NoTrt_F |
| CISD3 | 0.00716 | 0.38274 | 0.097 | 0.19 | 1 | MS.Trt_vs_MS.NoTrt_F |
| EMC6 | 0.00717 | 0.67227 | 0.086 | 0.17 | 1 | MS.Trt_vs_MS.NoTrt_F |
| HS2ST1 | 0.00718 | -0.7797 | 0.097 | 0.18 | 1 | MS.Trt_vs_MS.NoTrt_F |
| CEP290 | 0.0072 | 0.38241 | 0.112 | 0.21 | 1 | MS.Trt_vs_MS.NoTrt_F |
| AKT2 | 0.0072 | -0.3869 | 0.953 | 0.96 | 1 | MS.Trt_vs_MS.NoTrt_F |
| RGS2 | 0.00724 | 0.88419 | 0.137 | 0.24 | 1 | MS.Trt_vs_MS.NoTrt_F |

| | | | | | | |
|---|---|---|---|---|---|---|
| CXCR3 | 0.00725 | -0.3407 | 0.183 | 0.3 | 1 | MS.Trt_vs_MS.NoTrt_F |
| BACH2 | 0.00726 | -0.3557 | 0.155 | 0.26 | 1 | MS.Trt_vs_MS.NoTrt_F |
| EBLN2 | 0.00726 | -0.7892 | 0.104 | 0.19 | 1 | MS.Trt_vs_MS.NoTrt_F |
| UBE2G1 | 0.00729 | -0.4389 | 0.219 | 0.34 | 1 | MS.Trt_vs_MS.NoTrt_F |
| TELO2 | 0.00733 | 0.68991 | 0.115 | 0.22 | 1 | MS.Trt_vs_MS.NoTrt_F |
| VAMP4 | 0.00734 | 0.45596 | 0.219 | 0.34 | 1 | MS.Trt_vs_MS.NoTrt_F |
| ZC3H15 | 0.00734 | 0.46568 | 0.198 | 0.33 | 1 | MS.Trt_vs_MS.NoTrt_F |
| ANKRD36B | 0.00735 | 0.56068 | 0.644 | 0.77 | 1 | MS.Trt_vs_MS.NoTrt_F |
| NDUFAF4 | 0.00736 | 0.26668 | 0.094 | 0.18 | 1 | MS.Trt_vs_MS.NoTrt_F |
| PPP3R1 | 0.00738 | -0.5663 | 0.147 | 0.24 | 1 | MS.Trt_vs_MS.NoTrt_F |
| USP13 | 0.00741 | -0.3664 | 0.302 | 0.43 | 1 | MS.Trt_vs_MS.NoTrt_F |
| LRRC8A | 0.00741 | -0.6392 | 0.115 | 0.21 | 1 | MS.Trt_vs_MS.NoTrt_F |
| VPS37B | 0.00741 | -0.5752 | 0.615 | 0.74 | 1 | MS.Trt_vs_MS.NoTrt_F |
| TRPS1 | 0.00742 | -1.1908 | 0.104 | 0.19 | 1 | MS.Trt_vs_MS.NoTrt_F |
| SLAMF6 | 0.00743 | -0.4652 | 0.468 | 0.56 | 1 | MS.Trt_vs_MS.NoTrt_F |
| PIK3CD | 0.00743 | -0.3778 | 0.626 | 0.71 | 1 | MS.Trt_vs_MS.NoTrt_F |
| FKBP5 | 0.00746 | 0.40386 | 0.745 | 0.88 | 1 | MS.Trt_vs_MS.NoTrt_F |
| GREB1 | 0.00748 | 0.30996 | 0.827 | 0.82 | 1 | MS.Trt_vs_MS.NoTrt_F |
| AKAP17A | 0.00749 | -0.8926 | 0.101 | 0.19 | 1 | MS.Trt_vs_MS.NoTrt_F |
| UBAC2 | 0.00752 | -0.2647 | 0.576 | 0.67 | 1 | MS.Trt_vs_MS.NoTrt_F |
| CES3 | 0.00753 | -0.2918 | 0.259 | 0.39 | 1 | MS.Trt_vs_MS.NoTrt_F |
| SFPQ | 0.00758 | 0.46046 | 0.54 | 0.69 | 1 | MS.Trt_vs_MS.NoTrt_F |
| ESYT2 | 0.0076 | 0.31481 | 0.561 | 0.71 | 1 | MS.Trt_vs_MS.NoTrt_F |
| ZNF319 | 0.0076 | 0.26172 | 0.076 | 0.16 | 1 | MS.Trt_vs_MS.NoTrt_F |
| ANKRD46 | 0.00763 | -0.6885 | 0.198 | 0.3 | 1 | MS.Trt_vs_MS.NoTrt_F |
| CYB5D2 | 0.00763 | 0.31597 | 0.378 | 0.53 | 1 | MS.Trt_vs_MS.NoTrt_F |
| CCDC191 | 0.00767 | -0.6877 | 0.201 | 0.31 | 1 | MS.Trt_vs_MS.NoTrt_F |
| TSPAN32 | 0.0077 | 0.5906 | 0.227 | 0.36 | 1 | MS.Trt_vs_MS.NoTrt_F |
| ENTPD1 | 0.00771 | -0.3367 | 0.935 | 0.98 | 1 | MS.Trt_vs_MS.NoTrt_F |
| SLC22A23 | 0.00771 | -0.9009 | 0.079 | 0.16 | 1 | MS.Trt_vs_MS.NoTrt_F |
| MTPN | 0.00772 | 0.62804 | 0.198 | 0.33 | 1 | MS.Trt_vs_MS.NoTrt_F |
| QARS | 0.00773 | -0.4331 | 0.237 | 0.36 | 1 | MS.Trt_vs_MS.NoTrt_F |
| TTF2 | 0.00773 | 0.25534 | 0.709 | 0.82 | 1 | MS.Trt_vs_MS.NoTrt_F |
| PLEKHF1 | 0.00776 | 0.25623 | 0.158 | 0.27 | 1 | MS.Trt_vs_MS.NoTrt_F |
| IFT122 | 0.00777 | -0.312 | 0.849 | 0.93 | 1 | MS.Trt_vs_MS.NoTrt_F |
| TMEM104 | 0.00782 | -0.6043 | 0.155 | 0.26 | 1 | MS.Trt_vs_MS.NoTrt_F |
| SHROOM1 | 0.00782 | -0.2753 | 0.342 | 0.46 | 1 | MS.Trt_vs_MS.NoTrt_F |
| NECAP2 | 0.00783 | 0.37644 | 0.457 | 0.61 | 1 | MS.Trt_vs_MS.NoTrt_F |
| GPR171 | 0.00783 | -1.1139 | 0.058 | 0.13 | 1 | MS.Trt_vs_MS.NoTrt_F |
| NDRG3 | 0.00783 | 0.40397 | 0.381 | 0.53 | 1 | MS.Trt_vs_MS.NoTrt_F |

| | | | | | | |
|---|---|---|---|---|---|---|
| ARL6IP4 | 0.00784 | 0.34599 | 0.313 | 0.49 | 1 | MS.Trt_vs_MS.NoTrt_F |
| PARG | 0.00786 | 0.8502 | 0.155 | 0.27 | 1 | MS.Trt_vs_MS.NoTrt_F |
| RPGR | 0.0079 | -0.8729 | 0.068 | 0.14 | 1 | MS.Trt_vs_MS.NoTrt_F |
| VARS | 0.0079 | -0.4187 | 0.083 | 0.17 | 1 | MS.Trt_vs_MS.NoTrt_F |
| ACTR6 | 0.00791 | -0.45 | 0.547 | 0.65 | 1 | MS.Trt_vs_MS.NoTrt_F |
| NSUN2 | 0.00796 | -0.2667 | 0.594 | 0.74 | 1 | MS.Trt_vs_MS.NoTrt_F |
| SNX27 | 0.00797 | 0.34881 | 0.676 | 0.81 | 1 | MS.Trt_vs_MS.NoTrt_F |
| GCC1 | 0.00798 | -1.6026 | 0.327 | 0.45 | 1 | MS.Trt_vs_MS.NoTrt_F |
| PSAT1 | 0.00799 | -0.7825 | 0.076 | 0.16 | 1 | MS.Trt_vs_MS.NoTrt_F |
| PDRG1 | 0.00799 | -0.495 | 0.097 | 0.18 | 1 | MS.Trt_vs_MS.NoTrt_F |
| PANK4 | 0.008 | -0.3932 | 0.079 | 0.16 | 1 | MS.Trt_vs_MS.NoTrt_F |
| STK39 | 0.00802 | -0.2652 | 0.144 | 0.25 | 1 | MS.Trt_vs_MS.NoTrt_F |
| ADNP | 0.00803 | 0.78279 | 0.428 | 0.6 | 1 | MS.Trt_vs_MS.NoTrt_F |
| ADSL | 0.00803 | -0.4939 | 0.827 | 0.89 | 1 | MS.Trt_vs_MS.NoTrt_F |
| RNASET2 | 0.00804 | 0.26832 | 0.435 | 0.61 | 1 | MS.Trt_vs_MS.NoTrt_F |
| SPEF2 | 0.00806 | -0.8357 | 0.536 | 0.65 | 1 | MS.Trt_vs_MS.NoTrt_F |
| BTBD19 | 0.00809 | 0.3027 | 0.766 | 0.72 | 1 | MS.Trt_vs_MS.NoTrt_F |
| COPS4 | 0.00809 | 0.59346 | 0.453 | 0.57 | 1 | MS.Trt_vs_MS.NoTrt_F |
| CCDC93 | 0.0081 | 0.38981 | 0.151 | 0.26 | 1 | MS.Trt_vs_MS.NoTrt_F |
| BIN3 | 0.00812 | 0.50022 | 0.345 | 0.49 | 1 | MS.Trt_vs_MS.NoTrt_F |
| MCM2 | 0.00812 | -0.6936 | 0.058 | 0.13 | 1 | MS.Trt_vs_MS.NoTrt_F |
| TMEM164 | 0.00813 | 0.52752 | 0.079 | 0.16 | 1 | MS.Trt_vs_MS.NoTrt_F |
| RNF181 | 0.00817 | 0.31037 | 0.169 | 0.29 | 1 | MS.Trt_vs_MS.NoTrt_F |
| EAF1 | 0.00818 | 0.68912 | 0.331 | 0.49 | 1 | MS.Trt_vs_MS.NoTrt_F |
| PRDM4 | 0.00819 | 1.17586 | 0.086 | 0.17 | 1 | MS.Trt_vs_MS.NoTrt_F |
| FDX2 | 0.0082 | 0.54103 | 0.068 | 0.14 | 1 | MS.Trt_vs_MS.NoTrt_F |
| PSMB1 | 0.0082 | 0.37001 | 0.313 | 0.47 | 1 | MS.Trt_vs_MS.NoTrt_F |
| MAGOHB | 0.00821 | -0.3223 | 0.104 | 0.19 | 1 | MS.Trt_vs_MS.NoTrt_F |
| RPAP3 | 0.00821 | 0.28733 | 0.586 | 0.76 | 1 | MS.Trt_vs_MS.NoTrt_F |
| ARF1 | 0.00821 | 0.25232 | 0.46 | 0.65 | 1 | MS.Trt_vs_MS.NoTrt_F |
| SPOPL | 0.00824 | 0.33016 | 0.079 | 0.16 | 1 | MS.Trt_vs_MS.NoTrt_F |
| ZNF775 | 0.00825 | -0.4574 | 0.072 | 0.15 | 1 | MS.Trt_vs_MS.NoTrt_F |
| TRAF4 | 0.00827 | -1.0151 | 0.061 | 0.13 | 1 | MS.Trt_vs_MS.NoTrt_F |
| ABHD5 | 0.0083 | -0.5602 | 0.126 | 0.22 | 1 | MS.Trt_vs_MS.NoTrt_F |
| WIPF1 | 0.00831 | -0.3479 | 0.694 | 0.8 | 1 | MS.Trt_vs_MS.NoTrt_F |
| ISOC1 | 0.00836 | 0.34318 | 0.119 | 0.22 | 1 | MS.Trt_vs_MS.NoTrt_F |
| EMC9 | 0.00837 | -0.567 | 0.065 | 0.14 | 1 | MS.Trt_vs_MS.NoTrt_F |
| PDZD11 | 0.00837 | -0.305 | 0.112 | 0.2 | 1 | MS.Trt_vs_MS.NoTrt_F |
| NFXL1 | 0.00838 | -0.7506 | 0.112 | 0.2 | 1 | MS.Trt_vs_MS.NoTrt_F |
| ZNF624 | 0.00842 | -0.7996 | 0.378 | 0.52 | 1 | MS.Trt_vs_MS.NoTrt_F |

| | | | | | | |
|---|---|---|---|---|---|---|
| HDAC6 | 0.00845 | -0.5 | 0.86 | 0.92 | 1 | MS.Trt_vs_MS.NoTrt_F |
| CEP89 | 0.00847 | -0.3156 | 0.842 | 0.93 | 1 | MS.Trt_vs_MS.NoTrt_F |
| PLCXD2 | 0.00849 | -1.4772 | 0.09 | 0.17 | 1 | MS.Trt_vs_MS.NoTrt_F |
| FMO3 | 0.00853 | 0.35627 | 0.687 | 0.61 | 1 | MS.Trt_vs_MS.NoTrt_F |
| RHOQ | 0.00855 | 0.56989 | 0.086 | 0.17 | 1 | MS.Trt_vs_MS.NoTrt_F |
| POTEC | 0.00855 | 0.26316 | 0.356 | 0.51 | 1 | MS.Trt_vs_MS.NoTrt_F |
| ZNF155 | 0.00856 | -1.0261 | 0.054 | 0.12 | 1 | MS.Trt_vs_MS.NoTrt_F |
| EEF2KMT | 0.00857 | 0.25648 | 0.094 | 0.18 | 1 | MS.Trt_vs_MS.NoTrt_F |
| ATG4A | 0.00857 | 0.66005 | 0.083 | 0.17 | 1 | MS.Trt_vs_MS.NoTrt_F |
| FBXO22 | 0.00857 | -0.4446 | 0.748 | 0.82 | 1 | MS.Trt_vs_MS.NoTrt_F |
| IRF1 | 0.00858 | -0.5161 | 0.849 | 0.92 | 1 | MS.Trt_vs_MS.NoTrt_F |
| WDR35 | 0.00859 | 0.84921 | 0.083 | 0.17 | 1 | MS.Trt_vs_MS.NoTrt_F |
| CTSA | 0.00861 | -0.7721 | 0.644 | 0.74 | 1 | MS.Trt_vs_MS.NoTrt_F |
| ZNF514 | 0.00865 | -1.2963 | 0.586 | 0.69 | 1 | MS.Trt_vs_MS.NoTrt_F |
| ACVR2B | 0.00868 | -0.7078 | 0.396 | 0.53 | 1 | MS.Trt_vs_MS.NoTrt_F |
| RMI1 | 0.00869 | 0.54553 | 0.245 | 0.38 | 1 | MS.Trt_vs_MS.NoTrt_F |
| IPPK | 0.00872 | 0.41012 | 0.083 | 0.17 | 1 | MS.Trt_vs_MS.NoTrt_F |
| TMED9 | 0.00872 | 0.32594 | 0.32 | 0.47 | 1 | MS.Trt_vs_MS.NoTrt_F |
| RIC8B | 0.00875 | -0.3122 | 0.162 | 0.26 | 1 | MS.Trt_vs_MS.NoTrt_F |
| TULP3 | 0.00877 | -1.0386 | 0.058 | 0.13 | 1 | MS.Trt_vs_MS.NoTrt_F |
| GSTCD | 0.00877 | -0.3723 | 0.619 | 0.74 | 1 | MS.Trt_vs_MS.NoTrt_F |
| CNOT8 | 0.00879 | 0.39899 | 0.216 | 0.34 | 1 | MS.Trt_vs_MS.NoTrt_F |
| TYMP | 0.0088 | -0.4302 | 0.324 | 0.45 | 1 | MS.Trt_vs_MS.NoTrt_F |
| KANSL2 | 0.00882 | 0.58228 | 0.155 | 0.27 | 1 | MS.Trt_vs_MS.NoTrt_F |
| CDC42BPB | 0.00885 | 0.52777 | 0.061 | 0.13 | 1 | MS.Trt_vs_MS.NoTrt_F |
| ROBO4 | 0.00886 | -1.0911 | 0.169 | 0.27 | 1 | MS.Trt_vs_MS.NoTrt_F |
| TMEM186 | 0.00886 | -0.7939 | 0.478 | 0.61 | 1 | MS.Trt_vs_MS.NoTrt_F |
| PHPT1 | 0.00889 | 0.81757 | 0.144 | 0.25 | 1 | MS.Trt_vs_MS.NoTrt_F |
| FAM65A | 0.0089 | -0.4648 | 0.126 | 0.22 | 1 | MS.Trt_vs_MS.NoTrt_F |
| FOXP3 | 0.00891 | -0.2697 | 0.417 | 0.56 | 1 | MS.Trt_vs_MS.NoTrt_F |
| RGP1 | 0.00893 | -0.4165 | 0.162 | 0.27 | 1 | MS.Trt_vs_MS.NoTrt_F |
| EMG1 | 0.00897 | 0.37142 | 0.126 | 0.23 | 1 | MS.Trt_vs_MS.NoTrt_F |
| CCDC82 | 0.00897 | 0.48682 | 0.165 | 0.28 | 1 | MS.Trt_vs_MS.NoTrt_F |
| RPUSD1 | 0.00901 | 0.50135 | 0.115 | 0.21 | 1 | MS.Trt_vs_MS.NoTrt_F |
| ZCWPW1 | 0.00901 | 0.35551 | 0.086 | 0.17 | 1 | MS.Trt_vs_MS.NoTrt_F |
| RALGAPA2 | 0.00902 | 0.53807 | 0.14 | 0.25 | 1 | MS.Trt_vs_MS.NoTrt_F |
| PFKFB3 | 0.00903 | -0.2748 | 0.281 | 0.4 | 1 | MS.Trt_vs_MS.NoTrt_F |
| UQCRQ | 0.00903 | 0.32104 | 0.572 | 0.7 | 1 | MS.Trt_vs_MS.NoTrt_F |
| C12orf40 | 0.00904 | -0.925 | 0.237 | 0.36 | 1 | MS.Trt_vs_MS.NoTrt_F |
| LRIF1 | 0.00907 | -0.6475 | 0.101 | 0.19 | 1 | MS.Trt_vs_MS.NoTrt_F |

| | | | | | | |
|---|---|---|---|---|---|---|
| ZFYVE16 | 0.00908 | 0.30591 | 0.957 | 0.9 | 1 | MS.Trt_vs_MS.NoTrt_F |
| RABGAP1 | 0.00908 | -0.4608 | 0.363 | 0.5 | 1 | MS.Trt_vs_MS.NoTrt_F |
| PLCL2 | 0.00911 | 0.31118 | 0.072 | 0.15 | 1 | MS.Trt_vs_MS.NoTrt_F |
| ZNF518B | 0.00914 | 0.6442 | 0.094 | 0.18 | 1 | MS.Trt_vs_MS.NoTrt_F |
| EIF2B4 | 0.00915 | 0.42903 | 0.442 | 0.61 | 1 | MS.Trt_vs_MS.NoTrt_F |
| RAB5B | 0.00919 | -0.443 | 0.888 | 0.91 | 1 | MS.Trt_vs_MS.NoTrt_F |
| GXYLT1 | 0.0092 | -0.5229 | 0.417 | 0.53 | 1 | MS.Trt_vs_MS.NoTrt_F |
| SAP30L | 0.00922 | -1.2029 | 0.047 | 0.11 | 1 | MS.Trt_vs_MS.NoTrt_F |
| WRB | 0.00922 | 0.29602 | 0.083 | 0.17 | 1 | MS.Trt_vs_MS.NoTrt_F |
| RAD52 | 0.00922 | -0.323 | 0.266 | 0.39 | 1 | MS.Trt_vs_MS.NoTrt_F |
| HSBP1L1 | 0.00923 | -0.4344 | 0.345 | 0.47 | 1 | MS.Trt_vs_MS.NoTrt_F |
| SIRT6 | 0.00925 | 0.3768 | 0.083 | 0.17 | 1 | MS.Trt_vs_MS.NoTrt_F |
| PI16 | 0.00926 | 0.59305 | 0.158 | 0.27 | 1 | MS.Trt_vs_MS.NoTrt_F |
| TARBP2 | 0.00928 | -0.5367 | 0.097 | 0.18 | 1 | MS.Trt_vs_MS.NoTrt_F |
| FBRSL1 | 0.0093 | 0.30739 | 0.101 | 0.19 | 1 | MS.Trt_vs_MS.NoTrt_F |
| CHD9 | 0.00935 | -0.6794 | 0.41 | 0.54 | 1 | MS.Trt_vs_MS.NoTrt_F |
| SOWAHC | 0.0094 | -0.4413 | 0.457 | 0.55 | 1 | MS.Trt_vs_MS.NoTrt_F |
| TF | 0.00941 | -0.3013 | 0.892 | 0.91 | 1 | MS.Trt_vs_MS.NoTrt_F |
| NLRP2 | 0.00941 | 0.78224 | 0.824 | 0.8 | 1 | MS.Trt_vs_MS.NoTrt_F |
| ATPIF1 | 0.00942 | 0.49129 | 0.245 | 0.4 | 1 | MS.Trt_vs_MS.NoTrt_F |
| RUNDC1 | 0.00942 | 0.85245 | 0.752 | 0.71 | 1 | MS.Trt_vs_MS.NoTrt_F |
| EPB41L2 | 0.00943 | -1.0942 | 0.122 | 0.22 | 1 | MS.Trt_vs_MS.NoTrt_F |
| ZNF622 | 0.00946 | 0.3536 | 0.144 | 0.25 | 1 | MS.Trt_vs_MS.NoTrt_F |
| UTP3 | 0.00946 | -0.2854 | 0.05 | 0.12 | 1 | MS.Trt_vs_MS.NoTrt_F |
| MALSU1 | 0.00951 | 0.47692 | 0.126 | 0.22 | 1 | MS.Trt_vs_MS.NoTrt_F |
| FBXO3 | 0.00951 | 0.63342 | 0.266 | 0.39 | 1 | MS.Trt_vs_MS.NoTrt_F |
| TOP2B | 0.00956 | 0.30346 | 0.237 | 0.37 | 1 | MS.Trt_vs_MS.NoTrt_F |
| CCNE2 | 0.00957 | -0.5154 | 0.05 | 0.12 | 1 | MS.Trt_vs_MS.NoTrt_F |
| DIS3 | 0.00958 | -0.3743 | 0.651 | 0.76 | 1 | MS.Trt_vs_MS.NoTrt_F |
| RP11-77K12 | 0.00958 | -0.3054 | 0.061 | 0.13 | 1 | MS.Trt_vs_MS.NoTrt_F |
| TAF1 | 0.00963 | -0.2782 | 0.745 | 0.85 | 1 | MS.Trt_vs_MS.NoTrt_F |
| ARMC7 | 0.00964 | -0.2751 | 0.097 | 0.18 | 1 | MS.Trt_vs_MS.NoTrt_F |
| ZDHHC6 | 0.00967 | 0.40294 | 0.165 | 0.28 | 1 | MS.Trt_vs_MS.NoTrt_F |
| CCS | 0.0097 | -0.6439 | 0.183 | 0.29 | 1 | MS.Trt_vs_MS.NoTrt_F |
| BTAF1 | 0.0097 | 0.34612 | 0.248 | 0.37 | 1 | MS.Trt_vs_MS.NoTrt_F |
| TMTC4 | 0.00972 | -0.7702 | 0.086 | 0.17 | 1 | MS.Trt_vs_MS.NoTrt_F |
| ZNF620 | 0.00976 | -0.3094 | 0.665 | 0.81 | 1 | MS.Trt_vs_MS.NoTrt_F |
| SLC6A1 | 0.00976 | -0.3066 | 0.522 | 0.64 | 1 | MS.Trt_vs_MS.NoTrt_F |
| OR52A1 | 0.00978 | 0.39732 | 0.799 | 0.75 | 1 | MS.Trt_vs_MS.NoTrt_F |
| COX5B | 0.00978 | 0.65221 | 0.791 | 0.88 | 1 | MS.Trt_vs_MS.NoTrt_F |

| | | | | | | |
|---|---|---|---|---|---|---|
| EPS15L1 | 0.00979 | -0.56 | 0.799 | 0.9 | 1 | MS.Trt_vs_MS.NoTrt_F |
| PSMF1 | 0.0098 | 0.4376 | 0.227 | 0.36 | 1 | MS.Trt_vs_MS.NoTrt_F |
| EPHX1 | 0.00982 | -0.3299 | 0.097 | 0.18 | 1 | MS.Trt_vs_MS.NoTrt_F |
| USP8 | 0.00983 | 0.35043 | 0.986 | 0.99 | 1 | MS.Trt_vs_MS.NoTrt_F |
| NHP2 | 0.00985 | 0.55666 | 0.187 | 0.31 | 1 | MS.Trt_vs_MS.NoTrt_F |
| LIN54 | 0.00985 | -0.3132 | 0.565 | 0.66 | 1 | MS.Trt_vs_MS.NoTrt_F |
| CMC1 | 0.00986 | 0.42526 | 0.191 | 0.31 | 1 | MS.Trt_vs_MS.NoTrt_F |
| TBC1D2B | 0.00986 | 0.33713 | 0.295 | 0.41 | 1 | MS.Trt_vs_MS.NoTrt_F |
| SSSCA1 | 0.00988 | 0.64801 | 0.097 | 0.18 | 1 | MS.Trt_vs_MS.NoTrt_F |
| MBNL3 | 0.00991 | 0.32787 | 0.108 | 0.2 | 1 | MS.Trt_vs_MS.NoTrt_F |
| NRF1 | 0.00991 | 0.27264 | 0.306 | 0.44 | 1 | MS.Trt_vs_MS.NoTrt_F |
| NOG | 0.00993 | 1.55009 | 0.129 | 0.24 | 1 | MS.Trt_vs_MS.NoTrt_F |
| TTI1 | 0.00994 | 0.64168 | 0.234 | 0.37 | 1 | MS.Trt_vs_MS.NoTrt_F |
| ZNF671 | 0.00994 | -0.6748 | 0.097 | 0.18 | 1 | MS.Trt_vs_MS.NoTrt_F |
| MEI1 | 0.00994 | -0.3161 | 0.687 | 0.77 | 1 | MS.Trt_vs_MS.NoTrt_F |
| TBP | 0.01 | 0.56203 | 0.335 | 0.48 | 1 | MS.Trt_vs_MS.NoTrt_F |
| NCBP2-AS2 | 0.01002 | 0.39351 | 0.119 | 0.22 | 1 | MS.Trt_vs_MS.NoTrt_F |
| AL592183.1 | 0.01003 | -0.2845 | 0.119 | 0.21 | 1 | MS.Trt_vs_MS.NoTrt_F |
| PIGG | 0.01003 | -0.3277 | 0.511 | 0.62 | 1 | MS.Trt_vs_MS.NoTrt_F |
| CNPPD1 | 0.01005 | 0.34635 | 0.281 | 0.43 | 1 | MS.Trt_vs_MS.NoTrt_F |
| SLC25A32 | 0.01007 | 0.75777 | 0.338 | 0.47 | 1 | MS.Trt_vs_MS.NoTrt_F |
| FNDC3B | 0.01007 | 0.57367 | 0.745 | 0.65 | 1 | MS.Trt_vs_MS.NoTrt_F |
| MUT | 0.0101 | 0.26284 | 0.09 | 0.17 | 1 | MS.Trt_vs_MS.NoTrt_F |
| HSD17B8 | 0.01011 | 0.79816 | 0.079 | 0.16 | 1 | MS.Trt_vs_MS.NoTrt_F |
| TMEM221 | 0.01011 | -0.3537 | 0.183 | 0.28 | 1 | MS.Trt_vs_MS.NoTrt_F |
| UQCC2 | 0.01013 | 0.80353 | 0.112 | 0.2 | 1 | MS.Trt_vs_MS.NoTrt_F |
| SPICE1 | 0.01014 | -0.3552 | 0.234 | 0.34 | 1 | MS.Trt_vs_MS.NoTrt_F |
| MYSM1 | 0.01016 | 0.41907 | 0.227 | 0.35 | 1 | MS.Trt_vs_MS.NoTrt_F |
| STK11IP | 0.01016 | 0.28971 | 0.097 | 0.18 | 1 | MS.Trt_vs_MS.NoTrt_F |
| ADAM19 | 0.01017 | 0.56824 | 0.36 | 0.55 | 1 | MS.Trt_vs_MS.NoTrt_F |
| GANAB | 0.01024 | -0.8664 | 0.446 | 0.53 | 1 | MS.Trt_vs_MS.NoTrt_F |
| MDC1 | 0.01026 | 0.26353 | 0.115 | 0.2 | 1 | MS.Trt_vs_MS.NoTrt_F |
| TRIB3 | 0.01028 | 0.34722 | 0.072 | 0.15 | 1 | MS.Trt_vs_MS.NoTrt_F |
| ATP8B2 | 0.0103 | -0.3084 | 0.802 | 0.86 | 1 | MS.Trt_vs_MS.NoTrt_F |
| TRMT10C | 0.01036 | 0.93285 | 0.104 | 0.19 | 1 | MS.Trt_vs_MS.NoTrt_F |
| GALNT7 | 0.01039 | 0.46896 | 0.101 | 0.19 | 1 | MS.Trt_vs_MS.NoTrt_F |
| GS1-114I9.3 | 0.01039 | 1.06673 | 0.058 | 0.13 | 1 | MS.Trt_vs_MS.NoTrt_F |
| EIF2S2 | 0.0104 | 0.30124 | 0.241 | 0.38 | 1 | MS.Trt_vs_MS.NoTrt_F |
| C1orf146 | 0.01042 | -0.2912 | 0.151 | 0.25 | 1 | MS.Trt_vs_MS.NoTrt_F |
| OTUD7A | 0.01042 | 0.32972 | 0.158 | 0.26 | 1 | MS.Trt_vs_MS.NoTrt_F |

| | | | | | | |
|---|---|---|---|---|---|---|
| SMN2 | 0.01042 | 0.31899 | 0.878 | 0.73 | 1 | MS.Trt_vs_MS.NoTrt_F |
| FAM162A | 0.01046 | -0.431 | 0.403 | 0.5 | 1 | MS.Trt_vs_MS.NoTrt_F |
| B4GALT1 | 0.01047 | 0.76877 | 0.712 | 0.74 | 1 | MS.Trt_vs_MS.NoTrt_F |
| PARP9 | 0.01047 | 0.57946 | 0.331 | 0.46 | 1 | MS.Trt_vs_MS.NoTrt_F |
| ARL4C | 0.01053 | 0.26905 | 0.399 | 0.58 | 1 | MS.Trt_vs_MS.NoTrt_F |
| AK1 | 0.01055 | 0.74818 | 0.104 | 0.19 | 1 | MS.Trt_vs_MS.NoTrt_F |
| LRRC28 | 0.01056 | -0.7675 | 0.54 | 0.67 | 1 | MS.Trt_vs_MS.NoTrt_F |
| NDUFB10 | 0.01057 | 0.58912 | 0.248 | 0.39 | 1 | MS.Trt_vs_MS.NoTrt_F |
| MTAP | 0.01058 | 0.53211 | 0.658 | 0.87 | 1 | MS.Trt_vs_MS.NoTrt_F |
| QSER1 | 0.01061 | 0.48921 | 0.097 | 0.18 | 1 | MS.Trt_vs_MS.NoTrt_F |
| IBTK | 0.01062 | -0.5907 | 0.522 | 0.62 | 1 | MS.Trt_vs_MS.NoTrt_F |
| INTS12 | 0.01062 | 0.28825 | 0.104 | 0.19 | 1 | MS.Trt_vs_MS.NoTrt_F |
| AMPD3 | 0.01071 | -1.3494 | 0.331 | 0.47 | 1 | MS.Trt_vs_MS.NoTrt_F |
| IFT172 | 0.01072 | -0.9388 | 0.59 | 0.7 | 1 | MS.Trt_vs_MS.NoTrt_F |
| SOX6 | 0.01072 | -0.5072 | 0.378 | 0.5 | 1 | MS.Trt_vs_MS.NoTrt_F |
| RNASE10 | 0.01073 | -0.2614 | 0.173 | 0.27 | 1 | MS.Trt_vs_MS.NoTrt_F |
| ELMO1 | 0.01081 | -0.4114 | 0.788 | 0.86 | 1 | MS.Trt_vs_MS.NoTrt_F |
| ANKRD37 | 0.01082 | -0.8303 | 0.104 | 0.19 | 1 | MS.Trt_vs_MS.NoTrt_F |
| LRRC40 | 0.01084 | 0.64074 | 0.144 | 0.24 | 1 | MS.Trt_vs_MS.NoTrt_F |
| ANKRD11 | 0.01085 | 0.63206 | 0.263 | 0.4 | 1 | MS.Trt_vs_MS.NoTrt_F |
| PSMB5 | 0.01087 | -0.5499 | 0.133 | 0.22 | 1 | MS.Trt_vs_MS.NoTrt_F |
| NOL10 | 0.01091 | -1.001 | 0.165 | 0.27 | 1 | MS.Trt_vs_MS.NoTrt_F |
| FAM19A2 | 0.01092 | -0.2858 | 0.212 | 0.32 | 1 | MS.Trt_vs_MS.NoTrt_F |
| PCNX3 | 0.01093 | 0.36108 | 0.133 | 0.23 | 1 | MS.Trt_vs_MS.NoTrt_F |
| ARMT1 | 0.01095 | -0.5072 | 0.086 | 0.17 | 1 | MS.Trt_vs_MS.NoTrt_F |
| RASGRP2 | 0.01096 | -0.3022 | 0.313 | 0.44 | 1 | MS.Trt_vs_MS.NoTrt_F |
| EP300 | 0.011 | -0.5258 | 0.658 | 0.79 | 1 | MS.Trt_vs_MS.NoTrt_F |
| GNPDA2 | 0.01101 | -0.6101 | 0.079 | 0.16 | 1 | MS.Trt_vs_MS.NoTrt_F |
| UBAP1L | 0.01104 | -0.3155 | 0.324 | 0.45 | 1 | MS.Trt_vs_MS.NoTrt_F |
| PHF6 | 0.01108 | -0.887 | 0.647 | 0.74 | 1 | MS.Trt_vs_MS.NoTrt_F |
| PNMA1 | 0.0111 | 0.28254 | 0.097 | 0.18 | 1 | MS.Trt_vs_MS.NoTrt_F |
| TXNDC9 | 0.01111 | 0.39445 | 0.137 | 0.24 | 1 | MS.Trt_vs_MS.NoTrt_F |
| FAM206A | 0.01118 | -0.3651 | 0.083 | 0.16 | 1 | MS.Trt_vs_MS.NoTrt_F |
| PHF19 | 0.01119 | 0.72506 | 0.158 | 0.27 | 1 | MS.Trt_vs_MS.NoTrt_F |
| SF3A2 | 0.01125 | -0.3382 | 0.399 | 0.48 | 1 | MS.Trt_vs_MS.NoTrt_F |
| SCML4 | 0.01127 | 0.44351 | 0.572 | 0.75 | 1 | MS.Trt_vs_MS.NoTrt_F |
| NOMO1 | 0.0113 | 0.85187 | 0.165 | 0.27 | 1 | MS.Trt_vs_MS.NoTrt_F |
| MBOAT7 | 0.01134 | 0.59751 | 0.097 | 0.18 | 1 | MS.Trt_vs_MS.NoTrt_F |
| CD58 | 0.01135 | 0.28383 | 0.263 | 0.39 | 1 | MS.Trt_vs_MS.NoTrt_F |
| STK19 | 0.0114 | -0.3029 | 0.09 | 0.17 | 1 | MS.Trt_vs_MS.NoTrt_F |

| | | | | | | |
|---|---|---|---|---|---|---|
| ERCC3 | 0.0114 | -0.3507 | 0.353 | 0.48 | 1 | MS.Trt_vs_MS.NoTrt_F |
| AURKA | 0.0114 | 0.65111 | 0.306 | 0.43 | 1 | MS.Trt_vs_MS.NoTrt_F |
| IMPA2 | 0.01142 | 0.52332 | 0.874 | 0.82 | 1 | MS.Trt_vs_MS.NoTrt_F |
| HIRIP3 | 0.01143 | 0.3232 | 0.101 | 0.19 | 1 | MS.Trt_vs_MS.NoTrt_F |
| ZNF366 | 0.01144 | -0.3096 | 0.165 | 0.26 | 1 | MS.Trt_vs_MS.NoTrt_F |
| CARMIL2 | 0.01144 | 0.2581 | 0.086 | 0.17 | 1 | MS.Trt_vs_MS.NoTrt_F |
| TRMT2A | 0.01146 | 0.37758 | 0.158 | 0.27 | 1 | MS.Trt_vs_MS.NoTrt_F |
| POLL | 0.01147 | 0.37267 | 0.392 | 0.53 | 1 | MS.Trt_vs_MS.NoTrt_F |
| BCAS1 | 0.01149 | -1.9067 | 0.349 | 0.46 | 1 | MS.Trt_vs_MS.NoTrt_F |
| WNT2B | 0.0115 | 0.39598 | 0.453 | 0.56 | 1 | MS.Trt_vs_MS.NoTrt_F |
| DMXL1 | 0.01155 | 1.03494 | 0.187 | 0.31 | 1 | MS.Trt_vs_MS.NoTrt_F |
| ABCE1 | 0.01157 | -0.2702 | 0.36 | 0.46 | 1 | MS.Trt_vs_MS.NoTrt_F |
| ACTR1A | 0.01161 | -0.3555 | 0.471 | 0.58 | 1 | MS.Trt_vs_MS.NoTrt_F |
| AP1AR | 0.01162 | 0.89714 | 0.086 | 0.17 | 1 | MS.Trt_vs_MS.NoTrt_F |
| RAB10 | 0.01162 | 0.33357 | 0.241 | 0.37 | 1 | MS.Trt_vs_MS.NoTrt_F |
| NPIPB9 | 0.01165 | 0.45251 | 0.137 | 0.23 | 1 | MS.Trt_vs_MS.NoTrt_F |
| RPS14 | 0.01167 | 0.37597 | 0.964 | 1 | 1 | MS.Trt_vs_MS.NoTrt_F |
| PATZ1 | 0.01168 | 0.40871 | 0.112 | 0.2 | 1 | MS.Trt_vs_MS.NoTrt_F |
| MTMR3 | 0.01173 | -1.0194 | 0.241 | 0.35 | 1 | MS.Trt_vs_MS.NoTrt_F |
| ZNF692 | 0.01174 | 0.26836 | 0.119 | 0.21 | 1 | MS.Trt_vs_MS.NoTrt_F |
| TDG | 0.01177 | -0.5542 | 0.327 | 0.42 | 1 | MS.Trt_vs_MS.NoTrt_F |
| TARDBP | 0.01179 | -0.3884 | 0.586 | 0.69 | 1 | MS.Trt_vs_MS.NoTrt_F |
| KDM6B | 0.01179 | -0.3052 | 0.183 | 0.29 | 1 | MS.Trt_vs_MS.NoTrt_F |
| ADCY3 | 0.0118 | 0.28589 | 0.09 | 0.17 | 1 | MS.Trt_vs_MS.NoTrt_F |
| GRAP | 0.01185 | 0.40717 | 0.169 | 0.28 | 1 | MS.Trt_vs_MS.NoTrt_F |
| ZNF212 | 0.01185 | -0.8874 | 0.371 | 0.48 | 1 | MS.Trt_vs_MS.NoTrt_F |
| PEX19 | 0.01189 | 0.26987 | 0.14 | 0.24 | 1 | MS.Trt_vs_MS.NoTrt_F |
| ZDHHC24 | 0.01189 | 0.28722 | 0.91 | 0.87 | 1 | MS.Trt_vs_MS.NoTrt_F |
| STYX | 0.01192 | 0.45418 | 0.504 | 0.64 | 1 | MS.Trt_vs_MS.NoTrt_F |
| KDM6A | 0.01193 | -0.4988 | 0.245 | 0.36 | 1 | MS.Trt_vs_MS.NoTrt_F |
| PMPCA | 0.01194 | -0.2521 | 0.705 | 0.8 | 1 | MS.Trt_vs_MS.NoTrt_F |
| TNRC6C | 0.01194 | 0.46537 | 0.266 | 0.39 | 1 | MS.Trt_vs_MS.NoTrt_F |
| NSUN5 | 0.01194 | 0.4304 | 0.277 | 0.41 | 1 | MS.Trt_vs_MS.NoTrt_F |
| DPP8 | 0.01203 | -0.4931 | 0.363 | 0.44 | 1 | MS.Trt_vs_MS.NoTrt_F |
| DNAJC9 | 0.01207 | 0.44126 | 0.169 | 0.28 | 1 | MS.Trt_vs_MS.NoTrt_F |
| MLH3 | 0.0121 | 0.37038 | 0.122 | 0.22 | 1 | MS.Trt_vs_MS.NoTrt_F |
| SEL1L3 | 0.0121 | 0.36363 | 0.27 | 0.4 | 1 | MS.Trt_vs_MS.NoTrt_F |
| ATG2A | 0.0121 | -0.4604 | 0.705 | 0.75 | 1 | MS.Trt_vs_MS.NoTrt_F |
| MESDC2 | 0.01214 | 0.76524 | 0.504 | 0.66 | 1 | MS.Trt_vs_MS.NoTrt_F |
| NXF1 | 0.01215 | -0.3286 | 0.558 | 0.67 | 1 | MS.Trt_vs_MS.NoTrt_F |

| | | | | | | |
|---|---|---|---|---|---|---|
| LSR | 0.01217 | -0.7366 | 0.165 | 0.26 | 1 | MS.Trt_vs_MS.NoTrt_F |
| PGD | 0.01218 | -0.5187 | 0.104 | 0.19 | 1 | MS.Trt_vs_MS.NoTrt_F |
| GRSF1 | 0.01219 | 0.2613 | 0.68 | 0.77 | 1 | MS.Trt_vs_MS.NoTrt_F |
| F8A1 | 0.0122 | -0.3676 | 0.079 | 0.16 | 1 | MS.Trt_vs_MS.NoTrt_F |
| SURF1 | 0.01226 | 0.37995 | 0.126 | 0.22 | 1 | MS.Trt_vs_MS.NoTrt_F |
| ANAPC1 | 0.01228 | -0.5448 | 0.5 | 0.66 | 1 | MS.Trt_vs_MS.NoTrt_F |
| NEK9 | 0.01228 | -0.4182 | 0.899 | 0.95 | 1 | MS.Trt_vs_MS.NoTrt_F |
| PIAS4 | 0.01228 | 0.36875 | 0.205 | 0.32 | 1 | MS.Trt_vs_MS.NoTrt_F |
| CD2AP | 0.01235 | -0.3276 | 0.162 | 0.27 | 1 | MS.Trt_vs_MS.NoTrt_F |
| DYNLT3 | 0.01235 | -0.6568 | 0.615 | 0.74 | 1 | MS.Trt_vs_MS.NoTrt_F |
| FAM173A | 0.01236 | 0.58388 | 0.094 | 0.18 | 1 | MS.Trt_vs_MS.NoTrt_F |
| C14orf93 | 0.01237 | 0.4691 | 0.284 | 0.39 | 1 | MS.Trt_vs_MS.NoTrt_F |
| KIF18A | 0.01238 | 0.63765 | 0.291 | 0.42 | 1 | MS.Trt_vs_MS.NoTrt_F |
| CCDC36 | 0.01244 | -0.293 | 0.32 | 0.45 | 1 | MS.Trt_vs_MS.NoTrt_F |
| ZC3HAV1L | 0.01249 | -0.4495 | 0.396 | 0.52 | 1 | MS.Trt_vs_MS.NoTrt_F |
| NOC3L | 0.0125 | 0.40437 | 0.079 | 0.16 | 1 | MS.Trt_vs_MS.NoTrt_F |
| DCAF16 | 0.01251 | -0.6368 | 0.748 | 0.82 | 1 | MS.Trt_vs_MS.NoTrt_F |
| IPO8 | 0.01255 | 0.27591 | 0.489 | 0.62 | 1 | MS.Trt_vs_MS.NoTrt_F |
| PCNP | 0.01256 | 0.5949 | 0.241 | 0.38 | 1 | MS.Trt_vs_MS.NoTrt_F |
| PJA1 | 0.01262 | -0.2693 | 0.083 | 0.16 | 1 | MS.Trt_vs_MS.NoTrt_F |
| NUAK2 | 0.01265 | -0.8732 | 0.331 | 0.45 | 1 | MS.Trt_vs_MS.NoTrt_F |
| RAD18 | 0.01265 | -0.3316 | 0.076 | 0.15 | 1 | MS.Trt_vs_MS.NoTrt_F |
| RAP1GAP2 | 0.01266 | 0.60146 | 0.27 | 0.4 | 1 | MS.Trt_vs_MS.NoTrt_F |
| CEP104 | 0.01271 | -0.6876 | 0.709 | 0.8 | 1 | MS.Trt_vs_MS.NoTrt_F |
| MOV10 | 0.01272 | -0.4989 | 0.424 | 0.53 | 1 | MS.Trt_vs_MS.NoTrt_F |
| TNIP3 | 0.01273 | 0.47047 | 0.478 | 0.64 | 1 | MS.Trt_vs_MS.NoTrt_F |
| RGPD2 | 0.01275 | 0.79661 | 0.155 | 0.27 | 1 | MS.Trt_vs_MS.NoTrt_F |
| ANAPC11 | 0.01277 | 1.08317 | 0.403 | 0.54 | 1 | MS.Trt_vs_MS.NoTrt_F |
| UBAC1 | 0.01279 | 0.50168 | 0.076 | 0.15 | 1 | MS.Trt_vs_MS.NoTrt_F |
| ZNF609 | 0.01282 | -1.0707 | 0.565 | 0.66 | 1 | MS.Trt_vs_MS.NoTrt_F |
| RBFA | 0.01283 | -0.6856 | 0.295 | 0.4 | 1 | MS.Trt_vs_MS.NoTrt_F |
| USP10 | 0.01284 | 0.33105 | 0.23 | 0.35 | 1 | MS.Trt_vs_MS.NoTrt_F |
| CHRNA6 | 0.0129 | 0.56587 | 0.374 | 0.28 | 1 | MS.Trt_vs_MS.NoTrt_F |
| NLRP12 | 0.01292 | 0.29412 | 0.809 | 0.76 | 1 | MS.Trt_vs_MS.NoTrt_F |
| GYG1 | 0.01292 | -0.6948 | 0.554 | 0.64 | 1 | MS.Trt_vs_MS.NoTrt_F |
| PBLD | 0.01293 | -0.4627 | 0.572 | 0.67 | 1 | MS.Trt_vs_MS.NoTrt_F |
| WISP2 | 0.01295 | 0.94452 | 0.511 | 0.61 | 1 | MS.Trt_vs_MS.NoTrt_F |
| SLC39A6 | 0.01295 | -0.4726 | 0.097 | 0.18 | 1 | MS.Trt_vs_MS.NoTrt_F |
| UBE4B | 0.01297 | 0.30515 | 0.547 | 0.68 | 1 | MS.Trt_vs_MS.NoTrt_F |
| VPS45 | 0.013 | -0.5035 | 0.277 | 0.38 | 1 | MS.Trt_vs_MS.NoTrt_F |

| | | | | | | |
|---|---|---|---|---|---|---|
| GPR174 | 0.01301 | 0.43573 | 0.155 | 0.26 | 1 | MS.Trt_vs_MS.NoTrt_F |
| TBCK | 0.01307 | -0.4674 | 0.615 | 0.76 | 1 | MS.Trt_vs_MS.NoTrt_F |
| MCC | 0.01309 | -0.5259 | 0.432 | 0.53 | 1 | MS.Trt_vs_MS.NoTrt_F |
| AKAP1 | 0.01312 | -0.3114 | 0.104 | 0.19 | 1 | MS.Trt_vs_MS.NoTrt_F |
| GON7 | 0.01313 | -1.0943 | 0.058 | 0.12 | 1 | MS.Trt_vs_MS.NoTrt_F |
| ENGASE | 0.01315 | 1.42322 | 0.079 | 0.16 | 1 | MS.Trt_vs_MS.NoTrt_F |
| TMEM110 | 0.01315 | -0.4727 | 0.079 | 0.16 | 1 | MS.Trt_vs_MS.NoTrt_F |
| MED30 | 0.01319 | 0.60446 | 0.137 | 0.24 | 1 | MS.Trt_vs_MS.NoTrt_F |
| HTATSF1 | 0.01319 | 0.62916 | 0.169 | 0.28 | 1 | MS.Trt_vs_MS.NoTrt_F |
| TOP1 | 0.01322 | 0.30692 | 0.173 | 0.28 | 1 | MS.Trt_vs_MS.NoTrt_F |
| ZNF343 | 0.01324 | -0.8195 | 0.432 | 0.54 | 1 | MS.Trt_vs_MS.NoTrt_F |
| PDCD2L | 0.01329 | -0.5412 | 0.065 | 0.13 | 1 | MS.Trt_vs_MS.NoTrt_F |
| HDGF | 0.01336 | -0.3643 | 0.094 | 0.17 | 1 | MS.Trt_vs_MS.NoTrt_F |
| SESN2 | 0.01344 | -1.0904 | 0.061 | 0.13 | 1 | MS.Trt_vs_MS.NoTrt_F |
| CCSAP | 0.01346 | 0.94795 | 0.144 | 0.24 | 1 | MS.Trt_vs_MS.NoTrt_F |
| DAD1 | 0.01353 | 0.51352 | 0.324 | 0.53 | 1 | MS.Trt_vs_MS.NoTrt_F |
| 11-Sep | 0.01354 | 0.37144 | 0.392 | 0.51 | 1 | MS.Trt_vs_MS.NoTrt_F |
| BRCC3 | 0.01355 | 0.54402 | 0.248 | 0.37 | 1 | MS.Trt_vs_MS.NoTrt_F |
| KBTBD6 | 0.01356 | -0.3602 | 0.353 | 0.46 | 1 | MS.Trt_vs_MS.NoTrt_F |
| SUMO1 | 0.01358 | 0.31312 | 0.496 | 0.61 | 1 | MS.Trt_vs_MS.NoTrt_F |
| UBN1 | 0.01363 | 0.34949 | 0.147 | 0.24 | 1 | MS.Trt_vs_MS.NoTrt_F |
| ZNF626 | 0.01364 | 0.31355 | 0.763 | 0.72 | 1 | MS.Trt_vs_MS.NoTrt_F |
| CLEC18A | 0.01367 | 0.67374 | 0.32 | 0.43 | 1 | MS.Trt_vs_MS.NoTrt_F |
| FLCN | 0.01373 | 0.29203 | 0.881 | 0.82 | 1 | MS.Trt_vs_MS.NoTrt_F |
| SMC6 | 0.01376 | -0.3898 | 0.442 | 0.56 | 1 | MS.Trt_vs_MS.NoTrt_F |
| MC1R | 0.01377 | -0.9061 | 0.072 | 0.14 | 1 | MS.Trt_vs_MS.NoTrt_F |
| SAMD3 | 0.01379 | 0.29598 | 0.284 | 0.41 | 1 | MS.Trt_vs_MS.NoTrt_F |
| FLAD1 | 0.01382 | -0.3509 | 0.554 | 0.68 | 1 | MS.Trt_vs_MS.NoTrt_F |
| TXNRD2 | 0.01384 | -0.5734 | 0.827 | 0.91 | 1 | MS.Trt_vs_MS.NoTrt_F |
| KIF16B | 0.01385 | -1.2826 | 0.068 | 0.14 | 1 | MS.Trt_vs_MS.NoTrt_F |
| DAPP1 | 0.01386 | 0.39397 | 0.245 | 0.37 | 1 | MS.Trt_vs_MS.NoTrt_F |
| TNFRSF18 | 0.01387 | 0.7275 | 0.198 | 0.33 | 1 | MS.Trt_vs_MS.NoTrt_F |
| NME1 | 0.01388 | 0.32543 | 0.734 | 0.56 | 1 | MS.Trt_vs_MS.NoTrt_F |
| NR4A3 | 0.01394 | -0.691 | 0.133 | 0.22 | 1 | MS.Trt_vs_MS.NoTrt_F |
| PEX6 | 0.01395 | -0.4014 | 0.061 | 0.13 | 1 | MS.Trt_vs_MS.NoTrt_F |
| ABO | 0.01398 | -0.4101 | 0.331 | 0.45 | 1 | MS.Trt_vs_MS.NoTrt_F |
| ENOPH1 | 0.01399 | 0.32475 | 0.158 | 0.26 | 1 | MS.Trt_vs_MS.NoTrt_F |
| SCO2 | 0.01403 | 0.49523 | 0.076 | 0.15 | 1 | MS.Trt_vs_MS.NoTrt_F |
| CENPJ | 0.01405 | -0.9549 | 0.061 | 0.13 | 1 | MS.Trt_vs_MS.NoTrt_F |
| TRAPPC3 | 0.01405 | 0.32754 | 0.392 | 0.5 | 1 | MS.Trt_vs_MS.NoTrt_F |

| | | | | | | |
|---|---|---|---|---|---|---|
| METTL22 | 0.01407 | -0.3619 | 0.615 | 0.7 | 1 | MS.Trt_vs_MS.NoTrt_F |
| PCYT2 | 0.0141 | -0.2537 | 0.065 | 0.13 | 1 | MS.Trt_vs_MS.NoTrt_F |
| SBNO2 | 0.01415 | -0.3877 | 0.194 | 0.3 | 1 | MS.Trt_vs_MS.NoTrt_F |
| LDOC1L | 0.01415 | -0.6768 | 0.068 | 0.14 | 1 | MS.Trt_vs_MS.NoTrt_F |
| TAF3 | 0.01416 | 0.41326 | 0.165 | 0.27 | 1 | MS.Trt_vs_MS.NoTrt_F |
| GRAMD3 | 0.01424 | -0.4668 | 0.676 | 0.86 | 1 | MS.Trt_vs_MS.NoTrt_F |
| PAIP1 | 0.01424 | 0.38634 | 0.162 | 0.27 | 1 | MS.Trt_vs_MS.NoTrt_F |
| MPP6 | 0.01426 | -0.2511 | 0.255 | 0.38 | 1 | MS.Trt_vs_MS.NoTrt_F |
| HLA-C | 0.01426 | -0.4064 | 0.853 | 0.91 | 1 | MS.Trt_vs_MS.NoTrt_F |
| RTEL1 | 0.01432 | -0.964 | 0.115 | 0.2 | 1 | MS.Trt_vs_MS.NoTrt_F |
| ABCF1 | 0.01434 | -0.3444 | 0.817 | 0.89 | 1 | MS.Trt_vs_MS.NoTrt_F |
| PSD | 0.01437 | -0.2987 | 0.108 | 0.19 | 1 | MS.Trt_vs_MS.NoTrt_F |
| TRPM8 | 0.01438 | 0.26103 | 0.777 | 0.61 | 1 | MS.Trt_vs_MS.NoTrt_F |
| FAF2 | 0.01439 | 0.79136 | 0.122 | 0.22 | 1 | MS.Trt_vs_MS.NoTrt_F |
| REXO2 | 0.01439 | 0.53036 | 0.209 | 0.33 | 1 | MS.Trt_vs_MS.NoTrt_F |
| SNRPD2 | 0.01442 | 0.34144 | 0.529 | 0.71 | 1 | MS.Trt_vs_MS.NoTrt_F |
| TEX261 | 0.01442 | -0.3263 | 0.09 | 0.17 | 1 | MS.Trt_vs_MS.NoTrt_F |
| TOLLIP | 0.01442 | 0.53332 | 0.133 | 0.23 | 1 | MS.Trt_vs_MS.NoTrt_F |
| HDAC2 | 0.01445 | -0.2639 | 0.655 | 0.81 | 1 | MS.Trt_vs_MS.NoTrt_F |
| C1QBP | 0.01447 | 0.52434 | 0.273 | 0.4 | 1 | MS.Trt_vs_MS.NoTrt_F |
| GOLGA8N | 0.01448 | 1.05662 | 0.076 | 0.15 | 1 | MS.Trt_vs_MS.NoTrt_F |
| SGIP1 | 0.01449 | -0.3206 | 0.245 | 0.34 | 1 | MS.Trt_vs_MS.NoTrt_F |
| KLHL5 | 0.0145 | 0.59241 | 0.428 | 0.54 | 1 | MS.Trt_vs_MS.NoTrt_F |
| ZC4H2 | 0.01451 | -0.6733 | 0.094 | 0.17 | 1 | MS.Trt_vs_MS.NoTrt_F |
| SENP1 | 0.01452 | 0.82057 | 0.191 | 0.3 | 1 | MS.Trt_vs_MS.NoTrt_F |
| ZNF485 | 0.01452 | -0.3592 | 0.183 | 0.28 | 1 | MS.Trt_vs_MS.NoTrt_F |
| C2CD4A | 0.01454 | -0.4219 | 0.288 | 0.39 | 1 | MS.Trt_vs_MS.NoTrt_F |
| NUDT18 | 0.01455 | -1.9548 | 0.05 | 0.11 | 1 | MS.Trt_vs_MS.NoTrt_F |
| CCDC71 | 0.01459 | 0.5951 | 0.079 | 0.16 | 1 | MS.Trt_vs_MS.NoTrt_F |
| NSG1 | 0.01465 | -0.3088 | 0.615 | 0.74 | 1 | MS.Trt_vs_MS.NoTrt_F |
| HIBADH | 0.01467 | 0.62467 | 0.097 | 0.18 | 1 | MS.Trt_vs_MS.NoTrt_F |
| C2orf47 | 0.01473 | -0.7241 | 0.05 | 0.11 | 1 | MS.Trt_vs_MS.NoTrt_F |
| ETFDH | 0.01475 | 0.59963 | 0.619 | 0.74 | 1 | MS.Trt_vs_MS.NoTrt_F |
| SMC2 | 0.01477 | 0.57382 | 0.083 | 0.16 | 1 | MS.Trt_vs_MS.NoTrt_F |
| ZNRF3 | 0.0148 | -0.4017 | 0.23 | 0.34 | 1 | MS.Trt_vs_MS.NoTrt_F |
| BCL2 | 0.01482 | 0.4363 | 0.263 | 0.38 | 1 | MS.Trt_vs_MS.NoTrt_F |
| ATAD5 | 0.01485 | -0.5819 | 0.5 | 0.64 | 1 | MS.Trt_vs_MS.NoTrt_F |
| KIAA0391 | 0.01487 | -0.4449 | 0.691 | 0.8 | 1 | MS.Trt_vs_MS.NoTrt_F |
| BCL10 | 0.01487 | 0.41583 | 0.079 | 0.16 | 1 | MS.Trt_vs_MS.NoTrt_F |
| ABLIM3 | 0.01491 | -0.6337 | 0.065 | 0.13 | 1 | MS.Trt_vs_MS.NoTrt_F |

| | | | | | | |
|---|---|---|---|---|---|---|
| UBQLN4 | 0.01497 | 0.46718 | 0.065 | 0.13 | 1 | MS.Trt_vs_MS.NoTrt_F |
| BLK | 0.015 | 0.96836 | 0.065 | 0.13 | 1 | MS.Trt_vs_MS.NoTrt_F |
| N4BP2 | 0.01501 | -0.3782 | 0.313 | 0.44 | 1 | MS.Trt_vs_MS.NoTrt_F |
| DHRS13 | 0.01501 | -0.3803 | 0.065 | 0.13 | 1 | MS.Trt_vs_MS.NoTrt_F |
| PSKH1 | 0.01504 | 0.62928 | 0.097 | 0.18 | 1 | MS.Trt_vs_MS.NoTrt_F |
| SIGLEC10 | 0.01506 | -0.3466 | 0.579 | 0.73 | 1 | MS.Trt_vs_MS.NoTrt_F |
| ITM2A | 0.01506 | 0.47323 | 0.234 | 0.37 | 1 | MS.Trt_vs_MS.NoTrt_F |
| PLAGL2 | 0.01507 | -1.0114 | 0.068 | 0.14 | 1 | MS.Trt_vs_MS.NoTrt_F |
| LPCAT1 | 0.01508 | 0.30158 | 0.076 | 0.15 | 1 | MS.Trt_vs_MS.NoTrt_F |
| VPS13A | 0.01509 | 0.30928 | 0.27 | 0.38 | 1 | MS.Trt_vs_MS.NoTrt_F |
| PPARG | 0.0151 | -0.3361 | 0.424 | 0.52 | 1 | MS.Trt_vs_MS.NoTrt_F |
| PIGO | 0.01513 | -0.5108 | 0.381 | 0.48 | 1 | MS.Trt_vs_MS.NoTrt_F |
| PLEKHO1 | 0.01514 | 0.43648 | 0.076 | 0.15 | 1 | MS.Trt_vs_MS.NoTrt_F |
| ZNF429 | 0.01519 | 0.77918 | 0.896 | 0.91 | 1 | MS.Trt_vs_MS.NoTrt_F |
| MIB1 | 0.01521 | -0.8877 | 0.119 | 0.2 | 1 | MS.Trt_vs_MS.NoTrt_F |
| PUDP | 0.01522 | -0.4699 | 0.108 | 0.19 | 1 | MS.Trt_vs_MS.NoTrt_F |
| IFIT1 | 0.01524 | -0.7338 | 0.212 | 0.32 | 1 | MS.Trt_vs_MS.NoTrt_F |
| GPAM | 0.01525 | -1.2498 | 0.058 | 0.12 | 1 | MS.Trt_vs_MS.NoTrt_F |
| ADNP2 | 0.01525 | 0.54493 | 0.072 | 0.14 | 1 | MS.Trt_vs_MS.NoTrt_F |
| KLHDC7B | 0.01529 | -0.2991 | 0.061 | 0.13 | 1 | MS.Trt_vs_MS.NoTrt_F |
| LMAN2L | 0.01529 | -0.5441 | 0.05 | 0.11 | 1 | MS.Trt_vs_MS.NoTrt_F |
| PTRF | 0.01532 | 0.62805 | 0.353 | 0.46 | 1 | MS.Trt_vs_MS.NoTrt_F |
| TBC1D3L | 0.01536 | 1.58578 | 0.094 | 0.17 | 1 | MS.Trt_vs_MS.NoTrt_F |
| TMX4 | 0.01537 | -0.4008 | 0.565 | 0.62 | 1 | MS.Trt_vs_MS.NoTrt_F |
| ICK | 0.01538 | -1.442 | 0.201 | 0.29 | 1 | MS.Trt_vs_MS.NoTrt_F |
| ZNF581 | 0.01544 | -0.3013 | 0.09 | 0.17 | 1 | MS.Trt_vs_MS.NoTrt_F |
| RECQL5 | 0.01547 | -0.9357 | 0.191 | 0.29 | 1 | MS.Trt_vs_MS.NoTrt_F |
| OSGEPL1 | 0.0155 | 0.46988 | 0.252 | 0.39 | 1 | MS.Trt_vs_MS.NoTrt_F |
| ARHGAP5 | 0.0155 | -0.5472 | 0.414 | 0.53 | 1 | MS.Trt_vs_MS.NoTrt_F |
| BHLHE40 | 0.01553 | 0.48201 | 0.112 | 0.2 | 1 | MS.Trt_vs_MS.NoTrt_F |
| PRDX4 | 0.01559 | -0.3345 | 0.09 | 0.17 | 1 | MS.Trt_vs_MS.NoTrt_F |
| PLCG2 | 0.01565 | 1.13485 | 0.104 | 0.19 | 1 | MS.Trt_vs_MS.NoTrt_F |
| IGSF9B | 0.01571 | -1.1578 | 0.054 | 0.12 | 1 | MS.Trt_vs_MS.NoTrt_F |
| CCR2 | 0.01572 | -0.734 | 0.115 | 0.2 | 1 | MS.Trt_vs_MS.NoTrt_F |
| SLC17A5 | 0.01577 | -0.3461 | 0.424 | 0.54 | 1 | MS.Trt_vs_MS.NoTrt_F |
| AGO4 | 0.01584 | -0.9173 | 0.309 | 0.4 | 1 | MS.Trt_vs_MS.NoTrt_F |
| BRD3 | 0.01585 | 0.49648 | 0.18 | 0.29 | 1 | MS.Trt_vs_MS.NoTrt_F |
| CC2D2B | 0.01585 | 1.02766 | 0.119 | 0.21 | 1 | MS.Trt_vs_MS.NoTrt_F |
| VPS9D1 | 0.01589 | -0.3745 | 0.223 | 0.33 | 1 | MS.Trt_vs_MS.NoTrt_F |
| CCDC107 | 0.01596 | 0.32672 | 0.198 | 0.32 | 1 | MS.Trt_vs_MS.NoTrt_F |

| | | | | | | |
|---|---|---|---|---|---|---|
| SLC37A4 | 0.01598 | -0.4836 | 0.05 | 0.11 | 1 | MS.Trt_vs_MS.NoTrt_F |
| ZNF701 | 0.01602 | 0.90971 | 0.442 | 0.57 | 1 | MS.Trt_vs_MS.NoTrt_F |
| LUZP1 | 0.01606 | -0.5699 | 0.55 | 0.63 | 1 | MS.Trt_vs_MS.NoTrt_F |
| SEMA3G | 0.01607 | -0.6061 | 0.047 | 0.11 | 1 | MS.Trt_vs_MS.NoTrt_F |
| MRPL39 | 0.0161 | 0.32296 | 0.137 | 0.23 | 1 | MS.Trt_vs_MS.NoTrt_F |
| ECHS1 | 0.01611 | 0.57341 | 0.155 | 0.25 | 1 | MS.Trt_vs_MS.NoTrt_F |
| PRPF4 | 0.01612 | 0.37008 | 0.302 | 0.42 | 1 | MS.Trt_vs_MS.NoTrt_F |
| BRF2 | 0.01614 | -0.4266 | 0.147 | 0.24 | 1 | MS.Trt_vs_MS.NoTrt_F |
| POLM | 0.01617 | 0.36938 | 0.518 | 0.64 | 1 | MS.Trt_vs_MS.NoTrt_F |
| ANKS1A | 0.01625 | 0.76997 | 0.054 | 0.12 | 1 | MS.Trt_vs_MS.NoTrt_F |
| ARSG | 0.01628 | -0.3006 | 0.932 | 0.97 | 1 | MS.Trt_vs_MS.NoTrt_F |
| GGNBP2 | 0.01634 | 0.31691 | 0.676 | 0.83 | 1 | MS.Trt_vs_MS.NoTrt_F |
| RNF168 | 0.01639 | 0.6018 | 0.568 | 0.74 | 1 | MS.Trt_vs_MS.NoTrt_F |
| AQP11 | 0.0164 | -0.2917 | 0.05 | 0.11 | 1 | MS.Trt_vs_MS.NoTrt_F |
| AKT1 | 0.01642 | 0.41114 | 0.209 | 0.31 | 1 | MS.Trt_vs_MS.NoTrt_F |
| COMMD6 | 0.01643 | 0.46327 | 0.363 | 0.55 | 1 | MS.Trt_vs_MS.NoTrt_F |
| GOSR1 | 0.01646 | 0.52956 | 0.968 | 0.97 | 1 | MS.Trt_vs_MS.NoTrt_F |
| ZNF282 | 0.01649 | 0.82391 | 0.065 | 0.13 | 1 | MS.Trt_vs_MS.NoTrt_F |
| ARPC5 | 0.01649 | 0.26309 | 0.892 | 0.96 | 1 | MS.Trt_vs_MS.NoTrt_F |
| DPH2 | 0.01651 | 1.51377 | 0.083 | 0.16 | 1 | MS.Trt_vs_MS.NoTrt_F |
| FBXO42 | 0.01655 | -0.2821 | 0.486 | 0.58 | 1 | MS.Trt_vs_MS.NoTrt_F |
| CLDN5 | 0.01655 | -0.8988 | 0.047 | 0.11 | 1 | MS.Trt_vs_MS.NoTrt_F |
| SLC26A1 | 0.01658 | -0.53 | 0.047 | 0.11 | 1 | MS.Trt_vs_MS.NoTrt_F |
| AKT1S1 | 0.01659 | -0.5115 | 0.18 | 0.28 | 1 | MS.Trt_vs_MS.NoTrt_F |
| ZBTB16 | 0.01663 | 0.62816 | 0.327 | 0.44 | 1 | MS.Trt_vs_MS.NoTrt_F |
| ERP27 | 0.01663 | 0.56615 | 0.219 | 0.34 | 1 | MS.Trt_vs_MS.NoTrt_F |
| POC5 | 0.01669 | -0.2933 | 0.112 | 0.2 | 1 | MS.Trt_vs_MS.NoTrt_F |
| GPR160 | 0.01671 | 0.33777 | 0.054 | 0.12 | 1 | MS.Trt_vs_MS.NoTrt_F |
| ACTR10 | 0.01673 | -0.3551 | 0.741 | 0.82 | 1 | MS.Trt_vs_MS.NoTrt_F |
| CEP72 | 0.0168 | 0.64271 | 0.065 | 0.13 | 1 | MS.Trt_vs_MS.NoTrt_F |
| BSCL2 | 0.01682 | -0.641 | 0.201 | 0.3 | 1 | MS.Trt_vs_MS.NoTrt_F |
| FBXO38 | 0.01682 | -0.5058 | 0.507 | 0.6 | 1 | MS.Trt_vs_MS.NoTrt_F |
| CAMK2D | 0.01683 | 1.06885 | 0.227 | 0.35 | 1 | MS.Trt_vs_MS.NoTrt_F |
| PLPP5 | 0.01685 | 0.41188 | 0.878 | 0.93 | 1 | MS.Trt_vs_MS.NoTrt_F |
| SNAPC1 | 0.01689 | -0.2826 | 0.683 | 0.82 | 1 | MS.Trt_vs_MS.NoTrt_F |
| DDX41 | 0.0169 | 0.33062 | 0.129 | 0.22 | 1 | MS.Trt_vs_MS.NoTrt_F |
| TAPBPL | 0.01693 | 0.25916 | 0.216 | 0.33 | 1 | MS.Trt_vs_MS.NoTrt_F |
| LYAR | 0.01693 | 0.93488 | 0.112 | 0.2 | 1 | MS.Trt_vs_MS.NoTrt_F |
| PCNA | 0.01696 | 0.44513 | 0.097 | 0.18 | 1 | MS.Trt_vs_MS.NoTrt_F |
| DDX43 | 0.01698 | -0.7696 | 0.147 | 0.24 | 1 | MS.Trt_vs_MS.NoTrt_F |

| | | | | | | |
|---|---|---|---|---|---|---|
| FDXACB1 | 0.017 | -0.3816 | 0.388 | 0.51 | 1 | MS.Trt_vs_MS.NoTrt_F |
| ITGA11 | 0.01708 | -0.278 | 0.561 | 0.7 | 1 | MS.Trt_vs_MS.NoTrt_F |
| APOL3 | 0.01709 | -0.3397 | 0.176 | 0.27 | 1 | MS.Trt_vs_MS.NoTrt_F |
| APH1B | 0.01718 | 0.31451 | 0.094 | 0.17 | 1 | MS.Trt_vs_MS.NoTrt_F |
| CNNM4 | 0.0172 | 0.83636 | 0.05 | 0.11 | 1 | MS.Trt_vs_MS.NoTrt_F |
| SMARCA5 | 0.01721 | -0.3121 | 0.363 | 0.49 | 1 | MS.Trt_vs_MS.NoTrt_F |
| APEX1 | 0.01721 | 0.57736 | 0.155 | 0.26 | 1 | MS.Trt_vs_MS.NoTrt_F |
| ZBTB7A | 0.01722 | 1.02949 | 0.126 | 0.22 | 1 | MS.Trt_vs_MS.NoTrt_F |
| C2orf49 | 0.01726 | -0.3113 | 0.399 | 0.52 | 1 | MS.Trt_vs_MS.NoTrt_F |
| CHSY1 | 0.01729 | -0.4264 | 0.255 | 0.36 | 1 | MS.Trt_vs_MS.NoTrt_F |
| DDX18 | 0.01732 | 0.30988 | 0.694 | 0.8 | 1 | MS.Trt_vs_MS.NoTrt_F |
| PTK2 | 0.01738 | 0.37576 | 0.914 | 0.96 | 1 | MS.Trt_vs_MS.NoTrt_F |
| IKBKAP | 0.0174 | -0.2918 | 0.259 | 0.35 | 1 | MS.Trt_vs_MS.NoTrt_F |
| ETHE1 | 0.0174 | -0.3405 | 0.09 | 0.17 | 1 | MS.Trt_vs_MS.NoTrt_F |
| MRPL17 | 0.01742 | 0.74801 | 0.09 | 0.17 | 1 | MS.Trt_vs_MS.NoTrt_F |
| ZDHHC7 | 0.01743 | -0.3679 | 0.212 | 0.31 | 1 | MS.Trt_vs_MS.NoTrt_F |
| TSTA3 | 0.01745 | 0.3789 | 0.162 | 0.27 | 1 | MS.Trt_vs_MS.NoTrt_F |
| QSOX2 | 0.01747 | 0.27465 | 0.144 | 0.24 | 1 | MS.Trt_vs_MS.NoTrt_F |
| MARS | 0.01753 | -0.4945 | 0.414 | 0.5 | 1 | MS.Trt_vs_MS.NoTrt_F |
| TMEM30A | 0.01762 | 0.41476 | 0.219 | 0.33 | 1 | MS.Trt_vs_MS.NoTrt_F |
| DHX16 | 0.01762 | 0.36871 | 0.147 | 0.24 | 1 | MS.Trt_vs_MS.NoTrt_F |
| CES2 | 0.01769 | 0.9083 | 0.309 | 0.43 | 1 | MS.Trt_vs_MS.NoTrt_F |
| BIVM | 0.01775 | 0.56298 | 0.309 | 0.44 | 1 | MS.Trt_vs_MS.NoTrt_F |
| TARBP1 | 0.01775 | -0.9287 | 0.371 | 0.49 | 1 | MS.Trt_vs_MS.NoTrt_F |
| ABHD13 | 0.01779 | -0.3212 | 0.112 | 0.19 | 1 | MS.Trt_vs_MS.NoTrt_F |
| OGFR | 0.01789 | -0.4494 | 0.201 | 0.3 | 1 | MS.Trt_vs_MS.NoTrt_F |
| STXBP4 | 0.01794 | 0.43395 | 0.45 | 0.59 | 1 | MS.Trt_vs_MS.NoTrt_F |
| ST8SIA6 | 0.01796 | 0.68545 | 0.367 | 0.49 | 1 | MS.Trt_vs_MS.NoTrt_F |
| IDH1 | 0.01797 | -0.9368 | 0.216 | 0.32 | 1 | MS.Trt_vs_MS.NoTrt_F |
| DAP3 | 0.01802 | -0.4699 | 0.903 | 0.93 | 1 | MS.Trt_vs_MS.NoTrt_F |
| BACE1 | 0.01806 | 0.52473 | 0.536 | 0.7 | 1 | MS.Trt_vs_MS.NoTrt_F |
| TMEM223 | 0.0181 | 0.30787 | 0.176 | 0.28 | 1 | MS.Trt_vs_MS.NoTrt_F |
| ADPRHL1 | 0.01818 | -0.3569 | 0.277 | 0.38 | 1 | MS.Trt_vs_MS.NoTrt_F |
| PTP4A3 | 0.01819 | 0.31656 | 0.09 | 0.17 | 1 | MS.Trt_vs_MS.NoTrt_F |
| ARL17A | 0.01825 | 0.2662 | 0.583 | 0.77 | 1 | MS.Trt_vs_MS.NoTrt_F |
| BMF | 0.01832 | 0.43729 | 0.101 | 0.18 | 1 | MS.Trt_vs_MS.NoTrt_F |
| ZMYM6 | 0.01836 | 0.63708 | 0.824 | 0.79 | 1 | MS.Trt_vs_MS.NoTrt_F |
| SULT2B1 | 0.01836 | -0.5968 | 0.464 | 0.6 | 1 | MS.Trt_vs_MS.NoTrt_F |
| VWA5A | 0.01839 | -0.311 | 0.086 | 0.16 | 1 | MS.Trt_vs_MS.NoTrt_F |
| ULK4 | 0.01841 | -0.6216 | 0.291 | 0.4 | 1 | MS.Trt_vs_MS.NoTrt_F |

| | | | | | | |
|---|---|---|---|---|---|---|
| COL6A3 | 0.01842 | 0.36987 | 0.396 | 0.54 | 1 | MS.Trt_vs_MS.NoTrt_F |
| FBXO24 | 0.01843 | -0.3558 | 0.374 | 0.51 | 1 | MS.Trt_vs_MS.NoTrt_F |
| CASP7 | 0.01843 | -1.737 | 0.169 | 0.26 | 1 | MS.Trt_vs_MS.NoTrt_F |
| MEIS3 | 0.01847 | 1.28909 | 0.104 | 0.18 | 1 | MS.Trt_vs_MS.NoTrt_F |
| ARL1 | 0.01848 | 0.29848 | 0.327 | 0.45 | 1 | MS.Trt_vs_MS.NoTrt_F |
| USP21 | 0.01849 | -0.4656 | 0.137 | 0.23 | 1 | MS.Trt_vs_MS.NoTrt_F |
| PRKACA | 0.01849 | -0.3366 | 0.216 | 0.33 | 1 | MS.Trt_vs_MS.NoTrt_F |
| C19orf12 | 0.01858 | -0.5305 | 0.363 | 0.48 | 1 | MS.Trt_vs_MS.NoTrt_F |
| VASP | 0.01862 | -0.4281 | 0.572 | 0.68 | 1 | MS.Trt_vs_MS.NoTrt_F |
| PKIB | 0.01862 | -0.3611 | 0.701 | 0.79 | 1 | MS.Trt_vs_MS.NoTrt_F |
| EMILIN2 | 0.01864 | 0.99908 | 0.417 | 0.53 | 1 | MS.Trt_vs_MS.NoTrt_F |
| PGGT1B | 0.01869 | 0.87853 | 0.183 | 0.29 | 1 | MS.Trt_vs_MS.NoTrt_F |
| MED14 | 0.01869 | 0.25391 | 0.162 | 0.27 | 1 | MS.Trt_vs_MS.NoTrt_F |
| ELMOD2 | 0.01871 | 0.78385 | 0.097 | 0.18 | 1 | MS.Trt_vs_MS.NoTrt_F |
| RANBP3 | 0.01871 | -0.4258 | 0.755 | 0.86 | 1 | MS.Trt_vs_MS.NoTrt_F |
| SLC25A46 | 0.01871 | 0.64978 | 0.14 | 0.23 | 1 | MS.Trt_vs_MS.NoTrt_F |
| MAN2A1 | 0.01874 | 0.29271 | 0.245 | 0.37 | 1 | MS.Trt_vs_MS.NoTrt_F |
| ACTG2 | 0.01876 | -0.3728 | 0.259 | 0.35 | 1 | MS.Trt_vs_MS.NoTrt_F |
| ZFAT | 0.01886 | -1.0352 | 0.331 | 0.46 | 1 | MS.Trt_vs_MS.NoTrt_F |
| SLC10A7 | 0.01891 | -0.7683 | 0.097 | 0.17 | 1 | MS.Trt_vs_MS.NoTrt_F |
| ARMCX4 | 0.01891 | 0.49154 | 0.187 | 0.29 | 1 | MS.Trt_vs_MS.NoTrt_F |
| SSR4 | 0.01894 | 0.7491 | 0.317 | 0.45 | 1 | MS.Trt_vs_MS.NoTrt_F |
| CCR4 | 0.01905 | -0.2577 | 0.91 | 0.94 | 1 | MS.Trt_vs_MS.NoTrt_F |
| KAT8 | 0.01907 | -0.5427 | 0.101 | 0.18 | 1 | MS.Trt_vs_MS.NoTrt_F |
| GTF2A1 | 0.01909 | 0.31249 | 0.09 | 0.17 | 1 | MS.Trt_vs_MS.NoTrt_F |
| SLC4A2 | 0.0191 | -0.5648 | 0.083 | 0.16 | 1 | MS.Trt_vs_MS.NoTrt_F |
| GPN1 | 0.01913 | 0.39794 | 0.165 | 0.26 | 1 | MS.Trt_vs_MS.NoTrt_F |
| TRRAP | 0.01914 | 0.40601 | 0.14 | 0.24 | 1 | MS.Trt_vs_MS.NoTrt_F |
| CPT1B | 0.01923 | 0.27416 | 0.194 | 0.29 | 1 | MS.Trt_vs_MS.NoTrt_F |
| LDOC1 | 0.01925 | 0.48636 | 0.09 | 0.17 | 1 | MS.Trt_vs_MS.NoTrt_F |
| CALU | 0.01926 | 0.38143 | 0.353 | 0.49 | 1 | MS.Trt_vs_MS.NoTrt_F |
| EFCAB7 | 0.01935 | -0.4853 | 0.076 | 0.14 | 1 | MS.Trt_vs_MS.NoTrt_F |
| SEC11A | 0.0194 | -0.3262 | 0.766 | 0.88 | 1 | MS.Trt_vs_MS.NoTrt_F |
| RPL39 | 0.01941 | 0.26655 | 0.597 | 0.89 | 1 | MS.Trt_vs_MS.NoTrt_F |
| EHBP1L1 | 0.01942 | 0.87307 | 0.201 | 0.31 | 1 | MS.Trt_vs_MS.NoTrt_F |
| ZC3H12B | 0.01944 | 0.65571 | 0.504 | 0.62 | 1 | MS.Trt_vs_MS.NoTrt_F |
| NCOA1 | 0.01949 | 0.63261 | 0.187 | 0.28 | 1 | MS.Trt_vs_MS.NoTrt_F |
| RP11-47I22. | 0.01952 | -0.2904 | 0.478 | 0.61 | 1 | MS.Trt_vs_MS.NoTrt_F |
| C3orf58 | 0.01954 | 0.45088 | 0.133 | 0.22 | 1 | MS.Trt_vs_MS.NoTrt_F |
| MRPS33 | 0.01955 | -0.333 | 0.924 | 0.95 | 1 | MS.Trt_vs_MS.NoTrt_F |

| | | | | | | |
|---|---|---|---|---|---|---|
| RBBP9 | 0.01962 | -1.1163 | 0.378 | 0.5 | 1 | MS.Trt_vs_MS.NoTrt_F |
| TRIM58 | 0.01965 | 0.35617 | 0.662 | 0.61 | 1 | MS.Trt_vs_MS.NoTrt_F |
| IFFO2 | 0.01966 | -0.6151 | 0.165 | 0.25 | 1 | MS.Trt_vs_MS.NoTrt_F |
| ATP13A3 | 0.01974 | 0.49791 | 0.129 | 0.22 | 1 | MS.Trt_vs_MS.NoTrt_F |
| KATNA1 | 0.01981 | 0.8849 | 0.122 | 0.21 | 1 | MS.Trt_vs_MS.NoTrt_F |
| RBBP8 | 0.01992 | -0.2533 | 0.101 | 0.18 | 1 | MS.Trt_vs_MS.NoTrt_F |
| NVL | 0.01992 | 0.53484 | 0.741 | 0.88 | 1 | MS.Trt_vs_MS.NoTrt_F |
| SETD4 | 0.01997 | 0.6437 | 0.511 | 0.61 | 1 | MS.Trt_vs_MS.NoTrt_F |
| RP11-613M1 | 0.02003 | -0.3072 | 0.162 | 0.24 | 1 | MS.Trt_vs_MS.NoTrt_F |
| ALG12 | 0.02011 | -0.2548 | 0.155 | 0.25 | 1 | MS.Trt_vs_MS.NoTrt_F |
| PCDH9 | 0.02012 | -0.3142 | 0.669 | 0.75 | 1 | MS.Trt_vs_MS.NoTrt_F |
| MOB1A | 0.02013 | 0.32951 | 0.82 | 0.91 | 1 | MS.Trt_vs_MS.NoTrt_F |
| ZNF232 | 0.02013 | -0.5792 | 0.065 | 0.13 | 1 | MS.Trt_vs_MS.NoTrt_F |
| LTB4R | 0.02014 | -0.3008 | 0.349 | 0.45 | 1 | MS.Trt_vs_MS.NoTrt_F |
| ELL | 0.02018 | 0.79014 | 0.162 | 0.27 | 1 | MS.Trt_vs_MS.NoTrt_F |
| IRAK4 | 0.02021 | -0.3495 | 0.579 | 0.68 | 1 | MS.Trt_vs_MS.NoTrt_F |
| NCLN | 0.02026 | 0.27645 | 0.151 | 0.24 | 1 | MS.Trt_vs_MS.NoTrt_F |
| AEN | 0.02028 | 0.39591 | 0.137 | 0.23 | 1 | MS.Trt_vs_MS.NoTrt_F |
| SLC15A3 | 0.02028 | -0.3077 | 0.076 | 0.14 | 1 | MS.Trt_vs_MS.NoTrt_F |
| UBE2N | 0.02035 | 0.46578 | 0.536 | 0.66 | 1 | MS.Trt_vs_MS.NoTrt_F |
| GTF2H5 | 0.0204 | 0.30796 | 0.853 | 0.78 | 1 | MS.Trt_vs_MS.NoTrt_F |
| IGFLR1 | 0.02052 | 0.52898 | 0.543 | 0.71 | 1 | MS.Trt_vs_MS.NoTrt_F |
| TMEM2 | 0.02055 | 0.27167 | 0.608 | 0.71 | 1 | MS.Trt_vs_MS.NoTrt_F |
| CCNYL1 | 0.02059 | 0.41789 | 0.353 | 0.5 | 1 | MS.Trt_vs_MS.NoTrt_F |
| FBXL15 | 0.0206 | 0.51393 | 0.144 | 0.24 | 1 | MS.Trt_vs_MS.NoTrt_F |
| CDR2 | 0.02068 | 0.31451 | 0.147 | 0.24 | 1 | MS.Trt_vs_MS.NoTrt_F |
| ZC3H7B | 0.02068 | -0.6064 | 0.766 | 0.83 | 1 | MS.Trt_vs_MS.NoTrt_F |
| FXR1 | 0.0207 | 0.38957 | 0.205 | 0.31 | 1 | MS.Trt_vs_MS.NoTrt_F |
| WHSC1 | 0.02074 | 0.31766 | 0.683 | 0.74 | 1 | MS.Trt_vs_MS.NoTrt_F |
| OTUB1 | 0.02077 | 0.37028 | 0.881 | 0.98 | 1 | MS.Trt_vs_MS.NoTrt_F |
| IDI1 | 0.02079 | 0.62342 | 0.212 | 0.33 | 1 | MS.Trt_vs_MS.NoTrt_F |
| CASZ1 | 0.02085 | -0.8162 | 0.255 | 0.36 | 1 | MS.Trt_vs_MS.NoTrt_F |
| NISCH | 0.021 | 0.25833 | 0.399 | 0.49 | 1 | MS.Trt_vs_MS.NoTrt_F |
| ZNF197 | 0.02102 | -0.3028 | 0.471 | 0.59 | 1 | MS.Trt_vs_MS.NoTrt_F |
| C9orf72 | 0.02103 | 0.72179 | 0.169 | 0.27 | 1 | MS.Trt_vs_MS.NoTrt_F |
| ATXN1L | 0.02103 | -0.5812 | 0.097 | 0.17 | 1 | MS.Trt_vs_MS.NoTrt_F |
| DYNLRB1 | 0.02108 | 0.43263 | 0.198 | 0.31 | 1 | MS.Trt_vs_MS.NoTrt_F |
| DCTPP1 | 0.0211 | 0.40369 | 0.112 | 0.19 | 1 | MS.Trt_vs_MS.NoTrt_F |
| TMEM8A | 0.02117 | 0.29931 | 0.076 | 0.14 | 1 | MS.Trt_vs_MS.NoTrt_F |
| CEP55 | 0.02122 | -0.6461 | 0.14 | 0.23 | 1 | MS.Trt_vs_MS.NoTrt_F |

| | | | | | | |
|---|---|---|---|---|---|---|
| CRACR2A | 0.02122 | -0.2821 | 0.629 | 0.71 | 1 | MS.Trt_vs_MS.NoTrt_F |
| SLC2A8 | 0.02123 | -1.368 | 0.266 | 0.38 | 1 | MS.Trt_vs_MS.NoTrt_F |
| HARS2 | 0.02124 | 0.82185 | 0.129 | 0.22 | 1 | MS.Trt_vs_MS.NoTrt_F |
| SPRED3 | 0.02125 | 0.29193 | 0.676 | 0.62 | 1 | MS.Trt_vs_MS.NoTrt_F |
| CLUL1 | 0.02126 | -0.3945 | 0.09 | 0.16 | 1 | MS.Trt_vs_MS.NoTrt_F |
| ACADS | 0.02127 | 0.36813 | 0.112 | 0.19 | 1 | MS.Trt_vs_MS.NoTrt_F |
| HIST1H1C | 0.0213 | 0.53261 | 0.129 | 0.22 | 1 | MS.Trt_vs_MS.NoTrt_F |
| CCDC126 | 0.02131 | 0.62474 | 0.104 | 0.18 | 1 | MS.Trt_vs_MS.NoTrt_F |
| USP11 | 0.02133 | -0.3466 | 0.191 | 0.29 | 1 | MS.Trt_vs_MS.NoTrt_F |
| FNTA | 0.02134 | -0.3216 | 0.824 | 0.91 | 1 | MS.Trt_vs_MS.NoTrt_F |
| TNXB | 0.02135 | 0.31141 | 0.104 | 0.18 | 1 | MS.Trt_vs_MS.NoTrt_F |
| SCAND1 | 0.02136 | 0.43509 | 0.187 | 0.29 | 1 | MS.Trt_vs_MS.NoTrt_F |
| USP30 | 0.02141 | -0.4928 | 0.302 | 0.38 | 1 | MS.Trt_vs_MS.NoTrt_F |
| SEMA6A | 0.02142 | -0.4944 | 0.065 | 0.13 | 1 | MS.Trt_vs_MS.NoTrt_F |
| CHMP6 | 0.02142 | 0.39272 | 0.122 | 0.21 | 1 | MS.Trt_vs_MS.NoTrt_F |
| TRANK1 | 0.02142 | -0.3005 | 0.216 | 0.32 | 1 | MS.Trt_vs_MS.NoTrt_F |
| RP11-507K1 | 0.02143 | 0.32898 | 0.068 | 0.13 | 1 | MS.Trt_vs_MS.NoTrt_F |
| FAM199X | 0.02153 | 0.81596 | 0.072 | 0.14 | 1 | MS.Trt_vs_MS.NoTrt_F |
| HIST1H4C | 0.02155 | 0.74863 | 0.331 | 0.51 | 1 | MS.Trt_vs_MS.NoTrt_F |
| DNTTIP1 | 0.02155 | 0.7968 | 0.079 | 0.15 | 1 | MS.Trt_vs_MS.NoTrt_F |
| BAZ2A | 0.02156 | 0.44063 | 0.594 | 0.71 | 1 | MS.Trt_vs_MS.NoTrt_F |
| LMOD3 | 0.02157 | -0.6072 | 0.403 | 0.51 | 1 | MS.Trt_vs_MS.NoTrt_F |
| PELP1 | 0.02158 | -0.3643 | 0.263 | 0.36 | 1 | MS.Trt_vs_MS.NoTrt_F |
| GPATCH4 | 0.0216 | 0.56613 | 0.248 | 0.35 | 1 | MS.Trt_vs_MS.NoTrt_F |
| RPF1 | 0.02162 | 0.45851 | 0.245 | 0.35 | 1 | MS.Trt_vs_MS.NoTrt_F |
| SAMM50 | 0.02166 | 0.33125 | 0.453 | 0.56 | 1 | MS.Trt_vs_MS.NoTrt_F |
| GZMA | 0.02167 | 0.56978 | 0.112 | 0.19 | 1 | MS.Trt_vs_MS.NoTrt_F |
| CDK17 | 0.02167 | 0.7208 | 0.198 | 0.31 | 1 | MS.Trt_vs_MS.NoTrt_F |
| GOSR2 | 0.02168 | 0.7618 | 0.129 | 0.22 | 1 | MS.Trt_vs_MS.NoTrt_F |
| SLC5A3 | 0.02169 | 0.4968 | 0.126 | 0.21 | 1 | MS.Trt_vs_MS.NoTrt_F |
| WNT10A | 0.02173 | 0.36023 | 0.162 | 0.25 | 1 | MS.Trt_vs_MS.NoTrt_F |
| LOXL1 | 0.02174 | 0.28086 | 0.086 | 0.16 | 1 | MS.Trt_vs_MS.NoTrt_F |
| TBX19 | 0.02176 | 0.37824 | 0.09 | 0.16 | 1 | MS.Trt_vs_MS.NoTrt_F |
| MAP4K3 | 0.02177 | -0.9514 | 0.068 | 0.13 | 1 | MS.Trt_vs_MS.NoTrt_F |
| MEN1 | 0.02178 | -1.0492 | 0.094 | 0.17 | 1 | MS.Trt_vs_MS.NoTrt_F |
| MAP3K7CL | 0.02179 | -0.5956 | 0.424 | 0.53 | 1 | MS.Trt_vs_MS.NoTrt_F |
| APOBEC3C | 0.02181 | 0.75744 | 0.788 | 0.85 | 1 | MS.Trt_vs_MS.NoTrt_F |
| MSL1 | 0.02183 | 0.47914 | 0.525 | 0.66 | 1 | MS.Trt_vs_MS.NoTrt_F |
| PDCD2 | 0.02184 | 0.66607 | 0.119 | 0.2 | 1 | MS.Trt_vs_MS.NoTrt_F |
| COASY | 0.02184 | -0.6538 | 0.442 | 0.53 | 1 | MS.Trt_vs_MS.NoTrt_F |

| | | | | | | |
|---|---|---|---|---|---|---|
| MLLT3 | 0.02186 | 0.27463 | 0.371 | 0.5 | 1 | MS.Trt_vs_MS.NoTrt_F |
| DCTN6 | 0.02191 | -0.2883 | 0.511 | 0.62 | 1 | MS.Trt_vs_MS.NoTrt_F |
| VRK1 | 0.02191 | -0.6806 | 0.201 | 0.28 | 1 | MS.Trt_vs_MS.NoTrt_F |
| XKR6 | 0.02191 | 0.3471 | 0.079 | 0.15 | 1 | MS.Trt_vs_MS.NoTrt_F |
| PPIL2 | 0.02191 | 0.2985 | 0.392 | 0.49 | 1 | MS.Trt_vs_MS.NoTrt_F |
| CCDC12 | 0.02196 | 0.25274 | 0.281 | 0.4 | 1 | MS.Trt_vs_MS.NoTrt_F |
| EIF2AK3 | 0.02207 | 1.262 | 0.079 | 0.15 | 1 | MS.Trt_vs_MS.NoTrt_F |
| IFIT5 | 0.02208 | 0.99641 | 0.083 | 0.16 | 1 | MS.Trt_vs_MS.NoTrt_F |
| AL356053.1 | 0.02214 | -0.6549 | 0.061 | 0.12 | 1 | MS.Trt_vs_MS.NoTrt_F |
| ADCK1 | 0.02216 | -1.4668 | 0.234 | 0.33 | 1 | MS.Trt_vs_MS.NoTrt_F |
| MELK | 0.02217 | 0.47701 | 0.586 | 0.53 | 1 | MS.Trt_vs_MS.NoTrt_F |
| ANKRD6 | 0.02218 | -0.4291 | 0.14 | 0.22 | 1 | MS.Trt_vs_MS.NoTrt_F |
| SREBF2 | 0.02219 | 0.52254 | 0.165 | 0.27 | 1 | MS.Trt_vs_MS.NoTrt_F |
| CYP2D6 | 0.0223 | -0.7091 | 0.076 | 0.14 | 1 | MS.Trt_vs_MS.NoTrt_F |
| EPS8L2 | 0.02239 | -0.3436 | 0.065 | 0.13 | 1 | MS.Trt_vs_MS.NoTrt_F |
| CD226 | 0.02241 | -0.67 | 0.518 | 0.61 | 1 | MS.Trt_vs_MS.NoTrt_F |
| GSPT2 | 0.02245 | 0.34641 | 0.072 | 0.14 | 1 | MS.Trt_vs_MS.NoTrt_F |
| SLC39A13 | 0.02246 | 0.78885 | 0.14 | 0.23 | 1 | MS.Trt_vs_MS.NoTrt_F |
| ARV1 | 0.0225 | -0.6808 | 0.277 | 0.39 | 1 | MS.Trt_vs_MS.NoTrt_F |
| CNNM2 | 0.02251 | 0.47307 | 0.317 | 0.43 | 1 | MS.Trt_vs_MS.NoTrt_F |
| CALCOCO2 | 0.02252 | -0.327 | 0.982 | 1 | 1 | MS.Trt_vs_MS.NoTrt_F |
| CHD4 | 0.02256 | 0.34358 | 0.536 | 0.66 | 1 | MS.Trt_vs_MS.NoTrt_F |
| MRPL22 | 0.02259 | 0.59409 | 0.212 | 0.32 | 1 | MS.Trt_vs_MS.NoTrt_F |
| FUCA1 | 0.02261 | 0.62108 | 0.079 | 0.15 | 1 | MS.Trt_vs_MS.NoTrt_F |
| PACS2 | 0.02264 | 0.66966 | 0.248 | 0.35 | 1 | MS.Trt_vs_MS.NoTrt_F |
| ZC3HC1 | 0.02266 | 0.37907 | 0.414 | 0.56 | 1 | MS.Trt_vs_MS.NoTrt_F |
| NDUFB3 | 0.02272 | 0.80406 | 0.216 | 0.34 | 1 | MS.Trt_vs_MS.NoTrt_F |
| EIF4G1 | 0.02273 | -0.7094 | 0.482 | 0.58 | 1 | MS.Trt_vs_MS.NoTrt_F |
| ARHGEF39 | 0.02278 | -0.54 | 0.622 | 0.76 | 1 | MS.Trt_vs_MS.NoTrt_F |
| ASB7 | 0.02278 | 0.82652 | 0.097 | 0.17 | 1 | MS.Trt_vs_MS.NoTrt_F |
| SET | 0.02279 | 0.72599 | 0.874 | 0.96 | 1 | MS.Trt_vs_MS.NoTrt_F |
| ADAM28 | 0.02282 | 1.27062 | 0.144 | 0.23 | 1 | MS.Trt_vs_MS.NoTrt_F |
| SCOC | 0.02298 | 0.58318 | 0.112 | 0.19 | 1 | MS.Trt_vs_MS.NoTrt_F |
| CCDC120 | 0.02299 | 0.26541 | 0.507 | 0.65 | 1 | MS.Trt_vs_MS.NoTrt_F |
| VNN3 | 0.023 | 0.64013 | 0.32 | 0.47 | 1 | MS.Trt_vs_MS.NoTrt_F |
| MAP3K9 | 0.02306 | 0.58158 | 0.424 | 0.54 | 1 | MS.Trt_vs_MS.NoTrt_F |
| ZNF432 | 0.02312 | -0.9542 | 0.622 | 0.7 | 1 | MS.Trt_vs_MS.NoTrt_F |
| NDUFAF5 | 0.02312 | -0.4926 | 0.478 | 0.6 | 1 | MS.Trt_vs_MS.NoTrt_F |
| HERC6 | 0.02314 | 0.43169 | 0.302 | 0.4 | 1 | MS.Trt_vs_MS.NoTrt_F |
| TMPPE | 0.02315 | 0.43731 | 0.072 | 0.14 | 1 | MS.Trt_vs_MS.NoTrt_F |

| | | | | | | |
|---|---|---|---|---|---|---|
| CNOT6L | 0.02317 | 0.67341 | 0.241 | 0.35 | 1 | MS.Trt_vs_MS.NoTrt_F |
| ARMCX6 | 0.02318 | 0.30992 | 0.079 | 0.15 | 1 | MS.Trt_vs_MS.NoTrt_F |
| DISP1 | 0.02318 | -0.7052 | 0.097 | 0.17 | 1 | MS.Trt_vs_MS.NoTrt_F |
| ARL5A | 0.0232 | 0.66904 | 0.173 | 0.28 | 1 | MS.Trt_vs_MS.NoTrt_F |
| ZFP3 | 0.02322 | -1.3607 | 0.115 | 0.19 | 1 | MS.Trt_vs_MS.NoTrt_F |
| C9orf78 | 0.02323 | -0.2867 | 0.964 | 0.98 | 1 | MS.Trt_vs_MS.NoTrt_F |
| LZTR1 | 0.02324 | -0.5468 | 0.252 | 0.34 | 1 | MS.Trt_vs_MS.NoTrt_F |
| URI1 | 0.02331 | 0.32209 | 0.187 | 0.29 | 1 | MS.Trt_vs_MS.NoTrt_F |
| MRPL44 | 0.02332 | -0.7371 | 0.529 | 0.64 | 1 | MS.Trt_vs_MS.NoTrt_F |
| CD59 | 0.02334 | -0.4174 | 0.655 | 0.79 | 1 | MS.Trt_vs_MS.NoTrt_F |
| ATXN7L3 | 0.02337 | 0.99958 | 0.076 | 0.14 | 1 | MS.Trt_vs_MS.NoTrt_F |
| MRPL2 | 0.02337 | 0.33122 | 0.288 | 0.4 | 1 | MS.Trt_vs_MS.NoTrt_F |
| ZHX2 | 0.02359 | 0.54757 | 0.137 | 0.23 | 1 | MS.Trt_vs_MS.NoTrt_F |
| NUDT14 | 0.0236 | 0.56396 | 0.428 | 0.54 | 1 | MS.Trt_vs_MS.NoTrt_F |
| SLC25A43 | 0.02362 | -0.2656 | 0.219 | 0.31 | 1 | MS.Trt_vs_MS.NoTrt_F |
| ATPAF2 | 0.02363 | 0.28989 | 0.439 | 0.56 | 1 | MS.Trt_vs_MS.NoTrt_F |
| NSFL1C | 0.02363 | 0.54586 | 0.155 | 0.25 | 1 | MS.Trt_vs_MS.NoTrt_F |
| ITGAL | 0.02365 | 0.27691 | 0.752 | 0.84 | 1 | MS.Trt_vs_MS.NoTrt_F |
| HSF5 | 0.02368 | -0.2745 | 0.09 | 0.16 | 1 | MS.Trt_vs_MS.NoTrt_F |
| HIKESHI | 0.02368 | 0.30571 | 0.442 | 0.57 | 1 | MS.Trt_vs_MS.NoTrt_F |
| SESN1 | 0.02369 | 0.73781 | 0.183 | 0.29 | 1 | MS.Trt_vs_MS.NoTrt_F |
| ZNF48 | 0.02374 | -0.5322 | 0.058 | 0.12 | 1 | MS.Trt_vs_MS.NoTrt_F |
| TTC33 | 0.02388 | 0.28094 | 0.259 | 0.36 | 1 | MS.Trt_vs_MS.NoTrt_F |
| ZNF10 | 0.024 | 1.08885 | 0.126 | 0.21 | 1 | MS.Trt_vs_MS.NoTrt_F |
| HINT2 | 0.02407 | 0.71848 | 0.101 | 0.18 | 1 | MS.Trt_vs_MS.NoTrt_F |
| MED19 | 0.0242 | 0.25717 | 0.086 | 0.16 | 1 | MS.Trt_vs_MS.NoTrt_F |
| DDX10 | 0.02422 | -0.7917 | 0.658 | 0.75 | 1 | MS.Trt_vs_MS.NoTrt_F |
| NBPF1 | 0.02435 | 0.25717 | 0.219 | 0.33 | 1 | MS.Trt_vs_MS.NoTrt_F |
| GEMIN5 | 0.02438 | -2.2503 | 0.05 | 0.11 | 1 | MS.Trt_vs_MS.NoTrt_F |
| BRAF | 0.0244 | 0.37937 | 0.147 | 0.24 | 1 | MS.Trt_vs_MS.NoTrt_F |
| SPECC1L-AD | 0.02446 | -1.148 | 0.187 | 0.28 | 1 | MS.Trt_vs_MS.NoTrt_F |
| CLP1 | 0.02447 | -1.3381 | 0.05 | 0.11 | 1 | MS.Trt_vs_MS.NoTrt_F |
| OSBPL9 | 0.02451 | -0.5229 | 0.309 | 0.41 | 1 | MS.Trt_vs_MS.NoTrt_F |
| BAHD1 | 0.02455 | 0.79769 | 0.072 | 0.14 | 1 | MS.Trt_vs_MS.NoTrt_F |
| IDNK | 0.02456 | -0.3793 | 0.367 | 0.48 | 1 | MS.Trt_vs_MS.NoTrt_F |
| HARS | 0.02457 | 0.28833 | 0.871 | 0.87 | 1 | MS.Trt_vs_MS.NoTrt_F |
| AASDHPPT | 0.02462 | 0.4914 | 0.205 | 0.31 | 1 | MS.Trt_vs_MS.NoTrt_F |
| FEM1B | 0.02463 | -0.5941 | 0.09 | 0.16 | 1 | MS.Trt_vs_MS.NoTrt_F |
| ERLIN1 | 0.02463 | -1.8533 | 0.054 | 0.11 | 1 | MS.Trt_vs_MS.NoTrt_F |
| FANCM | 0.02463 | -0.6268 | 0.104 | 0.18 | 1 | MS.Trt_vs_MS.NoTrt_F |

| | | | | | | |
|---|---|---|---|---|---|---|
| ZFYVE1 | 0.02466 | -0.3212 | 0.108 | 0.18 | 1 | MS.Trt_vs_MS.NoTrt_F |
| RP11-449H3 | 0.02469 | -0.9919 | 0.234 | 0.33 | 1 | MS.Trt_vs_MS.NoTrt_F |
| PEX14 | 0.02482 | -0.5469 | 0.119 | 0.19 | 1 | MS.Trt_vs_MS.NoTrt_F |
| PARPBP | 0.02484 | 0.59105 | 0.338 | 0.45 | 1 | MS.Trt_vs_MS.NoTrt_F |
| ATP5J2-PTCI | 0.02487 | -0.5081 | 0.058 | 0.12 | 1 | MS.Trt_vs_MS.NoTrt_F |
| KYAT1 | 0.02488 | 0.50801 | 0.162 | 0.25 | 1 | MS.Trt_vs_MS.NoTrt_F |
| TOMM34 | 0.02491 | 0.8242 | 0.083 | 0.16 | 1 | MS.Trt_vs_MS.NoTrt_F |
| MRPL45 | 0.025 | -0.5422 | 0.076 | 0.14 | 1 | MS.Trt_vs_MS.NoTrt_F |
| TMEM184B | 0.02502 | -0.5384 | 0.09 | 0.16 | 1 | MS.Trt_vs_MS.NoTrt_F |
| SOCS4 | 0.02514 | -0.4267 | 0.299 | 0.41 | 1 | MS.Trt_vs_MS.NoTrt_F |
| LRTOMT | 0.02521 | 0.50641 | 0.849 | 0.83 | 1 | MS.Trt_vs_MS.NoTrt_F |
| FAM45A | 0.02521 | -0.476 | 0.673 | 0.76 | 1 | MS.Trt_vs_MS.NoTrt_F |
| BCDIN3D | 0.02523 | -1.5482 | 0.385 | 0.46 | 1 | MS.Trt_vs_MS.NoTrt_F |
| SCRIB | 0.02523 | -0.4501 | 0.054 | 0.11 | 1 | MS.Trt_vs_MS.NoTrt_F |
| MMRN2 | 0.02527 | -0.4126 | 0.162 | 0.26 | 1 | MS.Trt_vs_MS.NoTrt_F |
| IP6K1 | 0.02529 | 0.45069 | 0.151 | 0.24 | 1 | MS.Trt_vs_MS.NoTrt_F |
| MRPS34 | 0.02532 | 0.66071 | 0.18 | 0.29 | 1 | MS.Trt_vs_MS.NoTrt_F |
| WDR60 | 0.0255 | 0.4163 | 0.144 | 0.23 | 1 | MS.Trt_vs_MS.NoTrt_F |
| YDJC | 0.02553 | 0.38461 | 0.162 | 0.26 | 1 | MS.Trt_vs_MS.NoTrt_F |
| CCDC86 | 0.02554 | 0.56567 | 0.068 | 0.13 | 1 | MS.Trt_vs_MS.NoTrt_F |
| TRMT61B | 0.02572 | 1.11367 | 0.129 | 0.22 | 1 | MS.Trt_vs_MS.NoTrt_F |
| KLF11 | 0.02572 | -0.3631 | 0.054 | 0.11 | 1 | MS.Trt_vs_MS.NoTrt_F |
| FKBP4 | 0.02574 | -0.4021 | 0.108 | 0.18 | 1 | MS.Trt_vs_MS.NoTrt_F |
| MON2 | 0.02577 | 0.52428 | 0.36 | 0.49 | 1 | MS.Trt_vs_MS.NoTrt_F |
| FAM185A | 0.02578 | -0.8371 | 0.126 | 0.2 | 1 | MS.Trt_vs_MS.NoTrt_F |
| C7 | 0.02582 | -0.3618 | 0.381 | 0.51 | 1 | MS.Trt_vs_MS.NoTrt_F |
| CCDC146 | 0.02582 | -0.7162 | 0.09 | 0.16 | 1 | MS.Trt_vs_MS.NoTrt_F |
| SLC35A1 | 0.02589 | 0.30744 | 0.137 | 0.22 | 1 | MS.Trt_vs_MS.NoTrt_F |
| CCDC69 | 0.02597 | -0.6082 | 0.827 | 0.88 | 1 | MS.Trt_vs_MS.NoTrt_F |
| EVA1C | 0.02599 | -0.4939 | 0.144 | 0.23 | 1 | MS.Trt_vs_MS.NoTrt_F |
| SLC9B2 | 0.026 | 0.69078 | 0.079 | 0.15 | 1 | MS.Trt_vs_MS.NoTrt_F |
| MAMLD1 | 0.02604 | 0.99473 | 0.187 | 0.29 | 1 | MS.Trt_vs_MS.NoTrt_F |
| USP31 | 0.02607 | 0.84783 | 0.112 | 0.19 | 1 | MS.Trt_vs_MS.NoTrt_F |
| NR4A1 | 0.02608 | 0.88826 | 0.435 | 0.57 | 1 | MS.Trt_vs_MS.NoTrt_F |
| PPTC7 | 0.02615 | -0.2755 | 0.094 | 0.17 | 1 | MS.Trt_vs_MS.NoTrt_F |
| ABCD3 | 0.02619 | -0.4414 | 0.144 | 0.23 | 1 | MS.Trt_vs_MS.NoTrt_F |
| CNTROB | 0.02631 | -0.4864 | 0.201 | 0.29 | 1 | MS.Trt_vs_MS.NoTrt_F |
| ZNF235 | 0.02636 | 0.31503 | 0.133 | 0.22 | 1 | MS.Trt_vs_MS.NoTrt_F |
| DDIT4 | 0.0264 | 0.34192 | 0.342 | 0.5 | 1 | MS.Trt_vs_MS.NoTrt_F |
| RNF207 | 0.02641 | 0.7633 | 0.737 | 0.7 | 1 | MS.Trt_vs_MS.NoTrt_F |

| | | | | | | |
|---|---|---|---|---|---|---|
| FZR1 | 0.02644 | -0.3703 | 0.129 | 0.22 | 1 | MS.Trt_vs_MS.NoTrt_F |
| ZNF335 | 0.02645 | -0.4241 | 0.331 | 0.41 | 1 | MS.Trt_vs_MS.NoTrt_F |
| DYRK2 | 0.02645 | 0.91141 | 0.155 | 0.25 | 1 | MS.Trt_vs_MS.NoTrt_F |
| RAB33A | 0.02648 | 0.53155 | 0.097 | 0.17 | 1 | MS.Trt_vs_MS.NoTrt_F |
| SETD1A | 0.02652 | 1.03983 | 0.068 | 0.13 | 1 | MS.Trt_vs_MS.NoTrt_F |
| SLC26A11 | 0.0266 | -0.4955 | 0.597 | 0.72 | 1 | MS.Trt_vs_MS.NoTrt_F |
| ZFAND1 | 0.02661 | 0.74758 | 0.14 | 0.23 | 1 | MS.Trt_vs_MS.NoTrt_F |
| LIN52 | 0.02665 | 0.6958 | 0.676 | 0.78 | 1 | MS.Trt_vs_MS.NoTrt_F |
| DHX40 | 0.02668 | 0.92519 | 0.122 | 0.2 | 1 | MS.Trt_vs_MS.NoTrt_F |
| FBXO31 | 0.02677 | -0.3733 | 0.335 | 0.44 | 1 | MS.Trt_vs_MS.NoTrt_F |
| SUCLA2 | 0.02679 | 0.49468 | 0.126 | 0.21 | 1 | MS.Trt_vs_MS.NoTrt_F |
| ICOSLG | 0.02687 | -0.4602 | 0.709 | 0.85 | 1 | MS.Trt_vs_MS.NoTrt_F |
| PRIMPOL | 0.02691 | -0.3379 | 0.112 | 0.19 | 1 | MS.Trt_vs_MS.NoTrt_F |
| RFX3 | 0.02704 | -0.4877 | 0.187 | 0.28 | 1 | MS.Trt_vs_MS.NoTrt_F |
| L3MBTL4 | 0.02705 | -0.7966 | 0.432 | 0.59 | 1 | MS.Trt_vs_MS.NoTrt_F |
| SEC23IP | 0.02712 | -0.4543 | 0.385 | 0.5 | 1 | MS.Trt_vs_MS.NoTrt_F |
| CISD2 | 0.02721 | 0.64533 | 0.446 | 0.55 | 1 | MS.Trt_vs_MS.NoTrt_F |
| IL17RA | 0.02721 | -0.3261 | 0.658 | 0.76 | 1 | MS.Trt_vs_MS.NoTrt_F |
| PYGB | 0.02725 | 0.42616 | 0.144 | 0.23 | 1 | MS.Trt_vs_MS.NoTrt_F |
| PABPC1L | 0.0273 | -0.3682 | 0.288 | 0.39 | 1 | MS.Trt_vs_MS.NoTrt_F |
| KDM1B | 0.02734 | -0.807 | 0.086 | 0.16 | 1 | MS.Trt_vs_MS.NoTrt_F |
| OSBP | 0.02736 | 0.26665 | 0.09 | 0.16 | 1 | MS.Trt_vs_MS.NoTrt_F |
| CRADD | 0.0275 | 0.41125 | 0.094 | 0.17 | 1 | MS.Trt_vs_MS.NoTrt_F |
| TRIM11 | 0.02754 | -0.6106 | 0.554 | 0.64 | 1 | MS.Trt_vs_MS.NoTrt_F |
| ACOT13 | 0.02755 | -0.6361 | 0.374 | 0.46 | 1 | MS.Trt_vs_MS.NoTrt_F |
| EDA | 0.02755 | -0.6303 | 0.367 | 0.48 | 1 | MS.Trt_vs_MS.NoTrt_F |
| KCNMB3 | 0.02757 | 0.42663 | 0.378 | 0.47 | 1 | MS.Trt_vs_MS.NoTrt_F |
| AGA | 0.0276 | 0.33025 | 0.137 | 0.22 | 1 | MS.Trt_vs_MS.NoTrt_F |
| CLDN11 | 0.02764 | 0.57945 | 0.568 | 0.69 | 1 | MS.Trt_vs_MS.NoTrt_F |
| CENPP | 0.02769 | -0.5971 | 0.05 | 0.11 | 1 | MS.Trt_vs_MS.NoTrt_F |
| TMPRSS13 | 0.0277 | 1.04501 | 0.162 | 0.24 | 1 | MS.Trt_vs_MS.NoTrt_F |
| KTI12 | 0.02786 | 0.32775 | 0.09 | 0.16 | 1 | MS.Trt_vs_MS.NoTrt_F |
| THTPA | 0.02789 | 0.41461 | 0.115 | 0.19 | 1 | MS.Trt_vs_MS.NoTrt_F |
| MYDGF | 0.02789 | 0.4906 | 0.162 | 0.26 | 1 | MS.Trt_vs_MS.NoTrt_F |
| RNF215 | 0.02791 | -0.6309 | 0.468 | 0.54 | 1 | MS.Trt_vs_MS.NoTrt_F |
| ZNF181 | 0.02795 | -0.4573 | 0.169 | 0.27 | 1 | MS.Trt_vs_MS.NoTrt_F |
| PML | 0.02803 | -0.3842 | 0.601 | 0.73 | 1 | MS.Trt_vs_MS.NoTrt_F |
| COX8A | 0.02803 | 0.63447 | 0.23 | 0.36 | 1 | MS.Trt_vs_MS.NoTrt_F |
| FICD | 0.02822 | 0.31439 | 0.083 | 0.15 | 1 | MS.Trt_vs_MS.NoTrt_F |
| ATG7 | 0.02831 | 0.26187 | 0.536 | 0.69 | 1 | MS.Trt_vs_MS.NoTrt_F |

| | | | | | | |
|---|---|---|---|---|---|---|
| ATP13A2 | 0.02831 | 0.2805 | 0.421 | 0.53 | 1 | MS.Trt_vs_MS.NoTrt_F |
| GPR68 | 0.02838 | 0.52003 | 0.133 | 0.22 | 1 | MS.Trt_vs_MS.NoTrt_F |
| CEPT1 | 0.02847 | 0.42436 | 0.338 | 0.45 | 1 | MS.Trt_vs_MS.NoTrt_F |
| IFT57 | 0.02849 | 0.38665 | 0.112 | 0.19 | 1 | MS.Trt_vs_MS.NoTrt_F |
| POLR1E | 0.02852 | 0.71034 | 0.129 | 0.21 | 1 | MS.Trt_vs_MS.NoTrt_F |
| STK32C | 0.02853 | -0.5916 | 0.072 | 0.13 | 1 | MS.Trt_vs_MS.NoTrt_F |
| SACS | 0.02858 | -0.2807 | 0.108 | 0.18 | 1 | MS.Trt_vs_MS.NoTrt_F |
| SNRPG | 0.02858 | 0.70523 | 0.313 | 0.45 | 1 | MS.Trt_vs_MS.NoTrt_F |
| RAPGEF6 | 0.0286 | -0.4707 | 0.442 | 0.55 | 1 | MS.Trt_vs_MS.NoTrt_F |
| RPS26 | 0.02865 | 0.44722 | 0.691 | 0.77 | 1 | MS.Trt_vs_MS.NoTrt_F |
| C10orf54 | 0.0287 | 0.35077 | 0.338 | 0.49 | 1 | MS.Trt_vs_MS.NoTrt_F |
| SLC25A24 | 0.02897 | 0.50597 | 0.133 | 0.22 | 1 | MS.Trt_vs_MS.NoTrt_F |
| AP3M1 | 0.02899 | -0.311 | 0.468 | 0.55 | 1 | MS.Trt_vs_MS.NoTrt_F |
| NOL6 | 0.02899 | 0.90876 | 0.381 | 0.5 | 1 | MS.Trt_vs_MS.NoTrt_F |
| NRG4 | 0.02907 | -0.4368 | 0.586 | 0.7 | 1 | MS.Trt_vs_MS.NoTrt_F |
| ZNF180 | 0.0291 | 1.04966 | 0.353 | 0.43 | 1 | MS.Trt_vs_MS.NoTrt_F |
| IRF2BP2 | 0.02921 | 0.38291 | 0.216 | 0.32 | 1 | MS.Trt_vs_MS.NoTrt_F |
| CSE1L | 0.02921 | 0.31546 | 0.223 | 0.33 | 1 | MS.Trt_vs_MS.NoTrt_F |
| GFER | 0.02922 | 0.423 | 0.083 | 0.15 | 1 | MS.Trt_vs_MS.NoTrt_F |
| ARFGAP3 | 0.02926 | -0.2835 | 0.327 | 0.43 | 1 | MS.Trt_vs_MS.NoTrt_F |
| HLA-DMB | 0.02928 | -1.1508 | 0.101 | 0.17 | 1 | MS.Trt_vs_MS.NoTrt_F |
| THSD4 | 0.02939 | 0.2812 | 0.324 | 0.44 | 1 | MS.Trt_vs_MS.NoTrt_F |
| CNKSR1 | 0.02939 | -0.6048 | 0.173 | 0.26 | 1 | MS.Trt_vs_MS.NoTrt_F |
| TXN2 | 0.02943 | 0.44063 | 0.273 | 0.38 | 1 | MS.Trt_vs_MS.NoTrt_F |
| GLIPR1 | 0.02946 | 0.34755 | 0.309 | 0.43 | 1 | MS.Trt_vs_MS.NoTrt_F |
| INTS1 | 0.02946 | -0.5757 | 0.367 | 0.46 | 1 | MS.Trt_vs_MS.NoTrt_F |
| DCBLD2 | 0.02946 | 0.40731 | 0.64 | 0.77 | 1 | MS.Trt_vs_MS.NoTrt_F |
| UBXN6 | 0.02946 | 0.93975 | 0.101 | 0.18 | 1 | MS.Trt_vs_MS.NoTrt_F |
| NHSL2 | 0.02952 | 0.31094 | 0.633 | 0.74 | 1 | MS.Trt_vs_MS.NoTrt_F |
| BHLHB9 | 0.02954 | 0.4948 | 0.345 | 0.48 | 1 | MS.Trt_vs_MS.NoTrt_F |
| PLA2G6 | 0.02962 | -0.4239 | 0.428 | 0.52 | 1 | MS.Trt_vs_MS.NoTrt_F |
| CCDC24 | 0.02964 | 0.39302 | 0.745 | 0.66 | 1 | MS.Trt_vs_MS.NoTrt_F |
| MFSD14B | 0.02966 | 0.51032 | 0.122 | 0.2 | 1 | MS.Trt_vs_MS.NoTrt_F |
| TUSC2 | 0.02969 | 0.28691 | 0.115 | 0.19 | 1 | MS.Trt_vs_MS.NoTrt_F |
| ADRB2 | 0.02973 | 0.88376 | 0.205 | 0.31 | 1 | MS.Trt_vs_MS.NoTrt_F |
| ABCC9 | 0.02978 | 0.34352 | 0.924 | 0.96 | 1 | MS.Trt_vs_MS.NoTrt_F |
| HAUS4 | 0.02979 | 0.91741 | 0.147 | 0.24 | 1 | MS.Trt_vs_MS.NoTrt_F |
| FBXO48 | 0.02984 | -0.5884 | 0.655 | 0.8 | 1 | MS.Trt_vs_MS.NoTrt_F |
| ELOVL6 | 0.02986 | 0.43577 | 0.342 | 0.45 | 1 | MS.Trt_vs_MS.NoTrt_F |
| FIGNL1 | 0.0299 | 0.26478 | 0.151 | 0.23 | 1 | MS.Trt_vs_MS.NoTrt_F |

| | | | | | | |
|---|---|---|---|---|---|---|
| HACL1 | 0.02993 | 0.4571 | 0.137 | 0.22 | 1 | MS.Trt_vs_MS.NoTrt_F |
| EXOSC8 | 0.02995 | -0.6069 | 0.518 | 0.56 | 1 | MS.Trt_vs_MS.NoTrt_F |
| ZNF616 | 0.02996 | 0.2767 | 0.665 | 0.83 | 1 | MS.Trt_vs_MS.NoTrt_F |
| BTD | 0.03 | 0.4935 | 0.097 | 0.17 | 1 | MS.Trt_vs_MS.NoTrt_F |
| TPRG1L | 0.03001 | -0.8055 | 0.255 | 0.34 | 1 | MS.Trt_vs_MS.NoTrt_F |
| FLNB | 0.03002 | 0.50592 | 0.353 | 0.43 | 1 | MS.Trt_vs_MS.NoTrt_F |
| AIFM2 | 0.03019 | 1.03527 | 0.371 | 0.49 | 1 | MS.Trt_vs_MS.NoTrt_F |
| SMG6 | 0.03029 | -0.6016 | 0.237 | 0.33 | 1 | MS.Trt_vs_MS.NoTrt_F |
| BAZ2B | 0.0303 | -0.8906 | 0.806 | 0.87 | 1 | MS.Trt_vs_MS.NoTrt_F |
| XRCC4 | 0.03032 | 0.33071 | 0.079 | 0.14 | 1 | MS.Trt_vs_MS.NoTrt_F |
| ROCK2 | 0.03036 | -0.4479 | 0.464 | 0.59 | 1 | MS.Trt_vs_MS.NoTrt_F |
| IFI44L | 0.03039 | 2.11549 | 0.209 | 0.32 | 1 | MS.Trt_vs_MS.NoTrt_F |
| C8orf33 | 0.03048 | 0.87359 | 0.245 | 0.37 | 1 | MS.Trt_vs_MS.NoTrt_F |
| RUSC1 | 0.0305 | 0.55947 | 0.083 | 0.15 | 1 | MS.Trt_vs_MS.NoTrt_F |
| ENOSF1 | 0.03051 | 0.3626 | 0.813 | 0.9 | 1 | MS.Trt_vs_MS.NoTrt_F |
| NOL11 | 0.03063 | 0.47044 | 0.475 | 0.56 | 1 | MS.Trt_vs_MS.NoTrt_F |
| ANKAR | 0.03072 | -0.3543 | 0.137 | 0.22 | 1 | MS.Trt_vs_MS.NoTrt_F |
| BTBD2 | 0.03078 | 0.68072 | 0.345 | 0.44 | 1 | MS.Trt_vs_MS.NoTrt_F |
| SOCS5 | 0.03079 | 0.36977 | 0.09 | 0.16 | 1 | MS.Trt_vs_MS.NoTrt_F |
| ALS2 | 0.03084 | -0.4219 | 0.403 | 0.52 | 1 | MS.Trt_vs_MS.NoTrt_F |
| ZNF799 | 0.03089 | 1.48124 | 0.367 | 0.48 | 1 | MS.Trt_vs_MS.NoTrt_F |
| INSIG2 | 0.03096 | 0.84965 | 0.144 | 0.24 | 1 | MS.Trt_vs_MS.NoTrt_F |
| TMED1 | 0.03109 | -0.4404 | 0.083 | 0.15 | 1 | MS.Trt_vs_MS.NoTrt_F |
| FOPNL | 0.03114 | -0.3036 | 0.561 | 0.64 | 1 | MS.Trt_vs_MS.NoTrt_F |
| AP000275.6 | 0.03123 | 0.35972 | 0.094 | 0.17 | 1 | MS.Trt_vs_MS.NoTrt_F |
| HECTD3 | 0.03125 | 0.31577 | 0.151 | 0.24 | 1 | MS.Trt_vs_MS.NoTrt_F |
| CYP4F8 | 0.03128 | 0.32829 | 0.827 | 0.8 | 1 | MS.Trt_vs_MS.NoTrt_F |
| GALK2 | 0.03137 | -0.4691 | 0.892 | 0.94 | 1 | MS.Trt_vs_MS.NoTrt_F |
| TCF4 | 0.03139 | 0.29969 | 0.669 | 0.76 | 1 | MS.Trt_vs_MS.NoTrt_F |
| C9orf84 | 0.03145 | -0.7193 | 0.072 | 0.13 | 1 | MS.Trt_vs_MS.NoTrt_F |
| STARD6 | 0.03145 | 0.52159 | 0.129 | 0.2 | 1 | MS.Trt_vs_MS.NoTrt_F |
| CYP4V2 | 0.03145 | -0.327 | 0.752 | 0.82 | 1 | MS.Trt_vs_MS.NoTrt_F |
| FAM58A | 0.03148 | 0.59494 | 0.09 | 0.16 | 1 | MS.Trt_vs_MS.NoTrt_F |
| MRPL27 | 0.0315 | 0.49302 | 0.176 | 0.27 | 1 | MS.Trt_vs_MS.NoTrt_F |
| NDUFB5 | 0.0315 | -0.3056 | 0.691 | 0.75 | 1 | MS.Trt_vs_MS.NoTrt_F |
| NIFK | 0.03159 | -0.5369 | 0.496 | 0.59 | 1 | MS.Trt_vs_MS.NoTrt_F |
| MCFD2 | 0.0316 | 0.34363 | 0.824 | 0.95 | 1 | MS.Trt_vs_MS.NoTrt_F |
| TFPT | 0.03167 | 0.54557 | 0.112 | 0.19 | 1 | MS.Trt_vs_MS.NoTrt_F |
| TRPM1 | 0.03172 | -0.4348 | 0.112 | 0.18 | 1 | MS.Trt_vs_MS.NoTrt_F |
| FMNL1 | 0.03178 | 0.33587 | 0.683 | 0.85 | 1 | MS.Trt_vs_MS.NoTrt_F |

| | | | | | | |
|---|---|---|---|---|---|---|
| ENG | 0.03179 | 0.61233 | 0.076 | 0.14 | 1 | MS.Trt_vs_MS.NoTrt_F |
| ZMIZ2 | 0.03183 | 0.51388 | 0.183 | 0.28 | 1 | MS.Trt_vs_MS.NoTrt_F |
| MRPL32 | 0.03195 | 0.51683 | 0.18 | 0.28 | 1 | MS.Trt_vs_MS.NoTrt_F |
| DENND1B | 0.03195 | -0.5051 | 0.162 | 0.25 | 1 | MS.Trt_vs_MS.NoTrt_F |
| PHACTR4 | 0.03207 | 0.39807 | 0.95 | 0.98 | 1 | MS.Trt_vs_MS.NoTrt_F |
| UBASH3B | 0.03215 | -0.5735 | 0.421 | 0.51 | 1 | MS.Trt_vs_MS.NoTrt_F |
| SNX8 | 0.03215 | -0.3934 | 0.14 | 0.22 | 1 | MS.Trt_vs_MS.NoTrt_F |
| STIL | 0.03219 | -0.7697 | 0.554 | 0.63 | 1 | MS.Trt_vs_MS.NoTrt_F |
| RNF146 | 0.0322 | 0.69385 | 0.126 | 0.21 | 1 | MS.Trt_vs_MS.NoTrt_F |
| STX1A | 0.03235 | -1.2833 | 0.065 | 0.12 | 1 | MS.Trt_vs_MS.NoTrt_F |
| SKAP2 | 0.03236 | -0.3101 | 0.219 | 0.31 | 1 | MS.Trt_vs_MS.NoTrt_F |
| LYRM9 | 0.03238 | -0.3611 | 0.176 | 0.27 | 1 | MS.Trt_vs_MS.NoTrt_F |
| COPS9 | 0.03243 | 0.79206 | 0.187 | 0.29 | 1 | MS.Trt_vs_MS.NoTrt_F |
| DPY19L4 | 0.03256 | -1.0599 | 0.525 | 0.62 | 1 | MS.Trt_vs_MS.NoTrt_F |
| ELAVL1 | 0.03257 | 0.4524 | 0.964 | 0.97 | 1 | MS.Trt_vs_MS.NoTrt_F |
| F5 | 0.03266 | 0.72012 | 0.198 | 0.3 | 1 | MS.Trt_vs_MS.NoTrt_F |
| GPR65 | 0.03268 | 0.86282 | 0.273 | 0.37 | 1 | MS.Trt_vs_MS.NoTrt_F |
| SSFA2 | 0.03271 | -1.0694 | 0.144 | 0.22 | 1 | MS.Trt_vs_MS.NoTrt_F |
| ATG4C | 0.03275 | 0.31168 | 0.072 | 0.13 | 1 | MS.Trt_vs_MS.NoTrt_F |
| RIMS2 | 0.03287 | 0.2736 | 0.917 | 0.92 | 1 | MS.Trt_vs_MS.NoTrt_F |
| NUTM2G | 0.03291 | 0.26133 | 0.076 | 0.14 | 1 | MS.Trt_vs_MS.NoTrt_F |
| OSBPL7 | 0.03295 | 0.34183 | 0.09 | 0.16 | 1 | MS.Trt_vs_MS.NoTrt_F |
| GLCCI1 | 0.03295 | -0.3849 | 0.266 | 0.36 | 1 | MS.Trt_vs_MS.NoTrt_F |
| GLCE | 0.03299 | -0.6206 | 0.155 | 0.23 | 1 | MS.Trt_vs_MS.NoTrt_F |
| FUNDC1 | 0.03304 | -0.3983 | 0.076 | 0.14 | 1 | MS.Trt_vs_MS.NoTrt_F |
| CXCR2 | 0.03305 | 0.72452 | 0.302 | 0.38 | 1 | MS.Trt_vs_MS.NoTrt_F |
| ARID2 | 0.03306 | 0.27335 | 0.694 | 0.81 | 1 | MS.Trt_vs_MS.NoTrt_F |
| ENPP3 | 0.03325 | 1.25171 | 0.122 | 0.2 | 1 | MS.Trt_vs_MS.NoTrt_F |
| FHL1 | 0.03331 | -0.5266 | 0.072 | 0.13 | 1 | MS.Trt_vs_MS.NoTrt_F |
| PIK3CA | 0.03336 | -0.275 | 0.072 | 0.13 | 1 | MS.Trt_vs_MS.NoTrt_F |
| JOSD2 | 0.0334 | 0.28909 | 0.108 | 0.18 | 1 | MS.Trt_vs_MS.NoTrt_F |
| CYB5R4 | 0.03343 | -0.3782 | 0.888 | 0.95 | 1 | MS.Trt_vs_MS.NoTrt_F |
| BCKDHB | 0.0335 | -0.5103 | 0.568 | 0.65 | 1 | MS.Trt_vs_MS.NoTrt_F |
| MMAB | 0.03351 | -0.6891 | 0.64 | 0.71 | 1 | MS.Trt_vs_MS.NoTrt_F |
| ATP6AP1 | 0.03356 | -0.3916 | 0.464 | 0.54 | 1 | MS.Trt_vs_MS.NoTrt_F |
| CHGB | 0.03359 | -0.562 | 0.399 | 0.49 | 1 | MS.Trt_vs_MS.NoTrt_F |
| MANBAL | 0.03361 | -1.362 | 0.083 | 0.14 | 1 | MS.Trt_vs_MS.NoTrt_F |
| DHPS | 0.03362 | 0.42325 | 0.284 | 0.39 | 1 | MS.Trt_vs_MS.NoTrt_F |
| PARP3 | 0.03368 | -1.2609 | 0.076 | 0.14 | 1 | MS.Trt_vs_MS.NoTrt_F |
| MKS1 | 0.03371 | 0.60552 | 0.079 | 0.14 | 1 | MS.Trt_vs_MS.NoTrt_F |

| | | | | | | |
|---|---|---|---|---|---|---|
| SEC61G | 0.03381 | 0.41558 | 0.32 | 0.42 | 1 | MS.Trt_vs_MS.NoTrt_F |
| KIDINS220 | 0.03389 | 0.25659 | 0.255 | 0.35 | 1 | MS.Trt_vs_MS.NoTrt_F |
| PLCD1 | 0.03393 | 1.30375 | 0.097 | 0.17 | 1 | MS.Trt_vs_MS.NoTrt_F |
| ERI1 | 0.03393 | 0.30478 | 0.428 | 0.54 | 1 | MS.Trt_vs_MS.NoTrt_F |
| RTN2 | 0.03397 | 0.3393 | 0.345 | 0.26 | 1 | MS.Trt_vs_MS.NoTrt_F |
| MTMR1 | 0.034 | 0.7985 | 0.18 | 0.27 | 1 | MS.Trt_vs_MS.NoTrt_F |
| SREBF1 | 0.03406 | -0.6064 | 0.068 | 0.13 | 1 | MS.Trt_vs_MS.NoTrt_F |
| BRD1 | 0.03419 | 0.84538 | 0.115 | 0.19 | 1 | MS.Trt_vs_MS.NoTrt_F |
| NDUFC2 | 0.0343 | 0.46426 | 0.572 | 0.67 | 1 | MS.Trt_vs_MS.NoTrt_F |
| FAM102B | 0.03434 | 0.34492 | 0.223 | 0.32 | 1 | MS.Trt_vs_MS.NoTrt_F |
| PXK | 0.03434 | 0.84412 | 0.119 | 0.2 | 1 | MS.Trt_vs_MS.NoTrt_F |
| COX17 | 0.03435 | 0.37271 | 0.317 | 0.43 | 1 | MS.Trt_vs_MS.NoTrt_F |
| ZNF821 | 0.03437 | 0.78727 | 0.086 | 0.16 | 1 | MS.Trt_vs_MS.NoTrt_F |
| PKN2 | 0.03441 | 1.22963 | 0.313 | 0.43 | 1 | MS.Trt_vs_MS.NoTrt_F |
| CRAT | 0.03443 | -0.458 | 0.068 | 0.13 | 1 | MS.Trt_vs_MS.NoTrt_F |
| GPM6B | 0.03453 | 0.47845 | 0.129 | 0.21 | 1 | MS.Trt_vs_MS.NoTrt_F |
| WDR55 | 0.03455 | -0.3506 | 0.676 | 0.76 | 1 | MS.Trt_vs_MS.NoTrt_F |
| PLCH2 | 0.03456 | 1.23596 | 0.083 | 0.15 | 1 | MS.Trt_vs_MS.NoTrt_F |
| FCHSD1 | 0.03473 | 0.29992 | 0.299 | 0.43 | 1 | MS.Trt_vs_MS.NoTrt_F |
| KIF21B | 0.03475 | 0.41892 | 0.432 | 0.55 | 1 | MS.Trt_vs_MS.NoTrt_F |
| ATP11C | 0.03479 | 0.68155 | 0.072 | 0.13 | 1 | MS.Trt_vs_MS.NoTrt_F |
| ZNF362 | 0.03493 | 0.65207 | 0.147 | 0.23 | 1 | MS.Trt_vs_MS.NoTrt_F |
| DDX42 | 0.03495 | 0.29417 | 0.378 | 0.49 | 1 | MS.Trt_vs_MS.NoTrt_F |
| STK24 | 0.03495 | 0.6552 | 0.288 | 0.41 | 1 | MS.Trt_vs_MS.NoTrt_F |
| DBNDD2 | 0.03498 | -0.5183 | 0.065 | 0.12 | 1 | MS.Trt_vs_MS.NoTrt_F |
| HDAC1 | 0.03498 | 0.29173 | 0.288 | 0.41 | 1 | MS.Trt_vs_MS.NoTrt_F |
| CDC42SE1 | 0.03507 | -0.4956 | 0.511 | 0.57 | 1 | MS.Trt_vs_MS.NoTrt_F |
| FSD1L | 0.03511 | -0.4752 | 0.191 | 0.27 | 1 | MS.Trt_vs_MS.NoTrt_F |
| CTD-2568A1 | 0.03521 | -0.2504 | 0.388 | 0.47 | 1 | MS.Trt_vs_MS.NoTrt_F |
| PHRF1 | 0.03523 | 0.54264 | 0.158 | 0.25 | 1 | MS.Trt_vs_MS.NoTrt_F |
| SRR | 0.03536 | 1.75542 | 0.068 | 0.13 | 1 | MS.Trt_vs_MS.NoTrt_F |
| FAM92A1 | 0.03556 | -0.2795 | 0.27 | 0.39 | 1 | MS.Trt_vs_MS.NoTrt_F |
| ODF3B | 0.03557 | -0.3122 | 0.068 | 0.13 | 1 | MS.Trt_vs_MS.NoTrt_F |
| PSMB3 | 0.03557 | 0.2617 | 0.302 | 0.44 | 1 | MS.Trt_vs_MS.NoTrt_F |
| CDC16 | 0.03562 | 0.78824 | 0.18 | 0.27 | 1 | MS.Trt_vs_MS.NoTrt_F |
| CAPZA1 | 0.03564 | 0.73096 | 0.856 | 0.97 | 1 | MS.Trt_vs_MS.NoTrt_F |
| MTHFD2 | 0.03566 | -0.7148 | 0.683 | 0.75 | 1 | MS.Trt_vs_MS.NoTrt_F |
| NIPAL3 | 0.0358 | 0.25716 | 0.755 | 0.83 | 1 | MS.Trt_vs_MS.NoTrt_F |
| TNFRSF10D | 0.03587 | 0.33135 | 0.133 | 0.21 | 1 | MS.Trt_vs_MS.NoTrt_F |
| TRMT1L | 0.03597 | 0.37846 | 0.101 | 0.17 | 1 | MS.Trt_vs_MS.NoTrt_F |

| | | | | | | |
|---|---|---|---|---|---|---|
| ZXDB | 0.03607 | 0.4398 | 0.072 | 0.13 | 1 | MS.Trt_vs_MS.NoTrt_F |
| PDZD7 | 0.03607 | -0.3044 | 0.41 | 0.54 | 1 | MS.Trt_vs_MS.NoTrt_F |
| FAM120A | 0.03612 | 0.69343 | 0.162 | 0.25 | 1 | MS.Trt_vs_MS.NoTrt_F |
| DDX20 | 0.03624 | 1.07514 | 0.101 | 0.17 | 1 | MS.Trt_vs_MS.NoTrt_F |
| PDLIM7 | 0.03642 | -0.5297 | 0.068 | 0.13 | 1 | MS.Trt_vs_MS.NoTrt_F |
| ZNF707 | 0.03647 | 0.4003 | 0.727 | 0.79 | 1 | MS.Trt_vs_MS.NoTrt_F |
| GFM2 | 0.03693 | -0.9729 | 0.259 | 0.34 | 1 | MS.Trt_vs_MS.NoTrt_F |
| ALG11 | 0.03693 | 0.49009 | 0.813 | 0.89 | 1 | MS.Trt_vs_MS.NoTrt_F |
| C3orf18 | 0.03694 | 0.41615 | 0.266 | 0.35 | 1 | MS.Trt_vs_MS.NoTrt_F |
| UNC50 | 0.03698 | 0.65617 | 0.126 | 0.2 | 1 | MS.Trt_vs_MS.NoTrt_F |
| SLC25A34 | 0.03701 | 0.77205 | 0.554 | 0.67 | 1 | MS.Trt_vs_MS.NoTrt_F |
| CRYBG3 | 0.03705 | 0.80614 | 0.097 | 0.17 | 1 | MS.Trt_vs_MS.NoTrt_F |
| TSPAN4 | 0.03706 | -0.7413 | 0.058 | 0.11 | 1 | MS.Trt_vs_MS.NoTrt_F |
| APOBEC2 | 0.03709 | 0.74081 | 0.076 | 0.14 | 1 | MS.Trt_vs_MS.NoTrt_F |
| IER2 | 0.0371 | 0.33424 | 0.381 | 0.55 | 1 | MS.Trt_vs_MS.NoTrt_F |
| ZNF211 | 0.03715 | 0.95552 | 0.205 | 0.3 | 1 | MS.Trt_vs_MS.NoTrt_F |
| NLGN3 | 0.03731 | 0.4304 | 0.086 | 0.15 | 1 | MS.Trt_vs_MS.NoTrt_F |
| SNRNP48 | 0.03734 | -0.9475 | 0.223 | 0.3 | 1 | MS.Trt_vs_MS.NoTrt_F |
| SRPK2 | 0.03745 | 0.29225 | 0.317 | 0.44 | 1 | MS.Trt_vs_MS.NoTrt_F |
| CCT3 | 0.03746 | -0.313 | 0.547 | 0.63 | 1 | MS.Trt_vs_MS.NoTrt_F |
| SLC25A29 | 0.03746 | -0.4576 | 0.353 | 0.45 | 1 | MS.Trt_vs_MS.NoTrt_F |
| DUS3L | 0.03758 | 0.90486 | 0.151 | 0.24 | 1 | MS.Trt_vs_MS.NoTrt_F |
| NDUFS4 | 0.0376 | 0.88385 | 0.302 | 0.4 | 1 | MS.Trt_vs_MS.NoTrt_F |
| FAM83G | 0.03769 | 0.44003 | 0.41 | 0.52 | 1 | MS.Trt_vs_MS.NoTrt_F |
| BLVRB | 0.0377 | 0.55709 | 0.414 | 0.31 | 1 | MS.Trt_vs_MS.NoTrt_F |
| GOLPH3L | 0.03774 | -0.3762 | 0.201 | 0.28 | 1 | MS.Trt_vs_MS.NoTrt_F |
| NR3C1 | 0.03779 | 0.35921 | 0.209 | 0.3 | 1 | MS.Trt_vs_MS.NoTrt_F |
| CD320 | 0.03788 | 0.69702 | 0.662 | 0.8 | 1 | MS.Trt_vs_MS.NoTrt_F |
| BCO2 | 0.03789 | -0.3109 | 0.511 | 0.58 | 1 | MS.Trt_vs_MS.NoTrt_F |
| KDELR1 | 0.03791 | -0.2963 | 0.252 | 0.35 | 1 | MS.Trt_vs_MS.NoTrt_F |
| MTA2 | 0.03796 | -1.1555 | 0.302 | 0.43 | 1 | MS.Trt_vs_MS.NoTrt_F |
| GSDMB | 0.03801 | 0.4995 | 0.899 | 0.91 | 1 | MS.Trt_vs_MS.NoTrt_F |
| TSEN15 | 0.03803 | 0.55705 | 0.209 | 0.31 | 1 | MS.Trt_vs_MS.NoTrt_F |
| HLA-DRA | 0.03804 | 0.3091 | 0.173 | 0.26 | 1 | MS.Trt_vs_MS.NoTrt_F |
| ALS2CL | 0.03823 | 0.2676 | 0.151 | 0.23 | 1 | MS.Trt_vs_MS.NoTrt_F |
| GPX7 | 0.03832 | 0.50358 | 0.09 | 0.16 | 1 | MS.Trt_vs_MS.NoTrt_F |
| MAPKBP1 | 0.03833 | 0.42168 | 0.414 | 0.51 | 1 | MS.Trt_vs_MS.NoTrt_F |
| KLHL12 | 0.03839 | 0.45787 | 0.108 | 0.18 | 1 | MS.Trt_vs_MS.NoTrt_F |
| HABP4 | 0.03844 | 0.97784 | 0.165 | 0.24 | 1 | MS.Trt_vs_MS.NoTrt_F |
| ZNF492 | 0.0385 | 0.3262 | 0.622 | 0.5 | 1 | MS.Trt_vs_MS.NoTrt_F |

| | | | | | | |
|---|---|---|---|---|---|---|
| ZNF571 | 0.03856 | -0.5229 | 0.41 | 0.54 | 1 | MS.Trt_vs_MS.NoTrt_F |
| NPHS1 | 0.03858 | 0.28464 | 0.464 | 0.4 | 1 | MS.Trt_vs_MS.NoTrt_F |
| PNN | 0.03878 | 0.54307 | 0.252 | 0.37 | 1 | MS.Trt_vs_MS.NoTrt_F |
| ANKEF1 | 0.03884 | -0.3568 | 0.201 | 0.29 | 1 | MS.Trt_vs_MS.NoTrt_F |
| PPIL4 | 0.03886 | 0.41947 | 0.234 | 0.34 | 1 | MS.Trt_vs_MS.NoTrt_F |
| PRDX5 | 0.03887 | 0.52686 | 0.245 | 0.36 | 1 | MS.Trt_vs_MS.NoTrt_F |
| FAM219B | 0.03893 | 0.85655 | 0.421 | 0.53 | 1 | MS.Trt_vs_MS.NoTrt_F |
| C11orf71 | 0.03895 | 0.96532 | 0.094 | 0.16 | 1 | MS.Trt_vs_MS.NoTrt_F |
| RNPEPL1 | 0.03898 | 0.91171 | 0.162 | 0.26 | 1 | MS.Trt_vs_MS.NoTrt_F |
| TGDS | 0.03904 | 0.42645 | 0.119 | 0.19 | 1 | MS.Trt_vs_MS.NoTrt_F |
| CCDC153 | 0.0391 | 1.28956 | 0.245 | 0.35 | 1 | MS.Trt_vs_MS.NoTrt_F |
| TMEM5 | 0.03915 | 1.25046 | 0.496 | 0.6 | 1 | MS.Trt_vs_MS.NoTrt_F |
| LYRM7 | 0.03917 | 0.48909 | 0.917 | 0.9 | 1 | MS.Trt_vs_MS.NoTrt_F |
| NRP1 | 0.03917 | 0.90424 | 0.302 | 0.41 | 1 | MS.Trt_vs_MS.NoTrt_F |
| PILRB | 0.03926 | 0.43597 | 0.45 | 0.54 | 1 | MS.Trt_vs_MS.NoTrt_F |
| OSGEP | 0.03931 | -0.4145 | 0.576 | 0.68 | 1 | MS.Trt_vs_MS.NoTrt_F |
| SF3B6 | 0.03939 | 0.49217 | 0.198 | 0.3 | 1 | MS.Trt_vs_MS.NoTrt_F |
| RBAK | 0.03957 | -1.6884 | 0.054 | 0.11 | 1 | MS.Trt_vs_MS.NoTrt_F |
| ZSCAN16 | 0.03977 | 0.48592 | 0.065 | 0.12 | 1 | MS.Trt_vs_MS.NoTrt_F |
| SMAD9 | 0.03985 | 0.70698 | 0.331 | 0.44 | 1 | MS.Trt_vs_MS.NoTrt_F |
| UBXN8 | 0.03991 | 0.73263 | 0.198 | 0.28 | 1 | MS.Trt_vs_MS.NoTrt_F |
| IFITM2 | 0.03997 | 0.65777 | 0.849 | 0.81 | 1 | MS.Trt_vs_MS.NoTrt_F |
| TFB2M | 0.03998 | 1.30173 | 0.079 | 0.14 | 1 | MS.Trt_vs_MS.NoTrt_F |
| ZMYM1 | 0.04001 | 0.4567 | 0.547 | 0.66 | 1 | MS.Trt_vs_MS.NoTrt_F |
| CCZ1 | 0.04001 | -0.8721 | 0.673 | 0.77 | 1 | MS.Trt_vs_MS.NoTrt_F |
| ISOC2 | 0.04002 | 0.40783 | 0.068 | 0.13 | 1 | MS.Trt_vs_MS.NoTrt_F |
| AC013461.1 | 0.04004 | 0.57891 | 0.831 | 0.77 | 1 | MS.Trt_vs_MS.NoTrt_F |
| RP11-234B2 | 0.04005 | -0.3008 | 0.058 | 0.11 | 1 | MS.Trt_vs_MS.NoTrt_F |
| STX6 | 0.04005 | 0.26704 | 0.367 | 0.46 | 1 | MS.Trt_vs_MS.NoTrt_F |
| LRRC56 | 0.04006 | -0.5007 | 0.525 | 0.38 | 1 | MS.Trt_vs_MS.NoTrt_F |
| CBWD7 | 0.04012 | 0.34975 | 0.478 | 0.59 | 1 | MS.Trt_vs_MS.NoTrt_F |
| ARMC1 | 0.0402 | 0.53727 | 0.144 | 0.22 | 1 | MS.Trt_vs_MS.NoTrt_F |
| LUC7L3 | 0.04023 | 0.29108 | 0.529 | 0.7 | 1 | MS.Trt_vs_MS.NoTrt_F |
| ISG20 | 0.04023 | 0.26527 | 0.612 | 0.67 | 1 | MS.Trt_vs_MS.NoTrt_F |
| ANAPC15 | 0.04024 | 0.35572 | 0.763 | 0.85 | 1 | MS.Trt_vs_MS.NoTrt_F |
| TICAM1 | 0.04028 | -0.479 | 0.09 | 0.16 | 1 | MS.Trt_vs_MS.NoTrt_F |
| TMEM167B | 0.04042 | -0.6065 | 0.773 | 0.82 | 1 | MS.Trt_vs_MS.NoTrt_F |
| RPS21 | 0.04048 | 0.48408 | 0.435 | 0.64 | 1 | MS.Trt_vs_MS.NoTrt_F |
| ZNF790 | 0.04049 | -0.2572 | 0.468 | 0.6 | 1 | MS.Trt_vs_MS.NoTrt_F |
| PLEKHA4 | 0.04049 | -0.3193 | 0.086 | 0.15 | 1 | MS.Trt_vs_MS.NoTrt_F |

| | | | | | | |
|---|---|---|---|---|---|---|
| SPCS1 | 0.04049 | 0.28489 | 0.903 | 0.93 | 1 | MS.Trt_vs_MS.NoTrt_F |
| EMP1 | 0.0405 | 0.61423 | 0.162 | 0.25 | 1 | MS.Trt_vs_MS.NoTrt_F |
| C12orf29 | 0.04052 | 0.42902 | 0.223 | 0.32 | 1 | MS.Trt_vs_MS.NoTrt_F |
| STON1 | 0.04054 | -0.9005 | 0.335 | 0.43 | 1 | MS.Trt_vs_MS.NoTrt_F |
| TOP3B | 0.04055 | 0.32618 | 0.151 | 0.24 | 1 | MS.Trt_vs_MS.NoTrt_F |
| MTBP | 0.0406 | -0.6714 | 0.55 | 0.64 | 1 | MS.Trt_vs_MS.NoTrt_F |
| BNIP1 | 0.04069 | 1.46392 | 0.104 | 0.18 | 1 | MS.Trt_vs_MS.NoTrt_F |
| RAB15 | 0.04083 | 0.83489 | 0.058 | 0.11 | 1 | MS.Trt_vs_MS.NoTrt_F |
| KIF1B | 0.04096 | -0.7892 | 0.741 | 0.84 | 1 | MS.Trt_vs_MS.NoTrt_F |
| TMEM38A | 0.04097 | 0.2659 | 0.601 | 0.49 | 1 | MS.Trt_vs_MS.NoTrt_F |
| POP1 | 0.04099 | -0.2539 | 0.061 | 0.12 | 1 | MS.Trt_vs_MS.NoTrt_F |
| CUL4B | 0.041 | 0.67259 | 0.129 | 0.2 | 1 | MS.Trt_vs_MS.NoTrt_F |
| PRH1 | 0.04104 | 0.37467 | 0.115 | 0.19 | 1 | MS.Trt_vs_MS.NoTrt_F |
| CDK5 | 0.04115 | 0.35698 | 0.065 | 0.12 | 1 | MS.Trt_vs_MS.NoTrt_F |
| DHX30 | 0.04116 | -0.5023 | 0.633 | 0.69 | 1 | MS.Trt_vs_MS.NoTrt_F |
| TMEM209 | 0.04131 | 0.40345 | 0.752 | 0.81 | 1 | MS.Trt_vs_MS.NoTrt_F |
| ALKBH8 | 0.0414 | -0.5444 | 0.054 | 0.11 | 1 | MS.Trt_vs_MS.NoTrt_F |
| SLC38A9 | 0.04153 | 0.95459 | 0.331 | 0.45 | 1 | MS.Trt_vs_MS.NoTrt_F |
| MINPP1 | 0.04162 | -0.9367 | 0.097 | 0.16 | 1 | MS.Trt_vs_MS.NoTrt_F |
| PPME1 | 0.04165 | 0.32097 | 0.144 | 0.22 | 1 | MS.Trt_vs_MS.NoTrt_F |
| CBX4 | 0.04167 | -0.3944 | 0.083 | 0.14 | 1 | MS.Trt_vs_MS.NoTrt_F |
| ZBTB38 | 0.04167 | -0.2529 | 0.59 | 0.7 | 1 | MS.Trt_vs_MS.NoTrt_F |
| BMP8A | 0.04174 | -0.5354 | 0.601 | 0.71 | 1 | MS.Trt_vs_MS.NoTrt_F |
| FRMD5 | 0.04175 | 0.33243 | 0.061 | 0.12 | 1 | MS.Trt_vs_MS.NoTrt_F |
| MMD | 0.04177 | 0.32005 | 0.09 | 0.16 | 1 | MS.Trt_vs_MS.NoTrt_F |
| ADCK5 | 0.04182 | 0.64161 | 0.086 | 0.15 | 1 | MS.Trt_vs_MS.NoTrt_F |
| HAGH | 0.04186 | -0.3739 | 0.741 | 0.79 | 1 | MS.Trt_vs_MS.NoTrt_F |
| MTHFSD | 0.04204 | -0.509 | 0.09 | 0.16 | 1 | MS.Trt_vs_MS.NoTrt_F |
| BUB1 | 0.04206 | -0.5656 | 0.054 | 0.11 | 1 | MS.Trt_vs_MS.NoTrt_F |
| SLC39A9 | 0.04215 | -0.31 | 0.191 | 0.28 | 1 | MS.Trt_vs_MS.NoTrt_F |
| ZUFSP | 0.0422 | 0.65165 | 0.09 | 0.16 | 1 | MS.Trt_vs_MS.NoTrt_F |
| BNIP3L | 0.0422 | 0.4672 | 0.187 | 0.28 | 1 | MS.Trt_vs_MS.NoTrt_F |
| MAP7D1 | 0.04227 | -0.3843 | 0.46 | 0.58 | 1 | MS.Trt_vs_MS.NoTrt_F |
| SEC22A | 0.04234 | 0.30966 | 0.637 | 0.74 | 1 | MS.Trt_vs_MS.NoTrt_F |
| GABRP | 0.04237 | 0.36773 | 0.259 | 0.38 | 1 | MS.Trt_vs_MS.NoTrt_F |
| CKS2 | 0.0424 | -0.3493 | 0.076 | 0.13 | 1 | MS.Trt_vs_MS.NoTrt_F |
| COMT | 0.0424 | 0.32205 | 0.381 | 0.5 | 1 | MS.Trt_vs_MS.NoTrt_F |
| SCO1 | 0.04245 | -0.2638 | 0.788 | 0.85 | 1 | MS.Trt_vs_MS.NoTrt_F |
| RASSF3 | 0.04245 | 0.32641 | 0.583 | 0.78 | 1 | MS.Trt_vs_MS.NoTrt_F |
| ZNF550 | 0.04246 | 0.42551 | 0.288 | 0.38 | 1 | MS.Trt_vs_MS.NoTrt_F |

| | | | | | | |
|---|---|---|---|---|---|---|
| SMG8 | 0.04249 | 0.34695 | 0.133 | 0.21 | 1 | MS.Trt_vs_MS.NoTrt_F |
| SYTL2 | 0.04254 | 0.43495 | 0.737 | 0.74 | 1 | MS.Trt_vs_MS.NoTrt_F |
| TYK2 | 0.04262 | 0.51479 | 0.626 | 0.74 | 1 | MS.Trt_vs_MS.NoTrt_F |
| TSPAN5 | 0.04262 | -0.6312 | 0.615 | 0.67 | 1 | MS.Trt_vs_MS.NoTrt_F |
| EIF5B | 0.04267 | 0.46129 | 0.291 | 0.41 | 1 | MS.Trt_vs_MS.NoTrt_F |
| GSTP1 | 0.04271 | -0.6481 | 0.076 | 0.13 | 1 | MS.Trt_vs_MS.NoTrt_F |
| DUS4L | 0.04272 | 0.51148 | 0.122 | 0.2 | 1 | MS.Trt_vs_MS.NoTrt_F |
| SELM | 0.04281 | 0.39676 | 0.281 | 0.38 | 1 | MS.Trt_vs_MS.NoTrt_F |
| ABCC4 | 0.04284 | 0.51773 | 0.784 | 0.81 | 1 | MS.Trt_vs_MS.NoTrt_F |
| SLC25A23 | 0.04289 | 0.26402 | 0.486 | 0.6 | 1 | MS.Trt_vs_MS.NoTrt_F |
| KLHDC3 | 0.04295 | 0.6371 | 0.169 | 0.26 | 1 | MS.Trt_vs_MS.NoTrt_F |
| ZNF84 | 0.04296 | 0.84485 | 0.248 | 0.34 | 1 | MS.Trt_vs_MS.NoTrt_F |
| CEP57L1 | 0.04298 | -0.4615 | 0.529 | 0.61 | 1 | MS.Trt_vs_MS.NoTrt_F |
| GSTM3 | 0.04306 | 0.30943 | 0.471 | 0.6 | 1 | MS.Trt_vs_MS.NoTrt_F |
| LARP1B | 0.04306 | 0.41259 | 0.101 | 0.17 | 1 | MS.Trt_vs_MS.NoTrt_F |
| EIF3I | 0.04308 | -0.2912 | 0.536 | 0.6 | 1 | MS.Trt_vs_MS.NoTrt_F |
| IL18BP | 0.04321 | 0.75806 | 0.126 | 0.2 | 1 | MS.Trt_vs_MS.NoTrt_F |
| NFKB2 | 0.04334 | -1.5537 | 0.403 | 0.49 | 1 | MS.Trt_vs_MS.NoTrt_F |
| GPD2 | 0.04338 | 0.95298 | 0.086 | 0.15 | 1 | MS.Trt_vs_MS.NoTrt_F |
| MBTD1 | 0.04346 | 0.5941 | 0.421 | 0.53 | 1 | MS.Trt_vs_MS.NoTrt_F |
| RABGGTB | 0.04351 | -0.5338 | 0.277 | 0.38 | 1 | MS.Trt_vs_MS.NoTrt_F |
| LONP2 | 0.04355 | 0.33778 | 0.73 | 0.85 | 1 | MS.Trt_vs_MS.NoTrt_F |
| GARS | 0.04368 | 0.74657 | 0.187 | 0.29 | 1 | MS.Trt_vs_MS.NoTrt_F |
| PKNOX1 | 0.04372 | -0.3891 | 0.705 | 0.8 | 1 | MS.Trt_vs_MS.NoTrt_F |
| CLHC1 | 0.04376 | -0.4073 | 0.478 | 0.59 | 1 | MS.Trt_vs_MS.NoTrt_F |
| SLC2A1 | 0.04378 | 0.59118 | 0.133 | 0.21 | 1 | MS.Trt_vs_MS.NoTrt_F |
| C5orf22 | 0.04381 | 0.45762 | 0.288 | 0.39 | 1 | MS.Trt_vs_MS.NoTrt_F |
| SETDB2 | 0.04387 | 0.28409 | 0.849 | 0.92 | 1 | MS.Trt_vs_MS.NoTrt_F |
| PPP1R3E | 0.0439 | -1.01 | 0.248 | 0.33 | 1 | MS.Trt_vs_MS.NoTrt_F |
| LMBR1L | 0.04398 | -0.4284 | 0.705 | 0.82 | 1 | MS.Trt_vs_MS.NoTrt_F |
| FAM222B | 0.04407 | 0.74552 | 0.169 | 0.26 | 1 | MS.Trt_vs_MS.NoTrt_F |
| TAF11 | 0.04447 | 0.28826 | 0.453 | 0.58 | 1 | MS.Trt_vs_MS.NoTrt_F |
| TTPAL | 0.0445 | -0.2875 | 0.813 | 0.86 | 1 | MS.Trt_vs_MS.NoTrt_F |
| ERF | 0.04463 | 0.60372 | 0.104 | 0.18 | 1 | MS.Trt_vs_MS.NoTrt_F |
| PIK3CG | 0.04464 | 0.92047 | 0.295 | 0.41 | 1 | MS.Trt_vs_MS.NoTrt_F |
| PDK2 | 0.04466 | 0.32054 | 0.281 | 0.39 | 1 | MS.Trt_vs_MS.NoTrt_F |
| RP11-458D2 | 0.04471 | -0.3325 | 0.126 | 0.2 | 1 | MS.Trt_vs_MS.NoTrt_F |
| ZNF148 | 0.04486 | 0.40708 | 0.302 | 0.42 | 1 | MS.Trt_vs_MS.NoTrt_F |
| FRG1 | 0.04492 | 0.45292 | 0.396 | 0.48 | 1 | MS.Trt_vs_MS.NoTrt_F |
| NAA30 | 0.04508 | -1.1416 | 0.464 | 0.54 | 1 | MS.Trt_vs_MS.NoTrt_F |

| | | | | | | |
|---|---|---|---|---|---|---|
| SELP | 0.04508 | 0.95137 | 0.09 | 0.16 | 1 | MS.Trt_vs_MS.NoTrt_F |
| DDX47 | 0.04514 | 0.33742 | 0.471 | 0.56 | 1 | MS.Trt_vs_MS.NoTrt_F |
| RAB43 | 0.04526 | 0.33621 | 0.219 | 0.31 | 1 | MS.Trt_vs_MS.NoTrt_F |
| PGAP3 | 0.04527 | -0.5338 | 0.126 | 0.2 | 1 | MS.Trt_vs_MS.NoTrt_F |
| ZBED6 | 0.0453 | 0.47484 | 0.079 | 0.14 | 1 | MS.Trt_vs_MS.NoTrt_F |
| YIPF5 | 0.04531 | -0.9614 | 0.496 | 0.55 | 1 | MS.Trt_vs_MS.NoTrt_F |
| PGAP1 | 0.04533 | 0.29606 | 0.367 | 0.46 | 1 | MS.Trt_vs_MS.NoTrt_F |
| HBP1 | 0.04541 | -0.3117 | 0.799 | 0.87 | 1 | MS.Trt_vs_MS.NoTrt_F |
| NUDCD1 | 0.04541 | 0.44258 | 0.083 | 0.14 | 1 | MS.Trt_vs_MS.NoTrt_F |
| NDST2 | 0.04542 | 0.74198 | 0.094 | 0.16 | 1 | MS.Trt_vs_MS.NoTrt_F |
| WDR45 | 0.04551 | -0.4278 | 0.579 | 0.69 | 1 | MS.Trt_vs_MS.NoTrt_F |
| KRBA2 | 0.04557 | -1.1968 | 0.363 | 0.44 | 1 | MS.Trt_vs_MS.NoTrt_F |
| MAD2L2 | 0.04557 | 0.40469 | 0.137 | 0.22 | 1 | MS.Trt_vs_MS.NoTrt_F |
| SHTN1 | 0.04564 | 0.35196 | 0.119 | 0.19 | 1 | MS.Trt_vs_MS.NoTrt_F |
| MAST4 | 0.0458 | -0.5892 | 0.572 | 0.65 | 1 | MS.Trt_vs_MS.NoTrt_F |
| USP38 | 0.04593 | -0.3326 | 0.288 | 0.39 | 1 | MS.Trt_vs_MS.NoTrt_F |
| ALPK1 | 0.0461 | 0.34547 | 0.698 | 0.77 | 1 | MS.Trt_vs_MS.NoTrt_F |
| PIGU | 0.04617 | 0.76321 | 0.115 | 0.19 | 1 | MS.Trt_vs_MS.NoTrt_F |
| ANKRD49 | 0.04618 | 0.63697 | 0.183 | 0.27 | 1 | MS.Trt_vs_MS.NoTrt_F |
| DIP2C | 0.04621 | -0.45 | 0.086 | 0.15 | 1 | MS.Trt_vs_MS.NoTrt_F |
| PTEN | 0.04621 | 0.72628 | 0.518 | 0.66 | 1 | MS.Trt_vs_MS.NoTrt_F |
| AOAH | 0.04621 | 1.15084 | 0.363 | 0.47 | 1 | MS.Trt_vs_MS.NoTrt_F |
| PSMD11 | 0.04629 | 0.29245 | 0.482 | 0.59 | 1 | MS.Trt_vs_MS.NoTrt_F |
| CRELD1 | 0.04648 | 0.31226 | 0.126 | 0.19 | 1 | MS.Trt_vs_MS.NoTrt_F |
| ATXN1 | 0.04652 | 0.4144 | 0.309 | 0.42 | 1 | MS.Trt_vs_MS.NoTrt_F |
| CDADC1 | 0.04658 | 0.27165 | 0.083 | 0.14 | 1 | MS.Trt_vs_MS.NoTrt_F |
| C12orf66 | 0.04666 | -0.6357 | 0.353 | 0.44 | 1 | MS.Trt_vs_MS.NoTrt_F |
| PRKX | 0.04672 | 0.51489 | 0.54 | 0.68 | 1 | MS.Trt_vs_MS.NoTrt_F |
| DNAH17 | 0.04675 | 0.32771 | 0.313 | 0.4 | 1 | MS.Trt_vs_MS.NoTrt_F |
| CHST7 | 0.04686 | 1.18981 | 0.09 | 0.16 | 1 | MS.Trt_vs_MS.NoTrt_F |
| MUTYH | 0.04688 | -0.8201 | 0.507 | 0.57 | 1 | MS.Trt_vs_MS.NoTrt_F |
| BMI1 | 0.0469 | 1.24854 | 0.083 | 0.14 | 1 | MS.Trt_vs_MS.NoTrt_F |
| AHSA2 | 0.0469 | 0.45209 | 0.237 | 0.34 | 1 | MS.Trt_vs_MS.NoTrt_F |
| TNF | 0.04699 | 0.36934 | 0.151 | 0.23 | 1 | MS.Trt_vs_MS.NoTrt_F |
| FAM114A2 | 0.04705 | 0.70067 | 0.108 | 0.18 | 1 | MS.Trt_vs_MS.NoTrt_F |
| STK17B | 0.04708 | 0.5719 | 0.942 | 0.97 | 1 | MS.Trt_vs_MS.NoTrt_F |
| MRPS7 | 0.04713 | -0.4342 | 0.522 | 0.6 | 1 | MS.Trt_vs_MS.NoTrt_F |
| SHANK1 | 0.04715 | -0.2743 | 0.302 | 0.38 | 1 | MS.Trt_vs_MS.NoTrt_F |
| SREK1 | 0.04743 | 0.43443 | 0.525 | 0.62 | 1 | MS.Trt_vs_MS.NoTrt_F |
| COX6C | 0.04746 | 0.53413 | 0.486 | 0.62 | 1 | MS.Trt_vs_MS.NoTrt_F |

| | | | | | | |
|---|---|---|---|---|---|---|
| SFT2D1 | 0.04751 | 0.57038 | 0.414 | 0.49 | 1 | MS.Trt_vs_MS.NoTrt_F |
| PRMT5 | 0.04756 | 0.94913 | 0.09 | 0.16 | 1 | MS.Trt_vs_MS.NoTrt_F |
| WNT7A | 0.04762 | 0.92782 | 0.072 | 0.13 | 1 | MS.Trt_vs_MS.NoTrt_F |
| XPA | 0.0479 | 0.64606 | 0.147 | 0.23 | 1 | MS.Trt_vs_MS.NoTrt_F |
| STX11 | 0.04791 | 0.75097 | 0.079 | 0.14 | 1 | MS.Trt_vs_MS.NoTrt_F |
| POLN | 0.04811 | -0.4082 | 0.615 | 0.72 | 1 | MS.Trt_vs_MS.NoTrt_F |
| STARD3NL | 0.0482 | -0.2666 | 0.673 | 0.74 | 1 | MS.Trt_vs_MS.NoTrt_F |
| ABCD2 | 0.04837 | 0.54878 | 0.119 | 0.19 | 1 | MS.Trt_vs_MS.NoTrt_F |
| CYBA | 0.04837 | 0.35718 | 0.299 | 0.39 | 1 | MS.Trt_vs_MS.NoTrt_F |
| LIPT1 | 0.04843 | -0.35 | 0.676 | 0.78 | 1 | MS.Trt_vs_MS.NoTrt_F |
| BUB3 | 0.04851 | 0.30272 | 0.662 | 0.8 | 1 | MS.Trt_vs_MS.NoTrt_F |
| BAG6 | 0.04854 | 0.31518 | 0.554 | 0.59 | 1 | MS.Trt_vs_MS.NoTrt_F |
| ZNF596 | 0.04861 | 0.63426 | 0.45 | 0.52 | 1 | MS.Trt_vs_MS.NoTrt_F |
| SNAPC2 | 0.04865 | 0.285 | 0.083 | 0.14 | 1 | MS.Trt_vs_MS.NoTrt_F |
| ZBED1 | 0.04876 | -0.8472 | 0.468 | 0.55 | 1 | MS.Trt_vs_MS.NoTrt_F |
| USP9Y | 0.04878 | -0.9307 | 0.27 | 0.35 | 1 | MS.Trt_vs_MS.NoTrt_F |
| PSMB8 | 0.04886 | 0.33797 | 0.388 | 0.52 | 1 | MS.Trt_vs_MS.NoTrt_F |
| ZBTB21 | 0.04887 | 0.64254 | 0.147 | 0.23 | 1 | MS.Trt_vs_MS.NoTrt_F |
| MMP24 | 0.04892 | 0.70001 | 0.101 | 0.17 | 1 | MS.Trt_vs_MS.NoTrt_F |
| STK35 | 0.04893 | 1.50147 | 0.09 | 0.16 | 1 | MS.Trt_vs_MS.NoTrt_F |
| BTBD9 | 0.04918 | 0.63245 | 0.568 | 0.63 | 1 | MS.Trt_vs_MS.NoTrt_F |
| PAK1 | 0.04919 | 0.61698 | 0.173 | 0.26 | 1 | MS.Trt_vs_MS.NoTrt_F |
| C10orf67 | 0.04926 | 0.31834 | 0.288 | 0.38 | 1 | MS.Trt_vs_MS.NoTrt_F |
| CCR3 | 0.04938 | 0.61407 | 0.083 | 0.14 | 1 | MS.Trt_vs_MS.NoTrt_F |
| FAM188A | 0.04942 | 1.03622 | 0.115 | 0.18 | 1 | MS.Trt_vs_MS.NoTrt_F |
| UCK1 | 0.0495 | 0.79172 | 0.076 | 0.13 | 1 | MS.Trt_vs_MS.NoTrt_F |
| TTC5 | 0.0495 | -0.5282 | 0.511 | 0.61 | 1 | MS.Trt_vs_MS.NoTrt_F |
| TMEM179B | 0.0498 | 0.39221 | 0.216 | 0.32 | 1 | MS.Trt_vs_MS.NoTrt_F |
| RAP2A | 0.04987 | -0.6743 | 0.068 | 0.12 | 1 | MS.Trt_vs_MS.NoTrt_F |
| KPTN | 0.04987 | -0.6221 | 0.198 | 0.28 | 1 | MS.Trt_vs_MS.NoTrt_F |
| CNEP1R1 | 0.04991 | 0.73512 | 0.583 | 0.66 | 1 | MS.Trt_vs_MS.NoTrt_F |
| GLUD1 | 0.05005 | 0.58163 | 0.248 | 0.35 | 1 | MS.Trt_vs_MS.NoTrt_F |
| USF1 | 0.05008 | -0.747 | 0.18 | 0.25 | 1 | MS.Trt_vs_MS.NoTrt_F |
| AP1B1 | 0.05014 | 0.54585 | 0.162 | 0.25 | 1 | MS.Trt_vs_MS.NoTrt_F |
| CBX6 | 0.05017 | 0.96397 | 0.601 | 0.65 | 1 | MS.Trt_vs_MS.NoTrt_F |
| TSPAN3 | 0.05018 | -0.4721 | 0.349 | 0.43 | 1 | MS.Trt_vs_MS.NoTrt_F |
| RSPH3 | 0.05021 | 0.4027 | 0.799 | 0.82 | 1 | MS.Trt_vs_MS.NoTrt_F |
| TMEM234 | 0.05022 | 0.66837 | 0.112 | 0.18 | 1 | MS.Trt_vs_MS.NoTrt_F |
| ADPRM | 0.05022 | 0.41598 | 0.101 | 0.17 | 1 | MS.Trt_vs_MS.NoTrt_F |
| USP1 | 0.05023 | 1.1491 | 0.151 | 0.23 | 1 | MS.Trt_vs_MS.NoTrt_F |

| | | | | | | |
|---|---|---|---|---|---|---|
| INPP1 | 0.05026 | 0.58063 | 0.079 | 0.14 | 1 | MS.Trt_vs_MS.NoTrt_F |
| ERMN | 0.05031 | 0.58981 | 0.23 | 0.32 | 1 | MS.Trt_vs_MS.NoTrt_F |
| MRPL49 | 0.05033 | -0.3807 | 0.45 | 0.53 | 1 | MS.Trt_vs_MS.NoTrt_F |
| FAH | 0.05036 | 0.30226 | 0.122 | 0.19 | 1 | MS.Trt_vs_MS.NoTrt_F |
| NEIL2 | 0.05044 | 0.9637 | 0.119 | 0.19 | 1 | MS.Trt_vs_MS.NoTrt_F |
| EXOC3 | 0.05054 | -0.2744 | 0.216 | 0.3 | 1 | MS.Trt_vs_MS.NoTrt_F |
| GPAA1 | 0.05054 | 0.59382 | 0.158 | 0.24 | 1 | MS.Trt_vs_MS.NoTrt_F |
| ATMIN | 0.05058 | 0.2691 | 0.27 | 0.35 | 1 | MS.Trt_vs_MS.NoTrt_F |
| 8-Mar | 0.0507 | 0.48139 | 0.342 | 0.46 | 1 | MS.Trt_vs_MS.NoTrt_F |
| TMEM14C | 0.05076 | 0.61254 | 0.129 | 0.2 | 1 | MS.Trt_vs_MS.NoTrt_F |
| PLPP1 | 0.05085 | 0.62705 | 0.101 | 0.17 | 1 | MS.Trt_vs_MS.NoTrt_F |
| SNRNP27 | 0.05091 | 0.79286 | 0.234 | 0.34 | 1 | MS.Trt_vs_MS.NoTrt_F |
| SBK1 | 0.05092 | -0.7929 | 0.169 | 0.24 | 1 | MS.Trt_vs_MS.NoTrt_F |
| CD99L2 | 0.05099 | 0.70112 | 0.129 | 0.2 | 1 | MS.Trt_vs_MS.NoTrt_F |
| TMEM11 | 0.05099 | 0.81888 | 0.129 | 0.21 | 1 | MS.Trt_vs_MS.NoTrt_F |
| CCNG2 | 0.05106 | 0.30135 | 0.338 | 0.41 | 1 | MS.Trt_vs_MS.NoTrt_F |
| UPRT | 0.05113 | 0.91065 | 0.083 | 0.14 | 1 | MS.Trt_vs_MS.NoTrt_F |
| RAB6A | 0.05115 | 0.35174 | 0.36 | 0.49 | 1 | MS.Trt_vs_MS.NoTrt_F |
| TRMT6 | 0.05115 | 0.92252 | 0.137 | 0.22 | 1 | MS.Trt_vs_MS.NoTrt_F |
| KLHDC1 | 0.0514 | -0.2545 | 0.309 | 0.4 | 1 | MS.Trt_vs_MS.NoTrt_F |
| LMO7 | 0.05145 | -0.389 | 0.277 | 0.37 | 1 | MS.Trt_vs_MS.NoTrt_F |
| UBA5 | 0.0515 | 1.0706 | 0.147 | 0.23 | 1 | MS.Trt_vs_MS.NoTrt_F |
| CD63 | 0.05161 | 0.30365 | 0.187 | 0.27 | 1 | MS.Trt_vs_MS.NoTrt_F |
| MEMO1 | 0.05181 | 0.46477 | 0.36 | 0.45 | 1 | MS.Trt_vs_MS.NoTrt_F |
| LAPTM4A | 0.05181 | 0.33461 | 0.198 | 0.28 | 1 | MS.Trt_vs_MS.NoTrt_F |
| NDC80 | 0.05199 | 0.26059 | 0.662 | 0.77 | 1 | MS.Trt_vs_MS.NoTrt_F |
| BBS4 | 0.05219 | -0.6442 | 0.126 | 0.19 | 1 | MS.Trt_vs_MS.NoTrt_F |
| DNAJC5 | 0.05221 | 0.88441 | 0.097 | 0.16 | 1 | MS.Trt_vs_MS.NoTrt_F |
| NR2C2 | 0.05234 | 0.4535 | 0.622 | 0.72 | 1 | MS.Trt_vs_MS.NoTrt_F |
| ZNF570 | 0.05236 | 0.59435 | 0.791 | 0.88 | 1 | MS.Trt_vs_MS.NoTrt_F |
| FBRS | 0.05236 | 0.45511 | 0.176 | 0.27 | 1 | MS.Trt_vs_MS.NoTrt_F |
| ADTRP | 0.05238 | -0.4821 | 0.342 | 0.43 | 1 | MS.Trt_vs_MS.NoTrt_F |
| BLM | 0.05239 | 0.30841 | 0.27 | 0.36 | 1 | MS.Trt_vs_MS.NoTrt_F |
| ATP6AP1L | 0.05241 | -0.4384 | 0.453 | 0.55 | 1 | MS.Trt_vs_MS.NoTrt_F |
| FN3KRP | 0.05244 | 0.86047 | 0.112 | 0.18 | 1 | MS.Trt_vs_MS.NoTrt_F |
| CRTC1 | 0.05248 | 0.38135 | 0.245 | 0.35 | 1 | MS.Trt_vs_MS.NoTrt_F |
| SLC10A3 | 0.0525 | 0.88479 | 0.097 | 0.16 | 1 | MS.Trt_vs_MS.NoTrt_F |
| TP53INP1 | 0.05252 | 0.43396 | 0.353 | 0.48 | 1 | MS.Trt_vs_MS.NoTrt_F |
| TOMM5 | 0.05257 | 0.51109 | 0.219 | 0.32 | 1 | MS.Trt_vs_MS.NoTrt_F |
| CAD | 0.0526 | 0.77157 | 0.734 | 0.7 | 1 | MS.Trt_vs_MS.NoTrt_F |

| | | | | | | |
|---|---|---|---|---|---|---|
| PIGH | 0.0527 | 0.77557 | 0.561 | 0.68 | 1 | MS.Trt_vs_MS.NoTrt_F |
| PPT1 | 0.05285 | 0.50861 | 0.234 | 0.34 | 1 | MS.Trt_vs_MS.NoTrt_F |
| DSTYK | 0.053 | 0.5569 | 0.799 | 0.9 | 1 | MS.Trt_vs_MS.NoTrt_F |
| ZNF473 | 0.05302 | 1.34771 | 0.079 | 0.14 | 1 | MS.Trt_vs_MS.NoTrt_F |
| PLEKHA1 | 0.05303 | -0.3196 | 0.896 | 0.95 | 1 | MS.Trt_vs_MS.NoTrt_F |
| ORAI1 | 0.05304 | 0.60982 | 0.097 | 0.16 | 1 | MS.Trt_vs_MS.NoTrt_F |
| ACBD3 | 0.05305 | -0.3994 | 0.072 | 0.13 | 1 | MS.Trt_vs_MS.NoTrt_F |
| SMAD4 | 0.05307 | -0.3717 | 0.442 | 0.53 | 1 | MS.Trt_vs_MS.NoTrt_F |
| GNLY | 0.0532 | 2.38036 | 0.252 | 0.34 | 1 | MS.Trt_vs_MS.NoTrt_F |
| RHOT2 | 0.05333 | 0.31413 | 0.281 | 0.4 | 1 | MS.Trt_vs_MS.NoTrt_F |
| PSMD1 | 0.0535 | 0.49217 | 0.209 | 0.31 | 1 | MS.Trt_vs_MS.NoTrt_F |
| ZNF277 | 0.05351 | 0.45009 | 0.504 | 0.61 | 1 | MS.Trt_vs_MS.NoTrt_F |
| ZNF75D | 0.05355 | 0.64176 | 0.09 | 0.15 | 1 | MS.Trt_vs_MS.NoTrt_F |
| SLC5A5 | 0.05355 | -0.8786 | 0.432 | 0.5 | 1 | MS.Trt_vs_MS.NoTrt_F |
| C9orf142 | 0.05366 | 0.2615 | 0.263 | 0.38 | 1 | MS.Trt_vs_MS.NoTrt_F |
| ZNF816 | 0.0537 | 0.88635 | 0.428 | 0.52 | 1 | MS.Trt_vs_MS.NoTrt_F |
| NRROS | 0.05372 | 0.26402 | 0.126 | 0.19 | 1 | MS.Trt_vs_MS.NoTrt_F |
| ACAT1 | 0.05377 | 0.41948 | 0.604 | 0.72 | 1 | MS.Trt_vs_MS.NoTrt_F |
| COL4A3BP | 0.05378 | 0.62018 | 0.155 | 0.24 | 1 | MS.Trt_vs_MS.NoTrt_F |
| FUBP1 | 0.05384 | 0.61915 | 0.41 | 0.54 | 1 | MS.Trt_vs_MS.NoTrt_F |
| ZNF721 | 0.05393 | 0.348 | 0.371 | 0.46 | 1 | MS.Trt_vs_MS.NoTrt_F |
| ZNF100 | 0.05397 | 0.62906 | 0.838 | 0.83 | 1 | MS.Trt_vs_MS.NoTrt_F |
| ARHGAP19-! | 0.05402 | -0.3556 | 0.065 | 0.12 | 1 | MS.Trt_vs_MS.NoTrt_F |
| RORC | 0.05411 | 1.03833 | 0.122 | 0.19 | 1 | MS.Trt_vs_MS.NoTrt_F |
| ZNF880 | 0.05416 | 0.97446 | 0.5 | 0.56 | 1 | MS.Trt_vs_MS.NoTrt_F |
| C14orf169 | 0.05421 | 0.40236 | 0.108 | 0.18 | 1 | MS.Trt_vs_MS.NoTrt_F |
| NDUFV1 | 0.05424 | 0.32545 | 0.277 | 0.4 | 1 | MS.Trt_vs_MS.NoTrt_F |
| AHSA1 | 0.05425 | -0.4432 | 0.547 | 0.61 | 1 | MS.Trt_vs_MS.NoTrt_F |
| SHQ1 | 0.05425 | -0.4769 | 0.68 | 0.76 | 1 | MS.Trt_vs_MS.NoTrt_F |
| ZBED2 | 0.05427 | -0.2503 | 0.068 | 0.12 | 1 | MS.Trt_vs_MS.NoTrt_F |
| PLK1 | 0.05427 | 0.36466 | 0.457 | 0.53 | 1 | MS.Trt_vs_MS.NoTrt_F |
| ADGRL1 | 0.05437 | 0.90588 | 0.076 | 0.13 | 1 | MS.Trt_vs_MS.NoTrt_F |
| ARSD | 0.05447 | 0.51186 | 0.77 | 0.83 | 1 | MS.Trt_vs_MS.NoTrt_F |
| GBE1 | 0.05449 | 0.83723 | 0.076 | 0.13 | 1 | MS.Trt_vs_MS.NoTrt_F |
| MALT1 | 0.05453 | 0.70197 | 0.129 | 0.2 | 1 | MS.Trt_vs_MS.NoTrt_F |
| FAM110C | 0.05459 | 0.40625 | 0.259 | 0.35 | 1 | MS.Trt_vs_MS.NoTrt_F |
| ZNRF2 | 0.05481 | 0.68774 | 0.083 | 0.14 | 1 | MS.Trt_vs_MS.NoTrt_F |
| TM9SF1 | 0.0549 | 1.04957 | 0.255 | 0.34 | 1 | MS.Trt_vs_MS.NoTrt_F |
| AKAP10 | 0.05506 | 0.30449 | 0.471 | 0.57 | 1 | MS.Trt_vs_MS.NoTrt_F |
| PDIA4 | 0.05516 | 0.92366 | 0.097 | 0.16 | 1 | MS.Trt_vs_MS.NoTrt_F |

| | | | | | | |
|---|---|---|---|---|---|---|
| MED23 | 0.05516 | 0.27927 | 0.212 | 0.3 | 1 | MS.Trt_vs_MS.NoTrt_F |
| TREML2 | 0.0552 | 1.50611 | 0.129 | 0.2 | 1 | MS.Trt_vs_MS.NoTrt_F |
| UPF1 | 0.05526 | 0.37736 | 0.137 | 0.21 | 1 | MS.Trt_vs_MS.NoTrt_F |
| BABAM1 | 0.05528 | 0.498 | 0.619 | 0.7 | 1 | MS.Trt_vs_MS.NoTrt_F |
| SIPA1 | 0.05529 | 0.57779 | 0.532 | 0.68 | 1 | MS.Trt_vs_MS.NoTrt_F |
| IL17RD | 0.05538 | 0.71937 | 0.586 | 0.7 | 1 | MS.Trt_vs_MS.NoTrt_F |
| PAK3 | 0.05548 | -0.6551 | 0.421 | 0.55 | 1 | MS.Trt_vs_MS.NoTrt_F |
| APAF1 | 0.0555 | -0.3169 | 0.597 | 0.68 | 1 | MS.Trt_vs_MS.NoTrt_F |
| BROX | 0.05554 | 0.65167 | 0.644 | 0.71 | 1 | MS.Trt_vs_MS.NoTrt_F |
| AAR2 | 0.05557 | 0.86525 | 0.23 | 0.32 | 1 | MS.Trt_vs_MS.NoTrt_F |
| GORASP1 | 0.05569 | -0.5522 | 0.101 | 0.16 | 1 | MS.Trt_vs_MS.NoTrt_F |
| HSF1 | 0.05569 | -0.3787 | 0.424 | 0.51 | 1 | MS.Trt_vs_MS.NoTrt_F |
| GIPC1 | 0.05583 | 0.4061 | 0.457 | 0.59 | 1 | MS.Trt_vs_MS.NoTrt_F |
| MEGF11 | 0.05584 | -1.0298 | 0.065 | 0.12 | 1 | MS.Trt_vs_MS.NoTrt_F |
| SEC61A2 | 0.05585 | 1.05113 | 0.108 | 0.18 | 1 | MS.Trt_vs_MS.NoTrt_F |
| DNAH9 | 0.05597 | 0.52062 | 0.565 | 0.49 | 1 | MS.Trt_vs_MS.NoTrt_F |
| SLC4A7 | 0.05603 | 0.2885 | 0.342 | 0.44 | 1 | MS.Trt_vs_MS.NoTrt_F |
| ALG3 | 0.05606 | 1.12992 | 0.173 | 0.27 | 1 | MS.Trt_vs_MS.NoTrt_F |
| LNPK | 0.0561 | 0.25005 | 0.547 | 0.62 | 1 | MS.Trt_vs_MS.NoTrt_F |
| FAM50A | 0.05625 | 0.60022 | 0.147 | 0.22 | 1 | MS.Trt_vs_MS.NoTrt_F |
| NATD1 | 0.05628 | 0.3752 | 0.432 | 0.54 | 1 | MS.Trt_vs_MS.NoTrt_F |
| NCKIPSD | 0.05637 | -1.801 | 0.058 | 0.11 | 1 | MS.Trt_vs_MS.NoTrt_F |
| ANXA1 | 0.05639 | -0.3534 | 0.266 | 0.36 | 1 | MS.Trt_vs_MS.NoTrt_F |
| CAMKMT | 0.05642 | 0.7398 | 0.306 | 0.4 | 1 | MS.Trt_vs_MS.NoTrt_F |
| KLHL3 | 0.0565 | 0.34025 | 0.209 | 0.29 | 1 | MS.Trt_vs_MS.NoTrt_F |
| VDAC1 | 0.05683 | -0.3146 | 0.676 | 0.84 | 1 | MS.Trt_vs_MS.NoTrt_F |
| NBPF11 | 0.0569 | 0.6444 | 0.288 | 0.4 | 1 | MS.Trt_vs_MS.NoTrt_F |
| ITGA3 | 0.05691 | -0.3812 | 0.719 | 0.81 | 1 | MS.Trt_vs_MS.NoTrt_F |
| POLG2 | 0.05697 | -0.346 | 0.493 | 0.56 | 1 | MS.Trt_vs_MS.NoTrt_F |
| SORD | 0.05699 | 0.46418 | 0.18 | 0.25 | 1 | MS.Trt_vs_MS.NoTrt_F |
| DDX51 | 0.05703 | 0.63573 | 0.683 | 0.66 | 1 | MS.Trt_vs_MS.NoTrt_F |
| NEO1 | 0.05707 | 0.75345 | 0.241 | 0.36 | 1 | MS.Trt_vs_MS.NoTrt_F |
| EIF1AY | 0.05711 | -0.8168 | 0.615 | 0.67 | 1 | MS.Trt_vs_MS.NoTrt_F |
| TMEM187 | 0.05737 | 0.88169 | 0.068 | 0.12 | 1 | MS.Trt_vs_MS.NoTrt_F |
| ULBP3 | 0.05751 | 0.77485 | 0.543 | 0.6 | 1 | MS.Trt_vs_MS.NoTrt_F |
| USP42 | 0.05758 | -0.4099 | 0.435 | 0.51 | 1 | MS.Trt_vs_MS.NoTrt_F |
| SLC9A7 | 0.0576 | 0.57549 | 0.957 | 0.92 | 1 | MS.Trt_vs_MS.NoTrt_F |
| NUDT7 | 0.05786 | 0.61109 | 0.212 | 0.29 | 1 | MS.Trt_vs_MS.NoTrt_F |
| FAM104B | 0.05795 | 0.58552 | 0.086 | 0.14 | 1 | MS.Trt_vs_MS.NoTrt_F |
| MPDU1 | 0.05797 | -0.3702 | 0.791 | 0.89 | 1 | MS.Trt_vs_MS.NoTrt_F |

| | | | | | | |
|---|---|---|---|---|---|---|
| AMDHD1 | 0.05799 | 0.50162 | 0.191 | 0.28 | 1 | MS.Trt_vs_MS.NoTrt_F |
| GIPR | 0.058 | 0.51121 | 0.651 | 0.71 | 1 | MS.Trt_vs_MS.NoTrt_F |
| OSTF1 | 0.05804 | 0.43346 | 0.291 | 0.39 | 1 | MS.Trt_vs_MS.NoTrt_F |
| RP11-544M: | 0.05809 | 0.52248 | 0.227 | 0.31 | 1 | MS.Trt_vs_MS.NoTrt_F |
| GAA | 0.05819 | 0.33703 | 0.094 | 0.16 | 1 | MS.Trt_vs_MS.NoTrt_F |
| RABIF | 0.05841 | 0.46237 | 0.356 | 0.45 | 1 | MS.Trt_vs_MS.NoTrt_F |
| PAFAH1B2 | 0.05843 | 0.42999 | 0.82 | 0.95 | 1 | MS.Trt_vs_MS.NoTrt_F |
| C10orf95 | 0.05848 | 0.55388 | 0.169 | 0.25 | 1 | MS.Trt_vs_MS.NoTrt_F |
| KDM7A | 0.05859 | -0.3447 | 0.263 | 0.35 | 1 | MS.Trt_vs_MS.NoTrt_F |
| FAM91A1 | 0.0586 | -0.983 | 0.338 | 0.45 | 1 | MS.Trt_vs_MS.NoTrt_F |
| RPS6KA3 | 0.05866 | -0.4091 | 0.471 | 0.6 | 1 | MS.Trt_vs_MS.NoTrt_F |
| CALR | 0.0587 | -0.3272 | 0.806 | 0.86 | 1 | MS.Trt_vs_MS.NoTrt_F |
| DROSHA | 0.05875 | 0.4887 | 0.324 | 0.43 | 1 | MS.Trt_vs_MS.NoTrt_F |
| OSBPL3 | 0.05902 | 0.25506 | 0.45 | 0.57 | 1 | MS.Trt_vs_MS.NoTrt_F |
| CTCF | 0.05913 | 0.79305 | 0.151 | 0.23 | 1 | MS.Trt_vs_MS.NoTrt_F |
| MST1 | 0.05914 | -1.0698 | 0.058 | 0.11 | 1 | MS.Trt_vs_MS.NoTrt_F |
| CCR7 | 0.05925 | 0.33999 | 0.421 | 0.57 | 1 | MS.Trt_vs_MS.NoTrt_F |
| DEPDC7 | 0.05927 | 0.3956 | 0.079 | 0.13 | 1 | MS.Trt_vs_MS.NoTrt_F |
| RGL4 | 0.05934 | 0.38019 | 0.255 | 0.36 | 1 | MS.Trt_vs_MS.NoTrt_F |
| ZNF551 | 0.05941 | 0.70559 | 0.489 | 0.56 | 1 | MS.Trt_vs_MS.NoTrt_F |
| ZCCHC4 | 0.05943 | -0.3786 | 0.615 | 0.7 | 1 | MS.Trt_vs_MS.NoTrt_F |
| ZNF830 | 0.05944 | 0.5635 | 0.108 | 0.17 | 1 | MS.Trt_vs_MS.NoTrt_F |
| URB2 | 0.05946 | -1.1254 | 0.058 | 0.11 | 1 | MS.Trt_vs_MS.NoTrt_F |
| NAGPA | 0.0595 | -0.3257 | 0.669 | 0.71 | 1 | MS.Trt_vs_MS.NoTrt_F |
| GPCPD1 | 0.05951 | 0.30654 | 0.356 | 0.48 | 1 | MS.Trt_vs_MS.NoTrt_F |
| C1orf216 | 0.05956 | 0.78181 | 0.73 | 0.84 | 1 | MS.Trt_vs_MS.NoTrt_F |
| LCAT | 0.05965 | 0.4365 | 0.205 | 0.3 | 1 | MS.Trt_vs_MS.NoTrt_F |
| GIMAP5 | 0.05966 | 0.37121 | 0.338 | 0.45 | 1 | MS.Trt_vs_MS.NoTrt_F |
| SARNP | 0.05989 | 0.28173 | 0.579 | 0.65 | 1 | MS.Trt_vs_MS.NoTrt_F |
| PHB2 | 0.05991 | 0.40006 | 0.176 | 0.26 | 1 | MS.Trt_vs_MS.NoTrt_F |
| SLC25A30 | 0.05997 | -0.3259 | 0.644 | 0.72 | 1 | MS.Trt_vs_MS.NoTrt_F |
| SLC8A1 | 0.06 | 0.60923 | 0.378 | 0.29 | 1 | MS.Trt_vs_MS.NoTrt_F |
| RAB11FIP2 | 0.06002 | -0.695 | 0.086 | 0.14 | 1 | MS.Trt_vs_MS.NoTrt_F |
| ERCC6 | 0.06002 | -0.4679 | 0.266 | 0.34 | 1 | MS.Trt_vs_MS.NoTrt_F |
| CEP97 | 0.06007 | 0.53368 | 0.64 | 0.78 | 1 | MS.Trt_vs_MS.NoTrt_F |
| SNIP1 | 0.06008 | 0.52786 | 0.619 | 0.74 | 1 | MS.Trt_vs_MS.NoTrt_F |
| MINA | 0.06024 | 0.61541 | 0.126 | 0.2 | 1 | MS.Trt_vs_MS.NoTrt_F |
| AP1S2 | 0.06031 | 0.78192 | 0.201 | 0.3 | 1 | MS.Trt_vs_MS.NoTrt_F |
| NDRG2 | 0.06045 | -0.2671 | 0.835 | 0.88 | 1 | MS.Trt_vs_MS.NoTrt_F |
| CREB3 | 0.0606 | 0.46928 | 0.108 | 0.18 | 1 | MS.Trt_vs_MS.NoTrt_F |

| | | | | | | |
|---|---|---|---|---|---|---|
| RHOBTB3 | 0.06061 | -0.2593 | 0.572 | 0.69 | 1 | MS.Trt_vs_MS.NoTrt_F |
| STOM | 0.06072 | 0.8176 | 0.655 | 0.64 | 1 | MS.Trt_vs_MS.NoTrt_F |
| DHRS1 | 0.06076 | -0.365 | 0.608 | 0.66 | 1 | MS.Trt_vs_MS.NoTrt_F |
| UBE3B | 0.06089 | -0.5309 | 0.331 | 0.4 | 1 | MS.Trt_vs_MS.NoTrt_F |
| USE1 | 0.06093 | 0.69105 | 0.917 | 0.86 | 1 | MS.Trt_vs_MS.NoTrt_F |
| ARPC4 | 0.06102 | 0.62003 | 0.683 | 0.71 | 1 | MS.Trt_vs_MS.NoTrt_F |
| APLF | 0.06105 | 0.67849 | 0.176 | 0.25 | 1 | MS.Trt_vs_MS.NoTrt_F |
| MBIP | 0.06106 | 0.95542 | 0.097 | 0.16 | 1 | MS.Trt_vs_MS.NoTrt_F |
| ACOT7 | 0.06109 | 0.45814 | 0.058 | 0.11 | 1 | MS.Trt_vs_MS.NoTrt_F |
| TNS2 | 0.06116 | 0.26278 | 0.342 | 0.41 | 1 | MS.Trt_vs_MS.NoTrt_F |
| TOPORS-AS1 | 0.06116 | 0.89966 | 0.115 | 0.18 | 1 | MS.Trt_vs_MS.NoTrt_F |
| CTD-2021K4 | 0.06126 | 0.28425 | 0.115 | 0.18 | 1 | MS.Trt_vs_MS.NoTrt_F |
| GATAD2B | 0.0615 | 0.32097 | 0.514 | 0.65 | 1 | MS.Trt_vs_MS.NoTrt_F |
| C1QTNF6 | 0.06152 | 0.30066 | 0.547 | 0.67 | 1 | MS.Trt_vs_MS.NoTrt_F |
| ACADL | 0.06165 | -0.3153 | 0.288 | 0.37 | 1 | MS.Trt_vs_MS.NoTrt_F |
| ATRN | 0.06165 | 0.43168 | 0.119 | 0.19 | 1 | MS.Trt_vs_MS.NoTrt_F |
| MAP3K12 | 0.06203 | 0.6392 | 0.701 | 0.79 | 1 | MS.Trt_vs_MS.NoTrt_F |
| ZNF43 | 0.06212 | 1.08604 | 0.773 | 0.79 | 1 | MS.Trt_vs_MS.NoTrt_F |
| ANKRD26 | 0.06227 | 1.3373 | 0.144 | 0.22 | 1 | MS.Trt_vs_MS.NoTrt_F |
| AARSD1 | 0.0623 | -0.4026 | 0.676 | 0.77 | 1 | MS.Trt_vs_MS.NoTrt_F |
| RPS7 | 0.06234 | -0.3574 | 0.514 | 0.59 | 1 | MS.Trt_vs_MS.NoTrt_F |
| CTBS | 0.06241 | 0.57654 | 0.147 | 0.22 | 1 | MS.Trt_vs_MS.NoTrt_F |
| TFDP1 | 0.06241 | -0.4064 | 0.061 | 0.11 | 1 | MS.Trt_vs_MS.NoTrt_F |
| ADAT2 | 0.06248 | 0.51086 | 0.381 | 0.45 | 1 | MS.Trt_vs_MS.NoTrt_F |
| PLIN3 | 0.06265 | -0.3888 | 0.54 | 0.6 | 1 | MS.Trt_vs_MS.NoTrt_F |
| CAAP1 | 0.06269 | 0.41594 | 0.158 | 0.23 | 1 | MS.Trt_vs_MS.NoTrt_F |
| LRG1 | 0.06272 | 0.40345 | 0.813 | 0.76 | 1 | MS.Trt_vs_MS.NoTrt_F |
| EEF1E1 | 0.06286 | 0.29499 | 0.475 | 0.56 | 1 | MS.Trt_vs_MS.NoTrt_F |
| RELB | 0.06295 | 0.42183 | 0.126 | 0.19 | 1 | MS.Trt_vs_MS.NoTrt_F |
| TBRG1 | 0.06296 | 0.26155 | 0.651 | 0.74 | 1 | MS.Trt_vs_MS.NoTrt_F |
| USP32 | 0.063 | 1.80856 | 0.277 | 0.37 | 1 | MS.Trt_vs_MS.NoTrt_F |
| HSD17B1 | 0.06336 | -0.4525 | 0.766 | 0.82 | 1 | MS.Trt_vs_MS.NoTrt_F |
| ESCO1 | 0.06337 | 0.69724 | 0.378 | 0.51 | 1 | MS.Trt_vs_MS.NoTrt_F |
| ZEB2 | 0.06343 | 2.39712 | 0.694 | 0.61 | 1 | MS.Trt_vs_MS.NoTrt_F |
| ITGB1BP1 | 0.06355 | 0.72392 | 0.219 | 0.31 | 1 | MS.Trt_vs_MS.NoTrt_F |
| PALLD | 0.06376 | 0.25259 | 0.317 | 0.41 | 1 | MS.Trt_vs_MS.NoTrt_F |
| SPACA6 | 0.064 | 0.36857 | 0.881 | 0.91 | 1 | MS.Trt_vs_MS.NoTrt_F |
| RRM2B | 0.06401 | -0.9429 | 0.165 | 0.24 | 1 | MS.Trt_vs_MS.NoTrt_F |
| CCNK | 0.06407 | 0.48904 | 0.259 | 0.35 | 1 | MS.Trt_vs_MS.NoTrt_F |
| CXorf23 | 0.06408 | -0.3236 | 0.698 | 0.79 | 1 | MS.Trt_vs_MS.NoTrt_F |

| | | | | | | |
|---|---|---|---|---|---|---|
| ZSCAN12 | 0.06409 | 0.91056 | 0.493 | 0.4 | 1 | MS.Trt_vs_MS.NoTrt_F |
| AARS | 0.06422 | -0.5044 | 0.403 | 0.5 | 1 | MS.Trt_vs_MS.NoTrt_F |
| TRNT1 | 0.06434 | -0.3629 | 0.281 | 0.36 | 1 | MS.Trt_vs_MS.NoTrt_F |
| FITM2 | 0.0644 | -0.4432 | 0.673 | 0.75 | 1 | MS.Trt_vs_MS.NoTrt_F |
| SLAIN2 | 0.06445 | 0.31384 | 0.212 | 0.3 | 1 | MS.Trt_vs_MS.NoTrt_F |
| MIIP | 0.06446 | 0.5064 | 0.187 | 0.28 | 1 | MS.Trt_vs_MS.NoTrt_F |
| SLC10A1 | 0.06448 | 0.39844 | 0.061 | 0.11 | 1 | MS.Trt_vs_MS.NoTrt_F |
| ZNF552 | 0.06471 | -0.6353 | 0.432 | 0.5 | 1 | MS.Trt_vs_MS.NoTrt_F |
| SNURF | 0.06496 | 0.42311 | 0.176 | 0.25 | 1 | MS.Trt_vs_MS.NoTrt_F |
| SARM1 | 0.06516 | 0.9 | 0.417 | 0.51 | 1 | MS.Trt_vs_MS.NoTrt_F |
| MIEF2 | 0.06531 | -0.4049 | 0.457 | 0.56 | 1 | MS.Trt_vs_MS.NoTrt_F |
| VIPR1 | 0.06531 | 0.50841 | 0.086 | 0.14 | 1 | MS.Trt_vs_MS.NoTrt_F |
| THNSL1 | 0.06548 | 1.42816 | 0.119 | 0.19 | 1 | MS.Trt_vs_MS.NoTrt_F |
| IGF2R | 0.06554 | 0.49035 | 0.41 | 0.51 | 1 | MS.Trt_vs_MS.NoTrt_F |
| USP37 | 0.06555 | -0.9163 | 0.442 | 0.51 | 1 | MS.Trt_vs_MS.NoTrt_F |
| FOXRED2 | 0.06557 | 0.61444 | 0.504 | 0.6 | 1 | MS.Trt_vs_MS.NoTrt_F |
| FAM103A1 | 0.06558 | 0.70345 | 0.18 | 0.26 | 1 | MS.Trt_vs_MS.NoTrt_F |
| GRINA | 0.06562 | 0.49099 | 0.094 | 0.16 | 1 | MS.Trt_vs_MS.NoTrt_F |
| NMI | 0.06569 | -0.8887 | 0.306 | 0.38 | 1 | MS.Trt_vs_MS.NoTrt_F |
| PDE6D | 0.06577 | 0.38329 | 0.155 | 0.23 | 1 | MS.Trt_vs_MS.NoTrt_F |
| TBPL1 | 0.0658 | 0.36981 | 0.245 | 0.33 | 1 | MS.Trt_vs_MS.NoTrt_F |
| NT5C3B | 0.06592 | -0.8163 | 0.612 | 0.7 | 1 | MS.Trt_vs_MS.NoTrt_F |
| RINL | 0.06598 | -0.6946 | 0.741 | 0.77 | 1 | MS.Trt_vs_MS.NoTrt_F |
| EXOSC3 | 0.06629 | -0.469 | 0.392 | 0.46 | 1 | MS.Trt_vs_MS.NoTrt_F |
| CBFB | 0.06631 | 1.05832 | 0.173 | 0.26 | 1 | MS.Trt_vs_MS.NoTrt_F |
| TET1 | 0.06638 | 0.5379 | 0.097 | 0.16 | 1 | MS.Trt_vs_MS.NoTrt_F |
| DRC3 | 0.06646 | -0.39 | 0.565 | 0.65 | 1 | MS.Trt_vs_MS.NoTrt_F |
| ERGIC2 | 0.06651 | -0.7273 | 0.471 | 0.55 | 1 | MS.Trt_vs_MS.NoTrt_F |
| COMMD10 | 0.06657 | 0.27196 | 0.324 | 0.41 | 1 | MS.Trt_vs_MS.NoTrt_F |
| ZNF665 | 0.06668 | 0.68262 | 0.917 | 0.95 | 1 | MS.Trt_vs_MS.NoTrt_F |
| TRIM26 | 0.06673 | 0.54752 | 0.191 | 0.28 | 1 | MS.Trt_vs_MS.NoTrt_F |
| GORAB | 0.06676 | 0.32032 | 0.083 | 0.14 | 1 | MS.Trt_vs_MS.NoTrt_F |
| SNX21 | 0.06683 | 0.76921 | 0.421 | 0.56 | 1 | MS.Trt_vs_MS.NoTrt_F |
| DPYSL2 | 0.06689 | 0.36883 | 0.572 | 0.49 | 1 | MS.Trt_vs_MS.NoTrt_F |
| RBM48 | 0.06698 | 0.57152 | 0.719 | 0.85 | 1 | MS.Trt_vs_MS.NoTrt_F |
| RAB40B | 0.06704 | 0.26291 | 0.525 | 0.62 | 1 | MS.Trt_vs_MS.NoTrt_F |
| MTRF1L | 0.06709 | 1.02486 | 0.094 | 0.16 | 1 | MS.Trt_vs_MS.NoTrt_F |
| CLSTN3 | 0.06712 | 0.27702 | 0.183 | 0.27 | 1 | MS.Trt_vs_MS.NoTrt_F |
| MAP4K5 | 0.06714 | -0.6613 | 0.482 | 0.57 | 1 | MS.Trt_vs_MS.NoTrt_F |
| LIN7C | 0.06731 | -0.2998 | 0.388 | 0.45 | 1 | MS.Trt_vs_MS.NoTrt_F |

| | | | | | | |
|---|---|---|---|---|---|---|
| UQCR10 | 0.06734 | 0.84009 | 0.342 | 0.48 | 1 | MS.Trt_vs_MS.NoTrt_F |
| WDR46 | 0.06735 | 0.95118 | 0.122 | 0.19 | 1 | MS.Trt_vs_MS.NoTrt_F |
| ARF4 | 0.06736 | 0.54053 | 0.209 | 0.29 | 1 | MS.Trt_vs_MS.NoTrt_F |
| CASD1 | 0.0674 | 0.73964 | 0.083 | 0.14 | 1 | MS.Trt_vs_MS.NoTrt_F |
| CYP20A1 | 0.06743 | -0.2908 | 0.896 | 0.92 | 1 | MS.Trt_vs_MS.NoTrt_F |
| ZNF333 | 0.0677 | 0.71904 | 0.885 | 0.94 | 1 | MS.Trt_vs_MS.NoTrt_F |
| TRIM45 | 0.06778 | -0.6088 | 0.112 | 0.17 | 1 | MS.Trt_vs_MS.NoTrt_F |
| ACOX3 | 0.06799 | 0.40594 | 0.133 | 0.2 | 1 | MS.Trt_vs_MS.NoTrt_F |
| SCP2 | 0.06801 | 0.79147 | 0.277 | 0.39 | 1 | MS.Trt_vs_MS.NoTrt_F |
| RPS6KB2 | 0.06803 | -0.5591 | 0.853 | 0.87 | 1 | MS.Trt_vs_MS.NoTrt_F |
| PLEKHA3 | 0.06808 | -0.395 | 0.763 | 0.81 | 1 | MS.Trt_vs_MS.NoTrt_F |
| SMIM12 | 0.0681 | -0.3376 | 0.737 | 0.79 | 1 | MS.Trt_vs_MS.NoTrt_F |
| SH3PXD2A | 0.06814 | -0.3086 | 0.309 | 0.39 | 1 | MS.Trt_vs_MS.NoTrt_F |
| DNPH1 | 0.06818 | 0.99831 | 0.212 | 0.32 | 1 | MS.Trt_vs_MS.NoTrt_F |
| EDC3 | 0.06831 | 0.35438 | 0.371 | 0.46 | 1 | MS.Trt_vs_MS.NoTrt_F |
| MRPL46 | 0.06834 | -0.4318 | 0.252 | 0.33 | 1 | MS.Trt_vs_MS.NoTrt_F |
| NOP56 | 0.06841 | 0.64525 | 0.453 | 0.55 | 1 | MS.Trt_vs_MS.NoTrt_F |
| RASAL3 | 0.06846 | 0.45417 | 0.219 | 0.32 | 1 | MS.Trt_vs_MS.NoTrt_F |
| STEAP3 | 0.06847 | -0.7891 | 0.162 | 0.23 | 1 | MS.Trt_vs_MS.NoTrt_F |
| CLCN5 | 0.0685 | -0.3486 | 0.277 | 0.37 | 1 | MS.Trt_vs_MS.NoTrt_F |
| MYO10 | 0.0685 | -0.2901 | 0.619 | 0.71 | 1 | MS.Trt_vs_MS.NoTrt_F |
| DNMBP | 0.06865 | 1.09817 | 0.09 | 0.14 | 1 | MS.Trt_vs_MS.NoTrt_F |
| SLC25A14 | 0.06868 | -0.49 | 0.263 | 0.35 | 1 | MS.Trt_vs_MS.NoTrt_F |
| TMEM138 | 0.06868 | -0.5965 | 0.806 | 0.86 | 1 | MS.Trt_vs_MS.NoTrt_F |
| RP11-244H3 | 0.06876 | 0.86753 | 0.108 | 0.17 | 1 | MS.Trt_vs_MS.NoTrt_F |
| PCGF6 | 0.06895 | -0.4182 | 0.644 | 0.79 | 1 | MS.Trt_vs_MS.NoTrt_F |
| AC138645.2 | 0.06899 | -0.3001 | 0.151 | 0.22 | 1 | MS.Trt_vs_MS.NoTrt_F |
| MOB4 | 0.06899 | -0.4007 | 0.378 | 0.44 | 1 | MS.Trt_vs_MS.NoTrt_F |
| HRH2 | 0.06911 | -1.0327 | 0.115 | 0.17 | 1 | MS.Trt_vs_MS.NoTrt_F |
| CEP83 | 0.06913 | -0.4512 | 0.647 | 0.69 | 1 | MS.Trt_vs_MS.NoTrt_F |
| EIF2AK2 | 0.06921 | 0.30336 | 0.773 | 0.85 | 1 | MS.Trt_vs_MS.NoTrt_F |
| PDZD2 | 0.06924 | 0.64793 | 0.705 | 0.66 | 1 | MS.Trt_vs_MS.NoTrt_F |
| MYLIP | 0.06925 | 0.75536 | 0.104 | 0.17 | 1 | MS.Trt_vs_MS.NoTrt_F |
| AMACR | 0.06936 | -0.4537 | 0.101 | 0.16 | 1 | MS.Trt_vs_MS.NoTrt_F |
| DIMT1 | 0.06938 | 0.80276 | 0.147 | 0.23 | 1 | MS.Trt_vs_MS.NoTrt_F |
| DARS | 0.06942 | 0.51293 | 0.331 | 0.44 | 1 | MS.Trt_vs_MS.NoTrt_F |
| NUP54 | 0.06942 | 0.3562 | 0.194 | 0.27 | 1 | MS.Trt_vs_MS.NoTrt_F |
| METRN | 0.06959 | 1.09929 | 0.119 | 0.18 | 1 | MS.Trt_vs_MS.NoTrt_F |
| TMEM255B | 0.0699 | 0.66382 | 0.176 | 0.25 | 1 | MS.Trt_vs_MS.NoTrt_F |
| GATAD2A | 0.07005 | 0.48734 | 0.655 | 0.77 | 1 | MS.Trt_vs_MS.NoTrt_F |

| | | | | | | |
|---|---|---|---|---|---|---|
| FAM179B | 0.07014 | 0.4635 | 0.755 | 0.87 | 1 | MS.Trt_vs_MS.NoTrt_F |
| MYCBPAP | 0.0702 | -0.417 | 0.791 | 0.89 | 1 | MS.Trt_vs_MS.NoTrt_F |
| NCAPH2 | 0.07023 | 0.34265 | 0.712 | 0.81 | 1 | MS.Trt_vs_MS.NoTrt_F |
| CPNE8 | 0.07023 | 0.62009 | 0.151 | 0.23 | 1 | MS.Trt_vs_MS.NoTrt_F |
| TMEM116 | 0.07033 | -0.5656 | 0.871 | 0.91 | 1 | MS.Trt_vs_MS.NoTrt_F |
| MPI | 0.0704 | 0.52734 | 0.468 | 0.56 | 1 | MS.Trt_vs_MS.NoTrt_F |
| TNS1 | 0.07046 | -1.1693 | 0.23 | 0.31 | 1 | MS.Trt_vs_MS.NoTrt_F |
| YTHDC2 | 0.07046 | 0.52021 | 0.406 | 0.51 | 1 | MS.Trt_vs_MS.NoTrt_F |
| FRMD8 | 0.07052 | 0.73642 | 0.644 | 0.58 | 1 | MS.Trt_vs_MS.NoTrt_F |
| KIAA2013 | 0.07054 | 0.51302 | 0.284 | 0.37 | 1 | MS.Trt_vs_MS.NoTrt_F |
| ZNF91 | 0.07073 | 0.36562 | 0.727 | 0.82 | 1 | MS.Trt_vs_MS.NoTrt_F |
| FBXW4 | 0.07081 | 0.52688 | 0.446 | 0.56 | 1 | MS.Trt_vs_MS.NoTrt_F |
| MIEN1 | 0.071 | 0.79205 | 0.169 | 0.25 | 1 | MS.Trt_vs_MS.NoTrt_F |
| SMARCC1 | 0.07109 | -0.4873 | 0.442 | 0.51 | 1 | MS.Trt_vs_MS.NoTrt_F |
| VWF | 0.07127 | 0.35477 | 0.079 | 0.13 | 1 | MS.Trt_vs_MS.NoTrt_F |
| FAM101B | 0.07128 | 1.19368 | 0.288 | 0.4 | 1 | MS.Trt_vs_MS.NoTrt_F |
| LYZ | 0.0713 | 0.97044 | 0.371 | 0.48 | 1 | MS.Trt_vs_MS.NoTrt_F |
| VCL | 0.07152 | 0.45041 | 0.144 | 0.22 | 1 | MS.Trt_vs_MS.NoTrt_F |
| TEN1 | 0.07161 | -0.7201 | 0.327 | 0.41 | 1 | MS.Trt_vs_MS.NoTrt_F |
| PAPD5 | 0.07168 | -0.5355 | 0.205 | 0.28 | 1 | MS.Trt_vs_MS.NoTrt_F |
| NFYA | 0.07185 | 0.71579 | 0.144 | 0.22 | 1 | MS.Trt_vs_MS.NoTrt_F |
| KSR1 | 0.07203 | 0.83847 | 0.288 | 0.38 | 1 | MS.Trt_vs_MS.NoTrt_F |
| RDH14 | 0.07205 | 1.0108 | 0.079 | 0.13 | 1 | MS.Trt_vs_MS.NoTrt_F |
| RMND1 | 0.07205 | -0.3703 | 0.068 | 0.12 | 1 | MS.Trt_vs_MS.NoTrt_F |
| IL3RA | 0.07222 | 0.51934 | 0.101 | 0.16 | 1 | MS.Trt_vs_MS.NoTrt_F |
| NCMAP | 0.07233 | 0.32937 | 0.687 | 0.64 | 1 | MS.Trt_vs_MS.NoTrt_F |
| POT1 | 0.07255 | 0.51382 | 0.119 | 0.18 | 1 | MS.Trt_vs_MS.NoTrt_F |
| SYNGR2 | 0.07316 | 0.2637 | 0.525 | 0.62 | 1 | MS.Trt_vs_MS.NoTrt_F |
| UQCC1 | 0.07328 | -0.3435 | 0.496 | 0.6 | 1 | MS.Trt_vs_MS.NoTrt_F |
| INSR | 0.07345 | 0.42481 | 0.439 | 0.51 | 1 | MS.Trt_vs_MS.NoTrt_F |
| LAMB1 | 0.07351 | 0.34328 | 0.284 | 0.38 | 1 | MS.Trt_vs_MS.NoTrt_F |
| RPA1 | 0.07359 | 0.30366 | 0.263 | 0.35 | 1 | MS.Trt_vs_MS.NoTrt_F |
| CTD-3148I1( | 0.0737 | 0.55949 | 0.115 | 0.18 | 1 | MS.Trt_vs_MS.NoTrt_F |
| WDR5 | 0.07374 | 0.27386 | 0.14 | 0.21 | 1 | MS.Trt_vs_MS.NoTrt_F |
| B4GALT4 | 0.07379 | -0.3527 | 0.647 | 0.68 | 1 | MS.Trt_vs_MS.NoTrt_F |
| ATF5 | 0.07395 | 0.41971 | 0.263 | 0.36 | 1 | MS.Trt_vs_MS.NoTrt_F |
| GATS | 0.07399 | -0.3757 | 0.32 | 0.41 | 1 | MS.Trt_vs_MS.NoTrt_F |
| GSTM1 | 0.07402 | 1.1576 | 0.183 | 0.27 | 1 | MS.Trt_vs_MS.NoTrt_F |
| DPM3 | 0.07429 | 0.73018 | 0.079 | 0.13 | 1 | MS.Trt_vs_MS.NoTrt_F |
| KRBA1 | 0.07444 | -0.8521 | 0.068 | 0.12 | 1 | MS.Trt_vs_MS.NoTrt_F |

| | | | | | | |
|---|---|---|---|---|---|---|
| SEC24D | 0.07446 | 0.41744 | 0.522 | 0.6 | 1 | MS.Trt_vs_MS.NoTrt_F |
| CDYL | 0.07459 | -0.4447 | 0.097 | 0.16 | 1 | MS.Trt_vs_MS.NoTrt_F |
| SLC44A1 | 0.07467 | -1.4273 | 0.658 | 0.71 | 1 | MS.Trt_vs_MS.NoTrt_F |
| MEX3C | 0.07481 | 0.64909 | 0.104 | 0.17 | 1 | MS.Trt_vs_MS.NoTrt_F |
| C6orf163 | 0.07487 | 0.52956 | 0.421 | 0.54 | 1 | MS.Trt_vs_MS.NoTrt_F |
| ZBTB40 | 0.07488 | -0.3557 | 0.403 | 0.48 | 1 | MS.Trt_vs_MS.NoTrt_F |
| FAM213B | 0.07496 | 0.38717 | 0.122 | 0.19 | 1 | MS.Trt_vs_MS.NoTrt_F |
| FBXO6 | 0.07497 | 0.62734 | 0.716 | 0.65 | 1 | MS.Trt_vs_MS.NoTrt_F |
| CSNK1E | 0.07503 | -0.3711 | 0.676 | 0.78 | 1 | MS.Trt_vs_MS.NoTrt_F |
| ZXDC | 0.07511 | -0.5858 | 0.108 | 0.17 | 1 | MS.Trt_vs_MS.NoTrt_F |
| KIF2A | 0.07527 | 0.7431 | 0.252 | 0.35 | 1 | MS.Trt_vs_MS.NoTrt_F |
| LMNB1 | 0.07529 | 0.26425 | 0.126 | 0.19 | 1 | MS.Trt_vs_MS.NoTrt_F |
| TLK1 | 0.07538 | 0.55434 | 0.788 | 0.86 | 1 | MS.Trt_vs_MS.NoTrt_F |
| RNASEL | 0.07544 | 0.54214 | 0.345 | 0.45 | 1 | MS.Trt_vs_MS.NoTrt_F |
| NCR1 | 0.07547 | 0.76799 | 0.205 | 0.28 | 1 | MS.Trt_vs_MS.NoTrt_F |
| MOB3C | 0.07547 | 2.46341 | 0.079 | 0.13 | 1 | MS.Trt_vs_MS.NoTrt_F |
| CMAS | 0.07552 | 0.35693 | 0.119 | 0.18 | 1 | MS.Trt_vs_MS.NoTrt_F |
| FBXO8 | 0.07553 | 0.9535 | 0.104 | 0.17 | 1 | MS.Trt_vs_MS.NoTrt_F |
| NDUFB1 | 0.07561 | 0.39362 | 0.719 | 0.85 | 1 | MS.Trt_vs_MS.NoTrt_F |
| C1orf52 | 0.07564 | -0.6378 | 0.295 | 0.38 | 1 | MS.Trt_vs_MS.NoTrt_F |
| TRIM52 | 0.07571 | 0.54173 | 0.86 | 0.81 | 1 | MS.Trt_vs_MS.NoTrt_F |
| NUDT19 | 0.07591 | 0.62969 | 0.892 | 0.93 | 1 | MS.Trt_vs_MS.NoTrt_F |
| CPT2 | 0.07597 | -0.2756 | 0.727 | 0.78 | 1 | MS.Trt_vs_MS.NoTrt_F |
| FAM208B | 0.07615 | 0.60652 | 0.277 | 0.38 | 1 | MS.Trt_vs_MS.NoTrt_F |
| MS4A15 | 0.07616 | -0.2662 | 0.068 | 0.12 | 1 | MS.Trt_vs_MS.NoTrt_F |
| AKR1C3 | 0.07628 | -0.9273 | 0.065 | 0.11 | 1 | MS.Trt_vs_MS.NoTrt_F |
| CSNK1D | 0.07648 | -0.2967 | 0.921 | 0.96 | 1 | MS.Trt_vs_MS.NoTrt_F |
| ABHD10 | 0.07651 | 0.27508 | 0.137 | 0.2 | 1 | MS.Trt_vs_MS.NoTrt_F |
| PPP1R8 | 0.07658 | 0.49101 | 0.076 | 0.13 | 1 | MS.Trt_vs_MS.NoTrt_F |
| ICA1 | 0.07661 | 0.8874 | 0.612 | 0.56 | 1 | MS.Trt_vs_MS.NoTrt_F |
| SLC7A6OS | 0.07665 | 0.41387 | 0.216 | 0.3 | 1 | MS.Trt_vs_MS.NoTrt_F |
| DHX35 | 0.07678 | -0.8707 | 0.385 | 0.45 | 1 | MS.Trt_vs_MS.NoTrt_F |
| ALG13 | 0.07684 | 0.29049 | 0.338 | 0.43 | 1 | MS.Trt_vs_MS.NoTrt_F |
| ARHGEF40 | 0.07688 | 0.32017 | 0.18 | 0.25 | 1 | MS.Trt_vs_MS.NoTrt_F |
| IMPDH1 | 0.07693 | 0.38183 | 0.165 | 0.24 | 1 | MS.Trt_vs_MS.NoTrt_F |
| CLTC | 0.0771 | -0.3766 | 0.374 | 0.44 | 1 | MS.Trt_vs_MS.NoTrt_F |
| TRIM5 | 0.07714 | 0.57917 | 0.468 | 0.56 | 1 | MS.Trt_vs_MS.NoTrt_F |
| HIST1H2AC | 0.07746 | 0.88571 | 0.079 | 0.13 | 1 | MS.Trt_vs_MS.NoTrt_F |
| CTNS | 0.07754 | -0.4635 | 0.155 | 0.22 | 1 | MS.Trt_vs_MS.NoTrt_F |
| GALNT15 | 0.07769 | 0.70049 | 0.277 | 0.38 | 1 | MS.Trt_vs_MS.NoTrt_F |

| | | | | | | |
|---|---|---|---|---|---|---|
| KCNAB1 | 0.0778 | 1.33446 | 0.079 | 0.13 | 1 | MS.Trt_vs_MS.NoTrt_F |
| NR1H3 | 0.07781 | 0.70431 | 0.151 | 0.22 | 1 | MS.Trt_vs_MS.NoTrt_F |
| ADHFE1 | 0.07793 | 0.57899 | 0.158 | 0.24 | 1 | MS.Trt_vs_MS.NoTrt_F |
| MFGE8 | 0.07808 | -0.8546 | 0.547 | 0.62 | 1 | MS.Trt_vs_MS.NoTrt_F |
| PIH1D3 | 0.0782 | -0.3071 | 0.32 | 0.4 | 1 | MS.Trt_vs_MS.NoTrt_F |
| ARHGEF12 | 0.0783 | 0.38501 | 0.96 | 0.98 | 1 | MS.Trt_vs_MS.NoTrt_F |
| RGS13 | 0.07839 | 0.45259 | 0.399 | 0.57 | 1 | MS.Trt_vs_MS.NoTrt_F |
| ZSCAN21 | 0.07844 | -0.5495 | 0.227 | 0.31 | 1 | MS.Trt_vs_MS.NoTrt_F |
| CH507-9B2.! | 0.07846 | -0.5606 | 0.112 | 0.17 | 1 | MS.Trt_vs_MS.NoTrt_F |
| MCM5 | 0.07861 | 0.34682 | 0.216 | 0.29 | 1 | MS.Trt_vs_MS.NoTrt_F |
| ELOVL1 | 0.07876 | 0.63047 | 0.18 | 0.26 | 1 | MS.Trt_vs_MS.NoTrt_F |
| SMAD3 | 0.07892 | 0.56007 | 0.169 | 0.25 | 1 | MS.Trt_vs_MS.NoTrt_F |
| ETV7 | 0.07896 | -0.2604 | 0.068 | 0.12 | 1 | MS.Trt_vs_MS.NoTrt_F |
| SOX4 | 0.07913 | 1.50229 | 0.097 | 0.16 | 1 | MS.Trt_vs_MS.NoTrt_F |
| OSTM1 | 0.07946 | -0.4666 | 0.82 | 0.85 | 1 | MS.Trt_vs_MS.NoTrt_F |
| EGFL8 | 0.07958 | -0.4459 | 0.547 | 0.65 | 1 | MS.Trt_vs_MS.NoTrt_F |
| PAXBP1 | 0.07961 | 0.3641 | 0.205 | 0.29 | 1 | MS.Trt_vs_MS.NoTrt_F |
| SLC9A6 | 0.07961 | -0.3828 | 0.428 | 0.54 | 1 | MS.Trt_vs_MS.NoTrt_F |
| PRPF19 | 0.07973 | -0.4133 | 0.094 | 0.15 | 1 | MS.Trt_vs_MS.NoTrt_F |
| NPIPA8 | 0.07984 | -0.3152 | 0.068 | 0.12 | 1 | MS.Trt_vs_MS.NoTrt_F |
| FBXO9 | 0.08033 | -0.5148 | 0.809 | 0.84 | 1 | MS.Trt_vs_MS.NoTrt_F |
| KMT5B | 0.08033 | 0.63787 | 0.324 | 0.4 | 1 | MS.Trt_vs_MS.NoTrt_F |
| ADCY9 | 0.0804 | 0.74057 | 0.288 | 0.38 | 1 | MS.Trt_vs_MS.NoTrt_F |
| MTIF3 | 0.08058 | 0.63761 | 0.223 | 0.32 | 1 | MS.Trt_vs_MS.NoTrt_F |
| BRD7 | 0.08076 | 0.75122 | 0.54 | 0.63 | 1 | MS.Trt_vs_MS.NoTrt_F |
| ACAP3 | 0.08084 | -0.2507 | 0.133 | 0.19 | 1 | MS.Trt_vs_MS.NoTrt_F |
| OSTC | 0.08088 | 0.57444 | 0.266 | 0.36 | 1 | MS.Trt_vs_MS.NoTrt_F |
| WIZ | 0.08101 | -0.3532 | 0.313 | 0.4 | 1 | MS.Trt_vs_MS.NoTrt_F |
| CDRT4 | 0.08106 | 0.37253 | 0.129 | 0.19 | 1 | MS.Trt_vs_MS.NoTrt_F |
| ZC3H12C | 0.08117 | -0.9244 | 0.097 | 0.15 | 1 | MS.Trt_vs_MS.NoTrt_F |
| FAXDC2 | 0.08124 | 0.31279 | 0.849 | 0.82 | 1 | MS.Trt_vs_MS.NoTrt_F |
| ITGAV | 0.08142 | 0.89055 | 0.115 | 0.18 | 1 | MS.Trt_vs_MS.NoTrt_F |
| ZNF223 | 0.08147 | 1.45805 | 0.119 | 0.18 | 1 | MS.Trt_vs_MS.NoTrt_F |
| RNF170 | 0.08149 | -0.6884 | 0.55 | 0.64 | 1 | MS.Trt_vs_MS.NoTrt_F |
| C11orf80 | 0.08154 | 0.52659 | 0.482 | 0.56 | 1 | MS.Trt_vs_MS.NoTrt_F |
| MRPL19 | 0.08162 | 0.54111 | 0.273 | 0.36 | 1 | MS.Trt_vs_MS.NoTrt_F |
| KLC2 | 0.08169 | 0.32413 | 0.108 | 0.17 | 1 | MS.Trt_vs_MS.NoTrt_F |
| AD000671.6 | 0.08169 | 0.31544 | 0.126 | 0.19 | 1 | MS.Trt_vs_MS.NoTrt_F |
| ERCC1 | 0.08174 | 0.36434 | 0.777 | 0.81 | 1 | MS.Trt_vs_MS.NoTrt_F |
| ZFHX3 | 0.08182 | 0.28139 | 0.158 | 0.23 | 1 | MS.Trt_vs_MS.NoTrt_F |

| | | | | | | |
|---|---|---|---|---|---|---|
| WRAP53 | 0.08187 | 0.3827 | 0.072 | 0.12 | 1 | MS.Trt_vs_MS.NoTrt_F |
| EDF1 | 0.08198 | 0.39307 | 0.417 | 0.58 | 1 | MS.Trt_vs_MS.NoTrt_F |
| ARFGAP2 | 0.08217 | -0.3665 | 0.522 | 0.64 | 1 | MS.Trt_vs_MS.NoTrt_F |
| CDK2AP2 | 0.08218 | 0.31961 | 0.245 | 0.34 | 1 | MS.Trt_vs_MS.NoTrt_F |
| FGF1 | 0.08226 | 1.14669 | 0.086 | 0.14 | 1 | MS.Trt_vs_MS.NoTrt_F |
| SIL1 | 0.08238 | 0.36959 | 0.122 | 0.18 | 1 | MS.Trt_vs_MS.NoTrt_F |
| CRELD2 | 0.0825 | 0.38186 | 0.601 | 0.67 | 1 | MS.Trt_vs_MS.NoTrt_F |
| RNPC3 | 0.0827 | 0.63113 | 0.302 | 0.4 | 1 | MS.Trt_vs_MS.NoTrt_F |
| TMEM106A | 0.08278 | -0.3246 | 0.651 | 0.7 | 1 | MS.Trt_vs_MS.NoTrt_F |
| ELMO2 | 0.08279 | 0.47318 | 0.212 | 0.29 | 1 | MS.Trt_vs_MS.NoTrt_F |
| ILKAP | 0.08305 | 0.25149 | 0.597 | 0.72 | 1 | MS.Trt_vs_MS.NoTrt_F |
| INTS2 | 0.0832 | -0.5428 | 0.266 | 0.34 | 1 | MS.Trt_vs_MS.NoTrt_F |
| KIF5C | 0.08346 | -1.0081 | 0.068 | 0.12 | 1 | MS.Trt_vs_MS.NoTrt_F |
| CTPS2 | 0.08349 | 0.75657 | 0.108 | 0.17 | 1 | MS.Trt_vs_MS.NoTrt_F |
| CTSW | 0.08357 | 1.0813 | 0.669 | 0.61 | 1 | MS.Trt_vs_MS.NoTrt_F |
| GALNT8 | 0.08358 | 0.98109 | 0.065 | 0.11 | 1 | MS.Trt_vs_MS.NoTrt_F |
| GDAP1 | 0.08366 | 0.66404 | 0.108 | 0.17 | 1 | MS.Trt_vs_MS.NoTrt_F |
| TRAPPC6B | 0.08367 | -0.5747 | 0.518 | 0.59 | 1 | MS.Trt_vs_MS.NoTrt_F |
| ERCC5 | 0.08372 | 0.46843 | 0.392 | 0.48 | 1 | MS.Trt_vs_MS.NoTrt_F |
| MRPL21 | 0.0839 | 0.53465 | 0.126 | 0.19 | 1 | MS.Trt_vs_MS.NoTrt_F |
| MORN4 | 0.08391 | -0.3941 | 0.525 | 0.65 | 1 | MS.Trt_vs_MS.NoTrt_F |
| STX12 | 0.08401 | 0.59739 | 0.266 | 0.35 | 1 | MS.Trt_vs_MS.NoTrt_F |
| AKR1C2 | 0.0841 | -0.688 | 0.104 | 0.16 | 1 | MS.Trt_vs_MS.NoTrt_F |
| TMEM44 | 0.08419 | -0.4621 | 0.55 | 0.64 | 1 | MS.Trt_vs_MS.NoTrt_F |
| ZCRB1 | 0.08439 | 0.8532 | 0.18 | 0.26 | 1 | MS.Trt_vs_MS.NoTrt_F |
| BUD13 | 0.08441 | 1.0421 | 0.112 | 0.17 | 1 | MS.Trt_vs_MS.NoTrt_F |
| CPM | 0.08448 | 0.56287 | 0.799 | 0.77 | 1 | MS.Trt_vs_MS.NoTrt_F |
| MGME1 | 0.08453 | 0.30394 | 0.112 | 0.17 | 1 | MS.Trt_vs_MS.NoTrt_F |
| TMEM117 | 0.08454 | 0.77616 | 0.097 | 0.16 | 1 | MS.Trt_vs_MS.NoTrt_F |
| ENO3 | 0.08458 | 0.36659 | 0.5 | 0.59 | 1 | MS.Trt_vs_MS.NoTrt_F |
| CDC7 | 0.08474 | 0.26849 | 0.068 | 0.12 | 1 | MS.Trt_vs_MS.NoTrt_F |
| ZNF589 | 0.08494 | 0.69866 | 0.45 | 0.55 | 1 | MS.Trt_vs_MS.NoTrt_F |
| TCAF1 | 0.08505 | -0.937 | 0.338 | 0.41 | 1 | MS.Trt_vs_MS.NoTrt_F |
| SLC26A6 | 0.08507 | 0.6607 | 0.108 | 0.17 | 1 | MS.Trt_vs_MS.NoTrt_F |
| MSTO1 | 0.08526 | 1.39429 | 0.112 | 0.18 | 1 | MS.Trt_vs_MS.NoTrt_F |
| C2orf69 | 0.08536 | 0.56644 | 0.259 | 0.35 | 1 | MS.Trt_vs_MS.NoTrt_F |
| RNF25 | 0.08539 | 0.29029 | 0.112 | 0.17 | 1 | MS.Trt_vs_MS.NoTrt_F |
| SBF2 | 0.08567 | 0.3482 | 0.946 | 0.97 | 1 | MS.Trt_vs_MS.NoTrt_F |
| NDFIP2 | 0.08585 | 0.33562 | 0.205 | 0.28 | 1 | MS.Trt_vs_MS.NoTrt_F |
| RP11-295P9 | 0.08594 | 0.27151 | 0.126 | 0.19 | 1 | MS.Trt_vs_MS.NoTrt_F |

| | | | | | | |
|---|---|---|---|---|---|---|
| IFI44 | 0.08615 | 0.79449 | 0.09 | 0.14 | 1 | MS.Trt_vs_MS.NoTrt_F |
| ZNF394 | 0.08621 | -0.7822 | 0.547 | 0.61 | 1 | MS.Trt_vs_MS.NoTrt_F |
| FAHD2A | 0.08621 | 0.7459 | 0.165 | 0.24 | 1 | MS.Trt_vs_MS.NoTrt_F |
| DOCK11 | 0.08622 | 0.44014 | 0.543 | 0.73 | 1 | MS.Trt_vs_MS.NoTrt_F |
| SCNM1 | 0.08627 | -0.4382 | 0.723 | 0.78 | 1 | MS.Trt_vs_MS.NoTrt_F |
| KIAA0232 | 0.08633 | 0.6955 | 0.119 | 0.18 | 1 | MS.Trt_vs_MS.NoTrt_F |
| CFAP70 | 0.08639 | -0.8527 | 0.281 | 0.35 | 1 | MS.Trt_vs_MS.NoTrt_F |
| CHKA | 0.08663 | 0.92679 | 0.072 | 0.12 | 1 | MS.Trt_vs_MS.NoTrt_F |
| H2AFJ | 0.08679 | 0.40104 | 0.129 | 0.19 | 1 | MS.Trt_vs_MS.NoTrt_F |
| CARHSP1 | 0.08686 | -0.5702 | 0.705 | 0.72 | 1 | MS.Trt_vs_MS.NoTrt_F |
| TMBIM4 | 0.0869 | 0.36185 | 0.82 | 0.87 | 1 | MS.Trt_vs_MS.NoTrt_F |
| STAMBPL1 | 0.08693 | -0.2667 | 0.5 | 0.59 | 1 | MS.Trt_vs_MS.NoTrt_F |
| FNTB | 0.087 | -0.398 | 0.169 | 0.24 | 1 | MS.Trt_vs_MS.NoTrt_F |
| RCCD1 | 0.08706 | -0.9733 | 0.198 | 0.27 | 1 | MS.Trt_vs_MS.NoTrt_F |
| DECR1 | 0.08713 | 0.29336 | 0.478 | 0.59 | 1 | MS.Trt_vs_MS.NoTrt_F |
| GTPBP4 | 0.08714 | -1.004 | 0.489 | 0.57 | 1 | MS.Trt_vs_MS.NoTrt_F |
| LINC00371 | 0.08718 | -0.2929 | 0.144 | 0.21 | 1 | MS.Trt_vs_MS.NoTrt_F |
| PSMA2 | 0.08722 | 0.53594 | 0.586 | 0.66 | 1 | MS.Trt_vs_MS.NoTrt_F |
| GALNS | 0.08734 | 0.50041 | 0.532 | 0.61 | 1 | MS.Trt_vs_MS.NoTrt_F |
| CCDC117 | 0.08735 | 0.66106 | 0.367 | 0.45 | 1 | MS.Trt_vs_MS.NoTrt_F |
| RCOR3 | 0.08738 | -0.253 | 0.403 | 0.48 | 1 | MS.Trt_vs_MS.NoTrt_F |
| COQ3 | 0.08744 | 0.59956 | 0.252 | 0.33 | 1 | MS.Trt_vs_MS.NoTrt_F |
| ZNF165 | 0.0875 | 1.77211 | 0.068 | 0.12 | 1 | MS.Trt_vs_MS.NoTrt_F |
| EGLN1 | 0.08754 | 0.3134 | 0.374 | 0.45 | 1 | MS.Trt_vs_MS.NoTrt_F |
| MYO1D | 0.08769 | 0.38663 | 0.165 | 0.23 | 1 | MS.Trt_vs_MS.NoTrt_F |
| ACSL1 | 0.0879 | 0.59353 | 0.23 | 0.29 | 1 | MS.Trt_vs_MS.NoTrt_F |
| ARHGEF11 | 0.08794 | 0.71764 | 0.09 | 0.14 | 1 | MS.Trt_vs_MS.NoTrt_F |
| NOS3 | 0.08807 | -0.7895 | 0.335 | 0.43 | 1 | MS.Trt_vs_MS.NoTrt_F |
| KLHL9 | 0.08813 | -0.4458 | 0.381 | 0.46 | 1 | MS.Trt_vs_MS.NoTrt_F |
| IFIT2 | 0.08832 | -0.4547 | 0.094 | 0.15 | 1 | MS.Trt_vs_MS.NoTrt_F |
| JADE2 | 0.08835 | 0.37185 | 0.187 | 0.27 | 1 | MS.Trt_vs_MS.NoTrt_F |
| CAPN3 | 0.08838 | -0.662 | 0.083 | 0.13 | 1 | MS.Trt_vs_MS.NoTrt_F |
| TMEM263 | 0.08839 | 0.50166 | 0.388 | 0.46 | 1 | MS.Trt_vs_MS.NoTrt_F |
| FOXN3 | 0.08848 | -0.374 | 0.881 | 0.95 | 1 | MS.Trt_vs_MS.NoTrt_F |
| TSN | 0.08873 | 0.2531 | 0.36 | 0.44 | 1 | MS.Trt_vs_MS.NoTrt_F |
| LSM12 | 0.08881 | 0.44637 | 0.669 | 0.76 | 1 | MS.Trt_vs_MS.NoTrt_F |
| NUP85 | 0.08902 | 0.27865 | 0.428 | 0.53 | 1 | MS.Trt_vs_MS.NoTrt_F |
| LINC01125 | 0.08907 | 0.35213 | 0.96 | 0.96 | 1 | MS.Trt_vs_MS.NoTrt_F |
| WDR26 | 0.0891 | 0.39252 | 0.626 | 0.72 | 1 | MS.Trt_vs_MS.NoTrt_F |
| ETV4 | 0.08915 | -0.6535 | 0.097 | 0.15 | 1 | MS.Trt_vs_MS.NoTrt_F |

| | | | | | | |
|---|---|---|---|---|---|---|
| PPP3CB | 0.08929 | 0.92183 | 0.129 | 0.19 | 1 | MS.Trt_vs_MS.NoTrt_F |
| PTPRJ | 0.08929 | 0.3423 | 0.371 | 0.45 | 1 | MS.Trt_vs_MS.NoTrt_F |
| GALNT9 | 0.08942 | -0.3238 | 0.201 | 0.27 | 1 | MS.Trt_vs_MS.NoTrt_F |
| LSM7 | 0.08961 | 0.31485 | 0.637 | 0.71 | 1 | MS.Trt_vs_MS.NoTrt_F |
| RNF2 | 0.08964 | -0.3679 | 0.399 | 0.51 | 1 | MS.Trt_vs_MS.NoTrt_F |
| FLOT1 | 0.0897 | 0.51948 | 0.144 | 0.22 | 1 | MS.Trt_vs_MS.NoTrt_F |
| SLC20A1 | 0.08971 | 0.46345 | 0.237 | 0.34 | 1 | MS.Trt_vs_MS.NoTrt_F |
| BPGM | 0.08972 | 0.49259 | 0.094 | 0.15 | 1 | MS.Trt_vs_MS.NoTrt_F |
| SULT1A2 | 0.08977 | 0.25516 | 0.201 | 0.28 | 1 | MS.Trt_vs_MS.NoTrt_F |
| TRAF7 | 0.08988 | 0.88949 | 0.165 | 0.24 | 1 | MS.Trt_vs_MS.NoTrt_F |
| HIST1H2AG | 0.08993 | 0.55621 | 0.338 | 0.42 | 1 | MS.Trt_vs_MS.NoTrt_F |
| NPL | 0.09006 | -1.3931 | 0.266 | 0.34 | 1 | MS.Trt_vs_MS.NoTrt_F |
| FOXJ3 | 0.09019 | 0.33426 | 0.536 | 0.61 | 1 | MS.Trt_vs_MS.NoTrt_F |
| UTP14C | 0.09053 | 0.38581 | 0.205 | 0.28 | 1 | MS.Trt_vs_MS.NoTrt_F |
| SPECC1 | 0.09061 | 0.48088 | 0.932 | 0.93 | 1 | MS.Trt_vs_MS.NoTrt_F |
| RNF135 | 0.09061 | 0.8745 | 0.281 | 0.37 | 1 | MS.Trt_vs_MS.NoTrt_F |
| SPEG | 0.09076 | 0.46822 | 0.237 | 0.32 | 1 | MS.Trt_vs_MS.NoTrt_F |
| ABCG1 | 0.09083 | -0.266 | 0.432 | 0.5 | 1 | MS.Trt_vs_MS.NoTrt_F |
| RASSF2 | 0.09096 | 0.30411 | 0.356 | 0.45 | 1 | MS.Trt_vs_MS.NoTrt_F |
| FKBP11 | 0.09121 | -0.3341 | 0.701 | 0.76 | 1 | MS.Trt_vs_MS.NoTrt_F |
| STK11 | 0.0913 | -0.6022 | 0.381 | 0.47 | 1 | MS.Trt_vs_MS.NoTrt_F |
| ZNF512B | 0.09137 | -0.8551 | 0.061 | 0.11 | 1 | MS.Trt_vs_MS.NoTrt_F |
| METTL9 | 0.09138 | 0.62872 | 0.586 | 0.7 | 1 | MS.Trt_vs_MS.NoTrt_F |
| HSD17B4 | 0.0914 | 0.35044 | 0.594 | 0.7 | 1 | MS.Trt_vs_MS.NoTrt_F |
| TCEA2 | 0.09143 | -0.57 | 0.061 | 0.11 | 1 | MS.Trt_vs_MS.NoTrt_F |
| LRP10 | 0.09156 | 0.31762 | 0.759 | 0.82 | 1 | MS.Trt_vs_MS.NoTrt_F |
| CLOCK | 0.09165 | 0.84294 | 0.183 | 0.26 | 1 | MS.Trt_vs_MS.NoTrt_F |
| LCOR | 0.09199 | 0.46483 | 0.342 | 0.44 | 1 | MS.Trt_vs_MS.NoTrt_F |
| BRAT1 | 0.09202 | 0.74925 | 0.291 | 0.37 | 1 | MS.Trt_vs_MS.NoTrt_F |
| MTMR10 | 0.09218 | -1.2957 | 0.176 | 0.24 | 1 | MS.Trt_vs_MS.NoTrt_F |
| TSACC | 0.09219 | -0.3366 | 0.129 | 0.19 | 1 | MS.Trt_vs_MS.NoTrt_F |
| ADIPOR1 | 0.0925 | 0.35904 | 0.266 | 0.35 | 1 | MS.Trt_vs_MS.NoTrt_F |
| GPR25 | 0.0926 | 1.13573 | 0.065 | 0.11 | 1 | MS.Trt_vs_MS.NoTrt_F |
| C3orf67 | 0.09293 | 0.93192 | 0.442 | 0.37 | 1 | MS.Trt_vs_MS.NoTrt_F |
| CDC14B | 0.09302 | 0.55001 | 0.302 | 0.37 | 1 | MS.Trt_vs_MS.NoTrt_F |
| NEMP2 | 0.09316 | 0.77685 | 0.122 | 0.18 | 1 | MS.Trt_vs_MS.NoTrt_F |
| KLF9 | 0.09334 | 1.58858 | 0.068 | 0.12 | 1 | MS.Trt_vs_MS.NoTrt_F |
| SART3 | 0.09336 | 0.50469 | 0.176 | 0.25 | 1 | MS.Trt_vs_MS.NoTrt_F |
| RPL7L1 | 0.09337 | 0.26249 | 0.996 | 1 | 1 | MS.Trt_vs_MS.NoTrt_F |
| CCNC | 0.09341 | 0.35625 | 0.561 | 0.69 | 1 | MS.Trt_vs_MS.NoTrt_F |

| | | | | | | |
|---|---|---|---|---|---|---|
| MSC | 0.09343 | 0.99808 | 0.061 | 0.11 | 1 | MS.Trt_vs_MS.NoTrt_F |
| PTP4A2 | 0.09345 | 0.25194 | 0.525 | 0.61 | 1 | MS.Trt_vs_MS.NoTrt_F |
| USP45 | 0.09374 | 0.69787 | 0.18 | 0.25 | 1 | MS.Trt_vs_MS.NoTrt_F |
| SLC35B1 | 0.09429 | -0.2855 | 0.662 | 0.72 | 1 | MS.Trt_vs_MS.NoTrt_F |
| PSEN2 | 0.09438 | -1.0967 | 0.162 | 0.23 | 1 | MS.Trt_vs_MS.NoTrt_F |
| ST20-MTHFS | 0.09448 | 0.51121 | 0.151 | 0.22 | 1 | MS.Trt_vs_MS.NoTrt_F |
| DCAF6 | 0.09472 | 0.27489 | 0.223 | 0.31 | 1 | MS.Trt_vs_MS.NoTrt_F |
| MED6 | 0.09485 | -0.3113 | 0.504 | 0.56 | 1 | MS.Trt_vs_MS.NoTrt_F |
| DEAF1 | 0.09492 | -0.4465 | 0.331 | 0.42 | 1 | MS.Trt_vs_MS.NoTrt_F |
| C1orf109 | 0.09495 | -0.5554 | 0.921 | 0.86 | 1 | MS.Trt_vs_MS.NoTrt_F |
| ANGEL2 | 0.09499 | -0.3764 | 0.662 | 0.71 | 1 | MS.Trt_vs_MS.NoTrt_F |
| PTGIS | 0.09536 | 0.55423 | 0.752 | 0.75 | 1 | MS.Trt_vs_MS.NoTrt_F |
| DUSP23 | 0.0955 | 1.16947 | 0.122 | 0.19 | 1 | MS.Trt_vs_MS.NoTrt_F |
| ERBB3 | 0.09563 | 0.62155 | 0.723 | 0.82 | 1 | MS.Trt_vs_MS.NoTrt_F |
| ZMAT1 | 0.09577 | -0.4237 | 0.338 | 0.42 | 1 | MS.Trt_vs_MS.NoTrt_F |
| ABAT | 0.09608 | -0.6452 | 0.608 | 0.67 | 1 | MS.Trt_vs_MS.NoTrt_F |
| TPGS1 | 0.09612 | 0.51722 | 0.388 | 0.29 | 1 | MS.Trt_vs_MS.NoTrt_F |
| CGRRF1 | 0.09614 | 1.23626 | 0.245 | 0.33 | 1 | MS.Trt_vs_MS.NoTrt_F |
| TRIM61 | 0.09627 | -0.3152 | 0.234 | 0.3 | 1 | MS.Trt_vs_MS.NoTrt_F |
| SPG11 | 0.09633 | -0.3172 | 0.82 | 0.83 | 1 | MS.Trt_vs_MS.NoTrt_F |
| P2RX3 | 0.09636 | -0.2894 | 0.302 | 0.38 | 1 | MS.Trt_vs_MS.NoTrt_F |
| ACADSB | 0.09641 | 0.84173 | 0.612 | 0.72 | 1 | MS.Trt_vs_MS.NoTrt_F |
| NEURL1 | 0.09651 | -0.5876 | 0.23 | 0.29 | 1 | MS.Trt_vs_MS.NoTrt_F |
| TPH1 | 0.09653 | 0.62741 | 0.428 | 0.52 | 1 | MS.Trt_vs_MS.NoTrt_F |
| ADARB2 | 0.09666 | 0.56993 | 0.327 | 0.41 | 1 | MS.Trt_vs_MS.NoTrt_F |
| TTLL1 | 0.0968 | 0.53261 | 0.065 | 0.11 | 1 | MS.Trt_vs_MS.NoTrt_F |
| DLEC1 | 0.09688 | -0.9951 | 0.464 | 0.54 | 1 | MS.Trt_vs_MS.NoTrt_F |
| PPIP5K1 | 0.09694 | -0.4742 | 0.511 | 0.61 | 1 | MS.Trt_vs_MS.NoTrt_F |
| COG2 | 0.09697 | 1.30822 | 0.353 | 0.44 | 1 | MS.Trt_vs_MS.NoTrt_F |
| IFI27L1 | 0.09702 | 0.52401 | 0.076 | 0.12 | 1 | MS.Trt_vs_MS.NoTrt_F |
| RANBP6 | 0.09711 | 0.39119 | 0.101 | 0.16 | 1 | MS.Trt_vs_MS.NoTrt_F |
| SRD5A3 | 0.09718 | 1.03406 | 0.392 | 0.49 | 1 | MS.Trt_vs_MS.NoTrt_F |
| GPAT2 | 0.09718 | 0.51743 | 0.061 | 0.11 | 1 | MS.Trt_vs_MS.NoTrt_F |
| GTF2H2 | 0.09724 | -0.419 | 0.921 | 0.9 | 1 | MS.Trt_vs_MS.NoTrt_F |
| TIMM44 | 0.0973 | 0.72306 | 0.662 | 0.77 | 1 | MS.Trt_vs_MS.NoTrt_F |
| SPTY2D1 | 0.09751 | 0.51097 | 0.363 | 0.44 | 1 | MS.Trt_vs_MS.NoTrt_F |
| MYNN | 0.0976 | 0.421 | 0.277 | 0.35 | 1 | MS.Trt_vs_MS.NoTrt_F |
| LST1 | 0.09766 | 0.43574 | 0.255 | 0.33 | 1 | MS.Trt_vs_MS.NoTrt_F |
| SMN1 | 0.09768 | 0.29145 | 0.363 | 0.48 | 1 | MS.Trt_vs_MS.NoTrt_F |
| FO538757.1 | 0.09787 | 0.3583 | 0.191 | 0.27 | 1 | MS.Trt_vs_MS.NoTrt_F |

| | | | | | | |
|---|---|---|---|---|---|---|
| MRPL42 | 0.09802 | 0.29916 | 0.964 | 0.99 | 1 | MS.Trt_vs_MS.NoTrt_F |
| PIK3CB | 0.09804 | -0.8052 | 0.428 | 0.49 | 1 | MS.Trt_vs_MS.NoTrt_F |
| ZSCAN25 | 0.09854 | 0.27338 | 0.46 | 0.52 | 1 | MS.Trt_vs_MS.NoTrt_F |
| CGNL1 | 0.09866 | -0.535 | 0.335 | 0.41 | 1 | MS.Trt_vs_MS.NoTrt_F |
| DHX58 | 0.09868 | 1.59576 | 0.061 | 0.11 | 1 | MS.Trt_vs_MS.NoTrt_F |
| TSEN34 | 0.09872 | -0.5768 | 0.205 | 0.26 | 1 | MS.Trt_vs_MS.NoTrt_F |
| SERHL2 | 0.09877 | 0.59028 | 0.338 | 0.42 | 1 | MS.Trt_vs_MS.NoTrt_F |
| GPR35 | 0.0988 | -1.8272 | 0.27 | 0.33 | 1 | MS.Trt_vs_MS.NoTrt_F |
| ADAMTS17 | 0.0988 | 1.16926 | 0.183 | 0.25 | 1 | MS.Trt_vs_MS.NoTrt_F |
| STK40 | 0.09888 | 1.03436 | 0.112 | 0.17 | 1 | MS.Trt_vs_MS.NoTrt_F |
| CHD3 | 0.09888 | 0.31447 | 0.281 | 0.38 | 1 | MS.Trt_vs_MS.NoTrt_F |
| RPL27 | 0.09917 | 0.27237 | 0.586 | 0.77 | 1 | MS.Trt_vs_MS.NoTrt_F |
| JAGN1 | 0.09937 | 0.42729 | 0.129 | 0.19 | 1 | MS.Trt_vs_MS.NoTrt_F |
| PCBP4 | 0.09939 | 0.35349 | 0.119 | 0.18 | 1 | MS.Trt_vs_MS.NoTrt_F |
| ZNF74 | 0.09946 | -0.7171 | 0.446 | 0.51 | 1 | MS.Trt_vs_MS.NoTrt_F |
| CBX1 | 0.09958 | 0.35541 | 0.27 | 0.35 | 1 | MS.Trt_vs_MS.NoTrt_F |
| MIER1 | 0.09982 | 0.33112 | 0.658 | 0.78 | 1 | MS.Trt_vs_MS.NoTrt_F |
| RASGRF2 | 0.09984 | 0.39923 | 0.165 | 0.23 | 1 | MS.Trt_vs_MS.NoTrt_F |
| SLC12A9 | 0.09985 | -0.5516 | 0.867 | 0.9 | 1 | MS.Trt_vs_MS.NoTrt_F |
| TMEM39B | 0.09987 | -0.3979 | 0.155 | 0.21 | 1 | MS.Trt_vs_MS.NoTrt_F |
| SLC25A39 | 0.10001 | 0.36697 | 0.676 | 0.77 | 1 | MS.Trt_vs_MS.NoTrt_F |
| NEXN | 0.10012 | 0.25715 | 0.435 | 0.51 | 1 | MS.Trt_vs_MS.NoTrt_F |
| B3GNT5 | 0.10077 | -0.885 | 0.5 | 0.62 | 1 | MS.Trt_vs_MS.NoTrt_F |
| SRPK3 | 0.10083 | -0.3249 | 0.259 | 0.34 | 1 | MS.Trt_vs_MS.NoTrt_F |
| KLRK1 | 0.10106 | 1.78393 | 0.655 | 0.64 | 1 | MS.Trt_vs_MS.NoTrt_F |
| ASPH | 0.10111 | 0.60155 | 0.468 | 0.56 | 1 | MS.Trt_vs_MS.NoTrt_F |
| SPAG16 | 0.10118 | -0.7314 | 0.471 | 0.54 | 1 | MS.Trt_vs_MS.NoTrt_F |
| CAMK1D | 0.1012 | -0.2986 | 0.745 | 0.84 | 1 | MS.Trt_vs_MS.NoTrt_F |
| KCNJ6 | 0.10146 | 0.28482 | 0.827 | 0.83 | 1 | MS.Trt_vs_MS.NoTrt_F |
| MAD2L1BP | 0.10167 | 0.2886 | 0.18 | 0.25 | 1 | MS.Trt_vs_MS.NoTrt_F |
| BMPR2 | 0.10174 | -0.3961 | 0.54 | 0.61 | 1 | MS.Trt_vs_MS.NoTrt_F |
| PRKCB | 0.10179 | 0.51249 | 0.223 | 0.3 | 1 | MS.Trt_vs_MS.NoTrt_F |
| VWA2 | 0.1018 | -0.4062 | 0.73 | 0.79 | 1 | MS.Trt_vs_MS.NoTrt_F |
| THOC7 | 0.1019 | 0.5312 | 0.219 | 0.3 | 1 | MS.Trt_vs_MS.NoTrt_F |
| SLC25A19 | 0.10218 | -0.46 | 0.378 | 0.46 | 1 | MS.Trt_vs_MS.NoTrt_F |
| LAPTM4B | 0.10222 | -0.7884 | 0.183 | 0.24 | 1 | MS.Trt_vs_MS.NoTrt_F |
| BORCS8 | 0.10232 | 0.32441 | 0.906 | 0.95 | 1 | MS.Trt_vs_MS.NoTrt_F |
| ENOX2 | 0.10242 | 1.09221 | 0.129 | 0.19 | 1 | MS.Trt_vs_MS.NoTrt_F |
| CTNND1 | 0.10256 | 0.29273 | 0.356 | 0.45 | 1 | MS.Trt_vs_MS.NoTrt_F |
| JAK3 | 0.10265 | 0.63856 | 0.809 | 0.83 | 1 | MS.Trt_vs_MS.NoTrt_F |

| | | | | | | |
|---|---|---|---|---|---|---|
| MAP1LC3A | 0.10292 | 0.51252 | 0.083 | 0.13 | 1 | MS.Trt_vs_MS.NoTrt_F |
| AXL | 0.10305 | -0.8847 | 0.716 | 0.79 | 1 | MS.Trt_vs_MS.NoTrt_F |
| CD3EAP | 0.10336 | 0.42453 | 0.881 | 0.77 | 1 | MS.Trt_vs_MS.NoTrt_F |
| PTPN4 | 0.10345 | 0.32865 | 0.619 | 0.71 | 1 | MS.Trt_vs_MS.NoTrt_F |
| ZNF276 | 0.1035 | 0.27777 | 0.529 | 0.6 | 1 | MS.Trt_vs_MS.NoTrt_F |
| ZGRF1 | 0.1036 | 0.66506 | 0.705 | 0.83 | 1 | MS.Trt_vs_MS.NoTrt_F |
| CTDP1 | 0.10366 | 1.61401 | 0.086 | 0.14 | 1 | MS.Trt_vs_MS.NoTrt_F |
| KAT14 | 0.10376 | 0.49982 | 0.108 | 0.17 | 1 | MS.Trt_vs_MS.NoTrt_F |
| LANCL2 | 0.10414 | 0.38571 | 0.137 | 0.2 | 1 | MS.Trt_vs_MS.NoTrt_F |
| PROSER1 | 0.1042 | 0.44701 | 0.194 | 0.27 | 1 | MS.Trt_vs_MS.NoTrt_F |
| CHMP5 | 0.10434 | 0.59175 | 0.14 | 0.2 | 1 | MS.Trt_vs_MS.NoTrt_F |
| HOXB7 | 0.10447 | -0.3212 | 0.23 | 0.32 | 1 | MS.Trt_vs_MS.NoTrt_F |
| CSF1 | 0.10476 | 0.51397 | 0.129 | 0.19 | 1 | MS.Trt_vs_MS.NoTrt_F |
| ZNF700 | 0.10515 | 0.48507 | 0.191 | 0.26 | 1 | MS.Trt_vs_MS.NoTrt_F |
| WSB2 | 0.10518 | 0.61284 | 0.831 | 0.85 | 1 | MS.Trt_vs_MS.NoTrt_F |
| SEMA4F | 0.10546 | -0.5602 | 0.518 | 0.59 | 1 | MS.Trt_vs_MS.NoTrt_F |
| PLA2G15 | 0.10575 | -0.693 | 0.745 | 0.82 | 1 | MS.Trt_vs_MS.NoTrt_F |
| UNC45B | 0.10576 | -0.5145 | 0.151 | 0.21 | 1 | MS.Trt_vs_MS.NoTrt_F |
| WASL | 0.1058 | 0.57582 | 0.119 | 0.18 | 1 | MS.Trt_vs_MS.NoTrt_F |
| RP11-1035H | 0.10586 | 1.23619 | 0.126 | 0.19 | 1 | MS.Trt_vs_MS.NoTrt_F |
| C21orf33 | 0.10592 | 0.56207 | 0.374 | 0.46 | 1 | MS.Trt_vs_MS.NoTrt_F |
| GOLGA2 | 0.10595 | 0.28244 | 0.558 | 0.69 | 1 | MS.Trt_vs_MS.NoTrt_F |
| ATP5J | 0.10624 | 0.87141 | 0.237 | 0.34 | 1 | MS.Trt_vs_MS.NoTrt_F |
| ACSL3 | 0.1063 | -0.319 | 0.115 | 0.17 | 1 | MS.Trt_vs_MS.NoTrt_F |
| BSDC1 | 0.1063 | 0.30815 | 0.45 | 0.55 | 1 | MS.Trt_vs_MS.NoTrt_F |
| EME2 | 0.10644 | -0.4582 | 0.281 | 0.37 | 1 | MS.Trt_vs_MS.NoTrt_F |
| ZNF121 | 0.1065 | 0.43426 | 0.899 | 0.94 | 1 | MS.Trt_vs_MS.NoTrt_F |
| GMNN | 0.10657 | 0.25442 | 0.072 | 0.12 | 1 | MS.Trt_vs_MS.NoTrt_F |
| GMEB1 | 0.10663 | -0.9183 | 0.791 | 0.83 | 1 | MS.Trt_vs_MS.NoTrt_F |
| TPRG1 | 0.10725 | 0.75006 | 0.219 | 0.3 | 1 | MS.Trt_vs_MS.NoTrt_F |
| CYHR1 | 0.10737 | 0.43299 | 0.27 | 0.35 | 1 | MS.Trt_vs_MS.NoTrt_F |
| XRCC2 | 0.10743 | -0.4089 | 0.734 | 0.81 | 1 | MS.Trt_vs_MS.NoTrt_F |
| ERCC6L | 0.10745 | 1.39401 | 0.144 | 0.21 | 1 | MS.Trt_vs_MS.NoTrt_F |
| SH2D1A | 0.10752 | 0.71886 | 0.241 | 0.34 | 1 | MS.Trt_vs_MS.NoTrt_F |
| MED29 | 0.10765 | 0.41304 | 0.586 | 0.67 | 1 | MS.Trt_vs_MS.NoTrt_F |
| CHN1 | 0.10772 | 0.85021 | 0.079 | 0.13 | 1 | MS.Trt_vs_MS.NoTrt_F |
| GGA3 | 0.10782 | 0.25972 | 0.212 | 0.29 | 1 | MS.Trt_vs_MS.NoTrt_F |
| ZNF540 | 0.10814 | 0.97142 | 0.493 | 0.57 | 1 | MS.Trt_vs_MS.NoTrt_F |
| GTF2H2C.2 | 0.10819 | 0.67957 | 0.255 | 0.34 | 1 | MS.Trt_vs_MS.NoTrt_F |
| ATP5L | 0.10822 | 0.59536 | 0.421 | 0.58 | 1 | MS.Trt_vs_MS.NoTrt_F |

| | | | | | | |
|---|---|---|---|---|---|---|
| GARNL3 | 0.10823 | 0.78566 | 0.187 | 0.25 | 1 | MS.Trt_vs_MS.NoTrt_F |
| PAK1IP1 | 0.10849 | 0.85515 | 0.083 | 0.13 | 1 | MS.Trt_vs_MS.NoTrt_F |
| TUBD1 | 0.1086 | -0.7891 | 0.755 | 0.82 | 1 | MS.Trt_vs_MS.NoTrt_F |
| PDS5B | 0.10867 | 0.55168 | 0.363 | 0.44 | 1 | MS.Trt_vs_MS.NoTrt_F |
| SOS1 | 0.10875 | 0.39283 | 0.191 | 0.26 | 1 | MS.Trt_vs_MS.NoTrt_F |
| THOC2 | 0.10882 | 0.26423 | 0.532 | 0.61 | 1 | MS.Trt_vs_MS.NoTrt_F |
| CHRM5 | 0.109 | 0.31596 | 0.281 | 0.36 | 1 | MS.Trt_vs_MS.NoTrt_F |
| SORBS3 | 0.109 | 1.06742 | 0.083 | 0.13 | 1 | MS.Trt_vs_MS.NoTrt_F |
| POLRMT | 0.10914 | 1.16754 | 0.119 | 0.18 | 1 | MS.Trt_vs_MS.NoTrt_F |
| RP11-286N2 | 0.10915 | 0.32903 | 0.957 | 0.97 | 1 | MS.Trt_vs_MS.NoTrt_F |
| AIFM1 | 0.10918 | -0.5473 | 0.572 | 0.63 | 1 | MS.Trt_vs_MS.NoTrt_F |
| DHX29 | 0.10928 | 0.25486 | 0.18 | 0.25 | 1 | MS.Trt_vs_MS.NoTrt_F |
| TANC2 | 0.10928 | 1.06991 | 0.223 | 0.3 | 1 | MS.Trt_vs_MS.NoTrt_F |
| SETMAR | 0.1093 | 0.60287 | 0.331 | 0.45 | 1 | MS.Trt_vs_MS.NoTrt_F |
| GRK5 | 0.10962 | 0.68731 | 0.187 | 0.26 | 1 | MS.Trt_vs_MS.NoTrt_F |
| FOXK1 | 0.10968 | 0.32385 | 0.853 | 0.9 | 1 | MS.Trt_vs_MS.NoTrt_F |
| ZNF267 | 0.1097 | 0.38239 | 0.201 | 0.28 | 1 | MS.Trt_vs_MS.NoTrt_F |
| PLXNC1 | 0.10972 | -0.6324 | 0.101 | 0.16 | 1 | MS.Trt_vs_MS.NoTrt_F |
| EFCAB5 | 0.10976 | -0.6248 | 0.442 | 0.53 | 1 | MS.Trt_vs_MS.NoTrt_F |
| GPR132 | 0.11033 | -0.3853 | 0.378 | 0.46 | 1 | MS.Trt_vs_MS.NoTrt_F |
| GLA | 0.11052 | 0.92998 | 0.079 | 0.13 | 1 | MS.Trt_vs_MS.NoTrt_F |
| MSL3 | 0.11071 | 1.10168 | 0.18 | 0.26 | 1 | MS.Trt_vs_MS.NoTrt_F |
| FASTKD2 | 0.1109 | 0.68893 | 0.529 | 0.62 | 1 | MS.Trt_vs_MS.NoTrt_F |
| NAA15 | 0.11099 | 0.43399 | 0.788 | 0.79 | 1 | MS.Trt_vs_MS.NoTrt_F |
| MELTF | 0.11101 | 0.28562 | 0.112 | 0.17 | 1 | MS.Trt_vs_MS.NoTrt_F |
| TAF13 | 0.11126 | 0.39 | 0.165 | 0.23 | 1 | MS.Trt_vs_MS.NoTrt_F |
| CEP78 | 0.11148 | -0.5573 | 0.522 | 0.6 | 1 | MS.Trt_vs_MS.NoTrt_F |
| KIAA0408 | 0.11156 | -0.4917 | 0.291 | 0.36 | 1 | MS.Trt_vs_MS.NoTrt_F |
| MCF2L2 | 0.11189 | -0.5348 | 0.935 | 0.95 | 1 | MS.Trt_vs_MS.NoTrt_F |
| LYG2 | 0.11189 | 1.01873 | 0.083 | 0.13 | 1 | MS.Trt_vs_MS.NoTrt_F |
| RLF | 0.11192 | 0.4242 | 0.23 | 0.32 | 1 | MS.Trt_vs_MS.NoTrt_F |
| ZNF512 | 0.11207 | 0.54535 | 0.151 | 0.22 | 1 | MS.Trt_vs_MS.NoTrt_F |
| UBR1 | 0.11207 | 0.33409 | 0.691 | 0.72 | 1 | MS.Trt_vs_MS.NoTrt_F |
| NTPCR | 0.11224 | 0.56361 | 0.417 | 0.5 | 1 | MS.Trt_vs_MS.NoTrt_F |
| DHX32 | 0.11229 | 0.41675 | 0.068 | 0.11 | 1 | MS.Trt_vs_MS.NoTrt_F |
| UBE2E3 | 0.11236 | 1.19827 | 0.079 | 0.13 | 1 | MS.Trt_vs_MS.NoTrt_F |
| RNF41 | 0.11261 | -0.2526 | 0.719 | 0.82 | 1 | MS.Trt_vs_MS.NoTrt_F |
| AHCYL1 | 0.11265 | -0.3441 | 0.097 | 0.15 | 1 | MS.Trt_vs_MS.NoTrt_F |
| ILF3 | 0.11266 | 0.2502 | 0.723 | 0.83 | 1 | MS.Trt_vs_MS.NoTrt_F |
| PLPP6 | 0.1128 | 0.68361 | 0.094 | 0.14 | 1 | MS.Trt_vs_MS.NoTrt_F |

| | | | | | | |
|---|---|---|---|---|---|---|
| TNFSF8 | 0.11292 | 0.80483 | 0.151 | 0.22 | 1 | MS.Trt_vs_MS.NoTrt_F |
| MCTP2 | 0.11316 | 0.91147 | 0.378 | 0.46 | 1 | MS.Trt_vs_MS.NoTrt_F |
| AP2A1 | 0.1133 | -0.4306 | 0.716 | 0.82 | 1 | MS.Trt_vs_MS.NoTrt_F |
| NOL4L | 0.11333 | 0.26699 | 0.511 | 0.69 | 1 | MS.Trt_vs_MS.NoTrt_F |
| ALAS1 | 0.11348 | -0.404 | 0.108 | 0.16 | 1 | MS.Trt_vs_MS.NoTrt_F |
| COPB2 | 0.11366 | -0.3599 | 0.896 | 0.96 | 1 | MS.Trt_vs_MS.NoTrt_F |
| BTBD11 | 0.11391 | 0.7327 | 0.442 | 0.59 | 1 | MS.Trt_vs_MS.NoTrt_F |
| ZCCHC8 | 0.11403 | 0.3768 | 0.853 | 0.88 | 1 | MS.Trt_vs_MS.NoTrt_F |
| ALG6 | 0.11418 | 0.38316 | 0.183 | 0.26 | 1 | MS.Trt_vs_MS.NoTrt_F |
| CLEC2D | 0.11445 | 0.59717 | 0.827 | 0.88 | 1 | MS.Trt_vs_MS.NoTrt_F |
| MCCC1 | 0.11456 | -0.3233 | 0.493 | 0.58 | 1 | MS.Trt_vs_MS.NoTrt_F |
| SLC38A6 | 0.11472 | 0.68779 | 0.561 | 0.67 | 1 | MS.Trt_vs_MS.NoTrt_F |
| CENPA | 0.11493 | -0.2601 | 0.086 | 0.13 | 1 | MS.Trt_vs_MS.NoTrt_F |
| WDR3 | 0.11505 | -1.1051 | 0.741 | 0.79 | 1 | MS.Trt_vs_MS.NoTrt_F |
| EI24 | 0.11528 | 0.42876 | 0.716 | 0.84 | 1 | MS.Trt_vs_MS.NoTrt_F |
| RNF26 | 0.11546 | 1.59454 | 0.129 | 0.19 | 1 | MS.Trt_vs_MS.NoTrt_F |
| POLR2E | 0.11566 | 0.27771 | 0.583 | 0.66 | 1 | MS.Trt_vs_MS.NoTrt_F |
| SCAF1 | 0.11568 | 0.2652 | 0.09 | 0.14 | 1 | MS.Trt_vs_MS.NoTrt_F |
| MUM1 | 0.11576 | 0.25227 | 0.633 | 0.69 | 1 | MS.Trt_vs_MS.NoTrt_F |
| TRIM39 | 0.11586 | 0.3571 | 0.165 | 0.23 | 1 | MS.Trt_vs_MS.NoTrt_F |
| SRBD1 | 0.11604 | 1.10976 | 0.356 | 0.45 | 1 | MS.Trt_vs_MS.NoTrt_F |
| FAM86B1 | 0.11608 | -0.934 | 0.115 | 0.17 | 1 | MS.Trt_vs_MS.NoTrt_F |
| PFDN4 | 0.11629 | 1.13048 | 0.119 | 0.18 | 1 | MS.Trt_vs_MS.NoTrt_F |
| SC5D | 0.1163 | 0.26257 | 0.583 | 0.63 | 1 | MS.Trt_vs_MS.NoTrt_F |
| ARRB1 | 0.11662 | 0.54338 | 0.737 | 0.81 | 1 | MS.Trt_vs_MS.NoTrt_F |
| CDK18 | 0.11684 | -0.2872 | 0.205 | 0.27 | 1 | MS.Trt_vs_MS.NoTrt_F |
| MCRIP1 | 0.11686 | 0.39887 | 0.41 | 0.48 | 1 | MS.Trt_vs_MS.NoTrt_F |
| TANGO6 | 0.11692 | -0.4566 | 0.763 | 0.83 | 1 | MS.Trt_vs_MS.NoTrt_F |
| THUMPD2 | 0.1171 | 1.01253 | 0.417 | 0.5 | 1 | MS.Trt_vs_MS.NoTrt_F |
| SKIL | 0.11718 | 0.28106 | 0.374 | 0.47 | 1 | MS.Trt_vs_MS.NoTrt_F |
| ALG5 | 0.11722 | 0.47662 | 0.478 | 0.38 | 1 | MS.Trt_vs_MS.NoTrt_F |
| SNAP29 | 0.11722 | -0.371 | 0.683 | 0.77 | 1 | MS.Trt_vs_MS.NoTrt_F |
| SLC52A2 | 0.11734 | -0.2557 | 0.068 | 0.11 | 1 | MS.Trt_vs_MS.NoTrt_F |
| REL | 0.11767 | 0.33188 | 0.96 | 0.97 | 1 | MS.Trt_vs_MS.NoTrt_F |
| ANKHD1-EIF | 0.11769 | 1.56874 | 0.079 | 0.13 | 1 | MS.Trt_vs_MS.NoTrt_F |
| C15orf38-AF | 0.11777 | -0.3306 | 0.374 | 0.43 | 1 | MS.Trt_vs_MS.NoTrt_F |
| LILRB3 | 0.11778 | -0.7427 | 0.144 | 0.2 | 1 | MS.Trt_vs_MS.NoTrt_F |
| BAG5 | 0.11817 | -0.8469 | 0.331 | 0.4 | 1 | MS.Trt_vs_MS.NoTrt_F |
| SLC7A5 | 0.11836 | 0.4701 | 0.133 | 0.19 | 1 | MS.Trt_vs_MS.NoTrt_F |
| KLRG1 | 0.11839 | 0.80687 | 0.371 | 0.48 | 1 | MS.Trt_vs_MS.NoTrt_F |

| | | | | | | |
|---|---|---|---|---|---|---|
| B3GNTL1 | 0.11846 | 0.41757 | 0.288 | 0.35 | 1 | MS.Trt_vs_MS.NoTrt_F |
| MS4A2 | 0.11861 | 0.3569 | 0.568 | 0.59 | 1 | MS.Trt_vs_MS.NoTrt_F |
| SNX24 | 0.11871 | 0.63446 | 0.068 | 0.11 | 1 | MS.Trt_vs_MS.NoTrt_F |
| TBC1D32 | 0.11872 | 0.70143 | 0.46 | 0.53 | 1 | MS.Trt_vs_MS.NoTrt_F |
| RPS15 | 0.11894 | 0.29655 | 0.881 | 0.99 | 1 | MS.Trt_vs_MS.NoTrt_F |
| DTX3L | 0.11901 | 0.80694 | 0.658 | 0.79 | 1 | MS.Trt_vs_MS.NoTrt_F |
| YIF1B | 0.11916 | 0.36896 | 0.608 | 0.72 | 1 | MS.Trt_vs_MS.NoTrt_F |
| GCNT4 | 0.11936 | 0.46273 | 0.439 | 0.57 | 1 | MS.Trt_vs_MS.NoTrt_F |
| ABHD18 | 0.11974 | 0.33279 | 0.867 | 0.85 | 1 | MS.Trt_vs_MS.NoTrt_F |
| CCDC102B | 0.11977 | 0.2626 | 0.191 | 0.25 | 1 | MS.Trt_vs_MS.NoTrt_F |
| NSMF | 0.11998 | 1.94461 | 0.09 | 0.14 | 1 | MS.Trt_vs_MS.NoTrt_F |
| CHD1L | 0.12028 | 0.91063 | 0.14 | 0.2 | 1 | MS.Trt_vs_MS.NoTrt_F |
| HGS | 0.12035 | 0.40788 | 0.626 | 0.7 | 1 | MS.Trt_vs_MS.NoTrt_F |
| SERPINF1 | 0.12051 | 0.31849 | 0.791 | 0.88 | 1 | MS.Trt_vs_MS.NoTrt_F |
| ABCD4 | 0.12058 | 0.70877 | 0.799 | 0.83 | 1 | MS.Trt_vs_MS.NoTrt_F |
| RP11-15H20 | 0.12061 | -0.4463 | 0.151 | 0.21 | 1 | MS.Trt_vs_MS.NoTrt_F |
| GOLGA6L9 | 0.12082 | 0.65689 | 0.068 | 0.11 | 1 | MS.Trt_vs_MS.NoTrt_F |
| NKIRAS2 | 0.12086 | 0.92444 | 0.835 | 0.86 | 1 | MS.Trt_vs_MS.NoTrt_F |
| HLA-DOA | 0.121 | -0.9882 | 0.604 | 0.67 | 1 | MS.Trt_vs_MS.NoTrt_F |
| POU2F1 | 0.12137 | 0.2936 | 0.399 | 0.48 | 1 | MS.Trt_vs_MS.NoTrt_F |
| LY75-CD302 | 0.12149 | 1.76879 | 0.097 | 0.15 | 1 | MS.Trt_vs_MS.NoTrt_F |
| DCAF17 | 0.12166 | -0.5611 | 0.381 | 0.45 | 1 | MS.Trt_vs_MS.NoTrt_F |
| ATP5SL | 0.12169 | -0.3109 | 0.504 | 0.59 | 1 | MS.Trt_vs_MS.NoTrt_F |
| RAI14 | 0.12181 | 0.38465 | 0.101 | 0.15 | 1 | MS.Trt_vs_MS.NoTrt_F |
| RPS25 | 0.12222 | 0.25441 | 0.694 | 0.91 | 1 | MS.Trt_vs_MS.NoTrt_F |
| HIST1H4I | 0.12226 | -0.8774 | 0.068 | 0.11 | 1 | MS.Trt_vs_MS.NoTrt_F |
| WDR75 | 0.12239 | -0.2946 | 0.353 | 0.43 | 1 | MS.Trt_vs_MS.NoTrt_F |
| PTCD1 | 0.12278 | -0.5225 | 0.633 | 0.69 | 1 | MS.Trt_vs_MS.NoTrt_F |
| DNAJB7 | 0.12281 | 0.31441 | 0.227 | 0.31 | 1 | MS.Trt_vs_MS.NoTrt_F |
| STK16 | 0.12295 | 0.29319 | 0.115 | 0.17 | 1 | MS.Trt_vs_MS.NoTrt_F |
| BCS1L | 0.12295 | 0.7529 | 0.108 | 0.16 | 1 | MS.Trt_vs_MS.NoTrt_F |
| NOM1 | 0.12303 | -0.9726 | 0.619 | 0.69 | 1 | MS.Trt_vs_MS.NoTrt_F |
| TTC22 | 0.12305 | -1.1038 | 0.432 | 0.52 | 1 | MS.Trt_vs_MS.NoTrt_F |
| MAP2K2 | 0.1232 | 0.88811 | 0.705 | 0.78 | 1 | MS.Trt_vs_MS.NoTrt_F |
| ATP5H | 0.12322 | 0.55376 | 0.748 | 0.81 | 1 | MS.Trt_vs_MS.NoTrt_F |
| CNOT9 | 0.12339 | 0.34911 | 0.752 | 0.84 | 1 | MS.Trt_vs_MS.NoTrt_F |
| GIN1 | 0.12345 | 0.49762 | 0.367 | 0.46 | 1 | MS.Trt_vs_MS.NoTrt_F |
| SLC35A5 | 0.12356 | 0.4565 | 0.489 | 0.56 | 1 | MS.Trt_vs_MS.NoTrt_F |
| TMEM177 | 0.12356 | 0.2958 | 0.072 | 0.12 | 1 | MS.Trt_vs_MS.NoTrt_F |
| TCEA3 | 0.12357 | 1.43443 | 0.076 | 0.12 | 1 | MS.Trt_vs_MS.NoTrt_F |

| | | | | | | |
|---|---|---|---|---|---|---|
| TXNL4B | 0.1236 | 0.33732 | 0.353 | 0.43 | 1 | MS.Trt_vs_MS.NoTrt_F |
| SMTN | 0.12377 | 0.76227 | 0.399 | 0.32 | 1 | MS.Trt_vs_MS.NoTrt_F |
| KLB | 0.12379 | 0.26511 | 0.777 | 0.76 | 1 | MS.Trt_vs_MS.NoTrt_F |
| SEC13 | 0.12395 | 0.41802 | 0.187 | 0.26 | 1 | MS.Trt_vs_MS.NoTrt_F |
| NT5DC3 | 0.12404 | 0.57976 | 0.827 | 0.83 | 1 | MS.Trt_vs_MS.NoTrt_F |
| NDUFB9 | 0.12448 | 0.37122 | 0.237 | 0.32 | 1 | MS.Trt_vs_MS.NoTrt_F |
| PASK | 0.12481 | -0.265 | 0.586 | 0.65 | 1 | MS.Trt_vs_MS.NoTrt_F |
| TTC7A | 0.12488 | 0.4138 | 0.223 | 0.3 | 1 | MS.Trt_vs_MS.NoTrt_F |
| COQ2 | 0.12497 | -0.5475 | 0.065 | 0.11 | 1 | MS.Trt_vs_MS.NoTrt_F |
| DNAJA4 | 0.12506 | 0.4198 | 0.176 | 0.24 | 1 | MS.Trt_vs_MS.NoTrt_F |
| ANAPC16 | 0.12507 | 0.29811 | 0.946 | 0.97 | 1 | MS.Trt_vs_MS.NoTrt_F |
| HERPUD1 | 0.12515 | 0.28352 | 0.745 | 0.78 | 1 | MS.Trt_vs_MS.NoTrt_F |
| BRIP1 | 0.12517 | -0.4998 | 0.809 | 0.87 | 1 | MS.Trt_vs_MS.NoTrt_F |
| SLC2A11 | 0.12527 | -0.2592 | 0.863 | 0.83 | 1 | MS.Trt_vs_MS.NoTrt_F |
| SMURF1 | 0.12528 | 0.3072 | 0.191 | 0.27 | 1 | MS.Trt_vs_MS.NoTrt_F |
| MT1X | 0.12529 | 0.33177 | 0.155 | 0.22 | 1 | MS.Trt_vs_MS.NoTrt_F |
| NDUFB7 | 0.12531 | 0.84694 | 0.245 | 0.35 | 1 | MS.Trt_vs_MS.NoTrt_F |
| CDV3 | 0.12535 | 0.71572 | 0.263 | 0.36 | 1 | MS.Trt_vs_MS.NoTrt_F |
| ENTPD7 | 0.12549 | -0.4935 | 0.353 | 0.44 | 1 | MS.Trt_vs_MS.NoTrt_F |
| ZNF28 | 0.12549 | 0.82214 | 0.777 | 0.87 | 1 | MS.Trt_vs_MS.NoTrt_F |
| CDCA7L | 0.12573 | 0.53724 | 0.108 | 0.16 | 1 | MS.Trt_vs_MS.NoTrt_F |
| KLHL21 | 0.12575 | -0.3032 | 0.727 | 0.81 | 1 | MS.Trt_vs_MS.NoTrt_F |
| NR1D2 | 0.12604 | 0.50639 | 0.173 | 0.24 | 1 | MS.Trt_vs_MS.NoTrt_F |
| NR2C2AP | 0.12639 | 0.54625 | 0.122 | 0.18 | 1 | MS.Trt_vs_MS.NoTrt_F |
| UQCC3 | 0.12642 | 0.8418 | 0.428 | 0.5 | 1 | MS.Trt_vs_MS.NoTrt_F |
| HCCS | 0.12649 | 1.34812 | 0.097 | 0.15 | 1 | MS.Trt_vs_MS.NoTrt_F |
| TIPRL | 0.12652 | 0.3235 | 0.302 | 0.37 | 1 | MS.Trt_vs_MS.NoTrt_F |
| GOLGA7B | 0.12653 | 0.82253 | 0.086 | 0.13 | 1 | MS.Trt_vs_MS.NoTrt_F |
| CROCC | 0.12654 | 0.29535 | 0.554 | 0.48 | 1 | MS.Trt_vs_MS.NoTrt_F |
| CAMK2N1 | 0.12672 | -0.5323 | 0.108 | 0.16 | 1 | MS.Trt_vs_MS.NoTrt_F |
| TMEM126B | 0.12693 | 0.80322 | 0.205 | 0.27 | 1 | MS.Trt_vs_MS.NoTrt_F |
| RAB21 | 0.12694 | 0.27976 | 0.968 | 0.98 | 1 | MS.Trt_vs_MS.NoTrt_F |
| ZKSCAN1 | 0.12702 | 0.4047 | 0.96 | 0.96 | 1 | MS.Trt_vs_MS.NoTrt_F |
| RPS6KA2 | 0.12712 | 0.88752 | 0.104 | 0.16 | 1 | MS.Trt_vs_MS.NoTrt_F |
| ZACN | 0.12719 | 0.70674 | 0.086 | 0.13 | 1 | MS.Trt_vs_MS.NoTrt_F |
| DUSP7 | 0.12734 | 0.58847 | 0.147 | 0.21 | 1 | MS.Trt_vs_MS.NoTrt_F |
| IL27RA | 0.12772 | 0.61846 | 0.385 | 0.49 | 1 | MS.Trt_vs_MS.NoTrt_F |
| TBC1D9B | 0.12782 | 0.65684 | 0.446 | 0.55 | 1 | MS.Trt_vs_MS.NoTrt_F |
| MT-ND5 | 0.12798 | 0.47546 | 0.813 | 0.88 | 1 | MS.Trt_vs_MS.NoTrt_F |
| NFAT5 | 0.12798 | -0.5367 | 0.838 | 0.89 | 1 | MS.Trt_vs_MS.NoTrt_F |

| | | | | | | |
|---|---|---|---|---|---|---|
| ACTL6A | 0.12838 | 0.35332 | 0.259 | 0.34 | 1 | MS.Trt_vs_MS.NoTrt_F |
| LAP3 | 0.12853 | 0.71574 | 0.18 | 0.25 | 1 | MS.Trt_vs_MS.NoTrt_F |
| DPH6 | 0.12853 | 0.47233 | 0.363 | 0.45 | 1 | MS.Trt_vs_MS.NoTrt_F |
| RAP1GDS1 | 0.12857 | 0.60704 | 0.363 | 0.5 | 1 | MS.Trt_vs_MS.NoTrt_F |
| BCL2L14 | 0.12868 | 0.29407 | 0.716 | 0.7 | 1 | MS.Trt_vs_MS.NoTrt_F |
| PRG4 | 0.12871 | -0.3045 | 0.065 | 0.11 | 1 | MS.Trt_vs_MS.NoTrt_F |
| MKL2 | 0.12875 | -0.8305 | 0.705 | 0.8 | 1 | MS.Trt_vs_MS.NoTrt_F |
| SPTBN1 | 0.12907 | 0.36837 | 0.288 | 0.38 | 1 | MS.Trt_vs_MS.NoTrt_F |
| TESPA1 | 0.1291 | 0.56381 | 0.41 | 0.52 | 1 | MS.Trt_vs_MS.NoTrt_F |
| FREM2 | 0.12919 | 0.26664 | 0.673 | 0.62 | 1 | MS.Trt_vs_MS.NoTrt_F |
| DTD1 | 0.12946 | -0.7673 | 0.349 | 0.4 | 1 | MS.Trt_vs_MS.NoTrt_F |
| TMED3 | 0.12947 | 0.35269 | 0.558 | 0.65 | 1 | MS.Trt_vs_MS.NoTrt_F |
| PAQR8 | 0.12953 | 0.75721 | 0.514 | 0.61 | 1 | MS.Trt_vs_MS.NoTrt_F |
| KSR2 | 0.12954 | -0.5572 | 0.237 | 0.3 | 1 | MS.Trt_vs_MS.NoTrt_F |
| COA1 | 0.13013 | 0.49541 | 0.748 | 0.87 | 1 | MS.Trt_vs_MS.NoTrt_F |
| TGFBR1 | 0.1302 | 1.52705 | 0.122 | 0.18 | 1 | MS.Trt_vs_MS.NoTrt_F |
| FAM20A | 0.13025 | -0.7874 | 0.18 | 0.24 | 1 | MS.Trt_vs_MS.NoTrt_F |
| SHC1 | 0.13039 | 0.34239 | 0.601 | 0.67 | 1 | MS.Trt_vs_MS.NoTrt_F |
| CUL3 | 0.13078 | 0.28585 | 0.309 | 0.4 | 1 | MS.Trt_vs_MS.NoTrt_F |
| MAEA | 0.13111 | -0.2943 | 0.633 | 0.71 | 1 | MS.Trt_vs_MS.NoTrt_F |
| WBP4 | 0.13135 | 0.88056 | 0.112 | 0.17 | 1 | MS.Trt_vs_MS.NoTrt_F |
| FAM193B | 0.13198 | 0.45668 | 0.471 | 0.56 | 1 | MS.Trt_vs_MS.NoTrt_F |
| SURF4 | 0.13212 | 0.54582 | 0.317 | 0.44 | 1 | MS.Trt_vs_MS.NoTrt_F |
| APBA1 | 0.13214 | -1.1791 | 0.385 | 0.45 | 1 | MS.Trt_vs_MS.NoTrt_F |
| NEK6 | 0.13232 | 0.96095 | 0.119 | 0.18 | 1 | MS.Trt_vs_MS.NoTrt_F |
| PYGO1 | 0.13238 | 0.33801 | 0.381 | 0.34 | 1 | MS.Trt_vs_MS.NoTrt_F |
| GANC | 0.13249 | 0.83716 | 0.647 | 0.7 | 1 | MS.Trt_vs_MS.NoTrt_F |
| EXD3 | 0.13259 | 0.26464 | 0.478 | 0.56 | 1 | MS.Trt_vs_MS.NoTrt_F |
| MYBL1 | 0.13263 | 0.5731 | 0.065 | 0.11 | 1 | MS.Trt_vs_MS.NoTrt_F |
| FANCG | 0.13268 | 0.59398 | 0.104 | 0.16 | 1 | MS.Trt_vs_MS.NoTrt_F |
| SDF2L1 | 0.13319 | 0.56493 | 0.173 | 0.24 | 1 | MS.Trt_vs_MS.NoTrt_F |
| RNF180 | 0.13323 | 0.66879 | 0.306 | 0.4 | 1 | MS.Trt_vs_MS.NoTrt_F |
| NSMCE4A | 0.13357 | 0.47917 | 0.349 | 0.42 | 1 | MS.Trt_vs_MS.NoTrt_F |
| ST13 | 0.13376 | 0.51747 | 0.345 | 0.44 | 1 | MS.Trt_vs_MS.NoTrt_F |
| DPAGT1 | 0.13408 | -0.5963 | 0.112 | 0.16 | 1 | MS.Trt_vs_MS.NoTrt_F |
| SGSM2 | 0.13422 | 0.26637 | 0.263 | 0.33 | 1 | MS.Trt_vs_MS.NoTrt_F |
| PPAT | 0.13433 | 1.64413 | 0.281 | 0.34 | 1 | MS.Trt_vs_MS.NoTrt_F |
| TFDP2 | 0.13435 | 0.6306 | 0.932 | 0.94 | 1 | MS.Trt_vs_MS.NoTrt_F |
| NEDD8 | 0.13447 | 0.40985 | 0.788 | 0.88 | 1 | MS.Trt_vs_MS.NoTrt_F |
| DNAJC16 | 0.13456 | 0.44881 | 0.723 | 0.82 | 1 | MS.Trt_vs_MS.NoTrt_F |

| | | | | | | |
|---|---|---|---|---|---|---|
| PHKB | 0.1347 | -0.4648 | 0.694 | 0.77 | 1 | MS.Trt_vs_MS.NoTrt_F |
| FRS2 | 0.13495 | -0.36 | 0.129 | 0.18 | 1 | MS.Trt_vs_MS.NoTrt_F |
| EPN2 | 0.13505 | 0.47975 | 0.835 | 0.9 | 1 | MS.Trt_vs_MS.NoTrt_F |
| MSI2 | 0.13506 | -0.5548 | 0.863 | 0.91 | 1 | MS.Trt_vs_MS.NoTrt_F |
| VPS29 | 0.13519 | 0.57559 | 0.482 | 0.56 | 1 | MS.Trt_vs_MS.NoTrt_F |
| GID4 | 0.13521 | -0.422 | 0.673 | 0.67 | 1 | MS.Trt_vs_MS.NoTrt_F |
| TSSC4 | 0.13525 | 0.26061 | 0.191 | 0.26 | 1 | MS.Trt_vs_MS.NoTrt_F |
| LHFP | 0.13536 | 0.75673 | 0.173 | 0.23 | 1 | MS.Trt_vs_MS.NoTrt_F |
| PRR14 | 0.13539 | 0.35898 | 0.158 | 0.22 | 1 | MS.Trt_vs_MS.NoTrt_F |
| UHRF1BP1L | 0.1354 | -0.8153 | 0.41 | 0.47 | 1 | MS.Trt_vs_MS.NoTrt_F |
| APOBR | 0.13559 | 0.57508 | 0.165 | 0.23 | 1 | MS.Trt_vs_MS.NoTrt_F |
| RP11-894J14 | 0.13573 | -0.5906 | 0.14 | 0.19 | 1 | MS.Trt_vs_MS.NoTrt_F |
| DHRS11 | 0.1359 | -0.4056 | 0.291 | 0.37 | 1 | MS.Trt_vs_MS.NoTrt_F |
| XKR9 | 0.13598 | 0.35572 | 0.522 | 0.62 | 1 | MS.Trt_vs_MS.NoTrt_F |
| GAB1 | 0.13604 | 0.54534 | 0.471 | 0.6 | 1 | MS.Trt_vs_MS.NoTrt_F |
| C11orf1 | 0.13605 | 0.34732 | 0.799 | 0.82 | 1 | MS.Trt_vs_MS.NoTrt_F |
| WDR73 | 0.13617 | 0.50662 | 0.968 | 0.98 | 1 | MS.Trt_vs_MS.NoTrt_F |
| TLE3 | 0.13619 | 1.1304 | 0.234 | 0.34 | 1 | MS.Trt_vs_MS.NoTrt_F |
| XYLT1 | 0.13635 | -0.3846 | 0.144 | 0.2 | 1 | MS.Trt_vs_MS.NoTrt_F |
| SPATA7 | 0.13636 | 0.51759 | 0.295 | 0.37 | 1 | MS.Trt_vs_MS.NoTrt_F |
| UBL4A | 0.13637 | 0.75264 | 0.122 | 0.18 | 1 | MS.Trt_vs_MS.NoTrt_F |
| RHOC | 0.1364 | 0.42624 | 0.155 | 0.22 | 1 | MS.Trt_vs_MS.NoTrt_F |
| NPIPA5 | 0.13641 | -0.3542 | 0.112 | 0.16 | 1 | MS.Trt_vs_MS.NoTrt_F |
| MTF2 | 0.13656 | 0.32246 | 0.302 | 0.38 | 1 | MS.Trt_vs_MS.NoTrt_F |
| CWC25 | 0.13659 | 0.34545 | 0.396 | 0.46 | 1 | MS.Trt_vs_MS.NoTrt_F |
| SERTAD1 | 0.13667 | 0.6585 | 0.367 | 0.45 | 1 | MS.Trt_vs_MS.NoTrt_F |
| DUSP14 | 0.13682 | 0.99324 | 0.086 | 0.13 | 1 | MS.Trt_vs_MS.NoTrt_F |
| MVD | 0.13696 | 0.63193 | 0.5 | 0.58 | 1 | MS.Trt_vs_MS.NoTrt_F |
| NPIPB3 | 0.13704 | 0.72163 | 0.662 | 0.83 | 1 | MS.Trt_vs_MS.NoTrt_F |
| FAM213A | 0.1371 | 0.56011 | 0.773 | 0.87 | 1 | MS.Trt_vs_MS.NoTrt_F |
| UBA2 | 0.13735 | -0.533 | 0.568 | 0.62 | 1 | MS.Trt_vs_MS.NoTrt_F |
| PPP1R13B | 0.13736 | -0.3619 | 0.468 | 0.57 | 1 | MS.Trt_vs_MS.NoTrt_F |
| TCIRG1 | 0.13736 | 0.39733 | 0.331 | 0.45 | 1 | MS.Trt_vs_MS.NoTrt_F |
| CTSK | 0.13743 | 1.40779 | 0.068 | 0.11 | 1 | MS.Trt_vs_MS.NoTrt_F |
| CD101 | 0.1376 | 1.0392 | 0.104 | 0.16 | 1 | MS.Trt_vs_MS.NoTrt_F |
| TRIM16L | 0.13773 | 0.29687 | 0.777 | 0.8 | 1 | MS.Trt_vs_MS.NoTrt_F |
| TBPL2 | 0.13782 | 0.29277 | 0.097 | 0.14 | 1 | MS.Trt_vs_MS.NoTrt_F |
| MMACHC | 0.13803 | 0.27154 | 0.906 | 0.89 | 1 | MS.Trt_vs_MS.NoTrt_F |
| NBPF20 | 0.13825 | 0.70506 | 0.335 | 0.41 | 1 | MS.Trt_vs_MS.NoTrt_F |
| METTL17 | 0.13832 | 0.54895 | 0.629 | 0.73 | 1 | MS.Trt_vs_MS.NoTrt_F |

| | | | | | | |
|---|---|---|---|---|---|---|
| ATP8A1 | 0.13885 | 0.74496 | 0.165 | 0.23 | 1 | MS.Trt_vs_MS.NoTrt_F |
| CTDSP2 | 0.13888 | 0.28571 | 0.191 | 0.27 | 1 | MS.Trt_vs_MS.NoTrt_F |
| PTPN23 | 0.13903 | 0.33656 | 0.144 | 0.2 | 1 | MS.Trt_vs_MS.NoTrt_F |
| ZNF561 | 0.13926 | 0.84864 | 0.514 | 0.61 | 1 | MS.Trt_vs_MS.NoTrt_F |
| FBXL17 | 0.13932 | -1.484 | 0.237 | 0.3 | 1 | MS.Trt_vs_MS.NoTrt_F |
| PINK1 | 0.13954 | 0.25221 | 0.255 | 0.33 | 1 | MS.Trt_vs_MS.NoTrt_F |
| CTD-3105H1 | 0.13957 | -0.5171 | 0.306 | 0.4 | 1 | MS.Trt_vs_MS.NoTrt_F |
| KIAA1462 | 0.14022 | -0.3227 | 0.216 | 0.28 | 1 | MS.Trt_vs_MS.NoTrt_F |
| C11orf57 | 0.1408 | -0.4697 | 0.745 | 0.8 | 1 | MS.Trt_vs_MS.NoTrt_F |
| SOCS2 | 0.141 | 0.43668 | 0.468 | 0.38 | 1 | MS.Trt_vs_MS.NoTrt_F |
| GALNT3 | 0.14107 | 0.75569 | 0.108 | 0.16 | 1 | MS.Trt_vs_MS.NoTrt_F |
| PANK2 | 0.1411 | -0.2787 | 0.522 | 0.56 | 1 | MS.Trt_vs_MS.NoTrt_F |
| GEMIN8 | 0.14128 | 0.26824 | 0.086 | 0.13 | 1 | MS.Trt_vs_MS.NoTrt_F |
| SLC25A11 | 0.14148 | 0.34817 | 0.183 | 0.25 | 1 | MS.Trt_vs_MS.NoTrt_F |
| ITGB4 | 0.14166 | 0.88325 | 0.252 | 0.33 | 1 | MS.Trt_vs_MS.NoTrt_F |
| LETM1 | 0.14179 | 0.62208 | 0.888 | 0.87 | 1 | MS.Trt_vs_MS.NoTrt_F |
| PBX3 | 0.14202 | 0.27383 | 0.086 | 0.13 | 1 | MS.Trt_vs_MS.NoTrt_F |
| RNF220 | 0.14208 | 0.3583 | 0.601 | 0.73 | 1 | MS.Trt_vs_MS.NoTrt_F |
| KNOP1 | 0.14235 | 0.34855 | 0.705 | 0.77 | 1 | MS.Trt_vs_MS.NoTrt_F |
| NFKBID | 0.14249 | 0.85343 | 0.532 | 0.64 | 1 | MS.Trt_vs_MS.NoTrt_F |
| CES4A | 0.14289 | -0.5286 | 0.597 | 0.53 | 1 | MS.Trt_vs_MS.NoTrt_F |
| SYCP2 | 0.14294 | 0.59983 | 0.083 | 0.13 | 1 | MS.Trt_vs_MS.NoTrt_F |
| BTBD7 | 0.14299 | 0.61566 | 0.788 | 0.77 | 1 | MS.Trt_vs_MS.NoTrt_F |
| AKAP5 | 0.14309 | -0.4031 | 0.655 | 0.69 | 1 | MS.Trt_vs_MS.NoTrt_F |
| SGCB | 0.14335 | 0.51528 | 0.302 | 0.37 | 1 | MS.Trt_vs_MS.NoTrt_F |
| ZNF691 | 0.1434 | 0.31361 | 0.083 | 0.13 | 1 | MS.Trt_vs_MS.NoTrt_F |
| BICDL1 | 0.14344 | -0.5107 | 0.583 | 0.66 | 1 | MS.Trt_vs_MS.NoTrt_F |
| NFU1 | 0.14355 | -0.4089 | 0.169 | 0.23 | 1 | MS.Trt_vs_MS.NoTrt_F |
| CARMIL1 | 0.14411 | 0.70476 | 0.259 | 0.32 | 1 | MS.Trt_vs_MS.NoTrt_F |
| GTF2H1 | 0.14411 | -0.5608 | 0.414 | 0.48 | 1 | MS.Trt_vs_MS.NoTrt_F |
| VPS39 | 0.14435 | 0.34328 | 0.493 | 0.56 | 1 | MS.Trt_vs_MS.NoTrt_F |
| KPNA4 | 0.1445 | 0.30009 | 0.59 | 0.69 | 1 | MS.Trt_vs_MS.NoTrt_F |
| CD40 | 0.1448 | -0.7744 | 0.151 | 0.21 | 1 | MS.Trt_vs_MS.NoTrt_F |
| SNUPN | 0.14481 | 0.46549 | 0.791 | 0.92 | 1 | MS.Trt_vs_MS.NoTrt_F |
| COMMD2 | 0.14482 | 0.86873 | 0.345 | 0.44 | 1 | MS.Trt_vs_MS.NoTrt_F |
| TOR1B | 0.14486 | 0.6575 | 0.255 | 0.32 | 1 | MS.Trt_vs_MS.NoTrt_F |
| ITGA2 | 0.14486 | -0.5267 | 0.392 | 0.45 | 1 | MS.Trt_vs_MS.NoTrt_F |
| SEC61A1 | 0.14504 | 0.6097 | 0.396 | 0.5 | 1 | MS.Trt_vs_MS.NoTrt_F |
| ADH5 | 0.14516 | 0.49747 | 0.453 | 0.53 | 1 | MS.Trt_vs_MS.NoTrt_F |
| ZNF30 | 0.14544 | 1.66072 | 0.101 | 0.15 | 1 | MS.Trt_vs_MS.NoTrt_F |

| | | | | | | |
|---|---|---|---|---|---|---|
| ZNF557 | 0.14549 | 0.47854 | 0.867 | 0.91 | 1 | MS.Trt_vs_MS.NoTrt_F |
| MSH3 | 0.14575 | -0.4716 | 0.45 | 0.51 | 1 | MS.Trt_vs_MS.NoTrt_F |
| PLA2G4B | 0.1459 | 0.79866 | 0.468 | 0.54 | 1 | MS.Trt_vs_MS.NoTrt_F |
| CTC-454I21. | 0.14593 | 0.44844 | 0.205 | 0.28 | 1 | MS.Trt_vs_MS.NoTrt_F |
| TRNAU1AP | 0.14614 | -0.3925 | 0.878 | 0.92 | 1 | MS.Trt_vs_MS.NoTrt_F |
| DHRS4L2 | 0.1462 | 0.60045 | 0.212 | 0.29 | 1 | MS.Trt_vs_MS.NoTrt_F |
| SIK3 | 0.14641 | 0.41947 | 0.313 | 0.39 | 1 | MS.Trt_vs_MS.NoTrt_F |
| ERCC8 | 0.14662 | 0.5791 | 0.356 | 0.44 | 1 | MS.Trt_vs_MS.NoTrt_F |
| HIST1H1D | 0.1467 | 0.55101 | 0.104 | 0.16 | 1 | MS.Trt_vs_MS.NoTrt_F |
| ZCCHC7 | 0.14674 | 0.28082 | 0.302 | 0.39 | 1 | MS.Trt_vs_MS.NoTrt_F |
| ZNF763 | 0.14744 | 1.29418 | 0.183 | 0.24 | 1 | MS.Trt_vs_MS.NoTrt_F |
| ACACA | 0.14755 | -0.9173 | 0.586 | 0.68 | 1 | MS.Trt_vs_MS.NoTrt_F |
| TRMT1 | 0.14766 | 0.44548 | 0.169 | 0.23 | 1 | MS.Trt_vs_MS.NoTrt_F |
| HIF1AN | 0.1477 | 0.68572 | 0.806 | 0.87 | 1 | MS.Trt_vs_MS.NoTrt_F |
| NARFL | 0.14775 | 0.50501 | 0.759 | 0.83 | 1 | MS.Trt_vs_MS.NoTrt_F |
| TMEM199 | 0.14819 | -0.5753 | 0.658 | 0.7 | 1 | MS.Trt_vs_MS.NoTrt_F |
| C19orf48 | 0.14832 | -0.4335 | 0.374 | 0.44 | 1 | MS.Trt_vs_MS.NoTrt_F |
| BFSP2 | 0.14835 | 0.58069 | 0.083 | 0.13 | 1 | MS.Trt_vs_MS.NoTrt_F |
| PTGER2 | 0.14837 | 0.64774 | 0.219 | 0.29 | 1 | MS.Trt_vs_MS.NoTrt_F |
| LAMC3 | 0.14846 | 1.66325 | 0.302 | 0.39 | 1 | MS.Trt_vs_MS.NoTrt_F |
| NRIP2 | 0.14853 | 0.50971 | 0.331 | 0.4 | 1 | MS.Trt_vs_MS.NoTrt_F |
| RNF144A | 0.14861 | 1.67869 | 0.191 | 0.25 | 1 | MS.Trt_vs_MS.NoTrt_F |
| MAP3K3 | 0.14868 | 0.84778 | 0.137 | 0.2 | 1 | MS.Trt_vs_MS.NoTrt_F |
| HNRNPUL1 | 0.14878 | -0.6477 | 0.662 | 0.7 | 1 | MS.Trt_vs_MS.NoTrt_F |
| MRPL30 | 0.14896 | -0.7331 | 0.755 | 0.88 | 1 | MS.Trt_vs_MS.NoTrt_F |
| PNPLA4 | 0.14919 | 0.42767 | 0.252 | 0.33 | 1 | MS.Trt_vs_MS.NoTrt_F |
| IRAK1BP1 | 0.14929 | -0.5193 | 0.608 | 0.7 | 1 | MS.Trt_vs_MS.NoTrt_F |
| CTD-2192J1( | 0.14959 | 0.41033 | 0.194 | 0.26 | 1 | MS.Trt_vs_MS.NoTrt_F |
| SCARB2 | 0.14965 | 0.62165 | 0.324 | 0.4 | 1 | MS.Trt_vs_MS.NoTrt_F |
| LIMS2 | 0.14988 | 1.21006 | 0.083 | 0.13 | 1 | MS.Trt_vs_MS.NoTrt_F |
| MRPS15 | 0.15009 | -0.2892 | 0.572 | 0.62 | 1 | MS.Trt_vs_MS.NoTrt_F |
| CDON | 0.15024 | -0.3211 | 0.464 | 0.54 | 1 | MS.Trt_vs_MS.NoTrt_F |
| LENG1 | 0.15028 | 0.61016 | 0.133 | 0.19 | 1 | MS.Trt_vs_MS.NoTrt_F |
| SEZ6 | 0.15041 | 1.13469 | 0.112 | 0.16 | 1 | MS.Trt_vs_MS.NoTrt_F |
| E2F5 | 0.15044 | 0.45782 | 0.403 | 0.46 | 1 | MS.Trt_vs_MS.NoTrt_F |
| RPP21 | 0.15068 | 0.77513 | 0.183 | 0.25 | 1 | MS.Trt_vs_MS.NoTrt_F |
| LRP2BP | 0.15073 | -0.6082 | 0.421 | 0.49 | 1 | MS.Trt_vs_MS.NoTrt_F |
| METTL3 | 0.15077 | 0.47586 | 0.604 | 0.66 | 1 | MS.Trt_vs_MS.NoTrt_F |
| DNASE2 | 0.15086 | 0.52913 | 0.453 | 0.54 | 1 | MS.Trt_vs_MS.NoTrt_F |
| RUFY2 | 0.15106 | 0.59324 | 0.475 | 0.55 | 1 | MS.Trt_vs_MS.NoTrt_F |

| | | | | | | |
|---|---|---|---|---|---|---|
| S1PR1 | 0.15107 | 0.47279 | 0.432 | 0.6 | 1 | MS.Trt_vs_MS.NoTrt_F |
| TSC2 | 0.15109 | -0.2964 | 0.662 | 0.74 | 1 | MS.Trt_vs_MS.NoTrt_F |
| THBS1 | 0.15123 | -1.1012 | 0.248 | 0.3 | 1 | MS.Trt_vs_MS.NoTrt_F |
| ZNF425 | 0.15166 | -0.8044 | 0.439 | 0.51 | 1 | MS.Trt_vs_MS.NoTrt_F |
| TIMM9 | 0.15168 | 1.0865 | 0.09 | 0.14 | 1 | MS.Trt_vs_MS.NoTrt_F |
| DOCK9 | 0.15177 | 0.4375 | 0.302 | 0.37 | 1 | MS.Trt_vs_MS.NoTrt_F |
| MBD4 | 0.15209 | 0.61607 | 0.259 | 0.33 | 1 | MS.Trt_vs_MS.NoTrt_F |
| GCNT2 | 0.15229 | 0.27686 | 0.745 | 0.7 | 1 | MS.Trt_vs_MS.NoTrt_F |
| GALNT1 | 0.15238 | 2.50247 | 0.101 | 0.15 | 1 | MS.Trt_vs_MS.NoTrt_F |
| CACFD1 | 0.15285 | 0.64303 | 0.079 | 0.12 | 1 | MS.Trt_vs_MS.NoTrt_F |
| DDX27 | 0.15308 | 0.29319 | 0.644 | 0.72 | 1 | MS.Trt_vs_MS.NoTrt_F |
| ZNF33B | 0.15313 | 0.92889 | 0.507 | 0.45 | 1 | MS.Trt_vs_MS.NoTrt_F |
| MED24 | 0.15319 | -1.0169 | 0.608 | 0.64 | 1 | MS.Trt_vs_MS.NoTrt_F |
| ZBTB41 | 0.15499 | 1.34278 | 0.083 | 0.13 | 1 | MS.Trt_vs_MS.NoTrt_F |
| SNRNP200 | 0.15549 | 0.51598 | 0.759 | 0.83 | 1 | MS.Trt_vs_MS.NoTrt_F |
| RBMXL1 | 0.15557 | -0.7049 | 0.129 | 0.18 | 1 | MS.Trt_vs_MS.NoTrt_F |
| SLC39A3 | 0.1558 | 0.45546 | 0.576 | 0.68 | 1 | MS.Trt_vs_MS.NoTrt_F |
| GBP4 | 0.15592 | -0.3632 | 0.723 | 0.78 | 1 | MS.Trt_vs_MS.NoTrt_F |
| NCR3 | 0.15594 | 0.91889 | 0.115 | 0.17 | 1 | MS.Trt_vs_MS.NoTrt_F |
| SULT1A3 | 0.15597 | 0.38803 | 0.629 | 0.73 | 1 | MS.Trt_vs_MS.NoTrt_F |
| GSDMD | 0.1561 | 0.25019 | 0.802 | 0.86 | 1 | MS.Trt_vs_MS.NoTrt_F |
| RNFT1 | 0.15612 | 0.74032 | 0.388 | 0.45 | 1 | MS.Trt_vs_MS.NoTrt_F |
| AGO1 | 0.15647 | 0.27141 | 0.442 | 0.53 | 1 | MS.Trt_vs_MS.NoTrt_F |
| MT-CYB | 0.15692 | 0.54466 | 0.719 | 0.79 | 1 | MS.Trt_vs_MS.NoTrt_F |
| QRICH2 | 0.15718 | 0.47053 | 0.198 | 0.27 | 1 | MS.Trt_vs_MS.NoTrt_F |
| DYDC1 | 0.15722 | -0.3512 | 0.489 | 0.58 | 1 | MS.Trt_vs_MS.NoTrt_F |
| PPM1N | 0.15725 | 0.6614 | 0.137 | 0.19 | 1 | MS.Trt_vs_MS.NoTrt_F |
| MBTPS2 | 0.15737 | -0.5993 | 0.32 | 0.38 | 1 | MS.Trt_vs_MS.NoTrt_F |
| C19orf52 | 0.15748 | 0.45695 | 0.313 | 0.38 | 1 | MS.Trt_vs_MS.NoTrt_F |
| ANKRD42 | 0.15752 | -0.5488 | 0.388 | 0.48 | 1 | MS.Trt_vs_MS.NoTrt_F |
| UBE2V1 | 0.15762 | 0.48569 | 0.424 | 0.54 | 1 | MS.Trt_vs_MS.NoTrt_F |
| RAE1 | 0.15782 | 1.06477 | 0.133 | 0.19 | 1 | MS.Trt_vs_MS.NoTrt_F |
| LRMP | 0.15791 | 0.3296 | 0.122 | 0.18 | 1 | MS.Trt_vs_MS.NoTrt_F |
| DET1 | 0.15806 | 0.3452 | 0.543 | 0.59 | 1 | MS.Trt_vs_MS.NoTrt_F |
| KLHL2 | 0.15813 | 1.21009 | 0.335 | 0.39 | 1 | MS.Trt_vs_MS.NoTrt_F |
| PIGP | 0.15815 | -0.4751 | 0.781 | 0.81 | 1 | MS.Trt_vs_MS.NoTrt_F |
| LIX1L | 0.1584 | 0.36107 | 0.824 | 0.79 | 1 | MS.Trt_vs_MS.NoTrt_F |
| ICE2 | 0.1585 | -0.4278 | 0.842 | 0.85 | 1 | MS.Trt_vs_MS.NoTrt_F |
| RFX2 | 0.1585 | 0.63324 | 0.475 | 0.53 | 1 | MS.Trt_vs_MS.NoTrt_F |
| ZNF652 | 0.15853 | 0.51478 | 0.532 | 0.61 | 1 | MS.Trt_vs_MS.NoTrt_F |

| | | | | | | |
|---|---|---|---|---|---|---|
| ATG12 | 0.15864 | 0.39746 | 0.77 | 0.82 | 1 | MS.Trt_vs_MS.NoTrt_F |
| DRAXIN | 0.15883 | 0.33145 | 0.835 | 0.85 | 1 | MS.Trt_vs_MS.NoTrt_F |
| VAT1 | 0.15888 | 0.5274 | 0.137 | 0.19 | 1 | MS.Trt_vs_MS.NoTrt_F |
| SLX4 | 0.15903 | -0.7355 | 0.496 | 0.56 | 1 | MS.Trt_vs_MS.NoTrt_F |
| TMEM256-P | 0.15905 | 0.79887 | 0.489 | 0.53 | 1 | MS.Trt_vs_MS.NoTrt_F |
| NANS | 0.15925 | 0.67926 | 0.766 | 0.81 | 1 | MS.Trt_vs_MS.NoTrt_F |
| LLGL1 | 0.15926 | -0.7956 | 0.295 | 0.35 | 1 | MS.Trt_vs_MS.NoTrt_F |
| USP6 | 0.15974 | -0.3633 | 0.147 | 0.2 | 1 | MS.Trt_vs_MS.NoTrt_F |
| POLR3H | 0.15993 | -0.4657 | 0.295 | 0.35 | 1 | MS.Trt_vs_MS.NoTrt_F |
| FASTKD3 | 0.16001 | 0.82828 | 0.104 | 0.15 | 1 | MS.Trt_vs_MS.NoTrt_F |
| RARS | 0.16013 | 0.34531 | 0.493 | 0.54 | 1 | MS.Trt_vs_MS.NoTrt_F |
| BRI3 | 0.16016 | 0.37968 | 0.928 | 0.91 | 1 | MS.Trt_vs_MS.NoTrt_F |
| PPP2R3A | 0.16028 | 0.28023 | 0.705 | 0.81 | 1 | MS.Trt_vs_MS.NoTrt_F |
| C19orf44 | 0.16033 | -0.6669 | 0.349 | 0.43 | 1 | MS.Trt_vs_MS.NoTrt_F |
| PPP3CA | 0.16036 | 0.64669 | 0.561 | 0.69 | 1 | MS.Trt_vs_MS.NoTrt_F |
| PKD1 | 0.16058 | -0.6955 | 0.353 | 0.43 | 1 | MS.Trt_vs_MS.NoTrt_F |
| WBP11 | 0.16067 | 0.38926 | 0.299 | 0.39 | 1 | MS.Trt_vs_MS.NoTrt_F |
| PGAP2 | 0.16088 | 0.91981 | 0.094 | 0.14 | 1 | MS.Trt_vs_MS.NoTrt_F |
| MOB3A | 0.16105 | 0.32591 | 0.835 | 0.86 | 1 | MS.Trt_vs_MS.NoTrt_F |
| TEX14 | 0.1613 | 1.5643 | 0.112 | 0.16 | 1 | MS.Trt_vs_MS.NoTrt_F |
| COQ6 | 0.16161 | 0.39234 | 0.755 | 0.75 | 1 | MS.Trt_vs_MS.NoTrt_F |
| SSBP1 | 0.16215 | 0.51363 | 0.687 | 0.75 | 1 | MS.Trt_vs_MS.NoTrt_F |
| YIPF1 | 0.16228 | 1.77272 | 0.112 | 0.16 | 1 | MS.Trt_vs_MS.NoTrt_F |
| SAP30BP | 0.16231 | 0.36545 | 0.234 | 0.31 | 1 | MS.Trt_vs_MS.NoTrt_F |
| NECAB1 | 0.16264 | 0.26608 | 0.144 | 0.19 | 1 | MS.Trt_vs_MS.NoTrt_F |
| SPA17 | 0.16372 | 0.29865 | 0.252 | 0.32 | 1 | MS.Trt_vs_MS.NoTrt_F |
| USP33 | 0.16384 | 0.76451 | 0.867 | 0.95 | 1 | MS.Trt_vs_MS.NoTrt_F |
| AP5B1 | 0.16411 | -1.0033 | 0.288 | 0.37 | 1 | MS.Trt_vs_MS.NoTrt_F |
| SLC16A3 | 0.16421 | 0.55849 | 0.622 | 0.55 | 1 | MS.Trt_vs_MS.NoTrt_F |
| CWF19L2 | 0.16425 | 0.27452 | 0.299 | 0.38 | 1 | MS.Trt_vs_MS.NoTrt_F |
| SIRT5 | 0.16463 | 0.31494 | 0.791 | 0.86 | 1 | MS.Trt_vs_MS.NoTrt_F |
| TMEM185B | 0.16501 | -1.5171 | 0.338 | 0.39 | 1 | MS.Trt_vs_MS.NoTrt_F |
| CFL2 | 0.16509 | 0.3196 | 0.626 | 0.74 | 1 | MS.Trt_vs_MS.NoTrt_F |
| GPATCH2 | 0.1651 | 0.98805 | 0.604 | 0.5 | 1 | MS.Trt_vs_MS.NoTrt_F |
| CCDC18 | 0.16576 | 0.2649 | 0.198 | 0.26 | 1 | MS.Trt_vs_MS.NoTrt_F |
| GDE1 | 0.16581 | -0.9378 | 0.277 | 0.34 | 1 | MS.Trt_vs_MS.NoTrt_F |
| COPS6 | 0.16591 | 0.60757 | 0.723 | 0.66 | 1 | MS.Trt_vs_MS.NoTrt_F |
| SUMO4 | 0.16636 | 0.68562 | 0.068 | 0.11 | 1 | MS.Trt_vs_MS.NoTrt_F |
| ZNF845 | 0.16695 | 0.82552 | 0.651 | 0.59 | 1 | MS.Trt_vs_MS.NoTrt_F |
| SLC35C1 | 0.16706 | 0.79963 | 0.09 | 0.13 | 1 | MS.Trt_vs_MS.NoTrt_F |

| | | | | | | |
|---|---|---|---|---|---|---|
| CARD10 | 0.16716 | 0.30626 | 0.108 | 0.16 | 1 | MS.Trt_vs_MS.NoTrt_F |
| MRFAP1 | 0.16751 | 0.4519 | 0.338 | 0.41 | 1 | MS.Trt_vs_MS.NoTrt_F |
| MS4A6A | 0.16752 | 0.38535 | 0.176 | 0.23 | 1 | MS.Trt_vs_MS.NoTrt_F |
| DGKQ | 0.16772 | 2.2955 | 0.094 | 0.14 | 1 | MS.Trt_vs_MS.NoTrt_F |
| ADAT1 | 0.16829 | -0.6799 | 0.705 | 0.73 | 1 | MS.Trt_vs_MS.NoTrt_F |
| TBL3 | 0.16842 | 0.57097 | 0.27 | 0.34 | 1 | MS.Trt_vs_MS.NoTrt_F |
| WNT9A | 0.16856 | 0.50362 | 0.108 | 0.15 | 1 | MS.Trt_vs_MS.NoTrt_F |
| DUSP3 | 0.16871 | -0.3577 | 0.77 | 0.69 | 1 | MS.Trt_vs_MS.NoTrt_F |
| ANXA4 | 0.16902 | 0.58897 | 0.86 | 0.81 | 1 | MS.Trt_vs_MS.NoTrt_F |
| RP11-435I1( | 0.16922 | -0.427 | 0.36 | 0.45 | 1 | MS.Trt_vs_MS.NoTrt_F |
| HEMK1 | 0.16929 | 0.72746 | 0.96 | 0.97 | 1 | MS.Trt_vs_MS.NoTrt_F |
| SGF29 | 0.16967 | 0.61866 | 0.309 | 0.37 | 1 | MS.Trt_vs_MS.NoTrt_F |
| MDFIC | 0.16972 | -0.5785 | 0.356 | 0.41 | 1 | MS.Trt_vs_MS.NoTrt_F |
| MRM2 | 0.17024 | 0.36011 | 0.32 | 0.39 | 1 | MS.Trt_vs_MS.NoTrt_F |
| NSUN4 | 0.17043 | 0.27108 | 0.73 | 0.81 | 1 | MS.Trt_vs_MS.NoTrt_F |
| ZNF2 | 0.17054 | 0.9067 | 0.313 | 0.38 | 1 | MS.Trt_vs_MS.NoTrt_F |
| TMEM260 | 0.17075 | -0.5258 | 0.374 | 0.45 | 1 | MS.Trt_vs_MS.NoTrt_F |
| TRPV1 | 0.17078 | 0.48348 | 0.446 | 0.54 | 1 | MS.Trt_vs_MS.NoTrt_F |
| PLEKHM2 | 0.17088 | -0.72 | 0.068 | 0.11 | 1 | MS.Trt_vs_MS.NoTrt_F |
| PDCL3 | 0.17113 | 0.73249 | 0.147 | 0.21 | 1 | MS.Trt_vs_MS.NoTrt_F |
| SSB | 0.17118 | 0.56778 | 0.727 | 0.83 | 1 | MS.Trt_vs_MS.NoTrt_F |
| LPAR5 | 0.17118 | 1.93722 | 0.097 | 0.14 | 1 | MS.Trt_vs_MS.NoTrt_F |
| GSTT1 | 0.1713 | 0.66542 | 0.687 | 0.81 | 1 | MS.Trt_vs_MS.NoTrt_F |
| FBXO25 | 0.17143 | 0.76357 | 0.356 | 0.41 | 1 | MS.Trt_vs_MS.NoTrt_F |
| KIAA1683 | 0.17147 | -0.3129 | 0.604 | 0.67 | 1 | MS.Trt_vs_MS.NoTrt_F |
| SMAD5 | 0.17166 | 0.65285 | 0.471 | 0.45 | 1 | MS.Trt_vs_MS.NoTrt_F |
| DNAH6 | 0.17175 | 0.56114 | 0.219 | 0.29 | 1 | MS.Trt_vs_MS.NoTrt_F |
| SERP1 | 0.17186 | 0.60313 | 0.594 | 0.71 | 1 | MS.Trt_vs_MS.NoTrt_F |
| EDEM1 | 0.17188 | 1.01743 | 0.317 | 0.43 | 1 | MS.Trt_vs_MS.NoTrt_F |
| SWAP70 | 0.17189 | 0.34134 | 0.399 | 0.49 | 1 | MS.Trt_vs_MS.NoTrt_F |
| CDK2 | 0.17218 | -0.4199 | 0.104 | 0.15 | 1 | MS.Trt_vs_MS.NoTrt_F |
| B3GALNT2 | 0.17252 | -0.4191 | 0.626 | 0.72 | 1 | MS.Trt_vs_MS.NoTrt_F |
| EIF2A | 0.17275 | 0.33843 | 0.608 | 0.68 | 1 | MS.Trt_vs_MS.NoTrt_F |
| KIF19 | 0.17275 | -0.5794 | 0.392 | 0.45 | 1 | MS.Trt_vs_MS.NoTrt_F |
| MLLT1 | 0.17306 | 1.09756 | 0.097 | 0.14 | 1 | MS.Trt_vs_MS.NoTrt_F |
| ZNF689 | 0.17315 | 1.43359 | 0.076 | 0.12 | 1 | MS.Trt_vs_MS.NoTrt_F |
| POLR3B | 0.17322 | -0.6172 | 0.137 | 0.19 | 1 | MS.Trt_vs_MS.NoTrt_F |
| SLC25A4 | 0.17342 | 0.80478 | 0.306 | 0.38 | 1 | MS.Trt_vs_MS.NoTrt_F |
| FGGY | 0.1735 | 0.50519 | 0.442 | 0.53 | 1 | MS.Trt_vs_MS.NoTrt_F |
| ZNF566 | 0.17353 | -0.2858 | 0.878 | 0.87 | 1 | MS.Trt_vs_MS.NoTrt_F |

| | | | | | | |
|---|---|---|---|---|---|---|
| TP53BP2 | 0.17365 | 1.3157 | 0.18 | 0.24 | 1 | MS.Trt_vs_MS.NoTrt_F |
| IL12RB2 | 0.17384 | 0.9887 | 0.108 | 0.16 | 1 | MS.Trt_vs_MS.NoTrt_F |
| SLC25A17 | 0.17392 | 0.3442 | 0.324 | 0.4 | 1 | MS.Trt_vs_MS.NoTrt_F |
| PTPA | 0.17431 | -0.3339 | 0.727 | 0.82 | 1 | MS.Trt_vs_MS.NoTrt_F |
| MAN2A2 | 0.17431 | 1.12752 | 0.309 | 0.37 | 1 | MS.Trt_vs_MS.NoTrt_F |
| GSTM4 | 0.17439 | 2.19047 | 0.115 | 0.17 | 1 | MS.Trt_vs_MS.NoTrt_F |
| CCL5 | 0.17476 | 1.28259 | 0.899 | 0.94 | 1 | MS.Trt_vs_MS.NoTrt_F |
| COL15A1 | 0.17488 | 0.69925 | 0.363 | 0.32 | 1 | MS.Trt_vs_MS.NoTrt_F |
| ANKRD39 | 0.17539 | 0.89645 | 0.252 | 0.33 | 1 | MS.Trt_vs_MS.NoTrt_F |
| C20orf194 | 0.17543 | 0.90649 | 0.331 | 0.41 | 1 | MS.Trt_vs_MS.NoTrt_F |
| ZNF678 | 0.1756 | 0.97595 | 0.194 | 0.25 | 1 | MS.Trt_vs_MS.NoTrt_F |
| ANKRD12 | 0.17595 | 0.47674 | 0.791 | 0.87 | 1 | MS.Trt_vs_MS.NoTrt_F |
| AL928654.7 | 0.1761 | 0.27794 | 0.072 | 0.11 | 1 | MS.Trt_vs_MS.NoTrt_F |
| RPP38 | 0.17674 | -0.449 | 0.428 | 0.5 | 1 | MS.Trt_vs_MS.NoTrt_F |
| CAPS | 0.17698 | -0.4479 | 0.953 | 0.94 | 1 | MS.Trt_vs_MS.NoTrt_F |
| RFK | 0.17758 | 0.58733 | 0.496 | 0.53 | 1 | MS.Trt_vs_MS.NoTrt_F |
| SYPL1 | 0.17781 | 0.34219 | 0.414 | 0.48 | 1 | MS.Trt_vs_MS.NoTrt_F |
| CA13 | 0.17792 | 0.62403 | 0.536 | 0.65 | 1 | MS.Trt_vs_MS.NoTrt_F |
| FUOM | 0.17805 | 0.44525 | 0.068 | 0.11 | 1 | MS.Trt_vs_MS.NoTrt_F |
| ZDHHC14 | 0.17807 | 0.26402 | 0.356 | 0.44 | 1 | MS.Trt_vs_MS.NoTrt_F |
| BOLA2B | 0.17813 | 0.43261 | 0.158 | 0.22 | 1 | MS.Trt_vs_MS.NoTrt_F |
| PTDSS2 | 0.1782 | 1.46422 | 0.094 | 0.14 | 1 | MS.Trt_vs_MS.NoTrt_F |
| ZMIZ1 | 0.1785 | 1.13193 | 0.14 | 0.19 | 1 | MS.Trt_vs_MS.NoTrt_F |
| PUS3 | 0.17872 | 0.88377 | 0.309 | 0.39 | 1 | MS.Trt_vs_MS.NoTrt_F |
| LRFN3 | 0.17926 | -0.3999 | 0.597 | 0.67 | 1 | MS.Trt_vs_MS.NoTrt_F |
| JAKMIP2 | 0.17966 | 0.87961 | 0.442 | 0.51 | 1 | MS.Trt_vs_MS.NoTrt_F |
| IL1R1 | 0.17984 | 2.21598 | 0.324 | 0.39 | 1 | MS.Trt_vs_MS.NoTrt_F |
| PARP2 | 0.18015 | -0.4933 | 0.342 | 0.42 | 1 | MS.Trt_vs_MS.NoTrt_F |
| ZNF202 | 0.18024 | 0.84975 | 0.072 | 0.11 | 1 | MS.Trt_vs_MS.NoTrt_F |
| NDUFA3 | 0.18047 | 0.35997 | 0.946 | 0.95 | 1 | MS.Trt_vs_MS.NoTrt_F |
| RP11-111H1 | 0.18068 | -0.3965 | 0.137 | 0.18 | 1 | MS.Trt_vs_MS.NoTrt_F |
| TOB2 | 0.18095 | 0.6916 | 0.165 | 0.23 | 1 | MS.Trt_vs_MS.NoTrt_F |
| PKNOX2 | 0.18109 | 0.56495 | 0.097 | 0.14 | 1 | MS.Trt_vs_MS.NoTrt_F |
| PPP6R1 | 0.18119 | 0.40887 | 0.309 | 0.39 | 1 | MS.Trt_vs_MS.NoTrt_F |
| PSME1 | 0.18123 | 0.25912 | 0.576 | 0.73 | 1 | MS.Trt_vs_MS.NoTrt_F |
| NDUFV3 | 0.18143 | 0.27072 | 0.982 | 0.99 | 1 | MS.Trt_vs_MS.NoTrt_F |
| UBE2Z | 0.18167 | 0.64101 | 0.288 | 0.34 | 1 | MS.Trt_vs_MS.NoTrt_F |
| DDX54 | 0.18174 | 0.75418 | 0.158 | 0.22 | 1 | MS.Trt_vs_MS.NoTrt_F |
| JCHAIN | 0.18236 | 4.37604 | 0.133 | 0.18 | 1 | MS.Trt_vs_MS.NoTrt_F |
| SEPP1 | 0.1825 | 0.83962 | 0.162 | 0.22 | 1 | MS.Trt_vs_MS.NoTrt_F |

| | | | | | | |
|---|---|---|---|---|---|---|
| FBXL2 | 0.18292 | -0.3195 | 0.665 | 0.61 | 1 | MS.Trt_vs_MS.NoTrt_F |
| ENDOV | 0.18293 | -0.8085 | 0.604 | 0.66 | 1 | MS.Trt_vs_MS.NoTrt_F |
| ZNF124 | 0.18314 | 1.19336 | 0.665 | 0.67 | 1 | MS.Trt_vs_MS.NoTrt_F |
| FBXO36 | 0.18317 | 0.29203 | 0.687 | 0.66 | 1 | MS.Trt_vs_MS.NoTrt_F |
| DACT3 | 0.18336 | 0.25202 | 0.95 | 0.95 | 1 | MS.Trt_vs_MS.NoTrt_F |
| ZNF354A | 0.18363 | 0.49748 | 0.216 | 0.28 | 1 | MS.Trt_vs_MS.NoTrt_F |
| CTTN | 0.18368 | 0.88343 | 0.112 | 0.16 | 1 | MS.Trt_vs_MS.NoTrt_F |
| PEA15 | 0.18375 | 0.91167 | 0.133 | 0.19 | 1 | MS.Trt_vs_MS.NoTrt_F |
| GAREM1 | 0.18378 | 0.5335 | 0.101 | 0.07 | 1 | MS.Trt_vs_MS.NoTrt_F |
| QKI | 0.18388 | 0.32288 | 0.327 | 0.42 | 1 | MS.Trt_vs_MS.NoTrt_F |
| ZNF16 | 0.18422 | 0.2864 | 0.77 | 0.83 | 1 | MS.Trt_vs_MS.NoTrt_F |
| NSA2 | 0.18428 | 0.71817 | 0.367 | 0.44 | 1 | MS.Trt_vs_MS.NoTrt_F |
| ZNF254 | 0.18467 | 0.26223 | 0.45 | 0.48 | 1 | MS.Trt_vs_MS.NoTrt_F |
| SRRD | 0.18488 | 0.6349 | 0.453 | 0.54 | 1 | MS.Trt_vs_MS.NoTrt_F |
| ERO1A | 0.1849 | 0.56916 | 0.899 | 0.9 | 1 | MS.Trt_vs_MS.NoTrt_F |
| CHP1 | 0.18496 | 0.91268 | 0.781 | 0.82 | 1 | MS.Trt_vs_MS.NoTrt_F |
| DNAJA2 | 0.18521 | 0.35359 | 0.428 | 0.49 | 1 | MS.Trt_vs_MS.NoTrt_F |
| NUP35 | 0.18532 | 0.64301 | 0.504 | 0.61 | 1 | MS.Trt_vs_MS.NoTrt_F |
| GMEB2 | 0.18535 | 0.68765 | 0.122 | 0.17 | 1 | MS.Trt_vs_MS.NoTrt_F |
| SGK3 | 0.18547 | -0.2527 | 0.874 | 0.96 | 1 | MS.Trt_vs_MS.NoTrt_F |
| WDR20 | 0.18557 | 0.49469 | 0.432 | 0.53 | 1 | MS.Trt_vs_MS.NoTrt_F |
| ITIH4 | 0.18574 | 0.68096 | 0.248 | 0.32 | 1 | MS.Trt_vs_MS.NoTrt_F |
| ETNK1 | 0.18628 | 0.38348 | 0.514 | 0.63 | 1 | MS.Trt_vs_MS.NoTrt_F |
| CRTAM | 0.18629 | 0.80952 | 0.381 | 0.46 | 1 | MS.Trt_vs_MS.NoTrt_F |
| KTN1 | 0.18653 | 0.40344 | 0.565 | 0.71 | 1 | MS.Trt_vs_MS.NoTrt_F |
| B3GALNT1 | 0.18653 | 0.52169 | 0.205 | 0.27 | 1 | MS.Trt_vs_MS.NoTrt_F |
| OXCT1 | 0.18669 | 0.81763 | 0.36 | 0.46 | 1 | MS.Trt_vs_MS.NoTrt_F |
| RBM43 | 0.18705 | -0.5862 | 0.601 | 0.66 | 1 | MS.Trt_vs_MS.NoTrt_F |
| SELENBP1 | 0.18715 | 1.14806 | 0.273 | 0.33 | 1 | MS.Trt_vs_MS.NoTrt_F |
| FUT2 | 0.18731 | 0.28947 | 0.759 | 0.77 | 1 | MS.Trt_vs_MS.NoTrt_F |
| MSMO1 | 0.18735 | 0.4512 | 0.543 | 0.6 | 1 | MS.Trt_vs_MS.NoTrt_F |
| GATC | 0.18748 | -0.4045 | 0.532 | 0.6 | 1 | MS.Trt_vs_MS.NoTrt_F |
| SERPINB8 | 0.18761 | -0.3751 | 0.079 | 0.12 | 1 | MS.Trt_vs_MS.NoTrt_F |
| TBK1 | 0.18789 | 0.60553 | 0.122 | 0.17 | 1 | MS.Trt_vs_MS.NoTrt_F |
| HCST | 0.18792 | 0.66219 | 0.374 | 0.54 | 1 | MS.Trt_vs_MS.NoTrt_F |
| ZNF226 | 0.18801 | 0.26667 | 0.504 | 0.57 | 1 | MS.Trt_vs_MS.NoTrt_F |
| NRL | 0.18853 | 0.62444 | 0.09 | 0.13 | 1 | MS.Trt_vs_MS.NoTrt_F |
| RER1 | 0.18858 | 0.68979 | 0.227 | 0.3 | 1 | MS.Trt_vs_MS.NoTrt_F |
| PDK3 | 0.18862 | 0.85985 | 0.331 | 0.41 | 1 | MS.Trt_vs_MS.NoTrt_F |
| SMAD1 | 0.1889 | 1.64793 | 0.331 | 0.4 | 1 | MS.Trt_vs_MS.NoTrt_F |

| | | | | | | |
|---|---|---|---|---|---|---|
| FABP5 | 0.18926 | 0.28766 | 0.101 | 0.14 | 1 | MS.Trt_vs_MS.NoTrt_F |
| TMPRSS11B | 0.18938 | 0.50485 | 0.374 | 0.46 | 1 | MS.Trt_vs_MS.NoTrt_F |
| NNMT | 0.19043 | 0.56659 | 0.119 | 0.17 | 1 | MS.Trt_vs_MS.NoTrt_F |
| CYSTM1 | 0.19047 | 0.29403 | 0.331 | 0.4 | 1 | MS.Trt_vs_MS.NoTrt_F |
| NCF2 | 0.19057 | 1.27303 | 0.086 | 0.13 | 1 | MS.Trt_vs_MS.NoTrt_F |
| RARS2 | 0.19075 | -0.2936 | 0.594 | 0.63 | 1 | MS.Trt_vs_MS.NoTrt_F |
| KIF3C | 0.19091 | -0.8004 | 0.094 | 0.13 | 1 | MS.Trt_vs_MS.NoTrt_F |
| INTS4 | 0.19123 | -0.3768 | 0.396 | 0.44 | 1 | MS.Trt_vs_MS.NoTrt_F |
| INCA1 | 0.1916 | 0.73698 | 0.216 | 0.28 | 1 | MS.Trt_vs_MS.NoTrt_F |
| SSX2IP | 0.19175 | 0.7859 | 0.097 | 0.14 | 1 | MS.Trt_vs_MS.NoTrt_F |
| GPLD1 | 0.19176 | 0.69318 | 0.723 | 0.71 | 1 | MS.Trt_vs_MS.NoTrt_F |
| ADD1 | 0.1919 | 0.26674 | 0.784 | 0.81 | 1 | MS.Trt_vs_MS.NoTrt_F |
| LAMA4 | 0.19257 | 0.95285 | 0.234 | 0.3 | 1 | MS.Trt_vs_MS.NoTrt_F |
| ALKBH6 | 0.1928 | -0.8712 | 0.572 | 0.6 | 1 | MS.Trt_vs_MS.NoTrt_F |
| TTF1 | 0.19304 | -0.3313 | 0.45 | 0.51 | 1 | MS.Trt_vs_MS.NoTrt_F |
| CXCL14 | 0.19307 | 1.50284 | 0.147 | 0.2 | 1 | MS.Trt_vs_MS.NoTrt_F |
| SMARCD2 | 0.19309 | 1.04074 | 0.147 | 0.2 | 1 | MS.Trt_vs_MS.NoTrt_F |
| CDK7 | 0.19309 | 1.24205 | 0.216 | 0.29 | 1 | MS.Trt_vs_MS.NoTrt_F |
| EPS8 | 0.19361 | 0.56257 | 0.464 | 0.54 | 1 | MS.Trt_vs_MS.NoTrt_F |
| TIMELESS | 0.19363 | 0.3976 | 0.371 | 0.44 | 1 | MS.Trt_vs_MS.NoTrt_F |
| ZNF486 | 0.19432 | -0.2918 | 0.698 | 0.76 | 1 | MS.Trt_vs_MS.NoTrt_F |
| GEN1 | 0.19442 | 0.2657 | 0.755 | 0.75 | 1 | MS.Trt_vs_MS.NoTrt_F |
| WDR41 | 0.19449 | -0.3698 | 0.399 | 0.48 | 1 | MS.Trt_vs_MS.NoTrt_F |
| IL15 | 0.1946 | -0.9399 | 0.356 | 0.41 | 1 | MS.Trt_vs_MS.NoTrt_F |
| CYTH3 | 0.19479 | -0.9046 | 0.076 | 0.11 | 1 | MS.Trt_vs_MS.NoTrt_F |
| DCTN2 | 0.19565 | -0.4454 | 0.565 | 0.65 | 1 | MS.Trt_vs_MS.NoTrt_F |
| PRKRA | 0.19584 | 0.40666 | 0.259 | 0.33 | 1 | MS.Trt_vs_MS.NoTrt_F |
| NCF1 | 0.19628 | 0.45171 | 0.496 | 0.6 | 1 | MS.Trt_vs_MS.NoTrt_F |
| NME6 | 0.19636 | -0.3381 | 0.874 | 0.92 | 1 | MS.Trt_vs_MS.NoTrt_F |
| GEMIN6 | 0.19642 | 0.5371 | 0.917 | 0.93 | 1 | MS.Trt_vs_MS.NoTrt_F |
| XPO6 | 0.19647 | 0.37776 | 0.324 | 0.42 | 1 | MS.Trt_vs_MS.NoTrt_F |
| SVIL | 0.19661 | 0.8261 | 0.191 | 0.26 | 1 | MS.Trt_vs_MS.NoTrt_F |
| C2orf61 | 0.19703 | 0.28124 | 0.14 | 0.19 | 1 | MS.Trt_vs_MS.NoTrt_F |
| REEP3 | 0.19735 | 1.34922 | 0.532 | 0.53 | 1 | MS.Trt_vs_MS.NoTrt_F |
| SNAP23 | 0.19775 | 0.28551 | 0.655 | 0.74 | 1 | MS.Trt_vs_MS.NoTrt_F |
| DAZAP2 | 0.19782 | 0.32433 | 0.806 | 0.92 | 1 | MS.Trt_vs_MS.NoTrt_F |
| GTF2F1 | 0.19796 | 0.35214 | 0.309 | 0.39 | 1 | MS.Trt_vs_MS.NoTrt_F |
| RALB | 0.198 | 0.90672 | 0.133 | 0.18 | 1 | MS.Trt_vs_MS.NoTrt_F |
| FRMD4B | 0.1984 | -0.9718 | 0.96 | 0.96 | 1 | MS.Trt_vs_MS.NoTrt_F |
| CHI3L2 | 0.19845 | 1.15142 | 0.219 | 0.28 | 1 | MS.Trt_vs_MS.NoTrt_F |

| | | | | | | |
|---|---|---|---|---|---|---|
| P4HTM | 0.19902 | -0.4259 | 0.396 | 0.44 | 1 | MS.Trt_vs_MS.NoTrt_F |
| ATP10A | 0.19925 | 1.86202 | 0.115 | 0.17 | 1 | MS.Trt_vs_MS.NoTrt_F |
| ARHGAP44 | 0.19937 | 0.98503 | 0.313 | 0.36 | 1 | MS.Trt_vs_MS.NoTrt_F |
| TMEM108 | 0.19953 | -0.407 | 0.309 | 0.37 | 1 | MS.Trt_vs_MS.NoTrt_F |
| WDR78 | 0.20017 | -0.2832 | 0.486 | 0.54 | 1 | MS.Trt_vs_MS.NoTrt_F |
| CHFR | 0.20037 | 0.53871 | 0.942 | 0.96 | 1 | MS.Trt_vs_MS.NoTrt_F |
| GALE | 0.20044 | 0.34945 | 0.342 | 0.4 | 1 | MS.Trt_vs_MS.NoTrt_F |
| IFI27L2 | 0.20064 | 0.37517 | 0.263 | 0.32 | 1 | MS.Trt_vs_MS.NoTrt_F |
| ZSCAN22 | 0.20065 | -0.7382 | 0.486 | 0.57 | 1 | MS.Trt_vs_MS.NoTrt_F |
| SIKE1 | 0.20079 | 0.28803 | 0.453 | 0.5 | 1 | MS.Trt_vs_MS.NoTrt_F |
| DPEP2 | 0.20081 | 0.46775 | 0.227 | 0.29 | 1 | MS.Trt_vs_MS.NoTrt_F |
| NFIC | 0.20111 | 0.27795 | 0.432 | 0.49 | 1 | MS.Trt_vs_MS.NoTrt_F |
| NINJ1 | 0.20143 | 0.65933 | 0.543 | 0.65 | 1 | MS.Trt_vs_MS.NoTrt_F |
| ZNF23 | 0.2015 | -0.5363 | 0.371 | 0.42 | 1 | MS.Trt_vs_MS.NoTrt_F |
| LYPLA1 | 0.20165 | 0.51315 | 0.522 | 0.62 | 1 | MS.Trt_vs_MS.NoTrt_F |
| MRPS27 | 0.20207 | -0.765 | 0.543 | 0.56 | 1 | MS.Trt_vs_MS.NoTrt_F |
| ZNF562 | 0.2021 | -0.5737 | 0.986 | 0.97 | 1 | MS.Trt_vs_MS.NoTrt_F |
| CENPO | 0.2022 | 0.47271 | 0.079 | 0.12 | 1 | MS.Trt_vs_MS.NoTrt_F |
| PRKDC | 0.20232 | 0.39983 | 0.583 | 0.65 | 1 | MS.Trt_vs_MS.NoTrt_F |
| SNRPB2 | 0.20353 | 0.77153 | 0.223 | 0.3 | 1 | MS.Trt_vs_MS.NoTrt_F |
| BANK1 | 0.20369 | 0.82161 | 0.191 | 0.25 | 1 | MS.Trt_vs_MS.NoTrt_F |
| NR1H2 | 0.20401 | 0.41605 | 0.507 | 0.61 | 1 | MS.Trt_vs_MS.NoTrt_F |
| C15orf41 | 0.20422 | 0.76612 | 0.601 | 0.71 | 1 | MS.Trt_vs_MS.NoTrt_F |
| KIAA1324 | 0.20457 | -0.6184 | 0.752 | 0.81 | 1 | MS.Trt_vs_MS.NoTrt_F |
| C14orf80 | 0.20481 | 0.41375 | 0.284 | 0.35 | 1 | MS.Trt_vs_MS.NoTrt_F |
| B3GAT3 | 0.20541 | 0.68232 | 0.543 | 0.6 | 1 | MS.Trt_vs_MS.NoTrt_F |
| LGR4 | 0.20617 | -0.3432 | 0.104 | 0.14 | 1 | MS.Trt_vs_MS.NoTrt_F |
| MT-ATP8 | 0.20653 | 0.26428 | 0.662 | 0.8 | 1 | MS.Trt_vs_MS.NoTrt_F |
| RCC1L | 0.20677 | -0.4259 | 0.299 | 0.35 | 1 | MS.Trt_vs_MS.NoTrt_F |
| C14orf2 | 0.20678 | 0.3268 | 0.799 | 0.9 | 1 | MS.Trt_vs_MS.NoTrt_F |
| CEP68 | 0.20697 | -0.2767 | 0.511 | 0.55 | 1 | MS.Trt_vs_MS.NoTrt_F |
| ENY2 | 0.20726 | 0.58264 | 0.504 | 0.55 | 1 | MS.Trt_vs_MS.NoTrt_F |
| GSTM2 | 0.20733 | 0.25764 | 0.356 | 0.43 | 1 | MS.Trt_vs_MS.NoTrt_F |
| MFSD14C | 0.20783 | -0.3414 | 0.615 | 0.73 | 1 | MS.Trt_vs_MS.NoTrt_F |
| ZNF587B | 0.2082 | 0.80093 | 0.309 | 0.38 | 1 | MS.Trt_vs_MS.NoTrt_F |
| FAM96B | 0.20824 | 0.46378 | 0.237 | 0.31 | 1 | MS.Trt_vs_MS.NoTrt_F |
| SPDYE5 | 0.20844 | 0.30622 | 0.586 | 0.57 | 1 | MS.Trt_vs_MS.NoTrt_F |
| TMEM94 | 0.20875 | 0.31115 | 0.216 | 0.28 | 1 | MS.Trt_vs_MS.NoTrt_F |
| ZFYVE27 | 0.20875 | 0.5592 | 0.194 | 0.26 | 1 | MS.Trt_vs_MS.NoTrt_F |
| SGPP2 | 0.20915 | 0.29316 | 0.342 | 0.41 | 1 | MS.Trt_vs_MS.NoTrt_F |

| | | | | | | |
|---|---|---|---|---|---|---|
| NTN5 | 0.20935 | 0.28745 | 0.14 | 0.19 | 1 | MS.Trt_vs_MS.NoTrt_F |
| HVCN1 | 0.20967 | 0.36456 | 0.518 | 0.6 | 1 | MS.Trt_vs_MS.NoTrt_F |
| ING1 | 0.20983 | 0.48075 | 0.597 | 0.63 | 1 | MS.Trt_vs_MS.NoTrt_F |
| RBM7 | 0.21048 | 0.33304 | 0.759 | 0.68 | 1 | MS.Trt_vs_MS.NoTrt_F |
| TCP11L1 | 0.21053 | -0.6584 | 0.234 | 0.28 | 1 | MS.Trt_vs_MS.NoTrt_F |
| RRP1 | 0.21078 | 0.88123 | 0.647 | 0.71 | 1 | MS.Trt_vs_MS.NoTrt_F |
| PIP5K1C | 0.21091 | 0.77682 | 0.101 | 0.14 | 1 | MS.Trt_vs_MS.NoTrt_F |
| IFITM3 | 0.211 | 0.28733 | 0.529 | 0.66 | 1 | MS.Trt_vs_MS.NoTrt_F |
| RPS6KL1 | 0.21116 | -0.2516 | 0.655 | 0.76 | 1 | MS.Trt_vs_MS.NoTrt_F |
| LIMA1 | 0.21182 | 0.44203 | 0.45 | 0.48 | 1 | MS.Trt_vs_MS.NoTrt_F |
| POLR1D | 0.21194 | 0.31972 | 0.464 | 0.53 | 1 | MS.Trt_vs_MS.NoTrt_F |
| TCTEX1D2 | 0.21205 | 0.44081 | 0.335 | 0.39 | 1 | MS.Trt_vs_MS.NoTrt_F |
| WASH1 | 0.2121 | 0.46532 | 0.216 | 0.28 | 1 | MS.Trt_vs_MS.NoTrt_F |
| ZBTB2 | 0.21211 | 0.67319 | 0.126 | 0.18 | 1 | MS.Trt_vs_MS.NoTrt_F |
| BCL11B | 0.21258 | 0.46208 | 0.45 | 0.58 | 1 | MS.Trt_vs_MS.NoTrt_F |
| PRDX3 | 0.21276 | 0.5055 | 0.457 | 0.53 | 1 | MS.Trt_vs_MS.NoTrt_F |
| HIST2H2AA3 | 0.21303 | 0.69909 | 0.108 | 0.16 | 1 | MS.Trt_vs_MS.NoTrt_F |
| PDPN | 0.2134 | -0.5646 | 0.5 | 0.48 | 1 | MS.Trt_vs_MS.NoTrt_F |
| SCRN2 | 0.21353 | 1.23285 | 0.133 | 0.18 | 1 | MS.Trt_vs_MS.NoTrt_F |
| GUF1 | 0.2137 | 0.89487 | 0.371 | 0.45 | 1 | MS.Trt_vs_MS.NoTrt_F |
| C3 | 0.21396 | 0.25843 | 0.924 | 0.93 | 1 | MS.Trt_vs_MS.NoTrt_F |
| MPRIP | 0.21429 | -0.4146 | 0.892 | 0.92 | 1 | MS.Trt_vs_MS.NoTrt_F |
| CREBRF | 0.21492 | 0.29515 | 0.687 | 0.81 | 1 | MS.Trt_vs_MS.NoTrt_F |
| SUMF2 | 0.21505 | 0.59731 | 0.874 | 0.91 | 1 | MS.Trt_vs_MS.NoTrt_F |
| GSK3A | 0.21524 | -0.5247 | 0.324 | 0.37 | 1 | MS.Trt_vs_MS.NoTrt_F |
| NBPF19 | 0.21552 | 0.44242 | 0.381 | 0.46 | 1 | MS.Trt_vs_MS.NoTrt_F |
| ZNF324 | 0.21621 | 1.01256 | 0.101 | 0.14 | 1 | MS.Trt_vs_MS.NoTrt_F |
| FASLG | 0.21635 | 1.38779 | 0.104 | 0.14 | 1 | MS.Trt_vs_MS.NoTrt_F |
| ANP32E | 0.21715 | 1.09338 | 0.478 | 0.65 | 1 | MS.Trt_vs_MS.NoTrt_F |
| TANGO2 | 0.21719 | 0.66851 | 0.871 | 0.87 | 1 | MS.Trt_vs_MS.NoTrt_F |
| TBXAS1 | 0.21725 | 0.80846 | 0.392 | 0.46 | 1 | MS.Trt_vs_MS.NoTrt_F |
| BAG2 | 0.21736 | 0.42355 | 0.245 | 0.29 | 1 | MS.Trt_vs_MS.NoTrt_F |
| NCKAP1 | 0.21761 | 0.67588 | 0.68 | 0.65 | 1 | MS.Trt_vs_MS.NoTrt_F |
| BICD2 | 0.21788 | 1.1372 | 0.119 | 0.17 | 1 | MS.Trt_vs_MS.NoTrt_F |
| CARNS1 | 0.21799 | 0.75225 | 0.205 | 0.26 | 1 | MS.Trt_vs_MS.NoTrt_F |
| CFAP20 | 0.21804 | 0.53785 | 0.183 | 0.24 | 1 | MS.Trt_vs_MS.NoTrt_F |
| CLK2 | 0.21825 | 0.33349 | 0.122 | 0.17 | 1 | MS.Trt_vs_MS.NoTrt_F |
| RFC4 | 0.21835 | 0.25084 | 0.309 | 0.37 | 1 | MS.Trt_vs_MS.NoTrt_F |
| SYT2 | 0.21857 | 0.84633 | 0.072 | 0.11 | 1 | MS.Trt_vs_MS.NoTrt_F |
| VPS50 | 0.21911 | -1.0123 | 0.604 | 0.64 | 1 | MS.Trt_vs_MS.NoTrt_F |

| | | | | | | |
|---|---|---|---|---|---|---|
| RBM34 | 0.21942 | 0.59427 | 0.619 | 0.72 | 1 | MS.Trt_vs_MS.NoTrt_F |
| C16orf72 | 0.21989 | 0.46532 | 0.86 | 0.85 | 1 | MS.Trt_vs_MS.NoTrt_F |
| PLEKHA5 | 0.22018 | -0.5271 | 0.917 | 0.95 | 1 | MS.Trt_vs_MS.NoTrt_F |
| YRDC | 0.22035 | 0.54765 | 0.108 | 0.15 | 1 | MS.Trt_vs_MS.NoTrt_F |
| MXRA7 | 0.22038 | -0.3072 | 0.755 | 0.84 | 1 | MS.Trt_vs_MS.NoTrt_F |
| SH2B3 | 0.2206 | 0.30823 | 0.187 | 0.24 | 1 | MS.Trt_vs_MS.NoTrt_F |
| PRKD2 | 0.22069 | 0.81839 | 0.748 | 0.79 | 1 | MS.Trt_vs_MS.NoTrt_F |
| ATP1B1 | 0.22134 | 0.85051 | 0.309 | 0.39 | 1 | MS.Trt_vs_MS.NoTrt_F |
| ARHGAP8 | 0.22136 | 0.25439 | 0.683 | 0.72 | 1 | MS.Trt_vs_MS.NoTrt_F |
| TRPM7 | 0.22147 | 0.30474 | 0.906 | 0.94 | 1 | MS.Trt_vs_MS.NoTrt_F |
| RP11-514P8 | 0.2215 | -0.3996 | 0.072 | 0.11 | 1 | MS.Trt_vs_MS.NoTrt_F |
| RNF103 | 0.22158 | 0.44697 | 0.626 | 0.67 | 1 | MS.Trt_vs_MS.NoTrt_F |
| THOC3 | 0.22179 | 0.2747 | 0.532 | 0.65 | 1 | MS.Trt_vs_MS.NoTrt_F |
| NR2C1 | 0.22269 | 0.52493 | 0.658 | 0.76 | 1 | MS.Trt_vs_MS.NoTrt_F |
| BBS1 | 0.22288 | 0.25619 | 0.608 | 0.67 | 1 | MS.Trt_vs_MS.NoTrt_F |
| HLA-DRB3 | 0.22316 | 0.84042 | 0.18 | 0.24 | 1 | MS.Trt_vs_MS.NoTrt_F |
| CTD-2132N1 | 0.2239 | 0.58069 | 0.097 | 0.14 | 1 | MS.Trt_vs_MS.NoTrt_F |
| MCF2L | 0.22398 | -0.5688 | 0.295 | 0.35 | 1 | MS.Trt_vs_MS.NoTrt_F |
| CPNE1 | 0.22448 | 0.43377 | 0.522 | 0.69 | 1 | MS.Trt_vs_MS.NoTrt_F |
| EID2B | 0.22456 | 0.62854 | 0.514 | 0.57 | 1 | MS.Trt_vs_MS.NoTrt_F |
| USP12 | 0.22513 | 0.70899 | 0.115 | 0.16 | 1 | MS.Trt_vs_MS.NoTrt_F |
| TMEM41A | 0.22514 | 0.31682 | 0.799 | 0.79 | 1 | MS.Trt_vs_MS.NoTrt_F |
| G6PD | 0.22522 | 0.37564 | 0.115 | 0.16 | 1 | MS.Trt_vs_MS.NoTrt_F |
| BCL2L13 | 0.22538 | -0.8568 | 0.514 | 0.6 | 1 | MS.Trt_vs_MS.NoTrt_F |
| UVSSA | 0.22611 | -0.4903 | 0.741 | 0.81 | 1 | MS.Trt_vs_MS.NoTrt_F |
| TMEM127 | 0.22631 | 0.32599 | 0.687 | 0.71 | 1 | MS.Trt_vs_MS.NoTrt_F |
| ZNF563 | 0.22648 | 1.35616 | 0.55 | 0.62 | 1 | MS.Trt_vs_MS.NoTrt_F |
| ITGAX | 0.22663 | 0.35315 | 0.482 | 0.56 | 1 | MS.Trt_vs_MS.NoTrt_F |
| TEX2 | 0.2274 | 0.42438 | 0.633 | 0.73 | 1 | MS.Trt_vs_MS.NoTrt_F |
| TADA2B | 0.22758 | 1.07613 | 0.302 | 0.36 | 1 | MS.Trt_vs_MS.NoTrt_F |
| PZP | 0.22771 | 0.52611 | 0.083 | 0.12 | 1 | MS.Trt_vs_MS.NoTrt_F |
| HBA2 | 0.22807 | 0.65375 | 0.072 | 0.11 | 1 | MS.Trt_vs_MS.NoTrt_F |
| ZNF234 | 0.22834 | 0.69176 | 0.709 | 0.79 | 1 | MS.Trt_vs_MS.NoTrt_F |
| RRP12 | 0.22851 | 0.2918 | 0.906 | 0.87 | 1 | MS.Trt_vs_MS.NoTrt_F |
| UBE2V2 | 0.22854 | 0.38063 | 0.683 | 0.77 | 1 | MS.Trt_vs_MS.NoTrt_F |
| ERCC2 | 0.22917 | -0.7125 | 0.522 | 0.59 | 1 | MS.Trt_vs_MS.NoTrt_F |
| RGS5 | 0.2292 | 1.42784 | 0.104 | 0.14 | 1 | MS.Trt_vs_MS.NoTrt_F |
| IFT88 | 0.22921 | 0.9138 | 0.496 | 0.58 | 1 | MS.Trt_vs_MS.NoTrt_F |
| SLC38A10 | 0.22953 | 0.3674 | 0.388 | 0.43 | 1 | MS.Trt_vs_MS.NoTrt_F |
| XKR8 | 0.23012 | -0.3501 | 0.194 | 0.24 | 1 | MS.Trt_vs_MS.NoTrt_F |

| | | | | | | |
|---|---|---|---|---|---|---|
| STXBP5 | 0.23015 | 0.9034 | 0.176 | 0.23 | 1 | MS.Trt_vs_MS.NoTrt_F |
| RP11-343C2 | 0.23024 | 0.49745 | 0.072 | 0.11 | 1 | MS.Trt_vs_MS.NoTrt_F |
| NFE2L1 | 0.23045 | -0.3252 | 0.112 | 0.16 | 1 | MS.Trt_vs_MS.NoTrt_F |
| GPAT4 | 0.23057 | -0.611 | 0.73 | 0.85 | 1 | MS.Trt_vs_MS.NoTrt_F |
| LY6E | 0.23062 | 0.31842 | 0.522 | 0.63 | 1 | MS.Trt_vs_MS.NoTrt_F |
| IL17RE | 0.23069 | 0.81992 | 0.263 | 0.33 | 1 | MS.Trt_vs_MS.NoTrt_F |
| PPP2R5C | 0.23087 | 0.7143 | 0.924 | 0.96 | 1 | MS.Trt_vs_MS.NoTrt_F |
| DERA | 0.23108 | 1.63403 | 0.112 | 0.16 | 1 | MS.Trt_vs_MS.NoTrt_F |
| CEP120 | 0.23111 | 0.63731 | 0.165 | 0.22 | 1 | MS.Trt_vs_MS.NoTrt_F |
| PPIL3 | 0.2315 | 0.57739 | 0.734 | 0.85 | 1 | MS.Trt_vs_MS.NoTrt_F |
| NQO2 | 0.23158 | -0.2569 | 0.763 | 0.8 | 1 | MS.Trt_vs_MS.NoTrt_F |
| TRAF3IP1 | 0.23179 | 0.29647 | 0.349 | 0.4 | 1 | MS.Trt_vs_MS.NoTrt_F |
| IGF1R | 0.2326 | 1.47779 | 0.288 | 0.37 | 1 | MS.Trt_vs_MS.NoTrt_F |
| ORC5 | 0.23416 | 0.38109 | 0.263 | 0.34 | 1 | MS.Trt_vs_MS.NoTrt_F |
| ZBP1 | 0.23435 | 0.2816 | 0.547 | 0.61 | 1 | MS.Trt_vs_MS.NoTrt_F |
| NDUFA12 | 0.23442 | 0.85871 | 0.241 | 0.32 | 1 | MS.Trt_vs_MS.NoTrt_F |
| TUFT1 | 0.23469 | -0.3751 | 0.367 | 0.44 | 1 | MS.Trt_vs_MS.NoTrt_F |
| FSCN1 | 0.23494 | 0.616 | 0.094 | 0.13 | 1 | MS.Trt_vs_MS.NoTrt_F |
| ATAD3C | 0.23517 | 0.386 | 0.856 | 0.77 | 1 | MS.Trt_vs_MS.NoTrt_F |
| ZFP41 | 0.23522 | 0.8769 | 0.147 | 0.19 | 1 | MS.Trt_vs_MS.NoTrt_F |
| GPR155 | 0.23525 | -0.6452 | 0.899 | 0.91 | 1 | MS.Trt_vs_MS.NoTrt_F |
| OGFOD1 | 0.23546 | -0.2743 | 0.64 | 0.69 | 1 | MS.Trt_vs_MS.NoTrt_F |
| LMBR1 | 0.23568 | 0.68913 | 0.46 | 0.54 | 1 | MS.Trt_vs_MS.NoTrt_F |
| MATN1 | 0.23574 | 1.00125 | 0.09 | 0.13 | 1 | MS.Trt_vs_MS.NoTrt_F |
| UBOX5 | 0.23581 | -0.338 | 0.637 | 0.66 | 1 | MS.Trt_vs_MS.NoTrt_F |
| SPATA6 | 0.23586 | 0.3474 | 0.133 | 0.18 | 1 | MS.Trt_vs_MS.NoTrt_F |
| PEX11A | 0.23604 | 0.41771 | 0.565 | 0.61 | 1 | MS.Trt_vs_MS.NoTrt_F |
| LIPE | 0.23618 | 1.5981 | 0.342 | 0.44 | 1 | MS.Trt_vs_MS.NoTrt_F |
| SNX11 | 0.23631 | 0.35699 | 0.601 | 0.67 | 1 | MS.Trt_vs_MS.NoTrt_F |
| TRAF2 | 0.23638 | 0.32251 | 0.255 | 0.33 | 1 | MS.Trt_vs_MS.NoTrt_F |
| ISY1 | 0.23675 | 0.56629 | 0.597 | 0.69 | 1 | MS.Trt_vs_MS.NoTrt_F |
| POMZP3 | 0.23696 | 0.75178 | 0.439 | 0.54 | 1 | MS.Trt_vs_MS.NoTrt_F |
| CIPC | 0.23717 | 0.37484 | 0.45 | 0.5 | 1 | MS.Trt_vs_MS.NoTrt_F |
| AP5M1 | 0.23731 | 0.28953 | 0.176 | 0.23 | 1 | MS.Trt_vs_MS.NoTrt_F |
| DNPEP | 0.23765 | -0.9829 | 0.489 | 0.52 | 1 | MS.Trt_vs_MS.NoTrt_F |
| TAOK1 | 0.23795 | 0.4568 | 0.928 | 0.95 | 1 | MS.Trt_vs_MS.NoTrt_F |
| LRRN3 | 0.23816 | 1.30989 | 0.137 | 0.18 | 1 | MS.Trt_vs_MS.NoTrt_F |
| SPRYD7 | 0.23831 | 0.26402 | 0.691 | 0.69 | 1 | MS.Trt_vs_MS.NoTrt_F |
| PPP1R3B | 0.23849 | 0.90834 | 0.277 | 0.34 | 1 | MS.Trt_vs_MS.NoTrt_F |
| SLC16A7 | 0.23853 | -0.6383 | 0.608 | 0.66 | 1 | MS.Trt_vs_MS.NoTrt_F |

| | | | | | | |
|---|---|---|---|---|---|---|
| CLUAP1 | 0.23883 | 0.26741 | 0.817 | 0.79 | 1 | MS.Trt_vs_MS.NoTrt_F |
| ATP6AP2 | 0.23891 | 0.37417 | 0.813 | 0.87 | 1 | MS.Trt_vs_MS.NoTrt_F |
| ALMS1 | 0.2394 | -0.5443 | 0.446 | 0.48 | 1 | MS.Trt_vs_MS.NoTrt_F |
| HIP1 | 0.23986 | 0.37125 | 0.655 | 0.69 | 1 | MS.Trt_vs_MS.NoTrt_F |
| C16orf74 | 0.24014 | -0.3342 | 0.245 | 0.29 | 1 | MS.Trt_vs_MS.NoTrt_F |
| ATP13A4 | 0.24032 | -0.3105 | 0.424 | 0.48 | 1 | MS.Trt_vs_MS.NoTrt_F |
| MR1 | 0.24039 | -0.3061 | 0.896 | 0.91 | 1 | MS.Trt_vs_MS.NoTrt_F |
| RPP14 | 0.24045 | 0.50509 | 0.914 | 0.93 | 1 | MS.Trt_vs_MS.NoTrt_F |
| CCDC92 | 0.24057 | 0.61623 | 0.09 | 0.13 | 1 | MS.Trt_vs_MS.NoTrt_F |
| SH2D3A | 0.24062 | -0.2544 | 0.95 | 0.98 | 1 | MS.Trt_vs_MS.NoTrt_F |
| KIAA1429 | 0.24092 | 0.43045 | 0.835 | 0.88 | 1 | MS.Trt_vs_MS.NoTrt_F |
| CCDC34 | 0.24094 | 0.42488 | 0.137 | 0.18 | 1 | MS.Trt_vs_MS.NoTrt_F |
| HIST1H2BG | 0.24111 | 0.28526 | 0.486 | 0.43 | 1 | MS.Trt_vs_MS.NoTrt_F |
| LCLAT1 | 0.24114 | 0.61921 | 0.799 | 0.85 | 1 | MS.Trt_vs_MS.NoTrt_F |
| NQO1 | 0.24126 | 0.5621 | 0.579 | 0.64 | 1 | MS.Trt_vs_MS.NoTrt_F |
| CUBN | 0.24155 | -0.6881 | 0.392 | 0.44 | 1 | MS.Trt_vs_MS.NoTrt_F |
| BORCS7 | 0.24178 | -0.6479 | 0.331 | 0.39 | 1 | MS.Trt_vs_MS.NoTrt_F |
| ACRC | 0.24179 | 0.26985 | 0.439 | 0.49 | 1 | MS.Trt_vs_MS.NoTrt_F |
| ZNF623 | 0.24197 | 0.46805 | 0.629 | 0.72 | 1 | MS.Trt_vs_MS.NoTrt_F |
| ADSSL1 | 0.24219 | 0.39327 | 0.324 | 0.39 | 1 | MS.Trt_vs_MS.NoTrt_F |
| SHKBP1 | 0.24242 | 0.25822 | 0.734 | 0.83 | 1 | MS.Trt_vs_MS.NoTrt_F |
| ZNF680 | 0.24248 | 1.21632 | 0.173 | 0.22 | 1 | MS.Trt_vs_MS.NoTrt_F |
| UQCRH | 0.24268 | 0.48138 | 0.486 | 0.59 | 1 | MS.Trt_vs_MS.NoTrt_F |
| SLC16A5 | 0.24287 | 0.69502 | 0.176 | 0.23 | 1 | MS.Trt_vs_MS.NoTrt_F |
| C9orf64 | 0.24287 | -0.3668 | 0.583 | 0.62 | 1 | MS.Trt_vs_MS.NoTrt_F |
| ATP7A | 0.24305 | 0.76173 | 0.399 | 0.45 | 1 | MS.Trt_vs_MS.NoTrt_F |
| C9orf3 | 0.24307 | 0.39278 | 0.698 | 0.79 | 1 | MS.Trt_vs_MS.NoTrt_F |
| C9orf85 | 0.24333 | 0.28034 | 0.903 | 0.93 | 1 | MS.Trt_vs_MS.NoTrt_F |
| PRF1 | 0.24336 | 0.48955 | 0.263 | 0.33 | 1 | MS.Trt_vs_MS.NoTrt_F |
| TYRO3 | 0.24389 | 0.27308 | 0.968 | 0.96 | 1 | MS.Trt_vs_MS.NoTrt_F |
| HIST1H1E | 0.24424 | 1.51712 | 0.115 | 0.16 | 1 | MS.Trt_vs_MS.NoTrt_F |
| ALG10B | 0.2444 | -0.3627 | 0.748 | 0.67 | 1 | MS.Trt_vs_MS.NoTrt_F |
| FHDC1 | 0.24477 | 0.70678 | 0.486 | 0.49 | 1 | MS.Trt_vs_MS.NoTrt_F |
| POU2AF1 | 0.24498 | 0.43547 | 0.705 | 0.73 | 1 | MS.Trt_vs_MS.NoTrt_F |
| ZNF844 | 0.24526 | 0.27652 | 0.924 | 0.92 | 1 | MS.Trt_vs_MS.NoTrt_F |
| EXOSC9 | 0.2453 | 0.45637 | 0.227 | 0.28 | 1 | MS.Trt_vs_MS.NoTrt_F |
| STX7 | 0.24543 | 0.59769 | 0.719 | 0.76 | 1 | MS.Trt_vs_MS.NoTrt_F |
| ZNF532 | 0.24568 | -0.5676 | 0.935 | 0.97 | 1 | MS.Trt_vs_MS.NoTrt_F |
| ZNF77 | 0.24587 | -1.4561 | 0.083 | 0.12 | 1 | MS.Trt_vs_MS.NoTrt_F |
| CCDC159 | 0.24619 | -0.6019 | 0.759 | 0.79 | 1 | MS.Trt_vs_MS.NoTrt_F |

| | | | | | | |
|---|---|---|---|---|---|---|
| IGHMBP2 | 0.24666 | 1.26065 | 0.392 | 0.43 | 1 | MS.Trt_vs_MS.NoTrt_F |
| SGSM1 | 0.24671 | -0.437 | 0.558 | 0.58 | 1 | MS.Trt_vs_MS.NoTrt_F |
| ISPD | 0.24673 | 0.3282 | 0.791 | 0.82 | 1 | MS.Trt_vs_MS.NoTrt_F |
| DHCR7 | 0.24677 | 1.14586 | 0.094 | 0.13 | 1 | MS.Trt_vs_MS.NoTrt_F |
| CD38 | 0.2472 | -0.6029 | 0.356 | 0.41 | 1 | MS.Trt_vs_MS.NoTrt_F |
| BAIAP2L1 | 0.24754 | 0.42608 | 0.525 | 0.57 | 1 | MS.Trt_vs_MS.NoTrt_F |
| GOLGA6L1 | 0.24766 | 0.3826 | 0.457 | 0.43 | 1 | MS.Trt_vs_MS.NoTrt_F |
| THAP12 | 0.24771 | -0.5637 | 0.604 | 0.66 | 1 | MS.Trt_vs_MS.NoTrt_F |
| PNPT1 | 0.24822 | 0.5662 | 0.881 | 0.92 | 1 | MS.Trt_vs_MS.NoTrt_F |
| P3H3 | 0.24835 | 0.36764 | 0.079 | 0.11 | 1 | MS.Trt_vs_MS.NoTrt_F |
| PROSC | 0.24859 | 0.3505 | 0.392 | 0.44 | 1 | MS.Trt_vs_MS.NoTrt_F |
| RRBP1 | 0.24872 | -0.7123 | 0.32 | 0.39 | 1 | MS.Trt_vs_MS.NoTrt_F |
| B4GALNT2 | 0.2488 | 0.46141 | 0.353 | 0.44 | 1 | MS.Trt_vs_MS.NoTrt_F |
| TTI2 | 0.24885 | -0.3991 | 0.399 | 0.48 | 1 | MS.Trt_vs_MS.NoTrt_F |
| USP53 | 0.24902 | 0.52392 | 0.565 | 0.66 | 1 | MS.Trt_vs_MS.NoTrt_F |
| QPRT | 0.24919 | 0.3534 | 0.903 | 0.88 | 1 | MS.Trt_vs_MS.NoTrt_F |
| NAPB | 0.24938 | 0.89008 | 0.439 | 0.51 | 1 | MS.Trt_vs_MS.NoTrt_F |
| ZBED6CL | 0.24942 | 2.13484 | 0.094 | 0.13 | 1 | MS.Trt_vs_MS.NoTrt_F |
| PPARD | 0.2495 | -0.4399 | 0.561 | 0.65 | 1 | MS.Trt_vs_MS.NoTrt_F |
| ITM2B | 0.24961 | 0.39363 | 0.604 | 0.77 | 1 | MS.Trt_vs_MS.NoTrt_F |
| GRPEL1 | 0.24965 | 0.71833 | 0.18 | 0.24 | 1 | MS.Trt_vs_MS.NoTrt_F |
| NTAN1 | 0.24979 | 0.26016 | 0.662 | 0.77 | 1 | MS.Trt_vs_MS.NoTrt_F |
| SDK2 | 0.2506 | 0.25212 | 0.54 | 0.62 | 1 | MS.Trt_vs_MS.NoTrt_F |
| RP11-793H1 | 0.25079 | 0.45667 | 0.317 | 0.37 | 1 | MS.Trt_vs_MS.NoTrt_F |
| PRPSAP1 | 0.25138 | 0.41464 | 0.77 | 0.75 | 1 | MS.Trt_vs_MS.NoTrt_F |
| RPL26L1 | 0.25151 | 0.43701 | 0.165 | 0.21 | 1 | MS.Trt_vs_MS.NoTrt_F |
| IL11RA | 0.25164 | 0.25107 | 0.183 | 0.24 | 1 | MS.Trt_vs_MS.NoTrt_F |
| CIITA | 0.25185 | 0.33784 | 0.91 | 0.9 | 1 | MS.Trt_vs_MS.NoTrt_F |
| CLCN3 | 0.25212 | -0.3013 | 0.396 | 0.44 | 1 | MS.Trt_vs_MS.NoTrt_F |
| BEND7 | 0.25215 | 0.4704 | 0.363 | 0.43 | 1 | MS.Trt_vs_MS.NoTrt_F |
| 9-Mar | 0.25284 | -0.7553 | 0.59 | 0.63 | 1 | MS.Trt_vs_MS.NoTrt_F |
| UAP1 | 0.25306 | 1.34528 | 0.108 | 0.15 | 1 | MS.Trt_vs_MS.NoTrt_F |
| ZNF417 | 0.25331 | 0.68555 | 0.763 | 0.75 | 1 | MS.Trt_vs_MS.NoTrt_F |
| PIGS | 0.25383 | 0.49835 | 0.532 | 0.57 | 1 | MS.Trt_vs_MS.NoTrt_F |
| KIAA1456 | 0.25384 | 0.35888 | 0.914 | 0.93 | 1 | MS.Trt_vs_MS.NoTrt_F |
| SUOX | 0.25391 | -0.5629 | 0.781 | 0.84 | 1 | MS.Trt_vs_MS.NoTrt_F |
| PTPN18 | 0.25399 | 0.28581 | 0.406 | 0.48 | 1 | MS.Trt_vs_MS.NoTrt_F |
| ERC1 | 0.25404 | -0.4327 | 0.892 | 0.92 | 1 | MS.Trt_vs_MS.NoTrt_F |
| AARD | 0.25412 | 0.31275 | 0.755 | 0.7 | 1 | MS.Trt_vs_MS.NoTrt_F |
| ADCY2 | 0.25418 | 0.2833 | 0.727 | 0.8 | 1 | MS.Trt_vs_MS.NoTrt_F |

| | | | | | | |
|---|---|---|---|---|---|---|
| CDKL5 | 0.25422 | 0.29902 | 0.939 | 0.93 | 1 | MS.Trt_vs_MS.NoTrt_F |
| CNOT7 | 0.25454 | 0.49244 | 0.543 | 0.62 | 1 | MS.Trt_vs_MS.NoTrt_F |
| CLCN2 | 0.25477 | 0.50497 | 0.626 | 0.65 | 1 | MS.Trt_vs_MS.NoTrt_F |
| MIF | 0.25491 | 0.73241 | 0.46 | 0.64 | 1 | MS.Trt_vs_MS.NoTrt_F |
| RNF169 | 0.25525 | 1.09603 | 0.104 | 0.14 | 1 | MS.Trt_vs_MS.NoTrt_F |
| RHBDD3 | 0.25552 | -0.8155 | 0.554 | 0.58 | 1 | MS.Trt_vs_MS.NoTrt_F |
| RBM33 | 0.25555 | 0.65972 | 0.255 | 0.33 | 1 | MS.Trt_vs_MS.NoTrt_F |
| CD300LD | 0.25556 | 1.35939 | 0.392 | 0.46 | 1 | MS.Trt_vs_MS.NoTrt_F |
| ZNF699 | 0.2557 | -0.5198 | 0.306 | 0.36 | 1 | MS.Trt_vs_MS.NoTrt_F |
| ZSWIM8 | 0.25584 | 0.46405 | 0.831 | 0.86 | 1 | MS.Trt_vs_MS.NoTrt_F |
| MMP25 | 0.25615 | 0.59213 | 0.158 | 0.21 | 1 | MS.Trt_vs_MS.NoTrt_F |
| LGALS4 | 0.25628 | 0.36476 | 0.572 | 0.64 | 1 | MS.Trt_vs_MS.NoTrt_F |
| RRP15 | 0.25642 | 0.54735 | 0.532 | 0.58 | 1 | MS.Trt_vs_MS.NoTrt_F |
| LANCL3 | 0.25703 | 1.01638 | 0.309 | 0.35 | 1 | MS.Trt_vs_MS.NoTrt_F |
| ZNF749 | 0.25786 | 0.80897 | 0.46 | 0.41 | 1 | MS.Trt_vs_MS.NoTrt_F |
| LGR6 | 0.25798 | 1.16056 | 0.137 | 0.18 | 1 | MS.Trt_vs_MS.NoTrt_F |
| CTDSPL2 | 0.25805 | -0.6771 | 0.68 | 0.76 | 1 | MS.Trt_vs_MS.NoTrt_F |
| DKC1 | 0.25818 | 0.4248 | 0.385 | 0.42 | 1 | MS.Trt_vs_MS.NoTrt_F |
| SFT2D3 | 0.25848 | 0.77553 | 0.45 | 0.53 | 1 | MS.Trt_vs_MS.NoTrt_F |
| RBP1 | 0.25865 | 0.36048 | 0.399 | 0.48 | 1 | MS.Trt_vs_MS.NoTrt_F |
| SH3BGRL3 | 0.25879 | 0.26249 | 0.853 | 0.97 | 1 | MS.Trt_vs_MS.NoTrt_F |
| IRAK3 | 0.25921 | -0.3553 | 0.396 | 0.48 | 1 | MS.Trt_vs_MS.NoTrt_F |
| MT-ND1 | 0.25927 | 0.43734 | 0.878 | 0.94 | 1 | MS.Trt_vs_MS.NoTrt_F |
| HYPK | 0.26014 | -0.4148 | 0.896 | 0.95 | 1 | MS.Trt_vs_MS.NoTrt_F |
| MKNK1 | 0.26043 | 0.46334 | 0.917 | 0.93 | 1 | MS.Trt_vs_MS.NoTrt_F |
| ABL2 | 0.26055 | -0.5022 | 0.73 | 0.81 | 1 | MS.Trt_vs_MS.NoTrt_F |
| CLCN6 | 0.26059 | 0.93769 | 0.532 | 0.53 | 1 | MS.Trt_vs_MS.NoTrt_F |
| UROS | 0.26113 | -0.3987 | 0.446 | 0.5 | 1 | MS.Trt_vs_MS.NoTrt_F |
| CCDC25 | 0.26121 | 1.02098 | 0.309 | 0.37 | 1 | MS.Trt_vs_MS.NoTrt_F |
| ESRP2 | 0.26123 | 0.30412 | 0.094 | 0.13 | 1 | MS.Trt_vs_MS.NoTrt_F |
| KANSL3 | 0.26127 | -0.3359 | 0.932 | 0.97 | 1 | MS.Trt_vs_MS.NoTrt_F |
| EFCAB10 | 0.2613 | -0.3274 | 0.201 | 0.24 | 1 | MS.Trt_vs_MS.NoTrt_F |
| P2RY11 | 0.26154 | 0.5214 | 0.147 | 0.19 | 1 | MS.Trt_vs_MS.NoTrt_F |
| FAM134A | 0.26182 | 0.80883 | 0.421 | 0.5 | 1 | MS.Trt_vs_MS.NoTrt_F |
| RIMKLB | 0.26186 | 1.2715 | 0.119 | 0.16 | 1 | MS.Trt_vs_MS.NoTrt_F |
| ZNF273 | 0.26204 | 0.38973 | 0.892 | 0.9 | 1 | MS.Trt_vs_MS.NoTrt_F |
| CBX5 | 0.26215 | 0.34954 | 0.928 | 0.93 | 1 | MS.Trt_vs_MS.NoTrt_F |
| RCOR1 | 0.26267 | 0.36084 | 0.169 | 0.22 | 1 | MS.Trt_vs_MS.NoTrt_F |
| OSBPL2 | 0.26273 | -0.6177 | 0.486 | 0.55 | 1 | MS.Trt_vs_MS.NoTrt_F |
| LRRC8B | 0.26371 | 0.55563 | 0.421 | 0.48 | 1 | MS.Trt_vs_MS.NoTrt_F |

| | | | | | | |
|---|---|---|---|---|---|---|
| ATP7B | 0.26372 | 1.1477 | 0.191 | 0.24 | 1 | MS.Trt_vs_MS.NoTrt_F |
| CALM1 | 0.26389 | 0.393 | 0.799 | 0.91 | 1 | MS.Trt_vs_MS.NoTrt_F |
| VSIG1 | 0.26399 | 0.32559 | 0.55 | 0.65 | 1 | MS.Trt_vs_MS.NoTrt_F |
| AGAP3 | 0.264 | 0.6752 | 0.079 | 0.11 | 1 | MS.Trt_vs_MS.NoTrt_F |
| ZSCAN2 | 0.26422 | -0.6249 | 0.763 | 0.83 | 1 | MS.Trt_vs_MS.NoTrt_F |
| TCF7 | 0.26465 | 0.83504 | 0.766 | 0.76 | 1 | MS.Trt_vs_MS.NoTrt_F |
| NOP2 | 0.26504 | -0.4428 | 0.403 | 0.45 | 1 | MS.Trt_vs_MS.NoTrt_F |
| DIAPH3 | 0.26521 | 0.28257 | 0.403 | 0.38 | 1 | MS.Trt_vs_MS.NoTrt_F |
| MAPK1 | 0.26577 | 0.57821 | 0.54 | 0.64 | 1 | MS.Trt_vs_MS.NoTrt_F |
| CD8B | 0.26585 | 1.38308 | 0.115 | 0.16 | 1 | MS.Trt_vs_MS.NoTrt_F |
| MTO1 | 0.26611 | 0.26282 | 0.978 | 0.98 | 1 | MS.Trt_vs_MS.NoTrt_F |
| EIF1AD | 0.26675 | 0.77088 | 0.734 | 0.73 | 1 | MS.Trt_vs_MS.NoTrt_F |
| ASAH1 | 0.2669 | 0.32412 | 0.734 | 0.78 | 1 | MS.Trt_vs_MS.NoTrt_F |
| TM2D1 | 0.26729 | 0.53095 | 0.885 | 0.92 | 1 | MS.Trt_vs_MS.NoTrt_F |
| ACBD5 | 0.26762 | -0.2862 | 0.5 | 0.54 | 1 | MS.Trt_vs_MS.NoTrt_F |
| MAB21L3 | 0.26785 | 0.47203 | 0.327 | 0.4 | 1 | MS.Trt_vs_MS.NoTrt_F |
| EFHD1 | 0.26786 | -0.5149 | 0.281 | 0.34 | 1 | MS.Trt_vs_MS.NoTrt_F |
| RAB11FIP3 | 0.26835 | 0.58599 | 0.205 | 0.26 | 1 | MS.Trt_vs_MS.NoTrt_F |
| LMAN1 | 0.26871 | 0.37091 | 0.209 | 0.27 | 1 | MS.Trt_vs_MS.NoTrt_F |
| SIRPB1 | 0.26886 | 0.73309 | 0.248 | 0.31 | 1 | MS.Trt_vs_MS.NoTrt_F |
| MYOM1 | 0.26896 | -0.4099 | 0.335 | 0.39 | 1 | MS.Trt_vs_MS.NoTrt_F |
| KCNN3 | 0.26913 | 0.51575 | 0.504 | 0.45 | 1 | MS.Trt_vs_MS.NoTrt_F |
| UBE3C | 0.26926 | 0.69496 | 0.576 | 0.66 | 1 | MS.Trt_vs_MS.NoTrt_F |
| TMSB15B | 0.2694 | -0.6758 | 0.299 | 0.34 | 1 | MS.Trt_vs_MS.NoTrt_F |
| LATS1 | 0.2697 | 0.27192 | 0.665 | 0.72 | 1 | MS.Trt_vs_MS.NoTrt_F |
| SDC4 | 0.2699 | 1.20399 | 0.162 | 0.21 | 1 | MS.Trt_vs_MS.NoTrt_F |
| DEF8 | 0.27012 | 0.44453 | 0.46 | 0.53 | 1 | MS.Trt_vs_MS.NoTrt_F |
| TOMM70 | 0.27018 | 1.5151 | 0.176 | 0.22 | 1 | MS.Trt_vs_MS.NoTrt_F |
| ZFP36L2 | 0.27099 | 0.65322 | 0.493 | 0.64 | 1 | MS.Trt_vs_MS.NoTrt_F |
| CBWD1 | 0.27108 | 0.55123 | 0.665 | 0.74 | 1 | MS.Trt_vs_MS.NoTrt_F |
| MCTP1 | 0.27128 | 0.82851 | 0.083 | 0.12 | 1 | MS.Trt_vs_MS.NoTrt_F |
| LPCAT3 | 0.27133 | 0.36082 | 0.442 | 0.51 | 1 | MS.Trt_vs_MS.NoTrt_F |
| NFIB | 0.27149 | 1.95964 | 0.137 | 0.18 | 1 | MS.Trt_vs_MS.NoTrt_F |
| TSTD2 | 0.27153 | 0.80189 | 0.126 | 0.17 | 1 | MS.Trt_vs_MS.NoTrt_F |
| ZNF584 | 0.27156 | -0.4661 | 0.687 | 0.75 | 1 | MS.Trt_vs_MS.NoTrt_F |
| ASCC1 | 0.27167 | -0.3686 | 0.612 | 0.73 | 1 | MS.Trt_vs_MS.NoTrt_F |
| ALDH5A1 | 0.27216 | -0.2901 | 0.482 | 0.57 | 1 | MS.Trt_vs_MS.NoTrt_F |
| BPHL | 0.27217 | 0.73408 | 0.633 | 0.75 | 1 | MS.Trt_vs_MS.NoTrt_F |
| BEAN1 | 0.2723 | 0.29392 | 0.489 | 0.55 | 1 | MS.Trt_vs_MS.NoTrt_F |
| TNFRSF11A | 0.27249 | -0.2688 | 0.281 | 0.24 | 1 | MS.Trt_vs_MS.NoTrt_F |

| | | | | | | |
|---|---|---|---|---|---|---|
| G6PC3 | 0.27265 | 0.46781 | 0.856 | 0.83 | 1 | MS.Trt_vs_MS.NoTrt_F |
| DPP6 | 0.27277 | 0.60019 | 0.514 | 0.53 | 1 | MS.Trt_vs_MS.NoTrt_F |
| PLCD4 | 0.27315 | -0.3938 | 0.295 | 0.35 | 1 | MS.Trt_vs_MS.NoTrt_F |
| PSMG3 | 0.27316 | 0.92 | 0.594 | 0.7 | 1 | MS.Trt_vs_MS.NoTrt_F |
| PBRM1 | 0.27378 | 0.63126 | 0.683 | 0.79 | 1 | MS.Trt_vs_MS.NoTrt_F |
| ZNF98 | 0.27412 | -0.275 | 0.115 | 0.08 | 1 | MS.Trt_vs_MS.NoTrt_F |
| HSD17B12 | 0.27416 | -0.3307 | 0.457 | 0.52 | 1 | MS.Trt_vs_MS.NoTrt_F |
| ATAD1 | 0.27419 | 0.56408 | 0.176 | 0.23 | 1 | MS.Trt_vs_MS.NoTrt_F |
| PSMB9 | 0.27425 | 0.49536 | 0.748 | 0.8 | 1 | MS.Trt_vs_MS.NoTrt_F |
| TALDO1 | 0.2744 | 0.49538 | 0.669 | 0.76 | 1 | MS.Trt_vs_MS.NoTrt_F |
| WDR92 | 0.27457 | -0.4433 | 0.705 | 0.73 | 1 | MS.Trt_vs_MS.NoTrt_F |
| GABPB2 | 0.2746 | 0.51054 | 0.982 | 0.99 | 1 | MS.Trt_vs_MS.NoTrt_F |
| AAED1 | 0.27498 | 0.71007 | 0.777 | 0.83 | 1 | MS.Trt_vs_MS.NoTrt_F |
| ARHGAP27 | 0.27592 | 0.257 | 0.543 | 0.61 | 1 | MS.Trt_vs_MS.NoTrt_F |
| DCAKD | 0.27641 | 0.6703 | 0.374 | 0.44 | 1 | MS.Trt_vs_MS.NoTrt_F |
| CCND3 | 0.27688 | 0.45496 | 0.54 | 0.64 | 1 | MS.Trt_vs_MS.NoTrt_F |
| CHP2 | 0.27713 | 0.33258 | 0.406 | 0.53 | 1 | MS.Trt_vs_MS.NoTrt_F |
| TTC28 | 0.27721 | 0.46608 | 0.378 | 0.44 | 1 | MS.Trt_vs_MS.NoTrt_F |
| LRCH1 | 0.27771 | 0.74148 | 0.112 | 0.15 | 1 | MS.Trt_vs_MS.NoTrt_F |
| SAR1A | 0.27796 | 0.35148 | 0.745 | 0.74 | 1 | MS.Trt_vs_MS.NoTrt_F |
| CD79B | 0.27808 | 1.18518 | 0.151 | 0.2 | 1 | MS.Trt_vs_MS.NoTrt_F |
| GLRX5 | 0.27813 | 0.59582 | 0.119 | 0.16 | 1 | MS.Trt_vs_MS.NoTrt_F |
| ZNF549 | 0.27831 | 0.94314 | 0.673 | 0.73 | 1 | MS.Trt_vs_MS.NoTrt_F |
| SMCR8 | 0.27877 | 0.6557 | 0.302 | 0.36 | 1 | MS.Trt_vs_MS.NoTrt_F |
| ICE1 | 0.27901 | 1.24581 | 0.14 | 0.19 | 1 | MS.Trt_vs_MS.NoTrt_F |
| NBPF10 | 0.27915 | -0.3938 | 0.252 | 0.3 | 1 | MS.Trt_vs_MS.NoTrt_F |
| SFXN2 | 0.27942 | -0.3981 | 0.917 | 0.93 | 1 | MS.Trt_vs_MS.NoTrt_F |
| MBD5 | 0.27953 | -0.4602 | 0.835 | 0.82 | 1 | MS.Trt_vs_MS.NoTrt_F |
| DRG1 | 0.27957 | 1.04314 | 0.191 | 0.25 | 1 | MS.Trt_vs_MS.NoTrt_F |
| GOLGA6L22 | 0.27982 | 0.3787 | 0.421 | 0.41 | 1 | MS.Trt_vs_MS.NoTrt_F |
| PRAMEF1 | 0.27989 | 0.26402 | 0.108 | 0.14 | 1 | MS.Trt_vs_MS.NoTrt_F |
| COQ8A | 0.27996 | 0.42833 | 0.644 | 0.7 | 1 | MS.Trt_vs_MS.NoTrt_F |
| CREB3L4 | 0.28033 | 0.61139 | 0.374 | 0.44 | 1 | MS.Trt_vs_MS.NoTrt_F |
| DNAJC1 | 0.28051 | 1.5661 | 0.137 | 0.18 | 1 | MS.Trt_vs_MS.NoTrt_F |
| GGT5 | 0.28086 | 0.44213 | 0.737 | 0.75 | 1 | MS.Trt_vs_MS.NoTrt_F |
| ATP5G1 | 0.28107 | 0.52707 | 0.317 | 0.38 | 1 | MS.Trt_vs_MS.NoTrt_F |
| ZNF213 | 0.28129 | 0.90208 | 0.5 | 0.48 | 1 | MS.Trt_vs_MS.NoTrt_F |
| IFIT3 | 0.28132 | 1.076 | 0.558 | 0.55 | 1 | MS.Trt_vs_MS.NoTrt_F |
| TRIM66 | 0.28165 | 0.6262 | 0.489 | 0.55 | 1 | MS.Trt_vs_MS.NoTrt_F |
| ZNF675 | 0.28207 | 0.59865 | 0.903 | 0.96 | 1 | MS.Trt_vs_MS.NoTrt_F |

| | | | | | | |
|---|---|---|---|---|---|---|
| ZNF578 | 0.28329 | 0.4238 | 0.946 | 0.97 | 1 | MS.Trt_vs_MS.NoTrt_F |
| NFS1 | 0.28353 | 0.53693 | 0.906 | 0.92 | 1 | MS.Trt_vs_MS.NoTrt_F |
| FBXW8 | 0.28482 | -0.5451 | 0.331 | 0.38 | 1 | MS.Trt_vs_MS.NoTrt_F |
| 7-Mar | 0.28495 | 1.08659 | 0.273 | 0.33 | 1 | MS.Trt_vs_MS.NoTrt_F |
| ZNF559-ZNF | 0.28504 | 0.28368 | 0.076 | 0.11 | 1 | MS.Trt_vs_MS.NoTrt_F |
| PCBP1 | 0.28515 | 0.71357 | 0.464 | 0.56 | 1 | MS.Trt_vs_MS.NoTrt_F |
| TTBK2 | 0.28515 | -1.1502 | 0.529 | 0.55 | 1 | MS.Trt_vs_MS.NoTrt_F |
| AKAP6 | 0.28528 | 0.71285 | 0.662 | 0.76 | 1 | MS.Trt_vs_MS.NoTrt_F |
| RP11-80H18 | 0.28536 | 0.44233 | 0.306 | 0.37 | 1 | MS.Trt_vs_MS.NoTrt_F |
| SYNC | 0.28543 | 0.96605 | 0.191 | 0.25 | 1 | MS.Trt_vs_MS.NoTrt_F |
| SUSD3 | 0.28637 | 0.27994 | 0.396 | 0.48 | 1 | MS.Trt_vs_MS.NoTrt_F |
| FLG2 | 0.28644 | -0.6448 | 0.457 | 0.5 | 1 | MS.Trt_vs_MS.NoTrt_F |
| UBAP2 | 0.28659 | -0.5058 | 0.683 | 0.75 | 1 | MS.Trt_vs_MS.NoTrt_F |
| STXBP1 | 0.28703 | -0.4125 | 0.076 | 0.11 | 1 | MS.Trt_vs_MS.NoTrt_F |
| PMPCB | 0.28766 | 0.51494 | 0.554 | 0.65 | 1 | MS.Trt_vs_MS.NoTrt_F |
| PAPD4 | 0.28771 | 0.36572 | 0.299 | 0.36 | 1 | MS.Trt_vs_MS.NoTrt_F |
| KMT5A | 0.28817 | 0.58659 | 0.245 | 0.32 | 1 | MS.Trt_vs_MS.NoTrt_F |
| STRADA | 0.28823 | 0.37581 | 0.54 | 0.62 | 1 | MS.Trt_vs_MS.NoTrt_F |
| SPIN1 | 0.28836 | 0.72219 | 0.194 | 0.24 | 1 | MS.Trt_vs_MS.NoTrt_F |
| GPR89A | 0.28886 | 0.89108 | 0.432 | 0.52 | 1 | MS.Trt_vs_MS.NoTrt_F |
| ABR | 0.28909 | -0.3146 | 0.748 | 0.79 | 1 | MS.Trt_vs_MS.NoTrt_F |
| RSAD2 | 0.28991 | -0.6414 | 0.327 | 0.39 | 1 | MS.Trt_vs_MS.NoTrt_F |
| AASS | 0.28999 | 1.21058 | 0.353 | 0.44 | 1 | MS.Trt_vs_MS.NoTrt_F |
| EPPK1 | 0.29012 | 1.15618 | 0.101 | 0.14 | 1 | MS.Trt_vs_MS.NoTrt_F |
| PMS2 | 0.29017 | 0.34256 | 0.622 | 0.73 | 1 | MS.Trt_vs_MS.NoTrt_F |
| ZNF516 | 0.2902 | 1.99253 | 0.09 | 0.12 | 1 | MS.Trt_vs_MS.NoTrt_F |
| KLLN | 0.29057 | 0.8572 | 0.432 | 0.52 | 1 | MS.Trt_vs_MS.NoTrt_F |
| NUP50 | 0.2906 | 0.25986 | 0.673 | 0.75 | 1 | MS.Trt_vs_MS.NoTrt_F |
| HERC2 | 0.29074 | 0.25826 | 0.856 | 0.91 | 1 | MS.Trt_vs_MS.NoTrt_F |
| TMIGD2 | 0.29104 | 0.54433 | 0.252 | 0.3 | 1 | MS.Trt_vs_MS.NoTrt_F |
| ARHGAP18 | 0.29107 | 0.40008 | 0.151 | 0.19 | 1 | MS.Trt_vs_MS.NoTrt_F |
| SCYL3 | 0.29131 | 0.50394 | 0.712 | 0.72 | 1 | MS.Trt_vs_MS.NoTrt_F |
| ZBTB11 | 0.29155 | 0.5598 | 0.529 | 0.61 | 1 | MS.Trt_vs_MS.NoTrt_F |
| KIF3A | 0.29158 | 0.51058 | 0.694 | 0.72 | 1 | MS.Trt_vs_MS.NoTrt_F |
| TMEM25 | 0.29171 | 1.25316 | 0.396 | 0.48 | 1 | MS.Trt_vs_MS.NoTrt_F |
| USP22 | 0.29211 | 0.65319 | 0.817 | 0.84 | 1 | MS.Trt_vs_MS.NoTrt_F |
| ADAR | 0.2923 | -0.5877 | 0.809 | 0.8 | 1 | MS.Trt_vs_MS.NoTrt_F |
| GM2A | 0.29249 | -0.3391 | 0.572 | 0.58 | 1 | MS.Trt_vs_MS.NoTrt_F |
| ZFAND2A | 0.29267 | 0.45264 | 0.122 | 0.16 | 1 | MS.Trt_vs_MS.NoTrt_F |
| CEP250 | 0.29356 | 0.52415 | 0.784 | 0.87 | 1 | MS.Trt_vs_MS.NoTrt_F |

| | | | | | | |
|---|---|---|---|---|---|---|
| BECN1 | 0.29387 | 0.43701 | 0.727 | 0.79 | 1 | MS.Trt_vs_MS.NoTrt_F |
| NOVA1 | 0.29414 | -0.348 | 0.09 | 0.12 | 1 | MS.Trt_vs_MS.NoTrt_F |
| GPX3 | 0.29441 | -0.2747 | 0.104 | 0.14 | 1 | MS.Trt_vs_MS.NoTrt_F |
| KIAA1671 | 0.2949 | 0.43419 | 0.507 | 0.62 | 1 | MS.Trt_vs_MS.NoTrt_F |
| CNIH3 | 0.29528 | -0.3734 | 0.191 | 0.23 | 1 | MS.Trt_vs_MS.NoTrt_F |
| PMM2 | 0.29532 | -0.5084 | 0.745 | 0.79 | 1 | MS.Trt_vs_MS.NoTrt_F |
| NAGA | 0.29577 | 1.2694 | 0.144 | 0.19 | 1 | MS.Trt_vs_MS.NoTrt_F |
| MAPKAPK2 | 0.29652 | 0.31942 | 0.094 | 0.13 | 1 | MS.Trt_vs_MS.NoTrt_F |
| MAGT1 | 0.29672 | 0.43886 | 0.82 | 0.84 | 1 | MS.Trt_vs_MS.NoTrt_F |
| CDC26 | 0.29743 | 0.82266 | 0.248 | 0.33 | 1 | MS.Trt_vs_MS.NoTrt_F |
| EFL1 | 0.29803 | 1.09203 | 0.151 | 0.19 | 1 | MS.Trt_vs_MS.NoTrt_F |
| CEP70 | 0.29813 | 0.73504 | 0.162 | 0.2 | 1 | MS.Trt_vs_MS.NoTrt_F |
| HEXA | 0.2988 | 0.42447 | 0.701 | 0.77 | 1 | MS.Trt_vs_MS.NoTrt_F |
| PPP1CC | 0.29892 | -0.2976 | 0.763 | 0.79 | 1 | MS.Trt_vs_MS.NoTrt_F |
| CECR5 | 0.29946 | 0.46621 | 0.187 | 0.23 | 1 | MS.Trt_vs_MS.NoTrt_F |
| SLC15A4 | 0.29982 | 0.36195 | 0.755 | 0.8 | 1 | MS.Trt_vs_MS.NoTrt_F |
| SDR42E1 | 0.29983 | -0.3153 | 0.378 | 0.45 | 1 | MS.Trt_vs_MS.NoTrt_F |
| LPP | 0.30023 | -0.4103 | 0.957 | 0.96 | 1 | MS.Trt_vs_MS.NoTrt_F |
| CARM1 | 0.30047 | -0.2678 | 0.169 | 0.22 | 1 | MS.Trt_vs_MS.NoTrt_F |
| PDGFRA | 0.30049 | -0.2802 | 0.219 | 0.28 | 1 | MS.Trt_vs_MS.NoTrt_F |
| ST20 | 0.30055 | 0.2707 | 0.248 | 0.22 | 1 | MS.Trt_vs_MS.NoTrt_F |
| ZHX1 | 0.30105 | 0.28587 | 0.565 | 0.61 | 1 | MS.Trt_vs_MS.NoTrt_F |
| NUDCD2 | 0.30186 | 0.28774 | 0.77 | 0.76 | 1 | MS.Trt_vs_MS.NoTrt_F |
| ZFYVE26 | 0.30255 | 0.34115 | 0.871 | 0.86 | 1 | MS.Trt_vs_MS.NoTrt_F |
| VPS26A | 0.30261 | 0.59359 | 0.378 | 0.42 | 1 | MS.Trt_vs_MS.NoTrt_F |
| ZNF764 | 0.30264 | 0.76852 | 0.331 | 0.4 | 1 | MS.Trt_vs_MS.NoTrt_F |
| CREG1 | 0.30266 | 0.76612 | 0.313 | 0.38 | 1 | MS.Trt_vs_MS.NoTrt_F |
| TPMT | 0.30266 | 0.25879 | 0.737 | 0.8 | 1 | MS.Trt_vs_MS.NoTrt_F |
| SPOP | 0.30295 | -0.2981 | 0.946 | 0.98 | 1 | MS.Trt_vs_MS.NoTrt_F |
| MOSPD1 | 0.30347 | -0.8931 | 0.219 | 0.27 | 1 | MS.Trt_vs_MS.NoTrt_F |
| STARD9 | 0.30359 | 0.35936 | 0.896 | 0.91 | 1 | MS.Trt_vs_MS.NoTrt_F |
| FUBP3 | 0.30458 | 0.36294 | 0.115 | 0.16 | 1 | MS.Trt_vs_MS.NoTrt_F |
| GATSL3 | 0.30504 | 0.44927 | 0.076 | 0.11 | 1 | MS.Trt_vs_MS.NoTrt_F |
| B3GALT1 | 0.30542 | 0.27372 | 0.547 | 0.5 | 1 | MS.Trt_vs_MS.NoTrt_F |
| CTSH | 0.30543 | 0.74 | 0.108 | 0.14 | 1 | MS.Trt_vs_MS.NoTrt_F |
| AEBP2 | 0.30577 | 1.28741 | 0.691 | 0.77 | 1 | MS.Trt_vs_MS.NoTrt_F |
| ZNF524 | 0.3058 | 0.57078 | 0.147 | 0.19 | 1 | MS.Trt_vs_MS.NoTrt_F |
| PLD3 | 0.30589 | 0.45669 | 0.723 | 0.82 | 1 | MS.Trt_vs_MS.NoTrt_F |
| HINFP | 0.3059 | -0.3539 | 0.996 | 0.99 | 1 | MS.Trt_vs_MS.NoTrt_F |
| TBC1D3I | 0.30643 | 0.29723 | 0.245 | 0.29 | 1 | MS.Trt_vs_MS.NoTrt_F |

| | | | | | | |
|---|---|---|---|---|---|---|
| GTF2E2 | 0.30647 | 0.53797 | 0.604 | 0.7 | 1 | MS.Trt_vs_MS.NoTrt_F |
| ELL2 | 0.3065 | -0.5953 | 0.086 | 0.12 | 1 | MS.Trt_vs_MS.NoTrt_F |
| FAM60A | 0.30687 | 0.27254 | 0.504 | 0.56 | 1 | MS.Trt_vs_MS.NoTrt_F |
| ABHD6 | 0.30701 | 0.80577 | 0.493 | 0.56 | 1 | MS.Trt_vs_MS.NoTrt_F |
| CDH13 | 0.30704 | 0.32954 | 0.755 | 0.74 | 1 | MS.Trt_vs_MS.NoTrt_F |
| RFX5 | 0.30729 | 0.78604 | 0.327 | 0.38 | 1 | MS.Trt_vs_MS.NoTrt_F |
| SNAPC3 | 0.30758 | 0.50961 | 0.586 | 0.64 | 1 | MS.Trt_vs_MS.NoTrt_F |
| LPAR4 | 0.30776 | -0.8256 | 0.173 | 0.2 | 1 | MS.Trt_vs_MS.NoTrt_F |
| HIST1H3E | 0.30851 | -0.4365 | 0.158 | 0.2 | 1 | MS.Trt_vs_MS.NoTrt_F |
| GP1BA | 0.3086 | 0.68347 | 0.076 | 0.11 | 1 | MS.Trt_vs_MS.NoTrt_F |
| POLR1C | 0.30897 | 0.84344 | 0.432 | 0.49 | 1 | MS.Trt_vs_MS.NoTrt_F |
| HBB | 0.30976 | 0.37265 | 0.104 | 0.14 | 1 | MS.Trt_vs_MS.NoTrt_F |
| RABL6 | 0.31015 | 0.57961 | 0.572 | 0.6 | 1 | MS.Trt_vs_MS.NoTrt_F |
| ADRM1 | 0.31028 | 0.51707 | 0.273 | 0.34 | 1 | MS.Trt_vs_MS.NoTrt_F |
| CTSC | 0.31029 | 0.30012 | 0.853 | 0.91 | 1 | MS.Trt_vs_MS.NoTrt_F |
| ARIH1 | 0.31031 | 0.40301 | 0.96 | 0.97 | 1 | MS.Trt_vs_MS.NoTrt_F |
| XRCC3 | 0.31048 | -0.661 | 0.45 | 0.5 | 1 | MS.Trt_vs_MS.NoTrt_F |
| INPP5F | 0.31109 | 0.56767 | 0.317 | 0.38 | 1 | MS.Trt_vs_MS.NoTrt_F |
| CORIN | 0.3113 | -0.3211 | 0.597 | 0.63 | 1 | MS.Trt_vs_MS.NoTrt_F |
| PRKAB1 | 0.31225 | -0.2744 | 0.788 | 0.79 | 1 | MS.Trt_vs_MS.NoTrt_F |
| TRIM34 | 0.31237 | 1.6774 | 0.313 | 0.37 | 1 | MS.Trt_vs_MS.NoTrt_F |
| SLC16A4 | 0.31243 | 0.57965 | 0.691 | 0.69 | 1 | MS.Trt_vs_MS.NoTrt_F |
| EXTL2 | 0.31248 | 0.78061 | 0.094 | 0.13 | 1 | MS.Trt_vs_MS.NoTrt_F |
| ACOX1 | 0.313 | -0.3091 | 0.73 | 0.74 | 1 | MS.Trt_vs_MS.NoTrt_F |
| PUS10 | 0.31301 | -0.278 | 0.719 | 0.7 | 1 | MS.Trt_vs_MS.NoTrt_F |
| SMPD4 | 0.31395 | 0.49605 | 0.302 | 0.34 | 1 | MS.Trt_vs_MS.NoTrt_F |
| ZNF433 | 0.31433 | 1.09033 | 0.162 | 0.2 | 1 | MS.Trt_vs_MS.NoTrt_F |
| MAP7D3 | 0.31436 | 0.79883 | 0.597 | 0.65 | 1 | MS.Trt_vs_MS.NoTrt_F |
| PLOD3 | 0.31524 | -0.4586 | 0.669 | 0.78 | 1 | MS.Trt_vs_MS.NoTrt_F |
| RRM2 | 0.31563 | 0.27134 | 0.478 | 0.56 | 1 | MS.Trt_vs_MS.NoTrt_F |
| HLTF | 0.31566 | 1.17425 | 0.439 | 0.51 | 1 | MS.Trt_vs_MS.NoTrt_F |
| LCA5L | 0.31592 | 0.36739 | 0.41 | 0.5 | 1 | MS.Trt_vs_MS.NoTrt_F |
| PMF1-BGLA | 0.31595 | 1.49057 | 0.122 | 0.17 | 1 | MS.Trt_vs_MS.NoTrt_F |
| CRIM1 | 0.31667 | 1.00834 | 0.209 | 0.27 | 1 | MS.Trt_vs_MS.NoTrt_F |
| XAF1 | 0.31677 | 0.52192 | 0.723 | 0.84 | 1 | MS.Trt_vs_MS.NoTrt_F |
| FAAP20 | 0.31698 | 0.50784 | 0.899 | 0.87 | 1 | MS.Trt_vs_MS.NoTrt_F |
| C17orf89 | 0.31699 | 1.51432 | 0.119 | 0.16 | 1 | MS.Trt_vs_MS.NoTrt_F |
| PRPF40B | 0.31713 | 0.78042 | 0.288 | 0.36 | 1 | MS.Trt_vs_MS.NoTrt_F |
| MLST8 | 0.31737 | -0.6846 | 0.421 | 0.48 | 1 | MS.Trt_vs_MS.NoTrt_F |
| C1orf50 | 0.31742 | 0.35925 | 0.705 | 0.75 | 1 | MS.Trt_vs_MS.NoTrt_F |

| | | | | | | |
|---|---|---|---|---|---|---|
| ZNF687 | 0.3175 | -0.343 | 0.644 | 0.7 | 1 | MS.Trt_vs_MS.NoTrt_F |
| FSD1 | 0.31781 | 0.47704 | 0.432 | 0.42 | 1 | MS.Trt_vs_MS.NoTrt_F |
| HSF2 | 0.31785 | 1.30321 | 0.115 | 0.16 | 1 | MS.Trt_vs_MS.NoTrt_F |
| SP110 | 0.31814 | 0.50316 | 0.676 | 0.74 | 1 | MS.Trt_vs_MS.NoTrt_F |
| RPS27L | 0.3182 | 0.26928 | 0.982 | 0.99 | 1 | MS.Trt_vs_MS.NoTrt_F |
| RTCA | 0.31832 | 0.68502 | 0.576 | 0.64 | 1 | MS.Trt_vs_MS.NoTrt_F |
| MASP2 | 0.31897 | 0.3068 | 0.295 | 0.34 | 1 | MS.Trt_vs_MS.NoTrt_F |
| MAK | 0.31963 | 0.37936 | 0.187 | 0.23 | 1 | MS.Trt_vs_MS.NoTrt_F |
| PHLDA3 | 0.3206 | 0.27601 | 0.345 | 0.4 | 1 | MS.Trt_vs_MS.NoTrt_F |
| FAM98B | 0.32072 | 0.37106 | 0.432 | 0.5 | 1 | MS.Trt_vs_MS.NoTrt_F |
| FAM129C | 0.32103 | 0.2928 | 0.831 | 0.85 | 1 | MS.Trt_vs_MS.NoTrt_F |
| COIL | 0.32122 | 0.67157 | 0.151 | 0.19 | 1 | MS.Trt_vs_MS.NoTrt_F |
| IMPDH2 | 0.32126 | 0.63607 | 0.223 | 0.28 | 1 | MS.Trt_vs_MS.NoTrt_F |
| FPR2 | 0.32163 | -0.5349 | 0.482 | 0.43 | 1 | MS.Trt_vs_MS.NoTrt_F |
| LRRFIP1 | 0.32176 | 0.33945 | 0.989 | 1 | 1 | MS.Trt_vs_MS.NoTrt_F |
| SPDYE2B | 0.3222 | 0.77443 | 0.299 | 0.4 | 1 | MS.Trt_vs_MS.NoTrt_F |
| ZNF17 | 0.32252 | 1.28231 | 0.324 | 0.39 | 1 | MS.Trt_vs_MS.NoTrt_F |
| SKA2 | 0.32253 | 0.53541 | 0.594 | 0.63 | 1 | MS.Trt_vs_MS.NoTrt_F |
| LZIC | 0.32497 | -0.3911 | 0.817 | 0.85 | 1 | MS.Trt_vs_MS.NoTrt_F |
| SMG9 | 0.32545 | 0.69763 | 0.594 | 0.66 | 1 | MS.Trt_vs_MS.NoTrt_F |
| UBE2R2 | 0.3262 | 0.70586 | 0.108 | 0.14 | 1 | MS.Trt_vs_MS.NoTrt_F |
| ARSB | 0.32658 | 0.83825 | 0.615 | 0.59 | 1 | MS.Trt_vs_MS.NoTrt_F |
| GDPGP1 | 0.32665 | 0.44438 | 0.924 | 0.96 | 1 | MS.Trt_vs_MS.NoTrt_F |
| ABCB8 | 0.32672 | 0.60551 | 0.633 | 0.56 | 1 | MS.Trt_vs_MS.NoTrt_F |
| SPHK2 | 0.3275 | -0.3043 | 0.324 | 0.37 | 1 | MS.Trt_vs_MS.NoTrt_F |
| PRKCSH | 0.32754 | 0.5547 | 0.716 | 0.72 | 1 | MS.Trt_vs_MS.NoTrt_F |
| CASP4 | 0.32756 | 0.31625 | 0.396 | 0.46 | 1 | MS.Trt_vs_MS.NoTrt_F |
| ZHX3 | 0.32768 | 0.53772 | 0.36 | 0.41 | 1 | MS.Trt_vs_MS.NoTrt_F |
| CLN8 | 0.32775 | 0.55334 | 0.799 | 0.8 | 1 | MS.Trt_vs_MS.NoTrt_F |
| ELMOD3 | 0.32791 | 0.39954 | 0.86 | 0.93 | 1 | MS.Trt_vs_MS.NoTrt_F |
| SLC11A1 | 0.32792 | 0.26984 | 0.95 | 0.93 | 1 | MS.Trt_vs_MS.NoTrt_F |
| ANGPT2 | 0.32916 | 0.44485 | 0.795 | 0.76 | 1 | MS.Trt_vs_MS.NoTrt_F |
| RIT1 | 0.32936 | 1.08554 | 0.424 | 0.5 | 1 | MS.Trt_vs_MS.NoTrt_F |
| SLC25A42 | 0.3294 | 0.93226 | 0.536 | 0.61 | 1 | MS.Trt_vs_MS.NoTrt_F |
| CTD-2370N5 | 0.32952 | 0.39898 | 0.165 | 0.21 | 1 | MS.Trt_vs_MS.NoTrt_F |
| AK3 | 0.32955 | -0.5067 | 0.5 | 0.51 | 1 | MS.Trt_vs_MS.NoTrt_F |
| FUT4 | 0.32968 | 0.4009 | 0.353 | 0.4 | 1 | MS.Trt_vs_MS.NoTrt_F |
| TOE1 | 0.3305 | 0.5289 | 0.101 | 0.13 | 1 | MS.Trt_vs_MS.NoTrt_F |
| RDH11 | 0.33052 | 0.2557 | 0.504 | 0.55 | 1 | MS.Trt_vs_MS.NoTrt_F |
| PRC1 | 0.33069 | 0.34468 | 0.554 | 0.65 | 1 | MS.Trt_vs_MS.NoTrt_F |

| | | | | | | |
|---|---|---|---|---|---|---|
| KNTC1 | 0.33121 | 1.23489 | 0.813 | 0.8 | 1 | MS.Trt_vs_MS.NoTrt_F |
| ARID5B | 0.33205 | 0.64681 | 0.773 | 0.84 | 1 | MS.Trt_vs_MS.NoTrt_F |
| SUDS3 | 0.33256 | -0.8788 | 0.813 | 0.81 | 1 | MS.Trt_vs_MS.NoTrt_F |
| RPS17 | 0.33275 | 0.72508 | 0.417 | 0.46 | 1 | MS.Trt_vs_MS.NoTrt_F |
| XPOT | 0.33293 | 0.38923 | 0.457 | 0.5 | 1 | MS.Trt_vs_MS.NoTrt_F |
| NARF | 0.33304 | -0.2717 | 0.946 | 0.97 | 1 | MS.Trt_vs_MS.NoTrt_F |
| ZNF487 | 0.33332 | -0.6401 | 0.482 | 0.53 | 1 | MS.Trt_vs_MS.NoTrt_F |
| ZNF493 | 0.33334 | 0.7278 | 0.827 | 0.8 | 1 | MS.Trt_vs_MS.NoTrt_F |
| CXCL16 | 0.33348 | 0.33302 | 0.622 | 0.61 | 1 | MS.Trt_vs_MS.NoTrt_F |
| LIMD1 | 0.33369 | -0.5462 | 0.827 | 0.87 | 1 | MS.Trt_vs_MS.NoTrt_F |
| FBXO46 | 0.33473 | 0.51837 | 0.295 | 0.34 | 1 | MS.Trt_vs_MS.NoTrt_F |
| RINT1 | 0.33506 | 0.37521 | 0.55 | 0.62 | 1 | MS.Trt_vs_MS.NoTrt_F |
| PHACTR2 | 0.33526 | 0.68095 | 0.637 | 0.69 | 1 | MS.Trt_vs_MS.NoTrt_F |
| NLRP3 | 0.33536 | 1.78738 | 0.23 | 0.29 | 1 | MS.Trt_vs_MS.NoTrt_F |
| TOR1AIP2 | 0.33601 | 0.5543 | 0.982 | 0.99 | 1 | MS.Trt_vs_MS.NoTrt_F |
| COLGALT1 | 0.3361 | 0.59893 | 0.468 | 0.54 | 1 | MS.Trt_vs_MS.NoTrt_F |
| RPGRIP1L | 0.33614 | 0.50592 | 0.755 | 0.74 | 1 | MS.Trt_vs_MS.NoTrt_F |
| NIN | 0.33659 | 0.34331 | 0.219 | 0.27 | 1 | MS.Trt_vs_MS.NoTrt_F |
| CTSL | 0.337 | 1.62772 | 0.137 | 0.11 | 1 | MS.Trt_vs_MS.NoTrt_F |
| CYP46A1 | 0.33736 | 0.7456 | 0.55 | 0.6 | 1 | MS.Trt_vs_MS.NoTrt_F |
| GCDH | 0.33788 | 0.62332 | 0.194 | 0.24 | 1 | MS.Trt_vs_MS.NoTrt_F |
| PTRH1 | 0.33837 | 0.75552 | 0.712 | 0.74 | 1 | MS.Trt_vs_MS.NoTrt_F |
| RBM19 | 0.33847 | 0.53566 | 0.45 | 0.53 | 1 | MS.Trt_vs_MS.NoTrt_F |
| NEPRO | 0.33859 | -0.4946 | 0.219 | 0.25 | 1 | MS.Trt_vs_MS.NoTrt_F |
| NBR1 | 0.33918 | -0.3228 | 0.781 | 0.81 | 1 | MS.Trt_vs_MS.NoTrt_F |
| KCNC4 | 0.33953 | 0.79767 | 0.701 | 0.82 | 1 | MS.Trt_vs_MS.NoTrt_F |
| HN1L | 0.33965 | 0.59113 | 0.327 | 0.39 | 1 | MS.Trt_vs_MS.NoTrt_F |
| SELL | 0.34012 | 0.32072 | 0.55 | 0.69 | 1 | MS.Trt_vs_MS.NoTrt_F |
| COG6 | 0.34036 | 0.89272 | 0.176 | 0.22 | 1 | MS.Trt_vs_MS.NoTrt_F |
| CNIH4 | 0.34039 | 0.2933 | 0.615 | 0.65 | 1 | MS.Trt_vs_MS.NoTrt_F |
| PCIF1 | 0.34121 | 0.54381 | 0.32 | 0.4 | 1 | MS.Trt_vs_MS.NoTrt_F |
| KCTD10 | 0.34124 | 1.20299 | 0.414 | 0.37 | 1 | MS.Trt_vs_MS.NoTrt_F |
| TCOF1 | 0.34132 | 0.3927 | 0.773 | 0.88 | 1 | MS.Trt_vs_MS.NoTrt_F |
| SLC25A51 | 0.34142 | -0.6638 | 0.59 | 0.64 | 1 | MS.Trt_vs_MS.NoTrt_F |
| APMAP | 0.34166 | 0.76916 | 0.209 | 0.26 | 1 | MS.Trt_vs_MS.NoTrt_F |
| ZNF93 | 0.34214 | 0.41741 | 0.525 | 0.6 | 1 | MS.Trt_vs_MS.NoTrt_F |
| ATM | 0.3422 | 0.56317 | 0.957 | 0.95 | 1 | MS.Trt_vs_MS.NoTrt_F |
| APOE | 0.34303 | 0.30049 | 0.719 | 0.81 | 1 | MS.Trt_vs_MS.NoTrt_F |
| SCRG1 | 0.34305 | 0.30358 | 0.763 | 0.75 | 1 | MS.Trt_vs_MS.NoTrt_F |
| H1FX | 0.34373 | 1.15868 | 0.09 | 0.12 | 1 | MS.Trt_vs_MS.NoTrt_F |

| | | | | | | |
|---|---|---|---|---|---|---|
| CERKL | 0.34426 | 0.5034 | 0.36 | 0.45 | 1 | MS.Trt_vs_MS.NoTrt_F |
| STT3B | 0.34434 | 0.72235 | 0.604 | 0.58 | 1 | MS.Trt_vs_MS.NoTrt_F |
| PDCD7 | 0.34459 | 0.34913 | 0.442 | 0.48 | 1 | MS.Trt_vs_MS.NoTrt_F |
| FANCC | 0.3446 | -0.3305 | 0.313 | 0.34 | 1 | MS.Trt_vs_MS.NoTrt_F |
| DPY19L3 | 0.34481 | -0.4065 | 0.903 | 0.93 | 1 | MS.Trt_vs_MS.NoTrt_F |
| RP11-577H5 | 0.34489 | 1.04477 | 0.133 | 0.17 | 1 | MS.Trt_vs_MS.NoTrt_F |
| MYL6 | 0.345 | 0.4821 | 0.712 | 0.88 | 1 | MS.Trt_vs_MS.NoTrt_F |
| THBS4 | 0.3452 | 1.21903 | 0.086 | 0.12 | 1 | MS.Trt_vs_MS.NoTrt_F |
| POMT2 | 0.34554 | 0.26946 | 0.942 | 0.95 | 1 | MS.Trt_vs_MS.NoTrt_F |
| PFAS | 0.34607 | 0.34376 | 0.601 | 0.58 | 1 | MS.Trt_vs_MS.NoTrt_F |
| LAMP3 | 0.34726 | 0.68647 | 0.446 | 0.55 | 1 | MS.Trt_vs_MS.NoTrt_F |
| IST1 | 0.34738 | 0.28699 | 0.644 | 0.72 | 1 | MS.Trt_vs_MS.NoTrt_F |
| FUT3 | 0.34781 | -0.3137 | 0.219 | 0.27 | 1 | MS.Trt_vs_MS.NoTrt_F |
| PSTPIP1 | 0.34836 | 0.37649 | 0.46 | 0.51 | 1 | MS.Trt_vs_MS.NoTrt_F |
| CSF1R | 0.34838 | -0.8322 | 0.223 | 0.27 | 1 | MS.Trt_vs_MS.NoTrt_F |
| RP11-552F3 | 0.34869 | -0.4056 | 0.201 | 0.24 | 1 | MS.Trt_vs_MS.NoTrt_F |
| CCNG1 | 0.34915 | 0.81088 | 0.27 | 0.34 | 1 | MS.Trt_vs_MS.NoTrt_F |
| C1orf159 | 0.34922 | 1.25269 | 0.183 | 0.23 | 1 | MS.Trt_vs_MS.NoTrt_F |
| HDDC3 | 0.34948 | -0.9843 | 0.385 | 0.43 | 1 | MS.Trt_vs_MS.NoTrt_F |
| FAM110D | 0.34979 | 0.39124 | 0.464 | 0.47 | 1 | MS.Trt_vs_MS.NoTrt_F |
| IL10RB | 0.34985 | 1.15819 | 0.457 | 0.51 | 1 | MS.Trt_vs_MS.NoTrt_F |
| ABHD2 | 0.35025 | 0.61853 | 0.651 | 0.69 | 1 | MS.Trt_vs_MS.NoTrt_F |
| GFOD1 | 0.35115 | 1.16694 | 0.626 | 0.66 | 1 | MS.Trt_vs_MS.NoTrt_F |
| ZNF805 | 0.3513 | 1.16238 | 0.766 | 0.79 | 1 | MS.Trt_vs_MS.NoTrt_F |
| RPL36AL | 0.35141 | 0.57565 | 0.525 | 0.72 | 1 | MS.Trt_vs_MS.NoTrt_F |
| IL4R | 0.35144 | -0.7472 | 0.478 | 0.49 | 1 | MS.Trt_vs_MS.NoTrt_F |
| IREB2 | 0.35167 | -0.316 | 0.748 | 0.83 | 1 | MS.Trt_vs_MS.NoTrt_F |
| XYLT2 | 0.35224 | 0.72709 | 0.647 | 0.63 | 1 | MS.Trt_vs_MS.NoTrt_F |
| BHMT | 0.35235 | 0.33462 | 0.522 | 0.57 | 1 | MS.Trt_vs_MS.NoTrt_F |
| MICAL2 | 0.35311 | 0.72096 | 0.795 | 0.86 | 1 | MS.Trt_vs_MS.NoTrt_F |
| ZNF257 | 0.35316 | 0.29899 | 0.572 | 0.51 | 1 | MS.Trt_vs_MS.NoTrt_F |
| OR7A5 | 0.35317 | 0.25885 | 0.428 | 0.5 | 1 | MS.Trt_vs_MS.NoTrt_F |
| ABHD11 | 0.35325 | -0.4529 | 0.59 | 0.69 | 1 | MS.Trt_vs_MS.NoTrt_F |
| IGFBP2 | 0.3533 | 0.25642 | 0.637 | 0.64 | 1 | MS.Trt_vs_MS.NoTrt_F |
| ANKMY1 | 0.35377 | 0.26679 | 0.374 | 0.44 | 1 | MS.Trt_vs_MS.NoTrt_F |
| TEX101 | 0.3538 | 1.17642 | 0.086 | 0.12 | 1 | MS.Trt_vs_MS.NoTrt_F |
| GVQW1 | 0.35481 | 0.31541 | 0.532 | 0.5 | 1 | MS.Trt_vs_MS.NoTrt_F |
| ZNF264 | 0.3549 | 0.87147 | 0.953 | 0.95 | 1 | MS.Trt_vs_MS.NoTrt_F |
| DNAJC22 | 0.35495 | -0.4331 | 0.349 | 0.4 | 1 | MS.Trt_vs_MS.NoTrt_F |
| MAEL | 0.35498 | 0.26857 | 0.572 | 0.54 | 1 | MS.Trt_vs_MS.NoTrt_F |

| | | | | | | |
|---|---|---|---|---|---|---|
| GTF2H4 | 0.35504 | 0.31851 | 0.137 | 0.18 | 1 | MS.Trt_vs_MS.NoTrt_F |
| TMED10 | 0.35516 | 0.33507 | 0.669 | 0.71 | 1 | MS.Trt_vs_MS.NoTrt_F |
| WDR33 | 0.35542 | 0.56006 | 0.277 | 0.33 | 1 | MS.Trt_vs_MS.NoTrt_F |
| PDP2 | 0.35592 | -0.6498 | 0.795 | 0.8 | 1 | MS.Trt_vs_MS.NoTrt_F |
| VAV3 | 0.35625 | -0.5831 | 0.604 | 0.6 | 1 | MS.Trt_vs_MS.NoTrt_F |
| PURB | 0.35665 | -0.4354 | 0.478 | 0.45 | 1 | MS.Trt_vs_MS.NoTrt_F |
| CDNF | 0.35665 | 0.76032 | 0.227 | 0.28 | 1 | MS.Trt_vs_MS.NoTrt_F |
| ZNF414 | 0.3574 | 0.25888 | 0.687 | 0.67 | 1 | MS.Trt_vs_MS.NoTrt_F |
| DLGAP1 | 0.35818 | 0.69895 | 0.381 | 0.38 | 1 | MS.Trt_vs_MS.NoTrt_F |
| FCER1G | 0.35866 | 0.86382 | 0.705 | 0.72 | 1 | MS.Trt_vs_MS.NoTrt_F |
| TRIM68 | 0.3587 | -0.5265 | 0.561 | 0.62 | 1 | MS.Trt_vs_MS.NoTrt_F |
| ZRANB3 | 0.35879 | 0.33764 | 0.698 | 0.76 | 1 | MS.Trt_vs_MS.NoTrt_F |
| NDUFA10 | 0.35965 | 0.47541 | 0.335 | 0.39 | 1 | MS.Trt_vs_MS.NoTrt_F |
| TMEM64 | 0.36022 | 0.3513 | 0.385 | 0.46 | 1 | MS.Trt_vs_MS.NoTrt_F |
| FMNL3 | 0.36042 | 0.54878 | 0.327 | 0.39 | 1 | MS.Trt_vs_MS.NoTrt_F |
| UBE2J2 | 0.36188 | 0.64982 | 0.338 | 0.39 | 1 | MS.Trt_vs_MS.NoTrt_F |
| ACSF2 | 0.36209 | 0.69908 | 0.446 | 0.54 | 1 | MS.Trt_vs_MS.NoTrt_F |
| RUBCN | 0.36254 | 0.27904 | 0.867 | 0.9 | 1 | MS.Trt_vs_MS.NoTrt_F |
| SIMC1 | 0.36274 | 1.03258 | 0.439 | 0.53 | 1 | MS.Trt_vs_MS.NoTrt_F |
| KLHL11 | 0.36284 | 0.7969 | 0.504 | 0.59 | 1 | MS.Trt_vs_MS.NoTrt_F |
| AMT | 0.36347 | 0.63419 | 0.424 | 0.46 | 1 | MS.Trt_vs_MS.NoTrt_F |
| CCDC138 | 0.3637 | -0.291 | 0.187 | 0.22 | 1 | MS.Trt_vs_MS.NoTrt_F |
| POLH | 0.36392 | -0.5193 | 0.586 | 0.61 | 1 | MS.Trt_vs_MS.NoTrt_F |
| EPS15 | 0.36397 | 0.29422 | 0.687 | 0.72 | 1 | MS.Trt_vs_MS.NoTrt_F |
| MFAP1 | 0.36502 | 0.45648 | 0.14 | 0.18 | 1 | MS.Trt_vs_MS.NoTrt_F |
| GHRL | 0.36557 | 0.95018 | 0.129 | 0.17 | 1 | MS.Trt_vs_MS.NoTrt_F |
| BCL2L1 | 0.36592 | 0.41749 | 0.23 | 0.28 | 1 | MS.Trt_vs_MS.NoTrt_F |
| ZNF225 | 0.3661 | -0.5215 | 0.745 | 0.79 | 1 | MS.Trt_vs_MS.NoTrt_F |
| COQ9 | 0.36797 | 1.22672 | 0.133 | 0.17 | 1 | MS.Trt_vs_MS.NoTrt_F |
| MRPL12 | 0.36802 | 1.87237 | 0.09 | 0.12 | 1 | MS.Trt_vs_MS.NoTrt_F |
| TKFC | 0.36834 | 0.54884 | 0.525 | 0.61 | 1 | MS.Trt_vs_MS.NoTrt_F |
| BCL2L11 | 0.36875 | 0.67717 | 0.504 | 0.55 | 1 | MS.Trt_vs_MS.NoTrt_F |
| HSD11B1L | 0.36898 | 0.28865 | 0.709 | 0.7 | 1 | MS.Trt_vs_MS.NoTrt_F |
| MRPS11 | 0.36905 | -0.415 | 0.698 | 0.76 | 1 | MS.Trt_vs_MS.NoTrt_F |
| STRN4 | 0.36929 | 0.63133 | 0.478 | 0.55 | 1 | MS.Trt_vs_MS.NoTrt_F |
| MCAM | 0.36993 | -0.5303 | 0.471 | 0.53 | 1 | MS.Trt_vs_MS.NoTrt_F |
| AP003419.1 | 0.37037 | 0.6116 | 0.219 | 0.28 | 1 | MS.Trt_vs_MS.NoTrt_F |
| PITPNM1 | 0.37104 | 1.07852 | 0.27 | 0.32 | 1 | MS.Trt_vs_MS.NoTrt_F |
| USP28 | 0.37212 | 0.59691 | 0.392 | 0.44 | 1 | MS.Trt_vs_MS.NoTrt_F |
| STXBP3 | 0.37291 | -0.3913 | 0.68 | 0.65 | 1 | MS.Trt_vs_MS.NoTrt_F |

| | | | | | | |
|---|---|---|---|---|---|---|
| GIT2 | 0.37344 | 0.29117 | 0.694 | 0.76 | 1 | MS.Trt_vs_MS.NoTrt_F |
| EIF3B | 0.37383 | 0.84025 | 0.353 | 0.4 | 1 | MS.Trt_vs_MS.NoTrt_F |
| CACNB1 | 0.37418 | 0.66358 | 0.147 | 0.18 | 1 | MS.Trt_vs_MS.NoTrt_F |
| RNF130 | 0.37457 | 0.65379 | 0.676 | 0.69 | 1 | MS.Trt_vs_MS.NoTrt_F |
| KCTD18 | 0.37638 | 0.31463 | 0.755 | 0.86 | 1 | MS.Trt_vs_MS.NoTrt_F |
| LINS1 | 0.37651 | 0.355 | 0.586 | 0.69 | 1 | MS.Trt_vs_MS.NoTrt_F |
| RNASEH2B | 0.37704 | 0.36155 | 0.719 | 0.79 | 1 | MS.Trt_vs_MS.NoTrt_F |
| TAF6 | 0.3772 | -0.3198 | 0.532 | 0.58 | 1 | MS.Trt_vs_MS.NoTrt_F |
| DNAJC3 | 0.3772 | 0.72838 | 0.532 | 0.63 | 1 | MS.Trt_vs_MS.NoTrt_F |
| QSOX1 | 0.37725 | 0.4295 | 0.385 | 0.43 | 1 | MS.Trt_vs_MS.NoTrt_F |
| MANF | 0.37771 | 1.14602 | 0.083 | 0.11 | 1 | MS.Trt_vs_MS.NoTrt_F |
| STMN1 | 0.37807 | -0.3457 | 0.727 | 0.73 | 1 | MS.Trt_vs_MS.NoTrt_F |
| ZNF326 | 0.37817 | -0.5087 | 0.608 | 0.62 | 1 | MS.Trt_vs_MS.NoTrt_F |
| HMGCS1 | 0.37854 | -0.4342 | 0.558 | 0.62 | 1 | MS.Trt_vs_MS.NoTrt_F |
| FKBP2 | 0.37879 | 1.14461 | 0.137 | 0.18 | 1 | MS.Trt_vs_MS.NoTrt_F |
| UST | 0.37897 | 0.37408 | 0.277 | 0.33 | 1 | MS.Trt_vs_MS.NoTrt_F |
| MEGF6 | 0.37914 | 1.3975 | 0.144 | 0.18 | 1 | MS.Trt_vs_MS.NoTrt_F |
| LRRC58 | 0.37938 | 0.59234 | 0.712 | 0.74 | 1 | MS.Trt_vs_MS.NoTrt_F |
| CTNNB1 | 0.38105 | 0.3378 | 0.626 | 0.72 | 1 | MS.Trt_vs_MS.NoTrt_F |
| CNN3 | 0.38134 | 0.8152 | 0.255 | 0.3 | 1 | MS.Trt_vs_MS.NoTrt_F |
| TM9SF3 | 0.38164 | 0.99025 | 0.198 | 0.24 | 1 | MS.Trt_vs_MS.NoTrt_F |
| GPR89B | 0.38169 | 0.26254 | 0.396 | 0.44 | 1 | MS.Trt_vs_MS.NoTrt_F |
| DFFA | 0.38207 | 0.3451 | 0.946 | 0.97 | 1 | MS.Trt_vs_MS.NoTrt_F |
| PLLP | 0.38217 | 0.2704 | 0.939 | 0.94 | 1 | MS.Trt_vs_MS.NoTrt_F |
| GEMIN2 | 0.38247 | 0.80423 | 0.68 | 0.71 | 1 | MS.Trt_vs_MS.NoTrt_F |
| MAPKAPK5 | 0.3825 | -0.388 | 0.932 | 0.94 | 1 | MS.Trt_vs_MS.NoTrt_F |
| ZNF274 | 0.38303 | -0.2615 | 0.64 | 0.7 | 1 | MS.Trt_vs_MS.NoTrt_F |
| ALG2 | 0.38339 | 1.12961 | 0.133 | 0.17 | 1 | MS.Trt_vs_MS.NoTrt_F |
| MATN2 | 0.38351 | 0.45425 | 0.518 | 0.56 | 1 | MS.Trt_vs_MS.NoTrt_F |
| ZNF619 | 0.38387 | 0.29175 | 0.723 | 0.76 | 1 | MS.Trt_vs_MS.NoTrt_F |
| ESPL1 | 0.38391 | 0.84215 | 0.698 | 0.69 | 1 | MS.Trt_vs_MS.NoTrt_F |
| ELK4 | 0.3841 | 0.32338 | 0.543 | 0.6 | 1 | MS.Trt_vs_MS.NoTrt_F |
| CHRAC1 | 0.38433 | -0.4834 | 0.511 | 0.54 | 1 | MS.Trt_vs_MS.NoTrt_F |
| TMEM218 | 0.38439 | 0.42728 | 0.162 | 0.2 | 1 | MS.Trt_vs_MS.NoTrt_F |
| TBX6 | 0.38445 | 0.29069 | 0.46 | 0.51 | 1 | MS.Trt_vs_MS.NoTrt_F |
| TNFAIP2 | 0.38477 | 0.83134 | 0.331 | 0.39 | 1 | MS.Trt_vs_MS.NoTrt_F |
| RABL2B | 0.38544 | -0.4059 | 0.896 | 0.91 | 1 | MS.Trt_vs_MS.NoTrt_F |
| TRIQK | 0.38566 | 0.53685 | 0.547 | 0.5 | 1 | MS.Trt_vs_MS.NoTrt_F |
| SATB1 | 0.38577 | 1.13258 | 0.194 | 0.25 | 1 | MS.Trt_vs_MS.NoTrt_F |
| RMND5A | 0.38578 | 1.53468 | 0.331 | 0.32 | 1 | MS.Trt_vs_MS.NoTrt_F |

| | | | | | | |
|---|---|---|---|---|---|---|
| NDUFB2 | 0.3858 | 0.52815 | 0.932 | 0.95 | 1 | MS.Trt_vs_MS.NoTrt_F |
| LINC00116 | 0.38604 | 1.24485 | 0.169 | 0.21 | 1 | MS.Trt_vs_MS.NoTrt_F |
| C1orf112 | 0.38681 | 0.70964 | 0.468 | 0.55 | 1 | MS.Trt_vs_MS.NoTrt_F |
| ZFX | 0.38689 | 0.4666 | 0.622 | 0.72 | 1 | MS.Trt_vs_MS.NoTrt_F |
| CANX | 0.38721 | 0.28047 | 0.705 | 0.74 | 1 | MS.Trt_vs_MS.NoTrt_F |
| CYP51A1 | 0.38729 | 0.90941 | 0.46 | 0.54 | 1 | MS.Trt_vs_MS.NoTrt_F |
| C2orf88 | 0.38831 | -0.4291 | 0.335 | 0.38 | 1 | MS.Trt_vs_MS.NoTrt_F |
| CDKL1 | 0.38845 | 0.54038 | 0.651 | 0.73 | 1 | MS.Trt_vs_MS.NoTrt_F |
| EGR1 | 0.38961 | 2.57181 | 0.14 | 0.12 | 1 | MS.Trt_vs_MS.NoTrt_F |
| RP4-635E18 | 0.38972 | 0.85623 | 0.234 | 0.29 | 1 | MS.Trt_vs_MS.NoTrt_F |
| CTIF | 0.39041 | 1.21838 | 0.248 | 0.29 | 1 | MS.Trt_vs_MS.NoTrt_F |
| USP48 | 0.39048 | 0.53838 | 0.568 | 0.62 | 1 | MS.Trt_vs_MS.NoTrt_F |
| UBE2D4 | 0.39085 | 1.13929 | 0.719 | 0.64 | 1 | MS.Trt_vs_MS.NoTrt_F |
| MAPK9 | 0.39103 | 1.33227 | 0.457 | 0.5 | 1 | MS.Trt_vs_MS.NoTrt_F |
| SLC5A10 | 0.3914 | 0.50621 | 0.101 | 0.13 | 1 | MS.Trt_vs_MS.NoTrt_F |
| APC | 0.39154 | -0.279 | 0.522 | 0.46 | 1 | MS.Trt_vs_MS.NoTrt_F |
| SENP2 | 0.39182 | 0.39884 | 0.813 | 0.88 | 1 | MS.Trt_vs_MS.NoTrt_F |
| RABL2A | 0.3922 | -0.3771 | 0.561 | 0.56 | 1 | MS.Trt_vs_MS.NoTrt_F |
| KIF3B | 0.39237 | 0.35425 | 0.45 | 0.49 | 1 | MS.Trt_vs_MS.NoTrt_F |
| SOCS7 | 0.3929 | 0.66949 | 0.45 | 0.51 | 1 | MS.Trt_vs_MS.NoTrt_F |
| TMEM68 | 0.39338 | -0.3113 | 0.892 | 0.93 | 1 | MS.Trt_vs_MS.NoTrt_F |
| TMA7 | 0.39413 | 0.68531 | 0.576 | 0.79 | 1 | MS.Trt_vs_MS.NoTrt_F |
| ITGA4 | 0.39532 | 0.55865 | 0.252 | 0.3 | 1 | MS.Trt_vs_MS.NoTrt_F |
| NKAP | 0.39558 | 0.32616 | 0.946 | 0.94 | 1 | MS.Trt_vs_MS.NoTrt_F |
| TIGAR | 0.39563 | 0.43611 | 0.896 | 0.91 | 1 | MS.Trt_vs_MS.NoTrt_F |
| CHRNB1 | 0.39614 | 0.56513 | 0.586 | 0.6 | 1 | MS.Trt_vs_MS.NoTrt_F |
| TTC19 | 0.3963 | 0.26258 | 0.788 | 0.79 | 1 | MS.Trt_vs_MS.NoTrt_F |
| NABP2 | 0.39697 | 0.44874 | 0.529 | 0.48 | 1 | MS.Trt_vs_MS.NoTrt_F |
| TMEM144 | 0.39721 | 0.33477 | 0.209 | 0.24 | 1 | MS.Trt_vs_MS.NoTrt_F |
| ZC2HC1C | 0.3976 | 0.29718 | 0.14 | 0.17 | 1 | MS.Trt_vs_MS.NoTrt_F |
| TP53BP1 | 0.39777 | 0.6501 | 0.32 | 0.37 | 1 | MS.Trt_vs_MS.NoTrt_F |
| FCRL3 | 0.39792 | 0.98763 | 0.291 | 0.38 | 1 | MS.Trt_vs_MS.NoTrt_F |
| SIRT3 | 0.39796 | -0.3871 | 0.367 | 0.41 | 1 | MS.Trt_vs_MS.NoTrt_F |
| MVK | 0.3987 | 0.9632 | 0.097 | 0.13 | 1 | MS.Trt_vs_MS.NoTrt_F |
| ATP6V0A2 | 0.39911 | -0.3374 | 0.95 | 0.97 | 1 | MS.Trt_vs_MS.NoTrt_F |
| GPRIN3 | 0.39914 | 0.32226 | 0.504 | 0.56 | 1 | MS.Trt_vs_MS.NoTrt_F |
| DCAF13 | 0.39951 | 0.9696 | 0.662 | 0.72 | 1 | MS.Trt_vs_MS.NoTrt_F |
| ANK1 | 0.40006 | 1.15945 | 0.234 | 0.27 | 1 | MS.Trt_vs_MS.NoTrt_F |
| ALG8 | 0.40054 | -0.4725 | 0.489 | 0.51 | 1 | MS.Trt_vs_MS.NoTrt_F |
| PRKCD | 0.40078 | 0.40555 | 0.489 | 0.55 | 1 | MS.Trt_vs_MS.NoTrt_F |

| | | | | | | |
|---|---|---|---|---|---|---|
| EPAS1 | 0.40356 | 2.09482 | 0.108 | 0.14 | 1 | MS.Trt_vs_MS.NoTrt_F |
| RBM6 | 0.40399 | 0.30078 | 0.871 | 0.99 | 1 | MS.Trt_vs_MS.NoTrt_F |
| SH3RF2 | 0.40447 | 0.70949 | 0.216 | 0.19 | 1 | MS.Trt_vs_MS.NoTrt_F |
| KIAA0319L | 0.40449 | -0.4389 | 0.683 | 0.75 | 1 | MS.Trt_vs_MS.NoTrt_F |
| POU5F1B | 0.40484 | 0.36876 | 0.385 | 0.45 | 1 | MS.Trt_vs_MS.NoTrt_F |
| SGO1 | 0.40485 | 0.27561 | 0.522 | 0.59 | 1 | MS.Trt_vs_MS.NoTrt_F |
| PECR | 0.40491 | 0.30674 | 0.691 | 0.72 | 1 | MS.Trt_vs_MS.NoTrt_F |
| EIF5 | 0.40528 | 0.55457 | 0.583 | 0.6 | 1 | MS.Trt_vs_MS.NoTrt_F |
| TUBE1 | 0.40549 | 0.66421 | 0.514 | 0.55 | 1 | MS.Trt_vs_MS.NoTrt_F |
| B9D2 | 0.4059 | 0.54369 | 0.381 | 0.43 | 1 | MS.Trt_vs_MS.NoTrt_F |
| ZNF292 | 0.40592 | 0.4998 | 0.781 | 0.83 | 1 | MS.Trt_vs_MS.NoTrt_F |
| RASAL2 | 0.40596 | 0.35728 | 0.511 | 0.56 | 1 | MS.Trt_vs_MS.NoTrt_F |
| DZANK1 | 0.40611 | 0.25767 | 0.701 | 0.7 | 1 | MS.Trt_vs_MS.NoTrt_F |
| NIPAL1 | 0.40621 | -0.4704 | 0.432 | 0.49 | 1 | MS.Trt_vs_MS.NoTrt_F |
| PQLC1 | 0.40638 | 0.50373 | 0.453 | 0.54 | 1 | MS.Trt_vs_MS.NoTrt_F |
| HPS5 | 0.40641 | -0.6559 | 0.86 | 0.82 | 1 | MS.Trt_vs_MS.NoTrt_F |
| ZNF215 | 0.40701 | 0.36364 | 0.421 | 0.5 | 1 | MS.Trt_vs_MS.NoTrt_F |
| STX2 | 0.4073 | 0.49431 | 0.478 | 0.52 | 1 | MS.Trt_vs_MS.NoTrt_F |
| CCDC157 | 0.4074 | 0.48035 | 0.665 | 0.7 | 1 | MS.Trt_vs_MS.NoTrt_F |
| SYF2 | 0.40766 | 0.37674 | 0.414 | 0.48 | 1 | MS.Trt_vs_MS.NoTrt_F |
| SERPINB9 | 0.40874 | 0.57141 | 0.799 | 0.84 | 1 | MS.Trt_vs_MS.NoTrt_F |
| RFC5 | 0.409 | 0.31116 | 0.176 | 0.21 | 1 | MS.Trt_vs_MS.NoTrt_F |
| MAP2K6 | 0.40939 | 0.4763 | 0.378 | 0.44 | 1 | MS.Trt_vs_MS.NoTrt_F |
| ZNF140 | 0.40974 | 0.84216 | 0.187 | 0.23 | 1 | MS.Trt_vs_MS.NoTrt_F |
| ANO10 | 0.41001 | -0.4779 | 0.615 | 0.66 | 1 | MS.Trt_vs_MS.NoTrt_F |
| THUMPD1 | 0.41002 | 0.25524 | 0.788 | 0.81 | 1 | MS.Trt_vs_MS.NoTrt_F |
| FAM98C | 0.41037 | -0.2903 | 0.968 | 0.99 | 1 | MS.Trt_vs_MS.NoTrt_F |
| CASP2 | 0.41106 | 0.45868 | 0.806 | 0.89 | 1 | MS.Trt_vs_MS.NoTrt_F |
| SIPA1L1 | 0.41137 | 0.66651 | 0.576 | 0.6 | 1 | MS.Trt_vs_MS.NoTrt_F |
| MRE11A | 0.41158 | 0.3979 | 0.752 | 0.82 | 1 | MS.Trt_vs_MS.NoTrt_F |
| EXD2 | 0.41197 | 0.44283 | 0.212 | 0.25 | 1 | MS.Trt_vs_MS.NoTrt_F |
| LAT2 | 0.41201 | 0.35418 | 0.813 | 0.88 | 1 | MS.Trt_vs_MS.NoTrt_F |
| FGF23 | 0.41208 | 0.2686 | 0.817 | 0.85 | 1 | MS.Trt_vs_MS.NoTrt_F |
| MPV17L2 | 0.41224 | 0.81233 | 0.234 | 0.28 | 1 | MS.Trt_vs_MS.NoTrt_F |
| PLXDC2 | 0.41363 | 0.3966 | 0.698 | 0.69 | 1 | MS.Trt_vs_MS.NoTrt_F |
| C14orf119 | 0.41389 | 0.31907 | 0.529 | 0.56 | 1 | MS.Trt_vs_MS.NoTrt_F |
| CHPT1 | 0.41463 | 0.80181 | 0.626 | 0.61 | 1 | MS.Trt_vs_MS.NoTrt_F |
| ZNF785 | 0.41529 | -0.339 | 0.881 | 0.93 | 1 | MS.Trt_vs_MS.NoTrt_F |
| ZNF133 | 0.4165 | -0.6665 | 0.464 | 0.43 | 1 | MS.Trt_vs_MS.NoTrt_F |
| OLFML1 | 0.41669 | 0.32118 | 0.241 | 0.28 | 1 | MS.Trt_vs_MS.NoTrt_F |

| | | | | | | |
|---|---|---|---|---|---|---|
| WARS | 0.41743 | 0.58515 | 0.295 | 0.34 | 1 | MS.Trt_vs_MS.NoTrt_F |
| ZBTB25 | 0.41824 | 0.50437 | 0.975 | 0.98 | 1 | MS.Trt_vs_MS.NoTrt_F |
| KCMF1 | 0.41836 | 1.34881 | 0.518 | 0.55 | 1 | MS.Trt_vs_MS.NoTrt_F |
| SYT11 | 0.41847 | 0.509 | 0.604 | 0.68 | 1 | MS.Trt_vs_MS.NoTrt_F |
| TAF8 | 0.41894 | -0.5097 | 0.968 | 0.97 | 1 | MS.Trt_vs_MS.NoTrt_F |
| CD302 | 0.41903 | 1.10663 | 0.396 | 0.35 | 1 | MS.Trt_vs_MS.NoTrt_F |
| TRMT5 | 0.42011 | -0.7106 | 0.209 | 0.23 | 1 | MS.Trt_vs_MS.NoTrt_F |
| CCDC163 | 0.42047 | 0.86047 | 0.586 | 0.55 | 1 | MS.Trt_vs_MS.NoTrt_F |
| HSPBAP1 | 0.42058 | 0.49936 | 0.525 | 0.57 | 1 | MS.Trt_vs_MS.NoTrt_F |
| DCP1A | 0.42088 | 0.39279 | 0.367 | 0.4 | 1 | MS.Trt_vs_MS.NoTrt_F |
| ZNF613 | 0.42093 | 1.22009 | 0.36 | 0.42 | 1 | MS.Trt_vs_MS.NoTrt_F |
| PPIE | 0.42106 | 0.73592 | 0.781 | 0.82 | 1 | MS.Trt_vs_MS.NoTrt_F |
| KIF22 | 0.42175 | 0.44187 | 0.198 | 0.24 | 1 | MS.Trt_vs_MS.NoTrt_F |
| MTHFR | 0.42178 | 0.72084 | 0.799 | 0.9 | 1 | MS.Trt_vs_MS.NoTrt_F |
| PAAF1 | 0.42299 | -0.5029 | 0.95 | 0.99 | 1 | MS.Trt_vs_MS.NoTrt_F |
| ZNF345 | 0.42303 | -0.7408 | 0.795 | 0.85 | 1 | MS.Trt_vs_MS.NoTrt_F |
| CCDC152 | 0.42327 | -1.1778 | 0.237 | 0.27 | 1 | MS.Trt_vs_MS.NoTrt_F |
| FAM20B | 0.42394 | 0.35773 | 0.594 | 0.62 | 1 | MS.Trt_vs_MS.NoTrt_F |
| SASH1 | 0.42424 | 0.79825 | 0.309 | 0.34 | 1 | MS.Trt_vs_MS.NoTrt_F |
| PSPC1 | 0.42473 | 0.32577 | 0.335 | 0.38 | 1 | MS.Trt_vs_MS.NoTrt_F |
| APOD | 0.42509 | 1.63718 | 0.227 | 0.26 | 1 | MS.Trt_vs_MS.NoTrt_F |
| F2R | 0.42522 | 1.14925 | 0.558 | 0.56 | 1 | MS.Trt_vs_MS.NoTrt_F |
| RAPGEF2 | 0.42541 | -0.7307 | 0.371 | 0.41 | 1 | MS.Trt_vs_MS.NoTrt_F |
| MYO1C | 0.42548 | 1.86467 | 0.201 | 0.25 | 1 | MS.Trt_vs_MS.NoTrt_F |
| NOSTRIN | 0.42655 | 0.48841 | 0.374 | 0.35 | 1 | MS.Trt_vs_MS.NoTrt_F |
| TMEM268 | 0.42678 | -0.3915 | 0.496 | 0.54 | 1 | MS.Trt_vs_MS.NoTrt_F |
| ESAM | 0.42678 | 1.20627 | 0.165 | 0.2 | 1 | MS.Trt_vs_MS.NoTrt_F |
| GLRX3 | 0.42802 | 0.50681 | 0.827 | 0.85 | 1 | MS.Trt_vs_MS.NoTrt_F |
| IFI6 | 0.42851 | 0.65917 | 0.374 | 0.44 | 1 | MS.Trt_vs_MS.NoTrt_F |
| ZNF835 | 0.42863 | 0.44772 | 0.586 | 0.64 | 1 | MS.Trt_vs_MS.NoTrt_F |
| COPS5 | 0.42958 | -0.6109 | 0.824 | 0.88 | 1 | MS.Trt_vs_MS.NoTrt_F |
| CCDC88A | 0.42996 | 0.38194 | 0.658 | 0.67 | 1 | MS.Trt_vs_MS.NoTrt_F |
| ZNF284 | 0.43007 | 0.5381 | 0.576 | 0.61 | 1 | MS.Trt_vs_MS.NoTrt_F |
| CKAP4 | 0.43012 | 1.58578 | 0.126 | 0.16 | 1 | MS.Trt_vs_MS.NoTrt_F |
| EHBP1 | 0.43015 | 0.36991 | 0.435 | 0.46 | 1 | MS.Trt_vs_MS.NoTrt_F |
| NME9 | 0.43065 | 1.48373 | 0.241 | 0.29 | 1 | MS.Trt_vs_MS.NoTrt_F |
| RFC2 | 0.43136 | 0.38816 | 0.665 | 0.77 | 1 | MS.Trt_vs_MS.NoTrt_F |
| WIPI1 | 0.43178 | 0.26154 | 0.824 | 0.89 | 1 | MS.Trt_vs_MS.NoTrt_F |
| DCBLD1 | 0.43209 | -0.5549 | 0.543 | 0.58 | 1 | MS.Trt_vs_MS.NoTrt_F |
| SCFD1 | 0.43247 | 0.39585 | 0.741 | 0.76 | 1 | MS.Trt_vs_MS.NoTrt_F |

| | | | | | | |
|---|---|---|---|---|---|---|
| ZNF208 | 0.43256 | 0.25444 | 0.845 | 0.86 | 1 | MS.Trt_vs_MS.NoTrt_F |
| ING3 | 0.43262 | 0.44398 | 0.205 | 0.24 | 1 | MS.Trt_vs_MS.NoTrt_F |
| VASH2 | 0.43492 | 0.26908 | 0.77 | 0.77 | 1 | MS.Trt_vs_MS.NoTrt_F |
| METTL25 | 0.43511 | 0.87281 | 0.205 | 0.24 | 1 | MS.Trt_vs_MS.NoTrt_F |
| PRKCE | 0.43531 | -0.307 | 0.173 | 0.2 | 1 | MS.Trt_vs_MS.NoTrt_F |
| CINP | 0.43582 | 0.54704 | 0.896 | 0.9 | 1 | MS.Trt_vs_MS.NoTrt_F |
| ZNF80 | 0.43646 | 0.45444 | 0.219 | 0.27 | 1 | MS.Trt_vs_MS.NoTrt_F |
| KNSTRN | 0.43647 | 0.31033 | 0.525 | 0.5 | 1 | MS.Trt_vs_MS.NoTrt_F |
| UTP6 | 0.43884 | 0.25854 | 0.867 | 0.91 | 1 | MS.Trt_vs_MS.NoTrt_F |
| ALOX12 | 0.43906 | 0.44548 | 0.112 | 0.14 | 1 | MS.Trt_vs_MS.NoTrt_F |
| WBP2NL | 0.43914 | 0.64758 | 0.371 | 0.43 | 1 | MS.Trt_vs_MS.NoTrt_F |
| ATG9B | 0.43955 | 0.60328 | 0.158 | 0.19 | 1 | MS.Trt_vs_MS.NoTrt_F |
| MRS2 | 0.44018 | -0.3649 | 0.856 | 0.82 | 1 | MS.Trt_vs_MS.NoTrt_F |
| GNPNAT1 | 0.44032 | 0.84948 | 0.529 | 0.52 | 1 | MS.Trt_vs_MS.NoTrt_F |
| AVPR1A | 0.44041 | 0.42695 | 0.223 | 0.2 | 1 | MS.Trt_vs_MS.NoTrt_F |
| P2RX5 | 0.44046 | 0.70553 | 0.133 | 0.16 | 1 | MS.Trt_vs_MS.NoTrt_F |
| UPF3A | 0.44046 | -0.324 | 0.734 | 0.79 | 1 | MS.Trt_vs_MS.NoTrt_F |
| SRFBP1 | 0.4415 | 1.11538 | 0.464 | 0.45 | 1 | MS.Trt_vs_MS.NoTrt_F |
| AK6 | 0.44349 | 0.81553 | 0.241 | 0.28 | 1 | MS.Trt_vs_MS.NoTrt_F |
| YWHAE | 0.44376 | 0.53177 | 0.471 | 0.61 | 1 | MS.Trt_vs_MS.NoTrt_F |
| TP53I11 | 0.44383 | 1.19578 | 0.129 | 0.16 | 1 | MS.Trt_vs_MS.NoTrt_F |
| C11orf31 | 0.44454 | 0.39145 | 0.511 | 0.57 | 1 | MS.Trt_vs_MS.NoTrt_F |
| PDXK | 0.44541 | 0.79859 | 0.529 | 0.56 | 1 | MS.Trt_vs_MS.NoTrt_F |
| APBB2 | 0.44555 | 0.73131 | 0.335 | 0.39 | 1 | MS.Trt_vs_MS.NoTrt_F |
| ABCB10 | 0.44566 | 0.35499 | 0.129 | 0.16 | 1 | MS.Trt_vs_MS.NoTrt_F |
| CD177 | 0.44582 | 2.2992 | 0.09 | 0.12 | 1 | MS.Trt_vs_MS.NoTrt_F |
| LDHC | 0.44725 | -0.4944 | 0.209 | 0.24 | 1 | MS.Trt_vs_MS.NoTrt_F |
| SUMF1 | 0.44822 | -0.8364 | 0.752 | 0.71 | 1 | MS.Trt_vs_MS.NoTrt_F |
| GHDC | 0.44907 | 0.80759 | 0.284 | 0.32 | 1 | MS.Trt_vs_MS.NoTrt_F |
| UTP11 | 0.4491 | 0.76815 | 0.547 | 0.63 | 1 | MS.Trt_vs_MS.NoTrt_F |
| FAM13A | 0.44958 | 0.72293 | 0.237 | 0.28 | 1 | MS.Trt_vs_MS.NoTrt_F |
| VTA1 | 0.44987 | 0.91417 | 0.41 | 0.46 | 1 | MS.Trt_vs_MS.NoTrt_F |
| RPL37 | 0.44996 | 0.25193 | 0.91 | 0.98 | 1 | MS.Trt_vs_MS.NoTrt_F |
| NANOS1 | 0.45021 | 0.34608 | 0.572 | 0.55 | 1 | MS.Trt_vs_MS.NoTrt_F |
| GTF3C4 | 0.45099 | 0.31272 | 0.349 | 0.4 | 1 | MS.Trt_vs_MS.NoTrt_F |
| SLC30A2 | 0.45105 | 0.29201 | 0.701 | 0.74 | 1 | MS.Trt_vs_MS.NoTrt_F |
| CCDC15 | 0.45161 | 1.05626 | 0.248 | 0.29 | 1 | MS.Trt_vs_MS.NoTrt_F |
| MTF1 | 0.45194 | -0.3082 | 0.309 | 0.35 | 1 | MS.Trt_vs_MS.NoTrt_F |
| TSPOAP1 | 0.45206 | 0.25555 | 0.255 | 0.29 | 1 | MS.Trt_vs_MS.NoTrt_F |
| EML4 | 0.45239 | 0.30408 | 0.683 | 0.82 | 1 | MS.Trt_vs_MS.NoTrt_F |

| | | | | | | |
|---|---|---|---|---|---|---|
| ARID5A | 0.45242 | 0.84153 | 0.234 | 0.29 | 1 | MS.Trt_vs_MS.NoTrt_F |
| WDR76 | 0.45274 | 0.68144 | 0.687 | 0.74 | 1 | MS.Trt_vs_MS.NoTrt_F |
| KLF3 | 0.45403 | 0.53141 | 0.619 | 0.66 | 1 | MS.Trt_vs_MS.NoTrt_F |
| UMPS | 0.45409 | -0.3345 | 0.676 | 0.7 | 1 | MS.Trt_vs_MS.NoTrt_F |
| CEP95 | 0.45472 | 0.61611 | 0.453 | 0.43 | 1 | MS.Trt_vs_MS.NoTrt_F |
| LINC00854 | 0.45539 | -0.67 | 0.658 | 0.73 | 1 | MS.Trt_vs_MS.NoTrt_F |
| MOCS3 | 0.45578 | -0.477 | 0.831 | 0.88 | 1 | MS.Trt_vs_MS.NoTrt_F |
| APOBEC3D | 0.45663 | 0.38906 | 0.651 | 0.75 | 1 | MS.Trt_vs_MS.NoTrt_F |
| SPARC | 0.45797 | 1.68624 | 0.101 | 0.13 | 1 | MS.Trt_vs_MS.NoTrt_F |
| LETM2 | 0.45844 | 0.45088 | 0.709 | 0.81 | 1 | MS.Trt_vs_MS.NoTrt_F |
| HLA-DRB4 | 0.45846 | -0.6791 | 0.119 | 0.14 | 1 | MS.Trt_vs_MS.NoTrt_F |
| THUMPD3 | 0.45846 | 0.64995 | 0.41 | 0.45 | 1 | MS.Trt_vs_MS.NoTrt_F |
| PTPN7 | 0.45847 | 0.82058 | 0.745 | 0.75 | 1 | MS.Trt_vs_MS.NoTrt_F |
| GATB | 0.45856 | -0.5267 | 0.241 | 0.27 | 1 | MS.Trt_vs_MS.NoTrt_F |
| AKAP12 | 0.4591 | 0.29301 | 0.133 | 0.12 | 1 | MS.Trt_vs_MS.NoTrt_F |
| DAB2 | 0.45958 | 1.36043 | 0.234 | 0.27 | 1 | MS.Trt_vs_MS.NoTrt_F |
| NUDT22 | 0.45967 | 0.82115 | 0.716 | 0.71 | 1 | MS.Trt_vs_MS.NoTrt_F |
| SKP2 | 0.45968 | 0.38618 | 0.493 | 0.53 | 1 | MS.Trt_vs_MS.NoTrt_F |
| SECISBP2 | 0.4613 | 0.2724 | 0.629 | 0.66 | 1 | MS.Trt_vs_MS.NoTrt_F |
| GCM1 | 0.46152 | 0.72726 | 0.576 | 0.54 | 1 | MS.Trt_vs_MS.NoTrt_F |
| ANKRD10 | 0.46158 | 0.70956 | 0.73 | 0.8 | 1 | MS.Trt_vs_MS.NoTrt_F |
| B3GNT7 | 0.46159 | 0.71478 | 0.245 | 0.21 | 1 | MS.Trt_vs_MS.NoTrt_F |
| SENP6 | 0.46185 | 0.63675 | 0.863 | 0.93 | 1 | MS.Trt_vs_MS.NoTrt_F |
| ZPBP2 | 0.46214 | 0.25275 | 0.151 | 0.13 | 1 | MS.Trt_vs_MS.NoTrt_F |
| ZNF559 | 0.46215 | 0.33242 | 0.676 | 0.71 | 1 | MS.Trt_vs_MS.NoTrt_F |
| CD47 | 0.46236 | 0.45158 | 0.878 | 0.95 | 1 | MS.Trt_vs_MS.NoTrt_F |
| SLC25A15 | 0.46322 | -0.5071 | 0.669 | 0.7 | 1 | MS.Trt_vs_MS.NoTrt_F |
| MAF | 0.4638 | 1.03152 | 0.385 | 0.46 | 1 | MS.Trt_vs_MS.NoTrt_F |
| NUSAP1 | 0.4667 | -0.5409 | 0.734 | 0.79 | 1 | MS.Trt_vs_MS.NoTrt_F |
| TIMM50 | 0.46734 | 0.3047 | 0.982 | 0.99 | 1 | MS.Trt_vs_MS.NoTrt_F |
| C20orf196 | 0.46735 | 0.75469 | 0.331 | 0.38 | 1 | MS.Trt_vs_MS.NoTrt_F |
| STX17 | 0.46889 | 0.27927 | 0.468 | 0.55 | 1 | MS.Trt_vs_MS.NoTrt_F |
| SPN | 0.46901 | -0.3076 | 0.964 | 0.99 | 1 | MS.Trt_vs_MS.NoTrt_F |
| EP400NL | 0.46991 | -0.303 | 0.766 | 0.79 | 1 | MS.Trt_vs_MS.NoTrt_F |
| ZNF780A | 0.47023 | 0.66728 | 0.342 | 0.39 | 1 | MS.Trt_vs_MS.NoTrt_F |
| FIBP | 0.4705 | 0.55546 | 0.471 | 0.53 | 1 | MS.Trt_vs_MS.NoTrt_F |
| ANKRD40 | 0.47234 | 0.86063 | 0.817 | 0.89 | 1 | MS.Trt_vs_MS.NoTrt_F |
| TRIM59 | 0.47263 | 0.6308 | 0.82 | 0.9 | 1 | MS.Trt_vs_MS.NoTrt_F |
| VLDLR | 0.47273 | 1.66243 | 0.446 | 0.54 | 1 | MS.Trt_vs_MS.NoTrt_F |
| LIPT2 | 0.47282 | 0.72974 | 0.169 | 0.2 | 1 | MS.Trt_vs_MS.NoTrt_F |

| | | | | | | |
|---|---|---|---|---|---|---|
| PON2 | 0.47359 | 1.76649 | 0.417 | 0.46 | 1 | MS.Trt_vs_MS.NoTrt_F |
| P3H1 | 0.4737 | 0.32213 | 0.95 | 0.98 | 1 | MS.Trt_vs_MS.NoTrt_F |
| SCAF4 | 0.47385 | 0.62624 | 0.237 | 0.29 | 1 | MS.Trt_vs_MS.NoTrt_F |
| PARP12 | 0.47463 | 1.21834 | 0.252 | 0.29 | 1 | MS.Trt_vs_MS.NoTrt_F |
| POLE | 0.47495 | 0.3212 | 0.831 | 0.88 | 1 | MS.Trt_vs_MS.NoTrt_F |
| OCIAD1 | 0.47503 | 0.50984 | 0.676 | 0.67 | 1 | MS.Trt_vs_MS.NoTrt_F |
| MGP | 0.47558 | 1.01938 | 0.597 | 0.58 | 1 | MS.Trt_vs_MS.NoTrt_F |
| NUMA1 | 0.47596 | 0.26914 | 0.853 | 0.87 | 1 | MS.Trt_vs_MS.NoTrt_F |
| TMC6 | 0.47656 | 0.32813 | 0.809 | 0.86 | 1 | MS.Trt_vs_MS.NoTrt_F |
| FADS2 | 0.47701 | 0.49174 | 0.525 | 0.58 | 1 | MS.Trt_vs_MS.NoTrt_F |
| THAP5 | 0.4774 | 0.8211 | 0.903 | 0.93 | 1 | MS.Trt_vs_MS.NoTrt_F |
| LETMD1 | 0.4791 | 0.37784 | 0.831 | 0.89 | 1 | MS.Trt_vs_MS.NoTrt_F |
| STYXL1 | 0.48046 | 0.46749 | 0.324 | 0.35 | 1 | MS.Trt_vs_MS.NoTrt_F |
| PER2 | 0.48056 | 0.67269 | 0.507 | 0.58 | 1 | MS.Trt_vs_MS.NoTrt_F |
| LDLR | 0.48076 | 0.28022 | 0.921 | 0.96 | 1 | MS.Trt_vs_MS.NoTrt_F |
| SLC12A2 | 0.48107 | 1.43198 | 0.565 | 0.59 | 1 | MS.Trt_vs_MS.NoTrt_F |
| DNAH12 | 0.48132 | 0.67638 | 0.162 | 0.14 | 1 | MS.Trt_vs_MS.NoTrt_F |
| CGN | 0.48162 | -0.3686 | 0.216 | 0.25 | 1 | MS.Trt_vs_MS.NoTrt_F |
| TVP23B | 0.48248 | -0.5155 | 0.565 | 0.58 | 1 | MS.Trt_vs_MS.NoTrt_F |
| BOLA3 | 0.48261 | 0.63892 | 0.518 | 0.58 | 1 | MS.Trt_vs_MS.NoTrt_F |
| PCOLCE | 0.48272 | -0.2877 | 0.104 | 0.13 | 1 | MS.Trt_vs_MS.NoTrt_F |
| HINT3 | 0.48355 | -0.2607 | 0.309 | 0.35 | 1 | MS.Trt_vs_MS.NoTrt_F |
| ARHGEF26 | 0.48384 | 0.63812 | 0.486 | 0.53 | 1 | MS.Trt_vs_MS.NoTrt_F |
| AK9 | 0.48417 | 0.78056 | 0.658 | 0.76 | 1 | MS.Trt_vs_MS.NoTrt_F |
| UAP1L1 | 0.48444 | 0.95325 | 0.374 | 0.36 | 1 | MS.Trt_vs_MS.NoTrt_F |
| DCAF11 | 0.48522 | -0.4324 | 0.737 | 0.85 | 1 | MS.Trt_vs_MS.NoTrt_F |
| ZFP14 | 0.48625 | 0.29036 | 0.827 | 0.9 | 1 | MS.Trt_vs_MS.NoTrt_F |
| ZNF827 | 0.4865 | 1.32181 | 0.083 | 0.11 | 1 | MS.Trt_vs_MS.NoTrt_F |
| CH507-9B2.! | 0.48688 | 0.35356 | 0.464 | 0.54 | 1 | MS.Trt_vs_MS.NoTrt_F |
| PPIC | 0.48703 | -0.5114 | 0.086 | 0.11 | 1 | MS.Trt_vs_MS.NoTrt_F |
| VASH1 | 0.48708 | 0.50062 | 0.094 | 0.12 | 1 | MS.Trt_vs_MS.NoTrt_F |
| LINC00961 | 0.48769 | 0.31017 | 0.36 | 0.39 | 1 | MS.Trt_vs_MS.NoTrt_F |
| ARL2 | 0.48776 | 0.30377 | 0.788 | 0.83 | 1 | MS.Trt_vs_MS.NoTrt_F |
| MCRIP2 | 0.48861 | 0.26106 | 0.561 | 0.59 | 1 | MS.Trt_vs_MS.NoTrt_F |
| TRAPPC4 | 0.48879 | 0.43362 | 0.899 | 0.9 | 1 | MS.Trt_vs_MS.NoTrt_F |
| SIDT1 | 0.48912 | 0.32149 | 0.741 | 0.8 | 1 | MS.Trt_vs_MS.NoTrt_F |
| MRPS16 | 0.48941 | 0.25178 | 0.802 | 0.83 | 1 | MS.Trt_vs_MS.NoTrt_F |
| PDCD6 | 0.48993 | 0.29081 | 0.673 | 0.69 | 1 | MS.Trt_vs_MS.NoTrt_F |
| GALK1 | 0.49015 | 0.88187 | 0.284 | 0.33 | 1 | MS.Trt_vs_MS.NoTrt_F |
| CRB1 | 0.49056 | 0.5788 | 0.162 | 0.19 | 1 | MS.Trt_vs_MS.NoTrt_F |

| | | | | | | |
|---|---|---|---|---|---|---|
| TMEM97 | 0.491 | 1.23217 | 0.281 | 0.31 | 1 | MS.Trt_vs_MS.NoTrt_F |
| ESRP1 | 0.4914 | 0.75086 | 0.223 | 0.2 | 1 | MS.Trt_vs_MS.NoTrt_F |
| ICMT | 0.49197 | -1.1248 | 0.651 | 0.66 | 1 | MS.Trt_vs_MS.NoTrt_F |
| STARD4 | 0.49218 | 1.37128 | 0.165 | 0.19 | 1 | MS.Trt_vs_MS.NoTrt_F |
| FKRP | 0.49224 | 0.31286 | 0.651 | 0.69 | 1 | MS.Trt_vs_MS.NoTrt_F |
| MRPL40 | 0.49326 | 0.45094 | 0.453 | 0.47 | 1 | MS.Trt_vs_MS.NoTrt_F |
| PPM1F | 0.49352 | -0.5543 | 0.763 | 0.8 | 1 | MS.Trt_vs_MS.NoTrt_F |
| ARHGAP26 | 0.4936 | 0.51297 | 0.68 | 0.68 | 1 | MS.Trt_vs_MS.NoTrt_F |
| ACAA2 | 0.4943 | 0.32702 | 0.558 | 0.6 | 1 | MS.Trt_vs_MS.NoTrt_F |
| ZNF286B | 0.49455 | 1.32001 | 0.468 | 0.53 | 1 | MS.Trt_vs_MS.NoTrt_F |
| FAM210A | 0.49468 | -0.2585 | 0.327 | 0.4 | 1 | MS.Trt_vs_MS.NoTrt_F |
| FOXRED1 | 0.49505 | -0.544 | 0.482 | 0.52 | 1 | MS.Trt_vs_MS.NoTrt_F |
| ZADH2 | 0.49523 | 1.3178 | 0.817 | 0.83 | 1 | MS.Trt_vs_MS.NoTrt_F |
| ANKRD54 | 0.49644 | 0.66141 | 0.676 | 0.8 | 1 | MS.Trt_vs_MS.NoTrt_F |
| SYNE4 | 0.49656 | 0.78494 | 0.5 | 0.53 | 1 | MS.Trt_vs_MS.NoTrt_F |
| QTRT2 | 0.49695 | 0.52635 | 0.73 | 0.75 | 1 | MS.Trt_vs_MS.NoTrt_F |
| MPST | 0.49705 | 0.43764 | 0.82 | 0.9 | 1 | MS.Trt_vs_MS.NoTrt_F |
| CNBD2 | 0.49736 | 0.38834 | 0.439 | 0.49 | 1 | MS.Trt_vs_MS.NoTrt_F |
| KIAA0513 | 0.49743 | 0.55644 | 0.608 | 0.64 | 1 | MS.Trt_vs_MS.NoTrt_F |
| SERPINB6 | 0.49774 | 1.32778 | 0.486 | 0.56 | 1 | MS.Trt_vs_MS.NoTrt_F |
| ZNF250 | 0.49839 | 1.12628 | 0.59 | 0.64 | 1 | MS.Trt_vs_MS.NoTrt_F |
| BRD8 | 0.49905 | -0.3196 | 0.763 | 0.77 | 1 | MS.Trt_vs_MS.NoTrt_F |
| VMAC | 0.49918 | -0.4141 | 0.626 | 0.65 | 1 | MS.Trt_vs_MS.NoTrt_F |
| GTPBP2 | 0.49936 | 0.75416 | 0.626 | 0.61 | 1 | MS.Trt_vs_MS.NoTrt_F |
| SLC15A2 | 0.4997 | 0.53017 | 0.486 | 0.55 | 1 | MS.Trt_vs_MS.NoTrt_F |
| MFAP5 | 0.4997 | -0.2853 | 0.32 | 0.34 | 1 | MS.Trt_vs_MS.NoTrt_F |
| EVI5 | 0.50004 | -0.285 | 0.791 | 0.87 | 1 | MS.Trt_vs_MS.NoTrt_F |
| RDH10 | 0.50178 | -0.6489 | 0.223 | 0.25 | 1 | MS.Trt_vs_MS.NoTrt_F |
| NEDD1 | 0.50232 | 0.97421 | 0.162 | 0.2 | 1 | MS.Trt_vs_MS.NoTrt_F |
| PLEKHG5 | 0.50298 | 1.48619 | 0.162 | 0.19 | 1 | MS.Trt_vs_MS.NoTrt_F |
| VWA5B1 | 0.50351 | 0.30743 | 0.594 | 0.62 | 1 | MS.Trt_vs_MS.NoTrt_F |
| TMEM143 | 0.50369 | 0.84922 | 0.482 | 0.55 | 1 | MS.Trt_vs_MS.NoTrt_F |
| TTC38 | 0.50389 | 0.59817 | 0.529 | 0.52 | 1 | MS.Trt_vs_MS.NoTrt_F |
| ZNF157 | 0.50509 | -0.2984 | 0.752 | 0.83 | 1 | MS.Trt_vs_MS.NoTrt_F |
| ZNF761 | 0.50557 | 1.02001 | 0.73 | 0.78 | 1 | MS.Trt_vs_MS.NoTrt_F |
| SYK | 0.50608 | 0.41402 | 0.442 | 0.45 | 1 | MS.Trt_vs_MS.NoTrt_F |
| CD160 | 0.50678 | 1.29639 | 0.266 | 0.32 | 1 | MS.Trt_vs_MS.NoTrt_F |
| FAM217B | 0.50701 | 0.52604 | 0.385 | 0.43 | 1 | MS.Trt_vs_MS.NoTrt_F |
| CBS | 0.50724 | 0.63189 | 0.241 | 0.23 | 1 | MS.Trt_vs_MS.NoTrt_F |
| ZKSCAN3 | 0.50782 | 0.56909 | 0.838 | 0.89 | 1 | MS.Trt_vs_MS.NoTrt_F |

| | | | | | | |
|---|---|---|---|---|---|---|
| PAX5 | 0.50856 | 0.74886 | 0.133 | 0.16 | 1 | MS.Trt_vs_MS.NoTrt_F |
| SLC39A7 | 0.50903 | 0.75192 | 0.133 | 0.16 | 1 | MS.Trt_vs_MS.NoTrt_F |
| CCHCR1 | 0.50908 | -1.0928 | 0.349 | 0.37 | 1 | MS.Trt_vs_MS.NoTrt_F |
| FABP4 | 0.50954 | 1.37628 | 0.169 | 0.2 | 1 | MS.Trt_vs_MS.NoTrt_F |
| SAMD4A | 0.5099 | 0.39006 | 0.59 | 0.65 | 1 | MS.Trt_vs_MS.NoTrt_F |
| DYNC1H1 | 0.51019 | 0.55433 | 0.942 | 0.93 | 1 | MS.Trt_vs_MS.NoTrt_F |
| EIF4G2 | 0.51036 | 0.29707 | 0.601 | 0.71 | 1 | MS.Trt_vs_MS.NoTrt_F |
| ZNF280C | 0.51076 | -0.5489 | 0.291 | 0.32 | 1 | MS.Trt_vs_MS.NoTrt_F |
| CBLB | 0.51123 | 0.89647 | 0.414 | 0.4 | 1 | MS.Trt_vs_MS.NoTrt_F |
| TNFRSF10B | 0.51133 | 0.63243 | 0.46 | 0.51 | 1 | MS.Trt_vs_MS.NoTrt_F |
| R3HDM2 | 0.51138 | 0.71253 | 0.763 | 0.81 | 1 | MS.Trt_vs_MS.NoTrt_F |
| RBMS1 | 0.51151 | 0.97997 | 0.608 | 0.71 | 1 | MS.Trt_vs_MS.NoTrt_F |
| TPX2 | 0.51209 | 0.79679 | 0.23 | 0.27 | 1 | MS.Trt_vs_MS.NoTrt_F |
| TBC1D10B | 0.5127 | 0.45169 | 0.723 | 0.73 | 1 | MS.Trt_vs_MS.NoTrt_F |
| WNK3 | 0.51278 | -0.4569 | 0.302 | 0.34 | 1 | MS.Trt_vs_MS.NoTrt_F |
| SYS1 | 0.51285 | 0.74516 | 0.475 | 0.46 | 1 | MS.Trt_vs_MS.NoTrt_F |
| FAM220A | 0.51291 | -0.7475 | 0.342 | 0.38 | 1 | MS.Trt_vs_MS.NoTrt_F |
| STX8 | 0.5136 | 0.40428 | 0.835 | 0.87 | 1 | MS.Trt_vs_MS.NoTrt_F |
| GNB4 | 0.51423 | 0.29289 | 0.809 | 0.84 | 1 | MS.Trt_vs_MS.NoTrt_F |
| GMCL1 | 0.51444 | 0.94923 | 0.356 | 0.38 | 1 | MS.Trt_vs_MS.NoTrt_F |
| GOLGA3 | 0.51459 | 0.50539 | 0.799 | 0.81 | 1 | MS.Trt_vs_MS.NoTrt_F |
| GFRAL | 0.5148 | 0.33683 | 0.392 | 0.39 | 1 | MS.Trt_vs_MS.NoTrt_F |
| PICK1 | 0.5156 | 0.27833 | 0.46 | 0.5 | 1 | MS.Trt_vs_MS.NoTrt_F |
| GAN | 0.51569 | 0.28405 | 0.838 | 0.85 | 1 | MS.Trt_vs_MS.NoTrt_F |
| PPP5D1 | 0.51631 | 0.49512 | 0.748 | 0.75 | 1 | MS.Trt_vs_MS.NoTrt_F |
| IFNAR1 | 0.51684 | 1.03139 | 0.504 | 0.51 | 1 | MS.Trt_vs_MS.NoTrt_F |
| UTP15 | 0.51755 | -0.8778 | 0.363 | 0.39 | 1 | MS.Trt_vs_MS.NoTrt_F |
| HSPA13 | 0.51758 | 0.41218 | 0.09 | 0.11 | 1 | MS.Trt_vs_MS.NoTrt_F |
| PEX2 | 0.51795 | -0.2531 | 0.601 | 0.6 | 1 | MS.Trt_vs_MS.NoTrt_F |
| VAPA | 0.51925 | 0.29801 | 0.716 | 0.77 | 1 | MS.Trt_vs_MS.NoTrt_F |
| C16orf52 | 0.52289 | 0.73254 | 0.094 | 0.12 | 1 | MS.Trt_vs_MS.NoTrt_F |
| SLC36A4 | 0.52317 | -0.5605 | 0.104 | 0.13 | 1 | MS.Trt_vs_MS.NoTrt_F |
| RP11-505K9 | 0.52327 | 0.34253 | 0.108 | 0.13 | 1 | MS.Trt_vs_MS.NoTrt_F |
| CCDC142 | 0.52354 | 0.32643 | 0.579 | 0.61 | 1 | MS.Trt_vs_MS.NoTrt_F |
| SERINC5 | 0.5244 | 0.47422 | 0.896 | 0.95 | 1 | MS.Trt_vs_MS.NoTrt_F |
| RAB11FIP1 | 0.5246 | 0.50486 | 0.86 | 0.83 | 1 | MS.Trt_vs_MS.NoTrt_F |
| ASRGL1 | 0.52496 | 0.4859 | 0.54 | 0.63 | 1 | MS.Trt_vs_MS.NoTrt_F |
| LRRC37A | 0.52558 | 0.38371 | 0.399 | 0.45 | 1 | MS.Trt_vs_MS.NoTrt_F |
| DENR | 0.52574 | 0.40636 | 0.73 | 0.77 | 1 | MS.Trt_vs_MS.NoTrt_F |
| LYRM4 | 0.52607 | -0.3174 | 0.942 | 0.98 | 1 | MS.Trt_vs_MS.NoTrt_F |

| | | | | | | |
|---|---|---|---|---|---|---|
| RBM45 | 0.52658 | 1.04847 | 0.126 | 0.16 | 1 | MS.Trt_vs_MS.NoTrt_F |
| DLK2 | 0.52713 | 0.57093 | 0.363 | 0.4 | 1 | MS.Trt_vs_MS.NoTrt_F |
| GBA2 | 0.52729 | 0.3606 | 0.694 | 0.61 | 1 | MS.Trt_vs_MS.NoTrt_F |
| ST8SIA1 | 0.52756 | 0.82592 | 0.604 | 0.64 | 1 | MS.Trt_vs_MS.NoTrt_F |
| COL1A2 | 0.52826 | 0.26071 | 0.392 | 0.44 | 1 | MS.Trt_vs_MS.NoTrt_F |
| ZNF574 | 0.52844 | 0.25063 | 0.392 | 0.39 | 1 | MS.Trt_vs_MS.NoTrt_F |
| IL23R | 0.52954 | -1.4898 | 0.388 | 0.4 | 1 | MS.Trt_vs_MS.NoTrt_F |
| CLU | 0.53072 | 1.29462 | 0.547 | 0.6 | 1 | MS.Trt_vs_MS.NoTrt_F |
| TMEM231 | 0.5311 | 0.56842 | 0.766 | 0.82 | 1 | MS.Trt_vs_MS.NoTrt_F |
| HEXDC | 0.53167 | 0.55348 | 0.371 | 0.4 | 1 | MS.Trt_vs_MS.NoTrt_F |
| IDO1 | 0.53239 | 0.29442 | 0.532 | 0.54 | 1 | MS.Trt_vs_MS.NoTrt_F |
| CDK4 | 0.53246 | 1.04788 | 0.558 | 0.64 | 1 | MS.Trt_vs_MS.NoTrt_F |
| GALNT11 | 0.53254 | 0.32352 | 0.468 | 0.44 | 1 | MS.Trt_vs_MS.NoTrt_F |
| ZNF547 | 0.53276 | 1.47398 | 0.295 | 0.29 | 1 | MS.Trt_vs_MS.NoTrt_F |
| ERLIN2 | 0.53352 | 0.34175 | 0.385 | 0.42 | 1 | MS.Trt_vs_MS.NoTrt_F |
| SLC35A3 | 0.53374 | -0.2586 | 0.82 | 0.82 | 1 | MS.Trt_vs_MS.NoTrt_F |
| METTL15 | 0.53394 | -0.2714 | 0.414 | 0.42 | 1 | MS.Trt_vs_MS.NoTrt_F |
| NFKBIA | 0.53399 | 0.84208 | 0.309 | 0.38 | 1 | MS.Trt_vs_MS.NoTrt_F |
| PSIP1 | 0.5341 | 0.34331 | 0.759 | 0.82 | 1 | MS.Trt_vs_MS.NoTrt_F |
| C12orf57 | 0.53431 | 0.38498 | 0.403 | 0.41 | 1 | MS.Trt_vs_MS.NoTrt_F |
| L3MBTL1 | 0.53438 | 0.41842 | 0.651 | 0.65 | 1 | MS.Trt_vs_MS.NoTrt_F |
| SLAMF7 | 0.5346 | 1.29653 | 0.381 | 0.46 | 1 | MS.Trt_vs_MS.NoTrt_F |
| POFUT1 | 0.53513 | 0.67371 | 0.554 | 0.6 | 1 | MS.Trt_vs_MS.NoTrt_F |
| TRIM16 | 0.53573 | 0.26581 | 0.468 | 0.49 | 1 | MS.Trt_vs_MS.NoTrt_F |
| SEPSECS | 0.53577 | -0.4184 | 0.759 | 0.79 | 1 | MS.Trt_vs_MS.NoTrt_F |
| SPAG9 | 0.53652 | -0.4287 | 0.878 | 0.91 | 1 | MS.Trt_vs_MS.NoTrt_F |
| E2F6 | 0.53761 | -1.0754 | 0.309 | 0.34 | 1 | MS.Trt_vs_MS.NoTrt_F |
| ELAC1 | 0.53835 | -0.5981 | 0.547 | 0.58 | 1 | MS.Trt_vs_MS.NoTrt_F |
| PRCP | 0.53955 | 0.84021 | 0.845 | 0.79 | 1 | MS.Trt_vs_MS.NoTrt_F |
| PIGA | 0.5398 | -0.2544 | 0.457 | 0.51 | 1 | MS.Trt_vs_MS.NoTrt_F |
| PLPP2 | 0.54085 | 0.61322 | 0.914 | 0.87 | 1 | MS.Trt_vs_MS.NoTrt_F |
| TONSL | 0.54143 | 0.7427 | 0.622 | 0.66 | 1 | MS.Trt_vs_MS.NoTrt_F |
| FAM3A | 0.54196 | 0.44614 | 0.791 | 0.8 | 1 | MS.Trt_vs_MS.NoTrt_F |
| SLC35G2 | 0.54248 | 0.7875 | 0.219 | 0.25 | 1 | MS.Trt_vs_MS.NoTrt_F |
| KIFC3 | 0.54255 | 0.2955 | 0.576 | 0.67 | 1 | MS.Trt_vs_MS.NoTrt_F |
| RBPMS | 0.54257 | 1.4488 | 0.155 | 0.14 | 1 | MS.Trt_vs_MS.NoTrt_F |
| PITPNM2 | 0.54355 | 0.7755 | 0.414 | 0.48 | 1 | MS.Trt_vs_MS.NoTrt_F |
| RPUSD2 | 0.54377 | 0.72263 | 0.155 | 0.18 | 1 | MS.Trt_vs_MS.NoTrt_F |
| CEP41 | 0.54394 | -0.261 | 0.781 | 0.82 | 1 | MS.Trt_vs_MS.NoTrt_F |
| SPRYD4 | 0.54437 | 0.50307 | 0.856 | 0.93 | 1 | MS.Trt_vs_MS.NoTrt_F |

| | | | | | | |
|---|---|---|---|---|---|---|
| CFD | 0.5445 | 0.53298 | 0.223 | 0.24 | 1 | MS.Trt_vs_MS.NoTrt_F |
| GOT2 | 0.54453 | 0.3327 | 0.86 | 0.81 | 1 | MS.Trt_vs_MS.NoTrt_F |
| OCLN | 0.54516 | 0.55336 | 0.781 | 0.82 | 1 | MS.Trt_vs_MS.NoTrt_F |
| VNN2 | 0.5453 | 1.80533 | 0.223 | 0.2 | 1 | MS.Trt_vs_MS.NoTrt_F |
| BLOC1S4 | 0.54693 | 0.40538 | 0.205 | 0.23 | 1 | MS.Trt_vs_MS.NoTrt_F |
| PTPRF | 0.54749 | 0.31146 | 0.356 | 0.36 | 1 | MS.Trt_vs_MS.NoTrt_F |
| COMMD4 | 0.5476 | 0.37996 | 0.906 | 0.95 | 1 | MS.Trt_vs_MS.NoTrt_F |
| AC016577.1 | 0.54769 | 0.27216 | 0.417 | 0.5 | 1 | MS.Trt_vs_MS.NoTrt_F |
| WWTR1 | 0.5477 | 0.57429 | 0.468 | 0.52 | 1 | MS.Trt_vs_MS.NoTrt_F |
| NUCKS1 | 0.54835 | 0.89499 | 0.522 | 0.6 | 1 | MS.Trt_vs_MS.NoTrt_F |
| ALDH1B1 | 0.54883 | 0.28582 | 0.626 | 0.67 | 1 | MS.Trt_vs_MS.NoTrt_F |
| GLT8D1 | 0.54965 | 0.3127 | 0.302 | 0.33 | 1 | MS.Trt_vs_MS.NoTrt_F |
| MUC20 | 0.5497 | -0.3662 | 0.543 | 0.59 | 1 | MS.Trt_vs_MS.NoTrt_F |
| CCDC90B | 0.55065 | 0.39574 | 0.906 | 0.95 | 1 | MS.Trt_vs_MS.NoTrt_F |
| ST7L | 0.55114 | 0.43652 | 0.457 | 0.48 | 1 | MS.Trt_vs_MS.NoTrt_F |
| COG1 | 0.55173 | -0.3374 | 0.853 | 0.85 | 1 | MS.Trt_vs_MS.NoTrt_F |
| PRKAR2A | 0.55208 | -0.5709 | 0.867 | 0.93 | 1 | MS.Trt_vs_MS.NoTrt_F |
| IVD | 0.55218 | 0.32395 | 0.935 | 0.93 | 1 | MS.Trt_vs_MS.NoTrt_F |
| YES1 | 0.55269 | 0.26138 | 0.367 | 0.41 | 1 | MS.Trt_vs_MS.NoTrt_F |
| FABP2 | 0.55273 | 0.35629 | 0.421 | 0.44 | 1 | MS.Trt_vs_MS.NoTrt_F |
| TIE1 | 0.55327 | 0.7335 | 0.615 | 0.59 | 1 | MS.Trt_vs_MS.NoTrt_F |
| ORC2 | 0.55439 | 0.39415 | 0.673 | 0.73 | 1 | MS.Trt_vs_MS.NoTrt_F |
| TCEANC | 0.55449 | -0.3741 | 0.651 | 0.67 | 1 | MS.Trt_vs_MS.NoTrt_F |
| NME4 | 0.55512 | 0.29151 | 0.176 | 0.2 | 1 | MS.Trt_vs_MS.NoTrt_F |
| COQ10A | 0.5555 | -0.8386 | 0.543 | 0.54 | 1 | MS.Trt_vs_MS.NoTrt_F |
| TMEM71 | 0.55678 | 1.00179 | 0.119 | 0.14 | 1 | MS.Trt_vs_MS.NoTrt_F |
| KATNBL1 | 0.55679 | 0.57668 | 0.917 | 0.92 | 1 | MS.Trt_vs_MS.NoTrt_F |
| SEC31B | 0.55744 | 0.45667 | 0.435 | 0.49 | 1 | MS.Trt_vs_MS.NoTrt_F |
| WDR13 | 0.55864 | 0.32554 | 0.813 | 0.87 | 1 | MS.Trt_vs_MS.NoTrt_F |
| ASB2 | 0.55881 | 1.46826 | 0.104 | 0.13 | 1 | MS.Trt_vs_MS.NoTrt_F |
| ZNF160 | 0.55885 | 0.58293 | 0.536 | 0.59 | 1 | MS.Trt_vs_MS.NoTrt_F |
| LIN7B | 0.55937 | -0.2981 | 0.327 | 0.37 | 1 | MS.Trt_vs_MS.NoTrt_F |
| PAPOLG | 0.55951 | 1.05683 | 0.73 | 0.8 | 1 | MS.Trt_vs_MS.NoTrt_F |
| AGPAT3 | 0.55967 | 0.34021 | 0.655 | 0.68 | 1 | MS.Trt_vs_MS.NoTrt_F |
| TNRC6B | 0.55969 | 0.26207 | 0.96 | 0.98 | 1 | MS.Trt_vs_MS.NoTrt_F |
| GBA | 0.55972 | -0.3679 | 0.482 | 0.5 | 1 | MS.Trt_vs_MS.NoTrt_F |
| GPR107 | 0.55987 | 0.40782 | 0.209 | 0.23 | 1 | MS.Trt_vs_MS.NoTrt_F |
| CTD-3088G3 | 0.56079 | -0.4169 | 0.5 | 0.56 | 1 | MS.Trt_vs_MS.NoTrt_F |
| EXOC8 | 0.56165 | 0.25809 | 0.406 | 0.46 | 1 | MS.Trt_vs_MS.NoTrt_F |
| CFI | 0.56172 | 0.25358 | 0.561 | 0.61 | 1 | MS.Trt_vs_MS.NoTrt_F |

| | | | | | | |
|---|---|---|---|---|---|---|
| LRRC1 | 0.56184 | 0.57448 | 0.317 | 0.35 | 1 | MS.Trt_vs_MS.NoTrt_F |
| SLC39A11 | 0.56227 | 1.0907 | 0.32 | 0.38 | 1 | MS.Trt_vs_MS.NoTrt_F |
| CPT1A | 0.56328 | 0.61837 | 0.68 | 0.7 | 1 | MS.Trt_vs_MS.NoTrt_F |
| CHRNA3 | 0.56358 | -0.3113 | 0.223 | 0.24 | 1 | MS.Trt_vs_MS.NoTrt_F |
| RBBP5 | 0.56376 | 0.8746 | 0.568 | 0.63 | 1 | MS.Trt_vs_MS.NoTrt_F |
| STRN3 | 0.5669 | 0.61111 | 0.831 | 0.83 | 1 | MS.Trt_vs_MS.NoTrt_F |
| SLC31A2 | 0.5676 | 1.73928 | 0.133 | 0.16 | 1 | MS.Trt_vs_MS.NoTrt_F |
| CHST14 | 0.56863 | 0.92 | 0.245 | 0.28 | 1 | MS.Trt_vs_MS.NoTrt_F |
| GNE | 0.56922 | 0.69642 | 0.227 | 0.27 | 1 | MS.Trt_vs_MS.NoTrt_F |
| LRP12 | 0.56993 | 0.38777 | 0.604 | 0.67 | 1 | MS.Trt_vs_MS.NoTrt_F |
| ZSCAN20 | 0.57004 | 0.97959 | 0.302 | 0.32 | 1 | MS.Trt_vs_MS.NoTrt_F |
| TMTC3 | 0.5715 | 1.17103 | 0.241 | 0.28 | 1 | MS.Trt_vs_MS.NoTrt_F |
| SLFN13 | 0.57215 | 1.5951 | 0.626 | 0.67 | 1 | MS.Trt_vs_MS.NoTrt_F |
| PXMP4 | 0.5722 | 0.27525 | 0.849 | 0.87 | 1 | MS.Trt_vs_MS.NoTrt_F |
| THEMIS | 0.57329 | 1.04835 | 0.277 | 0.31 | 1 | MS.Trt_vs_MS.NoTrt_F |
| DHRS3 | 0.57429 | 2.01865 | 0.252 | 0.25 | 1 | MS.Trt_vs_MS.NoTrt_F |
| ZNF230 | 0.57494 | 0.44572 | 0.572 | 0.62 | 1 | MS.Trt_vs_MS.NoTrt_F |
| CA5B | 0.57538 | 0.39397 | 0.838 | 0.9 | 1 | MS.Trt_vs_MS.NoTrt_F |
| DIEXF | 0.57568 | 0.55841 | 0.525 | 0.49 | 1 | MS.Trt_vs_MS.NoTrt_F |
| ACTR3B | 0.57606 | 0.29791 | 0.496 | 0.5 | 1 | MS.Trt_vs_MS.NoTrt_F |
| FAM184B | 0.57618 | -0.3514 | 0.155 | 0.18 | 1 | MS.Trt_vs_MS.NoTrt_F |
| BCAS4 | 0.57659 | -0.3547 | 0.935 | 0.96 | 1 | MS.Trt_vs_MS.NoTrt_F |
| DRAM2 | 0.5766 | 0.97664 | 0.209 | 0.2 | 1 | MS.Trt_vs_MS.NoTrt_F |
| ZNF174 | 0.57681 | 2.52733 | 0.086 | 0.11 | 1 | MS.Trt_vs_MS.NoTrt_F |
| CASP9 | 0.57789 | 0.28943 | 0.374 | 0.4 | 1 | MS.Trt_vs_MS.NoTrt_F |
| PQLC2 | 0.57805 | -0.3677 | 0.403 | 0.45 | 1 | MS.Trt_vs_MS.NoTrt_F |
| CYB5D1 | 0.57841 | -1.2978 | 0.187 | 0.21 | 1 | MS.Trt_vs_MS.NoTrt_F |
| NEDD4L | 0.57848 | -0.9385 | 0.532 | 0.46 | 1 | MS.Trt_vs_MS.NoTrt_F |
| ABCB7 | 0.57918 | -0.316 | 0.705 | 0.72 | 1 | MS.Trt_vs_MS.NoTrt_F |
| SGPL1 | 0.57973 | 0.36159 | 0.511 | 0.52 | 1 | MS.Trt_vs_MS.NoTrt_F |
| MKLN1 | 0.58046 | 0.76218 | 0.576 | 0.66 | 1 | MS.Trt_vs_MS.NoTrt_F |
| TBL1X | 0.5807 | -0.6263 | 0.576 | 0.6 | 1 | MS.Trt_vs_MS.NoTrt_F |
| APP | 0.58193 | 0.89627 | 0.629 | 0.72 | 1 | MS.Trt_vs_MS.NoTrt_F |
| MTSS1 | 0.58239 | 1.13458 | 0.59 | 0.67 | 1 | MS.Trt_vs_MS.NoTrt_F |
| CORO2A | 0.585 | 0.36659 | 0.795 | 0.78 | 1 | MS.Trt_vs_MS.NoTrt_F |
| C21orf59 | 0.58508 | 0.48426 | 0.442 | 0.45 | 1 | MS.Trt_vs_MS.NoTrt_F |
| ZNF813 | 0.58514 | 0.97575 | 0.773 | 0.77 | 1 | MS.Trt_vs_MS.NoTrt_F |
| CD24 | 0.58519 | 0.29246 | 0.41 | 0.41 | 1 | MS.Trt_vs_MS.NoTrt_F |
| CRK | 0.58534 | 0.46047 | 0.626 | 0.69 | 1 | MS.Trt_vs_MS.NoTrt_F |
| PAN2 | 0.58591 | 0.78065 | 0.367 | 0.4 | 1 | MS.Trt_vs_MS.NoTrt_F |

| | | | | | | |
|---|---|---|---|---|---|---|
| IRAK2 | 0.58601 | -0.3783 | 0.353 | 0.38 | 1 | MS.Trt_vs_MS.NoTrt_F |
| FBXL22 | 0.58646 | 0.61936 | 0.14 | 0.16 | 1 | MS.Trt_vs_MS.NoTrt_F |
| BTBD8 | 0.58647 | 0.49453 | 0.32 | 0.36 | 1 | MS.Trt_vs_MS.NoTrt_F |
| EVI5L | 0.58659 | 0.2753 | 0.306 | 0.34 | 1 | MS.Trt_vs_MS.NoTrt_F |
| FAM208A | 0.58704 | 0.74607 | 0.737 | 0.73 | 1 | MS.Trt_vs_MS.NoTrt_F |
| BRPF3 | 0.58724 | 2.24241 | 0.101 | 0.12 | 1 | MS.Trt_vs_MS.NoTrt_F |
| CASC10 | 0.58731 | 0.25049 | 0.446 | 0.42 | 1 | MS.Trt_vs_MS.NoTrt_F |
| UIMC1 | 0.58778 | -0.4335 | 0.835 | 0.86 | 1 | MS.Trt_vs_MS.NoTrt_F |
| GNAS | 0.58943 | 0.75802 | 0.511 | 0.57 | 1 | MS.Trt_vs_MS.NoTrt_F |
| AKT3 | 0.58954 | 0.36178 | 0.712 | 0.76 | 1 | MS.Trt_vs_MS.NoTrt_F |
| IGFL2 | 0.58956 | 0.42512 | 0.399 | 0.47 | 1 | MS.Trt_vs_MS.NoTrt_F |
| ZNF778 | 0.5901 | 0.36564 | 0.637 | 0.66 | 1 | MS.Trt_vs_MS.NoTrt_F |
| POGZ | 0.59096 | 0.51509 | 0.856 | 0.86 | 1 | MS.Trt_vs_MS.NoTrt_F |
| LAMA5 | 0.59136 | 1.06148 | 0.277 | 0.32 | 1 | MS.Trt_vs_MS.NoTrt_F |
| FAU | 0.59239 | 0.26109 | 0.727 | 0.94 | 1 | MS.Trt_vs_MS.NoTrt_F |
| C11orf63 | 0.59259 | 0.51069 | 0.475 | 0.51 | 1 | MS.Trt_vs_MS.NoTrt_F |
| KLHDC2 | 0.59295 | 1.11961 | 0.27 | 0.3 | 1 | MS.Trt_vs_MS.NoTrt_F |
| GTF2IRD1 | 0.59435 | 0.25599 | 0.701 | 0.72 | 1 | MS.Trt_vs_MS.NoTrt_F |
| ZNF577 | 0.59447 | 0.88398 | 0.381 | 0.42 | 1 | MS.Trt_vs_MS.NoTrt_F |
| TOR1A | 0.59455 | -0.4176 | 0.752 | 0.78 | 1 | MS.Trt_vs_MS.NoTrt_F |
| CD300LG | 0.59467 | 0.33905 | 0.435 | 0.46 | 1 | MS.Trt_vs_MS.NoTrt_F |
| GRN | 0.59498 | 1.53246 | 0.165 | 0.19 | 1 | MS.Trt_vs_MS.NoTrt_F |
| KLHDC10 | 0.59683 | -0.3732 | 0.41 | 0.4 | 1 | MS.Trt_vs_MS.NoTrt_F |
| MT-ND2 | 0.59687 | 0.37428 | 0.838 | 0.89 | 1 | MS.Trt_vs_MS.NoTrt_F |
| MACROD2 | 0.59707 | 0.31011 | 0.662 | 0.68 | 1 | MS.Trt_vs_MS.NoTrt_F |
| ADGRA3 | 0.59774 | 0.37433 | 0.428 | 0.41 | 1 | MS.Trt_vs_MS.NoTrt_F |
| POU5F1 | 0.59781 | 0.35423 | 0.554 | 0.55 | 1 | MS.Trt_vs_MS.NoTrt_F |
| PIN1 | 0.59843 | 0.65641 | 0.781 | 0.79 | 1 | MS.Trt_vs_MS.NoTrt_F |
| GSR | 0.59957 | 0.72081 | 0.277 | 0.32 | 1 | MS.Trt_vs_MS.NoTrt_F |
| TSPYL1 | 0.60018 | 0.63604 | 0.622 | 0.69 | 1 | MS.Trt_vs_MS.NoTrt_F |
| ZNF780B | 0.60056 | -0.4469 | 0.644 | 0.65 | 1 | MS.Trt_vs_MS.NoTrt_F |
| SCARB1 | 0.60075 | -0.2854 | 0.478 | 0.51 | 1 | MS.Trt_vs_MS.NoTrt_F |
| ZC3H4 | 0.60119 | 0.61355 | 0.371 | 0.46 | 1 | MS.Trt_vs_MS.NoTrt_F |
| SLC30A7 | 0.6014 | 0.81517 | 0.68 | 0.74 | 1 | MS.Trt_vs_MS.NoTrt_F |
| IMPACT | 0.60193 | 1.37012 | 0.097 | 0.12 | 1 | MS.Trt_vs_MS.NoTrt_F |
| CTPS1 | 0.60285 | -0.3247 | 0.658 | 0.66 | 1 | MS.Trt_vs_MS.NoTrt_F |
| GALNT10 | 0.60401 | -0.3607 | 0.55 | 0.6 | 1 | MS.Trt_vs_MS.NoTrt_F |
| HOMEZ | 0.6042 | 0.56188 | 0.295 | 0.32 | 1 | MS.Trt_vs_MS.NoTrt_F |
| GGCX | 0.60476 | 0.53027 | 0.899 | 0.95 | 1 | MS.Trt_vs_MS.NoTrt_F |
| ZNF26 | 0.60482 | 0.35233 | 0.989 | 0.99 | 1 | MS.Trt_vs_MS.NoTrt_F |

| | | | | | | |
|---|---|---|---|---|---|---|
| TNFRSF25 | 0.60502 | 0.69496 | 0.676 | 0.75 | 1 | MS.Trt_vs_MS.NoTrt_F |
| PCCA | 0.60507 | 0.38094 | 0.608 | 0.66 | 1 | MS.Trt_vs_MS.NoTrt_F |
| SBNO1 | 0.60605 | 0.6088 | 0.655 | 0.7 | 1 | MS.Trt_vs_MS.NoTrt_F |
| CAPN12 | 0.60607 | 0.83183 | 0.493 | 0.54 | 1 | MS.Trt_vs_MS.NoTrt_F |
| NPIPB15 | 0.60687 | 1.08357 | 0.331 | 0.33 | 1 | MS.Trt_vs_MS.NoTrt_F |
| DDX52 | 0.60853 | 0.33894 | 0.896 | 0.89 | 1 | MS.Trt_vs_MS.NoTrt_F |
| FAM200A | 0.60878 | -0.4455 | 0.241 | 0.26 | 1 | MS.Trt_vs_MS.NoTrt_F |
| FAM83H | 0.60888 | -0.3159 | 0.144 | 0.16 | 1 | MS.Trt_vs_MS.NoTrt_F |
| MEF2A | 0.61067 | 0.39072 | 0.5 | 0.55 | 1 | MS.Trt_vs_MS.NoTrt_F |
| HAUS2 | 0.61092 | 0.2524 | 0.824 | 0.82 | 1 | MS.Trt_vs_MS.NoTrt_F |
| CCDC114 | 0.61131 | -0.7992 | 0.504 | 0.49 | 1 | MS.Trt_vs_MS.NoTrt_F |
| BRD4 | 0.6141 | 1.07163 | 0.317 | 0.37 | 1 | MS.Trt_vs_MS.NoTrt_F |
| ZNF418 | 0.61419 | 0.51375 | 0.205 | 0.23 | 1 | MS.Trt_vs_MS.NoTrt_F |
| EPN1 | 0.61523 | 0.26066 | 0.921 | 0.96 | 1 | MS.Trt_vs_MS.NoTrt_F |
| EXOG | 0.61539 | 0.43028 | 0.594 | 0.58 | 1 | MS.Trt_vs_MS.NoTrt_F |
| DFFB | 0.61544 | 0.39015 | 0.827 | 0.79 | 1 | MS.Trt_vs_MS.NoTrt_F |
| RPL35A | 0.61579 | 0.38286 | 0.842 | 0.98 | 1 | MS.Trt_vs_MS.NoTrt_F |
| IFI27 | 0.61583 | 0.68742 | 0.442 | 0.46 | 1 | MS.Trt_vs_MS.NoTrt_F |
| ZFP82 | 0.61612 | 0.29051 | 0.712 | 0.75 | 1 | MS.Trt_vs_MS.NoTrt_F |
| SOS2 | 0.61682 | 1.52744 | 0.115 | 0.14 | 1 | MS.Trt_vs_MS.NoTrt_F |
| SPTSSA | 0.61708 | -0.3297 | 0.363 | 0.41 | 1 | MS.Trt_vs_MS.NoTrt_F |
| CCNF | 0.61733 | -0.7736 | 0.504 | 0.53 | 1 | MS.Trt_vs_MS.NoTrt_F |
| STAT2 | 0.61791 | 0.36456 | 0.892 | 0.93 | 1 | MS.Trt_vs_MS.NoTrt_F |
| TMEM33 | 0.61839 | 0.26885 | 0.914 | 0.88 | 1 | MS.Trt_vs_MS.NoTrt_F |
| MPP5 | 0.6196 | -0.4322 | 0.342 | 0.39 | 1 | MS.Trt_vs_MS.NoTrt_F |
| ZCWPW2 | 0.62084 | 0.35162 | 0.493 | 0.5 | 1 | MS.Trt_vs_MS.NoTrt_F |
| HEG1 | 0.62096 | 1.00787 | 0.126 | 0.14 | 1 | MS.Trt_vs_MS.NoTrt_F |
| MBD3 | 0.62174 | 0.70206 | 0.712 | 0.77 | 1 | MS.Trt_vs_MS.NoTrt_F |
| COMMD9 | 0.62246 | 0.4596 | 0.486 | 0.46 | 1 | MS.Trt_vs_MS.NoTrt_F |
| KCNN2 | 0.62353 | 0.34094 | 0.317 | 0.32 | 1 | MS.Trt_vs_MS.NoTrt_F |
| ZNF526 | 0.62372 | -0.2582 | 0.504 | 0.56 | 1 | MS.Trt_vs_MS.NoTrt_F |
| CACNB2 | 0.62405 | 0.65384 | 0.471 | 0.49 | 1 | MS.Trt_vs_MS.NoTrt_F |
| RNF14 | 0.62606 | -0.665 | 0.576 | 0.62 | 1 | MS.Trt_vs_MS.NoTrt_F |
| GPR161 | 0.62626 | 0.86463 | 0.406 | 0.46 | 1 | MS.Trt_vs_MS.NoTrt_F |
| PRICKLE3 | 0.62631 | 0.26163 | 0.504 | 0.53 | 1 | MS.Trt_vs_MS.NoTrt_F |
| SSR3 | 0.62867 | 0.38537 | 0.601 | 0.66 | 1 | MS.Trt_vs_MS.NoTrt_F |
| ENTPD5 | 0.62945 | 1.06989 | 0.543 | 0.53 | 1 | MS.Trt_vs_MS.NoTrt_F |
| ZNF169 | 0.63031 | 0.29354 | 0.849 | 0.87 | 1 | MS.Trt_vs_MS.NoTrt_F |
| SLC37A2 | 0.63068 | 0.33552 | 0.511 | 0.55 | 1 | MS.Trt_vs_MS.NoTrt_F |
| RAD1 | 0.63079 | 0.55188 | 0.701 | 0.74 | 1 | MS.Trt_vs_MS.NoTrt_F |

| | | | | | | |
|---|---|---|---|---|---|---|
| PLOD1 | 0.6311 | 0.95158 | 0.68 | 0.66 | 1 | MS.Trt_vs_MS.NoTrt_F |
| CMC2 | 0.63144 | 0.53917 | 0.881 | 0.89 | 1 | MS.Trt_vs_MS.NoTrt_F |
| SLC14A1 | 0.63182 | 1.06785 | 0.504 | 0.54 | 1 | MS.Trt_vs_MS.NoTrt_F |
| RABAC1 | 0.63191 | 1.06131 | 0.273 | 0.33 | 1 | MS.Trt_vs_MS.NoTrt_F |
| SMARCB1 | 0.63268 | -0.3385 | 0.55 | 0.54 | 1 | MS.Trt_vs_MS.NoTrt_F |
| EXD1 | 0.63297 | 0.27875 | 0.54 | 0.55 | 1 | MS.Trt_vs_MS.NoTrt_F |
| MAVS | 0.63334 | -0.4183 | 0.95 | 0.94 | 1 | MS.Trt_vs_MS.NoTrt_F |
| IFNLR1 | 0.63452 | -1.8914 | 0.144 | 0.17 | 1 | MS.Trt_vs_MS.NoTrt_F |
| MED18 | 0.63566 | 0.62896 | 0.615 | 0.66 | 1 | MS.Trt_vs_MS.NoTrt_F |
| DCTN3 | 0.63602 | 0.32192 | 0.446 | 0.49 | 1 | MS.Trt_vs_MS.NoTrt_F |
| PTCD2 | 0.63612 | 1.0455 | 0.608 | 0.61 | 1 | MS.Trt_vs_MS.NoTrt_F |
| CDCA3 | 0.63726 | 0.50045 | 0.565 | 0.63 | 1 | MS.Trt_vs_MS.NoTrt_F |
| PIGM | 0.63741 | 0.48815 | 0.622 | 0.59 | 1 | MS.Trt_vs_MS.NoTrt_F |
| MRPS22 | 0.63792 | 0.28111 | 0.565 | 0.62 | 1 | MS.Trt_vs_MS.NoTrt_F |
| SPARCL1 | 0.63825 | 1.80326 | 0.18 | 0.17 | 1 | MS.Trt_vs_MS.NoTrt_F |
| NCAPG2 | 0.63858 | 2.57816 | 0.273 | 0.28 | 1 | MS.Trt_vs_MS.NoTrt_F |
| C17orf62 | 0.63955 | 0.42247 | 0.745 | 0.76 | 1 | MS.Trt_vs_MS.NoTrt_F |
| NUP155 | 0.64065 | 0.63961 | 0.917 | 0.91 | 1 | MS.Trt_vs_MS.NoTrt_F |
| P2RX5-TAX1 | 0.64203 | 0.46882 | 0.845 | 0.87 | 1 | MS.Trt_vs_MS.NoTrt_F |
| ST3GAL2 | 0.64214 | 0.54592 | 0.914 | 0.99 | 1 | MS.Trt_vs_MS.NoTrt_F |
| ANKRD16 | 0.64281 | 1.2142 | 0.46 | 0.49 | 1 | MS.Trt_vs_MS.NoTrt_F |
| SPTB | 0.64375 | 0.39342 | 0.453 | 0.5 | 1 | MS.Trt_vs_MS.NoTrt_F |
| TEN1-CDK3 | 0.64646 | -0.3569 | 0.583 | 0.56 | 1 | MS.Trt_vs_MS.NoTrt_F |
| TM2D2 | 0.647 | 1.37759 | 0.392 | 0.4 | 1 | MS.Trt_vs_MS.NoTrt_F |
| ZNF639 | 0.64701 | -0.3682 | 0.871 | 0.93 | 1 | MS.Trt_vs_MS.NoTrt_F |
| DNAJB4 | 0.6488 | 0.36602 | 0.396 | 0.41 | 1 | MS.Trt_vs_MS.NoTrt_F |
| SLX1B | 0.64885 | 0.70745 | 0.496 | 0.49 | 1 | MS.Trt_vs_MS.NoTrt_F |
| CORO1C | 0.65169 | 1.18311 | 0.094 | 0.11 | 1 | MS.Trt_vs_MS.NoTrt_F |
| TRAT1 | 0.65254 | 1.14138 | 0.237 | 0.24 | 1 | MS.Trt_vs_MS.NoTrt_F |
| POLR2J3 | 0.65346 | 0.37039 | 0.986 | 0.99 | 1 | MS.Trt_vs_MS.NoTrt_F |
| BIRC5 | 0.65528 | 0.48541 | 0.655 | 0.72 | 1 | MS.Trt_vs_MS.NoTrt_F |
| FBXL4 | 0.65536 | 0.8486 | 0.525 | 0.58 | 1 | MS.Trt_vs_MS.NoTrt_F |
| D2HGDH | 0.65843 | 0.28962 | 0.482 | 0.45 | 1 | MS.Trt_vs_MS.NoTrt_F |
| COX19 | 0.65972 | 0.43258 | 0.601 | 0.66 | 1 | MS.Trt_vs_MS.NoTrt_F |
| RWDD2A | 0.66065 | 0.45713 | 0.5 | 0.56 | 1 | MS.Trt_vs_MS.NoTrt_F |
| ZNF737 | 0.66152 | -0.3303 | 0.849 | 0.88 | 1 | MS.Trt_vs_MS.NoTrt_F |
| MT-CO3 | 0.66176 | 0.5106 | 0.608 | 0.67 | 1 | MS.Trt_vs_MS.NoTrt_F |
| RBM44 | 0.66225 | 0.8082 | 0.27 | 0.29 | 1 | MS.Trt_vs_MS.NoTrt_F |
| ZSWIM1 | 0.66306 | 0.36565 | 0.737 | 0.72 | 1 | MS.Trt_vs_MS.NoTrt_F |
| PECAM1 | 0.66334 | 0.34389 | 0.745 | 0.76 | 1 | MS.Trt_vs_MS.NoTrt_F |

| | | | | | | |
|---|---|---|---|---|---|---|
| KDM5B | 0.66388 | 0.63613 | 0.371 | 0.35 | 1 | MS.Trt_vs_MS.NoTrt_F |
| SMU1 | 0.66444 | 0.64587 | 0.773 | 0.82 | 1 | MS.Trt_vs_MS.NoTrt_F |
| TGM2 | 0.66453 | 0.27651 | 0.608 | 0.58 | 1 | MS.Trt_vs_MS.NoTrt_F |
| LGALS1 | 0.66468 | 0.97439 | 0.378 | 0.47 | 1 | MS.Trt_vs_MS.NoTrt_F |
| EIF1AX | 0.66521 | 0.99501 | 0.252 | 0.29 | 1 | MS.Trt_vs_MS.NoTrt_F |
| RPS29 | 0.66523 | 0.43423 | 0.896 | 0.94 | 1 | MS.Trt_vs_MS.NoTrt_F |
| CCDC180 | 0.66576 | -0.5266 | 0.709 | 0.78 | 1 | MS.Trt_vs_MS.NoTrt_F |
| SLCO4C1 | 0.66598 | 1.1815 | 0.194 | 0.22 | 1 | MS.Trt_vs_MS.NoTrt_F |
| PPM1L | 0.66629 | -0.8253 | 0.313 | 0.33 | 1 | MS.Trt_vs_MS.NoTrt_F |
| ZNF682 | 0.66728 | -0.822 | 0.403 | 0.38 | 1 | MS.Trt_vs_MS.NoTrt_F |
| RGS11 | 0.66833 | 0.28406 | 0.18 | 0.2 | 1 | MS.Trt_vs_MS.NoTrt_F |
| C14orf79 | 0.66859 | 0.60353 | 0.594 | 0.63 | 1 | MS.Trt_vs_MS.NoTrt_F |
| CDC42BPG | 0.67063 | 0.25492 | 0.543 | 0.56 | 1 | MS.Trt_vs_MS.NoTrt_F |
| MTA3 | 0.67073 | 0.72546 | 0.644 | 0.7 | 1 | MS.Trt_vs_MS.NoTrt_F |
| MRPS28 | 0.67098 | -0.3594 | 0.59 | 0.63 | 1 | MS.Trt_vs_MS.NoTrt_F |
| SHFM1 | 0.67102 | 0.468 | 0.91 | 0.92 | 1 | MS.Trt_vs_MS.NoTrt_F |
| C19orf60 | 0.67133 | 0.31812 | 0.888 | 0.87 | 1 | MS.Trt_vs_MS.NoTrt_F |
| F11R | 0.67139 | 0.82625 | 0.615 | 0.66 | 1 | MS.Trt_vs_MS.NoTrt_F |
| TPST2 | 0.67208 | 0.67753 | 0.619 | 0.65 | 1 | MS.Trt_vs_MS.NoTrt_F |
| CLNK | 0.67273 | 0.49836 | 0.687 | 0.69 | 1 | MS.Trt_vs_MS.NoTrt_F |
| COA7 | 0.67421 | -0.5786 | 0.906 | 0.9 | 1 | MS.Trt_vs_MS.NoTrt_F |
| TVP23A | 0.67488 | 0.4341 | 0.507 | 0.54 | 1 | MS.Trt_vs_MS.NoTrt_F |
| VPS33A | 0.67493 | 0.31847 | 0.878 | 0.86 | 1 | MS.Trt_vs_MS.NoTrt_F |
| CPSF4 | 0.67555 | 0.43108 | 0.417 | 0.46 | 1 | MS.Trt_vs_MS.NoTrt_F |
| DLG5 | 0.67601 | 0.5482 | 0.45 | 0.45 | 1 | MS.Trt_vs_MS.NoTrt_F |
| SERF2 | 0.67655 | 0.25378 | 0.827 | 0.9 | 1 | MS.Trt_vs_MS.NoTrt_F |
| NPIPB11 | 0.67688 | 0.30467 | 0.522 | 0.62 | 1 | MS.Trt_vs_MS.NoTrt_F |
| FBXL20 | 0.6771 | 0.39655 | 0.939 | 0.94 | 1 | MS.Trt_vs_MS.NoTrt_F |
| SFMBT2 | 0.67738 | 1.66744 | 0.338 | 0.38 | 1 | MS.Trt_vs_MS.NoTrt_F |
| PSMB4 | 0.67762 | 0.81969 | 0.371 | 0.45 | 1 | MS.Trt_vs_MS.NoTrt_F |
| LBHD1 | 0.67783 | 0.72513 | 0.529 | 0.56 | 1 | MS.Trt_vs_MS.NoTrt_F |
| NFKB1 | 0.678 | 3.05932 | 0.133 | 0.16 | 1 | MS.Trt_vs_MS.NoTrt_F |
| CDK13 | 0.6788 | 0.45247 | 0.835 | 0.89 | 1 | MS.Trt_vs_MS.NoTrt_F |
| EFHC1 | 0.6791 | 0.41535 | 0.906 | 0.87 | 1 | MS.Trt_vs_MS.NoTrt_F |
| GSE1 | 0.67933 | 0.85783 | 0.399 | 0.46 | 1 | MS.Trt_vs_MS.NoTrt_F |
| TMEM59 | 0.67967 | 0.26995 | 0.896 | 0.87 | 1 | MS.Trt_vs_MS.NoTrt_F |
| VRK3 | 0.68106 | 0.28293 | 0.788 | 0.83 | 1 | MS.Trt_vs_MS.NoTrt_F |
| ZNF66 | 0.68109 | -0.5421 | 0.626 | 0.62 | 1 | MS.Trt_vs_MS.NoTrt_F |
| SLC6A16 | 0.68203 | 1.08624 | 0.252 | 0.29 | 1 | MS.Trt_vs_MS.NoTrt_F |
| MCM4 | 0.68335 | -0.3761 | 0.604 | 0.6 | 1 | MS.Trt_vs_MS.NoTrt_F |

| | | | | | | |
|---|---|---|---|---|---|---|
| ZNF573 | 0.68337 | -0.7237 | 0.421 | 0.43 | 1 | MS.Trt_vs_MS.NoTrt_F |
| SMPD3 | 0.68422 | 0.61272 | 0.705 | 0.74 | 1 | MS.Trt_vs_MS.NoTrt_F |
| DDX3X | 0.68501 | 0.58607 | 0.381 | 0.4 | 1 | MS.Trt_vs_MS.NoTrt_F |
| COX20 | 0.68513 | 0.75284 | 0.622 | 0.65 | 1 | MS.Trt_vs_MS.NoTrt_F |
| IL17RC | 0.68585 | -0.6508 | 0.428 | 0.37 | 1 | MS.Trt_vs_MS.NoTrt_F |
| MIB2 | 0.68793 | 1.00963 | 0.245 | 0.26 | 1 | MS.Trt_vs_MS.NoTrt_F |
| ASPSCR1 | 0.68849 | 0.28077 | 0.64 | 0.66 | 1 | MS.Trt_vs_MS.NoTrt_F |
| GFOD2 | 0.68962 | 0.295 | 0.888 | 0.92 | 1 | MS.Trt_vs_MS.NoTrt_F |
| POMT1 | 0.69003 | 0.59253 | 0.547 | 0.58 | 1 | MS.Trt_vs_MS.NoTrt_F |
| TCEANC2 | 0.69073 | -0.3746 | 0.91 | 0.93 | 1 | MS.Trt_vs_MS.NoTrt_F |
| TCTN2 | 0.69093 | 0.43611 | 0.763 | 0.78 | 1 | MS.Trt_vs_MS.NoTrt_F |
| STX3 | 0.69133 | 0.4699 | 0.446 | 0.49 | 1 | MS.Trt_vs_MS.NoTrt_F |
| NFATC2IP | 0.69279 | 0.35481 | 0.737 | 0.73 | 1 | MS.Trt_vs_MS.NoTrt_F |
| PSMB6 | 0.69288 | 0.64953 | 0.629 | 0.62 | 1 | MS.Trt_vs_MS.NoTrt_F |
| INMT | 0.69288 | 0.31743 | 0.683 | 0.74 | 1 | MS.Trt_vs_MS.NoTrt_F |
| NUDT15 | 0.69525 | 1.1934 | 0.594 | 0.62 | 1 | MS.Trt_vs_MS.NoTrt_F |
| ZNF696 | 0.69555 | 0.32085 | 0.363 | 0.35 | 1 | MS.Trt_vs_MS.NoTrt_F |
| PDE6B | 0.69559 | 0.42585 | 0.615 | 0.66 | 1 | MS.Trt_vs_MS.NoTrt_F |
| TTC39C | 0.69626 | 0.47934 | 0.817 | 0.9 | 1 | MS.Trt_vs_MS.NoTrt_F |
| ATP1A3 | 0.69847 | -0.265 | 0.371 | 0.4 | 1 | MS.Trt_vs_MS.NoTrt_F |
| LSM5 | 0.69941 | 0.97955 | 0.5 | 0.5 | 1 | MS.Trt_vs_MS.NoTrt_F |
| HOPX | 0.7017 | 0.49317 | 0.781 | 0.85 | 1 | MS.Trt_vs_MS.NoTrt_F |
| NBPF12 | 0.70357 | 0.9542 | 0.371 | 0.38 | 1 | MS.Trt_vs_MS.NoTrt_F |
| PSMD12 | 0.70442 | 0.83505 | 0.723 | 0.79 | 1 | MS.Trt_vs_MS.NoTrt_F |
| ING5 | 0.70789 | -0.3065 | 0.95 | 0.94 | 1 | MS.Trt_vs_MS.NoTrt_F |
| DCLRE1C | 0.70806 | 0.66103 | 0.691 | 0.69 | 1 | MS.Trt_vs_MS.NoTrt_F |
| PLEKHH3 | 0.70823 | 0.2701 | 0.561 | 0.6 | 1 | MS.Trt_vs_MS.NoTrt_F |
| ACTR3C | 0.70946 | 0.42517 | 0.525 | 0.56 | 1 | MS.Trt_vs_MS.NoTrt_F |
| GCLC | 0.70951 | -0.7547 | 0.568 | 0.59 | 1 | MS.Trt_vs_MS.NoTrt_F |
| SETDB1 | 0.70961 | 0.3164 | 0.604 | 0.55 | 1 | MS.Trt_vs_MS.NoTrt_F |
| C4orf19 | 0.71106 | 0.33654 | 0.759 | 0.83 | 1 | MS.Trt_vs_MS.NoTrt_F |
| PHEX | 0.71152 | -1.1692 | 0.101 | 0.11 | 1 | MS.Trt_vs_MS.NoTrt_F |
| DHX57 | 0.71185 | -0.3077 | 0.597 | 0.57 | 1 | MS.Trt_vs_MS.NoTrt_F |
| FOSB | 0.71217 | 0.85817 | 0.36 | 0.4 | 1 | MS.Trt_vs_MS.NoTrt_F |
| SUCLG1 | 0.71232 | 0.28777 | 0.442 | 0.46 | 1 | MS.Trt_vs_MS.NoTrt_F |
| ZNF441 | 0.71322 | 1.69504 | 0.205 | 0.23 | 1 | MS.Trt_vs_MS.NoTrt_F |
| MORN1 | 0.71379 | 0.77774 | 0.568 | 0.55 | 1 | MS.Trt_vs_MS.NoTrt_F |
| ZNF625 | 0.71671 | 0.36923 | 0.284 | 0.28 | 1 | MS.Trt_vs_MS.NoTrt_F |
| PARVA | 0.71678 | -0.2804 | 0.719 | 0.79 | 1 | MS.Trt_vs_MS.NoTrt_F |
| AGPAT5 | 0.7175 | 1.73489 | 0.147 | 0.17 | 1 | MS.Trt_vs_MS.NoTrt_F |

| | | | | | | |
|---|---|---|---|---|---|---|
| PIPOX | 0.7179 | -0.4378 | 0.615 | 0.6 | 1 | MS.Trt_vs_MS.NoTrt_F |
| SPG7 | 0.719 | 0.50315 | 0.957 | 0.97 | 1 | MS.Trt_vs_MS.NoTrt_F |
| NCAPD2 | 0.71945 | 0.3805 | 0.45 | 0.46 | 1 | MS.Trt_vs_MS.NoTrt_F |
| KCTD20 | 0.71997 | 0.84682 | 0.701 | 0.72 | 1 | MS.Trt_vs_MS.NoTrt_F |
| P4HA2 | 0.72122 | 0.63621 | 0.464 | 0.53 | 1 | MS.Trt_vs_MS.NoTrt_F |
| TRAPPC12 | 0.72128 | 0.36829 | 0.594 | 0.6 | 1 | MS.Trt_vs_MS.NoTrt_F |
| SH2D1B | 0.7216 | 0.92568 | 0.234 | 0.24 | 1 | MS.Trt_vs_MS.NoTrt_F |
| CXorf57 | 0.72279 | -0.6896 | 0.147 | 0.16 | 1 | MS.Trt_vs_MS.NoTrt_F |
| PPP6R3 | 0.72285 | 0.41392 | 0.522 | 0.57 | 1 | MS.Trt_vs_MS.NoTrt_F |
| RGS16 | 0.72472 | 0.71206 | 0.112 | 0.13 | 1 | MS.Trt_vs_MS.NoTrt_F |
| MICU1 | 0.72537 | 0.26947 | 0.719 | 0.73 | 1 | MS.Trt_vs_MS.NoTrt_F |
| SAMD1 | 0.72579 | 0.97321 | 0.097 | 0.11 | 1 | MS.Trt_vs_MS.NoTrt_F |
| ORAI3 | 0.72675 | 0.87569 | 0.137 | 0.16 | 1 | MS.Trt_vs_MS.NoTrt_F |
| NEUROD2 | 0.72741 | 0.52758 | 0.212 | 0.22 | 1 | MS.Trt_vs_MS.NoTrt_F |
| CLMN | 0.72764 | 0.40996 | 0.633 | 0.7 | 1 | MS.Trt_vs_MS.NoTrt_F |
| SUN1 | 0.72781 | 0.40363 | 0.612 | 0.7 | 1 | MS.Trt_vs_MS.NoTrt_F |
| TMEM168 | 0.72829 | 1.04906 | 0.536 | 0.51 | 1 | MS.Trt_vs_MS.NoTrt_F |
| ZNF836 | 0.72878 | -0.6107 | 0.223 | 0.24 | 1 | MS.Trt_vs_MS.NoTrt_F |
| TBCEL | 0.729 | -0.3956 | 0.162 | 0.18 | 1 | MS.Trt_vs_MS.NoTrt_F |
| C15orf61 | 0.72932 | 0.61153 | 0.576 | 0.63 | 1 | MS.Trt_vs_MS.NoTrt_F |
| SSH1 | 0.7296 | 0.47771 | 0.86 | 0.89 | 1 | MS.Trt_vs_MS.NoTrt_F |
| ZNF525 | 0.73036 | -0.8711 | 0.59 | 0.59 | 1 | MS.Trt_vs_MS.NoTrt_F |
| CENPL | 0.73037 | -0.3013 | 0.719 | 0.75 | 1 | MS.Trt_vs_MS.NoTrt_F |
| ULK2 | 0.73092 | -0.333 | 0.727 | 0.72 | 1 | MS.Trt_vs_MS.NoTrt_F |
| C7orf50 | 0.73116 | 0.45135 | 0.371 | 0.39 | 1 | MS.Trt_vs_MS.NoTrt_F |
| ZNF346 | 0.73142 | -0.5463 | 0.799 | 0.8 | 1 | MS.Trt_vs_MS.NoTrt_F |
| TYMS | 0.7317 | 1.12546 | 0.237 | 0.26 | 1 | MS.Trt_vs_MS.NoTrt_F |
| USF3 | 0.73238 | 0.42622 | 0.727 | 0.77 | 1 | MS.Trt_vs_MS.NoTrt_F |
| FAM131A | 0.73395 | 0.51775 | 0.795 | 0.82 | 1 | MS.Trt_vs_MS.NoTrt_F |
| CFP | 0.7343 | 2.3235 | 0.363 | 0.38 | 1 | MS.Trt_vs_MS.NoTrt_F |
| ZNF385C | 0.73502 | 0.36043 | 0.572 | 0.61 | 1 | MS.Trt_vs_MS.NoTrt_F |
| MED13 | 0.73559 | 0.53675 | 0.554 | 0.56 | 1 | MS.Trt_vs_MS.NoTrt_F |
| RUVBL1 | 0.73561 | 0.45868 | 0.734 | 0.71 | 1 | MS.Trt_vs_MS.NoTrt_F |
| SRGAP2 | 0.73578 | 0.26545 | 0.543 | 0.59 | 1 | MS.Trt_vs_MS.NoTrt_F |
| ZNF683 | 0.73609 | 1.09757 | 0.112 | 0.11 | 1 | MS.Trt_vs_MS.NoTrt_F |
| IFI35 | 0.73738 | 0.30749 | 0.799 | 0.83 | 1 | MS.Trt_vs_MS.NoTrt_F |
| HMGCL | 0.73757 | -1.0167 | 0.601 | 0.6 | 1 | MS.Trt_vs_MS.NoTrt_F |
| TRHDE | 0.738 | 0.26791 | 0.234 | 0.26 | 1 | MS.Trt_vs_MS.NoTrt_F |
| MRPS24 | 0.73845 | 0.32774 | 0.547 | 0.55 | 1 | MS.Trt_vs_MS.NoTrt_F |
| RREB1 | 0.73851 | 0.72663 | 0.752 | 0.8 | 1 | MS.Trt_vs_MS.NoTrt_F |

| | | | | | | |
|---|---|---|---|---|---|---|
| PTAFR | 0.73868 | -0.3326 | 0.64 | 0.65 | 1 | MS.Trt_vs_MS.NoTrt_F |
| ACTN1 | 0.73986 | 1.18652 | 0.601 | 0.66 | 1 | MS.Trt_vs_MS.NoTrt_F |
| PPT2 | 0.73991 | -0.3377 | 0.576 | 0.56 | 1 | MS.Trt_vs_MS.NoTrt_F |
| FAM135A | 0.74156 | -0.3727 | 0.471 | 0.46 | 1 | MS.Trt_vs_MS.NoTrt_F |
| AC006449.1 | 0.7432 | 0.47133 | 0.18 | 0.2 | 1 | MS.Trt_vs_MS.NoTrt_F |
| LPIN3 | 0.7433 | -0.3115 | 0.471 | 0.49 | 1 | MS.Trt_vs_MS.NoTrt_F |
| RPH3AL | 0.74373 | 0.44707 | 0.363 | 0.38 | 1 | MS.Trt_vs_MS.NoTrt_F |
| APBA2 | 0.74412 | 0.33385 | 0.529 | 0.61 | 1 | MS.Trt_vs_MS.NoTrt_F |
| NHLRC2 | 0.74473 | 0.56542 | 0.763 | 0.79 | 1 | MS.Trt_vs_MS.NoTrt_F |
| FOXO3 | 0.74511 | 0.98226 | 0.421 | 0.43 | 1 | MS.Trt_vs_MS.NoTrt_F |
| SPOCD1 | 0.74564 | 0.5018 | 0.601 | 0.64 | 1 | MS.Trt_vs_MS.NoTrt_F |
| ZNF530 | 0.74621 | -1.1815 | 0.511 | 0.52 | 1 | MS.Trt_vs_MS.NoTrt_F |
| CACUL1 | 0.74659 | 1.03248 | 0.766 | 0.79 | 1 | MS.Trt_vs_MS.NoTrt_F |
| TMEM136 | 0.74667 | 1.14409 | 0.54 | 0.53 | 1 | MS.Trt_vs_MS.NoTrt_F |
| ZSWIM7 | 0.74698 | 0.52634 | 0.781 | 0.84 | 1 | MS.Trt_vs_MS.NoTrt_F |
| BTNL9 | 0.74722 | 0.41899 | 0.853 | 0.88 | 1 | MS.Trt_vs_MS.NoTrt_F |
| ZSCAN31 | 0.74731 | 0.29584 | 0.108 | 0.1 | 1 | MS.Trt_vs_MS.NoTrt_F |
| AFAP1L1 | 0.74833 | 0.28727 | 0.54 | 0.48 | 1 | MS.Trt_vs_MS.NoTrt_F |
| YARS | 0.74866 | -0.4078 | 0.871 | 0.9 | 1 | MS.Trt_vs_MS.NoTrt_F |
| LRRC57 | 0.74915 | 0.29937 | 0.968 | 0.98 | 1 | MS.Trt_vs_MS.NoTrt_F |
| CTSB | 0.74961 | 0.30962 | 0.917 | 0.96 | 1 | MS.Trt_vs_MS.NoTrt_F |
| ASB16 | 0.74996 | 0.3335 | 0.396 | 0.41 | 1 | MS.Trt_vs_MS.NoTrt_F |
| LARP4B | 0.75025 | 1.6878 | 0.306 | 0.36 | 1 | MS.Trt_vs_MS.NoTrt_F |
| LIG3 | 0.75083 | -0.6199 | 0.273 | 0.28 | 1 | MS.Trt_vs_MS.NoTrt_F |
| DDX59 | 0.7514 | 0.40791 | 0.46 | 0.45 | 1 | MS.Trt_vs_MS.NoTrt_F |
| RAC1 | 0.75251 | 1.06497 | 0.23 | 0.27 | 1 | MS.Trt_vs_MS.NoTrt_F |
| C3orf62 | 0.75259 | -0.2856 | 0.773 | 0.86 | 1 | MS.Trt_vs_MS.NoTrt_F |
| SHPK | 0.75359 | -0.597 | 0.486 | 0.49 | 1 | MS.Trt_vs_MS.NoTrt_F |
| GTSF1 | 0.7539 | -0.3572 | 0.273 | 0.27 | 1 | MS.Trt_vs_MS.NoTrt_F |
| BCR | 0.75409 | 1.95306 | 0.291 | 0.32 | 1 | MS.Trt_vs_MS.NoTrt_F |
| BAP1 | 0.75499 | 0.64504 | 0.299 | 0.28 | 1 | MS.Trt_vs_MS.NoTrt_F |
| TRIM22 | 0.75533 | 0.41258 | 0.885 | 0.91 | 1 | MS.Trt_vs_MS.NoTrt_F |
| MT-ND6 | 0.75552 | 0.50178 | 0.651 | 0.8 | 1 | MS.Trt_vs_MS.NoTrt_F |
| KLHL23 | 0.75565 | 0.26232 | 0.223 | 0.24 | 1 | MS.Trt_vs_MS.NoTrt_F |
| IRF2BPL | 0.75578 | 0.78096 | 0.489 | 0.51 | 1 | MS.Trt_vs_MS.NoTrt_F |
| ZNF630 | 0.75603 | -0.899 | 0.453 | 0.47 | 1 | MS.Trt_vs_MS.NoTrt_F |
| SYNJ2 | 0.75617 | -0.4088 | 0.899 | 0.91 | 1 | MS.Trt_vs_MS.NoTrt_F |
| CKAP2L | 0.75637 | 0.34035 | 0.522 | 0.57 | 1 | MS.Trt_vs_MS.NoTrt_F |
| ZBTB3 | 0.75674 | 0.27895 | 0.781 | 0.84 | 1 | MS.Trt_vs_MS.NoTrt_F |
| TUBB4A | 0.75701 | -0.6121 | 0.201 | 0.22 | 1 | MS.Trt_vs_MS.NoTrt_F |

| | | | | | | |
|---|---|---|---|---|---|---|
| GPATCH2L | 0.75708 | 0.29259 | 0.701 | 0.72 | 1 | MS.Trt_vs_MS.NoTrt_F |
| ZNF461 | 0.75715 | 0.50319 | 0.655 | 0.66 | 1 | MS.Trt_vs_MS.NoTrt_F |
| STPG1 | 0.75785 | -0.7302 | 0.791 | 0.81 | 1 | MS.Trt_vs_MS.NoTrt_F |
| BOP1 | 0.7582 | 0.37588 | 0.241 | 0.26 | 1 | MS.Trt_vs_MS.NoTrt_F |
| TSC22D2 | 0.75841 | 0.82608 | 0.435 | 0.48 | 1 | MS.Trt_vs_MS.NoTrt_F |
| SNRPD3 | 0.75843 | 0.64419 | 0.716 | 0.76 | 1 | MS.Trt_vs_MS.NoTrt_F |
| EML2 | 0.75884 | 0.26078 | 0.694 | 0.68 | 1 | MS.Trt_vs_MS.NoTrt_F |
| SLC39A8 | 0.75909 | -0.3457 | 0.385 | 0.39 | 1 | MS.Trt_vs_MS.NoTrt_F |
| FBXO43 | 0.75973 | 0.2742 | 0.442 | 0.47 | 1 | MS.Trt_vs_MS.NoTrt_F |
| EFCAB13 | 0.76 | -0.4159 | 0.342 | 0.35 | 1 | MS.Trt_vs_MS.NoTrt_F |
| DOPEY2 | 0.76055 | -0.3772 | 0.799 | 0.83 | 1 | MS.Trt_vs_MS.NoTrt_F |
| EBPL | 0.76084 | 0.34689 | 0.647 | 0.71 | 1 | MS.Trt_vs_MS.NoTrt_F |
| AC025335.1 | 0.76151 | -0.4513 | 0.615 | 0.58 | 1 | MS.Trt_vs_MS.NoTrt_F |
| C5 | 0.76199 | -0.2572 | 0.457 | 0.48 | 1 | MS.Trt_vs_MS.NoTrt_F |
| ARMC10 | 0.76279 | -0.3696 | 0.601 | 0.64 | 1 | MS.Trt_vs_MS.NoTrt_F |
| MTHFD1L | 0.7631 | 2.0431 | 0.144 | 0.16 | 1 | MS.Trt_vs_MS.NoTrt_F |
| SRSF6 | 0.76334 | 0.7984 | 0.723 | 0.8 | 1 | MS.Trt_vs_MS.NoTrt_F |
| COX15 | 0.7637 | -0.5729 | 0.633 | 0.62 | 1 | MS.Trt_vs_MS.NoTrt_F |
| TIMM10B | 0.76379 | 0.45974 | 0.604 | 0.58 | 1 | MS.Trt_vs_MS.NoTrt_F |
| DGKD | 0.76409 | 0.33507 | 0.622 | 0.64 | 1 | MS.Trt_vs_MS.NoTrt_F |
| KANK1 | 0.76438 | 0.3653 | 0.734 | 0.76 | 1 | MS.Trt_vs_MS.NoTrt_F |
| SCUBE3 | 0.76494 | -0.2774 | 0.626 | 0.66 | 1 | MS.Trt_vs_MS.NoTrt_F |
| C6orf25 | 0.76508 | 0.61631 | 0.518 | 0.55 | 1 | MS.Trt_vs_MS.NoTrt_F |
| POLK | 0.76514 | 0.50093 | 0.82 | 0.87 | 1 | MS.Trt_vs_MS.NoTrt_F |
| EBNA1BP2 | 0.76566 | 0.35005 | 0.543 | 0.51 | 1 | MS.Trt_vs_MS.NoTrt_F |
| ZCCHC9 | 0.76621 | 0.25528 | 0.673 | 0.68 | 1 | MS.Trt_vs_MS.NoTrt_F |
| SMIM8 | 0.7663 | 1.29044 | 0.604 | 0.63 | 1 | MS.Trt_vs_MS.NoTrt_F |
| CASS4 | 0.76691 | 0.48643 | 0.482 | 0.49 | 1 | MS.Trt_vs_MS.NoTrt_F |
| GDAP2 | 0.76884 | 0.69363 | 0.633 | 0.69 | 1 | MS.Trt_vs_MS.NoTrt_F |
| ZNF70 | 0.77039 | 0.62618 | 0.871 | 0.87 | 1 | MS.Trt_vs_MS.NoTrt_F |
| KIAA1586 | 0.77121 | 1.23002 | 0.155 | 0.17 | 1 | MS.Trt_vs_MS.NoTrt_F |
| PIGW | 0.77294 | -0.6871 | 0.439 | 0.43 | 1 | MS.Trt_vs_MS.NoTrt_F |
| PHTF1 | 0.77306 | 0.38722 | 0.536 | 0.57 | 1 | MS.Trt_vs_MS.NoTrt_F |
| NTHL1 | 0.77312 | -0.4772 | 0.119 | 0.11 | 1 | MS.Trt_vs_MS.NoTrt_F |
| EIF4E | 0.77356 | 0.45081 | 0.986 | 0.99 | 1 | MS.Trt_vs_MS.NoTrt_F |
| ATP5S | 0.77632 | 0.70208 | 0.91 | 0.93 | 1 | MS.Trt_vs_MS.NoTrt_F |
| CCT2 | 0.77967 | 0.31464 | 0.802 | 0.83 | 1 | MS.Trt_vs_MS.NoTrt_F |
| EEF1D | 0.77976 | 0.30134 | 0.842 | 0.94 | 1 | MS.Trt_vs_MS.NoTrt_F |
| PRMT2 | 0.78019 | 0.49431 | 0.417 | 0.46 | 1 | MS.Trt_vs_MS.NoTrt_F |
| ZNF135 | 0.7806 | 1.42292 | 0.432 | 0.49 | 1 | MS.Trt_vs_MS.NoTrt_F |

| | | | | | | |
|---|---|---|---|---|---|---|
| MIER2 | 0.7807 | 0.91743 | 0.532 | 0.54 | 1 | MS.Trt_vs_MS.NoTrt_F |
| TUBG1 | 0.78075 | 0.37338 | 0.615 | 0.61 | 1 | MS.Trt_vs_MS.NoTrt_F |
| RIN2 | 0.78105 | 0.76781 | 0.295 | 0.3 | 1 | MS.Trt_vs_MS.NoTrt_F |
| COQ5 | 0.7813 | 0.32867 | 0.817 | 0.9 | 1 | MS.Trt_vs_MS.NoTrt_F |
| YEATS2 | 0.78131 | 0.30259 | 0.698 | 0.7 | 1 | MS.Trt_vs_MS.NoTrt_F |
| LEO1 | 0.7824 | 0.42514 | 0.59 | 0.56 | 1 | MS.Trt_vs_MS.NoTrt_F |
| YPEL1 | 0.7831 | 1.34652 | 0.446 | 0.47 | 1 | MS.Trt_vs_MS.NoTrt_F |
| PPP2R1B | 0.78423 | 0.47229 | 0.442 | 0.47 | 1 | MS.Trt_vs_MS.NoTrt_F |
| COL1A1 | 0.7855 | 0.58832 | 0.525 | 0.55 | 1 | MS.Trt_vs_MS.NoTrt_F |
| ZNF610 | 0.78568 | 0.50258 | 0.468 | 0.45 | 1 | MS.Trt_vs_MS.NoTrt_F |
| GAL3ST4 | 0.78663 | 0.8601 | 0.453 | 0.45 | 1 | MS.Trt_vs_MS.NoTrt_F |
| PRPF39 | 0.78692 | 0.33292 | 0.378 | 0.4 | 1 | MS.Trt_vs_MS.NoTrt_F |
| TNFRSF17 | 0.78739 | 0.34775 | 0.453 | 0.53 | 1 | MS.Trt_vs_MS.NoTrt_F |
| SULT1B1 | 0.78817 | 0.27035 | 0.899 | 0.96 | 1 | MS.Trt_vs_MS.NoTrt_F |
| DTWD1 | 0.78876 | 0.61023 | 0.619 | 0.64 | 1 | MS.Trt_vs_MS.NoTrt_F |
| LSAMP | 0.78905 | 0.49277 | 0.45 | 0.5 | 1 | MS.Trt_vs_MS.NoTrt_F |
| ZC3H8 | 0.78929 | 0.46322 | 0.781 | 0.8 | 1 | MS.Trt_vs_MS.NoTrt_F |
| ITPK1 | 0.78949 | -0.8093 | 0.547 | 0.58 | 1 | MS.Trt_vs_MS.NoTrt_F |
| TMPRSS3 | 0.7897 | -0.4633 | 0.367 | 0.38 | 1 | MS.Trt_vs_MS.NoTrt_F |
| WDR11 | 0.7899 | -0.5678 | 0.619 | 0.61 | 1 | MS.Trt_vs_MS.NoTrt_F |
| SLC31A1 | 0.79026 | -0.564 | 0.784 | 0.79 | 1 | MS.Trt_vs_MS.NoTrt_F |
| NAA16 | 0.79037 | 0.38471 | 0.417 | 0.43 | 1 | MS.Trt_vs_MS.NoTrt_F |
| MZB1 | 0.79049 | 0.25834 | 0.626 | 0.66 | 1 | MS.Trt_vs_MS.NoTrt_F |
| PTPN12 | 0.7905 | 0.38519 | 0.77 | 0.81 | 1 | MS.Trt_vs_MS.NoTrt_F |
| FOXP2 | 0.79103 | 0.39605 | 0.284 | 0.32 | 1 | MS.Trt_vs_MS.NoTrt_F |
| GTF2H3 | 0.79125 | 0.42111 | 0.694 | 0.76 | 1 | MS.Trt_vs_MS.NoTrt_F |
| CARS2 | 0.7915 | 0.56455 | 0.745 | 0.77 | 1 | MS.Trt_vs_MS.NoTrt_F |
| PDCD4 | 0.79174 | 0.65093 | 0.629 | 0.71 | 1 | MS.Trt_vs_MS.NoTrt_F |
| SLC30A4 | 0.79191 | 1.00439 | 0.845 | 0.89 | 1 | MS.Trt_vs_MS.NoTrt_F |
| ZNF724 | 0.79233 | -0.5869 | 0.863 | 0.92 | 1 | MS.Trt_vs_MS.NoTrt_F |
| ASAH2B | 0.79237 | 0.43127 | 0.421 | 0.41 | 1 | MS.Trt_vs_MS.NoTrt_F |
| DAG1 | 0.79343 | -0.4735 | 0.554 | 0.59 | 1 | MS.Trt_vs_MS.NoTrt_F |
| ARPC2 | 0.79352 | 0.25641 | 0.971 | 0.99 | 1 | MS.Trt_vs_MS.NoTrt_F |
| CRKL | 0.79435 | 0.61792 | 0.601 | 0.64 | 1 | MS.Trt_vs_MS.NoTrt_F |
| SOGA1 | 0.79511 | 0.40447 | 0.597 | 0.65 | 1 | MS.Trt_vs_MS.NoTrt_F |
| DENND2D | 0.79564 | 0.34457 | 0.392 | 0.44 | 1 | MS.Trt_vs_MS.NoTrt_F |
| ATAD2B | 0.79646 | -0.3049 | 0.741 | 0.74 | 1 | MS.Trt_vs_MS.NoTrt_F |
| ZNF37A | 0.79875 | 0.9933 | 0.863 | 0.92 | 1 | MS.Trt_vs_MS.NoTrt_F |
| ZNF583 | 0.79891 | 0.39989 | 0.781 | 0.85 | 1 | MS.Trt_vs_MS.NoTrt_F |
| DR1 | 0.79918 | 0.62107 | 0.795 | 0.83 | 1 | MS.Trt_vs_MS.NoTrt_F |

| | | | | | | |
|---|---|---|---|---|---|---|
| SIN3B | 0.7992 | 0.51762 | 0.788 | 0.78 | 1 | MS.Trt_vs_MS.NoTrt_F |
| ADGRG1 | 0.79937 | 1.45451 | 0.694 | 0.72 | 1 | MS.Trt_vs_MS.NoTrt_F |
| SMYD4 | 0.79945 | 0.54083 | 0.813 | 0.87 | 1 | MS.Trt_vs_MS.NoTrt_F |
| FAM76A | 0.79953 | 0.36693 | 0.86 | 0.85 | 1 | MS.Trt_vs_MS.NoTrt_F |
| RNFT2 | 0.80022 | -0.3512 | 0.881 | 0.93 | 1 | MS.Trt_vs_MS.NoTrt_F |
| TSR2 | 0.80079 | 0.30059 | 0.446 | 0.46 | 1 | MS.Trt_vs_MS.NoTrt_F |
| SDE2 | 0.80092 | 0.26907 | 0.655 | 0.72 | 1 | MS.Trt_vs_MS.NoTrt_F |
| BRCA2 | 0.80229 | -0.4203 | 0.424 | 0.43 | 1 | MS.Trt_vs_MS.NoTrt_F |
| MAP3K10 | 0.80305 | 0.49535 | 0.342 | 0.38 | 1 | MS.Trt_vs_MS.NoTrt_F |
| TXK | 0.80315 | 1.13554 | 0.626 | 0.64 | 1 | MS.Trt_vs_MS.NoTrt_F |
| ZFP90 | 0.80332 | -0.7676 | 0.471 | 0.44 | 1 | MS.Trt_vs_MS.NoTrt_F |
| SF3B3 | 0.80429 | 0.28477 | 0.942 | 0.97 | 1 | MS.Trt_vs_MS.NoTrt_F |
| NWD1 | 0.80472 | -0.3155 | 0.511 | 0.53 | 1 | MS.Trt_vs_MS.NoTrt_F |
| DGKE | 0.80502 | 0.42408 | 0.709 | 0.75 | 1 | MS.Trt_vs_MS.NoTrt_F |
| MOAP1 | 0.80552 | 1.09224 | 0.388 | 0.42 | 1 | MS.Trt_vs_MS.NoTrt_F |
| RHBDF2 | 0.80643 | 0.4685 | 0.914 | 0.93 | 1 | MS.Trt_vs_MS.NoTrt_F |
| ZFYVE21 | 0.80686 | 0.44927 | 0.662 | 0.73 | 1 | MS.Trt_vs_MS.NoTrt_F |
| PKIG | 0.80764 | 0.30433 | 0.572 | 0.71 | 1 | MS.Trt_vs_MS.NoTrt_F |
| GRID1 | 0.80897 | 0.46962 | 0.147 | 0.14 | 1 | MS.Trt_vs_MS.NoTrt_F |
| PSMD13 | 0.80917 | 0.41677 | 0.597 | 0.63 | 1 | MS.Trt_vs_MS.NoTrt_F |
| ANGPT4 | 0.80978 | 0.25915 | 0.493 | 0.48 | 1 | MS.Trt_vs_MS.NoTrt_F |
| IL20RB | 0.81036 | 0.77008 | 0.299 | 0.32 | 1 | MS.Trt_vs_MS.NoTrt_F |
| DNAJC14 | 0.81047 | -0.3064 | 0.608 | 0.62 | 1 | MS.Trt_vs_MS.NoTrt_F |
| KCNT1 | 0.81079 | 0.29592 | 0.327 | 0.35 | 1 | MS.Trt_vs_MS.NoTrt_F |
| SEC22B | 0.81214 | 0.7498 | 0.853 | 0.84 | 1 | MS.Trt_vs_MS.NoTrt_F |
| BLZF1 | 0.81216 | -0.4716 | 0.496 | 0.49 | 1 | MS.Trt_vs_MS.NoTrt_F |
| EPHB1 | 0.81326 | 1.26818 | 0.112 | 0.12 | 1 | MS.Trt_vs_MS.NoTrt_F |
| NCKAP5L | 0.81347 | 0.39154 | 0.647 | 0.67 | 1 | MS.Trt_vs_MS.NoTrt_F |
| OAZ3 | 0.81428 | -0.2991 | 0.468 | 0.45 | 1 | MS.Trt_vs_MS.NoTrt_F |
| DCST1 | 0.81485 | 0.75638 | 0.155 | 0.17 | 1 | MS.Trt_vs_MS.NoTrt_F |
| ME2 | 0.81498 | 0.56121 | 0.888 | 0.92 | 1 | MS.Trt_vs_MS.NoTrt_F |
| CDC20B | 0.81513 | 0.25568 | 0.612 | 0.67 | 1 | MS.Trt_vs_MS.NoTrt_F |
| F2RL1 | 0.81517 | 1.6094 | 0.223 | 0.24 | 1 | MS.Trt_vs_MS.NoTrt_F |
| C17orf75 | 0.81579 | 0.30036 | 0.827 | 0.85 | 1 | MS.Trt_vs_MS.NoTrt_F |
| SPDYE2 | 0.81628 | 0.28683 | 0.633 | 0.71 | 1 | MS.Trt_vs_MS.NoTrt_F |
| TAPBP | 0.81692 | 0.26082 | 0.942 | 0.98 | 1 | MS.Trt_vs_MS.NoTrt_F |
| C12orf49 | 0.81719 | 0.63209 | 0.737 | 0.78 | 1 | MS.Trt_vs_MS.NoTrt_F |
| PDZD8 | 0.81723 | 0.57804 | 0.317 | 0.32 | 1 | MS.Trt_vs_MS.NoTrt_F |
| SGK1 | 0.81728 | 1.71924 | 0.417 | 0.43 | 1 | MS.Trt_vs_MS.NoTrt_F |
| FN1 | 0.81734 | 0.2583 | 0.403 | 0.41 | 1 | MS.Trt_vs_MS.NoTrt_F |

| | | | | | | |
|---|---|---|---|---|---|---|
| ATG3 | 0.81752 | 0.53977 | 0.46 | 0.44 | 1 | MS.Trt_vs_MS.NoTrt_F |
| CAGE1 | 0.81787 | 0.45467 | 0.36 | 0.38 | 1 | MS.Trt_vs_MS.NoTrt_F |
| DBF4 | 0.8181 | 0.62339 | 0.45 | 0.45 | 1 | MS.Trt_vs_MS.NoTrt_F |
| BCL7B | 0.81869 | 1.00808 | 0.442 | 0.48 | 1 | MS.Trt_vs_MS.NoTrt_F |
| ZNF891 | 0.81905 | 0.33287 | 0.957 | 0.97 | 1 | MS.Trt_vs_MS.NoTrt_F |
| PHF7 | 0.82021 | 0.26457 | 0.468 | 0.49 | 1 | MS.Trt_vs_MS.NoTrt_F |
| DSE | 0.82064 | -0.5888 | 0.115 | 0.12 | 1 | MS.Trt_vs_MS.NoTrt_F |
| MEGF8 | 0.82172 | 1.15442 | 0.511 | 0.54 | 1 | MS.Trt_vs_MS.NoTrt_F |
| GNB1 | 0.82174 | 1.05793 | 0.353 | 0.35 | 1 | MS.Trt_vs_MS.NoTrt_F |
| POLA2 | 0.82206 | 0.25624 | 0.655 | 0.66 | 1 | MS.Trt_vs_MS.NoTrt_F |
| ZNF177 | 0.82226 | 0.50485 | 0.133 | 0.13 | 1 | MS.Trt_vs_MS.NoTrt_F |
| CEP152 | 0.82237 | -0.3181 | 0.709 | 0.7 | 1 | MS.Trt_vs_MS.NoTrt_F |
| RPL35 | 0.82386 | 0.30927 | 0.691 | 0.87 | 1 | MS.Trt_vs_MS.NoTrt_F |
| ZNRF1 | 0.82413 | 0.53148 | 0.295 | 0.31 | 1 | MS.Trt_vs_MS.NoTrt_F |
| ABCA5 | 0.82436 | 0.90034 | 0.619 | 0.67 | 1 | MS.Trt_vs_MS.NoTrt_F |
| CCND2 | 0.82495 | 1.04958 | 0.518 | 0.59 | 1 | MS.Trt_vs_MS.NoTrt_F |
| FAM111A | 0.82597 | 0.48431 | 0.525 | 0.54 | 1 | MS.Trt_vs_MS.NoTrt_F |
| HYLS1 | 0.82603 | 1.33467 | 0.424 | 0.45 | 1 | MS.Trt_vs_MS.NoTrt_F |
| ZFPM1 | 0.82629 | 0.59272 | 0.112 | 0.12 | 1 | MS.Trt_vs_MS.NoTrt_F |
| PRR5-ARHG/ | 0.8274 | -0.6179 | 0.129 | 0.14 | 1 | MS.Trt_vs_MS.NoTrt_F |
| PLXNA3 | 0.82855 | 0.64614 | 0.809 | 0.85 | 1 | MS.Trt_vs_MS.NoTrt_F |
| SLC28A2 | 0.82935 | -0.2869 | 0.763 | 0.8 | 1 | MS.Trt_vs_MS.NoTrt_F |
| ANKRD13D | 0.82963 | 0.74079 | 0.687 | 0.74 | 1 | MS.Trt_vs_MS.NoTrt_F |
| CARD11 | 0.83111 | 0.82333 | 0.266 | 0.29 | 1 | MS.Trt_vs_MS.NoTrt_F |
| C22orf46 | 0.83118 | 1.07419 | 0.36 | 0.38 | 1 | MS.Trt_vs_MS.NoTrt_F |
| SLC30A5 | 0.83119 | 0.64775 | 0.734 | 0.76 | 1 | MS.Trt_vs_MS.NoTrt_F |
| AMY2B | 0.83136 | -0.7352 | 0.658 | 0.64 | 1 | MS.Trt_vs_MS.NoTrt_F |
| ZNF662 | 0.83144 | 1.33843 | 0.576 | 0.58 | 1 | MS.Trt_vs_MS.NoTrt_F |
| HSBP1 | 0.8316 | 1.03512 | 0.637 | 0.7 | 1 | MS.Trt_vs_MS.NoTrt_F |
| ATG13 | 0.83165 | 0.71302 | 0.802 | 0.84 | 1 | MS.Trt_vs_MS.NoTrt_F |
| HSPA2 | 0.83243 | -0.5393 | 0.565 | 0.59 | 1 | MS.Trt_vs_MS.NoTrt_F |
| TIGD6 | 0.8326 | -0.674 | 0.227 | 0.24 | 1 | MS.Trt_vs_MS.NoTrt_F |
| PFKFB2 | 0.83373 | 0.71474 | 0.281 | 0.28 | 1 | MS.Trt_vs_MS.NoTrt_F |
| ERCC4 | 0.83488 | -0.6127 | 0.885 | 0.91 | 1 | MS.Trt_vs_MS.NoTrt_F |
| TMED4 | 0.83513 | 0.54477 | 0.946 | 0.95 | 1 | MS.Trt_vs_MS.NoTrt_F |
| RELL1 | 0.83585 | 0.68399 | 0.568 | 0.6 | 1 | MS.Trt_vs_MS.NoTrt_F |
| METTL8 | 0.83687 | 0.38852 | 0.903 | 0.93 | 1 | MS.Trt_vs_MS.NoTrt_F |
| USP49 | 0.83703 | 0.38396 | 0.745 | 0.73 | 1 | MS.Trt_vs_MS.NoTrt_F |
| ZNF470 | 0.83877 | 0.39543 | 0.737 | 0.78 | 1 | MS.Trt_vs_MS.NoTrt_F |
| FGFR4 | 0.83883 | 0.50592 | 0.421 | 0.44 | 1 | MS.Trt_vs_MS.NoTrt_F |

| | | | | | | |
|---|---|---|---|---|---|---|
| SENP5 | 0.83892 | 0.55829 | 0.921 | 0.93 | 1 | MS.Trt_vs_MS.NoTrt_F |
| LEPR | 0.83914 | 0.51736 | 0.356 | 0.33 | 1 | MS.Trt_vs_MS.NoTrt_F |
| AGBL4 | 0.83939 | 0.28302 | 0.385 | 0.4 | 1 | MS.Trt_vs_MS.NoTrt_F |
| SLX1A | 0.83941 | 0.54127 | 0.543 | 0.56 | 1 | MS.Trt_vs_MS.NoTrt_F |
| MXD1 | 0.84095 | 1.32586 | 0.173 | 0.19 | 1 | MS.Trt_vs_MS.NoTrt_F |
| APPL1 | 0.84273 | -0.5326 | 0.486 | 0.48 | 1 | MS.Trt_vs_MS.NoTrt_F |
| PGS1 | 0.84288 | -0.5811 | 0.651 | 0.61 | 1 | MS.Trt_vs_MS.NoTrt_F |
| SGK494 | 0.84291 | 0.69377 | 0.701 | 0.76 | 1 | MS.Trt_vs_MS.NoTrt_F |
| PPARGC1B | 0.84294 | 0.8628 | 0.727 | 0.76 | 1 | MS.Trt_vs_MS.NoTrt_F |
| HYKK | 0.84333 | 0.52702 | 0.831 | 0.89 | 1 | MS.Trt_vs_MS.NoTrt_F |
| USP4 | 0.8435 | 0.33843 | 0.928 | 0.94 | 1 | MS.Trt_vs_MS.NoTrt_F |
| PLIN5 | 0.84452 | 0.29635 | 0.806 | 0.8 | 1 | MS.Trt_vs_MS.NoTrt_F |
| SLF2 | 0.84494 | -0.3763 | 0.547 | 0.55 | 1 | MS.Trt_vs_MS.NoTrt_F |
| PFKM | 0.84534 | 0.34045 | 0.871 | 0.91 | 1 | MS.Trt_vs_MS.NoTrt_F |
| CAPZA2 | 0.84567 | 0.4262 | 0.896 | 0.94 | 1 | MS.Trt_vs_MS.NoTrt_F |
| WHAMM | 0.84685 | 0.34237 | 0.838 | 0.87 | 1 | MS.Trt_vs_MS.NoTrt_F |
| TMEM19 | 0.84694 | 0.25443 | 0.802 | 0.79 | 1 | MS.Trt_vs_MS.NoTrt_F |
| UGT1A1 | 0.84732 | 0.28218 | 0.565 | 0.61 | 1 | MS.Trt_vs_MS.NoTrt_F |
| CEP19 | 0.84735 | 0.75133 | 0.633 | 0.64 | 1 | MS.Trt_vs_MS.NoTrt_F |
| HS3ST1 | 0.84793 | 0.63137 | 0.327 | 0.33 | 1 | MS.Trt_vs_MS.NoTrt_F |
| ACSS2 | 0.84803 | 0.99638 | 0.54 | 0.55 | 1 | MS.Trt_vs_MS.NoTrt_F |
| AVL9 | 0.84862 | 0.42775 | 0.896 | 0.93 | 1 | MS.Trt_vs_MS.NoTrt_F |
| TMEM154 | 0.8497 | 0.40366 | 0.921 | 0.93 | 1 | MS.Trt_vs_MS.NoTrt_F |
| RYR3 | 0.85001 | 0.28087 | 0.482 | 0.5 | 1 | MS.Trt_vs_MS.NoTrt_F |
| RP11-1070B | 0.85039 | 1.00911 | 0.223 | 0.23 | 1 | MS.Trt_vs_MS.NoTrt_F |
| LRRC2 | 0.85138 | -0.3262 | 0.745 | 0.79 | 1 | MS.Trt_vs_MS.NoTrt_F |
| PHLDA1 | 0.85226 | 0.88769 | 0.525 | 0.56 | 1 | MS.Trt_vs_MS.NoTrt_F |
| EIF2AK4 | 0.85246 | 0.52921 | 0.619 | 0.64 | 1 | MS.Trt_vs_MS.NoTrt_F |
| TIMP3 | 0.85353 | 2.04749 | 0.097 | 0.11 | 1 | MS.Trt_vs_MS.NoTrt_F |
| CLIP2 | 0.85389 | -0.7025 | 0.36 | 0.37 | 1 | MS.Trt_vs_MS.NoTrt_F |
| CDT1 | 0.85465 | 0.27405 | 0.475 | 0.49 | 1 | MS.Trt_vs_MS.NoTrt_F |
| KBTBD2 | 0.85641 | 0.48748 | 0.658 | 0.69 | 1 | MS.Trt_vs_MS.NoTrt_F |
| RAB36 | 0.85646 | 0.52191 | 0.36 | 0.38 | 1 | MS.Trt_vs_MS.NoTrt_F |
| NCR3LG1 | 0.85702 | -1.2707 | 0.104 | 0.11 | 1 | MS.Trt_vs_MS.NoTrt_F |
| UNKL | 0.85715 | 0.71605 | 0.482 | 0.47 | 1 | MS.Trt_vs_MS.NoTrt_F |
| CXorf21 | 0.85815 | -1.2357 | 0.381 | 0.41 | 1 | MS.Trt_vs_MS.NoTrt_F |
| HENMT1 | 0.85823 | -0.604 | 0.324 | 0.35 | 1 | MS.Trt_vs_MS.NoTrt_F |
| BLOC1S3 | 0.85894 | 0.48158 | 0.597 | 0.6 | 1 | MS.Trt_vs_MS.NoTrt_F |
| PLEKHA7 | 0.86083 | 0.73832 | 0.248 | 0.25 | 1 | MS.Trt_vs_MS.NoTrt_F |
| IMP4 | 0.86189 | 0.52948 | 0.277 | 0.29 | 1 | MS.Trt_vs_MS.NoTrt_F |

| | | | | | | |
|---|---|---|---|---|---|---|
| ZNF316 | 0.86194 | -0.5957 | 0.532 | 0.53 | 1 | MS.Trt_vs_MS.NoTrt_F |
| NMD3 | 0.86339 | 0.44886 | 0.709 | 0.71 | 1 | MS.Trt_vs_MS.NoTrt_F |
| A2M | 0.86495 | 1.42761 | 0.18 | 0.19 | 1 | MS.Trt_vs_MS.NoTrt_F |
| AVIL | 0.86597 | -0.6109 | 0.388 | 0.39 | 1 | MS.Trt_vs_MS.NoTrt_F |
| PNP | 0.86627 | 0.54951 | 0.385 | 0.4 | 1 | MS.Trt_vs_MS.NoTrt_F |
| PEX13 | 0.8676 | 0.26911 | 0.795 | 0.77 | 1 | MS.Trt_vs_MS.NoTrt_F |
| ZNF789 | 0.86769 | 0.90886 | 0.795 | 0.81 | 1 | MS.Trt_vs_MS.NoTrt_F |
| MFSD9 | 0.86852 | 0.33716 | 0.478 | 0.5 | 1 | MS.Trt_vs_MS.NoTrt_F |
| IDH3A | 0.87063 | 0.54737 | 0.73 | 0.83 | 1 | MS.Trt_vs_MS.NoTrt_F |
| KLHL36 | 0.87145 | 0.59629 | 0.644 | 0.67 | 1 | MS.Trt_vs_MS.NoTrt_F |
| LRRC27 | 0.87472 | 0.42219 | 0.777 | 0.87 | 1 | MS.Trt_vs_MS.NoTrt_F |
| RNF19B | 0.87566 | 1.46238 | 0.112 | 0.11 | 1 | MS.Trt_vs_MS.NoTrt_F |
| ZSCAN29 | 0.87614 | 1.33358 | 0.482 | 0.48 | 1 | MS.Trt_vs_MS.NoTrt_F |
| SSC5D | 0.87616 | -0.3117 | 0.112 | 0.12 | 1 | MS.Trt_vs_MS.NoTrt_F |
| NSUN3 | 0.8764 | 0.50211 | 0.806 | 0.81 | 1 | MS.Trt_vs_MS.NoTrt_F |
| USP6NL | 0.87902 | 0.29563 | 0.759 | 0.8 | 1 | MS.Trt_vs_MS.NoTrt_F |
| STAM2 | 0.87906 | 0.93302 | 0.579 | 0.62 | 1 | MS.Trt_vs_MS.NoTrt_F |
| IFITM1 | 0.88027 | 0.57957 | 0.55 | 0.66 | 1 | MS.Trt_vs_MS.NoTrt_F |
| RSRC1 | 0.88186 | 0.55039 | 0.759 | 0.76 | 1 | MS.Trt_vs_MS.NoTrt_F |
| CXADR | 0.88226 | -0.5885 | 0.327 | 0.34 | 1 | MS.Trt_vs_MS.NoTrt_F |
| UBB | 0.88289 | 0.52965 | 0.572 | 0.73 | 1 | MS.Trt_vs_MS.NoTrt_F |
| CNPY4 | 0.88295 | -0.2537 | 0.716 | 0.72 | 1 | MS.Trt_vs_MS.NoTrt_F |
| METTL12 | 0.88301 | -0.7262 | 0.655 | 0.64 | 1 | MS.Trt_vs_MS.NoTrt_F |
| GP6 | 0.88611 | 0.59013 | 0.482 | 0.53 | 1 | MS.Trt_vs_MS.NoTrt_F |
| CCP110 | 0.88637 | -0.3196 | 0.532 | 0.52 | 1 | MS.Trt_vs_MS.NoTrt_F |
| MDM2 | 0.88747 | 0.57265 | 0.986 | 0.98 | 1 | MS.Trt_vs_MS.NoTrt_F |
| TMEM80 | 0.88782 | 0.47602 | 0.594 | 0.6 | 1 | MS.Trt_vs_MS.NoTrt_F |
| LIFR | 0.88893 | 1.24212 | 0.176 | 0.19 | 1 | MS.Trt_vs_MS.NoTrt_F |
| RIPK2 | 0.88937 | 0.41106 | 0.23 | 0.23 | 1 | MS.Trt_vs_MS.NoTrt_F |
| PLEK | 0.89028 | 2.94957 | 0.119 | 0.12 | 1 | MS.Trt_vs_MS.NoTrt_F |
| AC013264.1 | 0.89111 | 0.90771 | 0.446 | 0.53 | 1 | MS.Trt_vs_MS.NoTrt_F |
| DAAM2 | 0.89136 | 0.71601 | 0.421 | 0.44 | 1 | MS.Trt_vs_MS.NoTrt_F |
| ZNF718 | 0.89167 | 0.28356 | 0.306 | 0.33 | 1 | MS.Trt_vs_MS.NoTrt_F |
| LIMS1 | 0.89217 | 0.29631 | 0.874 | 0.9 | 1 | MS.Trt_vs_MS.NoTrt_F |
| TTC39B | 0.8928 | 0.92028 | 0.759 | 0.77 | 1 | MS.Trt_vs_MS.NoTrt_F |
| DHTKD1 | 0.89361 | 0.88697 | 0.827 | 0.85 | 1 | MS.Trt_vs_MS.NoTrt_F |
| MBTPS1 | 0.89543 | 0.43734 | 0.568 | 0.6 | 1 | MS.Trt_vs_MS.NoTrt_F |
| DUSP12 | 0.89548 | 0.83876 | 0.349 | 0.37 | 1 | MS.Trt_vs_MS.NoTrt_F |
| CD8A | 0.89573 | 1.76908 | 0.133 | 0.14 | 1 | MS.Trt_vs_MS.NoTrt_F |
| COQ8B | 0.89665 | -0.3442 | 0.669 | 0.67 | 1 | MS.Trt_vs_MS.NoTrt_F |

| | | | | | | |
|---|---|---|---|---|---|---|
| U2AF1L4 | 0.89699 | 0.35312 | 0.799 | 0.83 | 1 | MS.Trt_vs_MS.NoTrt_F |
| ATP10D | 0.89726 | 0.7702 | 0.392 | 0.4 | 1 | MS.Trt_vs_MS.NoTrt_F |
| SSR1 | 0.89765 | 0.48293 | 0.917 | 0.93 | 1 | MS.Trt_vs_MS.NoTrt_F |
| TOB1 | 0.89848 | 0.41354 | 0.626 | 0.74 | 1 | MS.Trt_vs_MS.NoTrt_F |
| RANGAP1 | 0.89917 | -0.3326 | 0.734 | 0.69 | 1 | MS.Trt_vs_MS.NoTrt_F |
| ZNF846 | 0.89971 | 0.3765 | 0.496 | 0.53 | 1 | MS.Trt_vs_MS.NoTrt_F |
| RALBP1 | 0.90019 | 0.86112 | 0.439 | 0.44 | 1 | MS.Trt_vs_MS.NoTrt_F |
| SSU72 | 0.90045 | 0.43605 | 0.914 | 0.95 | 1 | MS.Trt_vs_MS.NoTrt_F |
| SLC39A14 | 0.90055 | -0.974 | 0.46 | 0.45 | 1 | MS.Trt_vs_MS.NoTrt_F |
| TP53 | 0.90379 | 0.33751 | 0.745 | 0.81 | 1 | MS.Trt_vs_MS.NoTrt_F |
| DNAJB5 | 0.90472 | 0.94248 | 0.201 | 0.21 | 1 | MS.Trt_vs_MS.NoTrt_F |
| THEM4 | 0.905 | 0.29483 | 0.809 | 0.79 | 1 | MS.Trt_vs_MS.NoTrt_F |
| SRGAP3 | 0.90515 | -0.3168 | 0.835 | 0.88 | 1 | MS.Trt_vs_MS.NoTrt_F |
| APOBEC3F | 0.90587 | 0.35825 | 0.924 | 0.88 | 1 | MS.Trt_vs_MS.NoTrt_F |
| MAP3K11 | 0.90648 | 0.25222 | 0.691 | 0.7 | 1 | MS.Trt_vs_MS.NoTrt_F |
| KLHL35 | 0.90779 | 0.46157 | 0.471 | 0.46 | 1 | MS.Trt_vs_MS.NoTrt_F |
| ZNF576 | 0.90874 | 0.27705 | 0.536 | 0.56 | 1 | MS.Trt_vs_MS.NoTrt_F |
| SWSAP1 | 0.91009 | -0.3013 | 0.701 | 0.71 | 1 | MS.Trt_vs_MS.NoTrt_F |
| MEIOC | 0.91059 | -0.2625 | 0.655 | 0.69 | 1 | MS.Trt_vs_MS.NoTrt_F |
| ORC1 | 0.91121 | 0.31532 | 0.245 | 0.27 | 1 | MS.Trt_vs_MS.NoTrt_F |
| PNPO | 0.91132 | -0.3069 | 0.658 | 0.67 | 1 | MS.Trt_vs_MS.NoTrt_F |
| GSDMA | 0.91185 | 0.36721 | 0.662 | 0.68 | 1 | MS.Trt_vs_MS.NoTrt_F |
| KNL1 | 0.91287 | 0.3161 | 0.432 | 0.43 | 1 | MS.Trt_vs_MS.NoTrt_F |
| SPACA9 | 0.91349 | -0.3828 | 0.644 | 0.64 | 1 | MS.Trt_vs_MS.NoTrt_F |
| PRKCA | 0.91534 | 0.29908 | 0.745 | 0.76 | 1 | MS.Trt_vs_MS.NoTrt_F |
| TM2D3 | 0.91745 | -0.2534 | 0.777 | 0.79 | 1 | MS.Trt_vs_MS.NoTrt_F |
| CEP135 | 0.91753 | -0.6042 | 0.799 | 0.8 | 1 | MS.Trt_vs_MS.NoTrt_F |
| ARF6 | 0.91799 | 0.66364 | 0.701 | 0.76 | 1 | MS.Trt_vs_MS.NoTrt_F |
| ATP2A2 | 0.91939 | 0.71163 | 0.748 | 0.72 | 1 | MS.Trt_vs_MS.NoTrt_F |
| ARID3A | 0.91988 | -0.299 | 0.594 | 0.58 | 1 | MS.Trt_vs_MS.NoTrt_F |
| C12orf60 | 0.92196 | 0.38115 | 0.619 | 0.62 | 1 | MS.Trt_vs_MS.NoTrt_F |
| TXLNA | 0.92201 | 0.66096 | 0.385 | 0.44 | 1 | MS.Trt_vs_MS.NoTrt_F |
| RGS12 | 0.92284 | -0.9961 | 0.86 | 0.85 | 1 | MS.Trt_vs_MS.NoTrt_F |
| STAT4 | 0.92297 | 0.65639 | 0.363 | 0.39 | 1 | MS.Trt_vs_MS.NoTrt_F |
| CDC73 | 0.92317 | 0.49927 | 0.572 | 0.55 | 1 | MS.Trt_vs_MS.NoTrt_F |
| MYL6B | 0.92403 | 0.41547 | 0.572 | 0.62 | 1 | MS.Trt_vs_MS.NoTrt_F |
| TBCCD1 | 0.92453 | 0.49564 | 0.406 | 0.41 | 1 | MS.Trt_vs_MS.NoTrt_F |
| TTC37 | 0.92515 | 0.27787 | 0.414 | 0.43 | 1 | MS.Trt_vs_MS.NoTrt_F |
| TXNL1 | 0.9258 | 0.34663 | 0.662 | 0.71 | 1 | MS.Trt_vs_MS.NoTrt_F |
| TIMM23B | 0.92771 | 0.50606 | 0.608 | 0.66 | 1 | MS.Trt_vs_MS.NoTrt_F |

| | | | | | | |
|---|---|---|---|---|---|---|
| ACY1 | 0.92881 | 0.51648 | 0.644 | 0.62 | 1 | MS.Trt_vs_MS.NoTrt_F |
| WDR90 | 0.93016 | 0.63652 | 0.878 | 0.91 | 1 | MS.Trt_vs_MS.NoTrt_F |
| ZNF14 | 0.93024 | 1.67306 | 0.396 | 0.41 | 1 | MS.Trt_vs_MS.NoTrt_F |
| LASP1 | 0.93058 | -0.402 | 0.809 | 0.82 | 1 | MS.Trt_vs_MS.NoTrt_F |
| IDH2 | 0.93205 | 0.81469 | 0.511 | 0.56 | 1 | MS.Trt_vs_MS.NoTrt_F |
| UBE2W | 0.9327 | 0.35646 | 0.658 | 0.7 | 1 | MS.Trt_vs_MS.NoTrt_F |
| AGO2 | 0.93322 | 0.95595 | 0.392 | 0.45 | 1 | MS.Trt_vs_MS.NoTrt_F |
| TANC1 | 0.93468 | 0.2918 | 0.378 | 0.39 | 1 | MS.Trt_vs_MS.NoTrt_F |
| PIP4K2A | 0.93505 | 0.46298 | 0.824 | 0.91 | 1 | MS.Trt_vs_MS.NoTrt_F |
| AC012314.1 | 0.93531 | 0.31939 | 0.691 | 0.88 | 1 | MS.Trt_vs_MS.NoTrt_F |
| AC012314.1 | 0.93531 | 0.31939 | 0.691 | 0.88 | 1 | MS.Trt_vs_MS.NoTrt_F |
| AC012314.1 | 0.93531 | 0.31939 | 0.691 | 0.88 | 1 | MS.Trt_vs_MS.NoTrt_F |
| AC012314.2 | 0.93531 | 0.31939 | 0.691 | 0.88 | 1 | MS.Trt_vs_MS.NoTrt_F |
| AC012314.4 | 0.93531 | 0.31939 | 0.691 | 0.88 | 1 | MS.Trt_vs_MS.NoTrt_F |
| AC012314.5 | 0.93531 | 0.31939 | 0.691 | 0.88 | 1 | MS.Trt_vs_MS.NoTrt_F |
| CD36 | 0.9358 | 1.40256 | 0.55 | 0.53 | 1 | MS.Trt_vs_MS.NoTrt_F |
| EXT1 | 0.93604 | 0.49072 | 0.201 | 0.21 | 1 | MS.Trt_vs_MS.NoTrt_F |
| RP11-231C1 | 0.93624 | 0.54585 | 0.694 | 0.79 | 1 | MS.Trt_vs_MS.NoTrt_F |
| NALCN | 0.93661 | -0.3416 | 0.543 | 0.54 | 1 | MS.Trt_vs_MS.NoTrt_F |
| POC1B | 0.93735 | 0.6054 | 0.755 | 0.83 | 1 | MS.Trt_vs_MS.NoTrt_F |
| SPDYE3 | 0.93812 | 0.31256 | 0.59 | 0.56 | 1 | MS.Trt_vs_MS.NoTrt_F |
| S1PR5 | 0.9387 | 0.28414 | 0.129 | 0.13 | 1 | MS.Trt_vs_MS.NoTrt_F |
| TOMM40 | 0.93924 | -0.3736 | 0.619 | 0.65 | 1 | MS.Trt_vs_MS.NoTrt_F |
| METTL21B | 0.93975 | 0.83127 | 0.5 | 0.53 | 1 | MS.Trt_vs_MS.NoTrt_F |
| LPGAT1 | 0.94048 | 0.74743 | 0.529 | 0.51 | 1 | MS.Trt_vs_MS.NoTrt_F |
| ZSCAN5A | 0.94055 | -0.2706 | 0.784 | 0.82 | 1 | MS.Trt_vs_MS.NoTrt_F |
| PGAM5 | 0.94132 | -0.3221 | 0.77 | 0.79 | 1 | MS.Trt_vs_MS.NoTrt_F |
| SLC7A13 | 0.94177 | 0.44781 | 0.155 | 0.17 | 1 | MS.Trt_vs_MS.NoTrt_F |
| IFFO1 | 0.94178 | 0.82021 | 0.464 | 0.46 | 1 | MS.Trt_vs_MS.NoTrt_F |
| ITPKB | 0.94278 | 0.76697 | 0.597 | 0.62 | 1 | MS.Trt_vs_MS.NoTrt_F |
| ATP6V1D | 0.94289 | 0.31551 | 0.586 | 0.66 | 1 | MS.Trt_vs_MS.NoTrt_F |
| SLC30A6 | 0.94295 | 0.33771 | 0.604 | 0.65 | 1 | MS.Trt_vs_MS.NoTrt_F |
| GTPBP3 | 0.94337 | 0.26194 | 0.723 | 0.75 | 1 | MS.Trt_vs_MS.NoTrt_F |
| UGDH | 0.9436 | -0.344 | 0.64 | 0.67 | 1 | MS.Trt_vs_MS.NoTrt_F |
| EXOSC2 | 0.94474 | 0.31003 | 0.755 | 0.78 | 1 | MS.Trt_vs_MS.NoTrt_F |
| DCUN1D5 | 0.94548 | -0.2963 | 0.468 | 0.45 | 1 | MS.Trt_vs_MS.NoTrt_F |
| CNTLN | 0.94581 | 0.33823 | 0.349 | 0.36 | 1 | MS.Trt_vs_MS.NoTrt_F |
| FGD2 | 0.94582 | 0.68325 | 0.759 | 0.78 | 1 | MS.Trt_vs_MS.NoTrt_F |
| OASL | 0.94655 | 0.87529 | 0.266 | 0.27 | 1 | MS.Trt_vs_MS.NoTrt_F |
| JKAMP | 0.94668 | -0.4936 | 0.795 | 0.85 | 1 | MS.Trt_vs_MS.NoTrt_F |

| | | | | | | |
|---|---|---|---|---|---|---|
| ELP3 | 0.94715 | -0.3758 | 0.536 | 0.5 | 1 | MS.Trt_vs_MS.NoTrt_F |
| SNX30 | 0.94722 | 0.33275 | 0.482 | 0.5 | 1 | MS.Trt_vs_MS.NoTrt_F |
| TIPIN | 0.94728 | 0.721 | 0.612 | 0.62 | 1 | MS.Trt_vs_MS.NoTrt_F |
| DND1 | 0.94756 | 0.3337 | 0.104 | 0.11 | 1 | MS.Trt_vs_MS.NoTrt_F |
| RNF125 | 0.94773 | 0.33178 | 0.91 | 0.95 | 1 | MS.Trt_vs_MS.NoTrt_F |
| TGFBRAP1 | 0.94927 | 0.37738 | 0.55 | 0.55 | 1 | MS.Trt_vs_MS.NoTrt_F |
| MLF1 | 0.94969 | 0.72219 | 0.381 | 0.41 | 1 | MS.Trt_vs_MS.NoTrt_F |
| HARBI1 | 0.94989 | 0.75236 | 0.604 | 0.62 | 1 | MS.Trt_vs_MS.NoTrt_F |
| INPP4A | 0.95079 | 0.47381 | 0.453 | 0.46 | 1 | MS.Trt_vs_MS.NoTrt_F |
| ZNF658 | 0.95104 | 0.59798 | 0.216 | 0.22 | 1 | MS.Trt_vs_MS.NoTrt_F |
| DBN1 | 0.95229 | 0.6956 | 0.234 | 0.24 | 1 | MS.Trt_vs_MS.NoTrt_F |
| HEATR4 | 0.95396 | 0.9968 | 0.396 | 0.44 | 1 | MS.Trt_vs_MS.NoTrt_F |
| EXOSC6 | 0.9549 | 0.71976 | 0.899 | 0.96 | 1 | MS.Trt_vs_MS.NoTrt_F |
| ZSCAN18 | 0.95563 | 0.53481 | 0.694 | 0.69 | 1 | MS.Trt_vs_MS.NoTrt_F |
| FKTN | 0.95608 | 0.62049 | 0.209 | 0.22 | 1 | MS.Trt_vs_MS.NoTrt_F |
| CD180 | 0.957 | 0.93504 | 0.295 | 0.3 | 1 | MS.Trt_vs_MS.NoTrt_F |
| CCDC124 | 0.95776 | 0.26113 | 0.522 | 0.54 | 1 | MS.Trt_vs_MS.NoTrt_F |
| NELFCD | 0.95859 | 0.67042 | 0.594 | 0.59 | 1 | MS.Trt_vs_MS.NoTrt_F |
| MYO7A | 0.95954 | 0.47304 | 0.385 | 0.41 | 1 | MS.Trt_vs_MS.NoTrt_F |
| WDPCP | 0.96008 | 0.5776 | 0.95 | 0.96 | 1 | MS.Trt_vs_MS.NoTrt_F |
| PLAC8 | 0.96172 | 0.9526 | 0.723 | 0.79 | 1 | MS.Trt_vs_MS.NoTrt_F |
| IFT80 | 0.96178 | 0.91568 | 0.752 | 0.79 | 1 | MS.Trt_vs_MS.NoTrt_F |
| TNRC18 | 0.96217 | 0.44634 | 0.777 | 0.81 | 1 | MS.Trt_vs_MS.NoTrt_F |
| ADPRH | 0.96247 | 3.26719 | 0.104 | 0.11 | 1 | MS.Trt_vs_MS.NoTrt_F |
| ZBTB20 | 0.96271 | 0.44894 | 0.752 | 0.8 | 1 | MS.Trt_vs_MS.NoTrt_F |
| PANK3 | 0.96303 | 0.80527 | 0.424 | 0.44 | 1 | MS.Trt_vs_MS.NoTrt_F |
| PCCB | 0.96309 | 0.37215 | 0.626 | 0.66 | 1 | MS.Trt_vs_MS.NoTrt_F |
| ZNF772 | 0.96447 | 1.02642 | 0.507 | 0.52 | 1 | MS.Trt_vs_MS.NoTrt_F |
| CD81 | 0.9645 | 1.19898 | 0.101 | 0.11 | 1 | MS.Trt_vs_MS.NoTrt_F |
| GSN | 0.96485 | 0.77901 | 0.637 | 0.64 | 1 | MS.Trt_vs_MS.NoTrt_F |
| PLXDC1 | 0.96531 | 1.08824 | 0.694 | 0.75 | 1 | MS.Trt_vs_MS.NoTrt_F |
| SPTLC2 | 0.96537 | 0.3214 | 0.719 | 0.74 | 1 | MS.Trt_vs_MS.NoTrt_F |
| TDRD3 | 0.96556 | 0.72186 | 0.543 | 0.55 | 1 | MS.Trt_vs_MS.NoTrt_F |
| PIK3C2B | 0.96636 | -0.333 | 0.45 | 0.42 | 1 | MS.Trt_vs_MS.NoTrt_F |
| SLC23A3 | 0.9667 | 1.11766 | 0.46 | 0.49 | 1 | MS.Trt_vs_MS.NoTrt_F |
| FBXO45 | 0.96719 | 0.88296 | 0.471 | 0.46 | 1 | MS.Trt_vs_MS.NoTrt_F |
| OBSL1 | 0.9672 | 0.27222 | 0.187 | 0.19 | 1 | MS.Trt_vs_MS.NoTrt_F |
| SH3YL1 | 0.96724 | -0.5507 | 0.777 | 0.71 | 1 | MS.Trt_vs_MS.NoTrt_F |
| MGA | 0.96835 | -0.316 | 0.881 | 0.89 | 1 | MS.Trt_vs_MS.NoTrt_F |
| EXOC5 | 0.96915 | 0.97165 | 0.439 | 0.47 | 1 | MS.Trt_vs_MS.NoTrt_F |

| | | | | | | |
|---|---|---|---|---|---|---|
| MSS51 | 0.96918 | -0.8115 | 0.536 | 0.56 | 1 | MS.Trt_vs_MS.NoTrt_F |
| BRDT | 0.96939 | 0.25162 | 0.554 | 0.59 | 1 | MS.Trt_vs_MS.NoTrt_F |
| NBEAL1 | 0.96976 | 0.49088 | 0.856 | 0.91 | 1 | MS.Trt_vs_MS.NoTrt_F |
| FNIP1 | 0.97029 | 0.83489 | 0.633 | 0.67 | 1 | MS.Trt_vs_MS.NoTrt_F |
| DNAJA3 | 0.97155 | 0.33281 | 0.91 | 0.95 | 1 | MS.Trt_vs_MS.NoTrt_F |
| EIF4EBP2 | 0.97286 | 0.38358 | 0.687 | 0.69 | 1 | MS.Trt_vs_MS.NoTrt_F |
| ZNF554 | 0.97299 | 0.29946 | 0.932 | 0.96 | 1 | MS.Trt_vs_MS.NoTrt_F |
| CSNK1A1 | 0.97554 | 0.4359 | 0.885 | 0.85 | 1 | MS.Trt_vs_MS.NoTrt_F |
| KBTBD12 | 0.97677 | 0.25889 | 0.385 | 0.4 | 1 | MS.Trt_vs_MS.NoTrt_F |
| HAVCR2 | 0.97692 | 0.3114 | 0.615 | 0.64 | 1 | MS.Trt_vs_MS.NoTrt_F |
| FAM161B | 0.977 | 0.63346 | 0.396 | 0.41 | 1 | MS.Trt_vs_MS.NoTrt_F |
| TMEM107 | 0.97735 | 0.86645 | 0.532 | 0.58 | 1 | MS.Trt_vs_MS.NoTrt_F |
| ZNF808 | 0.97938 | 1.40555 | 0.536 | 0.54 | 1 | MS.Trt_vs_MS.NoTrt_F |
| FOCAD | 0.98017 | 0.69586 | 0.522 | 0.52 | 1 | MS.Trt_vs_MS.NoTrt_F |
| KIF27 | 0.98117 | 0.86657 | 0.353 | 0.36 | 1 | MS.Trt_vs_MS.NoTrt_F |
| ADM2 | 0.98117 | -0.3425 | 0.356 | 0.35 | 1 | MS.Trt_vs_MS.NoTrt_F |
| A1BG | 0.9813 | 0.32864 | 0.892 | 0.87 | 1 | MS.Trt_vs_MS.NoTrt_F |
| ELMSAN1 | 0.98171 | 0.83021 | 0.727 | 0.75 | 1 | MS.Trt_vs_MS.NoTrt_F |
| UGT8 | 0.98469 | 0.48717 | 0.46 | 0.48 | 1 | MS.Trt_vs_MS.NoTrt_F |
| RHOF | 0.98669 | 0.40913 | 0.741 | 0.78 | 1 | MS.Trt_vs_MS.NoTrt_F |
| IBA57 | 0.9876 | 0.56556 | 0.813 | 0.86 | 1 | MS.Trt_vs_MS.NoTrt_F |
| GLTSCR1 | 0.98787 | 0.40605 | 0.619 | 0.64 | 1 | MS.Trt_vs_MS.NoTrt_F |
| NEU1 | 0.98821 | 1.00233 | 0.651 | 0.62 | 1 | MS.Trt_vs_MS.NoTrt_F |
| GMPS | 0.98847 | 0.48491 | 0.835 | 0.87 | 1 | MS.Trt_vs_MS.NoTrt_F |
| PARD6G | 0.98905 | 0.62991 | 0.511 | 0.55 | 1 | MS.Trt_vs_MS.NoTrt_F |
| GMPR | 0.98908 | 0.33337 | 0.356 | 0.4 | 1 | MS.Trt_vs_MS.NoTrt_F |
| BFAR | 0.98947 | 0.57572 | 0.543 | 0.58 | 1 | MS.Trt_vs_MS.NoTrt_F |
| UNC45A | 0.98949 | 0.31206 | 0.687 | 0.72 | 1 | MS.Trt_vs_MS.NoTrt_F |
| RBP7 | 0.98963 | 1.30453 | 0.331 | 0.32 | 1 | MS.Trt_vs_MS.NoTrt_F |
| KLRD1 | 0.98965 | 0.54575 | 0.975 | 0.98 | 1 | MS.Trt_vs_MS.NoTrt_F |
| ARIH2 | 0.98966 | 0.51449 | 0.968 | 0.99 | 1 | MS.Trt_vs_MS.NoTrt_F |
| RGS17 | 0.98983 | 0.43439 | 0.496 | 0.51 | 1 | MS.Trt_vs_MS.NoTrt_F |
| DBR1 | 0.99017 | 0.42285 | 0.788 | 0.81 | 1 | MS.Trt_vs_MS.NoTrt_F |
| ADAM15 | 0.99202 | 0.32786 | 0.374 | 0.36 | 1 | MS.Trt_vs_MS.NoTrt_F |
| POLR2D | 0.99219 | -0.3574 | 0.784 | 0.77 | 1 | MS.Trt_vs_MS.NoTrt_F |
| GNG7 | 0.99241 | -0.51 | 0.435 | 0.46 | 1 | MS.Trt_vs_MS.NoTrt_F |
| MROH1 | 0.99298 | -0.6764 | 0.547 | 0.48 | 1 | MS.Trt_vs_MS.NoTrt_F |
| KIF1BP | 0.9936 | 0.4052 | 0.719 | 0.76 | 1 | MS.Trt_vs_MS.NoTrt_F |
| CCDC30 | 0.99396 | -0.3479 | 0.871 | 0.93 | 1 | MS.Trt_vs_MS.NoTrt_F |
| TCTA | 0.99436 | 0.43273 | 0.583 | 0.64 | 1 | MS.Trt_vs_MS.NoTrt_F |

| | | | | | | |
|---|---|---|---|---|---|---|
| AHCY | 0.9963 | 1.98788 | 0.147 | 0.16 | 1 | MS.Trt_vs_MS.NoTrt_F |
| RASGEF1B | 0.99651 | 0.49368 | 0.698 | 0.76 | 1 | MS.Trt_vs_MS.NoTrt_F |
| ADAMTS1 | 0.99654 | 0.472 | 0.439 | 0.48 | 1 | MS.Trt_vs_MS.NoTrt_F |
| UBE2G2 | 0.99709 | 0.41498 | 0.705 | 0.72 | 1 | MS.Trt_vs_MS.NoTrt_F |
| FKBP9 | 0.99719 | 0.53667 | 0.457 | 0.48 | 1 | MS.Trt_vs_MS.NoTrt_F |
| OPN3 | 0.99719 | -0.7263 | 0.313 | 0.3 | 1 | MS.Trt_vs_MS.NoTrt_F |
| PLBD1 | 0.99783 | -0.6164 | 0.482 | 0.48 | 1 | MS.Trt_vs_MS.NoTrt_F |
| SMARCAL1 | 0.99854 | 0.66325 | 0.691 | 0.74 | 1 | MS.Trt_vs_MS.NoTrt_F |
| ZIK1 | 0.99863 | 1.34723 | 0.335 | 0.33 | 1 | MS.Trt_vs_MS.NoTrt_F |
| PTRH2 | 1 | 1.04764 | 0.18 | 0.18 | 1 | MS.Trt_vs_MS.NoTrt_F |
| HLA-B | 9.08E-30 | -0.7671 | 0.83 | 0.98 | 1.45E-25 | MS.Trt_vs_MS.NoTrt_B |
| RPS4Y1 | 2.18E-23 | -4.7027 | 0.063 | 0.35 | 3.48E-19 | MS.Trt_vs_MS.NoTrt_B |
| RPL13A | 1.66E-22 | -0.6421 | 0.743 | 1 | 2.65E-18 | MS.Trt_vs_MS.NoTrt_B |
| CD99 | 3.88E-21 | -0.8095 | 0.288 | 0.71 | 6.19E-17 | MS.Trt_vs_MS.NoTrt_B |
| S100A4 | 2.04E-20 | -0.8267 | 0.388 | 0.8 | 3.26E-16 | MS.Trt_vs_MS.NoTrt_B |
| RPL27A | 2.07E-19 | -0.6265 | 0.76 | 0.98 | 3.30E-15 | MS.Trt_vs_MS.NoTrt_B |
| MT-ND4L | 8.66E-19 | 1.46742 | 0.985 | 0.99 | 1.38E-14 | MS.Trt_vs_MS.NoTrt_B |
| RPL36 | 9.59E-19 | -0.6113 | 0.458 | 0.87 | 1.53E-14 | MS.Trt_vs_MS.NoTrt_B |
| EEF2 | 1.22E-18 | -0.6186 | 0.773 | 0.99 | 1.96E-14 | MS.Trt_vs_MS.NoTrt_B |
| RPS27A | 2.36E-18 | -0.5547 | 0.59 | 0.97 | 3.76E-14 | MS.Trt_vs_MS.NoTrt_B |
| RPS16 | 3.82E-18 | -0.5884 | 0.601 | 0.92 | 6.11E-14 | MS.Trt_vs_MS.NoTrt_B |
| RPL7A | 7.51E-18 | -0.5782 | 0.608 | 0.97 | 1.20E-13 | MS.Trt_vs_MS.NoTrt_B |
| RPL37A | 2.01E-17 | -0.558 | 0.603 | 0.94 | 3.22E-13 | MS.Trt_vs_MS.NoTrt_B |
| FGF23 | 2.08E-17 | 0.51084 | 0.791 | 0.54 | 3.33E-13 | MS.Trt_vs_MS.NoTrt_B |
| RPL24 | 2.20E-17 | -0.5859 | 0.664 | 0.91 | 3.51E-13 | MS.Trt_vs_MS.NoTrt_B |
| RPL36A | 1.24E-16 | -0.5904 | 0.556 | 0.84 | 1.97E-12 | MS.Trt_vs_MS.NoTrt_B |
| RPS3 | 1.45E-16 | -0.5014 | 0.8 | 1 | 2.31E-12 | MS.Trt_vs_MS.NoTrt_B |
| FXYD5 | 2.84E-16 | -0.5862 | 0.54 | 0.89 | 4.54E-12 | MS.Trt_vs_MS.NoTrt_B |
| CTLA4 | 3.40E-16 | -1.1243 | 0.131 | 0.41 | 5.44E-12 | MS.Trt_vs_MS.NoTrt_B |
| TAC4 | 4.30E-16 | -0.788 | 0.488 | 0.74 | 6.86E-12 | MS.Trt_vs_MS.NoTrt_B |
| TMSB10 | 4.90E-16 | -0.5663 | 0.784 | 1 | 7.83E-12 | MS.Trt_vs_MS.NoTrt_B |
| RPL28 | 6.54E-16 | -0.4774 | 0.708 | 1 | 1.05E-11 | MS.Trt_vs_MS.NoTrt_B |
| RPL37 | 9.12E-16 | -0.3166 | 0.858 | 0.99 | 1.46E-11 | MS.Trt_vs_MS.NoTrt_B |
| COMMD1 | 1.66E-15 | -0.3082 | 0.42 | 0.8 | 2.65E-11 | MS.Trt_vs_MS.NoTrt_B |
| ANXA2 | 1.67E-15 | -0.7003 | 0.542 | 0.82 | 2.66E-11 | MS.Trt_vs_MS.NoTrt_B |
| RAN | 1.93E-15 | -0.5889 | 0.325 | 0.68 | 3.08E-11 | MS.Trt_vs_MS.NoTrt_B |
| CSGALNACT | 2.03E-15 | -0.7741 | 0.37 | 0.69 | 3.25E-11 | MS.Trt_vs_MS.NoTrt_B |
| RPS23 | 2.81E-15 | -0.4648 | 0.623 | 0.97 | 4.49E-11 | MS.Trt_vs_MS.NoTrt_B |
| HOXB2 | 3.63E-15 | -1.264 | 0.052 | 0.27 | 5.81E-11 | MS.Trt_vs_MS.NoTrt_B |
| RPS12 | 6.52E-15 | -0.4954 | 0.673 | 0.98 | 1.04E-10 | MS.Trt_vs_MS.NoTrt_B |

| | | | | | | |
|---|---|---|---|---|---|---|
| TOMM7 | 1.68E-14 | -0.3352 | 0.414 | 0.75 | 2.68E-10 | MS.Trt_vs_MS.NoTrt_B |
| RPLP2 | 1.80E-14 | -0.4846 | 0.571 | 0.96 | 2.87E-10 | MS.Trt_vs_MS.NoTrt_B |
| RPL30 | 1.94E-14 | -0.4353 | 0.699 | 0.97 | 3.10E-10 | MS.Trt_vs_MS.NoTrt_B |
| PKM | 3.18E-14 | -0.4513 | 0.412 | 0.76 | 5.07E-10 | MS.Trt_vs_MS.NoTrt_B |
| RPL22 | 4.02E-14 | -0.3828 | 0.451 | 0.8 | 6.42E-10 | MS.Trt_vs_MS.NoTrt_B |
| ANXA11 | 4.55E-14 | -0.624 | 0.656 | 0.88 | 7.27E-10 | MS.Trt_vs_MS.NoTrt_B |
| MT-CO2 | 5.19E-14 | 1.04422 | 0.983 | 1 | 8.30E-10 | MS.Trt_vs_MS.NoTrt_B |
| RPS28 | 7.17E-14 | -0.5258 | 0.569 | 0.94 | 1.15E-09 | MS.Trt_vs_MS.NoTrt_B |
| EHD1 | 7.34E-14 | -0.9308 | 0.2 | 0.49 | 1.17E-09 | MS.Trt_vs_MS.NoTrt_B |
| LSP1 | 8.96E-14 | -0.5224 | 0.54 | 0.87 | 1.43E-09 | MS.Trt_vs_MS.NoTrt_B |
| CSTB | 1.22E-13 | -0.3258 | 0.322 | 0.65 | 1.95E-09 | MS.Trt_vs_MS.NoTrt_B |
| TMBIM6 | 1.28E-13 | -0.4766 | 0.636 | 0.92 | 2.05E-09 | MS.Trt_vs_MS.NoTrt_B |
| XRCC6 | 1.58E-13 | -0.4347 | 0.427 | 0.76 | 2.53E-09 | MS.Trt_vs_MS.NoTrt_B |
| RPSA | 1.84E-13 | -0.5601 | 0.556 | 0.83 | 2.94E-09 | MS.Trt_vs_MS.NoTrt_B |
| NACA | 2.13E-13 | -0.3118 | 0.802 | 0.98 | 3.40E-09 | MS.Trt_vs_MS.NoTrt_B |
| RPL19 | 2.52E-13 | -0.4867 | 0.654 | 0.94 | 4.02E-09 | MS.Trt_vs_MS.NoTrt_B |
| RPL27 | 2.63E-13 | -0.286 | 0.512 | 0.89 | 4.20E-09 | MS.Trt_vs_MS.NoTrt_B |
| TMEM50A | 2.81E-13 | -0.5817 | 0.192 | 0.48 | 4.48E-09 | MS.Trt_vs_MS.NoTrt_B |
| FTH1 | 2.95E-13 | -0.4833 | 0.606 | 0.93 | 4.72E-09 | MS.Trt_vs_MS.NoTrt_B |
| CASP8 | 4.34E-13 | -0.6692 | 0.808 | 0.95 | 6.93E-09 | MS.Trt_vs_MS.NoTrt_B |
| GAPDH | 5.04E-13 | -0.5695 | 0.61 | 0.89 | 8.06E-09 | MS.Trt_vs_MS.NoTrt_B |
| WDR45 | 5.99E-13 | -0.7539 | 0.429 | 0.72 | 9.56E-09 | MS.Trt_vs_MS.NoTrt_B |
| MT-CO1 | 7.19E-13 | -0.3184 | 0.965 | 1 | 1.15E-08 | MS.Trt_vs_MS.NoTrt_B |
| ATXN10 | 7.51E-13 | -0.2685 | 0.344 | 0.65 | 1.20E-08 | MS.Trt_vs_MS.NoTrt_B |
| CD53 | 8.39E-13 | -0.4835 | 0.468 | 0.82 | 1.34E-08 | MS.Trt_vs_MS.NoTrt_B |
| NDUFA4 | 8.72E-13 | -0.2762 | 0.462 | 0.75 | 1.39E-08 | MS.Trt_vs_MS.NoTrt_B |
| RASA4 | 9.08E-13 | -0.2835 | 0.505 | 0.77 | 1.45E-08 | MS.Trt_vs_MS.NoTrt_B |
| RPL18 | 9.12E-13 | -0.4078 | 0.545 | 0.92 | 1.46E-08 | MS.Trt_vs_MS.NoTrt_B |
| CDC37 | 9.53E-13 | -0.5756 | 0.253 | 0.56 | 1.52E-08 | MS.Trt_vs_MS.NoTrt_B |
| GBP5 | 1.13E-12 | -0.5387 | 0.484 | 0.78 | 1.80E-08 | MS.Trt_vs_MS.NoTrt_B |
| TAGLN2 | 1.17E-12 | -0.6194 | 0.407 | 0.71 | 1.86E-08 | MS.Trt_vs_MS.NoTrt_B |
| RAC2 | 1.27E-12 | -0.4321 | 0.566 | 0.87 | 2.02E-08 | MS.Trt_vs_MS.NoTrt_B |
| RPL5 | 1.37E-12 | -0.42 | 0.649 | 0.96 | 2.19E-08 | MS.Trt_vs_MS.NoTrt_B |
| UQCR11 | 1.72E-12 | -0.4057 | 0.473 | 0.77 | 2.75E-08 | MS.Trt_vs_MS.NoTrt_B |
| NME2 | 1.93E-12 | -0.4514 | 0.344 | 0.65 | 3.08E-08 | MS.Trt_vs_MS.NoTrt_B |
| VIM | 2.00E-12 | -0.4296 | 0.641 | 0.92 | 3.19E-08 | MS.Trt_vs_MS.NoTrt_B |
| RPS13 | 2.03E-12 | -0.3239 | 0.688 | 0.96 | 3.24E-08 | MS.Trt_vs_MS.NoTrt_B |
| CTB-133G6.1 | 2.06E-12 | -0.6907 | 0.179 | 0.45 | 3.29E-08 | MS.Trt_vs_MS.NoTrt_B |
| PTPRCAP | 2.50E-12 | -0.516 | 0.462 | 0.78 | 3.99E-08 | MS.Trt_vs_MS.NoTrt_B |
| IKZF3 | 2.74E-12 | -0.4422 | 0.856 | 0.97 | 4.38E-08 | MS.Trt_vs_MS.NoTrt_B |

| | | | | | | |
|---|---|---|---|---|---|---|
| UBE2D2 | 2.85E-12 | -0.4827 | 0.29 | 0.6 | 4.56E-08 | MS.Trt_vs_MS.NoTrt_B |
| EEF1A1 | 3.79E-12 | -0.2996 | 0.959 | 1 | 6.06E-08 | MS.Trt_vs_MS.NoTrt_B |
| S100A10 | 4.09E-12 | -0.4443 | 0.431 | 0.76 | 6.53E-08 | MS.Trt_vs_MS.NoTrt_B |
| EIF4A1 | 4.26E-12 | -0.514 | 0.375 | 0.68 | 6.80E-08 | MS.Trt_vs_MS.NoTrt_B |
| LIME1 | 5.71E-12 | -0.4464 | 0.29 | 0.62 | 9.12E-08 | MS.Trt_vs_MS.NoTrt_B |
| RUNDC3B | 6.07E-12 | -0.5488 | 0.198 | 0.44 | 9.69E-08 | MS.Trt_vs_MS.NoTrt_B |
| HS3ST3B1 | 6.25E-12 | -0.2662 | 0.298 | 0.6 | 9.99E-08 | MS.Trt_vs_MS.NoTrt_B |
| IL32 | 6.90E-12 | -0.4147 | 0.782 | 0.97 | 1.10E-07 | MS.Trt_vs_MS.NoTrt_B |
| RPL13 | 6.93E-12 | -0.4034 | 0.791 | 0.99 | 1.11E-07 | MS.Trt_vs_MS.NoTrt_B |
| RPL23 | 7.28E-12 | -0.2952 | 0.658 | 0.9 | 1.16E-07 | MS.Trt_vs_MS.NoTrt_B |
| CD247 | 9.45E-12 | -0.41 | 0.329 | 0.63 | 1.51E-07 | MS.Trt_vs_MS.NoTrt_B |
| H3F3A | 9.72E-12 | -0.2995 | 0.508 | 0.86 | 1.55E-07 | MS.Trt_vs_MS.NoTrt_B |
| CCR6 | 9.92E-12 | -0.5003 | 0.257 | 0.52 | 1.58E-07 | MS.Trt_vs_MS.NoTrt_B |
| LAPTM5 | 1.00E-11 | -0.3884 | 0.508 | 0.83 | 1.60E-07 | MS.Trt_vs_MS.NoTrt_B |
| PRDX1 | 1.48E-11 | -0.5592 | 0.407 | 0.67 | 2.36E-07 | MS.Trt_vs_MS.NoTrt_B |
| RPS25 | 2.08E-11 | -0.3424 | 0.61 | 0.95 | 3.32E-07 | MS.Trt_vs_MS.NoTrt_B |
| SLC25A6 | 2.09E-11 | -0.4332 | 0.44 | 0.76 | 3.34E-07 | MS.Trt_vs_MS.NoTrt_B |
| EEF1B2 | 2.19E-11 | -0.4107 | 0.375 | 0.7 | 3.49E-07 | MS.Trt_vs_MS.NoTrt_B |
| C9orf16 | 2.34E-11 | -0.5231 | 0.148 | 0.4 | 3.74E-07 | MS.Trt_vs_MS.NoTrt_B |
| RPS8 | 2.41E-11 | -0.3908 | 0.854 | 1 | 3.85E-07 | MS.Trt_vs_MS.NoTrt_B |
| ANXA6 | 2.50E-11 | -0.6236 | 0.34 | 0.64 | 4.00E-07 | MS.Trt_vs_MS.NoTrt_B |
| LDHB | 2.89E-11 | -0.4215 | 0.575 | 0.91 | 4.61E-07 | MS.Trt_vs_MS.NoTrt_B |
| PSMD8 | 3.08E-11 | -0.5562 | 0.462 | 0.73 | 4.92E-07 | MS.Trt_vs_MS.NoTrt_B |
| HCST | 3.25E-11 | -0.6281 | 0.216 | 0.48 | 5.19E-07 | MS.Trt_vs_MS.NoTrt_B |
| STMN3 | 3.42E-11 | -0.4286 | 0.155 | 0.39 | 5.47E-07 | MS.Trt_vs_MS.NoTrt_B |
| MSN | 3.48E-11 | -0.367 | 0.453 | 0.74 | 5.55E-07 | MS.Trt_vs_MS.NoTrt_B |
| CD52 | 3.50E-11 | -0.3679 | 0.643 | 0.94 | 5.59E-07 | MS.Trt_vs_MS.NoTrt_B |
| RPL31 | 3.69E-11 | -0.2967 | 0.593 | 0.94 | 5.90E-07 | MS.Trt_vs_MS.NoTrt_B |
| NONO | 3.71E-11 | -0.4099 | 0.49 | 0.78 | 5.93E-07 | MS.Trt_vs_MS.NoTrt_B |
| RPL15 | 4.13E-11 | -0.2752 | 0.882 | 1 | 6.60E-07 | MS.Trt_vs_MS.NoTrt_B |
| CD5 | 4.33E-11 | -0.4254 | 0.338 | 0.65 | 6.92E-07 | MS.Trt_vs_MS.NoTrt_B |
| ARPC3 | 6.65E-11 | -0.3827 | 0.331 | 0.64 | 1.06E-06 | MS.Trt_vs_MS.NoTrt_B |
| TBC1D4 | 7.27E-11 | -0.4045 | 0.237 | 0.51 | 1.16E-06 | MS.Trt_vs_MS.NoTrt_B |
| PSME1 | 7.38E-11 | -0.3703 | 0.464 | 0.76 | 1.18E-06 | MS.Trt_vs_MS.NoTrt_B |
| PBX1 | 7.41E-11 | -0.3348 | 0.728 | 0.95 | 1.18E-06 | MS.Trt_vs_MS.NoTrt_B |
| ATP2B4 | 7.64E-11 | -0.3595 | 0.344 | 0.62 | 1.22E-06 | MS.Trt_vs_MS.NoTrt_B |
| TXNIP | 8.08E-11 | -0.3632 | 0.691 | 0.94 | 1.29E-06 | MS.Trt_vs_MS.NoTrt_B |
| CAPN2 | 1.01E-10 | -0.3311 | 0.617 | 0.85 | 1.61E-06 | MS.Trt_vs_MS.NoTrt_B |
| RPS11 | 1.27E-10 | -0.321 | 0.556 | 0.87 | 2.03E-06 | MS.Trt_vs_MS.NoTrt_B |
| HNRNPH1 | 1.45E-10 | 1.10158 | 0.989 | 1 | 2.31E-06 | MS.Trt_vs_MS.NoTrt_B |

| | | | | | | |
|---|---|---|---|---|---|---|
| CNN2 | 1.67E-10 | -0.5012 | 0.954 | 0.99 | 2.66E-06 | MS.Trt_vs_MS.NoTrt_B |
| RASGRP1 | 1.69E-10 | -0.6075 | 0.218 | 0.45 | 2.69E-06 | MS.Trt_vs_MS.NoTrt_B |
| TUBA1B | 1.79E-10 | -0.5688 | 0.203 | 0.45 | 2.86E-06 | MS.Trt_vs_MS.NoTrt_B |
| EPS15 | 2.23E-10 | -0.3974 | 0.449 | 0.7 | 3.57E-06 | MS.Trt_vs_MS.NoTrt_B |
| RPS24 | 2.46E-10 | -0.3593 | 0.776 | 0.98 | 3.93E-06 | MS.Trt_vs_MS.NoTrt_B |
| P2RY8 | 3.11E-10 | -0.6018 | 0.368 | 0.64 | 4.97E-06 | MS.Trt_vs_MS.NoTrt_B |
| CALCOCO1 | 3.35E-10 | -0.368 | 0.338 | 0.63 | 5.34E-06 | MS.Trt_vs_MS.NoTrt_B |
| VCP | 3.94E-10 | -0.5354 | 0.2 | 0.45 | 6.29E-06 | MS.Trt_vs_MS.NoTrt_B |
| ARHGAP30 | 4.67E-10 | -0.55 | 0.298 | 0.56 | 7.45E-06 | MS.Trt_vs_MS.NoTrt_B |
| TTLL5 | 4.86E-10 | -0.6942 | 0.499 | 0.74 | 7.76E-06 | MS.Trt_vs_MS.NoTrt_B |
| GPI | 5.58E-10 | -0.5181 | 0.795 | 0.92 | 8.92E-06 | MS.Trt_vs_MS.NoTrt_B |
| EDF1 | 5.80E-10 | -0.3296 | 0.288 | 0.58 | 9.26E-06 | MS.Trt_vs_MS.NoTrt_B |
| RELL2 | 6.10E-10 | -1.6892 | 0.026 | 0.16 | 9.74E-06 | MS.Trt_vs_MS.NoTrt_B |
| RHOH | 6.45E-10 | -0.3573 | 0.721 | 0.89 | 1.03E-05 | MS.Trt_vs_MS.NoTrt_B |
| FGD3 | 6.90E-10 | -0.4055 | 0.414 | 0.68 | 1.10E-05 | MS.Trt_vs_MS.NoTrt_B |
| GBP2 | 7.03E-10 | -0.56 | 0.316 | 0.58 | 1.12E-05 | MS.Trt_vs_MS.NoTrt_B |
| ACTG1 | 7.10E-10 | -0.4061 | 0.802 | 0.96 | 1.13E-05 | MS.Trt_vs_MS.NoTrt_B |
| LDLRAP1 | 7.12E-10 | -0.6412 | 0.316 | 0.55 | 1.14E-05 | MS.Trt_vs_MS.NoTrt_B |
| RPL35A | 7.37E-10 | -0.2594 | 0.747 | 0.97 | 1.18E-05 | MS.Trt_vs_MS.NoTrt_B |
| HMGN4 | 7.44E-10 | -0.4796 | 0.176 | 0.41 | 1.19E-05 | MS.Trt_vs_MS.NoTrt_B |
| CHTF8 | 8.03E-10 | -0.7149 | 0.144 | 0.35 | 1.28E-05 | MS.Trt_vs_MS.NoTrt_B |
| AQP3 | 8.34E-10 | -0.9674 | 0.19 | 0.41 | 1.33E-05 | MS.Trt_vs_MS.NoTrt_B |
| TMEM63A | 8.52E-10 | -0.4917 | 0.599 | 0.83 | 1.36E-05 | MS.Trt_vs_MS.NoTrt_B |
| PBXIP1 | 8.69E-10 | -0.5873 | 0.466 | 0.71 | 1.39E-05 | MS.Trt_vs_MS.NoTrt_B |
| COX7C | 8.83E-10 | -0.3303 | 0.288 | 0.56 | 1.41E-05 | MS.Trt_vs_MS.NoTrt_B |
| PPP2R1A | 9.01E-10 | -0.2771 | 0.547 | 0.77 | 1.44E-05 | MS.Trt_vs_MS.NoTrt_B |
| JAKMIP1 | 9.31E-10 | -0.6981 | 0.163 | 0.38 | 1.49E-05 | MS.Trt_vs_MS.NoTrt_B |
| GGA2 | 9.72E-10 | -0.4619 | 0.479 | 0.75 | 1.55E-05 | MS.Trt_vs_MS.NoTrt_B |
| IL2RA | 1.00E-09 | -0.7032 | 0.268 | 0.51 | 1.60E-05 | MS.Trt_vs_MS.NoTrt_B |
| ANK3 | 1.00E-09 | 0.85485 | 0.926 | 0.82 | 1.60E-05 | MS.Trt_vs_MS.NoTrt_B |
| ANXA5 | 1.03E-09 | -0.764 | 0.192 | 0.42 | 1.65E-05 | MS.Trt_vs_MS.NoTrt_B |
| ITGB1 | 1.10E-09 | -0.408 | 0.523 | 0.76 | 1.76E-05 | MS.Trt_vs_MS.NoTrt_B |
| TALDO1 | 1.10E-09 | -0.7461 | 0.538 | 0.78 | 1.76E-05 | MS.Trt_vs_MS.NoTrt_B |
| PRKCQ | 1.12E-09 | -0.3362 | 0.259 | 0.52 | 1.79E-05 | MS.Trt_vs_MS.NoTrt_B |
| ATP5B | 1.14E-09 | -0.6441 | 0.251 | 0.5 | 1.82E-05 | MS.Trt_vs_MS.NoTrt_B |
| MT-ND4 | 1.15E-09 | 0.75919 | 0.989 | 1 | 1.84E-05 | MS.Trt_vs_MS.NoTrt_B |
| RBM4 | 1.23E-09 | -0.2539 | 0.551 | 0.8 | 1.96E-05 | MS.Trt_vs_MS.NoTrt_B |
| DEF6 | 1.30E-09 | -0.368 | 0.331 | 0.6 | 2.07E-05 | MS.Trt_vs_MS.NoTrt_B |
| ZNF362 | 1.38E-09 | -1.6086 | 0.048 | 0.2 | 2.20E-05 | MS.Trt_vs_MS.NoTrt_B |
| RPS6KA1 | 1.43E-09 | -0.9985 | 0.122 | 0.32 | 2.28E-05 | MS.Trt_vs_MS.NoTrt_B |

| | | | | | | |
|---|---|---|---|---|---|---|
| ZAP70 | 1.50E-09 | -0.5435 | 0.359 | 0.62 | 2.39E-05 | MS.Trt_vs_MS.NoTrt_B |
| LDLRAD4 | 1.60E-09 | -0.4628 | 0.218 | 0.45 | 2.55E-05 | MS.Trt_vs_MS.NoTrt_B |
| FAM118A | 1.72E-09 | -1.2386 | 0.447 | 0.68 | 2.75E-05 | MS.Trt_vs_MS.NoTrt_B |
| TMEM181 | 1.77E-09 | -0.632 | 0.29 | 0.55 | 2.82E-05 | MS.Trt_vs_MS.NoTrt_B |
| RARRES3 | 1.78E-09 | -0.4642 | 0.214 | 0.46 | 2.85E-05 | MS.Trt_vs_MS.NoTrt_B |
| LITAF | 1.79E-09 | -0.5347 | 0.41 | 0.65 | 2.87E-05 | MS.Trt_vs_MS.NoTrt_B |
| CHCHD2 | 1.92E-09 | -0.3186 | 0.292 | 0.55 | 3.06E-05 | MS.Trt_vs_MS.NoTrt_B |
| HLA-DQB1 | 1.98E-09 | -1.0829 | 0.394 | 0.65 | 3.17E-05 | MS.Trt_vs_MS.NoTrt_B |
| NCOA4 | 2.06E-09 | -0.8032 | 0.131 | 0.34 | 3.29E-05 | MS.Trt_vs_MS.NoTrt_B |
| REEP5 | 2.25E-09 | -0.2962 | 0.214 | 0.45 | 3.59E-05 | MS.Trt_vs_MS.NoTrt_B |
| MBP | 2.31E-09 | -0.5014 | 0.275 | 0.51 | 3.70E-05 | MS.Trt_vs_MS.NoTrt_B |
| TADA3 | 2.62E-09 | -0.3589 | 0.442 | 0.71 | 4.18E-05 | MS.Trt_vs_MS.NoTrt_B |
| SEPW1 | 3.22E-09 | -0.2904 | 0.834 | 0.96 | 5.14E-05 | MS.Trt_vs_MS.NoTrt_B |
| GLRX | 3.25E-09 | -0.5039 | 0.58 | 0.78 | 5.20E-05 | MS.Trt_vs_MS.NoTrt_B |
| RALY | 3.40E-09 | -0.4434 | 0.259 | 0.51 | 5.42E-05 | MS.Trt_vs_MS.NoTrt_B |
| LGALS3 | 3.68E-09 | -0.6202 | 0.466 | 0.67 | 5.87E-05 | MS.Trt_vs_MS.NoTrt_B |
| UFD1L | 3.95E-09 | -0.5585 | 0.66 | 0.83 | 6.31E-05 | MS.Trt_vs_MS.NoTrt_B |
| ERCC1 | 4.10E-09 | -0.3086 | 0.621 | 0.88 | 6.54E-05 | MS.Trt_vs_MS.NoTrt_B |
| UBE2F | 4.14E-09 | -0.2854 | 0.492 | 0.76 | 6.61E-05 | MS.Trt_vs_MS.NoTrt_B |
| ATP5F1 | 4.31E-09 | -0.2533 | 0.359 | 0.59 | 6.89E-05 | MS.Trt_vs_MS.NoTrt_B |
| TSC22D3 | 4.47E-09 | -0.3978 | 0.475 | 0.75 | 7.14E-05 | MS.Trt_vs_MS.NoTrt_B |
| DBI | 5.01E-09 | -0.6025 | 0.131 | 0.33 | 8.01E-05 | MS.Trt_vs_MS.NoTrt_B |
| RPL4 | 5.14E-09 | -0.3645 | 0.837 | 0.98 | 8.21E-05 | MS.Trt_vs_MS.NoTrt_B |
| SRP14 | 5.32E-09 | -0.2756 | 0.42 | 0.72 | 8.50E-05 | MS.Trt_vs_MS.NoTrt_B |
| HLA-A | 5.35E-09 | -0.4314 | 0.747 | 0.94 | 8.55E-05 | MS.Trt_vs_MS.NoTrt_B |
| TRPC3 | 5.72E-09 | 0.4049 | 0.42 | 0.18 | 9.13E-05 | MS.Trt_vs_MS.NoTrt_B |
| IRF9 | 6.30E-09 | -0.427 | 0.756 | 0.94 | 1.01E-04 | MS.Trt_vs_MS.NoTrt_B |
| FLT3LG | 7.07E-09 | -0.3836 | 0.702 | 0.83 | 1.13E-04 | MS.Trt_vs_MS.NoTrt_B |
| PEBP1 | 7.28E-09 | -0.4295 | 0.279 | 0.52 | 1.16E-04 | MS.Trt_vs_MS.NoTrt_B |
| MGAT1 | 7.30E-09 | -0.5059 | 0.392 | 0.63 | 1.17E-04 | MS.Trt_vs_MS.NoTrt_B |
| MRI1 | 7.33E-09 | -0.4848 | 0.573 | 0.83 | 1.17E-04 | MS.Trt_vs_MS.NoTrt_B |
| ACLY | 7.72E-09 | -0.7475 | 0.109 | 0.3 | 1.23E-04 | MS.Trt_vs_MS.NoTrt_B |
| ATG7 | 8.16E-09 | -0.6106 | 0.359 | 0.66 | 1.30E-04 | MS.Trt_vs_MS.NoTrt_B |
| MYH9 | 8.18E-09 | -0.2759 | 0.656 | 0.89 | 1.31E-04 | MS.Trt_vs_MS.NoTrt_B |
| PCSK1N | 8.20E-09 | -0.4319 | 0.2 | 0.41 | 1.31E-04 | MS.Trt_vs_MS.NoTrt_B |
| CEP63 | 8.29E-09 | -0.2676 | 0.473 | 0.76 | 0.000132 | MS.Trt_vs_MS.NoTrt_B |
| CKLF | 8.48E-09 | -0.3264 | 0.144 | 0.34 | 1.35E-04 | MS.Trt_vs_MS.NoTrt_B |
| LCP1 | 8.69E-09 | -0.2994 | 0.51 | 0.81 | 1.39E-04 | MS.Trt_vs_MS.NoTrt_B |
| DIP2B | 9.05E-09 | -0.4311 | 0.305 | 0.54 | 1.45E-04 | MS.Trt_vs_MS.NoTrt_B |
| SMIM10L1 | 9.11E-09 | -1.2153 | 0.102 | 0.28 | 1.45E-04 | MS.Trt_vs_MS.NoTrt_B |

| | | | | | | |
|---|---|---|---|---|---|---|
| HNRNPA1 | 9.43E-09 | -0.3017 | 0.671 | 0.93 | 1.51E-04 | MS.Trt_vs_MS.NoTrt_B |
| LRRC75A | 9.78E-09 | -0.3235 | 0.394 | 0.68 | 1.56E-04 | MS.Trt_vs_MS.NoTrt_B |
| ACTN4 | 9.84E-09 | -0.6462 | 0.231 | 0.46 | 1.57E-04 | MS.Trt_vs_MS.NoTrt_B |
| ARL17A | 1.06E-08 | -0.5043 | 0.503 | 0.75 | 1.69E-04 | MS.Trt_vs_MS.NoTrt_B |
| VAMP5 | 1.13E-08 | -0.3633 | 0.096 | 0.28 | 1.81E-04 | MS.Trt_vs_MS.NoTrt_B |
| TMEM219 | 1.20E-08 | -0.5019 | 0.146 | 0.36 | 1.91E-04 | MS.Trt_vs_MS.NoTrt_B |
| NCKAP1L | 1.21E-08 | -0.5269 | 0.601 | 0.8 | 1.93E-04 | MS.Trt_vs_MS.NoTrt_B |
| PPIA | 1.33E-08 | -0.2801 | 0.754 | 0.94 | 2.12E-04 | MS.Trt_vs_MS.NoTrt_B |
| WDR1 | 1.46E-08 | -0.4126 | 0.277 | 0.53 | 2.33E-04 | MS.Trt_vs_MS.NoTrt_B |
| CSNK2A2 | 1.52E-08 | -1.0503 | 0.159 | 0.35 | 2.43E-04 | MS.Trt_vs_MS.NoTrt_B |
| CCM2 | 1.55E-08 | -0.5249 | 0.562 | 0.74 | 2.48E-04 | MS.Trt_vs_MS.NoTrt_B |
| BRK1 | 1.64E-08 | -0.3863 | 0.168 | 0.39 | 2.62E-04 | MS.Trt_vs_MS.NoTrt_B |
| CPQ | 1.64E-08 | -0.5681 | 0.24 | 0.47 | 2.63E-04 | MS.Trt_vs_MS.NoTrt_B |
| LPP | 1.67E-08 | -0.4159 | 0.867 | 0.95 | 2.67E-04 | MS.Trt_vs_MS.NoTrt_B |
| TNFRSF25 | 1.68E-08 | -0.2967 | 0.545 | 0.76 | 2.68E-04 | MS.Trt_vs_MS.NoTrt_B |
| TAP1 | 1.71E-08 | -0.5056 | 0.514 | 0.74 | 2.73E-04 | MS.Trt_vs_MS.NoTrt_B |
| GLIPR2 | 1.89E-08 | -0.3636 | 0.174 | 0.38 | 3.02E-04 | MS.Trt_vs_MS.NoTrt_B |
| RAD17 | 1.91E-08 | -0.6572 | 0.357 | 0.59 | 3.06E-04 | MS.Trt_vs_MS.NoTrt_B |
| IK | 2.07E-08 | -0.2682 | 0.68 | 0.87 | 3.31E-04 | MS.Trt_vs_MS.NoTrt_B |
| TRAPPC3 | 2.08E-08 | -0.5443 | 0.277 | 0.5 | 3.32E-04 | MS.Trt_vs_MS.NoTrt_B |
| CD6 | 2.17E-08 | -0.3246 | 0.514 | 0.77 | 3.46E-04 | MS.Trt_vs_MS.NoTrt_B |
| STX8 | 2.19E-08 | -0.31 | 0.712 | 0.91 | 3.50E-04 | MS.Trt_vs_MS.NoTrt_B |
| EIF3K | 2.31E-08 | -0.2943 | 0.538 | 0.78 | 3.69E-04 | MS.Trt_vs_MS.NoTrt_B |
| TBC1D5 | 2.32E-08 | -0.5415 | 0.44 | 0.68 | 3.70E-04 | MS.Trt_vs_MS.NoTrt_B |
| APRT | 2.39E-08 | -0.391 | 0.277 | 0.51 | 3.81E-04 | MS.Trt_vs_MS.NoTrt_B |
| SERINC3 | 2.41E-08 | -0.4184 | 0.536 | 0.74 | 3.85E-04 | MS.Trt_vs_MS.NoTrt_B |
| SNRPN | 2.42E-08 | -0.2773 | 0.416 | 0.64 | 3.87E-04 | MS.Trt_vs_MS.NoTrt_B |
| CFDP1 | 2.44E-08 | -0.9415 | 0.516 | 0.71 | 3.90E-04 | MS.Trt_vs_MS.NoTrt_B |
| HSPA8 | 2.65E-08 | -0.2802 | 0.656 | 0.92 | 4.22E-04 | MS.Trt_vs_MS.NoTrt_B |
| GPR141 | 2.78E-08 | -0.2644 | 0.312 | 0.57 | 4.43E-04 | MS.Trt_vs_MS.NoTrt_B |
| LIMK2 | 2.84E-08 | -0.9598 | 0.111 | 0.29 | 4.54E-04 | MS.Trt_vs_MS.NoTrt_B |
| CCR4 | 2.90E-08 | -1.0251 | 0.786 | 0.91 | 4.63E-04 | MS.Trt_vs_MS.NoTrt_B |
| CDC25B | 2.91E-08 | -0.2531 | 0.333 | 0.58 | 4.65E-04 | MS.Trt_vs_MS.NoTrt_B |
| EIF1AY | 2.93E-08 | -1.2529 | 0.451 | 0.65 | 4.69E-04 | MS.Trt_vs_MS.NoTrt_B |
| COQ10B | 3.02E-08 | 0.2637 | 0.451 | 0.7 | 4.82E-04 | MS.Trt_vs_MS.NoTrt_B |
| CAPZB | 3.09E-08 | -0.3725 | 0.458 | 0.7 | 4.94E-04 | MS.Trt_vs_MS.NoTrt_B |
| EIF4A2 | 3.15E-08 | -0.3607 | 0.401 | 0.64 | 5.04E-04 | MS.Trt_vs_MS.NoTrt_B |
| DHFR2 | 3.31E-08 | -1.1647 | 0.423 | 0.63 | 5.29E-04 | MS.Trt_vs_MS.NoTrt_B |
| RABGAP1L | 3.33E-08 | -0.6569 | 0.78 | 0.91 | 5.33E-04 | MS.Trt_vs_MS.NoTrt_B |
| DNMT3A | 3.49E-08 | 0.31858 | 0.364 | 0.63 | 5.58E-04 | MS.Trt_vs_MS.NoTrt_B |

| | | | | | | |
|---|---|---|---|---|---|---|
| PXN | 3.50E-08 | -0.2858 | 0.283 | 0.51 | 5.59E-04 | MS.Trt_vs_MS.NoTrt_B |
| SLFN5 | 3.69E-08 | -0.2891 | 0.763 | 0.89 | 5.90E-04 | MS.Trt_vs_MS.NoTrt_B |
| GBP3 | 3.74E-08 | -1.2329 | 0.122 | 0.29 | 5.97E-04 | MS.Trt_vs_MS.NoTrt_B |
| POLD4 | 3.88E-08 | -0.2944 | 0.654 | 0.85 | 6.20E-04 | MS.Trt_vs_MS.NoTrt_B |
| PSMA3 | 4.01E-08 | -0.4109 | 0.595 | 0.82 | 6.40E-04 | MS.Trt_vs_MS.NoTrt_B |
| NGLY1 | 4.09E-08 | -0.4772 | 0.501 | 0.76 | 6.53E-04 | MS.Trt_vs_MS.NoTrt_B |
| DSCR3 | 4.15E-08 | -0.4739 | 0.503 | 0.71 | 6.63E-04 | MS.Trt_vs_MS.NoTrt_B |
| TMEM14B | 4.53E-08 | -0.3458 | 0.725 | 0.9 | 7.23E-04 | MS.Trt_vs_MS.NoTrt_B |
| TMEM179B | 4.70E-08 | -0.599 | 0.118 | 0.3 | 7.51E-04 | MS.Trt_vs_MS.NoTrt_B |
| RPL17 | 4.90E-08 | -0.2662 | 0.756 | 1 | 7.83E-04 | MS.Trt_vs_MS.NoTrt_B |
| VTI1B | 4.98E-08 | -0.5808 | 0.142 | 0.32 | 7.96E-04 | MS.Trt_vs_MS.NoTrt_B |
| SQSTM1 | 5.58E-08 | -0.4779 | 0.736 | 0.84 | 8.91E-04 | MS.Trt_vs_MS.NoTrt_B |
| ARPC1B | 5.70E-08 | -0.2734 | 0.841 | 0.98 | 9.10E-04 | MS.Trt_vs_MS.NoTrt_B |
| PGK1 | 5.82E-08 | -0.3515 | 0.431 | 0.66 | 9.29E-04 | MS.Trt_vs_MS.NoTrt_B |
| FUBP1 | 6.00E-08 | -0.4386 | 0.27 | 0.49 | 9.58E-04 | MS.Trt_vs_MS.NoTrt_B |
| C11orf31 | 6.06E-08 | -0.4224 | 0.397 | 0.6 | 9.68E-04 | MS.Trt_vs_MS.NoTrt_B |
| ATP2C1 | 6.30E-08 | -0.3941 | 0.085 | 0.24 | 0.001006 | MS.Trt_vs_MS.NoTrt_B |
| SMIM5 | 6.34E-08 | -0.3937 | 0.185 | 0.39 | 0.001013 | MS.Trt_vs_MS.NoTrt_B |
| ADI1 | 6.37E-08 | 0.28825 | 0.651 | 0.87 | 0.001017 | MS.Trt_vs_MS.NoTrt_B |
| HLA-F | 6.68E-08 | -0.2631 | 0.451 | 0.65 | 0.001067 | MS.Trt_vs_MS.NoTrt_B |
| NDUFS1 | 7.04E-08 | -0.4632 | 0.832 | 0.98 | 0.001124 | MS.Trt_vs_MS.NoTrt_B |
| DDX58 | 7.24E-08 | -0.3197 | 0.203 | 0.4 | 0.001157 | MS.Trt_vs_MS.NoTrt_B |
| SCAP | 7.30E-08 | -0.3496 | 0.427 | 0.68 | 0.001166 | MS.Trt_vs_MS.NoTrt_B |
| PYM1 | 7.33E-08 | -0.4458 | 0.203 | 0.41 | 0.00117 | MS.Trt_vs_MS.NoTrt_B |
| EIF3G | 7.65E-08 | -0.3932 | 0.743 | 0.89 | 0.001222 | MS.Trt_vs_MS.NoTrt_B |
| TLK1 | 7.83E-08 | 0.32078 | 0.632 | 0.85 | 0.00125 | MS.Trt_vs_MS.NoTrt_B |
| CYTH2 | 7.93E-08 | -0.2853 | 0.475 | 0.75 | 0.001267 | MS.Trt_vs_MS.NoTrt_B |
| MAPK6 | 8.42E-08 | -0.2591 | 0.305 | 0.54 | 0.001345 | MS.Trt_vs_MS.NoTrt_B |
| DDB2 | 8.61E-08 | -0.3844 | 0.771 | 0.9 | 0.001375 | MS.Trt_vs_MS.NoTrt_B |
| C12orf75 | 8.86E-08 | -0.6042 | 0.092 | 0.25 | 0.001415 | MS.Trt_vs_MS.NoTrt_B |
| SPOCK2 | 9.11E-08 | -0.446 | 0.514 | 0.71 | 0.001456 | MS.Trt_vs_MS.NoTrt_B |
| EIF3C | 9.16E-08 | -0.2645 | 0.641 | 0.85 | 0.001464 | MS.Trt_vs_MS.NoTrt_B |
| RPL12 | 9.25E-08 | -0.2625 | 0.715 | 0.95 | 0.001478 | MS.Trt_vs_MS.NoTrt_B |
| ATP5O | 9.53E-08 | -0.362 | 0.283 | 0.52 | 0.001522 | MS.Trt_vs_MS.NoTrt_B |
| DDX24 | 9.61E-08 | -0.2839 | 0.405 | 0.64 | 0.001534 | MS.Trt_vs_MS.NoTrt_B |
| ATP5A1 | 9.73E-08 | -0.3241 | 0.976 | 1 | 0.001553 | MS.Trt_vs_MS.NoTrt_B |
| SHMT2 | 9.74E-08 | -0.557 | 0.198 | 0.41 | 0.001556 | MS.Trt_vs_MS.NoTrt_B |
| CIRBP | 1.08E-07 | -0.257 | 0.477 | 0.78 | 0.001724 | MS.Trt_vs_MS.NoTrt_B |
| ZC3HC1 | 1.13E-07 | -0.2638 | 0.272 | 0.5 | 0.001807 | MS.Trt_vs_MS.NoTrt_B |
| PPM1M | 1.13E-07 | -0.5409 | 0.094 | 0.25 | 0.001811 | MS.Trt_vs_MS.NoTrt_B |

| | | | | | | |
|---|---|---|---|---|---|---|
| ARFGAP3 | 1.17E-07 | -0.5099 | 0.211 | 0.42 | 0.001876 | MS.Trt_vs_MS.NoTrt_B |
| RNF114 | 1.22E-07 | -0.4535 | 0.593 | 0.83 | 0.001954 | MS.Trt_vs_MS.NoTrt_B |
| CBWD2 | 1.23E-07 | -0.6947 | 0.331 | 0.54 | 0.001971 | MS.Trt_vs_MS.NoTrt_B |
| NDEL1 | 1.26E-07 | -0.3324 | 0.444 | 0.66 | 0.002012 | MS.Trt_vs_MS.NoTrt_B |
| FDFT1 | 1.29E-07 | -0.5096 | 0.224 | 0.42 | 0.002053 | MS.Trt_vs_MS.NoTrt_B |
| UXS1 | 1.29E-07 | -0.3296 | 0.172 | 0.37 | 0.002056 | MS.Trt_vs_MS.NoTrt_B |
| GLB1 | 1.31E-07 | -0.619 | 0.153 | 0.34 | 0.002098 | MS.Trt_vs_MS.NoTrt_B |
| PIM2 | 1.32E-07 | -0.4401 | 0.728 | 0.85 | 0.002106 | MS.Trt_vs_MS.NoTrt_B |
| FBXO7 | 1.41E-07 | -0.3169 | 0.233 | 0.44 | 0.00225 | MS.Trt_vs_MS.NoTrt_B |
| HINT2 | 1.51E-07 | -0.8318 | 0.041 | 0.16 | 0.002404 | MS.Trt_vs_MS.NoTrt_B |
| XPR1 | 1.53E-07 | -0.6756 | 0.211 | 0.41 | 0.002441 | MS.Trt_vs_MS.NoTrt_B |
| RPL29 | 1.63E-07 | -0.2645 | 0.695 | 0.98 | 0.002597 | MS.Trt_vs_MS.NoTrt_B |
| WWC3 | 1.64E-07 | 1.16987 | 0.227 | 0.44 | 0.002618 | MS.Trt_vs_MS.NoTrt_B |
| PAX6 | 1.68E-07 | -0.5336 | 0.24 | 0.44 | 0.002679 | MS.Trt_vs_MS.NoTrt_B |
| UCP2 | 1.68E-07 | -0.4005 | 0.669 | 0.86 | 0.002681 | MS.Trt_vs_MS.NoTrt_B |
| FAM161A | 1.73E-07 | -0.9156 | 0.508 | 0.76 | 0.002761 | MS.Trt_vs_MS.NoTrt_B |
| ADGRE5 | 1.77E-07 | -0.3156 | 0.458 | 0.69 | 0.002824 | MS.Trt_vs_MS.NoTrt_B |
| FKBP8 | 1.77E-07 | -0.5798 | 0.129 | 0.31 | 0.002825 | MS.Trt_vs_MS.NoTrt_B |
| KARS | 1.77E-07 | -0.4313 | 0.196 | 0.39 | 0.002829 | MS.Trt_vs_MS.NoTrt_B |
| PLA2G7 | 1.80E-07 | -0.439 | 0.414 | 0.62 | 0.002868 | MS.Trt_vs_MS.NoTrt_B |
| ANAPC5 | 1.82E-07 | -0.3551 | 0.885 | 0.98 | 0.00291 | MS.Trt_vs_MS.NoTrt_B |
| CHMP7 | 1.91E-07 | -0.4117 | 0.449 | 0.67 | 0.003058 | MS.Trt_vs_MS.NoTrt_B |
| CSNK2B | 1.92E-07 | -0.5475 | 0.545 | 0.74 | 0.003073 | MS.Trt_vs_MS.NoTrt_B |
| SCRN3 | 1.93E-07 | -0.7521 | 0.294 | 0.51 | 0.003076 | MS.Trt_vs_MS.NoTrt_B |
| ADAT1 | 1.96E-07 | -1.0545 | 0.525 | 0.74 | 0.003131 | MS.Trt_vs_MS.NoTrt_B |
| SDHA | 1.98E-07 | -0.472 | 0.501 | 0.7 | 0.00316 | MS.Trt_vs_MS.NoTrt_B |
| PERP | 1.98E-07 | -2.183 | 0.022 | 0.12 | 0.003164 | MS.Trt_vs_MS.NoTrt_B |
| ZC3H7A | 1.99E-07 | -0.43 | 0.259 | 0.47 | 0.003172 | MS.Trt_vs_MS.NoTrt_B |
| URM1 | 2.06E-07 | -1.0405 | 0.081 | 0.23 | 0.00329 | MS.Trt_vs_MS.NoTrt_B |
| ARHGAP4 | 2.12E-07 | -0.302 | 0.37 | 0.58 | 0.003393 | MS.Trt_vs_MS.NoTrt_B |
| HBP1 | 2.29E-07 | -0.3422 | 0.669 | 0.86 | 0.003658 | MS.Trt_vs_MS.NoTrt_B |
| LSM7 | 2.31E-07 | 0.67578 | 0.486 | 0.74 | 0.003693 | MS.Trt_vs_MS.NoTrt_B |
| APOL6 | 2.37E-07 | -0.6695 | 0.725 | 0.88 | 0.003787 | MS.Trt_vs_MS.NoTrt_B |
| ALDH8A1 | 2.38E-07 | -0.3553 | 0.401 | 0.67 | 0.003806 | MS.Trt_vs_MS.NoTrt_B |
| PSMA6 | 2.41E-07 | -0.3523 | 0.782 | 0.92 | 0.00385 | MS.Trt_vs_MS.NoTrt_B |
| TMEM199 | 2.48E-07 | -1.2226 | 0.444 | 0.67 | 0.003969 | MS.Trt_vs_MS.NoTrt_B |
| MYO1G | 2.55E-07 | -0.4159 | 0.455 | 0.68 | 0.004073 | MS.Trt_vs_MS.NoTrt_B |
| EZR | 2.62E-07 | -0.3977 | 0.562 | 0.74 | 0.004179 | MS.Trt_vs_MS.NoTrt_B |
| UHRF1BP1 | 2.66E-07 | -1.4044 | 0.425 | 0.65 | 0.004247 | MS.Trt_vs_MS.NoTrt_B |
| LDHA | 2.67E-07 | -0.5396 | 0.582 | 0.76 | 0.004273 | MS.Trt_vs_MS.NoTrt_B |

| | | | | | | |
|---|---|---|---|---|---|---|
| FAS | 2.72E-07 | -0.477 | 0.244 | 0.44 | 0.004344 | MS.Trt_vs_MS.NoTrt_B |
| MAP3K1 | 2.72E-07 | 0.28228 | 0.386 | 0.64 | 0.004351 | MS.Trt_vs_MS.NoTrt_B |
| RP11-529K1 | 2.74E-07 | -0.4362 | 0.039 | 0.16 | 0.004373 | MS.Trt_vs_MS.NoTrt_B |
| IL2RB | 2.91E-07 | -0.3107 | 0.364 | 0.6 | 0.004643 | MS.Trt_vs_MS.NoTrt_B |
| TMBIM1 | 2.96E-07 | -0.8063 | 0.44 | 0.64 | 0.004729 | MS.Trt_vs_MS.NoTrt_B |
| C14orf159 | 2.98E-07 | -0.815 | 0.525 | 0.73 | 0.004768 | MS.Trt_vs_MS.NoTrt_B |
| SS18 | 2.99E-07 | -0.4234 | 0.429 | 0.65 | 0.004771 | MS.Trt_vs_MS.NoTrt_B |
| ARHGAP25 | 2.99E-07 | -0.4901 | 0.383 | 0.59 | 0.004779 | MS.Trt_vs_MS.NoTrt_B |
| SHMT1 | 3.05E-07 | -0.5326 | 0.272 | 0.48 | 0.004867 | MS.Trt_vs_MS.NoTrt_B |
| GSDMD | 3.09E-07 | -0.6436 | 0.621 | 0.82 | 0.004937 | MS.Trt_vs_MS.NoTrt_B |
| GRB2 | 3.10E-07 | -0.4769 | 0.534 | 0.74 | 0.004957 | MS.Trt_vs_MS.NoTrt_B |
| TXLNA | 3.12E-07 | -0.3337 | 0.2 | 0.4 | 0.004985 | MS.Trt_vs_MS.NoTrt_B |
| COX7A2 | 3.17E-07 | 0.28062 | 0.627 | 0.82 | 0.005065 | MS.Trt_vs_MS.NoTrt_B |
| TMEM173 | 3.22E-07 | -0.4824 | 0.192 | 0.38 | 0.00514 | MS.Trt_vs_MS.NoTrt_B |
| TSTA3 | 3.22E-07 | -0.4098 | 0.096 | 0.25 | 0.005146 | MS.Trt_vs_MS.NoTrt_B |
| SRBD1 | 3.29E-07 | 0.26105 | 0.218 | 0.42 | 0.005259 | MS.Trt_vs_MS.NoTrt_B |
| MICU2 | 3.31E-07 | -0.9419 | 0.34 | 0.54 | 0.005281 | MS.Trt_vs_MS.NoTrt_B |
| CEP95 | 3.45E-07 | 0.33391 | 0.292 | 0.52 | 0.005519 | MS.Trt_vs_MS.NoTrt_B |
| TNFRSF1B | 3.48E-07 | -0.3042 | 0.603 | 0.76 | 0.005553 | MS.Trt_vs_MS.NoTrt_B |
| EIF4H | 3.56E-07 | -0.334 | 0.322 | 0.55 | 0.005689 | MS.Trt_vs_MS.NoTrt_B |
| DYNLL1 | 3.61E-07 | -0.3952 | 0.216 | 0.41 | 0.005762 | MS.Trt_vs_MS.NoTrt_B |
| NRF1 | 3.79E-07 | -0.9001 | 0.203 | 0.38 | 0.006055 | MS.Trt_vs_MS.NoTrt_B |
| C10orf54 | 3.84E-07 | -0.5159 | 0.281 | 0.49 | 0.006128 | MS.Trt_vs_MS.NoTrt_B |
| SEC11A | 3.91E-07 | -0.4019 | 0.641 | 0.82 | 0.006253 | MS.Trt_vs_MS.NoTrt_B |
| GPR63 | 3.93E-07 | -0.2573 | 0.229 | 0.43 | 0.006283 | MS.Trt_vs_MS.NoTrt_B |
| B3GALNT1 | 4.03E-07 | -0.2793 | 0.15 | 0.32 | 0.00644 | MS.Trt_vs_MS.NoTrt_B |
| TAZ | 4.22E-07 | -0.4339 | 0.379 | 0.62 | 0.006738 | MS.Trt_vs_MS.NoTrt_B |
| DYNLRB1 | 4.29E-07 | -0.5329 | 0.122 | 0.29 | 0.006854 | MS.Trt_vs_MS.NoTrt_B |
| RHOG | 4.43E-07 | -0.2889 | 0.24 | 0.46 | 0.007078 | MS.Trt_vs_MS.NoTrt_B |
| SLA2 | 4.46E-07 | -1.3419 | 0.377 | 0.58 | 0.007119 | MS.Trt_vs_MS.NoTrt_B |
| CCZ1B | 4.49E-07 | -0.4984 | 0.527 | 0.75 | 0.007168 | MS.Trt_vs_MS.NoTrt_B |
| C1orf56 | 4.49E-07 | 1.02698 | 0.911 | 0.89 | 0.007172 | MS.Trt_vs_MS.NoTrt_B |
| EIF3I | 4.51E-07 | -0.8236 | 0.373 | 0.56 | 0.007205 | MS.Trt_vs_MS.NoTrt_B |
| ADK | 4.53E-07 | -0.2651 | 0.17 | 0.35 | 0.007231 | MS.Trt_vs_MS.NoTrt_B |
| TNFRSF14 | 4.54E-07 | -0.3564 | 0.222 | 0.43 | 0.007253 | MS.Trt_vs_MS.NoTrt_B |
| DDX46 | 4.57E-07 | -0.9125 | 0.386 | 0.55 | 0.007295 | MS.Trt_vs_MS.NoTrt_B |
| FECH | 4.57E-07 | -0.7715 | 0.508 | 0.74 | 0.007298 | MS.Trt_vs_MS.NoTrt_B |
| NDUFB4 | 4.59E-07 | -0.2863 | 0.179 | 0.36 | 0.007333 | MS.Trt_vs_MS.NoTrt_B |
| HAPLN3 | 4.59E-07 | -0.3336 | 0.111 | 0.27 | 0.007339 | MS.Trt_vs_MS.NoTrt_B |
| ZNF200 | 4.62E-07 | -0.4608 | 0.63 | 0.83 | 0.007376 | MS.Trt_vs_MS.NoTrt_B |

| | | | | | | |
|---|---|---|---|---|---|---|
| RPS20 | 4.92E-07 | -0.2622 | 0.756 | 0.92 | 0.007863 | MS.Trt_vs_MS.NoTrt_B |
| XAB2 | 4.98E-07 | -0.2798 | 0.144 | 0.32 | 0.007956 | MS.Trt_vs_MS.NoTrt_B |
| EXOC4 | 5.04E-07 | -0.4427 | 0.464 | 0.67 | 0.00805 | MS.Trt_vs_MS.NoTrt_B |
| RNF214 | 5.09E-07 | -0.337 | 0.146 | 0.32 | 0.008123 | MS.Trt_vs_MS.NoTrt_B |
| CLCN7 | 5.42E-07 | 0.28001 | 0.183 | 0.4 | 0.008654 | MS.Trt_vs_MS.NoTrt_B |
| TTC7A | 5.45E-07 | -0.7305 | 0.133 | 0.3 | 0.008706 | MS.Trt_vs_MS.NoTrt_B |
| ACBD6 | 5.46E-07 | -0.2999 | 0.336 | 0.55 | 0.00872 | MS.Trt_vs_MS.NoTrt_B |
| CLINT1 | 5.48E-07 | 0.37758 | 0.532 | 0.78 | 0.008752 | MS.Trt_vs_MS.NoTrt_B |
| UBA1 | 5.52E-07 | -0.4079 | 0.612 | 0.8 | 0.008824 | MS.Trt_vs_MS.NoTrt_B |
| TRMT44 | 5.78E-07 | -0.6016 | 0.12 | 0.28 | 0.009226 | MS.Trt_vs_MS.NoTrt_B |
| RPL3 | 5.87E-07 | 0.45073 | 1 | 1 | 0.00937 | MS.Trt_vs_MS.NoTrt_B |
| APBA3 | 6.07E-07 | -0.3635 | 0.562 | 0.75 | 0.009699 | MS.Trt_vs_MS.NoTrt_B |
| CD58 | 6.11E-07 | -0.2885 | 0.109 | 0.27 | 0.009758 | MS.Trt_vs_MS.NoTrt_B |
| ERGIC2 | 6.26E-07 | 0.77001 | 0.349 | 0.58 | 0.010002 | MS.Trt_vs_MS.NoTrt_B |
| DGCR2 | 6.37E-07 | -0.3781 | 0.157 | 0.34 | 0.010177 | MS.Trt_vs_MS.NoTrt_B |
| CCDC167 | 6.38E-07 | -0.7114 | 0.1 | 0.25 | 0.010183 | MS.Trt_vs_MS.NoTrt_B |
| EIF3CL | 6.38E-07 | -0.5077 | 0.142 | 0.31 | 0.010187 | MS.Trt_vs_MS.NoTrt_B |
| TNIK | 6.46E-07 | -0.9054 | 0.257 | 0.46 | 0.010322 | MS.Trt_vs_MS.NoTrt_B |
| AP000304.1 | 6.48E-07 | -0.6035 | 0.144 | 0.31 | 0.010353 | MS.Trt_vs_MS.NoTrt_B |
| BEX3 | 6.53E-07 | -0.4052 | 0.074 | 0.21 | 0.010431 | MS.Trt_vs_MS.NoTrt_B |
| ERAP2 | 6.53E-07 | -0.6629 | 0.595 | 0.72 | 0.010434 | MS.Trt_vs_MS.NoTrt_B |
| APH1A | 6.84E-07 | -0.8078 | 0.122 | 0.28 | 0.010928 | MS.Trt_vs_MS.NoTrt_B |
| FAM134B | 7.09E-07 | -0.6746 | 0.035 | 0.14 | 0.011329 | MS.Trt_vs_MS.NoTrt_B |
| CTSB | 7.27E-07 | -0.4583 | 0.858 | 0.97 | 0.011608 | MS.Trt_vs_MS.NoTrt_B |
| PRELID3B | 7.35E-07 | 0.36303 | 0.166 | 0.35 | 0.011738 | MS.Trt_vs_MS.NoTrt_B |
| LETMD1 | 7.42E-07 | -0.4999 | 0.669 | 0.89 | 0.011847 | MS.Trt_vs_MS.NoTrt_B |
| ABHD15 | 7.59E-07 | -0.7987 | 0.179 | 0.37 | 0.012121 | MS.Trt_vs_MS.NoTrt_B |
| R3HDM4 | 7.71E-07 | -0.4833 | 0.364 | 0.58 | 0.012323 | MS.Trt_vs_MS.NoTrt_B |
| HIF1AN | 8.12E-07 | -0.3932 | 0.66 | 0.88 | 0.012966 | MS.Trt_vs_MS.NoTrt_B |
| UQCRB | 8.24E-07 | 0.57885 | 0.58 | 0.8 | 0.013165 | MS.Trt_vs_MS.NoTrt_B |
| LRRC37A3 | 8.24E-07 | -0.4974 | 0.359 | 0.56 | 0.013168 | MS.Trt_vs_MS.NoTrt_B |
| IRF2 | 8.28E-07 | -0.359 | 0.24 | 0.43 | 0.013223 | MS.Trt_vs_MS.NoTrt_B |
| HNRNPF | 8.57E-07 | -0.3449 | 0.309 | 0.52 | 0.013688 | MS.Trt_vs_MS.NoTrt_B |
| CRLF3 | 8.62E-07 | -0.4826 | 0.78 | 0.93 | 0.013776 | MS.Trt_vs_MS.NoTrt_B |
| EFTUD2 | 8.64E-07 | -0.6877 | 0.702 | 0.86 | 0.013795 | MS.Trt_vs_MS.NoTrt_B |
| ZFP36 | 8.76E-07 | -0.3573 | 0.237 | 0.45 | 0.014 | MS.Trt_vs_MS.NoTrt_B |
| NDUFS2 | 8.97E-07 | -0.4739 | 0.312 | 0.5 | 0.014332 | MS.Trt_vs_MS.NoTrt_B |
| RALGDS | 9.26E-07 | -0.3108 | 0.255 | 0.47 | 0.014793 | MS.Trt_vs_MS.NoTrt_B |
| PARP4 | 9.26E-07 | -0.4465 | 0.19 | 0.37 | 0.014795 | MS.Trt_vs_MS.NoTrt_B |
| BUB3 | 9.43E-07 | 0.29222 | 0.495 | 0.76 | 0.015067 | MS.Trt_vs_MS.NoTrt_B |

| | | | | | | |
|---|---|---|---|---|---|---|
| NR2C1 | 9.51E-07 | -0.4529 | 0.475 | 0.74 | 0.015191 | MS.Trt_vs_MS.NoTrt_B |
| TRABD | 9.60E-07 | -0.5019 | 0.266 | 0.46 | 0.015328 | MS.Trt_vs_MS.NoTrt_B |
| SYT5 | 9.81E-07 | -0.4798 | 0.542 | 0.7 | 0.015665 | MS.Trt_vs_MS.NoTrt_B |
| COG5 | 9.83E-07 | 0.61248 | 0.137 | 0.3 | 0.015709 | MS.Trt_vs_MS.NoTrt_B |
| MAP4 | 9.84E-07 | -0.6137 | 0.66 | 0.81 | 0.015712 | MS.Trt_vs_MS.NoTrt_B |
| SLC3A2 | 9.97E-07 | -0.3968 | 0.375 | 0.56 | 0.015926 | MS.Trt_vs_MS.NoTrt_B |
| PSMG3 | 1.01E-06 | 0.32665 | 0.447 | 0.69 | 0.016196 | MS.Trt_vs_MS.NoTrt_B |
| SEC61G | 1.02E-06 | -0.3416 | 0.198 | 0.37 | 0.016279 | MS.Trt_vs_MS.NoTrt_B |
| DPEP2 | 1.02E-06 | -0.3945 | 0.157 | 0.32 | 0.016364 | MS.Trt_vs_MS.NoTrt_B |
| MCM9 | 1.04E-06 | 0.42336 | 0.227 | 0.43 | 0.016671 | MS.Trt_vs_MS.NoTrt_B |
| KIAA0319L | 1.05E-06 | -0.6635 | 0.534 | 0.71 | 0.016736 | MS.Trt_vs_MS.NoTrt_B |
| RAP1GAP2 | 1.09E-06 | -0.7205 | 0.137 | 0.3 | 0.017388 | MS.Trt_vs_MS.NoTrt_B |
| RASA3 | 1.11E-06 | -0.3655 | 0.37 | 0.6 | 0.017666 | MS.Trt_vs_MS.NoTrt_B |
| LMF2 | 1.11E-06 | -0.3684 | 0.133 | 0.29 | 0.017747 | MS.Trt_vs_MS.NoTrt_B |
| SH2B1 | 1.12E-06 | -0.2855 | 0.458 | 0.68 | 0.017823 | MS.Trt_vs_MS.NoTrt_B |
| PMF1 | 1.12E-06 | -0.5204 | 0.325 | 0.51 | 0.017872 | MS.Trt_vs_MS.NoTrt_B |
| SUPV3L1 | 1.13E-06 | -0.3569 | 0.488 | 0.72 | 0.018123 | MS.Trt_vs_MS.NoTrt_B |
| SP140 | 1.14E-06 | -0.5603 | 0.133 | 0.3 | 0.018248 | MS.Trt_vs_MS.NoTrt_B |
| IL17RA | 1.17E-06 | -0.5117 | 0.503 | 0.7 | 0.018707 | MS.Trt_vs_MS.NoTrt_B |
| EPB41 | 1.19E-06 | 0.47819 | 0.325 | 0.59 | 0.019009 | MS.Trt_vs_MS.NoTrt_B |
| PAN3 | 1.19E-06 | 0.91011 | 0.767 | 0.94 | 0.019042 | MS.Trt_vs_MS.NoTrt_B |
| NCAPH2 | 1.19E-06 | -0.6986 | 0.606 | 0.77 | 0.019051 | MS.Trt_vs_MS.NoTrt_B |
| MTMR9 | 1.20E-06 | -0.585 | 0.205 | 0.38 | 0.019223 | MS.Trt_vs_MS.NoTrt_B |
| C19orf60 | 1.22E-06 | -0.4414 | 0.767 | 0.91 | 0.019461 | MS.Trt_vs_MS.NoTrt_B |
| DDX10 | 1.22E-06 | -1.0693 | 0.479 | 0.71 | 0.019552 | MS.Trt_vs_MS.NoTrt_B |
| NCK1 | 1.25E-06 | 0.27268 | 0.251 | 0.48 | 0.019903 | MS.Trt_vs_MS.NoTrt_B |
| EBP | 1.25E-06 | -0.489 | 0.129 | 0.28 | 0.020014 | MS.Trt_vs_MS.NoTrt_B |
| CCR8 | 1.29E-06 | -0.9569 | 0.063 | 0.19 | 0.020624 | MS.Trt_vs_MS.NoTrt_B |
| STRADA | 1.30E-06 | -0.3179 | 0.405 | 0.63 | 0.020733 | MS.Trt_vs_MS.NoTrt_B |
| STXBP2 | 1.34E-06 | -0.6306 | 0.205 | 0.38 | 0.021418 | MS.Trt_vs_MS.NoTrt_B |
| USP20 | 1.34E-06 | -0.2622 | 0.168 | 0.35 | 0.021477 | MS.Trt_vs_MS.NoTrt_B |
| ACP5 | 1.35E-06 | -1.0013 | 0.063 | 0.19 | 0.021486 | MS.Trt_vs_MS.NoTrt_B |
| ITGB1BP1 | 1.37E-06 | -0.8277 | 0.107 | 0.25 | 0.021815 | MS.Trt_vs_MS.NoTrt_B |
| DRAP1 | 1.37E-06 | -0.4088 | 0.131 | 0.3 | 0.021903 | MS.Trt_vs_MS.NoTrt_B |
| ATP6V0E2 | 1.38E-06 | -0.4502 | 0.161 | 0.34 | 0.022015 | MS.Trt_vs_MS.NoTrt_B |
| TPCN1 | 1.39E-06 | 0.431 | 0.294 | 0.5 | 0.022132 | MS.Trt_vs_MS.NoTrt_B |
| UBE4B | 1.40E-06 | -0.3238 | 0.475 | 0.7 | 0.02231 | MS.Trt_vs_MS.NoTrt_B |
| COL8A1 | 1.44E-06 | 0.50076 | 0.876 | 0.74 | 0.023014 | MS.Trt_vs_MS.NoTrt_B |
| C12orf4 | 1.44E-06 | 0.36501 | 0.453 | 0.72 | 0.023075 | MS.Trt_vs_MS.NoTrt_B |
| DNAJC9 | 1.51E-06 | -0.6343 | 0.094 | 0.24 | 0.024046 | MS.Trt_vs_MS.NoTrt_B |

| | | | | | | |
|---|---|---|---|---|---|---|
| MARS | 1.53E-06 | -0.4651 | 0.292 | 0.49 | 0.024377 | MS.Trt_vs_MS.NoTrt_B |
| MT2A | 1.54E-06 | -0.3618 | 0.122 | 0.28 | 0.024533 | MS.Trt_vs_MS.NoTrt_B |
| TUBE1 | 1.55E-06 | 0.2956 | 0.281 | 0.48 | 0.024733 | MS.Trt_vs_MS.NoTrt_B |
| C14orf166 | 1.55E-06 | -0.2712 | 0.473 | 0.64 | 0.024828 | MS.Trt_vs_MS.NoTrt_B |
| UQCRH | 1.56E-06 | -0.321 | 0.427 | 0.62 | 0.024878 | MS.Trt_vs_MS.NoTrt_B |
| TSC1 | 1.56E-06 | -0.6548 | 0.333 | 0.55 | 0.024975 | MS.Trt_vs_MS.NoTrt_B |
| SLC33A1 | 1.57E-06 | -0.4761 | 0.795 | 0.89 | 0.025096 | MS.Trt_vs_MS.NoTrt_B |
| ZNF770 | 1.60E-06 | -0.7822 | 0.174 | 0.34 | 0.025543 | MS.Trt_vs_MS.NoTrt_B |
| CBX7 | 1.63E-06 | -0.4211 | 0.275 | 0.47 | 0.026 | MS.Trt_vs_MS.NoTrt_B |
| C9orf78 | 1.65E-06 | -0.3286 | 0.885 | 0.97 | 0.026366 | MS.Trt_vs_MS.NoTrt_B |
| COPZ1 | 1.66E-06 | -0.472 | 0.142 | 0.31 | 0.026556 | MS.Trt_vs_MS.NoTrt_B |
| ARSA | 1.68E-06 | -0.4019 | 0.569 | 0.73 | 0.026803 | MS.Trt_vs_MS.NoTrt_B |
| ATP6V0C | 1.68E-06 | -0.329 | 0.157 | 0.34 | 0.026848 | MS.Trt_vs_MS.NoTrt_B |
| MRPL28 | 1.77E-06 | -0.8148 | 0.074 | 0.21 | 0.028213 | MS.Trt_vs_MS.NoTrt_B |
| PRMT2 | 1.77E-06 | -0.3718 | 0.285 | 0.47 | 0.028262 | MS.Trt_vs_MS.NoTrt_B |
| FKBP1A | 1.77E-06 | -0.254 | 0.242 | 0.44 | 0.028316 | MS.Trt_vs_MS.NoTrt_B |
| CDK5RAP3 | 1.79E-06 | -0.2648 | 0.464 | 0.69 | 0.028614 | MS.Trt_vs_MS.NoTrt_B |
| FLII | 1.81E-06 | -0.3981 | 0.157 | 0.33 | 0.028973 | MS.Trt_vs_MS.NoTrt_B |
| FO538757.1 | 1.89E-06 | -0.6867 | 0.12 | 0.28 | 0.030145 | MS.Trt_vs_MS.NoTrt_B |
| NIP7 | 1.91E-06 | -0.4894 | 0.447 | 0.66 | 0.030531 | MS.Trt_vs_MS.NoTrt_B |
| CANX | 1.97E-06 | 0.42038 | 0.575 | 0.81 | 0.031406 | MS.Trt_vs_MS.NoTrt_B |
| ABT1 | 2.00E-06 | -0.4057 | 0.12 | 0.28 | 0.032003 | MS.Trt_vs_MS.NoTrt_B |
| HNRNPU | 2.04E-06 | 0.49253 | 0.338 | 0.56 | 0.032515 | MS.Trt_vs_MS.NoTrt_B |
| BTF3L4 | 2.04E-06 | -0.4758 | 0.336 | 0.51 | 0.03261 | MS.Trt_vs_MS.NoTrt_B |
| ZNF655 | 2.05E-06 | -0.2971 | 0.475 | 0.68 | 0.032739 | MS.Trt_vs_MS.NoTrt_B |
| ASB18 | 2.06E-06 | -0.3371 | 0.375 | 0.57 | 0.032967 | MS.Trt_vs_MS.NoTrt_B |
| COMMD10 | 2.11E-06 | 0.49585 | 0.216 | 0.41 | 0.033632 | MS.Trt_vs_MS.NoTrt_B |
| SAR1A | 2.11E-06 | -0.2574 | 0.542 | 0.69 | 0.033666 | MS.Trt_vs_MS.NoTrt_B |
| SIGIRR | 2.12E-06 | -0.4153 | 0.253 | 0.44 | 0.033788 | MS.Trt_vs_MS.NoTrt_B |
| PTK2B | 2.12E-06 | -0.3174 | 0.294 | 0.49 | 0.033895 | MS.Trt_vs_MS.NoTrt_B |
| PAXIP1 | 2.15E-06 | 0.29015 | 0.309 | 0.5 | 0.034291 | MS.Trt_vs_MS.NoTrt_B |
| TNR | 2.16E-06 | -0.288 | 0.185 | 0.36 | 0.034563 | MS.Trt_vs_MS.NoTrt_B |
| HLA-DPB1 | 2.18E-06 | -0.6643 | 0.368 | 0.53 | 0.034815 | MS.Trt_vs_MS.NoTrt_B |
| ABI2 | 2.23E-06 | -0.5702 | 0.688 | 0.84 | 0.03569 | MS.Trt_vs_MS.NoTrt_B |
| NRBP1 | 2.29E-06 | -0.6903 | 0.425 | 0.63 | 0.036623 | MS.Trt_vs_MS.NoTrt_B |
| HSP90AA1 | 2.32E-06 | -0.2949 | 0.505 | 0.68 | 0.037086 | MS.Trt_vs_MS.NoTrt_B |
| ARL6IP1 | 2.32E-06 | -0.2983 | 0.333 | 0.51 | 0.037135 | MS.Trt_vs_MS.NoTrt_B |
| STAG1 | 2.38E-06 | 0.33289 | 0.157 | 0.33 | 0.038051 | MS.Trt_vs_MS.NoTrt_B |
| SPICE1 | 2.40E-06 | -0.3829 | 0.168 | 0.35 | 0.038308 | MS.Trt_vs_MS.NoTrt_B |
| UBE2B | 2.46E-06 | 0.37649 | 0.549 | 0.76 | 0.039357 | MS.Trt_vs_MS.NoTrt_B |

| | | | | | | |
|---|---|---|---|---|---|---|
| CD164 | 2.48E-06 | 0.31583 | 0.486 | 0.69 | 0.039619 | MS.Trt_vs_MS.NoTrt_B |
| CSK | 2.52E-06 | -0.3139 | 0.216 | 0.41 | 0.040232 | MS.Trt_vs_MS.NoTrt_B |
| NF2 | 2.55E-06 | -0.2612 | 0.645 | 0.81 | 0.040675 | MS.Trt_vs_MS.NoTrt_B |
| HDLBP | 2.56E-06 | -0.4681 | 0.294 | 0.48 | 0.040955 | MS.Trt_vs_MS.NoTrt_B |
| RNASEK | 2.61E-06 | -0.494 | 0.667 | 0.73 | 0.041705 | MS.Trt_vs_MS.NoTrt_B |
| CPSF7 | 2.63E-06 | -0.3347 | 0.146 | 0.31 | 0.04193 | MS.Trt_vs_MS.NoTrt_B |
| HADHA | 2.63E-06 | -0.3458 | 0.436 | 0.63 | 0.041973 | MS.Trt_vs_MS.NoTrt_B |
| JMJD7-PLA2 | 2.70E-06 | 0.58065 | 0.246 | 0.43 | 0.043086 | MS.Trt_vs_MS.NoTrt_B |
| PRDM2 | 2.73E-06 | -0.3777 | 0.292 | 0.48 | 0.043618 | MS.Trt_vs_MS.NoTrt_B |
| TARBP1 | 3.00E-06 | 0.27986 | 0.268 | 0.46 | 0.047886 | MS.Trt_vs_MS.NoTrt_B |
| SCN1A | 3.01E-06 | -0.2671 | 0.255 | 0.47 | 0.04806 | MS.Trt_vs_MS.NoTrt_B |
| SNX20 | 3.07E-06 | -0.5472 | 0.32 | 0.5 | 0.048997 | MS.Trt_vs_MS.NoTrt_B |
| METTL23 | 3.07E-06 | -0.2598 | 0.124 | 0.28 | 0.049089 | MS.Trt_vs_MS.NoTrt_B |
| VDAC1 | 3.10E-06 | -0.4238 | 0.608 | 0.84 | 0.049549 | MS.Trt_vs_MS.NoTrt_B |
| C7orf73 | 3.12E-06 | -0.4271 | 0.477 | 0.67 | 0.049768 | MS.Trt_vs_MS.NoTrt_B |
| SPAG7 | 3.15E-06 | -0.3726 | 0.695 | 0.88 | 0.050342 | MS.Trt_vs_MS.NoTrt_B |
| PSMD12 | 3.17E-06 | 0.37471 | 0.606 | 0.79 | 0.050565 | MS.Trt_vs_MS.NoTrt_B |
| RAB13 | 3.19E-06 | -0.3228 | 0.403 | 0.6 | 0.051015 | MS.Trt_vs_MS.NoTrt_B |
| CCDC107 | 3.25E-06 | -0.6069 | 0.107 | 0.25 | 0.051984 | MS.Trt_vs_MS.NoTrt_B |
| NDE1 | 3.31E-06 | -0.4703 | 0.525 | 0.76 | 0.052938 | MS.Trt_vs_MS.NoTrt_B |
| RNF10 | 3.32E-06 | -0.4114 | 0.697 | 0.82 | 0.053027 | MS.Trt_vs_MS.NoTrt_B |
| GTPBP10 | 3.33E-06 | 0.34506 | 0.649 | 0.87 | 0.053113 | MS.Trt_vs_MS.NoTrt_B |
| ERVW-1 | 3.33E-06 | -0.4137 | 0.251 | 0.44 | 0.05324 | MS.Trt_vs_MS.NoTrt_B |
| SRP9 | 3.36E-06 | 0.31495 | 0.342 | 0.56 | 0.053617 | MS.Trt_vs_MS.NoTrt_B |
| GLYR1 | 3.45E-06 | -0.54 | 0.244 | 0.43 | 0.055175 | MS.Trt_vs_MS.NoTrt_B |
| ATP5I | 3.49E-06 | 0.62065 | 0.196 | 0.39 | 0.055753 | MS.Trt_vs_MS.NoTrt_B |
| ENTPD1 | 3.50E-06 | -0.5169 | 0.863 | 0.97 | 0.055917 | MS.Trt_vs_MS.NoTrt_B |
| ZNF589 | 3.63E-06 | -0.8221 | 0.327 | 0.52 | 0.057936 | MS.Trt_vs_MS.NoTrt_B |
| RAB11FIP4 | 3.69E-06 | -0.4233 | 0.538 | 0.75 | 0.058876 | MS.Trt_vs_MS.NoTrt_B |
| NLRC3 | 3.71E-06 | -0.2581 | 0.664 | 0.77 | 0.059293 | MS.Trt_vs_MS.NoTrt_B |
| PIAS1 | 3.82E-06 | -0.4131 | 0.283 | 0.45 | 0.061064 | MS.Trt_vs_MS.NoTrt_B |
| COX5A | 3.88E-06 | -0.5238 | 0.163 | 0.33 | 0.061951 | MS.Trt_vs_MS.NoTrt_B |
| CD48 | 3.90E-06 | -0.2895 | 0.203 | 0.38 | 0.062335 | MS.Trt_vs_MS.NoTrt_B |
| CYCS | 3.99E-06 | -0.3219 | 0.619 | 0.79 | 0.063783 | MS.Trt_vs_MS.NoTrt_B |
| FLAD1 | 4.01E-06 | -0.6041 | 0.46 | 0.65 | 0.064099 | MS.Trt_vs_MS.NoTrt_B |
| BLZF1 | 4.18E-06 | -0.9105 | 0.307 | 0.5 | 0.066761 | MS.Trt_vs_MS.NoTrt_B |
| CAPN1 | 4.26E-06 | -0.6137 | 0.399 | 0.59 | 0.06804 | MS.Trt_vs_MS.NoTrt_B |
| RPAIN | 4.35E-06 | -0.2995 | 0.852 | 0.97 | 0.069468 | MS.Trt_vs_MS.NoTrt_B |
| CCNYL1 | 4.43E-06 | -0.6726 | 0.272 | 0.48 | 0.070738 | MS.Trt_vs_MS.NoTrt_B |
| TBRG1 | 4.47E-06 | 0.29251 | 0.479 | 0.68 | 0.071338 | MS.Trt_vs_MS.NoTrt_B |

| | | | | | | |
|---|---|---|---|---|---|---|
| GPR180 | 4.48E-06 | -1.3555 | 0.026 | 0.11 | 0.071488 | MS.Trt_vs_MS.NoTrt_B |
| NIF3L1 | 4.48E-06 | -0.8421 | 0.046 | 0.15 | 0.071637 | MS.Trt_vs_MS.NoTrt_B |
| ARMC8 | 4.52E-06 | -0.2526 | 0.397 | 0.6 | 0.072166 | MS.Trt_vs_MS.NoTrt_B |
| PSTPIP2 | 4.58E-06 | -0.4224 | 0.24 | 0.44 | 0.073184 | MS.Trt_vs_MS.NoTrt_B |
| SERPINB6 | 4.65E-06 | 0.36055 | 0.37 | 0.58 | 0.074336 | MS.Trt_vs_MS.NoTrt_B |
| ALDH1L2 | 4.68E-06 | -0.4611 | 0.161 | 0.31 | 0.074754 | MS.Trt_vs_MS.NoTrt_B |
| STAG3 | 4.72E-06 | -0.9255 | 0.102 | 0.24 | 0.075448 | MS.Trt_vs_MS.NoTrt_B |
| NDUFAB1 | 4.77E-06 | -0.4802 | 0.124 | 0.27 | 0.076255 | MS.Trt_vs_MS.NoTrt_B |
| POLDIP3 | 4.85E-06 | -0.7297 | 0.072 | 0.2 | 0.077411 | MS.Trt_vs_MS.NoTrt_B |
| FBXL5 | 4.87E-06 | -0.3123 | 0.205 | 0.38 | 0.07775 | MS.Trt_vs_MS.NoTrt_B |
| TMEM175 | 4.95E-06 | -0.4532 | 0.22 | 0.4 | 0.079 | MS.Trt_vs_MS.NoTrt_B |
| PDCD5 | 5.01E-06 | -0.4851 | 0.137 | 0.29 | 0.080005 | MS.Trt_vs_MS.NoTrt_B |
| HDAC2 | 5.03E-06 | 0.25554 | 0.56 | 0.78 | 0.080355 | MS.Trt_vs_MS.NoTrt_B |
| DDX19B | 5.07E-06 | -0.3594 | 0.307 | 0.48 | 0.080908 | MS.Trt_vs_MS.NoTrt_B |
| SKA2 | 5.12E-06 | 0.65063 | 0.444 | 0.65 | 0.081706 | MS.Trt_vs_MS.NoTrt_B |
| PAN2 | 5.14E-06 | -1.1293 | 0.214 | 0.38 | 0.082117 | MS.Trt_vs_MS.NoTrt_B |
| FABP4 | 5.15E-06 | 0.60859 | 0.172 | 0.04 | 0.082296 | MS.Trt_vs_MS.NoTrt_B |
| PPP3CA | 5.16E-06 | 0.56727 | 0.423 | 0.67 | 0.082453 | MS.Trt_vs_MS.NoTrt_B |
| POM121C | 5.23E-06 | -0.2702 | 0.353 | 0.52 | 0.083462 | MS.Trt_vs_MS.NoTrt_B |
| CCDC12 | 5.27E-06 | -0.5502 | 0.216 | 0.4 | 0.08423 | MS.Trt_vs_MS.NoTrt_B |
| C17orf62 | 5.42E-06 | -0.4446 | 0.595 | 0.75 | 0.086613 | MS.Trt_vs_MS.NoTrt_B |
| HMGB2 | 5.43E-06 | -0.2837 | 0.209 | 0.39 | 0.086701 | MS.Trt_vs_MS.NoTrt_B |
| ABL2 | 5.54E-06 | 0.44118 | 0.573 | 0.83 | 0.088461 | MS.Trt_vs_MS.NoTrt_B |
| TCEANC2 | 5.55E-06 | -0.5439 | 0.752 | 0.92 | 0.088574 | MS.Trt_vs_MS.NoTrt_B |
| TRAF1 | 5.63E-06 | -0.3131 | 0.211 | 0.4 | 0.089955 | MS.Trt_vs_MS.NoTrt_B |
| HDAC3 | 5.73E-06 | -1.439 | 0.061 | 0.17 | 0.091585 | MS.Trt_vs_MS.NoTrt_B |
| RBM5 | 5.75E-06 | -0.3851 | 0.749 | 0.9 | 0.091873 | MS.Trt_vs_MS.NoTrt_B |
| FLI1 | 5.84E-06 | -0.2755 | 0.359 | 0.55 | 0.093354 | MS.Trt_vs_MS.NoTrt_B |
| HMBOX1 | 5.90E-06 | -0.503 | 0.39 | 0.58 | 0.094234 | MS.Trt_vs_MS.NoTrt_B |
| NFKBIZ | 6.11E-06 | -0.4214 | 0.231 | 0.41 | 0.097626 | MS.Trt_vs_MS.NoTrt_B |
| TP63 | 6.15E-06 | -0.307 | 0.303 | 0.49 | 0.098165 | MS.Trt_vs_MS.NoTrt_B |
| MAX | 6.19E-06 | 0.31325 | 0.373 | 0.62 | 0.098943 | MS.Trt_vs_MS.NoTrt_B |
| AMBRA1 | 6.26E-06 | -0.8992 | 0.124 | 0.27 | 0.099929 | MS.Trt_vs_MS.NoTrt_B |
| CSNK1G1 | 6.27E-06 | -0.4078 | 0.423 | 0.61 | 0.100221 | MS.Trt_vs_MS.NoTrt_B |
| DUSP28 | 6.31E-06 | -0.5806 | 0.584 | 0.73 | 0.100729 | MS.Trt_vs_MS.NoTrt_B |
| CCND2 | 6.31E-06 | -0.2854 | 0.257 | 0.44 | 0.100747 | MS.Trt_vs_MS.NoTrt_B |
| TUBA4A | 6.31E-06 | -0.6092 | 0.159 | 0.31 | 0.100824 | MS.Trt_vs_MS.NoTrt_B |
| ERLEC1 | 6.50E-06 | 0.29419 | 0.285 | 0.5 | 0.103836 | MS.Trt_vs_MS.NoTrt_B |
| DTX2 | 6.55E-06 | -0.3472 | 0.283 | 0.45 | 0.104619 | MS.Trt_vs_MS.NoTrt_B |
| SPNS1 | 6.57E-06 | -0.4682 | 0.667 | 0.86 | 0.104888 | MS.Trt_vs_MS.NoTrt_B |

| | | | | | | |
|---|---|---|---|---|---|---|
| CDK7 | 6.70E-06 | -1.0542 | 0.094 | 0.22 | 0.107034 | MS.Trt_vs_MS.NoTrt_B |
| ZKSCAN3 | 6.84E-06 | -0.3058 | 0.68 | 0.84 | 0.109189 | MS.Trt_vs_MS.NoTrt_B |
| MED28 | 6.92E-06 | -0.4759 | 0.743 | 0.86 | 0.110566 | MS.Trt_vs_MS.NoTrt_B |
| HEXA | 7.09E-06 | -0.5874 | 0.597 | 0.76 | 0.113192 | MS.Trt_vs_MS.NoTrt_B |
| ACVR2B | 7.20E-06 | -0.5634 | 0.298 | 0.49 | 0.115008 | MS.Trt_vs_MS.NoTrt_B |
| ZNF577 | 7.27E-06 | 0.39534 | 0.237 | 0.43 | 0.1162 | MS.Trt_vs_MS.NoTrt_B |
| SUPT4H1 | 7.33E-06 | -0.2974 | 0.131 | 0.28 | 0.11707 | MS.Trt_vs_MS.NoTrt_B |
| PLEKHG2 | 7.36E-06 | -0.4731 | 0.706 | 0.87 | 0.117517 | MS.Trt_vs_MS.NoTrt_B |
| XPNPEP1 | 7.45E-06 | -0.9896 | 0.192 | 0.35 | 0.119021 | MS.Trt_vs_MS.NoTrt_B |
| CD82 | 7.50E-06 | -0.58 | 0.497 | 0.68 | 0.119747 | MS.Trt_vs_MS.NoTrt_B |
| RNF7 | 7.57E-06 | -0.5225 | 0.549 | 0.72 | 0.120953 | MS.Trt_vs_MS.NoTrt_B |
| ELP5 | 7.63E-06 | -1.1621 | 0.048 | 0.15 | 0.121804 | MS.Trt_vs_MS.NoTrt_B |
| ATP6AP2 | 7.64E-06 | -0.2657 | 0.71 | 0.85 | 0.121972 | MS.Trt_vs_MS.NoTrt_B |
| KANSL1 | 7.67E-06 | -0.4752 | 0.562 | 0.73 | 0.122496 | MS.Trt_vs_MS.NoTrt_B |
| ANP32E | 7.68E-06 | 0.37976 | 0.37 | 0.58 | 0.122684 | MS.Trt_vs_MS.NoTrt_B |
| EIF3H | 7.70E-06 | -0.3501 | 0.484 | 0.66 | 0.122955 | MS.Trt_vs_MS.NoTrt_B |
| TTC31 | 7.76E-06 | -0.876 | 0.146 | 0.29 | 0.123885 | MS.Trt_vs_MS.NoTrt_B |
| DDRGK1 | 7.76E-06 | -1.2812 | 0.081 | 0.2 | 0.123956 | MS.Trt_vs_MS.NoTrt_B |
| SMAP1 | 7.77E-06 | -0.9174 | 0.109 | 0.24 | 0.124102 | MS.Trt_vs_MS.NoTrt_B |
| HELLS | 7.82E-06 | -0.315 | 0.651 | 0.84 | 0.124952 | MS.Trt_vs_MS.NoTrt_B |
| KDELR2 | 7.86E-06 | -0.6778 | 0.102 | 0.23 | 0.125485 | MS.Trt_vs_MS.NoTrt_B |
| LRRC28 | 7.86E-06 | -0.9503 | 0.381 | 0.56 | 0.125545 | MS.Trt_vs_MS.NoTrt_B |
| ACP7 | 7.87E-06 | -0.3388 | 0.305 | 0.49 | 0.125756 | MS.Trt_vs_MS.NoTrt_B |
| ZBTB25 | 7.93E-06 | 0.25313 | 0.891 | 0.95 | 0.1267 | MS.Trt_vs_MS.NoTrt_B |
| TMEM230 | 7.96E-06 | 0.3309 | 0.331 | 0.52 | 0.127134 | MS.Trt_vs_MS.NoTrt_B |
| SLBP | 8.02E-06 | -0.7388 | 0.078 | 0.2 | 0.128137 | MS.Trt_vs_MS.NoTrt_B |
| PRPF3 | 8.03E-06 | -0.6977 | 0.743 | 0.86 | 0.128194 | MS.Trt_vs_MS.NoTrt_B |
| NUMB | 8.08E-06 | -0.4666 | 0.601 | 0.76 | 0.128999 | MS.Trt_vs_MS.NoTrt_B |
| SNRPD2 | 8.13E-06 | 0.28046 | 0.477 | 0.7 | 0.129923 | MS.Trt_vs_MS.NoTrt_B |
| ABCF2 | 8.19E-06 | -0.8336 | 0.161 | 0.32 | 0.130843 | MS.Trt_vs_MS.NoTrt_B |
| BIRC3 | 8.23E-06 | 0.26352 | 0.481 | 0.69 | 0.131533 | MS.Trt_vs_MS.NoTrt_B |
| MAN2B1 | 8.25E-06 | -0.625 | 0.484 | 0.68 | 0.131734 | MS.Trt_vs_MS.NoTrt_B |
| BCAS1 | 8.28E-06 | -1.7904 | 0.251 | 0.42 | 0.132191 | MS.Trt_vs_MS.NoTrt_B |
| CAPRIN1 | 8.42E-06 | -0.3032 | 0.222 | 0.4 | 0.134425 | MS.Trt_vs_MS.NoTrt_B |
| GLE1 | 8.44E-06 | -0.7829 | 0.065 | 0.18 | 0.134818 | MS.Trt_vs_MS.NoTrt_B |
| PPM1G | 8.55E-06 | -0.2967 | 0.542 | 0.77 | 0.136554 | MS.Trt_vs_MS.NoTrt_B |
| DERL2 | 8.68E-06 | 0.64154 | 0.2 | 0.37 | 0.138607 | MS.Trt_vs_MS.NoTrt_B |
| DDX27 | 8.70E-06 | -0.268 | 0.51 | 0.68 | 0.138995 | MS.Trt_vs_MS.NoTrt_B |
| GFPT1 | 8.71E-06 | -0.5613 | 0.545 | 0.74 | 0.139114 | MS.Trt_vs_MS.NoTrt_B |
| ZYX | 8.78E-06 | -0.305 | 0.207 | 0.38 | 0.140251 | MS.Trt_vs_MS.NoTrt_B |

| | | | | | | |
|---|---|---|---|---|---|---|
| PPP3CC | 8.78E-06 | -0.3485 | 0.12 | 0.27 | 0.14032 | MS.Trt_vs_MS.NoTrt_B |
| ARV1 | 8.80E-06 | 0.40726 | 0.155 | 0.31 | 0.140499 | MS.Trt_vs_MS.NoTrt_B |
| SSNA1 | 8.89E-06 | -0.6545 | 0.096 | 0.23 | 0.141965 | MS.Trt_vs_MS.NoTrt_B |
| DPYD | 9.06E-06 | -0.6476 | 0.359 | 0.52 | 0.144646 | MS.Trt_vs_MS.NoTrt_B |
| NUDT9 | 9.19E-06 | 0.39733 | 0.373 | 0.59 | 0.146833 | MS.Trt_vs_MS.NoTrt_B |
| ALG14 | 9.20E-06 | -0.469 | 0.786 | 0.93 | 0.147001 | MS.Trt_vs_MS.NoTrt_B |
| RSRP1 | 9.25E-06 | -0.2637 | 0.839 | 0.95 | 0.147675 | MS.Trt_vs_MS.NoTrt_B |
| CCDC25 | 9.28E-06 | 0.70035 | 0.168 | 0.34 | 0.148265 | MS.Trt_vs_MS.NoTrt_B |
| SIPA1L3 | 9.49E-06 | -0.412 | 0.501 | 0.71 | 0.151643 | MS.Trt_vs_MS.NoTrt_B |
| CAMK4 | 9.66E-06 | 0.26121 | 0.503 | 0.7 | 0.154358 | MS.Trt_vs_MS.NoTrt_B |
| RP1-27O5.3 | 9.85E-06 | -0.2532 | 0.111 | 0.25 | 0.157277 | MS.Trt_vs_MS.NoTrt_B |
| DHX36 | 9.92E-06 | 0.32472 | 0.56 | 0.77 | 0.158448 | MS.Trt_vs_MS.NoTrt_B |
| TOP3B | 9.97E-06 | -0.8089 | 0.089 | 0.22 | 0.159279 | MS.Trt_vs_MS.NoTrt_B |
| TFB1M | 1.01E-05 | -0.338 | 0.301 | 0.48 | 0.161698 | MS.Trt_vs_MS.NoTrt_B |
| MAZ | 1.02E-05 | -0.8662 | 0.102 | 0.23 | 0.163279 | MS.Trt_vs_MS.NoTrt_B |
| COTL1 | 1.03E-05 | -0.2962 | 0.767 | 0.91 | 0.163967 | MS.Trt_vs_MS.NoTrt_B |
| PSMA1 | 1.03E-05 | 0.42957 | 0.261 | 0.47 | 0.16452 | MS.Trt_vs_MS.NoTrt_B |
| RNF26 | 1.04E-05 | -1.0187 | 0.076 | 0.2 | 0.166744 | MS.Trt_vs_MS.NoTrt_B |
| ARHGEF6 | 1.05E-05 | -0.3089 | 0.246 | 0.42 | 0.167418 | MS.Trt_vs_MS.NoTrt_B |
| ARHGEF39 | 1.05E-05 | -0.9427 | 0.497 | 0.7 | 0.167939 | MS.Trt_vs_MS.NoTrt_B |
| FIS1 | 1.07E-05 | -0.7007 | 0.172 | 0.32 | 0.171511 | MS.Trt_vs_MS.NoTrt_B |
| GATC | 1.08E-05 | 0.412 | 0.451 | 0.67 | 0.172691 | MS.Trt_vs_MS.NoTrt_B |
| CYBA | 1.08E-05 | -0.3555 | 0.159 | 0.31 | 0.173114 | MS.Trt_vs_MS.NoTrt_B |
| RNASET2 | 1.12E-05 | -0.2698 | 0.307 | 0.48 | 0.178309 | MS.Trt_vs_MS.NoTrt_B |
| ARL5A | 1.14E-05 | -0.3286 | 0.092 | 0.22 | 0.182141 | MS.Trt_vs_MS.NoTrt_B |
| ADCY7 | 1.17E-05 | -0.3334 | 0.617 | 0.77 | 0.187106 | MS.Trt_vs_MS.NoTrt_B |
| TINF2 | 1.19E-05 | -0.4474 | 0.688 | 0.87 | 0.189536 | MS.Trt_vs_MS.NoTrt_B |
| AIFM1 | 1.19E-05 | -0.5424 | 0.355 | 0.56 | 0.189793 | MS.Trt_vs_MS.NoTrt_B |
| NADK2 | 1.20E-05 | -0.2834 | 0.283 | 0.47 | 0.191467 | MS.Trt_vs_MS.NoTrt_B |
| MRPL3 | 1.23E-05 | -0.3007 | 0.375 | 0.58 | 0.195767 | MS.Trt_vs_MS.NoTrt_B |
| TSPAN32 | 1.28E-05 | -0.3653 | 0.17 | 0.33 | 0.203886 | MS.Trt_vs_MS.NoTrt_B |
| PSMC5 | 1.28E-05 | -0.4481 | 0.244 | 0.42 | 0.203951 | MS.Trt_vs_MS.NoTrt_B |
| NT5C2 | 1.28E-05 | -0.6481 | 0.59 | 0.77 | 0.204811 | MS.Trt_vs_MS.NoTrt_B |
| STOML1 | 1.29E-05 | -0.785 | 0.634 | 0.82 | 0.205936 | MS.Trt_vs_MS.NoTrt_B |
| CCR10 | 1.29E-05 | -1.1168 | 0.057 | 0.16 | 0.20611 | MS.Trt_vs_MS.NoTrt_B |
| GNL3L | 1.29E-05 | -0.2565 | 0.821 | 0.96 | 0.206356 | MS.Trt_vs_MS.NoTrt_B |
| TMEM116 | 1.30E-05 | 0.44309 | 0.732 | 0.9 | 0.207101 | MS.Trt_vs_MS.NoTrt_B |
| NAIP | 1.31E-05 | -0.7526 | 0.505 | 0.68 | 0.209103 | MS.Trt_vs_MS.NoTrt_B |
| LIPT1 | 1.32E-05 | -0.3347 | 0.523 | 0.72 | 0.210203 | MS.Trt_vs_MS.NoTrt_B |
| NOSIP | 1.34E-05 | -0.2705 | 0.438 | 0.61 | 0.213354 | MS.Trt_vs_MS.NoTrt_B |

| | | | | | | |
|---|---|---|---|---|---|---|
| SNAP23 | 1.34E-05 | -0.3236 | 0.501 | 0.7 | 0.214782 | MS.Trt_vs_MS.NoTrt_B |
| IGBP1 | 1.35E-05 | -0.7022 | 0.133 | 0.28 | 0.21514 | MS.Trt_vs_MS.NoTrt_B |
| NEB | 1.36E-05 | -0.5066 | 0.401 | 0.62 | 0.217019 | MS.Trt_vs_MS.NoTrt_B |
| HELZ2 | 1.38E-05 | -0.4696 | 0.499 | 0.7 | 0.219755 | MS.Trt_vs_MS.NoTrt_B |
| OAZ2 | 1.40E-05 | -0.5424 | 0.627 | 0.83 | 0.22336 | MS.Trt_vs_MS.NoTrt_B |
| SNRPB2 | 1.41E-05 | -0.5377 | 0.137 | 0.28 | 0.224962 | MS.Trt_vs_MS.NoTrt_B |
| RAP2C | 1.41E-05 | -0.5759 | 0.198 | 0.36 | 0.225341 | MS.Trt_vs_MS.NoTrt_B |
| RNASEH2B | 1.41E-05 | 0.31228 | 0.588 | 0.82 | 0.22569 | MS.Trt_vs_MS.NoTrt_B |
| MED4 | 1.42E-05 | 0.34933 | 0.564 | 0.72 | 0.226959 | MS.Trt_vs_MS.NoTrt_B |
| SART3 | 1.44E-05 | -1.011 | 0.089 | 0.21 | 0.22962 | MS.Trt_vs_MS.NoTrt_B |
| RNASEH2A | 1.44E-05 | -0.2804 | 0.795 | 0.95 | 0.23021 | MS.Trt_vs_MS.NoTrt_B |
| VAMP3 | 1.44E-05 | -0.2913 | 0.214 | 0.38 | 0.230412 | MS.Trt_vs_MS.NoTrt_B |
| ZNF721 | 1.44E-05 | -0.8084 | 0.248 | 0.41 | 0.230501 | MS.Trt_vs_MS.NoTrt_B |
| RANGAP1 | 1.46E-05 | -0.2661 | 0.508 | 0.71 | 0.23304 | MS.Trt_vs_MS.NoTrt_B |
| HERC3 | 1.47E-05 | -0.3045 | 0.218 | 0.38 | 0.234258 | MS.Trt_vs_MS.NoTrt_B |
| PQLC3 | 1.47E-05 | -0.4887 | 0.102 | 0.23 | 0.234346 | MS.Trt_vs_MS.NoTrt_B |
| PRNP | 1.47E-05 | -0.4876 | 0.126 | 0.27 | 0.234391 | MS.Trt_vs_MS.NoTrt_B |
| ATP8B1 | 1.47E-05 | -0.6121 | 0.153 | 0.3 | 0.234634 | MS.Trt_vs_MS.NoTrt_B |
| DHRSX | 1.48E-05 | -0.6341 | 0.584 | 0.77 | 0.235642 | MS.Trt_vs_MS.NoTrt_B |
| AIM1 | 1.48E-05 | -0.3874 | 0.359 | 0.55 | 0.235666 | MS.Trt_vs_MS.NoTrt_B |
| YBEY | 1.48E-05 | -1.0671 | 0.035 | 0.12 | 0.23585 | MS.Trt_vs_MS.NoTrt_B |
| ZKSCAN5 | 1.48E-05 | -0.2879 | 0.401 | 0.61 | 0.235911 | MS.Trt_vs_MS.NoTrt_B |
| CDKN1B | 1.49E-05 | -0.2885 | 0.216 | 0.38 | 0.238499 | MS.Trt_vs_MS.NoTrt_B |
| DNAJB12 | 1.52E-05 | -0.6483 | 0.063 | 0.17 | 0.243253 | MS.Trt_vs_MS.NoTrt_B |
| USF1 | 1.53E-05 | -2.5167 | 0.102 | 0.22 | 0.243693 | MS.Trt_vs_MS.NoTrt_B |
| MPI | 1.53E-05 | -0.8604 | 0.383 | 0.54 | 0.24411 | MS.Trt_vs_MS.NoTrt_B |
| TMEM165 | 1.53E-05 | -0.27 | 0.512 | 0.71 | 0.244429 | MS.Trt_vs_MS.NoTrt_B |
| TGM2 | 1.58E-05 | -0.2596 | 0.401 | 0.6 | 0.252403 | MS.Trt_vs_MS.NoTrt_B |
| FBXO11 | 1.60E-05 | 0.77328 | 0.301 | 0.49 | 0.254777 | MS.Trt_vs_MS.NoTrt_B |
| MRPS6 | 1.60E-05 | -0.5495 | 0.126 | 0.27 | 0.255573 | MS.Trt_vs_MS.NoTrt_B |
| USP45 | 1.60E-05 | -0.7354 | 0.115 | 0.25 | 0.255671 | MS.Trt_vs_MS.NoTrt_B |
| NDUFS4 | 1.62E-05 | -0.779 | 0.168 | 0.31 | 0.259526 | MS.Trt_vs_MS.NoTrt_B |
| GIN1 | 1.63E-05 | 0.94371 | 0.244 | 0.43 | 0.260001 | MS.Trt_vs_MS.NoTrt_B |
| RFTN1 | 1.63E-05 | -0.4843 | 0.277 | 0.45 | 0.260012 | MS.Trt_vs_MS.NoTrt_B |
| C15orf52 | 1.63E-05 | -0.9584 | 0.285 | 0.48 | 0.260943 | MS.Trt_vs_MS.NoTrt_B |
| PSMD9 | 1.63E-05 | -0.3119 | 0.904 | 0.98 | 0.260947 | MS.Trt_vs_MS.NoTrt_B |
| CAPRIN2 | 1.65E-05 | -0.6021 | 0.532 | 0.77 | 0.264002 | MS.Trt_vs_MS.NoTrt_B |
| FBXO40 | 1.69E-05 | 0.38012 | 0.176 | 0.34 | 0.270246 | MS.Trt_vs_MS.NoTrt_B |
| SUPT7L | 1.72E-05 | -0.4295 | 0.344 | 0.54 | 0.275163 | MS.Trt_vs_MS.NoTrt_B |
| ZNF783 | 1.75E-05 | -0.7183 | 0.344 | 0.54 | 0.279031 | MS.Trt_vs_MS.NoTrt_B |

| | | | | | | |
|---|---|---|---|---|---|---|
| TRIM4 | 1.75E-05 | 0.30436 | 0.414 | 0.62 | 0.279136 | MS.Trt_vs_MS.NoTrt_B |
| GMPR2 | 1.75E-05 | -1.1306 | 0.076 | 0.19 | 0.279248 | MS.Trt_vs_MS.NoTrt_B |
| PPP2R3B | 1.75E-05 | -1.1172 | 0.333 | 0.53 | 0.279399 | MS.Trt_vs_MS.NoTrt_B |
| RGL4 | 1.75E-05 | -0.3296 | 0.142 | 0.29 | 0.280303 | MS.Trt_vs_MS.NoTrt_B |
| PTCD3 | 1.77E-05 | -0.2904 | 0.767 | 0.94 | 0.282904 | MS.Trt_vs_MS.NoTrt_B |
| NSA2 | 1.82E-05 | -0.4479 | 0.242 | 0.41 | 0.290633 | MS.Trt_vs_MS.NoTrt_B |
| GRIPAP1 | 1.82E-05 | -0.4798 | 0.575 | 0.77 | 0.290895 | MS.Trt_vs_MS.NoTrt_B |
| PRPF6 | 1.86E-05 | -0.6314 | 0.124 | 0.26 | 0.297362 | MS.Trt_vs_MS.NoTrt_B |
| RALB | 1.87E-05 | -1.5339 | 0.031 | 0.11 | 0.298064 | MS.Trt_vs_MS.NoTrt_B |
| DDX42 | 1.91E-05 | 0.61554 | 0.257 | 0.45 | 0.304538 | MS.Trt_vs_MS.NoTrt_B |
| TCHP | 1.93E-05 | -0.6202 | 0.381 | 0.55 | 0.307584 | MS.Trt_vs_MS.NoTrt_B |
| NUP107 | 1.94E-05 | -0.5391 | 0.597 | 0.78 | 0.309738 | MS.Trt_vs_MS.NoTrt_B |
| ZNF576 | 1.94E-05 | -0.2583 | 0.397 | 0.6 | 0.309943 | MS.Trt_vs_MS.NoTrt_B |
| FAM126A | 1.96E-05 | 0.8013 | 0.497 | 0.73 | 0.31364 | MS.Trt_vs_MS.NoTrt_B |
| TUBA1A | 1.99E-05 | -0.4619 | 0.187 | 0.34 | 0.317964 | MS.Trt_vs_MS.NoTrt_B |
| RGS10 | 2.02E-05 | -0.3463 | 0.166 | 0.31 | 0.322506 | MS.Trt_vs_MS.NoTrt_B |
| USP34 | 2.05E-05 | 0.36679 | 0.481 | 0.68 | 0.32776 | MS.Trt_vs_MS.NoTrt_B |
| CTSS | 2.08E-05 | -0.3112 | 0.556 | 0.73 | 0.332008 | MS.Trt_vs_MS.NoTrt_B |
| KLHL5 | 2.09E-05 | -0.3808 | 0.331 | 0.53 | 0.334378 | MS.Trt_vs_MS.NoTrt_B |
| TPI1 | 2.11E-05 | -0.2917 | 0.403 | 0.6 | 0.337725 | MS.Trt_vs_MS.NoTrt_B |
| RANBP3 | 2.12E-05 | -0.2913 | 0.601 | 0.81 | 0.339029 | MS.Trt_vs_MS.NoTrt_B |
| GALE | 2.13E-05 | 0.27145 | 0.192 | 0.36 | 0.340483 | MS.Trt_vs_MS.NoTrt_B |
| MAST4 | 2.20E-05 | -0.4494 | 0.481 | 0.65 | 0.351058 | MS.Trt_vs_MS.NoTrt_B |
| GDAP2 | 2.20E-05 | -0.7427 | 0.549 | 0.74 | 0.351639 | MS.Trt_vs_MS.NoTrt_B |
| AHSA2 | 2.21E-05 | 0.37154 | 0.166 | 0.32 | 0.35274 | MS.Trt_vs_MS.NoTrt_B |
| DIS3L2 | 2.22E-05 | -0.7258 | 0.333 | 0.5 | 0.353981 | MS.Trt_vs_MS.NoTrt_B |
| DTWD1 | 2.22E-05 | 0.38223 | 0.436 | 0.65 | 0.354362 | MS.Trt_vs_MS.NoTrt_B |
| 9-Sep | 2.22E-05 | 0.45039 | 0.42 | 0.67 | 0.355138 | MS.Trt_vs_MS.NoTrt_B |
| RP11-449H3 | 2.24E-05 | -0.4906 | 0.172 | 0.32 | 0.357739 | MS.Trt_vs_MS.NoTrt_B |
| SLC2A8 | 2.26E-05 | -0.939 | 0.196 | 0.35 | 0.361597 | MS.Trt_vs_MS.NoTrt_B |
| DPP8 | 2.30E-05 | -0.5408 | 0.29 | 0.47 | 0.367304 | MS.Trt_vs_MS.NoTrt_B |
| RAD21 | 2.30E-05 | 0.38671 | 0.298 | 0.48 | 0.367572 | MS.Trt_vs_MS.NoTrt_B |
| DMPK | 2.30E-05 | 0.34045 | 0.46 | 0.68 | 0.367653 | MS.Trt_vs_MS.NoTrt_B |
| COX5B | 2.31E-05 | 0.32354 | 0.682 | 0.83 | 0.368212 | MS.Trt_vs_MS.NoTrt_B |
| SVBP | 2.31E-05 | -0.7279 | 0.076 | 0.19 | 0.368506 | MS.Trt_vs_MS.NoTrt_B |
| ARL16 | 2.38E-05 | -0.6527 | 0.614 | 0.8 | 0.380429 | MS.Trt_vs_MS.NoTrt_B |
| MRPL52 | 2.39E-05 | 0.71492 | 0.473 | 0.67 | 0.381226 | MS.Trt_vs_MS.NoTrt_B |
| PSPC1 | 2.39E-05 | -0.4493 | 0.218 | 0.37 | 0.381977 | MS.Trt_vs_MS.NoTrt_B |
| MIXL1 | 2.41E-05 | 0.37281 | 0.619 | 0.42 | 0.385605 | MS.Trt_vs_MS.NoTrt_B |
| ASNS | 2.42E-05 | -0.897 | 0.163 | 0.3 | 0.386455 | MS.Trt_vs_MS.NoTrt_B |

| | | | | | | |
|---|---|---|---|---|---|---|
| PNPLA4 | 2.43E-05 | 0.5433 | 0.176 | 0.34 | 0.387641 | MS.Trt_vs_MS.NoTrt_B |
| STAT3 | 2.54E-05 | -0.344 | 0.636 | 0.82 | 0.405174 | MS.Trt_vs_MS.NoTrt_B |
| DBNL | 2.54E-05 | -0.2672 | 0.351 | 0.51 | 0.405671 | MS.Trt_vs_MS.NoTrt_B |
| KDM4A | 2.56E-05 | -0.7689 | 0.061 | 0.17 | 0.408685 | MS.Trt_vs_MS.NoTrt_B |
| TBCC | 2.58E-05 | -0.3276 | 0.259 | 0.43 | 0.411685 | MS.Trt_vs_MS.NoTrt_B |
| BIRC2 | 2.59E-05 | -0.3722 | 0.089 | 0.21 | 0.413592 | MS.Trt_vs_MS.NoTrt_B |
| GMPPA | 2.61E-05 | -0.2674 | 0.161 | 0.31 | 0.416807 | MS.Trt_vs_MS.NoTrt_B |
| FBXO32 | 2.62E-05 | -0.6579 | 0.098 | 0.22 | 0.417999 | MS.Trt_vs_MS.NoTrt_B |
| SEC61A1 | 2.62E-05 | -0.3466 | 0.253 | 0.42 | 0.418785 | MS.Trt_vs_MS.NoTrt_B |
| HIBCH | 2.63E-05 | -0.6524 | 0.139 | 0.28 | 0.41954 | MS.Trt_vs_MS.NoTrt_B |
| KAT2B | 2.70E-05 | 0.60791 | 0.115 | 0.25 | 0.430956 | MS.Trt_vs_MS.NoTrt_B |
| LRR1 | 2.71E-05 | -2.2748 | 0.057 | 0.16 | 0.432969 | MS.Trt_vs_MS.NoTrt_B |
| HLA-C | 2.72E-05 | -0.4072 | 0.85 | 0.92 | 0.434901 | MS.Trt_vs_MS.NoTrt_B |
| ASAP3 | 2.74E-05 | -0.4257 | 0.379 | 0.58 | 0.438334 | MS.Trt_vs_MS.NoTrt_B |
| RNF6 | 2.75E-05 | -0.3194 | 0.098 | 0.22 | 0.438561 | MS.Trt_vs_MS.NoTrt_B |
| GARS | 2.77E-05 | -0.4109 | 0.146 | 0.29 | 0.441817 | MS.Trt_vs_MS.NoTrt_B |
| LAIR2 | 2.78E-05 | -0.7484 | 0.057 | 0.16 | 0.44355 | MS.Trt_vs_MS.NoTrt_B |
| MTIF3 | 2.82E-05 | -0.6221 | 0.102 | 0.23 | 0.450813 | MS.Trt_vs_MS.NoTrt_B |
| ZNF266 | 2.83E-05 | -0.4922 | 0.235 | 0.4 | 0.451314 | MS.Trt_vs_MS.NoTrt_B |
| ADH5 | 2.84E-05 | 0.28233 | 0.309 | 0.5 | 0.452941 | MS.Trt_vs_MS.NoTrt_B |
| TOR3A | 2.84E-05 | -0.894 | 0.719 | 0.85 | 0.453745 | MS.Trt_vs_MS.NoTrt_B |
| VLDLR | 2.97E-05 | -0.3109 | 0.316 | 0.5 | 0.473872 | MS.Trt_vs_MS.NoTrt_B |
| SLC2A3 | 3.02E-05 | -0.2899 | 0.632 | 0.81 | 0.481793 | MS.Trt_vs_MS.NoTrt_B |
| IDNK | 3.02E-05 | -0.3102 | 0.272 | 0.45 | 0.482174 | MS.Trt_vs_MS.NoTrt_B |
| DHRS4 | 3.05E-05 | -0.7789 | 0.063 | 0.17 | 0.487064 | MS.Trt_vs_MS.NoTrt_B |
| TNIP3 | 3.17E-05 | -0.6101 | 0.368 | 0.56 | 0.506402 | MS.Trt_vs_MS.NoTrt_B |
| MAP4K3 | 3.18E-05 | -0.6362 | 0.028 | 0.11 | 0.507278 | MS.Trt_vs_MS.NoTrt_B |
| AP1S1 | 3.18E-05 | -0.2812 | 0.031 | 0.11 | 0.508433 | MS.Trt_vs_MS.NoTrt_B |
| MRPS23 | 3.29E-05 | -0.5505 | 0.658 | 0.82 | 0.524878 | MS.Trt_vs_MS.NoTrt_B |
| UTP20 | 3.30E-05 | 0.78227 | 0.368 | 0.57 | 0.526387 | MS.Trt_vs_MS.NoTrt_B |
| THEMIS2 | 3.33E-05 | -0.4731 | 0.081 | 0.2 | 0.53224 | MS.Trt_vs_MS.NoTrt_B |
| BMS1 | 3.34E-05 | -0.3191 | 0.394 | 0.58 | 0.533443 | MS.Trt_vs_MS.NoTrt_B |
| SVIP | 3.35E-05 | -0.3733 | 0.39 | 0.57 | 0.535611 | MS.Trt_vs_MS.NoTrt_B |
| KIAA1614 | 3.36E-05 | 0.45208 | 0.181 | 0.34 | 0.53736 | MS.Trt_vs_MS.NoTrt_B |
| C11orf21 | 3.37E-05 | -0.8749 | 0.081 | 0.2 | 0.537974 | MS.Trt_vs_MS.NoTrt_B |
| ZRANB2 | 3.37E-05 | 0.49764 | 0.303 | 0.49 | 0.538118 | MS.Trt_vs_MS.NoTrt_B |
| NOP14 | 3.41E-05 | -0.4705 | 0.314 | 0.5 | 0.544559 | MS.Trt_vs_MS.NoTrt_B |
| IQCC | 3.41E-05 | 0.52064 | 0.427 | 0.67 | 0.545213 | MS.Trt_vs_MS.NoTrt_B |
| ZNF215 | 3.43E-05 | -0.2612 | 0.283 | 0.45 | 0.547852 | MS.Trt_vs_MS.NoTrt_B |
| SEMA3C | 3.48E-05 | -0.2969 | 0.503 | 0.7 | 0.555067 | MS.Trt_vs_MS.NoTrt_B |

| | | | | | | |
|---|---|---|---|---|---|---|
| ATP5S | 3.48E-05 | -0.3852 | 0.793 | 0.93 | 0.556082 | MS.Trt_vs_MS.NoTrt_B |
| KIAA0391 | 3.48E-05 | -0.2931 | 0.573 | 0.75 | 0.556336 | MS.Trt_vs_MS.NoTrt_B |
| PEX16 | 3.50E-05 | -0.511 | 0.582 | 0.8 | 0.559008 | MS.Trt_vs_MS.NoTrt_B |
| BSDC1 | 3.50E-05 | -0.5827 | 0.353 | 0.51 | 0.559548 | MS.Trt_vs_MS.NoTrt_B |
| SH3GL1 | 3.56E-05 | -0.3356 | 0.634 | 0.78 | 0.567969 | MS.Trt_vs_MS.NoTrt_B |
| SMYD2 | 3.56E-05 | -0.3595 | 0.094 | 0.21 | 0.568165 | MS.Trt_vs_MS.NoTrt_B |
| CCDC191 | 3.60E-05 | -0.3675 | 0.155 | 0.3 | 0.5755 | MS.Trt_vs_MS.NoTrt_B |
| PCBP1 | 3.62E-05 | 0.31277 | 0.342 | 0.53 | 0.578576 | MS.Trt_vs_MS.NoTrt_B |
| ZNF428 | 3.67E-05 | -0.5699 | 0.113 | 0.23 | 0.585832 | MS.Trt_vs_MS.NoTrt_B |
| C4orf3 | 3.68E-05 | 0.26299 | 0.447 | 0.62 | 0.587096 | MS.Trt_vs_MS.NoTrt_B |
| EMD | 3.69E-05 | -0.4074 | 0.187 | 0.34 | 0.589147 | MS.Trt_vs_MS.NoTrt_B |
| GLYCTK | 3.74E-05 | -0.4509 | 0.063 | 0.17 | 0.597541 | MS.Trt_vs_MS.NoTrt_B |
| CES4A | 3.76E-05 | -0.3496 | 0.416 | 0.59 | 0.601358 | MS.Trt_vs_MS.NoTrt_B |
| DHX16 | 3.77E-05 | -0.65 | 0.092 | 0.21 | 0.601733 | MS.Trt_vs_MS.NoTrt_B |
| ZMYM6 | 3.78E-05 | 0.49049 | 0.664 | 0.89 | 0.603003 | MS.Trt_vs_MS.NoTrt_B |
| QRICH1 | 3.81E-05 | -0.3363 | 0.407 | 0.6 | 0.607854 | MS.Trt_vs_MS.NoTrt_B |
| BAX | 3.81E-05 | -0.3281 | 0.248 | 0.4 | 0.608911 | MS.Trt_vs_MS.NoTrt_B |
| ZNF507 | 3.85E-05 | -0.2533 | 0.118 | 0.25 | 0.614453 | MS.Trt_vs_MS.NoTrt_B |
| RP11-977G1 | 3.86E-05 | -0.2552 | 0.044 | 0.14 | 0.616379 | MS.Trt_vs_MS.NoTrt_B |
| AP003419.1 | 3.89E-05 | -0.5089 | 0.098 | 0.22 | 0.621135 | MS.Trt_vs_MS.NoTrt_B |
| FAM105A | 3.94E-05 | 0.34598 | 0.181 | 0.33 | 0.628629 | MS.Trt_vs_MS.NoTrt_B |
| ZNF226 | 3.94E-05 | -0.7097 | 0.316 | 0.47 | 0.629195 | MS.Trt_vs_MS.NoTrt_B |
| PSMB7 | 3.97E-05 | -0.3357 | 0.198 | 0.35 | 0.63421 | MS.Trt_vs_MS.NoTrt_B |
| RNF130 | 3.97E-05 | 0.6419 | 0.488 | 0.67 | 0.634516 | MS.Trt_vs_MS.NoTrt_B |
| CHST11 | 3.99E-05 | -0.3678 | 0.109 | 0.23 | 0.63773 | MS.Trt_vs_MS.NoTrt_B |
| ZNF85 | 4.00E-05 | -0.4592 | 0.514 | 0.72 | 0.639056 | MS.Trt_vs_MS.NoTrt_B |
| SIVA1 | 4.00E-05 | -0.3077 | 0.303 | 0.47 | 0.639271 | MS.Trt_vs_MS.NoTrt_B |
| HTATIP2 | 4.02E-05 | -0.4241 | 0.087 | 0.21 | 0.6416 | MS.Trt_vs_MS.NoTrt_B |
| TIMM23B | 4.03E-05 | -0.8321 | 0.46 | 0.68 | 0.644118 | MS.Trt_vs_MS.NoTrt_B |
| C1orf159 | 4.05E-05 | -0.9796 | 0.076 | 0.19 | 0.646534 | MS.Trt_vs_MS.NoTrt_B |
| POLA2 | 4.09E-05 | -0.7945 | 0.534 | 0.7 | 0.652982 | MS.Trt_vs_MS.NoTrt_B |
| SLC12A2 | 4.09E-05 | 0.7141 | 0.412 | 0.63 | 0.654022 | MS.Trt_vs_MS.NoTrt_B |
| BEX2 | 4.15E-05 | -0.7579 | 0.041 | 0.13 | 0.663462 | MS.Trt_vs_MS.NoTrt_B |
| TSEN15 | 4.17E-05 | -0.6113 | 0.107 | 0.23 | 0.666672 | MS.Trt_vs_MS.NoTrt_B |
| ERI1 | 4.19E-05 | 0.35913 | 0.268 | 0.45 | 0.669631 | MS.Trt_vs_MS.NoTrt_B |
| MTR | 4.21E-05 | -0.287 | 0.414 | 0.56 | 0.671742 | MS.Trt_vs_MS.NoTrt_B |
| ZNF687 | 4.25E-05 | -0.6164 | 0.497 | 0.66 | 0.679601 | MS.Trt_vs_MS.NoTrt_B |
| ZNF335 | 4.36E-05 | -0.3368 | 0.248 | 0.4 | 0.696227 | MS.Trt_vs_MS.NoTrt_B |
| ATP2B1 | 4.38E-05 | 0.48856 | 0.325 | 0.54 | 0.699107 | MS.Trt_vs_MS.NoTrt_B |
| C1orf43 | 4.46E-05 | -0.3596 | 0.451 | 0.62 | 0.71286 | MS.Trt_vs_MS.NoTrt_B |

| | | | | | | |
|---|---|---|---|---|---|---|
| VPS26A | 4.47E-05 | 0.63732 | 0.244 | 0.42 | 0.713716 | MS.Trt_vs_MS.NoTrt_B |
| FAHD2B | 4.48E-05 | -0.694 | 0.098 | 0.22 | 0.714833 | MS.Trt_vs_MS.NoTrt_B |
| ACOT9 | 4.49E-05 | -0.8023 | 0.8 | 0.9 | 0.71661 | MS.Trt_vs_MS.NoTrt_B |
| ARHGAP11A | 4.53E-05 | -1.0132 | 0.279 | 0.44 | 0.723538 | MS.Trt_vs_MS.NoTrt_B |
| ETF1 | 4.56E-05 | 0.30866 | 0.2 | 0.36 | 0.727615 | MS.Trt_vs_MS.NoTrt_B |
| DDOST | 4.57E-05 | -0.5817 | 0.444 | 0.57 | 0.730446 | MS.Trt_vs_MS.NoTrt_B |
| CMTR1 | 4.58E-05 | -1.1321 | 0.122 | 0.24 | 0.732327 | MS.Trt_vs_MS.NoTrt_B |
| ONECUT3 | 4.59E-05 | -0.3327 | 0.312 | 0.48 | 0.733245 | MS.Trt_vs_MS.NoTrt_B |
| ASB7 | 4.62E-05 | -1.3188 | 0.033 | 0.11 | 0.738497 | MS.Trt_vs_MS.NoTrt_B |
| ELOF1 | 4.77E-05 | -0.5975 | 0.473 | 0.62 | 0.762699 | MS.Trt_vs_MS.NoTrt_B |
| CCDC141 | 4.84E-05 | -0.3224 | 0.505 | 0.7 | 0.77339 | MS.Trt_vs_MS.NoTrt_B |
| COPS4 | 4.86E-05 | 0.53971 | 0.349 | 0.55 | 0.776267 | MS.Trt_vs_MS.NoTrt_B |
| CTSD | 4.86E-05 | -0.5412 | 0.139 | 0.28 | 0.776592 | MS.Trt_vs_MS.NoTrt_B |
| KIAA2013 | 4.92E-05 | -0.3852 | 0.209 | 0.36 | 0.785745 | MS.Trt_vs_MS.NoTrt_B |
| PHF7 | 4.93E-05 | -1.1447 | 0.351 | 0.54 | 0.787569 | MS.Trt_vs_MS.NoTrt_B |
| WBSCR22 | 4.95E-05 | -0.4817 | 0.584 | 0.75 | 0.791294 | MS.Trt_vs_MS.NoTrt_B |
| MCOLN2 | 4.96E-05 | 0.34069 | 0.316 | 0.49 | 0.792331 | MS.Trt_vs_MS.NoTrt_B |
| NOL6 | 4.97E-05 | -1.4045 | 0.275 | 0.42 | 0.794654 | MS.Trt_vs_MS.NoTrt_B |
| GALNT10 | 4.99E-05 | -0.6229 | 0.403 | 0.57 | 0.797088 | MS.Trt_vs_MS.NoTrt_B |
| RTN3 | 5.09E-05 | -0.4297 | 0.098 | 0.22 | 0.813735 | MS.Trt_vs_MS.NoTrt_B |
| GABRE | 5.12E-05 | -0.3915 | 0.133 | 0.26 | 0.817806 | MS.Trt_vs_MS.NoTrt_B |
| SIMC1 | 5.19E-05 | -0.2604 | 0.296 | 0.48 | 0.828983 | MS.Trt_vs_MS.NoTrt_B |
| WDR55 | 5.30E-05 | -0.3324 | 0.484 | 0.67 | 0.846807 | MS.Trt_vs_MS.NoTrt_B |
| SEL1L | 5.33E-05 | -0.7812 | 0.115 | 0.24 | 0.851888 | MS.Trt_vs_MS.NoTrt_B |
| TRIB1 | 5.34E-05 | -0.3288 | 0.033 | 0.11 | 0.852383 | MS.Trt_vs_MS.NoTrt_B |
| TLDC1 | 5.42E-05 | -0.536 | 0.593 | 0.81 | 0.866054 | MS.Trt_vs_MS.NoTrt_B |
| TSPAN5 | 5.46E-05 | -0.3382 | 0.46 | 0.62 | 0.871823 | MS.Trt_vs_MS.NoTrt_B |
| RCAN3 | 5.48E-05 | 0.25812 | 0.662 | 0.81 | 0.874648 | MS.Trt_vs_MS.NoTrt_B |
| WDR89 | 5.54E-05 | -0.2564 | 0.231 | 0.41 | 0.884475 | MS.Trt_vs_MS.NoTrt_B |
| ZC3H12A | 5.60E-05 | -1.1958 | 0.057 | 0.15 | 0.894389 | MS.Trt_vs_MS.NoTrt_B |
| C20orf196 | 5.60E-05 | -1.1335 | 0.244 | 0.41 | 0.894729 | MS.Trt_vs_MS.NoTrt_B |
| HYKK | 5.61E-05 | 0.25191 | 0.684 | 0.86 | 0.895481 | MS.Trt_vs_MS.NoTrt_B |
| MBD3 | 5.63E-05 | 0.42893 | 0.547 | 0.8 | 0.899478 | MS.Trt_vs_MS.NoTrt_B |
| TRMT5 | 5.64E-05 | -1.0639 | 0.089 | 0.2 | 0.90087 | MS.Trt_vs_MS.NoTrt_B |
| NHSL2 | 5.65E-05 | 0.38511 | 0.444 | 0.64 | 0.902273 | MS.Trt_vs_MS.NoTrt_B |
| NUDT2 | 5.68E-05 | -1.3169 | 0.031 | 0.11 | 0.906505 | MS.Trt_vs_MS.NoTrt_B |
| RBMX2 | 5.68E-05 | -0.9851 | 0.052 | 0.15 | 0.908043 | MS.Trt_vs_MS.NoTrt_B |
| MRPL11 | 5.72E-05 | -0.7593 | 0.07 | 0.17 | 0.913241 | MS.Trt_vs_MS.NoTrt_B |
| MAF1 | 5.83E-05 | -0.4039 | 0.122 | 0.25 | 0.931231 | MS.Trt_vs_MS.NoTrt_B |
| MS4A2 | 5.84E-05 | -0.2566 | 0.486 | 0.62 | 0.932981 | MS.Trt_vs_MS.NoTrt_B |

| | | | | | | |
|---|---|---|---|---|---|---|
| FAN1 | 5.85E-05 | 0.97198 | 0.192 | 0.34 | 0.933773 | MS.Trt_vs_MS.NoTrt_B |
| TNFRSF9 | 5.88E-05 | -0.4609 | 0.754 | 0.91 | 0.939299 | MS.Trt_vs_MS.NoTrt_B |
| RNASEH2C | 5.89E-05 | -0.3155 | 0.072 | 0.18 | 0.940958 | MS.Trt_vs_MS.NoTrt_B |
| HM13 | 5.98E-05 | -0.5029 | 0.715 | 0.83 | 0.955445 | MS.Trt_vs_MS.NoTrt_B |
| ZNF468 | 6.07E-05 | 0.3797 | 0.416 | 0.62 | 0.96963 | MS.Trt_vs_MS.NoTrt_B |
| STX10 | 6.09E-05 | -0.5476 | 0.083 | 0.2 | 0.972763 | MS.Trt_vs_MS.NoTrt_B |
| WEE1 | 6.09E-05 | -0.3158 | 0.51 | 0.65 | 0.972998 | MS.Trt_vs_MS.NoTrt_B |
| ZCCHC9 | 6.16E-05 | -0.4092 | 0.525 | 0.73 | 0.983722 | MS.Trt_vs_MS.NoTrt_B |
| VASP | 6.17E-05 | -0.7021 | 0.407 | 0.57 | 0.985657 | MS.Trt_vs_MS.NoTrt_B |
| HGS | 6.18E-05 | 0.30032 | 0.549 | 0.72 | 0.987444 | MS.Trt_vs_MS.NoTrt_B |
| CCDC130 | 6.18E-05 | 0.26865 | 0.316 | 0.49 | 0.987871 | MS.Trt_vs_MS.NoTrt_B |
| XPO6 | 6.20E-05 | -0.3156 | 0.233 | 0.38 | 0.989849 | MS.Trt_vs_MS.NoTrt_B |
| FNDC3A | 6.20E-05 | -0.2999 | 0.37 | 0.55 | 0.99019 | MS.Trt_vs_MS.NoTrt_B |
| CHD6 | 6.25E-05 | -0.409 | 0.135 | 0.27 | 0.998542 | MS.Trt_vs_MS.NoTrt_B |
| RCHY1 | 6.41E-05 | -0.8131 | 0.192 | 0.34 | 1 | MS.Trt_vs_MS.NoTrt_B |
| RINL | 6.45E-05 | -0.6398 | 0.597 | 0.76 | 1 | MS.Trt_vs_MS.NoTrt_B |
| CCDC14 | 6.47E-05 | 0.4885 | 0.129 | 0.27 | 1 | MS.Trt_vs_MS.NoTrt_B |
| CLCN3 | 6.49E-05 | 1.0745 | 0.227 | 0.39 | 1 | MS.Trt_vs_MS.NoTrt_B |
| EIF1AD | 6.51E-05 | 0.54578 | 0.571 | 0.74 | 1 | MS.Trt_vs_MS.NoTrt_B |
| EGLN1 | 6.55E-05 | -0.2654 | 0.24 | 0.4 | 1 | MS.Trt_vs_MS.NoTrt_B |
| RC3H1 | 6.60E-05 | 0.33164 | 0.353 | 0.5 | 1 | MS.Trt_vs_MS.NoTrt_B |
| ZNF621 | 6.61E-05 | -0.2734 | 0.828 | 0.94 | 1 | MS.Trt_vs_MS.NoTrt_B |
| AP5Z1 | 6.62E-05 | -1.0119 | 0.109 | 0.23 | 1 | MS.Trt_vs_MS.NoTrt_B |
| ARNTL2 | 6.67E-05 | -0.3318 | 0.532 | 0.63 | 1 | MS.Trt_vs_MS.NoTrt_B |
| C8orf59 | 6.70E-05 | 0.33163 | 0.283 | 0.45 | 1 | MS.Trt_vs_MS.NoTrt_B |
| MEI1 | 6.73E-05 | -0.6392 | 0.601 | 0.76 | 1 | MS.Trt_vs_MS.NoTrt_B |
| IARS2 | 6.78E-05 | -0.6984 | 0.085 | 0.2 | 1 | MS.Trt_vs_MS.NoTrt_B |
| TCAF1 | 6.81E-05 | -0.9715 | 0.187 | 0.34 | 1 | MS.Trt_vs_MS.NoTrt_B |
| ANAPC7 | 6.81E-05 | -0.3479 | 0.423 | 0.6 | 1 | MS.Trt_vs_MS.NoTrt_B |
| PTPN2 | 6.82E-05 | -0.4611 | 0.858 | 0.98 | 1 | MS.Trt_vs_MS.NoTrt_B |
| UBE2K | 6.88E-05 | -0.7427 | 0.111 | 0.23 | 1 | MS.Trt_vs_MS.NoTrt_B |
| KIAA1429 | 6.98E-05 | 0.33102 | 0.651 | 0.84 | 1 | MS.Trt_vs_MS.NoTrt_B |
| CNPY2 | 7.00E-05 | -0.4078 | 0.852 | 0.96 | 1 | MS.Trt_vs_MS.NoTrt_B |
| YPEL5 | 7.01E-05 | 0.28082 | 0.144 | 0.28 | 1 | MS.Trt_vs_MS.NoTrt_B |
| RARA | 7.04E-05 | -0.9172 | 0.037 | 0.12 | 1 | MS.Trt_vs_MS.NoTrt_B |
| ARHGAP19 | 7.05E-05 | 0.37037 | 0.207 | 0.36 | 1 | MS.Trt_vs_MS.NoTrt_B |
| FLYWCH1 | 7.10E-05 | -0.4672 | 0.575 | 0.79 | 1 | MS.Trt_vs_MS.NoTrt_B |
| RASSF1 | 7.11E-05 | -0.9732 | 0.109 | 0.22 | 1 | MS.Trt_vs_MS.NoTrt_B |
| HSDL2 | 7.17E-05 | -0.4857 | 0.087 | 0.2 | 1 | MS.Trt_vs_MS.NoTrt_B |
| GRAP2 | 7.17E-05 | -0.5246 | 0.209 | 0.35 | 1 | MS.Trt_vs_MS.NoTrt_B |

| | | | | | | |
|---|---|---|---|---|---|---|
| REEP4 | 7.17E-05 | -1.51 | 0.028 | 0.1 | 1 | MS.Trt_vs_MS.NoTrt_B |
| FAM153A | 7.28E-05 | -0.4554 | 0.373 | 0.54 | 1 | MS.Trt_vs_MS.NoTrt_B |
| HSBP1L1 | 7.33E-05 | -0.319 | 0.251 | 0.41 | 1 | MS.Trt_vs_MS.NoTrt_B |
| SMURF2 | 7.36E-05 | 0.31091 | 0.586 | 0.77 | 1 | MS.Trt_vs_MS.NoTrt_B |
| UXT | 7.44E-05 | -0.3357 | 0.157 | 0.3 | 1 | MS.Trt_vs_MS.NoTrt_B |
| PRPF31 | 7.51E-05 | -0.711 | 0.096 | 0.21 | 1 | MS.Trt_vs_MS.NoTrt_B |
| AKT1S1 | 7.58E-05 | -0.571 | 0.129 | 0.25 | 1 | MS.Trt_vs_MS.NoTrt_B |
| NAGPA | 7.59E-05 | -0.2978 | 0.505 | 0.7 | 1 | MS.Trt_vs_MS.NoTrt_B |
| SV2C | 7.60E-05 | -0.7581 | 0.444 | 0.66 | 1 | MS.Trt_vs_MS.NoTrt_B |
| BCL2L1 | 7.61E-05 | -0.6013 | 0.096 | 0.21 | 1 | MS.Trt_vs_MS.NoTrt_B |
| GORASP2 | 7.64E-05 | -0.487 | 0.129 | 0.25 | 1 | MS.Trt_vs_MS.NoTrt_B |
| APEH | 7.71E-05 | -0.3822 | 0.133 | 0.27 | 1 | MS.Trt_vs_MS.NoTrt_B |
| AFF1 | 7.75E-05 | 0.47566 | 0.279 | 0.46 | 1 | MS.Trt_vs_MS.NoTrt_B |
| POLR2L | 7.75E-05 | -0.2969 | 0.179 | 0.32 | 1 | MS.Trt_vs_MS.NoTrt_B |
| KBTBD6 | 7.88E-05 | -1.4414 | 0.253 | 0.43 | 1 | MS.Trt_vs_MS.NoTrt_B |
| FNTA | 7.91E-05 | 0.58284 | 0.695 | 0.84 | 1 | MS.Trt_vs_MS.NoTrt_B |
| GLCE | 7.94E-05 | -0.9136 | 0.074 | 0.18 | 1 | MS.Trt_vs_MS.NoTrt_B |
| RP11-435I1( | 7.94E-05 | -0.2844 | 0.24 | 0.39 | 1 | MS.Trt_vs_MS.NoTrt_B |
| SGSM3 | 7.98E-05 | -0.438 | 0.131 | 0.26 | 1 | MS.Trt_vs_MS.NoTrt_B |
| ACTR3B | 8.09E-05 | -0.5801 | 0.377 | 0.55 | 1 | MS.Trt_vs_MS.NoTrt_B |
| MCCC2 | 8.18E-05 | 0.42178 | 0.473 | 0.67 | 1 | MS.Trt_vs_MS.NoTrt_B |
| CCDC155 | 8.20E-05 | -0.5684 | 0.059 | 0.16 | 1 | MS.Trt_vs_MS.NoTrt_B |
| SREBF2 | 8.28E-05 | -0.5427 | 0.085 | 0.2 | 1 | MS.Trt_vs_MS.NoTrt_B |
| GOSR2 | 8.34E-05 | -0.9526 | 0.068 | 0.17 | 1 | MS.Trt_vs_MS.NoTrt_B |
| CALM2 | 8.35E-05 | 0.27316 | 0.78 | 0.9 | 1 | MS.Trt_vs_MS.NoTrt_B |
| SMARCA4 | 8.48E-05 | 0.53151 | 0.664 | 0.81 | 1 | MS.Trt_vs_MS.NoTrt_B |
| RPN1 | 8.53E-05 | -0.3454 | 0.527 | 0.68 | 1 | MS.Trt_vs_MS.NoTrt_B |
| PI4KB | 8.55E-05 | -0.6618 | 0.122 | 0.24 | 1 | MS.Trt_vs_MS.NoTrt_B |
| VPS4A | 8.56E-05 | -0.265 | 0.499 | 0.67 | 1 | MS.Trt_vs_MS.NoTrt_B |
| DDB1 | 8.64E-05 | -0.5385 | 0.756 | 0.89 | 1 | MS.Trt_vs_MS.NoTrt_B |
| GPN3 | 8.81E-05 | -1.2357 | 0.268 | 0.42 | 1 | MS.Trt_vs_MS.NoTrt_B |
| FAM63B | 8.84E-05 | -0.7426 | 0.623 | 0.8 | 1 | MS.Trt_vs_MS.NoTrt_B |
| WHSC1L1 | 8.87E-05 | 0.26738 | 0.477 | 0.69 | 1 | MS.Trt_vs_MS.NoTrt_B |
| IMMP1L | 8.92E-05 | 0.86892 | 0.405 | 0.58 | 1 | MS.Trt_vs_MS.NoTrt_B |
| MYDGF | 8.93E-05 | -0.2762 | 0.1 | 0.22 | 1 | MS.Trt_vs_MS.NoTrt_B |
| NUB1 | 9.14E-05 | -0.3243 | 0.778 | 0.91 | 1 | MS.Trt_vs_MS.NoTrt_B |
| RAB3IP | 9.25E-05 | 0.84011 | 0.51 | 0.67 | 1 | MS.Trt_vs_MS.NoTrt_B |
| PHKB | 9.47E-05 | 0.37711 | 0.623 | 0.81 | 1 | MS.Trt_vs_MS.NoTrt_B |
| NFKBIA | 9.47E-05 | -0.4282 | 0.183 | 0.32 | 1 | MS.Trt_vs_MS.NoTrt_B |
| MRPL2 | 9.53E-05 | 0.36727 | 0.183 | 0.34 | 1 | MS.Trt_vs_MS.NoTrt_B |

| | | | | | | |
|---|---|---|---|---|---|---|
| NSMAF | 9.61E-05 | -0.4023 | 0.137 | 0.27 | 1 | MS.Trt_vs_MS.NoTrt_B |
| RAD23B | 9.64E-05 | -0.4042 | 0.344 | 0.53 | 1 | MS.Trt_vs_MS.NoTrt_B |
| RUVBL2 | 9.66E-05 | -0.3749 | 0.723 | 0.92 | 1 | MS.Trt_vs_MS.NoTrt_B |
| RAB11B | 9.76E-05 | -0.2843 | 0.322 | 0.47 | 1 | MS.Trt_vs_MS.NoTrt_B |
| POLL | 9.76E-05 | -1.1229 | 0.283 | 0.42 | 1 | MS.Trt_vs_MS.NoTrt_B |
| TTC12 | 9.87E-05 | -0.9141 | 0.076 | 0.18 | 1 | MS.Trt_vs_MS.NoTrt_B |
| SLC25A15 | 9.88E-05 | -0.4861 | 0.532 | 0.72 | 1 | MS.Trt_vs_MS.NoTrt_B |
| FLT4 | 1.00E-04 | -0.9087 | 0.05 | 0.14 | 1 | MS.Trt_vs_MS.NoTrt_B |
| ZNF880 | 1.02E-04 | 0.40494 | 0.373 | 0.55 | 1 | MS.Trt_vs_MS.NoTrt_B |
| ZNF45 | 1.03E-04 | 0.95538 | 0.19 | 0.34 | 1 | MS.Trt_vs_MS.NoTrt_B |
| SLC25A20 | 1.04E-04 | -1.9575 | 0.048 | 0.14 | 1 | MS.Trt_vs_MS.NoTrt_B |
| PCNX1 | 1.04E-04 | 0.35873 | 0.715 | 0.88 | 1 | MS.Trt_vs_MS.NoTrt_B |
| DYNC1I2 | 1.04E-04 | -0.8045 | 0.135 | 0.26 | 1 | MS.Trt_vs_MS.NoTrt_B |
| ZBTB7A | 1.05E-04 | -0.4876 | 0.072 | 0.17 | 1 | MS.Trt_vs_MS.NoTrt_B |
| CES3 | 1.07E-04 | -0.2729 | 0.139 | 0.27 | 1 | MS.Trt_vs_MS.NoTrt_B |
| TMEM184A | 1.08E-04 | 0.62644 | 0.27 | 0.43 | 1 | MS.Trt_vs_MS.NoTrt_B |
| CXXC1 | 1.09E-04 | -0.6895 | 0.105 | 0.22 | 1 | MS.Trt_vs_MS.NoTrt_B |
| C1orf27 | 1.10E-04 | -0.9465 | 0.083 | 0.19 | 1 | MS.Trt_vs_MS.NoTrt_B |
| SREK1IP1 | 1.10E-04 | -0.4211 | 0.285 | 0.45 | 1 | MS.Trt_vs_MS.NoTrt_B |
| ZDHHC3 | 1.10E-04 | -0.3844 | 0.338 | 0.48 | 1 | MS.Trt_vs_MS.NoTrt_B |
| C12orf57 | 1.11E-04 | -0.3492 | 0.296 | 0.45 | 1 | MS.Trt_vs_MS.NoTrt_B |
| TMEM243 | 1.11E-04 | 0.39149 | 0.198 | 0.36 | 1 | MS.Trt_vs_MS.NoTrt_B |
| RFX5 | 1.11E-04 | -0.4418 | 0.216 | 0.35 | 1 | MS.Trt_vs_MS.NoTrt_B |
| GSPT1 | 1.13E-04 | 0.30335 | 0.386 | 0.56 | 1 | MS.Trt_vs_MS.NoTrt_B |
| BZW2 | 1.13E-04 | -0.6639 | 0.083 | 0.19 | 1 | MS.Trt_vs_MS.NoTrt_B |
| GNAL | 1.14E-04 | 0.44626 | 0.318 | 0.48 | 1 | MS.Trt_vs_MS.NoTrt_B |
| HMGA1 | 1.15E-04 | -0.3195 | 0.142 | 0.27 | 1 | MS.Trt_vs_MS.NoTrt_B |
| FBXL4 | 1.16E-04 | -0.574 | 0.386 | 0.55 | 1 | MS.Trt_vs_MS.NoTrt_B |
| SYAP1 | 1.16E-04 | 0.34969 | 0.887 | 0.98 | 1 | MS.Trt_vs_MS.NoTrt_B |
| ST8SIA3 | 1.16E-04 | -0.3209 | 0.107 | 0.22 | 1 | MS.Trt_vs_MS.NoTrt_B |
| LAPTM4A | 1.17E-04 | -0.27 | 0.098 | 0.21 | 1 | MS.Trt_vs_MS.NoTrt_B |
| ZNF675 | 1.17E-04 | 0.87316 | 0.795 | 0.92 | 1 | MS.Trt_vs_MS.NoTrt_B |
| MAP2K2 | 1.19E-04 | 0.2753 | 0.547 | 0.72 | 1 | MS.Trt_vs_MS.NoTrt_B |
| ASAH1 | 1.20E-04 | -0.31 | 0.595 | 0.76 | 1 | MS.Trt_vs_MS.NoTrt_B |
| SMARCE1 | 1.20E-04 | 0.47236 | 0.373 | 0.55 | 1 | MS.Trt_vs_MS.NoTrt_B |
| BFAR | 1.20E-04 | -0.9003 | 0.431 | 0.58 | 1 | MS.Trt_vs_MS.NoTrt_B |
| SPATA20 | 1.20E-04 | -0.4657 | 0.048 | 0.14 | 1 | MS.Trt_vs_MS.NoTrt_B |
| VPS13D | 1.21E-04 | 0.44909 | 0.187 | 0.34 | 1 | MS.Trt_vs_MS.NoTrt_B |
| SRSF6 | 1.21E-04 | 0.56602 | 0.597 | 0.75 | 1 | MS.Trt_vs_MS.NoTrt_B |
| FMR1 | 1.21E-04 | -0.4519 | 0.063 | 0.16 | 1 | MS.Trt_vs_MS.NoTrt_B |

| | | | | | | |
|---|---|---|---|---|---|---|
| CLTA | 1.22E-04 | -0.6571 | 0.499 | 0.64 | 1 | MS.Trt_vs_MS.NoTrt_B |
| KPNA6 | 1.23E-04 | 0.33003 | 0.399 | 0.6 | 1 | MS.Trt_vs_MS.NoTrt_B |
| PTCD2 | 1.23E-04 | 0.63103 | 0.388 | 0.61 | 1 | MS.Trt_vs_MS.NoTrt_B |
| SSB | 1.23E-04 | 0.51471 | 0.641 | 0.81 | 1 | MS.Trt_vs_MS.NoTrt_B |
| QRSL1 | 1.24E-04 | 0.99242 | 0.407 | 0.6 | 1 | MS.Trt_vs_MS.NoTrt_B |
| CREG1 | 1.25E-04 | -0.2972 | 0.19 | 0.34 | 1 | MS.Trt_vs_MS.NoTrt_B |
| EPPIN | 1.26E-04 | 0.42415 | 0.277 | 0.47 | 1 | MS.Trt_vs_MS.NoTrt_B |
| SUPT5H | 1.27E-04 | -0.274 | 0.316 | 0.48 | 1 | MS.Trt_vs_MS.NoTrt_B |
| KAT6A | 1.27E-04 | -0.4704 | 0.614 | 0.82 | 1 | MS.Trt_vs_MS.NoTrt_B |
| ZNF587 | 1.28E-04 | 0.40264 | 0.837 | 0.95 | 1 | MS.Trt_vs_MS.NoTrt_B |
| CAPS2 | 1.28E-04 | -0.3539 | 0.288 | 0.46 | 1 | MS.Trt_vs_MS.NoTrt_B |
| TRIM16 | 1.28E-04 | -0.9621 | 0.353 | 0.48 | 1 | MS.Trt_vs_MS.NoTrt_B |
| POU5F2 | 1.28E-04 | 0.38682 | 0.512 | 0.68 | 1 | MS.Trt_vs_MS.NoTrt_B |
| HAUS1 | 1.29E-04 | -0.7993 | 0.455 | 0.62 | 1 | MS.Trt_vs_MS.NoTrt_B |
| GPN2 | 1.29E-04 | -0.3038 | 0.651 | 0.76 | 1 | MS.Trt_vs_MS.NoTrt_B |
| ALDH16A1 | 1.29E-04 | -0.297 | 0.516 | 0.71 | 1 | MS.Trt_vs_MS.NoTrt_B |
| SDE2 | 1.29E-04 | -0.2748 | 0.534 | 0.71 | 1 | MS.Trt_vs_MS.NoTrt_B |
| SYS1 | 1.29E-04 | -0.5987 | 0.351 | 0.52 | 1 | MS.Trt_vs_MS.NoTrt_B |
| ENTPD3 | 1.30E-04 | -0.34 | 0.242 | 0.4 | 1 | MS.Trt_vs_MS.NoTrt_B |
| STRBP | 1.32E-04 | -1.0736 | 0.449 | 0.61 | 1 | MS.Trt_vs_MS.NoTrt_B |
| ALKBH2 | 1.33E-04 | -0.8524 | 0.24 | 0.39 | 1 | MS.Trt_vs_MS.NoTrt_B |
| CTNS | 1.34E-04 | -1.422 | 0.083 | 0.18 | 1 | MS.Trt_vs_MS.NoTrt_B |
| SULT1A2 | 1.34E-04 | -0.3446 | 0.129 | 0.25 | 1 | MS.Trt_vs_MS.NoTrt_B |
| ITGA10 | 1.34E-04 | 0.46815 | 0.344 | 0.54 | 1 | MS.Trt_vs_MS.NoTrt_B |
| ELMO2 | 1.35E-04 | -0.6792 | 0.144 | 0.27 | 1 | MS.Trt_vs_MS.NoTrt_B |
| TMEM35B | 1.35E-04 | -0.2744 | 0.109 | 0.23 | 1 | MS.Trt_vs_MS.NoTrt_B |
| MPPE1 | 1.37E-04 | -0.4954 | 0.776 | 0.87 | 1 | MS.Trt_vs_MS.NoTrt_B |
| PLEKHB2 | 1.37E-04 | -0.4983 | 0.732 | 0.87 | 1 | MS.Trt_vs_MS.NoTrt_B |
| PMM2 | 1.38E-04 | -0.3066 | 0.573 | 0.73 | 1 | MS.Trt_vs_MS.NoTrt_B |
| TMEM169 | 1.38E-04 | -0.5305 | 0.412 | 0.6 | 1 | MS.Trt_vs_MS.NoTrt_B |
| C12orf40 | 1.38E-04 | -0.4062 | 0.163 | 0.3 | 1 | MS.Trt_vs_MS.NoTrt_B |
| VCPIP1 | 1.40E-04 | 0.39679 | 0.242 | 0.41 | 1 | MS.Trt_vs_MS.NoTrt_B |
| GIGYF2 | 1.41E-04 | -0.2546 | 0.797 | 0.92 | 1 | MS.Trt_vs_MS.NoTrt_B |
| NDC80 | 1.41E-04 | -0.7288 | 0.521 | 0.7 | 1 | MS.Trt_vs_MS.NoTrt_B |
| STX12 | 1.42E-04 | -0.5402 | 0.161 | 0.29 | 1 | MS.Trt_vs_MS.NoTrt_B |
| RNF207 | 1.45E-04 | -0.8256 | 0.505 | 0.73 | 1 | MS.Trt_vs_MS.NoTrt_B |
| ASNSD1 | 1.46E-04 | -0.5608 | 0.081 | 0.18 | 1 | MS.Trt_vs_MS.NoTrt_B |
| PHF20 | 1.46E-04 | 0.6935 | 0.381 | 0.56 | 1 | MS.Trt_vs_MS.NoTrt_B |
| KCNA3 | 1.49E-04 | -0.491 | 0.122 | 0.24 | 1 | MS.Trt_vs_MS.NoTrt_B |
| PSMB2 | 1.50E-04 | -0.4427 | 0.603 | 0.77 | 1 | MS.Trt_vs_MS.NoTrt_B |

| | | | | | | |
|---|---|---|---|---|---|---|
| HLA-DRB1 | 1.51E-04 | -0.7169 | 0.168 | 0.29 | 1 | MS.Trt_vs_MS.NoTrt_B |
| HARS2 | 1.51E-04 | -0.3108 | 0.102 | 0.22 | 1 | MS.Trt_vs_MS.NoTrt_B |
| YIPF1 | 1.51E-04 | -0.3293 | 0.039 | 0.12 | 1 | MS.Trt_vs_MS.NoTrt_B |
| POLR2C | 1.51E-04 | -0.5877 | 0.466 | 0.63 | 1 | MS.Trt_vs_MS.NoTrt_B |
| ZNF134 | 1.53E-04 | 0.53452 | 0.126 | 0.25 | 1 | MS.Trt_vs_MS.NoTrt_B |
| RP3-461F17 | 1.53E-04 | -0.2709 | 0.139 | 0.26 | 1 | MS.Trt_vs_MS.NoTrt_B |
| ZNF511 | 1.54E-04 | -1.0092 | 0.039 | 0.12 | 1 | MS.Trt_vs_MS.NoTrt_B |
| MIEN1 | 1.54E-04 | -0.3452 | 0.076 | 0.18 | 1 | MS.Trt_vs_MS.NoTrt_B |
| GCC2 | 1.54E-04 | 0.32542 | 0.636 | 0.83 | 1 | MS.Trt_vs_MS.NoTrt_B |
| TUSC3 | 1.56E-04 | -0.2723 | 0.292 | 0.43 | 1 | MS.Trt_vs_MS.NoTrt_B |
| ST7 | 1.57E-04 | -0.8246 | 0.031 | 0.1 | 1 | MS.Trt_vs_MS.NoTrt_B |
| TERF2IP | 1.58E-04 | -0.4802 | 0.255 | 0.4 | 1 | MS.Trt_vs_MS.NoTrt_B |
| CEP170 | 1.58E-04 | 0.59115 | 0.442 | 0.63 | 1 | MS.Trt_vs_MS.NoTrt_B |
| TRIM59 | 1.58E-04 | 0.2944 | 0.747 | 0.91 | 1 | MS.Trt_vs_MS.NoTrt_B |
| CMTM3 | 1.58E-04 | -0.4135 | 0.098 | 0.21 | 1 | MS.Trt_vs_MS.NoTrt_B |
| FOXO3 | 1.59E-04 | -0.7582 | 0.292 | 0.43 | 1 | MS.Trt_vs_MS.NoTrt_B |
| RCC1 | 1.61E-04 | -0.6662 | 0.109 | 0.23 | 1 | MS.Trt_vs_MS.NoTrt_B |
| MMAB | 1.61E-04 | 0.66416 | 0.473 | 0.67 | 1 | MS.Trt_vs_MS.NoTrt_B |
| RPS10 | 1.62E-04 | -0.5935 | 0.046 | 0.13 | 1 | MS.Trt_vs_MS.NoTrt_B |
| MUTYH | 1.63E-04 | -0.3844 | 0.425 | 0.58 | 1 | MS.Trt_vs_MS.NoTrt_B |
| H2AFJ | 1.63E-04 | -0.7574 | 0.076 | 0.17 | 1 | MS.Trt_vs_MS.NoTrt_B |
| NAA16 | 1.64E-04 | 0.28413 | 0.272 | 0.43 | 1 | MS.Trt_vs_MS.NoTrt_B |
| SUB1 | 1.67E-04 | 0.38681 | 0.447 | 0.65 | 1 | MS.Trt_vs_MS.NoTrt_B |
| SNX2 | 1.67E-04 | -0.2678 | 0.473 | 0.62 | 1 | MS.Trt_vs_MS.NoTrt_B |
| RABL2A | 1.67E-04 | -0.4257 | 0.418 | 0.56 | 1 | MS.Trt_vs_MS.NoTrt_B |
| MSI2 | 1.68E-04 | 0.28626 | 0.719 | 0.85 | 1 | MS.Trt_vs_MS.NoTrt_B |
| STAT1 | 1.69E-04 | -0.2701 | 0.349 | 0.5 | 1 | MS.Trt_vs_MS.NoTrt_B |
| SMPD3 | 1.70E-04 | -0.6462 | 0.532 | 0.76 | 1 | MS.Trt_vs_MS.NoTrt_B |
| SUV39H2 | 1.70E-04 | -0.4065 | 0.309 | 0.48 | 1 | MS.Trt_vs_MS.NoTrt_B |
| AREL1 | 1.71E-04 | -0.551 | 0.349 | 0.51 | 1 | MS.Trt_vs_MS.NoTrt_B |
| DSTYK | 1.72E-04 | -0.9073 | 0.656 | 0.85 | 1 | MS.Trt_vs_MS.NoTrt_B |
| PAM | 1.72E-04 | -0.9612 | 0.216 | 0.35 | 1 | MS.Trt_vs_MS.NoTrt_B |
| NDUFA13 | 1.75E-04 | 0.29314 | 0.412 | 0.62 | 1 | MS.Trt_vs_MS.NoTrt_B |
| ERCC3 | 1.75E-04 | -0.2776 | 0.288 | 0.43 | 1 | MS.Trt_vs_MS.NoTrt_B |
| TEP1 | 1.78E-04 | 0.40629 | 0.305 | 0.49 | 1 | MS.Trt_vs_MS.NoTrt_B |
| FAM216B | 1.78E-04 | -0.2794 | 0.115 | 0.23 | 1 | MS.Trt_vs_MS.NoTrt_B |
| SNAP47 | 1.79E-04 | -1.0824 | 0.076 | 0.17 | 1 | MS.Trt_vs_MS.NoTrt_B |
| MTHFSD | 1.79E-04 | -1.2298 | 0.037 | 0.11 | 1 | MS.Trt_vs_MS.NoTrt_B |
| LSM3 | 1.80E-04 | 0.32269 | 0.423 | 0.6 | 1 | MS.Trt_vs_MS.NoTrt_B |
| HNRNPDL | 1.81E-04 | 0.25902 | 0.495 | 0.71 | 1 | MS.Trt_vs_MS.NoTrt_B |

| | | | | | | |
|---|---|---|---|---|---|---|
| NARS | 1.85E-04 | -0.4014 | 0.118 | 0.23 | 1 | MS.Trt_vs_MS.NoTrt_B |
| LUZP1 | 1.86E-04 | -0.3228 | 0.344 | 0.49 | 1 | MS.Trt_vs_MS.NoTrt_B |
| C2orf74 | 1.88E-04 | -0.2929 | 0.32 | 0.5 | 1 | MS.Trt_vs_MS.NoTrt_B |
| IRF1 | 1.89E-04 | -0.386 | 0.791 | 0.93 | 1 | MS.Trt_vs_MS.NoTrt_B |
| LATS1 | 1.90E-04 | 0.33706 | 0.525 | 0.74 | 1 | MS.Trt_vs_MS.NoTrt_B |
| TBC1D9B | 1.92E-04 | -0.8392 | 0.351 | 0.51 | 1 | MS.Trt_vs_MS.NoTrt_B |
| SPAG4 | 1.92E-04 | -0.2594 | 0.163 | 0.3 | 1 | MS.Trt_vs_MS.NoTrt_B |
| SIK2 | 1.93E-04 | -0.2615 | 0.527 | 0.68 | 1 | MS.Trt_vs_MS.NoTrt_B |
| NSMCE2 | 1.94E-04 | -0.4107 | 0.414 | 0.61 | 1 | MS.Trt_vs_MS.NoTrt_B |
| SLC30A9 | 1.94E-04 | -0.4363 | 0.386 | 0.53 | 1 | MS.Trt_vs_MS.NoTrt_B |
| PTBP2 | 1.94E-04 | 0.65918 | 0.471 | 0.65 | 1 | MS.Trt_vs_MS.NoTrt_B |
| SAP30BP | 1.94E-04 | -0.5285 | 0.163 | 0.29 | 1 | MS.Trt_vs_MS.NoTrt_B |
| MAPK1IP1L | 1.95E-04 | 0.3952 | 0.684 | 0.87 | 1 | MS.Trt_vs_MS.NoTrt_B |
| ULBP3 | 1.95E-04 | -0.44 | 0.405 | 0.59 | 1 | MS.Trt_vs_MS.NoTrt_B |
| ACAN | 1.95E-04 | -0.5221 | 0.192 | 0.34 | 1 | MS.Trt_vs_MS.NoTrt_B |
| ANKRD36C | 1.96E-04 | 0.42511 | 0.477 | 0.65 | 1 | MS.Trt_vs_MS.NoTrt_B |
| ACYP2 | 1.98E-04 | -0.3732 | 0.466 | 0.61 | 1 | MS.Trt_vs_MS.NoTrt_B |
| TNKS | 1.99E-04 | -0.3426 | 0.66 | 0.82 | 1 | MS.Trt_vs_MS.NoTrt_B |
| SPTSSA | 1.99E-04 | -0.3504 | 0.244 | 0.39 | 1 | MS.Trt_vs_MS.NoTrt_B |
| PPIL2 | 1.99E-04 | -0.6443 | 0.257 | 0.4 | 1 | MS.Trt_vs_MS.NoTrt_B |
| RARS2 | 2.00E-04 | -0.4864 | 0.453 | 0.62 | 1 | MS.Trt_vs_MS.NoTrt_B |
| SLC20A1 | 2.00E-04 | -0.7844 | 0.107 | 0.22 | 1 | MS.Trt_vs_MS.NoTrt_B |
| HDHD3 | 2.02E-04 | -0.5933 | 0.115 | 0.23 | 1 | MS.Trt_vs_MS.NoTrt_B |
| STK16 | 2.03E-04 | -1.6114 | 0.035 | 0.11 | 1 | MS.Trt_vs_MS.NoTrt_B |
| INTS1 | 2.03E-04 | -0.7476 | 0.24 | 0.37 | 1 | MS.Trt_vs_MS.NoTrt_B |
| KLRG1 | 2.03E-04 | 0.48454 | 0.246 | 0.42 | 1 | MS.Trt_vs_MS.NoTrt_B |
| TSSC1 | 2.04E-04 | -0.5917 | 0.068 | 0.16 | 1 | MS.Trt_vs_MS.NoTrt_B |
| C1orf131 | 2.05E-04 | -0.2882 | 0.488 | 0.67 | 1 | MS.Trt_vs_MS.NoTrt_B |
| C21orf59 | 2.05E-04 | -0.509 | 0.283 | 0.42 | 1 | MS.Trt_vs_MS.NoTrt_B |
| ACOX3 | 2.05E-04 | -1.8325 | 0.041 | 0.12 | 1 | MS.Trt_vs_MS.NoTrt_B |
| UPK3B | 2.08E-04 | -0.2827 | 0.065 | 0.16 | 1 | MS.Trt_vs_MS.NoTrt_B |
| PMVK | 2.08E-04 | -0.5064 | 0.074 | 0.17 | 1 | MS.Trt_vs_MS.NoTrt_B |
| ACSL5 | 2.09E-04 | -0.3493 | 0.098 | 0.21 | 1 | MS.Trt_vs_MS.NoTrt_B |
| WDR3 | 2.09E-04 | 0.34163 | 0.56 | 0.73 | 1 | MS.Trt_vs_MS.NoTrt_B |
| DNAJB14 | 2.11E-04 | 0.33428 | 0.375 | 0.52 | 1 | MS.Trt_vs_MS.NoTrt_B |
| CCDC28A | 2.13E-04 | -0.2955 | 0.135 | 0.25 | 1 | MS.Trt_vs_MS.NoTrt_B |
| RTCB | 2.13E-04 | -0.4811 | 0.109 | 0.22 | 1 | MS.Trt_vs_MS.NoTrt_B |
| ZNF555 | 2.14E-04 | 0.95589 | 0.438 | 0.61 | 1 | MS.Trt_vs_MS.NoTrt_B |
| LA16c-431H | 2.15E-04 | -0.366 | 0.416 | 0.57 | 1 | MS.Trt_vs_MS.NoTrt_B |
| PIGQ | 2.15E-04 | -0.7694 | 0.309 | 0.44 | 1 | MS.Trt_vs_MS.NoTrt_B |

| | | | | | | |
|---|---|---|---|---|---|---|
| CDK5RAP1 | 2.16E-04 | -0.6052 | 0.246 | 0.4 | 1 | MS.Trt_vs_MS.NoTrt_B |
| CHDH | 2.17E-04 | -0.486 | 0.366 | 0.51 | 1 | MS.Trt_vs_MS.NoTrt_B |
| CNIH4 | 2.18E-04 | 0.72241 | 0.423 | 0.62 | 1 | MS.Trt_vs_MS.NoTrt_B |
| EEF2KMT | 2.19E-04 | -1.0011 | 0.041 | 0.12 | 1 | MS.Trt_vs_MS.NoTrt_B |
| GGNBP2 | 2.20E-04 | 0.47041 | 0.595 | 0.76 | 1 | MS.Trt_vs_MS.NoTrt_B |
| ANTXR2 | 2.20E-04 | -0.4906 | 0.553 | 0.67 | 1 | MS.Trt_vs_MS.NoTrt_B |
| SLC35F6 | 2.21E-04 | -0.3233 | 0.632 | 0.81 | 1 | MS.Trt_vs_MS.NoTrt_B |
| FBL | 2.21E-04 | -0.4071 | 0.342 | 0.5 | 1 | MS.Trt_vs_MS.NoTrt_B |
| ERO1A | 2.22E-04 | 0.34905 | 0.786 | 0.92 | 1 | MS.Trt_vs_MS.NoTrt_B |
| HNRNPH2 | 2.23E-04 | -0.6113 | 0.129 | 0.24 | 1 | MS.Trt_vs_MS.NoTrt_B |
| VAPB | 2.23E-04 | -0.4491 | 0.148 | 0.27 | 1 | MS.Trt_vs_MS.NoTrt_B |
| PDCL | 2.24E-04 | -0.4655 | 0.29 | 0.44 | 1 | MS.Trt_vs_MS.NoTrt_B |
| LPXN | 2.25E-04 | -0.4935 | 0.279 | 0.41 | 1 | MS.Trt_vs_MS.NoTrt_B |
| TRDMT1 | 2.25E-04 | 0.3446 | 0.538 | 0.74 | 1 | MS.Trt_vs_MS.NoTrt_B |
| MAP3K14 | 2.26E-04 | -0.4541 | 0.113 | 0.23 | 1 | MS.Trt_vs_MS.NoTrt_B |
| BCKDHB | 2.26E-04 | 0.68991 | 0.479 | 0.65 | 1 | MS.Trt_vs_MS.NoTrt_B |
| EREG | 2.26E-04 | 0.93321 | 0.599 | 0.78 | 1 | MS.Trt_vs_MS.NoTrt_B |
| TCTA | 2.27E-04 | -0.4839 | 0.434 | 0.61 | 1 | MS.Trt_vs_MS.NoTrt_B |
| AARS2 | 2.30E-04 | -1.525 | 0.275 | 0.42 | 1 | MS.Trt_vs_MS.NoTrt_B |
| TTC39C | 2.31E-04 | 0.30475 | 0.756 | 0.9 | 1 | MS.Trt_vs_MS.NoTrt_B |
| KLHDC3 | 2.33E-04 | -0.5304 | 0.105 | 0.22 | 1 | MS.Trt_vs_MS.NoTrt_B |
| TMEM241 | 2.33E-04 | -0.6663 | 0.479 | 0.67 | 1 | MS.Trt_vs_MS.NoTrt_B |
| DDX23 | 2.34E-04 | -0.537 | 0.519 | 0.68 | 1 | MS.Trt_vs_MS.NoTrt_B |
| CRY1 | 2.35E-04 | -0.6959 | 0.081 | 0.18 | 1 | MS.Trt_vs_MS.NoTrt_B |
| ZFAND3 | 2.36E-04 | -0.5333 | 0.503 | 0.67 | 1 | MS.Trt_vs_MS.NoTrt_B |
| STARD7 | 2.37E-04 | -0.5403 | 0.425 | 0.58 | 1 | MS.Trt_vs_MS.NoTrt_B |
| ICK | 2.38E-04 | 0.86509 | 0.142 | 0.27 | 1 | MS.Trt_vs_MS.NoTrt_B |
| PTBP1 | 2.39E-04 | -0.434 | 0.163 | 0.29 | 1 | MS.Trt_vs_MS.NoTrt_B |
| UQCR10 | 2.40E-04 | 0.49709 | 0.279 | 0.45 | 1 | MS.Trt_vs_MS.NoTrt_B |
| ZNF91 | 2.40E-04 | 0.53243 | 0.643 | 0.79 | 1 | MS.Trt_vs_MS.NoTrt_B |
| DHRS12 | 2.43E-04 | -0.2753 | 0.061 | 0.15 | 1 | MS.Trt_vs_MS.NoTrt_B |
| ZFP64 | 2.43E-04 | -0.5818 | 0.46 | 0.62 | 1 | MS.Trt_vs_MS.NoTrt_B |
| SNX8 | 2.45E-04 | 0.63902 | 0.063 | 0.16 | 1 | MS.Trt_vs_MS.NoTrt_B |
| SNRPG | 2.45E-04 | 0.56464 | 0.2 | 0.36 | 1 | MS.Trt_vs_MS.NoTrt_B |
| TNFRSF18 | 2.46E-04 | -0.7741 | 0.041 | 0.12 | 1 | MS.Trt_vs_MS.NoTrt_B |
| NARF | 2.47E-04 | -0.5907 | 0.871 | 0.97 | 1 | MS.Trt_vs_MS.NoTrt_B |
| FAM219B | 2.47E-04 | -0.55 | 0.349 | 0.52 | 1 | MS.Trt_vs_MS.NoTrt_B |
| PCNP | 2.47E-04 | 0.71591 | 0.2 | 0.35 | 1 | MS.Trt_vs_MS.NoTrt_B |
| RRAGC | 2.48E-04 | -0.272 | 0.386 | 0.58 | 1 | MS.Trt_vs_MS.NoTrt_B |
| TMEM161A | 2.48E-04 | -0.8716 | 0.107 | 0.22 | 1 | MS.Trt_vs_MS.NoTrt_B |

| | | | | | | |
|---|---|---|---|---|---|---|
| PRDM15 | 2.50E-04 | -0.4225 | 0.434 | 0.64 | 1 | MS.Trt_vs_MS.NoTrt_B |
| ECT2L | 2.51E-04 | -0.4051 | 0.216 | 0.36 | 1 | MS.Trt_vs_MS.NoTrt_B |
| CUL3 | 2.52E-04 | 0.40144 | 0.288 | 0.46 | 1 | MS.Trt_vs_MS.NoTrt_B |
| TRMT2B | 2.53E-04 | -0.3877 | 0.492 | 0.68 | 1 | MS.Trt_vs_MS.NoTrt_B |
| ZNF397 | 2.53E-04 | -0.4213 | 0.904 | 0.96 | 1 | MS.Trt_vs_MS.NoTrt_B |
| MIF4GD | 2.53E-04 | -1.028 | 0.063 | 0.15 | 1 | MS.Trt_vs_MS.NoTrt_B |
| TMX2 | 2.54E-04 | -0.374 | 0.065 | 0.16 | 1 | MS.Trt_vs_MS.NoTrt_B |
| FAM20B | 2.54E-04 | -1.1951 | 0.416 | 0.62 | 1 | MS.Trt_vs_MS.NoTrt_B |
| GBA | 2.55E-04 | -0.6729 | 0.336 | 0.5 | 1 | MS.Trt_vs_MS.NoTrt_B |
| PATZ1 | 2.55E-04 | 0.30064 | 0.092 | 0.2 | 1 | MS.Trt_vs_MS.NoTrt_B |
| SRRM1 | 2.55E-04 | 0.29309 | 0.747 | 0.93 | 1 | MS.Trt_vs_MS.NoTrt_B |
| POLR2G | 2.58E-04 | -0.3042 | 0.15 | 0.28 | 1 | MS.Trt_vs_MS.NoTrt_B |
| CPD | 2.58E-04 | 0.34645 | 0.536 | 0.74 | 1 | MS.Trt_vs_MS.NoTrt_B |
| ZBTB33 | 2.58E-04 | -0.5055 | 0.041 | 0.12 | 1 | MS.Trt_vs_MS.NoTrt_B |
| C17orf51 | 2.58E-04 | 0.32744 | 0.296 | 0.47 | 1 | MS.Trt_vs_MS.NoTrt_B |
| ATP6V1E1 | 2.61E-04 | -0.4526 | 0.471 | 0.66 | 1 | MS.Trt_vs_MS.NoTrt_B |
| TKT | 2.61E-04 | -0.3367 | 0.477 | 0.61 | 1 | MS.Trt_vs_MS.NoTrt_B |
| LPO | 2.61E-04 | -0.3849 | 0.455 | 0.62 | 1 | MS.Trt_vs_MS.NoTrt_B |
| TTC14 | 2.66E-04 | 0.71834 | 0.294 | 0.46 | 1 | MS.Trt_vs_MS.NoTrt_B |
| ZNF549 | 2.67E-04 | 0.72904 | 0.538 | 0.7 | 1 | MS.Trt_vs_MS.NoTrt_B |
| CGRRF1 | 2.69E-04 | 0.31896 | 0.216 | 0.35 | 1 | MS.Trt_vs_MS.NoTrt_B |
| ZNF66 | 2.70E-04 | 0.52855 | 0.488 | 0.69 | 1 | MS.Trt_vs_MS.NoTrt_B |
| HPRT1 | 2.70E-04 | -0.5057 | 0.087 | 0.19 | 1 | MS.Trt_vs_MS.NoTrt_B |
| DCAF12 | 2.72E-04 | -1.0465 | 0.057 | 0.14 | 1 | MS.Trt_vs_MS.NoTrt_B |
| NFYB | 2.75E-04 | 0.7574 | 0.207 | 0.34 | 1 | MS.Trt_vs_MS.NoTrt_B |
| UROS | 2.77E-04 | -1.0152 | 0.338 | 0.49 | 1 | MS.Trt_vs_MS.NoTrt_B |
| AKR7A2 | 2.78E-04 | -0.4054 | 0.098 | 0.21 | 1 | MS.Trt_vs_MS.NoTrt_B |
| RAB22A | 2.78E-04 | 0.27318 | 0.527 | 0.7 | 1 | MS.Trt_vs_MS.NoTrt_B |
| PWP1 | 2.79E-04 | -0.3259 | 0.124 | 0.24 | 1 | MS.Trt_vs_MS.NoTrt_B |
| DBT | 2.79E-04 | -0.3398 | 0.682 | 0.85 | 1 | MS.Trt_vs_MS.NoTrt_B |
| RFT1 | 2.83E-04 | -0.5287 | 0.338 | 0.52 | 1 | MS.Trt_vs_MS.NoTrt_B |
| NDUFC2 | 2.83E-04 | 0.51682 | 0.462 | 0.65 | 1 | MS.Trt_vs_MS.NoTrt_B |
| OCIAD2 | 2.84E-04 | 0.53493 | 0.538 | 0.72 | 1 | MS.Trt_vs_MS.NoTrt_B |
| ACP2 | 2.88E-04 | -1.1642 | 0.033 | 0.1 | 1 | MS.Trt_vs_MS.NoTrt_B |
| AP3S2 | 2.89E-04 | -0.617 | 0.551 | 0.68 | 1 | MS.Trt_vs_MS.NoTrt_B |
| GCAT | 2.93E-04 | 0.85863 | 0.153 | 0.28 | 1 | MS.Trt_vs_MS.NoTrt_B |
| DHX57 | 2.96E-04 | 0.5123 | 0.414 | 0.6 | 1 | MS.Trt_vs_MS.NoTrt_B |
| PIH1D1 | 2.99E-04 | -0.3519 | 0.2 | 0.32 | 1 | MS.Trt_vs_MS.NoTrt_B |
| LDHAL6A | 3.00E-04 | 0.25491 | 0.227 | 0.37 | 1 | MS.Trt_vs_MS.NoTrt_B |
| UPF2 | 3.01E-04 | -0.3275 | 0.183 | 0.32 | 1 | MS.Trt_vs_MS.NoTrt_B |

| | | | | | | |
|---|---|---|---|---|---|---|
| CHFR | 3.03E-04 | -0.3448 | 0.817 | 0.94 | 1 | MS.Trt_vs_MS.NoTrt_B |
| WDR70 | 3.03E-04 | -0.7845 | 0.214 | 0.35 | 1 | MS.Trt_vs_MS.NoTrt_B |
| ZNF106 | 3.03E-04 | 0.29207 | 0.717 | 0.84 | 1 | MS.Trt_vs_MS.NoTrt_B |
| DUSP22 | 3.04E-04 | -0.4972 | 0.063 | 0.15 | 1 | MS.Trt_vs_MS.NoTrt_B |
| FAM134C | 3.04E-04 | -0.3819 | 0.475 | 0.62 | 1 | MS.Trt_vs_MS.NoTrt_B |
| CEP68 | 3.05E-04 | -0.5961 | 0.431 | 0.58 | 1 | MS.Trt_vs_MS.NoTrt_B |
| ARRDC5 | 3.05E-04 | -0.6219 | 0.072 | 0.17 | 1 | MS.Trt_vs_MS.NoTrt_B |
| ZNF550 | 3.06E-04 | -0.3032 | 0.198 | 0.33 | 1 | MS.Trt_vs_MS.NoTrt_B |
| FKBP3 | 3.06E-04 | 0.72327 | 0.242 | 0.38 | 1 | MS.Trt_vs_MS.NoTrt_B |
| PEPD | 3.07E-04 | -1.0709 | 0.094 | 0.19 | 1 | MS.Trt_vs_MS.NoTrt_B |
| TMEM223 | 3.08E-04 | -0.7973 | 0.109 | 0.21 | 1 | MS.Trt_vs_MS.NoTrt_B |
| NSUN2 | 3.10E-04 | -0.4553 | 0.529 | 0.72 | 1 | MS.Trt_vs_MS.NoTrt_B |
| NDOR1 | 3.10E-04 | -0.3326 | 0.041 | 0.12 | 1 | MS.Trt_vs_MS.NoTrt_B |
| ADA | 3.12E-04 | -0.942 | 0.046 | 0.12 | 1 | MS.Trt_vs_MS.NoTrt_B |
| RP11-619A1 | 3.14E-04 | -0.3489 | 0.453 | 0.62 | 1 | MS.Trt_vs_MS.NoTrt_B |
| ERN1 | 3.14E-04 | 0.31267 | 0.595 | 0.77 | 1 | MS.Trt_vs_MS.NoTrt_B |
| HECA | 3.19E-04 | 0.46444 | 0.44 | 0.61 | 1 | MS.Trt_vs_MS.NoTrt_B |
| ACACA | 3.24E-04 | -0.6153 | 0.46 | 0.65 | 1 | MS.Trt_vs_MS.NoTrt_B |
| HEMK1 | 3.25E-04 | -0.6252 | 0.865 | 0.98 | 1 | MS.Trt_vs_MS.NoTrt_B |
| WDR7 | 3.26E-04 | -0.5254 | 0.266 | 0.41 | 1 | MS.Trt_vs_MS.NoTrt_B |
| BAK1 | 3.28E-04 | -0.9166 | 0.072 | 0.16 | 1 | MS.Trt_vs_MS.NoTrt_B |
| SLC25A4 | 3.29E-04 | 0.34312 | 0.205 | 0.35 | 1 | MS.Trt_vs_MS.NoTrt_B |
| PHLDA3 | 3.33E-04 | -0.3918 | 0.257 | 0.41 | 1 | MS.Trt_vs_MS.NoTrt_B |
| LY6G5B | 3.33E-04 | -0.4195 | 0.664 | 0.85 | 1 | MS.Trt_vs_MS.NoTrt_B |
| SSR4 | 3.34E-04 | -0.3111 | 0.246 | 0.38 | 1 | MS.Trt_vs_MS.NoTrt_B |
| CRBN | 3.35E-04 | 0.50998 | 0.375 | 0.55 | 1 | MS.Trt_vs_MS.NoTrt_B |
| GOLPH3L | 3.35E-04 | 0.27666 | 0.122 | 0.24 | 1 | MS.Trt_vs_MS.NoTrt_B |
| TESC | 3.38E-04 | -0.4734 | 0.129 | 0.24 | 1 | MS.Trt_vs_MS.NoTrt_B |
| ZDHHC4 | 3.39E-04 | -0.2765 | 0.294 | 0.45 | 1 | MS.Trt_vs_MS.NoTrt_B |
| SCAPER | 3.40E-04 | -0.3402 | 0.172 | 0.3 | 1 | MS.Trt_vs_MS.NoTrt_B |
| TMEM156 | 3.40E-04 | 0.50621 | 0.617 | 0.81 | 1 | MS.Trt_vs_MS.NoTrt_B |
| PDP2 | 3.41E-04 | -0.5151 | 0.682 | 0.87 | 1 | MS.Trt_vs_MS.NoTrt_B |
| CDC16 | 3.41E-04 | -0.3347 | 0.1 | 0.2 | 1 | MS.Trt_vs_MS.NoTrt_B |
| ITPK1 | 3.44E-04 | 0.26102 | 0.464 | 0.65 | 1 | MS.Trt_vs_MS.NoTrt_B |
| MYO1F | 3.44E-04 | -0.3143 | 0.745 | 0.85 | 1 | MS.Trt_vs_MS.NoTrt_B |
| RRN3 | 3.45E-04 | -0.6259 | 0.196 | 0.32 | 1 | MS.Trt_vs_MS.NoTrt_B |
| RBM25 | 3.46E-04 | 0.27714 | 0.462 | 0.64 | 1 | MS.Trt_vs_MS.NoTrt_B |
| BRWD1 | 3.47E-04 | 0.57644 | 0.68 | 0.85 | 1 | MS.Trt_vs_MS.NoTrt_B |
| ZBTB46 | 3.47E-04 | 0.49652 | 0.052 | 0.14 | 1 | MS.Trt_vs_MS.NoTrt_B |
| PRRT3 | 3.49E-04 | 0.47215 | 0.231 | 0.37 | 1 | MS.Trt_vs_MS.NoTrt_B |

| | | | | | | |
|---|---|---|---|---|---|---|
| NAMPT | 3.49E-04 | -0.4515 | 0.56 | 0.71 | 1 | MS.Trt_vs_MS.NoTrt_B |
| PET100 | 3.51E-04 | 0.40638 | 0.527 | 0.7 | 1 | MS.Trt_vs_MS.NoTrt_B |
| LRRC37A | 3.53E-04 | 0.50331 | 0.294 | 0.44 | 1 | MS.Trt_vs_MS.NoTrt_B |
| PCDHGA8 | 3.54E-04 | -0.7178 | 0.458 | 0.58 | 1 | MS.Trt_vs_MS.NoTrt_B |
| RHNO1 | 3.59E-04 | -0.8567 | 0.542 | 0.68 | 1 | MS.Trt_vs_MS.NoTrt_B |
| GPR89B | 3.61E-04 | -0.6495 | 0.251 | 0.38 | 1 | MS.Trt_vs_MS.NoTrt_B |
| CBR1 | 3.61E-04 | -0.4544 | 0.052 | 0.14 | 1 | MS.Trt_vs_MS.NoTrt_B |
| ESR2 | 3.62E-04 | -0.2931 | 0.418 | 0.59 | 1 | MS.Trt_vs_MS.NoTrt_B |
| HAGH | 3.64E-04 | -0.592 | 0.612 | 0.77 | 1 | MS.Trt_vs_MS.NoTrt_B |
| OXSR1 | 3.65E-04 | -0.4522 | 0.057 | 0.14 | 1 | MS.Trt_vs_MS.NoTrt_B |
| FAM50A | 3.68E-04 | -0.5585 | 0.068 | 0.16 | 1 | MS.Trt_vs_MS.NoTrt_B |
| FAM78A | 3.69E-04 | -0.3078 | 0.255 | 0.4 | 1 | MS.Trt_vs_MS.NoTrt_B |
| LRRC58 | 3.71E-04 | -0.3261 | 0.597 | 0.73 | 1 | MS.Trt_vs_MS.NoTrt_B |
| ABCB7 | 3.75E-04 | 0.53687 | 0.519 | 0.7 | 1 | MS.Trt_vs_MS.NoTrt_B |
| GLTSCR1L | 3.75E-04 | 0.36241 | 0.115 | 0.23 | 1 | MS.Trt_vs_MS.NoTrt_B |
| HBG2 | 3.76E-04 | 1.5617 | 0.318 | 0.48 | 1 | MS.Trt_vs_MS.NoTrt_B |
| SMAD9 | 3.79E-04 | -0.6493 | 0.246 | 0.37 | 1 | MS.Trt_vs_MS.NoTrt_B |
| SP2 | 3.80E-04 | 0.60419 | 0.22 | 0.37 | 1 | MS.Trt_vs_MS.NoTrt_B |
| HAUS6 | 3.80E-04 | 0.29234 | 0.22 | 0.36 | 1 | MS.Trt_vs_MS.NoTrt_B |
| SH3BGRL | 3.81E-04 | 0.34983 | 0.272 | 0.44 | 1 | MS.Trt_vs_MS.NoTrt_B |
| PPM1K | 3.83E-04 | -0.6113 | 0.819 | 0.89 | 1 | MS.Trt_vs_MS.NoTrt_B |
| ZNF561 | 3.83E-04 | -0.7292 | 0.399 | 0.55 | 1 | MS.Trt_vs_MS.NoTrt_B |
| ATP8B2 | 3.83E-04 | -0.2666 | 0.71 | 0.83 | 1 | MS.Trt_vs_MS.NoTrt_B |
| DUS3L | 3.87E-04 | -0.4694 | 0.083 | 0.18 | 1 | MS.Trt_vs_MS.NoTrt_B |
| FCRL5 | 3.89E-04 | 0.39463 | 0.288 | 0.46 | 1 | MS.Trt_vs_MS.NoTrt_B |
| C6orf1 | 3.89E-04 | -0.6823 | 0.061 | 0.15 | 1 | MS.Trt_vs_MS.NoTrt_B |
| MRPL16 | 3.91E-04 | 0.40098 | 0.107 | 0.22 | 1 | MS.Trt_vs_MS.NoTrt_B |
| GAB3 | 3.91E-04 | 1.53365 | 0.229 | 0.39 | 1 | MS.Trt_vs_MS.NoTrt_B |
| CLDN19 | 3.92E-04 | -0.3854 | 0.155 | 0.28 | 1 | MS.Trt_vs_MS.NoTrt_B |
| RPTOR | 3.95E-04 | 0.4242 | 0.253 | 0.4 | 1 | MS.Trt_vs_MS.NoTrt_B |
| NPEPL1 | 3.96E-04 | 0.3248 | 0.105 | 0.22 | 1 | MS.Trt_vs_MS.NoTrt_B |
| DAD1 | 3.98E-04 | 0.37906 | 0.268 | 0.44 | 1 | MS.Trt_vs_MS.NoTrt_B |
| BCL7B | 4.00E-04 | -0.556 | 0.37 | 0.51 | 1 | MS.Trt_vs_MS.NoTrt_B |
| QSOX2 | 4.03E-04 | 0.38543 | 0.085 | 0.18 | 1 | MS.Trt_vs_MS.NoTrt_B |
| SLC38A9 | 4.07E-04 | 0.755 | 0.285 | 0.45 | 1 | MS.Trt_vs_MS.NoTrt_B |
| PPP1R10 | 4.09E-04 | -0.4175 | 0.353 | 0.5 | 1 | MS.Trt_vs_MS.NoTrt_B |
| HPF1 | 4.10E-04 | 0.33686 | 0.063 | 0.15 | 1 | MS.Trt_vs_MS.NoTrt_B |
| HSP90AB1 | 4.11E-04 | -0.2807 | 0.412 | 0.56 | 1 | MS.Trt_vs_MS.NoTrt_B |
| USP25 | 4.15E-04 | 0.82299 | 0.357 | 0.54 | 1 | MS.Trt_vs_MS.NoTrt_B |
| SRD5A1 | 4.15E-04 | -0.7753 | 0.466 | 0.68 | 1 | MS.Trt_vs_MS.NoTrt_B |

| | | | | | | |
|---|---|---|---|---|---|---|
| TNFAIP8 | 4.15E-04 | 0.27799 | 0.732 | 0.88 | 1 | MS.Trt_vs_MS.NoTrt_B |
| GNG2 | 4.21E-04 | 0.26097 | 0.244 | 0.38 | 1 | MS.Trt_vs_MS.NoTrt_B |
| PPP4R2 | 4.22E-04 | 0.25962 | 0.283 | 0.44 | 1 | MS.Trt_vs_MS.NoTrt_B |
| CDADC1 | 4.23E-04 | -0.3418 | 0.05 | 0.13 | 1 | MS.Trt_vs_MS.NoTrt_B |
| ZNF395 | 4.26E-04 | 0.67289 | 0.425 | 0.61 | 1 | MS.Trt_vs_MS.NoTrt_B |
| GPR137 | 4.28E-04 | -0.7604 | 0.135 | 0.25 | 1 | MS.Trt_vs_MS.NoTrt_B |
| BEAN1 | 4.36E-04 | 0.75769 | 0.29 | 0.46 | 1 | MS.Trt_vs_MS.NoTrt_B |
| POLE3 | 4.39E-04 | -0.3403 | 0.179 | 0.3 | 1 | MS.Trt_vs_MS.NoTrt_B |
| LRP10 | 4.40E-04 | 0.30262 | 0.63 | 0.8 | 1 | MS.Trt_vs_MS.NoTrt_B |
| RASGRP4 | 4.42E-04 | -0.6315 | 0.351 | 0.5 | 1 | MS.Trt_vs_MS.NoTrt_B |
| RPL26 | 4.44E-04 | -0.2644 | 0.806 | 0.92 | 1 | MS.Trt_vs_MS.NoTrt_B |
| SLC25A53 | 4.44E-04 | 0.72408 | 0.32 | 0.47 | 1 | MS.Trt_vs_MS.NoTrt_B |
| PRAMEF1 | 4.46E-04 | 0.79202 | 0.041 | 0.12 | 1 | MS.Trt_vs_MS.NoTrt_B |
| IKBKE | 4.50E-04 | -0.6164 | 0.218 | 0.35 | 1 | MS.Trt_vs_MS.NoTrt_B |
| LRRC37A2 | 4.55E-04 | 0.25951 | 0.566 | 0.76 | 1 | MS.Trt_vs_MS.NoTrt_B |
| FAM84B | 4.56E-04 | -0.5743 | 0.044 | 0.12 | 1 | MS.Trt_vs_MS.NoTrt_B |
| PIP5K1A | 4.56E-04 | -0.7631 | 0.386 | 0.54 | 1 | MS.Trt_vs_MS.NoTrt_B |
| AFDN | 4.60E-04 | -0.5612 | 0.044 | 0.12 | 1 | MS.Trt_vs_MS.NoTrt_B |
| SPOP | 4.64E-04 | -0.2539 | 0.893 | 0.98 | 1 | MS.Trt_vs_MS.NoTrt_B |
| STAM2 | 4.64E-04 | -0.396 | 0.416 | 0.58 | 1 | MS.Trt_vs_MS.NoTrt_B |
| ARMC1 | 4.67E-04 | -0.3314 | 0.065 | 0.15 | 1 | MS.Trt_vs_MS.NoTrt_B |
| CHRNA6 | 4.68E-04 | -0.6664 | 0.318 | 0.5 | 1 | MS.Trt_vs_MS.NoTrt_B |
| C16orf62 | 4.69E-04 | -0.326 | 0.902 | 0.99 | 1 | MS.Trt_vs_MS.NoTrt_B |
| C9orf3 | 4.73E-04 | -0.3471 | 0.536 | 0.73 | 1 | MS.Trt_vs_MS.NoTrt_B |
| PLA2G12A | 4.75E-04 | 0.26127 | 0.473 | 0.65 | 1 | MS.Trt_vs_MS.NoTrt_B |
| IFT57 | 0.00048 | -0.8521 | 0.048 | 0.12 | 1 | MS.Trt_vs_MS.NoTrt_B |
| SFXN1 | 4.83E-04 | 0.33816 | 0.654 | 0.82 | 1 | MS.Trt_vs_MS.NoTrt_B |
| KCNN4 | 4.85E-04 | -0.3162 | 0.115 | 0.22 | 1 | MS.Trt_vs_MS.NoTrt_B |
| C5orf45 | 4.85E-04 | -0.3475 | 0.767 | 0.91 | 1 | MS.Trt_vs_MS.NoTrt_B |
| HUS1 | 4.88E-04 | 0.48474 | 0.368 | 0.52 | 1 | MS.Trt_vs_MS.NoTrt_B |
| RMDN1 | 4.89E-04 | 0.36017 | 0.584 | 0.74 | 1 | MS.Trt_vs_MS.NoTrt_B |
| ATP5H | 4.91E-04 | 0.58439 | 0.636 | 0.78 | 1 | MS.Trt_vs_MS.NoTrt_B |
| NNT | 4.96E-04 | -0.721 | 0.495 | 0.63 | 1 | MS.Trt_vs_MS.NoTrt_B |
| PLEKHM3 | 4.99E-04 | -0.4943 | 0.475 | 0.63 | 1 | MS.Trt_vs_MS.NoTrt_B |
| WWP1 | 5.01E-04 | 0.64246 | 0.322 | 0.51 | 1 | MS.Trt_vs_MS.NoTrt_B |
| RABGGTA | 5.05E-04 | -0.4508 | 0.072 | 0.16 | 1 | MS.Trt_vs_MS.NoTrt_B |
| HIVEP1 | 5.09E-04 | 0.79415 | 0.527 | 0.73 | 1 | MS.Trt_vs_MS.NoTrt_B |
| FBXO22 | 5.09E-04 | 0.51422 | 0.619 | 0.81 | 1 | MS.Trt_vs_MS.NoTrt_B |
| KRCC1 | 5.11E-04 | -0.4177 | 0.072 | 0.16 | 1 | MS.Trt_vs_MS.NoTrt_B |
| ULK4 | 5.12E-04 | -0.3896 | 0.211 | 0.35 | 1 | MS.Trt_vs_MS.NoTrt_B |

| | | | | | | |
|---|---|---|---|---|---|---|
| PARVB | 5.19E-04 | -0.4402 | 0.551 | 0.7 | 1 | MS.Trt_vs_MS.NoTrt_B |
| KLRD1 | 5.21E-04 | 0.30715 | 0.874 | 0.98 | 1 | MS.Trt_vs_MS.NoTrt_B |
| PIGN | 5.24E-04 | 0.63351 | 0.414 | 0.6 | 1 | MS.Trt_vs_MS.NoTrt_B |
| PTRHD1 | 5.25E-04 | -0.4011 | 0.198 | 0.32 | 1 | MS.Trt_vs_MS.NoTrt_B |
| RNGTT | 5.28E-04 | -0.3394 | 0.259 | 0.39 | 1 | MS.Trt_vs_MS.NoTrt_B |
| ZNF639 | 5.29E-04 | 0.30017 | 0.756 | 0.93 | 1 | MS.Trt_vs_MS.NoTrt_B |
| HEBP1 | 5.30E-04 | -0.5919 | 0.153 | 0.27 | 1 | MS.Trt_vs_MS.NoTrt_B |
| APOA2 | 5.30E-04 | -0.3549 | 0.176 | 0.29 | 1 | MS.Trt_vs_MS.NoTrt_B |
| ZNF74 | 5.31E-04 | -0.3306 | 0.316 | 0.47 | 1 | MS.Trt_vs_MS.NoTrt_B |
| EDC3 | 5.32E-04 | -0.2874 | 0.266 | 0.41 | 1 | MS.Trt_vs_MS.NoTrt_B |
| FCHSD1 | 5.35E-04 | -0.4878 | 0.222 | 0.35 | 1 | MS.Trt_vs_MS.NoTrt_B |
| MUC15 | 5.35E-04 | -0.3134 | 0.081 | 0.17 | 1 | MS.Trt_vs_MS.NoTrt_B |
| INPP5K | 5.37E-04 | -0.5711 | 0.412 | 0.55 | 1 | MS.Trt_vs_MS.NoTrt_B |
| LIG1 | 5.38E-04 | -0.5129 | 0.115 | 0.22 | 1 | MS.Trt_vs_MS.NoTrt_B |
| TMEM184C | 5.42E-04 | 0.29086 | 0.414 | 0.58 | 1 | MS.Trt_vs_MS.NoTrt_B |
| SPATA4 | 5.42E-04 | 0.27362 | 0.492 | 0.36 | 1 | MS.Trt_vs_MS.NoTrt_B |
| SLC30A7 | 5.44E-04 | 0.47325 | 0.473 | 0.64 | 1 | MS.Trt_vs_MS.NoTrt_B |
| DNAJB6 | 5.45E-04 | -0.3144 | 0.486 | 0.6 | 1 | MS.Trt_vs_MS.NoTrt_B |
| BCAP31 | 5.46E-04 | -0.2803 | 0.124 | 0.24 | 1 | MS.Trt_vs_MS.NoTrt_B |
| RABGAP1 | 5.48E-04 | -0.6092 | 0.292 | 0.44 | 1 | MS.Trt_vs_MS.NoTrt_B |
| BROX | 5.48E-04 | 0.43356 | 0.516 | 0.67 | 1 | MS.Trt_vs_MS.NoTrt_B |
| PRAF2 | 5.51E-04 | -0.5182 | 0.054 | 0.14 | 1 | MS.Trt_vs_MS.NoTrt_B |
| NMT2 | 5.53E-04 | -0.6611 | 0.815 | 0.92 | 1 | MS.Trt_vs_MS.NoTrt_B |
| UTY | 5.54E-04 | -2.3705 | 0.349 | 0.47 | 1 | MS.Trt_vs_MS.NoTrt_B |
| EIF4A3 | 5.57E-04 | -0.5828 | 0.564 | 0.71 | 1 | MS.Trt_vs_MS.NoTrt_B |
| FARSA | 5.62E-04 | -0.4842 | 0.092 | 0.19 | 1 | MS.Trt_vs_MS.NoTrt_B |
| FAM234A | 5.62E-04 | -0.6088 | 0.161 | 0.28 | 1 | MS.Trt_vs_MS.NoTrt_B |
| NRROS | 5.66E-04 | -1.043 | 0.07 | 0.16 | 1 | MS.Trt_vs_MS.NoTrt_B |
| ZNF682 | 5.66E-04 | -0.7642 | 0.283 | 0.42 | 1 | MS.Trt_vs_MS.NoTrt_B |
| PTTG1 | 5.68E-04 | -0.2803 | 0.072 | 0.16 | 1 | MS.Trt_vs_MS.NoTrt_B |
| PHRF1 | 5.68E-04 | -0.723 | 0.087 | 0.18 | 1 | MS.Trt_vs_MS.NoTrt_B |
| TMEM167A | 5.69E-04 | 0.28192 | 0.279 | 0.43 | 1 | MS.Trt_vs_MS.NoTrt_B |
| PRDX6 | 5.69E-04 | -0.4046 | 0.129 | 0.24 | 1 | MS.Trt_vs_MS.NoTrt_B |
| VPS16 | 5.69E-04 | -0.8383 | 0.074 | 0.16 | 1 | MS.Trt_vs_MS.NoTrt_B |
| MRPL21 | 5.69E-04 | -0.6542 | 0.074 | 0.16 | 1 | MS.Trt_vs_MS.NoTrt_B |
| C19orf47 | 5.75E-04 | -0.3614 | 0.355 | 0.51 | 1 | MS.Trt_vs_MS.NoTrt_B |
| MDH1 | 5.76E-04 | 0.48984 | 0.488 | 0.69 | 1 | MS.Trt_vs_MS.NoTrt_B |
| EME2 | 5.76E-04 | 0.27303 | 0.19 | 0.32 | 1 | MS.Trt_vs_MS.NoTrt_B |
| RGMB | 5.76E-04 | -1.1294 | 0.052 | 0.13 | 1 | MS.Trt_vs_MS.NoTrt_B |
| SMG5 | 5.80E-04 | -0.4177 | 0.085 | 0.18 | 1 | MS.Trt_vs_MS.NoTrt_B |

| | | | | | | |
|---|---|---|---|---|---|---|
| PIGK | 5.81E-04 | 0.33894 | 0.061 | 0.15 | 1 | MS.Trt_vs_MS.NoTrt_B |
| HEATR3 | 5.84E-04 | -0.5378 | 0.468 | 0.68 | 1 | MS.Trt_vs_MS.NoTrt_B |
| POLM | 5.84E-04 | -0.4621 | 0.397 | 0.55 | 1 | MS.Trt_vs_MS.NoTrt_B |
| SURF6 | 5.86E-04 | -0.422 | 0.379 | 0.56 | 1 | MS.Trt_vs_MS.NoTrt_B |
| PKNOX1 | 5.90E-04 | -0.2769 | 0.632 | 0.72 | 1 | MS.Trt_vs_MS.NoTrt_B |
| PEX11A | 5.90E-04 | 0.81995 | 0.447 | 0.63 | 1 | MS.Trt_vs_MS.NoTrt_B |
| ATG12 | 5.93E-04 | 0.34934 | 0.649 | 0.78 | 1 | MS.Trt_vs_MS.NoTrt_B |
| RNF4 | 5.94E-04 | 0.38924 | 0.673 | 0.77 | 1 | MS.Trt_vs_MS.NoTrt_B |
| PHB | 5.95E-04 | -0.3577 | 0.105 | 0.21 | 1 | MS.Trt_vs_MS.NoTrt_B |
| RP5-994D16 | 5.96E-04 | -0.5607 | 0.39 | 0.57 | 1 | MS.Trt_vs_MS.NoTrt_B |
| HLCS | 5.96E-04 | 0.39381 | 0.564 | 0.74 | 1 | MS.Trt_vs_MS.NoTrt_B |
| UBE3A | 5.99E-04 | 0.48521 | 0.394 | 0.55 | 1 | MS.Trt_vs_MS.NoTrt_B |
| YIF1B | 6.00E-04 | 0.27074 | 0.499 | 0.67 | 1 | MS.Trt_vs_MS.NoTrt_B |
| NARS2 | 6.03E-04 | -0.3927 | 0.281 | 0.42 | 1 | MS.Trt_vs_MS.NoTrt_B |
| PPARD | 6.05E-04 | -0.2879 | 0.405 | 0.56 | 1 | MS.Trt_vs_MS.NoTrt_B |
| GEMIN4 | 6.06E-04 | -0.2619 | 0.035 | 0.1 | 1 | MS.Trt_vs_MS.NoTrt_B |
| MTRNR2L10 | 6.08E-04 | 0.38051 | 0.871 | 1 | 1 | MS.Trt_vs_MS.NoTrt_B |
| HECTD4 | 6.09E-04 | 0.55056 | 0.521 | 0.69 | 1 | MS.Trt_vs_MS.NoTrt_B |
| ARPC5 | 6.11E-04 | 0.4826 | 0.813 | 0.94 | 1 | MS.Trt_vs_MS.NoTrt_B |
| KLHL32 | 6.16E-04 | -0.9975 | 0.052 | 0.13 | 1 | MS.Trt_vs_MS.NoTrt_B |
| GDI2 | 6.17E-04 | 0.3251 | 0.316 | 0.48 | 1 | MS.Trt_vs_MS.NoTrt_B |
| SIGLEC6 | 6.17E-04 | -0.2573 | 0.355 | 0.5 | 1 | MS.Trt_vs_MS.NoTrt_B |
| SLC47A2 | 6.18E-04 | 0.47922 | 0.146 | 0.27 | 1 | MS.Trt_vs_MS.NoTrt_B |
| CTNNA1 | 6.21E-04 | -0.2959 | 0.44 | 0.63 | 1 | MS.Trt_vs_MS.NoTrt_B |
| KLHL22 | 6.27E-04 | -0.2958 | 0.129 | 0.24 | 1 | MS.Trt_vs_MS.NoTrt_B |
| TOR1AIP2 | 6.30E-04 | 0.25304 | 0.926 | 0.98 | 1 | MS.Trt_vs_MS.NoTrt_B |
| CRELD2 | 6.31E-04 | -0.9285 | 0.466 | 0.61 | 1 | MS.Trt_vs_MS.NoTrt_B |
| NOP56 | 6.34E-04 | 0.30628 | 0.316 | 0.47 | 1 | MS.Trt_vs_MS.NoTrt_B |
| ACSS1 | 6.35E-04 | -0.9556 | 0.462 | 0.62 | 1 | MS.Trt_vs_MS.NoTrt_B |
| MICA | 6.36E-04 | 0.31184 | 0.632 | 0.82 | 1 | MS.Trt_vs_MS.NoTrt_B |
| ZNF346 | 6.38E-04 | -0.4473 | 0.645 | 0.77 | 1 | MS.Trt_vs_MS.NoTrt_B |
| CNDP2 | 6.38E-04 | -0.4757 | 0.553 | 0.71 | 1 | MS.Trt_vs_MS.NoTrt_B |
| ISY1 | 6.41E-04 | -0.6385 | 0.466 | 0.64 | 1 | MS.Trt_vs_MS.NoTrt_B |
| TFIP11 | 6.48E-04 | -0.8697 | 0.346 | 0.51 | 1 | MS.Trt_vs_MS.NoTrt_B |
| MTRNR2L12 | 6.49E-04 | 0.25455 | 0.985 | 1 | 1 | MS.Trt_vs_MS.NoTrt_B |
| P4HB | 6.50E-04 | -0.6472 | 0.523 | 0.65 | 1 | MS.Trt_vs_MS.NoTrt_B |
| LRCH3 | 6.50E-04 | -0.2957 | 0.919 | 0.99 | 1 | MS.Trt_vs_MS.NoTrt_B |
| SPON2 | 6.51E-04 | -0.6121 | 0.283 | 0.42 | 1 | MS.Trt_vs_MS.NoTrt_B |
| FEZ2 | 6.51E-04 | -0.598 | 0.34 | 0.5 | 1 | MS.Trt_vs_MS.NoTrt_B |
| SF3B5 | 6.55E-04 | -0.3852 | 0.076 | 0.17 | 1 | MS.Trt_vs_MS.NoTrt_B |

| | | | | | | |
|---|---|---|---|---|---|---|
| MBTPS1 | 6.56E-04 | -0.382 | 0.41 | 0.55 | 1 | MS.Trt_vs_MS.NoTrt_B |
| CCDC127 | 6.61E-04 | -0.5555 | 0.593 | 0.79 | 1 | MS.Trt_vs_MS.NoTrt_B |
| B4GALT3 | 6.61E-04 | -0.6203 | 0.113 | 0.22 | 1 | MS.Trt_vs_MS.NoTrt_B |
| CDC42SE2 | 6.74E-04 | 0.44564 | 0.758 | 0.95 | 1 | MS.Trt_vs_MS.NoTrt_B |
| TMEM126A | 6.78E-04 | -0.7289 | 0.046 | 0.12 | 1 | MS.Trt_vs_MS.NoTrt_B |
| RCL1 | 6.80E-04 | 0.67194 | 0.331 | 0.5 | 1 | MS.Trt_vs_MS.NoTrt_B |
| TWSG1 | 6.80E-04 | -0.56 | 0.277 | 0.43 | 1 | MS.Trt_vs_MS.NoTrt_B |
| PLEKHH2 | 6.80E-04 | 0.4591 | 0.473 | 0.68 | 1 | MS.Trt_vs_MS.NoTrt_B |
| LCMT1 | 6.81E-04 | 0.34135 | 0.107 | 0.21 | 1 | MS.Trt_vs_MS.NoTrt_B |
| PPCDC | 6.83E-04 | -0.7639 | 0.039 | 0.11 | 1 | MS.Trt_vs_MS.NoTrt_B |
| CXorf38 | 6.87E-04 | -0.5934 | 0.708 | 0.82 | 1 | MS.Trt_vs_MS.NoTrt_B |
| SNRPA1 | 6.90E-04 | 0.34074 | 0.569 | 0.73 | 1 | MS.Trt_vs_MS.NoTrt_B |
| EIF2AK1 | 6.91E-04 | -0.5209 | 0.331 | 0.48 | 1 | MS.Trt_vs_MS.NoTrt_B |
| ROCK2 | 6.91E-04 | 0.36033 | 0.336 | 0.48 | 1 | MS.Trt_vs_MS.NoTrt_B |
| TDP1 | 7.04E-04 | -0.3031 | 0.203 | 0.34 | 1 | MS.Trt_vs_MS.NoTrt_B |
| ETNK1 | 7.07E-04 | 0.51919 | 0.379 | 0.57 | 1 | MS.Trt_vs_MS.NoTrt_B |
| LRFN1 | 7.10E-04 | -0.4045 | 0.068 | 0.15 | 1 | MS.Trt_vs_MS.NoTrt_B |
| PTDSS1 | 7.11E-04 | -0.2721 | 0.27 | 0.4 | 1 | MS.Trt_vs_MS.NoTrt_B |
| GPATCH1 | 7.16E-04 | -0.6819 | 0.266 | 0.38 | 1 | MS.Trt_vs_MS.NoTrt_B |
| EIF2B4 | 7.21E-04 | 0.90062 | 0.366 | 0.52 | 1 | MS.Trt_vs_MS.NoTrt_B |
| RP11-577H5 | 7.22E-04 | -0.8644 | 0.061 | 0.14 | 1 | MS.Trt_vs_MS.NoTrt_B |
| DYM | 7.23E-04 | -0.3904 | 0.551 | 0.69 | 1 | MS.Trt_vs_MS.NoTrt_B |
| WDR82 | 7.23E-04 | -0.8127 | 0.309 | 0.44 | 1 | MS.Trt_vs_MS.NoTrt_B |
| RCE1 | 7.27E-04 | -0.3781 | 0.052 | 0.13 | 1 | MS.Trt_vs_MS.NoTrt_B |
| RABL2B | 7.28E-04 | -0.4422 | 0.791 | 0.9 | 1 | MS.Trt_vs_MS.NoTrt_B |
| NDRG3 | 7.31E-04 | -0.332 | 0.275 | 0.42 | 1 | MS.Trt_vs_MS.NoTrt_B |
| VPS39 | 7.33E-04 | 0.58817 | 0.399 | 0.55 | 1 | MS.Trt_vs_MS.NoTrt_B |
| NOP58 | 7.36E-04 | 0.29799 | 0.373 | 0.55 | 1 | MS.Trt_vs_MS.NoTrt_B |
| FZD3 | 7.39E-04 | 0.76825 | 0.451 | 0.63 | 1 | MS.Trt_vs_MS.NoTrt_B |
| ANP32A | 7.41E-04 | -0.3811 | 0.227 | 0.35 | 1 | MS.Trt_vs_MS.NoTrt_B |
| C5AR2 | 7.42E-04 | -0.2524 | 0.85 | 0.94 | 1 | MS.Trt_vs_MS.NoTrt_B |
| IKBKB | 7.45E-04 | 0.30002 | 0.739 | 0.87 | 1 | MS.Trt_vs_MS.NoTrt_B |
| MAP4K2 | 7.46E-04 | -0.53 | 0.564 | 0.67 | 1 | MS.Trt_vs_MS.NoTrt_B |
| STX7 | 7.52E-04 | 0.49578 | 0.575 | 0.74 | 1 | MS.Trt_vs_MS.NoTrt_B |
| KLHL36 | 7.55E-04 | -0.3269 | 0.49 | 0.65 | 1 | MS.Trt_vs_MS.NoTrt_B |
| MTRNR2L3 | 7.57E-04 | -0.3446 | 0.285 | 0.17 | 1 | MS.Trt_vs_MS.NoTrt_B |
| NUDT19 | 7.58E-04 | 0.58968 | 0.769 | 0.91 | 1 | MS.Trt_vs_MS.NoTrt_B |
| GATAD1 | 7.60E-04 | 0.6342 | 0.71 | 0.88 | 1 | MS.Trt_vs_MS.NoTrt_B |
| VCPKMT | 7.61E-04 | -0.3218 | 0.063 | 0.15 | 1 | MS.Trt_vs_MS.NoTrt_B |
| CTD-2207O2 | 7.64E-04 | -0.8524 | 0.096 | 0.19 | 1 | MS.Trt_vs_MS.NoTrt_B |

| | | | | | | |
|---|---|---|---|---|---|---|
| BMP8A | 7.66E-04 | -0.3016 | 0.453 | 0.62 | 1 | MS.Trt_vs_MS.NoTrt_B |
| FAM60A | 7.70E-04 | 0.73613 | 0.39 | 0.55 | 1 | MS.Trt_vs_MS.NoTrt_B |
| SEC24D | 7.73E-04 | -0.5502 | 0.375 | 0.53 | 1 | MS.Trt_vs_MS.NoTrt_B |
| MTPAP | 7.73E-04 | 0.45921 | 0.691 | 0.89 | 1 | MS.Trt_vs_MS.NoTrt_B |
| MTAP | 7.77E-04 | -0.3806 | 0.547 | 0.76 | 1 | MS.Trt_vs_MS.NoTrt_B |
| NAV1 | 7.82E-04 | -0.5873 | 0.747 | 0.85 | 1 | MS.Trt_vs_MS.NoTrt_B |
| GPATCH2L | 7.82E-04 | 0.26864 | 0.577 | 0.78 | 1 | MS.Trt_vs_MS.NoTrt_B |
| GMIP | 7.88E-04 | 0.68691 | 0.17 | 0.3 | 1 | MS.Trt_vs_MS.NoTrt_B |
| DIAPH2 | 7.89E-04 | 0.26647 | 0.2 | 0.33 | 1 | MS.Trt_vs_MS.NoTrt_B |
| DNMT1 | 7.91E-04 | -0.2918 | 0.9 | 0.95 | 1 | MS.Trt_vs_MS.NoTrt_B |
| MFHAS1 | 7.94E-04 | -0.7977 | 0.037 | 0.1 | 1 | MS.Trt_vs_MS.NoTrt_B |
| SETD6 | 7.95E-04 | -0.2524 | 0.296 | 0.45 | 1 | MS.Trt_vs_MS.NoTrt_B |
| SMYD4 | 7.96E-04 | -0.6074 | 0.667 | 0.82 | 1 | MS.Trt_vs_MS.NoTrt_B |
| GDE1 | 7.96E-04 | -1.2399 | 0.172 | 0.29 | 1 | MS.Trt_vs_MS.NoTrt_B |
| PQBP1 | 7.97E-04 | -0.5932 | 0.102 | 0.2 | 1 | MS.Trt_vs_MS.NoTrt_B |
| SNRNP70 | 7.99E-04 | -0.4127 | 0.161 | 0.28 | 1 | MS.Trt_vs_MS.NoTrt_B |
| FLOT2 | 8.01E-04 | -0.3286 | 0.285 | 0.42 | 1 | MS.Trt_vs_MS.NoTrt_B |
| TMEM109 | 8.08E-04 | -0.3952 | 0.12 | 0.23 | 1 | MS.Trt_vs_MS.NoTrt_B |
| CHMP2A | 8.10E-04 | -0.5847 | 0.076 | 0.16 | 1 | MS.Trt_vs_MS.NoTrt_B |
| PTP4A2 | 8.14E-04 | 0.3733 | 0.44 | 0.6 | 1 | MS.Trt_vs_MS.NoTrt_B |
| NUCKS1 | 8.19E-04 | 0.58189 | 0.388 | 0.55 | 1 | MS.Trt_vs_MS.NoTrt_B |
| GTF3C5 | 8.28E-04 | -0.9692 | 0.209 | 0.32 | 1 | MS.Trt_vs_MS.NoTrt_B |
| TCIRG1 | 8.29E-04 | -0.3491 | 0.218 | 0.34 | 1 | MS.Trt_vs_MS.NoTrt_B |
| SLC26A11 | 8.35E-04 | -0.6612 | 0.54 | 0.71 | 1 | MS.Trt_vs_MS.NoTrt_B |
| QDPR | 8.36E-04 | -0.8015 | 0.039 | 0.11 | 1 | MS.Trt_vs_MS.NoTrt_B |
| PTPN18 | 8.37E-04 | 0.91667 | 0.24 | 0.4 | 1 | MS.Trt_vs_MS.NoTrt_B |
| SPPL3 | 8.47E-04 | -0.4982 | 0.447 | 0.62 | 1 | MS.Trt_vs_MS.NoTrt_B |
| ZNF496 | 8.49E-04 | 0.44547 | 0.481 | 0.67 | 1 | MS.Trt_vs_MS.NoTrt_B |
| TSTD1 | 8.51E-04 | 0.3281 | 0.166 | 0.3 | 1 | MS.Trt_vs_MS.NoTrt_B |
| VAMP1 | 8.52E-04 | -0.2681 | 0.118 | 0.22 | 1 | MS.Trt_vs_MS.NoTrt_B |
| USP32 | 8.53E-04 | 0.41971 | 0.179 | 0.29 | 1 | MS.Trt_vs_MS.NoTrt_B |
| POR | 8.59E-04 | -0.3773 | 0.227 | 0.35 | 1 | MS.Trt_vs_MS.NoTrt_B |
| EIF2D | 8.59E-04 | 0.51479 | 0.571 | 0.75 | 1 | MS.Trt_vs_MS.NoTrt_B |
| CENPC | 8.60E-04 | 0.46535 | 0.468 | 0.6 | 1 | MS.Trt_vs_MS.NoTrt_B |
| RSBN1L | 8.60E-04 | 0.35523 | 0.697 | 0.84 | 1 | MS.Trt_vs_MS.NoTrt_B |
| RTKN2 | 8.60E-04 | 0.29708 | 0.44 | 0.58 | 1 | MS.Trt_vs_MS.NoTrt_B |
| GSTM2 | 8.63E-04 | 0.59523 | 0.277 | 0.44 | 1 | MS.Trt_vs_MS.NoTrt_B |
| TNFRSF10B | 8.70E-04 | -0.8113 | 0.34 | 0.48 | 1 | MS.Trt_vs_MS.NoTrt_B |
| PARD6B | 8.72E-04 | 0.32075 | 0.364 | 0.51 | 1 | MS.Trt_vs_MS.NoTrt_B |
| OLFM2 | 8.77E-04 | 0.30861 | 0.039 | 0.11 | 1 | MS.Trt_vs_MS.NoTrt_B |

| | | | | | | |
|---|---|---|---|---|---|---|
| PGAM1 | 8.80E-04 | -0.2777 | 0.187 | 0.3 | 1 | MS.Trt_vs_MS.NoTrt_B |
| USP14 | 8.82E-04 | -0.4662 | 0.473 | 0.65 | 1 | MS.Trt_vs_MS.NoTrt_B |
| GIPR | 8.84E-04 | 0.48649 | 0.527 | 0.69 | 1 | MS.Trt_vs_MS.NoTrt_B |
| EZH2 | 8.85E-04 | -0.695 | 0.085 | 0.17 | 1 | MS.Trt_vs_MS.NoTrt_B |
| CSRP2 | 8.90E-04 | -0.4638 | 0.379 | 0.56 | 1 | MS.Trt_vs_MS.NoTrt_B |
| CLUAP1 | 8.90E-04 | 0.3305 | 0.68 | 0.87 | 1 | MS.Trt_vs_MS.NoTrt_B |
| HSPA9 | 8.92E-04 | -0.5085 | 0.303 | 0.43 | 1 | MS.Trt_vs_MS.NoTrt_B |
| PUM2 | 8.93E-04 | -0.327 | 0.233 | 0.36 | 1 | MS.Trt_vs_MS.NoTrt_B |
| VPS13A | 8.97E-04 | 0.53839 | 0.218 | 0.36 | 1 | MS.Trt_vs_MS.NoTrt_B |
| NPAT | 9.01E-04 | 0.6319 | 0.619 | 0.79 | 1 | MS.Trt_vs_MS.NoTrt_B |
| SIRT2 | 9.07E-04 | -0.717 | 0.808 | 0.91 | 1 | MS.Trt_vs_MS.NoTrt_B |
| RPIA | 9.09E-04 | -0.6128 | 0.061 | 0.14 | 1 | MS.Trt_vs_MS.NoTrt_B |
| BCL7C | 9.15E-04 | -0.4219 | 0.39 | 0.56 | 1 | MS.Trt_vs_MS.NoTrt_B |
| FAM149B1 | 9.18E-04 | 0.65003 | 0.129 | 0.24 | 1 | MS.Trt_vs_MS.NoTrt_B |
| KIAA0930 | 9.24E-04 | -0.2828 | 0.481 | 0.65 | 1 | MS.Trt_vs_MS.NoTrt_B |
| ZSCAN2 | 9.25E-04 | -0.5532 | 0.614 | 0.82 | 1 | MS.Trt_vs_MS.NoTrt_B |
| TNFRSF4 | 9.26E-04 | -0.5267 | 0.076 | 0.16 | 1 | MS.Trt_vs_MS.NoTrt_B |
| KXD1 | 9.29E-04 | -0.3874 | 0.706 | 0.8 | 1 | MS.Trt_vs_MS.NoTrt_B |
| ZNF567 | 9.32E-04 | -0.4063 | 0.688 | 0.88 | 1 | MS.Trt_vs_MS.NoTrt_B |
| TTC37 | 9.34E-04 | -0.275 | 0.272 | 0.41 | 1 | MS.Trt_vs_MS.NoTrt_B |
| AP1S3 | 9.36E-04 | -0.2885 | 0.791 | 0.93 | 1 | MS.Trt_vs_MS.NoTrt_B |
| RORC | 9.40E-04 | -1.1597 | 0.044 | 0.11 | 1 | MS.Trt_vs_MS.NoTrt_B |
| CAPNS1 | 9.43E-04 | 0.53034 | 0.22 | 0.35 | 1 | MS.Trt_vs_MS.NoTrt_B |
| NUP153 | 9.48E-04 | 0.27502 | 0.129 | 0.24 | 1 | MS.Trt_vs_MS.NoTrt_B |
| C1orf162 | 9.51E-04 | -0.5056 | 0.061 | 0.14 | 1 | MS.Trt_vs_MS.NoTrt_B |
| ZNF442 | 9.53E-04 | -0.3339 | 0.573 | 0.76 | 1 | MS.Trt_vs_MS.NoTrt_B |
| SLC25A12 | 9.54E-04 | -0.9501 | 0.059 | 0.14 | 1 | MS.Trt_vs_MS.NoTrt_B |
| MUL1 | 9.56E-04 | -0.5868 | 0.037 | 0.1 | 1 | MS.Trt_vs_MS.NoTrt_B |
| HACD3 | 9.56E-04 | 0.48476 | 0.58 | 0.8 | 1 | MS.Trt_vs_MS.NoTrt_B |
| B4GALT4 | 9.59E-04 | -0.5549 | 0.484 | 0.63 | 1 | MS.Trt_vs_MS.NoTrt_B |
| CDON | 9.61E-04 | -0.2853 | 0.349 | 0.49 | 1 | MS.Trt_vs_MS.NoTrt_B |
| MPP7 | 9.62E-04 | -0.8412 | 0.161 | 0.27 | 1 | MS.Trt_vs_MS.NoTrt_B |
| ABHD14A | 9.67E-04 | -0.7082 | 0.401 | 0.51 | 1 | MS.Trt_vs_MS.NoTrt_B |
| CCDC94 | 9.68E-04 | -0.3885 | 0.1 | 0.2 | 1 | MS.Trt_vs_MS.NoTrt_B |
| BIRC6 | 9.69E-04 | 0.40407 | 0.316 | 0.48 | 1 | MS.Trt_vs_MS.NoTrt_B |
| ACAD10 | 9.86E-04 | 0.31317 | 0.344 | 0.49 | 1 | MS.Trt_vs_MS.NoTrt_B |
| BRPF1 | 9.86E-04 | -0.6215 | 0.044 | 0.11 | 1 | MS.Trt_vs_MS.NoTrt_B |
| SRRT | 9.87E-04 | -0.3548 | 0.135 | 0.24 | 1 | MS.Trt_vs_MS.NoTrt_B |
| USP37 | 9.88E-04 | 0.43616 | 0.312 | 0.45 | 1 | MS.Trt_vs_MS.NoTrt_B |
| ABHD5 | 9.91E-04 | 0.43103 | 0.078 | 0.17 | 1 | MS.Trt_vs_MS.NoTrt_B |

| | | | | | | |
|---|---|---|---|---|---|---|
| NAGK | 9.91E-04 | -0.3801 | 0.647 | 0.84 | 1 | MS.Trt_vs_MS.NoTrt_B |
| METTL9 | 9.93E-04 | -0.4139 | 0.427 | 0.58 | 1 | MS.Trt_vs_MS.NoTrt_B |
| PGM1 | 9.93E-04 | -0.869 | 0.054 | 0.13 | 1 | MS.Trt_vs_MS.NoTrt_B |
| C10orf67 | 9.97E-04 | -0.7169 | 0.15 | 0.26 | 1 | MS.Trt_vs_MS.NoTrt_B |
| FKBP15 | 9.99E-04 | -1.089 | 0.059 | 0.14 | 1 | MS.Trt_vs_MS.NoTrt_B |
| LAMP1 | 9.99E-04 | -0.424 | 0.251 | 0.37 | 1 | MS.Trt_vs_MS.NoTrt_B |
| BTBD11 | 0.001 | 0.9407 | 0.338 | 0.53 | 1 | MS.Trt_vs_MS.NoTrt_B |
| LIN52 | 0.00101 | -0.2801 | 0.566 | 0.7 | 1 | MS.Trt_vs_MS.NoTrt_B |
| CDC14A | 0.00101 | 0.8312 | 0.144 | 0.26 | 1 | MS.Trt_vs_MS.NoTrt_B |
| PRPF40A | 0.00101 | 0.49895 | 0.322 | 0.48 | 1 | MS.Trt_vs_MS.NoTrt_B |
| TRMT61B | 0.00102 | 0.2653 | 0.096 | 0.19 | 1 | MS.Trt_vs_MS.NoTrt_B |
| MAP4K5 | 0.00103 | 0.3417 | 0.364 | 0.53 | 1 | MS.Trt_vs_MS.NoTrt_B |
| SLC35G1 | 0.00104 | -0.3551 | 0.068 | 0.15 | 1 | MS.Trt_vs_MS.NoTrt_B |
| C8orf33 | 0.00104 | -1.1153 | 0.124 | 0.23 | 1 | MS.Trt_vs_MS.NoTrt_B |
| CARHSP1 | 0.00104 | -0.454 | 0.536 | 0.66 | 1 | MS.Trt_vs_MS.NoTrt_B |
| TROAP | 0.00104 | -0.4394 | 0.338 | 0.51 | 1 | MS.Trt_vs_MS.NoTrt_B |
| ING3 | 0.00104 | -0.2617 | 0.1 | 0.2 | 1 | MS.Trt_vs_MS.NoTrt_B |
| KRTCAP2 | 0.00105 | 0.27014 | 0.623 | 0.8 | 1 | MS.Trt_vs_MS.NoTrt_B |
| ANKEF1 | 0.00105 | 0.6184 | 0.137 | 0.25 | 1 | MS.Trt_vs_MS.NoTrt_B |
| TCFL5 | 0.00105 | -0.8576 | 0.416 | 0.54 | 1 | MS.Trt_vs_MS.NoTrt_B |
| AKAP2 | 0.00106 | -0.5391 | 0.198 | 0.32 | 1 | MS.Trt_vs_MS.NoTrt_B |
| A2M | 0.00106 | 1.38854 | 0.157 | 0.06 | 1 | MS.Trt_vs_MS.NoTrt_B |
| NT5C | 0.00107 | -0.3379 | 0.102 | 0.2 | 1 | MS.Trt_vs_MS.NoTrt_B |
| ALG9 | 0.00108 | -0.7128 | 0.712 | 0.86 | 1 | MS.Trt_vs_MS.NoTrt_B |
| MOB2 | 0.00108 | -0.779 | 0.048 | 0.12 | 1 | MS.Trt_vs_MS.NoTrt_B |
| TAB3 | 0.00108 | 0.45413 | 0.558 | 0.76 | 1 | MS.Trt_vs_MS.NoTrt_B |
| KRBA2 | 0.00108 | 0.78948 | 0.259 | 0.41 | 1 | MS.Trt_vs_MS.NoTrt_B |
| NPIPB3 | 0.00108 | 0.43671 | 0.516 | 0.7 | 1 | MS.Trt_vs_MS.NoTrt_B |
| PSMC1 | 0.00109 | 0.26482 | 0.316 | 0.44 | 1 | MS.Trt_vs_MS.NoTrt_B |
| RAD51C | 0.0011 | -0.2569 | 0.789 | 0.94 | 1 | MS.Trt_vs_MS.NoTrt_B |
| ANKIB1 | 0.0011 | 0.63994 | 0.24 | 0.37 | 1 | MS.Trt_vs_MS.NoTrt_B |
| TPP2 | 0.00111 | -0.3021 | 0.412 | 0.52 | 1 | MS.Trt_vs_MS.NoTrt_B |
| LNPK | 0.00111 | 0.26079 | 0.41 | 0.55 | 1 | MS.Trt_vs_MS.NoTrt_B |
| LCMT2 | 0.00111 | -0.8271 | 0.538 | 0.61 | 1 | MS.Trt_vs_MS.NoTrt_B |
| SPCS2 | 0.00111 | 0.26549 | 0.229 | 0.36 | 1 | MS.Trt_vs_MS.NoTrt_B |
| KRIT1 | 0.00112 | -0.4949 | 0.499 | 0.67 | 1 | MS.Trt_vs_MS.NoTrt_B |
| MRPL24 | 0.00112 | -0.3975 | 0.065 | 0.15 | 1 | MS.Trt_vs_MS.NoTrt_B |
| FHOD1 | 0.00113 | -0.594 | 0.076 | 0.16 | 1 | MS.Trt_vs_MS.NoTrt_B |
| EIF2S1 | 0.00114 | 0.29602 | 0.233 | 0.36 | 1 | MS.Trt_vs_MS.NoTrt_B |
| ACTR3C | 0.00114 | -0.5333 | 0.42 | 0.55 | 1 | MS.Trt_vs_MS.NoTrt_B |

| | | | | | | |
|---|---|---|---|---|---|---|
| TJAP1 | 0.00114 | 0.75502 | 0.253 | 0.38 | 1 | MS.Trt_vs_MS.NoTrt_B |
| HNRNPL | 0.00114 | 0.33602 | 0.739 | 0.78 | 1 | MS.Trt_vs_MS.NoTrt_B |
| RGPD6 | 0.00115 | 0.46162 | 0.383 | 0.56 | 1 | MS.Trt_vs_MS.NoTrt_B |
| DLST | 0.00115 | -0.3829 | 0.793 | 0.95 | 1 | MS.Trt_vs_MS.NoTrt_B |
| PLEKHA2 | 0.00115 | -0.2567 | 0.68 | 0.77 | 1 | MS.Trt_vs_MS.NoTrt_B |
| MFGE8 | 0.00116 | 0.33072 | 0.383 | 0.54 | 1 | MS.Trt_vs_MS.NoTrt_B |
| MAPK1 | 0.00116 | 0.70139 | 0.392 | 0.55 | 1 | MS.Trt_vs_MS.NoTrt_B |
| ARHGEF9 | 0.00116 | -0.5517 | 0.423 | 0.55 | 1 | MS.Trt_vs_MS.NoTrt_B |
| SIAE | 0.00116 | -0.5601 | 0.49 | 0.67 | 1 | MS.Trt_vs_MS.NoTrt_B |
| RRP7A | 0.00117 | 0.34345 | 0.39 | 0.55 | 1 | MS.Trt_vs_MS.NoTrt_B |
| CELF1 | 0.00117 | -0.3665 | 0.309 | 0.44 | 1 | MS.Trt_vs_MS.NoTrt_B |
| SERPINH1 | 0.00117 | -0.2854 | 0.224 | 0.36 | 1 | MS.Trt_vs_MS.NoTrt_B |
| GOLGA8A | 0.00118 | 0.62736 | 0.314 | 0.47 | 1 | MS.Trt_vs_MS.NoTrt_B |
| PIGT | 0.00119 | -0.9271 | 0.057 | 0.13 | 1 | MS.Trt_vs_MS.NoTrt_B |
| ABHD10 | 0.00119 | -0.5716 | 0.052 | 0.12 | 1 | MS.Trt_vs_MS.NoTrt_B |
| ATP6V1C1 | 0.00119 | -0.2602 | 0.309 | 0.45 | 1 | MS.Trt_vs_MS.NoTrt_B |
| RRNAD1 | 0.00119 | 0.26573 | 0.497 | 0.67 | 1 | MS.Trt_vs_MS.NoTrt_B |
| FBXO45 | 0.0012 | 0.89049 | 0.303 | 0.44 | 1 | MS.Trt_vs_MS.NoTrt_B |
| TBC1D20 | 0.0012 | 0.73685 | 0.447 | 0.63 | 1 | MS.Trt_vs_MS.NoTrt_B |
| MAL | 0.00122 | -0.2676 | 0.222 | 0.35 | 1 | MS.Trt_vs_MS.NoTrt_B |
| MRPS16 | 0.00122 | -0.2888 | 0.647 | 0.79 | 1 | MS.Trt_vs_MS.NoTrt_B |
| DET1 | 0.00122 | 0.59961 | 0.355 | 0.5 | 1 | MS.Trt_vs_MS.NoTrt_B |
| LUC7L2 | 0.00122 | 0.37807 | 0.418 | 0.56 | 1 | MS.Trt_vs_MS.NoTrt_B |
| ECHDC1 | 0.00122 | 0.85787 | 0.688 | 0.9 | 1 | MS.Trt_vs_MS.NoTrt_B |
| EIF4E3 | 0.00122 | 0.82846 | 0.144 | 0.26 | 1 | MS.Trt_vs_MS.NoTrt_B |
| N4BP2 | 0.00124 | 0.55866 | 0.259 | 0.4 | 1 | MS.Trt_vs_MS.NoTrt_B |
| KPNB1 | 0.00124 | 0.38656 | 0.523 | 0.69 | 1 | MS.Trt_vs_MS.NoTrt_B |
| ELMOD3 | 0.00124 | -0.2994 | 0.745 | 0.86 | 1 | MS.Trt_vs_MS.NoTrt_B |
| SCML4 | 0.00124 | 0.35118 | 0.46 | 0.64 | 1 | MS.Trt_vs_MS.NoTrt_B |
| UTP23 | 0.00124 | -0.3109 | 0.418 | 0.56 | 1 | MS.Trt_vs_MS.NoTrt_B |
| MSH6 | 0.00125 | 0.2907 | 0.704 | 0.86 | 1 | MS.Trt_vs_MS.NoTrt_B |
| ALKBH7 | 0.00125 | -0.4875 | 0.096 | 0.19 | 1 | MS.Trt_vs_MS.NoTrt_B |
| ADTRP | 0.00125 | 0.26563 | 0.272 | 0.4 | 1 | MS.Trt_vs_MS.NoTrt_B |
| NRL | 0.00126 | -0.4725 | 0.037 | 0.1 | 1 | MS.Trt_vs_MS.NoTrt_B |
| ABLIM3 | 0.00126 | 0.45722 | 0.102 | 0.03 | 1 | MS.Trt_vs_MS.NoTrt_B |
| SLC17A5 | 0.00127 | -0.7705 | 0.349 | 0.51 | 1 | MS.Trt_vs_MS.NoTrt_B |
| GIT1 | 0.00127 | -0.8284 | 0.126 | 0.22 | 1 | MS.Trt_vs_MS.NoTrt_B |
| RDH5 | 0.00127 | -0.5621 | 0.373 | 0.51 | 1 | MS.Trt_vs_MS.NoTrt_B |
| CD109 | 0.00127 | -0.4074 | 0.237 | 0.36 | 1 | MS.Trt_vs_MS.NoTrt_B |
| UBE3D | 0.00127 | 0.68392 | 0.442 | 0.63 | 1 | MS.Trt_vs_MS.NoTrt_B |

| | | | | | | |
|---|---|---|---|---|---|---|
| SLC25A46 | 0.00127 | -0.5394 | 0.052 | 0.12 | 1 | MS.Trt_vs_MS.NoTrt_B |
| SLC9A9 | 0.00127 | -0.4335 | 0.089 | 0.18 | 1 | MS.Trt_vs_MS.NoTrt_B |
| POP4 | 0.00129 | -0.3257 | 0.089 | 0.18 | 1 | MS.Trt_vs_MS.NoTrt_B |
| SLC6A6 | 0.00129 | -0.5563 | 0.096 | 0.19 | 1 | MS.Trt_vs_MS.NoTrt_B |
| ATP5L | 0.00129 | 0.37841 | 0.373 | 0.55 | 1 | MS.Trt_vs_MS.NoTrt_B |
| MLH1 | 0.00129 | -0.8398 | 0.246 | 0.38 | 1 | MS.Trt_vs_MS.NoTrt_B |
| MICAL1 | 0.00129 | -0.339 | 0.185 | 0.3 | 1 | MS.Trt_vs_MS.NoTrt_B |
| SETMAR | 0.0013 | -0.4605 | 0.251 | 0.38 | 1 | MS.Trt_vs_MS.NoTrt_B |
| ZNF699 | 0.0013 | 0.63801 | 0.181 | 0.29 | 1 | MS.Trt_vs_MS.NoTrt_B |
| NR2C2 | 0.0013 | 0.42675 | 0.521 | 0.71 | 1 | MS.Trt_vs_MS.NoTrt_B |
| TAF1D | 0.0013 | 0.26238 | 0.405 | 0.55 | 1 | MS.Trt_vs_MS.NoTrt_B |
| RPH3AL | 0.00131 | -0.9111 | 0.209 | 0.33 | 1 | MS.Trt_vs_MS.NoTrt_B |
| KALRN | 0.00131 | -0.2874 | 0.222 | 0.36 | 1 | MS.Trt_vs_MS.NoTrt_B |
| USP48 | 0.00132 | 0.27838 | 0.39 | 0.54 | 1 | MS.Trt_vs_MS.NoTrt_B |
| VPS29 | 0.00133 | 0.34176 | 0.41 | 0.54 | 1 | MS.Trt_vs_MS.NoTrt_B |
| IFT46 | 0.00133 | -0.7804 | 0.15 | 0.25 | 1 | MS.Trt_vs_MS.NoTrt_B |
| GEN1 | 0.00134 | 0.42432 | 0.58 | 0.77 | 1 | MS.Trt_vs_MS.NoTrt_B |
| SLC16A6 | 0.00134 | 0.9227 | 0.209 | 0.34 | 1 | MS.Trt_vs_MS.NoTrt_B |
| FARS2 | 0.00134 | -0.4544 | 0.074 | 0.16 | 1 | MS.Trt_vs_MS.NoTrt_B |
| ZNF497 | 0.00135 | -1.1871 | 0.205 | 0.33 | 1 | MS.Trt_vs_MS.NoTrt_B |
| ZNF790 | 0.00135 | -0.8289 | 0.381 | 0.52 | 1 | MS.Trt_vs_MS.NoTrt_B |
| EXPH5 | 0.00136 | 0.35958 | 0.556 | 0.71 | 1 | MS.Trt_vs_MS.NoTrt_B |
| TMEM25 | 0.00136 | -0.7794 | 0.253 | 0.41 | 1 | MS.Trt_vs_MS.NoTrt_B |
| MRPL42 | 0.00136 | 0.31302 | 0.891 | 0.98 | 1 | MS.Trt_vs_MS.NoTrt_B |
| OGFOD2 | 0.00136 | -0.4312 | 0.076 | 0.16 | 1 | MS.Trt_vs_MS.NoTrt_B |
| ZNF563 | 0.00137 | -0.8533 | 0.423 | 0.59 | 1 | MS.Trt_vs_MS.NoTrt_B |
| MICU1 | 0.00137 | -0.2734 | 0.529 | 0.64 | 1 | MS.Trt_vs_MS.NoTrt_B |
| PTPMT1 | 0.00137 | -0.396 | 0.085 | 0.17 | 1 | MS.Trt_vs_MS.NoTrt_B |
| TCAIM | 0.00137 | 1.40633 | 0.153 | 0.27 | 1 | MS.Trt_vs_MS.NoTrt_B |
| AMMECR1 | 0.00137 | -0.3451 | 0.092 | 0.18 | 1 | MS.Trt_vs_MS.NoTrt_B |
| PTEN | 0.00138 | 0.3993 | 0.447 | 0.61 | 1 | MS.Trt_vs_MS.NoTrt_B |
| SCAMP3 | 0.00139 | -0.5413 | 0.139 | 0.24 | 1 | MS.Trt_vs_MS.NoTrt_B |
| ZNF654 | 0.00139 | -0.8471 | 0.072 | 0.15 | 1 | MS.Trt_vs_MS.NoTrt_B |
| PWWP2A | 0.0014 | 0.36957 | 0.102 | 0.2 | 1 | MS.Trt_vs_MS.NoTrt_B |
| IMPA1 | 0.0014 | 0.46564 | 0.216 | 0.34 | 1 | MS.Trt_vs_MS.NoTrt_B |
| SGMS1 | 0.00141 | -0.2657 | 0.159 | 0.27 | 1 | MS.Trt_vs_MS.NoTrt_B |
| LUC7L3 | 0.00141 | 0.43057 | 0.514 | 0.72 | 1 | MS.Trt_vs_MS.NoTrt_B |
| TSTD2 | 0.00141 | -0.449 | 0.057 | 0.13 | 1 | MS.Trt_vs_MS.NoTrt_B |
| METTL6 | 0.00141 | -0.3883 | 0.778 | 0.93 | 1 | MS.Trt_vs_MS.NoTrt_B |
| VPS35 | 0.00141 | 0.29205 | 0.425 | 0.55 | 1 | MS.Trt_vs_MS.NoTrt_B |

| | | | | | | |
|---|---|---|---|---|---|---|
| MSANTD1 | 0.00142 | 0.75475 | 0.349 | 0.48 | 1 | MS.Trt_vs_MS.NoTrt_B |
| SMAD3 | 0.00142 | -0.4423 | 0.094 | 0.18 | 1 | MS.Trt_vs_MS.NoTrt_B |
| PDCD10 | 0.00142 | 0.25212 | 0.166 | 0.28 | 1 | MS.Trt_vs_MS.NoTrt_B |
| AEBP2 | 0.00142 | 0.7706 | 0.532 | 0.7 | 1 | MS.Trt_vs_MS.NoTrt_B |
| ZER1 | 0.00142 | -0.3271 | 0.098 | 0.19 | 1 | MS.Trt_vs_MS.NoTrt_B |
| NADSYN1 | 0.00142 | 0.43386 | 0.453 | 0.61 | 1 | MS.Trt_vs_MS.NoTrt_B |
| FAM96B | 0.00143 | 0.40577 | 0.161 | 0.29 | 1 | MS.Trt_vs_MS.NoTrt_B |
| GALK1 | 0.00145 | -0.3542 | 0.218 | 0.34 | 1 | MS.Trt_vs_MS.NoTrt_B |
| AP1S2 | 0.00146 | 0.33198 | 0.135 | 0.24 | 1 | MS.Trt_vs_MS.NoTrt_B |
| CXCR6 | 0.00146 | -1.4527 | 0.092 | 0.18 | 1 | MS.Trt_vs_MS.NoTrt_B |
| TEX261 | 0.00146 | -0.4497 | 0.039 | 0.1 | 1 | MS.Trt_vs_MS.NoTrt_B |
| PPCS | 0.00146 | 0.49844 | 0.255 | 0.38 | 1 | MS.Trt_vs_MS.NoTrt_B |
| SFXN3 | 0.00146 | -1.1382 | 0.237 | 0.35 | 1 | MS.Trt_vs_MS.NoTrt_B |
| LRP2BP | 0.00147 | -0.6422 | 0.353 | 0.47 | 1 | MS.Trt_vs_MS.NoTrt_B |
| GRSF1 | 0.00148 | -0.4375 | 0.617 | 0.74 | 1 | MS.Trt_vs_MS.NoTrt_B |
| RTEL1 | 0.00148 | -0.689 | 0.057 | 0.13 | 1 | MS.Trt_vs_MS.NoTrt_B |
| SYNGR2 | 0.00148 | 0.28015 | 0.455 | 0.6 | 1 | MS.Trt_vs_MS.NoTrt_B |
| EIF2AK4 | 0.00149 | 0.33302 | 0.495 | 0.67 | 1 | MS.Trt_vs_MS.NoTrt_B |
| CNPPD1 | 0.0015 | -0.3747 | 0.198 | 0.31 | 1 | MS.Trt_vs_MS.NoTrt_B |
| RP11-343C2 | 0.0015 | -0.4434 | 0.089 | 0.18 | 1 | MS.Trt_vs_MS.NoTrt_B |
| THOC6 | 0.00151 | -0.4914 | 0.046 | 0.11 | 1 | MS.Trt_vs_MS.NoTrt_B |
| TOX | 0.00151 | -0.4402 | 0.179 | 0.29 | 1 | MS.Trt_vs_MS.NoTrt_B |
| EIF3B | 0.00152 | 0.70198 | 0.281 | 0.43 | 1 | MS.Trt_vs_MS.NoTrt_B |
| ICE2 | 0.00153 | 0.44418 | 0.769 | 0.87 | 1 | MS.Trt_vs_MS.NoTrt_B |
| RDH10 | 0.00153 | 0.99338 | 0.131 | 0.24 | 1 | MS.Trt_vs_MS.NoTrt_B |
| MREG | 0.00154 | 0.40706 | 0.497 | 0.69 | 1 | MS.Trt_vs_MS.NoTrt_B |
| IMMT | 0.00154 | 0.38231 | 0.34 | 0.48 | 1 | MS.Trt_vs_MS.NoTrt_B |
| DCLRE1C | 0.00154 | 0.69647 | 0.51 | 0.66 | 1 | MS.Trt_vs_MS.NoTrt_B |
| C16orf70 | 0.00155 | 0.62535 | 0.508 | 0.68 | 1 | MS.Trt_vs_MS.NoTrt_B |
| RBM43 | 0.00156 | -0.3788 | 0.481 | 0.68 | 1 | MS.Trt_vs_MS.NoTrt_B |
| EXD3 | 0.00157 | -0.4156 | 0.373 | 0.51 | 1 | MS.Trt_vs_MS.NoTrt_B |
| PRDX3 | 0.00158 | 0.54355 | 0.37 | 0.51 | 1 | MS.Trt_vs_MS.NoTrt_B |
| C17orf89 | 0.00158 | -0.4416 | 0.059 | 0.14 | 1 | MS.Trt_vs_MS.NoTrt_B |
| ALG1 | 0.00158 | 0.25931 | 0.492 | 0.65 | 1 | MS.Trt_vs_MS.NoTrt_B |
| GPX7 | 0.00158 | -0.6893 | 0.039 | 0.1 | 1 | MS.Trt_vs_MS.NoTrt_B |
| TRIM5 | 0.00159 | -0.8203 | 0.364 | 0.51 | 1 | MS.Trt_vs_MS.NoTrt_B |
| RNPC3 | 0.0016 | 0.6319 | 0.301 | 0.48 | 1 | MS.Trt_vs_MS.NoTrt_B |
| BHLHB9 | 0.00161 | 0.35456 | 0.224 | 0.35 | 1 | MS.Trt_vs_MS.NoTrt_B |
| CUL5 | 0.00163 | 0.35237 | 0.161 | 0.27 | 1 | MS.Trt_vs_MS.NoTrt_B |
| VPS37A | 0.00164 | -0.8575 | 0.105 | 0.2 | 1 | MS.Trt_vs_MS.NoTrt_B |

| | | | | | | |
|---|---|---|---|---|---|---|
| KLF13 | 0.00164 | 0.43249 | 0.222 | 0.35 | 1 | MS.Trt_vs_MS.NoTrt_B |
| RBBP9 | 0.00164 | -1.0993 | 0.233 | 0.35 | 1 | MS.Trt_vs_MS.NoTrt_B |
| GNPAT | 0.00165 | -0.4388 | 0.1 | 0.19 | 1 | MS.Trt_vs_MS.NoTrt_B |
| XPO5 | 0.00165 | -0.4915 | 0.538 | 0.72 | 1 | MS.Trt_vs_MS.NoTrt_B |
| TRIM24 | 0.00165 | 0.57736 | 0.331 | 0.47 | 1 | MS.Trt_vs_MS.NoTrt_B |
| INTS9 | 0.00166 | -0.6885 | 0.061 | 0.14 | 1 | MS.Trt_vs_MS.NoTrt_B |
| TMEM65 | 0.00166 | 0.42975 | 0.536 | 0.72 | 1 | MS.Trt_vs_MS.NoTrt_B |
| PPP6R3 | 0.00167 | 0.25652 | 0.366 | 0.52 | 1 | MS.Trt_vs_MS.NoTrt_B |
| MFSD13A | 0.00167 | -0.4819 | 0.584 | 0.73 | 1 | MS.Trt_vs_MS.NoTrt_B |
| P2RX7 | 0.00167 | -0.2587 | 0.586 | 0.75 | 1 | MS.Trt_vs_MS.NoTrt_B |
| CASP2 | 0.00167 | 0.45871 | 0.732 | 0.87 | 1 | MS.Trt_vs_MS.NoTrt_B |
| RBM48 | 0.00168 | 0.52384 | 0.614 | 0.81 | 1 | MS.Trt_vs_MS.NoTrt_B |
| TANGO2 | 0.00169 | -0.949 | 0.765 | 0.85 | 1 | MS.Trt_vs_MS.NoTrt_B |
| DRG2 | 0.00169 | -0.2857 | 0.17 | 0.28 | 1 | MS.Trt_vs_MS.NoTrt_B |
| NEMP1 | 0.00169 | -0.7904 | 0.044 | 0.11 | 1 | MS.Trt_vs_MS.NoTrt_B |
| EPPK1 | 0.0017 | -0.3819 | 0.057 | 0.13 | 1 | MS.Trt_vs_MS.NoTrt_B |
| KLF8 | 0.0017 | -0.3994 | 0.619 | 0.72 | 1 | MS.Trt_vs_MS.NoTrt_B |
| ACTA2 | 0.0017 | -0.5252 | 0.477 | 0.6 | 1 | MS.Trt_vs_MS.NoTrt_B |
| GON4L | 0.00171 | 0.4775 | 0.71 | 0.89 | 1 | MS.Trt_vs_MS.NoTrt_B |
| BTG2 | 0.00171 | -0.5769 | 0.089 | 0.17 | 1 | MS.Trt_vs_MS.NoTrt_B |
| ABCF3 | 0.00171 | -0.8365 | 0.113 | 0.2 | 1 | MS.Trt_vs_MS.NoTrt_B |
| AKT3 | 0.00171 | 0.45059 | 0.521 | 0.7 | 1 | MS.Trt_vs_MS.NoTrt_B |
| FBXO44 | 0.00173 | -0.2716 | 0.473 | 0.63 | 1 | MS.Trt_vs_MS.NoTrt_B |
| HSD17B10 | 0.00173 | -0.4402 | 0.083 | 0.17 | 1 | MS.Trt_vs_MS.NoTrt_B |
| NBEAL1 | 0.00173 | 0.28145 | 0.782 | 0.89 | 1 | MS.Trt_vs_MS.NoTrt_B |
| TXNDC12 | 0.00173 | -0.3483 | 0.111 | 0.21 | 1 | MS.Trt_vs_MS.NoTrt_B |
| CYP4F11 | 0.00174 | -0.315 | 0.275 | 0.39 | 1 | MS.Trt_vs_MS.NoTrt_B |
| COG4 | 0.00174 | -0.3594 | 0.597 | 0.72 | 1 | MS.Trt_vs_MS.NoTrt_B |
| SH3GLB1 | 0.00174 | -0.308 | 0.394 | 0.5 | 1 | MS.Trt_vs_MS.NoTrt_B |
| QTRT1 | 0.00174 | 0.65056 | 0.656 | 0.82 | 1 | MS.Trt_vs_MS.NoTrt_B |
| NAT6 | 0.00174 | -0.4756 | 0.039 | 0.1 | 1 | MS.Trt_vs_MS.NoTrt_B |
| HSPA2 | 0.00174 | -0.3108 | 0.407 | 0.56 | 1 | MS.Trt_vs_MS.NoTrt_B |
| IFT80 | 0.00175 | -0.2905 | 0.627 | 0.75 | 1 | MS.Trt_vs_MS.NoTrt_B |
| ZNF391 | 0.00175 | -0.7171 | 0.122 | 0.22 | 1 | MS.Trt_vs_MS.NoTrt_B |
| KLHDC4 | 0.00176 | 0.37342 | 0.588 | 0.7 | 1 | MS.Trt_vs_MS.NoTrt_B |
| RBX1 | 0.00178 | 0.31262 | 0.237 | 0.37 | 1 | MS.Trt_vs_MS.NoTrt_B |
| YARS | 0.00178 | -0.5237 | 0.747 | 0.87 | 1 | MS.Trt_vs_MS.NoTrt_B |
| HIKESHI | 0.00179 | 0.25109 | 0.314 | 0.47 | 1 | MS.Trt_vs_MS.NoTrt_B |
| NMD3 | 0.00179 | 0.29059 | 0.586 | 0.7 | 1 | MS.Trt_vs_MS.NoTrt_B |
| STMN1 | 0.00179 | -0.7001 | 0.597 | 0.7 | 1 | MS.Trt_vs_MS.NoTrt_B |

| | | | | | | |
|---|---|---|---|---|---|---|
| 2-Mar | 0.00181 | -1.1512 | 0.399 | 0.54 | 1 | MS.Trt_vs_MS.NoTrt_B |
| PLEKHG3 | 0.00181 | -1.0051 | 0.325 | 0.47 | 1 | MS.Trt_vs_MS.NoTrt_B |
| ATP6V1A | 0.00182 | -0.8919 | 0.377 | 0.5 | 1 | MS.Trt_vs_MS.NoTrt_B |
| C5orf15 | 0.00183 | 0.4758 | 0.094 | 0.18 | 1 | MS.Trt_vs_MS.NoTrt_B |
| METTL10 | 0.00183 | 0.35336 | 0.253 | 0.37 | 1 | MS.Trt_vs_MS.NoTrt_B |
| PIGX | 0.00183 | -0.6679 | 0.298 | 0.41 | 1 | MS.Trt_vs_MS.NoTrt_B |
| HMGN1 | 0.00183 | 0.45599 | 0.377 | 0.55 | 1 | MS.Trt_vs_MS.NoTrt_B |
| AHSA1 | 0.00184 | -0.5532 | 0.39 | 0.52 | 1 | MS.Trt_vs_MS.NoTrt_B |
| ATE1 | 0.00184 | -0.3372 | 0.218 | 0.34 | 1 | MS.Trt_vs_MS.NoTrt_B |
| P3H1 | 0.00185 | -0.6058 | 0.832 | 0.94 | 1 | MS.Trt_vs_MS.NoTrt_B |
| SLC41A3 | 0.00185 | -1.0603 | 0.196 | 0.31 | 1 | MS.Trt_vs_MS.NoTrt_B |
| RECQL | 0.00187 | -0.4302 | 0.122 | 0.22 | 1 | MS.Trt_vs_MS.NoTrt_B |
| SUSD3 | 0.00187 | 0.26273 | 0.309 | 0.45 | 1 | MS.Trt_vs_MS.NoTrt_B |
| CBL | 0.00187 | -0.3212 | 0.455 | 0.6 | 1 | MS.Trt_vs_MS.NoTrt_B |
| TCEAL4 | 0.00187 | -0.5031 | 0.089 | 0.17 | 1 | MS.Trt_vs_MS.NoTrt_B |
| DOPEY2 | 0.00188 | -0.5575 | 0.654 | 0.8 | 1 | MS.Trt_vs_MS.NoTrt_B |
| DCBLD2 | 0.00188 | 0.36171 | 0.521 | 0.7 | 1 | MS.Trt_vs_MS.NoTrt_B |
| ABCC1 | 0.00188 | 0.39004 | 0.285 | 0.41 | 1 | MS.Trt_vs_MS.NoTrt_B |
| GNPNAT1 | 0.00188 | 0.64419 | 0.401 | 0.58 | 1 | MS.Trt_vs_MS.NoTrt_B |
| MAN2A1 | 0.00189 | 0.33433 | 0.179 | 0.29 | 1 | MS.Trt_vs_MS.NoTrt_B |
| DSN1 | 0.00189 | -1.0574 | 0.2 | 0.31 | 1 | MS.Trt_vs_MS.NoTrt_B |
| MAML2 | 0.00189 | 0.34672 | 0.15 | 0.26 | 1 | MS.Trt_vs_MS.NoTrt_B |
| NUDCD3 | 0.0019 | -0.6271 | 0.264 | 0.37 | 1 | MS.Trt_vs_MS.NoTrt_B |
| TYK2 | 0.0019 | -0.3107 | 0.525 | 0.68 | 1 | MS.Trt_vs_MS.NoTrt_B |
| LRRC59 | 0.0019 | -0.9852 | 0.078 | 0.16 | 1 | MS.Trt_vs_MS.NoTrt_B |
| SIGMAR1 | 0.0019 | -0.4875 | 0.137 | 0.24 | 1 | MS.Trt_vs_MS.NoTrt_B |
| GOLGA8R | 0.00191 | 0.3704 | 0.368 | 0.53 | 1 | MS.Trt_vs_MS.NoTrt_B |
| FASTKD1 | 0.00191 | -0.6792 | 0.161 | 0.27 | 1 | MS.Trt_vs_MS.NoTrt_B |
| AOAH | 0.00191 | 0.66712 | 0.24 | 0.36 | 1 | MS.Trt_vs_MS.NoTrt_B |
| CHPT1 | 0.00191 | -0.3466 | 0.46 | 0.61 | 1 | MS.Trt_vs_MS.NoTrt_B |
| SLC35D1 | 0.00191 | 0.60628 | 0.216 | 0.35 | 1 | MS.Trt_vs_MS.NoTrt_B |
| SUSD6 | 0.00192 | 0.35438 | 0.218 | 0.34 | 1 | MS.Trt_vs_MS.NoTrt_B |
| GBAS | 0.00194 | 0.32067 | 0.59 | 0.71 | 1 | MS.Trt_vs_MS.NoTrt_B |
| SLC39A14 | 0.00195 | -0.806 | 0.312 | 0.45 | 1 | MS.Trt_vs_MS.NoTrt_B |
| TUBGCP5 | 0.00195 | 0.29715 | 0.575 | 0.73 | 1 | MS.Trt_vs_MS.NoTrt_B |
| GNG5 | 0.00198 | 0.2853 | 0.185 | 0.32 | 1 | MS.Trt_vs_MS.NoTrt_B |
| DHX30 | 0.00199 | -0.7165 | 0.516 | 0.64 | 1 | MS.Trt_vs_MS.NoTrt_B |
| EFCAB13 | 0.00199 | -0.5053 | 0.209 | 0.33 | 1 | MS.Trt_vs_MS.NoTrt_B |
| CBX1 | 0.002 | -0.2622 | 0.179 | 0.29 | 1 | MS.Trt_vs_MS.NoTrt_B |
| ZBTB38 | 0.002 | 0.77388 | 0.471 | 0.61 | 1 | MS.Trt_vs_MS.NoTrt_B |

| | | | | | | |
|---|---|---|---|---|---|---|
| FAAP100 | 0.00201 | -0.3338 | 0.12 | 0.22 | 1 | MS.Trt_vs_MS.NoTrt_B |
| SEMA4F | 0.00202 | -0.9236 | 0.401 | 0.52 | 1 | MS.Trt_vs_MS.NoTrt_B |
| EMC3 | 0.00202 | -0.3237 | 0.73 | 0.9 | 1 | MS.Trt_vs_MS.NoTrt_B |
| BUD31 | 0.00203 | 0.28505 | 0.586 | 0.7 | 1 | MS.Trt_vs_MS.NoTrt_B |
| UST | 0.00203 | 0.39698 | 0.198 | 0.31 | 1 | MS.Trt_vs_MS.NoTrt_B |
| NUP205 | 0.00203 | 0.78802 | 0.105 | 0.2 | 1 | MS.Trt_vs_MS.NoTrt_B |
| PCBP4 | 0.00204 | -0.7026 | 0.044 | 0.11 | 1 | MS.Trt_vs_MS.NoTrt_B |
| PLAA | 0.00207 | -0.2959 | 0.081 | 0.16 | 1 | MS.Trt_vs_MS.NoTrt_B |
| DCTN4 | 0.00207 | -0.2953 | 0.227 | 0.34 | 1 | MS.Trt_vs_MS.NoTrt_B |
| LIMS1 | 0.0021 | -0.3233 | 0.78 | 0.87 | 1 | MS.Trt_vs_MS.NoTrt_B |
| LRRC37B | 0.0021 | -0.2503 | 0.442 | 0.59 | 1 | MS.Trt_vs_MS.NoTrt_B |
| BANP | 0.0021 | -0.3182 | 0.521 | 0.66 | 1 | MS.Trt_vs_MS.NoTrt_B |
| MYCBP | 0.00211 | -1.4166 | 0.048 | 0.11 | 1 | MS.Trt_vs_MS.NoTrt_B |
| DDHD2 | 0.00212 | 0.51929 | 0.66 | 0.81 | 1 | MS.Trt_vs_MS.NoTrt_B |
| SNAP29 | 0.00213 | -0.2816 | 0.577 | 0.73 | 1 | MS.Trt_vs_MS.NoTrt_B |
| CNOT7 | 0.00214 | 0.33016 | 0.499 | 0.61 | 1 | MS.Trt_vs_MS.NoTrt_B |
| ELF2 | 0.00214 | 0.26073 | 0.516 | 0.71 | 1 | MS.Trt_vs_MS.NoTrt_B |
| PPAN | 0.00215 | 0.43613 | 0.266 | 0.39 | 1 | MS.Trt_vs_MS.NoTrt_B |
| GNAQ | 0.00215 | 0.36354 | 0.135 | 0.23 | 1 | MS.Trt_vs_MS.NoTrt_B |
| JAZF1 | 0.00216 | 0.28841 | 0.054 | 0.12 | 1 | MS.Trt_vs_MS.NoTrt_B |
| ZBP1 | 0.00216 | -0.3752 | 0.46 | 0.56 | 1 | MS.Trt_vs_MS.NoTrt_B |
| SRI | 0.00217 | 0.29951 | 0.342 | 0.47 | 1 | MS.Trt_vs_MS.NoTrt_B |
| FAM192A | 0.00217 | 0.54281 | 0.34 | 0.49 | 1 | MS.Trt_vs_MS.NoTrt_B |
| FBXO4 | 0.00217 | -0.2755 | 0.174 | 0.28 | 1 | MS.Trt_vs_MS.NoTrt_B |
| PANK2 | 0.00217 | -0.7898 | 0.418 | 0.54 | 1 | MS.Trt_vs_MS.NoTrt_B |
| PDCD4 | 0.00218 | 0.52882 | 0.495 | 0.67 | 1 | MS.Trt_vs_MS.NoTrt_B |
| CYB5B | 0.00218 | 0.40011 | 0.388 | 0.53 | 1 | MS.Trt_vs_MS.NoTrt_B |
| CNTROB | 0.00219 | -0.7026 | 0.111 | 0.2 | 1 | MS.Trt_vs_MS.NoTrt_B |
| SCAI | 0.0022 | 0.49528 | 0.715 | 0.83 | 1 | MS.Trt_vs_MS.NoTrt_B |
| CD99L2 | 0.00223 | -1.0354 | 0.072 | 0.15 | 1 | MS.Trt_vs_MS.NoTrt_B |
| ENOPH1 | 0.00224 | -0.2802 | 0.089 | 0.17 | 1 | MS.Trt_vs_MS.NoTrt_B |
| VTA1 | 0.00225 | 0.54723 | 0.264 | 0.4 | 1 | MS.Trt_vs_MS.NoTrt_B |
| TSPYL4 | 0.00225 | -0.3505 | 0.105 | 0.2 | 1 | MS.Trt_vs_MS.NoTrt_B |
| GLMN | 0.00226 | 0.41229 | 0.065 | 0.14 | 1 | MS.Trt_vs_MS.NoTrt_B |
| CNIH1 | 0.00226 | 0.27586 | 0.098 | 0.18 | 1 | MS.Trt_vs_MS.NoTrt_B |
| ATPAF1 | 0.00226 | -1.6024 | 0.139 | 0.23 | 1 | MS.Trt_vs_MS.NoTrt_B |
| ARID2 | 0.00227 | 0.62706 | 0.608 | 0.74 | 1 | MS.Trt_vs_MS.NoTrt_B |
| GGA3 | 0.00227 | -0.2908 | 0.122 | 0.22 | 1 | MS.Trt_vs_MS.NoTrt_B |
| STIP1 | 0.00227 | 0.33842 | 0.414 | 0.57 | 1 | MS.Trt_vs_MS.NoTrt_B |
| SGTB | 0.00228 | 0.33107 | 0.527 | 0.65 | 1 | MS.Trt_vs_MS.NoTrt_B |

| | | | | | | |
|---|---|---|---|---|---|---|
| ZNF253 | 0.00228 | 0.27639 | 0.66 | 0.53 | 1 | MS.Trt_vs_MS.NoTrt_B |
| SREK1 | 0.00228 | 0.37271 | 0.39 | 0.52 | 1 | MS.Trt_vs_MS.NoTrt_B |
| PAPOLA | 0.0023 | 0.4064 | 0.721 | 0.85 | 1 | MS.Trt_vs_MS.NoTrt_B |
| WDFY2 | 0.0023 | -1.0661 | 0.471 | 0.58 | 1 | MS.Trt_vs_MS.NoTrt_B |
| KLHL21 | 0.0023 | -0.8096 | 0.571 | 0.7 | 1 | MS.Trt_vs_MS.NoTrt_B |
| PPIH | 0.00233 | -0.8339 | 0.048 | 0.11 | 1 | MS.Trt_vs_MS.NoTrt_B |
| EXOC3 | 0.00234 | -0.5978 | 0.142 | 0.24 | 1 | MS.Trt_vs_MS.NoTrt_B |
| FYCO1 | 0.00234 | -0.403 | 0.094 | 0.18 | 1 | MS.Trt_vs_MS.NoTrt_B |
| ATR | 0.00235 | 0.51553 | 0.377 | 0.52 | 1 | MS.Trt_vs_MS.NoTrt_B |
| SLC9A8 | 0.00235 | -0.3322 | 0.131 | 0.23 | 1 | MS.Trt_vs_MS.NoTrt_B |
| DCTN5 | 0.00236 | -0.3415 | 0.678 | 0.82 | 1 | MS.Trt_vs_MS.NoTrt_B |
| PPP1R21 | 0.00236 | -0.2569 | 0.168 | 0.27 | 1 | MS.Trt_vs_MS.NoTrt_B |
| AVL9 | 0.00238 | 0.35871 | 0.78 | 0.92 | 1 | MS.Trt_vs_MS.NoTrt_B |
| CCDC163 | 0.00238 | -0.8795 | 0.449 | 0.58 | 1 | MS.Trt_vs_MS.NoTrt_B |
| UTS2B | 0.00238 | 0.36427 | 0.155 | 0.27 | 1 | MS.Trt_vs_MS.NoTrt_B |
| EIF5A2 | 0.00241 | 0.56837 | 0.253 | 0.39 | 1 | MS.Trt_vs_MS.NoTrt_B |
| NPTN | 0.00242 | -0.4408 | 0.174 | 0.28 | 1 | MS.Trt_vs_MS.NoTrt_B |
| EP400NL | 0.00243 | -0.6768 | 0.627 | 0.72 | 1 | MS.Trt_vs_MS.NoTrt_B |
| FARP2 | 0.00243 | -0.8164 | 0.27 | 0.39 | 1 | MS.Trt_vs_MS.NoTrt_B |
| CGGBP1 | 0.00244 | 0.35029 | 0.124 | 0.22 | 1 | MS.Trt_vs_MS.NoTrt_B |
| KCNAB3 | 0.00245 | 0.60212 | 0.329 | 0.47 | 1 | MS.Trt_vs_MS.NoTrt_B |
| CCDC81 | 0.00245 | -0.556 | 0.338 | 0.47 | 1 | MS.Trt_vs_MS.NoTrt_B |
| ZNF350 | 0.00245 | -0.2756 | 0.466 | 0.61 | 1 | MS.Trt_vs_MS.NoTrt_B |
| KIAA1109 | 0.00246 | 0.42533 | 0.264 | 0.38 | 1 | MS.Trt_vs_MS.NoTrt_B |
| TMEM19 | 0.00246 | -0.5881 | 0.664 | 0.78 | 1 | MS.Trt_vs_MS.NoTrt_B |
| PLEKHA3 | 0.00246 | 0.45673 | 0.61 | 0.75 | 1 | MS.Trt_vs_MS.NoTrt_B |
| SLC38A3 | 0.00247 | -0.3778 | 0.163 | 0.27 | 1 | MS.Trt_vs_MS.NoTrt_B |
| MINK1 | 0.00247 | 0.41512 | 0.153 | 0.26 | 1 | MS.Trt_vs_MS.NoTrt_B |
| 8-Mar | 0.00247 | 0.61545 | 0.288 | 0.44 | 1 | MS.Trt_vs_MS.NoTrt_B |
| KIF27 | 0.00248 | 0.71893 | 0.24 | 0.38 | 1 | MS.Trt_vs_MS.NoTrt_B |
| CSTF2T | 0.00248 | -0.8201 | 0.052 | 0.12 | 1 | MS.Trt_vs_MS.NoTrt_B |
| MCTS1 | 0.00248 | -0.2778 | 0.756 | 0.86 | 1 | MS.Trt_vs_MS.NoTrt_B |
| ZBED1 | 0.00248 | -0.5661 | 0.381 | 0.5 | 1 | MS.Trt_vs_MS.NoTrt_B |
| IKZF1 | 0.00249 | 0.26901 | 0.292 | 0.43 | 1 | MS.Trt_vs_MS.NoTrt_B |
| ZNF33B | 0.00249 | -0.6196 | 0.403 | 0.55 | 1 | MS.Trt_vs_MS.NoTrt_B |
| KIF5B | 0.00249 | 0.33972 | 0.211 | 0.34 | 1 | MS.Trt_vs_MS.NoTrt_B |
| NEMF | 0.00251 | 0.76343 | 0.63 | 0.8 | 1 | MS.Trt_vs_MS.NoTrt_B |
| RAB40B | 0.00252 | 0.51924 | 0.32 | 0.45 | 1 | MS.Trt_vs_MS.NoTrt_B |
| C9orf142 | 0.00252 | -0.3065 | 0.176 | 0.28 | 1 | MS.Trt_vs_MS.NoTrt_B |
| RP11-345J4. | 0.00252 | 0.40324 | 0.617 | 0.75 | 1 | MS.Trt_vs_MS.NoTrt_B |

| | | | | | | |
|---|---|---|---|---|---|---|
| PCNX4 | 0.00253 | -0.3579 | 0.749 | 0.87 | 1 | MS.Trt_vs_MS.NoTrt_B |
| GRAP | 0.00253 | -0.4244 | 0.115 | 0.21 | 1 | MS.Trt_vs_MS.NoTrt_B |
| GALNT6 | 0.00255 | 0.4625 | 0.56 | 0.71 | 1 | MS.Trt_vs_MS.NoTrt_B |
| SLC30A6 | 0.00256 | 0.63634 | 0.497 | 0.63 | 1 | MS.Trt_vs_MS.NoTrt_B |
| SLC35A1 | 0.00257 | -0.4301 | 0.074 | 0.15 | 1 | MS.Trt_vs_MS.NoTrt_B |
| ZNF441 | 0.00257 | -1.2184 | 0.113 | 0.21 | 1 | MS.Trt_vs_MS.NoTrt_B |
| PPP1R3F | 0.00257 | 1.06159 | 0.094 | 0.18 | 1 | MS.Trt_vs_MS.NoTrt_B |
| RRP1 | 0.00257 | 0.53786 | 0.538 | 0.7 | 1 | MS.Trt_vs_MS.NoTrt_B |
| CYSTM1 | 0.00258 | -0.4293 | 0.192 | 0.3 | 1 | MS.Trt_vs_MS.NoTrt_B |
| COMTD1 | 0.00259 | -1.0713 | 0.057 | 0.12 | 1 | MS.Trt_vs_MS.NoTrt_B |
| SYNJ2BP | 0.00259 | 0.3274 | 0.728 | 0.82 | 1 | MS.Trt_vs_MS.NoTrt_B |
| POLD1 | 0.0026 | -0.4014 | 0.113 | 0.2 | 1 | MS.Trt_vs_MS.NoTrt_B |
| STAMBP | 0.0026 | -0.3223 | 0.094 | 0.18 | 1 | MS.Trt_vs_MS.NoTrt_B |
| PPP2R5D | 0.00261 | -0.5779 | 0.12 | 0.21 | 1 | MS.Trt_vs_MS.NoTrt_B |
| RABGEF1 | 0.00261 | -0.4224 | 0.214 | 0.32 | 1 | MS.Trt_vs_MS.NoTrt_B |
| KIAA0922 | 0.00262 | 0.34525 | 0.381 | 0.53 | 1 | MS.Trt_vs_MS.NoTrt_B |
| MTRF1 | 0.00263 | -0.3462 | 0.455 | 0.57 | 1 | MS.Trt_vs_MS.NoTrt_B |
| ARMC9 | 0.00264 | 0.43631 | 0.512 | 0.68 | 1 | MS.Trt_vs_MS.NoTrt_B |
| KCNV1 | 0.00264 | 0.40205 | 0.423 | 0.6 | 1 | MS.Trt_vs_MS.NoTrt_B |
| THTPA | 0.00264 | -1.028 | 0.057 | 0.12 | 1 | MS.Trt_vs_MS.NoTrt_B |
| UBE4A | 0.00266 | 0.38348 | 0.307 | 0.46 | 1 | MS.Trt_vs_MS.NoTrt_B |
| WDR76 | 0.00266 | -0.5996 | 0.547 | 0.69 | 1 | MS.Trt_vs_MS.NoTrt_B |
| ANGEL2 | 0.00266 | 0.68133 | 0.571 | 0.69 | 1 | MS.Trt_vs_MS.NoTrt_B |
| HIF1A | 0.00267 | -0.468 | 0.802 | 0.94 | 1 | MS.Trt_vs_MS.NoTrt_B |
| SCO1 | 0.00268 | -0.4821 | 0.66 | 0.8 | 1 | MS.Trt_vs_MS.NoTrt_B |
| ITGB7 | 0.00268 | -0.3815 | 0.264 | 0.37 | 1 | MS.Trt_vs_MS.NoTrt_B |
| BDP1 | 0.00269 | 0.43561 | 0.58 | 0.74 | 1 | MS.Trt_vs_MS.NoTrt_B |
| DARS | 0.0027 | 0.36309 | 0.235 | 0.37 | 1 | MS.Trt_vs_MS.NoTrt_B |
| BTLA | 0.0027 | -0.3191 | 0.098 | 0.18 | 1 | MS.Trt_vs_MS.NoTrt_B |
| THOC7 | 0.00271 | 0.25764 | 0.133 | 0.24 | 1 | MS.Trt_vs_MS.NoTrt_B |
| COX20 | 0.00271 | 0.43964 | 0.475 | 0.64 | 1 | MS.Trt_vs_MS.NoTrt_B |
| ATP11A | 0.00272 | 0.2974 | 0.187 | 0.3 | 1 | MS.Trt_vs_MS.NoTrt_B |
| KATNAL2 | 0.00273 | 0.6019 | 0.203 | 0.32 | 1 | MS.Trt_vs_MS.NoTrt_B |
| MYB | 0.00274 | -0.3135 | 0.209 | 0.33 | 1 | MS.Trt_vs_MS.NoTrt_B |
| CERKL | 0.00276 | 0.72184 | 0.275 | 0.43 | 1 | MS.Trt_vs_MS.NoTrt_B |
| TMEM120A | 0.00276 | -0.4225 | 0.087 | 0.17 | 1 | MS.Trt_vs_MS.NoTrt_B |
| MRPS11 | 0.00277 | -0.2745 | 0.562 | 0.7 | 1 | MS.Trt_vs_MS.NoTrt_B |
| KDM5B | 0.0028 | 0.64126 | 0.253 | 0.37 | 1 | MS.Trt_vs_MS.NoTrt_B |
| FAM160B1 | 0.00281 | -0.4256 | 0.142 | 0.24 | 1 | MS.Trt_vs_MS.NoTrt_B |
| NEBL | 0.00281 | 0.26669 | 0.196 | 0.32 | 1 | MS.Trt_vs_MS.NoTrt_B |

| | | | | | | |
|---|---|---|---|---|---|---|
| MAGED2 | 0.00283 | -0.5287 | 0.133 | 0.22 | 1 | MS.Trt_vs_MS.NoTrt_B |
| IL15RA | 0.00283 | -1.0395 | 0.061 | 0.13 | 1 | MS.Trt_vs_MS.NoTrt_B |
| MRTO4 | 0.00283 | -0.4719 | 0.392 | 0.55 | 1 | MS.Trt_vs_MS.NoTrt_B |
| GLCCI1 | 0.00283 | 0.31523 | 0.194 | 0.31 | 1 | MS.Trt_vs_MS.NoTrt_B |
| SLC43A1 | 0.00284 | -0.6853 | 0.068 | 0.14 | 1 | MS.Trt_vs_MS.NoTrt_B |
| RNF44 | 0.00284 | 0.62751 | 0.22 | 0.32 | 1 | MS.Trt_vs_MS.NoTrt_B |
| MTHFD2 | 0.00285 | 0.6384 | 0.527 | 0.66 | 1 | MS.Trt_vs_MS.NoTrt_B |
| GNL3 | 0.00285 | -0.8417 | 0.076 | 0.15 | 1 | MS.Trt_vs_MS.NoTrt_B |
| MORC2 | 0.00286 | -0.301 | 0.34 | 0.44 | 1 | MS.Trt_vs_MS.NoTrt_B |
| WDR73 | 0.00286 | -0.4454 | 0.917 | 0.98 | 1 | MS.Trt_vs_MS.NoTrt_B |
| FAM35A | 0.00287 | 0.36995 | 0.063 | 0.14 | 1 | MS.Trt_vs_MS.NoTrt_B |
| KYAT3 | 0.00287 | -0.3571 | 0.046 | 0.11 | 1 | MS.Trt_vs_MS.NoTrt_B |
| CBWD7 | 0.00288 | 0.41159 | 0.377 | 0.5 | 1 | MS.Trt_vs_MS.NoTrt_B |
| SLC35A5 | 0.00288 | 0.77982 | 0.301 | 0.43 | 1 | MS.Trt_vs_MS.NoTrt_B |
| NCOA3 | 0.00289 | 0.3681 | 0.632 | 0.78 | 1 | MS.Trt_vs_MS.NoTrt_B |
| HLA-DPA1 | 0.00289 | -0.325 | 0.166 | 0.27 | 1 | MS.Trt_vs_MS.NoTrt_B |
| AASDHPPT | 0.00289 | -0.4793 | 0.092 | 0.17 | 1 | MS.Trt_vs_MS.NoTrt_B |
| CASP3 | 0.00292 | -0.4782 | 0.072 | 0.15 | 1 | MS.Trt_vs_MS.NoTrt_B |
| ZNF148 | 0.00295 | 0.32676 | 0.19 | 0.3 | 1 | MS.Trt_vs_MS.NoTrt_B |
| PAPSS1 | 0.00296 | -0.318 | 0.102 | 0.19 | 1 | MS.Trt_vs_MS.NoTrt_B |
| SNX14 | 0.00296 | 0.26414 | 0.379 | 0.51 | 1 | MS.Trt_vs_MS.NoTrt_B |
| LAS1L | 0.00296 | -0.3874 | 0.109 | 0.2 | 1 | MS.Trt_vs_MS.NoTrt_B |
| IKZF4 | 0.00297 | 0.47397 | 0.12 | 0.21 | 1 | MS.Trt_vs_MS.NoTrt_B |
| CHCHD3 | 0.00297 | -0.4219 | 0.113 | 0.2 | 1 | MS.Trt_vs_MS.NoTrt_B |
| UBXN8 | 0.00298 | -0.6447 | 0.115 | 0.21 | 1 | MS.Trt_vs_MS.NoTrt_B |
| FKBP11 | 0.00302 | -0.336 | 0.553 | 0.7 | 1 | MS.Trt_vs_MS.NoTrt_B |
| GNL2 | 0.00305 | -0.7595 | 0.527 | 0.7 | 1 | MS.Trt_vs_MS.NoTrt_B |
| C15orf61 | 0.00305 | 0.53728 | 0.451 | 0.64 | 1 | MS.Trt_vs_MS.NoTrt_B |
| SIRT5 | 0.00307 | -0.4488 | 0.749 | 0.88 | 1 | MS.Trt_vs_MS.NoTrt_B |
| FKBP14 | 0.00309 | -0.3234 | 0.44 | 0.58 | 1 | MS.Trt_vs_MS.NoTrt_B |
| 5-Mar | 0.00311 | -0.3309 | 0.118 | 0.21 | 1 | MS.Trt_vs_MS.NoTrt_B |
| SKIDA1 | 0.00312 | -0.5351 | 0.277 | 0.41 | 1 | MS.Trt_vs_MS.NoTrt_B |
| UNC13D | 0.00313 | -0.3329 | 0.508 | 0.62 | 1 | MS.Trt_vs_MS.NoTrt_B |
| MAST3 | 0.00314 | 0.36876 | 0.617 | 0.75 | 1 | MS.Trt_vs_MS.NoTrt_B |
| DUSP10 | 0.00315 | -0.4058 | 0.1 | 0.18 | 1 | MS.Trt_vs_MS.NoTrt_B |
| R3HDM2 | 0.00315 | 0.52753 | 0.756 | 0.88 | 1 | MS.Trt_vs_MS.NoTrt_B |
| RNF24 | 0.00315 | -0.2915 | 0.423 | 0.56 | 1 | MS.Trt_vs_MS.NoTrt_B |
| AGAP4 | 0.00317 | 1.26549 | 0.174 | 0.29 | 1 | MS.Trt_vs_MS.NoTrt_B |
| MX2 | 0.00318 | -0.3491 | 0.773 | 0.92 | 1 | MS.Trt_vs_MS.NoTrt_B |
| ACBD4 | 0.00319 | -0.9427 | 0.246 | 0.35 | 1 | MS.Trt_vs_MS.NoTrt_B |

| | | | | | | |
|---|---|---|---|---|---|---|
| EXOSC3 | 0.00319 | 0.5299 | 0.298 | 0.44 | 1 | MS.Trt_vs_MS.NoTrt_B |
| CHMP1A | 0.0032 | -0.476 | 0.155 | 0.25 | 1 | MS.Trt_vs_MS.NoTrt_B |
| AMPD3 | 0.00322 | -1.1664 | 0.316 | 0.45 | 1 | MS.Trt_vs_MS.NoTrt_B |
| FANK1 | 0.00323 | -0.2513 | 0.107 | 0.19 | 1 | MS.Trt_vs_MS.NoTrt_B |
| ZDHHC12 | 0.00324 | -0.6841 | 0.085 | 0.16 | 1 | MS.Trt_vs_MS.NoTrt_B |
| ACAD9 | 0.00324 | -0.4216 | 0.235 | 0.34 | 1 | MS.Trt_vs_MS.NoTrt_B |
| ETHE1 | 0.00324 | -0.6863 | 0.05 | 0.11 | 1 | MS.Trt_vs_MS.NoTrt_B |
| CRYZ | 0.00326 | 0.33436 | 0.068 | 0.14 | 1 | MS.Trt_vs_MS.NoTrt_B |
| MLST8 | 0.00328 | -0.3923 | 0.277 | 0.4 | 1 | MS.Trt_vs_MS.NoTrt_B |
| SPAG16 | 0.00329 | 0.38357 | 0.309 | 0.44 | 1 | MS.Trt_vs_MS.NoTrt_B |
| SFT2D3 | 0.00329 | 0.26665 | 0.336 | 0.47 | 1 | MS.Trt_vs_MS.NoTrt_B |
| HABP4 | 0.0033 | 0.83232 | 0.068 | 0.14 | 1 | MS.Trt_vs_MS.NoTrt_B |
| TRIM11 | 0.00333 | -0.2775 | 0.425 | 0.55 | 1 | MS.Trt_vs_MS.NoTrt_B |
| MAT2A | 0.00334 | -0.2847 | 0.087 | 0.17 | 1 | MS.Trt_vs_MS.NoTrt_B |
| ZNF569 | 0.00334 | 1.10291 | 0.126 | 0.22 | 1 | MS.Trt_vs_MS.NoTrt_B |
| CNNM3 | 0.00334 | 0.46559 | 0.492 | 0.63 | 1 | MS.Trt_vs_MS.NoTrt_B |
| PTER | 0.00334 | -0.6247 | 0.107 | 0.19 | 1 | MS.Trt_vs_MS.NoTrt_B |
| PTBP3 | 0.00335 | 0.72029 | 0.235 | 0.36 | 1 | MS.Trt_vs_MS.NoTrt_B |
| PPIP5K2 | 0.00335 | 0.47146 | 0.671 | 0.86 | 1 | MS.Trt_vs_MS.NoTrt_B |
| RTEL1-TNFR | 0.0034 | -0.3564 | 0.046 | 0.11 | 1 | MS.Trt_vs_MS.NoTrt_B |
| ACADS | 0.00342 | -0.9398 | 0.05 | 0.11 | 1 | MS.Trt_vs_MS.NoTrt_B |
| NHEJ1 | 0.00342 | -1.7969 | 0.111 | 0.2 | 1 | MS.Trt_vs_MS.NoTrt_B |
| RBM14 | 0.00346 | -0.3136 | 0.135 | 0.23 | 1 | MS.Trt_vs_MS.NoTrt_B |
| CSNK1G3 | 0.00346 | -0.4409 | 0.046 | 0.11 | 1 | MS.Trt_vs_MS.NoTrt_B |
| KLHL18 | 0.00346 | -0.8697 | 0.279 | 0.39 | 1 | MS.Trt_vs_MS.NoTrt_B |
| USB1 | 0.00347 | -0.3618 | 0.072 | 0.15 | 1 | MS.Trt_vs_MS.NoTrt_B |
| DDIT4 | 0.00347 | -0.7217 | 0.12 | 0.21 | 1 | MS.Trt_vs_MS.NoTrt_B |
| AZIN2 | 0.00348 | 1.02812 | 0.183 | 0.29 | 1 | MS.Trt_vs_MS.NoTrt_B |
| SYNJ2 | 0.00349 | -0.3198 | 0.789 | 0.89 | 1 | MS.Trt_vs_MS.NoTrt_B |
| MRPL46 | 0.00351 | 0.32104 | 0.183 | 0.29 | 1 | MS.Trt_vs_MS.NoTrt_B |
| ATF7 | 0.00352 | 0.42967 | 0.353 | 0.5 | 1 | MS.Trt_vs_MS.NoTrt_B |
| TGFBR3 | 0.00352 | 0.25084 | 0.34 | 0.47 | 1 | MS.Trt_vs_MS.NoTrt_B |
| SEMA4D | 0.00354 | 0.31273 | 0.375 | 0.52 | 1 | MS.Trt_vs_MS.NoTrt_B |
| SEMA3B | 0.00354 | 0.32315 | 0.157 | 0.26 | 1 | MS.Trt_vs_MS.NoTrt_B |
| AD000671.6 | 0.00355 | -0.3693 | 0.061 | 0.13 | 1 | MS.Trt_vs_MS.NoTrt_B |
| INO80E | 0.00356 | 0.49196 | 0.131 | 0.23 | 1 | MS.Trt_vs_MS.NoTrt_B |
| CLU | 0.00356 | -0.6082 | 0.355 | 0.48 | 1 | MS.Trt_vs_MS.NoTrt_B |
| STX5 | 0.00356 | -0.9362 | 0.044 | 0.1 | 1 | MS.Trt_vs_MS.NoTrt_B |
| C10orf76 | 0.00356 | -0.3222 | 0.218 | 0.34 | 1 | MS.Trt_vs_MS.NoTrt_B |
| CEP164 | 0.00357 | -0.8836 | 0.092 | 0.17 | 1 | MS.Trt_vs_MS.NoTrt_B |

| | | | | | | |
|---|---|---|---|---|---|---|
| LEKR1 | 0.00358 | -0.3583 | 0.115 | 0.21 | 1 | MS.Trt_vs_MS.NoTrt_B |
| THAP4 | 0.00359 | -0.2789 | 0.085 | 0.16 | 1 | MS.Trt_vs_MS.NoTrt_B |
| CLK4 | 0.00359 | 0.40847 | 0.551 | 0.69 | 1 | MS.Trt_vs_MS.NoTrt_B |
| PIKFYVE | 0.00359 | 0.34813 | 0.303 | 0.42 | 1 | MS.Trt_vs_MS.NoTrt_B |
| QSOX1 | 0.0036 | -0.4731 | 0.253 | 0.36 | 1 | MS.Trt_vs_MS.NoTrt_B |
| CD2BP2 | 0.00361 | -0.3623 | 0.113 | 0.2 | 1 | MS.Trt_vs_MS.NoTrt_B |
| VOPP1 | 0.00361 | -0.4004 | 0.227 | 0.33 | 1 | MS.Trt_vs_MS.NoTrt_B |
| CA5B | 0.00361 | 0.37711 | 0.734 | 0.89 | 1 | MS.Trt_vs_MS.NoTrt_B |
| INSIG1 | 0.00362 | 0.35148 | 0.098 | 0.18 | 1 | MS.Trt_vs_MS.NoTrt_B |
| USP53 | 0.00362 | 0.28681 | 0.425 | 0.56 | 1 | MS.Trt_vs_MS.NoTrt_B |
| SMARCC1 | 0.00363 | -0.4113 | 0.364 | 0.48 | 1 | MS.Trt_vs_MS.NoTrt_B |
| UBLCP1 | 0.00365 | -0.5455 | 0.124 | 0.21 | 1 | MS.Trt_vs_MS.NoTrt_B |
| NCOR1 | 0.00366 | 0.25344 | 0.943 | 1 | 1 | MS.Trt_vs_MS.NoTrt_B |
| GTF3C2 | 0.00366 | -0.2741 | 0.41 | 0.56 | 1 | MS.Trt_vs_MS.NoTrt_B |
| POMT1 | 0.00366 | -0.8006 | 0.436 | 0.58 | 1 | MS.Trt_vs_MS.NoTrt_B |
| DEPDC1B | 0.00367 | -0.2831 | 0.146 | 0.24 | 1 | MS.Trt_vs_MS.NoTrt_B |
| LYRM1 | 0.00368 | 0.76692 | 0.416 | 0.57 | 1 | MS.Trt_vs_MS.NoTrt_B |
| CBLL1 | 0.00369 | 0.30949 | 0.246 | 0.38 | 1 | MS.Trt_vs_MS.NoTrt_B |
| AMZ2 | 0.0037 | 0.3756 | 0.41 | 0.55 | 1 | MS.Trt_vs_MS.NoTrt_B |
| BICDL1 | 0.0037 | 0.63707 | 0.464 | 0.57 | 1 | MS.Trt_vs_MS.NoTrt_B |
| ALMS1 | 0.00371 | -0.694 | 0.342 | 0.46 | 1 | MS.Trt_vs_MS.NoTrt_B |
| CXorf21 | 0.00371 | -1.1269 | 0.227 | 0.33 | 1 | MS.Trt_vs_MS.NoTrt_B |
| ITGAE | 0.00372 | 0.32072 | 0.434 | 0.6 | 1 | MS.Trt_vs_MS.NoTrt_B |
| ZNF524 | 0.00372 | -0.3196 | 0.061 | 0.13 | 1 | MS.Trt_vs_MS.NoTrt_B |
| SAP18 | 0.00372 | 0.27082 | 0.316 | 0.45 | 1 | MS.Trt_vs_MS.NoTrt_B |
| SETX | 0.00373 | 0.41706 | 0.22 | 0.35 | 1 | MS.Trt_vs_MS.NoTrt_B |
| KTN1 | 0.00378 | 0.84388 | 0.449 | 0.67 | 1 | MS.Trt_vs_MS.NoTrt_B |
| EEA1 | 0.00381 | -0.3365 | 0.155 | 0.25 | 1 | MS.Trt_vs_MS.NoTrt_B |
| MALSU1 | 0.00382 | -0.3927 | 0.085 | 0.16 | 1 | MS.Trt_vs_MS.NoTrt_B |
| XXYLT1 | 0.00382 | -0.7181 | 0.159 | 0.25 | 1 | MS.Trt_vs_MS.NoTrt_B |
| RCN1 | 0.00383 | -0.8964 | 0.078 | 0.15 | 1 | MS.Trt_vs_MS.NoTrt_B |
| NBPF26 | 0.00384 | 0.25021 | 0.218 | 0.32 | 1 | MS.Trt_vs_MS.NoTrt_B |
| PFKL | 0.00386 | -0.369 | 0.658 | 0.73 | 1 | MS.Trt_vs_MS.NoTrt_B |
| KDM7A | 0.00387 | 0.28344 | 0.139 | 0.23 | 1 | MS.Trt_vs_MS.NoTrt_B |
| NET1 | 0.00388 | -0.4166 | 0.054 | 0.12 | 1 | MS.Trt_vs_MS.NoTrt_B |
| MRPL39 | 0.0039 | -0.2593 | 0.078 | 0.15 | 1 | MS.Trt_vs_MS.NoTrt_B |
| PIGA | 0.00391 | 0.37949 | 0.316 | 0.43 | 1 | MS.Trt_vs_MS.NoTrt_B |
| RNF41 | 0.00391 | 0.34492 | 0.606 | 0.75 | 1 | MS.Trt_vs_MS.NoTrt_B |
| KLHL12 | 0.00395 | -1.0176 | 0.048 | 0.11 | 1 | MS.Trt_vs_MS.NoTrt_B |
| TTN | 0.00396 | 0.42963 | 0.532 | 0.69 | 1 | MS.Trt_vs_MS.NoTrt_B |

| | | | | | | |
|---|---|---|---|---|---|---|
| ZNF365 | 0.00396 | -0.5862 | 0.229 | 0.35 | 1 | MS.Trt_vs_MS.NoTrt_B |
| TMEM222 | 0.00397 | -0.8107 | 0.078 | 0.15 | 1 | MS.Trt_vs_MS.NoTrt_B |
| OGFOD1 | 0.00397 | 0.44983 | 0.529 | 0.68 | 1 | MS.Trt_vs_MS.NoTrt_B |
| RNF212 | 0.00397 | -0.3924 | 0.427 | 0.58 | 1 | MS.Trt_vs_MS.NoTrt_B |
| YTHDC1 | 0.00399 | 0.25921 | 0.359 | 0.48 | 1 | MS.Trt_vs_MS.NoTrt_B |
| LMNA | 0.004 | -0.3942 | 0.451 | 0.56 | 1 | MS.Trt_vs_MS.NoTrt_B |
| CNTNAP1 | 0.004 | -0.3559 | 0.351 | 0.48 | 1 | MS.Trt_vs_MS.NoTrt_B |
| KATNA1 | 0.004 | -0.707 | 0.059 | 0.12 | 1 | MS.Trt_vs_MS.NoTrt_B |
| CDCA3 | 0.004 | -0.8344 | 0.455 | 0.57 | 1 | MS.Trt_vs_MS.NoTrt_B |
| FBXO24 | 0.00401 | -0.3859 | 0.294 | 0.43 | 1 | MS.Trt_vs_MS.NoTrt_B |
| CNNM2 | 0.00401 | -0.3874 | 0.207 | 0.31 | 1 | MS.Trt_vs_MS.NoTrt_B |
| SLC41A2 | 0.00404 | -0.3763 | 0.179 | 0.28 | 1 | MS.Trt_vs_MS.NoTrt_B |
| EPHA4 | 0.00404 | 0.33529 | 0.133 | 0.23 | 1 | MS.Trt_vs_MS.NoTrt_B |
| BCAT2 | 0.00406 | 0.40552 | 0.118 | 0.21 | 1 | MS.Trt_vs_MS.NoTrt_B |
| RCAN1 | 0.00406 | 0.26823 | 0.211 | 0.32 | 1 | MS.Trt_vs_MS.NoTrt_B |
| UBR4 | 0.00406 | 0.26531 | 0.608 | 0.78 | 1 | MS.Trt_vs_MS.NoTrt_B |
| SEC24C | 0.00406 | -0.4531 | 0.17 | 0.26 | 1 | MS.Trt_vs_MS.NoTrt_B |
| YPEL2 | 0.00407 | -0.368 | 0.142 | 0.23 | 1 | MS.Trt_vs_MS.NoTrt_B |
| SMARCA5 | 0.00408 | 0.63769 | 0.27 | 0.4 | 1 | MS.Trt_vs_MS.NoTrt_B |
| FBXO15 | 0.00409 | 0.25491 | 0.458 | 0.58 | 1 | MS.Trt_vs_MS.NoTrt_B |
| RUSC1 | 0.00411 | -0.9794 | 0.044 | 0.1 | 1 | MS.Trt_vs_MS.NoTrt_B |
| ADARB2 | 0.00411 | 0.56308 | 0.216 | 0.35 | 1 | MS.Trt_vs_MS.NoTrt_B |
| TPK1 | 0.00413 | 0.43868 | 0.253 | 0.37 | 1 | MS.Trt_vs_MS.NoTrt_B |
| SLC16A10 | 0.00413 | 0.33311 | 0.704 | 0.81 | 1 | MS.Trt_vs_MS.NoTrt_B |
| GTPBP8 | 0.00416 | -0.3847 | 0.545 | 0.66 | 1 | MS.Trt_vs_MS.NoTrt_B |
| CHMP5 | 0.00416 | -0.5142 | 0.078 | 0.15 | 1 | MS.Trt_vs_MS.NoTrt_B |
| MAP1LC3B | 0.00416 | 0.47706 | 0.272 | 0.4 | 1 | MS.Trt_vs_MS.NoTrt_B |
| ST3GAL3 | 0.00417 | -0.4907 | 0.654 | 0.82 | 1 | MS.Trt_vs_MS.NoTrt_B |
| HERPUD2 | 0.00417 | 0.2826 | 0.375 | 0.49 | 1 | MS.Trt_vs_MS.NoTrt_B |
| TRAPPC6A | 0.00421 | -0.4121 | 0.126 | 0.22 | 1 | MS.Trt_vs_MS.NoTrt_B |
| JKAMP | 0.00421 | 0.2676 | 0.678 | 0.81 | 1 | MS.Trt_vs_MS.NoTrt_B |
| MAN1B1 | 0.00422 | -0.5304 | 0.264 | 0.37 | 1 | MS.Trt_vs_MS.NoTrt_B |
| VWA9 | 0.00422 | -0.4974 | 0.059 | 0.12 | 1 | MS.Trt_vs_MS.NoTrt_B |
| C10orf95 | 0.00422 | -0.3648 | 0.179 | 0.27 | 1 | MS.Trt_vs_MS.NoTrt_B |
| COPS3 | 0.00423 | -0.2547 | 0.198 | 0.29 | 1 | MS.Trt_vs_MS.NoTrt_B |
| KDM2A | 0.00423 | 0.29591 | 0.778 | 0.88 | 1 | MS.Trt_vs_MS.NoTrt_B |
| HSD17B4 | 0.00423 | 0.44535 | 0.477 | 0.63 | 1 | MS.Trt_vs_MS.NoTrt_B |
| SETD7 | 0.00424 | -0.391 | 0.728 | 0.84 | 1 | MS.Trt_vs_MS.NoTrt_B |
| CLDND1 | 0.00425 | 0.39646 | 0.27 | 0.4 | 1 | MS.Trt_vs_MS.NoTrt_B |
| ACAA1 | 0.00425 | -0.4751 | 0.325 | 0.45 | 1 | MS.Trt_vs_MS.NoTrt_B |

| | | | | | | |
|---|---|---|---|---|---|---|
| LSS | 0.00425 | -0.3614 | 0.07 | 0.14 | 1 | MS.Trt_vs_MS.NoTrt_B |
| KDELC2 | 0.00426 | -0.8088 | 0.442 | 0.55 | 1 | MS.Trt_vs_MS.NoTrt_B |
| NDUFAF6 | 0.00427 | 0.3562 | 0.484 | 0.63 | 1 | MS.Trt_vs_MS.NoTrt_B |
| KIAA0907 | 0.00427 | 0.63969 | 0.632 | 0.78 | 1 | MS.Trt_vs_MS.NoTrt_B |
| FRA10AC1 | 0.00427 | 1.0311 | 0.423 | 0.61 | 1 | MS.Trt_vs_MS.NoTrt_B |
| COG2 | 0.0043 | -0.349 | 0.275 | 0.38 | 1 | MS.Trt_vs_MS.NoTrt_B |
| NCOA6 | 0.00431 | 0.42789 | 0.551 | 0.69 | 1 | MS.Trt_vs_MS.NoTrt_B |
| AIMP2 | 0.00432 | -0.9793 | 0.257 | 0.36 | 1 | MS.Trt_vs_MS.NoTrt_B |
| KDM4B | 0.00432 | -0.2799 | 0.49 | 0.6 | 1 | MS.Trt_vs_MS.NoTrt_B |
| PSKH1 | 0.00434 | -0.4287 | 0.065 | 0.14 | 1 | MS.Trt_vs_MS.NoTrt_B |
| NAT10 | 0.00435 | -0.6614 | 0.251 | 0.36 | 1 | MS.Trt_vs_MS.NoTrt_B |
| C19orf48 | 0.00436 | -1.1296 | 0.281 | 0.4 | 1 | MS.Trt_vs_MS.NoTrt_B |
| TOR2A | 0.00436 | -0.735 | 0.044 | 0.1 | 1 | MS.Trt_vs_MS.NoTrt_B |
| POLR3E | 0.00442 | 0.40048 | 0.386 | 0.54 | 1 | MS.Trt_vs_MS.NoTrt_B |
| EXO1 | 0.00447 | -1.1756 | 0.109 | 0.2 | 1 | MS.Trt_vs_MS.NoTrt_B |
| MIPOL1 | 0.00447 | 0.35758 | 0.974 | 0.98 | 1 | MS.Trt_vs_MS.NoTrt_B |
| C1R | 0.00447 | 0.40779 | 0.242 | 0.38 | 1 | MS.Trt_vs_MS.NoTrt_B |
| ZNF557 | 0.00449 | 0.51511 | 0.758 | 0.86 | 1 | MS.Trt_vs_MS.NoTrt_B |
| ARNT | 0.00449 | -0.6147 | 0.63 | 0.77 | 1 | MS.Trt_vs_MS.NoTrt_B |
| ANKRD27 | 0.0045 | 0.36064 | 0.131 | 0.22 | 1 | MS.Trt_vs_MS.NoTrt_B |
| UTP18 | 0.00451 | -0.5913 | 0.115 | 0.2 | 1 | MS.Trt_vs_MS.NoTrt_B |
| HAAO | 0.00452 | -0.7116 | 0.054 | 0.12 | 1 | MS.Trt_vs_MS.NoTrt_B |
| PQLC2 | 0.00452 | -0.704 | 0.32 | 0.45 | 1 | MS.Trt_vs_MS.NoTrt_B |
| ARHGAP31 | 0.00453 | 0.30546 | 0.176 | 0.29 | 1 | MS.Trt_vs_MS.NoTrt_B |
| RPL7 | 0.00455 | 0.36094 | 0.904 | 1 | 1 | MS.Trt_vs_MS.NoTrt_B |
| PPIF | 0.00455 | -0.541 | 0.044 | 0.1 | 1 | MS.Trt_vs_MS.NoTrt_B |
| SEC14L1 | 0.00456 | -0.6642 | 0.527 | 0.67 | 1 | MS.Trt_vs_MS.NoTrt_B |
| CEP104 | 0.00458 | 0.2638 | 0.564 | 0.65 | 1 | MS.Trt_vs_MS.NoTrt_B |
| CDC14B | 0.00458 | -0.9446 | 0.179 | 0.29 | 1 | MS.Trt_vs_MS.NoTrt_B |
| PITPNA | 0.00459 | -0.7384 | 0.059 | 0.12 | 1 | MS.Trt_vs_MS.NoTrt_B |
| FBXO5 | 0.00459 | 0.57079 | 0.268 | 0.4 | 1 | MS.Trt_vs_MS.NoTrt_B |
| DTYMK | 0.00459 | 0.30102 | 0.429 | 0.55 | 1 | MS.Trt_vs_MS.NoTrt_B |
| JMJD6 | 0.0046 | -0.6401 | 0.322 | 0.44 | 1 | MS.Trt_vs_MS.NoTrt_B |
| TRERF1 | 0.0046 | -0.2628 | 0.135 | 0.23 | 1 | MS.Trt_vs_MS.NoTrt_B |
| MUT | 0.0046 | -0.4826 | 0.044 | 0.1 | 1 | MS.Trt_vs_MS.NoTrt_B |
| FCGRT | 0.00461 | -0.3874 | 0.139 | 0.23 | 1 | MS.Trt_vs_MS.NoTrt_B |
| NABP2 | 0.00462 | -0.8274 | 0.368 | 0.49 | 1 | MS.Trt_vs_MS.NoTrt_B |
| PARPBP | 0.00462 | 0.68429 | 0.257 | 0.38 | 1 | MS.Trt_vs_MS.NoTrt_B |
| SLC25A11 | 0.00466 | -0.3341 | 0.102 | 0.18 | 1 | MS.Trt_vs_MS.NoTrt_B |
| TRMT1L | 0.0047 | -0.4477 | 0.068 | 0.14 | 1 | MS.Trt_vs_MS.NoTrt_B |

| | | | | | | |
|---|---|---|---|---|---|---|
| SMG8 | 0.00471 | -0.5896 | 0.063 | 0.13 | 1 | MS.Trt_vs_MS.NoTrt_B |
| ACYP1 | 0.00472 | 0.26153 | 0.168 | 0.27 | 1 | MS.Trt_vs_MS.NoTrt_B |
| VPS72 | 0.00472 | 0.30137 | 0.089 | 0.17 | 1 | MS.Trt_vs_MS.NoTrt_B |
| WDR18 | 0.00472 | -0.4741 | 0.063 | 0.13 | 1 | MS.Trt_vs_MS.NoTrt_B |
| ALG13 | 0.00473 | 0.30973 | 0.19 | 0.3 | 1 | MS.Trt_vs_MS.NoTrt_B |
| ENO2 | 0.00473 | -0.4748 | 0.174 | 0.27 | 1 | MS.Trt_vs_MS.NoTrt_B |
| AP4E1 | 0.00475 | 0.7327 | 0.264 | 0.38 | 1 | MS.Trt_vs_MS.NoTrt_B |
| PRKAA1 | 0.00475 | 0.31705 | 0.719 | 0.9 | 1 | MS.Trt_vs_MS.NoTrt_B |
| ARFGAP2 | 0.00475 | -0.2552 | 0.455 | 0.6 | 1 | MS.Trt_vs_MS.NoTrt_B |
| ASB6 | 0.00476 | -0.8921 | 0.436 | 0.54 | 1 | MS.Trt_vs_MS.NoTrt_B |
| CLPX | 0.00476 | 0.56586 | 0.569 | 0.73 | 1 | MS.Trt_vs_MS.NoTrt_B |
| FAM47E-STE | 0.00476 | 0.49338 | 0.115 | 0.21 | 1 | MS.Trt_vs_MS.NoTrt_B |
| TBXAS1 | 0.00477 | -0.35 | 0.292 | 0.42 | 1 | MS.Trt_vs_MS.NoTrt_B |
| CARD19 | 0.00478 | -0.2549 | 0.205 | 0.31 | 1 | MS.Trt_vs_MS.NoTrt_B |
| DAGLB | 0.0048 | 0.26517 | 0.654 | 0.84 | 1 | MS.Trt_vs_MS.NoTrt_B |
| MAGED1 | 0.0048 | -0.4596 | 0.571 | 0.69 | 1 | MS.Trt_vs_MS.NoTrt_B |
| NCBP1 | 0.00484 | 0.58687 | 0.556 | 0.71 | 1 | MS.Trt_vs_MS.NoTrt_B |
| COMMD6 | 0.00484 | 0.67633 | 0.34 | 0.53 | 1 | MS.Trt_vs_MS.NoTrt_B |
| CWF19L1 | 0.00487 | -0.4484 | 0.699 | 0.83 | 1 | MS.Trt_vs_MS.NoTrt_B |
| PLS3 | 0.00489 | 0.4617 | 0.283 | 0.43 | 1 | MS.Trt_vs_MS.NoTrt_B |
| C5orf63 | 0.00489 | -0.2895 | 0.257 | 0.37 | 1 | MS.Trt_vs_MS.NoTrt_B |
| PRPSAP1 | 0.00491 | -0.3648 | 0.575 | 0.71 | 1 | MS.Trt_vs_MS.NoTrt_B |
| CCDC126 | 0.00492 | 1.20824 | 0.072 | 0.15 | 1 | MS.Trt_vs_MS.NoTrt_B |
| SULT1A1 | 0.00492 | -0.3196 | 0.464 | 0.6 | 1 | MS.Trt_vs_MS.NoTrt_B |
| FAM3A | 0.00495 | 0.41155 | 0.638 | 0.84 | 1 | MS.Trt_vs_MS.NoTrt_B |
| EHD4 | 0.00495 | -0.3529 | 0.309 | 0.41 | 1 | MS.Trt_vs_MS.NoTrt_B |
| SPA17 | 0.00496 | -0.4016 | 0.153 | 0.25 | 1 | MS.Trt_vs_MS.NoTrt_B |
| RNF135 | 0.00498 | -0.7332 | 0.174 | 0.28 | 1 | MS.Trt_vs_MS.NoTrt_B |
| PPT1 | 0.00502 | 0.25051 | 0.17 | 0.27 | 1 | MS.Trt_vs_MS.NoTrt_B |
| MED31 | 0.00502 | 0.3836 | 0.076 | 0.15 | 1 | MS.Trt_vs_MS.NoTrt_B |
| RBBP8 | 0.00503 | 1.25321 | 0.05 | 0.11 | 1 | MS.Trt_vs_MS.NoTrt_B |
| DBN1 | 0.00505 | 0.27586 | 0.209 | 0.12 | 1 | MS.Trt_vs_MS.NoTrt_B |
| ITPKB | 0.00506 | 0.61231 | 0.423 | 0.59 | 1 | MS.Trt_vs_MS.NoTrt_B |
| GRAMD1B | 0.00507 | 0.32358 | 0.292 | 0.41 | 1 | MS.Trt_vs_MS.NoTrt_B |
| CEBPZ | 0.00509 | 0.31657 | 0.17 | 0.28 | 1 | MS.Trt_vs_MS.NoTrt_B |
| RUNX2 | 0.00509 | -0.3369 | 0.124 | 0.21 | 1 | MS.Trt_vs_MS.NoTrt_B |
| HSPB1 | 0.00511 | 0.32384 | 0.257 | 0.38 | 1 | MS.Trt_vs_MS.NoTrt_B |
| STK38L | 0.00512 | -0.5102 | 0.593 | 0.7 | 1 | MS.Trt_vs_MS.NoTrt_B |
| C2orf69 | 0.00514 | -0.3187 | 0.163 | 0.26 | 1 | MS.Trt_vs_MS.NoTrt_B |
| EIF2S3 | 0.00514 | 0.67727 | 0.943 | 0.95 | 1 | MS.Trt_vs_MS.NoTrt_B |

| | | | | | | |
|---|---|---|---|---|---|---|
| GM2A | 0.00517 | -1.0239 | 0.462 | 0.57 | 1 | MS.Trt_vs_MS.NoTrt_B |
| NPHP3 | 0.00518 | 0.61188 | 0.754 | 0.91 | 1 | MS.Trt_vs_MS.NoTrt_B |
| WNK1 | 0.00519 | 0.38017 | 0.41 | 0.57 | 1 | MS.Trt_vs_MS.NoTrt_B |
| CARD17 | 0.00519 | -0.3729 | 0.048 | 0.11 | 1 | MS.Trt_vs_MS.NoTrt_B |
| GCNT4 | 0.00521 | 1.45505 | 0.501 | 0.43 | 1 | MS.Trt_vs_MS.NoTrt_B |
| PCMTD1 | 0.00522 | 0.49916 | 0.224 | 0.35 | 1 | MS.Trt_vs_MS.NoTrt_B |
| KIAA0368 | 0.00522 | -0.4798 | 0.148 | 0.24 | 1 | MS.Trt_vs_MS.NoTrt_B |
| RNF20 | 0.00524 | -0.7602 | 0.061 | 0.12 | 1 | MS.Trt_vs_MS.NoTrt_B |
| UBE2W | 0.00524 | 0.62832 | 0.499 | 0.64 | 1 | MS.Trt_vs_MS.NoTrt_B |
| BLOC1S3 | 0.00524 | 0.29066 | 0.464 | 0.61 | 1 | MS.Trt_vs_MS.NoTrt_B |
| WDR45B | 0.00525 | -0.4006 | 0.333 | 0.45 | 1 | MS.Trt_vs_MS.NoTrt_B |
| NIT2 | 0.00527 | 0.30068 | 0.516 | 0.69 | 1 | MS.Trt_vs_MS.NoTrt_B |
| TMEM265 | 0.00528 | -0.4223 | 0.048 | 0.11 | 1 | MS.Trt_vs_MS.NoTrt_B |
| ZNF292 | 0.00528 | 0.55874 | 0.728 | 0.86 | 1 | MS.Trt_vs_MS.NoTrt_B |
| HOMER2 | 0.00528 | 0.7955 | 0.455 | 0.58 | 1 | MS.Trt_vs_MS.NoTrt_B |
| ASXL2 | 0.00528 | 0.30126 | 0.113 | 0.2 | 1 | MS.Trt_vs_MS.NoTrt_B |
| RABIF | 0.0053 | -0.2818 | 0.257 | 0.37 | 1 | MS.Trt_vs_MS.NoTrt_B |
| CCDC66 | 0.00531 | 0.66883 | 0.505 | 0.65 | 1 | MS.Trt_vs_MS.NoTrt_B |
| RPF2 | 0.00532 | -0.3344 | 0.19 | 0.29 | 1 | MS.Trt_vs_MS.NoTrt_B |
| TTC1 | 0.00532 | 0.39185 | 0.159 | 0.25 | 1 | MS.Trt_vs_MS.NoTrt_B |
| ZNF141 | 0.00533 | -0.3393 | 0.919 | 0.98 | 1 | MS.Trt_vs_MS.NoTrt_B |
| DCAF13 | 0.00533 | 0.29015 | 0.503 | 0.62 | 1 | MS.Trt_vs_MS.NoTrt_B |
| UTP15 | 0.00533 | -0.3934 | 0.279 | 0.41 | 1 | MS.Trt_vs_MS.NoTrt_B |
| CRIP2 | 0.00534 | -0.7126 | 0.096 | 0.17 | 1 | MS.Trt_vs_MS.NoTrt_B |
| ALOX12 | 0.00534 | -0.4041 | 0.048 | 0.11 | 1 | MS.Trt_vs_MS.NoTrt_B |
| GRK3 | 0.00535 | -0.3038 | 0.246 | 0.34 | 1 | MS.Trt_vs_MS.NoTrt_B |
| RGP1 | 0.00535 | -0.5919 | 0.087 | 0.16 | 1 | MS.Trt_vs_MS.NoTrt_B |
| EIF3E | 0.00536 | 0.52587 | 0.455 | 0.62 | 1 | MS.Trt_vs_MS.NoTrt_B |
| TXNDC15 | 0.00538 | -0.2923 | 0.514 | 0.64 | 1 | MS.Trt_vs_MS.NoTrt_B |
| TRAF3IP2 | 0.0054 | -0.8346 | 0.767 | 0.86 | 1 | MS.Trt_vs_MS.NoTrt_B |
| CSRP1 | 0.0054 | -0.5331 | 0.42 | 0.52 | 1 | MS.Trt_vs_MS.NoTrt_B |
| BAG6 | 0.00543 | 0.43005 | 0.444 | 0.56 | 1 | MS.Trt_vs_MS.NoTrt_B |
| L3MBTL2 | 0.00545 | -0.2643 | 0.597 | 0.75 | 1 | MS.Trt_vs_MS.NoTrt_B |
| FBXO48 | 0.00547 | 0.52563 | 0.534 | 0.68 | 1 | MS.Trt_vs_MS.NoTrt_B |
| RBM27 | 0.00549 | 0.49589 | 0.087 | 0.16 | 1 | MS.Trt_vs_MS.NoTrt_B |
| IQCF3 | 0.00549 | 0.29201 | 0.2 | 0.33 | 1 | MS.Trt_vs_MS.NoTrt_B |
| RALGAPB | 0.00551 | -0.2841 | 0.192 | 0.29 | 1 | MS.Trt_vs_MS.NoTrt_B |
| CORO7-PAM | 0.00551 | -0.7456 | 0.076 | 0.15 | 1 | MS.Trt_vs_MS.NoTrt_B |
| AMY2B | 0.00552 | 0.53631 | 0.464 | 0.63 | 1 | MS.Trt_vs_MS.NoTrt_B |
| TAF11 | 0.00553 | -0.329 | 0.351 | 0.51 | 1 | MS.Trt_vs_MS.NoTrt_B |

| | | | | | | |
|---|---|---|---|---|---|---|
| NUDT15 | 0.00557 | -0.2787 | 0.436 | 0.59 | 1 | MS.Trt_vs_MS.NoTrt_B |
| SIDT2 | 0.00558 | 0.64892 | 0.654 | 0.79 | 1 | MS.Trt_vs_MS.NoTrt_B |
| CXorf40A | 0.00559 | 0.39634 | 0.052 | 0.11 | 1 | MS.Trt_vs_MS.NoTrt_B |
| CBX6 | 0.0056 | 0.48329 | 0.366 | 0.53 | 1 | MS.Trt_vs_MS.NoTrt_B |
| ZBTB8OS | 0.00562 | -0.2935 | 0.878 | 1 | 1 | MS.Trt_vs_MS.NoTrt_B |
| CABIN1 | 0.00565 | -0.3454 | 0.137 | 0.23 | 1 | MS.Trt_vs_MS.NoTrt_B |
| RPAP3 | 0.0057 | 0.40814 | 0.532 | 0.69 | 1 | MS.Trt_vs_MS.NoTrt_B |
| ZSWIM1 | 0.0057 | 0.28145 | 0.636 | 0.76 | 1 | MS.Trt_vs_MS.NoTrt_B |
| ORMDL3 | 0.00572 | -0.7834 | 0.085 | 0.16 | 1 | MS.Trt_vs_MS.NoTrt_B |
| CHTOP | 0.00573 | 0.32323 | 0.44 | 0.59 | 1 | MS.Trt_vs_MS.NoTrt_B |
| GPRIN3 | 0.00573 | 0.64684 | 0.346 | 0.49 | 1 | MS.Trt_vs_MS.NoTrt_B |
| LYRM2 | 0.00573 | -0.5178 | 0.673 | 0.81 | 1 | MS.Trt_vs_MS.NoTrt_B |
| NEDD1 | 0.00574 | -0.4434 | 0.057 | 0.12 | 1 | MS.Trt_vs_MS.NoTrt_B |
| MCM8 | 0.00574 | -0.3141 | 0.573 | 0.7 | 1 | MS.Trt_vs_MS.NoTrt_B |
| NIPBL | 0.00575 | 0.51462 | 0.416 | 0.57 | 1 | MS.Trt_vs_MS.NoTrt_B |
| VAMP8 | 0.00577 | 0.34068 | 0.227 | 0.37 | 1 | MS.Trt_vs_MS.NoTrt_B |
| C21orf58 | 0.00578 | -0.4617 | 0.346 | 0.49 | 1 | MS.Trt_vs_MS.NoTrt_B |
| SMIM19 | 0.00578 | 0.69112 | 0.355 | 0.47 | 1 | MS.Trt_vs_MS.NoTrt_B |
| ATF1 | 0.00578 | 0.54767 | 0.096 | 0.17 | 1 | MS.Trt_vs_MS.NoTrt_B |
| LPGAT1 | 0.00579 | -0.3195 | 0.346 | 0.47 | 1 | MS.Trt_vs_MS.NoTrt_B |
| LILRB3 | 0.00579 | -1.5863 | 0.085 | 0.16 | 1 | MS.Trt_vs_MS.NoTrt_B |
| DSTN | 0.0058 | 0.71841 | 0.425 | 0.56 | 1 | MS.Trt_vs_MS.NoTrt_B |
| ARL13B | 0.0058 | -0.6417 | 0.137 | 0.23 | 1 | MS.Trt_vs_MS.NoTrt_B |
| ACTR6 | 0.00582 | 0.30754 | 0.412 | 0.54 | 1 | MS.Trt_vs_MS.NoTrt_B |
| COQ5 | 0.00583 | -0.5257 | 0.693 | 0.89 | 1 | MS.Trt_vs_MS.NoTrt_B |
| SLC16A1 | 0.00584 | -0.5546 | 0.089 | 0.16 | 1 | MS.Trt_vs_MS.NoTrt_B |
| DCAF7 | 0.00584 | -0.3536 | 0.229 | 0.32 | 1 | MS.Trt_vs_MS.NoTrt_B |
| KCTD18 | 0.00585 | 0.37303 | 0.647 | 0.81 | 1 | MS.Trt_vs_MS.NoTrt_B |
| PPP1R11 | 0.00586 | -0.2544 | 0.129 | 0.22 | 1 | MS.Trt_vs_MS.NoTrt_B |
| COQ8A | 0.00586 | -0.9191 | 0.497 | 0.62 | 1 | MS.Trt_vs_MS.NoTrt_B |
| EPB41L2 | 0.00586 | -0.9768 | 0.074 | 0.15 | 1 | MS.Trt_vs_MS.NoTrt_B |
| HERC1 | 0.00586 | 0.37912 | 0.669 | 0.79 | 1 | MS.Trt_vs_MS.NoTrt_B |
| WDTC1 | 0.0059 | -0.5776 | 0.233 | 0.35 | 1 | MS.Trt_vs_MS.NoTrt_B |
| LIN54 | 0.00593 | 0.62998 | 0.444 | 0.63 | 1 | MS.Trt_vs_MS.NoTrt_B |
| CCDC117 | 0.00594 | 1.08131 | 0.205 | 0.31 | 1 | MS.Trt_vs_MS.NoTrt_B |
| NTMT1 | 0.00594 | 0.67677 | 0.129 | 0.22 | 1 | MS.Trt_vs_MS.NoTrt_B |
| SETD5 | 0.00596 | 0.38009 | 0.388 | 0.53 | 1 | MS.Trt_vs_MS.NoTrt_B |
| APLP2 | 0.00602 | -0.2832 | 0.819 | 0.93 | 1 | MS.Trt_vs_MS.NoTrt_B |
| ZNF831 | 0.00602 | 0.27789 | 0.137 | 0.23 | 1 | MS.Trt_vs_MS.NoTrt_B |
| SMURF1 | 0.00605 | 0.51306 | 0.074 | 0.15 | 1 | MS.Trt_vs_MS.NoTrt_B |

| | | | | | | |
|---|---|---|---|---|---|---|
| TAF15 | 0.00607 | -0.3948 | 0.144 | 0.23 | 1 | MS.Trt_vs_MS.NoTrt_B |
| ALDH5A1 | 0.00609 | 0.9365 | 0.392 | 0.51 | 1 | MS.Trt_vs_MS.NoTrt_B |
| VPS26B | 0.00611 | -0.3682 | 0.061 | 0.12 | 1 | MS.Trt_vs_MS.NoTrt_B |
| SURF2 | 0.00613 | -0.5325 | 0.052 | 0.11 | 1 | MS.Trt_vs_MS.NoTrt_B |
| RRBP1 | 0.00614 | 0.54542 | 0.19 | 0.3 | 1 | MS.Trt_vs_MS.NoTrt_B |
| ATXN1 | 0.00616 | 0.28695 | 0.288 | 0.41 | 1 | MS.Trt_vs_MS.NoTrt_B |
| TARS | 0.00617 | -0.2556 | 0.135 | 0.22 | 1 | MS.Trt_vs_MS.NoTrt_B |
| MECP2 | 0.00618 | 0.3474 | 0.789 | 0.92 | 1 | MS.Trt_vs_MS.NoTrt_B |
| NBPF9 | 0.00619 | 0.25154 | 0.575 | 0.72 | 1 | MS.Trt_vs_MS.NoTrt_B |
| OTUB1 | 0.0062 | 0.40486 | 0.795 | 0.91 | 1 | MS.Trt_vs_MS.NoTrt_B |
| CWC25 | 0.00622 | -0.6677 | 0.257 | 0.35 | 1 | MS.Trt_vs_MS.NoTrt_B |
| VPS52 | 0.00622 | -0.4402 | 0.094 | 0.17 | 1 | MS.Trt_vs_MS.NoTrt_B |
| THUMPD2 | 0.00625 | -0.3409 | 0.296 | 0.4 | 1 | MS.Trt_vs_MS.NoTrt_B |
| MTBP | 0.00625 | 0.30274 | 0.484 | 0.62 | 1 | MS.Trt_vs_MS.NoTrt_B |
| FPR1 | 0.00626 | 0.29078 | 0.32 | 0.45 | 1 | MS.Trt_vs_MS.NoTrt_B |
| RNF121 | 0.00631 | -0.3322 | 0.24 | 0.35 | 1 | MS.Trt_vs_MS.NoTrt_B |
| SYNRG | 0.00634 | 0.40051 | 0.871 | 0.96 | 1 | MS.Trt_vs_MS.NoTrt_B |
| NUDT3 | 0.00634 | 0.35489 | 0.46 | 0.57 | 1 | MS.Trt_vs_MS.NoTrt_B |
| SORL1 | 0.00635 | 0.31074 | 0.673 | 0.78 | 1 | MS.Trt_vs_MS.NoTrt_B |
| HHLA1 | 0.00635 | 0.27792 | 0.279 | 0.41 | 1 | MS.Trt_vs_MS.NoTrt_B |
| CNOT6L | 0.00636 | 0.76386 | 0.166 | 0.27 | 1 | MS.Trt_vs_MS.NoTrt_B |
| ASMTL | 0.00637 | -0.5913 | 0.065 | 0.13 | 1 | MS.Trt_vs_MS.NoTrt_B |
| NSUN6 | 0.00638 | 0.56747 | 0.549 | 0.69 | 1 | MS.Trt_vs_MS.NoTrt_B |
| TTC16 | 0.00642 | 0.85582 | 0.438 | 0.62 | 1 | MS.Trt_vs_MS.NoTrt_B |
| TIMELESS | 0.00642 | -0.3253 | 0.172 | 0.27 | 1 | MS.Trt_vs_MS.NoTrt_B |
| CKAP2 | 0.00643 | -0.644 | 0.085 | 0.16 | 1 | MS.Trt_vs_MS.NoTrt_B |
| ZNF419 | 0.00644 | 0.53611 | 0.656 | 0.81 | 1 | MS.Trt_vs_MS.NoTrt_B |
| NUTF2 | 0.00646 | -0.5663 | 0.111 | 0.19 | 1 | MS.Trt_vs_MS.NoTrt_B |
| TM9SF1 | 0.00647 | -0.592 | 0.19 | 0.29 | 1 | MS.Trt_vs_MS.NoTrt_B |
| ZNF615 | 0.00648 | 0.51312 | 0.331 | 0.46 | 1 | MS.Trt_vs_MS.NoTrt_B |
| SLC4A1AP | 0.00651 | 0.30719 | 0.667 | 0.81 | 1 | MS.Trt_vs_MS.NoTrt_B |
| C19orf44 | 0.00654 | 0.3664 | 0.255 | 0.37 | 1 | MS.Trt_vs_MS.NoTrt_B |
| MCL1 | 0.00654 | 0.26619 | 0.261 | 0.38 | 1 | MS.Trt_vs_MS.NoTrt_B |
| TTC22 | 0.00658 | -0.3081 | 0.32 | 0.45 | 1 | MS.Trt_vs_MS.NoTrt_B |
| PRKDC | 0.00659 | 0.48021 | 0.521 | 0.65 | 1 | MS.Trt_vs_MS.NoTrt_B |
| NSL1 | 0.00661 | 0.34631 | 0.593 | 0.76 | 1 | MS.Trt_vs_MS.NoTrt_B |
| CCDC186 | 0.00662 | 0.51635 | 0.468 | 0.57 | 1 | MS.Trt_vs_MS.NoTrt_B |
| C12orf10 | 0.00663 | -0.3522 | 0.113 | 0.2 | 1 | MS.Trt_vs_MS.NoTrt_B |
| NCAPD3 | 0.00663 | -0.5471 | 0.05 | 0.11 | 1 | MS.Trt_vs_MS.NoTrt_B |
| RNF141 | 0.00664 | -0.384 | 0.285 | 0.41 | 1 | MS.Trt_vs_MS.NoTrt_B |

| | | | | | | |
|---|---|---|---|---|---|---|
| FAM127A | 0.00664 | -0.2589 | 0.065 | 0.13 | 1 | MS.Trt_vs_MS.NoTrt_B |
| CDC23 | 0.00664 | -0.9472 | 0.054 | 0.11 | 1 | MS.Trt_vs_MS.NoTrt_B |
| PBX4 | 0.00668 | -0.4419 | 0.085 | 0.16 | 1 | MS.Trt_vs_MS.NoTrt_B |
| CTD-2192J1( | 0.00668 | -0.3375 | 0.346 | 0.48 | 1 | MS.Trt_vs_MS.NoTrt_B |
| NETO2 | 0.00669 | 0.65133 | 0.416 | 0.55 | 1 | MS.Trt_vs_MS.NoTrt_B |
| PSMD2 | 0.00669 | -0.5075 | 0.129 | 0.21 | 1 | MS.Trt_vs_MS.NoTrt_B |
| SLC15A1 | 0.0067 | 0.33817 | 0.791 | 0.74 | 1 | MS.Trt_vs_MS.NoTrt_B |
| RBM6 | 0.0067 | 0.33821 | 0.765 | 0.9 | 1 | MS.Trt_vs_MS.NoTrt_B |
| NBEAL2 | 0.00676 | -0.268 | 0.135 | 0.22 | 1 | MS.Trt_vs_MS.NoTrt_B |
| RBM39 | 0.00677 | 0.25615 | 0.789 | 0.95 | 1 | MS.Trt_vs_MS.NoTrt_B |
| DIXDC1 | 0.00678 | 0.26805 | 0.172 | 0.28 | 1 | MS.Trt_vs_MS.NoTrt_B |
| C15orf53 | 0.00681 | -0.3255 | 0.046 | 0.1 | 1 | MS.Trt_vs_MS.NoTrt_B |
| ARRDC1 | 0.00682 | -0.2841 | 0.172 | 0.27 | 1 | MS.Trt_vs_MS.NoTrt_B |
| BPHL | 0.00682 | 0.35919 | 0.444 | 0.6 | 1 | MS.Trt_vs_MS.NoTrt_B |
| SNX11 | 0.00683 | 0.42396 | 0.501 | 0.65 | 1 | MS.Trt_vs_MS.NoTrt_B |
| NUMBL | 0.00685 | -0.4678 | 0.466 | 0.58 | 1 | MS.Trt_vs_MS.NoTrt_B |
| NDUFB6 | 0.00688 | 0.34064 | 0.442 | 0.56 | 1 | MS.Trt_vs_MS.NoTrt_B |
| TMEM53 | 0.00688 | 0.85852 | 0.218 | 0.34 | 1 | MS.Trt_vs_MS.NoTrt_B |
| CYP51A1 | 0.0069 | -0.3123 | 0.362 | 0.5 | 1 | MS.Trt_vs_MS.NoTrt_B |
| ATP6V1H | 0.00691 | -0.5993 | 0.272 | 0.38 | 1 | MS.Trt_vs_MS.NoTrt_B |
| SLC10A7 | 0.00691 | -0.399 | 0.046 | 0.1 | 1 | MS.Trt_vs_MS.NoTrt_B |
| CENPM | 0.00691 | -0.2664 | 0.187 | 0.29 | 1 | MS.Trt_vs_MS.NoTrt_B |
| HTATSF1 | 0.00691 | 0.3534 | 0.107 | 0.19 | 1 | MS.Trt_vs_MS.NoTrt_B |
| RP11-295P9 | 0.00692 | -0.4105 | 0.05 | 0.11 | 1 | MS.Trt_vs_MS.NoTrt_B |
| RNF40 | 0.00693 | -0.4537 | 0.065 | 0.13 | 1 | MS.Trt_vs_MS.NoTrt_B |
| ATAD5 | 0.00693 | 0.55722 | 0.429 | 0.58 | 1 | MS.Trt_vs_MS.NoTrt_B |
| CD46 | 0.00693 | 0.28712 | 0.688 | 0.84 | 1 | MS.Trt_vs_MS.NoTrt_B |
| PABPC1L | 0.00697 | -0.9913 | 0.2 | 0.31 | 1 | MS.Trt_vs_MS.NoTrt_B |
| MYO10 | 0.00699 | -0.3277 | 0.586 | 0.77 | 1 | MS.Trt_vs_MS.NoTrt_B |
| PYGM | 0.00703 | 0.67633 | 0.536 | 0.69 | 1 | MS.Trt_vs_MS.NoTrt_B |
| KIAA0100 | 0.00707 | 0.53913 | 0.375 | 0.5 | 1 | MS.Trt_vs_MS.NoTrt_B |
| TPMT | 0.00707 | 0.67027 | 0.61 | 0.75 | 1 | MS.Trt_vs_MS.NoTrt_B |
| STK11 | 0.0071 | -0.3067 | 0.251 | 0.36 | 1 | MS.Trt_vs_MS.NoTrt_B |
| USP54 | 0.00711 | 0.40553 | 0.479 | 0.63 | 1 | MS.Trt_vs_MS.NoTrt_B |
| MRPL57 | 0.00712 | 0.29591 | 0.632 | 0.71 | 1 | MS.Trt_vs_MS.NoTrt_B |
| ZNF540 | 0.00715 | 0.87653 | 0.336 | 0.46 | 1 | MS.Trt_vs_MS.NoTrt_B |
| PPP1R16A | 0.0072 | 0.94853 | 0.331 | 0.45 | 1 | MS.Trt_vs_MS.NoTrt_B |
| MCM5 | 0.00723 | -0.299 | 0.146 | 0.23 | 1 | MS.Trt_vs_MS.NoTrt_B |
| TSPOAP1 | 0.00727 | -0.5459 | 0.131 | 0.22 | 1 | MS.Trt_vs_MS.NoTrt_B |
| ERAP1 | 0.00728 | 0.43354 | 0.451 | 0.62 | 1 | MS.Trt_vs_MS.NoTrt_B |

| | | | | | | |
|---|---|---|---|---|---|---|
| PRR14L | 0.00728 | 0.46269 | 0.301 | 0.42 | 1 | MS.Trt_vs_MS.NoTrt_B |
| NPHP1 | 0.00728 | 0.32123 | 0.549 | 0.67 | 1 | MS.Trt_vs_MS.NoTrt_B |
| PIK3AP1 | 0.00728 | 1.04413 | 0.076 | 0.15 | 1 | MS.Trt_vs_MS.NoTrt_B |
| EMC1 | 0.0073 | -0.4836 | 0.654 | 0.77 | 1 | MS.Trt_vs_MS.NoTrt_B |
| PRRC2C | 0.00731 | 0.31861 | 0.656 | 0.82 | 1 | MS.Trt_vs_MS.NoTrt_B |
| UQCC1 | 0.00731 | 0.28105 | 0.397 | 0.52 | 1 | MS.Trt_vs_MS.NoTrt_B |
| UPF3A | 0.00733 | 0.51364 | 0.617 | 0.77 | 1 | MS.Trt_vs_MS.NoTrt_B |
| NUBPL | 0.00736 | -0.2527 | 0.597 | 0.74 | 1 | MS.Trt_vs_MS.NoTrt_B |
| MYBBP1A | 0.00743 | -0.6249 | 0.092 | 0.16 | 1 | MS.Trt_vs_MS.NoTrt_B |
| ANAPC4 | 0.00743 | -0.5189 | 0.176 | 0.27 | 1 | MS.Trt_vs_MS.NoTrt_B |
| CCNF | 0.00743 | 0.60408 | 0.336 | 0.46 | 1 | MS.Trt_vs_MS.NoTrt_B |
| HSBP1 | 0.00743 | 0.69443 | 0.503 | 0.64 | 1 | MS.Trt_vs_MS.NoTrt_B |
| RAB1A | 0.00744 | 0.62219 | 0.096 | 0.17 | 1 | MS.Trt_vs_MS.NoTrt_B |
| SMN1 | 0.00745 | 0.49463 | 0.296 | 0.42 | 1 | MS.Trt_vs_MS.NoTrt_B |
| ZFYVE26 | 0.00746 | -0.3639 | 0.708 | 0.86 | 1 | MS.Trt_vs_MS.NoTrt_B |
| BDH1 | 0.00747 | 0.36936 | 0.789 | 0.94 | 1 | MS.Trt_vs_MS.NoTrt_B |
| ATG4B | 0.00747 | -0.4431 | 0.129 | 0.21 | 1 | MS.Trt_vs_MS.NoTrt_B |
| ADAM19 | 0.00747 | 0.29784 | 0.312 | 0.43 | 1 | MS.Trt_vs_MS.NoTrt_B |
| NUDC | 0.0075 | 0.45088 | 0.468 | 0.62 | 1 | MS.Trt_vs_MS.NoTrt_B |
| NR1D2 | 0.0075 | 0.43282 | 0.076 | 0.15 | 1 | MS.Trt_vs_MS.NoTrt_B |
| ATP9B | 0.00752 | 0.28863 | 0.8 | 0.93 | 1 | MS.Trt_vs_MS.NoTrt_B |
| CLMP | 0.00752 | 0.63841 | 0.268 | 0.4 | 1 | MS.Trt_vs_MS.NoTrt_B |
| SRGAP3 | 0.00753 | 0.40515 | 0.752 | 0.9 | 1 | MS.Trt_vs_MS.NoTrt_B |
| EIF1AX | 0.00753 | 0.56103 | 0.218 | 0.32 | 1 | MS.Trt_vs_MS.NoTrt_B |
| ATP6V1B2 | 0.00755 | 0.41811 | 0.521 | 0.67 | 1 | MS.Trt_vs_MS.NoTrt_B |
| PICK1 | 0.00755 | 0.79747 | 0.316 | 0.43 | 1 | MS.Trt_vs_MS.NoTrt_B |
| FGFR1OP | 0.00756 | 0.52334 | 0.704 | 0.81 | 1 | MS.Trt_vs_MS.NoTrt_B |
| BCAS3 | 0.00759 | -0.4305 | 0.743 | 0.85 | 1 | MS.Trt_vs_MS.NoTrt_B |
| MRS2 | 0.00761 | -0.7474 | 0.708 | 0.77 | 1 | MS.Trt_vs_MS.NoTrt_B |
| SETD3 | 0.00761 | -0.6729 | 0.059 | 0.12 | 1 | MS.Trt_vs_MS.NoTrt_B |
| FAM120C | 0.00763 | 0.46418 | 0.155 | 0.25 | 1 | MS.Trt_vs_MS.NoTrt_B |
| TXNL4B | 0.00765 | -0.2952 | 0.257 | 0.37 | 1 | MS.Trt_vs_MS.NoTrt_B |
| TP53BP2 | 0.00765 | 0.39899 | 0.102 | 0.18 | 1 | MS.Trt_vs_MS.NoTrt_B |
| PDPR | 0.00765 | -0.2696 | 0.312 | 0.42 | 1 | MS.Trt_vs_MS.NoTrt_B |
| NOC2L | 0.00768 | 0.27621 | 0.625 | 0.77 | 1 | MS.Trt_vs_MS.NoTrt_B |
| RANBP2 | 0.00769 | 0.43724 | 0.266 | 0.38 | 1 | MS.Trt_vs_MS.NoTrt_B |
| CYP2R1 | 0.00771 | -0.9995 | 0.068 | 0.13 | 1 | MS.Trt_vs_MS.NoTrt_B |
| PIGC | 0.00773 | -0.5593 | 0.092 | 0.16 | 1 | MS.Trt_vs_MS.NoTrt_B |
| NDUFV2 | 0.00774 | 0.3897 | 0.349 | 0.48 | 1 | MS.Trt_vs_MS.NoTrt_B |
| NEIL3 | 0.00775 | -0.5238 | 0.198 | 0.3 | 1 | MS.Trt_vs_MS.NoTrt_B |

| | | | | | | |
|---|---|---|---|---|---|---|
| EFNA5 | 0.00777 | -0.3283 | 0.281 | 0.18 | 1 | MS.Trt_vs_MS.NoTrt_B |
| NIPAL3 | 0.00778 | 0.30818 | 0.647 | 0.77 | 1 | MS.Trt_vs_MS.NoTrt_B |
| ZNF581 | 0.0078 | -0.7129 | 0.05 | 0.11 | 1 | MS.Trt_vs_MS.NoTrt_B |
| LACTB | 0.00784 | -0.4251 | 0.458 | 0.58 | 1 | MS.Trt_vs_MS.NoTrt_B |
| IGSF9 | 0.00787 | -0.2582 | 0.17 | 0.27 | 1 | MS.Trt_vs_MS.NoTrt_B |
| MKRN2 | 0.00787 | -0.3633 | 0.065 | 0.13 | 1 | MS.Trt_vs_MS.NoTrt_B |
| DGKE | 0.00788 | 0.35659 | 0.603 | 0.74 | 1 | MS.Trt_vs_MS.NoTrt_B |
| BIVM | 0.00788 | 0.79675 | 0.218 | 0.32 | 1 | MS.Trt_vs_MS.NoTrt_B |
| DDX41 | 0.00792 | -0.8782 | 0.068 | 0.13 | 1 | MS.Trt_vs_MS.NoTrt_B |
| STAMBPL1 | 0.00793 | 0.36098 | 0.346 | 0.48 | 1 | MS.Trt_vs_MS.NoTrt_B |
| POLR2J2 | 0.00794 | 0.32619 | 0.27 | 0.38 | 1 | MS.Trt_vs_MS.NoTrt_B |
| SHPRH | 0.00794 | 0.63458 | 0.501 | 0.67 | 1 | MS.Trt_vs_MS.NoTrt_B |
| FBXO46 | 0.00794 | -0.2532 | 0.168 | 0.26 | 1 | MS.Trt_vs_MS.NoTrt_B |
| ELOVL6 | 0.00796 | -1.4948 | 0.205 | 0.3 | 1 | MS.Trt_vs_MS.NoTrt_B |
| RANGRF | 0.00798 | -0.2836 | 0.065 | 0.13 | 1 | MS.Trt_vs_MS.NoTrt_B |
| GSTCD | 0.00798 | 0.28245 | 0.46 | 0.63 | 1 | MS.Trt_vs_MS.NoTrt_B |
| TET2 | 0.00799 | 0.94881 | 0.355 | 0.51 | 1 | MS.Trt_vs_MS.NoTrt_B |
| SOX6 | 0.00804 | -0.4593 | 0.29 | 0.39 | 1 | MS.Trt_vs_MS.NoTrt_B |
| ACAD8 | 0.00804 | 0.3651 | 0.403 | 0.54 | 1 | MS.Trt_vs_MS.NoTrt_B |
| ZFYVE28 | 0.00805 | 0.26337 | 0.081 | 0.15 | 1 | MS.Trt_vs_MS.NoTrt_B |
| ZNF383 | 0.00807 | -0.2773 | 0.941 | 0.97 | 1 | MS.Trt_vs_MS.NoTrt_B |
| LINC00854 | 0.00809 | -0.3337 | 0.512 | 0.63 | 1 | MS.Trt_vs_MS.NoTrt_B |
| ARHGAP8 | 0.00809 | -0.3868 | 0.549 | 0.71 | 1 | MS.Trt_vs_MS.NoTrt_B |
| AL589743.1 | 0.00813 | -0.5008 | 0.425 | 0.55 | 1 | MS.Trt_vs_MS.NoTrt_B |
| PNISR | 0.00814 | 0.34242 | 0.479 | 0.67 | 1 | MS.Trt_vs_MS.NoTrt_B |
| SULT1A4 | 0.00814 | -0.2703 | 0.205 | 0.3 | 1 | MS.Trt_vs_MS.NoTrt_B |
| MCTP2 | 0.00816 | 0.52704 | 0.27 | 0.41 | 1 | MS.Trt_vs_MS.NoTrt_B |
| RABL6 | 0.00818 | 1.0937 | 0.42 | 0.52 | 1 | MS.Trt_vs_MS.NoTrt_B |
| LHPP | 0.00821 | -0.2869 | 0.094 | 0.17 | 1 | MS.Trt_vs_MS.NoTrt_B |
| TPP1 | 0.00822 | 0.35864 | 0.27 | 0.38 | 1 | MS.Trt_vs_MS.NoTrt_B |
| ZNF71 | 0.00823 | 0.50553 | 0.157 | 0.25 | 1 | MS.Trt_vs_MS.NoTrt_B |
| CHURC1-FN | 0.00823 | 0.29164 | 0.161 | 0.26 | 1 | MS.Trt_vs_MS.NoTrt_B |
| FAM47E | 0.00824 | 0.31354 | 0.549 | 0.69 | 1 | MS.Trt_vs_MS.NoTrt_B |
| LEMD3 | 0.00829 | -0.2667 | 0.503 | 0.64 | 1 | MS.Trt_vs_MS.NoTrt_B |
| CALR | 0.00831 | -0.4049 | 0.702 | 0.82 | 1 | MS.Trt_vs_MS.NoTrt_B |
| ZNF76 | 0.00832 | -0.4269 | 0.109 | 0.18 | 1 | MS.Trt_vs_MS.NoTrt_B |
| CELF5 | 0.00833 | -0.5383 | 0.179 | 0.28 | 1 | MS.Trt_vs_MS.NoTrt_B |
| RCCD1 | 0.00835 | -0.3351 | 0.102 | 0.18 | 1 | MS.Trt_vs_MS.NoTrt_B |
| DNAJC15 | 0.00837 | -0.5751 | 0.126 | 0.21 | 1 | MS.Trt_vs_MS.NoTrt_B |
| ING4 | 0.00837 | -0.4403 | 0.214 | 0.31 | 1 | MS.Trt_vs_MS.NoTrt_B |

| | | | | | | |
|---|---|---|---|---|---|---|
| C1orf35 | 0.00839 | -0.3808 | 0.17 | 0.25 | 1 | MS.Trt_vs_MS.NoTrt_B |
| SHTN1 | 0.00839 | 0.39441 | 0.05 | 0.11 | 1 | MS.Trt_vs_MS.NoTrt_B |
| ZNF773 | 0.00845 | -1.0232 | 0.464 | 0.56 | 1 | MS.Trt_vs_MS.NoTrt_B |
| USP28 | 0.00847 | -0.4424 | 0.266 | 0.38 | 1 | MS.Trt_vs_MS.NoTrt_B |
| METRN | 0.00848 | 0.29242 | 0.059 | 0.12 | 1 | MS.Trt_vs_MS.NoTrt_B |
| MUS81 | 0.00848 | -0.4834 | 0.092 | 0.16 | 1 | MS.Trt_vs_MS.NoTrt_B |
| ARPC4-TTLL3 | 0.00853 | -0.4239 | 0.054 | 0.11 | 1 | MS.Trt_vs_MS.NoTrt_B |
| PRKCSH | 0.00854 | 0.33361 | 0.623 | 0.77 | 1 | MS.Trt_vs_MS.NoTrt_B |
| HMGN3 | 0.00858 | 0.38867 | 0.194 | 0.29 | 1 | MS.Trt_vs_MS.NoTrt_B |
| SPOUT1 | 0.00858 | -0.4401 | 0.098 | 0.17 | 1 | MS.Trt_vs_MS.NoTrt_B |
| SCYL2 | 0.0086 | -0.4624 | 0.074 | 0.14 | 1 | MS.Trt_vs_MS.NoTrt_B |
| NFATC2 | 0.00861 | 0.3016 | 0.094 | 0.17 | 1 | MS.Trt_vs_MS.NoTrt_B |
| SMARCA2 | 0.00862 | 0.5051 | 0.305 | 0.43 | 1 | MS.Trt_vs_MS.NoTrt_B |
| HJURP | 0.00863 | -0.9551 | 0.275 | 0.39 | 1 | MS.Trt_vs_MS.NoTrt_B |
| MED18 | 0.00863 | -0.3766 | 0.447 | 0.59 | 1 | MS.Trt_vs_MS.NoTrt_B |
| MAPRE1 | 0.00864 | -0.2585 | 0.118 | 0.2 | 1 | MS.Trt_vs_MS.NoTrt_B |
| ST3GAL5 | 0.00864 | 0.33231 | 0.229 | 0.33 | 1 | MS.Trt_vs_MS.NoTrt_B |
| MKL2 | 0.00864 | 0.35833 | 0.573 | 0.72 | 1 | MS.Trt_vs_MS.NoTrt_B |
| CARD8 | 0.00864 | 0.32461 | 0.786 | 0.89 | 1 | MS.Trt_vs_MS.NoTrt_B |
| RABL3 | 0.00864 | 0.27352 | 0.525 | 0.67 | 1 | MS.Trt_vs_MS.NoTrt_B |
| PHB2 | 0.00869 | -0.3327 | 0.135 | 0.22 | 1 | MS.Trt_vs_MS.NoTrt_B |
| NDUFB8 | 0.00871 | 0.51421 | 0.316 | 0.47 | 1 | MS.Trt_vs_MS.NoTrt_B |
| FBXO31 | 0.00871 | 0.41162 | 0.2 | 0.31 | 1 | MS.Trt_vs_MS.NoTrt_B |
| NUFIP2 | 0.00871 | 0.45943 | 0.325 | 0.46 | 1 | MS.Trt_vs_MS.NoTrt_B |
| DDHD1 | 0.00879 | 0.55476 | 0.786 | 0.95 | 1 | MS.Trt_vs_MS.NoTrt_B |
| LSM8 | 0.0088 | 0.63182 | 0.754 | 0.89 | 1 | MS.Trt_vs_MS.NoTrt_B |
| SH3PXD2A | 0.00883 | 0.62995 | 0.172 | 0.27 | 1 | MS.Trt_vs_MS.NoTrt_B |
| IRAK4 | 0.00883 | 0.47028 | 0.471 | 0.58 | 1 | MS.Trt_vs_MS.NoTrt_B |
| KDM5D | 0.00885 | -2.8482 | 0.22 | 0.31 | 1 | MS.Trt_vs_MS.NoTrt_B |
| RSL24D1 | 0.0089 | 0.45888 | 0.24 | 0.35 | 1 | MS.Trt_vs_MS.NoTrt_B |
| PDZD11 | 0.00892 | -0.3467 | 0.059 | 0.12 | 1 | MS.Trt_vs_MS.NoTrt_B |
| EFCAB5 | 0.00892 | 0.54816 | 0.312 | 0.44 | 1 | MS.Trt_vs_MS.NoTrt_B |
| MAP3K13 | 0.00893 | -0.252 | 0.725 | 0.85 | 1 | MS.Trt_vs_MS.NoTrt_B |
| RPP30 | 0.009 | 0.40643 | 0.514 | 0.65 | 1 | MS.Trt_vs_MS.NoTrt_B |
| MTMR6 | 0.00901 | -0.7329 | 0.061 | 0.12 | 1 | MS.Trt_vs_MS.NoTrt_B |
| PDS5B | 0.00906 | 0.55589 | 0.272 | 0.38 | 1 | MS.Trt_vs_MS.NoTrt_B |
| GPKOW | 0.00908 | -0.2899 | 0.054 | 0.11 | 1 | MS.Trt_vs_MS.NoTrt_B |
| NUDT6 | 0.00914 | 0.28038 | 0.39 | 0.55 | 1 | MS.Trt_vs_MS.NoTrt_B |
| PORCN | 0.00918 | -0.9867 | 0.048 | 0.1 | 1 | MS.Trt_vs_MS.NoTrt_B |
| STRN4 | 0.00922 | -0.4617 | 0.368 | 0.48 | 1 | MS.Trt_vs_MS.NoTrt_B |

| | | | | | | |
|---|---|---|---|---|---|---|
| ANKMY1 | 0.00924 | -0.3246 | 0.318 | 0.43 | 1 | MS.Trt_vs_MS.NoTrt_B |
| RNF2 | 0.00929 | -0.5717 | 0.327 | 0.45 | 1 | MS.Trt_vs_MS.NoTrt_B |
| C8orf44 | 0.0093 | 0.49604 | 0.407 | 0.57 | 1 | MS.Trt_vs_MS.NoTrt_B |
| CAPN7 | 0.0093 | 0.41669 | 0.429 | 0.58 | 1 | MS.Trt_vs_MS.NoTrt_B |
| SUGP1 | 0.00931 | -0.3013 | 0.351 | 0.48 | 1 | MS.Trt_vs_MS.NoTrt_B |
| ALPK1 | 0.00932 | 0.42757 | 0.571 | 0.76 | 1 | MS.Trt_vs_MS.NoTrt_B |
| COQ9 | 0.00933 | -0.6975 | 0.048 | 0.1 | 1 | MS.Trt_vs_MS.NoTrt_B |
| GADD45B | 0.00934 | -0.8378 | 0.065 | 0.12 | 1 | MS.Trt_vs_MS.NoTrt_B |
| SLC12A7 | 0.00935 | -0.4071 | 0.083 | 0.15 | 1 | MS.Trt_vs_MS.NoTrt_B |
| GSDMB | 0.00935 | -0.3491 | 0.756 | 0.89 | 1 | MS.Trt_vs_MS.NoTrt_B |
| SRSF10 | 0.00936 | 0.48139 | 0.643 | 0.79 | 1 | MS.Trt_vs_MS.NoTrt_B |
| TRAF5 | 0.00937 | 0.49004 | 0.562 | 0.72 | 1 | MS.Trt_vs_MS.NoTrt_B |
| PIP4K2A | 0.00937 | 0.49325 | 0.699 | 0.87 | 1 | MS.Trt_vs_MS.NoTrt_B |
| ATIC | 0.00943 | 0.29505 | 0.301 | 0.41 | 1 | MS.Trt_vs_MS.NoTrt_B |
| ADAMTSL4 | 0.00944 | 0.35475 | 0.196 | 0.29 | 1 | MS.Trt_vs_MS.NoTrt_B |
| PINLYP | 0.00946 | 0.27366 | 0.442 | 0.61 | 1 | MS.Trt_vs_MS.NoTrt_B |
| ZNF680 | 0.00954 | 1.03205 | 0.061 | 0.12 | 1 | MS.Trt_vs_MS.NoTrt_B |
| EFCAB11 | 0.00956 | 0.39089 | 0.643 | 0.8 | 1 | MS.Trt_vs_MS.NoTrt_B |
| TAF7 | 0.00956 | 0.33662 | 0.285 | 0.4 | 1 | MS.Trt_vs_MS.NoTrt_B |
| LYRM7 | 0.00958 | 0.47646 | 0.808 | 0.91 | 1 | MS.Trt_vs_MS.NoTrt_B |
| CHGB | 0.00959 | -0.5054 | 0.309 | 0.41 | 1 | MS.Trt_vs_MS.NoTrt_B |
| SLC25A19 | 0.00962 | -0.3174 | 0.305 | 0.41 | 1 | MS.Trt_vs_MS.NoTrt_B |
| SEC22C | 0.00965 | -0.4116 | 0.523 | 0.67 | 1 | MS.Trt_vs_MS.NoTrt_B |
| MEF2A | 0.00972 | -0.4471 | 0.359 | 0.48 | 1 | MS.Trt_vs_MS.NoTrt_B |
| NRAS | 0.00973 | -0.8176 | 0.207 | 0.31 | 1 | MS.Trt_vs_MS.NoTrt_B |
| GTF2IRD1 | 0.00976 | -0.5805 | 0.525 | 0.64 | 1 | MS.Trt_vs_MS.NoTrt_B |
| IL15 | 0.00978 | -0.3543 | 0.307 | 0.42 | 1 | MS.Trt_vs_MS.NoTrt_B |
| ZNF330 | 0.00982 | -0.3329 | 0.048 | 0.1 | 1 | MS.Trt_vs_MS.NoTrt_B |
| CPT1A | 0.00983 | -0.4772 | 0.551 | 0.62 | 1 | MS.Trt_vs_MS.NoTrt_B |
| ZDHHC13 | 0.00983 | 0.79939 | 0.595 | 0.75 | 1 | MS.Trt_vs_MS.NoTrt_B |
| ISCA2 | 0.00986 | 0.40367 | 0.412 | 0.54 | 1 | MS.Trt_vs_MS.NoTrt_B |
| AIG1 | 0.00988 | -0.4493 | 0.453 | 0.6 | 1 | MS.Trt_vs_MS.NoTrt_B |
| OR8H2 | 0.00989 | -0.4417 | 0.12 | 0.2 | 1 | MS.Trt_vs_MS.NoTrt_B |
| HNRNPAB | 0.00996 | -0.3782 | 0.096 | 0.17 | 1 | MS.Trt_vs_MS.NoTrt_B |
| TBC1D14 | 0.00996 | -0.2692 | 0.146 | 0.23 | 1 | MS.Trt_vs_MS.NoTrt_B |
| XRCC5 | 0.01001 | 0.27469 | 0.429 | 0.56 | 1 | MS.Trt_vs_MS.NoTrt_B |
| PNKD | 0.01002 | 0.34097 | 0.098 | 0.17 | 1 | MS.Trt_vs_MS.NoTrt_B |
| TGFB1I1 | 0.01003 | -1.0107 | 0.166 | 0.25 | 1 | MS.Trt_vs_MS.NoTrt_B |
| 3-Mar | 0.01008 | -0.4226 | 0.259 | 0.36 | 1 | MS.Trt_vs_MS.NoTrt_B |
| TEN1-CDK3 | 0.01009 | 0.80764 | 0.377 | 0.49 | 1 | MS.Trt_vs_MS.NoTrt_B |

| | | | | | | |
|---|---|---|---|---|---|---|
| AP2A2 | 0.01009 | 0.43638 | 0.49 | 0.64 | 1 | MS.Trt_vs_MS.NoTrt_B |
| ZNF410 | 0.01009 | -0.3833 | 0.386 | 0.5 | 1 | MS.Trt_vs_MS.NoTrt_B |
| HSD17B1 | 0.0101 | 0.39709 | 0.621 | 0.8 | 1 | MS.Trt_vs_MS.NoTrt_B |
| MOB3B | 0.0101 | 0.26005 | 0.42 | 0.56 | 1 | MS.Trt_vs_MS.NoTrt_B |
| HGSNAT | 0.01012 | -0.4806 | 0.558 | 0.67 | 1 | MS.Trt_vs_MS.NoTrt_B |
| OSTC | 0.01013 | 0.34433 | 0.176 | 0.27 | 1 | MS.Trt_vs_MS.NoTrt_B |
| ZNF652 | 0.01016 | 0.5816 | 0.418 | 0.53 | 1 | MS.Trt_vs_MS.NoTrt_B |
| NCKAP1 | 0.01028 | 0.38128 | 0.536 | 0.7 | 1 | MS.Trt_vs_MS.NoTrt_B |
| REC8 | 0.0103 | -0.3556 | 0.048 | 0.1 | 1 | MS.Trt_vs_MS.NoTrt_B |
| PROSER1 | 0.0103 | -0.3103 | 0.12 | 0.2 | 1 | MS.Trt_vs_MS.NoTrt_B |
| OXR1 | 0.01034 | 0.46127 | 0.211 | 0.32 | 1 | MS.Trt_vs_MS.NoTrt_B |
| DENND1B | 0.01035 | 0.31964 | 0.096 | 0.17 | 1 | MS.Trt_vs_MS.NoTrt_B |
| CHRNB1 | 0.01035 | 0.54266 | 0.475 | 0.64 | 1 | MS.Trt_vs_MS.NoTrt_B |
| MRPS26 | 0.01037 | -0.4187 | 0.068 | 0.13 | 1 | MS.Trt_vs_MS.NoTrt_B |
| SRP54 | 0.01041 | 0.51274 | 0.131 | 0.21 | 1 | MS.Trt_vs_MS.NoTrt_B |
| TIMD4 | 0.01042 | 0.26312 | 0.048 | 0.1 | 1 | MS.Trt_vs_MS.NoTrt_B |
| PLEKHG4 | 0.01053 | -0.4821 | 0.111 | 0.18 | 1 | MS.Trt_vs_MS.NoTrt_B |
| ZFP36L1 | 0.01054 | 0.85386 | 0.702 | 0.69 | 1 | MS.Trt_vs_MS.NoTrt_B |
| MIIP | 0.01054 | -0.2896 | 0.105 | 0.18 | 1 | MS.Trt_vs_MS.NoTrt_B |
| DCTN6 | 0.01056 | -0.4105 | 0.427 | 0.55 | 1 | MS.Trt_vs_MS.NoTrt_B |
| PPP2CB | 0.01057 | 0.71528 | 0.46 | 0.6 | 1 | MS.Trt_vs_MS.NoTrt_B |
| LRRC34 | 0.01058 | 0.26965 | 0.503 | 0.65 | 1 | MS.Trt_vs_MS.NoTrt_B |
| NUP54 | 0.01058 | -0.4472 | 0.133 | 0.21 | 1 | MS.Trt_vs_MS.NoTrt_B |
| PSMB6 | 0.01064 | 0.36496 | 0.582 | 0.7 | 1 | MS.Trt_vs_MS.NoTrt_B |
| LYPLAL1 | 0.01066 | 0.33828 | 0.063 | 0.12 | 1 | MS.Trt_vs_MS.NoTrt_B |
| TAF9B | 0.01069 | 0.36042 | 0.107 | 0.18 | 1 | MS.Trt_vs_MS.NoTrt_B |
| CTD-3088G3 | 0.01073 | 0.96828 | 0.37 | 0.5 | 1 | MS.Trt_vs_MS.NoTrt_B |
| MINPP1 | 0.01073 | -0.5044 | 0.068 | 0.13 | 1 | MS.Trt_vs_MS.NoTrt_B |
| MED24 | 0.01074 | 0.31899 | 0.458 | 0.57 | 1 | MS.Trt_vs_MS.NoTrt_B |
| E2F4 | 0.01075 | -0.8028 | 0.089 | 0.16 | 1 | MS.Trt_vs_MS.NoTrt_B |
| POLR3B | 0.01078 | -0.9783 | 0.065 | 0.12 | 1 | MS.Trt_vs_MS.NoTrt_B |
| TEX2 | 0.0108 | -0.579 | 0.573 | 0.68 | 1 | MS.Trt_vs_MS.NoTrt_B |
| ZNF558 | 0.01084 | 0.42228 | 0.545 | 0.69 | 1 | MS.Trt_vs_MS.NoTrt_B |
| FAM161B | 0.01089 | 0.42071 | 0.261 | 0.37 | 1 | MS.Trt_vs_MS.NoTrt_B |
| GPATCH11 | 0.01097 | 0.63455 | 0.346 | 0.47 | 1 | MS.Trt_vs_MS.NoTrt_B |
| LAMTOR3 | 0.011 | 1.36654 | 0.231 | 0.34 | 1 | MS.Trt_vs_MS.NoTrt_B |
| ASB13 | 0.01108 | 0.8311 | 0.105 | 0.18 | 1 | MS.Trt_vs_MS.NoTrt_B |
| ELMSAN1 | 0.01109 | 0.44032 | 0.551 | 0.68 | 1 | MS.Trt_vs_MS.NoTrt_B |
| PDE4B | 0.0111 | 0.38635 | 0.207 | 0.31 | 1 | MS.Trt_vs_MS.NoTrt_B |
| SMARCAL1 | 0.01112 | -0.4389 | 0.564 | 0.68 | 1 | MS.Trt_vs_MS.NoTrt_B |

| | | | | | | |
|---|---|---|---|---|---|---|
| CTDSP2 | 0.01116 | 0.41911 | 0.159 | 0.25 | 1 | MS.Trt_vs_MS.NoTrt_B |
| PGM2 | 0.01119 | -0.5423 | 0.061 | 0.12 | 1 | MS.Trt_vs_MS.NoTrt_B |
| ARFGEF1 | 0.0112 | 0.40969 | 0.527 | 0.64 | 1 | MS.Trt_vs_MS.NoTrt_B |
| USP24 | 0.01123 | 0.30214 | 0.251 | 0.36 | 1 | MS.Trt_vs_MS.NoTrt_B |
| ING1 | 0.01125 | 0.87416 | 0.499 | 0.61 | 1 | MS.Trt_vs_MS.NoTrt_B |
| STYX | 0.01126 | 0.5854 | 0.444 | 0.57 | 1 | MS.Trt_vs_MS.NoTrt_B |
| LZIC | 0.01127 | 0.4269 | 0.684 | 0.85 | 1 | MS.Trt_vs_MS.NoTrt_B |
| BRMS1 | 0.0113 | -0.2848 | 0.098 | 0.17 | 1 | MS.Trt_vs_MS.NoTrt_B |
| PSMD10 | 0.01132 | 0.36802 | 0.207 | 0.3 | 1 | MS.Trt_vs_MS.NoTrt_B |
| KLHL2 | 0.01132 | 0.25116 | 0.242 | 0.34 | 1 | MS.Trt_vs_MS.NoTrt_B |
| PPP4R1 | 0.01136 | -0.9325 | 0.166 | 0.25 | 1 | MS.Trt_vs_MS.NoTrt_B |
| CAMK2D | 0.0114 | 0.54593 | 0.196 | 0.3 | 1 | MS.Trt_vs_MS.NoTrt_B |
| SLC43A3 | 0.01147 | -0.4245 | 0.298 | 0.4 | 1 | MS.Trt_vs_MS.NoTrt_B |
| ZC3H4 | 0.01148 | 0.89637 | 0.292 | 0.42 | 1 | MS.Trt_vs_MS.NoTrt_B |
| DDT | 0.01156 | -0.2568 | 0.122 | 0.2 | 1 | MS.Trt_vs_MS.NoTrt_B |
| TIAL1 | 0.01157 | 1.00925 | 0.743 | 0.89 | 1 | MS.Trt_vs_MS.NoTrt_B |
| TTC34 | 0.01158 | 0.51647 | 0.124 | 0.21 | 1 | MS.Trt_vs_MS.NoTrt_B |
| DLEC1 | 0.01159 | -0.5681 | 0.377 | 0.48 | 1 | MS.Trt_vs_MS.NoTrt_B |
| ARPC1A | 0.0116 | -0.4759 | 0.216 | 0.3 | 1 | MS.Trt_vs_MS.NoTrt_B |
| KCTD21 | 0.01161 | -0.6259 | 0.322 | 0.42 | 1 | MS.Trt_vs_MS.NoTrt_B |
| NDUFAF1 | 0.01164 | 0.70247 | 0.305 | 0.42 | 1 | MS.Trt_vs_MS.NoTrt_B |
| SLC25A5 | 0.01169 | 0.26449 | 0.261 | 0.37 | 1 | MS.Trt_vs_MS.NoTrt_B |
| ZNF333 | 0.0117 | 0.39639 | 0.78 | 0.89 | 1 | MS.Trt_vs_MS.NoTrt_B |
| LENG1 | 0.0117 | -0.2573 | 0.052 | 0.11 | 1 | MS.Trt_vs_MS.NoTrt_B |
| FAM213A | 0.01171 | -0.31 | 0.643 | 0.79 | 1 | MS.Trt_vs_MS.NoTrt_B |
| FUNDC2 | 0.01173 | 0.40717 | 0.386 | 0.49 | 1 | MS.Trt_vs_MS.NoTrt_B |
| LRPPRC | 0.01174 | 0.56802 | 0.41 | 0.54 | 1 | MS.Trt_vs_MS.NoTrt_B |
| ATRX | 0.01174 | 0.41238 | 0.501 | 0.64 | 1 | MS.Trt_vs_MS.NoTrt_B |
| ATP5G1 | 0.01176 | 0.44493 | 0.227 | 0.33 | 1 | MS.Trt_vs_MS.NoTrt_B |
| ANKRD40 | 0.01177 | 0.71823 | 0.658 | 0.78 | 1 | MS.Trt_vs_MS.NoTrt_B |
| SFN | 0.01179 | -0.343 | 0.2 | 0.28 | 1 | MS.Trt_vs_MS.NoTrt_B |
| AMT | 0.0118 | -0.2682 | 0.325 | 0.42 | 1 | MS.Trt_vs_MS.NoTrt_B |
| CSTF3 | 0.01182 | 0.25838 | 0.412 | 0.55 | 1 | MS.Trt_vs_MS.NoTrt_B |
| SP3 | 0.01184 | 0.37491 | 0.438 | 0.58 | 1 | MS.Trt_vs_MS.NoTrt_B |
| VGLL4 | 0.01184 | 0.48212 | 0.874 | 0.95 | 1 | MS.Trt_vs_MS.NoTrt_B |
| RBM41 | 0.01185 | 0.38266 | 0.331 | 0.44 | 1 | MS.Trt_vs_MS.NoTrt_B |
| CDK13 | 0.01187 | -0.3024 | 0.732 | 0.83 | 1 | MS.Trt_vs_MS.NoTrt_B |
| SKAP2 | 0.01189 | -1.0236 | 0.12 | 0.19 | 1 | MS.Trt_vs_MS.NoTrt_B |
| TGOLN2 | 0.01189 | 0.26067 | 0.516 | 0.65 | 1 | MS.Trt_vs_MS.NoTrt_B |
| FAM136A | 0.0119 | -0.3179 | 0.246 | 0.35 | 1 | MS.Trt_vs_MS.NoTrt_B |

| | | | | | | |
|---|---|---|---|---|---|---|
| ABCB10 | 0.01194 | -0.4394 | 0.061 | 0.12 | 1 | MS.Trt_vs_MS.NoTrt_B |
| CTC-487M23 | 0.01198 | -0.3518 | 0.852 | 0.91 | 1 | MS.Trt_vs_MS.NoTrt_B |
| ZNF486 | 0.012 | -0.2984 | 0.558 | 0.65 | 1 | MS.Trt_vs_MS.NoTrt_B |
| SPTLC1 | 0.01201 | 0.37211 | 0.595 | 0.74 | 1 | MS.Trt_vs_MS.NoTrt_B |
| SNUPN | 0.01202 | -0.3408 | 0.712 | 0.83 | 1 | MS.Trt_vs_MS.NoTrt_B |
| PDCD7 | 0.01205 | -0.4693 | 0.34 | 0.43 | 1 | MS.Trt_vs_MS.NoTrt_B |
| VSNL1 | 0.01206 | 0.28638 | 0.641 | 0.57 | 1 | MS.Trt_vs_MS.NoTrt_B |
| BRCA2 | 0.01208 | 1.14422 | 0.268 | 0.38 | 1 | MS.Trt_vs_MS.NoTrt_B |
| AUH | 0.01211 | -0.6369 | 0.065 | 0.12 | 1 | MS.Trt_vs_MS.NoTrt_B |
| CYB561A3 | 0.01211 | -0.4166 | 0.932 | 0.99 | 1 | MS.Trt_vs_MS.NoTrt_B |
| MTRNR2L6 | 0.01211 | 0.2748 | 0.88 | 1 | 1 | MS.Trt_vs_MS.NoTrt_B |
| GPATCH2 | 0.01213 | 0.56121 | 0.484 | 0.36 | 1 | MS.Trt_vs_MS.NoTrt_B |
| ALS2 | 0.01213 | 0.54772 | 0.312 | 0.45 | 1 | MS.Trt_vs_MS.NoTrt_B |
| ZC3H14 | 0.01217 | 0.39093 | 0.828 | 0.94 | 1 | MS.Trt_vs_MS.NoTrt_B |
| DHX29 | 0.01218 | -0.3113 | 0.109 | 0.18 | 1 | MS.Trt_vs_MS.NoTrt_B |
| CERS4 | 0.01219 | -0.6437 | 0.484 | 0.6 | 1 | MS.Trt_vs_MS.NoTrt_B |
| STX17 | 0.0122 | 0.85509 | 0.37 | 0.5 | 1 | MS.Trt_vs_MS.NoTrt_B |
| COPS7B | 0.0122 | -0.2846 | 0.686 | 0.76 | 1 | MS.Trt_vs_MS.NoTrt_B |
| EHMT2 | 0.01224 | -0.3107 | 0.523 | 0.64 | 1 | MS.Trt_vs_MS.NoTrt_B |
| SDR42E1 | 0.01227 | 0.37824 | 0.312 | 0.43 | 1 | MS.Trt_vs_MS.NoTrt_B |
| POGK | 0.0123 | -0.6274 | 0.176 | 0.26 | 1 | MS.Trt_vs_MS.NoTrt_B |
| MYCBP2 | 0.0123 | 0.68805 | 0.327 | 0.43 | 1 | MS.Trt_vs_MS.NoTrt_B |
| ZNF774 | 0.01232 | 0.37918 | 0.824 | 0.91 | 1 | MS.Trt_vs_MS.NoTrt_B |
| SENP8 | 0.01232 | 0.38383 | 0.386 | 0.48 | 1 | MS.Trt_vs_MS.NoTrt_B |
| MED29 | 0.01235 | 0.31608 | 0.407 | 0.51 | 1 | MS.Trt_vs_MS.NoTrt_B |
| ZNF500 | 0.01235 | 0.62945 | 0.466 | 0.64 | 1 | MS.Trt_vs_MS.NoTrt_B |
| ECHS1 | 0.01236 | -0.3457 | 0.089 | 0.16 | 1 | MS.Trt_vs_MS.NoTrt_B |
| ZNF331 | 0.01239 | 0.34838 | 0.492 | 0.61 | 1 | MS.Trt_vs_MS.NoTrt_B |
| FOCAD | 0.01243 | 0.30179 | 0.397 | 0.55 | 1 | MS.Trt_vs_MS.NoTrt_B |
| CARD16 | 0.01244 | 0.48617 | 0.224 | 0.34 | 1 | MS.Trt_vs_MS.NoTrt_B |
| CDHR3 | 0.01244 | -0.6549 | 0.51 | 0.62 | 1 | MS.Trt_vs_MS.NoTrt_B |
| SLC35F2 | 0.01244 | 0.48028 | 0.203 | 0.28 | 1 | MS.Trt_vs_MS.NoTrt_B |
| MFSD1 | 0.01249 | -0.4591 | 0.07 | 0.13 | 1 | MS.Trt_vs_MS.NoTrt_B |
| KPNA2 | 0.01249 | -0.7727 | 0.07 | 0.13 | 1 | MS.Trt_vs_MS.NoTrt_B |
| MSL1 | 0.01252 | 0.6253 | 0.479 | 0.64 | 1 | MS.Trt_vs_MS.NoTrt_B |
| LRRFIP2 | 0.01255 | 0.35036 | 0.571 | 0.67 | 1 | MS.Trt_vs_MS.NoTrt_B |
| RPS17 | 0.01259 | 0.36233 | 0.32 | 0.42 | 1 | MS.Trt_vs_MS.NoTrt_B |
| FAM227B | 0.01261 | -0.4494 | 0.307 | 0.4 | 1 | MS.Trt_vs_MS.NoTrt_B |
| CASP7 | 0.01266 | -0.8484 | 0.1 | 0.17 | 1 | MS.Trt_vs_MS.NoTrt_B |
| RNMT | 0.01267 | 0.37452 | 0.233 | 0.34 | 1 | MS.Trt_vs_MS.NoTrt_B |

| | | | | | | |
|---|---|---|---|---|---|---|
| HTRA2 | 0.0127 | -0.347 | 0.085 | 0.15 | 1 | MS.Trt_vs_MS.NoTrt_B |
| SEPHS2 | 0.01275 | -0.5632 | 0.07 | 0.13 | 1 | MS.Trt_vs_MS.NoTrt_B |
| CHURC1 | 0.01277 | 0.41022 | 0.61 | 0.7 | 1 | MS.Trt_vs_MS.NoTrt_B |
| GPR83 | 0.01278 | 0.73145 | 0.362 | 0.48 | 1 | MS.Trt_vs_MS.NoTrt_B |
| BEX4 | 0.01285 | -0.3102 | 0.083 | 0.15 | 1 | MS.Trt_vs_MS.NoTrt_B |
| NT5DC3 | 0.01286 | 0.31662 | 0.654 | 0.77 | 1 | MS.Trt_vs_MS.NoTrt_B |
| TMEM42 | 0.01286 | 0.27064 | 0.072 | 0.14 | 1 | MS.Trt_vs_MS.NoTrt_B |
| MTX2 | 0.01286 | -0.5395 | 0.135 | 0.21 | 1 | MS.Trt_vs_MS.NoTrt_B |
| CLCC1 | 0.01287 | 0.25423 | 0.682 | 0.83 | 1 | MS.Trt_vs_MS.NoTrt_B |
| PRRG3 | 0.01292 | 0.54071 | 0.514 | 0.42 | 1 | MS.Trt_vs_MS.NoTrt_B |
| SHARPIN | 0.01293 | -0.322 | 0.094 | 0.16 | 1 | MS.Trt_vs_MS.NoTrt_B |
| LMNB1 | 0.01294 | -0.7569 | 0.05 | 0.1 | 1 | MS.Trt_vs_MS.NoTrt_B |
| MKLN1 | 0.01296 | 0.36765 | 0.473 | 0.6 | 1 | MS.Trt_vs_MS.NoTrt_B |
| XKR9 | 0.01306 | 0.26134 | 0.42 | 0.56 | 1 | MS.Trt_vs_MS.NoTrt_B |
| GPR132 | 0.01311 | -0.6551 | 0.27 | 0.37 | 1 | MS.Trt_vs_MS.NoTrt_B |
| POU6F1 | 0.0132 | -0.8363 | 0.373 | 0.47 | 1 | MS.Trt_vs_MS.NoTrt_B |
| DAP | 0.01326 | -0.2804 | 0.094 | 0.16 | 1 | MS.Trt_vs_MS.NoTrt_B |
| AGAP2 | 0.01326 | -0.2848 | 0.124 | 0.2 | 1 | MS.Trt_vs_MS.NoTrt_B |
| RAB11A | 0.01328 | 0.3024 | 0.562 | 0.65 | 1 | MS.Trt_vs_MS.NoTrt_B |
| INTS4 | 0.01333 | -0.6977 | 0.281 | 0.37 | 1 | MS.Trt_vs_MS.NoTrt_B |
| DDX11 | 0.01334 | -0.7582 | 0.301 | 0.38 | 1 | MS.Trt_vs_MS.NoTrt_B |
| L3MBTL1 | 0.01335 | 0.41276 | 0.488 | 0.6 | 1 | MS.Trt_vs_MS.NoTrt_B |
| NPL | 0.01335 | 1.02956 | 0.153 | 0.24 | 1 | MS.Trt_vs_MS.NoTrt_B |
| NXPE3 | 0.01335 | -0.483 | 0.821 | 0.92 | 1 | MS.Trt_vs_MS.NoTrt_B |
| PREB | 0.01336 | -0.6117 | 0.078 | 0.14 | 1 | MS.Trt_vs_MS.NoTrt_B |
| RALGPS1 | 0.01337 | 0.53572 | 0.301 | 0.43 | 1 | MS.Trt_vs_MS.NoTrt_B |
| PHYH | 0.01338 | -0.4721 | 0.118 | 0.19 | 1 | MS.Trt_vs_MS.NoTrt_B |
| ABHD13 | 0.0134 | -0.4147 | 0.054 | 0.11 | 1 | MS.Trt_vs_MS.NoTrt_B |
| NAA30 | 0.01343 | 0.50754 | 0.342 | 0.46 | 1 | MS.Trt_vs_MS.NoTrt_B |
| SLC14A1 | 0.01344 | 0.33265 | 0.397 | 0.53 | 1 | MS.Trt_vs_MS.NoTrt_B |
| NECTIN4 | 0.01345 | -0.4354 | 0.129 | 0.21 | 1 | MS.Trt_vs_MS.NoTrt_B |
| DNAJC19 | 0.01345 | 0.70521 | 0.185 | 0.28 | 1 | MS.Trt_vs_MS.NoTrt_B |
| FSD2 | 0.01345 | 0.431 | 0.375 | 0.49 | 1 | MS.Trt_vs_MS.NoTrt_B |
| C22orf46 | 0.01347 | -0.3078 | 0.246 | 0.35 | 1 | MS.Trt_vs_MS.NoTrt_B |
| UBALD2 | 0.01351 | 0.7597 | 0.083 | 0.15 | 1 | MS.Trt_vs_MS.NoTrt_B |
| CARNS1 | 0.01356 | -0.5272 | 0.144 | 0.22 | 1 | MS.Trt_vs_MS.NoTrt_B |
| FGGY | 0.01358 | -0.5576 | 0.32 | 0.44 | 1 | MS.Trt_vs_MS.NoTrt_B |
| ZNF227 | 0.01362 | 0.77233 | 0.375 | 0.5 | 1 | MS.Trt_vs_MS.NoTrt_B |
| TMEM119 | 0.01366 | -0.6252 | 0.455 | 0.57 | 1 | MS.Trt_vs_MS.NoTrt_B |
| PPP2R5E | 0.01369 | 0.79688 | 0.174 | 0.27 | 1 | MS.Trt_vs_MS.NoTrt_B |

| | | | | | | |
|---|---|---|---|---|---|---|
| TRIM66 | 0.0137 | -0.3072 | 0.322 | 0.42 | 1 | MS.Trt_vs_MS.NoTrt_B |
| CEP89 | 0.01373 | 0.42123 | 0.789 | 0.92 | 1 | MS.Trt_vs_MS.NoTrt_B |
| CENPF | 0.01375 | -0.6641 | 0.259 | 0.38 | 1 | MS.Trt_vs_MS.NoTrt_B |
| RP11-80H18 | 0.01384 | -0.5444 | 0.194 | 0.28 | 1 | MS.Trt_vs_MS.NoTrt_B |
| RBL1 | 0.01389 | 0.40292 | 0.479 | 0.57 | 1 | MS.Trt_vs_MS.NoTrt_B |
| ZNF382 | 0.0139 | -0.5154 | 0.839 | 0.94 | 1 | MS.Trt_vs_MS.NoTrt_B |
| AKAP17A | 0.0139 | -0.8292 | 0.05 | 0.1 | 1 | MS.Trt_vs_MS.NoTrt_B |
| TAP2 | 0.01396 | -0.3416 | 0.211 | 0.3 | 1 | MS.Trt_vs_MS.NoTrt_B |
| RIN3 | 0.01397 | 0.58595 | 0.248 | 0.36 | 1 | MS.Trt_vs_MS.NoTrt_B |
| WDR11 | 0.01398 | -0.411 | 0.503 | 0.62 | 1 | MS.Trt_vs_MS.NoTrt_B |
| HIST1H4C | 0.01398 | 0.56439 | 0.283 | 0.42 | 1 | MS.Trt_vs_MS.NoTrt_B |
| ZCCHC4 | 0.01403 | -0.569 | 0.477 | 0.61 | 1 | MS.Trt_vs_MS.NoTrt_B |
| RBM12B | 0.01404 | -0.8722 | 0.059 | 0.11 | 1 | MS.Trt_vs_MS.NoTrt_B |
| MIB1 | 0.01405 | 0.50544 | 0.094 | 0.16 | 1 | MS.Trt_vs_MS.NoTrt_B |
| TMEM170A | 0.01405 | -0.5349 | 0.706 | 0.77 | 1 | MS.Trt_vs_MS.NoTrt_B |
| NAIF1 | 0.01405 | 0.27212 | 0.42 | 0.54 | 1 | MS.Trt_vs_MS.NoTrt_B |
| SLAMF7 | 0.01407 | 0.51066 | 0.322 | 0.44 | 1 | MS.Trt_vs_MS.NoTrt_B |
| RP11-552F3 | 0.01413 | 0.33009 | 0.109 | 0.18 | 1 | MS.Trt_vs_MS.NoTrt_B |
| IPO9 | 0.01415 | 0.36553 | 0.717 | 0.83 | 1 | MS.Trt_vs_MS.NoTrt_B |
| ZNF154 | 0.01419 | 0.80256 | 0.532 | 0.69 | 1 | MS.Trt_vs_MS.NoTrt_B |
| PSD4 | 0.01423 | 0.36597 | 0.834 | 0.91 | 1 | MS.Trt_vs_MS.NoTrt_B |
| PTCD1 | 0.01426 | 0.3956 | 0.49 | 0.61 | 1 | MS.Trt_vs_MS.NoTrt_B |
| ITGAL | 0.0143 | 0.30016 | 0.654 | 0.75 | 1 | MS.Trt_vs_MS.NoTrt_B |
| TRIM68 | 0.0143 | -0.37 | 0.405 | 0.54 | 1 | MS.Trt_vs_MS.NoTrt_B |
| CRTAM | 0.01431 | 0.34821 | 0.283 | 0.41 | 1 | MS.Trt_vs_MS.NoTrt_B |
| GINM1 | 0.01433 | -0.6254 | 0.397 | 0.48 | 1 | MS.Trt_vs_MS.NoTrt_B |
| AMD1 | 0.01434 | -0.3294 | 0.623 | 0.74 | 1 | MS.Trt_vs_MS.NoTrt_B |
| DONSON | 0.01434 | -0.5558 | 0.122 | 0.2 | 1 | MS.Trt_vs_MS.NoTrt_B |
| UBAP2 | 0.01438 | 0.47264 | 0.534 | 0.69 | 1 | MS.Trt_vs_MS.NoTrt_B |
| ATP13A3 | 0.01441 | 0.90348 | 0.085 | 0.15 | 1 | MS.Trt_vs_MS.NoTrt_B |
| MED22 | 0.01443 | 0.30937 | 0.423 | 0.55 | 1 | MS.Trt_vs_MS.NoTrt_B |
| RPL34 | 0.01443 | 0.52452 | 0.941 | 0.99 | 1 | MS.Trt_vs_MS.NoTrt_B |
| PCYOX1L | 0.01444 | -0.418 | 0.054 | 0.11 | 1 | MS.Trt_vs_MS.NoTrt_B |
| ARL6IP5 | 0.01452 | 0.38469 | 0.603 | 0.74 | 1 | MS.Trt_vs_MS.NoTrt_B |
| TNIP2 | 0.01454 | -0.9181 | 0.063 | 0.12 | 1 | MS.Trt_vs_MS.NoTrt_B |
| AP1B1 | 0.01456 | -0.5028 | 0.144 | 0.22 | 1 | MS.Trt_vs_MS.NoTrt_B |
| PCGF1 | 0.01461 | -0.3503 | 0.336 | 0.44 | 1 | MS.Trt_vs_MS.NoTrt_B |
| SGK494 | 0.01463 | 0.28736 | 0.56 | 0.71 | 1 | MS.Trt_vs_MS.NoTrt_B |
| ZBTB37 | 0.01471 | -0.3067 | 0.553 | 0.69 | 1 | MS.Trt_vs_MS.NoTrt_B |
| STARD9 | 0.01472 | 0.41526 | 0.756 | 0.93 | 1 | MS.Trt_vs_MS.NoTrt_B |

| | | | | | | |
|---|---|---|---|---|---|---|
| PCGF5 | 0.01491 | -0.5014 | 0.298 | 0.38 | 1 | MS.Trt_vs_MS.NoTrt_B |
| SCYL1 | 0.01495 | -0.2821 | 0.203 | 0.29 | 1 | MS.Trt_vs_MS.NoTrt_B |
| CHEK1 | 0.01495 | 0.5348 | 0.597 | 0.77 | 1 | MS.Trt_vs_MS.NoTrt_B |
| CLK1 | 0.01496 | 0.45396 | 0.453 | 0.59 | 1 | MS.Trt_vs_MS.NoTrt_B |
| RRP15 | 0.01498 | 0.5796 | 0.44 | 0.57 | 1 | MS.Trt_vs_MS.NoTrt_B |
| KIF3A | 0.01499 | 0.66971 | 0.584 | 0.72 | 1 | MS.Trt_vs_MS.NoTrt_B |
| PYGB | 0.015 | -0.9216 | 0.081 | 0.14 | 1 | MS.Trt_vs_MS.NoTrt_B |
| OTUD3 | 0.01501 | 0.97993 | 0.375 | 0.49 | 1 | MS.Trt_vs_MS.NoTrt_B |
| RBM34 | 0.01504 | 0.44106 | 0.593 | 0.74 | 1 | MS.Trt_vs_MS.NoTrt_B |
| RDH16 | 0.01508 | 0.26096 | 0.529 | 0.73 | 1 | MS.Trt_vs_MS.NoTrt_B |
| WASH1 | 0.01509 | 0.64698 | 0.183 | 0.28 | 1 | MS.Trt_vs_MS.NoTrt_B |
| NHLRC1 | 0.01511 | -0.846 | 0.146 | 0.23 | 1 | MS.Trt_vs_MS.NoTrt_B |
| SURF1 | 0.01514 | -0.3852 | 0.059 | 0.11 | 1 | MS.Trt_vs_MS.NoTrt_B |
| NCAPH | 0.01515 | -0.6842 | 0.303 | 0.42 | 1 | MS.Trt_vs_MS.NoTrt_B |
| PLS1 | 0.01518 | -0.6202 | 0.185 | 0.28 | 1 | MS.Trt_vs_MS.NoTrt_B |
| PTPRA | 0.01519 | -0.9485 | 0.081 | 0.14 | 1 | MS.Trt_vs_MS.NoTrt_B |
| EXOSC9 | 0.01521 | -0.3638 | 0.161 | 0.24 | 1 | MS.Trt_vs_MS.NoTrt_B |
| SLC22A5 | 0.01521 | 0.79431 | 0.294 | 0.41 | 1 | MS.Trt_vs_MS.NoTrt_B |
| SRSF3 | 0.01522 | 0.34861 | 0.76 | 0.84 | 1 | MS.Trt_vs_MS.NoTrt_B |
| B3GNTL1 | 0.01531 | -0.2665 | 0.2 | 0.29 | 1 | MS.Trt_vs_MS.NoTrt_B |
| ARHGAP18 | 0.01533 | -0.875 | 0.059 | 0.11 | 1 | MS.Trt_vs_MS.NoTrt_B |
| BAG5 | 0.01535 | 0.36701 | 0.257 | 0.36 | 1 | MS.Trt_vs_MS.NoTrt_B |
| NOCT | 0.0154 | 0.25592 | 0.542 | 0.69 | 1 | MS.Trt_vs_MS.NoTrt_B |
| POLE | 0.01542 | 0.73332 | 0.682 | 0.83 | 1 | MS.Trt_vs_MS.NoTrt_B |
| GAL3ST4 | 0.01546 | -1.6175 | 0.351 | 0.45 | 1 | MS.Trt_vs_MS.NoTrt_B |
| AGMAT | 0.0155 | -0.3163 | 0.773 | 0.85 | 1 | MS.Trt_vs_MS.NoTrt_B |
| RPAP2 | 0.01555 | 0.34931 | 0.978 | 1 | 1 | MS.Trt_vs_MS.NoTrt_B |
| ESD | 0.0156 | -0.4756 | 0.076 | 0.14 | 1 | MS.Trt_vs_MS.NoTrt_B |
| SLC25A37 | 0.01569 | 0.35087 | 0.549 | 0.68 | 1 | MS.Trt_vs_MS.NoTrt_B |
| PPP2R5C | 0.0157 | 0.33024 | 0.837 | 0.92 | 1 | MS.Trt_vs_MS.NoTrt_B |
| MKL1 | 0.01574 | 0.28277 | 0.292 | 0.4 | 1 | MS.Trt_vs_MS.NoTrt_B |
| LDB1 | 0.01578 | -0.4536 | 0.068 | 0.12 | 1 | MS.Trt_vs_MS.NoTrt_B |
| KIF1BP | 0.01588 | 0.35732 | 0.521 | 0.63 | 1 | MS.Trt_vs_MS.NoTrt_B |
| C9orf131 | 0.01588 | 0.61556 | 0.081 | 0.15 | 1 | MS.Trt_vs_MS.NoTrt_B |
| NKTR | 0.0159 | 0.51179 | 0.732 | 0.83 | 1 | MS.Trt_vs_MS.NoTrt_B |
| ZNF460 | 0.01592 | -0.6615 | 0.137 | 0.21 | 1 | MS.Trt_vs_MS.NoTrt_B |
| SPG21 | 0.01592 | -0.2843 | 0.76 | 0.88 | 1 | MS.Trt_vs_MS.NoTrt_B |
| JOSD1 | 0.01599 | 0.41921 | 0.403 | 0.5 | 1 | MS.Trt_vs_MS.NoTrt_B |
| SOCS4 | 0.01602 | 0.32741 | 0.205 | 0.3 | 1 | MS.Trt_vs_MS.NoTrt_B |
| ZC3H12C | 0.01606 | 0.53369 | 0.072 | 0.13 | 1 | MS.Trt_vs_MS.NoTrt_B |

| | | | | | | |
|---|---|---|---|---|---|---|
| ITFG1 | 0.01608 | 0.64442 | 0.15 | 0.23 | 1 | MS.Trt_vs_MS.NoTrt_B |
| PRMT1 | 0.01622 | -0.2521 | 0.695 | 0.82 | 1 | MS.Trt_vs_MS.NoTrt_B |
| TFR2 | 0.01635 | 0.67214 | 0.577 | 0.71 | 1 | MS.Trt_vs_MS.NoTrt_B |
| TYSND1 | 0.01638 | -0.3465 | 0.179 | 0.26 | 1 | MS.Trt_vs_MS.NoTrt_B |
| SLC25A25 | 0.01641 | -0.3021 | 0.087 | 0.15 | 1 | MS.Trt_vs_MS.NoTrt_B |
| IFRD2 | 0.01654 | -0.4729 | 0.081 | 0.14 | 1 | MS.Trt_vs_MS.NoTrt_B |
| ANGEL1 | 0.01656 | -0.8708 | 0.329 | 0.42 | 1 | MS.Trt_vs_MS.NoTrt_B |
| CRIPT | 0.01657 | 0.3492 | 0.172 | 0.25 | 1 | MS.Trt_vs_MS.NoTrt_B |
| DTNB | 0.0166 | -0.4639 | 0.111 | 0.18 | 1 | MS.Trt_vs_MS.NoTrt_B |
| EPRS | 0.01661 | -0.323 | 0.148 | 0.22 | 1 | MS.Trt_vs_MS.NoTrt_B |
| PFKM | 0.01664 | -0.2811 | 0.715 | 0.87 | 1 | MS.Trt_vs_MS.NoTrt_B |
| ZNF286A | 0.01671 | 0.68688 | 0.27 | 0.38 | 1 | MS.Trt_vs_MS.NoTrt_B |
| STARD4 | 0.01677 | -0.9504 | 0.085 | 0.15 | 1 | MS.Trt_vs_MS.NoTrt_B |
| ZNF3 | 0.01678 | -0.2916 | 0.58 | 0.69 | 1 | MS.Trt_vs_MS.NoTrt_B |
| CAPG | 0.01679 | -1.2552 | 0.057 | 0.11 | 1 | MS.Trt_vs_MS.NoTrt_B |
| PROSC | 0.01679 | -0.3573 | 0.331 | 0.41 | 1 | MS.Trt_vs_MS.NoTrt_B |
| ZNF461 | 0.0168 | -0.5648 | 0.564 | 0.64 | 1 | MS.Trt_vs_MS.NoTrt_B |
| TRIM37 | 0.01683 | -0.6559 | 0.113 | 0.18 | 1 | MS.Trt_vs_MS.NoTrt_B |
| BTBD7 | 0.01684 | 0.32537 | 0.595 | 0.73 | 1 | MS.Trt_vs_MS.NoTrt_B |
| AKTIP | 0.01687 | -0.3413 | 0.521 | 0.63 | 1 | MS.Trt_vs_MS.NoTrt_B |
| FAM156A | 0.01688 | -0.9293 | 0.1 | 0.16 | 1 | MS.Trt_vs_MS.NoTrt_B |
| ZNF641 | 0.01689 | 0.26346 | 0.373 | 0.5 | 1 | MS.Trt_vs_MS.NoTrt_B |
| VAPA | 0.01691 | 0.36299 | 0.638 | 0.74 | 1 | MS.Trt_vs_MS.NoTrt_B |
| GCHFR | 0.01692 | 0.31169 | 0.081 | 0.15 | 1 | MS.Trt_vs_MS.NoTrt_B |
| COA3 | 0.01694 | 0.38611 | 0.102 | 0.17 | 1 | MS.Trt_vs_MS.NoTrt_B |
| ZNF121 | 0.01696 | 0.37054 | 0.773 | 0.9 | 1 | MS.Trt_vs_MS.NoTrt_B |
| C20orf194 | 0.01697 | 0.42356 | 0.179 | 0.27 | 1 | MS.Trt_vs_MS.NoTrt_B |
| HMGXB4 | 0.01697 | -0.2701 | 0.068 | 0.12 | 1 | MS.Trt_vs_MS.NoTrt_B |
| POLI | 0.017 | 0.26816 | 0.266 | 0.37 | 1 | MS.Trt_vs_MS.NoTrt_B |
| AGTRAP | 0.01705 | -0.6541 | 0.076 | 0.14 | 1 | MS.Trt_vs_MS.NoTrt_B |
| PVRIG | 0.01705 | 0.46708 | 0.253 | 0.36 | 1 | MS.Trt_vs_MS.NoTrt_B |
| JMJD7 | 0.01706 | -0.4554 | 0.089 | 0.15 | 1 | MS.Trt_vs_MS.NoTrt_B |
| TADA2A | 0.01708 | -0.8996 | 0.782 | 0.88 | 1 | MS.Trt_vs_MS.NoTrt_B |
| CUBN | 0.0171 | 0.36269 | 0.301 | 0.41 | 1 | MS.Trt_vs_MS.NoTrt_B |
| SLC25A29 | 0.01711 | 0.36253 | 0.268 | 0.39 | 1 | MS.Trt_vs_MS.NoTrt_B |
| PLGLB1 | 0.01714 | 0.60567 | 0.089 | 0.16 | 1 | MS.Trt_vs_MS.NoTrt_B |
| PSMD3 | 0.01723 | -0.4769 | 0.394 | 0.51 | 1 | MS.Trt_vs_MS.NoTrt_B |
| CASP8AP2 | 0.01723 | 0.4324 | 0.305 | 0.41 | 1 | MS.Trt_vs_MS.NoTrt_B |
| FGD2 | 0.01731 | 0.25414 | 0.569 | 0.71 | 1 | MS.Trt_vs_MS.NoTrt_B |
| MED7 | 0.01732 | 0.33978 | 0.059 | 0.11 | 1 | MS.Trt_vs_MS.NoTrt_B |

| | | | | | | |
|---|---|---|---|---|---|---|
| DLD | 0.01734 | 0.57189 | 0.1 | 0.17 | 1 | MS.Trt_vs_MS.NoTrt_B |
| PM20D2 | 0.01741 | 0.35341 | 0.619 | 0.78 | 1 | MS.Trt_vs_MS.NoTrt_B |
| UBE2V2 | 0.01748 | 1.04734 | 0.621 | 0.72 | 1 | MS.Trt_vs_MS.NoTrt_B |
| FILIP1L | 0.0175 | -0.4771 | 0.231 | 0.33 | 1 | MS.Trt_vs_MS.NoTrt_B |
| ERCC6L | 0.01752 | -0.6386 | 0.083 | 0.15 | 1 | MS.Trt_vs_MS.NoTrt_B |
| PINK1 | 0.01758 | -0.6909 | 0.194 | 0.28 | 1 | MS.Trt_vs_MS.NoTrt_B |
| RASGRF1 | 0.01758 | 0.25394 | 0.235 | 0.34 | 1 | MS.Trt_vs_MS.NoTrt_B |
| TRAPPC11 | 0.01758 | -0.4577 | 0.305 | 0.39 | 1 | MS.Trt_vs_MS.NoTrt_B |
| SLC5A6 | 0.0176 | -0.6708 | 0.052 | 0.1 | 1 | MS.Trt_vs_MS.NoTrt_B |
| ELK4 | 0.01764 | 0.65115 | 0.399 | 0.52 | 1 | MS.Trt_vs_MS.NoTrt_B |
| TRAPPC9 | 0.01765 | -0.3431 | 0.072 | 0.13 | 1 | MS.Trt_vs_MS.NoTrt_B |
| PDE8A | 0.01771 | 0.26895 | 0.325 | 0.44 | 1 | MS.Trt_vs_MS.NoTrt_B |
| WASF2 | 0.01782 | 0.30966 | 0.481 | 0.64 | 1 | MS.Trt_vs_MS.NoTrt_B |
| LSM12 | 0.01783 | 0.40709 | 0.577 | 0.71 | 1 | MS.Trt_vs_MS.NoTrt_B |
| ISOC1 | 0.01789 | -0.4672 | 0.052 | 0.1 | 1 | MS.Trt_vs_MS.NoTrt_B |
| BRIP1 | 0.01789 | 0.6007 | 0.695 | 0.89 | 1 | MS.Trt_vs_MS.NoTrt_B |
| MDM2 | 0.0179 | 0.27477 | 0.952 | 1 | 1 | MS.Trt_vs_MS.NoTrt_B |
| SLC23A3 | 0.01792 | -0.5949 | 0.305 | 0.42 | 1 | MS.Trt_vs_MS.NoTrt_B |
| VRK2 | 0.01796 | -0.4143 | 0.172 | 0.25 | 1 | MS.Trt_vs_MS.NoTrt_B |
| TUBD1 | 0.01799 | 0.42244 | 0.723 | 0.8 | 1 | MS.Trt_vs_MS.NoTrt_B |
| ZNF470 | 0.01802 | 0.40558 | 0.569 | 0.74 | 1 | MS.Trt_vs_MS.NoTrt_B |
| SMS | 0.01804 | -0.4306 | 0.096 | 0.16 | 1 | MS.Trt_vs_MS.NoTrt_B |
| NME4 | 0.01805 | -0.3641 | 0.068 | 0.12 | 1 | MS.Trt_vs_MS.NoTrt_B |
| TAB2 | 0.01806 | 0.82383 | 0.44 | 0.57 | 1 | MS.Trt_vs_MS.NoTrt_B |
| CAPZA2 | 0.01808 | 0.71783 | 0.821 | 0.85 | 1 | MS.Trt_vs_MS.NoTrt_B |
| GMPS | 0.0181 | -0.4293 | 0.756 | 0.87 | 1 | MS.Trt_vs_MS.NoTrt_B |
| NEIL1 | 0.01813 | -0.2713 | 0.654 | 0.81 | 1 | MS.Trt_vs_MS.NoTrt_B |
| DDX3X | 0.01826 | 0.56048 | 0.224 | 0.32 | 1 | MS.Trt_vs_MS.NoTrt_B |
| NUP210 | 0.0184 | 0.42255 | 0.379 | 0.5 | 1 | MS.Trt_vs_MS.NoTrt_B |
| FCRL1 | 0.01843 | -0.7112 | 0.163 | 0.24 | 1 | MS.Trt_vs_MS.NoTrt_B |
| GPR68 | 0.01849 | -0.8742 | 0.061 | 0.11 | 1 | MS.Trt_vs_MS.NoTrt_B |
| TP53INP1 | 0.01849 | 0.42001 | 0.261 | 0.37 | 1 | MS.Trt_vs_MS.NoTrt_B |
| FBXL3 | 0.01854 | 0.59509 | 0.096 | 0.16 | 1 | MS.Trt_vs_MS.NoTrt_B |
| ABHD11 | 0.01863 | 0.37066 | 0.479 | 0.62 | 1 | MS.Trt_vs_MS.NoTrt_B |
| CHML | 0.01866 | -0.722 | 0.15 | 0.23 | 1 | MS.Trt_vs_MS.NoTrt_B |
| PCBD1 | 0.0187 | 0.34527 | 0.059 | 0.11 | 1 | MS.Trt_vs_MS.NoTrt_B |
| MTHFD2L | 0.0187 | -0.3253 | 0.39 | 0.49 | 1 | MS.Trt_vs_MS.NoTrt_B |
| PKP2 | 0.01875 | 0.54702 | 0.547 | 0.47 | 1 | MS.Trt_vs_MS.NoTrt_B |
| CCDC171 | 0.01879 | -0.2501 | 0.34 | 0.45 | 1 | MS.Trt_vs_MS.NoTrt_B |
| PTPN7 | 0.01881 | 0.25849 | 0.571 | 0.72 | 1 | MS.Trt_vs_MS.NoTrt_B |

| | | | | | | |
|---|---|---|---|---|---|---|
| TSC22D2 | 0.01883 | 0.37916 | 0.285 | 0.38 | 1 | MS.Trt_vs_MS.NoTrt_B |
| FDPS | 0.01884 | -0.3609 | 0.401 | 0.48 | 1 | MS.Trt_vs_MS.NoTrt_B |
| ARHGAP27 | 0.01884 | 0.42138 | 0.46 | 0.59 | 1 | MS.Trt_vs_MS.NoTrt_B |
| AP4M1 | 0.01892 | -0.6106 | 0.447 | 0.61 | 1 | MS.Trt_vs_MS.NoTrt_B |
| CMC1 | 0.01894 | 1.32656 | 0.115 | 0.19 | 1 | MS.Trt_vs_MS.NoTrt_B |
| ABCG5 | 0.01895 | -0.3961 | 0.159 | 0.24 | 1 | MS.Trt_vs_MS.NoTrt_B |
| ZNF681 | 0.01897 | 0.63818 | 0.549 | 0.68 | 1 | MS.Trt_vs_MS.NoTrt_B |
| PIP4K2C | 0.01897 | -0.5581 | 0.115 | 0.18 | 1 | MS.Trt_vs_MS.NoTrt_B |
| PHYKPL | 0.01901 | 0.345 | 0.336 | 0.44 | 1 | MS.Trt_vs_MS.NoTrt_B |
| NUP58 | 0.01902 | -0.4589 | 0.242 | 0.33 | 1 | MS.Trt_vs_MS.NoTrt_B |
| CDK9 | 0.01903 | -0.5494 | 0.061 | 0.11 | 1 | MS.Trt_vs_MS.NoTrt_B |
| PAPLN | 0.01909 | -0.6583 | 0.621 | 0.72 | 1 | MS.Trt_vs_MS.NoTrt_B |
| PPP2R3C | 0.01909 | -0.4288 | 0.098 | 0.16 | 1 | MS.Trt_vs_MS.NoTrt_B |
| USP7 | 0.0191 | 0.51801 | 0.235 | 0.34 | 1 | MS.Trt_vs_MS.NoTrt_B |
| EXOSC2 | 0.01913 | 0.58115 | 0.625 | 0.78 | 1 | MS.Trt_vs_MS.NoTrt_B |
| CLNK | 0.01917 | 0.37481 | 0.488 | 0.64 | 1 | MS.Trt_vs_MS.NoTrt_B |
| SEC23IP | 0.01923 | 0.49907 | 0.316 | 0.42 | 1 | MS.Trt_vs_MS.NoTrt_B |
| CCDC58 | 0.01923 | 0.30949 | 0.253 | 0.36 | 1 | MS.Trt_vs_MS.NoTrt_B |
| MKKS | 0.01925 | 0.30398 | 0.059 | 0.11 | 1 | MS.Trt_vs_MS.NoTrt_B |
| PEF1 | 0.01933 | -0.5169 | 0.129 | 0.2 | 1 | MS.Trt_vs_MS.NoTrt_B |
| CECR1 | 0.01933 | -0.4362 | 0.355 | 0.48 | 1 | MS.Trt_vs_MS.NoTrt_B |
| VIMP | 0.01936 | -0.3327 | 0.085 | 0.15 | 1 | MS.Trt_vs_MS.NoTrt_B |
| FITM2 | 0.01938 | 0.56863 | 0.573 | 0.7 | 1 | MS.Trt_vs_MS.NoTrt_B |
| RERE | 0.01945 | -0.4201 | 0.253 | 0.35 | 1 | MS.Trt_vs_MS.NoTrt_B |
| RP11-111H1 | 0.01947 | 0.78256 | 0.083 | 0.15 | 1 | MS.Trt_vs_MS.NoTrt_B |
| TOP3A | 0.01947 | -0.408 | 0.73 | 0.85 | 1 | MS.Trt_vs_MS.NoTrt_B |
| DHRS1 | 0.01948 | -0.5373 | 0.51 | 0.63 | 1 | MS.Trt_vs_MS.NoTrt_B |
| CEP152 | 0.01952 | 0.60077 | 0.562 | 0.67 | 1 | MS.Trt_vs_MS.NoTrt_B |
| FPGT | 0.01956 | -0.9069 | 0.074 | 0.13 | 1 | MS.Trt_vs_MS.NoTrt_B |
| ATXN7L1 | 0.0196 | 0.38997 | 0.264 | 0.37 | 1 | MS.Trt_vs_MS.NoTrt_B |
| HENMT1 | 0.01961 | -0.5318 | 0.264 | 0.36 | 1 | MS.Trt_vs_MS.NoTrt_B |
| FAM210A | 0.01971 | 1.05934 | 0.205 | 0.3 | 1 | MS.Trt_vs_MS.NoTrt_B |
| LTBP3 | 0.01978 | 0.54156 | 0.479 | 0.6 | 1 | MS.Trt_vs_MS.NoTrt_B |
| AK5 | 0.0198 | -0.3545 | 0.074 | 0.14 | 1 | MS.Trt_vs_MS.NoTrt_B |
| TFDP2 | 0.01981 | 0.45167 | 0.863 | 0.91 | 1 | MS.Trt_vs_MS.NoTrt_B |
| BLOC1S4 | 0.01985 | 0.55352 | 0.113 | 0.18 | 1 | MS.Trt_vs_MS.NoTrt_B |
| CSNK1E | 0.01985 | 0.39153 | 0.545 | 0.68 | 1 | MS.Trt_vs_MS.NoTrt_B |
| USP15 | 0.01993 | 0.5699 | 0.743 | 0.87 | 1 | MS.Trt_vs_MS.NoTrt_B |
| MCRIP2 | 0.01995 | 0.34593 | 0.394 | 0.51 | 1 | MS.Trt_vs_MS.NoTrt_B |
| ZNF570 | 0.01995 | 0.49561 | 0.66 | 0.75 | 1 | MS.Trt_vs_MS.NoTrt_B |

| | | | | | | |
|---|---|---|---|---|---|---|
| IFITM2 | 0.01997 | 0.38035 | 0.721 | 0.87 | 1 | MS.Trt_vs_MS.NoTrt_B |
| PTPN6 | 0.02 | 0.34053 | 0.309 | 0.43 | 1 | MS.Trt_vs_MS.NoTrt_B |
| NHLRC2 | 0.02003 | 0.59981 | 0.573 | 0.72 | 1 | MS.Trt_vs_MS.NoTrt_B |
| SDCBP | 0.02008 | 0.57093 | 0.124 | 0.2 | 1 | MS.Trt_vs_MS.NoTrt_B |
| YDJC | 0.02011 | 0.55127 | 0.081 | 0.14 | 1 | MS.Trt_vs_MS.NoTrt_B |
| CARD6 | 0.02016 | -0.5714 | 0.405 | 0.55 | 1 | MS.Trt_vs_MS.NoTrt_B |
| CACUL1 | 0.02021 | 0.56707 | 0.56 | 0.71 | 1 | MS.Trt_vs_MS.NoTrt_B |
| MIS18BP1 | 0.02026 | 0.26452 | 0.475 | 0.6 | 1 | MS.Trt_vs_MS.NoTrt_B |
| RNFT2 | 0.02028 | -0.2745 | 0.756 | 0.9 | 1 | MS.Trt_vs_MS.NoTrt_B |
| ZNF415 | 0.02029 | 0.39939 | 0.205 | 0.3 | 1 | MS.Trt_vs_MS.NoTrt_B |
| GOLT1B | 0.02031 | 0.56956 | 0.089 | 0.15 | 1 | MS.Trt_vs_MS.NoTrt_B |
| SAYSD1 | 0.02034 | 0.55592 | 0.126 | 0.2 | 1 | MS.Trt_vs_MS.NoTrt_B |
| SDF2 | 0.02034 | -0.3981 | 0.131 | 0.2 | 1 | MS.Trt_vs_MS.NoTrt_B |
| FBXO9 | 0.02034 | 0.38263 | 0.667 | 0.77 | 1 | MS.Trt_vs_MS.NoTrt_B |
| GPN1 | 0.02038 | 0.2952 | 0.081 | 0.14 | 1 | MS.Trt_vs_MS.NoTrt_B |
| PRKCD | 0.0204 | 0.35196 | 0.377 | 0.48 | 1 | MS.Trt_vs_MS.NoTrt_B |
| IRF5 | 0.02041 | 0.57579 | 0.279 | 0.38 | 1 | MS.Trt_vs_MS.NoTrt_B |
| EMC8 | 0.02043 | -1.0632 | 0.096 | 0.16 | 1 | MS.Trt_vs_MS.NoTrt_B |
| ANKRD54 | 0.0205 | -0.4492 | 0.564 | 0.69 | 1 | MS.Trt_vs_MS.NoTrt_B |
| TOMM5 | 0.02051 | 0.30367 | 0.139 | 0.22 | 1 | MS.Trt_vs_MS.NoTrt_B |
| BTD | 0.02057 | -0.373 | 0.089 | 0.15 | 1 | MS.Trt_vs_MS.NoTrt_B |
| PREP | 0.02059 | -0.5471 | 0.061 | 0.11 | 1 | MS.Trt_vs_MS.NoTrt_B |
| ACY1 | 0.02074 | 0.49449 | 0.473 | 0.61 | 1 | MS.Trt_vs_MS.NoTrt_B |
| GDPD5 | 0.0208 | -0.9733 | 0.198 | 0.27 | 1 | MS.Trt_vs_MS.NoTrt_B |
| PTPN4 | 0.02082 | 0.59426 | 0.473 | 0.58 | 1 | MS.Trt_vs_MS.NoTrt_B |
| IQCG | 0.02083 | -0.706 | 0.654 | 0.71 | 1 | MS.Trt_vs_MS.NoTrt_B |
| SIRT3 | 0.02088 | 0.77362 | 0.229 | 0.32 | 1 | MS.Trt_vs_MS.NoTrt_B |
| DFNA5 | 0.02093 | -0.4623 | 0.083 | 0.15 | 1 | MS.Trt_vs_MS.NoTrt_B |
| ACAT2 | 0.02095 | 0.28594 | 0.209 | 0.29 | 1 | MS.Trt_vs_MS.NoTrt_B |
| TIA1 | 0.02101 | 0.55286 | 0.198 | 0.29 | 1 | MS.Trt_vs_MS.NoTrt_B |
| KDM5C | 0.02109 | 0.3367 | 0.298 | 0.42 | 1 | MS.Trt_vs_MS.NoTrt_B |
| INVS | 0.02112 | -0.5115 | 0.098 | 0.16 | 1 | MS.Trt_vs_MS.NoTrt_B |
| KCTD10 | 0.02118 | -0.4986 | 0.314 | 0.41 | 1 | MS.Trt_vs_MS.NoTrt_B |
| BID | 0.02122 | -0.4544 | 0.418 | 0.51 | 1 | MS.Trt_vs_MS.NoTrt_B |
| MTERF1 | 0.02127 | -0.4023 | 0.159 | 0.24 | 1 | MS.Trt_vs_MS.NoTrt_B |
| CARM1 | 0.02129 | -0.4168 | 0.076 | 0.14 | 1 | MS.Trt_vs_MS.NoTrt_B |
| CRAMP1 | 0.02134 | -0.7371 | 0.061 | 0.11 | 1 | MS.Trt_vs_MS.NoTrt_B |
| PSMD7 | 0.02144 | 0.35838 | 0.266 | 0.36 | 1 | MS.Trt_vs_MS.NoTrt_B |
| SMC4 | 0.02144 | 0.6508 | 0.362 | 0.49 | 1 | MS.Trt_vs_MS.NoTrt_B |
| ARHGAP11B | 0.02149 | -0.6598 | 0.159 | 0.23 | 1 | MS.Trt_vs_MS.NoTrt_B |

| | | | | | | |
|---|---|---|---|---|---|---|
| MAP1LC3B2 | 0.02159 | 0.67146 | 0.203 | 0.31 | 1 | MS.Trt_vs_MS.NoTrt_B |
| GLO1 | 0.02162 | -0.3435 | 0.057 | 0.11 | 1 | MS.Trt_vs_MS.NoTrt_B |
| ZNF720 | 0.02172 | 0.51674 | 0.379 | 0.51 | 1 | MS.Trt_vs_MS.NoTrt_B |
| LAPTM4B | 0.02172 | 0.46542 | 0.122 | 0.2 | 1 | MS.Trt_vs_MS.NoTrt_B |
| LGALS9 | 0.02172 | 0.32275 | 0.198 | 0.29 | 1 | MS.Trt_vs_MS.NoTrt_B |
| FAM120B | 0.02175 | 0.44692 | 0.257 | 0.38 | 1 | MS.Trt_vs_MS.NoTrt_B |
| FAM214A | 0.02182 | 0.98769 | 0.142 | 0.22 | 1 | MS.Trt_vs_MS.NoTrt_B |
| OAS1 | 0.02185 | -0.2782 | 0.405 | 0.49 | 1 | MS.Trt_vs_MS.NoTrt_B |
| FAM193A | 0.02185 | 0.78839 | 0.198 | 0.29 | 1 | MS.Trt_vs_MS.NoTrt_B |
| AKIP1 | 0.02194 | -0.3044 | 0.538 | 0.7 | 1 | MS.Trt_vs_MS.NoTrt_B |
| GPATCH4 | 0.02194 | 1.1046 | 0.207 | 0.31 | 1 | MS.Trt_vs_MS.NoTrt_B |
| USE1 | 0.02196 | -0.4192 | 0.771 | 0.86 | 1 | MS.Trt_vs_MS.NoTrt_B |
| SPG11 | 0.022 | 0.26948 | 0.715 | 0.84 | 1 | MS.Trt_vs_MS.NoTrt_B |
| ASF1A | 0.02204 | 0.86846 | 0.133 | 0.21 | 1 | MS.Trt_vs_MS.NoTrt_B |
| RASAL3 | 0.02222 | 0.31654 | 0.098 | 0.16 | 1 | MS.Trt_vs_MS.NoTrt_B |
| IDH2 | 0.02228 | -0.3595 | 0.427 | 0.51 | 1 | MS.Trt_vs_MS.NoTrt_B |
| KBTBD4 | 0.02232 | -0.4103 | 0.057 | 0.11 | 1 | MS.Trt_vs_MS.NoTrt_B |
| PARP11 | 0.02235 | 0.70195 | 0.362 | 0.49 | 1 | MS.Trt_vs_MS.NoTrt_B |
| MAPK9 | 0.02258 | 0.26095 | 0.37 | 0.48 | 1 | MS.Trt_vs_MS.NoTrt_B |
| GAN | 0.0226 | 0.43848 | 0.736 | 0.82 | 1 | MS.Trt_vs_MS.NoTrt_B |
| RELA | 0.02274 | 0.33173 | 0.109 | 0.18 | 1 | MS.Trt_vs_MS.NoTrt_B |
| VRTN | 0.02285 | 0.72643 | 0.092 | 0.16 | 1 | MS.Trt_vs_MS.NoTrt_B |
| LTV1 | 0.02287 | 0.27249 | 0.1 | 0.17 | 1 | MS.Trt_vs_MS.NoTrt_B |
| ARHGAP5 | 0.02294 | 0.56315 | 0.362 | 0.49 | 1 | MS.Trt_vs_MS.NoTrt_B |
| MCEE | 0.02296 | 1.12553 | 0.096 | 0.16 | 1 | MS.Trt_vs_MS.NoTrt_B |
| PALB2 | 0.02296 | -0.3767 | 0.148 | 0.22 | 1 | MS.Trt_vs_MS.NoTrt_B |
| SMCR8 | 0.02298 | 0.32487 | 0.198 | 0.29 | 1 | MS.Trt_vs_MS.NoTrt_B |
| JADE1 | 0.02301 | 0.4264 | 0.229 | 0.32 | 1 | MS.Trt_vs_MS.NoTrt_B |
| SMPD1 | 0.02309 | 0.27382 | 0.065 | 0.12 | 1 | MS.Trt_vs_MS.NoTrt_B |
| RABEPK | 0.0231 | -0.6112 | 0.656 | 0.75 | 1 | MS.Trt_vs_MS.NoTrt_B |
| NDUFB3 | 0.02311 | 0.50263 | 0.122 | 0.2 | 1 | MS.Trt_vs_MS.NoTrt_B |
| ERCC6 | 0.02316 | -0.2546 | 0.181 | 0.27 | 1 | MS.Trt_vs_MS.NoTrt_B |
| TRMT10B | 0.02317 | 0.57532 | 0.743 | 0.87 | 1 | MS.Trt_vs_MS.NoTrt_B |
| SLFN13 | 0.02318 | 0.65184 | 0.466 | 0.57 | 1 | MS.Trt_vs_MS.NoTrt_B |
| ZFP41 | 0.0232 | 0.49348 | 0.072 | 0.13 | 1 | MS.Trt_vs_MS.NoTrt_B |
| ZNF785 | 0.02323 | -0.3349 | 0.771 | 0.9 | 1 | MS.Trt_vs_MS.NoTrt_B |
| AMDHD2 | 0.02323 | -0.3383 | 0.061 | 0.11 | 1 | MS.Trt_vs_MS.NoTrt_B |
| FAM83G | 0.02334 | 0.78308 | 0.316 | 0.43 | 1 | MS.Trt_vs_MS.NoTrt_B |
| STOM | 0.02335 | 0.44507 | 0.525 | 0.7 | 1 | MS.Trt_vs_MS.NoTrt_B |
| RNF103 | 0.02335 | 0.43047 | 0.473 | 0.58 | 1 | MS.Trt_vs_MS.NoTrt_B |

| | | | | | | |
|---|---|---|---|---|---|---|
| KAT6B | 0.02336 | 0.43985 | 0.061 | 0.11 | 1 | MS.Trt_vs_MS.NoTrt_B |
| DHX15 | 0.0235 | 0.42828 | 0.375 | 0.49 | 1 | MS.Trt_vs_MS.NoTrt_B |
| TMEM208 | 0.02354 | -0.4437 | 0.083 | 0.14 | 1 | MS.Trt_vs_MS.NoTrt_B |
| AC012314.1 | 0.02355 | 0.27508 | 0.124 | 0.21 | 1 | MS.Trt_vs_MS.NoTrt_B |
| AC012314.1 | 0.02355 | 0.27508 | 0.124 | 0.21 | 1 | MS.Trt_vs_MS.NoTrt_B |
| AC012314.1 | 0.02355 | 0.27508 | 0.124 | 0.21 | 1 | MS.Trt_vs_MS.NoTrt_B |
| AC012314.1 | 0.02355 | 0.27508 | 0.124 | 0.21 | 1 | MS.Trt_vs_MS.NoTrt_B |
| AC012314.3 | 0.02355 | 0.27508 | 0.124 | 0.21 | 1 | MS.Trt_vs_MS.NoTrt_B |
| AC012314.6 | 0.02355 | 0.27508 | 0.124 | 0.21 | 1 | MS.Trt_vs_MS.NoTrt_B |
| AC012314.7 | 0.02355 | 0.27508 | 0.124 | 0.21 | 1 | MS.Trt_vs_MS.NoTrt_B |
| AC012314.9 | 0.02355 | 0.27508 | 0.124 | 0.21 | 1 | MS.Trt_vs_MS.NoTrt_B |
| ADAD2 | 0.02355 | 0.92943 | 0.185 | 0.28 | 1 | MS.Trt_vs_MS.NoTrt_B |
| PCBP3 | 0.02361 | -0.665 | 0.336 | 0.47 | 1 | MS.Trt_vs_MS.NoTrt_B |
| ARMC2 | 0.02367 | -0.2535 | 0.052 | 0.1 | 1 | MS.Trt_vs_MS.NoTrt_B |
| NDUFS3 | 0.0237 | 0.41901 | 0.434 | 0.55 | 1 | MS.Trt_vs_MS.NoTrt_B |
| DNAJB4 | 0.02371 | -1.2685 | 0.27 | 0.36 | 1 | MS.Trt_vs_MS.NoTrt_B |
| C15orf57 | 0.02374 | -0.3293 | 0.66 | 0.74 | 1 | MS.Trt_vs_MS.NoTrt_B |
| RAPGEF1 | 0.02388 | 0.2512 | 0.148 | 0.22 | 1 | MS.Trt_vs_MS.NoTrt_B |
| RP11-231C1 | 0.02389 | 0.58731 | 0.523 | 0.69 | 1 | MS.Trt_vs_MS.NoTrt_B |
| TBRG4 | 0.0239 | -0.5345 | 0.285 | 0.38 | 1 | MS.Trt_vs_MS.NoTrt_B |
| HEMGN | 0.02393 | 0.66081 | 0.078 | 0.14 | 1 | MS.Trt_vs_MS.NoTrt_B |
| ZNF599 | 0.02405 | 0.28474 | 0.46 | 0.6 | 1 | MS.Trt_vs_MS.NoTrt_B |
| TTC32 | 0.02411 | 0.37173 | 0.068 | 0.12 | 1 | MS.Trt_vs_MS.NoTrt_B |
| PPARGC1B | 0.02419 | -0.3846 | 0.558 | 0.67 | 1 | MS.Trt_vs_MS.NoTrt_B |
| ARHGAP33 | 0.02426 | -0.6333 | 0.059 | 0.11 | 1 | MS.Trt_vs_MS.NoTrt_B |
| NEDD9 | 0.02427 | -0.3876 | 0.331 | 0.42 | 1 | MS.Trt_vs_MS.NoTrt_B |
| SDHAF3 | 0.02427 | 0.41635 | 0.052 | 0.1 | 1 | MS.Trt_vs_MS.NoTrt_B |
| CYLD | 0.02429 | 0.37619 | 0.584 | 0.72 | 1 | MS.Trt_vs_MS.NoTrt_B |
| ARFGEF2 | 0.02434 | 0.4317 | 0.377 | 0.48 | 1 | MS.Trt_vs_MS.NoTrt_B |
| ATXN7 | 0.02444 | -0.4214 | 0.416 | 0.54 | 1 | MS.Trt_vs_MS.NoTrt_B |
| PHF1 | 0.02455 | -0.288 | 0.111 | 0.18 | 1 | MS.Trt_vs_MS.NoTrt_B |
| TMEM2 | 0.02456 | 0.68556 | 0.466 | 0.57 | 1 | MS.Trt_vs_MS.NoTrt_B |
| DENND6B | 0.02459 | -0.6302 | 0.218 | 0.3 | 1 | MS.Trt_vs_MS.NoTrt_B |
| TRIM33 | 0.02461 | 0.4165 | 0.266 | 0.34 | 1 | MS.Trt_vs_MS.NoTrt_B |
| RPS14 | 0.02467 | 0.5136 | 0.954 | 1 | 1 | MS.Trt_vs_MS.NoTrt_B |
| SLC22A18 | 0.02469 | 0.5768 | 0.453 | 0.6 | 1 | MS.Trt_vs_MS.NoTrt_B |
| ARID3B | 0.02469 | -0.2689 | 0.599 | 0.69 | 1 | MS.Trt_vs_MS.NoTrt_B |
| NAA15 | 0.02469 | 0.29122 | 0.63 | 0.74 | 1 | MS.Trt_vs_MS.NoTrt_B |
| PPP4R3B | 0.02472 | 0.53183 | 0.338 | 0.45 | 1 | MS.Trt_vs_MS.NoTrt_B |
| ACSM3 | 0.02477 | 0.36597 | 0.455 | 0.6 | 1 | MS.Trt_vs_MS.NoTrt_B |

| | | | | | | |
|---|---|---|---|---|---|---|
| RBM12 | 0.02481 | -0.3102 | 0.161 | 0.23 | 1 | MS.Trt_vs_MS.NoTrt_B |
| DHX33 | 0.02484 | -0.9232 | 0.272 | 0.36 | 1 | MS.Trt_vs_MS.NoTrt_B |
| CIB1 | 0.0249 | 0.34349 | 0.168 | 0.26 | 1 | MS.Trt_vs_MS.NoTrt_B |
| CNPY4 | 0.02492 | 0.42883 | 0.606 | 0.75 | 1 | MS.Trt_vs_MS.NoTrt_B |
| BLVRA | 0.02503 | -0.2997 | 0.113 | 0.18 | 1 | MS.Trt_vs_MS.NoTrt_B |
| AC138517.6 | 0.02508 | 1.02638 | 0.065 | 0.12 | 1 | MS.Trt_vs_MS.NoTrt_B |
| SIK3 | 0.02515 | -0.5325 | 0.19 | 0.27 | 1 | MS.Trt_vs_MS.NoTrt_B |
| XRN1 | 0.02519 | 0.51058 | 0.821 | 0.92 | 1 | MS.Trt_vs_MS.NoTrt_B |
| MTA3 | 0.0252 | 0.4973 | 0.516 | 0.65 | 1 | MS.Trt_vs_MS.NoTrt_B |
| ST3GAL1 | 0.02528 | -0.3273 | 0.179 | 0.25 | 1 | MS.Trt_vs_MS.NoTrt_B |
| RWDD3 | 0.02528 | 0.30872 | 0.085 | 0.15 | 1 | MS.Trt_vs_MS.NoTrt_B |
| ZNF670 | 0.02531 | 0.58277 | 0.102 | 0.17 | 1 | MS.Trt_vs_MS.NoTrt_B |
| ATP11B | 0.02532 | 0.88101 | 0.185 | 0.27 | 1 | MS.Trt_vs_MS.NoTrt_B |
| PPP2R3A | 0.02537 | 0.27519 | 0.59 | 0.76 | 1 | MS.Trt_vs_MS.NoTrt_B |
| GLMP | 0.02538 | 0.2946 | 0.617 | 0.76 | 1 | MS.Trt_vs_MS.NoTrt_B |
| SOS2 | 0.02541 | 0.42121 | 0.07 | 0.12 | 1 | MS.Trt_vs_MS.NoTrt_B |
| EXOSC8 | 0.02543 | 0.59956 | 0.425 | 0.54 | 1 | MS.Trt_vs_MS.NoTrt_B |
| ZRANB3 | 0.02544 | -0.2536 | 0.545 | 0.71 | 1 | MS.Trt_vs_MS.NoTrt_B |
| PTAR1 | 0.02551 | 0.44842 | 0.124 | 0.2 | 1 | MS.Trt_vs_MS.NoTrt_B |
| ATP8A1 | 0.02551 | -0.3139 | 0.109 | 0.17 | 1 | MS.Trt_vs_MS.NoTrt_B |
| TDG | 0.02562 | 0.29695 | 0.229 | 0.32 | 1 | MS.Trt_vs_MS.NoTrt_B |
| ASAH2B | 0.02563 | 0.28869 | 0.331 | 0.41 | 1 | MS.Trt_vs_MS.NoTrt_B |
| MUC20 | 0.02568 | 0.36217 | 0.407 | 0.56 | 1 | MS.Trt_vs_MS.NoTrt_B |
| GCNT1 | 0.0257 | 0.86657 | 0.179 | 0.27 | 1 | MS.Trt_vs_MS.NoTrt_B |
| MAP2K3 | 0.02572 | -0.7335 | 0.089 | 0.15 | 1 | MS.Trt_vs_MS.NoTrt_B |
| NEO1 | 0.02573 | 0.97513 | 0.216 | 0.33 | 1 | MS.Trt_vs_MS.NoTrt_B |
| DMGDH | 0.02582 | 0.26426 | 0.322 | 0.45 | 1 | MS.Trt_vs_MS.NoTrt_B |
| MMS22L | 0.02592 | 0.26037 | 0.817 | 0.92 | 1 | MS.Trt_vs_MS.NoTrt_B |
| MAMLD1 | 0.02592 | 0.71752 | 0.163 | 0.24 | 1 | MS.Trt_vs_MS.NoTrt_B |
| TTC26 | 0.02596 | -0.2717 | 0.292 | 0.39 | 1 | MS.Trt_vs_MS.NoTrt_B |
| ZDHHC6 | 0.02603 | 0.29734 | 0.122 | 0.19 | 1 | MS.Trt_vs_MS.NoTrt_B |
| TG | 0.02606 | 0.6035 | 0.052 | 0.1 | 1 | MS.Trt_vs_MS.NoTrt_B |
| FBXO6 | 0.02608 | 0.30569 | 0.545 | 0.71 | 1 | MS.Trt_vs_MS.NoTrt_B |
| MED30 | 0.02624 | 0.30182 | 0.087 | 0.15 | 1 | MS.Trt_vs_MS.NoTrt_B |
| POU2AF1 | 0.02633 | 0.45765 | 0.542 | 0.7 | 1 | MS.Trt_vs_MS.NoTrt_B |
| GYG1 | 0.02635 | -0.8105 | 0.434 | 0.52 | 1 | MS.Trt_vs_MS.NoTrt_B |
| EMC10 | 0.0264 | 0.30836 | 0.708 | 0.83 | 1 | MS.Trt_vs_MS.NoTrt_B |
| ATP6V0E1 | 0.02642 | 0.33109 | 0.244 | 0.35 | 1 | MS.Trt_vs_MS.NoTrt_B |
| ZFC3H1 | 0.02656 | 0.47977 | 0.29 | 0.4 | 1 | MS.Trt_vs_MS.NoTrt_B |
| QKI | 0.0266 | -0.9587 | 0.292 | 0.39 | 1 | MS.Trt_vs_MS.NoTrt_B |

| | | | | | | |
|---|---|---|---|---|---|---|
| ZNF595 | 0.02662 | 0.95944 | 0.671 | 0.81 | 1 | MS.Trt_vs_MS.NoTrt_B |
| MTIF2 | 0.02662 | -1.7896 | 0.322 | 0.4 | 1 | MS.Trt_vs_MS.NoTrt_B |
| TUBA1C | 0.02667 | -0.479 | 0.479 | 0.61 | 1 | MS.Trt_vs_MS.NoTrt_B |
| TRIP4 | 0.02668 | -0.4144 | 0.455 | 0.55 | 1 | MS.Trt_vs_MS.NoTrt_B |
| RP11-514P8 | 0.02687 | -0.3264 | 0.054 | 0.1 | 1 | MS.Trt_vs_MS.NoTrt_B |
| ANAPC2 | 0.02687 | -0.873 | 0.054 | 0.1 | 1 | MS.Trt_vs_MS.NoTrt_B |
| TTBK2 | 0.02693 | 1.10818 | 0.397 | 0.5 | 1 | MS.Trt_vs_MS.NoTrt_B |
| MLH3 | 0.02698 | -0.2853 | 0.068 | 0.12 | 1 | MS.Trt_vs_MS.NoTrt_B |
| ZNF778 | 0.02698 | 1.14934 | 0.495 | 0.61 | 1 | MS.Trt_vs_MS.NoTrt_B |
| AC016577.1 | 0.02709 | 0.28885 | 0.298 | 0.42 | 1 | MS.Trt_vs_MS.NoTrt_B |
| NUDCD2 | 0.02711 | 0.29773 | 0.645 | 0.72 | 1 | MS.Trt_vs_MS.NoTrt_B |
| XPNPEP3 | 0.02712 | 0.27301 | 0.81 | 0.94 | 1 | MS.Trt_vs_MS.NoTrt_B |
| PAIP2 | 0.02712 | 0.55769 | 0.542 | 0.63 | 1 | MS.Trt_vs_MS.NoTrt_B |
| FAR2 | 0.02714 | 0.39527 | 0.137 | 0.21 | 1 | MS.Trt_vs_MS.NoTrt_B |
| MYO5A | 0.02717 | -0.3526 | 0.647 | 0.73 | 1 | MS.Trt_vs_MS.NoTrt_B |
| C11orf63 | 0.0272 | 0.48935 | 0.364 | 0.47 | 1 | MS.Trt_vs_MS.NoTrt_B |
| SAV1 | 0.02723 | -0.3513 | 0.235 | 0.32 | 1 | MS.Trt_vs_MS.NoTrt_B |
| TAF5L | 0.02723 | -0.7637 | 0.083 | 0.14 | 1 | MS.Trt_vs_MS.NoTrt_B |
| LRRC8B | 0.02725 | 0.83889 | 0.349 | 0.47 | 1 | MS.Trt_vs_MS.NoTrt_B |
| RIPK2 | 0.02732 | -0.7804 | 0.118 | 0.18 | 1 | MS.Trt_vs_MS.NoTrt_B |
| PLD1 | 0.02735 | -0.5842 | 0.39 | 0.52 | 1 | MS.Trt_vs_MS.NoTrt_B |
| BIRC5 | 0.02749 | -0.3547 | 0.593 | 0.71 | 1 | MS.Trt_vs_MS.NoTrt_B |
| NCOA5 | 0.0275 | -0.4228 | 0.078 | 0.14 | 1 | MS.Trt_vs_MS.NoTrt_B |
| THAP9 | 0.02755 | 0.36961 | 0.453 | 0.6 | 1 | MS.Trt_vs_MS.NoTrt_B |
| TXNRD1 | 0.02757 | 0.31412 | 0.246 | 0.33 | 1 | MS.Trt_vs_MS.NoTrt_B |
| KIF6 | 0.02757 | 0.41572 | 0.736 | 0.9 | 1 | MS.Trt_vs_MS.NoTrt_B |
| GMCL1 | 0.02757 | 0.73303 | 0.283 | 0.38 | 1 | MS.Trt_vs_MS.NoTrt_B |
| SNRNP40 | 0.02764 | 0.29293 | 0.135 | 0.21 | 1 | MS.Trt_vs_MS.NoTrt_B |
| GTF2F1 | 0.02769 | 0.28105 | 0.246 | 0.34 | 1 | MS.Trt_vs_MS.NoTrt_B |
| PRRC1 | 0.0277 | 0.38325 | 0.292 | 0.38 | 1 | MS.Trt_vs_MS.NoTrt_B |
| ZNF280B | 0.0278 | 1.10441 | 0.076 | 0.13 | 1 | MS.Trt_vs_MS.NoTrt_B |
| ZNF169 | 0.0278 | -0.4022 | 0.699 | 0.81 | 1 | MS.Trt_vs_MS.NoTrt_B |
| RPAP1 | 0.02782 | -0.4316 | 0.693 | 0.83 | 1 | MS.Trt_vs_MS.NoTrt_B |
| HTT | 0.02794 | 0.34522 | 0.163 | 0.24 | 1 | MS.Trt_vs_MS.NoTrt_B |
| GMPPB | 0.02813 | -0.6314 | 0.32 | 0.41 | 1 | MS.Trt_vs_MS.NoTrt_B |
| SYNE2 | 0.02816 | 0.31879 | 0.854 | 0.96 | 1 | MS.Trt_vs_MS.NoTrt_B |
| IP6K1 | 0.02819 | 0.51127 | 0.078 | 0.14 | 1 | MS.Trt_vs_MS.NoTrt_B |
| PLEKHA1 | 0.02829 | 0.41383 | 0.854 | 0.94 | 1 | MS.Trt_vs_MS.NoTrt_B |
| STK39 | 0.02851 | 0.82574 | 0.07 | 0.12 | 1 | MS.Trt_vs_MS.NoTrt_B |
| KIAA1958 | 0.02854 | -0.5083 | 0.573 | 0.69 | 1 | MS.Trt_vs_MS.NoTrt_B |

| | | | | | | |
|---|---|---|---|---|---|---|
| GNG10 | 0.02862 | 0.77165 | 0.113 | 0.18 | 1 | MS.Trt_vs_MS.NoTrt_B |
| CLSTN1 | 0.02865 | 0.25755 | 0.161 | 0.24 | 1 | MS.Trt_vs_MS.NoTrt_B |
| CCDC69 | 0.02866 | 0.33665 | 0.695 | 0.83 | 1 | MS.Trt_vs_MS.NoTrt_B |
| NT5C3B | 0.02874 | 0.36866 | 0.51 | 0.64 | 1 | MS.Trt_vs_MS.NoTrt_B |
| THRAP3 | 0.02874 | 0.3337 | 0.322 | 0.42 | 1 | MS.Trt_vs_MS.NoTrt_B |
| PIGM | 0.02875 | -0.5169 | 0.458 | 0.54 | 1 | MS.Trt_vs_MS.NoTrt_B |
| HSD17B7 | 0.02879 | 0.48926 | 0.412 | 0.51 | 1 | MS.Trt_vs_MS.NoTrt_B |
| CWC27 | 0.02892 | -0.2575 | 0.087 | 0.15 | 1 | MS.Trt_vs_MS.NoTrt_B |
| ZNF764 | 0.02901 | -0.4148 | 0.235 | 0.32 | 1 | MS.Trt_vs_MS.NoTrt_B |
| GRIA1 | 0.02908 | 0.42811 | 0.076 | 0.14 | 1 | MS.Trt_vs_MS.NoTrt_B |
| ABCF1 | 0.02908 | 0.35432 | 0.712 | 0.82 | 1 | MS.Trt_vs_MS.NoTrt_B |
| ERCC4 | 0.02913 | 0.67062 | 0.793 | 0.89 | 1 | MS.Trt_vs_MS.NoTrt_B |
| TRUB2 | 0.02915 | -0.4299 | 0.878 | 0.98 | 1 | MS.Trt_vs_MS.NoTrt_B |
| PRKAB1 | 0.02922 | 0.49418 | 0.651 | 0.76 | 1 | MS.Trt_vs_MS.NoTrt_B |
| RGPD8 | 0.02922 | 0.91595 | 0.124 | 0.2 | 1 | MS.Trt_vs_MS.NoTrt_B |
| ATP13A4 | 0.02926 | 0.61405 | 0.32 | 0.42 | 1 | MS.Trt_vs_MS.NoTrt_B |
| ASXL1 | 0.02929 | 0.39388 | 0.146 | 0.22 | 1 | MS.Trt_vs_MS.NoTrt_B |
| ORAOV1 | 0.02936 | -0.5642 | 0.529 | 0.64 | 1 | MS.Trt_vs_MS.NoTrt_B |
| AMACR | 0.02943 | -1.5475 | 0.054 | 0.1 | 1 | MS.Trt_vs_MS.NoTrt_B |
| SRPRB | 0.02945 | -0.4255 | 0.115 | 0.18 | 1 | MS.Trt_vs_MS.NoTrt_B |
| BMP1 | 0.0296 | 0.70866 | 0.381 | 0.51 | 1 | MS.Trt_vs_MS.NoTrt_B |
| FAM91A1 | 0.02965 | -0.8691 | 0.233 | 0.32 | 1 | MS.Trt_vs_MS.NoTrt_B |
| EFL1 | 0.02967 | -0.4703 | 0.068 | 0.12 | 1 | MS.Trt_vs_MS.NoTrt_B |
| RFX3 | 0.02974 | 0.77697 | 0.098 | 0.16 | 1 | MS.Trt_vs_MS.NoTrt_B |
| POGZ | 0.02979 | 0.44667 | 0.684 | 0.8 | 1 | MS.Trt_vs_MS.NoTrt_B |
| FUT11 | 0.02981 | -0.5885 | 0.603 | 0.72 | 1 | MS.Trt_vs_MS.NoTrt_B |
| VBP1 | 0.02983 | 0.49398 | 0.092 | 0.15 | 1 | MS.Trt_vs_MS.NoTrt_B |
| ITIH5 | 0.0299 | 0.48156 | 0.688 | 0.83 | 1 | MS.Trt_vs_MS.NoTrt_B |
| BTG3 | 0.02996 | -0.4787 | 0.196 | 0.27 | 1 | MS.Trt_vs_MS.NoTrt_B |
| CCDC9 | 0.02999 | -0.2931 | 0.172 | 0.24 | 1 | MS.Trt_vs_MS.NoTrt_B |
| PGS1 | 0.03004 | -0.534 | 0.512 | 0.58 | 1 | MS.Trt_vs_MS.NoTrt_B |
| INTS6 | 0.03006 | 0.36414 | 0.824 | 0.94 | 1 | MS.Trt_vs_MS.NoTrt_B |
| PAQR8 | 0.03007 | -0.3252 | 0.375 | 0.47 | 1 | MS.Trt_vs_MS.NoTrt_B |
| SERINC1 | 0.03012 | 0.34718 | 0.505 | 0.63 | 1 | MS.Trt_vs_MS.NoTrt_B |
| NLRP2 | 0.03029 | 0.58912 | 0.706 | 0.81 | 1 | MS.Trt_vs_MS.NoTrt_B |
| WDR33 | 0.03037 | 0.54675 | 0.172 | 0.25 | 1 | MS.Trt_vs_MS.NoTrt_B |
| SPRYD3 | 0.0304 | -0.3711 | 0.098 | 0.16 | 1 | MS.Trt_vs_MS.NoTrt_B |
| AKAP8 | 0.03041 | -0.3478 | 0.275 | 0.37 | 1 | MS.Trt_vs_MS.NoTrt_B |
| MRPS21 | 0.03056 | 0.5186 | 0.42 | 0.52 | 1 | MS.Trt_vs_MS.NoTrt_B |
| PARN | 0.03057 | -0.5267 | 0.606 | 0.69 | 1 | MS.Trt_vs_MS.NoTrt_B |

| | | | | | | |
|---|---|---|---|---|---|---|
| APBA2 | 0.03061 | 0.5622 | 0.412 | 0.54 | 1 | MS.Trt_vs_MS.NoTrt_B |
| NUMA1 | 0.03061 | 0.27256 | 0.78 | 0.91 | 1 | MS.Trt_vs_MS.NoTrt_B |
| TIRAP | 0.03064 | -1.1561 | 0.355 | 0.45 | 1 | MS.Trt_vs_MS.NoTrt_B |
| SLC24A1 | 0.03068 | 1.17004 | 0.344 | 0.45 | 1 | MS.Trt_vs_MS.NoTrt_B |
| RAC1 | 0.0307 | 0.51074 | 0.102 | 0.17 | 1 | MS.Trt_vs_MS.NoTrt_B |
| ZNF772 | 0.03077 | 0.4489 | 0.373 | 0.45 | 1 | MS.Trt_vs_MS.NoTrt_B |
| COQ3 | 0.03078 | -0.2663 | 0.185 | 0.26 | 1 | MS.Trt_vs_MS.NoTrt_B |
| MTUS1 | 0.0308 | 0.52269 | 0.27 | 0.37 | 1 | MS.Trt_vs_MS.NoTrt_B |
| TTC23 | 0.03085 | 0.27156 | 0.442 | 0.56 | 1 | MS.Trt_vs_MS.NoTrt_B |
| SOAT1 | 0.03087 | -0.505 | 0.61 | 0.64 | 1 | MS.Trt_vs_MS.NoTrt_B |
| BCAT1 | 0.0309 | -0.6216 | 0.344 | 0.45 | 1 | MS.Trt_vs_MS.NoTrt_B |
| HERC6 | 0.0309 | -0.9833 | 0.237 | 0.3 | 1 | MS.Trt_vs_MS.NoTrt_B |
| RIT1 | 0.03098 | 0.86031 | 0.327 | 0.42 | 1 | MS.Trt_vs_MS.NoTrt_B |
| TP53I13 | 0.03098 | 0.38418 | 0.083 | 0.14 | 1 | MS.Trt_vs_MS.NoTrt_B |
| TYW5 | 0.03103 | 1.07531 | 0.78 | 0.91 | 1 | MS.Trt_vs_MS.NoTrt_B |
| CAPN10 | 0.03105 | 0.79747 | 0.37 | 0.48 | 1 | MS.Trt_vs_MS.NoTrt_B |
| NR1H3 | 0.03116 | 0.65325 | 0.092 | 0.15 | 1 | MS.Trt_vs_MS.NoTrt_B |
| CPT1B | 0.03117 | -0.9328 | 0.118 | 0.18 | 1 | MS.Trt_vs_MS.NoTrt_B |
| SLMAP | 0.03117 | 0.54497 | 0.641 | 0.77 | 1 | MS.Trt_vs_MS.NoTrt_B |
| CHD9 | 0.03122 | 0.64348 | 0.305 | 0.4 | 1 | MS.Trt_vs_MS.NoTrt_B |
| HTN1 | 0.03124 | 0.32742 | 0.586 | 0.75 | 1 | MS.Trt_vs_MS.NoTrt_B |
| STK17A | 0.03131 | 0.71682 | 0.196 | 0.29 | 1 | MS.Trt_vs_MS.NoTrt_B |
| ANAPC1 | 0.03131 | 0.77484 | 0.434 | 0.56 | 1 | MS.Trt_vs_MS.NoTrt_B |
| SBK1 | 0.03134 | 0.3276 | 0.118 | 0.18 | 1 | MS.Trt_vs_MS.NoTrt_B |
| ACSF2 | 0.0314 | -1.0887 | 0.336 | 0.44 | 1 | MS.Trt_vs_MS.NoTrt_B |
| BBS9 | 0.03146 | -0.6021 | 0.072 | 0.12 | 1 | MS.Trt_vs_MS.NoTrt_B |
| DNASE2 | 0.03148 | -0.2635 | 0.349 | 0.46 | 1 | MS.Trt_vs_MS.NoTrt_B |
| ATG5 | 0.03156 | 0.30477 | 0.126 | 0.2 | 1 | MS.Trt_vs_MS.NoTrt_B |
| UBAP1L | 0.03156 | 0.50938 | 0.288 | 0.38 | 1 | MS.Trt_vs_MS.NoTrt_B |
| CEP126 | 0.03174 | 0.61268 | 0.577 | 0.73 | 1 | MS.Trt_vs_MS.NoTrt_B |
| RP11-226E2 | 0.0318 | 0.6305 | 0.516 | 0.63 | 1 | MS.Trt_vs_MS.NoTrt_B |
| HAUS5 | 0.03188 | 0.56385 | 0.318 | 0.43 | 1 | MS.Trt_vs_MS.NoTrt_B |
| SPC24 | 0.03188 | -0.3878 | 0.793 | 0.82 | 1 | MS.Trt_vs_MS.NoTrt_B |
| ZNF566 | 0.03191 | -0.4193 | 0.752 | 0.88 | 1 | MS.Trt_vs_MS.NoTrt_B |
| PHLDA1 | 0.03201 | -0.9513 | 0.357 | 0.44 | 1 | MS.Trt_vs_MS.NoTrt_B |
| GLB1L2 | 0.03205 | 0.66197 | 0.272 | 0.36 | 1 | MS.Trt_vs_MS.NoTrt_B |
| COL4A3BP | 0.03218 | 0.50331 | 0.087 | 0.15 | 1 | MS.Trt_vs_MS.NoTrt_B |
| ABO | 0.03218 | -0.8791 | 0.266 | 0.36 | 1 | MS.Trt_vs_MS.NoTrt_B |
| MVD | 0.03219 | 0.32662 | 0.386 | 0.5 | 1 | MS.Trt_vs_MS.NoTrt_B |
| JAKMIP2 | 0.03228 | -1.2083 | 0.305 | 0.38 | 1 | MS.Trt_vs_MS.NoTrt_B |

| | | | | | | |
|---|---|---|---|---|---|---|
| CHIT1 | 0.03229 | -1.2327 | 0.063 | 0.11 | 1 | MS.Trt_vs_MS.NoTrt_B |
| ZC3H12B | 0.03233 | 0.7214 | 0.429 | 0.52 | 1 | MS.Trt_vs_MS.NoTrt_B |
| BRD3 | 0.0324 | 0.3045 | 0.118 | 0.18 | 1 | MS.Trt_vs_MS.NoTrt_B |
| CCDC91 | 0.03245 | 0.30105 | 0.34 | 0.43 | 1 | MS.Trt_vs_MS.NoTrt_B |
| ZNF70 | 0.03245 | -0.3418 | 0.71 | 0.84 | 1 | MS.Trt_vs_MS.NoTrt_B |
| SCOC | 0.03255 | 0.35599 | 0.063 | 0.11 | 1 | MS.Trt_vs_MS.NoTrt_B |
| TPRG1 | 0.03267 | -0.6814 | 0.124 | 0.19 | 1 | MS.Trt_vs_MS.NoTrt_B |
| RAB2A | 0.03271 | 0.27273 | 0.155 | 0.23 | 1 | MS.Trt_vs_MS.NoTrt_B |
| ADAT2 | 0.03272 | 0.60688 | 0.279 | 0.38 | 1 | MS.Trt_vs_MS.NoTrt_B |
| TMCO6 | 0.03296 | -0.5836 | 0.412 | 0.51 | 1 | MS.Trt_vs_MS.NoTrt_B |
| GNAS | 0.03301 | 0.44795 | 0.377 | 0.5 | 1 | MS.Trt_vs_MS.NoTrt_B |
| COA5 | 0.03305 | 0.46759 | 0.388 | 0.49 | 1 | MS.Trt_vs_MS.NoTrt_B |
| DYNC1H1 | 0.03307 | 0.3885 | 0.841 | 0.96 | 1 | MS.Trt_vs_MS.NoTrt_B |
| MRPS25 | 0.0331 | 0.44517 | 0.346 | 0.44 | 1 | MS.Trt_vs_MS.NoTrt_B |
| CHST14 | 0.03315 | -0.6503 | 0.157 | 0.23 | 1 | MS.Trt_vs_MS.NoTrt_B |
| ITGAX | 0.03324 | 0.55165 | 0.412 | 0.54 | 1 | MS.Trt_vs_MS.NoTrt_B |
| ENO3 | 0.03324 | 0.52912 | 0.364 | 0.47 | 1 | MS.Trt_vs_MS.NoTrt_B |
| ZNF816 | 0.03327 | 0.36969 | 0.403 | 0.5 | 1 | MS.Trt_vs_MS.NoTrt_B |
| SNX30 | 0.0334 | 1.05903 | 0.344 | 0.45 | 1 | MS.Trt_vs_MS.NoTrt_B |
| TIMM17B | 0.03357 | 0.38422 | 0.584 | 0.67 | 1 | MS.Trt_vs_MS.NoTrt_B |
| PGF | 0.03358 | 0.29335 | 0.185 | 0.25 | 1 | MS.Trt_vs_MS.NoTrt_B |
| SGK1 | 0.03363 | -0.8105 | 0.27 | 0.37 | 1 | MS.Trt_vs_MS.NoTrt_B |
| IL13RA1 | 0.03378 | 0.37357 | 0.196 | 0.28 | 1 | MS.Trt_vs_MS.NoTrt_B |
| C14orf93 | 0.03384 | -0.3785 | 0.229 | 0.3 | 1 | MS.Trt_vs_MS.NoTrt_B |
| CHSY1 | 0.03384 | -0.8155 | 0.181 | 0.25 | 1 | MS.Trt_vs_MS.NoTrt_B |
| FAAP20 | 0.03388 | 0.64494 | 0.78 | 0.86 | 1 | MS.Trt_vs_MS.NoTrt_B |
| C1GALT1 | 0.03388 | 0.29281 | 0.473 | 0.58 | 1 | MS.Trt_vs_MS.NoTrt_B |
| RFC4 | 0.03417 | -0.2565 | 0.192 | 0.27 | 1 | MS.Trt_vs_MS.NoTrt_B |
| ZFP36L2 | 0.03425 | 0.68762 | 0.327 | 0.45 | 1 | MS.Trt_vs_MS.NoTrt_B |
| TUBGCP6 | 0.03435 | 0.36218 | 0.545 | 0.63 | 1 | MS.Trt_vs_MS.NoTrt_B |
| ZHX3 | 0.03438 | 0.49915 | 0.29 | 0.4 | 1 | MS.Trt_vs_MS.NoTrt_B |
| ZFAND4 | 0.03442 | -0.694 | 0.303 | 0.41 | 1 | MS.Trt_vs_MS.NoTrt_B |
| FAM101B | 0.03447 | 0.58485 | 0.17 | 0.25 | 1 | MS.Trt_vs_MS.NoTrt_B |
| MYO15A | 0.0347 | 0.32022 | 0.684 | 0.82 | 1 | MS.Trt_vs_MS.NoTrt_B |
| YEATS2 | 0.03514 | 0.71668 | 0.553 | 0.66 | 1 | MS.Trt_vs_MS.NoTrt_B |
| COPB2 | 0.0352 | -0.5005 | 0.824 | 0.91 | 1 | MS.Trt_vs_MS.NoTrt_B |
| HDDC3 | 0.03526 | -0.5061 | 0.292 | 0.37 | 1 | MS.Trt_vs_MS.NoTrt_B |
| TIMM22 | 0.03539 | -0.3155 | 0.07 | 0.12 | 1 | MS.Trt_vs_MS.NoTrt_B |
| CTSA | 0.03541 | -0.6762 | 0.562 | 0.65 | 1 | MS.Trt_vs_MS.NoTrt_B |
| APOBEC3D | 0.03552 | -0.2566 | 0.51 | 0.6 | 1 | MS.Trt_vs_MS.NoTrt_B |

| | | | | | | |
|---|---|---|---|---|---|---|
| BAG2 | 0.03555 | 0.33787 | 0.174 | 0.25 | 1 | MS.Trt_vs_MS.NoTrt_B |
| PXYLP1 | 0.03569 | 0.28132 | 0.353 | 0.47 | 1 | MS.Trt_vs_MS.NoTrt_B |
| CLN8 | 0.0357 | -0.2956 | 0.721 | 0.83 | 1 | MS.Trt_vs_MS.NoTrt_B |
| ATF7IP | 0.03571 | 0.37257 | 0.749 | 0.83 | 1 | MS.Trt_vs_MS.NoTrt_B |
| GOT2 | 0.03572 | -0.3356 | 0.763 | 0.86 | 1 | MS.Trt_vs_MS.NoTrt_B |
| FAM149A | 0.03573 | -0.6206 | 0.222 | 0.3 | 1 | MS.Trt_vs_MS.NoTrt_B |
| RPS6KA3 | 0.03594 | 0.44594 | 0.425 | 0.55 | 1 | MS.Trt_vs_MS.NoTrt_B |
| UBA5 | 0.03596 | 0.52584 | 0.098 | 0.16 | 1 | MS.Trt_vs_MS.NoTrt_B |
| TMEM33 | 0.036 | 0.57463 | 0.8 | 0.84 | 1 | MS.Trt_vs_MS.NoTrt_B |
| VPS53 | 0.03603 | 0.32789 | 0.887 | 0.95 | 1 | MS.Trt_vs_MS.NoTrt_B |
| RP11-599B1 | 0.03616 | 0.27584 | 0.248 | 0.35 | 1 | MS.Trt_vs_MS.NoTrt_B |
| HACL1 | 0.03618 | -0.2631 | 0.085 | 0.14 | 1 | MS.Trt_vs_MS.NoTrt_B |
| ZRANB1 | 0.03625 | -0.6739 | 0.061 | 0.11 | 1 | MS.Trt_vs_MS.NoTrt_B |
| SLC3A1 | 0.03626 | 0.65733 | 0.538 | 0.7 | 1 | MS.Trt_vs_MS.NoTrt_B |
| APOBEC3F | 0.03635 | 0.31477 | 0.795 | 0.88 | 1 | MS.Trt_vs_MS.NoTrt_B |
| TXK | 0.03636 | 0.61831 | 0.51 | 0.63 | 1 | MS.Trt_vs_MS.NoTrt_B |
| ICAM1 | 0.0364 | 0.48592 | 0.379 | 0.46 | 1 | MS.Trt_vs_MS.NoTrt_B |
| EFCAB10 | 0.03645 | 0.62021 | 0.133 | 0.21 | 1 | MS.Trt_vs_MS.NoTrt_B |
| ATP10B | 0.03656 | 0.32879 | 0.538 | 0.69 | 1 | MS.Trt_vs_MS.NoTrt_B |
| CDC26 | 0.03659 | 0.41785 | 0.198 | 0.28 | 1 | MS.Trt_vs_MS.NoTrt_B |
| SPTY2D1 | 0.0366 | -0.282 | 0.261 | 0.34 | 1 | MS.Trt_vs_MS.NoTrt_B |
| CCS | 0.03665 | -0.5924 | 0.15 | 0.22 | 1 | MS.Trt_vs_MS.NoTrt_B |
| TATDN3 | 0.03671 | -0.4248 | 0.697 | 0.82 | 1 | MS.Trt_vs_MS.NoTrt_B |
| CDKN2AIP | 0.03679 | -0.2715 | 0.057 | 0.1 | 1 | MS.Trt_vs_MS.NoTrt_B |
| MLLT10 | 0.0368 | 0.78188 | 0.253 | 0.33 | 1 | MS.Trt_vs_MS.NoTrt_B |
| TRIM26 | 0.03688 | 0.35107 | 0.142 | 0.21 | 1 | MS.Trt_vs_MS.NoTrt_B |
| C1orf52 | 0.03698 | -0.2736 | 0.216 | 0.29 | 1 | MS.Trt_vs_MS.NoTrt_B |
| IL17RE | 0.03709 | -0.8675 | 0.174 | 0.24 | 1 | MS.Trt_vs_MS.NoTrt_B |
| ATP10D | 0.03725 | 0.7879 | 0.251 | 0.35 | 1 | MS.Trt_vs_MS.NoTrt_B |
| TANK | 0.03734 | -0.3779 | 0.172 | 0.24 | 1 | MS.Trt_vs_MS.NoTrt_B |
| LANCL2 | 0.03755 | 0.38898 | 0.068 | 0.12 | 1 | MS.Trt_vs_MS.NoTrt_B |
| CCNC | 0.03755 | 0.53185 | 0.492 | 0.62 | 1 | MS.Trt_vs_MS.NoTrt_B |
| PPP3R1 | 0.03757 | 0.34122 | 0.098 | 0.16 | 1 | MS.Trt_vs_MS.NoTrt_B |
| RBFA | 0.03758 | 0.89306 | 0.205 | 0.29 | 1 | MS.Trt_vs_MS.NoTrt_B |
| LAP3 | 0.03758 | -0.4021 | 0.092 | 0.15 | 1 | MS.Trt_vs_MS.NoTrt_B |
| SLC35A3 | 0.03761 | -0.466 | 0.678 | 0.76 | 1 | MS.Trt_vs_MS.NoTrt_B |
| MRPS31 | 0.03767 | 0.66898 | 0.346 | 0.47 | 1 | MS.Trt_vs_MS.NoTrt_B |
| DUT | 0.03771 | 0.41226 | 0.118 | 0.18 | 1 | MS.Trt_vs_MS.NoTrt_B |
| IPO8 | 0.03776 | 0.70127 | 0.383 | 0.5 | 1 | MS.Trt_vs_MS.NoTrt_B |
| MECR | 0.03782 | -1.0071 | 0.224 | 0.3 | 1 | MS.Trt_vs_MS.NoTrt_B |

| | | | | | | |
|---|---|---|---|---|---|---|
| C3orf33 | 0.03783 | 0.28938 | 0.373 | 0.48 | 1 | MS.Trt_vs_MS.NoTrt_B |
| APOBEC3G | 0.03785 | -0.4284 | 0.654 | 0.74 | 1 | MS.Trt_vs_MS.NoTrt_B |
| FAM63A | 0.03805 | 1.14278 | 0.431 | 0.58 | 1 | MS.Trt_vs_MS.NoTrt_B |
| ODF2 | 0.03812 | -0.297 | 0.397 | 0.48 | 1 | MS.Trt_vs_MS.NoTrt_B |
| ZNF492 | 0.03814 | 0.35153 | 0.486 | 0.4 | 1 | MS.Trt_vs_MS.NoTrt_B |
| RUVBL1 | 0.03818 | 0.95112 | 0.625 | 0.58 | 1 | MS.Trt_vs_MS.NoTrt_B |
| PCDHGA1 | 0.03819 | 0.54417 | 0.211 | 0.3 | 1 | MS.Trt_vs_MS.NoTrt_B |
| SPEN | 0.03821 | 0.32294 | 0.198 | 0.28 | 1 | MS.Trt_vs_MS.NoTrt_B |
| DENND5B | 0.03825 | 0.35351 | 0.436 | 0.54 | 1 | MS.Trt_vs_MS.NoTrt_B |
| IGHMBP2 | 0.03825 | -1.0429 | 0.285 | 0.37 | 1 | MS.Trt_vs_MS.NoTrt_B |
| IFT122 | 0.03831 | 0.5724 | 0.734 | 0.88 | 1 | MS.Trt_vs_MS.NoTrt_B |
| GSR | 0.03837 | -0.4927 | 0.17 | 0.24 | 1 | MS.Trt_vs_MS.NoTrt_B |
| TRIM44 | 0.03838 | 0.47972 | 0.187 | 0.27 | 1 | MS.Trt_vs_MS.NoTrt_B |
| SLC16A7 | 0.03849 | -0.5138 | 0.512 | 0.6 | 1 | MS.Trt_vs_MS.NoTrt_B |
| AURKA | 0.03856 | 1.72642 | 0.227 | 0.3 | 1 | MS.Trt_vs_MS.NoTrt_B |
| ARMT1 | 0.03874 | 0.37132 | 0.057 | 0.1 | 1 | MS.Trt_vs_MS.NoTrt_B |
| PHACTR2 | 0.03877 | 0.99664 | 0.41 | 0.5 | 1 | MS.Trt_vs_MS.NoTrt_B |
| TRAPPC10 | 0.03877 | 0.346 | 0.168 | 0.24 | 1 | MS.Trt_vs_MS.NoTrt_B |
| ZNF594 | 0.0388 | 1.26569 | 0.353 | 0.45 | 1 | MS.Trt_vs_MS.NoTrt_B |
| TMX3 | 0.03883 | 0.91285 | 0.301 | 0.41 | 1 | MS.Trt_vs_MS.NoTrt_B |
| NUDT22 | 0.03887 | 0.36301 | 0.582 | 0.65 | 1 | MS.Trt_vs_MS.NoTrt_B |
| PFKFB3 | 0.03889 | 0.42994 | 0.157 | 0.23 | 1 | MS.Trt_vs_MS.NoTrt_B |
| POLK | 0.03891 | 0.6924 | 0.678 | 0.79 | 1 | MS.Trt_vs_MS.NoTrt_B |
| TDRD3 | 0.03893 | -0.2994 | 0.473 | 0.55 | 1 | MS.Trt_vs_MS.NoTrt_B |
| SIPA1 | 0.039 | -0.3185 | 0.481 | 0.59 | 1 | MS.Trt_vs_MS.NoTrt_B |
| KIF3B | 0.03903 | -0.82 | 0.325 | 0.4 | 1 | MS.Trt_vs_MS.NoTrt_B |
| FAM220A | 0.03917 | -0.2949 | 0.24 | 0.31 | 1 | MS.Trt_vs_MS.NoTrt_B |
| FADS1 | 0.03932 | -0.3208 | 0.508 | 0.6 | 1 | MS.Trt_vs_MS.NoTrt_B |
| RNF14 | 0.03939 | 0.30449 | 0.477 | 0.63 | 1 | MS.Trt_vs_MS.NoTrt_B |
| TOMM70 | 0.03951 | 0.51731 | 0.087 | 0.14 | 1 | MS.Trt_vs_MS.NoTrt_B |
| TBC1D7 | 0.03955 | 0.31719 | 0.39 | 0.48 | 1 | MS.Trt_vs_MS.NoTrt_B |
| ATP6AP1 | 0.03958 | -0.4306 | 0.364 | 0.42 | 1 | MS.Trt_vs_MS.NoTrt_B |
| ESYT3 | 0.03968 | 0.56623 | 0.388 | 0.51 | 1 | MS.Trt_vs_MS.NoTrt_B |
| PARP14 | 0.03972 | 0.35832 | 0.32 | 0.39 | 1 | MS.Trt_vs_MS.NoTrt_B |
| LTBP4 | 0.03976 | -0.2603 | 0.423 | 0.49 | 1 | MS.Trt_vs_MS.NoTrt_B |
| GJB7 | 0.03988 | 0.62289 | 0.083 | 0.14 | 1 | MS.Trt_vs_MS.NoTrt_B |
| DCP1A | 0.04005 | -0.4453 | 0.292 | 0.37 | 1 | MS.Trt_vs_MS.NoTrt_B |
| NASP | 0.04021 | 0.67315 | 0.856 | 0.88 | 1 | MS.Trt_vs_MS.NoTrt_B |
| VPS54 | 0.04026 | 0.47147 | 0.327 | 0.43 | 1 | MS.Trt_vs_MS.NoTrt_B |
| HSPA4L | 0.04029 | 0.29655 | 0.834 | 0.86 | 1 | MS.Trt_vs_MS.NoTrt_B |

| | | | | | | |
|---|---|---|---|---|---|---|
| DPP9 | 0.0403 | -0.3524 | 0.577 | 0.66 | 1 | MS.Trt_vs_MS.NoTrt_B |
| FUT10 | 0.0404 | 0.9057 | 0.277 | 0.35 | 1 | MS.Trt_vs_MS.NoTrt_B |
| ANKS3 | 0.04041 | -0.4551 | 0.571 | 0.7 | 1 | MS.Trt_vs_MS.NoTrt_B |
| GGA1 | 0.04052 | 0.33819 | 0.895 | 0.97 | 1 | MS.Trt_vs_MS.NoTrt_B |
| PYROXD2 | 0.04059 | 0.25595 | 0.115 | 0.18 | 1 | MS.Trt_vs_MS.NoTrt_B |
| SEC22B | 0.04067 | 0.59062 | 0.739 | 0.84 | 1 | MS.Trt_vs_MS.NoTrt_B |
| JMJD4 | 0.0407 | -0.7163 | 0.2 | 0.28 | 1 | MS.Trt_vs_MS.NoTrt_B |
| ARMC10 | 0.04074 | 0.47806 | 0.503 | 0.64 | 1 | MS.Trt_vs_MS.NoTrt_B |
| C5orf42 | 0.04082 | 1.26031 | 0.392 | 0.5 | 1 | MS.Trt_vs_MS.NoTrt_B |
| CDV3 | 0.04102 | 0.33988 | 0.192 | 0.27 | 1 | MS.Trt_vs_MS.NoTrt_B |
| CARS | 0.04106 | -0.2993 | 0.654 | 0.75 | 1 | MS.Trt_vs_MS.NoTrt_B |
| ZNF81 | 0.04109 | 0.42118 | 0.19 | 0.27 | 1 | MS.Trt_vs_MS.NoTrt_B |
| SIL1 | 0.0411 | -0.3353 | 0.061 | 0.11 | 1 | MS.Trt_vs_MS.NoTrt_B |
| AIFM2 | 0.0411 | 0.63622 | 0.259 | 0.35 | 1 | MS.Trt_vs_MS.NoTrt_B |
| DYRK2 | 0.04115 | 0.37526 | 0.135 | 0.2 | 1 | MS.Trt_vs_MS.NoTrt_B |
| CSNK1A1 | 0.04128 | 0.34582 | 0.791 | 0.82 | 1 | MS.Trt_vs_MS.NoTrt_B |
| KIAA0586 | 0.04135 | 0.47307 | 0.71 | 0.81 | 1 | MS.Trt_vs_MS.NoTrt_B |
| PRRC2B | 0.04137 | 0.53802 | 0.179 | 0.25 | 1 | MS.Trt_vs_MS.NoTrt_B |
| E2F3 | 0.0414 | 0.60752 | 0.129 | 0.2 | 1 | MS.Trt_vs_MS.NoTrt_B |
| XRCC2 | 0.04148 | 0.27515 | 0.636 | 0.85 | 1 | MS.Trt_vs_MS.NoTrt_B |
| LXN | 0.04155 | 0.73642 | 0.113 | 0.17 | 1 | MS.Trt_vs_MS.NoTrt_B |
| TSPAN6 | 0.04159 | 0.29195 | 0.418 | 0.55 | 1 | MS.Trt_vs_MS.NoTrt_B |
| RAPGEF6 | 0.04164 | 0.46097 | 0.442 | 0.54 | 1 | MS.Trt_vs_MS.NoTrt_B |
| RASSF2 | 0.04165 | 0.25294 | 0.261 | 0.35 | 1 | MS.Trt_vs_MS.NoTrt_B |
| CTD-2331H1 | 0.04166 | -0.9462 | 0.137 | 0.21 | 1 | MS.Trt_vs_MS.NoTrt_B |
| DCUN1D4 | 0.04168 | 0.99652 | 0.227 | 0.31 | 1 | MS.Trt_vs_MS.NoTrt_B |
| PJA2 | 0.04171 | 0.51926 | 0.203 | 0.28 | 1 | MS.Trt_vs_MS.NoTrt_B |
| DNAJC11 | 0.04183 | -0.5412 | 0.135 | 0.2 | 1 | MS.Trt_vs_MS.NoTrt_B |
| C22orf29 | 0.04184 | -0.3568 | 0.346 | 0.42 | 1 | MS.Trt_vs_MS.NoTrt_B |
| ZFAT | 0.04194 | -0.6769 | 0.227 | 0.31 | 1 | MS.Trt_vs_MS.NoTrt_B |
| CD180 | 0.04216 | 0.29084 | 0.209 | 0.29 | 1 | MS.Trt_vs_MS.NoTrt_B |
| ZNF211 | 0.04221 | 0.46219 | 0.159 | 0.23 | 1 | MS.Trt_vs_MS.NoTrt_B |
| NMRAL1 | 0.04226 | 0.60993 | 0.373 | 0.48 | 1 | MS.Trt_vs_MS.NoTrt_B |
| SLC25A51 | 0.0423 | 0.38613 | 0.466 | 0.57 | 1 | MS.Trt_vs_MS.NoTrt_B |
| RB1 | 0.04251 | 0.47086 | 0.111 | 0.17 | 1 | MS.Trt_vs_MS.NoTrt_B |
| STX18 | 0.04259 | -0.4601 | 0.102 | 0.16 | 1 | MS.Trt_vs_MS.NoTrt_B |
| APIP | 0.04263 | -0.4358 | 0.407 | 0.48 | 1 | MS.Trt_vs_MS.NoTrt_B |
| NDUFS6 | 0.04263 | 0.51862 | 0.105 | 0.17 | 1 | MS.Trt_vs_MS.NoTrt_B |
| MAP1S | 0.04268 | -0.3654 | 0.521 | 0.6 | 1 | MS.Trt_vs_MS.NoTrt_B |
| MBD5 | 0.04273 | 0.78422 | 0.706 | 0.88 | 1 | MS.Trt_vs_MS.NoTrt_B |

| | | | | | | |
|---|---|---|---|---|---|---|
| ATMIN | 0.04295 | 0.77764 | 0.192 | 0.27 | 1 | MS.Trt_vs_MS.NoTrt_B |
| REXO2 | 0.04295 | 0.25709 | 0.148 | 0.22 | 1 | MS.Trt_vs_MS.NoTrt_B |
| RUFY3 | 0.04299 | 0.77723 | 0.397 | 0.51 | 1 | MS.Trt_vs_MS.NoTrt_B |
| PKIG | 0.0432 | 0.77966 | 0.523 | 0.54 | 1 | MS.Trt_vs_MS.NoTrt_B |
| TEX14 | 0.04321 | 1.23218 | 0.059 | 0.11 | 1 | MS.Trt_vs_MS.NoTrt_B |
| HSH2D | 0.04345 | -0.6902 | 0.514 | 0.63 | 1 | MS.Trt_vs_MS.NoTrt_B |
| ZNHIT6 | 0.04348 | -0.5458 | 0.292 | 0.38 | 1 | MS.Trt_vs_MS.NoTrt_B |
| MAD2L1BP | 0.0435 | -0.5418 | 0.109 | 0.16 | 1 | MS.Trt_vs_MS.NoTrt_B |
| ZEB1 | 0.04354 | 0.91748 | 0.438 | 0.56 | 1 | MS.Trt_vs_MS.NoTrt_B |
| TPCN2 | 0.04377 | 0.6281 | 0.449 | 0.57 | 1 | MS.Trt_vs_MS.NoTrt_B |
| MOCS3 | 0.04405 | 0.52704 | 0.688 | 0.81 | 1 | MS.Trt_vs_MS.NoTrt_B |
| KLHL20 | 0.04412 | -0.2589 | 0.094 | 0.15 | 1 | MS.Trt_vs_MS.NoTrt_B |
| RP9 | 0.04436 | -0.3 | 0.061 | 0.11 | 1 | MS.Trt_vs_MS.NoTrt_B |
| C2orf49 | 0.04438 | 0.74488 | 0.32 | 0.42 | 1 | MS.Trt_vs_MS.NoTrt_B |
| SON | 0.04439 | 0.28211 | 0.847 | 0.97 | 1 | MS.Trt_vs_MS.NoTrt_B |
| CEP85L | 0.04446 | 0.25973 | 0.222 | 0.3 | 1 | MS.Trt_vs_MS.NoTrt_B |
| HEBP2 | 0.04466 | 1.00943 | 0.292 | 0.37 | 1 | MS.Trt_vs_MS.NoTrt_B |
| TAB1 | 0.04472 | -0.2898 | 0.065 | 0.11 | 1 | MS.Trt_vs_MS.NoTrt_B |
| ZC3H18 | 0.04479 | 0.48117 | 0.582 | 0.71 | 1 | MS.Trt_vs_MS.NoTrt_B |
| MRFAP1 | 0.04489 | -0.4499 | 0.22 | 0.29 | 1 | MS.Trt_vs_MS.NoTrt_B |
| RFX2 | 0.04497 | 0.47032 | 0.401 | 0.49 | 1 | MS.Trt_vs_MS.NoTrt_B |
| ZNF568 | 0.04509 | -0.4517 | 0.706 | 0.81 | 1 | MS.Trt_vs_MS.NoTrt_B |
| RSRC1 | 0.0452 | 0.30308 | 0.599 | 0.71 | 1 | MS.Trt_vs_MS.NoTrt_B |
| EID2B | 0.04531 | -0.8172 | 0.414 | 0.5 | 1 | MS.Trt_vs_MS.NoTrt_B |
| ERCC2 | 0.04565 | 0.32354 | 0.425 | 0.51 | 1 | MS.Trt_vs_MS.NoTrt_B |
| NEIL2 | 0.04598 | 0.36785 | 0.072 | 0.12 | 1 | MS.Trt_vs_MS.NoTrt_B |
| ZNF429 | 0.04598 | -0.2847 | 0.817 | 0.92 | 1 | MS.Trt_vs_MS.NoTrt_B |
| AL592183.1 | 0.04621 | 0.30052 | 0.078 | 0.13 | 1 | MS.Trt_vs_MS.NoTrt_B |
| ADIPOQ | 0.04624 | -0.2601 | 0.473 | 0.61 | 1 | MS.Trt_vs_MS.NoTrt_B |
| STRC | 0.0466 | 0.94729 | 0.427 | 0.51 | 1 | MS.Trt_vs_MS.NoTrt_B |
| SPDYA | 0.04675 | 1.01121 | 0.061 | 0.11 | 1 | MS.Trt_vs_MS.NoTrt_B |
| TSPAN17 | 0.04681 | -0.4181 | 0.068 | 0.11 | 1 | MS.Trt_vs_MS.NoTrt_B |
| DNASE1L1 | 0.04685 | -0.4083 | 0.063 | 0.11 | 1 | MS.Trt_vs_MS.NoTrt_B |
| GPD1L | 0.04698 | 1.0049 | 0.078 | 0.13 | 1 | MS.Trt_vs_MS.NoTrt_B |
| SNRNP27 | 0.04699 | 0.53573 | 0.218 | 0.31 | 1 | MS.Trt_vs_MS.NoTrt_B |
| CCDC137 | 0.04709 | 0.85101 | 0.166 | 0.23 | 1 | MS.Trt_vs_MS.NoTrt_B |
| MRM3 | 0.04713 | 0.3054 | 0.2 | 0.28 | 1 | MS.Trt_vs_MS.NoTrt_B |
| PNN | 0.04725 | 0.25632 | 0.251 | 0.34 | 1 | MS.Trt_vs_MS.NoTrt_B |
| NBPF1 | 0.04742 | 0.29926 | 0.174 | 0.25 | 1 | MS.Trt_vs_MS.NoTrt_B |
| RNPEP | 0.04742 | -0.321 | 0.059 | 0.1 | 1 | MS.Trt_vs_MS.NoTrt_B |

| | | | | | | |
|---|---|---|---|---|---|---|
| NFE2L2 | 0.04743 | 0.28726 | 0.405 | 0.5 | 1 | MS.Trt_vs_MS.NoTrt_B |
| ABLIM1 | 0.04756 | 0.46155 | 0.322 | 0.43 | 1 | MS.Trt_vs_MS.NoTrt_B |
| AFF3 | 0.04759 | 0.63698 | 0.599 | 0.73 | 1 | MS.Trt_vs_MS.NoTrt_B |
| PLAC8 | 0.04777 | 0.32746 | 0.632 | 0.77 | 1 | MS.Trt_vs_MS.NoTrt_B |
| MYSM1 | 0.04778 | 0.55571 | 0.146 | 0.21 | 1 | MS.Trt_vs_MS.NoTrt_B |
| GALNS | 0.0478 | -0.7768 | 0.42 | 0.5 | 1 | MS.Trt_vs_MS.NoTrt_B |
| CTD-2370N5 | 0.04794 | 0.25123 | 0.092 | 0.15 | 1 | MS.Trt_vs_MS.NoTrt_B |
| NDUFB1 | 0.048 | 0.37827 | 0.649 | 0.77 | 1 | MS.Trt_vs_MS.NoTrt_B |
| RBCK1 | 0.04816 | 0.28904 | 0.161 | 0.23 | 1 | MS.Trt_vs_MS.NoTrt_B |
| QRICH2 | 0.0482 | -0.501 | 0.12 | 0.18 | 1 | MS.Trt_vs_MS.NoTrt_B |
| SEC23B | 0.04825 | -0.3656 | 0.085 | 0.14 | 1 | MS.Trt_vs_MS.NoTrt_B |
| ZKSCAN8 | 0.0483 | 1.6512 | 0.209 | 0.29 | 1 | MS.Trt_vs_MS.NoTrt_B |
| IRF7 | 0.04845 | -0.4623 | 0.072 | 0.12 | 1 | MS.Trt_vs_MS.NoTrt_B |
| C14orf132 | 0.04848 | 0.25406 | 0.205 | 0.28 | 1 | MS.Trt_vs_MS.NoTrt_B |
| BSCL2 | 0.04849 | -0.4663 | 0.176 | 0.24 | 1 | MS.Trt_vs_MS.NoTrt_B |
| ATM | 0.0485 | 0.50393 | 0.898 | 0.98 | 1 | MS.Trt_vs_MS.NoTrt_B |
| MRPS14 | 0.04851 | 0.86561 | 0.111 | 0.17 | 1 | MS.Trt_vs_MS.NoTrt_B |
| COA4 | 0.04851 | 0.41308 | 0.216 | 0.3 | 1 | MS.Trt_vs_MS.NoTrt_B |
| TGIF2 | 0.0486 | -0.349 | 0.255 | 0.32 | 1 | MS.Trt_vs_MS.NoTrt_B |
| ZNF286B | 0.04871 | 1.13043 | 0.357 | 0.48 | 1 | MS.Trt_vs_MS.NoTrt_B |
| SLC9A1 | 0.04872 | -0.5271 | 0.166 | 0.23 | 1 | MS.Trt_vs_MS.NoTrt_B |
| GOLGA1 | 0.04874 | 0.25986 | 0.081 | 0.13 | 1 | MS.Trt_vs_MS.NoTrt_B |
| TLK2 | 0.04876 | 0.54354 | 0.272 | 0.35 | 1 | MS.Trt_vs_MS.NoTrt_B |
| ZNF564 | 0.04879 | 0.69911 | 0.519 | 0.65 | 1 | MS.Trt_vs_MS.NoTrt_B |
| ZMYND8 | 0.04897 | 0.50474 | 0.224 | 0.3 | 1 | MS.Trt_vs_MS.NoTrt_B |
| ACER3 | 0.04906 | 0.3023 | 0.427 | 0.52 | 1 | MS.Trt_vs_MS.NoTrt_B |
| GOLGA2 | 0.04908 | -0.4138 | 0.503 | 0.65 | 1 | MS.Trt_vs_MS.NoTrt_B |
| KIF21A | 0.04909 | 0.82831 | 0.105 | 0.16 | 1 | MS.Trt_vs_MS.NoTrt_B |
| SNRNP48 | 0.04911 | 0.65982 | 0.139 | 0.21 | 1 | MS.Trt_vs_MS.NoTrt_B |
| DOT1L | 0.0492 | 0.58588 | 0.627 | 0.77 | 1 | MS.Trt_vs_MS.NoTrt_B |
| MCRS1 | 0.04922 | 0.32121 | 0.144 | 0.21 | 1 | MS.Trt_vs_MS.NoTrt_B |
| MYL6 | 0.0493 | 0.287 | 0.682 | 0.85 | 1 | MS.Trt_vs_MS.NoTrt_B |
| GRK5 | 0.04934 | -0.3486 | 0.111 | 0.17 | 1 | MS.Trt_vs_MS.NoTrt_B |
| FAM65B | 0.04957 | 0.53641 | 0.545 | 0.67 | 1 | MS.Trt_vs_MS.NoTrt_B |
| PMAIP1 | 0.04968 | -0.3922 | 0.068 | 0.11 | 1 | MS.Trt_vs_MS.NoTrt_B |
| KIF13A | 0.0497 | 0.76248 | 0.224 | 0.3 | 1 | MS.Trt_vs_MS.NoTrt_B |
| C16orf74 | 0.0497 | -0.3033 | 0.181 | 0.24 | 1 | MS.Trt_vs_MS.NoTrt_B |
| SPARCL1 | 0.04985 | 0.4121 | 0.131 | 0.08 | 1 | MS.Trt_vs_MS.NoTrt_B |
| FCER1G | 0.05 | 0.5169 | 0.505 | 0.62 | 1 | MS.Trt_vs_MS.NoTrt_B |
| PHF20L1 | 0.05031 | -0.5414 | 0.514 | 0.62 | 1 | MS.Trt_vs_MS.NoTrt_B |

| | | | | | | |
|---|---|---|---|---|---|---|
| ATP6V0D1 | 0.0505 | -0.7528 | 0.573 | 0.62 | 1 | MS.Trt_vs_MS.NoTrt_B |
| CSAD | 0.0505 | 0.29514 | 0.68 | 0.81 | 1 | MS.Trt_vs_MS.NoTrt_B |
| FDX1 | 0.05052 | 0.59155 | 0.211 | 0.29 | 1 | MS.Trt_vs_MS.NoTrt_B |
| ASCC3 | 0.05053 | 0.80728 | 0.203 | 0.29 | 1 | MS.Trt_vs_MS.NoTrt_B |
| RIC8A | 0.05062 | -0.2808 | 0.157 | 0.22 | 1 | MS.Trt_vs_MS.NoTrt_B |
| CD93 | 0.05068 | -0.6618 | 0.118 | 0.17 | 1 | MS.Trt_vs_MS.NoTrt_B |
| BOD1L1 | 0.0507 | 0.69986 | 0.593 | 0.73 | 1 | MS.Trt_vs_MS.NoTrt_B |
| PRR14 | 0.05092 | 0.43382 | 0.1 | 0.16 | 1 | MS.Trt_vs_MS.NoTrt_B |
| MITD1 | 0.05098 | 0.39382 | 0.865 | 0.96 | 1 | MS.Trt_vs_MS.NoTrt_B |
| SMIM15 | 0.05113 | 0.32392 | 0.063 | 0.11 | 1 | MS.Trt_vs_MS.NoTrt_B |
| RASGEF1B | 0.05123 | 0.28071 | 0.614 | 0.72 | 1 | MS.Trt_vs_MS.NoTrt_B |
| ATP1A4 | 0.05147 | 0.34352 | 0.734 | 0.75 | 1 | MS.Trt_vs_MS.NoTrt_B |
| PMS2 | 0.05162 | 0.50247 | 0.558 | 0.7 | 1 | MS.Trt_vs_MS.NoTrt_B |
| ZDHHC7 | 0.05191 | -0.3751 | 0.124 | 0.18 | 1 | MS.Trt_vs_MS.NoTrt_B |
| ZNF543 | 0.05207 | 1.02567 | 0.342 | 0.45 | 1 | MS.Trt_vs_MS.NoTrt_B |
| THAP12 | 0.05235 | 0.61135 | 0.464 | 0.59 | 1 | MS.Trt_vs_MS.NoTrt_B |
| WDR19 | 0.05236 | -0.5748 | 0.418 | 0.51 | 1 | MS.Trt_vs_MS.NoTrt_B |
| MRPL10 | 0.05244 | -0.4672 | 0.301 | 0.39 | 1 | MS.Trt_vs_MS.NoTrt_B |
| RRM2B | 0.05255 | 0.95509 | 0.087 | 0.14 | 1 | MS.Trt_vs_MS.NoTrt_B |
| COX7B | 0.05257 | -0.4245 | 0.42 | 0.5 | 1 | MS.Trt_vs_MS.NoTrt_B |
| RASA1 | 0.05269 | 0.77266 | 0.159 | 0.22 | 1 | MS.Trt_vs_MS.NoTrt_B |
| SSSCA1 | 0.05276 | -0.3384 | 0.059 | 0.1 | 1 | MS.Trt_vs_MS.NoTrt_B |
| MED1 | 0.05302 | -0.6752 | 0.268 | 0.34 | 1 | MS.Trt_vs_MS.NoTrt_B |
| CD40 | 0.05302 | -0.5669 | 0.094 | 0.15 | 1 | MS.Trt_vs_MS.NoTrt_B |
| ZNF649 | 0.05312 | -0.6373 | 0.135 | 0.2 | 1 | MS.Trt_vs_MS.NoTrt_B |
| GABPB2 | 0.05318 | 0.28868 | 0.917 | 0.97 | 1 | MS.Trt_vs_MS.NoTrt_B |
| ZNF565 | 0.05327 | 1.4914 | 0.063 | 0.11 | 1 | MS.Trt_vs_MS.NoTrt_B |
| RFWD2 | 0.05328 | 1.00333 | 0.268 | 0.34 | 1 | MS.Trt_vs_MS.NoTrt_B |
| BLMH | 0.05333 | -0.504 | 0.545 | 0.63 | 1 | MS.Trt_vs_MS.NoTrt_B |
| IQCB1 | 0.05348 | 0.48261 | 0.063 | 0.11 | 1 | MS.Trt_vs_MS.NoTrt_B |
| CASZ1 | 0.05366 | -0.5956 | 0.155 | 0.22 | 1 | MS.Trt_vs_MS.NoTrt_B |
| BTG1 | 0.05369 | 0.33475 | 0.516 | 0.63 | 1 | MS.Trt_vs_MS.NoTrt_B |
| LPIN2 | 0.05373 | 0.56413 | 0.087 | 0.14 | 1 | MS.Trt_vs_MS.NoTrt_B |
| EXOC1 | 0.05382 | 0.58984 | 0.135 | 0.2 | 1 | MS.Trt_vs_MS.NoTrt_B |
| SZRD1 | 0.05392 | 0.2889 | 0.135 | 0.2 | 1 | MS.Trt_vs_MS.NoTrt_B |
| TMEM108 | 0.05424 | -0.7916 | 0.244 | 0.31 | 1 | MS.Trt_vs_MS.NoTrt_B |
| CTF1 | 0.05424 | -0.4931 | 0.17 | 0.24 | 1 | MS.Trt_vs_MS.NoTrt_B |
| STAT2 | 0.05438 | 0.36346 | 0.843 | 0.92 | 1 | MS.Trt_vs_MS.NoTrt_B |
| NOL4L | 0.05439 | 0.60699 | 0.551 | 0.69 | 1 | MS.Trt_vs_MS.NoTrt_B |
| XYLT2 | 0.05454 | 0.43456 | 0.523 | 0.62 | 1 | MS.Trt_vs_MS.NoTrt_B |

| | | | | | | |
|---|---|---|---|---|---|---|
| PRKACA | 0.05456 | -0.4101 | 0.146 | 0.21 | 1 | MS.Trt_vs_MS.NoTrt_B |
| RPS26 | 0.05475 | 0.38041 | 0.68 | 0.75 | 1 | MS.Trt_vs_MS.NoTrt_B |
| MTX3 | 0.05495 | 0.38598 | 0.545 | 0.64 | 1 | MS.Trt_vs_MS.NoTrt_B |
| DEPDC5 | 0.05499 | -0.3277 | 0.68 | 0.77 | 1 | MS.Trt_vs_MS.NoTrt_B |
| DIMT1 | 0.05502 | 0.3004 | 0.113 | 0.17 | 1 | MS.Trt_vs_MS.NoTrt_B |
| GPCPD1 | 0.05518 | 0.97371 | 0.203 | 0.29 | 1 | MS.Trt_vs_MS.NoTrt_B |
| ZDHHC18 | 0.05531 | -0.3782 | 0.059 | 0.1 | 1 | MS.Trt_vs_MS.NoTrt_B |
| DBP | 0.05536 | -0.2529 | 0.54 | 0.65 | 1 | MS.Trt_vs_MS.NoTrt_B |
| ANKRD13A | 0.05547 | 0.4448 | 0.712 | 0.81 | 1 | MS.Trt_vs_MS.NoTrt_B |
| PLK3 | 0.05553 | -0.3542 | 0.577 | 0.68 | 1 | MS.Trt_vs_MS.NoTrt_B |
| PLEKHB1 | 0.05566 | -0.8725 | 0.065 | 0.11 | 1 | MS.Trt_vs_MS.NoTrt_B |
| H2AFV | 0.0557 | 0.44802 | 0.514 | 0.61 | 1 | MS.Trt_vs_MS.NoTrt_B |
| ZMYM4 | 0.05571 | 0.59795 | 0.078 | 0.12 | 1 | MS.Trt_vs_MS.NoTrt_B |
| NUP62 | 0.05573 | -0.2714 | 0.399 | 0.48 | 1 | MS.Trt_vs_MS.NoTrt_B |
| NFATC3 | 0.05577 | 0.5182 | 0.499 | 0.58 | 1 | MS.Trt_vs_MS.NoTrt_B |
| TRPV1 | 0.05587 | -1.3194 | 0.288 | 0.39 | 1 | MS.Trt_vs_MS.NoTrt_B |
| ZNF267 | 0.05601 | 0.64754 | 0.081 | 0.13 | 1 | MS.Trt_vs_MS.NoTrt_B |
| IRF2BP2 | 0.05606 | 1.46346 | 0.174 | 0.12 | 1 | MS.Trt_vs_MS.NoTrt_B |
| SH2D1A | 0.05619 | 0.7738 | 0.159 | 0.22 | 1 | MS.Trt_vs_MS.NoTrt_B |
| TIGAR | 0.05633 | 0.29843 | 0.808 | 0.91 | 1 | MS.Trt_vs_MS.NoTrt_B |
| IWS1 | 0.05643 | 0.25077 | 0.094 | 0.15 | 1 | MS.Trt_vs_MS.NoTrt_B |
| KMT2A | 0.05649 | 0.42007 | 0.856 | 0.93 | 1 | MS.Trt_vs_MS.NoTrt_B |
| ZNF782 | 0.05657 | 0.80815 | 0.085 | 0.14 | 1 | MS.Trt_vs_MS.NoTrt_B |
| SPDYE1 | 0.05659 | 0.32307 | 0.407 | 0.52 | 1 | MS.Trt_vs_MS.NoTrt_B |
| NCOA2 | 0.05661 | 0.37558 | 0.183 | 0.25 | 1 | MS.Trt_vs_MS.NoTrt_B |
| PSIP1 | 0.05688 | 0.39752 | 0.691 | 0.81 | 1 | MS.Trt_vs_MS.NoTrt_B |
| RPF1 | 0.05696 | 0.42485 | 0.214 | 0.29 | 1 | MS.Trt_vs_MS.NoTrt_B |
| TOR4A | 0.05706 | 0.54378 | 0.129 | 0.19 | 1 | MS.Trt_vs_MS.NoTrt_B |
| NPIPA7 | 0.05712 | 0.67892 | 0.124 | 0.19 | 1 | MS.Trt_vs_MS.NoTrt_B |
| MDFIC | 0.05721 | 0.49339 | 0.261 | 0.35 | 1 | MS.Trt_vs_MS.NoTrt_B |
| RBM26 | 0.05726 | 0.66102 | 0.392 | 0.5 | 1 | MS.Trt_vs_MS.NoTrt_B |
| TECPR2 | 0.05726 | 0.65906 | 0.248 | 0.34 | 1 | MS.Trt_vs_MS.NoTrt_B |
| APOOL | 0.05731 | 0.73973 | 0.647 | 0.78 | 1 | MS.Trt_vs_MS.NoTrt_B |
| ATP1B1 | 0.05745 | -1.064 | 0.172 | 0.23 | 1 | MS.Trt_vs_MS.NoTrt_B |
| NOL11 | 0.05755 | -0.4601 | 0.418 | 0.49 | 1 | MS.Trt_vs_MS.NoTrt_B |
| PCIF1 | 0.05792 | 0.32264 | 0.268 | 0.35 | 1 | MS.Trt_vs_MS.NoTrt_B |
| PTRH1 | 0.05793 | -0.4439 | 0.603 | 0.74 | 1 | MS.Trt_vs_MS.NoTrt_B |
| TPRG1L | 0.05803 | 1.8834 | 0.176 | 0.24 | 1 | MS.Trt_vs_MS.NoTrt_B |
| TTC19 | 0.0583 | 0.41851 | 0.684 | 0.75 | 1 | MS.Trt_vs_MS.NoTrt_B |
| YBX3 | 0.05849 | 0.48722 | 0.608 | 0.74 | 1 | MS.Trt_vs_MS.NoTrt_B |

| | | | | | | |
|---|---|---|---|---|---|---|
| MSH3 | 0.05888 | 0.45997 | 0.292 | 0.4 | 1 | MS.Trt_vs_MS.NoTrt_B |
| FCHO2 | 0.05908 | 0.62746 | 0.22 | 0.3 | 1 | MS.Trt_vs_MS.NoTrt_B |
| ARPC4 | 0.05922 | 0.34218 | 0.588 | 0.75 | 1 | MS.Trt_vs_MS.NoTrt_B |
| BTBD1 | 0.05931 | -0.3422 | 0.471 | 0.57 | 1 | MS.Trt_vs_MS.NoTrt_B |
| PREPL | 0.05932 | 0.376 | 0.551 | 0.67 | 1 | MS.Trt_vs_MS.NoTrt_B |
| GOLGB1 | 0.05966 | 0.26908 | 0.438 | 0.51 | 1 | MS.Trt_vs_MS.NoTrt_B |
| RAI1 | 0.05987 | 0.31901 | 0.146 | 0.21 | 1 | MS.Trt_vs_MS.NoTrt_B |
| ZNF2 | 0.06004 | 1.25368 | 0.266 | 0.36 | 1 | MS.Trt_vs_MS.NoTrt_B |
| DOPEY1 | 0.06011 | 1.22052 | 0.224 | 0.31 | 1 | MS.Trt_vs_MS.NoTrt_B |
| VNN3 | 0.06013 | 0.60777 | 0.257 | 0.37 | 1 | MS.Trt_vs_MS.NoTrt_B |
| ZDHHC20 | 0.06015 | 0.37343 | 0.359 | 0.44 | 1 | MS.Trt_vs_MS.NoTrt_B |
| MRPL30 | 0.06024 | 0.51531 | 0.658 | 0.77 | 1 | MS.Trt_vs_MS.NoTrt_B |
| FAM160A1 | 0.06049 | 0.76313 | 0.131 | 0.19 | 1 | MS.Trt_vs_MS.NoTrt_B |
| RYK | 0.06057 | 0.25391 | 0.063 | 0.11 | 1 | MS.Trt_vs_MS.NoTrt_B |
| WDFY1 | 0.0607 | 0.43443 | 0.242 | 0.33 | 1 | MS.Trt_vs_MS.NoTrt_B |
| NQO1 | 0.06087 | -0.5483 | 0.46 | 0.52 | 1 | MS.Trt_vs_MS.NoTrt_B |
| TCF7 | 0.06095 | 0.75628 | 0.758 | 0.73 | 1 | MS.Trt_vs_MS.NoTrt_B |
| L3MBTL3 | 0.06104 | -0.7803 | 0.1 | 0.15 | 1 | MS.Trt_vs_MS.NoTrt_B |
| ERH | 0.06114 | 0.33028 | 0.192 | 0.26 | 1 | MS.Trt_vs_MS.NoTrt_B |
| RAD9B | 0.06133 | 0.30993 | 0.288 | 0.37 | 1 | MS.Trt_vs_MS.NoTrt_B |
| MS4A10 | 0.06142 | 0.25245 | 0.617 | 0.56 | 1 | MS.Trt_vs_MS.NoTrt_B |
| ACO2 | 0.06196 | -0.2752 | 0.092 | 0.14 | 1 | MS.Trt_vs_MS.NoTrt_B |
| FBXO21 | 0.06198 | -0.3602 | 0.113 | 0.17 | 1 | MS.Trt_vs_MS.NoTrt_B |
| CDH26 | 0.06209 | -0.3297 | 0.096 | 0.15 | 1 | MS.Trt_vs_MS.NoTrt_B |
| COX19 | 0.0621 | -0.2956 | 0.503 | 0.61 | 1 | MS.Trt_vs_MS.NoTrt_B |
| PAPOLG | 0.06212 | 0.63594 | 0.647 | 0.78 | 1 | MS.Trt_vs_MS.NoTrt_B |
| PRPF39 | 0.06225 | 0.35424 | 0.235 | 0.3 | 1 | MS.Trt_vs_MS.NoTrt_B |
| STK24 | 0.06228 | 0.75708 | 0.196 | 0.27 | 1 | MS.Trt_vs_MS.NoTrt_B |
| SSRP1 | 0.06234 | -0.3219 | 0.087 | 0.14 | 1 | MS.Trt_vs_MS.NoTrt_B |
| MRPL41 | 0.06235 | 0.56734 | 0.131 | 0.2 | 1 | MS.Trt_vs_MS.NoTrt_B |
| ZNF417 | 0.06235 | 0.25057 | 0.647 | 0.75 | 1 | MS.Trt_vs_MS.NoTrt_B |
| GTSF1 | 0.06257 | -0.5727 | 0.153 | 0.22 | 1 | MS.Trt_vs_MS.NoTrt_B |
| F2RL2 | 0.06267 | -0.4267 | 0.198 | 0.26 | 1 | MS.Trt_vs_MS.NoTrt_B |
| DHX34 | 0.06287 | -0.6341 | 0.061 | 0.1 | 1 | MS.Trt_vs_MS.NoTrt_B |
| ZNF585A | 0.06297 | 0.68651 | 0.244 | 0.31 | 1 | MS.Trt_vs_MS.NoTrt_B |
| ZNF225 | 0.063 | -0.2749 | 0.678 | 0.74 | 1 | MS.Trt_vs_MS.NoTrt_B |
| SLC30A4 | 0.0632 | 0.42098 | 0.756 | 0.87 | 1 | MS.Trt_vs_MS.NoTrt_B |
| RNASEH1 | 0.06353 | -0.3218 | 0.645 | 0.74 | 1 | MS.Trt_vs_MS.NoTrt_B |
| GTF3C4 | 0.06376 | 0.65452 | 0.266 | 0.35 | 1 | MS.Trt_vs_MS.NoTrt_B |
| HDAC10 | 0.06393 | 0.32217 | 0.087 | 0.14 | 1 | MS.Trt_vs_MS.NoTrt_B |

| | | | | | | |
|---|---|---|---|---|---|---|
| RAB11FIP3 | 0.06395 | 0.25224 | 0.115 | 0.17 | 1 | MS.Trt_vs_MS.NoTrt_B |
| SFR1 | 0.06407 | 0.43289 | 0.2 | 0.27 | 1 | MS.Trt_vs_MS.NoTrt_B |
| TAF1B | 0.06427 | 0.61074 | 0.381 | 0.49 | 1 | MS.Trt_vs_MS.NoTrt_B |
| JAK2 | 0.0643 | 0.47221 | 0.159 | 0.22 | 1 | MS.Trt_vs_MS.NoTrt_B |
| MRE11A | 0.0643 | 0.64651 | 0.603 | 0.7 | 1 | MS.Trt_vs_MS.NoTrt_B |
| FCRL3 | 0.06432 | 0.90012 | 0.316 | 0.25 | 1 | MS.Trt_vs_MS.NoTrt_B |
| ABCC10 | 0.06469 | -0.4386 | 0.813 | 0.95 | 1 | MS.Trt_vs_MS.NoTrt_B |
| PPM1B | 0.06509 | -0.265 | 0.301 | 0.38 | 1 | MS.Trt_vs_MS.NoTrt_B |
| AGO2 | 0.06512 | 0.43706 | 0.292 | 0.37 | 1 | MS.Trt_vs_MS.NoTrt_B |
| IRAK1BP1 | 0.06539 | 0.34407 | 0.529 | 0.67 | 1 | MS.Trt_vs_MS.NoTrt_B |
| TTC5 | 0.06559 | -0.6593 | 0.412 | 0.48 | 1 | MS.Trt_vs_MS.NoTrt_B |
| TGIF1 | 0.06584 | 0.28367 | 0.161 | 0.23 | 1 | MS.Trt_vs_MS.NoTrt_B |
| HYDIN | 0.06607 | 0.93415 | 0.179 | 0.24 | 1 | MS.Trt_vs_MS.NoTrt_B |
| BNC2 | 0.06608 | 0.56094 | 0.094 | 0.15 | 1 | MS.Trt_vs_MS.NoTrt_B |
| CHKB | 0.06624 | 0.41646 | 0.187 | 0.25 | 1 | MS.Trt_vs_MS.NoTrt_B |
| SAMHD1 | 0.06655 | -0.2733 | 0.113 | 0.16 | 1 | MS.Trt_vs_MS.NoTrt_B |
| TMEM45B | 0.06663 | -0.2941 | 0.481 | 0.57 | 1 | MS.Trt_vs_MS.NoTrt_B |
| C5 | 0.06675 | -0.9702 | 0.322 | 0.39 | 1 | MS.Trt_vs_MS.NoTrt_B |
| ZNF320 | 0.06684 | -0.3162 | 0.917 | 0.97 | 1 | MS.Trt_vs_MS.NoTrt_B |
| BCAP29 | 0.06691 | -0.384 | 0.253 | 0.34 | 1 | MS.Trt_vs_MS.NoTrt_B |
| KEAP1 | 0.06713 | -0.3803 | 0.353 | 0.43 | 1 | MS.Trt_vs_MS.NoTrt_B |
| KIAA0556 | 0.06751 | -0.332 | 0.244 | 0.31 | 1 | MS.Trt_vs_MS.NoTrt_B |
| CHD8 | 0.0677 | 0.3049 | 0.242 | 0.31 | 1 | MS.Trt_vs_MS.NoTrt_B |
| ELK1 | 0.0678 | -0.9406 | 0.068 | 0.11 | 1 | MS.Trt_vs_MS.NoTrt_B |
| MDN1 | 0.06801 | 0.33469 | 0.176 | 0.24 | 1 | MS.Trt_vs_MS.NoTrt_B |
| HYLS1 | 0.06805 | -0.251 | 0.281 | 0.37 | 1 | MS.Trt_vs_MS.NoTrt_B |
| JARID2 | 0.06806 | 0.29669 | 0.157 | 0.22 | 1 | MS.Trt_vs_MS.NoTrt_B |
| LAMB1 | 0.06809 | 0.74055 | 0.198 | 0.27 | 1 | MS.Trt_vs_MS.NoTrt_B |
| PPIG | 0.06861 | 0.43397 | 0.251 | 0.34 | 1 | MS.Trt_vs_MS.NoTrt_B |
| RCC2 | 0.06882 | 0.30898 | 0.061 | 0.1 | 1 | MS.Trt_vs_MS.NoTrt_B |
| RC3H2 | 0.06888 | 0.61165 | 0.41 | 0.5 | 1 | MS.Trt_vs_MS.NoTrt_B |
| RNF145 | 0.06894 | 0.81813 | 0.102 | 0.16 | 1 | MS.Trt_vs_MS.NoTrt_B |
| IQSEC1 | 0.06906 | 1.19525 | 0.146 | 0.21 | 1 | MS.Trt_vs_MS.NoTrt_B |
| USP3 | 0.06936 | 0.39944 | 0.669 | 0.74 | 1 | MS.Trt_vs_MS.NoTrt_B |
| CEP97 | 0.06941 | 0.52152 | 0.538 | 0.65 | 1 | MS.Trt_vs_MS.NoTrt_B |
| CLIP4 | 0.06983 | -0.3796 | 0.085 | 0.13 | 1 | MS.Trt_vs_MS.NoTrt_B |
| BAZ1A | 0.07007 | 0.54647 | 0.259 | 0.34 | 1 | MS.Trt_vs_MS.NoTrt_B |
| ZNF23 | 0.0702 | 0.35016 | 0.259 | 0.35 | 1 | MS.Trt_vs_MS.NoTrt_B |
| SLC27A5 | 0.0703 | 0.27081 | 0.776 | 0.89 | 1 | MS.Trt_vs_MS.NoTrt_B |
| ZNF737 | 0.07046 | 0.29939 | 0.702 | 0.82 | 1 | MS.Trt_vs_MS.NoTrt_B |

| | | | | | | |
|---|---|---|---|---|---|---|
| ITCH | 0.07047 | 0.73385 | 0.414 | 0.5 | 1 | MS.Trt_vs_MS.NoTrt_B |
| DENND3 | 0.07049 | 0.31338 | 0.351 | 0.45 | 1 | MS.Trt_vs_MS.NoTrt_B |
| MIOS | 0.07073 | 0.52812 | 0.251 | 0.32 | 1 | MS.Trt_vs_MS.NoTrt_B |
| DVL2 | 0.07074 | -0.3176 | 0.338 | 0.41 | 1 | MS.Trt_vs_MS.NoTrt_B |
| TGFB1 | 0.07091 | 0.63941 | 0.2 | 0.27 | 1 | MS.Trt_vs_MS.NoTrt_B |
| ZNF146 | 0.07094 | 0.29831 | 0.15 | 0.21 | 1 | MS.Trt_vs_MS.NoTrt_B |
| ORC3 | 0.07109 | -0.2793 | 0.098 | 0.15 | 1 | MS.Trt_vs_MS.NoTrt_B |
| AP2A1 | 0.0712 | -1.004 | 0.63 | 0.76 | 1 | MS.Trt_vs_MS.NoTrt_B |
| MALL | 0.07125 | 0.28578 | 0.508 | 0.65 | 1 | MS.Trt_vs_MS.NoTrt_B |
| HNRNPUL1 | 0.07126 | 0.25234 | 0.575 | 0.68 | 1 | MS.Trt_vs_MS.NoTrt_B |
| TGFBRAP1 | 0.07135 | -0.4145 | 0.355 | 0.44 | 1 | MS.Trt_vs_MS.NoTrt_B |
| CD59 | 0.0714 | -0.4659 | 0.577 | 0.7 | 1 | MS.Trt_vs_MS.NoTrt_B |
| CNIH3 | 0.07148 | -0.9564 | 0.122 | 0.17 | 1 | MS.Trt_vs_MS.NoTrt_B |
| WDR59 | 0.07174 | 0.6017 | 0.854 | 0.94 | 1 | MS.Trt_vs_MS.NoTrt_B |
| GK | 0.07202 | 0.54204 | 0.133 | 0.19 | 1 | MS.Trt_vs_MS.NoTrt_B |
| KLHL3 | 0.07234 | 0.46329 | 0.187 | 0.25 | 1 | MS.Trt_vs_MS.NoTrt_B |
| NAV2 | 0.07244 | 0.7691 | 0.379 | 0.32 | 1 | MS.Trt_vs_MS.NoTrt_B |
| SPHK2 | 0.07283 | -0.2526 | 0.222 | 0.29 | 1 | MS.Trt_vs_MS.NoTrt_B |
| AL354822.1 | 0.07286 | 0.47479 | 0.155 | 0.21 | 1 | MS.Trt_vs_MS.NoTrt_B |
| PI4KA | 0.07298 | 0.58146 | 0.203 | 0.27 | 1 | MS.Trt_vs_MS.NoTrt_B |
| PLIN2 | 0.07305 | -0.2631 | 0.553 | 0.63 | 1 | MS.Trt_vs_MS.NoTrt_B |
| TIMM21 | 0.07331 | 0.38225 | 0.542 | 0.63 | 1 | MS.Trt_vs_MS.NoTrt_B |
| LAX1 | 0.07338 | -1.0141 | 0.381 | 0.42 | 1 | MS.Trt_vs_MS.NoTrt_B |
| UNC45B | 0.07345 | -0.4804 | 0.072 | 0.11 | 1 | MS.Trt_vs_MS.NoTrt_B |
| DALRD3 | 0.0737 | -0.3448 | 0.089 | 0.14 | 1 | MS.Trt_vs_MS.NoTrt_B |
| GLB1L3 | 0.07384 | 0.26265 | 0.586 | 0.69 | 1 | MS.Trt_vs_MS.NoTrt_B |
| SCARB2 | 0.07385 | 0.40264 | 0.209 | 0.29 | 1 | MS.Trt_vs_MS.NoTrt_B |
| FIBP | 0.07394 | -0.4105 | 0.449 | 0.51 | 1 | MS.Trt_vs_MS.NoTrt_B |
| ZCWPW2 | 0.07408 | 0.40255 | 0.375 | 0.48 | 1 | MS.Trt_vs_MS.NoTrt_B |
| MAEA | 0.07433 | 0.28114 | 0.51 | 0.61 | 1 | MS.Trt_vs_MS.NoTrt_B |
| GLT8D1 | 0.07437 | -0.332 | 0.185 | 0.24 | 1 | MS.Trt_vs_MS.NoTrt_B |
| ZNF398 | 0.07466 | -0.6623 | 0.083 | 0.13 | 1 | MS.Trt_vs_MS.NoTrt_B |
| AKIRIN1 | 0.07473 | 0.3103 | 0.41 | 0.49 | 1 | MS.Trt_vs_MS.NoTrt_B |
| ZNF525 | 0.07483 | 1.39833 | 0.447 | 0.54 | 1 | MS.Trt_vs_MS.NoTrt_B |
| OGFOD3 | 0.07487 | -1.1044 | 0.447 | 0.52 | 1 | MS.Trt_vs_MS.NoTrt_B |
| RFC5 | 0.07519 | -0.5822 | 0.089 | 0.14 | 1 | MS.Trt_vs_MS.NoTrt_B |
| ZNF766 | 0.07526 | 0.36087 | 0.946 | 1 | 1 | MS.Trt_vs_MS.NoTrt_B |
| YY1 | 0.0753 | 0.56615 | 0.251 | 0.32 | 1 | MS.Trt_vs_MS.NoTrt_B |
| FAF1 | 0.07547 | 0.53617 | 0.383 | 0.47 | 1 | MS.Trt_vs_MS.NoTrt_B |
| MYO1D | 0.0755 | 0.89519 | 0.109 | 0.16 | 1 | MS.Trt_vs_MS.NoTrt_B |

| | | | | | | |
|---|---|---|---|---|---|---|
| UBXN2B | 0.07565 | 0.35625 | 0.285 | 0.36 | 1 | MS.Trt_vs_MS.NoTrt_B |
| DPF2 | 0.07593 | 0.34676 | 0.76 | 0.87 | 1 | MS.Trt_vs_MS.NoTrt_B |
| CCDC50 | 0.07616 | 1.09831 | 0.331 | 0.42 | 1 | MS.Trt_vs_MS.NoTrt_B |
| ANKRD52 | 0.07619 | -0.3321 | 0.113 | 0.16 | 1 | MS.Trt_vs_MS.NoTrt_B |
| RALBP1 | 0.07625 | 0.27847 | 0.29 | 0.36 | 1 | MS.Trt_vs_MS.NoTrt_B |
| NPIPB11 | 0.07638 | 0.89912 | 0.359 | 0.49 | 1 | MS.Trt_vs_MS.NoTrt_B |
| THOP1 | 0.07641 | -0.2801 | 0.312 | 0.41 | 1 | MS.Trt_vs_MS.NoTrt_B |
| FBXO34 | 0.07662 | -0.6324 | 0.063 | 0.1 | 1 | MS.Trt_vs_MS.NoTrt_B |
| ABCD4 | 0.07664 | 0.53797 | 0.669 | 0.79 | 1 | MS.Trt_vs_MS.NoTrt_B |
| SUZ12 | 0.07667 | 0.37719 | 0.527 | 0.61 | 1 | MS.Trt_vs_MS.NoTrt_B |
| TLN1 | 0.0767 | 0.49119 | 0.481 | 0.57 | 1 | MS.Trt_vs_MS.NoTrt_B |
| UBE2H | 0.07703 | 0.59218 | 0.105 | 0.16 | 1 | MS.Trt_vs_MS.NoTrt_B |
| DCBLD1 | 0.07703 | 1.19389 | 0.405 | 0.52 | 1 | MS.Trt_vs_MS.NoTrt_B |
| WDR46 | 0.07729 | 0.48318 | 0.083 | 0.13 | 1 | MS.Trt_vs_MS.NoTrt_B |
| CCDC18 | 0.07752 | 0.47048 | 0.133 | 0.19 | 1 | MS.Trt_vs_MS.NoTrt_B |
| ZBTB16 | 0.07763 | 1.17708 | 0.298 | 0.4 | 1 | MS.Trt_vs_MS.NoTrt_B |
| RP11-347C1 | 0.07764 | 0.44555 | 0.527 | 0.7 | 1 | MS.Trt_vs_MS.NoTrt_B |
| MED19 | 0.07765 | 0.42951 | 0.07 | 0.11 | 1 | MS.Trt_vs_MS.NoTrt_B |
| UFL1 | 0.07767 | 0.60029 | 0.135 | 0.19 | 1 | MS.Trt_vs_MS.NoTrt_B |
| NUP43 | 0.07812 | 0.58664 | 0.717 | 0.8 | 1 | MS.Trt_vs_MS.NoTrt_B |
| MTMR10 | 0.07831 | 0.66674 | 0.12 | 0.18 | 1 | MS.Trt_vs_MS.NoTrt_B |
| TRAT1 | 0.07842 | 1.30075 | 0.224 | 0.17 | 1 | MS.Trt_vs_MS.NoTrt_B |
| UHMK1 | 0.07847 | 0.2773 | 0.187 | 0.25 | 1 | MS.Trt_vs_MS.NoTrt_B |
| TTC33 | 0.07857 | 0.69633 | 0.163 | 0.22 | 1 | MS.Trt_vs_MS.NoTrt_B |
| SHQ1 | 0.07862 | 1.01003 | 0.575 | 0.67 | 1 | MS.Trt_vs_MS.NoTrt_B |
| SUN1 | 0.07867 | 0.60071 | 0.516 | 0.61 | 1 | MS.Trt_vs_MS.NoTrt_B |
| RRAGD | 0.07867 | 0.37586 | 0.113 | 0.17 | 1 | MS.Trt_vs_MS.NoTrt_B |
| CCHCR1 | 0.07878 | -0.4248 | 0.29 | 0.36 | 1 | MS.Trt_vs_MS.NoTrt_B |
| CASC10 | 0.07891 | 0.4845 | 0.301 | 0.37 | 1 | MS.Trt_vs_MS.NoTrt_B |
| CLTC | 0.07916 | -0.3325 | 0.294 | 0.35 | 1 | MS.Trt_vs_MS.NoTrt_B |
| NR1D1 | 0.07938 | -0.3322 | 0.072 | 0.11 | 1 | MS.Trt_vs_MS.NoTrt_B |
| ECSIT | 0.07979 | 0.4468 | 0.346 | 0.42 | 1 | MS.Trt_vs_MS.NoTrt_B |
| IL1R1 | 0.07985 | -0.4426 | 0.218 | 0.29 | 1 | MS.Trt_vs_MS.NoTrt_B |
| GLUL | 0.08005 | 0.84668 | 0.124 | 0.18 | 1 | MS.Trt_vs_MS.NoTrt_B |
| TIAF1 | 0.08006 | -0.6404 | 0.423 | 0.49 | 1 | MS.Trt_vs_MS.NoTrt_B |
| TTC21B | 0.0802 | -0.5948 | 0.264 | 0.35 | 1 | MS.Trt_vs_MS.NoTrt_B |
| CDK2AP2 | 0.08022 | 0.42925 | 0.155 | 0.22 | 1 | MS.Trt_vs_MS.NoTrt_B |
| ENY2 | 0.08028 | 1.06341 | 0.484 | 0.59 | 1 | MS.Trt_vs_MS.NoTrt_B |
| MAPRE2 | 0.08049 | 0.26966 | 0.205 | 0.27 | 1 | MS.Trt_vs_MS.NoTrt_B |
| NIPA1 | 0.08054 | -0.8491 | 0.122 | 0.17 | 1 | MS.Trt_vs_MS.NoTrt_B |

| | | | | | | |
|---|---|---|---|---|---|---|
| MT-ND5 | 0.08084 | 0.47071 | 0.854 | 0.89 | 1 | MS.Trt_vs_MS.NoTrt_B |
| RPA3 | 0.08088 | 0.33745 | 0.157 | 0.21 | 1 | MS.Trt_vs_MS.NoTrt_B |
| FBXL22 | 0.08097 | 0.47667 | 0.063 | 0.1 | 1 | MS.Trt_vs_MS.NoTrt_B |
| C1orf186 | 0.08115 | 0.30756 | 0.431 | 0.57 | 1 | MS.Trt_vs_MS.NoTrt_B |
| CAD | 0.08122 | 0.2783 | 0.58 | 0.7 | 1 | MS.Trt_vs_MS.NoTrt_B |
| MOCS2 | 0.08144 | 0.29608 | 0.063 | 0.1 | 1 | MS.Trt_vs_MS.NoTrt_B |
| KPNA5 | 0.0815 | 0.28597 | 0.9 | 0.98 | 1 | MS.Trt_vs_MS.NoTrt_B |
| TAS2R4 | 0.08155 | -0.3729 | 0.257 | 0.34 | 1 | MS.Trt_vs_MS.NoTrt_B |
| MAP3K7 | 0.08177 | 0.26344 | 0.115 | 0.17 | 1 | MS.Trt_vs_MS.NoTrt_B |
| CCNK | 0.0819 | 0.33787 | 0.218 | 0.29 | 1 | MS.Trt_vs_MS.NoTrt_B |
| ZMIZ2 | 0.08213 | 0.47798 | 0.111 | 0.16 | 1 | MS.Trt_vs_MS.NoTrt_B |
| C16orf54 | 0.08232 | 1.04193 | 0.73 | 0.84 | 1 | MS.Trt_vs_MS.NoTrt_B |
| FAM184B | 0.08249 | 0.45293 | 0.094 | 0.14 | 1 | MS.Trt_vs_MS.NoTrt_B |
| RBM22 | 0.08275 | 0.42638 | 0.105 | 0.16 | 1 | MS.Trt_vs_MS.NoTrt_B |
| SRPRA | 0.08283 | 0.2827 | 0.214 | 0.29 | 1 | MS.Trt_vs_MS.NoTrt_B |
| TPGS1 | 0.08289 | 0.82173 | 0.288 | 0.22 | 1 | MS.Trt_vs_MS.NoTrt_B |
| TMEM205 | 0.08289 | -0.3072 | 0.063 | 0.1 | 1 | MS.Trt_vs_MS.NoTrt_B |
| ALDH3A2 | 0.08291 | -0.7368 | 0.224 | 0.29 | 1 | MS.Trt_vs_MS.NoTrt_B |
| RCBTB2 | 0.08293 | 0.29259 | 0.1 | 0.15 | 1 | MS.Trt_vs_MS.NoTrt_B |
| IKZF5 | 0.08297 | 0.92293 | 0.181 | 0.25 | 1 | MS.Trt_vs_MS.NoTrt_B |
| GHRL | 0.08301 | 0.72694 | 0.061 | 0.1 | 1 | MS.Trt_vs_MS.NoTrt_B |
| THAP5 | 0.08309 | 0.96622 | 0.804 | 0.89 | 1 | MS.Trt_vs_MS.NoTrt_B |
| R3HCC1L | 0.08316 | -0.2679 | 0.211 | 0.28 | 1 | MS.Trt_vs_MS.NoTrt_B |
| AHRR | 0.08327 | 0.98368 | 0.081 | 0.13 | 1 | MS.Trt_vs_MS.NoTrt_B |
| MTHFD1L | 0.0834 | 1.45321 | 0.065 | 0.11 | 1 | MS.Trt_vs_MS.NoTrt_B |
| KBTBD2 | 0.08369 | 0.3744 | 0.51 | 0.6 | 1 | MS.Trt_vs_MS.NoTrt_B |
| ZSCAN32 | 0.08384 | 0.45649 | 0.172 | 0.24 | 1 | MS.Trt_vs_MS.NoTrt_B |
| EIF1B | 0.08397 | 0.4565 | 0.137 | 0.2 | 1 | MS.Trt_vs_MS.NoTrt_B |
| NAA35 | 0.0841 | -0.3661 | 0.27 | 0.34 | 1 | MS.Trt_vs_MS.NoTrt_B |
| SMG6 | 0.08418 | -0.4043 | 0.148 | 0.2 | 1 | MS.Trt_vs_MS.NoTrt_B |
| FASTKD2 | 0.08422 | 0.43962 | 0.464 | 0.57 | 1 | MS.Trt_vs_MS.NoTrt_B |
| USP6 | 0.08437 | 0.28323 | 0.1 | 0.15 | 1 | MS.Trt_vs_MS.NoTrt_B |
| FKBP9 | 0.0844 | 1.06262 | 0.344 | 0.44 | 1 | MS.Trt_vs_MS.NoTrt_B |
| KIFAP3 | 0.08452 | -0.9041 | 0.115 | 0.16 | 1 | MS.Trt_vs_MS.NoTrt_B |
| CR1 | 0.08461 | 0.48167 | 0.444 | 0.55 | 1 | MS.Trt_vs_MS.NoTrt_B |
| ABCG1 | 0.0848 | -0.4204 | 0.375 | 0.47 | 1 | MS.Trt_vs_MS.NoTrt_B |
| ZNF487 | 0.08489 | -0.8247 | 0.351 | 0.43 | 1 | MS.Trt_vs_MS.NoTrt_B |
| BACE2 | 0.08523 | 0.27514 | 0.172 | 0.23 | 1 | MS.Trt_vs_MS.NoTrt_B |
| FNIP1 | 0.08524 | 0.83056 | 0.46 | 0.55 | 1 | MS.Trt_vs_MS.NoTrt_B |
| ZNF221 | 0.08527 | -0.2642 | 0.187 | 0.26 | 1 | MS.Trt_vs_MS.NoTrt_B |

| | | | | | | |
|---|---|---|---|---|---|---|
| C6orf106 | 0.08527 | 0.3127 | 0.089 | 0.14 | 1 | MS.Trt_vs_MS.NoTrt_B |
| ZNF526 | 0.08542 | 0.47412 | 0.373 | 0.48 | 1 | MS.Trt_vs_MS.NoTrt_B |
| EPHX1 | 0.08549 | 0.4067 | 0.074 | 0.12 | 1 | MS.Trt_vs_MS.NoTrt_B |
| SELK | 0.0855 | 0.47752 | 0.325 | 0.38 | 1 | MS.Trt_vs_MS.NoTrt_B |
| STXBP4 | 0.08572 | -0.707 | 0.377 | 0.47 | 1 | MS.Trt_vs_MS.NoTrt_B |
| DZIP3 | 0.08614 | 1.44253 | 0.174 | 0.24 | 1 | MS.Trt_vs_MS.NoTrt_B |
| FAM96A | 0.08632 | 0.48634 | 0.298 | 0.38 | 1 | MS.Trt_vs_MS.NoTrt_B |
| INTU | 0.08646 | 0.37414 | 0.922 | 0.97 | 1 | MS.Trt_vs_MS.NoTrt_B |
| PRKCI | 0.08661 | -0.3784 | 0.224 | 0.3 | 1 | MS.Trt_vs_MS.NoTrt_B |
| EIF4G2 | 0.0867 | 0.27376 | 0.479 | 0.56 | 1 | MS.Trt_vs_MS.NoTrt_B |
| ZNF83 | 0.08679 | 0.56892 | 0.747 | 0.83 | 1 | MS.Trt_vs_MS.NoTrt_B |
| SLC15A4 | 0.08717 | -0.4739 | 0.654 | 0.73 | 1 | MS.Trt_vs_MS.NoTrt_B |
| CBFA2T2 | 0.08736 | 0.4393 | 0.843 | 0.95 | 1 | MS.Trt_vs_MS.NoTrt_B |
| SAXO2 | 0.08755 | 0.55764 | 0.148 | 0.21 | 1 | MS.Trt_vs_MS.NoTrt_B |
| CUL9 | 0.08777 | 0.55748 | 0.12 | 0.17 | 1 | MS.Trt_vs_MS.NoTrt_B |
| DHX35 | 0.08801 | 0.26358 | 0.325 | 0.4 | 1 | MS.Trt_vs_MS.NoTrt_B |
| PDE7A | 0.08817 | 0.59987 | 0.244 | 0.32 | 1 | MS.Trt_vs_MS.NoTrt_B |
| POLN | 0.08824 | 0.32681 | 0.514 | 0.6 | 1 | MS.Trt_vs_MS.NoTrt_B |
| SRFBP1 | 0.08836 | 0.57243 | 0.32 | 0.4 | 1 | MS.Trt_vs_MS.NoTrt_B |
| UNC50 | 0.08841 | 0.41465 | 0.081 | 0.12 | 1 | MS.Trt_vs_MS.NoTrt_B |
| TMEM231 | 0.08849 | -0.4103 | 0.58 | 0.71 | 1 | MS.Trt_vs_MS.NoTrt_B |
| ZCCHC11 | 0.08853 | 0.40332 | 0.852 | 0.9 | 1 | MS.Trt_vs_MS.NoTrt_B |
| TAF1A | 0.08866 | -0.7296 | 0.309 | 0.39 | 1 | MS.Trt_vs_MS.NoTrt_B |
| HMGXB3 | 0.08868 | -0.4583 | 0.109 | 0.16 | 1 | MS.Trt_vs_MS.NoTrt_B |
| B3GNT2 | 0.08872 | 0.28651 | 0.085 | 0.13 | 1 | MS.Trt_vs_MS.NoTrt_B |
| ZNF432 | 0.08877 | -0.5328 | 0.479 | 0.6 | 1 | MS.Trt_vs_MS.NoTrt_B |
| SYPL1 | 0.08889 | 0.52384 | 0.316 | 0.39 | 1 | MS.Trt_vs_MS.NoTrt_B |
| PSMC2 | 0.08899 | 0.3604 | 0.168 | 0.23 | 1 | MS.Trt_vs_MS.NoTrt_B |
| ZNF552 | 0.08904 | 0.25706 | 0.325 | 0.42 | 1 | MS.Trt_vs_MS.NoTrt_B |
| BDH2 | 0.08923 | -0.6362 | 0.139 | 0.19 | 1 | MS.Trt_vs_MS.NoTrt_B |
| LRRK1 | 0.08924 | 0.40722 | 0.688 | 0.81 | 1 | MS.Trt_vs_MS.NoTrt_B |
| PPP2R2D | 0.08936 | 1.45332 | 0.529 | 0.64 | 1 | MS.Trt_vs_MS.NoTrt_B |
| XYLB | 0.08937 | 0.35108 | 0.366 | 0.45 | 1 | MS.Trt_vs_MS.NoTrt_B |
| ADAP2 | 0.08961 | 0.52128 | 0.107 | 0.16 | 1 | MS.Trt_vs_MS.NoTrt_B |
| PSENEN | 0.08993 | -0.5281 | 0.118 | 0.16 | 1 | MS.Trt_vs_MS.NoTrt_B |
| YTHDC2 | 0.09022 | 0.68683 | 0.303 | 0.39 | 1 | MS.Trt_vs_MS.NoTrt_B |
| NKIRAS1 | 0.09061 | -0.8092 | 0.198 | 0.25 | 1 | MS.Trt_vs_MS.NoTrt_B |
| PPP1CB | 0.09072 | 1.18465 | 0.719 | 0.78 | 1 | MS.Trt_vs_MS.NoTrt_B |
| LRRC3 | 0.09078 | 0.31411 | 0.092 | 0.14 | 1 | MS.Trt_vs_MS.NoTrt_B |
| C7orf50 | 0.09085 | 0.26611 | 0.294 | 0.36 | 1 | MS.Trt_vs_MS.NoTrt_B |

| | | | | | | |
|---|---|---|---|---|---|---|
| FAM234B | 0.09095 | -0.888 | 0.325 | 0.38 | 1 | MS.Trt_vs_MS.NoTrt_B |
| PRKAG1 | 0.09108 | -0.5251 | 0.468 | 0.55 | 1 | MS.Trt_vs_MS.NoTrt_B |
| ABCE1 | 0.09128 | 0.44838 | 0.261 | 0.33 | 1 | MS.Trt_vs_MS.NoTrt_B |
| OSMR | 0.09135 | 0.97417 | 0.497 | 0.46 | 1 | MS.Trt_vs_MS.NoTrt_B |
| S1PR1 | 0.09138 | 0.44903 | 0.479 | 0.62 | 1 | MS.Trt_vs_MS.NoTrt_B |
| APPBP2 | 0.0918 | -0.4063 | 0.763 | 0.83 | 1 | MS.Trt_vs_MS.NoTrt_B |
| HELQ | 0.0919 | -0.3296 | 0.1 | 0.15 | 1 | MS.Trt_vs_MS.NoTrt_B |
| TMOD2 | 0.092 | 0.30035 | 0.359 | 0.47 | 1 | MS.Trt_vs_MS.NoTrt_B |
| TRAF6 | 0.09215 | 0.29474 | 0.556 | 0.65 | 1 | MS.Trt_vs_MS.NoTrt_B |
| POLH | 0.09218 | 0.71087 | 0.438 | 0.54 | 1 | MS.Trt_vs_MS.NoTrt_B |
| CH507-9B2.! | 0.09242 | 0.34685 | 0.072 | 0.11 | 1 | MS.Trt_vs_MS.NoTrt_B |
| ADCY9 | 0.0925 | 1.02819 | 0.207 | 0.28 | 1 | MS.Trt_vs_MS.NoTrt_B |
| AURKB | 0.09261 | -0.3838 | 0.301 | 0.38 | 1 | MS.Trt_vs_MS.NoTrt_B |
| ATP6V0A1 | 0.09267 | -0.8095 | 0.734 | 0.81 | 1 | MS.Trt_vs_MS.NoTrt_B |
| SLC35G2 | 0.09271 | 1.13489 | 0.105 | 0.15 | 1 | MS.Trt_vs_MS.NoTrt_B |
| EFHB | 0.09295 | 0.2976 | 0.246 | 0.31 | 1 | MS.Trt_vs_MS.NoTrt_B |
| PITRM1 | 0.09333 | -0.3099 | 0.131 | 0.18 | 1 | MS.Trt_vs_MS.NoTrt_B |
| FGF22 | 0.09339 | 0.46687 | 0.351 | 0.29 | 1 | MS.Trt_vs_MS.NoTrt_B |
| JUND | 0.09348 | 0.29596 | 0.085 | 0.13 | 1 | MS.Trt_vs_MS.NoTrt_B |
| APOBEC3C | 0.0935 | 0.30199 | 0.612 | 0.74 | 1 | MS.Trt_vs_MS.NoTrt_B |
| MAP1LC3C | 0.09364 | 0.63368 | 0.394 | 0.36 | 1 | MS.Trt_vs_MS.NoTrt_B |
| HCFC1 | 0.09376 | 0.35541 | 0.22 | 0.29 | 1 | MS.Trt_vs_MS.NoTrt_B |
| SPINK9 | 0.09383 | 0.40871 | 0.558 | 0.75 | 1 | MS.Trt_vs_MS.NoTrt_B |
| TDRD12 | 0.09384 | 1.10035 | 0.479 | 0.42 | 1 | MS.Trt_vs_MS.NoTrt_B |
| WNT2B | 0.09399 | 0.69149 | 0.329 | 0.41 | 1 | MS.Trt_vs_MS.NoTrt_B |
| HUWE1 | 0.09399 | 0.31321 | 0.344 | 0.42 | 1 | MS.Trt_vs_MS.NoTrt_B |
| CD200 | 0.09404 | 0.69404 | 0.54 | 0.51 | 1 | MS.Trt_vs_MS.NoTrt_B |
| ZNF287 | 0.09419 | -0.7271 | 0.305 | 0.37 | 1 | MS.Trt_vs_MS.NoTrt_B |
| PDXK | 0.09424 | 0.7114 | 0.401 | 0.51 | 1 | MS.Trt_vs_MS.NoTrt_B |
| LSM11 | 0.09452 | 0.26243 | 0.479 | 0.55 | 1 | MS.Trt_vs_MS.NoTrt_B |
| LRRC40 | 0.09464 | 0.80113 | 0.065 | 0.1 | 1 | MS.Trt_vs_MS.NoTrt_B |
| NPM1 | 0.09499 | 0.35436 | 0.584 | 0.76 | 1 | MS.Trt_vs_MS.NoTrt_B |
| RBM38 | 0.09549 | 0.35851 | 0.098 | 0.15 | 1 | MS.Trt_vs_MS.NoTrt_B |
| EPS8 | 0.09557 | -0.7587 | 0.298 | 0.38 | 1 | MS.Trt_vs_MS.NoTrt_B |
| RNF220 | 0.09611 | 0.46622 | 0.525 | 0.65 | 1 | MS.Trt_vs_MS.NoTrt_B |
| PIP4K2B | 0.09614 | 0.85525 | 0.102 | 0.15 | 1 | MS.Trt_vs_MS.NoTrt_B |
| AC138969.4 | 0.09617 | 0.67294 | 0.373 | 0.47 | 1 | MS.Trt_vs_MS.NoTrt_B |
| PECAM1 | 0.09625 | 1.01719 | 0.641 | 0.74 | 1 | MS.Trt_vs_MS.NoTrt_B |
| VIPR2 | 0.0964 | 0.30288 | 0.253 | 0.2 | 1 | MS.Trt_vs_MS.NoTrt_B |
| NREP | 0.09647 | 0.5247 | 0.109 | 0.16 | 1 | MS.Trt_vs_MS.NoTrt_B |

| | | | | | | |
|---|---|---|---|---|---|---|
| PFKP | 0.0967 | -0.4049 | 0.22 | 0.27 | 1 | MS.Trt_vs_MS.NoTrt_B |
| YES1 | 0.09681 | 0.55058 | 0.294 | 0.37 | 1 | MS.Trt_vs_MS.NoTrt_B |
| KNG1 | 0.09695 | 0.90633 | 0.144 | 0.21 | 1 | MS.Trt_vs_MS.NoTrt_B |
| ZNF354A | 0.097 | 0.50234 | 0.135 | 0.19 | 1 | MS.Trt_vs_MS.NoTrt_B |
| XPO1 | 0.09739 | 0.54093 | 0.869 | 0.93 | 1 | MS.Trt_vs_MS.NoTrt_B |
| MUM1 | 0.09774 | -0.5891 | 0.484 | 0.58 | 1 | MS.Trt_vs_MS.NoTrt_B |
| NAPG | 0.09786 | 0.37294 | 0.475 | 0.58 | 1 | MS.Trt_vs_MS.NoTrt_B |
| ACSL6 | 0.09793 | 0.60491 | 0.549 | 0.49 | 1 | MS.Trt_vs_MS.NoTrt_B |
| WDPCP | 0.09822 | 0.48919 | 0.871 | 0.98 | 1 | MS.Trt_vs_MS.NoTrt_B |
| PGAP1 | 0.09828 | 0.2626 | 0.279 | 0.35 | 1 | MS.Trt_vs_MS.NoTrt_B |
| LYST | 0.09849 | 1.18639 | 0.222 | 0.3 | 1 | MS.Trt_vs_MS.NoTrt_B |
| ABCA5 | 0.09853 | 0.41072 | 0.547 | 0.68 | 1 | MS.Trt_vs_MS.NoTrt_B |
| VAV3 | 0.09855 | 1.07488 | 0.492 | 0.43 | 1 | MS.Trt_vs_MS.NoTrt_B |
| CTD-2192J1( | 0.09892 | 1.86799 | 0.168 | 0.12 | 1 | MS.Trt_vs_MS.NoTrt_B |
| INPP5F | 0.09894 | 0.59353 | 0.248 | 0.32 | 1 | MS.Trt_vs_MS.NoTrt_B |
| PLGRKT | 0.09897 | -0.9491 | 0.074 | 0.11 | 1 | MS.Trt_vs_MS.NoTrt_B |
| PADI4 | 0.09914 | 0.49364 | 0.388 | 0.49 | 1 | MS.Trt_vs_MS.NoTrt_B |
| NFASC | 0.09919 | 0.76865 | 0.479 | 0.59 | 1 | MS.Trt_vs_MS.NoTrt_B |
| PAK1 | 0.09974 | -0.8122 | 0.12 | 0.17 | 1 | MS.Trt_vs_MS.NoTrt_B |
| TMEM97 | 0.09976 | -0.4377 | 0.185 | 0.25 | 1 | MS.Trt_vs_MS.NoTrt_B |
| CD96 | 0.09982 | 0.63478 | 0.514 | 0.6 | 1 | MS.Trt_vs_MS.NoTrt_B |
| TTF1 | 0.10011 | -0.3066 | 0.359 | 0.42 | 1 | MS.Trt_vs_MS.NoTrt_B |
| TMEM30A | 0.1002 | 0.31473 | 0.153 | 0.21 | 1 | MS.Trt_vs_MS.NoTrt_B |
| TTC8 | 0.10028 | -0.2587 | 0.56 | 0.69 | 1 | MS.Trt_vs_MS.NoTrt_B |
| CCDC62 | 0.10028 | 0.32851 | 0.349 | 0.45 | 1 | MS.Trt_vs_MS.NoTrt_B |
| POLR2H | 0.10029 | 0.41053 | 0.401 | 0.48 | 1 | MS.Trt_vs_MS.NoTrt_B |
| PLD2 | 0.10045 | -0.6756 | 0.065 | 0.1 | 1 | MS.Trt_vs_MS.NoTrt_B |
| GINS1 | 0.10049 | 0.3095 | 0.436 | 0.57 | 1 | MS.Trt_vs_MS.NoTrt_B |
| ETFBKMT | 0.10054 | 0.27035 | 0.54 | 0.64 | 1 | MS.Trt_vs_MS.NoTrt_B |
| LGALS3BP | 0.10058 | -0.3 | 0.418 | 0.54 | 1 | MS.Trt_vs_MS.NoTrt_B |
| ZNF69 | 0.10077 | -0.618 | 0.368 | 0.43 | 1 | MS.Trt_vs_MS.NoTrt_B |
| NFS1 | 0.10092 | -0.9751 | 0.776 | 0.87 | 1 | MS.Trt_vs_MS.NoTrt_B |
| VIL1 | 0.10103 | 0.36873 | 0.122 | 0.17 | 1 | MS.Trt_vs_MS.NoTrt_B |
| BANK1 | 0.10118 | 1.02634 | 0.122 | 0.18 | 1 | MS.Trt_vs_MS.NoTrt_B |
| CUL4B | 0.10146 | 0.65866 | 0.078 | 0.12 | 1 | MS.Trt_vs_MS.NoTrt_B |
| LAIR1 | 0.10186 | -1.2099 | 0.773 | 0.83 | 1 | MS.Trt_vs_MS.NoTrt_B |
| CENPN | 0.10198 | 0.5316 | 0.098 | 0.15 | 1 | MS.Trt_vs_MS.NoTrt_B |
| TRIP11 | 0.10219 | 0.50532 | 0.431 | 0.5 | 1 | MS.Trt_vs_MS.NoTrt_B |
| IGFL2 | 0.10229 | 0.37887 | 0.214 | 0.28 | 1 | MS.Trt_vs_MS.NoTrt_B |
| SYNGAP1 | 0.10251 | 0.5353 | 0.102 | 0.15 | 1 | MS.Trt_vs_MS.NoTrt_B |

| | | | | | | |
|---|---|---|---|---|---|---|
| PATL2 | 0.10292 | 0.54555 | 0.279 | 0.37 | 1 | MS.Trt_vs_MS.NoTrt_B |
| F3 | 0.10317 | 0.38322 | 0.392 | 0.52 | 1 | MS.Trt_vs_MS.NoTrt_B |
| CCP110 | 0.10326 | -0.7835 | 0.425 | 0.48 | 1 | MS.Trt_vs_MS.NoTrt_B |
| FBXO42 | 0.1033 | 0.4197 | 0.394 | 0.49 | 1 | MS.Trt_vs_MS.NoTrt_B |
| ZNF573 | 0.10334 | -0.8322 | 0.359 | 0.42 | 1 | MS.Trt_vs_MS.NoTrt_B |
| TMEM170B | 0.10372 | 0.29849 | 0.187 | 0.24 | 1 | MS.Trt_vs_MS.NoTrt_B |
| OTUD4 | 0.10379 | 0.73917 | 0.556 | 0.63 | 1 | MS.Trt_vs_MS.NoTrt_B |
| DNAJC10 | 0.10411 | 0.33575 | 0.88 | 0.96 | 1 | MS.Trt_vs_MS.NoTrt_B |
| SNRK | 0.10415 | 0.33877 | 0.477 | 0.55 | 1 | MS.Trt_vs_MS.NoTrt_B |
| CDK19 | 0.10434 | 0.84968 | 0.209 | 0.28 | 1 | MS.Trt_vs_MS.NoTrt_B |
| SLC35E4 | 0.1044 | 0.27733 | 0.233 | 0.3 | 1 | MS.Trt_vs_MS.NoTrt_B |
| FAM222B | 0.10445 | 0.36429 | 0.122 | 0.17 | 1 | MS.Trt_vs_MS.NoTrt_B |
| NRBF2 | 0.10451 | 0.36991 | 0.07 | 0.11 | 1 | MS.Trt_vs_MS.NoTrt_B |
| ZNF664 | 0.10454 | 0.65448 | 0.179 | 0.24 | 1 | MS.Trt_vs_MS.NoTrt_B |
| CASC3 | 0.10476 | 0.3984 | 0.423 | 0.52 | 1 | MS.Trt_vs_MS.NoTrt_B |
| LAMTOR5 | 0.10485 | 0.37895 | 0.392 | 0.47 | 1 | MS.Trt_vs_MS.NoTrt_B |
| AGFG2 | 0.10491 | -0.2644 | 0.268 | 0.34 | 1 | MS.Trt_vs_MS.NoTrt_B |
| ST5 | 0.10505 | -0.2878 | 0.251 | 0.34 | 1 | MS.Trt_vs_MS.NoTrt_B |
| ZBTB43 | 0.10539 | 0.5946 | 0.678 | 0.62 | 1 | MS.Trt_vs_MS.NoTrt_B |
| ABCC5 | 0.10544 | -0.5157 | 0.105 | 0.15 | 1 | MS.Trt_vs_MS.NoTrt_B |
| MYH10 | 0.10545 | 0.37297 | 0.074 | 0.11 | 1 | MS.Trt_vs_MS.NoTrt_B |
| RAB4B | 0.10558 | 0.36017 | 0.279 | 0.35 | 1 | MS.Trt_vs_MS.NoTrt_B |
| UBB | 0.10574 | 0.29758 | 0.54 | 0.69 | 1 | MS.Trt_vs_MS.NoTrt_B |
| GRWD1 | 0.10574 | -0.497 | 0.383 | 0.44 | 1 | MS.Trt_vs_MS.NoTrt_B |
| USP16 | 0.10584 | 0.27487 | 0.407 | 0.46 | 1 | MS.Trt_vs_MS.NoTrt_B |
| CDKL1 | 0.10586 | 0.86434 | 0.586 | 0.68 | 1 | MS.Trt_vs_MS.NoTrt_B |
| MRPS18B | 0.10589 | 0.57357 | 0.584 | 0.69 | 1 | MS.Trt_vs_MS.NoTrt_B |
| OCLN | 0.10594 | 0.34046 | 0.63 | 0.77 | 1 | MS.Trt_vs_MS.NoTrt_B |
| KIF11 | 0.106 | 0.29879 | 0.316 | 0.4 | 1 | MS.Trt_vs_MS.NoTrt_B |
| BCKDHA | 0.10602 | -0.3814 | 0.148 | 0.2 | 1 | MS.Trt_vs_MS.NoTrt_B |
| SPATA7 | 0.10657 | -0.3593 | 0.231 | 0.3 | 1 | MS.Trt_vs_MS.NoTrt_B |
| DDX17 | 0.1066 | 0.32539 | 0.749 | 0.95 | 1 | MS.Trt_vs_MS.NoTrt_B |
| VWA7 | 0.10665 | 0.49351 | 0.242 | 0.31 | 1 | MS.Trt_vs_MS.NoTrt_B |
| TRA2B | 0.10693 | 0.40558 | 0.431 | 0.51 | 1 | MS.Trt_vs_MS.NoTrt_B |
| FAM204A | 0.10764 | 0.39528 | 0.407 | 0.47 | 1 | MS.Trt_vs_MS.NoTrt_B |
| CLYBL | 0.10774 | 0.91943 | 0.068 | 0.11 | 1 | MS.Trt_vs_MS.NoTrt_B |
| FAM124B | 0.10795 | -0.2881 | 0.065 | 0.1 | 1 | MS.Trt_vs_MS.NoTrt_B |
| LMAN1 | 0.10797 | 0.67238 | 0.124 | 0.18 | 1 | MS.Trt_vs_MS.NoTrt_B |
| ENTPD6 | 0.10811 | -0.2985 | 0.078 | 0.12 | 1 | MS.Trt_vs_MS.NoTrt_B |
| WAPL | 0.10815 | 0.26012 | 0.357 | 0.45 | 1 | MS.Trt_vs_MS.NoTrt_B |

| | | | | | | |
|---|---|---|---|---|---|---|
| FN3KRP | 0.10823 | 0.32165 | 0.065 | 0.1 | 1 | MS.Trt_vs_MS.NoTrt_B |
| SAMD4B | 0.10841 | 0.56971 | 0.717 | 0.85 | 1 | MS.Trt_vs_MS.NoTrt_B |
| AP4B1 | 0.10844 | 0.63993 | 0.505 | 0.61 | 1 | MS.Trt_vs_MS.NoTrt_B |
| POLR2K | 0.10847 | 0.57284 | 0.092 | 0.14 | 1 | MS.Trt_vs_MS.NoTrt_B |
| CCDC148 | 0.10899 | -0.4055 | 0.159 | 0.22 | 1 | MS.Trt_vs_MS.NoTrt_B |
| TLR5 | 0.10908 | 1.10163 | 0.488 | 0.61 | 1 | MS.Trt_vs_MS.NoTrt_B |
| RB1CC1 | 0.10917 | 0.42523 | 0.27 | 0.34 | 1 | MS.Trt_vs_MS.NoTrt_B |
| NEK9 | 0.10923 | 0.28166 | 0.758 | 0.87 | 1 | MS.Trt_vs_MS.NoTrt_B |
| TRAPPC12 | 0.10963 | 0.29035 | 0.46 | 0.57 | 1 | MS.Trt_vs_MS.NoTrt_B |
| UBA2 | 0.10981 | 0.77789 | 0.462 | 0.54 | 1 | MS.Trt_vs_MS.NoTrt_B |
| LYZ | 0.10981 | 0.60547 | 0.301 | 0.38 | 1 | MS.Trt_vs_MS.NoTrt_B |
| MRLN | 0.10998 | 0.256 | 0.481 | 0.63 | 1 | MS.Trt_vs_MS.NoTrt_B |
| ZNF846 | 0.11001 | 0.58354 | 0.364 | 0.44 | 1 | MS.Trt_vs_MS.NoTrt_B |
| NADK | 0.11002 | 0.28141 | 0.198 | 0.27 | 1 | MS.Trt_vs_MS.NoTrt_B |
| LAMP3 | 0.11052 | -0.8738 | 0.342 | 0.41 | 1 | MS.Trt_vs_MS.NoTrt_B |
| PROX2 | 0.11086 | -0.5375 | 0.07 | 0.11 | 1 | MS.Trt_vs_MS.NoTrt_B |
| TTLL4 | 0.11113 | 0.3268 | 0.251 | 0.32 | 1 | MS.Trt_vs_MS.NoTrt_B |
| TSN | 0.11161 | 0.70515 | 0.272 | 0.35 | 1 | MS.Trt_vs_MS.NoTrt_B |
| MAPK8 | 0.11206 | 0.28177 | 0.327 | 0.39 | 1 | MS.Trt_vs_MS.NoTrt_B |
| PGM3 | 0.1121 | 0.89198 | 0.181 | 0.25 | 1 | MS.Trt_vs_MS.NoTrt_B |
| AGBL5 | 0.11221 | -0.305 | 0.113 | 0.16 | 1 | MS.Trt_vs_MS.NoTrt_B |
| FAM208A | 0.11249 | 0.7385 | 0.651 | 0.72 | 1 | MS.Trt_vs_MS.NoTrt_B |
| HEATR1 | 0.11277 | 0.41535 | 0.098 | 0.14 | 1 | MS.Trt_vs_MS.NoTrt_B |
| WDR97 | 0.11284 | 0.26798 | 0.351 | 0.44 | 1 | MS.Trt_vs_MS.NoTrt_B |
| MRPS27 | 0.11288 | -0.6217 | 0.477 | 0.51 | 1 | MS.Trt_vs_MS.NoTrt_B |
| TRAK2 | 0.11307 | 0.63824 | 0.176 | 0.24 | 1 | MS.Trt_vs_MS.NoTrt_B |
| ENOX2 | 0.11314 | 0.7222 | 0.07 | 0.11 | 1 | MS.Trt_vs_MS.NoTrt_B |
| LINC00116 | 0.11323 | 0.80044 | 0.124 | 0.17 | 1 | MS.Trt_vs_MS.NoTrt_B |
| NEDD4L | 0.11325 | -0.3084 | 0.39 | 0.43 | 1 | MS.Trt_vs_MS.NoTrt_B |
| EPN2 | 0.11429 | 0.38226 | 0.643 | 0.81 | 1 | MS.Trt_vs_MS.NoTrt_B |
| LPAR2 | 0.11483 | -0.6436 | 0.072 | 0.11 | 1 | MS.Trt_vs_MS.NoTrt_B |
| TFAP4 | 0.11484 | 0.31275 | 0.551 | 0.67 | 1 | MS.Trt_vs_MS.NoTrt_B |
| RHOC | 0.11487 | -0.8521 | 0.083 | 0.12 | 1 | MS.Trt_vs_MS.NoTrt_B |
| ZNF107 | 0.11517 | 0.5142 | 0.355 | 0.42 | 1 | MS.Trt_vs_MS.NoTrt_B |
| TATDN1 | 0.11525 | -0.3361 | 0.59 | 0.63 | 1 | MS.Trt_vs_MS.NoTrt_B |
| SLX4 | 0.1156 | 1.60549 | 0.355 | 0.45 | 1 | MS.Trt_vs_MS.NoTrt_B |
| RMI1 | 0.11573 | 0.57384 | 0.181 | 0.24 | 1 | MS.Trt_vs_MS.NoTrt_B |
| KYAT1 | 0.11587 | 0.37967 | 0.105 | 0.15 | 1 | MS.Trt_vs_MS.NoTrt_B |
| COL6A3 | 0.1159 | -0.4773 | 0.364 | 0.45 | 1 | MS.Trt_vs_MS.NoTrt_B |
| PFAS | 0.11613 | 0.27362 | 0.497 | 0.47 | 1 | MS.Trt_vs_MS.NoTrt_B |

| | | | | | | |
|---|---|---|---|---|---|---|
| RP2 | 0.11623 | 0.63862 | 0.588 | 0.62 | 1 | MS.Trt_vs_MS.NoTrt_B |
| NUCB2 | 0.1165 | 0.45016 | 0.891 | 0.94 | 1 | MS.Trt_vs_MS.NoTrt_B |
| CD38 | 0.11675 | 0.59201 | 0.246 | 0.31 | 1 | MS.Trt_vs_MS.NoTrt_B |
| MRPL35 | 0.11684 | 0.43115 | 0.065 | 0.1 | 1 | MS.Trt_vs_MS.NoTrt_B |
| SMPDL3B | 0.11685 | -0.3486 | 0.078 | 0.12 | 1 | MS.Trt_vs_MS.NoTrt_B |
| RGSL1 | 0.11709 | 0.54087 | 0.174 | 0.13 | 1 | MS.Trt_vs_MS.NoTrt_B |
| LZTFL1 | 0.11723 | -0.5623 | 0.481 | 0.57 | 1 | MS.Trt_vs_MS.NoTrt_B |
| MFSD14B | 0.11731 | 0.72239 | 0.078 | 0.12 | 1 | MS.Trt_vs_MS.NoTrt_B |
| SLC25A23 | 0.11737 | 0.84565 | 0.405 | 0.52 | 1 | MS.Trt_vs_MS.NoTrt_B |
| BPNT1 | 0.11761 | -0.5727 | 0.569 | 0.64 | 1 | MS.Trt_vs_MS.NoTrt_B |
| URGCP | 0.11762 | 0.49073 | 0.386 | 0.49 | 1 | MS.Trt_vs_MS.NoTrt_B |
| NDST1 | 0.11773 | 0.31273 | 0.318 | 0.25 | 1 | MS.Trt_vs_MS.NoTrt_B |
| SSFA2 | 0.11783 | 0.78499 | 0.078 | 0.12 | 1 | MS.Trt_vs_MS.NoTrt_B |
| AVPR1A | 0.11812 | 0.38363 | 0.144 | 0.1 | 1 | MS.Trt_vs_MS.NoTrt_B |
| MED12 | 0.11817 | 0.64433 | 0.144 | 0.2 | 1 | MS.Trt_vs_MS.NoTrt_B |
| OSER1 | 0.11833 | 0.33159 | 0.155 | 0.21 | 1 | MS.Trt_vs_MS.NoTrt_B |
| NISCH | 0.11833 | 0.30706 | 0.355 | 0.42 | 1 | MS.Trt_vs_MS.NoTrt_B |
| SMC3 | 0.11855 | 0.28669 | 0.194 | 0.25 | 1 | MS.Trt_vs_MS.NoTrt_B |
| SS18L1 | 0.11861 | 0.28097 | 0.322 | 0.41 | 1 | MS.Trt_vs_MS.NoTrt_B |
| EBLN2 | 0.11871 | -0.2845 | 0.085 | 0.12 | 1 | MS.Trt_vs_MS.NoTrt_B |
| GOLGA7 | 0.11873 | 0.58653 | 0.218 | 0.28 | 1 | MS.Trt_vs_MS.NoTrt_B |
| OTOA | 0.11947 | 0.30885 | 0.264 | 0.33 | 1 | MS.Trt_vs_MS.NoTrt_B |
| CUL1 | 0.12024 | 0.5008 | 0.137 | 0.19 | 1 | MS.Trt_vs_MS.NoTrt_B |
| ZNHIT3 | 0.12062 | -0.299 | 0.773 | 0.84 | 1 | MS.Trt_vs_MS.NoTrt_B |
| IGSF1 | 0.12066 | 0.42974 | 0.366 | 0.31 | 1 | MS.Trt_vs_MS.NoTrt_B |
| RP11-166B2 | 0.12103 | 0.39207 | 0.377 | 0.48 | 1 | MS.Trt_vs_MS.NoTrt_B |
| GTPBP4 | 0.12139 | -0.725 | 0.379 | 0.44 | 1 | MS.Trt_vs_MS.NoTrt_B |
| PTGIR | 0.12157 | 0.34671 | 0.54 | 0.62 | 1 | MS.Trt_vs_MS.NoTrt_B |
| ARHGAP35 | 0.12178 | -0.5046 | 0.146 | 0.19 | 1 | MS.Trt_vs_MS.NoTrt_B |
| AGGF1 | 0.12179 | 0.38547 | 0.078 | 0.12 | 1 | MS.Trt_vs_MS.NoTrt_B |
| ZNF341 | 0.12181 | 0.67143 | 0.312 | 0.41 | 1 | MS.Trt_vs_MS.NoTrt_B |
| FARP1 | 0.12219 | 0.37394 | 0.878 | 0.95 | 1 | MS.Trt_vs_MS.NoTrt_B |
| ZNF485 | 0.12275 | 0.2978 | 0.12 | 0.17 | 1 | MS.Trt_vs_MS.NoTrt_B |
| EIF4E | 0.12306 | 0.35422 | 0.928 | 1 | 1 | MS.Trt_vs_MS.NoTrt_B |
| MCF2L | 0.12319 | -0.3896 | 0.211 | 0.28 | 1 | MS.Trt_vs_MS.NoTrt_B |
| TMEM236 | 0.12351 | 0.47669 | 0.702 | 0.71 | 1 | MS.Trt_vs_MS.NoTrt_B |
| AIDA | 0.12363 | 0.7484 | 0.089 | 0.13 | 1 | MS.Trt_vs_MS.NoTrt_B |
| CC2D1B | 0.12379 | 0.36901 | 0.523 | 0.6 | 1 | MS.Trt_vs_MS.NoTrt_B |
| FER | 0.12396 | 0.27733 | 0.359 | 0.46 | 1 | MS.Trt_vs_MS.NoTrt_B |
| AMDHD1 | 0.12397 | -0.4174 | 0.126 | 0.17 | 1 | MS.Trt_vs_MS.NoTrt_B |

| | | | | | | |
|---|---|---|---|---|---|---|
| RREB1 | 0.12413 | 0.66587 | 0.593 | 0.74 | 1 | MS.Trt_vs_MS.NoTrt_B |
| PPME1 | 0.1242 | -0.886 | 0.098 | 0.14 | 1 | MS.Trt_vs_MS.NoTrt_B |
| FPGS | 0.12427 | -0.4248 | 0.159 | 0.21 | 1 | MS.Trt_vs_MS.NoTrt_B |
| HIST1H2BG | 0.12466 | 0.38556 | 0.403 | 0.47 | 1 | MS.Trt_vs_MS.NoTrt_B |
| PRPS2 | 0.12515 | -0.277 | 0.081 | 0.12 | 1 | MS.Trt_vs_MS.NoTrt_B |
| LINC00961 | 0.12521 | 0.30313 | 0.305 | 0.25 | 1 | MS.Trt_vs_MS.NoTrt_B |
| CPNE2 | 0.12526 | 0.63389 | 0.305 | 0.37 | 1 | MS.Trt_vs_MS.NoTrt_B |
| KLF2 | 0.12536 | 0.47839 | 0.207 | 0.27 | 1 | MS.Trt_vs_MS.NoTrt_B |
| OPA3 | 0.12543 | -0.2621 | 0.902 | 0.97 | 1 | MS.Trt_vs_MS.NoTrt_B |
| PKD2 | 0.12608 | -0.6705 | 0.131 | 0.18 | 1 | MS.Trt_vs_MS.NoTrt_B |
| LTB4R | 0.1261 | -0.8323 | 0.255 | 0.31 | 1 | MS.Trt_vs_MS.NoTrt_B |
| ATP13A1 | 0.12632 | 0.52235 | 0.244 | 0.3 | 1 | MS.Trt_vs_MS.NoTrt_B |
| UBXN11 | 0.1265 | 0.45621 | 0.15 | 0.2 | 1 | MS.Trt_vs_MS.NoTrt_B |
| SEMA4C | 0.12667 | 1.05053 | 0.643 | 0.76 | 1 | MS.Trt_vs_MS.NoTrt_B |
| ADAM32 | 0.1267 | 0.47468 | 0.673 | 0.82 | 1 | MS.Trt_vs_MS.NoTrt_B |
| CHORDC1 | 0.12698 | 0.29174 | 0.344 | 0.42 | 1 | MS.Trt_vs_MS.NoTrt_B |
| SPPL2B | 0.12705 | -0.6088 | 0.312 | 0.37 | 1 | MS.Trt_vs_MS.NoTrt_B |
| IER3IP1 | 0.12706 | 0.92614 | 0.102 | 0.15 | 1 | MS.Trt_vs_MS.NoTrt_B |
| MGAT4A | 0.12727 | 0.73298 | 0.34 | 0.41 | 1 | MS.Trt_vs_MS.NoTrt_B |
| AIPL1 | 0.12756 | 0.26532 | 0.375 | 0.32 | 1 | MS.Trt_vs_MS.NoTrt_B |
| ADAM15 | 0.12771 | -0.9327 | 0.285 | 0.34 | 1 | MS.Trt_vs_MS.NoTrt_B |
| SLC38A2 | 0.12871 | 0.31671 | 0.161 | 0.21 | 1 | MS.Trt_vs_MS.NoTrt_B |
| GTF2H4 | 0.12893 | -1.2783 | 0.068 | 0.1 | 1 | MS.Trt_vs_MS.NoTrt_B |
| C11orf24 | 0.12905 | 0.82228 | 0.174 | 0.23 | 1 | MS.Trt_vs_MS.NoTrt_B |
| EFCAB3 | 0.12931 | 0.35318 | 0.451 | 0.61 | 1 | MS.Trt_vs_MS.NoTrt_B |
| SLC39A4 | 0.12984 | 0.71946 | 0.301 | 0.37 | 1 | MS.Trt_vs_MS.NoTrt_B |
| ZMYM3 | 0.12986 | 0.48798 | 0.307 | 0.4 | 1 | MS.Trt_vs_MS.NoTrt_B |
| UMAD1 | 0.13015 | 0.63275 | 0.115 | 0.16 | 1 | MS.Trt_vs_MS.NoTrt_B |
| VARS2 | 0.1303 | -0.8979 | 0.39 | 0.45 | 1 | MS.Trt_vs_MS.NoTrt_B |
| SPAG9 | 0.13048 | 0.35065 | 0.747 | 0.87 | 1 | MS.Trt_vs_MS.NoTrt_B |
| MT-ATP8 | 0.13074 | 0.65811 | 0.614 | 0.84 | 1 | MS.Trt_vs_MS.NoTrt_B |
| DYNLL2 | 0.13089 | 0.98575 | 0.1 | 0.14 | 1 | MS.Trt_vs_MS.NoTrt_B |
| PFDN4 | 0.13149 | 1.10696 | 0.078 | 0.12 | 1 | MS.Trt_vs_MS.NoTrt_B |
| SATB1 | 0.13155 | 0.50882 | 0.176 | 0.12 | 1 | MS.Trt_vs_MS.NoTrt_B |
| MAF | 0.13159 | 0.86447 | 0.224 | 0.29 | 1 | MS.Trt_vs_MS.NoTrt_B |
| TRNT1 | 0.13267 | -0.2562 | 0.233 | 0.28 | 1 | MS.Trt_vs_MS.NoTrt_B |
| PON2 | 0.13312 | 0.42095 | 0.329 | 0.41 | 1 | MS.Trt_vs_MS.NoTrt_B |
| ZNF140 | 0.13315 | -0.546 | 0.124 | 0.17 | 1 | MS.Trt_vs_MS.NoTrt_B |
| KIAA0408 | 0.1339 | 0.74748 | 0.277 | 0.22 | 1 | MS.Trt_vs_MS.NoTrt_B |
| DHX9 | 0.13405 | 0.34473 | 0.187 | 0.24 | 1 | MS.Trt_vs_MS.NoTrt_B |

| | | | | | | |
|---|---|---|---|---|---|---|
| CEP19 | 0.13406 | 1.20579 | 0.492 | 0.58 | 1 | MS.Trt_vs_MS.NoTrt_B |
| AICDA | 0.13415 | 0.25758 | 0.453 | 0.54 | 1 | MS.Trt_vs_MS.NoTrt_B |
| UBL3 | 0.13419 | 0.41806 | 0.179 | 0.24 | 1 | MS.Trt_vs_MS.NoTrt_B |
| KIAA1191 | 0.13431 | 0.29585 | 0.512 | 0.63 | 1 | MS.Trt_vs_MS.NoTrt_B |
| ZNF480 | 0.13489 | 1.42768 | 0.549 | 0.72 | 1 | MS.Trt_vs_MS.NoTrt_B |
| LLGL2 | 0.13492 | 0.27858 | 0.386 | 0.34 | 1 | MS.Trt_vs_MS.NoTrt_B |
| C1orf174 | 0.13508 | 0.40832 | 0.455 | 0.55 | 1 | MS.Trt_vs_MS.NoTrt_B |
| HNRNPUL2 | 0.13521 | 0.49239 | 0.072 | 0.11 | 1 | MS.Trt_vs_MS.NoTrt_B |
| ACSL4 | 0.13536 | -0.3455 | 0.105 | 0.15 | 1 | MS.Trt_vs_MS.NoTrt_B |
| KCNMB3 | 0.13546 | 1.09598 | 0.301 | 0.38 | 1 | MS.Trt_vs_MS.NoTrt_B |
| ACAP3 | 0.13568 | -0.7058 | 0.087 | 0.12 | 1 | MS.Trt_vs_MS.NoTrt_B |
| CROCC | 0.13579 | 0.60107 | 0.397 | 0.47 | 1 | MS.Trt_vs_MS.NoTrt_B |
| ZSCAN22 | 0.13588 | -1.1466 | 0.325 | 0.41 | 1 | MS.Trt_vs_MS.NoTrt_B |
| L3HYPDH | 0.13618 | 1.15399 | 0.144 | 0.2 | 1 | MS.Trt_vs_MS.NoTrt_B |
| CD302 | 0.13619 | 0.80819 | 0.235 | 0.29 | 1 | MS.Trt_vs_MS.NoTrt_B |
| OARD1 | 0.13623 | 0.66128 | 0.172 | 0.23 | 1 | MS.Trt_vs_MS.NoTrt_B |
| RP11-644F5 | 0.13745 | 0.47719 | 0.551 | 0.64 | 1 | MS.Trt_vs_MS.NoTrt_B |
| ZBTB11 | 0.13782 | 1.00552 | 0.42 | 0.51 | 1 | MS.Trt_vs_MS.NoTrt_B |
| C1S | 0.13866 | 0.31302 | 0.481 | 0.6 | 1 | MS.Trt_vs_MS.NoTrt_B |
| PLAG1 | 0.13879 | 1.02791 | 0.312 | 0.25 | 1 | MS.Trt_vs_MS.NoTrt_B |
| PRKCB | 0.13891 | 0.64276 | 0.259 | 0.32 | 1 | MS.Trt_vs_MS.NoTrt_B |
| AFAP1 | 0.13929 | 0.86358 | 0.279 | 0.35 | 1 | MS.Trt_vs_MS.NoTrt_B |
| KLHL26 | 0.13968 | 1.07798 | 0.434 | 0.57 | 1 | MS.Trt_vs_MS.NoTrt_B |
| SPDYE6 | 0.14004 | 0.73234 | 0.094 | 0.14 | 1 | MS.Trt_vs_MS.NoTrt_B |
| TPRA1 | 0.1405 | 0.86775 | 0.118 | 0.16 | 1 | MS.Trt_vs_MS.NoTrt_B |
| CEP70 | 0.14102 | 1.26345 | 0.126 | 0.17 | 1 | MS.Trt_vs_MS.NoTrt_B |
| LINS1 | 0.14111 | 0.75979 | 0.449 | 0.55 | 1 | MS.Trt_vs_MS.NoTrt_B |
| VDAC2 | 0.14133 | 0.45291 | 0.248 | 0.31 | 1 | MS.Trt_vs_MS.NoTrt_B |
| ATXN7L3B | 0.14138 | 0.30391 | 0.126 | 0.17 | 1 | MS.Trt_vs_MS.NoTrt_B |
| TMED4 | 0.1415 | 0.38532 | 0.861 | 0.9 | 1 | MS.Trt_vs_MS.NoTrt_B |
| URI1 | 0.14165 | 1.16272 | 0.174 | 0.13 | 1 | MS.Trt_vs_MS.NoTrt_B |
| OSBPL10 | 0.14247 | 0.65449 | 0.292 | 0.35 | 1 | MS.Trt_vs_MS.NoTrt_B |
| TPX2 | 0.14274 | -0.3649 | 0.148 | 0.2 | 1 | MS.Trt_vs_MS.NoTrt_B |
| DUS4L | 0.14291 | -0.3899 | 0.07 | 0.1 | 1 | MS.Trt_vs_MS.NoTrt_B |
| OCEL1 | 0.14297 | -0.483 | 0.41 | 0.49 | 1 | MS.Trt_vs_MS.NoTrt_B |
| ARHGEF40 | 0.14305 | 0.78573 | 0.107 | 0.15 | 1 | MS.Trt_vs_MS.NoTrt_B |
| DENND4B | 0.14357 | 0.30863 | 0.179 | 0.24 | 1 | MS.Trt_vs_MS.NoTrt_B |
| PEX19 | 0.1438 | 0.25341 | 0.105 | 0.15 | 1 | MS.Trt_vs_MS.NoTrt_B |
| IL7R | 0.1439 | 0.36376 | 0.791 | 0.83 | 1 | MS.Trt_vs_MS.NoTrt_B |
| TPH1 | 0.14403 | 0.90898 | 0.353 | 0.44 | 1 | MS.Trt_vs_MS.NoTrt_B |

| | | | | | | |
|---|---|---|---|---|---|---|
| OXNAD1 | 0.14404 | 0.7056 | 0.211 | 0.28 | 1 | MS.Trt_vs_MS.NoTrt_B |
| BRE | 0.14439 | -0.2831 | 0.135 | 0.18 | 1 | MS.Trt_vs_MS.NoTrt_B |
| ADAMTS17 | 0.14442 | 1.25627 | 0.115 | 0.16 | 1 | MS.Trt_vs_MS.NoTrt_B |
| CLOCK | 0.14449 | 0.25232 | 0.115 | 0.16 | 1 | MS.Trt_vs_MS.NoTrt_B |
| GTF2F2 | 0.14465 | 0.53324 | 0.148 | 0.2 | 1 | MS.Trt_vs_MS.NoTrt_B |
| FASTKD5 | 0.14495 | 0.94306 | 0.113 | 0.08 | 1 | MS.Trt_vs_MS.NoTrt_B |
| SMAD5 | 0.14503 | 0.4154 | 0.366 | 0.47 | 1 | MS.Trt_vs_MS.NoTrt_B |
| CPT2 | 0.14537 | 0.47782 | 0.634 | 0.75 | 1 | MS.Trt_vs_MS.NoTrt_B |
| SGTA | 0.14539 | -0.3003 | 0.403 | 0.47 | 1 | MS.Trt_vs_MS.NoTrt_B |
| RP11-603J24 | 0.14556 | 0.39041 | 0.144 | 0.2 | 1 | MS.Trt_vs_MS.NoTrt_B |
| IRGQ | 0.14564 | 0.3044 | 0.928 | 0.99 | 1 | MS.Trt_vs_MS.NoTrt_B |
| VPS9D1 | 0.14577 | 1.05824 | 0.187 | 0.24 | 1 | MS.Trt_vs_MS.NoTrt_B |
| ANKDD1A | 0.14607 | 0.7517 | 0.431 | 0.52 | 1 | MS.Trt_vs_MS.NoTrt_B |
| RAB21 | 0.14623 | 0.67484 | 0.917 | 0.97 | 1 | MS.Trt_vs_MS.NoTrt_B |
| UBE2D4 | 0.14652 | 0.30153 | 0.599 | 0.53 | 1 | MS.Trt_vs_MS.NoTrt_B |
| IFRD1 | 0.14666 | -0.3325 | 0.264 | 0.33 | 1 | MS.Trt_vs_MS.NoTrt_B |
| ZNF426 | 0.14705 | 0.80846 | 0.83 | 0.9 | 1 | MS.Trt_vs_MS.NoTrt_B |
| HERPUD1 | 0.14753 | 0.44818 | 0.625 | 0.69 | 1 | MS.Trt_vs_MS.NoTrt_B |
| GADD45GIP | 0.14856 | 0.40121 | 0.102 | 0.15 | 1 | MS.Trt_vs_MS.NoTrt_B |
| JUNB | 0.1486 | 0.59276 | 0.288 | 0.38 | 1 | MS.Trt_vs_MS.NoTrt_B |
| LIG3 | 0.14881 | -0.747 | 0.281 | 0.33 | 1 | MS.Trt_vs_MS.NoTrt_B |
| SERF1B | 0.14891 | 0.41086 | 0.307 | 0.39 | 1 | MS.Trt_vs_MS.NoTrt_B |
| COIL | 0.14893 | 1.53847 | 0.083 | 0.12 | 1 | MS.Trt_vs_MS.NoTrt_B |
| CEP128 | 0.149 | 0.62085 | 0.172 | 0.23 | 1 | MS.Trt_vs_MS.NoTrt_B |
| STYXL1 | 0.14938 | -0.3348 | 0.2 | 0.25 | 1 | MS.Trt_vs_MS.NoTrt_B |
| AAED1 | 0.14989 | 0.87015 | 0.691 | 0.78 | 1 | MS.Trt_vs_MS.NoTrt_B |
| TCTN2 | 0.15033 | 0.56793 | 0.617 | 0.76 | 1 | MS.Trt_vs_MS.NoTrt_B |
| MOSPD1 | 0.15036 | 0.76144 | 0.137 | 0.18 | 1 | MS.Trt_vs_MS.NoTrt_B |
| SLC2A1 | 0.15101 | -0.6086 | 0.07 | 0.1 | 1 | MS.Trt_vs_MS.NoTrt_B |
| ANKRD39 | 0.15105 | 0.26459 | 0.159 | 0.21 | 1 | MS.Trt_vs_MS.NoTrt_B |
| TCP11L1 | 0.1511 | 0.68035 | 0.159 | 0.2 | 1 | MS.Trt_vs_MS.NoTrt_B |
| EML2 | 0.15125 | -0.5702 | 0.575 | 0.67 | 1 | MS.Trt_vs_MS.NoTrt_B |
| TOX4 | 0.15127 | 0.319 | 0.641 | 0.71 | 1 | MS.Trt_vs_MS.NoTrt_B |
| FARSB | 0.15146 | 0.47303 | 0.521 | 0.57 | 1 | MS.Trt_vs_MS.NoTrt_B |
| TIPIN | 0.15205 | 0.25082 | 0.484 | 0.59 | 1 | MS.Trt_vs_MS.NoTrt_B |
| MESDC2 | 0.15235 | 0.53897 | 0.436 | 0.5 | 1 | MS.Trt_vs_MS.NoTrt_B |
| PLCE1 | 0.15276 | 0.5838 | 0.797 | 0.88 | 1 | MS.Trt_vs_MS.NoTrt_B |
| RLF | 0.15297 | 0.27225 | 0.122 | 0.16 | 1 | MS.Trt_vs_MS.NoTrt_B |
| MT1X | 0.15325 | 0.6398 | 0.096 | 0.14 | 1 | MS.Trt_vs_MS.NoTrt_B |
| ITGA6 | 0.15329 | 0.34633 | 0.161 | 0.21 | 1 | MS.Trt_vs_MS.NoTrt_B |

| | | | | | | |
|---|---|---|---|---|---|---|
| DNAAF5 | 0.15371 | 2.54238 | 0.166 | 0.12 | 1 | MS.Trt_vs_MS.NoTrt_B |
| SMC1A | 0.1539 | 0.44877 | 0.176 | 0.23 | 1 | MS.Trt_vs_MS.NoTrt_B |
| ATPAF2 | 0.15429 | 0.33639 | 0.357 | 0.45 | 1 | MS.Trt_vs_MS.NoTrt_B |
| GEMIN7 | 0.1543 | 0.55861 | 0.172 | 0.22 | 1 | MS.Trt_vs_MS.NoTrt_B |
| MRPS5 | 0.15458 | 0.33362 | 0.699 | 0.88 | 1 | MS.Trt_vs_MS.NoTrt_B |
| CDRT4 | 0.15496 | -0.3521 | 0.083 | 0.12 | 1 | MS.Trt_vs_MS.NoTrt_B |
| PHF6 | 0.15521 | 1.16196 | 0.573 | 0.59 | 1 | MS.Trt_vs_MS.NoTrt_B |
| HOPX | 0.15533 | 0.47809 | 0.691 | 0.85 | 1 | MS.Trt_vs_MS.NoTrt_B |
| SKIV2L2 | 0.15547 | 0.27344 | 0.723 | 0.85 | 1 | MS.Trt_vs_MS.NoTrt_B |
| GPM6B | 0.15551 | 0.42904 | 0.107 | 0.15 | 1 | MS.Trt_vs_MS.NoTrt_B |
| PECR | 0.15589 | 0.42542 | 0.641 | 0.71 | 1 | MS.Trt_vs_MS.NoTrt_B |
| MAPK8IP3 | 0.15616 | 0.44358 | 0.118 | 0.16 | 1 | MS.Trt_vs_MS.NoTrt_B |
| WDR43 | 0.15626 | 0.39713 | 0.357 | 0.43 | 1 | MS.Trt_vs_MS.NoTrt_B |
| DCAF11 | 0.15629 | -0.863 | 0.632 | 0.78 | 1 | MS.Trt_vs_MS.NoTrt_B |
| OAZ3 | 0.15641 | 0.85814 | 0.307 | 0.38 | 1 | MS.Trt_vs_MS.NoTrt_B |
| FAM168A | 0.15645 | 0.57866 | 0.078 | 0.11 | 1 | MS.Trt_vs_MS.NoTrt_B |
| PSMG4 | 0.15705 | 0.32882 | 0.691 | 0.77 | 1 | MS.Trt_vs_MS.NoTrt_B |
| PDIA6 | 0.15711 | 0.46151 | 0.17 | 0.23 | 1 | MS.Trt_vs_MS.NoTrt_B |
| PPP1R3E | 0.1573 | -0.5081 | 0.181 | 0.23 | 1 | MS.Trt_vs_MS.NoTrt_B |
| BBS1 | 0.1577 | -0.309 | 0.501 | 0.58 | 1 | MS.Trt_vs_MS.NoTrt_B |
| CCDC174 | 0.15784 | 1.34597 | 0.153 | 0.11 | 1 | MS.Trt_vs_MS.NoTrt_B |
| MBNL2 | 0.15842 | 0.28832 | 0.137 | 0.18 | 1 | MS.Trt_vs_MS.NoTrt_B |
| KLHL8 | 0.15861 | 0.35917 | 0.51 | 0.58 | 1 | MS.Trt_vs_MS.NoTrt_B |
| SLC4A8 | 0.15882 | 0.96572 | 0.784 | 0.9 | 1 | MS.Trt_vs_MS.NoTrt_B |
| TMEM135 | 0.15907 | 0.46454 | 0.621 | 0.74 | 1 | MS.Trt_vs_MS.NoTrt_B |
| HOMEZ | 0.15941 | 0.82105 | 0.192 | 0.24 | 1 | MS.Trt_vs_MS.NoTrt_B |
| FXN | 0.15975 | 0.39536 | 0.146 | 0.2 | 1 | MS.Trt_vs_MS.NoTrt_B |
| KLF7 | 0.15989 | -1.0611 | 0.124 | 0.16 | 1 | MS.Trt_vs_MS.NoTrt_B |
| SET | 0.15998 | 0.47838 | 0.854 | 0.98 | 1 | MS.Trt_vs_MS.NoTrt_B |
| RSL1D1 | 0.16017 | 0.57642 | 0.73 | 0.86 | 1 | MS.Trt_vs_MS.NoTrt_B |
| UBR3 | 0.16026 | 1.30885 | 0.096 | 0.14 | 1 | MS.Trt_vs_MS.NoTrt_B |
| TMEM185B | 0.16071 | -0.9974 | 0.272 | 0.33 | 1 | MS.Trt_vs_MS.NoTrt_B |
| RIPK1 | 0.1611 | -0.6175 | 0.12 | 0.16 | 1 | MS.Trt_vs_MS.NoTrt_B |
| YARS2 | 0.16136 | 0.79128 | 0.283 | 0.36 | 1 | MS.Trt_vs_MS.NoTrt_B |
| SFT2D1 | 0.16168 | 0.49754 | 0.309 | 0.35 | 1 | MS.Trt_vs_MS.NoTrt_B |
| TMEM168 | 0.16176 | 0.76949 | 0.412 | 0.48 | 1 | MS.Trt_vs_MS.NoTrt_B |
| NPTXR | 0.16224 | 0.48785 | 0.272 | 0.35 | 1 | MS.Trt_vs_MS.NoTrt_B |
| KCTD19 | 0.16241 | 0.25499 | 0.571 | 0.69 | 1 | MS.Trt_vs_MS.NoTrt_B |
| ZNF84 | 0.16266 | 1.00652 | 0.209 | 0.26 | 1 | MS.Trt_vs_MS.NoTrt_B |
| SMIM8 | 0.16354 | 0.73757 | 0.466 | 0.52 | 1 | MS.Trt_vs_MS.NoTrt_B |

| | | | | | | |
|---|---|---|---|---|---|---|
| PIR | 0.1636 | 0.3388 | 0.366 | 0.47 | 1 | MS.Trt_vs_MS.NoTrt_B |
| KLHDC10 | 0.16363 | 1.14099 | 0.314 | 0.4 | 1 | MS.Trt_vs_MS.NoTrt_B |
| USP1 | 0.16387 | 0.2714 | 0.115 | 0.16 | 1 | MS.Trt_vs_MS.NoTrt_B |
| DSC2 | 0.16432 | -1.2221 | 0.22 | 0.27 | 1 | MS.Trt_vs_MS.NoTrt_B |
| ZNF684 | 0.16442 | 0.83668 | 0.573 | 0.69 | 1 | MS.Trt_vs_MS.NoTrt_B |
| SYK | 0.16449 | 0.49002 | 0.288 | 0.35 | 1 | MS.Trt_vs_MS.NoTrt_B |
| CES2 | 0.16496 | -0.6757 | 0.224 | 0.27 | 1 | MS.Trt_vs_MS.NoTrt_B |
| PARP10 | 0.16506 | 0.29453 | 0.092 | 0.13 | 1 | MS.Trt_vs_MS.NoTrt_B |
| RBBP7 | 0.16562 | 0.30119 | 0.632 | 0.69 | 1 | MS.Trt_vs_MS.NoTrt_B |
| MZF1 | 0.16592 | 0.62298 | 0.083 | 0.12 | 1 | MS.Trt_vs_MS.NoTrt_B |
| ZKSCAN4 | 0.16615 | -0.381 | 0.349 | 0.42 | 1 | MS.Trt_vs_MS.NoTrt_B |
| YLPM1 | 0.16619 | 0.57828 | 0.473 | 0.54 | 1 | MS.Trt_vs_MS.NoTrt_B |
| RHBG | 0.16707 | 0.55662 | 0.634 | 0.75 | 1 | MS.Trt_vs_MS.NoTrt_B |
| ALG5 | 0.16744 | 0.58012 | 0.434 | 0.37 | 1 | MS.Trt_vs_MS.NoTrt_B |
| CRLF2 | 0.16756 | 0.45375 | 0.216 | 0.28 | 1 | MS.Trt_vs_MS.NoTrt_B |
| MBTPS2 | 0.16787 | 0.36101 | 0.224 | 0.28 | 1 | MS.Trt_vs_MS.NoTrt_B |
| MFSD8 | 0.16802 | -0.4457 | 0.29 | 0.34 | 1 | MS.Trt_vs_MS.NoTrt_B |
| FZR1 | 0.16831 | -0.8253 | 0.072 | 0.1 | 1 | MS.Trt_vs_MS.NoTrt_B |
| TRA2A | 0.16897 | 0.55877 | 0.688 | 0.81 | 1 | MS.Trt_vs_MS.NoTrt_B |
| WRAP73 | 0.16907 | 0.25544 | 0.261 | 0.31 | 1 | MS.Trt_vs_MS.NoTrt_B |
| HIGD1A | 0.16932 | 0.72185 | 0.115 | 0.16 | 1 | MS.Trt_vs_MS.NoTrt_B |
| LRRC69 | 0.17 | -0.3227 | 0.466 | 0.51 | 1 | MS.Trt_vs_MS.NoTrt_B |
| GTPBP2 | 0.1702 | 0.51846 | 0.462 | 0.52 | 1 | MS.Trt_vs_MS.NoTrt_B |
| TTC28 | 0.17073 | -0.5456 | 0.292 | 0.35 | 1 | MS.Trt_vs_MS.NoTrt_B |
| NFIC | 0.17081 | 1.45994 | 0.275 | 0.34 | 1 | MS.Trt_vs_MS.NoTrt_B |
| NBAS | 0.17128 | 0.83975 | 0.153 | 0.2 | 1 | MS.Trt_vs_MS.NoTrt_B |
| FOXRED1 | 0.17131 | -0.2538 | 0.434 | 0.51 | 1 | MS.Trt_vs_MS.NoTrt_B |
| ADORA2A | 0.17136 | -0.2569 | 0.089 | 0.12 | 1 | MS.Trt_vs_MS.NoTrt_B |
| IVNS1ABP | 0.17155 | 0.35459 | 0.251 | 0.3 | 1 | MS.Trt_vs_MS.NoTrt_B |
| GPR1 | 0.1717 | 0.25325 | 0.205 | 0.16 | 1 | MS.Trt_vs_MS.NoTrt_B |
| SNAPC5 | 0.17221 | 0.44615 | 0.501 | 0.57 | 1 | MS.Trt_vs_MS.NoTrt_B |
| C8orf76 | 0.17227 | -0.4357 | 0.503 | 0.58 | 1 | MS.Trt_vs_MS.NoTrt_B |
| ERMN | 0.17231 | 1.19154 | 0.131 | 0.17 | 1 | MS.Trt_vs_MS.NoTrt_B |
| FAM200A | 0.17282 | 0.42586 | 0.174 | 0.22 | 1 | MS.Trt_vs_MS.NoTrt_B |
| COPG1 | 0.17335 | -0.4084 | 0.109 | 0.15 | 1 | MS.Trt_vs_MS.NoTrt_B |
| KIF22 | 0.17378 | 0.28678 | 0.172 | 0.22 | 1 | MS.Trt_vs_MS.NoTrt_B |
| C16orf72 | 0.17379 | 0.5716 | 0.706 | 0.82 | 1 | MS.Trt_vs_MS.NoTrt_B |
| OGFR | 0.17403 | 0.33384 | 0.137 | 0.18 | 1 | MS.Trt_vs_MS.NoTrt_B |
| TIMM17A | 0.17418 | 0.44184 | 0.8 | 0.9 | 1 | MS.Trt_vs_MS.NoTrt_B |
| GSN | 0.17448 | -0.8155 | 0.453 | 0.56 | 1 | MS.Trt_vs_MS.NoTrt_B |

| | | | | | | |
|---|---|---|---|---|---|---|
| 9-Mar | 0.17468 | 0.29129 | 0.455 | 0.55 | 1 | MS.Trt_vs_MS.NoTrt_B |
| FAM173B | 0.17482 | 0.52792 | 0.126 | 0.17 | 1 | MS.Trt_vs_MS.NoTrt_B |
| CSPP1 | 0.17485 | 1.03044 | 0.32 | 0.4 | 1 | MS.Trt_vs_MS.NoTrt_B |
| ZHX1 | 0.17498 | -0.2988 | 0.492 | 0.57 | 1 | MS.Trt_vs_MS.NoTrt_B |
| SCPEP1 | 0.1755 | -0.2631 | 0.645 | 0.74 | 1 | MS.Trt_vs_MS.NoTrt_B |
| AKAP11 | 0.17609 | 0.50447 | 0.161 | 0.21 | 1 | MS.Trt_vs_MS.NoTrt_B |
| AGK | 0.17719 | 0.47475 | 0.176 | 0.23 | 1 | MS.Trt_vs_MS.NoTrt_B |
| ZC3H15 | 0.17747 | 0.48875 | 0.187 | 0.24 | 1 | MS.Trt_vs_MS.NoTrt_B |
| RAPGEF2 | 0.17753 | 0.56587 | 0.316 | 0.38 | 1 | MS.Trt_vs_MS.NoTrt_B |
| DNPEP | 0.17769 | -0.2793 | 0.403 | 0.47 | 1 | MS.Trt_vs_MS.NoTrt_B |
| C2 | 0.17835 | 0.33223 | 0.556 | 0.68 | 1 | MS.Trt_vs_MS.NoTrt_B |
| CCDC138 | 0.17877 | 0.93931 | 0.111 | 0.15 | 1 | MS.Trt_vs_MS.NoTrt_B |
| CCDC85C | 0.17908 | 0.79937 | 0.316 | 0.37 | 1 | MS.Trt_vs_MS.NoTrt_B |
| ALG12 | 0.17911 | 0.51148 | 0.087 | 0.12 | 1 | MS.Trt_vs_MS.NoTrt_B |
| LEPR | 0.17934 | 0.49723 | 0.24 | 0.3 | 1 | MS.Trt_vs_MS.NoTrt_B |
| ZC3H13 | 0.17936 | 0.45904 | 0.227 | 0.29 | 1 | MS.Trt_vs_MS.NoTrt_B |
| RAP1GDS1 | 0.17946 | 0.47627 | 0.336 | 0.42 | 1 | MS.Trt_vs_MS.NoTrt_B |
| EVI5L | 0.1801 | -0.4658 | 0.207 | 0.27 | 1 | MS.Trt_vs_MS.NoTrt_B |
| ANKRD46 | 0.18078 | 0.56825 | 0.137 | 0.18 | 1 | MS.Trt_vs_MS.NoTrt_B |
| STAT5A | 0.18123 | -0.5639 | 0.107 | 0.14 | 1 | MS.Trt_vs_MS.NoTrt_B |
| CHRM3 | 0.18123 | 0.4758 | 0.148 | 0.2 | 1 | MS.Trt_vs_MS.NoTrt_B |
| FAM159A | 0.18128 | 1.12779 | 0.468 | 0.56 | 1 | MS.Trt_vs_MS.NoTrt_B |
| SMG7 | 0.18162 | 0.42629 | 0.159 | 0.21 | 1 | MS.Trt_vs_MS.NoTrt_B |
| DNAJC22 | 0.18215 | 0.25005 | 0.377 | 0.32 | 1 | MS.Trt_vs_MS.NoTrt_B |
| DCAF10 | 0.18282 | 0.48629 | 0.68 | 0.79 | 1 | MS.Trt_vs_MS.NoTrt_B |
| ADCK1 | 0.18342 | 0.30575 | 0.176 | 0.22 | 1 | MS.Trt_vs_MS.NoTrt_B |
| PPM1F | 0.18347 | 0.86965 | 0.695 | 0.81 | 1 | MS.Trt_vs_MS.NoTrt_B |
| COASY | 0.18379 | -0.9954 | 0.362 | 0.41 | 1 | MS.Trt_vs_MS.NoTrt_B |
| VEZT | 0.18425 | 0.35457 | 0.593 | 0.7 | 1 | MS.Trt_vs_MS.NoTrt_B |
| LRP1 | 0.18429 | -0.3928 | 0.166 | 0.21 | 1 | MS.Trt_vs_MS.NoTrt_B |
| PDZK1 | 0.18451 | -0.5509 | 0.214 | 0.26 | 1 | MS.Trt_vs_MS.NoTrt_B |
| EFR3A | 0.18455 | -0.3962 | 0.096 | 0.13 | 1 | MS.Trt_vs_MS.NoTrt_B |
| ZNF763 | 0.18464 | -0.6848 | 0.109 | 0.15 | 1 | MS.Trt_vs_MS.NoTrt_B |
| FAM111B | 0.18558 | 0.30339 | 0.414 | 0.55 | 1 | MS.Trt_vs_MS.NoTrt_B |
| NCAPG2 | 0.18625 | 1.434 | 0.2 | 0.16 | 1 | MS.Trt_vs_MS.NoTrt_B |
| PIK3CG | 0.18645 | 0.65177 | 0.183 | 0.23 | 1 | MS.Trt_vs_MS.NoTrt_B |
| GSDMA | 0.18652 | 0.47631 | 0.497 | 0.58 | 1 | MS.Trt_vs_MS.NoTrt_B |
| GATAD2A | 0.18654 | 0.3568 | 0.577 | 0.63 | 1 | MS.Trt_vs_MS.NoTrt_B |
| WIZ | 0.18684 | -0.3675 | 0.22 | 0.27 | 1 | MS.Trt_vs_MS.NoTrt_B |
| GTF2H1 | 0.18728 | 0.26998 | 0.277 | 0.34 | 1 | MS.Trt_vs_MS.NoTrt_B |

| | | | | | | |
|---|---|---|---|---|---|---|
| PPM1A | 0.18753 | -0.5111 | 0.111 | 0.15 | 1 | MS.Trt_vs_MS.NoTrt_B |
| LYRM9 | 0.18785 | -0.8987 | 0.176 | 0.22 | 1 | MS.Trt_vs_MS.NoTrt_B |
| FAM120A | 0.18808 | 0.90119 | 0.129 | 0.09 | 1 | MS.Trt_vs_MS.NoTrt_B |
| GDPGP1 | 0.18811 | 0.36266 | 0.841 | 0.94 | 1 | MS.Trt_vs_MS.NoTrt_B |
| RASEF | 0.18863 | 0.27756 | 0.745 | 0.96 | 1 | MS.Trt_vs_MS.NoTrt_B |
| MRPL32 | 0.18877 | 0.85831 | 0.098 | 0.14 | 1 | MS.Trt_vs_MS.NoTrt_B |
| MAP3K4 | 0.18917 | -0.4062 | 0.122 | 0.16 | 1 | MS.Trt_vs_MS.NoTrt_B |
| NT5DC1 | 0.1892 | 0.53243 | 0.146 | 0.19 | 1 | MS.Trt_vs_MS.NoTrt_B |
| PUS7 | 0.18968 | -0.6191 | 0.244 | 0.3 | 1 | MS.Trt_vs_MS.NoTrt_B |
| B4GALT1 | 0.18994 | 0.25758 | 0.566 | 0.63 | 1 | MS.Trt_vs_MS.NoTrt_B |
| GLS | 0.18996 | 0.68331 | 0.556 | 0.64 | 1 | MS.Trt_vs_MS.NoTrt_B |
| HID1 | 0.19002 | 1.10896 | 0.1 | 0.14 | 1 | MS.Trt_vs_MS.NoTrt_B |
| STRN | 0.19031 | 0.86715 | 0.126 | 0.17 | 1 | MS.Trt_vs_MS.NoTrt_B |
| IDH1 | 0.19045 | -0.5483 | 0.146 | 0.19 | 1 | MS.Trt_vs_MS.NoTrt_B |
| IRAK2 | 0.19048 | 1.00391 | 0.242 | 0.3 | 1 | MS.Trt_vs_MS.NoTrt_B |
| MEGF9 | 0.19054 | 0.50076 | 0.392 | 0.44 | 1 | MS.Trt_vs_MS.NoTrt_B |
| RPS6KB1 | 0.19057 | 0.51865 | 0.758 | 0.82 | 1 | MS.Trt_vs_MS.NoTrt_B |
| SLC25A40 | 0.19072 | -0.4674 | 0.222 | 0.29 | 1 | MS.Trt_vs_MS.NoTrt_B |
| MRPL13 | 0.19105 | 0.7866 | 0.102 | 0.14 | 1 | MS.Trt_vs_MS.NoTrt_B |
| GALNT2 | 0.19113 | 0.25231 | 0.137 | 0.18 | 1 | MS.Trt_vs_MS.NoTrt_B |
| SLC37A3 | 0.19143 | -0.4646 | 0.854 | 0.91 | 1 | MS.Trt_vs_MS.NoTrt_B |
| HMG20A | 0.19157 | 0.84361 | 0.222 | 0.28 | 1 | MS.Trt_vs_MS.NoTrt_B |
| ATAD3C | 0.19231 | 0.30668 | 0.741 | 0.69 | 1 | MS.Trt_vs_MS.NoTrt_B |
| ZCRB1 | 0.19299 | 0.43405 | 0.142 | 0.18 | 1 | MS.Trt_vs_MS.NoTrt_B |
| GTF2I | 0.19328 | 0.51903 | 0.8 | 0.88 | 1 | MS.Trt_vs_MS.NoTrt_B |
| SWAP70 | 0.19329 | -0.3043 | 0.399 | 0.48 | 1 | MS.Trt_vs_MS.NoTrt_B |
| MLXIP | 0.19386 | 0.32001 | 0.854 | 0.91 | 1 | MS.Trt_vs_MS.NoTrt_B |
| ZC3H6 | 0.19419 | 0.52378 | 0.695 | 0.83 | 1 | MS.Trt_vs_MS.NoTrt_B |
| ZSCAN21 | 0.19452 | 1.3538 | 0.187 | 0.15 | 1 | MS.Trt_vs_MS.NoTrt_B |
| PLCD4 | 0.19482 | -0.4569 | 0.19 | 0.24 | 1 | MS.Trt_vs_MS.NoTrt_B |
| C2orf42 | 0.19483 | 0.61797 | 0.072 | 0.1 | 1 | MS.Trt_vs_MS.NoTrt_B |
| RAD23A | 0.19529 | 0.28215 | 0.682 | 0.71 | 1 | MS.Trt_vs_MS.NoTrt_B |
| HPCAL1 | 0.19555 | 0.6947 | 0.305 | 0.36 | 1 | MS.Trt_vs_MS.NoTrt_B |
| PRKAR1A | 0.19566 | 0.27212 | 0.207 | 0.26 | 1 | MS.Trt_vs_MS.NoTrt_B |
| SCAMP1 | 0.19574 | 0.85281 | 0.399 | 0.44 | 1 | MS.Trt_vs_MS.NoTrt_B |
| ZNF92 | 0.19646 | -0.6485 | 0.107 | 0.14 | 1 | MS.Trt_vs_MS.NoTrt_B |
| NPC2 | 0.19676 | 0.5171 | 0.098 | 0.14 | 1 | MS.Trt_vs_MS.NoTrt_B |
| RP11-9J18.1 | 0.19702 | 0.27117 | 0.17 | 0.22 | 1 | MS.Trt_vs_MS.NoTrt_B |
| TTPAL | 0.19727 | 0.63349 | 0.691 | 0.8 | 1 | MS.Trt_vs_MS.NoTrt_B |
| CEP295 | 0.19733 | 0.75267 | 0.484 | 0.56 | 1 | MS.Trt_vs_MS.NoTrt_B |

| | | | | | | |
|---|---|---|---|---|---|---|
| SLCO2B1 | 0.19791 | 0.48774 | 0.246 | 0.31 | 1 | MS.Trt_vs_MS.NoTrt_B |
| ATP1A3 | 0.19853 | -0.4477 | 0.255 | 0.31 | 1 | MS.Trt_vs_MS.NoTrt_B |
| FEM1A | 0.19854 | 0.3516 | 0.867 | 0.92 | 1 | MS.Trt_vs_MS.NoTrt_B |
| OR11A1 | 0.19856 | 0.54012 | 0.183 | 0.24 | 1 | MS.Trt_vs_MS.NoTrt_B |
| TADA2B | 0.19861 | 1.64964 | 0.24 | 0.3 | 1 | MS.Trt_vs_MS.NoTrt_B |
| ZNF280C | 0.19941 | -0.4749 | 0.181 | 0.22 | 1 | MS.Trt_vs_MS.NoTrt_B |
| NEDD4 | 0.19966 | 0.41148 | 0.871 | 0.97 | 1 | MS.Trt_vs_MS.NoTrt_B |
| ZFP37 | 0.19975 | 0.54511 | 0.298 | 0.36 | 1 | MS.Trt_vs_MS.NoTrt_B |
| MIER2 | 0.20013 | -0.8812 | 0.39 | 0.45 | 1 | MS.Trt_vs_MS.NoTrt_B |
| PPP2R2A | 0.2003 | 0.98451 | 0.107 | 0.14 | 1 | MS.Trt_vs_MS.NoTrt_B |
| ZNF264 | 0.20036 | 0.28586 | 0.854 | 0.91 | 1 | MS.Trt_vs_MS.NoTrt_B |
| C2CD2 | 0.20081 | 0.84243 | 0.159 | 0.21 | 1 | MS.Trt_vs_MS.NoTrt_B |
| VPS18 | 0.20097 | -0.8137 | 0.174 | 0.21 | 1 | MS.Trt_vs_MS.NoTrt_B |
| AGAP6 | 0.20168 | 1.54113 | 0.089 | 0.12 | 1 | MS.Trt_vs_MS.NoTrt_B |
| RNF168 | 0.20175 | 0.49508 | 0.475 | 0.57 | 1 | MS.Trt_vs_MS.NoTrt_B |
| EIF5A | 0.20225 | 1.11836 | 0.61 | 0.67 | 1 | MS.Trt_vs_MS.NoTrt_B |
| LIPT2 | 0.2024 | -0.7268 | 0.092 | 0.12 | 1 | MS.Trt_vs_MS.NoTrt_B |
| CREBL2 | 0.20261 | 0.61942 | 0.072 | 0.1 | 1 | MS.Trt_vs_MS.NoTrt_B |
| UQCRC1 | 0.2033 | 0.47066 | 0.214 | 0.27 | 1 | MS.Trt_vs_MS.NoTrt_B |
| TBC1D23 | 0.20355 | 0.39412 | 0.092 | 0.12 | 1 | MS.Trt_vs_MS.NoTrt_B |
| CPNE8 | 0.2036 | 1.35965 | 0.109 | 0.08 | 1 | MS.Trt_vs_MS.NoTrt_B |
| MYO6 | 0.20361 | 1.64721 | 0.373 | 0.45 | 1 | MS.Trt_vs_MS.NoTrt_B |
| DENND2D | 0.2037 | 0.49618 | 0.322 | 0.28 | 1 | MS.Trt_vs_MS.NoTrt_B |
| MAN1C1 | 0.20412 | 0.84842 | 0.83 | 0.85 | 1 | MS.Trt_vs_MS.NoTrt_B |
| RBM7 | 0.20415 | 1.05648 | 0.601 | 0.67 | 1 | MS.Trt_vs_MS.NoTrt_B |
| TRIM34 | 0.20422 | -0.2653 | 0.246 | 0.3 | 1 | MS.Trt_vs_MS.NoTrt_B |
| STAG2 | 0.20428 | 0.47612 | 0.671 | 0.73 | 1 | MS.Trt_vs_MS.NoTrt_B |
| CLDN11 | 0.20473 | 0.49123 | 0.512 | 0.47 | 1 | MS.Trt_vs_MS.NoTrt_B |
| SFXN2 | 0.20486 | 0.38443 | 0.871 | 0.9 | 1 | MS.Trt_vs_MS.NoTrt_B |
| GLUD1 | 0.20493 | 0.5377 | 0.124 | 0.16 | 1 | MS.Trt_vs_MS.NoTrt_B |
| LINC00959 | 0.20531 | -0.941 | 0.083 | 0.11 | 1 | MS.Trt_vs_MS.NoTrt_B |
| SPATS2L | 0.20541 | -0.2991 | 0.856 | 0.92 | 1 | MS.Trt_vs_MS.NoTrt_B |
| USO1 | 0.20572 | 0.50707 | 0.102 | 0.14 | 1 | MS.Trt_vs_MS.NoTrt_B |
| PRKRA | 0.2059 | 0.63964 | 0.196 | 0.24 | 1 | MS.Trt_vs_MS.NoTrt_B |
| UPK1B | 0.2061 | 0.39634 | 0.412 | 0.37 | 1 | MS.Trt_vs_MS.NoTrt_B |
| CAMKMT | 0.20623 | 0.63175 | 0.231 | 0.28 | 1 | MS.Trt_vs_MS.NoTrt_B |
| TRABD2A | 0.20624 | 0.25758 | 0.741 | 0.79 | 1 | MS.Trt_vs_MS.NoTrt_B |
| LSR | 0.20651 | -0.6796 | 0.133 | 0.17 | 1 | MS.Trt_vs_MS.NoTrt_B |
| ZNF37A | 0.20695 | 0.40093 | 0.749 | 0.85 | 1 | MS.Trt_vs_MS.NoTrt_B |
| NANP | 0.20697 | -0.7252 | 0.244 | 0.3 | 1 | MS.Trt_vs_MS.NoTrt_B |

| | | | | | | |
|---|---|---|---|---|---|---|
| ITGA2B | 0.20762 | 0.31365 | 0.312 | 0.27 | 1 | MS.Trt_vs_MS.NoTrt_B |
| NOP16 | 0.20764 | 0.27463 | 0.15 | 0.2 | 1 | MS.Trt_vs_MS.NoTrt_B |
| UEVLD | 0.20766 | -0.274 | 0.828 | 0.9 | 1 | MS.Trt_vs_MS.NoTrt_B |
| INCA1 | 0.20786 | 0.27999 | 0.157 | 0.21 | 1 | MS.Trt_vs_MS.NoTrt_B |
| METTL25 | 0.20843 | 0.72804 | 0.124 | 0.16 | 1 | MS.Trt_vs_MS.NoTrt_B |
| ZNF12 | 0.20898 | -0.9904 | 0.237 | 0.28 | 1 | MS.Trt_vs_MS.NoTrt_B |
| ZNF407 | 0.20934 | 1.02852 | 0.076 | 0.11 | 1 | MS.Trt_vs_MS.NoTrt_B |
| TAF8 | 0.21012 | -0.283 | 0.882 | 0.96 | 1 | MS.Trt_vs_MS.NoTrt_B |
| TMEM127 | 0.21052 | 0.2524 | 0.571 | 0.63 | 1 | MS.Trt_vs_MS.NoTrt_B |
| ACACB | 0.21055 | 0.61003 | 0.828 | 0.94 | 1 | MS.Trt_vs_MS.NoTrt_B |
| LPCAT3 | 0.21066 | -0.9599 | 0.392 | 0.41 | 1 | MS.Trt_vs_MS.NoTrt_B |
| NIT1 | 0.21069 | -0.6789 | 0.471 | 0.56 | 1 | MS.Trt_vs_MS.NoTrt_B |
| DOK6 | 0.21136 | -0.3543 | 0.146 | 0.19 | 1 | MS.Trt_vs_MS.NoTrt_B |
| PHTF1 | 0.21155 | 0.39036 | 0.397 | 0.45 | 1 | MS.Trt_vs_MS.NoTrt_B |
| HIST1H2AG | 0.21169 | 0.52814 | 0.279 | 0.34 | 1 | MS.Trt_vs_MS.NoTrt_B |
| BCR | 0.21183 | 0.48223 | 0.216 | 0.27 | 1 | MS.Trt_vs_MS.NoTrt_B |
| ZP3 | 0.21197 | 0.41229 | 0.512 | 0.61 | 1 | MS.Trt_vs_MS.NoTrt_B |
| ASPH | 0.21233 | -0.3052 | 0.351 | 0.44 | 1 | MS.Trt_vs_MS.NoTrt_B |
| MAML1 | 0.21247 | -0.2576 | 0.098 | 0.13 | 1 | MS.Trt_vs_MS.NoTrt_B |
| TRIM39 | 0.21272 | -0.3255 | 0.089 | 0.12 | 1 | MS.Trt_vs_MS.NoTrt_B |
| RNF144B | 0.2128 | 0.4858 | 0.248 | 0.31 | 1 | MS.Trt_vs_MS.NoTrt_B |
| DGKZ | 0.21311 | 0.38788 | 0.407 | 0.46 | 1 | MS.Trt_vs_MS.NoTrt_B |
| WDR77 | 0.21326 | 0.35672 | 0.092 | 0.12 | 1 | MS.Trt_vs_MS.NoTrt_B |
| GMPR | 0.21335 | 0.67024 | 0.322 | 0.41 | 1 | MS.Trt_vs_MS.NoTrt_B |
| MFSD4A | 0.21343 | 1.05951 | 0.423 | 0.48 | 1 | MS.Trt_vs_MS.NoTrt_B |
| BET1L | 0.21349 | 0.31184 | 0.157 | 0.2 | 1 | MS.Trt_vs_MS.NoTrt_B |
| MTHFR | 0.2135 | 0.34969 | 0.699 | 0.84 | 1 | MS.Trt_vs_MS.NoTrt_B |
| ZNF845 | 0.21462 | 0.45968 | 0.514 | 0.6 | 1 | MS.Trt_vs_MS.NoTrt_B |
| RBMS3 | 0.21498 | 0.53 | 0.562 | 0.67 | 1 | MS.Trt_vs_MS.NoTrt_B |
| BEST1 | 0.21503 | 0.33259 | 0.708 | 0.8 | 1 | MS.Trt_vs_MS.NoTrt_B |
| TEFM | 0.21528 | 0.58204 | 0.595 | 0.7 | 1 | MS.Trt_vs_MS.NoTrt_B |
| SYTL2 | 0.21549 | 0.48411 | 0.736 | 0.8 | 1 | MS.Trt_vs_MS.NoTrt_B |
| HNRNPM | 0.21567 | 0.3753 | 0.242 | 0.3 | 1 | MS.Trt_vs_MS.NoTrt_B |
| ACOT4 | 0.21601 | -1.428 | 0.092 | 0.12 | 1 | MS.Trt_vs_MS.NoTrt_B |
| DNAJA4 | 0.21624 | 1.6615 | 0.076 | 0.11 | 1 | MS.Trt_vs_MS.NoTrt_B |
| VPS50 | 0.21658 | 0.30511 | 0.473 | 0.54 | 1 | MS.Trt_vs_MS.NoTrt_B |
| ZDHHC14 | 0.21689 | 0.43669 | 0.27 | 0.32 | 1 | MS.Trt_vs_MS.NoTrt_B |
| KLHL11 | 0.21694 | -0.2832 | 0.405 | 0.46 | 1 | MS.Trt_vs_MS.NoTrt_B |
| ENDOV | 0.21712 | 0.48768 | 0.484 | 0.54 | 1 | MS.Trt_vs_MS.NoTrt_B |
| PTPRC | 0.21723 | 0.48757 | 0.876 | 0.98 | 1 | MS.Trt_vs_MS.NoTrt_B |

| | | | | | | |
|---|---|---|---|---|---|---|
| AMFR | 0.21742 | -0.841 | 0.503 | 0.42 | 1 | MS.Trt_vs_MS.NoTrt_B |
| CD160 | 0.21761 | 0.64058 | 0.214 | 0.27 | 1 | MS.Trt_vs_MS.NoTrt_B |
| TSEN34 | 0.21798 | 0.28424 | 0.094 | 0.12 | 1 | MS.Trt_vs_MS.NoTrt_B |
| ABCC4 | 0.21916 | 0.76431 | 0.58 | 0.75 | 1 | MS.Trt_vs_MS.NoTrt_B |
| DDX47 | 0.21949 | 0.6712 | 0.41 | 0.47 | 1 | MS.Trt_vs_MS.NoTrt_B |
| APC | 0.21979 | 0.41569 | 0.416 | 0.46 | 1 | MS.Trt_vs_MS.NoTrt_B |
| EMC7 | 0.22 | 0.38523 | 0.148 | 0.19 | 1 | MS.Trt_vs_MS.NoTrt_B |
| PCGF6 | 0.22019 | 0.35837 | 0.59 | 0.75 | 1 | MS.Trt_vs_MS.NoTrt_B |
| UNKL | 0.22046 | 0.42019 | 0.353 | 0.4 | 1 | MS.Trt_vs_MS.NoTrt_B |
| PPIL3 | 0.22062 | 0.27031 | 0.643 | 0.78 | 1 | MS.Trt_vs_MS.NoTrt_B |
| LAYN | 0.22066 | 0.99569 | 0.166 | 0.21 | 1 | MS.Trt_vs_MS.NoTrt_B |
| KIAA1468 | 0.22149 | 0.52404 | 0.227 | 0.27 | 1 | MS.Trt_vs_MS.NoTrt_B |
| RBMS1 | 0.22165 | 0.58624 | 0.505 | 0.61 | 1 | MS.Trt_vs_MS.NoTrt_B |
| MACF1 | 0.2218 | 0.74319 | 0.789 | 0.84 | 1 | MS.Trt_vs_MS.NoTrt_B |
| FYN | 0.22187 | 0.44077 | 0.451 | 0.54 | 1 | MS.Trt_vs_MS.NoTrt_B |
| PRR23C | 0.22279 | 0.29413 | 0.338 | 0.42 | 1 | MS.Trt_vs_MS.NoTrt_B |
| SLK | 0.22322 | 0.49495 | 0.115 | 0.15 | 1 | MS.Trt_vs_MS.NoTrt_B |
| UNC13C | 0.22335 | 0.30926 | 0.479 | 0.62 | 1 | MS.Trt_vs_MS.NoTrt_B |
| USP6NL | 0.22461 | 0.67643 | 0.667 | 0.82 | 1 | MS.Trt_vs_MS.NoTrt_B |
| CKS1B | 0.22476 | 0.86709 | 0.15 | 0.19 | 1 | MS.Trt_vs_MS.NoTrt_B |
| PI16 | 0.22509 | -0.3714 | 0.133 | 0.17 | 1 | MS.Trt_vs_MS.NoTrt_B |
| DNAJC3 | 0.22524 | 0.91838 | 0.42 | 0.5 | 1 | MS.Trt_vs_MS.NoTrt_B |
| PRKRIP1 | 0.22532 | 0.75889 | 0.074 | 0.1 | 1 | MS.Trt_vs_MS.NoTrt_B |
| TTC3 | 0.22564 | 0.49126 | 0.645 | 0.73 | 1 | MS.Trt_vs_MS.NoTrt_B |
| SPN | 0.22601 | -0.3327 | 0.932 | 0.98 | 1 | MS.Trt_vs_MS.NoTrt_B |
| ZNF518A | 0.22604 | 0.27592 | 0.092 | 0.12 | 1 | MS.Trt_vs_MS.NoTrt_B |
| PPP1R1C | 0.22614 | 0.26579 | 0.203 | 0.25 | 1 | MS.Trt_vs_MS.NoTrt_B |
| SLC25A30 | 0.22628 | 0.28992 | 0.51 | 0.57 | 1 | MS.Trt_vs_MS.NoTrt_B |
| SIRT1 | 0.22665 | 1.01052 | 0.074 | 0.1 | 1 | MS.Trt_vs_MS.NoTrt_B |
| RABEP1 | 0.22715 | 0.57883 | 0.148 | 0.2 | 1 | MS.Trt_vs_MS.NoTrt_B |
| LPAR6 | 0.22717 | 0.62051 | 0.338 | 0.38 | 1 | MS.Trt_vs_MS.NoTrt_B |
| MIGA1 | 0.22719 | 0.62338 | 0.377 | 0.45 | 1 | MS.Trt_vs_MS.NoTrt_B |
| POLR2B | 0.22741 | 0.44738 | 0.438 | 0.48 | 1 | MS.Trt_vs_MS.NoTrt_B |
| C16orf45 | 0.22754 | 0.50955 | 0.146 | 0.18 | 1 | MS.Trt_vs_MS.NoTrt_B |
| TNFSF13B | 0.2276 | 0.46621 | 0.449 | 0.52 | 1 | MS.Trt_vs_MS.NoTrt_B |
| TBC1D3I | 0.22871 | -0.3156 | 0.157 | 0.2 | 1 | MS.Trt_vs_MS.NoTrt_B |
| MTFR1 | 0.22953 | 0.37363 | 0.111 | 0.15 | 1 | MS.Trt_vs_MS.NoTrt_B |
| MCMBP | 0.22974 | 1.22233 | 0.355 | 0.41 | 1 | MS.Trt_vs_MS.NoTrt_B |
| ADAMTS16 | 0.23002 | 0.34757 | 0.222 | 0.29 | 1 | MS.Trt_vs_MS.NoTrt_B |
| ZNF316 | 0.23019 | 1.08545 | 0.427 | 0.38 | 1 | MS.Trt_vs_MS.NoTrt_B |

| | | | | | | |
|---|---|---|---|---|---|---|
| CHST3 | 0.23021 | -0.4167 | 0.118 | 0.16 | 1 | MS.Trt_vs_MS.NoTrt_B |
| DHX37 | 0.23064 | -0.4431 | 0.349 | 0.41 | 1 | MS.Trt_vs_MS.NoTrt_B |
| SKP2 | 0.23133 | 0.34541 | 0.41 | 0.47 | 1 | MS.Trt_vs_MS.NoTrt_B |
| MORC3 | 0.23186 | 0.63747 | 0.403 | 0.44 | 1 | MS.Trt_vs_MS.NoTrt_B |
| FLOT1 | 0.23228 | 0.66515 | 0.12 | 0.09 | 1 | MS.Trt_vs_MS.NoTrt_B |
| IBA57 | 0.23284 | 1.12879 | 0.647 | 0.73 | 1 | MS.Trt_vs_MS.NoTrt_B |
| TMEM104 | 0.23373 | 0.35431 | 0.107 | 0.14 | 1 | MS.Trt_vs_MS.NoTrt_B |
| TNS2 | 0.23382 | 0.25392 | 0.257 | 0.31 | 1 | MS.Trt_vs_MS.NoTrt_B |
| SAP130 | 0.23392 | 0.66508 | 0.118 | 0.16 | 1 | MS.Trt_vs_MS.NoTrt_B |
| TMEM11 | 0.23403 | -0.2957 | 0.085 | 0.11 | 1 | MS.Trt_vs_MS.NoTrt_B |
| RNF13 | 0.23444 | 0.79226 | 0.131 | 0.17 | 1 | MS.Trt_vs_MS.NoTrt_B |
| AP3M1 | 0.23525 | -0.3562 | 0.407 | 0.44 | 1 | MS.Trt_vs_MS.NoTrt_B |
| TRIM61 | 0.23534 | -0.6524 | 0.174 | 0.21 | 1 | MS.Trt_vs_MS.NoTrt_B |
| AGO4 | 0.23631 | -0.5483 | 0.24 | 0.28 | 1 | MS.Trt_vs_MS.NoTrt_B |
| TLE2 | 0.23638 | 0.76969 | 0.375 | 0.44 | 1 | MS.Trt_vs_MS.NoTrt_B |
| ERBB3 | 0.23705 | -0.4448 | 0.654 | 0.68 | 1 | MS.Trt_vs_MS.NoTrt_B |
| UGT1A1 | 0.2373 | 0.39819 | 0.473 | 0.57 | 1 | MS.Trt_vs_MS.NoTrt_B |
| DNAH2 | 0.23764 | 0.33372 | 0.51 | 0.63 | 1 | MS.Trt_vs_MS.NoTrt_B |
| DEDD | 0.23799 | 0.32035 | 0.135 | 0.17 | 1 | MS.Trt_vs_MS.NoTrt_B |
| APOBR | 0.238 | 1.3632 | 0.083 | 0.11 | 1 | MS.Trt_vs_MS.NoTrt_B |
| C11orf54 | 0.23828 | 0.3656 | 0.732 | 0.76 | 1 | MS.Trt_vs_MS.NoTrt_B |
| KIAA1210 | 0.23907 | -0.3061 | 0.198 | 0.23 | 1 | MS.Trt_vs_MS.NoTrt_B |
| KIF20B | 0.23916 | 0.55811 | 0.111 | 0.15 | 1 | MS.Trt_vs_MS.NoTrt_B |
| IFITM1 | 0.23919 | 0.28173 | 0.54 | 0.63 | 1 | MS.Trt_vs_MS.NoTrt_B |
| AHI1 | 0.24012 | 0.38957 | 0.27 | 0.32 | 1 | MS.Trt_vs_MS.NoTrt_B |
| RP11-111M: | 0.24042 | 0.27456 | 0.778 | 0.88 | 1 | MS.Trt_vs_MS.NoTrt_B |
| GPR108 | 0.24047 | 0.55429 | 0.157 | 0.2 | 1 | MS.Trt_vs_MS.NoTrt_B |
| RAB43 | 0.24085 | 0.40069 | 0.157 | 0.2 | 1 | MS.Trt_vs_MS.NoTrt_B |
| TMEM185A | 0.24262 | 0.86757 | 0.386 | 0.46 | 1 | MS.Trt_vs_MS.NoTrt_B |
| GFOD1 | 0.24319 | -0.3365 | 0.497 | 0.53 | 1 | MS.Trt_vs_MS.NoTrt_B |
| NECAP2 | 0.24446 | 0.30739 | 0.399 | 0.47 | 1 | MS.Trt_vs_MS.NoTrt_B |
| MSMO1 | 0.24479 | 0.47024 | 0.442 | 0.52 | 1 | MS.Trt_vs_MS.NoTrt_B |
| SLC31A2 | 0.24488 | -0.4812 | 0.085 | 0.11 | 1 | MS.Trt_vs_MS.NoTrt_B |
| SERPINB9 | 0.2453 | 0.61293 | 0.693 | 0.8 | 1 | MS.Trt_vs_MS.NoTrt_B |
| RWDD2A | 0.24587 | -0.5969 | 0.349 | 0.41 | 1 | MS.Trt_vs_MS.NoTrt_B |
| TMA7 | 0.2459 | 0.76557 | 0.527 | 0.73 | 1 | MS.Trt_vs_MS.NoTrt_B |
| ZNF532 | 0.24623 | -0.2787 | 0.874 | 0.96 | 1 | MS.Trt_vs_MS.NoTrt_B |
| CD163 | 0.24732 | 0.36148 | 0.562 | 0.74 | 1 | MS.Trt_vs_MS.NoTrt_B |
| TMEM216 | 0.24734 | 0.45916 | 0.288 | 0.34 | 1 | MS.Trt_vs_MS.NoTrt_B |
| CD24 | 0.24735 | 0.26504 | 0.253 | 0.3 | 1 | MS.Trt_vs_MS.NoTrt_B |

| | | | | | | |
|---|---|---|---|---|---|---|
| DGCR8 | 0.24776 | -0.7752 | 0.081 | 0.11 | 1 | MS.Trt_vs_MS.NoTrt_B |
| TP53 | 0.24782 | 0.54775 | 0.741 | 0.84 | 1 | MS.Trt_vs_MS.NoTrt_B |
| AXIN1 | 0.24828 | 0.52645 | 0.074 | 0.1 | 1 | MS.Trt_vs_MS.NoTrt_B |
| KMT5B | 0.2491 | -0.4867 | 0.205 | 0.24 | 1 | MS.Trt_vs_MS.NoTrt_B |
| MALT1 | 0.24963 | 0.34794 | 0.078 | 0.11 | 1 | MS.Trt_vs_MS.NoTrt_B |
| CD69 | 0.24979 | 0.28913 | 0.148 | 0.18 | 1 | MS.Trt_vs_MS.NoTrt_B |
| ENPP3 | 0.24983 | 0.43467 | 0.098 | 0.13 | 1 | MS.Trt_vs_MS.NoTrt_B |
| BIN3 | 0.24984 | 0.57821 | 0.301 | 0.37 | 1 | MS.Trt_vs_MS.NoTrt_B |
| SCLY | 0.24994 | 0.4005 | 0.885 | 0.96 | 1 | MS.Trt_vs_MS.NoTrt_B |
| LCOR | 0.2501 | 0.35693 | 0.322 | 0.37 | 1 | MS.Trt_vs_MS.NoTrt_B |
| CLIP2 | 0.25038 | 0.45981 | 0.255 | 0.32 | 1 | MS.Trt_vs_MS.NoTrt_B |
| WNK3 | 0.25157 | 0.85963 | 0.222 | 0.27 | 1 | MS.Trt_vs_MS.NoTrt_B |
| TELO2 | 0.2516 | 0.33164 | 0.078 | 0.11 | 1 | MS.Trt_vs_MS.NoTrt_B |
| CRNKL1 | 0.25167 | -0.3255 | 0.081 | 0.11 | 1 | MS.Trt_vs_MS.NoTrt_B |
| PGD | 0.25172 | -0.2996 | 0.1 | 0.13 | 1 | MS.Trt_vs_MS.NoTrt_B |
| CDC73 | 0.25235 | 0.73703 | 0.473 | 0.54 | 1 | MS.Trt_vs_MS.NoTrt_B |
| CREBZF | 0.25337 | 0.63803 | 0.153 | 0.19 | 1 | MS.Trt_vs_MS.NoTrt_B |
| NOG | 0.25398 | 2.1113 | 0.216 | 0.19 | 1 | MS.Trt_vs_MS.NoTrt_B |
| ELF4 | 0.25484 | 0.54624 | 0.111 | 0.15 | 1 | MS.Trt_vs_MS.NoTrt_B |
| ARFGAP1 | 0.25495 | -0.2678 | 0.092 | 0.12 | 1 | MS.Trt_vs_MS.NoTrt_B |
| MFSD9 | 0.25575 | -1.0135 | 0.418 | 0.47 | 1 | MS.Trt_vs_MS.NoTrt_B |
| HSPG2 | 0.25584 | -0.7849 | 0.307 | 0.26 | 1 | MS.Trt_vs_MS.NoTrt_B |
| F5 | 0.25645 | 0.49404 | 0.192 | 0.24 | 1 | MS.Trt_vs_MS.NoTrt_B |
| OVCA2 | 0.25677 | 0.61636 | 0.092 | 0.12 | 1 | MS.Trt_vs_MS.NoTrt_B |
| PARP2 | 0.25683 | -0.2602 | 0.292 | 0.34 | 1 | MS.Trt_vs_MS.NoTrt_B |
| HTN3 | 0.25761 | 0.35096 | 0.486 | 0.58 | 1 | MS.Trt_vs_MS.NoTrt_B |
| 6-Mar | 0.25786 | 0.75458 | 0.732 | 0.85 | 1 | MS.Trt_vs_MS.NoTrt_B |
| CD47 | 0.25824 | 0.56556 | 0.854 | 0.91 | 1 | MS.Trt_vs_MS.NoTrt_B |
| ZSWIM7 | 0.25827 | 0.32227 | 0.682 | 0.76 | 1 | MS.Trt_vs_MS.NoTrt_B |
| CAMK1 | 0.25882 | 0.37011 | 0.475 | 0.56 | 1 | MS.Trt_vs_MS.NoTrt_B |
| SEC62 | 0.25981 | 0.50728 | 0.379 | 0.44 | 1 | MS.Trt_vs_MS.NoTrt_B |
| CTD-2006C1 | 0.26062 | -0.3571 | 0.22 | 0.26 | 1 | MS.Trt_vs_MS.NoTrt_B |
| CCDC90B | 0.26104 | 0.35204 | 0.865 | 0.97 | 1 | MS.Trt_vs_MS.NoTrt_B |
| LCAT | 0.26134 | 0.30177 | 0.157 | 0.2 | 1 | MS.Trt_vs_MS.NoTrt_B |
| TMEM259 | 0.26203 | 0.56309 | 0.081 | 0.11 | 1 | MS.Trt_vs_MS.NoTrt_B |
| RP1L1 | 0.26246 | 0.25358 | 0.597 | 0.52 | 1 | MS.Trt_vs_MS.NoTrt_B |
| IFT74 | 0.2626 | 0.47001 | 0.314 | 0.36 | 1 | MS.Trt_vs_MS.NoTrt_B |
| CALD1 | 0.26361 | -0.9368 | 0.139 | 0.18 | 1 | MS.Trt_vs_MS.NoTrt_B |
| U2AF2 | 0.26467 | 0.25065 | 0.133 | 0.17 | 1 | MS.Trt_vs_MS.NoTrt_B |
| RAB5A | 0.26524 | 0.86196 | 0.109 | 0.14 | 1 | MS.Trt_vs_MS.NoTrt_B |

| | | | | | | |
|---|---|---|---|---|---|---|
| CHD7 | 0.26545 | 2.36033 | 0.397 | 0.51 | 1 | MS.Trt_vs_MS.NoTrt_B |
| SDHB | 0.26572 | 0.73301 | 0.12 | 0.16 | 1 | MS.Trt_vs_MS.NoTrt_B |
| PRDX2 | 0.26623 | 0.36098 | 0.216 | 0.26 | 1 | MS.Trt_vs_MS.NoTrt_B |
| CPNE3 | 0.26651 | 0.36043 | 0.196 | 0.23 | 1 | MS.Trt_vs_MS.NoTrt_B |
| TBPL1 | 0.26661 | 0.74238 | 0.179 | 0.22 | 1 | MS.Trt_vs_MS.NoTrt_B |
| NBPF3 | 0.26677 | 0.85191 | 0.377 | 0.44 | 1 | MS.Trt_vs_MS.NoTrt_B |
| RPUSD3 | 0.26757 | 0.31585 | 0.695 | 0.77 | 1 | MS.Trt_vs_MS.NoTrt_B |
| C9orf85 | 0.26781 | 0.28013 | 0.797 | 0.84 | 1 | MS.Trt_vs_MS.NoTrt_B |
| ARF3 | 0.26782 | 0.27448 | 0.126 | 0.16 | 1 | MS.Trt_vs_MS.NoTrt_B |
| TRAF3IP1 | 0.26817 | 0.40123 | 0.292 | 0.35 | 1 | MS.Trt_vs_MS.NoTrt_B |
| MRPS30 | 0.26817 | 0.66528 | 0.259 | 0.32 | 1 | MS.Trt_vs_MS.NoTrt_B |
| AFMID | 0.26935 | 0.28971 | 0.869 | 0.9 | 1 | MS.Trt_vs_MS.NoTrt_B |
| SMN2 | 0.26955 | 0.73239 | 0.804 | 0.7 | 1 | MS.Trt_vs_MS.NoTrt_B |
| TAF6 | 0.26962 | -0.4378 | 0.473 | 0.54 | 1 | MS.Trt_vs_MS.NoTrt_B |
| NDRG4 | 0.26988 | 0.34998 | 0.105 | 0.14 | 1 | MS.Trt_vs_MS.NoTrt_B |
| MEGF8 | 0.27013 | 0.54587 | 0.318 | 0.38 | 1 | MS.Trt_vs_MS.NoTrt_B |
| ZNF133 | 0.27027 | -1.1438 | 0.344 | 0.41 | 1 | MS.Trt_vs_MS.NoTrt_B |
| SLF2 | 0.27031 | 0.308 | 0.434 | 0.49 | 1 | MS.Trt_vs_MS.NoTrt_B |
| FAM69A | 0.27129 | 0.97751 | 0.094 | 0.12 | 1 | MS.Trt_vs_MS.NoTrt_B |
| CHM | 0.27133 | 0.53178 | 0.41 | 0.44 | 1 | MS.Trt_vs_MS.NoTrt_B |
| H6PD | 0.27234 | 0.86174 | 0.684 | 0.83 | 1 | MS.Trt_vs_MS.NoTrt_B |
| ZMYM1 | 0.27386 | 0.54037 | 0.453 | 0.51 | 1 | MS.Trt_vs_MS.NoTrt_B |
| SRRD | 0.27398 | 0.48769 | 0.322 | 0.4 | 1 | MS.Trt_vs_MS.NoTrt_B |
| UBR5 | 0.27403 | 0.35991 | 0.697 | 0.81 | 1 | MS.Trt_vs_MS.NoTrt_B |
| FGFR4 | 0.2742 | 0.25535 | 0.312 | 0.36 | 1 | MS.Trt_vs_MS.NoTrt_B |
| CFAP36 | 0.27506 | 0.45454 | 0.322 | 0.38 | 1 | MS.Trt_vs_MS.NoTrt_B |
| LARP7 | 0.27563 | 0.8535 | 0.185 | 0.23 | 1 | MS.Trt_vs_MS.NoTrt_B |
| GPAA1 | 0.27602 | 0.45538 | 0.131 | 0.17 | 1 | MS.Trt_vs_MS.NoTrt_B |
| UTP14C | 0.27617 | 0.35557 | 0.133 | 0.17 | 1 | MS.Trt_vs_MS.NoTrt_B |
| NLK | 0.27631 | 1.02182 | 0.231 | 0.27 | 1 | MS.Trt_vs_MS.NoTrt_B |
| SETD4 | 0.27637 | 0.6775 | 0.423 | 0.48 | 1 | MS.Trt_vs_MS.NoTrt_B |
| PALLD | 0.27695 | -0.5339 | 0.257 | 0.3 | 1 | MS.Trt_vs_MS.NoTrt_B |
| BLOC1S5 | 0.27704 | 0.34236 | 0.523 | 0.63 | 1 | MS.Trt_vs_MS.NoTrt_B |
| FAM134A | 0.27707 | 0.71425 | 0.386 | 0.44 | 1 | MS.Trt_vs_MS.NoTrt_B |
| MT-ATP6 | 0.27715 | 0.28695 | 0.797 | 0.93 | 1 | MS.Trt_vs_MS.NoTrt_B |
| APOL1 | 0.27716 | 0.29023 | 0.588 | 0.65 | 1 | MS.Trt_vs_MS.NoTrt_B |
| HOOK2 | 0.27721 | 0.43745 | 0.911 | 0.96 | 1 | MS.Trt_vs_MS.NoTrt_B |
| CHP1 | 0.27724 | 0.72516 | 0.675 | 0.77 | 1 | MS.Trt_vs_MS.NoTrt_B |
| PITPNB | 0.27764 | 0.6346 | 0.166 | 0.21 | 1 | MS.Trt_vs_MS.NoTrt_B |
| IFI44L | 0.27851 | -0.5344 | 0.133 | 0.16 | 1 | MS.Trt_vs_MS.NoTrt_B |

| | | | | | | |
|---|---|---|---|---|---|---|
| SLC9A6 | 0.27865 | 0.44696 | 0.294 | 0.34 | 1 | MS.Trt_vs_MS.NoTrt_B |
| MEMO1 | 0.27904 | 0.53761 | 0.27 | 0.3 | 1 | MS.Trt_vs_MS.NoTrt_B |
| GNB1L | 0.27914 | 0.4471 | 0.15 | 0.18 | 1 | MS.Trt_vs_MS.NoTrt_B |
| GOLPH3 | 0.27928 | 0.30008 | 0.124 | 0.16 | 1 | MS.Trt_vs_MS.NoTrt_B |
| FLNB | 0.27933 | -0.5031 | 0.41 | 0.47 | 1 | MS.Trt_vs_MS.NoTrt_B |
| PIPOX | 0.27948 | 0.31879 | 0.451 | 0.52 | 1 | MS.Trt_vs_MS.NoTrt_B |
| SIM1 | 0.28034 | 0.67073 | 0.148 | 0.18 | 1 | MS.Trt_vs_MS.NoTrt_B |
| LRP8 | 0.28038 | 0.41559 | 0.381 | 0.47 | 1 | MS.Trt_vs_MS.NoTrt_B |
| FIGNL1 | 0.28042 | -0.8171 | 0.105 | 0.14 | 1 | MS.Trt_vs_MS.NoTrt_B |
| DUSP7 | 0.2806 | 0.25438 | 0.087 | 0.11 | 1 | MS.Trt_vs_MS.NoTrt_B |
| ZNF625 | 0.28087 | -0.3046 | 0.216 | 0.25 | 1 | MS.Trt_vs_MS.NoTrt_B |
| CD3EAP | 0.28094 | 0.50793 | 0.784 | 0.71 | 1 | MS.Trt_vs_MS.NoTrt_B |
| PSMG1 | 0.28129 | 0.52078 | 0.113 | 0.15 | 1 | MS.Trt_vs_MS.NoTrt_B |
| NOP2 | 0.28231 | 0.63386 | 0.331 | 0.38 | 1 | MS.Trt_vs_MS.NoTrt_B |
| KLHL9 | 0.28299 | 0.92786 | 0.277 | 0.34 | 1 | MS.Trt_vs_MS.NoTrt_B |
| AGPAT1 | 0.28361 | -0.2648 | 0.083 | 0.11 | 1 | MS.Trt_vs_MS.NoTrt_B |
| SLC39A11 | 0.28369 | 0.65938 | 0.209 | 0.25 | 1 | MS.Trt_vs_MS.NoTrt_B |
| CYC1 | 0.28429 | 0.49935 | 0.113 | 0.15 | 1 | MS.Trt_vs_MS.NoTrt_B |
| TCEB3 | 0.28606 | 0.68448 | 0.076 | 0.1 | 1 | MS.Trt_vs_MS.NoTrt_B |
| TMSB15B | 0.28631 | -0.4326 | 0.22 | 0.27 | 1 | MS.Trt_vs_MS.NoTrt_B |
| EXOSC6 | 0.28736 | 0.4349 | 0.815 | 0.88 | 1 | MS.Trt_vs_MS.NoTrt_B |
| NSUN5 | 0.28807 | 1.33203 | 0.279 | 0.35 | 1 | MS.Trt_vs_MS.NoTrt_B |
| CTRL | 0.28811 | 0.49693 | 0.096 | 0.12 | 1 | MS.Trt_vs_MS.NoTrt_B |
| PSMD1 | 0.29048 | 0.32001 | 0.15 | 0.18 | 1 | MS.Trt_vs_MS.NoTrt_B |
| GGT1 | 0.29061 | -0.7112 | 0.083 | 0.11 | 1 | MS.Trt_vs_MS.NoTrt_B |
| FUK | 0.29145 | -0.3411 | 0.434 | 0.49 | 1 | MS.Trt_vs_MS.NoTrt_B |
| ZNF302 | 0.29166 | 0.76205 | 0.418 | 0.35 | 1 | MS.Trt_vs_MS.NoTrt_B |
| PIGW | 0.29222 | -0.5453 | 0.41 | 0.38 | 1 | MS.Trt_vs_MS.NoTrt_B |
| ZNF708 | 0.29224 | 0.72607 | 0.68 | 0.75 | 1 | MS.Trt_vs_MS.NoTrt_B |
| HAVCR2 | 0.29382 | 0.5387 | 0.429 | 0.52 | 1 | MS.Trt_vs_MS.NoTrt_B |
| PIAS3 | 0.29429 | 0.29687 | 0.111 | 0.14 | 1 | MS.Trt_vs_MS.NoTrt_B |
| NEK7 | 0.29503 | 0.62233 | 0.087 | 0.11 | 1 | MS.Trt_vs_MS.NoTrt_B |
| RUNDC1 | 0.29506 | 0.38903 | 0.625 | 0.69 | 1 | MS.Trt_vs_MS.NoTrt_B |
| LSM1 | 0.29538 | 0.27319 | 0.092 | 0.12 | 1 | MS.Trt_vs_MS.NoTrt_B |
| PER2 | 0.29552 | 0.62231 | 0.401 | 0.47 | 1 | MS.Trt_vs_MS.NoTrt_B |
| ADAMTSL3 | 0.29574 | 0.29543 | 0.863 | 0.98 | 1 | MS.Trt_vs_MS.NoTrt_B |
| PSD | 0.29605 | 0.32846 | 0.078 | 0.11 | 1 | MS.Trt_vs_MS.NoTrt_B |
| KCTD9 | 0.29615 | -0.4792 | 0.146 | 0.18 | 1 | MS.Trt_vs_MS.NoTrt_B |
| TSPYL1 | 0.29623 | 0.64028 | 0.471 | 0.56 | 1 | MS.Trt_vs_MS.NoTrt_B |
| FAM107B | 0.29773 | 0.57986 | 0.272 | 0.32 | 1 | MS.Trt_vs_MS.NoTrt_B |

| | | | | | | |
|---|---|---|---|---|---|---|
| VAMP4 | 0.29795 | 0.93474 | 0.139 | 0.17 | 1 | MS.Trt_vs_MS.NoTrt_B |
| HIST2H4B | 0.29969 | 0.7075 | 0.118 | 0.15 | 1 | MS.Trt_vs_MS.NoTrt_B |
| TFEB | 0.29998 | -0.7781 | 0.388 | 0.43 | 1 | MS.Trt_vs_MS.NoTrt_B |
| GPX2 | 0.3018 | -0.6988 | 0.168 | 0.2 | 1 | MS.Trt_vs_MS.NoTrt_B |
| CAAP1 | 0.3035 | 0.93013 | 0.081 | 0.11 | 1 | MS.Trt_vs_MS.NoTrt_B |
| RP11-500M8 | 0.30416 | 1.11045 | 0.102 | 0.14 | 1 | MS.Trt_vs_MS.NoTrt_B |
| COX15 | 0.30506 | 0.30254 | 0.464 | 0.51 | 1 | MS.Trt_vs_MS.NoTrt_B |
| MYH7B | 0.30591 | -0.3385 | 0.678 | 0.78 | 1 | MS.Trt_vs_MS.NoTrt_B |
| FUT3 | 0.3072 | 0.28679 | 0.174 | 0.22 | 1 | MS.Trt_vs_MS.NoTrt_B |
| POP5 | 0.3075 | 0.5361 | 0.551 | 0.62 | 1 | MS.Trt_vs_MS.NoTrt_B |
| CTCF | 0.30763 | 0.29619 | 0.098 | 0.12 | 1 | MS.Trt_vs_MS.NoTrt_B |
| ITPRIPL2 | 0.30891 | 0.29058 | 0.492 | 0.55 | 1 | MS.Trt_vs_MS.NoTrt_B |
| NCOR2 | 0.30908 | 0.33323 | 0.264 | 0.31 | 1 | MS.Trt_vs_MS.NoTrt_B |
| BUB1B | 0.31018 | -0.3473 | 0.298 | 0.34 | 1 | MS.Trt_vs_MS.NoTrt_B |
| TMEM229B | 0.31032 | 0.91285 | 0.155 | 0.19 | 1 | MS.Trt_vs_MS.NoTrt_B |
| CYP17A1 | 0.3105 | 0.35141 | 0.207 | 0.18 | 1 | MS.Trt_vs_MS.NoTrt_B |
| INSIG2 | 0.31083 | 0.27888 | 0.078 | 0.1 | 1 | MS.Trt_vs_MS.NoTrt_B |
| ZNF718 | 0.31215 | 0.37649 | 0.198 | 0.24 | 1 | MS.Trt_vs_MS.NoTrt_B |
| ARFIP2 | 0.31307 | -0.5896 | 0.373 | 0.42 | 1 | MS.Trt_vs_MS.NoTrt_B |
| PAXBP1 | 0.31395 | 0.81603 | 0.137 | 0.17 | 1 | MS.Trt_vs_MS.NoTrt_B |
| ANKRD36B | 0.31405 | 0.44194 | 0.577 | 0.64 | 1 | MS.Trt_vs_MS.NoTrt_B |
| SNX3 | 0.31467 | 0.9487 | 0.109 | 0.14 | 1 | MS.Trt_vs_MS.NoTrt_B |
| PRIM1 | 0.31508 | 0.28496 | 0.12 | 0.15 | 1 | MS.Trt_vs_MS.NoTrt_B |
| PKN2 | 0.31518 | 0.42774 | 0.275 | 0.32 | 1 | MS.Trt_vs_MS.NoTrt_B |
| ZNF726 | 0.31584 | 0.56754 | 0.739 | 0.84 | 1 | MS.Trt_vs_MS.NoTrt_B |
| NIPSNAP1 | 0.31617 | -0.2845 | 0.316 | 0.35 | 1 | MS.Trt_vs_MS.NoTrt_B |
| SPP2 | 0.31678 | 0.31446 | 0.569 | 0.68 | 1 | MS.Trt_vs_MS.NoTrt_B |
| THEM4 | 0.31693 | 0.43641 | 0.717 | 0.81 | 1 | MS.Trt_vs_MS.NoTrt_B |
| RPP21 | 0.31731 | 0.50993 | 0.124 | 0.16 | 1 | MS.Trt_vs_MS.NoTrt_B |
| NCOA7 | 0.31754 | 0.32455 | 0.636 | 0.74 | 1 | MS.Trt_vs_MS.NoTrt_B |
| PRMT8 | 0.31785 | 0.81354 | 0.17 | 0.21 | 1 | MS.Trt_vs_MS.NoTrt_B |
| CXCR4 | 0.31789 | 0.83418 | 0.142 | 0.11 | 1 | MS.Trt_vs_MS.NoTrt_B |
| ATP7A | 0.31838 | 0.76443 | 0.314 | 0.28 | 1 | MS.Trt_vs_MS.NoTrt_B |
| NUDT14 | 0.32024 | 0.76988 | 0.379 | 0.48 | 1 | MS.Trt_vs_MS.NoTrt_B |
| PKD1 | 0.32025 | 1.45174 | 0.333 | 0.38 | 1 | MS.Trt_vs_MS.NoTrt_B |
| FN1 | 0.32121 | 0.89676 | 0.309 | 0.36 | 1 | MS.Trt_vs_MS.NoTrt_B |
| ZNF385D | 0.32134 | 0.26646 | 0.242 | 0.29 | 1 | MS.Trt_vs_MS.NoTrt_B |
| MSL2 | 0.3222 | 0.51934 | 0.109 | 0.14 | 1 | MS.Trt_vs_MS.NoTrt_B |
| SGPP2 | 0.32277 | 0.67585 | 0.253 | 0.28 | 1 | MS.Trt_vs_MS.NoTrt_B |
| DOCK6 | 0.32356 | 0.37642 | 0.451 | 0.55 | 1 | MS.Trt_vs_MS.NoTrt_B |

| | | | | | | |
|---|---|---|---|---|---|---|
| COQ10A | 0.32361 | 0.87358 | 0.401 | 0.47 | 1 | MS.Trt_vs_MS.NoTrt_B |
| TLE3 | 0.32391 | 0.88676 | 0.153 | 0.19 | 1 | MS.Trt_vs_MS.NoTrt_B |
| CCDC93 | 0.32472 | -0.5883 | 0.1 | 0.12 | 1 | MS.Trt_vs_MS.NoTrt_B |
| ZNF439 | 0.32542 | 0.69996 | 0.549 | 0.58 | 1 | MS.Trt_vs_MS.NoTrt_B |
| NPIPB6 | 0.32543 | -0.3166 | 0.166 | 0.2 | 1 | MS.Trt_vs_MS.NoTrt_B |
| MLF1 | 0.32712 | 0.82396 | 0.253 | 0.29 | 1 | MS.Trt_vs_MS.NoTrt_B |
| ZNF780B | 0.32811 | 0.65351 | 0.532 | 0.6 | 1 | MS.Trt_vs_MS.NoTrt_B |
| STK25 | 0.32838 | 0.40658 | 0.601 | 0.71 | 1 | MS.Trt_vs_MS.NoTrt_B |
| STAU1 | 0.32971 | 0.36595 | 0.325 | 0.36 | 1 | MS.Trt_vs_MS.NoTrt_B |
| ESF1 | 0.32972 | 0.49206 | 0.094 | 0.12 | 1 | MS.Trt_vs_MS.NoTrt_B |
| SLC39A9 | 0.33043 | -0.2883 | 0.124 | 0.15 | 1 | MS.Trt_vs_MS.NoTrt_B |
| CADM1 | 0.33057 | 1.29772 | 0.2 | 0.17 | 1 | MS.Trt_vs_MS.NoTrt_B |
| ATP5L2 | 0.33183 | 0.32185 | 0.255 | 0.23 | 1 | MS.Trt_vs_MS.NoTrt_B |
| DICER1 | 0.33193 | 1.29329 | 0.131 | 0.16 | 1 | MS.Trt_vs_MS.NoTrt_B |
| SPATA31A7 | 0.33353 | 0.28241 | 0.425 | 0.45 | 1 | MS.Trt_vs_MS.NoTrt_B |
| PHPT1 | 0.33427 | 0.41165 | 0.109 | 0.14 | 1 | MS.Trt_vs_MS.NoTrt_B |
| ITPR3 | 0.33459 | 0.55938 | 0.118 | 0.15 | 1 | MS.Trt_vs_MS.NoTrt_B |
| TOP2B | 0.33483 | 0.38616 | 0.198 | 0.23 | 1 | MS.Trt_vs_MS.NoTrt_B |
| TONSL | 0.33624 | 0.45531 | 0.54 | 0.62 | 1 | MS.Trt_vs_MS.NoTrt_B |
| SMUG1 | 0.33624 | 0.65606 | 0.582 | 0.7 | 1 | MS.Trt_vs_MS.NoTrt_B |
| SF3A2 | 0.33625 | 0.59421 | 0.296 | 0.34 | 1 | MS.Trt_vs_MS.NoTrt_B |
| MIER1 | 0.3364 | 1.03841 | 0.597 | 0.75 | 1 | MS.Trt_vs_MS.NoTrt_B |
| FRK | 0.33663 | 0.29573 | 0.946 | 1 | 1 | MS.Trt_vs_MS.NoTrt_B |
| TM9SF3 | 0.33796 | 0.9687 | 0.107 | 0.14 | 1 | MS.Trt_vs_MS.NoTrt_B |
| TOR1B | 0.33835 | 1.23549 | 0.144 | 0.11 | 1 | MS.Trt_vs_MS.NoTrt_B |
| NOTCH2NL | 0.33861 | 0.65237 | 0.146 | 0.18 | 1 | MS.Trt_vs_MS.NoTrt_B |
| HSPE1 | 0.33929 | 0.96845 | 0.242 | 0.29 | 1 | MS.Trt_vs_MS.NoTrt_B |
| ZNF688 | 0.33957 | 0.93685 | 0.479 | 0.42 | 1 | MS.Trt_vs_MS.NoTrt_B |
| NBPF10 | 0.33971 | 0.95003 | 0.2 | 0.17 | 1 | MS.Trt_vs_MS.NoTrt_B |
| ALDH1B1 | 0.34039 | 0.77745 | 0.495 | 0.59 | 1 | MS.Trt_vs_MS.NoTrt_B |
| COMMD8 | 0.34059 | 0.54068 | 0.083 | 0.11 | 1 | MS.Trt_vs_MS.NoTrt_B |
| DCAF1 | 0.34061 | 0.46319 | 0.17 | 0.21 | 1 | MS.Trt_vs_MS.NoTrt_B |
| AKR1B1 | 0.34062 | 0.38085 | 0.669 | 0.75 | 1 | MS.Trt_vs_MS.NoTrt_B |
| GTF2H2 | 0.34124 | -0.2621 | 0.817 | 0.89 | 1 | MS.Trt_vs_MS.NoTrt_B |
| OSGEPL1 | 0.34131 | 0.67227 | 0.257 | 0.31 | 1 | MS.Trt_vs_MS.NoTrt_B |
| TTC38 | 0.34169 | -1.2052 | 0.359 | 0.42 | 1 | MS.Trt_vs_MS.NoTrt_B |
| FAM21C | 0.34182 | 0.51184 | 0.183 | 0.22 | 1 | MS.Trt_vs_MS.NoTrt_B |
| TMPRSS3 | 0.34194 | 0.83798 | 0.292 | 0.33 | 1 | MS.Trt_vs_MS.NoTrt_B |
| SCYL3 | 0.34196 | 0.57276 | 0.575 | 0.64 | 1 | MS.Trt_vs_MS.NoTrt_B |
| KIAA1683 | 0.34227 | 0.33743 | 0.497 | 0.61 | 1 | MS.Trt_vs_MS.NoTrt_B |

| | | | | | | |
|---|---|---|---|---|---|---|
| SLFN11 | 0.34341 | -0.4405 | 0.207 | 0.24 | 1 | MS.Trt_vs_MS.NoTrt_B |
| KPTN | 0.34444 | -0.3823 | 0.174 | 0.22 | 1 | MS.Trt_vs_MS.NoTrt_B |
| CCNG2 | 0.34517 | 0.77685 | 0.246 | 0.29 | 1 | MS.Trt_vs_MS.NoTrt_B |
| ELP4 | 0.34541 | 0.58761 | 0.438 | 0.5 | 1 | MS.Trt_vs_MS.NoTrt_B |
| NUP35 | 0.34569 | -0.3641 | 0.427 | 0.5 | 1 | MS.Trt_vs_MS.NoTrt_B |
| NDUFV3 | 0.34656 | 0.42908 | 0.904 | 0.97 | 1 | MS.Trt_vs_MS.NoTrt_B |
| CCT5 | 0.34694 | 0.3681 | 0.155 | 0.19 | 1 | MS.Trt_vs_MS.NoTrt_B |
| ZC3H10 | 0.34697 | 0.35425 | 0.416 | 0.48 | 1 | MS.Trt_vs_MS.NoTrt_B |
| PDPN | 0.34725 | 0.28879 | 0.436 | 0.42 | 1 | MS.Trt_vs_MS.NoTrt_B |
| TBCD | 0.34769 | 0.32565 | 0.159 | 0.19 | 1 | MS.Trt_vs_MS.NoTrt_B |
| MAGEH1 | 0.3478 | 0.29385 | 0.089 | 0.11 | 1 | MS.Trt_vs_MS.NoTrt_B |
| ARMC6 | 0.34808 | 0.31231 | 0.24 | 0.28 | 1 | MS.Trt_vs_MS.NoTrt_B |
| SBNO1 | 0.3484 | 0.3208 | 0.558 | 0.6 | 1 | MS.Trt_vs_MS.NoTrt_B |
| TBP | 0.34938 | 0.43366 | 0.312 | 0.35 | 1 | MS.Trt_vs_MS.NoTrt_B |
| ACP1 | 0.34967 | 0.70596 | 0.285 | 0.35 | 1 | MS.Trt_vs_MS.NoTrt_B |
| MBD6 | 0.35042 | 0.33825 | 0.153 | 0.18 | 1 | MS.Trt_vs_MS.NoTrt_B |
| QTRT2 | 0.35105 | -0.4091 | 0.658 | 0.79 | 1 | MS.Trt_vs_MS.NoTrt_B |
| RDH11 | 0.35108 | 0.34518 | 0.442 | 0.48 | 1 | MS.Trt_vs_MS.NoTrt_B |
| KMT2D | 0.35145 | 0.41732 | 0.741 | 0.83 | 1 | MS.Trt_vs_MS.NoTrt_B |
| NR3C1 | 0.35211 | 0.25893 | 0.168 | 0.2 | 1 | MS.Trt_vs_MS.NoTrt_B |
| COX10 | 0.35265 | 0.72428 | 0.094 | 0.12 | 1 | MS.Trt_vs_MS.NoTrt_B |
| GTF2E1 | 0.35288 | 0.35835 | 0.192 | 0.22 | 1 | MS.Trt_vs_MS.NoTrt_B |
| PEX7 | 0.35339 | -0.7819 | 0.12 | 0.15 | 1 | MS.Trt_vs_MS.NoTrt_B |
| DNAH17 | 0.35367 | 0.28756 | 0.246 | 0.28 | 1 | MS.Trt_vs_MS.NoTrt_B |
| KCNE4 | 0.3546 | 0.33756 | 0.734 | 0.87 | 1 | MS.Trt_vs_MS.NoTrt_B |
| TRIM46 | 0.35463 | -0.3652 | 0.375 | 0.44 | 1 | MS.Trt_vs_MS.NoTrt_B |
| CREM | 0.3562 | 0.86119 | 0.157 | 0.13 | 1 | MS.Trt_vs_MS.NoTrt_B |
| C19orf54 | 0.3563 | 0.34953 | 0.394 | 0.46 | 1 | MS.Trt_vs_MS.NoTrt_B |
| PTPN23 | 0.35631 | 1.10486 | 0.102 | 0.08 | 1 | MS.Trt_vs_MS.NoTrt_B |
| INF2 | 0.35672 | 0.77289 | 0.179 | 0.22 | 1 | MS.Trt_vs_MS.NoTrt_B |
| ITSN1 | 0.35762 | 0.73627 | 0.767 | 0.85 | 1 | MS.Trt_vs_MS.NoTrt_B |
| SLC4A7 | 0.35829 | 0.38545 | 0.264 | 0.3 | 1 | MS.Trt_vs_MS.NoTrt_B |
| C3orf49 | 0.35889 | 0.69849 | 0.259 | 0.3 | 1 | MS.Trt_vs_MS.NoTrt_B |
| SNAPC3 | 0.35897 | 0.4648 | 0.495 | 0.55 | 1 | MS.Trt_vs_MS.NoTrt_B |
| CCDC144A | 0.35944 | 0.86804 | 0.749 | 0.84 | 1 | MS.Trt_vs_MS.NoTrt_B |
| GARNL3 | 0.35952 | 0.75174 | 0.131 | 0.16 | 1 | MS.Trt_vs_MS.NoTrt_B |
| FANCA | 0.35974 | -0.327 | 0.253 | 0.28 | 1 | MS.Trt_vs_MS.NoTrt_B |
| MYL6B | 0.35981 | 0.62817 | 0.458 | 0.51 | 1 | MS.Trt_vs_MS.NoTrt_B |
| GAPVD1 | 0.36032 | 0.45028 | 0.73 | 0.88 | 1 | MS.Trt_vs_MS.NoTrt_B |
| SGCB | 0.36044 | 0.27599 | 0.216 | 0.24 | 1 | MS.Trt_vs_MS.NoTrt_B |

| | | | | | | |
|---|---|---|---|---|---|---|
| TNK2 | 0.3608 | 0.58563 | 0.085 | 0.11 | 1 | MS.Trt_vs_MS.NoTrt_B |
| DENND5A | 0.36109 | 1.8391 | 0.203 | 0.17 | 1 | MS.Trt_vs_MS.NoTrt_B |
| OR7A5 | 0.3621 | 0.2981 | 0.359 | 0.42 | 1 | MS.Trt_vs_MS.NoTrt_B |
| FAM117B | 0.36252 | 0.76217 | 0.331 | 0.37 | 1 | MS.Trt_vs_MS.NoTrt_B |
| XRCC3 | 0.36329 | 0.86494 | 0.342 | 0.38 | 1 | MS.Trt_vs_MS.NoTrt_B |
| MANBA | 0.36344 | 0.31326 | 0.124 | 0.15 | 1 | MS.Trt_vs_MS.NoTrt_B |
| APPL1 | 0.36363 | 1.18085 | 0.355 | 0.41 | 1 | MS.Trt_vs_MS.NoTrt_B |
| ZNF124 | 0.36577 | 0.76756 | 0.51 | 0.63 | 1 | MS.Trt_vs_MS.NoTrt_B |
| FMO3 | 0.36591 | 0.43882 | 0.505 | 0.51 | 1 | MS.Trt_vs_MS.NoTrt_B |
| ABCA8 | 0.36624 | 0.29625 | 0.353 | 0.32 | 1 | MS.Trt_vs_MS.NoTrt_B |
| CYP7B1 | 0.36657 | 0.33107 | 0.359 | 0.42 | 1 | MS.Trt_vs_MS.NoTrt_B |
| CAT | 0.367 | 0.27974 | 0.109 | 0.14 | 1 | MS.Trt_vs_MS.NoTrt_B |
| FAM19A2 | 0.36754 | 0.80554 | 0.185 | 0.22 | 1 | MS.Trt_vs_MS.NoTrt_B |
| DNAH7 | 0.36803 | 0.31086 | 0.558 | 0.61 | 1 | MS.Trt_vs_MS.NoTrt_B |
| HLA-DRB3 | 0.36832 | 0.92003 | 0.109 | 0.14 | 1 | MS.Trt_vs_MS.NoTrt_B |
| MBD1 | 0.36866 | 0.71776 | 0.168 | 0.14 | 1 | MS.Trt_vs_MS.NoTrt_B |
| SNCAIP | 0.36869 | 0.25778 | 0.331 | 0.4 | 1 | MS.Trt_vs_MS.NoTrt_B |
| KIAA0825 | 0.3687 | 0.29079 | 0.222 | 0.27 | 1 | MS.Trt_vs_MS.NoTrt_B |
| MRPL14 | 0.36922 | 0.55672 | 0.085 | 0.11 | 1 | MS.Trt_vs_MS.NoTrt_B |
| COG6 | 0.36929 | 0.42954 | 0.096 | 0.12 | 1 | MS.Trt_vs_MS.NoTrt_B |
| PFKFB2 | 0.36938 | 0.52501 | 0.157 | 0.2 | 1 | MS.Trt_vs_MS.NoTrt_B |
| TMEM256-P | 0.36942 | 0.46867 | 0.481 | 0.62 | 1 | MS.Trt_vs_MS.NoTrt_B |
| TRADD | 0.37174 | 0.25695 | 0.168 | 0.2 | 1 | MS.Trt_vs_MS.NoTrt_B |
| ZNF138 | 0.37177 | 1.27876 | 0.081 | 0.1 | 1 | MS.Trt_vs_MS.NoTrt_B |
| FSIP2 | 0.3731 | 0.32092 | 0.645 | 0.76 | 1 | MS.Trt_vs_MS.NoTrt_B |
| UBE2A | 0.37401 | 0.636 | 0.288 | 0.32 | 1 | MS.Trt_vs_MS.NoTrt_B |
| RBM45 | 0.37436 | 0.80521 | 0.081 | 0.1 | 1 | MS.Trt_vs_MS.NoTrt_B |
| ZRSR2 | 0.37453 | 0.46978 | 0.63 | 0.76 | 1 | MS.Trt_vs_MS.NoTrt_B |
| CYB561D1 | 0.37495 | -0.2594 | 0.325 | 0.37 | 1 | MS.Trt_vs_MS.NoTrt_B |
| PTAFR | 0.37564 | 0.26566 | 0.523 | 0.61 | 1 | MS.Trt_vs_MS.NoTrt_B |
| TIMM8B | 0.37571 | 0.52452 | 0.089 | 0.11 | 1 | MS.Trt_vs_MS.NoTrt_B |
| ECI2 | 0.37608 | 0.2696 | 0.519 | 0.58 | 1 | MS.Trt_vs_MS.NoTrt_B |
| ZMAT3 | 0.37614 | 0.76589 | 0.499 | 0.6 | 1 | MS.Trt_vs_MS.NoTrt_B |
| GPR161 | 0.37714 | -0.2936 | 0.368 | 0.42 | 1 | MS.Trt_vs_MS.NoTrt_B |
| TBC1D24 | 0.37838 | 0.96245 | 0.599 | 0.68 | 1 | MS.Trt_vs_MS.NoTrt_B |
| SLC1A4 | 0.37847 | 0.49539 | 0.495 | 0.61 | 1 | MS.Trt_vs_MS.NoTrt_B |
| MFSD11 | 0.3795 | -0.316 | 0.377 | 0.42 | 1 | MS.Trt_vs_MS.NoTrt_B |
| EXT2 | 0.3799 | -0.2945 | 0.111 | 0.14 | 1 | MS.Trt_vs_MS.NoTrt_B |
| TBL3 | 0.37999 | 0.32593 | 0.233 | 0.27 | 1 | MS.Trt_vs_MS.NoTrt_B |
| ITGB8 | 0.38012 | 0.69905 | 0.636 | 0.71 | 1 | MS.Trt_vs_MS.NoTrt_B |

| | | | | | | |
|---|---|---|---|---|---|---|
| DBNDD1 | 0.38019 | -0.5857 | 0.122 | 0.15 | 1 | MS.Trt_vs_MS.NoTrt_B |
| PRKD3 | 0.38051 | 0.60996 | 0.294 | 0.32 | 1 | MS.Trt_vs_MS.NoTrt_B |
| PPIL4 | 0.38123 | 0.95487 | 0.185 | 0.22 | 1 | MS.Trt_vs_MS.NoTrt_B |
| SGF29 | 0.38129 | 0.34789 | 0.224 | 0.26 | 1 | MS.Trt_vs_MS.NoTrt_B |
| BAIAP2L1 | 0.38162 | 0.54066 | 0.405 | 0.46 | 1 | MS.Trt_vs_MS.NoTrt_B |
| ZMYM5 | 0.38171 | 0.3401 | 0.179 | 0.21 | 1 | MS.Trt_vs_MS.NoTrt_B |
| DHRS7B | 0.38206 | 0.43393 | 0.416 | 0.47 | 1 | MS.Trt_vs_MS.NoTrt_B |
| SYNC | 0.38228 | 0.85986 | 0.12 | 0.15 | 1 | MS.Trt_vs_MS.NoTrt_B |
| BRAF | 0.38254 | 0.79457 | 0.107 | 0.14 | 1 | MS.Trt_vs_MS.NoTrt_B |
| LINC00094 | 0.38373 | 1.15596 | 0.399 | 0.46 | 1 | MS.Trt_vs_MS.NoTrt_B |
| FAIM | 0.38424 | 1.06078 | 0.32 | 0.37 | 1 | MS.Trt_vs_MS.NoTrt_B |
| MRPS10 | 0.38488 | 0.28349 | 0.407 | 0.43 | 1 | MS.Trt_vs_MS.NoTrt_B |
| CAGE1 | 0.38521 | 0.47411 | 0.253 | 0.3 | 1 | MS.Trt_vs_MS.NoTrt_B |
| TCF20 | 0.38674 | 0.32057 | 0.109 | 0.14 | 1 | MS.Trt_vs_MS.NoTrt_B |
| ZNF347 | 0.38737 | -0.3312 | 0.817 | 0.9 | 1 | MS.Trt_vs_MS.NoTrt_B |
| KIAA0513 | 0.3877 | -1.0127 | 0.521 | 0.56 | 1 | MS.Trt_vs_MS.NoTrt_B |
| ZFHX3 | 0.38881 | 0.26597 | 0.1 | 0.12 | 1 | MS.Trt_vs_MS.NoTrt_B |
| ASPSCR1 | 0.38888 | -0.6892 | 0.558 | 0.61 | 1 | MS.Trt_vs_MS.NoTrt_B |
| USP9Y | 0.38945 | -0.6349 | 0.222 | 0.25 | 1 | MS.Trt_vs_MS.NoTrt_B |
| JUN | 0.39026 | 0.82168 | 0.142 | 0.12 | 1 | MS.Trt_vs_MS.NoTrt_B |
| DYRK1A | 0.39032 | 0.36355 | 0.601 | 0.68 | 1 | MS.Trt_vs_MS.NoTrt_B |
| PITPNC1 | 0.39099 | 0.49146 | 0.17 | 0.2 | 1 | MS.Trt_vs_MS.NoTrt_B |
| PAEP | 0.39121 | 1.10711 | 0.349 | 0.34 | 1 | MS.Trt_vs_MS.NoTrt_B |
| SULT1B1 | 0.39134 | 0.3002 | 0.852 | 0.96 | 1 | MS.Trt_vs_MS.NoTrt_B |
| PDCL3 | 0.39257 | 0.33222 | 0.092 | 0.11 | 1 | MS.Trt_vs_MS.NoTrt_B |
| PRKCE | 0.39259 | 0.40296 | 0.094 | 0.12 | 1 | MS.Trt_vs_MS.NoTrt_B |
| NR4A2 | 0.39315 | 1.38497 | 0.089 | 0.11 | 1 | MS.Trt_vs_MS.NoTrt_B |
| ZNF559 | 0.39346 | 0.43711 | 0.584 | 0.69 | 1 | MS.Trt_vs_MS.NoTrt_B |
| PPP1R15B | 0.39512 | -0.6616 | 0.261 | 0.28 | 1 | MS.Trt_vs_MS.NoTrt_B |
| ZNF451 | 0.39592 | 0.39611 | 0.673 | 0.71 | 1 | MS.Trt_vs_MS.NoTrt_B |
| SESN1 | 0.39718 | 0.33102 | 0.15 | 0.18 | 1 | MS.Trt_vs_MS.NoTrt_B |
| ZNF574 | 0.39776 | 0.66629 | 0.292 | 0.27 | 1 | MS.Trt_vs_MS.NoTrt_B |
| PNPLA1 | 0.39838 | 0.25184 | 0.248 | 0.23 | 1 | MS.Trt_vs_MS.NoTrt_B |
| MTMR4 | 0.39899 | 0.48561 | 0.196 | 0.23 | 1 | MS.Trt_vs_MS.NoTrt_B |
| FAM217B | 0.39928 | 0.5932 | 0.288 | 0.35 | 1 | MS.Trt_vs_MS.NoTrt_B |
| CTNND1 | 0.39992 | 0.47135 | 0.246 | 0.3 | 1 | MS.Trt_vs_MS.NoTrt_B |
| STIL | 0.40073 | 1.10427 | 0.503 | 0.55 | 1 | MS.Trt_vs_MS.NoTrt_B |
| ABCD3 | 0.40116 | -0.4278 | 0.102 | 0.12 | 1 | MS.Trt_vs_MS.NoTrt_B |
| THEMIS | 0.40123 | 0.63694 | 0.183 | 0.16 | 1 | MS.Trt_vs_MS.NoTrt_B |
| CAPZA1 | 0.40401 | 0.81123 | 0.813 | 0.9 | 1 | MS.Trt_vs_MS.NoTrt_B |

| | | | | | | |
|---|---|---|---|---|---|---|
| ITGA4 | 0.40454 | 0.97094 | 0.192 | 0.17 | 1 | MS.Trt_vs_MS.NoTrt_B |
| SERINC5 | 0.40465 | 0.50888 | 0.852 | 0.89 | 1 | MS.Trt_vs_MS.NoTrt_B |
| ASTN2 | 0.40507 | 0.39442 | 0.834 | 0.95 | 1 | MS.Trt_vs_MS.NoTrt_B |
| PTGES3L | 0.40525 | 0.32169 | 0.743 | 0.82 | 1 | MS.Trt_vs_MS.NoTrt_B |
| EHBP1 | 0.4054 | 1.13273 | 0.34 | 0.38 | 1 | MS.Trt_vs_MS.NoTrt_B |
| PLEKHG5 | 0.40771 | 0.76021 | 0.115 | 0.14 | 1 | MS.Trt_vs_MS.NoTrt_B |
| RNF157 | 0.40778 | 0.93811 | 0.501 | 0.56 | 1 | MS.Trt_vs_MS.NoTrt_B |
| WDR31 | 0.40802 | 0.34123 | 0.717 | 0.81 | 1 | MS.Trt_vs_MS.NoTrt_B |
| PCYT1A | 0.40812 | -0.256 | 0.438 | 0.5 | 1 | MS.Trt_vs_MS.NoTrt_B |
| GNLY | 0.40833 | 0.53801 | 0.166 | 0.2 | 1 | MS.Trt_vs_MS.NoTrt_B |
| FCGR2A | 0.40864 | 0.70155 | 0.137 | 0.16 | 1 | MS.Trt_vs_MS.NoTrt_B |
| CDNF | 0.40923 | 1.51182 | 0.118 | 0.14 | 1 | MS.Trt_vs_MS.NoTrt_B |
| FKTN | 0.4096 | 0.80533 | 0.118 | 0.14 | 1 | MS.Trt_vs_MS.NoTrt_B |
| ZIK1 | 0.40961 | -0.6287 | 0.253 | 0.28 | 1 | MS.Trt_vs_MS.NoTrt_B |
| LECT2 | 0.41 | 0.30143 | 0.094 | 0.12 | 1 | MS.Trt_vs_MS.NoTrt_B |
| GCNT2 | 0.41176 | 0.31899 | 0.575 | 0.67 | 1 | MS.Trt_vs_MS.NoTrt_B |
| LRRC8D | 0.41227 | 0.75915 | 0.113 | 0.14 | 1 | MS.Trt_vs_MS.NoTrt_B |
| RP11-849H4 | 0.41296 | 0.48146 | 0.479 | 0.51 | 1 | MS.Trt_vs_MS.NoTrt_B |
| TMEM64 | 0.41309 | 0.33961 | 0.281 | 0.32 | 1 | MS.Trt_vs_MS.NoTrt_B |
| ACRC | 0.41323 | -0.7665 | 0.336 | 0.37 | 1 | MS.Trt_vs_MS.NoTrt_B |
| HOMER3 | 0.4134 | -0.4 | 0.237 | 0.28 | 1 | MS.Trt_vs_MS.NoTrt_B |
| PRDM5 | 0.41361 | 0.33229 | 0.556 | 0.67 | 1 | MS.Trt_vs_MS.NoTrt_B |
| CARS2 | 0.41369 | 0.3315 | 0.647 | 0.77 | 1 | MS.Trt_vs_MS.NoTrt_B |
| AP1G2 | 0.41391 | 0.49929 | 0.612 | 0.72 | 1 | MS.Trt_vs_MS.NoTrt_B |
| RIC8B | 0.41429 | 0.79088 | 0.111 | 0.09 | 1 | MS.Trt_vs_MS.NoTrt_B |
| COLGALT1 | 0.41485 | 0.45276 | 0.403 | 0.47 | 1 | MS.Trt_vs_MS.NoTrt_B |
| PTOV1 | 0.41579 | 0.42344 | 0.096 | 0.12 | 1 | MS.Trt_vs_MS.NoTrt_B |
| FNTB | 0.41583 | -0.5435 | 0.118 | 0.14 | 1 | MS.Trt_vs_MS.NoTrt_B |
| SLC2A13 | 0.41743 | 0.43689 | 0.27 | 0.31 | 1 | MS.Trt_vs_MS.NoTrt_B |
| ZBTB20 | 0.41754 | 0.62786 | 0.678 | 0.74 | 1 | MS.Trt_vs_MS.NoTrt_B |
| MRPL44 | 0.41824 | -0.396 | 0.46 | 0.49 | 1 | MS.Trt_vs_MS.NoTrt_B |
| IMP4 | 0.41829 | 0.3862 | 0.172 | 0.2 | 1 | MS.Trt_vs_MS.NoTrt_B |
| GTF2H2C | 0.41866 | -0.2791 | 0.351 | 0.38 | 1 | MS.Trt_vs_MS.NoTrt_B |
| ZMAT2 | 0.41933 | 0.27722 | 0.087 | 0.11 | 1 | MS.Trt_vs_MS.NoTrt_B |
| SLC2A6 | 0.41939 | -0.6663 | 0.19 | 0.22 | 1 | MS.Trt_vs_MS.NoTrt_B |
| ZNF157 | 0.41959 | 0.28906 | 0.686 | 0.72 | 1 | MS.Trt_vs_MS.NoTrt_B |
| MARK2 | 0.4196 | 0.30423 | 0.083 | 0.1 | 1 | MS.Trt_vs_MS.NoTrt_B |
| MTF1 | 0.42011 | 0.84319 | 0.198 | 0.23 | 1 | MS.Trt_vs_MS.NoTrt_B |
| ZFP30 | 0.42064 | 0.38855 | 0.771 | 0.81 | 1 | MS.Trt_vs_MS.NoTrt_B |
| TMEM44 | 0.4214 | -0.3318 | 0.447 | 0.49 | 1 | MS.Trt_vs_MS.NoTrt_B |

| | | | | | | |
|---|---|---|---|---|---|---|
| ERCC6L2 | 0.42145 | 0.26967 | 0.157 | 0.18 | 1 | MS.Trt_vs_MS.NoTrt_B |
| ARL8B | 0.42234 | 0.45542 | 0.318 | 0.35 | 1 | MS.Trt_vs_MS.NoTrt_B |
| ZNF836 | 0.4235 | 1.24487 | 0.155 | 0.18 | 1 | MS.Trt_vs_MS.NoTrt_B |
| PARK2 | 0.42376 | 0.26251 | 0.527 | 0.57 | 1 | MS.Trt_vs_MS.NoTrt_B |
| FBLN5 | 0.42624 | 0.37663 | 0.133 | 0.16 | 1 | MS.Trt_vs_MS.NoTrt_B |
| CAMTA2 | 0.42714 | 0.46108 | 0.425 | 0.45 | 1 | MS.Trt_vs_MS.NoTrt_B |
| ADSS | 0.42777 | 0.39831 | 0.139 | 0.17 | 1 | MS.Trt_vs_MS.NoTrt_B |
| BLOC1S6 | 0.42805 | 0.30495 | 0.928 | 1 | 1 | MS.Trt_vs_MS.NoTrt_B |
| PTDSS2 | 0.42815 | -0.3572 | 0.092 | 0.11 | 1 | MS.Trt_vs_MS.NoTrt_B |
| XYLT1 | 0.42835 | -0.2885 | 0.096 | 0.11 | 1 | MS.Trt_vs_MS.NoTrt_B |
| TMEM191C | 0.42842 | 0.32837 | 0.617 | 0.72 | 1 | MS.Trt_vs_MS.NoTrt_B |
| NPIPB15 | 0.42896 | 0.35303 | 0.214 | 0.24 | 1 | MS.Trt_vs_MS.NoTrt_B |
| TMEM5 | 0.42919 | 1.31936 | 0.386 | 0.45 | 1 | MS.Trt_vs_MS.NoTrt_B |
| NRP1 | 0.42976 | 0.28961 | 0.322 | 0.37 | 1 | MS.Trt_vs_MS.NoTrt_B |
| MT-ND6 | 0.43055 | 0.39976 | 0.68 | 0.76 | 1 | MS.Trt_vs_MS.NoTrt_B |
| ARHGEF10 | 0.43119 | 0.4023 | 0.643 | 0.58 | 1 | MS.Trt_vs_MS.NoTrt_B |
| CCSAP | 0.43122 | 0.69833 | 0.098 | 0.12 | 1 | MS.Trt_vs_MS.NoTrt_B |
| TMEM39A | 0.43145 | 0.90264 | 0.087 | 0.11 | 1 | MS.Trt_vs_MS.NoTrt_B |
| TFPI | 0.43214 | 0.29355 | 0.843 | 0.98 | 1 | MS.Trt_vs_MS.NoTrt_B |
| BCDIN3D | 0.43223 | -0.3543 | 0.275 | 0.24 | 1 | MS.Trt_vs_MS.NoTrt_B |
| SMAD1 | 0.43328 | 0.4776 | 0.285 | 0.27 | 1 | MS.Trt_vs_MS.NoTrt_B |
| ZNF100 | 0.43352 | 0.57638 | 0.745 | 0.82 | 1 | MS.Trt_vs_MS.NoTrt_B |
| CAMKK1 | 0.43401 | 0.91468 | 0.161 | 0.2 | 1 | MS.Trt_vs_MS.NoTrt_B |
| IPO7 | 0.43484 | 0.96905 | 0.318 | 0.36 | 1 | MS.Trt_vs_MS.NoTrt_B |
| DNAJC24 | 0.43579 | 0.30716 | 0.612 | 0.68 | 1 | MS.Trt_vs_MS.NoTrt_B |
| CCDC170 | 0.43758 | 0.53862 | 0.307 | 0.28 | 1 | MS.Trt_vs_MS.NoTrt_B |
| SPTLC2 | 0.43874 | -0.5927 | 0.586 | 0.6 | 1 | MS.Trt_vs_MS.NoTrt_B |
| AGXT | 0.43935 | 0.3648 | 0.527 | 0.64 | 1 | MS.Trt_vs_MS.NoTrt_B |
| B3GAT2 | 0.43966 | 0.27385 | 0.15 | 0.12 | 1 | MS.Trt_vs_MS.NoTrt_B |
| TMEM45A | 0.44073 | 0.26492 | 0.95 | 1 | 1 | MS.Trt_vs_MS.NoTrt_B |
| STXBP3 | 0.44106 | 0.31063 | 0.542 | 0.52 | 1 | MS.Trt_vs_MS.NoTrt_B |
| METTL16 | 0.44168 | 0.44189 | 0.662 | 0.69 | 1 | MS.Trt_vs_MS.NoTrt_B |
| STON1 | 0.44241 | 0.35632 | 0.277 | 0.31 | 1 | MS.Trt_vs_MS.NoTrt_B |
| CD300LG | 0.44282 | 0.45439 | 0.359 | 0.36 | 1 | MS.Trt_vs_MS.NoTrt_B |
| CACNB3 | 0.44339 | 0.6979 | 0.458 | 0.56 | 1 | MS.Trt_vs_MS.NoTrt_B |
| TIMP2 | 0.44361 | 0.6663 | 0.131 | 0.16 | 1 | MS.Trt_vs_MS.NoTrt_B |
| GNG7 | 0.44392 | 0.41085 | 0.29 | 0.35 | 1 | MS.Trt_vs_MS.NoTrt_B |
| MED6 | 0.44434 | 0.36421 | 0.442 | 0.48 | 1 | MS.Trt_vs_MS.NoTrt_B |
| C6orf25 | 0.44449 | 0.25191 | 0.366 | 0.42 | 1 | MS.Trt_vs_MS.NoTrt_B |
| MRPL27 | 0.44536 | 0.62316 | 0.142 | 0.17 | 1 | MS.Trt_vs_MS.NoTrt_B |

| | | | | | | |
|---|---|---|---|---|---|---|
| PLCL1 | 0.4454 | 0.47962 | 0.111 | 0.13 | 1 | MS.Trt_vs_MS.NoTrt_B |
| MAN2A2 | 0.4462 | 0.70403 | 0.214 | 0.25 | 1 | MS.Trt_vs_MS.NoTrt_B |
| FGFR1 | 0.44621 | 0.64047 | 0.71 | 0.78 | 1 | MS.Trt_vs_MS.NoTrt_B |
| RAD52 | 0.44757 | 0.27029 | 0.244 | 0.27 | 1 | MS.Trt_vs_MS.NoTrt_B |
| XKR8 | 0.44759 | -0.7553 | 0.122 | 0.14 | 1 | MS.Trt_vs_MS.NoTrt_B |
| IFI27 | 0.44887 | 0.26685 | 0.292 | 0.34 | 1 | MS.Trt_vs_MS.NoTrt_B |
| ZFAND6 | 0.44931 | 0.42402 | 0.155 | 0.18 | 1 | MS.Trt_vs_MS.NoTrt_B |
| RHEB | 0.45246 | 0.6169 | 0.122 | 0.15 | 1 | MS.Trt_vs_MS.NoTrt_B |
| TERF2 | 0.45254 | 0.52529 | 0.782 | 0.84 | 1 | MS.Trt_vs_MS.NoTrt_B |
| CREB3L4 | 0.45255 | 1.52125 | 0.331 | 0.4 | 1 | MS.Trt_vs_MS.NoTrt_B |
| AP5S1 | 0.45259 | -0.6595 | 0.747 | 0.81 | 1 | MS.Trt_vs_MS.NoTrt_B |
| RP11-437B1 | 0.45266 | 0.88779 | 0.438 | 0.5 | 1 | MS.Trt_vs_MS.NoTrt_B |
| CCDC36 | 0.45357 | 0.96926 | 0.251 | 0.29 | 1 | MS.Trt_vs_MS.NoTrt_B |
| PARG | 0.45485 | -0.3801 | 0.107 | 0.12 | 1 | MS.Trt_vs_MS.NoTrt_B |
| MON1B | 0.455 | 0.31881 | 0.577 | 0.61 | 1 | MS.Trt_vs_MS.NoTrt_B |
| TBCEL | 0.45615 | 0.58104 | 0.092 | 0.11 | 1 | MS.Trt_vs_MS.NoTrt_B |
| TNRC6B | 0.45826 | 0.50588 | 0.891 | 1 | 1 | MS.Trt_vs_MS.NoTrt_B |
| ZYG11B | 0.4591 | -0.2986 | 0.608 | 0.62 | 1 | MS.Trt_vs_MS.NoTrt_B |
| DTD1 | 0.45929 | 0.45963 | 0.253 | 0.28 | 1 | MS.Trt_vs_MS.NoTrt_B |
| CDCP1 | 0.46019 | 0.50665 | 0.366 | 0.38 | 1 | MS.Trt_vs_MS.NoTrt_B |
| ADAMTS1 | 0.46064 | 0.25726 | 0.401 | 0.45 | 1 | MS.Trt_vs_MS.NoTrt_B |
| TMEM38A | 0.4607 | 0.67306 | 0.455 | 0.45 | 1 | MS.Trt_vs_MS.NoTrt_B |
| ST3GAL2 | 0.46253 | 0.44007 | 0.863 | 0.97 | 1 | MS.Trt_vs_MS.NoTrt_B |
| CYP4V2 | 0.46285 | 0.78709 | 0.601 | 0.69 | 1 | MS.Trt_vs_MS.NoTrt_B |
| ACSL1 | 0.46302 | 0.34804 | 0.185 | 0.21 | 1 | MS.Trt_vs_MS.NoTrt_B |
| POLR3D | 0.46315 | -0.4899 | 0.298 | 0.34 | 1 | MS.Trt_vs_MS.NoTrt_B |
| MAP7D3 | 0.46393 | -0.4053 | 0.429 | 0.49 | 1 | MS.Trt_vs_MS.NoTrt_B |
| SSBP2 | 0.46439 | 0.27932 | 0.773 | 0.92 | 1 | MS.Trt_vs_MS.NoTrt_B |
| SYMPK | 0.46526 | 0.35219 | 0.8 | 0.89 | 1 | MS.Trt_vs_MS.NoTrt_B |
| LY75 | 0.46552 | -0.3258 | 0.477 | 0.47 | 1 | MS.Trt_vs_MS.NoTrt_B |
| PNPLA2 | 0.46569 | 0.65137 | 0.133 | 0.16 | 1 | MS.Trt_vs_MS.NoTrt_B |
| BTBD2 | 0.46635 | 1.0541 | 0.288 | 0.34 | 1 | MS.Trt_vs_MS.NoTrt_B |
| TRIM35 | 0.46977 | -0.3151 | 0.166 | 0.18 | 1 | MS.Trt_vs_MS.NoTrt_B |
| OR6K3 | 0.47297 | 0.29709 | 0.614 | 0.78 | 1 | MS.Trt_vs_MS.NoTrt_B |
| BLM | 0.4733 | 0.32488 | 0.218 | 0.2 | 1 | MS.Trt_vs_MS.NoTrt_B |
| RFC2 | 0.47515 | 0.52993 | 0.584 | 0.65 | 1 | MS.Trt_vs_MS.NoTrt_B |
| DDX49 | 0.47544 | -0.4568 | 0.12 | 0.14 | 1 | MS.Trt_vs_MS.NoTrt_B |
| BTBD19 | 0.47643 | 0.47584 | 0.627 | 0.61 | 1 | MS.Trt_vs_MS.NoTrt_B |
| H3F3B | 0.4771 | 0.29341 | 0.786 | 0.96 | 1 | MS.Trt_vs_MS.NoTrt_B |
| DAZAP2 | 0.48048 | 0.37226 | 0.817 | 0.92 | 1 | MS.Trt_vs_MS.NoTrt_B |

| | | | | | | |
|---|---|---|---|---|---|---|
| CNOT6 | 0.48089 | 1.33651 | 0.089 | 0.11 | 1 | MS.Trt_vs_MS.NoTrt_B |
| TAF1 | 0.48137 | 0.57884 | 0.682 | 0.8 | 1 | MS.Trt_vs_MS.NoTrt_B |
| RALGPS2 | 0.4814 | 0.92929 | 0.303 | 0.32 | 1 | MS.Trt_vs_MS.NoTrt_B |
| HLTF | 0.48256 | 1.12269 | 0.399 | 0.44 | 1 | MS.Trt_vs_MS.NoTrt_B |
| JCHAIN | 0.48297 | 0.49723 | 0.1 | 0.12 | 1 | MS.Trt_vs_MS.NoTrt_B |
| STK17B | 0.48365 | 0.75335 | 0.867 | 0.96 | 1 | MS.Trt_vs_MS.NoTrt_B |
| TMEM17 | 0.4848 | 0.37061 | 0.416 | 0.49 | 1 | MS.Trt_vs_MS.NoTrt_B |
| SEC61A2 | 0.48516 | 0.55616 | 0.085 | 0.1 | 1 | MS.Trt_vs_MS.NoTrt_B |
| ZNF547 | 0.48583 | 1.41482 | 0.174 | 0.21 | 1 | MS.Trt_vs_MS.NoTrt_B |
| UGCG | 0.48646 | 0.3519 | 0.15 | 0.17 | 1 | MS.Trt_vs_MS.NoTrt_B |
| NFU1 | 0.48702 | -0.291 | 0.111 | 0.13 | 1 | MS.Trt_vs_MS.NoTrt_B |
| TRIOBP | 0.48732 | 0.96256 | 0.285 | 0.27 | 1 | MS.Trt_vs_MS.NoTrt_B |
| NOMO2 | 0.48932 | 0.96327 | 0.153 | 0.18 | 1 | MS.Trt_vs_MS.NoTrt_B |
| ARF6 | 0.48997 | 0.9823 | 0.614 | 0.64 | 1 | MS.Trt_vs_MS.NoTrt_B |
| VRK1 | 0.49074 | 0.41683 | 0.179 | 0.2 | 1 | MS.Trt_vs_MS.NoTrt_B |
| CYB5D1 | 0.49092 | -0.772 | 0.111 | 0.13 | 1 | MS.Trt_vs_MS.NoTrt_B |
| XPO4 | 0.49153 | 1.4607 | 0.102 | 0.12 | 1 | MS.Trt_vs_MS.NoTrt_B |
| ZNF248 | 0.49189 | 0.63708 | 0.096 | 0.11 | 1 | MS.Trt_vs_MS.NoTrt_B |
| RBMX | 0.49408 | 0.56161 | 0.664 | 0.71 | 1 | MS.Trt_vs_MS.NoTrt_B |
| GSE1 | 0.49452 | 0.62296 | 0.279 | 0.32 | 1 | MS.Trt_vs_MS.NoTrt_B |
| TTLL9 | 0.49561 | 0.29591 | 0.27 | 0.24 | 1 | MS.Trt_vs_MS.NoTrt_B |
| G6PC3 | 0.49626 | 0.46764 | 0.736 | 0.77 | 1 | MS.Trt_vs_MS.NoTrt_B |
| SEPSECS | 0.49806 | 0.27603 | 0.68 | 0.8 | 1 | MS.Trt_vs_MS.NoTrt_B |
| CTD-2369P2 | 0.49863 | 0.46762 | 0.251 | 0.24 | 1 | MS.Trt_vs_MS.NoTrt_B |
| SYF2 | 0.49954 | 0.62668 | 0.375 | 0.42 | 1 | MS.Trt_vs_MS.NoTrt_B |
| GAS7 | 0.49999 | 0.40933 | 0.505 | 0.56 | 1 | MS.Trt_vs_MS.NoTrt_B |
| CFAP54 | 0.50006 | 0.28553 | 0.802 | 0.96 | 1 | MS.Trt_vs_MS.NoTrt_B |
| PHLDB2 | 0.50021 | 1.44714 | 0.366 | 0.41 | 1 | MS.Trt_vs_MS.NoTrt_B |
| 1-Mar | 0.50172 | 1.42505 | 0.264 | 0.25 | 1 | MS.Trt_vs_MS.NoTrt_B |
| FOXM1 | 0.50204 | -0.2558 | 0.606 | 0.68 | 1 | MS.Trt_vs_MS.NoTrt_B |
| CCDC192 | 0.50206 | 0.71643 | 0.096 | 0.11 | 1 | MS.Trt_vs_MS.NoTrt_B |
| PLA2G4E | 0.50244 | 0.42441 | 0.109 | 0.09 | 1 | MS.Trt_vs_MS.NoTrt_B |
| ZNF257 | 0.50251 | 1.00142 | 0.451 | 0.5 | 1 | MS.Trt_vs_MS.NoTrt_B |
| BAP1 | 0.5028 | 0.43084 | 0.275 | 0.24 | 1 | MS.Trt_vs_MS.NoTrt_B |
| PLA2G4B | 0.50402 | 0.77198 | 0.379 | 0.43 | 1 | MS.Trt_vs_MS.NoTrt_B |
| MCM10 | 0.50402 | 0.28953 | 0.314 | 0.37 | 1 | MS.Trt_vs_MS.NoTrt_B |
| FANCI | 0.50416 | -0.6289 | 0.497 | 0.51 | 1 | MS.Trt_vs_MS.NoTrt_B |
| SRPK1 | 0.50432 | 0.32248 | 0.17 | 0.2 | 1 | MS.Trt_vs_MS.NoTrt_B |
| CASS4 | 0.50446 | 0.51372 | 0.353 | 0.39 | 1 | MS.Trt_vs_MS.NoTrt_B |
| SCN2B | 0.5052 | -0.7552 | 0.246 | 0.27 | 1 | MS.Trt_vs_MS.NoTrt_B |

| | | | | | | |
|---|---|---|---|---|---|---|
| TMC2 | 0.50608 | 0.30931 | 0.481 | 0.57 | 1 | MS.Trt_vs_MS.NoTrt_B |
| IFI27L2 | 0.50623 | 1.15906 | 0.129 | 0.15 | 1 | MS.Trt_vs_MS.NoTrt_B |
| CCDC40 | 0.50731 | 0.25476 | 0.741 | 0.75 | 1 | MS.Trt_vs_MS.NoTrt_B |
| SLC30A5 | 0.50848 | 0.62422 | 0.643 | 0.71 | 1 | MS.Trt_vs_MS.NoTrt_B |
| GABBR1 | 0.50968 | 0.35276 | 0.144 | 0.17 | 1 | MS.Trt_vs_MS.NoTrt_B |
| ZBTB44 | 0.51067 | 0.67178 | 0.246 | 0.28 | 1 | MS.Trt_vs_MS.NoTrt_B |
| MSH5 | 0.51112 | 0.36885 | 0.871 | 0.93 | 1 | MS.Trt_vs_MS.NoTrt_B |
| CKAP5 | 0.51127 | 0.76418 | 0.105 | 0.12 | 1 | MS.Trt_vs_MS.NoTrt_B |
| GPLD1 | 0.51141 | -0.2594 | 0.634 | 0.63 | 1 | MS.Trt_vs_MS.NoTrt_B |
| SH3RF2 | 0.51187 | -0.8092 | 0.155 | 0.14 | 1 | MS.Trt_vs_MS.NoTrt_B |
| SPTLC3 | 0.51282 | 0.51061 | 0.42 | 0.51 | 1 | MS.Trt_vs_MS.NoTrt_B |
| ITIH4 | 0.51338 | -0.8924 | 0.168 | 0.19 | 1 | MS.Trt_vs_MS.NoTrt_B |
| METTL21A | 0.51364 | 0.59876 | 0.863 | 0.97 | 1 | MS.Trt_vs_MS.NoTrt_B |
| ICA1 | 0.51384 | 1.00581 | 0.444 | 0.54 | 1 | MS.Trt_vs_MS.NoTrt_B |
| NAA20 | 0.51547 | 0.48483 | 0.135 | 0.16 | 1 | MS.Trt_vs_MS.NoTrt_B |
| NDUFA12 | 0.51674 | 0.9463 | 0.22 | 0.27 | 1 | MS.Trt_vs_MS.NoTrt_B |
| RP11-1070B | 0.51732 | 0.66289 | 0.111 | 0.13 | 1 | MS.Trt_vs_MS.NoTrt_B |
| TRAPPC3L | 0.51826 | 0.40909 | 0.464 | 0.58 | 1 | MS.Trt_vs_MS.NoTrt_B |
| FBN1 | 0.51902 | 0.26278 | 0.298 | 0.35 | 1 | MS.Trt_vs_MS.NoTrt_B |
| TRIM56 | 0.51988 | 0.65433 | 0.889 | 0.98 | 1 | MS.Trt_vs_MS.NoTrt_B |
| SENP1 | 0.52005 | 0.68827 | 0.174 | 0.15 | 1 | MS.Trt_vs_MS.NoTrt_B |
| TPRKB | 0.52117 | 0.37123 | 0.397 | 0.43 | 1 | MS.Trt_vs_MS.NoTrt_B |
| CCR7 | 0.52137 | 0.3959 | 0.499 | 0.57 | 1 | MS.Trt_vs_MS.NoTrt_B |
| HECTD3 | 0.52139 | 1.11239 | 0.118 | 0.1 | 1 | MS.Trt_vs_MS.NoTrt_B |
| ST3GAL6 | 0.52289 | 0.32225 | 0.233 | 0.23 | 1 | MS.Trt_vs_MS.NoTrt_B |
| TERB1 | 0.52402 | 0.27763 | 0.551 | 0.6 | 1 | MS.Trt_vs_MS.NoTrt_B |
| GALNT11 | 0.52483 | 0.28542 | 0.37 | 0.38 | 1 | MS.Trt_vs_MS.NoTrt_B |
| UIMC1 | 0.52486 | 0.40203 | 0.725 | 0.82 | 1 | MS.Trt_vs_MS.NoTrt_B |
| LARS2 | 0.52519 | -0.7591 | 0.403 | 0.42 | 1 | MS.Trt_vs_MS.NoTrt_B |
| SAMD9 | 0.52522 | 0.40631 | 0.157 | 0.18 | 1 | MS.Trt_vs_MS.NoTrt_B |
| ZCCHC10 | 0.52526 | 1.0214 | 0.486 | 0.56 | 1 | MS.Trt_vs_MS.NoTrt_B |
| VPS11 | 0.5259 | 0.64064 | 0.712 | 0.79 | 1 | MS.Trt_vs_MS.NoTrt_B |
| LONP1 | 0.52617 | 0.2888 | 0.497 | 0.46 | 1 | MS.Trt_vs_MS.NoTrt_B |
| PITPNM2 | 0.52689 | 0.80644 | 0.322 | 0.37 | 1 | MS.Trt_vs_MS.NoTrt_B |
| FAM135A | 0.52696 | 0.58825 | 0.351 | 0.4 | 1 | MS.Trt_vs_MS.NoTrt_B |
| RBFOX3 | 0.52702 | 0.27436 | 0.715 | 0.78 | 1 | MS.Trt_vs_MS.NoTrt_B |
| CRY2 | 0.52726 | 0.47487 | 0.41 | 0.38 | 1 | MS.Trt_vs_MS.NoTrt_B |
| DGKD | 0.52761 | 0.44496 | 0.527 | 0.57 | 1 | MS.Trt_vs_MS.NoTrt_B |
| CACNA1F | 0.52907 | 0.27538 | 0.192 | 0.22 | 1 | MS.Trt_vs_MS.NoTrt_B |
| ZNF736 | 0.52984 | -0.5752 | 0.423 | 0.47 | 1 | MS.Trt_vs_MS.NoTrt_B |

| | | | | | | |
|---|---|---|---|---|---|---|
| CXorf57 | 0.53016 | 1.14679 | 0.109 | 0.09 | 1 | MS.Trt_vs_MS.NoTrt_B |
| TENM1 | 0.5323 | 1.59715 | 0.085 | 0.1 | 1 | MS.Trt_vs_MS.NoTrt_B |
| PRKCZ | 0.53261 | -0.3461 | 0.187 | 0.21 | 1 | MS.Trt_vs_MS.NoTrt_B |
| C1orf50 | 0.53386 | 0.26774 | 0.577 | 0.64 | 1 | MS.Trt_vs_MS.NoTrt_B |
| RNF144A | 0.53396 | 0.49514 | 0.133 | 0.15 | 1 | MS.Trt_vs_MS.NoTrt_B |
| U2SURP | 0.53442 | 0.51002 | 0.292 | 0.34 | 1 | MS.Trt_vs_MS.NoTrt_B |
| THRA | 0.53701 | 0.69958 | 0.087 | 0.1 | 1 | MS.Trt_vs_MS.NoTrt_B |
| ADHFE1 | 0.53714 | 0.46492 | 0.12 | 0.14 | 1 | MS.Trt_vs_MS.NoTrt_B |
| ST20-MTHFS | 0.53819 | 0.59285 | 0.085 | 0.1 | 1 | MS.Trt_vs_MS.NoTrt_B |
| NUP155 | 0.53842 | 0.63662 | 0.804 | 0.89 | 1 | MS.Trt_vs_MS.NoTrt_B |
| OXCT1 | 0.53949 | -0.2987 | 0.312 | 0.35 | 1 | MS.Trt_vs_MS.NoTrt_B |
| HAVCR1 | 0.54088 | 0.50681 | 0.301 | 0.33 | 1 | MS.Trt_vs_MS.NoTrt_B |
| SMARCAD1 | 0.54192 | 0.66867 | 0.1 | 0.12 | 1 | MS.Trt_vs_MS.NoTrt_B |
| NMUR1 | 0.54331 | 0.28689 | 0.434 | 0.42 | 1 | MS.Trt_vs_MS.NoTrt_B |
| SLC16A5 | 0.54348 | 0.63923 | 0.107 | 0.12 | 1 | MS.Trt_vs_MS.NoTrt_B |
| BTBD8 | 0.54374 | 1.26915 | 0.288 | 0.27 | 1 | MS.Trt_vs_MS.NoTrt_B |
| TIGD5 | 0.54579 | 1.0604 | 0.155 | 0.17 | 1 | MS.Trt_vs_MS.NoTrt_B |
| ZNF114 | 0.54731 | 0.26858 | 0.377 | 0.43 | 1 | MS.Trt_vs_MS.NoTrt_B |
| ZNF622 | 0.54743 | 0.72736 | 0.102 | 0.12 | 1 | MS.Trt_vs_MS.NoTrt_B |
| DHPS | 0.54829 | 0.2822 | 0.224 | 0.25 | 1 | MS.Trt_vs_MS.NoTrt_B |
| DNAJB5 | 0.54871 | -1.3256 | 0.129 | 0.15 | 1 | MS.Trt_vs_MS.NoTrt_B |
| PRR5 | 0.54924 | 0.67121 | 0.436 | 0.48 | 1 | MS.Trt_vs_MS.NoTrt_B |
| ANKLE1 | 0.54969 | 0.48643 | 0.157 | 0.14 | 1 | MS.Trt_vs_MS.NoTrt_B |
| MELK | 0.55011 | 0.36854 | 0.423 | 0.48 | 1 | MS.Trt_vs_MS.NoTrt_B |
| ITM2C | 0.55125 | 0.51185 | 0.092 | 0.11 | 1 | MS.Trt_vs_MS.NoTrt_B |
| PUM3 | 0.55329 | 0.35394 | 0.137 | 0.12 | 1 | MS.Trt_vs_MS.NoTrt_B |
| TWISTNB | 0.55411 | 0.66079 | 0.087 | 0.1 | 1 | MS.Trt_vs_MS.NoTrt_B |
| G2E3 | 0.55424 | 0.68448 | 0.46 | 0.5 | 1 | MS.Trt_vs_MS.NoTrt_B |
| RARS | 0.55542 | 0.29649 | 0.414 | 0.43 | 1 | MS.Trt_vs_MS.NoTrt_B |
| CYB5D2 | 0.5557 | 0.31271 | 0.346 | 0.4 | 1 | MS.Trt_vs_MS.NoTrt_B |
| DPH3 | 0.55589 | -0.2953 | 0.174 | 0.19 | 1 | MS.Trt_vs_MS.NoTrt_B |
| EYS | 0.55735 | 0.40714 | 0.634 | 0.77 | 1 | MS.Trt_vs_MS.NoTrt_B |
| SLC12A4 | 0.55749 | -0.5575 | 0.142 | 0.12 | 1 | MS.Trt_vs_MS.NoTrt_B |
| ATG3 | 0.55817 | 0.25401 | 0.364 | 0.36 | 1 | MS.Trt_vs_MS.NoTrt_B |
| SLC25A42 | 0.5588 | 0.88498 | 0.471 | 0.52 | 1 | MS.Trt_vs_MS.NoTrt_B |
| TRAPPC4 | 0.55898 | 0.3594 | 0.837 | 0.92 | 1 | MS.Trt_vs_MS.NoTrt_B |
| KCTD6 | 0.5592 | 0.37304 | 0.471 | 0.52 | 1 | MS.Trt_vs_MS.NoTrt_B |
| NCDN | 0.56249 | 0.43172 | 0.17 | 0.19 | 1 | MS.Trt_vs_MS.NoTrt_B |
| NRDE2 | 0.56264 | 0.45205 | 0.695 | 0.72 | 1 | MS.Trt_vs_MS.NoTrt_B |
| WDR75 | 0.56328 | 0.73609 | 0.255 | 0.27 | 1 | MS.Trt_vs_MS.NoTrt_B |

| | | | | | | |
|---|---|---|---|---|---|---|
| REEP3 | 0.56476 | 1.29932 | 0.381 | 0.43 | 1 | MS.Trt_vs_MS.NoTrt_B |
| C4orf19 | 0.56512 | 0.30396 | 0.654 | 0.76 | 1 | MS.Trt_vs_MS.NoTrt_B |
| DVL3 | 0.56644 | 1.30105 | 0.229 | 0.26 | 1 | MS.Trt_vs_MS.NoTrt_B |
| PHF19 | 0.56668 | 0.33713 | 0.118 | 0.14 | 1 | MS.Trt_vs_MS.NoTrt_B |
| BCOR | 0.56734 | 0.76097 | 0.087 | 0.1 | 1 | MS.Trt_vs_MS.NoTrt_B |
| MAP2K6 | 0.56785 | 0.94443 | 0.285 | 0.32 | 1 | MS.Trt_vs_MS.NoTrt_B |
| ITM2A | 0.56845 | 0.5543 | 0.237 | 0.22 | 1 | MS.Trt_vs_MS.NoTrt_B |
| E2F6 | 0.56974 | -0.8679 | 0.216 | 0.24 | 1 | MS.Trt_vs_MS.NoTrt_B |
| STRN3 | 0.57006 | 1.03389 | 0.645 | 0.71 | 1 | MS.Trt_vs_MS.NoTrt_B |
| PTRH2 | 0.5746 | 0.74765 | 0.107 | 0.12 | 1 | MS.Trt_vs_MS.NoTrt_B |
| OFD1 | 0.57582 | 0.2687 | 0.216 | 0.24 | 1 | MS.Trt_vs_MS.NoTrt_B |
| LSM10 | 0.57773 | 0.54727 | 0.111 | 0.1 | 1 | MS.Trt_vs_MS.NoTrt_B |
| IFT88 | 0.57865 | 0.52646 | 0.425 | 0.47 | 1 | MS.Trt_vs_MS.NoTrt_B |
| BOLL | 0.57927 | 0.43324 | 0.142 | 0.13 | 1 | MS.Trt_vs_MS.NoTrt_B |
| NCOA1 | 0.57961 | 0.35206 | 0.207 | 0.18 | 1 | MS.Trt_vs_MS.NoTrt_B |
| ZNF696 | 0.58076 | 0.75 | 0.222 | 0.25 | 1 | MS.Trt_vs_MS.NoTrt_B |
| CD226 | 0.58154 | 0.42659 | 0.429 | 0.45 | 1 | MS.Trt_vs_MS.NoTrt_B |
| PLEKHA7 | 0.58172 | 0.95298 | 0.118 | 0.14 | 1 | MS.Trt_vs_MS.NoTrt_B |
| FRMPD2 | 0.58341 | 0.45307 | 0.098 | 0.11 | 1 | MS.Trt_vs_MS.NoTrt_B |
| IL17RC | 0.5841 | 0.67955 | 0.327 | 0.3 | 1 | MS.Trt_vs_MS.NoTrt_B |
| FAF2 | 0.58625 | 0.86289 | 0.098 | 0.11 | 1 | MS.Trt_vs_MS.NoTrt_B |
| CRYGS | 0.58702 | 1.21362 | 0.203 | 0.23 | 1 | MS.Trt_vs_MS.NoTrt_B |
| ANO10 | 0.58723 | -0.8153 | 0.447 | 0.51 | 1 | MS.Trt_vs_MS.NoTrt_B |
| PDZD8 | 0.58783 | 0.79306 | 0.246 | 0.22 | 1 | MS.Trt_vs_MS.NoTrt_B |
| SPG7 | 0.58792 | 0.56594 | 0.95 | 0.98 | 1 | MS.Trt_vs_MS.NoTrt_B |
| SVIL | 0.5882 | 0.32024 | 0.102 | 0.12 | 1 | MS.Trt_vs_MS.NoTrt_B |
| CCDC15 | 0.58826 | -1.2227 | 0.187 | 0.21 | 1 | MS.Trt_vs_MS.NoTrt_B |
| REPS1 | 0.58892 | 0.65374 | 0.547 | 0.62 | 1 | MS.Trt_vs_MS.NoTrt_B |
| GPANK1 | 0.58942 | 0.34574 | 0.257 | 0.25 | 1 | MS.Trt_vs_MS.NoTrt_B |
| YPEL1 | 0.58952 | 0.30288 | 0.366 | 0.41 | 1 | MS.Trt_vs_MS.NoTrt_B |
| FMN1 | 0.58986 | 0.53366 | 0.17 | 0.2 | 1 | MS.Trt_vs_MS.NoTrt_B |
| MYH3 | 0.5899 | 0.30361 | 0.584 | 0.63 | 1 | MS.Trt_vs_MS.NoTrt_B |
| PIK3R1 | 0.59033 | 0.54986 | 0.259 | 0.24 | 1 | MS.Trt_vs_MS.NoTrt_B |
| TNFRSF10D | 0.59065 | 0.45588 | 0.087 | 0.1 | 1 | MS.Trt_vs_MS.NoTrt_B |
| IFIT3 | 0.59163 | 0.36329 | 0.466 | 0.48 | 1 | MS.Trt_vs_MS.NoTrt_B |
| SC5D | 0.59457 | 0.60985 | 0.468 | 0.51 | 1 | MS.Trt_vs_MS.NoTrt_B |
| LOXL2 | 0.59481 | 0.33463 | 0.632 | 0.73 | 1 | MS.Trt_vs_MS.NoTrt_B |
| CDC42 | 0.59669 | 0.51864 | 0.806 | 0.95 | 1 | MS.Trt_vs_MS.NoTrt_B |
| DPY19L4 | 0.59804 | 1.3106 | 0.427 | 0.45 | 1 | MS.Trt_vs_MS.NoTrt_B |
| ADRB2 | 0.59814 | 1.11585 | 0.166 | 0.16 | 1 | MS.Trt_vs_MS.NoTrt_B |

| | | | | | | |
|---|---|---|---|---|---|---|
| SLC13A3 | 0.59938 | 0.51067 | 0.285 | 0.28 | 1 | MS.Trt_vs_MS.NoTrt_B |
| ANKRD36 | 0.59984 | 0.6261 | 0.673 | 0.66 | 1 | MS.Trt_vs_MS.NoTrt_B |
| ATP6V0B | 0.60163 | 0.38794 | 0.148 | 0.17 | 1 | MS.Trt_vs_MS.NoTrt_B |
| ORC4 | 0.60213 | 0.27225 | 0.904 | 1 | 1 | MS.Trt_vs_MS.NoTrt_B |
| E4F1 | 0.60226 | 0.5545 | 0.107 | 0.09 | 1 | MS.Trt_vs_MS.NoTrt_B |
| PNP | 0.6026 | -0.6713 | 0.244 | 0.27 | 1 | MS.Trt_vs_MS.NoTrt_B |
| GTF2H2C.2 | 0.60264 | 0.37705 | 0.248 | 0.27 | 1 | MS.Trt_vs_MS.NoTrt_B |
| RPGRIP1L | 0.60288 | 0.4628 | 0.59 | 0.7 | 1 | MS.Trt_vs_MS.NoTrt_B |
| TCEAL8 | 0.6029 | 0.66555 | 0.105 | 0.12 | 1 | MS.Trt_vs_MS.NoTrt_B |
| RPS19 | 0.60413 | 0.2588 | 0.913 | 1 | 1 | MS.Trt_vs_MS.NoTrt_B |
| SIRPB1 | 0.60447 | 1.48362 | 0.176 | 0.21 | 1 | MS.Trt_vs_MS.NoTrt_B |
| FNDC3B | 0.60586 | 0.38153 | 0.662 | 0.64 | 1 | MS.Trt_vs_MS.NoTrt_B |
| KHDRBS1 | 0.60605 | 0.62818 | 0.139 | 0.16 | 1 | MS.Trt_vs_MS.NoTrt_B |
| EDEM3 | 0.61295 | 0.52147 | 0.27 | 0.29 | 1 | MS.Trt_vs_MS.NoTrt_B |
| HAUS7 | 0.61299 | 0.93578 | 0.094 | 0.11 | 1 | MS.Trt_vs_MS.NoTrt_B |
| FAM200B | 0.61356 | 0.46497 | 0.307 | 0.34 | 1 | MS.Trt_vs_MS.NoTrt_B |
| MED14 | 0.61488 | 0.78069 | 0.126 | 0.11 | 1 | MS.Trt_vs_MS.NoTrt_B |
| AGBL2 | 0.61545 | 1.09508 | 0.218 | 0.25 | 1 | MS.Trt_vs_MS.NoTrt_B |
| LRRC1 | 0.61586 | -0.4053 | 0.209 | 0.22 | 1 | MS.Trt_vs_MS.NoTrt_B |
| C19orf35 | 0.61588 | 0.32091 | 0.255 | 0.28 | 1 | MS.Trt_vs_MS.NoTrt_B |
| MTSS1 | 0.61676 | 0.89025 | 0.492 | 0.56 | 1 | MS.Trt_vs_MS.NoTrt_B |
| RP4-635E18 | 0.61819 | 0.92441 | 0.259 | 0.3 | 1 | MS.Trt_vs_MS.NoTrt_B |
| DPH7 | 0.61879 | 0.38967 | 0.279 | 0.3 | 1 | MS.Trt_vs_MS.NoTrt_B |
| FAM168B | 0.61913 | 1.05798 | 0.089 | 0.1 | 1 | MS.Trt_vs_MS.NoTrt_B |
| TOB1 | 0.61935 | 0.68771 | 0.458 | 0.48 | 1 | MS.Trt_vs_MS.NoTrt_B |
| FAM160A2 | 0.62173 | 0.46584 | 0.092 | 0.1 | 1 | MS.Trt_vs_MS.NoTrt_B |
| CRIM1 | 0.62218 | 0.9648 | 0.115 | 0.13 | 1 | MS.Trt_vs_MS.NoTrt_B |
| LMNTD1 | 0.62257 | 0.25289 | 0.329 | 0.31 | 1 | MS.Trt_vs_MS.NoTrt_B |
| DSEL | 0.62267 | 0.27623 | 0.54 | 0.6 | 1 | MS.Trt_vs_MS.NoTrt_B |
| CTBS | 0.62277 | 1.09202 | 0.126 | 0.14 | 1 | MS.Trt_vs_MS.NoTrt_B |
| DNAH1 | 0.62568 | 0.79659 | 0.12 | 0.11 | 1 | MS.Trt_vs_MS.NoTrt_B |
| NEURL1 | 0.62635 | 0.42656 | 0.203 | 0.22 | 1 | MS.Trt_vs_MS.NoTrt_B |
| CNKSR1 | 0.62734 | -1.2978 | 0.137 | 0.15 | 1 | MS.Trt_vs_MS.NoTrt_B |
| ICMT | 0.62827 | 0.98198 | 0.527 | 0.55 | 1 | MS.Trt_vs_MS.NoTrt_B |
| PIGU | 0.62892 | 0.28606 | 0.092 | 0.1 | 1 | MS.Trt_vs_MS.NoTrt_B |
| NBPF14 | 0.62937 | 0.62456 | 0.346 | 0.33 | 1 | MS.Trt_vs_MS.NoTrt_B |
| TMEM182 | 0.62977 | 0.63043 | 0.625 | 0.79 | 1 | MS.Trt_vs_MS.NoTrt_B |
| LENG8 | 0.62989 | 0.73266 | 0.603 | 0.75 | 1 | MS.Trt_vs_MS.NoTrt_B |
| HAUS3 | 0.63192 | 0.29756 | 0.741 | 0.85 | 1 | MS.Trt_vs_MS.NoTrt_B |
| UTP6 | 0.63303 | 0.41003 | 0.789 | 0.88 | 1 | MS.Trt_vs_MS.NoTrt_B |

| | | | | | | |
|---|---|---|---|---|---|---|
| TCERG1 | 0.63317 | 0.57357 | 0.246 | 0.28 | 1 | MS.Trt_vs_MS.NoTrt_B |
| PTH2R | 0.63427 | 0.33909 | 0.144 | 0.14 | 1 | MS.Trt_vs_MS.NoTrt_B |
| LIAS | 0.63436 | -0.4635 | 0.105 | 0.09 | 1 | MS.Trt_vs_MS.NoTrt_B |
| PEX11B | 0.63466 | 0.44493 | 0.105 | 0.12 | 1 | MS.Trt_vs_MS.NoTrt_B |
| PELP1 | 0.63492 | 0.44558 | 0.222 | 0.24 | 1 | MS.Trt_vs_MS.NoTrt_B |
| CALHM2 | 0.63715 | 0.28091 | 0.131 | 0.11 | 1 | MS.Trt_vs_MS.NoTrt_B |
| USP31 | 0.63738 | 0.68105 | 0.094 | 0.11 | 1 | MS.Trt_vs_MS.NoTrt_B |
| RYR3 | 0.63795 | -0.3059 | 0.388 | 0.38 | 1 | MS.Trt_vs_MS.NoTrt_B |
| INTS7 | 0.63832 | 1.08711 | 0.377 | 0.36 | 1 | MS.Trt_vs_MS.NoTrt_B |
| SBNO2 | 0.6427 | -0.6219 | 0.133 | 0.12 | 1 | MS.Trt_vs_MS.NoTrt_B |
| RTTN | 0.64289 | -0.4003 | 0.192 | 0.22 | 1 | MS.Trt_vs_MS.NoTrt_B |
| CC2D2B | 0.64325 | 0.48333 | 0.094 | 0.11 | 1 | MS.Trt_vs_MS.NoTrt_B |
| USP8 | 0.64336 | 0.42898 | 0.963 | 0.99 | 1 | MS.Trt_vs_MS.NoTrt_B |
| ZNF236 | 0.6435 | 0.59915 | 0.111 | 0.12 | 1 | MS.Trt_vs_MS.NoTrt_B |
| NBPF19 | 0.64452 | 0.40461 | 0.344 | 0.36 | 1 | MS.Trt_vs_MS.NoTrt_B |
| FRMD8 | 0.64508 | -0.2531 | 0.501 | 0.58 | 1 | MS.Trt_vs_MS.NoTrt_B |
| FANCC | 0.64616 | -0.8816 | 0.214 | 0.22 | 1 | MS.Trt_vs_MS.NoTrt_B |
| IL20RB | 0.6468 | 0.43972 | 0.194 | 0.22 | 1 | MS.Trt_vs_MS.NoTrt_B |
| FAM174B | 0.64683 | -0.4612 | 0.109 | 0.1 | 1 | MS.Trt_vs_MS.NoTrt_B |
| STXBP5 | 0.64795 | 0.77498 | 0.142 | 0.16 | 1 | MS.Trt_vs_MS.NoTrt_B |
| GATM | 0.64909 | 0.28559 | 0.192 | 0.18 | 1 | MS.Trt_vs_MS.NoTrt_B |
| ZNF658 | 0.64935 | 1.31162 | 0.163 | 0.18 | 1 | MS.Trt_vs_MS.NoTrt_B |
| SCN7A | 0.65026 | 0.29789 | 0.438 | 0.44 | 1 | MS.Trt_vs_MS.NoTrt_B |
| ZNF584 | 0.65068 | 0.48974 | 0.58 | 0.65 | 1 | MS.Trt_vs_MS.NoTrt_B |
| H2AFY | 0.65083 | 0.29128 | 0.2 | 0.22 | 1 | MS.Trt_vs_MS.NoTrt_B |
| FAM110D | 0.65087 | 0.31035 | 0.39 | 0.44 | 1 | MS.Trt_vs_MS.NoTrt_B |
| ZNF610 | 0.65109 | -0.3741 | 0.312 | 0.33 | 1 | MS.Trt_vs_MS.NoTrt_B |
| EFCAB14 | 0.65296 | 0.36164 | 0.22 | 0.24 | 1 | MS.Trt_vs_MS.NoTrt_B |
| OSBP2 | 0.65385 | -0.9307 | 0.32 | 0.35 | 1 | MS.Trt_vs_MS.NoTrt_B |
| R3HDM1 | 0.65423 | 0.57749 | 0.763 | 0.84 | 1 | MS.Trt_vs_MS.NoTrt_B |
| LBHD1 | 0.6544 | 0.91538 | 0.383 | 0.39 | 1 | MS.Trt_vs_MS.NoTrt_B |
| FGF13 | 0.6544 | 0.2538 | 0.817 | 0.95 | 1 | MS.Trt_vs_MS.NoTrt_B |
| SH2D2A | 0.65559 | 0.7924 | 0.155 | 0.15 | 1 | MS.Trt_vs_MS.NoTrt_B |
| PPM1L | 0.65703 | -0.5725 | 0.187 | 0.2 | 1 | MS.Trt_vs_MS.NoTrt_B |
| TTLL7 | 0.65842 | 0.32214 | 0.693 | 0.77 | 1 | MS.Trt_vs_MS.NoTrt_B |
| ZNF660 | 0.65974 | 0.30909 | 0.481 | 0.51 | 1 | MS.Trt_vs_MS.NoTrt_B |
| IFIT1 | 0.65994 | 0.4214 | 0.192 | 0.18 | 1 | MS.Trt_vs_MS.NoTrt_B |
| ZNF678 | 0.66011 | -0.9034 | 0.124 | 0.14 | 1 | MS.Trt_vs_MS.NoTrt_B |
| NIPAL2 | 0.66025 | 0.65915 | 0.257 | 0.28 | 1 | MS.Trt_vs_MS.NoTrt_B |
| BRD7 | 0.6615 | 1.81169 | 0.447 | 0.52 | 1 | MS.Trt_vs_MS.NoTrt_B |

| | | | | | | |
|---|---|---|---|---|---|---|
| OGG1 | 0.66231 | 0.47675 | 0.32 | 0.34 | 1 | MS.Trt_vs_MS.NoTrt_B |
| DDX51 | 0.66255 | -0.2947 | 0.553 | 0.6 | 1 | MS.Trt_vs_MS.NoTrt_B |
| MAP3K3 | 0.66416 | 0.93422 | 0.148 | 0.14 | 1 | MS.Trt_vs_MS.NoTrt_B |
| ZFYVE21 | 0.66591 | 0.7511 | 0.599 | 0.66 | 1 | MS.Trt_vs_MS.NoTrt_B |
| IGF2R | 0.66607 | 0.87844 | 0.338 | 0.32 | 1 | MS.Trt_vs_MS.NoTrt_B |
| CTD-3105H1 | 0.66694 | 0.25329 | 0.222 | 0.24 | 1 | MS.Trt_vs_MS.NoTrt_B |
| PABPN1 | 0.66703 | 0.91306 | 0.168 | 0.16 | 1 | MS.Trt_vs_MS.NoTrt_B |
| PPT2 | 0.66996 | -0.4116 | 0.427 | 0.51 | 1 | MS.Trt_vs_MS.NoTrt_B |
| UBAP1 | 0.67193 | 0.44487 | 0.111 | 0.12 | 1 | MS.Trt_vs_MS.NoTrt_B |
| TCF7L1 | 0.67196 | 0.45936 | 0.556 | 0.61 | 1 | MS.Trt_vs_MS.NoTrt_B |
| ADGRG1 | 0.67208 | 0.97888 | 0.58 | 0.65 | 1 | MS.Trt_vs_MS.NoTrt_B |
| KLRK1 | 0.67362 | 1.66856 | 0.582 | 0.62 | 1 | MS.Trt_vs_MS.NoTrt_B |
| KMT2B | 0.67372 | 0.61924 | 0.196 | 0.22 | 1 | MS.Trt_vs_MS.NoTrt_B |
| RBPMS | 0.67637 | 1.19329 | 0.092 | 0.1 | 1 | MS.Trt_vs_MS.NoTrt_B |
| SDSL | 0.67719 | 0.25201 | 0.259 | 0.29 | 1 | MS.Trt_vs_MS.NoTrt_B |
| VSTM1 | 0.67853 | 0.31151 | 0.231 | 0.25 | 1 | MS.Trt_vs_MS.NoTrt_B |
| CSMD2 | 0.67966 | -0.2571 | 0.261 | 0.25 | 1 | MS.Trt_vs_MS.NoTrt_B |
| C5orf24 | 0.67974 | -0.6344 | 0.303 | 0.32 | 1 | MS.Trt_vs_MS.NoTrt_B |
| UTP14A | 0.68077 | -0.352 | 0.15 | 0.16 | 1 | MS.Trt_vs_MS.NoTrt_B |
| ADAM28 | 0.68212 | 0.58883 | 0.115 | 0.13 | 1 | MS.Trt_vs_MS.NoTrt_B |
| FAM216A | 0.68302 | 0.55465 | 0.449 | 0.51 | 1 | MS.Trt_vs_MS.NoTrt_B |
| TP53BP1 | 0.68332 | 0.61779 | 0.222 | 0.24 | 1 | MS.Trt_vs_MS.NoTrt_B |
| NPIPA5 | 0.68413 | 0.43615 | 0.107 | 0.1 | 1 | MS.Trt_vs_MS.NoTrt_B |
| ABC7-42404 | 0.68464 | -0.3225 | 0.59 | 0.63 | 1 | MS.Trt_vs_MS.NoTrt_B |
| COL15A1 | 0.68591 | 0.30149 | 0.261 | 0.26 | 1 | MS.Trt_vs_MS.NoTrt_B |
| TRIM36 | 0.68625 | 0.27396 | 0.316 | 0.31 | 1 | MS.Trt_vs_MS.NoTrt_B |
| HDAC5 | 0.68668 | 0.35675 | 0.22 | 0.24 | 1 | MS.Trt_vs_MS.NoTrt_B |
| ABCB8 | 0.69001 | -0.7411 | 0.505 | 0.5 | 1 | MS.Trt_vs_MS.NoTrt_B |
| DIRC1 | 0.69127 | 0.57553 | 0.139 | 0.13 | 1 | MS.Trt_vs_MS.NoTrt_B |
| ZNF528 | 0.6914 | 0.26315 | 0.471 | 0.51 | 1 | MS.Trt_vs_MS.NoTrt_B |
| RBBP6 | 0.69256 | 0.79175 | 0.211 | 0.23 | 1 | MS.Trt_vs_MS.NoTrt_B |
| SLC5A5 | 0.69312 | 1.38984 | 0.364 | 0.36 | 1 | MS.Trt_vs_MS.NoTrt_B |
| EI24 | 0.69343 | 0.51152 | 0.632 | 0.72 | 1 | MS.Trt_vs_MS.NoTrt_B |
| FADS6 | 0.69347 | 0.3028 | 0.492 | 0.52 | 1 | MS.Trt_vs_MS.NoTrt_B |
| UBE2S | 0.6947 | 0.41924 | 0.571 | 0.66 | 1 | MS.Trt_vs_MS.NoTrt_B |
| CALB1 | 0.69488 | 0.44084 | 0.1 | 0.11 | 1 | MS.Trt_vs_MS.NoTrt_B |
| RNF43 | 0.69514 | 1.03536 | 0.279 | 0.27 | 1 | MS.Trt_vs_MS.NoTrt_B |
| MT-CO3 | 0.69568 | 0.79005 | 0.59 | 0.72 | 1 | MS.Trt_vs_MS.NoTrt_B |
| OLR1 | 0.69725 | 0.2812 | 0.333 | 0.37 | 1 | MS.Trt_vs_MS.NoTrt_B |
| UBTF | 0.6973 | 0.59257 | 0.209 | 0.2 | 1 | MS.Trt_vs_MS.NoTrt_B |

| | | | | | | |
|---|---|---|---|---|---|---|
| MTHFS | 0.69752 | 1.2853 | 0.124 | 0.11 | 1 | MS.Trt_vs_MS.NoTrt_B |
| XPOT | 0.69804 | 0.74182 | 0.381 | 0.41 | 1 | MS.Trt_vs_MS.NoTrt_B |
| RFX7 | 0.70109 | 0.55014 | 0.148 | 0.16 | 1 | MS.Trt_vs_MS.NoTrt_B |
| ATP2A2 | 0.70129 | 0.28758 | 0.632 | 0.67 | 1 | MS.Trt_vs_MS.NoTrt_B |
| ATXN2L | 0.70245 | 0.44807 | 0.214 | 0.23 | 1 | MS.Trt_vs_MS.NoTrt_B |
| TMEM143 | 0.70671 | -0.7096 | 0.438 | 0.48 | 1 | MS.Trt_vs_MS.NoTrt_B |
| GIPC1 | 0.70682 | 0.38328 | 0.407 | 0.43 | 1 | MS.Trt_vs_MS.NoTrt_B |
| TUT1 | 0.70725 | -0.4378 | 0.122 | 0.11 | 1 | MS.Trt_vs_MS.NoTrt_B |
| DDI2 | 0.70764 | 0.4053 | 0.708 | 0.7 | 1 | MS.Trt_vs_MS.NoTrt_B |
| RIOK2 | 0.7084 | 0.84507 | 0.272 | 0.27 | 1 | MS.Trt_vs_MS.NoTrt_B |
| N4BP1 | 0.71178 | 0.77758 | 0.092 | 0.1 | 1 | MS.Trt_vs_MS.NoTrt_B |
| TMEM204 | 0.71281 | 0.83447 | 0.139 | 0.13 | 1 | MS.Trt_vs_MS.NoTrt_B |
| DNAJC5B | 0.71284 | 0.2528 | 0.438 | 0.45 | 1 | MS.Trt_vs_MS.NoTrt_B |
| MTMR2 | 0.71331 | -0.3428 | 0.133 | 0.15 | 1 | MS.Trt_vs_MS.NoTrt_B |
| ABCC2 | 0.7139 | 0.5666 | 0.444 | 0.51 | 1 | MS.Trt_vs_MS.NoTrt_B |
| ANKRD26 | 0.71422 | -0.757 | 0.105 | 0.11 | 1 | MS.Trt_vs_MS.NoTrt_B |
| DNAJC4 | 0.71584 | 0.29143 | 0.105 | 0.11 | 1 | MS.Trt_vs_MS.NoTrt_B |
| TIGD6 | 0.72012 | 1.17715 | 0.15 | 0.14 | 1 | MS.Trt_vs_MS.NoTrt_B |
| FAM65C | 0.72048 | 0.28988 | 0.301 | 0.34 | 1 | MS.Trt_vs_MS.NoTrt_B |
| STEAP2 | 0.72516 | 0.37267 | 0.601 | 0.69 | 1 | MS.Trt_vs_MS.NoTrt_B |
| MAB21L3 | 0.72615 | 0.39573 | 0.275 | 0.28 | 1 | MS.Trt_vs_MS.NoTrt_B |
| CCDC82 | 0.72653 | 0.949 | 0.129 | 0.14 | 1 | MS.Trt_vs_MS.NoTrt_B |
| HMGCR | 0.72661 | 0.58607 | 0.573 | 0.61 | 1 | MS.Trt_vs_MS.NoTrt_B |
| GRK7 | 0.72765 | 0.27962 | 0.261 | 0.27 | 1 | MS.Trt_vs_MS.NoTrt_B |
| ZNF850 | 0.72862 | 0.36389 | 0.834 | 0.88 | 1 | MS.Trt_vs_MS.NoTrt_B |
| TNFRSF13C | 0.72903 | 0.95607 | 0.538 | 0.58 | 1 | MS.Trt_vs_MS.NoTrt_B |
| ASB3 | 0.72969 | 0.39186 | 0.821 | 0.94 | 1 | MS.Trt_vs_MS.NoTrt_B |
| SLC25A45 | 0.7299 | 0.25577 | 0.939 | 0.99 | 1 | MS.Trt_vs_MS.NoTrt_B |
| PDE7B | 0.73045 | 0.3044 | 0.309 | 0.34 | 1 | MS.Trt_vs_MS.NoTrt_B |
| ROBO4 | 0.73113 | 0.29687 | 0.146 | 0.14 | 1 | MS.Trt_vs_MS.NoTrt_B |
| ANKRA2 | 0.73197 | 1.02461 | 0.098 | 0.11 | 1 | MS.Trt_vs_MS.NoTrt_B |
| ODF2L | 0.73407 | 0.37212 | 0.865 | 0.92 | 1 | MS.Trt_vs_MS.NoTrt_B |
| LSM2 | 0.73423 | 0.47833 | 0.118 | 0.13 | 1 | MS.Trt_vs_MS.NoTrt_B |
| SPDYE2B | 0.73432 | 0.96081 | 0.235 | 0.27 | 1 | MS.Trt_vs_MS.NoTrt_B |
| KCTD13 | 0.73604 | 0.45205 | 0.109 | 0.12 | 1 | MS.Trt_vs_MS.NoTrt_B |
| ST20 | 0.73651 | 0.431 | 0.159 | 0.17 | 1 | MS.Trt_vs_MS.NoTrt_B |
| CASP9 | 0.73794 | 0.68668 | 0.264 | 0.28 | 1 | MS.Trt_vs_MS.NoTrt_B |
| CTTNBP2NL | 0.74 | 0.25296 | 0.336 | 0.34 | 1 | MS.Trt_vs_MS.NoTrt_B |
| LLGL1 | 0.74082 | -1.0642 | 0.209 | 0.2 | 1 | MS.Trt_vs_MS.NoTrt_B |
| ZKSCAN1 | 0.74087 | 0.31884 | 0.863 | 0.92 | 1 | MS.Trt_vs_MS.NoTrt_B |

| | | | | | | |
|---|---|---|---|---|---|---|
| CACNB2 | 0.7409 | 0.62669 | 0.366 | 0.4 | 1 | MS.Trt_vs_MS.NoTrt_B |
| TDGF1 | 0.7414 | 0.33026 | 0.464 | 0.5 | 1 | MS.Trt_vs_MS.NoTrt_B |
| FAM178B | 0.74207 | 0.28515 | 0.586 | 0.6 | 1 | MS.Trt_vs_MS.NoTrt_B |
| MOSPD2 | 0.74217 | 0.51943 | 0.179 | 0.19 | 1 | MS.Trt_vs_MS.NoTrt_B |
| PUS3 | 0.74227 | -0.2717 | 0.301 | 0.32 | 1 | MS.Trt_vs_MS.NoTrt_B |
| MCM4 | 0.74235 | -0.2981 | 0.508 | 0.54 | 1 | MS.Trt_vs_MS.NoTrt_B |
| ZFYVE16 | 0.74241 | 1.05478 | 0.826 | 0.88 | 1 | MS.Trt_vs_MS.NoTrt_B |
| RBM44 | 0.74274 | 0.46114 | 0.174 | 0.2 | 1 | MS.Trt_vs_MS.NoTrt_B |
| ORC6 | 0.74542 | 0.47326 | 0.911 | 0.97 | 1 | MS.Trt_vs_MS.NoTrt_B |
| ABCA2 | 0.74853 | 1.17059 | 0.207 | 0.23 | 1 | MS.Trt_vs_MS.NoTrt_B |
| TMEM221 | 0.75071 | 0.54064 | 0.159 | 0.17 | 1 | MS.Trt_vs_MS.NoTrt_B |
| NOMO1 | 0.75076 | -0.6236 | 0.111 | 0.12 | 1 | MS.Trt_vs_MS.NoTrt_B |
| CPSF6 | 0.75366 | 0.69903 | 0.39 | 0.42 | 1 | MS.Trt_vs_MS.NoTrt_B |
| MED11 | 0.75766 | 0.77454 | 0.599 | 0.62 | 1 | MS.Trt_vs_MS.NoTrt_B |
| 11-Sep | 0.75955 | 0.71952 | 0.303 | 0.32 | 1 | MS.Trt_vs_MS.NoTrt_B |
| TRAPPC13 | 0.76091 | -0.372 | 0.148 | 0.16 | 1 | MS.Trt_vs_MS.NoTrt_B |
| PIK3R4 | 0.76176 | 0.29321 | 0.111 | 0.12 | 1 | MS.Trt_vs_MS.NoTrt_B |
| TBCCD1 | 0.76234 | 0.32609 | 0.279 | 0.28 | 1 | MS.Trt_vs_MS.NoTrt_B |
| GIMAP5 | 0.76246 | 0.42361 | 0.407 | 0.41 | 1 | MS.Trt_vs_MS.NoTrt_B |
| TMEM120B | 0.76332 | 0.2514 | 0.889 | 0.97 | 1 | MS.Trt_vs_MS.NoTrt_B |
| MRPL37 | 0.76514 | 0.51962 | 0.159 | 0.17 | 1 | MS.Trt_vs_MS.NoTrt_B |
| ZDHHC2 | 0.76516 | 0.7913 | 0.144 | 0.14 | 1 | MS.Trt_vs_MS.NoTrt_B |
| ZNF14 | 0.76692 | 2.74509 | 0.307 | 0.34 | 1 | MS.Trt_vs_MS.NoTrt_B |
| TANC2 | 0.76765 | 1.0413 | 0.183 | 0.2 | 1 | MS.Trt_vs_MS.NoTrt_B |
| FRS3 | 0.76814 | 1.76625 | 0.266 | 0.27 | 1 | MS.Trt_vs_MS.NoTrt_B |
| ZKSCAN7 | 0.76851 | -0.2521 | 0.516 | 0.53 | 1 | MS.Trt_vs_MS.NoTrt_B |
| PPFIBP2 | 0.77079 | 0.47563 | 0.187 | 0.18 | 1 | MS.Trt_vs_MS.NoTrt_B |
| IRAK1 | 0.77133 | 0.27661 | 0.166 | 0.17 | 1 | MS.Trt_vs_MS.NoTrt_B |
| HVCN1 | 0.77223 | 0.62822 | 0.386 | 0.42 | 1 | MS.Trt_vs_MS.NoTrt_B |
| RASSF3 | 0.77248 | 0.85284 | 0.645 | 0.81 | 1 | MS.Trt_vs_MS.NoTrt_B |
| OSBPL6 | 0.77406 | 0.25227 | 0.486 | 0.55 | 1 | MS.Trt_vs_MS.NoTrt_B |
| DIEXF | 0.77475 | -0.3161 | 0.431 | 0.41 | 1 | MS.Trt_vs_MS.NoTrt_B |
| RRP8 | 0.77498 | 0.50582 | 0.458 | 0.47 | 1 | MS.Trt_vs_MS.NoTrt_B |
| ZNF578 | 0.77562 | 0.36415 | 0.88 | 0.94 | 1 | MS.Trt_vs_MS.NoTrt_B |
| WBP2NL | 0.77581 | 0.75655 | 0.322 | 0.34 | 1 | MS.Trt_vs_MS.NoTrt_B |
| KLHL7 | 0.77993 | 0.73149 | 0.599 | 0.67 | 1 | MS.Trt_vs_MS.NoTrt_B |
| ATOX1 | 0.78006 | 0.43806 | 0.146 | 0.14 | 1 | MS.Trt_vs_MS.NoTrt_B |
| WSB2 | 0.78033 | 0.64498 | 0.688 | 0.81 | 1 | MS.Trt_vs_MS.NoTrt_B |
| ATN1 | 0.78053 | 0.38108 | 0.115 | 0.12 | 1 | MS.Trt_vs_MS.NoTrt_B |
| MFSD12 | 0.78221 | 1.03298 | 0.22 | 0.23 | 1 | MS.Trt_vs_MS.NoTrt_B |

| | | | | | | |
|---|---|---|---|---|---|---|
| DAXX | 0.78373 | 0.51393 | 0.19 | 0.2 | 1 | MS.Trt_vs_MS.NoTrt_B |
| ZNF430 | 0.78395 | 0.61117 | 0.832 | 0.92 | 1 | MS.Trt_vs_MS.NoTrt_B |
| GNE | 0.78458 | 0.34138 | 0.163 | 0.17 | 1 | MS.Trt_vs_MS.NoTrt_B |
| LRRC2 | 0.78466 | 1.05151 | 0.632 | 0.68 | 1 | MS.Trt_vs_MS.NoTrt_B |
| FAXC | 0.78613 | 0.3126 | 0.246 | 0.28 | 1 | MS.Trt_vs_MS.NoTrt_B |
| DSG2 | 0.78688 | 0.26222 | 0.362 | 0.38 | 1 | MS.Trt_vs_MS.NoTrt_B |
| CHID1 | 0.79009 | 0.4482 | 0.231 | 0.24 | 1 | MS.Trt_vs_MS.NoTrt_B |
| TACO1 | 0.79016 | 0.49091 | 0.309 | 0.32 | 1 | MS.Trt_vs_MS.NoTrt_B |
| GBA2 | 0.79092 | 0.453 | 0.564 | 0.59 | 1 | MS.Trt_vs_MS.NoTrt_B |
| DUSP11 | 0.79171 | 0.43191 | 0.523 | 0.48 | 1 | MS.Trt_vs_MS.NoTrt_B |
| SESN3 | 0.79195 | 1.11228 | 0.233 | 0.25 | 1 | MS.Trt_vs_MS.NoTrt_B |
| ERV3-1 | 0.79422 | -0.4345 | 0.438 | 0.45 | 1 | MS.Trt_vs_MS.NoTrt_B |
| NDFIP2 | 0.7956 | 0.60003 | 0.111 | 0.12 | 1 | MS.Trt_vs_MS.NoTrt_B |
| ZNF93 | 0.79578 | 0.93801 | 0.501 | 0.55 | 1 | MS.Trt_vs_MS.NoTrt_B |
| SGSM2 | 0.79717 | 1.21824 | 0.187 | 0.2 | 1 | MS.Trt_vs_MS.NoTrt_B |
| NUDT7 | 0.79987 | 0.44816 | 0.181 | 0.17 | 1 | MS.Trt_vs_MS.NoTrt_B |
| ZNF776 | 0.80066 | 0.43068 | 0.603 | 0.68 | 1 | MS.Trt_vs_MS.NoTrt_B |
| RUNX3 | 0.80154 | 0.46727 | 0.12 | 0.13 | 1 | MS.Trt_vs_MS.NoTrt_B |
| EIF4G3 | 0.80182 | 1.00525 | 0.179 | 0.19 | 1 | MS.Trt_vs_MS.NoTrt_B |
| SULF2 | 0.80192 | 0.31648 | 0.153 | 0.15 | 1 | MS.Trt_vs_MS.NoTrt_B |
| LRSAM1 | 0.80274 | 0.409 | 0.115 | 0.12 | 1 | MS.Trt_vs_MS.NoTrt_B |
| RP11-544M1 | 0.80319 | -0.4717 | 0.168 | 0.18 | 1 | MS.Trt_vs_MS.NoTrt_B |
| PGGT1B | 0.80323 | 1.11204 | 0.122 | 0.13 | 1 | MS.Trt_vs_MS.NoTrt_B |
| SAMD9L | 0.80524 | 0.90239 | 0.153 | 0.16 | 1 | MS.Trt_vs_MS.NoTrt_B |
| TMIGD2 | 0.80531 | 0.92061 | 0.174 | 0.19 | 1 | MS.Trt_vs_MS.NoTrt_B |
| PLCB2 | 0.80705 | 0.47363 | 0.218 | 0.22 | 1 | MS.Trt_vs_MS.NoTrt_B |
| BOP1 | 0.80929 | -0.5673 | 0.185 | 0.17 | 1 | MS.Trt_vs_MS.NoTrt_B |
| WDR60 | 0.80947 | 0.72257 | 0.096 | 0.1 | 1 | MS.Trt_vs_MS.NoTrt_B |
| THOC1 | 0.81457 | -0.3462 | 0.187 | 0.2 | 1 | MS.Trt_vs_MS.NoTrt_B |
| ID3 | 0.8146 | 0.87616 | 0.166 | 0.16 | 1 | MS.Trt_vs_MS.NoTrt_B |
| PTPN9 | 0.81575 | -0.6854 | 0.231 | 0.24 | 1 | MS.Trt_vs_MS.NoTrt_B |
| DHRS3 | 0.81602 | 0.65833 | 0.181 | 0.18 | 1 | MS.Trt_vs_MS.NoTrt_B |
| MPV17L2 | 0.81649 | 0.74241 | 0.163 | 0.18 | 1 | MS.Trt_vs_MS.NoTrt_B |
| CPEB1 | 0.8188 | -0.5637 | 0.118 | 0.12 | 1 | MS.Trt_vs_MS.NoTrt_B |
| BAZ1B | 0.8201 | 0.40749 | 0.161 | 0.17 | 1 | MS.Trt_vs_MS.NoTrt_B |
| SNF8 | 0.82076 | 0.3529 | 0.734 | 0.82 | 1 | MS.Trt_vs_MS.NoTrt_B |
| PACRGL | 0.822 | -0.4669 | 0.251 | 0.27 | 1 | MS.Trt_vs_MS.NoTrt_B |
| BTBD9 | 0.8227 | 0.39179 | 0.444 | 0.46 | 1 | MS.Trt_vs_MS.NoTrt_B |
| ZNF606 | 0.82306 | -0.2697 | 0.688 | 0.68 | 1 | MS.Trt_vs_MS.NoTrt_B |
| GTF3C6 | 0.82352 | 0.5623 | 0.12 | 0.13 | 1 | MS.Trt_vs_MS.NoTrt_B |

| | | | | | | |
|---|---|---|---|---|---|---|
| EVA1C | 0.82565 | 0.29982 | 0.102 | 0.1 | 1 | MS.Trt_vs_MS.NoTrt_B |
| SRPK3 | 0.82722 | 0.50659 | 0.214 | 0.21 | 1 | MS.Trt_vs_MS.NoTrt_B |
| ZNF614 | 0.82812 | 0.61337 | 0.19 | 0.18 | 1 | MS.Trt_vs_MS.NoTrt_B |
| AHCTF1 | 0.83009 | 1.0825 | 0.153 | 0.15 | 1 | MS.Trt_vs_MS.NoTrt_B |
| PPID | 0.83039 | 0.32355 | 0.388 | 0.37 | 1 | MS.Trt_vs_MS.NoTrt_B |
| PPP2R5A | 0.8311 | 0.9477 | 0.194 | 0.2 | 1 | MS.Trt_vs_MS.NoTrt_B |
| DUSP1 | 0.83168 | 0.59833 | 0.139 | 0.15 | 1 | MS.Trt_vs_MS.NoTrt_B |
| TSPYL2 | 0.83413 | 0.45799 | 0.168 | 0.16 | 1 | MS.Trt_vs_MS.NoTrt_B |
| BCL9L | 0.83428 | 0.3873 | 0.107 | 0.11 | 1 | MS.Trt_vs_MS.NoTrt_B |
| LIN7B | 0.83693 | 0.44047 | 0.233 | 0.25 | 1 | MS.Trt_vs_MS.NoTrt_B |
| LARP1 | 0.83711 | 0.80454 | 0.153 | 0.15 | 1 | MS.Trt_vs_MS.NoTrt_B |
| PTPRH | 0.83946 | -0.4312 | 0.115 | 0.12 | 1 | MS.Trt_vs_MS.NoTrt_B |
| ADH1B | 0.83963 | 0.30836 | 0.227 | 0.25 | 1 | MS.Trt_vs_MS.NoTrt_B |
| RAP1B | 0.83991 | 0.91245 | 0.961 | 0.99 | 1 | MS.Trt_vs_MS.NoTrt_B |
| CRKL | 0.84025 | 0.68405 | 0.503 | 0.57 | 1 | MS.Trt_vs_MS.NoTrt_B |
| CYB561 | 0.84046 | 0.62132 | 0.235 | 0.26 | 1 | MS.Trt_vs_MS.NoTrt_B |
| CTNNB1 | 0.84206 | 0.63261 | 0.503 | 0.56 | 1 | MS.Trt_vs_MS.NoTrt_B |
| FSTL4 | 0.84244 | 0.37204 | 0.131 | 0.13 | 1 | MS.Trt_vs_MS.NoTrt_B |
| USP39 | 0.84254 | -0.3031 | 0.416 | 0.39 | 1 | MS.Trt_vs_MS.NoTrt_B |
| CXorf36 | 0.84294 | 0.40778 | 0.309 | 0.34 | 1 | MS.Trt_vs_MS.NoTrt_B |
| ZNF638 | 0.84394 | 1.01878 | 0.303 | 0.31 | 1 | MS.Trt_vs_MS.NoTrt_B |
| CNN3 | 0.84423 | 0.81707 | 0.163 | 0.17 | 1 | MS.Trt_vs_MS.NoTrt_B |
| RANBP9 | 0.84519 | 0.9214 | 0.168 | 0.18 | 1 | MS.Trt_vs_MS.NoTrt_B |
| TNNI1 | 0.84558 | 0.30265 | 0.366 | 0.4 | 1 | MS.Trt_vs_MS.NoTrt_B |
| B3GALNT2 | 0.84815 | 0.48134 | 0.525 | 0.58 | 1 | MS.Trt_vs_MS.NoTrt_B |
| RPE | 0.84959 | 0.68458 | 0.275 | 0.28 | 1 | MS.Trt_vs_MS.NoTrt_B |
| ADARB1 | 0.85155 | 1.16536 | 0.176 | 0.17 | 1 | MS.Trt_vs_MS.NoTrt_B |
| ZNF669 | 0.85256 | -0.7844 | 0.466 | 0.5 | 1 | MS.Trt_vs_MS.NoTrt_B |
| MED16 | 0.85341 | 0.42034 | 0.185 | 0.2 | 1 | MS.Trt_vs_MS.NoTrt_B |
| POGLUT1 | 0.85362 | 0.76194 | 0.248 | 0.25 | 1 | MS.Trt_vs_MS.NoTrt_B |
| TMEM106A | 0.8543 | 0.73018 | 0.599 | 0.66 | 1 | MS.Trt_vs_MS.NoTrt_B |
| FAM21A | 0.85489 | 0.43654 | 0.19 | 0.2 | 1 | MS.Trt_vs_MS.NoTrt_B |
| SPIN1 | 0.85835 | 0.54523 | 0.137 | 0.13 | 1 | MS.Trt_vs_MS.NoTrt_B |
| FBXL2 | 0.86067 | 0.47171 | 0.538 | 0.58 | 1 | MS.Trt_vs_MS.NoTrt_B |
| CKAP2L | 0.86117 | 0.35914 | 0.373 | 0.41 | 1 | MS.Trt_vs_MS.NoTrt_B |
| TAF2 | 0.86229 | 1.23459 | 0.379 | 0.42 | 1 | MS.Trt_vs_MS.NoTrt_B |
| BMPR1B | 0.86589 | 1.19035 | 0.109 | 0.11 | 1 | MS.Trt_vs_MS.NoTrt_B |
| LRRD1 | 0.86637 | 0.31184 | 0.449 | 0.48 | 1 | MS.Trt_vs_MS.NoTrt_B |
| AGL | 0.86702 | 0.70426 | 0.111 | 0.11 | 1 | MS.Trt_vs_MS.NoTrt_B |
| HDAC7 | 0.86771 | 0.62827 | 0.2 | 0.21 | 1 | MS.Trt_vs_MS.NoTrt_B |

| | | | | | | |
|---|---|---|---|---|---|---|
| MT-CYB | 0.86775 | 0.52801 | 0.68 | 0.76 | 1 | MS.Trt_vs_MS.NoTrt_B |
| MFN1 | 0.86847 | 0.59802 | 0.383 | 0.38 | 1 | MS.Trt_vs_MS.NoTrt_B |
| CDAN1 | 0.86918 | 0.61836 | 0.375 | 0.41 | 1 | MS.Trt_vs_MS.NoTrt_B |
| VNN2 | 0.86941 | 0.53919 | 0.15 | 0.15 | 1 | MS.Trt_vs_MS.NoTrt_B |
| ADNP | 0.87217 | 1.36039 | 0.42 | 0.47 | 1 | MS.Trt_vs_MS.NoTrt_B |
| ZNF677 | 0.87233 | -0.2859 | 0.741 | 0.76 | 1 | MS.Trt_vs_MS.NoTrt_B |
| SBF1 | 0.87249 | 0.54411 | 0.148 | 0.15 | 1 | MS.Trt_vs_MS.NoTrt_B |
| PITPNM1 | 0.87655 | 0.55895 | 0.222 | 0.23 | 1 | MS.Trt_vs_MS.NoTrt_B |
| FAM13A | 0.87886 | 0.39811 | 0.153 | 0.15 | 1 | MS.Trt_vs_MS.NoTrt_B |
| FAM76A | 0.87916 | 0.81886 | 0.728 | 0.73 | 1 | MS.Trt_vs_MS.NoTrt_B |
| MORN1 | 0.87927 | 0.25778 | 0.464 | 0.49 | 1 | MS.Trt_vs_MS.NoTrt_B |
| MMACHC | 0.88046 | 0.45714 | 0.789 | 0.82 | 1 | MS.Trt_vs_MS.NoTrt_B |
| STT3A | 0.8814 | 0.50927 | 0.523 | 0.5 | 1 | MS.Trt_vs_MS.NoTrt_B |
| INTS2 | 0.88154 | 0.58875 | 0.216 | 0.22 | 1 | MS.Trt_vs_MS.NoTrt_B |
| MPP4 | 0.88192 | 0.27462 | 0.447 | 0.49 | 1 | MS.Trt_vs_MS.NoTrt_B |
| MCU | 0.88204 | 0.34193 | 0.196 | 0.2 | 1 | MS.Trt_vs_MS.NoTrt_B |
| CD3G | 0.88388 | 0.59626 | 0.917 | 1 | 1 | MS.Trt_vs_MS.NoTrt_B |
| PPIAL4G | 0.88544 | 0.55883 | 0.558 | 0.58 | 1 | MS.Trt_vs_MS.NoTrt_B |
| RPRD1A | 0.88553 | 0.31238 | 0.144 | 0.15 | 1 | MS.Trt_vs_MS.NoTrt_B |
| RAPH1 | 0.88559 | 0.34378 | 0.124 | 0.14 | 1 | MS.Trt_vs_MS.NoTrt_B |
| FBXW7 | 0.88685 | 0.86925 | 0.135 | 0.14 | 1 | MS.Trt_vs_MS.NoTrt_B |
| ZNF175 | 0.889 | -1.3709 | 0.176 | 0.17 | 1 | MS.Trt_vs_MS.NoTrt_B |
| C3orf18 | 0.88912 | 0.25965 | 0.185 | 0.18 | 1 | MS.Trt_vs_MS.NoTrt_B |
| NUDT16L1 | 0.88992 | 0.53295 | 0.098 | 0.1 | 1 | MS.Trt_vs_MS.NoTrt_B |
| DCUN1D3 | 0.89027 | -0.3827 | 0.129 | 0.12 | 1 | MS.Trt_vs_MS.NoTrt_B |
| UBA6 | 0.89094 | 0.56974 | 0.468 | 0.49 | 1 | MS.Trt_vs_MS.NoTrt_B |
| SGO2 | 0.89137 | 0.30994 | 0.715 | 0.8 | 1 | MS.Trt_vs_MS.NoTrt_B |
| TRPM4 | 0.89307 | 0.33579 | 0.721 | 0.76 | 1 | MS.Trt_vs_MS.NoTrt_B |
| ANKRD18A | 0.89366 | 0.36791 | 0.218 | 0.22 | 1 | MS.Trt_vs_MS.NoTrt_B |
| RNF31 | 0.8939 | 0.29966 | 0.253 | 0.25 | 1 | MS.Trt_vs_MS.NoTrt_B |
| WDR48 | 0.8969 | 0.75316 | 0.129 | 0.14 | 1 | MS.Trt_vs_MS.NoTrt_B |
| SPDYE3 | 0.89733 | 0.58628 | 0.542 | 0.55 | 1 | MS.Trt_vs_MS.NoTrt_B |
| FAM46C | 0.89806 | 0.62707 | 0.129 | 0.12 | 1 | MS.Trt_vs_MS.NoTrt_B |
| TIMM23 | 0.89855 | 0.82651 | 0.216 | 0.22 | 1 | MS.Trt_vs_MS.NoTrt_B |
| ZBTB1 | 0.89878 | 0.74841 | 0.155 | 0.16 | 1 | MS.Trt_vs_MS.NoTrt_B |
| RAB42 | 0.90165 | 0.35753 | 0.51 | 0.57 | 1 | MS.Trt_vs_MS.NoTrt_B |
| PVR | 0.90202 | 0.76328 | 0.392 | 0.4 | 1 | MS.Trt_vs_MS.NoTrt_B |
| GNB1 | 0.90234 | 0.42833 | 0.259 | 0.25 | 1 | MS.Trt_vs_MS.NoTrt_B |
| GFRAL | 0.90246 | 0.33664 | 0.331 | 0.34 | 1 | MS.Trt_vs_MS.NoTrt_B |
| C1orf228 | 0.90335 | 0.39606 | 0.207 | 0.22 | 1 | MS.Trt_vs_MS.NoTrt_B |

| | | | | | | |
|---|---|---|---|---|---|---|
| KLHDC1 | 0.90382 | 0.45599 | 0.272 | 0.3 | 1 | MS.Trt_vs_MS.NoTrt_B |
| AC025335.1 | 0.90423 | 0.39597 | 0.486 | 0.51 | 1 | MS.Trt_vs_MS.NoTrt_B |
| ZSCAN25 | 0.90765 | 0.3961 | 0.362 | 0.36 | 1 | MS.Trt_vs_MS.NoTrt_B |
| CSF1R | 0.90927 | 0.3109 | 0.109 | 0.11 | 1 | MS.Trt_vs_MS.NoTrt_B |
| ISPD | 0.91044 | 0.32308 | 0.704 | 0.78 | 1 | MS.Trt_vs_MS.NoTrt_B |
| KIF21B | 0.91222 | 1.06707 | 0.331 | 0.37 | 1 | MS.Trt_vs_MS.NoTrt_B |
| ARHGAP44 | 0.91248 | 0.72854 | 0.237 | 0.24 | 1 | MS.Trt_vs_MS.NoTrt_B |
| MCM3 | 0.9135 | 0.33974 | 0.133 | 0.13 | 1 | MS.Trt_vs_MS.NoTrt_B |
| WTAP | 0.91408 | 0.73242 | 0.664 | 0.75 | 1 | MS.Trt_vs_MS.NoTrt_B |
| ANKRD6 | 0.9143 | -0.5114 | 0.111 | 0.11 | 1 | MS.Trt_vs_MS.NoTrt_B |
| PLCXD1 | 0.91583 | 0.4586 | 0.874 | 0.91 | 1 | MS.Trt_vs_MS.NoTrt_B |
| ZNF17 | 0.91703 | 0.89923 | 0.235 | 0.24 | 1 | MS.Trt_vs_MS.NoTrt_B |
| ATRNL1 | 0.91778 | 0.28232 | 0.294 | 0.33 | 1 | MS.Trt_vs_MS.NoTrt_B |
| TMEM107 | 0.91975 | 1.14223 | 0.416 | 0.45 | 1 | MS.Trt_vs_MS.NoTrt_B |
| C6orf203 | 0.92038 | 0.78409 | 0.519 | 0.54 | 1 | MS.Trt_vs_MS.NoTrt_B |
| SLF1 | 0.92146 | -0.3803 | 0.118 | 0.12 | 1 | MS.Trt_vs_MS.NoTrt_B |
| DCAF17 | 0.92186 | 0.28911 | 0.307 | 0.32 | 1 | MS.Trt_vs_MS.NoTrt_B |
| SLC48A1 | 0.92558 | 0.70717 | 0.54 | 0.6 | 1 | MS.Trt_vs_MS.NoTrt_B |
| TCF4 | 0.92589 | 0.46594 | 0.641 | 0.67 | 1 | MS.Trt_vs_MS.NoTrt_B |
| BRAT1 | 0.92678 | 0.41064 | 0.266 | 0.27 | 1 | MS.Trt_vs_MS.NoTrt_B |
| NFATC1 | 0.92865 | 0.44499 | 0.146 | 0.15 | 1 | MS.Trt_vs_MS.NoTrt_B |
| GSTT1 | 0.92985 | 1.4096 | 0.577 | 0.64 | 1 | MS.Trt_vs_MS.NoTrt_B |
| SPINT2 | 0.92987 | -0.341 | 0.425 | 0.44 | 1 | MS.Trt_vs_MS.NoTrt_B |
| TERF1 | 0.93091 | 0.40088 | 0.61 | 0.66 | 1 | MS.Trt_vs_MS.NoTrt_B |
| CD36 | 0.93095 | 0.32579 | 0.486 | 0.55 | 1 | MS.Trt_vs_MS.NoTrt_B |
| ANXA1 | 0.93147 | 0.73757 | 0.131 | 0.13 | 1 | MS.Trt_vs_MS.NoTrt_B |
| MAP3K2 | 0.93298 | 0.35175 | 0.166 | 0.16 | 1 | MS.Trt_vs_MS.NoTrt_B |
| ZNF212 | 0.93353 | -0.6126 | 0.342 | 0.33 | 1 | MS.Trt_vs_MS.NoTrt_B |
| CD40LG | 0.93434 | 0.66287 | 0.131 | 0.14 | 1 | MS.Trt_vs_MS.NoTrt_B |
| PTCHD4 | 0.93554 | 0.25552 | 0.124 | 0.12 | 1 | MS.Trt_vs_MS.NoTrt_B |
| SLC1A5 | 0.93666 | -0.5748 | 0.115 | 0.11 | 1 | MS.Trt_vs_MS.NoTrt_B |
| RPS7 | 0.9382 | -0.4547 | 0.414 | 0.41 | 1 | MS.Trt_vs_MS.NoTrt_B |
| GALC | 0.93825 | 0.31271 | 0.804 | 0.89 | 1 | MS.Trt_vs_MS.NoTrt_B |
| CALM1 | 0.93865 | 0.4046 | 0.734 | 0.88 | 1 | MS.Trt_vs_MS.NoTrt_B |
| AAR2 | 0.93932 | 0.66296 | 0.2 | 0.21 | 1 | MS.Trt_vs_MS.NoTrt_B |
| BRD9 | 0.94102 | 1.46427 | 0.113 | 0.11 | 1 | MS.Trt_vs_MS.NoTrt_B |
| FAM229B | 0.94519 | 0.26169 | 0.264 | 0.28 | 1 | MS.Trt_vs_MS.NoTrt_B |
| POLE2 | 0.94559 | 0.39341 | 0.244 | 0.25 | 1 | MS.Trt_vs_MS.NoTrt_B |
| PPP1R16B | 0.94596 | 0.5428 | 0.19 | 0.19 | 1 | MS.Trt_vs_MS.NoTrt_B |
| POLR2M | 0.94602 | 0.63758 | 0.12 | 0.12 | 1 | MS.Trt_vs_MS.NoTrt_B |

| | | | | | | |
|---|---|---|---|---|---|---|
| ZNF789 | 0.9465 | 0.34846 | 0.649 | 0.71 | 1 | MS.Trt_vs_MS.NoTrt_B |
| C2orf68 | 0.94721 | 0.36547 | 0.767 | 0.77 | 1 | MS.Trt_vs_MS.NoTrt_B |
| PPIE | 0.94775 | 0.86755 | 0.673 | 0.77 | 1 | MS.Trt_vs_MS.NoTrt_B |
| GABRP | 0.94854 | 0.39737 | 0.216 | 0.22 | 1 | MS.Trt_vs_MS.NoTrt_B |
| TMTC3 | 0.94949 | 0.42762 | 0.159 | 0.16 | 1 | MS.Trt_vs_MS.NoTrt_B |
| PIAS2 | 0.95167 | 0.6867 | 0.678 | 0.7 | 1 | MS.Trt_vs_MS.NoTrt_B |
| LANCL1 | 0.95422 | 0.96021 | 0.133 | 0.14 | 1 | MS.Trt_vs_MS.NoTrt_B |
| NSUN7 | 0.95508 | 1.47136 | 0.203 | 0.21 | 1 | MS.Trt_vs_MS.NoTrt_B |
| DPP3 | 0.9557 | 0.30649 | 0.148 | 0.15 | 1 | MS.Trt_vs_MS.NoTrt_B |
| KLHL23 | 0.95733 | 0.4942 | 0.155 | 0.16 | 1 | MS.Trt_vs_MS.NoTrt_B |
| MBTD1 | 0.95852 | 0.83425 | 0.342 | 0.35 | 1 | MS.Trt_vs_MS.NoTrt_B |
| NDUFAF7 | 0.95935 | -0.6368 | 0.331 | 0.33 | 1 | MS.Trt_vs_MS.NoTrt_B |
| ABCC9 | 0.96002 | 0.34578 | 0.858 | 0.95 | 1 | MS.Trt_vs_MS.NoTrt_B |
| SLC13A4 | 0.9616 | -0.2952 | 0.27 | 0.28 | 1 | MS.Trt_vs_MS.NoTrt_B |
| TOR1A | 0.9617 | 0.75603 | 0.582 | 0.62 | 1 | MS.Trt_vs_MS.NoTrt_B |
| ZEB2 | 0.9632 | 1.12097 | 0.508 | 0.53 | 1 | MS.Trt_vs_MS.NoTrt_B |
| STT3B | 0.96359 | 0.37159 | 0.49 | 0.51 | 1 | MS.Trt_vs_MS.NoTrt_B |
| PLOD1 | 0.96402 | 0.56478 | 0.532 | 0.52 | 1 | MS.Trt_vs_MS.NoTrt_B |
| MKRN2OS | 0.96564 | 0.27087 | 0.15 | 0.16 | 1 | MS.Trt_vs_MS.NoTrt_B |
| TRAPPC2B | 0.96946 | 0.59439 | 0.259 | 0.26 | 1 | MS.Trt_vs_MS.NoTrt_B |
| MAP3K11 | 0.97281 | 0.67585 | 0.599 | 0.64 | 1 | MS.Trt_vs_MS.NoTrt_B |
| IRS2 | 0.97356 | 0.48797 | 0.292 | 0.3 | 1 | MS.Trt_vs_MS.NoTrt_B |
| ACADM | 0.97643 | 0.50771 | 0.135 | 0.14 | 1 | MS.Trt_vs_MS.NoTrt_B |
| FEZ1 | 0.97789 | 0.26077 | 0.514 | 0.57 | 1 | MS.Trt_vs_MS.NoTrt_B |
| THNSL1 | 0.97878 | 1.16527 | 0.098 | 0.1 | 1 | MS.Trt_vs_MS.NoTrt_B |
| ZSCAN5A | 0.9791 | 0.55992 | 0.678 | 0.7 | 1 | MS.Trt_vs_MS.NoTrt_B |
| SDCCAG8 | 0.97935 | 0.33427 | 0.678 | 0.74 | 1 | MS.Trt_vs_MS.NoTrt_B |
| FAM153C | 0.98038 | 0.38569 | 0.364 | 0.39 | 1 | MS.Trt_vs_MS.NoTrt_B |
| MORF4L2 | 0.98325 | 0.58893 | 0.159 | 0.16 | 1 | MS.Trt_vs_MS.NoTrt_B |
| PSD3 | 0.98382 | 0.46323 | 0.259 | 0.27 | 1 | MS.Trt_vs_MS.NoTrt_B |
| VDR | 0.98447 | -0.5908 | 0.261 | 0.25 | 1 | MS.Trt_vs_MS.NoTrt_B |
| IL24 | 0.98711 | 0.70315 | 0.135 | 0.14 | 1 | MS.Trt_vs_MS.NoTrt_B |
| MFAP5 | 0.98758 | 0.32064 | 0.281 | 0.3 | 1 | MS.Trt_vs_MS.NoTrt_B |
| CCL5 | 0.98876 | 0.75721 | 0.804 | 0.88 | 1 | MS.Trt_vs_MS.NoTrt_B |
| SRRM2 | 0.98892 | 0.77239 | 0.619 | 0.72 | 1 | MS.Trt_vs_MS.NoTrt_B |
| SFMBT2 | 0.98921 | 0.251 | 0.22 | 0.23 | 1 | MS.Trt_vs_MS.NoTrt_B |
| C19orf45 | 0.98938 | 0.36038 | 0.381 | 0.42 | 1 | MS.Trt_vs_MS.NoTrt_B |
| CHMP1B | 0.99016 | 0.34829 | 0.688 | 0.78 | 1 | MS.Trt_vs_MS.NoTrt_B |
| MFSD6 | 0.99101 | 0.5271 | 0.109 | 0.11 | 1 | MS.Trt_vs_MS.NoTrt_B |
| DISC1 | 0.99253 | 0.74828 | 0.42 | 0.44 | 1 | MS.Trt_vs_MS.NoTrt_B |

| | | | | | | |
|---|---|---|---|---|---|---|
| JMY | 0.99269 | 0.54546 | 0.211 | 0.22 | 1 | MS.Trt_vs_MS.NoTrt_B |
| SLC46A3 | 0.99464 | 1.095 | 0.1 | 0.1 | 1 | MS.Trt_vs_MS.NoTrt_B |
| ILVBL | 0.995 | -0.7125 | 0.209 | 0.21 | 1 | MS.Trt_vs_MS.NoTrt_B |
| ZNF626 | 0.99555 | 0.56033 | 0.593 | 0.63 | 1 | MS.Trt_vs_MS.NoTrt_B |
| CISH | 0.99752 | 0.33186 | 0.102 | 0.1 | 1 | MS.Trt_vs_MS.NoTrt_B |
| ZNF384 | 0.9987 | 0.3693 | 0.124 | 0.12 | 1 | MS.Trt_vs_MS.NoTrt_B |
| RTN4 | 0.99896 | 0.7052 | 0.211 | 0.22 | 1 | MS.Trt_vs_MS.NoTrt_B |
| ANKRD49 | 0.99973 | 1.47207 | 0.192 | 0.2 | 1 | MS.Trt_vs_MS.NoTrt_B |
| MAPK14 | 1 | 1.32611 | 0.107 | 0.11 | 1 | MS.Trt_vs_MS.NoTrt_B |

**Supplementary Table 4. Differentially expressed genes between healthy and MS donors.** Results of a Wilcoxon rank sum test, performed by Seurat's FindAllMarkers function with default parameter values as described in the text. The comparison column is formatted as "X_vs_Y_T" with X, Y being the two donor groups being compared, and T representing the tissue for whose cells the comparison is performed, B = peripheral blood, F = fat.

**Supplementary Table 5. Donor demographics for MS and healthy donors used for suppression assays.**

| | Age | Gender | Ethnicity | Diagnosis & Treatment |
|---|---|---|---|---|
| HC | 21 | M | White Non-Hispanic | N/A |
| | 31 | F | White Non-Hispanic | N/A |
| | 35 | M | White Non-Hispanic | N/A |
| | 22 | M | White Non-Hispanic | N/A |
| | 31 | F | White Non-Hispanic | N/A |
| | 34 | F | Asian | N/A |
| | 28 | F | White Non-Hispanic | N/A |
| | 30 | F | White Non-Hispanic | N/A |
| RRMS | 55 | F | White Non-Hispanic | New Onset; untreated RRMS |
| | 33 | F | White Non-Hispanic | New Onset; untreated RRMS |
| | 32 | F | Mixed Race, Hispanic | New Onset; untreated RRMS |
| | 44 | M | White Non-Hispanic | New Onset; untreated RRMS |
| | 50 | M | White Non-Hispanic | New Onset; untreated RRMS |
| | 34 | F | White, Hispanic | New Onset; untreated RRMS |
| | 32 | M | Black Non-Hispanic | Untreated since 12-2016 (copaxone) |
| | 35 | F | White Non-Hispanic | Untreated since 10/2017 (1 dose of rituxan) |

**Supplementary Table 5. Donor demographics for MS and healthy donors used for suppression assays.** Age, gender, ethnicity, and disease diagnosis and treatment information for peripheral blood $T_{regs}$ from healthy subjects or MS patients used for suppression assays.
